## Supplemental Table 5 for "Downregulation of Neurodevelopmental Gene Expression in iPSC-Derived Cerebral Organoids Upon Infection by Human Cytomegalovirus"

G-profiler Ontology Results based on list of most significantly differentially expressed genes as determined by padj-value for GFP (+) vs Mock

| source | term_name | term_id | adjusted_p_v | negative_log | term_size | query_size |
|---|---|---|---|---|---|---|
| **GO:MF** | RNA binding | GO:0003723 | 7.60E-86 | 85.1190743 | 1951 | 2622 |
| **GO:MF** | protein bindin | GO:0005515 | 7.63E-75 | 74.1171932 | 14767 | 2622 |
| **GO:MF** | enzyme bindir | GO:0019899 | 4.17E-65 | 64.3795408 | 2048 | 2622 |
| **GO:MF** | protein-contai | GO:0044877 | 1.76E-46 | 45.7555001 | 1304 | 2622 |
| **GO:MF** | cadherin bindi | GO:0045296 | 7.78E-38 | 37.109055 | 333 | 2622 |
| **GO:MF** | cell adhesion | GO:0050839 | 7.50E-37 | 36.124933 | 544 | 2622 |
| **GO:MF** | binding | GO:0005488 | 1.45E-34 | 33.8396045 | 17070 | 2622 |
| **GO:MF** | heterocyclic cc | GO:1901363 | 2.95E-32 | 31.5302343 | 6210 | 2622 |
| **GO:MF** | organic cyclic | GO:0097159 | 2.61E-31 | 30.5839167 | 6291 | 2622 |
| **GO:MF** | nucleic acid bi | GO:0003676 | 2.15E-29 | 28.6670225 | 4252 | 2622 |
| **GO:MF** | kinase binding | GO:0019900 | 2.73E-25 | 24.5632628 | 757 | 2622 |
| **GO:MF** | ubiquitin prote | GO:0031625 | 2.82E-22 | 21.5493455 | 298 | 2622 |
| **GO:MF** | ubiquitin-like | GO:0044389 | 4.51E-22 | 21.3461401 | 316 | 2622 |
| **GO:MF** | structural mol | GO:0005198 | 1.07E-21 | 20.9705081 | 701 | 2622 |
| **GO:MF** | protein kinase | GO:0019901 | 1.20E-21 | 20.9220187 | 677 | 2622 |
| **GO:MF** | structural con | GO:0003735 | 8.32E-21 | 20.079963 | 171 | 2622 |
| **GO:MF** | protein domai | GO:0019904 | 1.11E-19 | 18.9535784 | 706 | 2622 |
| **GO:MF** | cytoskeletal pr | GO:0008092 | 5.85E-19 | 18.232919 | 1001 | 2622 |
| **GO:MF** | transcription c | GO:0003712 | 1.50E-18 | 17.8237426 | 509 | 2622 |
| **GO:MF** | chromatin bin | GO:0003682 | 7.98E-15 | 14.0981075 | 573 | 2622 |
| **GO:MF** | nucleoside ph | GO:1901265 | 2.00E-14 | 13.6982647 | 2177 | 2622 |
| **GO:MF** | nucleotide bin | GO:0000166 | 3.06E-14 | 13.5135938 | 2176 | 2622 |
| **GO:MF** | carbohydrate | GO:0097367 | 3.79E-14 | 13.4218028 | 2282 | 2622 |
| **GO:MF** | purine ribonu | GO:0035639 | 4.61E-13 | 12.336587 | 1846 | 2622 |
| **GO:MF** | small molecul | GO:0036094 | 6.79E-13 | 12.1682048 | 2516 | 2622 |
| **GO:MF** | identical prote | GO:0042802 | 1.24E-12 | 11.9071042 | 2063 | 2622 |
| **GO:MF** | translation reg | GO:0045182 | 2.10E-12 | 11.6777418 | 154 | 2622 |
| **GO:MF** | purine nucleo | GO:0017076 | 4.07E-12 | 11.3903539 | 1932 | 2622 |
| **GO:MF** | purine ribonu | GO:0032555 | 5.27E-12 | 11.2779026 | 1917 | 2622 |
| **GO:MF** | transcription f | GO:0008134 | 5.42E-12 | 11.2662135 | 556 | 2622 |
| **GO:MF** | ribonucleotide | GO:0032553 | 7.85E-12 | 11.1049627 | 1934 | 2622 |
| **GO:MF** | anion binding | GO:0043168 | 3.05E-11 | 10.5150297 | 2413 | 2622 |
| **GO:MF** | ribonucleopro | GO:0043021 | 3.46E-11 | 10.4606332 | 150 | 2622 |
| **GO:MF** | transcription c | GO:0003713 | 4.60E-11 | 10.3368632 | 282 | 2622 |
| **GO:MF** | GTPase bindir | GO:0051020 | 2.08E-09 | 8.6809359 | 295 | 2622 |
| **GO:MF** | DNA-binding | GO:0140297 | 2.11E-09 | 8.67550426 | 234 | 2622 |
| **GO:MF** | chromatin DN | GO:0031490 | 2.96E-09 | 8.52886296 | 103 | 2622 |
| **GO:MF** | tau protein bir | GO:0048156 | 5.48E-09 | 8.26093801 | 45 | 2622 |
| **GO:MF** | unfolded prot | GO:0051082 | 6.74E-09 | 8.1710313 | 113 | 2622 |
| **GO:MF** | ATP binding | GO:0005524 | 1.26E-08 | 7.89845268 | 1500 | 2622 |
| **GO:MF** | translation reg | GO:0090079 | 1.33E-08 | 7.87565942 | 123 | 2622 |
| **GO:MF** | adenyl nucleo | GO:0030554 | 2.83E-08 | 7.54848338 | 1577 | 2622 |
| **GO:MF** | ATPase | GO:0016887 | 3.43E-08 | 7.46452064 | 553 | 2622 |
| **GO:MF** | RNA polymera | GO:0061629 | 3.88E-08 | 7.41156678 | 196 | 2622 |

| | | | | | | |
|---|---|---|---|---|---|---|
| **GO:MF** | adenyl ribonu | GO:0032559 | 4.20E-08 | 7.37634875 | 1564 | 2622 |
| **GO:MF** | nucleosome b | GO:0031491 | 8.99E-08 | 7.04609548 | 70 | 2622 |
| **GO:MF** | translation fac | GO:0008135 | 2.28E-07 | 6.64232341 | 99 | 2622 |
| **GO:MF** | small GTPase | GO:0031267 | 3.50E-07 | 6.45648008 | 263 | 2622 |
| **GO:MF** | nuclear recept | GO:0016922 | 1.011E-06 | 5.99526927 | 133 | 2622 |
| **GO:MF** | tubulin bindin | GO:0015631 | 1.037E-06 | 5.98421214 | 381 | 2622 |
| **GO:MF** | protein C-term | GO:0008022 | 2.7239E-06 | 5.56480237 | 201 | 2622 |
| **GO:MF** | nucleosomal I | GO:0031492 | 3.1546E-06 | 5.50105714 | 39 | 2622 |
| **GO:MF** | transcription c | GO:0003714 | 3.5074E-06 | 5.45501293 | 188 | 2622 |
| **GO:MF** | mRNA 5'-UTF | GO:0048027 | 3.567E-06 | 5.44769689 | 27 | 2622 |
| **GO:MF** | actin binding | GO:0003779 | 3.8723E-06 | 5.41203372 | 449 | 2622 |
| **GO:MF** | growth factor | GO:0019838 | 2.4928E-05 | 4.60331089 | 141 | 2622 |
| **GO:MF** | protein folding | GO:0044183 | 2.6085E-05 | 4.58361061 | 36 | 2622 |
| **GO:MF** | rRNA binding | GO:0019843 | 9.0911E-05 | 4.04138495 | 68 | 2622 |
| **GO:MF** | histone bindir | GO:0042393 | 9.7108E-05 | 4.01274708 | 240 | 2622 |
| **GO:MF** | protease bindi | GO:0002020 | 0.00017646 | 3.7533509 | 131 | 2622 |
| **GO:MF** | catalytic activit | GO:0140096 | 0.00022051 | 3.65657155 | 2336 | 2622 |
| **GO:MF** | molecular ada | GO:0060090 | 0.00025155 | 3.59937615 | 341 | 2622 |
| **GO:MF** | chaperone bir | GO:0051087 | 0.00030715 | 3.51264268 | 106 | 2622 |
| **GO:MF** | beta-tubulin b | GO:0048487 | 0.00036669 | 3.43570492 | 41 | 2622 |
| **GO:MF** | disordered do | GO:0097718 | 0.00036914 | 3.43281112 | 34 | 2622 |
| **GO:MF** | mRNA binding | GO:0003729 | 0.00045711 | 3.33998174 | 553 | 2622 |
| **GO:MF** | signal sequen | GO:0005048 | 0.0004789 | 3.31975441 | 49 | 2622 |
| **GO:MF** | catalytic activit | GO:0003824 | 0.00055127 | 3.25863797 | 5682 | 2622 |
| **GO:MF** | double-strand | GO:0003725 | 0.00062179 | 3.20635517 | 78 | 2622 |
| **GO:MF** | GTPase activit | GO:0003924 | 0.00066149 | 3.17947711 | 300 | 2622 |
| **GO:MF** | translation init | GO:0003743 | 0.00076236 | 3.11784022 | 58 | 2622 |
| **GO:MF** | actin filament | GO:0051015 | 0.0009085 | 3.04167737 | 213 | 2622 |
| **GO:MF** | low-density lip | GO:0050750 | 0.0013028 | 2.88512336 | 23 | 2622 |
| **GO:MF** | heat shock pr | GO:0031072 | 0.00160705 | 2.79397006 | 127 | 2622 |
| **GO:MF** | pre-mRNA bir | GO:0036002 | 0.00165286 | 2.78176343 | 37 | 2622 |
| **GO:MF** | structural con | GO:0005200 | 0.00173647 | 2.76033315 | 104 | 2622 |
| **GO:MF** | enzyme inhibi | GO:0004857 | 0.00190075 | 2.72107443 | 391 | 2622 |
| **GO:MF** | ion channel bi | GO:0044325 | 0.00214125 | 2.66933218 | 138 | 2622 |
| **GO:MF** | histone deace | GO:0042826 | 0.00255946 | 2.59185132 | 115 | 2622 |
| **GO:MF** | nucleoside-trip | GO:0017111 | 0.00315793 | 2.50059743 | 424 | 2622 |
| **GO:MF** | lipoprotein pa | GO:0070325 | 0.0036398 | 2.43892303 | 28 | 2622 |
| **GO:MF** | ion binding | GO:0043167 | 0.00384046 | 2.41561728 | 6056 | 2622 |
| **GO:MF** | GTP binding | GO:0005525 | 0.00397538 | 2.40062084 | 381 | 2622 |
| **GO:MF** | peroxiredoxin | GO:0051920 | 0.00425367 | 2.37123633 | 8 | 2622 |
| **GO:MF** | proteasome-a | GO:0036402 | 0.00432538 | 2.36397575 | 6 | 2622 |
| **GO:MF** | protein kinase | GO:0051018 | 0.00451533 | 2.34531092 | 51 | 2622 |
| **GO:MF** | protein-macro | GO:0030674 | 0.00471169 | 2.32682373 | 272 | 2622 |
| **GO:MF** | extracellular m | GO:0005201 | 0.00543024 | 2.26518099 | 173 | 2622 |
| **GO:MF** | proton transm | GO:0015078 | 0.00633721 | 2.19810213 | 129 | 2622 |

| | | | | | | |
|---|---|---|---|---|---|---|
| **GO:MF** | p53 binding | GO:0002039 | 0.00666576 | 2.17615021 | 69 | 2622 |
| **GO:MF** | nuclear recept | GO:0030374 | 0.00845392 | 2.07294202 | 57 | 2622 |
| **GO:MF** | single-strande | GO:0003697 | 0.00885561 | 2.05278137 | 121 | 2622 |
| **GO:MF** | extracellular m | GO:0030020 | 0.00900717 | 2.04541185 | 41 | 2622 |
| **GO:MF** | platelet-derive | GO:0048407 | 0.00940551 | 2.02661779 | 11 | 2622 |
| **GO:MF** | insulin recepto | GO:0005158 | 0.0096917 | 2.01359986 | 23 | 2622 |
| **GO:MF** | microtubule b | GO:0008017 | 0.01051661 | 1.97812419 | 273 | 2622 |
| **GO:MF** | enzyme regula | GO:0030234 | 0.01133128 | 1.94572103 | 911 | 2622 |
| **GO:MF** | protein kinase | GO:0019887 | 0.0131596 | 1.88075719 | 189 | 2622 |
| **GO:MF** | protein kinase | GO:0005080 | 0.01675993 | 1.77572784 | 55 | 2622 |
| **GO:MF** | ubiquitin ligas | GO:1990948 | 0.01680094 | 1.77466643 | 9 | 2622 |
| **GO:MF** | single-strande | GO:0003727 | 0.01786864 | 1.74790857 | 91 | 2622 |
| **GO:MF** | guanyl ribonu | GO:0032561 | 0.01863108 | 1.7297619 | 402 | 2622 |
| **GO:MF** | guanyl nucleo | GO:0019001 | 0.01863108 | 1.7297619 | 402 | 2622 |
| **GO:MF** | ligase activity, | GO:0016875 | 0.01893444 | 1.72274747 | 43 | 2622 |
| **GO:MF** | aminoacyl-tRI | GO:0004812 | 0.01893444 | 1.72274747 | 43 | 2622 |
| **GO:MF** | kinase inhibitc | GO:0019210 | 0.01943995 | 1.71130493 | 73 | 2622 |
| **GO:MF** | ubiquitin-spec | GO:1990381 | 0.02095225 | 1.67876925 | 21 | 2622 |
| **GO:MF** | kinase regulat | GO:0019207 | 0.0214615 | 1.66834 | 219 | 2622 |
| **GO:MF** | signaling rece | GO:0005102 | 0.02467298 | 1.60777839 | 1551 | 2622 |
| **GO:MF** | phosphatase | GO:0019902 | 0.02584964 | 1.58754552 | 199 | 2622 |
| **GO:MF** | molecular fun | GO:0098772 | 0.03028233 | 1.51881074 | 1115 | 2622 |
| **GO:MF** | protein phosp | GO:0019903 | 0.03042268 | 1.51680247 | 153 | 2622 |
| **GO:MF** | supercoiled D | GO:0097100 | 0.0308644 | 1.51054212 | 5 | 2622 |
| **GO:MF** | thioredoxin pe | GO:0008379 | 0.0308644 | 1.51054212 | 5 | 2622 |
| **GO:MF** | phosphatase | GO:0019208 | 0.03189092 | 1.49633295 | 75 | 2622 |
| **GO:MF** | ATPase-coupl | GO:0019829 | 0.0443712 | 1.35289878 | 54 | 2622 |
| **GO:MF** | nuclear localiz | GO:0008139 | 0.04596005 | 1.33761948 | 26 | 2622 |
| **GO:MF** | ubiquitin-prot | GO:0055105 | 0.04917048 | 1.30829556 | 10 | 2622 |

| intersection_ | effective_dor | intersections |
|---|---|---|
| 594 | 18679 | EEF1A1,PABPC1,LRP1,HSPA8,HNRNPA1,HSPA9,EEF2,HSP90AB1,HSP |
| 2403 | 18679 | EEF1A1,ORC4,IGFBP5,PABPC1,COL1A1,LRP1,HSPA8,GAPDH,MLLT1 |
| 569 | 18679 | EEF1A1,COL1A1,LRP1,HSPA8,HSPA9,ACTB,CTSC,USP22,ACTG1,EEF |
| 380 | 18679 | LRP1,HSPA8,ACTB,EEF2,CFL1,HSP90AB1,HSP90AA1,ACTR2,VIM,SPA |
| 146 | 18679 | HSPA8,NUDC,EEF2,HSP90AB1,EIF4G2,DBN1,ENO1,FLNA,RPL7A,ALD |
| 197 | 18679 | HSPA8,NUDC,EEF2,HSP90AB1,EIF4G2,DBN1,ENO1,FLNA,RPL7A,ALD |
| 2547 | 18679 | EEF1A1,ORC4,IGFBP5,PABPC1,COL1A1,LRP1,HSPA8,GAPDH,MLLT1 |
| 1152 | 18679 | EEF1A1,ORC4,PABPC1,LRP1,HSPA8,GAPDH,HNRNPA1,HSPA9,POLH, |
| 1160 | 18679 | EEF1A1,ORC4,PABPC1,LRP1,HSPA8,GAPDH,HNRNPA1,HSPA9,POLH, |
| 839 | 18679 | EEF1A1,ORC4,PABPC1,LRP1,HSPA8,HNRNPA1,HSPA9,POLH,ACTB,E |
| 220 | 18679 | EEF1A1,ACTB,EEF2,MAVS,HTT,HSP90AB1,HSP90AA1,DCX,RPS6,PIK3 |
| 113 | 18679 | HSPA8,HSPA9,ACTG1,HSP90AB1,HSP90AA1,TUBB,UBC,TUBA1B,CAL |
| 117 | 18679 | HSPA8,HSPA9,ACTG1,HSP90AB1,HSP90AA1,TUBB,UBC,TUBA1B,CAL |
| 200 | 18679 | COL1A1,COPA,ACTB,ACTG1,TUBA1A,ACTR2,VIM,SPARC,TUBB,NUM |
| 195 | 18679 | EEF1A1,ACTB,EEF2,MAVS,HSP90AB1,HSP90AA1,DCX,RPS6,PIK3R2,F |
| 78 | 18679 | RPL3,RPS6,RPL4,RPL7A,RPL8,RPLP0,RPL13A,RPL6,RPL13,RPS8,RP |
| 196 | 18679 | HSPA8,GAPDH,HNRNPA1,ATN1,DPYSL3,TUBA1A,MAVS,HSP90AB1,HS |
| 251 | 18679 | GAPDH,ACTB,EEF2,DPYSL3,CFL1,HTT,HSP90AB1,HSP90AA1,ACTR2, |
| 153 | 18679 | ATN1,USP22,MED18,TMF1,MAGED1,KAT6A,DDX17,NPM1,TRIM28,S |
| 157 | 18679 | ACTB,YBX1,H3-3B,DDX5,NPM1,TRIM28,NONO,SET,GTF2F1,HNRNPU |
| 439 | 18679 | EEF1A1,ORC4,HSPA8,GAPDH,HSPA9,ACTB,NRBP2,ACTG1,EEF2,TUB |
| 438 | 18679 | EEF1A1,ORC4,HSPA8,GAPDH,HSPA9,ACTB,NRBP2,ACTG1,EEF2,TUB |
| 455 | 18679 | EEF1A1,ORC4,LRP1,HSPA8,HSPA9,ACTB,NRBP2,ACTG1,EEF2,DPYSL |
| 378 | 18679 | EEF1A1,ORC4,HSPA8,HSPA9,ACTB,NRBP2,ACTG1,EEF2,TUBA1A,XR |
| 487 | 18679 | EEF1A1,ORC4,HSPA8,GAPDH,HSPA9,ACTB,NRBP2,ACTG1,EEF2,TUB |
| 412 | 18679 | COL1A1,GAPDH,HNRNPA1,ACTB,CTSC,ACTG1,DPYSL3,TUBA1A,HTT, |
| 61 | 18679 | EEF1A1,PABPC1,EEF2,EIF4G2,EIF3C,EIF4B,EIF1,EIF4A1,RPL10,LAR |
| 388 | 18679 | EEF1A1,ORC4,HSPA8,HSPA9,ACTB,NRBP2,ACTG1,EEF2,TUBA1A,XR |
| 385 | 18679 | EEF1A1,ORC4,HSPA8,HSPA9,ACTB,NRBP2,ACTG1,EEF2,TUBA1A,XR |
| 146 | 18679 | ACTB,FLNA,CALR,TMF1,DDX5,KAT6A,NPM1,RPS3,HMGB1,TPT1,DN |
| 387 | 18679 | EEF1A1,ORC4,HSPA8,HSPA9,ACTB,NRBP2,ACTG1,EEF2,TUBA1A,XR |
| 462 | 18679 | EEF1A1,ORC4,HSPA8,HSPA9,ACTB,CTSC,NRBP2,ACTG1,EEF2,TUBA |
| 58 | 18679 | EEF2,DDX5,RACK1,NPM1,EIF3C,EIF4B,EIF1,SEC61A1,RPLP1,HNRNP |
| 88 | 18679 | USP22,TMF1,MAGED1,KAT6A,DDX17,NPM1,TRIM28,HMGB1,ZMIZ1, |
| 87 | 18679 | HSP90AA1,ENO1,FLNA,YBX1,DVL3,SPTBN1,GNB1,GDI1,ANXA2,CDC |
| 74 | 18679 | ACTB,CALR,TMF1,DDX5,NPM1,DNAJA1,ATF4,JUND,ACTN4,CTNNB1, |
| 43 | 18679 | ACTB,H3-3B,HNRNPU,CHD4,HMGN2,RBBP4,HNRNPC,ACTN4,H1-0,H |
| 26 | 18679 | ACTB,HSP90AB1,HSP90AA1,CLU,MAP2,MAP1A,PPP2CA,FYN,DCTN1, |
| 45 | 18679 | HSPA8,HSPA9,NUDC,HSP90AB1,HSP90AA1,HSP90B1,CALR,HSPA1B, |
| 300 | 18679 | ORC4,HSPA8,HSPA9,ACTB,NRBP2,ACTG1,XRCC2,HSP90AB1,HSP90 |
| 47 | 18679 | EEF1A1,PABPC1,EEF2,EIF4G2,EIF3C,EIF4B,EIF1,EIF4A1,LARP1,EIF4 |
| 311 | 18679 | ORC4,HSPA8,HSPA9,ACTB,NRBP2,ACTG1,XRCC2,HSP90AB1,HSP90 |
| 134 | 18679 | ORC4,HSPA8,HSPA9,XRCC2,HSP90AB1,HSP90AA1,HSP90B1,ATP2B4 |
| 63 | 18679 | ACTB,CALR,TMF1,DDX5,NPM1,DNAJA1,JUND,ACTN4,CTNNB1,ARID1 |

| | | |
|---|---|---|
| 308 | 18679 | ORC4,HSPA8,HSPA9,ACTB,NRBP2,ACTG1,XRCC2,HSP90AB1,HSP90 |
| 32 | 18679 | ACTB,H3-3B,SMARCA5,CHD4,HMGN2,RBBP4,HNRNPC,H1-0,ARID1A |
| 39 | 18679 | EEF1A1,EEF2,EIF4G2,EIF3C,EIF4B,EIF1,EIF4A1,EIF4H,EIF5,EIF5A,EI |
| 75 | 18679 | FLNA,DVL3,GDI1,ANXA2,CDC42BPB,KPNB1,PFN1,XPO1,MYO5A,MT |
| 46 | 18679 | CALR,TMF1,DDX5,JUND,ACTN4,CTNNB1,ARID1A,PARP1,PKN1,ASXL |
| 97 | 18679 | GAPDH,HTT,NUMA1,DCX,MAP1B,ALDOA,NEFM,KIF1A,STMN2,MAPT, |
| 60 | 18679 | PABPC1,NUMA1,NCL,NEFL,FN1,YWHAQ,XRCC5,ATF4,ATP2A2,CCN2, |
| 21 | 18679 | ACTB,H3-3B,CHD4,HMGN2,RBBP4,HNRNPC,H1-0,H1-10,HDAC2,H3- |
| 57 | 18679 | ATN1,TRIM28,SPEN,BASP1,HNRNPU,TLE5,CTBP1,HSBP1,EID1,HDGF |
| 17 | 18679 | NCL,RPL5,RPS3A,LARP1,RPS14,RPL26,DDX3X,RPS7,MYH10,RPS13, |
| 108 | 18679 | EEF2,CFL1,ACTR2,TMSB4X,DBN1,FLNA,MAP1B,ALDOA,SPTBN1,MY |
| 45 | 18679 | IGFBP5,COL1A1,COL6A1,IGFBP2,COL5A1,COL4A1,HTRA4,PXDN,IG |
| 19 | 18679 | HSPA8,HSPA9,HSP90AB1,CALR,HSPA1B,CCT7,CCT6A,CCT2,TCP1,C |
| 27 | 18679 | EEF2,RPL8,RPLP0,RPL5,NPM1,RPL19,RPS3,RPS4X,RPL11,RPL23,R |
| 64 | 18679 | KAT6A,NPM1,SET,SMARCA5,BRD3,CBX5,RBBP4,TSPYL4,SSRP1,PKN |
| 41 | 18679 | COL1A1,LRP1,UBC,PSAP,DVL3,VCP,FN1,ITGB1,ANXA2,TIMP2,MARC |
| 408 | 18679 | GAPDH,NEDD4L,CTSC,NRBP2,USP22,MED18,PRPF8,UBA1,PGK1,FK |
| 82 | 18679 | HSPA8,MAVS,NEFL,RACK1,MYH9,SEPTIN2,FSCN1,MAPT,DDB1,SERI |
| 35 | 18679 | HSPA8,CTSC,CALR,CLU,HSPD1,MAPT,FN1,DNAJA1,SCARB2,ATP1A1, |
| 19 | 18679 | HTT,LRPPRC,APPL1,TBCD,GABARAP,CCT5,HDAC6,SMC3,TAOK1,VA |
| 17 | 18679 | GAPDH,HSP90AB1,HSP90AA1,NUMA1,CALM3,CALM1,FN1,AP2M1,CL |
| 119 | 18679 | PABPC1,EIF4G2,NCL,PKM,YBX1,CALR,DDX5,RPL13A,RPL7,RPL5,EIF |
| 21 | 18679 | AP2M1,SEC61A1,AP2B1,KPNB1,IPO5,TOMM20,TNPO1,SRP14,KDEL |
| 903 | 18679 | EEF1A1,HSPA8,GAPDH,ARSD,NEDD4L,POLH,CTSC,NRBP2,USP22,ME |
| 28 | 18679 | HSP90AB1,VIM,TUBA1B,HSPD1,EIF4B,EIF4A1,HMGB1,TUBB4B,HNR |
| 73 | 18679 | EEF1A1,EEF2,TUBA1A,TUBB,TUBA1B,TUBB2B,GNAS,GNB1,SEPTIN2 |
| 23 | 18679 | EIF4G2,EIF3C,EIF4B,EIF1,EIF4A1,EIF4H,EIF5,EIF3L,EIF3E,EIF3F,EIF |
| 56 | 18679 | EEF2,CFL1,ACTR2,DBN1,FLNA,MYH9,FSCN1,MARCKS,MARCKSL1,V |
| 13 | 18679 | HSP90B1,CLU,AP2M1,DNAJA1,CLTC,MESD,AP2A1,HSPG2,APBB1,SN |
| 38 | 18679 | HSPA8,HSPA9,HTT,HSP90AB1,HSPA1B,MAPT,RPS3,DNAJA1,KPNB1, |
| 17 | 18679 | HNRNPA1,PRPF8,DDX5,HNRNPU,TARDBP,U2AF2,HNRNPL,PTBP1,EP |
| 33 | 18679 | ACTB,ACTG1,TUBA1A,ACTR2,VIM,TUBB,TUBA1B,NEFL,NEFM,TUBB |
| 88 | 18679 | GAPDH,ATP2B4,RACK1,APP,RPL5,TMBIM6,NPM1,YWHAG,SET,RPL1 |
| 40 | 18679 | NEDD4L,HTT,HSP90AB1,FLNA,YWHAE,CALM3,CALM1,YWHAQ,PKD1, |
| 35 | 18679 | HSP90AB1,HSP90AA1,HSPA1B,YWHAE,CHD4,RBBP4,SFPQ,YWHAB,R |
| 93 | 18679 | EEF1A1,EEF2,TUBA1A,TUBB,TUBA1B,TUBB2B,KIF1A,GNAS,GNB1,M |
| 14 | 18679 | LRP1,HSP90B1,CLU,AP2M1,DNAJA1,CLTC,MESD,AP2A1,HSPG2,APB |
| 948 | 18679 | EEF1A1,ORC4,COL1A1,LRP1,HSPA8,ARSD,HSPA9,POLH,ACTB,CTSC, |
| 85 | 18679 | EEF1A1,EEF2,TUBA1A,HSP90AB1,TUBB,TUBA1B,TUBB2B,GNAS,SEP |
| 7 | 18679 | PRDX2,PRDX5,PRDX1,PRDX6,PRDX3,PARK7,PRDX4 |
| 6 | 18679 | PSMC3,PSMC1,PSMC5,PSMC2,PSMC4,PSMC6 |
| 20 | 18679 | RPS3,AKAP12,PJA2,PRKAR1A,ARFGEF2,PRKAR2A,AKAP9,RDX,WASF |
| 65 | 18679 | HSPA8,MAVS,NEFL,RACK1,MYH9,FSCN1,MAPT,DDB1,SERINC1,AP2 |
| 46 | 18679 | COL1A1,SPARC,COL6A1,COL5A1,COL4A1,COL18A1,COL4A2,FN1,A |
| 37 | 18679 | SLC9A7,ATP5F1B,ATP5F1A,ATP2A2,SLC25A3,ATP6V0C,ATP5PB,ATP |

| | | |
|---|---|---|
| 24 | 18679 | EEF2,HTT,HSPD1,PTTG1IP,NOP53,SMARCA4,KDM1A,USP7,TP53BP1 |
| 21 | 18679 | USP22,TMF1,ZMIZ1,ACTN4,PKN1,ACTN1,KDM1A,CCAR2,CARM1,ZM |
| 35 | 18679 | HNRNPA1,YBX1,HSPD1,MAPT,SSBP2,HMGB1,HNRNPU,LRPPRC,HNR |
| 17 | 18679 | COL1A1,COL6A1,COL5A1,COL4A1,COL18A1,COL4A2,COL11A1,CO |
| 8 | 18679 | COL1A1,COL6A1,COL5A1,COL4A1,COL1A2,COL2A1,PDGFRB,COL3 |
| 12 | 18679 | IGF2,PDPK1,PHIP,SNX2,PTPN11,IGF1R,PTPN1,SNX1,CCDC88A,IRS1, |
| 64 | 18679 | GAPDH,NUMA1,DCX,MAP1B,NEFM,KIF1A,MAPT,MAP9,RPS3,VAPA,PA |
| 172 | 18679 | GAPDH,CTSC,HSP90AB1,HSP90AA1,ATP2B4,RACK1,CALM3,APP,RPL |
| 48 | 18679 | HSP90AB1,KIDINS220,RACK1,CALM3,CALM1,NPM1,IGF2,YWHAG,H |
| 20 | 18679 | FLNA,RACK1,MARCKS,YWHAG,PRKCSH,TOP2B,SQSTM1,PKN1,ADA |
| 7 | 18679 | RPL5,RPL11,RPL23,RPS20,RPS15,RPS7,RPL37 |
| 28 | 18679 | PABPC1,HNRNPA1,EIF4B,HMGB1,HNRNPU,EIF4H,HNRNPC,KHDRBS |
| 86 | 18679 | EEF1A1,EEF2,TUBA1A,HSP90AB1,TUBB,TUBA1B,TUBB2B,GNAS,SEP |
| 86 | 18679 | EEF1A1,EEF2,TUBA1A,HSP90AB1,TUBB,TUBA1B,TUBB2B,GNAS,SEP |
| 17 | 18679 | NARS1,MARS1,QARS1,LARS1,SARS1,VARS1,GARS1,IARS1,IARS2,YA |
| 17 | 18679 | NARS1,MARS1,QARS1,LARS1,SARS1,VARS1,GARS1,IARS1,IARS2,YA |
| 24 | 18679 | RACK1,NPM1,YWHAG,PRKAR1A,YWHAB,TRIB2,CDKN1A,HSPB1,QAR |
| 11 | 18679 | VCP,MARCHF6,CLTC,BAG6,PTEN,RHBDD2,AMFR,SUMO1,PARK7,SA |
| 53 | 18679 | HSP90AB1,KIDINS220,PIK3R2,RACK1,CALM3,GSTP1,CALM1,NPM1, |
| 271 | 18679 | LRP1,HSPA8,COPA,MTRNR2L2,HSP90AB1,TUBB,PIK3R2,FLNA,MTR |
| 49 | 18679 | CTSC,PIK3R2,HSP90B1,ATP2B4,RACK1,VCP,MAPT,GTF2F1,CDH2,C |
| 202 | 18679 | GAPDH,NEDD4L,CTSC,MTRNR2L2,HSP90AB1,HSP90AA1,FLNA,MTR |
| 40 | 18679 | PIK3R2,HSP90B1,ATP2B4,RACK1,VCP,MAPT,GTF2F1,CDH2,CTNNB1 |
| 5 | 18679 | RPS3,HMGB1,TOP1,PSIP1,HMGB2 |
| 5 | 18679 | PRDX2,PRDX5,PRDX1,PRDX3,PRDX4 |
| 24 | 18679 | CALM3,CALM1,SET,SBF1,IGFBP3,PTN,PPP1R9B,PPP1R15A,PPP1R1 |
| 19 | 18679 | ATP2B4,ATP5F1B,ATP1A1,ATP2A2,ATP1A2,ATP6V0C,ATP6V1B2,ABC |
| 12 | 18679 | KPNB1,IPO5,TNPO1,KPNA2,POM121C,NUP153,KPNA6,NUP98,TNPO |
| 7 | 18679 | RPL5,RPL11,RPL23,RPS20,RPS15,RPS7,RPL37 |

'90AA1,VIM,UBC,TMSB4X,EIF4G2,RPL3,RPS6,RPL4,PRRC2B,ENO1,TUBA1B,FLNA,NCL,RPL7A,AL
,COPA,HNRNPA1,HSPA9,NEDD4L,POLH,ATN1,ACTB,CTSC,NUDC,USP22,MTRNR2L2,MED18,ACT(
2,MAVS,HTT,HSP90AB1,HSP90AA1,TUBB,UBC,TMSB4X,DCX,RPS6,ENO1,TUBA1B,PIK3R2,FLNA,N
RC,TUBB,NUMA1,DBN1,FLNA,MAP1B,PKM,COL6A1,NEFL,CALR,COL5A1,H3-3B,CLU,DDX5,YWH
OA,PKM,RPL6,YWHAE,RACK1,BSG,FASN,SPTBN1,MYH9,SEPTIN2,FSCN1,SPTAN1,DCHS1,RPL15,
OA,PKM,CALR,COL5A1,RPL6,YWHAE,RACK1,BSG,FASN,SPTBN1,MYH9,SEPTIN2,P4HB,FSCN1,SF
,COPA,HNRNPA1,ARSD,HSPA9,NEDD4L,POLH,ATN1,ACTB,CTSC,NUDC,NRBP2,USP22,MTRNR2L
ACTB,NRBP2,ACTG1,EEF2,TUBA1A,XRCC2,HSP90AB1,ZNF124,HSP90AA1,ACTR2,VIM,TUBB,UBC
ACTB,NRBP2,ACTG1,EEF2,TUBA1A,XRCC2,HSP90AB1,ZNF124,HSP90AA1,ACTR2,VIM,TUBB,UBC
EF2,XRCC2,HSP90AB1,ZNF124,HSP90AA1,VIM,UBC,TMSB4X,EIF4G2,RPL3,RPS6,ZNF316,RPL4,F
R2,FLNA,ATP2B4,RACK1,CALM3,GSTP1,MARCKS,CALM1,NPM1,YWHAG,MAPT,PKD1,RPS3,ITGE
R,HSPA1B,CLU,YWHAE,GPI,RPL5,TMBIM6,HSPD1,VCP,UBB,TRIM28,CKB,TPI1,UCHL1,RPL11,VC
R,HSPA1B,CLU,YWHAE,GPI,RPL5,TMBIM6,HSPD1,VCP,UBB,TRIM28,CKB,TPI1,UCHL1,RPL11,VC
A1,RPL3,RPS6,RPL4,TUBA1B,MAP1B,RPL7A,COL6A1,NEFL,NEFM,RPL8,TUBB2B,COL5A1,COL4
LNA,ATP2B4,RACK1,CALM3,GSTP1,MARCKS,CALM1,NPM1,YWHAG,MAPT,PKD1,RPS3,ITGB1,PG
L7,RPL5,RPL37A,RPL19,RPL15,RPS3,RPS4X,RPS3A,RPL18,RPL11,RPLP1,RPL10,RPS17,RPS16,F
SP90AA1,VIM,TUBB,KIDINS220,NUMA1,NEFL,ATP2B4,HSPA1B,DDX5,RACK1,CALM3,SHISA9,MYH
TMSB4X,NUMA1,DCX,DBN1,FLNA,MAP1B,ALDOA,NEFM,CLU,KIF1A,SPTBN1,CALM3,GNB1,STMN
PEN,BASP1,HMGB1,ZMIZ1,HNRNPU,NACA,TLE5,TADA2B,ACTN4,CTNNB1,ARID1A,SND1,CTBP1,B
J,SMARCA5,BRD3,CHD4,HMGN2,RBBP4,HNRNPC,TADA2B,SFPQ,ACTN4,H1-0,CTNNB1,ARID1A,H
A1A,XRCC2,HSP90AB1,HSP90AA1,ACTR2,TUBB,TUBA1B,PKM,HSP90B1,UBA1,ATP2B4,HSPA1B,T
A1A,XRCC2,HSP90AB1,HSP90AA1,ACTR2,TUBB,TUBA1B,PKM,HSP90B1,UBA1,ATP2B4,HSPA1B,T
3,TUBA1A,XRCC2,HSP90AB1,HSP90AA1,ACTR2,TUBB,TUBA1B,PKM,HSP90B1,UBA1,ATP2B4,FS
C2,HSP90AB1,HSP90AA1,ACTR2,TUBB,TUBA1B,PKM,HSP90B1,UBA1,ATP2B4,HSPA1B,TUBB2B,
A1A,XRCC2,HSP90AB1,HSP90AA1,ACTR2,TUBB,TUBA1B,ALDOA,PKM,HSP90B1,UBA1,ATP2B4,H
,HSP90AB1,HSP90AA1,VIM,ENO1,FLNA,NCL,ALDOA,FTL,NEFL,PSAP,MAGED1,YWHAE,KIF1A,HTR
'1,EIF4H,RPS9,RPL22,RPS14,EIF5,EIF5A,EIF3L,CIRBP,EEF1A2,MTRF1L,GCN1,EIF3E,EIF3F,EIF2A
C2,HSP90AB1,HSP90AA1,ACTR2,TUBB,TUBA1B,PKM,HSP90B1,UBA1,ATP2B4,HSPA1B,TUBB2B,
C2,HSP90AB1,HSP90AA1,ACTR2,TUBB,TUBA1B,PKM,HSP90B1,UBA1,ATP2B4,HSPA1B,TUBB2B,
JA1,GTF2F1,HNRNPU,RPL23,ATF4,CHD4,JUND,ACTN4,DDX3X,JUN,CTNNB1,ARID1A,CTBP1,HDA
C2,HSP90AB1,HSP90AA1,ACTR2,TUBB,TUBA1B,PKM,HSP90B1,UBA1,ATP2B4,HSPA1B,TUBB2B,
1A,XRCC2,HSP90AB1,HSP90AA1,ACTR2,TUBB,TUBA1B,PKM,HSP90B1,UBA1,ATP2B4,HSPA1B,TU
U,LARP1,RPSA,EIF4H,PTCD3,CBX5,EIF5A,DDX3X,CKAP5,SND1,SRP9,RPN2,GCN1,PRPF6,STRAP,E
NACA,TADA2B,ACTN4,CTNNB1,ARID1A,PKN1,ASXL1,PRPF6,SMARCA4,ACTN1,KDM1A,CCAR2,SM
2BPB,KPNB1,PFN1,XPO1,GNB2,MYO5A,MTSS2,CYFIP2,RAC1,ARHGDIA,IPO5,GDI2,PKN1,NCKAF
ARID1A,HDAC2,PARP1,PKN1,ASXL1,PRPF6,TAF7,NOTCH2,SMARCA4,HSPB1,KDM1A,SMARCE1,Y
1-10,HDAC2,H3-3A,KDM6B,SMARCA4,SMARCE1,SMARCC1,KDM3B,WBP2,STAT3,EP300,H2AZ1,
EP300,PPP2CB,HDAC6,BRSK1,TAOK1,FKBP4,GSK3A,APBB1,PPP5C,TAOK2,STUB1,GSK3B,ROCK
CANX,CLU,NPM1,HSPD1,TUBB4B,DNAJA1,NACA,CCT7,PTGES3,CCT6A,PPIB,TOMM20,CCT2,TCP
A1,ACTR2,PKM,HSP90B1,UBA1,ATP2B4,HSPA1B,PGK1,DDX5,KIF1A,DDX17,ATP5F1B,MYH9,SM(
H,EIF5,EIF5A,EIF3L,EEF1A2,MTRF1L,GCN1,EIF3E,EIF3F,EIF2AK1,GTPBP1,EIF3D,GSPT1,EIF5B,E
A1,ACTR2,PKM,HSP90B1,UBA1,ATP2B4,HSPA1B,PGK1,DDX5,KIF1A,DDX17,ALDH6A1,ATP5F1B,I
,HSPA1B,DDX5,KIF1A,DDX17,ATP5F1B,MYH9,HSPD1,SNRNP200,VCP,EIF4A1,DNAH8,CHD3,SLC4
A,HDAC2,PARP1,PKN1,ASXL1,PRPF6,TAF7,NOTCH2,SMARCA4,HSPB1,KDM1A,SMARCE1,YWHAH

A1,ACTR2,PKM,HSP90B1,UBA1,ATP2B4,HSPA1B,PGK1,DDX5,KIF1A,DDX17,ALDH6A1,ATP5F1B,I
,H1-10,HDAC2,SSRP1,H3-3A,SMARCA4,SMARCE1,SMARCC1,SUPT6H,NAP1L4,H2AZ1,ARID1B,NC
F3L,EEF1A2,MTRF1L,GCN1,EIF3E,EIF3F,EIF2AK1,GTPBP1,EIF3D,GSPT1,EIF5B,EIF2S2,EIF3H,EIF
S2,CYFIP2,RAC1,ARHGDIA,IPO5,GDI2,PKN1,NCKAP1,TNPO1,ATP6AP1,OPTN,IPO7,CORO1C,SOD
I,PRPF6,TAF7,SMARCA4,KDM1A,SMARCE1,YWHAH,NCOR2,WBP2,STAT3,EP300,PHB2,FUS,BAZ2
MAP9,RPS3,VAPA,STMN1,PAFAH1B1,MAP4,CNN3,KIF5B,LRPPRC,DDX3X,ARL4C,GJA1,MACF1,CK
CTNNB1,YWHAB,CTBP1,STIP1,TOP2B,VAMP2,OPTN,PPP2CA,MAPRE1,PFKM,FBLN1,USP7,SREBF
3A,SMARCA4,SMARCE1,SMARCC1,H2AZ1,MBD3,MTA2,MACROH2A1,GATAD2B,SMARCC2,SMAR
,CCND1,SMARCA4,TCF25,NCOR2,MTA1,RERE,KDM5B,SF1,APEX1,TLE1,SRSF2,NOC2L,CREBBP,
CCT5,IGF2BP1,SYNCRIP,DHX36,IGF2BP2,SHMT2,GNL3
H9,P4HB,FSCN1,SPTAN1,MARCKS,MARCKSL1,MAPT,TMSB10,ITGB1,VCL,HNRNPU,CTNNA1,CNN
FBP3,COL1A2,PTN,GLG1,CCN2,RPS19,GPC1,COL2A1,NRDC,IGFBP7,BMPR2,IL6ST,RPS2,A2M,C
CT3,CCT4,HSPB1,FKBP8,CCT8,CCT5,CCDC47,CD74,HSPA1A,DNAJB1
PS9,PTCD3,RPL9,RPS14,RPL12,RPS5,NGRN,CIRBP,RPL23A,NOP53,RPL37,RPS11,RPS13,MDM2,I
I1,YEATS2,CHD2,TAF7,SMARCA4,IPO7,SMARCC1,SUPT6H,PHC2,NAP1L4,TP53BP1,ANP32A,NAP1
HF6,COL1A2,CLTC,TIMP3,BAG6,OS9,FLOT1,CST3,PTEN,A2M,RHBDD2,LDLR,FLOT2,SH3PXD2A,I
BP14,KAT6A,HTRA4,P4HB,SMG1,TTC3,USP11,TRIM28,EGLN1,PPP2R1A,GTF3C4,PDIA3,CDKL1,U
NC1,AP2M1,COL11A1,AP2B1,COL1A2,CBX5,KHDRBS1,SEPTIN11,SEPTIN3,SKP1,MAP1A,DLG5,VA
PTGES3,ATP1A2,HLA-B,STIP1,TBCD,CDC37,HYOU1,SOD1,GRN,DNAJB2,AHSA1,BAG5,GAK,AMFF
PB,GABARAPL1,SYT11,ADNP,MAP1S,TBCA,ARL8A,GABARAPL2,FGF13,EMD
LTC,GJA1,CTNNB1,FKBP8,FYN,DFFB,AP2A2,HSPA1A,MDM2
4A1,SF3B1,RPS3,RPS3A,HNRNPU,LARP1,HNRNPA0,RPS14,RPL26,SRSF4,HNRNPC,KHDRBS1,HD
R2,KDELR1,KPNA2,POM121C,TOMM70,NUP153,KPNA6,NUP98,SRP54,TNPO2,KPNA3,NOLC1,NU
ED18,EEF2,DPYSL3,TUBA1A,XRCC2,HSP90AA1,TUBB,FTH1,ENO1,TUBA1B,ALDOA,PKM,GANAB,F
NPU,EIF4H,MSN,SLC3A2,DDX3X,CLTC,ILF3,ACTN1,DHX30,ILF2,ADAR,DHX15,ZFR,ELAVL1,LSM1
,TUBB4B,RHOA,RAB6B,RAB7A,DDX3X,ARL4C,GNB2,SEPTIN11,RHOB,RAB1A,SEPTIN3,RAC1,RAE
2AK1,EIF3D,EIF5B,EIF2S2,EIF3H,EIF4G3,EIF3G,EIF4G1,EIF3B,EIF3M,EIF1B,EIF2S1,EIF3J
CL,CTNNA1,TPM4,WDR1,LASP1,TPM1,ACTN4,TLN1,MYO5A,ANXA6,MACF1,TAGLN,MYH10,COTL
X17,APOE,SYT1,LANCL1
PTGES3,ARHGDIA,STIP1,HDAC2,CDC37,BAG6,CSNK2A1,HDAC6,HIF1A,MVD,FKBP4,DNAJB2,AHSA
300,SF1,SRSF6,TRA2B,HNRNPA2B1,SRSF2,RBM4,CELF3,SOX9
2B,SPTBN1,SPTAN1,INA,TUBB4B,AGRN,MSN,SPTBN2,TPM1,TLN1,TLN2,ACTR3,TUBB2A,ARPC2,F
1,ANXA2,TIMP2,RPL23,RPS20,OAZ1,PTN,PEBP1,GPC3,RPS15,RPS7,PRKAR1A,APLP2,YWHAB,IPO
AP2M1,ATP2A2,PRKCSH,ACTN4,CTNNB1,VAMP2,GRINA,ACTN1,FYN,YWHAH,PPP1R9B,VDAC1,AE
AC1,TOP2B,HDAC2,PARP1,PKN1,CCND1,NCOR2,HNRNPD,BEX4,MTA1,KPNA2,HDAC6,HIF1A,SMG
YH9,SEPTIN2,DNAH8,TUBB4B,RHOA,KIF5B,RAB6B,RAB7A,DDX3X,ARL4C,GNB2,MYO5A,MYL6,M
B1,SNX17,APOE,SYT1,LANCL1
,NRBP2,USP22,ACTG1,EEF2,TUBA1A,XRCC2,HSP90AB1,ZNF124,HSP90AA1,ACTR2,SPARC,TUBB
TIN2,TUBB4B,ARF1,RHOA,RAB6B,EIF5,PIP4K2B,RAB7A,ARL4C,ANXA6,SEPTIN11,RHOB,RAB1A,S

2,GSK3A,PRKACA,PKIA,AKAP8L,GSK3B,ACBD3,SOX9,WASF3,PRKAR2B,PRRC1,AKAP17A
11,COL11A1,AP2B1,COL1A2,CBX5,KHDRBS1,SKP1,MAP1A,DLG5,VAMP2,OPTN,VHL,G3BP2,GRB
GRN,COL11A1,PXDN,LAMA5,COL1A2,COL5A2,COL6A2,TNC,DCN,VCAN,TGFBI,COL2A1,LAMC1,A
V1B2,COX7C,SLC25A5,ATP6V0D1,COX6A1,ATP5MC3,ATP6V1G1,ATP5MC2,ATP5MG,TMCO3,ND

,EP300,BRD4,HIF1A,EHMT1,CREBBP,USP10,MDM2,EHMT2,PSME3,GSK3B,PRMT5,CHD8,RNF20,
IZ2,MED13,ATXN7L3,THRAP3,CALCOCO1,SLC30A9,HMGA1,RBM14,MED12,RNF4,NCOA6,MED1
NPDL,RAD23B,TERF2IP,TOP1,LONP1,PCBP2,HNRNPA2B1,SSBP3,NUCKS1,WBP11,SSBP4,CNBP,
L1A2,COL5A2,COL6A2,COL2A1,COL27A1,COL16A1,COL12A1,COL7A1,COL6A3,COL3A1
A1
SHC1,GRB10
FAH1B1,MAP4,CNN3,KIF5B,LRPPRC,MACF1,CKAP5,MAP2,MAP1A,CAMSAP1,DPYSL2,MAPRE1,G
.5,GDI1,CALM1,VCP,FN1,HACD3,SET,RPL11,ANXA2,TIMP2,SBF1,DNAJA1,RPL23,IGFBP3,LLGL1,
MGB1,RPLP1,CCNI,CCND2,DDX3X,PRKAR1A,YWHAB,TRIB2,CDKN1A,GCN1,CCND1,CCNL2,CDC3
M9,HSPB1,ADCY6,HINT1,SRSF2,ITGAV,NELL2,ABL1,FEZ1,CCDC88A,C1QBP,IRS1

1,DDX3X,HNRNPDL,HNRNPH1,U2AF2,PTBP1,ILF3,SSB,LONP1,ZFR,LSM14A,MSI1,EIF4A3,CNBP,T
TIN2,TUBB4B,ARF1,RHOA,RAB6B,EIF5,PIP4K2B,RAB7A,ARL4C,ANXA6,SEPTIN11,RHOB,RAB1A,S
TIN2,TUBB4B,ARF1,RHOA,RAB6B,EIF5,PIP4K2B,RAB7A,ARL4C,ANXA6,SEPTIN11,RHOB,RAB1A,S
RS1,DARS1,RARS1,WARS1,KARS1,FARSA,TARS1,HARS1
RS1,DARS1,RARS1,WARS1,KARS1,FARSA,TARS1,HARS1
S1,MAPK8IP1,PRKAR2A,WNK1,MACROH2A1,PKIA,TRIB3,WARS1,CDKN1B,SH3BP5,PRKAR2B,CA
RT3,RAD23A
GF2,YWHAG,HMGB1,RPLP1,CCNI,CCND2,DDX3X,PRKAR1A,YWHAB,TRIB2,CDKN1A,GCN1,CCND
NR2L8,HSP90B1,IGFBP2,CALR,HSPA1B,COL5A1,CANX,CLU,DVL3,GNAS,GPI,RACK1,ATP5F1B,CA
TNNB1,CADM4,HMGCR,TERF2IP,MAPK1,GRB2,SOD1,PPP6R1,PPP1R9B,STAT3,MASTL,PPP1CC,F
NR2L8,ATP2B4,YWHAE,RACK1,CALM3,APP,RPL5,GDI1,CALM1,VCP,FN1,HACD3,SET,RPL11,ANXA
,CADM4,HMGCR,GRB2,SOD1,PPP6R1,PPP1R9B,STAT3,MASTL,PPP1CC,PPP1R15A,STRN4,SMG5

),ARPP19,PHACTR4,MGAT5,ANP32E,PPP1R14B,PPME1,PPP2R5D,PPP1R37,PPP1R12A,PPP1R26,
C9,ATP6V0D1,ATP6V1G1,ATP13A1,ATP13A3,ATP13A2,ATP1B3,ATP6V0B,ATP6V0E2,ATP6V0A1,AT
2,KPNA3,NOLC1,NUP58

.DOA,PKM,GANAB,HSP90B1,PRPF8,RPL8,YBX1,UBA1,CALR,HSPA1B,RPLP0,CANX,DDX5,SOX4,RF
G1,EEF2,DPYSL3,TUBA1A,CFL1,BTBD9,XRCC2,MAVS,HTT,HSP90AB1,ZNF124,HSP90AA1,ACTR2,
NCL,HSP90B1,NEFL,YBX1,ATP2B4,CALR,HSPA1B,PSAP,CLU,DDX5,DVL3,YWHAE,GPI,RACK1,SPTE
AE,GNAS,RACK1,SPTBN1,GNB1,MYH9,P4HB,KLRD1,FSCN1,MARCKS,NPM1,HSPD1,IGF2,EIF3C,E
VAPA,ITGB1,ANXA2,VCL,LARP1,CTNNA1,LDHA,CNN3,RPL29,EIF4H,RTN4,RPL24,RPL14,KIF5B,EI
PTAN1,IGF2,FN1,DCHS1,RPL15,VAPA,HMGB1,ITGB1,ANXA2,TIMP2,VCL,LARP1,CTNNA1,LDHA,CF
2,MED18,ACTG1,EEF2,DPYSL3,TUBA1A,CFL1,BTBD9,XRCC2,MAVS,HTT,HSP90AB1,ZNF124,HSP9
C,TMSB4X,EIF4G2,RPL3,RPS6,ZNF316,RPL4,PRRC2B,ENO1,TUBA1B,FLNA,NCL,RPL7A,ALDOA,PK
C,TMSB4X,EIF4G2,RPL3,RPS6,ZNF316,RPL4,PRRC2B,ENO1,TUBA1B,FLNA,NCL,RPL7A,ALDOA,PK
PRRC2B,ENO1,TUBA1B,FLNA,NCL,RPL7A,ALDOA,PKM,GANAB,HSP90B1,PRPF8,RPL8,YBX1,UBA1
31,MIDN,PGAM1,HNRNPA0,LDHA,PJA2,MSN,LLGL1,NCS1,RHOA,ATF4,KIF5B,ATP1A1,NISCH,PRK(
L,DNAJA1,RPL23,RTN4,XRCC5,LRPPRC,PRKAR1A,JUN,ATP6V0C,LAPTM4B,TRIB2,CDKN1A,CCT2
L,DNAJA1,RPL23,RTN4,XRCC5,LRPPRC,PRKAR1A,JUN,ATP6V0C,LAPTM4B,TRIB2,CDKN1A,CCT2
A1,RPLP0,RPL13A,RPL6,RPL13,SPTBN1,RPS8,RPL7,RPL5,SPTAN1,COL18A1,MATR3,COL4A2,RPI
GAM1,HNRNPA0,PJA2,MSN,LLGL1,NCS1,RHOA,ATF4,KIF5B,ATP1A1,NISCH,PRKCSH,KHDRBS1,PT
RPL23,RPL29,RPSA,RPS24,RPS9,RPL24,RPL17,RPL14,RPL22,RPL9,RPS14,RPS20,RPL26,RPL31,I
9,APP,CALM1,VCP,TRIM28,YWHAG,MAPT,FN1,DDB1,YWHAQ,PKD1,BASP1,VAPA,AP2M1,DNAJA1,
I2,MYH9,P4HB,FSCN1,SPTAN1,MARCKS,CALM1,MARCKSL1,MAPT,MAP9,TMSB10,RPS3,VAPA,ST
CLAF1,HSBP1,EID1,TAF15,PKN1,ASXL1,HDGF,PRPF6,CCND1,SMARCA4,TCF25,ACTN1,KDM1A,C
1-10,CTBP1,TOP2B,HDAC2,TSPYL4,SSRP1,PKN1,RBMX,CBX1,ASXL1,H3-3A,KDM6B,TOP1,SMAR
TUBB2B,PGK1,DDX5,KIF1A,GNAS,DDX17,ALDH6A1,ATP5F1B,MYH9,SEPTIN2,SMG1,HSPD1,SNRN
TUBB2B,PGK1,DDX5,KIF1A,GNAS,DDX17,ALDH6A1,ATP5F1B,MYH9,SEPTIN2,SMG1,HSPD1,SNRN
TL1,HSPA1B,TUBB2B,COL5A1,PSAP,PGK1,DDX5,KIF1A,GNAS,DDX17,ALDH6A1,ATP5F1B,MYH9,S
,PGK1,DDX5,KIF1A,GNAS,DDX17,ATP5F1B,MYH9,SEPTIN2,SMG1,HSPD1,SNRNP200,ATP5F1A,VC
SPA1B,TUBB2B,PGK1,DDX5,KIF1A,GNAS,SLC2A3,DDX17,GPI,BSG,FASN,ALDH6A1,ATP5F1B,MYH
A4,RACK1,FASN,MYH9,RPL7,SEPTIN2,APP,MARCKS,NPM1,SNRNP200,VCP,MATR3,YWHAG,MAPT
K1,GTPBP1,EIF3D,GSPT1,EIF5B,EIF2S2,EIF3H,IREB2,EIF4G3,EEF1B2,ABCF1,IGF2BP1,EIF3G,M
,PGK1,DDX5,KIF1A,GNAS,DDX17,ALDH6A1,ATP5F1B,MYH9,SEPTIN2,SMG1,HSPD1,SNRNP200,AT
,PGK1,DDX5,KIF1A,GNAS,DDX17,ALDH6A1,ATP5F1B,MYH9,SEPTIN2,SMG1,HSPD1,SNRNP200,AT
C2,TRIB2,CHD6,PARP1,PKN1,YEATS2,ASXL1,HDGF,PRPF6,TAF7,NOTCH2,VHL,CCND1,MAPK1,SN
,PGK1,DDX5,KIF1A,GNAS,DDX17,ALDH6A1,ATP5F1B,MYH9,SEPTIN2,SMG1,HSPD1,SNRNP200,AT
IBB2B,PSAP,PGK1,DDX5,KIF1A,GNAS,DDX17,ATP5F1B,MYH9,SEPTIN2,GSTP1,SMG1,HSPD1,SNR
BAG6,SERBP1,NOMO1,TIMM50,SRP72,ABCF1,MTOR,CCDC47,EIF4A3,SRPRA,YTHDF3,MDM2,CD
ARCE1,SMARCC1,WBP2,BCL9L,MTA1,EP300,RERE,BRD4,ARID1B,NCOA4,APEX1,FUS,SSBP3,CAF
P1,TNPO1,ATP6AP1,OPTN,IPO7,CORO1C,SOD1,PPP6R1,XPOT,FARP1,RAB3GAP2,IARS1,FMNL2,U
WHAH,NCOR2,ARNT2,WBP2,STAT3,EP300,PHB2,APEX1,FUS,BAZ2A,GABARAPL1,TCF3,MED13,CF
APEX1,MACROH2A2,MBD3,CREBBP,MTA2,KDM3A,TOX4,MACROH2A1,RUVBL2,NOTCH1,SRF,HS
K2,MARK1,PIN1,APOE,PPP2R2A
1,CCT3,CCT4,CDC37,CCT8,HYOU1,NAP1L4,CCT5,LMAN1,SPG7,NUDCD3,TRAP1,DNAJB2,SCAP,T
G1,HSPD1,SNRNP200,ATP5F1A,VCP,CKB,EIF4A1,DNAH8,CHD3,CDKL1,DNAJA1,CDC42BPB,ITM2E
IF2S2,EIF3H,EIF4G3,EEF1B2,ABCF1,EIF3G,EIF4G1,TCEAL4,PAIP2,TCEA1,ETF1,C1QBP,EIF3B,DH
MYH9,SMG1,HSPD1,SNRNP200,ATP5F1A,VCP,CKB,EIF4A1,DNAH8,CHD3,CDKL1,DNAJA1,CYB5R3
A8,SMARCA5,XRCC5,KIF5B,CHD4,ATP1A1,ATP2A2,CCT7,RBBP4,ACSL6,DDX3X,MYO5A,MACF1,M
,NCOR2,ARNT2,WBP2,STAT3,EP300,PHB2,APEX1,FUS,BAZ2A,GABARAPL1,MED13,THRAP3,PAGF

MYH9,SMG1,HSPD1,SNRNP200,ATP5F1A,VCP,CKB,EIF4A1,DNAH8,CHD3,CDKL1,DNAJA1,CYB5R3
C2L,MBD3,MTA2,MACROH2A1,PWWP3A,HIRA,RNF4,CABIN1,GATAD2B,SMARCC2,SMARCB1,HM
4G3,EEF1B2,ABCF1,EIF3G,EIF4G1,TCEAL4,TCEA1,ETF1,EIF3B,EIF3M,ELOB,EIF1B,EIF2S1,TCEA
1,PPP6R1,XPOT,FARP1,RAB3GAP2,FMNL2,ULK1,BRK1,ABI2,LZTR1,CSE1L,EHD1,SH3BP4,EXOC
A,MED13,THRAP3,PAGR1,SLC30A9,TRIP12,HMGA1,KDM3A,PARK7,ASAH1,NSD1,TOB2,MED25,M
AP5,MAP2,APPL1,MAP1A,TBCD,HSPH1,CAMSAP1,HDGF,DPYSL2,MAPRE1,GABARAP,CLIP3,KIF3
2,PPP1R9B,EP300,AP2A1,PHB2,PPP2CB,PPP1CC,XRCC6,HSPG2,VPS4A,SH3GL1,FBLN5,DAPK3,
CB1,HMGA2
SAP18,MTA2,HSPA1A,RUVBL2,SIN3B,MECP2,DNAJB1,HIPK2,DRAP1,ATF7IP,TRIB3,ASAH1,C1QB

3,TPM4,MSN,ENC1,WDR1,LASP1,PFN1,SPTBN2,TPM1,ACTN4,TLN1,MYO5A,ANXA6,MACF1,MTSS
CN1,LTBP4,LTBP3,ITGAV,SORT1,FGFR1,NTRK2,IGFBP4,SDCBP,ACVR1B,IGF1R,HTRA1,PDGFRB

MRPS6,KDM2B,MRPS27
L1,KDM5B,BAZ1B,BRD4,BRD2,ATRX,HJURP,NAP1L3,ANP32B,MCM3AP,TNKS,BAZ2A,CARM1,KM
ITGAV,NTRK2,AMFR,PINK1,SUMO1,PARK7,SART3,ADRM1,GSK3B,LONP2,CFLAR,XBP1,FURIN,P
CHL1,MARCHF6,SBF1,CDC42BPB,CPE,CTSD,PJA2,PSMD2,KLHL9,CSNK1E,SCRN1,PDIA6,DDR1,
MP2,OPTN,VHL,G3BP2,GRB2,MAP2K2,MAPK8IP1,SGTA,HIP1,AP2A1,YKT6,SEPTIN7,AKAP9,SEPT
,DNAJB1,ERP29,STUB1,USP19,TBCA,CDKN1B,HSPE1,DNAJB6,OGDH,PRNP,GET4

LBP,DDX3X,RPS5,RPS7,RPL35,TARDBP,HNRNPK,CIRBP,MYH10,LUC7L3,NOVA2,SRSF5,CPSF1,H
P58
ADS1,PRPF8,UGT8,UBA1,ABHD2,TUBB2B,CES3,PGK1,DDX5,ENO2,FKBP14,KAT6A,KIF1A,GNAS,
A,DICER1,FXR1,DHX36,TFRC,DGCR8
27A,EEF1A2,GNAI3,TSR1,RAB5B,RAB11B,TUBB2A,ARF3,RAB11A,CDC42,GTPBP1,GSPT1,RAB2

1,GAS7,TLN2,ACTR3,ARPC2,DSTN,CAPZB,ACTN1,TPM3,CORO1C,MPRIP,PPP1R9B,HIP1,FMNL2,

1,BAG5,LMAN2,PPP5C,HSPA1A,DNAJB1,LIMK1,STUB1,UNC45A,HSF1,DNAJC8,USP19,BCOR,CD

LEC,HIP1,CROCC,TUBB4A,ANK2,ARPC3,TUBA1C,CCDC6,ARPC5,ADD1
5,TRIB2,CDKN1A,TIMP3,ANXA5,PRDX5,SKI,RPL37,PCSK1N,HSPB1,QARS1,CCAR2,MAPK8IP1,DY
CC9,CHERP,FKBP1A,AKAP9,TCAF1,SH3GL1,PACS1,PACS2,ANK2,RIMS3,USP10,NPTN,PHPT1,HE
5,WDTC1,MTA2,HSPA1A,KAT2A,AKAP8L,DHX36,SRF,BCOR,NACC1,PHB,SUDS3,SP1,RELA
YH10,SEPTIN11,RHOB,RAB1A,SEPTIN3,RAC1,RAB27A,EEF1A2,GNAI3,TSR1,KLC1,RAB5B,RAB11

,FTH1,ZNF316,ENO1,TUBA1B,PKM,FTL,HSP90B1,UBA1,ATP2B4,CALR,FSTL1,HSPA1B,ZNF99,TU
EPTIN3,RAC1,RAB27A,EEF1A2,GNAI3,TSR1,RAB5B,RAB11B,TUBB2A,ARF3,RABL6,ARF4,RAB11

2,MAP2K2,MAPK8IP1,HIP1,AP2A1,YKT6,FKBP4,ABI2,NCSTN,SPAG9,ATP1B2,AP2A2,AP1M1,ANK2
EBP1,FBLN1,COL27A1,IGFBP7,CCN1,HSPG2,LTBP4,FBLN5,BGN,COL16A1,COL12A1,ELN,MFA
UFA4,COX8A,COX7A2L,ATP6V0B,ATP5PO,CLCN4,ATP5F1C,COX6B1,ATP6V0E2,ATP5F1E,COX4I1

,TRIM24,SMARCB1
4
RTF1,TSNAX,SUB1,DHX36,HSF1,XPC,ANXA1,HMGB2,MCM7,MSH2,TSN,FUBP1,RAD23A,MYEF2,R

ABARAP,CLIP3,KIF3C,RAB11A,DCTN1,KIF5A,KIF21B,MAP6,HDAC6,EML1,MAPRE2,MAP1LC3B,VA
PSMD2,RPS20,EIF5,OAZ1,PFN1,PTN,PEBP1,GPC3,RPS15,RPS7,JUN,MTSS2,APLP2,ARHGDIA,IPO5
7,HSPB1,MAP2K2,QARS1,MAPK8IP1,PRKAR2A,TAOK1,DAZAP2,WNK1,MACROH2A1,PKIA,CAB39,

IA1,FXR1,PABPC4,EXOSC10,ANXA1,MSI2,CBX4
SEPTIN3,RAC1,RAB27A,EEF1A2,GNAI3,TSR1,RAB5B,RAB11B,TUBB2A,ARF3,RABL6,ARF4,RAB11A
SEPTIN3,RAC1,RAB27A,EEF1A2,GNAI3,TSR1,RAB5B,RAB11B,TUBB2A,ARF3,RABL6,ARF4,RAB11A

MK2N1,HEXIM1,GMFB,LRP6,IBTK

1,CCNL2,CDC37,HSPB1,MAP2K2,QARS1,MAPK8IP1,PRKAR2A,TAOK1,DAZAP2,WNK1,MACROH2A
LM3,MYH9,P4HB,APP,KLRD1,CALM1,IGF2,ATP5F1A,VCP,YWHAG,FN1,HMGB1,AP2M1,ITGB1,TU
PP1R15A,STRN4,SMG5,SH3GL1,KIFAP3,MTMR4,ARPP19,WNK1,SNX3,PHACTR4,MAST2,KAT2A,P
2,TIMP2,SBF1,DNAJA1,PXDN,RPL23,IGFBP3,LLGL1,RHOA,PSMD2,RPS20,EIF5,OAZ1,PFN1,PTN,P
,KIFAP3,MTMR4,ARPP19,SNX3,PHACTR4,KAT2A,PTPN1,PPP1CA,PPME1,JAK1,KIF3A,DLG4,CDKN

PPP2R5B,CABIN1,DMPK,PPP1R16A,PPP2R5C
P6V1F,ATP6V1D

PL13A,RPL6,YWHAE,RPL13,DDX17,RACK1,FASN,ALDH6A1,SPTBN1,RPS8,MYH9,RPL7,P4HB,RPL5
VIM,SPARC,TUBB,FTH1,UBC,KIDINS220,TMSB4X,NUMA1,DCX,EIF4G2,RPL3,RPS6,DBN1,RPL4,P
3N1,CALM3,GNB1,P4HB,APP,RPL5,GDI1,TMBIM6,GSTP1,MARCKS,CALM1,NPM1,HSPD1,VCP,UB
EIF4B,VCP,MARCKSL1,MAPT,FN1,DDB1,EIF1,MRC2,RPS3,HMGB1,SEC61A1,ITGB1,RPLP1,TIMP2
F5,LASP1,PFN1,SPTBN2,SLC3A2,HDLBP,CDH6,CDH2,DDX3X,TLN1,CTNNB1,MACF1,HNRNPK,H1-
E,CNN3,RPL29,RPSA,EIF4H,RTN4,LAMA5,MSN,RPL24,RPL14,KIF5B,NLGN2,CXADR,EIF5,CD81,L
0AA1,ACTR2,VIM,SPARC,TUBB,FTH1,UBC,KIDINS220,TMSB4X,NUMA1,DCX,EIF4G2,RPL3,RPS6,
KM,GANAB,HSP90B1,PRPF8,RPL8,YBX1,UBA1,ATP2B4,CALR,TMF1,HSPA1B,ZNF99,TUBB2B,PGK
KM,GANAB,HSP90B1,PRPF8,RPL8,YBX1,UBA1,ATP2B4,CALR,TMF1,HSPA1B,ZNF99,TUBB2B,PGK
,CALR,TMF1,HSPA1B,ZNF99,RPLP0,H3-3B,CANX,DDX5,SOX4,RPL13A,RPL6,KAT6A,YWHAE,RPL1
CSH,KHDRBS1,PTN,PFKL,PEBP1,CCND2,CDH2,CLTC,RPS7,RPS19,PRKAR1A,CTNNB1,CADM4,RH
,SQSTM1,TCP1,SKI,GABARAP,CASC3,USP7,PCBP2,BAG6,TXNIP,TRAF4,ARIH1,PRKAR2A,PSMD1,
,SQSTM1,TCP1,SKI,GABARAP,CASC3,USP7,HGS,PCBP2,BAG6,TXNIP,TRAF4,ARIH1,PRKAR2A,PS
L37A,FN1,INA,RPL19,RPL15,RPS3,RPS4X,RPS3A,RPL18,TUBB4B,RPL11,AGRN,RPLP1,COL11A1,
TN,PEBP1,CCND2,CDH2,CLTC,RPS7,RPS19,PRKAR1A,CTNNB1,CADM4,RHOB,RAC1,ARHGDIA,TO
RPS25,RPS15,RPL12,RPL30,RPS5,RPS7,RPS19,RPL35,RPS10,RPL35A,RPLP2,RPL36,RPS12,RPS
ARF1,GTF2F1,KPNB1,RHOA,ATF4,CXADR,ATP1A1,MCL1,KHDRBS1,XPO1,CLTC,PRKAR1A,TLN1,
MN1,ITGB1,ANXA2,VCL,PAFAH1B1,MAP4,HNRNPU,CTNNA1,CNN3,TPM4,MSN,LLGL1,RHOA,CLS
CAR2,SMARCE1,SMARCC1,NCOR2,KDM3B,WBP2,AKIRIN1,TP53BP1,BCL9L,MTA1,EP300,RERE,
CA4,PBX2,SUPT5H,KDM1A,POLR2B,MSL1,SMARCE1,SMARCC1,SUPT6H,DHX30,PHC2,NCOR2,H
P200,ATP5F1A,VCP,CKB,EIF4A1,DNAH8,CHD3,CDKL1,TUBB4B,DNAJA1,ARF1,CYB5R3,CDC42BP
P200,ATP5F1A,VCP,CKB,EIF4A1,DNAH8,CHD3,CDKL1,TUBB4B,DNAJA1,ARF1,CYB5R3,CDC42BP
EPTIN2,APP,SMG1,HSPD1,SNRNP200,ATP5F1A,VCP,CKB,FN1,EIF4A1,DNAH8,CHD3,HMGB1,CDK
P,CKB,EIF4A1,DNAH8,CHD3,CDKL1,TUBB4B,DNAJA1,ARF1,CDC42BPB,ITM2B,MAT2A,HNRNPU,
9,SEPTIN2,SMG1,HSPD1,SNRNP200,ATP5F1A,VCP,CKB,EGLN1,EIF4A1,DNAH8,CHD3,CDKL1,TU
,FN1,NONO,YWHAQ,TPI1,LDHB,PDIA3,NCAM1,ANXA2,PAFAH1B1,MAT2A,HNRNPU,CTNNA1,LDHA
TOR,EIF4A3,EIF4G1,CNBP,TCEAL4,FXR2,PAIP2,TCEA1,FXR1,CYFIP1,ETF1,C1QBP,EIF3B,IGF2BP
P5F1A,VCP,CKB,EIF4A1,DNAH8,CHD3,CDKL1,TUBB4B,DNAJA1,ARF1,CYB5R3,CDC42BPB,ITM2
P5F1A,VCP,CKB,EIF4A1,DNAH8,CHD3,CDKL1,TUBB4B,DNAJA1,ARF1,CYB5R3,CDC42BPB,ITM2
ARCA4,PBX2,PRKDC,CAND1,HSPB1,KDM1A,SMARCE1,YWHAH,USP7,NCOR2,HNRNPD,ARNT2,W
P5F1A,VCP,CKB,EIF4A1,DNAH8,CHD3,CDKL1,TUBB4B,DNAJA1,ARF1,CYB5R3,CDC42BPB,ITM2
NP200,ATP5F1A,VCP,CKB,EGLN1,EIF4A1,DNAH8,CHD3,CDKL1,TUBB4B,ANXA2,AGRN,DNAJA1,A
2BP2,ETF1,PPP1CA,CSNK2B,C1QBP,SNRPD2,PRMT5,SNRPB,SRP54,OLA1,PRPF31,YBX3,EIF1B,
RM1,ZMIZ2,NUCKS1,EDF1,MED13,UBE3A,SMARCD1,CREBBP,APBB1,ATXN7L3,TRIM8,THRAP3,C
JLK1,BRK1,ABI2,SH3GL1,FNBP1L,LZTR1,CSE1L,EHD1,SH3BP4,EXOC4,BNIP3,XPO6,DAPK3,MIC
REBBP,DAPK3,THRAP3,PAGR1,SLC30A9,TRIP12,HMGA1,USF2,MTA2,CGGBP1,KDM3A,PARK7,AS
F1,PER1,CTCF,THRA,XBP1,RELA,GATAD2B,SMARCC2,SMARCB1,HMGA2,LEMD2

APBP,HSPA1A,RUVBL2,DNAJB1,SRSF10,CLPX,NACAD,HSPE1,DNAJB6,DNAJB11
3,MAT2A,HNRNPU,SMARCA5,XRCC5,PAPOLA,HK1,KIF5B,CHD4,ATP1A1,ATP2A2,CCT7,CSNK1E,P
IFR,SHMT2,EIF3M,ELOB,PAIP1,EIF1B,EIF2S1,TCEA2,CTIF,EIF3J,ELOA
3,CDC42BPB,ITM2B,MAT2A,HNRNPU,SMARCA5,XRCC5,PAPOLA,HK1,KIF5B,CHD4,ATP1A1,ATP2
YH10,ATP1A2,CCT6A,TOP2B,SMCHD1,ATP6V0C,TMEM30A,DDX39B,CHD6,CCT2,HSPH1,ABCA3,
1,SLC30A9,TRIP12,HMGA1,KDM3A,PARK7,ASAH1,NSD1,TOB2,SRF,HSF1,GSK3B,MED25,MED12

3,CDC42BPB,ITM2B,MAT2A,HNRNPU,SMARCA5,XRCC5,PAPOLA,HK1,KIF5B,CHD4,ATP1A1,ATP2A
IGA2
2,EIF3J,ELOA
4,XPO6,DAPK3,MICAL1,RAB11FIP3,RANGAP1,RIMS3,DENND5A,SGSM2,RGPD5,SGSM3,TNFAIP1
ED12,RXRA,PRMT2,CTBP2,MED16,XBP1,PCNA,NCOA6,TRIM24,CNOT1,MED14
C,FYN,STMN3,RAB11A,CCT5,BEX4,DCTN1,KIF5A,KIF21B,MAP6,DIP2B,HDAC6,SMC3,EML1,MAPR
SAE1,PRDX3,CALCOCO1,ABL1,SDCBP,BANF1,RABAC1,MAPRE3,RAB3A,DLG4,ERBB2,MED12,PHE

P,PBXIP1,RBFOX2,NSD1,TOB2,PA2G4,RCOR2,BCOR,PRMT5,IRF2BP2,HIRA,PHB,CITED2,CTBP2,

2,TAGLN,MYH10,COTL1,MAP1A,CNN2,GAS7,TLN2,ACTR3,HDGF,ARPC2,DSTN,PLEC,CAPZB,ACTI
,ERBB2,NRP2,SHC1,TGFBR1,API5,FURIN,CRIM1,FGFBP3,COL3A1,EPHA2

T2E,SPTY2D1,MSH6,NOC2L,KDM4A,NASP,APBB1,PHIP,DEK,WDTC1,RSF1,TSPYL2,ANP32E,SAMD
RNP,COL3A1,RAD23A,CSTB
MMP14,B4GALNT4,CKAP5,UBE2H,HECTD4,CDK16,HDAC2,MMP2,TRIB2,PPIB,GLUL,CDKN1A,CTI
IN5,FKBP4,ABI2,SH3PXD2A,NCSTN,SPAG9,ATP1B2,AP2A2,AP1M1,SMARCD1,SH3PXD2B,EIF4G1,

NRNPL,HNRNPA3,RBMX,SNRNP70,G3BP2,SRSF1,NYNRIN,PTBP1,ILF3,SUPT5H,HNRNPM,SSB,SR

DX17,GPI,HTRA4,FASN,ALDH6A1,ATP5F1B,CALM3,GNB1,MYH9,SEPTIN2,P4HB,IDS,GSTP1,SMG

,EIF5B,SEPTIN7,SEPTIN5,RAB1B,RAB14,GNL1,GNA12,EFTUD2,RAB6A,RAB5C,GNA13,TUBB4A,S

CTTN,ANTXR1,CAPZA1,CLASP2,MICAL1,PLS3,ABL1,ARPC1A,HIP1R,CYFIP1,FLII,ARPC3,MAP1S,AF

N1B,DNAJB6,OGDH,PDXP

NLL1,TXNIP,PPP1R9B,PRKAR2A,CST3,OAZ2,SPINT2,SLIT2,A2M,PPP1R10,ARPP19,WNK1,PRDX3,
RPUD1,YWHAZ,FGF13,FHL1,PRNP,ID2

B,TUBB2A,ARF3,KIF3C,RAB11A,CDC42,KIF5A,GTPBP1,KIF21B,GSPT1,RAB2A,EIF5B,SMC3,SEPT

BB2B,COL5A1,PSAP,PGK1,CANX,DDX5,ENO2,FKBP14,KAT6A,KIF1A,GNAS,DDX17,ATP5F1B,CAL
,CDC42,GTPBP1,GSPT1,RAB2A,GLUD1,EIF5B,SEPTIN7,SEPTIN5,RAB1B,RAB14,GNL1,GNA12,EF

,AMFR,PTPN11,SUN1,SDCBP,HIP1R,OBSL1,ARRDC4,MAPK8IP2,ATP1B3,SUN2,NUP153,NCK2,ST
4,EMILIN1,PCOLCE,COL7A1,THBS2,ANOS1,NID1,COL6A3,MXRA5,MFGE8,COL3A1,POSTN,MFAF
,ATP6V0A1,COX5A,COX7A2,CLCN3,ATP6V1F,NNT,ATP6V1D,COX5B,ATP5PD

AD50

PB,VPS41,TUBGCP6,CLASP2,REEP2,REEP1,HDGFL3,PPP5C,CLIP2,KIFC2,MAST2,KIF26A,MAP4K
,TRIB2,TIMP3,EID1,NUCB1,TBCD,GDI2,HSPH1,PRDX5,TBC1D16,ACAP3,SKI,ABCA2,RAPGEF1,R
CCNY,CSNK2B,TRIB3,CCNG1,CDKN1B,SH3BP5,PRKAR2B,CAMK2N1,HEXIM1,GMFB,FGF13,TPX

,CDC42,GTPBP1,GSPT1,RAB2A,GLUD1,EIF5B,SEPTIN7,SEPTIN5,RAB1B,RAB14,GNL1,GNA12,E
,CDC42,GTPBP1,GSPT1,RAB2A,GLUD1,EIF5B,SEPTIN7,SEPTIN5,RAB1B,RAB14,GNL1,GNA12,E

A1,PKIA,CAB39,CCNY,CSNK2B,TRIB3,CCNG1,WARS1,CDKN1B,SH3BP5,PRKAR2B,CAMK2N1,HE
BB4B,UCHL1,TIMP2,DNAJA1,PXDN,CPE,LAMA5,MSN,NLGN2,CXADR,CD81,ATP2A2,BEX3,CLEC7
TPN1,PPP1CA,PPME1,JAK1,KIF3A,DLG4,CDKN1B,STAT1,ERBB2,CSRNP2,MAPK3,TRAF3,FER,TR
EBP1,GPC3,RAB7A,RPS15,RPS7,JUN,MTSS2,APLP2,RAC1,ARHGDIA,IPO5,TRIB2,TIMP3,EID1,NU
1B,STAT1,ERBB2,TRAF3,FER,PPP2R2A,PPP1R16A

,FSCN1,SMG1,NPM1,HSPD1,SNRNP200,CSDE1,EIF3C,ATP5F1A,EIF4B,VCP,TRIM28,MATR3,YWH
RRC2B,ENO1,TUBA1B,PIK3R2,FLNA,MAP1B,NCL,RPL7A,SLC9A7,ALDOA,MTRNR2L8,PKM,GANA
B,TRIM28,YWHAG,CKB,MAPT,FN1,HACD3,EGLN1,TPI1,PKD1,SERINC1,RPS3,HMGB1,ITGB1,UCH
,VCL,PAFAH1B1,CDC42BPB,AP2B1,GTF2F1,HNRNPU,SMARCA5,LARP1,CTNNA1,RPSA,TPM4,EIF
-10,YWHAB,CKAP5,SND1,RAB1A,RPL23A,CNN2,CDH11,GCN1,RAB11B,MAPRE1,PLEC,PRDX1,AR
ASP1,PFN1,NISCH,SPTBN2,SLC3A2,PTN,HDLBP,CDH6,CDH2,CCN2,ACTN4,MMP14,DDX3X,TLN1,
ZNF316,DBN1,RPL4,PRRC2B,ENO1,TUBA1B,PIK3R2,FLNA,MAP1B,NCL,RPL7A,SLC9A7,ALDOA,M
1,RPLP0,H3-3B,CANX,DDX5,SOX4,RPL13A,RPL6,KAT6A,YWHAE,KIF1A,RPL13,GNAS,DDX17,RAC
1,RPLP0,H3-3B,CANX,DDX5,SOX4,RPL13A,RPL6,KAT6A,YWHAE,KIF1A,RPL13,GNAS,DDX17,RAC
3,DDX17,RACK1,FASN,ALDH6A1,SPTBN1,RPS8,MYH9,RPL7,P4HB,APP,RPL5,FSCN1,SMG1,NPM1
IOB,RAC1,ARHGDIA,TOP2B,APPL1,LAPTM4B,TRIB2,CDKN1A,SRSF5,EEF1A2,PPP1CB,SQSTM1,P
UBA52,UQCRC1,HDAC6,CUL3,RPS27A,NPLOC4,HIF1A,SLC25A5,MAP1LC3B,PRDX6,ABI2,SMG5,
MD1,UBA52,UQCRC1,HDAC6,CUL3,RPS27A,NPLOC4,HIF1A,SLC25A5,MAP1LC3B,PRDX6,ABI2,SM
RPL10,RPS17,VCL,MAP4,PXDN,CTNNA1,RPS16,RPL23,RPL29,RPSA,TPM4,LAMA5,RPS24,RPS9,M
P2B,APPL1,TRIB2,CDKN1A,SRSF5,EEF1A2,PPP1CB,SQSTM1,PARP1,NOX4,PKN1,GCN1,SKI,DPY
27,RPL23A,RPL27,RPL36A,RPS15A,RPL38,RPL37,RPS11,RPL28,RPS28,FAU,RPL39,RPS13,RPL3
JA1,JUN,CTNNB1,NDFIP1,HNRNPK,YWHAB,LAMP1,CCT6A,CTBP1,SKP1,L1CAM,RAB27A,U2AF2,
TN1,KIF5B,RAB6B,ENC1,ATP1A1,WDR1,CRMP1,LASP1,PFN1,LRPPRC,SPTBN2,TPM1,ACTN4,DDX
DM5B,SF1,BRD4,ARID1B,NCOA4,APEX1,FUS,SSBP3,CARM1,RALY,ZMIZ2,TLE1,HDGFL2,SRSF2,
NRNPD,KDM3B,WBP2,NAP1L4,CIC,STAT3,NAP1L1,MTA1,EP300,RERE,H2AZ1,BRD4,BRD2,ATRX,
B,ITM2B,MAT2A,HNRNPU,SMARCA5,LDHA,XRCC5,PAPOLA,HK1,RHOA,KIF5B,RAB6B,CHD4,ATP1
B,ITM2B,MAT2A,HNRNPU,SMARCA5,LDHA,XRCC5,PAPOLA,HK1,RHOA,KIF5B,RAB6B,CHD4,ATP1
L1,TUBB4B,AGRN,COL11A1,DNAJA1,ARF1,CYB5R3,PAFAH1B1,CDC42BPB,ITM2B,MAT2A,HNRN
SMARCA5,XRCC5,PAPOLA,HK1,RHOA,KIF5B,RAB6B,CHD4,ATP1A1,EIF5,ATP2A2,CCT7,CSNK1E,
B4B,DNAJA1,ARF1,CYB5R3,CDC42BPB,ITM2B,MAT2A,HNRNPU,SMARCA5,LDHA,SCARB2,XRCC
,TPM4,SCARB2,RTN4,COL1A2,RPL22,CBX5,KIF5B,NLGN2,CXADR,CRMP1,BEX3,CCT7,MCL1,TL
2,DHFR,SHMT2,EIF3M,ELOB,PAIP1,EIF1B,EIF2S1,TCEA2,CTIF,EIF3J,ELOA
3,MAT2A,HNRNPU,SMARCA5,XRCC5,PAPOLA,HK1,RHOA,KIF5B,RAB6B,CHD4,ATP1A1,EIF5,ATP2
3,MAT2A,HNRNPU,SMARCA5,XRCC5,PAPOLA,HK1,RHOA,KIF5B,RAB6B,CHD4,ATP1A1,EIF5,ATP2
BP2,STAT3,EP300,PBX1,PHB2,CXXC5,APEX1,HIF1A,FUS,BAZ2A,PELP1,GABARAPL1,TLE1,NUCKS
3,MAT2A,HNRNPU,SMARCA5,XRCC5,PAPOLA,HK1,RHOA,KIF5B,RAB6B,CHD4,ATP1A1,EIF5,ATP2
RF1,CYB5R3,CDC42BPB,ITM2B,MAT2A,HNRNPU,SMARCA5,XRCC5,PAPOLA,GAP43,HK1,RHOA,K
ABCE1,NOLC1,TMCO1,EIF2S1,PES1,SECISBP2L,MRPS27,NEMF,PRMT7
ALCOCO1,SLC30A9,UBE2L3,HMGA1,ABL1,MTA2,NIBAN2,NOTCH1,PARK7,SUB1,PDLIM1,KAT2A,
L1,AMBRA1,RAB11FIP3,RANGAP1,RIMS3,DENND5A,SGSM2,CLTA,RGPD5,SGSM3,TNFAIP1,EXO
AH1,LDB1,NSD1,TOB2,SRF,HSF1,GSK3B,CRTC1,SOX9,MED25,MED12,RXRA,PRMT2,CTBP2,PS

FKL,PEBP1,DDR1,PIP4K2B,ACSL6,DDX3X,ITM2C,MYO5A,MYH10,NARS1,UBE2H,ATP1A2,CCT6A,

2,CCT7,PFN1,CSNK1E,PFKL,PEBP1,DDR1,PIP4K2B,ACSL6,DDX3X,ITM2C,PRKAR1A,MYO5A,MY
HSPA4,TCP1,CCT3,DDX42,CHD2,CCT4,ABCA2,SMARCA4,KIF3C,DHX30,ATP6V1B2,CCT8,LONP1,
,RXRA,PRMT2,CTBP2,PSMA6,MED16,XBP1,PCNA,RELA,NCOA6,TRIM24,SMARCB1,CDK5RAP3,H

A2,CCT7,CSNK1E,PFKL,PEBP1,DDR1,PIP4K2B,ACSL6,DDX3X,ITM2C,PRKAR1A,MYO5A,MYH10,NA

,EXOC5,CYFIP1,ANKFY1,MYCBP2,RNF41,MICAL3,DAAM1,NDRG1,RANBP9,GOLGA4,RCC2,KIF3A

E2,BRSK1,MAP1LC3B,NDN,TAOK1,VAPB,VPS41,GABARAPL1,TUBGCP6,CLASP2,REEP2,SYT11,R
3,SHANK1,KPNA3,AFDN,MDC1,PCNA,SP1,DYNLT1,SNX17,MSH2,SYT1,CENPF,DNM1

AJUBA,RYBP,CBX4,TLE4,HMGA2,DPF2,SIN3A,MYBBP1A

N1,FLNB,TPM3,CORO1C,YWHAH,MPRIP,PFN2,FLNC,PPP1R9B,PALLD,HIP1,EVL,HDAC6,FMNL2,C

D11,SART3,H2AX,STAT1,CHD8,RNF20,KDM5A,HIRA,WDR5,SPIN1,BRPF3,CBX4,TRIM24,SMARCC2

DSP2,CTSB,RNF187,PPP1CB,SQSTM1,PARP1,PTPRA,UBE4A,PKN1,PLOD1,PTK7,PSMB2,KDM6B,
ANK2,AMFR,PTPN11,SUN1,SDCBP,HIP1R,RUVBL2,OBSL1,ARRDC4,MAPK8IP2,ATP1B3,SUN2,NU

SF3,STRAP,PCBP2,HNRNPD,SERBP1,RPS13,CCT5,SF1,EIF3D,GRSF1,ZFP36L1,SRSF6,TRA2B,HN

1,CALM1,TTC3,PLD3,HSPD1,USP11,SNRNP200,ATP5F1A,VCP,TRIM28,CKB,HACD3,EGLN1,TPI1,

RPRA,ARF5,AGAP1,MFN2,RAN,RHEB,ARHGAP5,RAP2A,RAB18,TUBA1C,RAB3A,DIRAS1,ARL8A,NK

RPC5,AFDN,CAPZA2,ADD1,AJUBA,VPS16,GSN

WDTC1,MGAT5,MACROH2A1,ANP32E,PKIA,NOTCH1,PPP1R14B,USP14,FAF2,PPME1,TRIB3,SPOC

IN7,SEPTIN5,RAB1B,RAB14,GNL1,GNA12,EFTUD2,RAB6A,RAB5C,GNA13,TUBB4A,SRPRA,ARF5,

M3,MYH9,SEPTIN2,APP,IDS,SPTAN1,GSTP1,SMG1,CALM1,TTC3,HSPD1,COL18A1,SNRNP200,ATF
TUD2,RAB6A,RAB5C,EHD1,GNA13,TUBB4A,SRPRA,ARF5,AGAP1,MFN2,RAN,RHEB,ARHGAP5,RA

UB1,HERPUD1,NAPB,IRS1,NAPA,DGCR8,SRRT,SHC1,SHANK1,ZSWIM8,NOLC1,FBXO7,MAPK8IP3
2

4,SUN2,CCDC88A,CETN2,NDRG1,MAPRE3,RCC2,MAP1S,KIF3A,GABARAPL2,OPA1,FGF13,TPX2,F
PL37,AZIN1,PSMD3,IPO7,PCSK1N,LARS1,FBLN1,SEC23A,ARFGEF2,DYNLL1,PPP1R9B,BCR,TRIO,
2,CCNL1,CCNK,GPRC5B,IBTK

TUD2,RAB6A,RAB5C,EHD1,GNA13,TUBB4A,SRPRA,ARF5,AGAP1,MFN2,RAN,RHEB,ARHGAP5,RAI

TUD2,RAB6A,RAB5C,EHD1,GNA13,TUBB4A,SRPRA,ARF5,AGAP1,MFN2,RAN,RHEB,ARHGAP5,RAI

KIM1,GMFB,FGF13,TPX2,CCNL1,CCNK,GPRC5B,LRP6,PIK3R3,IBTK
,NES,NISCH,PTN,GLG1,CCN2,ACTN4,MMP14,CLTC,APLP1,MIF,TLN1,WLS,NARS1,CADM4,HLA-B
PC4AP,PPP2R2A,PPP1R16A
CB1,RAB27A,TBCD,GDI2,HSPH1,PRDX5,TBC1D16,ACAP3,SKI,ABCA2,RAPGEF1,RPL37,AZIN1,PSI

AG,RPL37A,MAPT,SPEN,NONO,EIF1,EIF4A1,RPL19,RPL15,SF3B1,PDIA3,RPS3,CHD3,HMGB1,TPT
3,COL6A1,FTL,HSP90B1,NEFL,NEFM,PRPF8,RPL8,FBXL4,YBX1,UBA1,ATP2B4,IGFBP2,CALR,FST
IL1,RPL11,ANXA2,TIMP2,MIDN,MARCHF6,VCL,DNAJA1,PAFAH1B1,AKAP12,CDC42BPB,GTF2F1,H
4H,LAMA5,XRCC5,PTCD3,CBX5,CXADR,CHD4,CD81,WDR1,HMGN2,LASP1,RBBP4,NES,NISCH,PR
GLU1,EIF3E,CAPZB,FLNB,CCT8,CSNK1D,ATXN2L,IDH1,MPRIP,SERBP1,BMPR2,RPL34,CTNND1,C
CTNNB1,MACF1,HNRNPK,H1-10,YWHAB,CKAP5,SND1,RAB1A,RPL23A,CNN2,CDH11,TLN2,TGFBI
MTRNR2L8,PKM,GANAB,COL6A1,FTL,HSP90B1,NEFL,NEFM,PRPF8,RPL8,FBXL4,YBX1,UBA1,ATP
K1,FASN,ALDH6A1,SPTBN1,ATP5F1B,RPS8,MYH9,RPL7,SEPTIN2,P4HB,APP,RPL5,FSCN1,SMG1,N
K1,FASN,ALDH6A1,SPTBN1,ATP5F1B,RPS8,MYH9,RPL7,SEPTIN2,P4HB,APP,RPL5,FSCN1,SMG1,N
,HSPD1,SNRNP200,CSDE1,EIF3C,ATP5F1A,EIF4B,VCP,TRIM28,MATR3,YWHAG,RPL37A,MAPT,SP
ARP1,NOX4,PKN1,GCN1,SKI,DPYSL2,CCND1,MAPK1,MAPRE1,SRSF1,ADAM9,GRB2,CDC37,HSPB
GABARAPL1,DNAJB2,ATXN3,MOAP1,SYT11,KDM4A,PRKACA,APBB1,BAG5,AMBRA1,UBE2K,RANG
MG5,GABARAPL1,DNAJB2,ATXN3,MOAP1,SYT11,KDM4A,PRKACA,APBB1,BAG5,AMBRA1,UBE2K,
ISN,RPL24,RPL17,COL1A2,LLGL1,RPL14,RPL22,RPL9,RPS14,RPS20,COL5A2,RPL26,PNN,RPL31
SL2,CCND1,MAPK1,MAPRE1,SRSF1,ADAM9,GRB2,CDC37,HSPB1,PLCG1,QARS1,DUSP4,PITPNM1
4,UBA52,RPL7L1,RPS2,RPS27A,RPS26,RPS21,RPL27A,MRPS6,RPS29,RPL36AL,RPL32,MRPL3,M
SQSTM1,GNAI3,RBMX,SKI,SCAF1,TOP1,RAPGEF1,SRSF9,ADAM9,GRB2,PRKDC,FKBP8,MAP2K2,
3X,ARL4C,TLN1,GJA1,MYO5A,ANXA6,MACF1,MTSS2,TAGLN,MYH10,CKAP5,MAP2,COTL1,APPL1
NUCKS1,EDF1,MED13,NOC2L,UBE3A,SMARCD1,CREBBP,SAP18,APBB1,ATXN7L3,TRIM8,THRAP
EP400,ARID1B,UPF1,SMC3,NAP1L3,MCM3AP,KLHDC3,APEX1,NDN,BAP1,FUS,PELP1,MACROH2A
A1,EIF5,ATP2A2,CCT7,PFN1,CSNK1E,PFKL,PEBP1,DDR1,PIP4K2B,ACSL6,RAB7A,DDX3X,ITM2C,
A1,EIF5,ATP2A2,CCT7,PFN1,CSNK1E,PFKL,PEBP1,DDR1,PIP4K2B,ACSL6,RAB7A,DDX3X,ITM2C,
IPU,SMARCA5,RPL29,XRCC5,PAPOLA,GAP43,HK1,RPL22,RHOA,KIF5B,RAB6B,CHD4,ATP1A1,EIF
PFKL,PEBP1,DDR1,PIP4K2B,ACSL6,RAB7A,DDX3X,ITM2C,ARL4C,MYO5A,ANXA6,MYH10,NARS1,S
5,PAPOLA,HK1,RHOA,KIF5B,RAB6B,CHD4,ATP1A1,EIF5,CD81,ATP2A2,CCT7,PFN1,ASTN2,XPR1,
E5,NISCH,HNRNPC,KHDRBS1,PFKL,TKT,SFPQ,TPM1,CDH2,PIP4K2B,ACSL6,ACTN4,APLP1,IK,RPS

A2,CCT7,PFN1,CSNK1E,PFKL,PEBP1,DDR1,PIP4K2B,ACSL6,RAB7A,DDX3X,ITM2C,ARL4C,PRKAR
A2,CCT7,CSNK1E,PFKL,PEBP1,DDR1,PIP4K2B,ACSL6,RAB7A,DDX3X,ITM2C,ARL4C,PRKAR1A,MY
1,TCF3,EDF1,PSMD4,RBPJ,MED13,MTOR,CREBBP,SAP18,APBB1,UBE2I,PSMC1,DAPK3,THRAP3,
A2,CCT7,CSNK1E,PFKL,PEBP1,DDR1,PIP4K2B,ACSL6,RAB7A,DDX3X,ITM2C,ARL4C,PRKAR1A,MY
IF5B,RAB6B,CHD4,ATP1A1,EIF5,ATP2A2,CCT7,PFN1,ASTN2,XPR1,CSNK1E,PFKL,TKT,PEBP1,DD

HIPK2,DRAP1,TRIM27,NME2,BCL9,PSIP1,NUP98,RRP1B,RNF20,KDM5A,RBM14,XPC,MED12,RN
C5,CYFIP1,ANKFY1,MYCBP2,RNF41,MICAL3,DAAM1,NDRG1,RANBP9,GOLGA4,RCC2,KIF3A,RAL
IA6,MED16,XBP1,PCNA,SP1,RELA,NCOA6,TRIM24,SMARCB1,CDK5RAP3,HES6,CNOT1,MED14

CDK16,TOP2B,SMCHD1,TRIB2,GLUL,DDX39B,CHD6,CCT2,HSPH1,ABCA3,ACTR3,PDXK,PKN1,HS

10,NARS1,UBE2H,ATP1A2,CCT6A,HMGCR,CDK16,TOP2B,SMCHD1,TRIB2,GLUL,DDX39B,CHD6,
HYOU1,PFN2,CCT5,YME1L1,DHX15,KIF5A,KIF21B,ATRX,EP400,UPF1,SMC3,ABCC9,SPG7,XRCC
ES6,CNOT1,MED14

\RS1,UBE2H,ATP1A2,CCT6A,HMGCR,CDK16,TOP2B,SMCHD1,TRIB2,GLUL,DDX39B,CHD6,CCT2,

\,RALBP1,CDC42EP4,CDC42SE1,INF2,RUSC2,TNPO2,AFDN,ROCK2,STRIP1,YBX3,TNPO3,MICALL

EEP1,HDGFL3,PPP5C,CLIP2,DAG1,KIFC2,MAST2,KIF26A,ADNP,GPAA1,MAP4K4,FEZ1,ATF5,SUN2

ALD1,CTTN,TMOD2,ANTXR1,RDX,CAPZA1,CLASP2,WASF2,CAP1,VASH1,MICAL1,CROCC,PLS3,AE

,BRD1,PRMT7,DPF2

NRDC,TOP1,RNF10,PPP2CA,CCND1,MAPK1,AEBP1,ADAM9,PRKDC,UBE2R2,NPEPPS,KDM1A,FKB
P153,NCK2,STUB1,EIF3B,HERPUD1,PDE4DIP,NAPB,IRS1,NAPA,DGCR8,SRRT,SHC1,SHANK1,ZSW

RNPA2B1,RPS2,HNRNPAB,PCBP4,FUS,EIF2S2,ELAVL1,IREB2,LSM14A,EIF4G3,LSM14B,RBM5,IG

CHSY1,EIF4A1,PPP2R1A,GTF3C4,LDHB,DNAH8,PDIA3,RPS3,CHD3,HMGB1,POLR2A,CDKL1,TUBE

KIRAS2,SRP54,RHOBTB2,GNAZ,RAB5A,SEPTIN9,RAB35,OPA1,ARF6,DNM1,GNAL,RAP1B,GNAO1,(

CK1,PSMF1,SERPINE2,WARS1,CDKN1B,SH3BP5,PPP1R37,PRKAR2B,CDC42SE1,COL7A1,ANOS1,I

AGAP1,KIFC2,MFN2,KIF26A,RAN,DYNLRB1,RHEB,ARHGAP5,RAP2A,RAB18,APPBP2,TUBA1C,RAB

P5F1A,VCP,TRIM28,MATR3,CKB,RPL37A,DCHS1,EGLN1,CHSY1,EIF4A1,DNAH8,ZNF43,CHD3,POLI
P2A,RTCB,RAB18,SEPHS1,TUBA1C,RAB3A,DIRAS1,ARL8A,NKIRAS2,SRP54,RHOBTB2,GNAZ,OLA

,PSMC6,GRB10,RAD50,STX12

PRNP,CENPF,DNM1,CCSER2,ZNF207
PSMD1,FLCN,CST3,FARP1,RAB3GAP2,TBC1D9B,OAZ2,SPINT2,GPSM1,PPP1R15A,RABEP1,SLIT2

P2A,RTCB,NME2,RAB18,SEPHS1,TUBA1C,RAB3A,DIRAS1,ARL8A,NKIRAS2,SRP54,RHOBTB2,GNA
P2A,RTCB,NME2,RAB18,SEPHS1,TUBA1C,RAB3A,DIRAS1,ARL8A,NKIRAS2,SRP54,RHOBTB2,GNA

,TTR,DNER,SQSTM1,GNAI3,TLN2,TGFBI,VPS35,COL2A1,MDK,ASXL1,HDGF,VEGFA,PPP2CA,ADAM

MD3,IPO7,PCSK1N,LARS1,YWHAH,MTRNR2L10,FBLN1,ID4,SEC23A,ARFGEF2,DYNLL1,CDC42,PP

1,RPS4X,RPS3A,RPL18,TUBB4B,RPL11,BTF3,ANXA2,RPL10,RPS17,ARF1,MAP4,SF3B2,GTF2F1,
L1,TMF1,HSPA1B,TUBB2B,COL5A1,COL4A1,PSAP,PGK1,RPLP0,H3-3B,CANX,CLU,PTMA,DDX5,E
NRNPU,PGAM1,HNRNPA0,RPL23,LDHA,SCARB2,RTN4,XRCC5,PJA2,MSN,KPNB1,COL1A2,LLGL1
KCSH,HNRNPC,KHDRBS1,PTN,TPM1,EIF5A,DDR1,RAB7A,CCN2,ACTN4,MMP14,DDX3X,GNB2,H1-
TNND2,PI4KA,UBFD1,TAGLN2,YKT6,CEMIP2,FMNL2,RPS2,CALD1,SEPTIN7,CTTN,LRRC59,PRDX
,PTK7,GCN1,RAB11B,MAPRE1,PLEC,PRDX1,ARGLU1,ADAM9,EIF3E,CAPZB,ACTN1,FLNB,CCT8,F
2B4,IGFBP2,CALR,FSTL1,TMF1,ABHD2,HSPA1B,ZNF99,TUBB2B,COL5A1,COL4A1,PSAP,PGK1,R
NPM1,HSPD1,SNRNP200,CSDE1,EIF3C,ATP5F1A,EIF4B,VCP,TRIM28,MATR3,YWHAG,CKB,RPL37
NPM1,HSPD1,SNRNP200,CSDE1,EIF3C,ATP5F1A,EIF4B,VCP,TRIM28,MATR3,YWHAG,CKB,RPL37
EN,NONO,DDB1,EIF1,SET,EIF4A1,RPL19,RPL15,GTF3C4,SF3B1,PDIA3,SSBP2,ZNF43,RPS3,CHD
1,PLCG1,QARS1,DUSP4,PFKM,PITPNM1,STXBP1,UBQLN1,MAPK8IP1,DGKD,BMPR2,CDC42,PPP
AP1,UBE2L3,PINK1,HSPA1A,ACVR1B,MFN2,MDM2,CUL1,SUMO1,FAF2,CUL4A,UBE2J1,TRIB3,JA
RANGAP1,UBE2L3,PINK1,HSPA1A,ACVR1B,MFN2,MDM2,CUL1,SUMO1,FAF2,CUL4A,UBE2J1,TRI
,SPTBN2,RPS25,TPM1,RPS15,RPL12,RPL30,CLTC,RPS5,RPS7,RPS19,RPL35,TLN1,COL6A2,MA
,STXBP1,MAPK8IP1,BMPR2,CDC42,CTNND1,VDAC1,TRAF4,DCTN1,STAT3,PRMT1,PRKAR2A,AP2
RPL28
,FYN,HNRNPM,YWHAH,STMN3,STXBP1,HGS,CDC42,SYP,TRAF4,DFFB,PRKAR2A,CNTNAP1,ATRX,
,MAP1A,DLG5,CNN2,RAB27A,TBCD,VAMP2,GAS7,TLN2,HSPH1,ACTR3,CAMSAP1,RAB11B,HDGF,
3,FBXL19,CALCOCO1,SLC30A9,UBE2L3,HMGA1,ABL1,RSF1,MTA2,KDM3A,HDGFL3,NIBAN2,HS
2,NUCKS1,MSH6,NSD2,RBPJ,PCGF2,NOC2L,MBD3,YY1,SMARCD1,CREBBP,APBB1,CENPB,SLC3
RL4C,PRKAR1A,MYO5A,ANXA6,MYH10,NARS1,SEPTIN11,RHOB,UBE2H,ATP1A2,RAB1A,SEPTIN3
RL4C,PRKAR1A,MYO5A,ANXA6,MYH10,NARS1,SEPTIN11,RHOB,UBE2H,ATP1A2,RAB1A,SEPTIN3
,ATP2A2,CCT7,PTN,CSNK1E,PFKL,PEBP1,DDR1,SULF1,PIP4K2B,ACSL6,RAB7A,CCN2,ACTN4,D
EPTIN11,RHOB,UBE2H,ATP1A2,RAB1A,SEPTIN3,CCT6A,RAC1,CDK16,TOP2B,SMCHD1,TRIB2,GL
SNK1E,PFKL,TKT,PEBP1,DDR1,PIP4K2B,ACSL6,RAB7A,ACTN4,DDX3X,ITM2C,ARL4C,HMGCS1,
19,MIF,HMGCS1,JUN,ANXA6,APLP2,TARDBP,ARL6IP1,ST3GAL2,WLS,HNRNPK,YWHAB,HNRNPH1

1A,MYO5A,ANXA6,MYH10,NARS1,SEPTIN11,RHOB,UBE2H,ATP1A2,RAB1A,SEPTIN3,CCT6A,HMG
O5A,ANXA6,MYH10,NARS1,SEPTIN11,RHOB,UBE2H,ATP1A2,RAB1A,SEPTIN3,CCT6A,HMGCR,RA
,PAGR1,SLC30A9,TRIP12,LMO4,HMGA1,USF2,MTA2,CGGBP1,KDM3A,RUVBL2,MECP2,SUMO1,
O5A,ANXA6,MYH10,NARS1,SEPTIN11,RHOB,UBE2H,ATP1A2,RAB1A,SEPTIN3,CCT6A,HMGCR,RA
1,PIP4K2B,ACSL6,RAB7A,DDX3X,ITM2C,ARL4C,PRKAR1A,MYO5A,ANXA6,SESN3,MTSS2,MYH10,

4,CITED2,PRMT2,SETD3,CTBP2,SS18L1,ARL2BP,MED16,HMGB2,NCOA6,MAML1,TRIM24,SMA
BP1,CDC42EP4,CDC42SE1,INF2,RUSC2,TNPO2,AFDN,ROCK2,STRIP1,YBX3,TNPO3,MICALL1,FG

A4,TCP1,ETNK1,CCT3,DDX42,PTK7,CHD2,CCT4,ABCA2,ATP6AP1,TOP1,MARS1,MAPK1,SMARC

CCT2,HSPH1,ABCA3,ACTR3,PDXK,PKN1,HSPA4,TCP1,ETNK1,CCT3,DDX42,PTK7,CHD2,CCT4,AB
,CLCN7,VPS4A,TRAP1,DICER1,ABCF1,MSH6,ATP6V0D1,PSMC3,DHX38,ATP6V1G1,PSMC1,EIF4A

HSPH1,ABCA3,ACTR3,PDXK,PKN1,HSPA4,TCP1,ETNK1,CCT3,DDX42,PTK7,CHD2,CCT4,ABCA2,AT

1,FGD1,AP3M1,NUTF2,UNC13B

,CCDC88A,CETN2,NDRG1,MAPRE3,RCC2,MAP1S,KIF3A,TBCA,STMN4,ARL8A,CACYBP,GABARAPI

3L1,PHACTR4,ARPC1A,DAG1,HIP1R,CYFIP1,PDLIM1,MAEA,FLII,CCDC88A,NCK2,MICAL3,ARPC3,D

P8,MAP2K2,DUSP4,XPNPEP3,FYN,NKTR,PTP4A2,EIF3F,TNKS2,LONP1,CSNK1D,USP7,HECTD1,PT
IM8,SEPTIN9,NOLC1,FBXO7,MAPK8IP3,PSMC6,GRB10,RAD50,STX12,RB1CC1,CNOT1

F2BP1,EIF4A3,EIF4G1,RPS26,CNBP,ELAVL3,PUM1,FXR2,LUC7L2,PAIP2,YTHDF3,RBM4,TIA1,RBM

34B,UCHL1,MARCHF6,SBF1,CYB5R3,CDC42BPB,MAT2A,PXDN,OGA,PGAM1,SMARCA5,FADS2,LD

GNAI1

PPP1R12A,CAMK2N1,GPS1,ANXA1,PPP1R26,COL6A3,HEXIM1,CABIN1,GMFB,ABCE1,FURIN,CRIM

3A,DIRAS1,KIF3A,ARL8A,NKIRAS2,SRP54,RHOBTB2,GNAZ,RAB5A,PIN1,SEPTIN9,RAB35,OPA1,AF

R2A,CDKL1,TPT1,ITGB1,TUBB4B,ANXA2,AGRN,ZMIZ1,TIMP2,XAF1,COL11A1,MARCHF6,DNAJA1,
1,RAB5A,SEPTIN9,RAB35,GNL3,OPA1,NOLC1,THG1L,ARF6,DNM1,GNAL,RAP1B,GNAO1,GNAI1

,A2M,PPP1R10,CYTH2,ARHGEF17,TBC1D9,DNAJB2,EEF1B2,RIC8A,AHSA1,ATP1B2,SH3BP4,COX

Z,OLA1,RAB5A,SEPTIN9,RAB35,GNL3,OPA1,NOLC1,THG1L,ARF6,DNM1,GNAL,RAP1B,GNAO1,GI

Z,OLA1,RAB5A,SEPTIN9,RAB35,GNL3,OPA1,NOLC1,THG1L,ARF6,DNM1,GNAL,RAP1B,GNAO1,GI

M9,GRB2,PCSK1N,GABARAP,ACTN1,PLCG1,ARF4,FYN,YWHAH,SLIT3,MTRNR2L10,FBLN1,PITPNM

P1R9B,BCR,TRIO,PSMD1,FLCN,CST3,FARP1,RAB3GAP2,TBC1D9B,OAZ2,ABCC9,SPINT2,GPSM1

HNRNPU,LARP1,HNRNPA0,CTNNA1,RPS16,RPL23,RPL29,RPSA,EIF4H,RTN4,RPS24,RPS9,XRCC5,I
NO2,SOX4,FKBP14,MAGED1,RPL6,KAT6A,DVL3,YWHAE,KIF1A,RPL13,GNAS,SLC2A3,DDX17,GPI,
,NCS1,RHOA,ATF4,KIF5B,CHD4,ATP1A1,JUND,ATP2A2,RBBP4,PFN1,NISCH,PRKCSH,LRPPRC,KH
-0,TLN1,JUN,MYO5A,COL6A2,ANXA6,MACF1,SLC25A3,ARID1A,TAGLN,H1-10,MYH10,YWHAB,CK/
6,VAPB,RDX,SH3GL1,STK24,CAPZA1,FNBP1L,AHSA1,WASF2,SCRIB,IST1,EHD1,CHMP4B,LYPLA2,
BLN1,CSNK1D,ATXN2L,IDH1,MPRIP,SERBP1,BMPR2,RPL34,CTNND1,PALLD,KIRREL1,CTNND2,F
PLP0,H3-3B,CANX,CLU,PTMA,DDX5,ENO2,SOX4,FKBP14,RPL13A,MAGED1,RPL6,KAT6A,DVL3,YV
A,MAPT,SPEN,NONO,DDB1,EIF1,SET,EIF4A1,RPL19,RPL15,GTF3C4,DNAH8,SF3B1,PDIA3,SSBP2,Z
A,MAPT,SPEN,NONO,DDB1,EIF1,SET,EIF4A1,RPL19,RPL15,GTF3C4,DNAH8,SF3B1,PDIA3,SSBP2,Z
3,BASP1,HMGB1,POLR2A,TPT1,RPS4X,RPS3A,RPL18,TUBB4B,RPL11,BTF3,ANXA2,RPL10,RPS17
1R9B,CTNND1,VDAC1,TRAF4,DCTN1,STAT3,PRMT1,PRKAR2A,AP2A1,ADCY6,PTEN,SV2A,PPP1CC,
K1,HM13,STUB1,YWHAZ,GSK3B,PA2G4,USP19,OTUB1,RNF20,PRKAR2B,UBXN1,PER1,CACYBP,U
33,JAK1,HM13,STUB1,STAT2,YWHAZ,GSK3B,PA2G4,USP19,OTUB1,STAT1,RNF20,PRKAR2B,UBX
CF1,RPS10,RPL35A,RPLP2,MYL6,RPL36,RPS12,MAP2,RPS27,TNC,RPL23A,DCN,MAP1A,VCAN,TLN
A1,ADCY6,PTEN,SV2A,PPP1CC,MAPRE2,BRSK1,HIF1A,PPP1R15A,SHC2,ELAVL1,HINT1,SRSF2,TR

EVL,TRA2B,PTEN,PPP1CC,CUL3,HIF1A,RABEP1,DAP,EHMT1,VAPB,GPX1,STRN4,ABI2,VPS4A,SRP7
DPYSL2,PPP2CA,MAPRE1,ARPC2,DSTN,PLEC,CAPZB,GABARAP,ACTN1,CLIP3,FLNB,TPM3,CORO
PA1A,RUVBL2,SIN3B,NOTCH1,MECP2,DNAJB1,PARK7,TRRAP,EWSR1,SUB1,PDLIM1,KAT2A,HIPK2
0A9,HMGA1,AUTS2,MTA2,KDM3A,TOX4,TSPYL2,MACROH2A1,RUVBL2,SIN3B,NOTCH1,MECP2,S
,CCT6A,CTBP1,HMGCR,RAC1,CDK16,TOP2B,SMCHD1,TRIB2,GLUL,SQLE,DDX39B,CHD6,RAB27
,CCT6A,CTBP1,HMGCR,RAC1,CDK16,TOP2B,SMCHD1,TRIB2,GLUL,SQLE,DDX39B,CHD6,RAB27
DX3X,ITM2C,ARL4C,APLP1,PRKAR1A,MYO5A,ANXA6,APLP2,MYH10,NARS1,SEPTIN11,RHOB,UBE2
UL,DDX39B,CHD6,RAB27A,EEF1A2,CCT2,GNAI3,TSR1,HSPH1,ABCA3,ACTR3,PDXK,PKN1,HSPA4,
PRKAR1A,MYO5A,ANXA6,MYH10,NARS1,SEPTIN11,RHOB,UBE2H,ATP1A2,RAB1A,SEPTIN3,CCT6A
I,TTR,SEPTIN3,CTBP1,HMGCR,APPL1,SMCHD1,GLUL,HSBP1,CRPPA,PGRMC1,DDX39B,SQSTM1,

CR,RAC1,CDK16,TOP2B,SMCHD1,TRIB2,GLUL,DDX39B,CHD6,RAB27A,EEF1A2,CCT2,GNAI3,TSR
C1,CDK16,TOP2B,SMCHD1,TRIB2,GLUL,DDX39B,CHD6,RAB27A,EEF1A2,CCT2,GNAI3,TSR1,HSPI
PARK7,KAT2A,DRAP1,CSNK2B,ASAH1,C1QBP,LDB1,CDK9,RBFOX2,PSMC5,PSMC2,NSD1,TOB2,S
C1,CDK16,TOP2B,SMCHD1,TRIB2,GLUL,DDX39B,CHD6,RAB27A,EEF1A2,CCT2,GNAI3,TSR1,HSPI
,NARS1,SEPTIN11,RHOB,UBE2H,ATP1A2,RAB1A,SEPTIN3,CCT6A,HMGCR,RAC1,ARHGDIA,CDK16

RCC2,SMARCB1,CEBPZ,SIN3A,MED14
D1,AP3M1,NUTF2,UNC13B

A4,DYNC1LI2,PRKDC,UBE2R2,MAP2K2,QARS1,KIF3C,DHX30,ATP6V1B2,FYN,LARS1,CCT8,PFKM,

CA2,ATP6AP1,TOP1,MARS1,MAPK1,SMARCA4,DYNC1LI2,ACADVL,PRKDC,UBE2R2,MAP2K2,QARS
3,DHX40,ATP13A1,TAPBP,KIFC2,ATP13A3,MTREX,HSPA1A,RUVBL2,ATP13A2,NSF,KIF26A,SUB1,A

TP6AP1,TOP1,MARS1,MAPK1,SMARCA4,DYNC1LI2,ACADVL,PRKDC,UBE2R2,MAP2K2,QARS1,KIF

L2,GIT1,OPA1,FGF13,TPX2,PRNP,CENPF,DNM1,CCSER2,ZNF207,EMD

AAM1,MAP1S,WASF3,INF2,RUSC1,ARPC5,SETD3,AFDN,EPB41L1,CAPZA2,ADD1,MIB2,AJUBA,GM

P4A1,PTPRS,P3H3,KDM3B,BMPR2,KDM4B,CDC42,PPP1R9B,YME1L1,PSMA7,TRAF7,ARIH1,EIF2

M25,MECP2,CLUH,FXR1,PARK7,SYNCRIP,PABPC4,ETF1,HNRNPR,DDX6,C1QBP,CELF3,DHX36,RE

HA,CPE,CTSD,LPCAT1,XRCC5,PJA2,PAPOLA,HK1,RHOA,PSMD2,KIF5B,RAB6B,CHD4,ATP1A1,CR

11,PRNP,LRP6,CRB2,CSTB,IBTK

RF6,DYNC1I2,DNM1,GNAL,RAP1B,GNAO1,GNAI1,MYL6B

,ARF1,CYB5R3,CDC42BPB,ITM2B,ZNF770,MAT2A,PXDN,HNRNPU,SMARCA5,CPE,TPM4,SCARB2,

6A1,ARPP19,DENND4B,BAG5,PRDX3,RANGAP1,DENND5A,UBE2L3,AGAP1,PINK1,SGSM2,PHACTF

NAI1

NAI1

M1,MESD,STRAP,BAG6,NCOR2,VGF,ARNT2,ARFGEF2,CDC42,PPP1R9B,CTNND1,TRAF4,STAT3,HI

,DKK3,PPP1R15A,RABEP1,SLIT2,A2M,AKAP9,PPP1R10,CYTH2,ARHGEF17,TBC1D9,DNAJB2,EEF1

MSN,RPL24,KPNB1,RPL17,PAPOLA,RPL14,RPL22,PTCD3,RPL9,RPS14,RPS20,EIF5,HMGN2,RPL2
HTRA4,RACK1,BSG,FASN,SPTBN1,ATP5F1B,CALM3,GNB1,SHISA9,STMN2,RPS8,MYH9,RPL7,SEF
IDRBS1,XPO1,PTN,PFKL,SFPQ,PEBP1,CCND2,CDH2,ACSL6,CLTC,GNB2,RPS7,RPS19,PRKAR1A,J
AP5,SND1,COTL1,RAC1,APPL1,HDAC2,PPIB,RPL23A,DCN,CDKN1A,DDX39B,SRP9,CTSB,RPN2,SQ
USO1,RPS26,RANGAP1,SNX2,PTPN11,KTN1,NOP56,NIBAN2,CLIC1,HSPA1A,PFKP,DNAJB1,PARK7
PI4KA,UBFD1,TAGLN2,YKT6,CEMIP2,FMNL2,PTPRF,ITGB5,RPS2,CALD1,SEPTIN7,CTTN,LRRC59,I
WHAE,KIF1A,RPL13,GNAS,SLC2A3,DDX17,GPI,HTRA4,RACK1,BSG,FASN,ALDH6A1,SPTBN1,ATP5F
ZNF43,RPS3,CHD3,BASP1,HMGB1,POLR2A,CDKL1,TPT1,RPS4X,RPS3A,RPL18,TUBB4B,RPL11,B
ZNF43,RPS3,CHD3,BASP1,HMGB1,POLR2A,CDKL1,TPT1,RPS4X,RPS3A,RPL18,TUBB4B,RPL11,B
,ARF1,MAP4,SF3B2,ZNF770,GTF2F1,HNRNPU,NFE2L1,SMARCA5,LARP1,HNRNPA0,CTNNA1,RPS
,MAPRE2,BRSK1,HIF1A,PPP1R15A,SLC2A1,SHC2,ABI2,ELAVL1,TAX1BP1,HINT1,SRSF2,TRAP1,SP
UBE2G1,ATF6,ELOB,GABARAPL2,TRAF3,XBP1,CUL7,RELA,CUL4B,FBXO7,NEDD8,FZD8,RNF40,A
KN1,PER1,CACYBP,UBE2G1,ATF6,ELOB,GABARAPL2,TRAF3,XBP1,CUL7,RELA,CUL4B,FBXO7,NEI
N2,TGFBI,RPL27,ACTR3,COL2A1,LAMC1,RPL36A,RPS15A,RPL38,TUBB2A,RPL37,ARPC2,PLEC,AE
AP1,SPAG9,CLASP2,TCF3,PDPK1,DAZAP2,ITGAV,ADAM10,FBXW5,GSK3A,MTOR,PRKACA,AP2A2,

72,SH3GL1,RAB6A,DICER1,WASF2,DAZAP2,WBP11,ADAM10,IST1,MTOR,PRKACA,AP2A2,EXOC4,U
1C,KIF3C,FYN,YWHAH,STMN3,MPRIP,MAPK8IP1,PFN2,RAB11A,FLNC,CCT5,PPP1R9B,PALLD,BEX
2,DRAP1,ATF7IP,TRIM27,TRIB3,NME2,ASAH1,C1QBP,PBXIP1,LDB1,BCL9,RBFOX2,PSIP1,NUP98,I
SAMD11,ADNP,KAT2A,ATF5,EHMT2,RAN,TASOR,CSNK2B,LDB1,CDK9,PWWP3A,PSIP1,NUP98,NSD
A,EEF1A2,CCT2,PGD,GNAI3,PARP1,TSR1,NOX4,HSPH1,ABCA3,ACTR3,PDXK,PKN1,HSPA4,TCP1,E
A,EEF1A2,CCT2,PGD,GNAI3,PARP1,TSR1,NOX4,HSPH1,ABCA3,ACTR3,PDXK,PKN1,HSPA4,TCP1,E
2H,ATP1A2,RAB1A,SEPTIN3,CCT6A,HMGCR,RAC1,CDK16,TOP2B,SMCHD1,TNC,TRIB2,GLUL,DCN
TCP1,ETNK1,CCT3,DDX42,PTK7,RAB5B,CHD2,CCT4,RAB11B,ABCA2,ATP6AP1,TOP1,MARS1,TUE
,CTBP1,HMGCR,RAC1,CDK16,TOP2B,SMCHD1,TRIB2,GLUL,SQLE,DDX39B,CHD6,RAB27A,EEF1A
,VAMP2,GAS7,NOP53,PARP1,TGFBI,LMNB2,PDXK,AHCYL1,COL2A1,RBMX,CBX1,SKI,DPYSL2,VEC

1,HSPH1,ABCA3,ACTR3,PDXK,PKN1,HSPA4,TCP1,ETNK1,CCT3,DDX42,PTK7,RAB5B,CHD2,CCT4,
H1,ABCA3,ACTR3,PDXK,PKN1,HSPA4,TCP1,ETNK1,CCT3,DDX42,PTK7,RAB5B,CHD2,CCT4,RAB1
RF,YWHAZ,HSF1,GSK3B,CRTC1,PDCD11,BCOR,SOX9,MED25,FBL,CEBPG,PER1,MED12,RXRA,RI
H1,ABCA3,ACTR3,PDXK,PKN1,HSPA4,TCP1,ETNK1,CCT3,DDX42,PTK7,RAB5B,CHD2,CCT4,RAB1
,TOP2B,SMCHD1,TRIB2,GLUL,SQLE,DDX39B,CHD6,RAB27A,EEF1A2,CCT2,GNAI3,TSR1,NOX4,H

LONP1,CSNK1D,PRPS1,HYOU1,SARS1,DGKD,BMPR2,CCT5,YME1L1,DHX15,KIF5A,EIF2AK1,BCR

S1,KIF3C,DHX30,ATP6V1B2,FYN,LARS1,CCT8,PFKM,LONP1,CSNK1D,PRPS1,HYOU1,SARS1,DGKI
ATP1B3,WRNIP1,DDX6,ATP6V0B,DHX36,PSMC5,CLPX,PSMC2,PSMC4,DDX41,ACSL4,SETX,KIF3A,(

3C,DHX30,ATP6V1B2,FYN,LARS1,CCT8,PFKM,LONP1,CSNK1D,PRPS1,HYOU1,SARS1,DGKD,BMF

IFB,VPS16,NF2,GSN,TLNRD1,EMD

AK1,PRMT1,BCR,CAPN2,TXNDC5,CPD,KDM5C,TRIO,EP300,KDM5B,PSMD1,BAZ1B,CSNK2A1,MM

3FOX2,NUP98,NCBP2,IGF2BP2,DHFR,SHMT2,RNF20,RBM14,THOC2,YBX3,FYTTD1,GNL3,FUBP1,

MP1,VAT1,RBBP4,KLHL9,SLC3A2,CSNK1E,PFKL,SCRN1,TKT,PDIA6,DDR1,SULF1,PIP4K2B,ACSL6

LPCAT1,XRCC5,PJA2,KPNB1,PAPOLA,IGFBP3,COL1A2,GAP43,HK1,NCS1,RHOA,ZNF865,CLSTN1

R4,MGAT5,RGL2,RGPD5,ANP32E,ARHGEF40,FZR1,SGSM3,DNAJB1,PPP1R14B,USP14,ACAP2,ATI

IP1,AP2A1,LRRC4B,FLOT1,PLXNB1,SFRP1,RER1,PTEN,FEM1B,CUL3,IL6ST,ITGB5,TIMM50,DKK3

1B2,RIC8A,AHSA1,ATP1B2,FXYD6,SH3BP4,COX6A1,ARPP19,DENND4B,WNK1,BAG5,PRDX3,RANG

26,SRSF4,PFN1,PNN,RPL31,PRKCSH,HNRNPC,LRPPRC,KHDRBS1,XPO1,SLC3A2,CSNK1E,HDLBP,

PTIN2,P4HB,APP,RPL5,KLRD1,GDI1,FSCN1,TMBIM6,SPTAN1,GSTP1,MARCKS,SMG1,CALM1,NPM

UN,MYO5A,CTNNB1,MTSS2,CYFIP2,YWHAB,PTGES3,CADM4,LAMP1,RHOB,COTL1,HMGCR,RAC

STM1,VAMP2,GNAI3,GAS7,TLN2,TGFBI,SSRP1,ACTR3,CAMSAP1,RAD23B,GCN1,H3-3A,PRPF6,TA

7,PDLIM1,GLOD4,PTPN1,RAN,KLC2,SNX1,PPP1CA,PPME1,DDX6,CRKL,NDRG1,SLK,UNC45A,YWH

PRDX6,VAPB,CCN1,LTBP4,RDX,SH3GL1,STK24,CAPZA1,FNBP1L,AHSA1,WASF2,ITGAV,SCRIB,ADA

F1B,CALM3,GNB1,SHISA9,STMN2,RPS8,MYH9,RPL7,SEPTIN2,P4HB,APP,IDS,RPL5,KLRD1,GDI1,F

TF3,ANXA2,RPL10,RPS17,DNAJA1,ARF1,CYB5R3,MAP4,SF3B2,CDC42BPB,ITM2B,ZNF770,MAT2A

TF3,ANXA2,RPL10,RPS17,DNAJA1,ARF1,CYB5R3,MAP4,SF3B2,CDC42BPB,ITM2B,ZNF770,MAT2A

16,RPL23,RPL29,RPSA,EIF4H,NACA,RTN4,RPS24,RPS9,XRCC5,MSN,RPL24,KPNB1,RPL17,PAPOL

AG9,CLASP2,TCF3,PDPK1,DAZAP2,ITGAV,ADAM10,FBXW5,GSK3A,MTOR,PRKACA,AP2A2,TTC28,

NAPC2,TOLLIP

OD8,FZD8,CDK5RAP3,RNF40,ANAPC2,TOLLIP

EBP1,RPS11,ACTN1,RPL28,RPS28,FAU,FBLN1,COL27A1,RPL39,RPS13,IGFBP7,RPL34,HIP1,UBA5

WNK1,BAG5,MICAL1,PRDX3,NELL2,PTPN23,ANK2,PTPN11,ABL1,PINK1,TAOK2,MACROH2A1,PKI

UBE2I,DAPK3,MICAL1,WBP1,SH3PXD2B,CALCOCO1,TCEAL4,USP47,PTPN11,ABL1,CHMP1A,PINK

K4,DCTN1,KIF5A,KCNQ2,CAPN2,KIRREL1,EP300,HIP1,KIF21B,MAP6,PPP2CB,DIP2B,FARP1,EVL,H

NSD1,TOB2,PA2G4,RCOR2,BCOR,PRMT5,RRP1B,IRF2BP2,KDM2A,RNF20,KDM5A,RBM14,XPC,N

01,MEIS1,SRF,WAC,HSF1,ZEB1,WDR82,SHMT2,ASH1L,SOX9,STAT1,CHD8,RNF20,PER1,MED12,HI

ETNK1,CCT3,DDX42,PTK7,RAB5B,CHD2,CCT4,RAB11B,HDGF,ABCA2,ATP6AP1,TOP1,MARS1,TUB

ETNK1,CCT3,DDX42,PTK7,RAB5B,CHD2,CCT4,RAB11B,HDGF,ABCA2,ATP6AP1,TOP1,MARS1,TUB

1,DDX39B,CTSB,CHD6,RAB27A,EEF1A2,CCT2,GNAI3,VCAN,TSR1,HSPH1,ABCA3,ACTR3,PDXK,PKN

B2A,ARF3,MAPK1,SMARCA4,RABL6,DYNC1LI2,PRKDC,UBE2R2,MAP2K2,QARS1,ARF4,KIF3C,DH

A2,CCT2,PGD,GNAI3,PARP1,TSR1,NOX4,HSPH1,ABCA3,ACTR3,PDXK,PKN1,HSPA4,TCP1,PLOD1,E

GFA,OPTN,MAPK1,MAPRE1,PRDX1,ODC1,DYNC1LI2,ACADVL,DYNLL2,GRB2,ATXN10,ACTN1,HSP

RAB11B,ABCA2,ATP6AP1,TOP1,MARS1,TUBB2A,ARF3,MAPK1,SMARCA4,RABL6,DYNC1LI2,ACAD

1B,ABCA2,ATP6AP1,TOP1,MARS1,TUBB2A,ARF3,MAPK1,SMARCA4,RABL6,DYNC1LI2,ACADVL,PF

NF4,PRMT2,CTCF,PSMD10,CTBP2,THRA,PSMA6,MED16,ADD1,XBP1,PCNA,TFDP1,SP1,HMGB2,R

1B,ABCA2,ATP6AP1,TOP1,MARS1,TUBB2A,ARF3,MAPK1,SMARCA4,RABL6,DYNC1LI2,ACADVL,PF

SPH1,ABCA3,ACTR3,PDXK,PKN1,HSPA4,TCP1,PLOD1,ETNK1,CCT3,DDX42,PTK7,RAB5B,CHD2,C

,ACTR1A,TRIO,VARS1,BAZ1B,CSNK2A1,KIF21B,CSNK1G2,ATRX,GARS1,EP400,PAPSS1,GLUD1,U

D,BMPR2,CCT5,YME1L1,DHX15,KIF5A,EIF2AK1,BCR,ACTR1A,TRIO,PRKAR2A,VARS1,BAZ1B,CSN

CLCN6,RALBP1,HSPE1,ATP5F1C,CHD8,ATP6V0E2,LONP2,SKIV2L,OLA1,ATP6V0A1,ANXA1,RAD54

PR2,CCT5,YME1L1,DHX15,KIF5A,EIF2AK1,BCR,ACTR1A,TRIO,PRKAR2A,VARS1,BAZ1B,CSNK2A1,

MP15,BRD4,MGAT4B,PPM1G,BRD2,PPP2CB,KLHL42,CSNK1G2,NXN,SFRP1,UBE2G2,FBXO11,MA

,RNPS1,SCAF8,MYEF2,MARF1,SECISBP2L,RNF40,JRK,MEX3D

,RAB7A,MMP14,PRDX2,DDX3X,ARL4C,GNB2,MIF,HMGCS1,GPAT4,PRKAR1A,CHPF,MYO5A,SESN

,NR2F1,KIF5B,RAB6B,CHD4,ATP1A1,EIF5,COL5A2,ATP2A2,LASP1,BEX3,CCT7,VAT1,PFN1,ASTN2

P1B3,RALGDS,RASA4,TBC1D10B,MYCBP2,ARHGAP5,IQSEC1,CCDC88A,PPME1,ARFGAP2,TBC1D

3,GALNT11,FKBP1A,RABEP1,SLIT2,A2M,AKAP9,GRK2,FUS,GRN,SLIT1,PALM,CCN1,YARS1,HSPG2

GAP1,DENND5A,UBE2L3,AGAP1,PINK1,SGSM2,PHACTR4,MGAT5,RGL2,RGPD5,ANP32E,ARHGEF4

RPS25,SFPQ,EIF5A,PEBP1,ACTN4,DDX3X,RPS15,RPL12,RPL30,CLTC,EIF3L,H1-0,RPS5,RPS7,RP

11,PLXNB2,TTC3,PLD3,HSPD1,USP11,COL18A1,SNRNP200,CSDE1,SMIM14,IGF2,EIF3C,ATP5F1A

1,ARHGDIA,TOP2B,APPL1,HDAC2,IPO5,ATP6V0C,LAPTM4B,TRIB2,PPIB,CDKN1A,TIMP3,EID1,SRS

AF7,CCND1,SMARCA4,ARPC2,DSTN,AEBP1,ADAM9,DYNLL2,GRB2,CAPZB,ACTN1,TPM3,PLTP,CO

HAZ,RARS1,STAT1,TMPO,OLA1,AFDN,CHMP5,ANXA1,SEPTIN9,ADD1,ZC3H15,MICALL1,OBSCN,PD

AM10,IST1,EHD1,FBLN5,CHMP4B,LYPLA2,USO1,RPS26,RANGAP1,SNX2,PTPN11,COL16A1,SDCB

SCN1,TMBIM6,SPTAN1,GSTP1,MARCKS,SMG1,CALM1,NPM1,PLXNB2,TTC3,PLD3,HSPD1,USP11

A,GTF2F1,PXDN,HNRNPU,NFE2L1,SMARCA5,LARP1,HNRNPA0,CTNNA1,RPS16,RPL23,LDHA,RPL

A,GTF2F1,PXDN,HNRNPU,NFE2L1,SMARCA5,LARP1,HNRNPA0,CTNNA1,RPS16,RPL23,LDHA,RPL

LA,RPL14,RPL22,PTCD3,ZNF865,RPL9,ATF4,NR2F1,RPS14,CHD4,JUND,RPS20,EIF5,HMGN2,RPL

WNK1,BAG5,MICAL1,PRDX3,NELL2,PTPN23,ANK2,PTPN11,ABL1,PINK1,TAOK2,MACROH2A1,PK

52,RPL7L1,POM121C,RPS2,RPS27A,SEPTIN7,SEPTIN5,CCN1,HSPG2,LTBP4,GNL1,FBLN5,CROC

IA,ACTA2,MAPK8IP2,STK39,NSF,FEZ1,CCNY,PTPN1,RHEB,PDGFRB,CCDC88A,PPME1,TBC1D14,

1,TRAK1,DAG1,HIP1R,HSPA1A,GRIK5,MDM2,MECP2,NSF,BANF1,EHMT2,SHISA5,ARHGAP5,CCD

IDAC6,SMC3,FMNL2,EML1,MAPRE2,BRSK1,CALD1,CTTN,MAP1LC3B,NDN,TMOD2,FUS,TAOK1,V

MED12,HIRA,PHB,RNF4,CITED2,PRMT2,SETD3,CTBP2,SS18L1,ARL2BP,MED16,AJUBA,HMGB2,N

IRA,RNF4,CITED2,CTCF,CTBP2,MEIS3,THRA,PBRM1,XBP1,PCNA,AJUBA,CABIN1,RELA,GATAD2B,

B2A,ARF3,MAPK1,SMARCA4,RABL6,DYNC1LI2,ACADVL,PRKDC,UBE2R2,KDM1A,MAP2K2,QARS

B2A,ARF3,MAPK1,SMARCA4,RABL6,DYNC1LI2,ACADVL,PRKDC,UBE2R2,KDM1A,MAP2K2,QARS

N1,HSPA4,TCP1,ETNK1,CCT3,COL2A1,MDK,DDX42,PTK7,RAB5B,CHD2,CCT4,RAB11B,HDGF,ABC

IX30,ATP6V1B2,FYN,LARS1,CCT8,PFKM,LONP1,CSNK1D,PRPS1,HYOU1,SARS1,DGKD,RAB11A,B

ETNK1,CCT3,DDX42,PTK7,RAB5B,CHD2,CCT4,RAB11B,HDGF,ABCA2,PYGB,ATP6AP1,TOP1,MAR

B1,BNIP3L,CASC3,FKBP8,FLNB,SDF4,XPNPEP3,FYN,PHC2,YWHAH,PFKM,FBLN1,EIF3F,B2M,STX

VL,PRKDC,UBE2R2,MAP2K2,QARS1,ARF4,KIF3C,DHX30,ATP6V1B2,FYN,LARS1,CCT8,PFKM,LO

KDC,UBE2R2,MAP2K2,QARS1,ARF4,KIF3C,DHX30,ATP6V1B2,FYN,LARS1,CCT8,PFKM,LONP1,C

ELA,NOLC1,NCOA6,SETD1A,TRIM24,CENPF,ID3,SMARCB1,CDK5RAP3,HMGA2,MAFB,NOP58,SIN

KDC,UBE2R2,MAP2K2,QARS1,ARF4,KIF3C,DHX30,ATP6V1B2,FYN,LARS1,CCT8,PFKM,LONP1,C

CT4,RAB11B,ABCA2,PYGB,ATP6AP1,TOP1,MARS1,TUBB2A,ARF3,MAPK1,SMARCA4,RABL6,DYN

PF1,PI4KA,UBE2G2,ADCY6,IARS1,MASTL,SMC3,ULK1,ABCC9,BRSK1,MVD,GRK2,STK35,SPG7,A

K2A1,KIF21B,CSNK1G2,ATRX,GARS1,EP400,PAPSS1,GLUD1,UPF1,PI4KA,UBE2G2,ADCY6,IARS1,

L2,ABCE1,MCM7,MSH2,ABCD3,ACIN1,PSMC6,DDX19A,ATP6V1F,DDX56,RAD50,ATP8A2,ATP6V1

KIF21B,CSNK1G2,ATRX,GARS1,EP400,PAPSS1,GLUD1,UPF1,PI4KA,UBE2G2,ADCY6,IARS1,MAST

STL,MBTPS1,HDAC6,PTEN,YKT6,ECEL1,PPP1CC,CUL3,PTPRF,ULK1,MCM3AP,PREPL,BRSK1,TIM

3,B4GALNT4,ST3GAL2,MYL6,MYH10,NARS1,PTGES3,CKAP5,SEPTIN11,SND1,RHOB,UBE2H,AT

,CLEC7A,PRKCSH,TADA2B,XPR1,CSNK1E,PFKL,TKT,CDH6,PEBP1,CDH2,DDR1,SULF1,PIP4K2B,

14,TRIB3,ADRM1,SYNGAP1,PPP2R5D,PSME3,CLPX,SPOCK1,PCOLCE,PSMF1,SERPINE2,RAB3A,F

2,SHC2,LINGO1,SEC14L1,LTBP4,GABARAPL1,GNA12,SH3GL1,TRAP1,FLOT2,NCSTN,SEMA3C,PD

0,FZR1,SGSM3,DNAJB1,PPP1R14B,SUMO1,USP14,ACAP2,ATP1B3,PHPT1,RALGDS,RASA4,TBC1

S19,RPL35,JUN,MYO5A,NGRN,MACF1,RPS10,RPL35A,HNRNPDL,TARDBP,HNRNPK,H1-10,CIRBP
,EIF4B,VCP,MARCKSL1,UBB,AFF4,TRIM28,MATR3,YWHAG,CKB,COL4A2,RPL37A,LRRC28,MAPT
SF5,U2AF2,EEF1A2,CCT2,CPSF1,PPP1CB,SQSTM1,PABPN1,GDI2,PARP1,NOX4,PKN1,TCP1,NCKA
RO1C,SMARCE1,SMARCC1,SUPT6H,FYN,FBLN1,STRAP,MPRIP,UQCRC2,BAG6,NCOR2,ARNT2,SE
XDC1,EPHA2,CC2D1A,MMP24,EMD
P,KTN1,NOP56,NRXN2,NIBAN2,CLIC1,HSPA1A,PFKP,NPTN,DNAJB1,JAM3,PARK7,PDLIM1,GLOD
,COL18A1,SNRNP200,CSDE1,SMIM14,IGF2,EIF3C,ATP5F1A,EIF4B,VCP,MARCKSL1,UBB,AFF4,T
29,RPSA,EIF4H,NACA,RTN4,RPS24,RPS9,XRCC5,MSN,RPL24,KPNB1,RPL17,PAPOLA,RPL14,HK1,
29,RPSA,EIF4H,SCARB2,NACA,RTN4,RPS24,RPS9,XRCC5,MSN,RPL24,KPNB1,RPL17,PAPOLA,RP
26,SRSF4,RBBP4,PFN1,PNN,RPL31,PRKCSH,HNRNPC,LRPPRC,KHDRBS1,XPO1,SLC3A2,CSNK1
IA,ACTA2,MAPK8IP2,STK39,NSF,CAB39,PARK7,FEZ1,ATF5,CCNY,PTPN1,RHEB,PDGFRB,CCDC88

,TUBB4A,RPS26,ANK2,BGN,HMGA1,COL16A1,CLTA,COL12A1,DAG1,ELN,MAPK8IP2,PSMD13,C
NF41,TRIM27,TRIB3,C1QBP,MAPKAPK2,CDK9,TOM1L2,MAPRE3,OXSR1,ZBTB4,LIMS1,YWHAZ,

88A,CSNK2B,SRSF10,RNF41,DDX6,PMEPA1,LSM4,SYNGAP1,LDB1,STUB1,SRSF7,YWHAZ,RCC2
APB,FKBP4,ANTXR1,VPS41,GABARAPL1,RAB14,RDX,KIFAP3,CAPZA1,TUBGCP6,RAB6A,SPAG9,P
OA6,RYBP,CBX4,KDM2B,MAML1,TRIM24,SMARCC2,AKIRIN2,SMARCB1,CEBPZ,TLE4,BTG1,HM
MSH2,MBD6,NCOA6,CBX4,TRIM24,SMARCC2,CENPF,PAF1,DNMT1,SMARCB1,TLE4,HMGA2,UB
1,ARF4,KIF3C,DHX30,ATP6V1B2,FYN,SDHA,LARS1,CCT8,PFKM,LONP1,CSNK1D,PRPS1,IDH1,HY
1,ARF4,KIF3C,DHX30,ATP6V1B2,FYN,SDHA,LARS1,CCT8,PFKM,LONP1,CSNK1D,PRPS1,IDH1,HY
A2,ATP6AP1,TOP1,VEGFA,MARS1,TUBB2A,ARF3,MAPK1,SMARCA4,RABL6,DYNC1LI2,ACADVL,P
MPR2,CCT5,CDC42,YME1L1,DHX15,KIF5A,EIF2AK1,BCR,ACTR1A,TRIO,VARS1,BAZ1B,GTPBP1,
S1,TUBB2A,ARF3,MAPK1,SMARCA4,DDAH2,RABL6,DYNC1LI2,ACADVL,RBP1,PRKDC,UBE2R2,KD
BP1,LONP1,UBQLN1,PRPS1,MESD,IDH1,SARS1,SOD1,DGKD,BAG6,FLNC,DYNLL1,CDC42,SYP,G

NP1,CSNK1D,PRPS1,HYOU1,SARS1,DGKD,RAB11A,BMPR2,CCT5,CDC42,YME1L1,DHX15,KIF5A,
SNK1D,PRPS1,HYOU1,SARS1,DGKD,RAB11A,BMPR2,CCT5,CDC42,YME1L1,DHX15,KIF5A,EIF2A
3A,HES6,CNOT1,MYBBP1A,MED14
SNK1D,PRPS1,HYOU1,SARS1,DGKD,RAB11A,BMPR2,CCT5,CDC42,YME1L1,DHX15,KIF5A,EIF2A
1LI2,ACADVL,PRKDC,UBE2R2,KDM1A,MAP2K2,QARS1,ARF4,PITPNA,PLTP,KIF3C,DHX30,ATP6V

CLY,XRCC6,CLCN7,TAOK1,IARS2,YARS1,UBE2M,PKDCC,UBE2D2,VPS4A,AKT3,DCLK1,KIAA0232

,MASTL,SMC3,ULK1,ABCC9,BRSK1,MVD,GRK2,STK35,SPG7,ACLY,XRCC6,CLCN7,TAOK1,IARS2,
,ABCA7

L,SMC3,ULK1,ABCC9,BRSK1,MVD,GRK2,STK35,SPG7,ACLY,XRCC6,CLCN7,TAOK1,IARS2,YARS

M50,TNKS,GALNT11,FKBP1A,LOX,FBXO21,RHBDD2,GRK2,BAP1,STK35,SPG7,PSMB4,TAOK1,E

1A2,RAB1A,PLPPR3,SEPTIN3,HECTD4,CTBP1,HMGCR,RAC1,CDK16,TOP2B,SMCHD1,HDAC2,AT

MEGF8,ACSL6,RAB7A,ACTN4,MMP14,DDX3X,ITM2C,ARL4C,RCN1,APLP1,PRKAR1A,CHPF,MYO5

RCC2,ARHGEF9,RALBP1,SH3BP5,PPP1R37,CDC42SE1,COL7A1,DNAJB6,ANOS1,SOS1,PPP1R12A

PK1,DAZAP2,ANKS1A,REEP2,ITGAV,ADAM10,ARPP19,GSK3A,FBLN5,SMARCD1,TMEFF1,APBB1,

D10B,MYCBP2,ARHGAP5,IQSEC1,CCDC88A,PPME1,ARFGAP2,RAP2A,TBC1D14,TRIB3,ADRM1,S

,MYH10,RPL36,SND1,HNRNPH1,CKAP4,RPS12,ARCN1,RPS27,CCT6A,CAPRIN1,STIP1,HDAC2,IPO
,FN1,SPEN,NONO,DDB1,PSMD5,YWHAQ,EIF1,HACD3,DCHS1,SET,EGLN1,MAP9,TPI1,TMSB10,IN
P1,MLEC,CBX1,GCN1,TNPO1,SKI,SCAF1,CNTNAP2,ATP6AP1,DPYSL2,TERF2IP,TAF7,NOTCH2,OP
RBP1,DYNLL1,NAP1L4,PPP1R9B,VDAC1,WDR45B,SGTA,CAPN2,HIP1,FLCN,AP2A1,H2AZ1,BRD4,

,PTPN1,RAN,KLC2,SNX1,PPP1CA,PPME1,NECTIN1,DDX6,EMILIN1,CRKL,NDRG1,SLK,UNC45A,S
IM28,MATR3,YWHAG,CKB,COL4A2,RPL37A,LRRC28,MAPT,FN1,SPEN,NONO,DDB1,PSMD5,YW
RPL22,RHOA,PTCD3,ZNF865,RPL9,ATF4,NR2F1,KIF5B,RAB6B,RPS14,CHD4,ATP1A1,JUND,RPS2
14,HK1,RPL22,RHOA,PTCD3,ZNF865,RPL9,ATF4,NR2F1,KIF5B,RAB6B,RPS14,CHD4,ATP1A1,JU
,HDLBP,RPS25,SFPQ,EIF5A,PEBP1,SOX12,ACTN4,DDX3X,RPS15,RPL12,RPL30,CLTC,EIF3L,NAN
A,PPME1,TBC1D14,RNF41,TRIM27,TRIB3,C1QBP,MAPKAPK2,CDK9,TOM1L2,STUB1,MAPRE3,O

OPE,RPS21,PRPH,RPL27A,NUP153,MFAP4,ARPC3,EMILIN1,NUP98,PCOLCE,MRPS6,TUBA1C,RPS
HSF1,GSK3B,RCC2,SOX9,TFRC,WARS1,CDKN1B,SLC12A7,DUSP3,DUSP8,IRS1,PRKAR2B,RHOB

,WARS1,CTR9,DLG4,SH3BP5,CHD8,IRS1,PRKAR2B,CCDC6,CACYBP,MED12,RXRA,SOS1,CITED2,
EKHM2,CLASP2,WASF2,REEP2,CAP1,SYT11,GSK3A,VASH1,APBB1,MICAL1,DPYSL4,REEP1,MYL
GA2,DPF2,SIN3A,MYBBP1A,MED14
TF,LEMD2,SKIL,SIN3A,CRAMP1
U1,SARS1,DGKD,RAB11A,BMPR2,CCT5,CDC42,YME1L1,DHX15,KIF5A,EIF2AK1,PRMT1,BCR,AC
U1,SARS1,DGKD,RAB11A,BMPR2,CCT5,CDC42,YME1L1,DHX15,KIF5A,EIF2AK1,PRMT1,BCR,AC
RKDC,UBE2R2,CLIP3,MAP2K2,QARS1,ARF4,POLR2B,KIF3C,DHX30,ATP6V1B2,FYN,LARS1,SLIT3
SNK2A1,KIF21B,GSPT1,CSNK1G2,ATRX,GARS1,RAB2A,EP400,PAPSS1,GLUD1,UPF1,PI4KA,UBE
M1A,MAP2K2,QARS1,ARF4,POLR2B,KIF3C,DHX30,ATP6V1B2,FYN,SDHA,LARS1,CCT8,PFKM,LO
IPAP1,PSMA7,VDAC1,TRAF4,DFFB,STAT3,EIF2AK1,PRMT1,SGTA,SF1,ALDH3A2,HIP1,UBQLN2,C

IF2AK1,BCR,ACTR1A,TRIO,PRKAR2A,VARS1,BAZ1B,GTPBP1,CSNK2A1,KIF21B,GSPT1,CSNK1G
1,BCR,ACTR1A,TRIO,PRKAR2A,VARS1,BAZ1B,GTPBP1,CSNK2A1,KIF21B,GSPT1,CSNK1G2,ATR

1,BCR,ACTR1A,TRIO,PRKAR2A,NDUFV1,VARS1,BAZ1B,GTPBP1,CSNK2A1,KIF21B,GSPT1,CSNK
1B2,FYN,SDHA,LARS1,CCT8,PFKM,PITPNM1,LONP1,CSNK1D,PRPS1,HYOU1,SARS1,DGKD,PFN2,

,STK24,TRAP1,PRPF4B,ACTR1B,DICER1,PDPK1,ABCF1,MSH6,AKT2,EHD1,GSK3A,PSMC3,MTOR

YARS1,UBE2M,PKDCC,UBE2D2,VPS4A,AKT3,DCLK1,KIAA0232,STK24,TRAP1,PRPF4B,ACTR1B,D

1,UBE2M,PKDCC,UBE2D2,VPS4A,AKT3,DCLK1,KIAA0232,STK24,TRAP1,PRPF4B,ACTR1B,DICER1

HMT1,FKBP4,EIF3H,CARM1,TPP1,UBR5,UBE2M,PKDCC,LTBP4,UBE2D2,AKT3,DCLK1,DTX4,STK2

P6V0C,MMP2,TRIB2,PPIB,GLUL,SQLE,CDKN1A,CRPPA,HS6ST1,CTDSP2,DDX39B,CTSB,MDH2,CH

A,ANXA6,MACF1,SESN3,MTSS2,APLP2,MYL6,MYH10,NARS1,SEPTIN11,RHOB,UBE2H,ATP1A2,RAE

,ARHGAP33,GPS1,PPP1R26,ARL2BP,COL6A3,ARHGEF18,VAV2,PPP2R5B,CABIN1,GIT1,THG1L,F[illegible]

[illegible]HIP,GNA13,CD74,REEP1,SMO,FGFR1,METRN,NSG1,SNX2,IGFBP4,TUB,PTPN11,TAPBP,HMGA1,

[illegible]NGAP1,PPP2R5D,PSME3,CLPX,MTRNR2L4,SPOCK1,PCOLCE,PSMF1,SERPINE2,RAB3A,RCC2,A[illegible]

5,BCLAF1,PPIB,RPL23A,LUC
A,EIF4A1,MRC2,RPL19,PPP2R1A,RPL15
TN,VHL,CCND1,MAPK1,SMARCA4,MAPRE1
ARID1B,CRTAP,UQCRC1,HDAC6,PTEN,SMC3,NOM

I PR3,YWHAZ,RARS1,STAT1,TMPO
HAQ,EIF1,HACD3,DCHS1,SET,EGLN1,MAP
0,EIF5,HMGN2,ATP2A2,RPL26,
ND,RPS20,EIF5,CD81,HMGN2,A
OG,H1-0,RPS5,RPS7,RPS19,
XSR1,ZBTB4,LIMS1,YWHAZ,HSF1,GSK3B,RCC

29,NUP205,CCDC6,LMNA,NUP133
TB2,PPP1R12A,ARHGAP33,SHC1,CAMK2N1,ADIP

SHANK1,MDC1,THRA,TFDP1,CLCN3,SP1,BCAR
12B,CROCC,ANK2,PLS3,ABL1,PHACTR4,KTN1,AR

TR1A,TRIO,PRKAR2A,NDUFV1,VARS1,BAZ1B
TR1A,TRIO,PRKAR2A,NDUFV1,VARS1,BAZ1B
,CCT8,PFKM,LONP1,CSNK1D,PRPS1,HYO
2G2,ADCY6,EIF5B,IARS1,MASTL,SMC3,
NP1,CSNK1D,PRPS1,IDH1,HYOU1,SARS1,DGK
SNK2A1,GPX4,PHB2,CST3,DECR1,HJURP,GAR

2,ATRX,GARS1,RAB2A,EP400,PAPSS1,GLUD
K,GARS1,RAB2A,EP400,PAPSS1,GLUD1,UPF

I G2,ATRX,GARS1,RAB2A,EP400,PAPSS1,GL
RAB11A,P3H3,BMPR2,CCT5,CDC42,YME1L

,DHX38,PRKACA,NDUFA13,UBE2I,UBE2D3,P

ICER1,PDPK1,ABCF1,MSH6,AKT2,EHD1,AHC

,PDPK1,ABCF1,MSH6,AKT2,EHD1,GSK3A,PS

24,GALNT1,NCSTN,ATXN3,PRPF4B,MTMR4,PDPK1,MKRN1,

ID6,RAB27A,RNF187,EEF1A2,FDFT1,RPN2,PPP

31A,CALU,SEPTIN3,ZNF454,RPS27,H

JRIN,CRIM1,PRNP,SRCIN1,OBSCN,ARH

ABL1,COL16A1,SDCBP,SEMA6B,HDGFL3,TA

RHGEF9,RALBP1,SH3BP5,PPP1R37,CDC42SE

| | | | | | | |
|---|---|---|---|---|---|---|
| **GO:BP** | cellular compo | GO:0071840 | 1.29E-85 | 84.8879839 | 6880 | 2574 |
| **GO:BP** | cellular localiza | GO:0051641 | 4.78E-81 | 80.3210143 | 3519 | 2574 |
| **GO:BP** | viral process | GO:0016032 | 2.51E-76 | 75.6006499 | 1011 | 2574 |
| **GO:BP** | cellular compo | GO:0016043 | 4.37E-75 | 74.3597231 | 6661 | 2574 |
| **GO:BP** | cellular macro | GO:0070727 | 5.53E-69 | 68.2569608 | 1856 | 2574 |
| **GO:BP** | cellular protein | GO:0034613 | 5.82E-68 | 67.2352358 | 1842 | 2574 |
| **GO:BP** | mRNA metabo | GO:0016071 | 2.94E-67 | 66.5321709 | 880 | 2574 |
| **GO:BP** | intracellular tra | GO:0046907 | 4.54E-66 | 65.3429903 | 1723 | 2574 |
| **GO:BP** | macromolecul | GO:0009057 | 3.52E-65 | 64.4537329 | 1479 | 2574 |
| **GO:BP** | establishment | GO:0051649 | 2.48E-63 | 62.6049939 | 2778 | 2574 |
| **GO:BP** | protein localiza | GO:0008104 | 1.83E-60 | 59.7368323 | 2670 | 2574 |
| **GO:BP** | regulation of c | GO:0051128 | 7.18E-60 | 59.1436738 | 2439 | 2574 |
| **GO:BP** | macromolecul | GO:0033036 | 4.23E-59 | 58.3737852 | 3148 | 2574 |
| **GO:BP** | cellular macro | GO:0044265 | 1.52E-58 | 57.8172896 | 1244 | 2574 |
| **GO:BP** | protein localiza | GO:0033365 | 1.65E-56 | 55.7823782 | 974 | 2574 |
| **GO:BP** | mRNA catabo | GO:0006402 | 4.83E-54 | 53.31644 | 388 | 2574 |
| **GO:BP** | localization | GO:0051179 | 2.84E-53 | 52.5472805 | 6938 | 2574 |
| **GO:BP** | cellular protein | GO:0044267 | 3.36E-53 | 52.4742573 | 5280 | 2574 |
| **GO:BP** | organelle orga | GO:0006996 | 8.60E-53 | 52.0656222 | 4136 | 2574 |
| **GO:BP** | cotranslationa | GO:0006613 | 5.32E-52 | 51.2740374 | 102 | 2574 |
| **GO:BP** | nuclear-transc | GO:0000184 | 2.18E-51 | 50.6623178 | 123 | 2574 |
| **GO:BP** | SRP-depende | GO:0006614 | 2.67E-50 | 49.5736733 | 96 | 2574 |
| **GO:BP** | RNA catabolic | GO:0006401 | 1.10E-49 | 48.9585274 | 431 | 2574 |
| **GO:BP** | cellular compo | GO:0044085 | 4.24E-49 | 48.3724409 | 3377 | 2574 |
| **GO:BP** | establishment | GO:0045184 | 4.68E-49 | 48.3297037 | 1866 | 2574 |
| **GO:BP** | intracellular pr | GO:0006886 | 2.46E-48 | 47.6083687 | 995 | 2574 |
| **GO:BP** | protein localiza | GO:0070972 | 2.91E-48 | 47.5357282 | 144 | 2574 |
| **GO:BP** | protein targeti | GO:0045047 | 1.11E-47 | 46.9544023 | 111 | 2574 |
| **GO:BP** | protein metab | GO:0019538 | 3.35E-47 | 46.4754023 | 5912 | 2574 |
| **GO:BP** | nitrogen comp | GO:0071705 | 7.09E-47 | 46.1493288 | 2216 | 2574 |
| **GO:BP** | viral gene expr | GO:0019080 | 8.11E-47 | 46.0909221 | 237 | 2574 |
| **GO:BP** | establishment | GO:0072599 | 9.58E-47 | 46.0186002 | 115 | 2574 |
| **GO:BP** | cytoplasmic tr | GO:0002181 | 4.20E-46 | 45.3765545 | 149 | 2574 |
| **GO:BP** | cellular metab | GO:0044237 | 4.30E-46 | 45.3663396 | 10941 | 2574 |
| **GO:BP** | catabolic proc | GO:0009056 | 4.97E-46 | 45.303964 | 2779 | 2574 |
| **GO:BP** | cellular catabo | GO:0044248 | 9.90E-46 | 45.0044354 | 2381 | 2574 |
| **GO:BP** | negative regul | GO:0048519 | 6.63E-45 | 44.1785623 | 5861 | 2574 |
| **GO:BP** | translational ir | GO:0006413 | 4.13E-44 | 43.3837519 | 203 | 2574 |
| **GO:BP** | regulation of c | GO:0009894 | 2.58E-43 | 42.5879658 | 1081 | 2574 |
| **GO:BP** | protein transp | GO:0015031 | 6.67E-43 | 42.1756937 | 1754 | 2574 |
| **GO:BP** | establishment | GO:0051234 | 4.06E-42 | 41.3916152 | 5364 | 2574 |
| **GO:BP** | establishment | GO:0072594 | 5.46E-42 | 41.262784 | 523 | 2574 |
| **GO:BP** | transport | GO:0006810 | 1.09E-41 | 40.9634025 | 5214 | 2574 |
| **GO:BP** | cellular compo | GO:0022607 | 2.32E-40 | 39.635153 | 3104 | 2574 |
| **GO:BP** | organic subst | GO:1901575 | 5.24E-40 | 39.2806682 | 2289 | 2574 |

| | | | | | | |
|---|---|---|---|---|---|---|
| **GO:BP** | organonitroge | GO:1901564 | 9.82E-40 | 39.0077327 | 6904 | 2574 |
| **GO:BP** | nervous syste | GO:0007399 | 1.03E-39 | 38.9883847 | 2484 | 2574 |
| **GO:BP** | protein localiz | GO:0072657 | 5.28E-39 | 38.2771742 | 698 | 2574 |
| **GO:BP** | positive regula | GO:0048522 | 1.21E-38 | 37.9178783 | 5863 | 2574 |
| **GO:BP** | neuron projec | GO:0031175 | 1.63E-38 | 37.7886141 | 1021 | 2574 |
| **GO:BP** | localization wi | GO:0051668 | 2.25E-38 | 37.6471019 | 775 | 2574 |
| **GO:BP** | regulation of c | GO:0031329 | 6.17E-38 | 37.2100262 | 913 | 2574 |
| **GO:BP** | viral transcripti | GO:0019083 | 7.57E-38 | 37.1207825 | 181 | 2574 |
| **GO:BP** | regulation of r | GO:0019222 | 1.07E-37 | 36.9713272 | 7175 | 2574 |
| **GO:BP** | nucleobase-co | GO:0034655 | 2.78E-37 | 36.5562555 | 573 | 2574 |
| **GO:BP** | negative regul | GO:0048523 | 5.05E-37 | 36.2968499 | 5153 | 2574 |
| **GO:BP** | negative regul | GO:0009892 | 1.23E-36 | 35.9114035 | 3498 | 2574 |
| **GO:BP** | regulation of p | GO:0051246 | 1.85E-36 | 35.7327775 | 2759 | 2574 |
| **GO:BP** | cellular macro | GO:0044260 | 2.38E-36 | 35.6233482 | 8430 | 2574 |
| **GO:BP** | negative regul | GO:0010605 | 3.29E-36 | 35.4833495 | 3262 | 2574 |
| **GO:BP** | cellular respor | GO:0033554 | 4.04E-36 | 35.3935771 | 2142 | 2574 |
| **GO:BP** | response to o | GO:0010033 | 1.89E-35 | 34.723084 | 3457 | 2574 |
| **GO:BP** | protein-contai | GO:0043933 | 2.51E-35 | 34.5997802 | 2017 | 2574 |
| **GO:BP** | neuron develo | GO:0048666 | 5.82E-35 | 34.2349279 | 1159 | 2574 |
| **GO:BP** | positive regula | GO:0051130 | 5.83E-35 | 34.2346175 | 1189 | 2574 |
| **GO:BP** | neurogenesis | GO:0022008 | 6.52E-35 | 34.1858594 | 1698 | 2574 |
| **GO:BP** | primary metab | GO:0044238 | 1.53E-34 | 33.8164813 | 10642 | 2574 |
| **GO:BP** | heterocycle ca | GO:0046700 | 1.83E-34 | 33.7370003 | 617 | 2574 |
| **GO:BP** | cellular nitroge | GO:0044270 | 2.85E-34 | 33.5454095 | 623 | 2574 |
| **GO:BP** | nuclear-transc | GO:0000956 | 4.19E-34 | 33.3780996 | 214 | 2574 |
| **GO:BP** | nitrogen comp | GO:0006807 | 1.03E-33 | 32.9877314 | 10148 | 2574 |
| **GO:BP** | neuron differe | GO:0030182 | 1.44E-33 | 32.8409103 | 1427 | 2574 |
| **GO:BP** | generation of | GO:0048699 | 4.44E-33 | 32.3527244 | 1576 | 2574 |
| **GO:BP** | organic substa | GO:0071702 | 1.03E-32 | 31.9887421 | 2657 | 2574 |
| **GO:BP** | regulation of r | GO:0060255 | 2.41E-32 | 31.6172605 | 6622 | 2574 |
| **GO:BP** | establishment | GO:0090150 | 2.48E-32 | 31.6052679 | 385 | 2574 |
| **GO:BP** | positive regula | GO:0048518 | 1.15E-31 | 30.9397782 | 6474 | 2574 |
| **GO:BP** | regulation of c | GO:0031323 | 1.32E-31 | 30.8785626 | 6309 | 2574 |
| **GO:BP** | peptide metab | GO:0006518 | 2.06E-31 | 30.686754 | 998 | 2574 |
| **GO:BP** | regulation of c | GO:0032268 | 2.36E-31 | 30.6264403 | 2591 | 2574 |
| **GO:BP** | protein targetir | GO:0006605 | 2.89E-31 | 30.5384789 | 400 | 2574 |
| **GO:BP** | aromatic comp | GO:0019439 | 3.08E-31 | 30.5111896 | 636 | 2574 |
| **GO:BP** | cell morphoge | GO:0000902 | 3.33E-31 | 30.4779673 | 1045 | 2574 |
| **GO:BP** | organic cyclic | GO:1901361 | 3.84E-31 | 30.4158327 | 664 | 2574 |
| **GO:BP** | cell projection | GO:0048858 | 5.14E-31 | 30.288785 | 688 | 2574 |
| **GO:BP** | plasma memb | GO:0120039 | 1.80E-30 | 29.7445425 | 684 | 2574 |
| **GO:BP** | cell part morpl | GO:0032990 | 3.33E-30 | 29.4777895 | 705 | 2574 |
| **GO:BP** | developmenta | GO:0032502 | 3.43E-30 | 29.4642695 | 6678 | 2574 |
| **GO:BP** | protein catabo | GO:0030163 | 3.53E-30 | 29.4516427 | 983 | 2574 |
| **GO:BP** | translation | GO:0006412 | 3.76E-30 | 29.4245234 | 804 | 2574 |

| | | | | | | |
|---|---|---|---|---|---|---|
| **GO:BP** | protein-contai | GO:0065003 | 4.02E-30 | 29.3956414 | 1724 | 2574 |
| **GO:BP** | metabolic pro | GO:0008152 | 4.23E-30 | 29.3740298 | 11915 | 2574 |
| **GO:BP** | plasma memb | GO:0120036 | 5.31E-30 | 29.2747333 | 1596 | 2574 |
| **GO:BP** | RNA processi | GO:0006396 | 5.44E-30 | 29.2646353 | 1010 | 2574 |
| **GO:BP** | cellular respor | GO:0070887 | 1.53E-29 | 28.8161064 | 3443 | 2574 |
| **GO:BP** | neuron projec | GO:0048812 | 1.56E-29 | 28.8073712 | 670 | 2574 |
| **GO:BP** | cellular proteir | GO:0034622 | 2.84E-29 | 28.5462703 | 1150 | 2574 |
| **GO:BP** | macromolecul | GO:0043170 | 4.09E-29 | 28.3880955 | 9867 | 2574 |
| **GO:BP** | regulation of p | GO:0080090 | 4.81E-29 | 28.3181486 | 6071 | 2574 |
| **GO:BP** | protein targeti | GO:0006612 | 4.90E-29 | 28.3099698 | 219 | 2574 |
| **GO:BP** | regulation of r | GO:1903311 | 7.85E-29 | 28.1052456 | 384 | 2574 |
| **GO:BP** | cell projection | GO:0030030 | 1.64E-28 | 27.7845654 | 1636 | 2574 |
| **GO:BP** | peptide biosyı | GO:0043043 | 2.84E-28 | 27.5471621 | 834 | 2574 |
| **GO:BP** | cellular respor | GO:0071310 | 3.79E-28 | 27.4216968 | 2821 | 2574 |
| **GO:BP** | anatomical str | GO:0048856 | 6.34E-28 | 27.1976513 | 6164 | 2574 |
| **GO:BP** | cell morphoge | GO:0000904 | 1.06E-27 | 26.9734414 | 754 | 2574 |
| **GO:BP** | anatomical str | GO:0009653 | 2.72E-27 | 26.5657826 | 2816 | 2574 |
| **GO:BP** | cellular compo | GO:0032989 | 2.88E-27 | 26.5401887 | 797 | 2574 |
| **GO:BP** | cell morphoge | GO:0048667 | 5.17E-27 | 26.2868243 | 607 | 2574 |
| **GO:BP** | regulation of k | GO:0065008 | 8.75E-27 | 26.0579049 | 4116 | 2574 |
| **GO:BP** | RNA splicing | GO:0008380 | 1.20E-26 | 25.9214729 | 459 | 2574 |
| **GO:BP** | multicellular o | GO:0007275 | 1.50E-26 | 25.825094 | 5670 | 2574 |
| **GO:BP** | regulation of p | GO:0042176 | 1.90E-26 | 25.7215522 | 401 | 2574 |
| **GO:BP** | regulation of r | GO:0051171 | 2.22E-26 | 25.6532587 | 5886 | 2574 |
| **GO:BP** | amide biosynt | GO:0043604 | 2.25E-26 | 25.648129 | 975 | 2574 |
| **GO:BP** | system develo | GO:0048731 | 3.71E-26 | 25.4307727 | 5095 | 2574 |
| **GO:BP** | organic subst | GO:0071704 | 7.67E-26 | 25.1151762 | 11397 | 2574 |
| **GO:BP** | mRNA splicing | GO:0000398 | 8.31E-26 | 25.0805885 | 360 | 2574 |
| **GO:BP** | RNA splicing, | GO:0000377 | 8.31E-26 | 25.0805885 | 360 | 2574 |
| **GO:BP** | RNA splicing, | GO:0000375 | 2.20E-25 | 24.6570826 | 363 | 2574 |
| **GO:BP** | cellular proteir | GO:0006464 | 5.63E-25 | 24.2497322 | 4108 | 2574 |
| **GO:BP** | protein modifi | GO:0036211 | 5.63E-25 | 24.2497322 | 4108 | 2574 |
| **GO:BP** | axon develop | GO:0061564 | 1.00E-24 | 23.9992107 | 524 | 2574 |
| **GO:BP** | positive regula | GO:0009893 | 2.10E-24 | 23.678609 | 3873 | 2574 |
| **GO:BP** | mRNA proces | GO:0006397 | 2.39E-24 | 23.621088 | 532 | 2574 |
| **GO:BP** | cellular proteir | GO:0044257 | 2.87E-24 | 23.5424411 | 822 | 2574 |
| **GO:BP** | cell developm | GO:0048468 | 3.05E-24 | 23.5161413 | 2122 | 2574 |
| **GO:BP** | regulation of c | GO:0033043 | 3.90E-24 | 23.4083956 | 1228 | 2574 |
| **GO:BP** | negative regul | GO:0010629 | 4.33E-24 | 23.3638097 | 1470 | 2574 |
| **GO:BP** | cellular amide | GO:0043603 | 6.07E-24 | 23.2168439 | 1284 | 2574 |
| **GO:BP** | ribonucleopro | GO:0022613 | 6.17E-24 | 23.2099271 | 473 | 2574 |
| **GO:BP** | positive regula | GO:0031325 | 1.98E-23 | 22.7041085 | 3342 | 2574 |
| **GO:BP** | regulation of k | GO:0050789 | 2.68E-23 | 22.5720373 | 12122 | 2574 |
| **GO:BP** | vesicle-mediat | GO:0016192 | 4.83E-23 | 22.3162035 | 2193 | 2574 |
| **GO:BP** | regulation of r | GO:0010975 | 5.90E-23 | 22.2293603 | 455 | 2574 |

| | | | | | | |
|---|---|---|---|---|---|---|
| **GO:BP** | positive regula | GO:0010604 | 8.42E-23 | 22.0744967 | 3568 | 2574 |
| **GO:BP** | negative regul | GO:0051172 | 9.15E-23 | 22.0385896 | 2592 | 2574 |
| **GO:BP** | axonogenesis | GO:0007409 | 9.22E-23 | 22.0352212 | 479 | 2574 |
| **GO:BP** | negative regul | GO:0031324 | 1.13E-22 | 21.9450832 | 2785 | 2574 |
| **GO:BP** | regulation of p | GO:0120035 | 1.20E-22 | 21.919593 | 642 | 2574 |
| **GO:BP** | response to e | GO:0009719 | 1.40E-22 | 21.8525143 | 1729 | 2574 |
| **GO:BP** | regulation of c | GO:0031344 | 2.60E-22 | 21.5846275 | 660 | 2574 |
| **GO:BP** | apoptotic proc | GO:0006915 | 3.60E-22 | 21.4435802 | 1998 | 2574 |
| **GO:BP** | positive regula | GO:0051173 | 3.91E-22 | 21.4075992 | 3143 | 2574 |
| **GO:BP** | gene expressi | GO:0010467 | 5.20E-22 | 21.2841771 | 6192 | 2574 |
| **GO:BP** | organonitroge | GO:1901566 | 2.39E-21 | 20.6212105 | 1889 | 2574 |
| **GO:BP** | cellular develo | GO:0048869 | 4.45E-21 | 20.35151 | 4468 | 2574 |
| **GO:BP** | response to c | GO:0042221 | 4.68E-21 | 20.3298009 | 4829 | 2574 |
| **GO:BP** | cellular nitroge | GO:0034641 | 5.33E-21 | 20.2729995 | 6576 | 2574 |
| **GO:BP** | cell differentiat | GO:0030154 | 1.23E-20 | 19.9097497 | 4392 | 2574 |
| **GO:BP** | RNA metaboli | GO:0016070 | 1.32E-20 | 19.8810128 | 4714 | 2574 |
| **GO:BP** | biosynthetic p | GO:0009058 | 1.63E-20 | 19.7877448 | 6391 | 2574 |
| **GO:BP** | organic subst | GO:1901576 | 1.63E-20 | 19.7872942 | 6297 | 2574 |
| **GO:BP** | macromolecul | GO:0043412 | 4.35E-20 | 19.3617803 | 4345 | 2574 |
| **GO:BP** | nucleobase-co | GO:0006139 | 4.46E-20 | 19.3510147 | 5752 | 2574 |
| **GO:BP** | growth | GO:0040007 | 8.27E-20 | 19.0823351 | 973 | 2574 |
| **GO:BP** | programmed c | GO:0012501 | 1.35E-19 | 18.8711757 | 2158 | 2574 |
| **GO:BP** | nucleic acid m | GO:0090304 | 2.02E-19 | 18.6944279 | 5234 | 2574 |
| **GO:BP** | regulation of c | GO:1903362 | 2.35E-19 | 18.6290895 | 258 | 2574 |
| **GO:BP** | macromolecul | GO:0009059 | 2.42E-19 | 18.6164392 | 5145 | 2574 |
| **GO:BP** | negative regul | GO:0051129 | 2.62E-19 | 18.5809973 | 760 | 2574 |
| **GO:BP** | regulation of c | GO:0044087 | 2.64E-19 | 18.5792135 | 991 | 2574 |
| **GO:BP** | organonitroge | GO:1901565 | 3.42E-19 | 18.4662786 | 1375 | 2574 |
| **GO:BP** | regulation of g | GO:0010468 | 4.04E-19 | 18.3934482 | 5218 | 2574 |
| **GO:BP** | regulation of c | GO:0050793 | 4.23E-19 | 18.3737531 | 2621 | 2574 |
| **GO:BP** | regulation of c | GO:1903827 | 5.39E-19 | 18.2684283 | 563 | 2574 |
| **GO:BP** | proteolysis inv | GO:0051603 | 6.33E-19 | 18.1986295 | 760 | 2574 |
| **GO:BP** | cellular macro | GO:0034645 | 6.47E-19 | 18.1893447 | 5081 | 2574 |
| **GO:BP** | cell death | GO:0008219 | 8.30E-19 | 18.0809085 | 2306 | 2574 |
| **GO:BP** | proteasomal p | GO:0010498 | 1.30E-18 | 17.8862548 | 501 | 2574 |
| **GO:BP** | cell growth | GO:0016049 | 1.51E-18 | 17.8215278 | 497 | 2574 |
| **GO:BP** | cellular respor | GO:0071495 | 1.99E-18 | 17.7003188 | 1467 | 2574 |
| **GO:BP** | biological regu | GO:0065007 | 2.74E-18 | 17.5615974 | 12802 | 2574 |
| **GO:BP** | regulation of s | GO:0023051 | 3.70E-18 | 17.4320699 | 3669 | 2574 |
| **GO:BP** | supramolecula | GO:0097435 | 4.09E-18 | 17.3887529 | 863 | 2574 |
| **GO:BP** | ribonucleopro | GO:0071826 | 4.36E-18 | 17.3603205 | 203 | 2574 |
| **GO:BP** | heterocycle m | GO:0046483 | 6.70E-18 | 17.1739131 | 5929 | 2574 |
| **GO:BP** | cellular biosyn | GO:0044249 | 1.53E-17 | 16.8159398 | 6210 | 2574 |
| **GO:BP** | regulation of p | GO:0043067 | 1.55E-17 | 16.8102056 | 1603 | 2574 |
| **GO:BP** | regulation of a | GO:0022603 | 2.22E-17 | 16.6529813 | 1048 | 2574 |

| | | | | | | |
|---|---|---|---|---|---|---|
| **GO:BP** | ribonucleopro | GO:0022618 | 2.52E-17 | 16.5991683 | 196 | 2574 |
| **GO:BP** | exocytosis | GO:0006887 | 2.53E-17 | 16.5965845 | 906 | 2574 |
| **GO:BP** | regulation of c | GO:0080135 | 3.03E-17 | 16.5179611 | 727 | 2574 |
| **GO:BP** | positive regula | GO:0051247 | 3.79E-17 | 16.421206 | 1577 | 2574 |
| **GO:BP** | regulated exoc | GO:0045055 | 4.16E-17 | 16.3808867 | 790 | 2574 |
| **GO:BP** | regulation of c | GO:0010646 | 4.81E-17 | 16.3181928 | 3634 | 2574 |
| **GO:BP** | cellular aroma | GO:0006725 | 6.77E-17 | 16.1696485 | 5973 | 2574 |
| **GO:BP** | cellular nitroge | GO:0044271 | 7.21E-17 | 16.1423407 | 5054 | 2574 |
| **GO:BP** | regulation of c | GO:0050794 | 1.11E-16 | 15.9532577 | 11517 | 2574 |
| **GO:BP** | proteasome-m | GO:0043161 | 1.17E-16 | 15.9309821 | 442 | 2574 |
| **GO:BP** | regulation of p | GO:0031399 | 1.24E-16 | 15.9052598 | 1685 | 2574 |
| **GO:BP** | modification-c | GO:0019941 | 1.30E-16 | 15.8871762 | 670 | 2574 |
| **GO:BP** | nuclear transp | GO:0051169 | 1.33E-16 | 15.877689 | 356 | 2574 |
| **GO:BP** | regulation of p | GO:0032880 | 1.35E-16 | 15.8698183 | 912 | 2574 |
| **GO:BP** | modification-c | GO:0043632 | 1.53E-16 | 15.8143266 | 681 | 2574 |
| **GO:BP** | response to e | GO:0034976 | 1.70E-16 | 15.7695569 | 300 | 2574 |
| **GO:BP** | regulation of D | GO:0043620 | 2.03E-16 | 15.6931014 | 117 | 2574 |
| **GO:BP** | ubiquitin-depe | GO:0006511 | 2.20E-16 | 15.657802 | 663 | 2574 |
| **GO:BP** | regulation of a | GO:0042981 | 2.49E-16 | 15.6040666 | 1560 | 2574 |
| **GO:BP** | positive regula | GO:0031346 | 2.83E-16 | 15.5479197 | 359 | 2574 |
| **GO:BP** | regulation of t | GO:0043618 | 3.02E-16 | 15.520298 | 111 | 2574 |
| **GO:BP** | regulation of c | GO:0010941 | 3.16E-16 | 15.5007773 | 1740 | 2574 |
| **GO:BP** | negative regul | GO:0051248 | 3.68E-16 | 15.4344168 | 1216 | 2574 |
| **GO:BP** | regulation of c | GO:0060341 | 3.93E-16 | 15.4054984 | 851 | 2574 |
| **GO:BP** | post-translatic | GO:0043687 | 5.98E-16 | 15.2234825 | 362 | 2574 |
| **GO:BP** | central nervou | GO:0007417 | 7.13E-16 | 15.1471146 | 1052 | 2574 |
| **GO:BP** | cell junction o | GO:0034330 | 8.94E-16 | 15.0486443 | 727 | 2574 |
| **GO:BP** | regulation of r | GO:0061013 | 9.72E-16 | 15.012312 | 255 | 2574 |
| **GO:BP** | regulation of r | GO:0050684 | 1.04E-15 | 14.9821628 | 145 | 2574 |
| **GO:BP** | organic cyclic | GO:1901360 | 1.06E-15 | 14.9761957 | 6227 | 2574 |
| **GO:BP** | regulation of r | GO:0043488 | 1.08E-15 | 14.9661952 | 247 | 2574 |
| **GO:BP** | autophagy | GO:0006914 | 1.37E-15 | 14.8642698 | 564 | 2574 |
| **GO:BP** | process utilizi | GO:0061919 | 1.37E-15 | 14.8642698 | 564 | 2574 |
| **GO:BP** | developmenta | GO:0048589 | 1.50E-15 | 14.8242735 | 654 | 2574 |
| **GO:BP** | nucleocytopla | GO:0006913 | 1.81E-15 | 14.7411237 | 353 | 2574 |
| **GO:BP** | regulation of p | GO:1903050 | 1.87E-15 | 14.7284084 | 224 | 2574 |
| **GO:BP** | response to a | GO:0009628 | 2.20E-15 | 14.6572774 | 1264 | 2574 |
| **GO:BP** | cytoskeleton c | GO:0007010 | 2.46E-15 | 14.6094831 | 1473 | 2574 |
| **GO:BP** | cell cycle | GO:0007049 | 2.90E-15 | 14.537103 | 1867 | 2574 |
| **GO:BP** | regulation of p | GO:0031647 | 3.23E-15 | 14.4903406 | 302 | 2574 |
| **GO:BP** | cell-cell signali | GO:0198738 | 3.68E-15 | 14.4337763 | 535 | 2574 |
| **GO:BP** | regulation of p | GO:0043254 | 3.72E-15 | 14.4295944 | 458 | 2574 |
| **GO:BP** | regulation of c | GO:0001558 | 4.62E-15 | 14.3350487 | 426 | 2574 |
| **GO:BP** | cellular compo | GO:0022411 | 6.02E-15 | 14.2202039 | 577 | 2574 |
| **GO:BP** | Wnt signaling | GO:0016055 | 6.36E-15 | 14.1963333 | 533 | 2574 |

| | | | | | | |
|---|---|---|---|---|---|---|
| **GO:BP** | positive regula | GO:0032270 | 6.71E-15 | 14.1732854 | 1488 | 2574 |
| **GO:BP** | response to o | GO:0010243 | 7.53E-15 | 14.1232868 | 1092 | 2574 |
| **GO:BP** | regulation of p | GO:0061136 | 1.10E-14 | 13.9568002 | 189 | 2574 |
| **GO:BP** | neutrophil deg | GO:0043312 | 1.23E-14 | 13.9115467 | 483 | 2574 |
| **GO:BP** | protein folding | GO:0006457 | 1.28E-14 | 13.8928793 | 230 | 2574 |
| **GO:BP** | RNA localizatio | GO:0006403 | 1.39E-14 | 13.8563926 | 222 | 2574 |
| **GO:BP** | protein modifi | GO:0032446 | 2.01E-14 | 13.6973142 | 980 | 2574 |
| **GO:BP** | regulation of I | GO:0043484 | 2.06E-14 | 13.6869422 | 152 | 2574 |
| **GO:BP** | negative regul | GO:0032269 | 2.75E-14 | 13.5611769 | 1141 | 2574 |
| **GO:BP** | neutrophil acti | GO:0002283 | 3.27E-14 | 13.4857528 | 488 | 2574 |
| **GO:BP** | regulation of I | GO:0043487 | 3.85E-14 | 13.4149342 | 259 | 2574 |
| **GO:BP** | response to n | GO:1901698 | 4.30E-14 | 13.3668721 | 1178 | 2574 |
| **GO:BP** | neutrophil me | GO:0002446 | 5.06E-14 | 13.2960374 | 500 | 2574 |
| **GO:BP** | response to g | GO:0070848 | 5.45E-14 | 13.2634121 | 769 | 2574 |
| **GO:BP** | protein modifi | GO:0070647 | 6.75E-14 | 13.1703952 | 1193 | 2574 |
| **GO:BP** | cellular proces | GO:0009987 | 7.38E-14 | 13.1319555 | 17186 | 2574 |
| **GO:BP** | granulocyte ac | GO:0036230 | 1.09E-13 | 12.9611857 | 509 | 2574 |
| **GO:BP** | regulation of s | GO:0009966 | 1.11E-13 | 12.9534748 | 3251 | 2574 |
| **GO:BP** | positive regula | GO:0010638 | 1.29E-13 | 12.8885193 | 630 | 2574 |
| **GO:BP** | cell cycle proc | GO:0022402 | 1.82E-13 | 12.739684 | 1388 | 2574 |
| **GO:BP** | neutrophil acti | GO:0042119 | 1.84E-13 | 12.7342682 | 502 | 2574 |
| **GO:BP** | protein ubiqui | GO:0016567 | 1.91E-13 | 12.7186624 | 900 | 2574 |
| **GO:BP** | mitotic cell cyc | GO:0000278 | 2.32E-13 | 12.6348444 | 1098 | 2574 |
| **GO:BP** | macroautopha | GO:0016236 | 2.38E-13 | 12.6238361 | 337 | 2574 |
| **GO:BP** | regulation of c | GO:0051726 | 4.36E-13 | 12.3602324 | 1120 | 2574 |
| **GO:BP** | secretion by c | GO:0032940 | 4.50E-13 | 12.3470517 | 1351 | 2574 |
| **GO:BP** | mitotic cell cyc | GO:1903047 | 4.94E-13 | 12.3063475 | 956 | 2574 |
| **GO:BP** | developmenta | GO:0060560 | 4.97E-13 | 12.3037764 | 238 | 2574 |
| **GO:BP** | regulation of t | GO:0061418 | 5.13E-13 | 12.2896906 | 78 | 2574 |
| **GO:BP** | positive regula | GO:0009891 | 5.30E-13 | 12.275562 | 2013 | 2574 |
| **GO:BP** | intracellular sig | GO:0035556 | 5.49E-13 | 12.2605325 | 2886 | 2574 |
| **GO:BP** | axon extensio | GO:0048675 | 6.31E-13 | 12.1999094 | 123 | 2574 |
| **GO:BP** | chromosome | GO:0051276 | 6.99E-13 | 12.1555418 | 1270 | 2574 |
| **GO:BP** | regulation of r | GO:0065009 | 7.76E-13 | 12.110198 | 3191 | 2574 |
| **GO:BP** | positive regula | GO:0031328 | 8.00E-13 | 12.0966629 | 1976 | 2574 |
| **GO:BP** | response to to | GO:0035966 | 8.26E-13 | 12.082872 | 206 | 2574 |
| **GO:BP** | cellular respor | GO:0071363 | 9.05E-13 | 12.0431551 | 741 | 2574 |
| **GO:BP** | membrane org | GO:0061024 | 9.53E-13 | 12.0210862 | 1016 | 2574 |
| **GO:BP** | head developr | GO:0060322 | 1.02E-12 | 11.9914692 | 816 | 2574 |
| **GO:BP** | protein localiz | GO:0034504 | 1.23E-12 | 11.911021 | 294 | 2574 |
| **GO:BP** | response to u | GO:0006986 | 1.39E-12 | 11.8563895 | 183 | 2574 |
| **GO:BP** | cell cycle G2/M | GO:0044839 | 1.70E-12 | 11.7691202 | 273 | 2574 |
| **GO:BP** | regulation of c | GO:0032535 | 1.76E-12 | 11.7537711 | 388 | 2574 |
| **GO:BP** | embryo develo | GO:0009790 | 1.92E-12 | 11.7159244 | 1055 | 2574 |
| **GO:BP** | regulation of c | GO:1902749 | 1.96E-12 | 11.7081786 | 213 | 2574 |

| | | | | | | |
|---|---|---|---|---|---|---|
| **GO:BP** | neuron projec | GO:1990138 | 2.01E-12 | 11.6961891 | 172 | 2574 |
| **GO:BP** | positive regula | GO:0010557 | 2.69E-12 | 11.5705242 | 1859 | 2574 |
| **GO:BP** | developmenta | GO:0048588 | 3.20E-12 | 11.4952222 | 236 | 2574 |
| **GO:BP** | regulation of c | GO:0010564 | 3.78E-12 | 11.4222353 | 815 | 2574 |
| **GO:BP** | positive regula | GO:0009896 | 4.56E-12 | 11.3409675 | 530 | 2574 |
| **GO:BP** | response to h | GO:0009725 | 4.86E-12 | 11.3134742 | 958 | 2574 |
| **GO:BP** | antigen proce | GO:0002478 | 5.56E-12 | 11.2545439 | 175 | 2574 |
| **GO:BP** | negative regul | GO:0031400 | 5.59E-12 | 11.2524465 | 587 | 2574 |
| **GO:BP** | regulation of ( | GO:0010389 | 5.79E-12 | 11.2373192 | 204 | 2574 |
| **GO:BP** | myeloid leuko | GO:0002444 | 7.21E-12 | 11.1422774 | 558 | 2574 |
| **GO:BP** | response to s | GO:0006950 | 7.23E-12 | 11.1409721 | 4321 | 2574 |
| **GO:BP** | export from ce | GO:0140352 | 7.49E-12 | 11.1255029 | 1413 | 2574 |
| **GO:BP** | leukocyte deg | GO:0043299 | 7.51E-12 | 11.1241552 | 538 | 2574 |
| **GO:BP** | cellular respor | GO:0051716 | 7.91E-12 | 11.1020281 | 7873 | 2574 |
| **GO:BP** | regulation of r | GO:0048024 | 8.31E-12 | 11.0802908 | 107 | 2574 |
| **GO:BP** | G2/M transitic | GO:0000086 | 9.57E-12 | 11.0190596 | 262 | 2574 |
| **GO:BP** | ribosome biog | GO:0042254 | 9.94E-12 | 11.0027287 | 335 | 2574 |
| **GO:BP** | regulation of c | GO:0045595 | 1.37E-11 | 10.8639656 | 1688 | 2574 |
| **GO:BP** | brain developr | GO:0007420 | 1.55E-11 | 10.8086767 | 772 | 2574 |
| **GO:BP** | protein stabiliz | GO:0050821 | 1.71E-11 | 10.7674101 | 195 | 2574 |
| **GO:BP** | negative regul | GO:0009895 | 1.91E-11 | 10.7198343 | 324 | 2574 |
| **GO:BP** | antigen proce | GO:0019884 | 1.98E-11 | 10.7036211 | 183 | 2574 |
| **GO:BP** | chromatin org | GO:0006325 | 2.13E-11 | 10.6719641 | 839 | 2574 |
| **GO:BP** | Golgi vesicle t | GO:0048193 | 2.16E-11 | 10.6660073 | 415 | 2574 |
| **GO:BP** | regulation of c | GO:0051493 | 2.44E-11 | 10.6125098 | 535 | 2574 |
| **GO:BP** | regulation of g | GO:0040008 | 2.52E-11 | 10.5979473 | 680 | 2574 |
| **GO:BP** | posttranscripti | GO:0010608 | 2.62E-11 | 10.5811646 | 928 | 2574 |
| **GO:BP** | regulation of c | GO:0034248 | 2.94E-11 | 10.5318112 | 577 | 2574 |
| **GO:BP** | myeloid cell ac | GO:0002275 | 3.23E-11 | 10.4904028 | 552 | 2574 |
| **GO:BP** | apoptotic sigr | GO:0097190 | 3.45E-11 | 10.4624219 | 609 | 2574 |
| **GO:BP** | nuclear export | GO:0051168 | 3.92E-11 | 10.4069061 | 202 | 2574 |
| **GO:BP** | negative regul | GO:0031330 | 4.84E-11 | 10.3152874 | 264 | 2574 |
| **GO:BP** | regulation of c | GO:0060284 | 5.02E-11 | 10.2994505 | 509 | 2574 |
| **GO:BP** | cell surface re | GO:1905114 | 6.20E-11 | 10.207572 | 639 | 2574 |
| **GO:BP** | regulation of k | GO:0009889 | 7.10E-11 | 10.1488698 | 4374 | 2574 |
| **GO:BP** | positive regula | GO:0045732 | 7.48E-11 | 10.1262475 | 239 | 2574 |
| **GO:BP** | secretion | GO:0046903 | 9.43E-11 | 10.0255914 | 1492 | 2574 |
| **GO:BP** | regulation of c | GO:0050790 | 1.11E-10 | 9.9553917 | 2452 | 2574 |
| **GO:BP** | proteolysis | GO:0006508 | 1.27E-10 | 9.89687797 | 1851 | 2574 |
| **GO:BP** | regulation of s | GO:1902903 | 1.45E-10 | 9.83801356 | 391 | 2574 |
| **GO:BP** | negative regul | GO:0050686 | 1.82E-10 | 9.74080676 | 31 | 2574 |
| **GO:BP** | rRNA processi | GO:0006364 | 1.84E-10 | 9.73521683 | 265 | 2574 |
| **GO:BP** | negative regul | GO:0043066 | 1.85E-10 | 9.73220091 | 928 | 2574 |
| **GO:BP** | negative regul | GO:0060548 | 2.26E-10 | 9.64544081 | 1047 | 2574 |
| **GO:BP** | rRNA metabol | GO:0016072 | 2.27E-10 | 9.64398728 | 275 | 2574 |

| | | | | | | |
|---|---|---|---|---|---|---|
| **GO:BP** | nucleic acid tr | GO:0050657 | 2.56E-10 | 9.59151274 | 183 | 2574 |
| **GO:BP** | RNA transport | GO:0050658 | 2.56E-10 | 9.59151274 | 183 | 2574 |
| **GO:BP** | cell cycle phas | GO:0044770 | 2.66E-10 | 9.57455148 | 681 | 2574 |
| **GO:BP** | regulation of t | GO:0006417 | 2.68E-10 | 9.57265175 | 504 | 2574 |
| **GO:BP** | positive regula | GO:0044089 | 3.16E-10 | 9.50054738 | 546 | 2574 |
| **GO:BP** | regulation of r | GO:0010556 | 3.26E-10 | 9.48727635 | 4134 | 2574 |
| **GO:BP** | enzyme linked | GO:0007167 | 3.29E-10 | 9.48335716 | 1107 | 2574 |
| **GO:BP** | cellular respor | GO:0071456 | 3.39E-10 | 9.47035997 | 214 | 2574 |
| **GO:BP** | regulation of a | GO:0050770 | 3.39E-10 | 9.46955544 | 159 | 2574 |
| **GO:BP** | negative regul | GO:0043069 | 4.28E-10 | 9.3682268 | 952 | 2574 |
| **GO:BP** | negative regul | GO:0033119 | 4.59E-10 | 9.33778887 | 27 | 2574 |
| **GO:BP** | positive regula | GO:0010720 | 4.65E-10 | 9.33276719 | 306 | 2574 |
| **GO:BP** | cellular respor | GO:0036294 | 4.68E-10 | 9.33001774 | 224 | 2574 |
| **GO:BP** | synapse orgar | GO:0050808 | 4.80E-10 | 9.31912807 | 432 | 2574 |
| **GO:BP** | regulation of r | GO:0050767 | 4.80E-10 | 9.31850357 | 373 | 2574 |
| **GO:BP** | positive regula | GO:0050769 | 4.84E-10 | 9.31482659 | 233 | 2574 |
| **GO:BP** | morphogenes | GO:0001738 | 4.94E-10 | 9.30612072 | 152 | 2574 |
| **GO:BP** | cellular respor | GO:0034620 | 5.02E-10 | 9.29937352 | 148 | 2574 |
| **GO:BP** | regulation of r | GO:0019219 | 5.22E-10 | 9.28197455 | 4123 | 2574 |
| **GO:BP** | regulation of p | GO:0032271 | 5.54E-10 | 9.25616097 | 238 | 2574 |
| **GO:BP** | regulation of c | GO:0031326 | 5.62E-10 | 9.25064767 | 4307 | 2574 |
| **GO:BP** | positive regula | GO:0050772 | 5.73E-10 | 9.24219898 | 84 | 2574 |
| **GO:BP** | cellular respor | GO:0035967 | 5.89E-10 | 9.22954511 | 169 | 2574 |
| **GO:BP** | establishment | GO:0051236 | 6.19E-10 | 9.20846126 | 186 | 2574 |
| **GO:BP** | cellular respor | GO:0071453 | 6.30E-10 | 9.20086884 | 243 | 2574 |
| **GO:BP** | negative regul | GO:1903312 | 6.70E-10 | 9.17388086 | 95 | 2574 |
| **GO:BP** | regulation of c | GO:2000112 | 6.90E-10 | 9.16129082 | 4102 | 2574 |
| **GO:BP** | mitotic cell cyc | GO:0044772 | 7.36E-10 | 9.1332413 | 588 | 2574 |
| **GO:BP** | Wnt signaling | GO:0060071 | 7.39E-10 | 9.13113531 | 110 | 2574 |
| **GO:BP** | negative regul | GO:0045934 | 7.90E-10 | 9.10248125 | 1569 | 2574 |
| **GO:BP** | regulation of u | GO:2000058 | 8.44E-10 | 9.07387536 | 166 | 2574 |
| **GO:BP** | viral life cycle | GO:0019058 | 8.81E-10 | 9.05482788 | 386 | 2574 |
| **GO:BP** | positive regula | GO:0045935 | 9.66E-10 | 9.01486748 | 1943 | 2574 |
| **GO:BP** | establishment | GO:0001736 | 9.83E-10 | 9.00752516 | 130 | 2574 |
| **GO:BP** | establishment | GO:0007164 | 9.83E-10 | 9.00752516 | 130 | 2574 |
| **GO:BP** | regulation of c | GO:0008361 | 1.19E-09 | 8.9255117 | 184 | 2574 |
| **GO:BP** | negative regul | GO:0042177 | 1.33E-09 | 8.87577471 | 123 | 2574 |
| **GO:BP** | antigen proce | GO:0048002 | 1.34E-09 | 8.87390174 | 193 | 2574 |
| **GO:BP** | regulation of p | GO:0032434 | 1.42E-09 | 8.84701404 | 135 | 2574 |
| **GO:BP** | positive regula | GO:1903829 | 1.55E-09 | 8.80913572 | 307 | 2574 |
| **GO:BP** | protein polym | GO:0051258 | 1.55E-09 | 8.80913572 | 307 | 2574 |
| **GO:BP** | biological proc | GO:0044403 | 1.62E-09 | 8.79122793 | 331 | 2574 |
| **GO:BP** | regulation of c | GO:0022604 | 1.64E-09 | 8.78524333 | 312 | 2574 |
| **GO:BP** | dendrite devel | GO:0016358 | 1.65E-09 | 8.78216603 | 247 | 2574 |
| **GO:BP** | positive regula | GO:0031331 | 1.83E-09 | 8.73770183 | 460 | 2574 |

| | | | | | | |
|---|---|---|---|---|---|---|
| **GO:BP** | response to o | GO:0070482 | 1.89E-09 | 8.72365939 | 410 | 2574 |
| **GO:BP** | response to o | GO:1901700 | 2.13E-09 | 8.67249546 | 1748 | 2574 |
| **GO:BP** | regulation of r | GO:0007346 | 2.36E-09 | 8.62751179 | 612 | 2574 |
| **GO:BP** | myeloid leuko | GO:0002274 | 2.49E-09 | 8.60371143 | 671 | 2574 |
| **GO:BP** | protein polyut | GO:0000209 | 2.57E-09 | 8.58946888 | 343 | 2574 |
| **GO:BP** | movement of | GO:0006928 | 2.82E-09 | 8.55026879 | 2319 | 2574 |
| **GO:BP** | phosphorylati | GO:0016310 | 3.35E-09 | 8.4751406 | 1996 | 2574 |
| **GO:BP** | regulation of e | GO:0090175 | 3.58E-09 | 8.44647796 | 114 | 2574 |
| **GO:BP** | regulation of F | GO:0051252 | 3.76E-09 | 8.42484092 | 3847 | 2574 |
| **GO:BP** | regulation of p | GO:0030162 | 4.04E-09 | 8.39371117 | 756 | 2574 |
| **GO:BP** | non-canonica | GO:0035567 | 5.04E-09 | 8.29766096 | 151 | 2574 |
| **GO:BP** | mRNA transp | GO:0051028 | 5.04E-09 | 8.29766096 | 151 | 2574 |
| **GO:BP** | regulation of a | GO:0010506 | 5.22E-09 | 8.28252118 | 337 | 2574 |
| **GO:BP** | positive regula | GO:0023056 | 5.32E-09 | 8.27385663 | 1814 | 2574 |
| **GO:BP** | negative regul | GO:0048025 | 5.41E-09 | 8.26650532 | 22 | 2574 |
| **GO:BP** | response to s | GO:0050896 | 5.49E-09 | 8.26067482 | 9487 | 2574 |
| **GO:BP** | regulation of p | GO:0019220 | 5.50E-09 | 8.25997373 | 1640 | 2574 |
| **GO:BP** | regulation of p | GO:0051174 | 5.50E-09 | 8.25997373 | 1640 | 2574 |
| **GO:BP** | ribonucleopro | GO:0071166 | 5.66E-09 | 8.24680374 | 131 | 2574 |
| **GO:BP** | regulation of p | GO:1903320 | 6.05E-09 | 8.2184623 | 248 | 2574 |
| **GO:BP** | positive regula | GO:0051962 | 6.74E-09 | 8.17124142 | 281 | 2574 |
| **GO:BP** | regulation of r | GO:1901990 | 6.94E-09 | 8.15882918 | 453 | 2574 |
| **GO:BP** | actin cytoskel | GO:0030036 | 7.48E-09 | 8.12607496 | 712 | 2574 |
| **GO:BP** | cell-cell signali | GO:0007267 | 8.12E-09 | 8.0904851 | 1735 | 2574 |
| **GO:BP** | regulation of b | GO:0051098 | 8.45E-09 | 8.07327553 | 364 | 2574 |
| **GO:BP** | negative regul | GO:0010648 | 1.06E-08 | 7.97528245 | 1499 | 2574 |
| **GO:BP** | regulation of r | GO:0048583 | 1.09E-08 | 7.96380248 | 4416 | 2574 |
| **GO:BP** | positive regula | GO:0051254 | 1.09E-08 | 7.96379623 | 1775 | 2574 |
| **GO:BP** | negative regul | GO:0051253 | 1.10E-08 | 7.95949179 | 1452 | 2574 |
| **GO:BP** | negative regul | GO:0009968 | 1.20E-08 | 7.91965319 | 1394 | 2574 |
| **GO:BP** | protein-contai | GO:0032984 | 1.31E-08 | 7.88277087 | 337 | 2574 |
| **GO:BP** | negative regul | GO:0023057 | 1.36E-08 | 7.86754107 | 1502 | 2574 |
| **GO:BP** | positive regula | GO:0010647 | 1.44E-08 | 7.84098908 | 1809 | 2574 |
| **GO:BP** | ribonucleopro | GO:0071426 | 1.58E-08 | 7.80066524 | 130 | 2574 |
| **GO:BP** | positive regula | GO:0034250 | 1.71E-08 | 7.7658888 | 172 | 2574 |
| **GO:BP** | regulation of c | GO:1901987 | 2.03E-08 | 7.69252453 | 543 | 2574 |
| **GO:BP** | covalent chror | GO:0016569 | 2.16E-08 | 7.6658837 | 486 | 2574 |
| **GO:BP** | positive regula | GO:0030177 | 2.24E-08 | 7.64986933 | 186 | 2574 |
| **GO:BP** | endoplasmic r | GO:0030968 | 2.24E-08 | 7.64932752 | 127 | 2574 |
| **GO:BP** | regulation of i | GO:0032386 | 2.44E-08 | 7.61263922 | 365 | 2574 |
| **GO:BP** | regulation of i | GO:1902531 | 2.45E-08 | 7.61155259 | 1883 | 2574 |
| **GO:BP** | actin filament- | GO:0030029 | 2.46E-08 | 7.60881507 | 810 | 2574 |
| **GO:BP** | regulation of \ | GO:0030111 | 2.47E-08 | 7.60667937 | 380 | 2574 |
| **GO:BP** | regulation of l | GO:0032879 | 2.68E-08 | 7.5711572 | 2906 | 2574 |
| **GO:BP** | histone modif | GO:0016570 | 2.71E-08 | 7.56695827 | 472 | 2574 |

| | | | | | | |
|---|---|---|---|---|---|---|
| **GO:BP** | protein-DNA c | GO:0071824 | 2.79E-08 | 7.55431207 | 346 | 2574 |
| **GO:BP** | endomembrai | GO:0010256 | 2.90E-08 | 7.53793022 | 572 | 2574 |
| **GO:BP** | response to o | GO:0014070 | 3.20E-08 | 7.49443633 | 1004 | 2574 |
| **GO:BP** | positive regula | GO:0090263 | 3.32E-08 | 7.47852254 | 153 | 2574 |
| **GO:BP** | regulation of r | GO:0051960 | 3.46E-08 | 7.46054121 | 453 | 2574 |
| **GO:BP** | response to h | GO:0001666 | 3.46E-08 | 7.46053439 | 367 | 2574 |
| **GO:BP** | response to d | GO:0036293 | 3.48E-08 | 7.45812024 | 382 | 2574 |
| **GO:BP** | antigen proces | GO:0002479 | 3.71E-08 | 7.43112246 | 75 | 2574 |
| **GO:BP** | RNA export fro | GO:0006405 | 4.37E-08 | 7.35981788 | 129 | 2574 |
| **GO:BP** | biological proc | GO:0051701 | 4.42E-08 | 7.35409955 | 243 | 2574 |
| **GO:BP** | cell activation | GO:0002263 | 4.58E-08 | 7.33925246 | 727 | 2574 |
| **GO:BP** | embryo develo | GO:0009792 | 4.81E-08 | 7.31818142 | 651 | 2574 |
| **GO:BP** | animal organ ( | GO:0048513 | 5.10E-08 | 7.29206968 | 3701 | 2574 |
| **GO:BP** | organelle local | GO:0051640 | 5.16E-08 | 7.28698099 | 657 | 2574 |
| **GO:BP** | chordate emb | GO:0043009 | 5.90E-08 | 7.22881785 | 631 | 2574 |
| **GO:BP** | retrograde ves | GO:0006890 | 6.11E-08 | 7.21395046 | 87 | 2574 |
| **GO:BP** | positive regula | GO:0044093 | 6.17E-08 | 7.20971215 | 1774 | 2574 |
| **GO:BP** | regulation of e | GO:0061387 | 6.21E-08 | 7.20689499 | 110 | 2574 |
| **GO:BP** | antigen proces | GO:0019882 | 7.32E-08 | 7.13565216 | 236 | 2574 |
| **GO:BP** | transmembrar | GO:0007169 | 7.62E-08 | 7.11781011 | 759 | 2574 |
| **GO:BP** | pre-replicative | GO:0036388 | 7.74E-08 | 7.11131143 | 66 | 2574 |
| **GO:BP** | regulation of c | GO:0006521 | 7.74E-08 | 7.11131143 | 66 | 2574 |
| **GO:BP** | regulation of r | GO:0080134 | 7.88E-08 | 7.1032424 | 1626 | 2574 |
| **GO:BP** | neuron projec | GO:0097485 | 8.71E-08 | 7.05977262 | 284 | 2574 |
| **GO:BP** | positive regula | GO:0031334 | 8.83E-08 | 7.05380942 | 265 | 2574 |
| **GO:BP** | leukocyte activ | GO:0002366 | 1.04E-07 | 6.98441889 | 723 | 2574 |
| **GO:BP** | positive regula | GO:1903364 | 1.06E-07 | 6.97559938 | 157 | 2574 |
| **GO:BP** | phosphate-co | GO:0006796 | 1.06E-07 | 6.97469151 | 2987 | 2574 |
| **GO:BP** | positive regula | GO:0009967 | 1.24E-07 | 6.9050009 | 1626 | 2574 |
| **GO:BP** | regulation of t | GO:0051338 | 1.35E-07 | 6.8700236 | 1036 | 2574 |
| **GO:BP** | cellular respor | GO:0071417 | 1.46E-07 | 6.83524379 | 671 | 2574 |
| **GO:BP** | regulation of p | GO:0042325 | 1.57E-07 | 6.80501399 | 1419 | 2574 |
| **GO:BP** | regulation of a | GO:0030516 | 1.62E-07 | 6.78983608 | 97 | 2574 |
| **GO:BP** | negative regul | GO:1902750 | 1.70E-07 | 6.76981847 | 101 | 2574 |
| **GO:BP** | regulation of c | GO:0060828 | 1.71E-07 | 6.76739195 | 302 | 2574 |
| **GO:BP** | axon guidanc | GO:0007411 | 1.84E-07 | 6.73585564 | 283 | 2574 |
| **GO:BP** | negative regul | GO:0010977 | 1.90E-07 | 6.72231966 | 142 | 2574 |
| **GO:BP** | regulation of r | GO:0031110 | 2.09E-07 | 6.67902333 | 90 | 2574 |
| **GO:BP** | organelle asse | GO:0070925 | 2.26E-07 | 6.64590234 | 921 | 2574 |
| **GO:BP** | protein-contai | GO:0031503 | 2.30E-07 | 6.63832916 | 289 | 2574 |
| **GO:BP** | regulation of a | GO:0090066 | 2.54E-07 | 6.59578631 | 519 | 2574 |
| **GO:BP** | negative regul | GO:1903363 | 3.06E-07 | 6.51475665 | 76 | 2574 |
| **GO:BP** | regulation of a | GO:2001233 | 3.15E-07 | 6.50203583 | 365 | 2574 |
| **GO:BP** | peptidyl-amin | GO:0018193 | 3.22E-07 | 6.49266848 | 1309 | 2574 |
| **GO:BP** | cell junction a | GO:0034329 | 3.24E-07 | 6.48890238 | 442 | 2574 |

| | | | | | | |
|---|---|---|---|---|---|---|
| **GO:BP** | cellular respor | GO:1901699 | 3.43E-07 | 6.46461415 | 728 | 2574 |
| **GO:BP** | antigen proce: | GO:0042590 | 3.51E-07 | 6.45444458 | 80 | 2574 |
| **GO:BP** | cell communic | GO:0007154 | 3.82E-07 | 6.41793543 | 6961 | 2574 |
| **GO:BP** | cell migration | GO:0016477 | 4.01E-07 | 6.39732293 | 1660 | 2574 |
| **GO:BP** | morphogenes | GO:0002009 | 4.43E-07 | 6.35340771 | 566 | 2574 |
| **GO:BP** | tissue morphc | GO:0048729 | 4.46E-07 | 6.35108487 | 675 | 2574 |
| **GO:BP** | phosphorus n | GO:0006793 | 4.75E-07 | 6.32361966 | 3014 | 2574 |
| **GO:BP** | positive regula | GO:0045727 | 4.89E-07 | 6.31093505 | 141 | 2574 |
| **GO:BP** | negative regul | GO:0009890 | 4.95E-07 | 6.30565194 | 1791 | 2574 |
| **GO:BP** | mRNA export | GO:0006406 | 4.99E-07 | 6.30190971 | 112 | 2574 |
| **GO:BP** | mRNA-contair | GO:0071427 | 4.99E-07 | 6.30190971 | 112 | 2574 |
| **GO:BP** | mRNA 3'-end | GO:0031124 | 5.02E-07 | 6.29923658 | 104 | 2574 |
| **GO:BP** | regulation of I | GO:0031056 | 5.05E-07 | 6.29706345 | 154 | 2574 |
| **GO:BP** | gliogenesis | GO:0042063 | 5.24E-07 | 6.28047482 | 318 | 2574 |
| **GO:BP** | regulation of r | GO:0070507 | 5.52E-07 | 6.25816425 | 150 | 2574 |
| **GO:BP** | locomotion | GO:0040011 | 5.53E-07 | 6.25688897 | 2039 | 2574 |
| **GO:BP** | regulation of c | GO:1902275 | 5.56E-07 | 6.25496199 | 222 | 2574 |
| **GO:BP** | canonical Wnt | GO:0060070 | 6.26E-07 | 6.20358379 | 349 | 2574 |
| **GO:BP** | hematopoietic | GO:0060218 | 6.33E-07 | 6.19857828 | 89 | 2574 |
| **GO:BP** | protein-DNA c | GO:0065004 | 7.15E-07 | 6.14544288 | 300 | 2574 |
| **GO:BP** | signaling | GO:0023052 | 7.65E-07 | 6.11656542 | 6937 | 2574 |
| **GO:BP** | cellular respor | GO:1901701 | 7.87E-07 | 6.10376547 | 1237 | 2574 |
| **GO:BP** | negative regul | GO:0010558 | 9.25E-07 | 6.03376412 | 1690 | 2574 |
| **GO:BP** | negative regul | GO:0031345 | 9.26E-07 | 6.03346396 | 192 | 2574 |
| **GO:BP** | intrinsic apopt | GO:0097193 | 1.0113E-06 | 5.99511298 | 297 | 2574 |
| **GO:BP** | regulation of r | GO:0016241 | 1.017E-06 | 5.99269142 | 161 | 2574 |
| **GO:BP** | ncRNA metab | GO:0034660 | 1.0366E-06 | 5.98440372 | 535 | 2574 |
| **GO:BP** | ATP metabolic | GO:0046034 | 1.0472E-06 | 5.97998254 | 317 | 2574 |
| **GO:BP** | postsynapse ( | GO:0099173 | 1.0536E-06 | 5.97733006 | 170 | 2574 |
| **GO:BP** | positive regula | GO:0051094 | 1.0901E-06 | 5.9625397 | 1354 | 2574 |
| **GO:BP** | regulation of e | GO:0070201 | 1.1012E-06 | 5.95813906 | 595 | 2574 |
| **GO:BP** | positive regula | GO:0010628 | 1.1387E-06 | 5.94357815 | 1206 | 2574 |
| **GO:BP** | negative regul | GO:0031333 | 1.1439E-06 | 5.94162528 | 144 | 2574 |
| **GO:BP** | activation of ir | GO:0002218 | 1.1439E-06 | 5.94162528 | 144 | 2574 |
| **GO:BP** | regulation of p | GO:0031396 | 1.144E-06 | 5.9415872 | 216 | 2574 |
| **GO:BP** | regulation of I | GO:1901532 | 1.3523E-06 | 5.86892065 | 91 | 2574 |
| **GO:BP** | positive regula | GO:0030307 | 1.3525E-06 | 5.86885777 | 171 | 2574 |
| **GO:BP** | negative regul | GO:0010972 | 1.4162E-06 | 5.84887602 | 99 | 2574 |
| **GO:BP** | nucleobase-cc | GO:0015931 | 1.6027E-06 | 5.79514066 | 246 | 2574 |
| **GO:BP** | cellular respor | GO:0032870 | 1.6452E-06 | 5.78378579 | 681 | 2574 |
| **GO:BP** | negative regul | GO:2001234 | 1.7896E-06 | 5.74723347 | 237 | 2574 |
| **GO:BP** | response to p | GO:1901652 | 2.0016E-06 | 5.69862487 | 551 | 2574 |
| **GO:BP** | positive regula | GO:0045773 | 2.0961E-06 | 5.67858285 | 42 | 2574 |
| **GO:BP** | negative regul | GO:2000113 | 2.3513E-06 | 5.62869883 | 1679 | 2574 |
| **GO:BP** | tissue develop | GO:0009888 | 2.3624E-06 | 5.62663875 | 2093 | 2574 |

| | | | | | | |
|---|---|---|---|---|---|---|
| **GO:BP** | positive regula | GO:0045862 | 2.415E-06 | 5.61708192 | 383 | 2574 |
| **GO:BP** | positive regula | GO:1902905 | 2.5164E-06 | 5.59922741 | 215 | 2574 |
| **GO:BP** | response to w | GO:0009611 | 2.7201E-06 | 5.56540814 | 680 | 2574 |
| **GO:BP** | regulation of h | GO:1902036 | 3.0789E-06 | 5.51160378 | 74 | 2574 |
| **GO:BP** | response to c | GO:0034097 | 3.0832E-06 | 5.51100014 | 1242 | 2574 |
| **GO:BP** | positive regula | GO:1903508 | 3.341E-06 | 5.47611997 | 1623 | 2574 |
| **GO:BP** | positive regula | GO:0045893 | 3.341E-06 | 5.47611997 | 1623 | 2574 |
| **GO:BP** | positive regula | GO:0045597 | 3.4729E-06 | 5.45930915 | 892 | 2574 |
| **GO:BP** | positive regula | GO:1902680 | 3.5801E-06 | 5.44610782 | 1624 | 2574 |
| **GO:BP** | animal organ | GO:0009887 | 3.7878E-06 | 5.42161649 | 1079 | 2574 |
| **GO:BP** | positive regula | GO:1904951 | 3.8421E-06 | 5.41542822 | 345 | 2574 |
| **GO:BP** | protein modifi | GO:0070646 | 4.0875E-06 | 5.38854672 | 310 | 2574 |
| **GO:BP** | modulation of | GO:0051817 | 4.298E-06 | 5.36673673 | 123 | 2574 |
| **GO:BP** | protein depoly | GO:0051261 | 4.5852E-06 | 5.33863815 | 119 | 2574 |
| **GO:BP** | response to p | GO:0043434 | 4.6792E-06 | 5.3298262 | 461 | 2574 |
| **GO:BP** | regulation of s | GO:2000736 | 4.8571E-06 | 5.3136191 | 115 | 2574 |
| **GO:BP** | regulation of s | GO:0050803 | 4.961E-06 | 5.30443186 | 223 | 2574 |
| **GO:BP** | regulation of r | GO:0031113 | 5.1624E-06 | 5.28714866 | 57 | 2574 |
| **GO:BP** | microtubule p | GO:0046785 | 5.3202E-06 | 5.27407022 | 83 | 2574 |
| **GO:BP** | antigen proces | GO:0002474 | 5.6061E-06 | 5.25134122 | 99 | 2574 |
| **GO:BP** | positive regula | GO:0031401 | 5.7713E-06 | 5.23872775 | 1084 | 2574 |
| **GO:BP** | negative regul | GO:0010563 | 6.2362E-06 | 5.20507949 | 544 | 2574 |
| **GO:BP** | positive regula | GO:1901800 | 6.5771E-06 | 5.18196711 | 116 | 2574 |
| **GO:BP** | regulation of p | GO:0001932 | 6.7324E-06 | 5.17182791 | 1246 | 2574 |
| **GO:BP** | alternative mR | GO:0000380 | 6.9318E-06 | 5.15915623 | 76 | 2574 |
| **GO:BP** | positive regula | GO:2001252 | 7.4442E-06 | 5.1281817 | 192 | 2574 |
| **GO:BP** | protein localiz | GO:1990778 | 7.4472E-06 | 5.12800944 | 339 | 2574 |
| **GO:BP** | protein phosp | GO:0006468 | 8.212E-06 | 5.08555171 | 1673 | 2574 |
| **GO:BP** | modulation of | GO:0050804 | 8.9756E-06 | 5.0469389 | 434 | 2574 |
| **GO:BP** | multicellular o | GO:0032501 | 9.2431E-06 | 5.0341836 | 8000 | 2574 |
| **GO:BP** | regulation of s | GO:0050807 | 9.6187E-06 | 5.01688444 | 212 | 2574 |
| **GO:BP** | regulation of t | GO:0099177 | 1.0273E-05 | 4.98829916 | 435 | 2574 |
| **GO:BP** | negative regul | GO:0044092 | 1.0611E-05 | 4.97425227 | 1228 | 2574 |
| **GO:BP** | negative regul | GO:0045936 | 1.0632E-05 | 4.97337531 | 543 | 2574 |
| **GO:BP** | positive regula | GO:0010976 | 1.0662E-05 | 4.97215866 | 166 | 2574 |
| **GO:BP** | regulation of i | GO:0033157 | 1.073E-05 | 4.96938359 | 256 | 2574 |
| **GO:BP** | response to o | GO:0006979 | 1.1277E-05 | 4.94780903 | 457 | 2574 |
| **GO:BP** | Ras protein si | GO:0007265 | 1.1357E-05 | 4.94474929 | 352 | 2574 |
| **GO:BP** | negative regul | GO:0031327 | 1.2098E-05 | 4.91729185 | 1759 | 2574 |
| **GO:BP** | endodermal c | GO:0035987 | 1.2527E-05 | 4.90214714 | 45 | 2574 |
| **GO:BP** | protein deubic | GO:0016579 | 1.3404E-05 | 4.87277275 | 292 | 2574 |
| **GO:BP** | protein localiz | GO:0072659 | 1.3426E-05 | 4.87205063 | 287 | 2574 |
| **GO:BP** | regulation of i | GO:2001242 | 1.3486E-05 | 4.8701026 | 167 | 2574 |
| **GO:BP** | dendrite morp | GO:0048813 | 1.3619E-05 | 4.86585042 | 149 | 2574 |
| **GO:BP** | cellular respor | GO:0071345 | 1.3857E-05 | 4.85834399 | 1148 | 2574 |

| | | | | | | |
|---|---|---|---|---|---|---|
| **GO:BP** | regulation of c | GO:0048638 | 1.4116E-05 | 4.8502983 | 338 | 2574 |
| **GO:BP** | microtubule p | GO:0031109 | 1.4653E-05 | 4.83406413 | 123 | 2574 |
| **GO:BP** | regulation of t | GO:0006446 | 1.5606E-05 | 4.80670563 | 82 | 2574 |
| **GO:BP** | regulation of t | GO:0032204 | 1.5606E-05 | 4.80670563 | 82 | 2574 |
| **GO:BP** | cellular respor | GO:0034605 | 1.5871E-05 | 4.79939949 | 119 | 2574 |
| **GO:BP** | negative regul | GO:1902532 | 1.7066E-05 | 4.76787722 | 558 | 2574 |
| **GO:BP** | ncRNA proces | GO:0034470 | 1.7499E-05 | 4.75697544 | 439 | 2574 |
| **GO:BP** | cell motility | GO:0048870 | 1.8432E-05 | 4.73443077 | 1840 | 2574 |
| **GO:BP** | localization of | GO:0051674 | 1.8432E-05 | 4.73443077 | 1840 | 2574 |
| **GO:BP** | cell-substrate | GO:0150115 | 1.9388E-05 | 4.71246468 | 107 | 2574 |
| **GO:BP** | regulation of p | GO:1900180 | 2.1999E-05 | 4.65759448 | 142 | 2574 |
| **GO:BP** | MAPK cascad | GO:0000165 | 2.2292E-05 | 4.65185715 | 965 | 2574 |
| **GO:BP** | anaphase-pro | GO:0031145 | 2.2572E-05 | 4.64642096 | 87 | 2574 |
| **GO:BP** | stem cell diffe | GO:0048863 | 2.6204E-05 | 4.58162821 | 261 | 2574 |
| **GO:BP** | negative regul | GO:2001243 | 2.9195E-05 | 4.53469892 | 100 | 2574 |
| **GO:BP** | external encap | GO:0045229 | 3.0925E-05 | 4.50969431 | 422 | 2574 |
| **GO:BP** | regulation of a | GO:2000027 | 3.3784E-05 | 4.47128691 | 185 | 2574 |
| **GO:BP** | Fc receptor si | GO:0038093 | 3.5172E-05 | 4.45380194 | 238 | 2574 |
| **GO:BP** | positive regula | GO:1903052 | 3.5428E-05 | 4.45065045 | 135 | 2574 |
| **GO:BP** | negative regul | GO:1903051 | 3.6649E-05 | 4.4359388 | 65 | 2574 |
| **GO:BP** | interleukin-1-r | GO:0070498 | 3.7631E-05 | 4.42445832 | 105 | 2574 |
| **GO:BP** | cytoskeleton-c | GO:0030705 | 3.9994E-05 | 4.39800119 | 200 | 2574 |
| **GO:BP** | cell activation | GO:0001775 | 4.0719E-05 | 4.39019904 | 1507 | 2574 |
| **GO:BP** | extracellular m | GO:0030198 | 4.2032E-05 | 4.37641647 | 419 | 2574 |
| **GO:BP** | neuron projec | GO:0106027 | 4.3338E-05 | 4.36313048 | 93 | 2574 |
| **GO:BP** | extracellular st | GO:0043062 | 4.791E-05 | 4.31957285 | 420 | 2574 |
| **GO:BP** | ameboidal-typ | GO:0001667 | 5.0206E-05 | 4.2992403 | 496 | 2574 |
| **GO:BP** | gastrulation | GO:0007369 | 5.1441E-05 | 4.28868748 | 187 | 2574 |
| **GO:BP** | dendritic spin | GO:0060996 | 5.3996E-05 | 4.26763549 | 102 | 2574 |
| **GO:BP** | cell surface re | GO:0007166 | 5.8149E-05 | 4.23545567 | 3226 | 2574 |
| **GO:BP** | establishment | GO:0051656 | 5.8383E-05 | 4.23371187 | 443 | 2574 |
| **GO:BP** | homeostatic p | GO:0042592 | 6.132E-05 | 4.21240039 | 1985 | 2574 |
| **GO:BP** | positive regula | GO:0032273 | 6.6833E-05 | 4.17500736 | 142 | 2574 |
| **GO:BP** | positive regula | GO:0032388 | 6.9108E-05 | 4.16046864 | 227 | 2574 |
| **GO:BP** | actin filament | GO:0007015 | 7.4982E-05 | 4.12504558 | 445 | 2574 |
| **GO:BP** | cytoplasmic tr | GO:0002183 | 7.5762E-05 | 4.12055 | 35 | 2574 |
| **GO:BP** | microtubule-b | GO:0007017 | 7.7487E-05 | 4.11076967 | 927 | 2574 |
| **GO:BP** | positive regula | GO:0051099 | 8.03E-05 | 4.09528222 | 175 | 2574 |
| **GO:BP** | peptidyl-lysine | GO:0018205 | 8.098E-05 | 4.09162351 | 408 | 2574 |
| **GO:BP** | regulation of c | GO:0033238 | 8.9758E-05 | 4.04692798 | 83 | 2574 |
| **GO:BP** | DNA metaboli | GO:0006259 | 9.0629E-05 | 4.04273493 | 976 | 2574 |
| **GO:BP** | anatomical str | GO:0048646 | 9.2622E-05 | 4.03328536 | 1209 | 2574 |
| **GO:BP** | negative regul | GO:2000059 | 9.9742E-05 | 4.00112405 | 49 | 2574 |
| **GO:BP** | negative regul | GO:0048585 | 0.00010549 | 3.97677524 | 1938 | 2574 |
| **GO:BP** | in utero embr | GO:0001701 | 0.00011959 | 3.92231136 | 379 | 2574 |

| | | | | | | |
|---|---|---|---|---|---|---|
| **GO:BP** | regulation of p | GO:0051223 | 0.00011967 | 3.92203084 | 570 | 2574 |
| **GO:BP** | regulation of r | GO:0032886 | 0.00012153 | 3.91532689 | 245 | 2574 |
| **GO:BP** | ephrin recepto | GO:0048013 | 0.0001218 | 3.9143674 | 88 | 2574 |
| **GO:BP** | positive regula | GO:0043085 | 0.0001236 | 3.90799678 | 1383 | 2574 |
| **GO:BP** | endoderm dev | GO:0007492 | 0.00012733 | 3.89507618 | 80 | 2574 |
| **GO:BP** | mitochondrion | GO:0007005 | 0.00012737 | 3.89492758 | 565 | 2574 |
| **GO:BP** | positive regula | GO:0051347 | 0.00013695 | 3.8634469 | 690 | 2574 |
| **GO:BP** | cell-substrate | GO:0007044 | 0.00014051 | 3.8522924 | 101 | 2574 |
| **GO:BP** | myeloid cell ho | GO:0002262 | 0.00014292 | 3.84490135 | 159 | 2574 |
| **GO:BP** | regulation of h | GO:0051336 | 0.00014979 | 3.82452084 | 1252 | 2574 |
| **GO:BP** | cellular respor | GO:0006974 | 0.00015214 | 3.81774261 | 894 | 2574 |
| **GO:BP** | cellular respor | GO:1901653 | 0.0001613 | 3.79235313 | 408 | 2574 |
| **GO:BP** | generation of | GO:0006091 | 0.00016317 | 3.78735622 | 545 | 2574 |
| **GO:BP** | cellular respor | GO:0071407 | 0.0001702 | 3.76903239 | 624 | 2574 |
| **GO:BP** | negative regul | GO:0045892 | 0.00017077 | 3.76758955 | 1339 | 2574 |
| **GO:BP** | formation of c | GO:0001732 | 0.00017149 | 3.76576354 | 16 | 2574 |
| **GO:BP** | positive regula | GO:0051222 | 0.00017284 | 3.7623627 | 329 | 2574 |
| **GO:BP** | organic cyclic | GO:1901362 | 0.00017297 | 3.76202207 | 4446 | 2574 |
| **GO:BP** | endoderm forn | GO:0001706 | 0.00018846 | 3.72477493 | 54 | 2574 |
| **GO:BP** | maintenance c | GO:0098727 | 0.00018896 | 3.72363285 | 151 | 2574 |
| **GO:BP** | negative regul | GO:1903507 | 0.00019532 | 3.7092424 | 1341 | 2574 |
| **GO:BP** | glial cell differe | GO:0010001 | 0.00020144 | 3.69585886 | 233 | 2574 |
| **GO:BP** | regulation of c | GO:0033044 | 0.00020464 | 3.68900547 | 299 | 2574 |
| **GO:BP** | histone ubiqu | GO:0016574 | 0.00020927 | 3.67928796 | 47 | 2574 |
| **GO:BP** | regulation of u | GO:0051438 | 0.00021262 | 3.6723962 | 58 | 2574 |
| **GO:BP** | regulation of r | GO:2000026 | 0.00021704 | 3.66346397 | 1459 | 2574 |
| **GO:BP** | negative regul | GO:1902679 | 0.00022325 | 3.65121389 | 1343 | 2574 |
| **GO:BP** | erythrocyte ho | GO:0034101 | 0.000226 | 3.64589617 | 129 | 2574 |
| **GO:BP** | negative regul | GO:0010948 | 0.0002286 | 3.64093068 | 400 | 2574 |
| **GO:BP** | vesicle buddir | GO:0006900 | 0.00022872 | 3.64070401 | 107 | 2574 |
| **GO:BP** | focal adhesior | GO:0048041 | 0.00023943 | 3.62082245 | 86 | 2574 |
| **GO:BP** | negative regul | GO:0043086 | 0.00024349 | 3.61351403 | 841 | 2574 |
| **GO:BP** | adherens junc | GO:0034332 | 0.00024482 | 3.61114483 | 66 | 2574 |
| **GO:BP** | IRE1-mediate | GO:0036498 | 0.00024482 | 3.61114483 | 66 | 2574 |
| **GO:BP** | signal transdu | GO:0007165 | 0.00024721 | 3.60693066 | 6447 | 2574 |
| **GO:BP** | regulation of c | GO:0150116 | 0.00025263 | 3.59750665 | 70 | 2574 |
| **GO:BP** | positive regula | GO:0010942 | 0.00025888 | 3.58690014 | 691 | 2574 |
| **GO:BP** | aerobic respira | GO:0009060 | 0.00026037 | 3.5844158 | 200 | 2574 |
| **GO:BP** | stem cell popu | GO:0019827 | 0.00027161 | 3.56605807 | 148 | 2574 |
| **GO:BP** | positive regula | GO:0045944 | 0.00027262 | 3.56444448 | 1200 | 2574 |
| **GO:BP** | microtubule cy | GO:0000226 | 0.00027461 | 3.56127917 | 663 | 2574 |
| **GO:BP** | negative regul | GO:0042326 | 0.00028659 | 3.54274319 | 418 | 2574 |
| **GO:BP** | nucleic acid-te | GO:0097659 | 0.00028715 | 3.54189556 | 3715 | 2574 |
| **GO:BP** | regulation of k | GO:0043549 | 0.00030023 | 3.52254584 | 920 | 2574 |
| **GO:BP** | regulation of p | GO:0043393 | 0.00030606 | 3.51419786 | 196 | 2574 |

| | | | | | | |
|---|---|---|---|---|---|---|
| **GO:BP** | nucleobase-co | GO:0034654 | 0.00032425 | 3.48911946 | 4202 | 2574 |
| **GO:BP** | transcription, | GO:0006351 | 0.00034704 | 3.45961694 | 3713 | 2574 |
| **GO:BP** | regulation of c | GO:1900034 | 0.00035363 | 3.45144792 | 79 | 2574 |
| **GO:BP** | cellular respor | GO:0062197 | 0.00039014 | 3.40878056 | 356 | 2574 |
| **GO:BP** | heterocycle bi | GO:0018130 | 0.00041529 | 3.38165303 | 4275 | 2574 |
| **GO:BP** | homeostasis c | GO:0048872 | 0.00042876 | 3.36778388 | 273 | 2574 |
| **GO:BP** | selective autoj | GO:0061912 | 0.00044506 | 3.35158519 | 88 | 2574 |
| **GO:BP** | dendritic spine | GO:0097061 | 0.00046933 | 3.32851829 | 84 | 2574 |
| **GO:BP** | regulation of a | GO:0000381 | 0.00047392 | 3.32429789 | 60 | 2574 |
| **GO:BP** | modulation by | GO:0044003 | 0.00047392 | 3.32429789 | 60 | 2574 |
| **GO:BP** | neural tube de | GO:0021915 | 0.00048362 | 3.31549179 | 160 | 2574 |
| **GO:BP** | positive regula | GO:0090316 | 0.00049555 | 3.30490986 | 184 | 2574 |
| **GO:BP** | embryonic mo | GO:0048598 | 0.00050777 | 3.29432977 | 601 | 2574 |
| **GO:BP** | regulation of p | GO:0045859 | 0.00051823 | 3.28547954 | 809 | 2574 |
| **GO:BP** | negative regul | GO:0032435 | 0.00052368 | 3.28093317 | 35 | 2574 |
| **GO:BP** | negative regul | GO:1901799 | 0.00052602 | 3.2790016 | 49 | 2574 |
| **GO:BP** | signal transdu | GO:0072331 | 0.00053892 | 3.26847872 | 290 | 2574 |
| **GO:BP** | RNA biosynth | GO:0032774 | 0.00055153 | 3.25843235 | 3731 | 2574 |
| **GO:BP** | lamellipodium | GO:0097581 | 0.00056011 | 3.2517232 | 93 | 2574 |
| **GO:BP** | hematopoietic | GO:0002244 | 0.00056023 | 3.25163207 | 175 | 2574 |
| **GO:BP** | formation of p | GO:0001704 | 0.0005996 | 3.2221363 | 124 | 2574 |
| **GO:BP** | regulation of s | GO:0062012 | 0.00062863 | 3.20160224 | 430 | 2574 |
| **GO:BP** | leukocyte acti | GO:0045321 | 0.00065061 | 3.18667889 | 1341 | 2574 |
| **GO:BP** | viral translatio | GO:0019081 | 0.00067614 | 3.16996383 | 20 | 2574 |
| **GO:BP** | RNA 3'-end p | GO:0031123 | 0.00069217 | 3.15978462 | 157 | 2574 |
| **GO:BP** | mitochondrion | GO:0061726 | 0.0006992 | 3.1553966 | 77 | 2574 |
| **GO:BP** | autophagy of | GO:0000422 | 0.0006992 | 3.1553966 | 77 | 2574 |
| **GO:BP** | transport alon | GO:0010970 | 0.00073065 | 3.13629079 | 162 | 2574 |
| **GO:BP** | regulation of a | GO:0032970 | 0.00073082 | 3.13618888 | 404 | 2574 |
| **GO:BP** | nucleus orgar | GO:0006997 | 0.00075356 | 3.12288287 | 148 | 2574 |
| **GO:BP** | nucleotide-exc | GO:0000715 | 0.00075658 | 3.12114309 | 23 | 2574 |
| **GO:BP** | positive regula | GO:0043068 | 0.00076136 | 3.118412 | 628 | 2574 |
| **GO:BP** | positive regula | GO:0043065 | 0.00077148 | 3.11267476 | 611 | 2574 |
| **GO:BP** | gland develop | GO:0048732 | 0.00079453 | 3.09989089 | 443 | 2574 |
| **GO:BP** | positive regula | GO:0045927 | 0.00080417 | 3.09465381 | 272 | 2574 |
| **GO:BP** | regulation of D | GO:2000142 | 0.00086415 | 3.06341286 | 86 | 2574 |
| **GO:BP** | positive regula | GO:0051495 | 0.00091923 | 3.03657407 | 232 | 2574 |
| **GO:BP** | modulation of | GO:0035821 | 0.00093276 | 3.03023232 | 149 | 2574 |
| **GO:BP** | protein import | GO:0006606 | 0.00099045 | 3.00416661 | 154 | 2574 |
| **GO:BP** | aromatic comp | GO:0019438 | 0.00100767 | 2.99668241 | 4285 | 2574 |
| **GO:BP** | ribosome asse | GO:0042255 | 0.00100842 | 2.99635926 | 62 | 2574 |
| **GO:BP** | synaptic signa | GO:0099536 | 0.00103169 | 2.98645185 | 747 | 2574 |
| **GO:BP** | histone methy | GO:0016571 | 0.00108068 | 2.96630091 | 145 | 2574 |
| **GO:BP** | interleukin-12- | GO:0035722 | 0.00109682 | 2.95986439 | 47 | 2574 |
| **GO:BP** | endoplasmic r | GO:0006888 | 0.0010995 | 2.95880544 | 208 | 2574 |

| | | | | | | |
|---|---|---|---|---|---|---|
| **GO:BP** | regulation of r | GO:0051239 | 0.00111097 | 2.95429837 | 2888 | 2574 |
| **GO:BP** | negative regul | GO:0001933 | 0.00112354 | 2.94941216 | 375 | 2574 |
| **GO:BP** | DNA conforma | GO:0071103 | 0.0011487 | 2.93979351 | 359 | 2574 |
| **GO:BP** | import into nu | GO:0051170 | 0.00117516 | 2.92990127 | 174 | 2574 |
| **GO:BP** | cellular respor | GO:0034599 | 0.00118642 | 2.92576331 | 306 | 2574 |
| **GO:BP** | neuron death | GO:0070997 | 0.0012407 | 2.90633266 | 365 | 2574 |
| **GO:BP** | regulation of a | GO:0032956 | 0.0012407 | 2.90633266 | 365 | 2574 |
| **GO:BP** | negative regul | GO:0032272 | 0.00124254 | 2.90568883 | 83 | 2574 |
| **GO:BP** | negative regul | GO:0090090 | 0.00126051 | 2.89945391 | 189 | 2574 |
| **GO:BP** | response to re | GO:0000302 | 0.0012948 | 2.88779883 | 224 | 2574 |
| **GO:BP** | protein import | GO:0017038 | 0.00130211 | 2.88535326 | 219 | 2574 |
| **GO:BP** | regeneration | GO:0031099 | 0.00130555 | 2.8842063 | 209 | 2574 |
| **GO:BP** | protein localiz | GO:0070198 | 0.00130881 | 2.88312343 | 30 | 2574 |
| **GO:BP** | cellular respor | GO:0071375 | 0.00138292 | 2.85920435 | 339 | 2574 |
| **GO:BP** | regulation of r | GO:1903506 | 0.00138657 | 2.8580586 | 3536 | 2574 |
| **GO:BP** | negative regul | GO:0045786 | 0.00140228 | 2.85316472 | 487 | 2574 |
| **GO:BP** | spliceosomal | GO:0000245 | 0.00141419 | 2.84949093 | 59 | 2574 |
| **GO:BP** | positive regula | GO:0032436 | 0.00142661 | 2.84569535 | 92 | 2574 |
| **GO:BP** | cognition | GO:0050890 | 0.00146538 | 2.83404937 | 297 | 2574 |
| **GO:BP** | protein methy | GO:0006479 | 0.00148639 | 2.82786808 | 185 | 2574 |
| **GO:BP** | protein alkylati | GO:0008213 | 0.00148639 | 2.82786808 | 185 | 2574 |
| **GO:BP** | inclusion body | GO:0070841 | 0.00157768 | 2.80198215 | 24 | 2574 |
| **GO:BP** | NADH regener | GO:0006735 | 0.00157768 | 2.80198215 | 24 | 2574 |
| **GO:BP** | glucose catab | GO:0061718 | 0.00157768 | 2.80198215 | 24 | 2574 |
| **GO:BP** | canonical glyc | GO:0061621 | 0.00157768 | 2.80198215 | 24 | 2574 |
| **GO:BP** | learning or me | GO:0007611 | 0.00161303 | 2.79235651 | 256 | 2574 |
| **GO:BP** | regulation of a | GO:0110053 | 0.00162314 | 2.78964473 | 282 | 2574 |
| **GO:BP** | immune effec | GO:0002252 | 0.00162421 | 2.78935851 | 1135 | 2574 |
| **GO:BP** | regulation of I | GO:0051052 | 0.00164355 | 2.78421763 | 389 | 2574 |
| **GO:BP** | regulation of t | GO:0006355 | 0.00166411 | 2.77881912 | 3534 | 2574 |
| **GO:BP** | regulation of F | GO:2001141 | 0.00169021 | 2.77205969 | 3541 | 2574 |
| **GO:BP** | negative regul | GO:0051348 | 0.00171488 | 2.76576665 | 314 | 2574 |
| **GO:BP** | response to ir | GO:0032868 | 0.00174526 | 2.75814022 | 293 | 2574 |
| **GO:BP** | ERAD pathwa | GO:0036503 | 0.00178787 | 2.74766411 | 106 | 2574 |
| **GO:BP** | cell-substrate | GO:0031589 | 0.00179642 | 2.74559169 | 368 | 2574 |
| **GO:BP** | positive regula | GO:0032206 | 0.00186134 | 2.73017371 | 52 | 2574 |
| **GO:BP** | response to s | GO:0048545 | 0.0019528 | 2.70934132 | 385 | 2574 |
| **GO:BP** | ossification | GO:0001503 | 0.00197919 | 2.70351273 | 418 | 2574 |
| **GO:BP** | vesicle organiz | GO:0016050 | 0.00202977 | 2.69255356 | 342 | 2574 |
| **GO:BP** | regulation of r | GO:0046822 | 0.00203623 | 2.69117391 | 111 | 2574 |
| **GO:BP** | cellular respor | GO:0032869 | 0.0020743 | 2.68312788 | 232 | 2574 |
| **GO:BP** | trans-synaptic | GO:0099537 | 0.00221727 | 2.65418239 | 721 | 2574 |
| **GO:BP** | regulation of r | GO:0010821 | 0.0022206 | 2.6535288 | 158 | 2574 |
| **GO:BP** | myeloid cell di | GO:0030099 | 0.00231986 | 2.63453792 | 436 | 2574 |
| **GO:BP** | cellular protein | GO:0043624 | 0.00246556 | 2.60808519 | 228 | 2574 |

| | | | | | | |
|---|---|---|---|---|---|---|
| **GO:BP** | telencephalon | GO:0021537 | 0.00251902 | 2.59876899 | 259 | 2574 |
| **GO:BP** | cellular respor | GO:0071349 | 0.00256235 | 2.59136075 | 49 | 2574 |
| **GO:BP** | stimulatory C- | GO:0002223 | 0.00259443 | 2.58595802 | 112 | 2574 |
| **GO:BP** | response to h | GO:0009408 | 0.00269964 | 2.56869438 | 159 | 2574 |
| **GO:BP** | axo-dendritic | GO:0008088 | 0.00270949 | 2.56711177 | 73 | 2574 |
| **GO:BP** | endosomal tra | GO:0016197 | 0.00271821 | 2.56571683 | 244 | 2574 |
| **GO:BP** | regulation of c | GO:0051270 | 0.00274398 | 2.56161858 | 1149 | 2574 |
| **GO:BP** | regulation of p | GO:0043244 | 0.00275695 | 2.55957054 | 126 | 2574 |
| **GO:BP** | ER-nucleus si | GO:0006984 | 0.00275705 | 2.5595555 | 53 | 2574 |
| **GO:BP** | regulation of c | GO:1902175 | 0.00276923 | 2.55764055 | 28 | 2574 |
| **GO:BP** | rhythmic proc | GO:0048511 | 0.00277788 | 2.55628629 | 307 | 2574 |
| **GO:BP** | energy derivat | GO:0015980 | 0.00287389 | 2.54152947 | 334 | 2574 |
| **GO:BP** | vascular endo | GO:0048010 | 0.0028893 | 2.53920778 | 99 | 2574 |
| **GO:BP** | regulation of c | GO:0090109 | 0.00289336 | 2.5385977 | 65 | 2574 |
| **GO:BP** | regulation of f | GO:0051893 | 0.00289336 | 2.5385977 | 65 | 2574 |
| **GO:BP** | pallium develo | GO:0021543 | 0.00292367 | 2.53407197 | 174 | 2574 |
| **GO:BP** | chemical syna | GO:0007268 | 0.00302208 | 2.51969395 | 713 | 2574 |
| **GO:BP** | anterograde tr | GO:0098916 | 0.00302208 | 2.51969395 | 713 | 2574 |
| **GO:BP** | biological adh | GO:0022610 | 0.00305339 | 2.51521725 | 1509 | 2574 |
| **GO:BP** | SCF-depende | GO:0031146 | 0.00321755 | 2.49247454 | 95 | 2574 |
| **GO:BP** | response to le | GO:1990840 | 0.00329195 | 2.48254676 | 113 | 2574 |
| **GO:BP** | cellular respor | GO:1990858 | 0.00329195 | 2.48254676 | 113 | 2574 |
| **GO:BP** | spindle organi | GO:0007051 | 0.00347077 | 2.45957463 | 256 | 2574 |
| **GO:BP** | negative regul | GO:0030178 | 0.00348286 | 2.45806392 | 220 | 2574 |
| **GO:BP** | regulation of a | GO:0030833 | 0.00350799 | 2.45494226 | 175 | 2574 |
| **GO:BP** | movement in | GO:0052126 | 0.00355548 | 2.44910226 | 180 | 2574 |
| **GO:BP** | response to ir | GO:0070671 | 0.00382866 | 2.4169537 | 50 | 2574 |
| **GO:BP** | regulation of h | GO:0031060 | 0.00388575 | 2.41052512 | 70 | 2574 |
| **GO:BP** | actin polymeri | GO:0008154 | 0.00407811 | 2.38954054 | 221 | 2574 |
| **GO:BP** | dendritic spine | GO:0060997 | 0.00411762 | 2.38535377 | 62 | 2574 |
| **GO:BP** | cell adhesion | GO:0007155 | 0.00416385 | 2.38050504 | 1502 | 2574 |
| **GO:BP** | protein localiz | GO:1902414 | 0.00417473 | 2.37937125 | 96 | 2574 |
| **GO:BP** | endoplasmic r | GO:0007029 | 0.00424272 | 2.37235554 | 105 | 2574 |
| **GO:BP** | NIK/NF-kappa | GO:0038061 | 0.00427901 | 2.36865709 | 196 | 2574 |
| **GO:BP** | regulation of a | GO:0008064 | 0.00428239 | 2.36831334 | 191 | 2574 |
| **GO:BP** | ubiquitin-depe | GO:0030433 | 0.00428744 | 2.3678018 | 83 | 2574 |
| **GO:BP** | wound healin | GO:0042060 | 0.00436744 | 2.35977281 | 555 | 2574 |
| **GO:BP** | regulation of v | GO:0060627 | 0.00455153 | 2.34184275 | 544 | 2574 |
| **GO:BP** | cerebral cortex | GO:0021987 | 0.00456142 | 2.34089977 | 119 | 2574 |
| **GO:BP** | regulation of t | GO:0060260 | 0.00464877 | 2.33266193 | 79 | 2574 |
| **GO:BP** | low-density lip | GO:0032802 | 0.00475108 | 2.32320728 | 14 | 2574 |
| **GO:BP** | low-density lip | GO:0032799 | 0.00475108 | 2.32320728 | 14 | 2574 |
| **GO:BP** | positive regula | GO:0045089 | 0.00484834 | 2.31440719 | 217 | 2574 |
| **GO:BP** | histone mono | GO:0010390 | 0.00498231 | 2.30256918 | 29 | 2574 |
| **GO:BP** | regulation of a | GO:0030832 | 0.00507318 | 2.29471988 | 192 | 2574 |

| | | | | | | |
|---|---|---|---|---|---|---|
| **GO:BP** | innate immun | GO:0002220 | 0.00523622 | 2.28098176 | 115 | 2574 |
| **GO:BP** | negative regul | GO:0031057 | 0.0052925 | 2.27633917 | 47 | 2574 |
| **GO:BP** | antigen proce: | GO:0019886 | 0.00538907 | 2.26848591 | 97 | 2574 |
| **GO:BP** | vesicle localiz | GO:0051648 | 0.00545001 | 2.26360309 | 228 | 2574 |
| **GO:BP** | negative regul | GO:0010639 | 0.00548552 | 2.26078258 | 361 | 2574 |
| **GO:BP** | erythrocyte dif | GO:0030218 | 0.00570188 | 2.24398197 | 120 | 2574 |
| **GO:BP** | chaperone-me | GO:0061077 | 0.00576083 | 2.23951484 | 63 | 2574 |
| **GO:BP** | glycolytic proc | GO:0061620 | 0.00586683 | 2.23159635 | 26 | 2574 |
| **GO:BP** | positive regula | GO:0048584 | 0.00587623 | 2.23090115 | 2368 | 2574 |
| **GO:BP** | regulation of p | GO:1905475 | 0.00591262 | 2.2282198 | 203 | 2574 |
| **GO:BP** | regulation of i | GO:0090083 | 0.00627573 | 2.20233546 | 17 | 2574 |
| **GO:BP** | muscle structu | GO:0061061 | 0.00641645 | 2.1927052 | 646 | 2574 |
| **GO:BP** | intracellular re | GO:0030522 | 0.00651419 | 2.1861397 | 357 | 2574 |
| **GO:BP** | innate immun | GO:0002758 | 0.00656519 | 2.18275297 | 116 | 2574 |
| **GO:BP** | cellular respor | GO:0031669 | 0.00656705 | 2.18262966 | 250 | 2574 |
| **GO:BP** | chromatin ass | GO:0006333 | 0.00661983 | 2.17915325 | 219 | 2574 |
| **GO:BP** | mRNA splice s | GO:0006376 | 0.00687044 | 2.16301555 | 33 | 2574 |
| **GO:BP** | response to ir | GO:0010035 | 0.00690264 | 2.16098504 | 583 | 2574 |
| **GO:BP** | transcription b | GO:0006366 | 0.00695104 | 2.15795014 | 2729 | 2574 |
| **GO:BP** | small GTPase | GO:0007264 | 0.00701002 | 2.1542808 | 520 | 2574 |
| **GO:BP** | vacuole organ | GO:0007033 | 0.00712785 | 2.14704122 | 184 | 2574 |
| **GO:BP** | establishment | GO:0007163 | 0.00771681 | 2.11256246 | 220 | 2574 |
| **GO:BP** | cellular respor | GO:0071496 | 0.00773517 | 2.11153041 | 353 | 2574 |
| **GO:BP** | axon regenera | GO:0031103 | 0.00820146 | 2.08610895 | 52 | 2574 |
| **GO:BP** | neuron projec | GO:0031102 | 0.00822263 | 2.08498937 | 60 | 2574 |
| **GO:BP** | synaptic vesic | GO:0099504 | 0.00840208 | 2.07561326 | 190 | 2574 |
| **GO:BP** | regulation of h | GO:0033182 | 0.00849583 | 2.07079422 | 12 | 2574 |
| **GO:BP** | adherens junc | GO:0034333 | 0.00849583 | 2.07079422 | 12 | 2574 |
| **GO:BP** | positive regula | GO:1904358 | 0.0085653 | 2.0672573 | 37 | 2574 |
| **GO:BP** | regulation of c | GO:0030334 | 0.00864583 | 2.0631933 | 1008 | 2574 |
| **GO:BP** | regulation of i | GO:0045088 | 0.00878791 | 2.05611417 | 316 | 2574 |
| **GO:BP** | negative regul | GO:1903321 | 0.00884823 | 2.0531438 | 99 | 2574 |
| **GO:BP** | regulation of l | GO:0040012 | 0.00932696 | 2.03025966 | 1113 | 2574 |
| **GO:BP** | vesicle-mediat | GO:0099003 | 0.00936785 | 2.02835989 | 211 | 2574 |
| **GO:BP** | response to h | GO:0042542 | 0.00939475 | 2.02711463 | 151 | 2574 |
| **GO:BP** | cellular respor | GO:0031668 | 0.00942538 | 2.02570131 | 279 | 2574 |
| **GO:BP** | actin filament | GO:0051017 | 0.00977359 | 2.009946 | 161 | 2574 |
| **GO:BP** | negative regul | GO:0030308 | 0.00990282 | 2.00424108 | 191 | 2574 |
| **GO:BP** | tube developm | GO:0035295 | 0.01055148 | 1.97668642 | 1158 | 2574 |
| **GO:BP** | glycolytic proc | GO:0061615 | 0.01059065 | 1.97507745 | 27 | 2574 |
| **GO:BP** | positive regula | GO:1903322 | 0.01072171 | 1.96973581 | 142 | 2574 |
| **GO:BP** | positive regula | GO:2000060 | 0.01086557 | 1.9639476 | 109 | 2574 |
| **GO:BP** | organelle disa: | GO:1903008 | 0.01086557 | 1.9639476 | 109 | 2574 |
| **GO:BP** | establishment | GO:0051650 | 0.01092639 | 1.96152319 | 212 | 2574 |
| **GO:BP** | protein K48-li | GO:0070936 | 0.01096304 | 1.96006896 | 65 | 2574 |

| | | | | | | |
|---|---|---|---|---|---|---|
| **GO:BP** | regulation of I | GO:0046578 | 0.01133891 | 1.94542852 | 202 | 2574 |
| **GO:BP** | regulation of s | GO:1901796 | 0.01133891 | 1.94542852 | 202 | 2574 |
| **GO:BP** | negative regul | GO:1901988 | 0.01140612 | 1.94286217 | 329 | 2574 |
| **GO:BP** | regulation of c | GO:0008360 | 0.01153656 | 1.93792363 | 157 | 2574 |
| **GO:BP** | response to n | GO:1990089 | 0.01172547 | 1.93086969 | 57 | 2574 |
| **GO:BP** | hemopoiesis | GO:0030097 | 0.01191423 | 1.92393392 | 946 | 2574 |
| **GO:BP** | circadian rhyth | GO:0007623 | 0.01242527 | 1.90569419 | 218 | 2574 |
| **GO:BP** | regulation of p | GO:1904814 | 0.01242799 | 1.90559919 | 15 | 2574 |
| **GO:BP** | positive regula | GO:0051240 | 0.01251691 | 1.9025028 | 1497 | 2574 |
| **GO:BP** | regulation of c | GO:1901888 | 0.01300164 | 1.8860017 | 208 | 2574 |
| **GO:BP** | hematopoietic | GO:0048534 | 0.01321814 | 1.87882953 | 984 | 2574 |
| **GO:BP** | regulation of r | GO:1901214 | 0.01355139 | 1.8680163 | 325 | 2574 |
| **GO:BP** | negative regul | GO:1902904 | 0.0140662 | 1.85182323 | 173 | 2574 |
| **GO:BP** | positive regula | GO:1904851 | 0.01410401 | 1.85065736 | 10 | 2574 |
| **GO:BP** | antigen proces | GO:0002495 | 0.01425547 | 1.84601857 | 101 | 2574 |
| **GO:BP** | regulation of p | GO:1901879 | 0.01446046 | 1.83981791 | 92 | 2574 |
| **GO:BP** | aggrephagy | GO:0035973 | 0.01456944 | 1.83655709 | 31 | 2574 |
| **GO:BP** | cellular respira | GO:0045333 | 0.0147317 | 1.8317471 | 240 | 2574 |
| **GO:BP** | microtubule cy | GO:1902850 | 0.01479232 | 1.82996361 | 214 | 2574 |
| **GO:BP** | forebrain deve | GO:0030900 | 0.01487659 | 1.82749665 | 392 | 2574 |
| **GO:BP** | epithelium dev | GO:0060429 | 0.0153878 | 1.8128236 | 1338 | 2574 |
| **GO:BP** | actin filament | GO:0030041 | 0.01603322 | 1.79497936 | 194 | 2574 |
| **GO:BP** | mitotic spindle | GO:0007052 | 0.01603322 | 1.79497936 | 194 | 2574 |
| **GO:BP** | regulation of e | GO:0045646 | 0.01613904 | 1.79212236 | 46 | 2574 |
| **GO:BP** | negative regul | GO:1903573 | 0.01613904 | 1.79212236 | 46 | 2574 |
| **GO:BP** | regulation of r | GO:2001020 | 0.01623022 | 1.78967566 | 225 | 2574 |
| **GO:BP** | membrane bio | GO:0044091 | 0.01625595 | 1.78898756 | 97 | 2574 |
| **GO:BP** | regulation of c | GO:0010769 | 0.01625595 | 1.78898756 | 97 | 2574 |
| **GO:BP** | regulation of t | GO:0006357 | 0.01625647 | 1.78897367 | 2567 | 2574 |
| **GO:BP** | regulation of s | GO:0048167 | 0.01625885 | 1.78891018 | 189 | 2574 |
| **GO:BP** | circulatory sys | GO:0072359 | 0.01662518 | 1.77923373 | 1215 | 2574 |
| **GO:BP** | actin filament | GO:0061572 | 0.0166278 | 1.77916511 | 164 | 2574 |
| **GO:BP** | positive regula | GO:0051050 | 0.01665026 | 1.77857901 | 951 | 2574 |
| **GO:BP** | cellular respor | GO:1990090 | 0.01670191 | 1.77723395 | 54 | 2574 |
| **GO:BP** | lysosomal trar | GO:0007041 | 0.01696123 | 1.77054264 | 111 | 2574 |
| **GO:BP** | lamellipodium | GO:0030032 | 0.01711422 | 1.7666428 | 75 | 2574 |
| **GO:BP** | immune syste | GO:0002376 | 0.01789353 | 1.74730408 | 3438 | 2574 |
| **GO:BP** | positive regula | GO:2000144 | 0.01802503 | 1.74412401 | 35 | 2574 |
| **GO:BP** | regulation of c | GO:0090287 | 0.01823933 | 1.73899119 | 333 | 2574 |
| **GO:BP** | organelle trans | GO:0072384 | 0.01825452 | 1.73862949 | 84 | 2574 |
| **GO:BP** | midbrain deve | GO:0030901 | 0.01847315 | 1.73345912 | 93 | 2574 |
| **GO:BP** | intracellular st | GO:0030518 | 0.01952571 | 1.70939308 | 155 | 2574 |
| **GO:BP** | proton transm | GO:1902600 | 0.01968543 | 1.7058551 | 160 | 2574 |
| **GO:BP** | circadian regu | GO:0032922 | 0.02014303 | 1.69587528 | 67 | 2574 |
| **GO:BP** | maintenance c | GO:0032507 | 0.02014303 | 1.69587528 | 67 | 2574 |

| | | | | | | |
|---|---|---|---|---|---|---|
| **GO:BP** | positive regula | GO:0090063 | 0.02028371 | 1.69285253 | 8 | 2574 |
| **GO:BP** | positive regula | GO:1904528 | 0.02041915 | 1.68996244 | 6 | 2574 |
| **GO:BP** | maintenance ( | GO:0045185 | 0.0205746 | 1.68666864 | 98 | 2574 |
| **GO:BP** | response to tr | GO:0071559 | 0.02065585 | 1.68495693 | 269 | 2574 |
| **GO:BP** | viral protein pr | GO:0019082 | 0.02070054 | 1.68401841 | 39 | 2574 |
| **GO:BP** | positive regula | GO:0033120 | 0.02070054 | 1.68401841 | 39 | 2574 |
| **GO:BP** | mesenchyme | GO:0060485 | 0.02089699 | 1.67991624 | 296 | 2574 |
| **GO:BP** | cellular respor | GO:0009267 | 0.02231378 | 1.65142681 | 191 | 2574 |
| **GO:BP** | NADH metabo | GO:0006734 | 0.02251363 | 1.64755439 | 43 | 2574 |
| **GO:BP** | maintenance ( | GO:0072595 | 0.02251363 | 1.64755439 | 43 | 2574 |
| **GO:BP** | regulation of t | GO:0043555 | 0.02255697 | 1.64671917 | 25 | 2574 |
| **GO:BP** | platelet-derive | GO:0048008 | 0.02262597 | 1.64539275 | 59 | 2574 |
| **GO:BP** | oxidative phos | GO:0006119 | 0.02308125 | 1.6367406 | 151 | 2574 |
| **GO:BP** | mitochondrial | GO:0007006 | 0.0232998 | 1.63264787 | 156 | 2574 |
| **GO:BP** | substantia nig | GO:0021762 | 0.02349919 | 1.62894713 | 47 | 2574 |
| **GO:BP** | nuclear envelo | GO:0006998 | 0.0235654 | 1.62772524 | 85 | 2574 |
| **GO:BP** | regulation of a | GO:1903146 | 0.02382784 | 1.6229154 | 32 | 2574 |
| **GO:BP** | positive regula | GO:1904816 | 0.02410795 | 1.61783963 | 13 | 2574 |
| **GO:BP** | positive regula | GO:0010508 | 0.0246487 | 1.60820596 | 127 | 2574 |
| **GO:BP** | immune syste | GO:0002520 | 0.02540829 | 1.5950246 | 1037 | 2574 |
| **GO:BP** | negative regul | GO:1901991 | 0.02571414 | 1.58982802 | 260 | 2574 |
| **GO:BP** | regulation of c | GO:0010810 | 0.02575897 | 1.58907147 | 223 | 2574 |
| **GO:BP** | regulation of r | GO:1902882 | 0.02592044 | 1.58635756 | 99 | 2574 |
| **GO:BP** | signal transdu | GO:0042770 | 0.02606874 | 1.58388002 | 192 | 2574 |
| **GO:BP** | response to a | GO:0048678 | 0.0264226 | 1.57802445 | 81 | 2574 |
| **GO:BP** | cell-cell junctio | GO:0045216 | 0.02660219 | 1.57508268 | 218 | 2574 |
| **GO:BP** | positive regula | GO:0048639 | 0.02695474 | 1.56936477 | 182 | 2574 |
| **GO:BP** | positive regula | GO:0042327 | 0.02713889 | 1.56640789 | 928 | 2574 |
| **GO:BP** | Fc-epsilon rec | GO:0038095 | 0.02771734 | 1.55724852 | 167 | 2574 |
| **GO:BP** | antigen proces | GO:0002504 | 0.02819205 | 1.54987333 | 104 | 2574 |
| **GO:BP** | ribosomal sma | GO:0000028 | 0.02908758 | 1.53629236 | 19 | 2574 |
| **GO:BP** | regulation of c | GO:2000145 | 0.0291875 | 1.53480304 | 1070 | 2574 |
| **GO:BP** | negative regul | GO:0000122 | 0.02977226 | 1.52618818 | 899 | 2574 |
| **GO:BP** | negative regul | GO:0031397 | 0.03025001 | 1.51927455 | 86 | 2574 |
| **GO:BP** | collagen fibril ( | GO:0030199 | 0.03026911 | 1.51900037 | 109 | 2574 |
| **GO:BP** | cell populatior | GO:0008283 | 0.03085827 | 1.51062848 | 2057 | 2574 |
| **GO:BP** | positive regula | GO:0010592 | 0.0309465 | 1.50938846 | 29 | 2574 |
| **GO:BP** | DNA geometri | GO:0032392 | 0.03213281 | 1.49305135 | 114 | 2574 |
| **GO:BP** | glial cell migra | GO:0008347 | 0.03249061 | 1.48824211 | 56 | 2574 |
| **GO:BP** | regulation of la | GO:0010591 | 0.03306297 | 1.48065818 | 44 | 2574 |
| **GO:BP** | cellular amino | GO:0006520 | 0.03383991 | 1.47057081 | 344 | 2574 |
| **GO:BP** | membrane ass | GO:0071709 | 0.0339284 | 1.46943658 | 91 | 2574 |
| **GO:BP** | positive regula | GO:1900182 | 0.0339284 | 1.46943658 | 91 | 2574 |
| **GO:BP** | aging | GO:0007568 | 0.03408618 | 1.46742172 | 333 | 2574 |
| **GO:BP** | cellular respor | GO:0071560 | 0.03780249 | 1.42247959 | 263 | 2574 |

| | | | | | | |
|---|---|---|---|---|---|---|
| **GO:BP** | negative regul | GO:0051494 | 0.03854423 | 1.41404058 | 169 | 2574 |
| **GO:BP** | post-Golgi ve | GO:0006892 | 0.03857234 | 1.41372398 | 144 | 2574 |
| **GO:BP** | negative regul | GO:1905268 | 0.03899566 | 1.4089837 | 65 | 2574 |
| **GO:BP** | vacuolar trans | GO:0007034 | 0.03905086 | 1.40836942 | 154 | 2574 |
| **GO:BP** | protein localiz | GO:0036010 | 0.03924712 | 1.40619226 | 26 | 2574 |
| **GO:BP** | response to a | GO:0043200 | 0.03946625 | 1.40377416 | 115 | 2574 |
| **GO:BP** | heart developi | GO:0007507 | 0.04050861 | 1.39245269 | 598 | 2574 |
| **GO:BP** | negative regul | GO:0045930 | 0.04054316 | 1.39208241 | 318 | 2574 |
| **GO:BP** | negative regul | GO:1903828 | 0.04124171 | 1.38466338 | 120 | 2574 |
| **GO:BP** | positive regula | GO:0033674 | 0.04126311 | 1.38443804 | 610 | 2574 |
| **GO:BP** | protein acylati | GO:0043543 | 0.04340557 | 1.36245457 | 248 | 2574 |
| **GO:BP** | positive regula | GO:0045937 | 0.04426191 | 1.35396988 | 1003 | 2574 |
| **GO:BP** | positive regula | GO:0010562 | 0.04426191 | 1.35396988 | 1003 | 2574 |
| **GO:BP** | intracellular tra | GO:0075733 | 0.04460361 | 1.35062998 | 57 | 2574 |
| **GO:BP** | response to h | GO:0052200 | 0.0452263 | 1.34460889 | 11 | 2574 |
| **GO:BP** | positive regula | GO:1904871 | 0.0452263 | 1.34460889 | 11 | 2574 |
| **GO:BP** | regulation of p | GO:1904869 | 0.0452263 | 1.34460889 | 11 | 2574 |
| **GO:BP** | regulation of I | GO:1905214 | 0.0452263 | 1.34460889 | 11 | 2574 |
| **GO:BP** | response to d | GO:0052173 | 0.0452263 | 1.34460889 | 11 | 2574 |
| **GO:BP** | regulation of e | GO:0070203 | 0.0452263 | 1.34460889 | 11 | 2574 |
| **GO:BP** | response to h | GO:0075136 | 0.0452263 | 1.34460889 | 11 | 2574 |
| **GO:BP** | insulin recepto | GO:0008286 | 0.04605653 | 1.33670877 | 145 | 2574 |
| **GO:BP** | regulation of p | GO:0035304 | 0.04605653 | 1.33670877 | 145 | 2574 |
| **GO:BP** | dopamine rec | GO:0007212 | 0.04635323 | 1.33391995 | 41 | 2574 |
| **GO:BP** | DNA biosynth | GO:0071897 | 0.04636875 | 1.33377458 | 201 | 2574 |
| **GO:BP** | chromatin rem | GO:0006338 | 0.04668847 | 1.33079038 | 222 | 2574 |
| **GO:BP** | negative regul | GO:0045926 | 0.04716117 | 1.32641545 | 254 | 2574 |
| **GO:BP** | NAD metaboli | GO:0019674 | 0.04781304 | 1.32045362 | 49 | 2574 |
| **GO:BP** | protein localiz | GO:0061462 | 0.04782394 | 1.32035461 | 45 | 2574 |
| **GO:BP** | leukocyte med | GO:0002443 | 0.04791241 | 1.31955202 | 888 | 2574 |
| **GO:BP** | protein refoldi | GO:0042026 | 0.04815609 | 1.31734875 | 23 | 2574 |
| **GO:BP** | regulation of p | GO:1901983 | 0.04939714 | 1.30629822 | 79 | 2574 |
| **GO:BP** | negative regul | GO:0033673 | 0.04965549 | 1.30403276 | 276 | 2574 |

| | | |
|---|---|---|
| 1440 | 18123 | ORC4,IGFBP5,COL1A1,LRP1,HSPA8,GAPDH,HNRNPA1,HSPA9,NEDD4 |
| 884 | 18123 | EEF1A1,COL1A1,LRP1,HSPA8,COPA,HNRNPA1,HSPA9,NEDD4L,ACTB |
| 377 | 18123 | PABPC1,HSPA8,HNRNPA1,NEDD4L,CFL1,MAVS,HSP90AB1,HSP90AA1 |
| 1378 | 18123 | ORC4,IGFBP5,COL1A1,LRP1,HSPA8,GAPDH,HNRNPA1,HSPA9,NEDD4 |
| 545 | 18123 | COL1A1,LRP1,HSPA8,COPA,HSPA9,NEDD4L,ACTB,MAVS,HSP90AB1,I |
| 540 | 18123 | COL1A1,LRP1,HSPA8,COPA,HSPA9,NEDD4L,ACTB,MAVS,HSP90AB1,I |
| 331 | 18123 | PABPC1,HSPA8,HNRNPA1,VIM,UBC,RPL3,RPS6,RPL4,FLNA,NCL,RPL |
| 512 | 18123 | HSPA8,COPA,HNRNPA1,HSPA9,CTSC,NRBP2,TUBA1A,MAVS,HTT,HSP |
| 460 | 18123 | EEF1A1,PABPC1,LRP1,HSPA8,NEDD4L,CTSC,USP22,HSP90AB1,HSP9 |
| 707 | 18123 | EEF1A1,LRP1,HSPA8,COPA,HNRNPA1,HSPA9,ACTB,CTSC,NUDC,NRB |
| 680 | 18123 | COL1A1,LRP1,HSPA8,COPA,HSPA9,NEDD4L,ACTB,ACTG1,MAVS,HSP |
| 636 | 18123 | IGFBP5,LRP1,HSPA8,HNRNPA1,NEDD4L,ODF2L,ACTG1,DPYSL3,TUB |
| 762 | 18123 | COL1A1,LRP1,HSPA8,COPA,HNRNPA1,HSPA9,NEDD4L,ACTB,ACTG1, |
| 397 | 18123 | PABPC1,LRP1,HSPA8,NEDD4L,CTSC,USP22,HSP90AB1,HSP90AA1,VI |
| 334 | 18123 | COL1A1,HSPA8,MAVS,HSP90AB1,HSP90AA1,NUMA1,RPL3,RPS6,RPL |
| 185 | 18123 | PABPC1,HSPA8,VIM,UBC,RPL3,RPS6,RPL4,RPL7A,RPL8,YBX1,HSPA1 |
| 1352 | 18123 | EEF1A1,IGFBP5,COL1A1,LRP1,HSPA8,COPA,HNRNPA1,HSPA9,NEDD |
| 1098 | 18123 | EEF1A1,IGFBP5,PABPC1,LRP1,GAPDH,MLLT1,NEDD4L,ACTB,CTSC,N |
| 912 | 18123 | LRP1,GAPDH,HNRNPA1,HSPA9,ACTB,CTSC,ODF2L,NUDC,USP22,ACT |
| 85 | 18123 | RPL3,RPS6,RPL4,RPL7A,RPL8,RPLP0,RPL13A,SEC62,RPL6,RPL13,R |
| 94 | 18123 | PABPC1,RPL3,RPS6,RPL4,RPL7A,RPL8,RPLP0,RPL13A,RPL6,RPL13, |
| 81 | 18123 | RPL3,RPS6,RPL4,RPL7A,RPL8,RPLP0,RPL13A,RPL6,RPL13,RPS8,RP |
| 191 | 18123 | PABPC1,HSPA8,VIM,UBC,RPL3,RPS6,RPL4,RPL7A,RPL8,YBX1,HSPA1 |
| 773 | 18123 | ORC4,LRP1,HSPA8,HSPA9,ACTB,CTSC,ODF2L,ACTG1,DPYSL3,TUBA |
| 501 | 18123 | COL1A1,LRP1,HSPA8,COPA,HSPA9,MAVS,HSP90AB1,HSP90AA1,RPL |
| 323 | 18123 | HSPA8,COPA,HSPA9,MAVS,HSP90AB1,HSP90AA1,RPL3,RPS6,RPL4,P |
| 100 | 18123 | RPL3,RPS6,RPL4,RPL7A,RPL8,RPLP0,RPL13A,SEC62,RPL6,RPL13,R |
| 86 | 18123 | RPL3,RPS6,RPL4,RPL7A,RPL8,RPLP0,RPL13A,SEC62,RPL6,RPL13,R |
| 1176 | 18123 | EEF1A1,IGFBP5,PABPC1,LRP1,HSPA8,GAPDH,MLLT1,NEDD4L,ACTB, |
| 561 | 18123 | COL1A1,LRP1,HSPA8,COPA,HNRNPA1,HSPA9,ACTB,MAVS,HSP90AB1 |
| 131 | 18123 | UBC,RPL3,RPS6,RPL4,RPL7A,GANAB,RPL8,RPLP0,CANX,RPL13A,RP |
| 87 | 18123 | RPL3,RPS6,RPL4,RPL7A,RPL8,RPLP0,RPL13A,SEC62,RPL6,RPL13,R |
| 100 | 18123 | EEF2,RPL3,RPS6,RPL4,RPL7A,PKM,RPL8,RPLP0,RPL13A,RPL6,RPL1 |
| 1886 | 18123 | EEF1A1,ORC4,IGFBP5,PABPC1,COL1A1,LRP1,HSPA8,GAPDH,MLLT1 |
| 660 | 18123 | EEF1A1,PABPC1,LRP1,HSPA8,GAPDH,NEDD4L,CTSC,NRBP2,USP22,H |
| 588 | 18123 | EEF1A1,PABPC1,LRP1,HSPA8,GAPDH,NEDD4L,CTSC,NRBP2,USP22,H |
| 1160 | 18123 | IGFBP5,PABPC1,COL1A1,LRP1,HSPA8,GAPDH,MLLT1,HNRNPA1,HSP |
| 117 | 18123 | PABPC1,EIF4G2,RPL3,RPS6,RPL4,RPL7A,RPL8,RPLP0,RPL13A,RPL6 |
| 331 | 18123 | EEF1A1,PABPC1,LRP1,HSPA8,GAPDH,NEDD4L,CTSC,NRBP2,HTT,HSI |
| 464 | 18123 | COL1A1,LRP1,HSPA8,COPA,HSPA9,MAVS,HSP90AB1,HSP90AA1,RPL |
| 1073 | 18123 | EEF1A1,COL1A1,LRP1,HSPA8,COPA,HNRNPA1,HSPA9,NEDD4L,ACTB |
| 203 | 18123 | HSPA8,MAVS,HSP90AB1,HSP90AA1,RPL3,RPS6,RPL4,PIK3R2,FLNA,F |
| 1048 | 18123 | EEF1A1,COL1A1,LRP1,HSPA8,COPA,HNRNPA1,HSPA9,NEDD4L,ACTB |
| 699 | 18123 | ORC4,LRP1,HSPA8,HSPA9,ACTB,CTSC,ODF2L,ACTG1,DPYSL3,TUBA |
| 555 | 18123 | EEF1A1,PABPC1,LRP1,HSPA8,GAPDH,NEDD4L,CTSC,USP22,HSP90AI |

| | | |
|---|---|---|
| 1298 | 18123 | EEF1A1,IGFBP5,PABPC1,LRP1,HSPA8,GAPDH,MLLT1,NEDD4L,ACTB, |
| 589 | 18123 | LRP1,NEDD4L,ATN1,ACTB,CTSC,NRBP2,EEF2,DPYSL3,CFL1,XRCC2, |
| 239 | 18123 | LRP1,ACTB,HSP90AA1,RPL3,RPS6,RPL4,FLNA,RPL7A,RPL8,ATP2B4, |
| 1137 | 18123 | IGFBP5,PABPC1,COL1A1,LRP1,HSPA8,HNRNPA1,NEDD4L,CTSC,USP |
| 308 | 18123 | LRP1,NEDD4L,ACTB,DPYSL3,CFL1,HSP90AB1,HSP90AA1,ACTR2,VIM |
| 255 | 18123 | LRP1,ACTB,HSP90AA1,RPL3,RPS6,RPL4,FLNA,RPL7A,RPL8,ATP2B4, |
| 284 | 18123 | EEF1A1,PABPC1,LRP1,HSPA8,GAPDH,CTSC,NRBP2,HTT,HSP90AB1,H |
| 103 | 18123 | RPL3,RPS6,RPL4,RPL7A,RPL8,RPLP0,RPL13A,RPL6,RPL13,RPS8,RP |
| 1330 | 18123 | EEF1A1,IGFBP5,PABPC1,COL1A1,LRP1,HSPA8,GAPDH,MLLT1,HNRN |
| 207 | 18123 | PABPC1,HSPA8,VIM,UBC,RPL3,RPS6,RPL4,RPL7A,RPL8,YBX1,HSPA1 |
| 1021 | 18123 | IGFBP5,PABPC1,COL1A1,LRP1,HSPA8,GAPDH,MLLT1,HNRNPA1,HSP |
| 753 | 18123 | IGFBP5,PABPC1,LRP1,HSPA8,GAPDH,MLLT1,HNRNPA1,NEDD4L,ATN |
| 627 | 18123 | EEF1A1,IGFBP5,PABPC1,LRP1,GAPDH,MLLT1,NEDD4L,ACTB,CTSC,E |
| 1506 | 18123 | EEF1A1,ORC4,IGFBP5,PABPC1,COL1A1,LRP1,HSPA8,GAPDH,MLLT1 |
| 712 | 18123 | IGFBP5,PABPC1,LRP1,HSPA8,GAPDH,MLLT1,HNRNPA1,NEDD4L,ATN |
| 517 | 18123 | LRP1,HSPA8,HNRNPA1,HSPA9,POLH,EEF2,XRCC2,HSP90AB1,HSP90 |
| 742 | 18123 | EEF1A1,IGFBP5,COL1A1,LRP1,HSPA8,GAPDH,HNRNPA1,HSPA9,ACT |
| 492 | 18123 | ORC4,HSPA8,CTSC,CFL1,HSP90AB1,HSP90AA1,ACTR2,UBC,TMSB4X |
| 328 | 18123 | LRP1,NEDD4L,ACTB,DPYSL3,CFL1,HSP90AB1,HSP90AA1,ACTR2,VIM |
| 334 | 18123 | LRP1,HNRNPA1,NEDD4L,DPYSL3,HTT,HSP90AA1,ACTR2,KIDINS220, |
| 432 | 18123 | LRP1,NEDD4L,ACTB,NRBP2,EEF2,DPYSL3,CFL1,XRCC2,HSP90AB1,H |
| 1804 | 18123 | EEF1A1,ORC4,IGFBP5,PABPC1,COL1A1,LRP1,HSPA8,GAPDH,MLLT1 |
| 212 | 18123 | PABPC1,HSPA8,VIM,UBC,RPL3,RPS6,RPL4,RPL7A,RPL8,YBX1,HSPA1 |
| 213 | 18123 | PABPC1,HSPA8,VIM,UBC,RPL3,RPS6,RPL4,RPL7A,RPL8,YBX1,HSPA1 |
| 109 | 18123 | PABPC1,RPL3,RPS6,RPL4,RPL7A,RPL8,RPLP0,DDX5,RPL13A,RPL6,F |
| 1734 | 18123 | EEF1A1,ORC4,IGFBP5,PABPC1,COL1A1,LRP1,HSPA8,GAPDH,MLLT1 |
| 377 | 18123 | LRP1,NEDD4L,ACTB,NRBP2,DPYSL3,CFL1,HSP90AB1,HSP90AA1,AC |
| 404 | 18123 | LRP1,NEDD4L,ACTB,NRBP2,DPYSL3,CFL1,XRCC2,HSP90AB1,HSP90 |
| 597 | 18123 | COL1A1,LRP1,HSPA8,COPA,HNRNPA1,HSPA9,ACTB,MAVS,HSP90AB1 |
| 1226 | 18123 | EEF1A1,IGFBP5,PABPC1,COL1A1,LRP1,HSPA8,GAPDH,MLLT1,HNRN |
| 153 | 18123 | HSP90AA1,RPL3,RPS6,RPL4,RPL7A,RPL8,RPLP0,RPL13A,SEC62,RPL |
| 1201 | 18123 | IGFBP5,PABPC1,COL1A1,LRP1,HSPA8,GAPDH,HNRNPA1,NEDD4L,AC |
| 1176 | 18123 | EEF1A1,IGFBP5,PABPC1,COL1A1,LRP1,HSPA8,GAPDH,MLLT1,HNRN |
| 287 | 18123 | EEF1A1,IGFBP5,PABPC1,GAPDH,EEF2,VIM,EIF4G2,RPL3,RPS6,RPL4 |
| 581 | 18123 | IGFBP5,PABPC1,LRP1,GAPDH,MLLT1,ACTB,CTSC,EEF2,MAVS,HTT,H |
| 155 | 18123 | HSPA8,HSP90AA1,RPL3,RPS6,RPL4,RPL7A,RPL8,RPLP0,CLU,RPL13 |
| 210 | 18123 | PABPC1,HSPA8,VIM,UBC,RPL3,RPS6,RPL4,RPL7A,RPL8,YBX1,HSPA1 |
| 296 | 18123 | LRP1,NEDD4L,ACTB,CFL1,HSP90AB1,HSP90AA1,ACTR2,SPARC,KIDI |
| 216 | 18123 | PABPC1,HSPA8,VIM,UBC,RPL3,RPS6,RPL4,RPL7A,RPL8,YBX1,HSPA1 |
| 221 | 18123 | LRP1,NEDD4L,ACTB,CFL1,HSP90AB1,HSP90AA1,ACTR2,KIDINS220, |
| 219 | 18123 | LRP1,NEDD4L,ACTB,CFL1,HSP90AB1,HSP90AA1,ACTR2,KIDINS220, |
| 223 | 18123 | LRP1,NEDD4L,ACTB,CFL1,HSP90AB1,HSP90AA1,ACTR2,KIDINS220, |
| 1225 | 18123 | IGFBP5,COL1A1,LRP1,HSPA9,NEDD4L,ATN1,ACTB,CTSC,NRBP2,USP |
| 281 | 18123 | EEF1A1,LRP1,HSPA8,NEDD4L,CTSC,USP22,HSP90AB1,HSP90AA1,UB |
| 244 | 18123 | EEF1A1,IGFBP5,PABPC1,GAPDH,EEF2,VIM,EIF4G2,RPL3,RPS6,RPL4 |

| | | |
|---|---|---|
| 423 | 18123 | ORC4,HSPA8,CTSC,HSP90AB1,HSP90AA1,ACTR2,UBC,TMSB4X,NUM |
| 1954 | 18123 | EEF1A1,ORC4,IGFBP5,PABPC1,COL1A1,LRP1,HSPA8,GAPDH,MLLT1 |
| 399 | 18123 | LRP1,NEDD4L,ACTB,ODF2L,DPYSL3,TUBA1A,CFL1,HTT,HSP90AB1, |
| 286 | 18123 | PABPC1,HSPA8,HNRNPA1,RPL3,RPS6,RPL4,NCL,RPL7A,PRPF8,RPL8 |
| 718 | 18123 | EEF1A1,IGFBP5,COL1A1,LRP1,HSPA8,GAPDH,HNRNPA1,HSPA9,ACT |
| 214 | 18123 | LRP1,NEDD4L,ACTB,CFL1,HSP90AB1,HSP90AA1,ACTR2,KIDINS220, |
| 312 | 18123 | ORC4,HSP90AB1,HSP90AA1,ACTR2,UBC,TMSB4X,NUMA1,RPL3,DBN |
| 1676 | 18123 | EEF1A1,ORC4,IGFBP5,PABPC1,COL1A1,LRP1,HSPA8,GAPDH,MLLT1 |
| 1129 | 18123 | EEF1A1,IGFBP5,PABPC1,COL1A1,LRP1,HSPA8,GAPDH,MLLT1,HNRN |
| 104 | 18123 | RPL3,RPS6,RPL4,RPL7A,RPL8,RPLP0,RPL13A,SEC62,RPL6,RPL13,R |
| 147 | 18123 | PABPC1,HSPA8,HNRNPA1,VIM,UBC,NCL,YBX1,DDX5,DDX17,NPM1,U |
| 402 | 18123 | LRP1,NEDD4L,ACTB,ODF2L,DPYSL3,TUBA1A,CFL1,HTT,HSP90AB1, |
| 246 | 18123 | EEF1A1,IGFBP5,PABPC1,GAPDH,EEF2,VIM,EIF4G2,RPL3,RPS6,RPL4 |
| 609 | 18123 | EEF1A1,IGFBP5,COL1A1,LRP1,HSPA8,GAPDH,HNRNPA1,HSPA9,ACT |
| 1138 | 18123 | IGFBP5,COL1A1,LRP1,HSPA9,NEDD4L,ATN1,ACTB,CTSC,NRBP2,US |
| 228 | 18123 | LRP1,NEDD4L,ACTB,CFL1,HSP90AB1,HSP90AA1,ACTR2,KIDINS220, |
| 605 | 18123 | IGFBP5,COL1A1,LRP1,NEDD4L,ACTB,ACTG1,CFL1,XRCC2,HSP90AB |
| 236 | 18123 | LRP1,NEDD4L,ACTB,ACTG1,CFL1,HSP90AB1,HSP90AA1,ACTR2,KIDI |
| 195 | 18123 | LRP1,NEDD4L,ACTB,CFL1,HSP90AB1,HSP90AA1,ACTR2,KIDINS220, |
| 816 | 18123 | IGFBP5,PABPC1,COL1A1,LRP1,HSPA8,GAPDH,COPA,HNRNPA1,HSP |
| 161 | 18123 | PABPC1,HSPA8,HNRNPA1,NCL,PRPF8,YBX1,DDX5,DDX17,TMBIM6,N |
| 1057 | 18123 | IGFBP5,COL1A1,LRP1,HSPA9,NEDD4L,ATN1,ACTB,CTSC,NRBP2,US |
| 147 | 18123 | EEF1A1,LRP1,NEDD4L,CTSC,HSP90AB1,HSP90AA1,FLNA,TMF1,HSP |
| 1089 | 18123 | EEF1A1,IGFBP5,PABPC1,COL1A1,LRP1,HSPA8,GAPDH,MLLT1,HNRN |
| 270 | 18123 | EEF1A1,IGFBP5,PABPC1,GAPDH,EEF2,VIM,EIF4G2,RPL3,RPS6,RPL4 |
| 967 | 18123 | IGFBP5,COL1A1,LRP1,HSPA9,NEDD4L,ATN1,ACTB,CTSC,NRBP2,ACT |
| 1868 | 18123 | EEF1A1,ORC4,IGFBP5,PABPC1,COL1A1,LRP1,HSPA8,GAPDH,MLLT1 |
| 136 | 18123 | PABPC1,HSPA8,HNRNPA1,NCL,PRPF8,YBX1,DDX5,DDX17,NPM1,SNR |
| 136 | 18123 | PABPC1,HSPA8,HNRNPA1,NCL,PRPF8,YBX1,DDX5,DDX17,NPM1,SNR |
| 136 | 18123 | PABPC1,HSPA8,HNRNPA1,NCL,PRPF8,YBX1,DDX5,DDX17,NPM1,SNR |
| 807 | 18123 | IGFBP5,GAPDH,MLLT1,NEDD4L,ACTB,NRBP2,USP22,MED18,MAVS, |
| 807 | 18123 | IGFBP5,GAPDH,MLLT1,NEDD4L,ACTB,NRBP2,USP22,MED18,MAVS, |
| 172 | 18123 | LRP1,ACTB,HSP90AB1,HSP90AA1,DCX,EIF4G2,DBN1,FLNA,MAP1B, |
| 767 | 18123 | PABPC1,COL1A1,LRP1,HSPA8,GAPDH,HNRNPA1,NEDD4L,ACTB,CTS |
| 173 | 18123 | PABPC1,HSPA8,HNRNPA1,NCL,PRPF8,YBX1,DDX5,DDX17,APP,NPM1, |
| 234 | 18123 | LRP1,NEDD4L,CTSC,USP22,HSP90AB1,HSP90AA1,UBC,HSP90B1,FB |
| 475 | 18123 | LRP1,NEDD4L,ACTB,ACTG1,DPYSL3,CFL1,XRCC2,HSP90AB1,HSP90 |
| 313 | 18123 | LRP1,HNRNPA1,ODF2L,ACTG1,MAVS,HTT,ACTR2,TMSB4X,NUMA1,D |
| 358 | 18123 | IGFBP5,PABPC1,HSPA8,GAPDH,MAVS,HSP90AB1,VIM,UBC,TMSB4X, |
| 323 | 18123 | EEF1A1,IGFBP5,PABPC1,GAPDH,EEF2,VIM,EIF4G2,RPL3,RPS6,RPL4 |
| 159 | 18123 | HSP90AB1,HSP90AA1,RPL3,RPS6,RPL4,RPL7A,PRPF8,RPL8,RPLP0, |
| 677 | 18123 | PABPC1,COL1A1,LRP1,HSPA8,HNRNPA1,CTSC,USP22,EEF2,MAVS,H |
| 1952 | 18123 | EEF1A1,IGFBP5,PABPC1,COL1A1,LRP1,HSPA8,GAPDH,MLLT1,COPA |
| 483 | 18123 | EEF1A1,LRP1,HSPA8,COPA,NEDD4L,ACTB,CTSC,NRBP2,ACTG1,EEF |
| 153 | 18123 | LRP1,NEDD4L,DPYSL3,CFL1,ACTR2,VIM,KIDINS220,EIF4G2,DBN1,F |

| | | |
|---|---|---|
| 711 | 18123 | PABPC1,COL1A1,LRP1,HSPA8,GAPDH,HNRNPA1,NEDD4L,ACTB,CTS |
| 550 | 18123 | IGFBP5,PABPC1,LRP1,HSPA8,GAPDH,MLLT1,HNRNPA1,NEDD4L,ATN |
| 158 | 18123 | LRP1,ACTB,HSP90AB1,HSP90AA1,DCX,EIF4G2,DBN1,MAP1B,NEFL,T |
| 582 | 18123 | IGFBP5,PABPC1,LRP1,HSPA8,GAPDH,MLLT1,HNRNPA1,NEDD4L,ATN |
| 193 | 18123 | LRP1,NEDD4L,ODF2L,DPYSL3,CFL1,HTT,ACTR2,VIM,KIDINS220,EIF |
| 400 | 18123 | EEF1A1,IGFBP5,COL1A1,LRP1,HNRNPA1,ACTB,EEF2,HSP90AB1,ACT |
| 196 | 18123 | LRP1,NEDD4L,ODF2L,DPYSL3,CFL1,HTT,ACTR2,VIM,KIDINS220,EIF |
| 446 | 18123 | LRP1,GAPDH,HSPA9,ATN1,CTSC,NRBP2,MTRNR2L2,CFL1,XRCC2,HT |
| 639 | 18123 | PABPC1,COL1A1,LRP1,HSPA8,HNRNPA1,NEDD4L,CTSC,USP22,EEF2 |
| 1114 | 18123 | EEF1A1,IGFBP5,PABPC1,COL1A1,LRP1,HSPA8,GAPDH,MLLT1,HNRN |
| 424 | 18123 | EEF1A1,IGFBP5,PABPC1,GAPDH,EEF2,VIM,TMSB4X,EIF4G2,RPL3,R |
| 846 | 18123 | IGFBP5,COL1A1,LRP1,HSPA9,NEDD4L,ACTB,NRBP2,ACTG1,EEF2,D |
| 902 | 18123 | EEF1A1,IGFBP5,COL1A1,LRP1,HSPA8,GAPDH,HNRNPA1,HSPA9,NED |
| 1166 | 18123 | EEF1A1,ORC4,IGFBP5,PABPC1,COL1A1,HSPA8,GAPDH,MLLT1,HNR |
| 832 | 18123 | IGFBP5,COL1A1,LRP1,HSPA9,NEDD4L,ACTB,NRBP2,ACTG1,EEF2,D |
| 882 | 18123 | PABPC1,COL1A1,HSPA8,MLLT1,HNRNPA1,NEDD4L,ATN1,USP22,ME |
| 1136 | 18123 | EEF1A1,ORC4,IGFBP5,PABPC1,COL1A1,HSPA8,GAPDH,MLLT1,HNR |
| 1122 | 18123 | EEF1A1,ORC4,IGFBP5,PABPC1,COL1A1,HSPA8,GAPDH,MLLT1,HNR |
| 822 | 18123 | IGFBP5,GAPDH,MLLT1,NEDD4L,ACTB,NRBP2,USP22,MED18,MAVS, |
| 1038 | 18123 | ORC4,PABPC1,COL1A1,HSPA8,GAPDH,MLLT1,HNRNPA1,NEDD4L,P |
| 252 | 18123 | IGFBP5,LRP1,NEDD4L,XRCC2,HSP90AB1,HSP90AA1,DCX,EIF4G2,D |
| 464 | 18123 | LRP1,GAPDH,HSPA9,ATN1,CTSC,NRBP2,MTRNR2L2,CFL1,XRCC2,HT |
| 956 | 18123 | ORC4,PABPC1,COL1A1,HSPA8,MLLT1,HNRNPA1,NEDD4L,POLH,ATN |
| 100 | 18123 | LRP1,CTSC,HSP90AB1,HSP90AA1,TMF1,HSPA1B,CLU,RACK1,RPL5, |
| 942 | 18123 | EEF1A1,ORC4,IGFBP5,PABPC1,COL1A1,HSPA8,GAPDH,MLLT1,HNR |
| 209 | 18123 | LRP1,HSPA8,HNRNPA1,ODF2L,DPYSL3,VIM,TMSB4X,FLNA,MAP1B,H |
| 254 | 18123 | LRP1,HSPA8,ODF2L,ACTG1,DPYSL3,HTT,HSP90AA1,ACTR2,TMSB4X |
| 325 | 18123 | EEF1A1,LRP1,HSPA8,NEDD4L,CTSC,USP22,HSP90AB1,HSP90AA1,U |
| 952 | 18123 | IGFBP5,PABPC1,COL1A1,LRP1,HSPA8,GAPDH,MLLT1,HNRNPA1,NE |
| 540 | 18123 | IGFBP5,COL1A1,LRP1,HSPA9,NEDD4L,CTSC,CFL1,XRCC2,HSP90AB |
| 168 | 18123 | LRP1,NEDD4L,ACTB,MAVS,HSP90AB1,HSP90AA1,NUMA1,PIK3R2,FL |
| 208 | 18123 | NEDD4L,CTSC,USP22,HSP90AB1,UBC,HSP90B1,FBXL4,UBA1,TMF1, |
| 930 | 18123 | EEF1A1,ORC4,IGFBP5,PABPC1,COL1A1,HSPA8,GAPDH,MLLT1,HNR |
| 486 | 18123 | LRP1,GAPDH,HSPA9,ATN1,CTSC,NRBP2,MTRNR2L2,CFL1,XRCC2,HT |
| 154 | 18123 | NEDD4L,HSP90AB1,UBC,HSP90B1,FBXL4,TMF1,HSPA1B,CLU,RACK |
| 153 | 18123 | IGFBP5,LRP1,NEDD4L,HSP90AB1,HSP90AA1,DCX,EIF4G2,DBN1,EN |
| 339 | 18123 | EEF1A1,IGFBP5,COL1A1,LRP1,HNRNPA1,ACTB,EEF2,HSP90AB1,ACT |
| 2017 | 18123 | EEF1A1,IGFBP5,PABPC1,COL1A1,LRP1,HSPA8,GAPDH,MLLT1,COPA |
| 706 | 18123 | IGFBP5,COL1A1,LRP1,CTSC,MTRNR2L2,BTBD9,MAVS,HTT,HSP90A |
| 226 | 18123 | COL1A1,HSPA8,ACTG1,DPYSL3,CFL1,HSP90AB1,ACTR2,VIM,UBC,T |
| 84 | 18123 | HSP90AB1,HSP90AA1,RPL3,PRPF8,RPLP0,RPL13A,RPL6,RPL5,SNRN |
| 1053 | 18123 | ORC4,PABPC1,COL1A1,HSPA8,GAPDH,MLLT1,HNRNPA1,NEDD4L,P |
| 1093 | 18123 | EEF1A1,ORC4,IGFBP5,PABPC1,COL1A1,HSPA8,GAPDH,MLLT1,HNR |
| 360 | 18123 | LRP1,HSPA9,CTSC,NRBP2,MTRNR2L2,CFL1,XRCC2,HTT,HSP90AB1, |
| 259 | 18123 | LRP1,NEDD4L,CFL1,ACTR2,SPARC,EIF4G2,DBN1,FLNA,MAP1B,ALD |

| | | |
|---|---|---|
| 81 | 18123 | HSP90AB1,HSP90AA1,RPL3,PRPF8,RPLP0,RPL13A,RPL6,RPL5,SNRN |
| 232 | 18123 | EEF1A1,HSPA8,CTSC,EEF2,HSP90AB1,HSP90AA1,ACTR2,SPARC,TUB |
| 197 | 18123 | LRP1,HSPA8,POLH,HSP90AB1,HSP90AA1,ACTR2,ENO1,FLNA,YBX1,H |
| 354 | 18123 | PABPC1,LRP1,NEDD4L,CTSC,EEF2,MAVS,HSP90AB1,HSP90AA1,VIM, |
| 209 | 18123 | EEF1A1,HSPA8,CTSC,EEF2,HSP90AB1,HSP90AA1,ACTR2,SPARC,TUB |
| 695 | 18123 | IGFBP5,COL1A1,LRP1,CTSC,MTRNR2L2,BTBD9,MAVS,HTT,HSP90AE |
| 1054 | 18123 | ORC4,PABPC1,COL1A1,HSPA8,GAPDH,MLLT1,HNRNPA1,NEDD4L,PC |
| 915 | 18123 | EEF1A1,IGFBP5,PABPC1,COL1A1,HSPA8,GAPDH,MLLT1,HNRNPA1,N |
| 1839 | 18123 | EEF1A1,IGFBP5,PABPC1,COL1A1,LRP1,HSPA8,GAPDH,MLLT1,COPA |
| 137 | 18123 | NEDD4L,HSP90AB1,UBC,HSP90B1,FBXL4,HSPA1B,CLU,RACK1,VCP, |
| 371 | 18123 | MLLT1,ACTB,MAVS,HTT,HSP90AB1,HSP90AA1,UBC,KIDINS220,HSP9 |
| 184 | 18123 | NEDD4L,USP22,HSP90AB1,UBC,HSP90B1,FBXL4,UBA1,HSPA1B,CLU |
| 118 | 18123 | HNRNPA1,HSPA9,MAVS,HSP90AB1,PIK3R2,FLNA,CALR,YWHAE,SMG |
| 231 | 18123 | LRP1,HSPA8,NEDD4L,ACTB,MAVS,HSP90AB1,HSP90AA1,NUMA1,PIK |
| 186 | 18123 | NEDD4L,USP22,HSP90AB1,UBC,HSP90B1,FBXL4,UBA1,HSPA1B,CLU |
| 105 | 18123 | EEF2,PIK3R2,HSP90B1,CALR,CANX,CLU,FKBP14,RACK1,P4HB,TMBI |
| 58 | 18123 | UBC,TMBIM6,UBB,PSMD5,EGLN1,ATF4,PSMD2,JUN,BCLAF1,CHD6, |
| 182 | 18123 | NEDD4L,USP22,HSP90AB1,UBC,HSP90B1,FBXL4,UBA1,HSPA1B,CLU |
| 348 | 18123 | LRP1,HSPA9,CTSC,NRBP2,MTRNR2L2,CFL1,XRCC2,HTT,HSP90AB1, |
| 118 | 18123 | LRP1,DPYSL3,HTT,ACTR2,KIDINS220,EIF4G2,DBN1,FLNA,MAP1B,NE |
| 56 | 18123 | UBC,TMBIM6,UBB,PSMD5,EGLN1,ATF4,PSMD2,JUN,CHD6,GCN1,PS |
| 379 | 18123 | LRP1,HSPA9,CTSC,NRBP2,MTRNR2L2,CFL1,XRCC2,HTT,HSP90AB1, |
| 286 | 18123 | IGFBP5,PABPC1,LRP1,GAPDH,MLLT1,HSP90AB1,FLNA,NCL,YBX1,AT |
| 218 | 18123 | LRP1,NEDD4L,ACTB,MAVS,HTT,HSP90AB1,HSP90AA1,NUMA1,PIK3R |
| 118 | 18123 | IGFBP5,HSP90B1,FBXL4,FSTL1,P4HB,APP,FN1,DDB1,PSMD5,WSB1, |
| 255 | 18123 | LRP1,ATN1,ACTB,HSP90AB1,HSP90AA1,VIM,DCX,FLNA,NEFL,UGT8, |
| 193 | 18123 | LRP1,HSPA8,ACTB,ACTG1,TUBA1A,CFL1,ACTR2,SPARC,TUBB,DBN1 |
| 93 | 18123 | PABPC1,HSPA8,VIM,UBC,YBX1,NPM1,UBB,PSMD5,SET,HNRNPU,LAR |
| 65 | 18123 | HSPA8,HNRNPA1,NCL,DDX5,DDX17,NPM1,HNRNPU,PAPOLA,SRSF4, |
| 1085 | 18123 | ORC4,PABPC1,COL1A1,HSPA8,GAPDH,MLLT1,HNRNPA1,NEDD4L,PC |
| 91 | 18123 | PABPC1,HSPA8,VIM,UBC,YBX1,NPM1,UBB,PSMD5,SET,HNRNPU,LAR |
| 160 | 18123 | EEF1A1,HSPA8,GAPDH,NRBP2,HTT,HSP90AA1,VIM,UBC,EIF4G2,PIK3 |
| 160 | 18123 | EEF1A1,HSPA8,GAPDH,NRBP2,HTT,HSP90AA1,VIM,UBC,EIF4G2,PIK3 |
| 178 | 18123 | LRP1,NEDD4L,XRCC2,HSP90AB1,HSP90AA1,DCX,EIF4G2,DBN1,MAP |
| 115 | 18123 | HNRNPA1,HSPA9,MAVS,HSP90AB1,PIK3R2,FLNA,CALR,YWHAE,SMG |
| 85 | 18123 | CTSC,HSP90AB1,TMF1,HSPA1B,CLU,RACK1,RPL5,VCP,UBB,PKD1,R |
| 292 | 18123 | COL1A1,HSPA8,POLH,XRCC2,HTT,HSP90AB1,HSP90AA1,SPARC,UBC |
| 329 | 18123 | LRP1,GAPDH,ACTB,NUDC,ACTG1,DPYSL3,TUBA1A,CFL1,XRCC2,HTT |
| 397 | 18123 | HSPA8,ACTB,NUDC,USP22,TUBA1A,CFL1,XRCC2,HTT,HSP90AB1,HSI |
| 103 | 18123 | HSPA8,GAPDH,NEDD4L,HTT,HSP90AB1,HSP90AA1,FLNA,CALR,HSPA |
| 153 | 18123 | COL1A1,LRP1,UBC,IGFBP2,SOX4,DVL3,RACK1,GNB1,APP,CALM1,V |
| 137 | 18123 | HSPA8,HSP90AA1,ACTR2,TMSB4X,NUMA1,DBN1,MAP1B,HSPA1B,CL |
| 130 | 18123 | IGFBP5,LRP1,NEDD4L,EIF4G2,DBN1,ENO1,MAP1B,HSPA1B,H3-3B,F |
| 161 | 18123 | LRP1,HSPA8,CFL1,HTT,MAP1B,DVL3,BSG,SPTBN1,STMN2,FSCN1,S |
| 152 | 18123 | COL1A1,LRP1,UBC,IGFBP2,SOX4,DVL3,RACK1,GNB1,APP,CALM1,V |

| | | |
|---|---|---|
| 330 | 18123 | PABPC1,LRP1,CTSC,EEF2,MAVS,HSP90AB1,HSP90AA1,VIM,UBC,KID |
| 259 | 18123 | IGFBP5,COL1A1,LRP1,ACTB,EEF2,HSP90AB1,ACTR2,VIM,SPARC,PIK |
| 75 | 18123 | HSP90AB1,TMF1,HSPA1B,CLU,RACK1,VCP,UBB,PKD1,RPL11,TMEM |
| 141 | 18123 | EEF1A1,HSPA8,CTSC,EEF2,HSP90AB1,HSP90AA1,ACTR2,TUBB,FTH1 |
| 85 | 18123 | HSPA8,HSPA9,NUDC,HSP90AB1,HSP90AA1,GANAB,HSP90B1,CALR,H |
| 83 | 18123 | HNRNPA1,YBX1,SMG1,NPM1,HNRNPU,CCT7,SRSF4,LRPPRC,KHDRB |
| 237 | 18123 | NEDD4L,USP22,MED18,HSP90AB1,HSP90AA1,UBC,FBXL4,UBA1,HSP |
| 65 | 18123 | HSPA8,HNRNPA1,NCL,DDX5,DDX17,TMBIM6,NPM1,HNRNPU,SRSF4, |
| 266 | 18123 | IGFBP5,PABPC1,LRP1,GAPDH,MLLT1,HSP90AB1,NCL,YBX1,ATP2B4 |
| 141 | 18123 | EEF1A1,HSPA8,CTSC,EEF2,HSP90AB1,HSP90AA1,ACTR2,TUBB,FTH1 |
| 91 | 18123 | PABPC1,HSPA8,VIM,UBC,YBX1,NPM1,UBB,PSMD5,SET,HNRNPU,LAR |
| 272 | 18123 | IGFBP5,COL1A1,LRP1,ACTB,EEF2,MAVS,HSP90AB1,ACTR2,VIM,SPA |
| 143 | 18123 | EEF1A1,HSPA8,CTSC,EEF2,HSP90AB1,HSP90AA1,ACTR2,TUBB,FTH1 |
| 196 | 18123 | EEF1A1,COL1A1,HNRNPA1,EEF2,HSP90AB1,UBC,KIDINS220,NUMA1 |
| 274 | 18123 | NEDD4L,ACTB,USP22,MED18,HSP90AB1,HSP90AA1,UBC,FBXL4,UB |
| 2520 | 18123 | EEF1A1,ORC4,IGFBP5,PABPC1,COL1A1,LRP1,HSPA8,GAPDH,MLLT1 |
| 144 | 18123 | EEF1A1,HSPA8,CTSC,EEF2,HSP90AB1,HSP90AA1,ACTR2,TUBB,FTH1 |
| 617 | 18123 | IGFBP5,COL1A1,LRP1,CTSC,MTRNR2L2,MAVS,HTT,HSP90AB1,HSP |
| 168 | 18123 | HNRNPA1,HTT,ACTR2,NUMA1,FLNA,MAP1B,HSPA1B,H3-3B,YWHAE, |
| 307 | 18123 | NUDC,TUBA1A,CFL1,XRCC2,HTT,HSP90AA1,ACTR2,TUBB,UBC,NUM |
| 142 | 18123 | EEF1A1,HSPA8,CTSC,EEF2,HSP90AB1,HSP90AA1,ACTR2,TUBB,FTH1 |
| 219 | 18123 | NEDD4L,USP22,MED18,HSP90AB1,HSP90AA1,UBC,FBXL4,UBA1,HSP |
| 255 | 18123 | NUDC,USP22,TUBA1A,XRCC2,HTT,HSP90AA1,TUBB,UBC,NUMA1,RP |
| 107 | 18123 | GAPDH,NRBP2,HTT,VIM,UBC,SMG1,CALM1,VCP,UBB,UCHL1,LARP1 |
| 258 | 18123 | HSPA8,ACTB,USP22,TUBA1A,HSP90AB1,HSP90AA1,TUBB,UBC,NUM |
| 299 | 18123 | EEF1A1,LRP1,HSPA8,CTSC,EEF2,HSP90AB1,HSP90AA1,ACTR2,SPAR |
| 228 | 18123 | NUDC,TUBA1A,HTT,HSP90AA1,TUBB,UBC,NUMA1,RPS6,FLNA,HSPA1 |
| 84 | 18123 | LRP1,NEDD4L,HSP90AB1,HSP90AA1,DCX,EIF4G2,DBN1,MAP1B,APP, |
| 42 | 18123 | UBC,UBB,PSMD5,EGLN1,PSMD2,PSMB2,VEGFA,VHL,PSMD3,PSMA7 |
| 412 | 18123 | PABPC1,COL1A1,HNRNPA1,USP22,EEF2,MAVS,HSP90AB1,HSP90AA1 |
| 555 | 18123 | IGFBP5,LRP1,NRBP2,CFL1,MAVS,HTT,HSP90AB1,HSP90AA1,UBC,T |
| 55 | 18123 | LRP1,HSP90AB1,HSP90AA1,DCX,EIF4G2,DBN1,MAP1B,GDI1,MAPT, |
| 284 | 18123 | HNRNPA1,ACTB,NUDC,USP22,HSP90AB1,HSP90AA1,UBC,NUMA1,H3 |
| 603 | 18123 | EEF1A1,LRP1,GAPDH,MLLT1,NEDD4L,ACTB,MTRNR2L2,MAVS,HTT, |
| 405 | 18123 | PABPC1,COL1A1,HNRNPA1,USP22,EEF2,MAVS,HSP90AB1,HSP90AA1 |
| 76 | 18123 | HSPA8,HSPA9,HSP90AB1,HSP90AA1,HSP90B1,CALR,HSPA1B,CANX, |
| 187 | 18123 | EEF1A1,COL1A1,HNRNPA1,EEF2,HSP90AB1,UBC,KIDINS220,NUMA1 |
| 238 | 18123 | HSPA8,HSPA9,ACTB,CTSC,ACTG1,HSP90AA1,ACTR2,UBC,CALR,COL |
| 201 | 18123 | COL1A1,LRP1,ACTB,DCX,FLNA,NEFL,ATP2B4,COL4A1,YWHAE,KIF1 |
| 96 | 18123 | COL1A1,MAVS,HSP90AB1,PIK3R2,FLNA,CALR,MAGED1,YWHAE,NPM |
| 70 | 18123 | HSPA8,HSPA9,HSP90AB1,HSP90AA1,HSP90B1,CALR,HSPA1B,CANX, |
| 91 | 18123 | TUBA1A,HSP90AA1,TUBB,YWHAE,CALM3,APP,CALM1,NPM1,YWHAG |
| 116 | 18123 | LRP1,HSP90AB1,ACTR2,TMSB4X,EIF4G2,DBN1,MAP1B,NEFL,SPTBN |
| 244 | 18123 | COL1A1,USP22,CFL1,XRCC2,KIDINS220,RPS6,DBN1,COL6A1,YBX1, |
| 77 | 18123 | TUBA1A,HSP90AA1,TUBB,YWHAE,APP,NPM1,YWHAG,PSMD5,PPP2R1 |

| | | |
|---|---|---|
| 67 | 18123 | LRP1,NEDD4L,HSP90AB1,HSP90AA1,DCX,EIF4G2,DBN1,MAP1B,GDI |
| 383 | 18123 | PABPC1,COL1A1,HNRNPA1,USP22,EEF2,MAVS,HSP90AB1,HSP90AA1 |
| 82 | 18123 | LRP1,NEDD4L,HSP90AB1,HSP90AA1,DCX,EIF4G2,DBN1,MAP1B,APP, |
| 199 | 18123 | TUBA1A,HSP90AA1,TUBB,UBC,NUMA1,ATP2B4,CALR,HSPA1B,SOX4, |
| 144 | 18123 | PABPC1,LRP1,NEDD4L,CTSC,HTT,HSP90AA1,HSPA1B,CLU,RACK1,A |
| 225 | 18123 | IGFBP5,COL1A1,SPARC,PIK3R2,MAP1B,PKM,FADS1,NEFL,IGFBP2,C |
| 67 | 18123 | CALR,CANX,PSMD5,HLA-C,PDIA3,AP2M1,ARF1,AP2B1,CTSD,PSMD2, |
| 155 | 18123 | MLLT1,ATP2B4,HSPA1B,SOX4,YWHAE,RACK1,CALM3,RPL5,GSTP1,C |
| 74 | 18123 | TUBA1A,HSP90AA1,TUBB,YWHAE,APP,YWHAG,PSMD5,PPP2R1A,TUB |
| 149 | 18123 | EEF1A1,HSPA8,CTSC,EEF2,MAVS,HSP90AB1,HSP90AA1,ACTR2,TUBB |
| 774 | 18123 | COL1A1,LRP1,HSPA8,GAPDH,HNRNPA1,HSPA9,POLH,ACTB,CTSC,AC |
| 305 | 18123 | EEF1A1,LRP1,HSPA8,CTSC,EEF2,HSP90AB1,HSP90AA1,ACTR2,SPAR |
| 145 | 18123 | EEF1A1,HSPA8,CTSC,EEF2,HSP90AB1,HSP90AA1,ACTR2,TUBB,FTH1 |
| 1301 | 18123 | EEF1A1,IGFBP5,COL1A1,LRP1,HSPA8,GAPDH,COPA,HNRNPA1,HSPA |
| 49 | 18123 | HSPA8,HNRNPA1,NCL,DDX5,DDX17,NPM1,HNRNPU,SRSF4,KHDRBS1 |
| 87 | 18123 | TUBA1A,HSP90AA1,TUBB,YWHAE,CALM3,APP,CALM1,YWHAG,PSMD |
| 103 | 18123 | RPL3,RPS6,RPL4,RPL7A,RPL8,RPLP0,RPL13A,RPL6,RPL13,DDX17,R |
| 351 | 18123 | IGFBP5,COL1A1,LRP1,HSPA9,XRCC2,HSP90AB1,ACTR2,NUMA1,EIF |
| 189 | 18123 | LRP1,ACTB,DCX,FLNA,NEFL,ATP2B4,COL4A1,YWHAE,KIF1A,CALM3, |
| 71 | 18123 | GAPDH,HSP90AB1,HSP90AA1,FLNA,CALR,HSPA1B,CLU,SOX4,DVL3,F |
| 100 | 18123 | PABPC1,NRBP2,HSP90AB1,EIF4G2,FLNA,YBX1,ATP2B4,RPL5,SMG1, |
| 68 | 18123 | CALR,CANX,PSMD5,HLA-C,PDIA3,AP2M1,AP3D1,ARF1,AP2B1,CTSD,F |
| 201 | 18123 | ACTB,USP22,H3-3B,KAT6A,NPM1,IGF2,TRIM28,DDB1,SET,GTF3C4,C |
| 119 | 18123 | HSPA8,COPA,CTSC,NRBP2,HTT,FTH1,FTL,RACK1,SPTBN1,APP,SPTA |
| 143 | 18123 | LRP1,ACTG1,ACTR2,TMSB4X,NUMA1,DBN1,FLNA,MAP1B,HSPA1B,S |
| 171 | 18123 | IGFBP5,LRP1,NEDD4L,EIF4G2,DBN1,ENO1,MAP1B,IGFBP2,HSPA1B, |
| 217 | 18123 | IGFBP5,PABPC1,HSPA8,GAPDH,EEF2,VIM,UBC,EIF4G2,NCL,PKM,YB |
| 151 | 18123 | IGFBP5,PABPC1,GAPDH,EEF2,VIM,EIF4G2,NCL,PKM,YBX1,CALR,CL |
| 146 | 18123 | EEF1A1,HSPA8,CTSC,EEF2,HSP90AB1,HSP90AA1,ACTR2,TUBB,FTH1 |
| 157 | 18123 | CTSC,HTT,ENO1,HSPA1B,CLU,DDX5,MAGED1,YWHAE,RACK1,P4HB, |
| 72 | 18123 | HNRNPA1,HSPA9,CALR,YWHAE,SMG1,NPM1,PKD1,SRSF4,KHDRBS1, |
| 86 | 18123 | PABPC1,NRBP2,HSP90AB1,EIF4G2,YBX1,ATP2B4,RPL5,SMG1,RPL1 |
| 137 | 18123 | LRP1,XRCC2,ACTR2,EIF4G2,DBN1,FLNA,MAP1B,NEFL,CALR,P4HB,G |
| 162 | 18123 | COL1A1,LRP1,UBC,IGFBP2,SOX4,DVL3,RACK1,GNB1,APP,CALM1,V |
| 776 | 18123 | IGFBP5,PABPC1,COL1A1,HSPA8,GAPDH,MLLT1,HNRNPA1,NEDD4L, |
| 80 | 18123 | LRP1,NEDD4L,CTSC,HSP90AA1,HSPA1B,CLU,RACK1,VCP,DDB1,LPC |
| 314 | 18123 | EEF1A1,LRP1,HSPA8,COPA,CTSC,EEF2,HSP90AB1,HSP90AA1,ACTR2 |
| 473 | 18123 | EEF1A1,LRP1,GAPDH,MLLT1,ACTB,HTT,HSP90AB1,HSP90AA1,UBC, |
| 374 | 18123 | LRP1,GAPDH,NEDD4L,ACTB,CTSC,USP22,HSP90AB1,UBC,ENO1,HSP |
| 112 | 18123 | HSPA8,ACTG1,ACTR2,TMSB4X,NUMA1,DBN1,FLNA,MAP1B,HSPA1B, |
| 23 | 18123 | NPM1,SRSF4,HNRNPK,U2AF2,RBMX,SRSF9,PTBP1,SUPT5H,SRSF6,H |
| 85 | 18123 | RPL3,RPS6,RPL4,RPL7A,RPL8,RPLP0,RPL13A,RPL6,RPL13,DDX17,R |
| 214 | 18123 | LRP1,HSPA9,NRBP2,MTRNR2L2,CFL1,XRCC2,HTT,HSP90AB1,UBC,F |
| 235 | 18123 | LRP1,HSPA9,NRBP2,MTRNR2L2,CFL1,XRCC2,HTT,HSP90AB1,UBC,F |
| 87 | 18123 | RPL3,RPS6,RPL4,RPL7A,RPL8,RPLP0,RPL13A,RPL6,RPL13,DDX17,R |

| | | |
|---|---|---|
| 66 | 18123 | HNRNPA1,YBX1,SMG1,NPM1,SRSF4,LRPPRC,KHDRBS1,XPO1,EIF5A, |
| 66 | 18123 | HNRNPA1,YBX1,SMG1,NPM1,SRSF4,LRPPRC,KHDRBS1,XPO1,EIF5A, |
| 168 | 18123 | TUBA1A,HSP90AA1,TUBB,UBC,RPS6,ATP2B4,SOX4,YWHAE,CALM3,A |
| 134 | 18123 | IGFBP5,PABPC1,GAPDH,EEF2,VIM,EIF4G2,NCL,PKM,YBX1,CALR,SO |
| 142 | 18123 | DPYSL3,HTT,HSP90AA1,ACTR2,NUMA1,FLNA,MAP1B,HSPA1B,CLU,R |
| 735 | 18123 | IGFBP5,PABPC1,COL1A1,HSPA8,GAPDH,MLLT1,HNRNPA1,NEDD4L,A |
| 245 | 18123 | IGFBP5,COL1A1,LRP1,HNRNPA1,ACTB,ACTG1,HSP90AB1,HSP90AA1 |
| 73 | 18123 | UBC,ENO1,PGK1,GNB1,P4HB,TMBIM6,UBB,PSMD5,EGLN1,RTN4,PS |
| 60 | 18123 | LRP1,EIF4G2,DBN1,MAP1B,NEFL,GDI1,PLXNB2,MAPT,FN1,PAFAH1E |
| 217 | 18123 | LRP1,HSPA9,NRBP2,MTRNR2L2,CFL1,XRCC2,HTT,HSP90AB1,UBC,F |
| 21 | 18123 | TMBIM6,NPM1,SRSF4,HNRNPK,U2AF2,RBMX,SRSF9,PTBP1,RPS13,S |
| 93 | 18123 | LRP1,XRCC2,ACTR2,EIF4G2,DBN1,FLNA,MAP1B,NEFL,CALR,P4HB,G |
| 75 | 18123 | UBC,ENO1,PGK1,GNB1,P4HB,TMBIM6,UBB,PSMD5,EGLN1,RTN4,AT |
| 119 | 18123 | HSPA8,ACTB,ACTG1,TUBA1A,CFL1,ACTR2,SPARC,TUBB,DBN1,FLNA |
| 107 | 18123 | LRP1,XRCC2,ACTR2,EIF4G2,DBN1,MAP1B,NEFL,GDI1,PLXNB2,MAPT |
| 77 | 18123 | LRP1,XRCC2,ACTR2,EIF4G2,DBN1,MAP1B,NEFL,GDI1,PLXNB2,MAPT |
| 58 | 18123 | ACTB,ACTG1,DVL3,PSMD5,AP2M1,PAFAH1B1,AP2B1,LAMA5,MSN,RI |
| 57 | 18123 | HSPA8,HSPA9,HSP90B1,CALR,HSPA1B,CANX,FKBP14,RACK1,TMBIM |
| 732 | 18123 | PABPC1,COL1A1,HSPA8,MLLT1,HNRNPA1,NEDD4L,POLH,ATN1,USP2 |
| 78 | 18123 | HSP90AA1,ACTR2,TMSB4X,NUMA1,DBN1,MAP1B,HSPA1B,SPTBN1,S |
| 760 | 18123 | IGFBP5,PABPC1,COL1A1,HSPA8,GAPDH,MLLT1,HNRNPA1,NEDD4L,A |
| 40 | 18123 | LRP1,EIF4G2,DBN1,MAP1B,NEFL,GDI1,PLXNB2,MAPT,FN1,PAFAH1E |
| 62 | 18123 | HSPA8,HSPA9,HSP90B1,CALR,HSPA1B,CANX,FKBP14,RACK1,TMBIM |
| 66 | 18123 | HNRNPA1,YBX1,SMG1,NPM1,SRSF4,LRPPRC,KHDRBS1,XPO1,EIF5A, |
| 79 | 18123 | UBC,ENO1,PGK1,GNB1,P4HB,TMBIM6,UBB,PSMD5,EGLN1,POLR2A |
| 43 | 18123 | PABPC1,YBX1,NPM1,HNRNPU,LARP1,HNRNPA0,SRSF4,HNRNPC,TAR |
| 728 | 18123 | IGFBP5,PABPC1,COL1A1,HSPA8,GAPDH,MLLT1,HNRNPA1,NEDD4L,A |
| 149 | 18123 | TUBA1A,HSP90AA1,TUBB,UBC,RPS6,SOX4,YWHAE,CALM3,APP,CALM |
| 47 | 18123 | DVL3,PSMD5,AP2M1,AP2B1,RHOA,PSMD2,PFN1,GPC3,CLTC,RAC1,F |
| 323 | 18123 | PABPC1,HSPA8,HNRNPA1,NEDD4L,ATN1,ZNF124,UBC,EIF4G2,ENO1 |
| 61 | 18123 | HSP90AB1,HSPA1B,CLU,RACK1,RPL5,VCP,RPL11,RPL23,XPO1,CSNK |
| 109 | 18123 | PABPC1,HSPA8,NEDD4L,MAVS,HSP90AB1,UBC,HSPA1B,DDX5,BSG,F |
| 385 | 18123 | PABPC1,COL1A1,HSPA8,HNRNPA1,USP22,MAVS,HSP90AB1,HSP90A |
| 52 | 18123 | DVL3,PSMD5,AP2M1,PAFAH1B1,AP2B1,RHOA,PSMD2,WDR1,PFN1,A |
| 52 | 18123 | DVL3,PSMD5,AP2M1,PAFAH1B1,AP2B1,RHOA,PSMD2,WDR1,PFN1,A |
| 65 | 18123 | LRP1,HSP90AB1,EIF4G2,DBN1,MAP1B,GDI1,NPM1,MAPT,FN1,PAFAH |
| 50 | 18123 | HSP90AB1,FLNA,RPL5,RPL11,ANXA2,TIMP2,RPL23,RPS7,HMGCR,LA |
| 67 | 18123 | CALR,CANX,PSMD5,HLA-C,PDIA3,AP2M1,ARF1,AP2B1,CTSD,PSMD2, |
| 53 | 18123 | HSP90AB1,HSPA1B,CLU,RACK1,VCP,RPL11,XPO1,CSNK1E,TRIB2,MA |
| 92 | 18123 | LRP1,MAVS,HSP90AB1,NUMA1,PIK3R2,FLNA,ATP2B4,YWHAE,RACK1 |
| 92 | 18123 | HSP90AA1,ACTR2,TMSB4X,NUMA1,DBN1,MAP1B,NEFL,HSPA1B,SPT |
| 97 | 18123 | HSPA8,GAPDH,CFL1,HSP90AB1,UBC,ENO1,HSPA1B,PGK1,BSG,P4HE |
| 93 | 18123 | NEDD4L,CFL1,ACTR2,SPARC,DBN1,FLNA,ALDOA,CALR,DVL3,MYH9, |
| 79 | 18123 | NEDD4L,CFL1,ACTR2,KIDINS220,DCX,EIF4G2,DBN1,MAP1B,KIF1A,A |
| 123 | 18123 | PABPC1,LRP1,CTSC,HTT,HSP90AA1,HSPA1B,CLU,RACK1,APP,VCP,HI |

| | | |
|---|---|---|
| 113 | 18123 | COL1A1,UBC,ENO1,HSP90B1,PGK1,GNB1,P4HB,TMBIM6,HSPD1,U |
| 351 | 18123 | IGFBP5,COL1A1,LRP1,ACTB,EEF2,HSP90AB1,ACTR2,VIM,SPARC,PI |
| 152 | 18123 | USP22,TUBA1A,HSP90AA1,TUBB,UBC,SOX4,YWHAE,APP,IGF2,UBB, |
| 163 | 18123 | EEF1A1,HSPA8,CTSC,EEF2,HSP90AB1,HSP90AA1,ACTR2,TUBB,FTH1 |
| 99 | 18123 | NEDD4L,UBC,FBXL4,TTC3,UBB,PSMD5,MARCHF6,WSB1,PSMD2,DD |
| 444 | 18123 | IGFBP5,COL1A1,LRP1,HSPA8,NEDD4L,ACTB,ACTG1,DPYSL3,CFL1, |
| 391 | 18123 | EEF1A1,GAPDH,MLLT1,ACTB,NRBP2,MAVS,HTT,HSP90AB1,HSP90A |
| 47 | 18123 | DVL3,PSMD5,AP2M1,AP2B1,RHOA,PSMD2,PFN1,GPC3,CLTC,RAC1, |
| 684 | 18123 | PABPC1,COL1A1,HSPA8,MLLT1,HNRNPA1,NEDD4L,ATN1,USP22,ME |
| 178 | 18123 | LRP1,GAPDH,CTSC,HSP90AB1,ENO1,TMF1,HSPA1B,CLU,YWHAE,GP |
| 56 | 18123 | DVL3,GNB1,CALM1,PSMD5,AP2M1,AP2B1,RHOA,PSMD2,PFN1,CSN |
| 56 | 18123 | HNRNPA1,SMG1,SRSF4,LRPPRC,XPO1,EIF5A,BCLAF1,CPSF2,DDX39 |
| 97 | 18123 | EEF1A1,GAPDH,NRBP2,HTT,EIF4G2,PIK3R2,PSAP,SMG1,MAPT,HMG |
| 360 | 18123 | IGFBP5,COL1A1,LRP1,CTSC,MAVS,HTT,HSP90AB1,HSP90AA1,UBC, |
| 18 | 18123 | NPM1,SRSF4,HNRNPK,U2AF2,RBMX,SRSF9,PTBP1,SRSF6,HNRNPA2 |
| 1510 | 18123 | EEF1A1,IGFBP5,COL1A1,LRP1,HSPA8,GAPDH,COPA,HNRNPA1,HSPA |
| 331 | 18123 | EEF1A1,MLLT1,ACTB,MAVS,HTT,HSP90AB1,HSP90AA1,UBC,KIDINS2 |
| 331 | 18123 | EEF1A1,MLLT1,ACTB,MAVS,HTT,HSP90AB1,HSP90AA1,UBC,KIDINS2 |
| 51 | 18123 | SMG1,NPM1,SRSF4,XPO1,EIF5A,RPS15,CPSF2,DDX39B,SRSF5,U2A |
| 78 | 18123 | HSP90AB1,HSP90AA1,HSPA1B,SOX4,RPL5,NPM1,VCP,UBB,RPS3,RP |
| 85 | 18123 | LRP1,XRCC2,ACTR2,EIF4G2,DBN1,MAP1B,NEFL,GDI1,PLXNB2,MAP |
| 120 | 18123 | TUBA1A,HSP90AA1,TUBB,UBC,SOX4,YWHAE,APP,UBB,YWHAG,DDB1 |
| 169 | 18123 | LRP1,ACTB,ACTG1,DPYSL3,CFL1,ACTR2,TMSB4X,DBN1,FLNA,ALDO |
| 346 | 18123 | COL1A1,LRP1,HSPA8,BTBD9,UBC,DBN1,FLNA,MAP1B,FADS1,IGFBP |
| 102 | 18123 | LRP1,ACTB,HSP90AB1,TMSB4X,RACK1,CALM3,APP,TMBIM6,CALM1 |
| 306 | 18123 | IGFBP5,LRP1,HTT,UBC,TMSB4X,ENO1,PIK3R2,ATP2B4,IGFBP2,CAL |
| 768 | 18123 | IGFBP5,COL1A1,LRP1,HSPA8,POLH,ACTB,CTSC,MTRNR2L2,ACTG1, |
| 352 | 18123 | PABPC1,COL1A1,HSPA8,USP22,MAVS,ACTR2,UBC,PIK3R2,NCL,YBX1 |
| 298 | 18123 | PABPC1,HSPA8,NEDD4L,ATN1,ZNF124,UBC,EIF4G2,ENO1,FLNA,YBX |
| 288 | 18123 | IGFBP5,LRP1,HTT,UBC,TMSB4X,ENO1,PIK3R2,ATP2B4,IGFBP2,CAL |
| 96 | 18123 | HSPA8,CFL1,MAP1B,DVL3,SPTBN1,STMN2,SPTAN1,CALM1,VCP,SET, |
| 306 | 18123 | IGFBP5,LRP1,HTT,UBC,TMSB4X,ENO1,PIK3R2,ATP2B4,IGFBP2,CAL |
| 357 | 18123 | IGFBP5,COL1A1,LRP1,CTSC,MAVS,HTT,HSP90AB1,HSP90AA1,UBC, |
| 50 | 18123 | SMG1,NPM1,SRSF4,XPO1,EIF5A,RPS15,CPSF2,DDX39B,SRSF5,U2A |
| 60 | 18123 | PABPC1,EEF2,VIM,EIF4G2,PKM,CLU,SOX4,RACK1,RPL5,NPM1,PLXN |
| 136 | 18123 | TUBA1A,HSP90AA1,TUBB,UBC,ATP2B4,SOX4,YWHAE,APP,NPM1,UB |
| 125 | 18123 | USP22,KAT6A,IGF2,TRIM28,DDB1,SET,GTF3C4,MORF4L1,TADA2B, |
| 63 | 18123 | COL1A1,SOX4,VCP,PSMD5,PSMD2,CSNK1E,GPC3,SULF1,DDX3X,MA |
| 49 | 18123 | HSP90B1,CALR,CANX,FKBP14,RACK1,VCP,ATF4,PDIA6,TLN1,CTDSP |
| 101 | 18123 | MAVS,HSP90AB1,NUMA1,PIK3R2,FLNA,MAP1B,YWHAE,GDI1,TRIM28 |
| 368 | 18123 | IGFBP5,LRP1,MAVS,HTT,HSP90AB1,HSP90AA1,UBC,TMSB4X,ENO1, |
| 185 | 18123 | LRP1,NEDD4L,ACTB,ACTG1,DPYSL3,CFL1,ACTR2,VIM,TMSB4X,DBN |
| 104 | 18123 | COL1A1,LRP1,IGFBP2,SOX4,DVL3,RACK1,APP,VCP,PSMD5,PPP2R1 |
| 532 | 18123 | IGFBP5,COL1A1,LRP1,HSPA8,NEDD4L,ACTB,ACTG1,DPYSL3,BTBD9 |
| 122 | 18123 | USP22,KAT6A,IGF2,DDB1,SET,GTF3C4,MORF4L1,TADA2B,SFPQ,CT |

| | | |
|---|---|---|
| 97 | 18123 | ORC4,UBC,H3-3B,KAT6A,NPM1,UBB,DDB1,PSMD5,SET,HMGB1,SM |
| 141 | 18123 | HTT,TMF1,COL5A1,CLU,SPTBN1,MYH9,APP,PPP2R1A,VAPA,SEC61A |
| 219 | 18123 | IGFBP5,COL1A1,ACTB,EEF2,MAVS,HSP90AB1,ACTR2,SPARC,FLNA, |
| 55 | 18123 | COL1A1,SOX4,VCP,PSMD5,PSMD2,CSNK1E,GPC3,DDX3X,WLS,PTK7 |
| 118 | 18123 | LRP1,CTSC,XRCC2,ACTR2,EIF4G2,DBN1,MAP1B,NEFL,GDI1,PLXNB |
| 101 | 18123 | UBC,ENO1,HSP90B1,PGK1,GNB1,P4HB,TMBIM6,HSPD1,UBB,PSMD |
| 104 | 18123 | UBC,ENO1,HSP90B1,PGK1,GNB1,P4HB,TMBIM6,HSPD1,UBB,PSMD |
| 35 | 18123 | CALR,PSMD5,HLA-C,PDIA3,PSMD2,HLA-B,PSMB2,PSMD3,B2M,PSM |
| 49 | 18123 | HNRNPA1,SMG1,NPM1,SRSF4,KHDRBS1,EIF5A,CPSF2,DDX39B,SRS |
| 75 | 18123 | UBC,ENO1,HSPA1B,PGK1,BSG,P4HB,UBB,TRIM28,DDB1,VAPA,ITGB |
| 169 | 18123 | EEF1A1,LRP1,HSPA8,CTSC,EEF2,HSP90AB1,HSP90AA1,ACTR2,TUBB |
| 155 | 18123 | COL1A1,USP22,XRCC2,KIDINS220,DBN1,YBX1,SOX4,DVL3,RPL13,G |
| 654 | 18123 | IGFBP5,COL1A1,LRP1,HSPA9,ACTB,EEF2,HSP90AB1,VIM,SPARC,NU |
| 156 | 18123 | CTSC,NUDC,TUBA1A,CFL1,HTT,HSP90AA1,ACTR2,TUBB,NUMA1,MA |
| 151 | 18123 | COL1A1,XRCC2,KIDINS220,DBN1,YBX1,SOX4,DVL3,RPL13,GPI,MYH |
| 38 | 18123 | COPA,HTT,RAB6B,RAB1A,ARCN1,KLC1,SURF4,TMED2,ARF3,KDELR |
| 348 | 18123 | LRP1,MAVS,HTT,HSP90AB1,HSP90AA1,UBC,KIDINS220,TMSB4X,AT |
| 44 | 18123 | LRP1,EIF4G2,DBN1,MAP1B,GDI1,MAPT,FN1,PAFAH1B1,RTN4,MEGF |
| 73 | 18123 | CALR,CANX,PSMD5,HLA-C,PDIA3,AP2M1,AP3D1,ARF1,AP2B1,CTSD, |
| 174 | 18123 | IGFBP5,COL1A1,LRP1,HNRNPA1,ACTB,ACTG1,HSP90AA1,ACTR2,PI |
| 32 | 18123 | ORC4,UBC,UBB,PSMD5,PSMD2,PSMB2,PSMD3,PSMA7,PSMD1,UBA |
| 32 | 18123 | ATP2B4,PSMD5,PSMD2,OAZ1,PSMB2,AZIN1,PSMD3,ODC1,PSMA7,P |
| 323 | 18123 | LRP1,HSPA8,POLH,CTSC,ACTG1,MAVS,HSP90AB1,HSP90AA1,ACTR2 |
| 83 | 18123 | LRP1,TUBB2B,BSG,SPTBN1,APP,SPTAN1,PLXNB2,NCAM1,LAMA5,R |
| 79 | 18123 | HSP90AA1,ACTR2,NUMA1,MAP1B,HSPA1B,CLU,RACK1,FSCN1,VCP, |
| 167 | 18123 | EEF1A1,HSPA8,CTSC,EEF2,HSP90AB1,HSP90AA1,ACTR2,TUBB,FTH1 |
| 55 | 18123 | LRP1,CTSC,HSP90AA1,HSPA1B,CLU,RACK1,VCP,MSN,TMEM259,CD |
| 541 | 18123 | EEF1A1,GAPDH,MLLT1,ACTB,NRBP2,MAVS,HTT,HSP90AB1,HSP90A |
| 322 | 18123 | IGFBP5,COL1A1,LRP1,CTSC,MAVS,HTT,HSP90AB1,HSP90AA1,UBC, |
| 222 | 18123 | EEF1A1,MLLT1,ACTB,HTT,HSP90AB1,HSP90AA1,UBC,KIDINS220,PIK |
| 157 | 18123 | IGFBP5,COL1A1,LRP1,ACTB,ACTR2,VIM,PIK3R2,FLNA,MAP1B,PKM, |
| 287 | 18123 | EEF1A1,MLLT1,ACTB,MAVS,HTT,HSP90AB1,HSP90AA1,UBC,KIDINS2 |
| 40 | 18123 | LRP1,EIF4G2,DBN1,MAP1B,GDI1,MAPT,FN1,PAFAH1B1,RTN4,MEGF |
| 41 | 18123 | PSMD5,PSMD2,SKP1,CDKN1A,NOP53,PSMB2,PSMD3,PSMA7,PSMD1 |
| 86 | 18123 | COL1A1,IGFBP2,SOX4,DVL3,VCP,PSMD5,PSMD2,TLE5,CSNK1E,CDH |
| 82 | 18123 | LRP1,TUBB2B,BSG,SPTBN1,APP,SPTAN1,PLXNB2,NCAM1,LAMA5,R |
| 51 | 18123 | LRP1,DPYSL3,VIM,FLNA,STMN2,GDI1,PAFAH1B1,RTN4,RHOA,NR2F1 |
| 38 | 18123 | NUMA1,MAP1B,HSPA1B,STMN2,MAPT,RPS3,STMN1,CKAP5,MAP2,R |
| 201 | 18123 | ODF2L,ACTG1,TUBA1A,HTT,HSP90AA1,ACTR2,TUBB,NUMA1,DCX,R |
| 83 | 18123 | DBN1,SMG1,NPM1,AP2M1,AP3D1,AP2B1,HNRNPU,CLSTN1,KIF5B,S |
| 128 | 18123 | LRP1,HSP90AB1,ACTR2,TMSB4X,EIF4G2,DBN1,MAP1B,NEFL,SPTBN |
| 34 | 18123 | HSP90AB1,RPL5,RPL11,ANXA2,RPL23,RPS7,LAPTM4B,MAP1A,NOP5 |
| 98 | 18123 | CTSC,HTT,ENO1,HSPA1B,CLU,MAGED1,RACK1,P4HB,TMBIM6,GSTF |
| 267 | 18123 | GAPDH,USP22,HSP90AB1,HSP90AA1,ATP2B4,SOX4,FKBP14,KAT6A, |
| 113 | 18123 | LRP1,ACTB,ACTG1,FLNA,MAP1B,UGT8,APP,FSCN1,PLXNB2,MAPT,F |

| | | |
|---|---|---|
| 166 | 18123 | IGFBP5,COL1A1,LRP1,ACTB,MAVS,ACTR2,VIM,PIK3R2,FLNA,MAP1B |
| 35 | 18123 | CALR,PSMD5,HLA-C,PDIA3,PSMD2,HLA-B,PSMB2,PSMD3,B2M,PSM |
| 1134 | 18123 | IGFBP5,COL1A1,LRP1,HSPA8,COPA,HNRNPA1,HSPA9,ACTB,CTSC,N |
| 325 | 18123 | IGFBP5,COL1A1,LRP1,ACTG1,DPYSL3,SPARC,TMSB4X,DCX,PIK3R2, |
| 136 | 18123 | IGFBP5,ACTB,ACTG1,FLNA,COL5A1,COL4A1,SOX4,MAGED1,DVL3,E |
| 156 | 18123 | IGFBP5,ACTB,ACTG1,FLNA,COL5A1,COL4A1,SOX4,MAGED1,DVL3, |
| 541 | 18123 | EEF1A1,GAPDH,MLLT1,ACTB,NRBP2,MAVS,HTT,HSP90AB1,HSP90A |
| 50 | 18123 | PABPC1,EEF2,VIM,EIF4G2,PKM,SOX4,RPL5,NPM1,PLXNB2,EIF3C,R |
| 346 | 18123 | IGFBP5,PABPC1,HSPA8,GAPDH,HNRNPA1,NEDD4L,ATN1,ZNF124,UB |
| 43 | 18123 | SMG1,SRSF4,EIF5A,CPSF2,DDX39B,SRSF5,U2AF2,CPSF1,PABPN1,A |
| 43 | 18123 | SMG1,SRSF4,EIF5A,CPSF2,DDX39B,SRSF5,U2AF2,CPSF1,PABPN1,A |
| 41 | 18123 | PABPC1,APP,PAPOLA,SRSF4,CPSF2,DDX39B,SRSF5,U2AF2,CPSF1,P |
| 53 | 18123 | IGF2,SET,TADA2B,CTNNB1,CTBP1,EID1,SKI,TAF7,VEGFA,KDM1A,SU |
| 88 | 18123 | LRP1,EEF2,VIM,DCX,CLU,SOX4,APP,GSTP1,MAPT,ZMIZ1,PAFAH1B1, |
| 52 | 18123 | NUMA1,MAP1B,HSPA1B,STMN2,MAPT,MAP9,PKD1,RPS3,STMN1,PA |
| 386 | 18123 | IGFBP5,COL1A1,LRP1,ACTB,ACTG1,DPYSL3,CFL1,SPARC,TMSB4X, |
| 68 | 18123 | H3-3B,IGF2,TRIM28,SET,HNRNPU,TADA2B,H1-0,CTNNB1,H1-10,CT |
| 94 | 18123 | COL1A1,IGFBP2,SOX4,DVL3,VCP,PSMD5,PSMD2,TLE5,CSNK1E,CDH |
| 37 | 18123 | HSPA9,PSMD5,XRCC5,PSMD2,CHD2,PSMB2,PSMD3,PRKDC,PSMA7, |
| 84 | 18123 | ORC4,UBC,H3-3B,KAT6A,NPM1,UBB,DDB1,PSMD5,SET,HMGB1,SM |
| 1128 | 18123 | IGFBP5,COL1A1,LRP1,HSPA8,COPA,HNRNPA1,HSPA9,ACTB,CTSC,N |
| 253 | 18123 | IGFBP5,COL1A1,LRP1,ACTB,ACTR2,VIM,PIK3R2,FLNA,MAP1B,PKM, |
| 328 | 18123 | IGFBP5,PABPC1,HSPA8,GAPDH,HNRNPA1,NEDD4L,ATN1,ZNF124,UB |
| 61 | 18123 | LRP1,ODF2L,DPYSL3,VIM,FLNA,STMN2,GDI1,PAFAH1B1,MAP4,RTN |
| 83 | 18123 | ENO1,CLU,DDX5,RACK1,P4HB,TMBIM6,UBB,NONO,RPS3,TPT1,RPL |
| 54 | 18123 | GAPDH,NRBP2,HTT,SMG1,UCHL1,LARP1,PIP4K2B,SESN3,ATP6V0C, |
| 129 | 18123 | RPL3,RPS6,RPL4,RPL7A,RPL8,RPLP0,RPL13A,RPL6,RPL13,DDX17,R |
| 87 | 18123 | HSPA8,GAPDH,TMSB4X,ENO1,ALDOA,PKM,HSPA1B,PGK1,ENO2,GPI |
| 56 | 18123 | HSPA8,ACTB,ACTG1,CFL1,ACTR2,DBN1,NEFL,KIF1A,INA,ARF1,PAFA |
| 272 | 18123 | COL1A1,LRP1,NEDD4L,CFL1,XRCC2,HSP90AB1,ACTR2,NUMA1,EIF4 |
| 140 | 18123 | LRP1,HSPA8,MAVS,HSP90AB1,PIK3R2,FLNA,SOX4,YWHAE,GNAS,RA |
| 247 | 18123 | PABPC1,LRP1,HSPA8,GAPDH,ACTB,ACTG1,EEF2,MAVS,HSP90AA1,V |
| 50 | 18123 | TMSB4X,CLU,RPL13A,SPTBN1,STMN2,SPTAN1,TMSB10,HMGB1,ST |
| 50 | 18123 | MAVS,HSP90AA1,KLRD1,MATR3,NONO,PSMD5,HMGB1,XRCC5,PSM |
| 66 | 18123 | HSP90AB1,HSP90AA1,HSPA1B,SOX4,RPL5,NPM1,UBB,RPS3,RPL11, |
| 37 | 18123 | HSPA9,PSMD5,PSMD2,PSMB2,PSMD3,PRKDC,PSMA7,PSMD1,FLCN, |
| 56 | 18123 | LRP1,NEDD4L,EIF4G2,DBN1,MAP1B,H3-3B,GDI1,MAPT,FN1,PAFAH1 |
| 39 | 18123 | PSMD5,PSMD2,SKP1,CDKN1A,NOP53,PSMB2,PSMD3,PSMA7,PSMD1 |
| 72 | 18123 | HNRNPA1,YBX1,SMG1,NPM1,SRSF4,LRPPRC,KHDRBS1,XPO1,EIF5A, |
| 155 | 18123 | PIK3R2,MAP1B,PKM,IGFBP2,CALR,TMF1,ABHD2,H3-3B,DDX5,GNAS |
| 70 | 18123 | HTT,ENO1,HSPA1B,CLU,TMBIM6,GSTP1,NONO,TPT1,DNAJA1,CTNN |
| 131 | 18123 | IGFBP5,COL1A1,LRP1,VIM,SPARC,PIK3R2,MAP1B,PKM,FADS1,NEFL |
| 23 | 18123 | LRP1,EIF4G2,DBN1,MAP1B,GDI1,MAPT,FN1,PAFAH1B1,MEGF8,MAC |
| 324 | 18123 | IGFBP5,PABPC1,HSPA8,GAPDH,HNRNPA1,NEDD4L,ATN1,ZNF124,UB |
| 391 | 18123 | IGFBP5,COL1A1,ACTB,ACTG1,EEF2,XRCC2,VIM,NUMA1,FLNA,COL6 |

| | | |
|---|---|---|
| 99 | 18123 | CTSC,ENO1,HSPA1B,CLU,RACK1,MYH9,APP,HSPD1,VCP,MAPT,FN1,F |
| 65 | 18123 | ACTR2,NUMA1,FLNA,MAP1B,HSPA1B,CLU,STMN2,APP,MAPT,RPS3,A |
| 154 | 18123 | COL1A1,LRP1,ACTB,ACTG1,DPYSL3,SPARC,FLNA,MAP1B,NEFL,ABH |
| 32 | 18123 | HSPA9,PSMD5,PSMD2,PSMB2,PSMD3,PRKDC,PSMA7,PSMD1,PSMD |
| 251 | 18123 | COL1A1,HSPA8,GAPDH,HSPA9,ACTG1,DPYSL3,CFL1,MAVS,HSP90A |
| 314 | 18123 | COL1A1,USP22,MAVS,ACTR2,UBC,PIK3R2,NCL,YBX1,TMF1,ZNF99,S |
| 314 | 18123 | COL1A1,USP22,MAVS,ACTR2,UBC,PIK3R2,NCL,YBX1,TMF1,ZNF99,S |
| 191 | 18123 | COL1A1,LRP1,XRCC2,HSP90AB1,ACTR2,NUMA1,EIF4G2,DBN1,FLN |
| 314 | 18123 | COL1A1,USP22,MAVS,ACTR2,UBC,PIK3R2,NCL,YBX1,TMF1,ZNF99,S |
| 223 | 18123 | IGFBP5,COL1A1,FLNA,COL5A1,SOX4,MAGED1,DVL3,BSG,COL18A1, |
| 91 | 18123 | LRP1,MAVS,HSP90AB1,PIK3R2,FLNA,SOX4,YWHAE,RACK1,BSG,TRI |
| 84 | 18123 | ACTB,USP22,UBC,USP11,VCP,UBB,PSMD5,UCHL1,RHOA,PSMD2,TA |
| 44 | 18123 | HSPA8,CFL1,UBC,ENO1,PGK1,UBB,VAPA,ANXA2,KPNB1,SFPQ,RAB7 |
| 43 | 18123 | HSPA8,CFL1,MAP1B,SPTBN1,STMN2,SPTAN1,STMN1,WDR1,NES,SP |
| 113 | 18123 | IGFBP5,COL1A1,SPARC,PIK3R2,MAP1B,PKM,FADS1,NEFL,IGFBP2,G |
| 42 | 18123 | HSPA9,PSMD5,HNRNPU,PSMD2,PTN,PSMB2,PSMD3,PRKDC,PSMA7, |
| 66 | 18123 | HSPA8,TUBA1A,CFL1,ACTR2,SPARC,TUBB,DBN1,KIF1A,APP,VCP,AG |
| 27 | 18123 | NUMA1,MAP1B,HSPA1B,STMN2,MAPT,RPS3,STMN1,CKAP5,MAP2,R |
| 34 | 18123 | NUMA1,MAP1B,HSPA1B,STMN2,MAPT,RPS3,STMN1,CKAP5,MAP2,R |
| 38 | 18123 | CALR,CANX,PSMD5,HLA-C,PDIA3,PSMD2,HLA-B,PSMB2,PSMD3,B2M |
| 223 | 18123 | MAVS,HSP90AB1,HSP90AA1,UBC,KIDINS220,ATP2B4,CLU,SOX4,MA |
| 128 | 18123 | MLLT1,YWHAE,RACK1,CALM3,GSTP1,CALM1,NPM1,YWHAG,MAPT,S |
| 42 | 18123 | HSPA1B,CLU,RACK1,VCP,TMEM259,CSNK1E,TRIB2,NOP53,GABARA |
| 250 | 18123 | MLLT1,ACTB,MAVS,HTT,HSP90AB1,HSP90AA1,UBC,KIDINS220,ATP2 |
| 32 | 18123 | HNRNPA1,DDX5,DDX17,HNRNPU,KHDRBS1,SFPQ,NOVA2,HNRNPL,R |
| 59 | 18123 | HNRNPA1,NUMA1,H3-3B,TRIM28,XRCC5,CCT7,TADA2B,SFPQ,CTNN |
| 89 | 18123 | LRP1,ACTB,NUMA1,FLNA,ATP2B4,RACK1,BSG,SPTBN1,DCHS1,AP2M |
| 320 | 18123 | MLLT1,ACTB,NRBP2,MAVS,HTT,HSP90AB1,HSP90AA1,UBC,KIDINS2 |
| 107 | 18123 | BTBD9,DBN1,MAP1B,TUBB2B,CALM3,SHISA9,APP,CALM1,YWHAG,M |
| 1272 | 18123 | IGFBP5,COL1A1,LRP1,GAPDH,COPA,HSPA9,NEDD4L,ATN1,ACTB,CT |
| 63 | 18123 | HSPA8,TUBA1A,CFL1,ACTR2,SPARC,TUBB,DBN1,KIF1A,APP,VCP,AG |
| 107 | 18123 | BTBD9,DBN1,MAP1B,TUBB2B,CALM3,SHISA9,APP,CALM1,YWHAG,M |
| 246 | 18123 | LRP1,GAPDH,MLLT1,NEDD4L,ACTB,MTRNR2L2,TMSB4X,FLNA,MTR |
| 127 | 18123 | MLLT1,YWHAE,RACK1,CALM3,GSTP1,CALM1,NPM1,YWHAG,MAPT,S |
| 53 | 18123 | LRP1,DPYSL3,ACTR2,KIDINS220,DBN1,FLNA,STMN2,PLXNB2,PAFAH |
| 72 | 18123 | MAVS,HSP90AB1,PIK3R2,FLNA,YWHAE,GDI1,TRIM28,YWHAG,YWHA |
| 111 | 18123 | COL1A1,EEF2,HSPA1B,RACK1,P4HB,APP,GSTP1,HSPD1,MAPT,NON |
| 91 | 18123 | CFL1,GNB1,GDI1,TRIM28,HACD3,STMN1,ITGB1,TIMP2,RTN4,KPNB |
| 333 | 18123 | IGFBP5,PABPC1,HSPA8,GAPDH,HNRNPA1,NEDD4L,ATN1,ZNF124,UB |
| 23 | 18123 | COL6A1,COL5A1,COL4A2,FN1,COL11A1,COL5A2,MMP14,NANOG,C |
| 79 | 18123 | ACTB,USP22,UBC,USP11,VCP,UBB,PSMD5,UCHL1,RHOA,PSMD2,TA |
| 78 | 18123 | LRP1,ACTB,FLNA,ATP2B4,RACK1,BSG,SPTBN1,DCHS1,AP2M1,ITGB |
| 53 | 18123 | ENO1,CLU,RACK1,P4HB,TMBIM6,UBB,NONO,RPS3,TPT1,RPL11,DN |
| 49 | 18123 | NEDD4L,CFL1,ACTR2,KIDINS220,DCX,DBN1,KIF1A,ITGB1,PAFAH1B1 |
| 232 | 18123 | COL1A1,HSPA8,GAPDH,HSPA9,ACTG1,DPYSL3,CFL1,MAVS,HSP90A |

| | | |
|---|---|---|
| 88 | 18123 | LRP1,NEDD4L,EIF4G2,DBN1,MAP1B,GNAS,APP,GDI1,IGF2,MAPT,FN |
| 43 | 18123 | NUMA1,MAP1B,HSPA1B,STMN2,MAPT,RPS3,STMN1,CKAP5,MAP2,R |
| 33 | 18123 | EIF4G2,RPL13A,NPM1,CSDE1,EIF4B,EIF1,LARP1,EIF4H,ATF4,EIF5,K |
| 33 | 18123 | HNRNPA1,SMG1,HNRNPU,XRCC5,CCT7,HNRNPC,CTNNB1,CCT6A,C |
| 42 | 18123 | HSPA8,HSP90AB1,HSP90AA1,HSPA1B,YWHAE,HSPD1,VCP,MAPT,ATP |
| 129 | 18123 | TMSB4X,ENO1,PIK3R2,ATP2B4,CLU,RACK1,TMBIM6,GSTP1,NONO, |
| 107 | 18123 | RPL3,RPS6,RPL4,RPL7A,RPL8,RPLP0,RPL13A,RPL6,RPL13,DDX17,R |
| 345 | 18123 | IGFBP5,COL1A1,LRP1,ACTB,ACTG1,DPYSL3,CFL1,SPARC,TMSB4X, |
| 345 | 18123 | IGFBP5,COL1A1,LRP1,ACTB,ACTG1,DPYSL3,CFL1,SPARC,TMSB4X, |
| 39 | 18123 | LRP1,ACTG1,FN1,VCL,RHOA,MMP14,TLN1,MACF1,RAC1,PTPRA,LA |
| 47 | 18123 | MAVS,HSP90AB1,PIK3R2,FLNA,MAGED1,NPM1,TRIM28,CTNNA1,CC |
| 200 | 18123 | LRP1,UBC,PIK3R2,NEFL,MAGED1,DVL3,YWHAE,SPTBN1,APP,SPTAN |
| 34 | 18123 | UBC,UBB,PSMD5,PSMD2,PSMB2,PSMD3,PSMA7,PSMD1,UBA52,CUL |
| 72 | 18123 | HSPA9,FN1,PSMD5,HNRNPU,LAMA5,XRCC5,PSMD2,PTN,CDH2,CTN |
| 37 | 18123 | ENO1,CLU,TMBIM6,NONO,TPT1,DNAJA1,MCL1,DDX3X,MIF,PTTG1I |
| 103 | 18123 | COL1A1,LRP1,SPARC,COL6A1,COL5A1,COL4A1,BSG,P4HB,APP,FSC |
| 56 | 18123 | MAGED1,DVL3,PSMD5,AP2M1,AP2B1,RHOA,PSMD2,PFN1,GPC3,SUL |
| 67 | 18123 | ACTB,ACTG1,HSP90AB1,HSP90AA1,ACTR2,PIK3R2,CALM1,PSMD5,P |
| 45 | 18123 | CTSC,HSPA1B,CLU,RACK1,VCP,TMEM259,CSNK1E,TRIB2,NOP53,G |
| 28 | 18123 | HSP90AB1,RPL5,RPL11,RPL23,RPS7,MAP1A,NOP53,CCAR2,SMARC |
| 38 | 18123 | UBC,UBB,PSMD5,PSMD2,SKP1,SQSTM1,PSMB2,PSMD3,PSMA7,PSM |
| 59 | 18123 | HSPA8,TUBA1A,HTT,TUBB,TUBA1B,MAP1B,NEFL,KIF1A,APP,UBB,M |
| 289 | 18123 | EEF1A1,COL1A1,LRP1,HSPA8,ACTB,CTSC,ACTG1,EEF2,HSP90AB1,H |
| 102 | 18123 | COL1A1,LRP1,SPARC,COL6A1,COL5A1,COL4A1,BSG,P4HB,APP,FSC |
| 35 | 18123 | CFL1,ACTR2,DBN1,KIF1A,APP,ARF1,PAFAH1B1,CAPRIN1,MAP1A,VP |
| 102 | 18123 | COL1A1,LRP1,SPARC,COL6A1,COL5A1,COL4A1,BSG,P4HB,APP,FSC |
| 116 | 18123 | SPARC,TMSB4X,ATP2B4,CALR,FSTL1,GPI,BSG,ATP5F1B,MYH9,ATP |
| 56 | 18123 | RPS6,COL6A1,COL5A1,GPI,RACK1,COL4A2,FN1,ITGB1,COL11A1,C |
| 37 | 18123 | CFL1,ACTR2,EIF4G2,DBN1,KIF1A,ARF1,PAFAH1B1,CAPRIN1,HDAC2, |
| 559 | 18123 | IGFBP5,COL1A1,LRP1,HSPA8,HNRNPA1,HSPA9,ACTB,ACTG1,CFL1, |
| 106 | 18123 | CTSC,NUDC,CFL1,HTT,ACTR2,NUMA1,MAP1B,NEFL,KIF1A,MYH9,NP |
| 365 | 18123 | IGFBP5,LRP1,HNRNPA1,HSPA9,NEDD4L,ACTB,ACTG1,BTBD9,HTT,H |
| 46 | 18123 | HSP90AA1,ACTR2,NUMA1,MAP1B,HSPA1B,MAPT,RPS3,ARF1,RHOA, |
| 64 | 18123 | MAVS,HSP90AB1,NUMA1,PIK3R2,FLNA,YWHAE,TRIM28,YWHAG,YW |
| 106 | 18123 | ACTG1,DPYSL3,CFL1,ACTR2,TMSB4X,DBN1,FLNA,ALDOA,HSP90B1, |
| 19 | 18123 | RPL13A,EIF3C,EIF4B,EIF4A1,EIF4H,EIF5,EIF3L,EIF3E,EIF3F,EIF3D,E |
| 191 | 18123 | HSPA8,GAPDH,NUDC,TUBA1A,CFL1,XRCC2,HTT,ACTR2,TUBB,NUMA |
| 53 | 18123 | LRP1,HSP90AB1,CALM3,APP,CALM1,EIF3C,TRIM28,PKD1,HMGB1,S |
| 99 | 18123 | USP22,SOX4,KAT6A,TRIM28,SET,GTF3C4,MORF4L1,ZMIZ1,TADA2B, |
| 32 | 18123 | ATP2B4,PSMD5,PSMD2,OAZ1,PSMB2,AZIN1,PSMD3,ODC1,PSMA7,P |
| 199 | 18123 | ORC4,HNRNPA1,POLH,XRCC2,HSP90AB1,HSP90AA1,ACTR2,UBC,SM |
| 238 | 18123 | COL1A1,ACTG1,XRCC2,SPARC,DCX,FLNA,NCL,PKM,COL6A1,UGT8, |
| 23 | 18123 | HSP90AB1,RPL5,RPL11,RPL23,RPS7,MAP1A,NOP53,CCAR2,SMARC |
| 356 | 18123 | IGFBP5,LRP1,HTT,UBC,TMSB4X,ENO1,PIK3R2,YBX1,ATP2B4,IGFBP |
| 93 | 18123 | XRCC2,KIDINS220,DBN1,YBX1,RPL13,GPI,MYH9,SMIM14,IGF2,TRI |

| | | |
|---|---|---|
| 128 | 18123 | LRP1,HSPA8,MAVS,HSP90AB1,PIK3R2,FLNA,SOX4,YWHAE,GNAS,RAC |
| 67 | 18123 | NUMA1,MAP1B,HSPA1B,STMN2,NPM1,MAPT,MAP9,PKD1,RPS3,STMI |
| 33 | 18123 | ACTB,ACTG1,ACTR2,AP2M1,AP2B1,RHOA,RAC1,MMP2,ACTR3,ARPC |
| 266 | 18123 | HSP90AB1,HSP90AA1,UBC,KIDINS220,TMSB4X,ATP2B4,PSAP,CLU,M |
| 31 | 18123 | COL6A1,COL5A1,COL4A2,FN1,COL11A1,COL5A2,MMP14,NANOG,C |
| 127 | 18123 | HSPA9,HTT,HSP90AA1,CLU,YWHAE,ATP5F1B,CALM3,CALM1,HSPD1, |
| 149 | 18123 | HSP90AB1,HSP90AA1,UBC,KIDINS220,ATP2B4,CLU,MAGED1,DVL3,C |
| 36 | 18123 | LRP1,ACTG1,FN1,VCL,RHOA,MMP14,TLN1,MACF1,RAC1,PTPRA,LAM |
| 49 | 18123 | HSPA9,RPS6,HSPA1B,GPI,HMGB1,RPS17,RPS24,RPS14,RPS19,VEGF |
| 244 | 18123 | LRP1,GAPDH,HTT,HSP90AB1,HSP90B1,PSAP,DVL3,YWHAE,GNAS,GP |
| 184 | 18123 | POLH,XRCC2,ACTR2,UBC,UBA1,CLU,DDX5,SOX4,SMG1,NPM1,VCP,U |
| 98 | 18123 | LRP1,VIM,PIK3R2,MAP1B,PKM,GNAS,APP,GSTP1,IGF2,YWHAG,RPL2 |
| 123 | 18123 | GAPDH,ENO1,ALDOA,PKM,PGK1,ENO2,GPI,ATP5F1B,APP,CALM1,IG |
| 137 | 18123 | IGFBP5,COL1A1,ACTB,MAVS,HSP90AB1,ACTR2,FLNA,HSP90B1,ATP2 |
| 258 | 18123 | HSPA8,NEDD4L,ATN1,ZNF124,UBC,EIF4G2,ENO1,FLNA,YBX1,CALR,I |
| 12 | 18123 | EIF3C,EIF5,EIF3L,EIF3E,EIF3F,EIF3D,EIF2S2,EIF3H,EIF3G,EIF3B,EIF3 |
| 83 | 18123 | LRP1,MAVS,HSP90AB1,PIK3R2,FLNA,SOX4,YWHAE,RACK1,BSG,TRIM |
| 740 | 18123 | COL1A1,HSPA8,MLLT1,HNRNPA1,NEDD4L,POLH,ATN1,USP22,MED1 |
| 24 | 18123 | COL6A1,COL5A1,COL4A2,FN1,COL11A1,COL5A2,MMP14,NANOG,C |
| 47 | 18123 | SOX4,TPT1,PTN,CDH2,NANOG,CTNNB1,SKI,NOTCH2,SELENON,STAT |
| 258 | 18123 | HSPA8,NEDD4L,ATN1,ZNF124,UBC,EIF4G2,ENO1,FLNA,YBX1,CALR,I |
| 64 | 18123 | LRP1,VIM,CLU,SOX4,APP,GSTP1,MAPT,GAP43,RHOA,WDR1,PTN,CDI |
| 77 | 18123 | HNRNPA1,NUMA1,H3-3B,SMG1,TRIM28,MAPT,HNRNPU,XRCC5,CCT |
| 22 | 18123 | USP22,DDB1,SKP1,KDM1A,UBR5,PCGF2,ATXN7L3,TRIP12,PARK7,PC |
| 25 | 18123 | RPL5,RPL11,RPL23,RPS20,RPS15,RPS7,RAB1A,SKP1,RPL37,PTEN,F |
| 277 | 18123 | LRP1,HSPA9,CTSC,CFL1,XRCC2,ACTR2,SPARC,NUMA1,EIF4G2,DBN |
| 258 | 18123 | HSPA8,NEDD4L,ATN1,ZNF124,UBC,EIF4G2,ENO1,FLNA,YBX1,CALR,I |
| 42 | 18123 | HSPA9,RPS6,HSPA1B,GPI,RPS17,RPS24,RPS14,RPS19,VEGFA,PRDX1 |
| 96 | 18123 | SOX4,NPM1,PSMD5,CBX5,PSMD2,RPL26,RBBP4,IK,PRKAR1A,DUX4, |
| 37 | 18123 | CTSC,VAPA,AP2M1,ANXA2,AP3D1,ARF1,RAB7A,RAB1A,TMED2,CSNK |
| 32 | 18123 | LRP1,ACTG1,VCL,RHOA,MMP14,MACF1,RAC1,PTPRA,VEGFA,ACTN1 |
| 174 | 18123 | LRP1,GAPDH,MLLT1,ATP2B4,YWHAE,GPI,RACK1,APP,RPL5,TMBIM6 |
| 27 | 18123 | ACTB,DCHS1,VCL,CTNNA1,CDH6,CDH2,CTNNB1,DLG5,CDH11,TBCD |
| 27 | 18123 | FKBP14,PDIA6,TLN1,CTDSP2,HDGF,ACADVL,HYOU1,SSR1,DCTN1,EX |
| 1034 | 18123 | IGFBP5,COL1A1,LRP1,HSPA8,COPA,HNRNPA1,HSPA9,ACTB,CTSC,NI |
| 28 | 18123 | LRP1,ACTG1,VCL,RHOA,MMP14,MACF1,RAC1,PTPRA,VEGFA,CORO |
| 148 | 18123 | LRP1,CTSC,HTT,UBC,RPS6,CLU,SOX4,MAGED1,RACK1,HSPD1,UBB, |
| 57 | 18123 | ATP5F1B,ATP5F1A,VCP,RHOA,MDH2,NOP53,ACO2,ATP5PB,SDHA,IDI |
| 46 | 18123 | SOX4,TPT1,PTN,CDH2,NANOG,CTNNB1,SKI,NOTCH2,STAT3,BCL9L,I |
| 234 | 18123 | MAVS,ACTR2,UBC,PIK3R2,NCL,YBX1,TMF1,ZNF99,SOX4,KAT6A,DVL |
| 143 | 18123 | GAPDH,NUDC,TUBA1A,CFL1,XRCC2,HTT,TUBB,NUMA1,DCX,TUBA1E |
| 99 | 18123 | MLLT1,RACK1,CALM3,GSTP1,CALM1,NPM1,YWHAG,MAPT,PPP2R1A |
| 628 | 18123 | COL1A1,HSPA8,MLLT1,NEDD4L,ATN1,USP22,MED18,MAVS,ZNF124, |
| 187 | 18123 | EEF1A1,MLLT1,ACTB,HTT,HSP90AB1,HSP90AA1,UBC,KIDINS220,PIK |
| 56 | 18123 | LRP1,ACTB,HSP90AB1,RACK1,APP,TMBIM6,PKD1,RPL11,ANXA2,GTI |

| | | |
|---|---|---|
| 701 | 18123 | COL1A1,HSPA8,MLLT1,HNRNPA1,NEDD4L,POLH,ATN1,USP22,MED1 |
| 627 | 18123 | COL1A1,HSPA8,MLLT1,NEDD4L,ATN1,USP22,MED18,MAVS,ZNF124, |
| 30 | 18123 | HSPA8,HSP90AB1,HSP90AA1,HSPA1B,YWHAE,MAPT,PTGES3,HSBP1, |
| 87 | 18123 | HSPA1B,RACK1,P4HB,GSTP1,MAPT,NONO,RPS3,DNAJA1,NFE2L1,XF |
| 711 | 18123 | COL1A1,HSPA8,MLLT1,HNRNPA1,NEDD4L,POLH,ATN1,USP22,MED1 |
| 71 | 18123 | HSPA9,RPS6,FSTL1,HSPA1B,SOX4,GPI,HMGB1,RPS17,RPS24,RPS14 |
| 32 | 18123 | HTT,VIM,UBC,UBB,RAB7A,SQSTM1,OPTN,ATG2B,DYNC1LI2,DYNLL2 |
| 31 | 18123 | CFL1,ACTR2,DBN1,KIF1A,ARF1,PAFAH1B1,CAPRIN1,VPS35,FYN,CDC |
| 25 | 18123 | HNRNPA1,DDX5,DDX17,HNRNPU,KHDRBS1,NOVA2,HNRNPL,RBMX,F |
| 25 | 18123 | UBC,ENO1,PGK1,UBB,KPNB1,SFPQ,RAB7A,PABPN1,MAPK1,FBLN1, |
| 48 | 18123 | SOX4,DVL3,PLXNB2,DCHS1,PKD1,PFN1,RPS7,PTK7,SKI,TMED2,FKB |
| 53 | 18123 | MAVS,HSP90AB1,PIK3R2,FLNA,YWHAE,TRIM28,YWHAG,YWHAQ,KIF5 |
| 131 | 18123 | RPS6,COL6A1,YBX1,COL5A1,SOX4,DVL3,GPI,RACK1,PLXNB2,IGF2,T |
| 167 | 18123 | MLLT1,ACTB,HTT,HSP90AB1,HSP90AA1,UBC,KIDINS220,ATP2B4,CL |
| 18 | 18123 | HSP90AB1,RPL11,MAP1A,NOP53,CCAR2,SMARCC1,USP7,BAG6,SGT |
| 22 | 18123 | HSP90AB1,RPL11,MAP1A,NOP53,CCAR2,SMARCC1,USP7,BAG6,SGT |
| 74 | 18123 | DDX5,SOX4,KAT6A,RPL5,NPM1,UBB,CHD3,RPL11,RPL23,CHD4,RPS |
| 628 | 18123 | COL1A1,HSPA8,MLLT1,NEDD4L,ATN1,USP22,MED18,MAVS,ZNF124, |
| 33 | 18123 | ACTR2,FSCN1,ITGB1,VCL,MTSS2,RAC1,ACTR3,NCKAP1,ARPC2,CAP |
| 51 | 18123 | HSPA9,EEF2,SOX4,PSMD5,XRCC5,PSMD2,CHD2,PSMB2,PSMD3,PRK |
| 40 | 18123 | COL6A1,COL5A1,GPI,COL4A2,FN1,ITGB1,COL11A1,COL5A2,MMP1 |
| 100 | 18123 | TMSB4X,ENO1,ATP2B4,APP,IGF2,VCP,DDB1,PSMD5,HMGB1,MIDN,F |
| 255 | 18123 | EEF1A1,HSPA8,CTSC,EEF2,HSP90AB1,HSP90AA1,ACTR2,TUBB,FTH1 |
| 13 | 18123 | UBC,CSDE1,UBB,EIF3L,PTBP1,SSB,EIF3F,PCBP2,EIF3D,UBA52,RPS2 |
| 47 | 18123 | PABPC1,APP,PAPOLA,SRSF4,CPSF2,DDX39B,SRSF5,U2AF2,CPSF1,P |
| 29 | 18123 | HTT,SQSTM1,OPTN,ATG2B,CDC37,GABARAP,BNIP3L,SREBF2,VDAC |
| 29 | 18123 | HTT,SQSTM1,OPTN,ATG2B,CDC37,GABARAP,BNIP3L,SREBF2,VDAC |
| 48 | 18123 | HSPA8,HTT,MAP1B,NEFL,KIF1A,APP,UBB,MAPT,UCHL1,AP3D1,PAFA |
| 95 | 18123 | LRP1,ACTG1,ACTR2,TMSB4X,DBN1,FLNA,SPTBN1,MYH9,FSCN1,SPT |
| 45 | 18123 | NUMA1,TMF1,PPP2R1A,PAFAH1B1,TMEM170A,RTN4,KPNB1,RPS19, |
| 14 | 18123 | UBC,UBB,DDB1,PARP1,RAD23B,UBA52,RPS27A,CUL4A,COPS6,CET |
| 135 | 18123 | CTSC,HTT,UBC,RPS6,CLU,SOX4,MAGED1,RACK1,HSPD1,UBB,PDIA3 |
| 132 | 18123 | CTSC,HTT,UBC,RPS6,CLU,SOX4,MAGED1,RACK1,HSPD1,UBB,PDIA3 |
| 102 | 18123 | IGFBP5,PSAP,FASN,IGF2,RPL19,PKD1,RPS16,RTN4,LAMA5,MSN,PR |
| 70 | 18123 | LRP1,NEDD4L,EIF4G2,DBN1,MAP1B,H3-3B,GDI1,IGF2,MAPT,FN1,B |
| 31 | 18123 | TRIM28,HMGB1,JUN,CTNNB1,BCLAF1,CDKN1A,CCND1,CAND1,PTE |
| 62 | 18123 | ACTR2,NUMA1,FLNA,MAP1B,HSPA1B,STMN2,MAPT,RPS3,ARF1,RHO |
| 45 | 18123 | HSPA8,CFL1,UBC,ENO1,PGK1,UBB,VAPA,ANXA2,KPNB1,CLEC7A,SF |
| 46 | 18123 | MAVS,HSP90AB1,PIK3R2,FLNA,TRIM28,RPL23,KPNB1,TARDBP,APPL |
| 709 | 18123 | COL1A1,HSPA8,MLLT1,HNRNPA1,NEDD4L,POLH,ATN1,USP22,MED1 |
| 25 | 18123 | RPL3,RPLP0,RPL6,RPL5,NPM1,RPL11,RPL10,RPSA,XRCC5,RPL24,R |
| 155 | 18123 | HSPA8,BTBD9,DBN1,MAP1B,TUBB2B,CALM3,GNB1,SHISA9,APP,CAL |
| 44 | 18123 | CTNNB1,KDM1A,SUPT6H,KDM4B,PRMT1,BRD4,ATRX,EHMT1,CARM |
| 21 | 18123 | HSPA9,CFL1,RPLP0,P4HB,ANXA2,ARF1,MSN,MIF,HNRNPDL,CNN2,TC |
| 57 | 18123 | COPA,CTSC,NRBP2,SPTBN1,SPTAN1,VCP,VAPA,ARF1,SPTBN2,RAB1 |

| | | |
|---|---|---|
| 497 | 18123 | IGFBP5,COL1A1,LRP1,GAPDH,HSPA9,CTSC,CFL1,XRCC2,MAVS,HSP[illegible] |
| 89 | 18123 | MLLT1,RACK1,CALM3,GSTP1,CALM1,NPM1,YWHAG,PPP2R1A,UCHL[illegible] |
| 86 | 18123 | UBC,H3-3B,KAT6A,NPM1,UBB,TRIM28,DDB1,SET,CHD3,HMGB1,SM[illegible] |
| 50 | 18123 | HNRNPA1,MAVS,HSP90AB1,PIK3R2,FLNA,TRIM28,RPL23,KPNB1,TA[illegible] |
| 76 | 18123 | HSPA1B,RACK1,P4HB,GSTP1,MAPT,NONO,RPS3,NFE2L1,ATF4,ATP2[illegible] |
| 87 | 18123 | LRP1,GAPDH,ATN1,NRBP2,XRCC2,HSP90AB1,NEFL,CLU,GPI,RACK1,[illegible] |
| 87 | 18123 | LRP1,ACTG1,ACTR2,TMSB4X,DBN1,FLNA,SPTBN1,FSCN1,SPTAN1,T[illegible] |
| 30 | 18123 | TMSB4X,SPTBN1,STMN2,SPTAN1,TMSB10,STMN1,PFN1,SPTBN2,M[illegible] |
| 53 | 18123 | IGFBP2,DVL3,PSMD5,PSMD2,TLE5,CDH2,GPC3,MDK,PSMB2,RAPGE[illegible] |
| 60 | 18123 | COL1A1,EEF2,RACK1,GSTP1,HSPD1,MAPT,RPS3,LDHA,TPM1,PRDX2[illegible] |
| 59 | 18123 | HSPA8,MAVS,HSP90AB1,HSP90AA1,PIK3R2,FLNA,CLU,HSPD1,TRIM2[illegible] |
| 57 | 18123 | LRP1,FLNA,MAP1B,NEFL,GSTP1,RPL19,CTNNA1,RPS16,NACA,GAP4[illegible] |
| 16 | 18123 | XRCC5,CCT7,CCT6A,CCT2,TCP1,CCT3,CCT4,TERF2IP,CCT8,TNKS2,[illegible] |
| 82 | 18123 | PIK3R2,MAP1B,PKM,GNAS,GSTP1,IGF2,YWHAG,PIP4K2B,PRKAR1A,[illegible] |
| 595 | 18123 | COL1A1,HSPA8,MLLT1,NEDD4L,ATN1,USP22,MED18,MAVS,ZNF124,[illegible] |
| 109 | 18123 | SOX4,NPM1,PSMD5,TIMP2,CBX5,PSMD2,RPL26,RBBP4,IK,PRKAR1A[illegible] |
| 24 | 18123 | SNRNP200,SF3B1,LUC7L3,DDX39B,SRSF5,PRPF6,SRSF9,SRSF1,SC[illegible] |
| 32 | 18123 | HSPA1B,CLU,RACK1,VCP,CSNK1E,TRIB2,NOP53,GABARAP,UBQLN1,[illegible] |
| 74 | 18123 | BTBD9,HTT,ACTR2,GNAS,GPI,APP,MAPT,ITGB1,PAFAH1B1,PJA2,PT[illegible] |
| 52 | 18123 | CTNNB1,KDM1A,SUPT6H,KDM4B,PRMT1,GSPT1,BRD4,ATRX,FBXO1[illegible] |
| 52 | 18123 | CTNNB1,KDM1A,SUPT6H,KDM4B,PRMT1,GSPT1,BRD4,ATRX,FBXO1[illegible] |
| 14 | 18123 | HSPA1B,CLU,VCP,MAPT,HDAC6,DNAJB2,BAG5,CDC34,HSPA1A,DNAJ[illegible] |
| 14 | 18123 | GAPDH,ENO1,ALDOA,PKM,PGK1,ENO2,GPI,TPI1,PGAM1,HK1,PFKL,[illegible] |
| 14 | 18123 | GAPDH,ENO1,ALDOA,PKM,PGK1,ENO2,GPI,TPI1,PGAM1,HK1,PFKL,[illegible] |
| 14 | 18123 | GAPDH,ENO1,ALDOA,PKM,PGK1,ENO2,GPI,TPI1,PGAM1,HK1,PFKL,[illegible] |
| 66 | 18123 | BTBD9,HTT,ACTR2,GPI,APP,MAPT,ITGB1,PAFAH1B1,PJA2,PTN,CCN[illegible] |
| 71 | 18123 | ACTG1,ACTR2,TMSB4X,DBN1,FLNA,SPTBN1,SPTAN1,TMSB10,STMN[illegible] |
| 219 | 18123 | EEF1A1,LRP1,HSPA8,ACTB,CTSC,ACTG1,EEF2,MAVS,HSP90AB1,HSP[illegible] |
| 91 | 18123 | HNRNPA1,POLH,HSP90AB1,HSP90AA1,ACTR2,SMG1,NPM1,TRIM28,[illegible] |
| 594 | 18123 | COL1A1,HSPA8,MLLT1,NEDD4L,ATN1,USP22,MED18,MAVS,ZNF124,[illegible] |
| 595 | 18123 | COL1A1,HSPA8,MLLT1,NEDD4L,ATN1,USP22,MED18,MAVS,ZNF124,[illegible] |
| 77 | 18123 | MLLT1,RACK1,RPL5,GSTP1,NPM1,YWHAG,MAPT,PPP2R1A,UCHL1,R[illegible] |
| 73 | 18123 | PIK3R2,PKM,FADS1,IGFBP2,GSTP1,IGF2,YWHAG,SRSF4,PIP4K2B,M[illegible] |
| 35 | 18123 | HSP90B1,VCP,MARCHF6,TMEM259,UBE4A,UBXN4,STT3B,UBQLN1,[illegible] |
| 87 | 18123 | COL1A1,LRP1,ACTG1,FLNA,CALR,P4HB,FN1,PKD1,ITGB1,VCL,LAM[illegible] |
| 22 | 18123 | HNRNPA1,XRCC5,CCT7,CTNNB1,CCT6A,CCT2,TCP1,CCT3,CCT4,MA[illegible] |
| 90 | 18123 | COL1A1,SPARC,NEFL,IGFBP2,CALR,TMF1,ABHD2,H3-3B,DDX5,DDX[illegible] |
| 96 | 18123 | IGFBP5,COL1A1,SPARC,COL6A1,DDX5,FASN,ATP5F1B,SNRNP200,I[illegible] |
| 82 | 18123 | CTSC,CALR,TMF1,VAPA,AP2M1,ANXA2,AP3D1,ARF1,PAFAH1B1,SCA[illegible] |
| 36 | 18123 | MAVS,HSP90AB1,PIK3R2,FLNA,YWHAE,TRIM28,KHDRBS1,XPO1,TAR[illegible] |
| 61 | 18123 | PIK3R2,PKM,GSTP1,IGF2,YWHAG,PIP4K2B,MYO5A,SESN3,APPL1,AT[illegible] |
| 149 | 18123 | HSPA8,BTBD9,DBN1,MAP1B,TUBB2B,CALM3,SHISA9,APP,CALM1,Y[illegible] |
| 46 | 18123 | HTT,CLU,YWHAE,YWHAG,MAPT,YWHAQ,VAT1,YWHAB,DCN,SQSTM1,[illegible] |
| 99 | 18123 | HSPA9,RPS6,HSPA1B,H3-3B,KAT6A,FASN,MYH9,APP,HMGB1,ANXA2,[illegible] |
| 60 | 18123 | HSPA8,CFL1,MAP1B,SPTBN1,STMN2,SPTAN1,STMN1,PTCD3,WDR1,[illegible] |

| | | |
|---|---|---|
| 66 | 18123 | LRP1,DCX,FLNA,NEFL,ATP2B4,ZMIZ1,PAFAH1B1,RTN4,RHOA,KIF5B, |
| 21 | 18123 | HSPA9,CFL1,RPLP0,P4HB,ANXA2,ARF1,MSN,MIF,HNRNPDL,CNN2,TC |
| 36 | 18123 | KLRD1,PSMD5,PSMD2,SKP1,PSMB2,PSMD3,FYN,PSMA7,EP300,PSM |
| 46 | 18123 | HSPA8,HSP90AB1,HSP90AA1,HSPA1B,YWHAE,HSPD1,VCP,MAPT,DNA |
| 27 | 18123 | HSPA8,NEFL,KIF1A,APP,MAPT,UCHL1,AP3D1,PAFAH1B1,HNRNPU,KII |
| 63 | 18123 | UBC,VCP,UBB,AP3D1,RAB6B,RAB7A,ARL4C,CLTC,LAPTM4B,SQSTM |
| 220 | 18123 | IGFBP5,COL1A1,LRP1,ACTG1,DPYSL3,SPARC,TMSB4X,FLNA,ATP2B |
| 39 | 18123 | MAP1B,SPTBN1,STMN2,SPTAN1,WDR1,NES,SPTBN2,EIF5A,MAP1A,C |
| 22 | 18123 | HSP90B1,CALR,ATF4,ATP2A2,TMED2,SREBF2,MBTPS1,PPP1R15A,C |
| 15 | 18123 | P4HB,NONO,MCL1,SFPQ,PARP1,HSPB1,UBQLN1,SOD1,HIF1A,GPX1, |
| 75 | 18123 | BTBD9,DDX5,MAGED1,RACK1,NONO,DDB1,HNRNPU,CIPC,ATF4,JUN |
| 80 | 18123 | ATP5F1B,CALM1,IGF2,ATP5F1A,VCP,HMGB1,RHOA,PTGES3,MDH2,F |
| 33 | 18123 | HSP90AA1,PIK3R2,RHOA,SULF1,CYFIP2,CADM4,RAC1,NCKAP1,VEGI |
| 25 | 18123 | LRP1,ACTG1,VCL,RHOA,MMP14,MACF1,RAC1,PTPRA,VEGFA,CORO |
| 25 | 18123 | LRP1,ACTG1,VCL,RHOA,MMP14,MACF1,RAC1,PTPRA,VEGFA,CORO |
| 49 | 18123 | LRP1,DCX,FLNA,NEFL,ATP2B4,ZMIZ1,PAFAH1B1,RTN4,RHOA,KIF5B, |
| 147 | 18123 | HSPA8,BTBD9,DBN1,MAP1B,TUBB2B,CALM3,SHISA9,APP,CALM1,YV |
| 147 | 18123 | HSPA8,BTBD9,DBN1,MAP1B,TUBB2B,CALM3,SHISA9,APP,CALM1,YV |
| 278 | 18123 | COL1A1,LRP1,ACTB,ACTG1,HSP90AB1,FLNA,COL6A1,IGFBP2,CALR |
| 32 | 18123 | FBXL4,PSMD5,PSMD2,SKP1,PSMB2,PSMD3,PSMA7,PSMD1,PSMD8,F |
| 36 | 18123 | KLRD1,PSMD5,PSMD2,SKP1,PSMB2,PSMD3,FYN,PSMA7,EP300,PSM |
| 36 | 18123 | KLRD1,PSMD5,PSMD2,SKP1,PSMB2,PSMD3,FYN,PSMA7,EP300,PSM |
| 65 | 18123 | NUDC,TUBB,NUMA1,FLNA,HSPA1B,MYH9,VCP,MAP9,PKD1,RPS3,CHI |
| 58 | 18123 | LRP1,IGFBP2,DVL3,RACK1,PSMD5,PSMD2,TLE5,CDH2,GPC3,MDK,F |
| 49 | 18123 | ACTR2,TMSB4X,DBN1,SPTBN1,SPTAN1,TMSB10,ARF1,RHOA,PFN1,S |
| 50 | 18123 | HSPA1B,BSG,P4HB,TRIM28,DDB1,VAPA,ITGB1,NCAM1,RPSA,SCARB |
| 21 | 18123 | HSPA9,CFL1,RPLP0,P4HB,ANXA2,ARF1,MSN,MIF,HNRNPDL,CNN2,TC |
| 26 | 18123 | CTNNB1,KDM1A,SUPT6H,KDM4B,BRD4,ATRX,KMT2E,OGT,KDM4A,A |
| 58 | 18123 | CFL1,ACTR2,TMSB4X,DBN1,SPTBN1,SPTAN1,TMSB10,ARF1,RHOA,W |
| 24 | 18123 | CFL1,ACTR2,DBN1,KIF1A,PAFAH1B1,CAPRIN1,CDC42,CTNND2,HDA |
| 276 | 18123 | COL1A1,LRP1,ACTB,ACTG1,FLNA,COL6A1,IGFBP2,CALR,COL5A1,S |
| 32 | 18123 | ACTB,ACTG1,FLNA,MAPT,VCL,CLSTN1,KIF5B,NLGN2,MAP1A,DLG5,V |
| 34 | 18123 | APP,VAPA,SEC61A1,TMEM170A,RTN4,RTN3,ARL6IP1,VAMP2,TMED2, |
| 53 | 18123 | TMSB4X,CALR,APP,PSMD5,RPS3,HMGB1,RHOA,PSMD2,ACTN4,DDX |
| 52 | 18123 | ACTR2,TMSB4X,DBN1,SPTBN1,SPTAN1,TMSB10,ARF1,RHOA,WDR1, |
| 29 | 18123 | HSP90B1,VCP,MARCHF6,UBE4A,UBXN4,STT3B,UBQLN1,BAG6,SGTA |
| 119 | 18123 | COL1A1,ACTB,ACTG1,SPARC,FLNA,COL5A1,H3-3B,GNAS,GNB1,MYI |
| 117 | 18123 | LRP1,NEDD4L,BTBD9,CALR,CLU,RACK1,CALM3,SEPTIN2,CALM1,HN |
| 37 | 18123 | LRP1,DCX,FLNA,NEFL,ZMIZ1,PAFAH1B1,RTN4,RHOA,CDH2,CTNNB1 |
| 28 | 18123 | TRIM28,HMGB1,CTNNB1,CDKN1A,CCND1,CAND1,PTEN,MED13,PSM |
| 10 | 18123 | AP2M1,ANXA2,AP2B1,CLTC,ABCA2,AP2A1,AP2A2,CLTA,APOE,FURIN |
| 10 | 18123 | AP2M1,ANXA2,AP2B1,CLTC,ABCA2,AP2A1,AP2A2,CLTA,APOE,FURIN |
| 57 | 18123 | MAVS,HSP90AA1,KLRD1,MATR3,NONO,PSMD5,HMGB1,XRCC5,PSMI |
| 15 | 18123 | DDB1,SKP1,PCGF2,ATXN7L3,PCGF3,WAC,BCOR,CTR9,RNF20,CUL4 |
| 52 | 18123 | ACTR2,TMSB4X,DBN1,SPTBN1,SPTAN1,TMSB10,ARF1,RHOA,WDR1, |

| | | |
|---|---|---|
| 36 | 18123 | KLRD1,PSMD5,PSMD2,SKP1,PSMB2,PSMD3,FYN,PSMA7,EP300,PSM |
| 20 | 18123 | SET,CTBP1,SKI,TAF7,KDM1A,SUPT6H,KDM4B,UBR5,NOC2L,KDM4A, |
| 32 | 18123 | CANX,AP2M1,ARF1,AP2B1,CTSD,SPTBN2,RAB7A,CLTC,KLC1,DYNC1 |
| 59 | 18123 | CTSC,HTT,KIF1A,AP3D1,ARF1,PAFAH1B1,KIF5B,NLGN2,CDH2,ACTN |
| 84 | 18123 | HNRNPA1,ODF2L,TMSB4X,MAP1B,CLU,SPTBN1,STMN2,SPTAN1,NP |
| 37 | 18123 | HSPA9,RPS6,HSPA1B,RPS14,RPS19,VEGFA,PRKDC,B2M,ADAR,STAT |
| 24 | 18123 | HSPA8,HSPA9,HSPA1B,CLU,DNAJC5,PTGES3,PPIB,CCT2,HSPH1,HSP |
| 14 | 18123 | GAPDH,ENO1,ALDOA,PKM,PGK1,ENO2,GPI,TPI1,PGAM1,HK1,PFKL, |
| 411 | 18123 | IGFBP5,COL1A1,LRP1,ACTB,CTSC,ACTG1,MAVS,HTT,HSP90AB1,HS |
| 54 | 18123 | LRP1,ACTB,ATP2B4,YWHAE,RACK1,SPTBN1,GDI1,YWHAG,YWHAQ,A |
| 11 | 18123 | HSPA1B,CLU,DNAJB2,BAG5,CDC34,HSPA1A,DNAJB1,PSMC5,HSF1,D |
| 134 | 18123 | IGFBP5,ACTG1,EEF2,YBX1,CALR,DDX5,DDX17,MYH9,PLD3,IGF2,BA |
| 83 | 18123 | UBC,CALR,TMF1,HSPA1B,H3-3B,DDX5,DDX17,FASN,UBB,ZMIZ1,DN |
| 36 | 18123 | KLRD1,PSMD5,PSMD2,SKP1,PSMB2,PSMD3,FYN,PSMA7,EP300,PSM |
| 63 | 18123 | COL1A1,HSPA8,HNRNPA1,FADS1,ATF4,XPR1,PTN,JUN,SESN3,TNC,A |
| 57 | 18123 | H3-3B,KAT6A,NPM1,TRIM28,SET,CHD3,HMGB1,SMARCA5,RBBP4,H |
| 16 | 18123 | LUC7L3,SRSF5,SRSF9,SRSF1,SF3A1,SF1,SRSF6,SF3A2,LUC7L2,SF3 |
| 123 | 18123 | COL1A1,HNRNPA1,NEDD4L,EEF2,SPARC,MAP1B,NEFL,ATP2B4,IGFB |
| 466 | 18123 | MLLT1,NEDD4L,ATN1,MED18,MAVS,ZNF124,ACTR2,UBC,ZNF316,E |
| 112 | 18123 | CFL1,PIK3R2,GNB1,GDI1,TRIM28,YWHAQ,HACD3,STMN1,ITGB1,TI |
| 50 | 18123 | ENO1,ENO2,ANXA2,ARF1,SCARB2,ATP2A2,PIP4K2B,RAB7A,RAB1A,L |
| 57 | 18123 | ACTB,HTT,HSP90AB1,HSP90AA1,ACTR2,NUMA1,MAP1B,RACK1,MYH |
| 82 | 18123 | COL1A1,HSPA8,HNRNPA1,FADS1,GSTP1,ATF4,ATP1A1,TLE5,XPR1,P |
| 21 | 18123 | LRP1,FLNA,MAP1B,NEFL,CTNNA1,GAP43,PTN,TNC,PTPRS,PTEN,PTP |
| 23 | 18123 | LRP1,FLNA,MAP1B,NEFL,CTNNA1,GAP43,PTN,TNC,PTPRS,PTEN,PTP |
| 51 | 18123 | ACTB,ACTG1,BTBD9,CANX,CALM3,CALM1,AP2M1,AP3D1,SLC4A8,N |
| 9 | 18123 | KDM1A,UBR5,TRIP12,PARK7,CDK9,CTR9,OTUB1,RNF20,RNF40 |
| 9 | 18123 | ACTB,VCL,CTNNB1,DLG5,TBCD,JAM3,PIP5K1C,ZNF703,HIPK1 |
| 17 | 18123 | HNRNPA1,XRCC5,CCT7,CTNNB1,CCT6A,CCT2,TCP1,CCT3,CCT4,MA |
| 194 | 18123 | IGFBP5,COL1A1,LRP1,ACTG1,DPYSL3,SPARC,TMSB4X,FLNA,ATP2B |
| 75 | 18123 | MAVS,HSP90AB1,HSP90AA1,KLRD1,MATR3,NONO,PSMD5,HMGB1,X |
| 32 | 18123 | HSPA1B,SOX4,RPL5,RPS3,RPL11,DNAJA1,RPL23,RPS20,RPS15,RPS |
| 211 | 18123 | IGFBP5,COL1A1,LRP1,ACTG1,DPYSL3,SPARC,TMSB4X,FLNA,ATP2B |
| 55 | 18123 | ACTB,ACTG1,BTBD9,CANX,CALM3,CALM1,AP2M1,AP3D1,AP2B1,SL |
| 43 | 18123 | COL1A1,EEF2,RACK1,HSPD1,RPS3,LDHA,JUN,RHOB,HDAC2,KDM6B |
| 68 | 18123 | COL1A1,HSPA8,HNRNPA1,FADS1,GSTP1,ATF4,TLE5,XPR1,PTN,JUN, |
| 45 | 18123 | ACTG1,DPYSL3,FLNA,HSP90B1,FSCN1,MARCKS,STMN1,RHOA,PFN1 |
| 51 | 18123 | ENO1,HSPA1B,RACK1,PPP2R1A,RTN4,DDX3X,GJA1,MAP2,CDKN1A, |
| 218 | 18123 | IGFBP5,LRP1,ACTG1,SPARC,FLNA,NCL,PKM,ATP2B4,COL4A1,PGK1 |
| 14 | 18123 | GAPDH,ENO1,ALDOA,PKM,PGK1,ENO2,GPI,TPI1,PGAM1,HK1,PFKL, |
| 41 | 18123 | NPM1,VCP,UBB,DDX3X,NDFIP1,RAB1A,SKP1,PTTG1IP,NOP53,GABA |
| 34 | 18123 | HSPA1B,CLU,RACK1,VCP,CSNK1E,TRIB2,NOP53,GABARAP,UBQLN1, |
| 34 | 18123 | HTT,VCP,KIF5B,SQSTM1,KLC1,OPTN,ATG2B,CDC37,GABARAP,BNIP |
| 55 | 18123 | CTSC,HTT,KIF1A,AP3D1,ARF1,PAFAH1B1,KIF5B,ACTN4,MYO5A,CTN |
| 24 | 18123 | NEDD4L,TTC3,MARCHF6,UBE2H,SKP1,RNF187,GABARAP,UBE2R2,U |

| | | |
|---|---|---|
| 53 | 18123 | STMN1,ITGB1,TIMP2,RTN4,CADM4,RAC1,ARHGDIA,SQSTM1,NOTCH |
| 53 | 18123 | DDX5,KAT6A,RPL5,NPM1,UBB,CHD3,RPL11,RPL23,CHD4,RPS20,RPI |
| 77 | 18123 | SOX4,PSMD5,PSMD2,RPL26,IK,SKP1,CDKN1A,CTDSP2,DDX39B,NOF |
| 44 | 18123 | ALDOA,MYH9,PLXNB2,FN1,MSN,RHOA,WDR1,PTN,TPM1,PLXNA1,MY |
| 22 | 18123 | EEF2,KIDINS220,STMN2,APP,MAPT,PTN,RAPGEF1,DYNC1LI2,CSNK1 |
| 183 | 18123 | HSPA9,EEF2,RPS6,HSPA1B,H3-3B,SOX4,KAT6A,FASN,MYH9,APP,PSM |
| 56 | 18123 | BTBD9,MAGED1,NONO,DDB1,HNRNPU,CIPC,ATF4,JUND,CSNK1E,SF |
| 10 | 18123 | CCT7,CCT6A,CCT2,TCP1,CCT3,CCT4,CCT8,CCT5,MACROH2A1,GNL |
| 272 | 18123 | COL1A1,LRP1,GAPDH,CFL1,XRCC2,MAVS,HSP90AA1,ACTR2,NUMA1 |
| 54 | 18123 | LRP1,ACTG1,APP,AGRN,VCL,RHOA,CLSTN1,NLGN2,MMP14,GJA1,M |
| 189 | 18123 | HSPA9,EEF2,RPS6,HSPA1B,H3-3B,SOX4,KAT6A,FASN,MYH9,APP,PSM |
| 76 | 18123 | LRP1,NRBP2,XRCC2,HSP90AB1,NEFL,CLU,GPI,RACK1,HSPD1,UBB,N |
| 47 | 18123 | HSPA8,TMSB4X,MAP1B,CLU,SPTBN1,STMN2,SPTAN1,TMSB10,STMN |
| 8 | 18123 | CCT7,CCT6A,CCT2,TCP1,CCT3,CCT4,CCT8,CCT5 |
| 32 | 18123 | CANX,AP2M1,ARF1,AP2B1,CTSD,SPTBN2,RAB7A,CLTC,KLC1,DYNC1 |
| 30 | 18123 | MAP1B,SPTBN1,STMN2,SPTAN1,WDR1,NES,SPTBN2,MAP1A,CAMSAF |
| 15 | 18123 | HTT,VIM,UBC,UBB,SQSTM1,DYNC1LI2,DYNLL2,UBQLN1,DYNLL1,UE |
| 60 | 18123 | ATP5F1B,ATP5F1A,VCP,RHOA,MDH2,NOP53,ACO2,ATP5PB,SDHA,IDI |
| 55 | 18123 | NUDC,HTT,NUMA1,FLNA,HSPA1B,VCP,MAP9,PKD1,STMN1,ITGB1,PA |
| 88 | 18123 | LRP1,DCX,FLNA,NEFL,ATP2B4,KIF1A,APP,UBB,ZMIZ1,PAFAH1B1,RTI |
| 246 | 18123 | IGFBP5,ACTB,ACTG1,XRCC2,VIM,NUMA1,FLNA,FSTL1,COL5A1,COL |
| 51 | 18123 | ACTR2,TMSB4X,DBN1,SPTBN1,SPTAN1,TMSB10,ARF1,RHOA,PFN1,S |
| 51 | 18123 | NUDC,NUMA1,FLNA,HSPA1B,VCP,MAP9,PKD1,STMN1,PAFAH1B1,MA |
| 19 | 18123 | HSPA9,HSPA1B,PRKDC,B2M,STAT3,PRMT1,ZFP36L1,HIF1A,FAM210I |
| 19 | 18123 | CLU,RACK1,TMBIM6,GRINA,TMED2,HYOU1,SGTA,PPP1R15A,HSPA1/ |
| 57 | 18123 | POLH,ACTR2,CLU,DDX5,SMG1,NPM1,TRIM28,MAPT,RPS3,HMGB1,T |
| 31 | 18123 | SPTBN1,PPP2R1A,SERINC1,ANXA2,KPNB1,NLGN2,CDH2,PPP2CA,CLI |
| 31 | 18123 | ACTR2,DBN1,FLNA,CALR,P4HB,PAFAH1B1,ACTN4,RAC1,CAPRIN1,MI |
| 438 | 18123 | NEDD4L,ATN1,MED18,MAVS,ZNF124,ACTR2,UBC,ZNF316,ENO1,PIK |
| 50 | 18123 | DBN1,MAP1B,SHISA9,APP,YWHAG,MAPT,ARF1,ATF4,PTN,MAP1A,SQ |
| 226 | 18123 | COL1A1,LRP1,ACTG1,SPARC,EIF4G2,FLNA,NCL,PKM,ATP2B4,CALR, |
| 45 | 18123 | ACTG1,DPYSL3,FLNA,HSP90B1,FSCN1,MARCKS,STMN1,RHOA,PFN1 |
| 183 | 18123 | LRP1,NEDD4L,ACTB,MAVS,HTT,HSP90AB1,TMSB4X,NUMA1,PIK3R2, |
| 21 | 18123 | EEF2,KIDINS220,STMN2,APP,MAPT,RAPGEF1,DYNC1LI2,CSNK1D,PT |
| 34 | 18123 | LRP1,HSPA8,PSAP,CLU,VCP,AP3D1,ARF1,SCARB2,RAB7A,LAMP1,RH |
| 26 | 18123 | ACTR2,FSCN1,ITGB1,VCL,RAC1,ACTR3,NCKAP1,ARPC2,CAPZB,AKIR |
| 570 | 18123 | EEF1A1,COL1A1,LRP1,HSPA8,GAPDH,HSPA9,ACTB,CTSC,ACTG1,EEI |
| 16 | 18123 | JUN,BCLAF1,CAND1,MED13,PSMC3,SUB1,DHX36,PSMC5,PSMC2,PS |
| 77 | 18123 | HSP90AB1,UBC,NUMA1,ATP2B4,FSTL1,HTRA4,UBB,GLG1,GPC3,SUI |
| 28 | 18123 | HTT,MAP1B,NEFL,KIF1A,UBB,MAPT,UCHL1,AP3D1,PAFAH1B1,KIF5B |
| 30 | 18123 | ACTB,YWHAE,CALM3,CALM1,CKB,YWHAQ,INA,BASP1,LDHA,RHOA,C |
| 43 | 18123 | CALR,TMF1,H3-3B,DDX5,DDX17,ZMIZ1,DNAJA1,RHOA,ARID1A,PTGE |
| 44 | 18123 | TMSB4X,SLC9A7,ATP5F1B,ATP5F1A,ATP1A1,ATP2A2,ACTN4,SLC25A |
| 24 | 18123 | MAGED1,HNRNPU,ATF4,CSNK1E,HDAC2,PPP1CB,TOP1,CSNK1D,ID4 |
| 24 | 18123 | TMSB4X,DBN1,TMSB10,HNRNPU,HK1,GJA1,SKP1,KDELR2,KDELR1, |

| | | |
|---|---|---|
| 7 | 18123 | HSPA1B,CKAP5,DCTN1,HSPA1A,MECP2,PDE4DIP,GIT1 |
| 6 | 18123 | PPP2CA,MAPRE1,PPP2CB,ABL1,MAPRE3,HSF1 |
| 31 | 18123 | TMSB4X,DBN1,FLNA,TMSB10,PKD1,HNRNPU,HK1,GJA1,YWHAB,SKP |
| 65 | 18123 | COL1A1,HSP90AB1,UBC,HTRA4,UBB,COL4A2,ZMIZ1,COL1A2,RHOA |
| 17 | 18123 | GANAB,CANX,PRKCSH,DDOST,ST3GAL2,RPN2,STT3A,MGAT4B,GALN |
| 17 | 18123 | HSPA8,NCL,CIRBP,SRSF5,U2AF2,RBMX,SNRNP70,SRSF1,TRA2B,PRD |
| 70 | 18123 | COL1A1,FLNA,DDX5,DDX17,TRIM28,FN1,DCHS1,BASP1,RTN4,LAMA |
| 50 | 18123 | HSPA8,HNRNPA1,FADS1,ATF4,XPR1,JUN,SESN3,ATP6V0C,GLUL,CDK |
| 18 | 18123 | GAPDH,ENO1,ALDOA,PKM,PGK1,ENO2,GPI,VCP,TPI1,PGAM1,HK1,PI |
| 18 | 18123 | HNRNPU,HK1,GJA1,SKP1,KDELR2,KDELR1,OS9,RER1,CIZ1,RANGAP |
| 13 | 18123 | NPM1,ATF4,DDX3X,TMED2,EIF2AK1,PPP1R15A,EIF4G1,RBM4,PPP1C |
| 22 | 18123 | LRP1,VEGFA,RAPGEF1,ZFAND5,HGS,TXNIP,BCR,HIP1,PTEN,LOX,CLA |
| 42 | 18123 | ATP5F1B,ATP5F1A,VCP,RHOA,ATP5PB,SDHA,UQCRC2,GHITM,NDUF |
| 43 | 18123 | HSPA9,HSP90AA1,YWHAE,ATP5F1B,CALM3,CALM1,ATP5F1A,YWHAG |
| 19 | 18123 | ACTB,YWHAE,CALM3,CALM1,CKB,YWHAQ,INA,BASP1,LDHA,RHOA,A |
| 28 | 18123 | PPP2R1A,PAFAH1B1,TMEM170A,KPNB1,TARDBP,PPP2CA,DCTN1,VP |
| 15 | 18123 | HTT,BNIP3L,SREBF2,VDAC1,HDAC6,HIF1A,CTTN,SLC25A5,GSK3A,BI |
| 9 | 18123 | CCT7,CCT6A,CCT2,TCP1,CCT3,CCT4,CCT8,CCT5,GNL3 |
| 37 | 18123 | HTT,HMGB1,LARP1,PIP4K2B,SESN3,DCN,GNAI3,OPTN,SUPT5H,BNIP |
| 196 | 18123 | HSPA9,EEF2,RPS6,HSPA1B,H3-3B,SOX4,KAT6A,FASN,MYH9,APP,HSP |
| 63 | 18123 | SOX4,PSMD5,PSMD2,RPL26,IK,SKP1,CDKN1A,CTDSP2,NOP53,PSME |
| 56 | 18123 | COL1A1,LRP1,ACTG1,FLNA,CALR,P4HB,FN1,VCL,RHOA,PTN,DDR1,A |
| 31 | 18123 | RACK1,P4HB,NONO,ATF4,MCL1,SFPQ,CTNNB1,SESN3,PARP1,SELEN |
| 50 | 18123 | DDX5,SOX4,NPM1,RPL26,MIF,PTTG1IP,CDKN1A,DDX39B,NOP53,CC |
| 27 | 18123 | LRP1,DPYSL3,FLNA,MAP1B,NEFL,CTNNA1,GAP43,PTN,TNC,ARF4,PT |
| 55 | 18123 | ACTB,ACTG1,FLNA,UGT8,FSCN1,DCHS1,VCL,CTNNA1,RHOA,NLGN2 |
| 48 | 18123 | LRP1,NEDD4L,EIF4G2,DBN1,MAP1B,GDI1,IGF2,MAPT,FN1,BASP1,A |
| 178 | 18123 | MAVS,HSP90AB1,HSP90AA1,UBC,KIDINS220,ATP2B4,CLU,MAGED1,I |
| 45 | 18123 | PIK3R2,CALM1,PSMD5,PSMD2,JUN,RAC1,SKP1,PSMB2,MAPK1,PSM |
| 32 | 18123 | CANX,AP2M1,ARF1,AP2B1,CTSD,SPTBN2,RAB7A,CLTC,KLC1,DYNC1 |
| 11 | 18123 | RPSA,XRCC5,RPS14,RPS15,RPS5,RPS19,RPS10,RPS27,RPL38,PRKD |
| 201 | 18123 | IGFBP5,COL1A1,LRP1,ACTG1,DPYSL3,SPARC,TMSB4X,FLNA,ATP2B |
| 173 | 18123 | NEDD4L,ATN1,ZNF124,UBC,ENO1,YBX1,CALR,DDX5,TMBIM6,IGF2,U |
| 28 | 18123 | HSPA1B,SOX4,RPL5,RPS3,RPL11,DNAJA1,RPL23,RPS20,RPS15,RPS |
| 33 | 18123 | COL1A1,COL6A1,COL5A1,COL4A1,P4HB,COL18A1,COL4A2,ANXA2, |
| 357 | 18123 | IGFBP5,LRP1,EEF2,SPARC,FTH1,RPS6,DBN1,FLNA,IGFBP2,CALR,HS |
| 14 | 18123 | ACTR2,FSCN1,RAC1,ACTR3,NCKAP1,ARPC2,AKIRIN1,CDC42,BRK1,A |
| 34 | 18123 | UBC,UBB,DDB1,CHD3,HMGB1,XRCC5,CHD4,DDX3X,CHD6,PARP1,R |
| 21 | 18123 | LRP1,DCX,ZMIZ1,PAFAH1B1,RTN4,MMP14,CTNNB1,VCAN,NDN,ATP1 |
| 18 | 18123 | ACTR2,FSCN1,RAC1,ACTR3,NCKAP1,ARPC2,CAPZB,AKIRIN1,CDC42, |
| 78 | 18123 | ATP2B4,ALDH6A1,PSMD5,ATF4,PSMD2,OAZ1,NARS1,GLUL,NOX4,PS |
| 29 | 18123 | SPTBN1,PPP2R1A,ANXA2,KPNB1,NLGN2,CDH2,PPP2CA,FLOT1,PTEN |
| 29 | 18123 | MAVS,HSP90AB1,PIK3R2,FLNA,NPM1,TRIM28,CCT7,TARDBP,CCT6A |
| 76 | 18123 | IGFBP5,LRP1,CTSC,EEF2,FADS1,YBX1,IGFBP2,CALR,CANX,KAT6A,A |
| 63 | 18123 | COL1A1,HSP90AB1,UBC,HTRA4,UBB,COL4A2,ZMIZ1,COL1A2,RHOA |

| | | |
|---|---|---|
| 45 | 18123 | TMSB4X,MAP1B,SPTBN1,STMN2,SPTAN1,NPM1,TMSB10,STMN1,PFI |
| 40 | 18123 | HSPA8,FTH1,FTL,RACK1,SPTBN1,APP,AP3D1,ARF1,LLGL1,CLTC,MY( |
| 23 | 18123 | SET,H1-0,H1-10,CTBP1,SKI,TAF7,KDM1A,SUPT6H,KDM4B,UBR5,NO |
| 42 | 18123 | LRP1,HSPA8,PSAP,CLU,VCP,AP3D1,ARF1,SCARB2,RAB7A,NDFIP1,LA |
| 13 | 18123 | MSN,TMEM30A,VPS35,VEGFA,RDX,SCRIB,MGAT3,ABHD17A,ROCK2, |
| 34 | 18123 | COL1A1,COL6A1,COL4A1,TMBIM6,GSTP1,COL1A2,RHOA,COL5A2,F |
| 122 | 18123 | SPARC,EIF4G2,FLNA,CALR,COL5A1,SOX4,MATR3,FN1,DCHS1,PKD1, |
| 73 | 18123 | SOX4,PSMD5,TIMP2,PSMD2,RPL26,IK,CTNNB1,SKP1,CDKN1A,CTDS |
| 35 | 18123 | NEDD4L,MAGED1,GDI1,MAPT,AP2M1,DNAJA1,CTNNA1,ASTN2,CLTC, |
| 124 | 18123 | HSP90AB1,HSP90AA1,UBC,KIDINS220,ATP2B4,CLU,MAGED1,DVL3,C |
| 60 | 18123 | USP22,SOX4,KAT6A,MAPT,SET,GTF3C4,MORF4L1,TADA2B,DDX3X,C |
| 189 | 18123 | MAVS,HSP90AB1,HSP90AA1,UBC,KIDINS220,TMSB4X,ENO1,ATP2B4 |
| 189 | 18123 | MAVS,HSP90AB1,HSP90AA1,UBC,KIDINS220,TMSB4X,ENO1,ATP2B4 |
| 21 | 18123 | UBC,UBB,KPNB1,XPO1,MVB12B,KPNA2,UBA52,POM121C,RPS27A,F |
| 8 | 18123 | UBC,UBB,RAB7A,B2M,UBA52,RPS27A,SCRIB,KPNA6 |
| 8 | 18123 | CCT7,CCT6A,CCT2,TCP1,CCT3,CCT4,CCT8,CCT5 |
| 8 | 18123 | CCT7,CCT6A,CCT2,TCP1,CCT3,CCT4,CCT8,CCT5 |
| 8 | 18123 | EIF3C,EIF3E,EIF3D,CARM1,NUCKS1,EIF4A3,EIF4G1,CDK9 |
| 8 | 18123 | UBC,UBB,RAB7A,B2M,UBA52,RPS27A,SCRIB,KPNA6 |
| 8 | 18123 | CCT7,CCT6A,CCT2,TCP1,CCT3,CCT4,CCT8,CCT5 |
| 8 | 18123 | UBC,UBB,RAB7A,B2M,UBA52,RPS27A,SCRIB,KPNA6 |
| 40 | 18123 | PIK3R2,IGF2,PIP4K2B,SESN3,APPL1,ATP6V0C,PTPRA,ATP6AP1,GRB: |
| 40 | 18123 | HTT,HSP90AB1,HSP90B1,YWHAE,CALM3,CALM1,SET,PPP2R1A,PTN,' |
| 17 | 18123 | FLNA,GNAS,GNB1,GNAI3,VPS35,NSG2,GNG2,ADCY6,PALM,NCSTN,A |
| 51 | 18123 | HNRNPA1,POLH,HSP90AB1,HSP90AA1,UBC,VCP,UBB,HNRNPU,XRCC |
| 55 | 18123 | ACTB,H3-3B,NPM1,TRIM28,CHD3,SMARCA5,CHD4,RBBP4,HNRNPC, |
| 61 | 18123 | IGFBP5,ENO1,HSPA1B,GNAS,RACK1,PPP2R1A,RTN4,GPC3,DDX3X,G |
| 19 | 18123 | GAPDH,ENO1,ALDOA,PKM,PGK1,ENO2,GPI,TPI1,PGAM1,LDHA,HK1,I |
| 18 | 18123 | HSPA8,CLU,AP3D1,SCARB2,RTN4,CD81,RAB7A,HGS,NCOA4,VPS4A,S |
| 170 | 18123 | EEF1A1,HSPA8,CTSC,EEF2,MAVS,HSP90AB1,HSP90AA1,ACTR2,TUBE |
| 12 | 18123 | HSPA8,HSPA9,HSP90AA1,HSPA1B,HSPD1,CCT7,SNRNP70,B2M,FKBF |
| 26 | 18123 | SOX4,SET,TADA2B,DDX3X,CTBP1,HDAC2,EID1,TERF2IP,TAF7,WBP2, |
| 65 | 18123 | MLLT1,RACK1,GSTP1,NPM1,YWHAG,MAPT,PPP2R1A,UCHL1,MIDN,D |

4L,ACTB,CTSC,ODF2L,NUDC,USP22,ACTG1,DPYSL3,TUBA1A,CFL1,BTBD9,XRCC2,MAVS,HTT,HS
,CTSC,NUDC,NRBP2,ACTG1,EEF2,TUBA1A,CFL1,BTBD9,MAVS,HTT,HSP90AB1,HSP90AA1,ACTR2
,VIM,UBC,RPL3,RPS6,RPL4,RPL7A,GANAB,RPL8,HSPA1B,RPLP0,CANX,DDX5,RPL13A,RPL6,YWH
4L,ACTB,CTSC,ODF2L,NUDC,USP22,ACTG1,DPYSL3,TUBA1A,CFL1,BTBD9,XRCC2,MAVS,HTT,HS
HSP90AA1,TMSB4X,NUMA1,RPL3,RPS6,DBN1,RPL4,PIK3R2,FLNA,RPL7A,HSP90B1,UGT8,RPL8,
HSP90AA1,TMSB4X,NUMA1,RPL3,RPS6,DBN1,RPL4,PIK3R2,FLNA,RPL7A,HSP90B1,UGT8,RPL8,
7A,PRPF8,RPL8,YBX1,HSPA1B,RPLP0,DDX5,RPL13A,RPL6,RPL13,DDX17,RPS8,RPL7,APP,RPL5,
90AB1,HSP90AA1,ACTR2,TUBB,UBC,NUMA1,RPL3,RPS6,RPL4,TUBA1B,PIK3R2,FLNA,MAP1B,RP
0AA1,VIM,UBC,RPL3,RPS6,RPL4,FLNA,RPL7A,HSP90B1,RPL8,FBXL4,YBX1,UBA1,TMF1,HSPA1B,
P2,ACTG1,EEF2,TUBA1A,CFL1,BTBD9,MAVS,HTT,HSP90AB1,HSP90AA1,ACTR2,SPARC,TUBB,FT
90AB1,HSP90AA1,UBC,TMSB4X,NUMA1,RPL3,RPS6,DBN1,RPL4,PIK3R2,FLNA,RPL7A,HSP90B1,
A1A,CFL1,BTBD9,MAVS,HTT,HSP90AA1,ACTR2,VIM,SPARC,TUBB,KIDINS220,TMSB4X,NUMA1,E
MAVS,HSP90AB1,HSP90AA1,UBC,TMSB4X,NUMA1,RPL3,RPS6,DBN1,RPL4,PIK3R2,FLNA,RPL7A,
M,UBC,RPL3,RPS6,RPL4,RPL7A,HSP90B1,RPL8,FBXL4,YBX1,UBA1,TMF1,HSPA1B,RPLP0,CLU,D
4,PIK3R2,FLNA,RPL7A,RPL8,YBX1,CALR,RPLP0,CLU,RPL13A,MAGED1,SEC62,RPL6,YWHAE,RPL
B,RPLP0,DDX5,RPL13A,RPL6,RPL13,RPS8,RPL7,RPL5,SMG1,NPM1,CSDE1,UBB,RPL37A,PSMD5
4L,ACTB,CTSC,NUDC,NRBP2,ACTG1,EEF2,DPYSL3,TUBA1A,CFL1,BTBD9,MAVS,HTT,HSP90AB1,
NRBP2,USP22,MED18,EEF2,MAVS,HTT,HSP90AB1,HSP90AA1,VIM,UBC,KIDINS220,EIF4G2,RPL3,
TG1,DPYSL3,TUBA1A,CFL1,XRCC2,MAVS,HTT,HSP90AB1,HSP90AA1,ACTR2,VIM,TUBB,UBC,TMS
PS8,RPL7,RPL5,RPL37A,RPL19,RPL15,RPS3,SEC61A1,RPS4X,RPS3A,RPL18,RPL11,RPLP1,RPL1
RPS8,RPL7,RPL5,SMG1,RPL37A,RPL19,PPP2R1A,RPL15,RPS3,RPS4X,RPS3A,RPL18,RPL11,RPL
L7,RPL5,RPL37A,RPL19,RPL15,RPS3,SEC61A1,RPS4X,RPS3A,RPL18,RPL11,RPLP1,RPL10,RPS1
B,RPLP0,DDX5,RPL13A,RPL6,RPL13,RPS8,RPL7,RPL5,SMG1,NPM1,CSDE1,UBB,RPL37A,PSMD5
1A,HTT,HSP90AB1,HSP90AA1,ACTR2,TUBB,UBC,TMSB4X,NUMA1,DCX,RPL3,RPS6,DBN1,RPL4,F
3,RPS6,RPL4,PIK3R2,FLNA,RPL7A,HSP90B1,RPL8,CALR,RPLP0,CANX,CLU,SOX4,RPL13A,SEC62,
IK3R2,FLNA,RPL7A,HSP90B1,RPL8,CALR,RPLP0,CLU,RPL13A,SEC62,RPL6,YWHAE,KIF1A,RPL13
PS8,RPL7,RPL5,RPL37A,RPL19,RPL15,RPS3,VAPA,SEC61A1,RPS4X,RPS3A,RPL18,RPL11,RPLP1,
PS8,RPL7,RPL5,RPL37A,RPL19,RPL15,RPS3,SEC61A1,RPS4X,RPS3A,RPL18,RPL11,RPLP1,RPL1
,CTSC,NRBP2,USP22,MED18,EEF2,MAVS,HTT,HSP90AB1,HSP90AA1,VIM,UBC,KIDINS220,EIF4G2
,HSP90AA1,RPL3,RPS6,RPL4,PIK3R2,FLNA,RPL7A,HSP90B1,RPL8,YBX1,CALR,PSAP,RPLP0,CAN
L6,RPL13,RPS8,RPL7,RPL5,CSDE1,UBB,RPL37A,RPL19,RPL15,RPS3,RPS4X,RPS3A,RPL18,RPL1
PS8,RPL7,RPL5,RPL37A,RPL19,RPL15,RPS3,SEC61A1,RPS4X,RPS3A,RPL18,RPL11,RPLP1,RPL1
3,RPS8,RPL7,RPL5,EIF3C,EIF4B,RPL37A,EIF4A1,RPL19,RPL15,RPS3,RPS4X,RPS3A,RPL18,RPL1
,HNRNPA1,HSPA9,NEDD4L,POLH,ATN1,ACTB,CTSC,NRBP2,USP22,MED18,EEF2,BTBD9,XRCC2,
HTT,HSP90AB1,HSP90AA1,VIM,UBC,EIF4G2,RPL3,RPS6,RPL4,ENO1,PIK3R2,FLNA,RPL7A,ALDOA,
HTT,HSP90AB1,HSP90AA1,VIM,UBC,EIF4G2,RPL3,RPS6,RPL4,PIK3R2,RPL7A,HSP90B1,RPL8,FB
A9,NEDD4L,ATN1,CTSC,ODF2L,NRBP2,MTRNR2L2,DPYSL3,CFL1,XRCC2,MAVS,HTT,HSP90AB1,
,RPL13,RPS8,RPL7,RPL5,NPM1,CSDE1,EIF3C,EIF4B,RPL37A,EIF1,EIF4A1,RPL19,RPL15,RPS3,R
90AB1,HSP90AA1,VIM,UBC,EIF4G2,PIK3R2,FLNA,YBX1,ATP2B4,TMF1,HSPA1B,PSAP,CLU,RACK
3,RPS6,RPL4,PIK3R2,FLNA,RPL7A,HSP90B1,RPL8,CALR,RPLP0,CANX,CLU,SOX4,RPL13A,SEC62,
,CTSC,NUDC,NRBP2,ACTG1,EEF2,TUBA1A,CFL1,BTBD9,MAVS,HTT,HSP90AB1,HSP90AA1,ACTR2
RPL7A,RPL8,RPLP0,CLU,RPL13A,SEC62,RPL6,YWHAE,RPL13,CALM3,RPS8,RPL7,RPL5,CALM1,H
,CTSC,NRBP2,ACTG1,EEF2,TUBA1A,BTBD9,MAVS,HTT,HSP90AB1,HSP90AA1,ACTR2,SPARC,TUB
1A,HTT,HSP90AB1,HSP90AA1,ACTR2,TUBB,UBC,TMSB4X,NUMA1,DCX,RPL3,DBN1,FLNA,MAP1B
31,HSP90AA1,VIM,UBC,RPL3,RPS6,RPL4,ENO1,FLNA,RPL7A,ALDOA,PKM,HSP90B1,RPL8,FBXL4,

,CTSC,NRBP2,USP22,MED18,EEF2,BTBD9,MAVS,HTT,HSP90AB1,HSP90AA1,VIM,UBC,KIDINS220,
HSP90AB1,HSP90AA1,ACTR2,VIM,KIDINS220,DCX,EIF4G2,DBN1,FLNA,MAP1B,NEFL,UGT8,ATP2
RPLP0,RPL13A,SEC62,RPL6,YWHAE,RPL13,RACK1,BSG,SPTBN1,CALM3,RPS8,RPL7,APP,RPL5,G
22,ACTG1,EEF2,DPYSL3,XRCC2,MAVS,HTT,HSP90AB1,HSP90AA1,ACTR2,VIM,SPARC,UBC,KIDIN
I,KIDINS220,DCX,EIF4G2,DBN1,FLNA,MAP1B,NEFL,UGT8,TUBB2B,DVL3,KIF1A,BSG,SPTBN1,STN
PSAP,RPLP0,RPL13A,SEC62,RPL6,YWHAE,RPL13,RACK1,BSG,SPTBN1,CALM3,RPS8,RPL7,APP,R
ISP90AA1,VIM,UBC,EIF4G2,PIK3R2,YBX1,ATP2B4,TMF1,HSPA1B,PSAP,CLU,RACK1,APP,RPL5,SM
L7,RPL5,RPL37A,RPL19,RPL15,RPS3,RPS4X,RPS3A,RPL18,RPL11,RPLP1,RPL10,RPS17,GTF2F1
IPA1,NEDD4L,POLH,ATN1,ACTB,CTSC,NRBP2,USP22,MED18,ACTG1,EEF2,MAVS,HTT,HSP90AB1,
B,RPLP0,DDX5,RPL13A,RPL6,RPL13,RPS8,RPL7,RPL5,SMG1,NPM1,CSDE1,VCP,UBB,RPL37A,PS
PA9,NEDD4L,ATN1,CTSC,ODF2L,NRBP2,MTRNR2L2,DPYSL3,CFL1,XRCC2,HTT,HSP90AB1,ZNF12
I1,NRBP2,MAVS,HSP90AB1,ZNF124,VIM,UBC,TMSB4X,EIF4G2,RPL3,RPS6,RPL4,ENO1,FLNA,NCL
EEF2,MAVS,HTT,HSP90AB1,HSP90AA1,VIM,UBC,KIDINS220,EIF4G2,ENO1,FLNA,NCL,PKM,HSP90
,HNRNPA1,NEDD4L,POLH,ATN1,ACTB,CTSC,NRBP2,USP22,MED18,EEF2,XRCC2,MAVS,HTT,HSP9
I1,MAVS,HSP90AB1,ZNF124,VIM,UBC,TMSB4X,EIF4G2,RPL3,RPS6,RPL4,ENO1,FLNA,NCL,RPL7A
AA1,ACTR2,UBC,ENO1,PIK3R2,FLNA,MAP1B,HSP90B1,FADS1,NEFL,YBX1,UBA1,CALR,HSPA1B,P
B,CTSC,ACTG1,EEF2,DPYSL3,CFL1,MAVS,HSP90AB1,HSP90AA1,ACTR2,VIM,SPARC,UBC,KIDINS2
X,NUMA1,RPL3,DBN1,MAP1B,ALDOA,NEFL,PRPF8,CALR,HSPA1B,RPLP0,H3-3B,CLU,RPL13A,RPL
I,KIDINS220,DCX,EIF4G2,DBN1,FLNA,MAP1B,NEFL,UGT8,TUBB2B,DVL3,KIF1A,BSG,SPTBN1,STN
NUMA1,EIF4G2,DBN1,FLNA,MAP1B,NEFL,HSPA1B,H3-3B,CLU,DVL3,YWHAE,RACK1,STMN2,APP,
ISP90AA1,ACTR2,VIM,KIDINS220,DCX,EIF4G2,DBN1,FLNA,MAP1B,NEFL,UGT8,CALR,TUBB2B,CL
,HNRNPA1,NEDD4L,POLH,ATN1,ACTB,CTSC,NRBP2,USP22,MED18,EEF2,XRCC2,MAVS,HTT,HSP9
B,RPLP0,DDX5,RPL13A,RPL6,RPL13,ALDH6A1,RPS8,RPL7,RPL5,SMG1,NPM1,CSDE1,VCP,UBB,F
B,RPLP0,DDX5,RPL13A,RPL6,RPL13,ALDH6A1,RPS8,RPL7,RPL5,SMG1,NPM1,CSDE1,VCP,UBB,F
RPL13,RPS8,RPL7,RPL5,SMG1,CSDE1,RPL37A,RPL19,PPP2R1A,RPL15,RPS3,RPS4X,RPS3A,RPL1
,HNRNPA1,NEDD4L,POLH,ATN1,ACTB,CTSC,NRBP2,USP22,MED18,EEF2,BTBD9,XRCC2,MAVS,H
TR2,VIM,KIDINS220,DCX,EIF4G2,DBN1,FLNA,MAP1B,NEFL,UGT8,CALR,TUBB2B,SOX4,DVL3,KIF
AA1,ACTR2,VIM,KIDINS220,DCX,EIF4G2,DBN1,FLNA,MAP1B,NEFL,UGT8,CALR,TUBB2B,SOX4,D
I,HSP90AA1,RPL3,RPS6,RPL4,PIK3R2,FLNA,RPL7A,HSP90B1,RPL8,YBX1,CALR,TMF1,PSAP,RPLF
IPA1,NEDD4L,POLH,ATN1,ACTB,CTSC,USP22,MED18,ACTG1,EEF2,MAVS,HTT,HSP90AB1,ZNF124
.6,YWHAE,RPL13,RACK1,SPTBN1,CALM3,RPS8,RPL7,APP,RPL5,GDI1,CALM1,YWHAG,RPL37A,YV
TB,CTSC,USP22,ACTG1,EEF2,DPYSL3,CFL1,XRCC2,MAVS,HTT,HSP90AB1,HSP90AA1,ACTR2,VIM
IPA1,NEDD4L,POLH,ATN1,ACTB,CTSC,NRBP2,USP22,MED18,EEF2,MAVS,HTT,HSP90AB1,ZNF124
,NCL,RPL7A,PKM,RPL8,YBX1,CALR,RPLP0,CLU,SOX4,RPL13A,RPL6,RPL13,RACK1,RPS8,RPL7,A
ISP90AB1,HSP90AA1,VIM,UBC,KIDINS220,EIF4G2,ENO1,NCL,PKM,HSP90B1,YBX1,ATP2B4,CALR
A,SEC62,RPL6,RPL13,RPS8,RPL7,RPL5,GDI1,HSPD1,YWHAG,RPL37A,YWHAQ,RPL19,RPL15,RPS
B,RPLP0,DDX5,RPL13A,RPL6,RPL13,RPS8,RPL7,RPL5,SMG1,NPM1,CSDE1,VCP,UBB,RPL37A,PS
NS220,DCX,EIF4G2,DBN1,FLNA,MAP1B,ALDOA,NEFL,UGT8,CALR,TUBB2B,CLU,DVL3,KIF1A,BSG
B,RPLP0,DDX5,RPL13A,RPL6,RPL13,ALDH6A1,RPS8,RPL7,RPL5,SMG1,NPM1,CSDE1,VCP,UBB,F
DCX,EIF4G2,DBN1,MAP1B,NEFL,UGT8,TUBB2B,DVL3,KIF1A,BSG,SPTBN1,APP,GDI1,SPTAN1,PLX
DCX,EIF4G2,DBN1,MAP1B,NEFL,UGT8,TUBB2B,DVL3,KIF1A,BSG,SPTBN1,APP,GDI1,SPTAN1,PLX
DCX,EIF4G2,DBN1,MAP1B,NEFL,UGT8,TUBB2B,DVL3,KIF1A,BSG,SPTBN1,APP,GDI1,SPTAN1,PLX
P22,ACTG1,EEF2,DPYSL3,CFL1,XRCC2,HSP90AB1,HSP90AA1,ACTR2,VIM,SPARC,KIDINS220,NUM
BC,FLNA,HSP90B1,FBXL4,UBA1,TMF1,HSPA1B,CLU,RACK1,RPL5,TTC3,USP11,VCP,UBB,DDB1,PS
,NCL,RPL7A,PKM,RPL8,YBX1,CALR,RPLP0,SOX4,RPL13A,RPL6,RPL13,RACK1,RPS8,RPL7,APP,R

A1,RPL3,DBN1,MAP1B,ALDOA,NEFL,PRPF8,CALR,HSPA1B,RPLP0,H3-3B,CLU,RPL13A,RPL6,KAT
,HNRNPA1,HSPA9,NEDD4L,POLH,ATN1,ACTB,CTSC,NRBP2,USP22,MED18,ACTG1,EEF2,BTBD9,X
ISP90AA1,ACTR2,VIM,TUBB,KIDINS220,DCX,EIF4G2,DBN1,FLNA,MAP1B,NEFL,UGT8,TUBB2B,DV
,YBX1,RPLP0,DDX5,RPL13A,RPL6,RPL13,DDX17,RPS8,RPL7,APP,RPL5,TMBIM6,NPM1,SNRNP20
3,ACTG1,EEF2,DPYSL3,CFL1,MAVS,HSP90AB1,HSP90AA1,ACTR2,VIM,UBC,KIDINS220,TMSB4X,N
DCX,EIF4G2,DBN1,MAP1B,NEFL,UGT8,TUBB2B,DVL3,KIF1A,BSG,SPTBN1,APP,GDI1,SPTAN1,PLX
I1,MAP1B,NEFL,PRPF8,CALR,HSPA1B,RPLP0,H3-3B,CLU,RPL13A,RPL6,KAT6A,SPTBN1,STMN2,F
,HNRNPA1,NEDD4L,POLH,ATN1,ACTB,CTSC,NRBP2,USP22,MED18,ACTG1,EEF2,XRCC2,MAVS,H
IPA1,NEDD4L,POLH,ATN1,ACTB,CTSC,USP22,MED18,EEF2,MAVS,HTT,HSP90AB1,ZNF124,HSP90
PS8,RPL7,RPL5,GDI1,RPL37A,RPL19,RPL15,RPS3,SEC61A1,RPS4X,RPS3A,RPL18,RPL11,RPLP1,
BB,PSMD5,SET,HNRNPU,LARP1,HNRNPA0,PAPOLA,PSMD2,SRSF4,HNRNPC,KHDRBS1,XPO1,TAR
ISP90AA1,ACTR2,VIM,TUBB,KIDINS220,DCX,EIF4G2,DBN1,FLNA,MAP1B,NEFL,UGT8,TUBB2B,DV
,NCL,RPL7A,PKM,RPL8,YBX1,CALR,RPLP0,SOX4,RPL13A,RPL6,RPL13,RACK1,RPS8,RPL7,APP,R
3,ACTG1,EEF2,DPYSL3,CFL1,MAVS,HSP90AB1,HSP90AA1,ACTR2,VIM,UBC,KIDINS220,TMSB4X,N
P22,ACTG1,EEF2,DPYSL3,CFL1,XRCC2,HSP90AB1,HSP90AA1,ACTR2,VIM,SPARC,KIDINS220,NUM
DCX,EIF4G2,DBN1,FLNA,MAP1B,NEFL,CALR,TUBB2B,KIF1A,BSG,SPTBN1,MYH9,P4HB,APP,GDI1
1,HSP90AA1,ACTR2,SPARC,KIDINS220,DCX,EIF4G2,RPS6,DBN1,FLNA,MAP1B,NCL,ALDOA,PKM,
NS220,DCX,EIF4G2,DBN1,MAP1B,NEFL,UGT8,TMF1,TUBB2B,DVL3,KIF1A,BSG,SPTBN1,APP,GDI
DCX,EIF4G2,DBN1,MAP1B,NEFL,TUBB2B,KIF1A,BSG,SPTBN1,APP,GDI1,SPTAN1,PLXNB2,MAPT,F
A9,NEDD4L,ACTB,ACTG1,TUBA1A,CFL1,BTBD9,HTT,HSP90AB1,HSP90AA1,ACTR2,VIM,SPARC,TU
PM1,SNRNP200,SPEN,NONO,PPP2R1A,SF3B1,SF3B2,GTF2F1,HNRNPU,HNRNPA0,PAPOLA,SRSF
P22,ACTG1,EEF2,DPYSL3,CFL1,XRCC2,HSP90AB1,HSP90AA1,ACTR2,VIM,SPARC,KIDINS220,NUM
A1B,CLU,RACK1,RPL5,VCP,UBB,DDB1,PKD1,RPL11,ANXA2,TIMP2,RPL23,LPCAT1,MSN,PSMD2,T
IPA1,NEDD4L,POLH,ATN1,ACTB,CTSC,USP22,MED18,EEF2,MAVS,HTT,HSP90AB1,ZNF124,HSP90
,NCL,RPL7A,PKM,UGT8,RPL8,YBX1,CALR,RPLP0,SOX4,RPL13A,RPL6,RPL13,RACK1,FASN,RPS8,
TG1,EEF2,DPYSL3,CFL1,XRCC2,HSP90AB1,HSP90AA1,ACTR2,VIM,SPARC,KIDINS220,NUMA1,DC
,HNRNPA1,NEDD4L,POLH,ATN1,ACTB,CTSC,NRBP2,USP22,MED18,ACTG1,EEF2,BTBD9,XRCC2,
NP200,SPEN,NONO,SF3B1,SF3B2,GTF2F1,HNRNPU,HNRNPA0,PAPOLA,SRSF4,PNN,HNRNPC,KH
NP200,SPEN,NONO,SF3B1,SF3B2,GTF2F1,HNRNPU,HNRNPA0,PAPOLA,SRSF4,PNN,HNRNPC,KH
NP200,SPEN,NONO,SF3B1,SF3B2,GTF2F1,HNRNPU,HNRNPA0,PAPOLA,SRSF4,PNN,HNRNPC,KH
HTT,HSP90AB1,HSP90AA1,UBC,KIDINS220,FLNA,HSP90B1,FBXL4,UBA1,ATP2B4,FSTL1,HSPA1B,
HTT,HSP90AB1,HSP90AA1,UBC,KIDINS220,FLNA,HSP90B1,FBXL4,UBA1,ATP2B4,FSTL1,HSPA1B,
IEFL,TUBB2B,BSG,SPTBN1,APP,GDI1,SPTAN1,PLXNB2,MAPT,FN1,STMN1,ITGB1,UCHL1,NCAM1,
C,USP22,ACTG1,EEF2,MAVS,HTT,HSP90AB1,HSP90AA1,ACTR2,VIM,UBC,KIDINS220,TMSB4X,EIF
,SNRNP200,SPEN,NONO,SF3B1,SF3B2,GTF2F1,HNRNPU,HNRNPA0,PAPOLA,SRSF4,PNN,HNRNP
XL4,UBA1,TMF1,HSPA1B,CLU,RACK1,RPL5,TTC3,USP11,VCP,UBB,DDB1,PSMD5,PKD1,AP2M1,U
AA1,ACTR2,VIM,KIDINS220,DCX,EIF4G2,RPS6,DBN1,FLNA,MAP1B,NEFL,UGT8,CALR,TMF1,ABH
BN1,ENO1,FLNA,MAP1B,CALR,HSPA1B,H3-3B,CLU,ENO2,YWHAE,SPTBN1,STMN2,FSCN1,SPTAN
RPL3,RPS6,RPL4,NCL,RPL7A,RPL8,YBX1,ATP2B4,CALR,TMF1,HSPA1B,RPLP0,H3-3B,DDX5,RPL1
,NCL,RPL7A,PKM,UGT8,RPL8,YBX1,CALR,RPLP0,CLU,SOX4,RPL13A,RPL6,RPL13,RACK1,FASN,
RPL13A,RPL6,RPL13,DDX17,RPS8,RPL7,RPL5,NPM1,SNRNP200,EIF3C,EIF4B,RPL37A,RPL19,RPL
TT,HSP90AB1,HSP90AA1,ACTR2,VIM,UBC,KIDINS220,TMSB4X,EIF4G2,ENO1,PIK3R2,NCL,PKM,Y
,HNRNPA1,HSPA9,NEDD4L,POLH,ATN1,ACTB,CTSC,ODF2L,NRBP2,USP22,MTRNR2L2,MED18,AC
2,BTBD9,HTT,HSP90AB1,HSP90AA1,ACTR2,SPARC,TUBB,FTH1,UBC,TMSB4X,PIK3R2,FLNA,ALD
LNA,MAP1B,NEFL,KIF1A,STMN2,GDI1,PLXNB2,MAPT,FN1,PAFAH1B1,SCARB2,RTN4,NCS1,RHOA

C,USP22,ACTG1,EEF2,MAVS,HSP90AB1,HSP90AA1,ACTR2,VIM,UBC,KIDINS220,EIF4G2,ENO1,PIK
11,HSP90AB1,ZNF124,UBC,EIF4G2,ENO1,FLNA,NCL,YBX1,ATP2B4,CALR,HSPA1B,H3-3B,CLU,DD
UBB2B,BSG,SPTBN1,APP,GDI1,SPTAN1,PLXNB2,MAPT,FN1,STMN1,ITGB1,UCHL1,NCAM1,VCL,F
11,NRBP2,HSP90AB1,ZNF124,UBC,EIF4G2,ENO1,FLNA,NCL,YBX1,ATP2B4,CALR,HSPA1B,H3-3B,
4G2,DBN1,FLNA,MAP1B,NEFL,KIF1A,STMN2,GDI1,FSCN1,PLXNB2,MAPT,FN1,CDKL1,AGRN,PAF
R2,VIM,SPARC,UBC,KIDINS220,NUMA1,PIK3R2,FLNA,MAP1B,NCL,PKM,COL6A1,HSP90B1,FADS
4G2,DBN1,FLNA,MAP1B,NEFL,DVL3,KIF1A,STMN2,GDI1,FSCN1,PLXNB2,MAPT,FN1,CDKL1,AGRI
TT,HSP90AB1,HSP90AA1,UBC,RPS6,ENO1,FLNA,MTRNR2L8,HSP90B1,NEFL,CALR,FSTL1,TMF1,H
,MAVS,HSP90AB1,HSP90AA1,ACTR2,VIM,UBC,KIDINS220,TMSB4X,EIF4G2,ENO1,PIK3R2,NCL,PK
IPA1,NEDD4L,ATN1,ACTB,USP22,MED18,ACTG1,EEF2,MAVS,HSP90AB1,ZNF124,HSP90AA1,ACTR
PS6,RPL4,ENO1,NCL,RPL7A,ALDOA,PKM,UGT8,RPL8,YBX1,ATP2B4,CALR,RPLP0,SOX4,RPL13A,F
PYSL3,CFL1,XRCC2,HSP90AB1,HSP90AA1,ACTR2,VIM,KIDINS220,NUMA1,DCX,EIF4G2,RPS6,DBN
D4L,ACTB,CTSC,ACTG1,EEF2,DPYSL3,CFL1,MAVS,HSP90AB1,HSP90AA1,ACTR2,VIM,SPARC,UBC
NPA1,NEDD4L,POLH,ATN1,USP22,MED18,EEF2,BTBD9,XRCC2,MAVS,HSP90AB1,ZNF124,HSP90A
PYSL3,CFL1,XRCC2,HSP90AB1,HSP90AA1,ACTR2,VIM,KIDINS220,NUMA1,DCX,EIF4G2,RPS6,DBN
D18,MAVS,ZNF124,ACTR2,VIM,UBC,TMSB4X,EIF4G2,RPL3,RPS6,ZNF316,RPL4,ENO1,PIK3R2,FL
NPA1,NEDD4L,POLH,ATN1,USP22,MED18,EEF2,MAVS,HSP90AB1,ZNF124,HSP90AA1,ACTR2,VIM,
NPA1,NEDD4L,POLH,ATN1,USP22,MED18,EEF2,MAVS,HSP90AB1,ZNF124,HSP90AA1,ACTR2,VIM,
HTT,HSP90AB1,HSP90AA1,UBC,KIDINS220,FLNA,HSP90B1,FBXL4,UBA1,ATP2B4,FSTL1,HSPA1B,
OLH,ATN1,USP22,MED18,XRCC2,MAVS,HSP90AB1,ZNF124,HSP90AA1,ACTR2,VIM,UBC,TMSB4X,E
3N1,ENO1,MAP1B,IGFBP2,HSPA1B,PSAP,H3-3B,GNAS,RACK1,APP,GDI1,NPM1,IGF2,TRIM28,MAF
TT,HSP90AB1,HSP90AA1,UBC,RPS6,ENO1,FLNA,MTRNR2L8,PKM,HSP90B1,NEFL,CALR,FSTL1,TM
11,USP22,MED18,XRCC2,MAVS,HSP90AB1,ZNF124,HSP90AA1,ACTR2,VIM,UBC,TMSB4X,EIF4G2,
/CP,UBB,PKD1,RPL11,ANXA2,RPL23,MSN,TMEM259,CD81,XPO1,CSNK1E,RPS7,LAPTM4B,TRIB2
NPA1,NEDD4L,POLH,ATN1,USP22,MED18,EEF2,MAVS,HSP90AB1,ZNF124,HSP90AA1,ACTR2,VIM,
SPA1B,CLU,RPL13A,SPTBN1,STMN2,GDI1,SPTAN1,NPM1,MAPT,SET,TMSB10,HMGB1,STMN1,PA
,NUMA1,DBN1,FLNA,MAP1B,HSPA1B,CLU,RPL13A,RACK1,SPTBN1,STMN2,APP,FSCN1,SPTAN1,N
3C,FLNA,HSP90B1,FBXL4,UBA1,ATP2B4,TMF1,HSPA1B,CLU,RACK1,ALDH6A1,IDS,RPL5,TTC3,US
DD4L,ATN1,ACTB,USP22,MED18,ACTG1,EEF2,MAVS,HSP90AB1,ZNF124,HSP90AA1,ACTR2,VIM,U
1,ACTR2,SPARC,NUMA1,EIF4G2,DBN1,FLNA,MAP1B,ALDOA,PKM,FADS1,NEFL,YBX1,ATP2B4,CA
NA,ATP2B4,MAGED1,DVL3,YWHAE,RACK1,SPTBN1,GDI1,NPM1,VCP,TRIM28,YWHAG,MAPT,YWH
HSPA1B,CLU,RACK1,RPL5,TTC3,USP11,VCP,UBB,DDB1,PSMD5,PKD1,UCHL1,RPL11,MARCHF6,F
NPA1,NEDD4L,POLH,ATN1,USP22,MED18,EEF2,MAVS,HSP90AB1,ZNF124,HSP90AA1,ACTR2,VIM,
TT,HSP90AB1,HSP90AA1,UBC,EIF4G2,RPS6,ENO1,FLNA,MTRNR2L8,PKM,HSP90B1,NEFL,CALR,F
1,VCP,UBB,DDB1,PSMD5,PKD1,UCHL1,RPL11,MARCHF6,PSMD2,ENC1,TMEM259,XPO1,CSNK1E
O1,MAP1B,HSPA1B,H3-3B,RACK1,APP,GDI1,NPM1,MAPT,FN1,PPP2R1A,ITGB1,VCL,PAFAH1B1,RT
R2,VIM,UBC,KIDINS220,NUMA1,PIK3R2,FLNA,MAP1B,NCL,PKM,COL6A1,HSP90B1,ATP2B4,IGFE
,HNRNPA1,HSPA9,NEDD4L,POLH,ATN1,ACTB,CTSC,ODF2L,NRBP2,USP22,MTRNR2L2,MED18,AC
31,HSP90AA1,UBC,TMSB4X,NUMA1,DBN1,ENO1,PIK3R2,FLNA,MAP1B,MTRNR2L8,NEFL,ATP2B4
MSB4X,NUMA1,DBN1,FLNA,MAP1B,ALDOA,COL6A1,HSP90B1,NEFL,NEFM,HSPA1B,COL5A1,COL
IP200,EIF3C,EIF4B,VCP,SF3B1,RPL11,RPL10,RPSA,EIF4H,XRCC5,RPL24,KIF5B,RPS14,EIF5,RPS1
OLH,ATN1,USP22,MED18,BTBD9,XRCC2,MAVS,HSP90AB1,ZNF124,HSP90AA1,ACTR2,VIM,UBC,TN
NPA1,NEDD4L,POLH,ATN1,USP22,MED18,EEF2,MAVS,HSP90AB1,ZNF124,HSP90AA1,ACTR2,VIM,
HSP90AA1,UBC,RPS6,ENO1,FLNA,MTRNR2L8,HSP90B1,NEFL,CALR,FSTL1,TMF1,HSPA1B,CLU,F
OA,PKM,NEFL,ATP2B4,CALR,PGK1,MAGED1,DVL3,KIF1A,RACK1,MYH9,P4HB,GDI1,PLXNB2,COL

IP200,EIF3C,EIF4B,SF3B1,RPL11,RPL10,RPSA,EIF4H,XRCC5,RPL24,RPS14,EIF5,RPS15,EIF3L,RF
B,FTH1,TMSB4X,FLNA,ALDOA,PKM,FTL,HSPA1B,PSAP,CLU,SLC2A3,GPI,CALM3,MYH9,SEPTIN2,
ISPA1B,CLU,DDX5,YWHAE,RACK1,P4HB,APP,TMBIM6,GSTP1,SMG1,NPM1,TRIM28,MAPT,NONO,
UBC,KIDINS220,EIF4G2,ENO1,PKM,ATP2B4,HSPA1B,CLU,SOX4,MAGED1,DVL3,RACK1,CALM3,M
B,FTH1,TMSB4X,FLNA,ALDOA,PKM,FTL,HSPA1B,PSAP,CLU,SLC2A3,GPI,CALM3,MYH9,APP,SPTA
31,HSP90AA1,UBC,TMSB4X,NUMA1,DBN1,ENO1,PIK3R2,FLNA,MAP1B,MTRNR2L8,NEFL,ATP2B4
OLH,ATN1,USP22,MED18,BTBD9,XRCC2,MAVS,HSP90AB1,ZNF124,HSP90AA1,ACTR2,VIM,UBC,TN
IEDD4L,POLH,ATN1,USP22,MED18,EEF2,MAVS,HSP90AB1,ZNF124,HSP90AA1,ACTR2,VIM,UBC,TI
,HNRNPA1,HSPA9,NEDD4L,POLH,ATN1,ACTB,CTSC,ODF2L,NRBP2,USP22,MTRNR2L2,MED18,AC
UBB,DDB1,PSMD5,UCHL1,RPL11,MARCHF6,PSMD2,XPO1,CSNK1E,CTNNB1,UBE2H,SKP1,TRIB2
0B1,ATP2B4,HSPA1B,CLU,SOX4,MAGED1,DVL3,YWHAE,RACK1,CALM3,APP,RPL5,GSTP1,CALM1
,RACK1,RPL5,TTC3,USP11,VCP,UBB,DDB1,PSMD5,UCHL1,RPL11,MARCHF6,RPL23,PSMD2,XPO
1,NPM1,TRIM28,PKD1,RPL23,KPNB1,SRSF4,KHDRBS1,XPO1,EIF5A,RPS15,TARDBP,APPL1,IPO5,
3R2,FLNA,ATP2B4,SOX4,MAGED1,DVL3,YWHAE,GNAS,RACK1,BSG,SPTBN1,GDI1,NPM1,VCP,TRI
,RACK1,RPL5,TTC3,USP11,VCP,UBB,DDB1,PSMD5,UCHL1,RPL11,MARCHF6,RPL23,PSMD2,XPO
M6,VCP,PDIA3,MARCHF6,ATF4,TMEM259,ATP2A2,PDIA6,DDX3X,TLN1,JUN,TARDBP,CTDSP2,UBI
GCN1,PSMB2,VEGFA,VHL,PSMD3,PSMA7,EP300,PSMD1,UBA52,RPS27A,PSMD8,HIF1A,PSMB4,PS
,RACK1,RPL5,TTC3,USP11,VCP,UBB,DDB1,PSMD5,UCHL1,RPL11,MARCHF6,RPL23,PSMD2,XPO
HSP90AA1,UBC,RPS6,ENO1,FLNA,MTRNR2L8,HSP90B1,NEFL,CALR,FSTL1,TMF1,HSPA1B,CLU,F
EFL,DVL3,STMN2,GDI1,FSCN1,PLXNB2,MAPT,FN1,AGRN,PAFAH1B1,SCARB2,RHOA,ENC1,PFN1,F
MB2,VEGFA,VHL,PSMD3,PSMA7,EP300,PSMD1,UBA52,RPS27A,PSMD8,HIF1A,PSMB4,PSMD4,RB
HSP90AA1,UBC,RPS6,ENO1,FLNA,MTRNR2L8,HSP90B1,NEFL,CALR,FSTL1,TMF1,HSPA1B,CLU,F
TP2B4,CALR,HSPA1B,CLU,SOX4,RPL13A,YWHAE,GPI,RACK1,CALM3,APP,RPL5,GSTP1,CALM1,NF
2,FLNA,MAP1B,ATP2B4,MAGED1,DVL3,YWHAE,RACK1,SPTBN1,CALM3,GDI1,TMBIM6,CALM1,NF
IGFBP3,PSMD2,PRKCSH,KLHL9,CDH2,PDIA6,GPC3,RCN1,GOLM1,APLP2,CKAP4,RAB1A,CALU,TI
ATP2B4,COL4A1,CLU,SOX4,YWHAE,KIF1A,CALM3,APP,GSTP1,MARCKS,CALM1,PLXNB2,MARCKS
,FLNA,MAP1B,NEFL,UGT8,COL4A1,KIF1A,APP,FSCN1,PLXNB2,VCP,MAPT,FN1,DCHS1,INA,AGRN,
P1,HNRNPA0,PSMD2,HNRNPC,XPO1,TARDBP,CIRBP,YWHAB,TAF15,ALKBH5,TNPO1,PSMB2,PSM
KHDRBS1,HNRNPK,CIRBP,NOVA2,U2AF2,PABPN1,HNRNPL,AHCYL1,RBMX,SNRNP70,SRSF9,PTBF
OLH,ATN1,USP22,MED18,BTBD9,XRCC2,MAVS,HSP90AB1,ZNF124,HSP90AA1,ACTR2,VIM,UBC,TN
P1,HNRNPA0,PSMD2,HNRNPC,XPO1,TARDBP,CIRBP,YWHAB,TAF15,ALKBH5,TNPO1,PSMB2,PSM
R2,PSAP,CLU,TMBIM6,SMG1,CALM1,VCP,UBB,MAPT,HMGB1,UCHL1,LARP1,ATP2A2,MCL1,PIP4
R2,PSAP,CLU,TMBIM6,SMG1,CALM1,VCP,UBB,MAPT,HMGB1,UCHL1,LARP1,ATP2A2,MCL1,PIP4
1B,PSAP,GNAS,APP,GDI1,IGF2,TRIM28,MAPT,FN1,BASP1,ITGB1,AGRN,ZMIZ1,VCL,PAFAH1B1,EI
1,NPM1,TRIM28,PKD1,RPL23,KPNB1,SRSF4,KHDRBS1,XPO1,EIF5A,RPS15,TARDBP,APPL1,IPO5,
PL11,RPL23,TMEM259,XPO1,CSNK1E,RPS7,TRIB2,MAP1A,NOP53,RAD23B,GABARAP,CCAR2,SM
,ENO1,MAP1B,HSP90B1,ATP2B4,IGFBP2,HSPA1B,PGK1,YWHAE,GPI,GNB1,P4HB,APP,TMBIM6,C
,ACTR2,VIM,TUBB,TMSB4X,NUMA1,DCX,DBN1,TUBA1B,FLNA,MAP1B,ALDOA,HSP90B1,NEFL,NE
P90AA1,ACTR2,TUBB,UBC,NUMA1,RPS6,TUBA1B,FLNA,ATP2B4,CALR,HSPA1B,TUBB2B,SOX4,YV
1B,CLU,SOX4,DVL3,RPL5,NPM1,HSPD1,RPL11,RPL23,RTN4,CD81,CCT7,PFN1,RPS7,TARDBP,DD
CP,UBB,DDB1,PSMD5,PPP2R1A,PKD1,AP2M1,AP2B1,CPE,RHOA,PSMD2,PFN1,TLE5,CSNK1E,CDH
U,RPL13A,RACK1,SPTBN1,STMN2,FSCN1,SPTAN1,VCP,MAPT,DDB1,TMSB10,RPS3,HMGB1,STMI
ACK1,GDI1,NPM1,MAPT,FN1,PPP2R1A,PAFAH1B1,RTN4,IGFBP3,RHOA,CLSTN1,DDR1,MEGF8,M
PTAN1,CALM1,VCP,SET,HMGB1,STMN1,TIMP2,PAFAH1B1,RPL23,KPNB1,PTCD3,KIF5B,WDR1,AT
CP,UBB,DDB1,PSMD5,PPP2R1A,PKD1,AP2M1,AP2B1,CPE,RHOA,PSMD2,PFN1,TLE5,CSNK1E,CDH

INS220,EIF4G2,ENO1,PKM,ATP2B4,HSPA1B,CLU,SOX4,MAGED1,DVL3,RACK1,CALM3,MYH9,APF
3R2,FLNA,MAP1B,PKM,COL6A1,HSP90B1,FADS1,NEFL,ATP2B4,IGFBP2,COL4A1,GNAS,GPI,BSG
259,XPO1,CSNK1E,TRIB2,MAP1A,NOP53,RAD23B,GABARAP,CCAR2,SMARCC1,UBQLN1,CSNK1D
,ALDOA,PKM,FTL,HSPA1B,PSAP,SLC2A3,GPI,SPTAN1,GSTP1,VCP,HLA-C,VAPA,HMGB1,TUBB4B,
SPA1B,CANX,CLU,GNB1,P4HB,HSPD1,VCP,PDIA3,DNAJA1,CCT7,PRKCSH,PDIA6,GNB2,DNAJC5,F
S1,XPO1,EIF5A,CKAP5,CCT6A,BCLAF1,CPSF2,DDX39B,SRSF5,TOMM20,U2AF2,CCT2,CPSF1,PAE
A1B,SOX4,RACK1,RPL5,NPM1,TTC3,VCP,UBB,TRIM28,DDB1,PSMD5,RPS3,RPL11,ZMIZ1,MARCH
KHDRBS1,HNRNPK,CIRBP,HNRNPH1,NOVA2,SRSF5,U2AF2,HNRNPL,RBMX,SNRNP70,SRSF9,SRS
,CALR,HSPA1B,SOX4,RPL13A,YWHAE,GPI,RACK1,CALM3,APP,RPL5,GSTP1,CALM1,NPM1,YWHAC
,ALDOA,PKM,FTL,HSPA1B,PSAP,SLC2A3,GPI,SPTAN1,GSTP1,VCP,HLA-C,VAPA,HMGB1,TUBB4B,
P1,HNRNPA0,PSMD2,HNRNPC,XPO1,TARDBP,CIRBP,YWHAB,TAF15,ALKBH5,TNPO1,PSMB2,PSM
RC,PIK3R2,FLNA,MAP1B,PKM,COL6A1,HSP90B1,FADS1,NEFL,ATP2B4,IGFBP2,COL4A1,GNAS,G
,ALDOA,PKM,FTL,HSPA1B,PSAP,SLC2A3,GPI,SPTAN1,GSTP1,VCP,HLA-C,VAPA,HMGB1,TUBB4B,
,MAP1B,NCL,ATP2B4,FSTL1,DDX5,HTRA4,RACK1,STMN2,APP,GSTP1,UBB,COL4A2,MAPT,ZMIZ1
1,HSPA1B,SOX4,RACK1,RPL5,NPM1,TTC3,USP11,VCP,UBB,TRIM28,DDB1,PSMD5,RPS3,UCHL1,
,COPA,HNRNPA1,HSPA9,NEDD4L,POLH,ATN1,ACTB,CTSC,ODF2L,NUDC,NRBP2,USP22,MTRNR2
,ALDOA,PKM,FTL,HSPA1B,PSAP,SLC2A3,GPI,SPTAN1,GSTP1,VCP,HLA-C,VAPA,HMGB1,TUBB4B,
0AA1,UBC,TMSB4X,NUMA1,ENO1,PIK3R2,FLNA,MTRNR2L8,NEFL,ATP2B4,IGFBP2,CALR,FSTL1,
STMN2,FSCN1,IGF2,TRIM28,YWHAG,MAPT,YWHAQ,RPS3,ANXA2,ARF1,XRCC5,MSN,RHOA,KIF5B,
A1,RPS6,FLNA,ATP2B4,CALR,HSPA1B,SOX4,YWHAE,SPTBN1,CALM3,MYH9,SEPTIN2,APP,CALM1,
,ALDOA,PKM,FTL,HSPA1B,PSAP,SLC2A3,GPI,SPTAN1,GSTP1,VCP,HLA-C,VAPA,HMGB1,TUBB4B,
A1B,SOX4,RACK1,RPL5,NPM1,TTC3,VCP,UBB,TRIM28,DDB1,PSMD5,RPS3,RPL11,MARCHF6,WS
S6,TUBA1B,FLNA,HSPA1B,TUBB2B,SOX4,YWHAE,SPTBN1,CALM3,APP,CALM1,IGF2,VCP,UBB,YW
,ATP2A2,PIP4K2B,RAB7A,SESN3,RAB1A,ATP6V0C,DCN,TOMM20,SQSTM1,GNAI3,VPS35,OPTN,AT
1,ATP2B4,CALR,HSPA1B,SOX4,YWHAE,RACK1,CALM3,APP,CALM1,NPM1,IGF2,UBB,YWHAG,DDI
C,TUBB,FTH1,TMSB4X,FLNA,ALDOA,PKM,FTL,TMF1,HSPA1B,PSAP,CANX,CLU,SOX4,GNAS,SLC2
B,SOX4,YWHAE,SPTBN1,CALM3,APP,CALM1,IGF2,VCP,UBB,YWHAG,DDB1,PSMD5,SET,MAP9,PP
GDI1,TRIM28,MAPT,FN1,ITGB1,VCL,PAFAH1B1,RTN4,DDR1,PLXNA1,MEGF8,CTNNB1,MACF1,CY
,EP300,PSMD1,UBA52,RPS27A,PSMD8,HIF1A,PSMB4,PSMD4,RBPJ,PSMC3,CREBBP,PSMC1,PSM
,ACTR2,VIM,UBC,TMSB4X,EIF4G2,ENO1,PIK3R2,NCL,PKM,YBX1,TMF1,ZNF99,CLU,SOX4,MAGE
MSB4X,DCX,RPS6,ENO1,PIK3R2,FLNA,NEFL,ATP2B4,CALR,HSPA1B,CLU,DDX5,SOX4,MAGED1,KA
N1,ITGB1,VCL,PAFAH1B1,RTN4,MEGF8,MACF1,MAP2,L1CAM,DPYSL2,VEGFA,SLIT3,PLXNA3,PTI
-3B,KAT6A,SMG1,NPM1,IGF2,UBB,TRIM28,MAPT,DDB1,SET,GTF3C4,CHD3,HMGB1,MORF4L1,Z
HSP90AB1,HSP90AA1,UBC,KIDINS220,TMSB4X,PIK3R2,FLNA,MTRNR2L8,HSP90B1,NEFL,ATP2B4
,ACTR2,VIM,UBC,TMSB4X,EIF4G2,ENO1,PIK3R2,NCL,PKM,YBX1,TMF1,ZNF99,CLU,SOX4,MAGE
LU,FKBP14,RACK1,TMBIM6,HSPD1,VCP,DNAJA1,ATF4,PDIA6,TLN1,CTDSP2,HSPH1,HSPA4,HDG
,MAP1B,NCL,ATP2B4,FSTL1,DDX5,HTRA4,RACK1,STMN2,APP,GSTP1,UBB,COL4A2,MAPT,ZMIZ1
5A1,CLU,YWHAE,SPTBN1,ATP5F1B,CALM3,MYH9,APP,CALM1,ATP5F1A,UBB,YWHAG,YWHAQ,PP
,CALM3,APP,CALM1,PLXNB2,UBB,CKB,YWHAQ,INA,BASP1,STMN1,ITGB1,ZMIZ1,PAFAH1B1,CTN
1,TRIM28,RPL11,HNRNPU,CTNNA1,RPL23,KPNB1,CCT7,XPO1,TARDBP,CCT6A,APPL1,IPO5,SKP
KBP14,RACK1,TMBIM6,HSPD1,VCP,DNAJA1,ATF4,PDIA6,TLN1,CTDSP2,HSPH1,HSPA4,HDGF,TM
,PSMD5,PPP2R1A,TUBB4B,PAFAH1B1,PSMD2,NES,KHDRBS1,CSNK1E,CKAP5,SKP1,CDKN1A,PPF
I1,GDI1,SPTAN1,NPM1,MAPT,FN1,TMSB10,AP2M1,ARF1,PAFAH1B1,RTN4,MSN,RHOA,WDR1,PFN
TUBB2B,COL5A1,SOX4,DVL3,RPL13,GPI,RACK1,MYH9,PLXNB2,SMIM14,IGF2,TRIM28,COL4A2,
A,TUBB4B,PAFAH1B1,PSMD2,CSNK1E,CKAP5,SKP1,CDKN1A,NOP53,PSMB2,CCND1,MAPRE1,PS

1,MAPT,FN1,ITGB1,VCL,PAFAH1B1,RTN4,DDR1,PLXNA1,MEGF8,CTNNB1,MACF1,CYFIP2,MAP2,L
,ACTR2,VIM,UBC,EIF4G2,PIK3R2,NCL,PKM,YBX1,TMF1,ZNF99,SOX4,MAGED1,KAT6A,DVL3,DD
GDI1,MAPT,FN1,ITGB1,VCL,PAFAH1B1,RTN4,DDR1,PLXNA1,MEGF8,CTNNB1,MACF1,CYFIP2,MA
YWHAE,CALM3,APP,CALM1,NPM1,IGF2,UBB,YWHAG,DDB1,PSMD5,MAP9,PPP2R1A,PKD1,TUBB
P,VCP,DDB1,HMGB1,LARP1,LPCAT1,MSN,TMEM259,CD81,OAZ1,CSNK1E,GPC3,PIP4K2B,RAB7
ALR,TMF1,ABHD2,H3-3B,DDX5,GNAS,DDX17,GPI,BSG,CALM3,GNB1,GSTP1,CALM1,HSPD1,IGF
SPTBN2,RAB7A,CLTC,HLA-B,KLC1,PSMB2,PSMD3,DYNC1LI2,DYNLL2,CAPZB,KIF3C,B2M,SEC23
ALM1,NPM1,YWHAG,MAPT,SET,PPP2R1A,RPS3,UCHL1,RPL11,DNAJA1,RPL23,IGFBP3,RPS20,PT
B4B,PAFAH1B1,PSMD2,CSNK1E,CKAP5,SKP1,CDKN1A,NOP53,PSMB2,CCND1,MAPRE1,PSMD3,
,FTH1,ALDOA,PKM,FTL,HSPA1B,PSAP,SLC2A3,GPI,SPTAN1,GSTP1,VCP,HLA-C,VAPA,HMGB1,TU
TG1,EEF2,DPYSL3,XRCC2,MAVS,HSP90AB1,HSP90AA1,ACTR2,VIM,SPARC,TUBB,UBC,TMSB4X,E
C,TUBB,FTH1,TMSB4X,FLNA,ALDOA,PKM,FTL,TMF1,HSPA1B,PSAP,CANX,CLU,SOX4,YWHAE,GN
,ALDOA,PKM,FTL,HSPA1B,PSAP,SLC2A3,GPI,SPTAN1,GSTP1,VCP,HLA-C,VAPA,HMGB1,TUBB4B,
9,POLH,ACTB,CTSC,NRBP2,MTRNR2L2,ACTG1,EEF2,DPYSL3,CFL1,XRCC2,MAVS,HTT,HSP90AB
,HNRNPK,CIRBP,NOVA2,U2AF2,HNRNPL,RBMX,SNRNP70,SRSF9,PTBP1,SON,SRSF3,RBM15B,S
5,PPP2R1A,TUBB4B,PAFAH1B1,PSMD2,NES,KHDRBS1,CSNK1E,CKAP5,SKP1,CDKN1A,PPP1CB,N
PS8,RPL7,RPL5,NPM1,RPL37A,RPL19,RPL15,RPL18,RPL11,RPLP1,RPL10,RPS17,RPS16,RPL23,
G2,DBN1,FLNA,MAP1B,FADS1,NEFL,YBX1,CALR,HSPA1B,COL5A1,H3-3B,DDX5,SOX4,DDX17,P4
APP,CALM1,PLXNB2,UBB,CKB,YWHAQ,INA,BASP1,STMN1,ITGB1,ZMIZ1,PAFAH1B1,CTNNA1,LD
RPL5,NPM1,HSPD1,RPL11,RPL23,RTN4,CCT7,PFN1,RPS7,PTGES3,LAMP1,CCT6A,PPIB,CDKN1A,
RPL11,ANXA2,TIMP2,HNRNPU,LARP1,HNRNPA0,RPL23,MCL1,HNRNPC,LRPPRC,RPS7,TARDBP,C
SMD2,SPTBN2,RAB7A,CLTC,HLA-B,KLC1,PSMB2,PSMD3,DYNC1LI2,DYNLL2,CAPZB,KIF3C,B2M
HD3,HMGB1,MORF4L1,ZMIZ1,HNRNPU,SMARCA5,BRD3,CHD4,HMGN2,RBBP4,HNRNPC,TADA2
N1,VCP,VAPA,AP3D1,ARF1,LLGL1,RAB6B,SPTBN2,CLTC,MYO5A,MACF1,LAMP1,RAB1A,ARCN1,V
PTBN1,STMN2,FSCN1,SPTAN1,NPM1,MAPT,MAP9,TMSB10,PKD1,RPS3,STMN1,ARF1,PAFAH1B1,
H3-3B,GNAS,RACK1,APP,GDI1,NPM1,IGF2,MAPT,FN1,PPP2R1A,BASP1,MORF4L1,AGRN,PAFAH1
X1,CALR,H3-3B,DDX5,SOX4,RPL13A,RACK1,APP,RPL5,NPM1,PLXNB2,CSDE1,EIF3C,EIF4B,UBB,
J,SOX4,RPL13A,RACK1,APP,RPL5,NPM1,PLXNB2,CSDE1,EIF3C,EIF4B,EIF1,RPS3,RPS4X,RPL10,
,ALDOA,PKM,FTL,HSPA1B,PSAP,SLC2A3,GPI,SPTAN1,GSTP1,VCP,HLA-C,VAPA,HMGB1,TUBB4B,
TMBIM6,GSTP1,UBB,YWHAG,NONO,YWHAQ,PDIA3,RPS3,TPT1,RPL11,DNAJA1,CTNNA1,ATF4,RP
,XPO1,EIF5A,RPS15,CPSF2,DDX39B,SRSF5,U2AF2,CPSF1,PABPN1,ALKBH5,AHCYL1,SRSF9,SRS
1,ANXA2,TIMP2,HNRNPU,LARP1,HNRNPA0,RPL23,MCL1,HNRNPC,LRPPRC,RPS7,TARDBP,CIRBP,
GDI1,PLXNB2,MAPT,FN1,SPEN,VCL,PAFAH1B1,CTNNA1,NACA,RTN4,XRCC5,RHOA,CLEC7A,PTN,P
CP,UBB,DDB1,PSMD5,PPP2R1A,PKD1,AP2M1,AP2B1,CPE,RHOA,PSMD2,NLGN2,PFN1,TLE5,CSN
TN1,USP22,MED18,EEF2,MAVS,HSP90AB1,ZNF124,HSP90AA1,ACTR2,VIM,UBC,TMSB4X,EIF4G2
AT1,MSN,TMEM259,CD81,OAZ1,CSNK1E,GPC3,RAB7A,GJA1,NDFIP1,TRIB2,PTTG1IP,NOP53,VP
,SPARC,TUBB,FTH1,TMSB4X,FLNA,ALDOA,PKM,FTL,TMF1,HSPA1B,PSAP,CANX,CLU,SOX4,GNAS
IDINS220,TMSB4X,PIK3R2,HSP90B1,ATP2B4,PSAP,CLU,MAGED1,DVL3,YWHAE,GNAS,GPI,RACK
90B1,PRPF8,FBXL4,UBA1,TMF1,HSPA1B,PGK1,CLU,YWHAE,GPI,HTRA4,RACK1,MYH9,APP,RPL5
CLU,SPTBN1,STMN2,APP,SPTAN1,MAPT,TMSB10,RPS3,STMN1,ARF1,RHOA,WDR1,PFN1,SPTBN2
INRNPA2B1,PCBP4,SAP18,SRSF10,C1QBP,CDK9,SRSF7,RBM10,CTR9,RNF20,SFSWAP,RNPS1,A
PS8,RPL7,RPL5,RPL37A,RPL19,RPL15,RPL18,RPL11,RPLP1,RPL10,RPS17,RPS16,RPL23,RPL29
RPS6,ENO1,FLNA,MTRNR2L8,HSP90B1,NEFL,FSTL1,TMF1,HSPA1B,CLU,PTMA,YWHAE,GPI,TMBI
RPS6,ENO1,FLNA,MTRNR2L8,HSP90B1,NEFL,FSTL1,TMF1,HSPA1B,CLU,PTMA,SOX4,YWHAE,GPI,
PS8,RPL7,RPL5,RPL37A,MAPT,RPL19,RPL15,RPL18,RPL11,RPLP1,RPL10,RPS17,RPS16,RPL23,

,CKAP5,BCLAF1,CPSF2,DDX39B,SRSF5,TOMM20,U2AF2,CPSF1,PABPN1,ALKBH5,HNRNPA3,G3B
,CKAP5,BCLAF1,CPSF2,DDX39B,SRSF5,TOMM20,U2AF2,CPSF1,PABPN1,ALKBH5,HNRNPA3,G3B
PP,CALM1,NPM1,UBB,YWHAG,DDB1,PSMD5,PPP2R1A,PKD1,ITGB1,TUBB4B,PAFAH1B1,RPL24,
X4,RPL13A,RACK1,APP,RPL5,NPM1,PLXNB2,CSDE1,EIF3C,EIF4B,EIF1,RPS3,RPS4X,RPL10,LARP
ACK1,FSCN1,VCP,TRIM28,MAPT,DDB1,RPS3,AGRN,ARF1,MSN,RHOA,CLSTN1,NLGN2,PFN1,CLE
TN1,USP22,MED18,EEF2,MAVS,HSP90AB1,ZNF124,HSP90AA1,ACTR2,VIM,UBC,TMSB4X,EIF4G2
,ACTR2,VIM,UBC,NUMA1,PIK3R2,IGFBP2,FSTL1,COL4A1,DDX5,HTRA4,SPTBN1,IGF2,UBB,COL4
MD2,PTN,NOP53,PSMB2,VEGFA,VHL,PSMD3,NPEPPS,BNIP3L,UBQLN1,HYOU1,PSMA7,EP300,PS
1,RTN4,RHOA,CDH2,PLXNA1,MEGF8,MACF1,MAP2,ARHGDIA,L1CAM,VEGFA,PLXNA3,PTPRS,RAE
PS6,ENO1,FLNA,MTRNR2L8,HSP90B1,NEFL,FSTL1,TMF1,HSPA1B,CLU,PTMA,YWHAE,GPI,TMBI
RSF6,HNRNPA2B1,PCBP4,SAP18,RPS26,SRSF10,C1QBP,SRSF7,RBM10,SFSWAP,RNPS1,ACIN1
DI1,PLXNB2,MAPT,FN1,SPEN,PAFAH1B1,XRCC5,RHOA,CLEC7A,PTN,PLXNA1,MEGF8,CTNNB1,M
F4,PSMD2,PTN,NOP53,PSMB2,VEGFA,VHL,PSMD3,NPEPPS,BNIP3L,UBQLN1,HYOU1,PSMA7,EP3
,MAP1B,NEFL,COL4A1,KIF1A,APP,PLXNB2,VCP,MAPT,INA,AGRN,ARF1,PAFAH1B1,GAP43,RHOA,
,FN1,SPEN,PAFAH1B1,CTNNA1,RTN4,XRCC5,RHOA,PTN,PLXNA1,MEGF8,CTNNB1,MACF1,MAP2
,FN1,SPEN,PAFAH1B1,XRCC5,RHOA,PTN,PLXNA1,MEGF8,CTNNB1,MACF1,ARHGDIA,CAPRIN1,H
OA,PSMD2,WDR1,PFN1,ASTN2,GPC3,CLTC,RAC1,DLG5,PTK7,PSMB2,PSMD3,VANGL2,CDC42,
6,HSPD1,VCP,ATF4,PDIA6,TLN1,CTDSP2,HDGF,TMED2,OPTN,CCND1,ACADVL,HYOU1,SSR1,DCT
2,MED18,MAVS,HSP90AB1,ZNF124,HSP90AA1,ACTR2,VIM,UBC,TMSB4X,EIF4G2,ZNF316,ENO1,
TMN2,SPTAN1,MAPT,TMSB10,RPS3,STMN1,ARF1,RHOA,PFN1,SPTBN2,CYFIP2,CKAP5,MAP2,CO
TN1,USP22,MED18,EEF2,MAVS,HSP90AB1,ZNF124,HSP90AA1,ACTR2,VIM,UBC,TMSB4X,EIF4G2
1,RHOA,PLXNA1,MEGF8,MACF1,ARHGDIA,L1CAM,VEGFA,PLXNA3,RAB11A,BMPR2,MAP6,PLXNE
6,HSPD1,VCP,ATF4,PDIA6,TLN1,CTDSP2,HDGF,TMED2,OPTN,CCND1,ACADVL,HYOU1,BAG6,SSF
,CKAP5,BCLAF1,CPSF2,DDX39B,SRSF5,TOMM20,U2AF2,CPSF1,PABPN1,ALKBH5,HNRNPA3,G3B
,RTN4,ATF4,PSMD2,PTN,NOP53,PSMB2,ATP6AP1,VEGFA,VHL,PSMD3,NPEPPS,BNIP3L,UBQLN1,
DBP,HNRNPK,CIRBP,U2AF2,TAF15,RBMX,SRSF9,PTBP1,SUPT5H,HNRNPD,SRSF6,HNRNPA2B1,P
TN1,USP22,MED18,EEF2,MAVS,HSP90AB1,ZNF124,HSP90AA1,ACTR2,VIM,UBC,TMSB4X,EIF4G2
1,UBB,YWHAG,DDB1,PSMD5,PPP2R1A,PKD1,ITGB1,TUBB4B,PAFAH1B1,RPL24,PSMD2,RPL26,
TK7,PSMB2,PSMD3,VANGL2,CDC42,PSMA7,PSMD1,AP2A1,SFRP1,PSMD8,GPC4,PSMB4,PSMD4,
,FLNA,YBX1,CALR,H3-3B,DDX5,MAGED1,KAT6A,TMBIM6,NPM1,IGF2,UBB,TRIM28,SPEN,NONO,
1E,RPS7,TRIB2,MAP1A,NOP53,RAD23B,GABARAP,CCAR2,SMARCC1,UBQLN1,CSNK1D,USP7,HE
4HB,VCP,UBB,TRIM28,DDB1,HACD3,VAPA,ITGB1,NCAM1,LARP1,RPSA,SCARB2,KPNB1,CXADR,C
1,ACTR2,UBC,TMSB4X,ENO1,PIK3R2,NCL,YBX1,TMF1,ZNF99,DDX5,SOX4,MAGED1,KAT6A,DVL
STN2,GPC3,CLTC,RAC1,PTK7,PSMB2,PSMD3,VANGL2,CDC42,PSMA7,PSMD1,AP2A1,SFRP1,BRS
STN2,GPC3,CLTC,RAC1,PTK7,PSMB2,PSMD3,VANGL2,CDC42,PSMA7,PSMD1,AP2A1,SFRP1,BRS
1B1,RTN4,MSN,RHOA,MEGF8,MACF1,MAP2,RAC1,L1CAM,DPYSL2,VEGFA,LUZP6,KDM1A,LARS
PTM4B,MAP1A,TIMP3,NOP53,SNRNP70,AZIN1,CCAR2,SMARCC1,FYN,USP7,BAG6,CYP51A1,SGT
SPTBN2,RAB7A,CLTC,HLA-B,KLC1,PSMB2,PSMD3,DYNC1LI2,DYNLL2,CAPZB,KIF3C,B2M,SEC23
P1A,NOP53,RAD23B,GABARAP,CCAR2,SMARCC1,UBQLN1,CSNK1D,USP7,HECTD1,BAG6,ARIH1
,SPTBN1,NPM1,TRIM28,YWHAG,MAPT,YWHAQ,ITGB1,AP2B1,RTN4,MSN,KIF5B,CD81,CCT7,PTN
BN1,STMN2,SPTAN1,MAPT,TMSB10,RPS3,STMN1,ARF1,RHOA,PFN1,SPTBN2,CYFIP2,CKAP5,MAF
,HSPD1,UBB,TRIM28,FN1,DDB1,VAPA,ITGB1,NCAM1,ANXA2,RPSA,SCARB2,KPNB1,CXADR,CD81
P4HB,PLXNB2,FN1,PAFAH1B1,MSN,RHOA,WDR1,PTN,TPM1,PLXNA1,ACTN4,MACF1,MYH10,RHO
PP,ITGB1,ARF1,PAFAH1B1,RHOA,PTN,MAP2,CAPRIN1,HDAC2,MAP1A,DLG5,ARF4,FYN,YWHAH,
MGB1,LARP1,MSN,TMEM259,CD81,CSNK1E,PIP4K2B,SESN3,TARDBP,TRIB2,PTTG1IP,DCN,GNA

3B,PSMD5,EGLN1,POLR2A,LDHA,RTN4,RHOA,ATF4,PSMD2,PTN,CCN2,MMP14,HDAC2,MMP2,CD
3R2,FLNA,MAP1B,PKM,COL6A1,HSP90B1,FADS1,NEFL,PRPF8,ATP2B4,IGFBP2,CALR,ABHD2,CC
WHAG,DDB1,PSMD5,PPP2R1A,PKD1,TUBB4B,TIMP2,PAFAH1B1,HNRNPU,PSMD2,RPL26,CSNK1
,ALDOA,PKM,FTL,HSPA1B,PSAP,CLU,SLC2A3,GPI,APP,SPTAN1,GSTP1,HSPD1,VCP,MAPT,HLA-C,
X3X,CTNNB1,UBE2H,SKP1,RNF187,UBE4A,PSMB2,PSMD3,GABARAP,UBE2R2,NPEPPS,TNKS2,HE
TT,ACTR2,VIM,SPARC,TMSB4X,DCX,PIK3R2,FLNA,MAP1B,NEFL,ATP2B4,CALR,FSTL1,TMF1,ABH
1,UBC,KIDINS220,ENO1,PIK3R2,ALDOA,PKM,ATP2B4,PGK1,CLU,ENO2,MAGED1,DVL3,GPI,RAC
TK7,PSMB2,PSMD3,VANGL2,CDC42,PSMA7,PSMD1,AP2A1,SFRP1,PSMD8,GPC4,PSMB4,PSMD4,
18,MAVS,ZNF124,ACTR2,VIM,UBC,TMSB4X,EIF4G2,ZNF316,ENO1,PIK3R2,FLNA,NCL,FADS1,YE
,RACK1,MYH9,APP,RPL5,HSPD1,VCP,UBB,MAPT,FN1,PKD1,RPS3,HMGB1,RPL11,ANXA2,TIMP2,
1E,GPC3,CLTC,CTNNB1,RAC1,PTK7,PSMB2,PSMD3,GNG2,CSNK1D,VANGL2,CDC42,PSMA7,PSM
B,SRSF5,U2AF2,CPSF1,PABPN1,ALKBH5,HNRNPA3,G3BP2,SRSF9,SRSF1,CASC3,SUPT6H,SRSF3
B1,UCHL1,LARP1,MCL1,PIP4K2B,SESN3,ATP6V0C,DCN,EEF1A2,GNAI3,VPS35,SNRNP70,OPTN,W
UMA1,PIK3R2,FLNA,CALR,TMF1,HSPA1B,DDX5,SOX4,MAGED1,DVL3,GNAS,RACK1,APP,IGF2,VC
B1,PCBP4,SAP18,SRSF10,C1QBP,SRSF7,RBM10,SFSWAP,RNPS1,ACIN1
9,NEDD4L,POLH,ACTB,CTSC,NRBP2,MTRNR2L2,ACTG1,EEF2,DPYSL3,CFL1,XRCC2,MAVS,HTT,
20,TMSB4X,ENO1,PIK3R2,HSP90B1,ATP2B4,CLU,MAGED1,DVL3,YWHAE,RACK1,CALM3,APP,GS
20,TMSB4X,ENO1,PIK3R2,HSP90B1,ATP2B4,CLU,MAGED1,DVL3,YWHAE,RACK1,CALM3,APP,GS
2,CPSF1,PABPN1,ALKBH5,SRSF9,SRSF1,CASC3,SUPT6H,SSB,SRSF3,RBM15B,XPOT,SRSF6,UPF
11,ZMIZ1,DNAJA1,RPL23,RPS20,DDX3X,RPS15,RPS7,CTNNB1,NDFIP1,RAB1A,SKP1,PTTG1IP,U
,FN1,SPEN,AGRN,PAFAH1B1,XRCC5,RHOA,CLSTN1,NLGN2,PTN,PLXNA1,MEGF8,CTNNB1,MAC
,PSMD5,PPP2R1A,PKD1,TUBB4B,PAFAH1B1,PSMD2,RPL26,CSNK1E,CCND2,DDX3X,IK,CKAP5,A
A,HSP90B1,CALR,SPTBN1,MYH9,FSCN1,SPTAN1,MARCKS,MARCKSL1,TMSB10,INA,STMN1,ITG
2,TMF1,TUBB2B,SOX4,DVL3,GNAS,RACK1,CALM3,GNB1,SHISA9,APP,CALM1,VCP,UBB,YWHAG,
,EIF3C,TRIM28,PKD1,HMGB1,STMN1,RPL11,ANXA2,GTF2F1,ATP2A2,CRMP1,TLE5,NES,CSNK1E
R,FSTL1,HSPA1B,CLU,DVL3,HTRA4,RACK1,TMBIM6,GSTP1,VCP,UBB,MAPT,NONO,PSMD5,PPP2
MAVS,HTT,HSP90AB1,HSP90AA1,ACTR2,UBC,TMSB4X,NUMA1,ENO1,PIK3R2,FLNA,MTRNR2L8,
,TMF1,ZNF99,DDX5,SOX4,MAGED1,KAT6A,DVL3,DDX17,APP,NPM1,IGF2,UBB,TRIM28,PKD1,SS
1,CALR,H3-3B,DDX5,MAGED1,KAT6A,TMBIM6,NPM1,IGF2,UBB,TRIM28,SPEN,NONO,YWHAQ,SE
R,FSTL1,HSPA1B,CLU,DVL3,HTRA4,RACK1,TMBIM6,GSTP1,VCP,UBB,NONO,PSMD5,PPP2R1A,T
,STMN1,RPL23,PTCD3,KIF5B,WDR1,ATP2A2,NES,SPTBN2,EIF5A,CTNNB1,ARID1A,CKAP5,MAP1A,
R,FSTL1,HSPA1B,CLU,DVL3,HTRA4,RACK1,TMBIM6,GSTP1,VCP,UBB,MAPT,NONO,PSMD5,PPP2
UMA1,PIK3R2,FLNA,CALR,TMF1,HSPA1B,DDX5,SOX4,MAGED1,DVL3,GNAS,RACK1,APP,IGF2,VC
2,CPSF1,PABPN1,ALKBH5,SRSF9,SRSF1,CASC3,SUPT6H,SSB,SRSF3,RBM15B,XPOT,SRSF6,UPF
B2,EIF3C,RPS4X,LARP1,RPS9,RHOA,RPL26,KHDRBS1,EIF5A,DDX3X,NGRN,CIRBP,DDX39B,ABC
,YWHAG,DDB1,PSMD5,PPP2R1A,PKD1,TUBB4B,PAFAH1B1,PSMD2,RPL26,CSNK1E,CCND2,DDX
FPQ,CTNNB1,CTBP1,HDAC2,SKP1,EID1,MORF4L2,PKN1,YEATS2,SKI,ASXL1,KDM6B,TAF7,VEGF
CF1,WLS,ATP6V0C,PTK7,SKI,PSMB2,SMARCA4,PSMD3,CCAR2,TNKS2,CSNK1D,PSMA7,SULF2,P
2,HDGF,TMED2,CCND1,ACADVL,HYOU1,SSR1,DCTN1,EP300,EXTL3,MBTPS1,KLHDC3,PPP1R15A
,YWHAG,YWHAQ,ANXA2,ARF1,MSN,KIF5B,CD81,OAZ1,KHDRBS1,XPO1,TARDBP,YWHAB,LAMP1,
IK3R2,FLNA,ATP2B4,CALR,HSPA1B,CLU,DDX5,MAGED1,KAT6A,DVL3,GNAS,RACK1,P4HB,APP,R
1,FLNA,ALDOA,HSP90B1,CALR,SPTBN1,MYH9,FSCN1,SPTAN1,MARCKS,MARCKSL1,TMSB10,IN
,PSMD2,TLE5,CSNK1E,CDH2,GPC3,SULF1,DDX3X,CTNNB1,MACF1,WLS,ATP6V0C,MDK,PTK7,SK
,MAVS,HTT,HSP90AB1,HSP90AA1,SPARC,TMSB4X,NUMA1,DBN1,PIK3R2,FLNA,MAP1B,NEFL,ATP
NB1,CTBP1,HDAC2,SKP1,EID1,MORF4L2,PKN1,YEATS2,SKI,ASXL1,KDM6B,TAF7,VEGFA,KDM1

RCA5,RPL23,PSMD2,RBBP4,H1-0,ARID1A,H1-10,PARP1,TSPYL4,RAD23B,PSMB2,H3-3A,TERF2I
,AP3D1,PAFAH1B1,TMEM170A,GOLGA8A,SCARB2,RTN4,KPNB1,ATP2A2,RTN3,RAB7A,ANXA6,M
AP1B,HSP90B1,FADS1,NEFL,ATP2B4,IGFBP2,CALR,TMF1,ABHD2,H3-3B,CLU,DDX5,GNAS,DDX1
,PSMB2,PSMD3,CCAR2,TNKS2,CSNK1D,PSMA7,SULF2,PSMD1,CSNK1G2,SFRP1,TNKS,PSMD8,A
,MAPT,FN1,SPEN,AGRN,PAFAH1B1,CTNNA1,RTN4,XRCC5,RHOA,CLSTN1,NLGN2,PTN,PLXNA1,M
,EGLN1,LDHA,RTN4,RHOA,PSMD2,PTN,MMP14,MMP2,NOP53,NOX4,ALKBH5,PLOD1,PSMB2,VE
,EGLN1,LDHA,RTN4,RHOA,ATF4,PSMD2,PTN,CCN2,MMP14,MMP2,NOP53,NOX4,ALKBH5,PLOD
7,PSMD1,ITGB5,PSMD8,PSMB4,PSMD4,ITGAV,PSMC3,PSMC1,TAPBP,PSMB1,PSMD13,HLA-A,PS
5,U2AF2,CPSF1,PABPN1,ALKBH5,SRSF9,SRSF1,CASC3,SUPT6H,SSB,SRSF3,RBM15B,XPOT,SRS
1,NCAM1,RPSA,SCARB2,KPNB1,CXADR,CD81,XPR1,SFPQ,RAB7A,CTNNB1,LAMP1,DDX39B,CTSE
,FTH1,RPS6,ALDOA,PKM,FTL,HSPA1B,PSAP,SLC2A3,GPI,APP,SPTAN1,GSTP1,HSPD1,VCP,HLA-C,
PI,MYH9,PLXNB2,SMIM14,IGF2,TRIM28,DCHS1,PKD1,ITGB1,BTF3,ZMIZ1,COL11A1,RTN4,PFN1,
MA1,DCX,EIF4G2,RPS6,FLNA,NEFL,ATP2B4,CALR,TMF1,HSPA1B,TUBB2B,COL5A1,COL4A1,PSA
1B,NEFL,YWHAE,KIF1A,MYH9,CALM1,NPM1,UBB,YWHAG,MAPT,PPP2R1A,ITGB1,TUBB4B,UCHL
9,PLXNB2,SMIM14,IGF2,TRIM28,DCHS1,PKD1,ITGB1,BTF3,ZMIZ1,COL11A1,RTN4,PFN1,PRKCS
,KDELR1,ARF4,KIF3C,TMED9,KIF5A,RER1,ERGIC1,RAB1B,KIFAP3,RAB6A,ERGIC3,COPZ1,ARF5
2B4,HSPA1B,PSAP,CLU,MAGED1,DVL3,GNAS,RACK1,CALM3,APP,GDI1,CALM1,NPM1,HSPD1,IG
8,MACF1,MAP2,L1CAM,DPYSL2,VEGFA,PLXNA3,PTPRS,RAB11A,BMPR2,DIP2B,CTTN,OLFM1,SE
SMD2,SPTBN2,RAB7A,CLTC,HLA-B,RAB27A,KLC1,RAB5B,PSMB2,PSMD3,DYNC1LI2,DYNLL2,C
3R2,IGFBP2,COL4A1,IGF2,COL4A2,AP2M1,STMN1,AP2B1,GTF2F1,RTN4,IGFBP3,RHOA,DDR1,
2,RPS27A,PSMD8,PSMB4,PSMD4,PSMC3,PSMC1,PSMB1,PSMD13,PSMD7,PSME3,PSMC5,PSMC
MD1,OAZ2,PSMD8,PSMB4,PSMD4,PSMC3,PSMC1,PSMB1,PSMD13,PARK7,PSMD7,PSME3,PSM
,TMSB4X,ENO1,FLNA,YBX1,ATP2B4,HSPA1B,CLU,DDX5,YWHAE,GNAS,RACK1,GNB1,MYH9,P4HE
L24,GAP43,KIF5B,CRMP1,SPTBN2,PLXNA1,MEGF8,RAC1,GPC1,L1CAM,CRPPA,PTPRA,PTK7,DP
APT,DDB1,RPS3,ARF1,MSN,RHOA,PFN1,CLEC7A,DDX3X,CKAP5,RAC1,PARP1,ABCA3,ACTR3,NC
,RPS6,ALDOA,PKM,FTL,HSPA1B,PSAP,SLC2A3,GPI,SPTAN1,GSTP1,HSPD1,VCP,HLA-C,VAPA,HM
1,CSNK1E,TRIB2,PTTG1IP,NOP53,VPS35,ABCA2,GABARAP,UBQLN1,CSNK1D,HECTD1,BAG6,AR
1,UBC,KIDINS220,TMSB4X,ENO1,PIK3R2,ALDOA,PKM,HSP90B1,FADS1,ATP2B4,PGK1,CLU,ENO
UMA1,PIK3R2,FLNA,CALR,HSPA1B,DDX5,SOX4,MAGED1,DVL3,GNAS,RACK1,APP,IGF2,VCP,UBE
3R2,ATP2B4,CLU,MAGED1,DVL3,RACK1,CALM3,APP,RPL5,GSTP1,CALM1,NPM1,IGF2,UBB,YW
COL6A1,HSP90B1,ATP2B4,COL4A1,GNAS,GNB1,APP,GSTP1,IGF2,YWHAG,LARP1,CTNNA1,RPL2
20,PIK3R2,ATP2B4,CLU,MAGED1,DVL3,RACK1,CALM3,APP,GSTP1,CALM1,NPM1,PLXNB2,IGF2,
8,MACF1,MAP2,L1CAM,DPYSL2,VEGFA,PLXNA3,PTPRS,RAB11A,BMPR2,DIP2B,CTTN,OLFM1,SE
,BRSK1,PSMD8,PSMB4,TAOK1,INTS3,VPS4A,PSMD4,PSMC3,PSMC1,USP47,PSMB1,TAOK2,MAC
2,GPC3,DDX3X,CTNNB1,WLS,MDK,PTK7,PSMB2,RAPGEF1,PSMD3,CCAR2,TNKS2,CSNK1D,CTN
L24,GAP43,KIF5B,CRMP1,SPTBN2,PLXNA1,MEGF8,RAC1,GPC1,L1CAM,CRPPA,PTPRA,PTK7,DP
,CRMP1,PRKCSH,ITM2C,MAP2,ARHGDIA,HDAC2,B2M,PLXNA3,PTPRS,DIP2B,ADCY6,PTEN,FKBP
C1,MAP1A,TBCD,CAMSAP1,MAPRE1,CLIP3,STMN3,DCTN1,HDAC6,MAPRE2,AKAP9,TAOK1,FKB
L3,FLNA,TMF1,HSPA1B,RPLP0,RPL6,YWHAE,SEPTIN2,RPL5,FSCN1,NPM1,CSDE1,YWHAG,MAPT
SF4,XPO1,SFPQ,EIF5A,RPS15,CPSF2,DDX39B,SRSF5,U2AF2,CPSF1,VAMP2,PABPN1,ALKBH5,V
1,GDI1,SPTAN1,NPM1,MAPT,FN1,TMSB10,AP2M1,ARF1,PAFAH1B1,RTN4,MSN,RHOA,WDR1,PFN
3,CCAR2,SMARCC1,USP7,BAG6,SGTA,CSNK2A1,EIF3H,OGT,BAG5,SDCBP,ATP13A2,MGAT3,PAR
1,UBB,NONO,PDIA3,RPS3,TPT1,RPL11,DNAJA1,CTNNA1,RPL26,MCL1,SFPQ,PRDX2,DDX3X,ITM2
ACK1,CALM3,P4HB,APP,SMG1,CALM1,IGF2,TRIM28,SET,EGLN1,PPP2R1A,GTF3C4,PKD1,MOR
1,DCHS1,AGRN,VCL,CTNNA1,GAP43,RHOA,CLSTN1,NLGN2,WDR1,SPTBN2,CDH6,CDH2,MMP1

,PKM,COL6A1,HSP90B1,ATP2B4,COL4A1,GNAS,GNB1,APP,GSTP1,NPM1,IGF2,YWHAG,AGRN,LA
A7,PSMD1,ITGB5,PSMD8,PSMB4,PSMD4,ITGAV,PSMC3,PSMC1,TAPBP,PSMB1,PSMD13,HLA-A,PS
RBP2,MTRNR2L2,ACTG1,CFL1,BTBD9,MAVS,HTT,HSP90AB1,HSP90AA1,ACTR2,VIM,UBC,KIDINS
FLNA,MAP1B,ATP2B4,CALR,FSTL1,ABHD2,TUBB2B,COL5A1,GPI,RACK1,BSG,ATP5F1B,MYH9,AF
SG,PLXNB2,TRIM28,PSMD5,DCHS1,PKD1,AP2M1,VCL,PAFAH1B1,AP2B1,LAMA5,MSN,RHOA,PSM
PI,BSG,PLXNB2,TRIM28,PSMD5,DCHS1,PKD1,AP2M1,ITGB1,COL11A1,VCL,PAFAH1B1,AP2B1,L
1,UBC,KIDINS220,TMSB4X,ENO1,PIK3R2,ALDOA,PKM,HSP90B1,FADS1,ATP2B4,PGK1,CLU,ENO
S4X,LARP1,RPS9,RHOA,RPL26,KHDRBS1,EIF5A,DDX3X,NGRN,CIRBP,DDX39B,MAPK1,EIF3E,PRK
C,EIF4G2,ENO1,FLNA,NCL,YBX1,ATP2B4,CALR,H3-3B,DDX5,RPL13A,MAGED1,KAT6A,RACK1,TM
KBH5,SRSF9,SRSF1,CASC3,SUPT6H,SRSF3,RBM15B,SRSF6,UPF1,HNRNPA2B1,MCM3AP,POM1
KBH5,SRSF9,SRSF1,CASC3,SUPT6H,SRSF3,RBM15B,SRSF6,UPF1,HNRNPA2B1,MCM3AP,POM1
ABPN1,AHCYL1,SRSF9,SRSF1,SUPT5H,CASC3,SRSF3,GRSF1,ZFP36L1,SRSF6,SYMPK,SRSF2,SRS
PT6H,WBP2,KDM4B,FLCN,BRD4,ATRX,UBR5,KMT2E,OGT,NOC2L,KDM4A,TRIP12,PINK1,AUTS2,
RTN4,GAP43,RHOA,WDR1,PTN,CDH2,MMP14,CTNNB1,GPC1,HDAC2,DNER,VCAN,MDK,SKI,ABCA
AH1B1,HNRNPU,RHOA,CLTC,CKAP5,MAP2,RAC1,MAP1A,TBCD,CAMSAP1,MAPRE1,CLIP3,STMN
DCX,PIK3R2,FLNA,MAP1B,NEFL,ATP2B4,CALR,FSTL1,TMF1,ABHD2,TUBB2B,COL5A1,GPI,RACK1
P1,EID1,SSRP1,SKI,H3-3A,TERF2IP,TAF7,VEGFA,KDM1A,SUPT6H,WBP2,KDM4B,FLCN,BRD4,AT
2,GPC3,DDX3X,CTNNB1,WLS,MDK,PTK7,PSMB2,RAPGEF1,PSMD3,CCAR2,TNKS2,CSNK1D,CTN
PSMD1,SFRP1,PSMD8,PSMB4,TCF3,PSMD4,PSMC3,PSMC1,ABL1,PSMB1,PSMD13,PSMD7,LDB1,
RCA5,PSMD2,RBBP4,H1-0,H1-10,PARP1,TSPYL4,RAD23B,PSMB2,H3-3A,TAF7,SMARCA4,PSMD
RBP2,MTRNR2L2,ACTG1,CFL1,BTBD9,MAVS,HTT,HSP90AB1,HSP90AA1,ACTR2,VIM,UBC,KIDINS
COL6A1,HSP90B1,PRPF8,ATP2B4,COL4A1,CLU,SOX4,GNAS,RACK1,GNB1,APP,GSTP1,IGF2,VCP,
C,EIF4G2,ENO1,FLNA,NCL,YBX1,CALR,H3-3B,DDX5,RPL13A,MAGED1,KAT6A,RACK1,TMBIM6,G
,RHOA,NR2F1,CRMP1,PRKCSH,ITM2C,MAP2,ARHGDIA,HDAC2,CAPZB,FYN,YWHAH,B2M,PLXNA
11,DNAJA1,ATF4,RPL26,MCL1,SFPQ,DDX3X,RPS7,MIF,HNRNPK,BCLAF1,PTTG1IP,CDKN1A,PARP
DCN,GNAI3,VPS35,OPTN,SUPT5H,BNIP3L,ATP6V1B2,UBQLN1,VDAC1,UBQLN2,RAB3GAP2,HDAC
PS8,RPL7,RPL5,RPL37A,MAPT,RPL19,RPL15,RPL18,RPL11,RPLP1,RPL10,RPS17,RPS16,RPL23,
,ATP5F1B,APP,ATP5F1A,VCP,TPI1,PGAM1,LDHA,HK1,RHOA,PFKL,ATP1A2,ATP6V0C,PARP1,ATP5
H1B1,GAP43,NLGN2,SPTBN2,PTN,CDH2,CAPRIN1,VPS35,ACTN1,ARF4,FYN,MESD,PTPRS,CDC42,
G2,DBN1,FLNA,MAP1B,PKM,NEFL,CALR,HSPA1B,SOX4,MAGED1,DVL3,RACK1,P4HB,GDI1,PLXN
CK1,BSG,GDI1,TRIM28,YWHAG,MAPT,YWHAQ,MIDN,DNAJA1,ARF1,CTSD,LLGL1,KIF5B,NLGN2,C
M,EIF4G2,ENO1,NCL,PKM,YBX1,CALR,TMF1,HSPA1B,CLU,DDX5,SOX4,GPI,BSG,SPTBN1,MYH9,
MN1,PFN1,SPTBN2,DDX3X,MAP2,TBCD,NOP53,MAPRE1,LUZP6,CAPZB,CLIP3,STXBP1,PFN2,CDC
D2,SFPQ,SKP1,PSMB2,PSMD3,PRKDC,FYN,PSMA7,EP300,PSMD1,PSMD8,XRCC6,PSMB4,PDPK1,
DNAJA1,RPL23,RPS20,DDX3X,RPS15,RPS7,NDFIP1,RAB1A,SKP1,PTTG1IP,U2AF2,SQSTM1,RPL37
PSMD8,PSMB4,TCF3,PSMD4,PSMC3,PSMC1,ABL1,PSMB1,NOTCH1,PSMD13,PSMD7,LDB1,PSME
B1,RHOA,MEGF8,MMP14,DDX3X,MACF1,L1CAM,DDX39B,H3-3A,VEGFA,LUZP6,RAB11A,BMPR2,
,BRSK1,PSMD8,PSMB4,TAOK1,INTS3,VPS4A,PSMD4,PSMC3,PSMC1,USP47,PSMB1,TAOK2,CUL
,GJA1,CKAP5,BCLAF1,CPSF2,DDX39B,SRSF5,TOMM20,U2AF2,CPSF1,PABPN1,ALKBH5,HNRNPA
,DDX17,GNB1,GSTP1,IGF2,YWHAG,ZMIZ1,DNAJA1,HNRNPU,RHOA,ATP1A1,PIP4K2B,HMGCS1,PI
A1,MCL1,PRDX2,DDX3X,MIF,CTNNB1,PEA15,PTTG1IP,GNAI3,GRINA,COL2A1,HSPB1,KDM1A,QAF
,IGFBP2,GNAS,BSG,APP,GSTP1,IGF2,YWHAG,RPL23,JUND,SRSF4,PIP4K2B,CCN2,PRKAR1A,GJA
F1,L1CAM,VEGFA,RAB11A,BMPR2,ADNP,CYFIP1,NTN1,LIMK1,ISLR2,GOLGA4,SRF,RUFY3,ANAP
C,EIF4G2,ENO1,FLNA,NCL,YBX1,CALR,H3-3B,DDX5,RPL13A,MAGED1,KAT6A,RACK1,TMBIM6,I
A1,YBX1,CALR,FSTL1,COL5A1,COL4A1,PSAP,PGK1,DDX5,SOX4,MAGED1,DVL3,GNAS,DDX17,GF

RPS3,HMGB1,CTSD,RHOA,TMEM259,BEX3,CLEC7A,CSNK1E,CCN2,MMP14,DDX3X,CYFIP2,HDAC
RF1,RHOA,WDR1,PFN1,TPM1,CCN2,CKAP5,RAC1,NOX4,ACTR3,NCKAP1,MAPRE1,ARPC2,DSTN,
D2,COL5A1,H3-3B,GNAS,GNB1,MYH9,FN1,HMGB1,ITGB1,ANXA2,VCL,CTNNA1,NACA,COL1A2,G
,PSMB4,TCF3,PSMD4,PSMC3,PSMC1,ABL1,PSMB1,PSMD13,PSMD7,LDB1,PSME3,PSMC5,PSMC
1,HSP90AA1,ACTR2,VIM,SPARC,UBC,TMSB4X,RPL3,TUBA1B,NCL,HSP90B1,PRPF8,YBX1,HSPA1
OX4,MAGED1,KAT6A,DVL3,DDX17,APP,NPM1,IGF2,UBB,TRIM28,PKD1,SSBP2,ZNF43,HMGB1,AC
OX4,MAGED1,KAT6A,DVL3,DDX17,APP,NPM1,IGF2,UBB,TRIM28,PKD1,SSBP2,ZNF43,HMGB1,AC
,MAP1B,NEFL,CALR,HSPA1B,SOX4,P4HB,GDI1,PLXNB2,MAPT,FN1,SPEN,HMGB1,ZMIZ1,TIMP2,
OX4,MAGED1,KAT6A,DVL3,DDX17,APP,NPM1,IGF2,UBB,TRIM28,PKD1,SSBP2,ZNF43,HMGB1,AC
,IGF2,PSMD5,DCHS1,CHSY1,PKD1,BASP1,AP2M1,AGRN,ZMIZ1,COL11A1,PAFAH1B1,AP2B1,CTN
M28,YWHAG,YWHAQ,ARF1,KIF5B,NLGN2,CD81,CCT7,OAZ1,GJA1,TARDBP,WLS,MYH10,YWHAB,
DA2B,TOMM20,NOP53,RAD23B,ASXL1,PSMB2,PSMD3,EIF3F,USP7,HGS,PSMA7,EP300,PSMD1,U
A,JUN,TARDBP,PPIB,PABPN1,MAPK1,SMARCA4,FBLN1,B2M,HGS,EP300,KPNA2,PHB2,BRD4,UBA
TBN2,CKAP5,MAP1A,CAMSAP1,DSTN,LUZP6,CAPZB,STMN3,PPP1R9B,HDAC6,TMOD2,TAOK1,VPS
NAS,BSG,GSTP1,IGF2,YWHAG,JUND,SRSF4,PIP4K2B,CCN2,PRKAR1A,GJA1,MYO5A,SESN3,APPL
STAT3,LBH,PSMD1,PSMD8,PSMB4,TCF3,LTBP3,PSMD4,PSMC3,PSMC1,ABL1,KDM3A,PSMB1,NO
RN,PAFAH1B1,RHOA,CLSTN1,NLGN2,PTN,CDH2,SEPTIN11,CAPRIN1,DLG5,VPS35,CLSTN3,ARF4,
AC1,TBCD,CAMSAP1,MAPRE1,CLIP3,DCTN1,AKAP9,FKBP4,CLASP2,TUBB4A,ABL1,HSPA1A,MECP
AC1,TBCD,CAMSAP1,MAPRE1,CLIP3,CSNK1D,DCTN1,AKAP9,FKBP4,TUBGCP6,CLASP2,TUBB4A,
,SEC23A,PSMA7,PSMD1,ITGB5,PSMD8,PSMB4,PSMD4,ITGAV,PSMC3,PSMC1,TAPBP,PSMB1,PSM
GED1,DVL3,RACK1,CALM3,APP,CALM1,NPM1,IGF2,VCP,UBB,HACD3,PKD1,RPS3,HMGB1,ANXA2,
ET,PPP2R1A,UCHL1,MIDN,DNAJA1,HNRNPU,LPCAT1,IGFBP3,PTN,PIP4K2B,PRKAR1A,TARDBP,Y
P,UBQLN1,CSNK1D,HECTD1,BAG6,ARIH1,SGTA,UBQLN2,DNAJB2,ATXN3,GSK3A,PSMC3,PSMC1
B4,CLU,MAGED1,DVL3,RACK1,CALM3,APP,GSTP1,CALM1,NPM1,PLXNB2,IGF2,UBB,YWHAG,FN
BMX,SRSF9,SRSF1,PTBP1,KDM1A,HNRNPM,STRAP,RBM15B,SRSF6,TRA2B,SRSF2,RBM5,SAP18,
B1,CCT6A,CTBP1,CCT2,TCP1,CCT3,CCT4,H3-3A,TERF2IP,VEGFA,MAPK1,KDM1A,CCT8,TNKS2,H
11,ITGB1,ANXA2,LAMA5,CLSTN1,KIF5B,CD81,CDH2,RAB7A,MMP14,CLTC,MYO5A,MACF1,APPL1
20,ATP2B4,CLU,MAGED1,DVL3,RACK1,SPTBN1,CALM3,APP,GSTP1,SMG1,CALM1,NPM1,PLXNB2
APT,ITGB1,ARF1,AKAP12,SLC4A8,NCS1,RPL22,CLSTN1,ATF4,KIF5B,NLGN2,ATP2A2,NISCH,PTN
SC,NRBP2,USP22,ACTG1,EEF2,DPYSL3,CFL1,BTBD9,XRCC2,MAVS,HTT,HSP90AB1,HSP90AA1,AC
RN,PAFAH1B1,RHOA,CLSTN1,NLGN2,PTN,CDH2,SEPTIN11,CAPRIN1,DLG5,VPS35,CLSTN3,ARF4,
APT,ITGB1,ARF1,AKAP12,SLC4A8,NCS1,RPL22,CLSTN1,ATF4,KIF5B,NLGN2,ATP2A2,NISCH,PTN
NR2L8,ATP2B4,YWHAE,GPI,RACK1,CALM3,APP,RPL5,TMBIM6,GSTP1,CALM1,NPM1,YWHAG,MA
ET,PPP2R1A,UCHL1,MIDN,DNAJA1,HNRNPU,LPCAT1,IGFBP3,PTN,PIP4K2B,PRKAR1A,TARDBP,Y
1B1,SCARB2,ENC1,PTN,CAPRIN1,TMEM30A,MDK,PTK7,RAPGEF1,KDM1A,FYN,EP300,AP2A1,GR
Q,KIF5B,CD81,OAZ1,XPO1,TARDBP,YWHAB,IPO5,TMEM30A,NUCB1,VAMP2,MAPK1,NPEPPS,BNI
,RPS3,PXDN,NFE2L1,LDHA,ATF4,ATP2A2,MCL1,SFPQ,TPM1,MMP14,PRDX2,JUN,CTNNB1,SESN
1,COL1A2,RHOA,NISCH,JUN,CADM4,RHOB,RAC1,ARHGDIA,CDKN1A,SQSTM1,NCKAP1,NOTCH2,
C,EIF4G2,ENO1,FLNA,NCL,YBX1,ATP2B4,CALR,H3-3B,DDX5,RPL13A,MAGED1,KAT6A,RACK1,T
TNNB1,MMP2,HSBP1,MMP15,ITGB5,SOX2,ITGAV,COL12A1,RTF1,MACROH2A1,CTR9,COL7A1,P
DA2B,TOMM20,NOP53,RAD23B,ASXL1,PSMB2,PSMD3,EIF3F,USP7,HGS,PSMA7,EP300,PSMD1,U
1,ANXA2,LAMA5,KIF5B,CD81,CDH2,RAB7A,MMP14,CLTC,MYO5A,MACF1,APPL1,SQSTM1,VAMP2
JA1,RPL26,MCL1,SFPQ,DDX3X,RPS7,MIF,HNRNPK,BCLAF1,PTTG1IP,PARP1,GRINA,HSPB1,KD
,PTN,MAP2,CAPRIN1,FYN,YWHAH,CDC42,RERE,MAP6,FARP1,CTNND2,HDAC6,PTEN,ABI2,DCLK1
1,HSP90AA1,ACTR2,VIM,UBC,TMSB4X,RPL3,TUBA1B,NCL,HSP90B1,PRPF8,YBX1,HSPA1B,RPLP

1,BASP1,AGRN,PAFAH1B1,NACA,RTN4,MEGF8,GPAT4,GJA1,MACF1,MAP2,L1CAM,PPIB,CDKN1A,
AC1,MAP1A,TBCD,CAMSAP1,MAPRE1,CLIP3,STMN3,CSNK1D,DCTN1,HDAC6,MAPRE2,AKAP9,TAC
HDRBS1,DDX3X,TMED2,EIF3E,HSPB1,EIF2AK1,EIF5B,PPP1R15A,EIF3H,EIF4G3,MTOR,EIF4G1,PA
CT2,PARP1,TCP1,CCT3,CCT4,TERF2IP,MAPK1,CCT8,TNKS2,USP7,HNRNPD,YLPM1,CCT5,ATRX,U
2A2,PTGES3,HDAC2,HSBP1,CDKN1A,HSPH1,MAPK1,CCAR2,RBBP7,EP300,POM121C,FKBP4,ATX
PPP2R1A,TPT1,STMN1,ITGB1,UCHL1,TIMP2,DNAJA1,PAFAH1B1,RHOA,MCL1,PEBP1,DDX3X,MIF,
PS8,RPL7,RPL5,RPL37A,RPL19,RPL15,RPL18,RPL11,RPLP1,RPL10,RPS17,RPS16,RPL23,RPL29
DCX,PIK3R2,FLNA,MAP1B,ATP2B4,CALR,FSTL1,TMF1,ABHD2,TUBB2B,COL5A1,GPI,RACK1,BSG,
DCX,PIK3R2,FLNA,MAP1B,ATP2B4,CALR,FSTL1,TMF1,ABHD2,TUBB2B,COL5A1,GPI,RACK1,BSG,
MC1,VEGFA,PLEC,ACTN1,CORO1C,BCR,SFRP1,PTEN,MAPRE2,CTTN,CLASP2,PDPK1,DAPK3,PTPN
T7,TARDBP,CCT6A,IPO5,GLUL,CCT2,PARP1,TCP1,CCT3,CCT4,MAPK1,FYN,CCT8,CCT5,FLCN,UB
1,GSTP1,CALM1,IGF2,UBB,FN1,PSMD5,HACD3,PPP2R1A,RPS3,HMGB1,UCHL1,NCAM1,TIMP2,DN
.3,RPS27A,PSMD8,PSMB4,PSMD4,PSMC3,PSMC1,PSMB1,FZR1,PSMD13,PSMD7,PSME3,PSMC5,
IB1,ANXA6,CHD2,PSMB2,MAPK1,PSMD3,PRKDC,CORO1C,PSMA7,STAT3,LBH,PSMD1,SFRP1,PSM
,GRINA,HSPB1,KDM1A,CCAR2,HYOU1,MAPK8IP1,HIF1A,GPX1,TRAP1,SOD2,NOC2L,NDUFA13,BA
N1,COL18A1,COL4A2,FN1,ITGB1,ANXA2,AGRN,TIMP2,COL11A1,PXDN,LAMA5,COL1A2,COL5A2,
F1,CLTC,CTNNB1,RAC1,PTK7,PSMB2,VEGFA,PSMD3,VANGL2,CDC42,PSMA7,PSMD1,AP2A1,PHE
SMD2,JUN,CYFIP2,RAC1,APPL1,SKP1,ACTR3,NCKAP1,PSMB2,RAPGEF1,MAPK1,ARPC2,PSMD3,
ABARAP,UBQLN1,CSNK1D,HECTD1,BAG6,ARIH1,SGTA,UBQLN2,PTEN,DNAJB2,ATXN3,GSK3A,PSI
C1,USP7,BAG6,SGTA,CSNK2A1,EIF3H,OGT,BAG5,SDCBP,PARK7,USP14,HIPK2,MARCHF7,PSMF1
ID1,UBA52,RPS27A,PSMD8,PSMB4,PSMD4,PSMC3,PSMC1,PSMB1,CUL1,PSMD13,PSMD7,PSME3
APT,UCHL1,AP3D1,PAFAH1B1,HNRNPU,KIF5B,LRPPRC,SFPQ,ACTN4,MYO5A,LAMP1,MAP2,RAB1
ISP90AA1,ACTR2,TUBB,FTH1,RPS6,FLNA,ALDOA,PKM,FTL,IGFBP2,HSPA1B,PSAP,CLU,SOX4,GNA
N1,COL18A1,COL4A2,FN1,ITGB1,ANXA2,AGRN,TIMP2,COL11A1,PXDN,LAMA5,COL1A2,COL5A2,
S35,FYN,CDC42,DCTN1,CTNND2,HDAC6,PTEN,CTTN,ABI2,ADAM10,UBE3A,TANC2,DIP2A,IGF1R,D
N1,COL18A1,COL4A2,FN1,ITGB1,ANXA2,AGRN,TIMP2,COL11A1,PXDN,LAMA5,COL1A2,COL5A2,
F1A,FN1,HMGB1,ITGB1,PAFAH1B1,AKAP12,RTN4,LAMA5,RHOA,PFN1,PTN,CDH2,GPC3,MEGF8,
DL5A2,GPC3,MEGF8,MMP14,NANOG,PRKAR1A,GJA1,CTNNB1,WLS,MMP2,HSBP1,KDM6B,DUSP
DLG5,ARF4,PTPRS,CDC42,CTNND2,HDAC6,PTEN,ABI2,ADAM10,MTOR,UBE3A,TANC2,DIP2A,IQS
MAVS,HTT,HSP90AB1,HSP90AA1,ACTR2,VIM,UBC,KIDINS220,TMSB4X,NUMA1,PIK3R2,FLNA,HSP
M1,UBB,MAPT,ITGB1,UCHL1,AP3D1,ARF1,PAFAH1B1,MAP4,HNRNPU,KPNB1,LLGL1,KIF5B,LRPF
SP90AB1,HSP90AA1,FTH1,RPS6,PIK3R2,SLC9A7,ALDOA,FTL,HSP90B1,ATP2B4,CALR,FSTL1,HSP
PFN1,CKAP5,RAC1,ACTR3,NCKAP1,MAPRE1,ARPC2,GRB2,PFN2,DCTN1,EVL,CTTN,BRK1,AKAP9,
IAQ,ANXA2,ARF1,MSN,KIF5B,CD81,OAZ1,KHDRBS1,TARDBP,YWHAB,MAP2,IPO5,TMEM30A,VAMI
SPTBN1,FSCN1,SPTAN1,MARCKS,MARCKSL1,TMSB10,STMN1,ARF1,CTNNA1,TPM4,RHOA,WDR1
F2S2,EIF3H,EIF3G,MTOR,EIF4G1,RBM4,EIF3B,EIF3M,EIF3J
.1,DCX,TUBA1B,FLNA,MAP1B,NEFL,ATP2B4,TMF1,HSPA1B,TUBB2B,KIF1A,STMN2,MYH9,APP,NPI
TMN1,RPL11,ANXA2,H1-0,CTNNB1,PARP1,SKI,PPP2CA,MAPRE1,EIF3E,KDM1A,B2M,EP300,FLOT
,CTNNB1,CTBP1,HDAC2,NNAT,EID1,MORF4L2,YEATS2,PLOD1,TAF7,KDM1A,MSL1,SUPT6H,BAG
SMD1,OAZ2,PSMD8,PSMB4,PSMD4,PSMC3,PSMC1,PSMB1,PSMD13,PARK7,PSMD7,PSME3,PSM
IG1,NPM1,PLD3,HSPD1,VCP,UBB,TRIM28,NONO,DDB1,RPS3,HMGB1,MORF4L1,HNRNPU,SMAR
ATP2B4,TMF1,COL5A1,COL4A1,PGK1,SOX4,DVL3,GNAS,GPI,BSG,ATP5F1B,MYH9,PLXNB2,COL1
C1,USP7,BAG6,SGTA,CSNK2A1,EIF3H,OGT,BAG5,SDCBP,PARK7,USP14,HIPK2,WAC,UBXN1,RYBF
2,CALR,FSTL1,HSPA1B,CLU,DVL3,GNAS,HTRA4,RACK1,GNB1,MYH9,KLRD1,TMBIM6,GSTP1,VCF
M28,PKD1,ITGB1,BTF3,ZMIZ1,RTN4,PRKCSH,CTNNB1,SLC35E2B,TMED2,NOTCH2,VEGFA,MAPK

:K1,BSG,GDI1,TRIM28,YWHAG,YWHAQ,MIDN,DNAJA1,ARF1,LLGL1,KIF5B,NLGN2,CD81,OAZ1,XI
N1,PAFAH1B1,HNRNPU,RHOA,XPO1,CLTC,MACF1,CKAP5,LAMP1,MAP2,RAC1,MAP1A,TBCD,CAM
:2,FYN,CDC42,AP2A1,NCSTN,ANKS1A,RBPJ,AP2A2,PTPN11,SDCBP,ARPC1A,EFNB2,EFNB1,NCK2,
IAGED1,DVL3,GNAS,RACK1,CALM3,APP,GDI1,CALM1,NPM1,HSPD1,IGF2,VCP,UBB,MAPT,FN1,HA
TNNB1,MMP2,HSBP1,LAMC1,DUSP4,MMP15,ZFP36L1,ITGB5,SSBP3,SOX2,ITGAV,COL12A1,RTF
ATP5F1A,YWHAG,MAPT,YWHAQ,CXADR,VAT1,JUN,YWHAB,DCN,TOMM20,SQSTM1,PARP1,HSPA4
:ALM3,CALM1,NPM1,IGF2,UBB,MAPT,HACD3,PKD1,SERINC1,RPS3,HMGB1,RPLP1,XRCC5,RHOA
MC1,VEGFA,PLEC,ACTN1,CORO1C,BCR,SFRP1,PTEN,CTTN,CLASP2,PDPK1,DAPK3,PTPN11,ABL1,
'A,PRDX1,PRKDC,B2M,SOD1,ADAR,STAT3,PRMT1,ZFP36L1,HIF1A,SLC25A5,BAP1,IREB2,KMT2E,
I,RACK1,CALM3,APP,GDI1,TMBIM6,CALM1,NPM1,PLXNB2,HSPD1,VCP,MAPT,FN1,HACD3,SET,EC
JBB,TRIM28,MAPT,NONO,DDB1,RPS3,HMGB1,TPT1,MORF4L1,DNAJA1,SMARCA5,XRCC5,CBX5,
!3,PIP4K2B,PRKAR1A,GJA1,MYO5A,SESN3,APPL1,ATP6V0C,SRSF5,VAMP2,PARP1,PTPRA,AHCYL1
F2,ATP5F1A,VCP,TPI1,HMGB1,PGAM1,LDHA,HK1,RHOA,PFKL,TKT,PTGES3,MDH2,PPP1CB,PGD,
2B4,CALR,TMF1,ABHD2,H3-3B,CLU,DDX5,GNAS,DDX17,GNB1,APP,GSTP1,NPM1,ZMIZ1,DNAJA1,
H3-3B,DDX5,MAGED1,KAT6A,TMBIM6,IGF2,UBB,TRIM28,SPEN,NONO,YWHAQ,SET,CHD3,BASP1,
3M,EIF3J
M28,YWHAG,YWHAQ,ARF1,KIF5B,NLGN2,CD81,OAZ1,GJA1,TARDBP,WLS,MYH10,YWHAB,RAC1,II
8,MAVS,HSP90AB1,ZNF124,HSP90AA1,ACTR2,UBC,TMSB4X,EIF4G2,ZNF316,ENO1,PIK3R2,FLNA
TNNB1,MMP2,HSBP1,DUSP4,MMP15,ITGB5,SOX2,ITGAV,COL12A1,RTF1,MACROH2A1,CTR9,CC
T3,BCL9L,LBH,PBX1,SFRP1,SMC3,SOX2,ELAVL1,IGF2BP1,RBPJ,TRIM8,SMO,KDM3A,RTF1,NOTC
H3-3B,DDX5,MAGED1,KAT6A,TMBIM6,IGF2,UBB,TRIM28,SPEN,NONO,YWHAQ,SET,CHD3,BASP1,
H2,CTNNB1,GPC1,HDAC2,DNER,MDK,SKI,ABCA2,MAPK1,SRSF1,SOD1,STAT3,CNTNAP1,PTEN,IL6
7,HNRNPC,TADA2B,SFPQ,IK,CTNNB1,CCT6A,CTBP1,CCT2,PARP1,TCP1,CCT3,CCT4,H3-3A,TERF
:GF3,CDK9,WAC,BCOR,CTR9,OTUB1,RNF20,CUL4B,RYBP,UBE2E1,KDM2B,PAF1,RNF40
EM1B,RPS2,DCUN1D2,BAG5,UBE2L3,ABL1,PINK1,ARRDC4,FZR1,PARK7,LIMK1,TRIB3,STUB1,PI
1,MAP1B,PKM,NEFL,ATP2B4,HSPA1B,PGK1,H3-3B,SOX4,RACK1,GDI1,PLXNB2,COL4A2,MAPT,FI
H3-3B,DDX5,MAGED1,KAT6A,TMBIM6,IGF2,UBB,TRIM28,SPEN,NONO,YWHAQ,SET,CHD3,BASP1,
I,PRKDC,B2M,ADAR,STAT3,PRMT1,ZFP36L1,HIF1A,SLC25A5,IREB2,KMT2E,FAM210B,KLF13,HSI
SKP1,CDKN1A,CTDSP2,DDX39B,NOP53,PSMB2,CCND1,PSMD3,CCAR2,TP53BP1,PSMA7,RBBP7,
1D,TMED9,PPP6R1,SEC23A,SEC16A,CUL3,LMAN1,VAPB,RAB1B,VPS4A,FNBP1L,CHMP4B,USO1,S
,CORO1C,BCR,SFRP1,PTEN,CTTN,CLASP2,PDPK1,DAPK3,PTPN11,ABL1,COL16A1,TAOK2,MAP4I
,GSTP1,NPM1,YWHAG,MAPT,SET,EGLN1,PPP2R1A,UCHL1,RPL11,ANXA2,TIMP2,MIDN,DNAJA1,HI
),VEGFA,CDC42,CTNND1,RDX,CADM3,ADAM10,PTPN23,JAM3,NECTIN1,PIP5K1C,ZNF703,NUMBL
KTL3,KLHDC3,SERP1,VAPB,TPP1,ATP6V0D1,GSK3A,SRPRA,TSPYL2,PTPN1,LMNA,SHC1,DNAJB11
RBP2,MTRNR2L2,ACTG1,CFL1,MAVS,HTT,HSP90AB1,HSP90AA1,ACTR2,VIM,UBC,KIDINS220,TM
1C,SFRP1,PTEN,MAPRE2,CLASP2,DAPK3,PTPN11,ABL1,COL16A1,MAP4K4,IQSEC1,LDB1,SLK,LII
MAPT,PDIA3,RPS3,HMGB1,ITGB1,RPL11,DNAJA1,AKAP12,CTNNA1,LDHA,CTSD,IGFBP3,RHOA,AT
H1,UQCRC2,GHITM,NDUFV1,UQCRC1,COX7C,HIF1A,IREB2,SLC25A23,COX6A1,ATP5MC3,NDUF
LBH,PBX1,SFRP1,SMC3,SOX2,ELAVL1,IGF2BP1,RBPJ,TRIM8,SMO,KDM3A,RTF1,NOTCH1,PCM1,
.3,DDX17,APP,NPM1,IGF2,UBB,PKD1,SSBP2,ZNF43,HMGB1,AGRN,ZMIZ1,AP3D1,ZNF770,GTF2F
3,FLNA,MAP1B,NEFL,HSPA1B,TUBB2B,STMN2,MYH9,NPM1,VCP,MAPT,MAP9,DNAH8,PKD1,RPS3,
,UCHL1,MIDN,DNAJA1,HNRNPU,IGFBP3,PIP4K2B,PRKAR1A,TARDBP,YWHAB,CADM4,HMGCR,IPC
ACTR2,UBC,TMSB4X,EIF4G2,ZNF316,ENO1,PIK3R2,FLNA,NCL,FADS1,YBX1,ATP2B4,CALR,TMF1
K3R2,ATP2B4,CLU,MAGED1,DVL3,RACK1,CALM3,APP,GSTP1,CALM1,NPM1,IGF2,UBB,YWHAG,M
F2F1,ATP2A2,CRMP1,TLE5,NES,CSNK1E,MAP2,CDKN1A,PPP2CA,MAPRE1,KDM1A,B2M,CDC42,EI

8,MAVS,HSP90AB1,ZNF124,HSP90AA1,ACTR2,UBC,TMSB4X,EIF4G2,ZNF316,ENO1,PIK3R2,FLN
ACTR2,UBC,TMSB4X,EIF4G2,ZNF316,ENO1,PIK3R2,FLNA,NCL,FADS1,YBX1,ATP2B4,CALR,TMF1
HSPH1,MAPK1,CCAR2,EP300,POM121C,FKBP4,MTOR,CREBBP,BAG5,HSPA1A,DNAJB1,NUP153,
CC5,ATF4,ATP2A2,MCL1,SFPQ,TPM1,PRDX2,DDX3X,JUN,CTNNB1,RHOB,HDAC2,MMP2,CHD6,P
8,MAVS,HSP90AB1,ZNF124,HSP90AA1,ACTR2,UBC,TMSB4X,EIF4G2,ZNF316,ENO1,PIK3R2,FLN
,CDH2,PRDX2,RPS19,PKN1,ASXL1,VEGFA,PRDX1,PRKDC,B2M,SOD1,ADAR,STAT3,PRMT1,ZFP36
,CDC37,UBQLN1,DYNLL1,VDAC1,PHB2,UBA52,HDAC6,RPS27A,SLC25A5,MAP1LC3B,AMBRA1,P
42,CTNND2,HDAC6,PTEN,CTTN,ABI2,ADAM10,UBE3A,TANC2,DIP2A,IGF1R,DHX36,DLG4,ARHGA
TBP1,RBM15B,SRSF6,TRA2B,SRSF2,RBM5,SAP18,THRAP3,FXR2,RBM4,RBM25,FXR1,CELF3,RE
2M,HGS,KPNA2,PHB2,UBA52,ULK1,RPS27A,VAPB,SCRIB,DAG1,HIPK2,TRIM27,RXRA,KPNA3,M
P8,HECTD1,VANGL2,TCTN1,ZFP36L1,BRD2,SFRP1,SPINT2,HIF1A,ATP6AP2,SSBP3,SEMA3C,PHG
B,CD81,OAZ1,TARDBP,YWHAB,IPO5,TMEM30A,VAMP2,MAPK1,NPEPPS,FYN,YWHAH,SREBF2,CD
RIM28,COL4A2,FN1,ITGB1,COL11A1,LAMA5,COL5A2,PFN1,GPC3,SULF1,MEGF8,MMP14,NANC
U,MAGED1,DVL3,RACK1,CALM3,APP,GSTP1,CALM1,NPM1,IGF2,UBB,YWHAG,HACD3,PPP2R1A,
A,EIF3H,OGT,BAG5,SDCBP,PARK7,USP14,WAC,UBXN1,RYBP
A,EIF3H,OGT,BAG5,SDCBP,PARK7,USP14,MARCHF7,PSMF1,WAC,USP19,UBXN1,GABARAPL2,R
20,RPL26,RBBP4,PIP4K2B,RPS15,RPS7,MIF,HNRNPK,HDAC2,PTTG1IP,CDKN1A,NOP53,TAF15,S
ACTR2,UBC,TMSB4X,EIF4G2,ZNF316,ENO1,PIK3R2,FLNA,NCL,FADS1,YBX1,ATP2B4,CALR,TMF1
ZB,CORO1C,AKIRIN1,CDC42,CTTN,SLIT2,BRK1,ABI2,WASF2,MTOR,SNX2,AUTS2,CYFIP1,CD44,G
DC,ADAR,PSMA7,PSMD1,FLCN,SFRP1,EML1,PSMD8,PSMB4,SSBP3,TCF3,PSMD4,PSMC3,PSMC1
,NANOG,PRKAR1A,GJA1,CTNNB1,WLS,MMP2,HSBP1,KDM6B,DUSP4,BMPR2,MMP15,ITGB5,SO
GAM1,KPNB1,IGFBP3,PSMD2,OAZ1,PPP1CB,PARP1,PSMB2,AZIN1,PSMD3,ODC1,ACADVL,USP7,
,RPS6,ALDOA,PKM,FTL,IGFBP2,HSPA1B,PSAP,CLU,SOX4,SLC2A3,GPI,MYH9,APP,SPTAN1,GSTP1
7A,EIF3G,EIF3B
BPN1,AHCYL1,SRSF9,SRSF1,SUPT5H,CASC3,SSB,SRSF3,GRSF1,ZFP36L1,SRSF6,SYMPK,SRSF2
1,WDR45B,PHB2,HDAC6,HIF1A,CTTN,SLC25A5,MAP1LC3B,GABARAPL1,GSK3A,BNIP3,AMBRA1,
1,WDR45B,PHB2,HDAC6,HIF1A,CTTN,SLC25A5,MAP1LC3B,GABARAPL1,GSK3A,BNIP3,AMBRA1,
1B1,HNRNPU,KIF5B,LRPPRC,SFPQ,LAMP1,MAP2,RAB1A,MAP1A,TNPO1,DYNLL2,HSPB1,CLIP3,
AN1,TMSB10,STMN1,ARF1,RHOA,ATP1A1,WDR1,ATP2A2,PFN1,SPTBN2,TPM1,CCN2,CYFIP2,RH
GOLM1,TARDBP,PPP2CA,SERBP1,DCTN1,SF1,VPS4A,IST1,CHMP4B,UBE2I,RANGAP1,SUN1,CHM
2,XPC,GPS1,CUL4B,COPS8
,RPS3,HMGB1,ITGB1,RPL11,DNAJA1,AKAP12,CTNNA1,LDHA,CTSD,IGFBP3,RHOA,ATF4,RPL26,
,RPS3,HMGB1,ITGB1,RPL11,DNAJA1,AKAP12,CTNNA1,LDHA,CTSD,IGFBP3,RHOA,ATF4,RPL26,
CSH,PTN,DDR1,SULF1,PLXNA1,PRDX2,RPS15,HMGCS1,GPAT4,CTNNB1,WLS,TNC,DDX39B,SRS
SP1,AGRN,PAFAH1B1,NACA,RHOA,MEGF8,MMP14,DDX3X,MACF1,L1CAM,PPIB,DDX39B,H3-3A,
,MED13,PSMC3,MECP2,SUB1,TNRC6A,ATF7IP,CDK4,DHX36,PSMC5,PSMC2,PSMC4,SRF,SETX,S
A,WDR1,PFN1,NES,TPM1,CCN2,CKAP5,RAC1,NOX4,ACTR3,NCKAP1,MAPRE1,ARPC2,DSTN,GRB
Q,RAB7A,JUN,TARDBP,PPIB,PABPN1,MAPK1,SMARCA4,FBLN1,B2M,HGS,EP300,KPNA2,PHB2,B
1,IPO5,PTTG1IP,CDKN1A,TNPO1,MAPK1,IPO7,TXNIP,ADAR,STAT3,KPNA2,PHB2,MBTPS1,POM1
8,MAVS,HSP90AB1,ZNF124,HSP90AA1,ACTR2,UBC,TMSB4X,EIF4G2,ZNF316,ENO1,PIK3R2,FLN
PS14,DDX3X,RPS15,RPS5,RPS19,RPS10,RPS27,RPL23A,NOP53,RPL38,PRKDC,DHX30,RPS28,TR
M1,YWHAG,MAPT,ITGB1,AGRN,ARF1,PAFAH1B1,AKAP12,SLC4A8,NCS1,RPL22,CLSTN1,ATF4,K
1,KMT2E,NSD2,OGT,KDM4A,CREBBP,PAGR1,AUTS2,KDM3A,RTF1,MACROH2A1,MECP2,TRMT11
P1,PITPNA,SOD1,CDC42,HNRNPA2B1,CAPZA1,SOD2,TYK2,JAK1,TALDO1,RAP1B
,ARCN1,TMED2,KDELR2,DYNC1LI2,DYNLL2,CAPZB,KDELR1,ARF4,CSNK1D,TMED9,HYOU1,PPP

90AB1,HSP90AA1,ACTR2,SPARC,TMSB4X,NUMA1,EIF4G2,DBN1,ENO1,FLNA,MAP1B,PKM,NEFL,
1,DNAJA1,IGFBP3,PRKAR1A,TARDBP,YWHAB,CADM4,HMGCR,IPO5,TRIB2,CDKN1A,CTDSP2,PKN
ARCA5,XRCC5,CHD4,RBBP4,DDX3X,H1-0,H1-10,TOP2B,SMCHD1,CHD6,PARP1,TSPYL4,RAD23B,
RDBP,APPL1,IPO5,PTTG1IP,CDKN1A,TNPO1,MAPK1,IPO7,TXNIP,ADAR,STAT3,KPNA2,PHB2,MBTF
A2,MCL1,SFPQ,TPM1,PRDX2,JUN,CTNNB1,RHOB,HDAC2,MMP2,CHD6,PARP1,NOX4,PRDX5,KDM
,APP,HSPD1,UBB,MAPT,NONO,SET,AP2B1,RHOA,ATF4,TMEM259,MCL1,NES,JUN,CTNNB1,DNAJ
MSB10,STMN1,ARF1,RHOA,WDR1,PFN1,SPTBN2,TPM1,CCN2,CYFIP2,RHOB,COTL1,RAC1,ARHG
P2,TBCD,MAPRE1,LUZP6,CAPZB,CLIP3,PFN2,SLIT2,TMOD2,FKBP4,RDX,CAPZA1,TUBB4A,HIP1R
F1,PSMD3,CTNND1,PSMA7,PSMD1,SFRP1,CUL3,PSMD8,DKK3,PSMB4,SOX2,TLE1,PSMD4,GSK3
,JUN,SESN3,RHOB,HDAC2,MMP2,PRDX5,KDM6B,MAPK1,PRDX1,ADAM9,FYN,SOD1,TXNIP,CAPN
8,RPL23,KPNB1,TARDBP,APPL1,IPO5,PTTG1IP,CDKN1A,TOMM20,HSPA4,TNPO1,MAPK1,IPO7,T
,CD81,PTN,RPS15,TNC,CDKN1A,SRSF5,MDK,SELENON,CCND1,SRSF1,LUZP6,PTPRS,AKIRIN1,S
CCT5,ATRX,TNKS,TPP1,MACROH2A1,GNL3
MYO5A,SESN3,APPL1,ATP6V0C,SRSF5,VAMP2,PARP1,PTPRA,AHCYL1,ATP6AP1,MARS1,GRB2,PR
ACTR2,UBC,TMSB4X,EIF4G2,ZNF316,ENO1,PIK3R2,FLNA,NCL,FADS1,YBX1,ATP2B4,CALR,TMF1
,CTNNB1,DUX4,RHOB,IPO5,SKP1,CDKN1A,CTDSP2,DDX39B,NOP53,PSMB2,CCND1,PSMD3,IPO7
AF11,SF3A1,SF1,SRSF6,RBM5,PRPF19,SF3A2,LUC7L2,SF3A3,SRSF10,CELF3,PSIP1,SNRPD2,SET
,CSNK1D,HECTD1,ARIH1,SGTA,UBQLN2,DNAJB2,GSK3A,ZER1,USP5,HSPA1A,MDM2,FZR1,SUMC
,CCND2,JUN,ATP1A2,HMGCR,MAP1A,MDK,CNTNAP2,MAPK1,ARF4,FYN,B2M,PPP1R9B,CIC,VDA
1,EHMT1,SNRPD3,CARM1,KMT2E,RAB6A,NSD2,OGT,KDM4A,CREBBP,PAGR1,AUTS2,KDM3A,RT
1,EHMT1,SNRPD3,CARM1,KMT2E,RAB6A,NSD2,OGT,KDM4A,CREBBP,PAGR1,AUTS2,KDM3A,RT
B1,PSMC5,HSF1,DNAJB6,PSMC6
PFKM,FOXK2,PFKP
PFKM,FOXK2,PFKP
PFKM,FOXK2,PFKP
2,JUN,ATP1A2,HMGCR,MAP1A,MDK,CNTNAP2,MAPK1,ARF4,FYN,B2M,PPP1R9B,CIC,VDAC1,EP3
1,ARF1,RHOA,WDR1,PFN1,SPTBN2,TPM1,CCN2,CYFIP2,COTL1,RAC1,NOX4,ACTR3,NCKAP1,AR
90AA1,ACTR2,TUBB,FTH1,RPS6,PIK3R2,ALDOA,PKM,FTL,HSPA1B,PSAP,CLU,SLC2A3,GPI,APP,K
RPS3,HMGB1,HNRNPU,XRCC5,CCT7,HNRNPC,H1-0,GJA1,CTNNB1,NDFIP1,H1-10,PTGES3,CCT6
ACTR2,UBC,TMSB4X,EIF4G2,ZNF316,ENO1,PIK3R2,FLNA,NCL,FADS1,YBX1,ATP2B4,CALR,TMF1
ACTR2,UBC,TMSB4X,EIF4G2,ZNF316,ENO1,PIK3R2,FLNA,NCL,FADS1,YBX1,ATP2B4,CALR,TMF1
PL11,MIDN,DNAJA1,HNRNPU,RPL23,RPS20,PIP4K2B,RPS15,RPS7,PRKAR1A,YWHAB,HMGCR,IP
YO5A,SESN3,APPL1,ATP6V0C,SRSF5,VAMP2,PARP1,PTPRA,ATP6AP1,MARS1,GRB2,PRKDC,SMA
BAG6,SGTA,OS9,UBQLN2,UBE2G2,NPLOC4,RHBDD2,DNAJB2,ATXN3,CCDC47,UBXN6,AMFR,UB
5,RHOA,PTN,DDR1,CCN2,ACTN4,MMP14,CTNNB1,CD63,MACF1,RAB1A,RAC1,L1CAM,TBCD,PT
PK1,CCT8,TNKS2,HNRNPD,CCT5,ATRX,HNRNPA2B1,TNKS,PPP1R10,DHX36,MAPK3,GNL3,RAD50
17,GPI,CALM3,GSTP1,CALM1,HSPD1,HMGB1,ZMIZ1,DNAJA1,HNRNPU,RHOA,ATP1A1,PTN,CCN2,
F2,DCHS1,CHSY1,MRC2,COL11A1,HNRNPU,TPM4,IGFBP3,COL1A2,RHOA,ATF4,JUND,COL5A2,
B2,KIF5B,PIP4K2B,RAB7A,RAB1A,LAPTM4B,RAB27A,SQSTM1,VAMP2,GNAI3,TMED2,CORO1C,
DBP,IPO5,MAPK1,SUPT6H,PPP1CC,ANP32B,UBR5,PRKACA,SMO,RANGAP1,PTPN11,RBM4,PKIA,
P6V0C,SRSF5,VAMP2,PARP1,PTPRA,ATP6AP1,MARS1,GRB2,PRKDC,SMARCC1,ATP6V1B2,ZFP36
VHAG,MAPT,ITGB1,ARF1,PAFAH1B1,AKAP12,SLC4A8,NCS1,RPL22,CLSTN1,ATF4,KIF5B,NLGN2,
VPS35,BNIP3L,YWHAH,GHITM,SREBF2,VDAC1,EP300,FLCN,HDAC6,HIF1A,CTTN,SLC25A5,GPX1,
PAFAH1B1,RPS14,CD81,WDR1,GPC3,RPS19,JUN,CTNNB1,NDFIP1,CDKN1A,PARP1,H3-3A,ATP6A
NES,SPTBN2,EIF5A,CKAP5,MAP1A,MTRF1L,CAMSAP1,DSTN,LUZP6,CAPZB,STMN3,PPP1R9B,GS

,CDH2,CTNNB1,ATP1A2,MDK,SKI,KDM6B,CNTNAP2,DPYSL2,KDM1A,PLXNA3,PTPRS,TCTN1,PPP1
P1,PITPNA,SOD1,CDC42,HNRNPA2B1,CAPZA1,SOD2,TYK2,JAK1,TALDO1,RAP1B
D1,PSMD8,PSMB4,PDPK1,PSMD4,PSMC3,PRKACA,CREBBP,PSMC1,PSMB1,CUL1,PSMD13,PSMD
JA1,ATP2A2,PTGES3,HDAC2,HSBP1,CDKN1A,HSPH1,MAPK1,CCAR2,SOD1,IGFBP7,RBBP7,EP300
5B,SFPQ,MAP2,MAP1A,HSPB1,SOD1,KIF5A,HIF1A,SPG7,FLOT2,TRAK1,FEZ1,KIF3A,ARL8A,SPG
1,VPS35,RAB11B,MVB12B,NSG2,CORO1C,USP7,HGS,RAB11A,GRIPAP1,DCTN1,UBA52,YKT6,PR
4,CALR,ABHD2,TUBB2B,GPI,RACK1,BSG,ATP5F1B,APP,GSTP1,PLXNB2,ATP5F1A,FN1,TMSB10,H
AMSAP1,DSTN,LUZP6,CAPZB,GSPT1,UPF1,HDAC6,TMOD2,TAOK1,RDX,CAPZA1,CLASP2,BNIP3,H
CDC47,SCAP,AMFR,PTPN1,NCK2,HERPUD1,GSK3B,ATF6B,ATF6,QRICH1,XBP1,TMCO1,EIF2S1,A
,TRAP1,SOD2,BAG5,PINK1,PARK7
ID,PTN,CSNK1E,SFPQ,JUN,TARDBP,HDAC2,PPP1CB,MDK,TOP1,PRKDC,CCAR2,SLIT3,CSNK1D,U
PP1CB,NOP53,ACO2,PYGB,ACADVL,ATP5PB,SDHA,PFKM,IDH1,UQCRC2,GHITM,NDUFV1,UQCR
A,HSPB1,FYN,HGS,CDC42,HIF1A,BRK1,SHC2,WASF2,ITGAV,NIBAN2,AAMP,CYFIP1,PTPN1,NCK2
1C,SFRP1,PTEN,CLASP2,DAPK3,PTPN11,ABL1,COL16A1,MAP4K4,LDB1,SLK,LIMS1,RCC2,DUSP3
1C,SFRP1,PTEN,CLASP2,DAPK3,PTPN11,ABL1,COL16A1,MAP4K4,LDB1,SLK,LIMS1,RCC2,DUSP3
,CDH2,CTNNB1,MDK,KDM6B,CNTNAP2,KDM1A,PLXNA3,PTPRS,PPP1R9B,TRA2B,PTEN,HIF1A,SLI
VHAG,MAPT,ITGB1,ARF1,PAFAH1B1,AKAP12,SLC4A8,NCS1,RPL22,CLSTN1,ATF4,KIF5B,NLGN2,
VHAG,MAPT,ITGB1,ARF1,PAFAH1B1,AKAP12,SLC4A8,NCS1,RPL22,CLSTN1,ATF4,KIF5B,NLGN2,
,COL5A1,SOX4,GNAS,BSG,MYH9,P4HB,APP,PLXNB2,HSPD1,COL18A1,IGF2,FN1,DCHS1,PPP2R1
SMB4,PSMD4,FBXW5,PSMC3,PSMC1,PSMB1,CUL1,PSMD13,PSMD7,CUL4A,PSME3,PSMC5,PSM
D1,PSMD8,PSMB4,PDPK1,PSMD4,PSMC3,PRKACA,CREBBP,PSMC1,PSMB1,CUL1,PSMD13,PSMD
D1,PSMD8,PSMB4,PDPK1,PSMD4,PSMC3,PRKACA,CREBBP,PSMC1,PSMB1,CUL1,PSMD13,PSMD
D3,STMN1,PAFAH1B1,MAP4,HNRNPU,KPNB1,RHOA,SPC25,XPO1,CLTC,CKAP5,RPS27,MAPRE1,D
SMB2,RAPGEF1,PSMD3,CTNND1,PSMA7,PSMD1,NXN,SFRP1,CUL3,PSMD8,DKK3,PSMB4,SOX2,T
PTBN2,CYFIP2,COTL1,ACTR3,NCKAP1,ARPC2,LUZP6,GRB2,CAPZB,PFN2,EVL,CTTN,SLIT2,BRK1
2,CXADR,CD81,XPR1,RAB7A,CTNNB1,LAMP1,CTSB,GRB2,IFITM3,CTNND1,ITGB5,GRK2,VAPB,V
P1,PITPNA,SOD1,CDC42,HNRNPA2B1,CAPZA1,SOD2,TYK2,JAK1,TALDO1,RAP1B
UTS2,KDM3A,RTF1,MACROH2A1,MECP2,EHMT2,NSD1,BCOR,CTR9,RNF20,LMNA,CTCF,WDR5,
VDR1,PFN1,SPTBN2,CYFIP2,COTL1,RAC1,GAS7,ACTR3,NCKAP1,ARPC2,DSTN,LUZP6,GRB2,CAP
C6,PTEN,ABI2,ADAM10,UBE3A,TANC2,DIP2A,DHX36,DLG4,ARHGAP33,SHANK1,OPA1,EPHB2,LRP
OX4,GNAS,BSG,MYH9,P4HB,APP,PLXNB2,HSPD1,COL18A1,IGF2,FN1,DCHS1,PPP2R1A,PKD1,RP
VAMP2,CLSTN3,HSPB1,RAB11A,GRIPAP1,KIF5A,GPC4,SCRIB,ADAM10,NSG1,DAG1,NRXN2,JAM3
,BAG6,SGTA,ALDH3A2,SEC16A,RAB3GAP2,LMAN1,SERP1,VAPB,TRAM1,REEP2,CCDC47,DOP1B,
3X,SKP1,PSMB2,TSPAN6,TERF2IP,PSMD3,PRDX1,PSMA7,TRAF4,EP300,PSMD1,PHB2,PSMD8,PSM
PFN1,SPTBN2,CYFIP2,COTL1,ACTR3,NCKAP1,ARPC2,DSTN,LUZP6,GRB2,CAPZB,PFN2,EVL,CTT
,OS9,UBQLN2,UBE2G2,NPLOC4,RHBDD2,DNAJB2,CCDC47,AMFR,UBE4B,USP14,FAF2,UBE2J1,
H9,FN1,HMGB1,ITGB1,ANXA2,VCL,NACA,COL1A2,RHOA,CBX5,CLEC7A,TPM1,DDR1,PRDX2,PRK
MGB1,AP2M1,ANXA2,ARF1,AP2B1,SLC4A8,MSN,NCS1,KIF5B,ATP2A2,CLEC7A,GPC3,RAB7A,CD63
,MDK,CNTNAP2,KDM1A,PTPRS,PPP1R9B,TRA2B,HIF1A,SLIT2,SLC2A1,BNIP3,SLC38A2,SMO,NTR
IC3,MECP2,SUB1,TNRC6A,ATF7IP,CDK4,DHX36,PSMC5,PSMC2,PSMC4,SRF,SOX9,MED25,MED1

D2,SFPQ,RPS19,LAMP1,SKP1,PSMB2,PSMD3,PRKDC,FYN,PSMA7,EP300,PSMD1,PSMD8,XRCC6,
B,RYBP,UBE2E1,KDM2B,PAF1,RNF40
PFN1,SPTBN2,CYFIP2,COTL1,ACTR3,NCKAP1,ARPC2,DSTN,LUZP6,GRB2,CAPZB,PFN2,EVL,CTT

D1,PSMD8,PSMB4,PDPK1,PSMD4,PSMC3,PRKACA,CREBBP,PSMC1,PSMB1,CUL1,PSMD13,PSMD
,TRIP12,KDM3A,MACROH2A1,BCOR,OTUB1,KDM5A,WDR5,DNMT1,SMARCB1,SIN3A
LI2,DYNLL2,CAPZB,KIF3C,SEC23A,DYNLL1,DCTN1,KIF5A,ACTR1A,AP2A1,KIFAP3,CAPZA1,ACTR1
4,MYO5A,CTNNB1,MAP2,RAB1A,RAB27A,RAB11B,DPYSL2,TMED2,CSNK1D,TMED9,RAB11A,PPP6
/I1,MAPT,TMSB10,STMN1,MAP4,HNRNPU,VAT1,PFN1,HNRNPC,SPTBN2,RAB7A,IK,PRKAR1A,MAP
3,PRMT1,ZFP36L1,HIF1A,SLC25A5,KMT2E,FAM210B,KLF13,HSPA1A,ACVR1B,MAEA,HIPK2,LDB1
B1,SGTA,CSNK2A1,CRTAP,FKBP1A,FKBP4,TRAP1,DNAJB2,CD74,GAK,HSPA1A,DNAJB1,UNC45A,I
PFKM,FOXK2,PFKP
P90AA1,ACTR2,UBC,TMSB4X,NUMA1,PIK3R2,FLNA,CALR,HSPA1B,CLU,DDX5,SOX4,MAGED1,DVL
P2M1,ITGB1,AP2B1,KIF5B,CD81,PTN,CDH2,GPC3,MMP14,CLTC,YWHAB,GPC1,APPL1,SQSTM1,C
NAJB6,PSMC6
SP1,ITGB1,COL11A1,HNRNPU,CTNNA1,NACA,LAMA5,IGFBP3,RHOA,CD81,WDR1,ATP2A2,TPM1,C
JA1,RHOA,NR2F1,ACTN4,ARL4C,ARID1A,PTGES3,NOP53,PARP1,PKN1,ASXL1,H3-3A,TSPAN6,TA
D1,PSMD8,PSMB4,PDPK1,PSMD4,PSMC3,PRKACA,CREBBP,PSMC1,PSMB1,CUL1,PSMD13,PSMD
TP6V0C,GLUL,CDKN1A,GCN1,MARS1,MAPK1,GABARAP,ATP6V1B2,LARS1,SREBF2,BMPR2,WDR4
1-0,ARID1A,H1-10,SMCHD1,TSPYL4,H3-3A,SMARCA4,SMARCE1,SMARCC1,NAP1L4,RBBP7,NAP1
A3,SRSF10,CELF3,PSIP1,SETX,SFSWAP,KHDC4
P2,CALR,GPI,RACK1,BSG,CALM3,APP,GDI1,CALM1,HSPD1,COL18A1,VCP,MAPT,EGLN1,RPS3,LD
IO1,PIK3R2,FLNA,NCL,YBX1,ATP2B4,CALR,TMF1,ZNF99,DDX5,SOX4,MAGED1,KAT6A,DVL3,DDX
/IP2,RTN4,KPNB1,COL1A2,RHOA,NISCH,JUN,CADM4,RHOB,RAC1,ARHGDIA,CDKN1A,NUCB1,SQS
APTM4B,ATG2B,GABARAP,UBQLN1,WDR45B,UBQLN2,RAB3GAP2,MBTPS1,ULK1,ATP6AP2,MAP
I9,FSCN1,PKD1,ITGB1,PAFAH1B1,MAP4,CDC42BPB,CTNNA1,MSN,LLGL1,RHOA,WDR1,GJA1,CK
TN,GJA1,JUN,SESN3,ATP1A2,RAC1,ARHGDIA,TNC,ATP6V0C,GLUL,CDKN1A,CNN2,GCN1,MARS1,I
RF,GRN,STK24,CERS2,DAG1,JAM3,DHFR,NREP,RGMA,EPHA4,MAPK8IP3
RF,ULK1,GRN,STK24,CERS2,DAG1,JAM3,MAP4K4,DHFR,NREP,RGMA,EPHA4,MAPK8IP3
CS1,NLGN2,ATP2A2,CDH2,CTNNB1,DNAJC5,VAMP2,STXBP1,PFN2,SYP,GRIPAP1,PTEN,SV2A,PREI

PK1,CCT8,TNKS2,CCT5,HNRNPA2B1,TNKS,DHX36,MAPK3
4,CALR,ABHD2,GPI,RACK1,BSG,ATP5F1B,APP,GSTP1,PLXNB2,ATP5F1A,FN1,TMSB10,HMGB1,IT
RCC5,PSMD2,SFPQ,RPS19,LAMP1,HLA-B,APPL1,SKP1,PSMB2,PSMD3,CDC37,PRKDC,FYN,ADAR
7,CTNNB1,U2AF2,SQSTM1,RPL37,FYN,NXN,UBR5,DNAJB2,OGT,MTOR,BAG5,TRIP12,ABL1,HSPA
4,CALR,ABHD2,TUBB2B,GPI,RACK1,BSG,ATP5F1B,APP,GSTP1,PLXNB2,ATP5F1A,FN1,TMSB10,H
C4A8,NCS1,CLSTN1,NLGN2,ATP2A2,CDH2,CTNNB1,DNAJC5,VAMP2,STXBP1,PFN2,RAB11A,SYP,C
,ADAM9,FYN,SOD1,TXNIP,CAPN2,PPP2CB,HDAC6,APEX1,GPX1,STK24,TRAP1,SOD2,PCGF2,UBE3
SESN3,TNC,ATP6V0C,GLUL,CDKN1A,GCN1,MARS1,MAPK1,GABARAP,ATP6V1B2,LARS1,SREBF2,I
,TPM1,CCN2,RAC1,NOX4,ACTN1,PFN2,CDC42,SFRP1,EVL,CUL3,ITGB5,CALD1,RDX,CLASP2,WAS
PP2CA,SMARCA4,CCAR2,SLIT3,PLXNA3,PTPRS,BMPR2,PPP1R9B,RBBP7,FLCN,DIP2B,GNG4,SFF
,SOX4,MAGED1,DVL3,BSG,ATP5F1B,MYH9,PLXNB2,COL18A1,UBB,COL4A2,FN1,DCHS1,EGLN1,
PFKM,FOXK2,PFKP
RAP,KDM1A,UBQLN1,MTA1,PTEN,CUL3,RPS2,FKBP1A,DNAJB2,DCUN1D2,GSK3A,UBE3A,SAE1,UI
,CSNK1D,HECTD1,ARIH1,SGTA,UBQLN2,PTEN,DNAJB2,GSK3A,ZER1,USP5,HSPA1A,MDM2,FZR1,
3L,SREBF2,VDAC1,WDR45B,PHB2,HDAC6,HIF1A,CTTN,SLC25A5,MAP1LC3B,GABARAPL1,GSK3A
NB1,MAP2,RAB1A,RAB27A,RAB11B,DPYSL2,TMED2,CSNK1D,TMED9,RAB11A,PPP6R1,SEC23A,K
BE2G2,CUL3,UBR5,UBE2D2,UBE3A,UBE2D3,UBE2K,AMFR,CDC34,CUL1,RNF216,ZNRF1,RNF4,U

I2,RAPGEF1,GRB2,GABARAP,STMN3,ARFGEF2,BCR,FLCN,FLOT1,PPP2CB,CUL3,MAPRE2,LRRC5
L26,RBBP4,PIP4K2B,RPS15,RPS7,MIF,HNRNPK,HDAC2,PTTG1IP,NOP53,TAF15,SSRP1,TAF7,RPL
P53,PSMB2,CCND1,PSMD3,CCAR2,TP53BP1,PSMA7,PRMT1,EP300,PSMD1,BRD4,ZFP36L1,PTEN,
H10,RHOB,RAC1,ARHGDIA,VEGFA,FYN,PLXNA3,PRPF40A,PLXNB1,FMNL2,SEPTIN7,PALM,RDX,P
D,PTEN,PDPK1,EHD1,UBE3A,SORT1,EIF4A3,NTRK2,ACAP2,KAT2A,ARPC3,RAB35,MICALL1,ARF6
MD5,HMGB1,ITGB1,ANXA2,ZMIZ1,AP3D1,PAFAH1B1,XRCC5,RPL22,RHOA,ATF4,PSMD2,RPS14,CI
PQ,JUN,TARDBP,HDAC2,PPP1CB,TOP1,PRKDC,CCAR2,CSNK1D,USP7,ID4,RAI1,HNRNPD,MTA1,E
3
,EIF4G2,DBN1,ENO1,MAP1B,PKM,NEFL,TMF1,HSPA1B,CLU,SOX4,MAGED1,GNAS,RACK1,BSG,S
ACF1,RAC1,DLG5,PTPRA,VPS35,CLSTN3,CNTNAP2,VEGFA,RAPGEF1,CORO1C,PTPRS,LRRC4B,F
MD5,HMGB1,ITGB1,ANXA2,ZMIZ1,AP3D1,PAFAH1B1,XRCC5,RPL22,RHOA,ATF4,PSMD2,RPS14,CI
MAPT,NONO,SET,AP2B1,RHOA,ATF4,TMEM259,MCL1,NES,JUN,CTNNB1,DNAJC5,PARP1,VPS35,M
N1,PFN1,SPTBN2,MAP2,TBCD,CAMSAP1,MAPRE1,LUZP6,CAPZB,CLIP3,PFN2,HDAC6,SLIT2,TMOD

LI2,DYNLL2,CAPZB,KIF3C,SEC23A,DYNLL1,DCTN1,KIF5A,ACTR1A,AP2A1,KIFAP3,CAPZA1,ACTR1
P1,DSTN,LUZP6,CAPZB,HDAC6,TMOD2,TAOK1,RDX,CAPZA1,CLASP2,HDGFL3,CRACD,FLII,NCK2,
3A52,HDAC6,RPS27A,PARK7,UBE2V1,DYNC1I2
H1,UQCRC2,GHITM,NDUFV1,UQCRC1,COX7C,HIF1A,IREB2,TRAP1,SLC25A23,COX6A1,ATP5MC3
FAH1B1,MAP4,HNRNPU,KPNB1,RHOA,SPC25,XPO1,CLTC,GJA1,CKAP5,RPS27,MAPRE1,DYNC1LI
N4,RHOA,KIF5B,PTN,CDH2,APLP1,CTNNB1,ATP1A2,TOP2B,MDK,SKI,KDM6B,CNTNAP2,DPYSL2,I
4A1,PSAP,PGK1,SOX4,MAGED1,DVL3,GNAS,BSG,FASN,PLXNB2,COL18A1,TRIM28,PSMD5,DCHS
SPTBN2,CYFIP2,COTL1,RAC1,GAS7,ACTR3,NCKAP1,ARPC2,LUZP6,GRB2,CAPZB,PFN2,EVL,CTTN
AP4,HNRNPU,KPNB1,RHOA,SPC25,XPO1,CLTC,CKAP5,RPS27,MAPRE1,DYNC1LI2,DYNLL2,RAB11
B,KLF13,HSPA1A,ACVR1B,LDB1,RBFOX2,HOXA5,STAT1,HMGB2,BRD1,MAFB
A,PARK7,USP14,PTPN1,NCK2,HERPUD1,UBXN1,ATF6B,XBP1,OPA1,ABCA7
PT1,RPL26,MCL1,MIF,HNRNPK,SMCHD1,BCLAF1,PTTG1IP,DDX39B,PARP1,TERF2IP,PRKDC,KDI
IP3,FLOT1,PTEN,VPS4A,FLOT2,SYT11,IST1,CHMP4B,UBE2I,GAK,RANGAP1,NRXN2,SUMO1,BANF
DK,ARPC2,PRKDC,CORO1C,FBLN1,CDC42,ABL1,OBSL1,C1QBP,DHX36,CRKL,LIMS1,RCC2,CNTI
3R2,NCL,YBX1,ATP2B4,CALR,TMF1,ZNF99,DDX5,SOX4,MAGED1,KAT6A,DVL3,DDX17,APP,TMBIN
STM1,VAMP2,MAPK1,YWHAH,STXBP1,VGF,RAB11A,SYP,PTEN,BRSK1,NCSTN,NSG1,NTRK2,RIMS
COL5A1,COL4A1,PGK1,SOX4,BSG,ATP5F1B,MYH9,COL18A1,MATR3,COL4A2,FN1,DCHS1,EGLN
,TPM1,CCN2,RAC1,NOX4,ACTN1,PFN2,CDC42,SFRP1,EVL,CUL3,ITGB5,CALD1,RDX,CLASP2,WAS
FLNA,CALR,TMF1,CLU,SOX4,YWHAE,RACK1,BSG,CALM3,GNB1,CALM1,TRIM28,YWHAG,YWHAQ
EN,PDPK1,EHD1,UBE3A,SORT1,EIF4A3,NTRK2,ACAP2,KAT2A,ARPC3,RAB35,MICALL1,ARF6
OB,VPS35,HGS,NCOA4,GRN,VPS41,VPS4A,UBXN6,SORT1,GAK,PINK1,TRAK1,HSPA1A,ATP13A2,F
IN1,CDC42,SLIT2,BRK1,ABI2,WASF2,MTOR,AUTS2,CYFIP1,GOLPH3,CCDC88A,NCK2,WASF3,AJU
F2,MAVS,HSP90AB1,HSP90AA1,ACTR2,VIM,TUBB,FTH1,RPS6,PIK3R2,FLNA,ALDOA,PKM,FTL,IGFI
MC4,SRF,SETX,MED12,MED16,PSMC6,MED14
LF1,CTNNB1,CD63,CYFIP2,CADM4,GPC1,DCN,SKI,NOTCH2,FKBP8,STRAP,HGS,BMPR2,SULF2,B
,LRPPRC,LAMP1,MAP2,RAB1A,KIF3C,CDC42,KIF5A,HIF1A,KIFAP3,RAB6A,SUN1,TRAK1,FEZ1,SUI
SNK1E,CTNNB1,WLS,DLG5,ATP5PB,YWHAH,CSNK1D,DYNLL1,CDC42,SEC16A,SFRP1,GLUD1,ND
S3,PARP1,PKN1,H3-3A,TAF7,SMARCA4,KDM1A,YWHAH,NCOR2,WBP2,EP300,LBH,PHB2,SFRP1,I
3,ATP1A2,ATP6V0C,ATP6AP1,ATP5PB,ATP6V1B2,PHB2,COX7C,SLC25A5,ATP6V0D1,COX6A1,ATP
,RAI1,MTA1,PPP1CC,OGT,MYCBP2,PPP1CA,LGR4,PRMT5,KDM2A,KDM5A,PER1,ID3,ID2,MYBBP1
OS9,RER1,CIZ1,RANGAP1,SUN1,PINK1,GRIK5,PARK7,GPAA1,SUN2,GOLPH3,CCDC88A,TXN,ARL

P1,G3BP2,KDELR2,KDELR1,OS9,RER1,DZIP1,CIZ1,RANGAP1,SUN1,PINK1,GRIK5,PARK7,GPAA1,S
,GLG1,JUN,APPL1,HDAC2,PARP1,NOX4,SKI,ADAM9,FYN,STRAP,BCL9L,EP300,FLCN,UBA52,ZFP3
NT1,UBE2I,MGAT5,SUMO1,DAD1,MOGS,GSK3B,MAN1B1,FURIN
DX6,PRPF19,THRAP3,SLC38A2,HSPA1A,CELF3,NUP98,SETX
5,CDH2,CTNNB1,ANXA6,HDAC2,DLG5,MDK,PTK7,PPP2CA,MAPK1,CORO1C,STRAP,BMPR2,BCL9
KN1A,GCN1,MARS1,MAPK1,GABARAP,ATP6V1B2,LARS1,SREBF2,BMPR2,WDR45B,FLCN,SFRP1,E
FKL,MDH2,PFKM,FOXK2,MDH1,PFKP,OGDH
P1,SUN1,PINK1,GRIK5,PARK7,GPAA1,SUN2,TXN,ARL2BP
CA,NCK2,SESN2,EIF2S1,SLC35A4
ASP2,PTPN11,ABL1,HIP1R,SMPD3,PTPN1,PDGFRB,NDRG4,SGPL1,FER,BCAR1,PTPN12
V1,UQCRC1,COX7C,SLC25A23,COX6A1,ATP5MC3,NDUFA13,ATP5MC2,ATP5MG,PINK1,NDUFA4,
,YWHAQ,YWHAB,HSPA4,ATP5PB,BNIP3L,YWHAH,GHITM,STAT3,TIMM50,SLC25A5,SPG7,CNP,NM
TP5PB,YWHAH,DYNLL1,CDC42,SEC16A,GLUD1,CNP,COX6B1,SUDS3
S4A,IST1,CHMP4B,UBE2I,RANGAP1,SUN1,SUMO1,BANF1,RAN,SUN2,AKAP8L,NUP98,TMEM43,NU
NIP3,PINK1,ATP13A2,PARK7,FBXO7,USP36

P3L,VDAC1,FLCN,PHB2,RAB3GAP2,ULK1,GPSM1,HIF1A,SLC25A5,SH3BP4,GSK3A,BNIP3,TRIM8,A
PD1,PSMD5,HMGB1,ITGB1,ANXA2,ZMIZ1,AP3D1,PAFAH1B1,XRCC5,RPL22,RHOA,ATF4,PSMD2,R
32,CCND1,PSMD3,PSMA7,PRMT1,EP300,PSMD1,ZFP36L1,PTEN,BRSK1,PSMD8,PCBP4,PSMB4,TA
ACTN4,MMP14,MACF1,RAC1,TBCD,PTPRA,MDK,VEGFA,ARPC2,CORO1C,FBLN1,CDC42,SFRP1,PT
NON,HSPB1,FYN,UBQLN1,SOD1,HDAC6,HIF1A,GPX1,TRAP1,SOD2,FBLN5,BAG5,PINK1,MACROH
ND1,GRB2,KDM1A,CCAR2,TP53BP1,PRMT1,EP300,BRD4,ATRX,FEM1B,BRSK1,PCBP4,PPP1R10,
PRS,SOD1,PTEN,PTPRF,GRN,STK24,BNIP3,CERS2,RANGAP1,LGALS1,DAG1,JAM3,DHFR,NREP,R
,CXADR,WDR1,CDH6,CDH2,TLN1,GJA1,CTNNB1,DLG5,CDH11,TBCD,TLN2,CNTNAP2,VEGFA,CDC
GRN,PAFAH1B1,NACA,MEGF8,MACF1,L1CAM,PPIB,DDX39B,VEGFA,MAPK1,PRKDC,RAB11A,BMP
DVL3,RACK1,CALM3,APP,CALM1,NPM1,IGF2,UBB,MAPT,HACD3,PKD1,RPS3,HMGB1,ANXA2,RPL
D3,GRB2,PLCG1,PSMA7,PSMD1,PSMD8,PSMB4,PDPK1,PSMD4,PSMC3,PSMC1,PSMB1,CUL1,PS
LI2,DYNLL2,CAPZB,KIF3C,SEC23A,DYNLL1,DCTN1,KIF5A,ACTR1A,AP2A1,KIFAP3,CAPZA1,ACTR1
C,RPS28
4,CALR,ABHD2,GPI,RACK1,BSG,ATP5F1B,APP,GSTP1,PLXNB2,ATP5F1A,FN1,TMSB10,HMGB1,IT
UBB,TRIM28,SPEN,CHD3,HMGB1,RPL10,HNRNPU,RPL23,NACA,CBX5,ATF4,NR2F1,RPS14,TLE5,K
7,U2AF2,SQSTM1,RPL37,FYN,NXN,UBR5,OGT,MTOR,BAG5,TRIP12,ABL1,HSPA1A,PARK7,LIMK1,
COL11A1,PXDN,COL1A2,COL5A2,COL6A2,PPIB,PLOD1,COL2A1,PLEC,AEBP1,COL27A1,CRTAP,L
SPA1B,CLU,SOX4,MAGED1,KIF1A,GNB1,APP,GSTP1,NPM1,PLXNB2,HSPD1,COL18A1,IGF2,ATP5F
ABI2,WASF2,MTOR,AUTS2,CYFIP1
AD23B,CHD2,UBA52,ATRX,HNRNPA2B1,RPS27A,XRCC6,HMGA1,RUVBL2,SUB1,HMGB3,CUL4A,C
B2,SCRIB,CERS2,SUN1,IDH2,SUN2,CRKL,MBOAT7,BMERB1,LRP8,TGFB2,EPHA4
SLIT2,BRK1,ABI2,WASF2,MTOR,AUTS2,CYFIP1,FER,EPHA2
SMB2,MARS1,AZIN1,DDAH2,PSMD3,ODC1,QARS1,LARS1,SARS1,SMS,PSMA7,PSMD1,VARS1,GARS
I,VPS4A,FLOT2,SYT11,IST1,CHMP4B,UBE2I,GAK,RANGAP1,NRXN2,SUMO1,BANF1,RAN,NUP98,N
,IPO5,CCT2,PARP1,TCP1,CCT3,CCT4,MAPK1,FYN,CCT8,CCT5,UBR5,TRIM8,SMO,PARK7,RAN,TF
PP,NPM1,COL4A2,ZMIZ1,TIMP2,CTNNA1,JUND,CCN2,MIF,JUN,HMGCR,DCN,CDKN1A,RPN2,NOX
,GLG1,JUN,APPL1,HDAC2,PARP1,NOX4,SKI,ADAM9,FYN,STRAP,BCL9L,EP300,FLCN,UBA52,ZFP3

N1,SPTBN2,MAP2,TBCD,CAMSAP1,MAPRE1,LUZP6,CAPZB,CLIP3,PFN2,HDAC6,SLIT2,TMOD2,TA(
O5A,MACF1,LAMP1,VAMP2,OPTN,HGS,ARFGEF2,TXNDC5,AP2A1,SEC16A,PREPL,RABEP1,PKDCC
C2L,KDM4A,TRIP12,HMGA1,KDM3A,MACROH2A1,BCOR,OTUB1,KDM5A,WDR5,DNMT1,SMARCE
MP1,RHOB,VPS35,HGS,NCOA4,GRN,VPS41,VPS4A,ATP6V0D1,CHMP4B,UBXN6,SORT1,GAK,CHM
RAB35,MICALL1,ARF6,TOLLIP
TN,CCN2,SESN3,IPO5,MMP2,TOMM20,FYN,LARS1,HNRNPD,PPP1R9B,CAPN2,SH3BP4,MTOR,NT
,BASP1,CDKL1,ITGB1,ZMIZ1,COL11A1,AP2B1,HNRNPU,CPE,NACA,RTN4,CXADR,PTN,TPM1,GPC
P2,NOP53,PSMB2,CCND1,PSMD3,PSMA7,PRMT1,EP300,PSMD1,ZFP36L1,PTEN,BRSK1,PSMD8,F
,MAP1A,HECTD1,OS9,FLCN,UBE2G2,LMAN1,PKDCC,DCLK1,RANGAP1,MACROH2A1,PKIA,PARK7
CALM3,CALM1,NPM1,IGF2,UBB,MAPT,HACD3,PKD1,RPS3,HMGB1,RPLP1,XRCC5,RHOA,CD81,CC
TBP1,HDAC2,GLUL,EID1,MORF4L2,YEATS2,TERF2IP,TAF7,CLIP3,MSL1,BAG6,WBP2,EP300,FLC
,CLU,MAGED1,DVL3,RACK1,CALM3,APP,CALM1,NPM1,IGF2,VCP,UBB,MAPT,HACD3,PKD1,RPS3,
,CLU,MAGED1,DVL3,RACK1,CALM3,APP,CALM1,NPM1,IGF2,VCP,UBB,MAPT,HACD3,PKD1,RPS3,
AN,NUP153,KPNA6,NUP98,NUP205,NUP133,KPNA3,DYNLT1,NUP50,NUP58,VPS28,VPS37B

2,SMARCC1,ATP6V1B2,NUCKS1,AKT2,OGT,ATP6V0D1,GSK3A,ATP6V1G1,PHIP,RABGAP1,PTPN1
YWHAB,PTBP1,PPP6R1,PPP1R9B,MASTL,PPP1R15A,FKBP1A,SYMPK,PPP1R10,ARPP19,MTOR,PI
RPP19,GSK3A,NSG1,GSK3B,PRMT5,GNAL,GNAO1
5,CCT7,HNRNPC,CCN2,GJA1,CTNNB1,PTGES3,CCT6A,CDKN1A,DDX39B,CCT2,TCP1,CCT3,CCT
TADA2B,ARID1A,SMCHD1,HDAC2,H3-3A,KDM6B,TERF2IP,TOP1,SMARCA4,SMARCE1,SMARCC1,
JA1,MAP2,CDKN1A,NOTCH2,PPP2CA,SMARCA4,CCAR2,SLIT3,PLXNA3,PTPRS,RAI1,BMPR2,PPP1
PFKL,PFKM,QPRT,FOXK2,NMNAT2,MDH1,PFKP,NADK
H3BP4,SORT1,ATP13A2,ROCK2,SZT2,CACNG7,AP3M1,M6PR
,FTH1,ALDOA,PKM,FTL,HSPA1B,PSAP,CLU,SLC2A3,GPI,KLRD1,SPTAN1,GSTP1,HSPD1,VCP,HLA
1A,DNAJB2,BAG5,HSPA1A
FLCN,DIP2B,TAOK1,NOC2L,DIP2A,AUTS2,RUVBL2,PARK7,KAT2A,GSK3B,SETD5,CTCF,MAPK3,X
NAJA1,HNRNPU,PIP4K2B,PRKAR1A,YWHAB,HMGCR,IPO5,TRIB2,CDKN1A,PKN1,TAF7,PPP2CA,IP

P90AB1,HSP90AA1,ACTR2,VIM,SPARC,TUBB,UBC,KIDINS220,TMSB4X,NUMA1,DCX,EIF4G2,RPL3
2,SPARC,TUBB,FTH1,UBC,TMSB4X,NUMA1,RPL3,RPS6,DBN1,RPL4,TUBA1B,PIK3R2,FLNA,MAP1B
IAE,RPL13,RACK1,BSG,RPS8,RPL7,P4HB,RPL5,CALM1,NPM1,HSPD1,USP11,CSDE1,VCP,UBB,TR
P90AB1,HSP90AA1,ACTR2,VIM,SPARC,TUBB,UBC,KIDINS220,TMSB4X,NUMA1,DCX,EIF4G2,RPL3
YBX1,ATP2B4,CALR,RPLP0,CLU,RPL13A,MAGED1,SEC62,RPL6,DVL3,YWHAE,KIF1A,RPL13,RACK
YBX1,ATP2B4,CALR,RPLP0,CLU,RPL13A,MAGED1,SEC62,RPL6,DVL3,YWHAE,KIF1A,RPL13,RACK
SMG1,NPM1,SNRNP200,CSDE1,UBB,RPL37A,SPEN,NONO,PSMD5,SET,RPL19,PPP2R1A,RPL15,SF
PL7A,HSP90B1,NEFL,RPL8,CALR,RPLP0,CLU,RPL13A,SEC62,RPL6,YWHAE,KIF1A,RPL13,GNAS,SF
,RPLP0,CLU,DDX5,RPL13A,RPL6,RPL13,RACK1,RPS8,RPL7,IDS,RPL5,SMG1,CALM1,NPM1,TTC3,
H1,UBC,TMSB4X,NUMA1,RPL3,RPS6,RPL4,TUBA1B,PIK3R2,FLNA,MAP1B,RPL7A,ALDOA,PKM,FT
UGT8,RPL8,YBX1,ATP2B4,CALR,RPLP0,CANX,CLU,SOX4,RPL13A,MAGED1,SEC62,RPL6,DVL3,YW
F4G2,DBN1,ENO1,FLNA,MAP1B,NEFL,CALR,HSPA1B,COL5A1,H3-3B,CLU,ENO2,RPL13A,DVL3,Y
HSP90B1,UGT8,RPL8,YBX1,ATP2B4,CALR,TMF1,PSAP,RPLP0,CANX,CLU,SOX4,RPL13A,MAGED1
DX5,RPL13A,RPL6,RPL13,RACK1,RPS8,RPL7,RPL5,SMG1,NPM1,TTC3,USP11,CSDE1,VCP,UBB,R
13,CALM3,RPS8,RPL7,APP,RPL5,CALM1,NPM1,HSPD1,VCP,TRIM28,YWHAG,RPL37A,MAPT,YWH
,SET,RPL19,PPP2R1A,RPL15,RPS3,RPS4X,RPS3A,RPL18,RPL11,RPLP1,RPL10,RPS17,HNRNPU,L
HSP90AA1,ACTR2,SPARC,TUBB,FTH1,UBC,TMSB4X,NUMA1,DCX,RPL3,RPS6,DBN1,RPL4,TUBA1B
RPS6,RPL4,ENO1,FLNA,NCL,RPL7A,PKM,HSP90B1,RPL8,FBXL4,YBX1,UBA1,ATP2B4,IGFBP2,CAL
B4X,NUMA1,DCX,RPL3,DBN1,ENO1,TUBA1B,FLNA,MAP1B,ALDOA,HSP90B1,NEFL,NEFM,UGT8,C
0,RPS17,RPS16,RPL23,RPL29,RPSA,RPS24,RPS9,RPL24,RPL17,RPL14,RPL22,RPL9,RPS14,RPS2
P1,RPL10,RPS17,RPS16,RPL23,RPL29,RPSA,RPS24,RPS9,RPL24,RPL17,RPL14,RPL22,RPL9,RPS
7,RPS16,RPL23,RPL29,RPSA,RPS24,RPS9,RPL24,RPL17,RPL14,RPL22,RPL9,RPS14,RPS20,RPL2
,SET,RPL19,PPP2R1A,RPL15,RPS3,RPS4X,RPS3A,RPL18,RPL11,RPLP1,RPL10,RPS17,HNRNPU,L
LNA,MAP1B,RPL7A,ALDOA,HSP90B1,NEFL,NEFM,PRPF8,UGT8,RPL8,CALR,TMF1,HSPA1B,RPLP0
,RPL6,YWHAE,KIF1A,RPL13,GNAS,RACK1,BSG,SPTBN1,CALM3,RPS8,MYH9,RPL7,APP,RPL5,GDI
,RPS8,RPL7,RPL5,GDI1,NPM1,HSPD1,VCP,TRIM28,YWHAG,RPL37A,YWHAQ,RPL19,RPL15,PKD1
RPL10,RPS17,RPS16,RPL23,RPL29,RPSA,RTN4,RPS24,RPS9,RPL24,RPL17,RPL14,RPL22,RPL9,
0,RPS17,RPS16,RPL23,RPL29,RPSA,RPS24,RPS9,RPL24,RPL17,RPL14,RPL22,RPL9,RPS14,RPS2
,RPL3,RPS6,RPL4,ENO1,FLNA,NCL,RPL7A,PKM,GANAB,HSP90B1,PRPF8,RPL8,FBXL4,YBX1,UBA
X,CLU,SOX4,RPL13A,SEC62,RPL6,YWHAE,KIF1A,RPL13,GNAS,RACK1,BSG,SPTBN1,RPS8,MYH9,
1,RPLP1,RPL10,RPS17,GTF2F1,RPS16,RPL23,RPL29,RPSA,RPS24,RPS9,RPL24,RPL17,RPL14,R
0,RPS17,RPS16,RPL23,RPL29,RPSA,RPS24,RPS9,RPL24,RPL17,RPL14,RPL22,RPL9,RPS14,RPS2
1,RPLP1,RPL10,RPS17,RPS16,RPL23,RPL29,RPSA,EIF4H,RPS24,RPS9,RPL24,RPL17,RPL14,RPL
MAVS,HTT,HSP90AB1,ZNF124,HSP90AA1,ACTR2,VIM,UBC,KIDINS220,TMSB4X,EIF4G2,RPL3,RPS
,PKM,HSP90B1,RPL8,FBXL4,YBX1,UBA1,ATP2B4,TMF1,ABHD2,HSPA1B,CES3,PSAP,PGK1,RPLP0
XL4,YBX1,UBA1,ATP2B4,TMF1,ABHD2,HSPA1B,PSAP,RPLP0,CLU,DDX5,RPL13A,RPL6,RPL13,RAC
ZNF124,VIM,SPARC,FTH1,UBC,TMSB4X,EIF4G2,RPL3,RPS6,RPL4,ENO1,PIK3R2,FLNA,MAP1B,N
PS4X,RPS3A,RPL18,RPL11,RPLP1,RPL10,RPS17,LARP1,RPS16,RPL23,RPL29,RPSA,EIF4H,RPS24
1,APP,RPL5,SMG1,NPM1,VCP,UBB,MAPT,DDB1,PSMD5,SET,PKD1,HMGB1,ITGB1,UCHL1,RPL11,
,RPL6,YWHAE,KIF1A,RPL13,GNAS,RACK1,BSG,SPTBN1,RPS8,MYH9,RPL7,RPL5,GDI1,NPM1,HSP
2,SPARC,TUBB,FTH1,UBC,TMSB4X,NUMA1,RPL3,RPS6,RPL4,TUBA1B,PIK3R2,FLNA,MAP1B,RPL7
SPD1,TRIM28,YWHAG,RPL37A,MAPT,YWHAQ,RPL19,RPL15,RPS3,SEC61A1,RPS4X,RPS3A,RPL18
B,FTH1,UBC,TMSB4X,NUMA1,RPL3,RPS6,RPL4,TUBA1B,PIK3R2,FLNA,MAP1B,RPL7A,SLC9A7,A
,ALDOA,HSP90B1,NEFL,NEFM,PRPF8,UGT8,CALR,TMF1,HSPA1B,RPLP0,H3-3B,CLU,RPL13A,RPL
,YBX1,UBA1,ATP2B4,TMF1,ABHD2,HSPA1B,CES3,PGK1,RPLP0,CLU,DDX5,ENO2,RPL13A,RPL6,R

,TMSB4X,EIF4G2,RPL3,RPS6,RPL4,ENO1,FLNA,NCL,RPL7A,ALDOA,PKM,GANAB,HSP90B1,PRPF8
34,CALR,TUBB2B,COL4A1,CLU,SOX4,DVL3,YWHAE,KIF1A,BSG,SPTBN1,CALM3,STMN2,SEPTIN2
GDI1,CALM1,YWHAG,RPL37A,YWHAQ,DCHS1,RPL19,RPL15,RPS3,VAPA,AP2M1,SEC61A1,RPS4X,R
S220,TMSB4X,NUMA1,EIF4G2,RPS6,DBN1,ENO1,PIK3R2,FLNA,MAP1B,NCL,PKM,NEFL,YBX1,ATP
MN2,SEPTIN2,APP,GDI1,SPTAN1,PLXNB2,UBB,MAPT,FN1,VAPA,HMGB1,STMN1,ITGB1,UCHL1,NC
PL5,GDI1,CALM1,YWHAG,RPL37A,YWHAQ,DCHS1,RPL19,RPL15,RPS3,VAPA,AP2M1,SEC61A1,RP
MG1,NPM1,VCP,UBB,MAPT,PSMD5,SET,PKD1,HMGB1,UCHL1,RPL11,ANXA2,TIMP2,HNRNPU,PGA
,RPS16,RPL23,RPL29,RPSA,RPS24,RPS9,RPL24,RPL17,RPL14,RPL22,RPL9,RPS14,RPS20,RPL26
ZNF124,HSP90AA1,ACTR2,VIM,UBC,KIDINS220,TMSB4X,EIF4G2,RPL3,RPS6,ZNF316,RPL4,ENO1
SMD5,SET,RPL19,PPP2R1A,RPL15,RPS3,HMGB1,RPS4X,RPS3A,RPL18,RPL11,RPLP1,RPL10,RPS1
4,VIM,SPARC,FTH1,UBC,TMSB4X,EIF4G2,RPS6,ENO1,PIK3R2,FLNA,MAP1B,NCL,MTRNR2L8,HSP
,RPL7A,RPL8,YBX1,ATP2B4,CALR,TMF1,HSPA1B,RPLP0,H3-3B,CLU,DDX5,SOX4,RPL13A,MAGED
B1,YBX1,ATP2B4,CALR,TMF1,HSPA1B,CLU,SOX4,RPL13A,MAGED1,DVL3,YWHAE,GPI,RACK1,CA
90AB1,ZNF124,HSP90AA1,ACTR2,VIM,UBC,KIDINS220,TMSB4X,EIF4G2,RPL3,RPS6,ZNF316,RPL4
,RPL8,YBX1,ATP2B4,CALR,TMF1,HSPA1B,RPLP0,H3-3B,CLU,DDX5,SOX4,RPL13A,MAGED1,RPL6
GK1,CANX,CLU,DDX5,SOX4,FKBP14,KAT6A,YWHAE,RACK1,GNB1,P4HB,APP,TMBIM6,GSTP1,SM
220,TMSB4X,NUMA1,RPL3,TUBA1B,PIK3R2,FLNA,MAP1B,NCL,PKM,COL6A1,HSP90B1,FADS1,NE
6,KAT6A,DVL3,RACK1,SPTBN1,STMN2,P4HB,RPL5,FSCN1,SPTAN1,CALM1,NPM1,HSPD1,SNRNP
MN2,SEPTIN2,APP,GDI1,SPTAN1,PLXNB2,UBB,MAPT,FN1,VAPA,HMGB1,STMN1,ITGB1,UCHL1,NC
GDI1,FSCN1,PLXNB2,IGF2,VCP,TRIM28,YWHAG,MAPT,FN1,DDB1,YWHAQ,RPS3,AP2M1,ANXA2,A
LU,SOX4,DVL3,KIF1A,BSG,SPTBN1,STMN2,SEPTIN2,APP,GDI1,SPTAN1,GSTP1,PLXNB2,UBB,YWH
90AB1,ZNF124,HSP90AA1,ACTR2,VIM,UBC,KIDINS220,TMSB4X,EIF4G2,RPL3,RPS6,ZNF316,RPL4
RPL37A,PSMD5,SET,RPL19,PPP2R1A,RPL15,RPS3,HMGB1,RPS4X,RPS3A,RPL18,RPL11,RPLP1,R
RPL37A,PSMD5,SET,RPL19,PPP2R1A,RPL15,RPS3,HMGB1,RPS4X,RPS3A,RPL18,RPL11,RPLP1,R
8,RPL11,RPLP1,RPL10,RPS17,RPS16,RPL23,RPL29,RPSA,RPS24,RPS9,RPL24,RPL17,RPL14,RPL
TT,HSP90AB1,ZNF124,HSP90AA1,ACTR2,VIM,UBC,KIDINS220,TMSB4X,EIF4G2,RPL3,RPS6,ZNF3
1A,BSG,SPTBN1,STMN2,SEPTIN2,APP,GDI1,SPTAN1,PLXNB2,UBB,YWHAG,MAPT,FN1,VAPA,HMG
VL3,KIF1A,BSG,SPTBN1,STMN2,SEPTIN2,APP,GDI1,SPTAN1,PLXNB2,UBB,YWHAG,MAPT,FN1,SPE
P0,CANX,CLU,SOX4,RPL13A,SEC62,RPL6,YWHAE,KIF1A,RPL13,GNAS,SLC2A3,RACK1,BSG,SPTBN
,HSP90AA1,ACTR2,VIM,UBC,KIDINS220,TMSB4X,EIF4G2,RPL3,RPS6,ZNF316,RPL4,ENO1,PIK3R
VHAQ,RPL19,RPL15,RPS3,VAPA,SEC61A1,RPS4X,RPS3A,RPL18,RPL11,RPLP1,RPL10,RPS17,RPS
1,SPARC,UBC,KIDINS220,TMSB4X,NUMA1,EIF4G2,RPS6,DBN1,ENO1,PIK3R2,FLNA,MAP1B,NCL,F
,HSP90AA1,ACTR2,VIM,UBC,KIDINS220,TMSB4X,EIF4G2,ZNF316,ENO1,PIK3R2,FLNA,NCL,PKM,
APP,RPL5,GSTP1,NPM1,PLXNB2,CSDE1,EIF3C,EIF4B,RPL37A,EIF1,EIF4A1,RPL19,RPL15,RPS3,RF
,TMF1,HSPA1B,CLU,SOX4,RPL13A,MAGED1,DVL3,YWHAE,GPI,RACK1,CALM3,MYH9,APP,RPL5,G
3,SEC61A1,RPS4X,RPS3A,RPL18,RPL11,RPLP1,RPL10,RPS17,AP3D1,RPS16,RPL23,RPL29,RPSA,
SMD5,SET,RPL19,PPP2R1A,RPL15,RPS3,HMGB1,RPS4X,RPS3A,RPL18,RPL11,RPLP1,RPL10,RPS1
,SPTBN1,MYH9,P4HB,APP,GDI1,SPTAN1,PLXNB2,COL18A1,UBB,MAPT,FN1,DCHS1,STMN1,ITGE
RPL37A,PSMD5,SET,RPL19,PPP2R1A,RPL15,RPS3,HMGB1,RPS4X,RPS3A,RPL18,RPL11,RPLP1,R
XNB2,UBB,MAPT,FN1,STMN1,ITGB1,UCHL1,NCAM1,VCL,PAFAH1B1,RTN4,LAMA5,RPL24,LLGL1,
XNB2,UBB,MAPT,FN1,STMN1,ITGB1,UCHL1,NCAM1,VCL,PAFAH1B1,RTN4,LAMA5,RPL24,LLGL1,
XNB2,UBB,MAPT,FN1,STMN1,ITGB1,UCHL1,NCAM1,VCL,PAFAH1B1,RTN4,LAMA5,RPL24,LLGL1,
1A1,DCX,EIF4G2,RPS6,DBN1,PRRC2B,FLNA,MAP1B,NCL,ALDOA,PKM,COL6A1,FADS1,NEFL,UGT
SMD5,PKD1,AP2M1,UCHL1,RPL11,ANXA2,TIMP2,MARCHF6,AP2B1,OGA,RPL23,CTSD,LPCAT1,MS
PL5,NPM1,PLXNB2,CSDE1,EIF3C,EIF4B,RPL37A,EIF1,EIF4A1,RPL19,RPL15,RPS3,RPS4X,RPS3A,

6A,RACK1,SPTBN1,STMN2,P4HB,RPL5,FSCN1,SPTAN1,NPM1,HSPD1,SNRNP200,EIF3C,EIF4B,VC
RCC2,MAVS,HTT,HSP90AB1,ZNF124,HSP90AA1,ACTR2,VIM,UBC,KIDINS220,TMSB4X,EIF4G2,RF
/L3,YWHAE,KIF1A,BSG,SPTBN1,STMN2,MYH9,SEPTIN2,APP,GDI1,FSCN1,SPTAN1,PLXNB2,UBB,Y
0,RPL37A,SPEN,NONO,RPL19,PPP2R1A,RPL15,SF3B1,RPL18,RPL11,RPLP1,RPL10,RPS17,SF3B
NUMA1,RPL3,ENO1,TUBA1B,PIK3R2,FLNA,MAP1B,NCL,PKM,COL6A1,HSP90B1,PRPF8,YBX1,ATP
XNB2,UBB,MAPT,FN1,STMN1,ITGB1,UCHL1,NCAM1,VCL,PAFAH1B1,RTN4,LAMA5,RPL24,LLGL1,
RPL5,SPTAN1,NPM1,HSPD1,SNRNP200,EIF3C,EIF4B,UBB,MAPT,DDB1,PSMD5,SET,TMSB10,DNAH
TT,HSP90AB1,ZNF124,HSP90AA1,ACTR2,VIM,UBC,KIDINS220,TMSB4X,EIF4G2,RPL3,RPS6,ZNF3
AA1,ACTR2,VIM,UBC,KIDINS220,TMSB4X,EIF4G2,ZNF316,ENO1,PIK3R2,FLNA,NCL,PKM,HSP90E
RPL10,RPS17,RPS16,RPL23,RPL29,RPSA,NACA,RPS24,RPS9,RPL24,RPL17,RPL14,RPL22,RPL9,
DBP,HNRNPK,CIRBP,YWHAB,NOVA2,U2AF2,PABPN1,HNRNPL,TAF15,ALKBH5,AHCYL1,RBMX,TN
/L3,YWHAE,KIF1A,BSG,SPTBN1,STMN2,MYH9,SEPTIN2,APP,GDI1,FSCN1,SPTAN1,PLXNB2,UBB,Y
PL5,NPM1,PLXNB2,CSDE1,EIF3C,EIF4B,RPL37A,EIF1,EIF4A1,RPL19,RPL15,RPS3,RPS4X,RPS3A,
NUMA1,RPL3,TUBA1B,PIK3R2,FLNA,MAP1B,NCL,PKM,COL6A1,HSP90B1,PRPF8,YBX1,ATP2B4,IG
MA1,DCX,EIF4G2,RPS6,DBN1,FLNA,MAP1B,NCL,ALDOA,PKM,COL6A1,NEFL,UGT8,YBX1,ATP2B4,
,SPTAN1,PLXNB2,COL18A1,MAPT,FN1,STMN1,ITGB1,UCHL1,NCAM1,VCL,PAFAH1B1,RTN4,LAM
COL6A1,NEFL,UGT8,YBX1,ATP2B4,CALR,TMF1,TUBB2B,COL5A1,COL4A1,PGK1,CLU,SOX4,MAG
1,SPTAN1,PLXNB2,UBB,MAPT,FN1,STMN1,ITGB1,UCHL1,NCAM1,VCL,PAFAH1B1,RTN4,LAMA5,F
FN1,STMN1,ITGB1,UCHL1,NCAM1,VCL,PAFAH1B1,RTN4,LAMA5,RPL24,LLGL1,GAP43,RHOA,KIF
BB,FTH1,UBC,TMSB4X,EIF4G2,RPS6,DBN1,PIK3R2,FLNA,MAP1B,SLC9A7,ALDOA,FTL,HSP90B1,
4,PNN,HNRNPC,KHDRBS1,SFPQ,IK,TARDBP,HNRNPK,CIRBP,HNRNPH1,BCLAF1,LUC7L3,CPSF2,D
MA1,DCX,EIF4G2,RPS6,DBN1,FLNA,MAP1B,NCL,PKM,COL6A1,NEFL,UGT8,YBX1,ATP2B4,CALR,F
MEM259,CD81,OAZ1,XPO1,CSNK1E,GPC3,RAB7A,RPS7,GJA1,NDFIP1,HMGCR,LAPTM4B,TRIB2,
AA1,ACTR2,VIM,UBC,KIDINS220,TMSB4X,EIF4G2,ZNF316,ENO1,PIK3R2,FLNA,NCL,PKM,HSP90E
,RPL7,APP,RPL5,NPM1,PLXNB2,CSDE1,EIF3C,EIF4B,RPL37A,EIF1,EIF4A1,RPL19,RPL15,RPS3,RF
X,EIF4G2,RPS6,DBN1,FLNA,MAP1B,NCL,PKM,NEFL,UGT8,ATP2B4,CALR,TMF1,HSPA1B,TUBB2B,
MAVS,HTT,HSP90AB1,ZNF124,HSP90AA1,ACTR2,VIM,UBC,KIDINS220,TMSB4X,EIF4G2,RPL3,RPS
DRBS1,SFPQ,IK,HNRNPK,CIRBP,HNRNPH1,LUC7L3,CPSF2,DDX39B,NOVA2,SRSF5,U2AF2,CPSF1
DRBS1,SFPQ,IK,HNRNPK,CIRBP,HNRNPH1,LUC7L3,CPSF2,DDX39B,NOVA2,SRSF5,U2AF2,CPSF1
DRBS1,SFPQ,IK,HNRNPK,CIRBP,HNRNPH1,LUC7L3,CPSF2,DDX39B,NOVA2,SRSF5,U2AF2,CPSF1
CLU,SOX4,FKBP14,MAGED1,KAT6A,DVL3,YWHAE,RACK1,SPTBN1,CALM3,P4HB,APP,RPL5,GSTP
CLU,SOX4,FKBP14,MAGED1,KAT6A,DVL3,YWHAE,RACK1,SPTBN1,CALM3,P4HB,APP,RPL5,GSTP
VCL,PAFAH1B1,CTNNA1,RTN4,LAMA5,RPL24,LLGL1,GAP43,RHOA,KIF5B,CRMP1,SPTBN2,PTN,C
4G2,ENO1,PIK3R2,NCL,PKM,YBX1,ATP2B4,CALR,TMF1,HSPA1B,ZNF99,CLU,DDX5,SOX4,MAGED
C,KHDRBS1,SFPQ,IK,TARDBP,HNRNPK,CIRBP,HNRNPH1,BCLAF1,LUC7L3,CPSF2,DDX39B,NOVA2
CHL1,RPL11,ANXA2,MARCHF6,AP2B1,RPL23,MSN,PSMD2,ENC1,TMEM259,CD81,XPO1,CSNK1E,
D2,TUBB2B,CLU,SOX4,DVL3,KIF1A,BSG,FASN,SPTBN1,STMN2,MYH9,SEPTIN2,P4HB,APP,GDI1,S
I1,SMG1,NPM1,IGF2,TRIM28,YWHAG,MAPT,YWHAQ,MAP9,TMSB10,PKD1,RPS3,CDKL1,STMN1,A
3A,RPL6,RPL13,DDX17,RACK1,RPS8,RPL7,APP,RPL5,TMBIM6,GSTP1,SMG1,NPM1,CSDE1,UBB,
RPS8,RPL7,APP,RPL5,GSTP1,NPM1,PLXNB2,CSDE1,EIF3C,EIF4B,RPL37A,EIF1,EIF4A1,RPL19,PPF
.15,SF3B1,RPL18,RPL11,RPLP1,RPL10,RPS17,RPS16,RPL23,RPL29,RPSA,EIF4H,RPS24,RPS9,XF
BX1,ATP2B4,TMF1,HSPA1B,ZNF99,CLU,DDX5,SOX4,MAGED1,KAT6A,DVL3,DDX17,RACK1,FASN,
CTG1,EEF2,DPYSL3,TUBA1A,CFL1,BTBD9,XRCC2,MAVS,HTT,HSP90AB1,ZNF124,HSP90AA1,ACTR
DA,PKM,FTL,HSP90B1,CALR,HSPA1B,PSAP,CANX,CLU,KIF1A,SLC2A3,GPI,RACK1,SPTBN1,CALM3
,NR2F1,ENC1,CRMP1,PRKCSH,PTN,CDH2,PLXNA1,MEGF8,ITM2C,MACF1,MAP2,ARHGDIA,CAPRI

3R2,NCL,PKM,YBX1,ATP2B4,CALR,TMF1,HSPA1B,ZNF99,CLU,DDX5,SOX4,MAGED1,KAT6A,DVL
X5,SOX4,RPL13A,MAGED1,KAT6A,YWHAE,GPI,RACK1,CALM3,APP,RPL5,TMBIM6,GSTP1,CALM1,
AFAH1B1,RTN4,LAMA5,RPL24,LLGL1,GAP43,RHOA,KIF5B,CRMP1,SPTBN2,CDH2,PLXNA1,MEG
CLU,DDX5,SOX4,RPL13A,MAGED1,KAT6A,YWHAE,GPI,RACK1,CALM3,APP,RPL5,TMBIM6,GSTP1,
H1B1,MAP4,SCARB2,RTN4,GAP43,NCS1,RHOA,NR2F1,ENC1,CRMP1,PFN1,PRKCSH,PTN,CDH2,
1,NEFL,ATP2B4,IGFBP2,CALR,FSTL1,TMF1,ABHD2,COL4A1,H3-3B,DDX5,GNAS,DDX17,GPI,HTR
,PAFAH1B1,MAP4,SCARB2,RTN4,GAP43,NCS1,RHOA,NR2F1,ENC1,CRMP1,PFN1,PRKCSH,PTN,
SPA1B,CLU,PTMA,DDX5,SOX4,MAGED1,YWHAE,GPI,RACK1,GNB1,P4HB,APP,TMBIM6,GSTP1,N
M,YBX1,ATP2B4,TMF1,HSPA1B,ZNF99,CLU,DDX5,SOX4,MAGED1,KAT6A,DVL3,DDX17,RACK1,C
2,VIM,UBC,TMSB4X,EIF4G2,RPL3,RPS6,ZNF316,RPL4,ENO1,PIK3R2,FLNA,NCL,RPL7A,PKM,FA
RPL6,RPL13,RACK1,FASN,ATP5F1B,RPS8,RPL7,APP,RPL5,GSTP1,NPM1,PLXNB2,CSDE1,EIF3C,A
1,PRRC2B,FLNA,MAP1B,COL6A1,FADS1,NEFL,UGT8,YBX1,CALR,FSTL1,TMF1,ABHD2,HSPA1B,T
,KIDINS220,TMSB4X,NUMA1,RPL3,ENO1,TUBA1B,PIK3R2,FLNA,MAP1B,NCL,PKM,COL6A1,HSP
A1,ACTR2,VIM,UBC,TMSB4X,EIF4G2,RPL3,RPS6,ZNF316,RPL4,ENO1,PIK3R2,FLNA,NCL,RPL7A,
1,PRRC2B,FLNA,MAP1B,COL6A1,FADS1,NEFL,UGT8,YBX1,CALR,FSTL1,TMF1,ABHD2,HSPA1B,T
NA,NCL,RPL7A,FADS1,PRPF8,RPL8,YBX1,ATP2B4,CALR,TMF1,HSPA1B,ZNF99,RPLP0,H3-3B,CL
UBC,TMSB4X,EIF4G2,RPL3,RPS6,ZNF316,RPL4,ENO1,PIK3R2,FLNA,NCL,RPL7A,ALDOA,PKM,FA
UBC,TMSB4X,EIF4G2,RPL3,RPS6,ZNF316,RPL4,ENO1,PIK3R2,FLNA,NCL,RPL7A,ALDOA,PKM,FA
CLU,SOX4,FKBP14,MAGED1,KAT6A,DVL3,YWHAE,RACK1,SPTBN1,CALM3,P4HB,APP,RPL5,GSTP
EIF4G2,RPL3,RPS6,ZNF316,RPL4,ENO1,PIK3R2,FLNA,NCL,RPL7A,ALDOA,PKM,FADS1,PRPF8,RP
T,FN1,PPP2R1A,BASP1,ITGB1,MORF4L1,AGRN,ZMIZ1,VCL,PAFAH1B1,EIF4H,NACA,RTN4,IGFBP
MF1,HSPA1B,CLU,PTMA,DDX5,SOX4,MAGED1,YWHAE,GPI,RACK1,GNB1,P4HB,APP,TMBIM6,GST
RPL3,RPS6,ZNF316,RPL4,ENO1,PIK3R2,FLNA,NCL,RPL7A,FADS1,PRPF8,RPL8,YBX1,ATP2B4,CA
,PTTG1IP,MAP1A,NOP53,VPS35,RAD23B,ABCA2,GABARAP,CCAR2,SMARCC1,UBQLN1,CSNK1D,
UBC,TMSB4X,EIF4G2,RPL3,RPS6,ZNF316,RPL4,ENO1,PIK3R2,FLNA,NCL,RPL7A,PKM,FADS1,RP
FAH1B1,MAP4,HNRNPU,RTN4,XRCC5,RHOA,NR2F1,CRMP1,VAT1,PFN1,PRKCSH,HNRNPC,SPTB
PM1,VCP,TRIM28,MAPT,DDB1,TMSB10,RPS3,HMGB1,CDKL1,STMN1,AGRN,VCL,ARF1,MAP4,H
P11,VCP,UBB,DDB1,PSMD5,PKD1,AP2M1,UCHL1,RPL11,ANXA2,AGRN,TIMP2,MARCHF6,AP2B1,
BC,TMSB4X,EIF4G2,RPL3,RPS6,ZNF316,RPL4,ENO1,PIK3R2,FLNA,NCL,RPL7A,PKM,FADS1,RPL
R,HSPA1B,COL5A1,PGK1,H3-3B,DDX5,SOX4,MAGED1,DVL3,KIF1A,GNAS,DDX17,RACK1,MYH9,
AQ,AP2M1,ITGB1,DNAJA1,AP2B1,CTNNA1,RTN4,MSN,KIF5B,CD81,CCT7,OAZ1,ASTN2,XPO1,PTN
RPL23,PSMD2,ENC1,TMEM259,XPO1,CSNK1E,RPS7,CTNNB1,NDFIP1,UBE2H,SKP1,TRIB2,MAP1
UBC,TMSB4X,EIF4G2,RPL3,RPS6,ZNF316,RPL4,ENO1,PIK3R2,FLNA,NCL,RPL7A,PKM,FADS1,RP
STL1,TMF1,HSPA1B,CLU,PTMA,DDX5,SOX4,MAGED1,YWHAE,GPI,RACK1,GNB1,P4HB,APP,TMB
,CTNNB1,UBE2H,SKP1,TRIB2,MAP1A,RNF187,NOP53,UBE4A,RAD23B,PSMB2,UBXN4,PSMD3,GA
N4,IGFBP3,RHOA,CLSTN1,DDR1,PLXNA1,MEGF8,MMP14,DDX3X,GJA1,CTNNB1,MACF1,CYFIP2,
P2,CALR,FSTL1,TMF1,ABHD2,COL4A1,H3-3B,DDX5,GNAS,DDX17,HTRA4,GNB1,STMN2,APP,GS
TG1,EEF2,DPYSL3,TUBA1A,CFL1,BTBD9,XRCC2,MAVS,HTT,HSP90AB1,ZNF124,HSP90AA1,ACTR
,IGFBP2,CALR,FSTL1,TMF1,HSPA1B,TUBB2B,H3-3B,CLU,DDX5,SOX4,MAGED1,KAT6A,DVL3,GN
4A1,H3-3B,CLU,SPTBN1,STMN2,P4HB,APP,FSCN1,SPTAN1,MARCKS,COL18A1,MARCKSL1,UBB,
15,EIF3L,RPS5,RPS19,RPS10,PTGES3,RPS27,RPL23A,LUC7L3,DDX39B,SRSF5,NOP53,KLC1,PRP
MSB4X,EIF4G2,RPL3,RPS6,ZNF316,RPL4,ENO1,PIK3R2,FLNA,NCL,RPL7A,ALDOA,PKM,FADS1,PR
UBC,TMSB4X,EIF4G2,RPL3,RPS6,ZNF316,RPL4,ENO1,PIK3R2,FLNA,NCL,RPL7A,ALDOA,PKM,FA
TMA,SOX4,MAGED1,YWHAE,GPI,RACK1,P4HB,TMBIM6,GSTP1,NPM1,HSPD1,VCP,UBB,NONO,D
4A2,MAPT,FN1,PSMD5,EGLN1,HMGB1,AP2M1,ITGB1,PAFAH1B1,AP2B1,RTN4,MSN,RHOA,PSMD

PS5,RPS19,RPS10,PTGES3,RPS27,RPL23A,LUC7L3,DDX39B,SRSF5,NOP53,PRPF6,RPL38,SRSF9,
APP,SPTAN1,GSTP1,CALM1,IGF2,VCP,FN1,HLA-C,VAPA,HMGB1,TUBB4B,ANXA2,TIMP2,VCL,ARF
RPS3,HMGB1,TPT1,DNAJA1,PAFAH1B1,PJA2,ATF4,TMEM259,RPL26,MCL1,CLEC7A,PTN,SFPQ,C
MYH9,APP,RPL5,CALM1,NPM1,PLXNB2,HSPD1,IGF2,EIF3C,VCP,UBB,MAPT,FN1,DDB1,HACD3,PK
N1,GSTP1,CALM1,IGF2,VCP,FN1,HLA-C,VAPA,HMGB1,TUBB4B,ANXA2,TIMP2,VCL,ARF1,CYB5R
,IGFBP2,CALR,FSTL1,TMF1,HSPA1B,TUBB2B,H3-3B,CLU,DDX5,SOX4,MAGED1,KAT6A,DVL3,GN
MSB4X,EIF4G2,RPL3,RPS6,ZNF316,RPL4,ENO1,PIK3R2,FLNA,NCL,RPL7A,ALDOA,PKM,FADS1,PR
MSB4X,EIF4G2,RPL3,RPS6,ZNF316,RPL4,ENO1,PIK3R2,FLNA,NCL,RPL7A,ALDOA,PKM,FADS1,UC
TG1,EEF2,DPYSL3,TUBA1A,CFL1,BTBD9,XRCC2,MAVS,HTT,HSP90AB1,ZNF124,HSP90AA1,ACTR
,MAP1A,RNF187,NOP53,UBE4A,RAD23B,PSMB2,UBXN4,PSMD3,GABARAP,CCAR2,SMARCC1,STT
,NPM1,PLXNB2,IGF2,VCP,UBB,YWHAG,MAPT,FN1,HACD3,SET,PPP2R1A,PKD1,RPS3,HMGB1,UC
1,CSNK1E,RPS7,CTNNB1,NDFIP1,UBE2H,SKP1,TRIB2,MAP1A,RNF187,SQSTM1,NOP53,UBE4A,R
PTTG1IP,CDKN1A,CPSF2,DDX39B,SRSF5,U2AF2,CPSF1,PABPN1,ALKBH5,AHCYL1,TNPO1,MAPK
M28,YWHAG,MAPT,YWHAQ,AP2M1,ITGB1,MIDN,VCL,DNAJA1,ARF1,AP2B1,CTNNA1,CTSD,RTN4,
1,CSNK1E,RPS7,CTNNB1,NDFIP1,UBE2H,SKP1,TRIB2,MAP1A,RNF187,SQSTM1,NOP53,UBE4A,R
E4A,GRINA,HDGF,TMED2,UBXN4,CCND1,ACADVL,STT3B,UBQLN1,HYOU1,BAG6,SSR1,DCTN1,SG
SMD4,RBPJ,PSMC3,CREBBP,PSMC1,PSMB1,HSPA1A,NOTCH1,DNAJB1,PSMD13,PSMD7,NCK2,PS
1,CSNK1E,RPS7,CTNNB1,NDFIP1,UBE2H,SKP1,TRIB2,MAP1A,RNF187,SQSTM1,NOP53,UBE4A,R
PTMA,SOX4,MAGED1,YWHAE,GPI,RACK1,P4HB,TMBIM6,GSTP1,NPM1,HSPD1,VCP,UBB,NONO,DD
PTN,PLXNA1,MEGF8,MACF1,RAC1,ARHGDIA,CAPRIN1,TMEM30A,L1CAM,ACTR3,NCKAP1,MDK,PT
PJ,PSMC3,CREBBP,PSMC1,PSMB1,HSPA1A,NOTCH1,DNAJB1,PSMD13,PSMD7,NCK2,PSME3,PSN
PTMA,SOX4,MAGED1,YWHAE,GPI,RACK1,P4HB,TMBIM6,GSTP1,NPM1,HSPD1,VCP,UBB,MAPT,NO
PM1,YWHAG,MAPT,SET,PPP2R1A,RPS3,UCHL1,RPL11,ANXA2,TIMP2,DNAJA1,ITM2B,LARP1,RPL2
PM1,VCP,TRIM28,YWHAG,MAPT,YWHAQ,AP2M1,ITGB1,ANXA2,DNAJA1,ARF1,AP2B1,HNRNPU,CT
NC,SKP1,TMEM132A,NUCB1,VCAN,LAMC1,PSMB2,RAB11B,ABCA2,VHL,PSMD3,CAND1,PTP4A2,
L1,UBB,CKB,MAPT,YWHAQ,INA,PKD1,BASP1,STMN1,ITGB1,ZMIZ1,TIMP2,PAFAH1B1,CTNNA1,L
VCL,ARF1,PAFAH1B1,CTNNA1,GAP43,RHOA,CLSTN1,NLGN2,CXADR,WDR1,PFN1,SPTBN2,PTN,C
ID3,FTO,HSPB1,HNRNPM,HNRNPD,SERBP1,PSMA7,ANP32A,EDC4,PSMD1,GTPBP1,UBA52,ZFP36
P1,SUPT5H,SON,SUPT6H,SRSF3,RBM15B,SF1,ZFP36L1,SRSF6,TRA2B,HNRNPA2B1,PCBP4,SAFB
MSB4X,EIF4G2,RPL3,RPS6,ZNF316,RPL4,ENO1,PIK3R2,FLNA,NCL,RPL7A,ALDOA,PKM,FADS1,PR
ID3,FTO,HSPB1,HNRNPM,HNRNPD,SERBP1,PSMA7,ANP32A,EDC4,PSMD1,UBA52,ZFP36L1,UPF1,
K2B,RAB7A,CLTC,SESN3,RAB1A,ATP6V0C,DCN,TOMM20,EEF1A2,SQSTM1,GNAI3,VPS35,SNRNP
K2B,RAB7A,CLTC,SESN3,RAB1A,ATP6V0C,DCN,TOMM20,EEF1A2,SQSTM1,GNAI3,VPS35,SNRNP
F4H,NACA,RTN4,GAP43,CD81,SPTBN2,PTN,DDR1,PLXNA1,MEGF8,GPAT4,PRKAR1A,GJA1,CTNN
PTTG1IP,CDKN1A,CPSF2,DDX39B,SRSF5,U2AF2,CPSF1,PABPN1,ALKBH5,AHCYL1,TNPO1,MAPK
ARCC1,UBQLN1,CSNK1D,USP7,HECTD1,BAG6,ARIH1,SGTA,UBQLN2,CSNK2A1,PTEN,GPX1,EIF3
ALM1,NPM1,HSPD1,COL18A1,VCP,UBB,MAPT,DDB1,PSMD5,EGLN1,PKD1,POLR2A,ITGB1,COL1
FM,UGT8,CALR,HSPA1B,TUBB2B,SPTBN1,STMN2,MYH9,FSCN1,SPTAN1,MARCKS,NPM1,VCP,M
WHAE,RACK1,SPTBN1,CALM3,MYH9,SEPTIN2,APP,CALM1,NPM1,IGF2,VCP,UBB,YWHAG,DDB1,PS
OST,PTGES3,LAMP1,CCT6A,PPIB,CDKN1A,SERF2,CCT2,NOP53,TCP1,CCT3,VPS35,CCT4,VHL,M
H2,GPC3,SULF1,DDX3X,CLTC,CTNNB1,MACF1,WLS,RAC1,ATP6V0C,SKP1,VPS35,MDK,PTK7,SKI,
N1,ARF1,MSN,RHOA,PFN1,CLEC7A,SPTBN2,DDX3X,CYFIP2,CKAP5,MAP2,COTL1,RAC1,TBCD,NO
IMP14,DDX3X,GJA1,MACF1,MAP2,L1CAM,CDKN1A,DDX39B,CLSTN3,H3-3A,DPYSL2,VEGFA,PPP2
P2A2,NES,SPTBN2,EIF5A,DDR1,MMP14,H1-0,CTNNB1,ARID1A,CKAP5,MMP2,MAP1A,SQSTM1,M
H2,GPC3,SULF1,DDX3X,CLTC,CTNNB1,MACF1,WLS,RAC1,ATP6V0C,SKP1,VPS35,MDK,PTK7,SKI,

?,RPL5,CALM1,NPM1,PLXNB2,HSPD1,IGF2,EIF3C,VCP,UBB,MAPT,FN1,HACD3,PKD1,RPS3,HMGB
i,CALM3,GNB1,APP,TMBIM6,GSTP1,CALM1,HSPD1,IGF2,VCP,YWHAG,ITGB1,MARCHF6,LARP1,C
),USP7,HECTD1,BAG6,ARIH1,SGTA,UBQLN2,GPX1,EIF3H,GNA12,DNAJB2,ATXN3,OGT,GSK3A,PSI
ANXA2,TIMP2,VCL,CYB5R3,PGAM1,CTSD,LPCAT1,XRCC5,KPNB1,RHOA,PSMD2,VAT1,PFKL,RAB7
'TGES3,CCT6A,PPIB,CCT2,TBCD,GNAI3,HSPH1,TCP1,CCT3,MLEC,RAD23B,CCT4,SNRNP70,CDC3
3PN1,ALKBH5,HNRNPA3,TCP1,CCT3,CCT4,PRPF6,G3BP2,SRSF9,SRSF1,CASC3,SUPT6H,SSB,SRS
HF6,WSB1,DNAJA1,RPL23,PJA2,PSMD2,ENC1,RPS20,KLHL9,DDX3X,RPS15,RPS7,CTNNB1,NDFIP
SF1,PTBP1,SON,SRSF3,RPS13,RBM15B,SF1,GRSF1,SRSF6,TRA2B,HNRNPA2B1,PCBP4,PRDX6,FL
G,MAPT,SET,PPP2R1A,RPS3,UCHL1,RPL11,ANXA2,TIMP2,DNAJA1,LARP1,RPL23,IGFBP3,ATF4,EN
ANXA2,TIMP2,VCL,CYB5R3,PGAM1,CTSD,LPCAT1,XRCC5,KPNB1,RHOA,PSMD2,VAT1,PFKL,RAB7
ID3,FTO,HSPB1,HNRNPM,HNRNPD,SERBP1,PSMA7,ANP32A,EDC4,PSMD1,UBA52,ZFP36L1,UPF1,
PI,BSG,CALM3,GNB1,APP,TMBIM6,GSTP1,CALM1,NPM1,HSPD1,IGF2,VCP,YWHAG,EGLN1,ITGB1
ANXA2,TIMP2,VCL,CYB5R3,PGAM1,CTSD,LPCAT1,XRCC5,KPNB1,RHOA,PSMD2,WDR1,VAT1,PFK
,GTF2F1,COL1A2,RHOA,PTN,GLG1,GPC3,SULF1,MEGF8,CCN2,JUN,CTNNB1,CD63,MTSS2,CYFI
RPL11,ZMIZ1,MARCHF6,WSB1,DNAJA1,RPL23,PJA2,RHOA,PSMD2,ENC1,RPS20,TADA2B,KLHL9,
L2,MED18,ACTG1,EEF2,DPYSL3,TUBA1A,CFL1,BTBD9,XRCC2,MAVS,HTT,HSP90AB1,ZNF124,HSI
ANXA2,TIMP2,VCL,CYB5R3,PGAM1,CTSD,LPCAT1,XRCC5,KPNB1,RHOA,PSMD2,VAT1,PFKL,RAB7
HSPA1B,H3-3B,CLU,DDX5,SOX4,MAGED1,KAT6A,DVL3,GNAS,HTRA4,RACK1,SPTBN1,SHISA9,P4I
,WDR1,CCT7,PFN1,NES,TADA2B,SFPQ,TPM1,PIP4K2B,CCN2,CTNNB1,YWHAB,CKAP5,CCT6A,CTE
NPM1,IGF2,VCP,UBB,YWHAG,DDB1,PSMD5,SET,MAP9,PPP2R1A,PKD1,RPS3,CHD3,STMN1,ITGB
ANXA2,TIMP2,VCL,CYB5R3,PGAM1,CTSD,LPCAT1,XRCC5,KPNB1,RHOA,PSMD2,VAT1,PFKL,RAB7
SB1,DNAJA1,RPL23,PJA2,PSMD2,ENC1,RPS20,KLHL9,DDX3X,RPS15,RPS7,CTNNB1,NDFIP1,UBE2
'HAG,DDB1,PSMD5,SET,MAP9,PPP2R1A,PKD1,STMN1,ITGB1,TUBB4B,TIMP2,ARF1,PAFAH1B1,M
TG2B,DYNC1LI2,DYNLL2,CDC37,GABARAP,SUPT5H,BNIP3L,ATP6V1B2,UBQLN1,HGS,DYNLL1,VD
31,PSMD5,MAP9,PPP2R1A,PKD1,CDKL1,ITGB1,TUBB4B,TIMP2,PAFAH1B1,HNRNPU,RHOA,CBX5
A3,GPI,BSG,CALM3,MYH9,SEPTIN2,APP,SPTAN1,GSTP1,CALM1,IGF2,VCP,FN1,HLA-C,VAPA,HMG
P2R1A,PKD1,STMN1,ITGB1,TUBB4B,ARF1,PAFAH1B1,MAP4,HNRNPU,RPL24,KPNB1,RHOA,PSMI
'FIP2,MAP2,TNC,L1CAM,PTK7,DPYSL2,VEGFA,SLIT3,PLXNA3,PTPRS,RAB11A,BMPR2,KDM5B,DII
B1,NOTCH1,PSMD13,PSMD7,PSME3,PSMC5,PSMC2,PSMC4,PSMF1,PSMB5,EGLN2,HIF3A,PSMA
D1,KAT6A,DVL3,DDX17,RACK1,APP,RPL5,NPM1,PLXNB2,IGF2,EIF3C,VCP,UBB,TRIM28,DDB1,PK
\T6A,DVL3,YWHAE,GNAS,RACK1,SPTBN1,CALM3,GNB1,P4HB,APP,RPL5,GDI1,TMBIM6,SPTAN1,G
PRS,RAB11A,BMPR2,DIP2B,ULK1,CTTN,SLIT2,NDN,SLIT1,OLFM1,DCLK1,SEMA3C,CLASP2,ABL1
ZMIZ1,HNRNPU,SMARCA5,BRD3,XRCC5,KPNB1,CHD4,HMGN2,CCT7,RBBP4,HNRNPC,TADA2B,SF
4,HSPA1B,PSAP,CLU,MAGED1,DVL3,YWHAE,GNAS,GPI,RACK1,CALM3,SHISA9,APP,RPL5,GDI1,TM
D1,KAT6A,DVL3,DDX17,RACK1,APP,RPL5,NPM1,PLXNB2,IGF2,EIF3C,VCP,UBB,TRIM28,DDB1,PK
F,TMED2,OPTN,UBXN4,CCND1,ACADVL,HSPB1,STT3B,HYOU1,BAG6,SSR1,DCTN1,EP300,EXTL3,
,GTF2F1,COL1A2,RHOA,PTN,GLG1,GPC3,SULF1,MEGF8,CCN2,JUN,CTNNB1,CD63,MTSS2,CYFI
P2R1A,VAPA,AP2M1,ANXA2,AGRN,AP3D1,ARF1,PAFAH1B1,AP2B1,TMEM170A,SCARB2,RTN4,KPN
INA1,LDHA,RTN4,XRCC5,RHOA,KIF5B,NES,SPTBN2,PTN,CSNK1E,CDH2,MMP14,APLP1,HMGCS1,
1,GLUL,PTTG1IP,CDKN1A,CCT2,NOP53,PARP1,TCP1,CCT3,CCT4,TNPO1,MAPK1,IPO7,FYN,CCT8
ED2,OPTN,UBXN4,CCND1,ACADVL,HSPB1,STT3B,HYOU1,SSR1,DCTN1,EP300,EXTL3,MBTPS1,KL
'1CB,NOP53,PSMB2,OPTN,CCND1,MAPRE1,PSMD3,CSNK1D,RAB11A,DYNLL1,PSMA7,DCTN1,AC
I1,SPTBN2,MEGF8,MACF1,CYFIP2,MAP2,COTL1,RAC1,L1CAM,ACTR3,NCKAP1,DPYSL2,VEGFA,AI
FN1,DCHS1,PKD1,BASP1,ITGB1,BTF3,ZMIZ1,COL11A1,RPL10,PAFAH1B1,RTN4,LAMA5,ATF4,CO
SMD3,CSNK1D,RAB11A,DYNLL1,PSMA7,DCTN1,ACTR1A,PSMD1,PBX1,BRD4,BRSK1,PSMD8,AKAF

.1CAM,DPYSL2,VEGFA,SLIT3,PLXNA3,PTPRS,RAB11A,BMPR2,DIP2B,ULK1,CTTN,SLIT2,NDN,SLIT
(17,APP,RPL5,NPM1,PLXNB2,IGF2,EIF3C,UBB,TRIM28,PKD1,SSBP2,ZNF43,HMGB1,RPS4X,AGRN
\P2,L1CAM,DDX39B,DPYSL2,VEGFA,SLIT3,PLXNA3,PTPRS,RAB11A,BMPR2,DIP2B,HDAC6,ULK1,C
4B,PAFAH1B1,HNRNPU,RHOA,CBX5,PSMD2,RPL26,RBBP4,KLHL9,XPO1,CSNK1E,SFPQ,CCND2,C
A,GJA1,SESN3,TARDBP,NDFIP1,TRIB2,PTTG1IP,DCN,GNAI3,NOP53,VPS35,ABCA2,NRDC,OPTN,A
2,YWHAG,HMGB1,ANXA2,ZMIZ1,TIMP2,DNAJA1,HNRNPU,CTNNA1,LDHA,RHOA,ATP1A1,JUND,SRS
A,DYNLL1,PSMA7,DCTN1,KIF5A,ACTR1A,PSMD1,AP2A1,ITGB5,PSMD8,PSMB4,KIFAP3,CAPZA1,A
N,RPS15,RPS7,PRKAR1A,CTNNB1,TARDBP,YWHAB,CADM4,CTBP1,HMGCR,HDAC2,IPO5,TRIB2,C
CSNK1D,RAB11A,DYNLL1,PSMA7,DCTN1,ACTR1A,PSMD1,PBX1,BRD4,BRSK1,PSMD8,AKAP9,PSM
BB4B,ANXA2,TIMP2,VCL,CYB5R3,PGAM1,CTSD,LPCAT1,XRCC5,KPNB1,RHOA,PSMD2,WDR1,VAT
ENO1,PIK3R2,FLNA,MAP1B,HSP90B1,FADS1,NEFL,YBX1,UBA1,ATP2B4,CALR,TMF1,ABHD2,HSPA
AS,SLC2A3,GPI,BSG,CALM3,MYH9,SEPTIN2,APP,SPTAN1,GSTP1,CALM1,IGF2,VCP,FN1,HLA-C,VA
ANXA2,TIMP2,VCL,CYB5R3,PGAM1,CTSD,LPCAT1,XRCC5,KPNB1,RHOA,PSMD2,VAT1,PFKL,RAB7
1,HSP90AA1,ACTR2,VIM,UBC,KIDINS220,TMSB4X,NUMA1,DCX,RPL3,RPS6,ENO1,TUBA1B,PIK3R
F1,SRSF6,TRA2B,HNRNPA2B1,PCBP4,PRDX6,SRSF2,RBM5,PRPF19,SAP18,THRAP3,FXR2,RBM4,
OP53,PSMB2,OPTN,CCND1,MAPRE1,PSMD3,CSNK1D,RAB11A,DYNLL1,PSMA7,DCTN1,ACTR1A,P
RPL29,RPSA,RPS24,RPS9,XRCC5,RPL24,RPL17,RPL14,RPL22,RPL9,RPS14,RPL26,RPL31,XPO1,
HB,APP,GDI1,PLXNB2,IGF2,YWHAG,MAPT,FN1,SPEN,PSMD5,PPP2R1A,HMGB1,ITGB1,ZMIZ1,TIM
IA,RTN4,XRCC5,RHOA,KIF5B,NES,SPTBN2,PTN,CSNK1E,CDH2,APLP1,HMGCS1,CTNNB1,WLS,AT
CCT2,NOP53,TCP1,CCT3,CCT4,VHL,CDC37,CCT8,STXBP1,USP7,BAG6,PFN2,CCT5,CTNND1,EP30
CIRBP,HMGCR,LAPTM4B,MAP1A,TIMP3,NOP53,TAF15,SNRNP70,AZIN1,WDR6,CCAR2,SMARCC1,
,SEC23A,DYNLL1,PSMA7,DCTN1,KIF5A,ACTR1A,PSMD1,AP2A1,ITGB5,PSMD8,PSMB4,KIFAP3,CA
B,SFPQ,H1-0,CTNNB1,ARID1A,H1-10,CTBP1,SMCHD1,HDAC2,SKP1,EID1,CHD6,TSPYL4,MORF4L
AMP2,KLC1,SURF4,TMED2,OPTN,ARF3,KDELR2,DYNC1LI2,DYNLL2,CAPZB,KDELR1,ARF4,KIF3C
HNRNPU,RHOA,WDR1,PFN1,NES,SPTBN2,TPM1,CCN2,CLTC,CYFIP2,CKAP5,RHOB,MAP2,COTL1,
B1,NACA,RTN4,IGFBP3,GAP43,RHOA,CLSTN1,TLE5,TKT,GPC3,DDR1,MEGF8,MMP14,DDX3X,GP
MATR3,PSMD5,EIF1,SET,RPS3,RPS4X,RPL10,HNRNPU,LARP1,HNRNPA0,EIF4H,RPS9,RPL22,RHO
PGAM1,LARP1,EIF4H,RTN4,RPS9,RPL22,RHOA,PTCD3,ATF4,ENC1,RPS14,EIF5,RPL26,RTN3,LRP
ANXA2,TIMP2,VCL,CYB5R3,PGAM1,CTSD,LPCAT1,XRCC5,KPNB1,RHOA,PSMD2,VAT1,PFKL,RAB7
_26,BEX3,MCL1,SFPQ,PRDX2,DDX3X,ITM2C,RPS7,MIF,JUN,CTNNB1,ANXA6,HNRNPK,YWHAB,AP
1,CASC3,SUPT6H,SSB,SRSF3,ADAR,RBM15B,XPOT,SRSF6,UPF1,HNRNPA2B1,ANP32B,MCM3AP,
LAPTM4B,MAP1A,TIMP3,NOP53,TAF15,SNRNP70,WDR6,CCAR2,SMARCC1,USP7,BAG6,HNRNPD,
LXNA1,MEGF8,ACTN4,RPS19,CTNNB1,MACF1,MAP2,RAC1,ARHGDIA,CAPRIN1,HDAC2,L1CAM,M
K1E,CDH2,GPC3,SULF1,DDX3X,CLTC,CTNNB1,MACF1,WLS,RAC1,ATP6V0C,SKP1,VPS35,MDK,PT
,ZNF316,ENO1,PIK3R2,FLNA,NCL,PKM,FADS1,YBX1,ATP2B4,CALR,TMF1,HSPA1B,ZNF99,H3-3B,
S35,ABCA2,NRDC,ADAM9,GABARAP,UBQLN1,CSNK1D,HECTD1,BAG6,ARIH1,SGTA,UBQLN2,CSN
,SLC2A3,GPI,BSG,CALM3,MYH9,SEPTIN2,APP,SPTAN1,GSTP1,CALM1,IGF2,VCP,FN1,HLA-C,VAP
1,CALM3,APP,RPL5,GDI1,TMBIM6,GSTP1,CALM1,NPM1,PLXNB2,HSPD1,IGF2,VCP,UBB,YWHAG,
,TTC3,HSPD1,USP11,VCP,UBB,MAPT,FN1,DDB1,PSMD5,PKD1,PDIA3,RPS3,HMGB1,UCHL1,RPL1
,TPM1,CCN2,CYFIP2,CKAP5,MAP2,COTL1,RAC1,MAP1A,TBCD,NOX4,ACTR3,CAMSAP1,NCKAP1,
CIN1,RNF40
,RPS24,RPS9,RPL24,RPL17,RPL14,RPL22,RPL9,RPS14,RPL26,RPL31,RPS25,RPS15,RPL12,RPL
M6,GSTP1,NPM1,HSPD1,UBB,NONO,DDB1,SET,TPT1,RPS3A,ITGB1,RPL10,DNAJA1,CTNNA1,NAC
RACK1,TMBIM6,GSTP1,NPM1,HSPD1,UBB,NONO,DDB1,SET,TPT1,RPS3A,ITGB1,RPL10,DNAJA1,
RPL29,RPS24,RPS9,RPL24,RPL17,RPL14,RPL22,RPL9,RPS14,RPL26,RPL31,RPS25,RPS15,RPL1

P2,SRSF9,SRSF1,CASC3,SUPT6H,SSB,SRSF3,RBM15B,XPOT,FLOT1,ZFP36L1,SRSF6,UPF1,HNRI
P2,SRSF9,SRSF1,CASC3,SUPT6H,SSB,SRSF3,RBM15B,XPOT,FLOT1,ZFP36L1,SRSF6,UPF1,HNRI
PSMD2,RPL26,NES,CCNI,KHDRBS1,CSNK1E,CCND2,DDX3X,IK,CKAP5,CDK16,APPL1,SKP1,CDKN
1,EIF4H,RPS9,RPL22,RHOA,PTCD3,ATF4,ENC1,RPS14,EIF5,RPL26,LRPPRC,KHDRBS1,EIF5A,DDX
C7A,TPM1,PIP4K2B,RAB7A,CCN2,DDX3X,CKAP5,RAC1,DLG5,PARP1,NOX4,ABCA3,ACTR3,NCKAF
,ZNF316,ENO1,PIK3R2,FLNA,NCL,PKM,FADS1,YBX1,ATP2B4,CALR,TMF1,HSPA1B,ZNF99,H3-3B,
A2,AP2M1,STMN1,ZMIZ1,AP2B1,GTF2F1,RTN4,IGFBP3,COL1A2,RHOA,PTN,GLG1,GPC3,DDR1,S
MD1,PHB2,UBA52,ZFP36L1,SFRP1,PTEN,RPS27A,PSMD8,HIF1A,PSMB4,PSMD4,RBPJ,PSMC3,M
311A,BMPR2,MAP6,PLXNB1,DIP2B,PTEN,BRSK1,SLIT2,LINGO1,SEMA3C,IST1,METRN,NTRK2,SE
M6,GSTP1,NPM1,HSPD1,UBB,NONO,DDB1,SET,TPT1,RPS3A,ITGB1,RPL10,DNAJA1,CTNNA1,NAC

ACF1,RAC1,ARHGDIA,CAPRIN1,HDAC2,L1CAM,MDK,ATP6AP1,VEGFA,ARPC2,PRKDC,KDM1A,PL
00,PSMD1,PHB2,UBA52,ZFP36L1,SFRP1,PTEN,RPS27A,PSMD8,HIF1A,PSMB4,PSMD4,RBPJ,PSM
CLSTN1,NLGN2,PFN1,SPTBN2,PTN,CDH2,CTNNB1,MYH10,SEPTIN11,CAPRIN1,TNC,DNER,L1CAM
,ARHGDIA,CAPRIN1,HDAC2,L1CAM,MDK,SKI,RNF10,VEGFA,KDM1A,YWHAH,B2M,PLXNA3,PTPRS
DAC2,L1CAM,MDK,VEGFA,KDM1A,PLXNA3,RAB11A,BMPR2,NAP1L1,MAP6,PLXNB1,PPP1CC,IL6S
PSMA7,PSMD1,AP2A1,SFRP1,BRSK1,PSMD8,GPC4,PSMB4,PSMD4,SCRIB,PSMC3,AP2A2,PSMC1,
N1,EP300,EXTL3,MBTPS1,KLHDC3,PPP1R15A,SERP1,RHBDD2,VAPB,TPP1,ATP6V0D1,GSK3A,SR
PIK3R2,FLNA,NCL,FADS1,YBX1,ATP2B4,CALR,TMF1,HSPA1B,ZNF99,H3-3B,CLU,DDX5,SOX4,MA
TL1,RAC1,TBCD,ACTR3,CAMSAP1,NCKAP1,MAPRE1,ARPC2,LUZP6,GRB2,CAPZB,CLIP3,PFN2,DC
,ZNF316,ENO1,PIK3R2,FLNA,NCL,PKM,FADS1,YBX1,ATP2B4,CALR,TMF1,HSPA1B,ZNF99,H3-3B,
1,SLIT2,IST1,METRN,NTRK2,TRAK1,PLXNA2,ADNP,CYFIP1,NTN1,LIMK1,ISLR2,GOLGA4,SRF,CR
1,DCTN1,EP300,EXTL3,MBTPS1,HDAC6,CUL3,KLHDC3,PPP1R15A,SERP1,RHBDD2,VAPB,TPP1,A
P2,SRSF9,SRSF1,CASC3,SUPT6H,SSB,SRSF3,RBM15B,XPOT,FLOT1,ZFP36L1,SRSF6,UPF1,HNRI
HYOU1,PSMA7,EP300,PSMD1,PHB2,UBA52,ZFP36L1,SFRP1,PTEN,RPS27A,PSMD8,HIF1A,PSMB4,
CBP4,FUS,ELAVL1,IGF2BP1,MTOR,SAP18,THRAP3,SYNCRIP,NBAS,SRSF10,C1QBP,MAPKAPK2,C
,ZNF316,ENO1,PIK3R2,FLNA,NCL,PKM,FADS1,YBX1,ATP2B4,CALR,TMF1,HSPA1B,ZNF99,H3-3B,
NES,CCNI,KHDRBS1,CSNK1E,CCND2,DDX3X,IK,CKAP5,CDK16,APPL1,SKP1,CDKN1A,CTDSP2,PPF
,PSMC3,AP2A2,PSMC1,CELSR1,ABL1,PSMB1,PSMD13,PSMD7,PSME3,DAAM1,PSMC5,PSMC2,PS
YWHAQ,SET,RPS3,CHD3,BASP1,HMGB1,MORF4L1,RPL10,HNRNPU,SMARCA5,LARP1,HNRNPA0,
ECTD1,BAG6,ARIH1,SGTA,UBQLN2,CSNK2A1,PTEN,EIF3H,GNA12,DNAJB2,OGT,GSK3A,UBE3A,B
CD81,RBBP4,XPR1,XPO1,RAB7A,DDX3X,HNRNPK,LAMP1,RAB1A,CTBP1,TOP2B,PPIB,CTSB,CBX1,
3,DDX17,APP,NPM1,IGF2,VCP,UBB,TRIM28,PKD1,SSBP2,ZNF43,RPS3,HMGB1,AGRN,ZMIZ1,AP3
K1,PSMD8,GPC4,PSMB4,PSMD4,PSMC3,AP2A2,PSMC1,CELSR1,ABL1,PSMB1,EXOC5,PSMD13,P
K1,PSMD8,GPC4,PSMB4,PSMD4,PSMC3,AP2A2,PSMC1,CELSR1,ABL1,PSMB1,EXOC5,PSMD13,P
1,PLXNA3,PTPRS,RAB11A,BMPR2,DIP2B,PTEN,CTTN,OLFM1,AKT3,RDX,SEMA3C,CLASP2,IST1,M
A,CSNK2A1,CST3,EIF3H,OGT,BAG5,SNX3,SDCBP,ATP13A2,MGAT3,PARK7,USP14,HIPK2,MYCBP
A,DYNLL1,PSMA7,DCTN1,KIF5A,ACTR1A,PSMD1,AP2A1,ITGB5,PSMD8,PSMB4,KIFAP3,CAPZA1,A
,SGTA,UBQLN2,EIF3H,GNA12,DNAJB2,OGT,GSK3A,BAG5,UBE2K,ZER1,USP5,SDCBP,HSPA1A,MI
,TARDBP,YWHAB,CCT6A,IPO5,MAP1A,CCT2,SQSTM1,PARP1,TCP1,CCT3,CCT4,VEGFA,MAPK1,NI
P2,COTL1,RAC1,TBCD,GAS7,ACTR3,CAMSAP1,NCKAP1,MAPRE1,ARPC2,LUZP6,GRB2,CAPZB,CLI
,XPR1,SFPQ,RAB7A,JUN,CTNNB1,TARDBP,LAMP1,PPIB,DDX39B,CTSB,PABPN1,TCP1,MAPK1,SM
B,RAC1,ARHGDIA,CAPRIN1,CAMSAP1,MDK,VEGFA,ARPC2,CAPZB,PRKDC,CORO1C,FYN,FBLN1,F
TPRS,CDC42,PPP1R9B,RERE,MAP6,FARP1,CTNND2,HDAC6,PTEN,CARM1,ABI2,DCLK1,ADAM10,I
I3,NOP53,VPS35,ABCA2,NRDC,OPTN,ADAM9,FTO,GABARAP,SUPT5H,BNIP3L,UBQLN1,CSNK1D,H

KN1A,NOP53,NOX4,ALKBH5,PLOD1,PSMB2,ATP6AP1,VEGFA,VHL,PSMD3,NPEPPS,BNIP3L,LONP
DL4A1,CLU,SOX4,GNAS,GPI,RACK1,BSG,CALM3,GNB1,APP,TMBIM6,GSTP1,CALM1,HSPD1,COL1
E,CCND2,DDX3X,IK,CTNNB1,CKAP5,APPL1,SKP1,CDKN1A,CTDSP2,NOP53,PSMB2,CCND1,MAPRE
VAPA,HMGB1,TUBB4B,ANXA2,TIMP2,VCL,CYB5R3,PGAM1,CTSD,LPCAT1,XRCC5,PJA2,KPNB1,RH
CTD1,PSMA7,TRAF4,ARIH1,PSMD1,UBA52,KLHL42,UBE2G2,FBXO11,HDAC6,CUL3,RPS27A,TNK
HD2,TUBB2B,COL5A1,KIF1A,GPI,RACK1,BSG,SPTBN1,ATP5F1B,MYH9,APP,FSCN1,SPTAN1,GSTP
K1,SPTBN1,CALM3,APP,GSTP1,SMG1,CALM1,NPM1,PLXNB2,IGF2,UBB,TRIM28,YWHAG,CKB,MA
,PSMC3,AP2A2,PSMC1,CELSR1,ABL1,PSMB1,PSMD13,PSMD7,PSME3,DAAM1,PSMC5,PSMC2,PS
3X1,ATP2B4,CALR,TMF1,HSPA1B,ZNF99,H3-3B,CLU,DDX5,SOX4,MAGED1,RPL6,KAT6A,DVL3,DD
RPL23,CTSD,RHOA,TMEM259,BEX3,CLEC7A,XPO1,CSNK1E,GLG1,PEBP1,GPC3,CCN2,MMP14,D
MD1,AP2A1,SFRP1,PSMD8,GPC4,PSMB4,PSMD4,PSMC3,AP2A2,PSMC1,CELSR1,ABL1,PSMB1,PS
,RBM15B,ZFP36L1,SRSF6,UPF1,HNRNPA2B1,MCM3AP,POM121C,TNKS,SYMPK,SMG5,SRSF2,SF
VDR6,SUPT5H,HSPB1,BNIP3L,ATP6V1B2,UBQLN1,SREBF2,VDAC1,STAT3,EP300,FLCN,UBQLN2,F
CP,UBB,FN1,PSMD5,HACD3,CHSY1,PDIA3,RPS3,VAPA,HMGB1,ITGB1,RPL11,ANXA2,ZMIZ1,TIMP2

HSP90AB1,HSP90AA1,ACTR2,VIM,SPARC,TUBB,FTH1,UBC,KIDINS220,TMSB4X,NUMA1,DCX,RPL3
TP1,CALM1,NPM1,PLXNB2,IGF2,VCP,UBB,YWHAG,MAPT,FN1,HACD3,SET,PPP2R1A,PKD1,RPS3,
TP1,CALM1,NPM1,PLXNB2,IGF2,VCP,UBB,YWHAG,MAPT,FN1,HACD3,SET,PPP2R1A,PKD1,RPS3,
1,HNRNPA2B1,MCM3AP,POM121C,SYMPK,SMG5,SRSF2,SRSF11,DHX38,EIF4A3,RAN,AKAP8L,NU
2AF2,SQSTM1,NOP53,RPL37,GABARAP,KDM1A,FYN,UBQLN1,MTA1,NXN,PTEN,CUL3,RPS2,HIF1A
F1,ARHGDIA,CAPRIN1,HDAC2,L1CAM,DLG5,CLSTN3,MDK,VEGFA,KDM1A,PLXNA3,RAB11A,BMPR
PPL1,SKP1,CDKN1A,CTDSP2,NOP53,PSMB2,CCND1,MAPRE1,PSMD3,CSNK1D,RAB11A,DYNLL1,F
B1,ARF1,PAFAH1B1,CDC42BPB,CTNNA1,CNN3,TPM4,LLGL1,RHOA,CXADR,WDR1,PFN1,NISCH,SF
MAPT,DDB1,PSMD5,PPP2R1A,PKD1,AP2M1,ITGB1,AGRN,MIDN,ARF1,PAFAH1B1,AKAP12,AP2B1,
,H1-0,JUN,CTNNB1,MAP2,HDAC2,CDKN1A,PARP1,C4ORF3,SKI,PPP2CA,MAPRE1,EIF3E,KDM1A,E
R1A,TPT1,STMN1,ITGB1,UCHL1,TIMP2,MIDN,DNAJA1,ARF1,PAFAH1B1,PXDN,CTNNA1,IGFBP3,RI
NEFL,YBX1,ATP2B4,IGFBP2,CALR,FSTL1,HSPA1B,TUBB2B,H3-3B,CLU,DDX5,SOX4,MAGED1,KAT
BP2,ZNF43,HMGB1,AGRN,ZMIZ1,AP3D1,ZNF770,GTF2F1,HNRNPU,SMARCA5,NACA,ATF4,NR2F1
ET,CHD3,BASP1,HMGB1,MORF4L1,RPL10,HNRNPU,SMARCA5,LARP1,HNRNPA0,RPL23,NACA,XR
PT1,STMN1,ITGB1,UCHL1,TIMP2,DNAJA1,PAFAH1B1,PXDN,CTNNA1,IGFBP3,RHOA,PSMD2,MCL1
MTRF1L,CAMSAP1,KLC1,SMARCA4,DSTN,LUZP6,CAPZB,GABARAP,SMARCE1,SMARCC1,STMN3,
R1A,TPT1,STMN1,ITGB1,UCHL1,TIMP2,MIDN,DNAJA1,ARF1,PAFAH1B1,PXDN,CTNNA1,IGFBP3,RI
CP,UBB,FN1,PSMD5,HACD3,CHSY1,PDIA3,RPS3,VAPA,HMGB1,ITGB1,RPL11,ZMIZ1,TIMP2,PAFAH
1,HNRNPA2B1,MCM3AP,POM121C,SYMPK,SMG5,SRSF2,SRSF11,DHX38,EIF4A3,RAN,AKAP8L,NU
A2,MAPK1,EIF3E,PRKDC,HNRNPD,EIF3D,PPP1R15A,SERP1,CCN1,ELAVL1,ABCF1,GSK3A,MTOR,E
3X,IK,CKAP5,APPL1,SKP1,CDKN1A,CTDSP2,DDX39B,NOP53,PSMB2,CCND1,MAPRE1,PSMD3,CC
A,KDM1A,MSL1,SUPT6H,USP7,KDM3B,WBP2,KDM4B,PRMT1,MTA1,KDM5C,EP300,KDM5B,FLC
SMD1,CSNK2A1,CSNK1G2,SFRP1,TNKS,PSMD8,ATP6AP2,PSMB4,UBR5,PSMD4,RBPJ,PSMC3,PS
,SERP1,RHBDD2,VAPB,TPP1,ATP6V0D1,GSK3A,SRPRA,AMFR,TSPYL2,PTPN1,NCK2,STUB1,HERP
MAP2,IPO5,TMEM30A,NUCB1,VAMP2,RAB11B,ABCA2,MAPK1,NPEPPS,BNIP3L,MAP2K2,SUPT6H,
PL5,TMBIM6,GSTP1,NPM1,IGF2,UBB,MAPT,FN1,NONO,HACD3,PPP2R1A,RPS3,CHD3,VAPA,HMG
,STMN1,ITGB1,ARF1,PAFAH1B1,CDC42BPB,CTNNA1,CNN3,TPM4,LLGL1,RHOA,CXADR,ATP1A1,
I,PSMB2,RAPGEF1,PPP2CA,SMARCA4,PSMD3,CCAR2,TNKS2,CSNK1D,CTNND1,PSMA7,SULF2,PS
2B4,CALR,TMF1,ABHD2,TUBB2B,CLU,SOX4,MAGED1,DVL3,YWHAE,GNAS,GPI,RACK1,BSG,SPTE
,MSL1,SUPT6H,USP7,KDM3B,WBP2,KDM4B,PRMT1,MTA1,KDM5C,EP300,KDM5B,FLCN,BAZ1B,

P,TAF7,SMARCA4,PSMD3,CAND1,SMARCE1,SMARCC1,SUPT6H,NAP1L4,PSMA7,RBBP7,NAP1L1,F
TSS2,TARDBP,ARL6IP1,RAB1A,LAPTM4B,RAB27A,SQSTM1,VAMP2,SURF4,TMED2,OPTN,PPP2CA
7,GPI,BSG,CALM3,GNB1,APP,GSTP1,CALM1,NPM1,HSPD1,HMGB1,POLR2A,ZMIZ1,DNAJA1,HNF
TP6AP2,PSMB4,UBR5,PSMD4,RBPJ,PSMC3,PSMC1,WNK1,DAPK3,USP47,PSMB1,PSMD13,PSMD7
IEGF8,CTNNB1,MACF1,MAP2,ARHGDIA,CAPRIN1,HDAC2,L1CAM,DLG5,CLSTN3,MDK,SKI,RNF10,
GFA,VHL,PSMD3,NPEPPS,BNIP3L,LONP1,UBQLN1,HYOU1,ARNT2,DDIT4,PSMA7,CAPN2,EP300,P
1,PSMB2,VEGFA,VHL,PSMD3,NPEPPS,BNIP3L,LONP1,UBQLN1,HYOU1,ARNT2,DDIT4,PSMA7,CAF
SMD7,PSME3,PSMC5,PSMC2,PSMC4,PSMF1,PSMB5,PSMA1,PSMD10,PSMA6,PSMD12,PSMA4,PSI
SF6,UPF1,HNRNPA2B1,MCM3AP,POM121C,SYMPK,SMG5,SRSF2,SRSF11,DHX38,EIF4A3,RAN,AK
3,PABPN1,TCP1,MAPK1,GRB2,FBLN1,B2M,IFITM3,HGS,CTNND1,KPNA2,PHB2,UBA52,ULK1,ITGE
VAPA,HMGB1,TUBB4B,ANXA2,TIMP2,VCL,CYB5R3,PGAM1,CTSD,LPCAT1,XRCC5,KPNB1,RHOA,F
PRKCSH,SULF1,MEGF8,MMP14,RPS7,CTNNB1,SLC35E2B,COL2A1,PTK7,SKI,TMED2,NOTCH2,V
P,H3-3B,DDX5,SOX4,MAGED1,KAT6A,DVL3,YWHAE,KIF1A,RPL13,GNAS,DDX17,BSG,FASN,CALM3
1,AP3D1,ARF1,PAFAH1B1,MAP4,HNRNPU,KPNB1,LLGL1,KIF5B,NLGN2,ATP2A2,LRPPRC,XPO1,C
H,SULF1,MEGF8,MMP14,RPS7,CTNNB1,SLC35E2B,COL2A1,PTK7,SKI,TMED2,NOTCH2,VEGFA,M
,TAPBP,LMAN2,NSF,COPE,KIF26A,KLC2,GOLPH3,NBAS,ARFGAP2,KIF3A,NAPA,COPB1,COG4,PLI
F2,EIF3C,VCP,UBB,TRIM28,MAPT,FN1,HACD3,PKD1,SERINC1,RPS3,HMGB1,STMN1,ITGB1,RPL1
MA3C,CLASP2,IST1,ABL1,SEMA6B,ADNP,CYFIP1,NTN1,LIMK1,ISLR2,GOLGA4,SEMA4C,SRF,GSK
APZB,KIF3C,B2M,SEC23A,DYNLL1,PSMA7,DCTN1,KIF5A,ACTR1A,PSMD1,AP2A1,ITGB5,PSMD8,PS
SULF1,PIP4K2B,RAB7A,CCN2,CTNNB1,CD63,SESN3,CYFIP2,CADM4,HNRNPH1,RAC1,GPC1,APPL
2,PSMC4,PSMF1,PSMB5,PSMA1,PSMD10,PSMA6,MCM7,PSMD12,PSMA4,PSMC6
C5,PSMC2,PSMC4,PSMF1,PSMB5,PSMA1,PSMD10,PSMA6,PSMD12,PSMA4,PSMC6
3,APP,KLRD1,TMBIM6,GSTP1,SMG1,NPM1,HSPD1,TRIM28,MATR3,MAPT,NONO,PSMD5,RPS3,HM
YSL2,NOTCH2,VEGFA,MAPK1,GRB2,PLCG1,FYN,SLIT3,PLXNA3,BMPR2,PALLD,KIF5A,TRIO,PLXN
KAP1,VEGFA,MAPRE1,ARPC2,GRB2,CAND1,PFN2,DCTN1,EVL,CTTN,BRK1,AKAP9,ABI2,WASF2,PS
GB1,TUBB4B,ANXA2,TIMP2,VCL,CYB5R3,PGAM1,CTSD,LPCAT1,XRCC5,KPNB1,RHOA,PSMD2,CD
IH1,SGTA,UBQLN2,PTEN,LDLR,RDX,DNAJB2,ATXN3,GSK3A,PSMC3,PSMC1,ZER1,USP5,HSPA1A,
2,MAGED1,DVL3,YWHAE,GPI,RACK1,FASN,SPTBN1,ATP5F1B,CALM3,APP,GSTP1,SMG1,CALM1,
3,FN1,PSMD5,HACD3,CHSY1,PDIA3,RPS3,VAPA,HMGB1,ITGB1,RPL11,ZMIZ1,TIMP2,PAFAH1B1,A
IAG,MAPT,HACD3,PPP2R1A,PKD1,SERINC1,RPS3,HMGB1,UCHL1,RPL11,RPLP1,MIDN,DNAJA1,HI
3,COL1A2,ATF4,COL5A2,PIP4K2B,APLP1,PRKAR1A,GJA1,MYO5A,CTNNB1,SESN3,APPL1,HDAC2,
UBB,YWHAG,MAPT,FN1,HACD3,PPP2R1A,PKD1,RPS3,HMGB1,UCHL1,ANXA2,RPLP1,MIDN,DNAJA
MA3C,CLASP2,ABL1,SEMA6B,ADNP,CYFIP1,NTN1,LIMK1,ISLR2,GOLGA4,SEMA4C,SRF,GSK3B,RU
ROH2A1,CUL1,PSMD13,ATF5,PSMD7,PSME3,PSMC5,PSMC2,PSMC4,PSMF1,PSMB5,PSMA1,PSM
ND1,PSMA7,SULF2,PSMD1,CSNK1G2,SFRP1,CTNND2,CUL3,TNKS,PSMD8,DKK3,ATP6AP2,PSMB4
YSL2,NOTCH2,VEGFA,MAPK1,GRB2,PLCG1,FYN,SLIT3,PLXNA3,BMPR2,PALLD,KIF5A,TRIO,PLXN
4,CARM1,LINGO1,SEMA3C,GSK3A,UBE3A,CERS2,BAG5,GAK,DENND5A,LGALS1,SEMA6B,MDM2,
P4,CLASP2,TUBB4A,ABL1,HDGFL3,HSPA1A,MECP2,GBA2,MAPRE3,PDE4DIP,MAP1S,STMN4,CDKN
,MAP9,PPP2R1A,DNAH8,PKD1,RPS3,CDKL1,ITGB1,TUBB4B,RPL11,RPL10,PAFAH1B1,MAP4,HNR
PS35,TNPO1,SRSF9,SRSF1,DYNLL2,CASC3,KIF3C,SUPT6H,SSB,SRSF3,RAB11A,DYNLL1,RBM15B
I1,SPTBN2,MEGF8,GJA1,MACF1,CYFIP2,ATP1A2,MAP2,COTL1,RAC1,L1CAM,ACTR3,NCKAP1,ASX
K7,USP14,HIPK2,MARCHF7,PSMF1,WAC,USP19,UBXN1,GABARAPL2,RYBP,FURIN,EPHA4,CDK5R
2C,RPS7,MIF,CTNNB1,HNRNPK,BCLAF1,PEA15,PTTG1IP,TIMP3,GNAI3,PARP1,GRINA,COL2A1,HS
F4L1,ZMIZ1,CDC42BPB,CD81,CLEC7A,TADA2B,CSNK1E,DDR1,MIF,CTNNB1,DDOST,CADM4,CTB
4,TLN1,GJA1,CTNNB1,MACF1,RAC1,DNER,DLG5,CDH11,TBCD,TLN2,PTPRA,VPS35,CLSTN3,LAM

ARP1,CTNNA1,RPL23,COL1A2,ATF4,COL5A2,PIP4K2B,APLP1,PRKAR1A,GJA1,MYO5A,CTNNB1,SE
SMD7,PSME3,PSMC5,PSMC2,PSMC4,PSMF1,PSMB5,PSMA1,PSMD10,PSMA6,PSMD12,PSMA4,PSI
220,TMSB4X,NUMA1,DCX,RPS6,DBN1,ENO1,PIK3R2,FLNA,MAP1B,MTRNR2L8,HSP90B1,FADS1,I
PP,FSCN1,GSTP1,PLXNB2,ATP5F1A,FN1,DCHS1,TMSB10,HMGB1,ITGB1,ZMIZ1,VCL,PAFAH1B1,A
MD2,WDR1,PFN1,ASTN2,GPC3,DDR1,SULF1,PLXNA1,MEGF8,MMP14,CLTC,RPS7,GJA1,CTNNB1,I
AMA5,MSN,RHOA,PSMD2,WDR1,PFN1,ASTN2,TPM1,GPC3,DDR1,SULF1,PLXNA1,MEGF8,MMP14,
2,MAGED1,DVL3,YWHAE,GPI,RACK1,FASN,SPTBN1,ATP5F1B,CALM3,APP,GSTP1,SMG1,CALM1,I
KDC,HNRNPD,EIF3D,PPP1R15A,SERP1,ELAVL1,ABCF1,MTOR,EIF4A3,EIF4G1,CNBP,FXR2,PINK1,
MBIM6,GSTP1,HSPD1,IGF2,UBB,TRIM28,SPEN,NONO,YWHAQ,SET,RPS3,CHD3,BASP1,HMGB1,M
21C,SYMPK,SMG5,SRSF2,SRSF11,DHX38,EIF4A3,AKAP8L,NUP153,SRSF7,NUP98,POLDIP3,NCBI
21C,SYMPK,SMG5,SRSF2,SRSF11,DHX38,EIF4A3,AKAP8L,NUP153,SRSF7,NUP98,POLDIP3,NCBI
SF11,DHX38,EIF4A3,SSU72,CDK9,DHX36,SRSF7,POLDIP3,NCBP2,HSF1,CTR9,RNF20,THOC2,THC
KDM3A,RTF1,MACROH2A1,RUVBL2,MECP2,PARK7,SART3,KAT2A,EHMT2,AKAP8L,CDK9,NSD1,B
A2,RNF10,MAPK1,SRSF1,SOD1,STAT3,CNTNAP1,PTEN,PPP1CC,IL6ST,NDN,GRN,SOX2,LDLR,CNP,
3,DCTN1,MAP6,HDAC6,MAPRE2,AKAP9,TAOK1,FKBP4,CLASP2,CHMP4B,TUBB4A,ABL1,CHMP1A,
,BSG,SPTBN1,ATP5F1B,MYH9,APP,FSCN1,SPTAN1,GSTP1,PLXNB2,ATP5F1A,FN1,DCHS1,TMSB1
RX,UBR5,KMT2E,SPTY2D1,OGT,NOC2L,KDM4A,TRIP12,HMGA1,PINK1,AUTS2,KDM3A,RTF1,MAC
ND1,PSMA7,SULF2,BCL9L,PSMD1,CSNK1G2,SFRP1,CTNND2,PTEN,CUL3,TNKS,PSMD8,DKK3,ATI
PSME3,PSMC5,PSMC2,PSMC4,PSMF1,SRF,PSMB5,PSMA1,PSMD10,PSMA6,EXT1,PSMD12,PSMA
3,CAND1,NAP1L4,PSMA7,RBBP7,NAP1L1,PSMD1,UBA52,BRD2,ATRX,HJURP,NAP1L3,ANP32B,RPS
220,TMSB4X,NUMA1,DCX,RPS6,DBN1,ENO1,PIK3R2,FLNA,MAP1B,MTRNR2L8,HSP90B1,FADS1,I
,YWHAG,MAPT,RPS3,HMGB1,AGRN,LARP1,CTNNA1,RPL23,XRCC5,MSN,COL1A2,RHOA,ATF4,KIF
STP1,IGF2,UBB,TRIM28,SPEN,NONO,YWHAQ,SET,RPS3,CHD3,BASP1,HMGB1,MORF4L1,RPL10,
3,PTPRS,PFN2,DIP2B,EVL,ADCY6,PTEN,SLIT2,FKBP4,CARM1,LINGO1,SEMA3C,GSK3A,UBE3A,CEI
1,GRINA,PRKDC,HSPB1,KDM1A,CCAR2,UBQLN1,HYOU1,MAPK8IP1,SOD1,BAG6,DDIT4,EP300,C
6,ULK1,GPSM1,HIF1A,SLC25A5,VPS26B,ATP6V0D1,CHMP4B,MTOR,PRKACA,EXOC4,BNIP3,ATP6
RPL29,RPS24,RPS9,RPL24,RPL17,RPL14,RPL22,RPL9,RPS14,RPL26,RPL31,RPS25,DDX3X,RPS1
PB,ATP6V1B2,SDHA,PFKM,UQCRC2,GHITM,DDIT4,STAT3,EP300,NDUFV1,FLCN,UQCRC1,COX7C
,LRRC4B,FARP1,CTNND2,RER1,HDAC6,PTEN,CTTN,ABI2,WASF2,ADAM10,UBE3A,TANC2,DIP2A,N
B2,IGF2,MAPT,FN1,SPEN,BASP1,HMGB1,ITGB1,AGRN,ZMIZ1,TIMP2,AP3D1,ARF1,PAFAH1B1,HNI
D81,CCT7,OAZ1,XPO1,PFKL,GJA1,TARDBP,NDFIP1,WLS,MYH10,YWHAB,CCT6A,HMGCR,RAC1,C
APP,RPL5,NPM1,PLXNB2,HSPD1,EIF3C,FN1,SF3B1,RPS3,HMGB1,RPS4X,RPL11,ZMIZ1,AKAP12,H
42,EP300,HDAC6,ULK1,SLIT2,TMOD2,FKBP4,RDX,CAPZA1,TUBB4A,LMO4,HIP1R,CRACD,JAM3,F
PSMD4,PSMC3,PRKACA,CREBBP,PSMC1,TOMM70,PSMB1,CUL1,PSMD13,PSMD7,ZCCHC3,PSMI
,GABARAP,KDM1A,FYN,UBQLN1,MTA1,NXN,PTEN,CUL3,RPS2,FKBP1A,UBR5,DNAJB2,DCUN1D2,
E3,DHX36,PSMC5,PSMC2,PSMC4,PSMF1,PSMB5,PSMA1,SOS1,PSMD10,PSMA6,PSMD12,PSMA4,S
CDC42,EXTL3,CSNK2A1,SFRP1,HDGFL2,ADAM10,IST1,MTOR,EIF4G1,USP47,SDCBP,ADNP,CYFI
1,PSMD13,PSMD7,PSME3,PSMC5,PSMC2,PSMC4,PSMF1,PSMB5,PSMA1,PSMD10,PSMA6,PSMD1
3,G3BP2,SRSF9,SRSF1,CASC3,SUPT6H,SSB,SRSF3,RBM15B,XPOT,FLOT1,ZFP36L1,SRSF6,UPF1
RKAR1A,GJA1,MYO5A,SESN3,ARID1A,PTGES3,ATP1A2,APPL1,TNC,ATP6V0C,CTSB,SRSF5,VAMP2
RS1,CCAR2,FYN,HYOU1,MAPK8IP1,GHITM,CSNK2A1,HIF1A,CTTN,SLC25A5,GPX1,MAZ,TRAP1,SO
A1,MYO5A,SESN3,APPL1,ATP6V0C,MMP2,SRSF5,VAMP2,PARP1,PTPRA,AHCYL1,ATP6AP1,MARS1,
C2
GF2,UBB,TRIM28,SPEN,NONO,YWHAQ,SET,RPS3,CHD3,BASP1,HMGB1,MORF4L1,RPL10,ITM2B,I
PI,BSG,FASN,PLXNB2,COL18A1,IGF2,UBB,TRIM28,COL4A2,FN1,PSMD5,DCHS1,CHSY1,PKD1,BAS

2,TRIB2,NOP53,NRDC,ADAM9,GABARAP,FYN,FBLN1,UBQLN1,CSNK1D,HECTD1,BAG6,STAT3,AR
GRB2,PFN2,CDC42,DCTN1,SFRP1,EVL,CTTN,BRK1,AKAP9,GPX1,ABI2,WASF2,MTOR,ABL1,ARPC
AP43,RHOA,CBX5,CLEC7A,PTN,TPM1,DDR1,CCN2,PRDX2,PRKAR1A,TLN1,GJA1,ANXA6,MACF1,C
C2,PSMC4,PSMF1,PSMB5,PSMA1,PSMD10,PSMA6,PSMD12,PSMA4,SETD1A,PSMC6
B,RPLP0,CANX,RPL13A,RACK1,FASN,P4HB,APP,FSCN1,GSTP1,HSPD1,UBB,FN1,PSMD5,INA,HLA-
GRN,ZMIZ1,AP3D1,ZNF770,GTF2F1,HNRNPU,SMARCA5,NACA,ATF4,NR2F1,JUND,CD81,TADA2B,S
GRN,ZMIZ1,AP3D1,ZNF770,GTF2F1,HNRNPU,SMARCA5,NACA,ATF4,NR2F1,JUND,CD81,TADA2B,S
AP3D1,PAFAH1B1,HNRNPU,CTNNA1,XRCC5,IGFBP3,RHOA,JUND,CLEC7A,PTN,CSNK1E,CDH2,PLX
GRN,ZMIZ1,AP3D1,ZNF770,GTF2F1,HNRNPU,SMARCA5,NACA,ATF4,NR2F1,JUND,CD81,TADA2B,S
NA1,CPE,NACA,RTN4,LAMA5,MSN,COL1A2,RHOA,ATF4,PSMD2,COL5A2,PFN1,TLE5,PTN,GLG1,T
CCT6A,RAC1,IPO5,TMEM30A,GLUL,NNAT,CCT2,VAMP2,TCP1,CCT3,CCT4,MAPK1,ADAM9,NPEPPS
BA52,PTEN,RPS27A,PSMD8,HIF1A,BAP1,PSMB4,DNAJB2,ATXN3,FOXK2,PSMD4,OGT,USP24,PSM
A52,ULK1,RPS27A,VAPB,NUCKS1,SCRIB,PSMC3,DAG1,HIPK2,TRIM27,RRP1B,RXRA,PHB,KPNA3,N
S4A,RDX,SH3GL1,CAPZA1,CLASP2,MICAL1,GAK,HDGFL3,CRACD,NCKAP5L,FLII,NCK2,MICAL3,MA
L1,ATP6V0C,SRSF5,VAMP2,PARP1,PTPRA,AHCYL1,ATP6AP1,MARS1,GRB2,PRKDC,SMARCC1,ATP
TCH1,PSMD13,PSMD7,LDB1,PSME3,DHX36,PSMC5,PSMC2,PSMC4,PSMF1,SOX9,PSMB5,PSMA1
FYN,PTPRS,CDC42,GRIPAP1,DCTN1,LRRC4B,FARP1,PTEN,GPC4,SLIT1,ABI2,ADAM10,UBE3A,TA
2,GBA2,MAPRE3,PDE4DIP,CDKN1B,GIT1
ABL1,HDGFL3,HSPA1A,MECP2,GBA2,MAPRE3,PDE4DIP,RANBP9,CDKN1B,GIT1,FGF13,TPX2,ZNF
MD13,HLA-A,PSMD7,PSME3,PSMC5,PSMC2,PSMC4,PSMF1,PSMB5,PSMA1,PSMD10,PSMA6,PSMD
RPLP1,XRCC5,RHOA,CD81,CLEC7A,TADA2B,CCND2,CCN2,PRDX2,DDX3X,MIF,PRKAR1A,CTNNB
WHAB,CADM4,HMGCR,IPO5,TRIB2,CDKN1A,CTDSP2,PARP1,PKN1,ABCA2,TERF2IP,TMED2,TAF7,
,ZER1,USP5,HSPA1A,MDM2,FZR1,SUMO1,TRIB3,STUB1,PSMC5,CSNK1A1,PSMC2,HERPUD1,PSM
1,HACD3,PPP2R1A,PKD1,RPS3,HMGB1,UCHL1,ANXA2,RPLP1,DNAJA1,XRCC5,IGFBP3,RHOA,ATF
,THRAP3,FXR2,RBM4,RBM25,FXR1,CELF3,RBFOX2,RBM10,RBM17,SFSWAP,RNPS1
HNRNPD,WBP2,CCT5,BRD4,ATRX,HNRNPA2B1,TNKS,PPP1R10,KMT2E,OGT,PINK1,AUTS2,RTF1,M
,SQSTM1,VAMP2,VPS35,ABCA2,TMED2,OPTN,CLIP3,STXBP1,HECTD1,RAB11A,SEC23A,PPP1R9E
2,IGF2,UBB,TRIM28,YWHAG,FN1,HACD3,PPP2R1A,PKD1,RPS3,HMGB1,CDKL1,UCHL1,ANXA2,RP
I,CDH2,ATP1A2,GLUL,MAP1A,CDH11,SQSTM1,VAMP2,PTPRA,CLSTN3,MAPK1,PLCG1,FYN,YWHA
CTR2,VIM,SPARC,KIDINS220,TMSB4X,NUMA1,DCX,EIF4G2,RPS6,DBN1,ENO1,FLNA,MAP1B,NCL,
FYN,PTPRS,CDC42,GRIPAP1,DCTN1,LRRC4B,FARP1,PTEN,GPC4,SLIT1,ABI2,ADAM10,UBE3A,TA
I,CDH2,ATP1A2,GLUL,MAP1A,CDH11,SQSTM1,VAMP2,PTPRA,CLSTN3,MAPK1,PLCG1,FYN,YWHA
PT,SET,EGLN1,PPP2R1A,PKD1,STMN1,UCHL1,RPL11,ANXA2,TIMP2,MIDN,DNAJA1,GTF2F1,HNRN
WHAB,CADM4,HMGCR,IPO5,TRIB2,CDKN1A,CTDSP2,PARP1,PKN1,ABCA2,TERF2IP,TMED2,TAF7,
N,GPC2,ATP1B2,EHD1,SF3A2,MTOR,APBB1,NTRK2,SNX3,NPTN,ADNP,CYFIP1,FEZ1,NME2,DHX3
P3L,FYN,YWHAH,SREBF2,CDC42,GRIPAP1,OS9,SEC16A,UBE2G2,OAZ2,ANP32B,LMAN1,BAP1,TC
3,RHOB,HDAC2,MMP2,CHD6,PARP1,NOX4,PRDX5,KDM6B,SELENON,MAPK1,PRDX1,ADAM9,HSPE
RAPGEF1,G3BP2,GRB2,GABARAP,STMN3,ARFGEF2,CDC42,BCR,FLCN,FLOT1,PPP2CB,CUL3,MA
MBIM6,IGF2,UBB,TRIM28,SPEN,NONO,YWHAQ,SET,RPS3,CHD3,BASP1,HMGB1,MORF4L1,RPL10
AF1,ITGA7,HMGA2
BA52,PTEN,RPS27A,PSMD8,HIF1A,BAP1,PSMB4,DNAJB2,ATXN3,FOXK2,PSMD4,OGT,USP24,PSM
2,VPS35,TMED2,OPTN,CLIP3,STXBP1,HECTD1,RAB11A,SEC23A,PPP1R9B,GRIPAP1,FLOT1,SEC16
I1A,CCAR2,UBQLN1,HYOU1,MAPK8IP1,SOD1,HIF1A,GPX1,TRAP1,SOD2,NOC2L,NDUFA13,BAG5,
,ADAM10,IGF2BP1,UBE3A,TANC2,DIP2A,TAOK2,TRAK1,OBSL1,MAPK8IP2,RAP2A,DHX36,RBFOX
0,CANX,RPL13A,RACK1,FASN,P4HB,FSCN1,GSTP1,HSPD1,UBB,FN1,PSMD5,INA,HLA-C,PDIA3,RP

,DDX39B,DPYSL2,VEGFA,MAPK1,FTO,PRKDC,TNKS2,PLXNA3,PTPRS,SOD1,RAI1,RAB11A,BMPR2
)K1,FKBP4,TUBGCP6,CLASP2,TUBB4A,ABL1,HDGFL3,HSPA1A,MECP2,NCKAP5L,GBA2,MAPRE3,I
IP2,YTHDF3,RBM4,PPP1CA,NCK2,EIF3B,NCBP2,PAIP1,EIF1B,EIF2S1,CTIF
PF1,HNRNPA2B1,TNKS,PPP1R10,SMG5,DHX36,EXOSC10,LMNA,MAPK3,GNL3,RAD50
N3,MTOR,PRKACA,CREBBP,BAG5,HSPA1A,DNAJB1,SUMO1,NUP153,MAPKAPK2,DHX36,STUB1,N
SESN3,HMGCR,PTTG1IP,DLG5,TIMP3,DDX39B,NOP53,GRINA,TSPAN6,RAPGEF1,OPTN,PPP2CA,H
,RPS24,RPS9,RPL24,RPL17,RPL14,RPL22,RPL9,RPS14,RPL26,RPL31,RPS25,DDX3X,RPS15,RPL
ATP5F1B,MYH9,APP,FSCN1,GSTP1,PLXNB2,ATP5F1A,FN1,DCHS1,TMSB10,DNAH8,HMGB1,ITGB
ATP5F1B,MYH9,APP,FSCN1,GSTP1,PLXNB2,ATP5F1A,FN1,DCHS1,TMSB10,DNAH8,HMGB1,ITGB
I11,ABL1,COL16A1,TAOK2,MAP4K4,IQSEC1,LDB1,SLK,LIMS1,RCC2,DUSP3,ROCK2,AJUBA,ARF6
R5,DCLK1,TRIM8,SMO,PKIA,ATP13A2,PARK7,RAN,TMEM98,GSK3B,ILRUN,TFRC,LMNA,LZTS2,PI
NAJA1,PAFAH1B1,AKAP12,PJA2,IGFBP3,PSMD2,CD81,CLEC7A,SPTBN2,PEBP1,CDH2,CCN2,PRDX
PSMC2,PSMC4,PSMF1,PSMB5,PSMA1,PSMD10,PSMA6,PSMD12,PSMA4,UBE2E1,PSMC6,ANAPC2
/ID8,HIF1A,A2M,PSMB4,SEMA3C,TCF3,LTBP3,PSMD4,RBPJ,PSMC3,PSMC1,SMO,ABL1,PUM1,SEI
AG5,CD74,USP47,PINK1,HSPA1A,MDM2,PARK7,CD44,PTPN1,MARCHF7,HERPUD1,VDAC2,XBP1,Y
,DDR1,SULF1,CCN2,MMP14,APLP1,COL6A2,TTR,HRNR,TNC,MMP2,PPIB,DCN,VCAN,TGFBI,PLOD
32,SFRP1,PSMD8,GPC4,PSMB4,PSMD4,PSMC3,AP2A2,PSMC1,SMO,CELSR1,ABL1,PSMB1,PSMD
GRB2,PLCG1,FYN,CDC42,PSMA7,PSMD1,PSMD8,BRK1,PSMB4,PDPK1,WASF2,PSMD4,PSMC3,PS
MC3,PSMC1,ZER1,USP5,HSPA1A,MDM2,FZR1,SUMO1,MAGEF1,TRIB3,STUB1,PSMC5,CSNK1A1,I
,WAC,USP19,UBXN1,GABARAPL2,RYBP,EPHA4
,PSMC5,PSMC2,PSMC4,PSMF1,UBE2V1,PSMB5,PSMA1,PSMD10,MAPK3,PSMA6,TAB3,RELA,PSN
A,MAP1A,TNPO1,DYNLL2,HSPB1,CLIP3,KIF3C,SOD1,DYNLL1,CDC42,KIF5A,MAP6,HIF1A,SPG7,KI
AS,SLC2A3,GPI,GNB1,MYH9,APP,SPTAN1,GSTP1,HSPD1,IGF2,VCP,MAPT,FN1,HLA-C,RPS3,VAPA,H
,DDR1,SULF1,CCN2,MMP14,APLP1,COL6A2,TTR,TNC,MMP2,PPIB,DCN,VCAN,TGFBI,PLOD1,COL
HX36,GSK3B,DLG4,ARHGAP33,SHANK1,OPA1,EPHB2,APOE,LRP8,PRNP,SRCIN1,EPHA4
,DDR1,SULF1,CCN2,MMP14,APLP1,COL6A2,TTR,TNC,MMP2,PPIB,DCN,VCAN,TGFBI,PLOD1,COL
GJA1,ANXA6,MACF1,RHOB,RAC1,APPL1,GLUL,DCN,PKN1,VEGFA,ADAM9,HSPB1,PLCG1,ZFAND5
4,BMPR2,MMP15,SFRP1,CUL3,ITGB5,SOX2,RIC8A,CLASP2,ITGAV,PRKACA,EXOC4,WNK1,COL12
EC1,NCK2,DHX36,PLK2,ACSL4,DLG4,ARHGAP33,SHANK1,OPA1,EPHB2,APOE,LRP8,ARF6,SRCIN
90B1,YBX1,IGFBP2,FSTL1,HSPA1B,COL4A1,RPLP0,CANX,DDX5,SOX4,DVL3,HTRA4,RACK1,BSG,S
PRC,XPO1,ACTN4,RPS15,GJA1,MYO5A,CTNNB1,LAMP1,MAP2,RAB1A,RAB27A,ACTR3,RAB11B,AT
A1B,SOX4,YWHAE,GNAS,GPI,RACK1,BSG,ATP5F1B,CALM3,GNB1,APP,TMBIM6,GSTP1,SMG1,CA
ABI2,WASF2,MTOR,PINK1,ARPC1A,HIP1R,HSPA1A,CRACD,MECP2,CYFIP1,TRIM27,NCK2,ARPC3,I
P2,MAPK1,NPEPPS,FYN,YWHAH,SREBF2,CDC42,SEC16A,OAZ2,ANP32B,BAP1,TCAF1,UBR5,RDX,
I,PFN1,SPTBN2,TPM1,CCN2,MYO5A,CYFIP2,RHOB,COTL1,RAC1,GAS7,NOX4,ACTR3,NCKAP1,AR

M1,VCP,UBB,MAPT,MAP9,DNAH8,PKD1,RPS3,CHD3,STMN1,ITGB1,TUBB4B,UCHL1,AP3D1,DNAJ
1,EIF3D,PPP2CB,GPSM1,FKBP1A,EDF1,EIF4G1,AMFR,ABL1,NIBAN2,HIP1R,PARK7,RAN,HIPK2,TI
,P3H3,WBP2,KDM4B,EP300,FLCN,BRD4,DIP2B,ATRX,EP400,HDAC6,MCM3AP,POM121C,LOX,EI
C5,PSMC2,PSMC4,PSMF1,PSMB5,PSMA1,PSMD10,PSMA6,PSMD12,PSMA4,PSMC6
CA5,XRCC5,KPNB1,CCT7,HNRNPC,CSNK1E,SFPQ,CCN2,H1-0,GJA1,CTNNB1,NDFIP1,H1-10,INIP,
8A1,COL4A2,FN1,DCHS1,EGLN1,HMGB1,ITGB1,ANXA2,COL11A1,PAFAH1B1,RTN4,RHOA,CD81,
[illegible]

P,UBB,NONO,PSMD5,PPP2R1A,RPS3,TPT1,STMN1,ITGB1,UCHL1,ANXA2,TIMP2,DNAJA1,PAFAH1E
1,GRB2,CMIP,PLCG1,ZFAND5,SUPT6H,STMN3,HECTD1,TCTN1,ARNT2,PRMT1,CAPN2,FLCN,ZFP

PO1,PFKL,GJA1,TARDBP,NDFIP1,WLS,MYH10,YWHAB,HMGCR,RAC1,CDK16,IPO5,TMEM30A,GLU
SAP1,MAPRE1,CLIP3,STMN3,DCTN1,MAP6,KLHL42,HDAC6,MAPRE2,AKAP9,TAOK1,FKBP4,CLAS
ARPC3,EFNB3,ARPC5,ROCK2,VAV2,GIT1,EPHB2,EPHA4,DNM1,EPHA2
ACD3,PKD1,SERINC1,RPS3,HMGB1,ITGB1,AGRN,RPLP1,TIMP2,HNRNPU,CTSD,XRCC5,LLGL1,RH
1,MACROH2A1,NOTCH1,LDB1,CTR9,COL7A1,MED12,EXT1,PAF1,ITGA7,HMGA2
,VPS35,SELENON,OPTN,ATG2B,ATP5PB,CDC37,GABARAP,NPEPPS,BNIP3L,CCAR2,YWHAH,LONP
A,CD81,CCND2,DDR1,PRDX2,DDX3X,PRKAR1A,CTNNB1,YWHAB,PTGES3,RAB1A,SKP1,PEA15,CDK
,COL16A1,TAOK2,MAP4K4,LDB1,SLK,LIMS1,RCC2,DUSP3,ROCK2,AJUBA,PIP5K1A,GPM6B
NCSTN,FAM210B,KLF13,HSPA1A,ACVR1B,JAM3,MAEA,HIPK2,LDB1,RBFOX2,SRF,HOXA5,STAT1,
GLN1,PPP2R1A,RPS3,HMGB1,STMN1,ITGB1,ANXA2,AGRN,TIMP2,SBF1,PAFAH1B1,CTSD,SCARB2
RPL26,MCL1,CSNK1E,SFPQ,MIF,HNRNPK,INIP,SMCHD1,BCLAF1,PTTG1IP,CDKN1A,DDX39B,NOI
I,ATP6AP1,MARS1,GRB2,PRKDC,SMARCC1,ATP6V1B2,FYN,PPP1R9B,STAT3,PRKAR2A,ZFP36L1,
NOP53,NOX4,ACO2,PYGB,ACADVL,ATP5PB,SDHA,PFKM,IDH1,UQCRC2,GHITM,VGF,DDIT4,STAT3
HNRNPU,LARP1,CTNNA1,MSN,RHOA,ATF4,ATP1A1,PTN,APLP1,HMGCS1,CTNNB1,ARID1A,PTGES
,HMGB1,MORF4L1,RPL10,HNRNPU,SMARCA5,RPL23,NACA,XRCC5,CIPC,CBX5,ATF4,NR2F1,RPS

PO5,TMEM30A,GLUL,NNAT,VAMP2,MAPK1,ADAM9,NPEPPS,FYN,YWHAH,PFKM,SREBF2,DYNLL1,
A,NCL,ALDOA,FADS1,YBX1,ATP2B4,CALR,TMF1,HSPA1B,ZNF99,H3-3B,CLU,PTMA,DDX5,SOX4,MA
DL7A1,PAF1,ITGA7,HMGA2
CH1,PCM1,KAT2A,CUL4A,DDX6,LDB1,BCL9,LOXL2,SOX9,CTR9,MED12,SRRT,ZNF358,CNOT3,GN
,HMGB1,MORF4L1,RPL10,HNRNPU,SMARCA5,RPL23,NACA,XRCC5,CIPC,CBX5,ATF4,NR2F1,RPS
SST,GRN,SOX2,LDLR,CNP,NCSTN,DICER1,PHGDH,AKT2,PRPF19,KDM4A,MTOR,BNIP3,SMO,MET
2IP,VEGFA,MAPK1,KDM1A,CCT8,TNKS2,USP7,HNRNPD,WBP2,YLPM1,CCT5,BRD4,ATRX,UPF1,HN

N1,ARRDC3
N1,SPEN,EGLN1,BASP1,HMGB1,ITGB1,AGRN,ZMIZ1,VCL,AP3D1,PAFAH1B1,CTNNA1,RTN4,LAMA
,HMGB1,MORF4L1,RPL10,HNRNPU,SMARCA5,RPL23,NACA,XRCC5,CIPC,CBX5,ATF4,NR2F1,RPS
PA1A,ACVR1B,MAEA,HIPK2,LDB1,RBFOX2,SRF,HOXA5,STAT1,CEBPG,TMEM14C,CITED2,THRA,A
PRMT1,EP300,PSMD1,BRD4,ZFP36L1,ATRX,PTEN,FEM1B,BRSK1,TNKS,PSMD8,PCBP4,PPP1R10
SCAP,SNX3,NSF,GOLPH3,ARFGAP2,MCFD2,NAPA,COL7A1,CHMP5,PRKCI,TRAPPC1,SEC23IP,SEC
K4,LDB1,SLK,LIMS1,RCC2,DUSP3,ROCK2,AJUBA,PIP5K1A,GPM6B
NRNPU,RPL23,RPS20,BEX3,OAZ1,NES,PTN,PEBP1,GPC3,PIP4K2B,DDX3X,RPS15,RPS7,PRKAR1A
.,AFDN,NECTIN2,ADD1,EPHA4,HIPK1
,ADD1,XBP1,CUL7,PPP2R5B,SEC31A
SB4X,NUMA1,DCX,RPS6,ENO1,PIK3R2,FLNA,MTRNR2L8,HSP90B1,NEFL,YBX1,ATP2B4,IGFBP2,C
MS1,RCC2,DUSP3,ROCK2,ARF6,GPM6B
TF4,RPL26,MCL1,TLE5,CLEC7A,PTN,SFPQ,PNMA1,CCN2,DDX3X,ITM2C,RPS7,JUN,CTNNB1,RHOI
A13,BNIP3,ATP5MC2,ATP5MG,MDH1,PINK1,PDHA1,NDUFA4,PARK7,IDH2,NDUFC2,COX8A,COX7A
KAT2A,CUL4A,DDX6,LDB1,BCL9,LOXL2,SOX9,CTR9,MED12,SRRT,ZNF358,CNOT3,GNL3,TUT4,K
1,HNRNPU,SMARCA5,NACA,ATF4,NR2F1,JUND,CD81,SFPQ,SOX12,DDX3X,NANOG,JUN,CTNNB1,
CHD3,STMN1,ITGB1,TUBB4B,PAFAH1B1,MAP4,HNRNPU,KPNB1,LLGL1,RHOA,CRMP1,SPC25,XF
D5,TRIB2,CDKN1A,CTDSP2,PKN1,TERF2IP,TMED2,TAF7,PPP2CA,IPO7,PRKDC,HSPB1,FKBP8,QAI
,HSPA1B,ZNF99,H3-3B,CLU,PTMA,DDX5,SOX4,MAGED1,RPL6,KAT6A,DVL3,DDX17,APP,TMBIM6
APT,HACD3,PPP2R1A,PKD1,RPS3,HMGB1,UCHL1,RPLP1,MIDN,DNAJA1,HNRNPU,XRCC5,RHOA,C
P300,FLOT1,PPP2CB,PTPRF,FKBP1A,SYMPK,LOX,DZIP1,DNAJB2,PRKACA,AMFR,ABL1,HIP1R,PAI

\,NCL,ALDOA,FADS1,YBX1,ATP2B4,CALR,TMF1,HSPA1B,ZNF99,H3-3B,CLU,PTMA,DDX5,SOX4,M/
,HSPA1B,ZNF99,H3-3B,CLU,PTMA,DDX5,SOX4,MAGED1,RPL6,KAT6A,DVL3,DDX17,APP,TMBIM6
MAPKAPK2,NUP98,HSF1,GSK3B,NUP205,NUP133,DNAJB6,MAPK3,NUP50,NUP58
ARP1,NOX4,PRDX5,KDM6B,SELENON,MAPK1,PRDX1,HSPB1,FYN,LONP1,UBQLN1,SOD1,ZFP36L
\,NCL,ALDOA,FADS1,YBX1,ATP2B4,CALR,TMF1,HSPA1B,ZNF99,H3-3B,CLU,PTMA,DDX5,SOX4,M/
L1,UBAP2L,HIF1A,SLC25A5,BAP1,AKT3,IREB2,KMT2E,NCSTN,CD74,SMO,PTPN11,FAM210B,ABL
NK1,MFN2,PARK7,UBE2V1,RETREG3,MAPK3,UFM1,DYNC1I2,NBR1,SESN2,RB1CC1
P33,SHANK1,OPA1,EPHB2,APOE,LRP8,PRNP,SRCIN1,EPHA4
FOX2,RBM10,RBM17,RNPS1
PK3
DH,SCRIB,PRKACA,AMBRA1,SMO,LMO4,CELSR1,ABL1,PHACTR4,PLXNA2,NOTCH1,KAT2A,SALL2
C42,SEC16A,OAZ2,ANP32B,BAP1,TCAF1,UBR5,TM9SF4,NMT1,AKT2,GSK3A,PRKACA,UBE2D3,S/
G,RPS7,PRKAR1A,GJA1,CTNNB1,WLS,HDAC2,MMP2,HSBP1,COL2A1,PTK7,SKI,KDM6B,TMED2,
KD1,RPS3,HMGB1,UCHL1,RPLP1,DNAJA1,XRCC5,RHOA,CD81,CCNI,CCND2,PRDX2,DDX3X,PRK/

BP
SRP1,TAF7,RPL37,KDM1A,USP7,BAG6,DDIT4,RBBP7,PRMT1,EP300,CSNK2A1,ATRX,PCBP4,EHM
,HSPA1B,ZNF99,H3-3B,CLU,PTMA,DDX5,SOX4,MAGED1,RPL6,KAT6A,DVL3,DDX17,APP,TMBIM6
OLPH3,SNX1,CCDC88A,NCK2,WASF3,ARPC5,AJUBA,VAV2,FER,EPHA2
,ABL1,PSMB1,SMPD3,NOTCH1,PSMD13,JAM3,PSMD7,LDB1,PSME3,DHX36,PSMC5,PSMC2,PSM
X2,ITGAV,PRKACA,EXOC4,COL12A1,RTF1,MACROH2A1,TXNRD1,EXT2,SRF,CTR9,COL7A1,PAF1,
SOD1,SREBF2,NCOR2,DDIT4,PSMA7,STAT3,EP300,PSMD1,FLCN,MBTPS1,OAZ2,POM121C,PSMI
,HSPD1,IGF2,VCP,MAPT,HLA-C,RPS3,VAPA,HMGB1,ITGB1,TUBB4B,ANXA2,ZMIZ1,TIMP2,VCL,AF

,SRSF11,DHX38,EIF4A3,RPS21,SSU72,CDK9,DHX36,SRSF7,POLDIP3,EXOSC10,NCBP2,HSF1,FBI
PINK1,ATP13A2,MFN2,PARK7,ATG9A,GABARAPL2,FBXO7,RB1CC1,USP36
PINK1,ATP13A2,MFN2,PARK7,ATG9A,GABARAPL2,FBXO7,RB1CC1,USP36
KIF3C,SOD1,DYNLL1,CDC42,KIF5A,HIF1A,SPG7,KIFAP3,FLOT2,RAB6A,TUB,SUN1,TRAK1,PCM1,
DB,ATP1A2,COTL1,RAC1,ARHGDIA,CNN2,NOX4,ACTR3,NCKAP1,MDK,NOTCH2,ARPC2,DSTN,LUZ
P1A,KDM3A,SUMO1,BANF1,RAN,SUN2,HIPK2,AKAP8L,NUP153,CELF3,NUP98,TMEM43,NUP205,

MCL1,TLE5,CLEC7A,PTN,SFPQ,PNMA1,DDX3X,ITM2C,RPS7,JUN,CTNNB1,RHOB,HMGCR,BCLAF1,
MCL1,TLE5,PTN,SFPQ,PNMA1,DDX3X,ITM2C,RPS7,JUN,CTNNB1,RHOB,HMGCR,BCLAF1,PEA15,T
F5,ACO2,MDK,ASXL1,NOTCH2,VEGFA,CCND1,MAPK1,SRSF1,PRKDC,SMARCC1,SOD1,NCOR2,HI
VEGFA,MAPK1,LUZP6,PRKDC,RAB11A,BMPR2,CDC42,EXTL3,CSNK2A1,SFRP1,SERP1,HDGFL2,A
OX9,MED25,MED12,THRA,MED16,RELA,CTNNBIP1,PSMC6,MED14
,PLXNA3,PFN2,CDC42,DCTN1,SFRP1,EVL,CTTN,BRK1,AKAP9,ABI2,WASF2,MTOR,ABL1,ARPC1A,
RD4,UBA52,ULK1,RPS27A,VAPB,NUCKS1,SCRIB,PSMC3,DAG1,HIPK2,TRIM27,RRP1B,RXRA,PHB,
21C,PPP1R10,UBR5,ELAVL1,CSE1L,SMO,RGPD5,PKIA,RAN,FAM53C,NUP153,KPNA6,NUP98,LMN
\,NCL,ALDOA,FADS1,YBX1,ATP2B4,CALR,TMF1,HSPA1B,ZNF99,H3-3B,CLU,PTMA,DDX5,SOX4,M/
AF7,C1QBP
F5B,NLGN2,ATP2A2,NISCH,PTN,CDH2,PRKAR1A,CTNNB1,DNAJC5,ATP1A2,GLUL,MAP1A,CDH11,
12,EHMT2,NSD1,WDR82,ASH1L,BCOR,PRMT5,FBL,SETD5,CTR9,RNF20,LMNA,PRMT2,CTCF,SETI

R1,SEC23A,DYNLL1,DCTN1,ACTR1A,SEC16A,RAB2A,YKT6,CUL3,LMAN1,VAPB,ERGIC1,RAB1B,(

ATP2B4,CALR,TMF1,HSPA1B,PGK1,H3-3B,CLU,SOX4,MAGED1,YWHAE,GNAS,GPI,RACK1,BSG,SF
N1,TERF2IP,TMED2,TAF7,PPP2CA,IPO7,PRKDC,HSPB1,FKBP8,QARS1,DUSP4,CORO1C,FBLN1,ST
CHD2,H3-3A,TOP1,NAP1L4,RBBP7,DFFB,NAP1L1,CSNK2A1,UBA52,BRD2,ATRX,HJURP,NAP1L3,H
PS1,POM121C,PPP1R10,SNRPD3,UBR5,ELAVL1,CSE1L,SMO,RGPD5,PKIA,RAN,FAM53C,NUP153,
M6B,SELENON,MAPK1,PRDX1,HSPB1,FYN,LONP1,UBQLN1,SOD1,HDAC6,APEX1,HIF1A,PRDX6,GF
C5,PARP1,VPS35,MDK,FYN,STXBP1,HYOU1,SOD1,CDC42,DDIT4,STAT3,CAPN2,HIF1A,GRN,UBE2I
DIA,NOX4,ACTR3,NCKAP1,MDK,NOTCH2,ARPC2,DSTN,LUZP6,GRB2,CAPZB,PFN2,CDC42,SFRP1
,CRACD,FLII,NCK2,VDAC2,CAPZA2,ADD1,MYADM,GSN
A,PSMC3,PSMC1,AMFR,IGFBP4,PSMB1,RUVBL2,NOTCH1,PSMD13,PSMD7,PSME3,PSMC5,CSNK
2,PPP2CB,TRA2B,HDAC6,APEX1,HIF1A,GPX1,STK24,TRAP1,SOD2,PCGF2,FBLN5,UBE3A,BNIP3,P
XNIP,ADAR,STAT3,ALDH3A2,KPNA2,PHB2,MBTPS1,POM121C,TIMM50,PPP1R10,UBR5,ELAVL1,C
ULF2,PTEN,PTPRF,ULK1,GRN,GPX1,SOX2,STK24,PTGFRN,CERS2,CAD,DAG1,NOTCH1,JAM3,MA

KDC,SMARCC1,ATP6V1B2,FYN,STAT3,PRKAR2A,ZFP36L1,ARID1B,ADCY6,PTEN,APEX1,NUCKS1,
,HSPA1B,ZNF99,H3-3B,CLU,DDX5,SOX4,MAGED1,RPL6,KAT6A,DVL3,DDX17,APP,TMBIM6,NPM1
7,CCAR2,TP53BP1,PSMA7,RBBP7,PRMT1,EP300,PSMD1,BRD4,ZFP36L1,ATRX,PTEN,FEM1B,BRS
X,SFSWAP,KHDC4
1,TRIB3,STUB1,CSNK1A1,HERPUD1,PLK2,GSK3B,PSMD10,ELOB,XBP1,RAD23A
C1,EP300,DGCR2,PTEN,BRSK1,HIF1A,TMOD2,LDLR,RIC8A,NCSTN,SYT11,MTOR,DOP1B,EIF4A3,
F1,MACROH2A1,MECP2,TRMT112,EHMT2,ETF1,NSD1,WDR82,ASH1L,BCOR,PRMT5,FBL,SETD5,
F1,MACROH2A1,MECP2,TRMT112,EHMT2,ETF1,NSD1,WDR82,ASH1L,BCOR,PRMT5,FBL,SETD5,

00,PTEN,BRSK1,HIF1A,TMOD2,LDLR,RIC8A,NCSTN,SYT11,MTOR,EIF4A3,NTRK2,AMFR,PAIP2,N
PC2,DSTN,LUZP6,GRB2,CAPZB,PFN2,CDC42,SFRP1,EVL,CTTN,SLIT2,BRK1,TMOD2,ABI2,RDX,CA
LRD1,TMBIM6,SPTAN1,GSTP1,HSPD1,VCP,HLA-C,VAPA,HMGB1,TUBB4B,ANXA2,TIMP2,VCL,CYB
A,SMCHD1,CDKN1A,DDX39B,CCT2,PARP1,TCP1,CCT3,CCT4,TERF2IP,VEGFA,MAPK1,PRKDC,KD
,HSPA1B,ZNF99,H3-3B,CLU,DDX5,SOX4,MAGED1,RPL6,KAT6A,DVL3,DDX17,APP,TMBIM6,NPM1
,HSPA1B,ZNF99,H3-3B,CLU,DDX5,SOX4,MAGED1,RPL6,KAT6A,DVL3,DDX17,APP,TMBIM6,NPM1
D5,TRIB2,CDKN1A,PKN1,TAF7,RPL37,PPP2CA,IPO7,HSPB1,QARS1,DUSP4,CORO1C,MAPK8IP1,A
RCC1,ATP6V1B2,VGF,ZFP36L1,SRSF6,PTEN,SLC2A1,NUCKS1,PDPK1,AKT2,OGT,ATP6V0D1,GSK
E4B,USP14,FAF2,UBE2J1,FAM8A1,HM13,STUB1,HERPUD1,USP19,UBXN1,XBP1,MAN1B1,PSMC6,
PRA,MDK,LAMC1,VEGFA,ARPC2,ADAM9,ACTN1,CORO1C,FBLN1,CDC42,BCR,SFRP1,PTEN,ITGB5
)
ARID1A,PTGES3,ATP1A2,CDKN1A,PARP1,ABCA3,RPL27,PKN1,MDK,H3-3A,ABCA2,TAF7,CCND1,S
HNRNPC,PTN,GPC3,CCN2,MMP14,RPS15,CLTC,CTNNB1,SND1,TNC,MMP2,CDH11,VCAN,COL2A1
STXBP1,CSNK1D,TMED9,HGS,RAB11A,PPP6R1,SEC23A,ARFGEF2,SYP,SEC16A,CUL3,LMAN1,VAP
MDM2,PARK7,RAN,AKAP8L,NUP153,NCBP2,GSK3B,TXN,PPP1R12A,XBP1,UFM1,NOLC1,NUP58,N
L1,PTEN,NUCKS1,PDPK1,AKT2,OGT,ATP6V0D1,GSK3A,ATP6V1G1,PHIP,USO1,RABGAP1,PTPN11
ATP2A2,NISCH,PTN,CDH2,PRKAR1A,CTNNB1,DNAJC5,ATP1A2,GLUL,MAP1A,CDH11,SQSTM1,VAM
NMT1,MOAP1,GSK3A,BNIP3,PINK1,HIP1R,HSPA1A,ATP13A2,PARK7,GOLPH3,STAT2,YWHAZ,GSK
P1,NOTCH2,VEGFA,PRKDC,ACTN1,B2M,CDC42,ADAR,STAT3,EIF2AK1,PRMT1,EP300,ZFP36L1,S
T1,UPF1,HDAC6,TMOD2,TAOK1,VPS4A,RDX,SH3GL1,CAPZA1,CLASP2,MICAL1,GAK,HDGFL3,CR

R9B,TRA2B,PTEN,HIF1A,SLIT2,SLC2A1,SLIT1,ATP1B2,IGF2BP1,BNIP3,SLC38A2,SMO,NTRK2,XF

)7,PSME3,PSMC5,PSMC2,PSMC4,PSMF1,PSMB5,PSMA1,PSMD10,PSMA6,TAB3,RELA,PSMD12,PS
),POM121C,FKBP4,ATXN3,MTOR,PRKACA,CREBBP,BAG5,HSPA1A,DNAJB1,SUMO1,NUP153,MAF
11,OPA1,MAPK8IP3,AP3M1
EPL,RPS27A,VPS26B,VPS4A,RAB14,DCLK1,RAB6A,SPAG9,SCRIB,EHD1,CHMP4B,UBXN6,SORT1,I
IMGB1,STMN1,ITGB1,VCL,AKAP12,CTNNA1,RTN4,LAMA5,MSN,IGFBP3,RHOA,ATP1A1,CD81,ATP
IDGFL3,CRACD,IGF1R,FLII,ETF1,NCK2,UBQLN4,SETX,MAP1S,BMERB1,CAPZA2,ADD1,FGF13,TP
BCA7

SP7,ID4,RAI1,NCOR2,HNRNPD,MTA1,KDM5C,EP300,KDM5B,CSNK2A1,PTEN,PPP1CC,SLIT2,IMPI
C1,COX7C,PPP1CC,IL6ST,HIF1A,IREB2,TRAP1,SLC25A23,AKT2,COX6A1,GSK3A,MTOR,ATP5MC3
,MAPKAPK2,EMILIN1,NRP2,ROCK2,XBP1,SHB,VAV2,BCAR1,GRB10,ELMO2
,ROCK2,GPM6B
,ROCK2,GPM6B
T2,SLC2A1,IGF2BP1,BNIP3,SLC38A2,SMO,NTRK2,XRN2,SUN1,SEMA6B,FEZ1,SUN2,H2AX,CRKL,
ATP2A2,NISCH,PTN,CDH2,PRKAR1A,CTNNB1,DNAJC5,ATP1A2,GLUL,MAP1A,CDH11,SQSTM1,VAN
ATP2A2,NISCH,PTN,CDH2,PRKAR1A,CTNNB1,DNAJC5,ATP1A2,GLUL,MAP1A,CDH11,SQSTM1,VAN
A,PKD1,RPS3,HMGB1,ITGB1,NCAM1,ANXA2,ZMIZ1,VCL,AP3D1,CTNNA1,CNN3,RPSA,RTN4,LAMA
IC2,PSMC4,PSMF1,PSMB5,PSMA1,PSMD10,PSMA6,CUL4B,PSMD12,PSMA4,PSMC6
)7,PSME3,PSMC5,PSMC2,PSMC4,PSMF1,PSMB5,PSMA1,PSMD10,PSMA6,TAB3,RELA,PSMD12,PS
)7,PSME3,PSMC5,PSMC2,PSMC4,PSMF1,PSMB5,PSMA1,PSMD10,PSMA6,TAB3,RELA,PSMD12,PS
)YNC1LI2,DYNLL2,CSNK1D,RAB11A,DYNLL1,DCTN1,ATRX,SMC3,PPP1CC,EML1,MAPRE2,TNKS,T
TLE1,PSMD4,GSK3A,PSMC3,PSMC1,AMFR,IGFBP4,PSMB1,RUVBL2,NOTCH1,PSMD13,PSMD7,PS
,TMOD2,ABI2,RDX,CAPZA1,WASF2,MTOR,ARPC1A,HIP1R,ELN,CRACD,CYFIP1,FLII,TRIM27,NCK
PS4A,LDLR,ITGAV,IST1,CHMP4B,TRIM8,CD74,SNX3,LGALS1,DAG1,HSPA1A,EFNB2,NECTIN1,TRII

KBP1,PAF1,DNMT1,SMARCB1
ZB,PFN2,PPP1R9B,EVL,CTTN,SLIT2,BRK1,TMOD2,ABI2,RDX,CAPZA1,WASF2,MTOR,MICAL1,ARP
'8,SRCIN1,EPHA4
S3,HMGB1,ITGB1,NCAM1,ANXA2,ZMIZ1,VCL,AP3D1,CTNNA1,CNN3,RPSA,RTN4,LAMA5,MSN,RHC
,NECTIN1,JAK1,KIF3A,DLG4,SHANK1,HEG1,ARHGEF18,MAPK8IP3,CACNG7
REEP1,GAK,LMAN2,ESYT1,REEP5,EMC1,RAB18,EMC10,GET1,SEC31A,PRNP,GET4,EMD
MB4,DICER1,PSMD4,PSMC3,PSMC1,PSMB1,CUL1,PSMD13,PSMD7,PSME3,PSMC5,PSMC2,TNFR
N,SLIT2,BRK1,TMOD2,ABI2,RDX,CAPZA1,WASF2,MTOR,ARPC1A,HIP1R,ELN,CRACD,CYFIP1,FLII,T
FAM8A1,STUB1,HERPUD1,USP19,XBP1,MAN1B1,PSMC6,RNF5
AR1A,TLN1,GJA1,ANXA6,MACF1,CADM4,HMGCR,RAC1,HDAC2,TNC,DCN,CDKN1A,CNN2,RAB27A
,DNAJC5,HNRNPK,LAMP1,RAC1,APPL1,CNN2,RAB27A,VAMP2,RAB5B,RAB11B,ABCA2,ATP6AP1,
K2,SUN1,SUN2,H2AX,CRKL,GSK3B,CCDC85C,MBOAT7,BMERB1,XAB2,LRP8,FGF13,COL3A1,PC
2,THRA,MED16,RELA,CTNNBIP1,PSMC6,MED14

PSMB4,LSM14A,PDPK1,PSMD4,PSMC3,PRKACA,CREBBP,PSMC1,TOMM70,UBE2K,PSMB1,CUL1

N,SLIT2,BRK1,TMOD2,ABI2,RDX,CAPZA1,WASF2,MTOR,ARPC1A,HIP1R,ELN,CRACD,CYFIP1,FLII,T

)7,PSME3,PSMC5,PSMC2,PSMC4,PSMF1,PSMB5,PSMA1,PSMD10,PSMA6,TAB3,RELA,PSMD12,PS

B,AP2A2,AP1M1,CD74,CLTA,KIF26A,KLC2,KIF3A,CAPZA2,SEC31A,DYNC1I2
SR1,SEC23A,KIF5A,SEC16A,PTEN,YKT6,CUL3,BRSK1,LMAN1,SEPTIN5,RAB1B,KIFAP3,FNBP1L,CI
2,TBCD,PARP1,CAMSAP1,TERF2IP,MAPRE1,LUZP6,CAPZB,CLIP3,TNKS2,GHITM,PFN2,FLCN,ATR
,RBFOX2,SRF,HOXA5,STAT1,CEBPG,TMEM14C,CITED2,THRA,ADD1,HMGB2,ACIN1,CDK5RAP3,II
P3H1,HSPE1

.3,GNAS,RACK1,APP,KLRD1,HSPD1,IGF2,VCP,UBB,TRIM28,MATR3,FN1,NONO,PSMD5,HACD3,CI
CLIP3,FYN,YWHAH,MESD,HECTD1,RAB11A,PPP1R9B,GRIPAP1,RER1,GPC4,TCAF1,PKDCC,GPC2,'

CDH2,MMP14,PRKAR1A,CTNNB1,TAGLN,MYL6,HMGCR,GPC1,DNER,DCN,DDX39B,MORF4L2,CHI
F7,SMARCA4,KDM1A,PLTP,YWHAH,NCOR2,WBP2,STAT3,EP300,LBH,PHB2,UBA52,SFRP1,HDAC6
)7,PSME3,PSMC5,PSMC2,PSMC4,PSMF1,PSMB5,PSMA1,PSMD10,PSMA6,TAB3,RELA,PSMD12,PS
45B,FLCN,SFRP1,ULK1,BRSK1,TNKS,MAP1LC3B,DAP,SLC2A1,VPS41,GABARAPL1,STK24,ATP6V(
L1,BAZ1B,BRD2,ATRX,HJURP,NAP1L3,ANP32B,BAZ2A,MACROH2A2,SPTY2D1,NOC2L,MBD3,NAS

)HA,ATF4,JUND,PRDX2,HMGCS1,JUN,RHOB,HDAC2,EEF1A2,PARP1,AHCYL1,KDM6B,PPP2CA,CCI
17,APP,TMBIM6,NPM1,HSPD1,IGF2,UBB,AFF4,TRIM28,SPEN,PSMD5,EGLN1,PKD1,SSBP2,ZNF43
STM1,GDI2,NCKAP1,NOTCH2,RAPGEF1,G3BP2,GRB2,GABARAP,STMN3,ARFGEF2,CDC42,BCR,T
1LC3B,GRN,TPP1,RAB1B,VPS4A,GABARAPL1,RAB14,CHMP4B,MTOR,PACS2,AMBRA1,PINK1,ATF
AP5,RHOB,RAC1,DLG5,ACTR3,PTK7,ARF4,AMOTL2,CDC42,DCTN1,SPINT2,BRSK1,FLOT2,CLASF
MAPK1,LUZP6,GABARAP,ATP6V1B2,LARS1,SREBF2,BMPR2,PPP1R9B,WDR45B,FLCN,SFRP1,ULK

PL,BRSK1,SEPTIN5,SH3GL1,SCRIB,SYT11,GAK,RIMS3,GRIK5,CYFIP1,ADCY1,PIP5K1C,GSK3B,R/

GB1,VCL,AKAP12,RTN4,LAMA5,MSN,IGFBP3,RHOA,CD81,PFN1,CLEC7A,NISCH,PTN,TPM1,SULF
,PSMA7,EP300,PSMD1,PSMD8,A2M,GRN,XRCC6,PSMB4,LSM14A,PDPK1,PSMD4,PSMC3,PRKAC/
1A,PARK7,LIMK1,MARCHF7,OTUB1,UBXN1,RELA,HMG20B,VPS28
IMGB1,ITGB1,VCL,AKAP12,CTNNA1,RTN4,LAMA5,MSN,IGFBP3,RHOA,CD81,PFN1,CLEC7A,NISCH
GRIPAP1,PTEN,SV2A,PREPL,BRSK1,SEPTIN5,SH3GL1,SCRIB,SYT11,GAK,NSG1,RIMS3,GRIK5,CYF
A,BNIP3,PRDX3,PTPN11,ABL1,PINK1,PPP5C,SMPD3,MDM2,PARK7,PDGFRB,HSF1,SETX,TXN,STA
BMPR2,PPP1R9B,WDR45B,FLCN,SFRP1,ULK1,BRSK1,TNKS,MAP1LC3B,DAP,SLC2A1,VPS41,GAE
SF2,MTOR,MICAL1,PLS3,ABL1,FAM171A1,ELN,TNFAIP1,PDLIM1,LIMK1,CCDC88A,NCK2,SRF,ZY
RP1,SLIT2,DNAJB2,SEMA3C,SPAG9,SH3BP4,GSK3A,YY1,APBB1,NDUFA13,SEMA6B,TSPYL2,HSPA
PKD1,BASP1,HMGB1,ITGB1,ANXA2,ZMIZ1,RTN4,LAMA5,RHOA,PFN1,PTN,CDH2,GPC3,DDR1,SUL

BE2L3,PINK1,ARRDC4,FZR1,SART3,MYCBP2,TRIB3,CDK9,STUB1,MARCHF7,CTR9,RNF20,PSMD
SUMO1,MAGEF1,TRIB3,STUB1,CSNK1A1,HERPUD1,PLK2,GSK3B,PSMD10,ELOB,XBP1,RAD23A
,BNIP3,AMBRA1,PINK1,ATP13A2,MFN2,PARK7,ATG9A,RETREG3,GABARAPL2,UFM1,FBXO7,RB1
IF5A,SEC16A,YKT6,CUL3,LMAN1,SEPTIN5,RAB1B,KIFAP3,FNBP1L,CLASP2,SCRIB,SYT11,EXOC4,
JBE2G1,UBE2E1,RNF5

ϡ9,CYTH2,RDX,LZTR1,OGT,ARPP19,DENND4B,ABL1,AUTS2,RGL2,NOTCH1,MFN2,SGSM3,TNFAIF
37,KDM1A,USP7,RBBP7,EP300,CSNK2A1,EHMT1,PDPK1,NOC2L,MBD3,MTOR,CD74,MTA2,MDM
FEM1B,BRSK1,PSMD8,PCBP4,PPP1R10,PSMB4,TAOK1,CARM1,INTS3,VPS4A,PSMD4,PSMC3,PSI
HIP,DAPK3,MYL12B,FAM171A1,TAOK2,DAG1,PLXNA2,CYFIP1,CFDP1,KIF3A,WASF3,MYL12A,RH(

D81,WDR1,PTN,GPC3,SOX12,MMP14,PRDX2,RPS19,JUN,CTNNB1,NDFIP1,HLA-B,CDKN1A,CNN2,
P300,PTEN,PPP1CC,IMPDH2,OGT,AHCY,MTOR,UBE3A,THRAP3,NTRK2,RBM4,ATF5,MYCBP2,PPF

PTBN1,APP,GDI1,PLXNB2,HSPD1,IGF2,MAPT,FN1,SPEN,RPS3,BASP1,HMGB1,ITGB1,ANXA2,AGR
LOT1,FARP1,SFRP1,PTEN,GPC4,SLIT1,CLASP2,PRKACA,DAPK3,EIF4G1,NTRK2,PTPN11,ABL1,CC
D81,WDR1,PTN,GPC3,SOX12,MMP14,PRDX2,RPS19,JUN,CTNNB1,NDFIP1,HLA-B,CDKN1A,CNN2,
IDK,FYN,STXBP1,HYOU1,SOD1,CDC42,DDIT4,STAT3,CAPN2,HIF1A,GRN,UBE2M,SOD2,PDPK1,GS
)2,TAOK1,FKBP4,LDLR,RDX,CAPZA1,CLASP2,WASF2,TUBB4A,HDGFL3,HIP1R,CRACD,FLII,NCK2,

B,AP2A2,AP1M1,CD74,CLTA,KIF26A,KLC2,KIF3A,CAPZA2,SEC31A,DYNC1I2
MAP1S,BMERB1,CAPZA2,ADD1,FGF13,TPX2,PDXP,GSN

,NDUFA13,BNIP3,ATP5MC2,ATP5MG,MDH1,PINK1,PDHA1,NDUFA4,PARK7,IDH2,NDUFC2,COX8A
2,DYNLL2,RAB11A,DYNLL1,DCTN1,SMC3,PPP1CC,EML1,TNKS,TAOK1,LSM14A,CLASP2,CHMP4I
KDM1A,FYN,B2M,PLXNA3,PTPRS,TCTN1,PPP1R9B,ATRX,TRA2B,PTEN,NDST1,HIF1A,SLIT2,SLC2A
1,PKD1,BASP1,AP2M1,VCL,PAFAH1B1,AP2B1,CNN3,RTN4,LAMA5,MSN,RHOA,ATF4,PSMD2,WDR
,SLIT2,BRK1,TMOD2,ABI2,RDX,CAPZA1,WASF2,MTOR,ARPC1A,HIP1R,ELN,CRACD,CYFIP1,FLII,T
A,DYNLL1,DCTN1,SMC3,PPP1CC,EML1,TNKS,TAOK1,LSM14A,CLASP2,CHMP4B,RANGAP1,CHMF

M1A,CCAR2,TP53BP1,BRD4,FEM1B,FUS,PPP1R10,UBR5,NSD2,APBB1,CD74,DEK,TRIP12,USP47,
1,RAN,NUP98,NUP205,LMNA,NUP133,EPHB2,PPP2R2A,LEMD2,EMD
N2,SS18L1,CUL7,OPA1,LRP8,EPHA4,MYADM,POSTN,ANAPC2
M6,NPM1,HSPD1,IGF2,UBB,TRIM28,SPEN,PSMD5,EGLN1,PKD1,SSBP2,ZNF43,CHD3,HMGB1,AGF
3,ABL1,PAIP2,NPTN,MECP2,KAT2A,ADCY1,SYNGAP1,PLK2,SRF,GSK3B,SYNGR1,SERPINE2,CRTC
I1,PKD1,BASP1,HMGB1,CDKL1,ITGB1,ANXA2,ZMIZ1,COL11A1,AP2B1,HNRNPU,CPE,NACA,RTN4,
F2,MTOR,MICAL1,PLS3,ABL1,FAM171A1,ELN,TNFAIP1,PDLIM1,LIMK1,CCDC88A,NCK2,SRF,ZY
,AP2M1,ITGB1,ANXA2,ARF1,AP2B1,SLC4A8,MSN,NCS1,ATF4,KIF5B,NLGN2,CD81,OAZ1,CLEC7A

RNF157,ANKFY1,VPS51,ARSB,MGRN1,CHMP5,VPS16,CACNG7,AP3M1,M6PR
JBA,VAV2,FER,EPHA2
BP2,CALR,TMF1,HSPA1B,PSAP,H3-3B,CANX,CLU,SOX4,RPL13A,KAT6A,GNAS,SLC2A3,DDX17,GP

CL9L,EP300,FLCN,UBA52,SFRP1,RPS27A,PEG10,DKK3,HIF1A,FKBP1A,LOX,SLIT2,CCN1,LTBP4,I
N2,MAP1S,KIF3A,OPA1,AP3M1
ST1,CNP,KAT2A,COX6B1,SOS1,SUDS3,KDM2B,LRP6
HDAC6,NCOA4,SAFB2,CARM1,UBR5,UBE3A,PAGR1,CALCOCO1,KDM3A,PPP5C,PARK7,PMEPA1,F
'5MC3,ATP6V1G1,ATP5MC2,ATP5MG,TMCO3,NDUFA4,PARK7,COX8A,COX7A2L,ATP6V0B,ATP5P
IA,NAMPT
2BP,GSN

SUN2,GOLPH3,CCDC88A,TXN,PSMD10,ARL2BP,GET4,GSN
36L1,SFRP1,ITGB5,RPS27A,PEG10,DKK3,FKBP1A,LOX,LTBP4,MTMR4,PDPK1,LTBP3,CREBBP,SD

L,ZFP36L1,SFRP1,PTEN,HNRNPAB,HIF1A,OLFM1,SEMA3C,CLASP2,RBPJ,MTOR,EXOC4,SMO,FG
3RSK1,MAP1LC3B,DAP,SLC2A1,VPS41,GABARAPL1,STK24,ATP6V0D1,MTOR,ATP6V1G1,AMBRA1

PARK7,NDUFC2,COX8A,COX7A2L,ATP5PO,DLD,NDUFB8,SHMT2,NDUFS1,ATP5F1C,COX6B1,UQC
T1,MOAP1,GSK3A,ATP5MC3,NDUFA13,BNIP3,TOMM70,ATP5MC2,ATP5MG,PINK1,HIP1R,HSPA1

JP205,LMNA,NUP133,CTDNEP1,PPP2R2A,LEMD2,DMPK,EMD

AMBRA1,PAFAH1B2,PINK1,PARK7,TRIM27,PLK2,WAC,GSK3B,ATF6,MAPK3,XBP1,TMEM59,FBXO
PS14,CD81,WDR1,PTN,GPC3,SOX12,MMP14,PRDX2,RPS19,JUN,CTNNB1,NDFIP1,HLA-B,CDKN1
AOK1,CARM1,INTS3,VPS4A,PSMD4,PSMC3,PSMC1,USP47,PSMB1,TAOK2,MDM2,CUL1,PSMD13,
TEN,CCN1,ATXN3,CLASP2,DAPK3,PTPN11,ABL1,COL16A1,LGALS1,DAG1,CCDC80,NOTCH1,MAP
2A1,PARK7,GGT7,DHFR,TXN,TLR6,FBXO7,LANCL1,SESN2
TAOK1,CARM1,CD74,PTPN11,ABL1,USP10,TAOK2,MDM2,CD44,HIPK2,CUL4A,H2AX,MARCHF7,N
GMA,EPHA4,MAPK8IP3
C42,CTNND1,CNTNAP1,FLCN,CTNND2,RDX,CADM3,ADAM10,PRKACA,PTPN23,ANK2,JAM3,NECTI
R2,SERP1,IST1,RBPJ,MTOR,SMO,RABGAP1,NOTCH1,ADNP,CYFIP1,RNF157,NTN1,LIMK1,ISLR2,
P1,XRCC5,RHOA,CD81,CLEC7A,CCND2,DDR1,CCN2,PRDX2,DDX3X,MIF,PRKAR1A,YWHAB,PTGES
MD13,PSMD7,PSME3,PSMC5,PSMC2,PSMC4,PSMF1,PSMB5,PSMA1,SOS1,SHC1,PSMD10,MAP2K
B,AP2A2,AP1M1,CD74,CLTA,KIF26A,KLC2,KIF3A,CAPZA2,SEC31A,DYNC1I2

GB1,VCL,AKAP12,CTNNA1,RTN4,LAMA5,MSN,IGFBP3,RHOA,CD81,PFN1,CLEC7A,NISCH,PTN,TP
KHDRBS1,XPO1,SFPQ,NANOG,H1-0,JUN,CTNNB1,MXD4,CTBP1,HDAC2,HSBP1,EID1,SQSTM1,NO
MARCHF7,OTUB1,UBXN1,VPS28
OX,COL16A1,COL12A1,EMILIN1,COLGALT1,LOXL2,P3H1,COL7A1,TGFBR1,COL6A3,EXT1,TGFB
1A,MARCKSL1,FN1,DCHS1,RPS3,HMGB1,RPS4X,ITGB1,UCHL1,ANXA2,ZMIZ1,TIMP2,PAFAH1B1,

ETN2,WRNIP1,DHX36,SETX,CHD8,XPC,ANXA1,RAD54L2,HMGB2,CUL4B,MCM7,RAD50

S1,GLUD1,IARS1,OAZ2,PSMD8,PSMB4,IARS2,YARS1,PHGDH,PSMD4,IVD,AHCY,PSMC3,GOT2,PS
UP205,LMNA,NUP133,EPHB2,PPP2R2A,LEMD2,EMD
RC,XBP1,SESN2,CDK5RAP3,NUTF2
4,MAPK1,GRB2,PRKDC,B2M,LONP1,SOD1,PPP1R9B,STAT3,GPX4,PTEN,APEX1,SERP1,CARM1,A
36L1,SFRP1,ITGB5,RPS27A,PEG10,DKK3,FKBP1A,LOX,LTBP4,MTMR4,PDPK1,LTBP3,CREBBP,SD

K1,FKBP4,RDX,CAPZA1,CLASP2,WASF2,TUBB4A,HDGFL3,HIP1R,CRACD,KAT2A,FLII,NCK2,RBM
,RAB14,AP1M1,EXOC4,SORT1,DOP1B,GAK,PUM1,CLTA,SCAMP1,EXOC5,NSF,ANKFY1,GOLPH3,
1,SIN3A
IP1A,PINK1,TRAK1,HSPA1A,ATP13A2,RNF157,ANKFY1,VPS51,ARSB,MGRN1,CHMP5,TMEM50A,V

RK2,COL16A1,ASNS,HSF1,ZEB1,CDKN1B,NSMF,XBP1,PCNA,RELA,OPA1,SESN2,DNMT1
,MEGF8,PRKAR1A,GJA1,CTNNB1,CDKN1A,DDX39B,COL2A1,PTK7,ASXL1,KDM6B,TMED2,NOTC
CBP4,PPP1R10,PSMB4,TAOK1,CARM1,INTS3,VPS4A,PSMD4,SCRIB,PSMC3,PSMC1,USP47,ABL1
,UBE2J1,TMEM98,GSK3B,ILRUN,ABHD17A,TXN,LZTS2,ROCK2,UFM1,TMEM59,ARF6,LEPROT,G
ND2,DDR1,PRDX2,DDX3X,PRKAR1A,YWHAB,PEA15,CDKN1A,EEF1A2,NOX4,PKN1,GCN1,VEGFA,
N,DIP2B,EP400,MCM3AP,TAOK1,NMT1,OGT,PCGF2,NOC2L,MBD3,CREBBP,APBB1,DIP2A,AUTS2
HMGB1,ANXA2,RPLP1,XRCC5,RHOA,CD81,CLEC7A,CCND2,DDR1,CCN2,PRDX2,DDX3X,MIF,PRK
HMGB1,ANXA2,RPLP1,XRCC5,RHOA,CD81,CLEC7A,CCND2,DDR1,CCN2,PRDX2,DDX3X,MIF,PRK

1,IGF1R,PTPN1,CDK4,ATP6V0B,GSK3B,IRS1,ATP6V0E2,SOS1,ATP6V0A1,SHC1,ADIPOR1,FER,BC
K1,PHACTR4,MGAT5,PPP1R14B,PDGFRB,PPME1,PPP2R5D,GSK3B,NSMF,PPP1R37,PPP1R12A,

,TERF2IP,VEGFA,MAPK1,CCT8,TNKS2,HNRNPD,BMPR2,CCT5,UBA52,HNRNPA2B1,RPS27A,TNK
KDM4B,RBBP7,KDM5C,RERE,KDM5B,BAZ1B,BRD4,ATRX,HJURP,ARID1B,BAZ2A,SPTY2D1,KDM4
R9B,RBBP7,FLCN,DIP2B,GNG4,SFRP1,PTEN,HIF1A,SLIT2,DNAJB2,SEMA3C,SPAG9,SH3BP4,GSK

-C,VAPA,HMGB1,TUBB4B,ANXA2,TIMP2,VCL,CYB5R3,PGAM1,CTSD,LPCAT1,XRCC5,KPNB1,RHO

BP1,SMARCB1,SIN3A
O7,HSPB1,QARS1,DUSP4,CORO1C,MAPK8IP1,ADAR,PRKAR2A,SFRP1,PTEN,PDPK1,WNK1,PRDX

3,RPS6,DBN1,RPL4,ENO1,TUBA1B,FLNA,MAP1B,RPL7A,ALDOA,COL6A1,HSP90B1,NEFL,NEFM,PF
3,RPL7A,ALDOA,PKM,FTL,HSP90B1,NEFL,UGT8,RPL8,YBX1,ATP2B4,CALR,HSPA1B,PSAP,RPLP0,
IM28,RPL37A,SPEN,DDB1,HACD3,SET,EIF4A1,RPL19,RPL15,HLA-C,RPS3,VAPA,HMGB1,AP2M1,F
3,DBN1,ENO1,TUBA1B,FLNA,MAP1B,ALDOA,COL6A1,HSP90B1,NEFL,NEFM,PRPF8,UGT8,CALR,T
1,BSG,SPTBN1,CALM3,RPS8,RPL7,SEPTIN2,APP,RPL5,GDI1,CALM1,NPM1,HSPD1,VCP,TRIM28,Y
1,BSG,SPTBN1,CALM3,RPS8,RPL7,SEPTIN2,APP,RPL5,GDI1,CALM1,NPM1,HSPD1,VCP,TRIM28,Y
F3B1,RPS3,RPS4X,RPS3A,RPL18,RPL11,RPLP1,RPL10,RPS17,SF3B2,GTF2F1,HNRNPU,LARP1,HN
PTBN1,ATP5F1B,RPS8,RPL7,APP,RPL5,GDI1,SPTAN1,SMG1,NPM1,HSPD1,ATP5F1A,VCP,UBB,TRI
,USP11,CSDE1,VCP,UBB,RPL37A,DDB1,PSMD5,SET,RPL19,PPP2R1A,RPL15,PKD1,RPS3,HMGB1,
L,HSP90B1,NEFL,RPL8,ATP2B4,CALR,HSPA1B,PSAP,RPLP0,CANX,CLU,RPL13A,SEC62,RPL6,YWH
VHAE,KIF1A,RPL13,GNAS,RACK1,BSG,SPTBN1,CALM3,RPS8,MYH9,RPL7,SEPTIN2,APP,RPL5,GDI
WHAE,KIF1A,RACK1,SPTBN1,CALM3,STMN2,MYH9,APP,GDI1,FSCN1,SPTAN1,SMG1,CALM1,NPM
,SEC62,RPL6,DVL3,YWHAE,KIF1A,RPL13,GNAS,RACK1,BSG,SPTBN1,CALM3,RPS8,MYH9,RPL7,S
RPL37A,DDB1,PSMD5,SET,RPL19,PPP2R1A,RPL15,PKD1,RPS3,HMGB1,AP2M1,RPS4X,RPS3A,RPL
AQ,RPL19,RPL15,RPS3,VAPA,SEC61A1,RPS4X,RPS3A,RPL18,RPL11,RPLP1,RPL10,RPS17,AP3D1,
LARP1,HNRNPA0,RPS16,RPL23,RPL29,RPSA,RPS24,RPS9,RPL24,RPL17,RPL14,RPL22,RPL9,PSM
3,PIK3R2,FLNA,MAP1B,RPL7A,SLC9A7,ALDOA,PKM,FTL,HSP90B1,NEFL,UGT8,RPL8,YBX1,ATP2E
LR,FSTL1,TMF1,HSPA1B,RPLP0,CLU,SOX4,FKBP14,RPL13A,MAGED1,RPL6,KAT6A,DVL3,YWHAE,
ALR,TMF1,HSPA1B,TUBB2B,RPLP0,H3-3B,CLU,ENO2,RPL6,KAT6A,YWHAE,SPTBN1,ATP5F1B,CA
20,RPL26,RPL31,RPS25,RPS15,RPL12,RPL30,RPS5,RPS7,RPS19,RPL35,RPS10,RPL35A,RPLP2,AI
S14,RPS20,RPL26,RPL31,RPS25,RPS15,RPL12,RPL30,RPS5,RPS7,RPS19,RPL35,RPS10,RPL35A,F
26,RPL31,RPS25,RPS15,RPL12,RPL30,RPS5,RPS7,RPS19,RPL35,RPS10,RPL35A,RPLP2,RPL36,RI
LARP1,HNRNPA0,RPS16,RPL23,RPL29,RPSA,RPS24,RPS9,RPL24,RPL17,RPL14,RPL22,RPL9,PSM
0,H3-3B,CLU,RPL13A,RPL6,KAT6A,YWHAE,RPL13,DDX17,RACK1,SPTBN1,STMN2,RPS8,RPL7,SEF
1,CALM1,NPM1,HSPD1,VCP,TRIM28,YWHAG,RPL37A,MAPT,YWHAQ,RPL19,RPL15,PKD1,RPS3,VA
,RPS3,AP2M1,SEC61A1,RPS4X,RPS3A,RPL18,RPL11,RPLP1,RPL10,RPS17,AP3D1,ARF1,AP2B1,RI
RPS14,RPS20,RPL26,RPL31,RPS25,RPS15,RPL12,RPL30,RPS5,RPS7,RPS19,RPL35,GJA1,RPS10,
20,RPL26,RPL31,RPS25,RPS15,RPL12,RPL30,RPS5,RPS7,RPS19,RPL35,RPS10,RPL35A,RPLP2,RI
A1,ATP2B4,IGFBP2,CALR,FSTL1,TMF1,HSPA1B,PGK1,RPLP0,H3-3B,CLU,SOX4,FKBP14,RPL13A,N
RPL7,RPL5,GDI1,SMG1,NPM1,HSPD1,VCP,TRIM28,YWHAG,RPL37A,YWHAQ,RPL19,RPL15,PKD1
PL22,RPL9,RPS14,RPS20,RPL26,RPL31,PRKCSH,RPS25,RPS15,RPL12,RPL30,EIF3L,RPS5,RPS7,
20,RPL26,RPL31,RPS25,RPS15,RPL12,RPL30,RPS5,RPS7,RPS19,RPL35,RPS10,RPL35A,RPLP2,RI
L22,RPL9,RPS14,RPS20,EIF5,RPL26,RPL31,RPS25,RPS15,RPL12,RPL30,EIF3L,RPS5,RPS7,RPS19
6,ZNF316,RPL4,ENO1,PIK3R2,FLNA,NCL,RPL7A,ALDOA,PKM,GANAB,HSP90B1,FADS1,PRPF8,UG
,CLU,DDX5,ENO2,RPL13A,RPL6,RPL13,GPI,RACK1,ALDH6A1,RPS8,RPL7,APP,IDS,RPL5,TMBIM6
K1,ALDH6A1,RPS8,RPL7,APP,IDS,RPL5,TMBIM6,SMG1,CALM1,NPM1,TTC3,USP11,CSDE1,VCP,U
CL,RPL7A,MTRNR2L8,HSP90B1,NEFL,RPL8,YBX1,ATP2B4,IGFBP2,CALR,FSTL1,TMF1,ABHD2,HS
,RPS9,RPL24,RPL17,RPL14,RPL22,RPL9,ATF4,RPS14,RPS20,EIF5,RPL26,RPL31,KHDRBS1,RPS2
,ANXA2,TIMP2,HNRNPU,PGAM1,LARP1,HNRNPA0,RPL23,SCARB2,LPCAT1,MSN,PSMD2,TMEM25
D1,VCP,TRIM28,YWHAG,RPL37A,YWHAQ,RPL19,RPL15,PKD1,RPS3,AP2M1,SEC61A1,RPS4X,RPS
7A,SLC9A7,ALDOA,PKM,FTL,HSP90B1,NEFL,RPL8,YBX1,ATP2B4,CALR,TMF1,HSPA1B,PSAP,RPLF
,RPL11,RPLP1,RPL10,RPS17,AP3D1,DNAJA1,RPS16,RPL23,RPL29,RPSA,SCARB2,RPS24,RPS9,RI
LDOA,PKM,FTL,HSP90B1,NEFL,RPL8,YBX1,ATP2B4,CALR,TMF1,HSPA1B,PSAP,RPLP0,CANX,CLU
L6,KAT6A,YWHAE,RACK1,SPTBN1,STMN2,SEPTIN2,P4HB,APP,RPL5,FSCN1,SPTAN1,MARCKS,NPM
PL13,GPI,RACK1,ALDH6A1,RPS8,RPL7,APP,IDS,RPL5,SMG1,CALM1,NPM1,TTC3,PLD3,USP11,CS

3,UGT8,RPL8,FBXL4,YBX1,UBA1,ATP2B4,IGFBP2,CALR,FSTL1,TMF1,HSPA1B,PSAP,PGK1,RPLP0
,APP,GDI1,SPTAN1,GSTP1,MARCKS,CALM1,PLXNB2,MARCKSL1,UBB,YWHAG,CKB,MAPT,FN1,SF
RPS3A,RPL18,ITGB1,RPL11,ANXA2,AGRN,RPLP1,RPL10,RPS17,AP3D1,AP2B1,RPS16,RPL23,CPE,
P2B4,IGFBP2,CALR,TMF1,HSPA1B,ZNF99,TUBB2B,H3-3B,CLU,DDX5,SOX4,MAGED1,KAT6A,DVL3
CAM1,VCL,ARF1,PAFAH1B1,MAP4,CTNNA1,SCARB2,RTN4,LAMA5,RPL24,LLGL1,GAP43,NCS1,RH
PS4X,RPS3A,RPL18,ITGB1,RPL11,ANXA2,AGRN,RPLP1,RPL10,RPS17,AP3D1,AP2B1,RPS16,RPL23
M1,LARP1,HNRNPA0,RPL23,SCARB2,MSN,PSMD2,TMEM259,CD81,MCL1,HNRNPC,LRPPRC,XPO
6,RPL31,RPS25,RPS15,RPL12,RPL30,RPS5,RPS7,RPS19,RPL35,JUN,RPS10,RPL35A,TARDBP,RPL
1,PIK3R2,FLNA,NCL,RPL7A,PKM,HSP90B1,FADS1,RPL8,YBX1,ATP2B4,CALR,TMF1,HSPA1B,ZNF9
17,HNRNPU,LARP1,HNRNPA0,RPS16,RPL23,RPL29,RPSA,RPS24,RPS9,RPL24,KPNB1,RPL17,RPL
P90B1,NEFL,YBX1,ATP2B4,IGFBP2,CALR,FSTL1,TMF1,ABHD2,HSPA1B,COL5A1,H3-3B,CLU,PTMA
D1,RPL6,KAT6A,YWHAE,RPL13,DDX17,GPI,RACK1,CALM3,RPS8,RPL7,APP,RPL5,TMBIM6,GSTP1
LM3,MYH9,APP,RPL5,GSTP1,CALM1,NPM1,PLXNB2,HSPD1,CSDE1,IGF2,EIF3C,EIF4B,VCP,UBB,Y
4,ENO1,PIK3R2,FLNA,NCL,RPL7A,PKM,GANAB,HSP90B1,FADS1,RPL8,FBXL4,YBX1,UBA1,ATP2B4
6,KAT6A,YWHAE,RPL13,DDX17,GPI,RACK1,CALM3,RPS8,RPL7,APP,RPL5,TMBIM6,GSTP1,SMG1,
G1,NPM1,HSPD1,VCP,UBB,TRIM28,MAPT,NONO,DDB1,PSMD5,HACD3,EGLN1,PDIA3,RPS3,HMG
FL,PRPF8,YBX1,ATP2B4,IGFBP2,CALR,FSTL1,TMF1,ABHD2,HSPA1B,COL4A1,RPLP0,H3-3B,CAN
200,EIF3C,EIF4B,VCP,UBB,MAPT,FN1,DDB1,PSMD5,SET,TMSB10,PPP2R1A,DNAH8,PKD1,SF3B1,
CAM1,AGRN,ZMIZ1,VCL,ARF1,PAFAH1B1,MAP4,CTNNA1,SCARB2,RTN4,LAMA5,RPL24,LLGL1,GA
GRN,ARF1,PAFAH1B1,AP2B1,SCARB2,XRCC5,MSN,RHOA,CLSTN1,KIF5B,ENC1,NLGN2,WDR1,CC
AG,MAPT,FN1,SPEN,DCHS1,VAPA,HMGB1,STMN1,ITGB1,UCHL1,NCAM1,AGRN,ZMIZ1,TIMP2,VC
4,ENO1,PIK3R2,FLNA,NCL,RPL7A,ALDOA,PKM,GANAB,HSP90B1,FADS1,PRPF8,UGT8,RPL8,FBXL
PL10,RPS17,HNRNPU,LARP1,HNRNPA0,RPS16,RPL23,RPL29,RPSA,RPS24,RPS9,RPL24,KPNB1,R
PL10,RPS17,HNRNPU,LARP1,HNRNPA0,RPS16,RPL23,RPL29,RPSA,RPS24,RPS9,RPL24,KPNB1,R
L22,RPL9,RPS14,RPS20,RPL26,RPL31,RPS25,RPS15,RPL12,RPL30,RPS5,RPS7,RPS19,RPL35,RP
16,RPL4,ENO1,PIK3R2,FLNA,NCL,RPL7A,ALDOA,PKM,GANAB,HSP90B1,FADS1,PRPF8,UGT8,RPL
B1,STMN1,ITGB1,UCHL1,NCAM1,AGRN,ZMIZ1,TIMP2,VCL,ARF1,PAFAH1B1,MAP4,CTNNA1,SCAR
EN,VAPA,HMGB1,STMN1,ITGB1,UCHL1,NCAM1,AGRN,ZMIZ1,TIMP2,VCL,ARF1,PAFAH1B1,MAP4,
I1,RPS8,MYH9,RPL7,RPL5,GDI1,SMG1,NPM1,HSPD1,VCP,TRIM28,YWHAG,RPL37A,YWHAQ,RPL1
2,FLNA,NCL,RPL7A,PKM,HSP90B1,FADS1,RPL8,YBX1,ATP2B4,CALR,TMF1,HSPA1B,ZNF99,RPLP
16,RPL23,RPL29,RPSA,NACA,RPS24,RPS9,RPL24,RPL17,RPL14,RPL22,RPL9,CLSTN1,RPS14,RPS
PKM,NEFL,YBX1,ATP2B4,IGFBP2,CALR,TMF1,HSPA1B,ZNF99,TUBB2B,H3-3B,CLU,DDX5,SOX4,M
HSP90B1,FADS1,YBX1,ATP2B4,CALR,TMF1,HSPA1B,ZNF99,PSAP,H3-3B,CLU,DDX5,SOX4,RPL13
PS4X,RPS3A,RPL18,RPL11,RPLP1,RPL10,RPS17,LARP1,RPS16,RPL23,CPE,RPL29,RPSA,EIF4H,NA
GSTP1,CALM1,NPM1,PLXNB2,HSPD1,CSDE1,IGF2,EIF3C,EIF4B,VCP,UBB,YWHAG,MAPT,FN1,EIF1
SCARB2,NACA,RPS24,RPS9,RPL24,RPL17,RPL14,RPL22,RPL9,RPS14,RPS20,RPL26,RPL31,RPS2
17,HNRNPU,LARP1,HNRNPA0,RPS16,RPL23,RPL29,RPSA,RPS24,RPS9,RPL24,KPNB1,RPL17,RPL
31,UCHL1,NCAM1,VCL,PAFAH1B1,RTN4,LAMA5,MSN,RPL24,LLGL1,GAP43,RHOA,ATF4,KIF5B,WI
PL10,RPS17,HNRNPU,LARP1,HNRNPA0,RPS16,RPL23,RPL29,RPSA,RPS24,RPS9,RPL24,KPNB1,R
GAP43,RHOA,KIF5B,CRMP1,NES,SPTBN2,PTN,CDH2,DDR1,PLXNA1,MEGF8,GJA1,CTNNB1,MACF
GAP43,RHOA,KIF5B,CRMP1,SPTBN2,PTN,CDH2,DDR1,PLXNA1,MEGF8,GJA1,CTNNB1,MACF1,CY
GAP43,RHOA,KIF5B,CRMP1,NES,SPTBN2,PTN,CDH2,DDR1,PLXNA1,MEGF8,GJA1,CTNNB1,MACF
8,YBX1,ATP2B4,IGFBP2,CALR,FSTL1,TMF1,ABHD2,HSPA1B,TUBB2B,COL5A1,COL4A1,PSAP,PGK
SN,PSMD2,ENC1,TMEM259,CD81,OAZ1,XPO1,CSNK1E,GPC3,RAB7A,CLTC,RPS7,GJA1,CTNNB1,N
RPL18,RPL11,RPLP1,RPL10,RPS17,LARP1,RPS16,RPL23,RPL29,RPSA,EIF4H,NACA,RPS24,RPS9,

P,UBB,MAPT,FN1,DDB1,PSMD5,SET,TMSB10,PPP2R1A,DNAH8,PKD1,SF3B1,RPS3,HMGB1,AP2M
L3,RPS6,ZNF316,RPL4,ENO1,PIK3R2,FLNA,NCL,RPL7A,ALDOA,PKM,GANAB,HSP90B1,FADS1,PF
WHAG,MAPT,FN1,PPP2R1A,DNAH8,PKD1,VAPA,HMGB1,CDKL1,STMN1,ITGB1,TUBB4B,UCHL1,N
2,GTF2F1,HNRNPU,HNRNPA0,RPS16,RPL23,RPL29,RPS24,RPS9,RPL24,RPL17,PAPOLA,RPL14,R
2B4,IGFBP2,CALR,FSTL1,TMF1,ABHD2,HSPA1B,COL4A1,CES3,PGK1,RPLP0,H3-3B,CANX,CLU,D
GAP43,RHOA,KIF5B,CRMP1,SPTBN2,PTN,CDH2,DDR1,PLXNA1,MEGF8,GJA1,CTNNB1,MACF1,CY
8,SF3B1,RPS3,HMGB1,AP2M1,STMN1,RPL11,RPL10,ARF1,AP2B1,TMEM170A,SMARCA5,RPSA,E
16,RPL4,ENO1,PIK3R2,FLNA,NCL,RPL7A,PKM,GANAB,HSP90B1,FADS1,PRPF8,RPL8,FBXL4,YBX
1,FADS1,YBX1,ATP2B4,CALR,TMF1,HSPA1B,ZNF99,PSAP,H3-3B,CLU,DDX5,SOX4,RPL13A,MAGE
RPS14,RPS20,RPL26,RPL31,RPS25,RPS15,RPL12,RPL30,RPS5,RPS7,RPS19,RPL35,RPS10,RPL3
O1,SNRNP70,PSMB2,SRSF9,PSMD3,PTBP1,FTO,SUPT5H,HSPB1,SON,SUPT6H,HNRNPM,SRSF3,
WHAG,MAPT,FN1,PPP2R1A,DNAH8,PKD1,VAPA,HMGB1,CDKL1,STMN1,ITGB1,TUBB4B,UCHL1,N
RPL18,RPL11,RPLP1,RPL10,RPS17,LARP1,RPS16,RPL23,RPL29,RPSA,EIF4H,NACA,RPS24,RPS9,
FBP2,CALR,FSTL1,TMF1,ABHD2,HSPA1B,COL4A1,RPLP0,H3-3B,CANX,CLU,DDX5,SOX4,FKBP14
CALR,FSTL1,TMF1,ABHD2,HSPA1B,TUBB2B,COL5A1,COL4A1,PSAP,PGK1,H3-3B,CLU,DDX5,SOX
5,RPL24,LLGL1,GAP43,RHOA,ATF4,KIF5B,WDR1,CRMP1,SPTBN2,PTN,CDH2,PLXNA1,MEGF8,A
ED1,DVL3,KIF1A,GNAS,GPI,RACK1,BSG,SPTBN1,ATP5F1B,MYH9,P4HB,APP,GDI1,SPTAN1,PLXN
PL24,LLGL1,GAP43,RHOA,KIF5B,WDR1,CRMP1,NES,SPTBN2,PTN,TPM1,CDH2,DDR1,PLXNA1,M
5B,CRMP1,SPTBN2,PTN,CDH2,PLXNA1,MEGF8,MACF1,MAP2,RAC1,ARHGDIA,CAPRIN1,GPC1,TO
NEFL,YBX1,ATP2B4,CALR,FSTL1,TMF1,HSPA1B,H3-3B,CLU,SOX4,DVL3,YWHAE,KIF1A,GNAS,GPI
DX39B,NOVA2,SRSF5,U2AF2,SUGP2,CPSF1,PABPN1,HNRNPL,SNRPN,HNRNPA3,RBMX,DDX42,S
STL1,TMF1,HSPA1B,TUBB2B,COL5A1,COL4A1,PSAP,PGK1,H3-3B,CLU,DDX5,SOX4,MAGED1,KA
PTTG1IP,MAP1A,TIMP3,EEF1A2,NOP53,VPS35,RAD23B,SNRNP70,ABCA2,NRDC,AZIN1,PSMD3,O
1,FADS1,YBX1,ATP2B4,CALR,TMF1,HSPA1B,ZNF99,H3-3B,CLU,DDX5,SOX4,RPL13A,MAGED1,R
S4X,RPS3A,RPL18,RPL11,RPLP1,RPL10,RPS17,LARP1,RPS16,RPL23,RPL29,RPSA,EIF4H,NACA,
,COL5A1,COL4A1,PSAP,PGK1,H3-3B,CLU,DDX5,SOX4,MAGED1,KAT6A,DVL3,YWHAE,KIF1A,RPL1
6,ZNF316,RPL4,ENO1,PIK3R2,FLNA,NCL,RPL7A,ALDOA,PKM,GANAB,HSP90B1,FADS1,PRPF8,U
,PABPN1,HNRNPL,SNRPN,HNRNPA3,RBMX,DDX42,SNRNP70,PRPF6,SRSF9,SRSF1,PTBP1,HNRN
,PABPN1,HNRNPL,SNRPN,HNRNPA3,RBMX,DDX42,SNRNP70,PRPF6,SRSF9,SRSF1,PTBP1,HNRN
,PABPN1,HNRNPL,SNRPN,HNRNPA3,RBMX,DDX42,SNRNP70,PRPF6,SRSF9,SRSF1,PTBP1,HNRN
1,SMG1,CALM1,NPM1,PLXNB2,TTC3,USP11,IGF2,VCP,UBB,TRIM28,YWHAG,MAPT,FN1,DDB1,PS
1,SMG1,CALM1,NPM1,PLXNB2,TTC3,USP11,IGF2,VCP,UBB,TRIM28,YWHAG,MAPT,FN1,DDB1,PS
DH2,DDR1,PLXNA1,MEGF8,MACF1,MAP2,RAC1,ARHGDIA,GPC1,TOP2B,TNC,L1CAM,MAP1A,CR
1,KAT6A,DVL3,GNAS,DDX17,GPI,RACK1,BSG,FASN,SPTBN1,CALM3,MYH9,APP,RPL5,GSTP1,CA
,SRSF5,U2AF2,SUGP2,CPSF1,PABPN1,HNRNPL,ALKBH5,SNRPN,HNRNPA3,AHCYL1,RBMX,DDX4
,RAB7A,CLTC,RPS7,CTNNB1,NDFIP1,UBE2H,SKP1,LAPTM4B,TRIB2,PTTG1IP,MAP1A,CTSB,RNF1
PTAN1,GSTP1,PLXNB2,COL18A1,UBB,AFF4,MAPT,FN1,SPEN,CHSY1,VAPA,HMGB1,STMN1,ITGB1
NXA2,ARF1,PAFAH1B1,MAP4,HNRNPU,SCARB2,XRCC5,MSN,GAP43,RHOA,KIF5B,WDR1,CCT7,V
RPL37A,MAPT,FN1,PSMD5,SET,RPL19,PPP2R1A,RPL15,RPS3,HMGB1,RPS4X,RPS3A,RPL18,RPL
2R1A,RPL15,RPS3,RPS4X,RPS3A,RPL18,RPL11,RPLP1,RPL10,RPS17,PGAM1,LARP1,RPS16,RPL
CC5,RPL24,RPL17,RPL14,RPL22,RPL9,RPS14,EIF5,RPL26,RPL31,XPO1,RPS25,DDX3X,RPS15,R
CALM3,MYH9,APP,RPL5,GSTP1,CALM1,NPM1,PLXNB2,HSPD1,IGF2,EIF3C,VCP,UBB,TRIM28,MA
2,VIM,SPARC,TUBB,FTH1,UBC,KIDINS220,TMSB4X,NUMA1,DCX,EIF4G2,RPL3,RPS6,ZNF316,DB
,MYH9,SEPTIN2,APP,GDI1,SPTAN1,GSTP1,CALM1,IGF2,VCP,UBB,FN1,MRC2,HLA-C,VAPA,HMGB
N1,HDAC2,TMEM30A,L1CAM,ACAP3,MDK,PTK7,DPYSL2,VEGFA,RAPGEF1,KDM1A,FYN,YWHAH,B

3,DDX17,GPI,RACK1,BSG,SPTBN1,CALM3,MYH9,APP,RPL5,CALM1,NPM1,PLXNB2,HSPD1,IGF2,E
,NPM1,IGF2,UBB,TRIM28,YWHAG,MAPT,SPEN,NONO,YWHAQ,SET,PPP2R1A,RPS3,CHD3,BASP1,H
F8,MACF1,MAP2,RAC1,ARHGDIA,GPC1,TOP2B,L1CAM,MAP1A,CRPPA,CDH11,PTPRA,PTK7,DPYSL
SMG1,CALM1,NPM1,HSPD1,IGF2,UBB,TRIM28,YWHAG,MAPT,SPEN,NONO,YWHAQ,SET,PPP2R1
PLXNA1,MEGF8,ITM2C,MACF1,MAP2,RAC1,ARHGDIA,CAPRIN1,HDAC2,TMEM30A,L1CAM,ACTR3,
A4,BSG,CALM3,GNB1,STMN2,APP,GSTP1,CALM1,HSPD1,IGF2,UBB,YWHAG,COL4A2,MAPT,HMG
CDH2,PLXNA1,MEGF8,ITM2C,MACF1,MAP2,RAC1,ARHGDIA,CAPRIN1,HDAC2,TMEM30A,L1CAM,A
PM1,HSPD1,VCP,UBB,YWHAG,NONO,DDB1,YWHAQ,SET,PPP2R1A,PDIA3,RPS3,HMGB1,TPT1,RPS
ALM3,MYH9,APP,RPL5,CALM1,NPM1,PLXNB2,HSPD1,IGF2,EIF3C,VCP,UBB,TRIM28,MAPT,FN1,DI
DS1,PRPF8,RPL8,YBX1,ATP2B4,CALR,TMF1,HSPA1B,ZNF99,PGK1,RPLP0,H3-3B,CLU,PTMA,DDX5
TP5F1A,EIF4B,VCP,CKB,RPL37A,EIF1,HACD3,CHSY1,EIF4A1,RPL19,RPL15,RPS3,VAPA,RPS4X,RP
UBB2B,COL5A1,COL4A1,PSAP,PGK1,H3-3B,CLU,DDX5,SOX4,KAT6A,DVL3,KIF1A,DDX17,BSG,FA
90B1,FADS1,NEFL,PRPF8,YBX1,ATP2B4,IGFBP2,CALR,FSTL1,TMF1,ABHD2,HSPA1B,TUBB2B,CO
ALDOA,PKM,FADS1,PRPF8,UGT8,RPL8,YBX1,ATP2B4,CALR,TMF1,HSPA1B,ZNF99,PGK1,RPLP0,H
UBB2B,COL5A1,COL4A1,PSAP,PGK1,H3-3B,CLU,DDX5,SOX4,KAT6A,DVL3,KIF1A,DDX17,BSG,FA
J,PTMA,DDX5,SOX4,RPL13A,MAGED1,RPL6,KAT6A,DVL3,RPL13,DDX17,RPS8,RPL7,APP,RPL5,TM
DS1,UGT8,RPL8,YBX1,ATP2B4,CALR,TMF1,ABHD2,HSPA1B,ZNF99,COL5A1,PGK1,RPLP0,H3-3B,
DS1,UGT8,RPL8,YBX1,ATP2B4,CALR,TMF1,ABHD2,HSPA1B,ZNF99,COL5A1,PGK1,RPLP0,H3-3B,
1,SMG1,CALM1,NPM1,PLXNB2,TTC3,USP11,IGF2,VCP,UBB,TRIM28,YWHAG,MAPT,FN1,DDB1,PS
L8,YBX1,ATP2B4,CALR,TMF1,HSPA1B,ZNF99,PGK1,RPLP0,H3-3B,CLU,PTMA,DDX5,ENO2,SOX4,
3,GAP43,RHOA,CLSTN1,CD81,TLE5,SPTBN2,PTN,TKT,GPC3,DDR1,PLXNA1,MEGF8,MMP14,DDX
P1,NPM1,HSPD1,VCP,UBB,YWHAG,NONO,DDB1,YWHAQ,SET,PPP2R1A,PDIA3,RPS3,HMGB1,TPT1
R,TMF1,HSPA1B,ZNF99,RPLP0,H3-3B,CLU,PTMA,DDX5,SOX4,RPL13A,MAGED1,RPL6,KAT6A,DV
JSP7,HECTD1,BAG6,ARIH1,SGTA,UBQLN2,CSNK2A1,PTEN,GPX1,EIF3H,LDLR,GNA12,RDX,DNAJE
L8,YBX1,ATP2B4,CALR,TMF1,HSPA1B,ZNF99,COL5A1,RPLP0,H3-3B,CLU,PTMA,DDX5,SOX4,RPL
N2,PTN,RAB7A,MMP14,DDX3X,ITM2C,H1-0,IK,PRKAR1A,H1-10,MAP2,CTBP1,RAC1,ARHGDIA,HDA
IRNPU,XRCC5,MSN,GAP43,RHOA,CLSTN1,NLGN2,PFN1,CLEC7A,SPTBN2,TPM1,PIP4K2B,RAB7A,
OGA,RPL23,CTSD,LPCAT1,MSN,PSMD2,ENC1,TMEM259,CD81,OAZ1,XPO1,CSNK1E,GPC3,RAB7A
3,YBX1,ATP2B4,CALR,TMF1,HSPA1B,ZNF99,RPLP0,H3-3B,CLU,DDX5,SOX4,RPL13A,MAGED1,RPI
P4HB,APP,GDI1,PLXNB2,IGF2,YWHAG,COL4A2,MAPT,FN1,SPEN,PSMD5,EGLN1,PPP2R1A,BASP1,
,CSNK1E,CDH2,GPC3,MMP14,CLTC,CTNNB1,TARDBP,YWHAB,MAP2,CCT6A,GPC1,APPL1,IPO5,T
A,CTSB,RNF187,SQSTM1,NOP53,UBE4A,RAD23B,PSMB2,UBXN4,MVB12B,PSMD3,GABARAP,UBE
L8,YBX1,ATP2B4,CALR,TMF1,HSPA1B,ZNF99,RPLP0,H3-3B,CLU,PTMA,DDX5,SOX4,RPL13A,MAG
M6,GSTP1,NPM1,HSPD1,VCP,UBB,YWHAG,MAPT,NONO,DDB1,YWHAQ,SET,PPP2R1A,PDIA3,RPS
ABARAP,CCAR2,SMARCC1,STT3B,UBQLN1,CSNK1D,USP7,HECTD1,PCBP2,BAG6,PSMA7,ARIH1,S
MAP2,L1CAM,CDKN1A,DDX39B,CLSTN3,H3-3A,DPYSL2,VEGFA,PPP2CA,SMARCA4,LUZP6,CCAR
TP1,IGF2,UBB,YWHAG,COL4A2,MAPT,ZMIZ1,DNAJA1,GTF2F1,HNRNPU,LARP1,CTNNA1,RPL23,C
2,VIM,SPARC,TUBB,FTH1,UBC,KIDINS220,TMSB4X,NUMA1,DCX,EIF4G2,RPL3,RPS6,ZNF316,DBI
AS,HTRA4,RACK1,SPTBN1,CALM3,SHISA9,P4HB,APP,RPL5,TMBIM6,GSTP1,CALM1,NPM1,IGF2,V
COL4A2,MAPT,TMSB10,RPS3,STMN1,ITGB1,ANXA2,COL11A1,ARF1,PAFAH1B1,ITM2B,PXDN,CTN
F6,RPL38,SRSF9,SRSF1,EIF3E,PRKDC,SCAF11,DHX30,RPS28,EIF3F,SF3A1,STRAP,ADAR,TRAF7,S
PF8,RPL8,YBX1,ATP2B4,CALR,TMF1,HSPA1B,ZNF99,PGK1,RPLP0,H3-3B,CLU,PTMA,DDX5,ENO2
DS1,UGT8,RPL8,YBX1,ATP2B4,CALR,TMF1,ABHD2,HSPA1B,ZNF99,RPLP0,H3-3B,CLU,PTMA,DDX
DB1,SET,PDIA3,RPS3,HMGB1,TPT1,RPS3A,ITGB1,RPL11,RPL10,DNAJA1,AKAP12,CTNNA1,LDHA,C
2,WDR1,VAT1,PFN1,PTN,TPM1,CDH2,GPC3,SULF1,PLXNA1,MEGF8,ACTN4,CLTC,GJA1,CTNNB1,

SRSF1,EIF3E,PRKDC,SCAF11,DHX30,RPS28,EIF3F,SF3A1,STRAP,ADAR,TRAF7,SF1,EIF3D,SRSF6,
1,CYB5R3,SLC4A8,PGAM1,CTSD,LPCAT1,XRCC5,KPNB1,LLGL1,NCS1,RHOA,PSMD2,WDR1,ATP2
CN2,MIF,CTNNB1,HNRNPK,PTGES3,HMGCR,SMCHD1,BCLAF1,HSBP1,PTTG1IP,DDX39B,PARP1,
D1,RPS3,HMGB1,RPS4X,ANXA2,RPLP1,LARP1,CTSD,LPCAT1,RPS9,XRCC5,MSN,RHOA,TMEM259
3,SLC4A8,PGAM1,CTSD,LPCAT1,XRCC5,KPNB1,NCS1,RHOA,PSMD2,WDR1,ATP2A2,VAT1,PFKL,R
AS,HTRA4,RACK1,SPTBN1,CALM3,SHISA9,P4HB,APP,RPL5,TMBIM6,GSTP1,CALM1,NPM1,IGF2,V
PF8,RPL8,YBX1,ATP2B4,CALR,TMF1,HSPA1B,ZNF99,PGK1,RPLP0,H3-3B,CLU,PTMA,DDX5,ENO2
GT8,RPL8,YBX1,ATP2B4,CALR,TMF1,HSPA1B,ZNF99,RPLP0,H3-3B,CLU,PTMA,DDX5,SOX4,RPL1
2,VIM,SPARC,TUBB,FTH1,UBC,KIDINS220,TMSB4X,NUMA1,DCX,EIF4G2,RPS6,ZNF316,DBN1,EN
T3B,UBQLN1,CSNK1D,USP7,HECTD1,PCBP2,BAG6,PSMA7,ARIH1,SGTA,OS9,MTA1,PSMD1,UBQL
HL1,RPL11,ANXA2,RPLP1,ZMIZ1,DNAJA1,RPL23,XRCC5,IGFBP3,RHOA,ATF4,RPS20,CD81,CLEC
AD23B,PSMB2,UBXN4,MVB12B,PSMD3,GABARAP,UBE2R2,CCAR2,SMARCC1,STT3B,LONP1,UBC
1,SRSF9,SRSF1,IPO7,CASC3,SUPT6H,SSB,SRSF3,TXNIP,ADAR,RBM15B,ANP32A,STAT3,XPOT,KP
MSN,LLGL1,KIF5B,NLGN2,CD81,CCT7,OAZ1,ASTN2,XPO1,PTN,CSNK1E,PFKL,CDH2,GPC3,MMP
AD23B,PSMB2,UBXN4,MVB12B,PSMD3,GABARAP,UBE2R2,CCAR2,SMARCC1,STT3B,LONP1,UBC
TA,OS9,EP300,EXTL3,UBQLN2,FLOT1,SEC16A,PPP2CB,UBE2G2,MBTPS1,KLHDC3,NPLOC4,PPP
ME3,PSMC5,PSMC2,PSMC4,PSMF1,HSF1,PSMB5,EGLN2,HIF3A,PSMA1,ATF6B,CITED2,PSMD10,
AD23B,PSMB2,UBXN4,MVB12B,PSMD3,GABARAP,UBE2R2,CCAR2,SMARCC1,STT3B,UBQLN1,CS
DB1,SET,PDIA3,RPS3,HMGB1,TPT1,RPS3A,ITGB1,RPL11,RPL10,DNAJA1,AKAP12,CTNNA1,LDHA,
K7,VEGFA,RAPGEF1,ARPC2,KDM1A,CORO1C,FYN,PLXNA3,RAB11A,AKIRIN1,BMPR2,CDC42,EP3
MC5,PSMC2,PSMC4,PSMF1,HSF1,PSMB5,EGLN2,HIF3A,PSMA1,ATF6B,CITED2,PSMD10,ATF6,EL
NO,DDB1,SET,PDIA3,RPS3,HMGB1,TPT1,RPS3A,ITGB1,RPL11,RPL10,DNAJA1,AKAP12,AP2B1,CT
3,RTN4,IGFBP3,ATF4,ENC1,RPS20,BEX3,RTN3,PTN,GLG1,PEBP1,GPC3,DDX3X,RPS15,ITM2C,RP
NNA1,RTN4,MSN,KIF5B,NLGN2,CD81,CCT7,OAZ1,ASTN2,KHDRBS1,XPO1,PTN,CSNK1E,CDH2,GP
STT3B,STT3A,VGF,RAB11A,IGFBP7,PSMA7,RBBP7,PSMD1,CST3,KLHL42,RAB2A,CRTAP,FBXO11,
DHA,RTN4,XRCC5,RHOA,KIF5B,WDR1,NES,SPTBN2,PTN,CSNK1E,CDH2,SOX12,APLP1,HMGCS1,
DH6,CDH2,MMP14,TLN1,GJA1,CTNNB1,MACF1,MYH10,SEPTIN11,RAC1,CAPRIN1,TNC,DNER,L1
L1,UPF1,RPS27A,PSMD8,APEX1,PCBP4,FUS,PSMB4,ELAVL1,TTC37,PSMD4,IGF2BP1,PSMC3,MT
2,PRDX6,SRSF2,RBM5,PRPF19,SAP18,THRAP3,FXR2,RBM4,TIA1,RBM25,FXR1,SRSF10,C1QBP,C
PF8,RPL8,YBX1,ATP2B4,CALR,TMF1,HSPA1B,ZNF99,PGK1,RPLP0,H3-3B,CLU,PTMA,DDX5,ENO2
,RPS27A,PSMD8,APEX1,PCBP4,FUS,PSMB4,ELAVL1,TTC37,PSMD4,IGF2BP1,PSMC3,MTOR,PSM
70,OPTN,ATG2B,DYNC1LI2,DYNLL2,CDC37,WDR6,GABARAP,SUPT5H,HSPB1,BNIP3L,ATP6V1B2
70,OPTN,ATG2B,DYNC1LI2,DYNLL2,CDC37,WDR6,GABARAP,SUPT5H,HSPB1,BNIP3L,ATP6V1B2
B1,ANXA6,MACF1,CYFIP2,MAP2,TNC,L1CAM,PPIB,CDKN1A,DDX39B,MDK,PTK7,DPYSL2,TMED2,
1,SRSF9,SRSF1,IPO7,CASC3,SUPT6H,SSB,SRSF3,TXNIP,ADAR,RBM15B,ANP32A,STAT3,XPOT,KP
H,GNA12,DNAJB2,ATXN3,OGT,GSK3A,PSMC3,UBE3A,PRKACA,PSMC1,BAG5,UBE2K,ZER1,USP5,
1A1,DNAJA1,AKAP12,LDHA,RTN4,XRCC5,RHOA,ATF4,PSMD2,ATP1A1,JUND,ATP2A2,RPL26,PTN,
ARCKSL1,MAPT,MAP9,TMSB10,INA,DNAH8,PKD1,RPS3,CHD3,STMN1,ITGB1,TUBB4B,AGRN,ARF
MD5,SET,MAP9,PPP2R1A,PKD1,RPS3,CHD3,CDKL1,STMN1,ITGB1,TUBB4B,TIMP2,ARF1,PAFAH1
APK1,CDC37,PRKDC,CCAR2,CCT8,LSS,STXBP1,USP7,BAG6,PFN2,CCT5,CTNND1,EP300,HIP1,FL
PSMB2,RAPGEF1,PPP2CA,CCND1,SMARCA4,PSMD3,CCAR2,GNG2,TNKS2,CSNK1D,MESD,VANGL
P53,PARP1,ABCA3,ACTR3,CAMSAP1,NCKAP1,VEGFA,MAPRE1,ARPC2,LUZP6,GRB2,CAPZB,CAND
CA,SMARCA4,LUZP6,CCAR2,SLIT3,PLXNA3,PTPRS,RAB11A,IGFBP7,BMPR2,CDC42,PPP1R9B,RB
TRF1L,CAMSAP1,KLC1,LAMC1,OPTN,SMARCA4,ATG2B,DSTN,LUZP6,CDC37,CAPZB,GABARAP,B
PSMB2,RAPGEF1,PPP2CA,CCND1,SMARCA4,PSMD3,CCAR2,GNG2,TNKS2,CSNK1D,MESD,VANGL

1,RPS4X,ANXA2,RPLP1,LARP1,CTSD,RPS9,XRCC5,MSN,RHOA,TMEM259,CD81,RPL26,BEX3,CLE
TNNA1,RPL23,LDHA,COL1A2,RHOA,ATF4,JUND,TMEM259,CD81,COL5A2,SRSF4,PTN,PIP4K2B,C
MC3,PRKACA,PSMC1,BAG5,UBE2K,ZER1,USP5,SDCBP,PINK1,HSPA1A,MDM2,FZR1,SUMO1,PARI
7A,DDX3X,MIF,CD63,DNAJC5,DDOST,LAMP1,CKAP4,HLA-B,TTR,COTL1,HRNR,RAC1,ATP6V0C,TM
7,HSPB1,FKBP8,GNG2,NKTR,CCT8,B2M,MESD,CCT5,SGTA,CSNK2A1,CRTAP,LMAN1,FKBP1A,G
F3,CCT8,CCT5,RBM15B,XPOT,FLOT1,ZFP36L1,SRSF6,UPF1,HNRNPA2B1,MCM3AP,POM121C,T
1,UBE2H,RAB1A,HECTD4,SKP1,PTTG1IP,RNF187,U2AF2,SQSTM1,KLHL24,UBE4A,PSMB2,RNF1C
JS,SRSF2,RBM12,RBM5,PRPF19,SAP18,THRAP3,SLC38A2,HNRNPH3,RPS26,FXR2,RBM4,HSPA1A
IC1,RPS20,BEX3,PTN,GLG1,PEBP1,GPC3,DDX3X,RPS15,RPS7,PRKAR1A,CTNNB1,APLP2,TARDBF
7A,DDX3X,MIF,CD63,DNAJC5,DDOST,LAMP1,CKAP4,HLA-B,TTR,COTL1,HRNR,RAC1,ATP6V0C,TM
,RPS27A,PSMD8,APEX1,PCBP4,FUS,PSMB4,ELAVL1,TTC37,PSMD4,IGF2BP1,PSMC3,MTOR,PSMC
,AGRN,MARCHF6,LARP1,CTNNA1,RPL23,LDHA,COL1A2,RHOA,ATF4,JUND,TMEM259,CD81,COL
L,RAB7A,DDX3X,MIF,CD63,DNAJC5,DDOST,LAMP1,CKAP4,HLA-B,TTR,COTL1,HRNR,RAC1,ATP6
P2,CADM4,HNRNPH1,GPC1,APPL1,HDAC2,TNC,DCN,PARP1,NOX4,COL2A1,SKI,NOTCH2,VEGFA,N
DDX3X,RPS15,RPS7,CTNNB1,NDFIP1,UBE2H,RAB1A,HECTD4,SKP1,PTTG1IP,TOMM20,RNF187,U
P90AA1,ACTR2,VIM,SPARC,TUBB,FTH1,UBC,KIDINS220,TMSB4X,NUMA1,DCX,EIF4G2,RPL3,RPS6
7A,DDX3X,MIF,CD63,DNAJC5,DDOST,LAMP1,CKAP4,HLA-B,TTR,COTL1,HRNR,RAC1,ATP6V0C,TM
HB,APP,RPL5,TMBIM6,GSTP1,CALM1,NPM1,IGF2,VCP,UBB,YWHAG,MAPT,FN1,NONO,PSMD5,HA
BP1,RAC1,DCN,CCT2,NOX4,ACTR3,TCP1,CCT3,NCKAP1,VPS35,SURF4,CCT4,H3-3A,TERF2IP,VEG
1,TUBB4B,ARF1,PAFAH1B1,MAP4,HNRNPU,RPL24,KPNB1,LLGL1,RHOA,CBX5,PSMD2,RPL26,RE
7A,DDX3X,MIF,CD63,DNAJC5,DDOST,LAMP1,CKAP4,HLA-B,TTR,COTL1,HRNR,RAC1,ATP6V0C,TM
H,RAB1A,HECTD4,SKP1,PTTG1IP,RNF187,U2AF2,SQSTM1,KLHL24,UBE4A,PSMB2,RNF10,RPL37
AP4,HNRNPU,RPL24,KPNB1,RHOA,PSMD2,RPL26,SPC25,NES,CCNI,KHDRBS1,XPO1,CSNK1E,CCI
AC1,WDR45B,UBQLN2,CSNK2A1,PHB2,UBA52,RAB3GAP2,HDAC6,ULK1,GPSM1,RPS27A,HIF1A,
,PSMD2,JUND,RPL26,RBBP4,CCNI,KLHL9,XPO1,CSNK1E,SFPQ,CCND2,CCN2,DDX3X,CLTC,IK,PF
B1,TUBB4B,ANXA2,TIMP2,MIDN,VCL,ARF1,CYB5R3,PAFAH1B1,SLC4A8,PGAM1,CTSD,LPCAT1,XI
D2,RPL26,SPC25,NES,CCNI,KHDRBS1,XPO1,CSNK1E,CCND2,DDX3X,CLTC,IK,GJA1,MYH10,CKAF
P2B,SFRP1,HDAC6,ULK1,CTTN,SLIT2,NDN,SLIT1,OLFM1,DCLK1,SEMA3C,SPAG9,CLASP2,IST1,AI
1,CITED2,PSMD10,ELOB,PSMA6,PSMD12,PSMA4,PSMC6
D1,SSBP2,ZNF43,HMGB1,RPS4X,AGRN,ZMIZ1,AP3D1,AKAP12,ZNF770,GTF2F1,HNRNPU,SMARC
STP1,SMG1,CALM1,NPM1,IGF2,UBB,TRIM28,MAPT,FN1,NONO,PSMD5,YWHAQ,HACD3,DCHS1,I
,SEMA6B,AUTS2,ADNP,CYFIP1,NTN1,LIMK1,ISLR2,GOLGA4,SEMA4C,SRF,GSK3B,ALCAM,NRP2,I
FPQ,DDX3X,H1-0,IK,CTNNB1,ARID1A,H1-10,PTGES3,CCT6A,CTBP1,TOP2B,SMCHD1,HDAC2,SKP
IBIM6,GSTP1,CALM1,NPM1,PLXNB2,HSPD1,IGF2,EIF3C,VCP,UBB,TRIM28,YWHAG,MAPT,FN1,HA
D1,SSBP2,ZNF43,HMGB1,RPS4X,AGRN,ZMIZ1,AP3D1,ZNF770,GTF2F1,HNRNPU,SMARCA5,LARF
MBTPS1,HDAC6,CUL3,KLHDC3,PPP1R15A,SERP1,RHBDD2,VAPB,TPP1,DNAJB2,ATXN3,ATP6V0D
P2,CADM4,HNRNPH1,GPC1,APPL1,HDAC2,DCN,PARP1,NOX4,COL2A1,SKI,NOTCH2,VEGFA,MARS
IB1,RHOA,KIF5B,NLGN2,ATP2A2,PTN,CDH2,PIP4K2B,RAB7A,CLTC,ANXA6,MTSS2,TARDBP,ARL6I
CTNNB1,WLS,ATP1A2,ARCN1,TOP2B,MMP2,NNAT,DLG5,GNAI3,COL2A1,MDK,SKI,KDM6B,CNTN
,TXNIP,CCT5,ADAR,STAT3,FLCN,KPNA2,PHB2,MBTPS1,POM121C,PPP1R10,UBR5,ELAVL1,DCLK
HDC3,PPP1R15A,SERP1,RHBDD2,VAPB,TPP1,DNAJB2,ATP6V0D1,GSK3A,SRPRA,AMFR,TSPYL2,I
TR1A,PSMD1,PBX1,BRD4,MASTL,BRSK1,PSMD8,AKAP9,PSMB4,TAOK1,INTS3,VPS4A,PSMD4,ARF
RPC2,DSTN,LUZP6,GRB2,CAPZB,KDM1A,LARS1,PLXNA3,PTPRS,PFN2,RAB11A,BMPR2,DIP2B,EVI
L5A2,PFN1,NES,PRKCSH,GPC3,SULF1,MEGF8,MMP14,NANOG,RPS7,PRKAR1A,GJA1,CTNNB1,W
P9,PSMB4,TAOK1,INTS3,VPS4A,PSMD4,PSMC3,PRKACA,PSMC1,TUBB4A,USP47,PSMB1,TAOK2,N

1,OLFM1,DCLK1,SEMA3C,SPAG9,CLASP2,ABL1,SEMA6B,AUTS2,ADNP,CYFIP1,RNF157,NTN1,LIN
I,ZMIZ1,AP3D1,AKAP12,ZNF770,GTF2F1,HNRNPU,SMARCA5,LARP1,NACA,RPS9,XRCC5,RHOA,AT
CTTN,SLIT2,NDN,SLIT1,OLFM1,DCLK1,SEMA3C,SPAG9,CLASP2,IST1,GSK3A,MTOR,YY1,ABL1,SE
CN2,DDX3X,CLTC,IK,PRKAR1A,CTNNB1,DUX4,CKAP5,SND1,APPL1,SKP1,CDKN1A,CTDSP2,DDX3
DAM9,FTO,GABARAP,SUPT5H,BNIP3L,UBQLN1,CSNK1D,HECTD1,BAG6,HNRNPD,VDAC1,ARIH1,E
F4,PTN,PIP4K2B,CCN2,MMP14,HMGCS1,PRKAR1A,GJA1,MYO5A,SESN3,ARID1A,PTGES3,ATP1A
CTR1B,PSMD4,ITGAV,PSMC3,AP2A2,AP1M1,PSMC1,CD74,TAPBP,CLTA,PSMB1,PSMD13,KIF26A,
CDKN1A,CTDSP2,U2AF2,SQSTM1,PKN1,SKI,TERF2IP,TMED2,TAF7,RPL37,PPP2CA,IPO7,PRKDC,H
B4,TAOK1,INTS3,VPS4A,PSMD4,PSMC3,PRKACA,PSMC1,TUBB4A,USP47,PSMB1,TAOK2,PKIA,PC
1,PFKL,RAB7A,DDX3X,MIF,CD63,DNAJC5,DDOST,LAMP1,CKAP4,HLA-B,TTR,COTL1,HRNR,RAC1
1B,COL5A1,PGK1,H3-3B,CANX,CLU,DDX5,SOX4,FKBP14,RPL13A,KAT6A,YWHAE,GNAS,DDX17,G
PA,HMGB1,TUBB4B,ANXA2,TIMP2,MIDN,VCL,ARF1,CYB5R3,PAFAH1B1,SLC4A8,PGAM1,CTSD,LP
A,DDX3X,MIF,CD63,DNAJC5,DDOST,LAMP1,CKAP4,HLA-B,TTR,COTL1,HRNR,RAC1,ATP6V0C,TM
2,FLNA,MAP1B,NCL,MTRNR2L8,PKM,COL6A1,HSP90B1,FADS1,NEFL,PRPF8,YBX1,UBA1,ATP2B4
TIA1,RBM25,FXR1,SRSF10,C1QBP,CELF3,RBFOX2,SRSF7,NUP98,RBM10,RBM17,SFSWAP,RNPS
SMD1,PBX1,BRD4,MASTL,BRSK1,PSMD8,AKAP9,PSMB4,TAOK1,INTS3,VPS4A,PSMD4,ARPP19,P
RPS25,DDX3X,RPS15,RPL12,RPL30,RPS5,RPS7,RPS19,RPL35,RPS10,RPL35A,RPLP2,TSR2,RPL3
P2,VCL,AP3D1,PAFAH1B1,HNRNPU,CTNNA1,NACA,RTN4,XRCC5,IGFBP3,RHOA,ATF4,PSMD2,JUN
P1A2,ARCN1,TOP2B,NNAT,DLG5,GNAI3,MDK,SKI,KDM6B,CNTNAP2,DPYSL2,PBX2,ATP5PB,LUZP
0,HIP1,FLOT1,SEC16A,PHB2,CRTAP,PTEN,GRN,FLOT2,ATP1B2,BAG5,CD74,SMO,ANK2,PINK1,H
FYN,USP7,BAG6,HNRNPD,CYP51A1,DDIT4,STAT3,SGTA,CSNK2A1,CST3,DAP,FUS,EIF3H,ELAVL1,I
PZA1,ACTR1B,PSMD4,ITGAV,PSMC3,AP2A2,AP1M1,PSMC1,CD74,TAPBP,CLTA,PSMB1,PSMD13,I
2,SSRP1,PKN1,YEATS2,CHD2,SKI,ASXL1,H3-3A,KDM6B,TERF2IP,TOP1,TAF7,VEGFA,SMARCA4,I
,CSNK1D,TMED9,HYOU1,HGS,PPP6R1,SEC23A,ARFGEF2,DYNLL1,DCTN1,KIF5A,TXNDC5,ACTR1
,RAC1,ARHGDIA,MAP1A,TBCD,NOX4,ACTR3,CAMSAP1,NCKAP1,MDK,NOTCH2,MAPK1,MAPRE1,A
AT4,GJA1,MACF1,MAP2,L1CAM,PPIB,CDKN1A,DDX39B,MORF4L2,CLSTN3,H3-3A,DPYSL2,NOTC
A,PTCD3,ATF4,PSMD2,ENC1,RPS14,EIF5,RPL26,HNRNPC,LRPPRC,KHDRBS1,XPO1,EIF5A,DDX3X
RC,KHDRBS1,EIF5A,DDX3X,NGRN,TARDBP,CIRBP,CAPRIN1,BCLAF1,DDX39B,SRP9,RTN1,GCN1
A,DDX3X,MIF,CD63,DNAJC5,DDOST,LAMP1,CKAP4,HLA-B,TTR,COTL1,HRNR,RAC1,ATP6V0C,TM
L1,BCLAF1,PEA15,PTTG1IP,CDKN1A,TIMP3,GNAI3,PARP1,GRINA,COL2A1,PRKDC,GABARAP,HS
POM121C,SYMPK,SMG5,SRSF2,SRSF11,CSE1L,DHX38,PRKACA,XPO6,EIF4A3,RANGAP1,PTPN11
STAT3,SGTA,CSNK2A1,CST3,DAP,FUS,EIF3H,ELAVL1,FOXK2,IGF2BP1,OGT,KDM4A,GSK3A,CHM
K,SKI,ATP6AP1,NOTCH2,RNF10,VEGFA,ARPC2,PRKDC,KDM1A,CORO1C,YWHAH,FBLN1,B2M,PI
K7,SKI,PSMB2,RAPGEF1,PPP2CA,CCND1,SMARCA4,PSMD3,CCAR2,GNG2,TNKS2,CSNK1D,MESD
,CLU,DDX5,SOX4,RPL13A,MAGED1,RPL6,KAT6A,DVL3,DDX17,RACK1,APP,RPL5,TMBIM6,GSTP1,
2A1,PTEN,OAZ2,LDLR,RDX,DNAJB2,ATXN3,GSK3A,PSMC3,NDUFA13,PSMC1,ATG4B,ZER1,USP5
A,HMGB1,TUBB4B,ANXA2,TIMP2,MIDN,VCL,ARF1,CYB5R3,PAFAH1B1,SLC4A8,PGAM1,CTSD,LPC
MAPT,FN1,HACD3,SET,EGLN1,PPP2R1A,PKD1,SERINC1,RPS3,HMGB1,STMN1,ITGB1,UCHL1,RPL
1,ANXA2,TIMP2,MARCHF6,RPL23,CPE,CTSD,RHOA,PSMD2,ENC1,TMEM259,BEX3,CLEC7A,TADA
MAPRE1,ARPC2,DSTN,AEBP1,LUZP6,GRB2,CAPZB,CLIP3,PFN2,CDC42,DCTN1,SFRP1,EVL,HDAC6

0,RPS7,RPS19,RPL35,RPL35A,RPLP2,TSR2,RPL36,RPS27,RPL23A,NOP53,TSR1,RPL27,RPL36A,
A,RHOA,BEX3,MCL1,NES,SFPQ,CCND2,PRDX2,DDX3X,MIF,CTNNB1,DNAJC5,ARL6IP1,HNRNPK,H
,AP2B1,CTNNA1,NACA,RHOA,ATF4,TMEM259,BEX3,MCL1,NES,SFPQ,CCND2,RAB7A,CCN2,PRDX
2,RPL30,RPS7,RPS19,RPL35,RPL35A,RPLP2,TSR2,RPL36,RPS27,RPL23A,NOP53,TSR1,RPL27,RI

NPA2B1,MCM3AP,POM121C,TNKS,SYMPK,SMG5,SRSF2,SRSF11,IGF2BP1,DHX38,EIF4A3,RAN,C
NPA2B1,MCM3AP,POM121C,TNKS,SYMPK,SMG5,SRSF2,SRSF11,IGF2BP1,DHX38,EIF4A3,RAN,C
1A,CTDSP2,DDX39B,PPP1CB,NOP53,PSMB2,OPTN,CCND1,MAPRE1,PSMD3,CCAR2,CSNK1D,RAE
3X,NGRN,TARDBP,CIRBP,CAPRIN1,BCLAF1,DDX39B,SRP9,GCN1,TMED2,RPL38,MAPK1,EIF3E,F
1,CLSTN3,CNTNAP2,VEGFA,G3BP2,MAPRE1,ARPC2,GRB2,CAND1,HGS,PFN2,AKIRIN1,CDC42,D
,CLU,DDX5,SOX4,RPL13A,MAGED1,RPL6,KAT6A,DVL3,DDX17,RACK1,APP,RPL5,TMBIM6,GSTP1,
SULF1,PIP4K2B,MEGF8,RAB7A,CCN2,JUN,CTNNB1,CD63,SESN3,CYFIP2,CADM4,HNRNPH1,RAC1
TOR,CREBBP,BNIP3,PSMC1,PINK1,PSMB1,NOTCH1,MDM2,PSMD13,PSMD7,HIPK2,PSME3,STUB
MA6B,TRAK1,PLXNA2,ADNP,CYFIP1,NTN1,LIMK1,SYNGAP1,ISLR2,GOLGA4,SEMA4C,SRF,GSK3E
A,RHOA,BEX3,MCL1,NES,SFPQ,CCND2,RAB7A,PRDX2,DDX3X,MIF,CTNNB1,DNAJC5,ARL6IP1,HN

NA3,RAB11A,BMPR2,CDC42,NAP1L1,MAP6,PLXNB1,PPP1CC,IL6ST,HIF1A,SLIT2,DICER1,IST1,P
C3,MTOR,CREBBP,BNIP3,PSMC1,PINK1,PSMB1,NOTCH1,MDM2,PSMD13,PSMD7,HIPK2,PSME3,
,DLG5,VPS35,CLSTN3,ACTN1,ARF4,FYN,MESD,PTPRS,PFN2,CDC42,GRIPAP1,DCTN1,LRRC4B,P
,RAB11A,BMPR2,NAP1L1,MAP6,PLXNB1,DIP2B,PTEN,PPP1CC,IL6ST,HIF1A,SLIT2,LINGO1,LDLR,
T,HIF1A,SLIT2,DICER1,IST1,PRPF19,MTOR,SMO,METRN,NTRK2,TRAK1,DAG1,PLXNA2,OBSL1,N
CELSR1,ABL1,PSMB1,EXOC5,PSMD13,PSMD7,PSME3,DAAM1,PSMC5,PSMC2,PSMC4,PSMF1,CE
PRA,AMFR,TSPYL2,HSPA1A,PTPN1,NCK2,STUB1,HERPUD1,HSF1,LMNA,ATF6B,SHC1,DNAJB11,A
GED1,RPL6,KAT6A,DVL3,DDX17,APP,TMBIM6,SMG1,NPM1,HSPD1,IGF2,VCP,UBB,TRIM28,SPEN
TN1,EVL,CTTN,SLIT2,BRK1,AKAP9,TMOD2,FKBP4,ABI2,RDX,CAPZA1,CLASP2,WASF2,MTOR,TUE
,CLU,DDX5,SOX4,RPL13A,MAGED1,RPL6,KAT6A,DVL3,DDX17,RACK1,APP,RPL5,TMBIM6,NPM1,
ABP2,RUFY3,PLXND1,ANAPC2,SKIL
TXN3,ATP6V0D1,GSK3A,SRPRA,AMFR,TSPYL2,HSPA1A,PTPN1,NCK2,STUB1,HERPUD1,HSF1,LM
NPA2B1,MCM3AP,POM121C,TNKS,SYMPK,SMG5,SRSF2,SRSF11,IGF2BP1,DHX38,EIF4A3,RAN,C
PSMD4,ATP6V0D1,RBPJ,PSMC3,MTOR,CREBBP,BNIP3,ATP6V1G1,PSMC1,PINK1,PSMB1,NOTC
DK9,SRSF7,RBM10,CTR9,RNF20,SFSWAP,PAIP1,YBX3,RNPS1,ACIN1,RNF40
,CLU,DDX5,SOX4,RPL13A,MAGED1,RPL6,KAT6A,DVL3,DDX17,RACK1,APP,RPL5,TMBIM6,NPM1,
1CB,NOP53,PSMB2,OPTN,CCND1,MAPRE1,PSMD3,CSNK1D,RAB11A,DYNLL1,PPP1R9B,PSMA7,
MC4,PSMF1,CELSR3,PSMB5,PSMA1,MED12,PSMD10,PSMA6,PSMD12,PSMA4,PSMC6,FZD7
RPL23,NACA,XRCC5,CIPC,CBX5,ATF4,NR2F1,RPS14,SRSF4,TLE5,HNRNPC,LRPPRC,KHDRBS1,X
AG5,UBE2K,ZER1,USP5,SDCBP,HSPA1A,MDM2,FZR1,SUMO1,PARK7,USP14,HIPK2,MAGEF1,TRIE
MVB12B,DYNC1LI2,DYNLL2,ILF3,IFITM3,PCBP2,DYNLL1,CDC42,ADAR,RBBP7,KPNA2,UBA52,P
1,ZNF770,GTF2F1,HNRNPU,SMARCA5,NACA,XRCC5,ATF4,NR2F1,JUND,CD81,CCT7,TADA2B,SF
SMD7,PSME3,DAAM1,PSMC5,PSMC2,PSMC4,PSMF1,CELSR3,PSMB5,PSMA1,MED12,PSMD10,P
SMD7,PSME3,DAAM1,PSMC5,PSMC2,PSMC4,PSMF1,CELSR3,PSMB5,PSMA1,MED12,PSMD10,P
TOR,WDTC1,ABL1,SEMA6B,ADNP,CYFIP1,NTN1,LIMK1,CDK4,ISLR2,GOLGA4,SEMA4C,SRF,GSK
2,MARCHF7,PSMF1,WAC,SERPINE2,USP19,UBXN1,PHB,GABARAPL2,PIN1,RELA,RYBP,FURIN,EP
CTR1B,PSMD4,ITGAV,PSMC3,AP2A2,AP1M1,PSMC1,CD74,TAPBP,CLTA,PSMB1,PSMD13,KIF26A,
M2,FZR1,SUMO1,PARK7,USP14,TRIB3,STUB1,CSNK1A1,HERPUD1,PLK2,WAC,GSK3B,UBQLN4,
EPPS,CLIP3,FYN,YWHAH,CCT8,MESD,SREBF2,RAB11A,CCT5,PPP1R9B,RER1,BAP1,TCAF1,UBR
P3,CSNK1D,PFN2,DCTN1,GPX4,EVL,CTTN,SLIT2,BRK1,AKAP9,TMOD2,FKBP4,ABI2,RDX,CAPZA1
MARCA4,GRB2,FBLN1,B2M,IFITM3,HGS,CTNND1,EP300,KPNA2,PHB2,BRD4,UBA52,ULK1,ITGB5,
LXNA3,PRPF40A,CDC42,BCL9L,PLXNB1,FMNL2,ZMYM4,SEPTIN7,ZMYM3,PALM,RDX,SPAG9,PH
GF2BP1,GSK3A,MTOR,UBE3A,TMEFF1,TANC2,DIP2A,TAOK2,TRAK1,OBSL1,MAPK8IP2,MECP2,
HECTD1,BAG6,HNRNPD,VDAC1,ARIH1,EDC4,SGTA,FLCN,GTPBP1,UBQLN2,GRSF1,PHB2,ZFP36L

1,UBQLN1,HYOU1,ARNT2,DDIT4,PSMA7,CAPN2,EP300,PSMD1,PHB2,UBA52,ZFP36L1,SFRP1,PTE
18A1,IGF2,VCP,YWHAG,MAPT,EGLN1,RPL15,RPS3,HMGB1,AGRN,AKAP12,LARP1,HNRNPA0,CTN
E1,PSMD3,CSNK1D,RAB11A,DYNLL1,CDC42,PPP1R9B,PSMA7,DCTN1,PRMT1,ACTR1A,EP300,PS
HOA,PSMD2,JUND,VAT1,PFKL,RAB7A,DDX3X,MIF,CD63,DNAJC5,DDOST,LAMP1,CKAP4,HLA-B,TT
S,PSMD8,FBXO21,PSMB4,UBR5,UBE2D2,MKRN1,PSMD4,PRPF19,FBXW5,PSMC3,UBE3A,UBE2D
1,PLXNB2,ATP5F1A,UBB,MAPT,FN1,DCHS1,TMSB10,DNAH8,HMGB1,STMN1,ITGB1,UCHL1,NCA
APT,FN1,HACD3,TPI1,PPP2R1A,PKD1,RPS3,HMGB1,CDKL1,UCHL1,ANXA2,RPLP1,MIDN,DNAJA1,
MC4,PSMF1,CELSR3,PSMB5,PSMA1,MED12,PSMD10,PSMA6,PSMD12,PSMA4,PSMC6,FZD7
X17,APP,TMBIM6,NPM1,HSPD1,IGF2,UBB,TRIM28,SPEN,NONO,PSMD5,YWHAQ,SET,EGLN1,PPP
DX3X,RPS7,CYFIP2,APLP2,ARL6IP1,HDAC2,TRIB2,MAP1A,TIMP3,NOP53,PRDX5,RAD23B,ABCA2,
MD13,PSMD7,TNRC6A,PSME3,DAAM1,PSMC5,PSMC2,PSMC4,PSMF1,CELSR3,PSMB5,PSMA1,M
RSF11,IGF2BP1,DHX38,EIF4A3,CETN2,AKAP8L,NUP153,SRSF7,NUP98,POLDIP3,NCBP2,HSF1,IG
PHB2,RAB3GAP2,HDAC6,ULK1,GPSM1,HIF1A,CTTN,SLC25A5,DAP,VPS26B,MTMR4,FOXK2,ATP6
,PAFAH1B1,AKAP12,SLC4A8,CTNNA1,RPL23,RTN4,PJA2,IGFBP3,RHOA,CLSTN1,PSMD2,KIF5B,N

,RPS6,ENO1,TUBA1B,PIK3R2,FLNA,MAP1B,NCL,ALDOA,MTRNR2L8,PKM,COL6A1,FTL,HSP90B1
HMGB1,UCHL1,ANXA2,RPLP1,MIDN,DNAJA1,ARF1,HNRNPU,PGAM1,LPCAT1,XRCC5,IGFBP3,RH
HMGB1,UCHL1,ANXA2,RPLP1,MIDN,DNAJA1,ARF1,HNRNPU,PGAM1,LPCAT1,XRCC5,IGFBP3,RH
JP153,SRSF7,NUP98,POLDIP3,NCBP2,NUP205,NUP133,THOC2,PRPF31,THOC3,FYTTD1,ABCE1,
,FKBP1A,UBR5,DNAJB2,DCUN1D2,OGT,GSK3A,MTOR,UBE3A,UBE2I,BAG5,SAE1,TRIP12,UBE2L
2,NAP1L1,LRRC4B,MAP6,PLXNB1,PPP1CC,IL6ST,HIF1A,SLIT2,DICER1,IST1,PRPF19,MTOR,SMO
PP1R9B,PSMA7,DCTN1,PRMT1,ACTR1A,EP300,PSMD1,PBX1,PHB2,BRD4,UBA52,ZFP36L1,PTEN
PTBN2,TPM1,CCN2,PRKAR1A,TLN1,MYO5A,CYFIP2,MYH10,RHOB,COTL1,RAC1,ARHGDIA,CNN2,
SLC4A8,CPE,NCS1,RPL22,RHOA,CLSTN1,ATF4,PSMD2,KIF5B,NLGN2,CXADR,ATP2A2,PFN1,TLE5
2M,CDC42,EP300,FLOT1,EIF3D,PPP2CB,HJURP,PTPRF,GPSM1,TNKS,FKBP1A,SYMPK,LOX,CAR
HOA,PSMD2,MCL1,TLE5,PFKL,GLG1,PEBP1,CDH2,GPC3,SULF1,PIP4K2B,MEGF8,RAB7A,MMP14
6A,DVL3,YWHAE,GNAS,HTRA4,RACK1,SPTBN1,GNB1,SHISA9,MYH9,P4HB,APP,RPL5,KLRD1,TME
,JUND,CD81,TADA2B,SFPQ,SOX12,ACTN4,DDX3X,NANOG,H1-0,JUN,CTNNB1,TARDBP,ARID1A,S
CC5,CIPC,CBX5,ATF4,NR2F1,RPS14,SRSF4,TLE5,HNRNPC,LRPPRC,KHDRBS1,XPO1,SFPQ,NANO
,TLE5,GLG1,PEBP1,CDH2,GPC3,SULF1,PIP4K2B,MEGF8,RAB7A,MMP14,PRDX2,DDX3X,APLP1,
,UBQLN1,DDIT4,PPP1R9B,GSPT1,UPF1,HDAC6,MAP1LC3B,TMOD2,TAOK1,VPS4A,GABARAPL1,R
HOA,PSMD2,MCL1,TLE5,PFKL,GLG1,PEBP1,CDH2,GPC3,SULF1,PIP4K2B,MEGF8,RAB7A,MMP14
1B1,AKAP12,SLC4A8,CTNNA1,RPL23,RTN4,PJA2,IGFBP3,RHOA,CLSTN1,PSMD2,KIF5B,NLGN2,
JP153,SRSF7,NUP98,POLDIP3,NCBP2,NUP205,NUP133,THOC2,THOC3,FYTTD1,ABCE1,NUP50,R
IF4A3,EIF4G1,CNBP,FXR2,PINK1,YTHDF3,SMPD3,FXR1,CDK4,NCK2,C1QBP,DHX36,POLDIP3,ER
AR2,CSNK1D,RAB11A,DYNLL1,TP53BP1,PPP1R9B,PSMA7,DCTN1,PRMT1,ACTR1A,EP300,PSMD1,
N,BAZ1B,BRD4,ATRX,EP400,HDAC6,MCM3AP,BAP1,BAZ2A,EHMT1,CARM1,UBR5,KMT2E,NSD2,O
MC1,WNK1,DAPK3,USP47,ABL1,PSMB1,PSMD13,PSMD7,GID8,PSME3,PSMC5,PSMC2,PSMC4,T
UD1,LMNA,ATF6B,SHC1,DNAJB11,ATF6,ADD1,QRICH1,XBP1,CUL7,PPP2R5B,SEC31A,EIF2S1,CD
FYN,YWHAH,USP7,SREBF2,CDC42,GRIPAP1,OS9,SEC16A,UBE2G2,PPP1CC,OAZ2,ANP32B,LMAN
B1,TPT1,STMN1,ITGB1,UCHL1,RPL11,TIMP2,DNAJA1,PAFAH1B1,AKAP12,RPL23,RTN4,PJA2,IG
WDR1,ATP2A2,PFN1,NISCH,SPTBN2,TPM1,CCN2,ACTN4,PRKAR1A,TLN1,GJA1,MYO5A,CYFIP2,
SMD1,CSNK2A1,CSNK1G2,NXN,SFRP1,CTNND2,CUL3,TNKS,PSMD8,DKK3,ATP6AP2,PSMB4,UBR
N1,ATP5F1B,CALM3,GNB1,SHISA9,SEPTIN2,APP,GDI1,TMBIM6,GSTP1,CALM1,NPM1,PLXNB2,A
,BRD4,ATRX,EP400,HDAC6,MCM3AP,BAP1,BAZ2A,EHMT1,CARM1,UBR5,KMT2E,NSD2,OGT,PCG

PSMD1,UBA52,BRD2,ATRX,HJURP,NAP1L3,ANP32B,MCM3AP,RPS27A,PSMD8,BAZ2A,PSMB4,MAC
,MAPK1,MAP2K2,CORO1C,STXBP1,CSNK1D,TMED9,HGS,SOD1,BAG6,RAB11A,ARFGEF2,CDC42
RNPU,LARP1,CTNNA1,LDHA,MSN,RHOA,ATF4,ATP1A1,JUND,SLC3A2,PTN,CCN2,MMP14,APLP1,HM
7,GID8,PSME3,PSMC5,PSMC2,PSMC4,TMEM9,PSMF1,LGR4,PSMB5,PSMA1,PSMD10,PSMA6,CTD
,VEGFA,KDM1A,YWHAH,B2M,PLXNA3,PTPRS,RAB11A,BMPR2,NAP1L1,LRRC4B,MAP6,PLXNB1,DI
SMD1,PHB2,UBA52,ZFP36L1,SFRP1,PTEN,RPS27A,PSMD8,HIF1A,SLC2A1,PSMB4,TM9SF4,SOD2
PN2,EP300,PSMD1,PHB2,UBA52,ZFP36L1,SFRP1,PTEN,RPS27A,PSMD8,HIF1A,SLC2A1,PSMB4,TM
MC6
AP8L,NUP153,SRSF7,NUP98,POLDIP3,NCBP2,NUP205,NUP133,THOC2,THOC3,FYTTD1,NUP50,R
35,RPS27A,GRK2,VAPB,VPS4A,LDLR,ITGAV,SCRIB,IST1,CHMP4B,TRIM8,CD74,SNX3,LGALS1,DAG
PSMD2,CD81,VAT1,CLEC7A,PFKL,RAB7A,DDX3X,MIF,CD63,DNAJC5,NDFIP1,DDOST,LAMP1,CKAP
EGFA,MAPK1,GRB2,PRKDC,CMIP,FKBP8,PLCG1,ZFAND5,SUPT6H,STMN3,HECTD1,VANGL2,TCT
3,GNB1,MYH9,APP,GSTP1,CALM1,PLXNB2,COL18A1,CSDE1,IGF2,UBB,TRIM28,MATR3,CKB,FN1
CSNK1E,CDH2,ACTN4,RPS15,GJA1,MYO5A,CTNNB1,CKAP5,LAMP1,MAP2,RAB1A,ARCN1,RAB27A
MAPK1,GRB2,PRKDC,CMIP,FKBP8,PLCG1,ZFAND5,SUPT6H,STMN3,HECTD1,VANGL2,TCTN1,ARN
PP3,ARFGAP3
1,ANXA2,AGRN,RPLP1,TIMP2,DNAJA1,HNRNPU,CTSD,XRCC5,LLGL1,RHOA,KIF5B,EIF5,CD81,BEX
3B,CRABP2,RGMA,RUFY3,SEMA6C,APOE,FGF13,ANAPC2,SIN3A
SMB4,KIFAP3,CAPZA1,RAB6A,ACTR1B,PSMD4,ITGAV,PSMC3,AP2A2,AP1M1,PSMC1,CD74,TAPBP,
1,ATP6V0C,MMP2,DCN,PTPRA,ACTR3,NCKAP1,ATP6AP1,VEGFA,RAPGEF1,MVB12B,MAPK1,ARPC

MGB1,TPT1,ITGB1,ANXA2,DNAJA1,PAFAH1B1,XRCC5,PJA2,ATF4,PSMD2,TMEM259,CD81,RPL26,
B1,EVL,SLIT2,SLIT1,SEMA3C,TMEFF1,DPYSL4,SMO,LYPLA2,PTPN11,SEMA6B,DAG1,NIBAN2,PLX
SMC3,MTOR,EIF4G1,PINK1,ARPC1A,HIP1R,HSPA1A,CRACD,MECP2,SUMO1,PARK7,CYFIP1,CUL4
81,VAT1,CLEC7A,PFKL,RAB7A,DDX3X,MIF,CD63,DNAJC5,NDFIP1,DDOST,LAMP1,CKAP4,HLA-B,T
MDM2,FZR1,SUMO1,MAGEF1,TRIB3,STUB1,PSMC5,CSNK1A1,PSMC2,HERPUD1,PSMC4,PLK2,G
NPM1,PLXNB2,IGF2,ATP5F1A,VCP,UBB,TRIM28,YWHAG,CKB,MAPT,FN1,HACD3,SET,TPI1,PPP2R
KAP12,CTNNA1,RPL23,RTN4,PJA2,IGFBP3,RHOA,PSMD2,NLGN2,RPS20,CD81,RPL26,MCL1,CLE
NRNPU,RPL23,XRCC5,RHOA,RPS20,CD81,CCNI,CCND2,DDR1,PIP4K2B,PRDX2,DDX3X,RPS15,RP
IPO5,ATP6V0C,MMP2,SRSF5,VAMP2,GNAI3,PARP1,NOX4,PTPRA,AHCYL1,VPS35,ATP6AP1,MARS
A1,HNRNPU,PGAM1,XRCC5,IGFBP3,RHOA,ATF4,CD81,CLEC7A,CCNI,CCND2,DDR1,PIP4K2B,CCN
JFY3,SEMA6C,APOE,ANAPC2,SIN3A
D10,PSMA6,PSMD12,PSMA4,FHL1,PSMC6,CDK5RAP3,HMGA2
4,UBR5,SOX2,TLE1,PSMD4,RBPJ,GSK3A,PSMC3,PSMC1,WNK1,DAPK3,AMFR,IGFBP4,USP47,PSM
B1,EVL,SLIT2,SLIT1,SEMA3C,TMEFF1,SMO,LYPLA2,PTPN11,SEMA6B,DAG1,NIBAN2,PLXNA2,NO
,EFNB2,MAP4K4,NTN1,SYNGAP1,SPOCK1,SEMA4C,THOC2,EFNB3,RGMA,RUFY3,SEMA6C,EPHB2
N1B,BMERB1,GIT1,FGF13,TPX2
NPU,RPSA,LAMA5,XRCC5,MSN,RPL24,KPNB1,GAP43,RHOA,NLGN2,RPS14,WDR1,ATP2A2,CSNK
,GRIPAP1,KIF5A,XPOT,HIP1,SRSF6,UPF1,HNRNPA2B1,MCM3AP,POM121C,SYMPK,GPC4,SMG5,
XL1,DPYSL2,VEGFA,ARPC2,DSTN,LUZP6,GRB2,CAPZB,KDM1A,LARS1,PLXNA3,PTPRS,SOD1,PFN2
AP3
PB1,KDM1A,QARS1,CCAR2,FYN,UBQLN1,HYOU1,MAPK8IP1,GHITM,SOD1,TRAF7,CSNK2A1,SFR
P1,HDAC2,PPIB,NNAT,DCN,EID1,RPN2,PARP1,MORF4L2,NOX4,PKN1,YEATS2,PLOD1,TERF2IP,TC
C1,CNTNAP2,VEGFA,RAPGEF1,PLEC,ACTN1,CORO1C,PTPRS,FLNC,CDC42,CTNND1,BCR,CNTNA

SN3,APPL1,HDAC2,IPO5,ATP6V0C,MMP2,SRSF5,VAMP2,GNAI3,PARP1,NOX4,PTPRA,AHCYL1,VPS
MC6
NEFL,YBX1,ATP2B4,IGFBP2,CALR,FSTL1,TMF1,ABHD2,HSPA1B,TUBB2B,COL4A1,PSAP,RPLP0,H
KAP12,CDC42BPB,CTNNA1,RTN4,LAMA5,MSN,IGFBP3,COL1A2,RHOA,SLC7A5,CXADR,CD81,WD
RHOB,RAC1,TNC,DLG5,MDK,PTK7,SKI,PSMB2,TMED2,NOTCH2,VEGFA,PSMD3,GRB2,HECTD1,V
,CLTC,RPS7,PRKAR1A,GJA1,CTNNB1,WLS,RHOB,RAC1,TNC,DLG5,MDK,PTK7,SKI,PSMB2,KDM6
NPM1,PLXNB2,IGF2,ATP5F1A,VCP,UBB,TRIM28,YWHAG,CKB,MAPT,FN1,HACD3,SET,TPI1,PPP2R
YTHDF3,FXR1,CDK4,NCK2,C1QBP,DHX36,POLDIP3,ERBB2,MAPK3,PAIP1,YBX3,CTIF,MRPS27,SL
ORF4L1,RPL10,ITM2B,HNRNPU,SMARCA5,LARP1,RPL23,NACA,LPCAT1,XRCC5,CIPC,RHOA,CBX
2,NUP205,NUP133,THOC2,THOC3,FYTTD1,NUP50,RNPS1,DDX19A,NUP58,WDR33
2,NUP205,NUP133,THOC2,THOC3,FYTTD1,NUP50,RNPS1,DDX19A,NUP58,WDR33
C3,RNPS1,PAF1,SCAF8,CPSF7,WDR33,RNF40
COR,SETD5,CTR9,OTUB1,RNF20,KDM5A,LMNA,CTCF,WDR5,MAPK3,XBP1,PAF1,DNMT1,SMARC
,NCSTN,DICER1,ATP1B2,PHGDH,SCRIB,AKT2,PRPF19,KDM4A,MTOR,BNIP3,CERS2,SMO,METRN
HDGFL3,HSPA1A,MECP2,SENP6,GBA2,MAPRE3,PDE4DIP,GSK3B,MAP1S,STMN4,CDKN1B,RNF4,
0,DNAH8,HMGB1,ITGB1,NCAM1,ZMIZ1,VCL,DNAJA1,PAFAH1B1,AKAP12,CDC42BPB,CTNNA1,RT
ROH2A1,RUVBL2,MECP2,PARK7,SART3,KAT2A,EHMT2,TASOR,ATF7IP,AKAP8L,CDK9,NSD1,BCC
P6AP2,PSMB4,UBR5,SOX2,TLE1,PSMD4,RBPJ,GSK3A,PSMC3,PSMC1,WNK1,DAPK3,AMFR,IGFBF
4,SETD1A,PSMC6
S27A,PSMD8,BAZ2A,PSMB4,MACROH2A2,PSMD4,SPTY2D1,PSMC3,NASP,PSMC1,RSF1,PSMB1,T
NEFL,YBX1,ATP2B4,IGFBP2,CALR,FSTL1,TMF1,ABHD2,HSPA1B,TUBB2B,COL4A1,PSAP,RPLP0,H
5B,COL5A2,CLEC7A,PTN,TPM1,PIP4K2B,PRDX2,APLP1,HMGCS1,PRKAR1A,GJA1,JUN,MYO5A,C
ITM2B,HNRNPU,SMARCA5,LARP1,RPL23,NACA,XRCC5,CIPC,CBX5,ATF4,NR2F1,ENC1,RPS14,TL
RS2,BAG5,GAK,DENND5A,LGALS1,SEMA6B,MDM2,EFNB2,MAP4K4,NTN1,SYNGAP1,SPOCK1,SEM
UL3,HIF1A,PPP1R15A,GPX1,HINT1,STK24,TRAP1,SOD2,MOAP1,MSH6,NOC2L,NDUFA13,BNIP3,B
V1G1,PAFAH1B2,POLDIP2,PINK1,ATP13A2,FEZ1,EHMT2,RHEB,CAPN1,ATP6V0B,EXOC7,WAC,UB
5,RPL12,RPL30,RPS7,RPS19,RPL35,RPL35A,RPLP2,TSR2,NARS1,RPL36,SND1,RPS27,RPL23A,C
,POM121C,HIF1A,SLC25A23,FOXK2,OGT,COX6A1,ATP5MC3,NDUFA13,ATP5MC2,ATP5MG,PINK
IRXN2,IGF1R,SYNGAP1,DHX36,CRKL,NDRG1,ABHD17A,DLG4,NRP2,ARHGAP33,SHANK1,OPA1,E
RNPU,CTNNA1,NACA,RTN4,XRCC5,IGFBP3,RHOA,CLSTN1,ATF4,NLGN2,JUND,CLEC7A,PTN,CSN
DK16,IPO5,TMEM30A,GLUL,NNAT,NUCB1,CCT2,VAMP2,TCP1,CCT3,CCT4,MAPK1,ADAM9,NPEP
HNRNPU,SMARCA5,LARP1,HNRNPA0,RPL23,RTN4,RPS9,XRCC5,MSN,HK1,RHOA,ATF4,SLC7A5,C
ARK7,FLII,NCK2,PMEPA1,HSF1,GSK3B,SOX9,VDAC2,CAPZA2,THRA,ADD1,CTNNBIP1,PPM1A,MY
E3,PSMC5,PSMC2,PSMC4,PSMF1,PSMB5,RBM14,PSMA1,PSMD10,PSMA6,TAB3,HEXIM1,RELA,P
,OGT,GSK3A,MTOR,UBE3A,BAG5,TRIP12,UBE2L3,ABL1,PINK1,HSPA1A,ARRDC4,FZR1,PARK7,LI
SETD1A,PSMC6,DPF2
P1,RNF157,NTN1,LIMK1,ISLR2,GOLGA4,SRF,ACSL4,CRABP2,ERBB2,UNC13A,PSMD10,TGFBR1,
2,PSMA4,FHL1,PSMC6,CDK5RAP3,HMGA2
,HNRNPA2B1,MCM3AP,POM121C,TNKS,SLC25A5,SYMPK,SMG5,SRSF2,SLC25A23,SRSF11,IGF2
,PARP1,PTPRA,PKN1,AHCYL1,H3-3A,ATP6AP1,TAF7,MARS1,SMARCA4,GRB2,PRKDC,KDM1A,GN
D2,ITGAV,PCGF2,NOC2L,NDUFA13,PHIP,BAG5,CD74,USP47,PINK1,HSPA1A,MDM2,MAPK8IP2,P
GRB2,PRKDC,SMARCC1,ATP6V1B2,FYN,VGF,PPP1R9B,STAT3,PRKAR2A,ZFP36L1,SRSF6,ARID1

HNRNPU,SMARCA5,LARP1,RPL23,NACA,XRCC5,CIPC,CBX5,ATF4,NR2F1,ENC1,RPS14,TLE5,HNR
P1,AP2M1,TPT1,ITGB1,AGRN,COL11A1,VCL,PAFAH1B1,AP2B1,HNRNPU,CNN3,NACA,RTN4,LAM

:IH1,SGTA,HIP1,UBQLN2,PTEN,ANP32B,DAP,MTCH1,GRN,CCN1,ANTXR1,DNAJB2,NCSTN,ATXN3,I
1A,HIP1R,HSPA1A,CRACD,MECP2,CYFIP1,LIMK1,CCDC88A,TRIM27,NCK2,ARPC3,COLGALT1,PDI
:ADM4,HMGCR,RAC1,HDAC2,TNC,DCN,CDKN1A,CNN2,RAB27A,ANXA5,GNAI3,MDK,PTK7,H3-3A,I

-C,PDIA3,RPS3,HMGB1,SEC61A1,ITGB1,NCAM1,ANXA2,TIMP2,XAF1,ARF1,PAFAH1B1,AKAP12,MA
SFPQ,SOX12,ACTN4,DDX3X,NANOG,H1-0,JUN,CTNNB1,ARID1A,SHOX,HNRNPK,H1-10,DUX4,HDA
SFPQ,SOX12,ACTN4,DDX3X,NANOG,H1-0,JUN,CTNNB1,ARID1A,SHOX,HNRNPK,H1-10,DUX4,HDA
KNA1,MEGF8,SOX12,CCN2,MMP14,GJA1,JUN,CTNNB1,MACF1,RAC1,ARHGDIA,CAPRIN1,GPC1,H
SFPQ,SOX12,ACTN4,DDX3X,NANOG,H1-0,JUN,CTNNB1,ARID1A,SHOX,HNRNPK,H1-10,DUX4,HDA
PM1,CDH2,GPC3,DDR1,SULF1,PLXNA1,MEGF8,CCN2,MMP14,CLTC,APLP1,CTNNB1,ANXA6,RAC
S,FYN,YWHAH,CCT8,PFKM,SREBF2,DYNLL1,CCT5,CDC42,SEC16A,GLUD1,OAZ2,ANP32B,HIF1A,S
C3,YY1,ATXN7L3,PSMC1,USP5,USP47,SNX3,USP10,PSMB1,MDM2,PSMD13,PARK7,USP14,TRRAI
MAPK3,SP1,APOE,CCNK,SMARCB1,HMGA2
AP1S,STMN4,BMERB1,CAPZA2,ADD1,FGF13,TPX2,PDXP,GSN
6V1B2,FYN,VGF,STAT3,PRKAR2A,ZFP36L1,SRSF6,ARID1B,ADCY6,PTEN,APEX1,SLC2A1,NUCKS1,
,PSMD10,PSMA6,PSMD12,TGFB2,PSMA4,SETD1A,PSMC6
NC2,EIF4G1,NTRK2,ABL1,DAG1,ADNP,CYFIP1,NTN1,NECTIN1,SYNGAP1,DHX36,YWHAZ,ABHD17A

207
D12,PSMA4,SEC31A,PSMC6
1,NDFIP1,YWHAB,RAB1A,CTBP1,RAC1,HDAC2,SKP1,PEA15,PTTG1IP,CDKN1A,SQSTM1,NOP53,N
PPP2CA,IPO7,PRKDC,HSPB1,FKBP8,QARS1,DUSP4,CORO1C,FBLN1,STRAP,MAPK8IP1,DYNLL1,D
MC4,PLK2,GSK3B,PSMD10,ELOB,XBP1,PSMC6,RAD23A,RNF40
4,CD81,CLEC7A,CCNI,CCND2,CCN2,PRDX2,DDX3X,MIF,PRKAR1A,TARDBP,YWHAB,CADM4,HMG

MACROH2A1,RUVBL2,MECP2,SART3,KAT2A,TASOR,ATF7IP,AKAP8L,CDK9,DHX36,CTR9,RNF20,LI
3,GRIPAP1,FLOT1,SEC16A,RER1,PREPL,RABEP1,GPC4,EFR3B,PALM,PKDCC,VPS4A,RDX,FLOT2,F
LP1,DNAJA1,CDC42BPB,XRCC5,IGFBP3,RHOA,ATF4,CD81,CLEC7A,CCNI,CSNK1E,CCND2,DDR1,
H,STXBP1,PTPRS,PFN2,VGF,RAB11A,PPP1R9B,SYP,STAT3,BCR,FLOT1,PTEN,PREPL,BRSK1,AKAF
ALDOA,PKM,COL6A1,NEFL,UGT8,YBX1,ATP2B4,IGFBP2,CALR,FSTL1,TMF1,ABHD2,HSPA1B,TUB
NC2,EIF4G1,NTRK2,ABL1,DAG1,ADNP,CYFIP1,NTN1,NECTIN1,DHX36,YWHAZ,ABHD17A,SETD5,TI
H,STXBP1,PTPRS,PFN2,VGF,RAB11A,PPP1R9B,SYP,STAT3,BCR,FLOT1,PTEN,PREPL,BRSK1,AKAF
IPU,RPL23,CNN3,RPS20,ATP2A2,CRMP1,BEX3,OAZ1,TLE5,NES,PTN,CSNK1E,PEBP1,GPC3,PIP4K
PPP2CA,IPO7,PRKDC,HSPB1,FKBP8,QARS1,DUSP4,CORO1C,FBLN1,STRAP,MAPK8IP1,DYNLL1,D
S,NDRG4,P3H1,CRTC1,SETX,CNTN1,RETREG3,ARSB,CFLAR,RGMA,PRKCI,DYNLT1,PPP2R5B,OPA
AF1,UBR5,TM9SF4,NMT1,AKT2,GSK3A,PRKACA,UBE2D3,SAE1,SMO,RANGAP1,UBE2L3,PTPN11,I
31,FYN,HNRNPM,LONP1,UBQLN1,IDH1,SOD1,TXNIP,CAPN2,GPX4,PPP2CB,TRA2B,HDAC6,APEX1,
APRE2,LRRC59,BRK1,CYTH2,SHC2,ABI2,GNA12,RDX,WASF2,LZTR1,OGT,ARPP19,DENND4B,GNA
),ITM2B,HNRNPU,SMARCA5,LARP1,RPL23,NACA,LPCAT1,XRCC5,CIPC,CBX5,ATF4,NR2F1,ENC1,F

C3,YY1,ATXN7L3,PSMC1,USP5,USP47,SNX3,USP10,PSMB1,MDM2,PSMD13,PARK7,USP14,TRRAI
SA,RER1,PREPL,EFR3B,PALM,PKDCC,VPS4A,RDX,FLOT2,PACS1,CLASP2,PDPK1,SCRIB,AKT2,EHD
CD74,USP47,PINK1,HSPA1A,MDM2,PARK7,CD44,ERP29,PTPN1,NCK2,MARCHF7,HERPUD1,VDAC
2,GSK3B,DLG4,NSMF,ARHGAP33,SHANK1,SS18L1,PPP3CA,CUL7,OPA1,EPHB2,LRP8,SRCIN1,EF
S3,HMGB1,ITGB1,NCAM1,ANXA2,XAF1,ARF1,PAFAH1B1,AKAP12,MAT2A,PXDN,HNRNPU,SMARCA

,STAT3,DIP2B,PTEN,CTTN,SERP1,OLFM1,SEMA3C,SPAG9,CLASP2,IST1,RBPJ,GSK3A,MTOR,YY1
PDE4DIP,RANBP9,MAP1S,STMN4,CDKN1B,BMERB1,GIT1,FGF13,TPX2,ZNF207

UP98,HSF1,GSK3B,NUP205,NUP133,DNAJB6,MAPK3,NUP50,EIF2S1,NUP58
HSPB1,KDM1A,QARS1,CCAR2,DUSP4,STMN3,FBLN1,HYOU1,MAPK8IP1,DGKD,DDIT4,FLCN,BRD
12,RPL30,RPS7,RPS19,RPL35,RPL35A,RPLP2,TSR2,RPL36,RPS27,RPL23A,CPSF1,NOP53,TSR1,
1,ZMIZ1,VCL,DNAJA1,PAFAH1B1,AKAP12,CDC42BPB,CTNNA1,RTN4,LAMA5,MSN,IGFBP3,COL1
1,ZMIZ1,VCL,DNAJA1,PAFAH1B1,AKAP12,CDC42BPB,CTNNA1,RTN4,LAMA5,MSN,IGFBP3,COL1
,PIP5K1A,GPM6B
IN1,XBP1,UFM1,NOLC1,SESN2,NUP58,NF2,CDK5RAP3,SIN3A,NUTF2
2,MIF,CTNNB1,YWHAB,HMGCR,SKP1,PEA15,TIMP3,PPP1CB,PABPN1,PTPRA,PKN1,PSMB2,ATP6

MA6B,KDM3A,PHACTR4,PSMB1,NOTCH1,PSMD13,PSMD7,EFNB1,LDB1,PSME3,DHX36,PSMC5,P
BX3,OPA1
1,COL2A1,LAMC1,PLEC,AEBP1,FBLN1,COL27A1,SULF2,CAPN2,MMP15,FLOT1,CST3,CRTAP,SP
13,PSMD7,PSME3,DAAM1,PSMC5,PSMC2,PSMC4,PSMF1,LGR4,CELSR3,SOX9,PSMB5,PSMA1,M
MC1,ABL1,ARPC1A,PSMB1,CUL1,PSMD13,CYFIP1,PSMD7,LIMK1,PSME3,ARPC3,PSMC5,PSMC2,
PSMC2,HERPUD1,PSMC4,PLK2,GSK3B,PSMD10,ELOB,XBP1,PSMC6,RAD23A,RNF40

D12,PSMA4,PSMC6,TOLLIP
FAP3,FLOT2,RAB6A,FNBP1L,TANC2,TUB,SUN1,TRAK1,PCM1,FEZ1,SUN2,DYNLRB1,CCDC88A,T
HMGB1,ITGB1,TUBB4B,ANXA2,ZMIZ1,TIMP2,VCL,AP3D1,CYB5R3,AKAP12,PGAM1,CTSD,LPCAT1
2A1,LAMC1,PLEC,AEBP1,FBLN1,COL27A1,SULF2,CAPN2,MMP15,FLOT1,CST3,CRTAP,SPINT2,IT

2A1,LAMC1,PLEC,AEBP1,FBLN1,COL27A1,SULF2,CAPN2,MMP15,FLOT1,CST3,CRTAP,SPINT2,IT
,CORO1C,STRAP,PFN2,RAB11A,BMPR2,AMOTL2,EVL,HDAC6,PTEN,MAPRE2,HIF1A,SLIT2,GRN,
A1,RTF1,DAG1,MACROH2A1,TXNRD1,TASOR,EXT2,LDB1,DLD,SRF,CTR9,COL7A1,EXT1,PAF1,M
1,EPHA4
SPTBN1,GNB1,MYH9,SEPTIN2,P4HB,APP,KLRD1,FSCN1,GSTP1,CALM1,PLXNB2,IGF2,VCP,UBB,
TP6AP1,DPYSL2,TMED2,KIF3C,STXBP1,CSNK1D,TMED9,RAB11A,PPP6R1,SEC23A,CDC42,DCTN1
LM1,NPM1,UBB,CKB,DDB1,HACD3,EGLN1,PKD1,HMGB1,TPT1,STMN1,ITGB1,RPS17,VCL,AP3D
PDE4DIP,CDKN1B,WASF3,CDC42EP4,ARPC5,FER,GMFB,GIT1,ARF6,GSN
TM9SF4,NMT1,AKT2,EHD1,GSK3A,PRKACA,UBE2D3,SAE1,SMO,PTPN23,UBE2L3,PINK1,MDM2,P
PC2,DSTN,LUZP6,GRB2,CAPZB,ACTN1,TPM3,CORO1C,MPRIP,PFN2,CDC42,PPP1R9B,HIP1,SFR

A1,PAFAH1B1,MAP4,HNRNPU,KPNB1,LLGL1,RHOA,KIF5B,CRMP1,SPC25,LRPPRC,XPO1,SFPQ,C
IB3,CDK9,MAPRE3,PLK2,HSF1,GSK3B,MED25,TXN,CEBPG,CTBP2,PIN1,ADD1,HMGB2,APOE,EP
HMT1,KMT2E,NSD2,OGT,PCGF2,NOC2L,KDM4A,MBD3,CREBBP,APBB1,UBE2I,SAE1,PAGR1,RAN

PTGES3,CCT6A,TOP2B,SMCHD1,CDKN1A,DDX39B,CCT2,NOP53,PARP1,MORF4L2,SSRP1,ALKB
COL5A2,WDR1,PFN1,PTN,TPM1,SULF1,CCN2,MMP14,NANOG,RPS7,PRKAR1A,GJA1,JUN,CTNN

1,PXDN,CTNNA1,IGFBP3,RHOA,PSMD2,MCL1,TLE5,GLG1,PEBP1,CDH2,GPC3,SULF1,PIP4K2B,
6L1,NXN,CUL3,SPINT2,RPL7L1,HIF1A,BRK1,EIF2S2,CCN1,RIC8A,SH3PXD2A,NMT1,KRT8,ADAM

UL,NNAT,NUCB1,VAMP2,MAPK1,ADAM9,NPEPPS,BNIP3L,FYN,YWHAH,PFKM,SREBF2,RAB11A,CY
P2,FBXW5,CHMP4B,TUBB4A,ABL1,CHMP1A,HDGFL3,HSPA1A,MECP2,FEZ1,KAT2A,ATF5,SENP6,(

IOA,EIF5,CD81,BEX3,CLEC7A,CCND2,DDR1,CCN2,PRDX2,DDX3X,PRKAR1A,JUN,CTNNB1,MTSS2,

'1,GHITM,SREBF2,YME1L1,VDAC1,STAT3,WDR45B,EP300,NDUFV1,FLCN,PHB2,PPP2CB,HDAC6,T
(N1A,EEF1A2,CCT2,PABPN1,NOX4,PKN1,TCP1,CCT4,GCN1,VEGFA,RAPGEF1,PPP2CA,CCND1,MA

CEBPG,TMEM14C,CITED2,ANXA1,THRA,ADD1,HMGB2,ACIN1,CDK5RAP3,ID2,BRD1,ITPKB,MAFB
,LLGL1,RHOA,EIF5,BEX3,CLEC7A,PTN,PEBP1,GPC3,PLXNA1,CCN2,DDX3X,JUN,MTSS2,CYFIP2,A
P53,PARP1,MORF4L2,SSRP1,ALKBH5,CBX1,RAD23B,CHD2,TERF2IP,CCND1,MAPK1,FTO,GRB2,F
ARID1B,ADCY6,PTEN,APEX1,NUCKS1,PDPK1,AKT2,OGT,ATP6V0D1,GSK3A,PRKACA,ATP6V1G1,PI
3,EP300,NDUFV1,PHB2,UQCRC1,COX7C,PPP1CC,ADH5,IL6ST,POM121C,HIF1A,IREB2,TRAP1,SL(
3,ATP1A2,ARHGDIA,HDAC2,TNC,GNAI3,PARP1,NOX4,PKN1,VPS35,H3-3A,TAF7,SELENON,RAPGE
14,TLE5,KHDRBS1,XPO1,SFPQ,NANOG,H1-0,JUN,CTNNB1,MXD4,H1-10,YWHAB,CTBP1,SMCHD

CDC42,SEC16A,GLUD1,OAZ2,ANP32B,HIF1A,SERP1,BAP1,TCAF1,UBR5,TM9SF4,NMT1,AKT2,GS
AGED1,RPL6,KAT6A,DVL3,DDX17,FASN,ATP5F1B,APP,TMBIM6,NPM1,HSPD1,IGF2,ATP5F1A,VCP,

IL3,TUT4,KDM2B,PAF1,FZD7,HMGA2,KLF10,CNOT1,MED14,MMP24
14,TLE5,KHDRBS1,XPO1,SFPQ,NANOG,H1-0,JUN,CTNNB1,MXD4,H1-10,YWHAB,CTBP1,SMCHD
RN,NTRK2,PTPN11,ABL1,DAG1,NOTCH1,B4GALT5,RHEB,TMEM98,NDRG1,SERPINE2,SOX9,PRM
IRNPA2B1,CUL3,TNKS,PPP1R10,SMG5,KMT2E,OGT,PINK1,AUTS2,RTF1,MACROH2A1,RUVBL2,N

5,XRCC5,RHOA,CLSTN1,ATF4,NLGN2,PTN,GLG1,SULF1,PLXNA1,MEGF8,SOX12,CCN2,MMP14,F
14,TLE5,KHDRBS1,XPO1,SFPQ,NANOG,H1-0,JUN,CTNNB1,MXD4,H1-10,YWHAB,CTBP1,SMCHD
DD1,HMGB2,ACIN1,CDK5RAP3,ID2,BRD1,MAFB
,PSMB4,TAOK1,EHMT1,CARM1,INTS3,VPS4A,PSMD4,PCGF2,PSMC3,PSMC1,USP47,PTPN11,PSM
31A,DNM1,ARFGAP3

,APLP2,ARL6IP1,YWHAB,MAP2,HMGCR,IPO5,TRIB2,CDKN1A,TIMP3,ANXA5,GNAI3,PRDX5,PKN1,

CALR,FSTL1,TMF1,ABHD2,HSPA1B,COL4A1,PSAP,RPLP0,H3-3B,CANX,CLU,DDX5,SOX4,FKBP14,N

3,HMGCR,BCLAF1,PEA15,CDKN1A,TIMP3,EEF1A2,SQSTM1,PARP1,NOX4,NOTCH2,PRKDC,BNIP3
2L,ATP5PO,DLD,NDUFB8,ME2,SHMT2,NDUFS1,ATP5F1C,COX6B1,UQCRFS1,ATP5F1E,COX4I1,C
(DM2B,PAF1,FZD7,HMGA2,KLF10,CNOT1,MED14,MMP24
SHOX,HNRNPK,DUX4,HDAC2,BCLAF1,DCN,CHD6,MLXIP,SQSTM1,PARP1,MORF4L2,ACTR3,PTOV
O1,CLTC,GJA1,CTNNB1,CKAP5,MAP2,RPS27,RAC1,MAP1A,TBCD,CAMSAP1,TUBB2A,MAPRE1,D
RS1,DUSP4,CORO1C,FBLN1,STRAP,MAPK8IP1,DYNLL1,DDIT4,ADAR,STAT3,PRKAR2A,SFRP1,PTE
,NPM1,HSPD1,IGF2,UBB,AFF4,TRIM28,COL4A2,SPEN,NONO,PSMD5,YWHAQ,SET,EGLN1,PPP2R
D81,CCNI,CCND2,DDR1,PIP4K2B,PRDX2,DDX3X,PRKAR1A,YWHAB,HMGCR,IPO5,TRIB2,PEA15,C
RK7,ADNP,RAN,HIPK2,PPP1CA,TRIB3,STUB1,MAPRE3,PLK2,HSF1,GSK3B,RGMA,TGFBR1,MAPK

AGED1,RPL6,KAT6A,DVL3,DDX17,FASN,ATP5F1B,APP,TMBIM6,NPM1,HSPD1,IGF2,ATP5F1A,VCP,
,NPM1,HSPD1,IGF2,UBB,AFF4,TRIM28,COL4A2,SPEN,NONO,PSMD5,YWHAQ,SET,EGLN1,PPP2R1

1,HDAC6,APEX1,HIF1A,SLC2A1,PRDX6,XRCC6,GPX1,STK24,TRAP1,SOD2,SLC25A23,PCGF2,FBLI
AGED1,RPL6,KAT6A,DVL3,DDX17,FASN,ATP5F1B,APP,TMBIM6,NPM1,HSPD1,IGF2,ATP5F1A,VCP,
L1,KLF13,HSPA1A,ACVR1B,NOTCH1,JAM3,SART3,MAEA,HIPK2,LDB1,RBFOX2,SRF,LGR4,SOX9,F

2,DEAF1,SEMA4C,NUP133,MED12,ITPK1,CITED2,RGMA,ZNF358,OPA1,TGFB2,KDM2B,LRP6,EPHA
AE1,SMO,UBE2L3,PINK1,MDM2,PCM1,RAN,YWHAZ,GSK3B,KIF3A,ERBB2,RUFY3,XBP1,TFDP1,PR
RPL38,NOTCH2,MAPK1,PBX2,GRB2,DUSP4,HECTD1,VANGL2,SOD1,TCTN1,BMPR2,PBX1,MMP15
AR1A,YWHAB,HMGCR,IPO5,TRIB2,PEA15,CDKN1A,NOX4,PKN1,GCN1,TAF7,VEGFA,RAPGEF1,PPF

T1,CARM1,HINT1,PDPK1,NOC2L,MBD3,MTOR,CD74,MTA2,USP10,MDM2,CD44,EHMT2,SHISA5,H
,NPM1,HSPD1,IGF2,UBB,AFF4,TRIM28,COL4A2,SPEN,NONO,PSMD5,YWHAQ,SET,EGLN1,PPP2R1

C4,PSMF1,SRF,PSMB5,PSMA1,SOS1,CITED2,PSMD10,PSMA6,EXT1,PSMD12,PSMA4,SETD1A,PSM
ITGA7,FZD7,NF2,CRB2,HMGA2,EPHA2
D8,DKK3,HIF1A,PSMB4,LDLR,FOXK2,PSMD4,AKT2,OGT,DHCR7,ARPP19,GSK3A,PSMC3,MTOR,P
P3D1,CYB5R3,PGAM1,CTSD,LPCAT1,XRCC5,PJA2,MSN,KPNB1,RPL22,RHOA,PSMD2,CXADR,JUND

L,CTR9,RNF20,THOC2,THOC3,TUT4,RNPS1,PAF1,SCAF8,ELAC2,CPSF7,WDR33,RNF40

FEZ1,SUN2,DYNLRB1,MAP1S,KIF3A,ARL8A,SPG11,OPA1,DYNC1I2,MAPK8IP3,AP3M1
ZP6,GRB2,CAPZB,PFN2,CDC42,SFRP1,EVL,CTTN,SLIT2,BRK1,AKAP9,TMOD2,TAOK1,ABI2,RDX,C
LMNA,NUP133,CHMP5,NECTIN2,CTDNEP1,HMGB2,NOLC1,PPP2R2A,LEMD2,DMPK,USP36,EMD

,PEA15,CDKN1A,TIMP3,EEF1A2,SQSTM1,NOX4,NOTCH2,PRKDC,BNIP3L,CLIP3,PLCG1,CCAR2,SO
IMP3,EEF1A2,SQSTM1,NOX4,NOTCH2,PRKDC,BNIP3L,CLIP3,PLCG1,CCAR2,SOD1,TXNIP,CDC42
NRNPD,CDC42,SULF2,LBH,KDM5B,PBX1,PHB2,SFRP1,PTEN,FEM1B,CUL3,DKK3,HIF1A,GPX1,SO
DAM10,IST1,RBPJ,MTOR,SMO,RABGAP1,EIF4G1,USP47,SDCBP,NOTCH1,ADNP,CYFIP1,RNF157,

,HIP1R,HSPA1A,CRACD,MECP2,CYFIP1,LIMK1,CCDC88A,TRIM27,NCK2,ARPC3,PDE4DIP,CDKN1B
,KPNA3,MAPK3,SP1,APOE,CCNK,SMARCB1,HMGA2
A,NUP133,TNPO2,KPNA3,PPP3CA,XBP1,UFM1,TNPO3,NOLC1,NUP50,NUP58,NUTF2
AGED1,RPL6,KAT6A,DVL3,DDX17,FASN,ATP5F1B,APP,TMBIM6,NPM1,HSPD1,IGF2,ATP5F1A,VCP,

SQSTM1,VAMP2,PTPRA,CLSTN3,MAPK1,GABARAP,PLCG1,FYN,YWHAH,STXBP1,PTPRS,PFN2,VGI
D3,WDR5,XBP1,TET3,SETD1A,PAF1,DNMT1,SMARCB1,SETD1B,PRMT7

CAPZA1,ERGIC3,COPZ1,USO1,SCAP,ANK2,LMAN2,NSF,COPE,ARFGAP2,MCFD2,NAPA,COL7A1,CA

TBN1,ATP5F1B,CALM3,SHISA9,APP,GDI1,GSTP1,CALM1,PLXNB2,PLD3,HSPD1,IGF2,ATP5F1A,C
RAP,MAPK8IP1,DDIT4,ADAR,PRKAR2A,SFRP1,PTEN,SLIT2,SPAG9,PDPK1,MTOR,WNK1,MICAL1,
HNRNPA2B1,ANP32B,RPS27A,BAZ2A,XRCC6,MACROH2A2,SPTY2D1,NOC2L,MBD3,NASP,HMGA1,
KPNA6,NUP98,SNRPD2,SNRPB,LMNA,NUP133,TNPO2,KPNA3,PPP3CA,XBP1,UFM1,TNPO3,NOLC
X1,STK24,TRAP1,SOD2,PCGF2,FBLN5,BNIP3,BAG5,PRDX3,PTPN11,ABL1,PINK1,PPP5C,HSPA1A,
,NCSTN,SOD2,PDPK1,AKT2,GSK3A,CHMP4B,MTOR,BNIP3,PSMC1,BAG5,DPYSL4,AMBRA1,EIF4
,EVL,CTTN,SLIT2,BRK1,TMOD2,TAOK1,ABI2,RDX,CAPZA1,CLASP2,WASF2,MTOR,DAPK3,CELSR

1A1,PSMC2,PSMC4,PSMF1,GSK3B,SOX9,PSMB5,CHD8,PSMA1,LZTS2,PSMD10,PSMA6,PSMD12,
RDX3,PTPN11,ABL1,PINK1,PPP5C,SMPD3,MDM2,PARK7,PDGFRB,HSF1,DHFR,SETX,TXN,STAT1,
SE1L,TOMM70,SMO,RGPD5,PKIA,RAN,FAM53C,NUP153,KPNA6,NUP98,LMNA,NUP133,LONP2,T
P4K4,CSNK2B,CDK4,BCL9,DHFR,NREP,PRMT5,CDKN1B,LARGE1,CFLAR,RGMA,ANXA1,PPP3CA,

DPK1,AKT2,OGT,ATP6V0D1,GSK3A,PRKACA,ATP6V1G1,PHIP,USO1,RABGAP1,RANGAP1,PTPN1
,HSPD1,IGF2,UBB,TRIM28,SPEN,NONO,PSMD5,YWHAQ,SET,EGLN1,PPP2R1A,PKD1,SSBP2,ZNF4
K1,TNKS,PSMD8,PCBP4,PPP1R10,PSMB4,TAOK1,EHMT1,CARM1,INTS3,VPS4A,PSMD4,SCRIB,P

NTRK2,AMFR,PAIP2,NRXN2,NPTN,MAPK8IP2,MGAT3,MECP2,ADNP,CYFIP1,ARL6IP5,KAT2A,EHM
CTR9,SNRPB,RNF20,LMNA,PRMT2,CTCF,SETD3,WDR5,XBP1,TET3,SETD1A,PAF1,DNMT1,SMAR
CTR9,SNRPB,RNF20,LMNA,PRMT2,CTCF,SETD3,WDR5,XBP1,TET3,SETD1A,PAF1,DNMT1,SMAR

RXN2,NPTN,MAPK8IP2,MECP2,ADNP,ARL6IP5,KAT2A,EHMT2,ADCY1,SYNGAP1,NDRG4,PLK2,SRF
PZA1,CLASP2,WASF2,MTOR,ABL1,ARPC1A,HIP1R,ELN,CRACD,CYFIP1,LIMK1,FLII,CCDC88A,TRI
5R3,PGAM1,CTSD,RTN4,LPCAT1,XRCC5,KPNB1,HK1,RHOA,PSMD2,KIF5B,SLC7A5,CD81,WDR1,
1A,POLR2B,SUPT6H,CCT8,TNKS2,USP7,HNRNPD,BMPR2,TP53BP1,YLPM1,CCT5,POLR2E,KPN
,HSPD1,IGF2,UBB,TRIM28,SPEN,NONO,PSMD5,YWHAQ,SET,EGLN1,PPP2R1A,PKD1,SSBP2,ZNF4
,HSPD1,IGF2,UBB,TRIM28,SPEN,NONO,PSMD5,YWHAQ,SET,EGLN1,PPP2R1A,PKD1,SSBP2,ZNF4
DAR,PRKAR2A,SFRP1,PTEN,PDPK1,GSK3A,WNK1,BAG5,PRDX3,ABL1,MACROH2A1,GGNBP2,PKI
A,MTOR,GOT2,ATP6V1G1,PHIP,SORT1,USO1,RABGAP1,SCAP,CAD,PTPN11,WDTC1,IGF1R,CYFI
RNF5,GET4
,CTTN,CCN1,ANTXR1,ATXN3,CLASP2,PDPK1,ATP1B2,ITGAV,FBLN5,DAPK3,PTPN11,ABL1,COL16

MARCA4,ADAM9,KDM1A,YWHAH,SLIT3,IDH1,NCOR2,WBP2,IGFBP7,TXNIP,DDIT4,PPP1R9B,RBB
,MDK,RBMX,SKI,ATP6AP1,RPL38,MAPK1,MESD,BMPR2,ADAR,PBX1,PLXNB1,SFRP1,IARS1,IL6S
B,RAB1B,VPS41,VPS4A,RAB14,FNBP1L,SYT11,AKT2,IST1,CHMP4B,AP1M1,SORT1,USO1,SCAP,T
UTF2,EMD
,WDTC1,IGF1R,CYFIP1,PTPN1,CDK4,TRIB3,ATP6V0B,GSK3B,STAT1,IRS1,ATP6V0E2,SOS1,ATP6
P2,PTPRA,CLSTN3,MAPK1,GABARAP,PLCG1,FYN,YWHAH,STXBP1,PTPRS,PFN2,VGF,RAB11A,PP
3B,TFRC,LMNA,INF2,DDHD2,PSMD10,TFDP1,OPA1,FBXO7,PRELID1,USP36
RP1,HIF1A,SLC25A5,LOX,GLO1,IREB2,KMT2E,SH3PXD2A,WASF2,RBPJ,MTOR,CREBBP,CD74,P
ACD,NSF,TRMT112,APEH,NCKAP5L,FLII,ETF1,NCK2,MICAL3,MRPS6,MAP1S,NAPB,STMN4,NAPA,

RN2,SUN1,SEMA6B,FEZ1,KAT2A,ATF5,SUN2,H2AX,CRKL,SRF,ZSWIM6,GSK3B,CCDC85C,MBOAT7

SMA4,PSMC6
PKAPK2,DHX36,STUB1,PSIP1,NUP98,HSF1,GSK3B,NUP205,NUP133,DNAJB6,MAPK3,NUP50,EIF2S

NSG1,PTPN23,RAB11FIP3,SNX2,DENND5A,SNX3,CHMP1A,SGSM2,ACAP2,TBC1D10B,ANKFY1,SN
2A2,PFN1,CLEC7A,NISCH,PTN,TPM1,SULF1,PLXNA1,ACTN4,MMP14,RPS19,MIF,GJA1,MACF1,CA
X2,PDXP,SCAF8,GSN

DH2,OGT,AHCY,MTOR,UBE3A,CREBBP,THRAP3,NTRK2,RBM4,ADNP,ATF5,CUL4A,MYCBP2,PPP1
,NDUFA13,BNIP3,ATP5MC2,ATP5MG,MDH1,PINK1,PDHA1,NDUFA4,PARK7,IDH2,NDUFC2,COX8A

SRF,GSK3B,CCDC85C,MBOAT7,BMERB1,OGDH,XAB2,LRP8,FGF13,COL3A1,NF2,POU3F3,LRP6
MP2,PTPRA,CLSTN3,MAPK1,GABARAP,PLCG1,FYN,YWHAH,STXBP1,PTPRS,PFN2,VGF,RAB11A,PP
MP2,PTPRA,CLSTN3,MAPK1,GABARAP,PLCG1,FYN,YWHAH,STXBP1,PTPRS,PFN2,VGF,RAB11A,PP
5,MSN,RHOA,CLSTN1,NLGN2,CXADR,CD81,ATP2A2,ASTN2,PNN,PTN,CDH6,TPM1,CDH2,DDR1,PL

SMA4,PSMC6
SMA4,PSMC6
AOK1,LSM14A,TUBGCP6,CLASP2,CHMP4B,RANGAP1,CHMP1A,KIFC2,HSPA1A,MECP2,RAN,SUN
ME3,PSMC5,CSNK1A1,PSMC2,PSMC4,PSMF1,GSK3B,SOX9,PSMB5,CHD8,PSMA1,LZTS2,PSMD1
,ARPC3,GBA2,WASF3,CDC42EP4,ARPC5,CAPZA2,ADD1,FER,GMFB,ARF6,MYADM,GSN
M27,NUP153,EXOC7,TFRC,PHB,EFNB3,KPNA3,NECTIN2,SLC1A5,DYNLT1,SCARB1,GSN,EPHA2,V

C1A,HIP1R,ELN,CRACD,CYFIP1,FLII,TRIM27,NCK2,MICAL3,ARPC3,GBA2,WASF3,CDC42EP4,ARP

A,CLSTN1,NLGN2,CXADR,CD81,ATP2A2,ASTN2,PNN,PTN,CDH6,TPM1,CDH2,DDR1,PLXNA1,SOX

SF10B,PSMC4,LIMS1,PSMF1,ZFP91,RASSF2,PSMB5,PSMA1,PHB,PSMD10,ADIPOR1,PSMA6,REL
TRIM27,NCK2,ARPC3,GBA2,WASF3,CDC42EP4,ARPC5,CAPZA2,ADD1,FER,GMFB,PDXP,ARF6,MY

,ANXA5,GNAI3,PTK7,H3-3A,NOTCH2,VEGFA,MAPK1,PLEC,CAPZB,HSPB1,KDM1A,GNG2,FYN,FBL
VEGFA,RAPGEF1,MAPK1,CLIP3,MAP2K2,B2M,STXBP1,USP7,HGS,SOD1,DGKD,PFN2,RAB11A,CD
U3F3,LRP6

,PSMD13,PSMD7,ZCCHC3,PSME3,PSMC5,PSMC2,PSMC4,PSMF1,PSMB5,RBM14,PSMA1,PSMD1

TRIM27,NCK2,ARPC3,GBA2,WASF3,CDC42EP4,ARPC5,CAPZA2,ADD1,FER,GMFB,PDXP,ARF6,MY

SMA4,PSMC6

LASP2,SCRIB,SYT11,EXOC4,TANC2,USO1,PINK1,TRAK1,NSF,ARFGAP2,WDR11,MCFD2,RAB3A,KI
X,HDAC6,TNKS,SLC25A5,SLIT2,TMOD2,TAOK1,GPX1,FKBP4,RDX,CAPZA1,CLASP2,WASF2,CHMF
D2,BRD1,MAFB

HSY1,PDIA3,RPS3,VAPA,HMGB1,ITGB1,RPL11,ZMIZ1,TIMP2,PAFAH1B1,AKAP12,CTNNA1,RPL23,
VPS4A,NMT1,PDPK1,ADAM10,AKT2,DAG1,CRKL,YWHAZ,ZDHHC5,ERBB2,TFDP1,PRKCI,TMEM59,

D2,SKI,KDM6B,VEGFA,SELENON,PTBP1,LUZP6,KDM1A,FLNB,SUPT6H,AKIRIN1,BMPR2,CDC42,B
,NCOA4,RPS27A,NPLOC4,SAFB2,CARM1,UBR5,SEC14L1,LSM14A,KDM4A,UBE3A,MN1,SCD,PAG
SMA4,PSMC6
D1,MTOR,MN1,ATP6V1G1,AMBRA1,SLC38A2,EIF4G1,USF2,MDM2,PIK3C2B,EHMT2,TNRC6A,RI
SP,SMARCD1,HMGA1,RSF1,MTA2,TSPYL2,MACROH2A1,SART3,TASOR,ATF7IP,H2AX,SOX9,MAP1

ND1,MAPK1,PRDX1,ADAM9,FYN,B2M,LONP1,SOD1,HNRNPD,TXNIP,CAPN2,PPP2CB,ATRX,HDAC6
,CHD3,HMGB1,POLR2A,BTF3,AGRN,ZMIZ1,RPL10,AP3D1,ZNF770,GTF2F1,HNRNPU,NFE2L1,SM
RIO,FLCN,FLOT1,PPP2CB,CUL3,MAPRE2,SLIT2,A2M,LRRC59,BRK1,CYTH2,SHC2,ABI2,ARHGEF
13A2,MFN2,PIK3C2B,FEZ1,EHMT2,TBC1D14,TMEM9,ATG9A,ARSB,CHMP5,RAB5A,GABARAPL2,
2,SCRIB,CAP1,JAM3,PDLIM1,FEZ1,GOLPH3,CRKL,GSK3B,KIF3A,LMNA,RHOBTB2,ARPC5,ROCK
1,BRSK1,TNKS,MAP1LC3B,DAP,SLC2A1,VPS41,GABARAPL1,STK24,ATP6V0D1,MTOR,BNIP3,MN

AB3A,NAPB,UNC13A,NAPA,BACE1,CTBP2,RAB5A,CLCN3,GIT1,ARF6,SYT1,DNM1,PPFIA3,RAP1B,C

1,PLXNA1,ACTN4,MMP14,MIF,GJA1,MACF1,RHOB,RAC1,ARHGDIA,APPL1,GLUL,DCN,DLG5,CNN
A,CREBBP,PSMC1,TOMM70,UBE2K,PTPN11,PSMB1,YTHDF3,CUL1,PSMD13,SUMO1,HLA-A,PTPN

H,PTN,TPM1,SULF1,PLXNA1,MEGF8,ACTN4,MMP14,MIF,GJA1,MACF1,CADM4,RHOB,RAC1,ARHG
FIP1,ADCY1,PIP5K1C,GSK3B,RAB3A,NAPB,UNC13A,NAPA,BACE1,CTBP2,RAB5A,CLCN3,GIT1,ARF
T1,ANXA1,PCNA,RELA,FBXO7,GNAO1
BARAPL1,STK24,ATP6V0D1,MTOR,MN1,ATP6V1G1,AMBRA1,SLC38A2,EIF4G1,USF2,MDM2,KIF26
,SHANK1,ROCK2,TGFBR1,ADD1,ARHGEF18,PDXP,NF2,ASAP3
1A,ACVR1B,NTN1,SEMA4C,SERPINE2,EI24,ADIPOR2,CDKN1B,PHB,ADIPOR1,RGMA,SEMA6C,TGF
F1,MEGF8,CCN2,MMP14,RPS7,GJA1,JUN,CTNNB1,PTGES3,RHOB,TNC,MMP2,GLUL,DCN,CDKN

10,PIN1,GNL3,CDK5RAP3,RNF40,ARRDC3,TOLLIP

CC1,USP36
,TANC2,USO1,PINK1,TRAK1,NSF,ARFGAP2,WDR11,MCFD2,RAB3A,KIF3A,STARD3,NAPA,COL7A1,

P1,MAP4K4,RASA4,IQSEC1,PDGFRB,SYNGAP1,CRKL,CDC42SE1,SHC1,HEG1,KCTD10,ARHGEF18
2,CD44,EHMT2,HIPK2,CSNK2B,CCNG1,MARCHF7,PRMT5,PSMD10,PIN1,HEXIM1,TPX2,TRIM24,
MC1,USP47,PTPN11,PSMB1,TAOK2,MACROH2A1,MDM2,CUL1,PSMD13,ATF5,PSMD7,CUL4A,CD
DBTB2,CDC42EP4,CDC42SE1,ANXA1,ARHGEF18,ITGA7,PLXND1,FGD1,SH3KBP1

PARP1,MDK,CHD2,ASXL1,PSMB2,H3-3A,ATP6AP1,NOTCH2,VEGFA,PSMD3,PRKDC,ACTN1,KDEL
P1CA,HNRNPR,CDK4,ADCY1,GSK3B,LGR4,CRTC1,SETX,PRMT5,KDM2A,KDM5A,PER1,NR2F6,RO

N,ZMIZ1,AP3D1,PAFAH1B1,AKAP12,NACA,RTN4,XRCC5,HK1,RHOA,CLSTN1,ATF4,SLC7A5,NLGN
OL16A1,ADNP,MAP4K4,NTN1,NECTIN1,LDB1,SLK,LIMS1,RCC2,SETD5,DUSP3,THBS2,ROCK2,LR
PARP1,PKN1,MDK,CHD2,ASXL1,PSMB2,H3-3A,ATP6AP1,NOTCH2,VEGFA,MAPK1,PSMD3,PRKDC,
SK3A,CHMP4B,MTOR,PSMC1,AMBRA1,EIF4G1,NTRK2,CDC34,ABL1,PINK1,PPP5C,GRIK5,ATP13
EMILIN1,BMERB1,SHANK1,CAPZA2,ADD1,ARHGEF18,GMFB,APOE,FGF13,TPX2,MYADM,GSN

,COX7A2L,ATP5PO,DLD,NDUFB8,ME2,SHMT2,NDUFS1,ATP5F1C,COX6B1,UQCRFS1,ATP5F1E,C
B,RANGAP1,CHMP1A,KIFC2,HSPA1A,MECP2,RAN,SUN2,TASOR,NUP98,PLK2,RCC2,NUP133,CHM
A1,SLIT1,SOX2,DCLK1,CNP,ATP1B2,IGF2BP1,RBPJ,BNIP3,SLC38A2,SMO,NTRK2,XRN2,SUN1,SE
1,PFN1,ASTN2,PTN,CDH2,GPC3,DDR1,SULF1,PLXNA1,MEGF8,MMP14,CLTC,RPS7,GPAT4,GJA1,
RIM27,NCK2,ARPC3,GBA2,WASF3,CDC42EP4,ARPC5,CAPZA2,ADD1,FER,GMFB,ARF6,MYADM,G
P1A,KIFC2,HSPA1A,MECP2,RAN,SUN2,TASOR,NUP98,PLK2,RCC2,NUP133,CHMP5,TPX2,DYNC1I2

ABL1,FXR2,MDM2,FXR1,CD44,CUL4A,WRNIP1,MAGEF1,FAM168A,H2AX,CDK9,MARCHF7,HSF1,

RN,ZMIZ1,RPL10,AP3D1,ZNF770,GTF2F1,HNRNPU,NFE2L1,SMARCA5,RPL23,NACA,BRD3,RHOA,
C1,RAB3A,DLG4,SORCS2,NSMF,CNTN2,SHANK1,RAB5A,EPHB2,APOE,PRNP,EPHA4,PPFIA3,ANAP
COL1A2,RHOA,CXADR,PTN,TPM1,CDH2,GPC3,SULF1,MEGF8,CCN2,MMP14,PRKAR1A,GJA1,JUN
K,SHANK1,ROCK2,TGFBR1,ADD1,ARHGEF18,PDXP,NF2,ASAP3
,KHDRBS1,CDH2,GPC3,RAB7A,ACTN4,GJA1,CD63,TARDBP,ARL6IP1,WLS,HNRNPK,MYH10,YWH

I,BSG,FASN,GNB1,MYH9,APP,KLRD1,TMBIM6,PTMS,SPTAN1,GSTP1,CALM1,PLD3,HSPD1,IGF2,

MTMR4,PDPK1,RBPJ,CREBBP,UBE2D3,FGFR1,ABL1,SDCBP,NIBAN2,HSPA1A,NOTCH1,NPTN,CY

RBFOX2,EGLN2,PER1,PHB,PRMT2,UFM1,HMGA2,CNOT1
O,CLCN4,ATP5F1C,COX6B1,ATP6V0E2,ATP5F1E,COX4I1,ATP6V0A1,COX5A,COX7A2,CLCN3,ATP

DCBP,HSPA1A,HTRA1,HIPK2,PMEPA1,STUB1,EMILIN1,CRKL,LIMS1,ZEB1,ZNF703,ZYX,NREP,SOX

FR1,SDCBP,SEMA6B,PHACTR4,DAG1,NOTCH1,ACTA2,MDM2,BNC2,EFNB1,TASOR,PDGFRB,LOX
,SLC38A2,PIK3C2B,EHMT2,TNRC6A,RHEB,ASNS,ATP6V0B,ATP6V0E2,GABARAPL2,MAPK3,XBP1,

CRFS1,ATP5F1E,COX4I1,COX5A,COX7A2,UQCRH,MSH2,NDUFA5,UQCR11,COX5B,NDUFA10,ATP5
A,MFN2,ATP5PO,YWHAZ,GSK3B,ATP5F1C,VDAC2,ATP5F1E,TFDP1,DYNLT1,OPA1,ATP5PD

7,SCOC,SESN2
A,CNN2,PARP1,PKN1,MDK,CHD2,ASXL1,PSMB2,H3-3A,ATP6AP1,NOTCH2,VEGFA,MAPK1,PSMD3,
PSMD7,CDK4,PSME3,PSMC5,PSMC2,PSMC4,PLK2,PSMF1,CDKN1B,PSMB5,PSMA1,PRMT2,PSMD
4K4,HSD17B12,C1QBP,LDB1,EMILIN1,CRKL,SLK,SPOCK1,LIMS1,GSK3B,RCC2,DUSP3,ROCK2,N

NDRG1,PLK2,CDKN1B,XPC,PSMD10,MDC1,CNOT3,PCNA,TFDP1,MSH2,SESN2,CDK5RAP3,HMGA2

N1,LIMS1,SRF,PIP5K1C,ZNF703,NUMBL,AFDN,ROCK2,NECTIN2,TGFBR1,HEG1,ADD1,EXT1,PRK
GOLGA4,SRF,CRABP2,UNC13A,RUFY3,PIN1,YBX3,SYT1,SYT4,ATP8A2,CACNG7,ANAPC2
S3,RAC1,HDAC2,PEA15,CDKN1A,EEF1A2,SQSTM1,NOX4,PKN1,GCN1,TERF2IP,VEGFA,RAPGEF1,F
K4,MAPK3,PPP3CA,PSMA6,VAV2,TAB3,FER,RELA,PSMD12,PSMA4,PSMC6

M1,SULF1,PLXNA1,ACTN4,MMP14,MIF,GJA1,MACF1,CADM4,RHOB,RAC1,ARHGDIA,APPL1,GLUL
P53,PARP1,YEATS2,SKI,HDGF,TAF7,NOTCH2,VEGFA,VHL,CCND1,SMARCA4,AEBP1,TCF25,SUPT5

2,COL3A1
HNRNPU,LARP1,CTNNA1,RPL23,NACA,RTN4,LAMA5,RPS9,XRCC5,MSN,IGFBP3,RHOA,JUND,CD8

MC1,CAD,PSAT1,AASDHPPT,PSMB1,MECP2,PSMD13,PARK7,ALDH18A1,PSMD7,ASNS,DARS1,ALD

KT3,CNP,SOD2,MSH6,MBD3,GSK3A,VASH1,MTOR,SCAP,AMFR,HMGA1,ABL1,DAG1,FZR1,PDGFR
DCBP,HSPA1A,HTRA1,HIPK2,PMEPA1,STUB1,EMILIN1,CRKL,LIMS1,ZEB1,ZNF703,ZYX,NREP,SOX

l14,BMERB1,SHANK1,CAPZA2,ADD1,ARHGEF18,GMFB,FGF13,TPX2,MYADM,GSN
GOLGA4,ACBD3,CCDC93,NAPA

PS16,VPS28,LEPROT,CACNG7,AP3M1,M6PR,VPS37B

H2,VEGFA,MAPK1,TCF25,PRKDC,SLIT3,HECTD1,VANGL2,BMPR2,CDC42,EP300,ZFP36L1,PTEN,
,PSMB1,TAOK2,MDM2,CUL1,PSMD13,PSMD7,CDK4,PSME3,TOM1L2,PSMC5,PSMC2,PSMC4,PLK
PM6B,SIN3A
RAPGEF1,PPP2CA,CCND1,MAPK1,ADAM9,MAP2K2,FYN,SOD1,CDC42,TRAF4,TRAF7,PRKAR2A,PH
,POR,RUVBL2,MECP2,PARK7,TRRAP,KAT2A,LDB1,DLD,GSK3B,SETD5,ZDHHC5,PER1,CTCF,OGD
AR1A,YWHAB,PTGES3,RAC1,HDAC2,PEA15,CDKN1A,EEF1A2,SQSTM1,NOX4,PKN1,GCN1,TERF2I
AR1A,YWHAB,PTGES3,RAC1,HDAC2,PEA15,CDKN1A,EEF1A2,SQSTM1,NOX4,PKN1,GCN1,TERF2I

AR1,RELA,OPA1,GRB10,ATP6V1F,ZNF106,PIK3R3,ATP6V1D
ROCK2,PPP1R26,PIN1,PPP2R5B,CABIN1,SLC39A10,PPP2R2A,PPP1R16A,PPP2R5C

S,NPLOC4,LOX,SMG5,POLDIP2,USP10,NIBAN2,SMPD3,PDGFRB,WRNIP1,POLD2,EXOSC10,PHB,N
A,MBD3,NASP,SMARCD1,HMGA1,RSF1,MTA2,ANP32E,RUVBL2,KAT2A,TASOR,ATF7IP,ASH1L,SO
3A,YY1,APBB1,NDUFA13,SEMA6B,TSPYL2,HSPA1A,ACVR1B,NOTCH1,NTN1,SEMA4C,MEIS1,SERF

A,PSMD2,KIF5B,CD81,WDR1,VAT1,CLEC7A,PFKL,RAB7A,DDX3X,MIF,CD63,DNAJC5,NDFIP1,DDO

3,ABL1,MACROH2A1,GGNBP2,PKIA,IGF1R,PARK7,PTPN1,TRIM27,TRIB3,WARS1,CDKN1B,SH3BF

RPF8,UGT8,RPL8,CALR,TMF1,HSPA1B,TUBB2B,COL5A1,COL4A1,RPLP0,H3-3B,CLU,ENO2,RPL1
CANX,CLU,RPL13A,MAGED1,SEC62,RPL6,DVL3,YWHAE,KIF1A,RPL13,GNAS,SLC2A3,GPI,RACK1,
RPS4X,RPS3A,RPL18,ITGB1,RPL11,NCAM1,ANXA2,RPLP1,RPL10,RPS17,ARF1,SF3B2,AP2B1,GTF
MF1,HSPA1B,TUBB2B,COL5A1,COL4A1,RPLP0,H3-3B,CLU,ENO2,RPL13A,RPL6,KAT6A,DVL3,YW
WHAG,RPL37A,MAPT,YWHAQ,DCHS1,TMSB10,RPL19,RPL15,PKD1,RPS3,VAPA,AP2M1,SEC61A1,
WHAG,RPL37A,MAPT,YWHAQ,DCHS1,TMSB10,RPL19,RPL15,PKD1,RPS3,VAPA,AP2M1,SEC61A1,
IRNPA0,RPS16,RPL23,RPL29,RPSA,RPS24,RPS9,RPL24,RPL17,PAPOLA,RPL14,RPL22,RPL9,ATF
M28,YWHAG,RPL37A,MAPT,YWHAQ,RPL19,RPL15,PKD1,RPS3,VAPA,AP2M1,SEC61A1,RPS4X,RPS
AP2M1,RPS4X,RPS3A,RPL18,UCHL1,RPL11,ANXA2,AGRN,RPLP1,TIMP2,RPL10,MARCHF6,RPS17
HAE,KIF1A,RPL13,GNAS,SLC2A3,GPI,SPTBN1,ATP5F1B,CALM3,RPS8,MYH9,RPL7,APP,RPL5,GDI1
1,CALM1,NPM1,HSPD1,VCP,UBB,TRIM28,YWHAG,RPL37A,MAPT,YWHAQ,DCHS1,TMSB10,RPL19
I1,PLXNB2,IGF2,VCP,TRIM28,YWHAG,MAPT,FN1,DDB1,YWHAQ,SET,MAP9,TMSB10,PPP2R1A,PKI
SEPTIN2,APP,RPL5,GDI1,SMG1,CALM1,NPM1,HSPD1,VCP,UBB,TRIM28,YWHAG,RPL37A,MAPT,YV
.18,UCHL1,RPL11,ANXA2,RPLP1,RPL10,MARCHF6,RPS17,AP2B1,HNRNPU,OGA,LARP1,HNRNPA0
DNAJA1,HNRNPU,CTNNA1,RPS16,RPL23,RPL29,RPSA,SCARB2,RTN4,RPS24,RPS9,XRCC5,MSN,F
D2,RPS14,RPS20,RPL26,RPL31,HNRNPC,XPO1,RPS25,RPS15,RPL12,RPL30,RPS5,RPS7,RPS19,F
34,CALR,FSTL1,TMF1,ABHD2,HSPA1B,TUBB2B,COL5A1,PSAP,RPLP0,CANX,CLU,SOX4,RPL13A,M
RPL13,GPI,RACK1,SPTBN1,CALM3,RPS8,MYH9,RPL7,P4HB,APP,RPL5,GSTP1,SMG1,CALM1,NPN
LM3,STMN2,MYH9,SEPTIN2,APP,RPL5,FSCN1,SPTAN1,MARCKS,SMG1,CALM1,NPM1,HSPD1,CSD
RL6IP1,RPL36,RPS12,RPS27,RPL23A,SRP9,RPL27,RPL36A,SRP14,RPS15A,RPL38,RPL37,RPS11,
RPLP2,RPL36,RPS12,RPS27,BCLAF1,RPL23A,RPL27,RPL36A,RPS15A,RPL38,RPL37,PPP2CA,EIF3
PS12,RPS27,RPL23A,SRP9,RPL27,RPL36A,SRP14,RPS15A,RPL38,RPL37,RPS11,RPL28,RPS28,F
D2,RPS14,RPS20,RPL26,RPL31,HNRNPC,LRPPRC,XPO1,RPS25,RPS15,RPL12,RPL30,RPS5,RPS7
PTIN2,P4HB,APP,RPL5,FSCN1,SPTAN1,MARCKS,NPM1,PLXNB2,HSPD1,SNRNP200,CSDE1,EIF3C,I
APA,AP2M1,SEC61A1,RPS4X,RPS3A,RPL18,RPL11,BTF3,RPLP1,MIDN,RPL10,RPS17,AP3D1,DNAJ
PS16,RPL23,RPL29,RPSA,SCARB2,NACA,RPS24,RPS9,RPL24,KPNB1,RPL17,RPL14,RPL22,RPL9,
,RPL35A,RPLP2,RPL36,RPS12,RPS27,RPL23A,SRP9,RPL27,RPL36A,SRP14,RPS15A,RPL38,RPL3
PL36,RPS12,RPS27,RPL23A,SRP9,RPL27,RPL36A,SRP14,RPS15A,RPL38,RPL37,RPS11,RPL28,RI
MAGED1,RPL6,KAT6A,DVL3,YWHAE,RPL13,GPI,HTRA4,RACK1,SPTBN1,CALM3,RPS8,MYH9,RPL7
,RPS3,AP2M1,SEC61A1,RPS4X,RPS3A,RPL18,ITGB1,RPL11,BTF3,RPLP1,MIDN,RPL10,RPS17,AP
RPS19,RPL35,JUN,RPS10,RPL35A,TARDBP,RPLP2,DDOST,ST3GAL2,RPL36,RPS12,RPS27,RPL23
PL36,RPS12,RPS27,RPL23A,SRP9,RPL27,RPL36A,SRP14,RPS15A,RPL38,RPL37,RPS11,RPL28,RI
),RPL35,RPS10,RPL35A,RPLP2,RPL36,RPS12,RPS27,RPL23A,RPL27,RPL36A,RPS15A,RPL38,RPL
GT8,RPL8,FBXL4,YBX1,UBA1,ATP2B4,IGFBP2,CALR,FSTL1,TMF1,ABHD2,HSPA1B,ZNF99,CES3,P
,SMG1,CALM1,NPM1,TTC3,PLD3,USP11,CSDE1,VCP,UBB,RPL37A,MAPT,DDB1,PSMD5,SET,TPI1,
UBB,RPL37A,MAPT,DDB1,PSMD5,SET,TPI1,RPL19,PPP2R1A,RPL15,PKD1,RPS3,HMGB1,AP2M1,R
PA1B,COL5A1,PGK1,RPLP0,H3-3B,CLU,PTMA,DDX5,SOX4,RPL13A,MAGED1,RPL6,KAT6A,DVL3,
25,DDX3X,RPS15,RPL12,RPL30,EIF3L,RPS5,RPS7,RPS19,RPL35,RPS10,RPL35A,RPLP2,RPL36,RF
9,CD81,OAZ1,MCL1,HNRNPC,LRPPRC,XPO1,CSNK1E,GPC3,PIP4K2B,RAB7A,RPS7,GJA1,SESN3,
3A,RPL18,RPL11,BTF3,RPLP1,MIDN,RPL10,RPS17,AP3D1,DNAJA1,ARF1,PAFAH1B1,AP2B1,RPS1
P0,CANX,CLU,SOX4,RPL13A,SEC62,RPL6,YWHAE,KIF1A,RPL13,GNAS,SLC2A3,GPI,RACK1,BSG,SF
PL24,KPNB1,RPL17,RPL14,HK1,RPL22,RPL9,RPS14,RPS20,RPL26,CCT7,RPL31,RPS25,RAB7A,R
,SOX4,RPL13A,SEC62,RPL6,YWHAE,KIF1A,RPL13,GNAS,SLC2A3,GPI,RACK1,BSG,SPTBN1,ATP5F
M1,PLXNB2,HSPD1,SNRNP200,CSDE1,EIF3C,EIF4B,VCP,UBB,TRIM28,YWHAG,MAPT,FN1,DDB1,P
SDE1,VCP,UBB,RPL37A,DDB1,PSMD5,SET,TPI1,RPL19,PPP2R1A,RPL15,PKD1,RPS3,HMGB1,AP2N

,H3-3B,CLU,ENO2,SOX4,FKBP14,RPL13A,MAGED1,RPL6,KAT6A,DVL3,YWHAE,RPL13,GPI,HTRA4
PEN,YWHAQ,DCHS1,INA,PKD1,BASP1,VAPA,HMGB1,STMN1,ITGB1,UCHL1,NCAM1,AGRN,ZMIZ1,T
,RPL29,RPSA,NACA,LAMA5,RPS24,RPS9,RPL24,RPL17,RPL14,RPL22,RPL9,CLSTN1,KIF5B,NLGN
3,YWHAE,GNAS,DDX17,GPI,RACK1,BSG,FASN,SPTBN1,ATP5F1B,CALM3,GNB1,STMN2,MYH9,P4H
OA,NR2F1,KIF5B,ENC1,CRMP1,PRKCSH,SPTBN2,PTN,CDH2,DDR1,PLXNA1,MEGF8,ITM2C,GJA1,
3,CPE,RPL29,RPSA,NACA,LAMA5,RPS24,RPS9,RPL24,RPL17,LLGL1,RPL14,RPL22,RPL9,CLSTN1
1,CSNK1E,PIP4K2B,RPS7,SESN3,TARDBP,CIRBP,YWHAB,ATP6V0C,LAPTM4B,TRIB2,PTTG1IP,DC
P2,RPL36,RPS12,RPS27,RPL23A,RPL27,RPL36A,RPS15A,RPL38,RPL37,SMARCA4,RPS11,SUPT5
9,PSAP,RPLP0,H3-3B,CLU,DDX5,SOX4,RPL13A,MAGED1,RPL6,KAT6A,DVL3,YWHAE,RPL13,GNAS
14,RPL22,RPL9,PSMD2,RPS14,RPS20,RPL26,RPL31,HNRNPC,LRPPRC,XPO1,RPS25,RPS15,RPL1
A,DDX5,SOX4,RPL13A,MAGED1,KAT6A,DVL3,YWHAE,GPI,HTRA4,RACK1,SPTBN1,CALM3,STMN2,
,SMG1,CALM1,NPM1,HSPD1,CSDE1,IGF2,UBB,TRIM28,YWHAG,RPL37A,MAPT,FN1,SPEN,NONO,
YWHAG,MAPT,FN1,DDB1,EIF1,HACD3,SET,PPP2R1A,PKD1,RPS3,HMGB1,RPS4X,UCHL1,RPL11,AN
4,IGFBP2,CALR,FSTL1,TMF1,HSPA1B,ZNF99,RPLP0,H3-3B,CLU,PTMA,DDX5,SOX4,FKBP14,RPL1
CALM1,NPM1,CSDE1,IGF2,UBB,TRIM28,YWHAG,RPL37A,MAPT,FN1,SPEN,NONO,PSMD5,YWHAQ
B1,TPT1,MORF4L1,MARCHF6,DNAJA1,PAFAH1B1,NFE2L1,SMARCA5,CTNNA1,RTN4,XRCC5,PJA2
X,CLU,DDX5,SOX4,FKBP14,RPL13A,GNAS,DDX17,GPI,HTRA4,RACK1,BSG,FASN,CALM3,GNB1,ST
RPS3,HMGB1,AP2M1,STMN1,RPL11,RPL10,ARF1,AP2B1,MAT2A,TMEM170A,SMARCA5,RPL23,R
P43,NCS1,RHOA,NR2F1,KIF5B,ENC1,CRMP1,PRKCSH,SPTBN2,PTN,CDH2,DDR1,PLXNA1,MEGF8
CT7,PFN1,CLEC7A,NES,TADA2B,PTN,SFPQ,TPM1,EIF5A,GPC3,PLXNA1,PIP4K2B,MEGF8,CCN2,DD
L,ARF1,PAFAH1B1,MAP4,CTNNA1,SCARB2,RTN4,LAMA5,XRCC5,RPL24,LLGL1,GAP43,NCS1,RHO
4,YBX1,UBA1,ATP2B4,IGFBP2,CALR,FSTL1,TMF1,ABHD2,HSPA1B,ZNF99,CES3,PSAP,PGK1,RPLI
PL17,RPL14,RPL22,RPL9,PSMD2,RPS14,RPS20,RPL26,RPL31,HNRNPC,LRPPRC,XPO1,RPS25,RI
PL17,RPL14,RPL22,RPL9,PSMD2,RPS14,RPS20,RPL26,RPL31,HNRNPC,LRPPRC,XPO1,RPS25,RI
S10,RPL35A,RPLP2,RPL36,RPS12,RPS27,BCLAF1,RPL23A,RPL27,RPL36A,RPS15A,RPL38,RPL37
8,FBXL4,YBX1,UBA1,ATP2B4,IGFBP2,CALR,FSTL1,TMF1,HSPA1B,ZNF99,PSAP,PGK1,RPLP0,H3-
B2,RTN4,LAMA5,RPL24,LLGL1,GAP43,NCS1,RHOA,ATF4,NR2F1,KIF5B,ENC1,CRMP1,PRKCSH,S
CTNNA1,SCARB2,RTN4,LAMA5,XRCC5,RPL24,LLGL1,GAP43,NCS1,RHOA,ATF4,NR2F1,KIF5B,EN
9,RPL15,PKD1,RPS3,AP2M1,SEC61A1,RPS4X,RPS3A,RPL18,ITGB1,RPL11,BTF3,ANXA2,RPLP1,M
0,H3-3B,CLU,DDX5,SOX4,RPL13A,MAGED1,RPL6,KAT6A,DVL3,YWHAE,RPL13,DDX17,GPI,RACK1
S20,RPL26,RPL31,RPS25,RAB7A,RPS15,RPL12,RPL30,RPS5,RPS7,RPS19,RPL35,MACF1,RPS10,I
IAGED1,KAT6A,DVL3,YWHAE,GNAS,DDX17,GPI,RACK1,BSG,FASN,SPTBN1,ATP5F1B,CALM3,GNB
A,MAGED1,RPL6,KAT6A,DVL3,YWHAE,DDX17,GPI,RACK1,FASN,CALM3,MYH9,APP,RPL5,TMBIM6
ACA,RTN4,RPS24,RPS9,RPL24,RPL17,RPL14,RPL22,RHOA,PTCD3,RPL9,ATF4,ENC1,RPS14,RPS2
,HACD3,SET,PPP2R1A,PKD1,RPS3,HMGB1,RPS4X,UCHL1,RPL11,ANXA2,RPLP1,ZMIZ1,TIMP2,RPI
25,RAB7A,RPS15,RPL12,RPL30,RPS5,RPS7,RPS19,RPL35,RPS10,RPL35A,RPLP2,ARL6IP1,RPL36
14,RPL22,RPL9,PSMD2,RPS14,RPS20,RPL26,RPL31,HNRNPC,LRPPRC,XPO1,RPS25,RPS15,RPL1
DR1,CRMP1,NES,SPTBN2,PTN,CDH6,TPM1,CDH2,DDR1,PLXNA1,MEGF8,ACTN4,GJA1,CTNNB1,M
PL17,RPL14,RPL22,RPL9,PSMD2,RPS14,RPS20,RPL26,RPL31,HNRNPC,LRPPRC,XPO1,RPS25,RI
F1,CYFIP2,MAP2,RAC1,ARHGDIA,CAPRIN1,GPC1,TOP2B,L1CAM,MAP1A,CRPPA,CDH11,GAS7,PTF
YFIP2,MAP2,RAC1,ARHGDIA,CAPRIN1,GPC1,TOP2B,L1CAM,MAP1A,CRPPA,CDH11,GAS7,PTPRA,N
F1,CYFIP2,MAP2,RAC1,ARHGDIA,CAPRIN1,GPC1,TOP2B,L1CAM,MAP1A,CRPPA,CDH11,GAS7,PTF
K1,H3-3B,CANX,CLU,DDX5,SOX4,MAGED1,KAT6A,DVL3,YWHAE,KIF1A,RPL13,GNAS,DDX17,GPI,F
NDFIP1,UBE2H,HMGCR,GPC1,SKP1,LAPTM4B,TRIB2,PTTG1IP,MAP1A,TIMP3,CTSB,RNF187,EEF1
RPL24,RPL17,RPL14,RPL22,RHOA,PTCD3,RPL9,ATF4,ENC1,RPS14,RPS20,EIF5,RPL26,RPL31,LI

11,STMN1,RPL11,RPL10,ARF1,AP2B1,MAT2A,TMEM170A,SMARCA5,RPSA,EIF4H,RTN4,XRCC5,M
RPF8,UGT8,RPL8,FBXL4,YBX1,UBA1,ATP2B4,IGFBP2,CALR,FSTL1,TMF1,ABHD2,HSPA1B,ZNF99,
ICAM1,AGRN,VCL,ARF1,PAFAH1B1,MAP4,CTNNA1,SCARB2,RTN4,LAMA5,RPL24,LLGL1,GAP43,N
PL22,RPL9,RPS14,RPL26,SRSF4,PNN,RPL31,HNRNPC,KHDRBS1,RPS25,SFPQ,DDX3X,RPS15,RPI
DX5,SOX4,FKBP14,RPL13A,GNAS,DDX17,HTRA4,RACK1,BSG,FASN,GNB1,STMN2,P4HB,APP,FSC
FIP2,MAP2,RAC1,ARHGDIA,CAPRIN1,GPC1,TOP2B,L1CAM,MAP1A,CRPPA,CDH11,GAS7,PTPRA,N
IF4H,RTN4,XRCC5,RPL24,RHOA,PSMD2,RPS14,EIF5,RBBP4,PFN1,SPTBN2,DDX3X,RPS15,CLTC,
1,UBA1,ATP2B4,IGFBP2,CALR,FSTL1,TMF1,HSPA1B,ZNF99,COL5A1,PGK1,RPLP0,H3-3B,CLU,PT
D1,RPL6,KAT6A,DVL3,YWHAE,DDX17,GPI,RACK1,FASN,CALM3,MYH9,APP,RPL5,TMBIM6,GSTP1
5A,RPLP2,ARL6IP1,RPL36,RPS12,RPS27,RPL23A,SRP9,RPL27,RPL36A,SRP14,RPS15A,RPL38,RF
HNRNPD,SERBP1,RBM15B,PSMA7,ANP32A,EDC4,PSMD1,SF1,GTPBP1,UBA52,ZFP36L1,SRSF6,U
ICAM1,AGRN,VCL,ARF1,PAFAH1B1,MAP4,CTNNA1,SCARB2,RTN4,LAMA5,RPL24,LLGL1,GAP43,N
RPL24,RPL17,RPL14,RPL22,RHOA,PTCD3,RPL9,ATF4,ENC1,RPS14,RPS20,EIF5,RPL26,RPL31,LF
,RPL13A,GNAS,DDX17,HTRA4,RACK1,FASN,GNB1,STMN2,P4HB,APP,FSCN1,TMBIM6,GSTP1,NPN
4,MAGED1,KAT6A,DVL3,YWHAE,KIF1A,RPL13,GNAS,DDX17,GPI,RACK1,BSG,FASN,SPTBN1,ATP5
CTN4,CTNNB1,MACF1,MAP2,RAB1A,RAC1,ARHGDIA,CAPRIN1,GPC1,TOP2B,L1CAM,MAP1A,CRPI
B2,COL18A1,IGF2,UBB,TRIM28,COL4A2,MAPT,FN1,PSMD5,DCHS1,EGLN1,CHSY1,PKD1,BASP1,I
IEGF8,PRKAR1A,GJA1,CTNNB1,MACF1,CYFIP2,MAP2,RAC1,ARHGDIA,CAPRIN1,GPC1,TOP2B,L1(
P2B,L1CAM,MAP1A,CRPPA,CDH11,TBCD,PTPRA,PTK7,DPYSL2,NOTCH2,VEGFA,MAPK1,GRB2,PL
,RACK1,BSG,SPTBN1,ATP5F1B,CALM3,GNB1,SHISA9,MYH9,APP,RPL5,GDI1,TMBIM6,SPTAN1,G
NRNP70,SCAF1,PRPF6,PPP2CA,SRSF9,SRSF1,PTBP1,HNRNPUL1,SCAF11,SON,KDM1A,CASC3,CC
T6A,DVL3,YWHAE,KIF1A,RPL13,GNAS,DDX17,GPI,RACK1,BSG,FASN,SPTBN1,ATP5F1B,CALM3,GI
DC1,ADAM9,GABARAP,CCAR2,SMARCC1,FYN,UBQLN1,CSNK1D,USP7,HECTD1,HGS,BAG6,CYP51
PL6,KAT6A,DVL3,YWHAE,DDX17,GPI,RACK1,CALM3,MYH9,APP,RPL5,TMBIM6,GSTP1,SMG1,CAL
RPS24,RPS9,RPL24,RPL17,RPL14,RPL22,RHOA,PTCD3,RPL9,ATF4,ENC1,RPS14,RPS20,EIF5,RPL
3,GNAS,DDX17,BSG,FASN,SPTBN1,ATP5F1B,CALM3,GNB1,STMN2,MYH9,SEPTIN2,APP,GDI1,SP
GT8,RPL8,FBXL4,YBX1,UBA1,ATP2B4,IGFBP2,CALR,FSTL1,TMF1,ABHD2,HSPA1B,ZNF99,COL5A
PUL1,SCAF11,SON,KDM1A,CASC3,POLR2B,HNRNPM,SRSF3,SF3A1,STRAP,PCBP2,HNRNPD,PRPF
PUL1,SCAF11,SON,KDM1A,CASC3,POLR2B,HNRNPM,SRSF3,SF3A1,STRAP,PCBP2,HNRNPD,PRPF
PUL1,SCAF11,SON,KDM1A,CASC3,POLR2B,HNRNPM,SRSF3,SF3A1,STRAP,PCBP2,HNRNPD,PRPF
MD5,HACD3,SET,EGLN1,PPP2R1A,GTF3C4,PKD1,PDIA3,RPS3,HMGB1,CDKL1,UCHL1,RPL11,MC
MD5,HACD3,SET,EGLN1,PPP2R1A,GTF3C4,PKD1,PDIA3,RPS3,HMGB1,CDKL1,UCHL1,RPL11,MC
PPA,CDH11,PTPRA,PTK7,DPYSL2,NOTCH2,VEGFA,MAPK1,GRB2,PLCG1,FYN,SLIT3,STXBP1,PLXN
M1,NPM1,PLXNB2,HSPD1,IGF2,EIF3C,VCP,UBB,TRIM28,MAPT,FN1,DDB1,HACD3,PKD1,SF3B1,S
2,SNRNP70,SCAF1,PRPF6,SRSF9,SRSF1,PTBP1,SUPT5H,HNRNPUL1,SCAF11,SON,KDM1A,CASC3
87,SQSTM1,NOP53,UBE4A,VPS35,RAD23B,PSMB2,ABCA2,UBXN4,MVB12B,PSMD3,GABARAP,UE
,UCHL1,NCAM1,ANXA2,AGRN,ZMIZ1,COL11A1,VCL,SBF1,ARF1,PAFAH1B1,MAP4,HNRNPU,CTNN
T1,PFN1,NES,HNRNPC,TADA2B,SPTBN2,SFPQ,TPM1,CDH2,PIP4K2B,RAB7A,CCN2,CLTC,IK,PRK
11,MORF4L1,RPLP1,RPL10,RPS17,HNRNPU,LARP1,HNRNPA0,RPS16,RPL23,RPL29,RPSA,RPS24,
23,CPE,RPL29,RPSA,EIF4H,NACA,RTN4,RPS24,RPS9,RPL24,RPL17,RPL14,RPL22,RHOA,PTCD3,I
PL12,RPL30,EIF3L,RPS5,RPS7,RPS19,RPL35,RPS10,RPL35A,RPLP2,TSR2,RPL36,PTGES3,RPS27
PT,FN1,DDB1,HACD3,PKD1,SSBP2,ZNF43,RPS3,HMGB1,AP2M1,RPS4X,ANXA2,AGRN,RPLP1,ZMI
N1,RPL4,ENO1,PIK3R2,FLNA,MAP1B,NCL,RPL7A,SLC9A7,ALDOA,MTRNR2L8,PKM,HSP90B1,FAD
1,AP2M1,ITGB1,TUBB4B,ANXA2,TIMP2,VCL,AP3D1,ARF1,CYB5R3,AP2B1,SLC4A8,PGAM1,CTSD,
2M,PLXNA3,PTPRS,RAB11A,BMPR2,EP300,AP2A1,MAP6,PLXNB1,DIP2B,SFRP1,ADCY6,PTEN,PTF

EIF3C,VCP,UBB,TRIM28,MAPT,FN1,DDB1,HACD3,PKD1,SF3B1,SSBP2,ZNF43,RPS3,HMGB1,AP2M

HMGB1,UCHL1,RPL11,MORF4L1,ANXA2,TIMP2,RPL10,DNAJA1,ITM2B,HNRNPU,SMARCA5,LARP1

L2,NOTCH2,VEGFA,MAPK1,GRB2,PLCG1,FYN,SLIT3,STXBP1,PLXNA3,PTPRS,RAB11A,TCTN1,BM

A,RPS3,CHD3,BASP1,HMGB1,UCHL1,RPL11,MORF4L1,ANXA2,TIMP2,MIDN,RPL10,DNAJA1,ITM2

,NCKAP1,VPS35,ACAP3,MDK,PTK7,DPYSL2,VEGFA,RAPGEF1,ARPC2,CAPZB,KDM1A,CORO1C,FY

B1,ITGB1,ANXA2,ZMIZ1,TIMP2,DNAJA1,GTF2F1,HNRNPU,LARP1,CTNNA1,RPL23,LDHA,COL1A2,

ACTR3,NCKAP1,VPS35,ACAP3,MDK,PTK7,DPYSL2,VEGFA,RAPGEF1,ARPC2,CAPZB,KDM1A,CORO

3A,ITGB1,RPL11,XAF1,RPL10,DNAJA1,AKAP12,CTNNA1,LDHA,CTSD,NACA,RTN4,KPNB1,IGFBP3,

DB1,HACD3,PKD1,SSBP2,ZNF43,RPS3,HMGB1,RPS4X,ANXA2,AGRN,RPLP1,ZMIZ1,AP3D1,ZNF77

,SOX4,RPL13A,MAGED1,RPL6,KAT6A,DVL3,RPL13,DDX17,GPI,RACK1,BSG,SPTBN1,RPS8,MYH9

S3A,RPL18,RPL11,AGRN,RPLP1,RPL10,RPS17,ITM2B,OGA,NFE2L1,LARP1,RPS16,RPL23,RPL29,

ASN,ALDH6A1,SPTBN1,ATP5F1B,STMN2,MYH9,SEPTIN2,P4HB,APP,GDI1,SPTAN1,GSTP1,NPM1,PL

L4A1,CES3,PGK1,RPLP0,H3-3B,CANX,CLU,DDX5,SOX4,FKBP14,RPL13A,DVL3,GNAS,DDX17,GPI,

H3-3B,CLU,PTMA,DDX5,ENO2,SOX4,RPL13A,MAGED1,RPL6,KAT6A,DVL3,RPL13,DDX17,GPI,RAC

ASN,ALDH6A1,SPTBN1,ATP5F1B,STMN2,MYH9,SEPTIN2,P4HB,APP,GDI1,SPTAN1,GSTP1,PLXNB2,

MBIM6,SMG1,NPM1,HSPD1,SNRNP200,CSDE1,IGF2,UBB,AFF4,TRIM28,COL4A2,RPL37A,MAPT,S

CLU,PTMA,DDX5,ENO2,SOX4,RPL13A,MAGED1,RPL6,KAT6A,DVL3,RPL13,DDX17,GPI,RACK1,FA

CLU,PTMA,DDX5,ENO2,SOX4,RPL13A,MAGED1,RPL6,KAT6A,DVL3,RPL13,DDX17,GPI,RACK1,FA

SMD5,HACD3,SET,EGLN1,PPP2R1A,GTF3C4,PKD1,PDIA3,RPS3,HMGB1,CDKL1,UCHL1,RPL11,MC

RPL13A,MAGED1,RPL6,KAT6A,DVL3,RPL13,DDX17,GPI,FASN,ALDH6A1,ATP5F1B,RPS8,RPL7,AP

3X,GPAT4,PRKAR1A,GJA1,CTNNB1,ANXA6,MACF1,CYFIP2,MAP2,TNC,L1CAM,PPIB,CDKN1A,DDX

1,RPS3A,ITGB1,RPL11,XAF1,RPL10,DNAJA1,AKAP12,CTNNA1,LDHA,CTSD,NACA,RTN4,KPNB1,IG

VL3,RPL13,DDX17,RPS8,RPL7,APP,RPL5,TMBIM6,SMG1,NPM1,PLD3,HSPD1,SNRNP200,CSDE1,I

B2,ATXN3,OGT,GSK3A,PSMC3,UBE3A,PRKACA,PSMC1,BAG5,UBE2K,ZER1,USP5,SDCBP,PINK1,H

13A,MAGED1,RPL6,KAT6A,DVL3,RPL13,DDX17,RACK1,RPS8,RPL7,APP,RPL5,TMBIM6,PTMS,GS

AC2,TBCD,NOP53,PARP1,CAMSAP1,SKI,ABCA2,TERF2IP,TAF7,VEGFA,MAPRE1,LUZP6,CAPZB,KD

CCN2,MMP14,DDX3X,GJA1,MACF1,CYFIP2,CKAP5,MAP2,COTL1,RAC1,DLG5,TBCD,NOP53,PAR

A,CLTC,RPS7,GJA1,CTNNB1,NDFIP1,UBE2H,HMGCR,GPC1,SKP1,LAPTM4B,TRIB2,GLUL,PTTG1I

L6,KAT6A,DVL3,RPL13,DDX17,GPI,RACK1,BSG,SPTBN1,RPS8,MYH9,RPL7,APP,RPL5,TMBIM6,G

,HMGB1,AP2M1,TPT1,ITGB1,AGRN,ZMIZ1,TIMP2,VCL,AP3D1,ARF1,PAFAH1B1,AP2B1,HNRNPU,C

TMEM30A,GLUL,MAP1A,NUCB1,CCT2,SQSTM1,VAMP2,PARP1,TCP1,CCT3,CCT4,RAB11B,ABCA2,

2R2,CCAR2,SMARCC1,STT3B,LONP1,UBQLN1,CSNK1D,USP7,HECTD1,PCBP2,BAG6,PCYOX1,Y

ED1,RPL6,KAT6A,DVL3,RPL13,DDX17,RACK1,RPS8,RPL7,APP,RPL5,TMBIM6,PTMS,NPM1,PLXN

B,VAPA,HMGB1,TPT1,RPS3A,ITGB1,RPL11,XAF1,RPL10,DNAJA1,AKAP12,AP2B1,CTNNA1,LDHA,C

GTA,OS9,MTA1,PSMD1,UBQLN2,UBA52,PPP2CB,KLHL42,UBE2G2,CUL3,RPS27A,PSMD8,NPLOC

2,SLIT3,PLXNA3,PTPRS,RAB11A,IGFBP7,BMPR2,CDC42,PPP1R9B,RBBP7,EXTL3,FLCN,CSNK2A1,

COL1A2,RHOA,ATF4,ATP1A1,COL5A2,PTN,GLG1,GPC3,SULF1,PIP4K2B,MEGF8,CCN2,APLP1,HM

N1,RPL4,ENO1,PIK3R2,FLNA,MAP1B,NCL,RPL7A,SLC9A7,ALDOA,MTRNR2L8,PKM,FTL,HSP90B1

VCP,UBB,YWHAG,MAPT,FN1,NONO,PSMD5,HACD3,CHSY1,PPP2R1A,PDIA3,RPS3,CHD3,VAPA,HM

NNA1,TPM4,KPNB1,COL1A2,RHOA,COL5A2,WDR1,PFN1,SPTBN2,TPM1,CCN2,PRKAR1A,MYO5A,

SF1,EIF3D,SRSF6,EIF2S2,EIF3H,SNRPD3,DICER1,RBM5,PRPF19,EIF3G,SF3A2,SART1,LUC7L2,RU

2,SOX4,RPL13A,MAGED1,RPL6,KAT6A,DVL3,RPL13,DDX17,GPI,FASN,ALDH6A1,ATP5F1B,RPS8,R

X5,SOX4,RPL13A,MAGED1,RPL6,KAT6A,DVL3,RPL13,DDX17,RACK1,FASN,ATP5F1B,RPS8,RPL7,A

CTSD,NACA,RTN4,IGFBP3,RHOA,ATF4,RPL26,BEX3,MCL1,TLE5,CLEC7A,NES,PTN,SFPQ,CCND2,

MACF1,MYH10,RHOB,MAP2,RAC1,ARHGDIA,CAPRIN1,L1CAM,GLUL,DCN,PKN1,CAMSAP1,VPS35

EIF2S2,EIF3H,SNRPD3,DICER1,RBM5,PRPF19,EIF3G,SF3A2,SART1,LUC7L2,RUVBL2,SF3A3,CD2
A2,VAT1,PFKL,SCRN1,RAB7A,ACTN4,DDX3X,MAGED2,MIF,TLN1,CD63,DNAJC5,APLP2,DDOST,L
HSPH1,GRINA,TERF2IP,TMED2,VEGFA,RAPGEF1,MAPK1,PRDX1,PRKDC,HSPB1,KDM1A,MAP2K2,
,CD81,RPL26,BEX3,OAZ1,CLEC7A,TADA2B,KHDRBS1,CSNK1E,EIF5A,CCND2,GPC3,RAB7A,CCN2
AB7A,ACTN4,DDX3X,MAGED2,MIF,TLN1,CD63,DNAJC5,APLP2,DDOST,LAMP1,LGALS3BP,CKAP4,
CP,UBB,YWHAG,MAPT,FN1,NONO,PSMD5,HACD3,CHSY1,PPP2R1A,PDIA3,RPS3,CHD3,VAPA,HM
,SOX4,RPL13A,MAGED1,RPL6,KAT6A,DVL3,RPL13,DDX17,GPI,FASN,ALDH6A1,ATP5F1B,RPS8,R
3A,MAGED1,RPL6,KAT6A,DVL3,RPL13,DDX17,RACK1,FASN,ATP5F1B,RPS8,RPL7,APP,RPL5,TMB
O1,PIK3R2,FLNA,MAP1B,NCL,SLC9A7,MTRNR2L8,PKM,HSP90B1,FADS1,NEFL,YBX1,ATP2B4,IG
N2,UBA52,PPP2CB,KLHL42,UBE2G2,CUL3,RPS27A,PSMD8,NPLOC4,RHBDD2,PSMB4,EIF3H,GN
7A,CCNI,TADA2B,PTN,CCND2,CCN2,PRDX2,DDX3X,RPS15,RPS7,MIF,PRKAR1A,CTNNB1,TARDBP,
LN1,CSNK1D,USP7,HECTD1,PCBP2,BAG6,PCYOX1,PSMA7,ARIH1,SGTA,OS9,MTA1,PSMD1,UBQ
A2,PHB2,SRSF6,UPF1,MBTPS1,HNRNPA2B1,PPP1CC,ANP32B,MCM3AP,POM121C,SYMPK,PPP1
14,CLTC,GJA1,CTNNB1,TARDBP,NDFIP1,WLS,MYH10,YWHAB,MAP2,CCT6A,HMGCR,RAC1,ARHG
LN1,CSNK1D,USP7,HECTD1,PCBP2,BAG6,PCYOX1,PSMA7,ARIH1,SGTA,OS9,MTA1,PSMD1,UBQ
1R15A,SERP1,RHBDD2,VAPB,TPP1,DNAJB2,ATXN3,ATP6V0D1,GSK3A,CCDC47,UBXN6,UBE2K,S
ATF6,ELOB,PSMA6,XBP1,RELA,PSMD12,PSMA4,SESN2,PSMC6,SIN3A
NK1D,USP7,HECTD1,PCBP2,BAG6,PSMA7,ARIH1,SGTA,OS9,MTA1,PSMD1,UBQLN2,CSNK2A1,U
TSD,NACA,RTN4,IGFBP3,RHOA,ATF4,RPL26,BEX3,MCL1,TLE5,NES,PTN,SFPQ,CCND2,PNMA1,A
00,AP2A1,MAP6,PLXNB1,SEPTIN7,SLIT2,BRK1,GRN,PALM,ABI2,GPC2,DZIP1,FNBP1L,WASF2,AT
OB,PSMA6,XBP1,PSMD12,PSMA4,SESN2,PSMC6,SIN3A
TNNA1,LDHA,CTSD,NACA,RTN4,IGFBP3,RHOA,ATF4,TMEM259,RPL26,BEX3,MCL1,TLE5,CLEC7A,
S7,PRKAR1A,CTNNB1,APLP2,TARDBP,ARL6IP1,CIRBP,YWHAB,CADM4,CTBP1,HMGCR,CAPRIN1,
PC3,MMP14,CLTC,CTNNB1,TARDBP,YWHAB,LAMP1,ATP1A2,MAP2,CCT6A,HMGCR,RAC1,GPC1,C
MBTPS1,FEM1B,CUL3,PSMD8,HIF1A,FBXO21,WSB2,PSMB4,CCN1,RAB1B,UBE2M,PSMD4,ADAM
TNNB1,WLS,ATP1A2,MAP2,ARCN1,TOP2B,HDAC2,DNER,NNAT,DLG5,CDH11,GNAI3,VCAN,NCKA
CAM,DLG5,CDH11,TBCD,TLN2,PTPRA,VPS35,CLSTN3,LAMC1,CNTNAP2,VEGFA,RAPGEF1,PLEC,
OR,PSMC1,THRAP3,EIF4G1,PUM1,FXR2,PSMB1,YTHDF3,HSPA1A,PSMD13,FXR1,SYNCRIP,PABP
DK9,CELF3,DHX36,RBFOX2,SRSF7,NUP98,NCBP2,RBM10,HSF1,CTR9,RNF20,RBM17,SFSWAP,
,SOX4,RPL13A,MAGED1,RPL6,KAT6A,DVL3,RPL13,DDX17,GPI,FASN,ALDH6A1,ATP5F1B,RPS8,R
C1,THRAP3,EIF4G1,PUM1,FXR2,PSMB1,YTHDF3,HSPA1A,PSMD13,FXR1,SYNCRIP,PABPC4,PSMD
,UBQLN1,HGS,SREBF2,DYNLL1,VDAC1,STAT3,WDR45B,EP300,FLCN,UBQLN2,CSNK2A1,SEC16
,UBQLN1,HGS,SREBF2,DYNLL1,VDAC1,STAT3,WDR45B,EP300,FLCN,UBQLN2,CSNK2A1,SEC16
NOTCH2,VEGFA,SELENON,MAPK1,FTO,LUZP6,PRKDC,FKBP8,SLIT3,TNKS2,PLXNA3,COL27A1,PT
A2,PHB2,SRSF6,UPF1,MBTPS1,HNRNPA2B1,PPP1CC,ANP32B,MCM3AP,POM121C,SYMPK,PPP1
SDCBP,PINK1,HSPA1A,MDM2,FZR1,SUMO1,PARK7,USP14,HIPK2,MAGEF1,TRIB3,PSME3,STUB1,
CCND2,CDH2,CCN2,MMP14,DDX3X,HMGCS1,GJA1,JUN,CTNNB1,INIP,CIRBP,PTGES3,RHOB,ATP
1,PAFAH1B1,MAP4,CDC42BPB,HNRNPU,CTNNA1,CNN3,TPM4,MSN,KPNB1,LLGL1,RHOA,CXADR,
B1,MAP4,HNRNPU,RPL24,KPNB1,LLGL1,RHOA,CBX5,PSMD2,JUND,RPL26,RBBP4,SPC25,NES,C
T1,SEC16A,PHB2,CRTAP,HDAC6,PTEN,CUL3,GRN,FLOT2,ATP1B2,CREBBP,BAG5,CD74,SMO,A
2,AMOTL2,CDC42,CTNND1,PSMA7,SULF2,BCL9L,PSMD1,AP2A1,CSNK2A1,UBA52,CSNK1G2,NX
1,CLIP3,STXBP1,PFN2,CDC42,DCTN1,EP300,HJURP,EVL,HDAC6,ULK1,CTTN,SLIT2,BRK1,AKAP9
BP7,EXTL3,FLCN,CSNK2A1,DIP2B,GNG4,SFRP1,CTTN,SLIT2,BAP1,OLFM1,LTBP4,HDGFL2,DNA
NIP3L,SMARCE1,SMARCC1,STMN3,UBQLN1,SREBF2,DDIT4,PPP1R9B,VDAC1,DCTN1,DFFB,WDR
2,AMOTL2,CDC42,CTNND1,PSMA7,SULF2,BCL9L,PSMD1,AP2A1,CSNK2A1,UBA52,CSNK1G2,NX

C7A,TADA2B,KHDRBS1,CSNK1E,EIF5A,CCND2,CCN2,MMP14,PRDX2,DDX3X,MIF,PRKAR1A,CTNN
CN2,APLP1,HMGCS1,PRKAR1A,GJA1,JUN,MYO5A,CTNNB1,SESN3,APPL1,HDAC2,IPO5,ATP6V0C
K7,USP14,TRIB3,PSME3,STUB1,MARCHF7,PSMC5,CSNK1A1,PSMC2,HERPUD1,PSMC4,PLK2,PSM
MEM30A,PGRMC1,CNN2,CTSB,RAB27A,CCT2,GDI2,PDXK,MLEC,SURF4,RAB5B,SRP14,PYGB,MAF
RN,FKBP4,LTBP4,NUDCD3,TRAP1,DNAJB2,AHSA1,FKBP9,PSMC1,CCDC47,BAG5,CD74,GAK,AMF
TNKS,SYMPK,SMG5,SRSF2,SRSF11,IGF2BP1,DHX38,YY1,EIF4A3,RUVBL2,RAN,CETN2,AKAP8L,NU
),RPL37,VHL,PSMD3,PRKDC,GABARAP,UBE2R2,CAND1,NPEPPS,KDM1A,FYN,TNKS2,UBQLN1,USI
A,TIA1,RBM25,FXR1,SRSF10,C1QBP,CELF3,RBFOX2,SRSF7,NUP98,RBM10,SETX,RRP1B,RBM17,
P,ARL6IP1,CIRBP,YWHAB,CADM4,CTBP1,HMGCR,CAPRIN1,HDAC2,IPO5,LAPTM4B,TRIB2,MAP1A,
MEM30A,PGRMC1,CNN2,CTSB,RAB27A,CCT2,GDI2,PDXK,MLEC,SURF4,RAB5B,SRP14,PYGB,MAF
C1,THRAP3,EIF4G1,PUM1,FXR2,PSMB1,YTHDF3,HSPA1A,PSMD13,FXR1,SYNCRIP,PABPC4,PSMD
5A2,SRSF4,SLC3A2,PTN,PIP4K2B,CCN2,APLP1,HMGCS1,PRKAR1A,GJA1,JUN,MYO5A,CTNNB1,S
V0C,TMEM30A,PGRMC1,CNN2,CTSB,RAB27A,CCT2,GDI2,PDXK,MLEC,SURF4,RAB5B,SRP14,PYG
MARS1,RAPGEF1,MAPK1,DYNC1LI2,ADAM9,PTBP1,GRB2,HSPB1,FKBP8,PLCG1,POLR2B,FYN,HN
U2AF2,SQSTM1,KLHL24,NOP53,UBE4A,RAD23B,ASXL1,PSMB2,RNF10,RPL37,VHL,PSMD3,PRKD
5,ZNF316,DBN1,RPL4,PRRC2B,ENO1,TUBA1B,PIK3R2,FLNA,MAP1B,NCL,RPL7A,SLC9A7,ALDOA,
MEM30A,PGRMC1,CNN2,CTSB,RAB27A,CCT2,VAMP2,GDI2,PDXK,MLEC,SURF4,RAB5B,SRP14,PY
ACD3,CHSY1,PPP2R1A,PDIA3,RPS3,CHD3,VAPA,HMGB1,TPT1,STMN1,ITGB1,UCHL1,RPL11,NCAM
FA,G3BP2,MAPK1,MAPRE1,ARPC2,DSTN,GRB2,KDM1A,YWHAH,CCT8,TNKS2,PLXNA3,TMED9,PF
BP4,SPC25,NES,CCNI,KHDRBS1,KLHL9,XPO1,CSNK1E,SFPQ,CCND2,CCN2,DDX3X,CLTC,IK,PRK
MEM30A,PGRMC1,CNN2,CTSB,RAB27A,CCT2,GDI2,PDXK,MLEC,SURF4,RAB5B,SRP14,PYGB,MAF
7,VHL,PSMD3,PRKDC,GABARAP,UBE2R2,CAND1,NPEPPS,KDM1A,FYN,TNKS2,UBQLN1,USP7,HEC
ND2,DDX3X,CLTC,IK,GJA1,CTNNB1,MYH10,CKAP5,RHOB,RPS27,CDK16,APPL1,SKP1,CDKN1A,C
SLC25A5,MAP1LC3B,VPS26B,RAB1B,VPS41,VPS4A,GABARAPL1,ATP6V0D1,CHMP4B,MTOR,PAC
RKAR1A,JUN,CTNNB1,TARDBP,DUX4,CKAP5,SND1,RHOB,CTBP1,CDK16,APPL1,IPO5,SKP1,CDKN
RCC5,KPNB1,LLGL1,NCS1,RHOA,PSMD2,KIF5B,NLGN2,WDR1,ATP2A2,VAT1,NISCH,PFKL,SCRN1
P5,RHOB,RPS27,CDK16,APPL1,SKP1,CDKN1A,CTDSP2,PPP1CB,NOP53,PSMB2,OPTN,PPP2CA,CC
BL1,SEMA6B,AUTS2,NOTCH1,ADNP,CYFIP1,RNF157,NTN1,LIMK1,ISLR2,GOLGA4,SEMA4C,SRF,

CA5,LARP1,NACA,RPS9,XRCC5,RHOA,ATF4,NR2F1,JUND,CD81,RPL26,CCT7,CLEC7A,TADA2B,KH
PPP2R1A,RPS3,CHD3,VAPA,HMGB1,TPT1,STMN1,ITGB1,UCHL1,RPL11,NCAM1,TIMP2,WSB1,DN
RUFY3,SPG11,SEMA6C,APOE,ANAPC2,SIN3A
1,EID1,CHD6,CCT2,PARP1,TSPYL4,MORF4L2,SSRP1,PKN1,YEATS2,TCP1,CCT3,RAD23B,CHD2,C
ACD3,SET,EGLN1,PPP2R1A,PKD1,SERINC1,RPS3,HMGB1,STMN1,ITGB1,UCHL1,RPL11,ANXA2,AG
P1,NACA,RPS9,XRCC5,RHOA,ATF4,NR2F1,JUND,CD81,RPL26,CCT7,CLEC7A,TADA2B,KHDRBS1,S
1,GSK3A,SRPRA,AMFR,TSPYL2,HSPA1A,MFN2,DNAJB1,PTPN1,FAF2,NCK2,STUB1,HERPUD1,HSF
S1,RAPGEF1,MAPK1,DYNC1LI2,ADAM9,PTBP1,GRB2,HSPB1,FKBP8,PLCG1,POLR2B,FYN,HNRNP
IP1,YWHAB,RAB1A,RAC1,TMEM30A,LAPTM4B,VAMP2,GNAI3,ABCA3,ACTR3,HSPA4,CNTNAP2,AB
IAP2,DPYSL2,MAPK1,PBX2,ATP5PB,LUZP6,PRKDC,KDM1A,QARS1,ARF4,ZFAND5,SDF4,FYN,YWH
1,CSE1L,CIZ1,TRIM8,SMO,RANGAP1,SUN1,RGPD5,PKIA,ATP13A2,MDM2,PARK7,RAN,SUN2,FAM
HSPA1A,MFN2,DNAJB1,PTPN1,FAF2,NCK2,STUB1,HERPUD1,HSF1,HSPE1,LMNA,ATF6B,SHC1,DN
PP19,PSMC3,PRKACA,PSMC1,TUBB4A,USP47,PSMB1,TAOK2,MACROH2A1,PKIA,PCM1,MECP2,C
L,PTEN,PPP1R15A,CTTN,SLIT2,BRK1,TMOD2,OLFM1,ABI2,AKT3,RDX,CAPZA1,SEMA3C,CLASP2,
YLS,SLC35E2B,HDAC2,MMP2,HSBP1,COL2A1,RAD23B,PTK7,SKI,KDM6B,TMED2,TOP1,RPL38,NC
MACROH2A1,PKIA,PCM1,MECP2,CUL1,PSMD13,ATF5,PSMD7,CETN2,CDK4,PSME3,PSMC5,PSMC

MK1,ISLR2,GOLGA4,SEMA4C,SRF,GSK3B,ALCAM,UNC13A,NRP2,RUFY3,SPG11,SEMA6C,APOE,S
TF4,NR2F1,JUND,CD81,RPL26,CCT7,TADA2B,KHDRBS1,SFPQ,EIF5A,SOX12,ACTN4,DDX3X,NANC
MA6B,AUTS2,ADNP,CYFIP1,RNF157,NTN1,LIMK1,ISLR2,GOLGA4,SEMA4C,SRF,GSK3B,MEX3C,A
9B,NOP53,PSMB2,CCND1,MAPRE1,PSMD3,CCAR2,CSNK1D,RAB11A,PRPF40A,DYNLL1,TP53BP1
EDC4,SGTA,FLCN,GTPBP1,UBQLN2,CSNK2A1,GRSF1,PHB2,ZFP36L1,UPF1,RAB3GAP2,PTEN,ULK
2,APPL1,TNC,ATP6V0C,CDKN1A,TIMP3,CTSB,SRSF5,VAMP2,PARP1,ABCA3,RPL27,SNRPN,PTPRA
HLA-A,PSMD7,KLC2,PSME3,PSMC5,PSMC2,PSMC4,PSMF1,KIF3A,PSMB5,PSMA1,PSMD10,CAPZ
HSPB1,KDM1A,FKBP8,QARS1,DUSP4,CORO1C,SUPT6H,FYN,FBLN1,STRAP,MAPK8IP1,KDM4B,DD
CM1,MECP2,CUL1,PSMD13,PSMD7,CETN2,CDK4,PSME3,PSMC5,PSMC2,PSMC4,PSMF1,RCC2,PS
,ATP6V0C,TMEM30A,PGRMC1,CNN2,CTSB,RAB27A,CCT2,VAMP2,GDI2,PDXK,MLEC,SURF4,RAB
PI,RACK1,GNB1,MYH9,P4HB,APP,KLRD1,TMBIM6,GSTP1,SMG1,NPM1,PLD3,HSPD1,COL18A1,V
PCAT1,XRCC5,KPNB1,LLGL1,NCS1,RHOA,PSMD2,KIF5B,NLGN2,ATP1A1,WDR1,ATP2A2,VAT1,NIS
MEM30A,PGRMC1,CNN2,CTSB,RAB27A,CCT2,VAMP2,GDI2,PDXK,MLEC,SURF4,RAB5B,SRP14,PY
,IGFBP2,CALR,FSTL1,TMF1,ABHD2,HSPA1B,COL4A1,CES3,PSAP,PGK1,RPLP0,H3-3B,CANX,CLU
1,ACIN1,RBM39
SMC3,PRKACA,PSMC1,TUBB4A,USP47,PSMB1,TAOK2,PKIA,PCM1,MECP2,CUL1,PSMD13,CCNY,
6,RPS27,RPL23A,GLUL,NOP53,TSR1,RPL27,RPL36A,RPL38,RPL37,PRKDC,RPL28,DHX30,RPS28
ND,COL5A2,WDR1,CLEC7A,PTN,CSNK1E,GLG1,CDH2,PLXNA1,MEGF8,SOX12,CCN2,ACTN4,MMF
6,PRKDC,KDM1A,QARS1,ARF4,SDF4,FYN,YWHAH,B2M,PITPNM1,PLXNA3,CSNK1D,PTPRS,BAG6,
HIP1R,HSPA1A,SUMO1,PARK7,ATP1B3,ATF7IP,WIZ,MARCHF7,P3H1,USP19,RASSF2,GET1,PHB,PIN
FOXK2,IGF2BP1,OGT,KDM4A,GSK3A,CHMP4B,MTOR,BAG5,THRAP3,EIF4G1,POLDIP2,SNX3,SDC
KIF26A,HLA-A,PSMD7,KLC2,PSME3,PSMC5,PSMC2,PSMC4,PSMF1,KIF3A,PSMB5,PSMA1,PSMD1
KDM1A,MSL1,SMARCE1,SMARCC1,SUPT6H,USP7,BAG6,KDM3B,WBP2,KDM4B,NAP1L4,RBBP7,F
A,AP2A1,SEC16A,RAB2A,RER1,YKT6,CUL3,PREPL,LMAN1,RABEP1,VAPB,ERGIC1,RAB1B,VPS41,
ARPC2,DSTN,LUZP6,GRB2,CAPZB,CLIP3,STMN3,PLXNA3,PFN2,CDC42,DCTN1,CAPN2,MAP6,PLXN
H2,VEGFA,PPP2CA,MAPK1,SMARCA4,FTO,LUZP6,PRKDC,CCAR2,SLIT3,TNKS2,PLXNA3,PTPRS,S
,NGRN,TARDBP,CIRBP,YWHAB,CAPRIN1,BCLAF1,DDX39B,SRP9,ZC3H7B,TAF15,ALKBH5,GCN1,T
,ABCA2,TMED2,RPL38,MAPK1,EIF3E,FTO,ILF3,LUZP6,PRKDC,HSPB1,CASC3,HNRNPD,STAT3,EIF
MEM30A,PGRMC1,CNN2,CTSB,RAB27A,CCT2,VAMP2,GDI2,PDXK,MLEC,SURF4,RAB5B,SRP14,PY
PB1,KDM1A,BNIP3L,QARS1,CCAR2,DIDO1,FYN,YWHAH,UBQLN1,HYOU1,MAPK8IP1,GHITM,SOD
,MDM2,PARK7,RAN,AKAP8L,NUP153,SRSF7,NUP98,POLDIP3,NCBP2,GSK3B,TXN,NUP205,NUP1
P4B,MTOR,BAG5,THRAP3,EIF4G1,POLDIP2,SDCBP,PINK1,ATP13A2,MGAT3,PARK7,USP14,SYNC
LXNA3,PTPRS,RAB11A,BMPR2,CDC42,NAP1L1,MAP6,PLXNB1,DIP2B,PTEN,PPP1CC,IL6ST,HIF1A,
),VANGL2,AMOTL2,CDC42,CTNND1,PSMA7,SULF2,BCL9L,PSMD1,AP2A1,CSNK2A1,UBA52,CSNK
NPM1,PLXNB2,HSPD1,CSDE1,IGF2,EIF3C,EIF4B,VCP,UBB,TRIM28,SPEN,NONO,DDB1,PSMD5,YW
,HSPA1A,MDM2,FZR1,SGSM3,NSF,SUMO1,SNX1,MAGEF1,AREL1,RNF41,TRIB3,STUB1,PSMC5,C
CAT1,XRCC5,KPNB1,LLGL1,NCS1,RHOA,PSMD2,KIF5B,NLGN2,WDR1,ATP2A2,VAT1,NISCH,PFKL,
L11,ANXA2,AGRN,RPLP1,TIMP2,MIDN,SBF1,DNAJA1,PAFAH1B1,HNRNPU,RPL23,CTSD,SCARB2,X
2B,XPO1,CSNK1E,SCRN1,GLG1,PEBP1,GPC3,CCN2,MMP14,DDX3X,RPS7,CTNNB1,CYFIP2,APLP
,CTTN,SLIT2,BRK1,AKAP9,TMOD2,TAOK1,GPX1,FKBP4,ABI2,LDLR,RDX,CAPZA1,CLASP2,WASF

RPL38,RPL37,PRKDC,RPL28,RPS28,RPL39,RPL34,UBA52,RPL7L1,RPS2,PELP1,WBP11,SART1,E
MGCR,ARHGDIA,HDAC2,PEA15,PTTG1IP,CDKN1A,ANXA5,SQSTM1,GNAI3,PRDX5,GRINA,COL2A1
2,DDX3X,MIF,CTNNB1,DNAJC5,ARL6IP1,HNRNPK,HMGCR,ARHGDIA,HDAC2,PEA15,PTTG1IP,CDK
PL36A,RPL38,RPL37,PRKDC,RPL28,RPS28,RPL39,RPL34,UBA52,RPL7L1,RPS2,TRIR,PELP1,WBF

ETN2,AKAP8L,NUP153,SRSF7,NUP98,POLDIP3,NCBP2,HSF1,IGF2BP2,NUP205,NUP133,THOC2,T
ETN2,AKAP8L,NUP153,SRSF7,NUP98,POLDIP3,NCBP2,HSF1,IGF2BP2,NUP205,NUP133,THOC2,T
311A,DYNLL1,TP53BP1,PPP1R9B,PSMA7,DCTN1,PRMT1,ACTR1A,EP300,PSMD1,PBX1,PHB2,GSP
TO,ILF3,LUZP6,PRKDC,HSPB1,CASC3,HNRNPD,STAT3,EIF2AK1,EDC4,EIF3D,GSPT1,ZFP36L1,UP
CTN1,LRRC4B,FLOT1,SFRP1,EVL,RAB3GAP2,ULK1,SEPTIN7,CTTN,BRK1,AKAP9,PALM,ABI2,VPS
NPM1,PLXNB2,HSPD1,CSDE1,IGF2,EIF3C,EIF4B,UBB,TRIM28,SPEN,NONO,PSMD5,YWHAQ,EIF1,
,GPC1,APPL1,ATP6V0C,MMP2,DCN,PARP1,PTPRA,ACTR3,NCKAP1,SKI,ATP6AP1,NOTCH2,VEGFA
1,PSMC5,NDRG1,PSMC2,PSMC4,PSMF1,DDAH1,USP19,STC2,PSMB5,EGLN2,HIF3A,LMNA,PSMA
,CRABP2,EFNB3,CNTN2,RGMA,RUFY3,SEMA6C,EPHB2,FGF13,EPHA4,PLXND1,ANAPC2,SKIL
RNPK,HMGCR,ARHGDIA,HDAC2,PEA15,PTTG1IP,CDKN1A,ANXA5,SQSTM1,GNAI3,PRDX5,GRINA,

RPF19,MTOR,SMO,METRN,NTRK2,ABL1,TRAK1,DAG1,NIBAN2,PLXNA2,OBSL1,NOTCH1,NPTN,AD
STUB1,PSMC5,NDRG1,PSMC2,PSMC4,PSMF1,DDAH1,USP19,STC2,PSMB5,EGLN2,HIF3A,LMNA,
LXNB1,FARP1,CTNND2,RER1,HDAC6,PTEN,PTPRF,CTTN,GPC4,SLIT1,PALM,ABI2,WASF2,ADAM10
SEMA3C,DICER1,IST1,PRPF19,KDM4A,MTOR,CERS2,SMO,METRN,NTRK2,SEMA6B,TRAK1,DAG1
IOTCH1,NPTN,ADNP,CYFIP1,NTN1,LIMK1,RHEB,ISLR2,DHX36,GOLGA4,SRF,SERPINE2,PRMT5,C
LSR3,PSMB5,PSMA1,MED12,PSMD10,PSMA6,PSMD12,PSMA4,PSMC6,FZD7
TF6,ADD1,QRICH1,XBP1,CUL7,PPP2R5B,SEC31A,EIF2S1,CDK5RAP3,ABCA7
,NONO,PSMD5,YWHAQ,SET,EGLN1,PPP2R1A,PKD1,SSBP2,ZNF43,RPS3,CHD3,BASP1,HMGB1,M
3B4A,ABL1,PINK1,ARPC1A,HIP1R,HSPA1A,ELN,CRACD,MECP2,CYFIP1,FLII,TRIM27,NCK2,ARPC3
PLXNB2,HSPD1,CSDE1,IGF2,EIF3C,EIF4B,VCP,UBB,TRIM28,SPEN,NONO,DDB1,PSMD5,YWHAQ,E

NA,ATF6B,SHC1,DNAJB11,ATF6,ADD1,QRICH1,XBP1,CUL7,PPP2R5B,SEC31A,EIF2S1,RNF5,CDK5
ETN2,AKAP8L,NUP153,SRSF7,NUP98,POLDIP3,NCBP2,HSF1,IGF2BP2,NUP205,NUP133,THOC2,T
11,MDM2,PSMD13,PSMD7,HIPK2,PSME3,STUB1,PSMC5,NDRG1,PSMC2,PSMC4,PSMF1,DDAH1,U

PLXNB2,HSPD1,CSDE1,IGF2,EIF3C,EIF4B,UBB,TRIM28,SPEN,NONO,PSMD5,YWHAQ,EIF1,SET,EG
DCTN1,PRMT1,ACTR1A,EP300,PSMD1,PBX1,PHB2,GSPT1,BRD4,UBA52,ZFP36L1,MASTL,PTEN,C

PO1,SFPQ,NANOG,H1-0,GJA1,JUN,CTNNB1,TARDBP,NDFIP1,MXD4,HNRNPK,H1-10,CIRBP,YWHA
33,STUB1,CSNK1A1,HERPUD1,PLK2,WAC,GSK3B,UBQLN4,UBXN1,PSMD10,ELOB,XBP1,RYBP,R
4KA,ITGB5,POM121C,RPS27A,GRK2,VAPB,RAB1B,VPS4A,LDLR,NUCKS1,ITGAV,IST1,CHMP4B,T
TPQ,SOX12,ACTN4,DDX3X,NANOG,H1-0,JUN,CTNNB1,TARDBP,ARID1A,SHOX,HNRNPK,H1-10,CIR
SMA6,PSMD12,PSMA4,PSMC6,FZD7
SMA6,PSMD12,PSMA4,PSMC6,FZD7
3B,CLCN6,CRABP2,SLC12A7,RGMA,RUFY3,ADD1,SLC12A4,CLCN3,VAV2,SEMA6C,APOE,FGF13,
HA4,CDK5RAP3,CTSA
HLA-A,PSMD7,KLC2,PSME3,PSMC5,PSMC2,PSMC4,PSMF1,KIF3A,PSMB5,PSMA1,PSMD10,CAPZ
UBXN1,PSMD10,ELOB,XBP1,RYBP,RAD23A
,RDX,TM9SF4,NMT1,PDPK1,SYT11,AKT2,GSK3A,UBE2D3,TRIM8,SAE1,SMO,CROCC,UBE2L3,PI
,TUBGCP6,CLASP2,WASF2,CHMP4B,MTOR,TUBB4A,UBE2K,ABL1,PINK1,ARPC1A,HDGFL3,HIP1
RPS27A,GRK2,VAPB,GPX1,VPS4A,LDLR,NUCKS1,ITGAV,SCRIB,IST1,PSMC3,CHMP4B,TRIM8,CD
P,DAPK3,MYL12B,ABL1,FAM171A1,TAOK2,DAG1,PLXNA2,OBSL1,CYFIP1,CD44,RNF157,CFDP1,
YFIP1,NTN1,IQSEC1,RAP2A,NCK2,SYNGAP1,DHX36,CRKL,RBFOX2,PLK2,GSK3B,ACSL4,CRTC1,
1,UPF1,RAB3GAP2,PTEN,ULK1,GPSM1,HIF1A,SLC25A5,LDLR,RDX,TTC37,DNAJB2,ATXN3,AKT2,

:N,RPS27A,PSMD8,HIF1A,SLC2A1,PSMB4,TM9SF4,SOD2,PSMD4,ATP6V0D1,RBPJ,AHCY,PSMC3,[illegible]
NA1,RPL23,LDHA,XRCC5,MSN,COL1A2,RHOA,ATF4,KIF5B,ATP1A1,JUND,COL5A2,ATP2A2,SRSF4
MD1,PBX1,PHB2,BRD4,UBA52,ZFP36L1,PTEN,CUL3,BRSK1,RPS27A,PSMD8,APEX1,PCBP4,AKAP[illegible]
R,COTL1,HRNR,RAC1,ATP6V0C,TMEM30A,PGRMC1,CNN2,CTSB,RAB27A,CCT2,VAMP2,GDI2,PD[illegible]
3,PSMC1,UBE2K,FBXL19,AMFR,TRIP12,UBE2L3,CDC34,PSMB1,MDM2,FZR1,UBE4B,TRIM2,CUL
/I1,ZMIZ1,VCL,AP3D1,DNAJA1,PAFAH1B1,AKAP12,MAP4,CDC42BPB,HNRNPU,CTNNA1,TPM4,RT[illegible]
CDC42BPB,HNRNPU,PGAM1,LDHA,XRCC5,IGFBP3,HK1,RHOA,ATF4,CD81,CLEC7A,CCNI,CSNK1E

2R1A,PKD1,SSBP2,ZNF43,RPS3,CHD3,BASP1,HMGB1,MORF4L1,AGRN,ZMIZ1,RPL10,AP3D1,ZNF
NRDC,VEGFA,ADAM9,PCSK1N,GABARAP,CCAR2,SMARCC1,FYN,FBLN1,UBQLN1,CSNK1D,USP7,[illegible]
ED12,PSMD10,PPP3CA,PSMA6,PSMD12,PSMA4,PSMC6,GNAO1,FZD7
F2BP2,NUP205,NUP133,THOC2,THOC3,FYTTD1,NUP50,RNPS1,DDX19A,NUP58,WDR33,NUTF2
/0D1,SH3BP4,KDM4A,GSK3A,CHMP4B,MTOR,PRKACA,EXOC4,BNIP3,ATP6V1G1,TRIM8,DAPK3,A[illegible]
LGN2,RPS20,CD81,RPL26,MCL1,CLEC7A,PTN,CSNK1E,SFPQ,CDH2,GPC3,SULF1,CCN2,ACTN4,[illegible]

,FADS1,NEFL,PRPF8,YBX1,UBA1,ATP2B4,IGFBP2,CALR,FSTL1,TMF1,ABHD2,HSPA1B,TUBB2B,C
)A,ATF4,CD81,CLEC7A,CCNI,PTN,CCND2,DDR1,PIP4K2B,CCN2,PRDX2,DDX3X,MIF,PRKAR1A,TAF
)A,ATF4,CD81,CLEC7A,CCNI,PTN,CCND2,DDR1,PIP4K2B,CCN2,PRDX2,DDX3X,MIF,PRKAR1A,TAF
NUP50,RNPS1,DDX19A,NUP58,WDR33
3,ABL1,PINK1,HSPA1A,ARRDC4,FZR1,PARK7,SART3,LIMK1,MYCBP2,TRIB3,CDK9,STUB1,MARC[illegible]
,METRN,NTRK2,TRAK1,DAG1,PLXNA2,OBSL1,NOTCH1,NPTN,ADNP,CYFIP1,NTN1,LIMK1,RHEB,I[illegible]
,CUL3,BRSK1,RPS27A,PSMD8,APEX1,PCBP4,AKAP9,PSMB4,TAOK1,CARM1,INTS3,VPS4A,RDX,K[illegible]
GAS7,NOX4,ACTR3,NCKAP1,MDK,PTK7,NOTCH2,ARPC2,DSTN,LUZP6,GRB2,CAPZB,ACTN1,FLNE
,NISCH,PTN,CSNK1E,PFKL,CDH2,GPC3,SULF1,DDX3X,CLTC,PRKAR1A,GJA1,CTNNB1,MACF1,D[illegible]
/I1,DZIP1,DNAJB2,NUCKS1,EDF1,PRKACA,SMO,EIF4A3,EIF4G1,AMFR,ABL1,RSF1,NIBAN2,HIP1R
,PRDX2,DDX3X,APLP1,MIF,GJA1,CTNNB1,SESN3,YWHAB,CADM4,HMGCR,GPC1,APPL1,PEA15,P[illegible]
3IM6,GSTP1,SMG1,CALM1,NPM1,HSPD1,IGF2,VCP,UBB,TRIM28,MATR3,YWHAG,MAPT,FN1,NON
HOX,HNRNPK,H1-10,CIRBP,DUX4,HDAC2,BCLAF1,DCN,DDX39B,SRSF5,CHD6,RNF187,U2AF2,ML
G,H1-0,JUN,CTNNB1,TARDBP,MXD4,HNRNPK,H1-10,CIRBP,YWHAB,CTBP1,SMCHD1,HDAC2,BCL
IIF,CTNNB1,SESN3,YWHAB,CADM4,HMGCR,GPC1,APPL1,PEA15,PTTG1IP,DCN,DLG5,TIMP3,DDX
DX,SH3GL1,CAPZA1,CLASP2,IST1,SMARCD1,BNIP3,MICAL1,GAK,HMGA1,CHMP1A,HDGFL3,CRA
,PRDX2,DDX3X,APLP1,MIF,GJA1,CTNNB1,SESN3,YWHAB,CADM4,HMGCR,GPC1,APPL1,PEA15,P[illegible]
RPS20,CD81,RPL26,MCL1,CLEC7A,PTN,CSNK1E,SFPQ,CDH2,GPC3,SULF1,CCN2,ACTN4,PRDX2,[illegible]
NPS1,DDX19A,NUP58,WDR33
BB2,ROCK2,MAPK3,PAIP1,YBX3,SP1,RELA,EPHA4,CTIF,MRPS27,SLC35A4
,PBX1,PHB2,BRD4,UBA52,ZFP36L1,PTEN,FEM1B,CUL3,BRSK1,RPS27A,PSMD8,APEX1,PCBP4,AK
GT,PCGF2,NOC2L,KDM4A,MBD3,CREBBP,APBB1,ATXN7L3,PAGR1,FBXL19,TRIP12,PINK1,AUTS
MEM9,PSMF1,LGR4,PSMB5,PSMA1,PSMD10,PSMA6,CTDNEP1,PIN1,SPIN1,PSMD12,PSMA4,PSMC
K5RAP3,ABCA7
I1,BAP1,TCAF1,UBR5,RDX,TM9SF4,NMT1,REEP2,AKT2,EHD1,GSK3A,PRKACA,UBE2D3,SAE1,SM[illegible]
FBP3,RHOA,CHD4,RPS20,CD81,RPL26,RBBP4,MCL1,CLEC7A,SFPQ,PEBP1,CDH2,PIP4K2B,CCN2,
YL6,MYH10,RHOB,ATP1A2,COTL1,RAC1,ARHGDIA,CNN2,GAS7,NOX4,ACTR3,NCKAP1,MDK,PTK[illegible]
5,SOX2,TLE1,PSMD4,RBPJ,GSK3A,PSMC3,PSMC1,WNK1,DAPK3,AMFR,IGFBP4,USP47,SNX3,ABL
TP5F1A,VCP,TRIM28,YWHAG,MAPT,FN1,YWHAQ,TMSB10,HMGB1,AP2M1,STMN1,ITGB1,ANXA2,[illegible]
F2,NOC2L,KDM4A,MBD3,CREBBP,APBB1,ATXN7L3,PAGR1,FBXL19,TRIP12,PINK1,AUTS2,MTA2,[illegible]

:ROH2A2,PSMD4,SPTY2D1,PSMC3,NASP,SMARCD1,PSMC1,HMGA1,RSF1,PSMB1,TSPYL2,MACR
,SYP,DCTN1,SGTA,ALDH3A2,FLOT1,SEC16A,RAB2A,RAB3GAP2,PTEN,LMAN1,NPLOC4,SERP1,AK
MGCS1,GJA1,JUN,CTNNB1,ARID1A,PTGES3,ATP1A2,HMGCR,ARHGDIA,HDAC2,TNC,PEA15,CDKN
)NEP1,PIN1,PSMD12,PSMA4,PSMC6,PPM1A,GPRC5B,JRK
P2B,PTEN,PPP1CC,IL6ST,HIF1A,SLIT2,SLIT1,LINGO1,LDLR,SEMA3C,DICER1,IST1,PRPF19,KDM4
,PSMD4,RBPJ,AHCY,PSMC3,MTOR,CREBBP,BNIP3,PSMC1,SCAP,PINK1,PSMB1,NOTCH1,MDM2,
/19SF4,SOD2,PSMD4,RBPJ,AHCY,PSMC3,MTOR,CREBBP,BNIP3,PSMC1,SCAP,PINK1,PSMB1,NOT

NPS1,DDX19A,NUP58,WDR33
G1,HSPA1A,EFNB2,HIPK2,NECTIN1,TRIM27,NUP153,KPNA6,EXOC7,TFRC,THOC2,RXRA,PHB,EFN
'4,HLA-B,TTR,COTL1,HRNR,RAC1,ATP6V0C,TMEM30A,PGRMC1,CNN2,CTSB,RAB27A,CCT2,VAM
N1,ARNT2,BMPR2,SULF2,PRMT1,CAPN2,EP300,PBX1,FLCN,ZFP36L1,BRD2,HOXB5,NXN,SFRP1,
,PSMD5,YWHAQ,DCHS1,CHSY1,INA,RPL19,PKD1,BASP1,HMGB1,CDKL1,AP2M1,SEC61A1,STMN1
,ACTR3,AHCYL1,RAB11B,ATP6AP1,DPYSL2,TMED2,MAPRE1,KIF3C,STXBP1,CSNK1D,TMED9,RAE
IT2,BMPR2,SULF2,PRMT1,CAPN2,EP300,PBX1,FLCN,ZFP36L1,BRD2,HOXB5,NXN,SFRP1,CUL3,N

X3,PFN1,CLEC7A,TPM1,CCND2,DDR1,CCN2,ACTN4,PRDX2,DDX3X,H1-0,PRKAR1A,JUN,CTNNB1,I

,CLTA,PSMB1,PSMD13,KIF26A,HLA-A,PSMD7,KLC2,PSME3,PSMC5,PSMC2,PSMC4,PSMF1,KIF3A
:2,PTBP1,GRB2,CDC37,HSPB1,PLCG1,ARF4,ZFAND5,POLR2B,SMARCC1,ATP6V1B2,FYN,HNRNP

MCL1,CLEC7A,PTN,SFPQ,RAB7A,CCN2,PRDX2,RPS19,MIF,PRKAR1A,GJA1,CTNNB1,SESN3,NDFI
XNA2,NOTCH1,NPTN,EFNB2,CYFIP1,FEZ1,NTN1,EFNB1,MYCBP2,NECTIN1,RANBP9,SEMA4C,ZSW
A,TRIM27,NCK2,ARPC3,STUB1,PSMC5,PSMC2,PDE4DIP,PSMC4,GSK3B,TFRC,MED25,WARS1,CD
TR,COTL1,HRNR,RAC1,ATP6V0C,TMEM30A,PGRMC1,CNN2,CTSB,RAB27A,CCT2,VAMP2,GDI2,P
SK3B,PSMD10,ELOB,XBP1,APOE,PSMC6,RAD23A,RNF40
1A,PKD1,SERINC1,RPS3,HMGB1,CDKL1,UCHL1,ANXA2,RPLP1,MIDN,SBF1,DNAJA1,ARF1,PAFAH1
C7A,CSNK1E,SFPQ,CDH2,GPC3,SULF1,CCN2,ACTN4,PRDX2,DDX3X,RPS15,ITM2C,RPS7,MIF,GJ
S7,PRKAR1A,CTNNB1,YWHAB,PTGES3,RAB1A,HMGCR,IPO5,SKP1,TRIB2,PEA15,CDKN1A,EEF1A
1,SELENON,RAPGEF1,MAPK1,GRB2,PRKDC,KDM1A,NSG2,GNG2,SMARCC1,ATP6V1B2,FYN,LAR
2,PRDX2,DDX3X,MIF,PRKAR1A,TARDBP,YWHAB,PTGES3,CADM4,HMGCR,RAC1,HDAC2,IPO5,TRI

/IB1,RUVBL2,NOTCH1,PSMD13,CCNY,PSMD7,PPP1CA,GID8,PSME3,PSMC5,CSNK1A1,PSMC2,PS
TCH1,NPTN,EFNB2,CYFIP1,FEZ1,NTN1,EFNB1,MYCBP2,NECTIN1,RANBP9,SEMA4C,ZSWIM6,ALC
!,APOE,FGF13,ARF6,EPHA4

1E,TPM1,PIP4K2B,RAB7A,DDX3X,RPS15,RPS5,RPS19,PRKAR1A,RPS10,CIRBP,CKAP5,RAB1A,RPS
KIFAP3,SRSF2,SRSF11,SCRIB,ADAM10,DHX38,NSG1,EIF4A3,TUB,DAG1,PCM1,NPTN,RAN,DYNLR
!,RAB11A,BMPR2,DIP2B,EVL,ADCY6,PTEN,PPP1R15A,CTTN,SLIT2,BRK1,TMOD2,GPX1,OLFM1,A

P1,PTEN,FEM1B,HIF1A,CTTN,SLC25A5,GPX1,MAZ,TRAP1,SOD2,ITGAV,PCGF2,NOC2L,NDUFA13,
)P1,TAF7,VEGFA,PPP2CA,MAPK1,PRKDC,KDM1A,CLIP3,FKBP8,MAP2K2,MSL1,SUPT6H,FYN,NKT
P1,FLCN,LRRC4B,FLOT1,PLXNB1,FARP1,SFRP1,CTNND2,PTEN,CTTN,GPC4,SLIT1,CAPZA1,CLAS

35,ATP6AP1,MARS1,SELENON,RAPGEF1,MAPK1,GRB2,PRKDC,KDM1A,NSG2,GNG2,SMARCC1,

3-3B,CANX,CLU,DDX5,SOX4,FKBP14,MAGED1,KAT6A,DVL3,YWHAE,GNAS,DDX17,GPI,HTRA4,RA
R1,PFN1,ASTN2,CLEC7A,NISCH,SLC3A2,PTN,GLG1,TPM1,CDH2,GPC3,DDR1,SULF1,PLXNA1,ME
ANGL2,TCTN1,CDC42,PSMA7,KDM5B,PSMD1,PBX1,AP2A1,GRSF1,PHB2,BRD2,SFRP1,PTEN,FEM
B,TMED2,NOTCH2,VEGFA,PSMD3,GRB2,HECTD1,VANGL2,TCTN1,BMPR2,CDC42,PSMA7,KDM5
1A,PKD1,SERINC1,RPS3,HMGB1,CDKL1,UCHL1,ANXA2,RPLP1,MIDN,SBF1,DNAJA1,ARF1,PAFAH1
C35A4
,ATF4,NR2F1,ENC1,RPS14,ATP1A1,TLE5,HNRNPC,KHDRBS1,XPO1,SFPQ,DDX3X,ITM2C,CLTC,N

B1,C6ORF89,RNF40,SIN3A
,NTRK2,PTPN11,ABL1,SUN1,DAG1,NOTCH1,IDH2,B4GALT5,SUN2,RHEB,TMEM98,CRKL,NDRG1,
BMERB1,CHMP5,GIT1,FGF13,TPX2,GNAI1
N4,LAMA5,MSN,RPL24,IGFBP3,COL1A2,GAP43,RHOA,KIF5B,SLC7A5,CXADR,CD81,WDR1,CRM
R,MAP1S,SETD5,CTR9,OTUB1,RNF20,KDM5A,LMNA,TLK1,CTCF,WDR5,MAPK3,XBP1,SETD1A,P
4,USP47,PSMB1,RUVBL2,NOTCH1,PSMD13,CCNY,PSMD7,PPP1CA,GID8,PSME3,BCL9,PSMC5,C

SPYL2,MACROH2A1,PSMD13,SART3,PSMD7,CUL4A,ATF7IP,CETN2,H2AX,PSME3,SENP6,PSMC5,
3-3B,CANX,CLU,DDX5,SOX4,FKBP14,MAGED1,KAT6A,DVL3,YWHAE,GNAS,DDX17,GPI,HTRA4,RA
TNNB1,SESN3,RHOB,HMGCR,RAC1,CDK16,APPL1,HDAC2,IPO5,TNC,ATP6V0C,MMP2,SRSF5,VA
E5,HNRNPC,KHDRBS1,XPO1,SFPQ,DDX3X,ITM2C,CLTC,NANOG,H1-0,GJA1,JUN,CTNNB1,TARDB
MA4C,THOC2,EFNB3,RGMA,PPP3CA,RUFY3,SEMA6C,EPHB2,APOE,FGF13,ARF6,PRNP,EPHA4
AG5,CD74,UBE2K,USP47,ABL1,PINK1,HSPA1A,MDM2,CUL1,PARK7,CD44,ARL6IP5,ERP29,PTPN1
QLN4,ATP6V0E2,ATP6V0A1,MAPK3,SCOC,SESN2,QSOX1,ATP6V1D,RB1CC1,USP36
PSF1,NOP53,TSR1,RPL27,RPL36A,RPL38,MARS1,RPL37,PRKDC,QARS1,RPL28,RPS28,SSB,LAR
1,HSPA1A,PFKP,NDUFA4,PARK7,NDUFC2,COX8A,COX7A2L,NUP153,NUP98,ATP5PO,CLPX,DLD,N
PHB2,APOE,LRP8,ARF6,PRNP,SRCIN1,EPHA4
1E,CDH2,PLXNA1,MEGF8,SOX12,CCN2,MMP14,GJA1,JUN,CTNNB1,MACF1,RHOB,HMGCR,RAC
PS,BNIP3L,FYN,YWHAH,CCT8,PFKM,SREBF2,RAB11A,CYP51A1,DYNLL1,CCT5,CDC42,GRIPAP1,
D81,RPL26,CLEC7A,HNRNPC,KHDRBS1,EIF5A,SULF1,CCN2,MMP14,DDX3X,H1-0,RPS7,MIF,GJA1
ADM,GSN
SMD12,PSMA4,PSMC6,SIN3A
MK1,MYCBP2,TRIB3,CDK9,STUB1,MARCHF7,HERPUD1,CTR9,OTUB1,RNF20,UBXN1,PSMD10,T

RUFY3,PIN1,TGFB2,SYT1,KDM2B,BRAT1,SYT4,CACNG7,ANAPC2

BP1,DHX38,EIF4A3,SLC25A36,SLC25A6,RAN,CETN2,AKAP8L,NUP153,SRSF7,NUP98,POLDIP3,N
G2,SMARCC1,ATP6V1B2,FYN,YWHAH,SLIT3,NCOR2,WBP2,IGFBP7,DDIT4,STAT3,EP300,LBH,PRK
RK7,CD44,PTPN1,PSME3,MARCHF7,HERPUD1,TNFRSF10B,SCG2,GSK3B,LMNA,VDAC2,CFLAR,
3,ADCY6,PTEN,APEX1,SLC2A1,NUCKS1,PDPK1,AKT2,OGT,ATP6V0D1,GSK3A,MTOR,PRKACA,GO

NPC,KHDRBS1,XPO1,SFPQ,DDX3X,ITM2C,NANOG,H1-0,GJA1,JUN,CTNNB1,TARDBP,MXD4,H1-1
A5,MSN,COL1A2,RHOA,ATF4,PSMD2,COL5A2,WDR1,ATP2A2,PFN1,ASTN2,PTN,GLG1,TPM1,CDH

IST1,OGT,GSK3A,PSMC3,NDUFA13,PSMC1,ZER1,USP5,PINK1,HIP1R,HSPA1A,MDM2,FZR1,SUMC
E4DIP,CDKN1B,WASF3,CDC42EP4,ARPC5,ROCK2,TGFBR1,FER,GMFB,GIT1,APOE,PDXP,ARF6,NF
NOTCH2,VEGFA,MAPK1,PLEC,CAPZB,HSPB1,KDM1A,ARF4,GNG2,FYN,FBLN1,STXBP1,PTPRS,SO

AT2A,PXDN,HNRNPU,SMARCA5,RPS16,LAMA5,XRCC5,MSN,COL1A2,RHOA,PSMD2,KIF5B,MCL1,T
C2,BCLAF1,DCN,DDX39B,CHD6,RNF187,MLXIP,SQSTM1,PARP1,MORF4L2,TAF15,ACTR3,PKN1,I
C2,BCLAF1,DCN,DDX39B,CHD6,RNF187,MLXIP,SQSTM1,PARP1,MORF4L2,TAF15,ACTR3,PKN1,I
DAC2,L1CAM,DDX39B,PARP1,MORF4L2,MDK,ATP6AP1,NOTCH2,VEGFA,VHL,ARPC2,PRKDC,KDN
C2,BCLAF1,DCN,DDX39B,CHD6,RNF187,MLXIP,SQSTM1,PARP1,MORF4L2,TAF15,ACTR3,PKN1,I
1,HDAC2,TNC,MMP2,DCN,DLG5,COL2A1,MDK,LAMC1,PTK7,SKI,ASXL1,PSMB2,TMED2,RPL38,N
SERP1,BAP1,TCAF1,UBR5,TM9SF4,NMT1,AKT2,GSK3A,PRKACA,APBB1,UBE2D3,SAE1,SMO,PTPN
P,SART3,KAT2A,PSMD7,COPS6,ADRM1,OTUD5,PSME3,SENP6,PSMC5,PSMC2,PSMC4,PSMF1,SHI

,PDPK1,AKT2,OGT,ATP6V0D1,GSK3A,MTOR,PRKACA,GOT2,ATP6V1G1,PHIP,SORT1,USO1,RABG

A,SETD5,THBS2,NRP2,ARHGAP33,LRRN1,ADGRL1,OPA1,EPHB2,APOE,LRP8,ARF6,PRNP,SRCIN1,

OX4,PKN1,TERF2IP,VEGFA,RAPGEF1,PPP2CA,CCND1,MAPK1,ADAM9,PTBP1,GABARAP,KDM1A,C
DDIT4,PPP1R9B,ADAR,STAT3,PRKAR2A,FLCN,FARP1,SFRP1,MASTL,PTEN,PPP1R15A,FKBP1A,SLI

CR,RAC1,HDAC2,IPO5,TRIB2,PEA15,CDKN1A,CTDSP2,SQSTM1,NOP53,NOX4,PKN1,GCN1,TERF2

MNA,WDR5,MAPK3,XBP1,GNL3,PAF1,DNMT1,SMARCB1,RAD50,C6ORF89,RNF40,SIN3A
PACS1,CLASP2,PDPK1,SCRIB,ADAM10,AKT2,EHD1,PACS2,GAK,NSG1,ANK2,TUB,DAG1,EXOC5,NS
CCN2,PRDX2,DDX3X,MIF,PRKAR1A,TARDBP,YWHAB,CADM4,CTBP1,HMGCR,RAC1,CDK16,HDAC
P9,SEPTIN5,SH3GL1,NCSTN,SYT11,MTOR,PRKACA,NSG1,EIF4A3,NTRK2,RIMS3,ABL1,PINK1,PAIF
B2B,COL5A1,COL4A1,CES3,PSAP,PGK1,H3-3B,CLU,DDX5,SOX4,MAGED1,KAT6A,DVL3,YWHAE,K
HBS2,NRP2,ARHGAP33,LRRN1,ADGRL1,OPA1,EPHB2,APOE,LRP8,ARF6,PRNP,SRCIN1,EPHA4,ANA
P9,SEPTIN5,SH3GL1,NCSTN,SYT11,MTOR,PRKACA,NSG1,EIF4A3,NTRK2,RIMS3,ABL1,PINK1,PAIF
2B,PRDX2,DDX3X,RPS15,RPS7,PRKAR1A,JUN,APLP2,ARL6IP1,NDFIP1,YWHAB,ATP1A2,MAP2,HM
DDIT4,PPP1R9B,ADAR,STAT3,PRKAR2A,FLCN,FARP1,SFRP1,MASTL,PTEN,PPP1R15A,FKBP1A,SLI
A1,APOE,LRP8,DDX56,ATP8A2
PINK1,PKIA,ATP13A2,MDM2,PCM1,PARK7,PTPN1,RAN,UBE2J1,YWHAZ,GSK3B,KIF3A,TXN,ERBB2
,HIF1A,PRDX6,GPX1,STK24,TRAP1,SOD2,PCGF2,FBLN5,UBE3A,BNIP3,BAG5,PRDX3,CYB5B,PTPN
A13,CELSR1,ABL1,SDCBP,AUTS2,PHACTR4,RGL2,NOTCH1,MFN2,SGSM3,TNFAIP1,PARK7,CYFIP1
RPS14,ATP1A1,TLE5,HNRNPC,KHDRBS1,XPO1,SFPQ,DDX3X,ITM2C,NANOG,H1-0,GJA1,JUN,CTNI

P,SART3,KAT2A,PSMD7,ADRM1,OTUD5,PSME3,PSMC5,PSMC2,PSMC4,PSMF1,SHMT2,USP48,USI
D1,PACS2,GAK,NSG1,ANK2,EXOC5,NSF,ARL6IP5,ATP1B3,TSPAN14,GOLPH3,CCDC88A,RAP2A,GC
C2,XBP1,YBX3,OPA1,SKIL
PHA4,ANAPC2
A5,RPS16,LAMA5,XRCC5,MSN,COL1A2,RHOA,PSMD2,KIF5B,MCL1,ACTN4,NANOG,MIF,HNRNPDL

,SMO,RABGAP1,ABL1,SEMA6B,NOTCH1,ADNP,CYFIP1,RNF157,NTN1,LIMK1,CDK4,ISLR2,GOLG[illegible]

[illegible],PPP2CB,SFRP1,PTEN,CUL3,NPLOC4,HIF1A,SLIT2,PPP1R10,GPX1,SEC14L1,TLE1,TRAP1,SOD2,[illegible]
[illegible]PL27,RPL36A,RPL38,RPL37,PRKDC,RPL28,RPS28,SSB,SARS1,RPL39,RPL34,ADAR,STAT3,GRSF[illegible]
[illegible]2,RHOA,SLC7A5,CXADR,CD81,WDR1,PFN1,ASTN2,CLEC7A,NISCH,SLC3A2,PTN,GLG1,TPM1,CD[illegible]
[illegible]2,RHOA,SLC7A5,CXADR,CD81,WDR1,PFN1,ASTN2,CLEC7A,NISCH,SLC3A2,PTN,GLG1,TPM1,CD[illegible]

[illegible]AP1,NOTCH2,VEGFA,RAPGEF1,PPP2CA,MAPK1,PSMD3,PRDX1,GRB2,MAP2K2,QARS1,DUSP4,FY[illegible]

SMC2,SEMA4C,PSMC4,PSMF1,SRF,SOX9,PSMB5,PSMA1,CITED2,NRP2,PSMD10,MAPK3,PSMA6,[illegible]

NT2,ITGB5,LOX,A2M,CCN1,HSPG2,ANTXR1,LTBP4,RIC8A,SH3PXD2A,CLASP2,LTBP3,ITGAV,ADA[illegible]
ED12,CITED2,PSMD10,PSMA6,PSMD12,PSMA4,PSMC6,FZD7
PSMC4,PSMF1,PSMB5,PSMA1,SOS1,ARPC5,SHC1,PSMD10,MAP2K4,MAPK3,PPP3CA,PSMA6,VA[illegible]

[illegible]BA1C,MAP1S,KIF3A,ARL8A,SPG11,OPA1,DYNC1I2,MAPK8IP3,SYT4,AP3M1
,XRCC5,PJA2,MSN,KPNB1,COL1A2,RPL22,RHOA,PSMD2,CXADR,JUND,CD81,VAT1,CLEC7A,PFKL
[illegible]GB5,LOX,A2M,CCN1,HSPG2,ANTXR1,LTBP4,RIC8A,SH3PXD2A,CLASP2,LTBP3,ITGAV,ADAM10,F[illegible]

[illegible]GB5,LOX,A2M,CCN1,HSPG2,ANTXR1,LTBP4,RIC8A,SH3PXD2A,CLASP2,LTBP3,ITGAV,ADAM10,F[illegible]
[illegible]PX1,AKT3,GNA12,RIC8A,SEMA3C,CLASP2,PDPK1,WASF2,CAP1,VASH1,MTOR,SMO,FGFR1,PTPN
[illegible]ADM,ITGA7,FZD7,NF2,LRP6,CRB2,HMGA2,EPHA2

[illegible]OL4A2,FN1,SPEN,DDB1,PSMD5,CHSY1,PPP2R1A,PKD1,HLA-C,PDIA3,RPS3,AP2M1,STMN1,ITGB[illegible]
,KIF5A,SEC16A,YKT6,CUL3,LMAN1,HIF1A,SEPTIN5,SLIT1,RAB1B,VPS4A,KIFAP3,RAB6A,FNBP1L,[illegible]
1,ARF1,SLC4A8,HNRNPU,RTN4,LPCAT1,RPS24,SLC22A17,XRCC5,HK1,NCS1,ATF4,KIF5B,RPS14,

CM1,FEZ1,RAN,NCBP2,YWHAZ,GSK3B,KIF3A,ERBB2,RUFY3,XBP1,TFDP1,PRNP,LEPROT,USP36,[illegible]
[illegible]1,EVL,CUL3,ITGB5,CALD1,CTTN,SLIT2,BRK1,TMOD2,ABI2,RDX,CAPZA1,CLASP2,WASF2,MTOR,

LTC,GJA1,CTNNB1,MACF1,CKAP5,LAMP1,MAP2,RAB1A,RPS27,RAC1,MAP1A,TBCD,ACTR3,CAM[illegible]
HA4,EIF2S1,PLXND1
GAP1,DIP2A,AUTS2,KDM3A,POR,RTF1,MACROH2A1,RUVBL2,MDM2,MECP2,TRMT112,SUMO1,[illegible]

[illegible]5,TCP1,CCT3,RAD23B,CCT4,TERF2IP,TOP1,VEGFA,MAPK1,NYNRIN,FTO,PRKDC,TDRD5,KDM1A
[illegible]1,WLS,RHOB,GPC1,HDAC2,MMP2,GLUL,HSBP1,DCN,TGFBI,PKN1,COL2A1,MDK,PTK7,SKI,KDM

[illegible]EGF8,RAB7A,MMP14,PRDX2,DDX3X,APLP1,RPS19,MIF,PRKAR1A,GJA1,CTNNB1,SESN3,NDFIP1,
10,RBPJ,PRPF19,PCGF2,MBD3,VASH1,NASP,TANC2,SMO,WDTC1,RTF1,ACVR1B,GGNBP2,NOTC[illegible]

P51A1,DYNLL1,CDC42,GRIPAP1,OS9,SEC16A,SFRP1,GLUD1,UBE2G2,OAZ2,ANP32B,LMAN1,HIF

GBA2,MAPRE3,PDE4DIP,PLK2,GSK3B,MAP1S,KIF3A,STMN4,CDKN1B,RBM14,ERBB2,RNF4,BME

CYFIP2,YWHAB,PTGES3,RAB1A,ARHGDIA,SKP1,PEA15,CDKN1A,EEF1A2,CCT2,TBCD,GNAI3,PAB

TIMM50,LMAN1,HIF1A,CTTN,SLC25A5,MAP1LC3B,BAP1,SPG7,GPX1,GABARAPL1,AKT3,CNP,NM

PK1,ADAM9,MAP2K2,FYN,SOD1,HNRNPD,CDC42,TRAF4,TRAF7,PRKAR2A,PHB2,UBA52,ADCY6,

PLP2,ARL6IP1,MAP2,ARHGDIA,IPO5,TIMP3,TBCD,GDI2,PRDX5,TBC1D16,ACAP3,ABCA2,NRDC,T

PRKDC,KDM1A,CCAR2,POLR2B,USP7,BAG6,KDM4B,TP53BP1,DDIT4,POLR2E,PRMT1,MTA1,EP3

HIP,USO1,RABGAP1,RANGAP1,PTPN11,WDTC1,POR,SMPD3,IGF1R,MDM2,CYFIP1,PTPN1,CDK4,

C25A23,PHGDH,FOXK2,AKT2,OGT,COX6A1,GSK3A,MTOR,ATP5MC3,NDUFA13,BNIP3,ATP5MC2,

F1,MAPK1,SMARCA4,KDM1A,NSG2,GNG2,YWHAH,SOD1,NCOR2,HNRNPD,WBP2,DDIT4,PPP1R9

1,HDAC2,BCLAF1,HSBP1,EID1,SQSTM1,NOP53,PARP1,MORF4L2,PRDX5,YEATS2,CBX1,SKI,H3-3

K3A,PRKACA,APBB1,UBE2D3,SAE1,SMO,PTPN23,UBE2L3,PINK1,ATP13A2,MDM2,PCM1,PHPT1,

UBB,AFF4,TRIM28,COL4A2,SPEN,NONO,PSMD5,YWHAQ,SET,EGLN1,PPP2R1A,GTF3C4,PKD1,S

1,HDAC2,BCLAF1,HSBP1,EID1,SQSTM1,NOP53,PARP1,MORF4L2,PRDX5,YEATS2,CBX1,SKI,H3-3

IT5,WASF3,ERBB2,MED12,CNTN2,MAPK3,RELA,EPHA4,NFIB,ID2,GSN,MXRA8,MMP24

MECP2,SART3,KAT2A,TASOR,ATF7IP,AKAP8L,CDK9,DHX36,SENP6,EXOSC10,CTR9,RNF20,LMNA,

RDX2,GJA1,JUN,CTNNB1,MACF1,NDFIP1,RHOB,HLA-B,MAP2,ARHGDIA,CAPRIN1,HDAC2,L1CAM

1,HDAC2,BCLAF1,HSBP1,EID1,SQSTM1,NOP53,PARP1,MORF4L2,PRDX5,YEATS2,CBX1,SKI,H3-3

IB1,TAOK2,MACROH2A1,MDM2,CUL1,PSMD13,KAT2A,ATF5,EHMT2,PSMD7,CUL4A,CDK4,H2AX,

TMED2,TAF7,VEGFA,RPL37,PPP2CA,IPO7,PCSK1N,CAND1,HSPB1,QARS1,CCAR2,DUSP4,CORO1

MAGED1,KAT6A,DVL3,YWHAE,GNAS,DDX17,GPI,HTRA4,RACK1,BSG,FASN,SPTBN1,CALM3,GNB1

L,CLIP3,PLCG1,CCAR2,FYN,SOD1,TXNIP,CDC42,DDIT4,TRAF7,CAPN2,TRIO,FLCN,UBA52,SFRP1

GDH,COX5A,COX7A2,UQCRH,FH,MSH2,PDHB,NNT,NDUFA5,UQCR11,COX5B,NDUFA10,ATP5PD

1,RBMX,GCN1,SKI,ASXL1,KDM6B,PRPF6,TAF7,RNF10,VEGFA,SMARCA4,PBX2,PRKDC,SUPT5H,

YNC1LI2,DYNLL2,GABARAP,SON,MAP7D1,CLIP3,STMN3,CSNK1D,RAB11A,DYNLL1,DCTN1,MAP6

N,SLIT2,SPAG9,PDPK1,MTOR,WNK1,MICAL1,PRDX3,EIF4G1,ABL1,PPP5C,MACROH2A1,GGNBP

1A,GTF3C4,PKD1,SSBP2,ZNF43,RPS3,CHD3,BASP1,HMGB1,POLR2A,MORF4L1,BTF3,AGRN,ZMI

DKN1A,EEF1A2,NOX4,PKN1,GCN1,TAF7,VEGFA,RAPGEF1,PPP2CA,CCND1,MAPK1,CCNL2,IPO7,

3,PIN1,ADD1,BTBD1,CTNNBIP1,APOE,PLXND1,PRMT7

UBB,AFF4,TRIM28,COL4A2,SPEN,NONO,PSMD5,YWHAQ,SET,EGLN1,PPP2R1A,GTF3C4,PKD1,SS
1A,GTF3C4,PKD1,SSBP2,ZNF43,RPS3,CHD3,BASP1,HMGB1,POLR2A,MORF4L1,BTF3,AGRN,ZMI

N5,BNIP3,BAG5,PRDX3,PTPN11,ABL1,PINK1,PPP5C,HSPA1A,TXNRD1,SMPD3,ATP13A2,MDM2,M
UBB,AFF4,TRIM28,COL4A2,SPEN,NONO,PSMD5,YWHAQ,SET,EGLN1,PPP2R1A,GTF3C4,PKD1,SS
ASSF2,HOXA5,STAT1,CEBPG,TMEM14C,SOS1,CITED2,ANXA1,THRA,ADD1,EXT1,HMGB2,FH,ACI

A2
NP,LEPROT,USP36,NUTF2,EMD
,GRSF1,ZFP36L1,BRD2,HOXB5,SFRP1,CUL3,NDST1,SPINT2,ITGB5,HIF1A,CCN1,SOX2,RIC8A,CL
2CA,CCND1,MAPK1,CCNL2,IPO7,ADAM9,CDC37,HSPB1,MAP2K2,QARS1,DUSP4,CORO1C,FYN,H

IPK2,CSNK2B,CCNG1,MARCHF7,NDRG1,PLK2,PRMT5,CDKN1B,PSMD10,PIN1,CNOT3,PCNA,TF
1A,GTF3C4,PKD1,SSBP2,ZNF43,RPS3,CHD3,BASP1,HMGB1,POLR2A,MORF4L1,BTF3,AGRN,ZMI

C6,DPF2,SIN3A

SMC1,CD74,ELOVL5,SCAP,IGFBP4,WDTC1,PINK1,POR,PSMB1,H6PD,PSMD13,PARK7,KAT2A,PS
,CD81,VAT1,CLEC7A,PFKL,SOX12,RAB7A,MMP14,PRDX2,DDX3X,MIF,PRKAR1A,CTNNB1,CD63,

APZA1,CLASP2,WASF2,MTOR,DAPK3,ANK2,CELSR1,ABL1,ARPC1A,TAOK2,HIP1R,ELN,CRACD,JA

D1,TXNIP,CDC42,TRAF7,CAPN2,TRIO,FLCN,UBA52,SFRP1,HDAC6,PTEN,RPS27A,SLIT2,MTCH1,
,TRAF7,CAPN2,TRIO,FLCN,UBA52,SFRP1,PTEN,RPS27A,SLIT2,MTCH1,GRN,CCN1,OLFM1,UBE2
X2,SEMA3C,SCRIB,AKT2,RBPJ,UBE3A,GOT2,SMO,CAD,LMO4,USF2,ABL1,DAG1,IGSF3,NOTCH1,
NTN1,LIMK1,ISLR2,GOLGA4,SRF,ACSL4,CRABP2,ERBB2,UNC13A,PSMD10,TGFBR1,RUFY3,PIN1

,WASF3,CDC42EP4,ARPC5,ROCK2,TGFBR1,FER,GMFB,GIT1,PDXP,ARF6,NF2,GSN

UBB,AFF4,TRIM28,COL4A2,SPEN,NONO,PSMD5,YWHAQ,SET,EGLN1,PPP2R1A,GTF3C4,PKD1,SS

F,RAB11A,PPP1R9B,SYP,VDAC1,KIF5A,STAT3,KCNQ2,BCR,PRKAR2A,FLOT1,KPNA2,FARP1,PTEN

PZA2,TRAPPC11,TRAPPC1,SEC23IP,COPB1,SEC31A,DYNC1I2,COG4,ARFGAP3,TMED5

DL4A2,MAPT,FN1,SPEN,EGLN1,CHSY1,RPS3,BASP1,HMGB1,ITGB1,ANXA2,AGRN,ZMIZ1,VCL,AP3
EIF4G1,ABL1,PPP5C,MACROH2A1,GGNBP2,PKIA,IGF1R,PARK7,PTPN1,TRIM27,NCK2,TRIB3,PME
,ABL1,RSF1,CHMP1A,TSPYL2,MACROH2A1,RUVBL2,SART3,SUB1,BANF1,HMGB3,CUL4A,TASOR,
1,NUP50,NUP58,NUTF2
TXNRD1,SMPD3,ATP13A2,MDM2,MGAT3,PARK7,ARL6IP5,GSR,NME2,HSF1,DHFR,SETX,TXN,VKC
G1,NTRK2,CDC34,ABL1,PINK1,PPP5C,GRIK5,ATP13A2,TRIM2,MECP2,EFNB2,PARK7,ADNP,ARL
,ABL1,ARPC1A,TAOK2,HIP1R,ELN,CRACD,JAM3,CYFIP1,PHPT1,LIMK1,FLII,PDGFRB,CCDC88A,T

,APOE,PSMA4,PSMC6,TLE4,LRP6,EMD
ANXA1,MAPK3,PCNA,FER,RELA,APOE,FBXO7,SESN2,GNAO1
NPO2,KPNA3,PPP3CA,XBP1,UFM1,TNPO3,NOLC1,APOE,NUP50,ABCD3,NUP58,TOMM40,NUTF2
PCNA,PTPN12,EPHA4,MAPK8IP3,POSTN,FZD7,GSN

1,WDTC1,POR,IGF1R,MDM2,CYFIP1,PTPN1,CDK4,TRIB3,ADCY1,ATP6V0B,HSF1,GSK3B,STAT1,IF
3,RPS3,CHD3,BASP1,HMGB1,MORF4L1,AGRN,ZMIZ1,RPL10,AP3D1,ZNF770,GTF2F1,HNRNPU,N
CGF2,PSMC3,PSMC1,USP47,PTPN11,ABL1,CHMP1A,PSMB1,TAOK2,TSPYL2,MACROH2A1,MDM2,

T2,SOBP,ADCY1,SYNGAP1,NDRG4,PLK2,SRF,CRTC1,SETD5,DLG4,PRKAR2B,DDHD2,CNTN2,SHA
CB1,SETD1B,BTG1,PCMTD2,PRMT7
CB1,SETD1B,BTG1,PCMTD2,PRMT7

,CRTC1,DLG4,PRKAR2B,DDHD2,CNTN2,SHANK1,THRA,B4GALT2,GIT1,EPHB2,APOE,FGF13,PRN
M27,NCK2,ARPC3,GBA2,WASF3,CDC42EP4,ARPC5,SHANK1,ROCK2,TGFBR1,CAPZA2,ADD1,ARH
VAT1,CLEC7A,PFKL,RAB7A,PRDX2,DDX3X,RPS19,MIF,CD63,CYFIP2,DNAJC5,NDFIP1,DDOST,LAM
A2,ATRX,UPF1,HNRNPA2B1,TNKS,FUS,PPP1R10,UBR5,SMG5,NUCKS1,MSH6,NSD2,CIZ1,MBD3,G
3,RPS3,CHD3,BASP1,HMGB1,MORF4L1,AGRN,ZMIZ1,RPL10,AP3D1,ZNF770,GTF2F1,HNRNPU,N
3,RPS3,CHD3,BASP1,HMGB1,MORF4L1,AGRN,ZMIZ1,RPL10,AP3D1,ZNF770,GTF2F1,HNRNPU,N
A,IGF1R,PARK7,PHPT1,PTPN1,LIMK1,TRIM27,TRIB3,GSK3B,WARS1,CDKN1B,SH3BP5,DUSP3,DU
P1,PTPN1,CDK4,TRIB3,ATP6V0B,GSK3B,STAT1,IRS1,ATP6V0E2,SOS1,ATP6V0A1,CFLAR,SHC1,AD

A1,LGALS1,TAOK2,DAG1,CCDC80,NOTCH1,JAM3,CD44,MAP4K4,HSD17B12,C1QBP,LDB1,EMIL

P7,EP300,LBH,PHB2,ZFP36L1,SFRP1,HDAC6,NCOA4,LOX,SLIT2,SAFB2,CARM1,UBR5,UBE3A,PA
T,HIF1A,LOX,CCN1,SOX2,PKDCC,LTBP3,HSD17B4,LRP3,RBPJ,PRKACA,MN1,CCDC47,SORT1,SM
APBP,SNX3,SDCBP,CHMP1A,GRIK5,ATP13A2,NSF,ANKFY1,GOLPH3,ARFGAP2,TMEM9,SERPINE2,

V0A1,CFLAR,SHC1,ADIPOR1,XBP1,PRKCI,SP1,FER,BCAR1,RELA,OPA1,GRB10,ATP6V1F,ZNF106,
P1R9B,SYP,VDAC1,KIF5A,STAT3,KCNQ2,BCR,PRKAR2A,FLOT1,FARP1,PTEN,SV2A,PREPL,BRSK1

RDX3,DEK,PTPN11,FAM210B,KLF13,HSPA1A,ACVR1B,PABPC4,MAEA,MTURN,HIPK2,CUL4A,TNR
BMERB1,CAPZA2,ADD1,ABCE1,FGF13,TPX2,PDXP,MRPL3,MRPS26,MRPS27,MRPL28,GSN

7,NUMBL,BMERB1,OGDH,CNTN2,EXT1,XAB2,EPHB2,LRP8,FGF13,SZT2,KDM2B,COL3A1,NF2,PO

S1,NUP58

X1,VPS51,TBC1D14,TRIM27,NDRG4,ANKRD50,CCDC93,CHMP5,TMEM50A,RAB35,MICALL1,SNX
DM4,LAMP1,RHOB,ATP1A2,MAP2,RAC1,ARHGDIA,APPL1,GLUL,DCN,DLG5,CNN2,NOX4,PKN1,MI

CA,HNRNPR,CDK4,ADCY1,GSK3B,LGR4,CRTC1,SETX,PRMT5,KDM2A,KDM5A,PER1,NR2F6,SGPL
,PPP1CA,COX7A2L,ATP5PO,DLD,GSK3B,NDUFB8,ME2,SHMT2,PHKB,NDUFS1,ATP5F1C,COX6B1,

P1R9B,SYP,VDAC1,KIF5A,STAT3,KCNQ2,BCR,PRKAR2A,FLOT1,PTEN,SV2A,PREPL,BRSK1,AKAP9
P1R9B,SYP,VDAC1,KIF5A,STAT3,KCNQ2,BCR,PRKAR2A,FLOT1,PTEN,SV2A,PREPL,BRSK1,AKAP9
XNA1,SOX12,CCN2,ACTN4,MMP14,PRDX2,APLP1,PRKAR1A,TLN1,CTNNB1,COL6A2,CD63,MACF

2,TASOR,SENP6,NUP98,GOLGA8B,MAPRE3,PDE4DIP,PLK2,RCC2,NUP133,RNF4,LZTS2,CHMP5,T
0,PSMA6,PSMD12,CTNNBIP1,APOE,PSMA4,PSMC6,GRB10,TLE4,LRP6,EMD

PS37B

C5,CAPZA2,ADD1,FER,GMFB,PDXP,ARF6,MYADM,GSN

12,CCN2,ACTN4,MMP14,PRDX2,APLP1,PRKAR1A,TLN1,CTNNB1,COL6A2,CD63,MACF1,CYFIP2,N

A,PSMD12,TLR6,PSMA4,COPS8,PSMC6,PPM1A
ADM,GSN

N1,STXBP1,DGKD,CDC42,PRKAR2A,SRSF6,PTEN,LMAN1,HIF1A,LOX,A2M,CCN1,GPX1,GNA12,CA
C42,SYP,GRIPAP1,HIP1,AP2A1,UBQLN2,FLOT1,MBTPS1,PTEN,PREPL,SEPTIN5,PKDCC,VPS4A,RD

0,NECTIN2,PSMA6,TAB3,HEXIM1,HMGB2,RELA,PSMD12,PSMA4,PSMC6,AKIRIN2,SIN3A

ADM,GSN

F3A,STARD3,NAPA,COL7A1,TRAPPC1,SPG11,SEC23IP,SEC31A,DNM1,SYT4,AP3M1,ARFGAP3
?4B,BNIP3,TUBB4A,PINK1,HDGFL3,HIP1R,HSPA1A,CRACD,FEZ1,KAT2A,EHMT2,FLII,NCK2,TOM1I

RTN4,XRCC5,PJA2,IGFBP3,RHOA,PSMD2,NLGN2,TMEM259,RPS20,CD81,RPL26,MCL1,CLEC7A,
,EPHB2,ARF6,PRNP,GSN,EPHA2

CL9L,CAPN2,EP300,FLOT1,ZFP36L1,CD164,FKBP1A,LOX,SERP1,TMOD2,GPX1,FLOT2,SOD2,SP
R1,CALCOCO1,PUM1,KDM3A,PPP5C,HSPA1A,AAMP,PARK7,TNRC6A,ZCCHC3,PMEPA1,C1QBP,CI

HEB,ASNS,ATP6V0B,SRF,ATP6V0E2,GABARAPL2,MAPK3,XBP1,SZT2,SESN2,TRIM24,EIF2S1,POST
S,HIRA,TLK1,CTCF,UBN1,CABIN1,HMGB2,SMARCC2,PAF1,DNMT1,SMARCB1,CENPT,HMGA2

,PTEN,SELENOW,APEX1,HIF1A,BAP1,FUS,GPX1,STK24,TRAP1,NCSTN,SOD2,SLC25A23,KRT8,SY
ARCA5,RPL23,NACA,BRD3,PAPOLA,RHOA,CBX5,ZNF865,ATF4,PSMD2,NR2F1,RPS14,CHD4,JUNI
17,GNA12,RDX,WASF2,LZTR1,OGT,ARPP19,DENND4B,GNA13,USO1,CELSR1,ABL1,SDCBP,AUTS
EXT1,SPG11,VPS16,PPT1,RNF5,STX12,RB1CC1
2,RUFY3,PRKCI,DYNLT1,FGF13,ARF6,RAP1B,CRB2,GSN,CYTH3
I1,ATP6V1G1,AMBRA1,SLC38A2,EIF4G1,PTPN11,USF2,DAG1,MDM2,KIF26A,ADNP,PIK3C2B,EHM

CASK,UNC13B

2,NOX4,MDK,VEGFA,MAPK1,MAPRE1,ADAM9,HSPB1,PLCG1,CORO1C,FBLN1,PLXNA3,STRAP,PTI
I1,PSMD7,ZCCHC3,JAK1,PSME3,PSMC5,PSMC2,PSMC4,STAT2,PSMF1,PSMB5,STAT1,RBM14,PSI

GDIA,APPL1,GLUL,DCN,DLG5,CNN2,NOX4,PKN1,MDK,VEGFA,MAPK1,MAPRE1,ADAM9,HSPB1,PL
F6,SYT1,DNM1,PPFIA3,RAP1B,CASK,UNC13B

6A,ADNP,PIK3C2B,EHMT2,TNRC6A,RHEB,ASNS,ATP6V0B,SRF,ATP6V0E2,GABARAPL2,MAPK3,XBI

FB2,FGF13,FHL1,PPT1,SESN2,BTG1
I1A,DLG5,TGFBI,ABCA3,COL2A1,MDK,PTK7,SKI,ASXL1,TMED2,NOTCH2,VEGFA,SELENON,MAPK

,TRAPPC1,SPG11,SEC23IP,SEC31A,DNM1,SYT4,AP3M1,ARFGAP3

3,DYNLT1,EPHB2,APOE,TGFB2,ARF6,COL3A1,ITPKB,CYTH3
CDK5RAP3,RAD50,HIPK1,PPP2R5C
K4,H2AX,PSME3,PSMC5,PSMC2,PSMC4,PLK2,PSMF1,CDKN1B,PSMB5,XPC,PSMA1,PRMT2,PSMD

R1,B2M,HECTD1,SOD1,CDC42,ADAR,PSMA7,STAT3,EIF2AK1,PRMT1,EP300,PSMD1,PBX1,FLCN,Z
CK2,ID3,ID2,KLF10,SIN3A,MYBBP1A,NAMPT

2,ATP1A1,CD81,ATP2A2,CLEC7A,PTN,TPM1,SULF1,PLXNA1,MEGF8,SOX12,CCN2,MMP14,PRDX2
RN1,EPHB2,ARF6,RAP1B,GPM6B,EPHA2
,ACTN1,KDELR1,B2M,HECTD1,SOD1,CDC42,ADAR,PSMA7,STAT3,EIF2AK1,PRMT1,EP300,PSMD1
A2,TRIM2,MECP2,EFNB2,PARK7,ADNP,HIPK2,SYNGAP1,HSF1,GSK3B,NSMF,EGLN2,BACE1,MAP2

DX4I1,OGDH,COX5A,COX7A2,UQCRH,FH,MSH2,PRELID1,PDHB,NNT,NDUFA5,UQCR11,COX5B,M
P5,DYNLT1,TPX2,DYNC1I2,CENPF,CENPT,ZNF207,GNAI1
MA6B,NOTCH1,PCM1,DNAJB1,FEZ1,KAT2A,ATF5,SUN2,H2AX,CRKL,SRF,ZSWIM6,GSK3B,CCDC8
CTNNB1,TAGLN,RHOB,HRNR,RAC1,HDAC2,TNC,CDKN1A,DLG5,CTSB,COL2A1,MDK,PTK7,SKI,AS
SN
,CENPF,CENPT,ZNF207,GNAI1

UBE2V1,UBQLN4,OTUB1,CEBPG,PSMD10,PCNA,HMGA2,SKIL

CBX5,ZNF865,ATF4,PSMD2,NR2F1,RPS14,CHD4,JUND,CD81,PFN1,TLE5,TADA2B,KHDRBS1,XPO
C2
N,CTNNB1,RHOB,MMP2,GLUL,DCN,CDKN1A,DDX39B,TGFBI,COL2A1,MDK,PTK7,ASXL1,KDM6B,

AB,LAMP1,MAP2,RAC1,APPL1,IPO5,TMEM30A,GLUL,NNAT,RAB27A,VAMP2,ABCA3,AHCYL1,ATP6

VCP,TRIM28,MATR3,MAPT,FN1,NONO,PSMD5,HLA-C,PDIA3,RPS3,VAPA,HMGB1,AP2M1,SEC61A1

FIP1,PTPN1,HTRA1,HIPK2,PMEPA1,STUB1,EMILIN1,ZEB1,ZNF703,NREP,NEO1,CITED2,CFLAR,DS

6V1F,NNT,ATP6V1D,COX5B,ATP5PD

9,GDF10,CITED2,CFLAR,ROCK2,TGFBR1,PIN1,XBP1,ARHGEF18,MXRA5,TGFB2,FURIN,COL3A1,[illegible]

[illegible]L2,SEMA4C,GSK3B,ZNF703,SOX9,HOXA5,STAT1,NUP133,CITED2,MCRIP1,NRP2,ROCK2,ADIPO[illegible]
SZT2,SESN2,EIF2S1,ATP6V1F,KLF10,ATP6V1D,MYBBP1A

[illegible]PD

PRKDC,ACTN1,KDELR1,SUPT6H,B2M,HECTD1,SOD1,TP53BP1,CDC42,ADAR,PSMA7,STAT3,EIF2[illegible]
[illegible]10,PSMA6,CNOT3,PCNA,TFDP1,PSMD12,PSMA4,FHL1,FBXO7,CENPF,PSMC6,CDK5RAP3,ZNF20[illegible]
[illegible]ID1,MYADM,POSTN,FZD7,NF2,GPM6B,CASK

[illegible],CNOT1,PPP2R5C

CI,TGFB2,EPHA4,GJC1,NF2,HIPK1,EPHA2

[illegible]PP2CA,CCND1,MAPK1,ADAM9,CLIP3,MAP2K2,FYN,CSNK1D,SOD1,PFN2,BMPR2,CDC42,TRAF4,[illegible]

[illegible],DCN,DLG5,CNN2,NOX4,PKN1,MDK,VEGFA,MAPK1,MAPRE1,ADAM9,HSPB1,PLCG1,CORO1C,FB[illegible]
[illegible]H,KDM1A,ID4,STRAP,SARS1,SREBF2,NCOR2,TXNIP,CIC,RBBP7,STAT3,MTA1,EP300,FLCN,UBA5

1,RBBP4,CLEC7A,NES,PTN,TPM1,EIF5A,CCND2,CDH2,GPC3,DDR1,SULF1,CCN2,MMP14,PRDX2,

[illegible]H7A1,SLC16A2,PSME3,PSMC5,PSMC2,GPT2,DLD,PSMC4,PSMF1,DDAH1,RIMKLB,SEPHS1,DHFR

[illegible]B,H2AX,LOXL2,DLD,LIMS1,PLK2,SRF,LMNA,CACYBP,CITED2,MAPK3,PPP3CA,RELA,OPA1,MSH2,
9,GDF10,CITED2,CFLAR,TGFBR1,PIN1,XBP1,ARHGEF18,TGFB2,FURIN,COL3A1,PPM1A,POSTN,[illegible]

NDST1,HIF1A,GALNT11,FKBP1A,LOX,SLIT2,GRK2,CCN1,OLFM1,SEMA3C,NSD2,RBPJ,MBD3,GS[illegible]
[illegible]2,PSMF1,CDKN1B,PSMB5,XPC,PSMA1,PRMT2,PSMD10,MDC1,PSMA6,CNOT3,PCNA,TFDP1,PS[illegible]

[illegible]B2,UBA52,ADCY6,RPS27A,XRCC6,TAOK1,CCN1,SHC2,SPAG9,PDPK1,DAZAP2,MTOR,PRKACA,W[illegible]
[illegible]H,ZDHHC9,WDR5,MAPK3,XBP1,BRPF3,ZDHHC17,PPM1A,SMARCB1,BRD1,SIN3A
P,VEGFA,RAPGEF1,PPP2CA,CCND1,MAPK1,ADAM9,PTBP1,CLIP3,MAP2K2,FYN,CSNK1D,SOD1,P[illegible]
P,VEGFA,RAPGEF1,PPP2CA,CCND1,MAPK1,ADAM9,PTBP1,CLIP3,MAP2K2,FYN,CSNK1D,SOD1,P[illegible]

[illegible]APK3,PCNA,TFDP1,CENPF,RAD50
X9,CHD8,KDM5A,PBRM1,GATAD2B,SMARCC2,DNMT1,SMARCB1,CENPT,HMGA2
[illegible]INE2,EI24,ADIPOR2,CDKN1B,PHB,CITED2,ADIPOR1,RGMA,SEMA6C,TGFB2,FGF13,FHL1,PPT1,[illegible]

[illegible]ST,LAMP1,CKAP4,HLA-B,TTR,COTL1,HRNR,RAC1,ATP6V0C,TMEM30A,PGRMC1,CNN2,CTSB,RA[illegible]

[illegible]5,DUSP3,DUSP8,PRKAR2B,NR2F2,CAMK2N1,GPS1,RTRAF,HEG1,AJUBA,HEXIM1,GMFB,EPHB2,[illegible]

3A,RPL6,KAT6A,DVL3,YWHAE,KIF1A,RPL13,DDX17,RACK1,BSG,SPTBN1,ATP5F1B,CALM3,STMN
BSG,SPTBN1,ATP5F1B,CALM3,RPS8,MYH9,RPL7,SEPTIN2,APP,RPL5,GDI1,TMBIM6,SPTAN1,GST
2F1,HNRNPU,OGA,LARP1,RPS16,RPL23,RPL29,RPSA,EIF4H,SCARB2,RPS24,RPS9,XRCC5,MSN,R
/HAE,KIF1A,RACK1,BSG,SPTBN1,ATP5F1B,CALM3,STMN2,MYH9,SEPTIN2,P4HB,APP,RPL5,GDI1,
RPS4X,RPS3A,RPL18,ITGB1,RPL11,ANXA2,AGRN,RPLP1,RPL10,RPS17,VCL,AP3D1,DNAJA1,ARF1
RPS4X,RPS3A,RPL18,ITGB1,RPL11,ANXA2,AGRN,RPLP1,RPL10,RPS17,VCL,AP3D1,DNAJA1,ARF1
,PSMD2,RPS14,RPS20,RPL26,SRSF4,PNN,RPL31,HNRNPC,KHDRBS1,XPO1,RPS25,SFPQ,RPS15,
3A,RPL18,UCHL1,RPL11,ANXA2,RPLP1,RPL10,RPS17,AP3D1,ARF1,PAFAH1B1,AP2B1,HNRNPU,F
,AP2B1,HNRNPU,OGA,LARP1,HNRNPA0,RPS16,RPL23,RPL29,RPSA,CTSD,LPCAT1,RPS24,RPS9,N
,TMBIM6,SPTAN1,GSTP1,SMG1,CALM1,NPM1,HSPD1,IGF2,ATP5F1A,VCP,UBB,TRIM28,YWHAG,
,RPL15,PKD1,RPS3,VAPA,AP2M1,SEC61A1,RPS4X,RPS3A,RPL18,ITGB1,RPL11,BTF3,ANXA2,AGR
D1,RPS3,HMGB1,CDKL1,AP2M1,STMN1,ANXA2,AGRN,VCL,ARF1,PAFAH1B1,MAP4,AP2B1,HNRNF
VHAQ,DCHS1,TMSB10,RPL19,RPL15,PKD1,RPS3,VAPA,AP2M1,SEC61A1,RPS4X,RPS3A,RPL18,IT
,RPS16,RPL23,RPL29,RPSA,RPS24,RPS9,MSN,RPL24,KPNB1,RPL17,RPL14,RPL22,RPL9,PSMD2
RPL24,KPNB1,RPL17,RPL14,HK1,RPL22,RPL9,RPS14,RPS20,CD81,RPL26,CCT7,RPL31,XPO1,RP
RPL35,RPS10,RPL35A,TARDBP,RPLP2,CIRBP,RPL36,YWHAB,RPS12,RPS27,BCLAF1,RPL23A,TAF1
AGED1,SEC62,RPL6,DVL3,YWHAE,KIF1A,RPL13,GNAS,SLC2A3,GPI,RACK1,BSG,SPTBN1,ATP5F1
/1,PLXNB2,TTC3,HSPD1,USP11,CSDE1,IGF2,EIF3C,EIF4B,VCP,UBB,TRIM28,YWHAG,RPL37A,MA
DE1,IGF2,ATP5F1A,VCP,MARCKSL1,UBB,TRIM28,YWHAG,MAPT,DDB1,YWHAQ,SET,MAP9,TMSB1
RPL28,RPS28,FAU,RPL39,RPS13,RPL34,SSR1,UBA52,RPS2,RPS27A,TRAM1,SRP72,SSR3,SSR2,S
E,RPS11,CASC3,RPL28,RPS28,FAU,RPL39,RPS13,RPL34,GSPT1,UBA52,UPF1,RPS2,RPS27A,SM
U,RPL39,RPS13,RPL34,UBA52,RPS2,RPS27A,TRAM1,SRP72,SSR3,SRPRA,RPS26,RPS21,RPL27
,RPS19,RPL35,RPS10,RPL35A,TARDBP,RPLP2,CIRBP,RPL36,YWHAB,SND1,RPS12,RPS27,BCLAF
EIF4B,VCP,UBB,TRIM28,YWHAG,RPL37A,MAPT,FN1,DDB1,PSMD5,DCHS1,SET,MAP9,TMSB10,RF
A1,ARF1,PAFAH1B1,AP2B1,RPS16,RPL23,RPL29,RPSA,CTSD,SCARB2,NACA,RPS24,RPS9,RPL24,
KIF5B,RAB6B,RPS14,RPS20,CD81,RPL26,OAZ1,RPL31,XPO1,RPS25,EIF5A,RAB7A,RPS15,RPL12
7,KDELR2,RPS11,KDELR1,RPL28,RPS28,FAU,RPL39,RPS13,RPL34,SGTA,OS9,SEC16A,UBA52,RE
PS28,FAU,RPL39,RPS13,RPL34,SGTA,UBA52,RPS2,RPS27A,TRAM1,SRP72,SSR3,CHMP4B,SRPR
,P4HB,APP,IDS,RPL5,GSTP1,SMG1,CALM1,NPM1,PLXNB2,TTC3,HSPD1,USP11,CSDE1,IGF2,EIF3
3D1,DNAJA1,ARF1,PAFAH1B1,AP2B1,RPS16,RPL23,RPL29,RPSA,SCARB2,NACA,RPS24,RPS9,RPL
A,RPN2,RPL27,RPL36A,RPS15A,RPL38,RPL37,SMARCA4,PTBP1,RPS11,SUPT5H,RPL28,POLR2B,
PS28,FAU,RPL39,RPS13,RPL34,SGTA,UBA52,RAB3GAP2,RPS2,RPS27A,TRAM1,SRP72,SSR3,CHM
37,EIF3E,RPS11,RPL28,RPS28,FAU,EIF3F,RPL39,RPS13,RPL34,EIF3D,GSPT1,UBA52,RPS2,RPS2
SAP,PGK1,RPLP0,H3-3B,CLU,PTMA,DDX5,ENO2,SOX4,FKBP14,RPL13A,MAGED1,RPL6,KAT6A,D
MRC2,RPL19,PPP2R1A,RPL15,PKD1,RPS3,HMGB1,AP2M1,RPS4X,RPS3A,RPL18,ITGB1,UCHL1,F
PS4X,RPS3A,RPL18,UCHL1,RPL11,ANXA2,RPLP1,TIMP2,RPL10,MARCHF6,RPS17,AP2B1,PXDN,H
YWHAE,RPL13,GNAS,DDX17,GPI,HTRA4,RACK1,SPTBN1,CALM3,GNB1,STMN2,RPS8,MYH9,RPL7
PS12,RPS27,RPL23A,RPL27,RPL36A,TMED2,RPS15A,RPL38,RPL37,EIF3E,RPS11,HSPB1,RPL28,R
TARDBP,NDFIP1,CIRBP,YWHAB,HMGCR,ATP6V0C,LAPTM4B,TRIB2,PTTG1IP,DCN,MAP1A,TIMP3,
6,RPL23,RPL29,RPSA,SCARB2,NACA,RPS24,RPS9,RPL24,KPNB1,RPL17,LLGL1,RPL14,RPL22,R
TBN1,ATP5F1B,CALM3,GNB1,SHISA9,RPS8,MYH9,RPL7,SEPTIN2,APP,RPL5,GDI1,TMBIM6,SPTA
PS15,RPL12,RPL30,RPS5,RPS7,RPS19,RPL35,RPS10,RPL35A,TARDBP,RPLP2,RPL36,YWHAB,LA
1B,CALM3,GNB1,SHISA9,RPS8,MYH9,RPL7,SEPTIN2,APP,RPL5,GDI1,TMBIM6,SPTAN1,GSTP1,S
SMD5,DCHS1,SET,MAP9,TMSB10,PPP2R1A,DNAH8,PKD1,SF3B1,RPS3,HMGB1,CDKL1,AP2M1,ST
/1,RPS4X,RPS3A,RPL18,UCHL1,RPL11,ANXA2,AGRN,RPLP1,TIMP2,RPL10,MARCHF6,RPS17,PAF

,RACK1,FASN,ALDH6A1,SPTBN1,ATP5F1B,CALM3,RPS8,MYH9,RPL7,P4HB,APP,IDS,RPL5,GSTP1
IMP2,VCL,ARF1,PAFAH1B1,MAP4,ITM2B,CTNNA1,LDHA,SCARB2,RTN4,LAMA5,XRCC5,RPL24,LL
2,RPS14,RPS20,CD81,RPL26,RPL31,PTN,RPS25,CDH2,GPC3,RAB7A,MMP14,RPS15,RPL12,RPL
B,APP,RPL5,KLRD1,GDI1,FSCN1,GSTP1,CALM1,NPM1,PLXNB2,HSPD1,IGF2,EIF3C,ATP5F1A,VC
,CTNNB1,MACF1,CYFIP2,MAP2,RAC1,ARHGDIA,CAPRIN1,GPC1,CDK16,TOP2B,HDAC2,TNC,TME
,KIF5B,NLGN2,RPS14,RPS20,CD81,RPL26,RPL31,PTN,RPS25,CDH2,GPC3,RAB7A,MMP14,RPS1
N,MAP1A,TIMP3,EEF1A2,PPP1CB,GNAI3,NOP53,TAF15,ALKBH5,VPS35,RAD23B,TNPO1,SNRNP7
H,RPL28,POLR2B,RPS28,FAU,IFITM3,RPL39,RPS13,RPL34,POLR2E,EP300,BRD4,UBA52,POM12
,DDX17,GPI,RACK1,BSG,FASN,SPTBN1,CALM3,RPS8,MYH9,RPL7,APP,RPL5,TMBIM6,GSTP1,SM
12,RPL30,H1-0,RPS5,RPS7,RPS19,RPL35,RPS10,RPL35A,TARDBP,RPLP2,CIRBP,RPL36,YWHAB,
MYH9,APP,RPL5,KLRD1,GDI1,TMBIM6,SPTAN1,GSTP1,SMG1,CALM1,NPM1,PLXNB2,HSPD1,CO
PSMD5,YWHAQ,SET,RPL19,PPP2R1A,RPL15,RPS3,CHD3,BASP1,HMGB1,RPS4X,RPS3A,RPL18,U
XA2,RPLP1,ZMIZ1,TIMP2,RPL10,DNAJA1,ITM2B,LARP1,RPL23,CTSD,EIF4H,RTN4,LPCAT1,RPS9
3A,MAGED1,RPL6,KAT6A,DVL3,YWHAE,RPL13,DDX17,GPI,RACK1,SPTBN1,CALM3,RPS8,MYH9,
,SET,RPL19,PPP2R1A,RPL15,RPS3,CHD3,BASP1,HMGB1,RPS4X,RPS3A,RPL18,UCHL1,RPL11,M
,GAP43,RHOA,CBX5,ATF4,PSMD2,TMEM259,ATP2A2,RPL26,MCL1,CLEC7A,XPR1,PTN,CSNK1E,
MN2,P4HB,APP,FSCN1,TMBIM6,GSTP1,CALM1,NPM1,HSPD1,IGF2,VCP,UBB,YWHAG,COL4A2,M
PSA,EIF4H,RTN4,XRCC5,MSN,RPL24,COL1A2,LLGL1,RHOA,PTCD3,PSMD2,KIF5B,RPS14,EIF5,W
,ITM2C,GJA1,CTNNB1,MACF1,CYFIP2,MAP2,RAC1,ARHGDIA,CAPRIN1,GPC1,CDK16,TOP2B,HDA
X3X,CTNNB1,CD63,MACF1,HNRNPK,YWHAB,CKAP5,CCT6A,CTBP1,RAC1,ARHGDIA,CAPRIN1,AP
A,ATF4,NR2F1,KIF5B,ENC1,WDR1,CRMP1,ASTN2,PRKCSH,SPTBN2,PTN,CSNK1E,CDH2,DDR1,P
0,H3-3B,CLU,PTMA,DDX5,ENO2,SOX4,FKBP14,RPL13A,MAGED1,RPL6,KAT6A,DVL3,YWHAE,RP
S15,RPL12,RPL30,H1-0,RPS5,RPS7,RPS19,RPL35,RPS10,RPL35A,TARDBP,RPLP2,CIRBP,RPL36
S15,RPL12,RPL30,H1-0,RPS5,RPS7,RPS19,RPL35,RPS10,RPL35A,TARDBP,RPLP2,CIRBP,RPL36
,PPP2CA,EIF3E,RPS11,CASC3,RPL28,RPS28,SSB,FAU,RPL39,RPS13,RPL34,EDC4,GSPT1,UBA52,
-3B,CLU,PTMA,DDX5,ENO2,SOX4,FKBP14,RPL13A,MAGED1,RPL6,KAT6A,DVL3,YWHAE,RPL13,D
PTBN2,PTN,CSNK1E,CDH2,DDR1,PLXNA1,MEGF8,ITM2C,GJA1,CTNNB1,NGRN,MACF1,CYFIP2,
C1,CRMP1,ASTN2,PRKCSH,SPTBN2,PTN,CSNK1E,CDH2,DDR1,PLXNA1,MEGF8,ITM2C,GJA1,CTN
IDN,RPL10,RPS17,AP3D1,DNAJA1,ARF1,PAFAH1B1,AP2B1,SLC4A8,RPS16,RPL23,RPL29,RPSA,S
,BSG,SPTBN1,CALM3,RPS8,MYH9,RPL7,APP,RPL5,TMBIM6,GSTP1,SMG1,CALM1,NPM1,PLXNB
RPL35A,RPLP2,ARL6IP1,RPL36,YWHAB,RPS12,RPS27,RPL23A,SRP9,VAMP2,RPL27,HSPA4,VPS35
1,STMN2,MYH9,P4HB,APP,RPL5,KLRD1,GDI1,FSCN1,GSTP1,CALM1,NPM1,PLXNB2,HSPD1,IGF2
,GSTP1,SMG1,CALM1,NPM1,PLXNB2,HSPD1,CSDE1,IGF2,EIF3C,EIF4B,VCP,UBB,TRIM28,YWHA
0,EIF5,RPL26,RTN3,RPL31,LRPPRC,KHDRBS1,RPS25,EIF5A,DDX3X,RPS15,RPL12,RPL30,EIF3L,
L10,DNAJA1,LARP1,RPL23,CTSD,EIF4H,RPS9,XRCC5,MSN,IGFBP3,RPL22,RHOA,PTCD3,ATF4,EN
,YWHAB,RPS12,RPS27,RPL23A,SRP9,NUCB1,TOMM20,RAB27A,RPL27,HSPA4,RPL36A,SRP14,R
12,RPL30,H1-0,RPS5,RPS7,RPS19,RPL35,RPS10,RPL35A,TARDBP,RPLP2,CIRBP,RPL36,YWHAB,
ACF1,CYFIP2,MYH10,RHOB,MAP2,RAB1A,RAC1,ARHGDIA,CAPRIN1,GPC1,TOP2B,L1CAM,MAP1
S15,RPL12,RPL30,H1-0,RPS5,RPS7,RPS19,RPL35,RPS10,RPL35A,TARDBP,RPLP2,CIRBP,RPL36
RA,NCKAP1,MDK,PTK7,CNTNAP2,DPYSL2,NOTCH2,VEGFA,MAPK1,GRB2,PLCG1,CORO1C,FYN,
CKAP1,PTK7,CNTNAP2,DPYSL2,NOTCH2,VEGFA,MAPK1,GRB2,PLCG1,CORO1C,FYN,YWHAH,SL
RA,NCKAP1,MDK,PTK7,CNTNAP2,DPYSL2,NOTCH2,VEGFA,MAPK1,GRB2,PLCG1,CORO1C,FYN,
ACK1,BSG,FASN,ALDH6A1,SPTBN1,ATP5F1B,CALM3,GNB1,STMN2,MYH9,SEPTIN2,P4HB,APP,G
A2,SQSTM1,NOP53,UBE4A,VPS35,RAD23B,SNRNP70,PSMB2,ABCA2,NRDC,UBXN4,MVB12B,AZI
PPRC,KHDRBS1,RPS25,EIF5A,DDX3X,RPS15,RPL12,RPL30,EIF3L,RPS5,RPS7,RPS19,RPL35,NG

SN,RPL24,COL1A2,LLGL1,RHOA,PSMD2,RPS14,EIF5,RBBP4,PFN1,CLEC7A,SPTBN2,DDX3X,RPS
COL5A1,CES3,PSAP,PGK1,RPLP0,H3-3B,CLU,PTMA,DDX5,ENO2,SOX4,FKBP14,RPL13A,MAGED1
ICS1,RHOA,NR2F1,KIF5B,ENC1,CRMP1,PFN1,PRKCSH,SPTBN2,PTN,CSNK1E,TPM1,CDH2,DDR1,
L12,RPL30,RPS7,IK,RPS19,RPL35,RPL35A,HNRNPDL,TARDBP,RPLP2,TSR2,HNRNPK,CIRBP,RPL
N1,TMBIM6,GSTP1,NPM1,HSPD1,IGF2,VCP,UBB,YWHAG,COL4A2,MAPT,FN1,NONO,PSMD5,EGL
ICKAP1,PTK7,CNTNAP2,DPYSL2,NOTCH2,VEGFA,MAPK1,GRB2,PLCG1,FYN,YWHAH,SLIT3,STXBF
EIF3L,H1-0,RPS5,RPS19,RPS10,CYFIP2,H1-10,PTGES3,CKAP5,MAP2,RPS27,COTL1,RAC1,RPL23
MA,DDX5,SOX4,FKBP14,RPL13A,MAGED1,RPL6,KAT6A,DVL3,YWHAE,RPL13,DDX17,GPI,HTRA4,
,SMG1,CALM1,NPM1,PLXNB2,HSPD1,CSDE1,IGF2,EIF3C,EIF4B,VCP,UBB,TRIM28,YWHAG,MAPT
L37,RPS11,RPL28,RPS28,FYN,FAU,RPL39,RPS13,RPL34,SSR1,SGTA,ALDH3A2,UBA52,RPS2,RPS
PF1,TRA2B,HNRNPA2B1,RPS27A,PSMD8,APEX1,PCBP4,SAFB2,PRDX6,FUS,PSMB4,ELAVL1,TTC3
ICS1,RHOA,NR2F1,KIF5B,ENC1,CRMP1,PFN1,NES,PRKCSH,SPTBN2,PTN,CSNK1E,TPM1,CDH2,DI
RPPRC,KHDRBS1,RPS25,EIF5A,DDX3X,RPS15,RPL12,RPL30,EIF3L,RPS5,RPS7,RPS19,RPL35,NG
M1,HSPD1,IGF2,VCP,UBB,YWHAG,COL4A2,MAPT,FN1,PSMD5,INA,HLA-C,PDIA3,RPS3,HMGB1,IT
F1B,CALM3,GNB1,STMN2,MYH9,SEPTIN2,P4HB,APP,GDI1,SPTAN1,GSTP1,MARCKS,CALM1,PLX
A,CDH11,TBCD,PTPRA,MDK,LAMC1,PTK7,DPYSL2,NOTCH2,VEGFA,MAPK1,ARPC2,GRB2,PRKD
HMGB1,AP2M1,STMN1,ITGB1,UCHL1,NCAM1,ANXA2,AGRN,ZMIZ1,COL11A1,VCL,PAFAH1B1,AP
AM,MAP1A,CRPPA,CDH11,GAS7,PTPRA,NCKAP1,MDK,PTK7,CNTNAP2,ABCA2,DPYSL2,NOTCH2,
LCG1,FYN,YWHAH,SLIT3,STXBP1,PLXNA3,PTPRS,SOD1,RAB11A,TCTN1,BMPR2,CDC42,PALLD,KI
STP1,SMG1,CALM1,NPM1,PLXNB2,HSPD1,VCP,UBB,YWHAG,CKB,MAPT,FN1,DDB1,PSMD5,YWH
AR2,POLR2B,SUPT6H,HNRNPM,SRSF3,SF3A1,STRAP,PCBP2,HNRNPD,RPS13,PRPF40A,PPP1R9
NB1,STMN2,MYH9,SEPTIN2,APP,GDI1,SPTAN1,GSTP1,MARCKS,CALM1,PLXNB2,HSPD1,COL18A1
A1,ARIH1,SGTA,PSMD1,UBQLN2,CSNK2A1,CST3,PTEN,OAZ2,GPX1,EIF3H,LDLR,GNA12,RDX,DN
M1,NPM1,PLXNB2,HSPD1,CSDE1,IGF2,EIF3C,EIF4B,VCP,UBB,TRIM28,YWHAG,MAPT,FN1,SPEN,
26,RPL31,LRPPRC,KHDRBS1,RPS25,EIF5A,ACSL6,DDX3X,RPS15,RPL12,RPL30,EIF3L,RPS5,RPS
TAN1,GSTP1,MARCKS,CALM1,PLXNB2,HSPD1,COL18A1,CSDE1,IGF2,MARCKSL1,UBB,TRIM28,
1,CES3,PSAP,PGK1,RPLP0,H3-3B,CLU,PTMA,DDX5,ENO2,SOX4,FKBP14,RPL13A,MAGED1,RPL6,
40A,RBM15B,DHX15,POLR2E,SF1,PRCC,SRSF6,TRA2B,HNRNPA2B1,CHERP,SYMPK,PCBP4,PRD
40A,RBM15B,DHX15,POLR2E,SF1,PRCC,SRSF6,TRA2B,HNRNPA2B1,CHERP,SYMPK,PCBP4,PRD
40A,RBM15B,DHX15,POLR2E,SF1,PRCC,SRSF6,TRA2B,HNRNPA2B1,CHERP,SYMPK,PCBP4,PRD
RF4L1,ANXA2,RPLP1,ZMIZ1,MARCHF6,WSB1,SBF1,DNAJA1,CDC42BPB,OGA,RPL23,CPE,XRCC
RF4L1,ANXA2,RPLP1,ZMIZ1,MARCHF6,WSB1,SBF1,DNAJA1,CDC42BPB,OGA,RPL23,CPE,XRCC
IA3,PTPRS,RAB11A,TCTN1,BMPR2,PALLD,KIF5A,TRIO,FLOT1,MAP6,PLXNB1,DIP2B,EVL,HDAC6,
SSBP2,ZNF43,RPS3,HMGB1,AP2M1,RPS4X,RPL11,ANXA2,AGRN,RPLP1,ZMIZ1,AP3D1,AKAP12,ZN
,CCAR2,POLR2B,SUPT6H,HNRNPM,SRSF3,SF3A1,STRAP,PCBP2,HNRNPD,PRPF40A,ADAR,RBM1
E2R2,BNIP3L,CCAR2,SMARCC1,STT3B,LONP1,UBQLN1,CSNK1D,USP7,HECTD1,PCBP2,BAG6,P
IA1,SCARB2,NACA,RTN4,LAMA5,XRCC5,MSN,RPL24,LLGL1,GAP43,NCS1,RHOA,ATF4,NR2F1,KIF
AR1A,CTNNB1,CYFIP2,ARL6IP1,YWHAB,CKAP5,RHOB,MAP2,COTL1,CCT6A,CTBP1,RAC1,ARHGD
,RPS9,RPL24,RPL17,RPL14,RPL22,RPL9,ATF4,PSMD2,SLC7A5,ENC1,RPS14,RPS20,RPL26,SRSF
RPL9,ATF4,ENC1,RPS14,RPS20,EIF5,RPL26,RTN3,RPL31,LRPPRC,KHDRBS1,RPS25,EIF5A,ACSL
,RPL23A,GLUL,LUC7L3,DDX39B,SRSF5,NOP53,TSR1,RPL27,RPL36A,PRPF6,RPL38,RPL37,SRSF
Z1,AP3D1,ZNF770,GTF2F1,HNRNPU,SMARCA5,LARP1,CTSD,NACA,RPS9,XRCC5,MSN,RHOA,ATF
S1,NEFL,RPL8,YBX1,ATP2B4,IGFBP2,CALR,FSTL1,TMF1,ABHD2,HSPA1B,ZNF99,TUBB2B,COL5
SCARB2,LPCAT1,XRCC5,MSN,KPNB1,C1ORF43,LLGL1,NCS1,RHOA,CLSTN1,PSMD2,KIF5B,RAB
RF,BRSK1,SLIT2,GRN,FKBP4,CARM1,LINGO1,ABI2,GPC2,STK24,SEMA3C,ATP1B2,ADAM10,IST

I1,RPS4X,RPL11,ANXA2,AGRN,RPLP1,ZMIZ1,AP3D1,AKAP12,ZNF770,GTF2F1,HNRNPU,SMARCA5
,HNRNPA0,RPL23,NACA,RTN4,LPCAT1,XRCC5,CIPC,IGFBP3,CBX5,ATF4,NR2F1,ENC1,RPS14,RP
PR2,PALLD,KIF5A,TRIO,FLOT1,MAP6,PLXNB1,DIP2B,EVL,HDAC6,PTEN,ULK1,BRSK1,CTTN,SLIT2
B,HNRNPU,SMARCA5,LARP1,HNRNPA0,RPL23,NACA,RTN4,LPCAT1,XRCC5,CIPC,IGFBP3,RHOA,C
N,YWHAH,B2M,PLXNA3,PTPRS,PFN2,RAB11A,AKIRIN1,BMPR2,CDC42,EP300,AP2A1,MAP6,PLXNE
RHOA,ATF4,ATP1A1,JUND,CD81,COL5A2,SRSF4,PTN,GLG1,GPC3,SULF1,PIP4K2B,MEGF8,CCN2
1C,FYN,YWHAH,B2M,PLXNA3,PTPRS,PFN2,RAB11A,AKIRIN1,BMPR2,CDC42,EP300,AP2A1,MAP6,
RHOA,ATF4,RPL26,BEX3,RTN3,MCL1,TLE5,NES,NISCH,PTN,SFPQ,EIF5A,CCND2,PNMA1,SULF1,
0,GTF2F1,HNRNPU,SMARCA5,LARP1,CTSD,NACA,LPCAT1,RPS9,XRCC5,MSN,RHOA,ATF4,NR2F1,
,RPL7,APP,RPL5,TMBIM6,GSTP1,SMG1,NPM1,PLXNB2,PLD3,HSPD1,SNRNP200,CSDE1,IGF2,EIF
,RPSA,EIF4H,NACA,LPCAT1,RPS24,RPS9,RPL24,RPL17,RPL14,RPL22,RHOA,PTCD3,RPL9,ATF4,E
LXNB2,PLD3,COL18A1,SNRNP200,IGF2,UBB,AFF4,TRIM28,YWHAG,COL4A2,MAPT,FN1,SPEN,PSN
HTRA4,RACK1,BSG,FASN,SPTBN1,CALM3,GNB1,STMN2,P4HB,APP,GDI1,FSCN1,TMBIM6,SPTAN
K1,FASN,ALDH6A1,ATP5F1B,RPS8,RPL7,APP,RPL5,TMBIM6,GSTP1,SMG1,NPM1,PLXNB2,PLD3,H
PLD3,COL18A1,SNRNP200,IGF2,UBB,AFF4,TRIM28,YWHAG,COL4A2,MAPT,FN1,SPEN,PSMD5,D
PEN,NONO,PSMD5,YWHAQ,SET,EGLN1,RPL19,PPP2R1A,RPL15,GTF3C4,PKD1,SF3B1,SSBP2,ZN
SN,ATP5F1B,RPS8,RPL7,APP,RPL5,TMBIM6,PTMS,GSTP1,SMG1,NPM1,PLXNB2,HSPD1,CSDE1,I
SN,ATP5F1B,RPS8,RPL7,APP,RPL5,TMBIM6,PTMS,GSTP1,SMG1,NPM1,PLXNB2,HSPD1,CSDE1,I
ORF4L1,ANXA2,RPLP1,ZMIZ1,MARCHF6,WSB1,SBF1,DNAJA1,CDC42BPB,OGA,RPL23,CPE,XRCC5
P,RPL5,TMBIM6,SMG1,NPM1,PLD3,HSPD1,SNRNP200,CSDE1,IGF2,ATP5F1A,VCP,UBB,AFF4,TRI
39B,MORF4L2,CLSTN3,MDK,PTK7,H3-3A,DPYSL2,TMED2,NOTCH2,VEGFA,SELENON,PPP2CA,M
FBP3,RHOA,ATF4,RPL26,BEX3,RTN3,MCL1,TLE5,CLEC7A,NES,NISCH,PTN,SFPQ,EIF5A,CCND2,P
GF2,VCP,UBB,AFF4,TRIM28,COL4A2,RPL37A,MAPT,SPEN,NONO,DDB1,PSMD5,YWHAQ,SET,EGL
ISPA1A,ATP13A2,MDM2,FZR1,MGAT3,SUMO1,PARK7,USP14,HIPK2,MAGEF1,TRIB3,PSME3,STUE
TP1,NPM1,PLXNB2,HSPD1,CSDE1,IGF2,EIF3C,EIF4B,VCP,UBB,AFF4,TRIM28,COL4A2,RPL37A,SP
M1A,CLIP3,CORO1C,SUPT6H,FYN,YWHAH,B2M,TNKS2,STXBP1,PLXNA3,PTPRS,GHITM,PFN2,KD
P1,NOX4,ABCA3,PTPRA,ACTR3,CAMSAP1,NCKAP1,VPS35,CLSTN3,CNTNAP2,ABCA2,VEGFA,RAPG
P,DCN,MAP1A,TIMP3,CTSB,RNF187,EEF1A2,SQSTM1,NOP53,VCAN,UBE4A,VPS35,RAD23B,SNRN
STP1,SMG1,NPM1,PLXNB2,PLD3,HSPD1,CSDE1,IGF2,EIF3C,EIF4B,UBB,AFF4,TRIM28,MATR3,R
TNNA1,NACA,RTN4,LAMA5,XRCC5,MSN,IGFBP3,RHOA,CLSTN1,ATF4,PSMD2,NLGN2,JUND,COL
VEGFA,MAPK1,NPEPPS,KDM1A,BNIP3L,CLIP3,FYN,YWHAH,CCT8,MESD,HECTD1,SREBF2,RAB11
ME1L1,PSMA7,ARIH1,SGTA,OS9,CAPN2,MTA1,PSMD1,UBQLN2,CSNK2A1,UBA52,PPP2CB,KLHL42
B2,HSPD1,CSDE1,IGF2,EIF3C,EIF4B,VCP,UBB,AFF4,TRIM28,COL4A2,RPL37A,SPEN,NONO,PSMD
TSD,NACA,RTN4,KPNB1,IGFBP3,RHOA,ATF4,TMEM259,RPL26,BEX3,RTN3,MCL1,TLE5,CLEC7A,
4,RHBDD2,PSMB4,GPX1,EIF3H,GNA12,DNAJB2,ATXN3,PSMD4,OGT,PRPF19,FBXW5,GSK3A,PSM
,DIP2B,GNG4,SFRP1,HDAC6,ULK1,CTTN,SLIT2,NDN,BAP1,SLIT1,OLFM1,LTBP4,HDGFL2,DCLK1,
GCS1,PRKAR1A,GJA1,JUN,MYO5A,CTNNB1,SESN3,MTSS2,ARID1A,PTGES3,HNRNPH1,ATP1A2,G
,FADS1,NEFL,RPL8,YBX1,ATP2B4,IGFBP2,CALR,FSTL1,TMF1,ABHD2,HSPA1B,ZNF99,TUBB2B,C
GB1,TPT1,STMN1,ITGB1,UCHL1,RPL11,NCAM1,ANXA2,ZMIZ1,TIMP2,MIDN,DNAJA1,ARF1,PAFAH
COL6A2,CYFIP2,CKAP5,RHOB,MAP2,TTR,COTL1,RAC1,PPIB,MAP1A,TBCD,KLHL24,GAS7,NOX4,T
JVBL2,SF3A3,CD2BP2,SART3,SRSF10,LSM4,CELF3,EIF3B,PSIP1,SNRPD2,SETX,PRMT5,EIF3M,SN
PL7,APP,RPL5,TMBIM6,SMG1,CALM1,NPM1,PLD3,HSPD1,SNRNP200,CSDE1,IGF2,ATP5F1A,VCP
APP,RPL5,TMBIM6,PTMS,GSTP1,SMG1,NPM1,PLXNB2,HSPD1,CSDE1,IGF2,EIF3C,ATP5F1A,EIF4B
PNMA1,RAB7A,ACTN4,PRDX2,DDX3X,ITM2C,RPS7,MIF,GJA1,JUN,CTNNB1,DNAJC5,TARDBP,ARL
,MDK,PTK7,PSMB2,VEGFA,ARPC2,PSMD3,CAPZB,PRKDC,HSPB1,PLCG1,CORO1C,FYN,YWHAH,

BP2,SART3,SRSF10,LSM4,CELF3,EIF3B,PSIP1,SNRPD2,SETX,PRMT5,EIF3M,SNRPB,SFSWAP,PRF
AMP1,LGALS3BP,CKAP4,HLA-B,TTR,COTL1,HRNR,RAC1,CDK16,ATP6V0C,TMEM30A,TIMP3,PGRI
,QARS1,CCAR2,FYN,B2M,UBQLN1,HYOU1,PTPRS,MAPK8IP1,SOD1,BAG6,TP53BP1,CDC42,TRAF
,MMP14,PRDX2,DDX3X,MIF,PRKAR1A,GJA1,CTNNB1,NGRN,CYFIP2,NDFIP1,CIRBP,YWHAB,RAB1
,HLA-B,TTR,COTL1,HRNR,RAC1,ATP6V0C,TMEM30A,TIMP3,PGRMC1,CNN2,CTSB,RAB27A,CCT2
GB1,TPT1,STMN1,ITGB1,UCHL1,RPL11,NCAM1,ZMIZ1,TIMP2,MIDN,DNAJA1,ARF1,PAFAH1B1,AK
PL7,APP,RPL5,TMBIM6,SMG1,CALM1,NPM1,PLD3,HSPD1,SNRNP200,CSDE1,IGF2,ATP5F1A,VCP
IM6,NPM1,PLXNB2,HSPD1,CSDE1,IGF2,EIF3C,ATP5F1A,EIF4B,VCP,UBB,AFF4,TRIM28,COL4A2,F
FBP2,CALR,FSTL1,TMF1,ABHD2,HSPA1B,ZNF99,TUBB2B,COL5A1,COL4A1,PSAP,RPLP0,H3-3B,C
A12,DNAJB2,ATXN3,PSMD4,OGT,FBXW5,GSK3A,PSMC3,UBE2D3,PSMC1,CCDC47,BAG5,UBE2K,
NDFIP1,YWHAB,CADM4,RAB1A,CTBP1,HMGCR,RAC1,HDAC2,IPO5,SKP1,TRIB2,PEA15,PTTG1IP,
LN2,CSNK2A1,UBA52,PPP2CB,KLHL42,UBE2G2,FBXO11,HDAC6,PTEN,CUL3,RPS27A,PSMD8,NP
R10,SNRPD3,UBR5,ELAVL1,SMG5,SRSF2,SRSF11,CSE1L,DHX38,PRKACA,XPO6,SMO,EIF4A3,RA
DIA,GPC1,CDK16,APPL1,IPO5,TMEM30A,GLUL,NNAT,MAP1A,NUCB1,CCT2,SQSTM1,VAMP2,PAR
LN2,CSNK2A1,UBA52,PPP2CB,KLHL42,UBE2G2,FBXO11,HDAC6,PTEN,CUL3,RPS27A,PSMD8,NP
RPRA,EIF4G1,AMFR,SCAMP5,TSPYL2,HSPA1A,TMX1,UBE4B,PARK7,USP14,ERP29,PTPN1,FAF2,U

BA52,PPP2CB,KLHL42,UBE2G2,FBXO11,HDAC6,PTEN,CUL3,RPS27A,PSMD8,NPLOC4,FBXO21,R
CTN4,PRDX2,DDX3X,ITM2C,RPS7,MIF,GJA1,JUN,CTNNB1,DNAJC5,TARDBP,ARL6IP1,HNRNPK,RH
P1B2,IST1,EHD1,SF3A2,MTOR,APBB1,METRN,CROCC,NTRK2,SNX3,AUTS2,TRAK1,PLXNA2,OBSL

,NES,PTN,SFPQ,CCND2,PNMA1,RAB7A,CCN2,ACTN4,PRDX2,DDX3X,ITM2C,RPS7,MIF,GJA1,JUN,
HDAC2,IPO5,LAPTM4B,TRIB2,MAP1A,CDKN1A,TIMP3,CTDSP2,SRP9,U2AF2,SQSTM1,NOP53,PRD
CDK16,APPL1,IPO5,TMEM30A,GLUL,MAP1A,NUCB1,CCT2,SQSTM1,VAMP2,PARP1,TCP1,CCT3,CC
I10,LRRC41,FBXW5,PSMC3,TGOLN2,PSMC1,FBXL19,IGFBP4,WDTC1,LGALS1,KTN1,PSMB1,OBS
P1,COL2A1,MDK,SKI,KDM6B,CNTNAP2,ABCA2,DPYSL2,MAPK1,PBX2,SRSF1,ATP5PB,LUZP6,PRK
ACTN1,ARF4,CORO1C,FYN,MESD,PTPRS,PFN2,FLNC,CDC42,CTNND1,GRIPAP1,DCTN1,BCR,CNT
C4,PSMD7,TNRC6A,NBAS,HNRNPR,DDX6,LSM4,MAPKAPK2,PSME3,DHX36,PSMC5,PSMC2,RBM1
RNPS1,ACIN1,PAF1,CPSF7,RNF40,RBM39
PL7,APP,RPL5,TMBIM6,SMG1,CALM1,NPM1,PLD3,HSPD1,SNRNP200,CSDE1,IGF2,ATP5F1A,VCP
7,TNRC6A,HNRNPR,DDX6,LSM4,MAPKAPK2,PSME3,DHX36,PSMC5,PSMC2,RBM10,PSMC4,PSMF
A,PHB2,UBA52,RAB3GAP2,HDAC6,ULK1,GPSM1,RPS27A,HIF1A,CTTN,SLC25A5,MAP1LC3B,DAP,V
A,PHB2,UBA52,RAB3GAP2,HDAC6,ULK1,GPSM1,RPS27A,HIF1A,CTTN,SLC25A5,MAP1LC3B,DAP,V
TPRS,SOD1,RAI1,RAB11A,AKIRIN1,BMPR2,STAT3,EP300,KDM5B,ZFP36L1,DIP2B,ATRX,SFRP1,HD
R10,SNRPD3,UBR5,ELAVL1,SMG5,SRSF2,SRSF11,CSE1L,DHX38,PRKACA,XPO6,SMO,EIF4A3,RA
,MARCHF7,PSMC5,CSNK1A1,PSMC2,HERPUD1,PSMC4,PLK2,PSMF1,WAC,GSK3B,USP19,UBQLN
1A2,HMGCR,RAC1,ARHGDIA,HDAC2,TNC,MMP2,HSBP1,DCN,CDKN1A,CNN2,EEF1A2,PPP1CB,NC
WDR1,CRMP1,PFN1,SPC25,NES,NISCH,XPO1,SPTBN2,TPM1,CCN2,CLTC,PRKAR1A,TLN1,GJA1,
CNI,KHDRBS1,KLHL9,XPO1,CSNK1E,SFPQ,CCND2,CCN2,DDX3X,CLTC,IK,PRKAR1A,GJA1,JUN,C
NK2,USP47,PINK1,HIP1R,HSPA1A,MDM2,SUMO1,PARK7,ATP1B3,KAT2A,B4GALT5,ATF7IP,DAD1,
N,SFRP1,CTNND2,PTEN,CUL3,RPS27A,TNKS,PSMD8,DKK3,ATP6AP2,GPC4,PSMB4,UBR5,SOX2,T
,TMOD2,FKBP4,ABI2,RDX,CAPZA1,CLASP2,WASF2,PSMC3,MTOR,TUBB4A,EIF4G1,LMO4,PTPN1
JB2,SEMA3C,SPAG9,CLASP2,ADAM10,IST1,SH3BP4,GSK3A,FBLN5,MTOR,YY1,APBB1,NDUFA13,
45B,CAPN2,MMP15,FLOT1,PHB2,GSPT1,CST3,UPF1,HDAC6,MAPRE2,HIF1A,CTTN,SLC25A5,A2M
N,SFRP1,CTNND2,PTEN,CUL3,RPS27A,TNKS,PSMD8,DKK3,ATP6AP2,GPC4,PSMB4,UBR5,SOX2,T

NB1,NGRN,CYFIP2,NDFIP1,CIRBP,YWHAB,RAB1A,CTBP1,RAC1,HDAC2,SKP1,TRIB2,PEA15,PTTG1
,MMP2,PEA15,CDKN1A,SRSF5,TOMM20,VAMP2,GNAI3,PARP1,NOX4,PTPRA,UBE4A,AHCYL1,VPS
MF1,WAC,GSK3B,USP19,UBQLN4,UBXN1,PSMD10,ELOB,GABARAPL2,XBP1,APOE,RYBP,PSMC6,
PK1,PSMD3,CAND1,ILF2,CCT8,B2M,IDH1,DYNLL1,TXNDC5,PSMD1,PDAP1,CST3,ATP6AP2,PRDX6,
FR,HSPA1A,RUVBL2,DNAJB1,ERP29,MOGS,CSNK2B,CLPX,UNC45A,P3H1,TBCA,HSPE1,UBXN1,DN
UP153,DHX36,SRSF7,NUP98,POLDIP3,EXOSC10,NCBP2,HSF1,IGF2BP2,FBL,NUP205,NUP133,TH
P7,HECTD1,CDC42,PSMA7,TRAF4,TRAF7,ARIH1,OS9,MTA1,PSMD1,UBA52,KLHL42,NXN,UBE2G2
,SFSWAP,RNPS1,ACIN1,AKAP17A,ZNF326,RBM39
CDKN1A,TIMP3,CTDSP2,SRP9,U2AF2,SQSTM1,NOP53,PRDX5,PKN1,SKI,TERF2IP,TMED2,TAF7,V
PK1,PSMD3,CAND1,ILF2,CCT8,B2M,IDH1,DYNLL1,TXNDC5,PSMD1,PDAP1,CST3,ATP6AP2,PRDX6,
7,TNRC6A,HNRNPR,DDX6,LSM4,MAPKAPK2,PSME3,DHX36,PSMC5,PSMC2,RBM10,PSMC4,PSMF
SESN3,APPL1,HDAC2,IPO5,ATP6V0C,MMP2,PEA15,CDKN1A,SRSF5,TOMM20,VAMP2,GNAI3,PARP
GB,MAPK1,PSMD3,CAND1,ILF2,CCT8,B2M,IDH1,DYNLL1,TXNDC5,PSMD1,PDAP1,CST3,ATP6AP2,
IRNPM,CSNK1D,STRAP,HGS,BMPR2,DDIT4,PPP1R9B,POLR2E,SULF2,BCL9L,EP300,KDM5B,FLC
C,GABARAP,UBE2R2,CAND1,NPEPPS,KDM1A,FYN,EIF3F,TNKS2,UBQLN1,USP7,HECTD1,HGS,CD
MTRNR2L8,PKM,GANAB,COL6A1,FTL,HSP90B1,FADS1,NEFL,NEFM,PRPF8,UGT8,RPL8,FBXL4,Y
GB,MAPK1,PSMD3,CAND1,ILF2,CCT8,B2M,IDH1,DYNLL1,TXNDC5,PSMD1,PDAP1,CST3,ATP6AP2
M1,ZMIZ1,TIMP2,DNAJA1,PAFAH1B1,AKAP12,PXDN,CTNNA1,RPL23,RTN4,PJA2,IGFBP3,LLGL1,R
N2,HNRNPD,WBP2,CCT5,CDC42,VDAC1,DCTN1,BRD4,ATRX,SFRP1,EVL,RAB3GAP2,HNRNPA2B1
AR1A,GJA1,CTNNB1,MYH10,DUX4,CKAP5,SEPTIN11,SND1,RHOB,SEPTIN3,RPS27,CDK16,TOP2E
PK1,PSMD3,CAND1,ILF2,CCT8,B2M,IDH1,DYNLL1,TXNDC5,PSMD1,PDAP1,CST3,ATP6AP2,PRDX6,
TD1,CDC42,PSMA7,TRAF4,TRAF7,ARIH1,OS9,MTA1,PSMD1,UBA52,KLHL42,NXN,UBE2G2,FBXO
TDSP2,TBCD,PPP1CB,NOP53,PSMB2,TUBB2A,OPTN,PPP2CA,CCND1,MAPRE1,PSMD3,DYNC1LI2
S2,PRKACA,EXOC4,BNIP3,ATP6V1G1,TOMM70,UBXN6,AMBRA1,ATG4B,PAFAH1B2,POLDIP2,PIN
I1A,CTDSP2,DDX39B,SRSF5,NOP53,PSMB2,RPS15A,CCND1,MAPRE1,CCNL2,PSMD3,IPO7,CDC37
I,RAB7A,CCN2,ACTN4,DDX3X,MAGED2,MIF,TLN1,GJA1,CD63,DNAJC5,APLP2,TARDBP,DDOST,W
ND1,MAPRE1,PSMD3,DYNC1LI2,DYNLL2,SON,CSNK1D,RAB11A,DYNLL1,CDC42,PPP1R9B,PSMA
GSK3B,ALCAM,SOX9,CRABP2,UNC13A,MED12,LZTS2,NRP2,RGMA,RUFY3,EXT1,SPG11,SEMA6C,

DRBS1,SFPQ,EIF5A,SOX12,CCN2,ACTN4,DDX3X,NANOG,H1-0,JUN,CTNNB1,NGRN,ARID1A,SHOX
AJA1,PAFAH1B1,AKAP12,CDC42BPB,LARP1,RPL23,TNS3,RTN4,PJA2,KPNB1,IGFBP3,COL1A2,NC

CT4,SKI,ASXL1,H3-3A,KDM6B,TERF2IP,TOP1,TAF7,VEGFA,MAPK1,SMARCA4,PRKDC,KDM1A,MS
GRN,RPLP1,TIMP2,MIDN,SBF1,DNAJA1,PAFAH1B1,GTF2F1,HNRNPU,RPL23,CNN3,CTSD,SCARB2,
FPQ,EIF5A,SOX12,ACTN4,DDX3X,NANOG,H1-0,JUN,CTNNB1,NGRN,ARID1A,SHOX,HNRNPK,H1-1
1,HSPE1,LMNA,ATF6B,SHC1,DNAJB11,ATF6,ADD1,QRICH1,XBP1,CUL7,PPP2R5B,SEC31A,EIF2S1
M,CSNK1D,STRAP,HGS,BMPR2,DDIT4,PPP1R9B,POLR2E,SULF2,BCL9L,EP300,KDM5B,FLCN,UB
CA2,TMED2,PPP2CA,ATG2B,ARPC2,ATP5PB,GRB2,BNIP3L,PITPNA,PLTP,CORO1C,YWHAH,PITPN
AH,B2M,PITPNM1,PLXNA3,CSNK1D,PTPRS,BAG6,HNRNPD,TCTN1,ARNT2,BMPR2,DYNLL1,CDC4
53C,TMEM98,NUP153,KPNA6,NUP98,GSK3B,ILRUN,SOX9,TFRC,TXN,LMNA,NUP133,DNAJB6,LZ
AJB11,ATF6,ADD1,QRICH1,XBP1,CUL7,PPP2R5B,SEC31A,EIF2S1,CDK5RAP3,ABCA7
UL1,PSMD13,ATF5,CCNY,PSMD7,CETN2,PPME1,CDK4,AKAP8L,PSME3,PSMC5,PSMC2,PSMC4,P
WASF2,IST1,MTOR,WDTC1,ABL1,SEMA6B,ARPC1A,HIP1R,ELN,ATP13A2,CRACD,ADNP,CYFIP1,NT
TCH2,VEGFA,MAPK1,PBX2,GRB2,PRKDC,TDRD5,CMIP,FKBP8,PLCG1,ZFAND5,DUSP4,SUPT6H,
2,PSMC4,PSMF1,RCC2,PSMB5,PRKAR2B,PSMA1,PSMD10,PSMA6,PSMD12,PSMA4,TPX2,FHL1,D

YT1,SYT4,POSTN,CACNG7,ANAPC2,SIN3A

G,H1-0,JUN,CTNNB1,NGRN,ARID1A,SHOX,HNRNPK,H1-10,CIRBP,DUX4,PTGES3,CCT6A,RAC1,H

LCAM,SOX9,CRABP2,UNC13A,PRMT2,NRP2,MAP2K4,RGMA,RUFY3,PIN1,SPG11,SEMA6C,APOE,

,CDC42,PPP1R9B,PSMA7,RBBP7,DCTN1,PRMT1,ACTR1A,EP300,PSMD1,PBX1,FLCN,CSNK2A1,P

1,OAZ2,GPSM1,HIF1A,SLC25A5,LDLR,RDX,TTC37,DNAJB2,ATXN3,AKT2,SH3BP4,GSK3A,PSMC3

,PKN1,AHCYL1,MDK,H3-3A,ABCA2,ATP6AP1,TAF7,MARS1,CCND1,MAPK1,SMARCA4,ADAM9,GR

A2,PSMA6,PSMD12,PSMA4,SEC31A,DYNC1I2,PSMC6

DIT4,PPP1R9B,ADAR,BEX4,PRKAR2A,NXN,SFRP1,CRTAP,MASTL,PTEN,PPP1R15A,FKBP1A,SLIT2,

MB5,PRKAR2B,PSMA1,PSMD10,PSMA6,PSMD12,PSMA4,TPX2,FHL1,DYNC1I2,CENPF,PSMC6,CD

5B,SRP14,PYGB,MAPK1,PSMD3,CAND1,ILF2,CCT8,B2M,STXBP1,IDH1,DYNLL1,TXNDC5,PSMD1,

CP,UBB,TRIM28,MATR3,MAPT,FN1,NONO,DDB1,PSMD5,HACD3,EGLN1,PKD1,HLA-C,PDIA3,RPS

CH,PFKL,SCRN1,RAB7A,CCN2,ACTN4,DDX3X,MAGED2,MIF,TLN1,GJA1,CD63,DNAJC5,APLP2,TA

GB,MAPK1,PSMD3,CAND1,ILF2,CCT8,B2M,STXBP1,IDH1,DYNLL1,TXNDC5,PSMD1,PDAP1,CST3,

,DDX5,SOX4,FKBP14,RPL13A,MAGED1,KAT6A,DVL3,YWHAE,GNAS,DDX17,GPI,HTRA4,RACK1,BS

SMD7,CETN2,PPME1,CDK4,PSME3,PSMC5,PSMC2,PSMC4,PSMF1,RCC2,PSMB5,PRKAR2B,PSM

,RPL39,RPL34,TRAF7,UBA52,PTEN,RPL7L1,RPS2,PELP1,WBP11,NOC2L,SART1,EIF4A3,XRN2,N

14,PRDX2,NANOG,RPS19,GJA1,JUN,CTNNB1,MACF1,NDFIP1,HLA-B,MAP2,RAC1,ARHGDIA,CAP

HNRNPD,TCTN1,ARNT2,BMPR2,DYNLL1,CDC42,DDIT4,PPP1R9B,CTNND1,CIC,RERE,PBX1,SEC1

1,EPHA4,PLPP3,ZNF207,STX12,USP36

BP,PINK1,ATP13A2,MGAT3,PARK7,USP14,BSCL2,SYNCRIP,FEZ1,EHMT2,HIPK2,NBAS,MYCBP2,

0,CAPZA2,PSMA6,PSMD12,PSMA4,SEC31A,DYNC1I2,PSMC6

RMT1,NAP1L1,MTA1,KDM5C,EP300,RERE,KDM5B,FLCN,BAZ1B,H2AZ1,GPX4,BRD4,BRD2,ATRX

KDCC,RAB14,KIFAP3,CAPZA1,RAB6A,ERGIC3,COPZ1,AP1M1,EXOC4,SORT1,DOP1B,GAK,USO1

B1,SFRP1,EVL,HDAC6,MAPRE2,CTTN,SLIT2,BRK1,AKAP9,TMOD2,TAOK1,FKBP4,ABI2,RDX,CAP

OD1,RAI1,RAB11A,IGFBP7,BMPR2,CDC42,PPP1R9B,RBBP7,STAT3,EXTL3,FLCN,CSNK2A1,DIP2

TNPO1,PSMB2,H3-3A,TMED2,RPL38,MAPK1,PSMD3,EIF3E,FTO,ILF3,LUZP6,CDC37,PRKDC,HSP

2AK1,EDC4,EIF3D,GSPT1,ZFP36L1,UPF1,EIF5B,PPP1R15A,SERP1,CCN1,EIF3H,ELAVL1,IREB2,L

GB,MAPK1,PSMD3,CAND1,ILF2,CCT8,B2M,STXBP1,IDH1,DYNLL1,TXNDC5,PSMD1,PDAP1,CST3,

1,BAG6,DDIT4,TRAF7,EP300,HIP1,CSNK2A1,SFRP1,PTEN,FEM1B,CUL3,TIMM50,HIF1A,PPP1R15

33,THOC2,LZTS2,THOC3,FYTTD1,ABCE1,NUP50,RNPS1,DDX19A,NUP58,NEMF,WDR33,NUTF2,E

RIP,FEZ1,EHMT2,HIPK2,NBAS,TBC1D14,MAPKAPK2,DHX36,MARCHF7,RBM10,PSMF1,WAC,USP

SLIT2,GRN,LINGO1,LDLR,SEMA3C,DICER1,IST1,PRPF19,KDM4A,GSK3A,MTOR,CERS2,SMO,ME

1G2,NXN,SFRP1,CTNND2,PTEN,CUL3,RPS27A,TNKS,PSMD8,DKK3,ATP6AP2,GPC4,PSMB4,UBR5

HAQ,EIF1,SET,EGLN1,PPP2R1A,PKD1,SSBP2,ZNF43,RPS3,CHD3,BASP1,HMGB1,RPS4X,MORF4

SNK1A1,PSMC2,HERPUD1,PSMC4,PLK2,GSK3B,SOX9,CDKN1B,EGLN2,PSMD10,ELOB,RGMA,X

SCRN1,DDR1,RAB7A,CCN2,ACTN4,DDX3X,MAGED2,MIF,GPAT4,TLN1,GJA1,CD63,DNAJC5,APLP

RCC5,IGFBP3,LLGL1,RHOA,PSMD2,RPS20,EIF5,CD81,BEX3,OAZ1,CLEC7A,NES,CCNI,PTN,PEBP

2,ARL6IP1,NDFIP1,UBE2H,HDAC2,MMP2,SKP1,TRIB2,MAP1A,TIMP3,CTSB,TOMM20,RNF187,SQ

,MTOR,DAPK3,TUBB4A,ABL1,ARPC1A,HDGFL3,HIP1R,HSPA1A,ELN,CRACD,MECP2,PARK7,CYFI

IF4A3,XRN2,NOP56,MTREX,TRMT112,RPS21,RPL27A,EXOSC10,PDCD11,PA2G4,FBL,RRP1B,SE

,MDK,NOTCH2,VEGFA,VHL,MAPK1,DDAH2,PRKDC,HSPB1,SON,KDM1A,BNIP3L,FKBP8,QARS1,C

N1A,ANXA5,SQSTM1,GNAI3,PRDX5,GRINA,VPS35,COL2A1,MDK,NOTCH2,VEGFA,VHL,MAPK1,DD

11,SART1,EIF4A3,XRN2,NOP56,MTREX,TRMT112,RPS21,RPL27A,EXOSC10,PDCD11,PA2G4,FBL

HOC3,RTRAF,FYTTD1,NUP50,RNPS1,DDX19A,NUP58,WDR33,NUTF2
HOC3,RTRAF,FYTTD1,NUP50,RNPS1,DDX19A,NUP58,WDR33,NUTF2
T1,BRD4,UBA52,ZFP36L1,UPF1,MASTL,PTEN,FEM1B,CUL3,BRSK1,RPS27A,PSMD8,APEX1,PCBP
F1,EIF5B,PPP1R15A,SERP1,EIF3H,ELAVL1,IREB2,LSM14A,TTC37,EIF4G3,TRAP1,LSM14B,ABCF1
4A,DZIP1,FNBP1L,CLASP2,WASF2,PSMC3,MTOR,CROCC,EIF4G1,NTRK2,PTPN11,CDC34,ABL1,C
SET,EGLN1,PPP2R1A,PKD1,SSBP2,ZNF43,RPS3,CHD3,BASP1,HMGB1,RPS4X,MORF4L1,AGRN,Z
,RAPGEF1,MVB12B,MAPK1,ARPC2,ADAM9,PTBP1,GRB2,CDC37,HSPB1,FKBP8,PLCG1,ARF4,ZF
1,CITED2,CFLAR,PSMD10,ROCK2,ELOB,PSMA6,AJUBA,PSMD12,OPA1,PSMA4,PSMC6

COL2A1,MDK,NOTCH2,VEGFA,VHL,MAPK1,DDAH2,PRKDC,HSPB1,SON,KDM1A,BNIP3L,FKBP8,Q

NP,CYFIP1,NTN1,LIMK1,RHEB,C1QBP,ISLR2,DHX36,CRKL,GOLGA4,LIMS1,SRF,SERPINE2,PRM
PSMA1,CITED2,CFLAR,PSMD10,ROCK2,ELOB,PSMA6,AJUBA,PSMD12,OPA1,PSMA4,PSMC6,NAM
,UBE3A,TANC2,EIF4G1,NTRK2,DIP2A,ABL1,SDCBP,DAG1,NRXN2,OBSL1,IGF1R,NPTN,MECP2,EF
,PLXNA2,OBSL1,NOTCH1,PCM1,NPTN,ADNP,IDH2,CYFIP1,NTN1,LIMK1,RHEB,TMEM98,SYNGAP
RABP2,NUMBL,SRRT,SS18L1,RUFY3,CUL7,PRKCI,RELA,OPA1,EPHB2,LRP8,EPHA4,PLXND1,ID2,

RF4L1,AGRN,ZMIZ1,RPL10,AP3D1,ZNF770,GTF2F1,HNRNPU,PGAM1,NFE2L1,SMARCA5,LARP1
,GBA2,MAPRE3,PDE4DIP,CDKN1B,WASF3,CDC42EP4,VDAC2,ARPC5,CAPZA2,ADD1,FER,GMFB,G
IF1,SET,EGLN1,PPP2R1A,PKD1,SSBP2,ZNF43,RPS3,CHD3,BASP1,HMGB1,RPS4X,MORF4L1,AGR

RAP3,ABCA7
HOC3,RTRAF,FYTTD1,NUP50,RNPS1,DDX19A,NUP58,WDR33,NUTF2
SP19,STC2,PSMB5,EGLN2,HIF3A,LMNA,PSMA1,CITED2,CFLAR,PSMD10,ROCK2,ELOB,PSMA6,A

LN1,PPP2R1A,PKD1,SSBP2,ZNF43,RPS3,CHD3,BASP1,HMGB1,RPS4X,MORF4L1,AGRN,ZMIZ1,R
UL3,BRSK1,RPS27A,PSMD8,APEX1,PCBP4,AKAP9,PSMB4,TAOK1,CARM1,INTS3,VPS4A,RDX,KMT

B,CTBP1,SMCHD1,HDAC2,BCLAF1,HSBP1,CDKN1A,EID1,U2AF2,SQSTM1,NOP53,PARP1,MORF4
D23A
IM8,CD74,DEK,SNX3,HTATSF1,LGALS1,CHMP1A,DAG1,HSPA1A,NOTCH1,EFNB2,BANF1,RAN,TA
BP,DUX4,PTGES3,CCT6A,SMCHD1,HDAC2,BCLAF1,DCN,DDX39B,SRSF5,CHD6,RNF187,U2AF2,C

NAPC2,RB1CC1,SIN3A

A2,PSMA6,PSMD12,PSMA4,SEC31A,DYNC1I2,PSMC6

K1,NPTN,MGAT3,PARK7,RAN,CCDC88A,YWHAZ,GSK3B,ABHD17A,TFRC,ZDHHC5,ERBB2,ROCK
,HSPA1A,ELN,CRACD,MECP2,CYFIP1,FLII,TRIM27,NCK2,ARPC3,GBA2,MAPRE3,PDE4DIP,RANBP
74,SNX3,LGALS1,DAG1,HSPA1A,EFNB2,HIPK2,NECTIN1,TRIM27,NUP153,KPNA6,EXOC7,TFRC,R
C1QBP,DHX36,CRKL,LIMS1,RCC2,KIF3A,DLG4,WASF3,MYL12A,RHOBTB2,CDC42EP4,CDC42SE
MAP1S,DLG4,NSMF,ARHGAP33,SHANK1,SS18L1,PPP3CA,MARK1,CUL7,DYNLT1,OPA1,EPHB2,A
H3BP4,GSK3A,PSMC3,BNIP3,PSMC1,TRIM8,AMBRA1,ZER1,USP5,PAFAH1B2,PUM1,PINK1,YTHD

MTOR,CREBBP,BNIP3,ATP6V1G1,PSMC1,SCAP,PINK1,PSMB1,NOTCH1,MDM2,MECP2,PSMD13,P
,CLEC7A,PTN,PFKL,TPM1,PIP4K2B,CCN2,PRDX2,APLP1,HMGCS1,PRKAR1A,GJA1,JUN,MYO5A,C
9,PPP1R10,PSMB4,TAOK1,CARM1,INTS3,VPS4A,RDX,KMT2E,TCF3,PSMD4,SCRIB,FBXW5,PSMC3
XK,MLEC,SURF4,RAB5B,SRP14,PYGB,NOTCH2,MAPK1,PSMD3,ADAM9,CAND1,ILF2,CCT8,B2M,S
1,PSMD13,PSMD7,UBE2J1,AREL1,RNF41,TRIM27,PSME3,UBE2Q1,STUB1,MARCHF7,PSMC5,PSI
N4,LAMA5,MSN,RPL24,KPNB1,IGFBP3,COL1A2,GAP43,RHOA,KIF5B,SLC7A5,CXADR,ATP1A1,CD8
E,PFKL,CCND2,DDR1,PIP4K2B,CCN2,PRDX2,DDX3X,MIF,PRKAR1A,TARDBP,YWHAB,PTGES3,CADI

770,GTF2F1,HNRNPU,NFE2L1,SMARCA5,LARP1,HNRNPA0,RPL23,NACA,BRD3,XRCC5,CIPC,PAP
HECTD1,BAG6,STAT3,ARIH1,SGTA,HIP1,UBQLN2,CSNK2A1,CST3,HDAC6,PTEN,ANP32B,SPINT2,A

MBRA1,EIF4G1,PAFAH1B2,POLDIP2,ABL1,PINK1,USP10,ATP13A2,PARK7,FEZ1,EHMT2,RHEB,CA
PRDX2,DDX3X,RPS15,ITM2C,RPS7,MIF,GJA1,CTNNB1,CD63,MACF1,CYFIP2,TARDBP,NDFIP1,WLS

OL5A1,COL4A1,CES3,PSAP,PGK1,RPLP0,H3-3B,CANX,CLU,DDX5,SOX4,FKBP14,RPL13A,MAGED
RDBP,YWHAB,PTGES3,CADM4,HMGCR,RAC1,HDAC2,IPO5,TRIB2,PEA15,CDKN1A,CTDSP2,EEF1A
RDBP,YWHAB,PTGES3,CADM4,HMGCR,RAC1,HDAC2,IPO5,TRIB2,PEA15,CDKN1A,CTDSP2,EEF1A

HF7,HERPUD1,CTR9,OTUB1,RNF20,UBXN1,PSMD10,TGFBR1,PIN1,GNL3,RELA,HMG20B,VPS28,
SLR2,DHX36,GOLGA4,SRF,SERPINE2,PRMT5,CRABP2,NUMBL,THBS2,SRRT,SS18L1,RUFY3,CUL
MT2E,TCF3,PSMD4,PSMC3,PRKACA,PSMC1,TUBB4A,EIF4G1,USP47,PSMB1,TAOK2,PKIA,MDM2,
3,TPM3,CORO1C,MPRIP,PFN2,AMOTL2,CDC42,PPP1R9B,PALLD,HIP1,FARP1,SFRP1,EVL,CUL3,F
NAJC5,TARDBP,WLS,ATP1A2,RAB1A,HMGCR,RAC1,CDK16,TNC,ATP6V0C,SKP1,GLUL,NNAT,MAP1
,SUMO1,PARK7,ADNP,RAN,ZNF462,HIPK2,PPP1CA,CSNK2B,TRIB3,PBXIP1,CDK9,STUB1,CSNK1
TTG1IP,DCN,DLG5,TIMP3,DDX39B,GNAI3,NOP53,GRINA,COL2A1,MDK,SKI,ASXL1,PSMB2,TSPAN
O,PSMD5,HACD3,CHSY1,PPP2R1A,HLA-C,PDIA3,RPS3,CHD3,VAPA,HMGB1,TPT1,STMN1,ITGB1,
XIP,SQSTM1,PARP1,MORF4L2,TAF15,ACTR3,PKN1,PTOV1,MDK,RBMX,GCN1,SKI,SNRNP70,ASX
AF1,HSBP1,EID1,U2AF2,SQSTM1,NOP53,PARP1,MORF4L2,TAF15,PRDX5,YEATS2,RBMX,CBX1,S
39B,GNAI3,NOP53,GRINA,COL2A1,MDK,SKI,ASXL1,PSMB2,TSPAN6,RAPGEF1,OPTN,VHL,PPP2C
CD,IGF1R,NSF,TRMT112,APEH,NCKAP5L,FLII,ETF1,PPP1CA,NCK2,MICAL3,CSNK1A1,GSK3B,MF
TTG1IP,DCN,DLG5,TIMP3,DDX39B,GNAI3,NOP53,GRINA,COL2A1,MDK,SKI,ASXL1,PSMB2,TSPAN
DX3X,RPS15,ITM2C,RPS7,MIF,GJA1,CTNNB1,CD63,MACF1,CYFIP2,TARDBP,NDFIP1,WLS,HMGC

AP9,PPP1R10,PSMB4,TAOK1,CARM1,INTS3,VPS4A,RDX,KMT2E,TCF3,PSMD4,PSMC3,PRKACA,M
2,MTA2,KDM3A,PPP5C,RTF1,MACROH2A1,RUVBL2,SIN3B,MECP2,TRMT112,PARK7,TRRAP,SAR
6,PPM1A,GPRC5B,JRK

,PTPN23,RANGAP1,UBE2L3,PTPN11,SNX3,PINK1,RBM4,PKIA,ATP13A2,REEP5,MDM2,PCM1,PAF
,ACTN4,PRDX2,DDX3X,RPS15,RPS7,MIF,PRKAR1A,GJA1,CTNNB1,SESN3,NDFIP1,WLS,HNRNPK,C
7,NOTCH2,ARPC2,DSTN,LUZP6,GRB2,CAPZB,ACTN1,FLNB,TPM3,CORO1C,MPRIP,PFN2,AMOTL
1,PSMB1,RUVBL2,NOTCH1,PSMD13,CCNY,PSMD7,PPP1CA,GID8,PSME3,PSMC5,CSNK1A1,PSM
SARAF,MIDN,VCL,AP3D1,DNAJA1,ARF1,AKAP12,AP2B1,SLC4A8,HNRNPU,CTNNA1,CTSD,RTN4,LA
KDM3A,PPP5C,RTF1,MACROH2A1,RUVBL2,SIN3B,MECP2,TRMT112,PARK7,TRRAP,SART3,KAT2

OH2A1,ANP32E,PSMD13,SART3,PSMD7,CUL4A,ATF7IP,CETN2,H2AX,PSME3,SENP6,PSMC5,PSMC
AP9,VAPB,TRAM1,VPS4A,FNBP1L,PLEKHM2,CLASP2,REEP2,SYT11,AKT2,IST1,CHMP4B,UBE2I,C
1A,TOMM20,GNAI3,PARP1,NOX4,ABCA3,RPL27,PKN1,VPS35,MDK,PSMB2,H3-3A,ABCA2,DPYSL2

A,MTOR,CERS2,SMO,METRN,NTRK2,SEMA6B,TRAK1,DAG1,PLXNA2,OBSL1,NOTCH1,PCM1,NPT
MECP2,PSMD13,PDLIM1,PSMD7,HIPK2,PSME3,STUB1,LOXL2,PSMC5,NDRG1,PSMC2,PSMC4,PS
CH1,MDM2,MECP2,PSMD13,PDLIM1,PSMD7,HIPK2,PSME3,STUB1,LOXL2,PSMC5,NDRG1,PSMC

B3,KPNA3,THOC3,NECTIN2,MAPK3,SLC1A5,DYNLT1,SCARB1,GSN,EPHA2,VPS37B
P2,GDI2,PDXK,MLEC,MDK,SURF4,RAB5B,SRP14,PYGB,NOTCH2,MAPK1,PSMD3,CAND1,SUPT6H
CUL3,NDST1,SPINT2,RPL7L1,HIF1A,ATP6AP2,BRK1,EIF2S2,CCN1,SSBP3,RIC8A,SH3PXD2A,SEM
,ITGB1,ANXA2,AGRN,ZMIZ1,COL11A1,RPL10,AP3D1,PAFAH1B1,AP2B1,HNRNPU,CTNNA1,RPS16
311A,PPP6R1,SEC23A,TCTN1,DYNLL1,CDC42,DCTN1,KIF5A,ACTR1A,FLCN,SEC16A,MAP6,HDAC6
DST1,SPINT2,RPL7L1,HIF1A,ATP6AP2,BRK1,EIF2S2,CCN1,SSBP3,RIC8A,SH3PXD2A,SEMA3C,NM

MTSS2,CYFIP2,HNRNPK,YWHAB,PTGES3,RAB1A,ARHGDIA,HDAC2,SKP1,PEA15,CDKN1A,EEF1A2,

,PSMB5,PSMA1,PSMD10,CAPZA2,PSMA6,EXT1,RAB35,PSMD12,PSMA4,SEC31A,DYNC1I2,PSMC6
M,GIGYF1,HGS,DGKD,TXNIP,CDC42,DDIT4,POLR2E,SULF2,STAT3,BCR,HIP1,AP2A1,PTEN,NDST1

P1,HNRNPK,PTGES3,CADM4,LAMP1,HLA-B,HMGCR,APPL1,SMCHD1,SKP1,BCLAF1,HSBP1,PTTG
IM6,ALCAM,ERBB2,ANOS1,EFNB3,SOS1,NEO1,CNTN2,NRP2,SHC1,ROBO3,MAPK3,ZSWIM8,EXT
KN1B,WASF3,CDC42EP4,ARPC5,AJUBA,PPP2R5B,FER,GMFB,CUL4B,GIT1,TLR6,ARF6,PSMC6,R
DXK,MLEC,MDK,SURF4,RAB5B,SRP14,PYGB,NOTCH2,MAPK1,PSMD3,CAND1,SUPT6H,ILF2,CCT

B1,CDC42BPB,HNRNPU,PGAM1,LDHA,LPCAT1,XRCC5,IGFBP3,HK1,RHOA,ATF4,ATP1A1,CD81,C
1,CTNNB1,CD63,MACF1,CYFIP2,NDFIP1,WLS,HMGCR,RAC1,ATP6V0C,BCLAF1,PEA15,DCN,DLG
,CCT2,PABPN1,NOX4,PKN1,TCP1,CCT4,GCN1,TAF7,VEGFA,RAPGEF1,RPL37,PPP2CA,CCND1,M
S1,SOD1,HNRNPD,PPP1R9B,STAT3,CAPN2,PRKAR2A,ZFP36L1,SFRP1,ARID1B,ADCY6,PTEN,APEX
B2,PEA15,CDKN1A,CTDSP2,EEF1A2,SQSTM1,NOP53,NOX4,PKN1,GCN1,TERF2IP,TMED2,TAF7,V

MC4,TMEM9,PSMF1,GSK3B,ZNF703,LGR4,SOX9,PSMB5,CHD8,PSMA1,LZTS2,PSMD10,PSMA6,C
AM,ERBB2,ANOS1,EFNB3,SOS1,NEO1,CNTN2,NRP2,SHC1,ROBO3,MAPK3,ZSWIM8,EXT1,SEMA6

S27,RPL23A,SQLE,NOP53,ABCA3,ACTR3,TNPO1,RPL38,G3BP2,ATG2B,MAPRE1,DYNLL2,PRKDC,
B1,AKAP8L,NUP153,SRSF7,NUP98,POLDIP3,NCBP2,RCC2,KIF3A,DLG4,NUP205,NUP133,THOC2,
BI2,AKT3,RDX,CAPZA1,SEMA3C,SOD2,CLASP2,WASF2,IST1,MTOR,WDTC1,ABL1,SEMA6B,ARPC1

PHIP,BAG5,DAPK3,CD74,FGFR1,USP47,PINK1,HSPA1A,MDM2,MAPK8IP2,PARK7,CD44,ERP29,PT
R,STT3B,STT3A,CSNK1D,BAG6,PFN2,P3H3,WBP2,KDM4B,DDIT4,STAT3,PRMT1,EP300,FLCN,BA
P2,PDPK1,PRKACA,DAPK3,EIF4G1,NTRK2,ANK2,PTPN11,ABL1,COL16A1,SDCBP,TAOK2,NRXN2,

ATP6V1B2,FYN,LARS1,SOD1,HNRNPD,PPP1R9B,STAT3,CAPN2,PRKAR2A,FLOT1,ZFP36L1,ATRX,S

CK1,BSG,FASN,SPTBN1,CALM3,GNB1,SHISA9,MYH9,SEPTIN2,P4HB,APP,RPL5,KLRD1,GDI1,FSC
GF8,CCN2,ACTN4,MMP14,RPS19,MIF,GJA1,CTNNB1,CD63,ANXA6,MACF1,NARS1,RHOB,RAB1A,
11B,SPINT2,BRSK1,PSMD8,HIF1A,SLIT2,GPC4,PSMB4,CCN1,SEMA3C,CLASP2,PSMD4,SCRIB,RB
3,PSMD1,PBX1,AP2A1,GRSF1,PHB2,BRD2,SFRP1,PTEN,FEM1B,SPINT2,BRSK1,PSMD8,HIF1A,FK
B1,CDC42BPB,HNRNPU,PGAM1,LDHA,LPCAT1,XRCC5,IGFBP3,HK1,RHOA,ATF4,ATP1A1,CD81,C

ANOG,H1-0,GJA1,JUN,CTNNB1,TARDBP,MXD4,H1-10,CIRBP,YWHAB,CTBP1,CAPRIN1,SMCHD1,

SERPINE2,SOX9,PRMT5,MBOAT7,WASF3,ERBB2,MED12,BMERB1,CNTN2,ANXA1,MAPK3,PRKCI,

1,PFN1,ASTN2,CLEC7A,NISCH,SPTBN2,SLC3A2,PTN,GLG1,TPM1,CDH2,GPC3,DDR1,SULF1,PLX
F1,DNMT1,SMARCB1,C6ORF89,RNF40,SIN3A
SNK1A1,PSMC2,PSMC4,TMEM9,PSMF1,GSK3B,ZNF703,LGR4,SOX9,PSMB5,CHD8,PSMA1,MED1

PSMC2,PSMC4,PSMF1,SOX9,PSMB5,XPC,PSMA1,HIRA,RNF4,PSMD10,UBN1,THRA,PSMA6,CABIN
CK1,BSG,FASN,SPTBN1,CALM3,GNB1,SHISA9,MYH9,SEPTIN2,P4HB,APP,RPL5,KLRD1,GDI1,FSC
IP2,GNAI3,PABPN1,PARP1,NOX4,PTPRA,PRDX5,AHCYL1,VPS35,PTK7,KDM6B,RAB11B,ATP6AP1,
P,MXD4,H1-10,CIRBP,YWHAB,CTBP1,CAPRIN1,SMCHD1,HDAC2,BCLAF1,HSBP1,CDKN1A,EID1,S

,SHISA5,HIPK2,CUL4A,NCK2,TRIB3,C1QBP,MARCHF7,HERPUD1,TNFRSF10B,VDAC2,QRICH1,X

1,SARS1,RPL39,RPL34,ADAR,STAT3,VARS1,GRSF1,UBA52,GARS1,IARS1,HNRNPA2B1,RPL7L1,R
DUFB8,SHMT2,NDUFS1,ATP5F1C,COX6B1,NUP205,UQCRFS1,NUP133,ATP5F1E,COX4I1,OGDH,

1,ARHGDIA,CAPRIN1,GPC1,HDAC2,L1CAM,PPIB,DCN,DLG5,DDX39B,PARP1,MORF4L2,VPS35,CL
S9,SEC16A,SFRP1,GLUD1,UBE2G2,HDAC6,OAZ2,ANP32B,LMAN1,HIF1A,SLC2A1,SERP1,BAP1,T
,CTNNB1,NGRN,TARDBP,ARID1A,H1-10,CIRBP,RAB1A,APPL1,HDAC2,TNC,DDX39B,CNN2,SRSF5,

GFBR1,PIN1,VPS28,CDK5RAP3,RNF40,ARRDC3

CBP2,HSF1,IGF2BP2,NUP205,NUP133,THOC2,THOC3,RTRAF,FYTTD1,NUP50,RNPS1,DDX19A,NU
AR2A,PHB2,ZFP36L1,SFRP1,ARID1B,ADCY6,HDAC6,PTEN,NCOA4,APEX1,SLIT2,SAFB2,PELP1,CA
PSMD10,MEIS3,TGFBR1,XBP1,YBX3,HMGB2,RELA,OPA1,PRELID1,RB1CC1
T2,ATP6V1G1,PHIP,SORT1,USO1,RABGAP1,SCAP,RANGAP1,CAD,PTPN11,WDTC1,POR,DAG1,SM

,CIRBP,YWHAB,CTBP1,CAPRIN1,SMCHD1,HDAC2,BCLAF1,HSBP1,CDKN1A,EID1,SRP9,SQSTM1,
2,GPC3,DDR1,SULF1,PLXNA1,MEGF8,CCN2,MMP14,CLTC,NANOG,RPS7,GPAT4,PRKAR1A,GJA1

1,SART3,ARL6IP5,MAGEF1,TRIB3,ADRM1,PSME3,STUB1,PSMC5,CSNK1A1,CLPX,PSMC2,HERPU
2,GSN
D1,DGKD,CDC42,SULF2,PRKAR2A,PHB2,ZFP36L1,SRSF6,PTEN,PTPRF,LMAN1,HIF1A,LOX,A2M,

LE5,ACTN4,NANOG,MIF,JUN,HNRNPDL,DDOST,HLA-B,STIP1,APPL1,MMP2,SKP1,CDKN1A,TIMP3
PTOV1,MDK,RBMX,GCN1,SKI,ASXL1,KDM6B,PRPF6,TAF7,RNF10,VEGFA,MARS1,VHL,MAPK1,SM
PTOV1,MDK,RBMX,GCN1,SKI,ASXL1,KDM6B,PRPF6,TAF7,RNF10,VEGFA,MARS1,VHL,MAPK1,SM
11A,PLXNA3,CSNK1D,RAB11A,AKIRIN1,BMPR2,CDC42,STAT3,BCL9L,PRMT1,NAP1L1,ZFP36L1,
PTOV1,MDK,RBMX,GCN1,SKI,ASXL1,KDM6B,PRPF6,TAF7,RNF10,VEGFA,MARS1,VHL,MAPK1,SM
OTCH2,VEGFA,MAPK1,PBX2,PSMD3,ZFAND5,SMARCC1,SLIT3,COL27A1,VANGL2,SOD1,BMPR2,
23,UBE2L3,PINK1,ATP13A2,MDM2,PCM1,PHPT1,RAN,GOLPH3,YWHAZ,GSK3B,ACSL4,KIF3A,ERE
MT2,USP48,USP19,PSMB5,OTUB1,UBXN1,PSMA1,PSMD10,SENP3,SUDS3,GPS1,TGFBR1,PSMA6,

AP1,SCAP,RANGAP1,CAD,PTPN11,WDTC1,POR,DAG1,IGF1R,MDM2,CYFIP1,PTPN1,CDK4,TRIB3,

PPT1,EPHA4,DNM1,ANAPC2

LIP3,MAP2K2,FYN,UBQLN1,CSNK1D,SOD1,PFN2,WBP2,BMPR2,TRAF4,TRAF7,STAT3,MTA1,PRK
T2,PPP1R10,SPAG9,PDPK1,ARPP19,GSK3A,MTOR,WNK1,MICAL1,PRDX3,EIF4G1,ABL1,MGAT5,

IP,TMED2,TAF7,VEGFA,RAPGEF1,PPP2CA,CCND1,MAPK1,CCNL2,IPO7,ADAM9,CDC37,PRKDC,H

F,ARL6IP5,ATP1B3,TSPAN14,GOLPH3,CCDC88A,RAP2A,GOLGA4,RAB3A,DLG4,ZDHHC5,MYL12
2,IPO5,TRIB2,PEA15,CDKN1A,CTDSP2,SQSTM1,NOP53,NOX4,PTPRA,PKN1,PTK7,GCN1,TERF2IP
2,GRIK5,NPTN,MAPK8IP2,MECP2,ADNP,CYFIP1,KAT2A,ADCY1,SYNGAP1,PSMC5,PLK2,SRF,GSK
IF1A,RPL13,GNAS,SLC2A3,DDX17,GPI,RACK1,BSG,FASN,SPTBN1,ATP5F1B,CALM3,GNB1,SHISA
APC2
2,GRIK5,NPTN,MAPK8IP2,MECP2,ADNP,CYFIP1,KAT2A,ADCY1,SYNGAP1,PSMC5,PLK2,SRF,GSK
GCR,HDAC2,IPO5,TRIB2,CDKN1A,TIMP3,ANXA5,GNAI3,PRDX5,C4ORF3,PKN1,ABCA2,TMED2,TA
T2,PPP1R10,SPAG9,PDPK1,ARPP19,MTOR,WNK1,MICAL1,PRDX3,EIF4G1,ABL1,MGAT5,PPP5C,

,RUFY3,XBP1,TFDP1,UFM1,NOLC1,PRNP,NUP58,LEPROT,USP36,NUTF2,EMD
11,ABL1,PINK1,PPP5C,MACROH2A1,HSPA1A,TXNRD1,SMPD3,ATP13A2,MDM2,MGAT3,PARK7,G
,MAP4K4,RALGDS,RASA4,NTN1,LIMK1,ARHGAP5,IQSEC1,PDGFRB,RAP2A,SYNGAP1,CRKL,PLK
NB1,TARDBP,MXD4,H1-10,CIRBP,YWHAB,CTBP1,CAPRIN1,SMCHD1,HDAC2,BCLAF1,HSBP1,CDK

P19,PSMB5,OTUB1,UBXN1,PSMA1,PSMD10,SUDS3,TGFBR1,PSMA6,TRAF3,PSMD12,MBD6,PSM
LGA4,RAB3A,ZDHHC5,MYL12A,AFDN,ROCK2,PRKCI,TMEM59,EPHB2,FGF13,ARF6,NFASC,PRN

,HLA-B,STIP1,APPL1,MMP2,SKP1,CDKN1A,CNN2,SQSTM1,ACTR3,TCP1,RBMX,RAD23B,PSMB2,

A4,SEMA4C,MEIS1,SRF,GSK3B,ATRN,CRABP2,CDKN1B,UNC13A,CITED2,SGPL1,RGMA,TGFBR1,

LZTR1,SH3BP4,NOC2L,GSK3A,MTOR,NDUFA13,BAG5,CD74,USP47,CDC34,ABL1,PINK1,USP10,D
1,UBA52,HNRNPA2B1,RPL7L1,RPS2,PELP1,INTS3,ELP1,INTS11,DICER1,WBP11,SART1,EIF4A3,X
H2,GPC3,DDR1,SULF1,PLXNA1,MEGF8,CCN2,ACTN4,MMP14,RPS19,MIF,GJA1,CTNNB1,CD63,AN
H2,GPC3,DDR1,SULF1,PLXNA1,MEGF8,CCN2,ACTN4,MMP14,RPS19,MIF,GJA1,CTNNB1,CD63,AN

N,FBLN1,MAPK8IP1,SOD1,CDC42,PSMA7,TRAF4,TRAF7,PRMT1,LBH,PSMD1,FLCN,PHB2,UBA52,

EXT1,SEMA6C,PSMD12,NOLC1,MSI2,TGFB2,PSMA4,SETD1A,PSMC6,LRP6,HMGA2

M10,FBLN5,SH3PXD2B,BGN,ABL1,COL16A1,COL12A1,DAG1,ELN,CCDC80,SMPD3,NOTCH1,JAM

/2,TAB3,FER,RELA,PSMD12,PSMA4,PSMC6,ELMO2

.,SOX12,RAB7A,CCN2,MMP14,PRDX2,DDX3X,MIF,PRKAR1A,TLN1,CTNNB1,CD63,DNAJC5,NDFIP
BLN5,SH3PXD2B,BGN,ABL1,COL16A1,COL12A1,DAG1,ELN,CCDC80,SMPD3,NOTCH1,JAM3,CD4

BLN5,SH3PXD2B,BGN,ABL1,COL16A1,COL12A1,DAG1,ELN,CCDC80,SMPD3,NOTCH1,JAM3,CD4
I23,PTPN11,ABL1,SEMA6B,MTA2,PHACTR4,YTHDF3,ACVR1B,NOTCH1,MECP2,EFNB2,AAMP,PDLI

1,NCAM1,ANXA2,ZMIZ1,XAF1,ARF1,PAFAH1B1,AP2B1,GTF2F1,PXDN,CTNNA1,CPE,RTN4,LAMA5,
CLASP2,PDPK1,SCRIB,SYT11,CHMP4B,EXOC4,TANC2,CROCC,USO1,SUN1,CHMP1A,PINK1,TRAK
CXADR,ATP1A1,ATP2A2,CCT7,MCL1,HNRNPC,XPR1,CDH2,RAB7A,CCN2,PRDX2,DDX3X,RPS19,P

NUTF2,EMD
MICAL1,PLS3,ABL1,ARPC1A,FAM171A1,HIP1R,ELN,CRACD,TNFAIP1,CYFIP1,PDLIM1,LIMK1,FLII,

SAP1,KLC1,TNPO1,TUBB2A,MAPRE1,DYNC1LI2,DYNLL2,GABARAP,HSPB1,SON,MAP7D1,CLIP3,P

PARK7,TRRAP,KAT2A,EHMT2,NUP153,LDB1,LOXL2,SENP6,NUP98,NSD1,WDR82,ASH1L,BCOR,SE

,POLR2B,SUPT6H,CCT8,TNKS2,LONP1,USP7,HNRNPD,BMPR2,KDM4B,TP53BP1,YLPM1,CCT5,P
6B,ABCA2,TMED2,NOTCH2,VEGFA,MAPK1,ADAM9,LUZP6,GRB2,PRKDC,ACTN1,HSPB1,PLCG1,D

,YWHAB,CADM4,HLA-B,HMGCR,GPC1,APPL1,SMCHD1,PEA15,PTTG1IP,DCN,DLG5,TIMP3,DDX39
H1,MFN2,KAT2A,CUL4A,TASOR,DAD1,EPN1,HM13,PSMC4,SRF,BCOR,CTR9,RRM2,CHD8,NR2F2,

1A,SLC2A1,SERP1,BAP1,TCAF1,UBR5,PKDCC,TM9SF4,NMT1,AKT2,GSK3A,PRKACA,APBB1,UBE
B1,ROCK2,CHMP5,GIT1,FGF13,TPX2,CFAP298,GNAI1

N1,GDI2,NOX4,PKN1,TBC1D16,TCP1,ACAP3,CCT4,GCN1,VEGFA,RAPGEF1,PPP2CA,CCND1,MA

1,SOD2,MOAP1,GSK3A,ATP5MC3,NDUFA13,BNIP3,UBE2D3,TOMM70,ATP5MC2,SAE1,PRDX3,A
TEN,HNRNPA2B1,RPS2,RPS27A,TNKS,XRCC6,TAOK1,CCN1,SHC2,SPAG9,PDPK1,DAZAP2,DCUN1

ED2,VEGFA,RAPGEF1,DYNLL2,PCSK1N,PLCG1,ARF4,CORO1C,FYN,LARS1,STMN3,FBLN1,PLX
0,BAZ1B,KPNA2,BRD4,UBA52,ATRX,UPF1,MASTL,SMC3,FEM1B,BRSK1,RPS27A,APEX1,NPLOC
TRIB3,ADCY1,ATP6V0B,HSF1,GSK3B,STAT1,IRS1,PRKAR2B,ATP6V0E2,BACE1,SOS1,GPR173,AT
CYB5B,ATP5MG,MDH1,PINK1,POR,H6PD,PDHA1,PFKP,NDUFA4,PARK7,IDH2,NDUFC2,COX8A,GS
,STAT3,EP300,LBH,H2AZ1,FLOT1,PHB2,ZFP36L1,SFRP1,ADCY6,HDAC6,NCOA4,APEX1,SAFB2,A
A,HDGF,TAF7,NOTCH2,VEGFA,VHL,CCND1,SMARCA4,AEBP1,TCF25,ILF3,SUPT5H,KDM1A,QARS

AN,GOLPH3,YWHAZ,GSK3B,ACSL4,KIF3A,ERBB2,NADK,CAMK2N1,RUFY3,XBP1,TFDP1,TGFB2,
BP2,ZNF43,RPS3,CHD3,BASP1,HMGB1,POLR2A,MORF4L1,BTF3,AGRN,ZMIZ1,RPL10,AP3D1,CY

A,HDGF,TAF7,NOTCH2,VEGFA,VHL,CCND1,SMARCA4,AEBP1,TCF25,ILF3,SUPT5H,KDM1A,QARS

RNF4,CTCF,WDR5,MAPK3,XBP1,GNL3,CENPF,PAF1,DNMT1,SMARCB1,ZNF207,RAD50,C6ORF8

,GLUL,DCN,DLG5,DDX39B,CLSTN3,MDK,SKI,H3-3A,ATP6AP1,NOTCH2,RNF10,VEGFA,CCND1,M
A,HDGF,TAF7,NOTCH2,VEGFA,VHL,CCND1,SMARCA4,AEBP1,TCF25,ILF3,SUPT5H,KDM1A,QARS

SME3,TOM1L2,PSMC5,PSMC2,PSMC4,PLK2,PSMF1,CDKN1B,PSMB5,RRM2,RBM14,XPC,PSMA

,MAPK8IP1,DYNLL1,TXNIP,PPP1R9B,ADAR,PRKAR2A,CSNK2A1,CST3,FARP1,SFRP1,MASTL,HDA

,SHISA9,MYH9,SEPTIN2,P4HB,APP,RPL5,KLRD1,GDI1,FSCN1,TMBIM6,SPTAN1,GSTP1,SMG1,CA

,HDAC6,PTEN,RPS27A,SLIT2,MTCH1,GRN,CCN1,OLFM1,UBE2M,ARHGEF17,NCSTN,SOD2,MOA

AND1,KDM1A,ARF4,SMARCC1,SUPT6H,PFKM,SREBF2,HNRNPD,ARNT2,WBP2,AKIRIN1,BMPR2,
,ATRX,HDAC6,SMC3,PPP1CC,EML1,MAPRE2,BRSK1,TNKS,AKAP9,TAOK1,FKBP4,LSM14A,CNP,T
2,PKIA,IGF1R,PARK7,PTPN1,TRIM27,NCK2,TRIB3,PMEPA1,PPP2R5D,EMILIN1,CRKL,WARS1,RAS
Z1,RPL10,AP3D1,ZNF770,GTF2F1,HNRNPU,NFE2L1,SMARCA5,RPL23,NACA,BRD3,XRCC5,CIPC,
DAM9,CDC37,HSPB1,MAP2K2,QARS1,DUSP4,CORO1C,FYN,HGS,MAPK8IP1,SOD1,CDC42,ADAR

SBP2,ZNF43,RPS3,CHD3,BASP1,HMGB1,POLR2A,MORF4L1,BTF3,AGRN,ZMIZ1,RPL10,AP3D1,ZN
Z1,RPL10,AP3D1,ZNF770,GTF2F1,HNRNPU,NFE2L1,SMARCA5,RPL23,NACA,BRD3,XRCC5,CIPC,F

GAT3,STK39,CAB39,PARK7,ARL6IP5,GSR,NME2,OXSR1,HSF1,DHFR,SETX,TXN,VKORC1L1,ANXA1
SBP2,ZNF43,RPS3,CHD3,BASP1,HMGB1,POLR2A,MORF4L1,BTF3,AGRN,ZMIZ1,RPL10,AP3D1,ZN
N1,NBEA,CDK5RAP3,ID2,BRD1,ITPKB,MAFB,SKIL

ASP2,ITGAV,SCRIB,PCGF2,PRKACA,EXOC4,CREBBP,WNK1,SMO,EIF4A3,LMO4,CELSR1,ABL1,LF
IGS,MAPK8IP1,SOD1,ADAR,TRAF4,TRAF7,PRKAR2A,PHB2,UBA52,SFRP1,ADCY6,PTEN,RPS27A,XI

DP1,HEXIM1,MSH2,TPX2,SESN2,TRIM24,CDK5RAP3,RAD50,HIPK1,CNOT1,PPP2R5C,MYBBP1A
Z1,RPL10,AP3D1,ZNF770,GTF2F1,HNRNPU,NFE2L1,SMARCA5,RPL23,NACA,BRD3,XRCC5,CIPC,F

MD7,MAEA,RAN,PPP1CA,NUP153,TRIB3,PSME3,NUP98,PSMC5,PSMC2,PSMC4,PSMF1,GSK3B,SC
DNAJC5,NDFIP1,DDOST,LAMP1,CKAP4,HLA-B,TTR,COTL1,HRNR,RAC1,ATP6V0C,TMEM30A,CDKI

M3,CYFIP1,PHPT1,LIMK1,FLII,PDGFRB,CCDC88A,TRIM27,NCK2,ARPC3,GBA2,WASF3,RHOBTB2

GRN,CCN1,OLFM1,UBE2M,ARHGEF17,NCSTN,SOD2,MOAP1,RBM5,SCRIB,ADAM10,GSK3A,SAP1
M,ARHGEF17,NCSTN,SOD2,MOAP1,RBM5,SCRIB,ADAM10,GSK3A,SAP18,APBB1,BNIP3,DAPK3,S
,NTN1,ARHGAP5,ASNS,CSNK2B,CAPN1,DEAF1,CRKL,SRF,ZNF703,ASH1L,SERPINE2,SOX9,PRMT
I,YBX3,TGFB2,SYT1,KDM2B,BRAT1,SYT4,ATP8A2,CACNG7,ANAPC2

SBP2,ZNF43,RPS3,CHD3,BASP1,HMGB1,POLR2A,MORF4L1,BTF3,AGRN,ZMIZ1,RPL10,AP3D1,ZN

,SV2A,PREPL,BRSK1,AKAP9,TMOD2,GRK2,SEPTIN5,SH3GL1,CNP,NCSTN,ATXN3,SCRIB,SYT11,G

D1,PAFAH1B1,AKAP12,CTNNA1,NACA,RTN4,LAMA5,XRCC5,HK1,RHOA,CLSTN1,ATF4,SLC7A5,NL
EPA1,PPP2R5D,EMILIN1,CRKL,WARS1,RASSF2,CDKN1B,SH3BP5,DUSP3,DUSP8,PRKAR2B,NR2F2
,ATF7IP,CETN2,CSNK2B,WRNIP1,AKAP8L,H2AX,DHX36,SETX,SOX9,CHD8,XPC,HIRA,CTCF,UBN1,A

ORC1L1,ANXA1,MAPK3,XBP1,PCNA,FER,RELA,TLR6,FBXO7,LANCL1,SESN2,EIF2S1,SIN3A
GIP5,HIPK2,SYNGAP1,HSF1,GSK3B,NSMF,EGLN2,BACE1,MAP2K4,PIN1,PRKCI,MSH2,APOE,TLR6
TRIM27,NCK2,ARPC3,GBA2,WASF3,RHOBTB2,CDC42EP4,ARPC5,SHANK1,ROCK2,TGFBR1,CAPZ

RS1,PRKAR2B,ATP6V0E2,SOS1,GPR173,ATP6V0A1,CFLAR,SHC1,ROCK2,ADIPOR1,XBP1,PRKCI,S
NFE2L1,SMARCA5,RPL23,NACA,BRD3,XRCC5,CIPC,RHOA,CBX5,ZNF865,ATF4,PSMD2,NR2F1,RPS
,CUL1,PSMD13,KAT2A,ATF5,EHMT2,PSMD7,CUL4A,CDK4,H2AX,PSME3,TOM1L2,PSMC5,PSMC2,

NK1,THRA,B4GALT2,SLC1A4,GIT1,EPHB2,APOE,FGF13,PRNP,PPT1,ABCA7

IP,PPT1,ABCA7
GEF18,FER,GMFB,PDXP,ARF6,MYADM,NF2,ASAP3,GSN
MP1,CKAP4,HLA-B,TTR,COTL1,HRNR,RAC1,APPL1,ATP6V0C,TMEM30A,PGRMC1,CNN2,CTSB,RAB
GSK3A,APBB1,DEK,TRIP12,NIBAN2,MECP2,SUB1,EHMT2,CUL4A,PDGFRB,WRNIP1,MAGEF1,FAM
NFE2L1,SMARCA5,RPL23,NACA,BRD3,XRCC5,CIPC,RHOA,CBX5,ZNF865,ATF4,PSMD2,NR2F1,RPS
NFE2L1,SMARCA5,RPL23,NACA,BRD3,XRCC5,CIPC,RHOA,CBX5,ZNF865,ATF4,PSMD2,NR2F1,RPS
USP8,PRKAR2B,NR2F2,CAMK2N1,GPS1,RTRAF,HEG1,AJUBA,HEXIM1,GMFB,EPHB2,APOE,FBXO7
DIPOR1,XBP1,PRKCI,SP1,FER,BCAR1,RELA,OPA1,SESN2,GRB10,ATP6V1F,ZNF106,PIK3R3,ATP6V

IN1,CRKL,SLK,SPOCK1,LIMS1,SRF,GSK3B,ZYX,RCC2,ATRN,DUSP3,ROCK2,NID1,AJUBA,FER,PIP

GR1,CALCOCO1,CAD,UBE2L3,KDM3A,PPP5C,NOTCH1,MDM2,PARK7,PMEPA1,RBFOX2,PGRMC2
O,PTPN11,SUN1,CLIC1,SMPD3,NOTCH1,SYNCRIP,EXT2,DHX36,PSMC2,TOB2,LGR4,BCOR,SOX9
,MCFD2,RAB3A,DLG4,UNC13A,NAPA,COL7A1,CHMP5,NECTIN2,ANXA1,PRKCI,CLCN3,TRAPPC1,S

PIK3R3,ATP6V1D
,AKAP9,TMOD2,SEPTIN5,SH3GL1,CNP,NCSTN,ATXN3,SCRIB,SYT11,GSK3A,MTOR,PRKACA,EXOC

C6A,CDK4,PRXL2A,NME2,LDB1,RBFOX2,TOB2,MEIS1,SRF,HSF1,TFRC,RASSF2,HOXA5,CTR9,STA

U3F3,LRP6,NFIB,ID2

17,VPS16,ARF6,VPS28,LEPROT,VPS37B
DK,VEGFA,MAPK1,MAPRE1,ADAM9,ACTN1,HSPB1,PLCG1,CORO1C,FBLN1,PLXNA3,STRAP,PTP4

1,ROCK2,ANXA1,PCNA,SP1,TGFB2,ID3,ID2,KLF10,PAM,SIN3A,MYBBP1A,NAMPT
,IRS1,UQCRFS1,ATP5F1E,COX4I1,OGDH,COX5A,COX7A2,UQCRH,FH,GYS1,MSH2,GRB10,PRELI

,TMOD2,SEPTIN5,SH3GL1,CNP,NCSTN,ATXN3,SCRIB,SYT11,GSK3A,MTOR,PRKACA,EXOC4,NSG
,TMOD2,SEPTIN5,SH3GL1,CNP,NCSTN,ATXN3,SCRIB,SYT11,GSK3A,MTOR,PRKACA,EXOC4,NSG
1,CYFIP2,NDFIP1,MYH10,CADM4,RHOB,LGALS3BP,RAB1A,RAC1,ARHGDIA,TNC,L1CAM,DLG5,C

PX2,DYNC1I2,CENPF,CENPT,ZNF207,GNAI1

IDFIP1,MYH10,CADM4,RHOB,LGALS3BP,RAB1A,RAC1,ARHGDIA,TNC,L1CAM,DLG5,CDH11,TBC

PZA1,CLASP2,PDPK1,SCRIB,SYT11,EHD1,MTOR,PRKACA,GNA13,MICAL1,PTPN11,CELSR1,DAG1
X,SH3GL1,RAB5C,CLASP2,PDPK1,ITGAV,SCRIB,SYT11,AKT2,EHD1,MICAL1,PTPN23,RIMS3,TUB,

L2,EXOSC10,TFRC,RBM14,LMNA,BMERB1,PSMD10,SHANK1,RPS6KA2,CAPZA2,ADD1,ARHGEF18

KHDRBS1,PTN,CSNK1E,SFPQ,CDH2,GPC3,SULF1,MEGF8,CCN2,ACTN4,PRDX2,DDX3X,RPS15,ITI

AG9,KRT8,TCF3,RBPJ,EHD1,GSK3A,PTGFRN,MTOR,YY1,SORT1,SMO,ANK2,OLFM2,ABL1,LGALS
RKL,RBFOX2,NR2C2,UBE2V1,EGLN2,PER1,NR2F2,RXRA,NR2F6,PHB,CITED2,PRMT2,CTBP2,THF

N,ATP6V1F,KLF10,ATP6V1D,MYBBP1A,NAMPT

T11,PCGF2,GSK3A,FBLN5,UBE3A,APBB1,BNIP3,UBE2D3,WNK1,PRDX3,SLC38A2,EIF4A3,PTPN11
D,CD81,PFN1,TLE5,TADA2B,KHDRBS1,XPO1,SFPQ,SOX12,CCN2,DDX3X,NANOG,H1-0,PRKAR1A,J
2,PHACTR4,RGL2,NOTCH1,MFN2,SGSM3,TNFAIP1,PARK7,CYFIP1,MAP4K4,RALGDS,RASA4,NTN1

T2,TNRC6A,RHEB,ASNS,ATP6V0B,TNFRSF10B,SRF,SOX9,ATP6V0E2,MAP2K4,GABARAPL2,MAPK

P4A1,PFN2,RAB11A,AKIRIN1,BMPR2,AMOTL2,BEX4,STAT3,FLCN,CD99,PLXNB1,SFRP1,EVL,HDA
MA1,PSMD10,NECTIN2,PSMA6,TAB3,HEXIM1,HMGB2,RELA,ABCE1,PSMD12,APOE,LRP8,PSMA4,I

CG1,CORO1C,FBLN1,PLXNA3,STRAP,PTP4A1,PFN2,RAB11A,AKIRIN1,BMPR2,AMOTL2,BEX4,STA

P1,SZT2,SESN2,TRIM24,EIF2S1,POSTN,ATP6V1F,KLF10,ATP6V1D,MYBBP1A,NAMPT

1,HSPB1,FKBP8,PLCG1,HECTD1,SARS1,VANGL2,HGS,BAG6,TCTN1,BMPR2,AMOTL2,CDC42,CI

)10,MDC1,PSMA6,CNOT3,PCNA,TFDP1,PSMD12,MSH2,PSMA4,FHL1,FBXO7,CENPF,PSMC6,CDK

ZFP36L1,SFRP1,CD164,EML1,PSMD8,HIF1A,SLC25A5,LOX,GLO1,XRCC6,PSMB4,CLPTM1,SSBP3

,DDX3X,RPS19,MIF,GJA1,JUN,CTNNB1,MACF1,RHOB,ATP1A2,RAB1A,HMGCR,ARHGDIA,CAPRIN

,PBX1,FLCN,ZFP36L1,SFRP1,CD164,EML1,PSMD8,HIF1A,SLC25A5,LOX,GLO1,XRCC6,PSMB4,C
K4,PIN1,PRKCI,MSH2,APOE,TLR6,TGFB2,FBXO7,PRNP,PPT1,KDM2B,LANCL1,EIF2S1

YBBP1A,NDUFA10,ATP5PD

5C,MBOAT7,NR2F2,NUMBL,BMERB1,OGDH,CNTN2,NRP2,ARPC5,EXT1,XAB2,EPHB2,LRP8,FGF1
SXL1,PSMB2,KDM6B,TMED2,NOTCH2,VEGFA,RAPGEF1,CCND1,MAPK1,PSMD3,ACADVL,ADAM9,

1,SFPQ,SOX12,DDX3X,NANOG,H1-0,PRKAR1A,JUN,CTNNB1,ARID1A,MXD4,SHOX,HNRNPK,DUX4

TMED2,NOTCH2,VEGFA,RAPGEF1,MAPK1,TCF25,PRKDC,HSPB1,PLCG1,ZFAND5,SLIT3,HECTD1,

AP1,DPYSL2,VEGFA,RAPGEF1,MAPK1,AZIN1,ADAM9,NPEPPS,CLIP3,PLCG1,GNG2,PLTP,FYN,YW

,ITGB1,TUBB4B,NCAM1,ANXA2,ZMIZ1,TIMP2,XAF1,RPS17,VCL,AP3D1,ARF1,CYB5R3,PAFAH1B1

TYK,RGMA,TGFBR1,PIN1,XBP1,PPP2R5B,ZDHHC17,CRIM1,FGFBP3,GRB10,PPM1A,CRB2,LEMD

PPM1A,POSTN,SKIL

R1,TGFBR1,MAPK3,CUL7,EXT1,SEMA6C,NOLC1,TGFB2,LRP6,CRB2,HMGA2

AK1,PRMT1,EP300,PSMD1,PBX1,FLCN,ZFP36L1,SFRP1,CD164,EML1,MCM3AP,PSMD8,HIF1A,SL
7,HMGA2,CNOT1,PPP2R5C

TRAF7,STAT3,PRKAR2A,FLCN,FLOT1,PHB2,UBA52,ADCY6,HDAC6,IL6ST,RPS27A,HIF1A,AKAP9,X

LN1,PLXNA3,STRAP,PTP4A1,PFN2,RAB11A,AKIRIN1,BMPR2,AMOTL2,BEX4,STAT3,FLCN,CD99,P
2,CXXC5,ATRX,CBX6,HNRNPA2B1,CUL3,RPS27A,NDN,EHMT1,SOX2,MAZ,MACROH2A2,DICER1,T

NANOG,MIF,PRKAR1A,GJA1,JUN,CTNNB1,NDFIP1,DUX4,PTGES3,CADM4,CTBP1,HMGCR,RAC1,

,SHMT2,OAT,RARS1,WARS1,PSMB5,KARS1,FARSA,PSMA1,OGDH,PSMD10,SLC1A3,PSMA6,TARS

,PRNP,EIF2S1,GNAO1,ID2,HMGA2,ATP8A2,GSN,SIN3A,NAMPT
KIL

K3A,MTOR,YY1,SMO,SH3PXD2B,ANK2,CAD,LMO4,PTPN11,ELN,OBSL1,NOTCH1,MDM2,EFNB2,P

ID12,MSH2,PSMA4,FHL1,FBXO7,CENPF,PSMC6,CDK5RAP3,BTG1,ZNF207,HMGA2,CNOT1,PPP2

NK1,CD74,AMBRA1,FGFR1,NTRK2,LMO4,PTPN11,ABL1,PINK1,TAOK2,DAG1,IGF1R,STK39,CAB3

FN2,BMPR2,CDC42,TRAF4,TRAF7,STAT3,CAPN2,PRKAR2A,FLCN,FLOT1,PHB2,UBA52,ADCY6,HD

FN2,BMPR2,CDC42,TRAF4,TRAF7,STAT3,CAPN2,PRKAR2A,FLCN,FLOT1,PHB2,UBA52,ADCY6,HD

ESN2,BTG1

327A,CCT2,VAMP2,GDI2,PDXK,MLEC,SURF4,RAB5B,SRP14,PYGB,MAPK1,PSMD3,PRDX1,CAND1

APOE,FBXO7,SRCIN1,NF2,CDK5RAP3,LRP6,IBTK

2,RPS8,MYH9,RPL7,SEPTIN2,P4HB,APP,RPL5,GDI1,FSCN1,SPTAN1,MARCKS,SMG1,CALM1,NPM

P1,SMG1,CALM1,NPM1,HSPD1,IGF2,ATP5F1A,VCP,UBB,TRIM28,YWHAG,RPL37A,MAPT,FN1,YW

PL24,KPNB1,RPL17,RPL14,RPL22,RHOA,CBX5,RPL9,RPS14,CXADR,RPS20,CD81,RPL26,RBBP4,

FSCN1,SPTAN1,MARCKS,SMG1,CALM1,NPM1,PLXNB2,HSPD1,COL18A1,SNRNP200,CSDE1,IGF2

,AP2B1,HNRNPU,CTNNA1,RPS16,RPL23,CPE,RPL29,RPSA,SCARB2,NACA,RTN4,LAMA5,RPS24,R

,AP2B1,HNRNPU,CTNNA1,RPS16,RPL23,CPE,RPL29,RPSA,SCARB2,NACA,RTN4,LAMA5,RPS24,R

RPL12,RPL30,RPS5,RPS7,IK,RPS19,RPL35,RPS10,RPL35A,TARDBP,RPLP2,HNRNPK,CIRBP,RPL

RPS16,RPL23,RPL29,RPSA,SCARB2,NACA,RPS24,RPS9,MSN,RPL24,KPNB1,RPL17,RPL14,RPL22,

MSN,RPL24,KPNB1,RPL17,RPL14,RPL22,RPL9,PSMD2,ENC1,RPS14,TMEM259,RPS20,CD81,RPL

RPL37A,MAPT,FN1,YWHAQ,RPL19,RPL15,PKD1,HLA-C,RPS3,VAPA,HMGB1,AP2M1,SEC61A1,RPS

N,RPLP1,MIDN,RPL10,RPS17,VCL,AP3D1,DNAJA1,ARF1,PAFAH1B1,AP2B1,HNRNPU,CTNNA1,RPS

U,SCARB2,RTN4,XRCC5,MSN,IGFBP3,GAP43,NCS1,RHOA,CLSTN1,NR2F1,KIF5B,ENC1,NLGN2,

GB1,RPL11,BTF3,ANXA2,AGRN,RPLP1,MIDN,RPL10,RPS17,VCL,AP3D1,DNAJA1,ARF1,PAFAH1B1,

,ENC1,RPS14,TMEM259,RPS20,CD81,RPL26,RPL31,HNRNPC,LRPPRC,XPO1,CSNK1E,RPS25,RA

S25,RAB7A,DDX3X,RPS15,RPL12,RPL30,RPS5,RPS7,IK,RPS19,RPL35,GJA1,RPS10,RPL35A,TAR

15,ALKBH5,RPL27,TNPO1,PSMB2,RPL36A,RPS15A,RPL38,RPL37,PPP2CA,PSMD3,EIF3E,FTO,RPS

B,CALM3,GNB1,SHISA9,RPS8,MYH9,RPL7,SEPTIN2,APP,RPL5,GDI1,FSCN1,TMBIM6,SPTAN1,GS

PT,FN1,DDB1,PSMD5,EIF1,HACD3,SET,EGLN1,EIF4A1,RPL19,PPP2R1A,RPL15,GTF3C4,PKD1,PD

0,INA,PPP2R1A,GTF3C4,DNAH8,PKD1,RPS3,CHD3,VAPA,HMGB1,CDKL1,AP2M1,SEC61A1,STMN

RPRA,RPS26,RPS21,RPL27A,RPS29,SRP54,RPL32

G5,EIF4A3,EIF4G1,RPS26,RPS21,PNRC2,ETF1,NBAS,RPL27A,EXOSC10,NCBP2,RPS29,SKIV2L,RF

,RPS29,SRP54,RPL32

1,RPL23A,TAF15,ALKBH5,RPL27,TNPO1,PSMB2,RPL36A,RPS15A,RPL38,RPL37,PPP2CA,PSMD3,

L19,PPP2R1A,RPL15,DNAH8,PKD1,SF3B1,SERINC1,RPS3,HMGB1,CDKL1,AP2M1,RPL18,STMN1,

KPNB1,RPL17,LLGL1,RPL14,HK1,RPL22,RPL9,CLSTN1,KIF5B,RAB6B,NLGN2,RPS14,RPS20,CD

,RPL30,ARL4C,CLTC,RPS5,RPS7,RPS19,RPL35,RPS10,RPL35A,TARDBP,RPLP2,ARL6IP1,WLS,RP

R1,RAB3GAP2,RPS2,RPS27A,PPP1R15A,TRAM1,SRP72,SSR3,CHMP4B,SRPRA,RPS26,ANK2,GRI

,RPS26,RPS21,RPL27A,HERPUD1,RPS29,SPCS2,SRP54,RPL32

C,EIF4B,VCP,UBB,TRIM28,YWHAG,RPL37A,MAPT,FN1,DDB1,PSMD5,EIF1,HACD3,SET,EGLN1,C

24,KPNB1,RPL17,LLGL1,RPL14,RPL22,RPL9,CLSTN1,KIF5B,RAB6B,SLC7A5,NLGN2,RPS14,RPS

,RPS28,SSB,FAU,EIF3F,STT3A,IFITM3,PCBP2,RPL39,RPS13,RPL34,POLR2E,EP300,EIF3D,BRD4,

P4B,SRPRA,RPS26,RPS21,RPL27A,HERPUD1,RPS29,SPCS2,SRP54,RPL32

7A,EIF2S2,EIF3H,EIF3G,MTOR,EIF4G1,RPS26,CNBP,RBM4,TMA7,RPS21,ETF1,RPL27A,DHX36,E

L3,YWHAE,RPL13,SLC2A3,DDX17,GPI,RACK1,BSG,FASN,ALDH6A1,SPTBN1,ATP5F1B,CALM3,RF

RPL11,ANXA2,AGRN,RPLP1,TIMP2,RPL10,MARCHF6,RPS17,PAFAH1B1,AP2B1,PXDN,HNRNPU,OG

NRNPU,OGA,PGAM1,LARP1,HNRNPA0,RPS16,RPL23,RPL29,RPSA,SCARB2,RPS24,RPS9,MSN,RP

,APP,RPL5,KLRD1,GDI1,TMBIM6,SPTAN1,GSTP1,SMG1,CALM1,NPM1,PLXNB2,HSPD1,COL18A1,

PS28,FAU,EIF3F,RPL39,RPS13,RPL34,EIF2AK1,EIF3D,UBA52,EIF5B,RPS2,RPS27A,PPP1R15A,EIF

EEF1A2,PPP1CB,GNAI3,NOP53,TAF15,ALKBH5,VPS35,RAD23B,TNPO1,SNRNP70,PSMB2,ABCA2,

PL9,CLSTN1,KIF5B,RAB6B,NLGN2,RPS14,RPS20,CD81,RPL26,OAZ1,ASTN2,MCL1,RPL31,XPO1,

N1,GSTP1,SMG1,CALM1,NPM1,HSPD1,IGF2,ATP5F1A,VCP,UBB,TRIM28,YWHAG,RPL37A,MAPT,

MP1,RPS12,RPS27,CCT6A,APPL1,IPO5,RPL23A,PTTG1IP,CDKN1A,SRP9,TOMM20,CCT2,RPL27,

MG1,CALM1,NPM1,HSPD1,IGF2,ATP5F1A,VCP,UBB,TRIM28,YWHAG,RPL37A,MAPT,FN1,PSMD5,

MN1,ITGB1,TUBB4B,RPL11,ANXA2,AGRN,RPL10,VCL,ARF1,PAFAH1B1,MAP4,AP2B1,MAT2A,TM

AH1B1,AP2B1,HNRNPU,OGA,PGAM1,LARP1,HNRNPA0,RPS16,RPL23,LDHA,RPL29,RPSA,CTSD,S

,SMG1,CALM1,NPM1,PLXNB2,TTC3,HSPD1,USP11,CSDE1,IGF2,EIF3C,ATP5F1A,EIF4B,VCP,UBB,
GL1,GAP43,NCS1,RHOA,CLSTN1,ATF4,NR2F1,KIF5B,ENC1,NLGN2,WDR1,CRMP1,PFN1,ASTN2,N
30,CLTC,RPS5,RPS7,RPS19,RPL35,MYO5A,MACF1,RPS10,RPL35A,RPLP2,ARL6IP1,RPL36,YWHA
P,MARCKSL1,UBB,TRIM28,YWHAG,MAPT,FN1,SPEN,DDB1,PSMD5,YWHAQ,HACD3,CHSY1,PKD1,
M30A,L1CAM,MAP1A,DLG5,CRPPA,CDH11,GAS7,PTPRA,CAMSAP1,NCKAP1,ACAP3,MDK,PTK7,CN
,RPL12,RPL30,ARL4C,CLTC,RPS5,RPS7,RPS19,RPL35,MYO5A,MACF1,RPS10,RPL35A,RPLP2,AF
0,PSMB2,ABCA2,NRDC,OPTN,PSMD3,ADAM9,FTO,WDR6,GABARAP,SUPT5H,HSPB1,BNIP3L,CCA
21C,RPS2,RPS27A,NUCKS1,PSMC3,TRIM8,RPS26,USF2,RSF1,RPS21,RPL27A,TRIM27,NUP153,CD
G1,CALM1,NPM1,PLXNB2,PLD3,HSPD1,CSDE1,IGF2,EIF3C,EIF4B,VCP,UBB,AFF4,TRIM28,MATR
SND1,RPS12,RPS27,BCLAF1,RPL23A,TAF15,ALKBH5,RPL27,TNPO1,PSMB2,RPL36A,RPS15A,RPL
L18A1,IGF2,ATP5F1A,VCP,UBB,TRIM28,YWHAG,MAPT,SPEN,NONO,DDB1,PSMD5,YWHAQ,SET,TN
CHL1,RPL11,MORF4L1,ANXA2,RPLP1,TIMP2,MIDN,RPL10,RPS17,DNAJA1,ITM2B,HNRNPU,SMAR
,XRCC5,MSN,IGFBP3,RPL22,RHOA,PTCD3,ATF4,PSMD2,ENC1,RPS14,TMEM259,RPS20,EIF5,CD
RPL7,P4HB,APP,IDS,RPL5,TMBIM6,PTMS,GSTP1,SMG1,CALM1,NPM1,PLXNB2,TTC3,PLD3,HSPD1
ORF4L1,ANXA2,RPLP1,TIMP2,RPL10,RPS17,DNAJA1,ITM2B,HNRNPU,SMARCA5,LARP1,HNRNPA0
SFPQ,TPM1,PDIA6,CCN2,PRDX2,DDX3X,MIF,TLN1,JUN,CTNNB1,SESN3,TARDBP,HNRNPK,INIP,PT
APT,FN1,PSMD5,INA,PPP2R1A,RPL15,HLA-C,PDIA3,RPS3,HMGB1,POLR2A,SEC61A1,ITGB1,NCA
DR1,ATP2A2,RBBP4,PFN1,CLEC7A,NES,SPTBN2,EIF5A,DDX3X,RPS15,CLTC,EIF3L,H1-0,RPS5,RF
C2,TNC,TMEM30A,L1CAM,MAP1A,DLG5,CRPPA,HS6ST1,CDH11,TBCD,GAS7,PTPRA,CAMSAP1,N
PL1,TMEM30A,L1CAM,DCN,DLG5,CCT2,PARP1,NOX4,ABCA3,ACTR3,TCP1,CCT3,NCKAP1,VPS35,
LXNA1,MEGF8,MMP14,ITM2C,GJA1,CTNNB1,NGRN,MACF1,CYFIP2,MAP2,RAC1,ARHGDIA,CAPRI
L13,SLC2A3,DDX17,GPI,HTRA4,RACK1,FASN,ALDH6A1,SPTBN1,ATP5F1B,CALM3,RPS8,MYH9,RF
,YWHAB,SND1,RPS12,RPS27,BCLAF1,RPL23A,TAF15,ALKBH5,RPL27,TNPO1,PSMB2,RPL36A,RF
,YWHAB,SND1,RPS12,RPS27,BCLAF1,RPL23A,TAF15,ALKBH5,RPL27,TNPO1,PSMB2,RPL36A,RF
,ZFP36L1,UPF1,RPS2,RPS27A,SMG5,TTC37,THRAP3,EIF4A3,EIF4G1,RPS26,XRN2,RPS21,PNRC2,
DX17,GPI,HTRA4,RACK1,FASN,ALDH6A1,SPTBN1,ATP5F1B,CALM3,RPS8,MYH9,RPL7,P4HB,APP,I
MAP2,RAC1,ARHGDIA,CAPRIN1,GPC1,CDK16,TOP2B,HDAC2,TNC,TMEM30A,L1CAM,NNAT,MAP1A
NB1,NGRN,MACF1,CYFIP2,MAP2,RAC1,ARHGDIA,CAPRIN1,GPC1,CDK16,TOP2B,HDAC2,TNC,TM
CARB2,NACA,RPS24,RPS9,RPL24,SLC16A7,KPNB1,RPL17,LLGL1,RPL14,RPL22,RPL9,CLSTN1,K
2,PLD3,HSPD1,CSDE1,IGF2,EIF3C,EIF4B,VCP,UBB,AFF4,TRIM28,MATR3,YWHAG,RPL37A,MAPT,F
,RPL36A,RAB11B,SRP14,RPS15A,RPL38,OPTN,RPL37,RPS11,RPL28,RPS28,FYN,YWHAH,FAU,BA
,EIF3C,ATP5F1A,VCP,MARCKSL1,UBB,TRIM28,MATR3,YWHAG,MAPT,FN1,SPEN,NONO,DDB1,PS
G,MAPT,FN1,SPEN,NONO,DDB1,PSMD5,YWHAQ,EIF1,HACD3,SET,EGLN1,PPP2R1A,PKD1,SSBP2,
RPS5,RPS7,RPS19,RPL35,NGRN,RPS10,RPL35A,TARDBP,RPLP2,CIRBP,NARS1,RPL36,RPS12,RPS
IC1,RPS14,TMEM259,RPS20,EIF5,CD81,RPL26,BEX3,CLEC7A,CCNI,LRPPRC,TADA2B,KHDRBS1,X
PS15A,RPL38,RPL37,CDC37,RPS11,GABARAP,NPEPPS,BNIP3L,RPL28,RPS28,FYN,FAU,HGS,SREB
SND1,RPS12,RPS27,BCLAF1,RPL23A,TAF15,ALKBH5,RPL27,TNPO1,PSMB2,RPL36A,RPS15A,RPL
A,CRPPA,CDH11,TBCD,GAS7,NOX4,PTPRA,CAMSAP1,NCKAP1,MDK,LAMC1,PTK7,ASXL1,CNTNAP
,YWHAB,SND1,RPS12,RPS27,BCLAF1,RPL23A,TAF15,ALKBH5,RPL27,TNPO1,PSMB2,RPL36A,RF
YWHAH,SLIT3,STXBP1,PLXNA3,PTPRS,RAB11A,TCTN1,BMPR2,CDC42,PALLD,KIF5A,TRIO,RERE,C
IT3,STXBP1,PLXNA3,PTPRS,RAB11A,TCTN1,BMPR2,CDC42,PALLD,KIF5A,TRIO,RERE,CNTNAP1,F
YWHAH,SLIT3,STXBP1,PLXNA3,PTPRS,RAB11A,TCTN1,BMPR2,CDC42,PALLD,KIF5A,TRIO,RERE,C
DI1,SPTAN1,GSTP1,MARCKS,CALM1,NPM1,PLXNB2,PLD3,HSPD1,COL18A1,SNRNP200,CSDE1,SI
N1,PSMD3,ODC1,ADAM9,GABARAP,UBE2R2,BNIP3L,CCAR2,SMARCC1,FYN,STT3B,LONP1,UBQL
RN,RPS10,RPL35A,TARDBP,RPLP2,CIRBP,NARS1,RPL36,RPS12,RPS27,CAPRIN1,BCLAF1,RPL23

15,CLTC,EIF3L,H1-0,RPS5,RPS19,MIF,TLN1,CTNNB1,RPS10,CYFIP2,H1-10,PTGES3,CKAP5,MAP2
1,RPL6,KAT6A,DVL3,YWHAE,RPL13,GNAS,SLC2A3,DDX17,GPI,HTRA4,RACK1,BSG,FASN,ALDH6A1
PLXNA1,MEGF8,ITM2C,GJA1,CTNNB1,MACF1,MTSS2,CYFIP2,CKAP5,MAP2,RAC1,ARHGDIA,CAPI
36,HNRNPH1,RPS27,BCLAF1,RPL23A,LUC7L3,CPSF2,DDX39B,NOVA2,SRSF5,ZC3H7B,U2AF2,SU
LN1,INA,HLA-C,PDIA3,RPS3,HMGB1,POLR2A,ITGB1,UCHL1,NCAM1,ANXA2,AGRN,ZMIZ1,XAF1,DN
P1,PLXNA3,PTPRS,RAB11A,TCTN1,BMPR2,CDC42,PALLD,KIF5A,TRIO,RERE,CNTNAP1,FLOT1,MA
3A,LUC7L3,DDX39B,SRSF5,CCT2,TBCD,VAMP2,GAS7,NOP53,PARP1,TSPYL4,ACTR3,HSPA4,TCP1
RACK1,BSG,SPTBN1,CALM3,RPS8,MYH9,RPL7,P4HB,APP,IDS,RPL5,TMBIM6,PTMS,GSTP1,SMG
,FN1,SPEN,NONO,DDB1,PSMD5,YWHAQ,EIF1,HACD3,SET,EGLN1,PPP2R1A,PKD1,SSBP2,ZNF43,I
S27A,TCAF1,TRAM1,SRP72,SSR3,AKT2,CHMP4B,SSR2,EXOC4,SRPRA,RPS26,SDCBP,TAOK2,RPS2
7,SRSF2,RBM5,PSMD4,IGF2BP1,PRPF19,PSMC3,MTOR,SAP18,PSMC1,THRAP3,EIF4G1,PUM1,F
DR1,PLXNA1,MEGF8,ITM2C,GJA1,CTNNB1,MACF1,MTSS2,CYFIP2,CKAP5,MAP2,RAC1,ARHGDIA,
RN,RPS10,RPL35A,TARDBP,RPLP2,CIRBP,NARS1,RPL36,RPS12,RPS27,CAPRIN1,BCLAF1,RPL23
GB1,NCAM1,ANXA2,ZMIZ1,XAF1,DNAJA1,ARF1,PAFAH1B1,AKAP12,MAT2A,GTF2F1,PXDN,HNRNP
NB2,PLD3,HSPD1,COL18A1,CSDE1,SMIM14,IGF2,MARCKSL1,UBB,AFF4,TRIM28,MATR3,YWHA
C,ACTN1,PLCG1,FLNB,CORO1C,FYN,YWHAH,SLIT3,FBLN1,STXBP1,PLXNA3,PTPRS,SOD1,RAB11
2B1,NFE2L1,CTNNA1,CPE,NACA,RTN4,LAMA5,MSN,RPL24,COL1A2,LLGL1,GAP43,RHOA,ATF4,P
VEGFA,MAPK1,GRB2,PLCG1,CORO1C,FYN,YWHAH,SLIT3,STXBP1,PLXNA3,PTPRS,RAB11A,TCT
F5A,TRIO,RERE,FLOT1,MAP6,PLXNB1,DIP2B,FARP1,CTNND2,EVL,HDAC6,PTEN,ULK1,BRSK1,CT
AQ,HACD3,SET,EGLN1,TMSB10,PKD1,HMGB1,AP2M1,TPT1,STMN1,ITGB1,RPL11,ANXA2,AGRN,
3,RBM15B,DHX15,POLR2E,SF1,PRCC,GRSF1,SRSF6,TRA2B,HNRNPA2B1,CHERP,SYMPK,PCBP4,
1,CSDE1,SMIM14,IGF2,MARCKSL1,UBB,TRIM28,MATR3,YWHAG,CKB,COL4A2,MAPT,FN1,SPEN,
AJB2,ATXN3,OGT,GSK3A,PSMC3,UBE3A,PRKACA,NDUFA13,PSMC1,BAG5,ATG4B,UBE2K,ZER1,
NONO,DDB1,PSMD5,YWHAQ,EIF1,HACD3,SET,EGLN1,PPP2R1A,PKD1,SSBP2,ZNF43,RPS3,CHD3,
7,RPS19,RPL35,NGRN,RPS10,RPL35A,TARDBP,RPLP2,ST3GAL2,CIRBP,NARS1,RPL36,RPS12,RP
MATR3,YWHAG,CKB,COL4A2,MAPT,FN1,SPEN,PSMD5,YWHAQ,DCHS1,EGLN1,CHSY1,INA,RPL19,
KAT6A,DVL3,YWHAE,RPL13,SLC2A3,DDX17,GPI,HTRA4,RACK1,BSG,FASN,ALDH6A1,SPTBN1,ATI
X6,FUS,SNRPD3,RALY,ELAVL1,EFTUD2,SRSF2,PRPF4B,RBM5,WBP11,SRSF11,PRPF19,SF3A2,SA
X6,FUS,SNRPD3,RALY,ELAVL1,EFTUD2,SRSF2,PRPF4B,RBM5,WBP11,SRSF11,PRPF19,SF3A2,SA
X6,FUS,SNRPD3,RALY,ELAVL1,EFTUD2,SRSF2,PRPF4B,RBM5,WBP11,SRSF11,PRPF19,SF3A2,SA
5,PJA2,IGFBP3,RHOA,ATF4,PSMD2,ENC1,RPS20,CD81,CLEC7A,PRKCSH,CCNI,TADA2B,KLHL9,PT
5,PJA2,IGFBP3,RHOA,ATF4,PSMD2,ENC1,RPS20,CD81,CLEC7A,PRKCSH,CCNI,TADA2B,KLHL9,PT
PTEN,PTPRF,ULK1,BRSK1,CTTN,SLIT2,NDN,GRN,SLIT1,OLFM1,LINGO1,DCLK1,CNP,STK24,SEM
IF770,GTF2F1,HNRNPU,SMARCA5,LARP1,HNRNPA0,RPL23,CTSD,NACA,RTN4,LPCAT1,RPS9,XRC
5B,DHX15,POLR2E,SF1,PRCC,GRSF1,ZFP36L1,SRSF6,TRA2B,HNRNPA2B1,CHERP,SYMPK,PCBP
CYOX1,YME1L1,PSMA7,ARIH1,SGTA,OS9,CAPN2,MTA1,PSMD1,AP2A1,UBQLN2,CSNK2A1,UBA52
75B,ENC1,JUND,WDR1,ATP2A2,CRMP1,CLEC7A,PRKCSH,SPTBN2,PTN,TPM1,CDH2,DDR1,SULF1,
IA,LAPTM4B,DCN,MAP1A,CCT2,TBCD,SQSTM1,PARP1,NOX4,ACTR3,TCP1,CAMSAP1,CCT3,NCKA
4,RBBP4,TLE5,RPL31,HNRNPC,XPO1,RPS25,GLG1,CCN2,DDX3X,RPS15,RPL12,RPL30,H1-0,RPS
5,DDX3X,RPS15,RPL12,RPL30,EIF3L,RPS5,RPS7,RPS19,RPL35,HMGCS1,GPAT4,NGRN,RPS10,R
79,SRSF1,EIF3E,PRKDC,SCAF11,RPL28,DHX30,RPS28,EIF3F,SF3A1,STRAP,RPL39,RPL34,ADAR,T
4,NR2F1,JUND,TMEM259,CD81,RPL26,BEX3,CCT7,CLEC7A,TADA2B,KHDRBS1,CSNK1E,SFPQ,EI
A1,COL4A1,CES3,PSAP,PGK1,RPLP0,H3-3B,CANX,CLU,PTMA,DDX5,ENO2,SOX4,FKBP14,RPL13A
6B,NLGN2,CD81,WDR1,ATP2A2,VAT1,RTN3,CLEC7A,SPTBN2,CSNK1E,PFKL,SCRN1,CDH2,GPC3,
1,EHD1,GSK3A,SF3A2,MTOR,UBE3A,APBB1,TANC2,CERS2,BAG5,METRN,GAK,NTRK2,DENND5A,

,LARP1,HNRNPA0,RPL23,CTSD,NACA,RTN4,LPCAT1,RPS9,XRCC5,MSN,HK1,RHOA,ATF4,NR2F1,
S20,BEX3,SRSF4,RTN3,TLE5,HNRNPC,LRPPRC,KHDRBS1,XPO1,PTN,GLG1,SFPQ,PEBP1,GPC3,D
,NDN,SLIT1,OLFM1,LINGO1,DCLK1,CNP,SEMA3C,CLASP2,IST1,TMEFF1,APBB1,SMO,METRN,LY
BX5,ATF4,NR2F1,ENC1,RPS14,ATP1A1,RPS20,BEX3,SRSF4,RTN3,MCL1,TLE5,HNRNPC,LRPPRC,
31,DIP2B,SFRP1,EVL,ADCY6,PTEN,PTPRF,BRSK1,SEPTIN7,SLIT2,BRK1,GRN,PALM,FKBP4,CARM
,MMP14,APLP1,HMGCS1,PRKAR1A,GJA1,JUN,MYO5A,CTNNB1,SESN3,MTSS2,ARID1A,PTGES3,
PLXNB1,DIP2B,SFRP1,EVL,ADCY6,PTEN,PTPRF,BRSK1,SEPTIN7,SLIT2,BRK1,GRN,PALM,FKBP4,
ACTN4,PRDX2,DDX3X,ITM2C,H1-0,APLP1,RPS7,MIF,GJA1,JUN,CTNNB1,ANXA6,CYFIP2,DNAJC5,T
,JUND,TMEM259,CD81,RPL26,BEX3,CCT7,OAZ1,CLEC7A,TADA2B,KHDRBS1,CSNK1E,SFPQ,EIF5
3C,EIF4B,UBB,AFF4,TRIM28,MATR3,COL4A2,RPL37A,MAPT,FN1,SPEN,NONO,PSMD5,YWHAQ,EI
NC1,RPS14,RPS20,EIF5,RPL26,OAZ1,RPL31,LRPPRC,KHDRBS1,RPS25,EIF5A,GPC3,ACSL6,DDX
MD5,DCHS1,CHSY1,INA,MRC2,PPP2R1A,BASP1,VAPA,HMGB1,RPS3A,STMN1,ITGB1,UCHL1,NCAM
1,GSTP1,CALM1,NPM1,PLXNB2,HSPD1,COL18A1,IGF2,VCP,UBB,YWHAG,COL4A2,MAPT,FN1,NC
HSPD1,SNRNP200,CSDE1,IGF2,EIF3C,ATP5F1A,EIF4B,VCP,UBB,AFF4,TRIM28,COL4A2,RPL37A,M
CHS1,CHSY1,INA,MRC2,PPP2R1A,BASP1,VAPA,HMGB1,RPS3A,STMN1,ITGB1,UCHL1,NCAM1,ANX
F43,RPS3,CHD3,BASP1,HMGB1,POLR2A,RPS4X,RPS3A,RPL18,RPL11,MORF4L1,BTF3,AGRN,RPL
GF2,EIF3C,ATP5F1A,EIF4B,VCP,UBB,AFF4,TRIM28,CKB,COL4A2,RPL37A,SPEN,NONO,DDB1,PSM
GF2,EIF3C,ATP5F1A,EIF4B,VCP,UBB,AFF4,TRIM28,CKB,COL4A2,RPL37A,SPEN,NONO,DDB1,PSM
,PJA2,IGFBP3,RHOA,ATF4,PSMD2,ENC1,RPS20,CD81,CLEC7A,PRKCSH,CCNI,TADA2B,KLHL9,PT
M28,COL4A2,RPL37A,MAPT,SPEN,NONO,DDB1,PSMD5,YWHAQ,SET,EGLN1,TPI1,RPL19,PPP2R1
APK1,SMARCA4,FTO,LUZP6,PRKDC,FKBP8,CCAR2,SLIT3,TNKS2,PLXNA3,COL27A1,PTPRS,SOD1
NMA1,SULF1,RAB7A,ACTN4,PRDX2,DDX3X,ITM2C,H1-0,APLP1,RPS7,MIF,GJA1,JUN,CTNNB1,ANX
N1,RPL19,PPP2R1A,RPL15,GTF3C4,PKD1,SF3B1,SSBP2,ZNF43,RPS3,CHD3,BASP1,HMGB1,POLI
31,MARCHF7,PSMC5,CSNK1A1,PSMC2,HERPUD1,PSMC4,PLK2,PSMF1,WAC,GSK3B,USP19,UBQL
EN,NONO,PSMD5,YWHAQ,EIF1,SET,EGLN1,CHSY1,EIF4A1,RPL19,PPP2R1A,RPL15,GTF3C4,PKD
M4B,CDC42,EP300,FLCN,UBQLN2,CST3,DIP2B,ATRX,EVL,ADCY6,HDAC6,PTEN,ULK1,TNKS,SLC2
EF1,G3BP2,MAPRE1,ARPC2,LUZP6,GRB2,CAPZB,CAND1,CLIP3,CORO1C,STXBP1,ATXN2L,PTPR
P70,PSMB2,ABCA2,NRDC,UBXN4,MVB12B,AZIN1,DDAH2,PDE7A,PSMD3,ODC1,ADAM9,GABARAF
L37A,MAPT,FN1,SPEN,NONO,PSMD5,YWHAQ,EIF1,SET,EGLN1,RPL19,PPP2R1A,RPL15,PKD1,SF
5A2,WDR1,VAT1,PFN1,CLEC7A,PTN,CSNK1E,GLG1,TPM1,CDH2,GPC3,SULF1,PLXNA1,MEGF8,SC
A,CCT5,CDC42,PPP1R9B,GRIPAP1,OS9,FLCN,SEC16A,RER1,UBE2G2,OAZ2,ANP32B,LMAN1,GPC
,UBE2G2,FBXO11,HDAC6,PTEN,CUL3,RPS27A,PSMD8,NPLOC4,FBXO21,RHBDD2,BAP1,PSMB4,
5,YWHAQ,EIF1,SET,EGLN1,CHSY1,EIF4A1,RPL19,PPP2R1A,RPL15,GTF3C4,PKD1,SSBP2,ZNF43,
NES,NISCH,PTN,SFPQ,EIF5A,CCND2,PNMA1,SULF1,RAB7A,CCN2,ACTN4,PRDX2,DDX3X,ITM2C,H
IC3,PRKACA,UBE2D3,PSMC1,CCDC47,BAG5,UBXN6,UBE2K,ZER1,FBXL19,USP5,AMFR,CDC34,SI
,DNAJB2,SEMA3C,SPAG9,CLASP2,ADAM10,IST1,SH3BP4,GSK3A,FBLN5,MTOR,YY1,APBB1,NDUF
PC1,APPL1,HDAC2,IPO5,TNC,ATP6V0C,MMP2,CTSB,SRSF5,VAMP2,GNAI3,PARP1,NOX4,PTPRA,P
OL5A1,COL4A1,CES3,PSAP,PGK1,RPLP0,H3-3B,CANX,CLU,PTMA,DDX5,ENO2,SOX4,FKBP14,RPL
1B1,AKAP12,SLC4A8,PXDN,CTNNA1,RPL23,RTN4,PJA2,IGFBP3,LLGL1,NCS1,RPL22,RHOA,CLST
TGFBI,ACTR3,PLOD1,CAMSAP1,NCKAP1,COL2A1,H3-3A,MAPRE1,ARPC2,DSTN,PLEC,AEBP1,LUZ
IRPB,SFSWAP,PRPF31,XAB2,KHDC4,CPSF7,PRMT7,EIF3J,SNU13
,UBB,AFF4,TRIM28,COL4A2,RPL37A,MAPT,SPEN,NONO,DDB1,PSMD5,YWHAQ,SET,EGLN1,TPI1,
,VCP,UBB,AFF4,TRIM28,COL4A2,RPL37A,SPEN,NONO,DDB1,PSMD5,YWHAQ,EIF1,HACD3,SET,E
IP1,HNRNPK,RHOB,HMGCR,ARHGDIA,HDAC2,BCLAF1,PEA15,PTTG1IP,CDKN1A,DLG5,TIMP3,C
FBLN1,PLXNA3,SARS1,PTPRS,VANGL2,HGS,RAB11A,PRPF40A,BMPR2,CDC42,PSMA7,STAT3,BCL

F31,XAB2,KHDC4,CPSF7,PRMT7,EIF3J,SNU13
MC1,CNN2,CTSB,RAB27A,CCT2,ANXA5,VAMP2,GDI2,PDXK,MLEC,SURF4,RAB5B,RAB11B,SRP14,
4,EIF2AK1,PRMT1,SGTA,EP300,UBQLN2,BRD4,SFRP1,HDAC6,PTEN,FEM1B,PTPRF,POM121C,HI
A,CTBP1,RAC1,HDAC2,SKP1,TRIB2,PEA15,PTTG1IP,CDKN1A,DDX39B,SQSTM1,NOP53,NOX4,PK
,ANXA5,VAMP2,GDI2,PDXK,MLEC,SURF4,RAB5B,RAB11B,SRP14,PYGB,VEGFA,MAPK1,PSMD3,A
AP12,SLC4A8,PXDN,CTNNA1,RPL23,RTN4,PJA2,IGFBP3,LLGL1,NCS1,RPL22,RHOA,CLSTN1,ATF
,UBB,AFF4,TRIM28,COL4A2,RPL37A,MAPT,SPEN,NONO,DDB1,PSMD5,YWHAQ,SET,EGLN1,TPI1,
RPL37A,SPEN,NONO,PSMD5,YWHAQ,EIF1,SET,EGLN1,EIF4A1,RPL19,PPP2R1A,RPL15,GTF3C4,P
ANX,CLU,PTMA,DDX5,ENO2,SOX4,FKBP14,RPL13A,MAGED1,RPL6,KAT6A,DVL3,YWHAE,KIF1A,G
ZER1,FBXL19,USP5,AMFR,CDC34,SDCBP,PSMB1,HSPA1A,MDM2,FZR1,UBE4B,TNFAIP1,TRIM2,C
CDKN1A,CTDSP2,EID1,U2AF2,SQSTM1,NOP53,NOX4,PKN1,GCN1,SKI,ABCA2,TERF2IP,TMED2,T
LOC4,FBXO21,RHBDD2,BAP1,PSMB4,EIF3H,UBE2D2,VPS4A,GNA12,DNAJB2,ATXN3,PSMD4,OGT
NGAP1,PTPN11,HMGA1,RBM4,RGPD5,PKIA,MDM2,PARK7,BANF1,RAN,FAM53C,AKAP8L,NUP153
P1,TCP1,CCT3,CLSTN3,CCT4,RAB11B,ABCA2,VEGFA,MAPK1,ADAM9,NPEPPS,KDM1A,BNIP3L,CL
LOC4,FBXO21,RHBDD2,BAP1,PSMB4,EIF3H,UBE2D2,VPS4A,GNA12,DNAJB2,ATXN3,PSMD4,OGT
BE2J1,FAM8A1,NCK2,TRIB3,HM13,STUB1,HERPUD1,TNFRSF10B,GSK3B,USP19,LMNA,UBXN1,A

HBDD2,BAP1,PSMB4,EIF3H,UBE2D2,VPS4A,GNA12,DNAJB2,ATXN3,PSMD4,OGT,USP24,FBXW5,
HOB,HMGCR,ARHGDIA,HDAC2,BCLAF1,PEA15,PTTG1IP,CDKN1A,DLG5,TIMP3,CTSB,EEF1A2,ANX
1,NPTN,ADNP,CYFIP1,FEZ1,KAT2A,NTN1,LIMK1,CCDC88A,NME2,ISLR2,DHX36,NDRG4,GOLGA4

TNNB1,DNAJC5,TARDBP,ARL6IP1,HNRNPK,RHOB,HMGCR,ARHGDIA,HDAC2,BCLAF1,PEA15,PT
X5,RTN1,PKN1,SKI,SNRNP70,TERF2IP,TMED2,TAF7,VEGFA,RPL37,PPP2CA,AZIN1,IPO7,EIF3E,FT
T4,RAB11B,ABCA2,VEGFA,SELENON,MAPK1,NPEPPS,KDM1A,BNIP3L,CLIP3,PLCG1,MAP2K2,SU
SL1,CUL1,PSMD13,DCAF5,ANO8,PSMD7,SHISA5,CUL4A,COPS6,PSME3,PSMC5,PSMC2,PSMC4,S
DC,KDM1A,QARS1,ARF4,SDF4,DHX30,FYN,YWHAH,B2M,PITPNM1,PLXNA3,CSNK1D,PTPRS,BAG
NAP1,FLCN,LRRC4B,FLOT1,PLXNB1,FARP1,SFRP1,CTNND2,RER1,HDAC6,PTEN,PTPRF,MAPRE2,
0,PSMC4,PSMF1,YWHAZ,IGF2BP2,PSMB5,PSMA1,SKIV2L,PSMD10,ROCK2,PSMA6,CNOT3,PAIP1,

,UBB,AFF4,TRIM28,COL4A2,RPL37A,MAPT,SPEN,NONO,DDB1,PSMD5,YWHAQ,SET,EGLN1,TPI1,
1,YWHAZ,IGF2BP2,PSMB5,PSMA1,SKIV2L,PSMD10,ROCK2,PSMA6,CNOT3,PAIP1,YBX3,PSMD12,
PS26B,RAB1B,VPS41,VPS4A,GABARAPL1,FNBP1L,MTMR4,FOXK2,SYT11,ATP6V0D1,SH3BP4,KD
PS26B,RAB1B,VPS41,VPS4A,GABARAPL1,FNBP1L,MTMR4,FOXK2,SYT11,ATP6V0D1,SH3BP4,KD
AC6,PTEN,ULK1,CTTN,SLIT2,SERP1,NDN,SLIT1,GPX1,OLFM1,SOX2,PKDCC,DCLK1,SEMA3C,SP
NGAP1,PTPN11,RBM4,RGPD5,PKIA,MDM2,PARK7,RAN,FAM53C,AKAP8L,NUP153,KPNA6,SRSF7,
4,UBXN1,PSMD10,ELOB,GABARAPL2,XBP1,APOE,RYBP,EPHA4,PSMC6,RAD23A,PITHD1,RNF40
P53,PARP1,NOX4,HSPH1,ALKBH5,PKN1,PLOD1,MDK,PSMB2,CNTNAP2,RAB11B,ATP6AP1,VEGF
YO5A,CTNNB1,MACF1,CYFIP2,MYH10,CKAP5,RHOB,MAP2,RPS27,COTL1,RAC1,ARHGDIA,MAP1
TNNB1,TARDBP,MYH10,DUX4,CKAP5,SEPTIN11,SND1,RHOB,SEPTIN3,RPS27,CTBP1,CDK16,TO
WIZ,STUB1,MARCHF7,P3H1,USP19,RASSF2,LMNA,GET1,PHB,PIN1,XBP1,FBXO7,PRNP,EPHA4,TR
LE1,PSMD4,RBPJ,GSK3A,PSMC3,AP2A2,PSMC1,WNK1,DAPK3,PTPN23,CALCOCO1,AMFR,IGFB
1,ABL1,PINK1,ARPC1A,HIP1R,HSPA1A,ELN,CRACD,MECP2,JAM3,SUMO1,PARK7,CYFIP1,FLII,CU
EIF4G1,IGFBP4,USP47,OGFR,ABL1,SDCBP,SEMA6B,TAOK2,TSPYL2,HSPA1A,ACVR1B,KIF26A,AD
,MAP1LC3B,TMOD2,TAOK1,VPS4A,GABARAPL1,RDX,SH3GL1,CAPZA1,BLCAP,DICER1,CLASP2,A
LE1,PSMD4,RBPJ,GSK3A,PSMC3,AP2A2,PSMC1,WNK1,DAPK3,CALCOCO1,AMFR,IGFBP4,USP47

IP,CDKN1A,DDX39B,SQSTM1,NOP53,NOX4,PKN1,VPS35,ABCA2,NRDC,TERF2IP,VEGFA,RAPGEF
35,PSMB2,ATP6AP1,DPYSL2,MARS1,SELENON,RAPGEF1,UBXN4,CCND1,MAPK1,SRSF9,GRB2,PF
RAD23A,PITHD1,RNF40
,GRN,ACLY,XRCC6,SLC44A2,IMPDH2,RAB14,NCSTN,RAB6A,RAB5C,ACTR1B,ITGAV,ADAM10,CAF
NAJB6,GNAZ,DNAJB11,ATF6,SELENOF,GNAO1,PRDX4,QSOX1,DNAJC21,GNAI1
OC2,THOC3,RTRAF,FYTTD1,NUP50,RNPS1,DDX19A,NUP58,WDR33,NOP58,MEX3D,NUTF2
2,FBXO11,HDAC6,PTEN,FEM1B,CUL3,POM121C,RPS2,RPS27A,TNKS,PSMD8,HIF1A,FKBP1A,FBX

VEGFA,RPL37,PPP2CA,IPO7,EIF3E,FTO,ILF3,PCSK1N,PRKDC,HSPB1,KDM1A,FKBP8,QARS1,CCAF
,GRN,ACLY,XRCC6,SLC44A2,IMPDH2,RAB14,NCSTN,RAB6A,RAB5C,ACTR1B,ITGAV,ADAM10,CAF
1,YWHAZ,IGF2BP2,PSMB5,PSMA1,SKIV2L,PSMD10,ROCK2,PSMA6,CNOT3,PAIP1,YBX3,PSMD12,
1,NOX4,PTPRA,UBE4A,AHCYL1,VPS35,PSMB2,ATP6AP1,DPYSL2,MARS1,SELENON,RAPGEF1,UBX
PRDX6,GRN,ACLY,XRCC6,SLC44A2,IMPDH2,RAB14,KMT2E,NCSTN,RAB6A,RAB5C,ACTR1B,ITGA
N,UBA52,ZFP36L1,SFRP1,HDAC6,PTEN,RGMB,NDST1,SPINT2,ITGB5,RPS27A,PEG10,DKK3,HIF1
C42,PSMA7,TRAF4,TRAF7,ARIH1,OS9,MTA1,EP300,PSMD1,UBA52,KLHL42,NXN,UBE2G2,FBXO1
BX1,UBA1,ATP2B4,IGFBP2,CALR,FSTL1,TMF1,ABHD2,HSPA1B,ZNF99,TUBB2B,COL5A1,COL4A1
,PRDX6,GRN,ACLY,XRCC6,SLC44A2,IMPDH2,RAB14,KMT2E,NCSTN,RAB6A,RAB5C,ACTR1B,ITG
HOA,PSMD2,NLGN2,CHD4,RPS20,CD81,RPL26,RBBP4,MCL1,TLE5,CLEC7A,CSNK1E,GLG1,SFPQ
,CUL3,ULK1,LMAN1,SEPTIN7,TNKS,HIF1A,CTTN,SLC25A5,BRK1,AKAP9,GRN,PPP1R10,ABI2,DZIF
3,APPL1,SKP1,CDKN1A,CTDSP2,DDX39B,PPP1CB,NOP53,ACTR3,PSMB2,TERF2IP,OPTN,PPP2CA,
,GRN,ACLY,XRCC6,SLC44A2,IMPDH2,RAB14,KMT2E,NCSTN,RAB6A,RAB5C,ACTR1B,ITGAV,ADAM
11,HDAC6,PTEN,FEM1B,CUL3,RPS2,RPS27A,TNKS,PSMD8,HIF1A,FKBP1A,FBXO21,WSB2,PSMB4
,DYNLL2,SON,CSNK1D,RAB11A,DYNLL1,CDC42,PPP1R9B,PSMA7,DCTN1,PRMT1,ACTR1A,EP300
K1,ATP13A2,MFN2,PARK7,PIK3C2B,FEZ1,EHMT2,RHEB,CSNK2B,CAPN1,TBC1D14,ATP6V0B,EXO
7,SON,CCAR2,CSNK1D,RAB11A,PRPF40A,DYNLL1,TP53BP1,CDC42,PPP1R9B,PSMA7,RBBP7,DCT
LS,MYH10,LAMP1,LGALS3BP,CKAP4,HLA-B,RAB1A,TTR,COTL1,HRNR,HMGCR,RAC1,CDK16,ATF
7,DCTN1,PRMT1,ACTR1A,EP300,PSMD1,PBX1,CSNK2A1,PHB2,GSPT1,BRD4,UBA52,ZFP36L1,AT
APOE,FGF13,SYT1,SYT4,POSTN,LRP6,CACNG7,ANAPC2,SIN3A

,HNRNPK,H1-10,CIRBP,DUX4,PTGES3,CCT6A,RAC1,APPL1,HDAC2,BCLAF1,DCN,DDX39B,CHD6,
S1,RHOA,ATF4,PSMD2,CHD4,RPS20,CD81,ATP2A2,RPL26,RBBP4,MCL1,CLEC7A,NISCH,PRKCSH

L1,SMARCE1,SMARCC1,SUPT6H,CCT8,TNKS2,USP7,BAG6,HNRNPD,RAB11A,KDM3B,WBP2,KDM
,XRCC5,IGFBP3,LLGL1,RHOA,PSMD2,KIF5B,NLGN2,RPS20,EIF5,CD81,ATP2A2,CRMP1,BEX3,OA
0,CIRBP,DUX4,PTGES3,CCT6A,RAC1,APPL1,HDAC2,BCLAF1,DCN,DDX39B,CHD6,RNF187,CCT2,
I,RNF5,CDK5RAP3,ABCA7
A52,ZFP36L1,SFRP1,PTEN,RGMB,NDST1,SPINT2,ITGB5,RPS27A,PEG10,DKK3,HIF1A,FKBP1A,LC
M1,STXBP1,CSNK1D,TMED9,MESD,HGS,GHITM,SOD1,BAG6,PPP6R1,SEC23A,CDC42,SYP,DCTN1
2,DDIT4,PPP1R9B,CTNND1,CIC,EP300,RERE,PBX1,SEC16A,ATRX,SFRP1,GLUD1,TRA2B,PTEN,ND
TS2,TNPO2,KPNA3,PPP3CA,CTDNEP1,ARL2BP,PIN1,XBP1,UFM1,TNPO3,NOLC1,NUP50,SESN2,PA

SMF1,RCC2,PSMB5,PRKAR2B,PSMA1,PPP1R12A,PSMD10,PSMA6,AJUBA,PSMD12,PSMA4,TPX2,F
N1,LIMK1,FLII,PPP1CA,CDK4,TRIM27,NCK2,ISLR2,ARPC3,GBA2,GOLGA4,SEMA4C,SRF,GSK3B,
STMN3,HECTD1,VANGL2,SOD1,BAG6,TCTN1,ARNT2,BMPR2,SULF2,PRMT1,CAPN2,EP300,PBX1,
YNC1I2,CENPF,PSMC6,CDK5RAP3,NINL,HMGA2,SIN3A

DAC2,BCLAF1,DCN,DDX39B,CHD6,RNF187,CCT2,MLXIP,SQSTM1,PARP1,MORF4L2,TAF15,HSPH
FGF13,SYT1,SYT4,POSTN,CACNG7,ANAPC2,SIN3A
HB2,BRD4,UBA52,ZFP36L1,ATRX,SFRP1,PTEN,FEM1B,CUL3,BRSK1,RPS27A,TNKS,PSMD8,APEX
,NDUFA13,BNIP3,PSMC1,TRIM8,AMBRA1,ATG4B,ZER1,USP5,PAFAH1B2,PUM1,PINK1,YTHDF3,H
32,PRKDC,KDM1A,GNG2,SMARCC1,ATP6V1B2,FYN,YWHAH,SLIT3,LONP1,IDH1,NCOR2,VGF,WBI

PPP1R10,UBR5,DNAJB2,SPAG9,PDPK1,OGT,NOC2L,KDM4A,ARPP19,MTOR,WNK1,BAG5,MICAL1
K5RAP3,NINL,HMGA2,SIN3A
PDAP1,CST3,ATP6AP2,PRDX6,GRN,ACLY,XRCC6,SLC44A2,IMPDH2,RAB14,KMT2E,NCSTN,RAB6A
3,HMGB1,TPT1,SEC61A1,ITGB1,TUBB4B,UCHL1,MORF4L1,NCAM1,ANXA2,XAF1,MARCHF6,VCL,
RDBP,DDOST,WLS,MYH10,LAMP1,LGALS3BP,ATP1A2,CKAP4,HLA-B,RAB1A,TTR,COTL1,HRNR,H
ATP6AP2,PRDX6,GRN,ACLY,XRCC6,SLC44A2,IMPDH2,RAB14,NCSTN,RAB6A,RAB5C,ACTR1B,PDF
G,FASN,SPTBN1,CALM3,GNB1,SHISA9,STMN2,MYH9,SEPTIN2,P4HB,APP,RPL5,KLRD1,GDI1,FSC

A1,PPP1R12A,PSMD10,PSMA6,AJUBA,PSMD12,PSMA4,TPX2,FHL1,DYNC1I2,CENPF,PSMC6,PPP2
OP56,MTREX,TRMT112,RPS21,RAN,CUL4A,RPL27A,C1QBP,EXOSC10,PDCD11,PA2G4,FBL,RRP1I
RIN1,GPC1,HDAC2,L1CAM,TRIB2,DDX39B,EID1,PARP1,MORF4L2,MDK,SKI,ASXL1,PSMB2,H3-3A,
6A,ATRX,SFRP1,GLUD1,TRA2B,PTEN,NDST1,EML1,ANP32B,HIF1A,SLIT2,SLC2A1,XRCC6,SLIT1,H

TBC1D14,MAPKAPK2,DHX36,MARCHF7,RBM10,PSMF1,WAC,SERPINE2,USP19,UBQLN4,UBXN1,F

,HJURP,EP400,ARID1B,HDAC6,NAP1L3,CBX6,ANP32B,MCM3AP,BAP1,BAZ2A,EHMT1,CARM1,UBI
,ARF5,SCAP,SNX2,ANK2,TAPBP,SNX3,PUM1,LMAN2,SGSM2,CLTA,SCAMP1,EXOC5,NSF,COPE,KII
ZA1,CLASP2,WASF2,CHMP4B,MTOR,DAPK3,TUBB4A,CELSR1,ABL1,CHMP1A,ARPC1A,HDGFL3,T
3,GNG4,SFRP1,PTEN,HIF1A,CTTN,SLIT2,SERP1,NDN,BAP1,OLFM1,LTBP4,HDGFL2,DNAJB2,SEM
31,CASC3,MAP2K2,POLR2B,HNRNPM,NCOR2,HNRNPD,SERBP1,ADAR,POLR2E,PSMA7,ANP32A,S
SM14A,TTC37,EIF4G3,TRAP1,LSM14B,ABCF1,IGF2BP1,AKT2,GSK3A,MTOR,DAPK3,EIF4A3,EIF4
ATP6AP2,PRDX6,GRN,ACLY,XRCC6,SLC44A2,IMPDH2,RAB14,NCSTN,RAB6A,RAB5C,ACTR1B,PDF
A,CTTN,SLC25A5,DAP,GPX1,HINT1,MAZ,STK24,TRAP1,NMT1,SOD2,MOAP1,KRT8,PDPK1,ITGAV,I
MD
19,UBQLN4,UBXN1,GABARAPL2,PAIP1,YBX3,TAB3,RYBP,FURIN,EPHA4,RNF5,CDK5RAP3,QSOX1,
TRN,NTRK2,ABL1,SEMA6B,TRAK1,DAG1,NIBAN2,PLXNA2,OBSL1,NOTCH1,PCM1,NPTN,ADNP,IDH
,SOX2,TLE1,SH3GL1,PSMD4,RBPJ,GSK3A,PSMC3,AP2A2,PSMC1,WNK1,DAPK3,SMO,EIF4A3,CAL
L1,AGRN,ZMIZ1,RPL10,AP3D1,AKAP12,ITM2B,ZNF770,GTF2F1,HNRNPU,NFE2L1,SMARCA5,LAR
BP1,APOE,FURIN,PSMC6,RAD23A,VPS28,RNF40
2,TARDBP,DDOST,WLS,MYH10,LAMP1,LGALS3BP,CKAP4,HLA-B,RAB1A,TTR,COTL1,HRNR,HMG
1,CCND2,GPC3,DDR1,PLXNA1,PIP4K2B,CCN2,PRDX2,DDX3X,RPS15,RPS7,PRKAR1A,JUN,CTNNB
STM1,NOP53,PARP1,PRDX5,UBE4A,RBMX,RAD23B,ASXL1,PSMB2,ABCA2,NRDC,VEGFA,VHL,UBX
P1,LIMK1,FLII,CCDC88A,TRIM27,NCK2,ARPC3,EMILIN1,COLGALT1,GBA2,MAPRE3,PDE4DIP,MA

IP3,NOLC1,RPL32,PES1,KRI1,DDX56,NOP58,USP36,SNU13
CAR2,ARF4,FYN,MTRNR2L10,STXBP1,HYOU1,MAPK8IP1,GHITM,SOD1,BAG6,ARNT2,YME1L1,AD
DAH2,PRKDC,HSPB1,SON,KDM1A,BNIP3L,FKBP8,QARS1,CCAR2,ARF4,FYN,MTRNR2L10,STXBP1,
,RRP1B,SENP3,NOLC1,RPL32,PES1,KRI1,DDX56,NOP58,USP36,SNU13

4,AKAP9,PPP1R10,PSMB4,TAOK1,CARM1,INTS3,VPS4A,RDX,KMT2E,CLASP2,TCF3,PSMD4,ARPP
,IGF2BP1,AKT2,MTOR,DAPK3,EIF4A3,EIF4G1,CNBP,PUM1,FXR2,PINK1,PAIP2,YTHDF3,RBM4,MI
OL16A1,SDCBP,PINK1,AUTS2,ARPC1A,DAG1,HIP1R,HSPA1A,SMPD3,ATP13A2,CRACD,MECP2,SU
MIZ1,RPL10,AP3D1,AKAP12,ITM2B,ZNF770,GTF2F1,HNRNPU,NFE2L1,SMARCA5,LARP1,RPL23,E
AND5,POLR2B,SMARCC1,ATP6V1B2,FYN,HNRNPM,GIGYF1,STRAP,HGS,DGKD,BMPR2,TXNIP,CD

ARS1,CCAR2,ARF4,FYN,MTRNR2L10,STXBP1,HYOU1,MAPK8IP1,GHITM,SOD1,BAG6,ARNT2,YME

T5,CRABP2,NUMBL,SRRT,SS18L1,RUFY3,ADD1,CUL7,PRKCI,RELA,OPA1,EPHB2,LRP8,EPHA4,MY
PT
NB2,ADNP,CYFIP1,NTN1,NECTIN1,SYNGAP1,DHX36,CRKL,NDRG1,YWHAZ,SEZ6L2,ABHD17A,RA
1,ISLR2,DHX36,GOLGA4,SEMA4C,SRF,SERPINE2,PRMT5,CRABP2,NUMBL,EFNB3,SRRT,SS18L1,
ANAPC2,SKIL

,HNRNPA0,RPL23,NACA,BRD3,XRCC5,CIPC,PAPOLA,RHOA,CBX5,ZNF865,ATF4,PSMD2,NR2F1,R
IT1,ARF6,MYADM,GSN
N,ZMIZ1,RPL10,AP3D1,ITM2B,ZNF770,GTF2F1,HNRNPU,NFE2L1,SMARCA5,LARP1,RPL23,EIF4H

JUBA,PSMD12,OPA1,PSMA4,PSMC6,NAMPT

PL10,AP3D1,ITM2B,ZNF770,GTF2F1,HNRNPU,NFE2L1,SMARCA5,LARP1,RPL23,EIF4H,NACA,BRD
2E,CLASP2,TCF3,PSMD4,ARPP19,PSMC3,CHMP4B,PRKACA,NASP,PSMC1,TUBB4A,EIF4G1,USP4

L2,TAF15,PRDX5,YEATS2,RBMX,CBX1,SKI,H3-3A,HDGF,TERF2IP,TAF7,NOTCH2,VEGFA,VHL,CCN

SOR,NECTIN1,DDX6,TRIM27,NUP153,KPNA6,NUP98,TFRC,NUP205,NUP133,PHB,EFNB3,KPNA3,
CT2,MLXIP,SQSTM1,PARP1,MORF4L2,TAF15,ACTR3,PKN1,TCP1,CCT3,PTOV1,MDK,RBMX,CCT

2,XBP1,TFDP1,PRKCI,GNL3,EPHB2,ARF6,PRNP,SESN2,VPS28,LEPROT,CDK5RAP3,EPHA2,USP36,
9,CDKN1B,WASF3,CDC42EP4,VDAC2,ARPC5,CAPZA2,ADD1,FER,GMFB,GIT1,OPA1,FGF13,TPX2,
RP1B,THOC2,RXRA,PHB,EFNB3,KPNA3,THOC3,NECTIN2,MAPK3,SLC1A5,SP1,DYNLT1,APOE,SCA
1,UNC13A,CNTN2,SS18L1,ANXA1,STRIP1,CUL7,ARHGEF18,OPA1,LRP8,SYT1,EPHA4,SYT4,MYAD
POE,LRP8,ARF6,SRCIN1,EPHA4,ANAPC2
F3,HSPA1A,MDM2,FZR1,SUMO1,PARK7,PTPN1,TNRC6A,HNRNPR,MAGEF1,DDX6,TRIM27,TRIB3,

'DLIM1,PSMD7,HIPK2,PDGFRB,CDK4,PSME3,STUB1,LOXL2,PSMC5,NDRG1,PSMC2,PSMC4,PSMI
TNNB1,COL6A2,SESN3,RHOB,ATP1A2,HMGCR,RAC1,CDK16,APPL1,HDAC2,IPO5,TNC,ATP6V0C,I
,PRKACA,PHIP,TTC28,PSMC1,DAPK3,TUBB4A,EIF4G1,POLDIP2,USP47,PTPN11,ABL1,SDCBP,PSI
TXBP1,IDH1,DYNLL1,TXNDC5,PSMD1,PDAP1,CST3,ATP6AP2,PRDX6,GRN,ACLY,XRCC6,SLC44A2
MC2,PSMC4,PSMF1,UBE3C,UBE2V1,RNF216,ZFP91,PSMB5,OTUB1,RNF20,PSMA1,ZNRF1,RNF4,
1,WDR1,ATP2A2,CRMP1,PFN1,ASTN2,CLEC7A,NISCH,LRPPRC,SPTBN2,SLC3A2,PTN,GLG1,SFPC
M4,CTBP1,HMGCR,RAC1,CDK16,HDAC2,IPO5,TRIB2,PEA15,CDKN1A,CTDSP2,EEF1A2,SQSTM1,N

OLA,RHOA,CBX5,ZNF865,ATF4,PSMD2,NR2F1,RPS14,CHD4,JUND,CD81,SRSF4,PFN1,TLE5,HNRI
2M,DAP,MTCH1,GRN,CCN1,GPX1,EIF3H,ANTXR1,SOX2,LTBP4,GNA12,DNAJB2,NCSTN,ATXN3,IST

PN1,TBC1D14,TRIM27,TRIB3,ATP6V0B,EXOC7,PLK2,WAC,GSK3B,UBQLN4,ATP6V0E2,ATP6V0A1
,HMGCR,RAC1,ATP6V0C,BCLAF1,GLUL,PEA15,NNAT,DCN,DLG5,TIMP3,SQSTM1,VAMP2,PARP1,

1,KAT6A,DVL3,YWHAE,GNAS,SLC2A3,DDX17,GPI,HTRA4,RACK1,BSG,FASN,SPTBN1,CALM3,GNE
2,SQSTM1,NOP53,PARP1,NOX4,ABCA3,PKN1,GCN1,ABCA2,TERF2IP,TMED2,TAF7,VEGFA,RAPGE
2,SQSTM1,NOP53,PARP1,NOX4,ABCA3,PKN1,GCN1,ABCA2,TERF2IP,TMED2,TAF7,VEGFA,RAPGE

CDK5RAP3,RNF40,ARRDC3,TOLLIP
7,PRKCI,LRRN1,RELA,OPA1,EPHB2,LRP8,EPHA4,PLXND1,ID2,ANAPC2,SKIL
FZR1,PCM1,MECP2,CUL1,PSMD13,PSMD7,CUL4A,CETN2,CDK4,PSME3,PSMC5,PSMC2,PSMC4,P
MNL2,ITGB5,CALD1,CTTN,SLIT2,BRK1,TMOD2,TAOK1,CYTH2,ANTXR1,ABI2,ARHGEF17,VPS4A,F
1A,CDH11,SQSTM1,VAMP2,PTPRA,VPS35,CLSTN3,MDK,PTK7,SKI,PSMB2,RAB11B,RAPGEF1,PPP
A1,MAPRE3,NSD1,PLK2,HSF1,GSK3B,ILRUN,MED25,TXN,CEBPG,CTBP2,RGMA,TGFBR1,MAPK3,I
I6,RAPGEF1,OPTN,VHL,PPP2CA,SMARCA4,PSMD3,GRB2,HSPB1,KDM1A,QARS1,CCAR2,DUSP4,D
UCHL1,RPL11,NCAM1,ANXA2,ZMIZ1,TIMP2,DNAJA1,PAFAH1B1,AKAP12,PXDN,CTNNA1,RPL23,RT
L1,KDM6B,PRPF6,TAF7,RNF10,VEGFA,MARS1,VHL,MAPK1,SMARCA4,PBX2,SRSF1,FTO,ILF3,PRI
KI,H3-3A,HDGF,TAF7,NOTCH2,VEGFA,VHL,CCND1,SMARCA4,SRSF9,AEBP1,TCF25,PTBP1,ILF3,S
A,SMARCA4,PSMD3,GRB2,HSPB1,KDM1A,QARS1,CCAR2,DUSP4,DLK1,FYN,SLIT3,STMN3,FBLN1
RPS6,UBQLN4,SETX,MAP1S,NAPB,STMN4,NAPA,BMERB1,CHMP5,GABARAPL2,CAPZA2,ADD1,ABC
I6,RAPGEF1,OPTN,VHL,PPP2CA,SMARCA4,PSMD3,GRB2,HSPB1,KDM1A,QARS1,CCAR2,DUSP4,D
CR,RAC1,ATP6V0C,BCLAF1,GLUL,PEA15,NNAT,DCN,DLG5,TIMP3,SQSTM1,VAMP2,PARP1,PKN1,V

IN1,PSMC1,TUBB4A,PAGR1,EIF4G1,USP47,PTPN11,PSMB1,TAOK2,MACROH2A1,PKIA,MDM2,FZI
T3,KAT2A,EHMT2,PCGF3,TASOR,ATF7IP,AKAP8L,LDB1,CDK9,NSD1,WAC,WDR82,ASH1L,RCOR2,

RK7,FEZ1,PTPN1,RAN,UBE2J1,AKAP8L,NUP153,NDRG4,NCBP2,YWHAZ,GSK3B,NUS1,KIF3A,TXN,I
ADM4,RHOB,HMGCR,RAC1,ARHGDIA,HDAC2,BCLAF1,PEA15,PTTG1IP,DCN,DLG5,TIMP3,DDX39I
2,CDC42,PPP1R9B,PALLD,HIP1,FARP1,SFRP1,EVL,CUL3,FMNL2,ITGB5,CALD1,CTTN,SLIT2,BRK1
C2,PSMC4,TMEM9,PSMF1,GSK3B,ZNF703,LGR4,SOX9,PSMB5,CHD8,PSMA1,LZTS2,PSMD10,PSI
MA5,MSN,IGFBP3,LLGL1,NCS1,RHOA,ATF4,KIF5B,NLGN2,ATP1A1,CD81,WDR1,ATP2A2,CCT7,O
A,EHMT2,PCGF3,AKAP8L,LDB1,CDK9,NSD1,WAC,WDR82,ASH1L,RCOR2,BCOR,PRMT5,FBL,SETI

C2,PSMC4,PSMF1,SOX9,PSMB5,XPC,PSMA1,HIRA,RNF4,CTCF,PSMD10,UBN1,THRA,PSMA6,CABI
CDC47,BAG5,DOP1B,REEP1,GAK,USO1,RANGAP1,ANK2,SNX3,SUN1,LMAN2,CHMP1A,ESYT1,GR
2,TAF7,SELENON,RAPGEF1,CCND1,MAPK1,SMARCA4,ADAM9,KDM1A,NSG2,GNG2,YWHAH,SLIT3

TN,ADNP,IDH2,CYFIP1,NTN1,LIMK1,RHEB,TMEM98,SYNGAP1,ISLR2,DHX36,GOLGA4,SEMA4C,SF
SMF1,SRF,DDAH1,HSF1,USP19,STC2,CDKN1B,PSMB5,EGLN2,HIF3A,LMNA,PSMA1,CITED2,CFLAR
2,PSMC4,PSMF1,SRF,DDAH1,HSF1,USP19,STC2,CDKN1B,PSMB5,EGLN2,HIF3A,LMNA,PSMA1,CI

,ILF2,CCT8,B2M,STXBP1,IDH1,DYNLL1,TP53BP1,STAT3,TXNDC5,PSMD1,PDAP1,CST3,ATP6AP2,P
A3C,NMT1,KRT8,PHGDH,SCRIB,ADAM10,RBPJ,PRPF19,PCGF2,MBD3,VASH1,PRKACA,NASP,TANC
5,LDHA,CPE,NACA,RTN4,LPCAT1,LAMA5,XRCC5,MSN,RPL24,COL1A2,RPL22,RHOA,ATF4,PSMD2,
,PTEN,YKT6,CUL3,BRSK1,LMAN1,HIF1A,AKAP9,SEPTIN5,VAPB,SLIT1,RAB1B,VPS4A,KIFAP3,RAB
T1,KRT8,PHGDH,SCRIB,ADAM10,RBPJ,PRPF19,PCGF2,MBD3,VASH1,PRKACA,NASP,TANC2,AMB

CCT2,TBCD,GNAI3,PABPN1,GDI2,PARP1,NOX4,PKN1,TBC1D16,TCP1,ACAP3,CCT4,GCN1,SKI,TE

,HIF1A,LOX,BRK1,NDN,SHC2,RAB14,NCSTN,NUCKS1,CLASP2,PDPK1,WASF2,ANKS1A,ITGAV,AKT

1IP,DDX39B,PARP1,HSPH1,GRINA,VPS35,MDK,PSMB2,TSPAN6,TERF2IP,TMED2,VEGFA,SELENO
1,SEMA6C,EPHB2,NFASC,EPHA4,GRB10,PLXND1,NFIB,SH3KBP1,EPHA2
AP1B,PIEZO1,GSN,UNC13B
3,B2M,STXBP1,IDH1,DYNLL1,TP53BP1,STAT3,TXNDC5,PSMD1,PDAP1,CST3,ATP6AP2,PRDX6,GR

CLEC7A,CCNI,PTN,CSNK1E,PFKL,TKT,CCND2,DDR1,PIP4K2B,ACSL6,CCN2,PRDX2,DDX3X,MIF,HM
5,TIMP3,PARP1,PKN1,VPS35,PTK7,SKI,ASXL1,PSMB2,TSPAN6,ATP6AP1,TERF2IP,NOTCH2,VEGF
APK1,CCNL2,IPO7,ADAM9,CDC37,HSPB1,MAP2K2,QARS1,DUSP4,CORO1C,FYN,HGS,MAPK8IP1,
1,SLIT2,AKAP9,PALM,NCSTN,NUCKS1,PDPK1,AKT2,OGT,ATP6V0D1,SH3BP4,ARPP19,GSK3A,MT
EGFA,RAPGEF1,PPP2CA,CCND1,MAPK1,CCNL2,IPO7,ADAM9,CDC37,PRKDC,HSPB1,CLIP3,FKBP

TDNEP1,PIN1,PSMD12,APOE,PSMA4,PSMC6,PPM1A,TLE4,FZD7,GPRC5B,LRP6,JRK,EMD
C,EPHB2,NFASC,EPHA4,GRB10,PLXND1,NFIB,SH3KBP1,EPHA2

GABARAP,ATXN10,ARF4,KIF3C,DHX30,RPS28,UBQLN1,CSNK1D,ATXN2L,PTPRS,VANGL2,HGS,R
,THOC3,FYTTD1,ABCE1,NUP50,RNPS1,DNM1,DDX19A,NUP58,WDR33,CACNG7
A,HIP1R,ELN,ATP13A2,CRACD,ACTA2,ADNP,CYFIP1,NTN1,LIMK1,FLII,PPP1CA,EXT2,CDK4,TBXA2

TPN1,PPP1CA,NCK2,PSME3,MARCHF7,HERPUD1,TNFRSF10B,SCG2,GSK3B,LMNA,VDAC2,CFLAR
Z1B,CSNK2A1,BRD4,CSNK1G2,DIP2B,ATRX,EP400,SFRP1,CRTAP,FBXO11,MASTL,HDAC6,PTEN,
OBSL1,NPTN,MECP2,EFNB2,JAM3,ADNP,MAP4K4,NTN1,NECTIN1,LDB1,CRKL,SLK,LIMS1,SRF,P

FRP1,ARID1B,ADCY6,PTEN,APEX1,SLIT2,AKAP9,GRK2,PALM,NCSTN,NUCKS1,PDPK1,AKT2,OGT,

N1,TMBIM6,SPTAN1,GSTP1,SMG1,CALM1,NPM1,PLXNB2,HSPD1,IGF2,VCP,UBB,TRIM28,YWHAG
RAC1,ARHGDIA,GPC1,TOP2B,APPL1,DNER,L1CAM,PPIB,GLUL,DCN,DLG5,CNN2,VCAN,NOX4,PKI
PJ,PSMC3,MTOR,PRKACA,AP2A2,PSMC1,SMO,LMO4,CELSR1,ABL1,PHACTR4,PSMB1,DAG1,NOT
BP1A,SLIT2,GPC4,PSMB4,CCN1,SEMA3C,CLASP2,PSMD4,SCRIB,RBPJ,PSMC3,MTOR,PRKACA,AI
LEC7A,CCNI,PTN,CSNK1E,PFKL,TKT,CCND2,DDR1,PIP4K2B,ACSL6,CCN2,PRDX2,DDX3X,MIF,HN

HDAC2,BCLAF1,HSBP1,CDKN1A,EID1,SRP9,SQSTM1,NOP53,PARP1,MORF4L2,PRDX5,YEATS2,CE

RELA,LRP8,TGFB2,EPHA4,NF2,NFIB,ID2,GSN,MXRA8,MMP24

IA1,MEGF8,CCN2,ACTN4,MMP14,RPS19,MIF,GJA1,CTNNB1,CD63,ANXA6,MACF1,NARS1,CADM

2,LZTS2,PSMD10,RAB5A,PSMA6,CTDNEP1,PIN1,EXT1,PSMD12,APOE,PSMA4,PSMC6,PLPP3,PPM

1,HMGB2,CUL4B,MCM7,PSMD12,PSMA4,CENPF,POGZ,PSMC6,SMARCB1,CENPT,UBTF
N1,TMBIM6,SPTAN1,GSTP1,SMG1,CALM1,NPM1,PLXNB2,HSPD1,IGF2,VCP,UBB,TRIM28,YWHAG
MARS1,RAPGEF1,MAPK1,SMARCA4,PRDX1,ADAM9,GRB2,PRKDC,KDM1A,NSG2,GNG2,SMARCC
RP9,SQSTM1,NOP53,PARP1,MORF4L2,PRDX5,YEATS2,CBX1,SKI,H3-3A,HDGF,TAF7,NOTCH2,VE

BP1,YBX3,OPA1,MSH2,HIPK1,EPHA2,SKIL,PPP2R5C,MYBBP1A

PS2,TRIR,PELP1,IARS2,YARS1,INTS3,ELP1,INTS11,DICER1,WBP11,SART1,EIF4A3,XRN2,NOP56,I
LA1,NADK,GUK1,COX5A,COX7A2,UQCRH,FAM20C,GIT1,MSH2,NUP50,NUP58,PFKFB3,NDUFA5,

STN3,MDK,ATP6AP1,NOTCH2,VEGFA,VHL,CCND1,MAPK1,ARPC2,ADAM9,PRKDC,HSPB1,KDM1A,
CAF1,UBR5,PKDCC,TM9SF4,NMT1,NSD2,AKT2,GSK3A,PRKACA,APBB1,UBE2D3,SAE1,SMO,PTPN
,RAB27A,U2AF2,TAF15,VPS35,MDK,RBMX,SNRNP70,TMED2,VEGFA,MAPK1,SRSF1,EIF3E,PRKDC

IP58,SLC35A4,WDR33,NUTF2
RM1,UBR5,RDX,NUCKS1,PDPK1,AKT2,OGT,ATP6V0D1,GSK3A,UBE3A,PRKACA,ATP6V1G1,PHIP,

PD3,NOTCH1,IGF1R,MDM2,CYFIP1,PTPN1,CDK4,TRIB3,NME2,ADCY1,ATP6V0B,STAT2,HSF1,GSK

NOP53,PARP1,MORF4L2,PRDX5,YEATS2,CBX1,SKI,H3-3A,HDGF,TAF7,NOTCH2,VEGFA,VHL,CCN
,CTNNB1,ANXA6,TAGLN,MYL6,WLS,RHOB,HRNR,HMGCR,RAC1,GPC1,HDAC2,TNC,MMP2,DNER,

JD1,TNFRSF10B,PCOLCE,PSMC4,PLK2,HSF1,GSK3B,HSPE1,CNTN2,CFLAR,PSMD10,ROCK2,ELC

GRN,CCN1,GPX1,SOX2,GNA12,STK24,CAPZA1,CLASP2,PDPK1,SCRIB,SYT11,EHD1,VASH1,MTOR,

,CNN2,SQSTM1,ACTR3,TCP1,RBMX,RAD23B,PSMB2,HDGF,VEGFA,RAPGEF1,CCND1,MAPK1,SM
ARCA4,PBX2,ILF3,PRKDC,SUPT5H,ACTN1,CAND1,KDM1A,MAP2K2,CCAR2,ARF4,SMARCE1,SMAF
ARCA4,PBX2,ILF3,PRKDC,SUPT5H,ACTN1,CAND1,KDM1A,MAP2K2,CCAR2,ARF4,SMARCE1,SMAF
MAP6,PLXNB1,SFRP1,PTEN,PPP1CC,IL6ST,HIF1A,SLIT2,XRCC6,CCN1,OLFM1,CARM1,SOX2,PKDC
ARCA4,PBX2,ILF3,PRKDC,SUPT5H,ACTN1,CAND1,KDM1A,MAP2K2,CCAR2,ARF4,SMARCE1,SMAF
CDC42,PSMA7,SULF2,STAT3,EP300,DGCR2,KDM5B,PSMD1,PBX1,AP2A1,PHB2,HOXB5,SFRP1,SF
3B2,NADK,CAMK2N1,RUFY3,XBP1,TFDP1,TGFB2,ARF6,PRNP,VPS28,LEPROT,USP36,NUTF2,EMD
TRAF3,PSMD12,MBD6,PSMA4,COPS8,PSMC6,RAD23A,NEDD8,USP36

NME2,ADCY1,ATP6V0B,STAT2,HSF1,GSK3B,STC2,CDKN1B,STAT1,IRS1,PRKAR2B,ATP6V0E2,CAC

AR2A,FLCN,FLOT1,PHB2,BRD4,UBA52,DIP2B,RAB3GAP2,ADCY6,HDAC6,PTEN,CUL3,IL6ST,RPS2
PPP5C,MACROH2A1,GGNBP2,PKIA,IGF1R,PPP1R14B,CD2BP2,PARK7,PTPN1,PPME1,TRIM27,NCK

SPB1,CLIP3,FKBP8,MAP2K2,QARS1,DUSP4,CORO1C,FYN,FBLN1,CSNK1D,STRAP,HGS,MAPK8IP

A,AFDN,ROCK2,PRKCI,TMEM59,EPHB2,FGF13,ARF6,NFASC,PRNP,PIP5K1A,MYADM,LRP6,CACN
,TMED2,TOP1,TAF7,VEGFA,RAPGEF1,PPP2CA,CCND1,MAPK1,CCNL2,IPO7,ADAM9,ILF3,CDC37,
3B,SYNGR1,SERPINE2,CRTC1,RAB3A,DLG4,SORCS2,NSMF,PRKAR2B,UNC13A,BACE1,CNTN2,SH
A9,STMN2,MYH9,SEPTIN2,P4HB,APP,GDI1,SPTAN1,GSTP1,MARCKS,CALM1,PLXNB2,PLD3,HSPD1

3B,SYNGR1,SERPINE2,CRTC1,RAB3A,DLG4,SORCS2,NSMF,PRKAR2B,UNC13A,BACE1,CNTN2,SH
F7,VEGFA,RPL37,PPP2CA,G3BP2,IPO7,PCSK1N,CAND1,HSPB1,KDM1A,QARS1,CCAR2,DUSP4,CC
MACROH2A1,GGNBP2,PKIA,IGF1R,PPP1R14B,CD2BP2,PARK7,PTPN1,PPME1,TRIM27,NCK2,TRIB

GT7,ARL6IP5,PDLIM1,GSR,PDGFRB,NME2,PSIP1,OXSR1,HSF1,DHFR,ATRN,SETX,TXN,PSMB5,ST
2,CDC42EP4,CDC42SE1,SOS1,SHC1,ROCK2,HEG1,KCTD10,ARHGEF18,RAB35,DYNLT1,EPHB2,A
N1A,EID1,SRP9,SQSTM1,NOP53,PARP1,MORF4L2,PRDX5,YEATS2,CBX1,SKI,H3-3A,HDGF,ABCA2

A4,PSMC6,RAD23A,NEDD8,USP36
,PIP5K1A,MYADM,LRP6,EPHA2

HDGF,VEGFA,RAPGEF1,CCND1,MAPK1,SMARCA4,PSMD3,GRB2,CDC37,CLIP3,FLNB,PITPNA,FYN

RUFY3,PIN1,YBX3,SEMA6C,APOE,FGF13,SYT1,SYT4,ATP8A2,CACNG7,ANAPC2,SIN3A

AG1,HSPA1A,MFN2,IGF1R,MDM2,TNFAIP1,PARK7,CD44,RNF157,RASA4,PTPN1,TRIB3,C1QBP,S
RN2,NOP56,MTREX,TRMT112,RPS21,RTCB,RPL27A,EXOSC10,PDCD11,PA2G4,FBL,RRP1B,KAR
XA6,MACF1,NARS1,CADM4,RHOB,RAB1A,RAC1,ARHGDIA,GPC1,TOP2B,APPL1,DNER,L1CAM,PP
XA6,MACF1,NARS1,CADM4,RHOB,RAB1A,RAC1,ARHGDIA,GPC1,TOP2B,APPL1,DNER,L1CAM,PP

ZFP36L1,SFRP1,PTEN,PPP1CC,CUL3,NDST1,RPS27A,PSMD8,ATP6AP2,PSMB4,TAOK1,CCN1,SHC

I3,CD44,HTRA1,CAPN1,HSD17B12,MFAP4,EMILIN1,COLGALT1,LOXL2,P3H1,SOX9,COL7A1,ADA

1,DDOST,LAMP1,CKAP4,HLA-B,TTR,COTL1,HRNR,RAC1,ATP6V0C,TMEM30A,CDKN1A,DLG5,PGF
4,HTRA1,CAPN1,HSD17B12,MFAP4,EMILIN1,COLGALT1,LOXL2,P3H1,SOX9,COL7A1,ADAMTS7,

4,HTRA1,CAPN1,HSD17B12,MFAP4,EMILIN1,COLGALT1,LOXL2,P3H1,SOX9,COL7A1,ADAMTS7,
M1,MAP4K4,EFNB1,IQSEC1,CSNK2B,TBXA2R,C1QBP,LOXL2,EMC10,SEMA4C,PLK2,SCG2,SRF,

MSN,IGFBP3,COL1A2,LLGL1,RHOA,PSMD2,NLGN2,CD81,BEX3,PFN1,MCL1,TLE5,KHDRBS1,PTN
1,PCM1,NSF,FEZ1,NTN1,RAN,SUN2,ARFGAP2,WDR11,YWHAZ,MCFD2,RAB3A,MAP1S,KIF3A,STA
KAR1A,GJA1,CTNNB1,ANXA6,SESN3,NDFIP1,PTGES3,ATP1A2,HRNR,CCT6A,HMGCR,RAC1,CDK

CCDC88A,TRIM27,NCK2,MICAL3,ARPC3,GBA2,SRF,ZYX,WASF3,RHOBTB2,CDC42EP4,ARPC5,S

TP,KIF3C,STMN3,CSNK1D,SOD1,RAB11A,DYNLL1,CDC42,DCTN1,KIF5A,KIF21B,MAP6,KLHL42,

TD5,CTR9,NUP205,LMNA,NUP133,PER1,CTCF,SETD3,WDR5,SENP3,MAPK3,TET3,GNL3,RELA,B

OLR2E,DFFB,MTA1,EP300,BAZ1B,KPNA2,UBA52,ATRX,UPF1,SMC3,HNRNPA2B1,RPS27A,KLHDC
USP4,SUPT6H,HECTD1,SARS1,VANGL2,HGS,TCTN1,BMPR2,AMOTL2,CDC42,STAT3,CAPN2,EP3

B,GNAI3,NOP53,GRINA,VPS35,COL2A1,MDK,SKI,ASXL1,PSMB2,TSPAN6,TMED2,RAPGEF1,OPTN
SLC5A7,DNAJB6,CITED2,TGFBR1,PLCD3,HEG1,CNOT3,ADD1,YBX3,XAB2,MSH2,FURIN,SMARCB

2D3,SAE1,SMO,PTPN23,RAB11FIP3,RANGAP1,UBE2L3,PTPN11,SNX3,PINK1,PKIA,ATP13A2,MDM2

PK1,ADAM9,DYNLL2,PLCG1,MAP2K2,ARF4,CORO1C,FYN,LARS1,FBLN1,SOD1,HNRNPD,SEC23A,

MBRA1,ATP5MG,POLDIP2,SLC25A36,UBE2L3,PINK1,TRAK1,HIP1R,HSPA1A,ATP13A2,MFN2,SLC2
D2,GSK3A,MTOR,PRKACA,UBE2I,WNK1,CD74,AMBRA1,FGFR1,NTRK2,LMO4,UBE2L3,PTPN11,A

NA3,SOD1,PPP6R1,SEC23A,DYNLL1,PPP1R9B,STAT3,BCR,HIP1,FLCN,CSNK2A1,PHB2,CST3,PLXN
,PPP1R15A,PCBP4,BAP1,FUS,PPP1R10,XRCC6,TAOK1,CARM1,INTS3,UBR5,GNL1,GTF2I,NUCKS
P6V0A1,CFLAR,SHC1,ROCK2,ADIPOR1,CHMP5,XBP1,PRKCI,SP1,FER,BCAR1,RELA,OPA1,TLR6,P
R,ALDH2,PPP1CA,COX7A2L,NUP153,LOXL2,NUP98,ATP5PO,DEGS1,DLD,GSK3B,NDUFB8,ME2,S
KAP9,PALM,CARM1,UBR5,NCSTN,ARPP19,GSK3A,UBE3A,PRKACA,MN1,SMO,NSG1,PAGR1,CALC
I,CCAR2,SMARCE1,PHC2,ID4,STRAP,SARS1,SREBF2,NCOR2,TXNIP,CIC,RBBP7,STAT3,MTA1,KDM

ARF6,PRNP,VPS28,LEPROT,USP36,NUTF2,EMD,UNC13B
B5R3,ZNF770,GTF2F1,HNRNPU,NFE2L1,SMARCA5,RPL23,NACA,BRD3,XRCC5,CIPC,KPNB1,PAP

I,CCAR2,SMARCE1,PHC2,ID4,STRAP,SARS1,SREBF2,NCOR2,TXNIP,CIC,RBBP7,STAT3,MTA1,KDM

),RNF40,WAPL,SIN3A

APK1,PRKDC,HSPB1,KDM1A,PLCG1,YWHAH,B2M,PLXNA3,SARS1,PTPRS,HGS,SOD1,BAG6,RAB1
I,CCAR2,SMARCE1,PHC2,ID4,STRAP,SARS1,SREBF2,NCOR2,TXNIP,CIC,RBBP7,STAT3,MTA1,KDM

1,PRMT2,PSMD10,RPS6KA2,MDC1,PSMA6,CNOT3,PCNA,TFDP1,PPP2R5B,PSMD12,MSH2,RYBP,

C6,PTEN,OAZ2,SPINT2,GPSM1,PPP1R15A,FKBP1A,SLIT2,A2M,AKAP9,PPP1R10,PALM,GPX1,RDX

LM1,NPM1,PLXNB2,HSPD1,IGF2,VCP,UBB,TRIM28,YWHAG,COL4A2,MAPT,FN1,SPEN,NONO,DDB

1,RBM5,SCRIB,ADAM10,GSK3A,MTOR,SAP18,APBB1,BNIP3,DAPK3,SORT1,GNA13,EIF4G1,CDC

TP53BP1,STAT3,BCL9L,EP300,PBX1,FLCN,H2AZ1,BRD4,UBA52,HOXB5,ATRX,RPS27A,TNKS,APE
UBGCP6,ATXN3,CLASP2,FBXW5,CHMP4B,CROCC,TUBB4A,RANGAP1,ABL1,SUN1,CHMP1A,HDG
SF2,CDKN1B,SH3BP5,DUSP3,DUSP8,PRKAR2B,NR2F2,CAMK2N1,GPS1,RTRAF,HEG1,XBP1,AJU
APOLA,RHOA,CBX5,ZNF865,ATF4,PSMD2,NR2F1,RPS14,CHD4,JUND,CD81,PFN1,TLE5,TADA2B,
,TRAF4,TRAF7,PRKAR2A,PHB2,UBA52,SFRP1,ADCY6,PTEN,RPS27A,XRCC6,TAOK1,CCN1,SHC2,

IF770,GTF2F1,HNRNPU,NFE2L1,SMARCA5,RPL23,NACA,BRD3,XRCC5,CIPC,PAPOLA,RHOA,CBX5
APOLA,RHOA,CBX5,ZNF865,ATF4,PSMD2,NR2F1,RPS14,CHD4,JUND,CD81,PFN1,TLE5,TADA2B,

,MAPK3,XBP1,PCNA,YBX3,FER,RELA,TLR6,FBXO7,LANCL1,SESN2,EIF2S1,SIN3A
IF770,GTF2F1,HNRNPU,NFE2L1,SMARCA5,RPL23,NACA,BRD3,XRCC5,CIPC,PAPOLA,RHOA,CBX5

IG1,PHACTR4,COL12A1,RTF1,DAG1,MACROH2A1,TXNRD1,NOTCH1,KAT2A,NTN1,HIPK2,TASOF
CC6,TAOK1,CCN1,SHC2,SPAG9,PDPK1,DAZAP2,MTOR,PRKACA,WNK1,CD74,FGFR1,NTRK2,PTI

APOLA,RHOA,CBX5,ZNF865,ATF4,PSMD2,NR2F1,RPS14,CHD4,JUND,CD81,PFN1,TLE5,TADA2B,

X9,ADIPOR2,PSMB5,IRS1,NUP205,NUP133,PSMA1,LONP2,PSMD10,ADIPOR1,WDR5,ANXA1,PSM
N1A,DLG5,PGRMC1,CNN2,CTSB,RAB27A,CCT2,VAMP2,GDI2,HSPH1,PDXK,PKN1,MLEC,MDK,SUI

,CDC42EP4,ARPC5,SHANK1,ROCK2,TGFBR1,CAPZA2,ADD1,ARHGEF18,FER,GMFB,FGF13,PDXP,

8,APBB1,BNIP3,DAPK3,SORT1,GNA13,CDC34,ABL1,HIP1R,GRIK5,TIA1,NOTCH1,MFN2,PARK7,A
ORT1,GNA13,CDC34,ABL1,HIP1R,GRIK5,TIA1,NOTCH1,MFN2,PARK7,ARL6IP5,PPP1CA,PDGFRB,
,HOXA5,CDKN1B,CEBPG,SRP54,CITED2,CFLAR,ANXA1,TGFBR1,MAPK3,THRA,XBP1,PCNA,RELA

IF770,GTF2F1,HNRNPU,NFE2L1,SMARCA5,RPL23,NACA,BRD3,XRCC5,CIPC,PAPOLA,RHOA,CBX5

SK3A,MTOR,PRKACA,EXOC4,NSG1,EIF4A3,NTRK2,RIMS3,ABL1,SDCBP,PINK1,DAG1,PAIP2,NRXI

.GN2,CXADR,ATP1A1,CD81,ATP2A2,PFN1,CLEC7A,PTN,GLG1,TPM1,SULF1,PLXNA1,MEGF8,SOX
,CAMK2N1,GPS1,RTRAF,HEG1,XBP1,HEXIM1,GMFB,EPHB2,APOE,FBXO7,PRNP,SRCIN1,PLPP3,M
ANXA1,RAD54L2,CABIN1,HMGB2,CUL4B,MCM7,ACIN1,PAF1,DNMT1,CENPT,HMGA2,RAD50,WAP

,TGFB2,PIGT,FBXO7,PRNP,PPT1,KDM2B,LANCL1,EIF2S1,NAMPT
A2,ADD1,ARHGEF18,FER,GMFB,PDXP,ARF6,MYADM,NF2,ASAP3,GSN,GPM6B,PAM

P1,FER,BCAR1,RELA,OPA1,GRB10,RAP1B,ATP6V1F,LEPROT,ZNF106,PIK3R3,ATP6V1D
S14,CHD4,JUND,CD81,PFN1,TLE5,TADA2B,KHDRBS1,XPO1,SFPQ,SOX12,ACTN4,PRDX2,DDX3X,N
PSMC4,PLK2,PSMF1,CDKN1B,PSMB5,RRM2,RBM14,XPC,PSMA1,NR2F2,PRMT2,PSMD10,CAMK

327A,CCT2,VAMP2,GDI2,ACTR3,PDXK,PKN1,NCKAP1,MLEC,MDK,SURF4,RAB5B,SRP14,PYGB,NC
168A,H2AX,CDK9,DHX36,EXOSC10,HSF1,UBE2V1,UBQLN4,TFRC,PRMT5,OTUB1,CEBPG,LMNA,M
S14,CHD4,JUND,CD81,PFN1,TLE5,TADA2B,KHDRBS1,XPO1,SFPQ,SOX12,ACTN4,PRDX2,DDX3X,N
S14,CHD4,JUND,CD81,PFN1,TLE5,TADA2B,KHDRBS1,XPO1,SFPQ,SOX12,ACTN4,PRDX2,DDX3X,N
',SRCIN1,NF2,CDK5RAP3,LRP6,IBTK
/1D

5K1A,SRCIN1,COL3A1,MYADM,POSTN,ITGA7,FZD7,NF2,GPM6B,CASK

',EGLN2,PER1,RXRA,PHB,PRMT2,CFLAR,ANXA1,PCNA,UFM1,HMGB2,RELA,TGFB2,HMGA2,PAM,
,FBL,RASSF2,HSPE1,RRBP1,PHB,GDF10,MAPK3,THRA,EXT1,GIT1,CTNNBIP1,TGFB2,CRIM1,NBF
PG11,SEC23IP,SEC31A,SYT1,DNM1,SYT4,VPS28,STX12,ARFGAP3,VPS37B,UNC13B

C4,NSG1,EIF4A3,NTRK2,RIMS3,ABL1,SDCBP,PINK1,DAG1,PAIP2,NRXN2,GRIK5,NPTN,MAPK8IP2,

AT1,CEBPG,SRP54,RXRA,TMEM14C,CITED2,WDR5,THRA,ADD1,HMGB2,CTNNBIP1,NCOA6,SETD1

A1,PFN2,RAB11A,AKIRIN1,BMPR2,AMOTL2,BEX4,STAT3,FLCN,CD99,PLXNB1,SFRP1,EVL,HDAC6,

D1,PDHB,NNT,NDUFA5,UQCR11,RB1CC1,COX5B,MYBBP1A,NDUFA10,ATP5PD

1,EIF4A3,NTRK2,RIMS3,ABL1,SDCBP,PINK1,PAIP2,NRXN2,GRIK5,NPTN,MAPK8IP2,MECP2,PARK
1,EIF4A3,NTRK2,RIMS3,ABL1,SDCBP,PINK1,PAIP2,NRXN2,GRIK5,NPTN,MAPK8IP2,MECP2,PARK
DH11,TBCD,PPP1CB,VCAN,TLN2,TGFBI,HSPH1,PTPRA,CLSTN3,MDK,LAMC1,PTK7,PCDHGB6,CN

),PPP1CB,VCAN,TLN2,TGFBI,HSPH1,PTPRA,CLSTN3,MDK,LAMC1,PTK7,PCDHGB6,CNTNAP2,VEG

,CLIC1,MFN2,PABPC4,CD44,PDGFRB,TBXA2R,C1QBP,SRF,YWHAZ,SERPINE2,RAB3A,ADIPOR2,F
SNX3,SCAMP5,ABL1,LMAN2,SDCBP,HIP1R,GRIK5,SMPD3,NOTCH1,ATP13A2,NSF,PTPN1,ANKFY1

3,GMFB,OPA1,FGF13,TPX2,CENPF,PRELID1,SYT4,MYADM,ZNF207,RAD50,GSN,WAPL

M2C,RPS7,RPS19,MIF,GJA1,CTNNB1,CD63,MACF1,CYFIP2,NDFIP1,WLS,LAMP1,HLA-B,HMGCR,F

S1,DAG1,NIBAN2,RBM4,ELN,OBSL1,NOTCH1,MDM2,MECP2,EFNB2,FXR1,PDLIM1,KAT2A,FLII,PD
RA,TAB3,UFM1,RELA,APOE,TRIM24,PRCP,HMGA2,CNOT1

,ABL1,PINK1,PPP5C,SMPD3,ATP13A2,MDM2,SUMO1,PARK7,ADNP,RASA4,SYT15,PPP1CA,PDGFF
JUN,CTNNB1,ARID1A,MXD4,SHOX,HNRNPK,DUX4,ZNF454,CTBP1,HDAC2,BCLAF1,HSBP1,DCN,CI
1,LIMK1,RHEB,ARHGAP5,IQSEC1,PDGFRB,CCDC88A,RAP2A,RAB18,SYNGAP1,CRKL,PLK2,ARHG

.3,XBP1,SZT2,SESN2,TRIM24,EIF2S1,POSTN,ATP6V1F,PIEZO1,KLF10,ATP6V1D,MYBBP1A,NAMPT

C6,PTEN,MAPRE2,APEX1,HIF1A,SLIT2,GRN,CCN1,TCAF1,AKT3,GNA12,RDX,MAZ,STK24,SEMA3C,
PSMC6,AKIRIN2,SIN3A

.T3,FLCN,CD99,PLXNB1,SFRP1,EVL,HDAC6,PTEN,SPINT2,MAPRE2,APEX1,HIF1A,SLIT2,GRN,CCN

C,TRAF4,STAT3,EP300,KDM5B,PBX1,PHB2,ZFP36L1,BRD2,ATRX,SFRP1,SRSF6,PTEN,CEMIP2,IL6 ϟRAP3,ZNF207,HMGA2,CNOT1,PPP2R5C

,IREB2,KMT2E,SH3PXD2A,TCF3,WASF2,PSMD4,RBPJ,PSMC3,SART1,MTOR,YY1,CREBBP,PSMC1

1,APPL1,HDAC2,L1CAM,PPIB,DLG5,DDX39B,PARP1,CLSTN3,MDK,KDM6B,ATP6AP1,NOTCH2,RN

LPTM1,SSBP3,IREB2,KMT2E,SH3PXD2A,TCF3,WASF2,PSMD4,RBPJ,PSMC3,SART1,MTOR,YY1,C

3,SZT2,KDM2B,COL3A1,GNAO1,NF2,POU3F3,LRP6,NFIB,ID2,SIN3A
GRB2,PRKDC,FKBP8,FLNB,HECTD1,PTPRS,VANGL2,SOD1,TCTN1,BMPR2,TXNIP,CDC42,PALLD,

,ZNF454,CTBP1,HDAC2,BCLAF1,HSBP1,DCN,CDKN1A,EID1,CHD6,MLXIP,SQSTM1,NOP53,PARP1

SARS1,VANGL2,HGS,BMPR2,AMOTL2,CDC42,STAT3,EP300,ZFP36L1,NXN,SFRP1,PTEN,CEMIP2,

HAH,PFKM,B2M,STXBP1,HGS,SOD1,DGKD,SREBF2,DYNLL1,CDC42,HIP1,AP2A1,FLOT1,SEC16A,

,AP2B1,PXDN,PGAM1,CTSD,RTN4,LPCAT1,RPS24,XRCC5,PJA2,MSN,KPNB1,COL1A2,HK1,RPL22,

2,SKIL,CASK

C25A5,LOX,GLO1,XRCC6,PSMB4,CLPTM1,SSBP3,IREB2,KMT2E,SH3PXD2A,TCF3,WASF2,PSMD4

RCC6,TAOK1,CCN1,SHC2,SPAG9,PDPK1,DAZAP2,AKT2,GSK3A,MTOR,PRKACA,WNK1,CD74,AMB

LXNB1,SFRP1,EVL,HDAC6,PTEN,SPINT2,MAPRE2,APEX1,HIF1A,SLIT2,GRN,CCN1,TCAF1,AKT3,GN
CF3,FOXK2,NSD2,RBPJ,PCGF2,NOC2L,MBD3,YY1,CREBBP,APBB1,UBE2I,UBE2D3,SMO,CNBP,W

HDAC2,TNC,MMP2,GLUL,SQLE,CDKN1A,DLG5,DDX39B,CNN2,RNF187,GNAI3,NOX4,TGFBI,PKN1

1,PSMD12,PSMA4,PSMC6,HARS1

DLIM1,KAT2A,PDGFRB,DHX36,EMILIN1,CRKL,WDR11,NDRG4,MEIS1,SRF,BCOR,SOX9,ADIPOR2,
R5C

9,PARK7,ADNP,CD44,MAP4K4,ERP29,CCNY,PTPN1,PDGFRB,CCDC88A,ADCY1,MAPKAPK2,CRKL

AC6,IL6ST,RPS27A,HIF1A,PPP1R15A,SYMPK,AKAP9,XRCC6,TAOK1,CCN1,SHC2,SPAG9,PDPK1,DA

AC6,IL6ST,RPS27A,HIF1A,PPP1R15A,SYMPK,AKAP9,XRCC6,TAOK1,CCN1,SHC2,SPAG9,PDPK1,DA

,SUPT6H,ILF2,CCT8,B2M,STXBP1,IDH1,DYNLL1,TP53BP1,TXNDC5,PSMD1,PDAP1,CST3,ATP6AP2

1,PLXNB2,HSPD1,COL18A1,SNRNP200,CSDE1,IGF2,EIF3C,ATP5F1A,EIF4B,VCP,MARCKSL1,UBB,
HAQ,DCHS1,TMSB10,RPL19,PPP2R1A,RPL15,PKD1,HLA-C,RPS3,VAPA,HMGB1,AP2M1,SEC61A1,
,RTN3,RPL31,PRKCSH,XPR1,XPO1,RPS25,RAB7A,DDX3X,RPS15,RPL12,RPL30,EIF3L,RPS5,RPS7,
,EIF3C,ATP5F1A,EIF4B,VCP,MARCKSL1,UBB,TRIM28,YWHAG,COL4A2,MAPT,FN1,DDB1,PSMD5,
PS9,XRCC5,MSN,RPL24,KPNB1,RPL17,RPL14,HK1,RPL22,RPL9,CLSTN1,KIF5B,RAB6B,NLGN2,F
PS9,XRCC5,MSN,RPL24,KPNB1,RPL17,RPL14,HK1,RPL22,RPL9,CLSTN1,KIF5B,RAB6B,NLGN2,F
36,YWHAB,HNRNPH1,RPS12,RPS27,BCLAF1,RPL23A,LUC7L3,CPSF2,DDX39B,NOVA2,SRSF5,U2A
RPL9,KIF5B,RAB6B,RPS14,RPS20,CD81,ATP2A2,RPL26,OAZ1,SRSF4,RPL31,LRPPRC,KHDRBS1,
26,OAZ1,RPL31,HNRNPC,LRPPRC,XPO1,CSNK1E,RPS25,GPC3,RAB7A,RPS15,RPL12,RPL30,CLT
S4X,RPS3A,RPL18,ITGB1,TUBB4B,UCHL1,RPL11,ANXA2,RPLP1,TIMP2,RPL10,RPS17,VCL,AP3D1,
S16,RPL23,CPE,RPL29,RPSA,CTSD,SCARB2,NACA,RTN4,LAMA5,RPS24,RPS9,XRCC5,MSN,RPL24
WDR1,CRMP1,CCT7,VAT1,PFN1,CLEC7A,NES,PRKCSH,HNRNPC,TADA2B,XPO1,SPTBN2,PTN,SFP
,AP2B1,HNRNPU,CTNNA1,RPS16,RPL23,CPE,RPL29,RPSA,CTSD,SCARB2,NACA,RTN4,LAMA5,RPS
37A,RPS15,RPL12,RPL30,CLTC,H1-0,RPS5,RPS7,RPS19,RPL35,CTNNB1,RPS10,RPL35A,TARDBF
DBP,RPLP2,RPL36,YWHAB,LAMP1,RPS12,RPS27,CCT6A,APPL1,IPO5,TMEM30A,SKP1,RPL23A,GL
S11,HSPB1,CASC3,RPL28,RPS28,HNRNPM,SSB,FAU,HNRNPD,RPL39,SERBP1,RPS13,RPL34,PSM
TP1,SMG1,CALM1,NPM1,PLXNB2,HSPD1,IGF2,ATP5F1A,VCP,UBB,TRIM28,YWHAG,RPL37A,MAP
IA3,RPS3,HMGB1,CDKL1,AP2M1,RPS4X,RPS3A,RPL18,UCHL1,RPL11,MORF4L1,ANXA2,RPLP1,Z
1,ITGB1,TUBB4B,RPL11,MORF4L1,ANXA2,AGRN,ZMIZ1,RPL10,AP3D1,ARF1,PAFAH1B1,MAP4,CD

PL32,RNPS1,PPP2R2A,CTIF

,EIF3E,FTO,RPS11,HSPB1,CASC3,RPL28,RPS28,HNRNPM,SSB,FAU,HNRNPD,RPL39,SERBP1,RPS
,ITGB1,TUBB4B,RPL11,ANXA2,AGRN,RPLP1,RPL10,RPS17,VCL,ARF1,PAFAH1B1,MAP4,AP2B1,M
31,RPL26,CCT7,OAZ1,ASTN2,MCL1,RPL31,XPO1,PFKL,RPS25,EIF5A,RAB7A,ACTN4,RPS15,RPL1
L36,YWHAB,RHOB,RPS12,RAB1A,ARCN1,RPS27,APPL1,IPO5,TMEM30A,RPL23A,PTTG1IP,MAP1A
K5,RPS21,GPAA1,RPL27A,HERPUD1,RPS29,SPCS2,SRP54,GABARAPL2,RPL32

ISY1,EIF4A1,RPL19,PPP2R1A,RPL15,GTF3C4,PKD1,PDIA3,RPS3,HMGB1,CDKL1,AP2M1,RPS4X,R
S20,CD81,RPL26,OAZ1,SRSF4,ASTN2,MCL1,RPL31,NISCH,LRPPRC,KHDRBS1,XPO1,SLC3A2,PFK
JBA52,MGAT4B,POM121C,RPS2,RPS27A,GALNT1,NUCKS1,EIF3G,PSMC3,UBE2I,TRIM8,RPS26,U

IF3B,EIF3M,RPS29,YBX3,ZC3H15,RPL32,EIF3J
PS8,MYH9,RPL7,P4HB,APP,IDS,RPL5,TMBIM6,PTMS,GSTP1,SMG1,CALM1,NPM1,PLXNB2,TTC3,P
A,PGAM1,LARP1,HNRNPA0,RPS16,RPL23,LDHA,RPL29,RPSA,CTSD,SCARB2,LPCAT1,RPS24,RPS
L24,KPNB1,RPL17,RPL14,RPL22,RPL9,PSMD2,ENC1,RPS14,TMEM259,RPS20,CD81,ATP2A2,RPL
,CSDE1,IGF2,ATP5F1A,VCP,UBB,TRIM28,YWHAG,COL4A2,RPL37A,MAPT,FN1,SPEN,NONO,DDB1
F2S2,EIF3H,EIF4G3,ABCF1,EIF3G,MTOR,EIF4G1,RPS26,PAIP2,YTHDF3,RBM4,RPS21,PPP1CA,RP
NRDC,OPTN,AZIN1,PSMD3,ODC1,ADAM9,FTO,WDR6,GABARAP,SUPT5H,HSPB1,BNIP3L,CCAR2,S
PFKL,RPS25,EIF5A,RAB7A,ACTN4,RPS15,RPL12,RPL30,ARL4C,CLTC,RPS5,RPS7,RPS19,RPL35,C
FN1,PSMD5,YWHAQ,MRC2,RPL19,RPL15,PKD1,HLA-C,RPS3,VAPA,HMGB1,AP2M1,TPT1,SEC61A1
HSPA4,TCP1,CCT3,CCT4,TNPO1,RPL36A,SRP14,RPS15A,RPL38,RPL37,MAPK1,IPO7,RPS11,NPEP
YWHAQ,MRC2,RPL19,RPL15,PKD1,HLA-C,RPS3,VAPA,HMGB1,AP2M1,TPT1,SEC61A1,RPS4X,RPS
EM170A,PXDN,HNRNPU,SMARCA5,CTNNA1,RPSA,EIF4H,RTN4,LAMA5,XRCC5,MSN,RPL24,KPNB1
CARB2,LPCAT1,RPS24,RPS9,MSN,RPL24,KPNB1,RPL17,RPL14,HK1,RPL22,RPL9,PSMD2,ENC1,F

TRIM28,YWHAG,CKB,RPL37A,MAPT,FN1,DDB1,PSMD5,EIF1,HACD3,SET,EGLN1,TPI1,CHSY1,EIF
NES,PRKCSH,SPTBN2,PTN,CSNK1E,CDH2,DDR1,SULF1,PLXNA1,MEGF8,SOX12,MMP14,ITM2C,A
3,RPS12,RPS27,GPC1,APPL1,RPL23A,SRP9,TM9SF3,SQSTM1,VAMP2,TM9SF2,RPL27,HSPA4,VP
PDIA3,SSBP2,ZNF43,RPS3,VAPA,HMGB1,AP2M1,RPS4X,STMN1,ITGB1,RPL11,ANXA2,AGRN,RPL
ITNAP2,DPYSL2,NOTCH2,VEGFA,RAPGEF1,MAPK1,GRB2,ATXN10,KDM1A,PLCG1,ARF4,FYN,YW
RL6IP1,RPL36,YWHAB,RPS12,RPS27,RAC1,GPC1,APPL1,RPL23A,SRP9,TM9SF3,SQSTM1,VAMP2,
R2,SMARCC1,ATP6V1B2,HNRNPM,UBQLN1,CSNK1D,USP7,IDH1,HECTD1,BAG6,SREBF2,HNRNP
K9,NUP98,RRP1B,RPS29,NUP205,NUP133,NELFCD,SP1,HEXIM1,RPL32,NUP50,SMARCB1,NUP5
3,YWHAG,RPL37A,MAPT,FN1,SPEN,NONO,DDB1,PSMD5,YWHAQ,EIF1,HACD3,SET,EGLN1,RPL19,
38,RPL37,PPP2CA,PDE7A,PSMD3,EIF3E,FTO,RPS11,HSPB1,CASC3,RPL28,RPS28,HNRNPM,SSB,
MSB10,PPP2R1A,RPS3,CHD3,BASP1,HMGB1,AP2M1,TPT1,RPS3A,STMN1,ITGB1,UCHL1,RPL11,M
CA5,LARP1,HNRNPA0,RPS16,RPL23,RPL29,RPSA,NACA,RTN4,LPCAT1,RPS24,RPS9,XRCC5,CIPC,
31,RPL26,BEX3,OAZ1,RTN3,CLEC7A,CCNI,LRPPRC,TADA2B,KHDRBS1,XPO1,PTN,CSNK1E,GLG1
,USP11,CSDE1,IGF2,EIF3C,EIF4B,VCP,UBB,AFF4,TRIM28,YWHAG,COL4A2,RPL37A,MAPT,FN1,S
,RPS16,RPL23,RPL29,RPSA,NACA,RTN4,RPS24,RPS9,XRCC5,CIPC,RPL24,RPL17,IGFBP3,RPL14
GES3,RHOB,HMGCR,SMCHD1,HDAC2,TNC,ATP6V0C,MMP2,SKP1,BCLAF1,GLUL,HSBP1,PTTG1I
M1,ANXA2,AGRN,ZMIZ1,TIMP2,XAF1,MARCHF6,DNAJA1,ARF1,PAFAH1B1,AKAP12,MAT2A,GTF2F
S19,MIF,TLN1,CTNNB1,RPS10,CYFIP2,ARID1A,H1-10,PTGES3,CKAP5,MAP2,RAB1A,RPS27,COTL
CKAP1,ACAP3,MDK,PTK7,CNTNAP2,DPYSL2,NOTCH2,VEGFA,RAPGEF1,MAPK1,PBX2,GRB2,ATX
CLSTN3,MDK,SURF4,PTK7,CCT4,H3-3A,CNTNAP2,TERF2IP,VEGFA,RAPGEF1,G3BP2,MAPK1,MA
N1,GPC1,CDK16,TOP2B,HDAC2,TNC,TMEM30A,DNER,L1CAM,NNAT,MAP1A,DLG5,CRPPA,HS6ST
L7,P4HB,APP,IDS,RPL5,TMBIM6,GSTP1,SMG1,CALM1,NPM1,PLXNB2,TTC3,PLD3,HSPD1,USP11,
S15A,RPL38,RPL37,PPP2CA,PDE7A,PSMD3,EIF3E,FTO,RPS11,HSPB1,CASC3,RPL28,RPS28,HNR
S15A,RPL38,RPL37,PPP2CA,PDE7A,PSMD3,EIF3E,FTO,RPS11,HSPB1,CASC3,RPL28,RPS28,HNR
,ETF1,TNRC6A,NBAS,RPL27A,DDX6,LSM4,DHX36,EXOSC10,NCBP2,RPS29,SKIV2L,CNOT3,RPL32
DS,RPL5,TMBIM6,GSTP1,SMG1,CALM1,NPM1,PLXNB2,TTC3,PLD3,HSPD1,USP11,SNRNP200,CS
,DLG5,CRPPA,HS6ST1,CDH11,TBCD,GAS7,PTPRA,RTN1,CAMSAP1,NCKAP1,ACAP3,MDK,PTK7,C
EM30A,DNER,L1CAM,NNAT,MAP1A,DLG5,CRPPA,HS6ST1,CDH11,TBCD,GAS7,PTPRA,RTN1,CAM
IF5B,RAB6B,SLC7A5,NLGN2,RPS14,RPS20,CD81,RPL26,OAZ1,SRSF4,ASTN2,MCL1,RPL31,NISC
N1,SPEN,NONO,DDB1,PSMD5,YWHAQ,EIF1,HACD3,SET,EGLN1,RPL19,PPP2R1A,RPL15,PKD1,S
G6,RAB11A,RPL39,RPS13,RPL34,GRIPAP1,SSR1,SGTA,ALDH3A2,SEC16A,UBA52,RAB3GAP2,PR
MD5,YWHAQ,HACD3,CHSY1,PKD1,SF3B1,PDIA3,SSBP2,ZNF43,RPS3,BASP1,VAPA,HMGB1,AP2M1
ZNF43,RPS3,CHD3,BASP1,HMGB1,AP2M1,RPS4X,UCHL1,RPL11,MORF4L1,ANXA2,AGRN,RPLP1,
27,CAPRIN1,BCLAF1,RPL23A,DDX39B,SRP9,EEF1A2,MTRF1L,RPL27,RTN1,GCN1,RPL36A,ABC
PO1,PTN,CSNK1E,GLG1,EIF5A,PEBP1,CCND2,GPC3,CCN2,MMP14,PRDX2,DDX3X,RPS15,RPS7,
F2,RPL39,RPS13,RPL34,SSR1,SGTA,OS9,ALDH3A2,UBA52,NCOA4,RPS2,RPS27A,TIMM50,LMAN
38,RPL37,PPP2CA,PDE7A,PSMD3,EIF3E,FTO,RPS11,HSPB1,CASC3,RPL28,RPS28,HNRNPM,SSB,
2,DPYSL2,NOTCH2,VEGFA,VHL,MAPK1,ARPC2,GRB2,CAPZB,PRKDC,ACTN1,PLCG1,FLNB,CORO
S15A,RPL38,RPL37,PPP2CA,PDE7A,PSMD3,EIF3E,FTO,RPS11,HSPB1,CASC3,RPL28,RPS28,HNR
NTNAP1,FLOT1,MAP6,PLXNB1,DIP2B,FARP1,CTNND2,EVL,HDAC6,PTEN,ULK1,BRSK1,CTTN,SLI
LOT1,MAP6,PLXNB1,DIP2B,FARP1,CTNND2,EVL,HDAC6,PTEN,ULK1,BRSK1,CTTN,SLIT2,NDN,TA
NTNAP1,FLOT1,MAP6,PLXNB1,DIP2B,FARP1,CTNND2,EVL,HDAC6,PTEN,ULK1,BRSK1,CTTN,SLI
MIM14,IGF2,MARCKSL1,UBB,AFF4,TRIM28,MATR3,YWHAG,CKB,COL4A2,MAPT,FN1,SPEN,PSM
N1,CSNK1D,USP7,HECTD1,HGS,PCBP2,BAG6,CYP51A1,PCYOX1,YME1L1,PSMA7,ARIH1,SGTA,O
,DDX39B,SRP9,EEF1A2,MTRF1L,RPL27,GCN1,RPL36A,TMED2,RPS15A,RPL38,MARS1,RPL37,M

2,RAB1A,RPS27,COTL1,RAC1,SKP1,RPL23A,LUC7L3,DLG5,DDX39B,SRSF5,CCT2,TBCD,VAMP2,

1,SPTBN1,ATP5F1B,CALM3,RPS8,MYH9,RPL7,P4HB,APP,IDS,RPL5,TMBIM6,PTMS,GSTP1,SMG1,

RIN1,GPC1,CDK16,TOP2B,HDAC2,TNC,TMEM30A,L1CAM,MAP1A,DLG5,CRPPA,CDH11,GAS7,PTP

GP2,CPSF1,NOP53,PABPN1,HNRNPL,TSR1,ALKBH5,RPL27,SNRPN,HNRNPA3,AHCYL1,RBMX,DD

JAJA1,ARF1,CYB5R3,PAFAH1B1,AKAP12,MAT2A,GTF2F1,PXDN,HNRNPU,NFE2L1,SMARCA5,LARP

P6,PLXNB1,DIP2B,FARP1,CTNND2,EVL,HDAC6,PTEN,ULK1,BRSK1,CTTN,SLIT2,NDN,TAOK1,SLIT

,CAMSAP1,NCKAP1,RAD23B,PSMB2,H3-3A,PRPF6,TAF7,RPL38,SMARCA4,MAPRE1,SRSF9,ARPC

1,CALM1,NPM1,PLXNB2,TTC3,PLD3,HSPD1,USP11,SNRNP200,CSDE1,IGF2,EIF3C,EIF4B,VCP,UB

RPS3,CHD3,BASP1,HMGB1,RPS4X,UCHL1,RPL11,MORF4L1,ANXA2,AGRN,RPLP1,ZMIZ1,TIMP2,M

21,RPL27A,NACAD,RPS29,ERBB2,SRP54,ZDHHC9,PRKCI,MICALL1,RPL32,ABCD3,PRNP,VPS37B

XR2,PSMB1,YTHDF3,RBM4,HSPA1A,TIA1,RBM25,PSMD13,FXR1,SYNCRIP,PABPC4,PSMD7,TNRC

CAPRIN1,GPC1,CDK16,TOP2B,HDAC2,TNC,TMEM30A,L1CAM,MAP1A,DLG5,CRPPA,CDH11,GAS7

A,DDX39B,SRP9,EEF1A2,MTRF1L,RPL27,GCN1,RPL36A,TMED2,RPS15A,RPL38,MARS1,RPL37,M

U,SMARCA5,LARP1,CTNNA1,RPS16,RPL23,LAMA5,XRCC5,MSN,COL1A2,RHOA,ATF4,PSMD2,KIF

G,CKB,COL4A2,MAPT,FN1,SPEN,PSMD5,YWHAQ,DCHS1,EGLN1,TPI1,CHSY1,INA,RPL19,PKD1,BA

A,TCTN1,BMPR2,CDC42,PALLD,KIF5A,TRIO,EP300,RERE,FLOT1,MAP6,PLXNB1,DIP2B,FARP1,CT

SMD2,KIF5B,CD81,COL5A2,WDR1,CRMP1,VAT1,PFN1,ASTN2,TLE5,NES,SPTBN2,PTN,GLG1,CDH

N1,BMPR2,CDC42,PALLD,KIF5A,TRIO,RERE,CNTNAP1,FLOT1,MAP6,PLXNB1,DIP2B,FARP1,CTNN

TN,SLIT2,NDN,SLIT1,OLFM1,LINGO1,ABI2,DCLK1,CNP,SEMA3C,CLASP2,SCRIB,ADAM10,IGF2B

MIDN,RPS17,VCL,AP3D1,ARF1,PAFAH1B1,AKAP12,AP2B1,SLC4A8,HNRNPU,LARP1,HNRNPA0,RPL

PRDX6,FUS,SNRPD3,RALY,ELAVL1,EFTUD2,SRSF2,PRPF4B,RBM12,RBM5,WBP11,SRSF11,PRPF

PSMD5,YWHAQ,DCHS1,EGLN1,TPI1,CHSY1,INA,RPL19,PKD1,BASP1,VAPA,HMGB1,CDKL1,AP2M1

USP5,SNX3,SDCBP,PINK1,HSPA1A,ATP13A2,MDM2,FZR1,SGSM3,MGAT3,NSF,SUMO1,PARK7,US

BASP1,HMGB1,RPS4X,UCHL1,RPL11,MORF4L1,ANXA2,AGRN,RPLP1,ZMIZ1,TIMP2,RPL10,AP3D

S27,CAPRIN1,BCLAF1,RPL23A,DDX39B,SRP9,EEF1A2,MTRF1L,RPL27,GCN1,RPL36A,TMED2,RP

PKD1,BASP1,VAPA,HMGB1,CDKL1,AP2M1,SEC61A1,STMN1,ITGB1,UCHL1,NCAM1,ANXA2,AGRN,

P5F1B,CALM3,RPS8,MYH9,RPL7,P4HB,APP,IDS,RPL5,TMBIM6,PTMS,GSTP1,SMG1,CALM1,NPM1

RT1,DHX38,RBM6,SAP18,THRAP3,HNRNPH3,EIF4A3,HTATSF1,FXR2,LUC7L2,RBM4,MTREX,TIA1

RT1,DHX38,RBM6,SAP18,THRAP3,HNRNPH3,EIF4A3,HTATSF1,FXR2,LUC7L2,RBM4,MTREX,TIA1

RT1,DHX38,RBM6,SAP18,THRAP3,HNRNPH3,EIF4A3,HTATSF1,FXR2,LUC7L2,RBM4,MTREX,TIA1

N,CSNK1E,SFPQ,CCND2,CDH2,PDIA6,GPC3,DDR1,CCN2,PRDX2,DDX3X,RPS15,RCN1,RPS7,MIF,

N,CSNK1E,SFPQ,CCND2,CDH2,PDIA6,GPC3,DDR1,CCN2,PRDX2,DDX3X,RPS15,RCN1,RPS7,MIF,

A3C,CLASP2,IST1,TMEFF1,APBB1,CERS2,SMO,METRN,LYPLA2,NTRK2,PTPN11,ABL1,SEMA6B,A

C5,MSN,HK1,RHOA,ATF4,NR2F1,SLC7A5,JUND,TMEM259,CD81,RPL26,BEX3,CCT7,OAZ1,CLEC

4,SAFB2,PRDX6,FUS,SNRPD3,RALY,ELAVL1,EFTUD2,SRSF2,PRPF4B,RBM5,WBP11,SRSF11,PRP

,PPP2CB,KLHL42,UBE2G2,FBXO11,HDAC6,PTEN,CUL3,RPS27A,PSMD8,NPLOC4,FBXO21,RHBD

PLXNA1,MEGF8,ACTN4,ITM2C,RPS19,GPAT4,PRKAR1A,GJA1,CTNNB1,ANXA6,MACF1,CYFIP2,M

P1,VPS35,MDK,SURF4,CCT4,H3-3A,TERF2IP,NOTCH2,VEGFA,G3BP2,MAPK1,MAPRE1,ARPC2,D

5,RPS7,RPS19,RPL35,MIF,CTNNB1,RPS10,RPL35A,TARDBP,RPLP2,NDFIP1,HNRNPK,H1-10,CIR

PL35A,TARDBP,RPLP2,ST3GAL2,CIRBP,NARS1,RPL36,RPS12,RPS27,CAPRIN1,BCLAF1,RPL23A,D

RAF7,SF1,EIF3D,UBA52,SRSF6,PTEN,RPL7L1,RPS2,PELP1,EIF2S2,EIF3H,SNRPD3,DICER1,RBM5

F5A,CCND2,DDR1,PIP4K2B,SOX12,CCN2,ACTN4,MMP14,PRDX2,DDX3X,NANOG,H1-0,MIF,PRKA

,MAGED1,RPL6,KAT6A,DVL3,YWHAE,KIF1A,RPL13,GNAS,DDX17,GPI,HTRA4,RACK1,BSG,FASN,S

PIP4K2B,RAB7A,ACTN4,DDX3X,ARL4C,CLTC,MAGED2,APLP1,MIF,TLN1,MYO5A,CTNNB1,CD63,

SNX3,LGALS1,SEMA6B,TRAK1,PLXNA2,OBSL1,MDM2,NPTN,EFNB2,ADNP,CYFIP1,MAP4K4,FEZ1,

SLC7A5,JUND,TMEM259,CD81,RPL26,BEX3,CCT7,OAZ1,CLEC7A,HNRNPC,TADA2B,KHDRBS1,CS
DX3X,RPS15,ITM2C,CLTC,NANOG,H1-0,RPS7,PRKAR1A,GJA1,JUN,CTNNB1,APLP2,TARDBP,ARL
PLA2,NTRK2,PTPN11,ABL1,SEMA6B,AUTS2,TAOK2,TRAK1,DAG1,NIBAN2,PLXNA2,NOTCH1,NPTN
,KHDRBS1,XPO1,PTN,GLG1,SFPQ,PEBP1,GPC3,PIP4K2B,DDX3X,RPS15,ITM2C,NANOG,H1-0,RPS
1,LINGO1,ABI2,GPC2,DZIP1,RDX,STK24,SEMA3C,FNBP1L,WASF2,ATP1B2,ADAM10,IST1,EHD1,G
HNRNPH1,ATP1A2,GPC1,APPL1,HDAC2,IPO5,TNC,ATP6V0C,MMP2,CDKN1A,TIMP3,CTSB,SRSF5,V
CARM1,LINGO1,ABI2,GPC2,DZIP1,RDX,STK24,SEMA3C,FNBP1L,WASF2,ATP1B2,ADAM10,IST1,E
ARDBP,ARL6IP1,HNRNPK,DUX4,YWHAB,RHOB,HMGCR,ARHGDIA,APPL1,HDAC2,BCLAF1,PEA15,
A,CCND2,GPC3,SOX12,RAB7A,CCN2,ACTN4,MMP14,PRDX2,DDX3X,NANOG,H1-0,MIF,PRKAR1A,
F1,SET,EGLN1,EIF4A1,RPL19,PPP2R1A,RPL15,GTF3C4,PKD1,SF3B1,SSBP2,ZNF43,RPS3,CHD3,
3X,RPS15,RPL12,ITM2C,RPL30,CLTC,EIF3L,RPS5,RPS7,RPS19,RPL35,GPAT4,CHPF,CTNNB1,NG
M1,ANXA2,AGRN,ZMIZ1,TIMP2,COL11A1,VCL,AP3D1,SBF1,ARF1,PAFAH1B1,MAP4,HNRNPU,CTNN
NO,PSMD5,EGLN1,INA,PPP2R1A,RPL15,HLA-C,PDIA3,RPS3,HMGB1,POLR2A,SEC61A1,OR8K1,I
IAPT,SPEN,NONO,DDB1,PSMD5,YWHAQ,EIF1,SET,EGLN1,TPI1,EIF4A1,RPL19,PPP2R1A,RPL15,G
A2,AGRN,ZMIZ1,TIMP2,COL11A1,VCL,AP3D1,SBF1,ARF1,PAFAH1B1,MAP4,HNRNPU,CTNNA1,CN
P1,ZMIZ1,RPL10,RPS17,AP3D1,SF3B2,ZNF770,GTF2F1,HNRNPU,NFE2L1,SMARCA5,LARP1,HNR
ID5,YWHAQ,EIF1,HACD3,SET,EGLN1,TPI1,CHSY1,EIF4A1,RPL19,PPP2R1A,RPL15,GTF3C4,PKD1,
ID5,YWHAQ,EIF1,HACD3,SET,EGLN1,TPI1,CHSY1,EIF4A1,RPL19,PPP2R1A,RPL15,GTF3C4,PKD1,
N,CSNK1E,SFPQ,CCND2,CDH2,PDIA6,GPC3,DDR1,CCN2,PRDX2,DDX3X,RPS15,RCN1,RPS7,MIF,
,RPL15,GTF3C4,PKD1,SF3B1,SSBP2,ZNF43,RPS3,CHD3,BASP1,HMGB1,POLR2A,RPS4X,RPS3A,
,RAI1,RAB11A,AKIRIN1,IGFBP7,BMPR2,CDC42,PPP1R9B,RBBP7,STAT3,EP300,EXTL3,KDM5B,FL
A6,CYFIP2,DNAJC5,TARDBP,ARL6IP1,HNRNPK,DUX4,YWHAB,LAMP1,RHOB,HMGCR,ARHGDIA,A
2A,RPS4X,RPS3A,RPL18,RPL11,MORF4L1,BTF3,AGRN,RPLP1,ZMIZ1,RPL10,RPS17,AP3D1,SF3
N4,UBXN1,PSMD10,ELOB,GABARAPL2,XBP1,APOE,RYBP,FURIN,EPHA4,PSMC6,RAD23A,CDK5R
1,SSBP2,ZNF43,RPS3,CHD3,BASP1,HMGB1,POLR2A,RPS4X,RPS3A,RPL18,RPL11,MORF4L1,BTF
5A5,SLIT2,TMOD2,TAOK1,SLIT1,GPX1,FKBP4,CARM1,ANTXR1,LINGO1,UBR5,LDLR,RDX,CAPZA
S,HGS,PFN2,AKIRIN1,CDC42,DCTN1,EP300,LRRC4B,UBQLN2,FLOT1,HJURP,FARP1,SFRP1,EVL,
,UBE2R2,NPEPPS,BNIP3L,CCAR2,SMARCC1,FYN,STT3B,LONP1,UBQLN1,CSNK1D,USP7,HECTD1
B1,SSBP2,ZNF43,RPS3,CHD3,BASP1,HMGB1,RPS4X,RPS3A,RPL18,RPL11,MORF4L1,AGRN,RPL
X12,CCN2,ACTN4,MMP14,PRDX2,CLTC,NANOG,RPS19,MIF,GPAT4,GJA1,JUN,CTNNB1,MACF1,
4,BAP1,TCAF1,UBR5,PKDCC,GPC2,VPS4A,RDX,TM9SF4,DCLK1,NMT1,PDPK1,ADAM10,SYT11,A
GPX1,EIF3H,UBE2D2,VPS4A,GNA12,DNAJB2,ATXN3,PSMD4,OGT,PRPF19,USP24,FBXW5,GSK3A,
RPS3,CHD3,BASP1,HMGB1,POLR2A,RPS4X,RPS3A,RPL18,RPL11,MORF4L1,BTF3,AGRN,RPLP1,
1-0,APLP1,RPS7,MIF,GJA1,JUN,CTNNB1,ANXA6,CYFIP2,DNAJC5,TARDBP,ARL6IP1,HNRNPK,DUX
CBP,PINK1,PSMB1,HSPA1A,MDM2,FZR1,UBE4B,TNFAIP1,TRIM2,CUL1,PSMD13,SUMO1,PARK7
A13,EIF4G1,IGFBP4,USP47,OGFR,ABL1,SDCBP,SEMA6B,AUTS2,TAOK2,TSPYL2,HSPA1A,ACVR1
N1,AHCYL1,VPS35,COL2A1,SKI,H3-3A,ATP6AP1,TAF7,NOTCH2,MARS1,SELENON,RAPGEF1,MA
13A,MAGED1,RPL6,KAT6A,DVL3,YWHAE,KIF1A,RPL13,GNAS,DDX17,GPI,HTRA4,RACK1,BSG,FA
N1,ATF4,PSMD2,KIF5B,NLGN2,CHD4,ATP1A1,RPS20,CD81,ATP2A2,RPL26,RBBP4,MCL1,TLE5,C
P6,GRB2,CAPZB,ACTN1,CLIP3,TPM3,CORO1C,STMN3,B2M,CSNK1D,MPRIP,COL27A1,PFN2,RAB

RPL19,PPP2R1A,RPL15,GTF3C4,PKD1,SF3B1,SSBP2,ZNF43,RPS3,CHD3,BASP1,HMGB1,POLR2
GLN1,TPI1,CHSY1,EIF4A1,RPL19,PPP2R1A,RPL15,GTF3C4,PKD1,SERINC1,SSBP2,ZNF43,RPS3,C
TSB,EEF1A2,ANXA5,SQSTM1,GNAI3,NOP53,PARP1,NOX4,PRDX5,TCP1,GRINA,COL2A1,MDK,DDX
9L,PSMD1,AP2A1,PHB2,CST3,MAP6,PLXNB1,DIP2B,SFRP1,PTEN,CEMIP2,FMNL2,BRSK1,ZMYM4

,PYGB,ATP6AP1,VEGFA,MAPK1,PSMD3,ACTN1,CAND1,SDF4,ILF2,CCT8,B2M,STXBP1,IDH1,HGS,
F1A,PPP1R15A,FUS,GRN,PPP1R10,TAOK1,GPX1,FKBP4,UBR5,AKT3,STK24,TRAP1,ATXN3,SOD2,
N1,VPS35,ABCA2,NRDC,TERF2IP,VEGFA,RAPGEF1,PPP2CA,CCND1,MAPK1,ADAM9,PTBP1,EIF3E,
CTN1,CAND1,SDF4,ILF2,CCT8,B2M,STXBP1,IDH1,SOD1,PFN2,DYNLL1,SYP,TXNDC5,PSMD1,PDA
4,PSMD2,KIF5B,NLGN2,CXADR,CHD4,RPS20,CD81,ATP2A2,RPL26,RBBP4,MCL1,TLE5,CLEC7A,
RPL19,PPP2R1A,RPL15,GTF3C4,PKD1,SF3B1,SSBP2,ZNF43,RPS3,CHD3,BASP1,HMGB1,POLR2
D1,SSBP2,ZNF43,RPS3,CHD3,BASP1,HMGB1,POLR2A,RPS4X,RPS3A,RPL18,RPL11,MORF4L1,B
NAS,DDX17,GPI,HTRA4,RACK1,BSG,FASN,SPTBN1,ATP5F1B,CALM3,GNB1,SHISA9,STMN2,MYH
UL1,PSMD13,SUMO1,PARK7,USP14,PSMD7,FAF2,MAEA,CUL4A,UBE2J1,AREL1,FAM8A1,TRIB3,
AF7,VEGFA,RAPGEF1,RPL37,PPP2CA,CCND1,MAPK1,CCNL2,IPO7,ADAM9,PTBP1,CDC37,PRKDC,
,USP24,FBXW5,GSK3A,PSMC3,UBE3A,UBE2I,UBE2D3,PSMC1,CCDC47,BAG5,UBE2K,PTPN23,ZE
,KPNA6,SRSF7,PSIP1,NUP98,POLDIP3,NCBP2,GSK3B,SNRPD2,TXN,SNRPB,NUP205,LMNA,NUP1
IP3,FYN,YWHAH,CCT8,PFKM,MESD,HECTD1,SREBF2,RAB11A,CYP51A1,DYNLL1,CCT5,CDC42,P
,USP24,FBXW5,GSK3A,PSMC3,UBE3A,UBE2I,UBE2D3,PSMC1,CCDC47,BAG5,UBE2K,PTPN23,ZE
TF6B,SHC1,DNAJB11,ATF6,ADD1,QRICH1,XBP1,CUL7,MAN1B1,PPP2R5B,UFM1,OPA1,SEC31A,S

GSK3A,PSMC3,UBE3A,UBE2I,UBE2D3,PSMC1,CCDC47,BAG5,UBE2K,PTPN23,ZER1,FBXL19,USP
A5,SQSTM1,GNAI3,NOP53,PARP1,NOX4,PRDX5,TCP1,GRINA,COL2A1,MDK,DDX42,NOTCH2,VEG
,P3H1,SRF,CRTC1,SETX,CRABP2,DLG4,ATMIN,CDC42EP4,CNTN1,RETREG3,ARSB,CFLAR,SS18L

TG1IP,CDKN1A,DLG5,TIMP3,CTSB,EEF1A2,ANXA5,SQSTM1,GNAI3,NOP53,PARP1,NOX4,PRDX5,T
O,ILF3,PCSK1N,PRKDC,HSPB1,KDM1A,FKBP8,QARS1,CCAR2,DUSP4,CORO1C,SMARCC1,SUPT
PT6H,FYN,YWHAH,CCT8,STXBP1,MESD,USP7,HECTD1,SREBF2,RAB11A,DYNLL1,CCT5,CDC42,P
CG2,PSMF1,P3H1,RAB3A,STC2,PSMB5,HIF3A,PSMA1,CHGB,PSMD10,ELOB,ASB6,RAB5A,WDR5,
5,HNRNPD,TCTN1,ARNT2,BMPR2,DYNLL1,CDC42,DDIT4,PPP1R9B,CTNND1,CIC,DCTN1,STAT3,T
CTTN,GPC4,SLIT1,PALM,ABI2,RDX,CAPZA1,CADM3,CLASP2,PDPK1,WASF2,ADAM10,UBE3A,PRK
,YBX3,PSMD12,TUT4,PSMA4,PSMC6,MYEF2,CACNG7,MEX3D,CNOT1

RPL19,PPP2R1A,RPL15,GTF3C4,PKD1,SF3B1,SSBP2,ZNF43,RPS3,CHD3,BASP1,HMGB1,POLR2
,TUT4,PSMA4,PSMC6,MYEF2,CACNG7,MEX3D,CNOT1
M4A,GSK3A,CHMP4B,MTOR,PACS2,PRKACA,EXOC4,BNIP3,ATP6V1G1,TOMM70,TRIM8,UBXN6,
M4A,GSK3A,CHMP4B,MTOR,PACS2,PRKACA,EXOC4,BNIP3,ATP6V1G1,TOMM70,TRIM8,UBXN6,
G9,CLASP2,IST1,RBPJ,PRPF19,GSK3A,PTGFRN,MTOR,UBE3A,YY1,SMO,RABGAP1,PTPN11,WD
NUP98,POLDIP3,NCBP2,GSK3B,SNRPD2,TXN,SNRPB,NUP205,LMNA,NUP133,THOC2,PPP1R12A,

A,VHL,CCND1,MAPK1,PSMD3,ACADVL,LUZP6,GRB2,PCSK1N,PRKDC,NPEPPS,KDM1A,BNIP3L,CC
A,CNN2,TBCD,KLHL24,GAS7,TLN2,NOX4,ACTR3,CAMSAP1,NCKAP1,MDK,PTK7,DPYSL2,NOTCH
2B,APPL1,IPO5,SKP1,CDKN1A,CTDSP2,DDX39B,EID1,SRSF5,TBCD,PPP1CB,GNAI3,NOP53,ACTR
IM24,RNF5,PLPP3,RAD23A,NF2,ZNF207,GET4,GSN,STX12,USP36,CTSA
4,USP47,CELSR1,SNX3,ABL1,PSMB1,RTF1,RUVBL2,NOTCH1,CUL1,PSMD13,CCNY,PSMD7,TNR
4A,ATF7IP,TRIM27,NCK2,PMEPA1,ARPC3,STUB1,SENP6,PSMC5,GBA2,MAPRE3,PSMC2,PDE4DI
NP,CYFIP1,RNF157,NTN1,LIMK1,ISLR2,CRKL,SPOCK1,GOLGA4,SEMA4C,SRF,GSK3B,SERPINE2
DAM10,IST1,GSK3A,SMARCD1,BNIP3,MICAL1,AMBRA1,GAK,SH3PXD2B,HMGA1,CHMP1A,PINK1,
,CELSR1,SNX3,ABL1,PSMB1,RTF1,RUVBL2,NOTCH1,CUL1,PSMD13,CCNY,PSMD7,TNRC6A,PPP

1,PPP2CA,CCND1,MAPK1,ADAM9,PTBP1,EIF3E,PRKDC,GABARAP,KDM1A,CLIP3,MAP2K2,FYN,FB
RKDC,KDM1A,NSG2,GNG2,SMARCC1,ATP6V1B2,FYN,LARS1,STT3B,UBQLN1,SOD1,BAG6,SREBF

P1,IST1,PSMC3,AP2A2,AP1M1,PAFAH1B2,SDCBP,SCAMP1,PSMB1,HSPA1A,PSMD13,CAB39,NDUF

O21,WSB2,PSMB4,UBR5,UBE2M,UBE2D2,DTX4,DNAJB2,MKRN1,DCUN1D2,LZTR1,RNF145,PSMI

R2,DUSP4,CORO1C,SMARCC1,SUPT6H,FYN,FBLN1,USP7,STRAP,MAPK8IP1,BAG6,HNRNPD,KDM4
P1,IST1,PSMC3,AP2A2,AP1M1,PAFAH1B2,SDCBP,SCAMP1,PSMB1,HSPA1A,PSMD13,CAB39,NDUF
,TUT4,PSMA4,PSMC6,MYEF2,CACNG7,MEX3D,CNOT1
KN4,CCND1,MAPK1,SRSF9,GRB2,PRKDC,KDM1A,NSG2,GNG2,SMARCC1,ATP6V1B2,FYN,LARS1,S
V,ADAM10,CAP1,IST1,PSMC3,AP2A2,AP1M1,PAFAH1B2,SDCBP,SCAMP1,PSMB1,HSPA1A,PSMD13
A,FKBP1A,LOX,SLIT2,NDN,CCN1,SOX2,LTBP4,RAB14,RDX,MTMR4,PDPK1,LTBP3,RBPJ,EHD1,UB
1,HDAC6,PTEN,FEM1B,CUL3,POM121C,RPS2,RPS27A,TNKS,PSMD8,HIF1A,FKBP1A,FBXO21,BAF
,CES3,PSAP,PGK1,RPLP0,H3-3B,CANX,CLU,PTMA,DDX5,ENO2,SOX4,FKBP14,RPL13A,MAGED1,S
AV,ADAM10,CAP1,IST1,PSMC3,AP2A2,AP1M1,PAFAH1B2,SDCBP,SCAMP1,PSMB1,HSPA1A,PSMD1
,PEBP1,CDH2,GPC3,SULF1,PIP4K2B,MEGF8,RAB7A,CCN2,ACTN4,MMP14,PRDX2,DDX3X,RPS15,
P1,KMT2E,NMT1,MOAP1,WASF2,AKT2,OGT,GSK3A,MTOR,BNIP3,PHIP,CROCC,ABL1,SDCBP,PINK
CCND1,MAPRE1,PSMD3,DYNC1LI2,DYNLL2,SON,CCAR2,CSNK1D,BAG6,RAB11A,PRPF40A,DYNLI
M10,CAP1,IST1,PSMC3,AP2A2,AP1M1,PAFAH1B2,SDCBP,SCAMP1,PSMB1,HSPA1A,PSMD13,CAB3
4,UBR5,UBE2M,UBE2D2,DTX4,DNAJB2,MKRN1,DCUN1D2,LZTR1,RNF145,PSMD4,LRRC41,OGT,F
,PSMD1,PBX1,CSNK2A1,PHB2,GSPT1,BRD4,UBA52,ZFP36L1,ATRX,MASTL,PTEN,SMC3,PPP1CC,
C7,WAC,ILRUN,UBE2V1,UBQLN4,EI24,ATP6V0E2,ATG9A,RETREG3,ATP6V0A1,RAB5A,GABARAPL
TN1,STAT3,PRMT1,ACTR1A,EP300,PSMD1,PBX1,FLCN,PRCC,CSNK2A1,PHB2,BRD4,UBA52,ZFP36
P6V0C,TMEM30A,GLUL,NNAT,TIMP3,PGRMC1,CNN2,CTSB,RAB27A,CCT2,ANXA5,VAMP2,GDI2,PD
RX,MASTL,PTEN,SMC3,PPP1CC,CUL3,EML1,BRSK1,RPS27A,TNKS,PSMD8,APEX1,PCBP4,AKAP9,

RAB27A,RNF187,CCT2,MLXIP,SQSTM1,PARP1,MORF4L2,TAF15,HSPH1,ACTR3,PKN1,TCP1,CCT3
I,SPTBN2,SFPQ,PEBP1,CDH2,PIP4K2B,CCN2,ACTN4,PRDX2,DDX3X,RPS15,RPS7,IK,MIF,PRKAR1A

4B,NAP1L4,YLPM1,CCT5,RBBP7,DFFB,PRMT1,NAP1L1,MTA1,KDM5C,EP300,RERE,KDM5B,FLCI
Z1,PFN1,TLE5,CLEC7A,NES,CCNI,PTN,CSNK1E,TPM1,PEBP1,CCND2,GPC3,DDR1,PLXNA1,PIP4K2
MLXIP,SQSTM1,PARP1,MORF4L2,TAF15,ACTR3,PKN1,TCP1,CCT3,PTOV1,MDK,RBMX,CCT4,GCN

DX,SLIT2,NDN,CCN1,LTBP4,RAB14,RDX,MTMR4,PDPK1,LTBP3,RBPJ,EHD1,UBE3A,CREBBP,UBE2
,STAT3,SGTA,ALDH3A2,HIP1,AP2A1,FLOT1,SEC16A,UBA52,RER1,PI4KA,PTEN,YKT6,CUL3,RPS27
ST1,EML1,ANP32B,HIF1A,ATP6AP2,SLIT2,SLC2A1,XRCC6,SLIT1,HSPG2,SSBP3,SOX2,AKT3,DCLK
AF1,NUP58,NF2,CDK5RAP3,PLRG1,SIN3A,NUTF2,ABCA7

HL1,DYNC1I2,CENPF,PSMC6,PPP2R2A,CDK5RAP3,NINL,HMGA2,SIN3A
CLCN6,CRABP2,SLC12A7,WASF3,CDC42EP4,CNTN2,ARPC5,RAB5A,RGMA,CAPZA2,RUFY3,ADD1,
FLCN,MMP15,GRSF1,ZFP36L1,BRD2,HOXB5,NXN,SFRP1,TRA2B,CUL3,NDST1,SPINT2,RPL7L1,IT

l1,ACTR3,PKN1,TCP1,CCT3,PTOV1,MDK,RBMX,CCT4,GCN1,SKI,ASXL1,KDM6B,ABCA2,PRPF6,TA

1,PCBP4,AKAP9,PPP1R10,PSMB4,TAOK1,EHMT1,CARM1,INTS3,VPS4A,RDX,KMT2E,TCF3,PSMD4
SPA1A,MDM2,FZR1,SGSM3,NSF,SUMO1,PARK7,PTPN1,SNX1,TNRC6A,HNRNPR,MAGEF1,AREL1,
P2,IGFBP7,TXNIP,DDIT4,PPP1R9B,RBBP7,STAT3,EP300,LBH,PRKAR2A,PHB2,ZFP36L1,SFRP1,SR

,EIF4G1,TRIP12,ABL1,KDM3A,MGAT5,PPP5C,MACROH2A1,HSPA1A,GGNBP2,PKIA,IGF1R,PPP1F

,RAB5C,ACTR1B,PDPK1,ITGAV,ADAM10,CAP1,IST1,PSMC3,AP2A2,AP1M1,PAFAH1B2,SDCBP,SC
DNAJA1,PAFAH1B1,PXDN,NFE2L1,SMARCA5,LARP1,HNRNPA0,CTNNA1,LDHA,NACA,RTN4,XRCC5
MGCR,RAC1,CDK16,ATP6V0C,TMEM30A,GLUL,NNAT,TIMP3,PGRMC1,CNN2,CTSB,RAB27A,CCT
K1,ITGAV,ADAM10,CAP1,IST1,PSMC3,AP2A2,AP1M1,PAFAH1B2,SDCBP,SCAMP1,PSMB1,HSPA1
N1,TMBIM6,SPTAN1,GSTP1,SMG1,CALM1,NPM1,PLXNB2,HSPD1,IGF2,VCP,UBB,TRIM28,YWHA

R2A,CDK5RAP3,NINL,HMGA2,SIN3A
B,SENP3,CUL4B,ABCE1,NOLC1,RPL32,PES1,KRI1,DDX56,NOP58,USP36,MYBBP1A,SNU13
ATP6AP1,DPYSL2,NOTCH2,RNF10,VEGFA,VHL,PPP2CA,CCND1,MAPK1,ARPC2,PSMD3,PTBP1,FT
SPG2,SSBP3,SOX2,AKT3,DCLK1,CNP,MACROH2A2,NCSTN,ATP1B2,PHGDH,SCRIB,IGF2BP1,ATP

HB,GABARAPL2,PIN1,PAIP1,YBX3,TAB3,RELA,GIT1,RYBP,FURIN,EPHA4,RNF5,CDK5RAP3,QSOX1

R5,SOX2,ZMIZ2,KMT2E,MACROH2A2,NUCKS1,SPTY2D1,NSD2,OGT,PCGF2,NOC2L,KDM4A,MBD
F26A,ANKFY1,KLC2,GOLPH3,SNX1,NBAS,VPS51,ARFGAP2,TBC1D14,GOLGA4,ACBD3,MCFD2,KI
AOK2,HIP1R,HSPA1A,ELN,CRACD,MECP2,JAM3,CYFIP1,PHPT1,KAT2A,LIMK1,FLII,MYCBP2,PDGF
A3C,SPAG9,CLASP2,ADAM10,IST1,RBPJ,SH3BP4,GSK3A,FBLN5,MTOR,YY1,APBB1,NDUFA13,SM
TAT3,EIF2AK1,EDC4,PSMD1,EIF3D,GSPT1,UBA52,ZFP36L1,UPF1,EIF5B,HNRNPA2B1,POM121C,
G1,CNBP,NTRK2,PUM1,FXR2,PINK1,PAIP2,YTHDF3,RBM4,MKNK2,SMPD3,TIA1,FXR1,CYFIP1,SY
K1,ITGAV,ADAM10,CAP1,IST1,PSMC3,AP2A2,AP1M1,PAFAH1B2,SDCBP,SCAMP1,PSMB1,HSPA1
MSH6,PCGF2,NOC2L,GSK3A,NDUFA13,BNIP3,PHIP,BAG5,DAPK3,SORT1,CD74,FGFR1,UBE2K,U

,USP36,CTSA,NAMPT
H2,CYFIP1,NTN1,LIMK1,RHEB,TMEM98,C1QBP,SYNGAP1,ISLR2,DHX36,CRKL,S1PR3,GOLGA4,S
COCO1,AMFR,IGFBP4,USP47,CELSR1,SNX3,ABL1,PSMB1,RTF1,RUVBL2,GRIK5,NOTCH1,MAPK
P1,RPL23,EIF4H,NACA,LPCAT1,BRD3,RPS9,XRCC5,CIPC,KPNB1,RPL22,RHOA,PTCD3,CBX5,ZNF

CR,RAC1,CDK16,ATP6V0C,TMEM30A,GLUL,NNAT,TIMP3,PGRMC1,CNN2,CTSB,RAB27A,CCT2,AN
1,MTSS2,CYFIP2,APLP2,ARL6IP1,YWHAB,PTGES3,MAP2,RAB1A,HMGCR,ARHGDIA,IPO5,SKP1,TF
N4,MVB12B,PSMD3,AEBP1,ADAM9,PCSK1N,GABARAP,UBE2R2,NPEPPS,CCAR2,XPNPEP3,SMAR
P1S,CDKN1B,WASF3,CDC42EP4,BMERB1,ARPC5,SHANK1,ROCK2,TGFBR1,CAPZA2,ADD1,ARHG

AR,VDAC1,TXNDC5,CSNK2A1,PHB2,UBA52,SFRP1,SRSF6,PTEN,IL6ST,RPS27A,HIF1A,CTTN,SLC2
HYOU1,MAPK8IP1,GHITM,SOD1,BAG6,ARNT2,YME1L1,ADAR,VDAC1,STAT3,TXNDC5,CSNK2A1,

19,PSMC3,CHMP4B,PRKACA,NASP,MN1,PSMC1,TUBB4A,PAGR1,EIF4G1,USP47,UBE2L3,PTPN11
KNK2,TIA1,FXR1,CYFIP1,SYNCRIP,ETF1,TNRC6A,PPP1CA,HNRNPR,CDK4,DDX6,NCK2,C1QBP,LS
MO1,PARK7,ADNP,CYFIP1,MAP4K4,LIMK1,CUL4A,TASOR,ATF7IP,CCDC88A,TRIM27,NCK2,ARPC
EIF4H,NACA,BRD3,RPS9,XRCC5,CIPC,RPL22,RHOA,PTCD3,CBX5,ZNF865,ATF4,PSMD2,NR2F1,E
C42,DDIT4,POLR2E,SULF2,STAT3,BCL9L,BCR,TRIO,EP300,HIP1,FLCN,AP2A1,UBA52,SFRP1,PTE

E1L1,ADAR,VDAC1,TXNDC5,CSNK2A1,GPX4,PHB2,UBA52,SFRP1,SRSF6,PTEN,IL6ST,RPS27A,HIF

YADM,PLXND1,ID2,ANAPC2,SKIL

B3A,SETD5,DLG4,WASF3,ERBB2,UNC13A,THBS2,CNTN2,NRP2,ARHGAP33,SHANK1,CTBP2,PCD
RGMA,PPP3CA,RUFY3,CUL7,PRKCI,DYNLT1,SEMA6C,HMGB2,RELA,OPA1,EPHB2,LRP8,FGF13,E

PS14,CHD4,JUND,CD81,CCT7,SRSF4,PFN1,TLE5,HNRNPC,LRPPRC,TADA2B,KHDRBS1,XPO1,SF

I,NACA,LPCAT1,BRD3,RPS9,XRCC5,CIPC,RPL22,RHOA,PTCD3,CBX5,ZNF865,ATF4,PSMD2,NR2F

D3,RPS9,XRCC5,CIPC,RPL22,RHOA,PTCD3,CBX5,ZNF865,ATF4,PSMD2,NR2F1,ENC1,RPS14,CHD
7,CDC34,PSMB1,TAOK2,ACVR1B,PKIA,MDM2,FZR1,PCM1,MECP2,CUL1,PSMD13,CCNY,PSMD7,

D1,SMARCA4,SRSF9,AEBP1,TCF25,PTBP1,ILF3,SUPT5H,KDM1A,QARS1,CCAR2,POLR2B,SMARC

ROCK2,CTBP2,CHMP5,RAB5A,NECTIN2,SLC1A5,DYNLT1,APOE,FURIN,NUP50,DYNC1I2,SCARB1,
I,GCN1,SKI,SNRNP70,ASXL1,KDM6B,PRPF6,TAF7,RNF10,VEGFA,MARS1,VHL,MAPK1,SMARCA4,

NUTF2,GNAI1,ABCA7
ARF6,DNM1,MYADM,ZNF207,GSN
RB1,CCNK,SMARCB1,HMGA2,GSN,EPHA2,VPS37B
M,POSTN,ITGA7,PLXND1,FGD1,SH3KBP1,CACNG7,ANAPC2

LSM4,DHX36,STUB1,PSMC5,CSNK1A1,PSMC2,HERPUD1,PSMC4,PLK2,WAC,HSF1,GSK3B,IRS1,

F1,SRF,DDAH1,HSF1,USP19,STC2,CDKN1B,PSMB5,EGLN2,HIF3A,LMNA,PSMA1,CITED2,CFLAR,P
MMP2,GLUL,DCN,CDKN1A,SRSF5,TOMM20,VAMP2,GNAI3,PABPN1,PARP1,NOX4,RPL27,PTPRA,F
MB1,TAOK2,SMPD3,OBSL1,PKIA,MDM2,FZR1,PCM1,MECP2,CUL1,PSMD13,PSMD7,MAEA,CUL4
,IMPDH2,RAB14,LDLR,KMT2E,NCSTN,RAB6A,RAB5C,ACTR1B,PDPK1,ITGAV,ADAM10,CAP1,SYT1
RUSC1,MGRN1,PSMD10,UBE2G1,PSMA6,TRAF3,MIB2,PSMD12,PSMA4,FBXO7,UBE2E1,PSMC6,
,TPM1,CDH2,GPC3,DDR1,SULF1,PLXNA1,MEGF8,CCN2,ACTN4,MMP14,RPS19,MIF,GJA1,MYO5
NOP53,NOX4,PTPRA,PDXK,PKN1,ETNK1,PTK7,GCN1,TERF2IP,TMED2,TOP1,TAF7,VEGFA,RAPGEF

NPC,LRPPRC,TADA2B,KHDRBS1,XPO1,SFPQ,SOX12,ACTN4,PRDX2,DDX3X,NANOG,H1-0,PRKAR1
T1,OGT,GSK3A,PSMC3,UBE3A,PRKACA,NDUFA13,PSMC1,BAG5,MICAL1,PRDX3,UBE2K,ZER1,US

,ATF6,MAPK3,XBP1,TAB3,TMEM59,FBXO7,SCOC,SESN2,RNF5,QSOX1,ATP6V1D,RB1CC1,USP36
PKN1,VPS35,CLSTN3,PTK7,SKI,ASXL1,PSMB2,TSPAN6,ATP6AP1,TERF2IP,NOTCH2,VEGFA,RAPG

31,SHISA9,STMN2,MYH9,SEPTIN2,P4HB,APP,RPL5,KLRD1,GDI1,FSCN1,TMBIM6,SPTAN1,GSTP1,
EF1,PPP2CA,CCND1,MAPK1,CCNL2,IPO7,ADAM9,PTBP1,CDC37,PRKDC,HSPB1,CLIP3,FKBP8,MAF
EF1,PPP2CA,CCND1,MAPK1,CCNL2,IPO7,ADAM9,PTBP1,CDC37,PRKDC,HSPB1,CLIP3,FKBP8,MAF

PLK2,PSMF1,RCC2,CDKN1B,PSMB5,PRKAR2B,XPC,PSMA1,PRMT2,PSMD10,ANXA1,PSMA6,CNOT
RDX,CAPZA1,ATXN3,KRT8,CLASP2,PDPK1,WASF2,CAP1,MTOR,DAPK3,MICAL1,PLS3,CELSR1,ABL
2CA,CCND1,MAPK1,SMARCA4,PSMD3,GABARAP,PLCG1,CCAR2,GNG2,FYN,YWHAH,PFKM,TNKS2
PPP3CA,PIN1,ADD1,BTBD1,HMGB2,CTNNBIP1,APOE,EPHA4,EIF2S1,PLXND1,HMGA2,PRMT7,WAF
LK1,FYN,SLIT3,STMN3,FBLN1,STXBP1,HYOU1,STRAP,PTPRS,HGS,MAPK8IP1,GHITM,DGKD,NCO
N4,XRCC5,PJA2,IGFBP3,COL1A2,LLGL1,RHOA,ATF4,PSMD2,NLGN2,CXADR,CHD4,TMEM259,RF
KDC,SUPT5H,ACTN1,CAND1,KDM1A,MAP2K2,CCAR2,ARF4,SMARCE1,SMARCC1,SUPT6H,YWHAH
SUPT5H,KDM1A,QARS1,CCAR2,SMARCE1,PHC2,ID4,STRAP,SARS1,SREBF2,NCOR2,HNRNPD,RPS
,HYOU1,STRAP,PTPRS,HGS,MAPK8IP1,GHITM,DGKD,NCOR2,BMPR2,CDC42,DDIT4,ADAR,CTNND
CE1,VPS16,FGF13,TPX2,PDXP,SMARCC2,MRPL3,SMARCB1,SCAF8,MRPS26,LRP6,MRPS27,MRPL
LK1,FYN,SLIT3,STMN3,FBLN1,STXBP1,HYOU1,STRAP,PTPRS,HGS,MAPK8IP1,GHITM,DGKD,NCO
VPS35,CLSTN3,PTK7,SKI,ASXL1,PSMB2,TSPAN6,ATP6AP1,TERF2IP,NOTCH2,VEGFA,RAPGEF1,RP

R1,PCM1,MECP2,CUL1,PSMD13,ATF5,PSMD7,CUL4A,CETN2,CDK4,H2AX,PSME3,PSMC5,PSMC2,
BCOR,PRMT5,FBL,SETD5,CTR9,OTUB1,KDM2A,RNF20,KDM5A,RBM14,LMNA,PER1,PHB,PRMT2

ERBB2,PPP1R12A,MAPK3,RUFY3,XBP1,TFDP1,UFM1,NOLC1,PRNP,NUP58,LEPROT,USP36,NUTF
B,SQSTM1,NOP53,PARP1,TAF15,SSRP1,PKN1,GRINA,TSPAN6,ATP6AP1,TERF2IP,TAF7,NOTCH2,V
,AKAP9,TMOD2,TAOK1,CYTH2,ANTXR1,ABI2,ARHGEF17,VPS4A,RDX,CAPZA1,ATXN3,FNBP1L,KR
MA6,CTDNEP1,PIN1,SPIN1,PSMD12,CTNNBIP1,APOE,PSMA4,PSMC6,PLPP3,GRB10,PPM1A,TLE4,
AZ1,PFN1,ASTN2,CLEC7A,NISCH,KHDRBS1,XPO1,PTN,CSNK1E,PFKL,TPM1,CDH2,GPC3,SULF1,F
D5,CTR9,OTUB1,KDM2A,RNF20,KDM5A,RBM14,LMNA,PER1,PHB,PRMT2,CTCF,SETD3,WDR5,SU

N1,HMGB2,CUL4B,MCM7,PSMD12,PSMA4,SMARCC2,CENPF,POGZ,PSMC6,PAF1,SMARCB1,CEN
IK5,OBSL1,REEP5,OSBPL2,NSF,EMC1,SUMO1,BANF1,RAN,SUN2,GOLPH3,VPS51,AKAP8L,RAB18
,STXBP1,IDH1,SOD1,NCOR2,HNRNPD,VGF,ARNT2,WBP2,IGFBP7,TXNIP,DDIT4,PPP1R9B,RBBP7,

RF,SERPINE2,PRMT5,CRABP2,WASF3,NUMBL,THBS2,EFNB3,SRRT,SS18L1,RGMA,PPP3CA,RUFY3
R,PSMD10,ROCK2,ELOB,PSMA6,PIN1,AJUBA,PSMD12,OPA1,TGFB2,PSMA4,PSMC6,POSTN,PAM
TED2,CFLAR,PSMD10,ROCK2,ELOB,PSMA6,PIN1,AJUBA,PSMD12,OPA1,TGFB2,PSMA4,PSMC6,P

PRDX6,GRN,ACLY,XRCC6,SLC44A2,IMPDH2,RAB14,NCSTN,RAB6A,RAB5C,ACTR1B,PDPK1,ITGAV
C2,AMBRA1,SMO,FGFR1,EIF4A3,LMO4,CELSR1,WDTC1,ABL1,PHACTR4,RTF1,PLXNA2,ACVR1B,G
KIF5B,RPS14,CXADR,CD81,COL5A2,WDR1,PFN1,TLE5,NES,PRKCSH,SPTBN2,PTN,CSNK1E,GLG1
6A,FNBP1L,SPAG9,PLEKHM2,CLASP2,PDPK1,SCRIB,SYT11,CHMP4B,MTOR,PACS2,PRKACA,EXO
RA1,SMO,FGFR1,EIF4A3,LMO4,CELSR1,WDTC1,ABL1,PHACTR4,RTF1,PLXNA2,ACVR1B,GGNBP2

RF2IP,VEGFA,RAPGEF1,PPP2CA,CCND1,MAPK1,SMARCA4,MAPRE1,ADAM9,EIF3E,DYNLL2,LUZP6

T2,OGT,ATP6V0D1,RBPJ,GSK3A,AP2A2,ATP6V1G1,PHIP,SORT1,FGFR1,RABGAP1,NTRK2,IGFBP4

N,RAPGEF1,OPTN,MAPK1,PSMD3,PRDX1,CDC37,PRKDC,HSPB1,KDM1A,PLCG1,MAP2K2,QARS1,

N,ACLY,XRCC6,SLC44A2,IMPDH2,RAB14,NCSTN,RAB6A,RAB5C,ACTR1B,PDPK1,ITGAV,ADAM10,

MGCS1,GPAT4,PRKAR1A,TARDBP,YWHAB,PTGES3,CADM4,ATP1A2,PLPPR3,CTBP1,HMGCR,RAC1
A,RAPGEF1,RPL37,MAPK1,SMARCA4,PSMD3,PTBP1,GRB2,FKBP8,MAP2K2,CCAR2,FYN,LARS1,TN
SOD1,HNRNPD,CDC42,ADAR,TRAF4,TRAF7,PRKAR2A,PHB2,UBA52,SFRP1,ADCY6,PTEN,FEM1B,H
OR,PRKACA,ATP6V1G1,PHIP,NSG1,USO1,RABGAP1,NTRK2,RANGAP1,PTPN11,WDTC1,ABL1,COL
8,MAP2K2,QARS1,DUSP4,CORO1C,FYN,FBLN1,CSNK1D,STRAP,HGS,MAPK8IP1,SOD1,PFN2,BMP

AB11A,TCTN1,DYNLL1,TRAF7,DCTN1,WDR45B,ACTR1A,LRRC4B,UBQLN2,UBAP2L,RAB3GAP2,HI

2R,TRIM27,NCK2,ISLR2,ARPC3,GBA2,GOLGA4,SEMA4C,SRF,GSK3B,CLCN6,CRABP2,SLC12A7,W

,PSMD10,MEIS3,TGFBR1,XBP1,YBX3,HMGB2,RELA,OPA1,PRELID1,SKIL,RB1CC1
ADH5,ULK1,IL6ST,MCM3AP,BRSK1,POM121C,LMAN1,TNKS,GALNT11,FKBP1A,LOX,AKAP9,GRK2
P5K1C,ZNF703,RCC2,SETD5,DUSP3,THBS2,SHANK1,AFDN,ROCK2,HEG1,PCDHB2,PRKCI,AJUBA

ATP6V0D1,SH3BP4,ARPP19,GSK3A,MTOR,PRKACA,ATP6V1G1,PHIP,NSG1,USO1,RABGAP1,NTRK

,COL4A2,MAPT,FN1,SPEN,NONO,DDB1,PSMD5,YWHAQ,HACD3,DCHS1,CHSY1,PPP2R1A,PKD1,H
N1,NCKAP1,ACAP3,MDK,LAMC1,PTK7,VEGFA,TUBB2A,MAPK1,MAPRE1,ADAM9,GRB2,HSPB1,PL
TCH1,EXOC5,PSMD13,CD44,KAT2A,NTN1,PSMD7,PPP1CA,CSNK2B,PSME3,DEAF1,DAAM1,PSMC
P2A2,EXOC4,PSMC1,SMO,LMO4,CELSR1,ABL1,PHACTR4,PSMB1,DAG1,TXNRD1,NOTCH1,ACTA2
MGCS1,GPAT4,PRKAR1A,TARDBP,YWHAB,PTGES3,CADM4,ATP1A2,PLPPR3,CTBP1,HMGCR,RAC1

X1,SKI,H3-3A,HDGF,ABCA2,TAF7,NOTCH2,VEGFA,VHL,CCND1,SMARCA4,AEBP1,ACADVL,TCF25

,RHOB,ATP1A2,RAB1A,RAC1,ARHGDIA,GPC1,TOP2B,APPL1,DNER,L1CAM,PPIB,GLUL,DCN,DLG

1A,TLE4,FZD7,GPRC5B,FZD8,LRP6,JRK,EMD

,COL4A2,MAPT,FN1,SPEN,NONO,DDB1,PSMD5,YWHAQ,HACD3,DCHS1,CHSY1,PPP2R1A,PKD1,H
1,ATP6V1B2,FYN,LARS1,SOD1,HNRNPD,DYNLL1,DDIT4,PPP1R9B,STAT3,CAPN2,PRKAR2A,H2AZ1
GFA,VHL,CCND1,SMARCA4,AEBP1,TCF25,EIF3E,FTO,ILF3,SUPT5H,KDM1A,QARS1,CCAR2,SMAR

MTREX,TRMT112,RPS21,RAN,RTCB,RPL27A,DARS1,EXOSC10,PDCD11,PA2G4,RARS1,FBL,WARS
,UQCR11,COX5B,NDUFA10,ATP5PD

PLCG1,CORO1C,B2M,PLXNA3,CSNK1D,RAB11A,AKIRIN1,BMPR2,TP53BP1,CDC42,STAT3,BCL9
N23,RAB11FIP3,RANGAP1,UBE2L3,PTPN11,SNX3,PINK1,PKIA,ATP13A2,MDM2,PCM1,PARK7,IDH2
,HSPB1,MAP2K2,FBLN1,B2M,HGS,SOD1,NCOR2,HNRNPD,ARFGEF2,WBP2,TP53BP1,ADAR,POL

JSO1,PAGR1,RABGAP1,CALCOCO1,RANGAP1,UBE2L3,PTPN11,WDTC1,KDM3A,POR,PPP5C,NOT

K3B,STC2,CDKN1B,STAT1,IRS1,PRKAR2B,ATP6V0E2,CACYBP,BACE1,SOS1,GPR173,ATP6V0A1,C

D1,SMARCA4,AEBP1,TCF25,EIF3E,FTO,ILF3,SUPT5H,KDM1A,QARS1,CCAR2,SMARCE1,PHC2,ID4
HSBP1,DCN,CDKN1A,DLG5,DDX39B,CTSB,MEST,TGFBI,PLOD1,COL2A1,MDK,LAMC1,PTK7,SKI,

)B,RGMA,XBP1,APOE,FURIN,EPHA4,PSMC6,PRELID1,AKIRIN2,RAD23A,RNF40,GSN

PRKACA,BNIP3,CERS2,GNA13,MICAL1,RANGAP1,PTPN11,CELSR1,LGALS1,DAG1,CLIC1,MFN2,J

ARCA4,PSMD3,ADAM9,GRB2,CDC37,CLIP3,FLNB,PITPNA,FYN,SRSF3,SLIT3,B2M,STXBP1,IFITM3
RCC1,SUPT6H,YWHAH,ILF2,PFKM,RAI1,SREBF2,HNRNPD,ARNT2,WBP2,AKIRIN1,BMPR2,TP53BP
RCC1,SUPT6H,YWHAH,ILF2,PFKM,RAI1,SREBF2,HNRNPD,ARNT2,WBP2,AKIRIN1,BMPR2,TP53BP
C,MACROH2A2,SOD2,SPAG9,DICER1,TCF3,LTBP3,LRP3,IST1,PRPF19,KDM4A,SART1,MTOR,CD
RCC1,SUPT6H,YWHAH,ILF2,PFKM,RAI1,SREBF2,HNRNPD,ARNT2,WBP2,AKIRIN1,BMPR2,TP53BP
RSF6,EVL,PTEN,FEM1B,CUL3,NDST1,PSMD8,HIF1A,FKBP1A,SLIT2,GPC4,SERP1,PSMB4,SLIT1,CC
),UNC13B

CYBP,SOS1,GPR173,ATP6V0A1,CFLAR,SHC1,ROCK2,ADIPOR1,ANXA1,XBP1,PRKCI,SP1,FER,BCA

,RPS27A,PPP1R15A,FKBP1A,SYMPK,NDN,AKAP9,XRCC6,TAOK1,CCN1,SHC2,KMT2E,DNAJB2,SP
K2,TRIB3,PMEPA1,PPP2R5D,EMILIN1,CRKL,GSK3B,WARS1,RRP1B,RASSF2,CDKN1B,SH3BP5,DU

1,SOD1,PFN2,BMPR2,DDIT4,PPP1R9B,ADAR,TRAF4,TRAF7,STAT3,EIF2AK1,PRKAR2A,FLCN,FLO

G7,EPHA2,GNAI1
PRKDC,HSPB1,CLIP3,FKBP8,MAP2K2,QARS1,DUSP4,CORO1C,FYN,FBLN1,CSNK1D,STRAP,HGS,
HANK1,RAB5A,PPP3CA,SLC1A3,GIT1,EPHB2,APOE,LRP8,SYT1,PRNP,EPHA4,DNM1,PPFIA3,SYT4,F
,COL18A1,SNRNP200,CSDE1,SMIM14,IGF2,ATP5F1A,MARCKSL1,UBB,AFF4,TRIM28,MATR3,YW

HANK1,RAB5A,PPP3CA,SLC1A3,GIT1,EPHB2,APOE,LRP8,SYT1,PRNP,EPHA4,DNM1,PPFIA3,SYT4,F
RO1C,MTRNR2L10,B2M,UBQLN1,USP7,MAPK8IP1,DYNLL1,TXNIP,PPP1R9B,ADAR,PRKAR2A,PB
3,PMEPA1,PPP2R5D,EMILIN1,CRKL,GSK3B,WARS1,RRP1B,RASSF2,CDKN1B,SH3BP5,DUSP3,DU

TAT1,VKORC1L1,ANXA1,MAPK3,XBP1,PCNA,SP1,FER,RELA,THG1L,APOE,TLR6,FBXO7,PRNP,LAN
POE,TGFB2,ARF6,COL3A1,RAP1B,DNMT1,ITPKB,CYTH3
,TAF7,NOTCH2,VEGFA,VHL,CCND1,SMARCA4,AEBP1,ACADVL,TCF25,EIF3E,FTO,ILF3,SUPT5H,K

,SRSF3,SLIT3,B2M,STXBP1,IFITM3,SOD1,CDC42,ADAR,PSMA7,TRAF4,ST3GAL6,STAT3,TXLNA,C

YNGAP1,MARCHF7,EMILIN1,HERPUD1,RANBP9,GSK3B,ASH1L,SERPINE2,RASSF2,STAT1,DUSP3,
31,TYW5,DGCR8,SRRT,SENP3,RTRAF,NOLC1,THG1L,RPL32,TUT4,PES1,KRI1,DDX56,ELAC2,DUS
IB,GLUL,DCN,DLG5,CNN2,VCAN,NOX4,ACTR3,PKN1,NCKAP1,ACAP3,MDK,LAMC1,PTK7,SKI,VEG
IB,GLUL,DCN,DLG5,CNN2,VCAN,NOX4,ACTR3,PKN1,NCKAP1,ACAP3,MDK,LAMC1,PTK7,SKI,VEG

2,SOX2,SPAG9,PSMD4,ITGAV,PSMC3,KLHDC10,PSMC1,CD74,FGFR1,NTRK2,IGFBP4,PTPN11,AE

MTS7,DNAJB6,CFLAR,TGFBR1,NID1,COL6A3,EXT1,TGFB2,FURIN,COL3A1,POSTN,ITGA7,PRDX4,

RMC1,CNN2,CTSB,RAB27A,CCT2,VAMP2,GNAI3,GDI2,HSPH1,PDXK,PKN1,MLEC,MDK,SURF4,RAE
DNAJB6,CFLAR,TGFBR1,NID1,COL6A3,EXT1,TGFB2,FURIN,COL3A1,POSTN,ITGA7,PRDX4,MFAP2

DNAJB6,CFLAR,TGFBR1,NID1,COL6A3,EXT1,TGFB2,FURIN,COL3A1,POSTN,ITGA7,PRDX4,MFAP2
NUS1,RCC2,SOX9,ADIPOR2,NR2F2,ARSB,NRP2,SGPL1,ROCK2,ADIPOR1,ANXA1,TGFBR1,SP1,FEI

I,CSNK1E,GLG1,CDH6,CDH2,GPC3,DDR1,SULF1,PLXNA1,PIP4K2B,MEGF8,RAB7A,CCN2,ACTN4,I
RD3,NAPA,COL7A1,CHMP5,NECTIN2,MARK1,DYNLT1,FER,TRAPPC1,SPG11,SEC23IP,ABCE1,OPA
16,ATP6V0C,SKP1,DCN,MAP1A,NOVA2,CCT2,PARP1,ABCA3,PRDX5,PKN1,TCP1,CCT3,GRINA,COL

HANK1,ROCK2,TGFBR1,CAPZA2,RUFY3,ADD1,ARHGEF18,PRKCI,FER,GMFB,BCAR1,PDXP,ARF6,N

ATRX,HDAC6,SMC3,PPP1CC,EML1,MAPRE2,BRSK1,TNKS,HIF1A,AKAP9,SPG7,TAOK1,FKBP4,LSN

RPF3,CBX4,NUP50,SETD1A,HMG20B,DNMT1,SMARCB1,NUP58,SETD1B,BRD1,SIN3A,TOLLIP

3,TNKS,APEX1,NPLOC4,LOX,BAP1,FUS,PPP1R10,BAZ2A,XRCC6,TAOK1,EHMT1,INTS3,UBR5,SMC
)0,CNTNAP1,MMP15,FLOT1,BRD2,SFRP1,PTEN,CEMIP2,CUL3,SPINT2,RPL7L1,ITGB5,CALD1,HIF

I,VHL,PPP2CA,SMARCA4,PSMD3,GRB2,PRKDC,HSPB1,KDM1A,PLCG1,QARS1,CCAR2,DUSP4,DLK
1,SLC39A1,SKIL,RBBP6,SIN3A,CNOT1

2,PCM1,PARK7,IDH2,MAP4K4,PHPT1,ERP29,PTPN1,RAN,GOLPH3,UBE2J1,P3H1,YWHAZ,GSK3B,/

DYNLL1,CDC42,TRAF4,TRAF7,STAT3,BCR,PRKAR2A,HIP1,FLCN,PHB2,UBA52,PLXNB1,SFRP1,RA

5A6,CLUH,PARK7,NDUFC2,FEZ1,KAT2A,PITRM1,GOLPH3,COX7A2L,ATP5PO,STAT2,YWHAZ,GSK
\BL1,PINK1,TAOK2,DAG1,ARRDC4,IGF1R,FZR1,STK39,CAB39,PARK7,ADNP,CD44,MAP4K4,ERP2

IB1,FARP1,SFRP1,RAB3GAP2,MASTL,TBC1D9B,ANP32B,SPINT2,MAPRE2,GPSM1,PPP1R15A,FKE
1,ATXN3,MOAP1,MSH6,NSD2,PRPF19,KDM4A,YY1,APBB1,UBE2D3,CD74,DEK,PAGR1,SLC30A9,F
RNP,EPHA4,KLF3,GRB10,RAP1B,ATP6V1F,LEPROT,ZNF106,KLF10,LDOC1,PIK3R3,ATP6V1D,NAN
HMT2,PHKB,NDUFS1,ATP5F1C,COX6B1,IRS1,NUP205,UQCRFS1,NUP133,ATP5F1E,COX4I1,OGD
COCO1,UBE2L3,FAM210B,ABL1,LGALS1,KDM3A,PPP5C,DAG1,RUVBL2,MDM2,PARK7,ARL6IP5,E
/I5C,EP300,LBH,RERE,KDM5B,SF1,FLCN,H2AZ1,PHB2,UBA52,CXXC5,ATRX,SFRP1,HDAC6,CBX6,

OLA,RHOA,CBX5,ZNF865,ATF4,PSMD2,NR2F1,RPS14,CHD4,ATP1A1,JUND,CD81,CCT7,PFN1,TLE

/I5C,EP300,LBH,RERE,KDM5B,SF1,FLCN,H2AZ1,PHB2,UBA52,CXXC5,ATRX,SFRP1,HDAC6,CBX6,

1A,BMPR2,STAT3,PRMT1,NAP1L1,EP300,LRRC4B,ZFP36L1,MAP6,PLXNB1,DIP2B,SFRP1,SRSF6,I
/I5C,EP300,LBH,RERE,KDM5B,SF1,FLCN,H2AZ1,PHB2,UBA52,CXXC5,ATRX,SFRP1,HDAC6,CBX6,

PSMA4,FHL1,FBXO7,CENPF,PSMC6,CDK5RAP3,ZNF207,HMGA2,WAPL,CNOT1,PPP2R5C

,PDPK1,SH3BP4,ARPP19,GSK3A,WNK1,BAG5,SORT1,MICAL1,PRDX3,USP47,WDTC1,ABL1,MGAT

1,PSMD5,YWHAQ,HACD3,DCHS1,CHSY1,PPP2R1A,PKD1,HLA-C,PDIA3,RPS3,CHD3,VAPA,HMGB1,

34,ABL1,HIP1R,GRIK5,TIA1,NOTCH1,MFN2,EFNB2,PARK7,ARL6IP5,PPP1CA,PDGFRB,CDK4,NCK

K1,HIF1A,NDN,XRCC6,PELP1,CCN1,SSBP3,SOX2,ZMIZ2,GTF2I,NUCKS1,TCF3,FOXK2,OGT,RBPJ,I
FL3,CLIP2,DAG1,KIFC2,HSPA1A,OBSL1,PCM1,MECP2,KAT2A,ATF5,NCKAP5L,RAN,SUN2,TASOR,C
3A,HEXIM1,GMFB,GIT1,EPHB2,APOE,FBXO7,PRNP,SRCIN1,PLPP3,GRB10,MYADM,NF2,CDK5RAF
KHDRBS1,XPO1,SFPQ,SOX12,CCN2,ACTN4,PRDX2,DDX3X,NANOG,H1-0,PRKAR1A,JUN,CTNNB1,
SPAG9,PDPK1,DAZAP2,MTOR,PRKACA,WNK1,CD74,PRDX3,AMBRA1,FGFR1,NTRK2,LMO4,PTPN1

,ZNF865,ATF4,PSMD2,NR2F1,RPS14,CHD4,JUND,CD81,CCT7,PFN1,TLE5,HNRNPC,TADA2B,KHD
KHDRBS1,XPO1,SFPQ,SOX12,CCN2,ACTN4,PRDX2,DDX3X,NANOG,H1-0,PRKAR1A,JUN,CTNNB1,

,ZNF865,ATF4,PSMD2,NR2F1,RPS14,CHD4,JUND,CD81,CCT7,PFN1,TLE5,HNRNPC,TADA2B,KHD

,EXT2,SOBP,LDB1,DEAF1,NDRG4,DLD,SEMA4C,SRF,ZEB1,SOX9,HOXA5,CRABP2,CTR9,COL7A1,
PN11,ABL1,PINK1,TAOK2,DAG1,TSPYL2,MACROH2A1,GGNBP2,PKIA,IGF1R,STK39,CAB39,PARK

KHDRBS1,XPO1,SFPQ,SOX12,CCN2,ACTN4,PRDX2,DDX3X,NANOG,H1-0,PRKAR1A,JUN,CTNNB1,

A6,GIT1,PSMD12,APOE,PSMA4,NUP50,SESN2,FAM3C,PSMC6,GRB10,NUP58,PFKFB3,PPP4R3B
RF4,RAB5B,SRP14,PYGB,NOTCH2,MAPK1,PSMD3,PRDX1,ADAM9,GRB2,PRKDC,CAND1,KDELR1,

,ARF6,MYADM,NF2,ASAP3,GSN,GPM6B,PAM

RL6IP5,PPP1CA,PDGFRB,CDK4,NCK2,C1QBP,TNFRSF10B,RBM10,ARHGEF9,RRP1B,RASSF2,HO
CDK4,NCK2,C1QBP,TNFRSF10B,RBM10,ARHGEF9,RRP1B,RASSF2,HOXA5,BACE1,SOS1,PRMT2
,TGFB2,ARF6,PLXND1,CDK5RAP3,LRP6,NFIB,ID2,MAFB,EPHA2,PAM,RB1CC1

,ZNF865,ATF4,PSMD2,NR2F1,RPS14,CHD4,JUND,CD81,CCT7,PFN1,TLE5,HNRNPC,TADA2B,KHD

N2,GRIK5,NPTN,MAPK8IP2,MECP2,PARK7,ADNP,CYFIP1,KAT2A,ADCY1,SYNGAP1,CRKL,PSMC5,F

12,CCN2,MMP14,PRDX2,DDX3X,RPS19,MIF,GPAT4,GJA1,JUN,CTNNB1,ANXA6,MACF1,NDFIP1,HN
/IYADM,NF2,CDK5RAP3,LRP6,NNT,IBTK
L

JANOG,H1-0,PRKAR1A,JUN,CTNNB1,ARID1A,MXD4,SHOX,HNRNPK,H1-10,DUX4,YWHAB,ZNF454,
2N1,RPS6KA2,MDC1,PSMA6,CNOT3,PCNA,TFDP1,HEXIM1,PPP2R5B,PSMD12,MSH2,RYBP,PSMA

)TCH2,RAPGEF1,MAPK1,ARPC2,PSMD3,PRDX1,GRB2,PRKDC,CAND1,PLCG1,SUPT6H,FYN,ILF2,(
/IAPK3,PCNA,TFDP1,GNL3,MSH2,HMGA2,RAD50
JANOG,H1-0,PRKAR1A,JUN,CTNNB1,ARID1A,MXD4,SHOX,HNRNPK,H1-10,DUX4,YWHAB,ZNF454,
JANOG,H1-0,PRKAR1A,JUN,CTNNB1,ARID1A,MXD4,SHOX,HNRNPK,H1-10,DUX4,YWHAB,ZNF454,

GLB1,CNOT1
31,ID3,LRP6,KLF10,GPM6B,EPHA2,MYBBP1A

MECP2,PARK7,ADNP,CYFIP1,KAT2A,ADCY1,SYNGAP1,PSMC5,PLK2,SRF,PIP5K1C,GSK3B,SYNGF

1A,ACIN1,PAF1,MYL9,CDK5RAP3,ID2,BRD1,KLF10,PITHD1,ITPKB,MAFB,EPHA2,SIN3A

PTEN,SPINT2,MAPRE2,APEX1,HIF1A,SLIT2,AKAP9,GRN,CCN1,TCAF1,AKT3,GNA12,RDX,MAZ,STK

7,ADNP,CYFIP1,KAT2A,ADCY1,SYNGAP1,PSMC5,PLK2,SRF,PIP5K1C,GSK3B,SYNGR1,SERPINE2,(
7,ADNP,CYFIP1,KAT2A,ADCY1,SYNGAP1,PSMC5,PLK2,SRF,PIP5K1C,GSK3B,SYNGR1,SERPINE2,(
TNAP2,VEGFA,RAPGEF1,PPP2CA,ARPC2,ADAM9,GRB2,ACTN1,HSPB1,CORO1C,FYN,FBLN1,STXE

FA,RAPGEF1,PPP2CA,ARPC2,ADAM9,GRB2,ACTN1,HSPB1,CORO1C,FYN,FBLN1,STXBP1,PLXNA3

'RKAR2B,MYL12A,ERBB2,ITPK1,CFLAR,RAB5A,ANXA1,TGFBR1,MAPK3,PPP3CA,CAPZA2,XBP1,EX
,SYT15,ADCY1,NDRG4,GSK3B,CACNA1H,RAB3A,KIF3A,DLG4,BACE1,RAB5A,ANXA1,MAPK3,PPP3

RAC1,APPL1,SMCHD1,ATP6V0C,SKP1,BCLAF1,PEA15,DCN,DLG5,TIMP3,PARP1,ACTR3,PKN1,NCK

DGFRB,CDK9,BCL9,UNC45A,SEMA4C,SRF,ZEB1,CACNA1H,USP19,SOX9,LARGE1,LMNA,NR2F2,C

RB,CDK4,ADCY1,DHX36,RTF2,LOXL2,NDRG1,HSF1,DHFR,CACNA1H,SETX,TXN,DLG4,CDKN1B,ST
DKN1A,CPSF2,EID1,CHD6,MLXIP,CPSF1,SQSTM1,NOP53,PABPN1,PARP1,MORF4L2,TAF15,SSRP
EF9,RALBP1,RHOBTB2,CDC42EP4,CDC42SE1,SOS1,ARHGAP33,SHC1,ROCK2,HEG1,KCTD10,AF

T

,SOD2,SPAG9,CLASP2,PDPK1,ATP1B2,ITGAV,ADAM10,AKT2,VASH1,MTOR,WNK1,CERS2,DAPK3,

1,TCAF1,AKT3,GNA12,RDX,MAZ,STK24,SEMA3C,SOD2,SPAG9,CLASP2,PDPK1,ATP1B2,ITGAV,AD

ST,SPINT2,CALD1,HIF1A,ATP6AP2,LOX,SLIT2,GRN,CCN1,GPX1,HSPG2,SSBP3,PKDCC,AKT3,LDL

,CD74,PRDX3,DEK,PTPN11,FAM210B,ABL1,LGALS1,PSMB1,KLF13,HSPA1A,MKNK2,ACVR1B,SM

F10,VEGFA,CCND1,MAPK1,LUZP6,PRKDC,HSPB1,KDM1A,PLCG1,B2M,PLXNA3,SOD1,RAB11A,AF

REBBP,PSMC1,CD74,PRDX3,DEK,LMO4,PTPN11,FAM210B,ABL1,LGALS1,PSMB1,KLF13,HSPA1A

PSMA7,VDAC1,BCR,EP300,LBH,KDM5B,PSMD1,PBX1,AP2A1,GRSF1,PHB2,ZFP36L1,BRD2,HOXB5

,MORF4L2,ACTR3,PKN1,YEATS2,PTOV1,ZNF302,RBMX,CHD2,GCN1,SKI,ASXL1,PSMB2,KDM6B,

NDST1,ANP32B,CALD1,HIF1A,GALNT11,FKBP1A,LOX,SLIT2,GRK2,GRN,CCN1,GPX1,HSPG2,OLF

GLUD1,OAZ2,ANP32B,HIF1A,SERP1,AKAP9,GRK2,BAP1,TCAF1,UBR5,VPS4A,RDX,TM9SF4,SH3GL

,RHOA,ATF4,PSMD2,KIF5B,SLC7A5,RPS14,CXADR,JUND,CD81,WDR1,HMGN2,VAT1,CLEC7A,KHD

,MSH6,NSD2,RBPJ,PSMC3,SART1,MTOR,YY1,CREBBP,PSMC1,CD74,PRDX3,DEK,LMO4,PTPN11

RA1,FGFR1,UBE2K,EIF4G1,NTRK2,LMO4,PTPN11,ABL1,FXR2,SDCBP,PINK1,TAOK2,DAG1,ACVR

A12,RDX,MAZ,STK24,SEMA3C,SOD2,SPAG9,CLASP2,PDPK1,ATP1B2,ITGAV,ADAM10,AKT2,VASH
DTC1,MTA2,CGGBP1,RTF1,MACROH2A1,HSPA1A,SIN3B,PKIA,NOTCH1,MDM2,FOXP4,MECP2,D

,MDK,LAMC1,SKI,RPS15A,NOTCH2,RNF10,VEGFA,SELENON,RAPGEF1,VHL,CCND1,MAPK1,PRD

,CDKN1B,ERBB2,LMNA,CACYBP,MED12,SOS1,CITED2,NRP2,SHC1,ROCK2,RPS6KA2,MAP2K4,TG

,MAPRE3,TNFRSF10B,UBE2V1,ZFP91,RASSF2,TXN,DLG4,CDKN1B,IRS1,PRKAR2B,ERBB2,PRRC

AZAP2,AKT2,GSK3A,MTOR,PRKACA,WNK1,CD74,AMBRA1,FGFR1,UBE2K,EIF4G1,NTRK2,LMO4,F
AZAP2,AKT2,GSK3A,MTOR,PRKACA,WNK1,CD74,AMBRA1,FGFR1,UBE2K,EIF4G1,NTRK2,LMO4,F

2,PRDX6,GRN,ACLY,XRCC6,SLC44A2,IMPDH2,RAB14,GNL1,KMT2E,NCSTN,RAB6A,RAB5C,PLEKH

,TRIM28,YWHAG,COL4A2,RPL37A,MAPT,FN1,DDB1,PSMD5,YWHAQ,DCHS1,SET,MAP9,TMSB10,I
RPS4X,RPS3A,RPL18,ITGB1,TUBB4B,UCHL1,RPL11,ANXA2,AGRN,RPLP1,TIMP2,RPL10,RPS17,V
,IK,RPS19,RPL35,TLN1,JUN,CTNNB1,RPS10,RPL35A,TARDBP,RPLP2,DDOST,ST3GAL2,HNRNPK,
YWHAQ,DCHS1,SET,MAP9,TMSB10,INA,PPP2R1A,GTF3C4,DNAH8,PKD1,SF3B1,RPS3,CHD3,VAPA
RPS14,RPS20,CD81,RPL26,CCT7,OAZ1,ASTN2,RPL31,XPO1,PTN,CSNK1E,RPS25,EIF5A,CDH2,GP
RPS14,RPS20,CD81,RPL26,CCT7,OAZ1,ASTN2,RPL31,XPO1,PTN,CSNK1E,RPS25,EIF5A,CDH2,GP
F2,SUGP2,CPSF1,PABPN1,HNRNPL,TAF15,ALKBH5,RPL27,SNRPN,HNRNPA3,AHCYL1,RBMX,DDX
XPO1,SPTBN2,RPS25,SFPQ,EIF5A,RAB7A,ACTN4,RPS15,RPL12,RPL30,ARL4C,CLTC,RPS5,RPS7,
C,H1-0,RPS5,RPS7,RPS19,RPL35,GJA1,CTNNB1,RPS10,RPL35A,TARDBP,RPLP2,NDFIP1,CIRBP,
ARF1,CYB5R3,PAFAH1B1,MAP4,AP2B1,SLC4A8,HNRNPU,PGAM1,RPS16,RPL23,RPL29,RPSA,CT
,KPNB1,RPL17,LLGL1,RPL14,HK1,RPL22,RPL9,CLSTN1,KIF5B,RAB6B,NLGN2,RPS14,RPS20,CD
Q,TPM1,EIF5A,CDH2,GPC3,DDR1,PLXNA1,PIP4K2B,MEGF8,RAB7A,CCN2,MMP14,DDX3X,ITM2C,
S24,RPS9,XRCC5,MSN,RPL24,KPNB1,RPL17,LLGL1,RPL14,HK1,RPL22,RPL9,CLSTN1,KIF5B,RAE
,RPLP2,NDFIP1,CIRBP,RPL36,YWHAB,SND1,UBE2H,RPS12,RPS27,GPC1,SKP1,BCLAF1,LAPTM4
UL,PTTG1IP,MAP1A,CDKN1A,SRP9,TOMM20,CCT2,VAMP2,NOP53,PARP1,RPL27,HSPA4,TCP1,C
A7,ANP32A,EDC4,PSMD1,GTPBP1,GSPT1,UBA52,ZFP36L1,UPF1,RPS2,RPS27A,PSMD8,APEX1,PC
T,FN1,PSMD5,YWHAQ,DCHS1,TMSB10,MRC2,RPL19,PPP2R1A,RPL15,DNAH8,PKD1,HLA-C,RPS3,
MIZ1,TIMP2,RPL10,MARCHF6,WSB1,RPS17,SBF1,DNAJA1,CDC42BPB,AP2B1,OGA,LARP1,RPS16
DC42BPB,TMEM170A,GOLGA8A,HNRNPU,SMARCA5,CTNNA1,CNN3,RPSA,TPM4,SCARB2,RTN4,L

13,RPL34,PSMA7,ANP32A,EDC4,PSMD1,GTPBP1,GRSF1,GSPT1,UBA52,ZFP36L1,UPF1,RPS2,RP
AT2A,TMEM170A,PXDN,HNRNPU,SMARCA5,CTNNA1,RPS16,RPL23,RPL29,RPSA,EIF4H,RTN4,LA
2,RPL30,ARL4C,CLTC,RPS5,RPS7,RPS19,RPL35,GJA1,MYO5A,CD63,MACF1,RPS10,RPL35A,TAR
A,CDKN1A,SRP9,NUCB1,TOMM20,RAB27A,VAMP2,RPL27,TBC1D16,HSPA4,AHCYL1,VPS35,RAB5

PS3A,RPL18,UCHL1,RPL11,MORF4L1,ANXA2,RPLP1,ZMIZ1,TIMP2,COL11A1,RPL10,MARCHF6,W
L,RPS25,EIF5A,RAB7A,ACTN4,RPS15,RPL12,RPL30,ARL4C,CLTC,RPS5,RPS7,RPS19,RPL35,GJA1
JSF2,RSF1,MGAT5,SUMO1,RPS21,DAD1,MOGS,RPL27A,TRIM27,NUP153,CDK9,EIF3B,NUP98,GS

LD3,HSPD1,USP11,SNRNP200,CSDE1,IGF2,EIF3C,ATP5F1A,EIF4B,VCP,UBB,AFF4,TRIM28,YWHA
9,MSN,RPL24,KPNB1,RPL17,RPL14,HK1,RPL22,RPL9,PSMD2,ENC1,RPS14,TMEM259,RPS20,CD
L26,MCL1,RPL31,HNRNPC,LRPPRC,XPO1,CSNK1E,RPS25,PIP4K2B,RAB7A,PRDX2,RPS15,RPL12
,PSMD5,YWHAQ,SET,CHSY1,TMSB10,RPL19,PPP2R1A,RPL15,RPS3,CHD3,BASP1,HMGB1,AP2M1
L27A,NCK2,EIF3B,NCBP2,EIF3M,RPS29,PAIP1,EIF1B,ABCE1,RPL32,EIF2S1,CTIF,EIF3J
MARCC1,ATP6V1B2,FYN,HNRNPM,UBQLN1,CSNK1D,USP7,IDH1,HECTD1,HGS,BAG6,SREBF2,H
GJA1,MYO5A,CD63,MACF1,RPS10,RPL35A,TARDBP,RPLP2,ARL6IP1,NDFIP1,WLS,MYH10,RPL36,
1,RPS4X,RPS3A,RPL18,ITGB1,TUBB4B,UCHL1,RPL11,BTF3,ANXA2,RPLP1,TIMP2,SARAF,MIDN,R
PS,BNIP3L,RPL28,RPS28,YWHAH,FAU,CCT8,HGS,SREBF2,RAB11A,RPL39,RPS13,TXNIP,RPL34,C
3A,RPL18,ITGB1,TUBB4B,UCHL1,RPL11,BTF3,ANXA2,RPLP1,TIMP2,SARAF,MIDN,RPL10,RPS17,
1,COL1A2,LLGL1,GAP43,RHOA,CLSTN1,PSMD2,NLGN2,RPS14,EIF5,WDR1,ATP2A2,RBBP4,RTN3
RPS14,TMEM259,RPS20,CD81,RPL26,OAZ1,RPL31,HNRNPC,LRPPRC,XPO1,CSNK1E,PFKL,RPS25

4A1,RPL19,PPP2R1A,RPL15,GTF3C4,PKD1,PDIA3,SERINC1,RPS3,VAPA,HMGB1,CDKL1,AP2M1,R
PLP1,RPS7,HMGCS1,GJA1,CTNNB1,NGRN,MACF1,CYFIP2,ARID1A,WLS,ATP1A2,MAP2,ARCN1,HM
S35,RPL36A,RAB11B,SRP14,TMED2,RPS15A,RPL38,OPTN,RPL37,RPS11,CLIP3,RPL28,RPS28,FYI
P1,ZMIZ1,TIMP2,AP3D1,DNAJA1,ARF1,PAFAH1B1,AKAP12,ZNF770,AP2B1,GTF2F1,SLC4A8,HNRN
HAH,SLIT3,STMN3,B2M,STXBP1,PLXNA3,PTPRS,SOD1,RAB11A,TCTN1,BMPR2,CDC42,PPP1R9B,F
TM9SF2,RPL27,HSPA4,VPS35,RPL36A,RAB11B,SRP14,TMED2,RPS15A,RPL38,OPTN,RPL37,RPS
),SERBP1,PSMA7,ANP32A,VDAC1,STAT3,ARIH1,EDC4,SGTA,EP300,PSMD1,FLCN,GTPBP1,UBQLN
8,HMGA2
,PPP2R1A,RPL15,PKD1,SF3B1,SSBP2,ZNF43,RPS3,CHD3,BASP1,HMGB1,AP2M1,RPS4X,RPS3A,R
FAU,HNRNPD,RPL39,SERBP1,RPS13,RPL34,PSMA7,ANP32A,DFFB,EDC4,PSMD1,GTPBP1,GRSF1,
IORF4L1,ANXA2,TIMP2,MIDN,RPL10,VCL,DNAJA1,ARF1,PAFAH1B1,MAP4,ITM2B,AP2B1,PXDN,HN
,RPL24,RPL17,IGFBP3,RPL14,RPL22,RHOA,CBX5,RPL9,ATF4,PSMD2,NR2F1,SLC7A5,ENC1,RPS1
,EIF5A,PEBP1,CCND2,GPC3,RAB7A,CCN2,MMP14,PRDX2,DDX3X,RPS15,ITM2C,RPS7,MIF,PRKAF
PEN,NONO,DDB1,PSMD5,YWHAQ,EIF1,HACD3,SET,EGLN1,CHSY1,EIF4A1,RPL19,PPP2R1A,RPL1
,RPL22,CBX5,RPL9,ATF4,PSMD2,NR2F1,SLC7A5,ENC1,RPS14,RPS20,RPL26,BEX3,SRSF4,RBBP
P,CDKN1A,CTDSP2,DDX39B,CHD6,SQSTM1,NOP53,PARP1,MORF4L2,NOX4,SSRP1,HSPH1,ALKB
1,PXDN,HNRNPU,SMARCA5,LARP1,HNRNPA0,CTNNA1,RPS16,RPL23,LDHA,LAMA5,XRCC5,MSN,C
.1,RAC1,SKP1,RPL23A,LUC7L3,MAP1A,DLG5,DDX39B,SRSF5,CCT2,TBCD,VAMP2,GAS7,NOP53,F
N10,KDM1A,PLCG1,ARF4,FYN,YWHAH,SLIT3,STMN3,B2M,STXBP1,PLXNA3,PTPRS,SOD1,RAB11A,
PRE1,ARPC2,DSTN,ADAM9,GRB2,CAND1,KDM1A,CLIP3,CORO1C,FYN,YWHAH,CCT8,B2M,TNKS2
T1,CDH11,TBCD,GAS7,VCAN,PTPRA,RTN1,CAMSAP1,NCKAP1,ACAP3,MDK,PTK7,SKI,CNTNAP2,AE
,SNRNP200,CSDE1,IGF2,EIF3C,ATP5F1A,EIF4B,VCP,UBB,AFF4,TRIM28,YWHAG,COL4A2,RPL37A
NPM,SSB,FAU,HNRNPD,RPL39,SERBP1,RPS13,RPL34,PSMA7,ANP32A,DFFB,EDC4,PSMD1,GTPBF
NPM,SSB,FAU,HNRNPD,RPL39,SERBP1,RPS13,RPL34,PSMA7,ANP32A,DFFB,EDC4,PSMD1,GTPBF
2,TUT4,RNPS1,PPP2R2A,CTIF,CNOT1
SDE1,IGF2,EIF3C,ATP5F1A,EIF4B,VCP,UBB,AFF4,TRIM28,YWHAG,CKB,COL4A2,RPL37A,MAPT,FN
NTNAP2,DPYSL2,NOTCH2,VEGFA,RAPGEF1,MAPK1,PBX2,LUZP6,GRB2,ATXN10,KDM1A,PLCG1,A
SAP1,NCKAP1,ACAP3,MDK,PTK7,SKI,CNTNAP2,DPYSL2,NOTCH2,RNF10,VEGFA,RAPGEF1,TUBB2
H,LRPPRC,KHDRBS1,XPO1,SLC3A2,PFKL,HDLBP,RPS25,EIF5A,GPC3,RAB7A,ACTN4,RPS15,RPL1
F3B1,SSBP2,ZNF43,RPS3,CHD3,BASP1,HMGB1,AP2M1,RPS4X,RPS3A,RPL18,UCHL1,RPL11,MOR
EPL,RPS2,RPS27A,SERP1,TCAF1,PKDCC,TRAM1,SRP72,SSR3,NMT1,MOAP1,REEP2,SCRIB,AKT2,
I,RPS4X,STMN1,ITGB1,RPL11,ANXA2,AGRN,RPLP1,ZMIZ1,TIMP2,AP3D1,DNAJA1,ARF1,PAFAH1B
ZMIZ1,TIMP2,MIDN,RPL10,AP3D1,DNAJA1,ARF1,ITM2B,ZNF770,GTF2F1,HNRNPU,PGAM1,NFE2I
A2,TMED2,RPS15A,RPL38,MARS1,RPL37,MAPK1,EIF3E,FTO,ILF3,LUZP6,PCSK1N,RPS11,PRKDC,I
MIF,PRKAR1A,CTNNB1,NGRN,CYFIP2,APLP2,TARDBP,ARL6IP1,NDFIP1,CIRBP,YWHAB,CADM4,RA
1,BAP1,TCAF1,VPS41,TRAM1,VPS4A,SRP72,SSR3,AKT2,GSK3A,CHMP4B,SSR2,EXOC4,NDUFA13,
FAU,HNRNPD,RPL39,SERBP1,RPS13,RPL34,PSMA7,ANP32A,DFFB,EDC4,PSMD1,GTPBP1,GRSF1,
1C,FYN,YWHAH,SLIT3,FBLN1,STXBP1,PLXNA3,PTPRS,SOD1,RAB11A,TCTN1,PRPF40A,BMPR2,CD
NPM,SSB,YWHAH,FAU,HNRNPD,RPL39,SERBP1,RPS13,RPL34,PSMA7,ANP32A,DFFB,EDC4,PSMD
T2,NDN,TAOK1,SLIT1,OLFM1,LINGO1,ABI2,DCLK1,CNP,SEMA3C,SPAG9,DICER1,CLASP2,WASF2
OK1,SLIT1,OLFM1,LINGO1,ABI2,DCLK1,CNP,SEMA3C,SPAG9,DICER1,CLASP2,WASF2,ADAM10,I
T2,NDN,TAOK1,SLIT1,OLFM1,LINGO1,ABI2,DCLK1,CNP,SEMA3C,SPAG9,DICER1,CLASP2,WASF2
D5,YWHAQ,DCHS1,EGLN1,TPI1,CHSY1,INA,MRC2,RPL19,PPP2R1A,PKD1,BASP1,VAPA,HMGB1,CD
S9,CAPN2,MTA1,PSMD1,AP2A1,UBQLN2,CSNK2A1,CST3,UBA52,PPP2CB,KLHL42,UBE2G2,FBXO
APK1,EIF3E,FTO,ILF3,LUZP6,RPS11,PRKDC,HSPB1,CASC3,QARS1,RPL28,RPS28,LARS1,FAU,EIF

AS7,NOP53,PARP1,TSPYL4,ABCA3,ACTR3,HSPA4,TCP1,CAMSAP1,CCT3,NCKAP1,RBMX,LAMC1,
CALM1,NPM1,PLXNB2,TTC3,PLD3,HSPD1,USP11,SNRNP200,CSDE1,IGF2,EIF3C,ATP5F1A,EIF4B,
RA,ACTR3,CAMSAP1,NCKAP1,VPS35,ACAP3,MDK,PTK7,TNPO1,CNTNAP2,DPYSL2,NOTCH2,VEG
42,SNRNP70,RPL36A,SCAF1,PRPF6,RPL38,RPL37,PPP2CA,SRSF9,SRSF1,PTBP1,PRKDC,SUPT5
1,CTNNA1,RPS16,RPL23,RTN4,LAMA5,XRCC5,MSN,COL1A2,RHOA,ATF4,PSMD2,KIF5B,CXADR,
1,OLFM1,LINGO1,ABI2,DCLK1,CNP,SEMA3C,SPAG9,DICER1,CLASP2,ADAM10,IGF2BP1,IST1,UB
2,PSMD3,SRSF1,EIF3E,LUZP6,GRB2,CAPZB,PRKDC,SCAF11,CAND1,NSG2,CLIP3,DHX30,RPS28,
B,AFF4,TRIM28,MATR3,YWHAG,COL4A2,RPL37A,MAPT,FN1,SPEN,NONO,DDB1,PSMD5,YWHAQ,
IDN,RPL10,AP3D1,DNAJA1,ARF1,ITM2B,ZNF770,GTF2F1,HNRNPU,PGAM1,NFE2L1,SMARCA5,L

6A,NBAS,HNRNPR,SRSF10,DDX6,C1QBP,LSM4,MAPKAPK2,CDK9,CELF3,PSME3,DHX36,RBFOX2
,PTPRA,ACTR3,CAMSAP1,NCKAP1,VPS35,ACAP3,MDK,PTK7,TNPO1,CNTNAP2,DPYSL2,NOTCH2,
APK1,EIF3E,FTO,ILF3,LUZP6,RPS11,PRKDC,HSPB1,CASC3,QARS1,RPL28,RPS28,LARS1,FAU,EIF
5B,ATP1A1,COL5A2,MCL1,CLEC7A,PTN,GLG1,PDIA6,GPC3,SULF1,PIP4K2B,MEGF8,CCN2,ACTN
SP1,VAPA,HMGB1,CDKL1,AP2M1,TPT1,SEC61A1,RPS4X,STMN1,ITGB1,UCHL1,BTF3,NCAM1,AN
NND2,EVL,HDAC6,PTEN,CUL3,ULK1,SPINT2,BRSK1,CTTN,SLIT2,NDN,SLIT1,OLFM1,ANTXR1,LIN
6,TPM1,CDH2,GPC3,DDR1,SULF1,PLXNA1,MEGF8,SOX12,CCN2,ACTN4,MMP14,CLTC,NANOG,A
D2,EVL,HDAC6,PTEN,ULK1,BRSK1,CTTN,SLIT2,NDN,TMOD2,TAOK1,SLIT1,OLFM1,LINGO1,ABI2,
1,IST1,UBE3A,TMEFF1,APBB1,TANC2,SMO,METRN,LYPLA2,NTRK2,DIP2A,PTPN11,ABL1,SEMA6
23,RTN4,LPCAT1,RPS24,SLC22A17,RPS9,XRCC5,MSN,COL1A2,HK1,NCS1,RHOA,CBX5,CLSTN1,
19,SF3A2,SART1,DHX38,RBM6,SAP18,THRAP3,SLC38A2,HNRNPH3,EIF4A3,RPS26,HTATSF1,FXR
,SEC61A1,RPS4X,STMN1,ITGB1,UCHL1,BTF3,NCAM1,ANXA2,AGRN,ZMIZ1,TIMP2,COL11A1,RPL
14,SNX1,HIPK2,MYCBP2,MAGEF1,AREL1,RNF41,TRIB3,PSME3,STUB1,MARCHF7,PSMC5,CSNK1
1,DNAJA1,ITM2B,ZNF770,GTF2F1,HNRNPU,PGAM1,NFE2L1,SMARCA5,LARP1,HNRNPA0,RPL23,
15A,RPL38,MARS1,RPL37,MAPK1,EIF3E,FTO,ILF3,LUZP6,RPS11,PRKDC,HSPB1,CASC3,QARS1,
ZMIZ1,TIMP2,COL11A1,RPL10,VCL,AP3D1,ARF1,PAFAH1B1,MAP4,ITM2B,AP2B1,HNRNPU,CTNN
,PLXNB2,TTC3,PLD3,HSPD1,USP11,SNRNP200,CSDE1,IGF2,EIF3C,ATP5F1A,EIF4B,VCP,UBB,AFF
,RBM25,SF3A3,CD2BP2,FXR1,SART3,SYNCRIP,HNRNPR,SRSF10,C1QBP,LSM4,CELF3,RBFOX2,S
,RBM25,SF3A3,CD2BP2,FXR1,SART3,SYNCRIP,HNRNPR,SRSF10,C1QBP,LSM4,CELF3,RBFOX2,S
,RBM25,SF3A3,CD2BP2,FXR1,SART3,SYNCRIP,HNRNPR,SRSF10,C1QBP,LSM4,CELF3,RBFOX2,S
PRKAR1A,CTNNB1,GOLM1,B4GALNT4,APLP2,TARDBP,NDFIP1,DDOST,ST3GAL2,YWHAB,CADM4
PRKAR1A,CTNNB1,GOLM1,B4GALNT4,APLP2,TARDBP,NDFIP1,DDOST,ST3GAL2,YWHAB,CADM4
UTS2,TAOK2,TRAK1,DAG1,NIBAN2,PLXNA2,NOTCH1,NPTN,EFNB2,JAM3,ADNP,CYFIP1,FEZ1,NT
A,HNRNPC,TADA2B,KHDRBS1,CSNK1E,SFPQ,EIF5A,CCND2,GPC3,DDR1,SULF1,PIP4K2B,SOX12
19,SF3A2,SART1,DHX38,PRKACA,RBM6,SAP18,THRAP3,HNRNPH3,EIF4A3,XRN2,HTATSF1,FXR2
D2,BAP1,PSMB4,GPX1,EIF3H,TPP1,UBE2D2,VPS4A,LDLR,GNA12,RDX,DNAJB2,ATXN3,PSMD4,OG
P2,RAB1A,RAC1,ARHGDIA,CAPRIN1,GPC1,CDK16,TOP2B,HDAC2,TNC,TMEM30A,DNER,L1CAM,
TN,LUZP6,GRB2,CAPZB,KDM1A,BNIP3L,CLIP3,MAP2K2,YWHAH,CCT8,STMN3,TNKS2,STXBP1,P
P,RPL36,YWHAB,SND1,RPS12,RPS27,CTBP1,RAC1,CAPRIN1,SMCHD1,HDAC2,BCLAF1,LAPTM4
DX39B,SRP9,EEF1A2,MTRF1L,RPL27,RTN1,GCN1,RPL36A,ABCA2,TMED2,RPS15A,RPL38,MARS
,WBP11,PRPF19,NOC2L,EIF3G,SF3A2,SART1,EIF4A3,XRN2,LUC7L2,NOP56,MTREX,RUVBL2,SF3
1A,JUN,CTNNB1,CD63,NGRN,SESN3,CYFIP2,TARDBP,ARID1A,NDFIP1,SHOX,HNRNPK,H1-10,CIR
PTBN1,ATP5F1B,CALM3,GNB1,SHISA9,STMN2,RPS8,MYH9,RPL7,SEPTIN2,P4HB,APP,RPL5,KLRD
ACF1,CYFIP2,DNAJC5,APLP2,DDOST,HNRNPK,LAMP1,RHOB,LGALS3BP,CKAP4,HLA-B,RAB1A,T
NTN1,LIMK1,CCDC88A,RAP2A,NME2,SYNGAP1,ISLR2,DHX36,CRKL,NDRG4,SPOCK1,GOLGA4,S

SNK1E,SFPQ,EIF5A,CCND2,GPC3,SULF1,SOX12,RAB7A,CCN2,ACTN4,MMP14,PRDX2,DDX3X,NAN
6IP1,NDFIP1,MXD4,HNRNPK,H1-10,CIRBP,YWHAB,CADM4,CTBP1,HMGCR,CAPRIN1,SMCHD1,HD
N,EFNB2,ADNP,CYFIP1,FEZ1,NTN1,LIMK1,B4GALT5,EFNB1,MYCBP2,NECTIN1,ADCY1,SYNGAP1,I
57,PRKAR1A,GJA1,JUN,CTNNB1,APLP2,TARDBP,ARL6IP1,NDFIP1,MXD4,HNRNPK,H1-10,CIRBP,Y
SK3A,SF3A2,MTOR,UBE3A,APBB1,TANC2,CERS2,BAG5,METRN,GAK,CROCC,NTRK2,DENND5A,S
AMP2,GNAI3,PARP1,NOX4,ABCA3,RPL27,SNRPN,PTPRA,PKN1,AHCYL1,VPS35,COL2A1,MDK,SKI
HD1,GSK3A,SF3A2,MTOR,UBE3A,APBB1,TANC2,CERS2,BAG5,METRN,GAK,CROCC,NTRK2,DENN
PTTG1IP,CDKN1A,DLG5,TIMP3,CTSB,EEF1A2,ANXA5,SQSTM1,GNAI3,NOP53,PARP1,NOX4,PRDX
GJA1,JUN,CTNNB1,NGRN,CYFIP2,TARDBP,ARID1A,NDFIP1,SHOX,HNRNPK,H1-10,CIRBP,DUX4,Y
BASP1,HMGB1,POLR2A,RPS4X,RPS3A,RPL18,RPL11,MORF4L1,BTF3,AGRN,RPLP1,ZMIZ1,RPL10
RN,RPS10,RPL35A,TARDBP,RPLP2,DDOST,ST3GAL2,CIRBP,NARS1,RPL36,RPS12,RPS27,CAPRIN
A1,CNN3,TPM4,SCARB2,NACA,RTN4,LAMA5,XRCC5,MSN,RPL24,IGFBP3,LLGL1,GAP43,NCS1,R
TGB1,UCHL1,NCAM1,ANXA2,AGRN,ZMIZ1,TIMP2,XAF1,MARCHF6,DNAJA1,ARF1,CYB5R3,PAFAH1
TF3C4,PKD1,SF3B1,SSBP2,ZNF43,RPS3,CHD3,BASP1,HMGB1,POLR2A,RPS4X,RPS3A,RPL18,RPL
N3,TPM4,SCARB2,NACA,RTN4,LAMA5,XRCC5,MSN,RPL24,IGFBP3,LLGL1,GAP43,NCS1,RPL22,R
NPA0,RPS16,RPL23,RPL29,RPSA,NACA,RPS24,BRD3,RPS9,XRCC5,CIPC,RPL24,RPL17,PAPOLA,R
SERINC1,SSBP2,ZNF43,RPS3,CHD3,BASP1,VAPA,HMGB1,POLR2A,RPS4X,RPS3A,RPL18,RPL11,M
SERINC1,SSBP2,ZNF43,RPS3,CHD3,BASP1,VAPA,HMGB1,POLR2A,RPS4X,RPS3A,RPL18,RPL11,M
PRKAR1A,CTNNB1,GOLM1,B4GALNT4,APLP2,TARDBP,NDFIP1,DDOST,ST3GAL2,YWHAB,CADM4
RPL18,RPL11,MORF4L1,BTF3,AGRN,RPLP1,ZMIZ1,RPL10,RPS17,AP3D1,SF3B2,ZNF770,GTF2F
CN,CSNK2A1,ZFP36L1,DIP2B,ATRX,GNG4,SFRP1,HDAC6,PTEN,ULK1,HIF1A,CTTN,SLIT2,SERP1,
PL1,HDAC2,BCLAF1,PEA15,PTTG1IP,CDKN1A,DLG5,TIMP3,CTSB,EEF1A2,ANXA5,SQSTM1,GNAI
32,ZNF770,GTF2F1,HNRNPU,NFE2L1,SMARCA5,LARP1,HNRNPA0,RPS16,RPL23,RPL29,RPSA,NA
AP3,PITHD1,RNF40
3,AGRN,RPLP1,ZMIZ1,RPL10,RPS17,AP3D1,AKAP12,ITM2B,ZNF770,GTF2F1,HNRNPU,OGA,NFE
1,DNAJB2,SEMA3C,CLASP2,WASF2,ITGAV,SYT11,NOC2L,KDM4A,GSK3A,CHMP4B,UBE3A,BNIP3,
RAB3GAP2,HDAC6,PTEN,ULK1,SEPTIN7,CTTN,SLIT2,GPC4,BRK1,AKAP9,TMOD2,SLIT1,PALM,CC
,HGS,PCBP2,BAG6,CYP51A1,PCYOX1,YME1L1,PSMA7,ARIH1,SGTA,OS9,CAPN2,MTA1,PSMD1,A
P1,ZMIZ1,RPL10,RPS17,AP3D1,AKAP12,ZNF770,GTF2F1,HNRNPU,NFE2L1,SMARCA5,LARP1,HN
NDFIP1,MYH10,RHOB,HLA-B,MAP2,HMGCR,RAC1,ARHGDIA,CAPRIN1,GPC1,HDAC2,L1CAM,TRIB
T2,GSK3A,PRKACA,UBE2D3,TRIM8,SAE1,SMO,CROCC,RANGAP1,UBE2L3,PTPN11,PINK1,DAG1,
PSMC3,CHMP4B,UBE3A,PRKACA,UBE2I,UBE2D3,PSMC1,CCDC47,BAG5,UBXN6,UBE2K,PTPN23,
ZMIZ1,RPL10,RPS17,AP3D1,ITM2B,ZNF770,GTF2F1,HNRNPU,OGA,NFE2L1,SMARCA5,LARP1,RP
4,YWHAB,LAMP1,RHOB,HMGCR,ARHGDIA,APPL1,HDAC2,BCLAF1,PEA15,PTTG1IP,CDKN1A,DLG
,USP14,PSMD7,FAF2,MAEA,CUL4A,UBE2J1,AREL1,RNF41,FAM8A1,TRIB3,GID8,PSME3,HM13,ST
3,KIF26A,ADNP,CYFIP1,RNF157,NTN1,LIMK1,ISLR2,CRKL,SPOCK1,GOLGA4,SEMA4C,SRF,GSK3
PK1,SMARCA4,DYNC1LI2,ADAM9,PTBP1,GRB2,PRKDC,KDM1A,NSG2,FKBP8,PLCG1,POLR2B,GN
SN,SPTBN1,ATP5F1B,CALM3,GNB1,SHISA9,STMN2,RPS8,MYH9,RPL7,SEPTIN2,P4HB,APP,RPL5,K
EC7A,NISCH,PTN,CSNK1E,PFKL,GLG1,SFPQ,PEBP1,CDH2,GPC3,SULF1,PIP4K2B,MEGF8,RAB7
11A,CDC42,PPP1R9B,DCTN1,HIP1,CST3,UBA52,SFRP1,EVL,CRTAP,HDAC6,CUL3,ITGB5,CALD1,F

,RPS4X,RPS3A,RPL18,RPL11,MORF4L1,BTF3,AGRN,RPLP1,ZMIZ1,RPL10,RPS17,AP3D1,SF3B2,
HD3,BASP1,VAPA,HMGB1,POLR2A,RPS4X,RPS3A,RPL18,RPL11,MORF4L1,BTF3,AGRN,RPLP1,Z
42,NOTCH2,VEGFA,VHL,MAPK1,DDAH2,CDC37,PRKDC,ACTN1,HSPB1,SON,KDM1A,BNIP3L,CLIP
,SEPTIN7,PSMD8,HIF1A,SLIT2,GPC4,ZMYM3,GRN,PSMB4,PALM,HSPG2,LINGO1,ABI2,AKT3,RDX

SOD1,PFN2,RAB11A,ARFGEF2,DYNLL1,SYP,TXLNA,TXNDC5,PSMD1,PDAP1,CST3,TAGLN2,YKT6,S
SLC25A23,NSD2,FBLN5,VASH1,MTOR,CREBBP,APBB1,KLHDC10,CERS2,BAG5,CD74,SLC38A2,D
,LUZP6,PRKDC,GABARAP,KDM1A,CLIP3,MAP2K2,FYN,FBLN1,UBQLN1,CSNK1D,HECTD1,SOD1,E
P1,CST3,TAGLN2,SV2A,PREPL,ATP6AP2,A2M,PRDX6,GRN,ACLY,SEPTIN5,XRCC6,SLC44A2,VPS41
IISCH,PTN,CSNK1E,PFKL,GLG1,SFPQ,PEBP1,CDH2,GPC3,SULF1,PIP4K2B,MEGF8,RAB7A,CCN2,
A,RPS4X,RPS3A,RPL18,RPL11,MORF4L1,BTF3,AGRN,RPLP1,ZMIZ1,RPL10,RPS17,AP3D1,SF3B2,
TF3,AGRN,RPLP1,ZMIZ1,RPL10,RPS17,AP3D1,CYB5R3,ZNF770,GTF2F1,HNRNPU,NFE2L1,SMAR
9,SEPTIN2,P4HB,APP,RPL5,KLRD1,GDI1,FSCN1,TMBIM6,SPTAN1,GSTP1,SMG1,CALM1,NPM1,PL
GID8,PSME3,STUB1,PSMC5,CSNK1A1,PSMC2,HERPUD1,PSMC4,PLK2,PSMF1,WAC,GSK3B,USP1
GABARAP,HSPB1,KDM1A,CLIP3,FKBP8,MAP2K2,QARS1,CCAR2,DUSP4,CORO1C,SUPT6H,FYN,F
R1,FBXL19,USP5,AMFR,TRIP12,USP47,UBE2L3,CDC34,RNF11,SDCBP,PINK1,USP10,PSMB1,HSF
33,THOC2,PPP1R12A,LZTS2,TNPO2,KPNA3,THOC3,PPP3CA,XBP1,FYTTD1,UFM1,TNPO3,ABCE1,
PP1R9B,GRIPAP1,OS9,FLCN,SEC16A,SFRP1,GLUD1,RER1,UBE2G2,HDAC6,OAZ2,ANP32B,LMAN
R1,FBXL19,USP5,AMFR,TRIP12,USP47,UBE2L3,CDC34,RNF11,SDCBP,PINK1,USP10,PSMB1,HSF
SESN2,TMCO1,EIF2S1,PSMC6,RNF5,CDK5RAP3,GET4,ABCA7

5,AMFR,TRIP12,USP47,UBE2L3,CDC34,RNF11,SDCBP,PINK1,USP10,PSMB1,HSPA1A,MDM2,FZR
FA,VHL,MAPK1,DDAH2,PRKDC,ACTN1,HSPB1,SON,KDM1A,BNIP3L,CLIP3,FKBP8,PLCG1,QARS1,
1,RGMA,TGFBR1,RUFY3,SEPTIN9,CUL7,PRKCI,DYNLT1,PPP2R5B,OPA1,APOE,LRP8,EPHA4,DDX

CP1,GRINA,VPS35,COL2A1,MDK,DDX42,NOTCH2,VEGFA,VHL,MAPK1,DDAH2,CDC37,PRKDC,ACT
H,FYN,FBLN1,USP7,STRAP,MAPK8IP1,BAG6,HNRNPD,CYP51A1,KDM4B,DDIT4,PPP1R9B,ADAR,E
PP1R9B,GRIPAP1,OS9,FLCN,SEC16A,ZFP36L1,RER1,UBE2G2,PTEN,PPP1CC,OAZ2,ANP32B,BRSK
GPS1,PSMA6,CUL7,BTBD1,CUL4B,FAM20C,PSMD12,APOE,PSMA4,MFGE8,FBXO7,COPS8,PSMC
RIO,RERE,PBX1,CNTNAP1,ALDH3A2,SEC16A,ATRX,SFRP1,GLUD1,TRA2B,PTEN,NDST1,EML1,IL6
ACA,TANC2,DAPK3,PTPN23,EIF4G1,NTRK2,ANK2,DIP2A,PTPN11,ABL1,COL16A1,SDCBP,TAOK2,

A,RPS4X,RPS3A,RPL18,RPL11,MORF4L1,BTF3,AGRN,RPLP1,ZMIZ1,RPL10,RPS17,AP3D1,CYB5R

DAPK3,AMBRA1,ATG4B,EIF4G1,PAFAH1B2,POLDIP2,ABL1,PINK1,USP10,ATP13A2,MFN2,PARK7,
DAPK3,AMBRA1,ATG4B,EIF4G1,PAFAH1B2,POLDIP2,ABL1,PINK1,USP10,ATP13A2,MFN2,PARK7,
C1,ABL1,SEMA6B,AUTS2,POR,RTF1,DAG1,SMPD3,NOTCH1,ADNP,CYFIP1,KAT2A,RNF157,BNC2,
LZTS2,TNPO2,KPNA3,THOC3,PPP3CA,XBP1,FYTTD1,UFM1,TNPO3,ABCE1,NOLC1,NUP50,RNPS1

AR2,SDF4,FYN,PITPNM1,LONP1,UBQLN1,HYOU1,SOD1,HNRNPD,VGF,ARNT2,IGFBP7,TXNIP,TP5
2,TUBB2A,MAPK1,MAPRE1,ARPC2,DSTN,PLEC,DYNC1LI2,DYNLL2,LUZP6,GRB2,CAPZB,GABARA
3,PSMB2,TERF2IP,RPS15A,TUBB2A,OPTN,PPP2CA,CCND1,MAPK1,MAPRE1,CCNL2,PSMD3,IPO7

C6A,PPP1CA,CCDC88A,CSNK2B,GID8,LDB1,PSME3,BCL9,DAAM1,PSMC5,CSNK1A1,PSMC2,PSMC
P,PSMC4,HSF1,GSK3B,RAB3A,SOX9,TFRC,MED25,WARS1,NAPB,CDKN1B,WASF3,CDC42EP4,VDA
,ACSL4,EI24,ADIPOR2,CRABP2,CDKN1B,EGLN2,ERBB2,UNC13A,PHB,PSMD10,ADIPOR1,RGMA,T
HDGFL3,ATP13A2,CRACD,MFN2,IGF1R,NSF,TRMT112,PARK7,APEH,CD44,MAP4K4,NCKAP5L,HT
1CA,CCDC88A,CSNK2B,GID8,LDB1,PSME3,BCL9,DAAM1,PSMC5,CSNK1A1,PSMC2,PSMC4,TMEM

LN1,UBQLN1,CSNK1D,HECTD1,SOD1,BAG6,PFN2,HNRNPD,WBP2,BMPR2,TRAF4,TRAF7,STAT3,

2,NCOR2,HNRNPD,VGF,PPP1R9B,DHX15,STAT3,SGTA,OS9,CAPN2,PRKAR2A,UBQLN2,ZFP36L1,

C2,APEH,CYFIP1,CD44,PSMD7,FAF2,TSPAN14,CSNK2B,CAPN1,NME2,ASAH1,RAB18,PSMC2,DEG

D4,LRRC41,OGT,PRPF19,PCGF2,FBXW5,GSK3A,PSMC3,MTOR,UBE3A,ATXN7L3,UBE2I,UBE2D3,

4B,DDIT4,PPP1R9B,ADAR,BEX4,STAT3,EIF2AK1,EDC4,SGTA,PRKAR2A,CSNK2A1,CST3,ZFP36L1,

C2,APEH,CYFIP1,CD44,PSMD7,FAF2,TSPAN14,CSNK2B,CAPN1,NME2,ASAH1,RAB18,PSMC2,DEG

STT3B,UBQLN1,SOD1,BAG6,SREBF2,NCOR2,HNRNPD,VGF,PPP1R9B,DHX15,STAT3,SGTA,OS9,C

3,CAB39,NDUFC2,APEH,CYFIP1,CD44,PSMD7,FAF2,TSPAN14,CSNK2B,CAPN1,NME2,ASAH1,RAB1

E3A,CREBBP,UBE2D3,SORT1,FGFR1,EIF4A3,IER2,NTRK2,CAD,PTPN11,CDC34,ABL1,SDCBP,NIB

1,WSB2,PSMB4,UBR5,UBE2M,UBE2D2,DTX4,DNAJB2,ATXN3,MKRN1,DCUN1D2,LZTR1,RNF145,

SEC62,RPL6,KAT6A,DVL3,YWHAE,KIF1A,RPL13,GNAS,SLC2A3,DDX17,GPI,HTRA4,RACK1,BSG,FA

3,CAB39,NDUFC2,APEH,CYFIP1,CD44,PSMD7,FAF2,TSPAN14,CSNK2B,CAPN1,NME2,ASAH1,RAB

,ITM2C,APLP1,RPS7,MIF,PRKAR1A,GJA1,CTNNB1,CD63,MACF1,SESN3,CYFIP2,NDFIP1,WLS,HNF

K1,AUTS2,ARPC1A,RTF1,HIP1R,MACROH2A1,HSPA1A,RUVBL2,SMPD3,CRACD,MECP2,PARK7,SA

L1,TP53BP1,CDC42,PPP1R9B,PSMA7,RBBP7,DCTN1,PRMT1,ACTR1A,EP300,PSMD1,PBX1,FLCN,

9,NDUFC2,APEH,CYFIP1,CD44,PSMD7,FAF2,TSPAN14,CSNK2B,CAPN1,NME2,ASAH1,RAB18,PSM

PRPF19,PCGF2,FBXW5,GSK3A,PSMC3,MTOR,UBE3A,ATXN7L3,UBE2D3,PSMC1,TRIM8,BAG5,SAE

CUL3,EML1,BRSK1,RPS27A,TNKS,PSMD8,APEX1,PCBP4,AKAP9,PPP1R10,PSMB4,TAOK1,CARM1

2,MAPK3,UFM1,VPS16,DYNC1I2,NBR1,SCOC,SESN2,RNF5,VPS28,QSOX1,TOMM40,ATP6V1D,ST

L1,ATRX,SFRP1,MASTL,PTEN,FEM1B,CUL3,BRSK1,RPS27A,TNKS,PSMD8,APEX1,PCBP4,AKAP9,

XK,MLEC,SURF4,RAB5B,RAB11B,SRP14,PYGB,ATP6AP1,DPYSL2,VEGFA,MAPK1,PSMD3,ADAM9

PPP1R10,PSMB4,TAOK1,CARM1,INTS3,VPS4A,RDX,LSM14A,KMT2E,CLASP2,TCF3,PSMD4,IST1,

,PTOV1,MDK,RBMX,CCT4,GCN1,SKI,ASXL1,KDM6B,ABCA2,PRPF6,TAF7,RNF10,VEGFA,MARS1,

A,GJA1,JUN,CTNNB1,SESN3,NDFIP1,WLS,HNRNPK,YWHAB,CADM4,RHOB,ATP1A2,HMGCR,RAC1,

N,BAZ1B,H2AZ1,CSNK2A1,GPX4,PHB2,BRD4,UBA52,BRD2,ATRX,HJURP,EP400,ARID1B,UPF1,HD

2B,CCN2,ACTN4,PRDX2,DDX3X,RPS15,H1-0,RPS7,PRKAR1A,GJA1,JUN,CTNNB1,MTSS2,CYFIP2,

N1,SKI,ASXL1,KDM6B,ABCA2,PRPF6,TAF7,RNF10,VEGFA,MARS1,VHL,MAPK1,SMARCA4,DDAH2,

2D3,SORT1,FGFR1,EIF4A3,NTRK2,CAD,PTPN11,ABL1,SDCBP,NIBAN2,HSPA1A,ACVR1B,SMPD3,T

A,TIMM50,LMAN1,CTTN,SLC25A5,RABEP1,MTCH1,SERP1,SPG7,VAPB,RAB1B,VPS41,TRAM1,VP

K1,CNP,MACROH2A2,NCSTN,ATP1B2,PHGDH,SCRIB,IGF2BP1,ATP6V0D1,RBPJ,MBD3,MTOR,UBE

SLC12A4,CLCN3,VAV2,FER,GMFB,SEMA6C,APOE,FGF13,PDXP,ARF6,MYADM,GSN,ANAPC2,RB1

GB5,SEPTIN7,HIF1A,ATP6AP2,BRK1,EIF2S2,EHMT1,CCN1,SSBP3,SOX2,PKDCC,RAB14,RIC8A,S

\F7,RNF10,VEGFA,MARS1,VHL,MAPK1,SMARCA4,PBX2,EIF3E,ILF3,LUZP6,PRKDC,SUPT5H,ACTN1

4,PCGF2,FBXW5,PSMC3,CHMP4B,PRKACA,MN1,PHIP,PSMC1,DAPK3,TUBB4A,PAGR1,RAB11FIP3
RNF41,DDX6,TRIM27,TRIB3,LSM4,DHX36,STUB1,PSMC5,CSNK1A1,PSMC2,HERPUD1,PSMC4,PL
SF6,ARID1B,ADCY6,HDAC6,PTEN,NCOA4,APEX1,LOX,SLIT2,SLC2A1,SAFB2,PELP1,CARM1,UBR5

R14B,PARK7,PTPN1,LIMK1,PPME1,TRIM27,NCK2,TRIB3,PMEPA1,PPP2R5D,MARCHF7,EMILIN1,C

AMP1,PSMB1,HSPA1A,PSMD13,CAB39,NDUFC2,APEH,CYFIP1,CD44,PSMD7,FAF2,TSPAN14,CSNK
5,PJA2,COL1A2,GAP43,HK1,RHOA,CBX5,ATF4,PSMD2,KIF5B,CXADR,TMEM259,CD81,ATP2A2,RP
2,ANXA5,VAMP2,GDI2,PDXK,MLEC,SURF4,RAB5B,RAB11B,SRP14,PYGB,ATP6AP1,DPYSL2,VEGF
A,PSMD13,CAB39,NDUFC2,APEH,CYFIP1,CD44,PSMD7,FAF2,TSPAN14,CSNK2B,CAPN1,NME2,ASA
G,COL4A2,MAPT,FN1,SPEN,NONO,DDB1,PSMD5,YWHAQ,HACD3,DCHS1,EGLN1,CHSY1,INA,PPP2

O,PRKDC,KDM1A,CORO1C,SUPT6H,YWHAH,FBLN1,B2M,PLXNA3,CSNK1D,STRAP,PTPRS,SOD1,F
6V0D1,RBPJ,MBD3,MTOR,UBE3A,BNIP3,SLC38A2,SMO,NTRK2,PAFAH1B2,XRN2,PTPN11,ABL1,S

1,USP36,CTSA,NAMPT

3,NASP,SMARCD1,CREBBP,APBB1,ATXN7L3,DAPK3,DEK,PAGR1,FBXL19,TRIP12,HMGA1,RSF1,P
F3A,CCDC93,NAPA,COL7A1,GABARAPL2,CAPZA2,TRAPPC11,PRKCI,TRAPPC1,SEC23IP,COPB1,SE
RB,CCDC88A,TRIM27,NCK2,ARPC3,SENP6,GBA2,MAPRE3,PDE4DIP,GSK3B,MAP1S,STMN4,CDKI
O,RABGAP1,EIF4G1,IGFBP4,USP47,OGFR,ABL1,SDCBP,SEMA6B,TAOK2,TSPYL2,HSPA1A,RUVBL
RPS27A,PSMD8,APEX1,PPP1R15A,PCBP4,SERP1,FUS,PSMB4,EIF3H,ELAVL1,IREB2,LSM14A,TTC3
ICRIP,ETF1,TNRC6A,PPP1CA,HNRNPR,CDK4,DDX6,NCK2,C1QBP,LSM4,DHX36,EIF3B,POLDIP3,N
A,PSMD13,CAB39,NDUFC2,APEH,CYFIP1,CD44,PSMD7,FAF2,TSPAN14,CSNK2B,CAPN1,NME2,ASA
SP47,ABL1,PINK1,HIP1R,HSPA1A,MKNK2,ACVR1B,MDM2,UBE4B,MAPK8IP2,CUL1,PARK7,CD44,A

EMA4C,LIMS1,SRF,GSK3B,SERPINE2,RCC2,PRMT5,CRABP2,NUMBL,EFNB3,SRRT,ROCK2,SS18L
8IP2,MECP2,CUL1,PSMD13,CCNY,PSMD7,TNRC6A,PPP1CA,CCDC88A,CSNK2B,GID8,LDB1,PSME
365,ATF4,PSMD2,NR2F1,ENC1,RPS14,CHD4,ATP1A1,JUND,EIF5,CD81,RPL26,CCT7,PFN1,TLE5,C

IXA5,VAMP2,GDI2,PDXK,MLEC,SURF4,RAB5B,RAB11B,SRP14,PYGB,ATP6AP1,DPYSL2,VEGFA,CC
RIB2,PEA15,CDKN1A,TIMP3,CTSB,NUCB1,EEF1A2,CCT2,TBCD,ANXA5,GNAI3,PABPN1,GDI2,PARP
CC1,FYN,FBLN1,STT3B,EIF3F,LONP1,UBQLN1,CSNK1D,USP7,HECTD1,UQCRC2,HGS,PCBP2,BAG
EF18,FER,GMFB,GIT1,APOE,FGF13,TPX2,PDXP,ARF6,MYADM,NF2,ASAP3,GSN

25A5,GLO1,GRN,PPP1R10,CCN1,GPX1,TLE1,TAX1BP1,MAZ,KIFAP3,TRAP1,SOD2,PDPK1,ITGAV,A
GPX4,PHB2,UBA52,SFRP1,SRSF6,PTEN,IL6ST,RPS27A,HIF1A,CTTN,SLC25A5,GLO1,GRN,PPP1R1

,CDC34,PSMB1,TAOK2,MACROH2A1,ACVR1B,PKIA,MDM2,FZR1,PCM1,MECP2,CUL1,PSMD13,AT
M4,DHX36,EIF3B,POLDIP3,NCBP2,IGF2BP2,DHFR,SHMT2,PA2G4,ERBB2,PER1,SKIV2L,ROCK2,M
3,STUB1,EMILIN1,PSMC5,PSMC2,PDE4DIP,PSMC4,LIMS1,SRF,HSF1,GSK3B,SOX9,TFRC,MED25,
NC1,RPS14,CHD4,JUND,EIF5,CD81,RPL26,CCT7,PFN1,TLE5,HNRNPC,LRPPRC,TADA2B,KHDRBS1
N,RGMB,PTPRF,NDST1,IL6ST,ITGB5,RPS27A,PEG10,DKK3,HIF1A,FKBP1A,LOX,BRK1,NDN,CCN1

1A,CTTN,SLC25A5,GLO1,GRN,PPP1R10,CCN1,GPX1,TLE1,TAX1BP1,MAZ,KIFAP3,TRAP1,SOD2,PD

IB2,PIN1,LRRN1,ADGRL1,OPA1,EPHB2,APOE,LRP8,FGF13,ARF6,NFASC,PRNP,SRCIN1,EPHA4,PP
PHA4,PLXND1,NF2,ID2,ANAPC2,SKIL,HES6

'Q,SOX12,ACTN4,PRDX2,DDX3X,NANOG,H1-0,PRKAR1A,GJA1,JUN,CTNNB1,TARDBP,ARID1A,NDF

1,ENC1,RPS14,CHD4,ATP1A1,JUND,EIF5,CD81,RPL26,CCT7,PFN1,TLE5,CLEC7A,HNRNPC,LRPPR

4,JUND,EIF5,CD81,RPL26,CCT7,PFN1,TLE5,HNRNPC,LRPPRC,TADA2B,KHDRBS1,XPO1,SFPQ,EIF
CUL4A,CETN2,PPME1,CDK4,CCNG1,PSME3,PSMC5,PSMC2,PSMC4,PLK2,PSMF1,DHFR,RCC2,CD

E1,PHC2,TNKS2,ID4,STRAP,SARS1,SREBF2,NCOR2,HNRNPD,RPS13,BMPR2,TXNIP,TP53BP1,DDI

CCNK,SMARCB1,RAD23A,NUP58,VPS28,HMGA2,GSN,EPHA2,VPS37B
PBX2,SRSF1,FTO,ILF3,PRKDC,SUPT5H,ACTN1,CAND1,KDM1A,MAP2K2,CCAR2,ARF4,SMARCE1,S

SKIV2L,PSMD10,ROCK2,ATF6,ELOB,RGMA,MAPK3,ZSWIM8,CNOT3,XBP1,TMEM59,APOE,TUT4,F

SMD10,ROCK2,ELOB,PSMA6,PIN1,AJUBA,PSMD12,OPA1,TGFB2,PSMA4,PSMC6,POSTN,PAM,NA
RDX5,AHCYL1,VPS35,PTK7,ASXL1,KDM6B,RAB11B,ABCA2,ATP6AP1,DPYSL2,MARS1,RAPGEF1,
,PDGFRB,CETN2,ASNS,CDK4,PSME3,TOM1L2,PSMC5,PSMC2,PSMC4,PLK2,PSMF1,HSF1,RCC2,
1,IST1,RBPJ,PSMC3,AP2A2,AP1M1,CD74,PAFAH1B2,SDCBP,SCAMP1,PSMB1,HSPA1A,PSMD13,C
RNF5,RNF40,ANAPC2,RBBP6
A,CTNNB1,CD63,ANXA6,MACF1,MYL6,MYH10,NARS1,CADM4,LAMP1,RHOB,ATP1A2,MAP2,RAB1
1,PPP2CA,CCND1,MAPK1,CCNL2,IPO7,ADAM9,ILF3,CDC37,PRKDC,HSPB1,CLIP3,FKBP8,MAP2K

A,JUN,CTNNB1,TARDBP,ARID1A,MXD4,SHOX,HNRNPK,H1-10,CIRBP,DUX4,YWHAB,HNRNPH1,ZN
P5,USP47,SDCBP,PINK1,POR,HIP1R,HSPA1A,ATP13A2,MDM2,FZR1,SUMO1,PARK7,USP14,SART

,CTSA,NAMPT
EF1,RPL37,MAPK1,SMARCA4,PSMD3,PTBP1,GRB2,FKBP8,MAP2K2,CCAR2,FYN,LARS1,PFKM,T

SMG1,CALM1,NPM1,PLXNB2,PLD3,HSPD1,COL18A1,IGF2,VCP,UBB,TRIM28,MATR3,YWHAG,CO
2K2,QARS1,DUSP4,CORO1C,FYN,FBLN1,CSNK1D,IDH1,STRAP,HGS,MAPK8IP1,SOD1,PFN2,PPP
2K2,QARS1,DUSP4,CORO1C,FYN,FBLN1,CSNK1D,IDH1,STRAP,HGS,MAPK8IP1,SOD1,PFN2,PPP

3,PCNA,TFDP1,PSMD12,PSMA4,TPX2,FHL1,DYNC1I2,FBXO7,UBE2E1,CENPF,PSMC6,CDK5RAP3,
1,SDCBP,AUTS2,PHACTR4,ARPC1A,FAM171A1,TAOK2,HIP1R,ELN,OBSL1,CRACD,TNFAIP1,JAM3
,STXBP1,CSNK1D,MESD,PTPRS,VANGL2,PFN2,VGF,RAB11A,DYNLL1,AMOTL2,CDC42,PPP1R9B,
L,SIN3A
R2,BMPR2,CDC42,DDIT4,ADAR,CTNND1,PSMA7,SULF2,BCL9L,LBH,PSMD1,FLCN,UBQLN2,CSN
S20,CD81,ATP2A2,RPL26,RBBP4,MCL1,TLE5,CLEC7A,KHDRBS1,PTN,CSNK1E,GLG1,SFPQ,PEB
,ILF2,PFKM,RAI1,SREBF2,HNRNPD,ARNT2,WBP2,AKIRIN1,BMPR2,TP53BP1,STAT3,EDC4,BCL9L
13,TXNIP,CIC,RBBP7,STAT3,MTA1,KDM5C,EP300,LBH,RERE,KDM5B,SF1,FLCN,H2AZ1,PHB2,UB
1,PSMA7,SULF2,BCL9L,LBH,PSMD1,FLCN,UBQLN2,CSNK2A1,PHB2,BRD4,UBA52,PPP2CB,NXN
28,GSN
R2,BMPR2,CDC42,DDIT4,ADAR,CTNND1,PSMA7,SULF2,BCL9L,LBH,PSMD1,FLCN,UBQLN2,CSN
L37,MAPK1,SMARCA4,PSMD3,PTBP1,GRB2,FKBP8,MAP2K2,CCAR2,FYN,LARS1,PFKM,TNKS2,ST

,PSMC4,PLK2,PSMF1,RCC2,CDKN1B,PSMB5,PRKAR2B,XPC,PSMA1,PRMT2,PSMD10,MDC1,ANX
,CTCF,SETD3,WDR5,SUDS3,MAPK3,XBP1,TET3,CUL4B,BRPF3,RYBP,SETD1A,UBE2E1,KDM2B,P

2,EMD
EGFA,RAPGEF1,OPTN,RPL37,PPP2CA,MAPK1,PRDX1,PTBP1,GRB2,GABARAP,HSPB1,KDM1A,M
T8,CLASP2,PDPK1,WASF2,CAP1,MTOR,DAPK3,MICAL1,ANK2,PLS3,CELSR1,ABL1,SDCBP,AUTS2
,FZD7,GPRC5B,LRP6,JRK,EMD
LXNA1,RAB7A,ACTN4,MMP14,CLTC,RPS19,MIF,GJA1,CTNNB1,CD63,ANXA6,MACF1,DNAJC5,TA
DS3,MAPK3,XBP1,TET3,CUL4B,BRPF3,RYBP,SETD1A,UBE2E1,KDM2B,PAF1,DNMT1,SMARCB1,

IPT,UBTF
3,NUP98,CSNK1A1,GOLGA8B,NDRG1,DEGS1,PDE4DIP,EMC10,TMEM9,YWHAZ,SYNGR1,SERPINE
STAT3,EP300,LBH,H2AZ1,MMP15,FLOT1,GPX4,PHB2,ZFP36L1,SFRP1,ADCY6,HDAC6,PTEN,NCO

3,CUL7,PRKCI,LRRN1,DYNLT1,SEMA6C,HMGB2,RELA,OPA1,EPHB2,LRP8,FGF13,EPHA4,PLXND1

POSTN,PAM,NAMPT

/,ADAM10,CAP1,MSH6,NSD2,IST1,PSMC3,MTOR,AP2A2,AP1M1,CD74,PAFAH1B2,ABL1,SDCBP,LC
GGNBP2,NOTCH1,MFN2,KAT2A,SALL2,CUL4A,TASOR,DAD1,EPN1,HM13,DEAF1,SEMA4C,PSMC4,
1,TPM1,CDH2,GPC3,DDR1,SULF1,PLXNA1,MEGF8,SOX12,CCN2,MMP14,PRDX2,RPS15,CLTC,API
C4,TANC2,CROCC,TUBB4A,USO1,SUN1,CHMP1A,PINK1,ESYT1,TRAK1,MFN2,PCM1,NSF,CLUH,F
2,NOTCH1,MFN2,KAT2A,SALL2,CUL4A,TASOR,DAD1,EPN1,HM13,DEAF1,SEMA4C,PSMC4,SRF,ZE

6,KDM1A,PLCG1,MAP2K2,ARF4,CORO1C,FYN,LARS1,FBLN1,B2M,SOD1,PFN2,HNRNPD,SEC23A,

4,PTPN11,ABL1,SDCBP,ARPC1A,NIBAN2,HIP1R,SMPD3,TIA1,IGF1R,NPTN,EFNB2,AAMP,CYFIP1,PT

CCAR2,GNG2,FYN,B2M,UBQLN1,HYOU1,PTPRS,PCBP2,MAPK8IP1,SOD1,BAG6,TP53BP1,CDC42

CAP1,MSH6,NSD2,IST1,PSMC3,MTOR,AP2A2,AP1M1,CD74,PAFAH1B2,ABL1,SDCBP,LGALS1,SCA

,CDK16,HDAC2,IPO5,ATP6V0C,TRIB2,PEA15,CDKN1A,CTDSP2,EEF1A2,FDFT1,PPP1CB,PGD,SQS
NKS2,CSNK1D,MAPK8IP1,SOD1,DGKD,WBP2,BMPR2,CDC42,PSMA7,TRAF4,TRAF7,SULF2,STAT3
HNRNPA2B1,RPS2,RPS27A,TNKS,XRCC6,TAOK1,CCN1,SHC2,SPAG9,PDPK1,DAZAP2,DCUN1D2,G
L16A1,POR,SMPD3,IGF1R,MDM2,CYFIP1,EHMT2,PTPN1,GOLPH3,CDK4,TRIB3,ADCY1,ATP6V0B,I
PR2,DYNLL1,CDC42,DDIT4,PPP1R9B,ADAR,TRAF4,TRAF7,STAT3,EIF2AK1,EP300,PRKAR2A,FLCN

DAC6,PTEN,SMC3,ULK1,MAPRE2,SEPTIN7,TNKS,GALNT11,MAP1LC3B,AKAP9,TMOD2,RAB1B,VP

WASF3,CDC42EP4,CNTN2,ARPC5,ROCK2,RAB5A,RGMA,CAPZA2,RUFY3,ADD1,EXT1,SLC12A4,CLC

2,EHMT1,FKBP4,CARM1,ABI2,PKDCC,AKT3,DCLK1,KMT2E,GALNT1,RAB6A,NMT1,PDPK1,NSD2,A
A,LRRN1,EPHB2,FGF13,ARF6,PIP5K1A,GJC1,RAP1B,PLXND1,HIPK1,GPM6B,EPHA2

K2,RANGAP1,PTPN11,WDTC1,ABL1,COL16A1,POR,SMPD3,IGF1R,MDM2,CYFIP1,EHMT2,PTPN1,(

LA-C,PDIA3,RPS3,CHD3,VAPA,HMGB1,AP2M1,TPT1,OR8K1,STMN1,ITGB1,UCHL1,RPL11,NCAM1
CG1,ARF4,ZFAND5,CORO1C,FYN,FBLN1,PLXNA3,STRAP,PTP4A1,VANGL2,PFN2,RAB11A,AKIRIN1
5,PSMC2,NDRG4,SEMA4C,PSMC4,PSMF1,SRF,LGR4,CELSR3,SOX9,HOXA5,PSMB5,STAT1,PSMA1
,MDM2,EXOC5,PSMD13,CD44,KAT2A,NTN1,PSMD7,PPP1CA,CSNK2B,EXT2,PSME3,DEAF1,DAAM
,CDK16,HDAC2,IPO5,ATP6V0C,TRIB2,PEA15,CDKN1A,CTDSP2,EEF1A2,FDFT1,PPP1CB,PGD,SQS

,EIF3E,FTO,ILF3,SUPT5H,KDM1A,QARS1,CCAR2,SMARCE1,FYN,PHC2,ID4,STRAP,SARS1,SOD1,S

5,CRPPA,CNN2,VCAN,NOX4,PTPRA,ACTR3,PKN1,NCKAP1,ACAP3,MDK,LAMC1,PTK7,SKI,DPYSL2,

LA-C,PDIA3,RPS3,CHD3,VAPA,HMGB1,AP2M1,TPT1,OR8K1,STMN1,ITGB1,UCHL1,RPL11,NCAM1
,PHB2,ZFP36L1,SFRP1,ARID1B,UPF1,ADCY6,TRA2B,HDAC6,PTEN,ADH5,NCOA4,APEX1,HIF1A,SL
CE1,PHC2,ID4,STRAP,SARS1,SREBF2,NCOR2,HNRNPD,BMPR2,TXNIP,CIC,RBBP7,STAT3,EIF2AK1

1,RRP1B,KARS1,FARSA,TYW5,DGCR8,SRRT,SENP3,RTRAF,ZSWIM8,TARS1,RELA,NOLC1,THG1L,

L,PRMT1,CAPN2,NAP1L1,LBH,KDM5B,LRRC4B,FLOT1,ZFP36L1,MAP6,PLXNB1,SFRP1,PTEN,PPP
2,MAP4K4,PHPT1,ERP29,PTPN1,RAN,GOLPH3,UBE2J1,P3H1,YWHAZ,GSK3B,ACSL4,KIF3A,TXN,IR
R2E,SULF2,STAT3,EP300,KDM5B,BAZ1B,FLOT1,EIF3D,PHB2,ARID1B,TRA2B,PTEN,AGPAT1,IL6ST

CH1,IGF1R,MDM2,PARK7,CYFIP1,PTPN1,ASNS,CSNK2B,CDK4,TRIB3,PMEPA1,ADCY1,ATP6V0B,

FLAR,SHC1,ROCK2,ADIPOR1,CHMP5,ANXA1,XBP1,PRKCI,SP1,FER,BCAR1,RELA,OPA1,TLR6,PRN

4,STRAP,SARS1,SREBF2,NCOR2,HNRNPD,BMPR2,TXNIP,CIC,RBBP7,STAT3,EIF2AK1,EDC4,MTA1,
ASXL1,PSMB2,KDM6B,TMED2,RPL38,NOTCH2,VEGFA,SELENON,RAPGEF1,PPP2CA,CCND1,MAP

AM3,PABPC4,CD44,PDGFRB,TBXA2R,C1QBP,SRF,YWHAZ,DHFR,SERPINE2,RAB3A,NREP,ADIPOR

,SOD1,CDC42,ADAR,PSMA7,TRAF4,ST3GAL6,STAT3,TXLNA,CAPN2,KDM5B,PSMD1,UBA52,ZFP3

1,STAT3,BCL9L,MTA1,EP300,LBH,RERE,PBX1,FLCN,H2AZ1,BRD4,UBA52,HOXB5,ATRX,SFRP1,AI

1,STAT3,BCL9L,MTA1,EP300,LBH,RERE,PBX1,FLCN,H2AZ1,BRD4,UBA52,HOXB5,ATRX,SFRP1,AI

74,SMO,FGFR1,METRN,EIF4G1,NTRK2,OLFM2,FAM210B,ABL1,SDCBP,POR,TRAK1,DAG1,NIBA

1,STAT3,BCL9L,MTA1,EP300,LBH,RERE,PBX1,FLCN,H2AZ1,BRD4,UBA52,HOXB5,ATRX,SFRP1,AI

N1,HSPG2,OLFM1,ABI2,TLE1,AKT3,SEMA3C,LTBP3,PSMD4,SCRIB,NSD2,RBPJ,PCGF2,GSK3A,P

R1,RELA,OPA1,SESN2,GRB10,RAP1B,ATP6V1F,LEPROT,ZNF106,LRP6,BTG1,RNF40,PIK3R3,ATP

G9,PDPK1,DAZAP2,DCUN1D2,AKT2,OGT,GSK3A,MTOR,UBE3A,PRKACA,WNK1,CD74,SAE1,FGF

SP3,DUSP8,PPP1R37,PRKAR2B,CSRNP2,NR2F2,LMTK3,ROCK2,CAMK2N1,GPS1,PPP1R26,RTRA

T1,PHB2,BRD4,UBA52,SFRP1,ADCY6,HDAC6,PTEN,IL6ST,RPS27A,LOX,SLIT2,AKAP9,XRCC6,TAO

MAPK8IP1,SOD1,PFN2,BMPR2,DDIT4,PPP1R9B,ADAR,TRAF4,TRAF7,STAT3,EIF2AK1,BCR,TRIO,F

AP1B,CACNG7,ABR,ANAPC2,CASK,GNAI1

HAG,CKB,COL4A2,MAPT,FN1,SPEN,PSMD5,YWHAQ,DCHS1,EGLN1,TPI1,CHSY1,INA,MRC2,RPL1

AP1B,CACNG7,ABR,ANAPC2,CASK,GNAI1

X1,CSNK2A1,PHB2,CST3,FARP1,SFRP1,MASTL,HDAC6,PTEN,PTPRF,OAZ2,SPINT2,GPSM1,TNKS,

SP8,PPP1R37,PRKAR2B,CSRNP2,NR2F2,LMTK3,ROCK2,CAMK2N1,GPS1,PPP1R26,RTRAF,HEG1

CL1,SESN2,EIF2S1,GNAO1,MYEF2,BTG1,PRDX4,SIN3A

DM1A,QARS1,CCAR2,SMARCE1,FYN,PHC2,ID4,STRAP,SARS1,SREBF2,NCOR2,HNRNPD,BMPR2,

APN2,KDM5B,PSMD1,UBA52,ZFP36L1,SFRP1,UPF1,UBE2G2,AGPAT1,HNRNPA2B1,IL6ST,RPS27

DUSP8,VDAC2,PER1,PHB,MGRN1,PSMD10,CAMK2N1,ADIPOR1,GPS1,HEG1,KCTD10,PIN1,XBP1,
1L,NOP58,USP36,SNU13
FA,TUBB2A,MAPK1,MAPRE1,ARPC2,DSTN,ADAM9,GRB2,HSPB1,PLCG1,ARF4,ZFAND5,PLTP,COI
FA,TUBB2A,MAPK1,MAPRE1,ARPC2,DSTN,ADAM9,GRB2,HSPB1,PLCG1,ARF4,ZFAND5,PLTP,COI

3L1,SDCBP,PINK1,PSMB1,PPP5C,TAOK2,DAG1,RGL2,NOTCH1,IGF1R,NPTN,MAPK8IP2,STK39,CI

MFAP2,QSOX1,GPM6B,MMP24

35B,SRP14,PYGB,NOTCH2,MAPK1,PSMD3,PRDX1,ADAM9,GRB2,PRKDC,CAND1,HSPB1,KDELR1,(
2,QSOX1,GPM6B,MMP24

2,QSOX1,GPM6B,MMP24
R,BCAR1,SEMA6C,APOE,TGFB2,ARF6,PIP5K1A,SCARB1,PDXDC1,PLPP3,PLXND1,PRCP,CRB2,PIK

MMP14,PRDX2,DDX3X,ITM2C,CLTC,NANOG,MIF,TLN1,JUN,CTNNB1,CD63,MACF1,SESN3,CYFIP2
1,SEC31A,CENPF,DNM1,SYT4,AP3M1,ARFGAP3
2A1,LAMC1,CCT4,ASXL1,KDM6B,RAB11B,ABCA2,NRDC,ATP6AP1,TERF2IP,VEGFA,SELENON,MA

MYADM,NF2,ELMO2,SH3KBP1,ASAP3,GSN

M14A,KIFAP3,CNP,FLOT2,TUBGCP6,ATXN3,RAB6A,CLASP2,FBXW5,CHMP4B,CROCC,TUBB4A,RA

G5,KMT2E,GTF2I,NUCKS1,ATXN3,DICER1,TCF3,MSH6,NSD2,PRPF19,KDM4A,CIZ1,MBD3,GSK3A
1A,FKBP1A,SLIT2,TMOD2,GRN,CCN1,GPX1,HSPG2,SSBP3,OLFM1,SOX2,AKT3,GTF2I,SH3PXD2A

K1,GNG2,FYN,SLIT3,STMN3,FBLN1,PLXNA3,HYOU1,STRAP,PTPRS,HGS,PCBP2,MAPK8IP1,GHITM

ACSL4,KIF3A,TXN,IRS1,SLC16A1,ERBB2,NADK,CAMK2N1,ANXA1,PPP3CA,RUFY3,XBP1,TFDP1,UF

B3GAP2,ADCY6,PTEN,TBC1D9B,HNRNPA2B1,ANP32B,MAPRE2,RPS2,RPS27A,TNKS,HIF1A,PPP1

3B,NDUFB8,RAB3A,TFRC,NDUFS1,ATP5F1C,LMNA,PMPCA,VDAC2,UQCRFS1,ATG9A,ATP5F1E,PH
9,CCNY,PTPN1,PDGFRB,CCDC88A,TRIB3,ADCY1,MAPKAPK2,STUB1,CRKL,MAPRE3,TNFRSF10B

P1A,RABEP1,SLIT2,A2M,DAP,MTCH1,GRN,PPP1R10,CCN1,GPX1,ANTXR1,SOX2,RDX,TBC1D9,RI
OLDIP2,TRIP12,USP47,PTPN11,HMGA1,ABL1,FXR2,USP10,PPP5C,TAOK2,MTREX,RUVBL2,TCEA
MPT
H,COX5A,TALDO1,COX7A2,UQCRH,GIT1,FH,GYS1,MSH2,NUP50,GRB10,PRELID1,NUP58,PDHB,
HMT2,GOLPH3,ZCCHC3,CDK4,PMEPA1,ADCY1,RBFOX2,HSF1,GSK3B,ZNF703,PGRMC2,PRMT5,
HNRNPA2B1,CUL3,RPS27A,APEX1,DKK3,DAP,NDN,BAP1,BAZ2A,XRCC6,EHMT1,BCL7A,SOX2,TLE

E5,HNRNPC,TADA2B,KHDRBS1,XPO1,SFPQ,ACSL6,SOX12,CCN2,ACTN4,PRDX2,DDX3X,NANOG,H

HNRNPA2B1,CUL3,RPS27A,APEX1,DKK3,DAP,NDN,BAP1,BAZ2A,XRCC6,EHMT1,BCL7A,SOX2,TLE

PTEN,CEMIP2,PPP1CC,IL6ST,SEPTIN7,HIF1A,LOX,SLIT2,GRN,XRCC6,EHMT1,SLIT1,CLPTM1,CCN
HNRNPA2B1,CUL3,RPS27A,APEX1,DKK3,DAP,NDN,BAP1,BAZ2A,XRCC6,EHMT1,BCL7A,SOX2,TLE

T5,POR,MACROH2A1,GGNBP2,PKIA,NOTCH1,IGF1R,MDM2,PPP1R14B,CD2BP2,PARK7,USP14,CI

,AP2M1,TPT1,OR8K1,STMN1,ITGB1,UCHL1,RPL11,NCAM1,ANXA2,AGRN,ZMIZ1,TIMP2,XAF1,WSI

2,C1QBP,TNFRSF10B,RBM10,GSK3B,ARHGEF9,RRP1B,RASSF2,HOXA5,CDKN1B,BACE1,PHB,SC

MED13,GSK3A,PSMC3,UBE3A,YY1,SMARCD1,CREBBP,APBB1,PHIP,SSBP4,THRAP3,SMO,TEAD1,
CDC88A,CETN2,SENP6,NUP98,GBA2,GOLGA8B,MAPRE3,PDE4DIP,SLK,RANBP9,PLK2,GSK3B,T
P3,LRP6,NNT,IBTK
ARID1A,MXD4,SHOX,HNRNPK,H1-10,DUX4,YWHAB,ZNF454,CTBP1,CDK16,SMCHD1,HDAC2,BCL
1,ABL1,PINK1,TAOK2,DAG1,TSPYL2,MACROH2A1,GGNBP2,PKIA,IGF1R,STK39,CAB39,PARK7,A

RBS1,XPO1,SFPQ,ACSL6,SOX12,CCN2,ACTN4,PRDX2,DDX3X,NANOG,H1-0,PRKAR1A,GJA1,JUN,

ARID1A,MXD4,SHOX,HNRNPK,H1-10,DUX4,YWHAB,ZNF454,CTBP1,CDK16,SMCHD1,HDAC2,BCL

RBS1,XPO1,SFPQ,ACSL6,SOX12,CCN2,ACTN4,PRDX2,DDX3X,NANOG,H1-0,PRKAR1A,GJA1,JUN,

,MED12,DNAJB6,SOS1,CITED2,ROCK2,RGMA,TGFBR1,MAPK3,ZNF358,EXT1,OPA1,EPHB2,TGFB

7,ADNP,MAP4K4,ERP29,CCNY,PTPN1,PDGFRB,CCDC88A,CSNK2B,TRIM27,NCK2,TRIB3,ADCY1,C

ARID1A,MXD4,SHOX,HNRNPK,H1-10,DUX4,YWHAB,ZNF454,CTBP1,CDK16,SMCHD1,HDAC2,BCL

SUPT6H,FYN,ILF2,CCT8,B2M,STXBP1,IDH1,HECTD1,MAPK8IP1,SOD1,BAG6,DYNLL1,TP53BP1,C

XA5,BACE1,SOS1,PRMT2,ATF6,RPS6KA2,MAP2K4,SUDS3,ANXA1,TGFBR1,PNMA2,PIN1,QRICH1,

,ATF6,RPS6KA2,MAP2K4,SUDS3,ANXA1,TGFBR1,PNMA2,PIN1,QRICH1,XBP1,ARHGEF18,PRKCI,V

RBS1,XPO1,SFPQ,ACSL6,SOX12,CCN2,ACTN4,PRDX2,DDX3X,NANOG,H1-0,PRKAR1A,GJA1,JUN,

LK2,SRF,PIP5K1C,GSK3B,SYNGR1,SERPINE2,CRTC1,RAB3A,RNF216,NAPB,DLG4,SLC12A7,SOF

NRNPK,RHOB,ATP1A2,HLA-B,MAP2,RAB1A,HMGCR,RAC1,ARHGDIA,CAPRIN1,APPL1,HDAC2,L1C

CTBP1,CDK16,SMCHD1,HDAC2,BCLAF1,HSBP1,DCN,CDKN1A,DDX39B,EID1,CHD6,RNF187,MLX
4,FHL1,FBXO7,CENPF,PSMC6,CDK5RAP3,BTG1,ZNF207,HMGA2,WAPL,CNOT1,PPP2R5C

CCT8,B2M,STXBP1,IDH1,DYNLL1,TP53BP1,CDC42,STAT3,TXNDC5,PSMD1,PDAP1,PHB2,CST3,AT

CTBP1,CDK16,SMCHD1,HDAC2,BCLAF1,HSBP1,DCN,CDKN1A,DDX39B,EID1,CHD6,RNF187,MLX
CTBP1,CDK16,SMCHD1,HDAC2,BCLAF1,HSBP1,DCN,CDKN1A,DDX39B,EID1,CHD6,RNF187,MLX

1,SERPINE2,CRTC1,RAB3A,RNF216,NAPB,DLG4,SLC12A7,SORCS2,NSMF,PRKAR2B,UNC13A,NA

24,SEMA3C,SOD2,SPAG9,CLASP2,PDPK1,ATP1B2,ITGAV,ADAM10,AKT2,VASH1,MTOR,WNK1,CE

CRTC1,RAB3A,RNF216,NAPB,DLG4,SLC12A7,SORCS2,NSMF,PRKAR2B,UNC13A,NAPA,BACE1,SL
CRTC1,RAB3A,RNF216,NAPB,DLG4,SLC12A7,SORCS2,NSMF,PRKAR2B,UNC13A,NAPA,BACE1,SL
BP1,PLXNA3,PTPRS,IGFBP7,CD99L2,CDC42,CTNND1,PALLD,BCR,KIRREL1,DGCR2,CNTNAP1,FL

,PTPRS,IGFBP7,CD99L2,CDC42,CTNND1,PALLD,BCR,KIRREL1,DGCR2,CNTNAP1,FLCN,LRRC4B

T1,AJUBA,VAV2,EPHB2,APOE,TGFB2,HMG20B,SCARB1,COL3A1,PLPP3,POSTN,MYL9,FZD7,PRC
CA,FER,SNX17,GIT1,APOE,ARF6,SYT1,MFGE8,SCARB1,PPT1,DNM1,ANKRD13B,SYT4,RAP1B,VP

KAP1,VPS35,MDK,PTK7,SKI,ASXL1,PSMB2,TSPAN6,ATP6AP1,TERF2IP,NOTCH2,VEGFA,SELENON,

ACYBP,CITED2,CFLAR,SETD3,MAP2K4,TGFBR1,PPP3CA,HEG1,PIN1,COL6A3,XBP1,YBX3,BTBD1,

AT1,CACYBP,BACE1,CFLAR,ANXA1,MAPK3,PPP3CA,ADD1,PCNA,RELA,SEC31A,SYT1,FBXO7,PRN
1,ACTR3,PKN1,YEATS2,PTOV1,ZNF302,RBMX,CHD2,GCN1,SKI,ASXL1,PSMB2,KDM6B,SCAF1,HD
HGEF18,RAB35,VAV2,DYNLT1,EPHB2,APOE,TGFB2,ARF6,OBSCN,ARHGAP23,COL3A1,RAP1B,DN

GNA13,CD74,SMO,FGFR1,PTPN23,LMO4,ABL1,SDCBP,SEMA6B,MTA2,MGAT5,DAG1,YTHDF3,PL

AM10,AKT2,VASH1,MTOR,WNK1,CERS2,DAPK3,GNA13,CD74,SMO,FGFR1,PTPN23,LMO4,ABL1,

R,GTF2I,SEMA3C,SLC4A2,PDPK1,WASF2,LTBP3,PHGDH,ITGAV,SCRIB,RBPJ,VASH1,PRKACA,AME

PD3,NOTCH1,PSMD13,CRIP2,JAM3,SART3,PABPC4,KAT2A,PSMD7,MAEA,MTURN,HIPK2,CUL4A,[illegible]

RFGEF2,BMPR2,TP53BP1,POLR2E,SULF2,STAT3,BCL9L,NAP1L1,EP300,LBH,KDM5B,LRRC4B,FL

,MKNK2,ACVR1B,SMPD3,NOTCH1,PSMD13,CRIP2,JAM3,SART3,PABPC4,KAT2A,PSMD7,MAEA,M[illegible]

5,ATRX,SFRP1,SRSF6,TAGLN2,PTEN,FEM1B,IL6ST,SPINT2,BRSK1,PSMD8,HIF1A,ATP6AP2,SLIT2,

HDGF,ABCA2,PRPF6,TAF7,NOTCH2,RNF10,VEGFA,VHL,CCND1,SMARCA4,CCNL2,PBX2,PSMD3,A

M1,ANTXR1,AKT3,LDLR,GTF2I,RIC8A,SEMA3C,SOD2,PDPK1,WASF2,ITGAV,ADAM10,NSD2,RBPJ,

.1,NMT1,CLASP2,PDPK1,ATP1B2,AKT2,EHD1,ARPP19,GSK3A,PRKACA,APBB1,UBE2D3,WNK1,SA[illegible]

ORBS1,SPTBN2,SLC3A2,PTN,PFKL,GLG1,SFPQ,PNMA1,GPC3,SOX12,RAB7A,MMP14,PRDX2,DDX[illegible]

,FAM210B,ABL1,LGALS1,PSMB1,KLF13,HSPA1A,MKNK2,ACVR1B,SMPD3,NOTCH1,PSMD13,CRI[illegible]

1B,IGF1R,NPTN,STK39,CAB39,FXR1,PARK7,ADNP,CD44,MAP4K4,ERP29,CCNY,PTPN1,KCTD20,[illegible]

[illegible]1,MTOR,WNK1,CERS2,DAPK3,GNA13,CD74,SMO,FGFR1,PTPN23,LMO4,ABL1,SDCBP,SEMA6B,[illegible]
NAJB1,SUMO1,EHMT2,SALL2,HIPK2,DRAP1,ATF7IP,TRIM27,NCK2,TRIB3,C1QBP,LDB1,DEAF1,L[illegible]

[illegible]1,FTO,WDR6,PRKDC,KDM1A,PLCG1,GNG2,FYN,SLIT3,FBLN1,B2M,STRAP,HGS,MAPK8IP1,BAG[illegible]

GFBR1,MAPK3,HEG1,PIN1,PCNA,EXT1,HEXIM1,SNX17,GYS1,TGFB2,NCOA6,MAML1,GJC1,COL3A

1,PHB,DSTYK,SHC1,PSMD10,MAP2K4,TGFBR1,MAPK3,AJUBA,VAV2,TAB3,EPHB2,TLR6,LRP8,TGI

PTPN11,ABL1,FXR2,SDCBP,PINK1,TAOK2,DAG1,ACVR1B,IGF1R,NPTN,STK39,CAB39,FXR1,PARK
PTPN11,ABL1,FXR2,SDCBP,PINK1,TAOK2,DAG1,ACVR1B,IGF1R,NPTN,STK39,CAB39,FXR1,PARK

HM2,ACTR1B,PDPK1,ITGAV,ADAM10,CAP1,MSH6,NSD2,IST1,PSMC3,AP2A2,AP1M1,CD74,PAFAH1

NA,RPL19,PPP2R1A,RPL15,GTF3C4,DNAH8,PKD1,SF3B1,SERINC1,RPS3,CHD3,VAPA,HMGB1,CDI
CL,AP3D1,DNAJA1,ARF1,CYB5R3,PAFAH1B1,MAP4,AP2B1,SLC4A8,HNRNPU,PGAM1,CTNNA1,RP
RPL36,YWHAB,SND1,LAMP1,RPS12,HLA-B,RAB1A,RPS27,CTBP1,RAC1,TOP2B,IPO5,ATP6V0C,SK
,HMGB1,CDKL1,AP2M1,SEC61A1,STMN1,ITGB1,TUBB4B,UCHL1,RPL11,MORF4L1,NCAM1,ANX
C3,RAB7A,MMP14,DDX3X,RPS15,RPL12,RPL30,ARL4C,CLTC,RPS5,RPS7,IK,RPS19,RPL35,GJA1,
C3,RAB7A,MMP14,DDX3X,RPS15,RPL12,RPL30,ARL4C,CLTC,RPS5,RPS7,IK,RPS19,RPL35,GJA1,
42,TNPO1,SNRNP70,PSMB2,RPL36A,SCAF1,PRPF6,RPS15A,RPL38,RPL37,PPP2CA,SRSF9,PSMD
RPS19,RPL35,MYO5A,CD63,RPS10,RPL35A,TARDBP,RPLP2,ARL6IP1,WLS,RPL36,YWHAB,LAMP1
RPL36,YWHAB,SND1,UBE2H,RPS12,RPS27,HMGCR,GPC1,SKP1,BCLAF1,LAPTM4B,TRIB2,RPL23
SD,SCARB2,NACA,LPCAT1,RPS24,RPS9,XRCC5,MSN,RPL24,KPNB1,RPL17,LLGL1,RPL14,NCS1,F
81,RPL26,CCT7,OAZ1,ASTN2,MCL1,RPL31,XPO1,PTN,CSNK1E,PFKL,RPS25,EIF5A,CDH2,GPC3,F
CLTC,H1-0,IK,PRKAR1A,GJA1,CTNNB1,CD63,MACF1,CYFIP2,ARL6IP1,HNRNPK,H1-10,YWHAB,C
36B,NLGN2,RPS14,RPS20,CD81,RPL26,CCT7,OAZ1,SRSF4,ASTN2,MCL1,RPL31,LRPPRC,KHDRB
3,TRIB2,RPL23A,PTTG1IP,MAP1A,CTSB,RNF187,SQSTM1,NOP53,TAF15,ALKBH5,RPL27,UBE4A,
CT3,VPS35,CCT4,TNPO1,RPL36A,SRP14,TERF2IP,RPS15A,RPL38,VEGFA,OPTN,RPL37,MAPK1,M
BP4,FUS,PSMB4,ELAVL1,SMG5,TTC37,PSMD4,IGF2BP1,PSMC3,MTOR,PSMC1,THRAP3,EIF4A3,
VAPA,HMGB1,AP2M1,TPT1,SEC61A1,RPS4X,RPS3A,RPL18,STMN1,ITGB1,TUBB4B,UCHL1,RPL1
,RPL23,CPE,RPL29,RPSA,CTSD,EIF4H,NACA,RPS24,RPS9,XRCC5,PJA2,MSN,RPL24,RPL17,IGFB
AMA5,BRD3,XRCC5,MSN,RPL24,KPNB1,LLGL1,GAP43,RHOA,KIF5B,NLGN2,RPS14,CXADR,CHD

S27A,PSMD8,APEX1,PCBP4,FUS,TRIR,PSMB4,ELAVL1,SMG5,TTC37,PSMD4,IGF2BP1,PSMC3,MT
MA5,RPS24,RPS9,XRCC5,MSN,RPL24,KPNB1,RPL17,COL1A2,LLGL1,GAP43,RPL14,RPL22,RHOA
DBP,RPLP2,ARL6IP1,NDFIP1,WLS,MYH10,RPL36,YWHAB,LAMP1,RHOB,RPS12,RAB1A,ARCN1,RF
3,TNPO1,RPL36A,SRP14,TMED2,RPS15A,RPL38,RPL37,ARF3,MAPK1,IPO7,CDC37,RPS11,GABAF

WSB1,RPS17,SBF1,DNAJA1,CDC42BPB,ITM2B,AP2B1,OGA,LARP1,RPS16,RPL23,CPE,RPL29,RPS
,MYO5A,CD63,MACF1,RPS10,RPL35A,TARDBP,RPLP2,ARL6IP1,NDFIP1,WLS,MYH10,RPL36,YW
K3B,RRP1B,RPS29,NUP205,NUP133,NELFCD,MAN1B1,SP1,HEXIM1,RPL32,FURIN,NUP50,SMAR

G,CKB,COL4A2,RPL37A,MAPT,FN1,SPEN,NONO,DDB1,PSMD5,YWHAQ,EIF1,HACD3,SET,EGLN1,
81,ATP2A2,RPL26,OAZ1,MCL1,RPL31,HNRNPC,LRPPRC,XPO1,CSNK1E,PFKL,RPS25,GPC3,PIP4
,RPL30,CLTC,H1-0,RPS5,RPS7,RPS19,RPL35,CTNNB1,RPS10,SESN3,RPL35A,TARDBP,RPLP2,ND
,TPT1,RPS4X,RPS3A,RPL18,STMN1,ITGB1,UCHL1,RPL11,MORF4L1,ANXA2,RPLP1,TIMP2,MIDN,

NRNPD,CYP51A1,SERBP1,DDIT4,PSMA7,ANP32A,VDAC1,STAT3,ARIH1,EDC4,SGTA,EP300,PSMD1,
YWHAB,RHOB,RPS12,RAB1A,ARCN1,RPS27,HMGCR,RAC1,CDK16,APPL1,IPO5,ATP6V0C,TMEM3
PL10,RPS17,VCL,AP3D1,DNAJA1,ARF1,CYB5R3,PAFAH1B1,MAP4,AP2B1,SLC4A8,HNRNPU,PGAM
CT5,ADAR,STAT3,SGTA,ALDH3A2,KPNA2,PHB2,UBA52,RAB3GAP2,MBTPS1,NCOA4,POM121C,R
VCL,AP3D1,DNAJA1,ARF1,CYB5R3,PAFAH1B1,AP2B1,SLC4A8,HNRNPU,PGAM1,RPS16,RPL23,RF
,PFN1,CLEC7A,SPTBN2,CSNK1E,CDH6,TPM1,CDH2,PIP4K2B,RAB7A,CCN2,MMP14,DDX3X,RPS1
,GPC3,RAB7A,RPS15,RPL12,RPL30,CLTC,H1-0,RPS5,RPS7,RPS19,RPL35,GJA1,CTNNB1,RPS10,

PS4X,RPS3A,RPL18,UCHL1,RPL11,MORF4L1,ANXA2,AGRN,RPLP1,ZMIZ1,TIMP2,COL11A1,RPL1
MGCR,RAC1,ARHGDIA,CAPRIN1,GPC1,CDK16,TOP2B,HDAC2,TNC,TMEM30A,DNER,L1CAM,NNAT,
N,YWHAH,FAU,STXBP1,MESD,HECTD1,HGS,BAG6,RAB11A,RPL39,SEC23A,RPS13,RPL34,PPP1R9
PU,SMARCA5,LARP1,CTNNA1,RPL23,LDHA,CTSD,SCARB2,NACA,RTN4,RPS9,XRCC5,PJA2,MSN,I
PALLD,KIF5A,PRMT1,TRIO,EP300,RERE,CNTNAP1,AP2A1,FLOT1,MAP6,PLXNB1,DIP2B,FARP1,SF
11,CLIP3,RPL28,RPS28,FYN,YWHAH,FAU,STXBP1,MESD,HECTD1,HGS,BAG6,RAB11A,RPL39,SEC
I2,CSNK2A1,GRSF1,PHB2,CST3,UBA52,ZFP36L1,UPF1,RAB3GAP2,HDAC6,PTEN,ULK1,GPSM1,R

RPL18,ITGB1,UCHL1,RPL11,MORF4L1,ANXA2,AGRN,RPLP1,ZMIZ1,TIMP2,MIDN,RPL10,RPS17,AP
GSPT1,UBA52,ZFP36L1,UPF1,RPS2,RPS27A,PSMD8,APEX1,PCBP4,FUS,TRIR,PSMB4,GPX1,ELAV
NRNPU,SMARCA5,LARP1,HNRNPA0,CTNNA1,RPL23,NACA,RTN4,LPCAT1,XRCC5,CIPC,IGFBP3,RH
14,ATP1A1,RPS20,RPL26,BEX3,SRSF4,RBBP4,RTN3,MCL1,TLE5,RPL31,HNRNPC,LRPPRC,KHDRE
R1A,GJA1,CTNNB1,NGRN,CYFIP2,APLP2,TARDBP,ARL6IP1,NDFIP1,CIRBP,YWHAB,CADM4,RAB1A
5,GTF3C4,PKD1,PDIA3,SSBP2,ZNF43,RPS3,CHD3,BASP1,HMGB1,POLR2A,CDKL1,AP2M1,RPS4X
4,RTN3,TLE5,RPL31,HNRNPC,LRPPRC,KHDRBS1,XPO1,PTN,RPS25,GLG1,SFPQ,PEBP1,GPC3,CC
H5,PRDX5,UBE4A,PKN1,GRINA,CBX1,RAD23B,CHD2,GCN1,PSMB2,KDM6B,HDGF,TERF2IP,TMED
COL1A2,RHOA,ATF4,PSMD2,KIF5B,ATP1A1,JUND,TMEM259,CD81,COL5A2,ATP2A2,SRSF4,MCL1
PARP1,TSPYL4,MTRF1L,ABCA3,ACTR3,HSPA4,TCP1,CAMSAP1,KLC1,CCT3,NCKAP1,RBMX,LAMC
,TCTN1,BMPR2,CDC42,PPP1R9B,PALLD,KIF5A,PRMT1,TRIO,EP300,RERE,PBX1,CNTNAP1,AP2A1
,PLXNA3,TMED9,DGKD,PFN2,HNRNPD,RAB11A,WBP2,AKIRIN1,BMPR2,CCT5,CDC42,VDAC1,DCT
3CA2,DPYSL2,NOTCH2,RNF10,VEGFA,RAPGEF1,TUBB2A,MAPK1,BTBD2,PBX2,SRSF1,LUZP6,GRE
,MAPT,FN1,SPEN,NONO,DDB1,PSMD5,YWHAQ,EIF1,HACD3,SET,EGLN1,TPI1,CHSY1,EIF4A1,RPL
P1,GRSF1,GSPT1,UBA52,ZFP36L1,UPF1,RPS2,RPS27A,PSMD8,APEX1,PCBP4,FUS,TRIR,PSMB4,G
P1,GRSF1,GSPT1,UBA52,ZFP36L1,UPF1,RPS2,RPS27A,PSMD8,APEX1,PCBP4,FUS,TRIR,PSMB4,G

I1,SPEN,NONO,DDB1,PSMD5,YWHAQ,EIF1,HACD3,SET,EGLN1,TPI1,CHSY1,EIF4A1,RPL19,PPP2R
RF4,FYN,YWHAH,SLIT3,STMN3,B2M,STXBP1,PLXNA3,CSNK1D,STRAP,PTPRS,SOD1,RAB11A,TCT
2A,MAPK1,PBX2,LUZP6,GRB2,ATXN10,KDM1A,PLCG1,ARF4,FYN,YWHAH,SLIT3,STMN3,B2M,STX
12,RPL30,ARL4C,CLTC,RPS5,RPS7,RPS19,RPL35,GJA1,MYO5A,CD63,MACF1,RPS10,RPL35A,TA
F4L1,ANXA2,AGRN,RPLP1,ZMIZ1,TIMP2,RPL10,RPS17,AP3D1,DNAJA1,ARF1,AKAP12,ITM2B,ZN
CHMP4B,SSR2,EXOC4,NDUFA13,TOMM70,CCDC47,REEP1,NSG1,ATP13A1,SRPRA,RPS26,SDCB
1,AKAP12,ZNF770,AP2B1,GTF2F1,SLC4A8,HNRNPU,SMARCA5,LARP1,HNRNPA0,CTNNA1,RPL23,
L1,SMARCA5,LARP1,HNRNPA0,RPL23,CTSD,EIF4H,SCARB2,NACA,RTN4,LPCAT1,BRD3,RPS9,XR
HSPB1,NPEPPS,CASC3,QARS1,RPL28,RPS28,LARS1,FAU,EIF3F,IDH1,SARS1,SOD1,HNRNPD,RPL3
AB1A,CTBP1,HMGCR,RAC1,CAPRIN1,HDAC2,IPO5,SKP1,BCLAF1,LAPTM4B,TRIB2,PEA15,PTTG1I
,UBE2D3,TOMM70,SORT1,SAE1,SRPRA,RPS26,UBE2L3,SDCBP,PINK1,TAOK2,TRAK1,MFN2,SLC
GSPT1,UBA52,ZFP36L1,UPF1,RPS2,RPS27A,PSMD8,APEX1,PCBP4,FUS,TRIR,PSMB4,GPX1,ELAV
C42,PALLD,KIF5A,BCL9L,TRIO,EP300,RERE,CNTNAP1,FLOT1,MAP6,PLXNB1,DIP2B,FARP1,CTN
1,GTPBP1,GRSF1,GSPT1,UBA52,ZFP36L1,UPF1,RPS2,RPS27A,PSMD8,APEX1,PCBP4,FUS,TRIR,F
,ADAM10,IGF2BP1,IST1,UBE3A,TMEFF1,APBB1,TANC2,DPYSL4,SMO,METRN,LYPLA2,NTRK2,SN
GF2BP1,IST1,UBE3A,TMEFF1,APBB1,TANC2,DPYSL4,SMO,METRN,LYPLA2,NTRK2,SNX2,DIP2A,F
,ADAM10,IGF2BP1,IST1,UBE3A,TMEFF1,APBB1,TANC2,DPYSL4,SMO,METRN,LYPLA2,NTRK2,SN
KL1,AP2M1,TPT1,SEC61A1,RPS4X,RPS3A,STMN1,ITGB1,UCHL1,BTF3,NCAM1,ANXA2,AGRN,ZMI
11,HDAC6,PTEN,CUL3,OAZ2,RPS27A,PSMD8,NPLOC4,FBXO21,RHBDD2,BAP1,PSMB4,GPX1,EIF
3F,SARS1,HNRNPD,RPL39,RPS13,RPL34,STAT3,EIF2AK1,EDC4,VARS1,GTPBP1,EIF3D,GSPT1,UB

RAD23B,PSMB2,H3-3A,PRPF6,TMED2,TAF7,RPL38,VEGFA,G3BP2,SMARCA4,MAPRE1,SRSF9,ARI
VCP,UBB,AFF4,TRIM28,MATR3,YWHAG,CKB,COL4A2,RPL37A,MAPT,FN1,SPEN,NONO,DDB1,PSN
=A,RAPGEF1,MAPK1,MAPRE1,ARPC2,DYNLL2,GRB2,CAPZB,ATXN10,KDM1A,PLCG1,ARF4,CORO
H,HNRNPUL1,SCAF11,SON,KDM1A,CASC3,MAP2K2,RPL28,CCAR2,POLR2B,SUPT6H,RPS28,HNRI
TP1A1,JUND,COL5A2,ATP2A2,MCL1,CLEC7A,PTN,GLG1,SFPQ,TPM1,PDIA6,GPC3,SULF1,PIP4K2
E3A,TMEFF1,APBB1,TANC2,DPYSL4,SMO,METRN,LYPLA2,NTRK2,DIP2A,PTPN11,ABL1,SEMA6B,
EIF3F,STXBP1,LONP1,CSNK1D,SF3A1,STRAP,PFN2,NAP1L4,ADAR,PSMA7,RBBP7,TRAF7,DCTN1,N
EIF1,HACD3,SET,EGLN1,CHSY1,EIF4A1,RPL19,PPP2R1A,RPL15,GTF3C4,PKD1,SF3B1,PDIA3,SSB
ARP1,HNRNPA0,RPL23,CTSD,EIF4H,SCARB2,NACA,RTN4,LPCAT1,BRD3,RPS9,XRCC5,CIPC,MSN,I

,SRSF7,NUP98,PSMC5,PSMC2,NCBP2,RBM10,PSMC4,PSMF1,YWHAZ,HSF1,IGF2BP2,CTR9,PSME
VEGFA,RAPGEF1,MAPK1,MAPRE1,ARPC2,DYNLL2,GRB2,CAPZB,ATXN10,KDM1A,PLCG1,ARF4,CO
3F,SARS1,HNRNPD,RPL39,RPS13,RPL34,STAT3,EIF2AK1,EDC4,VARS1,GTPBP1,EIF3D,GSPT1,UB
4,PRDX2,NANOG,APLP1,MIF,HMGCS1,PRKAR1A,TLN1,GJA1,JUN,MYO5A,CTNNB1,CD63,SESN3,N
(A2,AGRN,ZMIZ1,TIMP2,COL11A1,RPL10,VCL,AP3D1,SBF1,ARF1,PAFAH1B1,MAP4,ITM2B,AP2B1
IGO1,ABI2,DCLK1,CNP,SEMA3C,CLASP2,ITGAV,SCRIB,ADAM10,IGF2BP1,IST1,UBE3A,TMEFF1,AI
PLP1,RPS7,PRKAR1A,GJA1,JUN,CTNNB1,ANXA6,MACF1,CYFIP2,WLS,MYH10,RHOB,MAP2,RAB1A
DCLK1,CNP,SEMA3C,SPAG9,KRT8,DICER1,CLASP2,WASF2,ADAM10,IGF2BP1,IST1,UBE3A,TMEF
B,AUTS2,TAOK2,TRAK1,DAG1,NIBAN2,PLXNA2,OBSL1,NOTCH1,NPTN,MAPK8IP2,EFNB2,ADNP,C
ATF4,PSMD2,KIF5B,SLC7A5,NLGN2,RPS14,CXADR,ATP1A1,CD81,WDR1,ATP2A2,CCT7,PFN1,MC
2,LUC7L2,RBM4,MTREX,HSPA1A,TIA1,RBM25,SF3A3,CD2BP2,FXR1,SART3,SYNCRIP,HNRNPR,SF
10,VCL,AP3D1,ARF1,PAFAH1B1,MAP4,ITM2B,AP2B1,HNRNPU,CTNNA1,RPS16,LDHA,CPE,RPL29,
A1,PSMC2,HERPUD1,PSMC4,TMEM9,PLK2,PSMF1,WAC,GSK3B,SERPINE2,USP19,UBQLN4,SOX
CTSD,EIF4H,NACA,RTN4,LPCAT1,BRD3,RPS9,XRCC5,CIPC,MSN,PAPOLA,IGFBP3,RPL22,RHOA,PT
RPL28,RPS28,LARS1,FAU,EIF3F,SARS1,HNRNPD,RPL39,RPS13,RPL34,STAT3,EIF2AK1,EDC4,VAF
A1,RPS16,LDHA,CPE,SCARB2,NACA,RTN4,LPCAT1,LAMA5,XRCC5,MSN,RPL24,COL1A2,LLGL1,G
4,TRIM28,MATR3,YWHAG,CKB,COL4A2,RPL37A,MAPT,FN1,SPEN,NONO,DDB1,PSMD5,YWHAQ,E
RSF7,PSIP1,NUP98,NCBP2,RBM10,DNAJC8,DDX41,SNRPD2,SETX,PRMT5,RAVER1,SNRPB,RBM1
RSF7,PSIP1,NUP98,NCBP2,RBM10,DNAJC8,DDX41,SNRPD2,SETX,PRMT5,RAVER1,SNRPB,RBM1
RSF7,PSIP1,NUP98,NCBP2,RBM10,DNAJC8,DDX41,SNRPD2,SETX,PRMT5,RAVER1,SNRPB,RBM1
,UBE2H,CKAP4,RAB1A,CALU,HECTD4,CTBP1,HMGCR,RAC1,CDK16,HDAC2,IPO5,TNC,SKP1,TRIE
,UBE2H,CKAP4,RAB1A,CALU,HECTD4,CTBP1,HMGCR,RAC1,CDK16,HDAC2,IPO5,TNC,SKP1,TRIE
I1,LIMK1,B4GALT5,EFNB1,MYCBP2,NECTIN1,ADCY1,SYNGAP1,ISLR2,RANBP9,GOLGA4,SEMA4C
,RAB7A,CCN2,ACTN4,MMP14,PRDX2,DDX3X,NANOG,H1-0,RPS7,RPS19,MIF,PRKAR1A,GJA1,JUN
,LUC7L2,RBM4,MTREX,TIA1,RBM25,SF3A3,CD2BP2,FXR1,SART3,SYNCRIP,SSU72,HNRNPR,SRSI
iT,PRPF19,USP24,FBXW5,GSK3A,PSMC3,CHMP4B,UBE3A,PRKACA,AP2A2,BNIP3,UBE2I,UBE2D3,
MAP1A,CDKN1A,DLG5,CRPPA,HS6ST1,DDX39B,CDH11,TBCD,GAS7,PTPRA,CAMSAP1,NCKAP1,AC
LXNA3,ATXN2L,TMED9,USP7,PTPRS,GHITM,SREBF2,PFN2,HNRNPD,WBP2,YLPM1,CCT5,CDC42,\
3,TRIB2,RPL23A,CDKN1A,SRP9,ZC3H7B,U2AF2,MORF4L2,TAF15,ALKBH5,RPL27,RBMX,TNPO1,F
1,RPL37,PPP2CA,MAPK1,EIF3E,FTO,ILF3,LUZP6,PCSK1N,RPS11,PRKDC,HSPB1,NPEPPS,CASC3,C
A3,TRMT112,CD2BP2,RPS21,SART3,RAN,CUL4A,SRSF10,RPL27A,C1QBP,LSM4,CELF3,EIF3B,PS
BP,DUX4,YWHAB,PTGES3,RAB1A,CCT6A,CTBP1,RAC1,APPL1,SMCHD1,HDAC2,SKP1,BCLAF1,TR
)1,GDI1,FSCN1,TMBIM6,SPTAN1,GSTP1,SMG1,CALM1,NPM1,PLXNB2,PLD3,HSPD1,COL18A1,CS
TR,ARCN1,COTL1,HRNR,RAC1,CDK16,APPL1,ATP6V0C,TMEM30A,DNER,LAPTM4B,TIMP3,PGRM
EMA4C,P3H1,SRF,GSK3B,CRTC1,SETX,CRABP2,NSMF,THOC2,CNTN1,RETREG3,ARSB,EFNB3,CN

OG,H1-0,RPS7,MIF,PRKAR1A,GJA1,JUN,CTNNB1,CD63,NGRN,CYFIP2,TARDBP,ARID1A,NDFIP1,S
AC2,IPO5,BCLAF1,LAPTM4B,TRIB2,HSBP1,MAP1A,CDKN1A,TIMP3,CTDSP2,EID1,SRP9,U2AF2,S
SLR2,RANBP9,GOLGA4,SEMA4C,SRF,ZSWIM6,GSK3B,ALCAM,RAB3A,MAP1S,CRABP2,ERBB2,N
WHAB,CADM4,CTBP1,HMGCR,CAPRIN1,SMCHD1,HDAC2,IPO5,BCLAF1,LAPTM4B,TRIB2,HSBP1,
SNX3,FXR2,LGALS1,SEMA6B,AUTS2,TRAK1,PLXNA2,OBSL1,MDM2,NPTN,EFNB2,FXR1,ADNP,CYF
,H3-3A,ABCA2,ATP6AP1,TAF7,NOTCH2,MARS1,SELENON,RAPGEF1,CCND1,MAPK1,SMARCA4,DY
ND5A,SNX3,FXR2,LGALS1,SEMA6B,AUTS2,TRAK1,PLXNA2,OBSL1,MDM2,NPTN,EFNB2,FXR1,ADN
,PKN1,TCP1,AHCYL1,GRINA,NCKAP1,VPS35,COL2A1,MDK,DDX42,NOTCH2,VEGFA,VHL,PPP2CA
WHAB,PTGES3,RAB1A,CCT6A,CTBP1,RAC1,SMCHD1,HDAC2,SKP1,BCLAF1,TRIB2,PEA15,PTTG1I
,RPS17,AP3D1,AKAP12,SF3B2,ZNF770,GTF2F1,HNRNPU,NFE2L1,SMARCA5,LARP1,HNRNPA0,R
1,GPC1,ATP6V0C,BCLAF1,RPL23A,GLUL,DCN,CRPPA,HS6ST1,PGRMC1,DDX39B,SRP9,EEF1A2,F
PL22,RHOA,ATF4,PSMD2,NR2F1,KIF5B,ENC1,RPS14,JUND,CD81,COL5A2,WDR1,ATP2A2,CRMP1
B1,AKAP12,MAT2A,GTF2F1,PXDN,HNRNPU,NFE2L1,SMARCA5,LARP1,HNRNPA0,CTNNA1,RPS16,
11,MORF4L1,BTF3,AGRN,RPLP1,ZMIZ1,RPL10,RPS17,AP3D1,CYB5R3,SF3B2,ZNF770,GTF2F1,
RHOA,ATF4,PSMD2,NR2F1,KIF5B,ENC1,RPS14,JUND,CD81,COL5A2,WDR1,ATP2A2,CRMP1,ASTN
RPL14,RPL22,RHOA,CBX5,ZNF865,RPL9,ATF4,PSMD2,NR2F1,RPS14,CHD4,JUND,RPS20,CD81,R
ORF4L1,BTF3,AGRN,RPLP1,ZMIZ1,RPL10,RPS17,AP3D1,SBF1,ARF1,CYB5R3,AKAP12,ITM2B,Z
ORF4L1,BTF3,AGRN,RPLP1,ZMIZ1,RPL10,RPS17,AP3D1,SBF1,ARF1,CYB5R3,AKAP12,ITM2B,Z
,UBE2H,CKAP4,RAB1A,CALU,HECTD4,CTBP1,HMGCR,RAC1,CDK16,HDAC2,IPO5,TNC,SKP1,TRIE
1,HNRNPU,PGAM1,NFE2L1,SMARCA5,LARP1,HNRNPA0,RPS16,RPL23,LDHA,RPL29,RPSA,NACA,F
NDN,BAP1,SLIT1,GPX1,OLFM1,SOX2,PKDCC,LTBP4,HDGFL2,DCLK1,DNAJB2,SEMA3C,SPAG9,C
3,NOP53,PARP1,NOX4,PRDX5,PKN1,TCP1,AHCYL1,GRINA,NCKAP1,VPS35,COL2A1,MDK,DDX42,
CA,RPS24,BRD3,RPS9,XRCC5,CIPC,RPL24,KPNB1,RPL17,PAPOLA,RPL14,RPL22,RHOA,CBX5,ZN

L1,SMARCA5,LARP1,RPS16,RPL23,RPL29,RPSA,EIF4H,NACA,RPS24,BRD3,RPS9,XRCC5,CIPC,R
CERS2,BAG5,GAK,TUBB4A,DENND5A,TRIP12,LMO4,HMGA1,SCAMP5,SDCBP,LGALS1,PINK1,SE
N1,FKBP4,ANTXR1,ABI2,RAB1B,VPS4A,DZIP1,RDX,CAPZA1,DNAJB2,FNBP1L,CLASP2,WASF2,SPT
2A1,UBQLN2,CSNK2A1,CST3,UBA52,PPP2CB,KLHL42,GLUD1,UBE2G2,FBXO11,HDAC6,PTEN,C
RNPA0,RPS16,RPL23,RPL29,RPSA,EIF4H,NACA,RTN4,RPS24,BRD3,RPS9,XRCC5,CIPC,MSN,RPL
2,PPIB,GLUL,DCN,CDKN1A,DLG5,DDX39B,EID1,PARP1,MORF4L2,PKN1,CAMSAP1,VPS35,CLSTN
,MACROH2A1,PKIA,ATP13A2,MDM2,PCM1,NPTN,MGAT3,PARK7,PTPN1,RAN,UBE2J1,CCDC88A,T
ZER1,FBXL19,USP5,AMFR,TRIP12,USP47,UBE2L3,CDC34,RNF11,SDCBP,PINK1,USP10,PSMB1,H
16,RPL23,RPL29,RPSA,EIF4H,NACA,RPS24,BRD3,RPS9,XRCC5,CIPC,RPL24,RPL17,PAPOLA,RPL
,TIMP3,CTSB,EEF1A2,ANXA5,SQSTM1,GNAI3,NOP53,PARP1,NOX4,PRDX5,PKN1,TCP1,AHCYL1,
UB1,MARCHF7,PSMC5,CSNK1A1,PSMC2,HERPUD1,PSMC4,PLK2,PSMF1,WAC,GSK3B,USP19,UB
B,MEX3C,SERPINE2,ACSL4,ALCAM,SOX9,EI24,ADIPOR2,CRABP2,CDKN1B,EGLN2,ERBB2,UNC1
G2,SMARCC1,ATP6V1B2,FYN,HNRNPM,LARS1,YWHAH,SLIT3,CSNK1D,STRAP,SOD1,NCOR2,HN
LRD1,GDI1,FSCN1,TMBIM6,SPTAN1,GSTP1,SMG1,CALM1,NPM1,PLXNB2,PLD3,HSPD1,COL18A
,CCN2,ACTN4,MMP14,PRDX2,DDX3X,RPS15,ITM2C,APLP1,RPS7,MIF,PRKAR1A,GJA1,CTNNB1,C
RPS27A,FKBP1A,CTTN,LOX,SLIT2,BRK1,AKAP9,TMOD2,TAOK1,GPX1,HSPG2,FKBP4,ABI2,LTBP4

ZNF770,GTF2F1,HNRNPU,PGAM1,NFE2L1,SMARCA5,LARP1,HNRNPA0,RPS16,RPL23,LDHA,RPL
MIZ1,RPL10,RPS17,AP3D1,SBF1,ARF1,CYB5R3,ITM2B,ZNF770,MAT2A,GTF2F1,HNRNPU,OGA,N
,FKBP8,PLCG1,QARS1,CCAR2,ARF4,FYN,MTRNR2L10,STXBP1,UBQLN1,HYOU1,MAPK8IP1,GH
,GTF2I,SEMA3C,SPAG9,CLASP2,PDPK1,PSMD4,ADAM10,IST1,VASH1,PSMC3,UBE3A,AP2A2,BNIP

SV2A,PREPL,TMEM167A,ATP6AP2,A2M,PRDX6,GRN,ACLY,SEPTIN5,XRCC6,SLC44A2,VPS41,IMPD
DEK,EIF4G1,AMFR,TRIP12,USP47,HMGA1,ABL1,FXR2,SDCBP,PINK1,TAOK2,HSPA1A,IGF1R,MDM
BAG6,PFN2,HNRNPD,WBP2,BMPR2,TRAF4,TRAF7,STAT3,ARIH1,PRMT1,SGTA,MTA1,PRKAR2A,HI
,IMPDH2,RAB14,NCSTN,RAB6A,RAB5C,ACTR1B,PDPK1,ITGAV,SCRIB,ADAM10,CAP1,SYT11,IST1,
ACTN4,MMP14,PRDX2,DDX3X,RPS15,ITM2C,APLP1,RPS7,MIF,PRKAR1A,GJA1,CTNNB1,CD63,MA
ZNF770,GTF2F1,HNRNPU,PGAM1,NFE2L1,SMARCA5,LARP1,HNRNPA0,RPS16,RPL23,LDHA,RPL2
CA5,LARP1,RPS16,RPL23,RPL29,RPSA,EIF4H,NACA,RPS24,BRD3,RPS9,XRCC5,CIPC,RPL24,RPL
XNB2,HSPD1,COL18A1,CSDE1,IGF2,EIF3C,ATP5F1A,EIF4B,VCP,MARCKSL1,UBB,TRIM28,YWHA
9,UBQLN4,RNF216,PSMB5,UBXN1,PSMA1,ZNRF1,RNF4,PSMD10,UBE2G1,ELOB,ZSWIM8,PSMA6
BLN1,UBQLN1,CSNK1D,STRAP,HGS,MAPK8IP1,SOD1,PFN2,PPP6R1,WBP2,BMPR2,KDM4B,DDIT
PA1A,MDM2,FZR1,UBE4B,TNFAIP1,TRIM2,CUL1,PSMD13,SUMO1,PARK7,USP14,PSMD7,FAF2,MA
NOLC1,NUP50,RNPS1,DDX19A,NUP58,NEMF,WDR33,NUTF2,EMD
1,HIF1A,FKBP1A,GPC4,SLC2A1,SERP1,BAP1,TCAF1,UBR5,PKDCC,GPC2,VPS4A,RAB14,RDX,TM9
PA1A,MDM2,FZR1,UBE4B,TNFAIP1,TRIM2,CUL1,PSMD13,SUMO1,PARK7,USP14,PSMD7,FAF2,MA

1,UBE4B,TNFAIP1,TRIM2,CUL1,PSMD13,SUMO1,PARK7,USP14,PSMD7,FAF2,MAEA,HIPK2,CUL4
,CCAR2,ARF4,FYN,MTRNR2L10,STXBP1,UBQLN1,HYOU1,MAPK8IP1,GHITM,SOD1,BAG6,ARNT2,
56,PLXND1,ATP8A2,ANAPC2,SKIL

TN1,HSPB1,SON,KDM1A,BNIP3L,CLIP3,FKBP8,PLCG1,QARS1,CCAR2,ARF4,FYN,MTRNR2L10,ST
BEX4,STAT3,EIF2AK1,EDC4,SGTA,PRKAR2A,CSNK2A1,CST3,ZFP36L1,NXN,SFRP1,UPF1,CRTAP,M
K1,LMAN1,HIF1A,FKBP1A,GPC4,BAP1,TCAF1,UBR5,PKDCC,GPC2,VPS4A,RDX,TM9SF4,DCLK1,NM
6,NEDD8,SPSB3,QSOX1,PRSS23,MXRA8
ST,ANP32B,HIF1A,ATP6AP2,SLIT2,SLC2A1,NDN,GRN,XRCC6,SLIT1,HSPG2,SSBP3,TPP1,SOX2,AK
DAG1,NRXN2,OBSL1,IGF1R,NPTN,MECP2,EFNB2,JAM3,ADNP,CYFIP1,MAP4K4,NTN1,IQSEC1,NE

3,SF3B2,ZNF770,GTF2F1,HNRNPU,PGAM1,NFE2L1,SMARCA5,LARP1,HNRNPA0,RPS16,RPL23,LD

PIK3C2B,FEZ1,EHMT2,RHEB,VPS51,CSNK2B,CAPN1,TBC1D14,RNF41,TRIM27,TRIB3,ATP6V0B,S
PIK3C2B,FEZ1,EHMT2,RHEB,VPS51,CSNK2B,CAPN1,TBC1D14,RNF41,TRIM27,TRIB3,ATP6V0B,S
,ATF5,EHMT2,NTN1,LIMK1,PDGFRB,CDK4,ISLR2,BCL9,WDR11,NDRG4,GOLGA4,SEMA4C,MEIS1
,DDX19A,NUP58,NEMF,WDR33,NUTF2,EMD

3BP1,DDIT4,PSMA7,RBBP7,CAPN2,MTA1,EP300,PSMD1,PHB2,UBA52,ZFP36L1,SFRP1,PTEN,PPP
P,ACTN1,SON,MAP7D1,CLIP3,FLNB,TPM3,CORO1C,STMN3,PLXNA3,CSNK1D,MPRIP,SOD1,PFN2
,DYNC1LI2,DYNLL2,CDC37,WDR6,SON,CCAR2,CSNK1D,PTP4A1,BAG6,RAB11A,PRPF40A,DYNLL

C4,TMEM9,PSMF1,GSK3B,ZNF703,LGR4,CELSR3,SOX9,CTR9,PSMB5,CHD8,PSMA1,MED12,LZTS
AC2,NAPA,RNF4,ARPC5,CAPZA2,THRA,ADD1,AJUBA,PPP2R5B,FER,GMFB,CUL4B,GIT1,CTNNBIP1
TGFBR1,RUFY3,PIN1,XBP1,BCAR1,SEMA6C,APOE,TGFB2,CRIM1,FGF13,FHL1,SYT1,PPT1,KDM2
TRA1,FLII,ETF1,PPP1CA,IQSEC1,CAPN1,AKAP8L,NCK2,MICAL3,CSNK1A1,HSF1,GSK3B,MRPS6,U
M9,PSMF1,GSK3B,ZNF703,LGR4,CELSR3,SOX9,CTR9,PSMB5,CHD8,PSMA1,MED12,LZTS2,PSMD

ARIH1,SGTA,MTA1,PRKAR2A,HIP1,FLCN,UBQLN2,FLOT1,EIF3D,PHB2,BRD4,UBA52,DIP2B,RAB3
SFRP1,SRSF6,ARID1B,UBE2G2,ADCY6,UQCRC1,PTEN,ABCC9,APEX1,NPLOC4,SLIT2,SLC2A1,AK

S1,SYNGR1,PA2G4,RAB3A,ARL8A,MAN2B1,DPP7,BRI3,ARSB,ATP6V0A1,ARPC5,DYNLT1,TRAPPC

PSMC1,TRIM8,BAG5,SAE1,UBE2K,FBXL19,USP5,RANGAP1,AMFR,TRIP12,UBE2L3,CDC34,RNF1

NXN,SFRP1,UPF1,CRTAP,MASTL,HDAC6,PTEN,SPINT2,PPP1R15A,FKBP1A,SLIT2,A2M,PPP1R10,G
S1,SYNGR1,PA2G4,RAB3A,ARL8A,MAN2B1,DPP7,BRI3,ARSB,ATP6V0A1,ARPC5,DYNLT1,TRAPPC

CAPN2,PRKAR2A,UBQLN2,FLOT1,ZFP36L1,ATRX,SFRP1,SRSF6,ARID1B,UBE2G2,ADCY6,UQCRC1
8,PSMC2,DEGS1,SYNGR1,PA2G4,RAB3A,ARL8A,MAN2B1,DPP7,BRI3,ARSB,ATP6V0A1,ARPC5,DY
AN2,HSPA1A,ACVR1B,SMPD3,TIA1,NOTCH1,MDM2,NPTN,CYFIP1,CD44,FEZ1,ACAP2,KAT2A,PTP
FOXK2,PSMD4,LRRC41,OGT,PRPF19,USP24,PCGF2,FBXW5,GSK3A,PSMC3,MTOR,UBE3A,YY1,A
SN,ALDH6A1,SPTBN1,ATP5F1B,CALM3,GNB1,SHISA9,STMN2,RPS8,MYH9,RPL7,SEPTIN2,P4HB,
18,PSMC2,DEGS1,SYNGR1,PA2G4,RAB3A,ARL8A,MAN2B1,KARS1,DPP7,BRI3,ARSB,ATP6V0A1,A
NPK,YWHAB,CADM4,RHOB,HMGCR,RAC1,ARHGDIA,GPC1,APPL1,HDAC2,ATP6V0C,BCLAF1,PEA
RT3,CYFIP1,KAT2A,LIMK1,TASOR,ATF7IP,PDGFRB,CCDC88A,AKAP8L,TRIM27,NCK2,CDK9,DHX3
CSNK2A1,PHB2,GSPT1,BRD4,UBA52,ZFP36L1,ATRX,SFRP1,UPF1,MASTL,PTEN,SMC3,FEM1B,P
C2,DEGS1,SYNGR1,PA2G4,RAB3A,ARL8A,MAN2B1,DPP7,BRI3,ARSB,ATP6V0A1,ARPC5,DYNLT1,
E1,UBE2K,FBXL19,USP5,AMFR,TRIP12,UBE2L3,CDC34,RNF11,WDTC1,ABL1,PINK1,PSMB1,RTF1
,INTS3,VPS4A,RDX,LSM14A,KMT2E,SRSF2,TUBGCP6,CLASP2,TCF3,PSMD4,SCRIB,IST1,FBXW5,
X12,VPS37B,RB1CC1,USP36
BAP1,PPP1R10,PSMB4,TAOK1,EHMT1,CARM1,INTS3,VPS4A,RDX,KMT2E,TCF3,PSMD4,SCRIB,PC
,ACTN1,CAND1,SDF4,ILF2,CCT8,PFKM,B2M,STXBP1,IDH1,HGS,SOD1,PFN2,VGF,RAB11A,CYP51
FBXW5,ARPP19,PSMC3,CHMP4B,PRKACA,NASP,UBE2I,PHIP,PSMC1,DAPK3,TUBB4A,EIF4G1,RAN

VHL,MAPK1,SMARCA4,DDAH2,PBX2,EIF3E,ILF3,LUZP6,PRKDC,SUPT5H,ACTN1,CAND1,KDM1A,M
ARHGDIA,HDAC2,SKP1,BCLAF1,PEA15,PTTG1IP,DCN,CDKN1A,DLG5,TIMP3,DDX39B,NUCB1,PP

AC6,SMC3,NAP1L3,CBX6,HNRNPA2B1,CUL3,ANP32B,MCM3AP,RPS27A,TNKS,APEX1,BAP1,PPP1
APLP2,ARL6IP1,NDFIP1,HNRNPK,YWHAB,PTGES3,ATP1A2,MAP2,RAB1A,HMGCR,ARHGDIA,HDAC
PBX2,EIF3E,ILF3,PRKDC,SUPT5H,ACTN1,CAND1,KDM1A,MAP2K2,CCAR2,ARF4,SMARCE1,SMARC

IA1,NOTCH1,MDM2,NPTN,CYFIP1,CD44,FEZ1,ACAP2,KAT2A,PTPN1,HTRA1,HIPK2,PDGFRB,EXT2
S4A,LDLR,CNP,FLOT2,NMT1,FNBP1L,PLEKHM2,MOAP1,REEP2,SYT11,AKT2,IST1,GSK3A,CHMP4
E3A,BNIP3,SLC38A2,SMO,NTRK2,PAFAH1B2,XRN2,PTPN11,ABL1,SUN1,SEMA6B,PLXNA2,NOTCH

CC1,SIN3A
H3PXD2A,SEMA3C,NMT1,KRT8,CLASP2,PHGDH,ITGAV,SCRIB,ADAM10,RBPJ,PRPF19,PCGF2,MBI

l,CAND1,KDM1A,MAP2K2,CCAR2,ARF4,SMARCE1,SMARCC1,SUPT6H,YWHAH,ILF2,CCT8,PFKM,T

,EIF4G1,USP47,PTPN11,CHMP1A,PSMB1,TAOK2,MACROH2A1,HSPA1A,SMPD3,OBSL1,PKIA,MDI
K2,WAC,HSF1,GSK3B,SOX9,CDKN1B,IRS1,EGLN2,SKIV2L,PSMD10,ROCK2,ATF6,ELOB,RGMA,M
,RDX,NUCKS1,PDPK1,AKT2,OGT,ATP6V0D1,GSK3A,MTOR,UBE3A,PRKACA,GOT2,YY1,ATP6V1G1

RKL,P3H1,GSK3B,BCOR,WARS1,RASSF2,CDKN1B,SH3BP5,OTUB1,DUSP3,DUSP8,PPP1R37,KDM

2B,CAPN1,NME2,ASAH1,RAB18,DHX36,PSMC2,DEGS1,SYNGR1,PA2G4,RAB3A,ARL8A,MAN2B1,D
L26,MCL1,CLEC7A,XPR1,PTN,CSNK1E,SFPQ,TPM1,PNMA1,PDIA6,DDR1,RAB7A,CCN2,MMP14,PI
A,MAPK1,PSMD3,ADAM9,ACTN1,CAND1,SDF4,ILF2,CCT8,PFKM,B2M,STXBP1,IDH1,HGS,SOD1,PI
H1,RAB18,PSMC2,DEGS1,SYNGR1,PA2G4,RAB3A,ARL8A,MAN2B1,DPP7,BRI3,ARSB,ATP6V0A1,A
R1A,PKD1,HLA-C,PDIA3,RPS3,CHD3,VAPA,HMGB1,POLR2A,AP2M1,TPT1,OR8K1,STMN1,ITGB1,U

RAB11A,AKIRIN1,BMPR2,CDC42,PSMA7,STAT3,BCL9L,PRMT1,NAP1L1,TRIO,EP300,LBH,PSMD1,I
UN1,SEMA6B,PLXNA2,NOTCH1,PCM1,MECP2,DNAJB1,FEZ1,KAT2A,ATF5,SUN2,VPS51,H2AX,ADC

INK1,AUTS2,MTA2,KDM3A,PPP5C,RTF1,TSPYL2,MACROH2A1,ANP32E,RUVBL2,SIN3B,MECP2,TI
EC31A,DYNC1I2,COG4,PLPP3,CYTH3,ARFGAP3,TMED5
N1B,WASF3,RBM14,RHOBTB2,CDC42EP4,RNF4,BMERB1,ARPC5,SHANK1,ROCK2,CHMP5,TGFB
2,ACVR1B,NOTCH1,KIF26A,ADNP,CYFIP1,RNF157,NTN1,LIMK1,CDK4,ISLR2,CRKL,SPOCK1,GOI
37,EIF4G3,TRAP1,LSM14B,DICER1,ABCF1,PSMD4,IGF2BP1,AKT2,PSMC3,MTOR,PSMC1,DAPK3,T
CBP2,IGF2BP2,DHFR,SHMT2,PA2G4,ERBB2,PER1,SKIV2L,ROCK2,MAPK3,ZNF598,PIN1,CNOT3,I
H1,RAB18,PSMC2,DEGS1,SYNGR1,PA2G4,RAB3A,ARL8A,MAN2B1,KARS1,DPP7,BRI3,ARSB,ATP
RL6IP5,ERP29,PTPN1,SHISA5,HIPK2,CUL4A,PPP1CA,RNF41,NCK2,TRIB3,C1QBP,PSME3,MARCH

1,RGMA,PPP3CA,RUFY3,ADD1,CUL7,PRKCI,DYNLT1,SEMA6C,HMGB2,RELA,OPA1,EPHB2,LRP8,I
E3,BCL9,DAAM1,PSMC5,CSNK1A1,PSMC2,PSMC4,TMEM9,PSMF1,GSK3B,ZNF703,LGR4,CELSR3
LEC7A,HNRNPC,LRPPRC,TADA2B,KHDRBS1,XPO1,SFPQ,EIF5A,SOX12,CCN2,ACTN4,PRDX2,DDX

ND1,MAPK1,PSMD3,ADAM9,ACTN1,CAND1,SDF4,ILF2,CCT8,PFKM,B2M,STXBP1,IDH1,HGS,SOD
1,NOX4,HSPH1,PRDX5,PKN1,TBC1D16,TCP1,ACAP3,CCT4,GCN1,ABCA2,NRDC,TMED2,TAF7,VEG
36,PCYOX1,YME1L1,PSMA7,STAT3,ARIH1,SGTA,OS9,CAPN2,MTA1,CPD,EP300,PSMD1,HIP1,UBQ

DAM10,AKT2,PCGF2,NOC2L,MTOR,NDUFA13,BNIP3,PHIP,BAG5,CD74,MICAL1,PRDX3,AMBRA1,
),CCN1,GPX1,TLE1,TAX1BP1,MAZ,KIFAP3,TRAP1,SOD2,PDPK1,ITGAV,ADAM10,AKT2,PCGF2,NO

TF5,CCNY,PSMD7,CUL4A,CETN2,PPME1,CDK4,AKAP8L,H2AX,CCNG1,PSME3,PSMC5,PSMC2,PSM
IAPK3,ZNF598,CNOT3,PAIP1,YBX3,EIF1B,ABCE1,NOLC1,TUT4,SESN2,EIF2S1,SECISBP2L,CTIF,M
,WARS1,CDKN1B,WASF3,ATMIN,CDC42EP4,THBS2,ARPC5,ROCK2,TGFBR1,SEPTIN9,AJUBA,LRR
,XPO1,SFPQ,EIF5A,SOX12,ACTN4,PRDX2,DDX3X,ITM2C,CLTC,NANOG,H1-0,PRKAR1A,GJA1,JUN
,SHC2,LTBP4,RAB14,NCSTN,NUCKS1,CLASP2,MTMR4,PDPK1,WASF2,LTBP3,ANKS1A,ITGAV,AKT

DPK1,ITGAV,ADAM10,AKT2,PCGF2,NOC2L,MTOR,NDUFA13,BNIP3,PHIP,BAG5,CD74,MICAL1,PRD

FIA3,PLXND1,ANAPC2,UNC13B

FIP1,MXD4,SHOX,HNRNPK,H1-10,CIRBP,DUX4,YWHAB,PTGES3,HNRNPH1,ZNF454,CCT6A,CTBP1

C,TADA2B,KHDRBS1,XPO1,SFPQ,EIF5A,SOX12,ACTN4,PRDX2,DDX3X,ITM2C,NANOG,H1-0,PRKA

F5A,SOX12,ACTN4,PRDX2,DDX3X,ITM2C,NANOG,H1-0,PRKAR1A,GJA1,JUN,CTNNB1,NGRN,TARD
DKN1B,PSMB5,RRM2,PRKAR2B,XPC,PSMA1,PRMT2,PPP1R12A,PSMD10,ANXA1,PPP3CA,PSMA6,C

T4,POLR2E,CIC,RBBP7,STAT3,MTA1,KDM5C,EP300,LBH,RERE,KDM5B,SF1,FLCN,H2AZ1,PHB2,U

SMARCC1,SUPT6H,YWHAH,ILF2,CCT8,PFKM,TNKS2,USP7,RAI1,SREBF2,HNRNPD,ARNT2,WBP2,A

FURIN,FBXO7,SCOC,SESN2,PSMC6,RAD23A,NNT,PFKFB3,RNF40,MEX3D,CNOT1

MPT

CND1,MAPK1,SMARCA4,PRDX1,ADAM9,GRB2,PRKDC,KDM1A,NSG2,GNG2,SDF4,SMARCC1,AT

RPRM,PRMT5,CDKN1B,PSMB5,DUSP3,PRKAR2B,XPC,PSMA1,PRMT2,PSMD10,MDC1,ANXA1,PS

AB39,NDUFC2,APEH,CYFIP1,CD44,PSMD7,FAF2,TSPAN14,CSNK2B,CAPN1,NME2,ASAH1,RAB18,

A,RAC1,ARHGDIA,GPC1,TOP2B,APPL1,DNER,L1CAM,PPIB,GLUL,DCN,MAP1A,DLG5,CRPPA,CNN2

2,QARS1,DUSP4,CORO1C,FYN,PFKM,FBLN1,CSNK1D,PRPS1,STRAP,HGS,MAPK8IP1,SOD1,DGK

F454,CTBP1,CDK16,SMCHD1,HDAC2,BCLAF1,HSBP1,DCN,CDKN1A,DDX39B,EID1,NOVA2,SRSF5

,CD44,ARL6IP5,HIPK2,TMEM98,MAGEF1,TRIB3,ADRM1,PSME3,STUB1,MARCHF7,PSMC5,CSNK

IKS2,STXBP1,CSNK1D,MAPK8IP1,SOD1,DGKD,WBP2,BMPR2,DYNLL1,CDC42,PSMA7,TRAF4,TRA

4A2,MAPT,FN1,SPEN,NONO,DDB1,PSMD5,YWHAQ,HACD3,DCHS1,EGLN1,CHSY1,INA,PPP2R1A,

6R1,BMPR2,DYNLL1,CDC42,DDIT4,PPP1R9B,ADAR,TRAF4,TRAF7,STAT3,EIF2AK1,CAPN2,EP300,

6R1,BMPR2,DYNLL1,CDC42,DDIT4,PPP1R9B,ADAR,TRAF4,TRAF7,STAT3,EIF2AK1,CAPN2,EP300,

ZNF207,NINL,ID2,HMGA2,ANAPC2,PLRG1,SIN3A,CNOT1,PPP2R5C

,CYFIP1,PDLIM1,PHPT1,PTPN1,LIMK1,FLII,IQSEC1,PDGFRB,CCDC88A,RAP2A,TRIM27,NCK2,MI

SYP,CTNND1,PSMA7,VDAC1,KIF5A,SULF2,STAT3,KCNQ2,BCL9L,BCR,PRKAR2A,KDM5B,PSMD1,

K2A1,PHB2,BRD4,UBA52,PPP2CB,NXN,SFRP1,PTEN,CUL3,IL6ST,RPS27A,PSMD8,NPLOC4,PEG1

1,CDH2,GPC3,SULF1,PIP4K2B,MEGF8,RAB7A,CCN2,ACTN4,MMP14,PRDX2,DDX3X,RPS15,ITM2

,MTA1,EP300,LBH,RERE,PBX1,FLCN,H2AZ1,GTPBP1,GRSF1,BRD4,UBA52,ZFP36L1,HOXB5,ATR

A52,CXXC5,ATRX,SFRP1,SRSF6,HDAC6,CBX6,HNRNPA2B1,CUL3,RPS27A,APEX1,DKK3,DAP,PCB

,SFRP1,PTEN,CUL3,IL6ST,RPS27A,PSMD8,NPLOC4,PEG10,DKK3,HIF1A,PPP1R15A,FKBP1A,CTT

K2A1,PHB2,BRD4,UBA52,PPP2CB,NXN,SFRP1,PTEN,CUL3,IL6ST,RPS27A,PSMD8,NPLOC4,PEG1

XBP1,CSNK1D,MAPK8IP1,SOD1,DGKD,WBP2,BMPR2,DYNLL1,CDC42,PSMA7,TRAF4,TRAF7,SUL

1,PSMA6,CNOT3,PCNA,TFDP1,PSMD12,MSH2,PSMA4,TPX2,FHL1,DYNC1I2,FBXO7,UBE2E1,CEN

F1,DNMT1,SMARCB1,SETD1B,HMGA2,BRD1,C6ORF89,RNF40,PRMT7,SIN3A,USP36

P2K2,QARS1,CCAR2,DUSP4,FYN,LARS1,STMN3,FBLN1,UBQLN1,USP7,HYOU1,MAPK8IP1,SOD1,

,PHACTR4,ARPC1A,FAM171A1,TAOK2,HIP1R,ELN,OBSL1,CRACD,TNFAIP1,JAM3,CYFIP1,PDLIM1,

DBP,ARL6IP1,NDFIP1,WLS,HNRNPK,MYH10,YWHAB,CADM4,LAMP1,RHOB,ATP1A2,MAP2,CCT6

ETD1B,HMGA2,BRD1,C6ORF89,RNF40,PRMT7,SIN3A,USP36

2,TMEM43,RAB3A,PRMT5,NUP205,LMNA,NUP133,GET1,CHMP5,NECTIN2,MAPK3,CTDNEP1,TRA
A4,ABCC9,APEX1,LOX,SLIT2,SAFB2,AKAP9,PALM,CARM1,UBR5,EFTUD2,SRSF2,NCSTN,ARPP19,

,NF2,ID2,ANAPC2,SKIL,HES6

GALS1,SCAMP1,PSMB1,HSPA1A,PSMD13,CAB39,NDUFC2,APEH,CYFIP1,CD44,PSMD7,FAF2,TSPA
,SRF,ZEB1,ACSL4,BCOR,SOX9,HOXA5,CTR9,RRM2,CHD8,NUP133,NR2F2,MED12,SLC5A7,DNAJ
LP1,RPS19,HMGCS1,GPAT4,PRKAR1A,GJA1,JUN,CTNNB1,ANXA6,NDFIP1,TAGLN,MYL6,WLS,PTG
EZ1,KAT2A,NTN1,RAN,SUN2,KLC2,CETN2,ARFGAP2,WDR11,NDRG4,YWHAZ,MCFD2,RAB3A,MAP
B1,BCOR,SOX9,HOXA5,CTR9,RRM2,CHD8,NUP133,NR2F2,MED12,SLC5A7,DNAJB6,ITPK1,CITE

DYNLL1,TP53BP1,CDC42,TRAF4,TRAF7,STAT3,BCR,EP300,PRKAR2A,HIP1,FLCN,FLOT1,EIF3D,F

TPN1,EFNB1,PDGFRB,CCDC88A,CDK4,NCK2,MAPKAPK2,EPN1,ATP6V0B,ARPC3,STUB1,EMILIN1,

,ADAR,PSMA7,TRAF4,EIF2AK1,PRMT1,SGTA,EP300,PRKAR2A,PSMD1,UBQLN2,GPX4,BRD4,SFR

MP1,PSMB1,HSPA1A,PSMD13,CAB39,NDUFC2,APEH,CYFIP1,CD44,PSMD7,FAF2,TSPAN14,TMEM

TM1,GNAI3,NOP53,PARP1,NOX4,ABCA3,PTPRA,PDXK,PKN1,ETNK1,PTK7,GCN1,ABCA2,TERF2IP
,PRMT1,EP300,PSMD1,HIP1,FLCN,CSNK2A1,FLOT1,PHB2,BRD4,UBA52,CXXC5,PLXNB1,CSNK1
SK3A,MTOR,PRKACA,UBE2I,WNK1,BAG5,CD74,PRDX3,AMBRA1,FGFR1,NTRK2,LMO4,UBE2L3,F
HSF1,ZEB1,GSK3B,SOX9,PRMT5,STAT1,NSMF,IRS1,PRKAR2B,ATP6V0E2,BACE1,SOS1,GPR173,
,FLOT1,PHB2,BRD4,UBA52,SFRP1,ADCY6,HDAC6,PTEN,IL6ST,POM121C,RPS27A,HIF1A,LOX,SL

S4A,NUDCD3,GABARAPL1,RAB14,DZIP1,RDX,LSM14A,KIFAP3,TUBGCP6,FNBP1L,KRT8,CLASP2,

N3,VAV2,FER,GMFB,SEMA6C,APOE,FGF13,PDXP,ARF6,MYADM,GSN,ANAPC2,RB1CC1,SIN3A

KT2,OGT,FKBP9,PCGF2,NOC2L,KDM4A,MBD3,GSK3A,MTOR,PRKACA,CREBBP,APBB1,UBE2I,W

GOLPH3,ZCCHC3,CDK4,TRIB3,ADCY1,ATP6V0B,RTF2,CRKL,HSF1,ZEB1,GSK3B,SOX9,PRMT5,ST

,ANXA2,AGRN,ZMIZ1,TIMP2,XAF1,MIDN,WSB1,DNAJA1,ARF1,PAFAH1B1,AKAP12,CDC42BPB,AP2

,PRPF40A,BMPR2,AMOTL2,CDC42,DDIT4,PPP1R9B,PALLD,BEX4,STAT3,RERE,FLCN,CD99,PHB2

,MED12,SOS1,CITED2,LZTS2,PSMD10,RGMA,PSMA6,EXT1,AJUBA,PSMD12,OPA1,CTNNBIP1,TG

1,PSMC5,PSMC2,NDRG4,SEMA4C,PSMC4,PSMF1,SRF,LGR4,CELSR3,SOX9,HOXA5,PSMB5,STAT

TM1,GNAI3,NOP53,PARP1,NOX4,ABCA3,PTPRA,PDXK,PKN1,ETNK1,PTK7,GCN1,ABCA2,TERF2IP,

REBF2,NCOR2,HNRNPD,BMPR2,DYNLL1,TXNIP,CIC,RBBP7,STAT3,EIF2AK1,EDC4,MTA1,KDM5C

NOTCH2,VEGFA,TUBB2A,MAPK1,MAPRE1,ARPC2,DSTN,ADAM9,GRB2,HSPB1,PLCG1,ARF4,ZFA

,ANXA2,AGRN,ZMIZ1,TIMP2,XAF1,MIDN,WSB1,DNAJA1,ARF1,PAFAH1B1,AKAP12,CDC42BPB,AP2

IT2,AKAP9,GRK2,PALM,LDLR,TRAP1,NCSTN,NUCKS1,SOD2,PDPK1,AKT2,OGT,ATP6V0D1,PCGF2

,EDC4,MTA1,KDM5C,EP300,LBH,RERE,KDM5B,SF1,FLCN,AP2A1,H2AZ1,PHB2,UBA52,ZFP36L1,

,RPL32,TUT4,PES1,KRI1,HARS1,DDX56,ELAC2,DUS1L,NOP58,USP36,DTD1,SNU13

1CC,IL6ST,HIF1A,SLIT2,SERP1,GRN,XRCC6,CCN1,SSBP3,OLFM1,CARM1,SOX2,PKDCC,AKT3,M

S1,SLC16A1,ERBB2,NADK,CAMK2N1,ANXA1,PPP3CA,RUFY3,XBP1,TFDP1,UFM1,NOLC1,APOE,T

,HIF1A,PPP1R15A,ATP6AP2,SERP1,PRDX6,FUS,XRCC6,OLFM1,UBR5,ELAVL1,ZMIZ2,TLE1,LDLR,

RBFOX2,HSF1,GSK3B,LGR4,CACNA1H,PGRMC2,ADIPOR2,STAT1,NSMF,IRS1,EGLN2,PRKAR2B,

NP,EPHA4,SESN2,KLF3,DNM1,GRB10,RAP1B,ATP6V1F,LEPROT,ZNF106,LRP6,BTG1,KLF10,LDO

KDM5C,EP300,LBH,RERE,KDM5B,SF1,FLCN,H2AZ1,PHB2,UBA52,ZFP36L1,CXXC5,ATRX,SFRP1,

K1,PSMD3,ACADVL,ADAM9,FTO,LUZP6,GRB2,PRKDC,FKBP8,ZFAND5,DUSP4,FLNB,CORO1C,ME

2,PRKAR2B,MYL12A,ERBB2,ITPK1,CFLAR,MAP2K4,RAB5A,RGMA,ANXA1,TGFBR1,MAPK3,PPP3

6L1,SFRP1,UPF1,UBE2G2,AGPAT1,HNRNPA2B1,IL6ST,RPS27A,PSMD8,HIF1A,SLC25A5,LOX,SLIT

RID1B,RGMB,NCOA4,RPS27A,TNKS,APEX1,HNRNPAB,HIF1A,NDN,FUS,XRCC6,PELP1,CCN1,SSBP

RID1B,RGMB,NCOA4,RPS27A,TNKS,APEX1,HNRNPAB,HIF1A,NDN,FUS,XRCC6,PELP1,CCN1,SSBP

N2,MACROH2A1,RBM4,CLIC1,PLXNA2,HSPA1A,ACVR1B,OBSL1,NOTCH1,MDM2,NPTN,EFNB2,AD

RID1B,RGMB,NCOA4,RPS27A,TNKS,APEX1,HNRNPAB,HIF1A,NDN,FUS,XRCC6,PELP1,CCN1,SSBP

SMC3,MTOR,AP2A2,YY1,TMEFF1,EXOC4,PSMC1,FJX1,SMO,FGFR1,EIF4A3,NTRK2,CELSR1,ABL

6V1D,GNAI1

R1,UBE2K,EIF4G1,NTRK2,DIP2A,UBE2L3,PTPN11,ABL1,FXR2,SDCBP,PINK1,AUTS2,TAOK2,RTF1,

F,HEG1,XBP1,AJUBA,HEXIM1,CABIN1,GMFB,GIT1,EPHB2,APOE,TGFB2,FBXO7,PRNP,SRCIN1,PL

K1,CCN1,SHC2,SPAG9,PDPK1,DAZAP2,AKT2,GSK3A,MTOR,PRKACA,PHIP,WNK1,CD74,MICAL1,F

PRKAR2A,FLCN,BAZ1B,CSNK2A1,FLOT1,PHB2,BRD4,UBA52,BRD2,CSNK1G2,SFRP1,ADCY6,MAS

,PKD1,RPS3,BASP1,VAPA,HMGB1,CDKL1,AP2M1,TPT1,SEC61A1,RPS4X,OR8K1,STMN1,ITGB1,

DKK3,PPP1R15A,FKBP1A,SYMPK,SLIT2,A2M,DAP,NDN,AKAP9,PPP1R10,PALM,GPX1,TAX1BP1,RD

,XBP1,AJUBA,HEXIM1,CABIN1,GMFB,GIT1,EPHB2,APOE,TGFB2,FBXO7,PRNP,SRCIN1,PLPP3,GR

DYNLL1,TXNIP,CIC,RBBP7,STAT3,EIF2AK1,EDC4,MTA1,KDM5C,EP300,LBH,RERE,KDM5B,SF1,FL

A,PSMD8,HIF1A,SLC25A5,LOX,SLIT2,NDN,VPS26B,PSMB4,SOX2,IMPDH2,TAX1BP1,LSM14A,CAP

YBX3,AJUBA,PPP2R5B,RELA,OPA1,EPHB2,APOE,TGFB2,SZT2,PRNP,SESN2,PPM1A,MYADM,NF2,

RO1C,FYN,FBLN1,PLXNA3,STRAP,PTP4A1,VANGL2,PFN2,RAB11A,AKIRIN1,PRPF40A,BMPR2,AMC
RO1C,FYN,FBLN1,PLXNA3,STRAP,PTP4A1,VANGL2,PFN2,RAB11A,AKIRIN1,PRPF40A,BMPR2,AMC

JL1,PSMD13,CD44,ARL6IP5,MAP4K4,RALGDS,ERP29,RASA4,PTPN1,PSMD7,MTURN,HIPK2,PDGF

GNG2,SUPT6H,FYN,ILF2,CCT8,B2M,STXBP1,IDH1,HECTD1,MAPK8IP1,SOD1,DGKD,BAG6,DYNLL

3R3,EPHA2

,HNRNPDL,WLS,CADM4,HNRNPH1,HLA-B,RAC1,ARHGDIA,GPC1,APPL1,HDAC2,ATP6V0C,MMP2,

PK1,SMARCA4,PRDX1,ACADVL,RBP1,FTO,PRKDC,CAND1,HSPB1,KDM1A,PLCG1,ATP6V1B2,FYN,

NGAP1,TUB,ABL1,SUN1,CHMP1A,KTN1,HDGFL3,CLIP2,TRAK1,DAG1,KIFC2,HSPA1A,OBSL1,PCM

,YY1,APBB1,UBE2D3,DEK,PAGR1,CNBP,SLC30A9,POLDIP2,TRIP12,USP47,XRN2,CDC34,HMGA1,
,SEMA3C,KRT8,DICER1,CLASP2,PDPK1,WASF2,ITGAV,SCRIB,RBPJ,EHD1,VASH1,PTGFRN,PRKAC

,SOD1,DGKD,NCOR2,BMPR2,TP53BP1,CDC42,DDIT4,ADAR,CTNND1,PSMA7,SULF2,BCL9L,SGT

FM1,NOLC1,APOE,TGFB2,ARF6,PRNP,SRCIN1,NUP58,VPS28,LEPROT,PAM,USP36,NUTF2,EMD,UI

R15A,RABEP1,DAP,MTCH1,GRN,XRCC6,TAOK1,CCN1,SHC2,ANTXR1,TBC1D9,RIC8A,NCSTN,SPAC

IB,INF2,DDHD2,PSMD10,NECTIN2,COX7A2,GABARAPL2,TFDP1,DYNLT1,OPA1,THG1L,FBXO7,SES
,UBE2V1,ZFP91,RASSF2,TXN,DLG4,CDKN1B,IRS1,PRKAR2B,ERBB2,PRRC1,PHB,DSTYK,SHC1,PS

C8A,NCSTN,PDPK1,SCRIB,AKT2,SH3BP4,ARPP19,MTOR,NDUFA13,WNK1,SORT1,GNA13,MICAL1
1,MDM2,FZR1,SUMO1,FXR1,PARK7,TRRAP,CD44,SHISA5,HIPK2,CUL4A,COPS6,CETN2,WRNIP1,I

NNT,PFKFB3,NDUFA5,UQCR11,RB1CC1,COX5B,MYBBP1A,NDUFA10,ATP5PD
,CDKN1B,STAT1,EGLN2,SLC16A1,PER1,RXRA,PHB,PRMT2,CFLAR,ROCK2,ANXA1,MAPK3,SLC1A
E1,MAZ,SRSF2,MACROH2A2,DICER1,TCF3,FOXK2,NSD2,RBPJ,PCGF2,NOC2L,KDM4A,MBD3,YY1

1-0,HMGCS1,PRKAR1A,GJA1,JUN,CTNNB1,ARID1A,MXD4,SHOX,HNRNPK,H1-10,DUX4,YWHAB,PT

E1,MAZ,SRSF2,MACROH2A2,DICER1,TCF3,FOXK2,NSD2,RBPJ,PCGF2,NOC2L,KDM4A,MBD3,YY1

I1,HSPG2,LINGO1,PKDCC,AKT3,RAB14,LDLR,KMT2E,GTF2I,MACROH2A2,SEMA3C,DICER1,CLAS
E1,MAZ,SRSF2,MACROH2A2,DICER1,TCF3,FOXK2,NSD2,RBPJ,PCGF2,NOC2L,KDM4A,MBD3,YY1

D44,PHPT1,RASA4,PTPN1,FAF2,LIMK1,PPME1,HNRNPR,TRIM27,TRIB3,HERPUD1,SPOCK1,PSMF

B1,DNAJA1,ARF1,PAFAH1B1,AKAP12,CDC42BPB,AP2B1,GTF2F1,PXDN,LARP1,CTNNA1,RPL23,CF

DS1,PRMT2,ATF6,RPS6KA2,MAP2K4,SUDS3,ANXA1,TGFBR1,PNMA2,PIN1,QRICH1,XBP1,ARHGEF

PAGR1,IER2,CALCOCO1,CNBP,SLC30A9,LMO4,HMGA1,USF2,ABL1,RSF1,AUTS2,MTA2,KDM3A,K
UBA1C,RCC2,MAP1S,KIF3A,STMN4,CDKN1B,SLC16A1,RBM14,LMNA,NUP133,RNF4,BMERB1,PP

AF1,HSBP1,DCN,CDKN1A,CPSF2,DDX39B,EID1,CHD6,RNF187,MLXIP,CPSF1,SQSTM1,NOP53,PA
DNP,CD44,MAP4K4,ERP29,CCNY,PTPN1,PDGFRB,CCDC88A,CSNK2B,TRIM27,NCK2,TRIB3,ADCY

CTNNB1,ARID1A,MXD4,SHOX,HNRNPK,H1-10,DUX4,YWHAB,PTGES3,ZNF454,CCT6A,CTBP1,CDK
AF1,HSBP1,DCN,CDKN1A,CPSF2,DDX39B,EID1,CHD6,RNF187,MLXIP,CPSF1,SQSTM1,NOP53,PA

CTNNB1,ARID1A,MXD4,SHOX,HNRNPK,H1-10,DUX4,YWHAB,PTGES3,ZNF454,CCT6A,CTBP1,CDK

2,KDM2B,PAF1,MYADM,ITGA7,FZD7,NF2,LRIG3,LRP6,ID2,CRB2,HMGA2,MFAP2,ATP8A2,SLC39
CNG1,MAPKAPK2,CRKL,MAPRE3,TNFRSF10B,UBE2V1,ZFP91,WARS1,RASSF2,TXN,DLG4,CDKN

AF1,HSBP1,DCN,CDKN1A,CPSF2,DDX39B,EID1,CHD6,RNF187,MLXIP,CPSF1,SQSTM1,NOP53,PA

DC42,STAT3,TXLNA,TXNDC5,EP300,PSMD1,PBX1,PDAP1,PHB2,CST3,ZFP36L1,SFRP1,IL6ST,FKB

XBP1,ARHGEF18,PRKCI,VAV2,TGFB2,RYBP,PRNP,RNPS1,ACIN1,OBSCN,EIF2S1,ID3,PRELID1,BT
AV2,TGFB2,RYBP,PRNP,RNPS1,ACIN1,OBSCN,EIF2S1,ID3,PRELID1,BTG1,FGD1,HMGA2,GSN,AE

CTNNB1,ARID1A,MXD4,SHOX,HNRNPK,H1-10,DUX4,YWHAB,PTGES3,ZNF454,CCT6A,CTBP1,CDK

CS2,NSMF,PRKAR2B,UNC13A,NAPA,BACE1,SLC5A7,EFNB3,CNTN2,SHANK1,RPS6KA2,CTBP2,RA

AM,LAPTM4B,TRIB2,PPIB,GLUL,DCN,DLG5,DDX39B,NOVA2,ANXA5,PARP1,AHCYL1,CLSTN3,MDK

IP,SQSTM1,NOP53,PARP1,MORF4L2,TAF15,PRDX5,ACTR3,PKN1,YEATS2,PTOV1,MDK,ZNF302,R

P6AP2,A2M,BRK1,PRDX6,GRN,ACLY,XRCC6,SLC44A2,IMPDH2,RAB14,GNL1,KMT2E,NCSTN,RAB

IP,SQSTM1,NOP53,PARP1,MORF4L2,TAF15,PRDX5,ACTR3,PKN1,YEATS2,PTOV1,MDK,ZNF302,R
IP,SQSTM1,NOP53,PARP1,MORF4L2,TAF15,PRDX5,ACTR3,PKN1,YEATS2,PTOV1,MDK,ZNF302,R

APA,BACE1,SLC5A7,EFNB3,CNTN2,SHANK1,RPS6KA2,CTBP2,RAB5A,PPP3CA,SLC1A3,PCDHB2,EX

RS2,DAPK3,GNA13,CD74,SMO,FGFR1,PTPN23,ANK2,LMO4,ABL1,SDCBP,SEMA6B,MTA2,MGAT5

C5A7,EFNB3,CNTN2,SHANK1,RPS6KA2,CTBP2,RAB5A,PPP3CA,SLC1A3,PCDHB2,EXT1,SLC12A4,S

C5A7,EFNB3,CNTN2,SHANK1,RPS6KA2,CTBP2,RAB5A,PPP3CA,SLC1A3,PCDHB2,EXT1,SLC12A4,S

CN,LRRC4B,FLOT1,CD99,ZFP36L1,PLXNB1,SFRP1,CD164,CTNND2,PTEN,RGMB,PTPRF,IL6ST,SF

,FLOT1,CD99,ZFP36L1,PLXNB1,SFRP1,CD164,CTNND2,PTEN,RGMB,PTPRF,IL6ST,SPINT2,ITGB5

P,C6ORF89,GSN,CASK,GNAI1

S28,CASK,ABCA7

RAPGEF1,OPTN,RPL37,MAPK1,SMARCA4,ARPC2,PSMD3,PTBP1,GRB2,PRKDC,HSPB1,FKBP8,PL

,TGFB2,FHL1,HMG20B,OBSCN,MAML1,CENPF,ID3,DNMT1,ITGA7,MYEF2,FZD7,PIEZO1,BTG1,ID

P,EIF2S1,SYT4,GNAO1,CPNE2,ITPKB,GSN,PAM
GF,ABCA2,PRPF6,TAF7,NOTCH2,RNF10,VEGFA,VHL,CCND1,SMARCA4,CCNL2,PBX2,PSMD3,AEB
NMT1,FGD1,ITPKB,ABR,CYTH3

XNA2,ACVR1B,SMPD3,NOTCH1,IGF1R,MDM2,MGAT3,STK39,MECP2,JAM3,AAMP,IDH2,MAP4K4,

SDCBP,SEMA6B,MTA2,MGAT5,DAG1,YTHDF3,PLXNA2,ACVR1B,SMPD3,NOTCH1,IGF1R,MDM2,M

3RA1,SMO,NTRK2,LMO4,CELSR1,ABL1,PHACTR4,DAG1,NIBAN2,PLXNA2,SMPD3,NOTCH1,MECP

TNRC6A,TMEM98,CDK4,PRXL2A,NME2,LDB1,PSME3,DHX36,RBFOX2,PSMC5,PSMC2,PSMC4,TO[illegible]

[illegible]OT1,ZFP36L1,MAP6,PLXNB1,DECR1,AGPAT1,PPP1CC,IL6ST,HIF1A,CTTN,ATP6AP2,SLIT2,SERP1,

TURN,HIPK2,CUL4A,TNRC6A,TMEM98,CDK4,PRXL2A,NME2,LDB1,PSME3,DHX36,CRKL,RBFOX2,

GPC4,SAFB2,PSMB4,CCN1,GPX1,SSBP3,ABI2,TPP1,RDX,MACROH2A2,SEMA3C,KRT8,CLASP2,P[illegible]

[illegible]EBP1,TCF25,PRKDC,SUPT5H,CAND1,KDM1A,ARF4,TSC22D1,SMARCE1,SMARCC1,SUPT6H,PFK[illegible]

MBD3,GSK3A,VASH1,MTOR,YY1,SMO,SH3PXD2B,NTRK2,ANK2,CAD,LMO4,PTPN11,ABL1,DAG1,[illegible]

[illegible]E1,SMO,PTPN23,ANK2,TUB,UBE2L3,PTPN11,SCAMP5,ABL1,LMAN2,SDCBP,PINK1,HIP1R,SMPD3,

[illegible]3X,RPL30,CLTC,RPS19,MIF,PRKAR1A,JUN,CTNNB1,CD63,CYFIP2,DNAJC5,NDFIP1,DDOST,YWHA[illegible]

P2,JAM3,SART3,PABPC4,KAT2A,PSMD7,MAEA,MTURN,HIPK2,CUL4A,TNRC6A,TMEM98,CDK4,PR

HIPK2,PDGFRB,CCDC88A,CSNK2B,RAP2A,ADCY1,MAPKAPK2,CDK9,CRKL,MAPRE3,TNFRSF10B,

MTA2,MGAT5,DAG1,YTHDF3,PLXNA2,ACVR1B,SMPD3,NOTCH1,IGF1R,MDM2,MGAT3,STK39,ME

OXL2,NSD1,RBM10,ZBTB4,SRF,HSF1,ZEB1,NR2C2,BCOR,SOX9,MED25,TXN,IRF2BP2,CTR9,STA

6,P3H3,ARNT2,AKIRIN1,IGFBP7,BMPR2,TXNIP,CDC42,PPP1R9B,YME1L1,BEX4,SSR1,SULF2,STA

A1,ID3,PLXND1,FZD7,LRP6,ID2,LEMD2,PAM,RB1CC1

FB2,FGF13,TPX2,PRNP,SRCIN1,COPS8,CCNK,EPHA4,MAPK8IP3,GPRC5B,HMGA2,RAD50,EPHA2

7,ADNP,CD44,MAP4K4,ERP29,CCNY,PTPN1,KCTD20,HIPK2,PDGFRB,CCDC88A,CSNK2B,RAP2A,A
7,ADNP,CD44,MAP4K4,ERP29,CCNY,PTPN1,KCTD20,HIPK2,PDGFRB,CCDC88A,CSNK2B,RAP2A,A

B2,SDCBP,SCAMP1,PSMB1,HSPA1A,PSMD13,CAB39,NDUFC2,APEH,CYFIP1,CD44,HLA-A,PSMD7

KL1,AP2M1,SEC61A1,RPL18,STMN1,ITGB1,TUBB4B,UCHL1,RPL11,MORF4L1,NCAM1,ANXA2,AG
S16,RPL23,CPE,RPL29,RPSA,CTSD,SCARB2,NACA,RTN4,LPCAT1,LAMA5,RPS24,RPS9,XRCC5,MS
P1,PPIB,RPL23A,CTSB,ZC3H7B,CHD6,RPN2,NOP53,PABPN1,SSRP1,RPL27,KLC1,NCKAP1,VPS3
A2,AGRN,ZMIZ1,TIMP2,COL11A1,RPL10,VCL,AP3D1,ARF1,PAFAH1B1,MAP4,CDC42BPB,ITM2B,A
MYO5A,CTNNB1,CD63,MACF1,RPS10,RPL35A,TARDBP,RPLP2,ARL6IP1,WLS,RPL36,YWHAB,SEP
MYO5A,CTNNB1,CD63,MACF1,RPS10,RPL35A,TARDBP,RPLP2,ARL6IP1,WLS,RPL36,YWHAB,SEP
3,SRSF1,PTBP1,EIF3E,FTO,RPS11,SUPT5H,HNRNPUL1,SCAF11,HSPB1,SON,KDM1A,CASC3,RPL
,RHOB,RPS12,MAP2,RAB1A,ARCN1,RPS27,APPL1,IPO5,TMEM30A,LAPTM4B,RPL23A,PTTG1IP,
A,PTTG1IP,DCN,MAP1A,TIMP3,CTSB,RNF187,EEF1A2,PPP1CB,SQSTM1,NOP53,VCAN,TAF15,ALK
RPL22,RHOA,RPL9,PSMD2,KIF5B,RAB6B,NLGN2,RPS14,ATP1A1,RPS20,CD81,WDR1,ATP2A2,RPL
RAB7A,ACTN4,MMP14,DDX3X,RPS15,RPL12,RPL30,ARL4C,CLTC,RPS5,RPS7,IK,RPS19,RPL35,GJ
KAP5,SEPTIN11,RHOB,MAP2,COTL1,CCT6A,CTBP1,RAC1,ARHGDIA,CAPRIN1,APPL1,HDAC2,TME
S1,XPO1,PTN,CSNK1E,PFKL,HDLBP,RPS25,EIF5A,CDH2,GPC3,RAB7A,ACTN4,MMP14,DDX3X,RP
VPS35,RAD23B,TNPO1,PSMB2,RPL36A,ABCA2,RPS15A,RPL38,RPL37,UBXN4,MVB12B,PPP2CA,P
APRE1,KDELR2,IPO7,RPS11,NPEPPS,BNIP3L,KDELR1,RPL28,ARF4,RPS28,FYN,SSB,YWHAH,FAU,
EIF4G1,RPS26,XRN2,PUM1,FXR2,PSMB1,YTHDF3,HSPA1A,PSMD13,RPS21,FXR1,SYNCRIP,PABP
1,BTF3,ANXA2,AGRN,RPLP1,ZMIZ1,TIMP2,SARAF,MIDN,RPL10,RPS17,VCL,AP3D1,DNAJA1,ARF1,
P3,RPL14,RPL22,RHOA,PTCD3,RPL9,ATF4,PSMD2,ENC1,RPS14,TMEM259,RPS20,EIF5,CD81,RP
,WDR1,HMGN2,ATP2A2,CRMP1,CCT7,VAT1,RBBP4,RTN3,PFN1,SPC25,NES,NISCH,HNRNPC,TAD

OR,PSMC1,THRAP3,EIF4A3,EIF4G1,RPS26,XRN2,PUM1,FXR2,PSMB1,YTHDF3,MTREX,HSPA1A,P
,RPL9,CLSTN1,PSMD2,NLGN2,RPS14,EIF5,WDR1,ATP2A2,RPL26,RBBP4,RTN3,PFN1,CLEC7A,RP
S27,CCT6A,HMGCR,RAC1,CDK16,APPL1,IPO5,ATP6V0C,TMEM30A,RPL23A,GLUL,NNAT,PTTG1I
AP,HSPB1,NPEPPS,BNIP3L,CLIP3,RPL28,ARF4,RPS28,FYN,YWHAH,FAU,STXBP1,TMED9,HGS,SR

A,CTSD,EIF4H,NACA,RTN4,LPCAT1,RPS24,RPS9,XRCC5,PJA2,MSN,RPL24,RPL17,IGFBP3,RPL14,
AB,CKAP5,RHOB,ATP1A2,RPS12,RAB1A,ARCN1,RPS27,HMGCR,RAC1,CDK16,APPL1,IPO5,ATP6
B1,NUP58,HMGA2

PI1,CHSY1,EIF4A1,RPL19,PPP2R1A,RPL15,GTF3C4,LDHB,PKD1,SF3B1,PDIA3,SERINC1,SSBP2,
2B,RAB7A,MMP14,PRDX2,RPS15,RPL12,RPL30,CLTC,H1-0,RPS5,RPS7,RPS19,RPL35,GJA1,CT
FIP1,CIRBP,RPL36,YWHAB,SND1,UBE2H,RPS12,RAB1A,RPS27,GPC1,ATP6V0C,SKP1,BCLAF1,LA
RPL10,RPS17,VCL,DNAJA1,ARF1,PAFAH1B1,MAP4,ITM2B,AP2B1,PXDN,HNRNPU,SMARCA5,LAR

FLCN,GTPBP1,UBQLN2,CSNK2A1,GRSF1,PHB2,CST3,UBA52,ZFP36L1,UPF1,RAB3GAP2,HDAC6,
0A,RPL23A,GLUL,NNAT,PTTG1IP,MAP1A,CDKN1A,SRP9,NUCB1,TOMM20,RAB27A,SNX22,VAMP
1,RPS16,RPL23,RPL29,RPSA,CTSD,SCARB2,NACA,LPCAT1,RPS24,SLC22A17,RPS9,XRCC5,MSN,
S2,RPS27A,TIMM50,LMAN1,BAP1,PPP1R10,UBR5,VPS41,TRAM1,ELAVL1,VPS4A,SRP72,SSR3,M
L29,RPSA,CTSD,SCARB2,NACA,LPCAT1,RPS24,SLC22A17,RPS9,XRCC5,MSN,RPL24,SLC16A7,K
,CLTC,EIF3L,H1-0,RPS5,RPS19,MIF,PRKAR1A,TLN1,GJA1,CTNNB1,MACF1,RPS10,MTSS2,CYFI
RPL35A,TARDBP,RPLP2,NDFIP1,CIRBP,RPL36,YWHAB,SND1,UBE2H,RPS12,RPS27,HMGCR,GPC

0,MARCHF6,WSB1,RPS17,SBF1,DNAJA1,PAFAH1B1,CDC42BPB,ITM2B,AP2B1,OGA,PGAM1,NFE2
,MAP1A,DLG5,CRPPA,HS6ST1,CDH11,TBCD,GNAI3,GAS7,VCAN,PTPRA,RTN1,CAMSAP1,NCKAP1,
B,GRIPAP1,SSR1,SGTA,ALDH3A2,FLOT1,SEC16A,UBA52,RER1,RAB3GAP2,PREPL,RPS2,RPS27A,
GFBP3,NCS1,RHOA,CLSTN1,ATF4,PSMD2,NR2F1,KIF5B,ENC1,NLGN2,JUND,TMEM259,RPS20,CI
RP1,CTNND2,EVL,ADCY6,HDAC6,PTEN,PTPRF,ULK1,BRSK1,CTTN,SLIT2,NDN,GRN,TAOK1,SLIT1,
23A,RPS13,ARFGEF2,RPL34,PPP1R9B,GRIPAP1,SSR1,SGTA,ALDH3A2,FLOT1,SEC16A,UBA52,RE
PS27A,PSMD8,APEX1,HIF1A,CTTN,SLC25A5,DAP,PCBP4,VPS26B,FUS,PSMB4,GPX1,EIF3H,ELAVL

3D1,DNAJA1,ARF1,AKAP12,ITM2B,ZNF770,GTF2F1,HNRNPU,PGAM1,NFE2L1,SMARCA5,LARP1,
L1,SMG5,HINT1,CNP,TTC37,DICER1,PSMD4,IGF2BP1,AHCY,GSK3A,PSMC3,MTOR,PSMC1,THRA
OA,CBX5,ATF4,PSMD2,NR2F1,ENC1,RPS14,ATP1A1,TMEM259,RPS20,COL5A2,RPL26,CRMP1,B
3S1,XPO1,PTN,RPS25,GLG1,SFPQ,PEBP1,GPC3,PIP4K2B,CCN2,DDX3X,RPS15,RPL12,ITM2C,RPL
,CTBP1,HMGCR,RAC1,CAPRIN1,HDAC2,IPO5,SKP1,BCLAF1,LAPTM4B,TRIB2,PEA15,PTTG1IP,M
,RPS3A,RPL18,ITGB1,UCHL1,RPL11,MORF4L1,BTF3,ANXA2,AGRN,RPLP1,ZMIZ1,TIMP2,COL11
N2,DDX3X,RPS15,RPL12,ITM2C,RPL30,CLTC,NANOG,H1-0,RPS5,RPS7,RPS19,RPL35,MIF,PRKA
2,VEGFA,MARS1,SELENON,RAPGEF1,OPTN,VHL,UBXN4,CCND1,MAPK1,PSMD3,PRDX1,ACADVL,
,TLE5,CLEC7A,SLC3A2,PTN,PFKL,GLG1,PDIA6,GPC3,SULF1,PIP4K2B,MEGF8,CCN2,ACTN4,MM
1,RAD23B,PSMB2,H3-3A,PRPF6,TERF2IP,TMED2,TAF7,RPL38,VEGFA,G3BP2,SMARCA4,MAPRE1,
,FLOT1,MAP6,PLXNB1,DIP2B,FARP1,SFRP1,CTNND2,EVL,ADCY6,HDAC6,PTEN,PTPRF,ULK1,BRS
N1,CAPN2,EP300,HIP1,AP2A1,LRRC4B,FLOT1,BRD4,MAP6,PLXNB1,ATRX,SFRP1,EVL,RAB3GAP2
32,ATXN10,KDM1A,PLCG1,ARF4,SMARCE1,FYN,YWHAH,SLIT3,STMN3,B2M,STXBP1,PLXNA3,CS
19,PPP2R1A,RPL15,GTF3C4,LDHB,PKD1,SF3B1,PDIA3,SERINC1,SSBP2,ZNF43,RPS3,CHD3,BASP
PX1,ELAVL1,SMG5,HINT1,CNP,TTC37,SLCO1B3,QPRT,DICER1,PSMD4,IGF2BP1,AHCY,GSK3A,P
PX1,ELAVL1,SMG5,HINT1,CNP,TTC37,SLCO1B3,QPRT,DICER1,PSMD4,IGF2BP1,AHCY,GSK3A,P

1A,RPL15,GTF3C4,PKD1,SF3B1,PDIA3,SERINC1,SSBP2,ZNF43,RPS3,CHD3,BASP1,VAPA,HMGB1,
N1,BMPR2,CDC42,DDIT4,PPP1R9B,PALLD,KIF5A,STAT3,PRMT1,TRIO,EP300,RERE,PBX1,CNTN
BP1,PLXNA3,CSNK1D,STRAP,PTPRS,SOD1,RAB11A,TCTN1,BMPR2,CDC42,DDIT4,PPP1R9B,PALL
RDBP,RPLP2,ARL6IP1,NDFIP1,WLS,MYH10,RPL36,YWHAB,CKAP5,RHOB,ATP1A2,RPS12,RAB1A,
770,GTF2F1,HNRNPU,PGAM1,NFE2L1,SMARCA5,LARP1,HNRNPA0,RPS16,RPL23,RPL29,RPSA,C
,TAOK2,NSF,EMC1,RPS21,GOLPH3,RPL27A,NACAD,EMC10,GOLGA4,YWHAZ,RPS29,ERBB2,SR
,LDHA,CTSD,SCARB2,NACA,RTN4,LPCAT1,RPS9,XRCC5,PJA2,MSN,IGFBP3,HK1,NCS1,RHOA,CLS
C5,CIPC,MSN,PAPOLA,IGFBP3,RPL22,RHOA,PTCD3,CBX5,ZNF865,ATF4,PSMD2,NR2F1,ENC1,R
9,RPS13,RPL34,STAT3,EIF2AK1,EDC4,CPD,VARS1,GTPBP1,EIF3D,GPX4,GSPT1,UBA52,ZFP36L1
P,MAP1A,CDKN1A,TIMP3,CTDSP2,DDX39B,EID1,SRP9,U2AF2,SQSTM1,NOP53,NOX4,PRDX5,PKN
25A6,RPS21,PITRM1,RPL27A,HERPUD1,NACAD,YWHAZ,SYNGR1,RPS29,SPCS2,ERBB2,SRP54,P
L1,SMG5,HINT1,CNP,TTC37,SLCO1B3,DICER1,PSMD4,IGF2BP1,AHCY,GSK3A,PSMC3,MTOR,PS
ND2,EVL,HDAC6,PTEN,CUL3,FMNL2,ULK1,SPINT2,BRSK1,ZMYM4,SEPTIN7,CTTN,SLIT2,NDN,ZM
SMB4,GPX1,ELAVL1,SMG5,HINT1,CNP,TTC37,SLCO1B3,QPRT,DICER1,PSMD4,IGF2BP1,AHCY,
X2,DIP2A,PTPN11,ABL1,SEMA6B,AUTS2,TAOK2,TRAK1,DAG1,NIBAN2,PLXNA2,OBSL1,NOTCH1,
PTPN11,ABL1,SEMA6B,AUTS2,TAOK2,TRAK1,DAG1,NIBAN2,PLXNA2,OBSL1,NOTCH1,NPTN,MAP
X2,DIP2A,POLDIP2,PTPN11,ABL1,SEMA6B,AUTS2,TAOK2,TRAK1,DAG1,NIBAN2,PLXNA2,OBSL1,
Z1,TIMP2,COL11A1,RPL10,VCL,AP3D1,SBF1,DNAJA1,ARF1,PAFAH1B1,MAP4,ITM2B,AP2B1,HNR
3H,TPP1,UBE2D2,VPS4A,LDLR,GNA12,RDX,DNAJB2,ATXN3,PSMD4,OGT,PRPF19,USP24,FBXW5,
A52,ZFP36L1,GARS1,UPF1,EIF5B,IARS1,RPS2,RPS27A,PPP1R15A,SERP1,EIF2S2,IARS2,YARS1,E

PC2,PSMD3,SRSF1,EIF3E,LUZP6,GRB2,CAPZB,PRKDC,SCAF11,CAND1,NSG2,CLIP3,DHX30,RPS2
MD5,YWHAQ,EIF1,HACD3,SET,EGLN1,TPI1,CHSY1,EIF4A1,MRC2,RPL19,PPP2R1A,RPL15,GTF3C4
1C,KIF3C,FYN,YWHAH,SLIT3,STMN3,B2M,STXBP1,PLXNA3,CSNK1D,PTPRS,VANGL2,SOD1,PFN2,
NPM,SSB,SRSF3,SF3A1,STRAP,SARS1,PCBP2,NCOR2,HNRNPD,RPL39,RPS13,PRPF40A,RPL34,PF
2B,MEGF8,CCN2,ACTN4,PRDX2,DDX3X,NANOG,APLP1,RPS19,MIF,HMGCS1,PRKAR1A,TLN1,GJA
AUTS2,TAOK2,TRAK1,DAG1,NIBAN2,PLXNA2,OBSL1,NOTCH1,NPTN,MAPK8IP2,EFNB2,ADNP,CYF
NAP1L1,NDUFV1,PSMD1,SF1,HIP1,EIF3D,GPX4,UBA52,BRD2,ATRX,HJURP,SRSF6,EVL,SV2A,NAP1
P2,ZNF43,RPS3,CHD3,BASP1,HMGB1,POLR2A,CDKL1,AP2M1,RPS4X,RPS3A,RPL18,ITGB1,UCHL
KPNB1,PAPOLA,IGFBP3,RPL22,RHOA,PTCD3,CBX5,ZNF865,ATF4,PSMD2,NR2F1,ENC1,RPS14,CH

35,RNF20,PSMA1,SKIV2L,RBM17,PSMD10,ROCK2,SFSWAP,PSMA6,CNOT3,PAIP1,YBX3,PSMD12,
ORO1C,KIF3C,FYN,YWHAH,SLIT3,STMN3,B2M,STXBP1,PLXNA3,CSNK1D,PTPRS,VANGL2,SOD1,F
A52,ZFP36L1,GARS1,UPF1,EIF5B,IARS1,RPS2,RPS27A,PPP1R15A,SERP1,EIF2S2,IARS2,YARS1,EI
MTSS2,CYFIP2,HNRNPDL,ARID1A,PTGES3,CADM4,HNRNPH1,ATP1A2,HLA-B,HMGCR,RAC1,ARHG
,HNRNPU,NFE2L1,CTNNA1,RPS16,LDHA,CPE,CNN3,RPL29,SCARB2,NACA,RTN4,LPCAT1,LAMA5,
PBB1,TANC2,SMO,METRN,LYPLA2,NTRK2,DIP2A,PTPN11,ABL1,SEMA6B,AUTS2,TAOK2,TRAK1,D
,RAC1,ARHGDIA,CAPRIN1,GPC1,TOP2B,HDAC2,TNC,MMP2,L1CAM,GLUL,HSBP1,DCN,MAP1A,DI
F1,APBB1,TANC2,DPYSL4,SMO,METRN,LYPLA2,NTRK2,SNX2,ANK2,DIP2A,POLDIP2,PTPN11,ABL
YFIP1,FEZ1,NTN1,LIMK1,B4GALT5,EFNB1,MYCBP2,NECTIN1,RAP2A,ADCY1,SYNGAP1,ISLR2,DH
L1,NISCH,HNRNPC,XPR1,XPO1,SPTBN2,PTN,PFKL,TPM1,CDH2,DDR1,PLXNA1,MEGF8,RAB7A,CC
RSF10,RTCB,AKAP8L,C1QBP,LSM4,CELF3,ZRANB2,RBFOX2,SRSF7,PSIP1,NUP98,NCBP2,RBM10
SCARB2,NACA,RTN4,LPCAT1,LAMA5,XRCC5,MSN,RPL24,COL1A2,LLGL1,GAP43,NCS1,RPL22,RH
9,CDKN1B,EGLN2,UBXN1,PHB,PSMD10,ELOB,GABARAPL2,RGMA,PIN1,XBP1,RELA,APOE,RYBP,I
CD3,CBX5,ZNF865,ATF4,PSMD2,NR2F1,ENC1,RPS14,CHD4,JUND,TMEM259,RPS20,EIF5,CD81,F
RS1,GTPBP1,EIF3D,GSPT1,UBA52,ZFP36L1,GARS1,UPF1,EIF5B,IARS1,RPS2,RPS27A,PPP1R15A,S
AP43,NCS1,RPL22,RHOA,CLSTN1,ATF4,PSMD2,NR2F1,KIF5B,ENC1,NLGN2,RPS14,CXADR,CD81
EIF1,HACD3,SET,EGLN1,TPI1,CHSY1,EIF4A1,RPL19,PPP2R1A,RPL15,GTF3C4,LDHB,PKD1,SF3B1,
4,RBM17,SRRT,SFSWAP,PRPF31,XAB2,RNPS1,ACIN1,ZMAT2,KHDC4,SRSF8,CPSF7,WDR33,PRM
4,RBM17,SRRT,SFSWAP,PRPF31,XAB2,RNPS1,ACIN1,ZMAT2,KHDC4,SRSF8,CPSF7,WDR33,PRM
4,RBM17,SRRT,SFSWAP,PRPF31,XAB2,RNPS1,ACIN1,ZMAT2,KHDC4,SRSF8,CPSF7,WDR33,PRM
32,PPIB,GLUL,PEA15,NNAT,PTTG1IP,DCN,CDKN1A,CRPPA,TMEM132A,CTDSP2,EID1,NUCB1,TOM
32,PPIB,GLUL,PEA15,NNAT,PTTG1IP,DCN,CDKN1A,CRPPA,TMEM132A,CTDSP2,EID1,NUCB1,TOM
C,SRF,ZSWIM6,GSK3B,DHFR,ALCAM,RAB3A,NREP,MAP1S,CRABP2,ERBB2,NUMBL,ANOS1,EFNB
,CTNNB1,CD63,NGRN,SESN3,CYFIP2,TARDBP,ARID1A,NDFIP1,SHOX,HNRNPK,H1-10,CIRBP,DUX
F10,AKAP8L,C1QBP,LSM4,CDK9,CELF3,DHX36,ZRANB2,RBFOX2,SRSF7,PSIP1,NUP98,POLDIP3,
PSMC1,CCDC47,BAG5,UBXN6,UBE2K,PTPN23,ZER1,FBXL19,USP5,AMFR,TRIP12,USP47,UBE2L
CAP3,MDK,LAMC1,PTK7,SKI,ASXL1,CNTNAP2,ABCA2,ATP6AP1,DPYSL2,NOTCH2,RNF10,VEGFA,SI
VDAC1,DCTN1,CAPN2,EP300,FLCN,LRRC4B,UBQLN2,BRD4,MAP6,PLXNB1,ATRX,SFRP1,UPF1,EV
PSMB2,H3-3A,RPL36A,RPS15A,RPL38,NOTCH2,VEGFA,RPL37,VHL,PPP2CA,SRSF9,PSMD3,PTBP1
QARS1,RPL28,RPS28,LARS1,FAU,EIF3F,IDH1,SARS1,SOD1,HNRNPD,RPL39,RPS13,RPL34,STAT3,
IP1,EXOSC10,PDCD11,PA2G4,SNRPD2,SETX,PRMT5,EIF3M,FBL,RRP1B,SNRPB,SFSWAP,PRPF31
IB2,PEA15,PTTG1IP,DCN,CDKN1A,DDX39B,SRSF5,CHD6,RAB27A,RNF187,U2AF2,EEF1A2,CCT2,I
DE1,IGF2,EIF3C,ATP5F1A,EIF4B,VCP,MARCKSL1,UBB,AFF4,TRIM28,MATR3,YWHAG,COL4A2,RF
C1,CNN2,CTSB,RAB27A,CCT2,ANXA5,SQSTM1,VAMP2,GNAI3,GDI2,HSPH1,ACTR3,PDXK,KLC1,N
NTN2,ARHGAP33,CFLAR,SS18L1,RGMA,PPP3CA,MARK1,RUFY3,CUL7,PRKCI,DYNLT1,PPP2R5B,S

SHOX,HNRNPK,H1-10,CIRBP,DUX4,YWHAB,PTGES3,RAB1A,CCT6A,CTBP1,RAC1,APPL1,SMCHD1,
QSTM1,NOP53,PARP1,MORF4L2,TAF15,PRDX5,RTN1,PKN1,YEATS2,RBMX,CBX1,SKI,SNRNP70,H
UMBL,ANOS1,EFNB3,SOS1,NEO1,CNTN2,NRP2,SHC1,ROBO3,RGMA,MAPK3,ZSWIM8,RUFY3,EXT
MAP1A,CDKN1A,TIMP3,CTDSP2,EID1,SRP9,U2AF2,SQSTM1,NOP53,PARP1,MORF4L2,TAF15,PRD
IP1,CD44,MAP4K4,FEZ1,NTN1,LIMK1,CCDC88A,RAP2A,NME2,SYNGAP1,ISLR2,DHX36,CRKL,NDI
NC1LI2,ADAM9,PTBP1,GRB2,PRKDC,KDM1A,NSG2,FKBP8,PLCG1,POLR2B,GNG2,SMARCC1,AT
IP,CYFIP1,CD44,MAP4K4,FEZ1,KAT2A,NTN1,LIMK1,CCDC88A,RAP2A,NME2,SYNGAP1,ISLR2,DHX
,MAPK1,DDAH2,PRKDC,GABARAP,ACTN1,HSPB1,SON,KDM1A,BNIP3L,CLIP3,FKBP8,PLCG1,KDE
P,DCN,CDKN1A,DDX39B,SRSF5,CHD6,RNF187,U2AF2,CCT2,MLXIP,SQSTM1,NOP53,PARP1,MOR
PS16,RPL23,CPE,RPL29,RPSA,EIF4H,NACA,RTN4,RPS24,BRD3,RPS9,XRCC5,CIPC,MSN,RPL24,RI
RPN2,PARP1,VCAN,MTRF1L,RPL27,PDXK,GCN1,RPL36A,ABCA2,TMED2,RPS15A,RPL38,MARS1,RI
,ASTN2,MCL1,CLEC7A,NES,PRKCSH,HNRNPC,SPTBN2,PTN,CSNK1E,GLG1,TPM1,CDH2,GPC3,DI
,RPL23,LDHA,RTN4,LAMA5,XRCC5,MSN,RPL24,COL1A2,GAP43,RHOA,ATF4,PSMD2,KIF5B,CXAD
HNRNPU,PGAM1,NFE2L1,SMARCA5,LARP1,HNRNPA0,RPS16,RPL23,LDHA,CPE,RPL29,RPSA,EIF4
2,MCL1,CLEC7A,PRKCSH,HNRNPC,SPTBN2,PTN,CSNK1E,GLG1,TPM1,CDH2,GPC3,DDR1,SULF1
PL26,SRSF4,PFN1,TLE5,PNN,RPL31,HNRNPC,LRPPRC,TADA2B,KHDRBS1,XPO1,RPS25,SFPQ,SO
NF770,MAT2A,GTF2F1,HNRNPU,OGA,PGAM1,NFE2L1,SMARCA5,LARP1,FADS2,RPS16,RPL23,RPI
NF770,MAT2A,GTF2F1,HNRNPU,OGA,PGAM1,NFE2L1,SMARCA5,LARP1,FADS2,RPS16,RPL23,RPI
32,PPIB,GLUL,PEA15,NNAT,PTTG1IP,DCN,CDKN1A,CRPPA,TMEM132A,CTDSP2,EID1,NUCB1,TOM
RPS24,BRD3,RPS9,XRCC5,CIPC,RPL24,KPNB1,RPL17,PAPOLA,RPL14,HK1,RPL22,RHOA,CBX5,ZN
LASP2,ADAM10,IST1,RBPJ,PRPF19,SH3BP4,GSK3A,FBLN5,PTGFRN,MTOR,UBE3A,YY1,APBB1,NI
TOP1,NOTCH2,VEGFA,VHL,PPP2CA,MAPK1,DDAH2,CDC37,PRKDC,GABARAP,ACTN1,HSPB1,SON
F865,RPL9,ATF4,PSMD2,NR2F1,RPS14,CHD4,JUND,RPS20,CD81,RPL26,CCT7,SRSF4,PFN1,TLE5

PL24,RPL17,PAPOLA,RPL14,RPL22,RHOA,PTCD3,CBX5,ZNF865,RPL9,ATF4,PSMD2,NR2F1,ENC1
MA6B,KDM3A,HDGFL3,HIP1R,MACROH2A1,HSPA1A,NOTCH1,CRACD,MDM2,DNAJB1,EFNB2,JAM
Y2D1,SYT11,PSMC3,CHMP4B,MTOR,PRKACA,BAG5,DAPK3,CROCC,TUBB4A,EIF4G1,NTRK2,LM
EMIP2,CUL3,OAZ2,RPS27A,PSMD8,NPLOC4,GPC4,FBXO21,RHBDD2,BAP1,PSMB4,GPX1,HSPG2,
24,RPL17,RPL14,HK1,RPL22,RHOA,PTCD3,CBX5,ZNF865,RPL9,ATF4,PSMD2,NR2F1,SLC7A5,ENC
I3,MDK,PTK7,SKI,ASXL1,PSMB2,H3-3A,ATP6AP1,DPYSL2,NOTCH2,RNF10,VEGFA,VHL,PPP2CA,C
MEM98,CRKL,YWHAZ,GSK3B,ILRUN,ABHD17A,TFRC,KIF3A,TXN,ZDHHC5,ERBB2,LMNA,LZTS2,F
ISPA1A,MDM2,FZR1,UBE4B,TNFAIP1,TRIM2,CUL1,PSMD13,SUMO1,PARK7,USP14,PSMD7,FAF2,I
L14,RPL22,RHOA,PTCD3,CBX5,ZNF865,RPL9,ATF4,PSMD2,NR2F1,ENC1,RPS14,CHD4,JUND,RPS
GRINA,NCKAP1,VPS35,COL2A1,MDK,DDX42,TOP1,NOTCH2,VEGFA,OPTN,VHL,PPP2CA,MAPK1,DI
QLN4,RNF216,PSMB5,UBXN1,PSMA1,ZNRF1,RNF4,PSMD10,UBE2G1,ELOB,GABARAPL2,ZSWIM
3A,PHB,PRMT2,NRP2,PSMD10,ADIPOR1,MAP2K4,RGMA,TGFBR1,RUFY3,PIN1,XBP1,SPG11,BCAI
RNPD,WBP2,IGFBP7,BMPR2,DDIT4,PPP1R9B,POLR2E,SULF2,STAT3,BCL9L,CAPN2,EP300,LBH,PI
1,CSDE1,IGF2,EIF3C,ATP5F1A,EIF4B,VCP,MARCKSL1,UBB,AFF4,TRIM28,MATR3,YWHAG,CKB,C
D63,MACF1,SESN3,CYFIP2,TARDBP,NDFIP1,WLS,HNRNPK,YWHAB,CADM4,RHOB,ATP1A2,HMGC
,LDLR,RDX,CAPZA1,TUBGCP6,NCSTN,KRT8,CLASP2,WASF2,LTBP3,ADAM10,FBLN5,MTOR,DAPK

29,RPSA,NACA,RPS24,BRD3,RPS9,XRCC5,CIPC,RPL24,KPNB1,RPL17,PAPOLA,RPL14,HK1,RPL22
FE2L1,SMARCA5,LARP1,FADS2,RPS16,RPL23,RPL29,RPSA,EIF4H,NACA,LPCAT1,RPS24,BRD3,RP
TM,SOD1,BAG6,ARNT2,SERBP1,TXNIP,CDC42,YME1L1,ADAR,ANP32A,VDAC1,TRAF4,TRAF7,CAPI
3,PHIP,PSMC1,TANC2,DAPK3,SMO,METRN,MYL12B,NTRK2,CELSR1,ABL1,PINK1,SEMA6B,FAM1

H2,VPS4A,RAB14,NCSTN,RAB6A,RAB5C,ACTR1B,CLASP2,PDPK1,ITGAV,SCRIB,ADAM10,CAP1,SY
2,MAPK8IP2,DNAJB1,FXR1,PARK7,USP14,CD44,ARL6IP5,MAP4K4,ERP29,PTPN1,MTURN,HIPK2,
P1,FLCN,UBQLN2,CSNK2A1,FLOT1,EIF3D,PHB2,BRD4,UBA52,DIP2B,RAB3GAP2,ADCY6,HDAC6,F
PSMC3,AP2A2,AP1M1,MICAL1,RIMS3,PAFAH1B2,SCAMP5,SDCBP,SCAMP1,PSMB1,HSPA1A,GRIK
ACF1,SESN3,CYFIP2,TARDBP,NDFIP1,WLS,HNRNPK,YWHAB,CADM4,RHOB,ATP1A2,HMGCR,RAC1
29,RPSA,NACA,RPS24,BRD3,RPS9,XRCC5,CIPC,RPL24,KPNB1,RPL17,PAPOLA,RPL14,HK1,RPL22
17,PAPOLA,RPL14,RPL22,RHOA,PTCD3,CBX5,ZNF865,RPL9,ATF4,PSMD2,NR2F1,ENC1,RPS14,C
G,COL4A2,MAPT,FN1,SPEN,NONO,DDB1,PSMD5,YWHAQ,EIF1,HACD3,DCHS1,SET,EGLN1,MAP9,
,KCTD10,XBP1,MAN1B1,CUL4B,PSMD12,RYBP,PSMA4,UBE2E1,PSMC6,RNF5,RAD23A,SPSB3,NE
4,PPP1R9B,ADAR,BEX4,TRAF4,TRAF7,STAT3,EIF2AK1,MTA1,EP300,PRKAR2A,FLCN,FLOT1,PHB2
AEA,HIPK2,CUL4A,UBE2J1,MAGEF1,AREL1,FAM8A1,TRIB3,ADRM1,GID8,PSME3,STUB1,PSMC5,C

SF4,DCLK1,DNAJB2,NMT1,PLEKHM2,PDPK1,ADAM10,SYT11,NSD2,AKT2,GSK3A,PRKACA,APBB1
AEA,HIPK2,CUL4A,UBE2J1,MAGEF1,AREL1,FAM8A1,TRIB3,ADRM1,GID8,PSME3,STUB1,PSMC5,C

A,UBE2J1,MAGEF1,AREL1,FAM8A1,TRIB3,ADRM1,GID8,PSME3,STUB1,PSMC5,CSNK1A1,PSMC2,
SERBP1,TXNIP,CDC42,YME1L1,ADAR,ANP32A,VDAC1,TRAF4,TRAF7,CAPN2,TXNDC5,TRIO,HIP1,F

XBP1,UBQLN1,HYOU1,MAPK8IP1,GHITM,SOD1,BAG6,ARNT2,SERBP1,TXNIP,CDC42,DDIT4,YME1
IASTL,HDAC6,PTEN,SPINT2,PPP1R15A,FKBP1A,SLIT2,A2M,PPP1R10,GPX1,EIF3H,UBR5,LDLR,IRE
MT1,PDPK1,REEP2,ADAM10,SYT11,AKT2,EHD1,GSK3A,PRKACA,UBE2D3,TRIM8,SAE1,SMO,CROC

T3,LDLR,SH3GL1,DCLK1,CNP,MACROH2A2,NCSTN,ATP1B2,PHGDH,SCRIB,IGF2BP1,ATP6V0D1,
CTIN1,SYNGAP1,LDB1,DHX36,CRKL,NDRG1,SLK,LIMS1,SRF,YWHAZ,PIP5K1C,SEZ6L2,ZNF703,A

DHA,RPL29,RPSA,NACA,RPS24,BRD3,RPS9,XRCC5,CIPC,RPL24,KPNB1,RPL17,PAPOLA,RPL14,HK

STUB1,EXOC7,PLK2,WAC,GSK3B,ILRUN,UBE2V1,UBQLN4,MAP1S,EI24,ATP6V0E2,ATG9A,RETRE
STUB1,EXOC7,PLK2,WAC,GSK3B,ILRUN,UBE2V1,UBQLN4,MAP1S,EI24,ATP6V0E2,ATG9A,RETRE
,SRF,GSK3B,MEX3C,ALCAM,ATRN,SOX9,HOXA5,CRABP2,CTR9,CDKN1B,LARGE1,UNC13A,MED

P1CC,BRSK1,POM121C,RPS27A,PSMD8,HIF1A,SLC2A1,AKAP9,XRCC6,PSMB4,CLCN7,PALM,GPX1
,RAB11A,PRPF40A,DYNLL1,AMOTL2,CDC42,PPP1R9B,PALLD,DCTN1,CAPN2,CNTNAP1,HIP1,MA
1,TXNIP,TP53BP1,CDC42,PPP1R9B,PSMA7,RBBP7,DCTN1,STAT3,PRMT1,ACTR1A,EP300,PSMD1

S2,PSMD10,RAB5A,PPP3CA,MARK1,PSMA6,CTDNEP1,PIN1,EXT1,SPIN1,PSMD12,CTNNBIP1,APOE
,APOE,TLR6,ARF6,PSMC6,RAP1B,PPM1A,MYADM,PIEZO1,GSN,UNC13B
B,SESN2,BRAT1,SYT4,BTG1,CACNG7,ANAPC2,SIN3A
BQLN4,SETX,MAP1S,NAPB,STMN4,DUSP3,NAPA,ATG9A,RETREG3,BMERB1,CHMP5,GABARAPL2,
10,RAB5A,PPP3CA,MARK1,PSMA6,CTDNEP1,PIN1,EXT1,SPIN1,PSMD12,CTNNBIP1,APOE,PSMA4,

GAP2,ADCY6,HDAC6,PTEN,CUL3,IL6ST,ANP32B,RPS2,RPS27A,PPP1R15A,FKBP1A,SYMPK,DAP,M
AP9,RHBDD2,PALM,CARM1,LDLR,EFTUD2,DNAJB2,NCSTN,NUCKS1,ATXN3,PDPK1,AKT2,OGT,AT

1,PSMD12,COPB1,NFASC,RAP1B,PRDX4,PRCP,QSOX1,GSN,CSTB,ATP6V1D,GLB1,CTSA,ARSA,TC

,WDTC1,ABL1,PINK1,PSMB1,RTF1,HSPA1A,ARRDC4,MDM2,FZR1,UBE4B,TNFAIP1,TRIM2,CUL1,

PX1,EIF3H,UBR5,IREB2,LSM14A,TTC37,DNAJB2,SPAG9,PDPK1,IGF2BP1,OGT,NOC2L,KDM4A,AI
1,PSMD12,COPB1,NFASC,RAP1B,PRDX4,PRCP,QSOX1,GSN,CSTB,ATP6V1D,GLB1,CTSA,ARSA,TC

,PTEN,ABCC9,APEX1,NPLOC4,SLIT2,SLC2A1,AKAP9,RHBDD2,GRK2,PALM,CARM1,LDLR,EFTUD
NLT1,TRAPPC1,PSMD12,COPB1,NFASC,RAP1B,PRDX4,PRCP,QSOX1,GSN,CSTB,ATP6V1D,GLB1,
N1,HTRA1,HIPK2,PDGFRB,EXT2,PMEPA1,MAPKAPK2,ARPC3,STUB1,EMILIN1,CRKL,RBFOX2,NCB
TXN7L3,UBE2I,UBE2D3,PSMC1,TRIM8,BAG5,SAE1,UBE2K,FBXL19,USP5,RANGAP1,AMFR,TRIP1
APP,IDS,RPL5,KLRD1,GDI1,FSCN1,TMBIM6,PTMS,SPTAN1,GSTP1,MARCKS,SMG1,CALM1,NPM1,
RPC5,DYNLT1,TRAPPC1,PSMD12,COPB1,NFASC,RAP1B,PRDX4,PRCP,QSOX1,GSN,CSTB,ATP6V1
15,PTTG1IP,DCN,DLG5,TIMP3,DDX39B,SQSTM1,GNAI3,KLHL24,NOP53,PARP1,TAF15,SSRP1,PK
6,ARPC3,PDE4DIP,YWHAZ,GSK3B,CTR9,CDKN1B,WASF3,ATMIN,RNF20,LMNA,CDC42EP4,DDHD
P1CC,CUL3,EML1,MAPRE2,BRSK1,RPS27A,SEPTIN7,KLHDC3,TNKS,PSMD8,APEX1,PCBP4,AKAP
TRAPPC1,PSMD12,COPB1,NFASC,RAP1B,PRDX4,PRCP,QSOX1,GSN,CSTB,ATP6V1D,GLB1,CTSA,
,HSPA1A,ARRDC4,MDM2,FZR1,UBE4B,TNFAIP1,TRIM2,CUL1,PSMD13,DCAF5,PJA1,PARK7,RNF1
ARPP19,PSMC3,CHMP4B,PRKACA,NASP,UBE2I,PHIP,TTC28,PSMC1,DAPK3,TUBB4A,EIF4G1,RAN

GF2,FBXW5,PSMC3,CHMP4B,PRKACA,MN1,PHIP,TTC28,PSMC1,DAPK3,TUBB4A,PAGR1,RAB11
A1,ARFGEF2,DYNLL1,SYP,TXLNA,TXNDC5,KDM5B,PSMD1,PDAP1,CST3,SFRP1,GLUD1,TAGLN2,
GAP1,USP47,CDC34,CHMP1A,PSMB1,TAOK2,KIFC2,MACROH2A1,HSPA1A,ACVR1B,SMPD3,OBS

AP2K2,CCAR2,ARF4,SMARCE1,SMARCC1,SUPT6H,YWHAH,ILF2,CCT8,PFKM,TNKS2,RAI1,SREBF
P1CB,SQSTM1,NOP53,PABPN1,GDI2,PARP1,TAF15,SSRP1,PTPRA,PKN1,GRINA,NCKAP1,PSMB2,T

R10,BAZ2A,XRCC6,EHMT1,CARM1,UBR5,SOX2,VPS4A,SMG5,ZMIZ2,KMT2E,MACROH2A2,NUCK
,IPO5,SKP1,TRIB2,PEA15,CDKN1A,TIMP3,CTSB,NUCB1,EEF1A2,CCT2,TBCD,ANXA5,VAMP2,GNA
C1,SUPT6H,YWHAH,ILF2,CCT8,PFKM,TNKS2,RAI1,SREBF2,HNRNPD,ARNT2,WBP2,AKIRIN1,BMP

,PMEPA1,MAPKAPK2,ARPC3,STUB1,EMILIN1,CRKL,RBFOX2,NCBP2,LIMS1,ZEB1,GDF1,ZNF703,
B,ATP5MC3,AP2A2,TGOLN2,NDUFA13,BNIP3,UBE2I,TOMM70,ATP5MC2,CCDC47,REEP1,GAK,U
1,PCM1,MECP2,DNAJB1,FEZ1,KAT2A,ATF5,SUN2,VPS51,H2AX,ADCY1,RAB18,LDB1,CRKL,RBFO

)3,VASH1,PRKACA,NASP,EXOC4,CREBBP,TANC2,WNK1,AMBRA1,SMO,FGFR1,TEAD1,EIF4A3,TRI

'NKS2,RAI1,SREBF2,HNRNPD,ARNT2,WBP2,AKIRIN1,BMPR2,TP53BP1,CCT5,CDC42,STAT3,BCL9

M2,FZR1,PCM1,MECP2,CUL1,PSMD13,KAT2A,ATF5,EHMT2,PSMD7,CUL4A,PDGFRB,CETN2,WIZ,(
MAPK3,ZSWIM8,CNOT3,XBP1,TMEM59,APOE,TUT4,FURIN,FBXO7,SCOC,SESN2,PSMC6,RAD23A,\
,PHIP,SORT1,USO1,PAGR1,RABGAP1,SCAP,CALCOCO1,RANGAP1,CAD,TUB,UBE2L3,PTPN11,WD

15A,PRKAR2B,UBXN1,NR2F2,ROCK2,CAMK2N1,WDR5,GPS1,PPP1R26,RTRAF,HEG1,XBP1,HEXIM

)PP7,BRI3,ARSB,ATP6V0A1,ARPC5,DYNLT1,FER,TRAPPC1,PSMD12,COPB1,NFASC,RAP1B,PRDX4
RDX2,DDX3X,RPL30,RPS19,MIF,PRKAR1A,TLN1,GJA1,JUN,CTNNB1,ANXA6,MACF1,SESN3,TARDE
FN2,VGF,RAB11A,CYP51A1,ARFGEF2,DYNLL1,SYP,TXLNA,TXNDC5,KDM5B,PSMD1,PDAP1,CST3,S
ARPC5,DYNLT1,FER,TRAPPC1,PSMD12,COPB1,NFASC,RAP1B,PRDX4,PRCP,QSOX1,GSN,CSTB,AT
JCHL1,RPL11,MORF4L1,NCAM1,ANXA2,AGRN,ZMIZ1,TIMP2,XAF1,MARCHF6,WSB1,AP3D1,DNAJ

PBX1,FLCN,ZFP36L1,MAP6,PLXNB1,DIP2B,SFRP1,SRSF6,HDAC6,PTEN,PPP1CC,IL6ST,ANP32B,P
CY1,RAB18,LDB1,CRKL,RBFOX2,NDRG4,SEMA4C,MEIS1,SRF,ZSWIM6,GSK3B,SEZ6L2,SERPINE2

RMT112,FAM50A,PARK7,TRRAP,SART3,KAT2A,EHMT2,ZNF462,PCGF3,TASOR,ATF7IP,AKAP8L,H2

R1,MAPK3,CAPZA2,ADD1,ARHGEF18,DYNLT1,FER,GMFB,GIT1,FGF13,TPX2,PDXP,ARF6,MYADM,
LGA4,SEMA4C,MEIS1,SRF,GSK3B,SERPINE2,ACSL4,ATRN,EI24,ADIPOR2,CRABP2,CDKN1B,EGLN
THRAP3,EIF4A3,EIF4G1,CNBP,PUM1,FXR2,PINK1,PSMB1,TSNAX,PAIP2,YTHDF3,RBM4,HSPA1A,M
PAIP1,YBX3,SP1,EIF1B,RELA,ABCE1,NOLC1,APOE,TUT4,PRNP,EPHA4,SESN2,EIF2S1,SECISBP2L,
6V0A1,ARPC5,DYNLT1,FER,TRAPPC1,PSMD12,COPB1,NFASC,RAP1B,PRDX4,PRCP,QSOX1,GSN,C
HF7,HERPUD1,TNFRSF10B,SCG2,YWHAZ,GSK3B,LMNA,VDAC2,SGPL1,CFLAR,PSMD10,MEIS3,TG

FGF13,EPHA4,MYADM,POSTN,PLXND1,NF2,ID2,ANAPC2,SKIL,HES6
,SOX9,CTR9,DLG4,PSMB5,CHD8,PSMA1,MED12,LZTS2,PSMD10,SHANK1,RAB5A,PPP3CA,MARK
3X,ITM2C,CLTC,NANOG,H1-0,PRKAR1A,GJA1,JUN,CTNNB1,NGRN,TARDBP,ARID1A,MXD4,SHOX

1,PFN2,NCOR2,VGF,RAB11A,CYP51A1,ARFGEF2,DYNLL1,SYP,TXLNA,TXNDC5,KDM5B,PSMD1,PD
FA,RAPGEF1,RPL37,PPP2CA,CCND1,MAPK1,AZIN1,CCNL2,DDAH2,PSMD3,IPO7,ADAM9,DYNLL2,
LN2,CSNK2A1,MMP15,CST3,UBA52,PPP2CB,KLHL42,SFRP1,UBE2G2,FBXO11,MBTPS1,HDAC6,F

SMO,NTRK2,USP47,ABL1,PINK1,POR,NIBAN2,HIP1R,RGL2,HSPA1A,NOTCH1,IGF1R,MDM2,MAPK
C2L,KDM4A,CHMP4B,MTOR,NDUFA13,BNIP3,PHIP,PSMC1,BAG5,CD74,MICAL1,PRDX3,AMBRA1,

MC4,PLK2,PSMF1,DHFR,RCC2,CDKN1B,PSMB5,RRM2,PRKAR2B,XPC,PSMA1,PRMT2,PPP1R12A,F
RPS27,NEMF,SLC35A4,MEX3D,CNOT1
N1,PPP2R5B,FER,GMFB,CUL4B,GIT1,EPHB2,TLR6,ARF6,PSMC6,RAP1B,DNMT1,NF2,PIEZO1,GS
,CTNNB1,NGRN,TARDBP,ARID1A,MXD4,SHOX,HNRNPK,H1-10,CIRBP,DUX4,YWHAB,PTGES3,ZNF
T2,OGT,ATP6V0D1,RBPJ,GSK3A,AP2A2,CREBBP,ATP6V1G1,PHIP,UBE2D3,SORT1,FGFR1,RABGA

DX3,AMBRA1,SMO,NTRK2,USP47,ABL1,PINK1,POR,NIBAN2,HIP1R,RGL2,HSPA1A,NOTCH1,IGF1F

,CDK16,SMCHD1,HDAC2,BCLAF1,HSBP1,DCN,CDKN1A,DDX39B,EID1,NOVA2,SRSF5,CHD6,RNF1

R1A,GJA1,JUN,CTNNB1,NGRN,TARDBP,ARID1A,MXD4,SHOX,HNRNPK,H1-10,CIRBP,DUX4,YWHA

BP,ARID1A,MXD4,SHOX,HNRNPK,H1-10,CIRBP,DUX4,YWHAB,PTGES3,ZNF454,CCT6A,CTBP1,RA
CNOT3,PCNA,TFDP1,AJUBA,CUL4B,PSMD12,PSMA4,TPX2,FHL1,DYNC1I2,FBXO7,UBE2E1,CENPF

UBA52,CXXC5,ATRX,SFRP1,SRSF6,HDAC6,CBX6,HNRNPA2B1,CUL3,RPS27A,TNKS,APEX1,DKK3,D

KIRIN1,BMPR2,TP53BP1,CCT5,STAT3,EDC4,BCL9L,MTA1,EP300,LBH,RERE,PBX1,FLCN,H2AZ1,C

P6V1B2,FYN,LARS1,SLIT3,STXBP1,SOD1,NCOR2,HNRNPD,VGF,ARNT2,WBP2,IGFBP7,DYNLL1,TX
MA6,PIN1,CNOT3,PCNA,CUL7,TFDP1,PSMD12,MSH2,PSMA4,TPX2,PDXP,FHL1,DYNC1I2,FBXO7,U
NDRG1,PSMC2,DEGS1,SYNGR1,PA2G4,RAB3A,ARL8A,MAN2B1,KARS1,DPP7,BRI3,ARSB,ATP6V0

,VCAN,NOX4,PTPRA,ACTR3,PKN1,KLC1,NCKAP1,ACAP3,MDK,LAMC1,PTK7,TNPO1,SKI,DPYSL2,
D,PFN2,BMPR2,DYNLL1,CDC42,DDIT4,PPP1R9B,ADAR,TRAF4,TRAF7,STAT3,EIF2AK1,BCR,TRIO,

,CHD6,RNF187,U2AF2,MLXIP,SQSTM1,NOP53,PABPN1,HNRNPL,PARP1,MORF4L2,TAF15,ALKBH
1A1,CLPX,PSMC2,HERPUD1,TNFRSF10B,SPOCK1,PCOLCE,PSMC4,PLK2,PSMF1,WAC,HSF1,GSK

F7,SULF2,STAT3,PRMT1,EP300,PSMD1,HIP1,FLCN,CSNK2A1,FLOT1,PHB2,BRD4,UBA52,CXXC5,

RPL15,PKD1,HLA-C,PDIA3,RPS3,CHD3,VAPA,HMGB1,POLR2A,AP2M1,TPT1,SEC61A1,OR8K1,STM
PRKAR2A,FLCN,FLOT1,PHB2,BRD4,UBA52,FARP1,SFRP1,ADCY6,MASTL,HDAC6,PTEN,IL6ST,PO
PRKAR2A,FLCN,FLOT1,PHB2,BRD4,UBA52,FARP1,SFRP1,ADCY6,MASTL,HDAC6,PTEN,IL6ST,PO

CAL3,ARPC3,DAAM1,GBA2,SRF,PIP5K1C,ZYX,WASF3,RHOBTB2,CDC42EP4,DNAJB6,INF2,ARPC5,
AP2A1,CSNK2A1,FLOT1,KPNA2,UBA52,CSNK1G2,FARP1,NXN,SFRP1,GLUD1,CTNND2,PTEN,SV2

),DKK3,HIF1A,PPP1R15A,FKBP1A,CTTN,SLC25A5,SLIT2,GRK2,PPP1R10,PSMB4,PALM,GPX1,SO
C,APLP1,RPS7,RPS19,MIF,PRKAR1A,GJA1,JUN,CTNNB1,CD63,MACF1,SESN3,CYFIP2,NDFIP1,WL
,SFRP1,ARID1B,UPF1,TRA2B,RGMB,NCOA4,RPS27A,TNKS,APEX1,HNRNPAB,HIF1A,NDN,PRDX6,
P4,NDN,BAP1,FUS,BAZ2A,XRCC6,EHMT1,BCL7A,SOX2,ELAVL1,TLE1,MAZ,SRSF2,MACROH2A2,D
,SLC25A5,SLIT2,GRK2,PPP1R10,PSMB4,PALM,GPX1,SOX2,SEC14L1,TLE1,MAZ,TRAP1,SOD2,M

),DKK3,HIF1A,PPP1R15A,FKBP1A,CTTN,SLC25A5,SLIT2,GRK2,PPP1R10,PSMB4,PALM,GPX1,SO
F2,STAT3,PRMT1,EP300,PSMD1,HIP1,FLCN,CSNK2A1,FLOT1,PHB2,BRD4,UBA52,CXXC5,PLXNB

PF,PSMC6,PAF1,CDK5RAP3,ZNF207,NINL,ID2,HMGA2,ANAPC2,PLRG1,SIN3A,CNOT1,PPP2R5C

,DGKD,ARFGEF2,CDC42,DDIT4,TRAF4,RBBP7,TRAF7,PRMT1,BCR,TRIO,EP300,LBH,PRKAR2A,HI
,ACAP2,PHPT1,PTPN1,LIMK1,SUN2,FLII,IQSEC1,PDGFRB,CCDC88A,RAP2A,TRIM27,NCK2,MICAL

A,HMGCR,RAC1,ARHGDIA,GPC1,CDK16,APPL1,IPO5,TMEM30A,GLUL,PEA15,NNAT,DCN,MAP1A,

PPC11,CUL7,CLCN3,SEC23IP,MICALL1,TGFB2,SEC31A,SYT1,PRNP,COG4,DNM1,PPP2R2A,SYT4
MBD3,GSK3A,MTOR,UBE3A,PRKACA,NASP,GOT2,MN1,SMO,NSG1,EIF4A3,PAGR1,CALCOCO1,C

N14,TMEM98,CSNK2B,CAPN1,NME2,ASAH1,RAB18,PSMC2,DEGS1,SYNGR1,PA2G4,RAB3A,TFRC
36,ITPK1,CITED2,RGMA,TGFBR1,PLCD3,HEG1,ZNF358,CNOT3,ADD1,EXT1,YBX3,XAB2,OPA1,NO
ES3,ATP1A2,HLA-B,ARCN1,HRNR,HMGCR,RAC1,GPC1,TOP2B,SMCHD1,HDAC2,TNC,MMP2,DNE
1S,KIF3A,STARD3,PRKAR2B,LMNA,UNC13A,NAPA,COL7A1,CTBP2,CHMP5,NECTIN2,MARK1,DYN
D2,RGMA,TGFBR1,PLCD3,HEG1,ZNF358,CNOT3,ADD1,EXT1,YBX3,XAB2,OPA1,NOLC1,MSH2,TG

HB2,UBA52,PPP2CB,PLXNB1,SFRP1,RAB3GAP2,ADCY6,HDAC6,PTEN,TBC1D9B,HNRNPA2B1,AN

CRKL,RBFOX2,NDRG4,NCBP2,GSK3B,NUS1,SETX,SOX9,RASSF2,DUSP3,IRS1,ATP6V0E2,ERBB2

P1,SRSF6,ADCY6,HDAC6,PTEN,FEM1B,PTPRF,IL6ST,POM121C,PSMD8,NPLOC4,HIF1A,PPP1R15

98,CSNK2B,CAPN1,NME2,ASAH1,RAB18,PSMC2,DEGS1,SYNGR1,PA2G4,RAB3A,TFRC,ARL8A,M

,TMED2,TOP1,TAF7,VEGFA,RAPGEF1,ARF3,PPP2CA,CCND1,MAPK1,CCNL2,PDE7A,IPO7,ATP5PB,
G2,SFRP1,PTEN,AGPAT1,NDST1,IL6ST,MAPRE2,RPS27A,TNKS,PSMD8,HIF1A,CHERP,FKBP1A,ATP
TPN11,ABL1,PINK1,TAOK2,DAG1,TSPYL2,MACROH2A1,GGNBP2,PKIA,ARRDC4,IGF1R,FZR1,ST
TP6V0A1,CFLAR,SHC1,ROCK2,ADIPOR1,CHMP5,MAPK3,SLC1A3,XBP1,PRKCI,SP1,FER,BCAR1,
T2,AKAP9,XRCC6,TAOK1,CCN1,SHC2,SPAG9,PDPK1,DAZAP2,AKT2,OGT,GSK3A,MTOR,PRKACA,

IST1,ATP6V0D1,EHD1,CHMP4B,PACS2,PRKACA,EXOC4,AMBRA1,CROCC,TUBB4A,PTPN23,UNC1

NK1,CD74,SAE1,PRDX3,FGFR1,UBE2K,PAGR1,EIF4G1,NTRK2,RANGAP1,CAD,DIP2A,BGN,PTPN1

;AT1,NSMF,IRS1,PRKAR2B,ATP6V0E2,BACE1,SOS1,GPR173,ATP6V0A1,CFLAR,SHC1,ROCK2,ADII

2B1,GTF2F1,SLC4A8,PXDN,LARP1,CTNNA1,RPL23,CPE,TNS3,RTN4,LAMA5,PJA2,MSN,KPNB1,IGI
,PLXNB1,SFRP1,EVL,HDAC6,PTEN,CUL3,FMNL2,PTPRF,SLC7A6,MAPRE2,ITGB5,APEX1,HIF1A,PC
FB2,PSMA4,KDM2B,EPHA4,PSMC6,PLXND1,FZD7,LRP6,EPHA2
1,PSMA1,MED12,SOS1,CITED2,LZTS2,NRP2,PSMD10,RGMA,TGFBR1,HEG1,PSMA6,EXT1,AJUBA,
,TMED2,TOP1,TAF7,VEGFA,RAPGEF1,ARF3,PPP2CA,CCND1,MAPK1,CCNL2,PDE7A,IPO7,ATP5PB,

,EP300,LBH,RERE,KDM5B,SF1,FLCN,AP2A1,H2AZ1,PHB2,UBA52,ZFP36L1,CXXC5,ATRX,SFRP1,L

ND5,PLTP,CORO1C,FYN,SLIT3,FBLN1,PLXNA3,STRAP,PTP4A1,VANGL2,PFN2,RAB11A,AKIRIN1,PI

2B1,GTF2F1,SLC4A8,PXDN,LARP1,CTNNA1,RPL23,CPE,TNS3,RTN4,LAMA5,PJA2,MSN,KPNB1,IGI
2,SH3BP4,ARPP19,GSK3A,FBLN5,MTOR,PRKACA,SMARCD1,NDUFA13,BNIP3,MN1,ATP6V1G1,PH
CXXC5,ATRX,SFRP1,UPF1,HDAC6,CBX6,HNRNPA2B1,CUL3,RPS27A,APEX1,DKK3,DAP,NDN,BAP1,

ACROH2A2,SOD2,SPAG9,DICER1,CLASP2,TCF3,PDPK1,LTBP3,NSD2,LRP3,IST1,RBPJ,PRPF19,EH
GFB2,ARF6,PRNP,SRCIN1,NUP58,VPS28,LEPROT,PAM,USP36,NUTF2,EMD,UNC13B
,RDX,MAZ,ABCF1,SCRIB,IGF2BP1,NSD2,LRP3,RBPJ,PRPF19,KDM4A,GSK3A,MTOR,YY1,SMARCD

ATP6V0E2,PER1,RXRA,PHB,SOS1,PRMT2,GPR173,ATP6V0A1,CFLAR,SHC1,ROCK2,ADIPOR1,ANX

C1,RNF40,PIK3R3,ATP6V1D,GNAI1,NAMPT

UPF1,HDAC6,CBX6,HNRNPA2B1,CUL3,RPS27A,APEX1,DKK3,DAP,NDN,BAP1,BAZ2A,XRCC6,EHMT
SD,STRAP,HECTD1,COL27A1,PTPRS,VANGL2,SOD1,TCTN1,AKIRIN1,BMPR2,TXNIP,CDC42,PALLI

CA,CAPZA2,SLC1A3,XBP1,EXT1,AJUBA,VAV2,GIT1,EPHB2,APOE,TGFB2,HMG20B,SCARB1,EPHA4,

'2,NDN,VPS26B,PSMB4,SOX2,IMPDH2,TAX1BP1,LSM14A,CAPZA1,SOD2,KRT8,PSMD4,ADAM10,G
3,CARM1,SOX2,ZMIZ2,MAZ,KMT2E,GTF2I,NUCKS1,TCF3,FOXK2,EDF1,OGT,RBPJ,MED13,GSK3
3,CARM1,SOX2,ZMIZ2,MAZ,KMT2E,GTF2I,NUCKS1,TCF3,FOXK2,EDF1,OGT,RBPJ,MED13,GSK3
NP,CYFIP1,FEZ1,KAT2A,NTN1,MTURN,LIMK1,RHEB,NME2,C1QBP,ISLR2,DHX36,CRKL,LOXL2,G
3,CARM1,SOX2,ZMIZ2,MAZ,KMT2E,GTF2I,NUCKS1,TCF3,FOXK2,EDF1,OGT,RBPJ,MED13,GSK3
1,LRIG1,PHACTR4,POR,PSMB1,DAG1,ELN,SMPD3,OBSL1,NOTCH1,MFN2,ACTA2,MDM2,CUL1,P

,DAG1,RUVBL2,ACVR1B,ARRDC4,FZR1,NPTN,STK39,MECP2,CAB39,FXR1,PARK7,ADNP,SART3,C
PP3,GRB10,MYADM,MARF1,NF2,CDK5RAP3,LRP6,NNT,IBTK

GFR1,UBE2K,EIF4G1,NTRK2,PTPN11,ABL1,FXR2,SDCBP,PINK1,PPP5C,TAOK2,DAG1,TSPYL2,MA

TL,HDAC6,PTEN,ULK1,IL6ST,BRSK1,RPS27A,TNKS,LOX,SLIT2,AKAP9,GRK2,STK35,XRCC6,TAOK

JCHL1,BTF3,NCAM1,ANXA2,AGRN,ZMIZ1,TIMP2,COL11A1,RPL10,VCL,AP3D1,SBF1,DNAJA1,ARF

DX,DNAJB2,NUCKS1,PDPK1,ITGAV,SCRIB,SH3BP4,ARPP19,GSK3A,WNK1,BAG5,SORT1,MICAL1,F
B10,MYADM,MARF1,NF2,CDK5RAP3,LRP6,NNT,IBTK

CN,H2AZ1,PHB2,UBA52,ZFP36L1,CXXC5,ATRX,SFRP1,UPF1,HDAC6,CBX6,HNRNPA2B1,CUL3,RP

ZA1,SOD2,KRT8,PSMD4,GSK3A,PSMC3,PRKACA,YY1,NDUFA13,PSMC1,WNK1,TRIM8,DAPK3,CD7

CDK5RAP3,LEMD2,EPHA2,PPP2R5C,ABCA7

)TL2,CDC42,DDIT4,PPP1R9B,PALLD,BEX4,STAT3,RERE,FLCN,CD99,PHB2,PLXNB1,SFRP1,EVL,HI
)TL2,CDC42,DDIT4,PPP1R9B,PALLD,BEX4,STAT3,RERE,FLCN,CD99,PHB2,PLXNB1,SFRP1,EVL,HI

FRB,RAP2A,RNF41,MAPKAPK2,JAK1,SYNGAP1,PSME3,EMILIN1,CRKL,PSMC5,PSMC2,NDRG4,EM

1,TP53BP1,CDC42,STAT3,TXLNA,TXNDC5,EP300,PSMD1,PBX1,PDAP1,PHB2,CST3,ZFP36L1,SFRI

SKP1,DNER,PEA15,DCN,CDKN1A,DLG5,TIMP3,CNN2,SQSTM1,PARP1,PTPRA,ACTR3,TCP1,NCKAI

,CCT8,PFKM,B2M,TNKS2,USP7,STRAP,SOD1,SREBF2,HNRNPD,VGF,RAB11A,DYNLL1,YLPM1,CC

/I1,MECP2,KIF26A,FEZ1,KAT2A,ATF5,NCKAP5L,RAN,SUN2,KLC2,DYNLRB1,TASOR,CCDC88A,CET

ABL1,MTA2,USP10,PPP5C,NIBAN2,RUVBL2,SMPD3,TCEA1,FZR1,MECP2,SUMO1,PARK7,TRRAP,S
;A,EXOC4,FJX1,SMO,ANK2,LMO4,PTPN11,CELSR1,ABL1,PHACTR4,COL12A1,RTF1,DAG1,NIBAN2

A,LBH,PRKAR2A,PSMD1,FLCN,UBQLN2,CSNK2A1,PHB2,BRD4,UBA52,PPP2CB,NXN,SFRP1,ADCY

NC13B

G9,PDPK1,DAZAP2,DCUN1D2,SCRIB,MSH6,AKT2,GSK3A,MTOR,PRKACA,NDUFA13,UBE2I,WNK1,[illegible]

SN2,DNM1,PRELID1,LEPROT,TOMM40,NDUFA5,RB1CC1,USP36,NDUFA10,ATP5PD
SMD10,MAP2K4,TGFBR1,MAPK3,PIN1,PCNA,AJUBA,VAV2,TAB3,EPHB2,TLR6,LRP8,TGFB2,FGF1[illegible]

,PRDX3,FGFR1,NTRK2,RANGAP1,USP47,AGAP1,ABL1,PINK1,SGSM2,PHACTR4,MGAT5,POR,HIP1
MAGEF1,FAM168A,H2AX,MAPKAPK2,POLD2,CDK9,PWWP3A,STUB1,MARCHF7,NDRG1,ZBTB4,PL

3,UFM1,LRP8,PDXP,TRIM24,ID3,GNAL,RAP1B,GNAO1,LRP6,HMGA2,SIN3A,CNOT1,GNAI1
,CREBBP,SAP18,APBB1,NDUFA13,UBE2I,UBE2D3,SMO,NELFB,CNBP,USP47,HMGA1,WDTC1,RSF

TGES3,ZNF454,CCT6A,CTBP1,HMGCR,CDK16,APPL1,SMCHD1,HDAC2,ATP6V0C,BCLAF1,HSBP1,

,CREBBP,SAP18,APBB1,NDUFA13,UBE2I,UBE2D3,SMO,NELFB,CNBP,USP47,HMGA1,WDTC1,RSF

SP2,PDPK1,LTBP3,ADAM10,IST1,RBPJ,PRPF19,KDM4A,GSK3A,VASH1,SART1,MTOR,YY1,CREBBF
,CREBBP,SAP18,APBB1,NDUFA13,UBE2I,UBE2D3,SMO,NELFB,CNBP,USP47,HMGA1,WDTC1,RSF

1,GSK3B,SERPINE2,RCC2,WARS1,RRP1B,CDKN1B,SH3BP5,DUSP3,DUSP8,PPP1R37,KDM5A,PRK

PE,TNS3,RTN4,LAMA5,PJA2,MSN,KPNB1,IGFBP3,COL1A2,LLGL1,GAP43,NCS1,RHOA,ATF4,PSMD

18,PRKCI,VAV2,TLR6,TGFB2,RYBP,PRNP,RNPS1,ACIN1,OBSCN,EIF2S1,ID3,PRELID1,BTG1,FGD1

KLF13,RUVBL2,ACVR1B,NOTCH1,TCEA1,MDM2,MECP2,PARK7,SUB1,KAT2A,ATF5,ZNF462,SALL2
P1R12A,LZTS2,CNTN2,ROCK2,CHMP5,NAV1,MARK1,CUL7,DYNLT1,FER,GIT1,FGF13,TPX2,DYNC

BPN1,PARP1,MORF4L2,TAF15,SSRP1,PRDX5,ACTR3,PKN1,YEATS2,PTOV1,MDK,ZNF302,RBMX,[illegible]
[illegible]1,CCNG1,MAPKAPK2,LDB1,CRKL,MAPRE3,TNFRSF10B,UBE2V1,ZFP91,WARS1,RASSF2,TXN,DL[illegible]

:16,SMCHD1,HDAC2,ATP6V0C,BCLAF1,HSBP1,DCN,CDKN1A,CPSF2,DDX39B,EID1,CHD6,RNF187
BPN1,PARP1,MORF4L2,TAF15,SSRP1,PRDX5,ACTR3,PKN1,YEATS2,PTOV1,MDK,ZNF302,RBMX,C

:16,SMCHD1,HDAC2,ATP6V0C,BCLAF1,HSBP1,DCN,CDKN1A,CPSF2,PGRMC1,DDX39B,EID1,CHD

A1,HIPK1,MAFB,EPHA2
1B,SH3BP5,DUSP3,DUSP8,PRKAR2B,ERBB2,NR2F2,PRRC1,PHB,SHC1,PSMD10,CAMK2N1,MAP2

BPN1,PARP1,MORF4L2,TAF15,SSRP1,PRDX5,ACTR3,PKN1,YEATS2,PTOV1,MDK,ZNF302,RBMX,C

P1A,SLC25A5,ATP6AP2,PRDX6,GRN,ACLY,XRCC6,CLPTM1,SLC44A2,IMPDH2,RAB14,LDLR,KMT2

G1,FGD1,HMGA2,GSN,ABR,SKIL,MYBBP1A,UNC13B
3R,SKIL,MYBBP1A,UNC13B

:16,APPL1,SMCHD1,HDAC2,ATP6V0C,BCLAF1,HSBP1,DCN,CDKN1A,CPSF2,PGRMC1,DDX39B,EID

AB5A,PPP3CA,SLC1A3,PCDHB2,EXT1,SLC12A4,SLC1A4,CLCN3,SPG11,RELA,GIT1,EPHB2,APOE,L

,SKI,H3-3A,KDM6B,NRDC,ATP6AP1,NOTCH2,RNF10,VEGFA,SELENON,PPP2CA,CCND1,MAPK1,A

BMX,CBX1,CHD2,GCN1,SKI,ASXL1,PSMB2,H3-3A,KDM6B,HDGF,ABCA2,PRPF6,TERF2IP,TAF7,N

6A,RAB5C,PLEKHM2,ACTR1B,PDPK1,WASF2,ITGAV,ADAM10,CAP1,MSH6,NSD2,IST1,PSMC3,MT

BMX,CBX1,CHD2,GCN1,SKI,ASXL1,PSMB2,H3-3A,KDM6B,HDGF,ABCA2,PRPF6,TERF2IP,TAF7,N
BMX,CBX1,CHD2,GCN1,SKI,ASXL1,PSMB2,H3-3A,KDM6B,HDGF,ABCA2,PRPF6,TERF2IP,TAF7,N

T1,SLC12A4,SLC1A4,CLCN3,SPG11,GIT1,EPHB2,APOE,LRP8,SYT1,PRNP,PPT1,EPHA4,GJC1,DN

,DAG1,YTHDF3,PLXNA2,ACVR1B,SMPD3,NOTCH1,IGF1R,MDM2,MGAT3,STK39,MECP2,JAM3,AA

SLC1A4,CLCN3,SPG11,GIT1,EPHB2,APOE,LRP8,SYT1,PRNP,PPT1,EPHA4,GJC1,DNM1,PPFIA3,SY

SLC1A4,CLCN3,SPG11,GIT1,EPHB2,APOE,LRP8,SYT1,PRNP,PPT1,EPHA4,GJC1,DNM1,PPFIA3,SY

PINT2,ITGB5,PODXL2,CTTN,SYMPK,GPC4,CCN1,ANTXR1,SOX2,RDX,TM9SF4,KIFAP3,FLOT2,RIC

5,PODXL2,CTTN,SYMPK,GPC4,CCN1,ANTXR1,SOX2,RDX,TM9SF4,KIFAP3,FLOT2,RIC8A,CADM3,

CG1,MAP2K2,CCAR2,FYN,LARS1,B2M,TNKS2,STXBP1,UBQLN1,CSNK1D,MAPK8IP1,SOD1,DGK

2,LEMD2,SKIL,DMPK,MYL6B,EMD

P1,TCF25,PRKDC,SUPT5H,CAND1,KDM1A,ARF4,TSC22D1,POLR2B,SMARCE1,SMARCC1,SUPT6

NTN1,SUN2,IQSEC1,PDGFRB,CSNK2B,RAP2A,RNF41,TBXA2R,C1QBP,LDB1,EMILIN1,CRKL,NDR

GAT3,STK39,MECP2,EFNB2,JAM3,AAMP,USP14,IDH2,MAP4K4,PHPT1,NTN1,SUN2,MYCBP2,IQSE

2,EFNB2,JAM3,AAMP,ADGRB2,KAT2A,NTN1,SALL2,HIPK2,PPP1CA,PDGFRB,CSNK2B,TBXA2R,JA

B2,MEIS1,PSMF1,SRF,HSF1,ZEB1,TFRC,RASSF2,HOXA5,IRF2BP2,CTR9,PSMB5,STAT1,CEBPG,E

,GRN,XRCC6,EHMT1,CCN1,OLFM1,PKDCC,AKT3,MACROH2A2,DICER1,PDPK1,SCRIB,NSD2,IST1

,PSMC5,PSMC2,PSMC4,TOB2,MEIS1,PSMF1,SRF,HSF1,ZEB1,TFRC,RASSF2,HOXA5,IRF2BP2,CT

DPK1,PHGDH,EDF1,PSMD4,SCRIB,HSD17B4,AKT2,RBPJ,GSK3A,PSMC3,MTOR,PRKACA,AP2A2,P

,ID4,STRAP,SARS1,RAI1,SREBF2,NCOR2,HNRNPD,ZSCAN18,ARNT2,KDM3B,WBP2,AKIRIN1,BM

NIBAN2,ELN,GGNBP2,OBSL1,NOTCH1,ACTA2,MDM2,MECP2,EFNB2,JAM3,AAMP,ADGRB2,PDLI

,ATP13A2,MDM2,PCM1,STK39,NDUFA4,PARK7,FEZ1,ATP1B3,PHPT1,RAN,ANKFY1,GOLPH3,PDG

B,LAMP1,CKAP4,HLA-B,RAB1A,TTR,COTL1,HRNR,RAC1,GPC1,APPL1,ATP6V0C,TMEM30A,SKP1,

XL2A,NME2,LDB1,PSME3,DHX36,MARCHF7,CRKL,RBFOX2,PSMC5,PSMC2,PSMC4,TOB2,MEIS1,

HSF1,GDF1,UBE2V1,SOX9,TFRC,ADIPOR2,ZFP91,RASSF2,TXN,DLG4,CDKN1B,IRS1,PRKAR2B,ER

CP2,JAM3,AAMP,IDH2,MAP4K4,PHPT1,NTN1,SUN2,IQSEC1,PDGFRB,CSNK2B,RAP2A,RNF41,TBX

T1,CHD8,NACC1,PER1,NR2F2,RXRA,NR2F6,PHB,CITED2,CTCF,PSMD10,CTBP2,SUDS3,THRA,XB

T3,EIF2AK1,PRMT1,NAP1L1,LBH,RERE,KDM5B,SF1,PBX1,FLCN,CSNK2A1,PHB2,ZFP36L1,PLXNE

ADCY1,MAPKAPK2,CDK9,PPP2R5D,CRKL,MAPRE3,TNFRSF10B,HSF1,GDF1,UBE2V1,SOX9,TFRC,A
ADCY1,MAPKAPK2,CDK9,PPP2R5D,CRKL,MAPRE3,TNFRSF10B,HSF1,GDF1,UBE2V1,SOX9,TFRC,A

,FAF2,TSPAN14,CSNK2B,CAPN1,NME2,ASAH1,C1QBP,RAB18,DHX36,PSMC2,DEGS1,SYNGR1,PA

RN,RPLP1,ZMIZ1,TIMP2,COL11A1,RPL10,RPS17,VCL,AP3D1,ARF1,PAFAH1B1,MAP4,CDC42BPB
N,RPL24,KPNB1,RPL17,LLGL1,RPL14,HK1,NCS1,RPL22,RHOA,RPL9,CLSTN1,PSMD2,KIF5B,RA
,CBX1,TNPO1,PSMB2,RPL36A,TOP1,RPS15A,RPL38,OPTN,RPL37,MVB12B,G3BP2,MAPK1,SMA
P2B1,MAT2A,TMEM170A,GOLGA8A,PXDN,HNRNPU,SMARCA5,CTNNA1,RPL23,CNN3,RPSA,TPM4
TIN11,LAMP1,RHOB,RPS12,MAP2,RAB1A,ARCN1,SEPTIN3,RPS27,CCT6A,GPC1,APPL1,IPO5,TME
TIN11,LAMP1,RHOB,RPS12,MAP2,RAB1A,ARCN1,SEPTIN3,RPS27,CCT6A,GPC1,APPL1,IPO5,TME
8,CCAR2,POLR2B,SUPT6H,RPS28,HNRNPM,SSB,SRSF3,FAU,SF3A1,STRAP,PCBP2,HNRNPD,RPL
AP1A,CDKN1A,CPSF2,DDX39B,SRP9,NUCB1,SRSF5,TOMM20,RAB27A,U2AF2,CPSF1,SQSTM1,V
BH5,RPL27,UBE4A,VPS35,RAD23B,TNPO1,SNRNP70,PSMB2,RPL36A,ABCA2,NRDC,PYGB,RPS15
26,VAT1,OAZ1,SRSF4,RPL31,LRPPRC,KHDRBS1,XPO1,SPTBN2,PFKL,RPS25,SFPQ,EIF5A,CDH2,
A1,MYO5A,CTNNB1,CD63,MACF1,RPS10,RPL35A,TARDBP,RPLP2,ARL6IP1,NDFIP1,WLS,MYH10,
M30A,L1CAM,LAPTM4B,DCN,MAP1A,CDKN1A,DLG5,DDX39B,EID1,CCT2,TBCD,SQSTM1,NOP53,
15,RPL12,RPL30,ARL4C,CLTC,RPS5,RPS7,IK,RPS19,RPL35,GJA1,MYO5A,CTNNB1,CD63,MACF
SMD3,EIF3E,FTO,RPS11,GABARAP,UBE2R2,HSPB1,BNIP3L,CASC3,RPL28,CCAR2,SMARCC1,RPS
CCT8,TNKS2,CSNK1D,HGS,BAG6,SREBF2,RAB11A,RPL39,TCTN1,RPS13,TXNIP,RPL34,CCT5,PP
4,PNRC2,PSMD7,ETF1,TNRC6A,NBAS,HNRNPR,RPL27A,DDX6,LSM4,MAPKAPK2,PSME3,DHX36,
CYB5R3,PAFAH1B1,AKAP12,MAP4,CDC42BPB,AP2B1,SLC4A8,HNRNPU,PGAM1,CTNNA1,RPS16,
L26,BEX3,CLEC7A,RPL31,PRKCSH,CCNI,LRPPRC,TADA2B,KHDRBS1,KLHL9,XPO1,PTN,CSNK1E,
A2B,XPO1,SPTBN2,CSNK1E,SFPQ,TPM1,CDH2,PIP4K2B,RAB7A,CCN2,DDX3X,RPS15,CLTC,H1-0

MD13,RPS21,FXR1,SYNCRIP,PABPC4,PNRC2,PSMD7,ETF1,TNRC6A,NBAS,HNRNPR,RPL27A,DD
L31,XPO1,SPTBN2,CSNK1E,RPS25,CDH6,TPM1,CDH2,PIP4K2B,RAB7A,CCN2,MMP14,DDX3X,RP
,MAP1A,CDKN1A,SRP9,NUCB1,TOMM20,RAB27A,CCT2,SNX22,VAMP2,GDI2,RPL27,TBC1D16,H
EBF2,RAB11A,RPL39,SEC23A,RPS13,TXNIP,RPL34,CDC42,ADAR,GRIPAP1,SSR1,KIF5A,STAT3,SG

RPL22,RHOA,PTCD3,RPL9,ATF4,PSMD2,ENC1,RPS14,TMEM259,RPS20,EIF5,CD81,RPL26,BEX3,
V0C,TMEM30A,BCLAF1,RPL23A,GLUL,NNAT,PTTG1IP,MAP1A,CDKN1A,CPSF2,DDX39B,SRP9,NU

NF43,RPS3,CHD3,BASP1,VAPA,HMGB1,POLR2A,CDKL1,AP2M1,RPS4X,RPS3A,RPL18,ITGB1,UC
NNB1,RPS10,SESN3,RPL35A,TARDBP,RPLP2,NDFIP1,CIRBP,RPL36,YWHAB,SND1,UBE2H,RPS12,
PTM4B,TRIB2,RPL23A,GLUL,PTTG1IP,DCN,MAP1A,TIMP3,CTSB,TOMM20,RNF187,EEF1A2,PPP1
1,HNRNPA0,CTNNA1,RPS16,RPL23,RPL29,RPSA,NACA,RTN4,LPCAT1,RPS24,RPS9,XRCC5,CIPC,

PTEN,ULK1,OAZ2,GPSM1,POM121C,RPS27A,PSMD8,APEX1,HIF1A,CTTN,SLC25A5,DAP,PCBP4,V
,GDI2,RPL27,TBC1D16,HSPA4,TCP1,AHCYL1,VPS35,RAB5B,TNPO1,RPL36A,RAB11B,SRP14,ATF
RPL24,SLC16A7,KPNB1,RPL17,C1ORF43,LLGL1,RPL14,HK1,NCS1,RPL22,RHOA,RPL9,CLSTN1,
ACROH2A2,NMT1,MOAP1,SCRIB,CSE1L,GSK3A,CHMP4B,NDUFA13,UBE2D3,TOMM70,SORT1,S
NB1,RPL17,C1ORF43,LLGL1,RPL14,NCS1,RPL22,RHOA,RPL9,CLSTN1,ATF4,PSMD2,KIF5B,RA
2,ARL6IP1,H1-10,CIRBP,PTGES3,CKAP5,MAP2,RAB1A,RPS27,COTL1,RAC1,GPC1,SMCHD1,SKP
1,SKP1,BCLAF1,LAPTM4B,TRIB2,RPL23A,GLUL,PTTG1IP,DCN,MAP1A,TIMP3,CTSB,RNF187,EEF

?L1,LARP1,RPS16,RPL23,LDHA,CPE,RPL29,RPSA,CTSD,EIF4H,NACA,RTN4,LPCAT1,RPS24,RPS9,X
VPS35,COL2A1,CLSTN3,ACAP3,MDK,PTK7,SKI,KDM6B,CNTNAP2,ABCA2,DPYSL2,TMED2,NOTCH
RABEP1,GPC4,SERP1,EFR3B,PALM,TCAF1,PKDCC,TRAM1,GPC2,VPS4A,SRP72,SSR3,RDX,TM9SI
D81,WDR1,RPL26,BEX3,CCT7,OAZ1,PFN1,MCL1,TLE5,CLEC7A,NES,TADA2B,KHDRBS1,PTN,CSNI
FKBP4,OLFM1,CARM1,LINGO1,ABI2,GPC2,DCLK1,CNP,STK24,SEMA3C,SPAG9,DICER1,CLASP2,
ER1,RAB3GAP2,PREPL,RPS2,RPS27A,RABEP1,GPC4,SERP1,EFR3B,PALM,TCAF1,PKDCC,TRAM1,(
1,LDLR,GNA12,RDX,TTC37,DNAJB2,ATXN3,MTMR4,FOXK2,PSMD4,IGF2BP1,AKT2,OGT,ATP6V0D

HNRNPA0,RPS16,RPL23,RPL29,RPSA,CTSD,EIF4H,SCARB2,NACA,RTN4,LPCAT1,RPS24,BRD3,RPS
P3,EIF4A3,EIF4G1,RPS26,XRN2,PUM1,FXR2,PSMB1,YTHDF3,MTREX,HSPA1A,NUDT4,PSMD13,RI
EX3,VAT1,OAZ1,SRSF4,RBBP4,RTN3,PFN1,ASTN2,MCL1,TLE5,NES,NISCH,PRKCSH,HNRNPC,LRP
L30,CLTC,NANOG,H1-0,RPS5,RPS7,RPS19,RPL35,MIF,PRKAR1A,GJA1,JUN,CTNNB1,RPS10,RPL3
AP1A,CDKN1A,TIMP3,CTDSP2,DDX39B,EID1,SRP9,U2AF2,EEF1A2,SQSTM1,NOP53,NOX4,PRDX5,I
A1,RPL10,MARCHF6,WSB1,RPS17,AP3D1,SBF1,DNAJA1,ARF1,CDC42BPB,ITM2B,ZNF770,AP2B1,
R1A,GJA1,JUN,CTNNB1,RPS10,RPL35A,APLP2,TARDBP,RPLP2,ARL6IP1,NDFIP1,MXD4,HNRNPK,I
FTO,GRB2,CDC37,PRKDC,GABARAP,HSPB1,NPEPPS,KDM1A,BNIP3L,MAP2K2,QARS1,CCAR2,PC
P14,PRDX2,NANOG,APLP1,MIF,HMGCS1,PRKAR1A,TLN1,GJA1,JUN,MYO5A,CTNNB1,COL6A2,CD
,SRSF9,ARPC2,DSTN,PSMD3,SRSF1,EIF3E,LUZP6,GRB2,CAPZB,PRKDC,GABARAP,SCAF11,CAND
K1,CTTN,SLIT2,MTCH1,NDN,GRN,TAOK1,SLIT1,FKBP4,OLFM1,CARM1,LINGO1,ABI2,GPC2,DCL
?,HNRNPA2B1,CUL3,ULK1,MAPRE2,LMAN1,SEPTIN7,TNKS,HIF1A,PPP1R15A,CTTN,SLC25A5,SLIT
IK1D,STRAP,PTPRS,SOD1,RAB11A,TCTN1,BMPR2,CDC42,DDIT4,PPP1R9B,PALLD,KIF5A,STAT3,P
1,VAPA,HMGB1,POLR2A,CDKL1,AP2M1,RPS4X,RPS3A,RPL18,UCHL1,RPL11,MORF4L1,BTF3,AN
SMC3,MTOR,PSMC1,THRAP3,EIF4A3,EIF4G1,RPS26,XRN2,PUM1,FXR2,PSMB1,YTHDF3,MTREX,I
SMC3,MTOR,PSMC1,THRAP3,EIF4A3,EIF4G1,RPS26,XRN2,PUM1,FXR2,POR,PSMB1,YTHDF3,MT

,POLR2A,CDKL1,AP2M1,RPS4X,RPS3A,RPL18,UCHL1,RPL11,MORF4L1,BTF3,ANXA2,AGRN,RPLF
AP1,AP2A1,FLOT1,MAP6,PLXNB1,DIP2B,FARP1,SFRP1,CTNND2,EVL,ADCY6,HDAC6,PTEN,PPP1CC
D,KIF5A,STAT3,PRMT1,NAP1L1,TRIO,EP300,RERE,PBX1,CNTNAP1,AP2A1,FLOT1,MAP6,PLXNB1,
ARCN1,RPS27,HMGCR,RAC1,CDK16,APPL1,IPO5,ATP6V0C,TMEM30A,BCLAF1,RPL23A,GLUL,PEA
TSD,EIF4H,SCARB2,NACA,RTN4,LPCAT1,RPS24,BRD3,RPS9,XRCC5,CIPC,MSN,RPL24,RPL17,PAI
P54,GET1,AFDN,ZDHHC9,TFDP1,PRKCI,MICALL1,RPL32,ABCD3,PRNP,GET4,VPS37B,EMD
STN1,ATF4,PSMD2,NR2F1,KIF5B,SLC7A5,ENC1,NLGN2,ATP1A1,JUND,TMEM259,RPS20,CD81,WD
PS14,CHD4,ATP1A1,JUND,TMEM259,RPS20,EIF5,CD81,RPL26,BEX3,CCT7,OAZ1,SRSF4,RTN3,PF
,GARS1,UPF1,EIF5B,IARS1,RPS2,RPS27A,PPP1R15A,ATP6AP2,GLO1,SERP1,EIF2S2,IARS2,GPX1,
1,VPS35,RAD23B,GCN1,SKI,ABCA2,NRDC,TERF2IP,TMED2,TAF7,RPL38,VEGFA,RAPGEF1,RPL37
IPCA,LONP2,ZDHHC9,PRKCI,MICALL1,RPL32,ABCD3,FBXO7,PRNP,LEPROT,TOMM40,AP3M1,M6
MC1,THRAP3,EIF4A3,EIF4G1,RPS26,XRN2,PUM1,FXR2,PSMB1,YTHDF3,MTREX,HSPA1A,NUDT4,F
YM3,TAOK1,SLIT1,PALM,OLFM1,ANTXR1,LINGO1,ABI2,RDX,DCLK1,CNP,SEMA3C,SPAG9,DICER
GSK3A,PSMC3,MTOR,PSMC1,THRAP3,EIF4A3,EIF4G1,RPS26,XRN2,PUM1,FXR2,PSMB1,YTHDF3,I
IPTN,MAPK8IP2,EFNB2,ADNP,CYFIP1,CD44,MAP4K4,FEZ1,RNF157,NTN1,LIMK1,B4GALT5,EFNB
K8IP2,EFNB2,ADNP,CYFIP1,CD44,MAP4K4,FEZ1,RNF157,NTN1,LIMK1,B4GALT5,EFNB1,SNX1,MY
NOTCH1,NPTN,MAPK8IP2,EFNB2,ADNP,CYFIP1,CD44,MAP4K4,FEZ1,RNF157,NTN1,LIMK1,B4GAL
NPU,NFE2L1,CTNNA1,RPS16,LDHA,CPE,CNN3,RPL29,TPM4,EIF4H,SCARB2,NACA,RTN4,LPCAT1,
GSK3A,PSMC3,CHMP4B,MTOR,UBE3A,PRKACA,AP2A2,NDUFA13,BNIP3,UBE2I,UBE2D3,PSMC1,C
F3H,ELAVL1,IREB2,LSM14A,TTC37,EIF4G3,TRAP1,EEF1B2,LSM14B,ABCF1,IGF2BP1,AKT2,EIF3

8,CCT8,EIF3F,B2M,STXBP1,LONP1,CSNK1D,SF3A1,STRAP,PFN2,PPP6R1,SEC23A,NAP1L4,TP53E
,LDHB,PKD1,SF3B1,PDIA3,SERINC1,SSBP2,ZNF43,RPS3,CHD3,BASP1,VAPA,HMGB1,POLR2A,CD
RAB11A,TCTN1,AKIRIN1,BMPR2,DYNLL1,CDC42,PPP1R9B,PALLD,DCTN1,KIF5A,PRMT1,ACTR1
P1R9B,ADAR,RBM15B,DHX15,POLR2E,STAT3,SF1,PRCC,GRSF1,UBA52,ZFP36L1,SRSF6,TRA2B,
1,JUN,MYO5A,CTNNB1,CD63,SESN3,MTSS2,CYFIP2,HNRNPDL,ARID1A,PTGES3,CADM4,RHOB,H
IP1,MAP4K4,FEZ1,RNF157,NTN1,LIMK1,B4GALT5,EFNB1,MYCBP2,NECTIN1,RAP2A,ADCY1,SYN
L3,ANP32B,RPS27A,PSMD8,FKBP1A,CTTN,SLIT2,BRK1,AKAP9,TMOD2,BAZ2A,PSMB4,EIF2S2,FK
1,RPL11,MORF4L1,BTF3,ANXA2,AGRN,RPLP1,ZMIZ1,TIMP2,COL11A1,RPL10,MARCHF6,WSB1,
HD4,ATP1A1,JUND,TMEM259,RPS20,EIF5,CD81,RPL26,BEX3,CCT7,OAZ1,SRSF4,RTN3,PFN1,TLE

TUT4,PSMA4,RNPS1,ACIN1,PSMC6,PAF1,MYEF2,CPSF7,RNF40,CACNG7,MEX3D,RBM39,CNOT1
FN2,RAB11A,TCTN1,AKIRIN1,BMPR2,DYNLL1,CDC42,PPP1R9B,PALLD,DCTN1,KIF5A,PRMT1,AC
F3H,ELAVL1,IREB2,LSM14A,TTC37,EIF4G3,TRAP1,EEF1B2,LSM14B,ABCF1,IGF2BP1,AKT2,EIF3
DIA,STIP1,GPC1,CDK16,APPL1,HDAC2,IPO5,TNC,ATP6V0C,MMP2,SKP1,DCN,CDKN1A,CTDSP2,C
XRCC5,MSN,RPL24,IGFBP3,COL1A2,LLGL1,GAP43,NCS1,RPL22,RHOA,CLSTN1,ATF4,PSMD2,N
AG1,NIBAN2,PLXNA2,OBSL1,NOTCH1,NPTN,MAPK8IP2,EFNB2,ADNP,CYFIP1,FEZ1,NTN1,LIMK1,
G5,CRPPA,CDH11,TBCD,GAS7,NOX4,TGFBI,PTPRA,PKN1,CAMSAP1,NCKAP1,VPS35,COL2A1,MD
1,SEMA6B,AUTS2,TAOK2,TRAK1,DAG1,NIBAN2,PLXNA2,OBSL1,NOTCH1,NPTN,MAPK8IP2,EFNB
X36,RBFOX2,RANBP9,GOLGA4,SEMA4C,SRF,ZSWIM6,GSK3B,ALCAM,RAB3A,MAP1S,CRABP2,DL
N2,PRDX2,DDX3X,RPS5,RPS7,RPS19,GPAT4,PRKAR1A,TLN1,GJA1,JUN,CTNNB1,ANXA6,MACF1,
,DNAJC8,DDX41,SNRPD2,SETX,PRMT5,RRP1B,RAVER1,SNRPB,RBM14,THOC2,RBM17,SRRT,SF
HOA,CLSTN1,ATF4,PSMD2,NR2F1,KIF5B,ENC1,NLGN2,RPS14,CXADR,CD81,COL5A2,WDR1,CRM
FURIN,EPHA4,PSMC6,RAD23A,VPS28,CDK5RAP3,PITHD1,RNF40,CTSA
RPL26,BEX3,CCT7,OAZ1,SRSF4,RTN3,PFN1,TLE5,CLEC7A,HNRNPC,CCNI,LRPPRC,TADA2B,KHDF
ERP1,ACLY,EIF2S2,IARS2,CCN1,YARS1,EIF3H,ELAVL1,IREB2,LSM14A,TTC37,EIF4G3,TRAP1,EEF
,COL5A2,WDR1,CRMP1,PFN1,ASTN2,TLE5,NES,PRKCSH,SPTBN2,PTN,CSNK1E,GLG1,TPM1,CDH
,PDIA3,SERINC1,SSBP2,ZNF43,RPS3,CHD3,BASP1,VAPA,HMGB1,POLR2A,CDKL1,AP2M1,RPS4X,
T7,PLRG1,RBM39,SNU13
T7,PLRG1,RBM39,SNU13
T7,PLRG1,RBM39,SNU13
IM20,RNF187,U2AF2,RPN2,PPP1CB,SQSTM1,KLHL24,NOP53,PARP1,VCAN,MORF4L2,NOX4,PTP
IM20,RNF187,U2AF2,RPN2,PPP1CB,SQSTM1,KLHL24,NOP53,PARP1,VCAN,MORF4L2,NOX4,PTP
3,SOS1,NEO1,CNTN2,NRP2,SHC1,ROBO3,RGMA,MAPK3,ZSWIM8,RUFY3,EXT1,DYNLT1,SPG11,S
4,YWHAB,PTGES3,RAB1A,CCT6A,CTBP1,RAC1,APPL1,SMCHD1,HDAC2,TNC,SKP1,BCLAF1,TRIB2
NCBP2,RBM10,RNMT,HSF1,DNAJC8,DDX41,PDCD11,SNRPD2,SETX,PRMT5,RRP1B,RAVER1,CTR
3,CDC34,RNF11,SDCBP,PINK1,CLTA,USP10,PSMB1,HSPA1A,ATP13A2,MDM2,FZR1,UBE4B,MGAT
ELENON,RAPGEF1,MAPK1,ARPC2,PBX2,GRB2,PRKDC,TDRD5,ATXN10,ACTN1,KDM1A,PLCG1,AR
L,RAB3GAP2,HDAC6,HNRNPA2B1,CUL3,ULK1,MAPRE2,LMAN1,SEPTIN7,TNKS,HIF1A,CTTN,SLC2
,EIF3E,FTO,ILF3,RPS11,PRKDC,HSPB1,CASC3,MAP2K2,RPL28,POLR2B,RPS28,FYN,HNRNPM,SS
EIF2AK1,EDC4,CPD,VARS1,GTPBP1,EIF3D,GPX4,GSPT1,UBA52,ZFP36L1,GARS1,UPF1,EIF5B,IAF
,SENP3,XAB2,CUL4B,ABCE1,NOLC1,RPL32,PES1,KRI1,DDX56,KHDC4,CPSF7,PRMT7,NOP58,USF
MLXIP,SQSTM1,GNAI3,NOP53,PARP1,MORF4L2,NOX4,TAF15,ACTR3,PKN1,TCP1,CCT3,VPS35,PT
L37A,MAPT,FN1,SPEN,NONO,DDB1,PSMD5,YWHAQ,EIF1,HACD3,DCHS1,SET,EGLN1,MAP9,CHSY
CKAP1,VPS35,MLEC,SURF4,RAB5B,RAB11B,SRP14,ABCA2,PYGB,ATP6AP1,DPYSL2,TMED2,VEG
EMA6C,OPA1,EPHB2,APOE,LRP8,FGF13,ARF6,FBXO7,SRCIN1,EPHA4,DDX56,PLXND1,ATP8A2,A

HDAC2,TNC,SKP1,BCLAF1,TRIB2,PEA15,PTTG1IP,DCN,CDKN1A,DDX39B,CNN2,SRSF5,CHD6,RAI
I3-3A,HDGF,ABCA2,TERF2IP,TMED2,TAF7,NOTCH2,VEGFA,RPL37,VHL,PPP2CA,CCND1,SMARCA4
T1,SPG11,SEMA6C,ZDHHC17,EPHB2,APOE,FGF13,NFASC,SZT2,EPHA4,GRB10,PLXND1,NFIB,SH3
X5,RTN1,PKN1,YEATS2,RBMX,CBX1,SKI,SNRNP70,H3-3A,HDGF,ABCA2,TERF2IP,TMED2,TAF7,N
RG4,SPOCK1,GOLGA4,SEMA4C,P3H1,SRF,GSK3B,CRTC1,RCC2,SETX,CRABP2,ATMIN,NSMF,CD
P6V1B2,FYN,HNRNPM,LARS1,YWHAH,SLIT3,LONP1,CSNK1D,IDH1,STRAP,SOD1,SREBF2,NCOR2
36,CRKL,NDRG4,SPOCK1,GOLGA4,SEMA4C,P3H1,SRF,GSK3B,CRTC1,RCC2,SETX,CRABP2,DLG
LR1,QARS1,CCAR2,ARF4,DIDO1,FYN,YWHAH,SLIT3,MTRNR2L10,STXBP1,UBQLN1,HYOU1,MAPK
RF4L2,NOX4,TAF15,ACTR3,PKN1,TCP1,CCT3,VPS35,PTOV1,MDK,RBMX,CCT4,GCN1,SKI,SNRNP7
PL17,PAPOLA,RPL14,HK1,RPL22,RHOA,PTCD3,CBX5,ZNF865,RPL9,ATF4,PSMD2,NR2F1,SLC7A5
PL37,MAPK1,AZIN1,ODC1,ATP5PB,EIF3E,FTO,ILF3,LUZP6,RPS11,PRKDC,HSPB1,CASC3,CLIP3,Q
DR1,SULF1,PLXNA1,MEGF8,SOX12,CCN2,ACTN4,MMP14,PRDX2,DDX3X,RPS15,ITM2C,CLTC,NA
R,ATP1A1,JUND,TMEM259,CD81,COL5A2,ATP2A2,CRMP1,SRSF4,MCL1,TLE5,CLEC7A,XPO1,SPT
H,NACA,RTN4,RPS24,BRD3,RPS9,XRCC5,CIPC,RPL24,KPNB1,RPL17,PAPOLA,RPL14,HK1,RPL22
,PLXNA1,MEGF8,SOX12,CCN2,ACTN4,MMP14,PRDX2,DDX3X,RPS15,ITM2C,CLTC,NANOG,RPS7,
X12,CCN2,ACTN4,PRDX2,DDX3X,RPS15,RPL12,RPL30,NANOG,H1-0,RPS5,RPS7,IK,RPS19,RPL35
L29,RPSA,EIF4H,NACA,LPCAT1,RPS24,BRD3,RPS9,XRCC5,CIPC,RPL24,KPNB1,RPL17,PAPOLA,R
L29,RPSA,EIF4H,NACA,LPCAT1,RPS24,BRD3,RPS9,XRCC5,CIPC,RPL24,KPNB1,RPL17,PAPOLA,R
IM20,RNF187,U2AF2,RPN2,PPP1CB,SQSTM1,KLHL24,NOP53,PARP1,VCAN,MORF4L2,NOX4,ALK
NF865,RPL9,ATF4,PSMD2,NR2F1,RPS14,CHD4,JUND,RPS20,CD81,RPL26,CRMP1,CCT7,SRSF4,P
DUFA13,SMO,RABGAP1,EIF4G1,IGFBP4,USP47,PTPN11,WDTC1,OGFR,ABL1,SDCBP,SEMA6B,AU
,KDM1A,BNIP3L,CLIP3,FKBP8,PLCG1,KDELR1,QARS1,CCAR2,ARF4,DIDO1,FYN,YWHAH,SLIT3,M
,PNN,RPL31,HNRNPC,LRPPRC,TADA2B,KHDRBS1,XPO1,CSNK1E,RPS25,SFPQ,SOX12,CCN2,AC

,RPS14,CHD4,JUND,RPS20,EIF5,CD81,RPL26,CCT7,RBBP4,PFN1,TLE5,RPL31,HNRNPC,LRPPRC
I3,PARK7,MAP4K4,FEZ1,KAT2A,EHMT2,NTN1,FLII,NCK2,PMEPA1,SYNGAP1,TOM1L2,EMILIN1,EX
O4,PTPN11,CDC34,ABL1,COL16A1,FXR2,SDCBP,CHMP1A,PINK1,AUTS2,ARPC1A,DAG1,HIP1R,H
EIF3H,TPP1,GPC2,UBE2D2,VPS4A,HINT1,LDLR,GNA12,RDX,DNAJB2,ATXN3,SLCO1B3,QPRT,PSM
C1,RPS14,CHD4,JUND,RPS20,EIF5,CD81,ATXN7L3B,RPL26,SRSF4,RBBP4,PFN1,TLE5,CLEC7A,R
CND1,MAPK1,ARPC2,PSMD3,ADAM9,PTBP1,FTO,LUZP6,CAPZB,PRKDC,HSPB1,KDM1A,PLCG1,C
ROCK2,RUFY3,PIN1,XBP1,TFDP1,PRKCI,UFM1,TMEM59,GNL3,NOLC1,EPHB2,ARF6,PRNP,SESN2,
MAEA,HIPK2,CUL4A,UBE2J1,MAGEF1,AREL1,RNF41,FAM8A1,TRIB3,ADRM1,GID8,PSME3,HM13,
20,EIF5,CD81,RPL26,CCT7,RBBP4,PFN1,TLE5,RPL31,HNRNPC,LRPPRC,TADA2B,KHDRBS1,XPO1
DAH2,CDC37,PRKDC,GABARAP,ACTN1,HSPB1,SON,KDM1A,BNIP3L,CLIP3,FKBP8,PLCG1,KDELR
8,PSMA6,KCTD10,XBP1,MAN1B1,CUL4B,PSMD12,APOE,RYBP,PSMA4,UBE2E1,PSMC6,RNF5,RA
R1,SEMA6C,APOE,TGFB2,CRIM1,FGF13,FHL1,SYT1,PPT1,KDM2B,SESN2,BRAT1,SYT4,POSTN,B
RKAR2A,KDM5B,FLCN,PHB2,UBA52,ZFP36L1,SFRP1,ARID1B,ADCY6,HDAC6,PTEN,RGMB,NCOA
OL4A2,RPL37A,MAPT,FN1,SPEN,NONO,DDB1,PSMD5,YWHAQ,EIF1,HACD3,DCHS1,SET,EGLN1,M
CR,RAC1,ARHGDIA,GPC1,CDK16,APPL1,HDAC2,ATP6V0C,BCLAF1,GLUL,PEA15,NNAT,PTTG1IP,D
3,MICAL1,TUBB4A,PLS3,ABL1,COL16A1,ARPC1A,HDGFL3,COL12A1,FAM171A1,CLIP2,HIP1R,H

,RHOA,CBX5,ZNF865,RPL9,ATF4,PSMD2,NR2F1,RPS14,CHD4,JUND,RPS20,CD81,RPL26,CRMP1
S9,XRCC5,CIPC,RPL24,RPL17,PAPOLA,RPL14,RPL22,RHOA,PTCD3,CBX5,ZNF865,RPL9,ATF4,PS
N2,TXNDC5,TRIO,HIP1,FLCN,CSNK2A1,FLOT1,GPX4,PHB2,UBA52,ZFP36L1,SFRP1,SRSF6,FBXO
71A1,PSMB1,TAOK2,TRAK1,DAG1,NIBAN2,PLXNA2,OBSL1,NOTCH1,MECP2,PSMD13,ADNP,ADG

T11,IST1,PSMC3,AP2A2,AP1M1,EXOC4,MICAL1,RIMS3,PAFAH1B2,SCAMP5,SDCBP,SCAMP1,PSM
CUL4A,RAP2A,WRNIP1,MAGEF1,NUP153,NCK2,FAM168A,H2AX,MAPKAPK2,CDK9,MARCHF7,NUF
PTEN,CUL3,IL6ST,OAZ2,ANP32B,RPS2,RPS27A,PPP1R15A,FKBP1A,SYMPK,DAP,MTCH1,SERP1,NI
K5,NOTCH1,PSMD13,CAB39,NDUFC2,APEH,CYFIP1,CD44,PSMD7,FAF2,TSPAN14,SYT15,CSNK2B,
,ARHGDIA,GPC1,CDK16,APPL1,HDAC2,ATP6V0C,BCLAF1,GLUL,PEA15,NNAT,PTTG1IP,DCN,MAP
,RHOA,CBX5,ZNF865,RPL9,ATF4,PSMD2,NR2F1,RPS14,CHD4,JUND,RPS20,CD81,RPL26,CRMP1
HD4,JUND,RPS20,EIF5,CD81,RPL26,CCT7,OAZ1,PFN1,TLE5,CLEC7A,RPL31,HNRNPC,LRPPRC,T/
CHSY1,TMSB10,PPP2R1A,PKD1,HLA-C,PDIA3,SSBP2,ZNF43,RPS3,CHD3,BASP1,VAPA,HMGB1,CD
EMF,ANAPC2,PPP2R5C
,BRD4,UBA52,DIP2B,ATRX,NXN,SFRP1,RAB3GAP2,ADCY6,CRTAP,MASTL,HDAC6,PTEN,CUL3,IL6
SNK1A1,PSMC2,HERPUD1,PSMC4,PLK2,PSMF1,WAC,GSK3B,UBE3C,USP48,USP19,UBQLN4,RNI

,UBE2D3,TRIM8,SAE1,SMO,CROCC,PTPN23,RAB11FIP3,RANGAP1,UBE2L3,PTPN11,SNX3,PINK1
SNK1A1,PSMC2,HERPUD1,EXOSC10,PSMC4,PLK2,PSMF1,WAC,GSK3B,UBE3C,USP48,USP19,UE

HERPUD1,PSMC4,PLK2,PSMF1,WAC,GSK3B,UBE3C,USP48,USP19,UBQLN4,RNF216,PSMB5,RNI
FLCN,CSNK2A1,PHB2,UBA52,ZFP36L1,SFRP1,SRSF6,FBXO11,PTEN,FEM1B,IL6ST,ANP32B,RPS2

L1,ADAR,ANP32A,VDAC1,TRAF4,TRAF7,STAT3,CAPN2,TXNDC5,TRIO,HIP1,FLCN,CSNK2A1,FLOT
EB2,LSM14A,TTC37,FLOT2,DNAJB2,SPAG9,PDPK1,ITGAV,IGF2BP1,OGT,NOC2L,KDM4A,ARPP19
CC,PTPN23,RANGAP1,ANK2,UBE2L3,PTPN11,SNX3,ABL1,PINK1,DAG1,MACROH2A1,RBM4,PKIA,/

RBPJ,PRPF19,KDM4A,MBD3,MTOR,UBE3A,BNIP3,SLC38A2,SMO,NTRK2,PAFAH1B2,LMO4,XRN2
BHD17A,RAB3A,RCC2,SETD5,DLG4,WASF3,DUSP3,ERBB2,UNC13A,NUMBL,THBS2,CNTN2,NRP2

K1,RPL22,RHOA,CBX5,ZNF865,RPL9,ATF4,PSMD2,NR2F1,RPS14,CHD4,ATP1A1,JUND,RPS20,CD8

G3,ARSB,ATP6V0A1,ATF6,RAB5A,GABARAPL2,MAPK3,XBP1,TAB3,UFM1,TMEM59,VPS16,DYNC1I
G3,ARSB,ATP6V0A1,ATF6,RAB5A,GABARAPL2,MAPK3,XBP1,TAB3,UFM1,TMEM59,VPS16,DYNC1I
12,SOS1,CITED2,PRMT2,LZTS2,NRP2,SGPL1,CFLAR,SHANK1,MAP2K4,RGMA,ANXA1,TGFBR1,PF

I,FKBP4,INTS3,TM9SF4,RIC8A,NUCKS1,ATXN3,NMT1,SOD2,SLC25A23,KRT8,PDPK1,PSMD4,MSH
P6,PLXNB1,ATRX,FARP1,SFRP1,EVL,HDAC6,SMC3,PPP1CC,CUL3,FMNL2,EML1,MAPRE2,BRSK1,I
,PBX1,FLCN,PRCC,CSNK2A1,PHB2,GSPT1,BRD4,UBA52,ZFP36L1,KLHL42,ATRX,HJURP,SFRP1,U

E,PSMA4,PSMC6,PAF1,PLPP3,GRB10,PPM1A,GNAO1,TLE4,FZD7,GPRC5B,FZD8,LRP6,JRK,EMD

TGFBR1,CAPZA2,CTDNEP1,ADD1,UFM1,HMGB2,ABCE1,VPS16,FURIN,FGF13,TPX2,PDXP,ARF6,F
PSMC6,PAF1,PLPP3,GRB10,PPM1A,GNAO1,TLE4,FZD7,GPRC5B,FZD8,LRP6,JRK,EMD

TCH1,SERP1,NDN,AKAP9,GRN,XRCC6,TAOK1,CCN1,SHC2,ANTXR1,ELAVL1,LDLR,RDX,KMT2E,D
P6V0D1,SH3BP4,ARPP19,GSK3A,MTOR,UBE3A,PRKACA,GOT2,ATP6V1G1,PHIP,CCDC47,UBXN6,

)LLIP

PSMD13,DCAF5,SUMO1,PJA1,PARK7,RNF157,HLA-A,PSMD7,MAEA,LIMK1,PCGF3,CUL4A,MYCB

RPP19,MTOR,WNK1,BAG5,DAPK3,MICAL1,PRDX3,EIF4A3,EIF4G1,TRIP12,USP47,ABL1,PUM1,FX
)LLIP

2,DNAJB2,NCSTN,NUCKS1,ATXN3,PDPK1,AKT2,OGT,ATP6V0D1,SH3BP4,ARPP19,GSK3A,MTOR,
,CTSA,ARSA,TOLLIP
P2,LIMS1,ZEB1,GDF1,ZNF703,NUS1,ZYX,NREP,SETX,SOX9,DUSP3,ERBB2,ADAMTS7,ANOS1,GD
2,USP47,UBE2L3,CDC34,RNF11,SNX3,WDTC1,ABL1,PINK1,USP10,PSMB1,RTF1,HSPA1A,ARRDC
PLXNB2,TTC3,PLD3,HSPD1,USP11,COL18A1,SNRNP200,CSDE1,IGF2,EIF3C,ATP5F1A,EIF4B,VC
D,GLB1,CTSA,ARSA,TOLLIP
N1,GRINA,VPS35,COL2A1,MDK,PTK7,SKI,ASXL1,PSMB2,H3-3A,TSPAN6,ATP6AP1,TERF2IP,TMED
2,ARPC5,ROCK2,WDR5,ANXA1,TGFBR1,MAPK3,SEPTIN9,XBP1,TFDP1,FER,GNL3,GMFB,GIT1,O
9,PPP1R10,SEPTIN5,PSMB4,TAOK1,EHMT1,CARM1,INTS3,VPS4A,RDX,LSM14A,KMT2E,TUBGCP
ARSA,TOLLIP
57,HLA-A,PSMD7,MAEA,LIMK1,PCGF3,CUL4A,MYCBP2,UBE2J1,MAGEF1,AREL1,RNF41,TRIM27,
GAP1,POLDIP2,USP47,PTPN11,CDC34,ABL1,SDCBP,CHMP1A,PSMB1,PPP5C,TAOK2,KIFC2,MAC

FIP3,EIF4G1,POLDIP2,USP47,PTPN11,ABL1,PUM1,SDCBP,CHMP1A,PHACTR4,PSMB1,TAOK2,TS
KT6,SV2A,PREPL,BRSK1,TMEM167A,HIF1A,ATP6AP2,A2M,SLC2A1,SERP1,PRDX6,GRK2,GRN,A
L1,PKIA,MDM2,FZR1,PCM1,MECP2,CUL1,PSMD13,SUMO1,BANF1,CCNY,PSMD7,RAN,SUN2,CUL

2,HNRNPD,ARNT2,WBP2,AKIRIN1,BMPR2,TP53BP1,CCT5,CDC42,STAT3,BCL9L,PRMT1,CAPN2,
SPAN6,ATP6AP1,TERF2IP,TAF7,NOTCH2,VEGFA,RAPGEF1,OPTN,RPL37,PPP2CA,CCND1,G3BP2,

S1,SPTY2D1,NSD2,OGT,PCGF2,NOC2L,KDM4A,MBD3,CHMP4B,NASP,YY1,SMARCD1,CREBBP,A
I3,KLHL24,PABPN1,GDI2,PARP1,NOX4,HSPH1,PRDX5,C4ORF3,PKN1,TBC1D16,TCP1,AHCYL1,A
R2,TP53BP1,CCT5,CDC42,STAT3,BCL9L,PRMT1,CAPN2,MTA1,EP300,LBH,RERE,PBX1,FLCN,H2

NUS1,ZYX,NREP,SETX,SOX9,DUSP3,ERBB2,ADAMTS7,ANOS1,GDF10,SOS1,NEO1,CITED2,NRP2,
SO1,ATP5MG,SCAP,RANGAP1,ANK2,TAPBP,SNX3,SUN1,CHMP1A,PINK1,CLTA,ARPC1A,NRXN2,H
K2,WDR11,NDRG4,SEMA4C,MEIS1,SRF,ZSWIM6,GSK3B,SEZ6L2,SERPINE2,CCDC85C,ATRN,SOX

P12,LMO4,CELSR1,WDTC1,ABL1,LRIG1,PHACTR4,COL12A1,RTF1,DAG1,NIBAN2,MACROH2A1,

L,PRMT1,MTA1,EP300,LBH,RERE,PBX1,FLCN,H2AZ1,EIF3D,BRD4,UBA52,HOXB5,ATRX,SFRP1,A

CDK4,H2AX,PSME3,TOM1L2,SENP6,PSMC5,PSMC2,PSMC4,PLK2,PSMF1,USP19,RCC2,SOX9,PRM
/PS28,NNT,PFKFB3,RNF40,MEX3D,CNOT1
OTC1,KDM3A,POR,PPP5C,DAG1,NOTCH1,IGF1R,MDM2,CAB39,PARK7,CYFIP1,PTPN1,PDGFRB,A

11,TMEM59,CABIN1,GMFB,RELA,EPHB2,APOE,HMG20B,FBXO7,PRNP,SRCIN1,PLPP3,DNMT1,SM

,PRCP,QSOX1,GSN,CSTB,ATP6V1D,GLB1,CTSA,ARSA,TOLLIP
P,NDFIP1,HNRNPK,INIP,CIRBP,PTGES3,CADM4,LAMP1,RHOB,LGALS3BP,ATP1A2,HLA-B,RAB1A,
SFRP1,GLUD1,TAGLN2,YKT6,SV2A,PREPL,BRSK1,TMEM167A,HIF1A,ATP6AP2,A2M,SLC2A1,SER
P6V1D,GLB1,CTSA,ARSA,TOLLIP
A1,ARF1,CYB5R3,PAFAH1B1,AKAP12,CDC42BPB,AP2B1,MAT2A,GTF2F1,PXDN,HNRNPU,NFE2L1

SMD8,HIF1A,LOX,SLIT2,GRN,XRCC6,PSMB4,CLPTM1,CCN1,OLFM1,CARM1,LINGO1,SOX2,PKDC
,CCDC85C,ATRN,MAP1S,MBOAT7,SNRPB,COX6B1,CHD8,CNTN1,NR2F2,NAPA,NUMBL,SOS1,BM

AX,LDB1,CDK9,PWWP3A,LOXL2,NSD1,WAC,WDR82,ASH1L,RCOR2,BCOR,SOX9,MAP1S,PRMT5,

NF2,ASAP3,GSN,GPM6B,PAM,GNAI1
N2,ERBB2,UNC13A,PHB,CITED2,SGPL1,SHC1,PSMD10,ADIPOR1,RGMA,TGFBR1,RUFY3,PIN1,XB
KNK2,TIA1,PSMD13,FXR1,CYFIP1,SYNCRIP,PABPC4,PSMD7,RAN,ETF1,TNRC6A,PPP1CA,HNRNP
CTIF,MRPS27,NEMF,SLC35A4,MEX3D,CNOT1,ABCA7
CSTB,ATP6V1D,GLB1,CTSA,ARSA,TOLLIP
FBR1,QRICH1,XBP1,TFDP1,YBX3,HMGB2,RELA,OPA1,MSH2,TGFB2,PRELID1,HIPK1,DPF2,EPHA

1,PSMA6,CTDNEP1,PIN1,EXT1,SPIN1,PSMD12,CTNNBIP1,APOE,PSMA4,PSMC6,PAF1,PLPP3,GRB
,HNRNPK,H1-10,CIRBP,DUX4,YWHAB,PTGES3,ZNF454,CCT6A,CTBP1,RAC1,CAPRIN1,CDK16,APF

DAP1,CST3,SFRP1,GLUD1,TAGLN2,YKT6,SV2A,PREPL,BRSK1,TMEM167A,HIF1A,ATP6AP2,A2M,S
,CDC37,PCSK1N,CAND1,HSPB1,PLCG1,MAP2K2,QARS1,CCAR2,ARF4,DUSP4,CORO1C,FYN,LAR
PTEN,ECEL1,CUL3,ANP32B,SPINT2,PREPL,RPS27A,PSMD8,NPLOC4,HIF1A,ATP6AP2,A2M,DAP,FB

K8IP2,MECP2,PARK7,ADNP,CD44,MAP4K4,RNF157,ATF5,PTPN1,MAEA,CFDP1,GOLPH3,HIPK2,DA
SMO,EIF4G1,NTRK2,USP47,ABL1,PINK1,POR,PPP5C,NIBAN2,HIP1R,RGL2,HSPA1A,NOTCH1,ATF

PSMD10,MDC1,ANXA1,PPP3CA,PSMA6,CNOT3,PCNA,TFDP1,AJUBA,CUL4B,PSMD12,MSH2,PSMA4

N,EPHA2,CNOT1,UNC13B
454,CCT6A,CTBP1,RAC1,CAPRIN1,CDK16,SMCHD1,HDAC2,BCLAF1,HSBP1,DCN,CDKN1A,DDX39
P1,NTRK2,IGFBP4,PTPN11,ABL1,SDCBP,ARPC1A,NIBAN2,HIP1R,HSPA1A,ACVR1B,SMPD3,TIA1,N

R,MDM2,MAPK8IP2,MECP2,PARK7,ADNP,CD44,MAP4K4,RNF157,ATF5,PTPN1,MAEA,CFDP1,GOL

87,U2AF2,CCT2,MLXIP,SQSTM1,NOP53,PABPN1,HNRNPL,PARP1,MORF4L2,TAF15,ALKBH5,PRD

B,PTGES3,ZNF454,CCT6A,CTBP1,RAC1,CAPRIN1,CDK16,APPL1,SMCHD1,HDAC2,BCLAF1,HSBP1

AC1,CAPRIN1,CDK16,SMCHD1,HDAC2,BCLAF1,HSBP1,DCN,CDKN1A,DDX39B,EID1,SRP9,CHD6,R
,PSMC6,PPP2R2A,CDK5RAP3,ZNF207,NINL,ID2,HMGA2,ANAPC2,PLRG1,SIN3A,CNOT1,PPP2R5C

DAP,PCBP4,NDN,BAP1,FUS,BAZ2A,XRCC6,EHMT1,BCL7A,UBR5,SOX2,ELAVL1,TLE1,MAZ,SRSF2,N

GTPBP1,GRSF1,BRD4,UBA52,ZFP36L1,HOXB5,ATRX,SFRP1,ARID1B,UPF1,TRA2B,RGMB,HNRNP

(NIP,DDIT4,PPP1R9B,STAT3,CAPN2,PRKAR2A,H2AZ1,MMP15,GPX4,PHB2,ZFP36L1,PPP2CB,SFRF
BE2E1,CENPF,PSMC6,CDK5RAP3,BTG1,ZNF207,NINL,ID2,HMGA2,ANAPC2,PLRG1,SIN3A,CNOT1
A1,ARPC5,NECTIN2,DYNLT1,FER,TRAPPC1,PSMD12,TLR6,COPB1,NFASC,RAP1B,PRDX4,PRCP,Q

NOTCH2,VEGFA,TUBB2A,MAPK1,MAPRE1,ARPC2,DSTN,DYNC1LI2,ADAM9,DYNLL2,GRB2,ACTN1
EP300,PRKAR2A,FLCN,BAZ1B,CSNK2A1,FLOT1,PHB2,BRD4,UBA52,BRD2,CSNK1G2,SFRP1,PAP

I5,PRDX5,ACTR3,PKN1,YEATS2,AHCYL1,PTOV1,MDK,ZNF302,RBMX,CBX1,CHD2,GCN1,TNPO1,SI
3B,SERPINE2,USP19,UBQLN4,HSPE1,UBXN1,COL7A1,DNAJB6,ANOS1,CNTN2,CFLAR,PSMD10,R

PLXNB1,CSNK1G2,SFRP1,GLUD1,PTEN,AGPAT1,NDST1,IL6ST,MAPRE2,RPS27A,TNKS,PSMD8,HI

1N1,ITGB1,TUBB4B,UCHL1,RPL11,MORF4L1,NCAM1,ANXA2,AGRN,ZMIZ1,TIMP2,XAF1,COL11A1
M121C,RPS27A,HIF1A,PPP1R15A,FKBP1A,SYMPK,LOX,SLIT2,AKAP9,PPP1R10,XRCC6,TAOK1,CC
M121C,RPS27A,HIF1A,PPP1R15A,FKBP1A,SYMPK,LOX,SLIT2,AKAP9,PPP1R10,XRCC6,TAOK1,CC

CFLAR,SHC1,SETD3,SHANK1,ROCK2,ANXA1,TGFBR1,EPB41L1,CAPZA2,RUFY3,STRIP1,ADD1,AR
A,CUL3,PREPL,BRSK1,RPS27A,TNKS,PSMD8,DKK3,HIF1A,ATP6AP2,GPC4,SLC2A1,SERP1,AKAP9,

K2,SEC14L1,TLE1,MAZ,TRAP1,SOD2,MTMR4,PDPK1,LZTR1,PSMD4,ITGAV,SYT11,PCGF2,SH3BP4
LS,HNRNPK,YWHAB,PTGES3,CADM4,LAMP1,RHOB,HLA-B,HMGCR,RAC1,ARHGDIA,GPC1,APPL1,S
FUS,XRCC6,PELP1,CCN1,SSBP3,CARM1,SOX2,ZMIZ2,MAZ,KMT2E,GTF2I,TTC37,NUCKS1,TCF3,I
ICER1,TCF3,FOXK2,IGF2BP1,NSD2,RBPJ,PCGF2,NOC2L,KDM4A,MBD3,MTOR,YY1,CREBBP,SAF
TMR4,PDPK1,LZTR1,PSMD4,ITGAV,PCGF2,SH3BP4,NOC2L,GSK3A,PSMC3,MTOR,PRKACA,NDUF

K2,SEC14L1,TLE1,MAZ,TRAP1,SOD2,MTMR4,PDPK1,LZTR1,PSMD4,ITGAV,SYT11,PCGF2,SH3BP4
1,CSNK1G2,SFRP1,GLUD1,PTEN,AGPAT1,NDST1,IL6ST,MAPRE2,RPS27A,TNKS,PSMD8,HIF1A,CH

P1,FLCN,CSNK2A1,FLOT1,PHB2,BRD4,UBA52,CXXC5,PPP2CB,PLXNB1,SFRP1,PTEN,FEM1B,CUL
3,ARPC3,DAAM1,GBA2,SRF,PIP5K1C,ZYX,WASF3,RHOBTB2,CDC42EP4,DNAJB6,INF2,ARPC5,CFI

OLG5,CNN2,NUCB1,RAB27A,CCT2,SQSTM1,VAMP2,KLHL24,MEST,PARP1,NOX4,ABCA3,C4ORF3,

,VPS28,GET4,GSN,LEMD2,VPS37B,DMPK,TMED5,ABCA7,EMD
AD,UBE2L3,FAM210B,ABL1,LGALS1,KDM3A,PPP5C,DAG1,RUVBL2,NOTCH1,MDM2,PARK7,GGT[illegible]

;,ARL8A,MAN2B1,KARS1,DPP7,BRI3,ARSB,ATP6V0A1,ARPC5,ANXA1,XBP1,DYNLT1,FER,TRAPPC1
LC1,MSH2,TGFB2,FURIN,KDM2B,AKIRIN2,SMARCB1,LRP6,CRB2,STOX2,SLC39A1,EPHA2,SKIL,[illegible]
R,PPIB,NNAT,DCN,CDKN1A,DLG5,DDX39B,CNN2,CTSB,SRSF5,GNAI3,PARP1,TGFBI,ABCA3,PKN1
LT1,FER,TRAPPC1,SPG11,SEC23IP,ABCE1,OPA1,SEC31A,SYT1,DYNC1I2,CENPF,BRAT1,DNM1,P[illegible]
FB2,FURIN,KDM2B,SMARCB1,LRP6,CRB2,SLC39A1,EPHA2,SKIL,RBBP6,SIN3A,CNOT1

P32B,MAPRE2,GPSM1,RPS2,RPS27A,TNKS,HIF1A,PPP1R15A,FKBP1A,RABEP1,DAP,MTCH1,AKAP[illegible]

,ANOS1,EFNB3,SOS1,NRP2,ATP6V0A1,ARPC5,SGPL1,DSTYK,SHC1,ROCK2,ADIPOR1,MAPK3,XB[illegible]

A,A2M,BAP1,FUS,GRN,PPP1R10,XRCC6,PSMB4,TAOK1,GPX1,FKBP4,UBR5,SEC14L1,AKT3,LDLR,

AN2B1,KARS1,DPP7,BRI3,ARSB,ATP6V0A1,ARPC5,ANXA1,XBP1,DYNLT1,FER,TRAPPC1,PSMD12,[illegible]

ADAM9,PTBP1,ILF3,CDC37,PRKDC,HSPB1,CLIP3,FKBP8,PLCG1,MAP2K2,QARS1,DUSP4,CORO1
P6AP2,LRRC59,TMOD2,PSMB4,TAOK1,CCN1,GPX1,SLC44A2,SHC2,UBR5,SOX2,AKT3,MAZ,LSM1
X39,CAB39,PARK7,ADNP,CD44,MAP4K4,PHPT1,ERP29,CCNY,PTPN1,LIMK1,PDGFRB,CCDC88A,C
RELA,OPA1,TLR6,PDXP,PRNP,EPHA4,SESN2,KLF3,GNAL,GRB10,RAP1B,DNMT1,GNAO1,ATP6V1F,
PHIP,WNK1,CD74,MICAL1,PRDX3,AMBRA1,FGFR1,UBE2K,EIF4G1,NTRK2,LMO4,PTPN11,ABL1,[illegible]

19B,SH3PXD2B,RAB11FIP3,SDCBP,CHMP1A,PINK1,NRXN2,KIFC2,YTHDF3,HSPA1A,OBSL1,NOTC

11,ABL1,PINK1,AUTS2,KDM3A,MGAT5,POR,RTF1,MACROH2A1,MKNK2,RUVBL2,ACVR1B,GGNB[illegible]

POR1,CHMP5,MAPK3,SLC1A3,XBP1,PRKCI,SP1,FER,BCAR1,RELA,OPA1,TLR6,PDXP,PRNP,EPHA4

FBP3,COL1A2,LLGL1,GAP43,NCS1,RPL22,RHOA,CLSTN1,ATF4,PSMD2,NR2F1,KIF5B,NLGN2,CX
DXL2,CTTN,LOX,SLIT2,GPC4,NDN,GRN,SLIT1,CCN1,GPX1,TCAF1,ABI2,GPC2,AKT3,GNA12,RDX,

,PSMD12,OPA1,CTNNBIP1,TGFB2,PSMA4,KDM2B,EPHA4,PSMC6,PLXND1,FZD7,NF2,LRP6,CRB2,
ADAM9,PTBP1,ILF3,CDC37,PRKDC,HSPB1,CLIP3,FKBP8,PLCG1,MAP2K2,QARS1,DUSP4,CORO1

JPF1,HDAC6,CBX6,HNRNPA2B1,CUL3,RPS27A,APEX1,DKK3,DAP,NDN,BAP1,BAZ2A,XRCC6,EHMT

RPF40A,BMPR2,AMOTL2,CDC42,DDIT4,PPP1R9B,PALLD,BEX4,KIF5A,STAT3,TRIO,EP300,RERE,F

FBP3,COL1A2,LLGL1,GAP43,NCS1,RPL22,RHOA,CLSTN1,ATF4,PSMD2,NR2F1,KIF5B,NLGN2,CX
IP,PRDX3,SLC38A2,SMO,NSG1,USO1,RABGAP1,NTRK2,RANGAP1,PTPN11,FAM210B,WDTC1,ABI
,BAZ2A,XRCC6,EHMT1,BCL7A,SOX2,TLE1,MAZ,IREB2,LSM14A,TTC37,SRSF2,MACROH2A2,DICE

ID1,KDM4A,SART1,MTOR,BNIP3,CD74,SMO,FGFR1,METRN,RABGAP1,EIF4G1,NTRK2,OLFM2,FA

)1,CREBBP,TOMM70,THRAP3,CD74,SLC38A2,DEK,SMO,EIF4A3,EIF4G1,CALCOCO1,CNBP,NTRK

A1,THRA,XBP1,PRKCI,SP1,UFM1,FER,BCAR1,RELA,OPA1,TRIM24,GRB10,RAP1B,ATP6V1F,LEPR

T1,BCL7A,SOX2,TLE1,MAZ,IREB2,LSM14A,TTC37,SRSF2,MACROH2A2,DICER1,TCF3,FOXK2,IGF
),PSMA7,VDAC1,SULF2,BCL9L,BCR,EP300,LBH,KDM5B,PSMD1,PBX1,ALDH3A2,AP2A1,MMP15,F

,COL3A1,ID3,MAPK8IP3,PLPP3,POSTN,MYL9,FZD7,PRCP,C6ORF89,GSN,CASK,GNAI1

SK3A,PSMC3,PRKACA,YY1,NDUFA13,PSMC1,WNK1,TRIM8,DAPK3,CD74,UBE2K,PTPN23,PTPN11

A,PSMC3,MTOR,UBE3A,YY1,SMARCD1,CREBBP,APBB1,ATXN7L3,PHIP,TRIM8,SSBP4,THRAP3,CD

A,PSMC3,MTOR,UBE3A,YY1,SMARCD1,CREBBP,APBB1,ATXN7L3,PHIP,TRIM8,SSBP4,THRAP3,CD

DLGA4,LIMS1,SRF,HSF1,ZEB1,ZNF703,SERPINE2,PA2G4,SOX9,PRMT5,HOXA5,CRABP2,VEZF1,S

A,PSMC3,MTOR,UBE3A,YY1,SMARCD1,CREBBP,APBB1,ATXN7L3,PHIP,TRIM8,SSBP4,THRAP3,CD

SMD13,NTN1,PSMD7,HTRA1,HIPK2,PDGFRB,VPS51,CAPN1,NECTIN1,EXT2,SOBP,PSME3,CRKL,D

D44,MAP4K4,KAT2A,ERP29,CCNY,PTPN1,HIPK2,MYCBP2,PDGFRB,CCDC88A,CSNK2B,RAP2A,AK

CROH2A1,ACVR1B,GGNBP2,SMPD3,PKIA,IGF1R,NPTN,STK39,CAB39,FXR1,PARK7,ADNP,CD44,

1,CCN1,SHC2,ABI2,PKDCC,AKT3,DCLK1,STK24,PRPF4B,SPAG9,PDPK1,DAZAP2,ADAM10,AKT2,

1,CYB5R3,PAFAH1B1,AKAP12,MAP4,ITM2B,AP2B1,SLC4A8,HNRNPU,CTNNA1,RPS16,LDHA,CPE,

RDX3,SMO,EIF4A3,USP47,WDTC1,ABL1,RSF1,MGAT5,POR,MACROH2A1,GGNBP2,PKIA,NOTCH

S27A,APEX1,DKK3,DAP,NDN,BAP1,BAZ2A,XRCC6,EHMT1,BCL7A,SOX2,TLE1,MAZ,IREB2,LSM14A

4,UBE2K,PTPN23,PTPN11,CDC34,SPPL2B,USP10,KDM3A,PSMB1,YTHDF3,HSPA1A,SMPD3,PFKP

DAC6,PTEN,CUL3,FMNL2,PTPRF,SLC7A6,SPINT2,MAPRE2,ITGB5,APEX1,HIF1A,PODXL2,CTTN,LC
DAC6,PTEN,CUL3,FMNL2,PTPRF,SLC7A6,SPINT2,MAPRE2,ITGB5,APEX1,HIF1A,PODXL2,CTTN,LC

C10,RANBP9,SEMA4C,PSMC4,SCG2,PSMF1,YWHAZ,HSF1,ASH1L,ABHD17A,UBE2V1,SETX,SOX9,

P1,IL6ST,FKBP1A,SLC25A5,ATP6AP2,PRDX6,GRN,ACLY,XRCC6,CLPTM1,SLC44A2,IMPDH2,RAB1

P1,VPS35,COL2A1,MDK,PTK7,SKI,PSMB2,ATP6AP1,NOTCH2,VEGFA,RAPGEF1,VHL,MVB12B,PPP2

T5,ADAR,DCTN1,STAT3,EIF2AK1,PRMT1,PRKAR2A,FLCN,SLC39A6,ZFP36L1,SLC39A7,DECR1,AT

N2,CELF3,SENP6,NUP98,GBA2,GOLGA8B,MAPRE3,PDE4DIP,SLK,RANBP9,PLK2,GSK3B,ASH1L,

SUB1,EHMT2,HMGB3,RAN,CUL4A,COPS6,ATF7IP,PDGFRB,CETN2,WRNIP1,MAGEF1,FAM168A,H2
2,MACROH2A1,PLXNA2,TXNRD1,SMPD3,OBSL1,NOTCH1,MFN2,MECP2,EFNB2,JAM3,AAMP,ADG

6,PTEN,CUL3,IL6ST,RPS27A,PSMD8,NPLOC4,PEG10,DKK3,HIF1A,PPP1R15A,FKBP1A,CTTN,SLC2

GNA13,CD74,AMBRA1,FGFR1,NTRK2,RANGAP1,LMO4,UBE2L3,PTPN11,AGAP1,ABL1,PINK1,SGS

3,TPX2,PRNP,SRCIN1,COPS8,CCNK,EPHA4,MAPK8IP3,GPRC5B,HMGA2,RAD50,ARRDC3,EPHA2

IR,RGPD5,PLXNA2,HSPA1A,ATP13A2,TCEA1,MDM2,SGSM3,PPP1R14B,CD2BP2,PARK7,USP14,CI
K2,WAC,HSF1,UBE2V1,UBQLN4,SETX,CDKN1B,ATMIN,OTUB1,KDM2A,CEBPG,XPC,UBXN1,TLK1

F1,CHMP1A,MTA2,CGGBP1,RTF1,NIBAN2,MACROH2A1,HSPA1A,RUVBL2,SIN3B,PKIA,NOTCH1,MI

SQLE,DCN,CDKN1A,CPSF2,PGRMC1,DDX39B,EID1,CHD6,RNF187,FDFT1,CCT2,MLXIP,CPSF1,SC

F1,CHMP1A,MTA2,CGGBP1,RTF1,NIBAN2,MACROH2A1,HSPA1A,RUVBL2,SIN3B,PKIA,NOTCH1,MI

P,CERS2,CD74,SMO,METRN,NTRK2,TRIP12,PTPN11,FAM210B,ABL1,SEMA6B,POR,TRAK1,DAG1,
F1,CHMP1A,MTA2,CGGBP1,RTF1,NIBAN2,MACROH2A1,HSPA1A,RUVBL2,SIN3B,PKIA,NOTCH1,MI

AR2B,CDC42SE1,UBXN1,CSRNP2,NR2F2,COL7A1,DNAJB6,ANOS1,PPP1R12A,LMTK3,ROCK2,CA

D2,NR2F1,NLGN2,CHD4,RPS20,CD81,ATP2A2,RPL26,BEX3,RBBP4,PFN1,MCL1,TLE5,CLEC7A,NIS

,HMGA2,GSN,ABR,SKIL,MYBBP1A,UNC13B

2,HIPK2,DRAP1,AKAP8L,NCK2,NME2,KPNA6,LDB1,CDK9,DHX36,BCL9,PSIP1,PSMC5,PSMC2,PSM
C1I2,CENPF,CENPT,CCSER2,ZNF207,NINL,GNAI1

CBX1,CHD2,GCN1,SKI,ASXL1,PSMB2,H3-3A,KDM6B,SCAF1,HDGF,ABCA2,PRPF6,TERF2IP,TAF7,N
G4,CDKN1B,SH3BP5,DUSP3,DUSP8,IRS1,PRKAR2B,ERBB2,NR2F2,PRRC1,PHB,DSTYK,SHC1,PSM

,CCT2,MLXIP,CPSF1,SQSTM1,NOP53,PABPN1,PARP1,MORF4L2,TAF15,SSRP1,PRDX5,ACTR3,PK
CBX1,CHD2,GCN1,SKI,ASXL1,PSMB2,H3-3A,KDM6B,SCAF1,HDGF,ABCA2,PRPF6,TERF2IP,TAF7,N

6,RNF187,CCT2,MLXIP,CPSF1,SQSTM1,NOP53,PABPN1,PARP1,MORF4L2,TAF15,SSRP1,PRDX5,A

K4,GPS1,TGFBR1,MAPK3,RTRAF,HEG1,AJUBA,TAB3,HEXIM1,GMFB,EPHB2,APOE,TLR6,LRP8,TG

CBX1,CHD2,GCN1,SKI,ASXL1,PSMB2,H3-3A,KDM6B,SCAF1,HDGF,ABCA2,PRPF6,TERF2IP,TAF7,N

2E,FLOT2,NCSTN,RAB6A,RAB5C,SLC7A1,ACTR1B,TCF3,PDPK1,ITGAV,SCRIB,ADAM10,CAP1,MSH

D1,CHD6,RNF187,CCT2,MLXIP,CPSF1,SQSTM1,NOP53,PABPN1,PARP1,MORF4L2,TAF15,SSRP1,F

RP8,SYT1,PRNP,PPT1,EPHA4,GJC1,DNM1,PPFIA3,SYT4,RAP1B,SLC18A3,LRP6,CACNG7,ABR,ANA

DAM9,FTO,LUZP6,PRKDC,HSPB1,KDM1A,PLCG1,CORO1C,YWHAH,FBLN1,B2M,TNKS2,PLXNA3,S

OTCH2,RNF10,VEGFA,MARS1,VHL,PPP2CA,CCND1,G3BP2,MAPK1,SMARCA4,CCNL2,PBX2,PSMD

OR,AP2A2,AP1M1,CD74,PAFAH1B2,ABL1,SDCBP,LGALS1,ARPC1A,SCAMP1,PSMB1,HSPA1A,PSM

OTCH2,RNF10,VEGFA,MARS1,VHL,PPP2CA,CCND1,G3BP2,MAPK1,SMARCA4,CCNL2,PBX2,PSMD

OTCH2,RNF10,VEGFA,MARS1,VHL,PPP2CA,CCND1,G3BP2,MAPK1,SMARCA4,CCNL2,PBX2,PSMD

M1,PPFIA3,SYT4,RAP1B,SLC18A3,LRP6,CACNG7,ABR,ANAPC2,CASK,GNAI1,UNC13B

MP,IDH2,MAP4K4,FEZ1,PHPT1,NTN1,SUN2,MYCBP2,IQSEC1,PDGFRB,CSNK2B,RAP2A,RNF41,TE

T4,RAP1B,SLC18A3,LRP6,CACNG7,ABR,ANAPC2,CASK,GNAI1,UNC13B
T4,RAP1B,SLC18A3,LRP6,CACNG7,ABR,ANAPC2,CASK,GNAI1,UNC13B
3A,CADM3,ATXN3,SLC7A1,CLASP2,PDPK1,ATP1B2,ITGAV,SCRIB,ADAM10,FBLN5,SART1,WNK1,D

ATXN3,SLC7A1,CLASP2,PDPK1,ATP1B2,ITGAV,SCRIB,ADAM10,FBLN5,SART1,WNK1,DAPK3,CD74

,BAG6,WBP2,AKIRIN1,BMPR2,TP53BP1,CDC42,PSMA7,TRAF4,TRAF7,SULF2,STAT3,PRMT1,SG

,PFKM,ID4,STRAP,SARS1,RAI1,SREBF2,NCOR2,HNRNPD,ZSCAN18,ARNT2,KDM3B,WBP2,AKIRI

G4,OXSR1,SLK,EMC10,SEMA4C,PLK2,SRF,ZNF703,NUS1,SERPINE2,RCC2,SOX9,ADIPOR2,DUSP

C1,PDGFRB,CSNK2B,RAP2A,RNF41,TBXA2R,C1QBP,LDB1,EMILIN1,CRKL,NDRG4,OXSR1,SLK,

K1,EPN1,DEAF1,EMILIN1,CRKL,LOXL2,NDRG4,EMC10,SEMA4C,PLK2,SCG2,MEIS1,SRF,YWHAZ,

RBB2,SRP54,PSMA1,RXRA,TMEM14C,SOS1,CITED2,SGPL1,PSMD10,WDR5,ANXA1,THRA,PSMA6,

,OGT,RBPJ,PRPF19,GSK3A,SART1,MTOR,CREBBP,TOMM70,CD74,SMO,SCD,METRN,RABGAP1,I

R9,PSMB5,STAT1,CEBPG,ERBB2,SRP54,PSMA1,RXRA,TMEM14C,SOS1,CITED2,SGPL1,PSMD10,V

SMC1,AMBRA1,SMO,HNRNPH3,LMO4,CELSR1,ABL1,PHACTR4,PSMB1,DAG1,MACROH2A1,PLXN

PR2,TXNIP,TP53BP1,PSMA7,CIC,RBBP7,STAT3,BCL9L,MTA1,EP300,KDM5B,PSMD1,PBX1,FLCN,I

M1,KAT2A,HIPK2,PDGFRB,TBXA2R,JAK1,EPN1,DHX36,EMILIN1,CRKL,LOXL2,WDR11,NDRG4,EM(

FRB,NCBP2,YWHAZ,GSK3B,ACSL4,CACNA1H,RAB3A,ADIPOR2,KIF3A,IRS1,ERBB2,CNTN1,NADK,(

L1CAM,PPIB,CDKN1A,DLG5,PGRMC1,CNN2,CTSB,RAB27A,RNF187,CCT2,SQSTM1,VAMP2,NOP5

PSMF1,SRF,HSF1,ZEB1,TFRC,RASSF2,HOXA5,IRF2BP2,CTR9,PSMB5,STAT1,CEBPG,ERBB2,SRP[illegible]

[illegible]BB2,CNTN1,PRRC1,PHB,GDF10,DSTYK,SHC1,PSMD10,ROCK2,MAP2K4,TGFBR1,MAPK3,ARL2B

[illegible]A2R,C1QBP,LDB1,EMILIN1,CRKL,NDRG4,OXSR1,SLK,EMC10,SEMA4C,PLK2,SRF,ZNF703,NUS1,[illegible]
[illegible]P1,PCNA,YBX3,AJUBA,HEXIM1,RELA,GATAD2B,CTNNBIP1,RYBP,CBX4,KDM2B,KLF3,ID3,PAF1,AK[illegible]

[illegible]1,SFRP1,CD164,SRSF6,PTEN,PPP1CC,CUL3,NDST1,EML1,IL6ST,SPINT2,HIF1A,CHERP,SLC25A5,[illegible]

ADIPOR2,ZFP91,RASSF2,TXN,DLG4,CDKN1B,NSMF,IRS1,PRKAR2B,ERBB2,CNTN1,PRRC1,PHB,G
ADIPOR2,ZFP91,RASSF2,TXN,DLG4,CDKN1B,NSMF,IRS1,PRKAR2B,ERBB2,CNTN1,PRRC1,PHB,G

2G4,RAB3A,TFRC,ARL8A,MAN2B1,CEBPG,DPP7,BRI3,ARSB,ATP6V0A1,ARPC5,NECTIN2,DYNLT1,

,ITM2B,AP2B1,MAT2A,TMEM170A,GOLGA8A,PXDN,HNRNPU,SMARCA5,CTNNA1,RPS16,RPL23,C
36B,NLGN2,RPS14,ATP1A1,RPS20,CD81,WDR1,ATP2A2,RPL26,CCT7,VAT1,OAZ1,SRSF4,ASTN2,F
RCA4,IPO7,DYNC1LI2,PTBP1,DYNLL2,ILF3,GRB2,RPS11,SUPT5H,HSPB1,BNIP3L,FKBP8,PLCG1,I
,EIF4H,SCARB2,RTN4,LAMA5,BRD3,XRCC5,MSN,RPL24,KPNB1,IGFBP3,COL1A2,LLGL1,GAP43,I
M30A,SKP1,RPL23A,GLUL,PTTG1IP,MAP1A,CDKN1A,SRP9,NUCB1,TM9SF3,TOMM20,RAB27A,C
M30A,SKP1,RPL23A,GLUL,PTTG1IP,MAP1A,CDKN1A,SRP9,NUCB1,TM9SF3,TOMM20,RAB27A,C
39,SERBP1,RPS13,PRPF40A,RPL34,ADAR,RBM15B,DHX15,POLR2E,PSMA7,ANP32A,STAT3,EDC4
AMP2,PABPN1,ALKBH5,RPL27,TBC1D16,HSPA4,AHCYL1,VPS35,RAB5B,TNPO1,RPL36A,RAB11B,
A,RPL38,RPL37,UBXN4,MVB12B,PPP2CA,AZIN1,PSMD3,ODC1,ADAM9,EIF3E,FTO,RPS11,GABAR
RAB7A,ACTN4,DDX3X,RPS15,RPL12,RPL30,ARL4C,CLTC,MAGED2,RPS5,RPS7,RPS19,RPL35,MIF
RPL36,YWHAB,SEPTIN11,LAMP1,RHOB,RPS12,MAP2,RAB1A,ARCN1,SEPTIN3,RPS27,CCT6A,HM
PARP1,NOX4,SSRP1,ABCA3,PTPRA,ACTR3,TCP1,CAMSAP1,CCT3,NCKAP1,VPS35,CLSTN3,ACAP3,
1,RPS10,RPL35A,TARDBP,RPLP2,ARL6IP1,NDFIP1,WLS,MYH10,RPL36,YWHAB,CKAP5,SEPTIN11,
S28,HNRNPM,SSB,FAU,STT3B,LONP1,UBQLN1,CSNK1D,USP7,HECTD1,PCBP2,BAG6,HNRNPD,RF
P1R9B,ADAR,STAT3,SGTA,OS9,ALDH3A2,FLCN,SEC16A,KPNA2,PHB2,UBA52,ATRX,RER1,RAB3G
PSMC5,PSMC2,EXOSC10,NCBP2,RBM10,PSMC4,PSMF1,YWHAZ,IGF2BP2,RPS29,PSMB5,PSMA1,
RPL23,CPE,RPL29,RPSA,CTSD,SCARB2,NACA,RTN4,LPCAT1,LAMA5,RPS24,SLC22A17,RPS9,XRC
RPS25,GLG1,SFPQ,EIF5A,PEBP1,CCND2,CDH2,PDIA6,GPC3,DDR1,RAB7A,CCN2,MMP14,PRDX2,I
,RPS5,IK,RPS19,PRKAR1A,TLN1,GJA1,JUN,MYO5A,CTNNB1,MACF1,GOLM1,RPS10,CYFIP2,TARD

X6,LSM4,MAPKAPK2,PSME3,DHX36,PSMC5,PSMC2,EXOSC10,NCBP2,RBM10,PSMC4,PSMF1,YW
S15,RPL12,RPL30,CLTC,EIF3L,H1-0,RPS5,RPS7,RPS19,RPL35,MIF,PRKAR1A,TLN1,GJA1,CTNNB
SPA4,TCP1,AHCYL1,CCT3,VPS35,RAB5B,CCT4,TNPO1,RPL36A,RAB11B,SRP14,ATP6AP1,TMED2,
TA,XPOT,OS9,ALDH3A2,AP2A1,SEC16A,KPNA2,PHB2,UBA52,UBE2G2,RAB3GAP2,MBTPS1,HDAC

OAZ1,RTN3,CLEC7A,RPL31,PRKCSH,CCNI,LRPPRC,TADA2B,KHDRBS1,KLHL9,XPO1,PTN,CSNK1
CB1,SRSF5,TOMM20,RAB27A,U2AF2,CPSF1,SNX22,VAMP2,PABPN1,GDI2,ALKBH5,RPL27,HNRN

HL1,RPL11,MORF4L1,BTF3,ANXA2,AGRN,RPLP1,ZMIZ1,TIMP2,COL11A1,MIDN,RPL10,MARCHF6
RAB1A,RPS27,HMGCR,GPC1,ATP6V0C,MMP2,SKP1,BCLAF1,LAPTM4B,TRIB2,RPL23A,GLUL,PTT
CB,PGD,SQSTM1,GNAI3,NOP53,VCAN,TAF15,ALKBH5,RPL27,PRDX5,UBE4A,VPS35,RAD23B,TN
RPL24,RPL17,IGFBP3,RPL14,RPL22,RHOA,CBX5,RPL9,ATF4,PSMD2,NR2F1,SLC7A5,ENC1,RPS1

PS26B,FUS,PSMB4,GPX1,EIF3H,ELAVL1,LDLR,GNA12,RDX,TTC37,DNAJB2,ATXN3,MTMR4,FOXK
6AP1,TMED2,RPS15A,RPL38,OPTN,RPL37,ARF3,MVB12B,MAPK1,KDELR2,IPO7,ADAM9,CDC37,I
ATF4,PSMD2,KIF5B,RAB6B,SLC7A5,NLGN2,RPS14,CXADR,ATP1A1,RPS20,CD81,WDR1,ATP2A2,F
AE1,SMO,SRPRA,RPS26,UBE2L3,PINK1,MACROH2A1,RGPD5,RUVBL2,PKIA,MFN2,SLC25A6,RPS2
36B,SLC7A5,NLGN2,RPS14,CXADR,ATP1A1,RPS20,CD81,WDR1,ATP2A2,RPL26,LASP1,CCT7,VAT1
1,DNER,RPL23A,LUC7L3,SQLE,DLG5,DDX39B,SRSF5,CDH11,CCT2,TBCD,VAMP2,GAS7,NOP53,P
1A2,PPP1CB,PGD,SQSTM1,NOP53,VCAN,TAF15,ALKBH5,RPL27,UBE4A,VPS35,RAD23B,TNPO1,S

(RCC5,PJA2,MSN,RPL24,RPL17,IGFBP3,RPL14,HK1,RPL22,RHOA,PTCD3,RPL9,ATF4,PSMD2,EN(
I2,RNF10,VEGFA,RAPGEF1,TUBB2A,MAPK1,SMARCA4,BTBD2,PBX2,SRSF1,ATP5PB,LUZP6,GRB2
F4,FLOT2,NMT1,MOAP1,PACS1,CLASP2,PDPK1,REEP2,SCRIB,ADAM10,AKT2,EHD1,CHMP4B,PAC
K1E,SFPQ,TPM1,EIF5A,CCND2,CDH2,PNMA1,GPC3,DDR1,SULF1,PLXNA1,PIP4K2B,MEGF8,SOX1
,ATP1B2,PHGDH,SCRIB,ADAM10,IGF2BP1,IST1,EHD1,GSK3A,SF3A2,MTOR,UBE3A,TMEFF1,APBE
GPC2,VPS4A,RAB14,SRP72,SSR3,RDX,TM9SF4,FLOT2,NMT1,MOAP1,PACS1,CLASP2,PDPK1,REEI
)1,SH3BP4,KDM4A,GSK3A,PSMC3,CHMP4B,MTOR,UBE3A,PRKACA,EXOC4,BNIP3,ATP6V1G1,PSN

S9,XRCC5,CIPC,MSN,RPL24,KPNB1,RPL17,PAPOLA,IGFBP3,RPL14,HK1,RPL22,RHOA,PTCD3,CB
PS21,FXR1,SYNCRIP,PABPC4,PNRC2,PSMD7,ETF1,TNRC6A,NBAS,HNRNPR,RPL27A,DDX6,LSM4,N
PRC,KHDRBS1,XPO1,SPTBN2,PTN,PFKL,GLG1,SFPQ,TPM1,PEBP1,CCND2,CDH2,GPC3,DDR1,SU
35A,APLP2,TARDBP,RPLP2,ARL6IP1,NDFIP1,MXD4,HNRNPK,H1-10,CIRBP,RPL36,YWHAB,CADM4,
RTN1,PKN1,VPS35,RAD23B,GCN1,SKI,SNRNP70,ABCA2,NRDC,TERF2IP,TMED2,TAF7,RPL38,VEG
GTF2F1,HNRNPU,OGA,NFE2L1,SMARCA5,LARP1,HNRNPA0,RPS16,RPL23,CPE,RPL29,RPSA,CTSI
H1-10,CIRBP,RPL36,YWHAB,CADM4,SND1,RPS12,RPS27,CTBP1,HMGCR,RAC1,CAPRIN1,SMCHD
)LR2B,ATP6V1B2,FYN,HNRNPM,LARS1,STT3B,B2M,LONP1,UBQLN1,USP7,HYOU1,PTPRS,MAPK8
63,SESN3,MTSS2,CYFIP2,HNRNPDL,ARID1A,DDOST,PTGES3,CADM4,HNRNPH1,ATP1A2,HLA-B,H
1,NSG2,CLIP3,PLTP,SMARCE1,SMARCC1,SUPT6H,DHX30,RPS28,CCT8,STMN3,EIF3F,B2M,STXBI
K1,CNP,STK24,SEMA3C,SPAG9,DICER1,CLASP2,ATP1B2,ANKS1A,PHGDH,SCRIB,ADAM10,IGF2B
2,BRK1,AKAP9,GRN,PPP1R10,PALM,GPX1,ABI2,GPC2,DZIP1,SH3GL1,KMT2E,NMT1,FNBP1L,PLI
PRMT1,NAP1L1,TRIO,EP300,RERE,PBX1,CNTNAP1,AP2A1,FLOT1,MAP6,PLXNB1,DIP2B,FARP1,SFI
XA2,AGRN,RPLP1,ZMIZ1,TIMP2,COL11A1,MIDN,RPL10,MARCHF6,WSB1,RPS17,AP3D1,SBF1,DN/
HSPA1A,NUDT4,PSMD13,RPS21,FXR1,SYNCRIP,PABPC4,PNRC2,PSMD7,ETF1,TNRC6A,NBAS,HNR
REX,HSPA1A,NUDT4,PSMD13,RPS21,FXR1,SYNCRIP,PABPC4,PNRC2,PSMD7,ETF1,TNRC6A,NBAS

P1,ZMIZ1,TIMP2,COL11A1,RPL10,MARCHF6,WSB1,RPS17,AP3D1,SBF1,DNAJA1,CYB5R3,PAFAH1
;,PTPRF,ULK1,BRSK1,HIF1A,CTTN,SLIT2,MTCH1,NDN,GRN,TAOK1,SLIT1,FKBP4,OLFM1,CARM1,
DIP2B,FARP1,SFRP1,CTNND2,EVL,ADCY6,HDAC6,PTEN,PPP1CC,PTPRF,ULK1,EML1,IL6ST,BRSK
15,NNAT,PTTG1IP,MAP1A,CDKN1A,CPSF2,DDX39B,SRP9,NUCB1,SRSF5,TOMM20,RAB27A,U2AF
POLA,IGFBP3,RPL14,HK1,RPL22,RHOA,PTCD3,CBX5,ZNF865,RPL9,ATF4,PSMD2,NR2F1,SLC7A5

)R1,ATP2A2,RPL26,BEX3,CCT7,OAZ1,PFN1,MCL1,TLE5,CLEC7A,NES,HNRNPC,TADA2B,KHDRBS
N1,MCL1,TLE5,CLEC7A,HNRNPC,CCNI,LRPPRC,TADA2B,KHDRBS1,XPO1,PTN,CSNK1E,GLG1,SF
YARS1,EIF3H,TPP1,ELAVL1,IREB2,LSM14A,TTC37,EIF4G3,TRAP1,EEF1B2,NCSTN,LSM14B,ABCF
,PPP2CA,CCND1,MAPK1,CCNL2,IPO7,ADAM9,PTBP1,EIF3E,FTO,ILF3,LUZP6,CDC37,PCSK1N,PRI
PR,VPS37B,USP36
PSMD13,RPS21,FXR1,SYNCRIP,PABPC4,PNRC2,PSMD7,ETF1,TNRC6A,NBAS,HNRNPR,RPL27A,DD
1,CLASP2,WASF2,ITGAV,SCRIB,ADAM10,CAP1,IGF2BP1,IST1,UBE3A,TMEFF1,APBB1,PHIP,TANC
MTREX,HSPA1A,NUDT4,PSMD13,RPS21,FXR1,SYNCRIP,PABPC4,PNRC2,PSMD7,ETF1,TNRC6A,NE
1,SNX1,MYCBP2,NECTIN1,RAP2A,ADCY1,SYNGAP1,ISLR2,DHX36,RBFOX2,RANBP9,GOLGA4,SEN
YCBP2,NECTIN1,RAP2A,ADCY1,SYNGAP1,ISLR2,DHX36,RBFOX2,RANBP9,GOLGA4,SEMA4C,SRF,
T5,EFNB1,SNX1,MYCBP2,NECTIN1,RAP2A,ADCY1,SYNGAP1,ISLR2,DHX36,RBFOX2,RANBP9,GOI
LAMA5,XRCC5,MSN,RPL24,IGFBP3,COL1A2,LLGL1,GAP43,NCS1,RPL22,RHOA,CLSTN1,ATF4,PS
CDC47,BAG5,UBXN6,ATG4B,LYPLA2,UBE2K,PTPN23,ZER1,FBXL19,USP5,AMFR,TRIP12,USP47,U
G,MTOR,DAPK3,EIF4A3,EIF4G1,RPS26,CNBP,TCEAL4,PUM1,FXR2,PINK1,PAIP2,YTHDF3,RBM4,N

3P1,CDC42,YME1L1,ADAR,PSMA7,RBBP7,TRAF7,DCTN1,BCL9L,PRMT1,NAP1L1,EP300,NDUFV1,
OKL1,AP2M1,RPS4X,RPS3A,RPL18,ITGB1,UCHL1,RPL11,MORF4L1,BTF3,ANXA2,AGRN,RPLP1,ZM
,TRIO,EP300,RERE,CNTNAP1,AP2A1,FLOT1,MAP6,PLXNB1,DIP2B,FARP1,SFRP1,CTNND2,EVL,AD
HNRNPA2B1,RPL7L1,RPS2,CHERP,SYMPK,PCBP4,SAFB2,PRDX6,FUS,PELP1,SNRPD3,INTS3,RAL
NRNPH1,ATP1A2,HLA-B,HMGCR,RAC1,ARHGDIA,STIP1,GPC1,CDK16,APPL1,HDAC2,IPO5,TNC,AT
GAP1,ISLR2,DHX36,RBFOX2,RANBP9,GOLGA4,SEMA4C,SRF,ZSWIM6,GSK3B,ALCAM,RAB3A,MA
BP4,EIF3H,SNRPD3,ABI2,RDX,TM9SF4,CAPZA1,MACROH2A2,TUBGCP6,DICER1,CLASP2,WASF2,
RPS17,AP3D1,SBF1,DNAJA1,ARF1,AKAP12,SF3B2,CDC42BPB,ITM2B,ZNF770,AP2B1,GTF2F1,HN
5,CLEC7A,HNRNPC,CCNI,LRPPRC,TADA2B,KHDRBS1,XPO1,PTN,CSNK1E,GLG1,SFPQ,EIF5A,PEB

TR1A,TRIO,EP300,RERE,CNTNAP1,AP2A1,FLOT1,MAP6,PLXNB1,DIP2B,FARP1,SFRP1,CTNND2,E
G,MTOR,DAPK3,EIF4A3,EIF4G1,RPS26,CNBP,TCEAL4,PUM1,FXR2,PINK1,PAIP2,YTHDF3,RBM4,
NN2,CTSB,SRSF5,SQSTM1,VAMP2,GNAI3,PABPN1,PARP1,NOX4,PTPRA,ACTR3,PKN1,TCP1,AHC
R2F1,KIF5B,ENC1,NLGN2,RPS14,CXADR,JUND,CD81,COL5A2,WDR1,ATP2A2,CRMP1,BEX3,VAT1
B4GALT5,EFNB1,MYCBP2,NECTIN1,RAP2A,C1QBP,ADCY1,SYNGAP1,ISLR2,DHX36,CRKL,RBFOX
OK,LAMC1,PTK7,SKI,ASXL1,PSMB2,KDM6B,CNTNAP2,ABCA2,DPYSL2,TMED2,RPL38,NOTCH2,VE
2,ADNP,CYFIP1,CD44,MAP4K4,FEZ1,RNF157,NTN1,LIMK1,B4GALT5,EFNB1,SNX1,FLII,MYCBP2,
G4,NSMF,ERBB2,NUMBL,ANOS1,EFNB3,SOS1,NEO1,CNTN2,NRP2,ARHGAP33,SHC1,SHANK1,R
SESN3,CYFIP2,DNAJC5,TARDBP,NDFIP1,DDOST,CIRBP,MYH10,YWHAB,PTGES3,SEPTIN11,LAMP1
SWAP,PRPF31,THOC3,RTRAF,XAB2,RNPS1,ACIN1,AKAP17A,ZMAT2,KHDC4,SRSF8,CPSF7,WDR33
P1,BEX3,PFN1,ASTN2,MCL1,TLE5,NES,PRKCSH,SPTBN2,PTN,CSNK1E,GLG1,CDH6,TPM1,CDH2,

RBS1,XPO1,PTN,CSNK1E,GLG1,SFPQ,EIF5A,PEBP1,CCND2,GPC3,SOX12,RAB7A,CCN2,ACTN4,M
F1B2,LSM14B,ABCF1,IGF2BP1,AKT2,EIF3G,MTOR,CERS2,DAPK3,SCD,EIF4A3,ELOVL5,EIF4G1,R
2,GPC3,DDR1,SULF1,PLXNA1,MEGF8,SOX12,CCN2,MMP14,PRDX2,RPS15,ITM2C,CLTC,APLP1,
RPS3A,RPL18,ITGB1,UCHL1,RPL11,MORF4L1,BTF3,ANXA2,AGRN,RPLP1,ZMIZ1,TIMP2,COL11A1

RA,UBE4A,PKN1,YEATS2,PLOD1,LAMC1,RAD23B,PTK7,GCN1,SKI,ASXL1,PSMB2,KDM6B,RAB11
RA,UBE4A,PKN1,YEATS2,PLOD1,LAMC1,RAD23B,PTK7,GCN1,SKI,ASXL1,PSMB2,KDM6B,RAB11
EMA6C,ZDHHC17,EPHB2,APOE,FGF13,NFASC,SZT2,EPHA4,MAPK8IP3,GRB10,PLXND1,NFIB,SH3
,PEA15,PTTG1IP,DCN,CDKN1A,DDX39B,CNN2,SRSF5,CHD6,RAB27A,RNF187,U2AF2,EEF1A2,CC
9,SNRPB,RNF20,RBM14,THOC2,RBM17,SRRT,SFSWAP,PRPF31,THOC3,CMTR1,XAB2,RNPS1,AC
3,TNFAIP1,TRIM2,CUL1,PSMD13,SUMO1,PARK7,USP14,PSMD7,FAF2,MAEA,HIPK2,CUL4A,UBE2
F4,FLNB,CORO1C,FYN,YWHAH,SLIT3,STMN3,FBLN1,B2M,STXBP1,PLXNA3,COL27A1,PTPRS,SO
5A5,SLIT2,BRK1,AKAP9,TMOD2,GRN,PPP1R10,TAOK1,GPX1,FKBP4,ABI2,RAB1B,SMG5,DZIP1,
B,FAU,SLIT3,FBLN1,SARS1,PTPRS,PCBP2,NCOR2,HNRNPD,RPL39,SERBP1,RPS13,RPL34,ADAR,
RS1,RPS2,RPS27A,PPP1R15A,ATP6AP2,MVD,GLO1,SERP1,ACLY,EIF2S2,IARS2,CCN1,GPX1,YARS
36,EIF3J,MYBBP1A,SNU13
TOV1,MDK,RBMX,CCT4,GCN1,SKI,SNRNP70,ASXL1,KDM6B,ABCA2,NRDC,PRPF6,TERF2IP,TAF7,
1,TMSB10,RPL19,PPP2R1A,RPL15,PKD1,HLA-C,SF3B1,PDIA3,SSBP2,ZNF43,RPS3,CHD3,BASP1,
A,RAPGEF1,OPTN,ARF3,MVB12B,MAPK1,KDELR2,ARPC2,PSMD3,DYNC1LI2,DYNLL2,GRB2,CA
NAPC2,SKIL

327A,RNF187,U2AF2,CCT2,MLXIP,SQSTM1,NOP53,PARP1,MORF4L2,NOX4,TAF15,HSPH1,ACTR3
4,AZIN1,SRSF9,IPO7,AEBP1,TCF25,PTBP1,EIF3E,FTO,ILF3,PCSK1N,PRKDC,SUPT5H,HSPB1,KDM
KBP1,ATP8A2,ANAPC2,EPHA2,SKIL,SIN3A
OTCH2,VEGFA,OPTN,RPL37,VHL,PPP2CA,CCND1,SMARCA4,SRSF9,IPO7,AEBP1,ACADVL,TCF25,
C42EP4,THOC2,CNTN1,RETREG3,ARSB,EFNB3,CNTN2,ARHGAP33,CFLAR,RAB5A,SS18L1,RGMA
,HNRNPD,VGF,WBP2,IGFBP7,BMPR2,TXNIP,DDIT4,PPP1R9B,POLR2E,RBBP7,SULF2,STAT3,BCL
34,ATMIN,NSMF,CDC42EP4,THOC2,CNTN1,RETREG3,ARSB,EFNB3,CNTN2,ARHGAP33,CFLAR,RA
8IP1,GHITM,SOD1,BAG6,ARNT2,SERBP1,BMPR2,DYNLL1,TXNIP,CDC42,DDIT4,YME1L1,ADAR,A
'0,ASXL1,KDM6B,ABCA2,NRDC,PRPF6,TERF2IP,TAF7,RNF10,VEGFA,MARS1,RAPGEF1,VHL,PPP2
,ENC1,RPS14,CHD4,JUND,RPS20,EIF5,CD81,ATXN7L3B,RPL26,SRSF4,RBBP4,PFN1,TLE5,PNN,C
ARS1,RPL28,RPS28,LARS1,FAU,STT3B,EIF3F,STT3A,PRPS1,SARS1,HNRNPD,RPL39,RPS13,BMP
NOG,RPS7,RPS19,MIF,GPAT4,PRKAR1A,GJA1,JUN,CTNNB1,CD63,ANXA6,NGRN,MACF1,CYFIP2,
BN2,SLC3A2,PTN,PFKL,GLG1,SFPQ,TPM1,PDIA6,GPC3,SULF1,PLXNA1,PIP4K2B,MEGF8,CCN2,
,RHOA,PTCD3,CBX5,ZNF865,RPL9,ATF4,PSMD2,NR2F1,ENC1,RPS14,CHD4,JUND,RPS20,EIF5,C
RPS19,GPAT4,PRKAR1A,GJA1,JUN,CTNNB1,CD63,ANXA6,NGRN,MACF1,CYFIP2,NDFIP1,TAGLN,
,PRKAR1A,JUN,CTNNB1,RPS10,RPL35A,HNRNPDL,TARDBP,RPLP2,ARID1A,TSR2,MXD4,SHOX,H
PL14,RPL22,RHOA,PTCD3,CBX5,ZNF865,RPL9,ATF4,PSMD2,NR2F1,ENC1,RPS14,CHD4,ATP1A1,
PL14,RPL22,RHOA,PTCD3,CBX5,ZNF865,RPL9,ATF4,PSMD2,NR2F1,ENC1,RPS14,CHD4,ATP1A1,
BH5,PTPRA,UBE4A,PKN1,YEATS2,PLOD1,LAMC1,RAD23B,PTK7,GCN1,SKI,ASXL1,PSMB2,KDM6
FN1,TLE5,PNN,RPL31,HNRNPC,LRPPRC,TADA2B,KHDRBS1,XPO1,CSNK1E,PFKL,RPS25,SFPQ,AC
TS2,POR,TAOK2,RTF1,DAG1,TSPYL2,HSPA1A,RUVBL2,ACVR1B,SMPD3,NOTCH1,KIF26A,ADNP,C
TRNR2L10,STXBP1,UBQLN1,HYOU1,MAPK8IP1,GHITM,SOD1,BAG6,ARNT2,SERBP1,BMPR2,DY
TN4,PRDX2,DDX3X,RPS15,RPL12,RPL30,NANOG,H1-0,RPS5,RPS7,IK,RPS19,RPL35,PRKAR1A,GJ

,TADA2B,KHDRBS1,XPO1,RPS25,SFPQ,EIF5A,GPC3,SOX12,CCN2,ACTN4,PRDX2,DDX3X,RPS15,F
OSC10,SPOCK1,SEMA4C,HSF1,GSK3B,RCC2,UBQLN4,BCOR,SOX9,TFRC,DLG4,OTUB1,KDM5A
SPA1A,ELN,SMPD3,NOTCH1,ATP13A2,CRACD,MECP2,DNAJB1,JAM3,SUMO1,FXR1,PARK7,ADNP,
MD4,OGT,PRPF19,USP24,IVD,AHCY,FBXW5,GSK3A,PSMC3,CHMP4B,MTOR,UBE3A,PRKACA,GOT
PL31,HNRNPC,LRPPRC,TADA2B,KHDRBS1,XPO1,CSNK1E,RPS25,GLG1,SFPQ,EIF5A,SULF1,MEG
ORO1C,SUPT6H,FYN,YWHAH,FBLN1,B2M,TNKS2,PLXNA3,CSNK1D,STRAP,SARS1,PTPRS,VANGL2
,NUP58,VPS28,LEPROT,NF2,CDK5RAP3,GSN,GPM6B,EPHA2,WAPL,SIN3A,USP36,NUTF2,GNAI1,A
STUB1,MARCHF7,PSMC5,CSNK1A1,CLPX,PSMC2,HERPUD1,PSMC4,PLK2,PSMF1,WAC,GSK3B,U
I,RPS25,SFPQ,EIF5A,SOX12,CCN2,ACTN4,PRDX2,DDX3X,RPS15,RPL12,ITM2C,RPL30,EIF3L,NAN
1,QARS1,CCAR2,ARF4,DIDO1,FYN,YWHAH,SLIT3,MTRNR2L10,STXBP1,UBQLN1,HYOU1,MAPK8I
023A,SPSB3,GET4,NEMF,PITHD1,RNF40,ANAPC2,PPP2R5C
TG1,CACNG7,ANAPC2,SIN3A
4,NDST1,SPINT2,ITGB5,RPS27A,APEX1,PEG10,DKK3,FKBP1A,LOX,SLIT2,SAFB2,AKAP9,PELP1,PA
AP9,CHSY1,TMSB10,RPL19,PPP2R1A,RPL15,PKD1,HLA-C,SF3B1,PDIA3,SERINC1,SSBP2,ZNF43,
CN,MAP1A,DLG5,TIMP3,DDX39B,CDH11,SQSTM1,VAMP2,GNAI3,KLHL24,NOP53,PARP1,TAF15,S
SPA1A,ELN,OBSL1,CRACD,MDM2,PCM1,TNFAIP1,MECP2,PARK7,CYFIP1,PDLIM1,NCKAP5L,LIMK

,CCT7,SRSF4,PFN1,TLE5,PNN,RPL31,HNRNPC,LRPPRC,TADA2B,KHDRBS1,XPO1,CSNK1E,PFKL,
MD2,NR2F1,ENC1,RPS14,CHD4,ATP1A1,JUND,RPS20,EIF5,CD81,RPL26,CCT7,OAZ1,RBBP4,PFN
11,HDAC6,PTEN,FEM1B,IL6ST,ANP32B,RPS27A,APEX1,HIF1A,CTTN,SLC25A5,LOX,SLIT2,MTCH1,
RB2,CYFIP1,CD44,RNF157,NTN1,PSMD7,LIMK1,CFDP1,HIPK2,RAP2A,TBXA2R,C1QBP,JAK1,SYN

1B1,HSPA1A,GRIK5,SMPD3,NOTCH1,ATP13A2,EXOC5,NSF,PSMD13,CAB39,NDUFC2,APEH,CYFIP
'98,HERPUD1,EMC10,SEMA4C,PLK2,DDAH1,HSF1,GSK3B,DHFR,UBE2V1,USP19,UBQLN4,RASSF
DN,AKAP9,GRN,XRCC6,TAOK1,CCN1,SHC2,ANTXR1,RAB1B,ELAVL1,LDLR,RDX,KMT2E,DNAJB2,N
,CAPN1,NME2,ASAH1,ADCY1,RAB18,PSMC2,DEGS1,PIP5K1C,GSK3B,SYNGR1,CACNA1H,PA2G4,I
1A,DLG5,TIMP3,DDX39B,CDH11,SQSTM1,VAMP2,GNAI3,KLHL24,NOP53,PARP1,TAF15,SSRP1,P
,CCT7,SRSF4,PFN1,TLE5,PNN,RPL31,HNRNPC,LRPPRC,TADA2B,KHDRBS1,XPO1,CSNK1E,PFKL,I
ADA2B,KHDRBS1,XPO1,RPS25,SFPQ,EIF5A,ACSL6,SOX12,CCN2,ACTN4,PRDX2,DDX3X,RPS15,RF
KL1,AP2M1,TPT1,RPS4X,RPS3A,OR8K1,STMN1,ITGB1,TUBB4B,UCHL1,RPL11,MORF4L1,NCAM

ST,RPS2,RPS27A,HIF1A,PPP1R15A,FKBP1A,SYMPK,LOX,SLIT2,NDN,AKAP9,PPP1R10,XRCC6,TAC
=216,PSMB5,RNF20,UBXN1,PSMA1,ZNRF1,RNF4,PSMD10,UBE2G1,ELOB,ZSWIM8,PSMA6,KCTD

,DAG1,MACROH2A1,PKIA,ATP13A2,MDM2,PCM1,NPTN,MGAT3,SUMO1,PARK7,IDH2,MAP4K4,PH
QLN4,RNF216,PSMB5,RNF20,UBXN1,PSMA1,ZNRF1,RNF4,PSMD10,UBE2G1,ELOB,ZSWIM8,PSM

=20,UBXN1,PSMA1,ZNRF1,RNF4,PSMD10,UBE2G1,ELOB,ZSWIM8,PSMA6,KCTD10,XBP1,CUL7,M
7A,APEX1,HIF1A,CTTN,SLC25A5,LOX,SLIT2,MTCH1,GLO1,GRN,PPP1R10,CCN1,GPX1,OLFM1,UB

1,GPX4,PHB2,UBA52,ZFP36L1,SFRP1,SRSF6,FBXO11,HDAC6,PTEN,FEM1B,IL6ST,ANP32B,RPS2
,MTOR,WNK1,BAG5,DAPK3,MICAL1,PRDX3,EIF4A3,EIF4G1,NTRK2,TRIP12,USP47,SNX3,ABL1,PL
ATP13A2,REEP5,MDM2,PCM1,NPTN,MGAT3,PARK7,MAP4K4,FEZ1,PHPT1,PTPN1,RAN,UBE2J1,CC

,PTPN11,CELSR1,ABL1,SUN1,SEMA6B,DAG1,PLXNA2,ACVR1B,NOTCH1,PCM1,MECP2,DNAJB1,F
,ARHGAP33,SHANK1,AFDN,ROCK2,CTBP2,NECTIN2,TGFBR1,HEG1,PCDHB2,PIN1,ADD1,EXT1,PF

31,RPL26,CRMP1,CCT7,SRSF4,PFN1,TLE5,PNN,RPL31,HNRNPC,LRPPRC,TADA2B,KHDRBS1,XPO

2,NBR1,FBXO7,SCOC,SESN2,RNF5,VPS28,QSOX1,TOMM40,ATP6V1D,STX12,VPS37B,RB1CC1,U
2,NBR1,FBXO7,SCOC,SESN2,RNF5,VPS28,QSOX1,TOMM40,ATP6V1D,STX12,VPS37B,RB1CC1,U
'P3CA,HEG1,RUFY3,PIN1,ADD1,EXT1,YBX3,SPG11,SEMA6C,APOE,TGFB2,PTPN12,FGF13,SYT1,S

H6,AKT2,ATP6V0D1,RBPJ,AHCY,NOC2L,PSMC3,MTOR,PRKACA,YY1,CREBBP,BNIP3,ATP6V1G1,P
TGB5,CALD1,ZMYM4,TNKS,CTTN,SLIT2,BRK1,AKAP9,TMOD2,ZMYM3,TAOK1,PALM,CYTH2,FKB
IPF1,MASTL,PTEN,SMC3,FEM1B,PPP1CC,CUL3,EML1,MAPRE2,BRSK1,RPS27A,SEPTIN7,KLHDC3

BXO7,ACIN1,EPHA4,SMARCC2,MRPL3,SMARCB1,SCAF8,MRPS26,CDK5RAP3,LRP6,MRPS27,MR

NAJB2,NCSTN,ATXN3,SPAG9,PDPK1,ABCF1,DAZAP2,DCUN1D2,AKT2,IST1,OGT,GSK3A,PSMC3,M
SORT1,NSG1,USO1,RABGAP1,SCAP,NTRK2,RANGAP1,CAD,AMFR,PTPN11,WDTC1,ABL1,COL16A

P2,UBE2J1,MAGEF1,AREL1,RNF41,TRIM27,NUP153,TRIB3,CDK9,PSME3,UBE2Q1,STUB1,MARCH

R2,SDCBP,KDM3A,MGAT5,POR,PPP5C,PAIP2,MACROH2A1,YTHDF3,RBM4,HSPA1A,GGNBP2,TIA1

JBE3A,PRKACA,GOT2,ATP6V1G1,PHIP,CCDC47,UBXN6,SORT1,NSG1,USO1,RABGAP1,SCAP,NTR

F10,SOS1,NEO1,CITED2,NRP2,CFLAR,DSTYK,SHC1,ROCK2,RGMA,ANXA1,TGFBR1,MAPK3,PIN1,
4,MDM2,FZR1,UBE4B,TNFAIP1,TRIM2,CUL1,PSMD13,DCAF5,SUMO1,PJA1,PARK7,USP14,TRRAP
P,MARCKSL1,UBB,AFF4,TRIM28,YWHAG,CKB,COL4A2,RPL37A,MAPT,FN1,SPEN,NONO,DDB1,PSM

D2,TAF7,NOTCH2,VEGFA,RAPGEF1,OPTN,RPL37,VHL,MVB12B,PPP2CA,MAPK1,SMARCA4,PSMD3
PA1,PDXP,ARF6,SYT1,FBXO7,PAF1,DNMT1,SMARCB1,NF2,RAD50,C6ORF89,RNF40,GSN,SIN3A,C
5,CLASP2,TCF3,PHGDH,PSMD4,MSH6,IST1,PCGF2,FBXW5,ARPP19,PSMC3,CHMP4B,PRKACA,NA

,TRIB3,CDK9,PSME3,UBE2Q1,STUB1,MARCHF7,PSMC5,PSMC2,HERPUD1,PSMC4,PSMF1,WAC,U
ROH2A1,HSPA1A,ACVR1B,SMPD3,OBSL1,PKIA,MDM2,FZR1,PCM1,MECP2,CUL1,PSMD13,SUMO1

PYL2,MACROH2A1,CLIC1,HSPA1A,SMPD3,OBSL1,PKIA,MDM2,FZR1,PCM1,MECP2,CUL1,PSMD13
LY,SEPTIN5,XRCC6,SLC44A2,VPS41,LTBP4,IMPDH2,VPS4A,RAB14,NCSTN,RAB6A,RAB5C,ACTR1
4A,TASOR,PDGFRB,CETN2,PPME1,CSNK2B,CDK4,AKAP8L,CCNG1,PSME3,TOM1L2,RTF2,NUP98

ITA1,EP300,LBH,RERE,PBX1,FLCN,H2AZ1,EIF3D,BRD4,UBA52,HOXB5,ATRX,SFRP1,ARID1B,RAB
MAPK1,DDAH2,PDE7A,PSMD3,PRDX1,PTBP1,GRB2,PRKDC,GABARAP,HSPB1,KDM1A,FKBP8,PLC

PBB1,ATXN7L3,DAPK3,DEK,PAGR1,FBXL19,TRIP12,HMGA1,ABL1,RSF1,SUN1,CHMP1A,PINK1,AU
CAP3,CCT4,GCN1,SKI,SNRNP70,ABCA2,NRDC,TERF2IP,TMED2,TAF7,VEGFA,SELENON,RAPGEF1,
AZ1,EIF3D,BRD4,UBA52,HOXB5,ATRX,SFRP1,ARID1B,RAB3GAP2,RGMB,NCOA4,RPS27A,TNKS,A

CFLAR,DSTYK,SHC1,RGMA,ANXA1,TGFBR1,MAPK3,PIN1,XBP1,ARHGEF18,EXT1,RAB35,PPP2R5B
P1R,HSPA1A,GRIK5,ATP13A2,MFN2,OSBPL2,NSF,EMC1,SUMO1,BANF1,RAN,ANKFY1,SUN2,GOL
9,MAP1S,MBOAT7,SNRPB,COX6B1,CHD8,CNTN1,NR2F2,NAPA,NUMBL,SOS1,BMERB1,OGDH,CN

PLXNA2,ACVR1B,TXNRD1,GGNBP2,NOTCH1,MFN2,KAT2A,NTN1,B4GALT5,EFNB1,SALL2,HIPK2,C

RID1B,RGMB,NCOA4,RPS27A,TNKS,APEX1,HNRNPAB,HIF1A,PPP1R15A,SERP1,NDN,FUS,XRCC6,

MT5,CDKN1B,PSMB5,RRM2,PRKAR2B,RBM14,XPC,PSMA1,RNF4,PRMT2,CTCF,PSMD10,ROCK2,

SNS,CSNK2B,CDK4,TRIB3,PMEPA1,NME2,ADCY1,ATP6V0B,RBFOX2,STAT2,SRF,HSF1,GSK3B,LG

ARCB1,MYADM,VPS28,NF2,CDK5RAP3,LRP6,NNT,SIN3A,IBTK

COTL1,HMGCR,RAC1,APPL1,SMCHD1,HDAC2,TNC,ATP6V0C,MMP2,SKP1,BCLAF1,GLUL,HSBP1,
1,PRDX6,GRK2,GRN,ACLY,SEPTIN5,XRCC6,SLC44A2,VPS41,LTBP4,IMPDH2,VPS4A,RAB14,NCS

,SMARCA5,LARP1,CTNNA1,RPS16,RPL23,CPE,TNS3,RTN4,LAMA5,XRCC5,PJA2,MSN,KPNB1,IGF

C,LTBP4,LDLR,KMT2E,FLOT2,MACROH2A2,PLEKHB2,SEMA3C,SOD2,SPAG9,DICER1,CLASP2,T
ERB1,OGDH,CNTN2,NRP2,ARPC5,SHANK1,MEIS3,SUDS3,PPP3CA,B4GALT2,EXT1,XAB2,UFM1,G

FBL,SETD5,CTR9,OTUB1,KDM2A,CHD8,RNF20,KDM5A,RBM14,LMNA,PER1,HIRA,TLK1,PHB,PRM

1,YBX3,BCAR1,SEMA6C,APOE,TGFB2,CRIM1,FGF13,FHL1,SYT1,PPT1,KDM2B,SESN2,BRAT1,SY
,CDK4,DDX6,NUP153,NCK2,C1QBP,LSM4,MAPKAPK2,PSME3,DHX36,EIF3B,NUP98,PSMC5,POL

2,SKIL,RB1CC1,PPP2R5C,MYBBP1A

10,PPM1A,GNAO1,TLE4,FZD7,GPRC5B,FZD8,LRP6,JRK,EMD,UNC13B
L1,SMCHD1,HDAC2,BCLAF1,HSBP1,DCN,CDKN1A,DDX39B,EID1,SRP9,CHD6,RAB27A,RNF187,C

LC2A1,SERP1,PRDX6,GRK2,GRN,ACLY,SEPTIN5,XRCC6,SLC44A2,VPS41,LTBP4,IMPDH2,VPS4A,
1,STMN3,FBLN1,PLXNA3,HGS,MAPK8IP1,SOD1,HNRNPD,PPP6R1,SEC23A,ARFGEF2,DYNLL1,TX
XO21,MTCH1,RHBDD2,BAP1,GRN,SPG7,PSMB4,CCN1,GPX1,EIF3H,ANTXR1,TPP1,SOX2,LTBP4,

D1,PDGFRB,ASNS,AREL1,TRIM27,NME2,SYNGAP1,PSME3,MARCHF7,MTRNR2L4,HERPUD1,TNF
13A2,IGF1R,MDM2,MAPK8IP2,MECP2,PARK7,ADNP,CD44,MAP4K4,RNF157,ATF5,PTPN1,MAEA,

4,TPX2,FHL1,DYNC1I2,FBXO7,UBE2E1,CENPF,PSMC6,PAF1,PPP2R2A,CDK5RAP3,ZNF207,NINL,I[illegible]

9B,EID1,SRP9,CHD6,RNF187,CCT2,MLXIP,PPP1CB,SQSTM1,NOP53,PARP1,MORF4L2,TAF15,HSI[illegible]
IOTCH1,IGF1R,NPTN,EFNB2,AAMP,CYFIP1,SUB1,PTPN1,HTRA1,EFNB1,HIPK2,PDGFRB,CCDC88/[illegible]

PH3,HIPK2,DAD1,PDGFRB,ASNS,AREL1,TRIM27,NME2,SYNGAP1,PSME3,MARCHF7,PSMC5,MTR[illegible]

)X5,ACTR3,PKN1,YEATS2,TCP1,AHCYL1,CCT3,PTOV1,MDK,ZNF302,RBMX,CBX1,CHD2,CCT4,GC[illegible]

,DCN,CDKN1A,DDX39B,EID1,SRP9,CHD6,RNF187,CCT2,MLXIP,PPP1CB,SQSTM1,NOP53,PARP1,

NF187,CCT2,MLXIP,PPP1CB,SQSTM1,NOP53,PARP1,MORF4L2,TAF15,PRDX5,ACTR3,PKN1,YEAT

MACROH2A2,DICER1,TCF3,FOXK2,MSH6,IGF2BP1,NSD2,RBPJ,PCGF2,NOC2L,KDM4A,MBD3,GS

A2B1,NCOA4,RPS27A,TNKS,APEX1,HNRNPAB,HIF1A,NDN,PRDX6,FUS,PPP1R10,XRCC6,PELP1,CC

P1,SRSF6,ARID1B,UPF1,ADCY6,TRA2B,HDAC6,PTEN,ADH5,NCOA4,ABCC9,APEX1,HIF1A,SLIT2,SL
,PPP2R5C
SOX1,GSN,CSTB,ATP6V1D,GLB1,CTSA,NAMPT,ARSA,TOLLIP

,HSPB1,CLIP3,PLCG1,ARF4,ZFAND5,TPM3,PLTP,CORO1C,KIF3C,FYN,SLIT3,FBLN1,PLXNA3,STR
SS1,PI4KA,ADCY6,MASTL,HDAC6,PTEN,ULK1,IL6ST,BRSK1,POM121C,RPS27A,TNKS,HIF1A,LOX,

KI,SNRNP70,ASXL1,PSMB2,H3-3A,KDM6B,HDGF,ABCA2,PRPF6,TERF2IP,TAF7,NOTCH2,RNF10,V
OCK2,CAMK2N1,ELOB,RPS6KA2,GABARAPL2,RGMA,TRAF3,COL6A3,XBP1,TMEM59,APOE,RYBF

F1A,CHERP,FKBP1A,ATP6AP2,LRRC59,SERP1,TMOD2,PSMB4,TAOK1,CCN1,GPX1,SLC44A2,SHC

,MARCHF6,WSB1,VCL,AP3D1,DNAJA1,ARF1,CYB5R3,PAFAH1B1,AKAP12,CDC42BPB,AP2B1,MAT
N1,SHC2,SMG5,LDLR,SPAG9,MTMR4,PDPK1,DAZAP2,AKT2,OGT,ARPP19,GSK3A,MTOR,PRKACA
N1,SHC2,SMG5,LDLR,SPAG9,MTMR4,PDPK1,DAZAP2,AKT2,OGT,ARPP19,GSK3A,MTOR,PRKACA

HGEF18,PRKCI,FER,GMFB,BCAR1,PDXP,ARF6,PIP5K1A,OBSCN,CDC42BPA,MYADM,NF2,ELMO2
TMOD2,GRK2,SEPTIN5,PSMB4,UBR5,SOX2,LTBP4,TLE1,SH3GL1,CNP,NCSTN,ATXN3,PSMD4,SC

4,NOC2L,GSK3A,PSMC3,MTOR,PRKACA,NDUFA13,PHIP,UBE2D3,PSMC1,BAG5,CD74,EIF4A3,RAE
SMCHD1,HDAC2,ATP6V0C,SKP1,BCLAF1,PEA15,HSBP1,PTTG1IP,DCN,DLG5,TIMP3,DDX39B,SQS
FOXK2,EDF1,OGT,RBPJ,PRPF19,MED13,GSK3A,PSMC3,MTOR,UBE3A,YY1,SMARCD1,CREBBP,AI
P18,APBB1,NDUFA13,UBE2I,UBE2D3,THRAP3,SMO,NELFB,RPS26,CNBP,USP47,HMGA1,WDTC1,F
FA13,PHIP,UBE2D3,PSMC1,BAG5,CD74,EIF4A3,RABGAP1,AMFR,IGFBP4,USP47,CDC34,ABL1,PIN

4,NOC2L,GSK3A,PSMC3,MTOR,PRKACA,NDUFA13,PHIP,UBE2D3,PSMC1,BAG5,CD74,EIF4A3,RAE
HERP,FKBP1A,ATP6AP2,LRRC59,SERP1,TMOD2,PSMB4,TAOK1,CCN1,GPX1,SLC44A2,SHC2,UBR5

L3,NDST1,MAPRE2,RPS27A,NPLOC4,HIF1A,CHERP,FKBP1A,ATP6AP2,SLIT2,A2M,LRRC59,PPP1R1
LAR,SHC1,SETD3,SHANK1,ROCK2,ANXA1,TGFBR1,EPB41L1,CAPZA2,RUFY3,STRIP1,ADD1,ARHG

,PKN1,TCP1,AHCYL1,CCT3,VPS35,CLSTN3,MDK,RAB5B,CCT4,ASXL1,RAB11B,ABCA2,ATP6AP1,D

7,ARL6IP5,KAT2A,EHMT2,GOLPH3,PDGFRB,ASNS,CSNK2B,ZCCHC3,CDK4,TBXA2R,PMEPA1,ADC

,PSMD12,MSH2,COPB1,NFASC,RAP1B,PRDX4,PRCP,QSOX1,GSN,CSTB,ATP6V1D,GLB1,CTSA,AR
RBBP6,SIN3A,CNOT1
,ACO2,COL2A1,MDK,LAMC1,RAD23B,PTK7,CHD2,SKI,ASXL1,PSMB2,H3-3A,KDM6B,CNTNAP2,A
PFIA3,SYT4,NINL,AP3M1,STX12,ARFGAP3,UNC13B

9,GRN,XRCC6,TAOK1,CCN1,TCAF1,SHC2,ANTXR1,TBC1D9,FLOT2,DNAJB2,RIC8A,NCSTN,AHSA1

P1,EXT1,SHB,VAV2,PPP2R5B,FER,BCAR1,RELA,GIT1,ZDHHC17,OPA1,EPHB2,CRIM1,PTPN12,FGF

,LSM14A,STK24,TRAP1,ATXN3,SOD2,SLC25A23,CLASP2,PDPK1,PSMD4,SYT11,NSD2,FBLN5,VAS

MSH2,COPB1,NFASC,RAP1B,PRDX4,PRCP,QSOX1,GSN,CSTB,ATP6V1D,GLB1,CTSA,ARSA,TOLLIP

C,FYN,PTP4A2,PFKM,FBLN1,PITPNM1,ISYNA1,CSNK1D,PRPS1,IDH1,NT5DC2,STRAP,PTP4A1,PTF
4A,NUCKS1,SPAG9,PDPK1,PSMD4,AKT2,RBPJ,ARPP19,GSK3A,PSMC3,MTOR,UBE3A,CREBBP,UE
SNK2B,TRIM27,NCK2,TRIB3,ADCY1,CCNG1,MAPKAPK2,LDB1,STUB1,CRKL,MAPRE3,TNFRSF10
,LEPROT,ZNF106,KLF10,LDOC1,PIK3R3,ATP6V1D,SIN3A,NAMPT
FXR2,SDCBP,PINK1,PPP5C,TAOK2,DAG1,TSPYL2,MACROH2A1,ACVR1B,GGNBP2,SMPD3,PKIA,IG

H1,MFN2,PCM1,EXOC5,BSCL2,FEZ1,KAT2A,EHMT2,FLII,DYNLRB1,TASOR,PDGFRB,CCDC88A,C

P2,ATP13A2,MAST2,IGF1R,MDM2,STK39,MECP2,TRMT112,SUMO1,CAB39,PARK7,ADNP,TRRAP,

,SESN2,KLF3,GNAL,GRB10,RAP1B,DNMT1,GNAO1,ATP6V1F,LEPROT,ZNF106,KLF10,LDOC1,PIK

ADR,CHD4,ATP1A1,RPS20,CD81,ATP2A2,RPL26,BEX3,RBBP4,PFN1,MCL1,TLE5,CLEC7A,NISCH,P
,MAZ,DCLK1,STK24,RIC8A,SEMA3C,SOD2,SPAG9,CLASP2,PDPK1,WASF2,ATP1B2,ANKS1A,ITGAV

,HMGA2,EPHA2
C,FYN,PTP4A2,PFKM,FBLN1,PITPNM1,ISYNA1,CSNK1D,PRPS1,IDH1,NT5DC2,STRAP,PTP4A1,PTF

1,BCL7A,SOX2,TLE1,MAZ,IREB2,LSM14A,TTC37,SRSF2,TRAP1,MACROH2A2,DICER1,TCF3,FOXK

LCN,CD99,PHB2,PLXNB1,SFRP1,EVL,HDAC6,PTEN,CUL3,FMNL2,PTPRF,SLC7A6,SPINT2,MAPRE2

ADR,CHD4,ATP1A1,RPS20,CD81,ATP2A2,RPL26,BEX3,RBBP4,PFN1,MCL1,TLE5,CLEC7A,NISCH,P
L1,COL16A1,LGALS1,PINK1,POR,PPP5C,DAG1,RUVBL2,GRIK5,SMPD3,IGF1R,MDM2,PARK7,CYF
R1,TCF3,FOXK2,IGF2BP1,NSD2,RBPJ,PCGF2,NOC2L,KDM4A,MBD3,GSK3A,YY1,CREBBP,SAP18,

M210B,HMGA1,ABL1,SDCBP,PINK1,POR,TRAK1,DAG1,NIBAN2,MACROH2A1,RBM4,CLIC1,PLXN

2,ANK2,PTPN11,HMGA1,SCAMP5,ABL1,FXR2,PINK1,MACROH2A1,YTHDF3,HSPA1A,ACVR1B,NOT

OT,ZNF106,PRCP,HMGA2,PIK3R3,ATP6V1D,CNOT1

2BP1,NSD2,RBPJ,PCGF2,NOC2L,KDM4A,MBD3,GSK3A,YY1,CREBBP,SAP18,APBB1,NDUFA13,UE
LOT1,GRSF1,PHB2,ZFP36L1,BRD2,HOXB5,ATRX,SFRP1,SRSF6,TAGLN2,PTEN,FEM1B,IL6ST,SPI

,CDC34,SPPL2B,USP10,KDM3A,PSMB1,YTHDF3,HSPA1A,SMPD3,PFKP,STK39,CUL1,PSMD13,SU

74,SMO,TEAD1,PAGR1,IER2,CALCOCO1,CNBP,SLC30A9,LMO4,UBE2L3,HMGA1,USF2,ABL1,RS

74,SMO,TEAD1,PAGR1,IER2,CALCOCO1,CNBP,SLC30A9,LMO4,UBE2L3,HMGA1,USF2,ABL1,RS

TAT1,METRNL,NUMBL,SRRT,SETD3,SS18L1,ANXA1,TGFBR1,RUFY3,PIN1,ADD1,XBP1,CUL7,PRK

74,SMO,TEAD1,PAGR1,IER2,CALCOCO1,CNBP,SLC30A9,LMO4,UBE2L3,HMGA1,USF2,ABL1,RS

AAM1,PSMC5,PSMC2,NDRG4,PSMC4,MEIS1,PSMF1,SRF,ZEB1,ASH1L,LGR4,CELSR3,ATRN,BCO

AP8L,TRIB3,ADCY1,MAPKAPK2,CDK9,PPP2R5D,STUB1,MARCHF7,CRKL,MAPRE3,TNFRSF10B,H

MAP4K4,ERP29,CCNY,PTPN1,HIPK2,PDGFRB,CCDC88A,CSNK2B,RAP2A,AKAP8L,TRIM27,NCK2,T

GSK3A,MTOR,PRKACA,PHIP,WNK1,DAPK3,CD74,MICAL1,FGFR1,GAK,UBE2K,EIF4G1,NTRK2,CA

RPL29,TPM4,SCARB2,NACA,RTN4,LPCAT1,LAMA5,XRCC5,PJA2,MSN,RPL24,SLC16A7,IGFBP3,CC

1,IGF1R,MDM2,STK39,PPP1R14B,CD2BP2,SUMO1,CAB39,PARK7,ADNP,USP14,CD44,PHPT1,RA

,TTC37,SRSF2,MACROH2A2,DICER1,TCF3,FOXK2,IGF2BP1,NSD2,RBPJ,PCGF2,NOC2L,KDM4A,

,STK39,CUL1,PSMD13,SUMO1,SYNCRIP,CD44,KAT2A,HLA-A,PTPN1,PSMD7,CSNK2B,CDK4,ASAH

X,SLIT2,GPC4,BRK1,NDN,GRN,SLIT1,CCN1,GPX1,TCAF1,ABI2,GPC2,AKT3,GNA12,RDX,MAZ,DC
X,SLIT2,GPC4,BRK1,NDN,GRN,SLIT1,CCN1,GPX1,TCAF1,ABI2,GPC2,AKT3,GNA12,RDX,MAZ,DC

PRMT5,RASSF2,DLG4,PSMB5,KARS1,DUSP3,DUSP8,IRS1,ERBB2,PSMA1,PER1,PHB,SOS1,CFLA

4,LDLR,GNA12,KMT2E,FLOT2,NCSTN,RAB6A,RAB5C,SLC7A1,ACTR1B,TCF3,PDPK1,ITGAV,SCRI

CA,CCND1,MAPK1,SMARCA4,ARPC2,PSMD3,ADAM9,PTBP1,GRB2,CDC37,GABARAP,HSPB1,CLI

RX,UBAP2L,UPF1,ADCY6,TRA2B,SV2A,HNRNPA2B1,NCOA4,TNKS,APEX1,HIF1A,CHERP,FKBP1A,

TUBA1C,RCC2,MAP1S,KIF3A,STMN4,ARL8A,CDKN1B,SLC16A1,RBM14,ERBB2,LMNA,NUP133,R

AX,POLD2,CDK9,PWWP3A,DHX36,RTF2,STUB1,EXOSC10,HSF1,UBE2V1,UBQLN4,SETX,MAP1S,
RB2,KAT2A,HTRA1,HIPK2,FLII,PDGFRB,NECTIN1,EXT2,TBXA2R,JAK1,LDB1,EPN1,DEAF1,EMILIN

5A5,SLIT2,A2M,GRK2,GRN,PPP1R10,PSMB4,PALM,GPX1,UBR5,SOX2,SEC14L1,TLE1,AKT3,LDL ꞩM2,PHACTR4,POR,TAOK2,DAG1,HIP1R,RGPD5,HSPA1A,TCEA1,ARRDC4,IGF1R,FZR1,SGSM3,STI

Ɔ44,ARL6IP5,MAP4K4,ACAP2,RASA4,TBC1D10B,ARHGAP5,IQSEC1,PDGFRB,PPME1,ARFGAP2,TE
,RBM17,PSMD10,MDC1,GPS1,MAPK3,CNOT3,PCNA,TFDP1,XAB2,HMGB2,CUL4B,FH,MCM7,MSH

DM2,FOXP4,MECP2,DNAJB1,SAMD11,SUMO1,ATF5,EHMT2,SALL2,HIPK2,TASOR,DRAP1,ATF7IP,

ΩSTM1,NOP53,PABPN1,PARP1,MORF4L2,TAF15,SSRP1,PRDX5,ACTR3,PDXK,PKN1,YEATS2,TCP1

DM2,FOXP4,MECP2,DNAJB1,SAMD11,SUMO1,ATF5,EHMT2,SALL2,HIPK2,TASOR,DRAP1,ATF7IP,

NIBAN2,MACROH2A1,KLF13,PLXNA2,HSPA1A,ACVR1B,SMPD3,OBSL1,NOTCH1,PCM1,NPTN,MEC
DM2,FOXP4,MECP2,DNAJB1,SAMD11,SUMO1,ATF5,EHMT2,SALL2,HIPK2,TASOR,DRAP1,ATF7IP,

ⱭMK2N1,GPS1,ANXA1,PPP1R26,RTRAF,HEG1,COL6A3,AJUBA,HEXIM1,CABIN1,GMFB,ABCE1,EPH

ƧCH,PRKCSH,XPR1,KHDRBS1,SPTBN2,PTN,CSNK1E,GLG1,CDH6,SFPQ,PEBP1,CDH2,PDIA6,GPC3

IC4,MEIS1,SRF,HSF1,ZEB1,BEX1,NR2C2,ASH1L,CRTC1,SETX,SOX9,MED25,ZBED4,RRP1B,HOXA

IOTCH2,RNF10,VEGFA,MARS1,VHL,PPP2CA,CCND1,G3BP2,MAPK1,SMARCA4,CCNL2,PBX2,PSMI
ID10,CAMK2N1,MAP2K4,GPS1,TGFBR1,MAPK3,RTRAF,HEG1,AJUBA,VAV2,TAB3,HEXIM1,GMFB,

N1,YEATS2,TCP1,CCT3,PTOV1,MDK,ZNF302,RBMX,CBX1,CHD2,CCT4,GCN1,SKI,ASXL1,PSMB2,[illegible]
IOTCH2,RNF10,VEGFA,MARS1,VHL,PPP2CA,CCND1,G3BP2,MAPK1,SMARCA4,CCNL2,PBX2,PSM[illegible]

ACTR3,PDXK,PKN1,YEATS2,TCP1,CCT3,PTOV1,MDK,ZNF302,RBMX,CBX1,CHD2,CCT4,GCN1,SKI

GFB2,FGF13,TPX2,CCNL1,FBXO7,PRNP,SRCIN1,COPS8,CCNK,EPHA4,SESN2,MAPK8IP3,NF2,GP[illegible]

IOTCH2,RNF10,VEGFA,MARS1,VHL,PPP2CA,CCND1,G3BP2,MAPK1,SMARCA4,CCNL2,PBX2,PSM[illegible]

I6,SYT11,NSD2,IST1,RBPJ,PSMC3,SART1,MTOR,AP2A2,YY1,AP1M1,CD74,PAFAH1B2,PTPN11,AB[illegible]

PRDX5,ACTR3,PDXK,PKN1,YEATS2,TCP1,CCT3,PTOV1,MDK,ZNF302,RBMX,CBX1,CHD2,CCT4,GC

APC2,CASK,GNAI1,UNC13B

STRAP,SARS1,PTPRS,HGS,PCBP2,SOD1,BAG6,RAI1,PFN2,RAB11A,CYP51A1,ARFGEF2,BMPR2,TP

)3,AEBP1,TCF25,ILF3,LUZP6,PRKDC,SUPT5H,ACTN1,CAND1,KDM1A,MAP2K2,QARS1,CCAR2,ARF

D13,CAB39,NDUFC2,APEH,CYFIP1,CD44,HLA-A,PSMD7,FAF2,LIMK1,TSPAN14,TMEM98,CSNK2B

)3,AEBP1,TCF25,ILF3,LUZP6,PRKDC,SUPT5H,ACTN1,CAND1,KDM1A,MAP2K2,QARS1,CCAR2,ARF
)3,AEBP1,TCF25,ILF3,LUZP6,PRKDC,SUPT5H,ACTN1,CAND1,KDM1A,MAP2K2,QARS1,CCAR2,ARF

XA2R,C1QBP,LDB1,EMILIN1,CRKL,NDRG4,OXSR1,SLK,EMC10,SEMA4C,PLK2,SRF,ZSWIM6,ZN

APK3,CD74,TRO,PTPN23,PTPN11,CELSR1,ABL1,COL16A1,LGALS1,COL12A1,TAOK2,DAG1,NRX

,TRO,PTPN23,PTPN11,CELSR1,ABL1,COL16A1,LGALS1,COL12A1,TAOK2,DAG1,NRXN2,CLIC1,P

A,EP300,PSMD1,HIP1,FLCN,UBQLN2,CSNK2A1,FLOT1,PHB2,BRD4,UBA52,CXXC5,PLXNB1,CSNI

N1,BMPR2,TXNIP,TP53BP1,POLR2E,PSMA7,CIC,RBBP7,STAT3,BCL9L,MTA1,EP300,KDM5B,PSMI

3,NSMF,RNF20,LMNA,NR2F2,ARSB,CITED2,BMERB1,NRP2,AFDN,ROCK2,ADIPOR1,ANXA1,TGFB

MC10,SEMA4C,PLK2,SCG2,SRF,ZSWIM6,ZNF703,NUS1,SERPINE2,RCC2,SOX9,ADIPOR2,DUSP3,

DDAH1,ZEB1,NUS1,LGR4,SERPINE2,RAB3A,SOX9,ADIPOR2,WARS1,HOXA5,VEZF1,STAT1,CHD8,H

ADD1,XBP1,EXT1,HMGB2,PSMD12,MSH2,CTNNBIP1,TGFB2,NCOA6,PSMA4,SETD1A,FBXO7,ACIN

NTRK2,PTPN11,SCAMP5,ABL1,SDCBP,KDM3A,POR,TRAK1,DAG1,NIBAN2,MACROH2A1,PLXNA2,

WDR5,ANXA1,TGFBR1,MAPK3,THRA,PSMA6,ADD1,XBP1,EXT1,HMGB2,PSMD12,MSH2,CTNNBIP1

IA2,ACVR1B,NOTCH1,ACTA2,FZR1,EXOC5,PSMD13,CD44,KAT2A,NTN1,PSMD7,SALL2,RHEB,PPP

BAZ1B,H2AZ1,BRD4,UBA52,CXXC5,BRD2,HOXB5,ATRX,ARID1B,PTEN,CBX6,HNRNPA2B1,CUL3,R

C10,PLK2,SCG2,MEIS1,SRF,YWHAZ,DDAH1,NUS1,BCOR,SOX9,ADIPOR2,WARS1,HOXA5,VEZF1,C

CAMK2N1,RAB5A,ANXA1,PPP3CA,RUFY3,XBP1,TFDP1,PRKCI,EPHB2,APOE,TGFB2,FHL1,ARF6,SY

53,GDI2,PARP1,HSPH1,ACTR3,PDXK,PKN1,KLC1,NCKAP1,COL2A1,MLEC,MDK,SURF4,RAB5B,CH

54,PSMA1,RXRA,TMEM14C,SOS1,CITED2,SGPL1,PSMD10,WDR5,ANXA1,TGFBR1,MAPK3,THRA,F

P,PIN1,XBP1,AJUBA,VAV2,TAB3,EPHB2,TLR6,LRP8,TGFB2,FGF13,TPX2,PRNP,SRCIN1,COPS8,CC

SERPINE2,RCC2,SOX9,ADIPOR2,DUSP3,NSMF,RNF20,ERBB2,LMNA,NR2F2,ARSB,CITED2,BMERI
(IRIN2,PPM1A,DNMT1,SCAF8,TLE4,NFIB,ID2,HMGA2,KLF10,DPF2,SKIL,SIN3A,HES6,CNOT1,CC2I

,MVD,BRK1,NDN,BAP1,GRN,XRCC6,EIF2S2,CCN1,GPX1,SSBP3,CARM1,SOX2,IMPDH2,AKT3,GNA

GDF10,PPP1R12A,DSTYK,SHC1,PSMD10,ROCK2,MAP2K4,TGFBR1,MAPK3,ARL2BP,PIN1,XBP1,AJ
GDF10,PPP1R12A,DSTYK,SHC1,PSMD10,ROCK2,MAP2K4,TGFBR1,MAPK3,ARL2BP,PIN1,XBP1,AJ

FER,TRAPPC1,PSMD12,MSH2,COPB1,NFASC,RAP1B,PRDX4,PRCP,QSOX1,GSN,CSTB,ATP6V1D,[illegible]

NN3,RPL29,RPSA,TPM4,EIF4H,SCARB2,RTN4
RPL31,LRPPRC,KHDRBS1,XPO1,SPT
RPL28,POLR2B,SUPT6H,RPS
NCS1,RHOA,PTCD3,CLSTN1,PSMD2,NR2F1,KIF5B
CT2,SQSTM1,VAMP2,NOP53,PARP
CT2,SQSTM1,VAMP2,NOP53,PARP
,PSMD1,SF1,PRCC,GTPBP1,GRSF
,SRP14,ABCA2,TMED2,RPS15A,RP
AP,UBE2R2,HSPB1,BNIP3L,CASC3
,TLN1,GJA1,MYO5A,CTNNB1,CD
GCR,RAC1,ARHGDIA,GPC1,CDK16
MDK,SURF4,PTK7,RAB5B,CCT4,SKI,H3-3A,C
LAMP1,RHOB,RPS12,MAP2,RAB1A
L39,SERBP1,RPS13,RPL34,PCYOX1,
P2,MBTPS1,NCOA4,MAPRE2,
SKIV2L,PSMD10,ROCK2,PSMA6,CN
C5,MSN,RPL24,SLC16A7,KPNB1,RPL
DDX3X,RPS15,RPL12,RPL30
BP,ARL6IP1,ARID1A,H1-10,CIRBP,MYH10,YWHAB,P

HAZ,IGF2BP2,RPS29,PSMB5,PSMA1
1,MACF1,RPS10,MTSS2,RPL35A,CYF
RPS15A,RPL38,OPTN,RPL37,A
6,TBC1D9B,NCOA4,OAZ2

E,SCRN1,RPS25,GLG1,SFP
A3,TBC1D16,HSPA4,TCP1,AHC

,WSB1,RPS17,AP3D1,SBF1,DN
G1IP,DCN,MAP1A,TIMP3,CTSB,T
O1,SNRNP70,PSMB2,RPL36A,A
4,ATP1A1,TMEM259,RPS20,COL5A2,AT

2,PSMD4,IGF2BP1,AKT2,OGT,ATP6V0D1,SH3
RPS11,GABARAP,HSPB1,NPEPPS
RPL26,LASP1,CCT7,VAT1,OAZ1,S
1,RAN,PITRM1,FAM53C,RPL27A,
,OAZ1,SRSF4,RTN3,ASTN2,MCL
ARP1,TSPYL4,TLN2,NOX4,ABCA3,PTPRA,A
NRNP70,PSMB2,RPL36A,ABCA2,N

C1,RPS14,TMEM259,RPS20
2,PRKDC,ATXN10,KDM1A,FKBP8,PLCG1,QARS1,AR
CS2,SSR2,EXOC4,NDUFA13,TO
2,RAB7A,CCN2,ACTN4,MMP14,PRDX2,D
31,TANC2,CERS2,BAG5,DPYSL4,SMO,MET
P2,SCRIB,ADAM10,AKT2,EHD
MC1,TRIM8,BAG5,DAPK3,THRAP3,AMBRA1,UBE2K

X5,ZNF865,RPL9,ATF4,PSMD2,
MAPKAPK2,PSME3,DHX36,PSMC
JLF1,PLXNA1,PIP4K2B,MEGF8,RAB7A,CCN2,A
,SND1,RPS12,RPS27,CTBP1,HMGC
FA,RAPGEF1,RPL37,PPP2CA,
D,EIF4H,NACA,RPS24,BRD3,RPS
1,HDAC2,IPO5,BCLAF1,LAPTM4B,T
3IP1,SOD1,BAG6,SREBF2,BMPR2,KDM4B,TP53BP1,CDC42,
MGCR,RAC1,ARHGDIA,STIP1,GPC1,CDK16
P1,LONP1,UBQLN1,CSNK1D,SF3A1,S
P1,IST1,EHD1,GSK3A,SF3A2,MTOR,UBE3
EKHM2,MOAP1,CLASP2,WASF2,ATP1B2,AKT2
RP1,CTNND2,EVL,ADCY6,HDAC6,PTEN,PPP
AJA1,ARF1,CYB5R3,PAFAH1B1
NPR,RPL27A,DDX6,LSM4,MAPK
S,HNRNPR,RPL27A,DDX6,LSM4,

B1,SF3B2,CDC42BPB,ITM2B,Z
,LINGO1,ABI2,SOX2,GPC2,DCLK1,CNP,S
1,HIF1A,CTTN,SLIT2,MTCH1,NDN,GRN,
T2,CPSF1,SNX22,VAMP2,PABPN
,ENC1,RPS14,CHD4,JUND,TMEM

1,PTN,CSNK1E,SFPQ,TPM1,EIF5A,CCND
FPQ,EIF5A,PEBP1,CCND2,GPC3,DDR
T1,ADA
KDC,GABARAP,HSPB1,KDM1A,

DX6,LSM4,MAPKAPK2,PSME3,DH
2,DAPK3,DPYSL4,SMO,METRN,MYL12B,L
3AS,HNRNPR,RPL27A,DDX6,LSM
MA4C,SRF,ZSWIM6,GSK3B,ALCAM,RAB3
ZSWIM6,GSK3B,ALCAM,RAB3A,MAP1S,
LGA4,SEMA4C,SRF,ZSWIM6,GSK3B,AL
SMD2,NR2F1,KIF5B,ENC1,NLGN2,RPS1
UBE2L3,CDC34,RNF11,SNX3,SDCBP,PINK1,CLTA,
MKNK

PSMD1,SF1,HIP1,EIF3D,SEC16A,GP
MIZ1,TIMP2,COL11A1,MIDN,RPL
DCY6,HDAC6,PTEN,PTPRF,ULK1,BRSK1,SEPTI
Y,ELAVL1,EFTUD2,SRSF2,ELP1,INT
TP6V0C,MMP2,SKP1,PPIB,DCN,CDKN1A,CT
P1S,CRABP2,DLG4,NSMF,ERBB2,UNC13
,RBM5,PSMD4,SPTY2D1,PRPF19
RNPU,OGA,PGAM1,NFE2L1,SMARC
P1,CCND2,GPC3,PIP4K2B,SOX12,

VL,ADCY6,HDAC6,PTEN,CUL3,PTPRF,ULK1,BR
MKNK
VL1,VPS35,COL2A1,RBMX,RAD23B,PTK7,
,PFN1,ASTN2,MCL1,TLE5,CLEC7A,NES
2,RANBP9,GOLGA4,SEMA4C,LIMS1,SR
EGFA,VHL,MAPK1,ARPC2,PBX2,PSMD3,ADAM9,LU
PDGFRB,NECTIN1,RAP2A,ADCY1,SYNG
OBO3,SS18L1,RGMA,MAPK3,PPP3CA,ZSWI
I,RHOB,ATP1A2,MAP2,RAB1A,TTR,COTL
3,ZNF326,PRMT7,PLRG1,RBM39,SNU13
GPC3,DDR1,SULF1,PLXNA1,MEGF8,SOX12

MP14,PRDX2,DDX3X,RPS15,ITM2C,C
PS26
RPS7,RPS19,HMGCS1,GPAT4,PRKAR1A,GJA1,
I,MIDN,RPL10,MARCHF6,WSB1,

B,ABCA2,TERF2IP,TMED2,TOP1,TAF7,RNF10,VE
B,ABCA2,TERF2IP,TMED2,TOP1,TAF7,RNF10,VE
KBP1,ATP8A2,ANAPC2,EPHA2,SKIL,SIN3A
T2,MLXIP,SQSTM1,GNAI3,NOP53,PAR
IN1,AKAP17A,PAF1,SCAF8,ZMAT2,KHDC4,SRSF8,C
J1,MAGEF1,AREL1,RNF41,FAM8A1,TRIB3,ADRM1,
D1,RAB11A,TCTN1,BMPR2,DYNLL1,CDC42,PPP
RDX,KMT2E,CAPZA1,NMT1,MOAP1,CLASP2,WASF2,S
POLR2E,PSMA7,ANP32A,RBBP7
I,EIF3H,T

RNF10,VEGFA,MARS1,RAPGEF1,OPTN,
VAPA,HMGB1,CDKL1,AP2M1,TPT1,R
PZB,ACTN1,CAND1,NSG2,CLIP3,PLCG1,KDEL

3,PKN1,TCP1,CCT3,VPS35,PTOV1,MDK,
1A,FKBP8,QARS1,CCAR2,DUSP4,POLR2

PTBP1,EIF3E,FTO,ILF3,PCSK1N,WDR6,P
,TGFBR1,PPP3CA,MARK1,RUFY3,SEPTIN9,CUL
9L,CAPN2,EP300,LBH,PRKAR2A,KDM5B,FLCN,
B5A,SS18L1,RGMA,TGFBR1,PPP3CA,MARK1,R
NP32A,VDAC1,TRAF4,TRAF7,DFFB,CAPN2
CA,CCND1,MAPK1,SMARCA4,DDAH2,PBX2
LEC7A,RPL31,HNRNPC,LRPP
R2,SMS,RPL34,S
NDFIP1,TAGLN,SND1,RHOB,HLA-B,MAP2,
ACTN4,MMP14,PRDX2,DDX3X,NANOG,APLP
D81,RPL26,CRMP1,CCT7,OAZ
SND1,RHOB,HLA-B,MAP2,RAB1A,HRNR,CT
NRNPK,H1-10,CIRBP,NARS1,RPL3
JUND,RPS20,EIF5,CD81,RPL
JUND,RPS20,EIF5,CD81,RPL
3,RAB11B,ABCA2,TERF2IP,TMED2,TOP1,TAF7,R
CSL6,SOX12,CCN2,ACTN4,PRDX2,DD
CYFIP1,KAT2A,RNF157,BNC2,ATF5,EH
NLL1,TXNIP,CDC42,DDIT4,YME1L1,ADAR,A
A1,JUN,CTNNB1,RPS10,RPL35A,H

RPL12,ITM2C,RPL30,CLTC,E
,RBM14,LMNA,VDAC2,THOC2,DNAJB6,EFNB3,BMERB1
CYFIP1,MAP4K4,FEZ1,KAT2A,EHMT2,NTN1,LIM
2,AP2A2,NDUFA13,BNIP3,UBE2I,UBE2D3,PSMC1,C
F8,SOX12,CCN2,ACTN4,MMP14
2,HGS,SOD1,BAG6,RAI1,RAB11A,AKIRIN1,PRPF
BCA7,EMD
BE3C,USP48,USP19,UBQLN4,RNF216,PSMB5,RNF20,UBXN
OG,H1-0,RPS5,RPS7,RPS1
P1,GHITM,SOD1,BAG6,ARNT2,SERBP1,BMP

LM,CCN1,CARM1,UBR5,LTBP4,RAB14,RDX,
RPS3,CHD3,BASP1,VAPA,HMGB1,CD
SRP1,PTPRA,PKN1,TBC1D16,AHCYL1,GRI
1,TSPAN14,FLII,PDGFRB,CCDC88A,TRIM27,NC

RPS25,SFPQ,ACSL6,SOX12,CCN2,AC
I1,TLE5,CLEC7A,RPL31,HN
GLO1,GRN,PPP1R10,CCN1,GPX1,OLFM1
GAP1,EPN1,ISLR2,PSME3,DHX36,EMILIN1,CRKL

1,CD44,PSMD7,FAF2,TSPAN14,SYT15

2,TXN,OTUB1,DUSP3,CEBPG,NUP205,NUP133,UBXN

NCSTN,ATXN3,SPAG9,PDPK1,ABCF1,DAZAP

RAB3A,NAPB,ARL8A,MAN2B1,DPP7,UNC

TPRA,PKN1,GRINA,VPS35,COL2A1,CLSTN3

RPS25,SFPQ,ACSL6,SOX12,CCN2,AC

PL12,RPL30,EIF3L,NANOG,

1,ANXA2,AGRN,RPLP1,ZMIZ1,TIMP2,XAF

K1,CCN1,SHC2,UBR5,KMT2E,DNAJB2,

10,XBP1,CUL7,MAN1B1,CUL4B,PSMD12,TRPC4AP,RYBP,P

IPT1,ERP29,PTPN1,RAN,GOLPH3,MYCBP2,UBE2J1,C

MA6,KCTD10,XBP1,CUL7,MAN1B1,CUL4B,PSMD12,TUT4,T

IAN1B1,CUL4B,PSMD12,TRPC4AP,RYBP,PSMA4,FBXO7,UB

E2M,ARHGEF17,TLE1,TAX1BP1,MAZ,KIF

7A,APEX1,HIF1A,CTTN,SLC25A5,LOX,SL

JM1,FXR2,SDCBP,KDM3A,MGAT

CDC88A,TMEM98,AKAP8L,NUP153,DHX36,CRKL,NDRG4

EZ1,KAT2A,ATF5,B4GALT5,SUN2,MYCBP2,RH

RKCI,AJUBA,LRRN1,ADGRL1,OPA1,EPHB2,A

1,CSNK1E,PFKL,HDLBP,RPS25,SFPQ

SP36,CTSA,NAMPT,ARSA,TOLLIP

SP36,CTSA,NAMPT,ARSA,TOLLIP

YT4,POSTN,FZD7,LRP6,ATP8A2,CACNG7

SMC1,BAG5,SLC38A2,SCAP,USP47,PTPN11,SUN1,

P4,ANTXR1,ABI2,ARHGEF17,VPS4A,RDX,LSM14A,CNP,CAP

,TNKS,PSMD8,APEX1,PCBP4,AKAP9,BAP1,S

PL28,GSN,RB1CC1,USP36

ITOR,UBE3A,PRKACA,NDUFA13,PSMC1,WNK
\1,POR,PPP5C,DAG1,SMPD3,NOTCH1,IGF1R,MDM2,UB

HF7,SENP6,NUP98,PSMC5,PSMC2,HERPUD1,PSMC4,P

,PKIA,ATP13A2,IGF1R,MDM2,MGA

K2,RANGAP1,CAD,AMFR,PTPN11,WDTC1,ABL1,COL16

XBP1,ARHGEF18,EXT1,RAB35,MXRA5,PPP2
,SART3,KAT2A,RNF157,HLA-A,PSMD7,MAEA,LIM
MD5,YWHAQ,EIF1,HACD3,DCHS1,SET,E

,PRDX1,PTBP1,GRB2,CDC37,GABARAP,HSP
NOT1
SP,UBE2I,MN1,PHIP,PSMC1,DAPK3,CROCC,TUBB

JBE3C,MEX3C,UBE2V1,BCOR,RNF216,ZFP91,DTX3,CT
I,BANF1,CCNY,PSMD7,MAEA,RAN,SUN2,CUL4A,TASO

,KAT2A,ATF5,CCNY,EHMT2,PSMD7,MAEA,HIPK2,CU
B,CLASP2,PDPK1,ITGAV,SCRIB,ADAM10
,PSMC5,PSMC2,PSMC4,PLK2,PSMF1,DHFR,RCC2,PRMT5

3GAP2,RGMB,NCOA4,RPS27A,TNKS,APEX
G1,MAP2K2,QARS1,CCAR2,DUSP4,FYN,LARS1

TS2,MTA2,KDM3A,PPP5C,RTF1,TSPYL2,MACROH2A1,ANP3
RPL37,PPP2CA,CCND1,G3BP2,MAP
PEX1,HNRNPAB,HIF1A,PPP1R15A,SERP1,

,BCAR1,RELA,MICALL1,ZDHHC17,MCM7,LR
PH3,CCDC88A,ARFGAP2,AKAP8L,EPN1,ARPC3,
ITN2,NRP2,ARPC5,SGPL1,SHANK1,MEIS3,SUDS3,

CUL4A,TASOR,DAD1,EXT2,SOBP,LDB1,EPN1,DH

PELP1,CCN1,SSBP3,CARM1,SOX2,ELAV

RPS6KA2,MDC1,CHMP5,ANXA1,PSMA6,PIN1,CNOT3,PCNA,CUL

;R4,CACNA1H,PGRMC2,ADIPOR2,STC2,CDKN1B

,PTTG1IP,DCN,CDKN1A,CTDSP2,DDX39B,CNN2,
TN,RAB6A,RAB5C,ACTR1B,CLASP2,PDPK

3P3,COL1A2,LLGL1,GAP43,NCS1,RHOA,CBX

CF3,LTBP3,PSMD4,ITGAV,LRP3,IST1,RBPJ,
IT1,EPHB2,LRP8,NCOA6,FGF13,SETD1A,SZT2

1T2,CTCF,SETD3,UBN1,WDR5,SS18L1,SUDS3,MAPK3,PBRM

/T4,BTG1,HMGA2,ATP8A2,CACNG7,ANAP
DIP3,PSMC2,NCBP2,RBM10,PSMC4

;CT2,MLXIP,PPP1CB,SQSTM1,NOP53

RAB14,NCSTN,RAB6A,RAB5C,ACTR1B,CLA
NIP,CDC42,PPP1R9B,ADAR,TRAF4,TRA
JBE2D2,VPS4A,GNA12,DNAJB2,NCSTN,ATXN3,FO

RSF10B,PLK2,SCG2,YWHAZ,HSF1,GSK3B,PA2G4
;FDP1,GOLPH3,HIPK2,DAD1,PDGFRB,ASNS,A

D2,HMGA2,ANAPC2,PLRG1,SIN3A,CNOT1,PPP2R5C

PH1,PRDX5,ACTR3,PKN1,YEATS2,TCP
A,CSNK2B,CDK4,NCK2,PMEPA1,MAPKAPK

NR2L4,HERPUD1,TNFRSF10B,PLK2,SCG2,YWHAZ

N1,TNPO1,SKI,SNRNP70,ASXL1,PSMB2,H3-3A,

MORF4L2,TAF15,PRDX5,ACTR3,PKN

TS2,TCP1,CCT3,PTOV1,MDK,ZNF302,

K3A,MTOR,YY1,CREBBP,SAP18,APBB1,NDUFA

CN1,SSBP3,CARM1,SOX2,ZMIZ2,MAZ,KMT2E,

C2A1,AKAP9,GRK2,PALM,GPX1,CARM1,LDLR

AP,PTP4A1,VANGL2,SOD1,PFN2,RAB11A,AKI
SLIT2,AKAP9,GRK2,STK35,XRCC6,TAOK1,

'EGFA,MARS1,VHL,PPP2CA,CCND1,G3BP2,MAPK
',FURIN,CRIM1,PRNP,EPHA4,PSMC6,PRELID1,AKIR

:2,UBR5,SOX2,AKT3,MAZ,LSM14A,NCST

'2A,GTF2F1,PXDN,HNRNPU,PGAM1,NFE2L1,S
A,PHIP,WNK1,CD74,MICAL1,PRDX3,AMBRA1
A,PHIP,WNK1,CD74,MICAL1,PRDX3,AMBRA1

,FGD1,SH3KBP1,ASAP3,GSN,GPM6B,PAM
RIB,ADAM10,SYT11,RBPJ,GSK3A,PSMC3,MTOR,P

3GAP1,RAB11FIP3,AMFR,IGFBP4,USP47,PTP
TM1,GNAI3,KLHL24,NOP53,PARP1,TAF15,SSR
PBB1,ATXN7L3,PHIP,TRIM8,SSBP4,THRAP3,
3SF1,CHMP1A,MTA2,CGGBP1,RTF1,NIBAN2,MAC
K1,USP10,PHACTR4,PSMB1,DAG1,NIBAN2,YT

3GAP1,RAB11FIP3,AMFR,IGFBP4,USP47,PTP
5,SOX2,AKT3,MAZ,LSM14A,NCSTN,NUCK

10,TAOK1,EHMT1,CCN1,GPX1,CYTH2,SLC
EF18,PRKCI,FER,GMFB,BCAR1,FGF13,P

PYSL2,VEGFA,SELENON,RAPGEF1,MAPK1,AZIN1

:Y1,DHX36,RBFOX2,HSF1,GSK3B,DHFR,ZNF703,PGRM

SA,TOLLIP

TP6AP1,DPYSL2,TMED2,RPL38,NOTCH2,VEGFA

,SPAG9,TCF3,PDPK1,DAZAP2,DCUN1D2

BP3,EPHA4,DNM1,GRB10,ATP6V1F,LE

H1,PSMC3,MTOR,PRKACA,CREBBP,APBB1,KLHDC10,

,

'RS,HGS,MAPK8IP1,SOD1,DGKD,PFN2,P
3E2I,KLHDC10,MN1,PHIP,PSMC1,WNK1,T
B,GSK3B,UBE2V1,ZFP91,WARS1,RASSF2,TXN,DLG4

F1R,NPTN,STK39,PPP1R14B,CAB39,FXR1,P

ETN2,TBC1D14,DDX6,C1QBP,LSM4,EXOC7,SENP6,G

CD44,KAT2A,EHMT2,PTPN1,METAP2,HIPK2,UBE2J1,D

〈3R3,ATP6V1D,SIN3A,NAMPT

PRKCSH,XPR1,KHDRBS1,SPTBN2,PTN,CS
/,SCRIB,ADAM10,CAP1,AKT2,GSK3A,VAS

PRS,HGS,MAPK8IP1,SOD1,DGKD,PFN2,P

〈2,IGF2BP1,NSD2,RBPJ,PCGF2,NOC2L,

2,ITGB5,APEX1,HIF1A,PODXL2,CTTN,LO

PRKCSH,XPR1,KHDRBS1,SPTBN2,PTN,CS
IP1,MAP4K4,PHPT1,EHMT2,PTPN1,GOLPH3,CDK4,
,APBB1,NDUFA13,UBE2I,UBE2D3,DAPK3,

A2,HSPA1A,ACVR1B,OBSL1,NOTCH1,MDM2,NPTN,EFN

TCH1,ATP13A2,ACTA2,MDM2,FXR1,PARK7

3E2I,UBE2D3,DAPK3,SMO,EIF4A3,NELFB,
NT2,BRSK1,ITGB5,PSMD8,HNRNPAB,HIF1A,FKBP1A,A

MO1,GGT7,SYNCRIP,CD44,KAT2A,HLA-A,P
F1,AUTS2,MTA2,KDM3A,NIBAN2,KLF13,RUVBL2
F1,AUTS2,MTA2,KDM3A,NIBAN2,KLF13,RUVBL2
CI,HMGB2,RELA,OPA1,CTNNBIP1,EPHB2,LRP8,T
F1,AUTS2,MTA2,KDM3A,NIBAN2,KLF13,RUVBL2
R,SOX9,HOXA5,PSMB5,STAT1,PSMA1,MED12,SOS1,

SF1,GDF1,UBE2V1,SOX9,TFRC,ADIPOR2,ZFP91,

RIB3,PMEPA1,ADCY1,CCNG1,MAPKAPK2,CD

D,PTPN11,ABL1,FXR2,SDCBP,PINK1,PPP5C,

L1A2,LLGL1,GAP43,HK1,NCS1,RPL22

A4,PTPN1,FAF2,MTURN,LIMK1,ZNF462,PPP1

MBD3,GSK3A,YY1,CREBBP,SAP18,APBB1

1,TYK2,JAK1,CDK9,PSME3,CRKL,SRSF7,PS

_K1,STK24,RIC8A,SEMA3C,SOD2,SPAG9
_K1,STK24,RIC8A,SEMA3C,SOD2,SPAG9

R,DSTYK,SHC1,PSMD10,ROCK2,CAMK2N1,ZDHHC9,

B,ADAM10,CAP1,MSH6,SYT11,NSD2,IST

P3,FKBP8,PLCG1,CCAR2,ARF4,ZFAND5,PIT

SLC25A5,ATP6AP2,LOX,SLC2A1,NDN,PRDX6

NF4,BMERB1,PPP1R12A,LZTS2,CNTN2,ROCK2,CHMP

TFRC,PRMT5,OTUB1,KDM2A,CEBPG,LMNA,XPC,UBXN1,P
1,LOXL2,EMC10,SEMA4C,PLK2,SCG2,MEIS1,SRF,

R,MAZ,TRAP1,SEMA3C,SOD2,CLASP2,MTMR4,

K39,NDUFA4,CAB39,PARK7,ADNP,CD44,ARL6IP

BC1D14,ADRM1,PBXIP1,SYNGAP1,PPP2R5D
H2,NCOA6,COPS8,CCNK,SESN2,BRAT1,RAD23A,MRPS26,MAR

TRIM27,NCK2,TRIB3,ASAH1,H2AX,C1QBP,PBXI

,CCT3,PTOV1,MDK,ZNF302,RBMX,CBX1,CHD2,

TRIM27,NCK2,TRIB3,ASAH1,H2AX,C1QBP,PBXI

CP2,ADNP,IDH2,ADGRB2,CYFIP1,KAT2A,NTN1,
TRIM27,NCK2,TRIB3,ASAH1,H2AX,C1QBP,PBXI

B2,APOE,TGFB2,FURIN,CRIM1,FBXO7,PRNP,S

3,DDR1,SULF1,PLXNA1,PIP4K2B,MEGF8,

5,VEZF1,ATMIN,STAT1,CHD8,CEBPG,RBM14,P

D3,AEBP1,TCF25,ILF3,LUZP6,PRKDC,SUPT5H
EPHB2,APOE,TLR6,LRP8,TGFB2,FGF13,TPX2,CCNL1,

H3-3A,KDM6B,SCAF1,HDGF,ABCA2,PRPF6,T
D3,AEBP1,TCF25,ILF3,LUZP6,PRKDC,SUPT5H

,ASXL1,PSMB2,H3-3A,KDM6B,SCAF1,HDGF,A

RC5B,CDK5RAP3,LRP6,HMGA2,IBTK

D3,AEBP1,TCF25,ILF3,LUZP6,PRKDC,SUPT5H

L1,SDCBP,LGALS1,SCAMP1,PSMB1,HS

N1,SKI,ASXL1,PSMB2,H3-3A,KDM6B,SCAF1,

53BP1,POLR2E,SULF2,STAT3,BCL9L,PRMT1,

4,TSC22D1,SMARCE1,SMARCC1,SUPT6H,PHC2,

,CAPN1,NME2,ASAH1,C1QBP,MFAP4,RAB18

4,TSC22D1,SMARCE1,SMARCC1,SUPT6H,PHC2,
4,TSC22D1,SMARCE1,SMARCC1,SUPT6H,PHC2,

F703,NUS1,SERPINE2,RCC2,SOX9,ADIPOR2,KI

N2,CLIC1,PLXNA2,CCDC80,OBSL1,NOTCH1,NP

LXNA2,CCDC80,OBSL1,NOTCH1,NPTN,EFNB2,

K1G2,SFRP1,HDAC6,PTEN,AGPAT1,NDST1,IL

D1,PBX1,FLCN,BAZ1B,H2AZ1,BRD4,UBA52,CXX

R1,MAPK3,PPP3CA,RUFY3,XBP1,AJUBA,SP1,FE

NSMF,RNF20,ERBB2,LMNA,NR2F2,ARSB,CITED2

IIF3A,ERBB2,NUP133,NR2F2,MED12,THBS2,

N1,PSMC6,PAF1,PRELID1,MYL9,FZD7,CDK5RAP3,I

HSPA1A,ACVR1B,OBSL1,NOTCH1,MFN2,IGF1R,NPTN

,TGFB2,NCOA6,PSMA4,SETD1A,FBXO7,ACIN1,PSMC6

1CA,CSNK2B,CAPN1,NME2,ASAH1,LDB1,PSME3,DEAF1,

PS27A,TNKS,PSMD8,APEX1,HIF1A,SAFB2,GLO1

DKN1B,STAT1,HIF3A,ERBB2,LMNA,NR2F2,CACYBP,ME

T1,MFGE8,PRNP,PPT1,DNM1,PRELID1,SYT4,

D2,ASXL1,PSMB2,H3-3A,KDM6B,SRP14,

PSMA6,ADD1,XBP1,EXT1,HMGB2,PSMD12,MSH2,CTNNB

CNK,EPHA4,BRAT1,MAPK8IP3,GRB10,FZD7,GPR

B1,NRP2,AFDN,ROCK2,ADIPOR1,ANXA1,TGFBR1
D1A

A12,MAZ,KIFAP3,DNAJB2,NCSTN,SOD2,S

UBA,VAV2,TAB3,SLC39A10,EPHB2,TLR6,LRP8,T
UBA,VAV2,TAB3,SLC39A10,EPHB2,TLR6,LRP8,T

GLB1,CTSA,ARSA,TOLLIP

| | | | | | | |
|---|---|---|---|---|---|---|
| **GO:CC** | intracellular an | GO:0005622 | 4.06E-105 | 104.391344 | 14808 | 2630 |
| **GO:CC** | cytoplasm | GO:0005737 | 4.45E-103 | 102.351236 | 11951 | 2630 |
| **GO:CC** | intracellular or | GO:0043229 | 3.95E-95 | 94.4038696 | 13117 | 2630 |
| **GO:CC** | cytosol | GO:0005829 | 2.43E-90 | 89.6151455 | 5361 | 2630 |
| **GO:CC** | organelle | GO:0043226 | 1.05E-86 | 85.9773905 | 14027 | 2630 |
| **GO:CC** | intracellular m | GO:0043231 | 4.60E-85 | 84.3372723 | 12046 | 2630 |
| **GO:CC** | membrane-bo | GO:0043227 | 9.20E-81 | 80.0360405 | 13201 | 2630 |
| **GO:CC** | extracellular ex | GO:0070062 | 4.58E-79 | 78.3396005 | 2178 | 2630 |
| **GO:CC** | organelle lume | GO:0043233 | 2.96E-78 | 77.528804 | 5579 | 2630 |
| **GO:CC** | intracellular or | GO:0070013 | 2.96E-78 | 77.528804 | 5579 | 2630 |
| **GO:CC** | membrane-en | GO:0031974 | 2.96E-78 | 77.528804 | 5579 | 2630 |
| **GO:CC** | extracellular ve | GO:1903561 | 2.34E-74 | 73.6313181 | 2263 | 2630 |
| **GO:CC** | extracellular or | GO:0043230 | 3.46E-74 | 73.4609727 | 2265 | 2630 |
| **GO:CC** | extracellular m | GO:0065010 | 3.46E-74 | 73.4609727 | 2265 | 2630 |
| **GO:CC** | protein-contai | GO:0032991 | 4.26E-68 | 67.3707293 | 5541 | 2630 |
| **GO:CC** | vesicle | GO:0031982 | 5.88E-67 | 66.2306339 | 4058 | 2630 |
| **GO:CC** | nuclear lumen | GO:0031981 | 3.59E-62 | 61.4445926 | 4453 | 2630 |
| **GO:CC** | nucleoplasm | GO:0005654 | 2.54E-59 | 58.5948706 | 4112 | 2630 |
| **GO:CC** | nucleus | GO:0005634 | 3.27E-51 | 50.4861045 | 7607 | 2630 |
| **GO:CC** | focal adhesior | GO:0005925 | 6.27E-50 | 49.2027559 | 420 | 2630 |
| **GO:CC** | cell-substrate | GO:0030055 | 1.30E-48 | 47.8867005 | 427 | 2630 |
| **GO:CC** | endomembrar | GO:0012505 | 3.32E-46 | 45.478423 | 4651 | 2630 |
| **GO:CC** | ribonucleopro | GO:1990904 | 5.75E-46 | 45.2402515 | 694 | 2630 |
| **GO:CC** | cell junction | GO:0030054 | 1.20E-45 | 44.9197087 | 2107 | 2630 |
| **GO:CC** | cytosolic ribos | GO:0022626 | 1.52E-40 | 39.8185472 | 106 | 2630 |
| **GO:CC** | anchoring jun | GO:0070161 | 1.03E-39 | 38.9873523 | 838 | 2630 |
| **GO:CC** | non-membrar | GO:0043228 | 7.98E-37 | 36.0981305 | 5102 | 2630 |
| **GO:CC** | intracellular nc | GO:0043232 | 1.26E-36 | 35.9006457 | 5094 | 2630 |
| **GO:CC** | intracellular ve | GO:0097708 | 1.18E-33 | 32.9267339 | 2438 | 2630 |
| **GO:CC** | cytoplasmic v | GO:0031410 | 1.31E-33 | 32.8835043 | 2433 | 2630 |
| **GO:CC** | synapse | GO:0045202 | 4.52E-28 | 27.3451129 | 1348 | 2630 |
| **GO:CC** | extracellular sp | GO:0005615 | 1.45E-27 | 26.8375387 | 3595 | 2630 |
| **GO:CC** | cytosolic large | GO:0022625 | 3.06E-27 | 26.5144693 | 57 | 2630 |
| **GO:CC** | catalytic comp | GO:1902494 | 3.34E-25 | 24.475895 | 1377 | 2630 |
| **GO:CC** | melanosome | GO:0042470 | 1.94E-22 | 21.7117597 | 109 | 2630 |
| **GO:CC** | pigment granu | GO:0048770 | 1.94E-22 | 21.7117597 | 109 | 2630 |
| **GO:CC** | ribosomal sub | GO:0044391 | 8.90E-22 | 21.0505853 | 191 | 2630 |
| **GO:CC** | endoplasmic r | GO:0005783 | 1.08E-21 | 20.9668757 | 1995 | 2630 |
| **GO:CC** | nuclear protein | GO:0140513 | 1.79E-21 | 20.7480543 | 1236 | 2630 |
| **GO:CC** | ribosome | GO:0005840 | 1.86E-21 | 20.7305454 | 247 | 2630 |
| **GO:CC** | nuclear body | GO:0016604 | 4.50E-21 | 20.3467995 | 816 | 2630 |
| **GO:CC** | axon | GO:0030424 | 4.04E-20 | 19.393194 | 661 | 2630 |
| **GO:CC** | nuclear speck | GO:0016607 | 1.59E-19 | 18.7979187 | 413 | 2630 |
| **GO:CC** | neuron projec | GO:0043005 | 1.17E-17 | 16.9324879 | 1384 | 2630 |
| **GO:CC** | somatodendri | GO:0036477 | 2.32E-17 | 16.633737 | 875 | 2630 |

| | | | | | | |
|---|---|---|---|---|---|---|
| **GO:CC** | organelle men | GO:0031090 | 3.79E-17 | 16.4218377 | 3644 | 2630 |
| **GO:CC** | ficolin-1-rich g | GO:1904813 | 5.25E-17 | 16.2794487 | 124 | 2630 |
| **GO:CC** | cell body | GO:0044297 | 1.40E-16 | 15.8523352 | 595 | 2630 |
| **GO:CC** | endosome | GO:0005768 | 4.10E-16 | 15.3867118 | 993 | 2630 |
| **GO:CC** | supramolecula | GO:0099080 | 4.72E-16 | 15.3264807 | 1325 | 2630 |
| **GO:CC** | cell projection | GO:0042995 | 5.19E-16 | 15.284602 | 2337 | 2630 |
| **GO:CC** | Golgi apparatu | GO:0005794 | 1.03E-15 | 14.9891613 | 1593 | 2630 |
| **GO:CC** | cytosolic smal | GO:0022627 | 3.21E-15 | 14.4937036 | 46 | 2630 |
| **GO:CC** | extracellular re | GO:0005576 | 3.75E-15 | 14.4263073 | 4564 | 2630 |
| **GO:CC** | plasma memb | GO:0120025 | 4.56E-15 | 14.3414514 | 2234 | 2630 |
| **GO:CC** | ficolin-1-rich g | GO:0101002 | 4.61E-15 | 14.3358907 | 184 | 2630 |
| **GO:CC** | dendrite | GO:0030425 | 4.66E-15 | 14.3315154 | 641 | 2630 |
| **GO:CC** | postsynapse | GO:0098794 | 4.66E-15 | 14.3315154 | 641 | 2630 |
| **GO:CC** | dendritic tree | GO:0097447 | 6.45E-15 | 14.1902819 | 643 | 2630 |
| **GO:CC** | cytoplasmic ve | GO:0060205 | 2.15E-14 | 13.6681084 | 324 | 2630 |
| **GO:CC** | secretory gran | GO:0034774 | 2.48E-14 | 13.6053921 | 320 | 2630 |
| **GO:CC** | vesicle lumen | GO:0031983 | 3.45E-14 | 13.462249 | 326 | 2630 |
| **GO:CC** | polysome | GO:0005844 | 3.58E-14 | 13.4456045 | 72 | 2630 |
| **GO:CC** | bounding mer | GO:0098588 | 5.11E-14 | 13.2917138 | 2165 | 2630 |
| **GO:CC** | vacuole | GO:0005773 | 6.61E-14 | 13.1795345 | 818 | 2630 |
| **GO:CC** | endopeptidase | GO:1905369 | 6.87E-14 | 13.1629287 | 73 | 2630 |
| **GO:CC** | growth cone | GO:0030426 | 1.04E-13 | 12.9828068 | 185 | 2630 |
| **GO:CC** | nucleolus | GO:0005730 | 2.54E-13 | 12.5947835 | 955 | 2630 |
| **GO:CC** | spliceosomal c | GO:0005681 | 2.61E-13 | 12.5827298 | 192 | 2630 |
| **GO:CC** | secretory vesic | GO:0099503 | 4.46E-13 | 12.3508328 | 1027 | 2630 |
| **GO:CC** | peptidase com | GO:1905368 | 6.04E-13 | 12.2193017 | 97 | 2630 |
| **GO:CC** | proteasome co | GO:0000502 | 6.54E-13 | 12.1842365 | 67 | 2630 |
| **GO:CC** | site of polarize | GO:0030427 | 7.12E-13 | 12.1472611 | 191 | 2630 |
| **GO:CC** | lysosome | GO:0005764 | 8.50E-13 | 12.0706932 | 723 | 2630 |
| **GO:CC** | lytic vacuole | GO:0000323 | 8.50E-13 | 12.0706932 | 723 | 2630 |
| **GO:CC** | cytoskeleton | GO:0005856 | 1.00E-12 | 11.9997896 | 2339 | 2630 |
| **GO:CC** | lamellipodium | GO:0030027 | 2.05E-12 | 11.6883688 | 203 | 2630 |
| **GO:CC** | perinuclear reg | GO:0048471 | 3.91E-12 | 11.407498 | 745 | 2630 |
| **GO:CC** | secretory gran | GO:0030141 | 5.08E-12 | 11.2943592 | 857 | 2630 |
| **GO:CC** | organelle subc | GO:0031984 | 5.72E-12 | 11.2422681 | 1764 | 2630 |
| **GO:CC** | distal axon | GO:0150034 | 6.62E-12 | 11.1789966 | 307 | 2630 |
| **GO:CC** | glutamatergic | GO:0098978 | 1.50E-11 | 10.8226694 | 354 | 2630 |
| **GO:CC** | microtubule cy | GO:0015630 | 2.36E-11 | 10.6266185 | 1313 | 2630 |
| **GO:CC** | cell leading ed | GO:0031252 | 2.50E-11 | 10.6027869 | 426 | 2630 |
| **GO:CC** | cell cortex | GO:0005938 | 4.96E-11 | 10.3045652 | 326 | 2630 |
| **GO:CC** | neuronal cell b | GO:0043025 | 5.35E-11 | 10.2713045 | 523 | 2630 |
| **GO:CC** | large ribosoma | GO:0015934 | 6.82E-11 | 10.1662904 | 119 | 2630 |
| **GO:CC** | ribonucleopro | GO:0035770 | 6.92E-11 | 10.1601553 | 252 | 2630 |
| **GO:CC** | small ribosom | GO:0015935 | 1.22E-10 | 9.91484372 | 76 | 2630 |
| **GO:CC** | early endoson | GO:0005769 | 1.85E-10 | 9.73273969 | 387 | 2630 |

| | | | | | | |
|---|---|---|---|---|---|---|
| **GO:CC** | polysomal ribo | GO:0042788 | 3.40E-10 | 9.46793979 | 34 | 2630 |
| **GO:CC** | endosome me | GO:0010008 | 3.72E-10 | 9.42966373 | 520 | 2630 |
| **GO:CC** | endoplasmic r | GO:0098827 | 7.10E-10 | 9.14878404 | 1137 | 2630 |
| **GO:CC** | membrane | GO:0016020 | 9.49E-10 | 9.02264849 | 9829 | 2630 |
| **GO:CC** | nuclear outer | GO:0042175 | 1.17E-09 | 8.93256559 | 1154 | 2630 |
| **GO:CC** | endoplasmic r | GO:0005788 | 1.25E-09 | 8.90415744 | 313 | 2630 |
| **GO:CC** | catalytic step 2 | GO:0071013 | 1.64E-09 | 8.78392059 | 85 | 2630 |
| **GO:CC** | vacuolar mem | GO:0005774 | 3.09E-09 | 8.50984238 | 450 | 2630 |
| **GO:CC** | intracellular pr | GO:0140535 | 4.04E-09 | 8.39318382 | 768 | 2630 |
| **GO:CC** | endoplasmic r | GO:0005789 | 4.06E-09 | 8.39199228 | 1132 | 2630 |
| **GO:CC** | vacuolar lume | GO:0005775 | 4.84E-09 | 8.31539682 | 174 | 2630 |
| **GO:CC** | cytoplasmic ri | GO:0036464 | 4.92E-09 | 8.30814268 | 238 | 2630 |
| **GO:CC** | neuron to neu | GO:0098984 | 5.43E-09 | 8.26507868 | 371 | 2630 |
| **GO:CC** | cortical cytosk | GO:0030863 | 1.02E-08 | 7.99043501 | 121 | 2630 |
| **GO:CC** | proteasome re | GO:0005838 | 1.08E-08 | 7.9671934 | 22 | 2630 |
| **GO:CC** | vesicle membr | GO:0012506 | 1.62E-08 | 7.79151305 | 1183 | 2630 |
| **GO:CC** | microtubule | GO:0005874 | 1.70E-08 | 7.76969127 | 435 | 2630 |
| **GO:CC** | proteasome a | GO:0022624 | 2.05E-08 | 7.6884634 | 25 | 2630 |
| **GO:CC** | supramolecula | GO:0099081 | 2.64E-08 | 7.57761749 | 1005 | 2630 |
| **GO:CC** | azurophil gran | GO:0042582 | 2.76E-08 | 7.55874531 | 154 | 2630 |
| **GO:CC** | primary lysoso | GO:0005766 | 2.76E-08 | 7.55874531 | 154 | 2630 |
| **GO:CC** | asymmetric sy | GO:0032279 | 2.77E-08 | 7.55784312 | 345 | 2630 |
| **GO:CC** | supramolecula | GO:0099512 | 3.76E-08 | 7.42530171 | 997 | 2630 |
| **GO:CC** | postsynaptic c | GO:0014069 | 5.61E-08 | 7.25104172 | 339 | 2630 |
| **GO:CC** | postsynaptic s | GO:0099572 | 9.90E-08 | 7.00420138 | 363 | 2630 |
| **GO:CC** | lysosomal me | GO:0005765 | 1.11E-07 | 6.95285529 | 395 | 2630 |
| **GO:CC** | lytic vacuole m | GO:0098852 | 1.11E-07 | 6.95285529 | 395 | 2630 |
| **GO:CC** | inclusion body | GO:0016234 | 1.61E-07 | 6.79408439 | 77 | 2630 |
| **GO:CC** | cell-cell junctio | GO:0005911 | 2.18E-07 | 6.66060241 | 496 | 2630 |
| **GO:CC** | Golgi apparatu | GO:0098791 | 3.49E-07 | 6.45730506 | 845 | 2630 |
| **GO:CC** | presynapse | GO:0098793 | 6.08E-07 | 6.21579062 | 515 | 2630 |
| **GO:CC** | Golgi membra | GO:0000139 | 7.23E-07 | 6.140758 | 725 | 2630 |
| **GO:CC** | nuclear periph | GO:0034399 | 1.0106E-06 | 5.99542956 | 132 | 2630 |
| **GO:CC** | endoplasmic r | GO:0140534 | 1.1251E-06 | 5.94881404 | 128 | 2630 |
| **GO:CC** | proteasome re | GO:0008540 | 1.2439E-06 | 5.90519733 | 12 | 2630 |
| **GO:CC** | actin cytoskele | GO:0015629 | 1.524E-06 | 5.81701335 | 517 | 2630 |
| **GO:CC** | chaperone co | GO:0101031 | 1.7235E-06 | 5.76359374 | 27 | 2630 |
| **GO:CC** | adherens junc | GO:0005912 | 1.9574E-06 | 5.70832328 | 171 | 2630 |
| **GO:CC** | cortical actin c | GO:0030864 | 2.2236E-06 | 5.65294558 | 92 | 2630 |
| **GO:CC** | azurophil gran | GO:0035578 | 6.2079E-06 | 5.20705617 | 91 | 2630 |
| **GO:CC** | Schaffer collat | GO:0098685 | 9.3925E-06 | 5.02721703 | 84 | 2630 |
| **GO:CC** | translation pre | GO:0070993 | 9.7851E-06 | 5.00943547 | 18 | 2630 |
| **GO:CC** | spliceosomal s | GO:0097525 | 1.2597E-05 | 4.89971817 | 61 | 2630 |
| **GO:CC** | recycling endo | GO:0055038 | 1.3072E-05 | 4.88364202 | 85 | 2630 |
| **GO:CC** | coated vesicle | GO:0030135 | 3.0733E-05 | 4.51240015 | 296 | 2630 |

| | | | | | | |
|---|---|---|---|---|---|---|
| **GO:CC** | transport vesi | GO:0030133 | 3.617E-05 | 4.44165449 | 416 | 2630 |
| **GO:CC** | endocytic vesi | GO:0030139 | 3.9623E-05 | 4.40204877 | 335 | 2630 |
| **GO:CC** | ATPase comp | GO:1904949 | 4.2361E-05 | 4.37303615 | 115 | 2630 |
| **GO:CC** | eukaryotic 43 | GO:0016282 | 5.1378E-05 | 4.28922221 | 17 | 2630 |
| **GO:CC** | eukaryotic tra | GO:0005852 | 5.1378E-05 | 4.28922221 | 17 | 2630 |
| **GO:CC** | recycling end | GO:0055037 | 5.3923E-05 | 4.26822633 | 187 | 2630 |
| **GO:CC** | cytoplasmic s | GO:0010494 | 5.5389E-05 | 4.25657869 | 77 | 2630 |
| **GO:CC** | collagen-cont | GO:0062023 | 6.5813E-05 | 4.18168649 | 421 | 2630 |
| **GO:CC** | Golgi-associa | GO:0005798 | 7.504E-05 | 4.12470737 | 95 | 2630 |
| **GO:CC** | small nuclear | GO:0030532 | 8.3166E-05 | 4.08005629 | 70 | 2630 |
| **GO:CC** | eukaryotic 48 | GO:0033290 | 9.4332E-05 | 4.02533875 | 15 | 2630 |
| **GO:CC** | integral comp | GO:0030176 | 0.00010291 | 3.98753573 | 161 | 2630 |
| **GO:CC** | SWI/SNF sup | GO:0070603 | 0.00010503 | 3.97869238 | 79 | 2630 |
| **GO:CC** | dendritic spin | GO:0043197 | 0.00011265 | 3.94826802 | 186 | 2630 |
| **GO:CC** | nuclear matrix | GO:0016363 | 0.00011756 | 3.9297313 | 110 | 2630 |
| **GO:CC** | neuron projec | GO:0120111 | 0.00014559 | 3.83687635 | 93 | 2630 |
| **GO:CC** | transferase co | GO:1990234 | 0.00014696 | 3.83281354 | 795 | 2630 |
| **GO:CC** | heterochroma | GO:0000792 | 0.00015316 | 3.8148489 | 76 | 2630 |
| **GO:CC** | envelope | GO:0031975 | 0.00015978 | 3.79647785 | 1244 | 2630 |
| **GO:CC** | organelle enve | GO:0031967 | 0.00015978 | 3.79647785 | 1244 | 2630 |
| **GO:CC** | intrinsic comp | GO:0031227 | 0.00018945 | 3.72249718 | 169 | 2630 |
| **GO:CC** | neuron spine | GO:0044309 | 0.00019387 | 3.71248743 | 189 | 2630 |
| **GO:CC** | chromosome | GO:0005694 | 0.00021059 | 3.6765541 | 1795 | 2630 |
| **GO:CC** | PTW/PP1 pho | GO:0072357 | 0.00032448 | 3.48881597 | 7 | 2630 |
| **GO:CC** | intrinsic comp | GO:0031300 | 0.00035556 | 3.44908592 | 419 | 2630 |
| **GO:CC** | cytoplasmic v | GO:0030659 | 0.00038311 | 3.41667298 | 789 | 2630 |
| **GO:CC** | phagocytic ve | GO:0045335 | 0.0004246 | 3.37201752 | 139 | 2630 |
| **GO:CC** | U2-type splic | GO:0005684 | 0.00047971 | 3.31901974 | 93 | 2630 |
| **GO:CC** | coated vesicle | GO:0030662 | 0.00080848 | 3.09233308 | 177 | 2630 |
| **GO:CC** | chromosomal | GO:0098687 | 0.00082532 | 3.08337637 | 348 | 2630 |
| **GO:CC** | Sm-like protei | GO:0120114 | 0.00089167 | 3.04979355 | 82 | 2630 |
| **GO:CC** | banded collag | GO:0098643 | 0.00091754 | 3.03737291 | 12 | 2630 |
| **GO:CC** | fibrillar collage | GO:0005583 | 0.00091754 | 3.03737291 | 12 | 2630 |
| **GO:CC** | histone deace | GO:0000118 | 0.00099302 | 3.00304001 | 78 | 2630 |
| **GO:CC** | myelin sheath | GO:0043209 | 0.00105268 | 2.97770192 | 49 | 2630 |
| **GO:CC** | polymeric cyt | GO:0099513 | 0.00106452 | 2.97284587 | 766 | 2630 |
| **GO:CC** | filopodium | GO:0030175 | 0.00123577 | 2.90806323 | 106 | 2630 |
| **GO:CC** | lysosomal lur | GO:0043202 | 0.00131106 | 2.88237727 | 97 | 2630 |
| **GO:CC** | nBAF comple | GO:0071565 | 0.00131985 | 2.87947602 | 15 | 2630 |
| **GO:CC** | ruffle | GO:0001726 | 0.0013452 | 2.87121336 | 180 | 2630 |
| **GO:CC** | complex of co | GO:0098644 | 0.00141886 | 2.84806066 | 21 | 2630 |
| **GO:CC** | cytoplasmic re | GO:0099568 | 0.00141978 | 2.8477785 | 270 | 2630 |
| **GO:CC** | axonal growth | GO:0044295 | 0.00142916 | 2.84492013 | 31 | 2630 |
| **GO:CC** | proton-transp | GO:0016469 | 0.00151176 | 2.82051722 | 54 | 2630 |
| **GO:CC** | chaperonin-co | GO:0005832 | 0.00155445 | 2.80842361 | 10 | 2630 |

| | | | | | | |
|---|---|---|---|---|---|---|
| **GO:CC** | cell-cell conta | GO:0044291 | 0.00192767 | 2.71496734 | 76 | 2630 |
| **GO:CC** | nuclear envelo | GO:0005635 | 0.00215701 | 2.66614873 | 471 | 2630 |
| **GO:CC** | transcription r | GO:0005667 | 0.00226604 | 2.64473249 | 431 | 2630 |
| **GO:CC** | Lewy body | GO:0097413 | 0.00228154 | 2.64177172 | 8 | 2630 |
| **GO:CC** | presynaptic ac | GO:0048786 | 0.00252942 | 2.59697984 | 77 | 2630 |
| **GO:CC** | U2-type presp | GO:0071004 | 0.00307996 | 2.51145454 | 16 | 2630 |
| **GO:CC** | NuRD comple | GO:0016581 | 0.00307996 | 2.51145454 | 16 | 2630 |
| **GO:CC** | prespliceosom | GO:0071010 | 0.00307996 | 2.51145454 | 16 | 2630 |
| **GO:CC** | CHD-type con | GO:0090545 | 0.00307996 | 2.51145454 | 16 | 2630 |
| **GO:CC** | integral comp | GO:0031301 | 0.00336679 | 2.47278379 | 389 | 2630 |
| **GO:CC** | midbody | GO:0030496 | 0.0038345 | 2.416291 | 197 | 2630 |
| **GO:CC** | protein serine/ | GO:0008287 | 0.0044866 | 2.34808283 | 49 | 2630 |
| **GO:CC** | phosphatase | GO:1903293 | 0.0044866 | 2.34808283 | 49 | 2630 |
| **GO:CC** | host cellular c | GO:0018995 | 0.00556565 | 2.25448413 | 58 | 2630 |
| **GO:CC** | host cell | GO:0043657 | 0.00556565 | 2.25448413 | 58 | 2630 |
| **GO:CC** | membrane raf | GO:0045121 | 0.00699533 | 2.15519178 | 339 | 2630 |
| **GO:CC** | membrane mi | GO:0098857 | 0.00699533 | 2.15519178 | 339 | 2630 |
| **GO:CC** | mitochondrio | GO:0005739 | 0.00709385 | 2.14911818 | 1678 | 2630 |
| **GO:CC** | cell division si | GO:0032153 | 0.00715345 | 2.14548438 | 72 | 2630 |
| **GO:CC** | chromosome, | GO:0000781 | 0.00789572 | 2.10260847 | 160 | 2630 |
| **GO:CC** | late endosom | GO:0005770 | 0.00904399 | 2.04363976 | 285 | 2630 |
| **GO:CC** | rough endopl | GO:0005791 | 0.01221112 | 1.91324466 | 88 | 2630 |
| **GO:CC** | actin cortical p | GO:0030479 | 0.01404492 | 1.85248077 | 15 | 2630 |
| **GO:CC** | endocytic pat | GO:0061645 | 0.01404492 | 1.85248077 | 15 | 2630 |
| **GO:CC** | endoplasmic r | GO:0005793 | 0.01457667 | 1.83634169 | 133 | 2630 |
| **GO:CC** | centrosome | GO:0005813 | 0.01580832 | 1.80111437 | 620 | 2630 |
| **GO:CC** | axon cytoplas | GO:1904115 | 0.01758945 | 1.75474769 | 62 | 2630 |
| **GO:CC** | mitochondrial | GO:0042645 | 0.01872661 | 1.72754095 | 45 | 2630 |
| **GO:CC** | nucleoid | GO:0009295 | 0.01872661 | 1.72754095 | 45 | 2630 |
| **GO:CC** | clathrin-coate | GO:0005905 | 0.02247712 | 1.64825941 | 72 | 2630 |
| **GO:CC** | ubiquitin ligas | GO:0000151 | 0.0228375 | 1.64135139 | 299 | 2630 |
| **GO:CC** | U2-type preca | GO:0071005 | 0.023993 | 1.61991545 | 50 | 2630 |
| **GO:CC** | podosome | GO:0002102 | 0.02673581 | 1.57290672 | 30 | 2630 |
| **GO:CC** | dendrite cytop | GO:0032839 | 0.02813503 | 1.55075261 | 34 | 2630 |
| **GO:CC** | multivesicular | GO:0032585 | 0.02814272 | 1.55063394 | 16 | 2630 |
| **GO:CC** | multivesicular | GO:0005771 | 0.02975731 | 1.52640637 | 64 | 2630 |
| **GO:CC** | extracellular m | GO:0031012 | 0.03041165 | 1.51696006 | 562 | 2630 |
| **GO:CC** | actin filament | GO:0032432 | 0.03080212 | 1.51141939 | 78 | 2630 |
| **GO:CC** | presynaptic m | GO:0042734 | 0.0314562 | 1.50229381 | 148 | 2630 |
| **GO:CC** | external encap | GO:0030312 | 0.03274412 | 1.48486673 | 563 | 2630 |
| **GO:CC** | PcG protein c | GO:0031519 | 0.03536048 | 1.45148191 | 47 | 2630 |
| **GO:CC** | actomyosin | GO:0042641 | 0.03849 | 1.41465214 | 79 | 2630 |
| **GO:CC** | euchromatin | GO:0000791 | 0.04004349 | 1.39746812 | 39 | 2630 |
| **GO:CC** | COP9 signalo | GO:0008180 | 0.04093784 | 1.38787513 | 35 | 2630 |
| **GO:CC** | precatalytic sp | GO:0071011 | 0.043064 | 1.36588562 | 52 | 2630 |

| | | | | | | |
|---|---|---|---|---|---|---|
| **GO:CC** | Golgi-associa | GO:0030660 | 0.043064 | 1.36588562 | 52 | 2630 |

| | | |
|---|---|---|
| 2445 | 18964 | EEF1A1,ORC4,IGFBP5,PABPC1,COL1A1,LRP1,HSPA8,GAPDH,MLLT1 |
| 2137 | 18964 | EEF1A1,ORC4,IGFBP5,PABPC1,COL1A1,LRP1,HSPA8,GAPDH,MLLT1 |
| 2253 | 18964 | EEF1A1,ORC4,IGFBP5,PABPC1,COL1A1,LRP1,HSPA8,GAPDH,MLLT1 |
| 1198 | 18964 | EEF1A1,ORC4,PABPC1,HSPA8,GAPDH,MLLT1,COPA,NEDD4L,POLH, |
| 2334 | 18964 | EEF1A1,ORC4,IGFBP5,PABPC1,COL1A1,LRP1,HSPA8,GAPDH,MLLT1 |
| 2107 | 18964 | EEF1A1,ORC4,IGFBP5,PABPC1,COL1A1,LRP1,HSPA8,GAPDH,MLLT1 |
| 2231 | 18964 | EEF1A1,ORC4,IGFBP5,PABPC1,COL1A1,LRP1,HSPA8,GAPDH,MLLT1 |
| 620 | 18964 | EEF1A1,PABPC1,HSPA8,GAPDH,COPA,HNRNPA1,HSPA9,NEDD4L,ACT |
| 1200 | 18964 | EEF1A1,ORC4,IGFBP5,COL1A1,HSPA8,MLLT1,HNRNPA1,ARSD,HSPA |
| 1200 | 18964 | EEF1A1,ORC4,IGFBP5,COL1A1,HSPA8,MLLT1,HNRNPA1,ARSD,HSPA |
| 1200 | 18964 | EEF1A1,ORC4,IGFBP5,COL1A1,HSPA8,MLLT1,HNRNPA1,ARSD,HSPA |
| 626 | 18964 | EEF1A1,PABPC1,HSPA8,GAPDH,COPA,HNRNPA1,HSPA9,NEDD4L,ACT |
| 626 | 18964 | EEF1A1,PABPC1,HSPA8,GAPDH,COPA,HNRNPA1,HSPA9,NEDD4L,ACT |
| 626 | 18964 | EEF1A1,PABPC1,HSPA8,GAPDH,COPA,HNRNPA1,HSPA9,NEDD4L,ACT |
| 1165 | 18964 | EEF1A1,ORC4,IGFBP5,PABPC1,COL1A1,LRP1,HSPA8,GAPDH,MLLT1 |
| 924 | 18964 | EEF1A1,PABPC1,COL1A1,LRP1,HSPA8,GAPDH,COPA,HNRNPA1,HSPA |
| 975 | 18964 | EEF1A1,ORC4,HSPA8,MLLT1,HNRNPA1,HSPA9,NEDD4L,POLH,ATN1, |
| 911 | 18964 | ORC4,HSPA8,MLLT1,HNRNPA1,NEDD4L,POLH,ATN1,ACTB,CTSC,NU |
| 1418 | 18964 | EEF1A1,ORC4,PABPC1,LRP1,HSPA8,GAPDH,MLLT1,HNRNPA1,HSPA |
| 185 | 18964 | PABPC1,LRP1,HSPA8,HSPA9,ACTB,ACTG1,CFL1,ACTR2,VIM,RPL3,R |
| 185 | 18964 | PABPC1,LRP1,HSPA8,HSPA9,ACTB,ACTG1,CFL1,ACTR2,VIM,RPL3,R |
| 956 | 18964 | EEF1A1,IGFBP5,COL1A1,LRP1,HSPA8,GAPDH,COPA,ARSD,NEDD4L, |
| 246 | 18964 | PABPC1,HSPA8,GAPDH,HNRNPA1,ACTB,EEF2,VIM,RPL3,RPS6,RPL4, |
| 527 | 18964 | PABPC1,LRP1,HSPA8,HSPA9,ATN1,ACTB,ACTG1,EEF2,DPYSL3,TUBA |
| 77 | 18964 | RPL3,RPS6,RPL4,RPL7A,RPL8,RPLP0,RPL13A,RPL6,RPL13,RPS8,RP |
| 266 | 18964 | PABPC1,LRP1,HSPA8,HSPA9,ACTB,ACTG1,CFL1,ACTR2,VIM,EIF4G2, |
| 992 | 18964 | EEF1A1,ORC4,PABPC1,LRP1,HSPA8,GAPDH,MLLT1,HSPA9,POLH,AC |
| 990 | 18964 | EEF1A1,ORC4,PABPC1,LRP1,HSPA8,GAPDH,MLLT1,HSPA9,POLH,AC |
| 550 | 18964 | EEF1A1,COL1A1,LRP1,HSPA8,COPA,NEDD4L,CTSC,ACTG1,EEF2,DP |
| 549 | 18964 | EEF1A1,COL1A1,LRP1,HSPA8,COPA,NEDD4L,CTSC,ACTG1,EEF2,DP |
| 339 | 18964 | HSPA8,ACTB,ACTG1,EEF2,DPYSL3,TUBA1A,HTT,ACTR2,SPARC,DBN1 |
| 719 | 18964 | EEF1A1,IGFBP5,PABPC1,COL1A1,HSPA8,GAPDH,COPA,HNRNPA1,HS |
| 46 | 18964 | RPL3,RPL4,RPL7A,RPL8,RPLP0,RPL13A,RPL6,RPL13,RPL7,RPL5,RP |
| 336 | 18964 | PABPC1,HNRNPA1,ACTB,USP22,MED18,ENO1,PIK3R2,PRPF8,FBXL4 |
| 61 | 18964 | HSPA8,HSP90AB1,HSP90AA1,GANAB,HSP90B1,CANX,YWHAE,BSG,F |
| 61 | 18964 | HSPA8,HSP90AB1,HSP90AA1,GANAB,HSP90B1,CANX,YWHAE,BSG,F |
| 84 | 18964 | RPL3,RPS6,RPL4,RPL7A,RPL8,RPLP0,RPL13A,RPL6,RPL13,RACK1, |
| 433 | 18964 | IGFBP5,COL1A1,COPA,ARSD,CTSC,HTT,UBC,RPS6,RPL4,PKM,GANA |
| 299 | 18964 | ORC4,PABPC1,HSPA8,MLLT1,HNRNPA1,ACTB,USP22,MED18,XRCC2 |
| 98 | 18964 | EEF2,RPL3,RPS6,RPL4,RPL7A,RPL8,RPLP0,CANX,RPL13A,RPL6,RPL |
| 219 | 18964 | PRPF8,FBXL4,HSPA1B,KAT6A,DDX17,NPM1,MAPT,NONO,SF3B1,CH |
| 186 | 18964 | LRP1,HSPA8,COPA,ACTB,ACTG1,DPYSL3,HTT,HSP90AB1,HSP90AA1, |
| 133 | 18964 | PRPF8,FBXL4,HSPA1B,KAT6A,DDX17,NPM1,MAPT,NONO,SF3B1,BA |
| 313 | 18964 | LRP1,HSPA8,COPA,ACTB,ACTG1,DPYSL3,HTT,HSP90AB1,HSP90AA1, |
| 220 | 18964 | LRP1,HSPA8,HTT,HSP90AB1,HSP90AA1,NUMA1,RPS6,DBN1,FLNA,M |

| | | |
|---|---|---|
| 680 | 18964 | EEF1A1,LRP1,HSPA8,GAPDH,COPA,HSPA9,CTSC,MAVS,HTT,HSP90AE |
| 59 | 18964 | EEF1A1,HSPA8,EEF2,HSP90AB1,HSP90AA1,ACTR2,FTH1,ALDOA,PKM |
| 164 | 18964 | LRP1,DPYSL3,HSP90AB1,HSP90AA1,TUBB,NUMA1,RPS6,FLNA,MAP1 |
| 238 | 18964 | LRP1,HSPA8,NEDD4L,TUBA1A,HTT,UBC,KIDINS220,SLC9A7,UBA1,PS |
| 297 | 18964 | PABPC1,COL1A1,ACTB,NUDC,ACTG1,DPYSL3,TUBA1A,VIM,TUBB,NU |
| 467 | 18964 | EEF1A1,PABPC1,LRP1,HSPA8,COPA,ACTB,ODF2L,ACTG1,DPYSL3,CF |
| 342 | 18964 | COL1A1,LRP1,COPA,NEDD4L,CTSC,HTT,NUMA1,FLNA,SLC9A7,GANA |
| 32 | 18964 | RPS6,RPS8,RPS3,RPS4X,RPS3A,RPS17,RPS16,RPSA,RPS24,RPS9,RP |
| 810 | 18964 | EEF1A1,IGFBP5,PABPC1,COL1A1,HSPA8,GAPDH,COPA,HNRNPA1,HS |
| 446 | 18964 | EEF1A1,PABPC1,LRP1,HSPA8,COPA,ACTB,ODF2L,ACTG1,DPYSL3,CF |
| 72 | 18964 | EEF1A1,HSPA8,EEF2,HSP90AB1,HSP90AA1,ACTR2,FTH1,ALDOA,PKM |
| 169 | 18964 | LRP1,HSPA8,HTT,HSP90AB1,HSP90AA1,NUMA1,RPS6,DBN1,FLNA,MA |
| 169 | 18964 | HSPA8,ACTB,HTT,ACTR2,DBN1,FLNA,MAP1B,NEFL,RPL8,RPLP0,CAN |
| 169 | 18964 | LRP1,HSPA8,HTT,HSP90AB1,HSP90AA1,NUMA1,RPS6,DBN1,FLNA,MA |
| 103 | 18964 | EEF1A1,HSPA8,CTSC,EEF2,HSP90AB1,HSP90AA1,ACTR2,SPARC,TUB |
| 102 | 18964 | EEF1A1,HSPA8,CTSC,EEF2,HSP90AB1,HSP90AA1,ACTR2,SPARC,TUB |
| 103 | 18964 | EEF1A1,HSPA8,CTSC,EEF2,HSP90AB1,HSP90AA1,ACTR2,SPARC,TUB |
| 40 | 18964 | EEF2,VIM,RPS6,RPL7A,RPL8,CALR,RPL6,RPL19,RPS3,RPS4X,RPL18, |
| 430 | 18964 | EEF1A1,LRP1,HSPA8,COPA,HSPA9,CTSC,MAVS,HSP90AB1,SPARC,UE |
| 199 | 18964 | EEF1A1,LRP1,HSPA8,ARSD,CTSC,HTT,HSP90AB1,HSP90AA1,ACTR2,T |
| 40 | 18964 | VCP,PSMD5,PSMD2,RAD23B,PSMB2,PSMD3,HSPB1,UBQLN1,PSMA7 |
| 70 | 18964 | LRP1,COPA,DPYSL3,HSP90AB1,HSP90AA1,DBN1,FLNA,MAP1B,NEFL |
| 222 | 18964 | EEF1A1,ORC4,HSPA8,MLLT1,HSPA9,RPL3,RPS6,RPL4,FLNA,NCL,RP |
| 71 | 18964 | PABPC1,HSPA8,HNRNPA1,NCL,PRPF8,YBX1,DDX5,SNRNP200,SF3B1 |
| 234 | 18964 | EEF1A1,COL1A1,LRP1,HSPA8,CTSC,EEF2,DPYSL3,HSP90AB1,HSP90 |
| 46 | 18964 | USP22,VCP,PSMD5,PSMD2,RAD23B,PSMB2,PSMD3,HSPB1,UBQLN1, |
| 37 | 18964 | VCP,PSMD5,PSMD2,RAD23B,PSMB2,PSMD3,HSPB1,UBQLN1,PSMA7 |
| 70 | 18964 | LRP1,COPA,DPYSL3,HSP90AB1,HSP90AA1,DBN1,FLNA,MAP1B,NEFL |
| 178 | 18964 | EEF1A1,LRP1,HSPA8,ARSD,CTSC,HSP90AB1,HSP90AA1,ACTR2,TUBE |
| 178 | 18964 | EEF1A1,LRP1,HSPA8,ARSD,CTSC,HSP90AB1,HSP90AA1,ACTR2,TUBE |
| 452 | 18964 | EEF1A1,LRP1,GAPDH,ACTB,CTSC,ODF2L,NUDC,ACTG1,DPYSL3,TUE |
| 72 | 18964 | PABPC1,ACTB,DPYSL3,CFL1,ACTR2,DBN1,STMN2,P4HB,APP,FSCN1, |
| 180 | 18964 | HSPA8,GAPDH,ATN1,HTT,HSP90AB1,HSP90AA1,RPS6,FLNA,MAP1B,H |
| 200 | 18964 | EEF1A1,COL1A1,LRP1,HSPA8,CTSC,EEF2,HSP90AB1,HSP90AA1,ACT |
| 355 | 18964 | COPA,UBC,NUMA1,FLNA,SLC9A7,HSP90B1,FADS1,UBA1,CALR,TMF1 |
| 94 | 18964 | LRP1,HSPA8,COPA,ACTB,ACTG1,DPYSL3,HSP90AB1,HSP90AA1,DBN |
| 103 | 18964 | HSPA8,ACTB,SPARC,DBN1,FLNA,ATP2B4,CANX,YWHAE,SPTBN1,SHIS |
| 277 | 18964 | GAPDH,CTSC,ODF2L,NUDC,TUBA1A,XRCC2,HTT,VIM,TUBB,NUMA1,I |
| 117 | 18964 | EEF1A1,PABPC1,ACTB,DPYSL3,CFL1,ACTR2,VIM,DBN1,GNAS,SPTBN |
| 96 | 18964 | EEF1A1,ACTB,CFL1,ACTR2,NUMA1,DBN1,ENO1,FLNA,NCL,SPTBN1,I |
| 135 | 18964 | LRP1,HSP90AB1,HSP90AA1,NUMA1,FLNA,MAP1B,CANX,ENO2,KIF1A |
| 49 | 18964 | RPL3,RPL4,RPL7A,RPL8,RPLP0,RPL13A,RPL6,RPL13,RPL7,RPL5,NP |
| 80 | 18964 | PABPC1,ACTB,TUBA1A,TUBB,RPS6,NCL,YBX1,RPLP0,RPL6,CSDE1,V |
| 37 | 18964 | RPS6,RACK1,RPS8,NPM1,RPS3,RPS4X,RPS3A,RPS17,RPS16,RPSA,RI |
| 107 | 18964 | LRP1,HTT,APP,PLD3,HSPD1,HLA-C,HMGB1,ANXA2,LLGL1,ASTN2,NIS |

| | | |
|---|---|---|
| 23 | 18964 | EEF2,RPL7A,RPL8,RPL6,RPL19,RPL18,RPL11,BTF3,LARP1,RPL24,RI |
| 132 | 18964 | UBC,SLC9A7,UBA1,PLD3,UBB,HLA-C,PDIA3,AP2M1,ITGB1,ANXA2,AP |
| 241 | 18964 | COPA,UBC,HSP90B1,FADS1,UBA1,CALR,CANX,SEC62,BSG,TMBIM6, |
| 1527 | 18964 | EEF1A1,PABPC1,TMEM212,LRP1,HSPA8,GAPDH,COPA,HNRNPA1,AR: |
| 243 | 18964 | COPA,UBC,HSP90B1,FADS1,UBA1,CALR,CANX,SEC62,BSG,TMBIM6, |
| 90 | 18964 | IGFBP5,COL1A1,ARSD,CTSC,GANAB,COL6A1,HSP90B1,CALR,FSTL1 |
| 38 | 18964 | PABPC1,HNRNPA1,PRPF8,DDX5,SNRNP200,SF3B1,SF3B2,HNRNPU,F |
| 116 | 18964 | EEF1A1,LRP1,HSPA8,HSP90AB1,COL6A1,UBA1,PSAP,GNB1,PLD3,VA |
| 174 | 18964 | ACTB,USP22,MED18,HSP90AB1,FTH1,FTL,FBXL4,KAT6A,SMG1,VCP, |
| 237 | 18964 | COPA,UBC,HSP90B1,FADS1,UBA1,CALR,CANX,SEC62,BSG,TMBIM6, |
| 59 | 18964 | HSPA8,CTSC,HSP90AA1,ACTR2,TUBB,FTL,PSAP,IDS,PLD3,VCP,TUBB |
| 73 | 18964 | PABPC1,ACTB,TUBA1A,TUBB,RPS6,NCL,YBX1,RPLP0,RPL6,CSDE1,V |
| 100 | 18964 | ACTR2,DBN1,MAP1B,RPL8,RPLP0,RPL6,SPTBN1,SHISA9,RPL7,RPS3 |
| 46 | 18964 | EEF1A1,ACTB,CFL1,ACTR2,NUMA1,DBN1,FLNA,SPTBN1,MYH9,SPTA |
| 17 | 18964 | PSMD5,PSMD2,PSMD3,PSMD1,PSMD8,PSMD4,PSMC3,PSMC1,PSMD |
| 243 | 18964 | LRP1,HSPA8,COPA,HTT,SPARC,UBC,SLC9A7,UBA1,CALR,PSAP,KIF1/ |
| 111 | 18964 | NUDC,TUBA1A,TUBB,NUMA1,DCX,TUBA1B,MAP1B,TUBB2B,KIF1A,C |
| 18 | 18964 | PSMD5,PSMD2,PSMD3,PSMD1,PSMD8,PSMD4,PSMC3,PSMC1,PSMD |
| 212 | 18964 | COL1A1,ACTB,NUDC,ACTG1,DPYSL3,TUBA1A,VIM,TUBB,NUMA1,DC |
| 53 | 18964 | CTSC,ACTR2,TUBB,FTL,PSAP,VCP,VAPA,TUBB4B,ANXA2,CYB5R3,LP |
| 53 | 18964 | CTSC,ACTR2,TUBB,FTL,PSAP,VCP,VAPA,TUBB4B,ANXA2,CYB5R3,LP |
| 93 | 18964 | ACTR2,DBN1,MAP1B,RPL8,RPLP0,RPL6,SPTBN1,SHISA9,RPL7,RPS3 |
| 210 | 18964 | COL1A1,ACTB,NUDC,ACTG1,DPYSL3,TUBA1A,VIM,TUBB,NUMA1,DC |
| 91 | 18964 | ACTR2,DBN1,MAP1B,RPL8,RPLP0,RPL6,SPTBN1,SHISA9,RPL7,RPS3 |
| 95 | 18964 | HSPA8,ACTR2,DBN1,MAP1B,RPL8,RPLP0,RPL6,SPTBN1,SHISA9,RPL |
| 101 | 18964 | EEF1A1,LRP1,HSPA8,HSP90AB1,COL6A1,UBA1,PSAP,GNB1,PLD3,VA |
| 101 | 18964 | EEF1A1,LRP1,HSPA8,HSP90AB1,COL6A1,UBA1,PSAP,GNB1,PLD3,VA |
| 33 | 18964 | EEF2,HTT,HSP90AB1,NEFM,HSPA1B,CLU,MAPT,ATF4,SQSTM1,PABP |
| 119 | 18964 | ACTB,ACTG1,EIF4G2,DBN1,FLNA,UBA1,MYH9,APP,FSCN1,DCHS1,VA |
| 180 | 18964 | COPA,NUMA1,FLNA,SLC9A7,TMF1,GNAS,BSG,APP,PLD3,HSPD1,CHS |
| 121 | 18964 | HSPA8,ACTB,ACTG1,HTT,NEFL,ATP2B4,CANX,KIF1A,CALM3,SEPTIN2 |
| 158 | 18964 | COPA,NUMA1,SLC9A7,TMF1,GNAS,BSG,APP,PLD3,CHSY1,PKD1,HLA |
| 45 | 18964 | ATN1,CFL1,VIM,SPARC,NUMA1,MATR3,MAPT,NONO,BASP1,HNRNPU |
| 44 | 18964 | GANAB,HSP90B1,CALR,P4HB,VCP,PDIA3,SEC61A1,MARCHF6,PRKCS |
| 11 | 18964 | PSMD5,PSMD2,PSMD1,PSMD4,PSMC3,PSMC1,PSMC5,PSMC2,PSMC |
| 120 | 18964 | EEF1A1,ACTB,ACTG1,DPYSL3,CFL1,ACTR2,DBN1,FLNA,ALDOA,SPTE |
| 17 | 18964 | HSPA8,HSP90AB1,CCT7,PTGES3,CCT6A,STIP1,CCT2,TCP1,CCT3,CC |
| 53 | 18964 | ACTB,EIF4G2,MYH9,DCHS1,ANXA2,VCL,CTNNA1,CNN3,CXADR,CDH6 |
| 35 | 18964 | EEF1A1,CFL1,ACTR2,DBN1,SPTBN1,MYH9,SPTAN1,VCL,LLGL1,WDR |
| 34 | 18964 | CTSC,ACTR2,TUBB,FTL,VCP,TUBB4B,ANXA2,CYB5R3,VAT1,TTR,HRN |
| 32 | 18964 | ACTB,ACTG1,NEFL,TUBB2B,INA,ITGB1,AKAP12,CTNNB1,CDH11,PTF |
| 13 | 18964 | EIF3C,EIF1,EIF3L,EIF3E,EIF3F,EIF3D,EIF3H,EIF3G,EIF3B,EIF3M,EIF1 |
| 26 | 18964 | PRPF8,SNRNP200,SF3B1,SF3B2,LUC7L3,DDX39B,SNRPN,SNRNP70, |
| 32 | 18964 | SLC9A7,HLA-C,PDIA3,HLA-B,RAC1,RAB11B,OPTN,CLIP3,B2M,RAB11 |
| 75 | 18964 | LRP1,HSPA8,COPA,CTSC,APP,HSPD1,HLA-C,AP2M1,ARF1,AP2B1,SC/ |

| | | |
|---|---|---|
| 97 | 18964 | COPA,DPYSL3,CALR,KIF1A,GNAS,CALM3,SEPTIN2,APP,CALM1,HLA-( |
| 82 | 18964 | LRP1,ACTG1,HSP90AA1,VIM,SPARC,UBC,HSP90B1,CALR,UBB,HLA-C |
| 38 | 18964 | VCP,CHD3,SMARCA5,CHD4,ATP1A1,ATP2A2,RBBP4,ARID1A,ATP1A2, |
| 12 | 18964 | EIF3C,EIF1,EIF3L,EIF3E,EIF3F,EIF3D,EIF3H,EIF3G,EIF3B,EIF3M,EIF1 |
| 12 | 18964 | EIF3C,DDX3X,EIF3L,EIF3E,EIF3F,EIF3D,EIF3H,EIF3G,EIF3B,EIF3M,AE |
| 53 | 18964 | TUBA1A,SLC9A7,APP,HLA-C,PDIA3,ITGB1,NISCH,MYO5A,HLA-B,RAC |
| 29 | 18964 | PABPC1,YBX1,CSDE1,VCP,LARP1,KPNB1,DDX3X,TARDBP,HNRNPK,C |
| 97 | 18964 | COL1A1,CTSC,SPARC,PKM,COL6A1,HSP90B1,CALR,COL5A1,COL4A |
| 33 | 18964 | COPA,APP,PKD1,ARF1,ITM2B,CLTC,GJA1,ARCN1,NUCB1,TMED2,KD |
| 27 | 18964 | PRPF8,SNRNP200,SF3B1,SF3B2,LUC7L3,DDX39B,SNRPN,SNRNP70, |
| 11 | 18964 | EIF3C,EIF3L,EIF3E,EIF3F,EIF3D,EIF3H,EIF3G,EIF3B,EIF3M,EIF2S1,EI |
| 47 | 18964 | CALR,CANX,SEC62,HACD3,HLA-C,PDIA3,SEC61A1,SARAF,MARCHF6, |
| 29 | 18964 | CHD3,SMARCA5,CHD4,RBBP4,ARID1A,HDAC2,SMARCA4,SMARCE1,S |
| 52 | 18964 | MAP1B,CANX,SHISA9,APP,MAPT,ITGB1,CNN3,RHOA,CLSTN1,SEPTIN |
| 36 | 18964 | ATN1,CFL1,VIM,SPARC,NUMA1,MATR3,NONO,BASP1,HNRNPU,ENC1 |
| 32 | 18964 | NEFL,CANX,KIF1A,MAPT,UCHL1,AP3D1,PAFAH1B1,HNRNPU,KIF5B,S |
| 160 | 18964 | ACTB,USP22,MED18,PIK3R2,FBXL4,KAT6A,DDB1,MORF4L1,MARCH |
| 28 | 18964 | UBA1,TRIM28,CBX5,TOP2B,NOP53,TCP1,CBX1,PHC2,KDM4B,BAZ1E |
| 233 | 18964 | GAPDH,HSPA9,MAVS,TUBB,UBC,FBXL4,YBX1,CALR,CLU,ATP5F1B,C |
| 233 | 18964 | GAPDH,HSPA9,MAVS,TUBB,UBC,FBXL4,YBX1,CALR,CLU,ATP5F1B,C |
| 48 | 18964 | CALR,CANX,SEC62,HACD3,HLA-C,PDIA3,SEC61A1,SARAF,MARCHF6, |
| 52 | 18964 | MAP1B,CANX,SHISA9,APP,MAPT,ITGB1,CNN3,RHOA,CLSTN1,SEPTIN |
| 319 | 18964 | ORC4,POLH,ACTB,XRCC2,ACTR2,NUMA1,NCL,UBA1,H3-3B,SOX4,M |
| 7 | 18964 | PPP1CB,PPP1CC,PPP1R10,TOX4,PPP1CA,WDR82,PPP1R12A |
| 94 | 18964 | HSPA9,CALR,CANX,SEC62,HACD3,HLA-C,PDIA3,SEC61A1,SARAF,MA |
| 157 | 18964 | LRP1,HSPA8,COPA,HTT,SPARC,UBC,CALR,PSAP,KIF1A,SLC2A3,BSG, |
| 41 | 18964 | ACTG1,VIM,CALR,HLA-C,PDIA3,KIF5B,RAB7A,LAMP1,HLA-B,APPL1, |
| 31 | 18964 | PRPF8,SNRNP200,SF3B1,SF3B2,IK,LUC7L3,U2AF2,SNRPN,SNRNP70 |
| 48 | 18964 | HSPA8,COPA,HLA-C,AP2M1,AP2B1,SCARB2,CLTC,DNAJC5,HLA-B,AF |
| 80 | 18964 | ORC4,H3-3B,SEPTIN2,DDB1,PPP2R1A,PAFAH1B1,HNRNPU,XRCC5,C |
| 28 | 18964 | PRPF8,SNRNP200,SF3B1,SF3B2,LUC7L3,DDX39B,SNRPN,SNRNP70, |
| 9 | 18964 | COL1A1,COL5A1,COL11A1,COL1A2,COL5A2,COL2A1,COL27A1,COI |
| 9 | 18964 | COL1A1,COL5A1,COL11A1,COL1A2,COL5A2,COL2A1,COL27A1,COI |
| 27 | 18964 | CHD3,MORF4L1,CBX5,CHD4,RBBP4,HDAC2,NCOR2,KDM3B,RBBP7, |
| 20 | 18964 | TUBA1A,HSP90AA1,CALM3,GDI1,CALM1,ITGB1,ANXA2,GLUL,DPYSL |
| 151 | 18964 | ACTB,NUDC,ACTG1,DPYSL3,TUBA1A,VIM,TUBB,NUMA1,DCX,DBN1,T |
| 33 | 18964 | APP,FSCN1,ITGB1,MSN,GAP43,CXADR,ARL4C,MYO5A,MAP2,CDC42, |
| 31 | 18964 | HSPA8,HSP90AA1,PSAP,IDS,PLD3,AGRN,CTSD,SCARB2,GPC3,GPC1, |
| 10 | 18964 | ARID1A,SMARCA4,SMARCE1,SMARCC1,ARID1B,SMARCD1,SS18L1,S |
| 48 | 18964 | EEF1A1,CFL1,GNAS,MYH9,FSCN1,RPS3,ITGB1,ANXA2,RHOA,TPM1,T |
| 12 | 18964 | COL1A1,COL5A1,COL4A1,COL4A2,COL11A1,COL1A2,COL5A2,COL |
| 65 | 18964 | NUMA1,DCX,ENO1,NEFL,CANX,KIF1A,SEPTIN2,MAPT,DNAH8,UCHL1, |
| 15 | 18964 | LRP1,HSP90AB1,HSP90AA1,FLNA,KIF5B,L1CAM,MAPK8IP1,FKBP4,C |
| 21 | 18964 | ATP5F1B,ATP5F1A,ATP6V0C,ATP6AP1,ATP5PB,ATP6V1B2,ATP6AP2, |
| 8 | 18964 | CCT7,CCT6A,CCT2,TCP1,CCT3,CCT4,CCT8,CCT5 |

| | | |
|---|---|---|
| 26 | 18964 | ITGB1,VCL,CTNNA1,CXADR,ATP1A1,CDH2,GJA1,CTNNB1,CADM4,AT |
| 100 | 18964 | GAPDH,TUBB,YBX1,CALR,APP,MATR3,HACD3,VAPA,PAFAH1B1,TMEN |
| 93 | 18964 | MED18,SOX4,TRIM28,SPEN,NONO,GTF3C4,CHD3,HMGB1,GTF2F1,H |
| 7 | 18964 | ATF4,SQSTM1,ARIH1,GPX1,PINK1,HERPUD1,FBXO7 |
| 26 | 18964 | ATP2B4,CANX,APP,CDH2,CTNNB1,CTBP1,STXBP1,SYP,CTNND1,CNTI |
| 10 | 18964 | SF3B1,LUC7L3,U2AF2,SNRPN,SNRNP70,SF3A1,PRPF40A,SF3A2,LUC |
| 10 | 18964 | CHD3,CHD4,RBBP4,HDAC2,RBBP7,MTA1,CSNK2A1,MBD3,MTA2,GA |
| 10 | 18964 | SF3B1,LUC7L3,U2AF2,SNRPN,SNRNP70,SF3A1,PRPF40A,SF3A2,LUC |
| 10 | 18964 | CHD3,CHD4,RBBP4,HDAC2,RBBP7,MTA1,CSNK2A1,MBD3,MTA2,GA |
| 85 | 18964 | HSPA9,CALR,CANX,SEC62,HACD3,HLA-C,PDIA3,SEC61A1,SARAF,MA |
| 50 | 18964 | NUDC,HSP90B1,RACK1,SEPTIN2,GDI1,ANXA2,HNRNPU,RHOA,CRMP |
| 19 | 18964 | PPP2R1A,PPP1CB,PPP2CA,PPP1R9B,PPP2CB,PPP1CC,PPP1R15A,PPI |
| 19 | 18964 | PPP2R1A,PPP1CB,PPP2CA,PPP1R9B,PPP2CB,PPP1CC,PPP1R15A,PPI |
| 21 | 18964 | UBC,UBB,KPNB1,XPO1,MVB12B,KPNA2,UBA52,POM121C,RPS27A,F |
| 21 | 18964 | UBC,UBB,KPNB1,XPO1,MVB12B,KPNA2,UBA52,POM121C,RPS27A,F |
| 75 | 18964 | EEF2,TUBA1A,TUBB,TUBA1B,ATP2B4,PGK1,BSG,APP,HSPD1,MAPT,L |
| 75 | 18964 | EEF2,TUBA1A,TUBB,TUBA1B,ATP2B4,PGK1,BSG,APP,HSPD1,MAPT,L |
| 290 | 18964 | HSPA9,MAVS,HSP90AB1,HSP90AA1,UBC,PKM,FADS1,FBXL4,UBA1,H |
| 24 | 18964 | MYH9,SEPTIN2,ITGB1,RHOA,MYH10,SEPTIN11,RHOB,SEPTIN3,PKN1 |
| 42 | 18964 | ORC4,H3-3B,DDB1,XRCC5,CBX5,PTGES3,SMCHD1,PPP1CB,PARP1,( |
| 65 | 18964 | HSPA8,NEDD4L,HTT,KIDINS220,PSAP,PLD3,TPT1,ANXA2,ARF1,SCARI |
| 27 | 18964 | RPL4,PKM,UBA1,CANX,SEC62,RPL6,APP,HSPD1,SEC61A1,RPL18,ARI |
| 9 | 18964 | CFL1,ACTR2,ACTR3,DSTN,HIP1,CAPZA1,HIP1R,CAPZA2,GMFB |
| 9 | 18964 | CFL1,ACTR2,ACTR3,DSTN,HIP1,CAPZA1,HIP1R,CAPZA2,GMFB |
| 36 | 18964 | CTSC,CALR,P4HB,FN1,HMGB1,RAB6B,PDIA6,NUCB1,SURF4,ATP6AP |
| 121 | 18964 | CTSC,ODF2L,XRCC2,NUMA1,HSPA1B,CALM3,MARCKS,CALM1,NPM |
| 21 | 18964 | NEFL,KIF1A,MAPT,UCHL1,AP3D1,PAFAH1B1,KIF5B,MAP1A,HSPB1,S( |
| 17 | 18964 | HSPA9,ATP5F1B,LRPPRC,ACADVL,DHX30,LONP1,VDAC1,GRSF1,SLC |
| 17 | 18964 | HSPA9,ATP5F1B,LRPPRC,ACADVL,DHX30,LONP1,VDAC1,GRSF1,SLC |
| 23 | 18964 | LRP1,APP,HSPD1,AP2M1,AP2B1,CLTC,DGKD,BMPR2,AP2A1,CTTN,LI |
| 66 | 18964 | MED18,FBXL4,DDB1,MARCHF6,ENC1,KLHL9,SKP1,KLHL24,UBE4A,( |
| 18 | 18964 | PRPF8,SNRNP200,SF3B1,SF3B2,IK,PRPF6,SF3A1,SNRPD3,EFTUD2,S |
| 13 | 18964 | ACTR2,FSCN1,VCL,TPM4,WDR1,HNRNPK,PALLD,CTTN,SH3GL1,SH3I |
| 14 | 18964 | CANX,HNRNPU,KIF5B,SFPQ,WLS,MAP2,MAPK1,SOD1,KIF5A,GABAR/ |
| 9 | 18964 | CD63,LAPTM4B,RAB27A,ABCA3,NSG2,HGS,NSG1,ATP13A2,TMEM9 |
| 21 | 18964 | NEDD4L,TPT1,PRKAR1A,GJA1,CD63,LAMP1,LAPTM4B,RAB27A,ABC/ |
| 110 | 18964 | COL1A1,CTSC,SPARC,PKM,COL6A1,HSP90B1,CALR,COL5A1,COL4A |
| 24 | 18964 | FLNA,MYH9,FSCN1,MARCKS,TPM4,TPM1,MYH10,SEPTIN11,CNN2,N( |
| 38 | 18964 | ATP2B4,CANX,SLC4A8,NLGN2,CDH2,CTNNB1,STXBP1,PTPRS,SYP,CI |
| 110 | 18964 | COL1A1,CTSC,SPARC,PKM,COL6A1,HSP90B1,CALR,COL5A1,COL4A |
| 17 | 18964 | RBBP4,HDAC2,SKP1,ASXL1,PHC2,RBBP7,CSNK2A1,UBAP2L,CBX6,B |
| 24 | 18964 | DBN1,MYH9,FSCN1,CDC42BPB,TPM4,TPM1,MYH10,SEPTIN11,CNN2 |
| 15 | 18964 | TRIM28,POLR2A,H1-0,JUN,CTNNB1,RBMX,H2AZ1,TCF3,RUVBL2,HSI |
| 14 | 18964 | HSP90AB1,HSPA1B,MYH9,ATP5F1A,BASP1,GRB2,WDR6,PLCG1,DYN |
| 18 | 18964 | PRPF8,SNRNP200,SF3B1,SF3B2,IK,PRPF6,SF3A1,SNRPD3,EFTUD2,S |

| | | |
|---|---|---|
| 18 | 18964 | COPA,PKD1,ITM2B,CLTC,GJA1,ARCN1,TMED2,KDELR2,KDELR1,AP2 |

,COPA,HNRNPA1,ARSD,HSPA9,NEDD4L,POLH,ATN1,ACTB,CTSC,ODF2L,NUDC,NRBP2,USP22,MT
,COPA,HNRNPA1,ARSD,HSPA9,NEDD4L,POLH,ATN1,ACTB,CTSC,ODF2L,NUDC,NRBP2,MTRNR2L
,COPA,HNRNPA1,ARSD,HSPA9,NEDD4L,POLH,ATN1,ACTB,CTSC,ODF2L,NUDC,USP22,MED18,AC
ACTB,NUDC,ACTG1,EEF2,DPYSL3,TUBA1A,CFL1,HTT,HSP90AB1,HSP90AA1,ACTR2,VIM,TUBB,FT
,COPA,HNRNPA1,ARSD,HSPA9,NEDD4L,POLH,ATN1,ACTB,CTSC,ODF2L,NUDC,USP22,MED18,AC
,COPA,HNRNPA1,ARSD,HSPA9,NEDD4L,POLH,ATN1,ACTB,CTSC,NUDC,USP22,MED18,ACTG1,EE
,COPA,HNRNPA1,ARSD,HSPA9,NEDD4L,POLH,ATN1,ACTB,CTSC,ODF2L,NUDC,USP22,MED18,AC
TB,CTSC,ACTG1,EEF2,TUBA1A,CFL1,HSP90AB1,HSP90AA1,ACTR2,VIM,TUBB,FTH1,UBC,NUMA1,
A9,NEDD4L,POLH,ATN1,ACTB,CTSC,NUDC,USP22,EEF2,CFL1,XRCC2,HTT,HSP90AB1,HSP90AA1,A
A9,NEDD4L,POLH,ATN1,ACTB,CTSC,NUDC,USP22,EEF2,CFL1,XRCC2,HTT,HSP90AB1,HSP90AA1,A
A9,NEDD4L,POLH,ATN1,ACTB,CTSC,NUDC,USP22,EEF2,CFL1,XRCC2,HTT,HSP90AB1,HSP90AA1,A
TB,CTSC,ACTG1,EEF2,TUBA1A,CFL1,HSP90AB1,HSP90AA1,ACTR2,VIM,TUBB,FTH1,UBC,NUMA1,
TB,CTSC,ACTG1,EEF2,TUBA1A,CFL1,HSP90AB1,HSP90AA1,ACTR2,VIM,TUBB,FTH1,UBC,NUMA1,
TB,CTSC,ACTG1,EEF2,TUBA1A,CFL1,HSP90AB1,HSP90AA1,ACTR2,VIM,TUBB,FTH1,UBC,NUMA1,
,COPA,HNRNPA1,HSPA9,ACTB,USP22,MED18,ACTG1,EEF2,DPYSL3,XRCC2,HTT,HSP90AB1,HSP9
A9,NEDD4L,ACTB,CTSC,ACTG1,EEF2,DPYSL3,TUBA1A,CFL1,HTT,HSP90AB1,HSP90AA1,ACTR2,VI
ACTB,CTSC,NUDC,USP22,CFL1,XRCC2,HTT,HSP90AB1,HSP90AA1,VIM,SPARC,UBC,NUMA1,RPL3
DC,USP22,XRCC2,HTT,HSP90AB1,HSP90AA1,UBC,NUMA1,RPS6,NCL,PRPF8,FBXL4,YBX1,UBA1,H
),NEDD4L,POLH,ATN1,ACTB,CTSC,NUDC,USP22,MED18,ACTG1,EEF2,TUBA1A,CFL1,XRCC2,HTT,
PL4,FLNA,RPL7A,HSP90B1,RPL8,CALR,HSPA1B,RPLP0,RPL13A,RPL6,YWHAE,BSG,RPS8,MYH9,R
PL4,FLNA,RPL7A,HSP90B1,RPL8,CALR,HSPA1B,RPLP0,RPL13A,RPL6,YWHAE,BSG,RPS8,MYH9,R
CTSC,NRBP2,EEF2,DPYSL3,TUBA1A,HTT,HSP90AB1,HSP90AA1,ACTR2,SPARC,TUBB,FTH1,UBC,K
NCL,RPL7A,PRPF8,RPL8,YBX1,CALR,HSPA1B,RPLP0,DDX5,RPL13A,RPL6,RPL13,DDX17,RACK1,R
1A,CFL1,HTT,ACTR2,VIM,SPARC,EIF4G2,RPL3,DBN1,RPL4,FLNA,MAP1B,RPL7A,HSP90B1,NEFL,
L7,RPL5,RPL37A,RPL19,RPL15,RPS3,RPS4X,RPS3A,RPL18,RPL11,RPLP1,RPL10,RPS17,RPS16,R
RPL3,DBN1,RPL4,FLNA,RPL7A,HSP90B1,RPL8,UBA1,CALR,HSPA1B,RPLP0,RPL13A,RPL6,YWHAE
TB,CTSC,ODF2L,NUDC,ACTG1,EEF2,DPYSL3,TUBA1A,CFL1,XRCC2,HTT,HSP90AB1,ACTR2,VIM,T
TB,CTSC,ODF2L,NUDC,ACTG1,EEF2,DPYSL3,TUBA1A,CFL1,XRCC2,HTT,HSP90AB1,ACTR2,VIM,T
YSL3,TUBA1A,HTT,HSP90AB1,HSP90AA1,ACTR2,VIM,SPARC,TUBB,FTH1,UBC,KIDINS220,TMSB4X
YSL3,TUBA1A,HTT,HSP90AB1,HSP90AA1,ACTR2,VIM,SPARC,TUBB,FTH1,UBC,KIDINS220,TMSB4X
I,FLNA,MAP1B,NEFL,RPL8,ATP2B4,TUBB2B,RPLP0,CANX,CLU,RPL6,YWHAE,KIF1A,SPTBN1,CALM
SPA9,NEDD4L,ACTB,CTSC,ACTG1,EEF2,DPYSL3,TUBA1A,CFL1,HSP90AB1,HSP90AA1,ACTR2,VIM,
L37A,RPL19,RPL15,RPL18,RPL11,RPLP1,RPL10,RPL23,RPL29,RPL24,RPL17,RPL14,RPL22,RPL
,DDX5,ENO2,KAT6A,GNAS,CALM3,GNB1,P4HB,CALM1,SNRNP200,VCP,DDB1,PSMD5,PPP2R1A,D
ASN,P4HB,PDIA3,ITGB1,ANXA2,CTSD,ATP1A1,SLC3A2,PDIA6,RAB7A,MMP14,CLTC,MYO5A,CD63,
ASN,P4HB,PDIA3,ITGB1,ANXA2,CTSD,ATP1A1,SLC3A2,PDIA6,RAB7A,MMP14,CLTC,MYO5A,CD63,
RPS8,RPL7,RPL5,NPM1,RPL37A,RPL19,RPL15,RPS3,RPS4X,RPS3A,RPL18,RPL11,RPLP1,RPL10,R
B,COL6A1,HSP90B1,FADS1,UGT8,UBA1,CALR,FSTL1,TMF1,HSPA1B,COL5A1,COL4A1,CES3,RPL
,HSP90AB1,FLNA,NCL,PRPF8,YBX1,HSPA1B,DDX5,KAT6A,MYH9,SNRNP200,ATP5F1A,AFF4,TRIM
13,RACK1,RPS8,RPL7,RPL5,NPM1,RPL37A,RPL19,RPL15,RPS3,RPS4X,RPS3A,RPL18,RPL11,BTF
D3,BASP1,MORF4L1,SBF1,SF3B2,HNRNPU,CBX5,ATF4,SRSF4,PNN,XPO1,TKT,SFPQ,ACTN4,H1-0
EIF4G2,DBN1,FLNA,MAP1B,NEFL,NEFM,CANX,YWHAE,KIF1A,BSG,SPTBN1,CALM3,STMN2,APP,G
SP1,MORF4L1,SF3B2,HNRNPU,ATF4,SRSF4,PNN,SFPQ,IK,TARDBP,CKAP4,BCLAF1,LUC7L3,DDX3
VIM,NUMA1,DCX,EIF4G2,RPS6,DBN1,FLNA,MAP1B,NEFL,NEFM,ATP2B4,RPLP0,CANX,CLU,YWHA
AP1B,RPLP0,CANX,CLU,ENO2,KIF1A,SLC2A3,RACK1,GNB1,SHISA9,STMN2,APP,GDI1,EIF4B,UBB

31,SPARC,UBC,NUMA1,SLC9A7,COL6A1,HSP90B1,FADS1,YBX1,UBA1,CALR,TMF1,PSAP,CANX,CL
1,HSPA1B,GPI,GSTP1,VCP,HMGB1,TIMP2,VCL,PGAM1,CTSD,KPNB1,PSMD2,PFKL,DDX3X,MIF,CC
B,CANX,ENO2,KIF1A,SLC2A3,RACK1,GNB1,STMN2,APP,GDI1,EIF4B,UBB,CKB,MAPT,UCHL1,TIM
SAP,BSG,STMN2,APP,PLD3,HSPD1,UBB,HLA-C,PDIA3,HMGB1,AP2M1,TPT1,ITGB1,ANXA2,AP3D1,
JMA1,DCX,RPS6,DBN1,ENO1,TUBA1B,FLNA,MAP1B,NCL,ALDOA,COL6A1,NEFL,NEFM,YBX1,ATP2
FL1,HTT,HSP90AB1,HSP90AA1,ACTR2,VIM,NUMA1,DCX,EIF4G2,RPS6,DBN1,FLNA,MAP1B,PKM,N
AB,CALR,TMF1,CLU,GNAS,BSG,FASN,STMN2,APP,GDI1,TTC3,PLD3,HSPD1,CSDE1,CHSY1,PKD1,H
S14,RPS20,RPS25,DDX3X,RPS15,RPS5,RPS7,RPS19,RPS10,RPS12,RPS27,RPS15A,RPS11,RPS28
PA9,NEDD4L,ACTB,CTSC,MTRNR2L2,ACTG1,EEF2,DPYSL3,TUBA1A,CFL1,HSP90AB1,HSP90AA1,
FL1,HTT,HSP90AB1,HSP90AA1,ACTR2,VIM,NUMA1,DCX,EIF4G2,RPS6,DBN1,FLNA,MAP1B,PKM,N
1,HSPA1B,SLC2A3,GPI,GSTP1,VCP,HMGB1,TIMP2,VCL,PGAM1,CTSD,KPNB1,RHOA,PSMD2,PFKL
AP1B,RPLP0,CANX,CLU,KIF1A,RACK1,GNB1,SHISA9,APP,EIF4B,CKB,MAPT,PPP2R1A,ITGB1,SLC4
IX,RPL6,KIF1A,SPTBN1,SHISA9,RPL7,APP,EIF4B,MAPT,INA,RPS3,AP2M1,ITGB1,AP3D1,ARF1,AP2
AP1B,RPLP0,CANX,CLU,KIF1A,RACK1,GNB1,SHISA9,APP,EIF4B,CKB,MAPT,PPP2R1A,ITGB1,SLC4
B,TMSB4X,ALDOA,PKM,FTL,CLU,GPI,APP,SPTAN1,GSTP1,IGF2,VCP,FN1,HMGB1,TUBB4B,ANXA2
B,TMSB4X,ALDOA,PKM,FTL,CLU,GPI,APP,SPTAN1,GSTP1,IGF2,VCP,FN1,HMGB1,TUBB4B,ANXA2
B,TMSB4X,ALDOA,PKM,FTL,CLU,GPI,APP,SPTAN1,GSTP1,IGF2,VCP,FN1,HMGB1,TUBB4B,ANXA2
,RPL11,BTF3,LARP1,RPL24,RPL31,HDLBP,RPL30,MYH10,RPL36,GCN1,RPL36A,RPL38,RPS28,RI
3C,NUMA1,SLC9A7,COL6A1,UBA1,CALR,TMF1,PSAP,KIF1A,GNAS,SLC2A3,GPI,BSG,CALM3,GNB1
TUBB,FTH1,COL6A1,FTL,UBA1,PSAP,GNB1,IDS,PLD3,VCP,VAPA,AP2M1,TUBB4B,ANXA2,AGRN,AF
,PSMD1,PSMD8,PSMB4,DNAJB2,PSMD4,PSMC3,UBE3A,PSMC1,PSMB1,PSMD13,USP14,GPAA1,F
,YWHAE,CALM3,STMN2,APP,FSCN1,CALM1,MAPT,BASP1,TIMP2,PAFAH1B1,GAP43,KIF5B,CXADF
L7A,DDX5,RPL13A,KAT6A,RPL13,DDX17,SPTBN1,RPL7,RPL5,NPM1,TTC3,AFF4,NONO,RPL19,RP
,SF3B2,HNRNPU,PNN,HNRNPC,IK,HNRNPDL,HNRNPK,CIRBP,HNRNPH1,LUC7L3,DDX39B,U2AF2,
AA1,ACTR2,SPARC,TUBB,FTH1,TMSB4X,ALDOA,PKM,FTL,CALR,ABHD2,HSPA1B,PSAP,CLU,KIF1
,PSMA7,PSMD1,PSMD8,SPG7,PSMB4,DNAJB2,PSMD4,PSMC3,UBE3A,ATXN7L3,PSMC1,PSMB1,P
,PSMD1,PSMD8,PSMB4,DNAJB2,PSMD4,PSMC3,UBE3A,PSMC1,PSMB1,PSMD13,USP14,PSMD7,
,YWHAE,CALM3,STMN2,APP,FSCN1,CALM1,MAPT,BASP1,TIMP2,PAFAH1B1,GAP43,KIF5B,CXADF
3,FTH1,COL6A1,FTL,UBA1,PSAP,GNB1,IDS,PLD3,VCP,VAPA,AP2M1,TUBB4B,ANXA2,AGRN,AP3D1
3,FTH1,COL6A1,FTL,UBA1,PSAP,GNB1,IDS,PLD3,VCP,VAPA,AP2M1,TUBB4B,ANXA2,AGRN,AP3D1
3A1A,CFL1,XRCC2,HTT,ACTR2,VIM,TUBB,TMSB4X,NUMA1,DCX,DBN1,TUBA1B,FLNA,MAP1B,ALD
,ITGB1,CDC42BPB,CTNNA1,RHOA,CDH2,DDX3X,CTNNB1,MTSS2,MYH10,RAC1,CAPRIN1,ACTR3,
HSP90B1,CALR,HSPA1B,CLU,GNAS,RACK1,STMN2,SEPTIN2,APP,VCP,SET,VAPA,ITGB1,ANXA2,SBI
R2,SPARC,TUBB,FTH1,TMSB4X,ALDOA,PKM,FTL,CALR,ABHD2,HSPA1B,PSAP,CLU,KIF1A,SLC2A3
I,CANX,SEC62,GNAS,BSG,APP,TMBIM6,PLD3,HSPD1,SMIM14,VCP,UBB,HACD3,CHSY1,PKD1,HLA
1,FLNA,MAP1B,NEFL,YWHAE,CALM3,STMN2,APP,FSCN1,CALM1,MAPT,BASP1,TIMP2,AP3D1,PAF
SA9,VCP,AP2M1,ITGB1,AP3D1,ARF1,AP2B1,SLC4A8,NCS1,RPL22,RHOA,CLSTN1,PFN1,SPTBN2,PI
DCX,TUBA1B,MAP1B,HSPA1B,TUBB2B,YWHAE,KIF1A,CALM3,MYH9,SEPTIN2,APP,SPTAN1,MARCI
I1,SHISA9,STMN2,MYH9,P4HB,APP,FSCN1,MAPT,RPS3,ITGB1,ANXA2,ARF1,PAFAH1B1,CDC42BPI
MYH9,SEPTIN2,FSCN1,SPTAN1,MARCKS,ANXA2,VCL,PAFAH1B1,AKAP12,TPM4,LLGL1,RHOA,WDI
,SLC2A3,RACK1,STMN2,APP,GDI1,EIF4B,UBB,CKB,MAPT,UCHL1,TIMP2,PAFAH1B1,AKAP12,SLC
M1,RPL37A,RPL19,RPL15,RPL18,RPL11,RPLP1,RPL10,RPL23,RPL29,RPL24,RPL17,RPL14,RPL2
CP,MAPT,INA,RPS4X,HNRNPU,LARP1,KPNB1,DDX3X,TARDBP,HNRNPK,CIRBP,CKAP4,RAC1,CAPF
PS24,RPS9,RPS14,RPS20,RPS25,DDX3X,RPS15,RPS5,RPS7,RPS19,RPS10,RPS12,RPS27,RPS15A,
SCH,GJA1,MYO5A,WLS,RHOB,HLA-B,RAB1A,APPL1,DNER,LAPTM4B,NUCB1,TBC1D16,VPS35,RAI

PL31,RPL30,RPL36,RPL36A,RPL38,RPS28,RPL39,EIF3H,ABCF1,RPS26,RPS21,RPS29,RPL32
'3D1,ITM2B,AP2B1,CTSD,SCARB2,LLGL1,RAB7A,CLTC,CD63,ANXA6,NDFIP1,WLS,LAMP1,RHOB,I
PLD3,SMIM14,VCP,UBB,HACD3,HLA-C,PDIA3,SERINC1,VAPA,SEC61A1,UCHL1,SARAF,MARCHF6,
SD,MS4A10,HSPA9,NEDD4L,ACTB,CTSC,NUDC,ACTG1,EEF2,TUBA1A,CFL1,MAVS,HTT,HSP90AB1
PLD3,SMIM14,VCP,UBB,HACD3,HLA-C,PDIA3,SERINC1,VAPA,SEC61A1,UCHL1,SARAF,MARCHF6,
,COL5A1,COL4A1,CES3,CANX,CLU,FKBP14,P4HB,APP,COL18A1,COL4A2,FN1,PDIA3,COL11A1,I(
'NN,HNRNPC,HNRNPK,HNRNPH1,SNRPN,HNRNPA3,RBMX,PRPF6,SRSF1,HNRNPM,SF3A1,HNRNP
.PA,AP2M1,ANXA2,AP3D1,AP2B1,CTSD,SCARB2,LPCAT1,SLC22A17,SLC7A5,SLC3A2,RAB7A,ITM2
DDB1,PSMD5,PPP2R1A,MORF4L1,MARCHF6,MAT2A,GTF2F1,LARP1,XRCC5,PSMD2,ENC1,TADA:
PLD3,SMIM14,VCP,UBB,HACD3,HLA-C,PDIA3,SERINC1,VAPA,SEC61A1,UCHL1,SARAF,MARCHF6,
.4B,ANXA2,AGRN,CYB5R3,CTSD,SCARB2,VAT1,GPC3,TTR,HRNR,GPC1,DCN,CTSB,CCT2,GDI2,VC
CP,MAPT,INA,RPS4X,HNRNPU,LARP1,KPNB1,DDX3X,TARDBP,HNRNPK,CIRBP,CKAP4,RAC1,CAPF
,ARF1,SLC4A8,RPL23,CNN3,RTN4,PJA2,GAP43,RPL14,NCS1,CLSTN1,NLGN2,RPS14,ATP1A1,RPS
N1,VCL,TPM4,LLGL1,WDR1,LASP1,SPTBN2,CDH2,MTSS2,COTL1,ACTR3,MAPRE1,DSTN,CAPZB,I
)13,PSMD7,ADRM1,PSMC5,PSMC2,PSMC4,PSMD10,PSMD12,PSMC6
\,SLC2A3,BSG,CALM3,CALM1,PLD3,UBB,PKD1,HLA-C,PDIA3,VAPA,AP2M1,ITGB1,ANXA2,AP3D1,I
ALM3,CALM1,MAPT,MAP9,DNAH8,TPT1,STMN1,TUBB4B,PAFAH1B1,MAP4,HNRNPU,KIF5B,CCT7
)13,PSMD7,ADRM1,PSME3,PSMC5,PSMC2,PSMC4,PSMD10,PSMD12,PSMC6
X,DBN1,ENO1,TUBA1B,FLNA,MAP1B,ALDOA,COL6A1,NEFL,NEFM,ATP2B4,TUBB2B,COL5A1,COL
CAT1,VAT1,CD63,DNAJC5,DDOST,LAMP1,CKAP4,TTR,HRNR,ATP6V0C,TMEM30A,CCT2,GDI2,SUF
CAT1,VAT1,CD63,DNAJC5,DDOST,LAMP1,CKAP4,TTR,HRNR,ATP6V0C,TMEM30A,CCT2,GDI2,SUF
,ARF1,SLC4A8,RPL23,CNN3,RTN4,PJA2,GAP43,RPL14,NCS1,CLSTN1,RPS14,ATP1A1,RPS25,CDH
X,DBN1,ENO1,TUBA1B,FLNA,MAP1B,ALDOA,COL6A1,NEFL,NEFM,ATP2B4,TUBB2B,COL5A1,KIF
,ARF1,RPL23,CNN3,RTN4,PJA2,GAP43,RPL14,NCS1,CLSTN1,RPS14,ATP1A1,RPS25,CDH2,RPL12
7,RPS3,ARF1,RPL23,CNN3,RTN4,PJA2,GAP43,RPL14,NCS1,CLSTN1,NLGN2,RPS14,ATP1A1,RPS2
.PA,AP2M1,ANXA2,AP3D1,AP2B1,CTSD,SCARB2,LPCAT1,SLC7A5,SLC3A2,RAB7A,ITM2C,CLTC,GN
.PA,AP2M1,ANXA2,AP3D1,AP2B1,CTSD,SCARB2,LPCAT1,SLC7A5,SLC3A2,RAB7A,ITM2C,CLTC,GN
N1,UBQLN1,ARIH1,HDAC6,GPX1,DNAJB2,ATXN3,BAG5,RANGAP1,XRN2,PINK1,HSPA1A,NUP153,S
\PA,ITGB1,ANXA2,VCL,CDC42BPB,CTNNA1,CNN3,RHOA,CXADR,ATP1A1,WDR1,ATP2A2,PNN,CDH
;Y1,PKD1,HLA-C,NCAM1,AP3D1,ARF1,ITM2B,GOLGA8A,SCARB2,LPCAT1,PJA2,LLGL1,CLSTN1,R
2,APP,CALM1,YWHAG,AP2M1,AP3D1,SLC4A8,NCS1,RPL22,RAB6B,NLGN2,SPTBN2,PTN,CDH2,CT
\-C,NCAM1,AP3D1,ARF1,ITM2B,GOLGA8A,SCARB2,LPCAT1,PJA2,LLGL1,CLSTN1,RAB6B,RTN3,G
J,ATF4,ENC1,SFPQ,DDX39B,HNRNPM,NCOR2,PRPF40A,XPOT,PHB2,SMC3,HNRNPA2B1,ATXN3,KI
3H,PDIA6,ARL6IP1,DDOST,PPIB,SRP9,RPN2,STT3B,B2M,STT3A,HYOU1,SREBF2,OS9,NPLOC4,FKI
:4,PSMD10,PSMC6
3N1,MYH9,SEPTIN2,FSCN1,SPTAN1,MARCKS,VCL,CDC42BPB,CTNNA1,CNN3,TPM4,LLGL1,WDR1
T4,CDC37,CCT8,CCT5,PPP5C,STUB1,HSF1,DNAJB11
;,CDH2,TLN1,CTNNB1,DLG5,CDH11,TBCD,KLHL24,VEGFA,BMPR2,PPP1R9B,CTNND1,FLOT1,CTI
:1,LASP1,SPTBN2,CDH2,MTSS2,COTL1,ACTR3,DSTN,PPP1R9B,CAPN2,HIP1,FLOT1,CALD1,CTTN,
NR,CCT2,GDI2,PYGB,MAPK1,CCT8,TXNDC5,PSMD1,PRDX6,GRN,ACLY,CAP1,IST1,SDCBP,FAF2,P/
'RA,CAPZB,PLCG1,FYN,PTPRS,PFN2,RAB11A,CDC42,SYP,CTNND1,STAT3,BCR,SH3GL1,SYT11,NI
B,EIF2S1,EIF3J
PRPF6,SF3A1,PRPF40A,SNRPD3,EFTUD2,SF3A2,SART1,HTATSF1,LUC7L2,SF3A3,CD2BP2,SART3
IA,GRIPAP1,RAB14,PLEKHB2,LZTR1,SYT11,EHD1,NSG1,RAB11FIP3,SCAMP5,SCAMP1,SCAMP4,H
\RB2,ASTN2,CLTC,DNAJC5,HLA-B,ARCN1,RAB27A,VAMP2,TMED2,KDELR2,KDELR1,B2M,TMED9

C,SLC4A8,CPE,CLTC,DNAJC5,LAMP1,HLA-B,RAB1A,ARCN1,CDK16,TMEM30A,RAB27A,VAMP2,SL
C,PDIA3,AP2M1,AP2B1,SCARB2,KIF5B,RAB7A,CLTC,WLS,LAMP1,HLA-B,APPL1,ATP6V0C,VAMP2,I
HDAC2,TMEM30A,ABCA2,SMARCA4,SMARCE1,SMARCC1,RBBP7,MTA1,CSNK2A1,EP400,ARID1B
B,EIF3J
3CE1,EIF3J
1,RAB11B,OPTN,CLIP3,B2M,RAB11A,ARFGEF2,AMOTL2,GRIPAP1,ULK1,RABEP1,TPP1,RAB14,PI
IRBP,CAPRIN1,G3BP2,CASC3,ATXN2L,UBAP2L,ELAVL1,LSM14A,IGF2BP1,EIF4G1,PUM1,YTHDF3
1,PSAP,CLU,COL18A1,COL4A2,FN1,NCAM1,ANXA2,AGRN,TIMP2,COL11A1,PXDN,CTSD,LAMA5,C
ELR2,KDELR1,TMED9,ACTR1A,AP2A1,MAP6,PI4KA,VPS41,RAB14,PACS1,ADAM10,COPZ1,TGOLN
PRPF6,SF3A1,PRPF40A,SNRPD3,EFTUD2,SF3A2,SART1,HTATSF1,LUC7L2,SF3A3,CD2BP2,SART3
F3J
RTN4,ARL6IP1,HLA-B,RTN1,FKBP8,B2M,SREBF2,RHBDD2,ERGIC1,TRAM1,ERGIC3,DHCR7,CCD
SMARCC1,RBBP7,MTA1,CSNK2A1,EP400,ARID1B,MBD3,YY1,SMARCD1,RSF1,MTA2,ANP32E,RUV
11,ATP1A2,RAC1,ARF4,ARFGEF2,CDC42,PPP1R9B,CTNND1,BCR,FARP1,PTEN,PPP1CC,CTTN,ATI
,SFPQ,DDX39B,HNRNPM,NCOR2,PRPF40A,XPOT,PHB2,SMC3,HNRNPA2B1,ATXN3,KRT8,YY1,CAI
FPQ,WLS,MAP2,MAP1A,MAPK1,HSPB1,SOD1,DYNLL1,KIF5A,HIF1A,SPG7,GABARAPL1,FLOT2,RA
F6,MAT2A,GTF2F1,CBX5,ENC1,RBBP4,CCNI,TADA2B,KLHL9,PFKL,CCND2,PRKAR1A,DDOST,HD
3,H2AZ1,ATRX,BAZ2A,PCGF2,KDM4A,MBD3,CENPB,HMGA1,MACROH2A1,MECP2,TASOR,DDX6,F
ALM3,APP,TMBIM6,CALM1,HSPD1,ATP5F1A,UBB,MATR3,HACD3,RPS3,VAPA,CYB5R3,PAFAH1B1,
ALM3,APP,TMBIM6,CALM1,HSPD1,ATP5F1A,UBB,MATR3,HACD3,RPS3,VAPA,CYB5R3,PAFAH1B1,
RTN4,ARL6IP1,HLA-B,RTN1,FKBP8,B2M,SREBF2,RHBDD2,ERGIC1,TRAM1,DNAJB2,ERGIC3,DHC
11,ATP1A2,RAC1,ARF4,ARFGEF2,CDC42,PPP1R9B,CTNND1,BCR,FARP1,PTEN,PPP1CC,CTTN,ATI
AGED1,KAT6A,DVL3,SEPTIN2,USP11,VCP,AFF4,TRIM28,DDB1,SET,PPP2R1A,CHD3,BASP1,HMGB

RCHF6,ITM2B,RTN4,SLC22A17,CLSTN1,DNAJC5,SLC25A3,ARL6IP1,WLS,LAMP1,HLA-B,TOMM20
CALM3,CALM1,UBB,HLA-C,VAPA,AP2M1,AP2B1,CPE,SCARB2,LPCAT1,RHOA,RAB7A,CLTC,CD63,
TP6V0C,RAB11B,RAPGEF1,FLNB,B2M,STXBP1,HGS,RAB11A,CDC42,ITGB5,VPS26B,RAB14,ITG
),PRPF6,CASC3,SF3A1,PRPF40A,DHX15,SNRPD3,EFTUD2,PRPF19,SF3A2,SART1,EIF4A3,HTATSF1
RCN1,VAMP2,TMED2,KDELR2,KDELR1,B2M,SREBF2,SEC23A,HIP1,AP2A1,SEC16A,LMAN1,LDLR,S
BX5,SPC25,XPO1,PTGES3,CKAP5,SMCHD1,PPP1CB,PARP1,CBX1,H3-3A,TERF2IP,PPP2CA,DYNC1
PRPF6,SF3A1,STRAP,PRPF40A,SNRPD3,EFTUD2,SF3A2,SART1,HTATSF1,LUC7L2,SF3A3,CD2BP2
_16A1,COL3A1
_16A1,COL3A1
,MTA1,RERE,CSNK2A1,EP400,HDAC6,HINT1,MBD3,SAP18,MTA2,KDM3A,ANP32E,RUVBL2,SIN3B,
2,PTEN,CNP,SCRIB,EXOC4,TUBB4A,JAM3,ERBB2,CNTN2,PRKCI,GNAO1,GSN
UBA1B,FLNA,MAP1B,NEFL,NEFM,TUBB2B,KIF1A,CALM3,CALM1,MAPT,MAP9,INA,DNAH8,TPT1,S
PPP1R9B,AP2A1,FARP1,PALM,ANTXR1,ABI2,RDX,ITGAV,IGF2BP1,APBB1,UBE2K,FXR2,DAG1,AC
DCN,CTSB,VCAN,NSG2,TXNDC5,GPC4,HSPG2,TPP1,GPC2,CD74,NSG1,BGN,ATP13A2,PDGFRB,A
MARCC2,SMARCB1,DPF2
LN1,MYO5A,MACF1,MTSS2,RAC1,APPL1,TLN2,NCKAP1,ACTN1,PLCG1,ARF4,CORO1C,PPP1R9B
2A1,COL27A1,COL16A1,COL7A1,COL3A1
AP3D1,PAFAH1B1,MAP4,HNRNPU,KIF5B,SFPQ,PRKAR1A,CTNNB1,WLS,MAP2,CTBP1,MAP1A,MA
LFM1,CLASP2,TAOK2,TRAK1,CYFIP1,DYNLT1,EPHA4
TP6V0D1,ATP5MC3,ATP6V1G1,ATP5MC2,ATP5MG,ATP6V0B,ATP5PO,ATP5F1C,ATP6V0E2,ATP5F

P1A2,ACTN1,YWHAH,FLCN,FLOT1,SLC2A1,FLOT2,SCRIB,ANK2,OBSL1,JAM3,NECTIN1,AFDN,NE
/I170A,RTN4,KPNB1,CBX5,CCNI,LRPPRC,XPO1,SCRN1,EIF5A,CCND2,DUX4,IPO5,PARP1,LMNB2,
HNRNPU,CBX5,ATF4,CHD4,JUND,RBBP4,TLE5,SFPQ,H1-0,JUN,CTNNB1,YWHAB,CTBP1,HDAC2,P

NAP1,FLOT1,PHB2,SV2A,BRSK1,PNISR,FLOT2,SYT11,RIMS3,NPTN,NECTIN1,C1QBP,RAB3A,UNC
:7L2,SNRPB
TAD2B
:7L2,SNRPB
TAD2B
RCHF6,ITM2B,RTN4,SLC22A17,CLSTN1,SLC25A3,ARL6IP1,WLS,LAMP1,HLA-B,TOMM20,VAMP2,
1,KLHL9,MYH10,GNAI3,PKN1,PITPNM1,CDC42,CTNND1,FLCN,PPP1CC,SEPTIN7,SLC2A1,ERH,VP
P1R10,TOX4,PPP1CA,PPP2R5D,WDR82,PPP1R12A,PPP3CA,CTDNEP1,PPP2R5B,PPP2R2A,PPP4R3
P1R10,TOX4,PPP1CA,PPP2R5D,WDR82,PPP1R12A,PPP3CA,CTDNEP1,PPP2R5B,PPP2R2A,PPP4R3
RAN,NUP153,KPNA6,NUP98,NUP205,NUP133,KPNA3,DYNLT1,NUP50,NUP58,VPS28,VPS37B
RAN,NUP153,KPNA6,NUP98,NUP205,NUP133,KPNA3,DYNLT1,NUP50,NUP58,VPS28,VPS37B
LDHB,ITGB1,ANXA2,VCL,CTNNA1,CTSD,HK1,CXADR,ATP1A1,CDH2,SULF1,PRKAR1A,GJA1,CTNNE
LDHB,ITGB1,ANXA2,VCL,CTNNA1,CTSD,HK1,CXADR,ATP1A1,CDH2,SULF1,PRKAR1A,GJA1,CTNNE
SPA1B,CLU,SOX4,YWHAE,RACK1,BSG,ALDH6A1,ATP5F1B,CALM3,TMBIM6,GSTP1,CALM1,HSPD
,PITPNM1,RAB11A,MASTL,PPP1CC,SEPTIN7,SEPTIN5,RDX,TUBGCP6,RAB11FIP3,MAEA,PLCD3,S
CBX1,H3-3A,TERF2IP,PRKDC,KDM1A,POLR2B,SSB,TNKS2,TP53BP1,ATRX,UPF1,HNRNPA2B1,PPF
B2,ASTN2,RAB7A,PRKAR1A,GJA1,MYO5A,CD63,ANXA6,LAMP1,RHOB,LAPTM4B,RAB27A,SQSTM
L6IP1,CKAP4,GLUL,SRP9,NUCB1,RPN2,RPL27,PLOD1,RPS28,RAB14,CCDC47,SRPRA,RPS26,RPS


P1,TMED2,KDELR1,CSNK1D,TMED9,MGAT4B,RAB2A,RER1,YKT6,RGMB,LMAN1,ERGIC1,RAB1B,
1,CHD3,PAFAH1B1,HNRNPU,ATF4,KIF5B,CHD4,CRMP1,DDX3X,RPS7,PRKAR1A,CTNNB1,CKAP5,S
OD1,DYNLL1,KIF5A,HIF1A,SPG7,RANGAP1,TRAK1,KIF3A,ARL8A,OPA1,MAPK8IP3,AP3M1
:25A5,LRRC59,SOD2,POLDIP2,CLPX,SHMT2,VDAC2,HADHB,ELAC2
:25A5,LRRC59,SOD2,POLDIP2,CLPX,SHMT2,VDAC2,HADHB,ELAC2
RP10,LDLR,LRP3,SH3BP4,AP2A2,SORT1,CLTA,HIP1R,EPN1,TFRC,RAB35,SLC18A3,NECAP1
CAND1,PHC2,TRAF7,ARIH1,OS9,KLHL42,FBXO11,CUL3,FBXO21,RHBDD2,UBE2D2,DCUN1D2,LZ
SF3A2,SART1,SF3A3,LSM4,SNRPD2,SNRPB,PRPF31,ZMAT2,SNU13
PXD2A,SH3PXD2B,PTPN12,GSN
APL1,FLOT2,HIP1R,DLG4,MAP2K4

A3,NSG2,HGS,RAB11A,HDAC6,CHMP4B,CD74,NSG1,CHMP1A,ATP13A2,TMEM9,BACE1,CHMP5
A1,PSAP,CLU,COL18A1,COL4A2,FN1,NCAM1,ANXA2,AGRN,TIMP2,COL11A1,PXDN,CTSD,LAMA5,C
OX4,ACTN1,FLNB,TPM3,PALLD,SEPTIN7,MYL12B,PLS3,PDLIM1,DAAM1,ZYX,MYL12A,SEPTIN9,M
NTNAP1,HIP1,LRRC4B,KCTD12,GPC4,FLOT2,NCSTN,SCRIB,SYT11,FXYD6,APBB1,RIMS3,CLTA,G
A1,PSAP,CLU,COL18A1,COL4A2,FN1,NCAM1,ANXA2,AGRN,TIMP2,COL11A1,PXDN,CTSD,LAMA5,C
AP1,PCGF2,YY1,PCGF3,CSNK2B,RYBP,CBX4,KDM2B
2,NOX4,ACTN1,FLNB,TPM3,PALLD,SEPTIN7,MYL12B,PDLIM1,DAAM1,ZYX,MYL12A,SEPTIN9,CDC
F1,SETD5,NSMF,SP1,TRIM24,ID2
LL1,FLOT1,HSPA1A,COPS6,GPS1,COPS8
SF3A2,SART1,SF3A3,LSM4,SNRPD2,SNRPB,PRPF31,ZMAT2,SNU13

2A1,PI4KA,COPZ1,SPPL2B,CLTA,COPE,ZDHHC17,COPB1,SLC18A3

RNR2L2,MED18,ACTG1,EEF2,DPYSL3,TUBA1A,CFL1,XRCC2,MAVS,HTT,HSP90AB1,ZNF124,HSP9
2,ACTG1,EEF2,DPYSL3,TUBA1A,CFL1,XRCC2,MAVS,HTT,HSP90AB1,HSP90AA1,ACTR2,VIM,SPAR
TG1,EEF2,DPYSL3,TUBA1A,CFL1,XRCC2,MAVS,HTT,HSP90AB1,ZNF124,HSP90AA1,ACTR2,VIM,S
H1,UBC,KIDINS220,TMSB4X,NUMA1,DCX,EIF4G2,RPL3,RPS6,DBN1,RPL4,ENO1,PIK3R2,FLNA,M
TG1,EEF2,DPYSL3,TUBA1A,CFL1,XRCC2,MAVS,HTT,HSP90AB1,ZNF124,HSP90AA1,ACTR2,VIM,S
F2,DPYSL3,TUBA1A,CFL1,XRCC2,MAVS,HTT,HSP90AB1,ZNF124,HSP90AA1,ACTR2,VIM,SPARC,T
TG1,EEF2,DPYSL3,TUBA1A,CFL1,XRCC2,MAVS,HTT,HSP90AB1,ZNF124,HSP90AA1,ACTR2,VIM,S
RPL3,RPL4,ENO1,FLNA,NCL,ALDOA,PKM,GANAB,COL6A1,FTL,HSP90B1,YBX1,UBA1,IGFBP2,CA
ACTR2,VIM,SPARC,TUBB,FTH1,UBC,TMSB4X,NUMA1,RPL3,RPS6,RPL4,FLNA,NCL,RPL7A,ALDOA,
ACTR2,VIM,SPARC,TUBB,FTH1,UBC,TMSB4X,NUMA1,RPL3,RPS6,RPL4,FLNA,NCL,RPL7A,ALDOA,
ACTR2,VIM,SPARC,TUBB,FTH1,UBC,TMSB4X,NUMA1,RPL3,RPS6,RPL4,FLNA,NCL,RPL7A,ALDOA,
RPL3,RPL4,ENO1,FLNA,NCL,ALDOA,PKM,GANAB,COL6A1,FTL,HSP90B1,YBX1,UBA1,IGFBP2,CA
RPL3,RPL4,ENO1,FLNA,NCL,ALDOA,PKM,GANAB,COL6A1,FTL,HSP90B1,YBX1,UBA1,IGFBP2,CA
RPL3,RPL4,ENO1,FLNA,NCL,ALDOA,PKM,GANAB,COL6A1,FTL,HSP90B1,YBX1,UBA1,IGFBP2,CA
0AA1,ACTR2,VIM,TUBB,FTH1,KIDINS220,NUMA1,DCX,EIF4G2,RPL3,RPS6,RPL4,ENO1,PIK3R2,F
M,SPARC,TUBB,FTH1,UBC,KIDINS220,TMSB4X,NUMA1,RPL3,RPL4,ENO1,FLNA,NCL,SLC9A7,AL
,RPS6,RPL4,FLNA,NCL,RPL7A,PRPF8,FBXL4,YBX1,UBA1,HSPA1B,H3-3B,PTMA,DDX5,SOX4,RPL1
HSPA1B,H3-3B,PTMA,DDX5,SOX4,KAT6A,DDX17,GPI,RACK1,ALDH6A1,RPS8,SEPTIN2,RPL5,SMG1
HSP90AB1,ZNF124,HSP90AA1,ACTR2,VIM,SPARC,TUBB,FTH1,UBC,TMSB4X,NUMA1,RPL3,RPS6,
PL7,P4HB,RPL5,MARCKS,NPM1,YWHAG,RPL37A,YWHAQ,HACD3,MRC2,RPL19,PDIA3,RPS3,RPS
PL7,P4HB,RPL5,MARCKS,NPM1,YWHAG,RPL37A,YWHAQ,HACD3,MRC2,RPL19,PDIA3,RPS3,RPS
IDINS220,TMSB4X,NUMA1,RPS6,RPL4,FLNA,SLC9A7,ALDOA,PKM,GANAB,COL6A1,FTL,HSP90B1
RPS8,RPL7,RPL5,NPM1,SNRNP200,EIF3C,RPL37A,EIF1,RPL19,RPL15,SF3B1,RPS3,RPS4X,RPS3A
RPL8,UBA1,ATP2B4,CALR,HSPA1B,TUBB2B,RPLP0,CANX,CLU,RPL13A,RPL6,YWHAE,KIF1A,BSG,
RPL23,RPL29,RPSA,RPS24,RPS9,RPL24,RPL17,RPL14,RPL22,RPL9,RPS14,RPS20,RPL26,RPL31,
E,BSG,RPS8,MYH9,RPL7,P4HB,APP,RPL5,FSCN1,MARCKS,NPM1,YWHAG,RPL37A,YWHAQ,HACD
UBB,TMSB4X,NUMA1,DCX,RPL3,RPS6,DBN1,RPL4,ENO1,TUBA1B,FLNA,MAP1B,NCL,RPL7A,AL
UBB,TMSB4X,NUMA1,DCX,RPL3,RPS6,DBN1,RPL4,ENO1,TUBA1B,FLNA,MAP1B,NCL,RPL7A,AL
X,SLC9A7,ALDOA,PKM,GANAB,FTL,HSP90B1,UBA1,IGFBP2,CALR,ABHD2,HSPA1B,PSAP,CANX,CL
X,SLC9A7,ALDOA,PKM,GANAB,FTL,HSP90B1,UBA1,IGFBP2,CALR,ABHD2,HSPA1B,PSAP,CANX,CL
M3,GNB1,SHISA9,MYH9,RPL7,SEPTIN2,APP,RPL5,CALM1,EIF4B,VCP,YWHAG,MAPT,YWHAQ,INA,
SPARC,TUBB,FTH1,UBC,NUMA1,RPL3,RPL4,ENO1,FLNA,NCL,ALDOA,PKM,GANAB,COL6A1,FTL,
9,RPL26,RPL31,RPL12,RPL30,RPL35,RPL35A,RPLP2,RPL36,RPL23A,RPL27,RPL36A,RPL38,RPL
NAH8,SF3B1,CHD3,MORF4L1,MARCHF6,CYB5R3,PAFAH1B1,SF3B2,MAT2A,GTF2F1,HNRNPU,C
ANXA6,DNAJC5,YWHAB,SND1,LAMP1,RAB1A,CALU,RAC1,PPIB,CTSB,RAB27A,RAB5B,CCT4,PRDX
ANXA6,DNAJC5,YWHAB,SND1,LAMP1,RAB1A,CALU,RAC1,PPIB,CTSB,RAB27A,RAB5B,CCT4,PRDX
RPS17,RPS16,RPL23,RPL29,RPSA,RPS24,RPS9,RPL24,RPL17,RPL14,RPL22,RPL9,RPS14,RPS20,
P0,CANX,CLU,FKBP14,SEC62,RPL6,RPL13,BSG,RPS8,P4HB,APP,RPL5,TMBIM6,PLD3,HSPD1,CO
28,NONO,SF3B1,RPS3,CHD3,BASP1,MORF4L1,SF3B2,GTF2F1,HNRNPU,SMARCA5,XRCC5,KPNE
3,RPLP1,RPL10,RPS17,LARP1,RPS16,RPL23,RPL29,RPSA,RPS24,RPS9,RPL24,RPL17,RPL14,RP
,IK,NGRN,TARDBP,CKAP4,BCLAF1,LUC7L3,CDKN1A,DDX39B,SRSF5,U2AF2,RPN2,SUGP2,SQST
DI1,FSCN1,CALM1,MAPT,MAP9,BASP1,UCHL1,TIMP2,AP3D1,PAFAH1B1,MAP4,SLC4A8,LLGL1,G
9B,SRSF5,U2AF2,PABPN1,ALKBH5,DDX42,SNRNP70,PRPF6,SRSF9,CCNL2,SRSF1,FTO,SON,BN
E,KIF1A,GPI,RACK1,BSG,SPTBN1,CALM3,GNB1,SHISA9,STMN2,SEPTIN2,APP,GDI1,FSCN1,CAL
,CKB,MAPT,PPP2R1A,ITGB1,UCHL1,TIMP2,PAFAH1B1,AKAP12,SLC4A8,HNRNPU,CNN3,RTN4,GA

LU,SEC62,KIF1A,GNAS,SLC2A3,GPI,BSG,ATP5F1B,CALM3,GNB1,SEPTIN2,APP,TMBIM6,CALM1,F
OTL1,CTSB,SRP14,MAPK1,PSMD3,CAND1,ILF2,CCT8,IDH1,PDAP1,CST3,ACLY,XRCC6,IMPDH2,AC
P2,PAFAH1B1,AKAP12,SLC4A8,CNN3,RTN4,GAP43,ENC1,CXADR,CRMP1,CCT7,ASTN2,SPTBN2,S
ARF1,ITM2B,AP2B1,CTSD,SCARB2,LLGL1,RHOA,CLSTN1,ATP1A1,ASTN2,NISCH,RAB7A,MMP14,
B4,TUBB2B,COL5A1,COL4A1,RPLP0,RPL6,KIF1A,SPTBN1,CALM3,MYH9,SEPTIN2,CALM1,CSDE1
EFL,NEFM,ATP2B4,ABHD2,RPLP0,CANX,CLU,YWHAE,KIF1A,GNAS,SLC2A3,GPI,RACK1,BSG,SPTE
HLA-C,NCAM1,AGRN,AP3D1,ARF1,ITM2B,GOLGA8A,CPE,SCARB2,LPCAT1,PJA2,C1ORF43,LLGL1
,FAU,RPS13,UBA52,RPS2,RPS27A,RPS26,RPS21,RPS29
ACTR2,VIM,SPARC,TUBB,FTH1,UBC,TMSB4X,NUMA1,RPL3,RPL4,ENO1,FLNA,NCL,ALDOA,MTRN
EFL,NEFM,ATP2B4,ABHD2,RPLP0,CANX,CLU,YWHAE,KIF1A,GNAS,GPI,RACK1,BSG,SPTBN1,CALN
,DDX3X,MIF,LAMP1,COTL1,RAC1,ATP6V0C,CTSB,SRP14,MAPK1,PSMD3,CAND1,ILF2,CCT8,IDH1
4A8,HNRNPU,CNN3,GAP43,NCS1,RHOA,CLSTN1,ATF4,KIF5B,SFPQ,NDFIP1,WLS,SEPTIN11,LAMP
B1,RPL23,CNN3,RTN4,PJA2,GAP43,RPL14,NCS1,RHOA,CLSTN1,NLGN2,RPS14,ATP1A1,SPTBN2,
4A8,HNRNPU,CNN3,GAP43,NCS1,RHOA,CLSTN1,ATF4,KIF5B,SFPQ,NDFIP1,WLS,SEPTIN11,LAMP
2,TIMP2,VCL,CYB5R3,PGAM1,CTSD,XRCC5,KPNB1,PSMD2,VAT1,PFKL,ACTN4,DDX3X,MAGED2,M
2,TIMP2,VCL,CYB5R3,PGAM1,CTSD,XRCC5,KPNB1,PSMD2,VAT1,PFKL,ACTN4,DDX3X,MAGED2,M
2,TIMP2,VCL,CYB5R3,PGAM1,CTSD,XRCC5,KPNB1,PSMD2,VAT1,PFKL,ACTN4,DDX3X,MAGED2,M
PL39,FUS,EIF3H,ABCF1,MSI1,EIF4G1,RPS26,FXR2,RPS21,FXR1,RPS29,PSMA1,PSMA6,MSI2,RPL
I,SEPTIN2,APP,CALM1,PLD3,UBB,CHSY1,PKD1,HLA-C,PDIA3,VAPA,AP2M1,SEC61A1,ITGB1,NCAM
P3D1,CYB5R3,AP2B1,CTSD,SCARB2,LPCAT1,SLC22A17,SLC7A5,VAT1,SLC3A2,GPC3,PIP4K2B,RA
PSMD7,ADRM1,PSME3,PSMC5,PSMC2,PSMC4,PSMF1,UBE3C,UBQLN4,PSMB5,PSMA1,PSMD10,F
R,CRMP1,MYO5A,MAP2,L1CAM,KLC1,ACAP3,ORAI2,DPYSL2,STMN3,PTPRS,MAPK8IP1,PPP1R9B,
S3,CHD3,RPS3A,RPL18,RPL11,MIDN,SMARCA5,RPL23,RPS9,XRCC5,CIPC,CBX5,ENC1,RPS14,RP
SNRPN,HNRNPA3,RBMX,SNRNP70,PRPF6,SRSF1,CASC3,HNRNPM,SF3A1,PRPF40A,ADAR,DHX15,
A,SLC2A3,GPI,BSG,CALM3,MYH9,SEPTIN2,APP,SPTAN1,GSTP1,CALM1,HSPD1,IGF2,VCP,FN1,HLA
SMD13,USP14,TRRAP,GPAA1,PSMD7,ADRM1,PSME3,PSMC5,PSMC2,PSMC4,PSMF1,UBE3C,UBQ
ADRM1,PSME3,PSMC5,PSMC2,PSMC4,PSMF1,UBE3C,UBQLN4,PSMB5,PSMA1,PSMD10,PSMA6,F
R,CRMP1,MYO5A,MAP2,L1CAM,KLC1,ACAP3,ORAI2,DPYSL2,STMN3,PTPRS,MAPK8IP1,PPP1R9B,
,CYB5R3,AP2B1,CTSD,SCARB2,LPCAT1,SLC7A5,VAT1,SLC3A2,GPC3,RAB7A,ITM2C,CLTC,GNB2,
,CYB5R3,AP2B1,CTSD,SCARB2,LPCAT1,SLC7A5,VAT1,SLC3A2,GPC3,RAB7A,ITM2C,CLTC,GNB2,
OA,NEFL,NEFM,HSPA1B,TUBB2B,CLU,YWHAE,KIF1A,SPTBN1,CALM3,MYH9,SEPTIN2,P4HB,APP,
NCKAP1,ARPC2,CAPZB,PLCG1,CORO1C,PPP1R9B,CTNND1,PALLD,FLOT1,SLC39A6,EVL,CTTN,B
F1,DNAJA1,ARF1,PAFAH1B1,EIF4H,MSN,NCS1,KIF5B,ATP2A2,LRPPRC,PTN,CCN2,ACTN4,ITM2C,
3,GPI,BSG,MYH9,APP,SPTAN1,GSTP1,HSPD1,IGF2,VCP,FN1,HLA-C,VAPA,HMGB1,TUBB4B,ANXA2
A-C,PDIA3,SERINC1,VAPA,SEC61A1,UCHL1,NCAM1,SARAF,MARCHF6,AP3D1,SBF1,DNAJA1,ARF1,
AH1B1,SLC4A8,GAP43,NCS1,KIF5B,CXADR,CRMP1,MYO5A,MAP2,L1CAM,GLUL,KLC1,ACAP3,OF
RKAR1A,SEPTIN11,CTBP1,RAC1,CDH11,CLSTN3,ARPC2,DYNLL2,ACTN1,PLCG1,ARF4,FYN,STXBI
KS,CALM1,NPM1,MAPT,MAP9,PPP2R1A,DNAH8,RPS3,CHD3,VAPA,TPT1,STMN1,TUBB4B,DNAJA1,
3,CTNNA1,RHOA,ATF4,TPM1,CDH2,DDX3X,TLN1,MYO5A,CTNNB1,MACF1,MTSS2,WLS,MYH10,CA
R1,LASP1,PFN1,ASTN2,SPTBN2,CDH2,CCN2,CTNNB1,MTSS2,NDFIP1,MYH10,SEPTIN11,RHOB,SE
4A8,CNN3,RTN4,GAP43,ENC1,CRMP1,ASTN2,SPTBN2,SLC3A2,LAMP1,ATP1A2,MAP2,DNER,L1CA
22,RPL9,RPL26,RPL31,RPL12,RPL30,RPL35,RPL35A,RPLP2,RPL36,RPL23A,RPL27,RPL36A,RPL3
RIN1,EID1,SQSTM1,HNRNPL,HNRNPA3,TOP1,G3BP2,BTBD2,NYNRIN,TDRD5,CASC3,RPL28,DHX3
,RPS11,RPS28,FAU,RPS13,UBA52,RPS2,RPS27A,RPS26,RPS21,MRPS6,RPS29,MRPS26,MRPS27
35B,CNTNAP2,RAPGEF1,MVB12B,MAPK1,NSG2,CLIP3,MAP2K2,PTP4A2,B2M,GPR107,IFITM3,PT

HLA-B,RAC1,APPL1,ATP6V0C,LAPTM4B,RAB27A,TM9SF2,ABCA3,VPS35,RAB5B,RAB11B,ABCA2,[illegible]
SBF1,DNAJA1,CYB5R3,TMEM170A,NFE2L1,FADS2,SCARB2,RTN4,LPCAT1,PJA2,KPNB1,RHOA,CL[illegible]
,HSP90AA1,ACTR2,VIM,SPARC,TUBB,FTH1,UBC,KIDINS220,NUMA1,EIF4G2,RPS6,DBN1,RPL4,EN[illegible]
SBF1,DNAJA1,CYB5R3,TMEM170A,NFE2L1,FADS2,SCARB2,RTN4,LPCAT1,PJA2,RHOA,CLSTN1,T[illegible]
GFBP3,COL1A2,COL5A2,PRKCSH,CDH2,PDIA6,GPC3,RCN1,COL6A2,GOLM1,APLP2,CKAP4,CALU[illegible]
A2B1,SNRPD3,RALY,EFTUD2,PRPF4B,PRPF19,SF3A2,SART1,DHX38,EIF4A3,MTREX,SF3A3,SYNC[illegible]
C,CLTC,GNB2,CD63,ANXA6,DNAJC5,DDOST,YWHAB,LAMP1,CKAP4,ATP6V0C,TMEM30A,LAPTM4[illegible]
2B,KLHL9,PRKAR1A,SESN3,SKP1,KLHL24,MORF4L2,UBE4A,YEATS2,RAD23B,PSMB2,TAF7,PPP2[illegible]
SBF1,DNAJA1,CYB5R3,TMEM170A,NFE2L1,FADS2,SCARB2,RTN4,LPCAT1,PJA2,RHOA,CLSTN1,T[illegible]
CAN,PYGB,MAPK1,NSG2,CCT8,TXNDC5,PSMD1,GPC4,PRDX6,GRN,ACLY,HSPG2,TPP1,GPC2,CAP[illegible]
RIN1,EID1,SQSTM1,TOP1,G3BP2,BTBD2,NYNRIN,TDRD5,CASC3,RPL28,ATXN2L,MEX3A,CDC42,E[illegible]
S25,CDH2,RPL12,RPL30,RPS19,CTNNB1,SEPTIN11,RPS27,DLG5,HNRNPA3,CLSTN3,RPL38,MAPK[illegible]
PPP1R9B,CAPN2,HIP1,FLOT1,CALD1,CTTN,SLC2A1,RDX,FLOT2,CAPZA1,CLASP2,CAP1,RIMS3,HI[illegible]

TM2B,AP2B1,CPE,CTSD,SCARB2,LPCAT1,LLGL1,RHOA,ATP2A2,RAB7A,CLTC,GJA1,CD63,ANXA6
,LRPPRC,CLTC,MACF1,CKAP5,MAP2,CCT6A,MAP1A,CCT2,TBCD,HSPH1,TCP1,CAMSAP1,KLC1,C[illegible]

L4A1,KIF1A,SPTBN1,CALM3,MYH9,CALM1,COL4A2,MAPT,MAP9,INA,RPL15,DNAH8,TPT1,STMN1,
RF4,PYGB,MAPK1,CCT8,TXNDC5,PSMD1,PRDX6,GRN,ACLY,NCSTN,RAB5C,CAP1,IST1,SDCBP,ND
RF4,PYGB,MAPK1,CCT8,TXNDC5,PSMD1,PRDX6,GRN,ACLY,NCSTN,RAB5C,CAP1,IST1,SDCBP,ND
H2,RPL12,RPL30,RPS19,CTNNB1,RPS27,DLG5,HNRNPA3,CLSTN3,RPL38,MAPK1,DYNLL2,CAPZB,
1A,SPTBN1,CALM3,MYH9,CALM1,MAPT,MAP9,INA,RPL15,DNAH8,TPT1,STMN1,TUBB4B,COL11A[illegible]
2,RPL30,RPS19,CTNNB1,RPS27,DLG5,HNRNPA3,CLSTN3,RPL38,MAPK1,DYNLL2,CAPZB,FYN,PTF[illegible]
25,CDH2,RPL12,RPL30,RPS19,CTNNB1,SEPTIN11,RPS27,DLG5,HNRNPA3,CLSTN3,RPL38,MAPK1
IB2,CD63,ANXA6,DNAJC5,DDOST,LAMP1,CKAP4,ATP6V0C,TMEM30A,LAPTM4B,EEF1A2,GNAI3,A[illegible]
IB2,CD63,ANXA6,DNAJC5,DDOST,LAMP1,CKAP4,ATP6V0C,TMEM30A,LAPTM4B,EEF1A2,GNAI3,A[illegible]
STUB1,NUP98,PSMC5,HERPUD1,PSMC4,CABIN1,ZDHHC17,GYS1,FBXO7,PRNP,PSMC6
6,CDH2,TLN1,GJA1,CTNNB1,CADM4,ATP1A2,DLG5,CNN2,CDH11,TBCD,KLHL24,ACTR3,PTK7,C[illegible]
AB6B,RTN3,GLG1,SULF1,CLTC,CHPF,GJA1,B4GALNT4,NDFIP1,ST3GAL2,WLS,HLA-B,RAB1A,AR[illegible]
NNB1,DNAJC5,LAMP1,SEPTIN3,CTBP1,CDK16,GLUL,VAMP2,RAB11B,DPYSL2,STXBP1,TMED9,P[illegible]
GLG1,CLTC,CHPF,GJA1,B4GALNT4,NDFIP1,ST3GAL2,WLS,HLA-B,RAB1A,ARCN1,RAC1,HS6ST1,T[illegible]
RT8,YY1,CAD,CHMP1A,DAG1,RUVBL2,MAEA,CSNK2B,AKAP8L,NUP153,PSIP1,NUP98,ZNF703,NS[illegible]
BP1A,RHBDD2,SRPRA,AMFR,TAPBP,EMC1,GPAA1,HLA-A,FAF2,NBAS,DAD1,EXT2,FAM8A1,HSD17

,LASP1,HNRNPC,SPTBN2,TPM1,CDH2,ACTN4,H1-0,MYO5A,MACF1,MTSS2,MYL6,HNRNPK,MYH1

NND2,CYTH2,ABI2,RDX,FLOT2,SCRIB,ADAM10,SDCBP,DAG1,NIBAN2,NOTCH1,EFNB2,PARK7,PD
SLC2A1,RDX,FLOT2,CAPZA1,CAP1,HIP1R,MAEA,CAPZA2,GMFB,MYADM,NF2,GSN
A2G4,MAN2B1,DPP7,ARSB,TRAPPC1,GLB1,CTSA,ARSA,TOLLIP
PTN,ADCY1,RNF216,SHANK1,PPP3CA,PRKCI,EPHA4,ABR,CASK

,LSM4,SNRPD2,SNRPB,PRPF31,ZMAT2,SNU13
HLA-A,RAP2A,NDRG1,CLCN4,ABHD17A,SORCS2,RAB35,MICALL1,ARF6,FZD7,STX12
),GPR107,SREBF2,SEC23A,ACTR1A,HIP1,AP2A1,SEC16A,LMAN1,ERGIC1,VPS41,RAB14,LDLR,PA[illegible]

JRF4,RAB11B,TMED2,KDELR2,KDELR1,B2M,TMED9,PTPRS,SREBF2,VGF,RAB11A,SEC23A,SYP,V
HSPH1,RAB5B,RAB11B,RAPGEF1,FLNB,B2M,STXBP1,HYOU1,HGS,RAB11A,CDC42,AP2A1,UBA52
,ABCC9,ATP1B2,MBD3,YY1,SMARCD1,RSF1,MTA2,ANP32E,RUVBL2,TRRAP,ATP1B3,ASH1L,SS18

LEKHB2,LZTR1,SYT11,EHD1,NSG1,RAB11FIP3,ANK2,RNF11,SCAMP5,SCAMP1,SCAMP4,HLA-A,R
,RBM4,TIA1,PABPC4,DDX6,DHX36,MCRIP1,EIF2S1,DDX19A
COL1A2,COL5A2,PTN,CDH2,GPC3,SULF1,CCN2,APLP1,COL6A2,ANXA6,LGALS3BP,HRNR,GPC1,T
N2,SORT1,SPPL2B,CLTA,COPE,CCDC88A,BACE1,ZDHHC17,COPB1,FURIN,SLC18A3
,LSM4,SNRPD2,SNRPB,PRPF31,HEXIM1,ZMAT2,SNU13

C47,CD74,SCD,ELOVL5,AMFR,TECR,TAPBP,SPPL2B,KTN1,ESYT1,EMC1,GPAA1,BSCL2,HLA-A,TM
VBL2,TRRAP,ASH1L,SS18L1,PBRM1,GATAD2B,SMARCC2,SMARCB1,DPF2
P6AP2,FUS,PALM,STRN4,ABI2,ADAM10,IGF2BP1,SYT11,PRKACA,APBB1,TANC2,NTRK2,DIP2A,FX
D,CHMP1A,RUVBL2,MAEA,CSNK2B,AKAP8L,ZNF703,NSMF,LMNA,PSMA6,CENPF,SCAF8,ZNF326,
NGAP1,TRAK1,HIP1R,KIF3A,DLG4,ARL8A,MAP2K4,OPA1,MAPK8IP3,AP3M1
AC2,SKP1,CDKN1A,RPN2,KLHL24,MORF4L2,UBE4A,YEATS2,KDM6B,TAF7,CCND1,CAND1,POLR2
PSIP1,HSF1,RNF20,DNMT1,HMGA2,RNF40
,TMEM170A,RTN4,KPNB1,HK1,PTCD3,CBX5,VAT1,MCL1,CCNI,LRPPRC,XPO1,SCRN1,EIF5A,CCN
,TMEM170A,RTN4,KPNB1,HK1,PTCD3,CBX5,VAT1,MCL1,CCNI,LRPPRC,XPO1,SCRN1,EIF5A,CCN
CR7,CCDC47,CD74,SCD,ELOVL5,AMFR,TECR,TAPBP,SPPL2B,KTN1,ESYT1,EMC1,GPAA1,BSCL2,H
P6AP2,FUS,PALM,STRN4,ABI2,ADAM10,IGF2BP1,SYT11,PRKACA,APBB1,TANC2,NTRK2,DIP2A,FX
1,POLR2A,MORF4L1,PAFAH1B1,HNRNPU,NFE2L1,SMARCA5,XRCC5,CBX5,ATF4,ENC1,CHD4,JUN

),VAMP2,RTN1,RAB11B,FKBP8,B2M,PTPRS,GHITM,SREBF2,SYP,RER1,SV2A,B4GAT1,RHBDD2,S
,DNAJC5,APLP2,DDOST,WLS,LAMP1,CKAP4,HLA-B,RAB1A,ARCN1,RAC1,ATP6V0C,TMEM30A,LA
AV,SYT11,ATP6V0D1,MTOR,AMBRA1,TAPBP,SNX3,HLA-A,ATP6V0B,ZYX,ZDHHC5,ATP6V0E2,CDC4
,LUC7L2,SF3A3,LSM4,SNRPD2,SNRPB,PRPF31,XAB2,ZMAT2,PLRG1,SNU13
SYT11,COPZ1,AP2A2,TGOLN2,AP1M1,CD74,USO1,SCAP,CLTA,HIP1R,COPE,HLA-A,EPN1,MCFD2,
LI2,PRKDC,KDM1A,POLR2B,SSB,TNKS2,KDM4B,DYNLL1,TP53BP1,DCTN1,BAZ1B,PPP2CB,ATR
,SART3,LSM4,SNRPD2,SNRPB,PRPF31,HEXIM1,ZMAT2,SNU13

,TRRAP,RCOR2,SUDS3,GATAD2B,SIN3A

STMN1,TUBB4B,PAFAH1B1,MAP4,HNRNPU,TPM4,KIF5B,CCT7,PNN,NES,LRPPRC,TPM1,CLTC,MY
TA2,FXR1,CYFIP1,UBE2Q1,RUFY3,FGF13,ARF6,SRCIN1,EPHA4,NF2
SAH1,MAN2B1,ARSB,PPT1,GLB1,CTSA,ARSA

,PALLD,CTTN,CLASP2,WASF2,ITGAV,AKT2,HIP1R,CYFIP1,ACAP2,NME2,PIP5K1C,RAB5A,CLCN3,

PK1,GABARAP,HSPB1,SOD1,ARFGEF2,DYNLL1,CTNND1,DCTN1,KIF5A,ACTR1A,PRKAR2A,SEPTI

F1E,ATP6V0A1,ATP6V1F,ATP6V1D,ATP5PD

CTIN2,FGF13,OBSCN,GJC1
JPT1,UBXN4,CCND1,IPO7,BNIP3L,CASC3,TNKS2,AKIRIN1,RBM15B,DCTN1,XPOT,MTA1,GPX4,TR
ARP1,PTOV1,SKI,HDGF,TAF7,CCND1,PBX2,PRKDC,KDM1A,NCOR2,ARNT2,RBBP7,STAT3,BCL9L,

13A,CTBP2,PPFIA3,UNC13B

RTN1,FKBP8,B2M,PTPRS,GHITM,SREBF2,SYP,RER1,SV2A,B4GAT1,RHBDD2,SPG7,ERGIC1,TRAN
S4A,RDX,IST1,CHMP4B,EXOC4,TTC28,MICAL1,RAB11FIP3,EXOC5,RAN,WIZ,RAP2A,MICAL3,EXO(
B,PPP2R5C
B,PPP2R5C

31,ATP1A2,GPC1,GNAI3,PPP2CA,MAPK1,CLIP3,CORO1C,FYN,BMPR2,VDAC1,CAPN2,KIRREL1,PF
31,ATP1A2,GPC1,GNAI3,PPP2CA,MAPK1,CLIP3,CORO1C,FYN,BMPR2,VDAC1,CAPN2,KIRREL1,PF
1,ATP5F1A,UBB,YWHAG,MAPT,YWHAQ,PPP2R1A,GTF3C4,RPS3,XAF1,DNAJA1,CYB5R3,C1ORF43
EPTIN9,PDXP,ARF6,NF2
P1CC,TNKS,APEX1,PPP1R10,XRCC6,MACROH2A2,TOX4,MACROH2A1,SUN2,PPP1CA,WRNIP1,H2A
1,ABCA3,VPS35,MVB12B,MAPK1,LAPTM4A,DYNC1LI2,NSG2,MAP2K2,SDF4,IFITM3,HGS,RAB11A
S21,HM13,RPS29,ARSB

GALNT1,DICER1,ERGIC3,IST1,TAPBP,LMAN2,MCFD2,COL7A1,DDHD2,SEC23IP,COPB1,PLPP3,PII
KP1,TBCD,GNAI3,TCP1,CCT4,SKI,MAPRE1,RABL6,DYNC1LI2,DYNLL2,CCT8,TNKS2,CSNK1D,RAE

TR1,PCGF2,FBXW5,ZER1,FBXL19,AMFR,USP47,UBE2L3,RNF11,WDTC1,FZR1,UBE4B,TNFAIP1,CI

COL1A2,COL5A2,PTN,GLG1,CDH2,GPC3,SULF1,CCN2,MMP14,APLP1,COL6A2,ANXA6,LGALS3BP
YL9,TLNRD1
RIK5,NPTN,EFNB2,NECTIN1,UNC13A,CNTN1,EFNB3,ZDHHC17,ADGRL1,EPHB2,SYT1,EPHA4,DNM
COL1A2,COL5A2,PTN,GLG1,CDH2,GPC3,SULF1,CCN2,MMP14,APLP1,COL6A2,ANXA6,LGALS3BP

42BPA,MYL9,TLNRD1

90AA1,ACTR2,VIM,SPARC,TUBB,FTH1,UBC,KIDINS220,TMSB4X,NUMA1,DCX,EIF4G2,RPL3,RPS6,
C,TUBB,FTH1,UBC,KIDINS220,TMSB4X,NUMA1,DCX,EIF4G2,RPL3,RPS6,DBN1,RPL4,ENO1,TUBA
PARC,TUBB,FTH1,UBC,KIDINS220,TMSB4X,NUMA1,DCX,RPL3,RPS6,ZNF316,DBN1,RPL4,ENO1,T
AP1B,RPL7A,ALDOA,PKM,FTL,HSP90B1,NEFL,RPL8,FBXL4,YBX1,UBA1,CALR,TMF1,HSPA1B,CES
PARC,TUBB,FTH1,UBC,KIDINS220,TMSB4X,NUMA1,DCX,RPL3,RPS6,ZNF316,DBN1,RPL4,ENO1,T
UBB,FTH1,UBC,KIDINS220,TMSB4X,NUMA1,RPL3,RPS6,ZNF316,RPL4,ENO1,PIK3R2,FLNA,NCL,F
PARC,TUBB,FTH1,UBC,KIDINS220,TMSB4X,NUMA1,DCX,RPL3,RPS6,ZNF316,RPL4,ENO1,PIK3R2
LR,FSTL1,HSPA1B,CES3,PSAP,PGK1,RPLP0,H3-3B,CANX,CLU,DDX5,ENO2,YWHAE,GNAS,SLC2A3
PKM,GANAB,COL6A1,FTL,HSP90B1,PRPF8,FBXL4,YBX1,UBA1,CALR,FSTL1,HSPA1B,COL5A1,CC
PKM,GANAB,COL6A1,FTL,HSP90B1,PRPF8,FBXL4,YBX1,UBA1,CALR,FSTL1,HSPA1B,COL5A1,CC
PKM,GANAB,COL6A1,FTL,HSP90B1,PRPF8,FBXL4,YBX1,UBA1,CALR,FSTL1,HSPA1B,COL5A1,CC
LR,FSTL1,HSPA1B,CES3,PSAP,PGK1,RPLP0,H3-3B,CANX,CLU,DDX5,ENO2,YWHAE,GNAS,SLC2A3
LR,FSTL1,HSPA1B,CES3,PSAP,PGK1,RPLP0,H3-3B,CANX,CLU,DDX5,ENO2,YWHAE,GNAS,SLC2A3
LR,FSTL1,HSPA1B,CES3,PSAP,PGK1,RPLP0,H3-3B,CANX,CLU,DDX5,ENO2,YWHAE,GNAS,SLC2A3
LNA,MAP1B,NCL,RPL7A,GANAB,COL6A1,FTL,HSP90B1,PRPF8,RPL8,FBXL4,YBX1,ATP2B4,CALR,I
DOA,PKM,GANAB,COL6A1,FTL,HSP90B1,YBX1,UBA1,IGFBP2,CALR,FSTL1,ABHD2,HSPA1B,CES3,
13A,KAT6A,RPL13,DDX17,GPI,RACK1,ALDH6A1,SPTBN1,RPS8,RPL7,SEPTIN2,RPL5,SMG1,CALM1
1,CALM1,NPM1,TTC3,USP11,SNRNP200,VCP,UBB,AFF4,TRIM28,MAPT,SPEN,NONO,DDB1,PSMD5
ZNF316,RPL4,ENO1,PIK3R2,FLNA,NCL,RPL7A,ALDOA,PKM,HSP90B1,FADS1,PRPF8,FBXL4,YBX1
4X,RPS3A,RPL18,ITGB1,RPLP1,RPS17,VCL,ARF1,AKAP12,CTNNA1,RPS16,RPL23,CNN3,TPM4,TNS
4X,RPS3A,RPL18,ITGB1,RPLP1,RPS17,VCL,ARF1,AKAP12,CTNNA1,RPS16,RPL23,CNN3,TPM4,TNS
1,FADS1,UGT8,YBX1,UBA1,CALR,FSTL1,TMF1,ABHD2,HSPA1B,COL5A1,COL4A1,CES3,PSAP,RPLP
A,RPL18,RPL11,BTF3,RPLP1,RPL10,RPS17,SF3B2,HNRNPU,LARP1,RPS16,RPL23,RPL29,RPSA,RP
SPTBN1,CALM3,GNB1,SHISA9,RPS8,MYH9,RPL7,SEPTIN2,P4HB,APP,RPL5,FSCN1,MARCKS,CALI
RPS25,DDX3X,RPS15,RPL12,RPL30,RPS5,RPS7,RPS19,RPL35,RPS10,RPL35A,RPLP2,RPL36,RPS
3,DCHS1,MRC2,RPL19,PDIA3,RPS3,VAPA,RPS4X,RPS3A,RPL18,ITGB1,ANXA2,RPLP1,RPS17,VCL,A
DOA,NEFL,NEFM,RPL8,YBX1,UBA1,ATP2B4,HSPA1B,TUBB2B,RPLP0,H3-3B,CANX,CLU,DDX5,SOX
DOA,NEFL,NEFM,RPL8,YBX1,UBA1,ATP2B4,HSPA1B,TUBB2B,RPLP0,H3-3B,CANX,CLU,DDX5,SOX
LU,YWHAE,KIF1A,GNAS,SLC2A3,GPI,BSG,FASN,CALM3,STMN2,MYH9,SEPTIN2,P4HB,APP,SPTAN1
LU,YWHAE,KIF1A,GNAS,SLC2A3,GPI,BSG,FASN,CALM3,STMN2,MYH9,SEPTIN2,P4HB,APP,SPTAN1
RPL19,RPS3,AP2M1,RPS4X,ITGB1,AGRN,AP3D1,ARF1,PAFAH1B1,AKAP12,AP2B1,SLC4A8,RPL23,
HSP90B1,YBX1,UBA1,IGFBP2,CALR,FSTL1,HSPA1B,COL5A1,COL4A1,CES3,PSAP,PGK1,RPLP0,H3
37,RPL28,RPL39,RPL34,UBA52,RPL7L1,FXR2,RPL27A,RPL36AL,RPL32
BX5,PSMD2,ENC1,CHD4,RBBP4,PNN,HNRNPC,CCNI,TADA2B,KLHL9,PFKL,CCND2,GNB2,PRKAR1
1,ATP6V1B2,NAP1L1,FLOT1,RAB2A,SLC2A1,TPP1,CNP,NCSTN,RAB5C,AHCY,GNA13,SDCBP,SGS
1,ATP6V1B2,NAP1L1,FLOT1,RAB2A,SLC2A1,TPP1,CNP,NCSTN,RAB5C,AHCY,GNA13,SDCBP,SGS
RPL26,RPL31,RPS25,DDX3X,RPS15,RPL12,RPL30,RPS5,RPS7,RPS19,RPL35,RPS10,RPL35A,RPL
L18A1,SMIM14,VCP,UBB,COL4A2,FN1,HACD3,SET,PKD1,HLA-C,PDIA3,SERINC1,RPS3,VAPA,SEC
31,CBX5,ATF4,CHD4,JUND,RBBP4,PNN,HNRNPC,TADA2B,SFPQ,EIF5A,IK,JUN,CTNNB1,HNRNPDL
L22,PTCD3,RPL9,RPS14,RPS20,RPL26,RPL31,RPS25,DDX3X,RPS15,RPL12,RPL30,RPS5,RPS7,RP
M1,PABPN1,PARP1,ALKBH5,DDX42,SKI,SNRNP70,PRPF6,TERF2IP,SRSF9,CCNL2,SRSF1,EIF3E,FT
AP43,NCS1,RHOA,KIF5B,CXADR,CRMP1,MYO5A,SEPTIN11,MAP2,L1CAM,GLUL,MAP1A,KLHL24,
P3L,CASC3,MSL1,SRSF3,SF3A1,ATXN2L,PRPF40A,YLPM1,RBM15B,DHX15,PRCC,BRD2,EP400,SI
M1,EIF4B,UBB,CKB,MAPT,MAP9,PPP2R1A,BASP1,HMGB1,STMN1,ITGB1,UCHL1,NCAM1,TIMP2,A
P43,NCS1,RHOA,CLSTN1,ATF4,KIF5B,ENC1,CRMP1,ASTN2,SPTBN2,SLC3A2,SFPQ,NDFIP1,WLS,S

PLD3,HSPD1,SMIM14,ATP5F1A,VCP,UBB,MATR3,HACD3,CHSY1,PKD1,HLA-C,PDIA3,SERINC1,RPS
TR1B,PSMC3,PAFAH1B2,PSMB1,HSPA1A,PSMD13,CAB39,APEH,PSMD7,CSNK2B,CAPN1,NME2,A
LC3A2,GNB2,LAMP1,ATP1A2,MAP2,DNER,L1CAM,GLUL,MAP1A,PGRMC1,EEF1A2,CCT2,KLHL24
CLTC,PRKAR1A,GJA1,MYO5A,CD63,ANXA6,NDFIP1,WLS,LAMP1,RHOB,ATP1A2,HLA-B,RAB1A,RA
l,VCP,COL4A2,MAPT,MAP9,INA,RPL15,DNAH8,TPT1,RPS4X,STMN1,TUBB4B,COL11A1,VCL,ARF1
3N1,CALM3,GNB1,SHISA9,STMN2,MYH9,SEPTIN2,P4HB,APP,GDI1,FSCN1,CALM1,EIF4B,UBB,CKE
,NCS1,CLSTN1,RAB6B,ATP1A1,RTN3,XPR1,GLG1,GPC3,SULF1,CCN2,MMP14,ITM2C,CLTC,H1-0

IR2L8,PKM,GANAB,COL6A1,FTL,HSP90B1,YBX1,UBA1,IGFBP2,CALR,FSTL1,HSPA1B,COL5A1,CC
M3,GNB1,SHISA9,STMN2,MYH9,SEPTIN2,P4HB,APP,GDI1,FSCN1,CALM1,EIF4B,UBB,CKB,MAPT,M
,DYNLL1,PDAP1,CST3,ATP6AP2,ACLY,XRCC6,IMPDH2,ACTR1B,PSMC3,AP2A2,PAFAH1B2,PSMB1
1,ATP1A2,MAP2,RAC1,CAPRIN1,DNER,L1CAM,MAP1A,RAB27A,CLSTN3,CNTNAP2,DPYSL2,MAPK1
PTN,RPS25,CDH2,RPL12,RPL30,RPS19,CTNNB1,SEPTIN11,ATP1A2,RPS27,RAC1,DLG5,HNRNPA3
1,ATP1A2,MAP2,RAC1,CAPRIN1,DNER,L1CAM,MAP1A,RAB27A,CLSTN3,CNTNAP2,DPYSL2,MAPK1
MIF,LGALS3BP,TTR,COTL1,HRNR,TIMP3,CNN2,RAB27A,CCT2,GDI2,PDXK,SRP14,PYGB,VEGFA,M
MIF,LGALS3BP,TTR,COTL1,HRNR,TIMP3,CNN2,RAB27A,CCT2,GDI2,PDXK,SRP14,PYGB,VEGFA,M
MIF,LGALS3BP,TTR,COTL1,HRNR,TIMP3,CNN2,RAB27A,CCT2,GDI2,PDXK,SRP14,PYGB,VEGFA,M
32,EIF2S1
11,ANXA2,AP3D1,ARF1,CYB5R3,ITM2B,AP2B1,GOLGA8A,CPE,CTSD,SCARB2,LPCAT1,SLC22A17,I
AB7A,ITM2C,CLTC,GNB2,GJA1,MYO5A,CD63,ANXA6,DNAJC5,DDOST,YWHAB,LAMP1,CKAP4,TTR,
PSMA6,PIGS,PSMD12,PSMA4,PIGT,PSMC6,RAD23A
CTNND1,PALLD,KIF21B,CTTN,TMOD2,CYTH2,FKBP4,OLFM1,DICER1,CLASP2,IGF2BP1,EXOC4,A
L26,RPL31,XPO1,RPS25,CCND2,PNMA1,EIF3L,NANOG,MAGED2,RPS7,RPS19,RPL35,RPS10,H1-
,SF1,UPF1,TRA2B,HNRNPA2B1,SNRPD3,RALY,EFTUD2,SRSF2,PRPF4B,RBM5,WBP11,PRPF19,SF
A-C,VAPA,HMGB1,TUBB4B,ANXA2,TIMP2,VCL,CYB5R3,SLC4A8,PGAM1,CTNNA1,CTSD,LPCAT1,XI
LN4,PSMB5,SPCS2,PSMA1,PSMD10,PSMA6,PIGS,PSMD12,PSMA4,PIGT,PSMC6,RAD23A,SEC11A
PSMD12,PSMA4,PSMC6,RAD23A
CTNND1,PALLD,KIF21B,CTTN,TMOD2,CYTH2,FKBP4,OLFM1,DICER1,CLASP2,IGF2BP1,EXOC4,A
GJA1,MYO5A,CD63,ANXA6,DNAJC5,DDOST,LAMP1,CKAP4,TTR,HRNR,GPC1,ATP6V0C,TMEM30A
GJA1,MYO5A,CD63,ANXA6,DNAJC5,DDOST,LAMP1,CKAP4,TTR,HRNR,GPC1,ATP6V0C,TMEM30A
FSCN1,SPTAN1,MARCKS,CALM1,NPM1,MARCKSL1,MAPT,MAP9,TMSB10,INA,PPP2R1A,DNAH8,R
RK1,ANTXR1,ABI2,RDX,FLOT2,WASF2,ITGAV,SCRIB,IGF2BP1,APBB1,SNX2,PHACTR4,DAG1,ACT
MRFAP1,GNB2,APLP1,CTNNB1,ANXA6,CYFIP2,NDFIP1,YWHAB,LAMP1,CKAP4,HRNR,PPIB,CDKN1
,TIMP2,VCL,CYB5R3,PGAM1,CTNNA1,CTSD,LPCAT1,XRCC5,KPNB1,NCS1,RHOA,PSMD2,CXADR,
CYB5R3,ITM2B,TMEM170A,GOLGA8A,NFE2L1,FADS2,SCARB2,RTN4,LPCAT1,PJA2,KPNB1,LLGL
RAI2,DPYSL2,STMN3,PTPRS,MAPK8IP1,PPP1R9B,SYP,CTNND1,PALLD,KIF21B,BRSK1,CTTN,TMO
P1,PTPRS,PFN2,RAB11A,CTNND1,GRIPAP1,STAT3,BCR,HIP1,LRRC4B,FLOT1,KPNA2,PHB2,FARP
PAFAH1B1,MAP4,HNRNPU,ATF4,KIF5B,CHD4,CRMP1,CCT7,NISCH,LRPPRC,DDX3X,CLTC,RPS7,I
ADM4,RAC1,CAPRIN1,APPL1,TLN2,ACTR3,NCKAP1,CNTNAP2,ARPC2,CAPZB,ACTN1,PLCG1,ARF4
EPTIN3,COTL1,CTBP1,RAC1,ACTR3,MAPRE1,DSTN,CAPZB,FLNB,CORO1C,PPP1R9B,CTNND1,DC
AM,GLUL,MAP1A,PGRMC1,EEF1A2,KLHL24,CNTNAP2,DPYSL2,TOP1,MAPK1,ATXN10,FLNB,PTPR
8,RPL37,RPL28,RPL39,RPL34,UBA52,RPL7L1,FXR2,RPL27A,RPL36AL,RPL32,MRPL3,MRPL28
0,ATXN2L,MEX3A,CDC42,EDC4,GRSF1,ZFP36L1,UBAP2L,UPF1,SYMPK,ELAVL1,LSM14A,IGF2BP

P4A1,HGS,GRIPAP1,FLOT1,RABEP1,VPS26B,VPS41,VPS4A,RAB14,LDLR,TM9SF4,SH3GL1,NCSTI

ATP6AP1,OPTN,MVB12B,LAPTM4A,NSG2,CLIP3,CORO1C,B2M,IFITM3,HGS,RAB11A,GRIPAP1,AP
LSTN1,TMEM259,ATP2A2,RTN3,TKT,EIF5A,PDIA6,PIP4K2B,ACSL6,GPAT4,GJA1,ARL6IP1,DDOST,
NO1,TUBA1B,PIK3R2,FLNA,MAP1B,NCL,RPL7A,SLC9A7,ALDOA,GANAB,COL6A1,FTL,HSP90B1,FA
MEM259,ATP2A2,RTN3,LRPPRC,XPO1,TKT,EIF5A,PDIA6,PIP4K2B,ACSL6,GPAT4,GJA1,ARL6IP1,D
J,TNC,PPIB,TMEM132A,NUCB1,VCAN,COL2A1,LAMC1,MAPK1,B2M,HYOU1,COL27A1,VGF,IGFBP
RIP,HNRNPR,DDX41,SNRPD2,SNRPB,XAB2,PLRG1
B,EEF1A2,RPN2,GNAI3,ABCA3,VPS35,SURF4,ABCA2,LAPTM4A,GABARAP,ATP6V1B2,IFITM3,FLC
CA,PSMD3,PRKDC,CAND1,HSPB1,CCAR2,POLR2B,MSL1,PHC2,UBQLN1,ARNT2,POLR2E,PSMA7,
MEM259,ATP2A2,RTN3,TKT,EIF5A,PDIA6,PIP4K2B,ACSL6,GPAT4,GJA1,ARL6IP1,DDOST,WLS,CK
1,IST1,CD74,NSG1,BGN,SDCBP,ATP13A2,FAF2,PDGFRB,ASAH1,PA2G4,MAN2B1,DPP7,ARSB,TRA
DC4,ZFP36L1,UBAP2L,UPF1,ELAVL1,LSM14A,IGF2BP1,PSMC3,EIF4G1,PUM1,FXR2,YTHDF3,RBM
K1,DYNLL2,CAPZB,FYN,PTPRS,RPS13,ARFGEF2,BMPR2,PPP1R9B,CTNND1,STAT3,BCR,LRRC4B,
P1R,MAEA,PDE4DIP,DLG4,NSMF,CAPZA2,GMFB,MYADM,NF2,GSN

,DNAJC5,APLP2,NDFIP1,DDOST,WLS,LAMP1,RHOB,CKAP4,HLA-B,RAB1A,ARCN1,RAC1,APPL1,AT
CT3,CCT4,DPYSL2,TUBB2A,MAPRE1,DYNC1LI2,DYNLL2,GABARAP,CLIP3,MAP2K2,KIF3C,CCT8,C

TUBB4B,COL11A1,VCL,ARF1,PAFAH1B1,MAP4,HNRNPU,TPM4,COL1A2,KIF5B,COL5A2,CCT7,PN
UFC2,FAF2,SYNGR1,PA2G4,ARL8A,MAN2B1,DPP7,BRI3,ARSB,TRAPPC1,RAP1B,PRCP,GLB1,CTS
UFC2,FAF2,SYNGR1,PA2G4,ARL8A,MAN2B1,DPP7,BRI3,ARSB,TRAPPC1,RAP1B,PRCP,GLB1,CTS
,FYN,PTPRS,RPS13,ARFGEF2,BMPR2,PPP1R9B,CTNND1,STAT3,BCR,LRRC4B,PHB2,CTNND2,AK
1,VCL,ARF1,PAFAH1B1,MAP4,HNRNPU,TPM4,COL1A2,KIF5B,COL5A2,CCT7,PNN,NES,LRPPRC,T
PRS,RPS13,BMPR2,PPP1R9B,CTNND1,STAT3,BCR,LRRC4B,PHB2,CTNND2,AKAP9,PALM,SH3GL1
I,DYNLL2,CAPZB,FYN,PTPRS,RPS13,BMPR2,PPP1R9B,CTNND1,STAT3,BCR,LRRC4B,KPNA2,PHB
BCA3,VPS35,SURF4,ABCA2,LAPTM4A,ATP6V1B2,IFITM3,FLCN,AP2A1,FLOT1,UBA52,RAB2A,CD1
BCA3,VPS35,SURF4,ABCA2,LAPTM4A,ATP6V1B2,IFITM3,FLCN,AP2A1,FLOT1,UBA52,RAB2A,CD1

NTNAP2,VEGFA,CCND1,GRB2,ACTN1,PLCG1,MAP2K2,YWHAH,VANGL2,BMPR2,AMOTL2,CDC42,
CN1,RAC1,HS6ST1,TMEM132A,NUCB1,VAMP2,GNAI3,TGFBI,RTN1,CLSTN3,SURF4,TMED2,NOTC
TPRS,PFN2,SYP,CTNND1,VDAC1,SGTA,CNTNAP1,HIP1,LRRC4B,FLOT1,PHB2,KCTD12,SV2A,PPP1
MEM132A,NUCB1,GNAI3,RTN1,CLSTN3,SURF4,TMED2,NOTCH2,OPTN,ARF3,KDELR2,GABARAP,
MF,NUP205,LMNA,NARF,PSMA6,PCNA,CENPF,SCAF8,ZNF326,CASK
B12,HM13,EMC10,SPCS2,UBXN1,GET1,PIGS,PIGT,SEC11A

0,SEPTIN11,COTL1,RAC1,CNN2,GAS7,TLN2,NOX4,ACTR3,NCKAP1,ARPC2,DSTN,DYNLL2,LUZP6

LIM1,PPP1CA,NECTIN1,DDX6,NDRG1,PIP5K1C,CCDC85C,ZYX,CDC42EP4,AFDN,NECTIN2,ANXA1,

CS1,ERGIC3,ADAM10,SYT11,COPZ1,SH3BP4,AP2A2,TGOLN2,AP1M1,SORT1,CD74,USO1,SCAP,S

DAC1,SGTA,AP2A1,SEC16A,MAP6,YKT6,SV2A,BRSK1,LMAN1,SEPTIN5,RAB1B,RAB14,NCSTN,RA
,ITGB5,RPS27A,RABEP1,VPS26B,RAB14,LDLR,FLOT2,RAB5C,ITGAV,SYT11,ATP6V0D1,EHD1,MT
L1,PBRM1,GATAD2B,SMARCC2,SMARCB1,DPF2

AN,VPS51,RAP2A,TBC1D14,NDRG1,CLCN4,ABHD17A,TFRC,SORCS2,BACE1,ATG9A,RAB35,CLCN

NC,MMP2,L1CAM,DCN,TIMP3,CTSB,ANXA5,VCAN,TGFBI,COL2A1,MDK,LAMC1,HDGF,AEBP1,HN

7SF2,HM13,EMC10,RRBP1,ATF6B,SGPL1,ATF6,XBP1,EXT1,PIGS,PIGT,TMCO1

R2,HIP1R,DNAJB1,FXR1,CYFIP1,PPP1CA,PSMC2,ABHD17A,DLG4,SORCS2,PRKAR2B,ARHGAP33,
CASK

B,MSL1,PHC2,PFKM,STT3B,STT3A,PRPS1,RBM15B,POLR2E,RBBP7,TRAF7,ARIH1,PRMT1,OS9,

D2,ACSL6,GJA1,NGRN,SLC25A3,DUX4,IPO5,PGRMC1,TOMM20,MLXIP,PARP1,LMNB2,JPT1,UBX
D2,ACSL6,GJA1,NGRN,SLC25A3,DUX4,IPO5,PGRMC1,TOMM20,MLXIP,PARP1,LMNB2,JPT1,UBX
LA-A,TM7SF2,HM13,EMC10,RRBP1,ATF6B,SGPL1,ATF6,XBP1,EXT1,PIGS,PIGT,TMCO1
R2,HIP1R,DNAJB1,FXR1,CYFIP1,PPP1CA,PSMC2,ABHD17A,DLG4,SORCS2,PRKAR2B,ARHGAP33,
D,HMGN2,RBBP4,SPC25,HNRNPC,LRPPRC,XPO1,SFPQ,CCND2,SOX12,NANOG,H1-0,IK,JUN,CT

G7,ERGIC1,TRAM1,DNAJB2,SFXN1,ERGIC3,DHCR7,BNIP3,CCDC47,CD74,SCD,ELOVL5,AMFR,T
TM4B,PGRMC1,RAB27A,SNX22,VAMP2,ABCA3,MLEC,SURF4,RAB5B,RAB11B,TMED2,RAPGEF1,
2EP4,ATP6V0A1,RAB5A,CLCN3,TLR6,GSN,STX12

RAB3A,TFRC,RAB5A,RAB35,SEC23IP,APOE,COPB1,SEC31A,SYT1,SLC18A3,M6PR,NECAP1
,HJURP,UPF1,SMC3,HNRNPA2B1,PPP1CC,SEPTIN7,TNKS,APEX1,PPP1R10,XRCC6,MACROH2A2,

O5A,MACF1,CKAP5,MAP2,COTL1,CCT6A,RAC1,MAP1A,CCT2,TBCD,GAS7,HSPH1,LMNB2,TCP1,

CAR1,PDXP,ARF6,PIP5K1A,MYADM,NF2,FGD1,ASAP3,GSN,CYTH3,EPHA2,ABCA7

7,HIF1A,MAP1LC3B,SPG7,GABARAPL1,KIFAP3,FLOT2,CLASP2,PRKACA,AMBRA1,MYL12B,TUBB

A2B,MCM3AP,POM121C,TNKS,LRRC59,DNAJB2,ABCF1,IST1,DHCR7,CSE1L,TEX2,CHMP4B,MTC
MTA1,EP300,PBX1,CSNK2A1,APEX1,HNRNPAB,HIF1A,XRCC6,SSBP3,SOX2,TLE1,TCF3,RBPJ,MBI

M1,SFXN1,ERGIC3,DHCR7,BNIP3,CCDC47,CD74,SCD,ELOVL5,AMFR,TECR,TAPBP,SPPL2B,SUN1,
C7,MAPRE3,RCC2,ARL8A,LZTS2,ARL2BP,PIN1,GNL3,PDXP,ARF6,CENPF,C6ORF89,VPS37B,SCCF

RKAR2A,FLOT1,HDAC6,RGMB,SLC25A5,SLC2A1,TPP1,FLOT2,PRKACA,DAPK3,SMO,ANK2,PTPN1
RKAR2A,FLOT1,HDAC6,RGMB,SLC25A5,SLC2A1,TPP1,FLOT2,PRKACA,DAPK3,SMO,ANK2,PTPN1
,HK1,PTCD3,VAT1,MCL1,LRPPRC,ACSL6,CHPF,GJA1,ANXA6,NGRN,DNAJC5,SLC25A3,YWHAB,MI

AX,DHX36,WDR82,SETX,THOC2,THOC3,PCNA,MCM7,MSH2,RAD50
,HDAC6,VPS26B,GRN,VPS41,VPS4A,RAB14,LDLR,CHMP4B,NMNAT2,UBXN6,CD74,NSG1,CHMP1

EZO1,TMED5
311A,DYNLL1,CCT5,CDC42,DCTN1,ACTR1A,PRKAR2A,FLCN,FLOT1,MASTL,CUL3,BRSK1,TNKS,A

UL1,DCAF5,WDR26,MAEA,PCGF3,CUL4A,FAM8A1,GID8,STUB1,RANBP9,ZSWIM6,UBE2V1,BCOR,

,HRNR,GPC1,TNC,MMP2,L1CAM,DCN,TIMP3,CTSB,ANXA5,VCAN,TGFBI,COL2A1,MDK,LAMC1,HI

M1,ADAM23,CASK,UNC13B
,HRNR,GPC1,TNC,MMP2,L1CAM,DCN,TIMP3,CTSB,ANXA5,VCAN,TGFBI,COL2A1,MDK,LAMC1,HI

ZNF316,DBN1,RPL4,ENO1,TUBA1B,PIK3R2,FLNA,MAP1B,NCL,RPL7A,SLC9A7,ALDOA,MTRNR2L
A1B,PIK3R2,FLNA,MAP1B,NCL,RPL7A,SLC9A7,ALDOA,MTRNR2L8,PKM,GANAB,COL6A1,FTL,HSP
TUBA1B,PIK3R2,FLNA,MAP1B,NCL,RPL7A,SLC9A7,ALDOA,PKM,GANAB,COL6A1,FTL,HSP90B1,F
3,PGK1,RPLP0,CLU,PTMA,DDX5,ENO2,RPL13A,RPL6,KAT6A,DVL3,YWHAE,KIF1A,RPL13,GNAS,D
TUBA1B,PIK3R2,FLNA,MAP1B,NCL,RPL7A,SLC9A7,ALDOA,PKM,GANAB,COL6A1,FTL,HSP90B1,F
RPL7A,SLC9A7,ALDOA,PKM,GANAB,COL6A1,FTL,HSP90B1,FADS1,PRPF8,UGT8,FBXL4,YBX1,UB
2,FLNA,MAP1B,NCL,RPL7A,SLC9A7,ALDOA,PKM,GANAB,COL6A1,FTL,HSP90B1,FADS1,PRPF8,U
,GPI,RACK1,BSG,FASN,SPTBN1,ATP5F1B,GNB1,RPS8,MYH9,SEPTIN2,P4HB,APP,RPL5,FSCN1,S
L4A1,CES3,PSAP,H3-3B,CANX,CLU,PTMA,DDX5,SOX4,FKBP14,RPL13A,KAT6A,RPL13,DDX17,GP
L4A1,CES3,PSAP,H3-3B,CANX,CLU,PTMA,DDX5,SOX4,FKBP14,RPL13A,KAT6A,RPL13,DDX17,GP
L4A1,CES3,PSAP,H3-3B,CANX,CLU,PTMA,DDX5,SOX4,FKBP14,RPL13A,KAT6A,RPL13,DDX17,GP
,GPI,RACK1,BSG,FASN,SPTBN1,ATP5F1B,GNB1,RPS8,MYH9,SEPTIN2,P4HB,APP,RPL5,FSCN1,S
,GPI,RACK1,BSG,FASN,SPTBN1,ATP5F1B,GNB1,RPS8,MYH9,SEPTIN2,P4HB,APP,RPL5,FSCN1,S
,GPI,RACK1,BSG,FASN,SPTBN1,ATP5F1B,GNB1,RPS8,MYH9,SEPTIN2,P4HB,APP,RPL5,FSCN1,S
HSPA1B,COL5A1,COL4A1,RPLP0,H3-3B,CANX,CLU,DDX5,ENO2,SOX4,RPL13A,MAGED1,RPL6,K
PSAP,PGK1,RPLP0,H3-3B,CANX,CLU,DDX5,ENO2,YWHAE,KIF1A,GNAS,SLC2A3,GPI,RACK1,BSG,
,NPM1,TTC3,USP11,SNRNP200,VCP,UBB,AFF4,TRIM28,MATR3,MAPT,SPEN,NONO,DDB1,PSMD
,SET,GTF3C4,SF3B1,RPS3,CHD3,BASP1,HMGB1,CDKL1,TPT1,RPS4X,RPS3A,UCHL1,RPL11,MO
,UBA1,CALR,TMF1,HSPA1B,ZNF99,TUBB2B,RPLP0,H3-3B,CLU,PTMA,DDX5,SOX4,RPL13A,MAG
S3,SCARB2,RPS9,MSN,RPL22,RHOA,RPL9,RPS14,CD81,LASP1,PFN1,RPL31,CDH2,ACTN4,MMP1
S3,SCARB2,RPS9,MSN,RPL22,RHOA,RPL9,RPS14,CD81,LASP1,PFN1,RPL31,CDH2,ACTN4,MMP1
P0,CANX,CLU,FKBP14,SEC62,RPL6,KIF1A,RPL13,GNAS,SLC2A3,GPI,BSG,FASN,CALM3,STMN2,
PS24,RPS9,XRCC5,RPL24,RPL17,RPL14,RPL22,CBX5,RPL9,RPS14,RPS20,RPL26,PNN,RPL31,HN
M1,NPM1,EIF4B,VCP,YWHAG,RPL37A,MAPT,YWHAQ,HACD3,DCHS1,INA,MRC2,RPL19,PDIA3,RPS
12,RPS27,RPL23A,RPL27,RPL36A,RPS15A,RPL38,RPL37,RPS11,RPL28,RPS28,FAU,RPL39,RPS1
ARF1,AKAP12,CDC42BPB,CTNNA1,RPS16,RPL23,CNN3,TPM4,TNS3,SCARB2,RPS9,MSN,RPL22,R
4,RPL13A,MAGED1,RPL6,KAT6A,DVL3,YWHAE,KIF1A,RPL13,DDX17,RACK1,SPTBN1,ATP5F1B,C
4,RPL13A,MAGED1,RPL6,KAT6A,DVL3,YWHAE,KIF1A,RPL13,DDX17,RACK1,SPTBN1,ATP5F1B,C
,GSTP1,CALM1,PLD3,HSPD1,IGF2,VCP,UBB,FN1,PKD1,HLA-C,PDIA3,VAPA,HMGB1,AP2M1,TPT1,
,GSTP1,CALM1,PLD3,HSPD1,IGF2,VCP,UBB,FN1,PKD1,HLA-C,PDIA3,VAPA,HMGB1,AP2M1,TPT1,
CNN3,RTN4,LAMA5,RPS9,PJA2,RPL24,GAP43,RPL14,NCS1,RPL22,RHOA,CLSTN1,RAB6B,NLGN2
3-3B,CANX,CLU,DDX5,ENO2,YWHAE,GNAS,SLC2A3,GPI,RACK1,BSG,FASN,SPTBN1,ATP5F1B,GN

A,DDOST,HNRNPK,HNRNPH1,HDAC2,SKP1,CDKN1A,RPN2,PPP1CB,GNAI3,KLHL24,MORF4L2,N
M2,ATP1B3,ERP29,RAN,YWHAZ,SYNGR1,TFRC,ATP6V0A1,RAB5A,SLC1A5,SLC1A4,RAB35,GSN
M2,ATP1B3,ERP29,RAN,YWHAZ,SYNGR1,TFRC,ATP6V0A1,RAB5A,SLC1A5,SLC1A4,RAB35,GSN
P2,RPL36,RPS12,RPS27,RPL23A,RPL27,RPL36A,RPS15A,RPL38,RPL37,RPS11,RPL28,RPS28,FA
61A1,RPS3A,RPL18,UCHL1,SARAF,COL11A1,RPL10,MARCHF6,SBF1,DNAJA1,CYB5R3,TMEM170
,ARID1A,HNRNPK,INIP,CIRBP,PTGES3,HNRNPH1,HDAC2,IPO5,SKP1,BCLAF1,LUC7L3,CDKN1A,C
PS19,RPL35,RPS10,RPL35A,RPLP2,RPL36,RPS12,RPS27,RPL23A,RPL27,GCN1,RPL36A,RPS15A,
O,SCAF11,SON,BNIP3L,CASC3,MSL1,HNRNPM,SRSF3,LARS1,SF3A1,ATXN2L,USP7,NCOR2,PRP
KLC1,ACAP3,ORAI2,CNTNAP2,DPYSL2,MAPK1,FAM168B,LUZP6,HSPB1,STMN3,STXBP1,PTPRS,
RSF6,PPP1CC,TIMM50,APEX1,HIF1A,BAZ2A,KMT2E,EFTUD2,PNISR,SRSF2,PRPF4B,DAZAP2,SRS
P3D1,ARF1,PAFAH1B1,MAP4,SLC4A8,HNRNPU,CNN3,LLGL1,GAP43,NCS1,RHOA,CLSTN1,ATF4,
SEPTIN11,LAMP1,ATP1A2,MAP2,RAC1,CAPRIN1,DNER,L1CAM,GLUL,MAP1A,PGRMC1,RAB27A,E

3,VAPA,AP2M1,SEC61A1,ITGB1,UCHL1,NCAM1,ANXA2,SARAF,MARCHF6,AP3D1,SBF1,DNAJA1,A
SAH1,PSMC2,ARSB,ARPC5,DYNLT1,PSMD12,PRDX4,GSN,CSTB,GLB1
,TCP1,CCT3,CCT4,CNTNAP2,DPYSL2,TOP1,MAPK1,GABARAP,ATXN10,RPL28,FLNB,FYN,CCT8,P
C1,GPC1,APPL1,ATP6V0C,DNER,LAPTM4B,CTSB,NUCB1,RAB27A,SQSTM1,TM9SF2,ABCA3,PKN1
,PAFAH1B1,MAP4,HNRNPU,LARP1,TPM4,KPNB1,COL1A2,KIF5B,COL5A2,CCT7,PNN,SPC25,NES,
3,MAPT,MAP9,PPP2R1A,DNAH8,PKD1,RPS3,BASP1,HMGB1,CDKL1,STMN1,ITGB1,UCHL1,NCAM1
,APLP1,CHPF,GJA1,MACF1,GOLM1,B4GALNT4,DNAJC5,NDFIP1,ST3GAL2,WLS,DUX4,RPS12,HL

L4A1,CES3,PSAP,PGK1,RPLP0,H3-3B,CANX,CLU,DDX5,ENO2,YWHAE,GNAS,SLC2A3,GPI,HTRA4,
AP9,PPP2R1A,DNAH8,PKD1,RPS3,BASP1,HMGB1,CDKL1,STMN1,ITGB1,UCHL1,NCAM1,ANXA2,
,HSPA1A,PSMD13,CAB39,APEH,PSMD7,CSNK2B,CAPN1,NME2,ASAH1,PSMC2,ARL8A,ARSB,ATP6
,ATXN10,CASC3,NSG2,RPL28,ARF4,FYN,MAPK8IP1,SOD1,ARFGEF2,BMPR2,CDC42,PPP1R9B,C
3,CLSTN3,RPL38,MAPK1,DYNLL2,CAPZB,ARF4,FYN,STXBP1,PTPRS,PFN2,RAB11A,RPS13,ARFGE
,ATXN10,CASC3,NSG2,RPL28,ARF4,FYN,MAPK8IP1,SOD1,ARFGEF2,BMPR2,CDC42,PPP1R9B,C
APK1,PSMD3,ACTN1,CAND1,ILF2,CCT8,B2M,IDH1,TXNDC5,PSMD1,A2M,PRDX6,GRN,ACLY,XRC
APK1,PSMD3,ACTN1,CAND1,ILF2,CCT8,B2M,IDH1,TXNDC5,PSMD1,A2M,PRDX6,GRN,ACLY,XRC
APK1,PSMD3,ACTN1,CAND1,ILF2,CCT8,B2M,IDH1,TXNDC5,PSMD1,A2M,PRDX6,GRN,ACLY,XRC

PJA2,LLGL1,HK1,RHOA,CLSTN1,RAB6B,SLC7A5,ATP2A2,VAT1,RTN3,MCL1,LRPPRC,SLC3A2,GL
,HRNR,GPC1,ATP6V0C,TMEM30A,LAPTM4B,DCN,CTSB,RAB27A,EEF1A2,CCT2,RPN2,SQSTM1,G

PBB1,AMFR,FXR2,PINK1,AUTS2,TAOK2,TRAK1,FXR1,CYFIP1,FEZ1,HNRNPR,NECTIN1,ARPC3,EX
10,RPL36,DUX4,CKAP5,TOP2B,IPO5,RPL23A,CDKN1A,SRSF5,PPP1CB,NOP53,PARP1,TSR1,MOR
3A2,SART1,DHX38,HNRNPH3,EIF4A3,HTATSF1,LUC7L2,MTREX,SF3A3,SYNCRIP,RHEB,HNRNPR,
RCC5,KPNB1,NCS1,RHOA,PSMD2,CXADR,VAT1,PFKL,RAB7A,ACTN4,DDX3X,MAGED2,MIF,MYO5

PBB1,AMFR,FXR2,PINK1,AUTS2,TAOK2,TRAK1,FXR1,CYFIP1,FEZ1,HNRNPR,NECTIN1,ARPC3,EX
,LAPTM4B,DCN,CTSB,RAB27A,EEF1A2,CCT2,SQSTM1,GNAI3,GDI2,VCAN,ABCA3,VPS35,SURF4,
,LAPTM4B,DCN,CTSB,RAB27A,EEF1A2,CCT2,SQSTM1,GNAI3,GDI2,VCAN,ABCA3,VPS35,SURF4,
PS3,CHD3,BASP1,VAPA,TPT1,STMN1,TUBB4B,VCL,DNAJA1,PAFAH1B1,AKAP12,MAP4,CDC42BPB
2,CYFIP1,CD44,LIMK1,SNX1,CCDC88A,NME2,ARPC3,WASF3,ARPC5,RUFY3,AJUBA,DYNLT1,FER
A,CTSB,VAMP2,NOX4,PRDX5,PTOV1,RAPGEF1,OPTN,ARF3,PLEC,ODC1,FAM168B,LUZP6,GABA
VAT1,PFKL,RAB7A,ACTN4,DDX3X,MAGED2,MIF,MYO5A,CD63,DNAJC5,APLP2,DDOST,LAMP1,LG
1,RHOA,CLSTN1,RAB6B,TMEM259,ATP2A2,RTN3,TKT,GLG1,EIF5A,PDIA6,SULF1,PIP4K2B,ACSL
D2,CYTH2,FKBP4,OLFM1,DICER1,CLASP2,IGF2BP1,SYT11,ATP6V0D1,EXOC4,APBB1,MICAL1,NT
1,SV2A,PPP1CC,GPC4,AKAP9,ELAVL1,SH3GL1,FLOT2,SCRIB,ADAM10,FXYD6,MTOR,NSG1,EIF4A
K,PRKAR1A,CTNNB1,NGRN,MACF1,CKAP5,SEPTIN11,MAP2,SEPTIN3,CCT6A,CDK16,SKP1,PEA15
,CORO1C,CDC42,PPP1R9B,CTNND1,PALLD,FLOT1,SLC39A6,EVL,HDAC6,CTTN,ATP6AP2,BRK1,
TN1,CAPN2,ACTR1A,HIP1,FLOT1,CSNK1G2,GPSM1,CALD1,SEPTIN7,CTTN,SLC2A1,SEPTIN5,RDX
S,SOD1,BMPR2,CDC42,PPP1R9B,DCTN1,KIF5A,CAPN2,DIP2B,CTNND2,HDAC6,YKT6,SV2A,PTPR

1,PSMC3,EIF4G1,PUM1,FXR2,YTHDF3,RBM4,TIA1,SUMO1,FXR1,PABPC4,PNRC2,TNRC6A,DDX6,

N,RAB5C,MTMR4,WASF2,AKT2,ATP6V0D1,EHD1,UBXN6,SORT1,DOP1B,PRDX3,NSG1,PTPN23,NT

2A1,UBA52,CD164,RPS27A,ATP6AP2,RABEP1,ANTXR1,VPS41,VPS4A,RAB14,LDLR,SH3GL1,PLEK
WLS,CKAP4,HLA-B,CALU,ARCN1,HMGCR,RAC1,SQLE,TMEM132A,PGRMC1,SRP9,FDFT1,RPN2,G
ADS1,PRPF8,UGT8,RPL8,YBX1,UBA1,ATP2B4,IGFBP2,CALR,TMF1,ABHD2,HSPA1B,COL4A1,PSAP,
DDOST,WLS,CKAP4,HLA-B,CALU,ARCN1,HMGCR,RAC1,SQLE,TMEM132A,PGRMC1,SRP9,FDFT1,
?7,OS9,TXNDC5,CST3,CRTAP,MBTPS1,CCN1,ADAM10,TGOLN2,RCN2,IGFBP4,COL16A1,LGALS1,

N,AP2A1,FLOT1,UBA52,RAB2A,CD164,ULK1,ATP6AP2,MAP1LC3B,GRN,CLCN7,SLC44A2,VPS41,V
,TRAF7,ARIH1,OS9,EP300,PRKAR2A,PSMD1,CSNK2A1,PPP2CB,KLHL42,EP400,UBAP2L,FBXO11,
AP4,HLA-B,CALU,ARCN1,HMGCR,RAC1,SQLE,TMEM132A,PGRMC1,SRP9,FDFT1,RPN2,GNAI3,M
APPC1,PPT1,GLB1,CTSA,ARSA,TOLLIP
M4,TIA1,FXR1,PABPC4,PNRC2,TNRC6A,DDX6,LSM4,CDK9,DHX36,POLDIP3,PSMC2,MCRIP1,ROC
PHB2,CTNND2,AKAP9,PALM,SH3GL1,DCLK1,PRR12,PDPK1,ATP1B2,SCRIB,ADAM10,SYT11,BNIP

TP6V0C,TMEM30A,LAPTM4B,PGRMC1,RAB27A,SNX22,VAMP2,TM9SF2,ABCA3,VPS35,MLEC,SUR
CSNK1D,RAB11A,ARFGEF2,DYNLL1,CCT5,BEX4,DCTN1,KIF5A,MTA1,KIF21B,MAP6,HDAC6,CUL3,

IN,NES,LRPPRC,TPM1,CLTC,MYO5A,CTNNB1,MACF1,MYH10,CKAP5,MAP2,COTL1,CCT6A,RAC1,
A,ARSA,TOLLIP
A,ARSA,TOLLIP
AP9,PALM,SH3GL1,DCLK1,PRR12,PDPK1,SCRIB,ADAM10,SYT11,BNIP3,NTRK2,FXR2,HIP1R,GRIK
PM1,CLTC,MYO5A,CTNNB1,MACF1,MYH10,CKAP5,MAP2,COTL1,CCT6A,RAC1,MMP2,DCN,MAP1
,DCLK1,PRR12,PDPK1,SCRIB,ADAM10,SYT11,BNIP3,NTRK2,FXR2,HIP1R,GRIK5,NPTN,MAPK8IP2
2,CTNND2,AKAP9,PALM,SH3GL1,DCLK1,PRR12,PDPK1,SCRIB,ADAM10,SYT11,BNIP3,NTRK2,FX
64,ATP6AP2,GRN,CLCN7,SLC44A2,VPS41,PGAP6,RAB14,LDLR,NCSTN,RAB5C,SPAG9,PLEKHM2,
64,ATP6AP2,GRN,CLCN7,SLC44A2,VPS41,PGAP6,RAB14,LDLR,NCSTN,RAB5C,SPAG9,PLEKHM2,

PPP1R9B,CTNND1,TRAF4,KIRREL1,CNTNAP1,FLCN,FLOT1,CTNND2,SV2A,SYMPK,SLC2A1,CYTH2
H2,OPTN,ARF3,KDELR2,PCSK1N,GABARAP,NSG2,CLIP3,KDELR1,B2M,PITPNM1,TNKS2,TMED9,S
CC,BRSK1,GPC4,SLC2A1,SEPTIN5,SH3GL1,PNISR,FLOT2,NCSTN,SCRIB,ADAM10,SYT11,ATP6V0
NSG2,CLIP3,KDELR1,B2M,PITPNM1,TNKS2,TMED9,SREBF2,SEC23A,ARFGEF2,CDC42,ST3GAL6

,CAPZB,GABARAP,ACTN1,FLNB,TPM3,CORO1C,FYN,MPRIP,PPP1R9B,PALLD,CAPN2,ACTR1A,HIP

ADD1,AJUBA,EPHA4,PLPP3,NF2,CYTH3

NX3,LMAN2,CLTA,SCAMP1,HIP1R,COPE,HLA-A,CCDC88A,EPN1,MCFD2,RAB3A,TFRC,COL7A1,DI

36A,YIPF3,ADAM10,SYT11,ATP6V0D1,COPZ1,TGOLN2,SORT1,CD74,USO1,SCAP,BGN,SCAMP5,C
OR,AP2A2,TGOLN2,CD74,AMBRA1,SMO,RAB11FIP3,TAPBP,SNX3,CLTA,MDM2,PIK3C2B,HLA-A,A

I3,MICALL1,VPS16,ARF6,C1ORF210,FZD7,STX12

RNPM,FBLN1,COL27A1,IGFBP7,SFRP1,LMAN1,A2M,GPC4,CCN1,HSPG2,GPC2,LTBP4,LTBP3,FBI

SHANK1,PPP3CA,EPHA4,DNM1,ABR

EP300,PRKAR2A,KLHL42,EP400,UBAP2L,FBXO11,CUL3,ULK1,FBXO21,RHBDD2,ERH,PELP1,SNR

N4,CCND1,IPO7,ACADVL,ATP5PB,BNIP3L,CASC3,FKBP8,SDHA,TNKS2,UQCRC2,MAPK8IP1,GHITN
N4,CCND1,IPO7,ACADVL,ATP5PB,BNIP3L,CASC3,FKBP8,SDHA,TNKS2,UQCRC2,MAPK8IP1,GHITN

SHANK1,PPP3CA,EPHA4,DNM1,ABR
NNB1,TARDBP,ARID1A,MXD4,SHOX,HNRNPK,H1-10,INIP,PTGES3,CKAP5,TOP2B,SMCHD1,HDAC2

ECR,TAPBP,SPPL2B,SUN1,PINK1,KTN1,ESYT1,ATP13A2,MFN2,EMC1,GPAA1,BSCL2,HLA-A,SUN2,
KDELR2,NSG2,KDELR1,B2M,PTPRS,SREBF2,RAB11A,SEC23A,DYNLL1,SYP,SGTA,HIP1,AP2A1,SE

CLASP2,KDM4A,CENPB,RANGAP1,TOX4,MACROH2A1,SUN2,CFDP1,PPP1CA,WRNIP1,H2AX,DHX3

CAMSAP1,KLC1,CCT3,NCKAP1,CCT4,DPYSL2,TUBB2A,MAPRE1,PLEC,DYNC1LI2,DYNLL2,GABAR

4A,RANGAP1,RIMS3,CLTA,TRAK1,HIP1R,WDR11,KIF3A,DLG4,ARL8A,UNC13A,MAP2K4,CTBP2,SI

)R,BNIP3,UBE2I,SORT1,RANGAP1,ABL1,SUN1,SDCBP,RGPD5,CLIC1,FZR1,PCM1,SUMO1,APEH,B
)3,YY1,CREBBP,TEAD1,LMO4,HMGA1,MTA2,GTF3C1,TCEA1,ADNP,TRRAP,SUB1,PDLIM1,KAT2A,

PINK1,KTN1,ESYT1,ATP13A2,EMC1,GPAA1,BSCL2,HLA-A,SUN2,TM7SF2,HM13,EMC10,SYNGR1,
'DH,GNAI1,CCDC124

1,SDCBP,DAG1,PARK7,ATP1B3,EFNB1,ADCY1,IRS1,PRKAR2B,CNTN1,BACE1,CFLAR,RAB5A,TGFI
1,SDCBP,DAG1,PARK7,ATP1B3,EFNB1,ADCY1,IRS1,PRKAR2B,CNTN1,BACE1,CFLAR,RAB5A,TGFI
MP2,GLUL,PGRMC1,MDH2,TOMM20,MLXIP,SQSTM1,PARP1,NOX4,MTRF1L,PRDX5,ACO2,HSPA4

A,ATP13A2,TMEM9,CLCN4,ARL8A,STARD3,BACE1,ATG9A,CHMP5,MAPK3,CLCN3,TMEM59,MICAI

PEX1,NDN,AKAP9,CEP170,VPS4A,DZIP1,KIFAP3,TUBGCP6,SPAG9,ACTR1B,IST1,ATP6V0D1,PRKA

CDKN1B,RNF20,CACYBP,MED12,ELOB,ZSWIM8,KCTD10,MIB2,CUL7,CUL4B,TRPC4AP,CBX4,FB

)GF,VEGFA,AEBP1,HNRNPM,FBLN1,COL27A1,IGFBP7,MMP15,SFRP1,LMAN1,LOX,A2M,GPC4,CC

)GF,VEGFA,AEBP1,HNRNPM,FBLN1,COL27A1,IGFBP7,MMP15,SFRP1,LMAN1,LOX,A2M,GPC4,CC

8,PKM,GANAB,COL6A1,FTL,HSP90B1,FADS1,NEFL,NEFM,PRPF8,UGT8,RPL8,FBXL4,YBX1,UBA1,

90B1,FADS1,NEFL,NEFM,UGT8,RPL8,FBXL4,YBX1,UBA1,ATP2B4,IGFBP2,CALR,FSTL1,TMF1,AB

ADS1,NEFL,NEFM,PRPF8,UGT8,RPL8,FBXL4,YBX1,UBA1,ATP2B4,IGFBP2,CALR,FSTL1,TMF1,AB

DX17,GPI,RACK1,FASN,SPTBN1,GNB1,RPS8,MYH9,RPL7,P4HB,APP,RPL5,GDI1,FSCN1,SPTAN1,

ADS1,NEFL,NEFM,PRPF8,UGT8,RPL8,FBXL4,YBX1,UBA1,ATP2B4,IGFBP2,CALR,FSTL1,TMF1,AB

A1,ATP2B4,IGFBP2,CALR,FSTL1,TMF1,ABHD2,HSPA1B,ZNF99,TUBB2B,COL5A1,COL4A1,CES3,

T8,FBXL4,YBX1,UBA1,ATP2B4,IGFBP2,CALR,FSTL1,TMF1,ABHD2,HSPA1B,ZNF99,TUBB2B,COL

TAN1,GSTP1,MARCKS,PLXNB2,PLD3,HSPD1,COL18A1,ATP5F1A,VCP,MARCKSL1,UBB,YWHAG,C

I,RACK1,ALDH6A1,SPTBN1,ATP5F1B,RPS8,RPL7,SEPTIN2,P4HB,APP,IDS,RPL5,SPTAN1,GSTP1,S

I,RACK1,ALDH6A1,SPTBN1,ATP5F1B,RPS8,RPL7,SEPTIN2,P4HB,APP,IDS,RPL5,SPTAN1,GSTP1,S

I,RACK1,ALDH6A1,SPTBN1,ATP5F1B,RPS8,RPL7,SEPTIN2,P4HB,APP,IDS,RPL5,SPTAN1,GSTP1,S

TAN1,GSTP1,MARCKS,PLXNB2,PLD3,HSPD1,COL18A1,ATP5F1A,VCP,MARCKSL1,UBB,YWHAG,C

TAN1,GSTP1,MARCKS,PLXNB2,PLD3,HSPD1,COL18A1,ATP5F1A,VCP,MARCKSL1,UBB,YWHAG,C

TAN1,GSTP1,MARCKS,PLXNB2,PLD3,HSPD1,COL18A1,ATP5F1A,VCP,MARCKSL1,UBB,YWHAG,C

AT6A,YWHAE,KIF1A,RPL13,GNAS,DDX17,RACK1,SPTBN1,ATP5F1B,CALM3,GNB1,SHISA9,RPS8,M

FASN,SPTBN1,ATP5F1B,CALM3,GNB1,STMN2,RPS8,MYH9,SEPTIN2,P4HB,APP,RPL5,FSCN1,SPT

,SET,RPL19,GTF3C4,SF3B1,RPS3,CHD3,BASP1,HMGB1,CDKL1,TPT1,RPS4X,RPS3A,RPL18,UCH

RF4L1,ZMIZ1,XAF1,RPS17,SBF1,SF3B2,GTF2F1,HNRNPU,NFE2L1,SMARCA5,HNRNPA0,RPS16,R

ED1,RPL6,KAT6A,YWHAE,RPL13,GNAS,DDX17,GPI,RACK1,ALDH6A1,SPTBN1,ATP5F1B,CALM3,R

4,RPS15,RPL12,RPL30,CLTC,GNB2,RPS5,RPS7,RPS19,TLN1,GJA1,CTNNB1,ANXA6,RPS10,RPLP

4,RPS15,RPL12,RPL30,CLTC,GNB2,RPS5,RPS7,RPS19,TLN1,GJA1,CTNNB1,ANXA6,RPS10,RPLP

RPS8,MYH9,SEPTIN2,P4HB,APP,RPL5,GDI1,TMBIM6,SPTAN1,GSTP1,CALM1,TTC3,PLD3,HSPD1,C

RNPC,LRPPRC,XPO1,HDLBP,RPS25,ACTN4,DDX3X,RPS15,RPL12,RPL30,EIF3L,RPS5,RPS7,IK,RP

3,BASP1,VAPA,AP2M1,RPS4X,RPS3A,RPL18,ITGB1,ANXA2,AGRN,RPLP1,RPS17,VCL,AP3D1,ARF1

3,RPL34,UBA52,RPL7L1,RPS2,RPS27A,RPS26,FXR2,RPS21,RPL27A,RPS29,RPL36AL,RPL32

HOA,RPL9,RPS14,CXADR,ATP1A1,CD81,WDR1,ATP2A2,LASP1,PFN1,PNN,RPL31,CDH6,CDH2,AC

LM3,RPS8,MYH9,RPL7,SEPTIN2,P4HB,APP,RPL5,FSCN1,SPTAN1,MARCKS,CALM1,NPM1,TTC3,

LM3,RPS8,MYH9,RPL7,SEPTIN2,P4HB,APP,RPL5,FSCN1,SPTAN1,MARCKS,CALM1,NPM1,TTC3,

ITGB1,TUBB4B,ANXA2,TIMP2,VCL,AP3D1,ARF1,CYB5R3,ITM2B,AP2B1,SLC4A8,PGAM1,CTNNA1

ITGB1,TUBB4B,ANXA2,TIMP2,VCL,AP3D1,ARF1,CYB5R3,ITM2B,AP2B1,SLC4A8,PGAM1,CTNNA1

,RPS14,ATP1A1,RPS20,ATP2A2,RTN3,PFN1,SPTBN2,SLC3A2,PTN,RPS25,CDH2,RPS15,RPL12,R

31,RPS8,MYH9,SEPTIN2,P4HB,APP,RPL5,FSCN1,SPTAN1,GSTP1,MARCKS,PLXNB2,PLD3,HSPD1,

X4,SNRPN,HNRNPA3,UBE4A,YEATS2,PLOD1,RBMX,RAD23B,PSMB2,KDM6B,PRPF6,TAF7,PPP2C

,RPL39,RPS13,RPL34,UBA52,RPL7L1,RPS2,RPS27A,RPS26,FXR2,RPS21,RPL27A,MRPS6,RPS29

A,PXDN,NFE2L1,FADS2,SCARB2,RTN4,LPCAT1,RPS24,PJA2,RPL24,KPNB1,IGFBP3,COL1A2,RHC

PSF2,DDX39B,U2AF2,CPSF1,MORF4L2,SSRP1,SNRPN,HNRNPA3,YEATS2,RBMX,RAD23B,SNRNP

RPL38,RPL37,RPS11,RPL28,RPS28,FAU,RPL39,RPS13,RPL34,SF1,UBA52,RPL7L1,RPS2,RPS27A

40A,TP53BP1,YLPM1,YME1L1,RBM15B,DHX15,ARIH1,SF1,PRCC,BAZ1B,BRD2,ATRX,EP400,SRS

MAPK8IP1,SOD1,RAB11A,DYNLL1,PPP1R9B,SYP,CTNND1,PALLD,GRIPAP1,DCTN1,KIF5A,KCNQ2,

11,PRPF19,SF3A2,SART1,PRKACA,SAP18,APBB1,THRAP3,EIF4A3,TRIP12,CDC34,LUC7L2,RBM4

KIF5B,CXADR,CRMP1,SFPQ,CDH2,MYO5A,CYFIP2,NDFIP1,WLS,SEPTIN11,LAMP1,ATP1A2,MAP2,

EF1A2,KLHL24,CLSTN3,CNTNAP2,DPYSL2,TOP1,MAPK1,ATXN10,CASC3,NSG2,RPL28,ARF4,FLN

RF1,CYB5R3,PAFAH1B1,ITM2B,AP2B1,TMEM170A,GOLGA8A,NFE2L1,FADS2,CPE,CTSD,SCARB2

TPNM1,PTPRS,MAPK8IP1,SOD1,BMPR2,CCT5,CDC42,PPP1R9B,DCTN1,KIF5A,CAPN2,DIP2B,CTN
I,TBC1D16,VPS35,RAB5B,CNTNAP2,RAB11B,ABCA2,ATP6AP1,RAPGEF1,OPTN,MVB12B,MAPK1,L
LRPPRC,XPO1,TPM1,DDX3X,CLTC,MYO5A,CTNNB1,MACF1,TARDBP,HNRNPK,CIRBP,MYH10,CKA
,ANXA2,TIMP2,AP3D1,ARF1,PAFAH1B1,MAP4,CDC42BPB,SLC4A8,HNRNPU,CTNNA1,CNN3,RTN4
A-B,RAB1A,CALU,ARCN1,RAC1,STIP1,GPC1,SLC35E2B,TMEM30A,DCN,HS6ST1,TMEM132A,NUCI

RACK1,BSG,FASN,SPTBN1,ATP5F1B,GNB1,RPS8,MYH9,SEPTIN2,P4HB,APP,RPL5,FSCN1,SPTAN1
TIMP2,AP3D1,ARF1,PAFAH1B1,MAP4,CDC42BPB,SLC4A8,HNRNPU,CTNNA1,CNN3,MSN,LLGL1,G
V0A1,ARPC5,DYNLT1,PSMD12,COPB1,NFASC,PRDX4,PRCP,GSN,CSTB,GLB1
TNND1,GRIPAP1,KIF5A,BCR,CAPN2,KIF21B,MAP6,DIP2B,FARP1,CTNND2,HDAC6,PTEN,YKT6,SV2
F2,BMPR2,CDC42,PPP1R9B,CTNND1,GRIPAP1,PSMA7,STAT3,BCR,HIP1,LRRC4B,KPNA2,PHB2,F
TNND1,GRIPAP1,KIF5A,BCR,CAPN2,KIF21B,MAP6,DIP2B,FARP1,CTNND2,HDAC6,PTEN,YKT6,SV2
C6,IMPDH2,ACTR1B,CAP1,IST1,PSMC3,PAFAH1B2,SDCBP,PSMB1,PSMD13,CAB39,CYFIP1,PSMD
C6,IMPDH2,ACTR1B,CAP1,IST1,PSMC3,PAFAH1B2,SDCBP,PSMB1,PSMD13,CAB39,CYFIP1,PSMD
C6,IMPDH2,ACTR1B,CAP1,IST1,PSMC3,PAFAH1B2,SDCBP,PSMB1,PSMD13,CAB39,CYFIP1,PSMD

G1,ACSL6,RAB7A,ITM2C,CLTC,GNB2,CHPF,GJA1,CD63,ANXA6,B4GALNT4,DNAJC5,APLP2,ARL6I
NAI3,GDI2,VCAN,ABCA3,VPS35,SURF4,ABCA2,PYGB,OPTN,MAPK1,LAPTM4A,DYNC1LI2,GABARAI

OC7,SETX,STMN4,ARPC5,GPRIN1,RUFY3,DYNLT1,ADGRL1,FGF13,EPHA4,MAPK8IP3
F4L2,NOX4,SSRP1,RPL27,JPT1,TOP1,MARS1,SMARCA4,SRSF9,ACADVL,PTBP1,ILF3,GRB2,RPS1
LSM4,DDX41,SNRPD2,SNRPB,RBM17,PRPF31,API5,XAB2,AKAP17A,MYEF2,ZMAT2,KHDC4,PLRG1
A,CD63,DNAJC5,APLP2,DDOST,LAMP1,LGALS3BP,CKAP4,HLA-B,TTR,COTL1,HRNR,RAC1,CDK16

OC7,SETX,STMN4,ARPC5,GPRIN1,RUFY3,DYNLT1,ADGRL1,FGF13,EPHA4,MAPK8IP3
BCA2,PYGB,MAPK1,LAPTM4A,DYNC1LI2,GABARAP,NSG2,ATP6V1B2,LARS1,CCT8,IFITM3,HGS,S
BCA2,PYGB,MAPK1,LAPTM4A,DYNC1LI2,GABARAP,NSG2,ATP6V1B2,LARS1,CCT8,IFITM3,HGS,S
,HNRNPU,CTNNA1,CNN3,TPM4,PJA2,MSN,LLGL1,RHOA,ATF4,KIF5B,ENC1,CHD4,WDR1,CRMP1,
,BCAR1,PDXP,PIP5K1A,SRCIN1,CDC42BPA,PLXND1,NF2,PIEZO1,FGD1,GSN,EPHA2
AP,ATXN10,CASC3,MAP2K2,FYN,TNKS2,STXBP1,UBQLN1,CSNK1D,IFITM3,MAPK8IP1,RAB11A,S
ALS3BP,CKAP4,HLA-B,TTR,COTL1,HRNR,RAC1,ATP6V0C,TMEM30A,TIMP3,PGRMC1,CNN2,CTSE
6,CLTC,GPAT4,CHPF,GJA1,B4GALNT4,ARL6IP1,NDFIP1,DDOST,ST3GAL2,WLS,CKAP4,HLA-B,RA
RK2,CAD,AMFR,FXR2,PINK1,AUTS2,TAOK2,TRAK1,GRIK5,FXR1,CYFIP1,FEZ1,HNRNPR,NECTIN1
3,DAG1,GRIK5,NPTN,DNAJB1,EFNB2,FXR1,PPP1CA,C1QBP,ADCY1,SYNGAP1,NDRG1,YWHAZ,GS
,MAP1A,DLG5,CCT2,TBCD,GNAI3,HSPH1,YEATS2,TCP1,CAMSAP1,KLC1,CCT3,CBX1,CCT4,SKI,A
ALM,ANTXR1,ABI2,GABARAPL1,RDX,FLOT2,CLASP2,WASF2,ITGAV,SCRIB,IGF2BP1,AKT2,APBB1
,FLOT2,CAPZA1,FNBP1L,CLASP2,CAP1,AKT2,EXOC4,MYL12B,RIMS3,CLTA,TRAK1,HIP1R,EXOC5
,IL6ST,NDN,AKAP9,FUS,FKBP4,OLFM1,PDPK1,ADAM10,IGF2BP1,SYT11,GSK3A,MTOR,GOT2,AI

LSM4,CDK9,DHX36,POLDIP3,PSMC2,HSF1,MCRIP1,ROCK2,PSMA6,CNOT3,SHB,AJUBA,BTBD1,T

RK2,SNX2,ANK2,RNF11,SNX3,CHMP1A,USP10,TRAK1,ARRDC4,RNF157,HLA-A,PTPN1,ANKFY1,S

HB2,NCSTN,RAB5C,PLEKHM2,MTMR4,LZTR1,SYT11,ATP6V0D1,EHD1,CHMP4B,AP2A2,UBE2D3,

NAI3,MEST,NOX4,RPL27,C4ORF3,RTN1,PLOD1,AHCYL1,MLEC,CLSTN3,SURF4,ATP6AP1,TMED2

,PGK1,RPLP0,CANX,CLU,DDX5,ENO2,RPL13A,MAGED1,SEC62,RPL6,YWHAE,KIF1A,RPL13,GNAS,

RPN2,GNAI3,MEST,NOX4,RPL27,C4ORF3,RTN1,PLOD1,AHCYL1,MLEC,CLSTN3,SURF4,ATP6AP1,

KTN1,COL12A1,DAG1,H6PD,ANO8,ERP29,SHISA5,TSPAN14,COLGALT1,SCG2,P3H1,TMEM43,ST

PS4A,GABARAPL1,PGAP6,RAB14,LDLR,NCSTN,RAB5C,SPAG9,PLEKHM2,BORCS7,SYT11,ATP6V

CUL3,ULK1,PSMD8,FBXO21,RHBDD2,XRCC6,PSMB4,UBE2D2,DNAJB2,ELP1,DCUN1D2,LZTR1,P

EST,NOX4,RPL27,C4ORF3,RTN1,PLOD1,AHCYL1,MLEC,CLSTN3,SURF4,ATP6AP1,TMED2,NOTC

K2,PSMA6,CNOT3,SHB,AJUBA,BTBD1,TUT4,PSMA4,EIF2S1,DDX19A,MEX3B,CNOT1

3,NTRK2,FXR2,HIP1R,GRIK5,NPTN,MAPK8IP2,DNAJB1,NSF,FXR1,RHEB,IQSEC1,NECTIN1,NCK2,

F4,RAB5B,RAB11B,ABCA2,ATP6AP1,TMED2,RAPGEF1,OPTN,MVB12B,KDELR2,LAPTM4A,GRB2,

EML1,MAPRE2,MAP1LC3B,SERP1,CEP170,FKBP4,GABARAPL1,KIFAP3,CNP,TUBGCP6,CLASP2,

MMP2,DCN,MAP1A,CCT2,TBCD,SQSTM1,GAS7,HSPH1,LMNB2,TCP1,CAMSAP1,KLC1,CCT3,NCK

5,NPTN,MAPK8IP2,DNAJB1,NSF,FXR1,RHEB,IQSEC1,NCK2,ADCY1,SYNGAP1,SPOCK1,SEMA4C,

A,CCT2,TBCD,SQSTM1,GAS7,HSPH1,LMNB2,TCP1,CAMSAP1,KLC1,CCT3,NCKAP1,COL2A1,CCT

,DNAJB1,NSF,FXR1,RHEB,IQSEC1,NCK2,ADCY1,SYNGAP1,SPOCK1,SEMA4C,ABHD17A,CRTC1,A

R2,HIP1R,GRIK5,NPTN,MAPK8IP2,DNAJB1,NSF,FXR1,RHEB,IQSEC1,NCK2,ADCY1,SYNGAP1,SPO

BORCS7,SYT11,ATP6V0D1,MTOR,AP2A2,AP1M1,ATP6V1G1,UBXN6,SORT1,CD74,SPPL2B,CLTA,

BORCS7,SYT11,ATP6V0D1,MTOR,AP2A2,AP1M1,ATP6V1G1,UBXN6,SORT1,CD74,SPPL2B,CLTA,

,ABI2,RDX,FLOT2,CADM3,KRT8,WASF2,SCRIB,ADAM10,ANK2,SDCBP,DAG1,NIBAN2,OBSL1,KIA

SREBF2,RAB11A,SEC23A,ARFGEF2,CDC42,ST3GAL6,SULF2,SEC16A,KPNA2,MGAT4B,RAB2A,RE

D1,FXYD6,APBB1,MICAL1,GAK,NTRK2,RIMS3,CAD,SCAMP5,FXR2,CLTA,SCAMP1,NRXN2,GRIK5,

,SEC16A,KPNA2,MGAT4B,RAB2A,RER1,MBTPS1,YKT6,NDST1,B4GAT1,GPSM1,TMEM167A,LMA

1,FLOT1,CALD1,SEPTIN7,CTTN,SLC2A1,TMOD2,EFR3B,RDX,SH3GL1,FLOT2,CAPZA1,TMEM63B

HD2,RAB5A,RAB35,SEC23IP,APOE,COPB1,VPS16,FURIN,SEC31A,SYT1,SLC18A3,M6PR,TMED5,

LTA,SCAMP1,ATP13A2,COPE,ERP29,HLA-A,SYT15,IQSEC1,CCDC88A,RABAC1,SEMA4C,SYNGR1,
TP6V0B,ZYX,TFRC,DLG4,ZDHHC5,ATP6V0E2,CDC42EP4,ATP6V0A1,RAB5A,RAB35,CLCN3,APOE,

LN5,TMEFF1,BGN,COL16A1,LGALS1,COL12A1,DAG1,ELN,CCDC80,NTN1,HTRA1,MFAP4,EMILIN

PD3,UBE2D2,DCUN1D2,LZTR1,OGT,PCGF2,FBXW5,PRKACA,CREBBP,ATXN7L3,UBE2I,ZER1,PA

M,SOD1,AKIRIN1,TXNIP,YME1L1,RBM15B,VDAC1,DCTN1,STAT3,XPOT,CAPN2,MTA1,STARD7,ND

M,SOD1,AKIRIN1,TXNIP,YME1L1,RBM15B,VDAC1,DCTN1,STAT3,XPOT,CAPN2,MTA1,STARD7,ND

,MLXIP,PPP1CB,NOP53,PARP1,TSPYL4,SSRP1,ACTR3,TCP1,RBMX,CBX1,H3-3A,TERF2IP,TOP1,

,TM7SF2,HM13,EMC10,SYNGR1,ABHD17A,TMEM43,RAB3A,LARGE1,RRBP1,ATF6B,SGPL1,ZDH
C16A,UBA52,MAP6,YKT6,SV2A,RPS27A,LMAN1,ATP6AP2,SLC44A2,GABARAPL1,RAB14,LDLR,FL

6,NUP98,CSNK1A1,HSF1,WDR82,RCC2,SETX,RASSF2,NUP133,THOC2,CTCF,PPP1R12A,THOC3,

AP,ACTN1,CLIP3,MAP2K2,TPM3,KIF3C,FYN,CCT8,CSNK1D,RAB11A,ARFGEF2,DYNLL1,CCT5,PAL

EPTIN9,OPA1,CENPF,MAPK8IP3,PPFIA3,AP3M1,GNAI1,UNC13B

ANF1,SHISA5,RAN,SUN2,CETN2,TM7SF2,CDK4,TRIM27,NUP153,NUP98,SEPHS1,TMEM43,AGPAT
ATF5,HIPK2,DRAP1,ATF7IP,CDK4,LDB1,BCL9,MEIS1,BEX1,RCOR2,SOX9,MED25,RBM14,RXRA,A

TMEM43,LARGE1,RRBP1,ATF6B,SGPL1,ATF6,VOPP1,GALNT2,XBP1,EXT1,CLCN3,PIGS,POMGNT

3R1,MAPK3,TLR6,LRP8,FURIN,PRNP,SCARB1,PPT1,PLPP3,MYADM,LRP6,GPM6B,STX12,GNAI1
3R1,MAPK3,TLR6,LRP8,FURIN,PRNP,SCARB1,PPT1,PLPP3,MYADM,LRP6,GPM6B,STX12,GNAI1
,VPS35,NRDC,VHL,PPP2CA,MAPK1,DDAH2,ARGLU1,ACADVL,ATP5PB,ILF3,BNIP3L,FKBP8,MAP2K

L1,VPS16,NBR1,ANKRD13B,VPS28,M6PR,VPS37B

CA,CROCC,RAB11FIP3,MZT2B,MDH1,SPPL2B,HSPA1A,RUVBL2,OBSL1,PCM1,MECP2,GPAA1,FE

XO7,UBE2E1,SPSB3,KLHL7,RNF40,ANAPC2,PLRG1

N1,HSPG2,LINGO1,GPC2,LTBP4,LTBP3,FBLN5,TMEFF1,BGN,COL16A1,LGALS1,LRIG1,COL12A

N1,HSPG2,LINGO1,GPC2,LTBP4,LTBP3,FBLN5,TMEFF1,BGN,COL16A1,LGALS1,LRIG1,COL12A

ATP2B4,IGFBP2,CALR,FSTL1,TMF1,ABHD2,HSPA1B,ZNF99,TUBB2B,COL5A1,COL4A1,CES3,PSA
HD2,HSPA1B,TUBB2B,COL5A1,COL4A1,CES3,PSAP,PGK1,RPLP0,CANX,CLU,PTMA,DDX5,ENO2,S
HD2,HSPA1B,ZNF99,TUBB2B,COL5A1,COL4A1,CES3,PSAP,RPLP0,H3-3B,CANX,CLU,PTMA,DDX5,
GSTP1,SMG1,CALM1,NPM1,TTC3,HSPD1,USP11,CSDE1,EIF3C,EIF4B,VCP,UBB,YWHAG,CKB,RPL
HD2,HSPA1B,ZNF99,TUBB2B,COL5A1,COL4A1,CES3,PSAP,PGK1,RPLP0,H3-3B,CANX,CLU,PTMA,
PSAP,RPLP0,H3-3B,CANX,CLU,PTMA,DDX5,SOX4,FKBP14,RPL13A,MAGED1,SEC62,RPL6,KAT6A,Y
5A1,COL4A1,CES3,PSAP,PGK1,RPLP0,H3-3B,CANX,CLU,PTMA,DDX5,ENO2,SOX4,FKBP14,RPL13
KB,COL4A2,RPL37A,FN1,SPEN,DDB1,YWHAQ,TPI1,EIF4A1,PPP2R1A,LDHB,PKD1,HLA-C,PDIA3,R
MG1,CALM1,NPM1,TTC3,PLD3,HSPD1,USP11,COL18A1,SNRNP200,IGF2,ATP5F1A,VCP,UBB,AFF
MG1,CALM1,NPM1,TTC3,PLD3,HSPD1,USP11,COL18A1,SNRNP200,IGF2,ATP5F1A,VCP,UBB,AFF
MG1,CALM1,NPM1,TTC3,PLD3,HSPD1,USP11,COL18A1,SNRNP200,IGF2,ATP5F1A,VCP,UBB,AFF
KB,COL4A2,RPL37A,FN1,SPEN,DDB1,YWHAQ,TPI1,EIF4A1,PPP2R1A,LDHB,PKD1,HLA-C,PDIA3,R
KB,COL4A2,RPL37A,FN1,SPEN,DDB1,YWHAQ,TPI1,EIF4A1,PPP2R1A,LDHB,PKD1,HLA-C,PDIA3,R
KB,COL4A2,RPL37A,FN1,SPEN,DDB1,YWHAQ,TPI1,EIF4A1,PPP2R1A,LDHB,PKD1,HLA-C,PDIA3,R
YH9,RPL7,SEPTIN2,P4HB,APP,RPL5,KLRD1,GDI1,SPTAN1,GSTP1,SMG1,CALM1,NPM1,PLXNB2,H
AN1,GSTP1,MARCKS,CALM1,PLXNB2,PLD3,HSPD1,COL18A1,IGF2,ATP5F1A,VCP,MARCKSL1,UBB
L1,RPL11,MORF4L1,ZMIZ1,XAF1,MIDN,RPS17,SBF1,SF3B2,GTF2F1,HNRNPU,NFE2L1,SMARCA5
PL23,RPSA,RPS24,RPS9,XRCC5,CIPC,SLC16A7,KPNB1,PAPOLA,PTCD3,CBX5,ATF4,PSMD2,NR2F
PS8,MYH9,RPL7,SEPTIN2,APP,RPL5,TMBIM6,PTMS,GSTP1,MARCKS,SMG1,CALM1,NPM1,TTC3,U
2,HNRNPK,YWHAB,RHOB,RAC1,TNC,ATP6V0C,L1CAM,PPIB,CNN2,ANXA5,PPP1CB,GDI2,TLN2,NO
2,HNRNPK,YWHAB,RHOB,RAC1,TNC,ATP6V0C,L1CAM,PPIB,CNN2,ANXA5,PPP1CB,GDI2,TLN2,NO
OL18A1,CSDE1,SMIM14,IGF2,VCP,UBB,MATR3,COL4A2,FN1,HACD3,SET,CHSY1,PKD1,HLA-C,PD
S19,RPL35,RPS10,RPL35A,HNRNPDL,RPLP2,HNRNPK,CIRBP,MYH10,RPL36,PTGES3,SND1,HNRN
,PAFAH1B1,AKAP12,CDC42BPB,AP2B1,GTF2F1,SLC4A8,CTNNA1,RPS16,RPL23,CNN3,TPM4,TNS

TN4,MMP14,RPS15,RPL12,RPL30,CLTC,GNB2,RPS5,RPS7,RPS19,TLN1,GJA1,CTNNB1,ANXA6,RF
JSP11,CSDE1,VCP,MARCKSL1,AFF4,TRIM28,RPL37A,MAPT,NONO,DDB1,SET,MAP9,TMSB10,INA,
JSP11,CSDE1,VCP,MARCKSL1,AFF4,TRIM28,RPL37A,MAPT,NONO,DDB1,SET,MAP9,TMSB10,INA,
,CPE,CTSD,SCARB2,LPCAT1,XRCC5,KPNB1,LLGL1,NCS1,RHOA,CLSTN1,PSMD2,KIF5B,RAB6B,C
,CPE,CTSD,SCARB2,LPCAT1,XRCC5,KPNB1,LLGL1,NCS1,RHOA,CLSTN1,PSMD2,KIF5B,RAB6B,C
PL30,RPS7,RPS19,PRKAR1A,CTNNB1,CYFIP2,DNAJC5,NDFIP1,SEPTIN11,LAMP1,ATP1A2,SEPTIN3
COL18A1,IGF2,ATP5F1A,VCP,MARCKSL1,UBB,YWHAG,CKB,COL4A2,RPL37A,FN1,SPEN,DDB1,YW

A,CCND1,PSMD3,SRSF1,DYNC1LI2,DYNLL2,CAND1,HSPB1,CASC3,POLR2B,GNG2,MSL1,HNRNP

,RPL36AL,RPL32,MRPL3,MRPS26,MRPS27,MRPL28
A,CLSTN1,ATP1A1,TMEM259,COL5A2,ATP2A2,RTN3,PRKCSH,PTN,TKT,EIF5A,CDH2,PDIA6,GPC3
0,ASXL1,KDM6B,PRPF6,TERF2IP,TAF7,SMARCA4,SRSF1,IPO7,GRB2,WDR6,PRKDC,SUPT5H,CAS
,APEX1,SERP1,EIF3H,ABCF1,RPS26,FXR2,RPS21,RPL27A,MRPS6,RPS29,RRBP1,RPS6KA2,RPL36
F6,PTEN,HNRNPA2B1,PPP1CC,TIMM50,TNKS,APEX1,HIF1A,SYMPK,SAFB2,STK35,PPP1R10,BAZ2
BCR,CNTNAP1,LRRC4B,KIF21B,PHB2,MAP6,DIP2B,GARS1,HDAC6,ULK1,BRSK1,HIF1A,CTTN,ATI
,MTREX,HSPA1A,RBM25,SF3A3,CD2BP2,SUMO1,SART3,EHMT2,LIMK1,PPP4R3A,SRSF10,TBXA2I
SEPTIN3,RAC1,CAPRIN1,CDK16,DNER,L1CAM,GLUL,MAP1A,PGRMC1,RAB27A,VAMP2,KLHL24,V
B,FYN,PTPRS,MAPK8IP1,SOD1,ARFGEF2,BMPR2,CDC42,PPP1R9B,CTNND1,GRIPAP1,DCTN1,KIF

,RTN4,LPCAT1,SLC22A17,PJA2,KPNB1,LLGL1,HK1,RHOA,PTCD3,CLSTN1,RAB6B,SLC7A5,ATP1A

NND2,HDAC6,YKT6,SV2A,PTPRF,IL6ST,ATP6AP2,NDN,AKAP9,FUS,FKBP4,OLFM1,GABARAPL1,PDI
LAPTM4A,ARPC2,DYNC1LI2,GRB2,NSG2,CLIP3,MAP2K2,CORO1C,SDF4,FYN,PTP4A2,B2M,GPR1(
AP5,CKAP4,MAP2,COTL1,CCT6A,RAC1,CAPRIN1,MMP2,DCN,MAP1A,EID1,CCT2,TBCD,SQSTM1,G
,MSN,LLGL1,GAP43,NCS1,RHOA,CLSTN1,ATF4,KIF5B,SLC7A5,NLGN2,CXADR,ATP1A1,WDR1,ATI
31,RAB27A,VAMP2,GNAI3,VCAN,TM9SF2,TGFBI,ALKBH5,RTN1,TCP1,GRINA,CLSTN3,SURF4,CNT

I,GSTP1,MARCKS,CALM1,PLXNB2,PLD3,HSPD1,COL18A1,IGF2,ATP5F1A,VCP,MARCKSL1,UBB,Y
AP43,NCS1,RHOA,CLSTN1,ATF4,KIF5B,SLC7A5,CXADR,ATP1A1,ATP2A2,CRMP1,SFPQ,TPM1,CDI

A,PPP1CC,IL6ST,CTTN,ATP6AP2,AKAP9,FUS,PALM,STRN4,ABI2,GABARAPL1,FLOT2,DICER1,ADA
ARP1,CTNND2,KCTD12,PTEN,PPP1CC,CTTN,ATP6AP2,AKAP9,FUS,PALM,STRN4,ABI2,ELAVL1,SH
A,PPP1CC,IL6ST,CTTN,ATP6AP2,AKAP9,FUS,PALM,STRN4,ABI2,GABARAPL1,FLOT2,DICER1,ADA
7,FAF2,CSNK2B,NME2,PSMC2,PA2G4,MAN2B1,DPP7,BACE1,ARSB,OLA1,ARPC5,DYNLT1,TRAPP(
7,FAF2,CSNK2B,NME2,PSMC2,PA2G4,MAN2B1,DPP7,ARSB,OLA1,ARPC5,DYNLT1,TRAPPC1,PSM
7,FAF2,CSNK2B,NME2,PSMC2,PA2G4,MAN2B1,DPP7,BACE1,ARSB,OLA1,ARPC5,DYNLT1,TRAPP(

P1,NDFIP1,DDOST,ST3GAL2,WLS,YWHAB,LAMP1,RHOB,CKAP4,HLA-B,RAB1A,ARCN1,HMGCR,R
P,NSG2,ATP6V1B2,LARS1,CCT8,UBQLN1,IFITM3,HGS,SOD1,PCYOX1,CDC42,WDR45B,CAPN2,TX

1,PRKDC,SCAF11,HNRNPM,SDHA,ILF2,RPS13,ADAR,DHX15,POLR2E,TRAF4,TRAF7,DFFB,BCL9L,
I,SNU13
,ATP6V0C,TMEM30A,TIMP3,PGRMC1,CNN2,CTSB,RAB27A,CCT2,VAMP2,GNAI3,GDI2,ABCA3,PD

OD1,PCYOX1,WDR45B,CAPN2,TXNDC5,PSMD1,FLCN,AP2A1,FLOT1,UBA52,RAB2A,CD164,NCO
OD1,PCYOX1,WDR45B,CAPN2,TXNDC5,PSMD1,FLCN,AP2A1,FLOT1,UBA52,RAB2A,CD164,NCO
LASP1,CCT7,PFN1,PNN,NES,NISCH,HNRNPC,LRPPRC,SPTBN2,GLG1,TPM1,CDH2,ACTN4,MMP14

EC23A,SERBP1,ARFGEF2,SYP,ANP32A,TRAF4,KIF5A,CAPN2,KIRREL1,SEC16A,MAP6,TRA2B,HDA(
3,RAB27A,CCT2,VAMP2,GNAI3,GDI2,ABCA3,PDXK,TCP1,MLEC,SURF4,RAB5B,SRP14,PYGB,TMED
B1A,CALU,ARCN1,HMGCR,RAC1,SQLE,HS6ST1,TMEM132A,PGRMC1,SRP9,NUCB1,FDFT1,RPN2,
,ARPC3,EXOC7,RAB3A,SETX,STMN4,UNC13A,ARPC5,GPRIN1,RAB5A,RUFY3,CLCN3,DYNLT1,GIT
K3B,ABHD17A,CRTC1,RNF216,NAPB,DLG4,WASF3,PRKAR2B,NAPA,EFNB3,NRP2,SHANK1,CTBP
3CA2,DPYSL2,TUBB2A,PPP2CA,MAPK1,MAPRE1,RABL6,DYNC1LI2,DYNLL2,GABARAP,HSPB1,MA
,SNX2,LMO4,ABL1,PHACTR4,DAG1,HIP1R,ACTA2,CYFIP1,CD44,ACAP2,LIMK1,SNX1,CCDC88A,N
,CRIP2,MAEA,MICAL3,EXOC7,PDE4DIP,DLG4,NSMF,RHOBTB2,UNC13A,CTBP2,CAPZA2,SEPTIN9
PBB1,SORT1,SLC38A2,DMWD,TUBB4A,EIF4A3,ELOVL5,CAD,AMFR,ABL1,FXR2,PPP5C,HIP1R,GRI

UT4,PSMA4,EIF2S1,DDX19A,MEX3B,CNOT1

NX1,TRIM27,PMEPA1,WDFY1,CLCN4,RCC2,TFRC,CCDC93,SORCS2,ERBB2,BACE1,PHB,RUSC1,N

JBXN6,SORT1,DOP1B,NSG1,RAB11FIP3,NTRK2,SNX2,SNX3,SCAMP5,SPPL2B,CHMP1A,CLTA,SC/
,NOTCH2,SELENON,UBXN4,ATG2B,KDELR2,FKBP8,KDELR1,RPS28,LSS,STT3B,B2M,PITPNM1,S
SLC2A3,DDX17,GPI,RACK1,BSG,FASN,SPTBN1,ATP5F1B,CALM3,GNB1,SHISA9,STMN2,RPS8,MY
,TMED2,NOTCH2,SELENON,UBXN4,ATG2B,KDELR2,FKBP8,KDELR1,RPS28,LSS,STT3B,B2M,PIT
C2,BACE1,COL7A1,ADAMTS7,ARSB,CHGB,DNAJB11,MAPK3,COL6A3,FAM20C,APOE,MFGE8,SEL

0D1,MTOR,AP2A2,AP1M1,ATP6V1G1,UBXN6,SORT1,CD74,SPPL2B,CLTA,ATP13A2,NSF,NDUFC2,
SMD4,OGT,PCGF2,FBXW5,PSMC3,MTOR,UBE3A,PRKACA,CREBBP,ATXN7L3,PSMC1,ZER1,FBXL
I2,SELENON,UBXN4,ATG2B,KDELR2,FKBP8,KDELR1,RPS28,LSS,STT3B,B2M,PITPNM1,STT3A,TN

ADCY1,SYNGAP1,SPOCK1,SEMA4C,YWHAZ,ABHD17A,CRTC1,ARHGEF9,DLG4,SORCS2,NSMF,BA

NSG2,CLIP3,KDELR1,CORO1C,B2M,IFITM3,PTPRS,HGS,SREBF2,RAB11A,SEC23A,DYNLL1,SYP,G
REEP2,GSK3A,REEP1,TUBB4A,CLIP2,KIFC2,KIF26A,FEZ1,NCKAP5L,KLC2,DYNLRB1,PBXIP1,APPB

AP1,COL2A1,CCT4,DPYSL2,TUBB2A,MAPRE1,PLEC,DYNC1LI2,DYNLL2,CAPZB,GABARAP,ACTN1,

ABHD17A,CRTC1,ARHGEF9,DLG4,SORCS2,NSMF,EFNB3,RUSC1,SOS1,SHANK1,CAMK2N1,ADD1,
4,DPYSL2,TUBB2A,MAPRE1,PLEC,DYNC1LI2,DYNLL2,CAPZB,GABARAP,ACTN1,HSPB1,CLIP3,MA
RHGEF9,DLG4,SORCS2,NSMF,EFNB3,RUSC1,SOS1,SHANK1,CAMK2N1,ADD1,ADGRL1,LRP8,PR
CK1,SEMA4C,ABHD17A,CRTC1,ARHGEF9,DLG4,SORCS2,NSMF,EFNB3,RUSC1,SOS1,SHANK1,CA
ATP13A2,NSF,NDUFC2,ANKFY1,RHEB,ATP6V0B,WDR11,TMEM9,SYNGR1,CLCN4,CLCN6,ARL8A,S
ATP13A2,NSF,NDUFC2,ANKFY1,RHEB,ATP6V0B,WDR11,TMEM9,SYNGR1,CLCN4,CLCN6,ARL8A,S

A1210,NOTCH1,SGSM3,EFNB2,JAM3,PARK7,PDLIM1,PPP1CA,NECTIN1,CDK4,DDX6,NDRG1,LIM
R1,MBTPS1,YKT6,NDST1,PREPL,B4GAT1,GPSM1,TMEM167A,LMAN1,TNKS,GALNT11,SLC2A1,A
NPTN,EFNB2,FXR1,PARK7,CYFIP1,PPP1CA,IQSEC1,HNRNPR,NECTIN1,C1QBP,RABAC1,SEMA4C,
N1,TNKS,GALNT11,SLC2A1,RHBDD2,ERGIC1,CYTH2,RAB1B,RAB14,GALNT1,RAB6A,ERGIC3,ADA

,SH3PXD2A,ACTR1B,WASF2,CAP1,DAPK3,MICAL1,MYL12B,CROCC,SH3PXD2B,PLS3,ABL1,SPPL

NECAP1

CLCN4,MCFD2,RAB3A,DLG4,VDAC2,UNC13A,BACE1,ZNRF1,RAB5A,RAB35,CLCN3,SEC23IP,COP
TLR6,ARF6,SYT1,SCARB1,SLC18A3,SH3KBP1,GSN,STX12,M6PR

1,LOXL2,PCOLCE,P3H1,SERPINE2,ATRN,COL7A1,THBS2,GDF10,ANXA1,NID1,COL6A3,MXRA5,AF

GR1,FBXL19,AMFR,USP47,UBE2L3,RNF11,WDTC1,RTF1,RUVBL2,ACVR1B,TCEA1,PFKP,IGF1R,FZ

JFV1,GPX4,PHB2,UBA52,TRA2B,UQCRC1,COX7C,PPP1CC,ULK1,MCM3AP,POM121C,RPS27A,TIM

JFV1,GPX4,PHB2,UBA52,TRA2B,UQCRC1,COX7C,PPP1CC,ULK1,MCM3AP,POM121C,RPS27A,TIM

PP2CA,SMARCA4,ARPC2,PBX2,DYNC1LI2,EIF3E,PRKDC,KDM1A,CCAR2,POLR2B,SMARCE1,SMA

IC9,ATF6,VOPP1,RAB5A,GALNT2,XBP1,EXT1,RAB35,CLCN3,PIGS,POMGNT1,FURIN,SYT1,PIGT,T
DT2,NCSTN,RAB6A,RAB5C,ITGAV,ADAM10,SYT11,ATP6V0D1,COPZ1,AP2A2,TGOLN2,AP1M1,NMN

SS18L1,PCNA,MCM7,MSH2,CENPF,DNMT1,CENPT,ZNF207,RAD50,WAPL,SIN3A,PPP2R5C

LD,BEX4,DCTN1,KIF5A,MTA1,KIF21B,MAP6,HDAC6,CUL3,EML1,MAPRE2,CTTN,MAP1LC3B,SERF

T3,PGRMC2,EI24,NSMF,TMPO,NUP205,LMNA,NUP133,CACYBP,GNAZ,KPNA3,ATF6,MAPK3,CTDN
ATF6B,CTBP2,TFDP1,AJUBA,SP1,RELA,GATAD2B,NCOA6,AKIRIN2,TLE4,MAFB,SKIL,SIN3A,HES6,M

T1,FURIN,SYT1,PIGT,TMCO1,SYT4,SLC35A4,QSOX1,TOMM40,LEMD2,STX12,DMPK

K2,QARS1,CCAR2,XPNPEP3,DHX30,FYN,SDHA,YWHAH,SLIT3,STXBP1,LONP1,IDH1,UQCRC2,MAPK

Z1,KAT2A,ATF5,NCKAP5L,FLII,DYNLRB1,CCDC88A,CETN2,H2AX,MAPKAPK2,EXOC7,CSNK1A1,ND

A1,DAG1,ELN,CCDC80,NTN1,HTRA1,HSD17B12,MFAP4,EMILIN1,LOXL2,PCOLCE,P3H1,SERPINE2

A1,DAG1,ELN,CCDC80,NTN1,HTRA1,HSD17B12,MFAP4,EMILIN1,LOXL2,PCOLCE,P3H1,SERPINE2

\P,PGK1,RPLP0,H3-3B,CANX,CLU,PTMA,DDX5,ENO2,SOX4,FKBP14,RPL13A,MAGED1,SEC62,RPL
SOX4,FKBP14,RPL13A,MAGED1,SEC62,RPL6,KAT6A,DVL3,YWHAE,KIF1A,RPL13,GNAS,SLC2A3,DI
SOX4,FKBP14,RPL13A,MAGED1,SEC62,RPL6,KAT6A,DVL3,YWHAE,KIF1A,RPL13,GNAS,SLC2A3,D
37A,MAPT,PSMD5,YWHAQ,SET,EGLN1,TPI1,EIF4A1,RPL19,PPP2R1A,RPL15,LDHB,RPS3,VAPA,AP2
DDX5,ENO2,SOX4,FKBP14,RPL13A,MAGED1,SEC62,RPL6,KAT6A,DVL3,YWHAE,KIF1A,RPL13,GN
WHAE,KIF1A,RPL13,GNAS,SLC2A3,DDX17,GPI,RACK1,BSG,FASN,ALDH6A1,SPTBN1,ATP5F1B,C/
3A,MAGED1,SEC62,RPL6,KAT6A,YWHAE,KIF1A,RPL13,GNAS,SLC2A3,DDX17,GPI,RACK1,BSG,FAS
PS3,BASP1,CDKL1,AP2M1,TPT1,RPS4X,RPS3A,STMN1,ITGB1,TUBB4B,RPL11,ANXA2,AGRN,VCL,
4,TRIM28,MATR3,COL4A2,MAPT,FN1,SPEN,NONO,DDB1,PSMD5,SET,RPL19,GTF3C4,SF3B1,PDI/
4,TRIM28,MATR3,COL4A2,MAPT,FN1,SPEN,NONO,DDB1,PSMD5,SET,RPL19,GTF3C4,SF3B1,PDI/
4,TRIM28,MATR3,COL4A2,MAPT,FN1,SPEN,NONO,DDB1,PSMD5,SET,RPL19,GTF3C4,SF3B1,PDI/
PS3,BASP1,CDKL1,AP2M1,TPT1,RPS4X,RPS3A,STMN1,ITGB1,TUBB4B,RPL11,ANXA2,AGRN,VCL,
PS3,BASP1,CDKL1,AP2M1,TPT1,RPS4X,RPS3A,STMN1,ITGB1,TUBB4B,RPL11,ANXA2,AGRN,VCL,
PS3,BASP1,CDKL1,AP2M1,TPT1,RPS4X,RPS3A,STMN1,ITGB1,TUBB4B,RPL11,ANXA2,AGRN,VCL,
ISPD1,COL18A1,SNRNP200,CSDE1,EIF3C,ATP5F1A,EIF4B,VCP,AFF4,TRIM28,COL4A2,RPL37A,M/
3,YWHAG,CKB,COL4A2,RPL37A,FN1,SPEN,DDB1,YWHAQ,TPI1,EIF4A1,PPP2R1A,LDHB,PKD1,HLA
,HNRNPA0,RPS16,RPL23,RPSA,RPS24,RPS9,XRCC5,CIPC,SLC16A7,KPNB1,PAPOLA,PTCD3,CBX5
1,ENC1,RPS14,CXADR,CHD4,JUND,RPS20,RPL26,SRSF4,RBBP4,MCL1,PNN,RPL31,NISCH,HNRNI
ISP11,SNRNP200,ATP5F1A,VCP,UBB,AFF4,TRIM28,MATR3,CKB,RPL37A,MAPT,SPEN,NONO,DDB
X4,RPL27,PTPRA,ACTR3,NCKAP1,PTK7,RPL38,MAPK1,MAPRE1,ARPC2,PLEC,ADAM9,RPS11,ACT
X4,RPL27,PTPRA,ACTR3,NCKAP1,PTK7,RPL38,MAPK1,MAPRE1,ARPC2,PLEC,ADAM9,RPS11,ACT
)IA3,SERINC1,RPS3,VAPA,HMGB1,AP2M1,TPT1,SEC61A1,RPS3A,RPL18,ITGB1,TUBB4B,UCHL1,N
IPH1,RPS12,RPS27,TOP2B,RPL23A,LUC7L3,DDX39B,SRP9,U2AF2,PABPN1,HNRNPL,TSR1,RPL27
3,SCARB2,RTN4,LAMA5,RPS9,PJA2,MSN,RPL24,GAP43,RPL14,NCS1,RPL22,RHOA,RPL9,CLSTN

PS10,RPLP2,HNRNPK,YWHAB,CADM4,RHOB,ATP1A2,RAC1,TNC,ATP6V0C,L1CAM,PPIB,DLG5,CNI
RPL19,PPP2R1A,RPL15,DNAH8,RPS3,CHD3,BASP1,VAPA,HMGB1,POLR2A,TPT1,RPS4X,RPS3A,RI
RPL19,PPP2R1A,RPL15,DNAH8,RPS3,CHD3,BASP1,VAPA,HMGB1,POLR2A,TPT1,RPS4X,RPS3A,RI
XADR,ATP1A1,VAT1,ASTN2,NISCH,SLC3A2,PFKL,PDIA6,RAB7A,ACTN4,MMP14,DDX3X,CLTC,MAG
XADR,ATP1A1,VAT1,ASTN2,NISCH,SLC3A2,PFKL,PDIA6,RAB7A,ACTN4,MMP14,DDX3X,CLTC,MAG
,RPS27,CTBP1,RAC1,ARHGDIA,CAPRIN1,GPC1,CDK16,GLUL,MAP1A,DLG5,PGRMC1,CDH11,EEF
VHAQ,TPI1,INA,EIF4A1,PPP2R1A,LDHB,PKD1,HLA-C,PDIA3,RPS3,BASP1,HMGB1,CDKL1,AP2M1,T

M,PHC2,SDHA,PFKM,STT3B,STT3A,UBQLN1,SF3A1,PRPS1,UQCRC2,NCOR2,P3H3,KDM3B,DYNL

,SULF1,PIP4K2B,ACSL6,RCN1,GPAT4,GJA1,MYO5A,COL6A2,GOLM1,APLP2,ARL6IP1,NDFIP1,DD
C3,PLCG1,POLR2B,MSL1,SMARCE1,SMARCC1,SUPT6H,HNRNPM,PHC2,SF3A1,NCOR2,KDM3B,I
AL,RPL32,MRPL3,MRPS26,MRPS27,MRPL28,DNAJC21
2A,EHMT1,YARS1,SNRPD3,KMT2E,EFTUD2,PNISR,SRSF2,PRPF4B,DAZAP2,SRSF11,PRPF19,PCGF
P6AP2,TMOD2,SPG7,PALM,CYTH2,FKBP4,OLFM1,DICER1,CLASP2,ADAM10,IGF2BP1,SYT11,ATP
R,AKAP8L,H2AX,DHX36,SRSF7,CSNK1A1,POLDIP3,UNC45A,RBM10,WAC,ILRUN,ABHD17A,CTR9,I
CAN,HNRNPA3,KLC1,CLSTN3,ACAP3,ORAI2,CNTNAP2,DPYSL2,MAPK1,ARPC2,FAM168B,LUZP6,A
5A,BCR,CAPN2,KIF21B,MAP6,DIP2B,FARP1,CTNND2,HDAC6,PTEN,YKT6,SV2A,PPP1CC,PTPRF,IL

1,TMEM259,ATP2A2,VAT1,RTN3,MCL1,CCNI,LRPPRC,XPO1,SLC3A2,SCRN1,TKT,GLG1,EIF5A,C

K1,ATP1B2,ADAM10,IGF2BP1,SYT11,GSK3A,MTOR,GOT2,APBB1,SORT1,SLC38A2,DMWD,TUB
7,IFITM3,PTP4A1,HGS,RAB11A,ARFGEF2,AMOTL2,GRIPAP1,AP2A1,FLOT1,UBA52,CD164,HDAC
AS7,HNRNPL,HSPH1,LMNB2,HNRNPA3,TCP1,CAMSAP1,KLC1,CCT3,NCKAP1,COL2A1,CCT4,DPY
P2A2,CRMP1,SFPQ,TPM1,CDH2,ACTN4,DDX3X,ARL4C,PRKAR1A,TLN1,MYO5A,CTNNB1,MACF1,
NAP2,TMED2,NOTCH2,OPTN,ARF3,MAPK1,MAPRE1,KDELR2,LAPTM4A,GRB2,PCSK1N,GABARA

WHAG,CKB,COL4A2,RPL37A,MAPT,FN1,SPEN,DDB1,YWHAQ,TPI1,CHSY1,INA,EIF4A1,PPP2R1A,L
H2,ACTN4,DDX3X,ARL4C,PRKAR1A,TLN1,MYO5A,CTNNB1,MACF1,MTSS2,CYFIP2,NDFIP1,WLS,

M10,IGF2BP1,SYT11,GSK3A,MTOR,PRKACA,APBB1,BNIP3,TANC2,SORT1,SLC38A2,DMWD,SMO
GL1,DCLK1,PRR12,PDPK1,SCRIB,ADAM10,IGF2BP1,SYT11,FXYD6,GSK3A,MTOR,PRKACA,APB
M10,IGF2BP1,SYT11,GSK3A,MTOR,PRKACA,APBB1,BNIP3,TANC2,SORT1,SLC38A2,DMWD,SMO
1,PSMD12,TGFB2,FAM3C,PRDX4,QSOX1,GSN,CSTB,GLB1,SCCPDH,CTSA,ARSA,TOLLIP
D12,TGFB2,FAM3C,PRDX4,QSOX1,GSN,CSTB,GLB1,SCCPDH,CTSA,ARSA,TOLLIP
1,PSMD12,TGFB2,FAM3C,PRDX4,QSOX1,GSN,CSTB,GLB1,SCCPDH,CTSA,ARSA,TOLLIP

AC1,APPL1,ATP6V0C,TMEM30A,LAPTM4B,HS6ST1,TMEM132A,PGRMC1,SRP9,NUCB1,TOMM20,
NDC5,PSMD1,FLCN,AP2A1,UBQLN2,FLOT1,UBA52,RAB2A,CD164,NCOA4,ULK1,ATP6AP2,GPC4,

HOXB5,HJURP,FBXO11,PPP1CC,ANP32B,RPL7L1,RPS27A,APEX1,STK35,BAZ2A,XRCC6,PELP1,S

K,TCP1,MLEC,SURF4,RAB5B,RAB11B,SRP14,PYGB,TMED2,VEGFA,MAPK1,PSMD3,PCSK1N,ACT

4,ATP6AP2,GPC4,PRDX6,GRN,ACLY,CLCN7,SLC44A2,HSPG2,TPP1,VPS41,GPC2,VPS4A,PGAP6,
4,ATP6AP2,GPC4,PRDX6,GRN,ACLY,CLCN7,SLC44A2,HSPG2,TPP1,VPS41,GPC2,VPS4A,PGAP6,
,DDX3X,CLTC,H1-0,RPS7,IK,PRKAR1A,TLN1,MYO5A,CTNNB1,NGRN,MACF1,MTSS2,MYL6,HNRN

6,EML1,APEX1,CHERP,PRDX6,RHBDD2,FUS,TAOK1,FKBP4,RAB1B,UBR5,VPS4A,RAB14,CNP,FL
2,VEGFA,MAPK1,PSMD3,PCSK1N,ACTN1,CAND1,ILF2,CCT8,B2M,STXBP1,IDH1,SOD1,DYNLL1,C
VAMP2,GNAI3,MEST,NOX4,TGFBI,RPL27,C4ORF3,RTN1,PLOD1,AHCYL1,MLEC,CLSTN3,SURF4,
1,ADGRL1,FGF13,PRNP,EPHA4,MAPK8IP3,SLC18A3,RNF40,UNC13B
,PPP3CA,PIN1,PRKCI,CLCN3,RELA,EPHB2,APOE,ARF6,SYT1,EPHA4,DNM1,ADAM23,PPP2R2A,PP
P7D1,CLIP3,MAP2K2,CCAR2,DIDO1,KIF3C,CCT8,TNKS2,CSNK1D,PTP4A1,RAB11A,ARFGEF2,DY
ME2,LDB1,ARPC3,SLK,PIP5K1C,WASF3,ARPC5,RAB5A,RUFY3,PRKCI,AJUBA,CLCN3,DYNLT1,FE
,FER,GMFB,ARF6,PPFIA3,MYADM,MYL9,NF2,GSN,GNAI1,EMD,UNC13B
K5,ATP13A2,SLC38A1,MAPK8IP2,DNAJB1,FXR1,ADNP,CYFIP1,FEZ1,PRPH,PPP1CA,DHX36,SEZ6L

GRN1,RAB5A,NIPA1,ANXA1,MAPK3,MIB2,CLCN3,SNX17,APOE,VPS16,ARF6,C1ORF210,EPHA4,

AMP1,ATP13A2,SCAMP4,ACAP2,HLA-A,ANKFY1,SUN2,SNX1,RAP2A,PMEPA1,ATP6V0B,NDRG1,TM
TT3A,TMED9,MAPK8IP1,SREBF2,CYP51A1,SEC23A,CDC42,SSR1,MSMO1,OS9,EXTL3,ALDH3A2,S
H9,RPL7,SEPTIN2,P4HB,APP,ERVV-1,RPL5,KLRD1,FSCN1,TMBIM6,SPTAN1,GSTP1,MARCKS,CAL
NM1,STT3A,TMED9,MAPK8IP1,SREBF2,CYP51A1,SEC23A,CDC42,SSR1,MSMO1,OS9,EXTL3,ALD
ENOF,COL3A1,QSOX1,PRSS23,MXRA8,ARSA

ANKFY1,RHEB,ATP6V0B,WDR11,TMEM9,SYNGR1,CLCN4,CLCN6,NAPB,ARL8A,MAN2B1,STARD3
19,AMFR,USP47,UBE2L3,RNF11,WDTC1,PSMB1,RTF1,RUVBL2,TCEA1,FZR1,UBE4B,TNFAIP1,CU
MED9,MAPK8IP1,SREBF2,CYP51A1,SEC23A,CDC42,SSR1,MSMO1,OS9,EXTL3,ALDH3A2,SEC16A,

CE1,EFNB3,RUSC1,SOS1,SHANK1,CAMK2N1,ADD1,ADGRL1,LRP8,SYT1,PRNP,SRCIN1,EPHA4,DN

RIPAP1,SGTA,HIP1,AP2A1,SEC16A,UBA52,MAP6,CD164,PI4KA,YKT6,SV2A,RPS27A,LMAN1,ATP6
P2,NDRG1,MAPRE3,PDE4DIP,TUBA1C,RCC2,MAP1S,KIF3A,TBCA,WDR47,RNF4,RUSC1,LZTS2,N

HSPB1,CLIP3,MAP2K2,FLNB,TPM3,CORO1C,KIF3C,FYN,CCT8,FBLN1,CSNK1D,COL27A1,RAB11

,ADGRL1,LRP8,PRNP,SRCIN1,EPHA4,DNM1,CACNG7
2K2,FLNB,TPM3,CORO1C,KIF3C,FYN,CCT8,FBLN1,CSNK1D,COL27A1,RAB11A,FLNC,ARFGEF2
NP,SRCIN1,EPHA4,DNM1,CACNG7
MK2N1,ADD1,ADGRL1,LRP8,PRNP,SRCIN1,EPHA4,DNM1,CACNG7
TARD3,ATP6V0E2,BRI3,ATP6V0A1,SLC12A4,CLCN3,TMEM59,SPG11,VPS16,SZT2,SCARB1,RAP1
TARD3,ATP6V0E2,BRI3,ATP6V0A1,SLC12A4,CLCN3,TMEM59,SPG11,VPS16,SZT2,SCARB1,RAP1

1,PIP5K1C,ASH1L,CCDC85C,ZYX,CDC42EP4,AFDN,UBN1,NECTIN2,ANXA1,TGFBR1,PPP3CA,HEG
AP9,RHBDD2,GRN,ERGIC1,CYTH2,RAB1B,RAB14,GALNT1,RAB6A,CLASP2,ERGIC3,ADAM10,SY
IP5K1C,SYNGR1,CLCN4,RAB3A,DLG4,VDAC2,UNC13A,CNTN1,NAPA,BACE1,ZNRF1,EFNB3,CTB
M10,COPZ1,MTOR,AP1M1,NMNAT2,SORT1,DOP1B,CD74,NSG1,USO1,SCAP,DENND5A,HS2ST1,

B,AUTS2,ARPC1A,TAOK2,HIP1R,ACTA2,JAM3,PDLIM1,MAEA,FLII,ARPC3,DAAM1,ZYX,MYL12A,C

B1,FURIN,SEC31A,SYT1,PPT1,DNM1,SYT4,SLC18A3,PAM,STX12,M6PR,UNC13B

POE,TGFB2,FGFBP3,MFGE8,COL3A1,POSTN,MFAP2,MXRA7,CSTB,CASK

ZR1,UBE4B,TNFAIP1,CUL1,DCAF5,WDR26,CAB39,TRRAP,PIK3C2B,KAT2A,CCNY,MAEA,PCGF3,CU

MM50,TNKS,PPP1R15A,SLC25A5,LRRC59,MTCH1,SPG7,SNN,CNP,TRAP1,DNAJB2,SFXN1,ATXN3,

MM50,TNKS,PPP1R15A,SLC25A5,LRRC59,MTCH1,SPG7,SNN,CNP,TRAP1,DNAJB2,SFXN1,ATXN3,

RCC1,SUPT6H,SSB,PHC2,TNKS2,USP7,SREBF2,NCOR2,ARNT2,KDM3B,WBP2,AKIRIN1,KDM4B,D

MCO1,SYT4,SLC35A4,QSOX1,TOMM40,LEMD2,STX12,DMPK

NAT2,SORT1,CD74,SMO,NSG1,USO1,SCAP,TAPBP,SCAMP5,CLTA,SCAMP1,TAOK2,HIP1R,MDM2,

P1,CEP170,FKBP4,GABARAPL1,KIFAP3,CNP,TUBGCP6,KRT8,CLASP2,REEP2,GSK3A,DAPK3,MICA

IEP1,SPIN1,TNPO3,NUP50,PRNP,CENPF,NUP58,LEMD2,DMPK,RB1CC1,PLRG1,NUTF2,EMD
/IED14

K8IP1,GHITM,SOD1,DYNLL1,TXNIP,DDIT4,YME1L1,VDAC1,STAT3,CAPN2,STARD7,NDUFV1,GRSF1

DRG1,PDE4DIP,PLK2,HSF1,GSK3B,ILRUN,KIF3A,PSMB5,SLC16A1,PRKAR2B,PSMA1,DDHD2,PPP1

,ATRN,COL7A1,ADAMTS7,THBS2,ANOS1,GDF10,ANXA1,NID1,COL6A3,MXRA5,LRRN1,APOE,TGF

,ATRN,COL7A1,ADAMTS7,THBS2,ANOS1,GDF10,ANXA1,NID1,COL6A3,MXRA5,LRRN1,APOE,TGF

.6,KAT6A,DVL3,YWHAE,KIF1A,RPL13,GNAS,SLC2A3,DDX17,GPI,RACK1,BSG,FASN,ALDH6A1,SPTE
DX17,GPI,RACK1,BSG,FASN,ALDH6A1,SPTBN1,ATP5F1B,CALM3,GNB1,STMN2,RPS8,MYH9,RPL7
DDX17,GPI,RACK1,BSG,FASN,ALDH6A1,SPTBN1,ATP5F1B,CALM3,GNB1,STMN2,RPS8,MYH9,RPL
2M1,TPT1,RPS4X,RPS3A,RPL18,STMN1,TUBB4B,UCHL1,RPL11,BTF3,NCAM1,ANXA2,RPLP1,ZMI
IAS,SLC2A3,DDX17,GPI,RACK1,BSG,FASN,ALDH6A1,SPTBN1,ATP5F1B,CALM3,GNB1,SHISA9,STM
ALM3,GNB1,STMN2,RPS8,MYH9,RPL7,SEPTIN2,P4HB,APP,IDS,RPL5,GDI1,TMBIM6,PTMS,SPTAN
SN,ALDH6A1,SPTBN1,ATP5F1B,CALM3,GNB1,STMN2,RPS8,MYH9,RPL7,SEPTIN2,P4HB,APP,IDS,F
DNAJA1,ARF1,PAFAH1B1,CDC42BPB,ITM2B,PXDN,PGAM1,RPS16,RPL23,LDHA,CPE,RPSA,TPM4,
A3,RPS3,CHD3,BASP1,HMGB1,CDKL1,TPT1,RPS4X,RPS3A,RPL18,TUBB4B,UCHL1,RPL11,MORF
A3,RPS3,CHD3,BASP1,HMGB1,CDKL1,TPT1,RPS4X,RPS3A,RPL18,TUBB4B,UCHL1,RPL11,MORF
A3,RPS3,CHD3,BASP1,HMGB1,CDKL1,TPT1,RPS4X,RPS3A,RPL18,TUBB4B,UCHL1,RPL11,MORF
DNAJA1,ARF1,PAFAH1B1,CDC42BPB,ITM2B,PXDN,PGAM1,RPS16,RPL23,LDHA,CPE,RPSA,TPM4,
DNAJA1,ARF1,PAFAH1B1,CDC42BPB,ITM2B,PXDN,PGAM1,RPS16,RPL23,LDHA,CPE,RPSA,TPM4,
DNAJA1,ARF1,PAFAH1B1,CDC42BPB,ITM2B,PXDN,PGAM1,RPS16,RPL23,LDHA,CPE,RPSA,TPM4,
APT,FN1,SPEN,NONO,DDB1,PSMD5,YWHAQ,EIF1,DCHS1,SET,EIF4A1,RPL19,PPP2R1A,RPL15,GT
-C,PDIA3,RPS3,BASP1,VAPA,HMGB1,CDKL1,AP2M1,TPT1,RPS4X,RPS3A,STMN1,ITGB1,TUBB4B,
,ATF4,PSMD2,NR2F1,ENC1,RPS14,CXADR,CHD4,JUND,RPS20,RPL26,SRSF4,RBBP4,MCL1,PNN,
PC,LRPPRC,TADA2B,KHDRBS1,XPO1,SLC3A2,CSNK1E,RPS25,TKT,CDH6,SFPQ,CCND2,PNMA1,P
1,PSMD5,EIF1,HACD3,SET,EGLN1,TPI1,RPL19,PPP2R1A,RPL15,GTF3C4,PKD1,SF3B1,PDIA3,SSB
N1,HSPB1,MAP2K2,FLNB,CORO1C,B2M,HYOU1,MPRIP,PCBP2,FLNC,RPS13,CD99L2,CDC42,PA
N1,HSPB1,MAP2K2,FLNB,CORO1C,B2M,HYOU1,MPRIP,PCBP2,FLNC,RPS13,CD99L2,CDC42,PA
ICAM1,ANXA2,AGRN,TIMP2,SARAF,COL11A1,RPL10,MARCHF6,VCL,AP3D1,SBF1,DNAJA1,ARF1,C
,SNRPN,HNRNPA3,RBMX,GCN1,SNRNP70,RPL36A,SRP14,PRPF6,RPS15A,RPL38,RPL37,G3BP2,S
1,RAB6B,NLGN2,RPS14,CXADR,ATP1A1,RPS20,CD81,WDR1,ATP2A2,LASP1,RTN3,PFN1,PNN,RPL

N2,CDH11,TBCD,ANXA5,PPP1CB,KLHL24,GDI2,TLN2,NOX4,RPL27,PTPRA,ACTR3,NCKAP1,PTK7,
PL18,STMN1,TUBB4B,RPL11,MORF4L1,BTF3,ANXA2,RPLP1,MIDN,RPL10,RPS17,VCL,DNAJA1,A
PL18,STMN1,TUBB4B,RPL11,MORF4L1,BTF3,ANXA2,RPLP1,MIDN,RPL10,RPS17,VCL,DNAJA1,A
ED2,MIF,PRKAR1A,GJA1,MYO5A,CD63,ANXA6,DNAJC5,APLP2,NDFIP1,DDOST,WLS,YWHAB,SND
ED2,MIF,PRKAR1A,GJA1,MYO5A,CD63,ANXA6,DNAJC5,APLP2,NDFIP1,DDOST,WLS,YWHAB,SND
1A2,VAMP2,GDI2,TLN2,HNRNPA3,PTPRA,CLSTN3,RAB11B,DPYSL2,RPL38,RPL37,PPP2CA,MAPK
TPT1,RPS4X,RPS3A,STMN1,ITGB1,TUBB4B,RPL11,ANXA2,AGRN,TIMP2,COL11A1,VCL,DNAJA1,A

L1,PPP1R9B,RBM15B,POLR2E,PSMA7,RBBP7,TRAF7,DCTN1,ARIH1,PRMT1,OS9,MTA1,EP300,R

DOST,WLS,CKAP4,HLA-B,RAB1A,CALU,ARCN1,HMGCR,RAC1,TNC,TMEM30A,PPIB,GLUL,SQLE,T
PRPF40A,DYNLL1,ADAR,DHX15,POLR2E,RBBP7,STAT3,BCL9L,XPOT,MTA1,EP300,RERE,SF1,PB

2,SF3A2,SART1,MTOR,PRKACA,CREBBP,SAP18,APBB1,UBE2I,TRIM8,DAPK3,THRAP3,EIF4A3,PT
6V0D1,GSK3A,EXOC4,APBB1,TANC2,NMNAT2,MICAL1,SLC38A2,TUBB4A,NTRK2,RANGAP1,CAD,
RBM14,HIF3A,SRP54,LMNA,THOC2,ATP6V0A1,AFDN,ZNF395,PRPF31,THOC3,SLC2A4RG,PIN1,A
TXN10,HSPB1,CASC3,NSG2,RPL28,ARF4,FLNB,FYN,STMN3,STXBP1,CSNK1D,PTPRS,MAPK8IP1,
6ST,CTTN,ATP6AP2,NDN,AKAP9,FUS,PALM,FKBP4,OLFM1,STRN4,ABI2,GABARAPL1,LDLR,FLOT

CND2,PDIA6,PIP4K2B,ACSL6,RAB7A,ITM2C,CLTC,GNB2,GPAT4,CHPF,GJA1,CD63,ANXA6,NGRN,E

34A,EIF4A3,ELOVL5,CAD,AMFR,ABL1,FXR2,PINK1,PPP5C,HIP1R,GRIK5,ATP13A2,ACTA2,SLC38A
6,YKT6,ULK1,RPS27A,ATP6AP2,RABEP1,VPS26B,GRN,ANTXR1,TPP1,VPS41,VPS4A,RAB14,LDLR,
SL2,TOP1,TUBB2A,G3BP2,MAPRE1,PLEC,BTBD2,DYNC1LI2,NYNRIN,DYNLL2,CAPZB,GABARAP,T
MTSS2,CYFIP2,NDFIP1,WLS,HNRNPK,MYH10,SEPTIN11,LAMP1,RHOB,ATP1A2,MAP2,SEPTIN3,RA
P,CAND1,NSG2,CLIP3,KDELR1,MAP2K2,ARF4,SDF4,STMN3,B2M,PITPNM1,TNKS2,CSNK1D,TMED

DHB,PKD1,HLA-C,PDIA3,RPS3,BASP1,HMGB1,CDKL1,AP2M1,TPT1,RPS4X,RPS3A,STMN1,ITGB1,
MYH10,SEPTIN11,LAMP1,ATP1A2,MAP2,SEPTIN3,RAC1,CAPRIN1,CDK16,APPL1,DNER,L1CAM,GLU

,NSG1,EIF4A3,ELOVL5,NTRK2,RANGAP1,DIP2A,AMFR,ABL1,FXR2,PPP5C,TAOK2,TRAK1,HIP1R,G
31,BNIP3,TANC2,NSG1,EIF4A3,NTRK2,ANK2,DIP2A,ABL1,FXR2,CLTA,DAG1,HIP1R,GRIK5,NPTN,N
,NSG1,EIF4A3,ELOVL5,NTRK2,RANGAP1,DIP2A,AMFR,ABL1,FXR2,PPP5C,TAOK2,TRAK1,HIP1R,G

RAB27A,EEF1A2,RPN2,MLXIP,VAMP2,GNAI3,TM9SF2,ABCA3,RPL27,RTN1,PLOD1,VPS35,MLEC,C
MAP1LC3B,PRDX6,GRN,ACLY,CLCN7,SLC44A2,HSPG2,TPP1,VPS41,GPC2,VPS4A,GABARAPL1,PC

TK24,NUCKS1,ATXN3,EDF1,SPTY2D1,NSD2,RBPJ,NOC2L,KDM4A,XPO6,SCD,EIF4A3,USO1,RCN2,

N1,CAND1,ILF2,CCT8,B2M,STXBP1,TMED9,IDH1,PTPRS,SOD1,DYNLL1,CDC42,SYP,VDAC1,SGTA

RAB14,LDLR,NCSTN,RAB5C,SPAG9,PLEKHM2,BORCS7,CAP1,SYT11,IST1,ATP6V0D1,MTOR,AP2A
RAB14,LDLR,NCSTN,RAB5C,SPAG9,PLEKHM2,BORCS7,CAP1,SYT11,IST1,ATP6V0D1,MTOR,AP2A
IPK,MYH10,CKAP5,SEPTIN11,RHOB,CKAP4,MAP2,SEPTIN3,COTL1,CCT6A,RAC1,ARHGDIA,CDK16

OT2,GALNT1,SPAG9,DICER1,ADAM10,IGF2BP1,SYT11,EHD1,EIF3G,PRKACA,APBB1,BAG5,SORT
DC42,TXNDC5,PSMD1,PDAP1,CST3,GARS1,ATP6AP2,A2M,PRDX6,GRN,ACLY,XRCC6,SLC44A2,IM
ATP6AP1,TMED2,NOTCH2,SELENON,OPTN,ARF3,UBXN4,ATG2B,KDELR2,PCSK1N,GABARAP,NSG

FIA3,CACNG7,ABR
NLL1,CCT5,CDC42,BEX4,DCTN1,KIF5A,MTA1,ACTR1A,PRKAR2A,FLCN,FLOT1,KIF21B,MAP6,PPP
R,BCAR1,PDXP,ARF6,PIP5K1A,SRCIN1,CDC42BPA,MYADM,PLXND1,NF2,PIEZO1,FGD1,ASAP3,GSI

2,ACSL4,CRTC1,ALCAM,MAP1S,KIF3A,SORCS2,NSMF,PRKAR2B,UBXN1,BACE1,SLC5A7,SOS1,CN

NKRD13B,VPS28,NF2,LRP6,CACNG7,ARRDC3,STX12,ABCA7

EM9,PIP5K1C,CLCN4,ABHD17A,RCC2,TFRC,CLCN6,ARL8A,SORCS2,STARD3,ATP6V0E2,ERBB2,F
;EC16A,KPNA2,UBA52,SLC39A7,RAB2A,RAB3GAP2,MBTPS1,NOMO1,AGPAT1,ULK1,GPSM1,POM
.M1,NPM1,PLXNB2,PLD3,HSPD1,SNRNP200,CSDE1,SMIM14,ATP5F1A,VCP,MARCKSL1,UBB,MAT
)H3A2,SEC16A,KPNA2,UBA52,SLC39A7,RAB2A,RAB3GAP2,MBTPS1,NOMO1,AGPAT1,ULK1,GPSM

,ATP6V0E2,NAPA,ATG9A,BRI3,ATP6V0A1,GABARAPL2,SLC12A4,CLCN3,TMEM59,SPG11,VPS16,S
L1,PSMD13,DCAF5,WDR26,USP14,TRRAP,CYFIP1,KAT2A,PSMD7,MAEA,PCGF3,CUL4A,CSNK2B,F
KPNA2,UBA52,SLC39A7,RAB2A,RAB3GAP2,MBTPS1,NOMO1,AGPAT1,ULK1,GPSM1,POM121C,R

NM1,CACNG7

AP2,RABEP1,SLC44A2,ANTXR1,VPS41,VPS4A,GABARAPL1,RAB14,LDLR,SH3GL1,FLOT2,PLEKHB
V1,SEPTIN9,DYNLT1,FGF13,TPX2,DYNC1I2,HID1,DNM1,CDK5RAP3,ZNF207,NINL,TMEM214,TPC

A,FLNC,ARFGEF2,DYNLL1,CCT5,PALLD,BEX4,DCTN1,KIF5A,MTA1,KIF21B,MAP6,HDAC6,CUL3,EI

,DYNLL1,CCT5,PALLD,BEX4,DCTN1,KIF5A,MTA1,KIF21B,MAP6,HDAC6,CUL3,EML1,ABCC9,MAPF

B,PRCP,AP3M1,ATP6V1D,M6PR,GNAI1,CTSA
B,PRCP,AP3M1,ATP6V1D,M6PR,GNAI1,CTSA

;1,ADD1,PRKCI,YBX3,AJUBA,FGF13,PDXP,NFASC,OBSCN,EPHA4,GJC1,CDC42BPA,PLPP3,RAP1B
T11,COPZ1,MTOR,TGOLN2,AP1M1,NMNAT2,SORT1,DOP1B,CD74,LYPLA2,NSG1,USO1,SCAP,DEI
?2,RAB5A,RAB35,CLCN3,GIT1,ZDHHC17,ADGRL1,EPHB2,ARF6,SYT1,PRNP,SRCIN1,PPT1,EPHA4,
TAPBP,SCAMP5,SPPL2B,LMAN2,CLTA,MGAT5,SCAMP1,SMPD3,NOTCH1,MGAT3,COPE,SCAMP4,C

DC42EP4,PPP1R12A,ARPC5,RAB5A,ANXA1,CAPZA2,SEPTIN9,ADD1,FER,GMFB,BCAR1,PTPN12,PD

JL4A,DAD1,CDK4,FAM8A1,HSD17B12,CCNG1,POLD2,GID8,CDK9,STUB1,DLD,RANBP9,RNMT,ZS

SLC25A23,MOAP1,ABCF1,IST1,OGT,DHCR7,CSE1L,IVD,TEX2,COX6A1,CHMP4B,MTOR,ATP5MC3
SLC25A23,MOAP1,ABCF1,IST1,OGT,DHCR7,CSE1L,IVD,TEX2,COX6A1,CHMP4B,MTOR,ATP5MC3

YNLL1,NAP1L4,TP53BP1,CIC,RBBP7,DCTN1,DFFB,STAT3,CAPN2,NAP1L1,MTA1,EP300,PBX1,BA

COPE,NDUFC2,CD44,HLA-A,TSPAN14,RAB18,EPN1,APPBP2,ATP6V0B,DEGS1,SEMA4C,SYNGR1,M

L1,REEP1,TUBB4A,PLS3,CLIP2,KIFC2,KIF26A,JAM3,PDLIM1,FEZ1,NCKAP5L,KLC2,PRPH,DYNLR

1,GPX4,PHB2,UBA52,DECR1,HJURP,GARS1,GLUD1,UQCRC1,COX7C,PTEN,YKT6,PPP1CC,ADH5,U

R12A,LZTS2,OLA1,ROCK2,UBN1,RTRAF,ARL2BP,PCNA,CUL7,SLC1A4,GIT1,DYNC1I2,CENPF,TC

B2,FGFBP3,MFGE8,COL3A1,POSTN,LRIG3,MFAP2,MXRA7,CSTB,CASK,MMP24

B2,FGFBP3,MFGE8,COL3A1,POSTN,LRIG3,MFAP2,MXRA7,CSTB,CASK,MMP24

3N1,ATP5F1B,CALM3,GNB1,STMN2,RPS8,MYH9,RPL7,SEPTIN2,P4HB,APP,IDS,RPL5,GDI1,FSCN1

7,SEPTIN2,P4HB,APP,IDS,RPL5,GDI1,FSCN1,TMBIM6,SPTAN1,GSTP1,MARCKS,SMG1,CALM1,NPI

.7,SEPTIN2,P4HB,APP,IDS,RPL5,GDI1,FSCN1,TMBIM6,PTMS,SPTAN1,GSTP1,MARCKS,SMG1,CAL

Z1,XAF1,MIDN,RPL10,WSB1,RPS17,VCL,SBF1,DNAJA1,ARF1,CYB5R3,PAFAH1B1,AKAP12,MAP4,A

MN2,RPS8,MYH9,RPL7,SEPTIN2,P4HB,APP,IDS,RPL5,GDI1,FSCN1,TMBIM6,PTMS,SPTAN1,GSTP1,

1,GSTP1,MARCKS,SMG1,CALM1,NPM1,TTC3,PLD3,HSPD1,USP11,COL18A1,SNRNP200,CSDE1,S

RPL5,GDI1,FSCN1,TMBIM6,PTMS,SPTAN1,GSTP1,MARCKS,SMG1,CALM1,NPM1,PLXNB2,TTC3,PI

CTSD,SCARB2,NACA,TSPAN3,LAMA5,RPS9,MSN,RPL24,KPNB1,COL1A2,RPL14,RPL22,RHOA,PSN

4L1,ANXA2,AGRN,ZMIZ1,TIMP2,XAF1,COL11A1,MIDN,RPS17,VCL,SBF1,CYB5R3,SF3B2,GTF2F1,

4L1,ANXA2,AGRN,ZMIZ1,TIMP2,XAF1,COL11A1,MIDN,RPS17,VCL,SBF1,CYB5R3,SF3B2,GTF2F1,

4L1,ANXA2,AGRN,ZMIZ1,TIMP2,XAF1,COL11A1,MIDN,RPS17,VCL,SBF1,CYB5R3,SF3B2,GTF2F1,

CTSD,SCARB2,NACA,TSPAN3,LAMA5,RPS9,MSN,RPL24,KPNB1,COL1A2,RPL14,RPL22,RHOA,PSN

CTSD,SCARB2,NACA,TSPAN3,LAMA5,RPS9,MSN,RPL24,KPNB1,COL1A2,RPL14,RPL22,RHOA,PSN

CTSD,SCARB2,NACA,TSPAN3,LAMA5,RPS9,MSN,RPL24,KPNB1,COL1A2,RPL14,RPL22,RHOA,PSN

F3C4,DNAH8,PKD1,HLA-C,SF3B1,PDIA3,RPS3,CHD3,BASP1,HMGB1,AP2M1,SEC61A1,RPS4X,RPS

RPL11,ANXA2,AGRN,TIMP2,VCL,AP3D1,DNAJA1,ARF1,CYB5R3,PAFAH1B1,CDC42BPB,ITM2B,AP2

RPL31,NISCH,HNRNPC,LRPPRC,TADA2B,KHDRBS1,XPO1,SLC3A2,CSNK1E,RPS25,TKT,CDH6,SFI

LXNA1,PIP4K2B,SOX12,ACTN4,DDX3X,RPS15,EIF3L,NANOG,MRFAP1,MAGED2,H1-0,RPS5,APLP1

P2,ZNF43,RPS3,CHD3,BASP1,VAPA,HMGB1,POLR2A,CDKL1,TPT1,RPS4X,RPS3A,RPL18,TUBB4B,

LLD,CAPN2,PRKAR2A,FLOT1,CD99,EVL,PI4KA,PPP1CC,MAPRE2,ITGB5,RPS2,CTTN,HSPG2,GNA1

LLD,CAPN2,PRKAR2A,FLOT1,CD99,EVL,PI4KA,PPP1CC,MAPRE2,ITGB5,RPS2,CTTN,HSPG2,GNA1

CYB5R3,PAFAH1B1,ITM2B,AP2B1,TMEM170A,SLC4A8,GOLGA8A,PXDN,PGAM1,NFE2L1,CTNNA1,

SRSF1,EIF3E,ILF3,RPS11,PRKDC,CASC3,RPL28,RPS28,HNRNPM,SSB,FAU,ILF2,EIF3F,SF3A1,HNF

31,SPTBN2,SLC3A2,PTN,RPS25,CDH6,CDH2,ACTN4,MMP14,RPS15,RPL12,RPL30,CLTC,GNB2,R

CNTNAP2,RPL38,VEGFA,CCND1,MAPK1,MAPRE1,ARPC2,PLEC,ADAM9,GRB2,RPS11,ACTN1,HSPE

RF1,CYB5R3,PAFAH1B1,AKAP12,MAP4,CDC42BPB,HNRNPU,NFE2L1,SMARCA5,LARP1,CTNNA1,F

RF1,CYB5R3,PAFAH1B1,AKAP12,MAP4,CDC42BPB,HNRNPU,NFE2L1,SMARCA5,LARP1,CTNNA1,F

1,LAMP1,RHOB,LGALS3BP,ATP1A2,CKAP4,HLA-B,RAB1A,TTR,CALU,ARCN1,COTL1,HRNR,RAC1,

1,LAMP1,RHOB,LGALS3BP,ATP1A2,CKAP4,HLA-B,RAB1A,TTR,CALU,ARCN1,COTL1,HRNR,RAC1,

1,ARPC2,DYNLL2,CAPZB,GABARAP,ACTN1,PLCG1,ARF4,CORO1C,FYN,STXBP1,TMED9,PTPRS,M

RF1,PAFAH1B1,CDC42BPB,ITM2B,PXDN,PGAM1,RPS16,RPL23,LDHA,CPE,RPSA,TPM4,CTSD,SCAI

ERE,PRKAR2A,NDUFV1,PSMD1,GTPBP1,CSNK2A1,PPP2CB,KLHL42,DECR1,GNG4,EP400,UBAP2L

MEM132A,PGRMC1,SRP9,NUCB1,FDFT1,RPN2,SQSTM1,GNAI3,MEST,VCAN,NOX4,RPL27,C4ORF

K1,CSNK2A1,FLOT1,EP400,ARID1B,UBAP2L,UPF1,TRA2B,HDAC6,SMC3,CBX6,HNRNPA2B1,MCM3

PN23,CENPB,TRIP12,CDC34,ABL1,LUC7L2,RBM4,MTREX,HSPA1A,MKNK2,RBM25,MDM2,SF3A3,

AMFR,FXR2,PINK1,AUTS2,TAOK2,TRAK1,DAG1,GRIK5,IGF1R,NPTN,SLC38A1,FXR1,PARK7,ADNI

PI5,FYTTD1,GATAD2B,CBX4,SETD1A,CCNL1,PIP5K1A,RNPS1,ACIN1,MAML1,AKAP17A,FAM193B,Z

SOD1,RAB11A,ARFGEF2,BMPR2,DYNLL1,CDC42,PPP1R9B,SYP,CTNND1,PALLD,GRIPAP1,DCTN1

2,DICER1,PDPK1,ADAM10,IGF2BP1,SYT11,GSK3A,MTOR,PRKACA,GOT2,APBB1,BNIP3,TANC2,SC

34GALNT4,DNAJC5,APLP2,SLC25A3,ARL6IP1,NDFIP1,DDOST,ST3GAL2,WLS,DUX4,YWHAB,LAMP

1,MAPK8IP2,DNAJB1,FXR1,PARK7,ADNP,CYFIP1,FEZ1,RNF157,PRPH,PPP1CA,DHX36,SEZ6L2,AC
TM9SF4,SH3GL1,FLOT2,PLEKHB2,NCSTN,RAB5C,PLEKHM2,MTMR4,WASF2,LZTR1,SYT11,AKT2
TDRD5,ACTN1,HSPB1,CASC3,CLIP3,MAP2K2,RPL28,FLNB,TPM3,CORO1C,KIF3C,DHX30,FYN,CC
C1,CAPRIN1,CDK16,APPL1,DNER,L1CAM,LAPTM4B,GLUL,MAP1A,DLG5,PGRMC1,RAB27A,SQST
09,GPR107,SREBF2,VGF,RAB11A,SEC23A,ARFGEF2,CDC42,ST3GAL6,SULF2,CAPN2,EXTL3,HIP1,

TUBB4B,RPL11,NCAM1,ANXA2,AGRN,TIMP2,COL11A1,VCL,DNAJA1,ARF1,CYB5R3,PAFAH1B1,CI
JL,MAP1A,DLG5,PGRMC1,RAB27A,SQSTM1,VAMP2,KLHL24,VCAN,TLN2,HNRNPA3,ACTR3,KLC1,

GRIK5,NPTN,DNAJB1,NSF,FXR1,ADNP,CYFIP1,FEZ1,PPP1CA,HNRNPR,NECTIN1,SYNGAP1,DHX36,
MAPK8IP2,MECP2,DNAJB1,NSF,FXR1,CYFIP1,RHEB,PPP1CA,IQSEC1,NCK2,ADCY1,SYNGAP1,CRK
GRIK5,NPTN,DNAJB1,NSF,FXR1,ADNP,CYFIP1,FEZ1,PPP1CA,HNRNPR,NECTIN1,SYNGAP1,DHX36,

CLSTN3,SURF4,RAB5B,RAB11B,ABCA2,ATP6AP1,TMED2,NOTCH2,RAPGEF1,OPTN,ARF3,MVB12E
GAP6,RAB14,LDLR,NCSTN,RAB5C,SPAG9,PLEKHM2,BORCS7,CAP1,SYT11,IST1,ATP6V0D1,MTOF

,PAFAH1B2,XRN2,ABL1,TAOK2,RTF1,TSPYL2,NOP56,GTF3C1,MACROH2A1,RBM4,MTREX,TXNRI

A,TXNDC5,PSMD1,PDAP1,CST3,GARS1,SV2A,BRSK1,ATP6AP2,A2M,PRDX6,GRN,ACLY,SEPTIN5,XF

2,AP1M1,ATP6V1G1,UBXN6,SORT1,CD74,NSG1,USP5,SNX2,ANK2,BGN,SPPL2B,SDCBP,CLTA,ATI
2,AP1M1,ATP6V1G1,UBXN6,SORT1,CD74,NSG1,USP5,SNX2,ANK2,BGN,SPPL2B,SDCBP,CLTA,ATI
5,SKP1,TRIB2,PEA15,HSBP1,MAP1A,DLG5,CNN2,CCT2,TBCD,GNAI3,GAS7,TLN2,TM9SF2,NOX4,H

1,AMBRA1,GAK,USO1,ARF5,NTRK2,RANGAP1,AMFR,ABL1,PINK1,TRAK1,TSNAX,HIP1R,CLIC1,HS
IPDH2,RAB14,FLOT2,NCSTN,RAB6A,RAB5C,SPAG9,ACTR1B,ITGAV,ADAM10,CAP1,IST1,PSMC3,P
32,CLIP3,FKBP8,KDELR1,RPS28,LSS,STT3B,B2M,PITPNM1,TNKS2,STT3A,TMED9,MAPK8IP1,SRE

2CB,KLHL42,MASTL,HDAC6,SMC3,CUL3,EML1,MAPRE2,BRSK1,SEPTIN7,TNKS,APEX1,CTTN,SLC
N,CYTH3,EPHA2,ABCA7

NTN2,CAMK2N1,MAP2K4,RAB5A,TMEM50A,SLC1A3,RUFY3,SLC1A4,DYNLT1,EPHB2,APOE,LRP8,

RHOBTB2,BACE1,ATG9A,ATP6V0A1,CHMP5,RAB5A,ANXA1,RAB35,CLCN3,TAB3,TMEM59,MICALL
121C,RPS27A,LMAN1,NPLOC4,CHERP,PPP1R15A,FKBP1A,ATP6AP2,LRRC59,SERP1,RHBDD2,VA
R3,YWHAG,COL4A2,MAPT,FN1,NONO,YWHAQ,HACD3,DCHS1,CHSY1,EIF4A1,MRC2,RPL19,PPP2
M1,POM121C,RPS27A,LMAN1,NPLOC4,CHERP,PPP1R15A,FKBP1A,ATP6AP2,LRRC59,SERP1,RHB

ZT2,SCARB1,RAP1B,ATP6V1F,PRCP,AP3M1,ATP6V1D,M6PR,RB1CC1,GNAI1,CTSA
RTCB,FAM8A1,ADRM1,POLD2,GID8,PPP2R5D,PSME3,STUB1,PSMC5,PSMC2,NCBP2,RANBP9,RN
PS27A,LMAN1,NPLOC4,CHERP,PPP1R15A,FKBP1A,ATP6AP2,LRRC59,SERP1,RHBDD2,VAPB,ERG

2,NCSTN,RAB6A,RAB5C,PLEKHM2,MTMR4,LZTR1,ITGAV,ADAM10,SYT11,ATP6V0D1,EHD1,COPZ
GS2,TBCB,EMD

ML1,ABCC9,MAPRE2,CALD1,LMAN1,FKBP1A,CTTN,MAP1LC3B,SLC2A1,SERP1,TMOD2,CEP170,

RE2,CALD1,LMAN1,FKBP1A,CTTN,MAP1LC3B,SLC2A1,SERP1,TMOD2,CEP170,FKBP4,LTBP4,GA

,MYADM,NF2,SH3KBP1,CYTH3,EPHA2,CASK,MXRA8
NND5A,HS2ST1,TAPBP,SCAMP5,SPPL2B,LMAN2,CLTA,MGAT5,SCAMP1,SMPD3,NOTCH1,MGAT3,
DNM1,ADAM23,PPFIA3,SYT4,SLC18A3,RNF40,STX12,CASK,UNC13B
GALNT10,HLA-A,B4GALT5,GOLPH3,GALNT17,RHEB,VPS51,ARFGAP2,EXT2,PMEPA1,GBA2,GOLG

XP,SRCIN1,CDC42BPA,MYADM,MYL9,NF2,GSN,TLNRD1,CASK,MYL6B

WIM6,WDR82,NUS1,PHKB,UBE2V1,SNRPD2,BCOR,PRMT5,SETD5,CTR9,CDKN1B,SNRPB,CHD8,F

,GOT2,NDUFA13,BNIP3,UBE2I,TOMM70,ATP5MC2,SORT1,AMBRA1,REEP1,CYB5B,ATP5MG,RAN

,GOT2,NDUFA13,BNIP3,UBE2I,TOMM70,ATP5MC2,SORT1,AMBRA1,REEP1,CYB5B,ATP5MG,RAN

Z1B,H2AZ1,CSNK2A1,BRD4,PPP2CB,HOXB5,ATRX,HJURP,EP400,ARID1B,UPF1,FBXO11,ZFR,SM(

MCFD2,RAB3A,TFRC,DLG4,ARL8A,SH3BP5,SORCS2,ATP6V0E2,UNC13A,BACE1,ATG9A,ZNRF1,BF

B1,DDX6,NME2,PBXIP1,APPBP2,ARPC3,CSNK1A1,NDRG1,MAPRE3,PDE4DIP,TUBA1C,RCC2,MAP

JLK1,RPS27A,TIMM50,APEX1,PPP1R15A,SLC25A5,MAP1LC3B,LRRC59,MTCH1,PRDX6,SPG7,PSN

EA2,RAD23A,CDK5RAP3,NINL,PPP4R3B,HIPK1,DPF2,ATP6V1D,WAPL,RBBP6,RBM39,GNAI1,EMD

,TMBIM6,PTMS,SPTAN1,GSTP1,MARCKS,SMG1,CALM1,NPM1,TTC3,PLD3,HSPD1,USP11,COL18

M1,TTC3,PLD3,HSPD1,USP11,COL18A1,CSDE1,SMIM14,IGF2,EIF3C,ATP5F1A,EIF4B,VCP,MARCK

.M1,NPM1,TTC3,PLD3,HSPD1,USP11,COL18A1,SNRNP200,CSDE1,SMIM14,IGF2,ATP5F1A,VCP,M

P2B1,MAT2A,GOLGA8A,OGA,PGAM1,NFE2L1,SMARCA5,CTNNA1,RPS16,RPL23,LDHA,CNN3,RPL

,MARCKS,SMG1,CALM1,NPM1,PLXNB2,TTC3,PLD3,HSPD1,USP11,COL18A1,SNRNP200,CSDE1,S

MIM14,IGF2,ATP5F1A,VCP,UBB,AFF4,TRIM28,MATR3,YWHAG,CKB,COL4A2,RPL37A,MAPT,FN1,

LD3,HSPD1,USP11,COL18A1,SNRNP200,CSDE1,SMIM14,IGF2,ATP5F1A,VCP,MARCKSL1,UBB,AF

MD2,SLC7A5,RPS14,ATP1A1,RPS20,CD81,WDR1,RPL26,CCT7,VAT1,PFN1,RPL31,HNRNPC,SLC3A

HNRNPU,PGAM1,NFE2L1,SMARCA5,HNRNPA0,RPS16,RPL23,RPSA,CTSD,SCARB2,RPS24,RPS9,X

HNRNPU,PGAM1,NFE2L1,SMARCA5,HNRNPA0,RPS16,RPL23,RPSA,CTSD,SCARB2,RPS24,RPS9,X

HNRNPU,PGAM1,NFE2L1,SMARCA5,HNRNPA0,RPS16,RPL23,RPSA,CTSD,SCARB2,RPS24,RPS9,X

MD2,SLC7A5,RPS14,ATP1A1,RPS20,CD81,WDR1,RPL26,CCT7,VAT1,PFN1,RPL31,HNRNPC,SLC3A

MD2,SLC7A5,RPS14,ATP1A1,RPS20,CD81,WDR1,RPL26,CCT7,VAT1,PFN1,RPL31,HNRNPC,SLC3A

MD2,SLC7A5,RPS14,ATP1A1,RPS20,CD81,WDR1,RPL26,CCT7,VAT1,PFN1,RPL31,HNRNPC,SLC3A

3A,RPL18,ITGB1,RPL11,MORF4L1,BTF3,ANXA2,RPLP1,COL11A1,RPL10,MARCHF6,RPS17,VCL,

2B1,SLC4A8,PXDN,PGAM1,CTNNA1,RPS16,RPL23,LDHA,CPE,RPSA,TPM4,CTSD,SCARB2,NACA,TS

PQ,CCND2,PNMA1,PLXNA1,PIP4K2B,SOX12,ACTN4,DDX3X,RPS15,EIF3L,NANOG,MRFAP1,MAGE

,RPS7,IK,RPS19,MIF,GJA1,JUN,CTNNB1,CD63,NGRN,RPS10,HNRNPDL,TARDBP,ARID1A,MXD4,S

,UCHL1,RPL11,MORF4L1,BTF3,ANXA2,ZMIZ1,XAF1,MIDN,RPL10,RPS17,SBF1,DNAJA1,PAFAH1B

2,RDX,FLOT2,NCSTN,SLC4A2,CLASP2,PDPK1,ITGAV,ADAM10,CAP1,GNA13,GAK,HMGA1,SDCBP,

2,RDX,FLOT2,NCSTN,SLC4A2,CLASP2,PDPK1,ITGAV,ADAM10,CAP1,GNA13,GAK,HMGA1,SDCBP,

FADS2,CPE,CTSD,SCARB2,RTN4,LPCAT1,RPS24,XRCC5,PJA2,RPL24,KPNB1,C1ORF43,IGFBP3,C

RNPD,RPL39,RPS13,PRPF40A,RPL34,ADAR,DHX15,SF1,EIF3D,GRSF1,UBA52,ZFP36L1,UPF1,TRA2

PS5,RPS7,RPS19,PRKAR1A,TLN1,GJA1,CTNNB1,ANXA6,MACF1,RPS10,CYFIP2,DNAJC5,RPLP2,N

31,PLCG1,MAP2K2,FLNB,CORO1C,YWHAH,B2M,HYOU1,MPRIP,VANGL2,PCBP2,FLNC,RPS13,BM

RPS16,RPL23,CNN3,RPL29,RPSA,TPM4,LPCAT1,RPS24,RPS9,XRCC5,CIPC,PJA2,MSN,RPL24,KPN

RPS16,RPL23,CNN3,RPL29,RPSA,TPM4,LPCAT1,RPS24,RPS9,XRCC5,CIPC,PJA2,MSN,RPL24,KPN

GPC1,CDK16,APPL1,ATP6V0C,TMEM30A,DNER,LAPTM4B,PPIB,TIMP3,PGRMC1,CNN2,CTSB,NU

GPC1,CDK16,APPL1,ATP6V0C,TMEM30A,DNER,LAPTM4B,PPIB,TIMP3,PGRMC1,CNN2,CTSB,NU

APK8IP1,PFN2,HNRNPD,RAB11A,RPS13,ARFGEF2,BMPR2,CDC42,PPP1R9B,SYP,CTNND1,PALLD,

RB2,NACA,TSPAN3,LAMA5,RPS9,MSN,RPL24,KPNB1,IGFBP3,COL1A2,RPL14,RPL22,RHOA,PSMD

,FBXO11,UQCRC1,HDAC6,HNRNPA2B1,PPP1CC,CUL3,ULK1,PSMD8,PPP1R15A,FBXO21,RHBDD

3,RTN1,PLOD1,AHCYL1,GRINA,COL2A1,MLEC,CLSTN3,SURF4,LAMC1,RPL36A,ATP6AP1,TMED2

3AP,POM121C,TNKS,HNRNPAB,HIF1A,SYMPK,BAP1,BAZ2A,XRCC6,PELP1,SNRPD3,INTS3,RALY,S

,CD2BP2,SUMO1,PARK7,SART3,PHPT1,EHMT2,LIMK1,HIPK2,ATF7IP,PPP4R3A,SRSF10,TBXA2R,S

P,CYFIP1,FEZ1,PRPH,MYCBP2,HNRNPR,NECTIN1,DHX36,ARPC3,EXOC7,SPOCK1,GSK3B,ALCAM,

ZNF106,SETD1B,SRSF8,BRD1,PPP4R3B,HIPK1,ALYREF,RBBP6,PLRG1,USP36,RBM39,NAMPT

,KIF5A,KCNQ2,BCR,CAPN2,CNTNAP1,LRRC4B,KIF21B,PHB2,MAP6,DIP2B,GARS1,FARP1,CTNND

ORT1,SLC38A2,DMWD,SMO,NSG1,TUBB4A,EIF4A3,ELOVL5,NTRK2,RANGAP1,CAD,DIP2A,AMFR,

1,RHOB,ATP1A2,CKAP4,HLA-B,RAB1A,CALU,ARCN1,HMGCR,RAC1,APPL1,ATP6V0C,TMEM30A,L

SL4,CRTC1,ALCAM,MAP1S,KIF3A,SORCS2,NSMF,PRKAR2B,UBXN1,CACYBP,BACE1,SLC5A7,SOS
,ATP6V0D1,EHD1,CHMP4B,AP2A2,TGOLN2,UBE2D3,NMNAT2,UBXN6,SORT1,DOP1B,CD74,PRD
T8,FBLN1,CSNK1D,ATXN2L,COL27A1,MEX3A,RAB11A,FLNC,ARFGEF2,DYNLL1,TP53BP1,CCT5,C
M1,VAMP2,KLHL24,VCAN,TLN2,HNRNPA3,ACTR3,KLC1,NCKAP1,CLSTN3,ACAP3,MDK,CCT4,OR
,AP2A1,SEC16A,KPNA2,CST3,MGAT4B,SLC39A7,MAP6,RAB2A,RER1,MBTPS1,YKT6,CUL3,NDST1

C42BPB,ITM2B,PXDN,PGAM1,RPS16,RPL23,LDHA,CPE,RPSA,TPM4,CTSD,SCARB2,NACA,TSPAN
,NCKAP1,CLSTN3,ACAP3,ORAI2,TNPO1,CNTNAP2,DPYSL2,NOTCH2,MAPK1,MAPRE1,ARPC2,FAM

PSMC2,PLK2,GSK3B,ABHD17A,CRTC1,ALCAM,MAP1S,DLG4,STAT1,SORCS2,NSMF,PRKAR2B,ZD
L,PSMC5,PSMC2,SPOCK1,SEMA4C,GSK3B,ABHD17A,CRTC1,RNF216,ARHGEF9,DLG4,WASF3,S
PSMC2,PLK2,GSK3B,ABHD17A,CRTC1,ALCAM,MAP1S,DLG4,STAT1,SORCS2,NSMF,PRKAR2B,ZD

,KDELR2,LAPTM4A,GABARAP,BNIP3L,NSG2,CLIP3,KDELR1,MAP2K2,CORO1C,RPS28,ATP6V1B
,AP2A2,AP1M1,ATP6V1G1,UBXN6,SORT1,CD74,AMBRA1,NSG1,USP5,SNX2,ANK2,BGN,SPPL2B,

1,TCEA1,PDHA1,MDM2,DNAJB1,CMPK1,CD2BP2,SUMO1,FXR1,EWSR1,SUB1,RAN,IP6K1,PPP1C

CC6,SLC44A2,IMPDH2,RAB14,FLOT2,NCSTN,RAB6A,RAB5C,SPAG9,ACTR1B,ITGAV,ADAM10,CA

P13A2,NSF,NDUFC2,FAF2,ANKFY1,SNX1,RHEB,PDGFRB,CAPN1,ASAH1,ATP6V0B,WDR11,TMEM9
P13A2,NSF,NDUFC2,FAF2,ANKFY1,SNX1,RHEB,PDGFRB,CAPN1,ASAH1,ATP6V0B,WDR11,TMEM9
HSPH1,LMNB2,ACTR3,YEATS2,TCP1,CAMSAP1,KLC1,CCT3,NCKAP1,CBX1,CCT4,SKI,ABCA2,DPY

PA1A,OBSL1,TRMT112,FXR1,PARK7,CYFIP1,COPS6,WRNIP1,CDK4,RNF41,DDX6,NME2,NDRG1,
RKACA,AP2A2,AP1M1,PAFAH1B2,SDCBP,SCAMP1,PSMB1,HSPA1A,KIAA1210,NOTCH1,PSMD13,C
BF2,RAB11A,CYP51A1,SEC23A,ARFGEF2,CDC42,SSR1,ST3GAL6,MSMO1,SULF2,OS9,EXTL3,AL

25A5,MAP1LC3B,SERP1,NDN,AKAP9,SEPTIN5,PSMB4,CEP170,TAOK1,FKBP4,VPS41,VPS4A,NUD

VPS16,TGFB2,TPX2,SRCIN1,PPT1,EPHA4,SYT4,LRP6,BRD1,CACNG7,PAM

.1,SNX17,VPS16,FURIN,ARF6,EPHA4,VPS28,LEPROT,FZD7,LRP6,STX12,VPS37B,ABCA7
.PB,ERGIC1,RAB1B,TRAM1,SSR3,GALNT1,DNAJB2,ATXN3,RAB6A,ERGIC3,RNF145,REEP2,DHCR7
R1A,RPL15,LDHB,PKD1,HLA-C,PDIA3,SERINC1,RPS3,BASP1,VAPA,HMGB1,AP2M1,SEC61A1,RPS4
DD2,VAPB,ERGIC1,RAB1B,TRAM1,RAB14,SSR3,GALNT1,DNAJB2,ATXN3,RAB6A,ERGIC3,RNF145

MT,PSMC4,PSMF1,ZSWIM6,UBE3C,UBE2V1,UBQLN4,BCOR,SETD5,CTR9,CDKN1B,PSMB5,RNF2
IC1,RAB1B,TRAM1,SSR3,GALNT1,DNAJB2,ATXN3,RAB6A,ERGIC3,RNF145,REEP2,DHCR7,COPZ1

:1,CHMP4B,AP2A2,TGOLN2,AP1M1,UBE2D3,NMNAT2,UBXN6,SORT1,DOP1B,CD74,SMO,NSG1,U

FKBP4,LTBP4,GABARAPL1,KIFAP3,CNP,TUBGCP6,KRT8,CLASP2,REEP2,GSK3A,FBLN5,DAPK3,M

3ARAPL1,KIFAP3,CNP,TUBGCP6,KRT8,CLASP2,REEP2,GSK3A,FBLN5,DAPK3,MICAL1,DEK,REEP1

NSF,COPE,SCAMP4,GALNT10,HLA-A,B4GALT5,GOLPH3,GALNT17,RHEB,VPS51,ARFGAP2,EXT2,

iA8B,CHST3,GOLGA4,ACBD3,AGPAT3,LARGE1,FAM20B,LMTK3,ZDHHC9,ATF6,GALNT2,GABARA

RNF20,IRS1,CACYBP,MED12,OGDH,ZDHHC9,ELOB,PRPF31,WDR5,SENP3,TGFBR1,ZSWIM8,KCTD

IGAP1,SLC25A36,FAM210B,ABL1,SUN1,SDCBP,PINK1,TRAK1,RGPD5,CLIC1,MFN2,FZR1,PCM1,S

IGAP1,SLC25A36,FAM210B,ABL1,SUN1,SDCBP,PINK1,TRAK1,RGPD5,CLIC1,MFN2,FZR1,PCM1,S

C3,NAP1L3,CBX6,HNRNPA2B1,PPP1CC,MCM3AP,SEPTIN7,KLHDC3,TNKS,APEX1,HIF1A,PPP1R10,

RI3,ATP6V0A1,VOPP1,RAB5A,ANXA1,RAB35,CLCN3,SEC23IP,SNX17,ZDHHC17,APOE,TLR6,COPB1

1S,KIF3A,TBCA,WDR47,LMNA,RNF4,RUSC1,LZTS2,NARF,NAV1,ANXA1,SEPTIN9,SLC1A4,DYNLT1,

MB4,IARS2,GPX1,SNN,SLC44A2,FKBP4,ZMIZ2,GABARAPL1,IREB2,CNP,TRAP1,NCSTN,SFXN1,ATX

A1,SNRNP200,CSDE1,SMIM14,IGF2,EIF3C,ATP5F1A,EIF4B,VCP,MARCKSL1,UBB,AFF4,TRIM28,M
SL1,UBB,YWHAG,CKB,COL4A2,RPL37A,MAPT,FN1,DDB1,PSMD5,YWHAQ,EIF1,HACD3,SET,EGLN
ARCKSL1,UBB,AFF4,TRIM28,MATR3,YWHAG,CKB,COL4A2,RPL37A,MAPT,FN1,SPEN,NONO,DDB
29,RPSA,TPM4,TNS3,EIF4H,RPS24,RPS9,XRCC5,CIPC,MSN,RPL24,KPNB1,RPL17,RPL14,HK1,NC
MIM14,IGF2,ATP5F1A,VCP,MARCKSL1,UBB,AFF4,TRIM28,MATR3,YWHAG,CKB,COL4A2,RPL37A
SPEN,NONO,DDB1,PSMD5,YWHAQ,EIF1,HACD3,SET,EGLN1,TPI1,CHSY1,RPL19,PPP2R1A,RPL15,
F4,TRIM28,MATR3,YWHAG,CKB,COL4A2,RPL37A,MAPT,FN1,SPEN,NONO,DDB1,PSMD5,YWHAQ,
2,PFKL,RPS25,TKT,GLG1,PEBP1,PDIA6,DDR1,PLXNA1,MEGF8,RAB7A,ACTN4,PRDX2,DDX3X,RPL
RCC5,CIPC,SLC16A7,KPNB1,PAPOLA,IGFBP3,COL1A2,PTCD3,CBX5,ATF4,PSMD2,NR2F1,ENC1,
RCC5,CIPC,SLC16A7,KPNB1,PAPOLA,IGFBP3,COL1A2,PTCD3,CBX5,ATF4,PSMD2,NR2F1,ENC1,
RCC5,CIPC,SLC16A7,KPNB1,PAPOLA,IGFBP3,COL1A2,PTCD3,CBX5,ATF4,PSMD2,NR2F1,ENC1,
2,PFKL,RPS25,TKT,GLG1,PEBP1,PDIA6,DDR1,PLXNA1,MEGF8,RAB7A,ACTN4,PRDX2,DDX3X,RPL
2,PFKL,RPS25,TKT,GLG1,PEBP1,PDIA6,DDR1,PLXNA1,MEGF8,RAB7A,ACTN4,PRDX2,DDX3X,RPL
2,PFKL,RPS25,TKT,GLG1,PEBP1,PDIA6,DDR1,PLXNA1,MEGF8,RAB7A,ACTN4,PRDX2,DDX3X,RPL
AP3D1,ARF1,CYB5R3,PAFAH1B1,MAP4,SF3B2,AP2B1,MAT2A,GTF2F1,HNRNPU,SMARCA5,LARP1
PAN3,LPCAT1,LAMA5,RPS9,XRCC5,MSN,RPL24,KPNB1,COL1A2,LLGL1,RPL14,NCS1,RPL22,RH
2,H1-0,RPS5,APLP1,RPS7,IK,RPS19,RPL35,MIF,GJA1,JUN,CTNNB1,CD63,NGRN,RPS10,HNRNP
HOX,HNRNPK,H1-10,INIP,CIRBP,DUX4,PTGES3,HNRNPH1,CKAP4,RPS12,RPS27,CTBP1,GPC1,TC
1,SF3B2,ZNF770,TMEM170A,GTF2F1,HNRNPU,OGA,NFE2L1,SMARCA5,HNRNPA0,RPS16,RPL23,
,DAG1,HSPA1A,EFNB2,CYFIP1,CD44,PDLIM1,MAP4K4,LIMK1,FLII,PDGFRB,CAPN1,JAK1,ARPC3,L
,DAG1,HSPA1A,EFNB2,CYFIP1,CD44,PDLIM1,MAP4K4,LIMK1,FLII,PDGFRB,CAPN1,JAK1,ARPC3,L
OL1A2,LLGL1,NCS1,RHOA,CBX5,CLSTN1,PSMD2,RAB6B,CXADR,ATP1A1,TMEM259,COL5A2,AT
B,HNRNPA2B1,RPL7L1,RPS2,RPS27A,HNRNPAB,FUS,EIF3H,SNRPD3,RALY,ELAVL1,SMG5,SRP72
DFIP1,HNRNPK,YWHAB,SEPTIN11,CADM4,LAMP1,RHOB,ATP1A2,SEPTIN3,RPS27,CTBP1,RAC1,A

PR2,CD99L2,AMOTL2,CDC42,PPP1R9B,CTNND1,PALLD,TRAF4,CAPN2,KIRREL1,PRKAR2A,CNTN
B1,RPL17,LLGL1,RPL14,RPL22,RHOA,PTCD3,CBX5,RPL9,ATF4,PSMD2,KIF5B,ENC1,RPS14,CHD
B1,RPL17,LLGL1,RPL14,RPL22,RHOA,PTCD3,CBX5,RPL9,ATF4,PSMD2,KIF5B,ENC1,RPS14,CHD
B1,RAB27A,CCT2,SNX22,SQSTM1,VAMP2,GNAI3,GDI2,TM9SF2,HSPH1,ABCA3,PRDX5,PDXK,PK
B1,RAB27A,CCT2,SNX22,SQSTM1,VAMP2,GNAI3,GDI2,TM9SF2,HSPH1,ABCA3,PRDX5,PDXK,PK
,GRIPAP1,PSMA7,VDAC1,KIF5A,STAT3,KCNQ2,BCR,SGTA,PRKAR2A,CNTNAP1,HIP1,LRRC4B,FLC
2,SLC7A5,GLIPR1L2,RPS14,CXADR,ATP1A1,RPS20,CD81,COL5A2,WDR1,HMGN2,RPL26,CCT7,

2,PPP1R10,SPG7,ERH,PSMB4,PELP1,SNRPD3,RALY,UBE2D2,NUDCD3,HINT1,GNA12,EFTUD2,DN

,NOTCH2,SELENON,VHL,UBXN4,MAPK1,ATG2B,KDELR2,GABARAP,BNIP3L,FKBP8,KDELR1,MAP
MG5,TLE1,HINT1,EFTUD2,SRSF2,INTS11,PRPF4B,DICER1,TCF3,RBM5,WBP11,MSH6,OGT,PRPF

H3BGRL3,AKAP8L,TRIM27,H2AX,CDK9,CELF3,DHX36,SRSF7,NUP98,CSNK1A1,POLDIP3,UNC45
RAB3A,SETX,KIF3A,DLG4,STMN4,ARL8A,STAT1,UNC13A,BACE1,SLC5A7,CNTN2,NRP2,ARPC5,G

2,ADCY6,HDAC6,PTEN,YKT6,SV2A,PPP1CC,PTPRF,ULK1,IL6ST,BRSK1,HIF1A,CTTN,ATP6AP2,AK
BL1,FXR2,PPP5C,TAOK2,TRAK1,HIP1R,GRIK5,ATP13A2,NPTN,SLC38A1,MAPK8IP2,DNAJB1,NS

APTM4B,SQLE,HS6ST1,TMEM132A,PGRMC1,SRP9,NUCB1,TOMM20,RAB27A,EEF1A2,FDFT1,RP

1,CNTN2,CAMK2N1,MAP2K4,RAB5A,TMEM50A,SLC1A3,RUFY3,SLC1A4,DYNLT1,EPHB2,APOE,L
K3,NSG1,PTPN23,RAB11FIP3,NTRK2,SNX2,ANK2,RNF11,SNX3,SCAMP5,SPPL2B,CHMP1A,CLTA,U
DC42,PALLD,BEX4,DCTN1,KIF5A,EDC4,MTA1,GRSF1,KIF21B,ZFP36L1,MAP6,UBAP2L,UPF1,HDA
I2,TNPO1,CNTNAP2,DPYSL2,NOTCH2,MAPK1,MAPRE1,ARPC2,FAM168B,DYNLL2,LUZP6,CAPZB,
,PREPL,B4GAT1,GPSM1,TMEM167A,LMAN1,TNKS,PPP1R15A,PODXL2,GALNT11,CTTN,GPC4,SL

I3,LAMA5,RPS9,XRCC5,MSN,RPL24,KPNB1,IGFBP3,COL1A2,RPL14,RPL22,RHOA,CLSTN1,PSMD
168B,DYNLL2,LUZP6,CAPZB,GABARAP,ATXN10,ACTN1,HSPB1,CASC3,NSG2,PLCG1,RPL28,ARF

HHC5,UBXN1,BACE1,SLC5A7,ARHGAP33,LMTK3,SHANK1,CAMK2N1,MAP2K4,RAB5A,MAF1,PPP
RCS2,NSMF,PRKAR2B,CNTN1,NAPA,EFNB3,RUSC1,SOS1,CNTN2,NRP2,ARHGAP33,SHANK1,CA
HHC5,UBXN1,BACE1,SLC5A7,ARHGAP33,LMTK3,SHANK1,CAMK2N1,MAP2K4,RAB5A,MAF1,PPP

2,B2M,PITPNM1,TNKS2,CSNK1D,TMED9,IFITM3,PTPRS,HGS,SREBF2,RAB11A,SEC23A,ARFGEF2,
DCBP,CLTA,ATP13A2,NSF,NDUFC2,FAF2,ANKFY1,SNX1,RHEB,PDGFRB,CAPN1,TBC1D14,ASAH1

A,IQSEC1,VPS51,COX7A2L,CDK4,DDX6,TRIM27,NUP153,C1QBP,FAM32A,DEAF1,EXOSC10,RNM

P1,SYT11,IST1,ATP6V0D1,PSMC3,PRKACA,AP2A2,AP1M1,PAFAH1B2,SCAMP5,SDCBP,CLTA,SCA

,SYNGR1,CLCN4,PA2G4,RAB3A,CLCN6,ARL8A,MAN2B1,STARD3,DPP7,ATP6V0E2,BACE1,ZNRF1,
,SYNGR1,CLCN4,PA2G4,RAB3A,CLCN6,ARL8A,MAN2B1,STARD3,DPP7,ATP6V0E2,BACE1,ZNRF1,
L2,TUBB2A,PPP2CA,MAPK1,MAPRE1,ARPC2,DSTN,PLEC,RABL6,DYNC1LI2,DYNLL2,LUZP6,CAP

MAPRE3,SLK,UNC45A,LIMS1,PSMF1,HSF1,RAB3A,UBQLN4,MAP1S,TFRC,STC2,STAT1,PRKAR2B,
AB39,NDUFC2,APEH,CYFIP1,CD44,PSMD7,FAF2,TSPAN14,CSNK2B,CAPN1,TBXA2R,NME2,ASAH1
H3A2,SEC16A,KPNA2,UBA52,MGAT4B,SLC39A7,RAB2A,RER1,RAB3GAP2,MBTPS1,YKT6,NOMO

CD3,GABARAPL1,DZIP1,LSM14A,KMT2E,KIFAP3,CNP,TUBGCP6,SPAG9,ACTR1B,CLASP2,REEP2,

7,COPZ1,TEX2,PTGFRN,MTOR,SSR2,CERS2,CCDC47,SORT1,CD74,REEP1,SCD,NSG1,ELOVL5,AT
4X,OR8K1,RPL18,STMN1,ITGB1,UCHL1,RPL11,NCAM1,ANXA2,AGRN,SARAF,RPL10,MARCHF6,RI
5,REEP2,DHCR7,COPZ1,TEX2,PTGFRN,MTOR,SSR2,CERS2,CCDC47,SORT1,CD74,REEP1,SCD,NS

0,PRKAR2B,PSMA1,CACYBP,MED12,PSMD10,ELOB,WDR5,PPP3CA,RTRAF,ZSWIM8,PSMA6,TRAF
1,TEX2,PTGFRN,MTOR,SSR2,CERS2,CCDC47,SORT1,CD74,REEP1,SCD,NSG1,ELOVL5,ATP13A1,S

SO1,SCAP,RAB11FIP3,NTRK2,SNX2,TAPBP,SNX3,SCAMP5,SPPL2B,CHMP1A,CLTA,SCAMP1,TAOK

ICAL1,DEK,REEP1,MYL12B,TUBB4A,ANK2,PLS3,COL16A1,CLIP2,DAG1,KIFC2,ELN,OBSL1,ACTA2

,MYL12B,TUBB4A,ANK2,PLS3,COL16A1,CLIP2,DAG1,KIFC2,ELN,OBSL1,ACTA2,KIF26A,JAM3,CA

PMEPA1,GBA2,GOLGA8B,CHST3,WDR11,GOLGA4,ACBD3,AGPAT3,LARGE1,BACE1,ATG9A,FAM2

PL2,B4GALT2,EXT1,PRKCI,CHPF2,CLCN3,TMEM59,POMGNT1,ZDHHC17,COPB1,FURIN,HID1,CC

D10,MIB2,PCNA,CUL7,CUL4B,BRPF3,THG1L,TRPC4AP,NCOA6,CBX4,SETD1A,FBXO7,UBE2E1,CC

SLC25A6,NDUFA4,SUMO1,PARK7,NDUFC2,APEH,ALDH18A1,BANF1,PTPN1,COX8A,SHISA5,RAN,SU
SLC25A6,NDUFA4,SUMO1,PARK7,NDUFC2,APEH,ALDH18A1,BANF1,PTPN1,COX8A,SHISA5,RAN,SU

,BAZ2A,XRCC6,PELP1,EHMT1,INTS3,SOX2,ZMIZ2,KMT2E,KIFAP3,TTC37,MACROH2A2,NUCKS1,F

,NFASC,SEC31A,SYT1,MFGE8,SCARB1,SYT4,RAP1B,SLC18A3,GPRC5B,LRIG3,PRCP,SH3KBP1,A

FGF13,TPX2,DYNC1I2,HID1,DNM1,CDK5RAP3,ZNF207,NINL,TMEM214,TPGS2,TBCB,EMD

N3,NMT1,SOD2,SLC25A23,MOAP1,FOXK2,OGT,IVD,COX6A1,GSK3A,MTOR,PACS2,ATP5MC3,PR

ATR3,YWHAG,CKB,COL4A2,RPL37A,MAPT,FN1,SPEN,NONO,DDB1,PSMD5,YWHAQ,EIF1,HACD3,
1,MAP9,TPI1,CHSY1,TMSB10,INA,EIF4A1,RPL19,PPP2R1A,RPL15,GTF3C4,LDHB,DNAH8,PKD1,
1,PSMD5,YWHAQ,EIF1,HACD3,SET,EGLN1,MAP9,TPI1,CHSY1,TMSB10,INA,RPL19,PPP2R1A,RPL
S1,RPL22,RHOA,PTCD3,RPL9,PSMD2,NR2F1,KIF5B,RAB6B,SLC7A5,RPS14,RPS20,EIF5,WDR1,
,MAPT,FN1,SPEN,NONO,DDB1,PSMD5,YWHAQ,EIF1,HACD3,SET,EGLN1,MAP9,TPI1,CHSY1,TMS
GTF3C4,PKD1,HLA-C,SF3B1,PDIA3,SERINC1,SSBP2,ZNF43,RPS3,CHD3,BASP1,VAPA,HMGB1,PO
EIF1,HACD3,SET,EGLN1,TPI1,CHSY1,EIF4A1,RPL19,PPP2R1A,RPL15,GTF3C4,LDHB,DNAH8,PKD
12,ITM2C,RPL30,CLTC,GNB2,RPS5,RPS19,MIF,TLN1,MYO5A,CTNNB1,COL6A2,CD63,ANXA6,R
RPS14,CXADR,CHD4,JUND,RPS20,COL5A2,RPL26,VAT1,SRSF4,RBBP4,MCL1,PNN,RPL31,NISCH,
RPS14,CXADR,CHD4,JUND,RPS20,COL5A2,RPL26,VAT1,SRSF4,RBBP4,MCL1,PNN,RPL31,NISCH,
RPS14,CXADR,CHD4,JUND,RPS20,COL5A2,RPL26,VAT1,SRSF4,RBBP4,MCL1,PNN,RPL31,NISCH,
12,ITM2C,RPL30,CLTC,GNB2,RPS5,RPS19,MIF,TLN1,MYO5A,CTNNB1,COL6A2,CD63,ANXA6,R
12,ITM2C,RPL30,CLTC,GNB2,RPS5,RPS19,MIF,TLN1,MYO5A,CTNNB1,COL6A2,CD63,ANXA6,R
12,ITM2C,RPL30,CLTC,GNB2,RPS5,RPS19,MIF,TLN1,MYO5A,CTNNB1,COL6A2,CD63,ANXA6,R
,CTNNA1,RPS16,RPL23,RPL29,RPSA,TPM4,EIF4H,NACA,LAMA5,RPS24,RPS9,XRCC5,RPL24,KPN
A,CLSTN1,PSMD2,KIF5B,RAB6B,SLC7A5,RPS14,CXADR,ATP1A1,RPS20,CD81,WDR1,ATP2A2,R
L,TARDBP,ARID1A,MXD4,SHOX,HNRNPK,H1-10,INIP,CIRBP,RPL36,DUX4,PTGES3,CKAP5,HNRN
P2B,HDAC2,SKP1,BCLAF1,PPIB,PEA15,HSBP1,LUC7L3,PTTG1IP,CDKN1A,CTDSP2,CPSF2,DDX3
LDHA,RPSA,NACA,RTN4,LAMA5,RPS24,BRD3,RPS9,XRCC5,CIPC,MSN,SLC16A7,KPNB1,RPL17,P
IMS1,YWHAZ,PIP5K1C,ALCAM,ZYX,RPS29,PPP1R12A,ARPC5,MDC1,NECTIN2,ANXA1,MAPK3,AD
IMS1,YWHAZ,PIP5K1C,ALCAM,ZYX,RPS29,PPP1R12A,ARPC5,MDC1,NECTIN2,ANXA1,MAPK3,AD
P2A2,VAT1,RTN3,ASTN2,NISCH,PRKCSH,CCNI,LRPPRC,XPR1,XPO1,PTN,PFKL,SCRN1,TKT,GLG1
,LSM14A,EFTUD2,SRSF2,PRPF4B,DICER1,RBM12,RBM5,ABCF1,MSI1,WBP11,IGF2BP1,PRPF19,
HGDIA,CAPRIN1,GPC1,CDK16,TNC,ATP6V0C,L1CAM,PPIB,GLUL,MAP1A,DLG5,PGRMC1,CNN2,

AP1,FLCN,FLOT1,CD99,CTNND2,EVL,PI4KA,SV2A,PPP1CC,MAPRE2,ITGB5,RPS2,CTTN,SYMPK,S
4,JUND,RPS20,WDR1,HMGN2,RPL26,CRMP1,LASP1,CCT7,RBBP4,PFN1,PNN,SPC25,NES,RPL31,
4,JUND,RPS20,WDR1,HMGN2,RPL26,CRMP1,LASP1,CCT7,RBBP4,PFN1,PNN,SPC25,NES,RPL31,
1,TBC1D16,TCP1,KLC1,VPS35,MLEC,SURF4,RAB5B,CCT4,CNTNAP2,RAB11B,SRP14,ABCA2,PY
1,TBC1D16,TCP1,KLC1,VPS35,MLEC,SURF4,RAB5B,CCT4,CNTNAP2,RAB11B,SRP14,ABCA2,PY
T1,KPNA2,PHB2,FARP1,CTNND2,KCTD12,PTEN,SV2A,PPP1CC,BRSK1,CTTN,ATP6AP2,GPC4,SLC
AT1,RTN3,PFN1,RPL31,HNRNPC,SPTBN2,SLC3A2,PTN,PFKL,HDLBP,RPS25,TKT,GLG1,PEBP1,P

AJB2,PRPF4B,QPRT,DICER1,DCUN1D2,LZTR1,PSMD4,OGT,PRPF19,PCGF2,FBXW5,MBD3,SF3A

2K2,SDF4,RPS28,FYN,LARS1,LSS,STT3B,B2M,PITPNM1,STT3A,UBQLN1,TMED9,MESD,HYOU1,P
19,MED13,PCGF2,NOC2L,MBD3,SF3A2,SART1,DHX38,YY1,SMARCD1,CREBBP,SAP18,ATXN7L3,

A,RANBP9,RBM10,ZBTB4,WAC,HSF1,ILRUN,ABHD17A,CRTC1,SETX,FBL,CTR9,ATMIN,SH3BP5,R
RIN1,LMTK3,ROBO3,MAP2K4,RAB5A,NAV1,MAF1,RUFY3,CLCN3,DYNLT1,SPG11,BCAR1,GIT1,

P9,TMOD2,FUS,SPG7,PALM,CYTH2,FKBP4,OLFM1,STRN4,ABI2,GABARAPL1,KIFAP3,PRR12,FL
,FXR1,ADNP,CYFIP1,FEZ1,PRPH,PPP1CA,HNRNPR,NECTIN1,SYNGAP1,DHX36,PSMC2,PLK2,GSK

N2,MLXIP,SNX22,VAMP2,GNAI3,MEST,TM9SF2,NOX4,LMNB2,ABCA3,RPL27,C4ORF3,RTN1,PLOD

RP8,VPS16,TGFB2,TPX2,SRCIN1,PPT1,EPHA4,MAPK8IP3,SYT4,GNAO1,NF2,LRP6,BRD1,CACNG7,
ISP10,SCAMP1,TRAK1,ATP13A2,ARRDC4,TNFAIP1,SCAMP4,ACAP2,RNF157,HLA-A,PTPN1,RAN,AN
AC6,PPP1CC,CUL3,EML1,ABCC9,MAPRE2,CALD1,LMAN1,SEPTIN7,FKBP1A,CTTN,SYMPK,MAP1LC
,GABARAP,ATXN10,ACTN1,HSPB1,CASC3,NSG2,PLCG1,RPL28,ARF4,FLNB,CORO1C,SDF4,KIF3C
C2A1,AKAP9,RHBDD2,GRN,VAPB,ERGIC1,CYTH2,HSPG2,TPP1,RAB1B,VPS41,SEC14L1,PKDCC,G

2,SLC7A5,GLIPR1L2,RPS14,CXADR,ATP1A1,RPS20,CD81,COL5A2,WDR1,HMGN2,RPL26,CCT7,V
4,FLNB,CORO1C,SDF4,KIF3C,FYN,CCT8,STMN3,PFKM,STXBP1,CSNK1D,PTPRS,MAPK8IP1,SOD1

3CA,MARK1,RUFY3,SLC1A4,SPG11,OPA1,EPHB2,APOE,LRP8,FGF13,NFASC,PRNP,SRCIN1,PPT1,
MK2N1,RAB5A,PPP3CA,PIN1,ADD1,GIT1,ADGRL1,EPHB2,LRP8,PRNP,SRCIN1,EPHA4,DNM1,CACN
3CA,MARK1,RUFY3,SLC1A4,SPG11,OPA1,EPHB2,APOE,LRP8,FGF13,NFASC,PRNP,SRCIN1,PPT1,

DYNLL1,CDC42,SYP,GRIPAP1,VDAC1,ST3GAL6,SGTA,STARD7,ALDH3A2,HIP1,FLCN,AP2A1,FLOT
,ATP6V0B,WDR11,DLD,TMEM9,SYNGR1,CLCN4,PA2G4,RAB3A,UBQLN4,CLCN6,NAPB,ARL8A,MA

T,WDR82,PDCD11,PA2G4,RCC2,SETX,MAP1S,ZFP91,FBL,RRP1B,TBCA,STAT1,RNF20,KDM5A,RB

MP1,PSMB1,HSPA1A,KIAA1210,NOTCH1,PSMD13,CAB39,NDUFC2,APEH,CYFIP1,CD44,PSMD7,FA

,BRI3,ARSB,ATP6V0A1,ANXA1,SLC12A4,CLCN3,TMEM59,TRAPPC1,SPG11,VPS16,SZT2,NBR1,SC
,BRI3,ARSB,ATP6V0A1,ANXA1,SLC12A4,CLCN3,TMEM59,TRAPPC1,SPG11,VPS16,SZT2,NBR1,SC
ZB,GABARAP,ACTN1,HSPB1,MAP7D1,CLIP3,MAP2K2,CCAR2,DIDO1,FLNB,TPM3,CORO1C,KIF3C,

ERBB2,LMNA,DNAJB6,BRI3,INF2,ATP6V0A1,GALNT2,MAF1,RTRAF,SLC1A3,SEPTIN9,CUL7,YBX3
1,RAB18,PSMC2,DEGS1,DLD,SCG2,SYNGR1,SERPINE2,PA2G4,RAB3A,ARL8A,MAN2B1,DPP7,VDA
1,AGPAT1,NDST1,ULK1,PREPL,B4GAT1,GPSM1,POM121C,RPS27A,TMEM167A,LMAN1,TNKS,NP

IST1,ATP6V0D1,PRPF19,GSK3A,PRKACA,TTC28,UBXN6,REEP1,CROCC,TUBB4A,PTPN23,USO1,R

ˈP13A1,SRPRA,SCAP,RPS26,AMFR,TECR,TAPBP,SPPL2B,LMAN2,SDCBP,KTN1,ESYT1,POR,SC5D,
PS17,VCL,AP3D1,SBF1,DNAJA1,ARF1,CYB5R3,PAFAH1B1,AKAP12,MAP4,CDC42BPB,ITM2B,AP2E
G1,ELOVL5,ATP13A1,SRPRA,SCAP,RPS26,AMFR,TECR,TAPBP,SPPL2B,LMAN2,SDCBP,KTN1,ESY

PC11,KCTD10,CNOT3,MIB2,CUL7,PPP2R5B,TRAPPC1,CUL4B,BRPF3,PSMD12,TRPC4AP,PSMA4,
SRPRA,SCAP,RPS26,AMFR,TECR,TAPBP,SPPL2B,LMAN2,SDCBP,KTN1,ESYT1,POR,SC5D,TMX1,N

K2,HIP1R,ATP13A2,MDM2,COPE,SCAMP4,NDUFC2,CD44,ACAP2,HLA-A,ANKFY1,TSPAN14,SUN2,S

2,KIF26A,JAM3,CAB39,FXR1,PDLIM1,FEZ1,NCKAP5L,KLC2,PRPH,DYNLRB1,DDX6,NME2,PBXIP1,

AB39,FXR1,PDLIM1,FEZ1,NCKAP5L,KLC2,PRPH,DYNLRB1,DDX6,NME2,PBXIP1,MFAP4,APPBP2,A

20B,LMTK3,ZDHHC9,ATF6,GALNT2,GABARAPL2,B4GALT2,EXT1,PRKCI,CHPF2,CLCN3,TMEM59,

G4,SCOC,TMCO1,MAPK8IP3,PLPP3,SYT4,LEPROT,SLC35A4,C6ORF89,QSOX1,CYTH3,STX12,M

NK,SESN2,PAF1,SETD1B,SPSB3,KLHL7,BRD1,RNF40,PIK3R3,ANAPC2,RB1CC1,PLRG1

JN2,GOLPH3,CETN2,TM7SF2,COX7A2L,CDK4,TRIM27,NUP153,SLC25A1,NME2,NUP98,ATP5PO,
JN2,GOLPH3,CETN2,TM7SF2,COX7A2L,CDK4,TRIM27,NUP153,SLC25A1,NME2,NUP98,ATP5PO,

RPF4B,CLASP2,TCF3,FOXK2,MSH6,NSD2,IST1,RBPJ,PRPF19,PCGF2,NOC2L,KDM4A,MBD3,NAS

P3M1,PAM,ATP6V1D,STX12,M6PR,NECAP1,GDE1,UNC13B

KACA,GOT2,NDUFA13,BNIP3,TOMM70,ATP5MC2,BAG5,PRDX3,AMBRA1,REEP1,CYB5B,HSDL1,A

SET,EGLN1,MAP9,TPI1,CHSY1,TMSB10
ILA-C,PDIA3,SERINC1,SSBP2,RPS3,
15,GTF3C4,DNAH8,PKD1,HLA-C,SF3B
RPL26,CRMP1,BEX3,CCT7,OAZ
310,INA,EIF4A1,RPL19,PPP2R1A,RPL
LR2A,CDKL1,AP2M1,TPT1,SEC61A
1,HLA-C,SF3B1,PDIA3,SERINC1,SSB
L35A,CYFIP2,APLP2,SLC25A3,RP
PRKCSH,HNRNPC,LRPPRC,TADA2B
PRKCSH,HNRNPC,LRPPRC,TADA2B
PRKCSH,HNRNPC,LRPPRC,TADA2B
L35A,CYFIP2,APLP2,SLC25A3,RP
L35A,CYFIP2,APLP2,SLC25A3,RP
L35A,CYFIP2,APLP2,SLC25A3,RP
B1,RPL17,IGFBP3,COL1A2,RPL
L26,CCT7,VAT1,PFN1,ASTN2,RPL
H1,CKAP4,RPS12,RPS27,CTBP1,G
B,EID1,SRSF5,CHD6,RNF187,U2AF2
APOLA,IGFBP3,RPL22,RHOA,PTCD3,C
1,AJUBA,BCAR1,GIT1,PTPN12,FHL
1,AJUBA,BCAR1,GIT1,PTPN12,FHL
,EIF5A,CCND2,CDH2,PDIA6,GPC3,
,NOC2L,EIF3G,SF3A2,
CDH11,EEF1A2,TBCD,ANXA5,PP

LC2A1,CYTH2,HSPG2,ABI2,GNA12,RDX,F
NISCH,HNRNPC,LRPPRC,XPO1,SPTBN2
NISCH,HNRNPC,LRPPRC,XPO1,SPTBN2
GB,ATP6AP1,TMED2,VEGFA,RAPGEF1
GB,ATP6AP1,TMED2,VEGFA,RAPGEF1
2A1,AKAP9,TMOD2,GRK2,FUS,SE
DIA6,DDR1,SULF1,PLXNA1,MEGF8,

2,PSMC3,SART1,DHX38,UBE3A,PRKACA,YY1,CREBBP,SAP18,NDU

TP4A1,COL27A1,MAPK8IP1,SREBF2,VGF,P
THRAP3,SAE1,TEAD1,HNRNPH3,EIF4A3,PAGR1,R

BM14,HIF3A,SRP54,LMNA,THOC2,HI
DGRL1,OPA1,EPHB2,LRP8,VPS16,TGFB2,

DT2,DICER1,CLASP2,ANKS1A,ADAM10
3B,SEZ6L2,ABHD17A,ACSL4,CRTC

)1,AHCYL1,VPS35,MLEC,CLSTN3,SURF4,

,PAM
IKFY1,SUN2,GOLPH3,SNX1,VPS51,RAP2A,
C3B,SLC2A1,SERP1,TMOD2,CEP170,FKBP4,LTBP4
;,FYN,CCT8,STMN3,PFKM,STXBP1,CSNK1D,PTPR
PC2,GABARAPL1,RAB14,LDLR,TM9SF4,KIFAP3,S

AT1,RTN3,PFN1,RPL31,NISCH,HNRNPC,
,RAB11A,TCTN1,ARFGEF2,BMPR2,DYNLL1,CD

EPHA4,DNM1,MAPK8IP3,GNAO1,BRD1
NG7,ABR,STX12
EPHA4,DNM1,MAPK8IP3,GNAO1,BRD1

1,SEC16A,KPNA2,PHB2,UBA52,MGAT4B,MA
N2B1,STARD3,DPP7,ATP6V0E2,NAPA,BACE1,A

M14,XPC,DGCR8,PRMT2,CTCF,PPP1R12

F2,TSPAN14,SYT15,IQSEC1,CSNK2B,CA

ARB1,PPT1,RAP1B,PRCP,ARRDC3,AP3M1,
ARB1,PPT1,RAP1B,PRCP,ARRDC3,AP3M1,
FYN,CCT8,TNKS2,STXBP1,CSNK1D,MPRIP,PTP4

,HMGB2,SEC31A,COPS8,CENPF,MA
C2,THBS2,BRI3,ARSB,CHGB,OLA1,ATP6V0
LOC4,CHERP,PPP1R15A,GALNT11,FKBP1A,ATP6A

AB11FIP3,RANGAP1,MZT2B,MDH1,SPPL2B

TMX1,NOTCH1,NOMO2,REEP5,COPE,EMC1,GPAA1
31,TMEM170A,SLC4A8,GOLGA8A,HNRNPU,
T1,POR,SC5D,TMX1,NOTCH1,NOMO2,COPE,EMC1

CBX4,FBXO7,UBE2E1,SESN2,PSMC6,PAF1,PPP2R2A,RAD23A,S
OTCH1,NOMO2,COPE,EMC1,GPAA1,BSCL2,ARL6I

SNX1,RAP2A,NCK2,PMEPA1,RAB18,EPN1,A

MFAP4,APPBP2,ARPC3,STUB1,CSNK1A1,NDRG1,M

RPC3,STUB1,CSNK1A1,NDRG1,MAPRE3,PDE4DIP,

POMGNT1,ZDHHC17,COPB1,FURIN,HID1,COG4,SCOC,TMCO1,

SPR,ARFGAP3,ABCA7,MMP24

CLPX,DEGS1,SEPHS1,NDUFB8,MRPS6,SHMT2,ACSL4
CLPX,DEGS1,SEPHS1,NDUFB8,MRPS6,SHMT2,ACSL4

P,YY1,SMARCD1,CREBBP,UBE2I,TEAD1,CEN

ATP5MG,RAB11FIP3,ANK2,DIP2A,ACAT2,POLDIP2,SL

| | | | | | | |
|---|---|---|---|---|---|---|
| **KEGG** | Ribosome | KEGG:03010 | 4.01E-19 | 18.3971724 | 153 | 1382 |
| **KEGG** | Parkinson dise | KEGG:05012 | 4.60E-16 | 15.3377059 | 249 | 1382 |
| **KEGG** | Protein proces | KEGG:04141 | 4.66E-15 | 14.3315877 | 170 | 1382 |
| **KEGG** | Amyotrophic l | KEGG:05014 | 5.95E-14 | 13.2252422 | 363 | 1382 |
| **KEGG** | Coronavirus d | KEGG:05171 | 2.22E-13 | 12.6539245 | 231 | 1382 |
| **KEGG** | Pathways of r | KEGG:05022 | 7.80E-13 | 12.1077889 | 475 | 1382 |
| **KEGG** | Salmonella inf | KEGG:05132 | 1.17E-10 | 9.9317625 | 249 | 1382 |
| **KEGG** | Prion disease | KEGG:05020 | 3.24E-10 | 9.48904285 | 273 | 1382 |
| **KEGG** | Alzheimer dise | KEGG:05010 | 3.71E-10 | 9.43059624 | 369 | 1382 |
| **KEGG** | Huntington di | KEGG:05016 | 8.91E-10 | 9.05006277 | 306 | 1382 |
| **KEGG** | Spliceosome | KEGG:03040 | 3.50E-09 | 8.4563469 | 150 | 1382 |
| **KEGG** | Focal adhesio | KEGG:04510 | 9.85E-08 | 7.00659321 | 200 | 1382 |
| **KEGG** | mRNA surveill | KEGG:03015 | 1.0627E-06 | 5.97359833 | 98 | 1382 |
| **KEGG** | Endocytosis | KEGG:04144 | 1.1322E-06 | 5.94608039 | 252 | 1382 |
| **KEGG** | Proteasome | KEGG:03050 | 1.531E-06 | 5.81502781 | 46 | 1382 |
| **KEGG** | Renal cell carc | KEGG:05211 | 6.5165E-06 | 5.18598682 | 68 | 1382 |
| **KEGG** | RNA transport | KEGG:03013 | 1.242E-05 | 4.9058754 | 180 | 1382 |
| **KEGG** | Thyroid horm | KEGG:04919 | 1.4475E-05 | 4.83938762 | 121 | 1382 |
| **KEGG** | Diabetic cardi | KEGG:05415 | 2.0142E-05 | 4.69589172 | 203 | 1382 |
| **KEGG** | Adherens junc | KEGG:04520 | 2.2068E-05 | 4.6562421 | 71 | 1382 |
| **KEGG** | Proteoglycans | KEGG:05205 | 3.0212E-05 | 4.51981989 | 205 | 1382 |
| **KEGG** | HIF-1 signalin | KEGG:04066 | 3.7004E-05 | 4.43175107 | 109 | 1382 |
| **KEGG** | Shigellosis | KEGG:05131 | 4.7354E-05 | 4.32464333 | 246 | 1382 |
| **KEGG** | Bacterial invas | KEGG:05100 | 5.5468E-05 | 4.25595695 | 77 | 1382 |
| **KEGG** | Spinocerebella | KEGG:05017 | 0.00015052 | 3.82241658 | 142 | 1382 |
| **KEGG** | Carbon metab | KEGG:01200 | 0.00066654 | 3.17617339 | 116 | 1382 |
| **KEGG** | Central carbor | KEGG:05230 | 0.00071181 | 3.14763862 | 70 | 1382 |
| **KEGG** | Mitophagy - a | KEGG:04137 | 0.00122878 | 2.91052495 | 68 | 1382 |
| **KEGG** | Oxidative pho | KEGG:00190 | 0.00193065 | 2.71429711 | 133 | 1382 |
| **KEGG** | Human papillo | KEGG:05165 | 0.00226483 | 2.64496403 | 331 | 1382 |
| **KEGG** | Phagosome | KEGG:04145 | 0.00254791 | 2.59381599 | 147 | 1382 |
| **KEGG** | Ubiquitin med | KEGG:04120 | 0.00279469 | 2.55366695 | 139 | 1382 |
| **KEGG** | Regulation of | KEGG:04810 | 0.00440153 | 2.35639668 | 216 | 1382 |
| **KEGG** | Tight junction | KEGG:04530 | 0.00601736 | 2.22059389 | 169 | 1382 |
| **KEGG** | Neurotrophin | KEGG:04722 | 0.01871563 | 1.72779546 | 119 | 1382 |
| **KEGG** | Insulin signali | KEGG:04910 | 0.02148399 | 1.667885 | 137 | 1382 |
| **KEGG** | Glycolysis / G | KEGG:00010 | 0.02594661 | 1.58591933 | 67 | 1382 |
| **KEGG** | Biosynthesis o | KEGG:01230 | 0.02639242 | 1.57852083 | 75 | 1382 |
| **KEGG** | Chronic myelo | KEGG:05220 | 0.033634 | 1.47322149 | 76 | 1382 |
| **KEGG** | Prostate canc | KEGG:05215 | 0.03672948 | 1.43498521 | 97 | 1382 |
| **KEGG** | Axon guidanc | KEGG:04360 | 0.0393482 | 1.4050751 | 181 | 1382 |
| **KEGG** | Thermogenes | KEGG:04714 | 0.03989849 | 1.39904353 | 232 | 1382 |

| | | |
|---|---|---|
| 77 | 8000 | RPL3,RPS6,RPL4,RPL7A,RPL8,RPLP0,RPL13A,RPL6,RPL13,RPS8,RP |
| 100 | 8000 | TUBA1A,TUBB,UBC,TUBA1B,UBA1,TUBB2B,GNAS,ATP5F1B,CALM3, |
| 76 | 8000 | HSPA8,HSP90AB1,HSP90AA1,GANAB,HSP90B1,CALR,HSPA1B,CANX, |
| 125 | 8000 | HNRNPA1,ACTB,ACTG1,TUBA1A,TUBB,TUBA1B,NEFL,NEFM,TUBB2 |
| 90 | 8000 | MAVS,RPL3,RPS6,RPL4,PIK3R2,RPL7A,RPL8,RPLP0,RPL13A,RPL6,R |
| 149 | 8000 | TUBA1A,HTT,TUBB,UBC,TUBA1B,NEFL,NEFM,UBA1,TUBB2B,DVL3, |
| 89 | 8000 | GAPDH,ACTB,ACTG1,TUBA1A,HSP90AB1,HSP90AA1,ACTR2,TUBB,T |
| 94 | 8000 | HSPA8,TUBA1A,TUBB,TUBA1B,PIK3R2,HSPA1B,TUBB2B,ATP5F1B, |
| 117 | 8000 | LRP1,GAPDH,TUBA1A,TUBB,TUBA1B,PIK3R2,TUBB2B,DVL3,ATP5F1 |
| 101 | 8000 | TUBA1A,HTT,TUBB,TUBA1B,TUBB2B,ATP5F1B,ATP5F1A,DNAH8,POL |
| 60 | 8000 | HSPA8,HNRNPA1,PRPF8,HSPA1B,DDX5,SNRNP200,SF3B1,SF3B2,HN |
| 70 | 8000 | COL1A1,ACTB,ACTG1,PIK3R2,FLNA,COL6A1,COL4A1,COL4A2,FN1, |
| 41 | 8000 | PABPC1,SMG1,PPP2R1A,PAPOLA,PNN,TARDBP,CPSF2,DDX39B,CPS |
| 80 | 8000 | HSPA8,NEDD4L,ACTR2,HSPA1B,HLA-C,AP2M1,ARF1,AP2B1,RHOA,K |
| 25 | 8000 | PSMD2,PSMB2,PSMD3,PSMA7,PSMD1,PSMD8,PSMB4,PSMD4,PSMC |
| 31 | 8000 | PIK3R2,EGLN1,JUN,RAC1,CDKN1A,VEGFA,RAPGEF1,VHL,MAPK1,G |
| 60 | 8000 | EEF1A1,PABPC1,EIF4G2,EIF3C,EIF4B,EIF1,EIF4A1,KPNB1,EIF5,PNN, |
| 45 | 8000 | ACTB,ACTG1,PIK3R2,ATP1A1,ATP2A2,PFKL,CTNNB1,ATP1A2,HDAC2 |
| 65 | 8000 | COL1A1,GAPDH,PIK3R2,ATP5F1B,ATP5F1A,CTSD,COL1A2,ATP2A2, |
| 31 | 8000 | ACTB,ACTG1,VCL,CTNNA1,RHOA,ACTN4,CTNNB1,RAC1,MAPK1,ACT |
| 65 | 8000 | COL1A1,ACTB,ACTG1,RPS6,PIK3R2,FLNA,DDX5,IGF2,EIF4B,FN1,IT |
| 41 | 8000 | GAPDH,RPS6,ENO1,PIK3R2,ALDOA,PGK1,ENO2,EGLN1,LDHB,LDHA, |
| 74 | 8000 | ACTB,ACTG1,ACTR2,UBC,PIK3R2,H3-3B,SEPTIN2,UBB,ITGB1,VCL, |
| 32 | 8000 | ACTB,ACTG1,ACTR2,PIK3R2,SEPTIN2,FN1,ITGB1,VCL,CTNNA1,RHO |
| 48 | 8000 | PIK3R2,PSMD2,ATP2A2,SPTBN2,PSMB2,ATG2B,PSMD3,ATXN10,ATX |
| 40 | 8000 | GAPDH,ENO1,ALDOA,PKM,PGK1,ENO2,GPI,ALDH6A1,TPI1,PGAM1,H |
| 28 | 8000 | PIK3R2,PKM,LDHB,PGAM1,LDHA,HK1,SLC7A5,PFKL,MAPK1,MAP2K |
| 27 | 8000 | UBC,UBB,ATF4,RAB7A,JUN,SQSTM1,OPTN,GABARAP,BNIP3L,CSNK |
| 43 | 8000 | ATP5F1B,ATP5F1A,ATP6V0C,ATP6AP1,ATP5PB,ATP6V1B2,SDHA,UQC |
| 87 | 8000 | COL1A1,PIK3R2,PKM,COL6A1,COL4A1,DVL3,GNAS,COL4A2,FN1,PP |
| 46 | 8000 | ACTB,ACTG1,TUBA1A,TUBB,TUBA1B,CALR,TUBB2B,CANX,MRC2,HL |
| 44 | 8000 | NEDD4L,UBC,UBA1,UBB,DDB1,KLHL9,UBE2H,SKP1,UBE4A,VHL,UBE |
| 61 | 8000 | ACTB,ACTG1,CFL1,ACTR2,TMSB4X,PIK3R2,MYH9,FN1,ITGB1,VCL, |
| 50 | 8000 | NEDD4L,ACTB,ACTG1,TUBA1A,ACTR2,TUBA1B,MYH9,PPP2R1A,ITG |
| 37 | 8000 | KIDINS220,PIK3R2,MAGED1,YWHAE,CALM3,CALM1,RHOA,ATF4,BE |
| 41 | 8000 | RPS6,PIK3R2,FASN,CALM3,CALM1,HK1,PRKAR1A,PPP1CB,PYGB,RA |
| 24 | 8000 | GAPDH,ENO1,ALDOA,PKM,PGK1,ENO2,GPI,TPI1,LDHB,PGAM1,LDH |
| 26 | 8000 | GAPDH,ENO1,ALDOA,PKM,PGK1,ENO2,TPI1,MAT2A,PGAM1,PFKL,T |
| 26 | 8000 | PIK3R2,CTBP1,HDAC2,CDKN1A,CCND1,MAPK1,GRB2,MAP2K2,BCR, |
| 31 | 8000 | HSP90AB1,HSP90AA1,PIK3R2,HSP90B1,GSTP1,ATF4,CTNNB1,CDKN |
| 50 | 8000 | CFL1,PIK3R2,PLXNB2,ITGB1,RHOA,PLXNA1,RAC1,L1CAM,GNAI3,DP |
| 61 | 8000 | ACTB,ACTG1,RPS6,GNAS,ATP5F1B,ATP5F1A,ACSL6,ARID1A,SMARC |

L7,RPL5,RPL37A,RPL19,RPL15,RPS3,RPS4X,RPS3A,RPL18,RPL11,RPLP1,RPL10,RPS17,RPS16,F
CALM1,ATP5F1A,UBB,MAPT,TUBB4B,UCHL1,ATF4,PSMD2,KIF5B,GNAI3,KLC1,PSMB2,TUBB2A,F
SEC62,P4HB,VCP,PDIA3,SEC61A1,MARCHF6,DNAJA1,ATF4,PRKCSH,PDIA6,DNAJC5,DDOST,CKAF
3,ATP5F1B,ATP5F1A,VCP,DNAH8,TUBB4B,ATF4,PSMD2,KIF5B,PFN1,TARDBP,RAB1A,RAC1,SQSTI
PL13,RPS8,RPL7,RPL5,RPL37A,RPL19,RPL15,RPS3,RPS4X,RPS3A,RPL18,RPL11,RPLP1,RPL10,F
TP5F1B,CALM3,APP,CALM1,ATP5F1A,VCP,UBB,MAPT,DNAH8,TUBB4B,UCHL1,ATF4,PSMD2,KIF5
JBA1B,FLNA,HSP90B1,TUBB2B,RPS3,TUBB4B,ANXA2,ARF1,RHOA,KIF5B,PFN1,RAB7A,JUN,CTNN
TP5F1A,TUBB4B,NCAM1,ATF4,PSMD2,KIF5B,RAC1,STIP1,KLC1,LAMC1,PSMB2,TUBB2A,MAPK1,
B,CALM3,APP,CALM1,ATP5F1A,MAPT,TUBB4B,RTN4,ATF4,PSMD2,KIF5B,ATP2A2,RTN3,CSNK1E
R2A,AP2M1,TUBB4B,AP2B1,PSMD2,KIF5B,CLTC,HDAC2,KLC1,PSMB2,TUBB2A,ATG2B,PSMD3,
RNPU,SRSF4,HNRNPC,HNRNPK,DDX39B,SRSF5,U2AF2,HNRNPA3,RBMX,DDX42,SNRNP70,PRPF6
TGB1,VCL,LAMA5,COL1A2,RHOA,CCND2,ACTN4,TLN1,JUN,CTNNB1,COL6A2,RAC1,TNC,PPP1CE
-1,PPP1CB,PABPN1,PPP2CA,CASC3,GSPT1,PPP2CB,UPF1,PPP1CC,SYMPK,FUS,SMG5,MSI1,SAP1
F5B,RAB7A,CLTC,HLA-B,ACTR3,VPS35,ACAP3,RAB5B,RAB11B,ARF3,MVB12B,ARPC2,CAPZB,AR
3,PSMC1,PSMB1,PSMD13,PSMD7,ADRM1,PSME3,PSMC5,PSMC2,PSMC4,PSMF1,PSMB5,PSMA1,
B2,MAP2K2,ARNT2,CDC42,EP300,FLCN,PRCC,HIF1A,SLC2A1,AKT3,AKT2,CREBBP,PTPN11,CRK
XPO1,CYFIP2,TARDBP,DDX39B,EEF1A2,EIF3E,CASC3,EIF3F,STRAP,XPOT,EIF3D,UPF1,EIF5B,POM
,NOTCH2,CCND1,MAPK1,PLCG1,MAP2K2,PFKM,EP300,HIF1A,SLC2A1,AKT3,PDPK1,ATP1B2,ITG
AC1,MMP2,PPP1CB,PARP1,ATP5PB,SDHA,UQCRC2,VDAC1,NDUFV1,UQCRC1,COX7C,PTEN,PPP1
N1,FYN,CDC42,CTNND1,EP300,CSNK2A1,PTPRF,WASF2,CREBBP,FGFR1,IGF1R,PTPN1,CSNK2B,
B1,MSN,COL1A2,RHOA,GPC3,NANOG,CTNNB1,CD63,RAC1,GPC1,MMP2,DCN,CDKN1A,TIMP3,F
HK1,PFKL,CDKN1A,VEGFA,VHL,MAPK1,PLCG1,MAP2K2,PFKM,STAT3,EP300,HIF1A,SLC2A1,AKT
RF1,HK1,RHOA,PFN1,ACTN4,TLN1,JUN,SEPTIN11,SEPTIN3,RAC1,SKP1,SQSTM1,TLN2,ACTR3,H3
A,CLTC,CTNNB1,SEPTIN11,SEPTIN3,RAC1,ACTR3,ARPC2,CDC42,CTTN,SHC2,WASF2,CLTA,ARPC
N2L,PSMA7,CIC,VDAC1,PSMD1,ULK1,PSMD8,SLC25A5,PSMB4,AKT3,ATXN3,PSMD4,AKT2,RBPJ,
K1,PFKL,TKT,MDH2,PGD,ACO2,SDHA,PFKM,PRPS1,IDH1,GLUD1,ADH5,PHGDH,GOT2,ACAT2,MD
2,PFKM,IDH1,PTEN,HIF1A,SLC2A1,AKT3,AKT2,MTOR,FGFR1,PDHA1,PFKP,IDH2,PDGFRB,ERBB2
2A1,UBA52,ULK1,RPS27A,HIF1A,GABARAPL1,TAX1BP1,BNIP3,AMBRA1,PINK1,MFN2,CSNK2B,AT
RC2,NDUFV1,UQCRC1,COX7C,ATP6V0D1,COX6A1,ATP5MC3,NDUFA13,ATP6V1G1,ATP5MC2,AT
P2R1A,HLA-C,ITGB1,LAMA5,COL1A2,LLGL1,CHD4,CCND2,CTNNB1,COL6A2,HLA-B,HDAC2,TNC,
A-C,SEC61A1,ITGB1,TUBB4B,CLEC7A,RAB7A,LAMP1,HLA-B,RAC1,ATP6V0C,RAB5B,ATP6AP1,TU
2R2,UBA52,UBE2G2,CUL3,RPS27A,UBR5,UBE2M,UBE2D2,PRPF19,UBE3A,UBE2I,UBE2D3,SAE1,
ISN,RHOA,PFN1,PIP4K2B,ACTN4,CYFIP2,MYH10,RAC1,PPP1CB,ACTR3,NCKAP1,MAPK1,ARPC2,A
31,MSN,LLGL1,RHOA,ACTN4,JUN,MYL6,MYH10,RAC1,ACTR3,HSPA4,PPP2CA,CCND1,ARPC2,ACT
3,JUN,RAC1,ARHGDIA,RAPGEF1,MAPK1,GRB2,PLCG1,MAP2K2,CDC42,SHC2,AKT3,PDPK1,AKT2
PGEF1,MAPK1,GRB2,MAP2K2,PRKAR2A,FLOT1,PPP1CC,PTPRF,SHC2,AKT3,FLOT2,PDPK1,AKT2
,HK1,PFKL,PFKM,ALDH3A2,ADH5,PDHA1,PFKP,ALDH2,ALDH7A1,DLD,ALDH9A1,G6PC3,PDHB
T,GLUL,ACO2,PFKM,PRPS1,IDH1,PHGDH,GOT2,PSAT1,PFKP,IDH2,ALDH18A1,ASNS,GPT2,SHMT
SHC2,AKT3,AKT2,PTPN11,ABL1,MDM2,CDK4,CRKL,CDKN1B,SOS1,SHC1,CTBP2,TGFBR1,MAPK
1A,CCND1,MAPK1,GRB2,MAP2K2,EP300,PTEN,AKT3,PDPK1,AKT2,MTOR,CREBBP,FGFR1,IGF1F
YSL2,MAPK1,PLCG1,FYN,SLIT3,PLXNA3,BMPR2,CDC42,PLXNB1,SLIT2,SLIT1,SEMA3C,PDPK1,SN
A4,ATP5PB,GRB2,KDM1A,SMARCE1,SMARCC1,SDHA,UQCRC2,KDM3B,NDUFV1,ARID1B,ADCY6,U

RPL23,RPL29,RPSA,RPS24,RPS9,RPL24,RPL17,RPL14,RPL22,RPL9,RPS14,RPS20,RPL26,RPL31,
PSMD3,ATP5PB,PLCG1,SDHA,UQCRC2,PSMA7,VDAC1,KIF5A,NDUFV1,PSMD1,UBA52,UBE2G2,UQ
P4,SKP1,RPN2,HSPH1,RAD23B,UBXN4,STT3B,STT3A,UBQLN1,HYOU1,SEC23A,SSR1,EIF2AK1,OS
M1,HNRNPA3,KLC1,PSMB2,TUBB2A,OPTN,ATG2B,PSMD3,ATP5PB,GABARAP,SRSF3,SDHA,UBQL
RPS17,RPS16,RPL23,RPL29,RPSA,RPS24,RPS9,RPL24,RPL17,RPL14,RPL22,RPL9,RPS14,RPS20,
5B,ATP2A2,SPTBN2,CSNK1E,CTNNB1,TARDBP,RAB1A,RAC1,SQSTM1,NOX4,KLC1,PSMB2,TUBB2
NB1,CYFIP2,RHOB,RAC1,SKP1,ACTR3,PKN1,KLC1,NCKAP1,RAB5B,TUBB2A,MAPK1,ARPC2,DYNC
PSMD3,ATP5PB,FYN,SDHA,UQCRC2,SOD1,PSMA7,VDAC1,KIF5A,NDUFV1,PSMD1,CSNK2A1,UQC
,CTNNB1,NOX4,KLC1,PSMB2,TUBB2A,MAPK1,ATG2B,PSMD3,ATP5PB,MAP2K2,SDHA,UQCRC2,
ATP5PB,POLR2B,SDHA,UQCRC2,SOD1,POLR2E,PSMA7,VDAC1,DCTN1,KIF5A,ACTR1A,EP300,NDU
,SRSF9,SRSF1,HNRNPM,SRSF3,SF3A1,PRPF40A,DHX15,SRSF6,TRA2B,CHERP,FUS,SNRPD3,EFT
3,TLN2,COL2A1,LAMC1,VEGFA,RAPGEF1,CCND1,MAPK1,GRB2,ACTN1,FLNB,FYN,FLNC,CDC42,
8,EIF4A3,PABPC4,SSU72,ETF1,PPP1CA,PPP2R5D,NCBP2,RNMT,WDR82,PPP2R5B,MSI2,RNPS1,A
F4,HGS,RAB11A,ARFGEF2,CDC42,KIF5A,AP2A1,RABEP1,GRK2,VPS26B,CYTH2,VPS4A,LDLR,SH3
,PSMA6,PSMD12,PSMA4,PSMC6
L,EGLN2,SOS1,ELOB,MAPK3,FH,TGFB2,RAP1B,PIK3R3
M121C,FUS,EIF2S2,EIF3H,EIF4G3,EIF3G,SAP18,UBE2I,EIF4A3,EIF4G1,RANGAP1,FXR2,SUMO1,F
AV,AKT2,MED13,MTOR,PRKACA,CREBBP,NOTCH1,PFKP,MDM2,ATP1B3,KAT2A,RHEB,SLC16A2,
CC,SLC25A5,SLC2A1,AKT3,AKT2,COX6A1,MTOR,ATP5MC3,NDUFA13,ATP5MC2,PDHA1,SLC25A
,NECTIN1,WASF3,ERBB2,ACP1,AFDN,NECTIN2,TGFBR1,MAPK3,FER
PP1CB,VEGFA,CCND1,MAPK1,GRB2,PLCG1,MAP2K2,FLNB,FLNC,CDC42,STAT3,PPP1CC,ITGB5,
3,AKT2,MTOR,CREBBP,MKNK2,PDHA1,PFKP,IGF1R,TFRC,CDKN1B,EGLN2,ERBB2,ELOB,MAPK3
3-3A,MAPK1,ARPC2,GABARAP,ACTN1,PLCG1,PFN2,CDC42,VDAC1,CAPN2,UBA52,RPS27A,SEPTIN
1A,ARPC3,CRKL,ARPC5,SHC1,SEPTIN9,BCAR1,DNM1,ELMO2,PIK3R3
PSMC3,MTOR,PSMC1,AMBRA1,PUM1,PSMB1,NOP56,SLC25A6,PSMD13,PSMD7,ADRM1,PSMC5,
DH1,PSAT1,H6PD,PDHA1,PFKP,IDH2,ECHS1,GPT2,DLD,ME2,SHMT2,OGDH,TALDO1,FH,PDHB
,MAPK3,SLC1A5,PDHB,PIK3R3
G9A,CITED2,GABARAPL2,SP1,RELA,NBR1
P5MG,NDUFA4,NDUFC2,COX8A,COX7A2L,ATP6V0B,ATP5PO,NDUFB8,NDUFS1,ATP5F1C,COX6B1
,ATP6V0C,CDKN1A,COL2A1,LAMC1,ATP6AP1,NOTCH2,VEGFA,PPP2CA,CCND1,MAPK1,GRB2,MA
JBB2A,DYNC1LI2,ATP6V1B2,HGS,ITGB5,RAB5C,ITGAV,ATP6V0D1,ATP6V1G1,TUBB4A,HLA-A,ATP
,UBE2K,TRIP12,UBE2L3,CDC34,MDM2,FZR1,UBE4B,CUL1,CUL4A,UBE2J1,UBE2Q1,STUB1,UBE
ACTN1,MAP2K2,PFN2,CDC42,PPP1CC,ITGB5,BRK1,ABI2,GNA12,RDX,WASF2,ITGAV,GNA13,FGFR
N1,AMOTL2,CDC42,PPP2CB,CTTN,SYMPK,RDX,SCRIB,PRKACA,MYL12B,ARPC1A,JAM3,CDK4,AF
2,SORT1,NTRK2,PTPN11,ABL1,MAPKAPK2,CRKL,GSK3B,IRS1,SOS1,SHC1,RPS6KA2,MAPK3,REL
,MTOR,PRKACA,MKNK2,PTPN1,RHEB,PPP1CA,EXOC7,CRKL,GSK3B,PHKB,IRS1,PRKAR2B,SOS1

T2,TALDO1
3,RELA,TGFB2,PIK3R3
R,MDM2,PDGFRB,ZEB1,GSK3B,CDKN1B,ERBB2,SOS1,MAPK3,RELA,PIK3R3
MO,MYL12B,PTPN11,ABL1,SEMA6B,PLXNA2,EFNB2,NTN1,LIMK1,EFNB1,NCK2,SEMA4C,GSK3B,
JQCRC1,COX7C,COX6A1,MTOR,ATP5MC3,PRKACA,SMARCD1,NDUFA13,ATP5MC2,FGFR1,ATP5

RPS25,RPS15,RPL12,RPL30,RPS5,RPS7,RPS19,RPL35,RPS10,RPL35A,RPLP2,RPL36,RPS12,RPS
CRC1,COX7C,RPS27A,PSMD8,SLC25A5,SEPTIN5,PSMB4,TRAP1,PSMD4,COX6A1,PSMC3,ATP5M
9,CAPN2,TXNDC5,UBQLN2,UBE2G2,MBTPS1,LMAN1,NPLOC4,PPP1R15A,TRAM1,UBE2D2,SSR3,
N1,ATXN2L,UQCRC2,SOD1,PFN2,PSMA7,VDAC1,DCTN1,KIF5A,ACTR1A,NDUFV1,PSMD1,UBQLN
RPL26,RPL31,RPS25,RPS15,RPL12,RPL30,RPS5,RPS7,RPS19,RPL35,JUN,RPS10,RPL35A,RPLP2,
A,OPTN,MAPK1,ATG2B,PSMD3,ATP5PB,GABARAP,PLCG1,MAP2K2,SDHA,ATXN2L,UQCRC2,SOD
1LI2,DYNLL2,MAP2K2,FLNB,PFN2,FLNC,DYNLL1,CDC42,DCTN1,KIF5A,ACTR1A,BRK1,CYTH2,VI
RC1,COX7C,PSMD8,SLC25A5,PSMB4,PSMD4,COX6A1,PSMC3,ATP5MC3,PRKACA,NDUFA13,PSM
SMA7,VDAC1,KIF5A,CAPN2,NDUFV1,PSMD1,CSNK2A1,UQCRC1,COX7C,ULK1,PSMD8,SLC25A5,
JFV1,PSMD1,HIP1,AP2A1,UQCRC1,COX7C,ULK1,PSMD8,SLC25A5,PSMB4,GPX1,SOD2,ACTR1B,
UD2,SRSF2,WBP11,PRPF19,SF3A2,SART1,DHX38,EIF4A3,HSPA1A,RBM25,SF3A3,SRSF10,LSM4,
CAPN2,PTEN,PPP1CC,ITGB5,SHC2,AKT3,PDPK1,ITGAV,AKT2,MYL12B,IGF1R,PPP1CA,ARHGAP5,
CIN1,PPP2R2A,DDX19A,CPSF7,WDR33,ALYREF,PPP2R5C
GL1,CAPZA1,RAB5C,IST1,EHD1,CHMP4B,AP2A2,ARF5,RAB11FIP3,SNX2,SNX3,AGAP1,CHMP1A,C

XR1,CYFIP1,PABPC4,RAN,NUP153,EIF3B,NUP98,NCBP2,PRMT5,NUP205,NUP133,THOC2,THOC3,
GSK3B,STAT1,MED12,RXRA,MAPK3,PLCD3,THRA,MED16,PIK3R3,SIN3A,MED14
6,NDUFA4,NDUFC2,COX8A,GSR,PPP1CA,COX7A2L,ATP5PO,GSK3B,NDUFB8,NDUFS1,ATP5F1C,C

HIF1A,CTTN,HSPG2,AKT3,RDX,PDPK1,ITGAV,AKT2,MTOR,PRKACA,SMO,FGFR1,ANK2,PTPN11,I
,RELA,PDHB,PFKFB3,PIK3R3
I7,CTTN,CYTH2,UBE2D2,GABARAPL1,AKT3,FNBP1L,WASF2,AKT2,GSK3A,MTOR,BNIP3,UBE2D3,

PSMC2,PSMC4,PSMB5,VDAC2,PSMA1,PSMA6,XBP1,SP1,PSMD12,OPA1,PSMA4,PSMC6,PIK3R3,F

I,ATP6V0E2,UQCRFS1,ATP5F1E,COX4I1,ATP6V0A1,COX5A,COX7A2,UQCRH,ATP6V1F,NDUFA5,U
P2K2,ATP6V1B2,CDC42,EP300,PPP2CB,PTEN,ITGB5,AKT3,ITGAV,SCRIB,AKT2,ATP6V0D1,RBPJ,
6V0B,TUBA1C,TFRC,ATP6V0E2,THBS2,ATP6V0A1,RAB5A,TLR6,DYNC1I2,SCARB1,ATP6V1F,ATP6
3C,RHOBTB2,MGRN1,UBE2G1,ELOB,CUL7,CUL4B,UBE2E1,ANAPC2
1,MYL12B,ARPC1A,CYFIP1,LIMK1,PPP1CA,PDGFRB,ARPC3,CRKL,PIP5K1C,MYL12A,SOS1,PPP1
RPC3,TUBA1C,MYL12A,ERBB2,ARPC5,AFDN,ROCK2,PCNA,ARHGEF18,PRKCI,YBX3,PPP2R2A,MY
A,RAP1B,PIK3R3
,SHC1,MAPK3,G6PC3,PRKCI,GYS1,PIK3R3

MYL12A,EFNB3,NEO1,ROCK2,ROBO3,RGMA,MAPK3,PPP3CA,SEMA6C,EPHB2,EPHA4,MYL9,PIK3
MG,KDM3A,NDUFA4,NDUFC2,COX8A,RHEB,COX7A2L,ADCY1,ATP5PO,NDUFB8,ACSL4,NDUFS1,A

27,RPL23A,RPL27,RPL36A,RPS15A,RPL38,RPL37,RPS11,RPL28,RPS28,FAU,RPL39,RPS13,RPL3
C3,PRKACA,NDUFA13,PSMC1,ATP5MC2,TUBB4A,UBE2L3,PINK1,PSMB1,MFN2,SLC25A6,NDUFA
DNAJB2,ATXN3,SSR2,UBE2D3,UBXN6,AMFR,LMAN2,HSPA1A,UBE4B,CUL1,DNAJB1,ERP29,UBE2
2,UQCRC1,COX7C,HDAC6,HNRNPA2B1,ULK1,POM121C,PSMD8,FUS,PSMB4,VAPB,GPX1,GABAR
,RPL36,RPS12,RPS27,RPL23A,RPL27,RPL36A,RPS15A,RPL38,RPL37,MAPK1,RPS11,PLCG1,RPL2
1,PSMA7,VDAC1,DCTN1,KIF5A,CAPN2,ACTR1A,NDUFV1,PSMD1,HIP1,CSNK2A1,UBA52,UBE2G2,U
PS41,AKT3,RAB5C,PLEKHM2,ACTR1B,AKT2,CSE1L,EXOC4,MYL12B,TUBB4A,ARPC1A,EXOC5,CY
C1,ATP5MC2,TUBB4A,PSMB1,HSPA1A,NOTCH1,SLC25A6,NDUFA4,PSMD13,NDUFC2,PSMD7,CO
PSMB4,AKT3,NCSTN,PSMD4,ADAM10,AKT2,COX6A1,PSMC3,MTOR,ATP5MC3,APBB1,NDUFA13,F
PSMD4,COX6A1,PSMC3,MTOR,ATP5MC3,AP2A2,CREBBP,NDUFA13,PSMC1,ATP5MC2,AMBRA1,T
SRSF7,NCBP2,SNRPD2,SNRPB,THOC2,RBM17,PRPF31,THOC3,XAB2,ACIN1,ZMAT2,SRSF8,ALYRE
PDGFRB,CRKL,PIP5K1C,GSK3B,ZYX,MYL12A,ERBB2,THBS2,SOS1,PPP1R12A,SHC1,ROCK2,MAP

CLTA,ARPC1A,HSPA1A,IGF1R,MDM2,ACAP2,HLA-A,SNX1,IQSEC1,ARFGAP2,RNF41,EPN1,ARPC3,F

,PAIP1,EIF1B,NUP50,RNPS1,ACIN1,EIF2S1,DDX19A,NUP58,ELAC2,ALYREF,EIF3J

COX6B1,IRS1,VDAC2,UQCRFS1,ATP5F1E,COX4I1,COX5A,COX7A2,TGFBR1,UQCRH,SP1,RELA,GY

GF1R,MDM2,CD44,PPP1CA,ERBB2,SOS1,PPP1R12A,ROCK2,MAPK3,VAV2,TGFB2,FZD7,FZD8,PIK

MYL12B,ARPC1A,MDM2,CUL1,CD44,CAPN1,ARPC3,CRKL,GSK3B,UBE2V1,MYL12A,ARPC5,ROC

B1CC1

QCR11,ATP6V1D,COX5B,NDUFA10,ATP5PD
MTOR,UBE3A,PRKACA,CREBBP,ATP6V1G1,PSMC1,NOTCH1,MDM2,HLA-A,RHEB,PDGFRB,CDK4,
V1D,STX12,M6PR

R12A,ARPC5,ROCK2,MAPK3,VAV2,BCAR1,GIT1,PIP5K1A,ITGA7,MYL9,FGD1,PIK3R3,GSN
L9,NF2,MYL6B

R3,EPHA2,GNAI1
TP5F1C,COX6B1,UQCRFS1,ATP5F1E,SOS1,COX4I1,RPS6KA2,COX5A,COX7A2,SLC25A29,UQCR

4,UBA52,RPS2,RPS27A,RPS26,RPS21,RPL27A,MRPS6,RPS29,RPL36AL,RPL32,MRPL3,MRPL28
4,PSMD13,PARK7,NDUFC2,PSMD7,COX8A,KLC2,UBE2J1,COX7A2L,ADRM1,PSMC5,ATP5PO,PSM
J1,DAD1,MOGS,CAPN1,STUB1,HERPUD1,UBQLN4,RRBP1,UBXN1,ATF6B,DNAJB11,UBE2G1,ATF
APL1,ACTR1B,PSMD4,COX6A1,PSMC3,MTOR,ATP5MC3,NDUFA13,PSMC1,ATP5MC2,AMBRA1,TU
28,RPS28,FAU,RPL39,RPS13,RPL34,ADAR,STAT3,UBA52,IL6ST,RPS2,RPS27A,RPS26,RPS21,RPL2
UQCRC1,COX7C,ULK1,RPS27A,PSMD8,SLC25A5,FUS,SEPTIN5,PSMB4,VAPB,GPX1,GABARAPL1,T
FIP1,PIK3C2B,KLC2,DYNLRB1,ARPC3,EXOC7,TNFRSF10B,ACBD3,TUBA1C,TXN,ARL8A,WASF3,M
X8A,KLC2,CSNK2B,COX7A2L,ADRM1,PSMC5,ATP5PO,PSMC2,PSMC4,GSK3B,NDUFB8,TUBA1C,N
PSMC1,ATP5MC2,AMBRA1,TUBB4A,PSMB1,SLC25A6,NDUFA4,PSMD13,NDUFC2,PSMD7,COX8A,K
UBB4A,CLTA,PSMB1,SLC25A6,NDUFA4,PSMD13,NDUFC2,PSMD7,COX8A,KLC2,COX7A2L,ADRM
EF,PLRG1,SNU13
K3,COL6A3,VAV2,BCAR1,PIP5K1A,RAP1B,ITGA7,MYL9,PIK3R3

PIP5K1C,TFRC,ARPC5,CHMP5,RAB5A,TGFBR1,CAPZA2,PRKCI,RAB35,GIT1,ARF6,PIP5K1A,DNM1,

S1,TGFB2,COL3A1,PDHB,PIK3R3,NDUFA5,UQCR11,COX5B,NDUFA10,ATP5PD

K3R3

K2,GABARAPL2,MAPK3,PLCD3,SEPTIN9,TAB3,BCAR1,RELA,ARF6,MYL9,ELMO2,PIK3R3,CYTH3

TYK2,JAK1,PPP2R5D,ATP6V0B,CSNK1A1,STAT2,GSK3B,CDKN1B,STAT1,ATP6V0E2,THBS2,SOS1,

H,ZNF516,SMARCC2,SMARCB1,NDUFA5,UQCR11,COX5B,NDUFA10,ATP5PD

C2,PSMC4,NDUFB8,TUBA1C,TXN,NDUFS1,PSMB5,ATP5F1C,COX6B1,VDAC2,UQCRFS1,PSMA1,A
5,XBP1,MAN1B1,SEC31A,EIF2S1,RNF5,RAD23A
JBB4A,PINK1,PSMB1,NDUFA4,PSMD13,NDUFC2,PSMD7,COX8A,KLC2,PRPH,COX7A2L,NUP153,A
27A,TYK2,JAK1,STAT2,RPS29,STAT1,MAPK3,TRAF3,RPL36AL,RELA,RPL32,PIK3R3
TRAP1,ATXN3,ACTR1B,PSMD4,COX6A1,PSMC3,MTOR,ATP5MC3,NDUFA13,PSMC1,ATP5MC2,AMI
YL12A,ARPC5,KPNA3,ROCK2,MAP2K4,RAB5A,MAPK3,DYNLT1,TAB3,RELA,TLR6,VPS16,ARF6,DY
NDUFS1,PSMB5,ATP5F1C,COX6B1,VDAC2,UQCRFS1,PSMA1,ATP5F1E,ATF6B,COX4I1,COX5A,CO
KLC2,CSNK2B,CAPN1,COX7A2L,ADRM1,PSMC5,ATP5PO,CSNK1A1,PSMC2,PSMC4,GSK3B,NDUF
1,PSMC5,ATP5PO,PSMC2,PSMC4,NDUFB8,TUBA1C,NDUFS1,DLG4,PSMB5,ATP5F1C,COX6B1,VD

VPS28,SH3KBP1,ASAP3,CYTH3,ARFGAP3,VPS37B

ATP6V0A1,MAPK3,TRAF3,COL6A3,PRKCI,PPP2R5B,RELA,MAML1,PPP2R2A,ATP6V1F,ITGA7,FZD

ATP5F1E,COX4I1,UBE2G1,ATF6,COX5A,COX7A2,PSMA6,XBP1,UQCRH,PSMD12,PSMA4,EIF2S1,PS

DRM1,SRSF7,NUP98,PSMC5,ATP5PO,PSMC2,PSMC4,NDUFB8,TUBA1C,UBQLN4,SETX,NDUFS1,F

BRA1,TUBB4A,UBE2L3,PINK1,PSMB1,MFN2,SLC25A6,NDUFA4,PSMD13,PARK7,NDUFC2,PSMD7,
NC1I2,MYL9,ELMO2,CYTH3,M6PR
X7A2,MAPK3,PPP3CA,PSMA6,UQCRH,PSMD12,PSMA4,PRNP,EIF2S1,PSMC6,PIK3R3,NDUFA5,UQ
B8,TUBA1C,NDUFS1,PSMB5,ATP5F1C,COX6B1,IRS1,VDAC2,UQCRFS1,PSMA1,BACE1,ATP5F1E,C
DAC2,UQCRFS1,PSMA1,ATP5F1E,COX4I1,COX5A,COX7A2,SLC1A3,PSMA6,UQCRH,SP1,PSMD12,P

7,FZD8,PIK3R3,ATP6V1D,HES6,PPP2R5C

SMC6,GNAL,NDUFA5,UQCR11,COX5B,GNAI1,NDUFA10,ATP5PD

PSMB5,ATP5F1C,COX6B1,NUP205,UQCRFS1,NUP133,PSMA1,ATP5F1E,COX4I1,ATF6,COX5A,RAE

COX8A,KLC2,PRPH,UBE2J1,CSNK2B,CAPN1,COX7A2L,ADRM1,PSMC5,ATP5PO,CSNK1A1,PSMC2

CR11,COX5B,NDUFA10,ATP5PD
COX4I1,ATF6,COX5A,COX7A2,MAPK3,PPP3CA,PSMA6,XBP1,UQCRH,RELA,PSMD12,APOE,PSMA4,
PSMA4,PSMC6,NDUFA5,UQCR11,RB1CC1,SIN3A,COX5B,NDUFA10,ATP5PD

35A,COX7A2,GABARAPL2,PPP3CA,PSMA6,XBP1,UQCRH,SPG11,PSMD12,PSMA4,NUP50,EIF2S1,PS

2,PSMC4,GSK3B,NDUFB8,TUBA1C,NDUFS1,DLG4,PSMB5,ATP5F1C,COX6B1,VDAC2,UQCRFS1,P

,EIF2S1,PSMC6,FZD7,FZD8,LRP6,PIK3R3,NDUFA5,UQCR11,RB1CC1,COX5B,NDUFA10,ATP5PD

SMC6,NUP58,TOMM40,NDUFA5,UQCR11,ALYREF,RB1CC1,COX5B,NDUFA10,ATP5PD

SMA1,ATP5F1E,COX4I1,UBE2G1,ATF6,COX5A,RAB5A,COX7A2,GABARAPL2,MAPK3,PPP3CA,PSM

A6,XBP1,UQCRH,SPG11,RELA,PSMD12,PSMA4,PRNP,EIF2S1,PSMC6,FZD7,FZD8,LRP6,TOMM40

,NDUFA5,UQCR11,RB1CC1,COX5B,NDUFA10,ATP5PD

| | | | | | | |
|---|---|---|---|---|---|---|
| **REAC** | Axon guidanc | REAC:R-HSA- | 1.82E-53 | 52.7401737 | 549 | 1833 |
| **REAC** | Nervous syste | REAC:R-HSA- | 1.06E-52 | 51.9732271 | 574 | 1833 |
| **REAC** | GTP hydrolysi | REAC:R-HSA- | 3.89E-48 | 47.4096035 | 113 | 1833 |
| **REAC** | Regulation of | REAC:R-HSA- | 1.12E-47 | 46.9502395 | 171 | 1833 |
| **REAC** | L13a-mediate | REAC:R-HSA- | 1.91E-47 | 46.7181076 | 112 | 1833 |
| **REAC** | Signaling by F | REAC:R-HSA- | 5.82E-47 | 46.2351273 | 217 | 1833 |
| **REAC** | Eukaryotic Tra | REAC:R-HSA- | 3.33E-45 | 44.4769076 | 120 | 1833 |
| **REAC** | Cap-depender | REAC:R-HSA- | 3.33E-45 | 44.4769076 | 120 | 1833 |
| **REAC** | Formation of a | REAC:R-HSA- | 1.50E-44 | 43.8244698 | 102 | 1833 |
| **REAC** | Nonsense Me | REAC:R-HSA- | 4.09E-43 | 42.3878187 | 96 | 1833 |
| **REAC** | SRP-depende | REAC:R-HSA- | 5.56E-43 | 42.2547755 | 113 | 1833 |
| **REAC** | Nonsense Me | REAC:R-HSA- | 1.57E-42 | 41.8041144 | 116 | 1833 |
| **REAC** | Nonsense-Me | REAC:R-HSA- | 1.57E-42 | 41.8041144 | 116 | 1833 |
| **REAC** | Eukaryotic Tra | REAC:R-HSA- | 1.02E-41 | 40.990283 | 94 | 1833 |
| **REAC** | Eukaryotic Tra | REAC:R-HSA- | 1.02E-41 | 40.990283 | 94 | 1833 |
| **REAC** | Peptide chain | REAC:R-HSA- | 1.04E-41 | 40.983722 | 90 | 1833 |
| **REAC** | Cellular respor | REAC:R-HSA- | 1.31E-40 | 39.8833551 | 582 | 1833 |
| **REAC** | Response of I | REAC:R-HSA- | 1.36E-40 | 39.8661047 | 102 | 1833 |
| **REAC** | Viral mRNA Tr | REAC:R-HSA- | 2.63E-40 | 39.5793602 | 90 | 1833 |
| **REAC** | Cellular respor | REAC:R-HSA- | 1.57E-38 | 37.8029568 | 596 | 1833 |
| **REAC** | Metabolism o | REAC:R-HSA- | 8.36E-38 | 37.077832 | 662 | 1833 |
| **REAC** | Selenocystein | REAC:R-HSA- | 1.01E-37 | 36.9960335 | 94 | 1833 |
| **REAC** | Selenoamino a | REAC:R-HSA- | 2.39E-37 | 36.6220759 | 117 | 1833 |
| **REAC** | Influenza Infe | REAC:R-HSA- | 3.76E-36 | 35.4249121 | 152 | 1833 |
| **REAC** | Metabolism o | REAC:R-HSA- | 1.03E-34 | 33.9860337 | 2016 | 1833 |
| **REAC** | Cellular respor | REAC:R-HSA- | 1.18E-31 | 30.927739 | 157 | 1833 |
| **REAC** | Influenza Viral | REAC:R-HSA- | 1.19E-31 | 30.9241686 | 133 | 1833 |
| **REAC** | Translation | REAC:R-HSA- | 2.72E-31 | 30.5657122 | 292 | 1833 |
| **REAC** | Infectious dise | REAC:R-HSA- | 2.04E-30 | 29.6905346 | 909 | 1833 |
| **REAC** | Ribosomal sca | REAC:R-HSA- | 5.52E-23 | 22.2579922 | 59 | 1833 |
| **REAC** | Translation ini | REAC:R-HSA- | 5.52E-23 | 22.2579922 | 59 | 1833 |
| **REAC** | Disease | REAC:R-HSA- | 5.97E-23 | 22.2241631 | 1616 | 1833 |
| **REAC** | Activation of tl | REAC:R-HSA- | 2.13E-22 | 21.6722928 | 60 | 1833 |
| **REAC** | Major pathwa | REAC:R-HSA- | 6.95E-21 | 20.1577299 | 183 | 1833 |
| **REAC** | Formation of t | REAC:R-HSA- | 8.10E-21 | 20.0916014 | 52 | 1833 |
| **REAC** | Developmenta | REAC:R-HSA- | 1.48E-19 | 18.8284179 | 1116 | 1833 |
| **REAC** | rRNA processi | REAC:R-HSA- | 1.72E-19 | 18.7656335 | 193 | 1833 |
| **REAC** | Metabolism o | REAC:R-HSA- | 3.08E-19 | 18.5121144 | 370 | 1833 |
| **REAC** | rRNA processi | REAC:R-HSA- | 3.18E-18 | 17.4975095 | 203 | 1833 |
| **REAC** | mRNA Splicin | REAC:R-HSA- | 1.27E-16 | 15.8965023 | 178 | 1833 |
| **REAC** | mRNA Splicin | REAC:R-HSA- | 3.55E-15 | 14.4498147 | 186 | 1833 |
| **REAC** | Membrane Tra | REAC:R-HSA- | 6.20E-14 | 13.207518 | 625 | 1833 |
| **REAC** | Processing of | REAC:R-HSA- | 6.98E-13 | 12.1561538 | 237 | 1833 |
| **REAC** | Degradation o | REAC:R-HSA- | 1.66E-12 | 11.7802761 | 83 | 1833 |
| **REAC** | Cellular respor | REAC:R-HSA- | 2.54E-12 | 11.5951647 | 75 | 1833 |

| | | | | | | |
|---|---|---|---|---|---|---|
| **REAC** | Signaling by F | REAC:R-HSA- | 3.43E-12 | 11.4648347 | 486 | 1833 |
| **REAC** | AUF1 (hnRNF | REAC:R-HSA- | 4.61E-12 | 11.3358929 | 54 | 1833 |
| **REAC** | Oxygen-deper | REAC:R-HSA- | 1.20E-11 | 10.920772 | 66 | 1833 |
| **REAC** | PTEN Regulat | REAC:R-HSA- | 1.36E-11 | 10.8674829 | 138 | 1833 |
| **REAC** | Regulation of | REAC:R-HSA- | 1.37E-11 | 10.8630664 | 69 | 1833 |
| **REAC** | Host Interactic | REAC:R-HSA- | 1.63E-11 | 10.7888852 | 125 | 1833 |
| **REAC** | Vesicle-mediat | REAC:R-HSA- | 2.38E-11 | 10.6235142 | 719 | 1833 |
| **REAC** | SCF(Skp2)-m | REAC:R-HSA- | 6.24E-11 | 10.2049376 | 60 | 1833 |
| **REAC** | Ubiquitin-dep | REAC:R-HSA- | 6.67E-11 | 10.1761151 | 52 | 1833 |
| **REAC** | Signaling by N | REAC:R-HSA- | 1.01E-10 | 9.995413 | 81 | 1833 |
| **REAC** | Diseases of si | REAC:R-HSA- | 1.03E-10 | 9.9874365 | 380 | 1833 |
| **REAC** | SARS-CoV Inf | REAC:R-HSA- | 1.46E-10 | 9.83586677 | 144 | 1833 |
| **REAC** | Negative regul | REAC:R-HSA- | 1.48E-10 | 9.83101589 | 53 | 1833 |
| **REAC** | Regulation of | REAC:R-HSA- | 2.93E-10 | 9.53269758 | 86 | 1833 |
| **REAC** | Degradation o | REAC:R-HSA- | 4.43E-10 | 9.35393931 | 60 | 1833 |
| **REAC** | GLI3 is proce | REAC:R-HSA- | 4.43E-10 | 9.35393931 | 60 | 1833 |
| **REAC** | Golgi-to-ER re | REAC:R-HSA- | 6.54E-10 | 9.18431791 | 134 | 1833 |
| **REAC** | SCF-beta-TrC | REAC:R-HSA- | 6.62E-10 | 9.17916336 | 55 | 1833 |
| **REAC** | G2/M Transiti | REAC:R-HSA- | 7.23E-10 | 9.14082393 | 196 | 1833 |
| **REAC** | The role of GT | REAC:R-HSA- | 9.85E-10 | 9.00638123 | 79 | 1833 |
| **REAC** | Programmed | REAC:R-HSA- | 1.06E-09 | 8.97532027 | 186 | 1833 |
| **REAC** | PCP/CE pathv | REAC:R-HSA- | 1.25E-09 | 8.90482831 | 92 | 1833 |
| **REAC** | Mitotic G2-G2 | REAC:R-HSA- | 1.31E-09 | 8.88296698 | 198 | 1833 |
| **REAC** | Post-translatic | REAC:R-HSA- | 1.62E-09 | 8.79101804 | 1415 | 1833 |
| **REAC** | HIV Infection | REAC:R-HSA- | 1.78E-09 | 8.74863162 | 226 | 1833 |
| **REAC** | Regulation of | REAC:R-HSA- | 2.34E-09 | 8.63053382 | 54 | 1833 |
| **REAC** | Dectin-1 medi | REAC:R-HSA- | 2.94E-09 | 8.53218262 | 60 | 1833 |
| **REAC** | Intracellular si | REAC:R-HSA- | 2.99E-09 | 8.52365866 | 292 | 1833 |
| **REAC** | Neutrophil de | REAC:R-HSA- | 3.49E-09 | 8.45722934 | 476 | 1833 |
| **REAC** | Vpu mediated | REAC:R-HSA- | 4.06E-09 | 8.39122508 | 52 | 1833 |
| **REAC** | FBXL7 down- | REAC:R-HSA- | 4.68E-09 | 8.32995091 | 55 | 1833 |
| **REAC** | Degradation o | REAC:R-HSA- | 4.68E-09 | 8.32995091 | 55 | 1833 |
| **REAC** | Regulation of | REAC:R-HSA- | 7.03E-09 | 8.15300477 | 50 | 1833 |
| **REAC** | Vif-mediated c | REAC:R-HSA- | 8.18E-09 | 8.08736722 | 53 | 1833 |
| **REAC** | Regulation of | REAC:R-HSA- | 8.18E-09 | 8.08736722 | 53 | 1833 |
| **REAC** | Hedgehog 'of | REAC:R-HSA- | 8.71E-09 | 8.05991387 | 113 | 1833 |
| **REAC** | Hh mutants a | REAC:R-HSA- | 9.12E-09 | 8.04001369 | 56 | 1833 |
| **REAC** | NIK-->noncar | REAC:R-HSA- | 9.82E-09 | 8.00806465 | 59 | 1833 |
| **REAC** | Regulation of | REAC:R-HSA- | 1.43E-08 | 7.84558731 | 51 | 1833 |
| **REAC** | Degradation o | REAC:R-HSA- | 1.74E-08 | 7.76010579 | 57 | 1833 |
| **REAC** | Degradation o | REAC:R-HSA- | 1.82E-08 | 7.73934623 | 60 | 1833 |
| **REAC** | Cyclin A:Cdk2 | REAC:R-HSA- | 2.07E-08 | 7.68425983 | 85 | 1833 |
| **REAC** | PIP3 activates | REAC:R-HSA- | 2.28E-08 | 7.6420505 | 252 | 1833 |
| **REAC** | L1CAM intera | REAC:R-HSA- | 2.33E-08 | 7.63255958 | 119 | 1833 |
| **REAC** | p53-Indepenc | REAC:R-HSA- | 2.82E-08 | 7.54992531 | 52 | 1833 |

| | | | | | | |
|---|---|---|---|---|---|---|
| **REAC** | Autodegradati | REAC:R-HSA- | 2.82E-08 | 7.54992531 | 52 | 1833 |
| **REAC** | Ubiquitin Med | REAC:R-HSA- | 2.82E-08 | 7.54992531 | 52 | 1833 |
| **REAC** | p53-Independ | REAC:R-HSA- | 2.82E-08 | 7.54992531 | 52 | 1833 |
| **REAC** | Metabolism o | REAC:R-HSA- | 3.24E-08 | 7.48966521 | 58 | 1833 |
| **REAC** | Asymmetric lo | REAC:R-HSA- | 3.32E-08 | 7.47943663 | 64 | 1833 |
| **REAC** | Defective CFT | REAC:R-HSA- | 3.32E-08 | 7.47923147 | 61 | 1833 |
| **REAC** | Cyclin E asso | REAC:R-HSA- | 3.70E-08 | 7.43140641 | 83 | 1833 |
| **REAC** | Hh mutants a | REAC:R-HSA- | 5.91E-08 | 7.22817788 | 59 | 1833 |
| **REAC** | Transcriptiona | REAC:R-HSA- | 6.12E-08 | 7.21319191 | 94 | 1833 |
| **REAC** | Mitotic Anaph | REAC:R-HSA- | 6.17E-08 | 7.20985011 | 228 | 1833 |
| **REAC** | Apoptosis | REAC:R-HSA- | 7.63E-08 | 7.11722865 | 174 | 1833 |
| **REAC** | Mitotic Metap | REAC:R-HSA- | 7.86E-08 | 7.10462603 | 229 | 1833 |
| **REAC** | p53-Depende | REAC:R-HSA- | 1.00E-07 | 6.99965072 | 66 | 1833 |
| **REAC** | p53-Depende | REAC:R-HSA- | 1.00E-07 | 6.99965072 | 66 | 1833 |
| **REAC** | Stabilization o | REAC:R-HSA- | 1.05E-07 | 6.97724365 | 57 | 1833 |
| **REAC** | ER-Phagoson | REAC:R-HSA- | 1.66E-07 | 6.77906937 | 83 | 1833 |
| **REAC** | Autodegradati | REAC:R-HSA- | 1.79E-07 | 6.74629671 | 64 | 1833 |
| **REAC** | EPH-Ephrin si | REAC:R-HSA- | 1.94E-07 | 6.71308132 | 90 | 1833 |
| **REAC** | CLEC7A (Dec | REAC:R-HSA- | 2.12E-07 | 6.67406937 | 97 | 1833 |
| **REAC** | Regulation of | REAC:R-HSA- | 2.61E-07 | 6.58392377 | 71 | 1833 |
| **REAC** | G1/S DNA Da | REAC:R-HSA- | 2.84E-07 | 6.54717186 | 68 | 1833 |
| **REAC** | Hedgehog lig | REAC:R-HSA- | 3.04E-07 | 6.51653979 | 65 | 1833 |
| **REAC** | CDT1 associa | REAC:R-HSA- | 3.33E-07 | 6.4773204 | 59 | 1833 |
| **REAC** | RHO GTPase | REAC:R-HSA- | 3.46E-07 | 6.46036802 | 319 | 1833 |
| **REAC** | Signaling by V | REAC:R-HSA- | 3.75E-07 | 6.42552124 | 328 | 1833 |
| **REAC** | MAPK6/MAPK | REAC:R-HSA- | 5.22E-07 | 6.2822858 | 89 | 1833 |
| **REAC** | Activation of N | REAC:R-HSA- | 8.38E-07 | 6.07688667 | 67 | 1833 |
| **REAC** | Regulation of | REAC:R-HSA- | 8.38E-07 | 6.07688667 | 67 | 1833 |
| **REAC** | Signaling by H | REAC:R-HSA- | 1.1015E-06 | 5.95800876 | 149 | 1833 |
| **REAC** | Separation of | REAC:R-HSA- | 1.1359E-06 | 5.94467142 | 188 | 1833 |
| **REAC** | Orc1 removal | REAC:R-HSA- | 1.2173E-06 | 5.91461291 | 71 | 1833 |
| **REAC** | COPI-mediate | REAC:R-HSA- | 1.4306E-06 | 5.84447243 | 102 | 1833 |
| **REAC** | Antigen proce | REAC:R-HSA- | 1.736E-06 | 5.760439 | 99 | 1833 |
| **REAC** | Beta-catenin i | REAC:R-HSA- | 1.7498E-06 | 5.75701629 | 143 | 1833 |
| **REAC** | Cross-present | REAC:R-HSA- | 1.9264E-06 | 5.71525119 | 50 | 1833 |
| **REAC** | Potential thera | REAC:R-HSA- | 2.2538E-06 | 5.64707667 | 79 | 1833 |
| **REAC** | Transcriptiona | REAC:R-HSA- | 2.4685E-06 | 5.60755942 | 118 | 1833 |
| **REAC** | Downstream s | REAC:R-HSA- | 3.4237E-06 | 5.4654994 | 80 | 1833 |
| **REAC** | ABC transport | REAC:R-HSA- | 4.0077E-06 | 5.39710531 | 77 | 1833 |
| **REAC** | Intra-Golgi an | REAC:R-HSA- | 5.9282E-06 | 5.22707536 | 203 | 1833 |
| **REAC** | APC/C:Cdc20 | REAC:R-HSA- | 6.1565E-06 | 5.21066768 | 68 | 1833 |
| **REAC** | Assembly of t | REAC:R-HSA- | 6.1565E-06 | 5.21066768 | 68 | 1833 |
| **REAC** | Signaling by F | REAC:R-HSA- | 8.9319E-06 | 5.0490559 | 459 | 1833 |
| **REAC** | Signaling by F | REAC:R-HSA- | 8.9319E-06 | 5.0490559 | 459 | 1833 |
| **REAC** | TGF-beta rece | REAC:R-HSA- | 1.1824E-05 | 4.9272537 | 30 | 1833 |

| | | | | | | |
|---|---|---|---|---|---|---|
| **REAC** | Interleukin-1 s | REAC:R-HSA- | 1.2381E-05 | 4.90723744 | 101 | 1833 |
| **REAC** | UCH proteinas | REAC:R-HSA- | 1.2381E-05 | 4.90723744 | 101 | 1833 |
| **REAC** | CDK-mediated | REAC:R-HSA- | 1.266E-05 | 4.8975748 | 73 | 1833 |
| **REAC** | Selective auto | REAC:R-HSA- | 1.3539E-05 | 4.86839978 | 80 | 1833 |
| **REAC** | ER to Golgi A | REAC:R-HSA- | 1.6138E-05 | 4.79216001 | 155 | 1833 |
| **REAC** | Signaling by N | REAC:R-HSA- | 1.7579E-05 | 4.75500959 | 233 | 1833 |
| **REAC** | APC/C:Cdh1 r | REAC:R-HSA- | 1.9098E-05 | 4.71900379 | 74 | 1833 |
| **REAC** | Regulation of | REAC:R-HSA- | 1.9923E-05 | 4.70063606 | 81 | 1833 |
| **REAC** | Attenuation pl | REAC:R-HSA- | 2.2491E-05 | 4.64799018 | 14 | 1833 |
| **REAC** | Hedgehog 'or | REAC:R-HSA- | 2.3986E-05 | 4.62004179 | 85 | 1833 |
| **REAC** | ABC-family pr | REAC:R-HSA- | 2.3997E-05 | 4.61985208 | 103 | 1833 |
| **REAC** | MAPK family s | REAC:R-HSA- | 2.4265E-05 | 4.61501208 | 308 | 1833 |
| **REAC** | M Phase | REAC:R-HSA- | 2.5858E-05 | 4.58740044 | 407 | 1833 |
| **REAC** | Post-translatic | REAC:R-HSA- | 2.6541E-05 | 4.57608206 | 107 | 1833 |
| **REAC** | COPI-depend | REAC:R-HSA- | 2.9787E-05 | 4.52597024 | 100 | 1833 |
| **REAC** | Asparagine N- | REAC:R-HSA- | 3.0702E-05 | 4.51283312 | 305 | 1833 |
| **REAC** | G1/S Transitic | REAC:R-HSA- | 3.8792E-05 | 4.41125757 | 131 | 1833 |
| **REAC** | mRNA 3'-end | REAC:R-HSA- | 4.5112E-05 | 4.34570633 | 56 | 1833 |
| **REAC** | Cdc20:Phosp | REAC:R-HSA- | 5.0373E-05 | 4.29780516 | 73 | 1833 |
| **REAC** | Downstream T | REAC:R-HSA- | 5.123E-05 | 4.29047831 | 98 | 1833 |
| **REAC** | Innate Immur | REAC:R-HSA- | 5.4463E-05 | 4.263895 | 1090 | 1833 |
| **REAC** | Clathrin-media | REAC:R-HSA- | 5.4522E-05 | 4.26343011 | 144 | 1833 |
| **REAC** | Cytosolic tRN | REAC:R-HSA- | 5.4573E-05 | 4.26302544 | 24 | 1833 |
| **REAC** | MAPK1/MAPK | REAC:R-HSA- | 5.4693E-05 | 4.2620705 | 269 | 1833 |
| **REAC** | Switching of c | REAC:R-HSA- | 5.6401E-05 | 4.24871419 | 91 | 1833 |
| **REAC** | XBP1(S) activa | REAC:R-HSA- | 5.8907E-05 | 4.22983642 | 47 | 1833 |
| **REAC** | RHO GTPases | REAC:R-HSA- | 5.9008E-05 | 4.22909218 | 35 | 1833 |
| **REAC** | N-glycan trimr | REAC:R-HSA- | 5.9008E-05 | 4.22909218 | 35 | 1833 |
| **REAC** | APC:Cdc20 m | REAC:R-HSA- | 7.4201E-05 | 4.12959209 | 74 | 1833 |
| **REAC** | TCF depende | REAC:R-HSA- | 7.5602E-05 | 4.121464 | 232 | 1833 |
| **REAC** | Signaling by T | REAC:R-HSA- | 8.8956E-05 | 4.05082719 | 71 | 1833 |
| **REAC** | RAF/MAP kina | REAC:R-HSA- | 0.00010471 | 3.98000383 | 264 | 1833 |
| **REAC** | Signaling by V | REAC:R-HSA- | 0.000105 | 3.97880734 | 104 | 1833 |
| **REAC** | Cell Cycle, Mit | REAC:R-HSA- | 0.00011077 | 3.9555761 | 549 | 1833 |
| **REAC** | RNA Polymera | REAC:R-HSA- | 0.0001258 | 3.90031722 | 65 | 1833 |
| **REAC** | Regulation of | REAC:R-HSA- | 0.00013585 | 3.86692897 | 124 | 1833 |
| **REAC** | Mitotic G1 ph | REAC:R-HSA- | 0.00014678 | 3.83334584 | 148 | 1833 |
| **REAC** | VEGFA-VEGF | REAC:R-HSA- | 0.00015078 | 3.8216658 | 94 | 1833 |
| **REAC** | APC/C:Cdc20 | REAC:R-HSA- | 0.00015627 | 3.80612283 | 76 | 1833 |
| **REAC** | Signaling by N | REAC:R-HSA- | 0.00015627 | 3.80612283 | 76 | 1833 |
| **REAC** | IRE1alpha act | REAC:R-HSA- | 0.00016186 | 3.79086766 | 49 | 1833 |
| **REAC** | Protein folding | REAC:R-HSA- | 0.00017811 | 3.74931254 | 102 | 1833 |
| **REAC** | Autophagy | REAC:R-HSA- | 0.00018634 | 3.72969918 | 149 | 1833 |
| **REAC** | Transport to th | REAC:R-HSA- | 0.00019021 | 3.72077104 | 186 | 1833 |
| **REAC** | Response of I | REAC:R-HSA- | 0.00021349 | 3.67061353 | 23 | 1833 |

| | | | | | | |
|---|---|---|---|---|---|---|
| **REAC** | Activation of A | REAC:R-HSA- | 0.00022358 | 3.65055902 | 77 | 1833 |
| **REAC** | COPI-indeper | REAC:R-HSA- | 0.00023007 | 3.63813473 | 53 | 1833 |
| **REAC** | Immune Syste | REAC:R-HSA- | 0.00023013 | 3.63802269 | 2038 | 1833 |
| **REAC** | APC/C-mediat | REAC:R-HSA- | 0.00023376 | 3.63123254 | 88 | 1833 |
| **REAC** | Regulation of | REAC:R-HSA- | 0.00023376 | 3.63123254 | 88 | 1833 |
| **REAC** | MHC class II a | REAC:R-HSA- | 0.00029274 | 3.53351318 | 123 | 1833 |
| **REAC** | HSP90 chaper | REAC:R-HSA- | 0.00030879 | 3.51033403 | 57 | 1833 |
| **REAC** | Aggrephagy | REAC:R-HSA- | 0.00031935 | 3.49573042 | 44 | 1833 |
| **REAC** | Transcriptiona | REAC:R-HSA- | 0.00033105 | 3.4801057 | 361 | 1833 |
| **REAC** | Semaphorin ir | REAC:R-HSA- | 0.00033191 | 3.47898502 | 64 | 1833 |
| **REAC** | RHO GTPases | REAC:R-HSA- | 0.00035349 | 3.45162227 | 21 | 1833 |
| **REAC** | C-type lectin r | REAC:R-HSA- | 0.00038731 | 3.41194206 | 136 | 1833 |
| **REAC** | Neddylation | REAC:R-HSA- | 0.00043786 | 3.35866521 | 233 | 1833 |
| **REAC** | Downregulatic | REAC:R-HSA- | 0.00047836 | 3.32024146 | 24 | 1833 |
| **REAC** | Infection with | REAC:R-HSA- | 0.00057274 | 3.24204167 | 27 | 1833 |
| **REAC** | Folding of act | REAC:R-HSA- | 0.00058178 | 3.23524459 | 10 | 1833 |
| **REAC** | Class I MHC r | REAC:R-HSA- | 0.00060301 | 3.21967697 | 370 | 1833 |
| **REAC** | Macroautopha | REAC:R-HSA- | 0.00062782 | 3.20216429 | 134 | 1833 |
| **REAC** | Chaperonin-m | REAC:R-HSA- | 0.00064144 | 3.1928419 | 95 | 1833 |
| **REAC** | Signaling by N | REAC:R-HSA- | 0.00099103 | 3.00391447 | 112 | 1833 |
| **REAC** | RHO GTPases | REAC:R-HSA- | 0.00100407 | 2.9982354 | 136 | 1833 |
| **REAC** | Metabolism | REAC:R-HSA- | 0.00116614 | 2.93324803 | 2077 | 1833 |
| **REAC** | Cell Cycle | REAC:R-HSA- | 0.00122814 | 2.91075093 | 679 | 1833 |
| **REAC** | Interleukin-12 | REAC:R-HSA- | 0.00124188 | 2.90591928 | 57 | 1833 |
| **REAC** | ECM proteogl | REAC:R-HSA- | 0.0013295 | 2.8763104 | 75 | 1833 |
| **REAC** | Antigen proce | REAC:R-HSA- | 0.00155916 | 2.80710955 | 308 | 1833 |
| **REAC** | Signaling by N | REAC:R-HSA- | 0.00163421 | 2.78669235 | 130 | 1833 |
| **REAC** | Unfolded Prot | REAC:R-HSA- | 0.00182223 | 2.73939788 | 91 | 1833 |
| **REAC** | Formation of t | REAC:R-HSA- | 0.00199657 | 2.69971445 | 26 | 1833 |
| **REAC** | Recycling patl | REAC:R-HSA- | 0.00204919 | 2.68841739 | 48 | 1833 |
| **REAC** | TCR signaling | REAC:R-HSA- | 0.00268568 | 2.5709454 | 120 | 1833 |
| **REAC** | Regulation of | REAC:R-HSA- | 0.00290547 | 2.53678291 | 85 | 1833 |
| **REAC** | DNA Replicatic | REAC:R-HSA- | 0.00290547 | 2.53678291 | 85 | 1833 |
| **REAC** | Adaptive Imm | REAC:R-HSA- | 0.00297488 | 2.52653029 | 799 | 1833 |
| **REAC** | Transcriptiona | REAC:R-HSA- | 0.00313833 | 2.50330106 | 236 | 1833 |
| **REAC** | RUNX1 interac | REAC:R-HSA- | 0.00337755 | 2.47139793 | 36 | 1833 |
| **REAC** | SARS-CoV-2 | REAC:R-HSA- | 0.00353752 | 2.45130083 | 67 | 1833 |
| **REAC** | Mitophagy | REAC:R-HSA- | 0.00377637 | 2.42292586 | 27 | 1833 |
| **REAC** | Synthesis of a | REAC:R-HSA- | 0.00378676 | 2.42173221 | 30 | 1833 |
| **REAC** | Regulation of | REAC:R-HSA- | 0.00378676 | 2.42173221 | 30 | 1833 |
| **REAC** | Interleukin-1 f | REAC:R-HSA- | 0.0038883 | 2.4102402 | 138 | 1833 |
| **REAC** | EPHB-mediate | REAC:R-HSA- | 0.00489034 | 2.31066132 | 40 | 1833 |
| **REAC** | Cellular respor | REAC:R-HSA- | 0.00512203 | 2.29055769 | 87 | 1833 |
| **REAC** | Signaling by I | REAC:R-HSA- | 0.00527086 | 2.27811821 | 459 | 1833 |
| **REAC** | RHO GTPases | REAC:R-HSA- | 0.00527859 | 2.27748175 | 19 | 1833 |

| | | | | | | |
|---|---|---|---|---|---|---|
| **REAC** | RHO GTPases | REAC:R-HSA- | 0.00527859 | 2.27748175 | 19 | 1833 |
| **REAC** | Translocation | REAC:R-HSA- | 0.00539424 | 2.26806986 | 72 | 1833 |
| **REAC** | Interleukin-12 | REAC:R-HSA- | 0.00555399 | 2.25539492 | 47 | 1833 |
| **REAC** | Signaling by F | REAC:R-HSA- | 0.00567827 | 2.24578389 | 54 | 1833 |
| **REAC** | Spry regulatio | REAC:R-HSA- | 0.00601298 | 2.22090999 | 14 | 1833 |
| **REAC** | Translation of | REAC:R-HSA- | 0.00655294 | 2.1835638 | 44 | 1833 |
| **REAC** | Prefoldin med | REAC:R-HSA- | 0.00683858 | 2.16503437 | 28 | 1833 |
| **REAC** | S Phase | REAC:R-HSA- | 0.00699864 | 2.15498604 | 162 | 1833 |
| **REAC** | Signaling by T | REAC:R-HSA- | 0.00711404 | 2.1478835 | 100 | 1833 |
| **REAC** | Gene and pro | REAC:R-HSA- | 0.00880692 | 2.05517618 | 38 | 1833 |
| **REAC** | Association of | REAC:R-HSA- | 0.00880692 | 2.05517618 | 38 | 1833 |
| **REAC** | HSF1 activatio | REAC:R-HSA- | 0.00915475 | 2.03835333 | 12 | 1833 |
| **REAC** | Extracellular m | REAC:R-HSA- | 0.00916793 | 2.03772849 | 298 | 1833 |
| **REAC** | EPH-ephrin m | REAC:R-HSA- | 0.01215221 | 1.91534458 | 49 | 1833 |
| **REAC** | Calnexin/calre | REAC:R-HSA- | 0.01236418 | 1.90783466 | 26 | 1833 |
| **REAC** | Regulation of | REAC:R-HSA- | 0.01432702 | 1.84384411 | 53 | 1833 |
| **REAC** | Response of I | REAC:R-HSA- | 0.01523892 | 1.81704591 | 15 | 1833 |
| **REAC** | Iron uptake ar | REAC:R-HSA- | 0.01634872 | 1.78651629 | 57 | 1833 |
| **REAC** | Synthesis of D | REAC:R-HSA- | 0.01679526 | 1.77481325 | 120 | 1833 |
| **REAC** | Cooperation c | REAC:R-HSA- | 0.01803428 | 1.74390131 | 33 | 1833 |
| **REAC** | Transport of N | REAC:R-HSA- | 0.01808748 | 1.74262187 | 80 | 1833 |
| **REAC** | TP53 Regulat | REAC:R-HSA- | 0.01852238 | 1.73230324 | 84 | 1833 |
| **REAC** | TNFR2 non-c | REAC:R-HSA- | 0.01875778 | 1.72681847 | 100 | 1833 |
| **REAC** | HSF1-depend | REAC:R-HSA- | 0.02238383 | 1.65006565 | 24 | 1833 |
| **REAC** | Non-integrin r | REAC:R-HSA- | 0.02271188 | 1.64374694 | 58 | 1833 |
| **REAC** | Chromatin org | REAC:R-HSA- | 0.02428444 | 1.61467197 | 273 | 1833 |
| **REAC** | Chromatin mc | REAC:R-HSA- | 0.02428444 | 1.61467197 | 273 | 1833 |
| **REAC** | TBC/RABGAP | REAC:R-HSA- | 0.02604219 | 1.58432255 | 44 | 1833 |
| **REAC** | The citric acid | REAC:R-HSA- | 0.03189772 | 1.49624034 | 175 | 1833 |
| **REAC** | Collagen biosy | REAC:R-HSA- | 0.03561571 | 1.44835836 | 67 | 1833 |
| **REAC** | DNA Damage | REAC:R-HSA- | 0.03751671 | 1.42577531 | 38 | 1833 |
| **REAC** | Smooth Musc | REAC:R-HSA- | 0.03751671 | 1.42577531 | 38 | 1833 |
| **REAC** | RHO GTPases | REAC:R-HSA- | 0.04020207 | 1.3957516 | 11 | 1833 |
| **REAC** | PINK1-PRKN | REAC:R-HSA- | 0.04058352 | 1.39165026 | 22 | 1833 |
| **REAC** | Collagen degr | REAC:R-HSA- | 0.04511639 | 1.34566565 | 64 | 1833 |
| **REAC** | Negative regul | REAC:R-HSA- | 0.04614546 | 1.33587106 | 42 | 1833 |
| **REAC** | Cooperation c | REAC:R-HSA- | 0.04614546 | 1.33587106 | 42 | 1833 |
| **REAC** | Deactivation o | REAC:R-HSA- | 0.04614546 | 1.33587106 | 42 | 1833 |
| **REAC** | Deubiquitinati | REAC:R-HSA- | 0.04676908 | 1.33004122 | 297 | 1833 |
| **REAC** | Signaling by N | REAC:R-HSA- | 0.04871951 | 1.31229711 | 72 | 1833 |

| | | |
|---|---|---|
| 249 | 10622 | PABPC1,HSPA8,ACTB,ACTG1,DPYSL3,TUBA1A,CFL1,HSP90AB1,HSP[illegible] |
| 255 | 10622 | PABPC1,HSPA8,ACTB,ACTG1,DPYSL3,TUBA1A,CFL1,HSP90AB1,HSP[illegible] |
| 92 | 10622 | RPL3,RPS6,RPL4,RPL7A,RPL8,RPLP0,RPL13A,RPL6,RPL13,RPS8,RP |
| 117 | 10622 | PABPC1,UBC,RPL3,RPS6,RPL4,RPL7A,RPL8,RPLP0,RPL13A,RPL6,R[illegible] |
| 91 | 10622 | PABPC1,RPL3,RPS6,RPL4,RPL7A,RPL8,RPLP0,RPL13A,RPL6,RPL13, |
| 134 | 10622 | PABPC1,UBC,RPL3,RPS6,RPL4,RPL7A,RPL8,RPLP0,RPL13A,RPL6,R[illegible] |
| 93 | 10622 | PABPC1,RPL3,RPS6,RPL4,RPL7A,RPL8,RPLP0,RPL13A,RPL6,RPL13, |
| 93 | 10622 | PABPC1,RPL3,RPS6,RPL4,RPL7A,RPL8,RPLP0,RPL13A,RPL6,RPL13, |
| 84 | 10622 | RPL3,RPS6,RPL4,RPL7A,RPL8,RPLP0,RPL13A,RPL6,RPL13,RPS8,RP |
| 80 | 10622 | PABPC1,RPL3,RPS6,RPL4,RPL7A,RPL8,RPLP0,RPL13A,RPL6,RPL13, |
| 88 | 10622 | RPL3,RPS6,RPL4,RPL7A,RPL8,RPLP0,RPL13A,RPL6,RPL13,RPS8,RP |
| 89 | 10622 | PABPC1,RPL3,RPS6,RPL4,RPL7A,RPL8,RPLP0,RPL13A,RPL6,RPL13, |
| 89 | 10622 | PABPC1,RPL3,RPS6,RPL4,RPL7A,RPL8,RPLP0,RPL13A,RPL6,RPL13, |
| 78 | 10622 | EEF1A1,EEF2,RPL3,RPS6,RPL4,RPL7A,RPL8,RPLP0,RPL13A,RPL6,R[illegible] |
| 78 | 10622 | RPL3,RPS6,RPL4,RPL7A,RPL8,RPLP0,RPL13A,RPL6,RPL13,RPS8,RP |
| 76 | 10622 | EEF1A1,EEF2,RPL3,RPS6,RPL4,RPL7A,RPL8,RPLP0,RPL13A,RPL6,R[illegible] |
| 237 | 10622 | EEF1A1,HSPA8,HSPA9,TUBA1A,HSP90AB1,HSP90AA1,UBC,RPL3,RPS |
| 81 | 10622 | RPL3,RPS6,RPL4,RPL7A,RPL8,RPLP0,RPL13A,RPL6,RPL13,RPS8,RP |
| 75 | 10622 | RPL3,RPS6,RPL4,RPL7A,RPL8,RPLP0,RPL13A,RPL6,RPL13,RPS8,RP |
| 237 | 10622 | EEF1A1,HSPA8,HSPA9,TUBA1A,HSP90AB1,HSP90AA1,UBC,RPL3,RPS |
| 253 | 10622 | PABPC1,HSPA8,HNRNPA1,UBC,RPL3,RPS6,RPL4,NCL,RPL7A,PRPF8, |
| 75 | 10622 | RPL3,RPS6,RPL4,RPL7A,RPL8,RPLP0,RPL13A,RPL6,RPL13,RPS8,RP |
| 85 | 10622 | RPL3,RPS6,RPL4,RPL7A,RPL8,RPLP0,RPL13A,RPL6,RPL13,RPS8,RP |
| 98 | 10622 | HSP90AA1,RPL3,RPS6,RPL4,RPL7A,RPL8,CALR,RPLP0,CANX,RPL13[illegible] |
| 552 | 10622 | EEF1A1,IGFBP5,PABPC1,HSPA8,COPA,ARSD,ACTB,CTSC,USP22,EEF[illegible] |
| 95 | 10622 | RPL3,RPS6,RPL4,RPL7A,RPL8,RPLP0,RPL13A,RPL6,RPL13,RPS8,RP |
| 86 | 10622 | HSP90AA1,RPL3,RPS6,RPL4,RPL7A,RPL8,RPLP0,RPL13A,RPL6,RPL1 |
| 139 | 10622 | EEF1A1,PABPC1,EEF2,RPL3,RPS6,RPL4,RPL7A,RPL8,RPLP0,RPL13A |
| 298 | 10622 | NEDD4L,ACTB,ACTG1,EEF2,TUBA1A,HSP90AB1,HSP90AA1,ACTR2,T[illegible] |
| 47 | 10622 | RPS6,RPS8,EIF3C,EIF4B,EIF4A1,RPS3,RPS4X,RPS3A,RPS17,RPS16,[illegible] |
| 47 | 10622 | PABPC1,RPS6,RPS8,EIF3C,EIF4B,EIF4A1,RPS3,RPS4X,RPS3A,RPS17, |
| 432 | 10622 | NEDD4L,ACTB,ACTG1,EEF2,TUBA1A,HSP90AB1,HSP90AA1,ACTR2,T[illegible] |
| 47 | 10622 | PABPC1,RPS6,RPS8,EIF3C,EIF4B,EIF4A1,RPS3,RPS4X,RPS3A,RPS17, |
| 90 | 10622 | RPL3,RPS6,RPL4,NCL,RPL7A,RPL8,RPLP0,RPL13A,RPL6,RPL13,RPS |
| 42 | 10622 | RPS6,RPS8,EIF3C,RPS3,RPS4X,RPS3A,RPS17,RPS16,RPSA,RPS24,R[illegible] |
| 315 | 10622 | PABPC1,HSPA8,ACTB,MED18,ACTG1,DPYSL3,TUBA1A,CFL1,HSP90A |
| 91 | 10622 | RPL3,RPS6,RPL4,NCL,RPL7A,RPL8,RPLP0,RPL13A,RPL6,RPL13,RPS |
| 140 | 10622 | RPL3,RPS6,RPL4,RPL7A,RPL8,RPLP0,RPL13A,RPL6,RPL13,ALDH6A1 |
| 92 | 10622 | RPL3,RPS6,RPL4,NCL,RPL7A,RPL8,RPLP0,RPL13A,RPL6,RPL13,RPS |
| 82 | 10622 | HSPA8,HNRNPA1,PRPF8,YBX1,DDX5,SNRNP200,SF3B1,SF3B2,GTF2[illegible] |
| 82 | 10622 | HSPA8,HNRNPA1,PRPF8,YBX1,DDX5,SNRNP200,SF3B1,SF3B2,GTF2[illegible] |
| 189 | 10622 | HSPA8,COPA,ACTB,CTSC,ACTG1,TUBA1A,ACTR2,FTH1,UBC,TUBA1[illegible] |
| 92 | 10622 | HSPA8,HNRNPA1,PRPF8,YBX1,DDX5,SNRNP200,SF3B1,SF3B2,GTF2[illegible] |
| 46 | 10622 | UBC,UBB,PSMD5,PPP2R1A,PSMD2,TLE5,CTNNB1,CTBP1,SKP1,PSM[illegible] |
| 43 | 10622 | UBC,UBB,PSMD5,EGLN1,PSMD2,PSMB2,VEGFA,VHL,PSMD3,PSMA7, |

| | | |
|---|---|---|
| 152 | 10622 | COL1A1,HNRNPA1,ACTB,ACTG1,HSP90AA1,SPARC,UBC,KIDINS220, |
| 35 | 10622 | PABPC1,HSPA8,UBC,UBB,PSMD5,PSMD2,PSMB2,PSMD3,HSPB1,PSM |
| 39 | 10622 | UBC,UBB,PSMD5,EGLN1,PSMD2,PSMB2,VHL,PSMD3,PSMA7,PSMD1 |
| 62 | 10622 | ATN1,UBC,UBB,PSMD5,CHD3,PSMD2,CHD4,RBBP4,JUN,HDAC2,PSM |
| 40 | 10622 | UBC,UBB,PSMD5,PSMD2,PSMB2,PSMD3,TNKS2,PSMA7,PSMD1,CSN |
| 58 | 10622 | UBC,NPM1,UBB,PSMD5,AP2M1,ARF1,AP2B1,KPNB1,PSMD2,XPO1,R |
| 202 | 10622 | COL1A1,LRP1,HSPA8,COPA,ACTB,CTSC,ACTG1,TUBA1A,HSP90AA1, |
| 36 | 10622 | UBC,UBB,PSMD5,PSMD2,SKP1,CDKN1A,PSMB2,CCND1,PSMD3,PSM |
| 33 | 10622 | UBC,UBB,PSMD5,PSMD2,PSMB2,CCND1,PSMD3,PSMA7,PSMD1,UBA |
| 43 | 10622 | UBC,UBB,PSMD5,PSMD2,SKP1,PSMB2,NOTCH2,PSMD3,PSMA7,EP3 |
| 123 | 10622 | ACTB,ACTG1,HSP90AA1,UBC,PIK3R2,SPTBN1,CALM1,VCP,UBB,FN1, |
| 62 | 10622 | HSP90AB1,HSP90AA1,TUBB,UBC,GANAB,CANX,DDX5,VCP,UBB,CHD |
| 33 | 10622 | UBC,UBB,PSMD5,PSMD2,SKP1,PSMB2,PSMD3,PSMA7,PSMD1,UBA5 |
| 44 | 10622 | PABPC1,HSPA8,UBC,UBB,PSMD5,SET,PSMD2,XPO1,YWHAB,TNPO1, |
| 35 | 10622 | UBC,UBB,PSMD5,PSMD2,SKP1,PSMB2,PSMD3,PSMA7,PSMD1,UBA5 |
| 35 | 10622 | UBC,UBB,PSMD5,PSMD2,SKP1,PSMB2,PSMD3,PSMA7,PSMD1,UBA5 |
| 58 | 10622 | COPA,TUBA1A,TUBA1B,TUBB2B,KIF1A,TUBB4B,ARF1,PAFAH1B1,KI |
| 33 | 10622 | UBC,UBB,PSMD5,PSMD2,SKP1,PSMB2,PSMD3,PSMA7,PSMD1,UBA5 |
| 75 | 10622 | TUBA1A,HSP90AB1,HSP90AA1,TUBB,UBC,TUBA1B,TUBB2B,YWHAE, |
| 41 | 10622 | TUBA1A,HSP90AB1,HSP90AA1,UBC,TUBA1B,TUBB2B,UBB,PSMD5,T |
| 72 | 10622 | HSP90AA1,VIM,UBC,MAGED1,YWHAE,SPTAN1,UBB,YWHAG,MAPT,PS |
| 45 | 10622 | UBC,DVL3,UBB,PSMD5,AP2M1,AP2B1,RHOA,PSMD2,PFN1,CLTC,RA |
| 75 | 10622 | TUBA1A,HSP90AB1,HSP90AA1,TUBB,UBC,TUBA1B,TUBB2B,YWHAE, |
| 339 | 10622 | EEF1A1,IGFBP5,HSPA8,COPA,ARSD,ACTB,CTSC,USP22,EEF2,TUBA1 |
| 82 | 10622 | NEDD4L,UBC,NPM1,UBB,PSMD5,AP2M1,ARF1,AP2B1,GTF2F1,XRCC |
| 32 | 10622 | UBC,UBB,PSMD5,PSMD2,PSMB2,PSMD3,PSMA7,EP300,PSMD1,UBA |
| 34 | 10622 | UBC,UBB,PSMD5,PSMD2,SKP1,PSMB2,PSMD3,PSMA7,PSMD1,UBA5 |
| 98 | 10622 | ATN1,UBC,PIK3R2,CALM1,UBB,PSMD5,PPP2R1A,CHD3,PSMD2,CHD |
| 141 | 10622 | EEF1A1,HSPA8,CTSC,EEF2,HSP90AB1,HSP90AA1,ACTR2,TUBB,FTH1 |
| 31 | 10622 | UBC,UBB,PSMD5,PSMD2,SKP1,PSMB2,PSMD3,PSMA7,PSMD1,UBA5 |
| 32 | 10622 | UBC,UBB,PSMD5,PSMD2,SKP1,PSMB2,PSMD3,PSMA7,PSMD1,UBA5 |
| 32 | 10622 | UBC,UBB,PSMD5,PSMD2,PSMB2,PSMD3,TNKS2,PSMA7,PSMD1,UBA |
| 30 | 10622 | UBC,UBB,PSMD5,PSMD2,PSMB2,PSMD3,PSMA7,PSMD1,UBA52,RPS |
| 31 | 10622 | UBC,UBB,PSMD5,PSMD2,PSMB2,PSMD3,PSMA7,PSMD1,UBA52,RPS |
| 31 | 10622 | UBC,UBB,PSMD5,PSMD2,PSMB2,PSMD3,PSMA7,PSMD1,UBA52,RPS |
| 50 | 10622 | TUBA1A,UBC,TUBA1B,TUBB2B,GNAS,UBB,PSMD5,TUBB4B,PSMD2, |
| 32 | 10622 | UBC,VCP,UBB,PSMD5,PSMD2,PSMB2,PSMD3,PSMA7,OS9,PSMD1,U |
| 33 | 10622 | UBC,UBB,PSMD5,PSMD2,SKP1,PSMB2,PSMD3,PSMA7,PSMD1,UBA5 |
| 30 | 10622 | PSMD5,PSMD2,OAZ1,PSMB2,AZIN1,PSMD3,ODC1,PSMA7,PSMD1,O |
| 32 | 10622 | UBC,DVL3,UBB,PSMD5,PSMD2,PSMB2,PSMD3,PSMA7,PSMD1,UBA5 |
| 33 | 10622 | UBC,UBB,PSMD5,PSMD2,SKP1,PSMB2,PSMD3,PSMA7,PSMD1,UBA5 |
| 41 | 10622 | UBC,UBB,PSMD5,PSMD2,RBBP4,SKP1,CDKN1A,PSMB2,CCND1,PSM |
| 86 | 10622 | ATN1,UBC,PIK3R2,UBB,PSMD5,PPP2R1A,CHD3,PSMD2,CHD4,RBBP |
| 51 | 10622 | HSPA8,ACTB,ACTG1,TUBA1A,DCX,TUBA1B,TUBB2B,SPTBN1,SPTAN |
| 30 | 10622 | UBC,UBB,PSMD5,PSMD2,PSMB2,PSMD3,PSMA7,PSMD1,UBA52,RPS |

| | | |
|---|---|---|
| 30 | 10622 | UBC,UBB,PSMD5,PSMD2,PSMB2,PSMD3,PSMA7,PSMD1,UBA52,RPS |
| 30 | 10622 | UBC,UBB,PSMD5,PSMD2,PSMB2,PSMD3,PSMA7,PSMD1,UBA52,RPS |
| 30 | 10622 | UBC,UBB,PSMD5,PSMD2,PSMB2,PSMD3,PSMA7,PSMD1,UBA52,RPS |
| 32 | 10622 | PSMD5,PSMD2,OAZ1,PSMB2,AZIN1,PSMD3,ODC1,SMS,PSMA7,PSMD |
| 34 | 10622 | UBC,UBB,PSMD5,PSMD2,PSMB2,PSMD3,VANGL2,PSMA7,PSMD1,UE |
| 33 | 10622 | UBC,VCP,UBB,PSMD5,PSMD2,PSMB2,PSMD3,PSMA7,OS9,PSMD1,UI |
| 40 | 10622 | UBC,UBB,PSMD5,PSMD2,RBBP4,SKP1,CDKN1A,PSMB2,CCND1,PSM |
| 32 | 10622 | UBC,VCP,UBB,PSMD5,PSMD2,PSMB2,PSMD3,PSMA7,OS9,PSMD1,UI |
| 43 | 10622 | UBC,UBB,PSMD5,PSMD2,CCN2,CTNNB1,CDKN1A,PSMB2,CCND1,PS |
| 79 | 10622 | NUDC,TUBA1A,UBC,TUBA1B,TUBB2B,UBB,PSMD5,PPP2R1A,TUBB4I |
| 65 | 10622 | VIM,UBC,MAGED1,YWHAE,SPTAN1,UBB,YWHAG,MAPT,PSMD5,YWHA |
| 79 | 10622 | NUDC,TUBA1A,UBC,TUBA1B,TUBB2B,UBB,PSMD5,PPP2R1A,TUBB4I |
| 34 | 10622 | UBC,UBB,PSMD5,PSMD2,CDKN1A,PSMB2,PSMD3,PSMA7,PSMD1,UE |
| 34 | 10622 | UBC,UBB,PSMD5,PSMD2,CDKN1A,PSMB2,PSMD3,PSMA7,PSMD1,UE |
| 31 | 10622 | UBC,UBB,PSMD5,PSMD2,PSMB2,PSMD3,PSMA7,PSMD1,UBA52,RPS |
| 39 | 10622 | UBC,CALR,UBB,PSMD5,HLA-C,PDIA3,SEC61A1,PSMD2,HLA-B,PSMB |
| 33 | 10622 | UBC,UBB,PSMD5,PSMD2,PSMB2,PSMD3,PSMA7,PSMD1,UBA52,RPS |
| 41 | 10622 | ACTB,ACTG1,CFL1,ACTR2,MYH9,AP2M1,AP2B1,RHOA,CLTC,MYL6,N |
| 43 | 10622 | UBC,CALM1,UBB,PSMD5,PSMD2,CLEC7A,SKP1,AHCYL1,PSMB2,PSN |
| 35 | 10622 | UBC,UBB,PSMD5,PSMD2,SKP1,PSMB2,PSMD3,PSMA7,PSMD1,UBA5 |
| 34 | 10622 | UBC,UBB,PSMD5,PSMD2,CDKN1A,PSMB2,PSMD3,PSMA7,PSMD1,UE |
| 33 | 10622 | UBC,P4HB,VCP,UBB,PSMD5,PSMD2,PSMB2,PSMD3,PSMA7,OS9,PSN |
| 31 | 10622 | ORC4,UBC,UBB,PSMD5,PSMD2,PSMB2,PSMD3,PSMA7,PSMD1,UBA5 |
| 99 | 10622 | ACTB,NUDC,ACTG1,TUBA1A,CFL1,ACTR2,TUBA1B,FLNA,TUBB2B,H3 |
| 101 | 10622 | UBC,H3-3B,SOX4,DVL3,GNB1,CALM1,UBB,PSMD5,PPP2R1A,AP2M1, |
| 40 | 10622 | UBC,UBB,PSMD5,PSMD2,XPO1,JUN,RAC1,PSMB2,PSMD3,HSPB1,CD |
| 33 | 10622 | UBC,UBB,PSMD5,PSMD2,SKP1,PSMB2,PSMD3,PSMA7,PSMD1,UBA5 |
| 33 | 10622 | UBC,UBB,PSMD5,PSMD2,PSMB2,PSMD3,PSMA7,PSMD1,UBA52,CUL |
| 56 | 10622 | TUBA1A,UBC,TUBA1B,TUBB2B,GNAS,P4HB,VCP,UBB,PSMD5,TUBB4 |
| 66 | 10622 | NUDC,TUBA1A,UBC,TUBA1B,TUBB2B,UBB,PSMD5,PPP2R1A,TUBB4I |
| 34 | 10622 | ORC4,UBC,UBB,PSMD5,PSMD2,SKP1,PSMB2,PSMD3,PSMA7,PSMD1 |
| 43 | 10622 | COPA,TUBA1A,TUBA1B,TUBB2B,SPTBN1,SPTAN1,TUBB4B,ARF1,SPT |
| 42 | 10622 | UBC,CALR,UBB,PSMD5,MRC2,HLA-C,PDIA3,SEC61A1,PSMD2,HLA-B |
| 54 | 10622 | UBC,DVL3,GNB1,CALM1,UBB,PSMD5,AP2M1,AP2B1,RHOA,PSMD2,F |
| 27 | 10622 | PSMD5,MRC2,PSMD2,PSMB2,PSMD3,PSMA7,PSMD1,PSMD8,PSMB4 |
| 36 | 10622 | HSP90AB1,HSP90AA1,TUBB,CHD3,AP2M1,ITGB1,AP2B1,CHD4,ATP1, |
| 47 | 10622 | COL1A1,UBC,UBB,PSMD5,PSMD2,SKP1,CDKN1A,PSMB2,CCND1,MA |
| 36 | 10622 | UBC,CALM1,UBB,PSMD5,PSMD2,SKP1,PSMB2,PSMD3,PSMA7,PSMD |
| 35 | 10622 | UBC,VCP,UBB,PSMD5,PSMD2,ABCA3,PSMB2,PSMD3,PSMA7,OS9,PS |
| 68 | 10622 | COPA,TUBA1A,TUBA1B,TMF1,TUBB2B,KIF1A,TUBB4B,ARF1,PAFAH1 |
| 32 | 10622 | UBC,UBB,PSMD5,PSMD2,PSMB2,PSMD3,PSMA7,PSMD1,UBA52,RPS |
| 32 | 10622 | ORC4,UBC,UBB,PSMD5,PSMD2,PSMB2,PSMD3,PSMA7,PSMD1,UBA5 |
| 126 | 10622 | ACTB,NUDC,ACTG1,TUBA1A,CFL1,HSP90AB1,HSP90AA1,ACTR2,TUE |
| 126 | 10622 | ACTB,NUDC,ACTG1,TUBA1A,CFL1,HSP90AB1,HSP90AA1,ACTR2,TUE |
| 19 | 10622 | NEDD4L,UBC,UBB,XPO1,PPP1CB,STRAP,UBA52,PPP1CC,RPS27A,PPI |

| | | |
|---|---|---|
| 41 | 10622 | UBC,APP,UBB,PSMD5,HMGB1,PSMD2,SKP1,SQSTM1,PSMB2,PSMD |
| 41 | 10622 | ACTB,UBC,UBB,PSMD5,UCHL1,PSMD2,ASXL1,PSMB2,PSMD3,PSMA |
| 33 | 10622 | UBC,UBB,PSMD5,PSMD2,PSMB2,PSMD3,PSMA7,PSMD1,UBA52,RPS |
| 35 | 10622 | HSPA8,TUBA1A,HSP90AA1,VIM,UBC,TUBA1B,TUBB2B,VCP,UBB,TUB |
| 55 | 10622 | COPA,CTSC,TUBA1A,TUBA1B,TUBB2B,SPTBN1,SPTAN1,TUBB4B,AR |
| 74 | 10622 | UBC,YBX1,H3-3B,UBB,PSMD5,PSMD2,ATP2A2,JUN,HDAC2,SKP1,DN |
| 33 | 10622 | UBC,UBB,PSMD5,PSMD2,PSMB2,PSMD3,PSMA7,PSMD1,UBA52,RPS |
| 35 | 10622 | UBC,UBB,PSMD5,PSMD2,SKP1,PSMB2,PSMD3,PSMA7,PSMD1,UBA5 |
| 12 | 10622 | HSPA8,HSP90AB1,HSP90AA1,HSPA1B,PTGES3,HSBP1,EP300,FKBP4, |
| 36 | 10622 | UBC,UBB,PSMD5,PSMD2,PSMB2,PSMD3,PSMA7,PSMD1,UBA52,CUL |
| 41 | 10622 | UBC,VCP,UBB,PSMD5,PSMD2,ABCA3,PSMB2,ABCA2,PSMD3,PSMA7, |
| 91 | 10622 | ACTB,ACTG1,UBC,PIK3R2,NEFL,SPTBN1,SPTAN1,CALM1,UBB,FN1, |
| 113 | 10622 | NUDC,TUBA1A,HSP90AA1,TUBB,UBC,NUMA1,TUBA1B,TUBB2B,H3-3 |
| 42 | 10622 | IGFBP5,HSP90B1,FSTL1,P4HB,APP,FN1,IGFBP3,PRKCSH,CDH2,PDI |
| 40 | 10622 | COPA,TUBA1A,TUBA1B,TUBB2B,KIF1A,TUBB4B,ARF1,KIF5B,RAB1A, |
| 90 | 10622 | COPA,CTSC,TUBA1A,UBC,TUBA1B,GANAB,CALR,TUBB2B,CANX,SPT |
| 48 | 10622 | ORC4,UBC,UBB,PSMD5,PPP2R1A,PSMD2,RBBP4,SKP1,CDKN1A,PSM |
| 27 | 10622 | PAPOLA,SRSF4,CPSF2,DDX39B,SRSF5,U2AF2,CPSF1,PABPN1,SRSF |
| 32 | 10622 | UBC,UBB,PSMD5,PSMD2,PSMB2,PSMD3,PSMA7,PSMD1,UBA52,RPS |
| 39 | 10622 | UBC,PIK3R2,UBB,PSMD5,PSMD2,SKP1,PSMB2,PSMD3,PSMA7,PSMD |
| 252 | 10622 | EEF1A1,HSPA8,ACTB,CTSC,ACTG1,EEF2,CFL1,MAVS,HSP90AB1,HSP |
| 51 | 10622 | HSPA8,ACTB,ACTG1,ACTR2,UBC,UBB,AP2M1,AP2B1,SCARB2,CLTC, |
| 16 | 10622 | NARS1,MARS1,QARS1,LARS1,SARS1,VARS1,GARS1,IARS1,YARS1,DA |
| 81 | 10622 | ACTB,ACTG1,UBC,PIK3R2,NEFL,SPTBN1,SPTAN1,CALM1,UBB,FN1, |
| 37 | 10622 | ORC4,UBC,UBB,PSMD5,PSMD2,SKP1,PSMB2,PSMD3,PSMA7,PSMD1 |
| 24 | 10622 | FKBP14,PDIA6,TLN1,CTDSP2,HDGF,ACADVL,HYOU1,SSR1,DCTN1,E |
| 20 | 10622 | ACTB,ACTG1,ACTR2,CYFIP2,RAC1,ACTR3,NCKAP1,MAPK1,ARPC2,C |
| 20 | 10622 | UBC,GANAB,CALR,CANX,VCP,UBB,PDIA3,MARCHF6,PRKCSH,MLEC, |
| 32 | 10622 | UBC,UBB,PSMD5,PSMD2,PSMB2,PSMD3,PSMA7,PSMD1,UBA52,RPS |
| 72 | 10622 | UBC,H3-3B,SOX4,DVL3,UBB,PSMD5,PPP2R1A,PSMD2,XPO1,CSNK1 |
| 31 | 10622 | NEDD4L,UBC,UBB,RHOA,XPO1,PPP1CB,PARP1,SKI,STRAP,NCOR2,U |
| 79 | 10622 | ACTB,ACTG1,UBC,PIK3R2,NEFL,SPTBN1,SPTAN1,CALM1,UBB,FN1, |
| 40 | 10622 | ACTB,ACTG1,HSP90AA1,PIK3R2,CALM1,CTNNA1,RHOA,CTNNB1,CY |
| 141 | 10622 | ORC4,NUDC,TUBA1A,HSP90AB1,HSP90AA1,TUBB,UBC,NUMA1,TUB |
| 29 | 10622 | PAPOLA,SRSF4,CPSF2,DDX39B,SRSF5,U2AF2,CPSF1,PABPN1,SRSF |
| 45 | 10622 | IGFBP5,HSP90B1,IGFBP2,FSTL1,P4HB,APP,IGF2,FN1,IGFBP3,PRKC |
| 51 | 10622 | ORC4,UBC,UBB,PSMD5,PPP2R1A,PSMD2,RBBP4,CCND2,SKP1,CDK |
| 37 | 10622 | ACTB,ACTG1,HSP90AA1,PIK3R2,CALM1,CTNNA1,RHOA,CTNNB1,CY |
| 32 | 10622 | UBC,UBB,PSMD5,PSMD2,PSMB2,PSMD3,PSMA7,PSMD1,UBA52,RPS |
| 32 | 10622 | COL1A1,UBC,COL5A1,UBB,FN1,ITGB1,COL11A1,TNS3,LAMA5,COL1 |
| 24 | 10622 | FKBP14,PDIA6,TLN1,CTDSP2,HDGF,ACADVL,HYOU1,SSR1,DCTN1,E |
| 39 | 10622 | ACTB,TUBA1A,TUBA1B,TUBB2B,GNB1,USP11,TUBB4B,CCT7,GNB2, |
| 51 | 10622 | EEF1A1,HSPA8,TUBA1A,HSP90AB1,HSP90AA1,VIM,UBC,TUBA1B,TU |
| 60 | 10622 | COPA,CTSC,TUBA1A,TUBA1B,TUBB2B,SPTBN1,SPTAN1,TUBB4B,AR |
| 15 | 10622 | UBC,ENO1,PGK1,UBB,KPNB1,SFPQ,RAB7A,MAPK1,HGS,UBA52,RPS |

| | | |
|---|---|---|
| 32 | 10622 | UBC,UBB,PSMD5,PSMD2,PSMB2,PSMD3,PSMA7,PSMD1,UBA52,RPS |
| 25 | 10622 | TUBA1A,TUBA1B,TUBB2B,TUBB4B,PAFAH1B1,RAB6B,TUBB2A,DYN |
| 429 | 10622 | EEF1A1,COL1A1,HSPA8,HSPA9,NEDD4L,ACTB,CTSC,ACTG1,EEF2,TU |
| 35 | 10622 | UBC,UBB,PSMD5,PSMD2,SKP1,PSMB2,PSMD3,PSMA7,PSMD1,UBA5 |
| 35 | 10622 | UBC,UBB,PSMD5,PSMD2,SKP1,PSMB2,PSMD3,PSMA7,PSMD1,UBA5 |
| 44 | 10622 | CTSC,TUBA1A,TUBA1B,TUBB2B,CANX,AP2M1,TUBB4B,ARF1,AP2B1, |
| 26 | 10622 | HSPA8,TUBA1A,HSP90AB1,HSP90AA1,TUBA1B,HSPA1B,TUBB2B,TUB |
| 22 | 10622 | TUBA1A,HSP90AA1,VIM,UBC,TUBA1B,TUBB2B,VCP,UBB,TUBB4B,TU |
| 99 | 10622 | UBC,KAT6A,YWHAE,GPI,NPM1,UBB,YWHAG,YWHAQ,PPP2R1A,CHD3, |
| 28 | 10622 | DPYSL3,CFL1,HSP90AB1,HSP90AA1,MYH9,ITGB1,RHOA,CRMP1,PLX |
| 14 | 10622 | FLNA,MYH9,CALM1,MYL6,MYH10,RAC1,PPP1CB,CDC42,CTTN,MYL1 |
| 47 | 10622 | UBC,CALM1,UBB,PSMD5,PSMD2,CLEC7A,SKP1,AHCYL1,PSMB2,PSM |
| 70 | 10622 | UBC,FBXL4,UBB,DDB1,PSMD5,WSB1,PSMD2,KLHL9,SKP1,PSMB2,V |
| 15 | 10622 | NEDD4L,UBC,UBB,XPO1,PPP1CB,STRAP,UBA52,PPP1CC,RPS27A,PP |
| 16 | 10622 | UBC,ENO1,PGK1,UBB,KPNB1,SFPQ,RAB7A,MAPK1,B2M,HGS,UBA5 |
| 9 | 10622 | ACTB,CCT7,CCT6A,CCT2,TCP1,CCT3,CCT4,CCT8,CCT5 |
| 100 | 10622 | NEDD4L,UBC,FBXL4,UBA1,CALR,CANX,UBB,PSMD5,MRC2,HLA-C,PD |
| 46 | 10622 | HSPA8,TUBA1A,HSP90AA1,VIM,UBC,TUBA1B,TUBB2B,VCP,UBB,TUB |
| 36 | 10622 | ACTB,TUBA1A,TUBA1B,TUBB2B,GNB1,USP11,TUBB4B,CCT7,GNB2, |
| 40 | 10622 | KIDINS220,PIK3R2,PPP2R1A,AP2M1,AP2B1,RHOA,CHD4,JUND,CLTC |
| 46 | 10622 | ACTB,NUDC,ACTG1,TUBA1A,TUBA1B,TUBB2B,DVL3,PPP2R1A,ITGB1 |
| 431 | 10622 | LRP1,GAPDH,ARSD,MED18,HSP90AB1,HSP90AA1,UBC,RPL3,RPS6,R |
| 163 | 10622 | ORC4,NUDC,TUBA1A,HSP90AB1,HSP90AA1,TUBB,UBC,NUMA1,TUB |
| 25 | 10622 | HSPA9,CFL1,RPLP0,CANX,P4HB,ANXA2,ARF1,MSN,MIF,HNRNPDL,C |
| 30 | 10622 | COL1A1,SPARC,COL6A1,COL5A1,COL4A1,APP,COL4A2,FN1,ITGB1, |
| 85 | 10622 | NEDD4L,UBC,FBXL4,UBA1,UBB,PSMD5,WSB1,PJA2,PSMD2,KLHL9, |
| 44 | 10622 | KIDINS220,PIK3R2,PPP2R1A,AP2M1,AP2B1,RHOA,CHD4,JUND,CLTC |
| 34 | 10622 | HSP90B1,CALR,FKBP14,ATF4,PDIA6,TLN1,CTDSP2,HDGF,ACADVL,H |
| 15 | 10622 | TUBA1A,TUBA1B,TUBB2B,TUBB4B,CCT7,CCT6A,CCT2,TCP1,CCT3, |
| 22 | 10622 | ACTB,ACTG1,TUBA1A,TUBA1B,TUBB2B,AP2M1,TUBB4B,AP2B1,MS |
| 41 | 10622 | UBC,PIK3R2,UBB,PSMD5,PSMD2,SKP1,PSMB2,PSMD3,PLCG1,PSM |
| 32 | 10622 | TUBA1A,HSP90AA1,TUBB,UBC,YWHAE,UBB,YWHAG,PPP2R1A,TUBB |
| 32 | 10622 | ORC4,UBC,UBB,PSMD5,PSMD2,PSMB2,PSMD3,PSMA7,PSMD1,UBA |
| 185 | 10622 | COL1A1,NEDD4L,CTSC,TUBA1A,UBC,TUBA1B,PIK3R2,FBXL4,UBA1, |
| 68 | 10622 | UBC,H3-3B,UBB,PSMD5,PSMD2,CCND2,ARID1A,PSMB2,H3-3A,CCN |
| 18 | 10622 | ARID1A,SMARCA4,SMARCE1,SMARCC1,PHC2,EP300,CSNK2A1,ARID |
| 27 | 10622 | UBC,GANAB,CANX,DDX5,VCP,UBB,PRKCSH,DDOST,ST3GAL2,RPN2, |
| 15 | 10622 | UBC,UBB,TOMM20,SQSTM1,VDAC1,CSNK2A1,UBA52,ULK1,RPS27A |
| 16 | 10622 | UBC,UBA1,UBB,UBE2H,UBE2R2,USP7,UBA52,UBE2G2,RPS27A,UBE |
| 16 | 10622 | KAT6A,CHD3,CHD4,RBBP4,PIP4K2B,HDAC2,RBBP7,EP300,AKT3,AK |
| 45 | 10622 | UBC,APP,UBB,PSMD5,HMGB1,PSMD2,SKP1,SQSTM1,PSMB2,PSMD |
| 19 | 10622 | ACTB,ACTG1,CFL1,ACTR2,RHOA,RAC1,ACTR3,ARPC2,FYN,CDC42,A |
| 32 | 10622 | EEF1A1,HSPA8,HSPA9,HSP90AB1,HSP90AA1,HSPA1B,YWHAE,VCP,PT |
| 115 | 10622 | HSPA8,HSPA9,CFL1,HSP90AA1,VIM,UBC,PIK3R2,HSP90B1,RPLP0,C |
| 12 | 10622 | CFL1,MYH9,RHOA,MYL6,MYH10,RHOB,PPP1CB,MYL12B,LIMK1,PP |

| | | |
|---|---|---|
| 12 | 10622 | MYH9,RHOA,MYL6,MYH10,RHOB,RAC1,PPP1CB,MYL12B,DLG4,CDK |
| 28 | 10622 | ACTB,ACTG1,TUBA1A,TUBA1B,TUBB2B,YWHAE,MYH9,CALM1,YWHA |
| 21 | 10622 | HSPA9,CFL1,RPLP0,P4HB,ANXA2,ARF1,MSN,MIF,HNRNPDL,CNN2,TC |
| 23 | 10622 | PIK3R2,COL6A1,COL5A1,COL4A1,COL4A2,COL5A2,COL6A2,COL2A |
| 10 | 10622 | UBC,UBB,PPP2R1A,PPP2CA,MAPK1,UBA52,PPP2CB,RPS27A,PTPN11 |
| 20 | 10622 | UBC,GANAB,CANX,UBB,PRKCSH,DDOST,ST3GAL2,RPN2,STT3A,UBA |
| 15 | 10622 | ACTB,TUBA1A,TUBB2B,TUBB4B,CCT7,CCT6A,CCT2,TCP1,CCT3,CCT |
| 50 | 10622 | ORC4,UBC,UBB,PSMD5,PSMD2,RBBP4,SKP1,CDKN1A,PSMB2,CCND |
| 35 | 10622 | NEDD4L,UBC,FSTL1,UBB,RHOA,XPO1,PPP1CB,PARP1,SKI,STRAP,NC |
| 18 | 10622 | HSPA9,CFL1,RPLP0,ANXA2,ARF1,MSN,MIF,HNRNPDL,CNN2,TCP1,PI |
| 18 | 10622 | USP11,CCT7,CCT6A,CCT2,TCP1,CCT3,CCT4,CCT8,ARFGEF2,CCT5,S |
| 9 | 10622 | EEF1A1,HSP90AB1,HSP90AA1,YWHAE,VCP,PTGES3,HSBP1,HDAC6,H |
| 80 | 10622 | COL1A1,SPARC,COL6A1,COL5A1,COL4A1,BSG,P4HB,APP,COL18A1, |
| 21 | 10622 | ACTB,ACTG1,AP2M1,AP2B1,CLTC,RAC1,MMP2,FYN,AP2A1,NCSTN,A |
| 14 | 10622 | UBC,GANAB,CALR,CANX,UBB,PDIA3,MARCHF6,PRKCSH,OS9,UBA52 |
| 22 | 10622 | FASN,KPNB1,HMGCS1,SQLE,FDFT1,LSS,SREBF2,CYP51A1,SEC23A, |
| 10 | 10622 | ATF4,EIF2AK1,PPP1R15A,EIF2S2,ATF5,ASNS,TRIB3,CEBPG,EIF2S1,G |
| 23 | 10622 | FTH1,UBC,FTL,UBB,SLC22A17,ATP6V0C,SKP1,ATP6AP1,CAND1,ATP |
| 39 | 10622 | ORC4,UBC,UBB,PSMD5,PSMD2,SKP1,PSMB2,PSMD3,PSMA7,PSMD1 |
| 16 | 10622 | ACTB,TUBA1A,TUBA1B,TUBB2B,TUBB4B,CCT7,CCT6A,CCT2,TCP1,C |
| 29 | 10622 | SRSF4,CPSF2,DDX39B,SRSF5,U2AF2,CPSF1,SRSF9,SRSF1,CASC3,S |
| 30 | 10622 | YWHAE,GPI,YWHAG,YWHAQ,LRPPRC,PRDX2,SESN3,YWHAB,PRDX1,Y |
| 34 | 10622 | UBC,UBB,PSMD5,PSMD2,SKP1,PSMB2,PSMD3,PSMA7,PSMD1,UBA5 |
| 13 | 10622 | HSPA8,HSP90AB1,HSP90AA1,HSPA1B,PTGES3,HSBP1,EP300,FKBP4, |
| 23 | 10622 | COL1A1,COL5A1,COL4A1,COL4A2,FN1,ITGB1,AGRN,COL11A1,LAM |
| 73 | 10622 | ACTB,USP22,KAT6A,CHD3,MORF4L1,CHD4,RBBP4,TADA2B,ARID1A, |
| 73 | 10622 | ACTB,USP22,KAT6A,CHD3,MORF4L1,CHD4,RBBP4,TADA2B,ARID1A, |
| 19 | 10622 | RAB6B,RAB7A,TBC1D16,RAB5B,RAB11B,OPTN,GABARAP,RAB11A,U |
| 51 | 10622 | BSG,ATP5F1B,ATP5F1A,LDHB,LDHA,LRPPRC,MDH2,ACO2,ATP5PB,S |
| 25 | 10622 | COL1A1,COL6A1,COL5A1,COL4A1,P4HB,COL18A1,COL4A2,COL11 |
| 17 | 10622 | ACTB,UBC,UBB,DDB1,PARP1,RAD23B,UBA52,RPS27A,YY1,CUL4A,C |
| 17 | 10622 | CALM1,ANXA2,VCL,TPM4,TPM1,TLN1,ANXA6,MYL6,TPM3,ITGB5,CA |
| 8 | 10622 | RHOA,KIF5B,RAC1,KLC1,CDC42,KIF5A,KTN1,KLC2 |
| 12 | 10622 | UBC,UBB,TOMM20,SQSTM1,VDAC1,UBA52,RPS27A,MAP1LC3B,TO |
| 24 | 10622 | COL1A1,COL6A1,COL5A1,COL4A1,COL18A1,COL4A2,COL11A1,CTS |
| 18 | 10622 | UBC,UBB,PPP2R1A,PEBP1,YWHAB,PPP2CA,MAPK1,MAP2K2,DUSP4, |
| 18 | 10622 | GNB1,CCT7,GNB2,CCT6A,CCT2,GNAI3,TCP1,CCT3,CCT4,GNG2,CCT |
| 18 | 10622 | UBC,SOX4,UBB,XPO1,CTNNB1,CTBP1,BCL9L,UBA52,RPS27A,SOX2, |
| 77 | 10622 | ACTB,USP22,MAVS,UBC,USP11,VCP,UBB,PSMD5,UCHL1,RHOA,PSM |
| 26 | 10622 | UBC,UBB,HDAC2,SKP1,DNER,DLK1,NCOR2,EP300,UBA52,HDAC6,R |

90AA1,ACTR2,UBC,DCX,RPL3,RPS6,RPL4,TUBA1B,PIK3R2,RPL7A,COL6A1,RPL8,TUBB2B,COL5A
90AA1,ACTR2,UBC,DCX,RPL3,RPS6,RPL4,TUBA1B,PIK3R2,RPL7A,COL6A1,RPL8,TUBB2B,COL5A
L7,RPL5,EIF3C,EIF4B,RPL37A,EIF4A1,RPL19,RPL15,RPS3,RPS4X,RPS3A,RPL18,RPL11,RPLP1,RI
PL13,RPS8,RPL7,RPL5,UBB,RPL37A,PSMD5,RPL19,RPL15,RPS3,RPS4X,RPS3A,RPL18,RPL11,RPI
RPS8,RPL7,RPL5,EIF3C,EIF4B,RPL37A,EIF4A1,RPL19,RPL15,RPS3,RPS4X,RPS3A,RPL18,RPL11,I
PL13,RPS8,RPL7,RPL5,UBB,RPL37A,PSMD5,RPL19,RPL15,RPS3,RPS4X,RPS3A,RPL18,RPL11,RPI
RPS8,RPL7,RPL5,EIF3C,EIF4B,RPL37A,EIF4A1,RPL19,RPL15,RPS3,RPS4X,RPS3A,RPL18,RPL11,I
RPS8,RPL7,RPL5,EIF3C,EIF4B,RPL37A,EIF4A1,RPL19,RPL15,RPS3,RPS4X,RPS3A,RPL18,RPL11,I
L7,RPL5,EIF3C,RPL37A,RPL19,RPL15,RPS3,RPS4X,RPS3A,RPL18,RPL11,RPLP1,RPL10,RPS17,R
RPS8,RPL7,RPL5,RPL37A,RPL19,RPL15,RPS3,RPS4X,RPS3A,RPL18,RPL11,RPLP1,RPL10,RPS17
L7,RPL5,RPL37A,RPL19,RPL15,RPS3,SEC61A1,RPS4X,RPS3A,RPL18,RPL11,RPLP1,RPL10,RPS1
RPS8,RPL7,RPL5,SMG1,RPL37A,RPL19,PPP2R1A,RPL15,RPS3,RPS4X,RPS3A,RPL18,RPL11,RPLI
RPS8,RPL7,RPL5,SMG1,RPL37A,RPL19,PPP2R1A,RPL15,RPS3,RPS4X,RPS3A,RPL18,RPL11,RPLI
PL13,RPS8,RPL7,RPL5,RPL37A,RPL19,RPL15,RPS3,RPS4X,RPS3A,RPL18,RPL11,RPLP1,RPL10,R
L7,RPL5,RPL37A,RPL19,RPL15,RPS3,RPS4X,RPS3A,RPL18,RPL11,RPLP1,RPL10,RPS17,RPS16,F
PL13,RPS8,RPL7,RPL5,RPL37A,RPL19,RPL15,RPS3,RPS4X,RPS3A,RPL18,RPL11,RPLP1,RPL10,R
6,RPL4,TUBA1B,RPL7A,RPL8,HSPA1B,TUBB2B,RPLP0,H3-3B,RPL13A,RPL6,YWHAE,RPL13,RPS8
L7,RPL5,RPL37A,RPL19,RPL15,RPS3,RPS4X,RPS3A,RPL18,RPL11,RPLP1,RPL10,RPS17,RPS16,F
L7,RPL5,RPL37A,RPL19,RPL15,RPS3,RPS4X,RPS3A,RPL18,RPL11,RPLP1,RPL10,RPS17,RPS16,F
6,RPL4,TUBA1B,RPL7A,RPL8,HSPA1B,TUBB2B,RPLP0,H3-3B,RPL13A,RPL6,YWHAE,RPL13,RPS8
RPL8,YBX1,RPLP0,DDX5,RPL13A,RPL6,RPL13,RPS8,RPL7,RPL5,SMG1,SNRNP200,EIF4B,UBB,RI
L7,RPL5,RPL37A,RPL19,RPL15,RPS3,RPS4X,RPS3A,RPL18,RPL11,RPLP1,RPL10,RPS17,RPS16,F
L7,RPL5,RPL37A,RPL19,RPL15,RPS3,RPS4X,RPS3A,RPL18,RPL11,RPLP1,RPL10,RPS17,RPS16,F
A,RPL6,RPL13,RPS8,RPL7,RPL5,RPL37A,RPL19,RPL15,RPS3,RPS4X,RPS3A,RPL18,RPL11,RPLP1
2,TUBA1A,MAVS,UBC,RPL3,RPS6,RPL4,TUBA1B,RPL7A,GANAB,HSP90B1,RPL8,FBXL4,UBA1,IGF
L7,RPL5,RPL37A,RPL19,RPL15,RPS3,RPS4X,RPS3A,RPL18,RPL11,RPLP1,RPL10,RPS17,RPS16,F
3,RPS8,RPL7,RPL5,RPL37A,RPL19,RPL15,RPS3,RPS4X,RPS3A,RPL18,RPL11,RPLP1,RPL10,RPS
,RPL6,RPL13,RPS8,RPL7,RPL5,EIF3C,EIF4B,RPL37A,EIF4A1,RPL19,RPL15,RPS3,SEC61A1,RPS4
UBB,UBC,RPL3,RPS6,RPL4,ENO1,TUBA1B,RPL7A,GANAB,RPL8,CALR,TUBB2B,PGK1,RPLP0,CAN
RPSA,EIF4H,RPS24,RPS9,RPS14,RPS20,EIF5,RPS25,RPS15,EIF3L,RPS5,RPS7,RPS19,RPS10,RPS1
,RPS16,RPSA,EIF4H,RPS24,RPS9,RPS14,RPS20,RPS25,RPS15,EIF3L,RPS5,RPS7,RPS19,RPS10,RI
UBB,UBC,RPL3,RPS6,RPL4,ENO1,TUBA1B,PIK3R2,RPL7A,GANAB,RPL8,CALR,TUBB2B,PGK1,RPI
,RPS16,RPSA,EIF4H,RPS24,RPS9,RPS14,RPS20,RPS25,RPS15,EIF3L,RPS5,RPS7,RPS19,RPS10,RI
8,RPL7,RPL5,RPL37A,RPL19,RPL15,RPS3,RPS4X,RPS3A,RPL18,RPL11,RPLP1,RPL10,RPS17,RPS
PS9,RPS14,RPS20,RPS25,RPS15,EIF3L,RPS5,RPS7,RPS19,RPS10,RPS12,RPS27,RPS15A,EIF3E,RI
B1,HSP90AA1,ACTR2,UBC,DCX,RPL3,RPS6,RPL4,TUBA1B,PIK3R2,RPL7A,COL6A1,RPL8,TUBB2E
8,RPL7,RPL5,RPL37A,RPL19,RPL15,RPS3,RPS4X,RPS3A,RPL18,RPL11,RPLP1,RPL10,RPS17,RPS
,RPS8,RPL7,RPL5,CKB,RPL37A,PSMD5,RPL19,RPL15,SERINC1,RPS3,RPS4X,RPS3A,RPL18,RPL1
8,RPL7,RPL5,RPL37A,RPL19,RPL15,RPS3,RPS4X,RPS3A,RPL18,RPL11,RPLP1,RPL10,RPS17,RPS
F1,HNRNPU,HNRNPA0,PAPOLA,SRSF4,HNRNPC,HNRNPK,HNRNPH1,CPSF2,SRSF5,U2AF2,CPSF1,
F1,HNRNPU,HNRNPA0,PAPOLA,SRSF4,HNRNPC,HNRNPK,HNRNPH1,CPSF2,SRSF5,U2AF2,CPSF1,
,FTL,TMF1,TUBB2B,YWHAE,KIF1A,SPTBN1,MYH9,APP,GDI1,SPTAN1,CALM1,UBB,YWHAG,YWHA
F1,HNRNPU,HNRNPA0,PAPOLA,SRSF4,HNRNPC,HNRNPK,HNRNPH1,CPSF2,DDX39B,SRSF5,U2AF
B2,PPP2CA,PSMD3,PSMA7,PSMD1,UBA52,PPP2CB,RPS27A,PSMD8,PSMB4,TLE1,PSMD4,PSMC3,
,EP300,PSMD1,UBA52,RPS27A,PSMD8,HIF1A,PSMB4,UBE2D2,PSMD4,PSMC3,CREBBP,UBE2D3,I

PIK3R2,COL6A1,COL5A1,COL4A1,CALM1,IGF2,UBB,COL4A2,FN1,PPP2R1A,AP2M1,STMN1,ITGB
MA7,PSMD1,UBA52,RPS27A,PSMD8,PSMB4,PSMD4,PSMC3,PSMC1,EIF4G1,PSMB1,HSPA1A,PSMD
,UBA52,RPS27A,PSMD8,HIF1A,PSMB4,UBE2D2,PSMD4,PSMC3,UBE2D3,PSMC1,PSMB1,PSMD13
IB2,MAPK1,PSMD3,KDM1A,PHC2,TNKS2,USP7,PSMA7,RBBP7,MTA1,PSMD1,CSNK2A1,UBA52,PT
IK2A1,UBA52,PTEN,RPS27A,TNKS,PSMD8,PSMB4,AKT3,MKRN1,PSMD4,AKT2,PSMC3,PSMC1,PSM
AC1,SKP1,PSMB2,PSMD3,FYN,B2M,PSMA7,PSMD1,AP2A1,UBA52,POM121C,RPS27A,PSMD8,SLC
ACTR2,SPARC,FTH1,UBC,TUBA1B,FTL,HSP90B1,CALR,TMF1,TUBB2B,COL4A1,YWHAE,KIF1A,SPT
IA7,PSMD1,UBA52,RPS27A,PSMD8,PSMB4,PSMD4,PSMC3,PSMC1,PSMB1,CUL1,PSMD13,PSMD7
A52,RPS27A,PSMD8,PSMB4,PSMD4,PSMC3,PSMC1,PSMB1,PSMD13,PSMD7,CDK4,PSME3,PSMC
00,PSMD1,UBA52,RPS27A,PSMD8,PSMB4,NCSTN,PSMD4,ADAM10,RBPJ,PSMC3,CREBBP,PSMC1
,PSMD5,PPP2R1A,VCL,GTF2F1,PSMD2,PEBP1,TLN1,CTNNB1,YWHAB,SND1,CTBP1,RAC1,HDAC2,
3,AP2M1,ITGB1,AP2B1,CHD4,ATP1A1,RBBP4,PRKCSH,DDOST,ST3GAL2,PTGES3,ATP1A2,HDAC2
2,RPS27A,PSMD8,PSMB4,PSMD4,PSMC3,PSMC1,PSMB1,CUL1,PSMD13,PSMD7,PSME3,PSMC5,P
PSMB2,PSMD3,HSPB1,PSMA7,ANP32A,PSMD1,UBA52,ZFP36L1,RPS27A,PSMD8,PSMB4,ELAVL1,P
2,RPS27A,PSMD8,PSMB4,PSMD4,PSMC3,PRKACA,PSMC1,PSMB1,CUL1,PSMD13,PSMD7,PSME3
2,RPS27A,PSMD8,PSMB4,PSMD4,PSMC3,PRKACA,PSMC1,PSMB1,CUL1,PSMD13,PSMD7,PSME3
F5B,RAB6B,RAB1A,ARCN1,KLC1,SURF4,TMED2,TUBB2A,ARF3,KDELR2,DYNC1LI2,DYNLL2,CAPZ
2,RPS27A,PSMD8,PSMB4,PSMD4,PSMC3,PSMC1,PSMB1,FZR1,CUL1,PSMD13,PSMD7,PSME3,PS
UBB,YWHAG,PSMD5,PPP2R1A,TUBB4B,PAFAH1B1,PSMD2,RBBP4,XPO1,CSNK1E,CKAP5,SKP1,C
UBB4B,PSMD2,CDKN1A,PSMB2,TUBB2A,MAPRE1,PSMD3,PSMA7,PSMD1,UBA52,RPS27A,PSMD8
SMD5,YWHAQ,HMGB1,KPNB1,PSMD2,H1-0,CTNNB1,YWHAB,APPL1,PSMB2,MAPK1,PLEC,PSMD3,
C1,PSMB2,PSMD3,VANGL2,PSMA7,PSMD1,AP2A1,UBA52,RPS27A,PSMD8,PSMB4,PSMD4,SCRIB,
UBB,YWHAG,PSMD5,PPP2R1A,TUBB4B,PAFAH1B1,PSMD2,RBBP4,XPO1,CSNK1E,CKAP5,SKP1,C
A,MAVS,UBC,TUBA1B,GANAB,HSP90B1,FBXL4,UBA1,CALR,FSTL1,TUBB2B,CANX,DDX5,DDX17,S
5,KPNB1,PSMD2,XPO1,RAC1,SKP1,TAF15,SSRP1,PSMB2,TAF7,MVB12B,PSMD3,SUPT5H,POLR2
52,RPS27A,PSMD8,PSMB4,PSMD4,PSMC3,PSMC1,PSMB1,MDM2,PSMD13,PSMD7,PSME3,PSMC
2,RPS27A,PSMD8,PSMB4,UBE2M,PSMD4,PSMC3,PSMC1,PSMB1,CUL1,PSMD13,PSMD7,PSME3,
4,RBBP4,PIP4K2B,PRKAR1A,JUN,RAC1,HDAC2,CDKN1A,AHCYL1,PSMB2,PPP2CA,MAPK1,PSMD3
,ALDOA,PKM,FTL,HSPA1B,PSAP,SLC2A3,GPI,SPTAN1,GSTP1,VCP,HLA-C,VAPA,HMGB1,TUBB4B,
2,RPS27A,PSMD8,PSMB4,PSMD4,PSMC3,PSMC1,PSMB1,PSMD13,PSMD7,PSME3,PSMC5,PSMC2
2,RPS27A,PSMD8,PSMB4,PSMD4,PSMC3,PSMC1,PSMB1,CUL1,PSMD13,PSMD7,PSME3,PSMC5,P
52,RPS27A,TNKS,PSMD8,PSMB4,PSMD4,PSMC3,PSMC1,PSMB1,PSMD13,PSMD7,PSME3,PSMC5
27A,PSMD8,PSMB4,PSMD4,PSMC3,PSMC1,PSMB1,PSMD13,PSMD7,PSME3,PSMC5,PSMC2,PSM
27A,PSMD8,PSMB4,PSMD4,PSMC3,PSMC1,PSMB1,PSMD13,PSMD7,PSME3,PSMC5,PSMC2,PSM
27A,PSMD8,PSMB4,PSMD4,PSMC3,PSMC1,PSMB1,PSMD13,PSMD7,PSME3,PSMC5,PSMC2,PSM
PRKAR1A,SKP1,PSMB2,TUBB2A,PSMD3,PSMA7,PRKAR2A,PSMD1,UBA52,ADCY6,RPS27A,PSMD8
BA52,RPS27A,PSMD8,PSMB4,PSMD4,PSMC3,PSMC1,PSMB1,PSMD13,PSMD7,PSME3,PSMC5,PSM
2,RPS27A,PSMD8,PSMB4,UBE2M,PSMD4,PSMC3,PSMC1,PSMB1,CUL1,PSMD13,PSMD7,PSME3,
AZ2,PSMD8,PSMB4,PSMD4,PSMC3,PSMC1,PSMB1,PSMD13,PSMD7,PSME3,PSMC5,PSMC2,PSMC
2,CUL3,RPS27A,PSMD8,PSMB4,PSMD4,PSMC3,PSMC1,PSMB1,PSMD13,PSMD7,PSME3,PSMC5,P
2,RPS27A,PSMD8,PSMB4,PSMD4,PSMC3,PRKACA,PSMC1,PSMB1,CUL1,PSMD13,PSMD7,PSME3
D3,PSMA7,PSMD1,UBA52,RPS27A,PSMD8,PSMB4,AKT3,PSMD4,AKT2,PSMC3,PSMC1,PSMB1,FZ
4,PIP4K2B,JUN,RAC1,HDAC2,CDKN1A,PSMB2,PPP2CA,MAPK1,PSMD3,KDM1A,FYN,PHC2,TNKS2,
1,AP2M1,ITGB1,TUBB4B,NCAM1,AP2B1,MSN,GAP43,SPTBN2,CLTC,RAC1,L1CAM,LAMC1,DPYSL
27A,PSMD8,PSMB4,PSMD4,PSMC3,PSMC1,PSMB1,PSMD13,PSMD7,PSME3,PSMC5,PSMC2,PSM

27A,PSMD8,PSMB4,PSMD4,PSMC3,PSMC1,PSMB1,PSMD13,PSMD7,PSME3,PSMC5,PSMC2,PSM
27A,PSMD8,PSMB4,PSMD4,PSMC3,PSMC1,PSMB1,PSMD13,PSMD7,PSME3,PSMC5,PSMC2,PSM
27A,PSMD8,PSMB4,PSMD4,PSMC3,PSMC1,PSMB1,PSMD13,PSMD7,PSME3,PSMC5,PSMC2,PSM
)1,OAZ2,PSMD8,PSMB4,PSMD4,PSMC3,PSMC1,PSMB1,PSMD13,PSMD7,PSME3,PSMC5,PSMC2,
3A52,RPS27A,PSMD8,PSMB4,PSMD4,SCRIB,PSMC3,PSMC1,PSMB1,PSMD13,PSMD7,PSME3,PSM
BA52,RPS27A,PSMD8,PSMB4,PSMD4,PSMC3,PSMC1,PSMB1,PSMD13,PSMD7,PSME3,PSMC5,PS
D3,PSMA7,PSMD1,UBA52,RPS27A,PSMD8,PSMB4,AKT3,PSMD4,AKT2,PSMC3,PSMC1,PSMB1,CL
3A52,RPS27A,PSMD8,PSMB4,PSMD4,PSMC3,PSMC1,PSMB1,PSMD13,PSMD7,PSME3,PSMC5,PS
MD3,PSMA7,EP300,PSMD1,UBA52,BRD2,RPS27A,PSMD8,PSMB4,PSMD4,RBPJ,PSMC3,CREBBP,
3,PAFAH1B1,KPNB1,PSMD2,SPC25,XPO1,CKAP5,RPS27,TNPO1,PSMB2,TUBB2A,PPP2CA,MAPRE
AQ,HMGB1,KPNB1,PSMD2,H1-0,CTNNB1,YWHAB,APPL1,PSMB2,MAPK1,PLEC,PSMD3,DYNLL2,Y
3,PAFAH1B1,KPNB1,PSMD2,SPC25,XPO1,CKAP5,RPS27,TNPO1,PSMB2,TUBB2A,PPP2CA,MAPRE
3A52,RPS27A,PSMD8,PCBP4,PSMB4,PSMD4,PSMC3,PSMC1,PSMB1,MDM2,PSMD13,PSMD7,PSM
3A52,RPS27A,PSMD8,PCBP4,PSMB4,PSMD4,PSMC3,PSMC1,PSMB1,MDM2,PSMD13,PSMD7,PSM
27A,PSMD8,PSMB4,PSMD4,PSMC3,PSMC1,PSMB1,MDM2,PSMD13,PSMD7,PSME3,PSMC5,PSMC
2,PSMD3,B2M,PSMA7,PSMD1,UBA52,RPS27A,PSMD8,PSMB4,PSMD4,PSMC3,PSMC1,TAPBP,PSM
27A,PSMD8,PSMB4,PSMD4,PSMC3,PSMC1,PSMB1,FZR1,PSMD13,PSMD7,PSME3,PSMC5,PSMC2
MYH10,RAC1,MMP2,ACTR3,ARPC2,FYN,CDC42,AP2A1,NCSTN,ADAM10,AP2A2,MYL12B,SDCBP,C
MD3,PSMA7,PSMD1,UBA52,RPS27A,PSMD8,PSMB4,UBE2M,UBE2D2,PDPK1,PSMD4,PSMC3,PSM
2,RPS27A,PSMD8,PSMB4,PSMD4,PSMC3,PSMC1,PSMB1,CUL1,PSMD13,PSMD7,PSME3,STUB1,
3A52,RPS27A,PSMD8,PCBP4,PSMB4,PSMD4,PSMC3,PSMC1,PSMB1,MDM2,PSMD13,PSMD7,PSM
MD1,UBA52,RPS27A,PSMD8,PSMB4,PSMD4,PSMC3,PSMC1,PSMB1,PSMD13,PSMD7,PSME3,PSM
52,RPS27A,PSMD8,PSMB4,PSMD4,PSMC3,PSMC1,PSMB1,PSMD13,PSMD7,PSME3,PSMC5,PSMC
3-3B,DVL3,YWHAE,MYH9,CALM1,YWHAG,YWHAQ,PPP2R1A,ITGB1,TUBB4B,PAFAH1B1,CTNNA1,
,AP2B1,RHOA,PSMD2,PFN1,TLE5,XPO1,CSNK1E,CLTC,GNB2,CTNNB1,WLS,CTBP1,RAC1,SKP1,V
C42,PSMA7,PSMD1,UBA52,RPS27A,SEPTIN7,PSMD8,PSMB4,PSMD4,IGF2BP1,PSMC3,PRKACA,
2,RPS27A,PSMD8,PSMB4,PSMD4,PSMC3,PSMC1,PSMB1,CUL1,PSMD13,PSMD7,PSME3,PSMC5,
3,RPS27A,PSMD8,PSMB4,PSMD4,PSMC3,PSMC1,PSMB1,PSMD13,RASA4,PSMD7,SYNGAP1,PSM
B,PSMD2,PRKAR1A,SKP1,PSMB2,TUBB2A,PSMD3,PSMA7,OS9,PRKAR2A,PSMD1,UBA52,ADCY6,
3,PAFAH1B1,PSMD2,SPC25,XPO1,CKAP5,RPS27,PSMB2,TUBB2A,PPP2CA,MAPRE1,PSMD3,DYNC
,UBA52,RPS27A,PSMD8,PSMB4,PSMD4,PSMC3,PSMC1,PSMB1,CUL1,PSMD13,PSMD7,PSME3,P
TBN2,RAB1A,ARCN1,TMED2,TUBB2A,ARF3,KDELR2,DYNC1LI2,DYNLL2,CAPZB,KDELR1,ARF4,T
,PSMB2,PSMD3,B2M,PSMA7,PSMD1,UBA52,ITGB5,RPS27A,PSMD8,PSMB4,PSMD4,ITGAV,PSMC
FN1,CLTC,GNB2,CTNNB1,RAC1,PSMB2,PSMD3,GNG2,VANGL2,PSMA7,PSMD1,AP2A1,UBA52,G
,PSMD4,PSMC3,PSMC1,PSMB1,PSMD13,PSMD7,PSME3,PSMC5,PSMC2,PSMC4,PSMF1,PSMB5,
A1,RBBP4,PTGES3,ATP1A2,HDAC2,KDM1A,RBBP7,MTA1,AP2A1,BRD4,FKBP1A,FKBP4,IMPDH2,A
PK1,PSMD3,PSMA7,PSMD1,UBA52,HDAC6,RPS27A,PSMD8,PSMB4,AKT3,PSMD4,AKT2,PSMC3,P
)1,UBA52,RPS27A,PSMD8,FKBP1A,PSMB4,PSMD4,PSMC3,PSMC1,PSMB1,CUL1,PSMD13,PSMD7
MD1,UBA52,ABCC9,RPS27A,PSMD8,PSMB4,PSMD4,PSMC3,PSMC1,PSMB1,PSMD13,PSMD7,PSM
B1,KIF5B,RAB6B,RAB1A,ARCN1,KLC1,SURF4,TMED2,TUBB2A,ARF3,KDELR2,DYNC1LI2,DYNLL
27A,PSMD8,PSMB4,PSMD4,PSMC3,PSMC1,PSMB1,PSMD13,PSMD7,PSME3,PSMC5,PSMC2,PSM
52,RPS27A,PSMD8,PSMB4,PSMD4,PSMC3,PSMC1,PSMB1,PSMD13,PSMD7,PSME3,PSMC5,PSMC
3A1B,PIK3R2,FLNA,TUBB2B,H3-3B,DVL3,YWHAE,MYH9,CALM1,YWHAG,YWHAQ,PPP2R1A,ITGB1,
3A1B,PIK3R2,FLNA,TUBB2B,H3-3B,DVL3,YWHAE,MYH9,CALM1,YWHAG,YWHAQ,PPP2R1A,ITGB1,
P1R15A,FKBP1A,UBE2M,MTMR4,PPP1CA,PMEPA1,STUB1,TGFBR1,FURIN,NEDD8

3,PSMA7,PSMD1,UBA52,RPS27A,PSMD8,PSMB4,PSMD4,PSMC3,PSMC1,PSMB1,CUL1,PSMD13,PS
7,PSMD1,UBA52,RPS27A,PSMD8,BAP1,PSMB4,FOXK2,PSMD4,OGT,PSMC3,YY1,PSMC1,PSMB1,F
27A,PSMD8,PSMB4,PSMD4,PSMC3,PSMC1,PSMB1,FZR1,PSMD13,PSMD7,PSME3,PSMC5,PSMC2
B4B,TOMM20,SQSTM1,TUBB2A,DYNC1LI2,DYNLL2,DYNLL1,VDAC1,CSNK2A1,UBA52,HDAC6,UL
F1,SPTBN2,RAB1A,ARCN1,TMED2,TUBB2A,ARF3,KDELR2,DYNC1LI2,DYNLL2,CAPZB,KDELR1,AF
ER,MDK,PSMB2,H3-3A,TMED2,NOTCH2,CCND1,PSMD3,DLK1,NCOR2,PSMA7,ST3GAL6,EP300,PS
27A,PSMD8,PSMB4,PSMD4,PSMC3,PSMC1,PSMB1,FZR1,PSMD13,PSMD7,PSME3,PSMC5,PSMC2
2,RPS27A,PSMD8,PSMB4,PSMD4,PSMC3,PSMC1,PSMB1,FZR1,CUL1,PSMD13,PSMD7,PSME3,PS
CREBBP,HSPA1A,DNAJB1,HSF1
.3,RPS27A,PSMD8,GRK2,PSMB4,DZIP1,PSMD4,PSMC3,PSMC1,SMO,PSMB1,PSMD13,PSMD7,PSN
OS9,PSMD1,UBA52,ABCC9,RPS27A,PSMD8,PSMB4,EIF2S2,ABCF1,PSMD4,PSMC3,PSMC1,PSMB
PSMD5,PPP2R1A,NCAM1,VCL,PSMD2,XPO1,SPTBN2,PEBP1,TLN1,JUN,YWHAB,RAC1,PEA15,PPP1
B,YWHAE,UBB,YWHAG,PSMD5,SET,PPP2R1A,TUBB4B,PAFAH1B1,KPNB1,PSMD2,SPC25,XPO1,C
A6,GPC3,RCN1,GOLM1,APLP2,CKAP4,CALU,TNC,TMEM132A,NUCB1,VCAN,LAMC1,VGF,IGFBP7,
,ARCN1,KLC1,SURF4,TMED2,TUBB2A,ARF3,KDELR2,KDELR1,ARF4,KIF3C,TMED9,KIF5A,KIF21B
BN1,SPTAN1,VCP,UBB,PDIA3,TUBB4B,MARCHF6,ARF1,PRKCSH,SPTBN2,DDOST,ST3GAL2,RAB1
MB2,PPP2CA,CCND1,PSMD3,PSMA7,PSMD1,UBA52,PPP2CB,RPS27A,PSMD8,PSMB4,AKT3,PSMD
),SRSF1,CASC3,SRSF3,SRSF6,SYMPK,SRSF2,SRSF11,DHX38,EIF4A3,SRSF7,POLDIP3,NCBP2,THC
27A,PSMD8,PSMB4,PSMD4,PSMC3,PSMC1,PSMB1,PSMD13,PSMD7,PSME3,PSMC5,PSMC2,PSM
D1,UBA52,PTEN,RPS27A,PSMD8,PSMB4,UBE2D2,PDPK1,PSMD4,PSMC3,PSMC1,CDC34,PSMB1,C
P90AA1,ACTR2,TUBB,FTH1,UBC,PIK3R2,ALDOA,PKM,FTL,HSP90B1,HSPA1B,PSAP,CLU,SLC2A3,G
VAMP2,ACTR3,RAB5B,ARPC2,UBQLN1,HGS,HIP1,AP2A1,UBQLN2,UBA52,RPS27A,CTTN,GRK2,LI
ARS1,RARS1,WARS1,KARS1,FARSA,TARS1,HARS1
PSMD5,PPP2R1A,NCAM1,VCL,PSMD2,SPTBN2,PEBP1,TLN1,YWHAB,PEA15,PPP1CB,PTPRA,PSMB2
,UBA52,RPS27A,PSMD8,PSMB4,PSMD4,PSMC3,PSMC1,PSMB1,FZR1,CUL1,PSMD13,PSMD7,PSN
KTL3,KLHDC3,SERP1,TPP1,ATP6V0D1,GSK3A,SRPRA,TSPYL2,LMNA,SHC1,DNAJB11,ADD1,CUL7,
DC42,BRK1,ABI2,WASF2,ABL1,ARPC1A,CYFIP1,ARPC3,WASF3,ARPC5,MAPK3
RAD23B,OS9,UBA52,RPS27A,PSMC1,AMFR,MOGS,UBXN1,MAN1B1,RNF5
27A,PSMD8,PSMB4,PSMD4,PSMC3,PSMC1,PSMB1,PSMD13,PSMD7,PSME3,PSMC5,PSMC2,PSM
E,CTNNB1,CTBP1,PSMB2,H3-3A,PPP2CA,SMARCA4,PSMD3,TNKS2,PSMA7,BCL9L,EP300,PSMD1,
BA52,PPP1CC,RPS27A,PPP1R15A,FKBP1A,UBE2M,MTMR4,UBE2D3,PPP1CA,PMEPA1,CDK9,STUI
PSMD5,PPP2R1A,NCAM1,VCL,PSMD2,SPTBN2,PEBP1,TLN1,YWHAB,PEA15,PPP1CB,PTPRA,PSMB2
FIP2,RAC1,AHCYL1,NCKAP1,VEGFA,HSPB1,PLCG1,FYN,CDC42,CTNND1,BRK1,SHC2,ABI2,AKT3,
A1B,TUBB2B,H3-3B,YWHAE,UBB,YWHAG,PSMD5,SET,PPP2R1A,TUBB4B,PAFAH1B1,KPNB1,PSMD
),SRSF1,CASC3,SRSF3,SRSF6,SYMPK,SNRPD3,SRSF2,SRSF11,DHX38,EIF4A3,SRSF7,POLDIP3,NC
SH,CDH2,PDIA6,GPC3,RCN1,GOLM1,APLP2,CKAP4,CALU,TNC,MMP2,TMEM132A,NUCB1,VCAN,L
N1A,PSMB2,PPP2CA,CCND1,PSMD3,PSMA7,PSMD1,UBA52,PPP2CB,RPS27A,PSMD8,PSMB4,AKT
FIP2,RAC1,AHCYL1,NCKAP1,HSPB1,PLCG1,FYN,CDC42,CTNND1,BRK1,SHC2,ABI2,AKT3,PDPK1,
27A,PSMD8,PSMB4,PSMD4,PSMC3,PSMC1,PSMB1,PSMD13,PSMD7,PSME3,PSMC5,PSMC2,PSM
IA2,COL5A2,RAC1,COL2A1,LAMC1,RAPGEF1,COL27A1,HGS,STAT3,UBA52,SPINT2,RPS27A,SH3
KTL3,KLHDC3,SERP1,TPP1,ATP6V0D1,GSK3A,SRPRA,TSPYL2,LMNA,SHC1,DNAJB11,ADD1,CUL7,
CCT6A,CCT2,TBCD,GNAI3,TCP1,CCT3,CCT4,TUBB2A,GNG2,CCT8,ARFGEF2,CCT5,STAT3,CSNK
3B2B,VCP,UBB,TUBB4B,TOMM20,SQSTM1,TUBB2A,MVB12B,DYNC1LI2,DYNLL2,GABARAP,DYNI
F1,SPTBN2,RAB1A,ARCN1,TMED2,TUBB2A,ARF3,KDELR2,DYNC1LI2,DYNLL2,CAPZB,KDELR1,AF
27A,GSK3A,TRIM27,RAB5A,MAPK3

27A,PSMD8,PSMB4,PSMD4,PSMC3,PSMC1,PSMB1,PSMD13,PSMD7,PSME3,PSMC5,PSMC2,PSM
C1LI2,DYNLL2,CAPZB,DYNLL1,DCTN1,ACTR1A,RAB3GAP2,CAPZA1,GALNT1,RAB6A,TUBB4A,PAF
JBA1A,CFL1,MAVS,HSP90AB1,HSP90AA1,ACTR2,VIM,TUBB,FTH1,UBC,EIF4G2,TUBA1B,PIK3R2,F
2,RPS27A,PSMD8,PSMB4,PSMD4,PSMC3,PSMC1,PSMB1,FZR1,CUL1,PSMD13,PSMD7,PSME3,PS
2,RPS27A,PSMD8,PSMB4,PSMD4,PSMC3,PSMC1,PSMB1,FZR1,CUL1,PSMD13,PSMD7,PSME3,PS
,CTSD,KIF5B,SPTBN2,RAB7A,CLTC,CTSB,KLC1,TUBB2A,DYNC1LI2,DYNLL2,CAPZB,KIF3C,SEC23
3B4B,DNAJA1,PTGES3,STIP1,TUBB2A,DYNC1LI2,DYNLL2,CAPZB,DYNLL1,DCTN1,ACTR1A,FKBP4
JBB2A,DYNC1LI2,DYNLL2,DYNLL1,UBA52,HDAC6,RPS27A,TUBB4A,PARK7,HSF1,UBE2V1,TUBA1
GTF2F1,IGFBP3,CHD4,RBBP4,LRPPRC,PIP4K2B,PRDX2,JUN,SESN3,YWHAB,HDAC2,CDKN1A,TA
NA1,TLN1,MYL6,MYH10,RHOB,RAC1,DPYSL2,FYN,PLXNA3,PLXNB1,DPYSL4,MYL12B,PLXNA2,LI
2B,LIMK1,PPP1R12A,MYL9,NF2
ID3,FYN,PSMA7,EP300,PSMD1,UBA52,RPS27A,PSMD8,PSMB4,UBE2M,UBE2D2,PDPK1,PSMD4,P
HL,PSMD3,CAND1,PSMA7,RBBP7,PSMD1,UBA52,KLHL42,FBXO11,FEM1B,CUL3,RPS27A,PSMD8
P1R15A,MTMR4,PPP1CA,PMEPA1,STUB1,TGFBR1
2,RPS27A,GSK3A,TRIM27,RAB5A,MAPK3

DIA3,SEC61A1,WSB1,PJA2,PSMD2,KLHL9,UBE2H,HLA-B,SKP1,UBE4A,PSMB2,VHL,PSMD3,UBE2F
B4B,TOMM20,SQSTM1,TUBB2A,DYNC1LI2,DYNLL2,GABARAP,DYNLL1,VDAC1,WDR45B,CSNK2A
CCT6A,CCT2,GNAI3,TCP1,CCT3,CCT4,TUBB2A,GNG2,CCT8,ARFGEF2,CCT5,STAT3,CSNK2A1,GN
,YWHAB,RAPGEF1,PPP2CA,MAPK1,PLCG1,MAP2K2,DUSP4,ID4,VGF,STAT3,EP300,AP2A1,PPP2C
,TUBB4B,PAFAH1B1,RHOA,PFN1,SPC25,XPO1,CKAP5,RHOB,RPS27,RAC1,TUBB2A,PPP2CA,MAF
PL4,ENO1,PIK3R2,RPL7A,ALDOA,FADS1,UGT8,RPL8,CES3,PSAP,PGK1,RPLP0,ENO2,RPL13A,RP
A1B,TUBB2B,H3-3B,YWHAE,NPM1,UBB,YWHAG,PSMD5,YWHAQ,SET,PPP2R1A,TUBB4B,PAFAH1B
NN2,TCP1,PITPNA,SOD1,CDC42,STAT3,HNRNPA2B1,IL6ST,CAPZA1,SOD2,TYK2,JAK1,STAT1,TALD
NCAM1,AGRN,LAMA5,COL1A2,COL5A2,COL6A2,TNC,DCN,VCAN,COL2A1,LAMC1,PTPRS,ITGB5,H
JBE2H,SKP1,UBE4A,PSMB2,VHL,PSMD3,UBE2R2,NPEPPS,HECTD1,PSMA7,TRAF7,PSMD1,UBA52,
,YWHAB,RAC1,RAPGEF1,PPP2CA,MAPK1,PLCG1,MAP2K2,DUSP4,ID4,PTPRS,VGF,STAT3,EP300,
YOU1,SSR1,DCTN1,EXTL3,MBTPS1,KLHDC3,SERP1,EIF2S2,TPP1,ATP6V0D1,GSK3A,SRPRA,TSPY
CCT4,TUBB2A,CCT8,CCT5,TUBB4A,TUBA1C
,CLTC,L1CAM,DPYSL2,TUBB2A,MAPK1,AP2A1,RDX,AP2A2,TUBB4A,CLTA,TUBA1C,RPS6KA2,DN
A7,PSMD1,UBA52,EVL,PTEN,RPS27A,PSMD8,PSMB4,UBE2D2,PDPK1,PSMD4,PSMC3,PSMC1,CDC
4B,PAFAH1B1,CSNK1E,CKAP5,SKP1,PPP1CB,OPTN,MAPRE1,CSNK1D,DYNLL1,ACTR1A,UBA52,R
52,RPS27A,PSMD8,PSMB4,PSMD4,PSMC3,PSMC1,PSMB1,PSMD13,PSMD7,PSME3,PSMC5,PSMC
CALR,TUBB2B,CANX,KLRD1,CALM1,UBB,PSMD5,MRC2,PPP2R1A,HLA-C,PDIA3,AP2M1,SEC61A1,
D1,SMARCA4,PSMD3,SMARCE1,SMARCC1,PHC2,PSMA7,PRMT1,EP300,PSMD1,H2AZ1,CSNK2A1,
1B,CBX6,SMARCD1,AUTS2,HIPK2,CSNK2B,PBRM1,RYBP,CBX4,SMARCC2,SMARCB1
VHL,STT3A,UBA52,MGAT4B,RPS27A,MAP1LC3B,GALNT1,GSK3A,CHMP4B,UBE2I,MGAT5,SUMO1
,MAP1LC3B,TOMM70,PINK1,MFN2,CSNK2B,TOMM40
2D2,UBE2K,USP5,UBE2L3,CDC34,UBE2G1,UBE2E1
T2,MBD3,MTA2,PIN1,GATAD2B,BRPF3,BRD1
3,PSMA7,STAT3,PSMD1,UBA52,RPS27A,PSMD8,PSMB4,PSMD4,PSMC3,PSMC1,PTPN23,PTPN11,
RPC1A,EFNB2,LIMK1,EFNB1,ARPC3,EFNB3,ARPC5,ROCK2,EPHB2
GES3,HSBP1,HSPH1,HSPA4,MAPK1,CCAR2,EP300,HDAC6,POM121C,FKBP4,MTOR,CREBBP,BA
ANX,P4HB,APP,FSCN1,UBB,FN1,PSMD5,PPP2R1A,HMGB1,ITGB1,ANXA2,ARF1,LAMA5,MSN,COL1
P1R12A,ROCK2,MYL9

:N1B,PPP1R12A,MYL9
\G,YWHAQ,TUBB4B,MYO5A,YWHAB,RAC1,VAMP2,TUBB2A,YWHAH,RAB11A,RAB14,KIFAP3,AKT2,
:P1,PITPNA,SOD1,CDC42,HNRNPA2B1,CAPZA1,SOD2,TYK2,JAK1,TALDO1,RAP1B
.1,RAPGEF1,PLCG1,STAT3,PTPN11,PDGFRB,NCK2,CRKL,STAT1,THBS2,SOS1,COL6A3,BCAR1,FL
,MAPK3
52,MGAT4B,RPS27A,GALNT1,UBE2I,MGAT5,SUMO1,DAD1,MOGS,GSK3B,MAN1B1
T4,TUBB2A,CCT8,CCT5,TUBB4A,TUBA1C
)1,PSMD3,PSMA7,PSMD1,UBA52,SMC3,RPS27A,PSMD8,PSMB4,AKT3,PSMD4,AKT2,PSMC3,PSM(
:OR2,BMPR2,UBA52,PPP1CC,RPS27A,PPP1R15A,FKBP1A,UBE2M,MTMR4,UBE2D3,ACVR1B,DRA
TPNA,SOD1,CDC42,HNRNPA2B1,CAPZA1,SOD2,TALDO1,RAP1B
STAT3,FKBP9,FBXW5,XRN2,NOP56,LONP2,SKIV2L,AP3M1
SF1
,COL4A2,FN1,ITGB1,NCAM1,AGRN,TIMP2,COL11A1,PXDN,CTSD,LAMA5,COL1A2,COL5A2,DDR1,
DAM10,AP2A2,CLTA,EFNB2,EFNB1,EFNB3,VAV2,EPHB2,EPHA4,DNM1,EPHA2
,RPS27A,AMFR,MAN1B1,RNF5
MBTPS1,MVD,CARM1,DHCR7,CREBBP,SCD,SCAP,SC5D,RAN,TM7SF2,RXRA,SP1,NCOA6
RB10
6V1B2,UBA52,RPS27A,IREB2,ATP6V0D1,ATP6V1G1,CUL1,ATP6V0B,TFRC,ATP6V0E2,ATP6V0A1,A
,UBA52,RPS27A,PSMD8,PSMB4,PSMD4,PSMC3,PSMC1,PSMB1,FZR1,CUL1,PSMD13,PSMD7,POI
CT3,CCT4,TUBB2A,CCT8,CCT5,TUBB4A,TUBA1C
RSF3,SRSF6,POM121C,SYMPK,SRSF2,SRSF11,DHX38,EIF4A3,NUP153,SRSF7,POLDIP3,NCBP2,N
WHAH,DDIT4,COX7C,PTEN,AKT3,AKT2,COX6A1,MTOR,TXNRD1,NDUFA4,COX8A,TNRC6A,RHEB,
2,RPS27A,PSMD8,PSMB4,UBE2M,PSMD4,PSMC3,PSMC1,PSMB1,CUL1,PSMD13,PSMD7,PSME3,
MTOR,CREBBP,HSPA1A,DNAJB1,HSF1
A5,COL1A2,COL5A2,DDR1,TTR,TNC,COL2A1,LAMC1,ACTN1,ITGB5,HSPG2,ITGAV,DAG1,COL3A1
HDAC2,MORF4L2,YEATS2,KDM6B,CCND1,SMARCA4,KDM1A,MSL1,SMARCE1,SMARCC1,NCOR2
HDAC2,MORF4L2,YEATS2,KDM6B,CCND1,SMARCA4,KDM1A,MSL1,SMARCE1,SMARCC1,NCOR2
LK1,RABEP1,MAP1LC3B,RAB6A,RAB5C,TBC1D10B,TBC1D14,RAB5A,GABARAPL2,RAB35,ARF6
DHA,UQCRC2,VDAC1,NDUFV1,UQCRC1,COX7C,GLO1,TRAP1,COX6A1,ATP5MC3,NDUFA13,ATP5
\1,COL1A2,COL5A2,COL6A2,PPIB,PLOD1,COL2A1,COL27A1,P3H3,CRTAP,COL16A1,COL12A1,C
OPS6,CETN2,XPC,GPS1,CUL4B,COPS8,RAD23A
D1,MYL12B,ACTA2,MYL12A,ANXA1,MYL9,MYL6B

M70,PINK1,MFN2,TOMM40
D,COL1A2,COL5A2,MMP14,COL6A2,MMP2,CTSB,COL2A1,ADAM9,MMP15,ADAM10,COL16A1,C
JBA52,PPP2CB,RPS27A,PPP5C,PPP2R5D,DUSP8,MAPK3,PPP2R5B,PPP2R5C
T8,CCT5,CSNK2A1,GNG4,CSNK2B,GNAZ,GNAO1,GNAI1
TLE1,AKT2,BCL9,YWHAZ,SOX9,CHD8,CTNNBIP1,TLE4
D2,TADA2B,TOMM20,RAD23B,ASXL1,PSMB2,PSMD3,FKBP8,TNKS2,USP7,HGS,PSMA7,VDAC1,EF
PS27A,HIF1A,TLE1,DTX4,NCSTN,ADAM10,RBPJ,CREBBP,NOTCH1,CUL1,KAT2A,CNTN1,MIB2,MAN

\1,COL4A1,RPLP0,RPL13A,RPL6,RPL13,SPTBN1,RPS8,MYH9,RPL7,RPL5,SPTAN1,UBB,COL4A2,F
\1,COL4A1,RPLP0,RPL13A,RPL6,RPL13,SPTBN1,RPS8,MYH9,RPL7,RPL5,SPTAN1,UBB,COL4A2,F
PL10,RPS17,RPS16,RPL23,RPL29,RPSA,EIF4H,RPS24,RPS9,RPL24,RPL17,RPL14,RPL22,RPL9,RI
LP1,RPL10,RPS17,RPS16,RPL23,RPL29,RPSA,RPS24,RPS9,RPL24,RPL17,RPL14,RPL22,RPL9,PS
RPLP1,RPL10,RPS17,RPS16,RPL23,RPL29,RPSA,EIF4H,RPS24,RPS9,RPL24,RPL17,RPL14,RPL22
LP1,RPL10,RPS17,RPS16,RPL23,RPL29,RPSA,RPS24,RPS9,RPL24,RPL17,RPL14,RPL22,RHOA,RI
RPLP1,RPL10,RPS17,RPS16,RPL23,RPL29,RPSA,EIF4H,RPS24,RPS9,RPL24,RPL17,RPL14,RPL22
RPLP1,RPL10,RPS17,RPS16,RPL23,RPL29,RPSA,EIF4H,RPS24,RPS9,RPL24,RPL17,RPL14,RPL22
PS16,RPL23,RPL29,RPSA,RPS24,RPS9,RPL24,RPL17,RPL14,RPL22,RPL9,RPS14,RPS20,RPL26,F
,RPS16,RPL23,RPL29,RPSA,RPS24,RPS9,RPL24,RPL17,RPL14,RPL22,RPL9,RPS14,RPS20,RPL26
7,RPS16,RPL23,RPL29,RPSA,RPS24,RPS9,RPL24,RPL17,RPL14,RPL22,RPL9,RPS14,RPS20,RPL2
P1,RPL10,RPS17,RPS16,RPL23,RPL29,RPSA,RPS24,RPS9,RPL24,RPL17,RPL14,RPL22,RPL9,RPS
P1,RPL10,RPS17,RPS16,RPL23,RPL29,RPSA,RPS24,RPS9,RPL24,RPL17,RPL14,RPL22,RPL9,RPS
PS17,RPS16,RPL23,RPL29,RPSA,RPS24,RPS9,RPL24,RPL17,RPL14,RPL22,RPL9,RPS14,RPS20,F
RPL23,RPL29,RPSA,RPS24,RPS9,RPL24,RPL17,RPL14,RPL22,RPL9,RPS14,RPS20,RPL26,RPL31,I
PS17,RPS16,RPL23,RPL29,RPSA,RPS24,RPS9,RPL24,RPL17,RPL14,RPL22,RPL9,RPS14,RPS20,F
,RPL7,P4HB,RPL5,GSTP1,VCP,UBB,RPL37A,PSMD5,EGLN1,RPL19,RPL15,RPS3,RPS4X,RPS3A,R
RPL23,RPL29,RPSA,RPS24,RPS9,RPL24,RPL17,RPL14,RPL22,RPL9,ATF4,RPS14,RPS20,RPL26,RI
RPL23,RPL29,RPSA,RPS24,RPS9,RPL24,RPL17,RPL14,RPL22,RPL9,RPS14,RPS20,RPL26,RPL31,I
,RPL7,P4HB,RPL5,GSTP1,VCP,UBB,RPL37A,PSMD5,EGLN1,RPL19,RPL15,RPS3,RPS4X,RPS3A,R
PL37A,PSMD5,SET,EIF4A1,RPL19,PPP2R1A,RPL15,SF3B1,RPS3,RPS4X,RPS3A,RPL18,RPL11,RPL
RPL23,RPL29,RPSA,RPS24,RPS9,RPL24,RPL17,RPL14,RPL22,RPL9,RPS14,RPS20,RPL26,RPL31,I
RPL23,RPL29,RPSA,RPS24,RPS9,RPL24,RPL17,RPL14,RPL22,RPL9,RPS14,RPS20,RPL26,RPL31,I
,RPL10,RPS17,GTF2F1,RPS16,RPL23,RPL29,RPSA,RPS24,RPS9,RPL24,KPNB1,RPL17,RPL14,RF
BP2,CALR,FSTL1,TUBB2B,RPLP0,H3-3B,CANX,DDX5,FKBP14,RPL13A,RPL6,RPL13,DDX17,SPTB
RPL23,RPL29,RPSA,RPS24,RPS9,RPL24,RPL17,RPL14,RPL22,RPL9,ATF4,RPS14,RPS20,RPL26,RI
17,GTF2F1,RPS16,RPL23,RPL29,RPSA,RPS24,RPS9,RPL24,RPL17,RPL14,RPL22,RPL9,RPS14,RF
X,RPS3A,RPL18,RPL11,RPLP1,RPL10,RPS17,RPS16,RPL23,RPL29,RPSA,EIF4H,RPS24,RPS9,RPL
IX,DDX5,RPL13A,RPL6,DVL3,RPL13,GNAS,GNB1,RPS8,MYH9,RPL7,APP,RPL5,CALM1,NPM1,VCP,
2,RPS27,RPS15A,EIF3E,RPS11,RPS28,FAU,EIF3F,RPS13,EIF3D,RPS2,RPS27A,EIF2S2,EIF3H,EIF3
PS12,RPS27,RPS15A,EIF3E,RPS11,RPS28,FAU,EIF3F,RPS13,EIF3D,RPS2,RPS27A,EIF2S2,EIF3H,E
LP0,CANX,DDX5,RPL13A,RPL6,DVL3,YWHAE,RPL13,GNAS,BSG,SPTBN1,GNB1,RPS8,MYH9,RPL7
PS12,RPS27,RPS15A,EIF3E,RPS11,RPS28,FAU,EIF3F,RPS13,EIF3D,RPS2,RPS27A,EIF2S2,EIF3H,E
S16,RPL23,RPL29,RPSA,RPS24,RPS9,RPL24,RPL17,RPL14,RPL22,RPL9,RPS14,RPS20,RPL26,RF
PS11,RPS28,FAU,EIF3F,RPS13,EIF3D,RPS2,RPS27A,EIF2S2,EIF3H,EIF3G,RPS26,RPS21,EIF3B,EIF
,COL5A1,COL4A1,RPLP0,H3-3B,RPL13A,RPL6,RPL13,SPTBN1,RPS8,MYH9,RPL7,RPL5,SPTAN1,
S16,RPL23,RPL29,RPSA,RPS24,RPS9,RPL24,RPL17,RPL14,RPL22,RPL9,RPS14,RPS20,RPL26,RF
11,RPLP1,RPL10,RPS17,RPS16,RPL23,RPL29,RPSA,RPS24,RPS9,RPL24,RPL17,RPL14,RPL22,RF
S16,RPL23,RPL29,RPSA,RPS24,RPS9,RPL24,RPL17,RPL14,RPL22,RPL9,RPS14,RPS20,RPL26,RF
PABPN1,HNRNPL,SNRPN,HNRNPA3,RBMX,DDX42,SNRNP70,PRPF6,SRSF9,SRSF1,PTBP1,HNRNP
PABPN1,HNRNPL,SNRPN,HNRNPA3,RBMX,DDX42,SNRNP70,PRPF6,SRSF9,SRSF1,PTBP1,HNRNP
Q,AP2M1,TUBB4B,SBF1,ARF1,PAFAH1B1,AP2B1,SCARB2,KIF5B,RAB6B,SPTBN2,RAB7A,CLTC,G
2,CPSF1,PABPN1,HNRNPL,SNRPN,HNRNPA3,RBMX,DDX42,SNRNP70,PRPF6,SRSF9,SRSF1,PTBP
PSMC1,PSMB1,CUL1,PSMD13,PSMD7,PPP2R5D,PSME3,PSMC5,CSNK1A1,PSMC2,PSMC4,PSMF
PSMC1,PSMB1,PSMD13,PSMD7,PSME3,PSMC5,PSMC2,PSMC4,PSMF1,PSMB5,EGLN2,HIF3A,PSN

1,COL11A1,AP2B1,GTF2F1,CTNNA1,TNS3,LAMA5,COL1A2,RHOA,CHD4,JUND,COL5A2,CLTC,CT
D13,PSMD7,PSME3,PSMC5,PSMC2,PSMC4,PSMF1,PSMB5,PSMA1,PSMD10,PSMA6,PSMD12,PSM
,PSMD7,PSME3,PSMC5,PSMC2,PSMC4,PSMF1,PSMB5,EGLN2,HIF3A,PSMA1,PSMD10,ELOB,PSM
EN,CBX6,RPS27A,TNKS,PSMD8,PSMB4,AKT3,MKRN1,PSMD4,AKT2,MBD3,PSMC3,MTOR,PSMC1
MB1,PSMD13,PSMD7,CSNK2B,TRIM27,PSME3,STUB1,PSMC5,PSMC2,PSMC4,PSMF1,PSMB5,PSI
C25A5,PSMB4,PACS1,PSMD4,PSMC3,AP2A2,AP1M1,PSMC1,RANGAP1,HMGA1,PSMB1,SLC25A6,
TBN1,MYH9,APP,GDI1,SPTAN1,CALM1,UBB,YWHAG,COL4A2,YWHAQ,AP2M1,TUBB4B,SBF1,ARF
,CDK4,PSME3,PSMC5,PSMC2,PSMC4,PSMF1,CDKN1B,PSMB5,PSMA1,PSMD10,PSMA6,PSMD12,
5,PSMC2,PSMC4,PSMF1,GSK3B,PSMB5,PSMA1,PSMD10,PSMA6,PSMD12,PSMA4,PSMC6
,PSMB1,NOTCH1,ACTA2,CUL1,PSMD13,KAT2A,PSMD7,PSME3,PSMC5,PSMC2,PSMC4,PSMF1,Y
,SKP1,CDKN1A,PPP1CB,NOX4,PSMB2,PPP2CA,MAPK1,PSMD3,GRB2,CDC37,PLCG1,MAP2K2,PO
,RPN2,VHL,KDM1A,STT3A,RBBP7,MTA1,AP2A1,BRD4,UBA52,MGAT4B,RPS27A,FKBP1A,MAP1LC
PSMC2,PSMC4,PSMF1,YWHAZ,PSMB5,PSMA1,PSMD10,PSMA6,PSMD12,PSMA4,PSMC6
PSMD4,PSMC3,PSMC1,EIF4G1,PSMB1,HSPA1A,PSMD13,PSMD7,MAPKAPK2,PSME3,PSMC5,PSM
,PSMC5,CSNK1A1,PSMC2,PSMC4,PSMF1,GSK3B,PSMB5,PSMA1,PSMD10,PSMA6,PSMD12,PSMA
,PSMC5,CSNK1A1,PSMC2,PSMC4,PSMF1,GSK3B,PSMB5,PSMA1,PSMD10,PSMA6,PSMD12,PSMA
ZB,KDELR1,ARF4,KIF3C,TMED9,DYNLL1,DCTN1,KIF5A,ACTR1A,KIF21B,RAB3GAP2,RAB1B,KIFAP
MC5,PSMC2,PSMC4,PSMF1,PSMB5,PSMA1,PSMD10,PSMA6,PSMD12,PSMA4,PSMC6
DKN1A,PPP1CB,PSMB2,TUBB2A,OPTN,PPP2CA,MAPRE1,PSMD3,CSNK1D,DYNLL1,PSMA7,ACTR1
,PSMB4,PSMD4,PSMC3,PSMC1,TUBB4A,PSMB1,PSMD13,PSMD7,PSME3,PSMC5,PSMC2,PSMC4
,DYNLL2,CDC37,YWHAH,DYNLL1,PSMA7,DFFB,STAT3,PSMD1,FLOT1,UBA52,RPS27A,PSMD8,PSM
PSMC3,AP2A2,PSMC1,CLTA,PSMB1,PSMD13,PSMD7,PSME3,DAAM1,PSMC5,PSMC2,PSMC4,PSM
DKN1A,PPP1CB,PSMB2,TUBB2A,OPTN,PPP2CA,MAPRE1,PSMD3,CSNK1D,DYNLL1,PSMA7,ACTR1
PTBN1,P4HB,APP,SPTAN1,CALM1,NPM1,USP11,VCP,UBB,TRIM28,FN1,DDB1,PSMD5,PDIA3,CHD
B,FYN,B2M,POLR2E,PSMA7,PSMD1,AP2A1,UBA52,POM121C,RPS27A,PSMD8,SLC25A5,XRCC6,P
5,PSMC2,PSMC4,PSMF1,PSMB5,PSMA1,PSMD10,PSMA6,PSMD12,PSMA4,PSMC6
PSMC5,PSMC2,PSMC4,PSMF1,PSMB5,PSMA1,PSMD10,PSMA6,RELA,PSMD12,PSMA4,PSMC6
,KDM1A,PLCG1,FYN,PHC2,TNKS2,USP7,PSMA7,RBBP7,MTA1,PRKAR2A,PSMD1,CSNK2A1,KPNA
ANXA2,TIMP2,VCL,CYB5R3,PGAM1,CTSD,LPCAT1,XRCC5,KPNB1,RHOA,PSMD2,VAT1,PFKL,RAB
,PSMC4,PSMF1,PSMB5,PSMA1,PSMD10,PSMA6,PSMD12,PSMA4,PSMC6
PSMC2,PSMC4,PSMF1,PSMB5,PSMA1,PSMD10,PSMA6,PSMD12,PSMA4,PSMC6
,PSMC2,PSMC4,PSMF1,PSMB5,PSMA1,PSMD10,PSMA6,PSMD12,PSMA4,PSMC6
C4,PSMF1,PSMB5,PSMA1,PSMD10,PSMA6,PSMD12,PSMA4,PSMC6
C4,PSMF1,PSMB5,PSMA1,PSMD10,ELOB,PSMA6,PSMD12,PSMA4,PSMC6
C4,PSMF1,PSMB5,PSMA1,PSMD10,PSMA6,PSMD12,OPA1,PSMA4,PSMC6
,PSMB4,PSMD4,PSMC3,PRKACA,PSMC1,SMO,TUBB4A,PSMB1,CUL1,PSMD13,PSMD7,ADCY1,PS
MC2,PSMC4,PSMF1,PSMB5,PSMA1,PSMD10,PSMA6,PSMD12,PSMA4,PSMC6
PSMC5,PSMC2,PSMC4,PSMF1,PSMB5,PSMA1,PSMD10,PSMA6,PSMD12,PSMA4,PSMC6
C4,PSMF1,PSMB5,PSMA1,PSMD10,PSMA6,PSMD12,PSMA4,PSMC6
PSMC2,PSMC4,PSMF1,PSMB5,PSMA1,PSMD10,PSMA6,PSMD12,PSMA4,PSMC6
,PSMC5,PSMC2,PSMC4,PSMF1,PSMB5,PSMA1,PSMD10,PSMA6,PSMD12,PSMA4,PSMC6
R1,CUL1,PSMD13,PSMD7,CDK4,PSME3,PSMC5,PSMC2,PSMC4,PSMF1,CDKN1B,PSMB5,PSMA1,
USP7,PSMA7,RBBP7,MTA1,PSMD1,CSNK2A1,UBA52,PPP2CB,PTEN,CBX6,RPS27A,TNKS,PSMD8,
2,TUBB2A,MAPK1,MAP2K2,KCNQ2,CNTNAP1,AP2A1,CSNK2A1,RDX,ITGAV,AP2A2,LYPLA2,TUBB4
C4,PSMF1,PSMB5,PSMA1,PSMD10,PSMA6,PSMD12,PSMA4,PSMC6

C4,PSMF1,PSMB5,PSMA1,PSMD10,PSMA6,PSMD12,PSMA4,PSMC6
C4,PSMF1,PSMB5,PSMA1,PSMD10,PSMA6,PSMD12,PSMA4,PSMC6
C4,PSMF1,PSMB5,PSMA1,PSMD10,PSMA6,PSMD12,PSMA4,PSMC6
PSMC4,PSMF1,PSMB5,PSMA1,PSMD10,PSMA6,AMD1,PSMD12,PSMA4,PSMC6
C5,PSMC2,PSMC4,PSMF1,PSMB5,PSMA1,PSMD10,PSMA6,PSMD12,PSMA4,PSMC6,FZD7,FZD8
MC2,PSMC4,PSMF1,PSMB5,PSMA1,PSMD10,PSMA6,PSMD12,PSMA4,PSMC6,RNF5
JL1,PSMD13,PSMD7,CDK4,PSME3,PSMC5,PSMC2,PSMC4,PSMF1,CDKN1B,PSMB5,PSMA1,PSMD
MC2,PSMC4,PSMF1,PSMB5,PSMA1,PSMD10,PSMA6,PSMD12,PSMA4,PSMC6
PSMC1,TEAD1,PSMB1,NOTCH1,MDM2,PSMD13,KAT2A,PSMD7,PSME3,PSMC5,PSMC2,PSMC4,P
1,PSMD3,DYNC1LI2,DYNLL2,DYNLL1,PSMA7,PSMD1,UBA52,PPP2CB,SMC3,PPP1CC,RPS27A,PSM
WHAH,DYNLL1,PSMA7,DFFB,STAT3,PSMD1,UBA52,RPS27A,PSMD8,PSMB4,AKT3,STK24,NMT1,P
1,PSMD3,DYNC1LI2,DYNLL2,DYNLL1,PSMA7,PSMD1,UBA52,PPP2CB,SMC3,PPP1CC,RPS27A,PSM
E3,PSMC5,PSMC2,PSMC4,PSMF1,CDKN1B,PSMB5,PSMA1,PSMD10,PSMA6,PSMD12,PSMA4,PSM
E3,PSMC5,PSMC2,PSMC4,PSMF1,CDKN1B,PSMB5,PSMA1,PSMD10,PSMA6,PSMD12,PSMA4,PSM
2,PSMC4,PSMF1,PSMB5,PSMA1,PSMD10,PSMA6,PSMD12,PSMA4,PSMC6
IB1,PSMD13,HLA-A,PSMD7,PSME3,PSMC5,PSMC2,PSMC4,PSMF1,PSMB5,PSMA1,PSMD10,PSMA
2,PSMC4,PSMF1,PSMB5,PSMA1,PSMD10,PSMA6,PSMD12,PSMA4,UBE2E1,PSMC6,ANAPC2
LTA,ARPC1A,EFNB2,LIMK1,EFNB1,NCK2,ARPC3,MYL12A,EFNB3,ARPC5,ROCK2,VAV2,GIT1,EPHE
C1,CDC34,PSMB1,CUL1,PSMD13,PSMD7,PSME3,PSMC5,PSMC2,PSMC4,PSMF1,UBE2V1,PSMB5,
PSMC5,PSMC2,PSMC4,PSMF1,GSK3B,PSMB5,STAT1,PSMA1,PSMD10,PSMA6,PSMD12,PSMA4,PS
E3,PSMC5,PSMC2,PSMC4,PSMF1,CDKN1B,PSMB5,PSMA1,PSMD10,PSMA6,PSMD12,PSMA4,PSM
C5,PSMC2,PSMC4,PSMF1,PSMB5,PSMA1,PSMD10,PSMA6,PSMD12,PSMA4,PSMC6
2,PSMC4,PSMF1,PSMB5,PSMA1,PSMD10,PSMA6,PSMD12,PSMA4,PSMC6
RHOA,KIF5B,PFN1,SPC25,XPO1,CTNNB1,CYFIP2,MYL6,MYH10,YWHAB,CKAP5,RHOB,RPS27,RAC
PS35,PSMB2,H3-3A,PPP2CA,SMARCA4,PSMD3,GNG2,TNKS2,VANGL2,PSMA7,BCL9L,EP300,PSM
PSMC1,PSMB1,DNAJB1,PSMD13,PSMD7,TNRC6A,PSME3,PSMC5,PSMC2,PSMC4,PSMF1,PSMB5,F
PSMC2,PSMC4,PSMF1,PSMB5,PSMA1,PSMD10,PSMA6,RELA,PSMD12,PSMA4,PSMC6
E3,PSMC5,PSMC2,PSMC4,PSMF1,PSMB5,PSMA1,PSMD10,PSMA6,PSMD12,PSMA4,PSMC6
CUL3,RPS27A,PSMD8,GRK2,PSMB4,DZIP1,PSMD4,PSMC3,PRKACA,PSMC1,SMO,TUBB4A,PSMB
1LI2,DYNLL2,DYNLL1,PSMA7,PSMD1,UBA52,PPP2CB,SMC3,PPP1CC,RPS27A,PSMD8,PSMB4,TA
SMC5,PSMC2,PSMC4,PSMF1,PSMB5,PSMA1,PSMD10,PSMA6,MCM7,PSMD12,PSMA4,PSMC6
IED9,DYNLL1,DCTN1,ACTR1A,YKT6,RAB1B,CAPZA1,COPZ1,TUBB4A,USO1,ARF5,ANK2,NSF,COP
3,PSMC1,TAPBP,PSMB1,PSMD13,HLA-A,PSMD7,PSME3,PSMC5,PSMC2,PSMC4,PSMF1,PSMB5,P
NG4,RPS27A,PSMD8,PSMB4,PSMD4,SCRIB,PSMC3,AP2A2,PSMC1,CLTA,PSMB1,PSMD13,PSMD7
PSMA1,PSMD10,PSMA6,PSMD12,PSMA4,PSMC6
TP1B2,FXYD6,MBD3,AP2A2,SAP18,MTA2,ATP1B3,TYK2,JAK1,STAT2,ROCK2,SUDS3,GATAD2B,FU
SMC1,ABL1,PSMB1,CUL1,PSMD13,PSMD7,CDK4,PSME3,STUB1,PSMC5,PSMC2,PSMC4,PSMF1,C
,PSME3,PSMC5,PSMC2,PSMC4,PSMF1,PSMB5,PSMA1,PSMD10,PPP3CA,PSMA6,RELA,PSMD12,P
ME3,PSMC5,PSMC2,PSMC4,PSMF1,PSMB5,PSMA1,PSMD10,PSMA6,PSMD12,PSMA4,PSMC6,RNF
2,CAPZB,KDELR1,ARF4,KIF3C,TMED9,DYNLL1,DCTN1,KIF5A,ACTR1A,KIF21B,RAB3GAP2,YKT6,C
C4,PSMF1,PSMB5,PSMA1,PSMD10,PSMA6,PSMD12,PSMA4,UBE2E1,PSMC6,ANAPC2
2,PSMC4,PSMF1,PSMB5,PSMA1,PSMD10,PSMA6,MCM7,PSMD12,PSMA4,PSMC6
TUBB4B,PAFAH1B1,CTNNA1,RHOA,KIF5B,PFN1,SPC25,XPO1,CTNNB1,CYFIP2,MYL6,MYH10,YW
TUBB4B,PAFAH1B1,CTNNA1,RHOA,KIF5B,PFN1,SPC25,XPO1,CTNNB1,CYFIP2,MYL6,MYH10,YW

SMD7,PSME3,PSMC5,PSMC2,PSMC4,PSMF1,UBE2V1,PSMB5,NKIRAS2,PSMA1,PSMD10,MAP2K4
PSMD13,PSMD7,ADRM1,PSME3,PSMC5,PSMC2,PSMC4,PSMF1,PSMB5,PSMA1,PSMD10,TGFBR1,
2,PSMC4,PSMF1,PSMB5,PSMA1,PSMD10,PSMA6,PSMD12,PSMA4,UBE2E1,PSMC6,ANAPC2
K1,RPS27A,MAP1LC3B,TOMM70,TUBB4A,PINK1,MFN2,PARK7,CSNK2B,HSF1,UBE2V1,TUBA1C,
RF4,CSNK1D,TMED9,PPP6R1,SEC23A,DYNLL1,DCTN1,ACTR1A,SEC16A,YKT6,LMAN1,RAB1B,CAP
SMD1,PBX1,H2AZ1,UBA52,HDAC6,RPS27A,PSMD8,HIF1A,PSMB4,TLE1,DTX4,NCSTN,RAB6A,PSM
2,PSMC4,PSMF1,PSMB5,PSMA1,PSMD10,PSMA6,PSMD12,PSMA4,UBE2E1,PSMC6,ANAPC2
MC5,PSMC2,PSMC4,PSMF1,PSMB5,PSMA1,PSMD10,PSMA6,PSMD12,PSMA4,UBE2E1,PSMC6,AN

ME3,PSMC5,CSNK1A1,PSMC2,PSMC4,PSMF1,KIF3A,PSMB5,PSMA1,PSMD10,PSMA6,PSMD12,PS
1,PSMD13,PSMD7,PSME3,PSMC5,PSMC2,PSMC4,PSMF1,PSMB5,PSMA1,PSMD10,PSMA6,PSMD
CB,PTPRA,PSMB2,PPP2CA,MAPK1,PSMD3,HSPB1,MAP2K2,DUSP4,FYN,CDC42,PSMA7,PSMD1,UE
SNK1E,CKAP5,RAB1A,RPS27,TNPO1,PSMB2,H3-3A,TUBB2A,PPP2CA,MAPK1,MAPRE1,PSMD3,DY
CST3,MBTPS1,CCN1,ADAM10,TGOLN2,IGFBP4,LGALS1,KTN1,ANO8,SHISA5,SCG2,STC2,CHGB,
,RAB1B,KIFAP3,COPZ1,TUBB4A,ARF5,KIFC2,NSF,COPE,KIF26A,KLC2,NBAS,ARFGAP2,TUBA1C,
A,ARCN1,RPN2,MLEC,RAD23B,TMED2,TUBB2A,ARF3,KDELR2,DYNC1LI2,DYNLL2,CAPZB,KDELR
4,AKT2,PSMC3,PSMC1,PSMB1,CUL1,PSMD13,PSMD7,CDK4,PSME3,PSMC5,PSMC2,PSMC4,PSM
DC2,THOC3,FYTTD1,RNPS1,CPSF7,WDR33
C4,PSMF1,PSMB5,PSMA1,PSMD10,PSMA6,PSMD12,PSMA4,UBE2E1,PSMC6,ANAPC2
UL1,PSMD13,PSMD7,PSME3,PSMC5,PSMC2,PSMC4,PSMF1,UBE2V1,PSMB5,PSMA1,PSMD10,PS
PI,MYH9,APP,KLRD1,SPTAN1,GSTP1,CALM1,PLD3,VCP,UBB,PSMD5,PPP2R1A,HLA-C,VAPA,HMG
DLR,SH3GL1,FNBP1L,RAB5C,SYT11,AP2A2,TGOLN2,GAK,CLTA,ARPC1A,HIP1R,COPS6,EPN1,AR

2,PPP2CA,MAPK1,PSMD3,MAP2K2,DUSP4,FYN,PSMA7,PSMD1,UBA52,PPP2CB,PPP1CC,CUL3,RP
ME3,PSMC5,PSMC2,PSMC4,PSMF1,PSMB5,PSMA1,PSMD10,PSMA6,MCM7,PSMD12,PSMA4,UBE2
PPP2R5B,SEC31A

C4,PSMF1,PSMB5,PSMA1,PSMD10,PSMA6,PSMD12,PSMA4,UBE2E1,PSMC6,ANAPC2
H2AZ1,CSNK2A1,UBA52,PPP2CB,CSNK1G2,SFRP1,CUL3,RPS27A,TNKS,PSMD8,PSMB4,SOX2,TL
31,TGFBR1,ARHGEF18,TFDP1,SP1,FURIN,CCNK,PPM1A,NEDD8,SKIL
2,PPP2CA,MAPK1,PSMD3,MAP2K2,DUSP4,FYN,PSMA7,PSMD1,UBA52,PPP2CB,PPP1CC,CUL3,RP
PDPK1,WASF2,ITGAV,AKT2,MTOR,PRKACA,AAMP,CYFIP1,NCK2,TRIB3,MAPKAPK2,WASF3,NRP2,
D2,RBBP4,SPC25,XPO1,CSNK1E,CCND2,CKAP5,RAB1A,RPS27,SKP1,CDKN1A,PPP1CB,TNPO1,PS
CBP2,SNRPB,THOC2,THOC3,FYTTD1,RNPS1,CPSF7,WDR33
LAMC1,VGF,IGFBP7,CST3,MBTPS1,CCN1,ADAM10,TGOLN2,IGFBP4,LGALS1,KTN1,ANO8,SHISA5
3,PSMD4,AKT2,PSMC3,PSMC1,ABL1,PSMB1,CUL1,PSMD13,PSMD7,CDK4,PSME3,PSMC5,PSMC2
WASF2,ITGAV,AKT2,MTOR,PRKACA,CYFIP1,NCK2,TRIB3,MAPKAPK2,WASF3,ROCK2,SHB,VAV2,B
C4,PSMF1,PSMB5,PSMA1,PSMD10,PSMA6,PSMD12,PSMA4,UBE2E1,PSMC6,ANAPC2
GL1,PTPN11,LRIG1,PTPN1,CRKL,RANBP9,SOS1,ARF6,COL3A1,RAP1B,SH3KBP1
PPP2R5B,SEC31A
2A1,GNG4,FKBP9,FBXW5,TUBB4A,XRN2,NOP56,CSNK2B,TUBA1C,TBCA,LONP2,SKIV2L,GNAZ,G
L1,VDAC1,WDR45B,CSNK2A1,UBA52,HDAC6,ULK1,RPS27A,MAP1LC3B,GABARAPL1,CHMP4B,
RF4,CSNK1D,TMED9,PPP6R1,SEC23A,DYNLL1,DCTN1,ACTR1A,SEC16A,MGAT4B,YKT6,LMAN1,R

C4,PSMF1,PSMB5,PSMA1,PSMD10,PSMA6,PSMD12,PSMA4,UBE2E1,PSMC6,ANAPC2
AH1B2,RAB18,TUBA1C,AGPAT3,GALNT2,CAPZA2,DYNC1I2
LNA,ALDOA,PKM,FTL,HSP90B1,FBXL4,UBA1,CALR,HSPA1B,TUBB2B,PSAP,RPLP0,CANX,CLU,SL(
SMC5,PSMC2,PSMC4,PSMF1,PSMB5,PSMA1,PSMD10,PSMA6,PSMD12,PSMA4,UBE2E1,PSMC6,AN
SMC5,PSMC2,PSMC4,PSMF1,PSMB5,PSMA1,PSMD10,PSMA6,PSMD12,PSMA4,UBE2E1,PSMC6,AN
3A,DYNLL1,DCTN1,KIF5A,ACTR1A,AP2A1,KIFAP3,CAPZA1,ACTR1B,AP2A2,AP1M1,CD74,TUBB4A,(
,CAPZA1,TUBB4A,HSPA1A,DNAJB1,TUBA1C,CAPZA2,DYNC1I2
C,DYNC1I2
F15,SSRP1,TAF7,PPP2CA,PRDX1,SUPT5H,BNIP3L,POLR2B,YWHAH,USP7,DDIT4,POLR2E,RBBP7,I
MK1,PIP5K1C,GSK3B,ERBB2,ROCK2,PLXND1,MYL9

SMC3,PRKACA,CREBBP,PSMC1,CDC34,PSMB1,CUL1,PSMD13,PSMD7,PSME3,PSMC5,PSMC2,PS
,HIF1A,FBXO21,WSB2,PSMB4,UBE2M,UBE2D2,DCUN1D2,PSMD4,LRRC41,FBXW5,PSMC3,UBE2I

R2,NPEPPS,B2M,HECTD1,SEC23A,PSMA7,TRAF7,PSMD1,UBA52,KLHL42,UBE2G2,FBXO11,CUL3,
A1,UBA52,HDAC6,ULK1,RPS27A,MAP1LC3B,GABARAPL1,CHMP4B,MTOR,TOMM70,AMBRA1,ATG
NG4,FKBP9,FBXW5,TUBB4A,XRN2,NOP56,CSNK2B,TUBA1C,LONP2,SKIV2L,GNAZ,GNAO1,AP3M
3,SHC2,AP2A2,NTRK2,CLTA,RALGDS,MAPKAPK2,PPP2R5D,CRKL,SRF,DUSP3,IRS1,SOS1,SHC1,F
PRE1,DYNC1LI2,DYNLL2,PFN2,DYNLL1,CDC42,PPP2CB,EVL,PPP1CC,FMNL2,TAOK1,CLASP2,TUE
L6,RPL13,GNAS,SLC2A3,GPI,BSG,FASN,ALDH6A1,ATP5F1B,GNB1,RPS8,RPL7,IDS,RPL5,GSTP1,N
1,SMARCA5,KPNB1,PSMD2,RBBP4,SPC25,XPO1,CSNK1E,CCND2,YWHAB,CKAP5,RAB1A,RPS27,S
DO1,RAP1B
HSPG2,ITGAV,BGN,DAG1,COL6A3,TGFB2,COL3A1,ITGA7
,KLHL42,UBE2G2,FBXO11,CUL3,RPS27A,PSMD8,FBXO21,PSMB4,UBE2M,UBE2D2,MKRN1,PSMI
AP2A1,PPP2CB,SHC2,AP2A2,NTRK2,PTPN11,CLTA,RALGDS,MAPKAPK2,PPP2R5D,CRKL,SRF,DUS
YL2,ASNS,HERPUD1,CEBPG,LMNA,SHC1,DNAJB11,ATF6,ADD1,CUL7,PPP2R5B,SEC31A,EIF2S1

M1
34,PSMB1,CUL1,PSMD13,PSMD7,PSME3,PSMC5,PSMC2,PSMC4,PSMF1,UBE2V1,PSMB5,PSMA1
PS27A,AKAP9,PRKACA,TUBB4A,PCM1,CUL1,CETN2,PRKAR2B,PPP1R12A,AJUBA,DYNC1I2,NINL
2,PSMC4,PSMF1,PSMB5,PSMA1,PSMD10,PSMA6,MCM7,PSMD12,PSMA4,PSMC6
,ITGB1,TUBB4B,WSB1,ARF1,AP2B1,CTSD,PJA2,COL1A2,PSMD2,KIF5B,CXADR,CD81,KLHL9,SPT
UBA52,ARID1B,CBX6,RPS27A,PSMD8,PSMB4,KMT2E,TCF3,PSMD4,PSMC3,SMARCD1,CREBBP,F

I,DAD1,MOGS,GSK3B,MAN1B1,FURIN

PSMB1,CUL1,PSMD13,PSMD7,PSME3,PSMC5,PSMC2,PSMC4,PSMF1,UBE2V1,PSMB5,NKIRAS2,P

G5,HSPA1A,DNAJB1,NUP153,MAPKAPK2,HSF1,GSK3B,NUP205,NUP133,DNAJB6,MAPK3,NUP50
A2,PSMD2,MCL1,NANOG,MIF,JUN,HNRNPDL,MMP2,SKP1,CDKN1A,CNN2,SQSTM1,VAMP2,TCP1,

EXOC4,TUBB4A,EXOC5,EXOC7,YWHAZ,TUBA1C,KIF3A

JRIN,PTPN12,COL3A1

C1,PSMB1,FZR1,CUL1,PSMD13,PSMD7,CDK4,POLD2,PSME3,PSMC5,PSMC2,PSMC4,PSMF1,GSK
P1,PPP1CA,PMEPA1,CDK9,STUB1,TGFBR1,ARHGEF18,TFDP1,SP1,FURIN,CCNK,PPM1A,NEDD8,S

,MMP14,COL6A2,TTR,TNC,MMP2,PPIB,DCN,CTSB,VCAN,PLOD1,COL2A1,LAMC1,PLEC,ADAM9,A

ATP6V1F,NEDD8,ATP6V1D
LD2,PSME3,PSMC5,PSMC2,PSMC4,PSMF1,PSMB5,PSMA1,PSMD10,PSMA6,PCNA,MCM7,PSMD12

IUP205,NUP133,THOC2,THOC3,FYTTD1,NUP50,RNPS1,WDR33
COX7A2L,YWHAZ,TXN,COX6B1,COX4I1,COX5A,SESN2,COX5B
PSMC5,PSMC2,PSMC4,PSMF1,PSMB5,PSMA1,PSMD10,PSMA6,TRAF3,PSMD12,PSMA4,PSMC6

I,CASK
,KDM3B,KDM4B,RBBP7,PRMT1,MTA1,KDM5C,EP300,KDM5B,H2AZ1,EP400,ARID1B,RPS2,EHMT
,KDM3B,KDM4B,RBBP7,PRMT1,MTA1,KDM5C,EP300,KDM5B,H2AZ1,EP400,ARID1B,RPS2,EHMT

MC2,ATP5MG,PDHA1,NDUFA4,IDH2,NDUFC2,COX8A,COX7A2L,ATP5PO,DLD,NDUFB8,ME2,NDUF
OLGALT1,PCOLCE,P3H1,COL7A1,COL6A3,COL3A1

OL12A1,COL7A1,COL6A3,FURIN,COL3A1

P300,PSMD1,UBA52,PTEN,RPS27A,TNKS,PSMD8,HIF1A,BAP1,PSMB4,ATXN3,FOXK2,PSMD4,OGT,
ML1,NBEA,TLE4

RPL37A,PSMD5,RPL19,RPL15,RPS3,AP2M1,RPS4X,RPS3A,RPL18,ITGB1,TUBB4B,RPL11,NCAM1,
RPL37A,PSMD5,RPL19,RPL15,RPS3,AP2M1,RPS4X,RPS3A,RPL18,ITGB1,TUBB4B,RPL11,NCAM1,
PS14,RPS20,EIF5,RPL26,RPL31,RPS25,RPS15,RPL12,RPL30,EIF3L,RPS5,RPS7,RPS19,RPL35,RP
MD2,RPS14,RPS20,RPL26,RPL31,RPS25,RPS15,RPL12,RPL30,RPS5,RPS7,RPS19,RPL35,RPS10,
,RPL9,RPS14,RPS20,RPL26,RPL31,RPS25,RPS15,RPL12,RPL30,EIF3L,RPS5,RPS7,RPS19,RPL35
PL9,PSMD2,RPS14,RPS20,RPL26,PFN1,RPL31,RPS25,RPS15,RPL12,RPL30,RPS5,RPS7,RPS19,R
,RPL9,RPS14,RPS20,EIF5,RPL26,RPL31,RPS25,RPS15,RPL12,RPL30,EIF3L,RPS5,RPS7,RPS19,R
,RPL9,RPS14,RPS20,EIF5,RPL26,RPL31,RPS25,RPS15,RPL12,RPL30,EIF3L,RPS5,RPS7,RPS19,R
RPL31,RPS25,RPS15,RPL12,RPL30,EIF3L,RPS5,RPS7,RPS19,RPL35,RPS10,RPL35A,RPLP2,RPL3
,RPL31,RPS25,RPS15,RPL12,RPL30,RPS5,RPS7,RPS19,RPL35,RPS10,RPL35A,RPLP2,RPL36,RP
6,RPL31,RPS25,RPS15,RPL12,RPL30,RPS5,RPS7,RPS19,RPL35,RPS10,RPL35A,RPLP2,DDOST,F
14,RPS20,RPL26,RPL31,RPS25,RPS15,RPL12,RPL30,RPS5,RPS7,RPS19,RPL35,RPS10,RPL35A,F
14,RPS20,RPL26,RPL31,RPS25,RPS15,RPL12,RPL30,RPS5,RPS7,RPS19,RPL35,RPS10,RPL35A,F
RPL26,RPL31,RPS25,RPS15,RPL12,RPL30,RPS5,RPS7,RPS19,RPL35,RPS10,RPL35A,RPLP2,RPL3
RPS25,RPS15,RPL12,RPL30,RPS5,RPS7,RPS19,RPL35,RPS10,RPL35A,RPLP2,RPL36,RPS12,RPS
RPL26,RPL31,RPS25,RPS15,RPL12,RPL30,RPS5,RPS7,RPS19,RPL35,RPS10,RPL35A,RPLP2,RPL3
PL18,TUBB4B,RPL11,RPLP1,RPL10,RPS17,DNAJA1,RPS16,RPL23,RPL29,RPSA,RPS24,RPS9,RPL
PL31,RPS25,RPS15,RPL12,RPL30,RPS5,RPS7,RPS19,RPL35,RPS10,RPL35A,RPLP2,RPL36,RPS1
RPS25,RPS15,RPL12,RPL30,RPS5,RPS7,RPS19,RPL35,RPS10,RPL35A,RPLP2,RPL36,RPS12,RPS
PL18,TUBB4B,RPL11,RPLP1,RPL10,RPS17,DNAJA1,RPS16,RPL23,RPL29,RPSA,RPS24,RPS9,RPL
P1,RPL10,RPS17,SF3B2,GTF2F1,HNRNPU,HNRNPA0,RPS16,RPL23,RPL29,RPSA,RPS24,RPS9,RF
RPS25,RPS15,RPL12,RPL30,RPS5,RPS7,RPS19,RPL35,RPS10,RPL35A,RPLP2,RPL36,RPS12,RPS
RPS25,RPS15,RPL12,RPL30,RPS5,RPS7,RPS19,RPL35,RPS10,RPL35A,RPLP2,RPL36,RPS12,RPS
L22,RPL9,RPS14,RPS20,RPL26,RPL31,XPO1,RPS25,RPS15,RPL12,RPL30,CLTC,RPS5,RPS7,RPS
N1,GNB1,RPS8,RPL7,P4HB,APP,RPL5,SPTAN1,CALM1,NPM1,USP11,IGF2,EIF3C,EIF4B,VCP,UBB,
PL31,RPS25,RPS15,RPL12,RPL30,RPS5,RPS7,RPS19,RPL35,RPS10,RPL35A,RPLP2,RPL36,RPS1
PS20,RPL26,RPL31,RPS25,RPS15,RPL12,RPL30,RPS5,RPS7,RPS19,RPL35,RPS10,RPL35A,RPLP2
24,RPL17,RPL14,RPL22,PTCD3,RPL9,RPS14,RPS20,EIF5,RPL26,RPL31,RPS25,RPS15,RPL12,RP
,UBB,TRIM28,RPL37A,PSMD5,RPL19,RPL15,RPS3,CHD3,AP2M1,RPS4X,RPS3A,RPL18,ITGB1,TU
G,EIF4G1,RPS26,RPS21,EIF3B,EIF3M,RPS29,EIF2S1,EIF3J
IF3G,EIF4G1,RPS26,RPS21,EIF3B,EIF3M,RPS29,EIF2S1,EIF3J
,APP,IDS,RPL5,CALM1,NPM1,VCP,UBB,TRIM28,RPL37A,FN1,PSMD5,CHSY1,RPL19,PPP2R1A,RPL
IF3G,EIF4G1,RPS26,RPS21,EIF3B,EIF3M,RPS29,EIF2S1,EIF3J
L31,CSNK1E,RPS25,RPS15,RPL12,RPL30,RPS5,RPS7,RPS19,RPL35,RPS10,RPL35A,RPLP2,RPL3
3M,RPS29,EIF2S1,EIF3J
UBB,COL4A2,RPL37A,PSMD5,RPL19,RPL15,RPS3,AP2M1,RPS4X,RPS3A,RPL18,ITGB1,TUBB4B,
L31,CSNK1E,RPS25,RPS15,RPL12,RPL30,RPS5,RPS7,RPS19,RPL35,RPS10,RPL35A,RPLP2,RPL3
L9,PSMD2,SLC7A5,RPS14,RPS20,RPL26,OAZ1,RPL31,SLC3A2,RPS25,RPS15,RPL12,RPL30,RPS
L31,CSNK1E,RPS25,RPS15,RPL12,RPL30,RPS5,RPS7,RPS19,RPL35,RPS10,RPL35A,RPLP2,RPL3
UL1,CASC3,POLR2B,HNRNPM,SRSF3,SF3A1,PCBP2,HNRNPD,PRPF40A,DHX15,POLR2E,SF1,PRC
UL1,CASC3,POLR2B,HNRNPM,SRSF3,SF3A1,PCBP2,HNRNPD,PRPF40A,DHX15,POLR2E,SF1,PRC
JA1,MYO5A,YWHAB,RAB1A,ARCN1,RAC1,RAB27A,VAMP2,GDI2,ACTR3,TBC1D16,KLC1,SURF4,R
1,HNRNPUL1,CASC3,POLR2B,HNRNPM,SRSF3,SF3A1,PCBP2,HNRNPD,PRPF40A,DHX15,POLR2E,
1,GSK3B,PSMB5,PSMA1,PSMD10,CTBP2,PSMA6,PPP2R5B,PSMD12,PSMA4,PSMC6,TLE4,PPP2R5
IA1,CITED2,PSMD10,ELOB,PSMA6,AJUBA,PSMD12,PSMA4,PSMC6

NNB1,COL6A2,CYFIP2,MXD4,YWHAB,HNRNPH1,RAC1,ATP6V0C,AHCYL1,NCKAP1,COL2A1,LAMC
A4,PSMC6
1A6,AJUBA,PSMD12,PSMA4,PSMC6
,MTA2,PSMB1,PSMD13,PSMD7,TNRC6A,RHEB,CSNK2B,TRIM27,PSME3,STUB1,PSMC5,PSMC2,F
MA1,PSMD10,PSMA6,PSMD12,PSMA4,PSMC6
PSMD13,BANF1,PSMD7,RAN,NUP153,CDK9,PSME3,PSIP1,PSMC5,PSMC2,PSMC4,PSMF1,PSMB5,
1,PAFAH1B1,AP2B1,SCARB2,COL1A2,KIF5B,RAB6B,SPTBN2,RAB7A,CLTC,GJA1,MYO5A,YWHAB,
,PSMA4,PSMC6

WHAZ,PSMB5,PSMA1,PSMD10,PSMA6,PSMD12,PSMA4,MAML1,PSMC6
LR2B,FYN,TNKS2,MPRIP,NCOR2,POLR2E,PSMA7,STAT3,BCR,OS9,EP300,PSMD1,UBA52,PPP2CE
3B,FKBP4,IMPDH2,GALNT1,ATP1B2,FXYD6,MBD3,GSK3A,CHMP4B,AP2A2,SAP18,UBE2I,MTA2,M

C2,PSMC4,PSMF1,YWHAZ,PSMB5,PSMA1,PSMD10,PSMA6,PSMD12,PSMA4,PSMC6
A4,PSMC6
A4,PSMC6
P3,CAPZA1,GALNT1,RAB6A,COPZ1,TUBB4A,ARF5,PAFAH1B2,KIFC2,NSF,COPE,KIF26A,KLC2,NBA

1A,EP300,PSMD1,UBA52,PPP2CB,RPS27A,PSMD8,AKAP9,PSMB4,TUBGCP6,PSMD4,PSMC3,PRKA
,PSMF1,TUBA1C,PSMB5,PSMA1,PSMD10,PSMA6,PSMD12,PSMA4,PSMC6
MB4,AKT3,STK24,FLOT2,NMT1,PSMD4,AKT2,OGT,PSMC3,PSMC1,DAPK3,SDCBP,PSMB1,PSMD1
IF1,PSMB5,PSMA1,PSMD10,PSMA6,PSMD12,PSMA4,PSMC6,FZD7,FZD8
1A,EP300,PSMD1,UBA52,PPP2CB,RPS27A,PSMD8,AKAP9,PSMB4,TUBGCP6,PSMD4,PSMC3,PRKA
3,TUBB4B,UCHL1,MARCHF6,WSB1,ARF1,IGFBP3,RHOA,CBX5,PSMD2,RAB6B,PRKCSH,HNRNPC,
PSMB4,VPS4A,NMT1,PACS1,PSMD4,PSMC3,CHMP4B,AP2A2,AP1M1,PSMC1,RANGAP1,HMGA1,PS

2,UBA52,PPP2CB,ADCY6,PTEN,CBX6,RPS27A,TNKS,PSMD8,GRK2,PSMB4,AKT3,PDPK1,MKRN1,F
7A,DDX3X,MIF,CD63,DNAJC5,DDOST,LAMP1,CKAP4,HLA-B,TTR,COTL1,HRNR,RAC1,ATP6V0C,TM

ME3,PSMC5,CSNK1A1,PSMC2,PSMC4,PSMF1,GSK3B,TUBA1C,KIF3A,PSMB5,PRKAR2B,PSMA1,F

PSMD10,PSMA6,TFDP1,PSMD12,PSMA4,PSMC6
PSMB4,AKT3,PDPK1,MKRN1,PSMD4,AKT2,MBD3,GSK3A,PSMC3,MTOR,PSMC1,PTPN11,MTA2,P
4A,ANK2,SDCBP,CLTA,CSNK2B,RANBP9,ALCAM,TUBA1C,DLG4,CNTN1,CNTN2,NRP2,RPS6KA2,M

10,PSMA6,TFDP1,PSMD12,PSMA4,PSMC6

SMF1,PSMB5,PSMA1,PSMD10,PSMA6,PSMD12,PSMA4,MAML1,PSMC6
MD8,PSMB4,TAOK1,VPS4A,CLASP2,PSMD4,IST1,PSMC3,CHMP4B,UBE2I,PSMC1,TUBB4A,RANGA
SMD4,AKT2,PSMC3,PSMC1,DAPK3,PSMB1,PSMD13,PSMD7,C1QBP,PSME3,PSMC5,PSMC2,TNFR
MD8,PSMB4,TAOK1,VPS4A,CLASP2,PSMD4,IST1,PSMC3,CHMP4B,UBE2I,PSMC1,TUBB4A,RANGA
MC6
MC6

A6,PSMD12,TLR6,PSMA4,PSMC6

32,EPHA4,DNM1,MYL9,EPHA2
PSMA1,PSMD10,PPP3CA,PSMA6,TAB3,RELA,PSMD12,PSMA4,PSMC6
MC6
MC6

1,PPP1CB,ACTR3,PKN1,KLC1,NCKAP1,H3-3A,TUBB2A,PPP2CA,MAPK1,MAPRE1,ARPC2,DYNC1LI
ID1,AP2A1,H2AZ1,CSNK2A1,UBA52,PPP2CB,CSNK1G2,GNG4,SFRP1,CUL3,RPS27A,TNKS,PSMD8
PSMA1,PSMD10,PSMA6,PSMD12,PSMA4,PSMC6

1,CUL1,PSMD13,PSMD7,ADCY1,PSME3,PSMC5,CSNK1A1,PSMC2,PSMC4,PSMF1,GSK3B,TUBA1C
OK1,CLASP2,PSMD4,PSMC3,PSMC1,TUBB4A,RANGAP1,PSMB1,PSMD13,PSMD7,PPP2R5D,PSME

E,ARFGAP2,TUBA1C,NAPB,NAPA,CAPZA2,COPB1,DYNC1I2,COG4,ARFGAP3
SMA1,PSMD10,PSMA6,PSMD12,TLR6,PSMA4,PSMC6
,TNRC6A,PSME3,DAAM1,PSMC5,PSMC2,PSMC4,PSMF1,PSMB5,PSMA1,PSMD10,PPP3CA,PSMA6,

URIN,HMG20B
GSK3B,SOX9,PSMB5,STAT1,RBM14,PSMA1,PSMD10,MAPK3,PSMA6,PSMD12,PSMA4,PSMC6
SMA4,PSMC6
5
CYTH2,RAB1B,KIFAP3,CAPZA1,GALNT1,RAB6A,COPZ1,TGOLN2,TUBB4A,ARF5,PAFAH1B2,KIFC2,

HAB,CKAP5,RHOB,RPS27,RAC1,ARHGDIA,PPP1CB,ACTR3,PKN1,KLC1,NCKAP1,H3-3A,TUBB2A,V
HAB,CKAP5,RHOB,RPS27,RAC1,ARHGDIA,PPP1CB,ACTR3,PKN1,KLC1,NCKAP1,H3-3A,TUBB2A,V

,PSMA6,TAB3,RELA,PSMD12,PSMA4,PSMC6,TOLLIP
PSMA6,PSMD12,MBD6,PSMA4,PSMC6,NEDD8

DYNC1I2,NBR1,TOMM40
ZA1,COPZ1,TUBB4A,USO1,ARF5,ANK2,LMAN2,NSF,COPE,ARFGAP2,TUBA1C,MCFD2,NAPB,NAPA
D4,ADAM10,RBPJ,PSMC3,CREBBP,PSMC1,PSMB1,NOTCH1,ACTA2,CUL1,PSMD13,KAT2A,PSMD7

NAPC2

MA4,PSMC6
12,PSMA4,ABCD3,EIF2S1,PSMC6,RNF5,ABCA7
BA52,PPP2CB,PPP1CC,CUL3,RPS27A,SEPTIN7,PSMD8,PSMB4,SHC2,PSMD4,IGF2BP1,PSMC3,PR
NC1LI2,DYNLL2,CSNK1D,DYNLL1,PSMA7,ACTR1A,PSMD1,H2AZ1,CSNK2A1,UBA52,PPP2CB,RAB
FAM20C,APOE,MFGE8,QSOX1,PRSS23,MXRA8
KIF3A,NAPB,NAPA,COPB1,ARFGAP3
1,ARF4,STT3A,CSNK1D,TMED9,PPP6R1,SEC23A,DYNLL1,ST3GAL6,DCTN1,OS9,ACTR1A,SEC16A
F1,DHFR,CDKN1B,PSMB5,RRM2,PSMA1,PSMD10,PSMA6,PCNA,TFDP1,MCM7,PSMD12,PSMA4,PS

MA6,RELA,PSMD12,PSMA4,PSMC6
B1,TUBB4B,ANXA2,TIMP2,VCL,CYB5R3,PGAM1,CTSD,LPCAT1,XRCC5,KPNB1,RHOA,PSMD2,CD8
PC3,PIP5K1C,TFRC,ARPC5,RAB5A,GPS1,ARF6,SYT1,COPS8,DNM1,NEDD8,SLC18A3,SH3KBP1,M6

S27A,PSMD8,PSMB4,SHC2,PSMD4,PSMC3,PSMC1,PTPN11,PSMB1,PPP5C,RGL2,PSMD13,RALGD
2E1,PSMC6,ANAPC2

LE1,PSMD4,AKT2,PSMC3,CREBBP,PSMC1,PSMB1,PSMD13,TRRAP,PSMD7,CSNK2B,H2AX,PPP2R5

S27A,PSMD8,PSMB4,SHC2,PSMD4,PSMC3,PSMC1,PSMB1,PPP5C,RGL2,PSMD13,RALGDS,RASA4
ROCK2,SHB,VAV2,BCAR1,ELMO2
MB2,H3-3A,TUBB2A,OPTN,PPP2CA,CCND1,MAPK1,MAPRE1,PSMD3,DYNC1LI2,DYNLL2,CSNK1D

,SCG2,STC2,CHGB,FAM20C,APOE,MFGE8,QSOX1,PRSS23,MXRA8
2,PSMC4,PSMF1,DHFR,CDKN1B,PSMB5,RRM2,PSMA1,PSMD10,PSMA6,PCNA,TFDP1,MCM7,PSMI
CAR1,ELMO2

NAO1,AP3M1,GNAI1,TBCB
MTOR,TOMM70,AMBRA1,ATG4B,TUBB4A,PINK1,MFN2,PARK7,RHEB,CSNK2B,HSF1,UBE2V1,TUB
AB1B,CAPZA1,COPZ1,TUBB4A,USO1,ARF5,ANK2,LMAN2,MGAT5,MGAT3,NSF,COPE,B4GALT5,AF

C2A3,GPI,MYH9,P4HB,APP,KLRD1,FSCN1,SPTAN1,GSTP1,CALM1,PLD3,VCP,UBB,FN1,PSMD5,EIF
NAPC2
NAPC2
CLTA,KIF26A,KLC2,TUBA1C,KIF3A,CAPZA2,SEC31A,DYNC1I2,DNM1,CTSA

PRMT1,EP300,CSNK2A1,UBA52,PPP2CB,COX7C,PTEN,RPS27A,PCBP4,EHMT1,CARM1,AKT3,PDP

SMC4,PSMF1,UBE2V1,PSMB5,PSMA1,PSMD10,PPP3CA,PSMA6,TAB3,RELA,PSMD12,PSMA4,PSM
D3,PSMC1,FBXL19,WDTC1,PSMB1,OBSL1,CUL1,PSMD13,DCAF5,PSMD7,CUL4A,COPS6,PSME3,

ITGB5,RPS27A,PSMD8,FBXO21,PSMB4,UBE2M,UBE2D2,MKRN1,PSMD4,ITGAV,LRRC41,FBXW5,
34B,TUBB4A,PINK1,MFN2,PARK7,RHEB,CSNK2B,HSF1,UBE2V1,TUBA1C,ATG9A,GABARAPL2,DY
1,GNAI1
RPS6KA2,MAPK3,DNM1,ID3,ID2
3B4A,RANGAP1,PPP2R5D,DAAM1,SRF,TUBA1C,RCC2,NUP133,PPP2R5B,DYNC1I2,CENPF,CENPT,
MARCKS,CALM1,PLD3,ATP5F1A,UBB,CKB,RPL37A,PSMD5,HACD3,TPI1,CHSY1,RPL19,PPP2R1A,
SKP1,CDKN1A,PPP1CB,TNPO1,PSMB2,H3-3A,TERF2IP,TUBB2A,OPTN,PPP2CA,CCND1,MAPK1,M

D4,LRRC41,FBXW5,PSMC3,UBE3A,UBE2D3,PSMC1,UBE2K,FBXL19,TRIP12,UBE2L3,CDC34,PSM
P3,IRS1,SOS1,SHC1,RPS6KA2,MAPK3,FURIN,DNM1,ID3,ID2

,PSMD10,PSMA6,RELA,PSMD12,PSMA4,PSMC6

BN2,RAB7A,CLTC,YWHAB,UBE2H,HLA-B,RAC1,SKP1,CTSB,UBE4A,AHCYL1,KLC1,COL2A1,ORAI2
PSMC1,PTPN11,ABL1,AUTS2,PSMB1,SIN3B,PSMD13,PSMD7,HIPK2,TNRC6A,CSNK2B,H2AX,LDB1,

SMA1,PSMD10,MAP2K4,PSMA6,TAB3,RELA,PSMD12,PTPN12,PSMA4,PSMC6,TOLLIP

,PSMB2,VEGFA,RAPGEF1,PPP2CA,CCND1,MAPK1,SMARCA4,PSMD3,GRB2,DUSP4,PITPNA,FYN,S

(3B,CDKN1B,PSMB5,PSMA1,PSMD10,PSMA6,PCNA,TFDP1,MCM7,PSMD12,PSMA4,UBE2E1,PSM(
SKIL

CTN1,FBLN1,COL27A1,PTPRS,P3H3,CAPN2,MMP15,CRTAP,ITGB5,LOX,A2M,HSPG2,LTBP4,SH3F

2,PSMA4,UBE2E1,PSMC6,ANAPC2

T1,CARM1,KMT2E,ELP1,NSD2,OGT,KDM4A,MBD3,SMARCD1,CREBBP,SAP18,ATXN7L3,MTA2,KD
T1,CARM1,KMT2E,ELP1,NSD2,OGT,KDM4A,MBD3,SMARCD1,CREBBP,SAP18,ATXN7L3,MTA2,KD

FS1,ATP5F1C,COX6B1,SLC16A1,UQCRFS1,ATP5F1E,RXRA,COX4I1,OGDH,COX5A,UQCRH,FH,PDI

,USP24,PSMC3,YY1,PSMC1,TOMM70,USP5,USP47,SNX3,USP10,PSMB1,MDM2,PSMD13,USP14,T

AGRN,RPLP1,RPL10,RPS17,AP2B1,RPS16,RPL23,RPL29,RPSA,RPS24,RPS9,MSN,RPL24,RPL17,(
AGRN,RPLP1,RPL10,RPS17,AP2B1,RPS16,RPL23,RPL29,RPSA,RPS24,RPS9,MSN,RPL24,RPL17,(
S10,RPL35A,RPLP2,RPL36,RPS12,RPS27,RPL23A,RPL27,RPL36A,RPS15A,RPL38,RPL37,EIF3E,R
RPL35A,RPLP2,RPL36,RPS12,RPS27,RPL23A,RPL27,PSMB2,RPL36A,RPS15A,RPL38,RPL37,PSM
,RPS10,RPL35A,RPLP2,RPL36,RPS12,RPS27,RPL23A,RPL27,RPL36A,RPS15A,RPL38,RPL37,EIF3
PL35,RPS10,RPL35A,RPLP2,RPL36,RPS12,RPS27,RAC1,GPC1,RPL23A,RPL27,PSMB2,RPL36A,RI
PL35,RPS10,RPL35A,RPLP2,RPL36,RPS12,RPS27,RPL23A,RPL27,RPL36A,RPS15A,RPL38,RPL37
PL35,RPS10,RPL35A,RPLP2,RPL36,RPS12,RPS27,RPL23A,RPL27,RPL36A,RPS15A,RPL38,RPL37
,RPS12,RPS27,RPL23A,RPL27,RPL36A,RPS15A,RPL38,RPL37,EIF3E,RPS11,RPL28,RPS28,FAU,E
S12,RPS27,RPL23A,RPL27,RPL36A,RPS15A,RPL38,RPL37,RPS11,RPL28,RPS28,FAU,RPL39,RPS
RPL36,RPS12,RPS27,RPL23A,SRP9,RPN2,RPL27,RPL36A,SRP14,RPS15A,RPL38,RPL37,RPS11,RF
RPLP2,RPL36,RPS12,RPS27,RPL23A,RPL27,RPL36A,RPS15A,RPL38,RPL37,PPP2CA,RPS11,CASC
RPLP2,RPL36,RPS12,RPS27,RPL23A,RPL27,RPL36A,RPS15A,RPL38,RPL37,PPP2CA,RPS11,CASC
36,RPS12,RPS27,RPL23A,EEF1A2,RPL27,RPL36A,RPS15A,RPL38,RPL37,RPS11,RPL28,RPS28,FA
27,RPL23A,RPL27,RPL36A,RPS15A,RPL38,RPL37,RPS11,RPL28,RPS28,FAU,RPL39,RPS13,RPL3
36,RPS12,RPS27,RPL23A,RPL27,RPL36A,RPS15A,RPL38,RPL37,RPS11,RPL28,RPS28,FAU,RPL39
.24,RPL17,RPL14,RPL22,RPL9,ATF4,PSMD2,RPS14,RPS20,RPL26,RBBP4,RPL31,RPS25,PRDX2,F
2,RPS27,RPL23A,RPL27,GCN1,RPL36A,RPS15A,RPL38,RPL37,RPS11,RPL28,RPS28,FAU,RPL39,F
27,RPL23A,RPL27,RPL36A,RPS15A,RPL38,RPL37,RPS11,RPL28,RPS28,FAU,RPL39,RPS13,RPL3
.24,RPL17,RPL14,RPL22,RPL9,ATF4,PSMD2,RPS14,RPS20,RPL26,RBBP4,RPL31,RPS25,PRDX2,F
PL24,RPL17,PAPOLA,RPL14,RPL22,RPL9,PSMD2,RPS14,RPS20,RPL26,SRSF4,RPL31,HNRNPC,XF
27,RPL23A,RPL27,RPL36A,RPS15A,RPL38,RPL37,RPS11,RPL28,RPS28,FAU,SARS1,RPL39,RPS1
27,RPL23A,RPL27,RPL36A,RPS15A,RPL38,MARS1,RPL37,RPS11,QARS1,RPL28,RPS28,LARS1,F
19,RPL35,RPS10,RPL35A,RPLP2,RPL36,RPS12,RPS27,IPO5,RPL23A,PABPN1,PARP1,RPL27,RPL
,TRIM28,RPL37A,FN1,DDB1,PSMD5,EIF4A1,RPL19,RPL15,PDIA3,RPS3,CHD3,SEC61A1,RPS4X,RF
2,RPS27,ATP6V0C,RPL23A,RPL27,GCN1,RPL36A,RPS15A,RPL38,RPL37,RPS11,RPL28,RPS28,AT
,RPL36,RPS12,RPS27,IPO5,RPL23A,PARP1,RPL27,RPL36A,RPS15A,RPL38,RPL37,RPS11,RPL28,
L30,EIF3L,RPS5,RPS7,RPS19,RPL35,RPS10,RPL35A,RPLP2,DDOST,NARS1,RPL36,RPS12,RPS27,
B4B,RPL11,RPLP1,RPL10,RPS17,ARF1,AP2B1,GTF2F1,RPS16,RPL23,RPL29,RPSA,RPS24,RPS9

.15,RPS3,CHD3,AP2M1,RPS4X,RPS3A,RPL18,ITGB1,TUBB4B,RPL11,AGRN,RPLP1,RPL10,RPS17,

6,RPS12,RPS27,RPL23A,TSR1,RPL27,RPL36A,RPS15A,RPL38,RPL37,RPS11,RPL28,RPS28,FAU,(

RPL11,NCAM1,AGRN,RPLP1,RPL10,RPS17,AP2B1,CTNNA1,RPS16,RPL23,RPL29,RPSA,RPS24,RF
6,RPS12,RPS27,RPL23A,TSR1,RPL27,RPL36A,RPS15A,RPL38,RPL37,RPS11,RPL28,RPS28,FAU,(
,RPS7,RPS19,RPL35,RPS10,RPL35A,RPLP2,RPL36,RPS12,RPS27,RPL23A,GLUL,RPL27,PSMB2,I
6,RPS12,RPS27,RPL23A,TSR1,RPL27,RPL36A,RPS15A,RPL38,RPL37,RPS11,RPL28,RPS28,FAU,(
C,SRSF6,TRA2B,HNRNPA2B1,CHERP,SYMPK,FUS,SNRPD3,ELAVL1,EFTUD2,SRSF2,RBM5,WBP1
C,SRSF6,TRA2B,HNRNPA2B1,CHERP,SYMPK,FUS,SNRPD3,ELAVL1,EFTUD2,SRSF2,RBM5,WBP1
AB5B,RAB11B,TMED2,TUBB2A,OPTN,ARF3,MVB12B,KDELR2,ARPC2,DYNC1LI2,DYNLL2,CAPZB,
,SF1,PRCC,SRSF6,TRA2B,HNRNPA2B1,POM121C,CHERP,SYMPK,FUS,SNRPD3,ELAVL1,EFTUD2,S
C

:1,ATP6AP1,VEGFA,RAPGEF1,PPP2CA,MAPK1,PTBP1,CDC37,HSPB1,PLCG1,MAP2K2,DUSP4,POLI

PSMC4,PSMF1,PSMB5,PSMA1,PSMD10,MAPK3,MAF1,PSMA6,GATAD2B,PSMD12,PSMA4,CBX4,PS

,NUP205,NUP133,PSMA1,PSMD10,ELOB,PSMA6,PSMD12,PSMA4,NUP50,PSMC6
RAB1A,ARCN1,RAC1,RAB27A,VAMP2,GDI2,HSPH1,ACTR3,TBC1D16,KLC1,SURF4,RAB5B,RAB11

3,PAPSS1,HDAC6,PTEN,PPP1CC,RPS27A,TNKS,PSMD8,FKBP1A,AKAP9,PSMB4,AKT3,NCSTN,PDPK
1GAT5,SUMO1,ATP1B3,DAD1,MOGS,TYK2,JAK1,STAT2,GSK3B,ROCK2,SUDS3,MAN1B1,GATAD2

AS,ARFGAP2,RAB18,TUBA1C,AGPAT3,KIF3A,NAPB,NAPA,GALNT2,CAPZA2,COPB1,DYNC1I2,ARFG

CA,PSMC1,TUBB4A,MZT2B,PSMB1,PCM1,CUL1,PSMD13,PSMD7,CETN2,PPME1,PSME3,PSMC5,I

3,PSMD7,C1QBP,PSME3,STUB1,PSMC5,PSMC2,TNFRSF10B,PSMC4,PSMF1,YWHAZ,PSMB5,PSM

CA,PSMC1,TUBB4A,MZT2B,PSMB1,PCM1,CUL1,PSMD13,PSMD7,CETN2,PPME1,PSME3,PSMC5,I
TADA2B,KLHL9,SPTBN2,EIF5A,CDH2,PDIA6,GPC3,RAB7A,RCN1,GOLM1,APLP2,DDOST,ST3GAL2
MB1,TCEA1,SLC25A6,PSMD13,BANF1,PSMD7,RAN,NUP153,CDK9,PSME3,PSIP1,PSMC5,PSMC2,I

PSMD4,AKT2,MBD3,GSK3A,PSMC3,MTOR,PRKACA,PSMC1,PTPN11,MTA2,PSMB1,MDM2,PSMD13
MEM30A,PGRMC1,CNN2,CTSB,RAB27A,CCT2,GDI2,PDXK,MLEC,SURF4,RAB5B,SRP14,PYGB,MAF

PSMD10,PSMA6,PSMD12,PSMA4,PSMC6

SMB1,MDM2,PSMD13,PSMD7,TNRC6A,RHEB,PDGFRB,CSNK2B,TRIM27,TRIB3,PPP2R5D,PSME3
APK3,VAV2,EPHB2,NFASC,DNM1

AP1,PSMB1,PSMD13,SUMO1,BANF1,PSMD7,RAN,PPP2R5D,PSME3,PSMC5,PSMC2,PSMC4,PSMF1

SF10B,PSMC4,PSMF1,YWHAZ,PSMB5,PSMA1,PSMD10,MAPK3,PSMA6,ADD1,TFDP1,HMGB2,PSM

AP1,PSMB1,PSMD13,SUMO1,BANF1,PSMD7,RAN,PPP2R5D,PSME3,PSMC5,PSMC2,PSMC4,PSMF1

I2,DYNLL2,KDM1A,YWHAH,PFN2,DYNLL1,CDC42,KIF5A,H2AZ1,PPP2CB,EVL,PPP1CC,FMNL2,CTT

3,PSMB4,SOX2,TLE1,PSMD4,SCRIB,AKT2,PSMC3,AP2A2,CREBBP,PSMC1,SNX3,CLTA,PSMB1,CUL

C,KIF3A,PSMB5,PRKAR2B,PSMA1,PSMD10,PSMA6,PSMD12,PSMA4,PSMC6

E3,PSMC5,PSMC2,PSMC4,PSMF1,TUBA1C,RCC2,PSMB5,NUP133,PSMA1,PSMD10,PSMA6,PPP2R5

,PSMD12,PSMA4,PSMC6,GNAO1,FZD7,FZD8

NSF,COPE,KIF26A,KLC2,NBAS,VPS51,ARFGAP2,RAB18,GOLGA4,TUBA1C,AGPAT3,KIF3A,NAPB,N

VHL,PPP2CA,MAPK1,MAPRE1,ARPC2,DYNC1LI2,DYNLL2,CDC37,KDM1A,YWHAH,HGS,PFN2,DYNLL

VHL,PPP2CA,MAPK1,MAPRE1,ARPC2,DYNC1LI2,DYNLL2,CDC37,KDM1A,YWHAH,HGS,PFN2,DYNLL

A,COL7A1,CAPZA2,TRAPPC1,SEC23IP,COPB1,SEC31A,DYNC1I2,COG4,ARFGAP3
7,TNRC6A,H2AX,PSME3,PSMC5,PSMC2,PSMC4,PSMF1,YWHAZ,PSMB5,STAT1,PSMA1,CNTN1,PSM

KACA,PSMC1,PTPN11,PSMB1,PPP5C,RGL2,DNAJB1,PSMD13,RALGDS,RASA4,PSMD7,TNRC6A,PD
2A,MASTL,SMC3,PPP1CC,POM121C,RPS27A,PSMD8,AKAP9,PSMB4,TAOK1,RAB1B,VPS4A,TUBG

,UBA52,MGAT4B,YKT6,RPS27A,LMAN1,MVD,RAB1B,CAPZA1,COPZ1,PSMC1,TUBB4A,USO1,ARF
SMC6

1,VAT1,CLEC7A,PRKCSH,PFKL,RAB7A,DDX3X,MIF,JUN,MYO5A,CTNNB1,CD63,CYFIP2,DNAJC5,D
6PR,NECAP1

DS,RASA4,PSMD7,PDGFRB,TYK2,JAK1,SYNGAP1,PPP2R5D,PSME3,PSMC5,PSMC2,RANBP9,PSMC

5D,PSME3,BCL9,PSMC5,CSNK1A1,PSMC2,PSMC4,PSMF1,YWHAZ,GSK3B,LGR4,SOX9,PSMB5,CH

,PSMD7,PDGFRB,JAK1,SYNGAP1,PPP2R5D,PSME3,PSMC5,PSMC2,RANBP9,PSMC4,PSMF1,ABHI

,DYNLL1,PSMA7,ACTR1A,EP300,PSMD1,H2AZ1,CSNK2A1,UBA52,PPP2CB,RAB2A,MASTL,SMC3,P

D12,PSMA4,PSMC6,PPP2R2A

A1C,ATG9A,GABARAPL2,DYNC1I2,NBR1,VPS28,TOMM40,VPS37B,RB1CC1
RFGAP2,TUBA1C,MCFD2,NAPB,NAPA,COL7A1,CAPZA2,B4GALT2,TRAPPC1,SEC23IP,COPB1,SEC3

:4A1,MRC2,PPP2R1A,HLA-C,PDIA3,VAPA,HMGB1,AP2M1,SEC61A1,ITGB1,TUBB4B,NCAM1,ANXA2

K1,AKT2,NOC2L,COX6A1,MBD3,MTOR,CREBBP,MTA2,TXNRD1,TCEA1,MDM2,NDUFA4,EHMT2,C

C6
PSMC5,PSMC2,PSMC4,PSMF1,PSMB5,HIF3A,PSMA1,PSMD10,ELOB,ASB6,WDR5,GPS1,PSMA6,C

PSMC3,UBE3A,UBE2D3,PSMC1,UBE2K,FBXL19,TRIP12,UBE2L3,TAPBP,CDC34,PSMB1,FZR1,CU
NC1I2,NBR1,TOMM40,RB1CC1

PPP2R5C
RPL15,LDHB,SERINC1,RPS3,VAPA,RPS4X,RPS3A,RPL18,RPL11,AGRN,RPLP1,RPL10,RPS17,SBF1,
APRE1,PSMD3,DYNC1LI2,DYNLL2,POLR2B,YWHAH,CSNK1D,DYNLL1,TP53BP1,POLR2E,PSMA7,F

B1,FZR1,CUL1,PSMD13,PJA1,PSMD7,UBE2J1,AREL1,RNF41,PSME3,UBE2Q1,STUB1,PSMC5,PSM

2,PSMB2,TUBB2A,VHL,PPP2CA,PSMD3,DYNC1LI2,DYNLL2,CAPZB,UBE2R2,NPEPPS,PLCG1,KIF3C
,PSME3,PSMC5,PSMC2,PSMC4,PSMF1,PSMB5,PSMA1,PSMD10,WDR5,PSMA6,PBRM1,PSMD12,F

OD1,CDC42,PSMA7,STAT3,TXLNA,PSMD1,UBA52,PPP2CB,HNRNPA2B1,IL6ST,RPS27A,PSMD8,HI

C6,ANAPC2,WAPL

PXD2A,NCSTN,LTBP3,ITGAV,ADAM10,FBLN5,BGN,COL16A1,COL12A1,DAG1,ELN,JAM3,CD44,HT[illegible]

M3A,RUVBL2,TRRAP,KAT2A,EHMT2,ATF7IP,CDK4,H2AX,NSD1,ASH1L,PRMT5,KDM2A,KDM5A,SE[illegible]
M3A,RUVBL2,TRRAP,KAT2A,EHMT2,ATF7IP,CDK4,H2AX,NSD1,ASH1L,PRMT5,KDM2A,KDM5A,SE[illegible]

HB,NNT,NDUFA5,UQCR11,COX5B,NDUFA10,ATP5PD

TRRAP,KAT2A,PSMD7,ADRM1,OTUD5,PSME3,PSMC5,PSMC2,PSMC4,PSMF1,USP48,USP19,PSMB

GAP43,RPL14,RPL22,RHOA,RPL9,PSMD2,RPS14,RPS20,COL5A2,RPL26,CRMP1,PFN1,RPL31,SPT
GAP43,RPL14,RPL22,RHOA,RPL9,PSMD2,RPS14,RPS20,COL5A2,RPL26,CRMP1,PFN1,RPL31,SPT
PS11,RPL28,RPS28,FAU,EIF3F,RPL39,RPS13,RPL34,EIF3D,UBA52,EIF5B,RPS2,RPS27A,EIF2S2,E
ID3,RPS11,CASC3,RPL28,RPS28,FAU,RPL39,RPS13,RPL34,PSMA7,PSMD1,GSPT1,UBA52,RPS2,F
E,RPS11,RPL28,RPS28,FAU,EIF3F,RPL39,RPS13,RPL34,EIF3D,UBA52,RPS2,RPS27A,EIF2S2,EIF3
PS15A,RPL38,RPL37,PSMD3,RPS11,CASC3,RPL28,RPS28,FAU,SLIT3,PFN2,RPL39,RPS13,RPL34,
,EIF3E,RPS11,RPL28,RPS28,FAU,EIF3F,RPL39,RPS13,RPL34,EIF3D,UBA52,EIF5B,RPS2,RPS27A,
,EIF3E,RPS11,RPL28,RPS28,FAU,EIF3F,RPL39,RPS13,RPL34,EIF3D,UBA52,EIF5B,RPS2,RPS27A,
EIF3F,RPL39,RPS13,RPL34,EIF3D,UBA52,RPS2,RPS27A,EIF3H,EIF3G,RPS26,RPS21,RPL27A,EIF3
13,RPL34,GSPT1,UBA52,UPF1,RPS2,RPS27A,EIF4G1,RPS26,RPS21,ETF1,RPL27A,NCBP2,RPS29,
PL28,RPS28,FAU,RPL39,RPS13,RPL34,SSR1,UBA52,RPS2,RPS27A,TRAM1,SRP72,SSR3,SSR2,SRF
3,RPL28,RPS28,FAU,RPL39,RPS13,RPL34,GSPT1,UBA52,UPF1,RPS2,RPS27A,SMG5,EIF4A3,EIF4
3,RPL28,RPS28,FAU,RPL39,RPS13,RPL34,GSPT1,UBA52,UPF1,RPS2,RPS27A,SMG5,EIF4A3,EIF4
U,RPL39,RPS13,RPL34,UBA52,RPS2,RPS27A,EEF1B2,RPS26,RPS21,RPL27A,RPS29,RPL36AL,RF
4,GSPT1,UBA52,RPS2,RPS27A,RPS26,TRMT112,RPS21,APEH,ETF1,RPL27A,RPS29,RPL36AL,RPL
,RPS13,RPL34,UBA52,RPS2,RPS27A,RPS26,RPS21,RPL27A,RPS29,RPL36AL,RPL32
RPS15,RPL12,RPL30,H1-0,RPS5,RPS7,RPS19,RPL35,JUN,RPS10,RPL35A,RPLP2,RPL36,PTGES3,
RPS13,RPL34,UBA52,RPS2,RPS27A,EIF2S2,RPS26,RPS21,ASNS,RPL27A,TRIB3,RPS29,CEBPG,RF
4,GRSF1,UBA52,RPS2,RPS27A,RPS26,RPS21,RPL27A,RPS29,RPL36AL,RPL32
RPS15,RPL12,RPL30,H1-0,RPS5,RPS7,RPS19,RPL35,JUN,RPS10,RPL35A,RPLP2,RPL36,PTGES3,
PO1,CSNK1E,RPS25,RPS15,RPL12,RPL30,RPS5,RPS7,RPS19,RPL35,RPS10,RPL35A,RPLP2,HNRN
3,RPL34,UBA52,RPS2,RPS27A,RPS26,RPS21,RPL27A,RPS29,RPL36AL,RPL32
AU,SARS1,RPL39,RPS13,RPL34,UBA52,PAPSS1,IARS1,RPS2,RPS27A,AHCY,RPS26,TXNRD1,RPS21
36A,RPS15A,RPL38,RPL37,RPS11,RPL28,POLR2B,RPS28,FAU,RPL39,RPS13,RPL34,POLR2E,GRS
PS3A,RPL18,TUBB4B,UCHL1,RPL11,RPLP1,RPL10,MARCHF6,WSB1,RPS17,ARF1,ITM2B,RPS16,F
P6V1B2,FAU,RPL39,RPS13,RPL34,FLCN,UBA52,RPS2,RPS27A,EIF2S2,ATP6V0D1,SH3BP4,MTOR,
,POLR2B,RPS28,FAU,RPL39,RPS13,RPL34,POLR2E,GRSF1,UBA52,POM121C,RPS2,RPS27A,RPS
,RPL23A,SRP9,EEF1A2,RPN2,MTRF1L,RPL27,RPL36A,SRP14,RPS15A,RPL38,MARS1,RPL37,EIF3
,XRCC5,RPL24,KPNB1,RPL17,RPL14,RPL22,RPL9,PSMD2,RPS14,CHD4,ATP1A1,RPS20,RPL26,R

,VCL,ARF1,AP2B1,GTF2F1,RPS16,RPL23,RPL29,RPSA,RPS24,RPS9,XRCC5,RPL24,KPNB1,RPL17

CSNK1D,RPL39,RPS13,RPL34,UBA52,RPS2,RPS27A,PELP1,RPS26,XRN2,NOP56,MTREX,RPS21,R

PS9,MSN,RPL24,RPL17,GAP43,RPL14,RPL22,RHOA,RPL9,PSMD2,RPS14,RPS20,COL5A2,RPL26,C
CSNK1D,RPL39,RPS13,RPL34,UBA52,RPS2,RPS27A,PELP1,RPS26,XRN2,NOP56,MTREX,TRMT11
RPL36A,RPS15A,RPL38,MARS1,RPL37,AZIN1,PSMD3,ODC1,RPS11,QARS1,RPL28,RPS28,LARS1,
CSNK1D,RPL39,RPS13,RPL34,UBA52,RPS2,RPS27A,PELP1,RPS26,XRN2,NOP56,MTREX,TRMT11
1,SRSF11,PRPF19,SF3A2,SART1,DHX38,EIF4A3,MTREX,SF3A3,CD2BP2,HNRNPR,SRSF10,LSM4,S
1,SRSF11,PRPF19,SF3A2,SART1,DHX38,EIF4A3,MTREX,SF3A3,CD2BP2,HNRNPR,SRSF10,LSM4,S
GABARAP,KDELR1,ARF4,KIF3C,YWHAH,UBQLN1,CSNK1D,TMED9,HGS,RAB11A,PPP6R1,SEC23A
SRSF2,RBM5,WBP11,SRSF11,PRPF19,SF3A2,SART1,DHX38,EIF4A3,MTREX,SF3A3,CD2BP2,HNRN

R2B,ATP6V1B2,FYN,HNRNPM,ID4,COL27A1,PTPRS,HGS,VGF,CDC42,CTNND1,POLR2E,STAT3,EF

SMC6

B,TMED2,TUBB2A,OPTN,ARF3,MVB12B,KDELR2,ARPC2,DYNC1LI2,DYNLL2,CAPZB,GABARAP,KD

K1,PSMD4,ADAM10,AKT2,RBPJ,GSK3A,PSMC3,MTOR,CREBBP,PSMC1,FGFR1,PTPN11,PSMB1,TI
B,FURIN,HMG20B

AP3

PSMC2,PSMC4,PSMF1,TUBA1C,PSMB5,PRKAR2B,PSMA1,PPP1R12A,PSMD10,PSMA6,AJUBA,PSM

IA1,PSMD10,MAPK3,PSMA6,ADD1,TFDP1,HMGB2,PSMD12,OPA1,PSMA4,ACIN1,PSMC6,GSN

PSMC2,PSMC4,PSMF1,TUBA1C,PSMB5,PRKAR2B,PSMA1,PPP1R12A,PSMD10,PSMA6,AJUBA,PSM
2,HNRNPK,UBE2H,CKAP4,RAB1A,CALU,ARCN1,CTBP1,TOP2B,HDAC2,TNC,SKP1,TMEM132A,NUC
NCBP2,RNMT,PSMC4,PSMF1,PSMB5,NUP205,NUP133,PSMA1,NELFCD,PSMD10,ELOB,CHMP5,PS

3,PSMD7,TNRC6A,RHEB,PDGFRB,CSNK2B,TRIM27,TRIB3,ADCY1,PPP2R5D,PSME3,STUB1,PSMC
PK1,PSMD3,CAND1,ILF2,CCT8,B2M,IDH1,DYNLL1,TXNDC5,PSMD1,PDAP1,CST3,ATP6AP2,PRDX6,

,STUB1,PSMC5,PSMC2,PSMC4,PSMF1,PIP5K1C,GSK3B,CDKN1B,PSMB5,IRS1,ERBB2,PSMA1,PS

,TUBA1C,RCC2,PSMB5,NUP205,NUP133,PSMA1,PSMD10,PSMA6,PPP2R5B,PSMD12,PSMA4,DYN
1D12,OPA1,PSMA4,ACIN1,PSMC6,GSN
,TUBA1C,RCC2,PSMB5,NUP205,NUP133,PSMA1,PSMD10,PSMA6,PPP2R5B,PSMD12,PSMA4,DYN

N,BRK1,TAOK1,ABI2,CLASP2,PDPK1,WASF2,MYL12B,TUBB4A,RANGAP1,ABL1,KTN1,ARPC1A,C
_1,PSMD13,TRRAP,PSMD7,TNRC6A,CSNK2B,H2AX,PPP2R5D,PSME3,BCL9,DAAM1,PSMC5,CSNK

5B,PSMD12,PSMA4,DYNC1I2,UBE2E1,CENPF,PSMC6,CENPT,ANAPC2,WAPL,PPP2R5C

NAPA,GALNT2,CAPZA2,COPB1,DYNC1I2,COG4,SCOC,CYTH3,M6PR,ARFGAP3

L1,CDC42,KIF5A,BCR,TRIO,H2AZ1,PPP2CB,EVL,PPP1CC,CUL3,FMNL2,CTTN,A2M,BRK1,TAOK1,
L1,CDC42,KIF5A,BCR,TRIO,H2AZ1,PPP2CB,EVL,PPP1CC,CUL3,FMNL2,CTTN,A2M,BRK1,TAOK1,

ʼD10,PSMA6,MIB2,TFDP1,PRKCI,PSMD12,FURIN,PSMA4,MAML1,NBEA,PSMC6,PLXND1,TLE4

)GFRB,TYK2,JAK1,SYNGAP1,PPP2R5D,PSME3,PSMC5,PSMC2,RANBP9,PSMC4,PSMF1,ABHD17A
CP6,CLASP2,PSMD4,IST1,ARPP19,PSMC3,CHMP4B,PRKACA,UBE2I,PSMC1,TUBB4A,USO1,RANG

5,ANK2,AMFR,LMAN2,MGAT5,MGAT3,NSF,COPE,B4GALT5,DAD1,ARFGAP2,MOGS,NUS1,TUBA1(

)DOST,LAMP1,CKAP4,HLA-B,TTR,COTL1,HRNR,RAC1,ATP6V0C,TMEM30A,SKP1,PGRMC1,CNN2,

;4,PSMF1,ABHD17A,DLG4,PSMB5,DUSP8,IRS1,ERBB2,PSMA1,PHB,SOS1,SHC1,PSMD10,ZDHHC9

ID8,PSMA1,PSMD10,PSMA6,PPP2R5B,PSMD12,CTNNBIP1,PSMA4,PSMC6,TLE4,FZD8,LRP6,PPP2

)17A,DLG4,PSMB5,DUSP8,IRS1,ERBB2,PSMA1,PHB,SOS1,SHC1,PSMD10,ZDHHC9,MAPK3,PSMA

ʼPP1CC,POM121C,RPS27A,PSMD8,AKAP9,PSMB4,TAOK1,RAB1B,VPS4A,AKT3,TUBGCP6,CLASP2

1A,DYNC1I2,COG4,ARFGAP3

2,TIMP2,XAF1,WSB1,VCL,ARF1,CYB5R3,AP2B1,PGAM1,CTSD,LPCAT1,LAMA5,XRCC5,PJA2,MSN,

OX8A,HIPK2,TNRC6A,RHEB,CSNK2B,COX7A2L,CCNG1,CDK9,NDRG1,TNFRSF10B,PLK2,YWHAZ,

UL7,BTBD1,CUL4B,PSMD12,PSMA4,FBXO7,COPS8,PSMC6,NEDD8,SPSB3

L1,PSMD13,PJA1,HLA-A,PSMD7,UBE2J1,AREL1,RNF41,PSME3,UBE2Q1,STUB1,PSMC5,PSMC2,F

,ARF1,CYB5R3,MAT2A,PGAM1,FADS2,RPS16,RPL23,LDHA,RPL29,RPSA,LPCAT1,RPS24,RPS9,RP
BBP7,ACTR1A,EP300,PSMD1,H2AZ1,CSNK2A1,UBA52,PPP2CB,ATRX,HJURP,RAB2A,MASTL,SMC

IC2,PSMC4,PSMF1,UBE3C,MEX3C,UBE2V1,PSMB5,PSMA1,ZNRF1,RNF4,MGRN1,PSMD10,UBE2

,FYN,B2M,HECTD1,SEC23A,DYNLL1,CDC42,PSMA7,TRAF7,DCTN1,KIF5A,ACTR1A,PSMD1,AP2A1
YBP,PSMA4,CBX4,SETD1A,SMARCC2,PSMC6,SMARCB1,MYL9,SETD1B,SIN3A

F1A,NDN,PSMB4,SOX2,CAPZA1,SOD2,PSMD4,PSMC3,PRKACA,PSMC1,PTPN23,PTPN11,PSMB1,C

RA1,CAPN1,MFAP4,EMILIN1,COLGALT1,LOXL2,PCOLCE,P3H1,COL7A1,NID1,COL6A3,TGFB2,FU

TD3,WDR5,SUDS3,PBRM1,RELA,GATAD2B,BRPF3,SETD1A,HMG20B,KDM2B,SMARCC2,SMARCB
TD3,WDR5,SUDS3,PBRM1,RELA,GATAD2B,BRPF3,SETD1A,HMG20B,KDM2B,SMARCC2,SMARCB

5,OTUB1,VDAC2,PSMA1,PSMD10,SUDS3,TGFBR1,PSMA6,TRAF3,PSMD12,MBD6,PSMA4,PSMC6,

TBN2,RPS25,PLXNA1,RPS15,RPL12,RPL30,CLTC,RPS5,RPS7,RPS19,RPL35,TLN1,COL6A2,RPS10
TBN2,RPS25,PLXNA1,RPS15,RPL12,RPL30,CLTC,RPS5,RPS7,RPS19,RPL35,TLN1,COL6A2,RPS10
EIF3H,EIF3G,EIF4G1,RPS26,RPS21,RPL27A,EIF3B,EIF3M,RPS29,RPL36AL,RPL32,EIF2S1,EIF3J
RPS27A,PSMD8,SLIT2,PSMB4,SLIT1,MSI1,PSMD4,PSMC3,PSMC1,EIF4A3,EIF4G1,RPS26,PSMB1,[illegible]
3H,EIF3G,EIF4G1,RPS26,RPS21,RPL27A,EIF3B,EIF3M,RPS29,RPL36AL,RPL32,EIF2S1,EIF3J
CDC42,PSMA7,PRKAR2A,PSMD1,GSPT1,UBA52,EVL,RPS2,RPS27A,PSMD8,SLIT2,PSMB4,SLIT1,[illegible]
EIF2S2,EIF3H,EIF3G,EIF4G1,RPS26,RPS21,RPL27A,EIF3B,EIF3M,RPS29,RPL36AL,RPL32,EIF2S1
EIF2S2,EIF3H,EIF3G,EIF4G1,RPS26,RPS21,RPL27A,EIF3B,EIF3M,RPS29,RPL36AL,RPL32,EIF2S1
B,EIF3M,RPS29,RPL36AL,RPL32,EIF3J
,RPL36AL,RPL32
PRA,RPS26,RPS21,RPL27A,RPS29,SPCS2,SRP54,RPL36AL,RPL32,SEC11A
G1,RPS26,RPS21,PNRC2,ETF1,RPL27A,NCBP2,RPS29,RPL36AL,RPL32,RNPS1,PPP2R2A
G1,RPS26,RPS21,PNRC2,ETF1,RPL27A,NCBP2,RPS29,RPL36AL,RPL32,RNPS1,PPP2R2A
PL32
.32

RPS12,RPS27,STIP1,ATP6V0C,RPL23A,HSBP1,CDKN1A,NOX4,HSPH1,RPL27,HSPA4,GCN1,PSMB[illegible]
PL36AL,RPL32,EIF2S1

RPS12,RPS27,STIP1,ATP6V0C,RPL23A,HSBP1,CDKN1A,NOX4,HSPH1,RPL27,HSPA4,GCN1,PSMB[illegible]
IPK,RPL36,YWHAB,HNRNPH1,RPS12,RPS27,RPL23A,CPSF2,DDX39B,SRSF5,U2AF2,CPSF1,PABPN

,RPL27A,DARS1,RARS1,RPS29,KARS1,RPL36AL,RPL32
SF1,KPNA2,UBA52,POM121C,RPS2,RPS27A,RPS26,CLTA,HSPA1A,SLC25A6,RPS21,RAN,RPL27A,[illegible]
RPL23,CPE,RPL29,RPSA,CTSD,EIF4H,RPS24,RPS9,RPL24,RPL17,IGFBP3,RPL14,RPL22,RHOA,PT[illegible]
,ATP6V1G1,RPS26,RPS21,RHEB,ASNS,RPL27A,TRIB3,ATP6V0B,RPS29,CEBPG,ATP6V0E2,RPL36[illegible]
26,RPS21,RPL27A,NUP153,RPS29,NUP205,NUP133,RPL36AL,RPL32,NUP50
E,RPS11,QARS1,RPL28,RPS28,LARS1,FAU,EIF3F,SARS1,RPL39,RPS13,RPL34,SSR1,VARS1,EIF3[illegible]
BBP4,RPL31,PRKCSH,XPO1,RPS25,SFPQ,RAB7A,RPS15,RPL12,RPL30,CLTC,GNB2,RPS5,RPS7,[illegible]

,RPL14,HK1,RPL22,RPL9,PSMD2,RPS14,CHD4,ATP1A1,RPS20,RPL26,RBBP4,RPL31,PRKCSH,XP[illegible]

PL27A,EXOSC10,PDCD11,FBL,RPS29,SENP3,RPL36AL,GNL3,RPL32,PES1,NOP58,SNU13

CRMP1,RBBP4,PFN1,RPL31,SPTBN2,RPS25,CDH2,PLXNA1,RPS15,RPL12,RPL30,CLTC,NANOG,R[illegible]
2,RPS21,RPL27A,EXOSC10,PDCD11,FBL,RPS29,SENP3,RPL36AL,GNL3,RPL32,PES1,NOP58,SNU
FAU,SARS1,RPL39,RPS13,SMS,RPL34,PSMA7,PSMD1,UBA52,PAPSS1,GLUD1,IARS1,OAZ2,RPS2,[illegible]
2,RPS21,RPL27A,EXOSC10,PDCD11,FBL,RPS29,SENP3,RPL36AL,GNL3,RPL32,PES1,ELAC2,NOP
SRSF7,NCBP2,DNAJC8,SNRPD2,SNRPB,RBM17,SRRT,PRPF31,XAB2,RNPS1,CPSF7,WDR33,PLRG[illegible]
SRSF7,NCBP2,DNAJC8,SNRPD2,SNRPB,RBM17,SRRT,PRPF31,XAB2,RNPS1,CPSF7,WDR33,PLRG[illegible]
,DYNLL1,DCTN1,KIF5A,TXNDC5,CPD,ACTR1A,HIP1,AP2A1,UBQLN2,KIF21B,SEC16A,UBA52,RAB[illegible]
NPR,SRSF10,NUP153,LSM4,SRSF7,POLDIP3,NCBP2,DNAJC8,SNRPD2,SNRPB,NUP205,NUP133,T[illegible]

'300,AP2A1,UBA52,PPP2CB,SPINT2,RPS27A,BRK1,SHC2,ABI2,AKT3,SH3GL1,NCSTN,PDPK1,WASI

)ELR1,ARF4,KIF3C,YWHAH,UBQLN1,CSNK1D,TMED9,HYOU1,HGS,RAB11A,PPP6R1,SEC23A,DYN

RAK1,NOTCH1,MDM2,CUL1,PSMD13,FXR1,KAT2A,PSMD7,PDGFRB,PPP2R5D,PSME3,PSMC5,CSI

/ID12,PSMA4,TPX2,DYNC1I2,CENPF,PSMC6,PPP2R2A,NINL

/ID12,PSMA4,TPX2,DYNC1I2,CENPF,PSMC6,PPP2R2A,NINL
;B1,TOMM20,RAB27A,RPN2,PARP1,VCAN,MLEC,LAMC1,RAD23B,RAB5B,ASXL1,PSMB2,RAB11B,
SMA6,PSMD12,FURIN,PSMA4,NUP50,CCNK,PSMC6,VPS28,VPS37B,ELOA

5,PSMC2,PSMC4,PSMF1,PIP5K1C,GSK3B,CDKN1B,PSMB5,IRS1,PRKAR2B,ERBB2,PSMA1,PSMD
,GRN,ACLY,XRCC6,SLC44A2,IMPDH2,RAB14,NCSTN,RAB6A,RAB5C,ACTR1B,ITGAV,ADAM10,CAF

;MD10,MAPK3,MAF1,PSMA6,PPP2R5B,GATAD2B,PSMD12,PSMA4,CBX4,PIP5K1A,PSMC6,PIK3R3

IC1I2,UBE2E1,CENPF,PSMC6,PPP2R2A,CENPT,LEMD2,ANAPC2,WAPL,PPP2R5C,EMD

IC1I2,UBE2E1,CENPF,PSMC6,PPP2R2A,CENPT,LEMD2,ANAPC2,WAPL,PPP2R5C,EMD

YFIP1,LIMK1,KLC2,H2AX,PPP2R5D,ARPC3,DAAM1,SRF,YWHAZ,TUBA1C,RCC2,DLG4,CDKN1B,WA

1A1,PSMC2,PSMC4,PSMF1,YWHAZ,GSK3B,LGR4,SOX9,PSMB5,CHD8,PSMA1,PSMD10,CTBP2,PP

ABI2,ARHGEF17,CLASP2,PDPK1,WASF2,LRRC41,GNA13,MYL12B,TUBB4A,RANGAP1,ABL1,KTN1

ABI2,ARHGEF17,CLASP2,PDPK1,WASF2,LRRC41,GNA13,MYL12B,TUBB4A,RANGAP1,ABL1,KTN1

,DLG4,PSMB5,DUSP8,IRS1,ERBB2,PSMA1,PHB,SOS1,SHC1,PSMD10,ZDHHC9,MAPK3,PSMA6,PP[illegible]

AP1,MZT2B,PSMB1,PCM1,PSMD13,SUMO1,BANF1,PSMD7,RAN,CETN2,CSNK2B,NUP153,H2AX,[illegible]

[illegible],MCFD2,NAPB,UBXN1,NAPA,COL7A1,CAPZA2,B4GALT2,MAN1B1,TRAPPC1,SEC23IP,COPB1,SE[illegible]

CTSB,RAB27A,CCT2,GDI2,ACTR3,PDXK,AHCYL1,NCKAP1,MLEC,SURF4,RAB5B,PSMB2,SRP14,P[illegible]

[illegible],MAPK3,PSMA6,PPP2R5B,PSMD12,PSMA4,PSMC6,RAP1B,PPP2R5C

R5C

[illegible]6,PPP2R5B,PSMD12,PSMA4,PSMC6,RAP1B,PPP2R5C

,PSMD4,AKT2,IST1,ARPP19,PSMC3,CHMP4B,PRKACA,UBE2I,PSMC1,TUBB4A,USO1,RANGAP1,M[illegible]

KPNB1,COL1A2,RHOA,PSMD2,KIF5B,CXADR,CD81,VAT1,MCL1,CLEC7A,PRKCSH,KLHL9,SPTBN2

PRMT5,TXN,CDKN1B,COX6B1,NELFCD,COX4I1,ELOB,MDC1,COX5A,PIN1,CNOT3,PCNA,TFDP1,C

PSMC4,PSMF1,UBE3C,MEX3C,UBE2V1,PSMB5,PSMA1,ZNRF1,RNF4,MGRN1,PSMD10,UBE2G1,EL

L24,KPNB1,RPL17,RPL14,HK1,RPL22,RPL9,PSMD2,SLC7A5,RPS14,RPS20,RPL26,OAZ1,RPL31,L
3,PPP1CC,POM121C,RPS27A,PSMD8,PCBP4,AKAP9,PSMB4,TAOK1,RAB1B,VPS4A,AKT3,TUBGCP

G1,ELOB,ASB6,PSMA6,MIB2,CUL7,BTBD1,PSMD12,PSMA4,FBXO7,UBE2E1,PSMC6,ANAPC2,RBE

,CD99,UBA52,PPP2CB,KLHL42,EVL,UBE2G2,FBXO11,PTEN,CUL3,NPDC1,ITGB5,RPS27A,PSMD8

CUL1,PSMD13,PSMD7,MAPKAPK2,TYK2,JAK1,PPP2R5D,PSME3,CRKL,PSMC5,PSMC2,PSMC4,STA

RIN,COL3A1,ITGA7,MFAP2,CASK,MMP24

1,SETD1B,BRD1,PRMT7

1,SETD1B,BRD1,PRMT7

RAD23A,NEDD8

,RPL35A,RPLP2,MYL6,MYH10,RPL36,RHOB,RPS12,RPS27,RAC1,GPC1,MMP2,L1CAM,RPL23A,R
,RPL35A,RPLP2,MYL6,MYH10,RPL36,RHOB,RPS12,RPS27,HMGCR,RAC1,GPC1,HDAC2,MMP2,L

DAG1,PSMD13,RPS21,PSMD7,ETF1,RPL27A,LDB1,PSME3,PSMC5,PSMC2,NCBP2,PSMC4,PSMF1,

CLASP2,MSI1,PSMD4,CAP1,PSMC3,PRKACA,PSMC1,NELL2,EIF4A3,EIF4G1,RPS26,ABL1,PSMB1,D
,EIF3J
,EIF3J

2,H3-3A,RPL36A,KDM6B,TERF2IP,RPS15A,RPL38,VEGFA,TUBB2A,RPL37,VHL,MAPK1,PSMD3,PR

2,H3-3A,RPL36A,KDM6B,TERF2IP,RPS15A,RPL38,VEGFA,TUBB2A,RPL37,VHL,MAPK1,PSMD3,PR
N1,HNRNPL,TSR1,RPL27,SNRPN,HNRNPA3,RBMX,DDX42,TNPO1,SNRNP70,PSMB2,RPL36A,PRPF

NUP153,RPS29,NUP205,NUP133,KPNA3,RPL36AL,RPL32,NUP50
CD3,CBX5,RPL9,ATF4,PSMD2,KIF5B,RAB6B,RPS14,RPS20,EIF5,RPL26,CCT7,RPL31,PRKCSH,HN
AL,RPL32,SZT2,SESN2,EIF2S1,ATP6V1F,ATP6V1D

D,GSPT1,UBA52,GARS1,EIF5B,IARS1,RPS2,RPS27A,EIF2S2,IARS2,YARS1,EIF3H,TRAM1,SRP72,S
RPS19,RPL35,PRKAR1A,JUN,MYO5A,CTNNB1,RPS10,RPL35A,CYFIP2,RPLP2,DDOST,ST3GAL2,HN

PO1,SLC3A2,RPS25,SFPQ,PEBP1,CCND2,GPC3,RAB7A,PRDX2,RPS15,RPL12,RPL30,CLTC,GNB2,

PS5,RPS7,RPS19,RPL35,TLN1,JUN,CTNNB1,COL6A2,RPS10,RPL35A,RPLP2,MYL6,MYH10,RPL36
I13
RPS27A,PSMD8,PSMB4,SLC6A8,PHGDH,PSMD4,IVD,AHCY,PSMC3,GOT2,PSMC1,RPS26,PSAT1,P
58,SNU13
1,SNU13
1,SNU13
3GAP2,YKT6,ULK1,RPS27A,LMAN1,CTTN,RABEP1,MAP1LC3B,GRK2,CYTH2,RAB1B,VPS4A,AKT3,
HOC2,RBM17,SRRT,PRPF31,THOC3,FYTTD1,XAB2,NUP50,RNPS1,CPSF7,WDR33,PLRG1,SNU13

F2,ITGAV,ADAM10,AKT2,ATP6V0D1,MTOR,PRKACA,AP2A2,ATP6V1G1,NTRK2,PTPN11,CLTA,LRIG

LL1,DCTN1,KIF5A,TXNDC5,CPD,ACTR1A,HIP1,AP2A1,UBQLN2,KIF21B,SEC16A,UBA52,RAB3GAP

NK1A1,PSMC2,NCBP2,GOLGA4,PSMC4,PSMF1,GSK3B,CLCN6,CDKN1B,PSMB5,STAT1,DUSP8,IF

TMED2,TOP1,TUBB2A,ARF3,VHL,KDELR2,PSMD3,DYNC1LI2,DYNLL2,CAPZB,PRKDC,UBE2R2,CA

10,MAPK3,MAF1,PSMA6,PPP2R5B,GATAD2B,PSMD12,PSMA4,CBX4,PIP5K1A,NBEA,PSMC6,PIK3I
'1,IST1,PSMC3,AP2A2,AP1M1,PAFAH1B2,SDCBP,SCAMP1,PSMB1,HSPA1A,PSMD13,CAB39,NDUF

,PPP2R5C

ASF3,NUP133,PPP1R12A,ARPC5,ROCK2,MAPK3,PIN1,PPP2R5B,DYNC1I2,CENPF,CENPT,MYL9,NF
P3CA,PSMA6,PPP2R5B,PSMD12,CTNNBIP1,PSMA4,PSMC6,GNAO1,TLE4,FZD7,FZD8,LRP6,TMED

,ARPC1A,ARHGEF40,CYFIP1,LIMK1,KLC2,ARHGAP5,H2AX,PPP2R5D,ARPC3,DAAM1,SRF,YWHAZ,
,ARPC1A,ARHGEF40,CYFIP1,LIMK1,KLC2,ARHGAP5,H2AX,PPP2R5D,ARPC3,DAAM1,SRF,YWHAZ,

P2R5B,PSMD12,PSMA4,PSMC6,RAP1B,PPP2R5C
PPP2R5D,PSME3,PSMC5,PSMC2,PSMC4,PSMF1,TUBA1C,RCC2,PSMB5,PRKAR2B,NUP205,NUP13

C31A,DYNC1I2,COG4,RNF5,GLB1,ARFGAP3,CTSA

YGB,PPP2CA,MAPK1,ARPC2,PSMD3,PRKDC,CAND1,PLCG1,DUSP4,ATP6V1B2,FYN,ILF2,CCT8,B2

ZT2B,ABL1,PSMB1,FZR1,PCM1,CUL1,PSMD13,SUMO1,BANF1,PSMD7,RAN,CETN2,PPME1,CSNK

?,PFKL,RAB7A,DDX3X,CLTC,NANOG,MIF,JUN,MYO5A,CTNNB1,CD63,CYFIP2,DNAJC5,HNRNPDL,[illegible]

GATAD2B,BRPF3,MSH2,TPX2,CCNK,SESN2,PRELID1,BRD1,RAD50,HIPK1,COX5B,CNOT1,PPP2R5

LOB,ASB6,PSMA6,MIB2,CUL7,BTBD1,PSMD12,TLR6,PSMA4,SEC31A,FBXO7,UBE2E1,PSMC6,ANA

LRPPRC,SLC3A2,PFKL,RPS25,TKT,GPC3,PIP4K2B,ACSL6,RPS15,RPL12,RPL30,GNB2,RPS5,RPS7
P6,CLASP2,PSMD4,NSD2,AKT2,IST1,ARPP19,PSMC3,CHMP4B,PRKACA,UBE2I,PSMC1,TUBB4A,U[illegible]

3P6

,FKBP1A,FBXO21,PSMB4,UBE2M,UBE2D2,AKT3,KIFAP3,CAPZA1,ACTR1B,PDPK1,MKRN1,PSMD[illegible]

AT2,PSMF1,YWHAZ,ZEB1,UBE2V1,PSMB5,STAT1,DUSP3,IRS1,NKIRAS2,PSMA1,SOS1,SHC1,PSMD

PL27,PTPRA,ACTR3,COL2A1,LAMC1,PSMB2,RPL36A,DPYSL2,RPS15A,RPL38,TUBB2A,RPL37,MA
1CAM,RPL23A,RPL27,PTPRA,ACTR3,COL2A1,LAMC1,PSMB2,RPL36A,DPYSL2,RPS15A,RPL38,TU

RPS29,PSMB5,PSMA1,PSMD10,ELOB,ZSWIM8,PSMA6,RPL36AL,PSMD12,RPL32,PSMA4,RNPS1,

AG1,PSMD13,RPS21,NTN1,PSMD7,ETF1,RPL27A,NCK2,LDB1,PSME3,PSMC5,PSMC2,NCBP2,PS

DX1,DYNC1LI2,DYNLL2,CAPZB,RPS11,RPL28,CCAR2,RPS28,ATP6V1B2,PHC2,FAU,SOD1,RPL39,

DX1,DYNC1LI2,DYNLL2,CAPZB,RPS11,RPL28,CCAR2,RPS28,ATP6V1B2,PHC2,FAU,SOD1,RPL39,
6,RPS15A,RPL38,RPL37,PPP2CA,SRSF9,PSMD3,SRSF1,PTBP1,RPS11,SUPT5H,HNRNPUL1,HSPB

RNPC,TADA2B,KLHL9,SPTBN2,RPS25,EIF5A,CDH2,PDIA6,GPC3,RAB7A,RPS15,RPL12,RPL30,EIF

R3,EEF1B2,EIF3G,SSR2,SRPRA,EIF4G1,RPS26,TRMT112,RPS21,APEH,ETF1,RPL27A,DARS1,EIF
NRNPK,RPL36,PTGES3,ATP1A2,RPS12,RPS27,RAC1,HDAC2,IPO5,SKP1,RPL23A,RPN2,VAMP2,GN

RPS5,RPS7,RPS19,RPL35,PRKAR1A,TLN1,JUN,MYO5A,CTNNB1,RPS10,RPL35A,CYFIP2,RPLP2,D

,RHOB,RPS12,RPS27,HMGCR,RAC1,GPC1,HDAC2,MMP2,L1CAM,RPL23A,CDKN1A,RPL27,PTPR

SMB1,TXNRD1,PDHA1,PSMD13,RPS21,PSMD7,ASNS,ECHS1,RPL27A,DARS1,ALDH7A1,PSME3,PS

RAB14,LDLR,SH3GL1,KIFAP3,CAPZA1,GALNT1,RAB6A,FNBP1L,RAB5C,SYT11,AKT2,COPZ1,DEN

G1,TIA1,IGF1R,AAMP,CYFIP1,RALGDS,PTPN1,PDGFRB,RNF41,NCK2,TRIB3,MAPKAPK2,EPN1,PPP

2,YKT6,ULK1,RPS27A,LMAN1,CTTN,RABEP1,MAP1LC3B,GRK2,CYTH2,RAB1B,VPS4A,AKT3,RAB1

RS1,ERBB2,LMNA,PSMA1,PHB,SOS1,SHC1,PSMD10,CTBP2,TGFBR1,MAPK3,PSMA6,MIB2,PPP2R5

AND1,FKBP8,KDELR1,ARF4,PHC2,PTP4A2,TNKS2,STT3A,CSNK1D,TMED9,USP7,HGS,NCOR2,VGF

R3,PPP2R5C
C2,APEH,CYFIP1,CD44,PSMD7,FAF2,TSPAN14,CSNK2B,CAPN1,NME2,ASAH1,RAB18,PSMC2,DEG

:2,PPP2R5C
)5,PPP2R5C

TUBA1C,RCC2,ARHGEF9,DLG4,RALBP1,CDKN1B,WASF3,RHOBTB2,NUP133,SOS1,PPP1R12A,AF
TUBA1C,RCC2,ARHGEF9,DLG4,RALBP1,CDKN1B,WASF3,RHOBTB2,NUP133,SOS1,PPP1R12A,AF

3,PSMA1,PSMD10,PSMA6,CTDNEP1,PPP2R5B,PSMD12,PSMA4,NUP50,DYNC1I2,UBE2E1,CENPF,

M,IDH1,PCBP2,DYNLL1,TXNIP,CDC42,POLR2E,PSMA7,TXNDC5,EP300,PSMD1,PDAP1,CST3,UBA

2B,CDK4,NUP153,H2AX,POLD2,PPP2R5D,PSME3,PSMC5,PSMC2,PSMC4,PSMF1,GSK3B,DHFR,T

DDOST,YWHAB,LAMP1,UBE2H,CKAP4,HLA-B,TTR,COTL1,HRNR,RAC1,ATP6V0C,TMEM30A,MMP2

C,ELOA

APC2,RBBP6

,RPS19,RPL35,HMGCS1,GPAT4,PRKAR1A,CHPF,RPS10,RPL35A,RPLP2,ST3GAL2,RPL36,PTGES3
SO1,RANGAP1,MZT2B,ABL1,RSF1,SUN1,PSMB1,RUVBL2,MDM2,FZR1,PCM1,CUL1,PSMD13,SUM

4,ITGAV,LRRC41,AKT2,FBXW5,PSMC3,MTOR,UBE3A,PRKACA,AP2A2,AP1M1,UBE2D3,PSMC1,CD

10,RPS6KA2,MAP2K4,TALDO1,ANXA1,MAPK3,PSMA6,TAB3,RELA,PSMD12,PTPN12,PSMA4,PSMC

APK1,ARPC2,PSMD3,RPS11,CASC3,PLCG1,MAP2K2,RPL28,PITPNA,RPS28,FYN,FAU,SLIT3,PLXNA
IBB2A,RPL37,MAPK1,SMARCA4,ARPC2,PSMD3,RPS11,CASC3,PLCG1,MAP2K2,RPL28,PITPNA,RP

PSMC6

MC4,PSMF1,RPS29,PSMB5,PSMA1,SOS1,PSMD10,ELOB,ZSWIM8,PSMA6,RPL36AL,PSMD12,RPL

RPS13,IGFBP7,DYNLL1,RPL34,PSMA7,RBBP7,DCTN1,STAT3,EIF2AK1,ACTR1A,EP300,PSMD1,FL

RPS13,IGFBP7,DYNLL1,RPL34,PSMA7,RBBP7,DCTN1,STAT3,EIF2AK1,ACTR1A,EP300,PSMD1,FL
1,CASC3,RPL28,POLR2B,RPS28,HNRNPM,SRSF3,FAU,CSNK1D,SF3A1,PCBP2,HNRNPD,RPL39,RF

3L,GNB2,RCN1,RPS5,RPS7,RPS19,RPL35,TLN1,MYO5A,CTNNB1,GOLM1,RPS10,RPL35A,APLP2,

3B,MRPS6,EIF3M,RARS1,WARS1,RPS29,KARS1,SPCS2,FARSA,SRP54,RPL36AL,TARS1,RPL32,EIF
AI3,PABPN1,PARP1,TAF15,SSRP1,RPL27,ACTR3,AHCYL1,NCKAP1,CBX1,PSMB2,RPL36A,TAF7,RF

DOST,ST3GAL2,HNRNPK,RPL36,YWHAB,PTGES3,SND1,ATP1A2,RPS12,TTR,RPS27,CTBP1,RAC1,

A,ACTR3,COL2A1,LAMC1,PSMB2,H3-3A,RPL36A,DPYSL2,RPS15A,RPL38,TUBB2A,RPL37,MAPK1

MC5,PSMC2,DLD,PSMC4,PSMF1,RIMKLB,OAT,RARS1,ALDH9A1,RPS29,PSMB5,KARS1,PSMA1,GI

ND4B,CHMP4B,AP2A2,TGOLN2,AP1M1,EXOC4,SORT1,GAK,TUBB4A,USO1,ARF5,SNX2,ANK2,PAI

2R5D,ATP6V0B,STUB1,CRKL,RBFOX2,NCBP2,RANBP9,SRF,DLG4,WASF3,STAT1,DUSP3,IRS1,AT

4,LDLR,SH3GL1,KIFAP3,CAPZA1,GALNT1,RAB6A,FNBP1L,RAB5C,SYT11,AKT2,COPZ1,DENND4E

5B,PSMD12,PTPN12,PSMA4,MAML1,TRIM24,PSMC6,RAP1B,FZD8,LRP6,PIK3R3,PPP2R5C

,RAB11A,PPP6R1,SEC23A,IGFBP7,DYNLL1,TP53BP1,PSMA7,VDAC1,RBBP7,ST3GAL6,DCTN1,OS

S1,SYNGR1,PA2G4,RAB3A,ARL8A,MAN2B1,DPP7,BRI3,ARSB,ATP6V0A1,ARPC5,DYNLT1,TRAPPC

RHGAP33,ARPC5,ROCK2,MAPK3,PIN1,ARHGEF18,VAV2,PPP2R5B,DYNC1I2,OBSCN,ARHGAP23,CI
RHGAP33,ARPC5,ROCK2,MAPK3,PIN1,ARHGEF18,VAV2,PPP2R5B,DYNC1I2,OBSCN,ARHGAP23,CI

PSMC6,PPP2R2A,CENPT,NINL,LEMD2,ANAPC2,WAPL,PPP2R5C,EMD

52,PPP2CB,RPS27A,PSMD8,ATP6AP2,BRK1,PRDX6,GRN,ACLY,XRCC6,PSMB4,SLC44A2,ABI2,UBE

TUBA1C,RCC2,CDKN1B,PSMB5,RRM2,PRKAR2B,NUP205,NUP133,PSMA1,PPP1R12A,PSMD10,PS

!,SKP1,CDKN1A,PGRMC1,CNN2,CTSB,RAB27A,CCT2,SQSTM1,VAMP2,GDI2,ACTR3,PDXK,UBE4A,

3,RPS12,TTR,RPS27,GPC1,RPL23A,GLUL,SQLE,DCN,HS6ST1,MDH2,FDFT1,PPP1CB,PGD,VAMP2,
1O1,BANF1,PSMD7,RAN,SUN2,CETN2,PPME1,CSNK2B,CDK4,NUP153,H2AX,POLD2,PPP2R5D,PSN

74,TUBB4A,UBE2K,FBXL19,TRIP12,UBE2L3,PTPN11,TAPBP,CDC34,CLTA,PSMB1,FZR1,CUL1,PS

C6,RAP1B,PIK3R3,TOLLIP

3,PFN2,RPL39,RPS13,RPL34,CDC42,PSMA7,KCNQ2,TRIO,PRKAR2A,PSMD1,CNTNAP1,AP2A1,CS
S28,FYN,FAU,SLIT3,PLXNA3,SREBF2,PFN2,RPL39,CYP51A1,RPS13,RPL34,CDC42,PSMA7,KCNQ

32,PSMA4,RNPS1,PSMC6

CN,H2AZ1,UBA52,EP400,HDAC6,CBX6,POM121C,RPS2,RPS27A,PSMD8,HIF1A,PPP1R15A,PRDX6,

CN,H2AZ1,UBA52,EP400,HDAC6,CBX6,POM121C,RPS2,RPS27A,PSMD8,HIF1A,PPP1R15A,PRDX6,
PS13,PRPF40A,RPL34,ADAR,DHX15,POLR2E,PSMA7,ANP32A,EDC4,XPOT,PSMD1,SF1,PRCC,GSPT

RPLP2,DDOST,ST3GAL2,HNRNPK,NARS1,RPL36,UBE2H,CKAP4,RPS12,RAB1A,TTR,CALU,ARCN1

F2S1,HARS1,MRPL3,MRPS26,MRPS27,SEC11A,MRPL28,EIF3J
PS15A,RPL38,TUBB2A,RPL37,VHL,MVB12B,MAPK1,ARPC2,PSMD3,DYNC1LI2,DYNLL2,RPS11,SU

GPC1,HDAC2,IPO5,SKP1,RPL23A,DCN,CDKN1A,RPN2,PPP1CB,VAMP2,GNAI3,PABPN1,PARP1,VC

,SMARCA4,ARPC2,PSMD3,RPS11,CASC3,PLCG1,MAP2K2,RPL28,PITPNA,POLR2B,RPS28,FYN,FA

RHPR,OGDH,PSMD10,PSMA6,RPL36AL,AMD1,PSMD12,RPL32,PSMA4,PSMC6,PDHB

FAH1B2,DENND5A,PUM1,LMAN2,CLTA,ARPC1A,KIFC2,HIP1R,EXOC5,NSF,COPE,KIF26A,TBC1D1

P6V0E2,ERBB2,THBS2,ANOS1,SOS1,NRP2,ATP6V0A1,SHC1,ROCK2,RPS6KA2,MAPK3,COL6A3,S[illegible]

[illegible],CHMP4B,AP2A2,TGOLN2,AP1M1,EXOC4,SORT1,GAK,TUBB4A,USO1,ARF5,SNX2,ANK2,PAFAH1[illegible]

9,MTA1,ACTR1A,EP300,PSMD1,SEC16A,CST3,UBA52,MGAT4B,KLHL42,RAB2A,UBE2G2,FBXO11

1,PSMD12,COPB1,NFASC,RAP1B,PRDX4,PRCP,QSOX1,GSN,CSTB,ATP6V1D,GLB1,CTSA,ARSA,T[illegible]

ENPF,CENPT,MYL9,NF2,FGD1,ABR,PPP2R5C
ENPF,CENPT,MYL9,NF2,FGD1,ABR,PPP2R5C

E2M,UBE2D2,IMPDH2,TAX1BP1,RAB14,DTX4,CAPZA1,NCSTN,RAB6A,RAB5C,ACTR1B,PDPK1,WAS

MA6,CTDNEP1,PCNA,TFDP1,AJUBA,PPP2R5B,MCM7,PSMD12,PSMA4,TPX2,NUP50,DYNC1I2,UBE

,TCP1,AHCYL1,KLC1,NCKAP1,COL2A1,MLEC,SURF4,RAB5B,ORAI2,PSMB2,SRP14,PYGB,VEGFA,

VCAN,RPL27,PDXK,ACO2,AHCYL1,ETNK1,PSMB2,RPL36A,PYGB,RPS15A,RPL38,MARS1,RPL37,A
ME3,PSMC5,PSMC2,PSMC4,PSMF1,YWHAZ,GSK3B,DHFR,TUBA1C,RCC2,CDKN1B,PSMB5,RRM2,

MD13,KIF26A,PJA1,HLA-A,PSMD7,KLC2,UBE2J1,AREL1,RNF41,TRIB3,PPP2R5D,PSME3,UBE2Q1

NK2A1,GSPT1,UBA52,PLXNB1,EVL,RGMB,RPS2,RPS27A,PSMD8,SLIT2,PSMB4,SLIT1,RDX,NCSTN
2,TRIO,PRKAR2A,PSMD1,CNTNAP1,AP2A1,CSNK2A1,GSPT1,UBA52,PLXNB1,EVL,RGMB,RPS2,RP

,PSMB4,EIF2S2,EHMT1,GPX1,FKBP4,UBE2D2,CAPZA1,SOD2,PSMD4,ATP6V0D1,SH3BP4,PSMC3,

,PSMB4,EIF2S2,EHMT1,GPX1,FKBP4,UBE2D2,CAPZA1,SOD2,PSMD4,ATP6V0D1,SH3BP4,PSMC3,
T1,UBA52,ZFP36L1,SRSF6,UPF1,TRA2B,HNRNPA2B1,POM121C,RPS2,RPS27A,PSMD8,CHERP,SY

,RPS27,CCT6A,CTBP1,TOP2B,HDAC2,TNC,MMP2,SKP1,RPL23A,TMEM132A,CTDSP2,SRP9,NUCE

PT5H,KDM1A,PLCG1,MAP2K2,RPL28,POLR2B,GNG2,RPS28,FYN,FAU,B2M,STT3A,HGS,NCOR2,F

AN,NOX4,TAF15,SSRP1,ABCA3,RPL27,ACTR3,AHCYL1,NCKAP1,CBX1,PSMB2,RPL36A,TAF7,RPS1

U,SLIT3,PLXNA3,SREBF2,PFN2,NCOR2,RPL39,CYP51A1,RPS13,RPL34,CDC42,POLR2E,PSMA7,R

0B,KLC2,NBAS,COPS6,VPS51,ARFGAP2,TBC1D14,RAB18,EPN1,ARPC3,EXOC7,GOLGA4,YWHAZ,

HB,VAV2,FER,BCAR1,APOE,FURIN,PTPN12,ARF6,FGFBP3,COL3A1,DNM1,ID3,GRB10,RAP1B,ATF

B2,DENND5A,PUM1,LMAN2,CLTA,ARPC1A,KIFC2,HIP1R,EXOC5,NSF,COPE,KIF26A,TBC1D10B,KL

,MBTPS1,PTEN,SMC3,YKT6,FEM1B,CUL3,B4GAT1,POM121C,RPS2,RPS27A,LMAN1,TNKS,PSMD

)LLIP

SF2,PSMD4,ITGAV,ADAM10,CAP1,IST1,ATP6V0D1,PSMC3,PRKACA,AP2A2,AP1M1,CREBBP,ATP6V

2E1,CENPF,PSMC6,PPP2R2A,CENPT,NINL,LEMD2,ANAPC2,WAPL,PPP2R5C,EMD

RAPGEF1,TUBB2A,VHL,PPP2CA,CCND1,MAPK1,SMARCA4,ARPC2,PSMD3,DYNC1LI2,DYNLL2,GR

ARF3,PPP2CA,AZIN1,DDAH2,PSMD3,ODC1,ACADVL,RBP1,ATP5PB,RPS11,PLCG1,QARS1,RPL28,G
,PRKAR2B,NUP205,NUP133,PSMA1,PPP1R12A,PSMD10,MDC1,PSMA6,CTDNEP1,PCNA,TFDP1,AJ

,STUB1,PSMC5,PSMC2,CD300E,PSMC4,PSMF1,YWHAZ,UBE3C,MEX3C,UBE2V1,TUBA1C,KIF3A,F

,CLASP2,MSI1,PSMD4,ITGAV,ADAM10,CAP1,PSMC3,PRKACA,AP2A2,PSMC1,DPYSL4,MYL12B,LY
S27A,PSMD8,SLIT2,PSMB4,SLIT1,RDX,NCSTN,CLASP2,MSI1,PSMD4,ITGAV,ADAM10,CAP1,PSMC

MTOR,CREBBP,ATP6V1G1,UBE2D3,PSMC1,BAG5,PRDX3,TUBB4A,RPS26,HMGA1,PSMB1,HSPA1

MTOR,CREBBP,ATP6V1G1,UBE2D3,PSMC1,BAG5,PRDX3,TUBB4A,RPS26,HMGA1,PSMB1,HSPA1
MPK,FUS,PSMB4,PELP1,SNRPD3,ELAVL1,SMG5,EFTUD2,TTC37,SRSF2,RBM5,WBP11,PSMD4,SF

31,TOMM20,RAB27A,EEF1A2,CCT2,RPN2,TBCD,VAMP2,GNAI3,PARP1,VCAN,TGFBI,MTRF1L,ABC

RPL39,RPS13,DYNLL1,TXNIP,RPL34,CDC42,CTNND1,POLR2E,PSMA7,RBBP7,MTA1,PRKAR2A,PSM

5A,RPL38,NOTCH2,TUBB2A,RPL37,VHL,MVB12B,PPP2CA,CCND1,MAPK1,ARPC2,PSMD3,PRDX1

BBP7,STAT3,KCNQ2,TRIO,EP300,PRKAR2A,PSMD1,PBX1,CNTNAP1,AP2A1,H2AZ1,CSNK2A1,GSF

PIP5K1C,ACBD3,TUBA1C,MCFD2,RAB3A,AGPAT3,TFRC,KIF3A,NAPB,NAPA,COL7A1,ARPC5,CHMF [illegible]6V1F,ELMO2,ID2,SH3KBP1,PIK3R3,ATP6V1D

[illegible]C2,NBAS,COPS6,VPS51,ARFGAP2,TBC1D14,RAB18,EPN1,ARPC3,EXOC7,GOLGA4,YWHAZ,PIP5K[illegible]

8,HIF1A,GALNT11,MVD,FBXO21,BAP1,WSB2,PSMB4,CCN1,RAB1B,UBE2M,UBE2D2,RAB14,CAP[illegible]

1G1,UBE2D3,PSMC1,UBE2K,PAFAH1B2,PTPN11,CDC34,ABL1,SDCBP,ARPC1A,SCAMP1,PSMB1,[illegible]

B2,CAPZB,PRKDC,UBE2R2,CAND1,NPEPPS,PLCG1,DUSP4,FLNB,PITPNA,KIF3C,ATP6V1B2,FYN,IL

NG2,RPS28,SDHA,LARS1,FAU,PFKM,LSS,PITPNM1,STXBP1,ISYNA1,PRPS1,IDH1,SARS1,UQCRC2
UBA,PPP2R5B,MCM7,PSMD12,PSMA4,TPX2,NUP50,DYNC1I2,UBE2E1,CENPF,PSMC6,PPP2R2A,C

PSMB5,PSMA1,ZNRF1,RNF4,SOS1,MGRN1,PSMD10,UBE2G1,ELOB,ASB6,NECTIN2,PPP3CA,CAPZ

'PLA2,NELL2,TUBB4A,EIF4A3,EI
3,PRKACA,AP2A2,PSMC1,DPYSL4,T

A,TXNRD1,MDM2,FZR1,DNAJB

A,TXNRD1,MDM2,FZR1,DNAJB
RSF11,IGF2BP1,PRPF19,SF3A2,PSM

A3,RPL27,TCP1,CCT3,MLE

MD1,AP2A1,GRSF1,KPNA2,BRD4,UB

,DYNC1LI2,RBP1,DYNLL2,GRB2,C

PT1,UBA52,PLXNB1,EVL,RGMB,RPS2,R

P5,RAB5A,GALNT2,GABARAPL2,GPS1,CAPZA2,TRA

[illegible]1C,ACBD3,TUBA1C,MCFD2,RAB3A,AGPAT3,TFRC,

[illegible]A1,GALNT1,ATXN3,RAB6A,RAB5C,DCUN1D2,FOXK2,PS

HSPA1A,CUL1,PSMD13,AAMP,CAB39,NDUFC2

_F2,CCT8,B2M,IDH1,HECTD1,IFITM3,PCBP

,SREBF2,NCOR2,RPL39,CYP

ENPT,NINL,RAD50,LEMD2,ANAPC2,WAPL,PPP2R5C,EMD

'A2,PSMA6,MIB2,CUL7,BTBD1,PPP2R5B,RELA

| | | | | | | |
|---|---|---|---|---|---|---|
| **WP** | Cytoplasmic F | WP:WP477 | 1.56E-36 | 35.8079366 | 90 | 1436 |
| **WP** | VEGFA-VEGF | WP:WP3888 | 8.29E-25 | 24.0812571 | 437 | 1436 |
| **WP** | mRNA Proces | WP:WP411 | 1.75E-15 | 14.7569908 | 133 | 1436 |
| **WP** | Proteasome D | WP:WP183 | 1.07E-09 | 8.9712292 | 64 | 1436 |
| **WP** | Parkin-Ubiquit | WP:WP2359 | 7.52E-09 | 8.12358736 | 70 | 1436 |
| **WP** | Androgen rece | WP:WP138 | 4.9176E-06 | 5.30824623 | 90 | 1436 |
| **WP** | Metabolic repr | WP:WP4290 | 5.0693E-06 | 5.29505152 | 44 | 1436 |
| **WP** | Translation Fa | WP:WP107 | 6.4066E-06 | 5.19337316 | 53 | 1436 |
| **WP** | Pathways in c | WP:WP4018 | 1.9366E-05 | 4.71296834 | 87 | 1436 |
| **WP** | Computationa | WP:WP4629 | 2.2279E-05 | 4.65209709 | 12 | 1436 |
| **WP** | Focal Adhesio | WP:WP306 | 6.2142E-05 | 4.20661428 | 201 | 1436 |
| **WP** | Pathogenic Es | WP:WP2272 | 0.00033143 | 3.47960419 | 55 | 1436 |
| **WP** | 7q11.23 copy | WP:WP4932 | 0.00087771 | 3.05664801 | 106 | 1436 |
| **WP** | nsp1 from SA | WP:WP5027 | 0.00100989 | 2.99572514 | 17 | 1436 |
| **WP** | Glycolysis anc | WP:WP534 | 0.00104612 | 2.98041843 | 45 | 1436 |
| **WP** | Brain-Derived | WP:WP2380 | 0.00231842 | 2.63480819 | 144 | 1436 |
| **WP** | Fragile X Sync | WP:WP4549 | 0.0029486 | 2.53038449 | 122 | 1436 |
| **WP** | Ciliary landsca | WP:WP4352 | 0.00324875 | 2.48828394 | 218 | 1436 |
| **WP** | Association Be | WP:WP3680 | 0.00365108 | 2.43757898 | 68 | 1436 |
| **WP** | 16p11.2 prox | WP:WP4949 | 0.00496191 | 2.30435123 | 76 | 1436 |
| **WP** | Cori Cycle | WP:WP1946 | 0.00892808 | 2.04924201 | 17 | 1436 |
| **WP** | TGF-beta Sigr | WP:WP366 | 0.01381806 | 1.85955293 | 133 | 1436 |
| **WP** | Cholesterol Bi | WP:WP197 | 0.01397167 | 1.85475175 | 15 | 1436 |
| **WP** | Leptin signalir | WP:WP2034 | 0.0140704 | 1.85169362 | 76 | 1436 |
| **WP** | Prolactin Signa | WP:WP2037 | 0.0140704 | 1.85169362 | 76 | 1436 |
| **WP** | EGF/EGFR Si | WP:WP437 | 0.01972727 | 1.70493303 | 163 | 1436 |
| **WP** | PDGF Pathwa | WP:WP2526 | 0.02950656 | 1.53008142 | 40 | 1436 |
| **WP** | Human Thyro | WP:WP2032 | 0.03165326 | 1.49958153 | 68 | 1436 |
| **WP** | Photodynamic | WP:WP3614 | 0.03286176 | 1.48330915 | 37 | 1436 |
| **WP** | Cell migration | WP:WP4561 | 0.0389429 | 1.40957167 | 31 | 1436 |

| | | |
|---|---|---|
| 74 | 7562 | RPL3,RPS6,RPL4,RPL7A,RPL8,RPLP0,RPL13A,RPL6,RPL13,RPS8,RP |
| 177 | 7562 | PABPC1,GAPDH,ACTG1,CFL1,HSP90AA1,TMSB4X,EIF4G2,RPS6,PIK3 |
| 69 | 7562 | HNRNPA1,PRPF8,YBX1,NONO,SF3B1,POLR2A,SF3B2,HNRNPU,PAPO |
| 37 | 7562 | UBC,UBA1,UBB,PSMD5,HLA-C,UCHL1,PSMD2,HLA-B,RPN2,PSMB2, |
| 38 | 7562 | HSPA8,HSPA9,TUBA1A,TUBB,TUBA1B,UBA1,HSPA1B,TUBB2B,PSMD |
| 40 | 7562 | PIK3R2,FLNA,CALR,RACK1,ZMIZ1,RHOA,TLE5,JUN,CTNNB1,RHOB, |
| 25 | 7562 | GAPDH,ENO1,PKM,PGK1,GPI,FASN,PGAM1,LDHA,PFKL,TKT,MDH2, |
| 28 | 7562 | EEF1A1,PABPC1,EEF2,EIF3C,EIF4B,EIF1,EIF4A1,EIF4H,EIF5,EIF5A,E |
| 38 | 7562 | GAPDH,ENO1,ALDOA,PKM,PGK1,ENO2,GPI,FASN,TPI1,LDHB,LDHA, |
| 11 | 7562 | GAPDH,ENO1,ALDOA,PKM,PGK1,GPI,TPI1,LDHA,HK1,PFKM,SLC2A1 |
| 68 | 7562 | COL1A1,ACTB,ACTG1,PIK3R2,FLNA,COL4A1,COL4A2,FN1,ITGB1,VC |
| 26 | 7562 | ACTB,ACTG1,TUBA1A,TUBB,TUBA1B,NCL,TUBB2B,YWHAQ,ITGB1,T |
| 40 | 7562 | GAPDH,CFL1,ATP5F1B,ATP5F1A,SF3B1,SMARCA5,EIF4H,ATF4,CLTC |
| 12 | 7562 | EIF3C,EIF1,EIF5,EIF3E,EIF3F,EIF3D,EIF2S2,EIF3H,EIF3G,EIF3B,EIF2 |
| 22 | 7562 | GAPDH,ENO1,ALDOA,PKM,PGK1,ENO2,SLC2A3,GPI,TPI1,LDHB,PGA |
| 49 | 7562 | EEF2,CFL1,KIDINS220,RPS6,PIK3R2,YBX1,RACK1,MARCKS,MAPT,N |
| 43 | 7562 | EEF1A1,MAP1B,APP,EIF4A1,AP2M1,AP2B1,CLTC,PRKAR1A,CYFIP2, |
| 67 | 7562 | NUDC,VIM,NEFL,NEFM,DDX5,DVL3,GDI1,CALM1,PAFAH1B1,LRPPRC |
| 28 | 7562 | ACTR2,TMSB4X,NEFL,NEFM,FN1,RHOA,PFN1,JUN,CTNNB1,CDKN1A |
| 30 | 7562 | ALDOA,PPP2R1A,MSN,IGFBP3,CCT7,CCT6A,CCT2,TCP1,CCT3,CCT4, |
| 11 | 7562 | GAPDH,ALDOA,PGK1,GPI,TPI1,PGAM1,LDHA,HK1,SLC2A1,PFKP,TAL |
| 44 | 7562 | NEDD4L,PIK3R2,SPTBN1,APP,FN1,ITGB1,COL1A2,RHOA,JUND,JUN, |
| 10 | 7562 | HMGCS1,HMGCR,SQLE,FDFT1,LSS,CYP51A1,MSMO1,MVD,DHCR7, |
| 29 | 7562 | RPS6,PIK3R2,RHOA,KHDRBS1,RAC1,CCND1,MAPK1,GRB2,PLCG1, |
| 29 | 7562 | RPS6,PIK3R2,FLNA,YWHAG,ITGB1,CTSD,JUN,RAC1,PPIB,MAPK1,GR |
| 51 | 7562 | CFL1,PIK3R2,AP2M1,STMN1,AP2B1,PXDN,JUND,PEBP1,GJA1,JUN, |
| 18 | 7562 | RHOA,JUN,RAC1,MAPK1,GRB2,PLCG1,CDC42,STAT3,PTPN11,PDGF |
| 26 | 7562 | RPS6,PIK3R2,GNAS,GNB1,JUN,GNAI3,MAPK1,GNG2,STAT3,APEX1,G |
| 17 | 7562 | PKM,IGFBP2,PGK1,SLC2A3,EGLN1,LDHA,IGFBP3,HK1,MCL1,PFKL, |
| 15 | 7562 | KIDINS220,ARF1,RHOA,JUN,RAC1,MMP2,CDH11,STAT3,TRIO,AKT3, |

L7,RPL5,RPL37A,RPL19,RPL15,RPS3,RPS4X,RPS3A,RPL18,RPL11,RPLP1,RPL10,RPS17,RPS16,F
3R2,NCL,ALDOA,CALR,PGK1,PTMA,RPL13A,YWHAE,RACK1,BSG,MYH9,RPL7,P4HB,RPL5,FSCN1,I
LA,SRSF4,HNRNPC,SFPQ,HNRNPK,HNRNPH1,CPSF2,SRSF5,U2AF2,SUGP2,CPSF1,PABPN1,HNRN
PSMD3,PSMA7,PSMD1,H2AZ1,PSMD8,PSMB4,UBE2D2,PSMD4,PSMC3,UBE2D3,PSMC1,PSMB1,PS
5,TUBB4B,PSMD2,HSPA4,TUBB2A,PSMD3,PSMD1,UBE2G2,PSMD8,SEPTIN5,PSMD4,PSMC3,PSM
RAC1,CDKN1A,CCND1,DSTN,PRDX1,KDM1A,SMARCE1,NCOR2,CDC42,STAT3,EP300,PTEN,NCOA
PGD,ACO2,GLUD1,SLC2A1,ACLY,GOT2,PSAT1,PDHA1,IDH2,SHMT2,TALDO1,SLC1A5,FH,PDHB
EF1A2,EIF3E,EIF3F,EIF2AK1,EIF3D,EIF5B,EIF2S2,EIF3H,EIF4G3,EEF1B2,EIF3G,EIF4G1,CLUH,ETF
HK1,PFKL,SQSTM1,VEGFA,VHL,PFKM,STAT3,KDM5C,EP300,PTEN,HIF1A,SLC2A1,BAP1,ACLY,PH(

CL,LAMA5,COL1A2,RHOA,COL5A2,CCND2,ACTN4,TLN1,JUN,CTNNB1,COL6A2,RAC1,TNC,PPP1CE
UBB4B,RHOA,CTNNB1,TUBB2A,ARPC2,FYN,CDC42,CTTN,TUBB4A,ABL1,ARPC1A,NCK2,ARPC3,Y
;,CTNNB1,HDAC2,SQSTM1,VAMP2,ATP5PB,GRB2,BAZ1B,BRD4,HDAC6,ULK1,GTF2I,CLASP2,FBL
S1,EIF3J
M1,LDHA,HK1,PFKL,MDH2,PFKM,SLC2A1,GOT2,MDH1,PDHA1,PFKP,DLD
CAM1,CDH2,JUN,CTNNB1,RAC1,SQSTM1,DPYSL2,PPP2CA,MAPK1,GRB2,PLCG1,MAP2K2,FYN,CI
MAPK1,GRB2,PLCG1,MAP2K2,FYN,ALDH3A2,AP2A1,PTEN,SH3GL1,DICER1,MTOR,PRKACA,EIF4G
;,CTNNB1,MYL6,UBE2H,RAC1,ARHGDIA,HDAC2,H3-3A,DYNLL2,XPNPEP3,DYNLL1,APMAP,RAB2A,
,ACTR3,PPP2CA,MAPK1,GRB2,PLCG1,SOD1,PFN2,GPX1,DAAM1,GSK3B,CDKN1B,ERBB2,SOS1,S
,PPP2CA,CCT8,CCT5,PPP2CB,PTEN,MAZ,QPRT,PAGR1,TAOK2,PPP4R3A,PPP2R5D,SEZ6L2,CCDC
DO1
RAC1,TNC,SKP1,CDKN1A,SKI,CCND1,MAPK1,GRB2,MAP2K2,STRAP,HGS,CDC42,EP300,PRKAR2
SC5D
MAP2K2,FYN,CDC42,STAT3,PTEN,GSK3A,MTOR,PTPN11,PTPN1,JAK1,GSK3B,STAT1,IRS1,ERBB2,
B2,MAP2K2,FYN,STAT3,MTOR,PTPN11,PTPN1,JAK1,YWHAZ,GSK3B,STAT1,IRS1,ERBB2,SOS1,SH
AC1,MAPK1,GRB2,PLCG1,MAP2K2,STXBP1,HGS,CDC42,STAT3,AP2A1,PTEN,PDPK1,MTOR,NDUF
RB,JAK1,SRF,STAT1,SOS1,SHC1,MAP2K4,MAPK3,VAV2
NA12,PDPK1,SCRIB,MTOR,GNA13,MYL12B,IGF1R,RALGDS,CDK4,JAK1,CDKN1B,STAT1,MAPK3,
VEGFA,BNIP3L,HIF1A,SLC2A1,BNIP3,PDHA1,SLC16A1
AKT2,NTRK2,EFNB2,EFNB1,EFNB3

RPL23,RPL29,RPSA,RPS24,RPS9,RPL24,RPL17,RPL14,RPL22,RPL9,RPS14,RPS20,RPL26,RPL31,
FN1,SET,TMSB10,HMGB1,ITGB1,VCL,DNAJA1,CDC42BPB,TMEM170A,CTNNA1,LDHA,IGFBP3,RH
NPL,SNRPN,RBMX,SNRNP70,PRPF6,SRSF9,SRSF1,PTBP1,SUPT5H,HNRNPM,SRSF3,SF3A1,PCBP2
SMD13,HLA-A,PSMD7,H2AX,PSME3,PSMC5,PSMC2,PSMC4,PSMB5,PSMA1,PSMD10,PSMA6,PSMD
C1,TUBB4A,UBE2L3,HSPA1A,CUL1,PSMD13,PSMD7,UBE2J1,STUB1,PSMC5,PSMC2,PSMC4,TUB
4,FKBP4,CARM1,UBE3A,CREBBP,UBE2I,MDM2,SUMO1,PARK7,RAN,STUB1,GSK3B,NR2C2,RNF4

F1,EIF3B,PAIP1,EIF2S1,EIF3J
GDH,MTOR,CREBBP,MDH1,PSAT1,PFKP,RHEB,PDGFRB,ZEB1,SHMT2,PBRM1,TGFB2,GRB10

3,TLN2,COL2A1,LAMC1,VEGFA,RAPGEF1,CCND1,MAPK1,GRB2,ACTN1,FLNB,FYN,FLNC,CDC42,C
WHAZ,TUBA1C,ARPC5,ROCK2
N5,ATP5MC3,ATP5MC2,DEK,HMGA1,CLIP2,ELN,LIMK1,H2AX,ATP5PO,DLD,ATP5F1E,OGDH,MAP

DC42,STAT3,CSNK2A1,PTPRF,EIF2S2,SHC2,LINGO1,PDPK1,IGF2BP1,MTOR,SORT1,NTRK2,PTPN
1,NTRK2,TECR,PTPN11,CLTA,DAG1,MECP2,CYFIP1,RHEB,PPP1CA,SYNGAP1,DLG4,DUSP3,SLC1
EIF5B,MAPRE2,PSMD8,RHBDD2,CEP170,VAPB,UBE2D2,VPS4A,RAB14,EFTUD2,EXOC4,CREBBP,
SHC1,ROCK2,MAP2K4,FZD7,FZD8
6,UNC13A,HIRA,MAPK3,PCNA,CDIPT,PPP4R3B,UNC13B

A,TRAP1,CREBBP,UBE2I,CUL1,SUMO1,PJA1,NUP153,ZEB1,SOS1,SHC1,MAP2K4,TGFBR1,TFDP1

,SOS1,SHC1,ROCK2,MAPK3,SP1,RELA
HC1,RPS6KA2,MAPK3,VAV2,RELA
FA13,PTPN11,ABL1,PIK3C2B,RALGDS,IQSEC1,NCK2,JAK1,EPN1,CRKL,RALBP1,STAT1,ERBB2,SC

,RAP1B,GNAO1,GNAI1

RPS25,RPS15,RPL12,RPL30,RPS5,RPS7,RPS19,RPL35,RPS10,RPL35A,RPLP2,RPL36,RPS12,RPS[illegible]
[illegible]IOA,ATF4,RPL26,CCT7,PFN1,TKT,PDIA6,CCN2,MMP14,PRDX2,CLTC,RCN1,GJA1,JUN,CTNNB1,R[illegible]
,HNRNPD,PRPF40A,DHX15,PRMT1,PPM1G,SRSF6,TRA2B,HNRNPA2B1,HNRNPAB,FUS,SNRPD3,E[illegible]
[illegible]12,PSMA4,PSMC6
A1C,PSMD10,UBE2G1,PSMD12,PSMC6,CASK
,ROCK2,SP1,RELA,SIN3A

[illegible]APN2,PTEN,PPP1CC,ITGB5,SHC2,AKT3,PDPK1,ITGAV,AKT2,MYL12B,IGF1R,PPP1CA,ARHGAP5,[illegible]

K3,PCNA,UBE2E1,VPS28,VPS37B,MYBBP1A

[illegible]11,NSF,CYFIP1,NCK2,RANBP9,GSK3B,CRTC1,RAB3A,STAT1,IRS1,SHC1,MAPK3,VAV2,RELA,EIF2[illegible]
6A1,SOS1,SHC1,SHANK1,PPP3CA,PPP2R5B,ABCD3,EPHA4,DNM1
WNK1,EXOC5,PSMD13,WDR26,CD2BP2,PSMD7,MAEA,DYNLRB1,COPS6,ECHS1,LSM4,GID8,EXO[illegible]

,SP1,BCAR1,PPM1A,KLF10,SKIL,SIN3A,KLF6

[illegible]S1,SHC1,RPS6KA2,RAB5A,PCNA,PRKCI,VAV2,SP1,BCAR1,PTPN12,ARF6,DNM1,GRB10,NEDD8,S[illegible]

27,RPL23A,RPL27,RPL36A,RPS15A,RPL38,RPL37,RPS11,RPL28,RPS28,FAU,RPL39,RPS13,RPL3[illegible]
PLP2,SND1,CALU,RAC1,STIP1,GPC1,MMP2,PGD,NOX4,RPL27,VEGFA,RAPGEF1,PPP2CA,CCND1,[illegible]
FTUD2,SRSF2,PRPF4B,DICER1,RBM5,SF3A2,DHX38,XRN2,SF3A3,CD2BP2,HNRNPR,SRSF10,SRS[illegible]

PDGFRB,CRKL,PIP5K1C,GSK3B,ZYX,MYL12A,ERBB2,THBS2,SOS1,PPP1R12A,SHC1,ROCK2,MAP

2S1

C7,RANBP9,PSMC4,PGRMC2,RBM14,NUP133,CTBP2,DYNLT1,MCM7,PSMD12,MSH2,DYNC1I2,C

H3KBP1

4,UBA52,RPS2,RPS27A,RPS26,RPS21,RPL27A,RPS29,RPS6KA2,RPL32
MAPK1,ADAM9,GRB2,CAPZB,RPS11,HSPB1,PLCG1,MAP2K2,ARF4,FLNB,TPM3,FYN,EIF3F,HYOU
F7,NCBP2,RNMT,DNAJC8,SNRPD2,SNRPB,SRP54,RBM17,PRMT2,SFSWAP,RNPS1,RBM39,SNU1

K3,VAV2,BCAR1,RAP1B,ITGA7,MYL9,PIK3R3

OPS8,PSMC6,NINL,NDUFA5,GLB1,ARFGAP3,CNOT1,CTSA,MYL6B

1,SARS1,HGS,RAB11A,IGFBP7,TXNIP,CDC42,CTNND1,STAT3,CAPN2,NAP1L1,TXNDC5,AP2A1,EIF[illegible]
[illegible]

3D,GRSF1,BRD4,PPM1G,DECR1,UBAP2L,GLUD1,ITGB5,LMAN1,LRRC59,PRDX6,CCN1,GPX1,SH[illegible]

C2,EIF3H,TPP1,SSR3,CNP,SOD2,SLC7A1,PDPK1,PSMD4,ITGAV,ADAM10,ATP6V0D1,MTOR,CREBE

3P,FJX1,EIF4G1,PTPN11,ABL1,FXR2,SDCBP,USP10,TAOK2,CLIC1,HSPA1A,MDM2,CRIP2,IDH2,EW

'SR1,PTPN1,HTRA1,LIMK1,FLII,PPP1CA,SH3BGRL3,PBXIP1,MAPKAPK2,EPN1,DHX36,NDRG1,HEF

RPUD1,PDE4DIP,SRF,GSK3B,TUBA1C,TXN,TBCA,STAT1,SRP54,NAPA,ACP1,NRP2,SHC1,AFDN,ROC

CK2,ATF6,MAP2K4,RAB5A,ANXA1,MAPK3,PPP3CA,SHB,PRKCI,ZC3H15,VAV2,BCAR1,RELA,EPHB2

,ARF6,GRB10,RAP1B,EPHA2,RBM39,

| | | | | | | |
|---|---|---|---|---|---|---|
| **TF** | Factor: ZF5; n | TF:M10438_1 | 5.03E-140 | 139.298798 | 15001 | 2679 |
| **TF** | Factor: E2F-4 | TF:M09894_1 | 1.37E-139 | 138.86322 | 14041 | 2679 |
| **TF** | Factor: ZF5; n | TF:M00716_1 | 3.64E-133 | 132.439369 | 14230 | 2679 |
| **TF** | Factor: E2F-2 | TF:M11531 | 6.38E-133 | 132.19546 | 13379 | 2679 |
| **TF** | Factor: ETF; n | TF:M00695 | 4.41E-131 | 130.355318 | 10500 | 2679 |
| **TF** | Factor: E2F-2 | TF:M11529_1 | 6.77E-126 | 125.169389 | 14592 | 2679 |
| **TF** | Factor: E2F-3 | TF:M08526_1 | 1.15E-124 | 123.938441 | 15130 | 2679 |
| **TF** | Factor: ETF; n | TF:M00695_1 | 1.21E-124 | 123.917424 | 7067 | 2679 |
| **TF** | Factor: E2F-2 | TF:M11531_1 | 4.48E-122 | 121.348928 | 12431 | 2679 |
| **TF** | Factor: E2F; n | TF:M00803 | 1.13E-121 | 120.945464 | 13289 | 2679 |
| **TF** | Factor: E2F-3 | TF:M02089 | 1.79E-118 | 117.746064 | 13271 | 2679 |
| **TF** | Factor: E2F-3 | TF:M02089_1 | 3.29E-118 | 117.482297 | 9425 | 2679 |
| **TF** | Factor: Sp1; n | TF:M00196 | 5.09E-114 | 113.293622 | 10303 | 2679 |
| **TF** | Factor: Sp1; n | TF:M07395 | 1.20E-112 | 111.921924 | 10721 | 2679 |
| **TF** | Factor: Churcl | TF:M00986_1 | 3.45E-110 | 109.462105 | 10429 | 2679 |
| **TF** | Factor: TCF-1 | TF:M11603_1 | 4.05E-109 | 108.392988 | 13293 | 2679 |
| **TF** | Factor: Sp1; n | TF:M00932 | 1.27E-108 | 107.897306 | 10490 | 2679 |
| **TF** | Factor: sp4; n | TF:M10072 | 2.74E-107 | 106.561474 | 10571 | 2679 |
| **TF** | Factor: TIEG1 | TF:M12351_1 | 3.30E-107 | 106.480941 | 8242 | 2679 |
| **TF** | Factor: Sp1; n | TF:M00931 | 4.56E-107 | 106.341077 | 10344 | 2679 |
| **TF** | Factor: E2F; n | TF:M00803_1 | 2.35E-106 | 105.62932 | 10294 | 2679 |
| **TF** | Factor: Egr-1; | TF:M04869 | 1.26E-105 | 104.900667 | 11319 | 2679 |
| **TF** | Factor: Egr-1; | TF:M04869_1 | 8.01E-105 | 104.096234 | 10315 | 2679 |
| **TF** | Factor: ZF5; n | TF:M10438 | 4.72E-104 | 103.325745 | 16373 | 2679 |
| **TF** | Factor: E2F-1 | TF:M07206_1 | 3.30E-103 | 102.481475 | 10947 | 2679 |
| **TF** | Factor: ZF5; n | TF:M00333_1 | 5.54E-102 | 101.256737 | 12512 | 2679 |
| **TF** | Factor: E2F-1 | TF:M08205_1 | 2.90E-101 | 100.537721 | 10502 | 2679 |
| **TF** | Factor: MAZ; | TF:M09636_1 | 1.83E-100 | 99.7377753 | 14202 | 2679 |
| **TF** | Factor: ZF5; n | TF:M00716 | 2.85E-100 | 99.5445513 | 16245 | 2679 |
| **TF** | Factor: E2F-2 | TF:M11529 | 6.37E-100 | 99.1956397 | 16407 | 2679 |
| **TF** | Factor: E2F-1 | TF:M07206 | 6.45E-100 | 99.1905205 | 14705 | 2679 |
| **TF** | Factor: Sp1; n | TF:M00933 | 1.02E-99 | 98.9923261 | 9772 | 2679 |
| **TF** | Factor: MOVC | TF:M01104 | 5.56E-98 | 97.2549757 | 10336 | 2679 |
| **TF** | Factor: E2F-1 | TF:M08525_1 | 1.56E-97 | 96.8062398 | 15410 | 2679 |
| **TF** | Factor: E2F-4 | TF:M09894 | 2.94E-97 | 96.5317212 | 16345 | 2679 |
| **TF** | Factor: SP2; n | TF:M03807 | 4.17E-96 | 95.3798345 | 8460 | 2679 |
| **TF** | Factor: E2F-1 | TF:M08205 | 1.71E-95 | 94.7681787 | 14560 | 2679 |
| **TF** | Factor: TCF-1 | TF:M11601_1 | 3.50E-95 | 94.4555344 | 11153 | 2679 |
| **TF** | Factor: BTEB | TF:M09723 | 1.90E-94 | 93.7206885 | 10217 | 2679 |
| **TF** | Factor: E2F-1 | TF:M09892 | 2.69E-93 | 92.5696467 | 13661 | 2679 |
| **TF** | Factor: ZF5; n | TF:M00333 | 2.93E-93 | 92.5324788 | 15217 | 2679 |
| **TF** | Factor: TCF-1 | TF:M11601 | 3.93E-93 | 92.4059502 | 14964 | 2679 |
| **TF** | Factor: E2F-1 | TF:M07250 | 1.38E-92 | 91.8614101 | 10809 | 2679 |
| **TF** | Factor: TIEG1 | TF:M12351 | 1.45E-92 | 91.837282 | 12403 | 2679 |
| **TF** | Factor: ETF; n | TF:M07039_1 | 2.33E-92 | 91.6329744 | 13735 | 2679 |

| | | | | | | |
|---|---|---|---|---|---|---|
| **TF** | Factor: Churcl | TF:M00986 | 4.14E-92 | 91.3826162 | 14305 | 2679 |
| **TF** | Factor: BTEB3 | TF:M09826 | 4.20E-92 | 91.3771979 | 11560 | 2679 |
| **TF** | Factor: BTEB3 | TF:M09826_1 | 1.91E-91 | 90.7194034 | 7557 | 2679 |
| **TF** | Factor: E2F-2 | TF:M11530 | 3.68E-89 | 88.4336692 | 14722 | 2679 |
| **TF** | Factor: p300; | TF:M04826_1 | 6.16E-89 | 88.2102168 | 15413 | 2679 |
| **TF** | Factor: WT1; | TF:M02036 | 2.45E-88 | 87.6107068 | 9820 | 2679 |
| **TF** | Factor: Sp1; n | TF:M10071 | 3.31E-86 | 85.4806449 | 8550 | 2679 |
| **TF** | Factor: pax-6; | TF:M11882_1 | 1.05E-85 | 84.9778855 | 11519 | 2679 |
| **TF** | Factor: Sp1; n | TF:M00932_1 | 3.67E-85 | 84.4356636 | 6076 | 2679 |
| **TF** | Factor: E2F-4 | TF:M07380 | 5.22E-85 | 84.2826907 | 13143 | 2679 |
| **TF** | Factor: BTEB1 | TF:M09723_1 | 2.36E-83 | 82.6270237 | 6142 | 2679 |
| **TF** | Factor: sp4; n | TF:M10072_1 | 3.34E-83 | 82.4758158 | 6272 | 2679 |
| **TF** | Factor: WT1; | TF:M03893 | 4.58E-83 | 82.3386641 | 8336 | 2679 |
| **TF** | Factor: KLF3; | TF:M09970 | 5.77E-83 | 82.2387625 | 7770 | 2679 |
| **TF** | Factor: E2F-1 | TF:M04515_1 | 1.15E-82 | 81.9389044 | 12267 | 2679 |
| **TF** | Factor: MOVC | TF:M01104_1 | 2.11E-82 | 81.6758447 | 5650 | 2679 |
| **TF** | Factor: E2F-1 | TF:M04515 | 4.10E-82 | 81.3871994 | 13444 | 2679 |
| **TF** | Factor: Sp2; n | TF:M03567 | 5.02E-82 | 81.2995342 | 7746 | 2679 |
| **TF** | Factor: Sp1; n | TF:M00933_1 | 5.68E-82 | 81.2458124 | 5191 | 2679 |
| **TF** | Factor: SP1; n | TF:M07226 | 4.52E-81 | 80.3448015 | 8333 | 2679 |
| **TF** | Factor: BEN; r | TF:M01240_1 | 1.43E-80 | 79.8434616 | 13334 | 2679 |
| **TF** | Factor: Sp1; n | TF:M00931_1 | 1.58E-80 | 79.8025724 | 5934 | 2679 |
| **TF** | Factor: AP-2g | TF:M08441_1 | 1.61E-79 | 78.7936876 | 12848 | 2679 |
| **TF** | Factor: Egr-1; | TF:M07354 | 2.22E-79 | 78.6533893 | 7842 | 2679 |
| **TF** | Factor: Sp1; n | TF:M07395_1 | 4.50E-79 | 78.3472298 | 6376 | 2679 |
| **TF** | Factor: RNF96 | TF:M01199 | 1.05E-78 | 77.9805028 | 8535 | 2679 |
| **TF** | Factor: CPBP; | TF:M09973 | 4.46E-78 | 77.350425 | 10922 | 2679 |
| **TF** | Factor: ZNF14 | TF:M09834 | 5.08E-78 | 77.2939958 | 6998 | 2679 |
| **TF** | Factor: IRX-1; | TF:M11018_1 | 5.84E-78 | 77.2337237 | 16079 | 2679 |
| **TF** | Factor: E2F-4 | TF:M04823 | 3.00E-77 | 76.5226213 | 12880 | 2679 |
| **TF** | Factor: E2F-3 | TF:M08207_1 | 5.58E-77 | 76.2533417 | 15626 | 2679 |
| **TF** | Factor: KROX | TF:M00982 | 8.11E-77 | 76.0908069 | 7866 | 2679 |
| **TF** | Factor: KLF15 | TF:M12160 | 1.69E-76 | 75.772714 | 12836 | 2679 |
| **TF** | Factor: E2F-2 | TF:M11530_1 | 3.19E-76 | 75.4959493 | 13561 | 2679 |
| **TF** | Factor: GKLF; | TF:M12173 | 1.09E-75 | 74.9615624 | 10088 | 2679 |
| **TF** | Factor: KLF15 | TF:M12160_1 | 1.34E-75 | 74.8726562 | 8085 | 2679 |
| **TF** | Factor: Sp1; n | TF:M00196_1 | 1.96E-75 | 74.7088308 | 5949 | 2679 |
| **TF** | Factor: E2F-3 | TF:M08526 | 2.45E-75 | 74.6114655 | 17025 | 2679 |
| **TF** | Factor: PATZ; | TF:M10026 | 2.98E-75 | 74.5260225 | 8440 | 2679 |
| **TF** | Factor: Sp1; n | TF:M07063 | 3.49E-75 | 74.4565726 | 7736 | 2679 |
| **TF** | Factor: Sp2; n | TF:M10435 | 3.49E-75 | 74.4565726 | 7736 | 2679 |
| **TF** | Factor: EGR; | TF:M08878 | 2.48E-74 | 73.6048292 | 7527 | 2679 |
| **TF** | Factor: TCF-1 | TF:M11603 | 5.37E-74 | 73.2698642 | 16517 | 2679 |
| **TF** | Factor: AP-2al | TF:M01857 | 1.51E-73 | 72.8198738 | 9223 | 2679 |
| **TF** | Factor: E2F-4 | TF:M07380_1 | 2.31E-73 | 72.6364834 | 8046 | 2679 |

| | | | | | | |
|---|---|---|---|---|---|---|
| **TF** | Factor: BTEB2 | TF:M07277 | 1.09E-72 | 71.962094 | 8539 | 2679 |
| **TF** | Factor: Sp3; n | TF:M07615 | 1.45E-72 | 71.838248 | 6845 | 2679 |
| **TF** | Factor: SP1; n | TF:M02281 | 1.77E-72 | 71.751377 | 6518 | 2679 |
| **TF** | Factor: CPBP; | TF:M09973_1 | 1.84E-72 | 71.7345189 | 6648 | 2679 |
| **TF** | Factor: Sp1; n | TF:M04953 | 2.09E-72 | 71.6789045 | 8758 | 2679 |
| **TF** | Factor: E2F-1 | TF:M08525 | 1.83E-70 | 69.7366687 | 17224 | 2679 |
| **TF** | Factor: WT1; | TF:M02036_1 | 2.26E-70 | 69.64581 | 5308 | 2679 |
| **TF** | Factor: SP1; n | TF:M09765 | 4.61E-70 | 69.3366937 | 7518 | 2679 |
| **TF** | Factor: ETF; n | TF:M07039 | 4.61E-70 | 69.336067 | 16527 | 2679 |
| **TF** | Factor: E2F-1 | TF:M09892_1 | 9.69E-70 | 69.0136726 | 9248 | 2679 |
| **TF** | Factor: E2F-4 | TF:M07084 | 1.25E-69 | 68.9041007 | 10945 | 2679 |
| **TF** | Factor: sp4; n | TF:M10530 | 2.48E-69 | 68.6060899 | 8285 | 2679 |
| **TF** | Factor: Egr-1; | TF:M01873 | 1.06E-68 | 67.9763801 | 6773 | 2679 |
| **TF** | Factor: ZBP89 | TF:M07397 | 2.53E-68 | 67.5967461 | 7190 | 2679 |
| **TF** | Factor: MAZ; | TF:M09636 | 3.66E-68 | 67.4360716 | 16380 | 2679 |
| **TF** | Factor: E2F-4 | TF:M04823_1 | 4.21E-68 | 67.3752647 | 7916 | 2679 |
| **TF** | Factor: SP1; n | TF:M07226_1 | 4.94E-68 | 67.3064589 | 3931 | 2679 |
| **TF** | Factor: Sp1; n | TF:M00008 | 1.14E-66 | 65.9438749 | 10131 | 2679 |
| **TF** | Factor: E2F-1 | TF:M08523_1 | 4.39E-66 | 65.3575484 | 7463 | 2679 |
| **TF** | Factor: SP1:SI | TF:M01219 | 4.50E-66 | 65.3470703 | 6856 | 2679 |
| **TF** | Factor: GCMa | TF:M08487_1 | 6.42E-65 | 64.1923246 | 8853 | 2679 |
| **TF** | Factor: ZIC4; | TF:M12227_1 | 8.95E-65 | 64.0480859 | 5925 | 2679 |
| **TF** | Factor: PATZ; | TF:M10026_1 | 1.09E-64 | 63.9619038 | 4924 | 2679 |
| **TF** | Factor: AP-2al | TF:M01857_1 | 1.21E-64 | 63.9167049 | 4625 | 2679 |
| **TF** | Factor: KLF15 | TF:M12158 | 2.42E-64 | 63.6165437 | 11165 | 2679 |
| **TF** | Factor: RNF96 | TF:M01199_1 | 5.73E-64 | 63.2415402 | 4335 | 2679 |
| **TF** | Factor: AP-2; | TF:M00189 | 7.36E-64 | 63.1334075 | 10687 | 2679 |
| **TF** | Factor: ZIC4; | TF:M12227 | 1.33E-63 | 62.876011 | 11151 | 2679 |
| **TF** | Factor: Miz-1; | TF:M10112 | 1.66E-63 | 62.7786225 | 8768 | 2679 |
| **TF** | Factor: TAFII2 | TF:M10086 | 2.06E-63 | 62.6863364 | 9342 | 2679 |
| **TF** | Factor: pax-6; | TF:M11882 | 3.41E-63 | 62.4678747 | 15642 | 2679 |
| **TF** | Factor: Kaiso; | TF:M03876_1 | 1.40E-62 | 61.8546331 | 9160 | 2679 |
| **TF** | Factor: Sp1; n | TF:M04953_1 | 1.54E-62 | 61.8117959 | 4099 | 2679 |
| **TF** | Factor: SP1; n | TF:M03920 | 5.60E-62 | 61.2515163 | 7329 | 2679 |
| **TF** | Factor: MAZ; | TF:M10432 | 8.41E-62 | 61.0749807 | 9328 | 2679 |
| **TF** | Factor: E2F-4 | TF:M02090 | 1.97E-61 | 60.7059783 | 12068 | 2679 |
| **TF** | Factor: CNOT | TF:M01253 | 2.70E-61 | 60.5683588 | 3948 | 2679 |
| **TF** | Factor: ZNF46 | TF:M12313 | 5.89E-61 | 60.230213 | 5877 | 2679 |
| **TF** | Factor: WT1; | TF:M03893_1 | 2.86E-60 | 59.543357 | 3997 | 2679 |
| **TF** | Factor: WT1; | TF:M01118 | 4.26E-60 | 59.3704791 | 6869 | 2679 |
| **TF** | Factor: AP-2b | TF:M01858 | 1.25E-59 | 58.9042185 | 7460 | 2679 |
| **TF** | Factor: E2F-1 | TF:M07250_1 | 1.69E-59 | 58.7711734 | 5555 | 2679 |
| **TF** | Factor: SP2; n | TF:M01783 | 2.18E-59 | 58.6619999 | 9222 | 2679 |
| **TF** | Factor: Sp2; n | TF:M07129 | 2.75E-59 | 58.5604547 | 6297 | 2679 |
| **TF** | Factor: E2F-3 | TF:M08207 | 3.40E-59 | 58.4680167 | 17225 | 2679 |

| | | | | | | |
|---|---|---|---|---|---|---|
| **TF** | Factor: AP2; n | TF:M08867 | 3.69E-59 | 58.4329899 | 10120 | 2679 |
| **TF** | Factor: Zbtb3 | TF:M12345 | 4.98E-59 | 58.3025401 | 5578 | 2679 |
| **TF** | Factor: AP-2; | TF:M00189_1 | 1.14E-58 | 57.9436006 | 6114 | 2679 |
| **TF** | Factor: E2F-1 | TF:M00428 | 2.02E-58 | 57.6956951 | 9030 | 2679 |
| **TF** | Factor: E2F-1 | TF:M08523 | 2.47E-58 | 57.6075726 | 12268 | 2679 |
| **TF** | Factor: GKLF | TF:M07289_1 | 6.00E-58 | 57.2216064 | 10645 | 2679 |
| **TF** | Factor: KLF14 | TF:M12183 | 9.24E-58 | 57.0343792 | 6908 | 2679 |
| **TF** | Factor: E2F-7 | TF:M09896 | 4.33E-57 | 56.3631296 | 10034 | 2679 |
| **TF** | Factor: GCMa | TF:M08487 | 4.42E-57 | 56.3549276 | 14054 | 2679 |
| **TF** | Factor: BEN; r | TF:M01240 | 7.61E-57 | 56.118619 | 16656 | 2679 |
| **TF** | Factor: YY2; n | TF:M03925 | 1.44E-56 | 55.8425089 | 4690 | 2679 |
| **TF** | Factor: Sp1; n | TF:M10071_1 | 1.86E-56 | 55.7312148 | 4641 | 2679 |
| **TF** | Factor: sp4; n | TF:M10530_1 | 3.46E-56 | 55.4614872 | 3799 | 2679 |
| **TF** | Factor: RREB | TF:M05439 | 4.21E-56 | 55.3760099 | 8689 | 2679 |
| **TF** | Factor: AP-2; | TF:M00800 | 5.44E-56 | 55.2640435 | 9172 | 2679 |
| **TF** | Factor: E2F-1 | TF:M04516_1 | 1.03E-55 | 54.9884814 | 10299 | 2679 |
| **TF** | Factor: MAZ; | TF:M10432_1 | 1.25E-55 | 54.9033625 | 4390 | 2679 |
| **TF** | Factor: Miz-1; | TF:M10112_1 | 1.51E-55 | 54.8206205 | 4096 | 2679 |
| **TF** | Factor: Hey1; | TF:M11058 | 1.78E-55 | 54.7501169 | 13278 | 2679 |
| **TF** | Factor: E2F-1 | TF:M04516 | 2.09E-55 | 54.6792998 | 11467 | 2679 |
| **TF** | Factor: Kaiso; | TF:M03876 | 2.31E-55 | 54.6364785 | 13737 | 2679 |
| **TF** | Factor: SP1; n | TF:M01303 | 6.46E-55 | 54.1899497 | 7935 | 2679 |
| **TF** | Factor: RUNX | TF:M04106_1 | 7.05E-55 | 54.1519842 | 8449 | 2679 |
| **TF** | Factor: IRX-1; | TF:M11018 | 7.30E-55 | 54.1368519 | 17131 | 2679 |
| **TF** | Factor: Zfx; m | TF:M01593 | 8.54E-55 | 54.0687856 | 4198 | 2679 |
| **TF** | Factor: EGR1 | TF:M03896 | 1.82E-54 | 53.7410438 | 6153 | 2679 |
| **TF** | Factor: TAFII2 | TF:M10086_1 | 3.24E-54 | 53.489936 | 4184 | 2679 |
| **TF** | Factor: CHD2 | TF:M04710 | 2.73E-53 | 52.563096 | 14245 | 2679 |
| **TF** | Factor: Sp2; n | TF:M09658 | 5.49E-53 | 52.2604112 | 5297 | 2679 |
| **TF** | Factor: E2F; n | TF:M00426 | 1.14E-52 | 51.9442634 | 12013 | 2679 |
| **TF** | Factor: WT1; | TF:M01118_1 | 2.33E-52 | 51.6328889 | 2791 | 2679 |
| **TF** | Factor: SP4; n | TF:M03922 | 2.50E-52 | 51.6025364 | 6073 | 2679 |
| **TF** | Factor: E2F-1 | TF:M00938 | 3.44E-52 | 51.4631847 | 7376 | 2679 |
| **TF** | Factor: Sp1; n | TF:M10529 | 5.79E-52 | 51.2370308 | 6933 | 2679 |
| **TF** | Factor: CHD2 | TF:M04710_1 | 2.31E-51 | 50.6358044 | 11589 | 2679 |
| **TF** | Factor: ZBP89 | TF:M07397_1 | 2.69E-51 | 50.5700786 | 2973 | 2679 |
| **TF** | Factor: SP2; n | TF:M03807_1 | 3.40E-51 | 50.4681478 | 4218 | 2679 |
| **TF** | Factor: MAZ; | TF:M09984 | 7.88E-51 | 50.1034008 | 9627 | 2679 |
| **TF** | Factor: EGR1 | TF:M07208 | 1.23E-50 | 49.9101302 | 6221 | 2679 |
| **TF** | Factor: Sp1; n | TF:M12152 | 1.46E-50 | 49.8365397 | 7381 | 2679 |
| **TF** | Factor: E2F-1 | TF:M00736 | 2.67E-50 | 49.5736652 | 10495 | 2679 |
| **TF** | Factor: ZNF14 | TF:M09834_1 | 3.37E-50 | 49.4724865 | 2891 | 2679 |
| **TF** | Factor: Sp6; n | TF:M05361 | 4.14E-50 | 49.3826368 | 8651 | 2679 |
| **TF** | Factor: Sp2; n | TF:M05332 | 4.14E-50 | 49.3826368 | 8651 | 2679 |
| **TF** | Factor: CPBP; | TF:M05444 | 4.14E-50 | 49.3826368 | 8651 | 2679 |

| | | | | | | |
|---|---|---|---|---|---|---|
| **TF** | Factor: E2F; n | TF:M00427 | 5.61E-50 | 49.2510757 | 9597 | 2679 |
| **TF** | Factor: Osx; n | TF:M07329 | 6.66E-50 | 49.1765228 | 6851 | 2679 |
| **TF** | Factor: E2F; n | TF:M00425 | 1.26E-49 | 48.8983004 | 11168 | 2679 |
| **TF** | Factor: ZBP9 | TF:M12057 | 1.38E-49 | 48.8601763 | 5253 | 2679 |
| **TF** | Factor: SP1; n | TF:M02281_1 | 1.58E-49 | 48.8002191 | 2575 | 2679 |
| **TF** | Factor: GKLF | TF:M07289 | 1.81E-49 | 48.741303 | 14779 | 2679 |
| **TF** | Factor: KLF1 | TF:M12158_1 | 2.44E-49 | 48.6120325 | 6041 | 2679 |
| **TF** | Factor: E2F-6 | TF:M09895 | 3.55E-49 | 48.4495756 | 7857 | 2679 |
| **TF** | Factor: ctcf; n | TF:M10426 | 4.59E-49 | 48.3382891 | 9389 | 2679 |
| **TF** | Factor: BTEB | TF:M07277_1 | 6.44E-49 | 48.1908608 | 4169 | 2679 |
| **TF** | Factor: AP-2b | TF:M01858_1 | 7.15E-49 | 48.145849 | 3152 | 2679 |
| **TF** | Factor: Hey1; | TF:M11058_1 | 7.70E-49 | 48.1134023 | 11001 | 2679 |
| **TF** | Factor: E2F-3 | TF:M08208 | 1.84E-48 | 47.7361087 | 12690 | 2679 |
| **TF** | Factor: Egr-1; | TF:M07354_1 | 1.85E-48 | 47.7325868 | 3573 | 2679 |
| **TF** | Factor: Sp2; n | TF:M06948 | 3.83E-48 | 47.4164974 | 8931 | 2679 |
| **TF** | Factor: TEL1; | TF:M02070 | 5.30E-48 | 47.275768 | 10938 | 2679 |
| **TF** | Factor: AP-2g | TF:M08441 | 5.66E-48 | 47.2471383 | 16491 | 2679 |
| **TF** | Factor: E2F-4 | TF:M07084_1 | 6.63E-48 | 47.1781596 | 5470 | 2679 |
| **TF** | Factor: pax-6; | TF:M11883 | 7.58E-48 | 47.1203082 | 13587 | 2679 |
| **TF** | Factor: Sp1; n | TF:M12151 | 1.07E-47 | 46.9714646 | 6864 | 2679 |
| **TF** | Factor: SP1:S | TF:M01219_1 | 1.15E-47 | 46.9411499 | 2952 | 2679 |
| **TF** | Factor: E2F-4 | TF:M11535 | 1.49E-47 | 46.8275745 | 10966 | 2679 |
| **TF** | Factor: WT1; | TF:M07436 | 1.55E-47 | 46.8096156 | 6227 | 2679 |
| **TF** | Factor: GKLF | TF:M12173_1 | 1.92E-47 | 46.7166435 | 4992 | 2679 |
| **TF** | Factor: MAZ; | TF:M09984_1 | 5.98E-47 | 46.2229689 | 5599 | 2679 |
| **TF** | Factor: BTEB | TF:M12186 | 7.14E-47 | 46.146004 | 9195 | 2679 |
| **TF** | Factor: AP2; n | TF:M08867_1 | 1.38E-46 | 45.8602573 | 5202 | 2679 |
| **TF** | Factor: Egr-1; | TF:M04950 | 1.38E-46 | 45.8599777 | 11032 | 2679 |
| **TF** | Factor: AP-2; | TF:M00915 | 1.55E-46 | 45.8106941 | 9683 | 2679 |
| **TF** | Factor: CTCF; | TF:M08911 | 1.68E-46 | 45.7757715 | 9168 | 2679 |
| **TF** | Factor: E2F-1 | TF:M00940 | 4.21E-46 | 45.3761228 | 7873 | 2679 |
| **TF** | Factor: WT1; | TF:M10108 | 7.76E-46 | 45.1099998 | 6585 | 2679 |
| **TF** | Factor: AP-2a | TF:M00469 | 9.83E-46 | 45.0073264 | 5182 | 2679 |
| **TF** | Factor: E2F-4 | TF:M00738 | 4.25E-45 | 44.3712068 | 9912 | 2679 |
| **TF** | Factor: CKRO | TF:M01175 | 5.20E-45 | 44.2837636 | 7354 | 2679 |
| **TF** | Factor: p300; | TF:M04826 | 7.90E-45 | 44.1022492 | 17982 | 2679 |
| **TF** | Factor: RREB | TF:M05439_1 | 1.34E-44 | 43.8737048 | 3218 | 2679 |
| **TF** | Factor: KROX | TF:M00982_1 | 1.43E-44 | 43.8461503 | 3531 | 2679 |
| **TF** | Factor: EGR; | TF:M08878_1 | 1.06E-43 | 42.9765104 | 3235 | 2679 |
| **TF** | Factor: TR4; r | TF:M04934_1 | 1.12E-43 | 42.9504201 | 15594 | 2679 |
| **TF** | Factor: E2F-4 | TF:M02090_1 | 1.21E-43 | 42.9183256 | 6427 | 2679 |
| **TF** | Factor: IRX2a | TF:M11022 | 1.56E-43 | 42.8061534 | 9572 | 2679 |
| **TF** | Factor: E2F-1 | TF:M00431 | 2.07E-43 | 42.6840427 | 7333 | 2679 |
| **TF** | Factor: Sp3; n | TF:M12153 | 2.74E-43 | 42.5620875 | 6237 | 2679 |
| **TF** | Factor: ZNF6 | TF:M09734 | 4.78E-43 | 42.3207137 | 11788 | 2679 |

| | | | | | | |
|---|---|---|---|---|---|---|
| **TF** | Factor: TFAP2 | TF:M04148_1 | 7.14E-43 | 42.1460357 | 9666 | 2679 |
| **TF** | Factor: TFAP2 | TF:M04148 | 9.41E-43 | 42.0264972 | 10108 | 2679 |
| **TF** | Factor: WT1; | TF:M05327 | 1.37E-42 | 41.8641072 | 5543 | 2679 |
| **TF** | Factor: Sp1; n | TF:M00008_1 | 1.46E-42 | 41.8357691 | 4966 | 2679 |
| **TF** | Factor: E2F-3 | TF:M08209_1 | 3.09E-42 | 41.5104354 | 7849 | 2679 |
| **TF** | Factor: E2F-7 | TF:M09896_1 | 4.50E-42 | 41.346564 | 4566 | 2679 |
| **TF** | Factor: pax-6; | TF:M11883_1 | 8.22E-42 | 41.0849555 | 7825 | 2679 |
| **TF** | Factor: BTEB: | TF:M07409 | 8.70E-42 | 41.0606129 | 5229 | 2679 |
| **TF** | Factor: Sp2; n | TF:M10435_1 | 9.25E-42 | 41.0338853 | 3513 | 2679 |
| **TF** | Factor: Sp1; n | TF:M07063_1 | 9.25E-42 | 41.0338853 | 3513 | 2679 |
| **TF** | Factor: ZBP9 | TF:M12057_1 | 1.14E-41 | 40.9446268 | 1366 | 2679 |
| **TF** | Factor: NRF-1 | TF:M09641 | 1.64E-41 | 40.7845009 | 3253 | 2679 |
| **TF** | Factor: E2F-1 | TF:M11533 | 2.92E-41 | 40.5345155 | 13223 | 2679 |
| **TF** | Factor: NGFI- | TF:M00244 | 2.98E-41 | 40.5254005 | 6221 | 2679 |
| **TF** | Factor: Sp4; n | TF:M07617 | 3.62E-41 | 40.4414381 | 3606 | 2679 |
| **TF** | Factor: SP1; n | TF:M09765_1 | 3.92E-41 | 40.4062481 | 3328 | 2679 |
| **TF** | Factor: Sp2; n | TF:M03567_1 | 4.66E-41 | 40.3315442 | 3627 | 2679 |
| **TF** | Factor: AP-2; | TF:M00915_1 | 5.77E-41 | 40.238718 | 4572 | 2679 |
| **TF** | Factor: ER71: | TF:M08266_1 | 7.21E-41 | 40.1422432 | 9053 | 2679 |
| **TF** | Factor: Elk-1:I | TF:M08225_1 | 7.60E-41 | 40.1192894 | 8075 | 2679 |
| **TF** | Factor: YY1; n | TF:M03924 | 8.03E-41 | 40.0955356 | 10195 | 2679 |
| **TF** | Factor: BTEB: | TF:M01865 | 8.82E-41 | 40.0543888 | 9976 | 2679 |
| **TF** | Factor: Rb:E2 | TF:M00740 | 1.13E-40 | 39.945773 | 10536 | 2679 |
| **TF** | Factor: Sp1; n | TF:M10529_1 | 2.14E-40 | 39.6696081 | 2838 | 2679 |
| **TF** | Factor: EGR1; | TF:M09761 | 2.37E-40 | 39.6253337 | 5191 | 2679 |
| **TF** | Factor: GKLF; | TF:M07040_1 | 3.78E-40 | 39.4225443 | 8225 | 2679 |
| **TF** | Factor: AP-2g | TF:M00470 | 5.51E-40 | 39.2587009 | 9473 | 2679 |
| **TF** | Factor: E2F; n | TF:M00425_1 | 5.61E-40 | 39.250672 | 5407 | 2679 |
| **TF** | Factor: Osx; n | TF:M07329_1 | 5.64E-40 | 39.2488877 | 2519 | 2679 |
| **TF** | Factor: DP-1; | TF:M10098 | 5.91E-40 | 39.228077 | 7103 | 2679 |
| **TF** | Factor: RUNX: | TF:M04106 | 6.64E-40 | 39.1777995 | 13969 | 2679 |
| **TF** | Factor: KLF3; | TF:M09970_1 | 7.36E-40 | 39.133395 | 3803 | 2679 |
| **TF** | Factor: AP-2re | TF:M09967 | 9.54E-40 | 39.0204126 | 4040 | 2679 |
| **TF** | Factor: ZFX; n | TF:M10121 | 1.68E-39 | 38.7743542 | 4020 | 2679 |
| **TF** | Factor: AP-2; | TF:M00800_1 | 2.04E-39 | 38.6893221 | 4278 | 2679 |
| **TF** | Factor: YY1; n | TF:M10436 | 2.82E-39 | 38.5502166 | 8301 | 2679 |
| **TF** | Factor: PLAG | TF:M01973 | 3.48E-39 | 38.4580546 | 6207 | 2679 |
| **TF** | Factor: Egr-1; | TF:M09603 | 3.68E-39 | 38.4346703 | 4660 | 2679 |
| **TF** | Factor: Pax-5; | TF:M00144 | 5.06E-39 | 38.2954445 | 13330 | 2679 |
| **TF** | Factor: E2F-3 | TF:M08208_1 | 6.88E-39 | 38.1621665 | 7175 | 2679 |
| **TF** | Factor: Egr-3; | TF:M00245 | 8.29E-39 | 38.0813418 | 7361 | 2679 |
| **TF** | Factor: RUNX: | TF:M04110 | 1.53E-38 | 37.815195 | 9352 | 2679 |
| **TF** | Factor: AP-2al | TF:M00469_1 | 1.56E-38 | 37.8066444 | 3350 | 2679 |
| **TF** | Factor: EKLF; | TF:M09969 | 1.60E-38 | 37.7961068 | 4610 | 2679 |
| **TF** | Factor: Egr-2; | TF:M09898 | 3.35E-38 | 37.4745098 | 6551 | 2679 |

| | | | | | | |
|---|---|---|---|---|---|---|
| **TF** | Factor: CTCF; | TF:M03895 | 3.48E-38 | 37.4578375 | 9333 | 2679 |
| **TF** | Factor: Elk-1; | TF:M11435 | 3.71E-38 | 37.4307002 | 11838 | 2679 |
| **TF** | Factor: pax-6; | TF:M11880_1 | 3.98E-38 | 37.4000469 | 7112 | 2679 |
| **TF** | Factor: ELK1; | TF:M03971 | 4.23E-38 | 37.3737167 | 11390 | 2679 |
| **TF** | Factor: GKLF; | TF:M01588 | 8.07E-38 | 37.0931009 | 9003 | 2679 |
| **TF** | Factor: Sp2; n | TF:M12156 | 9.04E-38 | 37.0440636 | 4355 | 2679 |
| **TF** | Factor: Egr-1; | TF:M01873_1 | 9.49E-38 | 37.0225867 | 2678 | 2679 |
| **TF** | Factor: RUNX | TF:M04107_1 | 1.43E-37 | 36.8436753 | 6513 | 2679 |
| **TF** | Factor: E2F-3 | TF:M08209 | 1.54E-37 | 36.8114499 | 13784 | 2679 |
| **TF** | Factor: Sp2; n | TF:M06948_1 | 1.84E-37 | 36.734738 | 6233 | 2679 |
| **TF** | Factor: E2F; n | TF:M00426_1 | 2.03E-37 | 36.6931307 | 5892 | 2679 |
| **TF** | Factor: AP-2g | TF:M00470_1 | 2.58E-37 | 36.5886293 | 5637 | 2679 |
| **TF** | Factor: E2F; n | TF:M08875 | 3.14E-37 | 36.5027744 | 6260 | 2679 |
| **TF** | Factor: MAZ; | TF:M02023 | 4.03E-37 | 36.3950341 | 7183 | 2679 |
| **TF** | Factor: GABP | TF:M01660 | 4.06E-37 | 36.3911267 | 11217 | 2679 |
| **TF** | Factor: ctcf; m | TF:M10426_1 | 7.62E-37 | 36.1181014 | 4390 | 2679 |
| **TF** | Factor: E2F-1 | TF:M00430 | 8.13E-37 | 36.0899974 | 8113 | 2679 |
| **TF** | Factor: E2F1; | TF:M08874 | 1.17E-36 | 35.93342 | 7267 | 2679 |
| **TF** | Factor: GKLF; | TF:M07040 | 1.19E-36 | 35.924354 | 12809 | 2679 |
| **TF** | Factor: Egr-1; | TF:M00243 | 1.29E-36 | 35.889569 | 5386 | 2679 |
| **TF** | Factor: IRX2a | TF:M11022_1 | 1.33E-36 | 35.8754571 | 6298 | 2679 |
| **TF** | Factor: ER71: | TF:M08266 | 1.55E-36 | 35.8109254 | 14504 | 2679 |
| **TF** | Factor: Sp3; n | TF:M07615_1 | 1.93E-36 | 35.7147179 | 2902 | 2679 |
| **TF** | Factor: TF3C- | TF:M04863 | 1.97E-36 | 35.706585 | 13733 | 2679 |
| **TF** | Factor: RUNX | TF:M04110_1 | 3.55E-36 | 35.449866 | 3796 | 2679 |
| **TF** | Factor: KLF17 | TF:M05386 | 3.57E-36 | 35.4473796 | 7952 | 2679 |
| **TF** | Factor: LKLF; | TF:M05499 | 3.57E-36 | 35.4473796 | 7952 | 2679 |
| **TF** | Factor: Elk-1; | TF:M11435_1 | 4.04E-36 | 35.3935087 | 5791 | 2679 |
| **TF** | Factor: SP2; n | TF:M01783_1 | 4.23E-36 | 35.3738252 | 4369 | 2679 |
| **TF** | Factor: CKRO | TF:M01175_1 | 7.39E-36 | 35.1315365 | 2690 | 2679 |
| **TF** | Factor: WT1; | TF:M10108_1 | 1.15E-35 | 34.9407487 | 2881 | 2679 |
| **TF** | Factor: pax-6; | TF:M11881 | 1.16E-35 | 34.9366258 | 13307 | 2679 |
| **TF** | Factor: Egr-1; | TF:M04797 | 1.51E-35 | 34.8210617 | 5133 | 2679 |
| **TF** | Factor: MAZ; | TF:M00649_1 | 2.07E-35 | 34.684259 | 4185 | 2679 |
| **TF** | Factor: YY1; n | TF:M03924_1 | 2.14E-35 | 34.6690592 | 3744 | 2679 |
| **TF** | Factor: TF3C- | TF:M04863_1 | 3.32E-35 | 34.4788654 | 8620 | 2679 |
| **TF** | Factor: AhR,; | TF:M00976 | 4.27E-35 | 34.3693165 | 4387 | 2679 |
| **TF** | Factor: MAZ; | TF:M00649 | 4.60E-35 | 34.3368352 | 9407 | 2679 |
| **TF** | Factor: SP1; n | TF:M01303_1 | 4.95E-35 | 34.3049658 | 3525 | 2679 |
| **TF** | Factor: RUNX | TF:M04107 | 5.02E-35 | 34.2996873 | 12479 | 2679 |
| **TF** | Factor: Kaiso; | TF:M04691_1 | 5.11E-35 | 34.2913977 | 10595 | 2679 |
| **TF** | Factor: E2F-1 | TF:M00736_1 | 7.50E-35 | 34.1246691 | 4694 | 2679 |
| **TF** | Factor: YY1; n | TF:M10436_1 | 9.83E-35 | 34.0075556 | 2313 | 2679 |
| **TF** | Factor: Egr-1; | TF:M12140 | 1.17E-34 | 33.9310485 | 5391 | 2679 |
| **TF** | Factor: ZNF5( | TF:M12085 | 1.66E-34 | 33.7809538 | 8554 | 2679 |

| | | | | | | |
|---|---|---|---|---|---|---|
| **TF** | Factor: ER81; | TF:M02065 | 1.80E-34 | 33.7451854 | 12666 | 2679 |
| **TF** | Factor: ZNF37 | TF:M12354_1 | 3.76E-34 | 33.4244886 | 5648 | 2679 |
| **TF** | Factor: AP-2b | TF:M11478 | 9.98E-34 | 33.0008944 | 10020 | 2679 |
| **TF** | Factor: NRF-1 | TF:M02102 | 1.05E-33 | 32.9805835 | 2040 | 2679 |
| **TF** | Factor: E2F-1 | TF:M11533_1 | 1.05E-33 | 32.9770609 | 8777 | 2679 |
| **TF** | Factor: Hey2; | TF:M11052 | 1.15E-33 | 32.9395266 | 12619 | 2679 |
| **TF** | Factor: GABP | TF:M01660_1 | 1.17E-33 | 32.9312803 | 5163 | 2679 |
| **TF** | Factor: Elk-1; | TF:M11434 | 1.23E-33 | 32.9115237 | 8290 | 2679 |
| **TF** | Factor: WT1; | TF:M07436_1 | 1.58E-33 | 32.8009106 | 2424 | 2679 |
| **TF** | Factor: AP-2g | TF:M03811 | 2.12E-33 | 32.6743586 | 8919 | 2679 |
| **TF** | Factor: EGR-1 | TF:M01972 | 2.78E-33 | 32.5565972 | 5388 | 2679 |
| **TF** | Factor: ZNF46 | TF:M12313_1 | 4.05E-33 | 32.3925109 | 2225 | 2679 |
| **TF** | Factor: SP4; n | TF:M01273 | 4.16E-33 | 32.3809804 | 3311 | 2679 |
| **TF** | Factor: HES-7 | TF:M11066 | 4.40E-33 | 32.3567734 | 11596 | 2679 |
| **TF** | Factor: Elk-1:I | TF:M08225 | 7.15E-33 | 32.1454526 | 14162 | 2679 |
| **TF** | Factor: E2F-1 | TF:M00516 | 8.14E-33 | 32.0895594 | 6972 | 2679 |
| **TF** | Factor: CTCF; | TF:M08911_1 | 1.25E-32 | 31.9027998 | 3956 | 2679 |
| **TF** | Factor: AP-2al | TF:M07348 | 2.04E-32 | 31.6911256 | 8104 | 2679 |
| **TF** | Factor: Zbtb3 | TF:M12345_1 | 2.20E-32 | 31.6574659 | 1703 | 2679 |
| **TF** | Factor: ER81; | TF:M02065_1 | 2.65E-32 | 31.5767193 | 6798 | 2679 |
| **TF** | Factor: NRF-1 | TF:M07052 | 2.89E-32 | 31.5393722 | 2141 | 2679 |
| **TF** | Factor: BTEB | TF:M01865_1 | 2.90E-32 | 31.5376178 | 4243 | 2679 |
| **TF** | Factor: nerf; n | TF:M09606 | 4.33E-32 | 31.3638744 | 6310 | 2679 |
| **TF** | Factor: CPBP; | TF:M01822_1 | 4.53E-32 | 31.3439707 | 18835 | 2679 |
| **TF** | Factor: ETS1; | TF:M03978 | 1.09E-31 | 30.9634901 | 12179 | 2679 |
| **TF** | Factor: Egr-1; | TF:M09897 | 1.41E-31 | 30.8499323 | 4267 | 2679 |
| **TF** | Factor: RUNX | TF:M04109_1 | 1.52E-31 | 30.8190287 | 3899 | 2679 |
| **TF** | Factor: pax-6; | TF:M11881_1 | 1.55E-31 | 30.8104562 | 7216 | 2679 |
| **TF** | Factor: GKLF | TF:M01588_1 | 2.07E-31 | 30.6849185 | 3986 | 2679 |
| **TF** | Factor: MAZ; | TF:M02023_1 | 2.32E-31 | 30.6345305 | 3138 | 2679 |
| **TF** | Factor: Egr-1; | TF:M04950_1 | 2.36E-31 | 30.6274661 | 5591 | 2679 |
| **TF** | Factor: ETS1; | TF:M03977 | 5.33E-31 | 30.2735898 | 10612 | 2679 |
| **TF** | Factor: Pax-5; | TF:M00144_1 | 8.01E-31 | 30.0961501 | 7203 | 2679 |
| **TF** | Factor: Elk-1:I | TF:M08226 | 8.96E-31 | 30.0478094 | 12154 | 2679 |
| **TF** | Factor: AP-2b | TF:M11478_1 | 1.11E-30 | 29.9563136 | 8102 | 2679 |
| **TF** | Factor: RUNX | TF:M04109 | 1.20E-30 | 29.9207673 | 9858 | 2679 |
| **TF** | Factor: Kaiso; | TF:M04691 | 1.22E-30 | 29.913386 | 15263 | 2679 |
| **TF** | Factor: ZNF61 | TF:M09862 | 2.07E-30 | 29.6832714 | 9432 | 2679 |
| **TF** | Factor: PLAG | TF:M01973_1 | 2.09E-30 | 29.6803803 | 2413 | 2679 |
| **TF** | Factor: TR4; r | TF:M04934 | 2.63E-30 | 29.580095 | 18272 | 2679 |
| **TF** | Factor: GCMa | TF:M08313_1 | 2.65E-30 | 29.5775103 | 9802 | 2679 |
| **TF** | Factor: ELK1; | TF:M03971_1 | 2.76E-30 | 29.5594699 | 7220 | 2679 |
| **TF** | Factor: pax-2; | TF:M11877_1 | 2.94E-30 | 29.5321947 | 5338 | 2679 |
| **TF** | Factor: ZSCAI | TF:M05432 | 3.01E-30 | 29.5216051 | 4145 | 2679 |
| **TF** | Factor: BTEB | TF:M12186_1 | 5.16E-30 | 29.2874839 | 3976 | 2679 |

| | | | | | | |
|---|---|---|---|---|---|---|
| **TF** | Factor: Sp1; n | TF:M12152_1 | 7.29E-30 | 29.1370056 | 2874 | 2679 |
| **TF** | Factor: Sp2; n | TF:M07129_1 | 1.04E-29 | 28.9820943 | 2433 | 2679 |
| **TF** | Factor: Kaiso; | TF:M09632 | 1.66E-29 | 28.7810145 | 3018 | 2679 |
| **TF** | Factor: pax-6; | TF:M11880 | 2.54E-29 | 28.5952257 | 13224 | 2679 |
| **TF** | Factor: Fli-1; r | TF:M11401 | 3.10E-29 | 28.5080271 | 12056 | 2679 |
| **TF** | Factor: KLF14 | TF:M12183_1 | 3.51E-29 | 28.455126 | 2498 | 2679 |
| **TF** | Factor: SP1; n | TF:M03920_1 | 4.72E-29 | 28.3257128 | 3047 | 2679 |
| **TF** | Factor: Erg; m | TF:M11396 | 7.95E-29 | 28.0994789 | 10863 | 2679 |
| **TF** | Factor: Sp6; n | TF:M05361_1 | 9.83E-29 | 28.0074083 | 3343 | 2679 |
| **TF** | Factor: CPBP; | TF:M05444_1 | 9.83E-29 | 28.0074083 | 3343 | 2679 |
| **TF** | Factor: Sp2; n | TF:M05332_1 | 9.83E-29 | 28.0074083 | 3343 | 2679 |
| **TF** | Factor: LBP-1 | TF:M11490 | 1.52E-28 | 27.8179568 | 6803 | 2679 |
| **TF** | Factor: ZBTB7 | TF:M03806 | 3.07E-28 | 27.5135507 | 3047 | 2679 |
| **TF** | Factor: ZNF37 | TF:M12354 | 3.79E-28 | 27.4215738 | 11033 | 2679 |
| **TF** | Factor: Erg; m | TF:M11397_1 | 4.98E-28 | 27.3028427 | 11846 | 2679 |
| **TF** | Factor: ETV7; | TF:M02071 | 5.50E-28 | 27.2593731 | 10214 | 2679 |
| **TF** | Factor: Pet-1; | TF:M09918 | 6.13E-28 | 27.2123753 | 13738 | 2679 |
| **TF** | Factor: Sp2; n | TF:M09658_1 | 6.37E-28 | 27.1956393 | 2040 | 2679 |
| **TF** | Factor: E2F; n | TF:M00939 | 7.40E-28 | 27.1310349 | 6496 | 2679 |
| **TF** | Factor: ZNF61 | TF:M09862_1 | 1.02E-27 | 26.9916661 | 4212 | 2679 |
| **TF** | Factor: ZNF69 | TF:M09734_1 | 1.30E-27 | 26.8856545 | 6295 | 2679 |
| **TF** | Factor: E2F; n | TF:M00427_1 | 1.50E-27 | 26.8248111 | 3878 | 2679 |
| **TF** | Factor: BTEB2 | TF:M09972_1 | 2.16E-27 | 26.6658811 | 3947 | 2679 |
| **TF** | Factor: Fli-1; r | TF:M11401_1 | 2.80E-27 | 26.5525563 | 11904 | 2679 |
| **TF** | Factor: EGR1; | TF:M07208_1 | 3.68E-27 | 26.4343909 | 2203 | 2679 |
| **TF** | Factor: pax-2; | TF:M11877 | 5.26E-27 | 26.2787315 | 11632 | 2679 |
| **TF** | Factor: FKLF; | TF:M01837 | 5.39E-27 | 26.2685061 | 5489 | 2679 |
| **TF** | Factor: BTEB2 | TF:M03814 | 6.00E-27 | 26.2221601 | 5165 | 2679 |
| **TF** | Factor: Rb:E2 | TF:M00740_1 | 7.12E-27 | 26.1477243 | 4638 | 2679 |
| **TF** | Factor: PEA3; | TF:M11430 | 8.23E-27 | 26.0848536 | 10973 | 2679 |
| **TF** | Factor: AP-2al | TF:M07348_1 | 9.46E-27 | 26.0239968 | 2957 | 2679 |
| **TF** | Factor: PEA3; | TF:M11430_1 | 1.01E-26 | 25.9938957 | 4943 | 2679 |
| **TF** | Factor: DB1; r | TF:M10107 | 1.09E-26 | 25.961873 | 5875 | 2679 |
| **TF** | Factor: E2F; n | TF:M00919 | 1.21E-26 | 25.9183552 | 6208 | 2679 |
| **TF** | Factor: REX-1 | TF:M10119 | 1.53E-26 | 25.8150658 | 1902 | 2679 |
| **TF** | Factor: E2F-1 | TF:M00737 | 1.90E-26 | 25.7208159 | 8859 | 2679 |
| **TF** | Factor: Erm; n | TF:M11424 | 2.10E-26 | 25.6776476 | 8225 | 2679 |
| **TF** | Factor: EHF; r | TF:M02052 | 2.70E-26 | 25.5681109 | 10199 | 2679 |
| **TF** | Factor: ZF5; n | TF:M10111 | 3.01E-26 | 25.5209651 | 2665 | 2679 |
| **TF** | Factor: FLI1; | TF:M03989 | 3.67E-26 | 25.4355503 | 10530 | 2679 |
| **TF** | Factor: Hey2; | TF:M11052_1 | 3.70E-26 | 25.4322072 | 9588 | 2679 |
| **TF** | Factor: Foxn2 | TF:M11541 | 4.34E-26 | 25.3625609 | 5784 | 2679 |
| **TF** | Factor: Erg; m | TF:M11397 | 4.96E-26 | 25.304418 | 12122 | 2679 |
| **TF** | Factor: E2F-4 | TF:M00738_1 | 5.43E-26 | 25.2650016 | 4050 | 2679 |
| **TF** | Factor: Pet-1; | TF:M09918_1 | 5.57E-26 | 25.2540352 | 7213 | 2679 |

| | | | | | | |
|---|---|---|---|---|---|---|
| **TF** | Factor: c-ets-2 | TF:M02064_1 | 6.54E-26 | 25.1844682 | 4626 | 2679 |
| **TF** | Factor: DB1; r | TF:M10107_1 | 7.28E-26 | 25.1380139 | 2686 | 2679 |
| **TF** | Factor: E2F-1 | TF:M00428_1 | 7.75E-26 | 25.1105497 | 3652 | 2679 |
| **TF** | Factor: PEA3; | TF:M11427 | 8.93E-26 | 25.0491839 | 10689 | 2679 |
| **TF** | Factor: Elk-1; | TF:M00025 | 9.36E-26 | 25.0285541 | 10355 | 2679 |
| **TF** | Factor: LBP-1 | TF:M11490_1 | 1.01E-25 | 24.9959032 | 6037 | 2679 |
| **TF** | Factor: EGR1; | TF:M03896_1 | 1.29E-25 | 24.8896201 | 2109 | 2679 |
| **TF** | Factor: LRF; r | TF:M01100 | 1.33E-25 | 24.8760292 | 7597 | 2679 |
| **TF** | Factor: LKLF; | TF:M07261 | 1.73E-25 | 24.7608582 | 9867 | 2679 |
| **TF** | Factor: Foxn2 | TF:M11541_1 | 2.28E-25 | 24.6428783 | 5526 | 2679 |
| **TF** | Factor: Sp1; n | TF:M12151_1 | 2.30E-25 | 24.6380754 | 2562 | 2679 |
| **TF** | Factor: Sp2; n | TF:M12155 | 2.54E-25 | 24.59508 | 4630 | 2679 |
| **TF** | Factor: SP4; n | TF:M03922_1 | 2.90E-25 | 24.5379835 | 2158 | 2679 |
| **TF** | Factor: KLF15 | TF:M12157 | 4.07E-25 | 24.3907535 | 3626 | 2679 |
| **TF** | Factor: c-Myc | TF:M00322 | 4.92E-25 | 24.3076155 | 7919 | 2679 |
| **TF** | Factor: ER71: | TF:M08457_1 | 5.12E-25 | 24.2907677 | 5633 | 2679 |
| **TF** | Factor: AP-2g | TF:M11482 | 5.84E-25 | 24.2332764 | 6739 | 2679 |
| **TF** | Factor: Elf-1; | TF:M11448 | 1.03E-24 | 23.9882129 | 7657 | 2679 |
| **TF** | Factor: E2F-1 | TF:M00938_1 | 1.04E-24 | 23.9831027 | 2583 | 2679 |
| **TF** | Factor: Fli-1; r | TF:M11400 | 1.28E-24 | 23.8941743 | 10042 | 2679 |
| **TF** | Factor: BEN; r | TF:M01241 | 1.47E-24 | 23.8331625 | 7555 | 2679 |
| **TF** | Factor: YY1; n | TF:M02044 | 2.13E-24 | 23.6712208 | 6150 | 2679 |
| **TF** | Factor: ELK-1 | TF:M02059 | 2.25E-24 | 23.6486748 | 9512 | 2679 |
| **TF** | Factor: KLF15 | TF:M09968 | 2.54E-24 | 23.5947621 | 3180 | 2679 |
| **TF** | Factor: E2F-6 | TF:M09895_1 | 3.73E-24 | 23.4282103 | 2732 | 2679 |
| **TF** | Factor: PEA3; | TF:M11427_1 | 3.76E-24 | 23.424542 | 4645 | 2679 |
| **TF** | Factor: Erg; m | TF:M11396_1 | 4.18E-24 | 23.3791099 | 5009 | 2679 |
| **TF** | Factor: TEL1; | TF:M02070_1 | 7.83E-24 | 23.1060319 | 4876 | 2679 |
| **TF** | Factor: ZNF51 | TF:M09738 | 9.38E-24 | 23.0277423 | 10956 | 2679 |
| **TF** | Factor: SAP-1 | TF:M09905 | 1.23E-23 | 22.909501 | 8354 | 2679 |
| **TF** | Factor: NRF-1 | TF:M10018 | 1.79E-23 | 22.746398 | 1838 | 2679 |
| **TF** | Factor: BTEB2 | TF:M09972 | 2.36E-23 | 22.6272103 | 9283 | 2679 |
| **TF** | Factor: Sp3; n | TF:M12153_1 | 2.70E-23 | 22.5688077 | 1948 | 2679 |
| **TF** | Factor: E2F-4 | TF:M11535_1 | 2.74E-23 | 22.5621394 | 6579 | 2679 |
| **TF** | Factor: meis1 | TF:M08377 | 3.56E-23 | 22.4481562 | 11363 | 2679 |
| **TF** | Factor: c-Ets- | TF:M08759 | 4.27E-23 | 22.3698322 | 8396 | 2679 |
| **TF** | Factor: E2F; n | TF:M00050 | 4.38E-23 | 22.3582771 | 7130 | 2679 |
| **TF** | Factor: Egr-2; | TF:M09898_1 | 5.11E-23 | 22.2912468 | 2573 | 2679 |
| **TF** | Factor: Elk-1; | TF:M11434_1 | 1.44E-22 | 21.8431095 | 2959 | 2679 |
| **TF** | Factor: KLF16 | TF:M03914 | 1.52E-22 | 21.8195457 | 3380 | 2679 |
| **TF** | Factor: Fli-1; r | TF:M11399_1 | 1.62E-22 | 21.7914276 | 11613 | 2679 |
| **TF** | Factor: CTCF; | TF:M03895_1 | 1.62E-22 | 21.7905339 | 3529 | 2679 |
| **TF** | Factor: DPF2; | TF:M09760 | 1.67E-22 | 21.7764696 | 8251 | 2679 |
| **TF** | Factor: Egr-2; | TF:M00246 | 1.83E-22 | 21.7380209 | 6265 | 2679 |
| **TF** | Factor: NRF-1 | TF:M09641_1 | 2.08E-22 | 21.6825499 | 997 | 2679 |

| | | | | | | |
|---|---|---|---|---|---|---|
| **TF** | Factor: ERG; ı | TF:M03976_1 | 3.16E-22 | 21.5002623 | 7856 | 2679 |
| **TF** | Factor: EHF; r | TF:M02052_1 | 3.22E-22 | 21.4915383 | 4086 | 2679 |
| **TF** | Factor: PLAG1 | TF:M01778 | 3.40E-22 | 21.4688405 | 2761 | 2679 |
| **TF** | Factor: Zfx; m | TF:M01593_1 | 3.83E-22 | 21.4173109 | 974 | 2679 |
| **TF** | Factor: Erg; m | TF:M11392 | 4.04E-22 | 21.3931274 | 8777 | 2679 |
| **TF** | Factor: PEA3; | TF:M02066 | 4.44E-22 | 21.3525882 | 7124 | 2679 |
| **TF** | Factor: KLF; r | TF:M07461 | 4.72E-22 | 21.3264317 | 8831 | 2679 |
| **TF** | Factor: Erg; m | TF:M11393 | 7.25E-22 | 21.1397412 | 14242 | 2679 |
| **TF** | Factor: Egr-1; | TF:M04797_1 | 7.79E-22 | 21.1087144 | 1566 | 2679 |
| **TF** | Factor: Erg; m | TF:M11393_1 | 8.24E-22 | 21.0839597 | 13785 | 2679 |
| **TF** | Factor: E2F-1 | TF:M00940_1 | 9.43E-22 | 21.02547 | 2672 | 2679 |
| **TF** | Factor: Elf-1; ı | TF:M04714 | 1.07E-21 | 20.972283 | 7876 | 2679 |
| **TF** | Factor: Elf-1; ı | TF:M11447 | 1.34E-21 | 20.8741553 | 7786 | 2679 |
| **TF** | Factor: SALL2 | TF:M04595 | 1.37E-21 | 20.8637201 | 13123 | 2679 |
| **TF** | Factor: Elk-1:I | TF:M08226_1 | 1.41E-21 | 20.8507425 | 5665 | 2679 |
| **TF** | Factor: E2F; n | TF:M00918 | 1.77E-21 | 20.7512455 | 5634 | 2679 |
| **TF** | Factor: ZNF59 | TF:M12310 | 1.77E-21 | 20.7509631 | 438 | 2679 |
| **TF** | Factor: NGFI- | TF:M00244_1 | 1.84E-21 | 20.7354932 | 1778 | 2679 |
| **TF** | Factor: ER71: | TF:M08568_1 | 2.01E-21 | 20.6965715 | 15263 | 2679 |
| **TF** | Factor: E2F; n | TF:M00920 | 2.07E-21 | 20.6849808 | 5719 | 2679 |
| **TF** | Factor: GCMa | TF:M08313 | 2.32E-21 | 20.6347592 | 15556 | 2679 |
| **TF** | Factor: ERG; ı | TF:M03976 | 3.07E-21 | 20.5127274 | 12913 | 2679 |
| **TF** | Factor: ESE-1 | TF:M11381_1 | 3.37E-21 | 20.4729402 | 9844 | 2679 |
| **TF** | Factor: ZNF77 | TF:M08993 | 3.50E-21 | 20.4559747 | 10713 | 2679 |
| **TF** | Factor: meis1: | TF:M08377_1 | 3.72E-21 | 20.429371 | 5084 | 2679 |
| **TF** | Factor: FLI-1; | TF:M08913 | 4.58E-21 | 20.3390707 | 8401 | 2679 |
| **TF** | Factor: TFAP2 | TF:M04146 | 7.37E-21 | 20.1327423 | 6582 | 2679 |
| **TF** | Factor: NRF-1 | TF:M04694 | 8.14E-21 | 20.0895412 | 1568 | 2679 |
| **TF** | Factor: EZI; m | TF:M10151 | 9.62E-21 | 20.0169738 | 2123 | 2679 |
| **TF** | Factor: HES-1 | TF:M02011 | 9.94E-21 | 20.0026734 | 4284 | 2679 |
| **TF** | Factor: Pax-4; | TF:M00378_1 | 1.04E-20 | 19.9827138 | 14850 | 2679 |
| **TF** | Factor: Kaiso; | TF:M09966 | 1.17E-20 | 19.9309479 | 2376 | 2679 |
| **TF** | Factor: c-Ets-1 | TF:M01078 | 1.19E-20 | 19.924201 | 7621 | 2679 |
| **TF** | Factor: SALL2 | TF:M04595_1 | 1.24E-20 | 19.9077638 | 7428 | 2679 |
| **TF** | Factor: Fli-1; r | TF:M11399 | 1.36E-20 | 19.8674891 | 13090 | 2679 |
| **TF** | Factor: ATF-3 | TF:M04744 | 1.75E-20 | 19.7562822 | 10196 | 2679 |
| **TF** | Factor: ZFX; n | TF:M10121_1 | 2.17E-20 | 19.6643928 | 818 | 2679 |
| **TF** | Factor: GCMa | TF:M11595 | 2.85E-20 | 19.5447804 | 12217 | 2679 |
| **TF** | Factor: Erg; m | TF:M02062 | 3.14E-20 | 19.5028879 | 8143 | 2679 |
| **TF** | Factor: ELK-1 | TF:M02059_1 | 3.48E-20 | 19.4584251 | 3792 | 2679 |
| **TF** | Factor: Egr-1; | TF:M09603_1 | 3.73E-20 | 19.4282604 | 1327 | 2679 |
| **TF** | Factor: Fli-1; r | TF:M11400_1 | 4.01E-20 | 19.3963528 | 4230 | 2679 |
| **TF** | Factor: ER71: | TF:M08457 | 4.70E-20 | 19.328211 | 12171 | 2679 |
| **TF** | Factor: c-ets-2 | TF:M02064 | 4.97E-20 | 19.3036796 | 11050 | 2679 |
| **TF** | Factor: FLI1; ı | TF:M03988 | 5.80E-20 | 19.2362087 | 8401 | 2679 |

| | | | | | | |
|---|---|---|---|---|---|---|
| **TF** | Factor: ER-be | TF:M09910 | 5.89E-20 | 19.2298784 | 9603 | 2679 |
| **TF** | Factor: DP-1; | TF:M10098_1 | 6.02E-20 | 19.2200995 | 2184 | 2679 |
| **TF** | Factor: BTEB2 | TF:M03814_1 | 6.61E-20 | 19.1796127 | 1303 | 2679 |
| **TF** | Factor: CNOT | TF:M01253_1 | 7.38E-20 | 19.1319787 | 1030 | 2679 |
| **TF** | Factor: Erg; m | TF:M11391 | 9.27E-20 | 19.0326868 | 10173 | 2679 |
| **TF** | Factor: Zfp28 | TF:M01597 | 9.84E-20 | 19.0069529 | 1021 | 2679 |
| **TF** | Factor: c-Ets-2 | TF:M11417_1 | 1.06E-19 | 18.9730237 | 14160 | 2679 |
| **TF** | Factor: Egr-1; | TF:M04918 | 1.27E-19 | 18.8955863 | 5665 | 2679 |
| **TF** | Factor: AP-2g | TF:M03811_1 | 1.34E-19 | 18.8734207 | 3823 | 2679 |
| **TF** | Factor: ZNF50 | TF:M12085_1 | 2.08E-19 | 18.6829787 | 3005 | 2679 |
| **TF** | Factor: EGR1; | TF:M09761_1 | 2.53E-19 | 18.5966824 | 1574 | 2679 |
| **TF** | Factor: Erg; m | TF:M11395_1 | 2.53E-19 | 18.5963396 | 8027 | 2679 |
| **TF** | Factor: ZNF50 | TF:M12086 | 2.58E-19 | 18.5880892 | 1835 | 2679 |
| **TF** | Factor: Pax-4; | TF:M00378 | 3.23E-19 | 18.4905005 | 18032 | 2679 |
| **TF** | Factor: ERG; r | TF:M01752 | 3.40E-19 | 18.468791 | 9560 | 2679 |
| **TF** | Factor: GABP; | TF:M00341 | 3.89E-19 | 18.410394 | 8296 | 2679 |
| **TF** | Factor: PAX5; | TF:M09817_1 | 4.06E-19 | 18.3911013 | 6212 | 2679 |
| **TF** | Factor: DPF2; | TF:M09760_1 | 5.77E-19 | 18.2387225 | 3011 | 2679 |
| **TF** | Factor: nerf; m | TF:M09606_1 | 6.31E-19 | 18.2002909 | 1718 | 2679 |
| **TF** | Factor: Elk-1; | TF:M00025_1 | 7.43E-19 | 18.1287286 | 4674 | 2679 |
| **TF** | Factor: c-Ets-2 | TF:M11417 | 7.58E-19 | 18.120416 | 14456 | 2679 |
| **TF** | Factor: nerf; m | TF:M09901 | 1.01E-18 | 17.9971747 | 5408 | 2679 |
| **TF** | Factor: AP-2al | TF:M01047_1 | 1.07E-18 | 17.9708055 | 9306 | 2679 |
| **TF** | Factor: AP-2al | TF:M01047 | 1.07E-18 | 17.9708055 | 9306 | 2679 |
| **TF** | Factor: LKLF; | TF:M08819 | 1.18E-18 | 17.9283266 | 5645 | 2679 |
| **TF** | Factor: ER71; | TF:M02067 | 1.33E-18 | 17.8764361 | 6001 | 2679 |
| **TF** | Factor: ctcf; m | TF:M07249 | 1.64E-18 | 17.7850166 | 6749 | 2679 |
| **TF** | Factor: Fli-1; r | TF:M11404 | 1.77E-18 | 17.7530256 | 8828 | 2679 |
| **TF** | Factor: Egr-3; | TF:M00245_1 | 2.09E-18 | 17.6802458 | 2382 | 2679 |
| **TF** | Factor: ZNF51 | TF:M09738_1 | 2.64E-18 | 17.5778264 | 4781 | 2679 |
| **TF** | Factor: E2F-1 | TF:M00430_1 | 2.66E-18 | 17.5746085 | 2541 | 2679 |
| **TF** | Factor: E2F; n | TF:M08875_1 | 2.85E-18 | 17.5455177 | 1744 | 2679 |
| **TF** | Factor: ETS1; | TF:M03978_1 | 3.14E-18 | 17.5033488 | 6527 | 2679 |
| **TF** | Factor: ETV4; | TF:M03982 | 3.34E-18 | 17.475891 | 6903 | 2679 |
| **TF** | Factor: ZNF77 | TF:M08993_1 | 4.40E-18 | 17.3566697 | 4143 | 2679 |
| **TF** | Factor: E2F-1 | TF:M00431_1 | 4.40E-18 | 17.3562081 | 2297 | 2679 |
| **TF** | Factor: Elf-1; r | TF:M11448_1 | 4.62E-18 | 17.3354396 | 2341 | 2679 |
| **TF** | Factor: Elk-1; | TF:M09904 | 4.73E-18 | 17.3247306 | 8239 | 2679 |
| **TF** | Factor: FLI-1; | TF:M08913_1 | 5.12E-18 | 17.2905671 | 3083 | 2679 |
| **TF** | Factor: ETS1; | TF:M03977_1 | 5.21E-18 | 17.283142 | 4710 | 2679 |
| **TF** | Factor: ERG; r | TF:M03975_1 | 6.18E-18 | 17.2087323 | 4048 | 2679 |
| **TF** | Factor: LKLF; | TF:M05499_1 | 7.02E-18 | 17.1538169 | 2527 | 2679 |
| **TF** | Factor: KLF17 | TF:M05386_1 | 7.02E-18 | 17.1538169 | 2527 | 2679 |
| **TF** | Factor: c-ets-1 | TF:M02063 | 9.06E-18 | 17.042801 | 7686 | 2679 |
| **TF** | Factor: Erg; m | TF:M11391_1 | 9.87E-18 | 17.0058298 | 9162 | 2679 |

| | | | | | | |
|---|---|---|---|---|---|---|
| **TF** | Factor: AP-2g | TF:M11482_1 | 1.07E-17 | 16.9711529 | 2066 | 2679 |
| **TF** | Factor: KLF; r | TF:M07461_1 | 1.73E-17 | 16.7614512 | 3423 | 2679 |
| **TF** | Factor: Erg; m | TF:M11395 | 1.90E-17 | 16.7219217 | 8791 | 2679 |
| **TF** | Factor: YY2; n | TF:M03925_1 | 3.39E-17 | 16.4697833 | 778 | 2679 |
| **TF** | Factor: etv3; r | TF:M02068 | 3.54E-17 | 16.4507192 | 3485 | 2679 |
| **TF** | Factor: ATF-3 | TF:M04744_1 | 3.58E-17 | 16.4463387 | 4937 | 2679 |
| **TF** | Factor: LKLF; | TF:M07261_1 | 4.46E-17 | 16.3506596 | 4110 | 2679 |
| **TF** | Factor: Elf-1; | TF:M11451 | 4.54E-17 | 16.342754 | 6956 | 2679 |
| **TF** | Factor: GABP | TF:M09935 | 4.87E-17 | 16.3123481 | 4467 | 2679 |
| **TF** | Factor: PEA3; | TF:M11428 | 5.78E-17 | 16.2378758 | 7260 | 2679 |
| **TF** | Factor: NRF-1 | TF:M00652 | 6.09E-17 | 16.215524 | 1382 | 2679 |
| **TF** | Factor: TFAP2 | TF:M04152_1 | 6.17E-17 | 16.209607 | 5599 | 2679 |
| **TF** | Factor: Erg; m | TF:M02062_1 | 7.67E-17 | 16.1153867 | 2739 | 2679 |
| **TF** | Factor: Egr; m | TF:M00807 | 7.74E-17 | 16.1114533 | 4373 | 2679 |
| **TF** | Factor: SREBI | TF:M11081 | 7.74E-17 | 16.1113415 | 11156 | 2679 |
| **TF** | Factor: Erm; n | TF:M11424_1 | 8.07E-17 | 16.0929463 | 2861 | 2679 |
| **TF** | Factor: HES-1 | TF:M11061 | 9.32E-17 | 16.0305463 | 6318 | 2679 |
| **TF** | Factor: ESE-1 | TF:M11382 | 1.32E-16 | 15.8780347 | 10437 | 2679 |
| **TF** | Factor: HES-7 | TF:M11066_1 | 1.85E-16 | 15.7324924 | 8227 | 2679 |
| **TF** | Factor: TFAP2 | TF:M04152 | 2.23E-16 | 15.6507444 | 7781 | 2679 |
| **TF** | Factor: ERG; | TF:M03975 | 2.83E-16 | 15.5475614 | 10098 | 2679 |
| **TF** | Factor: ELF1; | TF:M02053 | 2.86E-16 | 15.5434778 | 6085 | 2679 |
| **TF** | Factor: Fli-1; r | TF:M11404_1 | 2.93E-16 | 15.5331095 | 3194 | 2679 |
| **TF** | Factor: ERG; | TF:M01752_1 | 3.23E-16 | 15.4902119 | 3740 | 2679 |
| **TF** | Factor: HES-1 | TF:M07042 | 3.73E-16 | 15.427912 | 3988 | 2679 |
| **TF** | Factor: LRF; r | TF:M10115 | 4.39E-16 | 15.3575164 | 7671 | 2679 |
| **TF** | Factor: FLI1; | TF:M03988_1 | 5.40E-16 | 15.2677003 | 2969 | 2679 |
| **TF** | Factor: E2F-2 | TF:M11528 | 5.80E-16 | 15.2362875 | 10466 | 2679 |
| **TF** | Factor: YY1; n | TF:M08982 | 5.84E-16 | 15.2338306 | 7188 | 2679 |
| **TF** | Factor: Sp4; n | TF:M07617_1 | 7.76E-16 | 15.1100072 | 902 | 2679 |
| **TF** | Factor: ER-be | TF:M09910_1 | 8.85E-16 | 15.0529366 | 4108 | 2679 |
| **TF** | Factor: GABP | TF:M03990 | 8.87E-16 | 15.0522747 | 5312 | 2679 |
| **TF** | Factor: ERF; r | TF:M02061 | 8.99E-16 | 15.0462171 | 2604 | 2679 |
| **TF** | Factor: BTEB | TF:M07409_1 | 9.56E-16 | 15.0195657 | 1838 | 2679 |
| **TF** | Factor: PAX5; | TF:M09817 | 1.08E-15 | 14.9670332 | 12624 | 2679 |
| **TF** | Factor: ctcf; m | TF:M09890 | 1.45E-15 | 14.8382368 | 8544 | 2679 |
| **TF** | Factor: ER81; | TF:M11419 | 1.70E-15 | 14.7698683 | 5760 | 2679 |
| **TF** | Factor: ZBP89 | TF:M01816 | 1.96E-15 | 14.7079565 | 2210 | 2679 |
| **TF** | Factor: PEA3; | TF:M02066_1 | 2.34E-15 | 14.6306265 | 2306 | 2679 |
| **TF** | Factor: NRF-1 | TF:M04059 | 3.64E-15 | 14.4394649 | 1478 | 2679 |
| **TF** | Factor: LRF; r | TF:M01100_1 | 4.19E-15 | 14.3773275 | 2777 | 2679 |
| **TF** | Factor: YY1; n | TF:M02044_1 | 5.07E-15 | 14.2951264 | 1211 | 2679 |
| **TF** | Factor: E2F-1 | TF:M00737_1 | 5.66E-15 | 14.2475366 | 2968 | 2679 |
| **TF** | Factor: ZBTB7 | TF:M03806_1 | 6.02E-15 | 14.2207601 | 547 | 2679 |
| **TF** | Factor: HIC1; | TF:M01072 | 8.38E-15 | 14.0767938 | 4190 | 2679 |

| | | | | | | |
|---|---|---|---|---|---|---|
| **TF** | Factor: HIC1; | TF:M01073 | 8.52E-15 | 14.0696114 | 4123 | 2679 |
| **TF** | Factor: LRF; r | TF:M12231_1 | 8.93E-15 | 14.0492372 | 5186 | 2679 |
| **TF** | Factor: XBP-1 | TF:M01770 | 8.96E-15 | 14.0474872 | 11113 | 2679 |
| **TF** | Factor: DATF1 | TF:M09724 | 8.97E-15 | 14.0471008 | 8463 | 2679 |
| **TF** | Factor: PEA3; | TF:M11425 | 9.33E-15 | 14.0303206 | 7499 | 2679 |
| **TF** | Factor: c-Myc | TF:M00322_1 | 1.14E-14 | 13.942921 | 2874 | 2679 |
| **TF** | Factor: ESE-1 | TF:M11382_1 | 1.31E-14 | 13.8822861 | 4150 | 2679 |
| **TF** | Factor: ETV7; | TF:M02071_1 | 1.34E-14 | 13.8737261 | 4495 | 2679 |
| **TF** | Factor: LRF; r | TF:M12231 | 1.42E-14 | 13.8480207 | 11597 | 2679 |
| **TF** | Factor: WT1; | TF:M05327_1 | 1.60E-14 | 13.7972086 | 1764 | 2679 |
| **TF** | Factor: Erg; m | TF:M11392_1 | 2.03E-14 | 13.6915116 | 3253 | 2679 |
| **TF** | Factor: E2F1; | TF:M08874_1 | 2.38E-14 | 13.6225274 | 2158 | 2679 |
| **TF** | Factor: E2F-4 | TF:M00739 | 2.43E-14 | 13.61499 | 5682 | 2679 |
| **TF** | Factor: Pax-3; | TF:M00327_1 | 2.60E-14 | 13.5844514 | 6960 | 2679 |
| **TF** | Factor: LRF; r | TF:M04617 | 2.79E-14 | 13.5539966 | 6210 | 2679 |
| **TF** | Factor: XBP-1 | TF:M01770_1 | 3.07E-14 | 13.5127341 | 4604 | 2679 |
| **TF** | Factor: Egr-1; | TF:M12140_1 | 3.17E-14 | 13.4994596 | 1565 | 2679 |
| **TF** | Factor: Egr-1; | TF:M00243_1 | 3.74E-14 | 13.427294 | 1287 | 2679 |
| **TF** | Factor: VDR; r | TF:M00444 | 3.78E-14 | 13.4220942 | 10400 | 2679 |
| **TF** | Factor: Elf-1; | TF:M11452 | 4.33E-14 | 13.3631784 | 6324 | 2679 |
| **TF** | Factor: E2F-3 | TF:M09893 | 4.63E-14 | 13.3341721 | 6128 | 2679 |
| **TF** | Factor: ZGPA | TF:M09739 | 5.70E-14 | 13.2444562 | 8274 | 2679 |
| **TF** | Factor: AP-2re | TF:M09967_1 | 6.06E-14 | 13.2174237 | 1386 | 2679 |
| **TF** | Factor: ELF4; | TF:M02056 | 6.14E-14 | 13.2117331 | 5379 | 2679 |
| **TF** | Factor: E2F-3 | TF:M08529_1 | 8.97E-14 | 13.0471795 | 4841 | 2679 |
| **TF** | Factor: CREM | TF:M01820 | 9.37E-14 | 13.0282002 | 6374 | 2679 |
| **TF** | Factor: EKLF; | TF:M09969_1 | 9.61E-14 | 13.0174488 | 1498 | 2679 |
| **TF** | Factor: YY1; n | TF:M04696_1 | 1.03E-13 | 12.9882387 | 1988 | 2679 |
| **TF** | Factor: A-Myb | TF:M08390_1 | 1.07E-13 | 12.9722905 | 4844 | 2679 |
| **TF** | Factor: Pax-3; | TF:M00360 | 1.10E-13 | 12.9581088 | 12366 | 2679 |
| **TF** | Factor: VDR; r | TF:M00444_1 | 1.14E-13 | 12.9441184 | 4558 | 2679 |
| **TF** | Factor: ATF-3 | TF:M09868 | 1.17E-13 | 12.9333554 | 6456 | 2679 |
| **TF** | Factor: Erm; n | TF:M02069 | 1.18E-13 | 12.9277909 | 6961 | 2679 |
| **TF** | Factor: Elf-1; | TF:M04714_1 | 1.61E-13 | 12.793887 | 2684 | 2679 |
| **TF** | Factor: SAP-1 | TF:M09905_1 | 1.91E-13 | 12.7196331 | 3017 | 2679 |
| **TF** | Factor: MAZ; | TF:M07297 | 2.29E-13 | 12.6405113 | 2987 | 2679 |
| **TF** | Factor: YY1; n | TF:M07370 | 2.94E-13 | 12.5312475 | 5854 | 2679 |
| **TF** | Factor: NRF-1 | TF:M07052_1 | 5.08E-13 | 12.2940758 | 739 | 2679 |
| **TF** | Factor: FLI1; | TF:M03989_1 | 5.80E-13 | 12.2365083 | 5657 | 2679 |
| **TF** | Factor: E2F-3 | TF:M08529 | 6.02E-13 | 12.220649 | 11553 | 2679 |
| **TF** | Factor: AhR:A | TF:M00237 | 6.04E-13 | 12.218832 | 6389 | 2679 |
| **TF** | Factor: Elk-1: | TF:M08213 | 6.55E-13 | 12.1837459 | 6334 | 2679 |
| **TF** | Factor: ETV5; | TF:M03983 | 8.00E-13 | 12.0971547 | 3169 | 2679 |
| **TF** | Factor: EGR-1 | TF:M01972_1 | 8.47E-13 | 12.0719846 | 1478 | 2679 |
| **TF** | Factor: GABP | TF:M02074 | 9.17E-13 | 12.0377857 | 4642 | 2679 |

| | | | | | | |
|---|---|---|---|---|---|---|
| **TF** | Factor: E2F-1 | TF:M00516_1 | 1.25E-12 | 11.9027115 | 2093 | 2679 |
| **TF** | Factor: Elk-1; | TF:M11433 | 1.29E-12 | 11.8884892 | 7772 | 2679 |
| **TF** | Factor: Kaiso; | TF:M09632_1 | 1.48E-12 | 11.8289304 | 681 | 2679 |
| **TF** | Factor: GKLF; | TF:M09971 | 2.02E-12 | 11.6947399 | 5583 | 2679 |
| **TF** | Factor: Elf-1; r | TF:M11447_1 | 2.08E-12 | 11.6815187 | 2616 | 2679 |
| **TF** | Factor: c-ets-1 | TF:M02063_1 | 2.85E-12 | 11.5458748 | 2421 | 2679 |
| **TF** | Factor: Elk-1:I | TF:M08224 | 2.93E-12 | 11.5332368 | 9010 | 2679 |
| **TF** | Factor: LRF; r | TF:M10115_1 | 2.95E-12 | 11.5307351 | 2332 | 2679 |
| **TF** | Factor: c-Ets-1 | TF:M01078_1 | 3.00E-12 | 11.5229985 | 2473 | 2679 |
| **TF** | Factor: NF90; | TF:M09859 | 3.35E-12 | 11.4748453 | 4775 | 2679 |
| **TF** | Factor: GABP- | TF:M04748_1 | 3.45E-12 | 11.4621904 | 10467 | 2679 |
| **TF** | Factor: FKLF; | TF:M01837_1 | 3.75E-12 | 11.4263018 | 1421 | 2679 |
| **TF** | Factor: Erm:H | TF:M08700 | 4.09E-12 | 11.3880503 | 2070 | 2679 |
| **TF** | Factor: CREB | TF:M11221 | 4.14E-12 | 11.3827032 | 9789 | 2679 |
| **TF** | Factor: Elf-1; r | TF:M07415 | 6.34E-12 | 11.1978225 | 8060 | 2679 |
| **TF** | Factor: GLI; m | TF:M03871 | 7.09E-12 | 11.1493891 | 3348 | 2679 |
| **TF** | Factor: YY1; n | TF:M08982_1 | 9.52E-12 | 11.0211664 | 1675 | 2679 |
| **TF** | Factor: AP-2al | TF:M11480 | 9.98E-12 | 11.0008976 | 6233 | 2679 |
| **TF** | Factor: E2F-2 | TF:M11528_1 | 1.01E-11 | 10.9976403 | 4809 | 2679 |
| **TF** | Factor: Elk-1:T | TF:M08231 | 1.02E-11 | 10.9909936 | 6702 | 2679 |
| **TF** | Factor: Elk-1; | TF:M01165 | 1.05E-11 | 10.9797928 | 7853 | 2679 |
| **TF** | Factor: A-Myb | TF:M08390 | 1.19E-11 | 10.9227032 | 11199 | 2679 |
| **TF** | Factor: YY1; n | TF:M04696 | 1.30E-11 | 10.8858243 | 7894 | 2679 |
| **TF** | Factor: ER71; | TF:M02067_1 | 1.38E-11 | 10.85935 | 1630 | 2679 |
| **TF** | Factor: PUR1; | TF:M01721 | 1.41E-11 | 10.8504029 | 9200 | 2679 |
| **TF** | Factor: ER71: | TF:M08568 | 1.47E-11 | 10.8323579 | 18420 | 2679 |
| **TF** | Factor: Elk-1:I | TF:M08224_1 | 1.55E-11 | 10.8100725 | 2781 | 2679 |
| **TF** | Factor: ELK1; | TF:M03970 | 1.57E-11 | 10.8041425 | 4838 | 2679 |
| **TF** | Factor: NRF-1 | TF:M10018_1 | 1.59E-11 | 10.7998126 | 854 | 2679 |
| **TF** | Factor: TFAP2 | TF:M04146_1 | 1.70E-11 | 10.7707321 | 3817 | 2679 |
| **TF** | Factor: ELF1; | TF:M02053_1 | 1.71E-11 | 10.7677337 | 1571 | 2679 |
| **TF** | Factor: Sp2; n | TF:M12156_1 | 1.78E-11 | 10.7490966 | 1004 | 2679 |
| **TF** | Factor: ETV4; | TF:M03982_1 | 1.84E-11 | 10.7340956 | 2026 | 2679 |
| **TF** | Factor: N-Myc | TF:M03834 | 1.92E-11 | 10.7166774 | 1420 | 2679 |
| **TF** | Factor: AP-2al | TF:M11480_1 | 2.16E-11 | 10.6661412 | 5889 | 2679 |
| **TF** | Factor: Kaiso; | TF:M09966_1 | 2.57E-11 | 10.5893719 | 450 | 2679 |
| **TF** | Factor: E2F; n | TF:M00939_1 | 2.69E-11 | 10.5702166 | 1687 | 2679 |
| **TF** | Factor: LRF; r | TF:M12230 | 2.70E-11 | 10.5692145 | 8640 | 2679 |
| **TF** | Factor: ER81; | TF:M09611 | 2.79E-11 | 10.5541542 | 2692 | 2679 |
| **TF** | Factor: AP-2g | TF:M08662 | 3.31E-11 | 10.4805478 | 1983 | 2679 |
| **TF** | Factor: Pax-3; | TF:M00360_1 | 3.32E-11 | 10.47868 | 5547 | 2679 |
| **TF** | Factor: Fli-1; r | TF:M02073 | 3.35E-11 | 10.4749656 | 5905 | 2679 |
| **TF** | Factor: E2F; n | TF:M00919_1 | 3.41E-11 | 10.4667812 | 1542 | 2679 |
| **TF** | Factor: REX1; | TF:M01744 | 3.47E-11 | 10.4595986 | 632 | 2679 |
| **TF** | Factor: EGR4; | TF:M03899 | 3.63E-11 | 10.4399956 | 2041 | 2679 |

| | | | | | | |
|---|---|---|---|---|---|---|
| **TF** | Factor: GABP· | TF:M11415 | 3.88E-11 | 10.4106967 | 3289 | 2679 |
| **TF** | Factor: FXR; r | TF:M10014 | 4.68E-11 | 10.329372 | 5687 | 2679 |
| **TF** | Factor: LRF; r | TF:M12230_1 | 4.81E-11 | 10.317901 | 2674 | 2679 |
| **TF** | Factor: ZBP9 | TF:M10137 | 4.85E-11 | 10.3145008 | 618 | 2679 |
| **TF** | Factor: Egr-1; | TF:M09897_1 | 5.14E-11 | 10.289419 | 996 | 2679 |
| **TF** | Factor: FPM3 | TF:M10134 | 6.89E-11 | 10.1615373 | 2517 | 2679 |
| **TF** | Factor: CSX; r | TF:M10008 | 6.90E-11 | 10.1608674 | 10227 | 2679 |
| **TF** | Factor: AP-2g | TF:M08440 | 7.00E-11 | 10.1546427 | 1205 | 2679 |
| **TF** | Factor: YY1; n | TF:M12275 | 7.46E-11 | 10.1273044 | 7095 | 2679 |
| **TF** | Factor: Sp2; n | TF:M04932 | 7.74E-11 | 10.1110751 | 2616 | 2679 |
| **TF** | Factor: PEA3; | TF:M11428_1 | 8.17E-11 | 10.0875315 | 2166 | 2679 |
| **TF** | Factor: BEN; r | TF:M01241_1 | 9.45E-11 | 10.0246464 | 2117 | 2679 |
| **TF** | Factor: FPM3 | TF:M07141 | 9.54E-11 | 10.0205524 | 2200 | 2679 |
| **TF** | Factor: ER81; | TF:M11419_1 | 1.01E-10 | 9.99412973 | 1438 | 2679 |
| **TF** | Factor: USF; r | TF:M00187 | 1.07E-10 | 9.97044328 | 3795 | 2679 |
| **TF** | Factor: CREB | TF:M00981 | 1.08E-10 | 9.9676673 | 8437 | 2679 |
| **TF** | Factor: ELF4; | TF:M02056_1 | 1.15E-10 | 9.93772149 | 1300 | 2679 |
| **TF** | Factor: YY1; n | TF:M12273 | 1.28E-10 | 9.89407499 | 6272 | 2679 |
| **TF** | Factor: p53; n | TF:M01652_1 | 1.30E-10 | 9.88762364 | 5302 | 2679 |
| **TF** | Factor: GABP | TF:M03791 | 1.31E-10 | 9.88220028 | 3004 | 2679 |
| **TF** | Factor: Tax/Cl | TF:M00114 | 1.40E-10 | 9.85525068 | 6608 | 2679 |
| **TF** | Factor: YY1; n | TF:M01035 | 1.56E-10 | 9.80793953 | 5958 | 2679 |
| **TF** | Factor: Spz1; | TF:M00446 | 1.64E-10 | 9.78641315 | 8740 | 2679 |
| **TF** | Factor: AhR,; | TF:M00976_1 | 1.95E-10 | 9.7093221 | 1014 | 2679 |
| **TF** | Factor: NF90; | TF:M09859_1 | 2.18E-10 | 9.66253323 | 1015 | 2679 |
| **TF** | Factor: Elf-1; | TF:M09900 | 2.21E-10 | 9.65566593 | 5264 | 2679 |
| **TF** | Factor: KLF1 | TF:M09968_1 | 2.27E-10 | 9.64412642 | 992 | 2679 |
| **TF** | Factor: CREB | TF:M00113 | 2.30E-10 | 9.63872062 | 4862 | 2679 |
| **TF** | Factor: SP4; n | TF:M01273_1 | 2.46E-10 | 9.6094222 | 1022 | 2679 |
| **TF** | Factor: ZF5; n | TF:M10111_1 | 2.47E-10 | 9.60667242 | 500 | 2679 |
| **TF** | Factor: NRF-1 | TF:M02102_1 | 2.50E-10 | 9.60278191 | 657 | 2679 |
| **TF** | Factor: YY1; n | TF:M07370_1 | 2.52E-10 | 9.59849688 | 1087 | 2679 |
| **TF** | Factor: HES-1 | TF:M08767 | 2.75E-10 | 9.56109339 | 7868 | 2679 |
| **TF** | Factor: Pax-3; | TF:M00327 | 2.91E-10 | 9.53680196 | 13445 | 2679 |
| **TF** | Factor: GABP | TF:M00341_1 | 3.33E-10 | 9.47786612 | 2333 | 2679 |
| **TF** | Factor: GABP· | TF:M08917 | 3.59E-10 | 9.44527448 | 2724 | 2679 |
| **TF** | Factor: YY1; n | TF:M00793 | 4.34E-10 | 9.36236554 | 5222 | 2679 |
| **TF** | Factor: LKLF; | TF:M08819_1 | 4.50E-10 | 9.34693313 | 1547 | 2679 |
| **TF** | Factor: E2F; n | TF:M00920_1 | 4.69E-10 | 9.32855763 | 1321 | 2679 |
| **TF** | Factor: ESE-1 | TF:M11381 | 4.72E-10 | 9.32615291 | 15871 | 2679 |
| **TF** | Factor: ATF; n | TF:M00017 | 5.01E-10 | 9.30052621 | 7303 | 2679 |
| **TF** | Factor: KLF1 | TF:M12159 | 5.33E-10 | 9.27310776 | 1426 | 2679 |
| **TF** | Factor: Erm; n | TF:M02069_1 | 5.34E-10 | 9.27232955 | 1975 | 2679 |
| **TF** | Factor: ctcf; m | TF:M09602 | 5.44E-10 | 9.2640795 | 7507 | 2679 |
| **TF** | Factor: Egr-2; | TF:M00246_1 | 5.83E-10 | 9.23443038 | 1612 | 2679 |

| | | | | | | |
|---|---|---|---|---|---|---|
| **TF** | Factor: GABP· | TF:M04698 | 5.93E-10 | 9.22699757 | 3221 | 2679 |
| **TF** | Factor: ATF-3 | TF:M09868_1 | 6.34E-10 | 9.19823633 | 2143 | 2679 |
| **TF** | Factor: LBP9; | TF:M11492 | 6.84E-10 | 9.1647468 | 7657 | 2679 |
| **TF** | Factor: Elf-1; ı | TF:M09605 | 7.05E-10 | 9.15194633 | 5835 | 2679 |
| **TF** | Factor: GABP, | TF:M03990_1 | 7.31E-10 | 9.13618579 | 1217 | 2679 |
| **TF** | Factor: ZNF3· | TF:M10144 | 7.35E-10 | 9.13371851 | 4809 | 2679 |
| **TF** | Factor: DRRS; | TF:M10056 | 7.49E-10 | 9.12535006 | 4844 | 2679 |
| **TF** | Factor: ETV3; | TF:M03981 | 8.15E-10 | 9.08860663 | 2346 | 2679 |
| **TF** | Factor: SAP-1 | TF:M01167 | 8.57E-10 | 9.06688363 | 1975 | 2679 |
| **TF** | Factor: ZGPA' | TF:M09739_1 | 8.95E-10 | 9.04801395 | 2758 | 2679 |
| **TF** | Factor: c-Ets-: | TF:M08759_1 | 9.23E-10 | 9.03460273 | 2943 | 2679 |
| **TF** | Factor: ESE-1 | TF:M01214 | 1.03E-09 | 8.98854881 | 7623 | 2679 |
| **TF** | Factor: ESE-1 | TF:M02055 | 1.03E-09 | 8.98854881 | 7623 | 2679 |
| **TF** | Factor: HES-1 | TF:M08767_1 | 1.08E-09 | 8.9647613 | 7762 | 2679 |
| **TF** | Factor: GABP; | TF:M01258 | 1.10E-09 | 8.95904472 | 2860 | 2679 |
| **TF** | Factor: PSF; n | TF:M07612 | 1.24E-09 | 8.90616117 | 5636 | 2679 |
| **TF** | Factor: TEF-3 | TF:M08423 | 1.33E-09 | 8.87474358 | 3888 | 2679 |
| **TF** | Factor: Elk-1:( | TF:M08223_1 | 1.45E-09 | 8.83877065 | 5900 | 2679 |
| **TF** | Factor: SAP-1 | TF:M11436 | 1.63E-09 | 8.78670145 | 1445 | 2679 |
| **TF** | Factor: Fli-1; r | TF:M02073_1 | 1.73E-09 | 8.76153777 | 1495 | 2679 |
| **TF** | Factor: GABP· | TF:M11414 | 2.27E-09 | 8.64316803 | 2938 | 2679 |
| **TF** | Factor: ctcf; m | TF:M07249_1 | 2.32E-09 | 8.63430737 | 2078 | 2679 |
| **TF** | Factor: NRF-1 | TF:M04059_1 | 2.40E-09 | 8.61936528 | 1038 | 2679 |
| **TF** | Factor: Elk-1:( | TF:M08223 | 2.65E-09 | 8.57607633 | 12856 | 2679 |
| **TF** | Factor: SAP-1 | TF:M11437 | 2.86E-09 | 8.54378439 | 2803 | 2679 |
| **TF** | Factor: EZI; m | TF:M10151_1 | 2.99E-09 | 8.52453383 | 576 | 2679 |
| **TF** | Factor: YY1; n | TF:M10104 | 3.73E-09 | 8.42796575 | 6112 | 2679 |
| **TF** | Factor: Erm; n | TF:M11422 | 3.83E-09 | 8.4164712 | 4711 | 2679 |
| **TF** | Factor: MAZR | TF:M00491 | 3.97E-09 | 8.40127382 | 1223 | 2679 |
| **TF** | Factor: MZF-1 | TF:M00084 | 4.10E-09 | 8.3877313 | 6598 | 2679 |
| **TF** | Factor: SP8; n | TF:M12148 | 4.11E-09 | 8.38637348 | 2928 | 2679 |
| **TF** | Factor: ZNF5ε | TF:M10491_1 | 4.25E-09 | 8.37162283 | 7780 | 2679 |
| **TF** | Factor: CREB, | TF:M00981_1 | 4.56E-09 | 8.34141336 | 2943 | 2679 |
| **TF** | Factor: YY2; n | TF:M08020 | 5.20E-09 | 8.28424139 | 1354 | 2679 |
| **TF** | Factor: ER81; | TF:M09913 | 5.58E-09 | 8.25370317 | 2900 | 2679 |
| **TF** | Factor: GABP· | TF:M04748 | 6.41E-09 | 8.19291855 | 16173 | 2679 |
| **TF** | Factor: PEA3; | TF:M11425_1 | 6.44E-09 | 8.19085436 | 2351 | 2679 |
| **TF** | Factor: Elf-1; ı | TF:M11450 | 7.47E-09 | 8.12668717 | 5640 | 2679 |
| **TF** | Factor: KLF8; | TF:M05318 | 7.64E-09 | 8.11706907 | 832 | 2679 |
| **TF** | Factor: KLF7; | TF:M05319 | 7.64E-09 | 8.11706907 | 832 | 2679 |
| **TF** | Factor: ELK1; | TF:M03970_1 | 8.08E-09 | 8.09259509 | 1104 | 2679 |
| **TF** | Factor: ZIC1; | TF:M12219 | 8.73E-09 | 8.05909432 | 839 | 2679 |
| **TF** | Factor: Elf-1; ı | TF:M11451_1 | 9.26E-09 | 8.03358657 | 2130 | 2679 |
| **TF** | Factor: SP9; n | TF:M12149 | 1.05E-08 | 7.98047719 | 2391 | 2679 |
| **TF** | Factor: MAZ; ı | TF:M07297_1 | 1.11E-08 | 7.95307931 | 692 | 2679 |

| | | | | | | |
|---|---|---|---|---|---|---|
| **TF** | Factor: Egr-1; | TF:M12139 | 1.14E-08 | 7.94447104 | 3813 | 2679 |
| **TF** | Factor: LBP9; | TF:M11492_1 | 1.27E-08 | 7.89534575 | 6069 | 2679 |
| **TF** | Factor: Sp3; n | TF:M00665_1 | 1.60E-08 | 7.79533635 | 2313 | 2679 |
| **TF** | Factor: Elk-1; | TF:M11433_1 | 1.65E-08 | 7.78157324 | 2281 | 2679 |
| **TF** | Factor: ZIC4; | TF:M03932 | 1.69E-08 | 7.77163779 | 2740 | 2679 |
| **TF** | Factor: CREM | TF:M01820_1 | 1.71E-08 | 7.76709631 | 1908 | 2679 |
| **TF** | Factor: nerf; n | TF:M09901_1 | 1.84E-08 | 7.73509442 | 1375 | 2679 |
| **TF** | Factor: ELF2; | TF:M02054 | 1.85E-08 | 7.73383661 | 3087 | 2679 |
| **TF** | Factor: MTF-1 | TF:M01243 | 2.04E-08 | 7.69031867 | 2869 | 2679 |
| **TF** | Factor: ESE-1 | TF:M01214_1 | 2.12E-08 | 7.67423444 | 2194 | 2679 |
| **TF** | Factor: ESE-1 | TF:M02055_1 | 2.12E-08 | 7.67423444 | 2194 | 2679 |
| **TF** | Factor: KLF3; | TF:M03553 | 2.20E-08 | 7.65849561 | 1886 | 2679 |
| **TF** | Factor: Erg; m | TF:M09608_1 | 2.47E-08 | 7.60674275 | 2398 | 2679 |
| **TF** | Factor: ASH-2 | TF:M11179 | 2.48E-08 | 7.6049434 | 2177 | 2679 |
| **TF** | Factor: GCMa | TF:M11595_1 | 2.52E-08 | 7.59883664 | 5851 | 2679 |
| **TF** | Factor: ATF; n | TF:M00017_1 | 2.77E-08 | 7.55698174 | 2250 | 2679 |
| **TF** | Factor: p53; n | TF:M01652 | 2.78E-08 | 7.55669877 | 8154 | 2679 |
| **TF** | Factor: E2F-1 | TF:M00024 | 2.91E-08 | 7.53573338 | 9014 | 2679 |
| **TF** | Factor: SP9; n | TF:M12150 | 2.92E-08 | 7.53432016 | 2368 | 2679 |
| **TF** | Factor: Egr-1; | TF:M04918_1 | 3.06E-08 | 7.51386892 | 1301 | 2679 |
| **TF** | Factor: Elf-1; | TF:M11452_1 | 3.71E-08 | 7.43078646 | 1709 | 2679 |
| **TF** | Factor: YY1; n | TF:M07234 | 3.90E-08 | 7.40877948 | 6088 | 2679 |
| **TF** | Factor: BTEB3 | TF:M12180 | 3.96E-08 | 7.40243996 | 2100 | 2679 |
| **TF** | Factor: ETV6; | TF:M03985 | 5.09E-08 | 7.29346101 | 1148 | 2679 |
| **TF** | Factor: ETV7; | TF:M08760 | 5.31E-08 | 7.27469088 | 2371 | 2679 |
| **TF** | Factor: ER81; | TF:M11420 | 5.54E-08 | 7.25636065 | 3241 | 2679 |
| **TF** | Factor: YY2; n | TF:M12277 | 5.83E-08 | 7.23444668 | 139 | 2679 |
| **TF** | Factor: Sp3; n | TF:M00665 | 6.06E-08 | 7.21765719 | 7795 | 2679 |
| **TF** | Factor: Elk-1; | TF:M09904_1 | 6.29E-08 | 7.20136619 | 2769 | 2679 |
| **TF** | Factor: LRF; r | TF:M04617_1 | 7.16E-08 | 7.14505306 | 1460 | 2679 |
| **TF** | Factor: Net; m | TF:M11439 | 7.41E-08 | 7.13042345 | 2580 | 2679 |
| **TF** | Factor: KLF15 | TF:M12157_1 | 7.46E-08 | 7.12718814 | 818 | 2679 |
| **TF** | Factor: BTEB2 | TF:M12165 | 8.84E-08 | 7.05373368 | 4931 | 2679 |
| **TF** | Factor: Elk-1; | TF:M11432_1 | 8.96E-08 | 7.04763648 | 1538 | 2679 |
| **TF** | Factor: FPM3 | TF:M01587 | 1.00E-07 | 6.999694 | 3279 | 2679 |
| **TF** | Factor: ATF-1 | TF:M01861_1 | 1.13E-07 | 6.94683806 | 1916 | 2679 |
| **TF** | Factor: ZNF34 | TF:M10144_1 | 1.15E-07 | 6.93859111 | 982 | 2679 |
| **TF** | Factor: AhR; r | TF:M00778 | 1.22E-07 | 6.91327807 | 3567 | 2679 |
| **TF** | Factor: HDAC | TF:M07041 | 1.48E-07 | 6.83096199 | 12632 | 2679 |
| **TF** | Factor: TCFL5 | TF:M11078 | 1.53E-07 | 6.81589056 | 1476 | 2679 |
| **TF** | Factor: Pet-1; | TF:M02072 | 1.54E-07 | 6.81281995 | 2408 | 2679 |
| **TF** | Factor: AhR:A | TF:M00237_1 | 1.92E-07 | 6.71610999 | 1479 | 2679 |
| **TF** | Factor: FPM3 | TF:M07141_1 | 1.95E-07 | 6.71047038 | 898 | 2679 |
| **TF** | Factor: CREB | TF:M11217 | 1.96E-07 | 6.70687892 | 2809 | 2679 |
| **TF** | Factor: NF-E4 | TF:M02105 | 2.05E-07 | 6.68720719 | 2657 | 2679 |

| | | | | | | |
|---|---|---|---|---|---|---|
| **TF** | Factor: CREB | TF:M11218_1 | 2.13E-07 | 6.67255341 | 2466 | 2679 |
| **TF** | Factor: BTEB | TF:M09974 | 2.29E-07 | 6.63981606 | 870 | 2679 |
| **TF** | Factor: CREB | TF:M00916_1 | 2.42E-07 | 6.61575147 | 1716 | 2679 |
| **TF** | Factor: GABP | TF:M09935_1 | 2.60E-07 | 6.58438013 | 1057 | 2679 |
| **TF** | Factor: PLAG | TF:M01778_1 | 2.66E-07 | 6.57489245 | 455 | 2679 |
| **TF** | Factor: ZNF4 | TF:M09837 | 2.70E-07 | 6.56810427 | 4296 | 2679 |
| **TF** | Factor: NRF-1 | TF:M04694_1 | 2.71E-07 | 6.56764195 | 466 | 2679 |
| **TF** | Factor: E2F-3 | TF:M08206 | 3.08E-07 | 6.51168684 | 11930 | 2679 |
| **TF** | Factor: Kaiso; | TF:M07140 | 3.34E-07 | 6.4760832 | 556 | 2679 |
| **TF** | Factor: Elk-1; | TF:M01163 | 3.36E-07 | 6.47429381 | 5326 | 2679 |
| **TF** | Factor: RREB | TF:M00257 | 3.39E-07 | 6.46956138 | 10599 | 2679 |
| **TF** | Factor: TFDP1 | TF:M09789 | 3.78E-07 | 6.42244975 | 2057 | 2679 |
| **TF** | Factor: E2F; n | TF:M00918_1 | 3.92E-07 | 6.4071286 | 1270 | 2679 |
| **TF** | Factor: SREBI | TF:M11081_1 | 4.09E-07 | 6.38836844 | 7954 | 2679 |
| **TF** | Factor: SREBI | TF:M01177 | 4.19E-07 | 6.37776612 | 8915 | 2679 |
| **TF** | Factor: Sp2; n | TF:M12155_1 | 4.66E-07 | 6.33118478 | 1009 | 2679 |
| **TF** | Factor: CREB | TF:M00917_1 | 5.07E-07 | 6.29488599 | 1612 | 2679 |
| **TF** | Factor: Elf-1; | TF:M07415_1 | 5.10E-07 | 6.2926185 | 2415 | 2679 |
| **TF** | Factor: ZSCAN | TF:M05432_1 | 5.18E-07 | 6.28552667 | 819 | 2679 |
| **TF** | Factor: SAP-1 | TF:M09607 | 5.24E-07 | 6.28025956 | 4762 | 2679 |
| **TF** | Factor: YY1; n | TF:M12275_1 | 5.31E-07 | 6.27469788 | 1619 | 2679 |
| **TF** | Factor: PDEF; | TF:M02075 | 5.38E-07 | 6.26894778 | 2694 | 2679 |
| **TF** | Factor: Elf-1; | TF:M09900_1 | 5.43E-07 | 6.26532585 | 1163 | 2679 |
| **TF** | Factor: Tax/CI | TF:M00114_1 | 5.59E-07 | 6.25239079 | 1588 | 2679 |
| **TF** | Factor: SP8; n | TF:M12147 | 5.74E-07 | 6.24132383 | 2483 | 2679 |
| **TF** | Factor: ZNF58 | TF:M10491 | 5.99E-07 | 6.22287739 | 13992 | 2679 |
| **TF** | Factor: CREB | TF:M03544 | 6.41E-07 | 6.19335615 | 4156 | 2679 |
| **TF** | Factor: ATF-1 | TF:M07034 | 6.96E-07 | 6.1574304 | 4296 | 2679 |
| **TF** | Factor: CPBP; | TF:M01822 | 7.03E-07 | 6.15300107 | 19532 | 2679 |
| **TF** | Factor: MZF-1 | TF:M00084_1 | 7.13E-07 | 6.14697532 | 1706 | 2679 |
| **TF** | Factor: SP3; n | TF:M03921 | 7.45E-07 | 6.1280664 | 3884 | 2679 |
| **TF** | Factor: ER71; | TF:M11413 | 8.51E-07 | 6.07014984 | 2122 | 2679 |
| **TF** | Factor: Sp3; n | TF:M12154 | 9.96E-07 | 6.00178152 | 2203 | 2679 |
| **TF** | Factor: Net; m | TF:M11442 | 1.1015E-06 | 5.958034 | 2022 | 2679 |
| **TF** | Factor: YY1; n | TF:M00069 | 1.1053E-06 | 5.95651807 | 8449 | 2679 |
| **TF** | Factor: YY2; n | TF:M08020_1 | 1.1794E-06 | 5.92834731 | 74 | 2679 |
| **TF** | Factor: pax-2; | TF:M11876 | 1.2096E-06 | 5.91736592 | 6258 | 2679 |
| **TF** | Factor: CREB | TF:M11218 | 1.2141E-06 | 5.91575185 | 8547 | 2679 |
| **TF** | Factor: Zic3; r | TF:M00450_1 | 1.2251E-06 | 5.91183419 | 7991 | 2679 |
| **TF** | Factor: Elk-1; | TF:M01165_1 | 1.3081E-06 | 5.88335358 | 2201 | 2679 |
| **TF** | Factor: MECP | TF:M01299 | 1.352E-06 | 5.86901387 | 5275 | 2679 |
| **TF** | Factor: DEAF1 | TF:M01001 | 1.4667E-06 | 5.83366487 | 2308 | 2679 |
| **TF** | Factor: KLF16 | TF:M03914_1 | 1.5141E-06 | 5.81983667 | 712 | 2679 |
| **TF** | Factor: CREB | TF:M09887 | 1.5557E-06 | 5.80808595 | 5483 | 2679 |
| **TF** | Factor: ER81; | TF:M11421 | 1.6357E-06 | 5.78628399 | 2356 | 2679 |

| | | | | | | |
|---|---|---|---|---|---|---|
| **TF** | Factor: ATF-1 | TF:M09866 | 1.7627E-06 | 5.7538183 | 3193 | 2679 |
| **TF** | Factor: CREB | TF:M07248 | 1.8572E-06 | 5.73113198 | 4325 | 2679 |
| **TF** | Factor: Elk-1: | TF:M08213_1 | 2.215E-06 | 5.65463279 | 1430 | 2679 |
| **TF** | Factor: ELF1; | TF:M03966 | 2.3382E-06 | 5.63112615 | 6662 | 2679 |
| **TF** | Factor: FPM3 | TF:M10134_1 | 2.3594E-06 | 5.62719012 | 589 | 2679 |
| **TF** | Factor: FPM3 | TF:M01587_1 | 2.5918E-06 | 5.58639468 | 550 | 2679 |
| **TF** | Factor: TFII-I; | TF:M00706_1 | 3.0733E-06 | 5.51239852 | 1606 | 2679 |
| **TF** | Factor: HIC1; | TF:M01072_1 | 3.0826E-06 | 5.51108715 | 897 | 2679 |
| **TF** | Factor: Erg; m | TF:M09608 | 3.1318E-06 | 5.50420943 | 8251 | 2679 |
| **TF** | Factor: AP-2b | TF:M11477 | 3.1788E-06 | 5.49773385 | 8310 | 2679 |
| **TF** | Factor: HDAC | TF:M07041_1 | 3.3222E-06 | 5.47856918 | 6406 | 2679 |
| **TF** | Factor: E2F-3 | TF:M09893_1 | 3.6216E-06 | 5.44109473 | 1336 | 2679 |
| **TF** | Factor: Elk-1; | TF:M11432 | 3.638E-06 | 5.43914036 | 6201 | 2679 |
| **TF** | Factor: p54NI | TF:M07611 | 4.3029E-06 | 5.36624248 | 7053 | 2679 |
| **TF** | Factor: E2F; n | TF:M00050_1 | 4.733E-06 | 5.3248623 | 1909 | 2679 |
| **TF** | Factor: GABP | TF:M02074_1 | 4.7533E-06 | 5.32300125 | 1011 | 2679 |
| **TF** | Factor: Elk-1: | TF:M08229 | 4.9257E-06 | 5.30753142 | 9636 | 2679 |
| **TF** | Factor: CREB | TF:M00177 | 4.9532E-06 | 5.30511388 | 5279 | 2679 |
| **TF** | Factor: ATF-1 | TF:M01861 | 5.2293E-06 | 5.28155971 | 6368 | 2679 |
| **TF** | Factor: Pax-5; | TF:M04064_1 | 5.3056E-06 | 5.2752653 | 3563 | 2679 |
| **TF** | Factor: DATF1 | TF:M09724_1 | 6.0277E-06 | 5.21985014 | 2907 | 2679 |
| **TF** | Factor: CREB | TF:M00039_1 | 6.8773E-06 | 5.16258282 | 1503 | 2679 |
| **TF** | Factor: TFII-I; | TF:M00706 | 6.9831E-06 | 5.15595423 | 5748 | 2679 |
| **TF** | Factor: CREM | TF:M08803 | 7.0327E-06 | 5.15287867 | 1145 | 2679 |
| **TF** | Factor: ZBP8 | TF:M01816_1 | 7.1954E-06 | 5.14294298 | 239 | 2679 |
| **TF** | Factor: ZNF6 | TF:M10550_1 | 1.0089E-05 | 4.99617336 | 9427 | 2679 |
| **TF** | Factor: EGR3 | TF:M08877 | 1.1706E-05 | 4.93159915 | 6168 | 2679 |
| **TF** | Factor: Pet-1; | TF:M11407 | 1.2293E-05 | 4.9103514 | 774 | 2679 |
| **TF** | Factor: LRF; r | TF:M07387_1 | 1.2388E-05 | 4.90700805 | 1047 | 2679 |
| **TF** | Factor: TFAP2 | TF:M04147 | 1.2868E-05 | 4.89049259 | 5127 | 2679 |
| **TF** | Factor: KLF17 | TF:M12188 | 1.3432E-05 | 4.87187359 | 3707 | 2679 |
| **TF** | Factor: BTEB | TF:M12165_1 | 1.366E-05 | 4.86454314 | 944 | 2679 |
| **TF** | Factor: RREB | TF:M00257_1 | 1.4673E-05 | 4.83349246 | 4513 | 2679 |
| **TF** | Factor: NR1B | TF:M11796 | 1.6215E-05 | 4.79009586 | 10778 | 2679 |
| **TF** | Factor: ATF-3 | TF:M07313 | 1.645E-05 | 4.78384671 | 2934 | 2679 |
| **TF** | Factor: ctcf; m | TF:M09890_1 | 1.6875E-05 | 4.77276094 | 2894 | 2679 |
| **TF** | Factor: DEAF1 | TF:M01002 | 1.7013E-05 | 4.76921786 | 1088 | 2679 |
| **TF** | Factor: CREB | TF:M00916 | 1.8421E-05 | 4.73468337 | 5959 | 2679 |
| **TF** | Factor: CREB | TF:M09600 | 1.8719E-05 | 4.7277109 | 5391 | 2679 |
| **TF** | Factor: KLF17 | TF:M12187 | 2.0764E-05 | 4.68268349 | 2905 | 2679 |
| **TF** | Factor: ATF2; | TF:M00179 | 2.0908E-05 | 4.67967947 | 6141 | 2679 |
| **TF** | Factor: Egr; m | TF:M00807_1 | 2.1683E-05 | 4.66387427 | 822 | 2679 |
| **TF** | Factor: TIEG1 | TF:M08905 | 2.1743E-05 | 4.66268972 | 5197 | 2679 |
| **TF** | Factor: Spz1; | TF:M00446_1 | 2.2082E-05 | 4.65595472 | 2886 | 2679 |
| **TF** | Factor: P73; n | TF:M03558 | 2.2927E-05 | 4.63965748 | 4734 | 2679 |

| | | | | | | |
|---|---|---|---|---|---|---|
| **TF** | Factor: E2F-3 | TF:M08206_1 | 2.3317E-05 | 4.63232013 | 5093 | 2679 |
| **TF** | Factor: LHX4; | TF:M11005 | 2.4148E-05 | 4.61711996 | 10745 | 2679 |
| **TF** | Factor: BTEB3 | TF:M12182 | 2.6989E-05 | 4.56881617 | 1749 | 2679 |
| **TF** | Factor: CREB; | TF:M00039 | 2.9694E-05 | 4.52733753 | 5290 | 2679 |
| **TF** | Factor: rfx3:SI | TF:M08623 | 3.3152E-05 | 4.47949279 | 14793 | 2679 |
| **TF** | Factor: ETV5; | TF:M03983_1 | 3.4846E-05 | 4.45784532 | 538 | 2679 |
| **TF** | Factor: GLIS2 | TF:M12217 | 3.5589E-05 | 4.44868622 | 567 | 2679 |
| **TF** | Factor: SAP1A | TF:M02058 | 3.6774E-05 | 4.43445971 | 846 | 2679 |
| **TF** | Factor: Elk-1:I | TF:M08451 | 3.816E-05 | 4.41838658 | 8515 | 2679 |
| **TF** | Factor: CREB | TF:M11217_1 | 3.9214E-05 | 4.40655679 | 465 | 2679 |
| **TF** | Factor: AhR; r | TF:M07347 | 3.9223E-05 | 4.40645509 | 1433 | 2679 |
| **TF** | Factor: CP2; r | TF:M11486 | 4.3722E-05 | 4.35930409 | 3679 | 2679 |
| **TF** | Factor: LRF; r | TF:M07387 | 4.5464E-05 | 4.34233696 | 5266 | 2679 |
| **TF** | Factor: LUMA | TF:M09729 | 4.9162E-05 | 4.30836971 | 10782 | 2679 |
| **TF** | Factor: GATA- | TF:M00075 | 4.9697E-05 | 4.30367062 | 7101 | 2679 |
| **TF** | Factor: GMEE | TF:M12005 | 5.4107E-05 | 4.2667505 | 1178 | 2679 |
| **TF** | Factor: Elf-1; i | TF:M09605_1 | 5.6081E-05 | 4.25118193 | 1311 | 2679 |
| **TF** | Factor: AhR; r | TF:M00139 | 5.6739E-05 | 4.24611789 | 3761 | 2679 |
| **TF** | Factor: Pasx-5 | TF:M04765 | 6.7795E-05 | 4.16880406 | 11048 | 2679 |
| **TF** | Factor: BTEB4 | TF:M12185 | 6.7945E-05 | 4.16784036 | 2298 | 2679 |
| **TF** | Factor: CP2; r | TF:M11486_1 | 6.9324E-05 | 4.15911465 | 2971 | 2679 |
| **TF** | Factor: Zic1; r | TF:M00448 | 7.1018E-05 | 4.14863007 | 11057 | 2679 |
| **TF** | Factor: ATF-2 | TF:M07312 | 8.7665E-05 | 4.05717277 | 3530 | 2679 |
| **TF** | Factor: FEV; n | TF:M03987 | 8.8533E-05 | 4.05289382 | 1324 | 2679 |
| **TF** | Factor: AP-2b | TF:M11477_1 | 9.0303E-05 | 4.04429804 | 7003 | 2679 |
| **TF** | Factor: Elk-1:7 | TF:M08231_1 | 9.217E-05 | 4.03541116 | 1607 | 2679 |
| **TF** | Factor: DEAF1 | TF:M01001_1 | 9.3118E-05 | 4.03096486 | 243 | 2679 |
| **TF** | Factor: etv3; r | TF:M11410 | 9.3333E-05 | 4.02996354 | 1054 | 2679 |
| **TF** | Factor: SREBI | TF:M01177_1 | 0.00010316 | 3.98647162 | 2871 | 2679 |
| **TF** | Factor: DEAF1 | TF:M01002_1 | 0.00011022 | 3.95775003 | 58 | 2679 |
| **TF** | Factor: GABP- | TF:M01876 | 0.000125 | 3.90309195 | 2170 | 2679 |
| **TF** | Factor: etv3; r | TF:M02068_1 | 0.00012556 | 3.90114872 | 614 | 2679 |
| **TF** | Factor: KLF14 | TF:M12184 | 0.00012992 | 3.8863267 | 2545 | 2679 |
| **TF** | Factor: NF-YA | TF:M02106 | 0.00013142 | 3.88132339 | 1769 | 2679 |
| **TF** | Factor: ATF-1 | TF:M07034_1 | 0.00013498 | 3.86973876 | 1197 | 2679 |
| **TF** | Factor: GLI; n | TF:M07290 | 0.00013524 | 3.86888697 | 7189 | 2679 |
| **TF** | Factor: ELK3; | TF:M03972 | 0.00013539 | 3.86841891 | 1216 | 2679 |
| **TF** | Factor: USF2; | TF:M09670 | 0.00015365 | 3.81348073 | 2374 | 2679 |
| **TF** | Factor: E2F-4 | TF:M00739_1 | 0.00015538 | 3.80860478 | 1199 | 2679 |
| **TF** | Factor: Kaiso; | TF:M07140_1 | 0.00015673 | 3.80483663 | 190 | 2679 |
| **TF** | Factor: ZNF44 | TF:M09837_1 | 0.00016278 | 3.7884079 | 766 | 2679 |
| **TF** | Factor: Zic3; r | TF:M00450 | 0.00016645 | 3.77872738 | 14133 | 2679 |
| **TF** | Factor: HES-1 | TF:M07042_1 | 0.000179 | 3.74713521 | 665 | 2679 |
| **TF** | Factor: ATF2; | TF:M00179_1 | 0.00018375 | 3.73578311 | 1801 | 2679 |
| **TF** | Factor: Cdx-2 | TF:M10849_1 | 0.00018743 | 3.7271526 | 4557 | 2679 |

| | | | | | | |
|---|---|---|---|---|---|---|
| **TF** | Factor: NF-YC | TF:M02107 | 0.00019365 | 3.71298465 | 1020 | 2679 |
| **TF** | Factor: Pax-5; | TF:M00143 | 0.00019793 | 3.70349515 | 11490 | 2679 |
| **TF** | Factor: Erm; n | TF:M11422_1 | 0.00020435 | 3.68962931 | 940 | 2679 |
| **TF** | Factor: c-Myb | TF:M00004 | 0.00020538 | 3.68744441 | 7638 | 2679 |
| **TF** | Factor: YY1; n | TF:M12273_1 | 0.00020798 | 3.68197563 | 1273 | 2679 |
| **TF** | Factor: Elf-1; | TF:M11450_1 | 0.00020801 | 3.68190843 | 1324 | 2679 |
| **TF** | Factor: BTEB | TF:M12181 | 0.00020876 | 3.68034347 | 1601 | 2679 |
| **TF** | Factor: Elk-1: | TF:M08229_1 | 0.00020988 | 3.67803865 | 3125 | 2679 |
| **TF** | Factor: ZNF71 | TF:M05938 | 0.00021837 | 3.66080526 | 697 | 2679 |
| **TF** | Factor: c-Krox | TF:M12232 | 0.00021979 | 3.65800107 | 3010 | 2679 |
| **TF** | Factor: NRF-1 | TF:M00652_1 | 0.00023851 | 3.62248993 | 318 | 2679 |
| **TF** | Factor: TEF-3 | TF:M08423_1 | 0.00025972 | 3.5854925 | 657 | 2679 |
| **TF** | Factor: CREB | TF:M00917 | 0.00031616 | 3.50009668 | 5647 | 2679 |
| **TF** | Factor: Pet-1; | TF:M11406 | 0.00032016 | 3.49462692 | 1305 | 2679 |
| **TF** | Factor: rfx3:SI | TF:M08623_1 | 0.00032833 | 3.48368881 | 8201 | 2679 |
| **TF** | Factor: LHX4; | TF:M11005_1 | 0.00037262 | 3.42873335 | 4253 | 2679 |
| **TF** | Factor: ZNF21 | TF:M01122 | 0.00037483 | 3.42616941 | 566 | 2679 |
| **TF** | Factor: CREB | TF:M09887_1 | 0.00039899 | 3.39904269 | 1638 | 2679 |
| **TF** | Factor: Pasx-5 | TF:M04765_1 | 0.00039992 | 3.39802729 | 4171 | 2679 |
| **TF** | Factor: CREB | TF:M11221_1 | 0.00040252 | 3.39520917 | 5754 | 2679 |
| **TF** | Factor: GABP | TF:M01258_1 | 0.00041257 | 3.38449773 | 428 | 2679 |
| **TF** | Factor: ZNF51 | TF:M01231 | 0.00041313 | 3.38391602 | 1431 | 2679 |
| **TF** | Factor: Cdx-2 | TF:M10849 | 0.00042689 | 3.36968234 | 11281 | 2679 |
| **TF** | Factor: CREB | TF:M07248_1 | 0.00043283 | 3.36368492 | 1233 | 2679 |
| **TF** | Factor: CREB | TF:M00177_1 | 0.00045467 | 3.34230669 | 1381 | 2679 |
| **TF** | Factor: E2F-6 | TF:M07085 | 0.00045937 | 3.33784159 | 1063 | 2679 |
| **TF** | Factor: NF-KA | TF:M08952 | 0.00048035 | 3.3184436 | 7811 | 2679 |
| **TF** | Factor: GKLF | TF:M09971_1 | 0.00049327 | 3.30691358 | 1363 | 2679 |
| **TF** | Factor: DREF; | TF:M12238 | 0.00050712 | 3.29489351 | 351 | 2679 |
| **TF** | Factor: ATF-2 | TF:M07312_1 | 0.00053429 | 3.27221959 | 1084 | 2679 |
| **TF** | Factor: batf3; | TF:M11303 | 0.00053994 | 3.26765407 | 623 | 2679 |
| **TF** | Factor: Kid3; | TF:M01160_1 | 0.00056346 | 3.24913555 | 19632 | 2679 |
| **TF** | Factor: ELK4; | TF:M03973 | 0.00064595 | 3.1897988 | 703 | 2679 |
| **TF** | Factor: ERF; r | TF:M03974 | 0.00068604 | 3.1636495 | 962 | 2679 |
| **TF** | Factor: ATF-1 | TF:M09866_1 | 0.0007257 | 3.13924221 | 969 | 2679 |
| **TF** | Factor: CSX; r | TF:M10008_1 | 0.00074387 | 3.12850368 | 3985 | 2679 |
| **TF** | Factor: SAP-1 | TF:M09607_1 | 0.00079476 | 3.09976613 | 989 | 2679 |
| **TF** | Factor: DRRS | TF:M10056_1 | 0.0008101 | 3.0914604 | 983 | 2679 |
| **TF** | Factor: BSX; r | TF:M10591 | 0.00083389 | 3.07888938 | 7851 | 2679 |
| **TF** | Factor: Elf-1; | TF:M11449 | 0.00087227 | 3.05934951 | 4339 | 2679 |
| **TF** | Factor: GABP | TF:M08917_1 | 0.00087919 | 3.05591532 | 389 | 2679 |
| **TF** | Factor: batf3; | TF:M11303_1 | 0.0009306 | 3.03123625 | 593 | 2679 |
| **TF** | Factor: CREB | TF:M03544_1 | 0.00094897 | 3.02274618 | 1213 | 2679 |
| **TF** | Factor: ZNF43 | TF:M10477 | 0.00094918 | 3.02264958 | 14655 | 2679 |
| **TF** | Factor: RREB | TF:M07062 | 0.00126686 | 2.89727234 | 1482 | 2679 |

| | | | | | | |
|---|---|---|---|---|---|---|
| **TF** | Factor: ELF2; | TF:M02054_1 | 0.00139365 | 2.85584675 | 456 | 2679 |
| **TF** | Factor: E2F-1 | TF:M00024_1 | 0.00157437 | 2.80289255 | 2721 | 2679 |
| **TF** | Factor: elf5; m | TF:M11389 | 0.00159386 | 2.79754925 | 1311 | 2679 |
| **TF** | Factor: GKLF; | TF:M01835 | 0.00166306 | 2.77909172 | 15737 | 2679 |
| **TF** | Factor: Nkx2-3 | TF:M10635 | 0.00167604 | 2.77571691 | 4660 | 2679 |
| **TF** | Factor: ELF1; | TF:M03966_1 | 0.00169673 | 2.77038666 | 1585 | 2679 |
| **TF** | Factor: AIBZIP | TF:M11256 | 0.00171428 | 2.76591791 | 1338 | 2679 |
| **TF** | Factor: YY1; n | TF:M01035_1 | 0.00171586 | 2.76551755 | 1120 | 2679 |
| **TF** | Factor: foxl2; | TF:M11578 | 0.00174712 | 2.75767612 | 5520 | 2679 |
| **TF** | Factor: HES-1 | TF:M11061_1 | 0.00175096 | 2.7567242 | 2479 | 2679 |
| **TF** | Factor: ELF3; | TF:M03967 | 0.00178324 | 2.74879137 | 7672 | 2679 |
| **TF** | Factor: GABP- | TF:M11415_1 | 0.00190526 | 2.72004546 | 547 | 2679 |
| **TF** | Factor: GABP- | TF:M04698_1 | 0.00195808 | 2.70817034 | 583 | 2679 |
| **TF** | Factor: FXR; r | TF:M10014_1 | 0.0020067 | 2.69751767 | 1503 | 2679 |
| **TF** | Factor: BTEB3 | TF:M12179 | 0.00206328 | 2.68544254 | 1354 | 2679 |
| **TF** | Factor: SP8; n | TF:M03923 | 0.00210568 | 2.67660692 | 2878 | 2679 |
| **TF** | Factor: CREB | TF:M09600_1 | 0.00225344 | 2.64715399 | 1577 | 2679 |
| **TF** | Factor: ATF-3 | TF:M07313_1 | 0.00230974 | 2.6364374 | 549 | 2679 |
| **TF** | Factor: LBP9; | TF:M01592 | 0.00240937 | 2.61809569 | 1428 | 2679 |
| **TF** | Factor: YY1; n | TF:M10104_1 | 0.00246032 | 2.60900869 | 1151 | 2679 |
| **TF** | Factor: Pax-5; | TF:M04064 | 0.00253122 | 2.59666931 | 10224 | 2679 |
| **TF** | Factor: c-Ets- | TF:M00032_1 | 0.00270818 | 2.56732208 | 2185 | 2679 |
| **TF** | Factor: ZNF56 | TF:M06534 | 0.00273805 | 2.56255848 | 227 | 2679 |
| **TF** | Factor: CREB; | TF:M00178 | 0.00273815 | 2.56254263 | 5320 | 2679 |
| **TF** | Factor: NF-Y; | TF:M00287 | 0.00273826 | 2.56252532 | 1666 | 2679 |
| **TF** | Factor: GCMa | TF:M08312 | 0.00273828 | 2.56252222 | 659 | 2679 |
| **TF** | Factor: ATF-1 | TF:M09592 | 0.00276006 | 2.55908221 | 4470 | 2679 |
| **TF** | Factor: Pax-5; | TF:M00143_1 | 0.00285945 | 2.54371761 | 5115 | 2679 |
| **TF** | Factor: ATF; n | TF:M00338_1 | 0.00322075 | 2.49204276 | 1136 | 2679 |
| **TF** | Factor: ZNF68 | TF:M10550 | 0.00339667 | 2.46894655 | 15402 | 2679 |
| **TF** | Factor: Sall1; | TF:M05464 | 0.00350045 | 2.4558757 | 928 | 2679 |
| **TF** | Factor: DREF; | TF:M12238_1 | 0.00363438 | 2.43956952 | 294 | 2679 |
| **TF** | Factor: Elk-1:I | TF:M08451_1 | 0.00405353 | 2.39216647 | 2600 | 2679 |
| **TF** | Factor: Fli-1; r | TF:M11402_1 | 0.00405647 | 2.39185145 | 603 | 2679 |
| **TF** | Factor: ELF3; | TF:M03967_1 | 0.00411322 | 2.38581769 | 1986 | 2679 |
| **TF** | Factor: ZNF74 | TF:M12237 | 0.00414799 | 2.38216203 | 484 | 2679 |
| **TF** | Factor: Pet-1; | TF:M02072_1 | 0.0042298 | 2.37367971 | 323 | 2679 |
| **TF** | Factor: TEF-3 | TF:M08415_1 | 0.0044433 | 2.35229456 | 2766 | 2679 |
| **TF** | Factor: CP2; r | TF:M11485 | 0.00446468 | 2.35020964 | 7041 | 2679 |
| **TF** | Factor: Elf5; n | TF:M02057 | 0.00450914 | 2.34590667 | 10484 | 2679 |
| **TF** | Factor: GABP- | TF:M11414_1 | 0.00460896 | 2.33639692 | 444 | 2679 |
| **TF** | Factor: NR1B | TF:M11796_1 | 0.00463538 | 2.33391462 | 4931 | 2679 |
| **TF** | Factor: batf3; | TF:M11304_1 | 0.0047245 | 2.32564385 | 739 | 2679 |
| **TF** | Factor: Lumar | TF:M11251_1 | 0.00507697 | 2.29439542 | 777 | 2679 |
| **TF** | Factor: ZNF32 | TF:M10139 | 0.00513235 | 2.28968351 | 697 | 2679 |

| | | | | | | |
|---|---|---|---|---|---|---|
| **TF** | Factor: Elk-1; | TF:M01163_1 | 0.00543522 | 2.26478278 | 1093 | 2679 |
| **TF** | Factor: ZNF51 | TF:M08835 | 0.00546081 | 2.26274333 | 3558 | 2679 |
| **TF** | Factor: XBP-1 | TF:M11277_1 | 0.00563696 | 2.24895474 | 897 | 2679 |
| **TF** | Factor: batf3; | TF:M11305_1 | 0.00581702 | 2.23529943 | 754 | 2679 |
| **TF** | Factor: batf3; | TF:M11305 | 0.00581702 | 2.23529943 | 754 | 2679 |
| **TF** | Factor: CREB | TF:M00113_1 | 0.00586696 | 2.23158702 | 885 | 2679 |
| **TF** | Factor: GEMII | TF:M09727 | 0.005978 | 2.22344434 | 8631 | 2679 |
| **TF** | Factor: BTEB2 | TF:M12164 | 0.00602168 | 2.22028203 | 3000 | 2679 |
| **TF** | Factor: CP2; r | TF:M11485_1 | 0.00609934 | 2.21471723 | 6256 | 2679 |
| **TF** | Factor: PRX-2 | TF:M10885 | 0.00621229 | 2.20674854 | 5775 | 2679 |
| **TF** | Factor: PRX-2 | TF:M10887 | 0.00621229 | 2.20674854 | 5775 | 2679 |
| **TF** | Factor: CTCF; | TF:M01200 | 0.00631575 | 2.1995751 | 3974 | 2679 |
| **TF** | Factor: ATF-4 | TF:M07246_1 | 0.00637218 | 2.19571192 | 590 | 2679 |
| **TF** | Factor: GKLF | TF:M01835_1 | 0.00660044 | 2.18042685 | 10214 | 2679 |
| **TF** | Factor: TCFL5 | TF:M11077 | 0.00703715 | 2.15260307 | 995 | 2679 |
| **TF** | Factor: XBP-1 | TF:M11276 | 0.00704529 | 2.15210107 | 561 | 2679 |
| **TF** | Factor: Thap1 | TF:M04723 | 0.00755058 | 2.12201988 | 12290 | 2679 |
| **TF** | Factor: ATF-2 | TF:M11309 | 0.00763292 | 2.1173092 | 6509 | 2679 |
| **TF** | Factor: hoxa9 | TF:M10785 | 0.00764604 | 2.1165633 | 5703 | 2679 |
| **TF** | Factor: ZBP9 | TF:M12058 | 0.00786825 | 2.10412175 | 234 | 2679 |
| **TF** | Factor: ELK4; | TF:M07209 | 0.00812746 | 2.09004504 | 1125 | 2679 |
| **TF** | Factor: Pax-6; | TF:M04065_1 | 0.00819278 | 2.08656854 | 3604 | 2679 |
| **TF** | Factor: REX-1 | TF:M10119_1 | 0.00852203 | 2.06945714 | 166 | 2679 |
| **TF** | Factor: NF-KA | TF:M08952_1 | 0.00857903 | 2.06656163 | 3473 | 2679 |
| **TF** | Factor: ERF; r | TF:M02061_1 | 0.00862637 | 2.06417179 | 363 | 2679 |
| **TF** | Factor: AML3 | TF:M12000 | 0.00881421 | 2.05481657 | 6876 | 2679 |
| **TF** | Factor: batf3; | TF:M11304 | 0.00892284 | 2.04949667 | 772 | 2679 |
| **TF** | Factor: YY1; n | TF:M07234_1 | 0.00896103 | 2.04764194 | 1191 | 2679 |
| **TF** | Factor: AHR; r | TF:M01855 | 0.00925251 | 2.03374052 | 8566 | 2679 |
| **TF** | Factor: NF-ka | TF:M00051 | 0.00929454 | 2.03177215 | 6458 | 2679 |
| **TF** | Factor: SAP-1 | TF:M11437_1 | 0.00953144 | 2.02084132 | 457 | 2679 |
| **TF** | Factor: LIM-1 | TF:M10999 | 0.0096993 | 2.01325969 | 5955 | 2679 |
| **TF** | Factor: jdp2; r | TF:M11298_1 | 0.01096829 | 1.95986092 | 875 | 2679 |
| **TF** | Factor: ATF-2 | TF:M11309_1 | 0.01136992 | 1.94424269 | 2325 | 2679 |
| **TF** | Factor: ER81; | TF:M11420_1 | 0.01205847 | 1.9187079 | 495 | 2679 |
| **TF** | Factor: ATF-3 | TF:M01863_1 | 0.01257366 | 1.90053812 | 448 | 2679 |
| **TF** | Factor: AIBZIF | TF:M11254 | 0.01257708 | 1.90042026 | 1145 | 2679 |
| **TF** | Factor: P73; n | TF:M03558_1 | 0.01289516 | 1.88957314 | 827 | 2679 |
| **TF** | Factor: ETV1; | TF:M03979 | 0.01306749 | 1.8838078 | 1030 | 2679 |
| **TF** | Factor: ATF-3 | TF:M01863 | 0.01312196 | 1.88200118 | 1910 | 2679 |
| **TF** | Factor: KLF15 | TF:M12159_1 | 0.01343068 | 1.87190184 | 179 | 2679 |
| **TF** | Factor: Lumar | TF:M11251 | 0.01344106 | 1.87156644 | 790 | 2679 |
| **TF** | Factor: CREB | TF:M00178_1 | 0.01376554 | 1.86120679 | 1545 | 2679 |
| **TF** | Factor: AIBZIF | TF:M11257_1 | 0.01387767 | 1.8576834 | 910 | 2679 |
| **TF** | Factor: ER81; | TF:M11421_1 | 0.01406375 | 1.8518988 | 299 | 2679 |

| | | | | | | |
|---|---|---|---|---|---|---|
| **TF** | Factor: ATF-6 | TF:M11226_1 | 0.01563186 | 1.80598921 | 836 | 2679 |
| **TF** | Factor: ZNF74 | TF:M03939 | 0.0156494 | 1.80550232 | 468 | 2679 |
| **TF** | Factor: jdp2; r | TF:M11294_1 | 0.01596302 | 1.79688503 | 357 | 2679 |
| **TF** | Factor: jdp2; r | TF:M11294 | 0.01596302 | 1.79688503 | 357 | 2679 |
| **TF** | Factor: XBP-1 | TF:M11277 | 0.01615611 | 1.79166333 | 944 | 2679 |
| **TF** | Factor: BSX; r | TF:M10591_1 | 0.01635766 | 1.78627884 | 2286 | 2679 |
| **TF** | Factor: ATF-1 | TF:M09592_1 | 0.01666781 | 1.77812151 | 1273 | 2679 |
| **TF** | Factor: USF; r | TF:M00121_1 | 0.01700008 | 1.76954908 | 2498 | 2679 |
| **TF** | Factor: USF; r | TF:M00121 | 0.01700008 | 1.76954908 | 2498 | 2679 |
| **TF** | Factor: GCMb | TF:M08488 | 0.01850903 | 1.73261635 | 1564 | 2679 |
| **TF** | Factor: Egr-1; | TF:M12139_1 | 0.01861774 | 1.7300731 | 701 | 2679 |
| **TF** | Factor: Pax-6; | TF:M04065 | 0.01880141 | 1.72580948 | 10276 | 2679 |
| **TF** | Factor: tcfec; | TF:M11088 | 0.01935906 | 1.71311566 | 96 | 2679 |
| **TF** | Factor: ATF-6 | TF:M11227 | 0.01992089 | 1.70069134 | 208 | 2679 |
| **TF** | Factor: Fli-1; r | TF:M11402 | 0.02021368 | 1.69435463 | 3924 | 2679 |
| **TF** | Factor: AIBZIF | TF:M11254_1 | 0.02042819 | 1.68977009 | 752 | 2679 |
| **TF** | Factor: CREBI | TF:M11231 | 0.02139808 | 1.66962522 | 1070 | 2679 |
| **TF** | Factor: ELFR; | TF:M11455 | 0.02159127 | 1.66572177 | 635 | 2679 |
| **TF** | Factor: Net; m | TF:M11439_1 | 0.02168327 | 1.6638752 | 383 | 2679 |
| **TF** | Factor: PDEF; | TF:M02075_1 | 0.02207979 | 1.65600499 | 269 | 2679 |
| **TF** | Factor: E2F2; | TF:M03958_1 | 0.02253838 | 1.64707737 | 4230 | 2679 |
| **TF** | Factor: E2F-8 | TF:M11537 | 0.02283301 | 1.64143682 | 526 | 2679 |
| **TF** | Factor: GCMa | TF:M08306 | 0.02331126 | 1.63243426 | 2794 | 2679 |
| **TF** | Factor: AP-2g | TF:M08662_1 | 0.02373302 | 1.6246469 | 220 | 2679 |
| **TF** | Factor: GKLF; | TF:M12174 | 0.02374867 | 1.62436069 | 2657 | 2679 |
| **TF** | Factor: CPBP; | TF:M12166 | 0.02478672 | 1.60578091 | 1339 | 2679 |
| **TF** | Factor: ZNF50 | TF:M12086_1 | 0.02759617 | 1.55915126 | 147 | 2679 |
| **TF** | Factor: HES-1 | TF:M02011_1 | 0.02783711 | 1.55537584 | 857 | 2679 |
| **TF** | Factor: CREB | TF:M11223 | 0.02792146 | 1.55406184 | 2847 | 2679 |
| **TF** | Factor: Elk-1:I | TF:M08222_1 | 0.02993816 | 1.52377495 | 2090 | 2679 |
| **TF** | Factor: BTEB2 | TF:M12164_1 | 0.03007154 | 1.52184426 | 392 | 2679 |
| **TF** | Factor: Egr-2; | TF:M05344 | 0.03066393 | 1.51337215 | 266 | 2679 |
| **TF** | Factor: NF-E4 | TF:M08826_1 | 0.03204556 | 1.49423214 | 4737 | 2679 |
| **TF** | Factor: CREB | TF:M11223_1 | 0.03220073 | 1.49213422 | 2713 | 2679 |
| **TF** | Factor: nerf; n | TF:M11457 | 0.03276879 | 1.48453966 | 1090 | 2679 |
| **TF** | Factor: E2F-8 | TF:M11536 | 0.03283155 | 1.48370868 | 387 | 2679 |
| **TF** | Factor: ETV3; | TF:M03981_1 | 0.03297993 | 1.48175032 | 329 | 2679 |
| **TF** | Factor: isx; m | TF:M10956_1 | 0.03952849 | 1.40308975 | 3370 | 2679 |
| **TF** | Factor: ATF-6 | TF:M11226 | 0.03977872 | 1.40034923 | 856 | 2679 |
| **TF** | Factor: KLF17 | TF:M12188_1 | 0.04194827 | 1.3772859 | 588 | 2679 |
| **TF** | Factor: AIBZIF | TF:M11257 | 0.04466414 | 1.35004101 | 992 | 2679 |
| **TF** | Factor: CREBI | TF:M11231_1 | 0.04646038 | 1.33291729 | 852 | 2679 |
| **TF** | Factor: ER81; | TF:M09913_1 | 0.04652127 | 1.33234848 | 420 | 2679 |
| **TF** | Factor: GABPa | TF:M03791_1 | 0.04702327 | 1.32768716 | 432 | 2679 |
| **TF** | Factor: AhR; r | TF:M07347_1 | 0.04740651 | 1.32416201 | 81 | 2679 |

| | | | | | | |
|---|---|---|---|---|---|---|
| **TF** | Factor: USF; r | TF:M00187_1 | 0.04783349 | 1.32026795 | 1530 | 2679 |
| **TF** | Factor: KLF14 | TF:M12184_1 | 0.04907244 | 1.30916236 | 350 | 2679 |

| | | |
|---|---|---|
| 2493 | 19917 | EEF1A1,ORC4,IGFBP5,PABPC1,COL1A1,LRP1,HSPA8,GAPDH,MLLT1 |
| 2403 | 19917 | EEF1A1,ORC4,PABPC1,COL1A1,LRP1,HSPA8,GAPDH,MLLT1,COPA,H |
| 2411 | 19917 | EEF1A1,ORC4,IGFBP5,PABPC1,COL1A1,LRP1,HSPA8,GAPDH,MLLT1 |
| 2325 | 19917 | EEF1A1,ORC4,PABPC1,COL1A1,LRP1,HSPA8,GAPDH,MLLT1,COPA,H |
| 1993 | 19917 | EEF1A1,PABPC1,LRP1,GAPDH,MLLT1,COPA,HNRNPA1,ARSD,HSPA9, |
| 2434 | 19917 | EEF1A1,ORC4,PABPC1,COL1A1,LRP1,HSPA8,GAPDH,MLLT1,COPA,H |
| 2482 | 19917 | EEF1A1,ORC4,IGFBP5,PABPC1,COL1A1,LRP1,HSPA8,GAPDH,MLLT1 |
| 1518 | 19917 | PABPC1,MLLT1,HNRNPA1,ARSD,NEDD4L,POLH,ACTB,NRBP2,USP22 |
| 2202 | 19917 | ORC4,PABPC1,COL1A1,HSPA8,GAPDH,MLLT1,COPA,HNRNPA1,ARSD |
| 2295 | 19917 | EEF1A1,PABPC1,COL1A1,LRP1,HSPA8,GAPDH,MLLT1,HNRNPA1,ARS |
| 2287 | 19917 | ORC4,PABPC1,COL1A1,LRP1,HSPA8,GAPDH,MLLT1,COPA,HNRNPA1 |
| 1828 | 19917 | ORC4,PABPC1,COL1A1,LRP1,HSPA8,GAPDH,MLLT1,COPA,HNRNPA1 |
| 1931 | 19917 | ORC4,PABPC1,COL1A1,LRP1,HSPA8,GAPDH,MLLT1,HNRNPA1,ARSD |
| 1980 | 19917 | ORC4,PABPC1,COL1A1,LRP1,HSPA8,GAPDH,MLLT1,HNRNPA1,ARSD |
| 1938 | 19917 | IGFBP5,PABPC1,LRP1,HSPA8,GAPDH,MLLT1,COPA,ARSD,HSPA9,NE |
| 2271 | 19917 | EEF1A1,IGFBP5,PABPC1,COL1A1,LRP1,HSPA8,GAPDH,MLLT1,COPA |
| 1942 | 19917 | EEF1A1,ORC4,PABPC1,COL1A1,LRP1,GAPDH,MLLT1,HNRNPA1,HSP |
| 1949 | 19917 | EEF1A1,ORC4,PABPC1,COL1A1,LRP1,GAPDH,MLLT1,HNRNPA1,ARS |
| 1643 | 19917 | EEF1A1,ORC4,IGFBP5,PABPC1,COL1A1,LRP1,HSPA8,GAPDH,MLLT1 |
| 1920 | 19917 | ORC4,PABPC1,COL1A1,LRP1,HSPA8,GAPDH,MLLT1,COPA,HNRNPA1 |
| 1912 | 19917 | EEF1A1,PABPC1,COL1A1,HSPA8,GAPDH,MLLT1,ARSD,HSPA9,NEDD4 |
| 2037 | 19917 | EEF1A1,ORC4,IGFBP5,PABPC1,COL1A1,LRP1,HSPA8,GAPDH,MLLT1 |
| 1911 | 19917 | EEF1A1,ORC4,PABPC1,COL1A1,LRP1,HSPA8,GAPDH,MLLT1,HNRNP |
| 2559 | 19917 | EEF1A1,ORC4,IGFBP5,PABPC1,COL1A1,LRP1,HSPA8,GAPDH,MLLT1 |
| 1986 | 19917 | EEF1A1,ORC4,PABPC1,COL1A1,LRP1,HSPA8,GAPDH,MLLT1,HNRNP |
| 2169 | 19917 | EEF1A1,PABPC1,TMEM212,COL1A1,LRP1,HSPA8,GAPDH,MLLT1,CO |
| 1926 | 19917 | ORC4,COL1A1,LRP1,HSPA8,MLLT1,HNRNPA1,ARSD,HSPA9,NEDD4L, |
| 2350 | 19917 | EEF1A1,ORC4,IGFBP5,PABPC1,COL1A1,LRP1,HSPA8,GAPDH,MLLT1 |
| 2543 | 19917 | EEF1A1,ORC4,IGFBP5,PABPC1,COL1A1,LRP1,HSPA8,GAPDH,MLLT1 |
| 2556 | 19917 | EEF1A1,ORC4,IGFBP5,PABPC1,COL1A1,LRP1,HSPA8,GAPDH,MLLT1 |
| 2400 | 19917 | EEF1A1,ORC4,PABPC1,COL1A1,LRP1,HSPA8,GAPDH,MLLT1,COPA,H |
| 1829 | 19917 | EEF1A1,ORC4,PABPC1,COL1A1,LRP1,HSPA8,GAPDH,MLLT1,HNRNP |
| 1897 | 19917 | ORC4,PABPC1,LRP1,HSPA8,GAPDH,MLLT1,COPA,HNRNPA1,ARSD,H |
| 2464 | 19917 | EEF1A1,IGFBP5,PABPC1,COL1A1,LRP1,HSPA8,GAPDH,MLLT1,COPA |
| 2547 | 19917 | EEF1A1,ORC4,PABPC1,COL1A1,LRP1,HSPA8,GAPDH,MLLT1,COPA,H |
| 1645 | 19917 | ORC4,PABPC1,COL1A1,LRP1,GAPDH,MLLT1,HNRNPA1,HSPA9,POLH |
| 2377 | 19917 | ORC4,IGFBP5,COL1A1,LRP1,HSPA8,GAPDH,MLLT1,COPA,HNRNPA1 |
| 1992 | 19917 | IGFBP5,PABPC1,COL1A1,HSPA8,GAPDH,MLLT1,COPA,ARSD,NEDD4L |
| 1873 | 19917 | ORC4,PABPC1,LRP1,HSPA8,GAPDH,MLLT1,ARSD,NEDD4L,POLH,NR |
| 2278 | 19917 | EEF1A1,ORC4,PABPC1,COL1A1,LRP1,HSPA8,GAPDH,MLLT1,COPA,H |
| 2438 | 19917 | EEF1A1,ORC4,PABPC1,TMEM212,COL1A1,LRP1,HSPA8,GAPDH,MLL |
| 2413 | 19917 | EEF1A1,ORC4,IGFBP5,PABPC1,COL1A1,LRP1,HSPA8,GAPDH,MLLT1 |
| 1943 | 19917 | EEF1A1,COL1A1,HSPA8,GAPDH,MLLT1,HNRNPA1,MS4A10,HSPA9,NE |
| 2135 | 19917 | EEF1A1,ORC4,IGFBP5,PABPC1,COL1A1,LRP1,HSPA8,GAPDH,MLLT1 |
| 2284 | 19917 | EEF1A1,ORC4,IGFBP5,PABPC1,COL1A1,LRP1,HSPA8,GAPDH,MLLT1 |

| | | |
|---|---|---|
| 2344 | 19917 | ORC4,IGFBP5,PABPC1,COL1A1,LRP1,HSPA8,GAPDH,MLLT1,COPA,H |
| 2034 | 19917 | EEF1A1,ORC4,PABPC1,COL1A1,LRP1,HSPA8,GAPDH,MLLT1,HNRNP/ |
| 1507 | 19917 | EEF1A1,ORC4,PABPC1,LRP1,GAPDH,MLLT1,NEDD4L,POLH,NRBP2,U |
| 2381 | 19917 | EEF1A1,ORC4,IGFBP5,PABPC1,COL1A1,LRP1,HSPA8,GAPDH,MLLT1 |
| 2449 | 19917 | EEF1A1,ORC4,IGFBP5,PABPC1,COL1A1,LRP1,HSPA8,GAPDH,MLLT1 |
| 1806 | 19917 | EEF1A1,ORC4,PABPC1,COL1A1,LRP1,HSPA8,GAPDH,MLLT1,HNRNP/ |
| 1631 | 19917 | ORC4,PABPC1,COL1A1,LRP1,HSPA8,GAPDH,MLLT1,NEDD4L,POLH,/ |
| 2013 | 19917 | EEF1A1,ORC4,PABPC1,LRP1,HSPA8,GAPDH,MLLT1,COPA,ARSD,HSP |
| 1274 | 19917 | ORC4,PABPC1,COL1A1,LRP1,GAPDH,MLLT1,ACTB,NRBP2,ACTG1,TU |
| 2202 | 19917 | EEF1A1,ORC4,PABPC1,COL1A1,LRP1,HSPA8,GAPDH,MLLT1,HNRNP/ |
| 1279 | 19917 | ORC4,PABPC1,LRP1,GAPDH,MLLT1,NEDD4L,NRBP2,USP22,ACTG1,T |
| 1298 | 19917 | ORC4,PABPC1,LRP1,GAPDH,MLLT1,HNRNPA1,NEDD4L,POLH,NRBP2 |
| 1593 | 19917 | EEF1A1,ORC4,PABPC1,LRP1,HSPA8,MLLT1,HNRNPA1,NEDD4L,NRBI |
| 1514 | 19917 | ORC4,PABPC1,COL1A1,LRP1,GAPDH,MLLT1,HSPA9,POLH,NRBP2,US |
| 2095 | 19917 | EEF1A1,ORC4,IGFBP5,PABPC1,COL1A1,LRP1,GAPDH,MLLT1,HNRNI |
| 1202 | 19917 | PABPC1,HSPA8,MLLT1,ARSD,NEDD4L,ACTB,NRBP2,USP22,ACTG1,D |
| 2229 | 19917 | EEF1A1,ORC4,IGFBP5,PABPC1,TMEM212,COL1A1,LRP1,GAPDH,ML |
| 1508 | 19917 | ORC4,PABPC1,COL1A1,LRP1,GAPDH,MLLT1,HNRNPA1,HSPA9,POLH |
| 1130 | 19917 | ORC4,PABPC1,COL1A1,LRP1,HSPA8,GAPDH,MLLT1,ACTB,NRBP2,US |
| 1587 | 19917 | EEF1A1,ORC4,PABPC1,COL1A1,LRP1,HSPA8,GAPDH,MLLT1,HNRNP/ |
| 2213 | 19917 | EEF1A1,IGFBP5,PABPC1,LRP1,HSPA8,GAPDH,MLLT1,HNRNPA1,ARSI |
| 1240 | 19917 | ORC4,PABPC1,COL1A1,LRP1,GAPDH,MLLT1,ACTB,NRBP2,ACTG1,TU |
| 2155 | 19917 | EEF1A1,PABPC1,COL1A1,LRP1,HSPA8,GAPDH,MLLT1,COPA,ARSD,M |
| 1514 | 19917 | EEF1A1,ORC4,PABPC1,COL1A1,LRP1,HSPA8,MLLT1,HNRNPA1,HSPA |
| 1302 | 19917 | ORC4,PABPC1,COL1A1,LRP1,HSPA8,MLLT1,ARSD,POLH,ACTB,NRBI |
| 1608 | 19917 | PABPC1,GAPDH,MLLT1,ARSD,NEDD4L,POLH,ACTB,NRBP2,USP22,AC |
| 1919 | 19917 | EEF1A1,ORC4,IGFBP5,PABPC1,LRP1,HSPA8,GAPDH,MLLT1,HSPA9,N |
| 1390 | 19917 | EEF1A1,ORC4,IGFBP5,PABPC1,LRP1,HSPA8,GAPDH,MLLT1,HNRNPA |
| 2492 | 19917 | EEF1A1,IGFBP5,PABPC1,COL1A1,LRP1,HSPA8,GAPDH,MLLT1,COPA |
| 2153 | 19917 | EEF1A1,ORC4,PABPC1,COL1A1,LRP1,HSPA8,GAPDH,MLLT1,COPA,H |
| 2447 | 19917 | EEF1A1,IGFBP5,PABPC1,LRP1,HSPA8,GAPDH,MLLT1,HNRNPA1,ARSI |
| 1510 | 19917 | EEF1A1,ORC4,PABPC1,COL1A1,LRP1,HSPA8,GAPDH,MLLT1,HNRNP/ |
| 2146 | 19917 | EEF1A1,ORC4,IGFBP5,PABPC1,COL1A1,LRP1,HSPA8,GAPDH,MLLT1 |
| 2228 | 19917 | EEF1A1,ORC4,PABPC1,COL1A1,LRP1,GAPDH,MLLT1,HNRNPA1,ARSI |
| 1806 | 19917 | EEF1A1,ORC4,PABPC1,COL1A1,LRP1,GAPDH,MLLT1,ARSD,MS4A10, |
| 1537 | 19917 | EEF1A1,ORC4,IGFBP5,PABPC1,LRP1,HSPA8,GAPDH,MLLT1,NEDD4L |
| 1228 | 19917 | ORC4,PABPC1,COL1A1,LRP1,MLLT1,HNRNPA1,ARSD,POLH,ACTB,NI |
| 2571 | 19917 | EEF1A1,ORC4,IGFBP5,PABPC1,COL1A1,LRP1,HSPA8,GAPDH,MLLT1 |
| 1585 | 19917 | ORC4,IGFBP5,PABPC1,COL1A1,LRP1,HSPA8,GAPDH,MLLT1,NEDD4I |
| 1487 | 19917 | ORC4,PABPC1,COL1A1,LRP1,MLLT1,HNRNPA1,HSPA9,ACTB,NRBP2 |
| 1487 | 19917 | ORC4,PABPC1,COL1A1,LRP1,MLLT1,HNRNPA1,HSPA9,ACTB,NRBP2 |
| 1455 | 19917 | EEF1A1,ORC4,PABPC1,COL1A1,LRP1,HSPA8,MLLT1,HNRNPA1,NEDI |
| 2525 | 19917 | EEF1A1,ORC4,IGFBP5,PABPC1,COL1A1,LRP1,HSPA8,GAPDH,MLLT1 |
| 1686 | 19917 | PABPC1,LRP1,HSPA8,MLLT1,ARSD,NEDD4L,ACTB,ODF2L,NUDC,NRE |
| 1525 | 19917 | EEF1A1,PABPC1,HSPA8,GAPDH,MLLT1,ARSD,HSPA9,NEDD4L,POLH,/ |

| | | |
|---|---|---|
| 1591 | 19917 | ORC4,PABPC1,COL1A1,LRP1,GAPDH,MLLT1,HNRNPA1,HSPA9,ACTB |
| 1352 | 19917 | ORC4,PABPC1,COL1A1,LRP1,MLLT1,HNRNPA1,HSPA9,ACTB,NRBP2 |
| 1304 | 19917 | EEF1A1,ORC4,PABPC1,LRP1,HSPA8,GAPDH,MLLT1,HNRNPA1,NEDD |
| 1323 | 19917 | ORC4,PABPC1,LRP1,HSPA8,GAPDH,MLLT1,NEDD4L,NRBP2,USP22, |
| 1620 | 19917 | EEF1A1,ORC4,IGFBP5,PABPC1,COL1A1,LRP1,HSPA8,GAPDH,MLLT1 |
| 2580 | 19917 | EEF1A1,ORC4,IGFBP5,PABPC1,COL1A1,LRP1,HSPA8,GAPDH,MLLT1 |
| 1116 | 19917 | LRP1,HSPA8,MLLT1,ARSD,NEDD4L,ACTB,NRBP2,USP22,ACTG1,TUB |
| 1441 | 19917 | ORC4,PABPC1,COL1A1,LRP1,GAPDH,MLLT1,HNRNPA1,POLH,CTSC, |
| 2519 | 19917 | EEF1A1,ORC4,IGFBP5,PABPC1,COL1A1,LRP1,HSPA8,GAPDH,MLLT1 |
| 1678 | 19917 | EEF1A1,ORC4,PABPC1,COL1A1,LRP1,HSPA8,GAPDH,MLLT1,COPA, |
| 1898 | 19917 | EEF1A1,ORC4,PABPC1,COL1A1,LRP1,HSPA8,GAPDH,MLLT1,COPA, |
| 1546 | 19917 | PABPC1,COL1A1,LRP1,GAPDH,MLLT1,HNRNPA1,HSPA9,NEDD4L,PO |
| 1330 | 19917 | EEF1A1,ORC4,PABPC1,COL1A1,LRP1,HSPA8,MLLT1,HNRNPA1,HSP |
| 1389 | 19917 | EEF1A1,ORC4,IGFBP5,PABPC1,LRP1,HSPA8,GAPDH,MLLT1,HNRNP |
| 2502 | 19917 | EEF1A1,ORC4,IGFBP5,PABPC1,COL1A1,LRP1,HSPA8,GAPDH,MLLT1 |
| 1491 | 19917 | EEF1A1,ORC4,PABPC1,COL1A1,LRP1,HSPA8,GAPDH,MLLT1,HNRNP |
| 891 | 19917 | ORC4,PABPC1,LRP1,HSPA8,GAPDH,MLLT1,NRBP2,USP22,ACTG1,TU |
| 1785 | 19917 | EEF1A1,ORC4,PABPC1,COL1A1,LRP1,HSPA8,GAPDH,MLLT1,COPA, |
| 1421 | 19917 | EEF1A1,HSPA8,MLLT1,ARSD,HSPA9,POLH,ACTB,CTSC,USP22,MED1 |
| 1334 | 19917 | EEF1A1,PABPC1,LRP1,HSPA8,MLLT1,HNRNPA1,ARSD,NEDD4L,NRB |
| 1610 | 19917 | EEF1A1,IGFBP5,PABPC1,LRP1,HSPA8,GAPDH,MLLT1,COPA,HNRNP |
| 1193 | 19917 | PABPC1,GAPDH,NEDD4L,POLH,ACTB,NRBP2,USP22,ACTG1,CFL1,M |
| 1040 | 19917 | ORC4,PABPC1,COL1A1,LRP1,GAPDH,MLLT1,NEDD4L,NRBP2,USP22 |
| 993 | 19917 | PABPC1,LRP1,MLLT1,ARSD,NEDD4L,ACTB,USP22,CFL1,MAVS,HTT, |
| 1910 | 19917 | EEF1A1,ORC4,IGFBP5,PABPC1,COL1A1,LRP1,MLLT1,HNRNPA1,ARS |
| 945 | 19917 | PABPC1,MLLT1,NEDD4L,POLH,ACTB,USP22,EEF2,TUBA1A,CFL1,XR |
| 1848 | 19917 | EEF1A1,PABPC1,LRP1,HSPA8,MLLT1,ARSD,HSPA9,NEDD4L,ATN1,AC |
| 1906 | 19917 | PABPC1,COL1A1,HSPA8,GAPDH,MLLT1,ARSD,NEDD4L,POLH,ACTB, |
| 1594 | 19917 | EEF1A1,ORC4,IGFBP5,PABPC1,COL1A1,LRP1,HSPA8,GAPDH,MLLT1 |
| 1671 | 19917 | EEF1A1,ORC4,IGFBP5,PABPC1,COL1A1,LRP1,HSPA8,GAPDH,MLLT1 |
| 2420 | 19917 | EEF1A1,ORC4,IGFBP5,PABPC1,COL1A1,LRP1,HSPA8,GAPDH,MLLT1 |
| 1644 | 19917 | PABPC1,LRP1,HSPA8,GAPDH,NEDD4L,POLH,ODF2L,NRBP2,USP22, |
| 903 | 19917 | ORC4,PABPC1,LRP1,HSPA8,GAPDH,MLLT1,HNRNPA1,ACTB,NRBP2, |
| 1389 | 19917 | ORC4,PABPC1,COL1A1,LRP1,MLLT1,MS4A10,POLH,CTSC,NRBP2,U |
| 1664 | 19917 | EEF1A1,ORC4,PABPC1,COL1A1,LRP1,HSPA8,GAPDH,MLLT1,COPA, |
| 2013 | 19917 | EEF1A1,ORC4,IGFBP5,PABPC1,COL1A1,LRP1,HSPA8,GAPDH,COPA, |
| 875 | 19917 | PABPC1,MLLT1,HSPA9,ACTB,NRBP2,USP22,ACTG1,BTBD9,XRCC2, |
| 1174 | 19917 | EEF1A1,ORC4,IGFBP5,PABPC1,LRP1,MLLT1,ARSD,HSPA9,NEDD4L, |
| 880 | 19917 | PABPC1,LRP1,HSPA8,MLLT1,NEDD4L,NRBP2,ACTG1,TUBA1A,CFL1, |
| 1317 | 19917 | EEF1A1,ORC4,IGFBP5,PABPC1,LRP1,HSPA8,GAPDH,MLLT1,ARSD,M |
| 1400 | 19917 | EEF1A1,PABPC1,GAPDH,MLLT1,ARSD,HSPA9,NEDD4L,POLH,CTSC,N |
| 1121 | 19917 | HSPA8,GAPDH,MLLT1,HNRNPA1,ACTB,NRBP2,USP22,DPYSL3,CFL1, |
| 1642 | 19917 | PABPC1,COL1A1,GAPDH,MLLT1,ARSD,MS4A10,HSPA9,NEDD4L,POL |
| 1231 | 19917 | EEF1A1,ORC4,PABPC1,COL1A1,LRP1,MLLT1,HNRNPA1,ARSD,NRBP |
| 2562 | 19917 | EEF1A1,ORC4,IGFBP5,PABPC1,LRP1,HSPA8,GAPDH,MLLT1,COPA,H |

| | | |
|---|---|---|
| 1760 | 19917 | EEF1A1,IGFBP5,PABPC1,LRP1,MLLT1,ARSD,HSPA9,NEDD4L,ATN1, |
| 1123 | 19917 | PABPC1,HSPA8,MLLT1,COPA,ARSD,POLH,ACTB,ACTG1,EEF2,BTBD9 |
| 1202 | 19917 | PABPC1,MLLT1,NEDD4L,ACTB,NRBP2,USP22,ACTG1,EEF2,CFL1,M |
| 1613 | 19917 | EEF1A1,COL1A1,GAPDH,MLLT1,HSPA9,NEDD4L,POLH,ACTB,CTSC, |
| 2028 | 19917 | EEF1A1,PABPC1,HSPA8,GAPDH,MLLT1,ARSD,HSPA9,NEDD4L,POLH, |
| 1824 | 19917 | EEF1A1,ORC4,IGFBP5,PABPC1,COL1A1,LRP1,HSPA8,GAPDH,MLLT1 |
| 1315 | 19917 | ORC4,PABPC1,LRP1,HSPA8,GAPDH,MLLT1,HNRNPA1,ARSD,NEDD4L |
| 1742 | 19917 | EEF1A1,ORC4,PABPC1,LRP1,GAPDH,MLLT1,HNRNPA1,ARSD,MS4A1 |
| 2234 | 19917 | EEF1A1,ORC4,IGFBP5,PABPC1,LRP1,HSPA8,GAPDH,MLLT1,COPA,H |
| 2506 | 19917 | EEF1A1,ORC4,IGFBP5,PABPC1,TMEM212,LRP1,HSPA8,GAPDH,MLL |
| 979 | 19917 | GAPDH,HNRNPA1,HSPA9,NUDC,NRBP2,MED18,EEF2,DPYSL3,TUBA1 |
| 971 | 19917 | PABPC1,LRP1,MLLT1,NRBP2,ACTG1,TUBA1A,BTBD9,HTT,KIDINS22 |
| 836 | 19917 | PABPC1,LRP1,MLLT1,NRBP2,USP22,ACTG1,BTBD9,MAVS,HTT,HSP9 |
| 1559 | 19917 | EEF1A1,IGFBP5,PABPC1,COL1A1,LRP1,HSPA8,MLLT1,ARSD,HSPA9, |
| 1624 | 19917 | EEF1A1,PABPC1,LRP1,GAPDH,MLLT1,HSPA9,NEDD4L,POLH,ACTB,O |
| 1772 | 19917 | ORC4,IGFBP5,COL1A1,LRP1,GAPDH,MLLT1,HNRNPA1,ARSD,HSPA9, |
| 929 | 19917 | COL1A1,LRP1,HSPA8,GAPDH,MLLT1,NEDD4L,ACTB,NRBP2,USP22,T |
| 882 | 19917 | ORC4,IGFBP5,PABPC1,COL1A1,LRP1,HSPA8,GAPDH,MLLT1,NEDD4 |
| 2140 | 19917 | EEF1A1,ORC4,PABPC1,COL1A1,LRP1,HSPA8,GAPDH,MLLT1,HNRNP |
| 1920 | 19917 | ORC4,IGFBP5,PABPC1,COL1A1,LRP1,GAPDH,MLLT1,HNRNPA1,ARS |
| 2193 | 19917 | ORC4,IGFBP5,PABPC1,LRP1,HSPA8,GAPDH,MLLT1,HNRNPA1,ARSD, |
| 1451 | 19917 | ORC4,IGFBP5,PABPC1,COL1A1,LRP1,MLLT1,HNRNPA1,MS4A10,AC |
| 1522 | 19917 | IGFBP5,PABPC1,COL1A1,GAPDH,MLLT1,COPA,HNRNPA1,HSPA9,NE |
| 2546 | 19917 | EEF1A1,IGFBP5,PABPC1,COL1A1,LRP1,HSPA8,GAPDH,MLLT1,COPA |
| 896 | 19917 | GAPDH,HSPA9,NEDD4L,ACTB,NRBP2,USP22,ACTG1,EEF2,CFL1,XRC |
| 1194 | 19917 | ORC4,PABPC1,COL1A1,HSPA8,MLLT1,HSPA9,POLH,ATN1,ACTB,OD |
| 892 | 19917 | ORC4,LRP1,HSPA8,HNRNPA1,NEDD4L,POLH,BTBD9,MAVS,HTT,VIM, |
| 2245 | 19917 | EEF1A1,ORC4,PABPC1,COL1A1,LRP1,HSPA8,GAPDH,MLLT1,ARSD, |
| 1061 | 19917 | ORC4,PABPC1,LRP1,MLLT1,ARSD,ACTB,NRBP2,ACTG1,EEF2,DPYSL |
| 1979 | 19917 | EEF1A1,PABPC1,TMEM212,COL1A1,LRP1,GAPDH,HNRNPA1,ARSD, |
| 657 | 19917 | ORC4,PABPC1,LRP1,HSPA8,GAPDH,MLLT1,NEDD4L,USP22,CFL1,HT |
| 1175 | 19917 | ORC4,PABPC1,LRP1,HSPA8,GAPDH,MLLT1,POLH,ACTB,NRBP2,USP |
| 1363 | 19917 | EEF1A1,ORC4,COL1A1,LRP1,GAPDH,COPA,NEDD4L,ACTB,CTSC,OD |
| 1299 | 19917 | EEF1A1,ORC4,IGFBP5,PABPC1,LRP1,HSPA8,MLLT1,NEDD4L,ACTB, |
| 1922 | 19917 | EEF1A1,PABPC1,HSPA8,GAPDH,MLLT1,ARSD,MS4A10,HSPA9,NEDD4 |
| 685 | 19917 | ORC4,PABPC1,LRP1,HSPA8,GAPDH,MLLT1,NRBP2,USP22,ACTG1,C |
| 888 | 19917 | PABPC1,LRP1,MLLT1,NRBP2,ACTG1,BTBD9,XRCC2,HTT,ZNF124,KI |
| 1667 | 19917 | ORC4,IGFBP5,PABPC1,COL1A1,LRP1,HSPA8,GAPDH,MLLT1,ARSD, |
| 1191 | 19917 | EEF1A1,ORC4,IGFBP5,PABPC1,LRP1,HSPA8,GAPDH,MLLT1,HNRNP |
| 1358 | 19917 | ORC4,PABPC1,LRP1,HSPA8,GAPDH,MLLT1,ARSD,NEDD4L,POLH,AC |
| 1779 | 19917 | EEF1A1,PABPC1,COL1A1,LRP1,HSPA8,GAPDH,ARSD,HSPA9,NEDD4L |
| 668 | 19917 | ORC4,PABPC1,LRP1,HSPA8,GAPDH,MLLT1,NRBP2,USP22,CFL1,HTT |
| 1533 | 19917 | COL1A1,LRP1,GAPDH,MLLT1,HNRNPA1,ARSD,ACTB,CTSC,ODF2L,N |
| 1533 | 19917 | COL1A1,LRP1,GAPDH,MLLT1,HNRNPA1,ARSD,ACTB,CTSC,ODF2L,N |
| 1533 | 19917 | COL1A1,LRP1,GAPDH,MLLT1,HNRNPA1,ARSD,ACTB,CTSC,ODF2L,N |

| | | |
|---|---|---|
| 1660 | 19917 | EEF1A1,PABPC1,COL1A1,GAPDH,HNRNPA1,MS4A10,HSPA9,NEDD4L |
| 1280 | 19917 | IGFBP5,PABPC1,LRP1,HSPA8,GAPDH,MLLT1,ARSD,NEDD4L,NRBP2,[illegible] |
| 1863 | 19917 | EEF1A1,IGFBP5,PABPC1,COL1A1,GAPDH,HNRNPA1,MS4A10,HSPA9, |
| 1043 | 19917 | EEF1A1,ORC4,IGFBP5,PABPC1,LRP1,HSPA8,GAPDH,MLLT1,NEDD4L |
| 612 | 19917 | ORC4,PABPC1,LRP1,HSPA8,GAPDH,MLLT1,NRBP2,USP22,ACTG1,C[illegible] |
| 2294 | 19917 | EEF1A1,ORC4,IGFBP5,PABPC1,COL1A1,LRP1,HSPA8,GAPDH,MLLT1 |
| 1160 | 19917 | EEF1A1,ORC4,PABPC1,LRP1,MLLT1,ARSD,NEDD4L,POLH,CTSC,NR[illegible] |
| 1420 | 19917 | EEF1A1,ORC4,PABPC1,LRP1,GAPDH,MLLT1,ARSD,MS4A10,HSPA9,N |
| 1629 | 19917 | PABPC1,COL1A1,LRP1,HSPA8,MLLT1,HNRNPA1,MS4A10,HSPA9,NE[illegible] |
| 873 | 19917 | PABPC1,LRP1,GAPDH,MLLT1,ACTB,NRBP2,USP22,ACTG1,TUBA1A,[illegible] |
| 708 | 19917 | PABPC1,GAPDH,MLLT1,USP22,ACTG1,DPYSL3,ZNF316,DBN1,NEFL,[illegible] |
| 1839 | 19917 | EEF1A1,ORC4,PABPC1,COL1A1,HSPA8,GAPDH,MLLT1,HNRNPA1,AR[illegible] |
| 2048 | 19917 | EEF1A1,ORC4,PABPC1,LRP1,HSPA8,GAPDH,MLLT1,COPA,ARSD,MS[illegible] |
| 776 | 19917 | LRP1,HSPA8,MLLT1,NEDD4L,NRBP2,ACTG1,TUBA1A,BTBD9,HTT,HS |
| 1564 | 19917 | ORC4,COL1A1,LRP1,HSPA8,MLLT1,HSPA9,NEDD4L,POLH,ACTB,NR[illegible] |
| 1828 | 19917 | EEF1A1,ORC4,IGFBP5,PABPC1,LRP1,HSPA8,GAPDH,MLLT1,COPA,N[illegible] |
| 2471 | 19917 | EEF1A1,ORC4,PABPC1,COL1A1,LRP1,HSPA8,GAPDH,MLLT1,COPA,[illegible] |
| 1070 | 19917 | EEF1A1,PABPC1,LRP1,HSPA8,GAPDH,MLLT1,HNRNPA1,ARSD,HSPA9 |
| 2153 | 19917 | EEF1A1,ORC4,IGFBP5,PABPC1,COL1A1,LRP1,HSPA8,GAPDH,MLLT1 |
| 1274 | 19917 | ORC4,PABPC1,LRP1,HSPA8,MLLT1,POLH,ACTB,CTSC,NRBP2,USP22 |
| 671 | 19917 | LRP1,MLLT1,NEDD4L,NRBP2,USP22,DPYSL3,HTT,ZNF124,NUMA1,Z[illegible] |
| 1830 | 19917 | EEF1A1,ORC4,TMEM212,COL1A1,LRP1,HSPA8,GAPDH,MLLT1,HNR[illegible] |
| 1181 | 19917 | EEF1A1,ORC4,LRP1,HSPA8,GAPDH,MLLT1,COPA,HNRNPA1,MS4A10 |
| 996 | 19917 | ORC4,PABPC1,LRP1,MLLT1,ARSD,HSPA9,NRBP2,ACTG1,TUBA1A,C[illegible] |
| 1086 | 19917 | ORC4,IGFBP5,PABPC1,LRP1,GAPDH,MLLT1,NEDD4L,NRBP2,USP22, |
| 1595 | 19917 | EEF1A1,ORC4,PABPC1,COL1A1,LRP1,MLLT1,ARSD,MS4A10,HSPA9,[illegible] |
| 1025 | 19917 | PABPC1,LRP1,NEDD4L,ACTB,CTSC,NRBP2,USP22,TUBA1A,CFL1,ZN[illegible] |
| 1835 | 19917 | EEF1A1,ORC4,IGFBP5,PABPC1,COL1A1,LRP1,HSPA8,GAPDH,MLLT1 |
| 1659 | 19917 | IGFBP5,PABPC1,LRP1,MLLT1,ARSD,MS4A10,HSPA9,NEDD4L,ACTB,[illegible] |
| 1590 | 19917 | IGFBP5,PABPC1,COL1A1,LRP1,HSPA8,MLLT1,HNRNPA1,MS4A10,NE |
| 1411 | 19917 | EEF1A1,COL1A1,GAPDH,NEDD4L,NUDC,USP22,MED18,DPYSL3,CFL[illegible] |
| 1227 | 19917 | ORC4,IGFBP5,COL1A1,LRP1,HSPA8,GAPDH,MLLT1,NEDD4L,POLH,[illegible] |
| 1019 | 19917 | LRP1,HSPA8,MLLT1,ARSD,HSPA9,NEDD4L,ACTB,NRBP2,USP22,ACT[illegible] |
| 1684 | 19917 | EEF1A1,PABPC1,COL1A1,LRP1,HSPA8,GAPDH,HNRNPA1,MS4A10,HS |
| 1334 | 19917 | EEF1A1,ORC4,IGFBP5,PABPC1,COL1A1,LRP1,HSPA8,GAPDH,MLLT1 |
| 2602 | 19917 | EEF1A1,ORC4,IGFBP5,PABPC1,TMEM212,COL1A1,LRP1,HSPA8,GA[illegible] |
| 706 | 19917 | EEF1A1,PABPC1,HSPA8,ARSD,NEDD4L,ACTB,USP22,ACTG1,HTT,ACT |
| 757 | 19917 | PABPC1,LRP1,HSPA8,MLLT1,ARSD,NEDD4L,POLH,NRBP2,ACTG1,HT |
| 706 | 19917 | LRP1,HSPA8,MLLT1,NEDD4L,NRBP2,TUBA1A,HTT,HSP90AB1,HSP90 |
| 2367 | 19917 | EEF1A1,ORC4,IGFBP5,PABPC1,COL1A1,LRP1,HSPA8,GAPDH,MLLT1 |
| 1196 | 19917 | EEF1A1,ORC4,PABPC1,COL1A1,LRP1,HSPA8,GAPDH,COPA,HNRNPA[illegible] |
| 1633 | 19917 | PABPC1,COL1A1,GAPDH,MLLT1,NEDD4L,POLH,ACTB,CTSC,ODF2L,[illegible] |
| 1325 | 19917 | EEF1A1,COL1A1,GAPDH,HNRNPA1,MS4A10,HSPA9,ACTB,NRBP2,US[illegible] |
| 1167 | 19917 | ORC4,PABPC1,LRP1,MLLT1,NEDD4L,POLH,ACTB,CTSC,NRBP2,USP[illegible] |
| 1918 | 19917 | EEF1A1,ORC4,IGFBP5,PABPC1,COL1A1,LRP1,HSPA8,GAPDH,MLLT1 |

| | | |
|---|---|---|
| 1643 | 19917 | IGFBP5,PABPC1,LRP1,HSPA8,GAPDH,MLLT1,NEDD4L,POLH,ACTB,O |
| 1701 | 19917 | EEF1A1,IGFBP5,PABPC1,LRP1,HSPA8,GAPDH,MLLT1,NEDD4L,POLH |
| 1062 | 19917 | PABPC1,COL1A1,LRP1,MLLT1,NEDD4L,NRBP2,USP22,ACTG1,EEF2, |
| 975 | 19917 | ORC4,PABPC1,COL1A1,LRP1,GAPDH,MLLT1,HNRNPA1,NRBP2,ACT |
| 1393 | 19917 | LRP1,GAPDH,COPA,ARSD,MS4A10,HSPA9,NEDD4L,POLH,NRBP2,ME |
| 912 | 19917 | ORC4,PABPC1,LRP1,GAPDH,MLLT1,HNRNPA1,ARSD,HSPA9,POLH,C |
| 1388 | 19917 | ORC4,PABPC1,MLLT1,ARSD,HSPA9,NEDD4L,POLH,NUDC,NRBP2,US |
| 1012 | 19917 | IGFBP5,PABPC1,COL1A1,MLLT1,ODF2L,NRBP2,USP22,ACTG1,TUB |
| 745 | 19917 | ORC4,PABPC1,COL1A1,LRP1,MLLT1,ACTB,NRBP2,ACTG1,TUBA1A, |
| 745 | 19917 | ORC4,PABPC1,COL1A1,LRP1,MLLT1,ACTB,NRBP2,ACTG1,TUBA1A, |
| 373 | 19917 | ORC4,LRP1,HSPA8,GAPDH,USP22,CFL1,VIM,FTH1,DBN1,PIK3R2,HS |
| 702 | 19917 | PABPC1,LRP1,MLLT1,HNRNPA1,NEDD4L,POLH,ACTB,CTSC,MED18, |
| 2088 | 19917 | EEF1A1,ORC4,IGFBP5,TMEM212,COL1A1,LRP1,HSPA8,GAPDH,MLL |
| 1157 | 19917 | ORC4,IGFBP5,PABPC1,COL1A1,LRP1,MLLT1,HSPA9,NEDD4L,POLH, |
| 758 | 19917 | PABPC1,COL1A1,MLLT1,HNRNPA1,NRBP2,ACTG1,DPYSL3,TUBA1A, |
| 713 | 19917 | PABPC1,LRP1,MLLT1,NRBP2,ACTG1,HTT,KIDINS220,NUMA1,ZNF31 |
| 761 | 19917 | ORC4,PABPC1,MLLT1,ACTB,NRBP2,ACTG1,TUBA1A,BTBD9,HTT,HS |
| 909 | 19917 | PABPC1,LRP1,MLLT1,ARSD,ACTB,NRBP2,USP22,CFL1,HTT,ZNF124, |
| 1553 | 19917 | PABPC1,COL1A1,HSPA8,GAPDH,COPA,ARSD,MS4A10,HSPA9,NEDD4 |
| 1419 | 19917 | EEF1A1,COL1A1,HSPA8,GAPDH,ARSD,MS4A10,NEDD4L,POLH,ATN1, |
| 1705 | 19917 | IGFBP5,COL1A1,HSPA8,GAPDH,MLLT1,HNRNPA1,HSPA9,NEDD4L,P |
| 1676 | 19917 | EEF1A1,ORC4,IGFBP5,PABPC1,COL1A1,LRP1,HSPA8,GAPDH,MLLT1 |
| 1749 | 19917 | EEF1A1,PABPC1,COL1A1,LRP1,HSPA8,GAPDH,HNRNPA1,MS4A10,H |
| 630 | 19917 | ORC4,PABPC1,LRP1,HSPA8,ACTB,NRBP2,USP22,TUBA1A,CFL1,HTT, |
| 1001 | 19917 | ORC4,PABPC1,COL1A1,LRP1,MLLT1,HNRNPA1,NEDD4L,POLH,NRB |
| 1437 | 19917 | ORC4,PABPC1,COL1A1,LRP1,HSPA8,GAPDH,MLLT1,ARSD,NEDD4L, |
| 1606 | 19917 | EEF1A1,COL1A1,LRP1,HSPA8,GAPDH,MLLT1,ARSD,MS4A10,HSPA9, |
| 1032 | 19917 | EEF1A1,PABPC1,GAPDH,HNRNPA1,MS4A10,HSPA9,NEDD4L,ACTB,C |
| 575 | 19917 | PABPC1,LRP1,MLLT1,NEDD4L,NRBP2,USP22,EEF2,HTT,HSP90AA1, |
| 1279 | 19917 | EEF1A1,PABPC1,GAPDH,MLLT1,HNRNPA1,ARSD,HSPA9,NEDD4L,PO |
| 2172 | 19917 | EEF1A1,IGFBP5,PABPC1,COL1A1,LRP1,HSPA8,GAPDH,MLLT1,COPA |
| 785 | 19917 | PABPC1,LRP1,MLLT1,NRBP2,ACTG1,XRCC2,HTT,HSP90AB1,ZNF124 |
| 822 | 19917 | PABPC1,COL1A1,MLLT1,NRBP2,USP22,ACTG1,TUBA1A,XRCC2,MAV |
| 818 | 19917 | EEF1A1,LRP1,GAPDH,HSPA9,ACTB,NRBP2,ACTG1,EEF2,CFL1,XRCC |
| 858 | 19917 | EEF1A1,PABPC1,LRP1,GAPDH,MLLT1,NEDD4L,ACTB,USP22,ACTG1, |
| 1444 | 19917 | PABPC1,HSPA8,GAPDH,COPA,HNRNPA1,HSPA9,ATN1,ACTB,NUDC,N |
| 1147 | 19917 | EEF1A1,PABPC1,LRP1,GAPDH,MLLT1,ARSD,NEDD4L,ACTB,NRBP2,U |
| 916 | 19917 | EEF1A1,ORC4,IGFBP5,PABPC1,COL1A1,LRP1,HSPA8,MLLT1,HNRN |
| 2093 | 19917 | EEF1A1,ORC4,PABPC1,COL1A1,LRP1,HSPA8,GAPDH,MLLT1,COPA, |
| 1285 | 19917 | EEF1A1,ORC4,PABPC1,LRP1,GAPDH,ARSD,MS4A10,POLH,ACTB,NU |
| 1311 | 19917 | ORC4,IGFBP5,PABPC1,COL1A1,LRP1,HSPA9,NEDD4L,POLH,ATN1,C |
| 1584 | 19917 | EEF1A1,ORC4,PABPC1,HSPA8,GAPDH,COPA,ARSD,HSPA9,NEDD4L,A |
| 708 | 19917 | ARSD,NEDD4L,USP22,ACTG1,CFL1,MAVS,ZNF124,FTH1,DBN1,ENO1 |
| 906 | 19917 | PABPC1,GAPDH,MLLT1,NRBP2,USP22,ACTG1,TUBA1A,XRCC2,HTT, |
| 1193 | 19917 | EEF1A1,ORC4,PABPC1,LRP1,HSPA8,GAPDH,MLLT1,HNRNPA1,NEDD |

| | | |
|---|---|---|
| 1580 | 19917 | ORC4,IGFBP5,PABPC1,COL1A1,HSPA8,GAPDH,MLLT1,ARSD,MS4A1 |
| 1906 | 19917 | EEF1A1,ORC4,PABPC1,LRP1,HSPA8,GAPDH,COPA,HNRNPA1,ARSD, |
| 1273 | 19917 | ORC4,IGFBP5,PABPC1,ARSD,HSPA9,NEDD4L,POLH,ACTB,NUDC,NR |
| 1849 | 19917 | IGFBP5,PABPC1,HSPA8,GAPDH,MLLT1,COPA,HNRNPA1,MS4A10,PO |
| 1534 | 19917 | IGFBP5,PABPC1,COL1A1,HSPA8,GAPDH,MLLT1,ARSD,POLH,ACTB, |
| 864 | 19917 | PABPC1,LRP1,HNRNPA1,ARSD,POLH,NRBP2,ACTG1,TUBA1A,XRCC2 |
| 595 | 19917 | LRP1,HSPA8,MLLT1,NEDD4L,NRBP2,TUBA1A,HTT,HSP90AA1,VIM,FT |
| 1185 | 19917 | PABPC1,HSPA8,GAPDH,NEDD4L,ACTB,ODF2L,NRBP2,USP22,ACTG1, |
| 2142 | 19917 | EEF1A1,IGFBP5,LRP1,HSPA8,GAPDH,MLLT1,COPA,ARSD,MS4A10,H |
| 1144 | 19917 | COL1A1,LRP1,HSPA8,MLLT1,HSPA9,NEDD4L,POLH,ACTB,NRBP2,US |
| 1094 | 19917 | EEF1A1,LRP1,GAPDH,HNRNPA1,NEDD4L,ACTB,NRBP2,MED18,DPYS |
| 1056 | 19917 | HSPA8,ARSD,HSPA9,NEDD4L,ACTB,NRBP2,USP22,ACTG1,DPYSL3,T |
| 1147 | 19917 | EEF1A1,COL1A1,GAPDH,NEDD4L,ACTB,USP22,MED18,DPYSL3,CFL1 |
| 1279 | 19917 | ORC4,IGFBP5,PABPC1,LRP1,HSPA8,GAPDH,MLLT1,NEDD4L,POLH, |
| 1823 | 19917 | EEF1A1,ORC4,PABPC1,LRP1,HSPA8,GAPDH,COPA,HNRNPA1,ARSD, |
| 866 | 19917 | PABPC1,LRP1,ACTB,USP22,ACTG1,DPYSL3,HTT,HSP90AB1,HSP90A |
| 1408 | 19917 | COL1A1,HSPA8,GAPDH,COPA,HNRNPA1,MS4A10,NEDD4L,ODF2L,N |
| 1289 | 19917 | EEF1A1,ORC4,COL1A1,GAPDH,COPA,POLH,ACTB,USP22,MED18,DP |
| 2021 | 19917 | EEF1A1,ORC4,PABPC1,COL1A1,LRP1,HSPA8,GAPDH,MLLT1,COPA, |
| 1016 | 19917 | PABPC1,COL1A1,HNRNPA1,HSPA9,NEDD4L,POLH,ODF2L,NRBP2,US |
| 1150 | 19917 | PABPC1,POLH,ACTB,CTSC,NRBP2,USP22,ACTG1,EEF2,XRCC2,HSP9 |
| 2223 | 19917 | EEF1A1,ORC4,IGFBP5,PABPC1,COL1A1,HSPA8,GAPDH,COPA,ARSD, |
| 628 | 19917 | ORC4,PABPC1,LRP1,MLLT1,ACTB,NRBP2,ACTG1,TUBA1A,BTBD9,X |
| 2132 | 19917 | EEF1A1,IGFBP5,PABPC1,TMEM212,COL1A1,LRP1,HSPA8,GAPDH,M |
| 771 | 19917 | PABPC1,HSPA8,GAPDH,NEDD4L,ACTB,USP22,EEF2,BTBD9,HTT,HSP |
| 1383 | 19917 | ORC4,PABPC1,COL1A1,LRP1,HNRNPA1,POLH,CTSC,NUDC,NRBP2,U |
| 1383 | 19917 | ORC4,PABPC1,COL1A1,LRP1,HNRNPA1,POLH,CTSC,NUDC,NRBP2,U |
| 1074 | 19917 | ORC4,PABPC1,HSPA8,GAPDH,COPA,ARSD,NEDD4L,POLH,ACTB,ODF |
| 860 | 19917 | PABPC1,MLLT1,ARSD,NRBP2,ACTG1,HTT,HSP90AB1,ZNF124,KIDIN |
| 591 | 19917 | ORC4,PABPC1,COL1A1,LRP1,HSPA8,GAPDH,MLLT1,ARSD,USP22,AC |
| 622 | 19917 | ORC4,COL1A1,LRP1,HSPA8,GAPDH,MLLT1,NEDD4L,USP22,DPYSL3, |
| 2078 | 19917 | EEF1A1,ORC4,IGFBP5,PABPC1,COL1A1,LRP1,HSPA8,GAPDH,HNRN |
| 974 | 19917 | ORC4,PABPC1,COL1A1,LRP1,HSPA8,GAPDH,MLLT1,POLH,ACTB,CT |
| 829 | 19917 | ORC4,PABPC1,COL1A1,LRP1,HSPA8,GAPDH,MLLT1,ARSD,NEDD4L, |
| 760 | 19917 | HSPA8,GAPDH,HNRNPA1,HSPA9,NEDD4L,ATN1,NUDC,NRBP2,MED18 |
| 1471 | 19917 | EEF1A1,IGFBP5,PABPC1,COL1A1,MLLT1,NEDD4L,POLH,ATN1,CTSC |
| 859 | 19917 | EEF1A1,ORC4,HNRNPA1,NEDD4L,POLH,ACTB,CTSC,NUDC,DPYSL3, |
| 1577 | 19917 | EEF1A1,ORC4,IGFBP5,PABPC1,COL1A1,LRP1,HSPA8,GAPDH,MLLT1 |
| 724 | 19917 | ORC4,PABPC1,COL1A1,LRP1,MLLT1,ACTB,NRBP2,ACTG1,TUBA1A, |
| 1974 | 19917 | EEF1A1,ORC4,PABPC1,COL1A1,HSPA8,GAPDH,COPA,ARSD,MS4A10, |
| 1734 | 19917 | EEF1A1,LRP1,HSPA8,MLLT1,ARSD,MS4A10,HSPA9,NEDD4L,POLH,A |
| 905 | 19917 | EEF1A1,PABPC1,COL1A1,GAPDH,ARSD,HSPA9,ODF2L,NUDC,DPYSL |
| 524 | 19917 | HSPA8,GAPDH,HNRNPA1,HSPA9,ATN1,NUDC,EEF2,MAVS,HTT,FTH1, |
| 1009 | 19917 | ORC4,PABPC1,COL1A1,LRP1,MLLT1,NEDD4L,POLH,ODF2L,NRBP2, |
| 1459 | 19917 | ORC4,PABPC1,TMEM212,COL1A1,LRP1,GAPDH,COPA,HNRNPA1,M |

| | | |
|---|---|---|
| 1995 | 19917 | EEF1A1,ORC4,PABPC1,LRP1,HSPA8,GAPDH,MLLT1,COPA,ARSD,HSP |
| 1045 | 19917 | IGFBP5,PABPC1,HSPA8,MLLT1,ARSD,NEDD4L,NRBP2,USP22,ACTG1 |
| 1653 | 19917 | PABPC1,LRP1,GAPDH,ARSD,HSPA9,ACTB,NUDC,NRBP2,USP22,DPYS |
| 474 | 19917 | LRP1,GAPDH,HNRNPA1,ACTB,MED18,EEF2,CFL1,XRCC2,HSP90AA1, |
| 1486 | 19917 | EEF1A1,ORC4,COL1A1,GAPDH,HSPA9,NEDD4L,ACTB,CTSC,ODF2L, |
| 1986 | 19917 | EEF1A1,ORC4,PABPC1,COL1A1,LRP1,HSPA8,GAPDH,MLLT1,HNRNP |
| 971 | 19917 | ORC4,PABPC1,HSPA8,COPA,ARSD,NEDD4L,POLH,ACTB,ODF2L,NUD |
| 1419 | 19917 | ORC4,PABPC1,LRP1,HSPA8,COPA,ARSD,HSPA9,POLH,ACTB,CTSC,O |
| 539 | 19917 | ORC4,LRP1,HSPA8,MLLT1,HNRNPA1,NEDD4L,USP22,CFL1,VIM,NU |
| 1504 | 19917 | EEF1A1,ORC4,PABPC1,LRP1,GAPDH,MLLT1,MS4A10,NEDD4L,ATN1, |
| 1003 | 19917 | ORC4,PABPC1,COL1A1,LRP1,MLLT1,NEDD4L,POLH,CTSC,ODF2L,N |
| 504 | 19917 | EEF1A1,ORC4,PABPC1,LRP1,ARSD,ACTB,USP22,ACTG1,HTT,ZNF12 |
| 683 | 19917 | PABPC1,COL1A1,MLLT1,NRBP2,USP22,ACTG1,TUBA1A,XRCC2,HTT, |
| 1855 | 19917 | EEF1A1,ORC4,PABPC1,COL1A1,LRP1,HSPA8,GAPDH,MLLT1,HNRNP |
| 2170 | 19917 | EEF1A1,ORC4,IGFBP5,COL1A1,HSPA8,GAPDH,COPA,ARSD,MS4A10, |
| 1231 | 19917 | EEF1A1,PABPC1,COL1A1,GAPDH,MLLT1,HNRNPA1,MS4A10,HSPA9, |
| 783 | 19917 | PABPC1,LRP1,MLLT1,NEDD4L,ACTB,NRBP2,USP22,ACTG1,DPYSL3, |
| 1388 | 19917 | PABPC1,LRP1,GAPDH,ARSD,HSPA9,NEDD4L,ACTB,USP22,ACTG1,DP |
| 411 | 19917 | ARSD,ACTB,ACTG1,EEF2,HTT,FTH1,UBC,ZNF316,PIK3R2,ALDOA,NE |
| 1204 | 19917 | ORC4,PABPC1,LRP1,HSPA8,COPA,ARSD,NEDD4L,POLH,ACTB,CTSC, |
| 487 | 19917 | PABPC1,LRP1,MLLT1,HNRNPA1,POLH,ACTB,CTSC,MED18,CFL1,BT |
| 826 | 19917 | ORC4,IGFBP5,COL1A1,LRP1,HSPA8,GAPDH,MLLT1,NEDD4L,ODF2L, |
| 1133 | 19917 | EEF1A1,ORC4,PABPC1,HSPA8,COPA,ARSD,POLH,ACTB,ODF2L,NUD |
| 2649 | 19917 | EEF1A1,ORC4,IGFBP5,PABPC1,COL1A1,LRP1,HSPA8,GAPDH,MLLT1 |
| 1923 | 19917 | ORC4,PABPC1,LRP1,GAPDH,MLLT1,HNRNPA1,MS4A10,HSPA9,NEDD |
| 827 | 19917 | EEF1A1,ORC4,IGFBP5,PABPC1,COL1A1,HSPA8,MLLT1,HNRNPA1,NE |
| 770 | 19917 | IGFBP5,PABPC1,HSPA8,GAPDH,HNRNPA1,MS4A10,HSPA9,NEDD4L,A |
| 1260 | 19917 | ORC4,IGFBP5,PABPC1,ARSD,NEDD4L,POLH,NUDC,NRBP2,USP22,M |
| 783 | 19917 | PABPC1,MLLT1,NRBP2,ACTG1,CFL1,HTT,HSP90AB1,FTH1,UBC,KID |
| 649 | 19917 | ORC4,PABPC1,LRP1,HSPA8,GAPDH,MLLT1,POLH,USP22,TUBA1A,X |
| 1025 | 19917 | EEF1A1,PABPC1,COL1A1,HSPA8,GAPDH,HNRNPA1,MS4A10,NEDD4L |
| 1719 | 19917 | ORC4,PABPC1,LRP1,GAPDH,MLLT1,COPA,NEDD4L,POLH,ATN1,ACT |
| 1255 | 19917 | EEF1A1,PABPC1,COL1A1,HSPA8,COPA,NEDD4L,ODF2L,NUDC,NRBP |
| 1916 | 19917 | EEF1A1,ORC4,COL1A1,HSPA8,GAPDH,COPA,HNRNPA1,ARSD,MS4A1 |
| 1380 | 19917 | LRP1,GAPDH,ARSD,HSPA9,ACTB,NRBP2,DPYSL3,TUBA1A,CFL1,BTB |
| 1618 | 19917 | EEF1A1,IGFBP5,PABPC1,COL1A1,HSPA8,GAPDH,MLLT1,COPA,HNR |
| 2290 | 19917 | EEF1A1,ORC4,PABPC1,COL1A1,LRP1,HSPA8,GAPDH,MLLT1,ARSD, |
| 1560 | 19917 | IGFBP5,PABPC1,COL1A1,LRP1,GAPDH,MLLT1,HNRNPA1,ATN1,ACT |
| 527 | 19917 | PABPC1,LRP1,MLLT1,ARSD,ACTB,NRBP2,TUBA1A,HTT,HSP90AA1,D |
| 2601 | 19917 | EEF1A1,ORC4,IGFBP5,PABPC1,COL1A1,LRP1,HSPA8,GAPDH,MLLT1 |
| 1609 | 19917 | PABPC1,COL1A1,LRP1,HSPA8,GAPDH,MLLT1,HNRNPA1,ARSD,NEDD |
| 1255 | 19917 | IGFBP5,PABPC1,HSPA8,HNRNPA1,ATN1,ACTB,NRBP2,USP22,MED18 |
| 983 | 19917 | PABPC1,GAPDH,COPA,ARSD,NEDD4L,POLH,ACTB,NUDC,BTBD9,MA |
| 803 | 19917 | EEF1A1,NUDC,HTT,HSP90AA1,UBC,EIF4G2,ZNF316,DBN1,TUBA1B, |
| 776 | 19917 | EEF1A1,ORC4,PABPC1,COL1A1,LRP1,MLLT1,ARSD,NRBP2,USP22,A |

| | | |
|---|---|---|
| 601 | 19917 | PABPC1,LRP1,NRBP2,USP22,ACTG1,HTT,KIDINS220,NUMA1,RPL4,F |
| 528 | 19917 | PABPC1,LRP1,MLLT1,ARSD,NRBP2,HTT,KIDINS220,NUMA1,DBN1,PI |
| 623 | 19917 | HSPA8,USP22,CFL1,HTT,HSP90AB1,VIM,NUMA1,EIF4G2,RPS6,ZNF3 |
| 2043 | 19917 | ORC4,IGFBP5,PABPC1,COL1A1,LRP1,GAPDH,MLLT1,HNRNPA1,ARS |
| 1897 | 19917 | EEF1A1,ORC4,IGFBP5,COL1A1,LRP1,HSPA8,GAPDH,MLLT1,COPA,A |
| 537 | 19917 | ORC4,PABPC1,LRP1,MLLT1,ACTB,NRBP2,ACTG1,HTT,KIDINS220,DE |
| 626 | 19917 | PABPC1,COL1A1,LRP1,MLLT1,CTSC,NRBP2,ACTG1,TUBA1A,HTT,HS |
| 1742 | 19917 | ORC4,PABPC1,COL1A1,LRP1,HSPA8,GAPDH,MLLT1,COPA,NEDD4L, |
| 672 | 19917 | GAPDH,MLLT1,ARSD,NUDC,NRBP2,MAVS,HTT,HSP90AB1,HSP90AA1 |
| 672 | 19917 | GAPDH,MLLT1,ARSD,NUDC,NRBP2,MAVS,HTT,HSP90AB1,HSP90AA1 |
| 672 | 19917 | GAPDH,MLLT1,ARSD,NUDC,NRBP2,MAVS,HTT,HSP90AB1,HSP90AA1 |
| 1188 | 19917 | EEF1A1,PABPC1,LRP1,HSPA8,MLLT1,COPA,NEDD4L,POLH,ACTB,CT |
| 623 | 19917 | GAPDH,MLLT1,MS4A10,NEDD4L,NUDC,CFL1,MAVS,HTT,HSP90AA1,Z |
| 1761 | 19917 | ORC4,IGFBP5,PABPC1,COL1A1,LRP1,HSPA8,GAPDH,MLLT1,COPA,A |
| 1865 | 19917 | EEF1A1,ORC4,IGFBP5,COL1A1,LRP1,HSPA8,GAPDH,MLLT1,COPA,A |
| 1653 | 19917 | ORC4,IGFBP5,PABPC1,LRP1,HSPA8,MLLT1,COPA,ARSD,NEDD4L,PO |
| 2100 | 19917 | ORC4,IGFBP5,PABPC1,TMEM212,COL1A1,LRP1,HSPA8,GAPDH,MLL |
| 456 | 19917 | PABPC1,MLLT1,NRBP2,TUBA1A,HTT,ZNF316,DBN1,FLNA,PKM,COL |
| 1141 | 19917 | EEF1A1,PABPC1,COL1A1,GAPDH,HNRNPA1,MS4A10,NEDD4L,ACTB, |
| 803 | 19917 | IGFBP5,PABPC1,MLLT1,ACTB,NRBP2,DPYSL3,TUBA1A,BTBD9,MAV |
| 1111 | 19917 | EEF1A1,ORC4,IGFBP5,PABPC1,LRP1,HSPA8,ARSD,ACTB,NRBP2,ACT |
| 751 | 19917 | EEF1A1,GAPDH,HNRNPA1,ACTB,NRBP2,USP22,MED18,DPYSL3,VIM, |
| 761 | 19917 | IGFBP5,PABPC1,LRP1,HSPA8,ACTB,CTSC,NRBP2,ACTG1,TUBA1A,C |
| 1869 | 19917 | EEF1A1,ORC4,IGFBP5,COL1A1,LRP1,HSPA8,GAPDH,MLLT1,COPA,A |
| 481 | 19917 | ORC4,LRP1,HSPA8,NEDD4L,NRBP2,USP22,ACTG1,DPYSL3,HTT,ZNF |
| 1833 | 19917 | EEF1A1,PABPC1,COL1A1,LRP1,GAPDH,MLLT1,COPA,HNRNPA1,ARS |
| 991 | 19917 | ORC4,IGFBP5,PABPC1,LRP1,GAPDH,ARSD,NEDD4L,ODF2L,USP22,C |
| 943 | 19917 | EEF1A1,ORC4,LRP1,GAPDH,HNRNPA1,ARSD,NEDD4L,CTSC,NRBP2, |
| 864 | 19917 | EEF1A1,PABPC1,COL1A1,GAPDH,HNRNPA1,HSPA9,ACTB,ODF2L,ME |
| 1747 | 19917 | ORC4,PABPC1,COL1A1,LRP1,HSPA8,GAPDH,MLLT1,COPA,ARSD,NE |
| 603 | 19917 | PABPC1,HSPA9,NEDD4L,ACTB,HTT,ZNF124,HSP90AA1,DBN1,ENO1, |
| 909 | 19917 | ORC4,PABPC1,GAPDH,COPA,ARSD,ACTB,ODF2L,NUDC,MED18,ACT |
| 1046 | 19917 | ORC4,IGFBP5,PABPC1,COL1A1,LRP1,HSPA8,GAPDH,MLLT1,MS4A1 |
| 1094 | 19917 | COL1A1,GAPDH,HNRNPA1,MS4A10,HSPA9,NEDD4L,ACTB,NRBP2,ME |
| 428 | 19917 | HSPA8,GAPDH,COPA,HNRNPA1,HSPA9,MED18,HSP90AB1,FTH1,UBC |
| 1464 | 19917 | EEF1A1,PABPC1,COL1A1,LRP1,HSPA8,GAPDH,HNRNPA1,ARSD,HSPA |
| 1377 | 19917 | EEF1A1,ORC4,PABPC1,LRP1,HSPA8,GAPDH,COPA,ARSD,NEDD4L,PC |
| 1643 | 19917 | EEF1A1,ORC4,PABPC1,TMEM212,LRP1,HSPA8,MLLT1,COPA,ARSD, |
| 554 | 19917 | COL1A1,COPA,NEDD4L,ACTB,CTSC,ACTG1,CFL1,MAVS,HSP90AA1,V |
| 1686 | 19917 | EEF1A1,ORC4,IGFBP5,COL1A1,LRP1,HSPA8,GAPDH,MLLT1,COPA,A |
| 1561 | 19917 | EEF1A1,ORC4,PABPC1,HSPA8,GAPDH,MLLT1,HNRNPA1,ARSD,HSPA |
| 1030 | 19917 | IGFBP5,LRP1,HSPA8,GAPDH,ARSD,NEDD4L,ACTB,NRBP2,TUBA1A,B |
| 1891 | 19917 | EEF1A1,ORC4,IGFBP5,COL1A1,LRP1,HSPA8,GAPDH,MLLT1,COPA,A |
| 771 | 19917 | EEF1A1,COL1A1,GAPDH,HNRNPA1,HSPA9,ACTB,CTSC,MED18,DPYS |
| 1234 | 19917 | IGFBP5,PABPC1,COL1A1,HSPA8,GAPDH,MLLT1,COPA,HNRNPA1,AR |

| | | |
|---|---|---|
| 858 | 19917 | PABPC1,GAPDH,MLLT1,COPA,ACTB,ODF2L,NUDC,MED18,MAVS,HS[illegible] |
| 556 | 19917 | ORC4,IGFBP5,PABPC1,COL1A1,LRP1,MLLT1,NEDD4L,USP22,DPYSL[illegible] |
| 709 | 19917 | EEF1A1,GAPDH,MLLT1,NEDD4L,ACTB,CTSC,ACTG1,DPYSL3,BTBD9,[illegible] |
| 1705 | 19917 | ORC4,PABPC1,COL1A1,LRP1,HSPA8,GAPDH,MLLT1,COPA,ARSD,HS[illegible] |
| 1661 | 19917 | ORC4,PABPC1,LRP1,HSPA8,MLLT1,COPA,NEDD4L,POLH,ATN1,ACT[illegible] |
| 1065 | 19917 | EEF1A1,PABPC1,LRP1,HSPA8,MLLT1,COPA,NEDD4L,POLH,ACTB,CT[illegible] |
| 460 | 19917 | ORC4,MLLT1,ACTG1,HTT,HSP90AA1,FTH1,ZNF316,DBN1,ENO1,FBX[illegible] |
| 1286 | 19917 | EEF1A1,IGFBP5,PABPC1,COL1A1,LRP1,GAPDH,MLLT1,NEDD4L,ACT[illegible] |
| 1595 | 19917 | EEF1A1,ORC4,IGFBP5,PABPC1,COL1A1,LRP1,GAPDH,COPA,HNRNP[illegible] |
| 989 | 19917 | IGFBP5,LRP1,HSPA8,ARSD,NEDD4L,ACTB,NRBP2,TUBA1A,BTBD9,X[illegible] |
| 534 | 19917 | PABPC1,LRP1,NRBP2,USP22,ACTG1,HTT,KIDINS220,NUMA1,RPL4,P[illegible] |
| 856 | 19917 | PABPC1,LRP1,MLLT1,HNRNPA1,ARSD,MS4A10,POLH,NRBP2,ACTG1[illegible] |
| 467 | 19917 | PABPC1,NRBP2,ACTG1,HTT,KIDINS220,NUMA1,PKM,FBXL4,ABHD2,[illegible] |
| 702 | 19917 | ORC4,PABPC1,LRP1,MLLT1,NEDD4L,NRBP2,ACTG1,TUBA1A,XRCC2[illegible] |
| 1328 | 19917 | PABPC1,COL1A1,LRP1,HSPA8,MLLT1,HNRNPA1,ARSD,MS4A10,HSP[illegible] |
| 1003 | 19917 | PABPC1,HSPA8,GAPDH,ARSD,HSPA9,POLH,ACTB,CFL1,MAVS,HTT,Z[illegible] |
| 1162 | 19917 | EEF1A1,PABPC1,LRP1,GAPDH,HSPA9,ATN1,ACTB,CTSC,USP22,ACT[illegible] |
| 1290 | 19917 | EEF1A1,ORC4,PABPC1,HSPA8,COPA,ARSD,NEDD4L,POLH,ATN1,ACT[illegible] |
| 535 | 19917 | COL1A1,NEDD4L,ACTB,ACTG1,BTBD9,XRCC2,MAVS,FTH1,EIF4G2,T[illegible] |
| 1614 | 19917 | ORC4,PABPC1,LRP1,GAPDH,MLLT1,COPA,NEDD4L,POLH,ACTB,CTS[illegible] |
| 1275 | 19917 | EEF1A1,LRP1,GAPDH,HSPA9,POLH,ACTB,CTSC,ODF2L,NRBP2,USP2[illegible] |
| 1075 | 19917 | COL1A1,HSPA8,GAPDH,MLLT1,COPA,HNRNPA1,HSPA9,ATN1,NRBP2[illegible] |
| 1542 | 19917 | ORC4,PABPC1,LRP1,HSPA8,MLLT1,COPA,HSPA9,NEDD4L,POLH,ACT[illegible] |
| 629 | 19917 | ORC4,PABPC1,COL1A1,LRP1,GAPDH,MLLT1,NEDD4L,USP22,TUBA1[illegible] |
| 557 | 19917 | PABPC1,GAPDH,MLLT1,ARSD,POLH,CFL1,XRCC2,HTT,TUBB,FTH1,N[illegible] |
| 853 | 19917 | ORC4,PABPC1,GAPDH,COPA,ARSD,ACTB,ODF2L,MED18,ACTG1,MA[illegible] |
| 907 | 19917 | ORC4,PABPC1,COPA,ACTB,ODF2L,NUDC,MED18,ACTG1,CFL1,MAV[illegible] |
| 886 | 19917 | ORC4,PABPC1,LRP1,HSPA8,COPA,POLH,ODF2L,MED18,ACTG1,DPY[illegible] |
| 1730 | 19917 | EEF1A1,ORC4,PABPC1,COL1A1,LRP1,HSPA8,GAPDH,MLLT1,HNRNP[illegible] |
| 1381 | 19917 | ORC4,PABPC1,LRP1,HSPA8,MLLT1,COPA,NEDD4L,POLH,ACTB,CTS[illegible] |
| 407 | 19917 | PABPC1,LRP1,HNRNPA1,POLH,ACTB,MED18,EEF2,CFL1,XRCC2,HSP[illegible] |
| 1506 | 19917 | EEF1A1,ORC4,IGFBP5,PABPC1,LRP1,HSPA8,GAPDH,MLLT1,POLH,A[illegible] |
| 425 | 19917 | PABPC1,LRP1,ACTG1,HTT,NUMA1,DBN1,PKM,RPL8,ABHD2,TUBB2B[illegible] |
| 1131 | 19917 | ORC4,COL1A1,GAPDH,MLLT1,ARSD,HSPA9,ACTB,CTSC,ODF2L,USP[illegible] |
| 1780 | 19917 | ORC4,PABPC1,HSPA8,GAPDH,COPA,HNRNPA1,ARSD,MS4A10,HSPA9[illegible] |
| 1384 | 19917 | ORC4,PABPC1,LRP1,HSPA8,GAPDH,COPA,POLH,ACTB,ODF2L,NUDC[illegible] |
| 1208 | 19917 | EEF1A1,PABPC1,COL1A1,LRP1,HSPA8,GAPDH,HNRNPA1,MS4A10,H[illegible] |
| 527 | 19917 | EEF1A1,ORC4,LRP1,HSPA8,GAPDH,USP22,ACTG1,HTT,ZNF124,HSP[illegible] |
| 587 | 19917 | PABPC1,COPA,ARSD,POLH,ACTB,ODF2L,NUDC,MED18,ACTG1,CFL1[illegible] |
| 653 | 19917 | PABPC1,LRP1,MLLT1,NRBP2,USP22,ACTG1,TUBA1A,XRCC2,HTT,ZN[illegible] |
| 1809 | 19917 | EEF1A1,ORC4,IGFBP5,LRP1,GAPDH,MLLT1,COPA,ARSD,HSPA9,NED[illegible] |
| 676 | 19917 | PABPC1,COL1A1,GAPDH,MLLT1,HSPA9,CTSC,ACTG1,HTT,HSP90AB[illegible] |
| 1361 | 19917 | ORC4,IGFBP5,PABPC1,COL1A1,LRP1,HSPA8,GAPDH,HNRNPA1,HSP[illegible] |
| 1082 | 19917 | ORC4,COL1A1,LRP1,HSPA8,NEDD4L,POLH,ATN1,ODF2L,NRBP2,US[illegible] |
| 255 | 19917 | LRP1,HNRNPA1,POLH,ACTB,MED18,CFL1,XRCC2,HSP90AA1,ZNF31[illegible] |

| | | |
|---|---|---|
| 1305 | 19917 | ORC4,IGFBP5,COL1A1,LRP1,GAPDH,COPA,ARSD,MS4A10,HSPA9,NE |
| 760 | 19917 | ORC4,HSPA8,COPA,ODF2L,NUDC,MED18,ACTG1,BTBD9,MAVS,ZNF1 |
| 554 | 19917 | PABPC1,MLLT1,ARSD,ACTB,NRBP2,USP22,HTT,HSP90AB1,UBC,DB |
| 250 | 19917 | NRBP2,ACTG1,XRCC2,HTT,HSP90AA1,ZNF316,SLC9A7,YBX1,CANX, |
| 1431 | 19917 | ORC4,PABPC1,LRP1,MLLT1,COPA,NEDD4L,POLH,ACTB,CTSC,ODF2 |
| 1202 | 19917 | ORC4,PABPC1,LRP1,HSPA8,MLLT1,COPA,ACTB,ODF2L,NUDC,USP2 |
| 1438 | 19917 | EEF1A1,ORC4,IGFBP5,PABPC1,COL1A1,LRP1,HSPA8,COPA,HNRNP |
| 2135 | 19917 | EEF1A1,ORC4,IGFBP5,PABPC1,COL1A1,LRP1,GAPDH,MLLT1,COPA, |
| 355 | 19917 | MLLT1,NRBP2,USP22,HTT,HSP90AA1,ZNF316,DBN1,PIK3R2,TUBB2 |
| 2079 | 19917 | EEF1A1,ORC4,IGFBP5,PABPC1,COL1A1,LRP1,GAPDH,MLLT1,COPA, |
| 538 | 19917 | EEF1A1,COL1A1,GAPDH,USP22,DPYSL3,CFL1,TUBB,UBC,EIF4G2,E |
| 1305 | 19917 | EEF1A1,ORC4,PABPC1,LRP1,HSPA8,COPA,NEDD4L,POLH,ACTB,OD |
| 1292 | 19917 | ORC4,PABPC1,LRP1,HSPA8,COPA,NEDD4L,POLH,ACTB,ODF2L,NUD |
| 1996 | 19917 | EEF1A1,ORC4,IGFBP5,PABPC1,COL1A1,LRP1,HSPA8,GAPDH,COPA, |
| 991 | 19917 | COL1A1,HSPA8,GAPDH,MS4A10,HSPA9,ATN1,ACTB,CTSC,ODF2L,NU |
| 986 | 19917 | COL1A1,HSPA8,GAPDH,HNRNPA1,MS4A10,NEDD4L,ACTB,CTSC,ME |
| 142 | 19917 | GAPDH,HNRNPA1,UBC,EIF4G2,GANAB,KAT6A,YWHAE,DDX17,RACK1 |
| 390 | 19917 | NEDD4L,NRBP2,MAVS,HSP90AB1,HSP90AA1,DBN1,PIK3R2,RPL7A,F |
| 2255 | 19917 | EEF1A1,ORC4,PABPC1,LRP1,GAPDH,MLLT1,HNRNPA1,ARSD,MS4A1 |
| 998 | 19917 | PABPC1,COL1A1,GAPDH,HNRNPA1,MS4A10,NEDD4L,ACTB,MED18, |
| 2289 | 19917 | EEF1A1,IGFBP5,PABPC1,COL1A1,LRP1,HSPA8,GAPDH,MLLT1,HNR |
| 1968 | 19917 | EEF1A1,ORC4,IGFBP5,COL1A1,LRP1,GAPDH,MLLT1,COPA,ARSD,M |
| 1570 | 19917 | ORC4,IGFBP5,PABPC1,GAPDH,MLLT1,ARSD,NEDD4L,ATN1,NUDC,U |
| 1685 | 19917 | EEF1A1,ORC4,PABPC1,TMEM212,COL1A1,MLLT1,COPA,HNRNPA1, |
| 904 | 19917 | PABPC1,HSPA8,GAPDH,COPA,ARSD,HSPA9,NEDD4L,ACTB,NUDC,ME |
| 1374 | 19917 | ORC4,PABPC1,LRP1,HSPA8,MLLT1,COPA,ACTB,CTSC,ODF2L,NUDC, |
| 1119 | 19917 | MLLT1,HNRNPA1,ARSD,ACTB,CTSC,NRBP2,USP22,ACTG1,TUBA1A, |
| 352 | 19917 | PABPC1,LRP1,HNRNPA1,POLH,ACTB,MED18,EEF2,XRCC2,HSP90AA |
| 445 | 19917 | PABPC1,COL1A1,LRP1,GAPDH,USP22,TUBA1A,VIM,KIDINS220,NUM |
| 783 | 19917 | EEF1A1,PABPC1,HSPA8,MLLT1,ARSD,HSPA9,NEDD4L,POLH,CTSC,A |
| 2203 | 19917 | EEF1A1,ORC4,IGFBP5,PABPC1,COL1A1,LRP1,HSPA8,GAPDH,COPA, |
| 486 | 19917 | NRBP2,USP22,TUBA1A,CFL1,HTT,EIF4G2,RPS6,ZNF316,RPL4,NCL, |
| 1264 | 19917 | ORC4,PABPC1,LRP1,HSPA8,GAPDH,COPA,ARSD,NEDD4L,POLH,ACT |
| 1237 | 19917 | EEF1A1,ORC4,IGFBP5,PABPC1,COL1A1,LRP1,HSPA8,COPA,MS4A10 |
| 1987 | 19917 | EEF1A1,ORC4,IGFBP5,COL1A1,LRP1,GAPDH,MLLT1,COPA,ARSD,M |
| 1613 | 19917 | EEF1A1,ORC4,PABPC1,GAPDH,MLLT1,COPA,HNRNPA1,ARSD,NEDD4 |
| 216 | 19917 | GAPDH,ACTB,NRBP2,ACTG1,XRCC2,ZNF316,KAT6A,MYH9,SMG1,N |
| 1875 | 19917 | EEF1A1,ORC4,IGFBP5,PABPC1,COL1A1,LRP1,HSPA8,GAPDH,HNRN |
| 1334 | 19917 | ORC4,PABPC1,LRP1,GAPDH,MLLT1,COPA,NEDD4L,ACTB,ODF2L,NU |
| 706 | 19917 | ORC4,PABPC1,COPA,NEDD4L,ACTB,ODF2L,NUDC,MED18,ACTG1,C |
| 308 | 19917 | PABPC1,LRP1,HSPA8,POLH,DPYSL3,HTT,VIM,DBN1,RPL7A,TUBB2B, |
| 772 | 19917 | ORC4,PABPC1,COPA,ACTB,ODF2L,NUDC,MED18,CFL1,MAVS,ZNF12 |
| 1868 | 19917 | PABPC1,LRP1,HSPA8,GAPDH,MLLT1,HNRNPA1,ARSD,HSPA9,NEDD4 |
| 1723 | 19917 | EEF1A1,ORC4,PABPC1,COL1A1,LRP1,GAPDH,MLLT1,COPA,HNRNP |
| 1368 | 19917 | ORC4,PABPC1,LRP1,MLLT1,COPA,ACTB,CTSC,ODF2L,NUDC,USP22, |

| | | |
|---|---|---|
| 1531 | 19917 | IGFBP5,PABPC1,HSPA8,GAPDH,COPA,HSPA9,POLH,ATN1,ACTB,USP2 |
| 452 | 19917 | PABPC1,MLLT1,ARSD,POLH,XRCC2,HTT,VIM,TUBB,FTH1,NUMA1,TU |
| 303 | 19917 | ORC4,LRP1,HNRNPA1,NEDD4L,USP22,HTT,MAP1B,PKM,IGFBP2,CO |
| 254 | 19917 | ACTG1,HSP90AB1,HSP90AA1,NCL,YBX1,TUBB2B,COL4A1,PTMA,DVL |
| 1606 | 19917 | EEF1A1,ORC4,IGFBP5,COL1A1,HSPA8,GAPDH,COPA,ARSD,HSPA9,N |
| 252 | 19917 | HSPA8,GAPDH,MLLT1,CFL1,FTH1,DCX,RPL3,ENO1,NEFM,DVL3,SEP |
| 2115 | 19917 | PABPC1,LRP1,GAPDH,MLLT1,ARSD,MS4A10,HSPA9,NEDD4L,ACTB,C |
| 981 | 19917 | PABPC1,COL1A1,GAPDH,HNRNPA1,MS4A10,NRBP2,USP22,MTRNR2 |
| 708 | 19917 | PABPC1,GAPDH,MLLT1,ATN1,ACTB,NRBP2,USP22,TUBA1A,CFL1,MA |
| 581 | 19917 | COL1A1,LRP1,GAPDH,COPA,ATN1,CTSC,NRBP2,DPYSL3,TUBA1A,HS |
| 348 | 19917 | LRP1,NEDD4L,NRBP2,DPYSL3,HTT,NUMA1,DBN1,YBX1,PTMA,YWHA |
| 1313 | 19917 | ORC4,IGFBP5,COL1A1,MLLT1,COPA,ARSD,MS4A10,HSPA9,NEDD4L, |
| 392 | 19917 | LRP1,GAPDH,NEDD4L,HTT,HSP90AA1,TUBA1B,NCL,RPL7A,NEFM,YB |
| 2553 | 19917 | EEF1A1,ORC4,IGFBP5,PABPC1,COL1A1,LRP1,HSPA8,GAPDH,MLLT1 |
| 1521 | 19917 | ORC4,PABPC1,LRP1,HSPA8,GAPDH,MLLT1,COPA,NEDD4L,POLH,AC |
| 1349 | 19917 | PABPC1,TMEM212,GAPDH,MLLT1,COPA,HNRNPA1,ARSD,HSPA9,NE |
| 1057 | 19917 | PABPC1,GAPDH,COPA,MS4A10,HSPA9,NEDD4L,ACTB,NUDC,NRBP2, |
| 580 | 19917 | ORC4,IGFBP5,PABPC1,COL1A1,LRP1,HSPA8,GAPDH,HSPA9,NEDD4L |
| 371 | 19917 | PABPC1,HSPA8,COPA,ARSD,POLH,ACTB,NUDC,MED18,ACTR2,FTH1, |
| 832 | 19917 | ORC4,PABPC1,COPA,ACTB,ODF2L,NUDC,MED18,CFL1,MAVS,ZNF12 |
| 2147 | 19917 | PABPC1,LRP1,GAPDH,MLLT1,ARSD,MS4A10,HSPA9,NEDD4L,ACTB,C |
| 939 | 19917 | ORC4,PABPC1,HSPA8,COPA,POLH,ACTB,ODF2L,NUDC,USP22,MED1 |
| 1484 | 19917 | PABPC1,LRP1,GAPDH,MLLT1,ARSD,HSPA9,ACTB,ACTG1,DPYSL3,TU |
| 1484 | 19917 | PABPC1,LRP1,GAPDH,MLLT1,ARSD,HSPA9,ACTB,ACTG1,DPYSL3,TU |
| 973 | 19917 | ORC4,IGFBP5,PABPC1,COL1A1,LRP1,HNRNPA1,MS4A10,ACTB,CTS |
| 1024 | 19917 | ORC4,PABPC1,LRP1,MLLT1,COPA,ACTB,CTSC,ODF2L,NUDC,USP22, |
| 1130 | 19917 | GAPDH,MLLT1,MS4A10,ACTB,NRBP2,USP22,ACTG1,EEF2,DPYSL3,C |
| 1418 | 19917 | ORC4,PABPC1,LRP1,GAPDH,MLLT1,COPA,NEDD4L,ACTB,CTSC,ODF |
| 478 | 19917 | ORC4,NEDD4L,CTSC,NRBP2,USP22,ACTG1,HSP90AB1,HSP90AA1,VI |
| 845 | 19917 | PABPC1,COL1A1,LRP1,GAPDH,MLLT1,HNRNPA1,ACTB,NRBP2,MED1 |
| 503 | 19917 | COL1A1,GAPDH,HNRNPA1,NRBP2,USP22,MAVS,ACTR2,VIM,UBC,KID |
| 373 | 19917 | EEF1A1,COL1A1,GAPDH,DPYSL3,UBC,KIDINS220,TUBA1B,FLNA,NCL |
| 1097 | 19917 | LRP1,GAPDH,MS4A10,NEDD4L,NRBP2,USP22,ACTG1,EEF2,TUBA1A, |
| 1150 | 19917 | ORC4,PABPC1,LRP1,GAPDH,MLLT1,COPA,NEDD4L,ACTB,ODF2L,NU |
| 749 | 19917 | EEF1A1,ORC4,ARSD,NEDD4L,POLH,ACTB,NRBP2,ACTG1,EEF2,CFL1 |
| 463 | 19917 | GAPDH,NRBP2,USP22,DPYSL3,UBC,KIDINS220,EIF4G2,TUBA1B,FLN |
| 470 | 19917 | ORC4,HSPA8,COPA,ARSD,POLH,NUDC,MED18,ACTG1,BTBD9,ZNF12 |
| 1335 | 19917 | ORC4,PABPC1,TMEM212,LRP1,HSPA8,MLLT1,COPA,POLH,ACTB,OD |
| 587 | 19917 | PABPC1,COPA,ACTB,ODF2L,NUDC,MED18,CFL1,MAVS,ZNF124,HSP |
| 833 | 19917 | ORC4,PABPC1,COPA,ACTB,ODF2L,NUDC,MED18,DPYSL3,CFL1,MAV |
| 734 | 19917 | ORC4,PABPC1,COPA,ACTB,ODF2L,NUDC,MED18,CFL1,MAVS,ZNF12 |
| 499 | 19917 | HNRNPA1,POLH,NRBP2,USP22,BTBD9,HSP90AB1,ZNF124,TUBB,FTH |
| 499 | 19917 | HNRNPA1,POLH,NRBP2,USP22,BTBD9,HSP90AB1,ZNF124,TUBB,FTH |
| 1257 | 19917 | ORC4,PABPC1,LRP1,HSPA8,GAPDH,MLLT1,COPA,POLH,ACTB,ODF2 |
| 1459 | 19917 | EEF1A1,ORC4,IGFBP5,GAPDH,COPA,ARSD,HSPA9,NEDD4L,ACTB,CT |

| | | |
|---|---|---|
| 424 | 19917 | LRP1,GAPDH,ACTB,USP22,HSP90AA1,FLNA,NCL,RPL8,FBXL4,COL5 |
| 637 | 19917 | ORC4,IGFBP5,PABPC1,COL1A1,LRP1,HSPA8,ACTB,NRBP2,CFL1,HT |
| 1407 | 19917 | EEF1A1,ORC4,IGFBP5,COL1A1,GAPDH,MLLT1,COPA,ARSD,MS4A10, |
| 200 | 19917 | EEF2,HTT,FTH1,UBC,RPL3,NCL,YBX1,RPL13A,YWHAE,DDX17,RACK1 |
| 645 | 19917 | ORC4,COPA,ACTB,ODF2L,MED18,ACTG1,HSP90AB1,ACTR2,FTH1,K |
| 862 | 19917 | EEF1A1,GAPDH,HNRNPA1,ODF2L,USP22,ACTG1,EEF2,BTBD9,XRCC |
| 739 | 19917 | ORC4,IGFBP5,PABPC1,COL1A1,LRP1,HNRNPA1,NEDD4L,ACTB,NRB |
| 1151 | 19917 | ORC4,PABPC1,LRP1,HSPA8,COPA,NEDD4L,POLH,ACTB,ODF2L,NUD |
| 792 | 19917 | ORC4,HSPA8,COPA,POLH,ACTB,ODF2L,NUDC,MED18,XRCC2,MAVS, |
| 1193 | 19917 | ORC4,PABPC1,LRP1,GAPDH,COPA,NEDD4L,ACTB,ODF2L,NUDC,USP |
| 307 | 19917 | LRP1,HNRNPA1,POLH,ACTB,CTSC,MED18,CFL1,BTBD9,XRCC2,TUB |
| 957 | 19917 | GAPDH,MLLT1,HNRNPA1,ARSD,ACTB,NRBP2,USP22,ACTG1,TUBA1A |
| 528 | 19917 | ORC4,PABPC1,COPA,ACTB,ODF2L,NUDC,MED18,MAVS,ZNF124,HSP |
| 777 | 19917 | LRP1,GAPDH,ACTB,ODF2L,NRBP2,USP22,TUBA1A,MAVS,HTT,ACTR2 |
| 1719 | 19917 | ORC4,PABPC1,HSPA8,GAPDH,MLLT1,HNRNPA1,ARSD,HSPA9,NEDD4 |
| 547 | 19917 | ORC4,COPA,ARSD,POLH,ACTB,ODF2L,MED18,ZNF124,HSP90AA1,A |
| 1059 | 19917 | EEF1A1,ORC4,PABPC1,HSPA8,GAPDH,ARSD,HSPA9,NEDD4L,POLH,C |
| 1623 | 19917 | EEF1A1,ORC4,PABPC1,LRP1,HSPA8,GAPDH,COPA,HNRNPA1,ARSD, |
| 1324 | 19917 | ORC4,PABPC1,HSPA8,GAPDH,MLLT1,HNRNPA1,ARSD,HSPA9,NEDD4 |
| 1262 | 19917 | GAPDH,MLLT1,HNRNPA1,ARSD,ACTB,NRBP2,USP22,ACTG1,TUBA1A |
| 1576 | 19917 | ORC4,PABPC1,LRP1,MLLT1,COPA,NEDD4L,ACTB,CTSC,ODF2L,NUD |
| 1023 | 19917 | PABPC1,LRP1,GAPDH,MLLT1,COPA,POLH,ACTB,CTSC,ODF2L,NUDC |
| 596 | 19917 | ORC4,PABPC1,COPA,ACTB,ODF2L,NUDC,MED18,CFL1,MAVS,ZNF12 |
| 679 | 19917 | ORC4,PABPC1,COPA,ACTB,ODF2L,NUDC,MED18,CFL1,MAVS,ZNF12 |
| 716 | 19917 | HSPA8,GAPDH,NEDD4L,POLH,CTSC,NUDC,CFL1,BTBD9,XRCC2,HSP |
| 1245 | 19917 | ORC4,PABPC1,LRP1,MLLT1,COPA,ARSD,HSPA9,NEDD4L,ACTB,ODF |
| 560 | 19917 | PABPC1,COPA,ACTB,ODF2L,NUDC,MED18,CFL1,MAVS,ZNF124,HSP |
| 1623 | 19917 | EEF1A1,IGFBP5,TMEM212,COL1A1,HSPA8,GAPDH,COPA,HNRNPA1, |
| 1177 | 19917 | COL1A1,HSPA8,GAPDH,MLLT1,COPA,HNRNPA1,HSPA9,ATN1,NUDC, |
| 219 | 19917 | HTT,DBN1,NCL,DVL3,TRIM28,YWHAQ,PKD1,ITGB1,UCHL1,AGRN,MI |
| 732 | 19917 | PABPC1,GAPDH,HSPA9,USP22,ACTG1,TUBA1A,XRCC2,MAVS,HTT,ZN |
| 909 | 19917 | PABPC1,LRP1,GAPDH,COPA,ACTB,ODF2L,NUDC,USP22,MED18,CFL |
| 502 | 19917 | ORC4,MLLT1,COPA,ACTB,MED18,ACTG1,HSP90AB1,HSP90AA1,ACT |
| 379 | 19917 | PABPC1,MLLT1,NRBP2,ACTG1,HTT,HSP90AB1,KIDINS220,TMSB4X, |
| 1902 | 19917 | ORC4,IGFBP5,PABPC1,TMEM212,COL1A1,LRP1,GAPDH,MLLT1,CO |
| 1362 | 19917 | EEF1A1,IGFBP5,PABPC1,COL1A1,GAPDH,MLLT1,HNRNPA1,ARSD,M |
| 972 | 19917 | ORC4,PABPC1,LRP1,GAPDH,COPA,ARSD,ACTB,ODF2L,NUDC,USP22 |
| 438 | 19917 | IGFBP5,HSPA8,GAPDH,MLLT1,CFL1,FTH1,DCX,RPL3,ENO1,FLNA,AL |
| 453 | 19917 | PABPC1,COPA,ACTB,ODF2L,NUDC,MED18,ZNF124,HSP90AA1,ACTR |
| 317 | 19917 | PABPC1,LRP1,HSPA8,HNRNPA1,POLH,MED18,XRCC2,HSP90AA1,TU |
| 526 | 19917 | IGFBP5,LRP1,GAPDH,MLLT1,ACTB,NRBP2,ACTG1,TUBA1A,UBC,ZN |
| 271 | 19917 | GAPDH,MLLT1,HNRNPA1,ATN1,NRBP2,FTH1,EIF4G2,RPS6,NCL,RPL |
| 555 | 19917 | EEF1A1,PABPC1,GAPDH,MED18,UBC,NUMA1,TUBA1B,NCL,GANAB, |
| 150 | 19917 | MLLT1,HSP90AA1,ZNF316,DBN1,RACK1,GNB1,IGF2,DCHS1,EGLN1, |
| 739 | 19917 | PABPC1,LRP1,POLH,ACTB,NRBP2,MTRNR2L2,EEF2,DPYSL3,TUBA1 |

| | | |
|---|---|---|
| 729 | 19917 | PABPC1,COL1A1,LRP1,HSPA9,NEDD4L,POLH,ACTB,ACTG1,DPYSL3, |
| 885 | 19917 | COL1A1,GAPDH,NEDD4L,ODF2L,NUDC,NRBP2,ACTG1,EEF2,TUBA1 |
| 1701 | 19917 | EEF1A1,PABPC1,COL1A1,HSPA8,GAPDH,MLLT1,COPA,ARSD,MS4A1 |
| 1346 | 19917 | ORC4,IGFBP5,PABPC1,HSPA8,GAPDH,MLLT1,HSPA9,NEDD4L,ACTB, |
| 1213 | 19917 | ORC4,PABPC1,LRP1,HSPA8,GAPDH,COPA,NEDD4L,ACTB,ODF2L,NU |
| 539 | 19917 | PABPC1,ARSD,HSPA9,POLH,ACTB,NUDC,ACTG1,MAVS,HTT,HSP90A |
| 732 | 19917 | ORC4,HSPA8,GAPDH,COPA,ARSD,NEDD4L,MED18,TUBA1A,CFL1,BT |
| 783 | 19917 | PABPC1,COPA,ACTB,ODF2L,NUDC,MED18,CFL1,MAVS,ZNF124,HSP |
| 1763 | 19917 | IGFBP5,PABPC1,COL1A1,GAPDH,COPA,HNRNPA1,ARSD,NEDD4L,PO |
| 362 | 19917 | MLLT1,USP22,ZNF124,FLNA,SLC9A7,PKM,YBX1,PGK1,H3-3B,ENO2, |
| 596 | 19917 | ORC4,PABPC1,COPA,ACTB,ODF2L,NUDC,MED18,CFL1,MAVS,ZNF12 |
| 425 | 19917 | EEF1A1,GAPDH,POLH,MED18,DPYSL3,XRCC2,HTT,UBC,KIDINS220, |
| 954 | 19917 | EEF1A1,COL1A1,GAPDH,NEDD4L,POLH,MED18,DPYSL3,BTBD9,XRC |
| 1135 | 19917 | HSPA8,GAPDH,ARSD,NEDD4L,ODF2L,NUDC,ACTG1,HTT,ZNF124,HSP |
| 1029 | 19917 | COL1A1,LRP1,HSPA8,GAPDH,COPA,HNRNPA1,ARSD,MS4A10,NEDD4 |
| 797 | 19917 | EEF1A1,PABPC1,ARSD,NEDD4L,CTSC,MED18,ACTG1,DPYSL3,TUBA1 |
| 328 | 19917 | NEDD4L,NRBP2,USP22,HTT,HSP90AA1,VIM,UBC,DBN1,PIK3R2,TUB |
| 281 | 19917 | HNRNPA1,HSPA9,NEDD4L,TUBA1A,MAVS,HTT,HSP90AA1,FTH1,PIK3 |
| 1603 | 19917 | EEF1A1,ORC4,IGFBP5,PABPC1,COL1A1,LRP1,HSPA8,GAPDH,MLLT1 |
| 1044 | 19917 | ORC4,PABPC1,HSPA8,COPA,ARSD,NEDD4L,POLH,ATN1,ACTB,CTSC, |
| 1016 | 19917 | EEF1A1,ORC4,PABPC1,COL1A1,GAPDH,ARSD,HSPA9,NEDD4L,POLH, |
| 1315 | 19917 | ORC4,IGFBP5,PABPC1,COL1A1,LRP1,GAPDH,MLLT1,HNRNPA1,MS4 |
| 297 | 19917 | PABPC1,MLLT1,NRBP2,ACTG1,HTT,KIDINS220,NUMA1,COL6A1,FBX |
| 908 | 19917 | PABPC1,MLLT1,COPA,POLH,ACTB,CTSC,ODF2L,NUDC,MED18,CFL1 |
| 829 | 19917 | PABPC1,LRP1,GAPDH,HNRNPA1,ARSD,POLH,ACTB,CTSC,EEF2,BTB |
| 1049 | 19917 | EEF1A1,PABPC1,COPA,HNRNPA1,HSPA9,POLH,ODF2L,USP22,MED1 |
| 315 | 19917 | NRBP2,ACTG1,HTT,KIDINS220,ZNF316,TUBA1B,PKM,COL6A1,FBXL |
| 395 | 19917 | GAPDH,HNRNPA1,ATN1,NUDC,MTRNR2L2,EEF2,HSP90AB1,FTH1,TM |
| 829 | 19917 | GAPDH,HNRNPA1,MS4A10,NEDD4L,POLH,CTSC,NUDC,ACTG1,TUBA |
| 1857 | 19917 | ORC4,IGFBP5,PABPC1,TMEM212,COL1A1,HSPA8,COPA,HNRNPA1,A |
| 787 | 19917 | ORC4,IGFBP5,PABPC1,LRP1,HSPA8,GAPDH,MLLT1,COPA,ATN1,NRB |
| 1060 | 19917 | EEF1A1,ORC4,PABPC1,GAPDH,HNRNPA1,NEDD4L,CTSC,ODF2L,NUD |
| 1131 | 19917 | ORC4,PABPC1,LRP1,GAPDH,MLLT1,COPA,NEDD4L,ACTB,ODF2L,NU |
| 504 | 19917 | ORC4,PABPC1,HSPA8,COPA,ACTB,ODF2L,MED18,MAVS,ZNF124,HS |
| 555 | 19917 | ORC4,PABPC1,COPA,ACTB,ODF2L,NUDC,MED18,CFL1,MAVS,ZNF12 |
| 550 | 19917 | ORC4,PABPC1,LRP1,HSPA8,MLLT1,NEDD4L,NRBP2,CFL1,HSP90AA1 |
| 972 | 19917 | LRP1,HSPA8,GAPDH,COPA,HNRNPA1,HSPA9,POLH,ATN1,MTRNR2L2 |
| 181 | 19917 | LRP1,HNRNPA1,POLH,ACTB,MED18,CFL1,TUBA1B,RPL8,TMF1,PSA |
| 942 | 19917 | ORC4,LRP1,COPA,ARSD,HSPA9,NEDD4L,ACTG1,DPYSL3,MAVS,HSP |
| 1747 | 19917 | PABPC1,LRP1,GAPDH,HNRNPA1,ARSD,POLH,ACTB,CTSC,NRBP2,US |
| 1046 | 19917 | COL1A1,GAPDH,HNRNPA1,ARSD,HSPA9,NEDD4L,ACTB,CTSC,DPYSL |
| 1038 | 19917 | IGFBP5,LRP1,GAPDH,MLLT1,COPA,ARSD,HSPA9,POLH,ATN1,ACTB, |
| 575 | 19917 | COPA,ACTB,ODF2L,MED18,HSP90AB1,HSP90AA1,ACTR2,FTH1,RPS |
| 308 | 19917 | NRBP2,USP22,DPYSL3,HTT,HSP90AA1,PIK3R2,YBX1,SOX4,SPTBN1, |
| 794 | 19917 | PABPC1,MLLT1,COPA,ACTB,CTSC,ODF2L,NUDC,MED18,CFL1,MAVS |

| | | |
|---|---|---|
| 407 | 19917 | PABPC1,GAPDH,HNRNPA1,ACTB,MED18,DPYSL3,VIM,EIF4G2,FLNA,N |
| 1237 | 19917 | ORC4,PABPC1,COL1A1,LRP1,GAPDH,COPA,ACTB,ODF2L,NUDC,USP |
| 169 | 19917 | HTT,EIF4G2,RPS6,RPL4,SEC62,NPM1,CSDE1,MATR3,PSMD5,EIF4A1 |
| 928 | 19917 | IGFBP5,PABPC1,HSPA8,GAPDH,POLH,ACTB,ODF2L,NRBP2,ACTG1,C |
| 488 | 19917 | ORC4,PABPC1,COPA,ACTB,ODF2L,MED18,ACTG1,MAVS,ZNF124,HS |
| 457 | 19917 | PABPC1,COPA,ACTB,ODF2L,MED18,ZNF124,HSP90AA1,ACTR2,RPS6 |
| 1404 | 19917 | ORC4,COL1A1,HSPA8,GAPDH,COPA,ARSD,MS4A10,NEDD4L,POLH,A |
| 443 | 19917 | PABPC1,MLLT1,ACTB,TUBA1A,ZNF124,VIM,ZNF316,MAP1B,COL6A1 |
| 465 | 19917 | GAPDH,COPA,ACTB,ODF2L,MED18,CFL1,MAVS,HSP90AB1,ZNF124,H |
| 810 | 19917 | PABPC1,COL1A1,LRP1,MLLT1,ACTB,NRBP2,USP22,DPYSL3,CFL1,M |
| 1599 | 19917 | EEF1A1,PABPC1,HSPA8,GAPDH,MLLT1,COPA,HNRNPA1,MS4A10,NE |
| 296 | 19917 | GAPDH,NEDD4L,DBN1,TUBA1B,FLNA,PTMA,KAT6A,FASN,SPTBN1,SH |
| 401 | 19917 | USP22,HSP90AA1,FLNA,MAP1B,ALDOA,PKM,FADS1,YBX1,RPL13,GP |
| 1508 | 19917 | EEF1A1,PABPC1,TMEM212,LRP1,HSPA8,GAPDH,MLLT1,HNRNPA1,A |
| 1272 | 19917 | ORC4,PABPC1,TMEM212,LRP1,MLLT1,COPA,NEDD4L,ACTB,ODF2L, |
| 597 | 19917 | PABPC1,LRP1,GAPDH,MS4A10,CTSC,NRBP2,USP22,ACTG1,DPYSL3, |
| 336 | 19917 | GAPDH,HNRNPA1,HSPA9,ATN1,NRBP2,MTRNR2L2,EEF2,HSP90AB1, |
| 1016 | 19917 | LRP1,GAPDH,HSPA9,ACTB,DPYSL3,BTBD9,HTT,ZNF124,HSP90AA1,F |
| 812 | 19917 | COL1A1,GAPDH,HSPA9,ACTB,MED18,ACTG1,EEF2,DPYSL3,TUBA1A, |
| 1082 | 19917 | LRP1,GAPDH,MLLT1,COPA,ARSD,HSPA9,POLH,ATN1,ACTB,ODF2L,N |
| 1242 | 19917 | EEF1A1,ORC4,PABPC1,HSPA8,MLLT1,COPA,NEDD4L,ACTB,CTSC,OD |
| 1692 | 19917 | EEF1A1,PABPC1,COL1A1,HSPA8,GAPDH,MLLT1,HNRNPA1,ARSD,MS |
| 1247 | 19917 | PABPC1,HSPA8,GAPDH,MLLT1,COPA,HNRNPA1,HSPA9,ATN1,CTSC,N |
| 328 | 19917 | COPA,ODF2L,NUDC,CFL1,ZNF124,HSP90AA1,ACTR2,RPS6,ZNF316, |
| 1425 | 19917 | IGFBP5,PABPC1,COL1A1,LRP1,GAPDH,HNRNPA1,ARSD,POLH,ACTB, |
| 2570 | 19917 | EEF1A1,ORC4,IGFBP5,PABPC1,TMEM212,LRP1,GAPDH,MLLT1,COP |
| 509 | 19917 | HSPA8,ATN1,CTSC,ODF2L,NUDC,MED18,CFL1,HSP90AA1,SPARC,EIF |
| 815 | 19917 | ORC4,PABPC1,LRP1,COPA,ACTB,ODF2L,NUDC,USP22,MED18,CFL1, |
| 197 | 19917 | LRP1,HNRNPA1,POLH,ENO1,TUBA1B,RPL13A,RPL5,TMBIM6,TRIM2 |
| 665 | 19917 | MLLT1,ACTB,NRBP2,USP22,ACTG1,CFL1,XRCC2,MAVS,TMSB4X,DB |
| 318 | 19917 | PABPC1,COPA,MED18,CFL1,ZNF124,HSP90AA1,RPS6,ZNF316,DBN1 |
| 223 | 19917 | PABPC1,LRP1,NRBP2,XRCC2,NUMA1,NCL,BSG,ALDH6A1,MYH9,IGF |
| 391 | 19917 | ORC4,PABPC1,COPA,ACTB,ODF2L,MED18,MAVS,ZNF124,HSP90AA1 |
| 293 | 19917 | POLH,ACTB,CTSC,ACTG1,CFL1,BTBD9,XRCC2,TUBB,ENO1,TUBA1B |
| 965 | 19917 | LRP1,GAPDH,HSPA9,ACTB,DPYSL3,BTBD9,HTT,ZNF124,HSP90AA1,F |
| 122 | 19917 | USP22,RPS6,RPL4,NCL,SEC62,NPM1,CSDE1,MATR3,PSMD5,GTF3C |
| 336 | 19917 | HNRNPA1,MED18,DPYSL3,VIM,EIF4G2,ENO1,FLNA,NCL,NEFL,RPL8, |
| 1347 | 19917 | PABPC1,GAPDH,COPA,ARSD,NEDD4L,POLH,ODF2L,NUDC,NRBP2,AC |
| 494 | 19917 | COPA,ACTB,NUDC,MED18,HSP90AB1,ACTR2,NUMA1,RPL3,ZNF316, |
| 383 | 19917 | HSPA9,ACTB,ACTG1,CFL1,ZNF316,FLNA,NCL,HSP90B1,PSAP,RPLP0 |
| 915 | 19917 | PABPC1,TMEM212,HNRNPA1,POLH,ODF2L,USP22,MED18,ACTG1,C |
| 966 | 19917 | PABPC1,LRP1,MLLT1,COPA,ACTB,ODF2L,NUDC,USP22,MED18,DPY |
| 312 | 19917 | HNRNPA1,DPYSL3,VIM,EIF4G2,ENO1,NCL,FBXL4,IGFBP2,TUBB2B,C |
| 156 | 19917 | HSPA8,GAPDH,MED18,EIF4G2,RPL3,RPL4,GANAB,KAT6A,DDX17,RA |
| 392 | 19917 | PABPC1,COL1A1,POLH,ODF2L,NRBP2,USP22,TUBA1A,HSP90AA1,FT |

| | | |
|---|---|---|
| 584 | 19917 | COPA,ACTB,ODF2L,NUDC,MED18,CFL1,HSP90AB1,HSP90AA1,ACTR |
| 934 | 19917 | PABPC1,LRP1,MLLT1,ARSD,POLH,ACTB,CTSC,NUDC,NRBP2,USP22, |
| 490 | 19917 | NRBP2,EEF2,ZNF124,ZNF316,RPL4,PIK3R2,MAP1B,RPL7A,GANAB,H |
| 153 | 19917 | ORC4,HSPA8,GAPDH,MLLT1,USP22,CFL1,DCX,ENO1,NCL,NEFM,GN |
| 220 | 19917 | DPYSL3,BTBD9,VIM,NUMA1,DBN1,TUBB2B,PTMA,YWHAE,SPTBN1,I |
| 465 | 19917 | ORC4,IGFBP5,COL1A1,LRP1,HSPA8,MLLT1,NEDD4L,XRCC2,KIDINS |
| 1558 | 19917 | EEF1A1,ORC4,IGFBP5,PABPC1,COL1A1,LRP1,GAPDH,COPA,HNRNP |
| 255 | 19917 | PABPC1,XRCC2,ZNF124,ZNF316,DBN1,NCL,ALDOA,GANAB,HSPA1B, |
| 1131 | 19917 | HSPA8,GAPDH,MLLT1,COPA,HNRNPA1,HSPA9,ATN1,NUDC,MTRNR2 |
| 480 | 19917 | PABPC1,LRP1,ARSD,ACTG1,CFL1,ACTR2,VIM,FTH1,KIDINS220,RPL |
| 410 | 19917 | ORC4,PABPC1,COPA,ACTB,ODF2L,MED18,ZNF124,HSP90AA1,ACTR |
| 402 | 19917 | POLH,ACTB,ODF2L,USP22,ACTG1,BTBD9,MAVS,HTT,RPL3,ENO1,NC |
| 415 | 19917 | ORC4,COL1A1,LRP1,MLLT1,NEDD4L,NRBP2,HSP90AB1,HSP90AA1, |
| 293 | 19917 | GAPDH,COPA,ACTB,ODF2L,ZNF124,HSP90AA1,ACTR2,RPS6,ZNF316 |
| 657 | 19917 | ORC4,HSPA8,GAPDH,MLLT1,ODF2L,ACTG1,TUBA1A,XRCC2,HTT,AC |
| 1315 | 19917 | EEF1A1,PABPC1,TMEM212,COPA,HNRNPA1,ARSD,HSPA9,NEDD4L,O |
| 270 | 19917 | COPA,MED18,CFL1,HSP90AA1,RPS6,ZNF316,PIK3R2,HSP90B1,FKB |
| 1014 | 19917 | TMEM212,HSPA8,GAPDH,MLLT1,COPA,HNRNPA1,HSPA9,ATN1,MTR |
| 876 | 19917 | LRP1,HSPA8,ARSD,HSPA9,ACTB,USP22,MED18,EEF2,BTBD9,XRCC2, |
| 538 | 19917 | COPA,ACTB,ODF2L,NUDC,MED18,CFL1,ACTR2,FTH1,RPL3,ZNF316, |
| 1061 | 19917 | EEF1A1,PABPC1,LRP1,GAPDH,ARSD,HSPA9,POLH,USP22,MTRNR2L |
| 969 | 19917 | HSPA8,GAPDH,MLLT1,COPA,HNRNPA1,HSPA9,POLH,ATN1,NUDC,MT |
| 1355 | 19917 | EEF1A1,ORC4,PABPC1,TMEM212,COL1A1,LRP1,HSPA8,COPA,HNRN |
| 221 | 19917 | EEF1A1,POLH,ACTB,DPYSL3,CFL1,BTBD9,HSP90AA1,NEFL,YBX1,IG |
| 221 | 19917 | PABPC1,LRP1,NRBP2,HTT,ZNF316,FSTL1,GNAS,PTMS,USP11,EIF3C, |
| 869 | 19917 | ORC4,PABPC1,TMEM212,HSPA8,MLLT1,COPA,POLH,ACTB,ODF2L, |
| 217 | 19917 | LRP1,MLLT1,NEDD4L,TUBA1A,BTBD9,HTT,KIDINS220,EIF4G2,COL6 |
| 811 | 19917 | EEF1A1,PABPC1,COPA,ARSD,POLH,USP22,MED18,ACTG1,EEF2,DPY |
| 222 | 19917 | MLLT1,NRBP2,ACTG1,HTT,FBXL4,DVL3,GDI1,MARCKSL1,TRIM28,L |
| 129 | 19917 | NEDD4L,ACTG1,SPARC,NUMA1,COL5A1,COL4A1,GNAS,SPTBN1,GN |
| 158 | 19917 | LRP1,HNRNPA1,ACTB,MED18,TUBA1B,RPL8,HSPD1,TRIM28,SPEN,S |
| 233 | 19917 | GAPDH,HNRNPA1,HSP90AB1,FTH1,TMSB4X,EIF4G2,RPLP0,H3-3B,C |
| 1234 | 19917 | EEF1A1,ORC4,PABPC1,HSPA8,GAPDH,MLLT1,HNRNPA1,ARSD,HSPA |
| 1973 | 19917 | ORC4,PABPC1,COL1A1,HSPA8,GAPDH,MLLT1,COPA,HNRNPA1,ARSD |
| 433 | 19917 | COPA,ARSD,ACTB,MAVS,ACTR2,RPL3,TUBA1B,GANAB,CANX,PTMA,E |
| 493 | 19917 | COPA,ACTB,ODF2L,MED18,MAVS,HSP90AB1,HSP90AA1,ACTR2,RPS |
| 861 | 19917 | HSPA8,GAPDH,MLLT1,COPA,HNRNPA1,HSPA9,ATN1,NUDC,MTRNR2 |
| 308 | 19917 | IGFBP5,PABPC1,LRP1,NRBP2,HTT,VIM,FTH1,TMSB4X,HSP90B1,CAL |
| 271 | 19917 | HNRNPA1,DPYSL3,VIM,EIF4G2,ENO1,FLNA,NCL,RPL7A,NEFL,FBXL4, |
| 2273 | 19917 | EEF1A1,ORC4,IGFBP5,PABPC1,TMEM212,LRP1,GAPDH,MLLT1,HNR |
| 1154 | 19917 | EEF1A1,ORC4,COPA,HNRNPA1,ARSD,HSPA9,NEDD4L,POLH,ODF2L, |
| 288 | 19917 | NRBP2,HTT,ZNF124,KIDINS220,NUMA1,PKM,GANAB,ALDH6A1,STM |
| 376 | 19917 | PABPC1,COPA,ACTB,ODF2L,MED18,ZNF124,HSP90AA1,ACTR2,ZNF |
| 1182 | 19917 | IGFBP5,PABPC1,COL1A1,GAPDH,MLLT1,HNRNPA1,MS4A10,ATN1,A |
| 318 | 19917 | NEDD4L,HSP90AA1,VIM,UBC,RPL7A,FTL,FSTL1,H3-3B,DDX5,SOX4, |

| | | |
|---|---|---|
| 567 | 19917 | ORC4,HSPA8,COPA,POLH,ACTB,ODF2L,MED18,XRCC2,HSP90AB1,Z |
| 402 | 19917 | HNRNPA1,MED18,EEF2,BTBD9,UBC,ENO1,PKM,NEFL,YBX1,TMF1,P |
| 1202 | 19917 | EEF1A1,IGFBP5,PABPC1,HSPA8,MLLT1,COPA,POLH,ACTB,CTSC,NU |
| 947 | 19917 | ORC4,PABPC1,TMEM212,LRP1,HSPA8,MLLT1,COPA,ACTB,ODF2L,N |
| 253 | 19917 | PABPC1,COPA,ACTB,ODF2L,ZNF124,HSP90AA1,ACTR2,ZNF316,PIK |
| 800 | 19917 | PABPC1,LRP1,NEDD4L,POLH,NUDC,ACTG1,TUBA1A,CFL1,BTBD9,M |
| 805 | 19917 | ORC4,PABPC1,COL1A1,LRP1,HSPA8,GAPDH,COPA,HNRNPA1,MS4A1 |
| 433 | 19917 | ORC4,COPA,ACTB,ODF2L,MED18,HSP90AB1,HSP90AA1,RPS6,ZNF3 |
| 375 | 19917 | COPA,ACTB,ODF2L,MED18,ACTR2,FTH1,RPS6,ZNF316,PIK3R2,NCL, |
| 496 | 19917 | IGFBP5,PABPC1,COL1A1,LRP1,ACTB,DPYSL3,BTBD9,HTT,ZNF124,E |
| 524 | 19917 | PABPC1,COPA,ACTB,ODF2L,MED18,ZNF124,HSP90AA1,ACTR2,KIDI |
| 1196 | 19917 | EEF1A1,ORC4,PABPC1,TMEM212,LRP1,HSPA8,MLLT1,COPA,POLH, |
| 1196 | 19917 | EEF1A1,ORC4,PABPC1,TMEM212,LRP1,HSPA8,MLLT1,COPA,POLH, |
| 1215 | 19917 | EEF1A1,ORC4,PABPC1,HSPA8,GAPDH,MLLT1,HNRNPA1,ARSD,HSPA |
| 511 | 19917 | COPA,ACTB,ODF2L,MED18,HSP90AB1,HSP90AA1,ACTR2,NUMA1,RP |
| 917 | 19917 | EEF1A1,TMEM212,LRP1,COPA,ARSD,HSPA9,NEDD4L,TUBA1A,CFL1, |
| 664 | 19917 | PABPC1,HSPA8,COPA,ACTB,ODF2L,MED18,MAVS,HSP90AB1,HSP90 |
| 954 | 19917 | EEF1A1,ORC4,PABPC1,TMEM212,GAPDH,HNRNPA1,ARSD,NEDD4L, |
| 289 | 19917 | COPA,ACTB,MED18,ACTR2,ZNF316,PIK3R2,NCL,ALDOA,DDX5,MAG |
| 297 | 19917 | PABPC1,COPA,ODF2L,ZNF124,HSP90AA1,ACTR2,ZNF316,PIK3R2,YE |
| 521 | 19917 | COPA,ACTB,ODF2L,NUDC,MED18,HSP90AB1,HSP90AA1,ACTR2,FTH |
| 389 | 19917 | HSP90AA1,EIF4G2,ZNF316,FADS1,COL4A1,CES3,PSAP,PTMA,KAT6A |
| 221 | 19917 | LRP1,HSPA8,HNRNPA1,POLH,MED18,HSP90AA1,TUBA1B,ALDOA,TM |
| 1891 | 19917 | EEF1A1,ORC4,PABPC1,TMEM212,LRP1,HSPA8,GAPDH,COPA,HNRN |
| 500 | 19917 | COPA,ACTB,ODF2L,NUDC,MED18,CFL1,HSP90AB1,HSP90AA1,ACTR |
| 140 | 19917 | PABPC1,LRP1,GAPDH,USP22,EIF4G2,COL6A1,FSTL1,CES3,PTMS,PL |
| 981 | 19917 | HSPA8,GAPDH,MLLT1,COPA,HNRNPA1,HSPA9,ATN1,NUDC,MTRNR2 |
| 781 | 19917 | ORC4,TMEM212,MLLT1,COPA,ACTB,ODF2L,NUDC,MED18,HSP90A |
| 251 | 19917 | USP22,HTT,ZNF124,FTH1,NUMA1,YBX1,DVL3,SPTBN1,SHISA9,FSCN |
| 1049 | 19917 | ORC4,IGFBP5,COL1A1,LRP1,HSPA8,GAPDH,HNRNPA1,ARSD,MS4A1 |
| 518 | 19917 | EEF1A1,ORC4,PABPC1,LRP1,NEDD4L,USP22,HTT,HSP90AA1,ACTR2, |
| 1213 | 19917 | EEF1A1,ORC4,PABPC1,COL1A1,LRP1,HSPA8,GAPDH,MS4A10,HSPA |
| 520 | 19917 | HNRNPA1,HSPA9,USP22,MED18,EEF2,DPYSL3,BTBD9,XRCC2,HSP90 |
| 272 | 19917 | HNRNPA1,HSPA9,NUDC,EEF2,TUBA1A,ACTR2,FTH1,UBC,EIF4G2,RPL |
| 513 | 19917 | COPA,ACTB,NUDC,USP22,MED18,HSP90AB1,HSP90AA1,ACTR2,NUM |
| 2303 | 19917 | EEF1A1,PABPC1,COL1A1,HSPA8,GAPDH,MLLT1,COPA,HNRNPA1,AR |
| 429 | 19917 | ORC4,PABPC1,COPA,ACTB,ODF2L,MED18,ZNF124,HSP90AA1,ACTR |
| 912 | 19917 | ORC4,PABPC1,HSPA8,GAPDH,COPA,ARSD,POLH,ACTB,ODF2L,MED1 |
| 184 | 19917 | ACTB,NRBP2,HTT,DBN1,SLC9A7,YBX1,KIF1A,RPL13,SPTBN1,SMG1, |
| 184 | 19917 | ACTB,NRBP2,HTT,DBN1,SLC9A7,YBX1,KIF1A,RPL13,SPTBN1,SMG1, |
| 230 | 19917 | COPA,ODF2L,HSP90AA1,ACTR2,RPS6,ZNF316,PIK3R2,RPL6,RACK1, |
| 185 | 19917 | ACTB,PKM,FASN,GDI1,TMSB10,GTF3C4,PKD1,VAPA,ITGB1,MARCH |
| 394 | 19917 | ORC4,COPA,ODF2L,MED18,MAVS,ZNF124,HSP90AA1,ACTR2,KIDINS |
| 434 | 19917 | ORC4,PABPC1,LRP1,NEDD4L,HTT,ACTR2,UBC,KIDINS220,NUMA1,M |
| 159 | 19917 | ORC4,LRP1,MLLT1,NEDD4L,DBN1,PKM,CES3,KAT6A,SLC2A3,MARC |

| | | |
|---|---|---|
| 647 | 19917 | IGFBP5,COL1A1,MLLT1,NEDD4L,POLH,NRBP2,BTBD9,XRCC2,HTT, |
| 971 | 19917 | EEF1A1,IGFBP5,PABPC1,HSPA8,MLLT1,COPA,POLH,ACTB,CTSC,NU |
| 421 | 19917 | PABPC1,LRP1,MLLT1,ARSD,CTSC,TUBA1A,CFL1,HTT,ZNF316,DBN1, |
| 416 | 19917 | PABPC1,COPA,ACTB,ODF2L,NUDC,ZNF124,HSP90AA1,ACTR2,RPS6, |
| 486 | 19917 | ACTB,ACTG1,ZNF124,TUBB,DBN1,ALDOA,CALR,GNAS,DDX17,FASN, |
| 358 | 19917 | USP22,MED18,EEF2,BTBD9,XRCC2,HSP90AB1,UBC,NUMA1,RPL3,R |
| 273 | 19917 | COPA,ACTB,ODF2L,MED18,ZNF124,HSP90AA1,ACTR2,RPS6,ZNF316 |
| 538 | 19917 | PABPC1,LRP1,COPA,ACTB,ODF2L,USP22,MED18,MAVS,HSP90AB1,Z |
| 505 | 19917 | LRP1,GAPDH,ARSD,HSPA9,NEDD4L,POLH,TUBA1A,XRCC2,HSP90AA1 |
| 402 | 19917 | PABPC1,HSPA8,COPA,ACTB,MED18,BTBD9,ZNF124,HSP90AA1,RPS6 |
| 402 | 19917 | PABPC1,HSPA8,COPA,ACTB,MED18,BTBD9,ZNF124,HSP90AA1,RPS6 |
| 354 | 19917 | IGFBP5,COL1A1,LRP1,HNRNPA1,BTBD9,VIM,FTH1,KIDINS220,TMSE |
| 433 | 19917 | COPA,ODF2L,MED18,MAVS,HSP90AA1,ACTR2,UBC,RPS6,ZNF316,PI |
| 399 | 19917 | PABPC1,COPA,CTSC,HTT,UBC,KIDINS220,NCL,FADS1,HSPA1B,ZNF9 |
| 938 | 19917 | IGFBP5,PABPC1,COL1A1,HSPA8,HNRNPA1,NEDD4L,NUDC,CFL1,HTT |
| 410 | 19917 | EEF1A1,USP22,MED18,EEF2,DPYSL3,BTBD9,XRCC2,HSP90AB1,UBC |
| 1258 | 19917 | LRP1,HSPA8,ARSD,HSPA9,NEDD4L,ACTB,NRBP2,USP22,MTRNR2L2, |
| 1375 | 19917 | EEF1A1,ORC4,IGFBP5,PABPC1,GAPDH,HNRNPA1,ARSD,HSPA9,NEDD |
| 428 | 19917 | ORC4,PABPC1,LRP1,NEDD4L,HTT,ACTR2,VIM,UBC,KIDINS220,MAP1 |
| 260 | 19917 | PABPC1,HNRNPA1,NRBP2,USP22,CFL1,BTBD9,ACTR2,NCL,SLC9A7, |
| 325 | 19917 | ORC4,HSPA8,COPA,POLH,MED18,ACTG1,ZNF124,ACTR2,KIDINS220 |
| 970 | 19917 | HSPA8,GAPDH,MLLT1,COPA,HNRNPA1,HSPA9,ATN1,NUDC,MTRNR2 |
| 386 | 19917 | ORC4,PABPC1,LRP1,NEDD4L,HTT,ACTR2,UBC,KIDINS220,MAP1B,A |
| 234 | 19917 | POLH,MED18,XRCC2,RPS6,ZNF316,PIK3R2,IGFBP2,DDX5,FKBP14,F |
| 427 | 19917 | ORC4,HSPA8,POLH,ACTB,ODF2L,MED18,HSP90AB1,RPS6,PIK3R2,S |
| 558 | 19917 | COPA,ACTB,ODF2L,MED18,HSP90AB1,HSP90AA1,ACTR2,FTH1,RPS6 |
| 50 | 19917 | UBC,EIF4G2,YWHAE,DDX17,WSB1,CIPC,SFPQ,RAB1A,ATP6V0C,LUC |
| 1206 | 19917 | IGFBP5,PABPC1,COL1A1,LRP1,GAPDH,MLLT1,COPA,HNRNPA1,ARS |
| 487 | 19917 | ORC4,PABPC1,COPA,ACTB,ODF2L,MED18,ZNF124,HSP90AA1,ACTR |
| 284 | 19917 | COL1A1,LRP1,COPA,ARSD,ACTG1,ZNF124,PIK3R2,MAP1B,CALR,TU |
| 458 | 19917 | COPA,ACTB,ODF2L,NUDC,MED18,HSP90AB1,HSP90AA1,ACTR2,RPS |
| 178 | 19917 | HTT,ABHD2,ALDH6A1,STMN2,TRIM28,PKD1,STMN1,UCHL1,AGRN,M |
| 803 | 19917 | EEF1A1,ORC4,PABPC1,LRP1,ARSD,MS4A10,HSPA9,NEDD4L,NUDC,N |
| 296 | 19917 | COPA,ACTB,ODF2L,HSP90AA1,ACTR2,FTH1,RPS6,ZNF316,PIK3R2,C |
| 562 | 19917 | ORC4,IGFBP5,PABPC1,COL1A1,LRP1,MLLT1,HSPA9,NEDD4L,NRBP |
| 355 | 19917 | USP22,MED18,EEF2,BTBD9,XRCC2,HSP90AB1,UBC,RPL3,RPL4,FLN |
| 205 | 19917 | POLH,NUDC,VIM,COL6A1,ATP2B4,IGFBP2,PTMA,SEC62,KAT6A,YWH |
| 604 | 19917 | EEF1A1,GAPDH,HNRNPA1,HSPA9,NEDD4L,POLH,ACTB,ODF2L,ACTG |
| 1849 | 19917 | EEF1A1,IGFBP5,PABPC1,COL1A1,LRP1,HSPA8,GAPDH,MLLT1,ARSD, |
| 285 | 19917 | HSPA9,POLH,NUDC,XRCC2,HSP90AA1,TUBB,NEFL,IGFBP2,COL4A1, |
| 430 | 19917 | COPA,ACTB,ODF2L,MED18,HSP90AB1,HSP90AA1,ACTR2,RPS6,ZNF |
| 285 | 19917 | GAPDH,ARSD,NEDD4L,HTT,KIDINS220,ZNF316,FLNA,PKM,YBX1,PTM |
| 190 | 19917 | ORC4,LRP1,NEDD4L,DBN1,MAP1B,UBA1,IGFBP2,COL5A1,GNAS,CA |
| 490 | 19917 | USP22,BTBD9,XRCC2,UBC,RPL4,ENO1,FLNA,HSP90B1,TMF1,DDX5, |
| 467 | 19917 | PABPC1,COL1A1,LRP1,HSPA8,ARSD,NEDD4L,NRBP2,EIF4G2,MAP1B |

| | | |
|---|---|---|
| 438 | 19917 | ORC4,PABPC1,LRP1,NUDC,CFL1,BTBD9,HSP90AA1,UBC,ZNF316,NE |
| 185 | 19917 | HTT,KIDINS220,MAP1B,PKM,ALDH6A1,FSCN1,LDHB,AGRN,ZMIZ1,AI |
| 322 | 19917 | EEF1A1,HSPA9,USP22,MED18,EEF2,BTBD9,XRCC2,HSP90AB1,UBC,N |
| 216 | 19917 | COPA,ACTB,ODF2L,MED18,HSP90AA1,ACTR2,RPS6,ZNF316,PIK3R2, |
| 112 | 19917 | MLLT1,ARSD,ACTB,NRBP2,HTT,UBC,PSAP,SPTBN1,FSCN1,IGF2,TRII |
| 708 | 19917 | EEF1A1,COL1A1,GAPDH,MLLT1,NRBP2,TUBA1A,MAVS,ZNF124,SPAF |
| 114 | 19917 | HNRNPA1,POLH,RPL13A,RPL5,TRIM28,POLR2A,DNAJA1,OGA,RPS9,I |
| 1755 | 19917 | EEF1A1,IGFBP5,TMEM212,LRP1,HSPA8,GAPDH,MLLT1,COPA,HNRNI |
| 130 | 19917 | CFL1,HTT,EIF4G2,RPS6,ZNF316,RPL4,NCL,RPL7A,YBX1,SEC62,P4HI |
| 855 | 19917 | ORC4,PABPC1,COPA,ACTB,ODF2L,NUDC,MED18,XRCC2,HSP90AB1, |
| 1579 | 19917 | EEF1A1,IGFBP5,PABPC1,COL1A1,LRP1,HSPA8,GAPDH,MLLT1,COPA |
| 374 | 19917 | EEF1A1,PABPC1,ARSD,HSPA9,CFL1,TUBB,NUMA1,TUBA1B,NCL,PKM |
| 250 | 19917 | HNRNPA1,MED18,DPYSL3,VIM,EIF4G2,ENO1,NCL,FBXL4,IGFBP2,TU |
| 1221 | 19917 | ORC4,PABPC1,HSPA8,GAPDH,MLLT1,HNRNPA1,HSPA9,NEDD4L,POL |
| 1352 | 19917 | COL1A1,LRP1,GAPDH,MS4A10,HSPA9,NEDD4L,ACTB,NUDC,NRBP2,U |
| 207 | 19917 | HNRNPA1,ARSD,NRBP2,HTT,HSP90AB1,ENO1,NCL,MYH9,TRIM28,YW |
| 304 | 19917 | USP22,MED18,EEF2,BTBD9,XRCC2,HSP90AB1,UBC,RPL3,RPL4,FLNA |
| 428 | 19917 | PABPC1,MLLT1,COPA,ACTB,ODF2L,MED18,MAVS,ZNF124,HSP90AA |
| 175 | 19917 | HSP90AA1,TUBA1B,HSPA1B,KIF1A,FSCN1,PTMS,CALM1,SPEN,YWHA |
| 773 | 19917 | ORC4,HSPA8,COPA,POLH,ACTB,ODF2L,MED18,ACTG1,XRCC2,HSP9 |
| 305 | 19917 | GAPDH,HNRNPA1,ATN1,NUDC,MTRNR2L2,HSP90AB1,FTH1,EIF4G2,( |
| 470 | 19917 | LRP1,GAPDH,ACTG1,DPYSL3,BTBD9,MAVS,HSP90AB1,HSP90AA1,EII |
| 232 | 19917 | COPA,ACTB,MED18,ZNF124,HSP90AA1,ACTR2,KIDINS220,RPS6,ZNF |
| 300 | 19917 | EEF1A1,PABPC1,LRP1,HSPA9,ACTG1,EEF2,SPARC,FTH1,NUMA1,TUE |
| 438 | 19917 | ORC4,PABPC1,LRP1,NEDD4L,USP22,HTT,UBC,KIDINS220,NUMA1,EN |
| 2019 | 19917 | EEF1A1,ORC4,IGFBP5,PABPC1,COL1A1,LRP1,HSPA8,GAPDH,COPA,I |
| 685 | 19917 | COPA,HNRNPA1,HSPA9,USP22,MED18,EEF2,DPYSL3,CFL1,BTBD9,XI |
| 705 | 19917 | EEF1A1,COPA,HNRNPA1,HSPA9,USP22,MED18,EEF2,DPYSL3,BTBD9 |
| 2664 | 19917 | EEF1A1,ORC4,IGFBP5,PABPC1,COL1A1,LRP1,HSPA8,GAPDH,MLLT1 |
| 318 | 19917 | ORC4,COL1A1,LRP1,GAPDH,USP22,TUBA1A,HSP90AB1,VIM,TUBB,N |
| 645 | 19917 | EEF1A1,ORC4,PABPC1,LRP1,NEDD4L,NRBP2,CFL1,HTT,ACTR2,VIM, |
| 382 | 19917 | COPA,ACTB,ODF2L,MED18,HSP90AB1,HSP90AA1,RPS6,ZNF316,PIK3 |
| 394 | 19917 | ORC4,PABPC1,LRP1,NEDD4L,HTT,UBC,KIDINS220,MAP1B,ABHD2,AI |
| 366 | 19917 | COPA,ACTB,MED18,HSP90AB1,HSP90AA1,ACTR2,RPS6,ZNF316,PIK3 |
| 1285 | 19917 | EEF1A1,IGFBP5,HSPA8,GAPDH,COPA,HSPA9,ATN1,NUDC,USP22,MTI |
| 32 | 19917 | FTH1,UBC,NCL,YWHAE,DDX17,PFN1,HNRNPDL,HNRNPK,H1-10,HNRI |
| 982 | 19917 | EEF1A1,PABPC1,COL1A1,GAPDH,COPA,HNRNPA1,ARSD,ACTB,NRBP |
| 1298 | 19917 | EEF1A1,ORC4,PABPC1,LRP1,HSPA8,GAPDH,MLLT1,COPA,ARSD,POL |
| 1222 | 19917 | ORC4,PABPC1,LRP1,MLLT1,MS4A10,NEDD4L,ODF2L,NUDC,NRBP2, |
| 393 | 19917 | PABPC1,COPA,ODF2L,NUDC,USP22,ZNF124,HSP90AA1,ACTR2,KIDIN |
| 843 | 19917 | IGFBP5,MS4A10,NEDD4L,ACTB,ODF2L,MED18,BTBD9,MAVS,HSP90A |
| 409 | 19917 | GAPDH,MLLT1,NEDD4L,NUDC,MED18,BTBD9,VIM,NUMA1,PIK3R2,H |
| 155 | 19917 | LRP1,HTT,NUMA1,TRIM28,PKD1,AGRN,MIDN,BRD3,NCS1,ZNF865,C |
| 872 | 19917 | EEF1A1,HNRNPA1,HSPA9,ODF2L,USP22,MED18,ACTG1,EEF2,BTBD9 |
| 416 | 19917 | COPA,ACTB,ODF2L,NUDC,MED18,HSP90AB1,HSP90AA1,ACTR2,RPS( |

| | | |
|---|---|---|
| 541 | 19917 | HNRNPA1,USP22,MED18,EEF2,BTBD9,XRCC2,HSP90AB1,UBC,RPL3, |
| 706 | 19917 | COPA,HNRNPA1,HSPA9,USP22,MED18,EEF2,DPYSL3,BTBD9,XRCC2, |
| 272 | 19917 | ZNF124,RPL3,ZNF316,PIK3R2,NCL,GANAB,H3-3B,RPL13A,MAGED1, |
| 1036 | 19917 | ORC4,PABPC1,TMEM212,HSPA8,GAPDH,MLLT1,COPA,POLH,ACTB,C |
| 133 | 19917 | NEDD4L,MAP1B,COL6A1,FSTL1,COL5A1,GNAS,SMG1,SET,GTF3C4, |
| 126 | 19917 | IGFBP5,ZNF316,ALDOA,GNAS,SPTBN1,MRC2,AGRN,PXDN,HK1,KIF5 |
| 299 | 19917 | ORC4,LRP1,MLLT1,NEDD4L,NUMA1,ZNF316,PKM,FADS1,NEFM,COL |
| 185 | 19917 | PABPC1,PKM,COL4A1,RPL13,GPI,SHISA9,MYH9,FSCN1,COL4A2,DCI |
| 1254 | 19917 | EEF1A1,ORC4,PABPC1,MLLT1,COPA,ACTB,ODF2L,NUDC,MED18,CF |
| 1262 | 19917 | EEF1A1,LRP1,GAPDH,HSPA9,ATN1,ACTB,CTSC,ODF2L,DPYSL3,CFL1 |
| 999 | 19917 | EEF1A1,PABPC1,GAPDH,MLLT1,NEDD4L,ACTB,USP22,ACTG1,DPYSL |
| 256 | 19917 | ORC4,PABPC1,CFL1,XRCC2,TUBB,TUBA1B,PKM,GANAB,PTMA,BSG, |
| 970 | 19917 | ORC4,LRP1,MLLT1,COPA,ACTB,ODF2L,NUDC,USP22,MED18,CFL1,H |
| 1088 | 19917 | ORC4,IGFBP5,PABPC1,LRP1,HSPA8,GAPDH,MLLT1,ARSD,POLH,ATN |
| 345 | 19917 | EEF1A1,GAPDH,HNRNPA1,MED18,VIM,UBC,KIDINS220,NUMA1,TUBA |
| 203 | 19917 | COPA,NUDC,CFL1,ACTR2,ZNF316,PIK3R2,RPL6,RACK1,SHISA9,P4HI |
| 1440 | 19917 | ORC4,COL1A1,MLLT1,COPA,HNRNPA1,MS4A10,HSPA9,NEDD4L,ACT |
| 839 | 19917 | COL1A1,HNRNPA1,HSPA9,USP22,MED18,EEF2,DPYSL3,BTBD9,XRCC |
| 992 | 19917 | EEF1A1,PABPC1,COPA,HNRNPA1,HSPA9,NEDD4L,USP22,MED18,EEF |
| 592 | 19917 | PABPC1,GAPDH,ARSD,NEDD4L,POLH,EEF2,BTBD9,HTT,HSP90AB1,Z |
| 495 | 19917 | ORC4,PABPC1,HSPA8,GAPDH,NEDD4L,HTT,NUMA1,ZNF316,DBN1,M |
| 281 | 19917 | USP22,MED18,EEF2,BTBD9,XRCC2,HSP90AB1,UBC,RPL3,RPL4,FLN |
| 904 | 19917 | ORC4,LRP1,HSPA8,MLLT1,HNRNPA1,NEDD4L,NRBP2,MAVS,HSP90A |
| 224 | 19917 | EEF1A1,USP22,UBC,RPL4,ENO1,FLNA,HSP90B1,KIF1A,FSCN1,PLD3, |
| 67 | 19917 | GAPDH,CFL1,NEFM,KAT6A,CALM3,SEPTIN2,EIF4A1,MRC2,LDHB,POI |
| 1409 | 19917 | ORC4,IGFBP5,PABPC1,COL1A1,LRP1,GAPDH,MLLT1,HNRNPA1,NED |
| 961 | 19917 | EEF1A1,ORC4,IGFBP5,PABPC1,COL1A1,LRP1,HSPA8,HNRNPA1,NED |
| 162 | 19917 | COPA,ACTB,MED18,ZNF316,PIK3R2,NCL,DDX5,MAGED1,RPL6,P4HE |
| 207 | 19917 | PABPC1,LRP1,ARSD,NUDC,NCL,RPL8,HSPA1B,COL4A1,ENO2,GNAS, |
| 814 | 19917 | LRP1,GAPDH,HSPA9,ACTB,DPYSL3,BTBD9,HTT,ZNF124,HSP90AA1,K |
| 610 | 19917 | LRP1,GAPDH,COPA,ARSD,ACTB,NRBP2,ACTG1,BTBD9,XRCC2,HTT,A |
| 190 | 19917 | LRP1,ARSD,HSPA9,HTT,KIDINS220,PIK3R2,ABHD2,ALDH6A1,CHSY1, |
| 726 | 19917 | EEF1A1,IGFBP5,COL1A1,LRP1,HSPA8,COPA,ARSD,NEDD4L,POLH,AT |
| 1588 | 19917 | EEF1A1,IGFBP5,PABPC1,TMEM212,LRP1,COPA,ARSD,MS4A10,HSPA |
| 496 | 19917 | EEF1A1,USP22,MED18,EEF2,DPYSL3,BTBD9,MAVS,HTT,HSP90AB1,U |
| 490 | 19917 | PABPC1,COL1A1,GAPDH,MS4A10,NRBP2,USP22,MAVS,FLNA,NCL,FA |
| 213 | 19917 | NEDD4L,ZNF124,PIK3R2,NCL,COL6A1,IGFBP2,YWHAE,DDX17,GNB1 |
| 930 | 19917 | EEF1A1,PABPC1,COPA,HNRNPA1,HSPA9,NEDD4L,ODF2L,USP22,MEI |
| 850 | 19917 | EEF1A1,PABPC1,HNRNPA1,ODF2L,USP22,MED18,EEF2,BTBD9,XRCC |
| 491 | 19917 | EEF1A1,LRP1,HSPA8,COPA,ARSD,MS4A10,ACTB,NRBP2,HTT,VIM,TU |
| 955 | 19917 | COPA,HNRNPA1,HSPA9,NEDD4L,ODF2L,USP22,MED18,EEF2,DPYSL3 |
| 169 | 19917 | LRP1,ACTB,HTT,ACTR2,VIM,PIK3R2,CALR,TUBB2B,DDX5,SPTBN1,G |
| 822 | 19917 | EEF1A1,ORC4,IGFBP5,PABPC1,COL1A1,LRP1,COPA,HSPA9,NEDD4L |
| 488 | 19917 | ORC4,LRP1,NEDD4L,NRBP2,HSP90AB1,HSP90AA1,DBN1,RPL7A,SLC |
| 756 | 19917 | LRP1,MLLT1,ACTB,NUDC,MTRNR2L2,CFL1,HTT,HSP90AA1,ENO1,TU |

| | | |
|---|---|---|
| 807 | 19917 | TMEM212,GAPDH,HSPA9,POLH,ACTB,DPYSL3,ACTR2,RPL3,ENO1, |
| 1582 | 19917 | EEF1A1,PABPC1,HSPA8,GAPDH,COPA,HNRNPA1,ARSD,HSPA9,POLH, |
| 316 | 19917 | ORC4,PABPC1,LRP1,NEDD4L,HTT,UBC,KIDINS220,MAP1B,ABHD2, |
| 834 | 19917 | EEF1A1,COPA,HNRNPA1,HSPA9,NEDD4L,USP22,MED18,EEF2,DPYSL |
| 2107 | 19917 | EEF1A1,ORC4,IGFBP5,PABPC1,HSPA8,GAPDH,MLLT1,HNRNPA1,AR |
| 120 | 19917 | ODF2L,ACTR2,RPS6,ZNF316,PIK3R2,RPL6,TMBIM6,RPL29,ZNF865, |
| 125 | 19917 | ACTB,HTT,KIF1A,GNAS,RACK1,APP,GSTP1,IGF2,SET,TMSB10,AGRN |
| 172 | 19917 | COPA,ACTB,MED18,ZNF316,PIK3R2,NCL,DDX5,MAGED1,RPL6,P4HE |
| 1280 | 19917 | EEF1A1,ORC4,PABPC1,LRP1,COPA,MS4A10,NEDD4L,ATN1,ACTB,CT |
| 107 | 19917 | BTBD9,UBC,HSP90B1,KIF1A,ALDH6A1,SEPTIN2,GSTP1,USP11,TRIM |
| 266 | 19917 | GAPDH,ACTG1,DPYSL3,TUBA1A,NUMA1,RPL8,ENO2,DDX17,BSG,RP |
| 602 | 19917 | EEF1A1,HSPA8,COPA,ARSD,POLH,CTSC,ODF2L,NUDC,USP22,MTRN |
| 829 | 19917 | IGFBP5,PABPC1,TMEM212,LRP1,MLLT1,COPA,ARSD,MS4A10,NEDD |
| 1584 | 19917 | EEF1A1,ORC4,IGFBP5,PABPC1,COL1A1,LRP1,HSPA8,GAPDH,MLLT1 |
| 1085 | 19917 | IGFBP5,LRP1,GAPDH,ATN1,ACTB,NUDC,EEF2,CFL1,HTT,SPARC,UBC |
| 225 | 19917 | GAPDH,COPA,ODF2L,BTBD9,FLNA,MTRNR2L8,NEFL,DDX5,SEC62,R |
| 246 | 19917 | COPA,ACTB,MED18,ZNF124,HSP90AA1,ACTR2,ZNF316,PIK3R2,NCL, |
| 613 | 19917 | COL1A1,HNRNPA1,POLH,ACTB,NUDC,ACTG1,TUBA1A,HSP90AB1,NU |
| 1618 | 19917 | ORC4,PABPC1,COL1A1,GAPDH,ARSD,MS4A10,HSPA9,NEDD4L,POL |
| 397 | 19917 | EEF1A1,LRP1,ARSD,NRBP2,CFL1,HTT,VIM,KIDINS220,NUMA1,PIK3R |
| 497 | 19917 | EEF1A1,POLH,CTSC,NUDC,USP22,MTRNR2L2,EEF2,CFL1,VIM,FTH1, |
| 1619 | 19917 | EEF1A1,ORC4,PABPC1,LRP1,GAPDH,MLLT1,MS4A10,HSPA9,NEDD4 |
| 578 | 19917 | HNRNPA1,HSPA9,USP22,MED18,EEF2,DPYSL3,BTBD9,XRCC2,HSP90 |
| 247 | 19917 | COPA,ACTB,MED18,HSP90AB1,HSP90AA1,ZNF316,PIK3R2,NCL,HSP |
| 1069 | 19917 | LRP1,GAPDH,HSPA9,ACTB,ODF2L,DPYSL3,CFL1,BTBD9,HTT,ZNF12 |
| 291 | 19917 | ZNF124,RPL3,ZNF316,RPL4,PIK3R2,NCL,GANAB,RPL13A,MAGED1, |
| 65 | 19917 | PIK3R2,HSP90B1,KAT6A,YWHAE,RPS8,P4HB,SPTAN1,UBB,TPT1,MT |
| 204 | 19917 | COPA,HNRNPA1,HSPA9,MED18,NCL,HSP90B1,FBXL4,FSTL1,SEPTIN |
| 481 | 19917 | LRP1,GAPDH,ACTB,XRCC2,MAVS,ACTR2,VIM,FTH1,EIF4G2,MAP1B, |
| 25 | 19917 | PIK3R2,SPTAN1,SF3B1,RPL18,NARS1,U2AF2,LMNB2,RAPGEF1,RPL |
| 376 | 19917 | COPA,ODF2L,MED18,FTH1,NUMA1,RPS6,ZNF316,TUBA1B,PIK3R2, |
| 131 | 19917 | COPA,ODF2L,ACTR2,RPS6,ZNF316,PIK3R2,RPL6,DVL3,ATP5F1B,US |
| 432 | 19917 | EEF1A1,ORC4,PABPC1,LRP1,NEDD4L,HTT,VIM,UBC,KIDINS220,NUM |
| 315 | 19917 | COL1A1,HNRNPA1,MS4A10,NEDD4L,ACTB,FTH1,NUMA1,FTL,HSPA1 |
| 226 | 19917 | USP22,MED18,EEF2,BTBD9,XRCC2,HSP90AB1,UBC,RPL3,RPL4,FLN |
| 1093 | 19917 | IGFBP5,PABPC1,LRP1,GAPDH,MLLT1,COPA,ARSD,MS4A10,NEDD4L, |
| 229 | 19917 | COPA,ACTB,MED18,HSP90AB1,ZNF316,PIK3R2,NCL,HSP90B1,DDX5 |
| 406 | 19917 | ORC4,GAPDH,ACTG1,EEF2,KIDINS220,ENO1,NCL,ALDOA,GANAB,R |
| 226 | 19917 | EEF1A1,GAPDH,DPYSL3,UBC,TUBA1B,NCL,GANAB,TMF1,PTMA,DDX |
| 54 | 19917 | RPS6,CSDE1,MATR3,PSMD5,RPS9,RPL17,EIF5,RPL26,DDX3X,RPS15 |
| 156 | 19917 | NRBP2,ZNF316,PIK3R2,KAT6A,CALM3,CALM1,HSPD1,TMSB10,MRC |
| 2017 | 19917 | EEF1A1,ORC4,IGFBP5,PABPC1,LRP1,HSPA8,GAPDH,MLLT1,HNRNP |
| 139 | 19917 | POLH,ZNF124,HSP90B1,RPL8,YBX1,TUBB2B,BSG,FASN,P4HB,NPM1 |
| 319 | 19917 | HSPA9,USP22,MED18,EEF2,DPYSL3,BTBD9,XRCC2,HSP90AB1,UBC, |
| 723 | 19917 | EEF1A1,TMEM212,HSPA8,GAPDH,ARSD,HSPA9,POLH,ODF2L,NUDC, |

| | | |
|---|---|---|
| 197 | 19917 | COL1A1,HNRNPA1,ACTB,ZNF124,FTH1,NUMA1,NCL,GANAB,PRPF8, |
| 1672 | 19917 | EEF1A1,ORC4,IGFBP5,PABPC1,COL1A1,MLLT1,HNRNPA1,ARSD,NE |
| 184 | 19917 | ACTB,ODF2L,ACTR2,FTH1,RPS6,ZNF316,PIK3R2,MAGED1,RPL6,GN |
| 1153 | 19917 | EEF1A1,PABPC1,HNRNPA1,HSPA9,NEDD4L,POLH,ACTB,CTSC,USP22 |
| 237 | 19917 | GAPDH,HNRNPA1,ATN1,MTRNR2L2,HSP90AB1,FTH1,TMSB4X,EIF4G |
| 245 | 19917 | PABPC1,POLH,MED18,ACTG1,ACTR2,KIDINS220,RPS6,ZNF316,PIK3 |
| 288 | 19917 | ORC4,PABPC1,LRP1,NEDD4L,HTT,FTH1,UBC,KIDINS220,MAP1B,AB |
| 516 | 19917 | COL1A1,COPA,HNRNPA1,ODF2L,MED18,ACTG1,TUBA1A,XRCC2,ZN |
| 144 | 19917 | ARSD,ODF2L,NUDC,BTBD9,MAVS,HSP90B1,PRPF8,CANX,GDI1,PTM |
| 499 | 19917 | NEDD4L,ACTB,ODF2L,NRBP2,EEF2,XRCC2,HTT,FTH1,TMSB4X,ZNF3 |
| 78 | 19917 | POLH,ACTB,CFL1,TUBA1B,NPM1,SERINC1,CDC42BPB,SMARCA5,R |
| 137 | 19917 | COPA,ZNF316,PIK3R2,RACK1,P4HB,AFF4,POLR2A,MORF4L1,SF3B2 |
| 875 | 19917 | EEF1A1,COPA,HNRNPA1,HSPA9,NEDD4L,USP22,MED18,EEF2,DPYSL |
| 241 | 19917 | COPA,ACTB,MED18,ZNF316,PIK3R2,NCL,DDX5,MAGED1,RPL6,P4HE |
| 1228 | 19917 | ORC4,HSPA8,GAPDH,MLLT1,ARSD,MS4A10,POLH,ACTB,USP22,MED |
| 677 | 19917 | EEF1A1,HSPA8,HNRNPA1,HSPA9,ACTB,CTSC,NUDC,USP22,ACTG1,E |
| 121 | 19917 | PABPC1,LRP1,TUBB,NEFM,KAT6A,GNAS,SPTBN1,IGF2,MORF4L1,A |
| 292 | 19917 | EEF1A1,HNRNPA1,USP22,MED18,EEF2,BTBD9,XRCC2,HSP90AB1,UE |
| 665 | 19917 | PABPC1,COL1A1,GAPDH,NEDD4L,POLH,ODF2L,USP22,ACTG1,HTT, |
| 889 | 19917 | EEF1A1,PABPC1,LRP1,GAPDH,MLLT1,HNRNPA1,ARSD,HSPA9,POLH, |
| 97 | 19917 | COPA,ACTB,HSP90AA1,ZNF316,PIK3R2,RHOA,ZNF865,WDR1,PFN1, |
| 260 | 19917 | PABPC1,LRP1,MLLT1,NEDD4L,ZNF124,VIM,RPL7A,SLC9A7,COL6A1, |
| 1641 | 19917 | EEF1A1,PABPC1,TMEM212,HSPA8,GAPDH,MLLT1,COPA,HNRNPA1,A |
| 229 | 19917 | USP22,MED18,EEF2,BTBD9,XRCC2,HSP90AB1,UBC,RPL3,RPL4,FLN |
| 252 | 19917 | USP22,EEF2,XRCC2,UBC,RPL3,RPL4,ENO1,HSP90B1,FADS1,FKBP1 |
| 202 | 19917 | EEF1A1,ARSD,TUBB,NUMA1,TUBA1B,PKM,GANAB,CANX,PTMA,DVL3 |
| 1173 | 19917 | IGFBP5,PABPC1,LRP1,HSPA8,MLLT1,COPA,HNRNPA1,HSPA9,NEDD4 |
| 249 | 19917 | ODF2L,NRBP2,CFL1,HTT,FTH1,UBC,TMSB4X,DBN1,MAP1B,PKM,AB |
| 83 | 19917 | RPS6,PTMA,CSDE1,EIF4B,MATR3,YWHAG,RPS9,RPL17,EIF5,RPL26, |
| 205 | 19917 | USP22,MED18,EEF2,BTBD9,XRCC2,HSP90AB1,UBC,RPL3,RPL4,FLN |
| 130 | 19917 | MED18,EEF2,HSP90AB1,UBC,RPL3,TMF1,KIF1A,FSCN1,USP11,TRIM |
| 2667 | 19917 | EEF1A1,ORC4,IGFBP5,PABPC1,TMEM212,COL1A1,LRP1,HSPA8,GA |
| 143 | 19917 | COPA,MED18,ZNF316,PIK3R2,NCL,DDX5,MAGED1,RPL6,RACK1,P4 |
| 185 | 19917 | PABPC1,COPA,MED18,ZNF316,PIK3R2,NCL,YBX1,PTMA,DDX5,MAGE |
| 186 | 19917 | EEF2,BTBD9,XRCC2,UBC,RPL3,FADS1,TMF1,KIF1A,ALDH6A1,SEPTI |
| 636 | 19917 | ORC4,IGFBP5,PABPC1,COL1A1,LRP1,COPA,NEDD4L,POLH,NRBP2, |
| 189 | 19917 | ACTB,MED18,XRCC2,RPS6,ZNF316,PIK3R2,HSP90B1,RPL13A,MAGE |
| 188 | 19917 | ORC4,LRP1,ACTB,ACTG1,VIM,UBC,DBN1,MAP1B,RPL7A,COL4A1,E |
| 1176 | 19917 | EEF1A1,PABPC1,HSPA8,COPA,HNRNPA1,HSPA9,POLH,ATN1,ACTB,C |
| 686 | 19917 | ORC4,PABPC1,COPA,ACTB,ODF2L,MED18,XRCC2,MAVS,HSP90AB1, |
| 89 | 19917 | COPA,RPS6,ZNF316,PIK3R2,RPL6,P4HB,POLR2A,PFN1,EIF5A,EIF3L, |
| 124 | 19917 | MED18,EEF2,HSP90AB1,UBC,RPL3,KIF1A,FSCN1,USP11,TRIM28,CK |
| 224 | 19917 | USP22,MED18,EEF2,BTBD9,XRCC2,HSP90AB1,UBC,RPL3,RPL4,FLN |
| 2077 | 19917 | EEF1A1,ORC4,IGFBP5,PABPC1,COL1A1,LRP1,HSPA8,GAPDH,MS4A1 |
| 265 | 19917 | EEF1A1,PABPC1,LRP1,MLLT1,RPL7A,FADS1,IGFBP2,SLC2A3,CALM |

| | | |
|---|---|---|
| 100 | 19917 | COPA,ZNF124,HSP90AA1,ZNF316,PIK3R2,RPL6,SHISA9,ZNF43,SF3B |
| 450 | 19917 | GAPDH,HNRNPA1,DPYSL3,TUBA1A,ACTR2,TUBB,NUMA1,TUBA1B,RF |
| 238 | 19917 | ORC4,PABPC1,MED18,XRCC2,ZNF124,ACTR2,HSP90B1,FBXL4,IGFE |
| 2212 | 19917 | EEF1A1,ORC4,IGFBP5,PABPC1,TMEM212,COL1A1,LRP1,HSPA8,GAF |
| 729 | 19917 | EEF1A1,PABPC1,GAPDH,MLLT1,HNRNPA1,POLH,ATN1,CFL1,ACTR2, |
| 280 | 19917 | COPA,MED18,ZNF124,ACTR2,KIDINS220,RPS6,ZNF316,PIK3R2,ALD( |
| 242 | 19917 | COPA,UBC,ALDOA,FTL,FADS1,RPLP0,DDX5,FKBP14,SEC62,FASN,AL |
| 208 | 19917 | GAPDH,COPA,HNRNPA1,ATN1,MTRNR2L2,HSP90AB1,FTH1,EIF4G2,F |
| 850 | 19917 | LRP1,HNRNPA1,HSPA9,NEDD4L,ODF2L,NUDC,ACTG1,EEF2,TUBA1A, |
| 414 | 19917 | GAPDH,POLH,ODF2L,ACTG1,RPL3,DBN1,ENO1,NCL,FTL,FADS1,UGT |
| 1148 | 19917 | ORC4,PABPC1,TMEM212,COL1A1,HSPA8,MLLT1,COPA,POLH,ACTB, |
| 115 | 19917 | ACTB,ODF2L,ACTR2,ZNF316,PIK3R2,RPL6,HMGB1,RPL29,RPS14,PF |
| 121 | 19917 | MED18,RPS6,ZNF316,PIK3R2,HSP90B1,PPP2R1A,DNAJA1,ZNF865,R |
| 267 | 19917 | NUDC,ACTR2,ALDOA,COL6A1,COL4A1,PTMA,GNB1,MYH9,RPL5,YWI |
| 244 | 19917 | LRP1,NEDD4L,HTT,UBC,KIDINS220,MAP1B,ALDH6A1,FSCN1,SMIM1 |
| 472 | 19917 | ORC4,IGFBP5,PABPC1,COL1A1,LRP1,NEDD4L,USP22,HTT,UBC,KIDI |
| 278 | 19917 | EEF1A1,HNRNPA1,MED18,EEF2,BTBD9,XRCC2,HSP90AB1,UBC,RPL3 |
| 115 | 19917 | EEF2,UBC,RPL3,DVL3,SEPTIN2,FSCN1,USP11,FN1,EIF1,TPT1,RPL18 |
| 255 | 19917 | GAPDH,USP22,EEF2,DBN1,UBA1,TUBB2B,KIF1A,NPM1,AFF4,RPL37A |
| 212 | 19917 | GAPDH,HNRNPA1,ATN1,MTRNR2L2,HSP90AB1,FTH1,EIF4G2,GANAB |
| 1492 | 19917 | PABPC1,TMEM212,COL1A1,GAPDH,HNRNPA1,ARSD,NEDD4L,POLH,I |
| 369 | 19917 | GAPDH,MLLT1,COPA,ACTB,ODF2L,HSP90AB1,ZNF124,HSP90AA1,AC |
| 58 | 19917 | DBN1,HTRA4,GSTP1,INA,ITGB1,LDHA,CPE,TSPAN3,TLE5,RPS25,HEC |
| 820 | 19917 | PABPC1,HNRNPA1,HSPA9,ODF2L,USP22,MED18,EEF2,DPYSL3,CFL1 |
| 291 | 19917 | COL1A1,HNRNPA1,MS4A10,NEDD4L,ACTB,ZNF124,FTH1,NUMA1,HS |
| 133 | 19917 | NUDC,TUBA1A,UBC,TMSB4X,HSP90B1,GNAS,FASN,PTMS,CALM1,RF |
| 700 | 19917 | PABPC1,TMEM212,HNRNPA1,ODF2L,USP22,MED18,EEF2,BTBD9,XR |
| 791 | 19917 | EEF1A1,MLLT1,ARSD,NEDD4L,ACTG1,DPYSL3,MAVS,HTT,HSP90AB1 |
| 209 | 19917 | USP22,MED18,EEF2,XRCC2,HSP90AB1,UBC,RPL3,RPL4,FADS1,TMF |
| 2167 | 19917 | EEF1A1,ORC4,IGFBP5,PABPC1,COL1A1,LRP1,HSPA8,GAPDH,MLLT1 |
| 176 | 19917 | MLLT1,HNRNPA1,NEDD4L,POLH,DPYSL3,HTT,RPS6,TUBA1B,PRPF8, |
| 70 | 19917 | RPS6,CSDE1,EIF4B,MATR3,RPS9,RPL17,EIF5,RPL26,DDX3X,RPS15,F |
| 429 | 19917 | COPA,NEDD4L,ACTB,ODF2L,MED18,HSP90AA1,KIDINS220,NUMA1,F |
| 123 | 19917 | ZNF316,PIK3R2,MAGED1,RPL6,SHISA9,P4HB,TMBIM6,POLR2A,SF3E |
| 338 | 19917 | COPA,MED18,ACTR2,KIDINS220,RPS6,ZNF316,PIK3R2,ALDOA,RPL8 |
| 103 | 19917 | LRP1,CALR,SHISA9,SEPTIN2,PLXNB2,EIF4B,PPP2R1A,RPLP1,DNAJA1 |
| 75 | 19917 | ZNF316,PIK3R2,RPL6,POLR2A,PFN1,EIF5A,EIF3L,RPLP2,DDOST,MD |
| 453 | 19917 | LRP1,HSPA8,GAPDH,ARSD,MED18,BTBD9,UBC,KIDINS220,TMSB4X,I |
| 1057 | 19917 | EEF1A1,ORC4,PABPC1,HSPA8,MLLT1,COPA,ARSD,MS4A10,POLH,CT |
| 1524 | 19917 | EEF1A1,ORC4,PABPC1,TMEM212,COL1A1,LRP1,HSPA8,MLLT1,COP |
| 96 | 19917 | ODF2L,ACTR2,ZNF316,PIK3R2,RPL6,HMGB1,POLR2A,RPL29,RPS14 |
| 763 | 19917 | PABPC1,TMEM212,MS4A10,NUDC,USP22,ACTG1,CFL1,HTT,EIF4G2, |
| 145 | 19917 | MED18,EEF2,XRCC2,HSP90AB1,UBC,RPL3,TMF1,KIF1A,FSCN1,USP1 |
| 151 | 19917 | MED18,EEF2,XRCC2,HSP90AB1,UBC,RPL3,FADS1,TMF1,KIF1A,FSCN |
| 138 | 19917 | ACTB,UBC,TUBA1B,HSP90B1,CES3,SPTBN1,GNB1,MAP9,TIMP2,MID |

| | | |
|---|---|---|
| 201 | 19917 | COPA,ODF2L,RPS6,ZNF316,PIK3R2,ALDOA,FKBP14,RPL13A,MAGED |
| 567 | 19917 | PABPC1,COL1A1,MLLT1,COPA,ARSD,MS4A10,NUDC,VIM,TUBB,TMS |
| 170 | 19917 | USP22,MED18,EEF2,XRCC2,HSP90AB1,UBC,RPL3,FADS1,TMF1,KIF1 |
| 147 | 19917 | MED18,XRCC2,HSP90AB1,UBC,FADS1,TMF1,KIF1A,FSCN1,USP11,TI |
| 147 | 19917 | MED18,XRCC2,HSP90AB1,UBC,FADS1,TMF1,KIF1A,FSCN1,USP11,TI |
| 168 | 19917 | EEF1A1,PABPC1,EEF2,NUMA1,RPL3,PSAP,PGK1,DVL3,KIF1A,SEPTIN |
| 1273 | 19917 | ORC4,IGFBP5,PABPC1,LRP1,HSPA8,MLLT1,COPA,MS4A10,NEDD4L, |
| 486 | 19917 | ORC4,ARSD,MS4A10,NEDD4L,NUDC,HTT,UBC,KIDINS220,FLNA,MAI |
| 947 | 19917 | EEF1A1,ORC4,PABPC1,HSPA8,MLLT1,COPA,ARSD,POLH,CTSC,ODF |
| 880 | 19917 | EEF1A1,PABPC1,HSPA8,HNRNPA1,POLH,ATN1,ACTB,CTSC,USP22,CI |
| 880 | 19917 | EEF1A1,PABPC1,HSPA8,HNRNPA1,POLH,ATN1,ACTB,CTSC,USP22,CI |
| 626 | 19917 | PABPC1,COL1A1,GAPDH,MLLT1,MS4A10,NEDD4L,ATN1,ACTB,USP2 |
| 120 | 19917 | EEF2,HSP90AB1,UBC,RPL3,MAP1B,FADS1,KIF1A,FSCN1,USP11,FN1, |
| 1486 | 19917 | ORC4,COL1A1,LRP1,HSPA8,GAPDH,MLLT1,COPA,HNRNPA1,ARSD,M |
| 185 | 19917 | HSPA9,POLH,HSP90AA1,GNB1,MYH9,APP,NPM1,CSDE1,EIF4B,TMSB |
| 115 | 19917 | EEF1A1,BTBD9,UBC,RPLP0,FKBP14,ALDH6A1,RPL5,SNRNP200,SMI |
| 1761 | 19917 | EEF1A1,ORC4,TMEM212,HSPA8,GAPDH,COPA,HNRNPA1,ARSD,MS4 |
| 981 | 19917 | EEF1A1,COPA,HNRNPA1,HSPA9,NEDD4L,USP22,MED18,ACTG1,EEF2 |
| 869 | 19917 | EEF1A1,HSPA8,HNRNPA1,ARSD,POLH,ATN1,ODF2L,USP22,MTRNR2I |
| 58 | 19917 | GAPDH,MLLT1,FTH1,NEFM,SEPTIN2,SERINC1,MORF4L1,VCL,PJA2,H |
| 205 | 19917 | ACTB,MED18,HSP90AB1,PIK3R2,DDX5,MAGED1,RPL6,GPI,USP11,EI |
| 572 | 19917 | HSPA9,POLH,NUDC,ACTG1,BTBD9,ACTR2,FTH1,UBC,KIDINS220,NU |
| 45 | 19917 | GAPDH,EIF4G2,YWHAE,SEPTIN2,RPSA,ATF4,NISCH,PRKCSH,HDLBP, |
| 553 | 19917 | IGFBP5,LRP1,ACTB,NRBP2,ACTG1,DPYSL3,TUBA1A,HTT,ZNF124,HS |
| 81 | 19917 | RPS6,ZNF316,PIK3R2,FKBP14,RPL6,RACK1,USP11,PPP2R1A,POLR2 |
| 1031 | 19917 | EEF1A1,ORC4,PABPC1,HSPA8,GAPDH,MS4A10,ATN1,ACTB,USP22,C |
| 149 | 19917 | MED18,EEF2,XRCC2,HSP90AB1,UBC,RPL3,TMF1,KIF1A,FSCN1,USP1 |
| 215 | 19917 | GAPDH,HNRNPA1,ATN1,NUDC,MTRNR2L2,HSP90AB1,FTH1,EIF4G2, |
| 1262 | 19917 | EEF1A1,ORC4,IGFBP5,COL1A1,GAPDH,HNRNPA1,MS4A10,NEDD4L, |
| 973 | 19917 | EEF1A1,IGFBP5,LRP1,MLLT1,COPA,NEDD4L,ACTB,NRBP2,MED18,A |
| 97 | 19917 | COPA,ACTR2,ZNF316,PIK3R2,RPL6,HMGB1,TPT1,RPS14,PFN1,EIF5 |
| 903 | 19917 | EEF1A1,PABPC1,HSPA8,HNRNPA1,POLH,ATN1,ACTB,CTSC,USP22,CI |
| 165 | 19917 | MED18,EEF2,XRCC2,HSP90AB1,UBC,RPL3,FADS1,TMF1,KIF1A,FSCN |
| 385 | 19917 | HNRNPA1,USP22,MED18,EEF2,BTBD9,XRCC2,HSP90AB1,UBC,RPL3, |
| 103 | 19917 | ODF2L,ACTR2,ZNF316,PIK3R2,RPL6,POLR2A,RPL29,RPS14,PFN1,P |
| 95 | 19917 | UBC,KIF1A,TRIM28,CKB,FN1,EIF1,RPL18,SARAF,WSB1,AP2B1,LDHA |
| 207 | 19917 | MED18,EEF2,XRCC2,HSP90AB1,UBC,TMSB4X,RPL3,MAP1B,TMF1,K |
| 157 | 19917 | HSP90AA1,ENO1,COL6A1,BSG,GNB1,GSTP1,PLXNB2,HSPD1,UBB,Y |
| 189 | 19917 | COPA,MED18,ACTR2,FTH1,ZNF316,PIK3R2,NCL,ALDOA,DDX5,MAGE |
| 323 | 19917 | EEF1A1,HSPA9,USP22,MED18,BTBD9,XRCC2,HSP90AB1,UBC,EIF4G |
| 47 | 19917 | AGRN,MIDN,ZNF865,CHD4,ATP2A2,OAZ1,SOX12,APLP1,CADM4,CHD |
| 151 | 19917 | MED18,EEF2,XRCC2,HSP90AB1,UBC,RPL3,FADS1,TMF1,KIF1A,FSCN |
| 268 | 19917 | HSPA9,USP22,MED18,EEF2,BTBD9,XRCC2,HSP90AB1,UBC,RPL3,RP |
| 170 | 19917 | USP22,MED18,EEF2,XRCC2,HSP90AB1,UBC,RPL3,FADS1,TMF1,KIF1 |
| 69 | 19917 | ZNF316,PIK3R2,RPL6,POLR2A,PFN1,EIF5A,EIF3L,RPLP2,DDOST,TM |

| | | |
|---|---|---|
| 158 | 19917 | USP22,MED18,EEF2,XRCC2,HSP90AB1,UBC,RPL3,FADS1,TMF1,KIF1 |
| 98 | 19917 | LRP1,CALR,SHISA9,SEPTIN2,PLXNB2,PPP2R1A,RPLP1,DNAJA1,CDC4 |
| 79 | 19917 | UBC,KIF1A,TRIM28,CKB,FN1,RPL18,WSB1,AP2B1,TMEM259,RPLP2, |
| 79 | 19917 | UBC,KIF1A,TRIM28,CKB,FN1,RPL18,WSB1,AP2B1,TMEM259,RPLP2, |
| 175 | 19917 | USP22,MED18,EEF2,XRCC2,HSP90AB1,UBC,RPL3,FADS1,TMF1,TUB |
| 378 | 19917 | HNRNPA1,HSPA9,ACTB,USP22,EEF2,CFL1,ACTR2,VIM,HSP90B1,RPL |
| 226 | 19917 | HNRNPA1,MED18,EEF2,BTBD9,XRCC2,HSP90AB1,UBC,RPL3,FADS1, |
| 409 | 19917 | ORC4,HSPA8,GAPDH,HNRNPA1,NUDC,KIDINS220,EIF4G2,DBN1,RPL |
| 409 | 19917 | ORC4,HSPA8,GAPDH,HNRNPA1,NUDC,KIDINS220,EIF4G2,DBN1,RPL |
| 270 | 19917 | LRP1,MLLT1,HNRNPA1,CFL1,HTT,NUMA1,NEFM,PRPF8,ENO2,SNRN |
| 136 | 19917 | NEDD4L,VIM,DBN1,RPL7A,FSTL1,KAT6A,GDI1,USP11,YWHAG,CKB,T |
| 1489 | 19917 | EEF1A1,IGFBP5,COL1A1,HSPA8,GAPDH,HNRNPA1,HSPA9,NEDD4L,P |
| 30 | 19917 | KAT6A,CTSD,HMGN2,LAMP1,SERF2,CTSB,MVB12B,PLTP,HNRNPM,V |
| 52 | 19917 | EEF2,UBC,ALDH6A1,VCP,POLR2A,TPT1,MORF4L1,MIDN,MARCHF6,A |
| 614 | 19917 | PABPC1,COPA,ACTB,ODF2L,MED18,HSP90AB1,HSP90AA1,ACTR2,D |
| 144 | 19917 | MED18,EEF2,XRCC2,HSP90AB1,UBC,RPL3,TMF1,KIF1A,FSCN1,USP1 |
| 194 | 19917 | MED18,EEF2,XRCC2,HSP90AB1,UBC,TMSB4X,RPL3,ENO1,HSP90B1, |
| 125 | 19917 | XRCC2,RPS6,HSP90B1,IGFBP2,FKBP14,RPL13A,RPS8,USP11,ATP5F |
| 83 | 19917 | ACTR2,ZNF316,PIK3R2,RPL6,HMGB1,RPS14,PFN1,EIF5A,EIF3L,RPL |
| 63 | 19917 | BTBD9,HSP90AB1,ALDH6A1,PTMS,AFF4,TPT1,XRCC5,RPS20,LASP1, |
| 657 | 19917 | IGFBP5,TMEM212,HSPA8,GAPDH,HNRNPA1,HSPA9,ACTB,MTRNR2L |
| 107 | 19917 | HNRNPA1,MS4A10,XRCC2,ALDOA,NPM1,EIF4B,MAPT,NFE2L1,TNS3, |
| 451 | 19917 | HSPA8,GAPDH,HSPA9,POLH,ACTB,TUBA1A,CFL1,XRCC2,MAVS,HSP9 |
| 54 | 19917 | ACTB,NCL,FASN,AGRN,ARF1,EIF5A,CDH2,TLN1,RPLP2,KLHL24,ACO |
| 431 | 19917 | ARSD,NEDD4L,NUDC,NRBP2,HTT,KIDINS220,PIK3R2,FLNA,ABHD2,P |
| 235 | 19917 | HTT,UBC,KIDINS220,MAP1B,ABHD2,CANX,ALDH6A1,FSCN1,SMIM14 |
| 40 | 19917 | GNAS,ATP5F1A,SPEN,CDC42BPB,JUND,PFKL,SOX12,MRFAP1,PLPPR |
| 160 | 19917 | POLH,FLNA,FTL,FADS1,PTMA,KIF1A,RPL13,GPI,RACK1,FASN,GNB1, |
| 458 | 19917 | HNRNPA1,USP22,MED18,ACTG1,EEF2,XRCC2,HSP90AB1,UBC,RPL3, |
| 347 | 19917 | ORC4,COPA,ACTB,ODF2L,MED18,ZNF124,NUMA1,ZNF316,RPL4,PIK |
| 84 | 19917 | ARSD,HTT,KIDINS220,ABHD2,ALDH6A1,STMN1,UCHL1,ANXA2,AGRN |
| 62 | 19917 | COL1A1,POLH,ZNF316,IDS,YWHAG,PKD1,VAPA,ATP1A1,HNRNPDL,T |
| 727 | 19917 | EEF1A1,COL1A1,HNRNPA1,HSPA9,ACTG1,DPYSL3,CFL1,XRCC2,MAV |
| 438 | 19917 | HNRNPA1,USP22,MED18,ACTG1,EEF2,XRCC2,HSP90AB1,UBC,RPL3, |
| 196 | 19917 | HSPA8,MED18,XRCC2,KIDINS220,RPS6,RPL4,HSP90B1,IGFBP2,PTM |
| 83 | 19917 | XRCC2,ENO2,AFF4,BASP1,GTF2F1,NFE2L1,RPS16,CNN3,TNS3,RPS2 |
| 73 | 19917 | RPS6,ZNF316,PIK3R2,TMBIM6,POLR2A,PFN1,CHPF,RPLP2,DDOST,Y |
| 532 | 19917 | EEF1A1,HSPA8,HNRNPA1,HSPA9,ACTB,CTSC,USP22,EEF2,CFL1,HSP |
| 159 | 19917 | USP22,MED18,EEF2,XRCC2,HSP90AB1,UBC,RPL3,FADS1,TMF1,KIF1 |
| 116 | 19917 | COPA,HTT,UBC,TUBA1B,PKM,ABHD2,PGK1,KAT6A,ALDH6A1,MYH9, |
| 180 | 19917 | USP22,MED18,EEF2,XRCC2,HSP90AB1,UBC,RPL3,FADS1,TMF1,KIF1 |
| 158 | 19917 | MED18,EEF2,XRCC2,HSP90AB1,UBC,RPL3,FADS1,TMF1,KIF1A,FSCN |
| 88 | 19917 | COPA,ACTB,ACTR2,ZNF316,RPL6,EIF4B,RHOA,ZNF865,WDR1,RPLP |
| 90 | 19917 | ACTB,ACTR2,ZNF316,RPL6,EIF4B,HMGB1,TUBB4B,RPL29,CIPC,PNN |
| 26 | 19917 | RPL5,PAPOLA,CCT7,KHDRBS1,HNRNPDL,DDOST,MAPRE1,BEX4,PSM |

| | | |
|---|---|---|
| 262 | 19917 | GAPDH,ODF2L,NCL,ALDOA,YBX1,IGFBP2,PSAP,CANX,DDX5,KAT6A,F |
| 76 | 19917 | LRP1,HTT,ABHD2,ALDH6A1,STMN1,GOLGA8A,CTNNA1,SCARB2,RPS |

,COPA,HNRNPA1,ARSD,HSPA9,NEDD4L,POLH,ATN1,ACTB,CTSC,ODF2L,NUDC,NRBP2,USP22,ME
INRNPA1,ARSD,HSPA9,NEDD4L,POLH,ACTB,CTSC,ODF2L,NUDC,NRBP2,USP22,MED18,ACTG1,E
,HNRNPA1,ARSD,HSPA9,NEDD4L,POLH,ACTB,CTSC,ODF2L,NUDC,NRBP2,USP22,MED18,ACTG1,
INRNPA1,ARSD,HSPA9,NEDD4L,POLH,ATN1,ACTB,CTSC,ODF2L,NUDC,NRBP2,USP22,MED18,ACT
NEDD4L,POLH,ACTB,ODF2L,NRBP2,USP22,ACTG1,EEF2,TUBA1A,CFL1,BTBD9,MAVS,HTT,HSP9
INRNPA1,ARSD,HSPA9,NEDD4L,POLH,ACTB,CTSC,ODF2L,NUDC,NRBP2,USP22,ACTG1,DPYSL3,T
,HNRNPA1,ARSD,HSPA9,NEDD4L,POLH,ACTB,CTSC,ODF2L,NUDC,NRBP2,USP22,MTRNR2L2,AC
,ACTG1,EEF2,TUBA1A,BTBD9,HTT,HSP90AA1,ACTR2,FTH1,TMSB4X,EIF4G2,ZNF316,DBN1,ENO1
),HSPA9,NEDD4L,POLH,ATN1,ACTB,CTSC,NUDC,NRBP2,USP22,MED18,ACTG1,DPYSL3,CFL1,BT
SD,HSPA9,NEDD4L,POLH,ACTB,CTSC,ODF2L,NUDC,NRBP2,USP22,MED18,ACTG1,EEF2,DPYSL3,
,ARSD,HSPA9,NEDD4L,POLH,ACTB,CTSC,ODF2L,NRBP2,USP22,MTRNR2L2,ACTG1,EEF2,DPYSL
,NEDD4L,POLH,ACTB,CTSC,NRBP2,USP22,ACTG1,EEF2,DPYSL3,TUBA1A,BTBD9,XRCC2,MAVS,
),HSPA9,POLH,ACTB,CTSC,NRBP2,USP22,ACTG1,DPYSL3,TUBA1A,CFL1,BTBD9,XRCC2,MAVS,HT
),HSPA9,NEDD4L,POLH,ACTB,NRBP2,USP22,ACTG1,EEF2,DPYSL3,TUBA1A,CFL1,BTBD9,XRCC2,
DD4L,ATN1,ACTB,NUDC,NRBP2,USP22,ACTG1,EEF2,DPYSL3,TUBA1A,CFL1,BTBD9,XRCC2,MAVS
,ARSD,HSPA9,NEDD4L,POLH,ACTB,CTSC,ODF2L,NUDC,NRBP2,USP22,MED18,ACTG1,EEF2,DPY
A9,POLH,ACTB,CTSC,NRBP2,USP22,ACTG1,EEF2,DPYSL3,TUBA1A,CFL1,BTBD9,XRCC2,MAVS,H
D,HSPA9,NEDD4L,POLH,ACTB,NRBP2,USP22,ACTG1,DPYSL3,TUBA1A,CFL1,BTBD9,XRCC2,MAV
,ARSD,NEDD4L,POLH,ACTB,NRBP2,USP22,ACTG1,EEF2,TUBA1A,CFL1,BTBD9,HTT,HSP90AB1,Z
,HSPA9,POLH,ACTB,CTSC,NRBP2,USP22,ACTG1,EEF2,DPYSL3,TUBA1A,CFL1,BTBD9,XRCC2,MA
L,POLH,ACTB,CTSC,NUDC,NRBP2,USP22,MED18,ACTG1,EEF2,DPYSL3,CFL1,BTBD9,XRCC2,MA
,HNRNPA1,HSPA9,NEDD4L,POLH,ACTB,CTSC,NUDC,NRBP2,USP22,MTRNR2L2,MED18,ACTG1,E
A1,HSPA9,NEDD4L,POLH,ACTB,CTSC,NUDC,NRBP2,USP22,MED18,ACTG1,EEF2,DPYSL3,CFL1,B
,COPA,HNRNPA1,ARSD,HSPA9,NEDD4L,POLH,ATN1,ACTB,CTSC,ODF2L,NUDC,NRBP2,USP22,ME
A1,ARSD,HSPA9,NEDD4L,POLH,ACTB,ODF2L,NUDC,NRBP2,USP22,MED18,DPYSL3,CFL1,BTBD9,
PA,HNRNPA1,ARSD,HSPA9,NEDD4L,POLH,ATN1,ACTB,CTSC,NUDC,NRBP2,USP22,ACTG1,EEF2,D
,POLH,ACTB,CTSC,NUDC,NRBP2,USP22,MED18,ACTG1,DPYSL3,TUBA1A,CFL1,BTBD9,MAVS,HT
,HNRNPA1,ARSD,HSPA9,NEDD4L,POLH,ACTB,CTSC,ODF2L,NRBP2,USP22,ACTG1,EEF2,DPYSL3,
,COPA,HNRNPA1,ARSD,HSPA9,NEDD4L,POLH,ACTB,CTSC,ODF2L,NUDC,NRBP2,USP22,MED18,A
,COPA,HNRNPA1,ARSD,MS4A10,HSPA9,NEDD4L,POLH,ATN1,ACTB,CTSC,ODF2L,NUDC,NRBP2,U
INRNPA1,ARSD,MS4A10,HSPA9,NEDD4L,POLH,ATN1,ACTB,CTSC,ODF2L,NUDC,NRBP2,USP22,M
A1,HSPA9,NEDD4L,POLH,ACTB,CTSC,NRBP2,USP22,ACTG1,DPYSL3,TUBA1A,CFL1,BTBD9,XRCC
SPA9,NEDD4L,POLH,ACTB,ODF2L,NRBP2,USP22,ACTG1,EEF2,DPYSL3,TUBA1A,CFL1,BTBD9,XR
,HNRNPA1,ARSD,HSPA9,NEDD4L,POLH,ATN1,ACTB,CTSC,NUDC,NRBP2,USP22,MED18,ACTG1,E
INRNPA1,ARSD,MS4A10,HSPA9,NEDD4L,POLH,ATN1,ACTB,CTSC,ODF2L,NUDC,NRBP2,USP22,M
,ACTB,NRBP2,USP22,ACTG1,DPYSL3,TUBA1A,BTBD9,XRCC2,MAVS,HTT,HSP90AB1,ZNF124,HSP
,ARSD,HSPA9,NEDD4L,POLH,ACTB,CTSC,NUDC,NRBP2,USP22,MED18,ACTG1,DPYSL3,TUBA1A,
,POLH,ACTB,ODF2L,NUDC,NRBP2,USP22,ACTG1,EEF2,DPYSL3,TUBA1A,CFL1,BTBD9,XRCC2,M
BP2,USP22,ACTG1,DPYSL3,TUBA1A,CFL1,BTBD9,XRCC2,MAVS,HTT,HSP90AB1,ZNF124,HSP90A
INRNPA1,ARSD,MS4A10,HSPA9,NEDD4L,POLH,ATN1,ACTB,CTSC,ODF2L,NUDC,NRBP2,MED18,A
T1,COPA,HNRNPA1,ARSD,HSPA9,NEDD4L,POLH,ATN1,ACTB,CTSC,NUDC,NRBP2,USP22,ACTG1,
,COPA,HNRNPA1,ARSD,HSPA9,NEDD4L,POLH,ACTB,CTSC,ODF2L,NUDC,NRBP2,USP22,MED18,A
EDD4L,POLH,ACTB,CTSC,ODF2L,NRBP2,USP22,MED18,DPYSL3,CFL1,BTBD9,MAVS,ZNF124,HSP
,HNRNPA1,ARSD,HSPA9,NEDD4L,POLH,ACTB,ODF2L,NRBP2,USP22,ACTG1,EEF2,DPYSL3,TUBA1
,HNRNPA1,ARSD,HSPA9,NEDD4L,POLH,ACTB,CTSC,ODF2L,NUDC,NRBP2,USP22,ACTG1,EEF2,D

HNRNPA1,ARSD,HSPA9,NEDD4L,POLH,ATN1,ACTB,ODF2L,NUDC,NRBP2,USP22,MTRNR2L2,MED1
A1,HSPA9,NEDD4L,POLH,NRBP2,USP22,ACTG1,DPYSL3,TUBA1A,CFL1,BTBD9,XRCC2,MAVS,HTT
JSP22,ACTG1,DPYSL3,TUBA1A,CFL1,BTBD9,HTT,HSP90AB1,ZNF124,KIDINS220,EIF4G2,RPL3,ZI
,HNRNPA1,ARSD,HSPA9,NEDD4L,POLH,ACTB,CTSC,ODF2L,NUDC,NRBP2,USP22,MTRNR2L2,ME
,HNRNPA1,ARSD,MS4A10,HSPA9,NEDD4L,POLH,ATN1,ACTB,CTSC,ODF2L,NUDC,NRBP2,USP22,I
A1,ARSD,NEDD4L,ACTB,ODF2L,NRBP2,USP22,ACTG1,EEF2,DPYSL3,TUBA1A,CFL1,BTBD9,MAVS
ACTB,NRBP2,USP22,ACTG1,EEF2,DPYSL3,TUBA1A,CFL1,BTBD9,XRCC2,MAVS,HTT,HSP90AB1,ZI
A9,NEDD4L,POLH,ACTB,NUDC,NRBP2,USP22,MED18,ACTG1,EEF2,DPYSL3,CFL1,BTBD9,XRCC2
UBA1A,BTBD9,XRCC2,MAVS,HTT,HSP90AB1,ZNF124,FTH1,KIDINS220,NUMA1,ZNF316,DBN1,EN
A1,ARSD,MS4A10,HSPA9,NEDD4L,POLH,ACTB,ODF2L,NUDC,NRBP2,USP22,MED18,ACTG1,EEF2,
TUBA1A,CFL1,BTBD9,HTT,ZNF124,FTH1,KIDINS220,NUMA1,RPL3,ZNF316,DBN1,TUBA1B,PIK3R
2,USP22,ACTG1,TUBA1A,BTBD9,XRCC2,MAVS,HTT,HSP90AB1,ZNF124,KIDINS220,NUMA1,ZNF31
P2,USP22,ACTG1,DPYSL3,TUBA1A,CFL1,BTBD9,XRCC2,MAVS,HTT,HSP90AB1,ZNF124,HSP90AA1
SP22,ACTG1,DPYSL3,TUBA1A,BTBD9,XRCC2,MAVS,HTT,HSP90AB1,ZNF124,TUBB,FTH1,KIDINS2
PA1,ARSD,HSPA9,NEDD4L,POLH,ACTB,CTSC,ODF2L,NUDC,NRBP2,USP22,MED18,ACTG1,EEF2,D
PYSL3,TUBA1A,CFL1,BTBD9,MAVS,HTT,HSP90AA1,FTH1,UBC,ZNF316,DBN1,ENO1,TUBA1B,FLN
LT1,HNRNPA1,ARSD,HSPA9,NEDD4L,POLH,ACTB,CTSC,ODF2L,NUDC,NRBP2,USP22,MTRNR2L2,
I,ACTB,NRBP2,USP22,ACTG1,DPYSL3,TUBA1A,CFL1,BTBD9,XRCC2,MAVS,HTT,HSP90AB1,ZNF12
SP22,ACTG1,TUBA1A,CFL1,BTBD9,XRCC2,MAVS,HTT,HSP90AB1,ZNF124,VIM,FTH1,KIDINS220,N
A1,HSPA9,NEDD4L,NRBP2,USP22,ACTG1,DPYSL3,TUBA1A,CFL1,BTBD9,XRCC2,MAVS,HTT,HSP9
D,HSPA9,NEDD4L,POLH,ACTB,CTSC,ODF2L,NUDC,NRBP2,USP22,ACTG1,EEF2,DPYSL3,TUBA1A,
UBA1A,BTBD9,XRCC2,MAVS,HTT,HSP90AB1,ZNF124,HSP90AA1,KIDINS220,TMSB4X,NUMA1,ZNF
IS4A10,HSPA9,NEDD4L,POLH,ATN1,ACTB,ODF2L,NUDC,NRBP2,USP22,MED18,ACTG1,EEF2,DPY
A9,NEDD4L,POLH,ACTB,ODF2L,NRBP2,USP22,ACTG1,DPYSL3,TUBA1A,CFL1,BTBD9,HTT,HSP90
P2,USP22,ACTG1,TUBA1A,BTBD9,XRCC2,MAVS,HTT,HSP90AB1,ZNF124,HSP90AA1,FTH1,KIDINS
CTG1,EEF2,DPYSL3,TUBA1A,CFL1,XRCC2,HTT,ZNF124,HSP90AA1,ACTR2,FTH1,UBC,KIDINS220,
NEDD4L,POLH,ACTB,ODF2L,NRBP2,USP22,ACTG1,DPYSL3,TUBA1A,CFL1,BTBD9,XRCC2,HTT,HS
A1,NEDD4L,NRBP2,USP22,ACTG1,EEF2,DPYSL3,CFL1,BTBD9,XRCC2,HTT,HSP90AB1,HSP90AA1,
,HNRNPA1,ARSD,MS4A10,HSPA9,NEDD4L,POLH,ATN1,ACTB,CTSC,NUDC,NRBP2,USP22,MED18,
HNRNPA1,ARSD,MS4A10,HSPA9,NEDD4L,POLH,ATN1,ACTB,CTSC,ODF2L,NUDC,NRBP2,MED18,A
D,HSPA9,NEDD4L,POLH,ACTB,CTSC,ODF2L,NUDC,NRBP2,USP22,MED18,ACTG1,EEF2,DPYSL3,C
A1,ARSD,NEDD4L,POLH,ACTB,ODF2L,NRBP2,USP22,ACTG1,DPYSL3,TUBA1A,CFL1,HTT,HSP90A
,HNRNPA1,ARSD,MS4A10,NEDD4L,POLH,ACTB,CTSC,NUDC,NRBP2,USP22,ACTG1,EEF2,DPYSL3
D,HSPA9,NEDD4L,POLH,ACTB,CTSC,ODF2L,NUDC,NRBP2,USP22,MED18,ACTG1,EEF2,DPYSL3,T
,HSPA9,NEDD4L,POLH,ACTB,CTSC,NUDC,NRBP2,USP22,ACTG1,EEF2,TUBA1A,CFL1,BTBD9,XRC
,POLH,ACTB,CTSC,NRBP2,USP22,ACTG1,DPYSL3,CFL1,MAVS,HTT,HSP90AB1,ZNF124,HSP90AA
RBP2,USP22,ACTG1,TUBA1A,BTBD9,XRCC2,MAVS,HTT,HSP90AB1,ZNF124,KIDINS220,NUMA1,ZI
,COPA,HNRNPA1,ARSD,HSPA9,NEDD4L,POLH,ATN1,ACTB,CTSC,ODF2L,NUDC,NRBP2,USP22,MT
L,ACTB,ODF2L,NRBP2,USP22,ACTG1,DPYSL3,TUBA1A,CFL1,BTBD9,HTT,HSP90AB1,ZNF124,ACT
,USP22,ACTG1,DPYSL3,TUBA1A,CFL1,BTBD9,XRCC2,MAVS,HTT,HSP90AB1,ZNF124,HSP90AA1,V
,USP22,ACTG1,DPYSL3,TUBA1A,CFL1,BTBD9,XRCC2,MAVS,HTT,HSP90AB1,ZNF124,HSP90AA1,V
D4L,POLH,ACTB,ODF2L,NRBP2,USP22,ACTG1,DPYSL3,TUBA1A,BTBD9,HTT,HSP90AB1,ZNF124,H
,COPA,HNRNPA1,ARSD,HSPA9,NEDD4L,POLH,ACTB,CTSC,ODF2L,NUDC,NRBP2,USP22,MTRNR2
BP2,USP22,ACTG1,TUBA1A,CFL1,BTBD9,MAVS,HTT,ZNF124,HSP90AA1,ACTR2,FTH1,UBC,ZNF31
ACTB,NUDC,NRBP2,USP22,MED18,ACTG1,EEF2,DPYSL3,CFL1,BTBD9,XRCC2,MAVS,HTT,HSP90

;,NRBP2,USP22,ACTG1,DPYSL3,TUBA1A,BTBD9,XRCC2,MAVS,HTT,HSP90AB1,ZNF124,VIM,FTH1,
,USP22,ACTG1,DPYSL3,TUBA1A,CFL1,BTBD9,XRCC2,MAVS,HTT,HSP90AB1,ZNF124,KIDINS220,I
4L,NRBP2,USP22,ACTG1,DPYSL3,TUBA1A,CFL1,BTBD9,XRCC2,HTT,HSP90AB1,ZNF124,HSP90A
ACTG1,TUBA1A,CFL1,BTBD9,HTT,HSP90AB1,ZNF124,FTH1,UBC,KIDINS220,TMSB4X,NUMA1,EIF
,HNRNPA1,ARSD,NEDD4L,POLH,ACTB,CTSC,NRBP2,USP22,ACTG1,EEF2,DPYSL3,TUBA1A,CFL1,I
,COPA,HNRNPA1,ARSD,MS4A10,HSPA9,NEDD4L,POLH,ATN1,ACTB,CTSC,ODF2L,NUDC,NRBP2,U
A1A,BTBD9,HTT,HSP90AB1,HSP90AA1,VIM,KIDINS220,NUMA1,ZNF316,DBN1,ENO1,PIK3R2,FLN
NRBP2,USP22,ACTG1,DPYSL3,TUBA1A,CFL1,BTBD9,XRCC2,MAVS,HTT,HSP90AB1,ZNF124,HSP9
,COPA,HNRNPA1,ARSD,MS4A10,HSPA9,NEDD4L,POLH,ACTB,CTSC,ODF2L,NUDC,NRBP2,USP22,
INRNPA1,ARSD,HSPA9,NEDD4L,POLH,ACTB,CTSC,ODF2L,NUDC,NRBP2,MED18,CFL1,XRCC2,MA
INRNPA1,ARSD,MS4A10,HSPA9,NEDD4L,POLH,ACTB,ODF2L,NUDC,NRBP2,MED18,ACTG1,DPYSL
LH,ACTB,NRBP2,USP22,ACTG1,DPYSL3,TUBA1A,CFL1,BTBD9,XRCC2,MAVS,HTT,HSP90AB1,ZNF
A9,NEDD4L,POLH,ACTB,ODF2L,NRBP2,USP22,ACTG1,DPYSL3,TUBA1A,BTBD9,HTT,HSP90AB1,ZN
A1,NEDD4L,NRBP2,USP22,ACTG1,DPYSL3,TUBA1A,CFL1,BTBD9,XRCC2,HTT,HSP90AB1,ZNF124,
,COPA,HNRNPA1,ARSD,HSPA9,NEDD4L,POLH,ACTB,CTSC,ODF2L,NUDC,NRBP2,USP22,MTRNR2
A1,ARSD,HSPA9,NEDD4L,POLH,ACTB,ODF2L,NUDC,NRBP2,MED18,DPYSL3,CFL1,XRCC2,MAVS,H
JBA1A,CFL1,BTBD9,HTT,HSP90AB1,VIM,FTH1,KIDINS220,NUMA1,RPL3,DBN1,ENO1,FLNA,NCL,F
INRNPA1,ARSD,NEDD4L,POLH,ACTB,CTSC,ODF2L,NRBP2,USP22,MTRNR2L2,ACTG1,EEF2,TUBA
8,TUBA1A,CFL1,BTBD9,XRCC2,MAVS,HSP90AA1,FTH1,NUMA1,RPL3,ENO1,TUBA1B,PIK3R2,FLN
P2,USP22,DPYSL3,BTBD9,HTT,HSP90AB1,ZNF124,ACTR2,VIM,NUMA1,EIF4G2,ZNF316,DBN1,PIK
A1,ARSD,NEDD4L,POLH,ATN1,NRBP2,USP22,ACTG1,CFL1,BTBD9,MAVS,HTT,HSP90AA1,ACTR2,N
AVS,HTT,ZNF124,HSP90AA1,VIM,EIF4G2,RPS6,ZNF316,DBN1,MAP1B,RPL7A,PKM,GANAB,RPL8,F
,DPYSL3,TUBA1A,HTT,FTH1,KIDINS220,EIF4G2,ZNF316,DBN1,TUBA1B,PIK3R2,MAP1B,RPL7A,A
ZNF124,HSP90AA1,DBN1,ENO1,PIK3R2,FLNA,MAP1B,NCL,RPL7A,SLC9A7,PKM,GANAB,HSP90B1,
SD,HSPA9,NEDD4L,POLH,ACTB,CTSC,NRBP2,USP22,ACTG1,DPYSL3,TUBA1A,CFL1,BTBD9,XRCC
CC2,HTT,HSP90AA1,UBC,ZNF316,DBN1,ENO1,MAP1B,SLC9A7,RPL8,FBXL4,YBX1,IGFBP2,FSTL1
TB,NUDC,NRBP2,USP22,ACTG1,EEF2,DPYSL3,TUBA1A,CFL1,XRCC2,MAVS,HTT,ZNF124,HSP90A
NUDC,NRBP2,USP22,MTRNR2L2,ACTG1,DPYSL3,TUBA1A,CFL1,BTBD9,XRCC2,MAVS,HTT,ZNF12
,MS4A10,NEDD4L,POLH,NRBP2,USP22,ACTG1,EEF2,DPYSL3,TUBA1A,CFL1,BTBD9,MAVS,HTT,H
,HNRNPA1,ARSD,NEDD4L,POLH,ATN1,ACTB,ODF2L,NUDC,NRBP2,MTRNR2L2,ACTG1,EEF2,DPY
,COPA,HNRNPA1,ARSD,HSPA9,NEDD4L,POLH,ATN1,ACTB,NUDC,NRBP2,USP22,MTRNR2L2,MED
MED18,ACTG1,DPYSL3,CFL1,MAVS,HTT,HSP90AA1,ACTR2,SPARC,FTH1,TMSB4X,NUMA1,RPS6,Z
JSP22,ACTG1,CFL1,HTT,ZNF124,VIM,TMSB4X,NUMA1,ZNF316,DBN1,FLNA,NCL,SLC9A7,PKM,F
SP22,ACTG1,TUBA1A,CFL1,XRCC2,MAVS,HTT,HSP90AB1,ZNF124,HSP90AA1,FTH1,KIDINS220,T
INRNPA1,ARSD,MS4A10,HSPA9,NEDD4L,ACTB,NRBP2,USP22,ACTG1,EEF2,DPYSL3,TUBA1A,CFL
HNRNPA1,ARSD,HSPA9,NEDD4L,POLH,ACTB,CTSC,ODF2L,NUDC,NRBP2,USP22,MED18,ACTG1,E
MAVS,HTT,HSP90AB1,ZNF124,HSP90AA1,DBN1,NCL,SLC9A7,ALDOA,COL6A1,FTL,FADS1,RPL8,Y
ACTB,NRBP2,USP22,ACTG1,TUBA1A,CFL1,BTBD9,HTT,ZNF124,VIM,TUBB,FTH1,TMSB4X,NUMA1
BTBD9,HTT,ZNF316,DBN1,PIK3R2,FLNA,MAP1B,NCL,SLC9A7,PKM,HSP90B1,FBXL4,YBX1,CALR,
S4A10,NEDD4L,ACTB,NRBP2,USP22,ACTG1,TUBA1A,CFL1,MAVS,HTT,ZNF124,HSP90AA1,ACTR2
IRBP2,USP22,ACTG1,DPYSL3,TUBA1A,HTT,ZNF124,HSP90AA1,FTH1,ZNF316,DBN1,SLC9A7,PKM
BTBD9,ZNF124,ACTR2,VIM,UBC,KIDINS220,NUMA1,EIF4G2,ZNF316,ENO1,TUBA1B,FLNA,NCL,R
H,NRBP2,USP22,ACTG1,TUBA1A,XRCC2,MAVS,HTT,HSP90AB1,ZNF124,HSP90AA1,VIM,SPARC,T
2,USP22,ACTG1,DPYSL3,XRCC2,MAVS,HTT,HSP90AB1,ZNF124,HSP90AA1,VIM,KIDINS220,NUMA
NRNPA1,ARSD,HSPA9,NEDD4L,POLH,ATN1,ACTB,CTSC,ODF2L,NUDC,NRBP2,USP22,MTRNR2L2,

CTB,CTSC,NUDC,NRBP2,USP22,DPYSL3,TUBA1A,CFL1,MAVS,HTT,ZNF124,HSP90AA1,FTH1,KIDI
),HTT,FTH1,UBC,TMSB4X,ZNF316,DBN1,PIK3R2,FLNA,NCL,ALDOA,PKM,COL6A1,FADS1,NEFL,R
VS,ZNF124,ACTR2,UBC,KIDINS220,ZNF316,DBN1,PIK3R2,FLNA,NCL,ALDOA,PKM,GANAB,HSP90
DF2L,NUDC,USP22,MED18,ACTG1,EEF2,DPYSL3,TUBA1A,CFL1,BTBD9,XRCC2,MAVS,HSP90AA1
ACTB,CTSC,ODF2L,NRBP2,USP22,MED18,ACTG1,DPYSL3,TUBA1A,CFL1,BTBD9,XRCC2,MAVS,H
,COPA,HNRNPA1,ARSD,MS4A10,HSPA9,NEDD4L,POLH,ACTB,CTSC,NUDC,NRBP2,USP22,EEF2,T
,POLH,ACTB,CTSC,NRBP2,USP22,ACTG1,DPYSL3,TUBA1A,CFL1,BTBD9,XRCC2,HTT,HSP90AB1,
0,HSPA9,NEDD4L,POLH,NUDC,NRBP2,MED18,DPYSL3,CFL1,XRCC2,MAVS,HTT,HSP90AB1,ZNF1
NRNPA1,ARSD,HSPA9,NEDD4L,POLH,ATN1,ACTB,CTSC,ODF2L,NRBP2,USP22,ACTG1,EEF2,TUB
T1,COPA,HNRNPA1,ARSD,HSPA9,NEDD4L,POLH,ACTB,CTSC,ODF2L,NUDC,NRBP2,USP22,MED18
A,HTT,HSP90AB1,ACTR2,FTH1,UBC,EIF4G2,RPL3,ZNF316,RPL4,NCL,GANAB,HSP90B1,NEFL,PR
),NUMA1,ZNF316,DBN1,TUBA1B,FLNA,NCL,PKM,COL6A1,HSP90B1,NEFM,RPL8,FBXL4,IGFBP2,
0AB1,KIDINS220,NUMA1,DBN1,FLNA,MAP1B,NCL,SLC9A7,PKM,COL6A1,FTL,NEFM,PRPF8,RPL
NEDD4L,ACTB,ODF2L,NUDC,NRBP2,USP22,MTRNR2L2,ACTG1,EEF2,XRCC2,HTT,HSP90AB1,HS
DF2L,NRBP2,USP22,MED18,ACTG1,EEF2,DPYSL3,CFL1,MAVS,HTT,ZNF124,ACTR2,VIM,SPARC,K
,POLH,ACTB,CTSC,ODF2L,NRBP2,USP22,ACTG1,EEF2,DPYSL3,TUBA1A,CFL1,BTBD9,MAVS,HTT,
UBA1A,CFL1,BTBD9,DBN1,ENO1,FLNA,MAP1B,NCL,SLC9A7,COL6A1,FTL,HSP90B1,NEFM,YBX1,
,POLH,USP22,CFL1,HTT,HSP90AB1,VIM,FTH1,DBN1,PKM,COL6A1,FTL,HSP90B1,NEFM,UGT8,Y
A1,ARSD,MS4A10,HSPA9,NEDD4L,POLH,ATN1,ACTB,CTSC,ODF2L,NUDC,NRBP2,USP22,MED18,
D,HSPA9,NEDD4L,POLH,ACTB,CTSC,ODF2L,NRBP2,USP22,MED18,ACTG1,EEF2,DPYSL3,TUBA1
NEDD4L,POLH,ACTB,CTSC,ODF2L,NUDC,NRBP2,USP22,MED18,ACTG1,DPYSL3,CFL1,BTBD9,M
TB,CTSC,NUDC,NRBP2,USP22,ACTG1,TUBA1A,CFL1,BTBD9,XRCC2,HTT,HSP90AB1,ZNF124,HSP
DD4L,POLH,ACTB,NUDC,USP22,EEF2,DPYSL3,CFL1,BTBD9,XRCC2,MAVS,HTT,ZNF124,HSP90AA
,HNRNPA1,ARSD,MS4A10,HSPA9,NEDD4L,POLH,ATN1,ACTB,CTSC,ODF2L,NUDC,NRBP2,USP22,
C2,HTT,HSP90AA1,ACTR2,VIM,EIF4G2,ZNF316,ENO1,FLNA,NCL,RPL7A,SLC9A7,COL6A1,RPL8,F
F2L,NRBP2,USP22,ACTG1,DPYSL3,TUBA1A,CFL1,BTBD9,HTT,HSP90AA1,FTH1,UBC,RPL3,ZNF31
FTH1,TMSB4X,EIF4G2,ZNF316,DBN1,MAP1B,SLC9A7,ALDOA,GANAB,NEFM,PRPF8,RPL8,UBA1,
IS4A10,HSPA9,NEDD4L,POLH,ACTB,CTSC,ODF2L,NUDC,NRBP2,USP22,MED18,ACTG1,EEF2,DPY
3,TUBA1A,XRCC2,HTT,ZNF124,HSP90AA1,VIM,KIDINS220,NUMA1,ZNF316,DBN1,ENO1,PIK3R2,
IS4A10,HSPA9,NEDD4L,ACTB,CTSC,NRBP2,USP22,MTRNR2L2,MED18,ACTG1,EEF2,DPYSL3,TUB
T,ZNF124,VIM,FTH1,NUMA1,DBN1,PIK3R2,SLC9A7,PKM,COL6A1,FTL,HSP90B1,NEFM,UGT8,YB
22,ACTG1,XRCC2,MAVS,HTT,ZNF124,UBC,KIDINS220,NUMA1,RPL3,ZNF316,DBN1,RPL4,MAP1B,
F2L,NRBP2,USP22,MED18,ACTG1,DPYSL3,TUBA1A,BTBD9,XRCC2,MAVS,HTT,HSP90AB1,ZNF12
NRBP2,USP22,ACTG1,TUBA1A,CFL1,BTBD9,HTT,ZNF124,VIM,FTH1,KIDINS220,TMSB4X,NUMA1,
L,POLH,ACTB,CTSC,NUDC,NRBP2,USP22,MED18,ACTG1,EEF2,DPYSL3,TUBA1A,CFL1,MAVS,HT
FL1,HTT,ZNF124,VIM,NUMA1,DBN1,FLNA,NCL,PKM,NEFM,IGFBP2,TUBB2B,PTMA,ENO2,KAT6A,
DINS220,NUMA1,ZNF316,DBN1,ENO1,FLNA,MAP1B,NCL,PKM,COL6A1,RPL8,FBXL4,TUBB2B,PG
IEDD4L,NRBP2,USP22,DPYSL3,TUBA1A,CFL1,BTBD9,XRCC2,HTT,HSP90AB1,VIM,TUBB,FTH1,UB
1,NEDD4L,POLH,NRBP2,USP22,ACTG1,DPYSL3,TUBA1A,BTBD9,HTT,HSP90AB1,ZNF124,HSP90
TB,CTSC,NRBP2,USP22,ACTG1,TUBA1A,XRCC2,MAVS,HTT,ZNF124,VIM,UBC,KIDINS220,NUMA1,
,POLH,ACTB,ODF2L,NUDC,NRBP2,MED18,DPYSL3,TUBA1A,CFL1,BTBD9,XRCC2,MAVS,HTT,HSP
,VIM,DBN1,FLNA,NCL,PKM,NEFM,IGFBP2,PTMA,KAT6A,DVL3,GNAS,SLC2A3,FASN,ALDH6A1,SPT
UDC,NRBP2,USP22,ACTG1,TUBA1A,BTBD9,MAVS,HTT,HSP90AB1,ZNF124,HSP90AA1,ACTR2,UB
UDC,NRBP2,USP22,ACTG1,TUBA1A,BTBD9,MAVS,HTT,HSP90AB1,ZNF124,HSP90AA1,ACTR2,UB
UDC,NRBP2,USP22,ACTG1,TUBA1A,BTBD9,MAVS,HTT,HSP90AB1,ZNF124,HSP90AA1,ACTR2,UB

,ACTB,CTSC,NRBP2,USP22,MED18,DPYSL3,TUBA1A,CFL1,BTBD9,XRCC2,MAVS,ACTR2,VIM,SPA
USP22,ACTG1,EEF2,DPYSL3,TUBA1A,CFL1,BTBD9,XRCC2,MAVS,HTT,ZNF124,HSP90AA1,FTH1,N
NEDD4L,ACTB,CTSC,ODF2L,NRBP2,USP22,MED18,DPYSL3,TUBA1A,CFL1,BTBD9,MAVS,HSP90A
,NRBP2,USP22,TUBA1A,CFL1,BTBD9,HTT,HSP90AB1,ACTR2,VIM,FTH1,DCX,RPL3,DBN1,ENO1,T
L1,HTT,ZNF124,VIM,FTH1,NUMA1,DBN1,FLNA,NCL,PKM,NEFM,IGFBP2,TUBB2B,PTMA,ENO2,K
,COPA,HNRNPA1,ARSD,MS4A10,HSPA9,NEDD4L,POLH,ATN1,ACTB,CTSC,ODF2L,NUDC,NRBP2,U
P2,USP22,ACTG1,DPYSL3,CFL1,XRCC2,MAVS,HTT,HSP90AB1,ZNF124,HSP90AA1,ACTR2,VIM,FT
EDD4L,POLH,NUDC,NRBP2,MED18,DPYSL3,CFL1,XRCC2,MAVS,HTT,VIM,TUBB,FTH1,NUMA1,EI
D4L,ACTB,NRBP2,USP22,ACTG1,EEF2,DPYSL3,CFL1,MAVS,HTT,HSP90AB1,ZNF124,HSP90AA1,
TBD9,XRCC2,HTT,HSP90AB1,ZNF124,KIDINS220,NUMA1,DBN1,ENO1,TUBA1B,FLNA,NCL,PKM,
FBXL4,YBX1,ABHD2,TUBB2B,COL5A1,KAT6A,KIF1A,GNAS,SPTBN1,GNB1,RPS8,RPL7,SEPTIN2,G
SD,HSPA9,POLH,ATN1,ACTB,CTSC,ODF2L,NUDC,USP22,ACTG1,DPYSL3,CFL1,BTBD9,XRCC2,HS
A10,HSPA9,POLH,ACTB,CTSC,NUDC,NRBP2,USP22,MED18,ACTG1,EEF2,DPYSL3,TUBA1A,CFL1,
P90AB1,HSP90AA1,VIM,FTH1,NUMA1,ZNF316,DBN1,ENO1,PIK3R2,NCL,HSP90B1,FBXL4,YBX1,
P2,USP22,ACTG1,DPYSL3,TUBA1A,CFL1,BTBD9,MAVS,HTT,HSP90AA1,VIM,TUBB,FTH1,UBC,TM
EDD4L,POLH,ATN1,ACTB,CTSC,ODF2L,NRBP2,USP22,MED18,ACTG1,EEF2,DPYSL3,CFL1,BTBD
RSD,MS4A10,HSPA9,NEDD4L,POLH,ATN1,ACTB,CTSC,ODF2L,NUDC,NRBP2,USP22,MTRNR2L2,
,POLH,NRBP2,MED18,CFL1,XRCC2,MAVS,HTT,VIM,TUBB,FTH1,UBC,NUMA1,TUBA1B,PIK3R2,N
,HNRNPA1,ARSD,HSPA9,NEDD4L,POLH,ACTB,NUDC,NRBP2,USP22,MTRNR2L2,MED18,ACTG1,E
,ACTG1,TUBA1A,XRCC2,MAVS,HTT,HSP90AB1,ZNF124,VIM,FTH1,UBC,KIDINS220,NUMA1,EIF4G
NF316,DBN1,PIK3R2,FLNA,MAP1B,GANAB,NEFM,YBX1,IGFBP2,PTMA,SEC62,KAT6A,YWHAE,GN
PA1,ARSD,HSPA9,NEDD4L,ACTB,CTSC,ODF2L,USP22,MED18,ACTG1,EEF2,DPYSL3,TUBA1A,CFL
,NEDD4L,ACTB,NUDC,NRBP2,USP22,DPYSL3,TUBA1A,CFL1,BTBD9,MAVS,ZNF124,VIM,NUMA1,D
L1,HTT,ZNF124,HSP90AA1,KIDINS220,NUMA1,ZNF316,DBN1,ENO1,TUBA1B,PIK3R2,MAP1B,SL
TUBA1A,CFL1,BTBD9,HTT,HSP90AB1,VIM,KIDINS220,NUMA1,EIF4G2,ZNF316,DBN1,TUBA1B,PI
NEDD4L,NUDC,NRBP2,USP22,ACTG1,TUBA1A,CFL1,XRCC2,MAVS,HTT,HSP90AB1,ZNF124,HSP9
124,HSP90AA1,DBN1,ENO1,PIK3R2,MAP1B,RPL7A,SLC9A7,PKM,GANAB,FBXL4,YBX1,TUBB2B,
,HNRNPA1,MS4A10,NEDD4L,POLH,ATN1,ACTB,CTSC,ODF2L,NUDC,NRBP2,ACTG1,EEF2,DPYSL3
DF2L,NUDC,NRBP2,USP22,ACTG1,DPYSL3,TUBA1A,CFL1,BTBD9,MAVS,HTT,ZNF124,HSP90AA1
DD4L,ACTB,NRBP2,USP22,ACTG1,EEF2,DPYSL3,HTT,HSP90AB1,HSP90AA1,ACTR2,UBC,NUMA1,
1,XRCC2,VIM,TUBB,UBC,KIDINS220,TMSB4X,NUMA1,EIF4G2,ENO1,TUBA1B,PIK3R2,FLNA,NCL,
UDC,USP22,DPYSL3,CFL1,BTBD9,HTT,HSP90AB1,HSP90AA1,VIM,SPARC,FTH1,UBC,NUMA1,EIF
1,CFL1,MAVS,ZNF124,FTH1,ZNF316,DBN1,ENO1,PIK3R2,MAP1B,RPL7A,SLC9A7,ALDOA,PKM,
PA9,NEDD4L,POLH,ACTB,CTSC,ODF2L,USP22,MTRNR2L2,MED18,DPYSL3,TUBA1A,CFL1,BTBD
,ARSD,NEDD4L,NRBP2,USP22,ACTG1,TUBA1A,CFL1,XRCC2,HTT,HSP90AB1,VIM,TUBB,FTH1,UB
DH,MLLT1,COPA,HNRNPA1,ARSD,MS4A10,HSPA9,NEDD4L,POLH,ATN1,ACTB,CTSC,ODF2L,NUD
R2,TUBB,KIDINS220,EIF4G2,RPS6,ZNF316,PIK3R2,NCL,SLC9A7,COL6A1,NEFM,RPL8,PTMA,RP
T,HSP90AA1,VIM,DBN1,ENO1,PIK3R2,NCL,HSP90B1,FBXL4,YBX1,TUBB2B,PTMA,KAT6A,YWHAE
AA1,VIM,NUMA1,ZNF316,DBN1,ENO1,PIK3R2,NCL,HSP90B1,FBXL4,YBX1,CALR,TUBB2B,PTMA,
,COPA,HNRNPA1,ARSD,MS4A10,HSPA9,NEDD4L,POLH,ATN1,ACTB,CTSC,NUDC,NRBP2,USP22,M
1,ARSD,NEDD4L,POLH,ACTB,CTSC,ODF2L,NRBP2,XRCC2,MAVS,HTT,HSP90AB1,VIM,TUBB,FTH1
NRBP2,USP22,MTRNR2L2,ACTG1,EEF2,DPYSL3,TUBA1A,XRCC2,HSP90AB1,HSP90AA1,VIM,TUB
22,MED18,DPYSL3,BTBD9,XRCC2,VIM,UBC,KIDINS220,TMSB4X,NUMA1,EIF4G2,DBN1,RPL4,E
22,ACTG1,TUBA1A,CFL1,XRCC2,HTT,ZNF124,HSP90AA1,ACTR2,FTH1,UBC,KIDINS220,NUMA1,D
,ARSD,NEDD4L,ACTB,CTSC,ODF2L,NUDC,NRBP2,ACTG1,EEF2,TUBA1A,CFL1,BTBD9,MAVS,HTT,

DF2L,NUDC,NRBP2,USP22,ACTG1,EEF2,DPYSL3,BTBD9,XRCC2,HTT,ACTR2,VIM,TUBB,FTH1,KID
,ACTB,ODF2L,NUDC,NRBP2,USP22,ACTG1,EEF2,DPYSL3,BTBD9,XRCC2,HTT,ACTR2,VIM,SPARC,
TUBA1A,CFL1,HTT,HSP90AB1,ZNF124,HSP90AA1,ACTR2,UBC,NUMA1,EIF4G2,ZNF316,DBN1,ENO
G1,TUBA1A,CFL1,XRCC2,MAVS,HTT,HSP90AB1,ZNF124,VIM,KIDINS220,TMSB4X,NUMA1,ZNF316
D18,ACTG1,EEF2,DPYSL3,TUBA1A,BTBD9,XRCC2,HTT,ZNF124,HSP90AA1,TUBB,FTH1,UBC,KIDI
FL1,XRCC2,HTT,HSP90AB1,TUBB,FTH1,NUMA1,ZNF316,DBN1,TUBA1B,RPL7A,PKM,GANAB,COL
P22,MED18,CFL1,BTBD9,MAVS,HTT,HSP90AA1,ACTR2,FTH1,UBC,KIDINS220,TMSB4X,NUMA1,E
A1A,CFL1,XRCC2,MAVS,HTT,HSP90AB1,ZNF124,FTH1,KIDINS220,TMSB4X,NUMA1,RPL3,ZNF316
BTBD9,XRCC2,HTT,HSP90AB1,ZNF124,KIDINS220,NUMA1,DBN1,ENO1,FLNA,NCL,GANAB,COL6A
BTBD9,XRCC2,HTT,HSP90AB1,ZNF124,KIDINS220,NUMA1,DBN1,ENO1,FLNA,NCL,GANAB,COL6A
P90B1,NEFM,YBX1,PTMA,ENO2,KAT6A,KIF1A,GNAS,SEPTIN2,PLXNB2,EIF4B,MARCKSL1,CKB,HA
EEF2,CFL1,BTBD9,XRCC2,HTT,HSP90AA1,TUBB,ZNF316,ENO1,TUBA1B,NCL,RPL8,TMF1,TUBB2
T1,COPA,HNRNPA1,ARSD,HSPA9,NEDD4L,POLH,ACTB,CTSC,ODF2L,NUDC,NRBP2,USP22,MTRNI
NRBP2,USP22,MTRNR2L2,MED18,ACTG1,DPYSL3,CFL1,BTBD9,XRCC2,MAVS,HTT,HSP90AB1,HS
BTBD9,XRCC2,HTT,ZNF124,NUMA1,DBN1,FLNA,MAP1B,NCL,SLC9A7,PKM,GANAB,FTL,HSP90B1
6,DBN1,FLNA,MAP1B,NCL,PKM,COL6A1,RPL8,FBXL4,ABHD2,TUBB2B,DDX5,ENO2,KAT6A,DVL3,
P90AB1,ZNF124,KIDINS220,ZNF316,DBN1,ENO1,TUBA1B,FLNA,MAP1B,PKM,COL6A1,FBXL4,ABI
ZNF316,DBN1,ENO1,PIK3R2,FLNA,SLC9A7,PKM,GANAB,HSP90B1,FADS1,PRPF8,FBXL4,IGFBP2,
L,POLH,ATN1,ACTB,CTSC,ODF2L,NUDC,MED18,ACTG1,DPYSL3,CFL1,BTBD9,MAVS,HSP90AA1,A
ACTB,CTSC,ODF2L,NUDC,MED18,ACTG1,DPYSL3,CFL1,MAVS,HSP90AB1,HSP90AA1,ACTR2,VIM,
OLH,ATN1,ACTB,CTSC,NUDC,NRBP2,USP22,MED18,ACTG1,EEF2,DPYSL3,TUBA1A,BTBD9,XRCC2
,HNRNPA1,HSPA9,NEDD4L,POLH,ODF2L,NRBP2,USP22,ACTG1,EEF2,DPYSL3,CFL1,BTBD9,XRC
SPA9,NEDD4L,POLH,ACTB,CTSC,ODF2L,USP22,MTRNR2L2,MED18,EEF2,DPYSL3,TUBA1A,CFL1,
ZNF124,NUMA1,EIF4G2,ZNF316,DBN1,PKM,HSP90B1,NEFM,UGT8,FBXL4,ATP2B4,FSTL1,ENO2,
P2,DPYSL3,TUBA1A,HTT,HSP90AB1,ZNF124,HSP90AA1,ACTR2,VIM,UBC,NUMA1,DCX,ZNF316,DB
ODF2L,NRBP2,USP22,ACTG1,TUBA1A,CFL1,XRCC2,HTT,HSP90AB1,ZNF124,VIM,FTH1,UBC,KIDIN
NEDD4L,ATN1,ACTB,CTSC,ODF2L,NRBP2,USP22,ACTG1,EEF2,DPYSL3,TUBA1A,CFL1,BTBD9,XR
TSC,NRBP2,USP22,MED18,DPYSL3,TUBA1A,ACTR2,VIM,UBC,KIDINS220,EIF4G2,RPL3,RPL4,ENO
TH1,NUMA1,ZNF316,DBN1,PIK3R2,NCL,HSP90B1,TUBB2B,COL5A1,H3-3B,PTMA,SOX4,KAT6A,G
LH,NUDC,NRBP2,CFL1,XRCC2,HTT,HSP90AB1,VIM,TUBB,FTH1,NUMA1,ZNF316,DBN1,ENO1,TUE
,HNRNPA1,ARSD,MS4A10,HSPA9,NEDD4L,POLH,ACTB,ODF2L,NUDC,NRBP2,USP22,EEF2,DPYSL
4,KIDINS220,NUMA1,ZNF316,DBN1,TUBA1B,FLNA,PKM,COL6A1,FBXL4,PGK1,DDX5,KAT6A,DVL3
S,HTT,ZNF124,KIDINS220,NUMA1,RPL3,ZNF316,RPL4,ENO1,TUBA1B,PIK3R2,NCL,PKM,GANAB,
2,HSP90AA1,VIM,NUMA1,ZNF316,NCL,RPL7A,SLC9A7,COL6A1,HSP90B1,NEFM,RPL8,FBXL4,YBX
DPYSL3,CFL1,MAVS,HTT,KIDINS220,ZNF316,DBN1,TUBA1B,PIK3R2,FLNA,SLC9A7,ALDOA,PKM,
RBP2,MTRNR2L2,MED18,EEF2,DPYSL3,MAVS,HTT,HSP90AB1,ACTR2,FTH1,UBC,KIDINS220,TMS
SP22,TUBA1A,MAVS,HTT,HSP90AB1,HSP90AA1,UBC,EIF4G2,ZNF316,DBN1,ENO1,TUBA1B,FLNA
A1,POLH,NRBP2,DPYSL3,BTBD9,HTT,HSP90AA1,VIM,NUMA1,ZNF316,DBN1,MAP1B,RPL7A,FADS
HNRNPA1,ARSD,MS4A10,NEDD4L,POLH,ATN1,ACTB,ODF2L,NUDC,NRBP2,USP22,MTRNR2L2,MEI
DC,NRBP2,USP22,MED18,ACTG1,DPYSL3,CFL1,BTBD9,MAVS,HTT,HSP90AA1,ACTR2,VIM,TUBB,F
TSC,ODF2L,NRBP2,USP22,MTRNR2L2,ACTG1,TUBA1A,CFL1,BTBD9,XRCC2,MAVS,HTT,HSP90AE
TN1,ACTB,ODF2L,USP22,EEF2,TUBA1A,BTBD9,XRCC2,HTT,HSP90AB1,ZNF124,HSP90AA1,ACTR
,MAP1B,RPL7A,SLC9A7,ALDOA,PKM,COL6A1,YBX1,COL5A1,PTMA,KAT6A,GNAS,HTRA4,BSG,FA
KIDINS220,ZNF316,RPL4,TUBA1B,PKM,COL6A1,FADS1,NEFM,RPL8,FBXL4,DDX5,FKBP14,KAT6A
4L,POLH,NRBP2,USP22,ACTG1,DPYSL3,CFL1,BTBD9,XRCC2,MAVS,HTT,ZNF124,HSP90AA1,ACT

0,HSPA9,ATN1,ACTB,CTSC,NRBP2,USP22,ACTG1,DPYSL3,TUBA1A,CFL1,MAVS,HTT,HSP90AB1,H
ISPA9,NEDD4L,POLH,ATN1,ACTB,CTSC,ODF2L,NUDC,USP22,MED18,ACTG1,EEF2,DPYSL3,TUBA
3P2,USP22,MED18,ACTG1,CFL1,BTBD9,HSP90AA1,ACTR2,FTH1,UBC,NUMA1,RPL3,ZNF316,TUB
LH,ATN1,ACTB,CTSC,NRBP2,USP22,MED18,ACTG1,EEF2,DPYSL3,TUBA1A,CFL1,BTBD9,XRCC2,N
DF2L,NRBP2,USP22,ACTG1,TUBA1A,CFL1,BTBD9,XRCC2,MAVS,HTT,HSP90AB1,ZNF124,VIM,TU
,MAVS,HSP90AB1,ZNF124,FTH1,KIDINS220,NUMA1,DBN1,ENO1,FLNA,NCL,SLC9A7,PKM,GANAB
H1,NUMA1,ZNF316,DBN1,ENO1,PIK3R2,HSP90B1,FBXL4,YBX1,CALR,PTMA,SEC62,YWHAE,GNAS
,EEF2,BTBD9,HTT,HSP90AB1,ACTR2,FTH1,UBC,ZNF316,PIK3R2,FLNA,MAP1B,ALDOA,COL6A1,F
SPA9,NEDD4L,POLH,ACTB,CTSC,NUDC,NRBP2,USP22,MED18,ACTG1,EEF2,DPYSL3,TUBA1A,BTE
P22,ACTG1,CFL1,MAVS,HTT,HSP90AA1,VIM,FTH1,UBC,TMSB4X,RPL3,ZNF316,DBN1,ENO1,PIK3
L3,TUBA1A,BTBD9,VIM,UBC,KIDINS220,NUMA1,EIF4G2,RPL3,DBN1,RPL4,ENO1,TUBA1B,PIK3R
JBA1A,CFL1,MAVS,ZNF124,ACTR2,VIM,FTH1,TMSB4X,ZNF316,DBN1,ENO1,PIK3R2,MAP1B,RPL7
,XRCC2,TUBB,UBC,KIDINS220,TMSB4X,NUMA1,EIF4G2,TUBA1B,PIK3R2,FLNA,NCL,RPL7A,SLC
TSC,NRBP2,USP22,ACTG1,TUBA1A,CFL1,BTBD9,XRCC2,MAVS,HTT,HSP90AB1,ZNF124,VIM,FTH
ISPA9,NEDD4L,POLH,ATN1,ACTB,CTSC,ODF2L,NUDC,USP22,MED18,ACTG1,EEF2,DPYSL3,TUBA
A1,UBC,EIF4G2,ZNF316,DBN1,ENO1,FLNA,NCL,FADS1,NEFL,YBX1,COL5A1,COL4A1,RPLP0,H3-3
JDC,NRBP2,USP22,MED18,EEF2,DPYSL3,CFL1,XRCC2,MAVS,HSP90AA1,ACTR2,VIM,SPARC,UBC,
YSL3,CFL1,BTBD9,XRCC2,HTT,TUBB,FTH1,UBC,KIDINS220,TMSB4X,NUMA1,EIF4G2,DBN1,TUB
RSD,NEDD4L,ACTB,CTSC,ODF2L,NUDC,NRBP2,USP22,ACTG1,EEF2,TUBA1A,CFL1,XRCC2,HTT,H
P22,MTRNR2L2,MED18,ACTG1,EEF2,TUBA1A,BTBD9,MAVS,HTT,HSP90AB1,HSP90AA1,VIM,FTH1
0AB1,HSP90AA1,TUBB,UBC,NUMA1,ENO1,TUBA1B,NCL,RPL7A,HSP90B1,FADS1,NEFL,NEFM,RP
MS4A10,HSPA9,NEDD4L,POLH,ATN1,ACTB,CTSC,ODF2L,NUDC,NRBP2,USP22,MED18,ACTG1,DP
RCC2,HTT,HSP90AB1,ZNF124,KIDINS220,NUMA1,DBN1,ENO1,FLNA,NCL,GANAB,COL6A1,FBXL4
LLT1,COPA,HSPA9,NEDD4L,POLH,ATN1,ACTB,CTSC,ODF2L,NUDC,NRBP2,USP22,ACTG1,EEF2,D
90AB1,ACTR2,FTH1,ZNF316,FLNA,MAP1B,ALDOA,FTL,NEFL,NEFM,PRPF8,YBX1,IGFBP2,ABHD2,
SP22,EEF2,TUBA1A,CFL1,BTBD9,XRCC2,HSP90AB1,ZNF124,HSP90AA1,ACTR2,VIM,SPARC,TUBE
SP22,EEF2,TUBA1A,CFL1,BTBD9,XRCC2,HSP90AB1,ZNF124,HSP90AA1,ACTR2,VIM,SPARC,TUBE
2L,NUDC,MED18,ACTG1,CFL1,BTBD9,MAVS,ZNF124,HSP90AA1,ACTR2,SPARC,FTH1,KIDINS220
S220,NUMA1,ZNF316,DBN1,ENO1,TUBA1B,PKM,COL6A1,FADS1,NEFM,PRPF8,FBXL4,YBX1,CAL
TG1,CFL1,VIM,KIDINS220,EIF4G2,ZNF316,DBN1,NCL,SLC9A7,COL6A1,FTL,NEFM,RPL8,YBX1,I
BTBD9,HTT,ZNF316,DBN1,MAP1B,SLC9A7,COL6A1,NEFM,FBXL4,UBA1,ATP2B4,FSTL1,COL5A1,
A1,ARSD,HSPA9,NEDD4L,POLH,ATN1,ACTB,NUDC,NRBP2,USP22,MTRNR2L2,MED18,ACTG1,EE
SC,ODF2L,NRBP2,USP22,EEF2,DPYSL3,TUBA1A,CFL1,BTBD9,HTT,HSP90AA1,VIM,UBC,KIDINS22
NRBP2,USP22,ACTG1,CFL1,VIM,UBC,KIDINS220,EIF4G2,ZNF316,DBN1,NCL,SLC9A7,ALDOA,COL
,ACTG1,EEF2,TUBA1A,XRCC2,MAVS,HTT,FTH1,UBC,NUMA1,EIF4G2,RPL3,RPS6,PIK3R2,NCL,SL
,ODF2L,NUDC,NRBP2,USP22,CFL1,XRCC2,HTT,HSP90AA1,ACTR2,FTH1,UBC,TMSB4X,EIF4G2,Z
CFL1,BTBD9,XRCC2,HSP90AA1,TUBB,NUMA1,RPL3,ENO1,TUBA1B,ALDOA,HSP90B1,NEFL,NEFM
,ARSD,NEDD4L,NRBP2,USP22,ACTG1,TUBA1A,CFL1,XRCC2,HTT,HSP90AB1,ZNF124,HSP90AA1,
CFL1,XRCC2,HTT,HSP90AB1,ZNF124,VIM,KIDINS220,DBN1,COL6A1,HSP90B1,RPL8,FBXL4,TUBE
,HSPA9,NEDD4L,ATN1,ACTB,CTSC,ODF2L,NUDC,NRBP2,USP22,ACTG1,EEF2,TUBA1A,CFL1,BTB
CTB,CTSC,NRBP2,USP22,MED18,ACTG1,EEF2,DPYSL3,TUBA1A,CFL1,MAVS,HTT,ZNF124,HSP90
,CFL1,HSP90AA1,VIM,TUBB,UBC,KIDINS220,NUMA1,TUBA1B,FLNA,NCL,PKM,GANAB,COL6A1,H
KIDINS220,EIF4G2,RPL3,RPL4,NCL,NEFM,YBX1,IGFBP2,DDX5,RPL13A,DVL3,YWHAE,GNAS,DDX1
USP22,DPYSL3,TUBA1A,CFL1,BTBD9,XRCC2,MAVS,HTT,HSP90AB1,HSP90AA1,ACTR2,VIM,UBC,N
4A10,NEDD4L,ATN1,ACTB,CTSC,NRBP2,USP22,ACTG1,DPYSL3,TUBA1A,CFL1,BTBD9,MAVS,HTT

PA9,NEDD4L,POLH,ATN1,ACTB,CTSC,ODF2L,NUDC,USP22,MED18,ACTG1,EEF2,DPYSL3,CFL1,BT
,HSP90AA1,ACTR2,ZNF316,DBN1,FLNA,MAP1B,ALDOA,PKM,RPL8,FBXL4,IGFBP2,FSTL1,TMF1,C
L3,TUBA1A,CFL1,BTBD9,MAVS,HTT,HSP90AA1,FTH1,KIDINS220,EIF4G2,DBN1,ENO1,PIK3R2,FL
TUBB,ENO1,TUBA1B,RPL8,TMF1,PSAP,RPL13A,BSG,RPL5,TMBIM6,PLD3,HSPD1,EIF4B,UBB,TRI
JSP22,MED18,ACTG1,EEF2,DPYSL3,TUBA1A,CFL1,BTBD9,HSP90AB1,HSP90AA1,VIM,FTH1,UBC,I
A1,ARSD,HSPA9,NEDD4L,POLH,ATN1,ACTB,CTSC,ODF2L,NUDC,ACTG1,DPYSL3,TUBA1A,CFL1,B
C,MED18,ACTG1,CFL1,BTBD9,ZNF124,HSP90AA1,ACTR2,SPARC,FTH1,KIDINS220,TMSB4X,NUM
DF2L,NUDC,USP22,MED18,ACTG1,EEF2,TUBA1A,CFL1,BTBD9,XRCC2,HSP90AB1,ZNF124,HSP9
1A1,DBN1,SLC9A7,HSP90B1,ENO2,YWHAE,KIF1A,GNAS,SPTBN1,CALM3,GNB1,SHISA9,MYH9,SP
ACTB,CTSC,ODF2L,NRBP2,USP22,EEF2,DPYSL3,TUBA1A,CFL1,MAVS,HTT,ZNF124,HSP90AA1,A
RBP2,USP22,ACTG1,DPYSL3,TUBA1A,CFL1,BTBD9,MAVS,HTT,HSP90AB1,HSP90AA1,ACTR2,VIM,
,VIM,FTH1,NUMA1,DBN1,FLNA,MAP1B,SLC9A7,HSP90B1,YBX1,KAT6A,DVL3,YWHAE,GNAS,FASN
ZNF124,KIDINS220,NUMA1,ZNF316,RPL4,NCL,PKM,GANAB,COL6A1,NEFM,RPL8,FBXL4,TUBB2
A1,ARSD,HSPA9,NEDD4L,POLH,ATN1,ACTB,CTSC,ODF2L,NUDC,ACTG1,DPYSL3,CFL1,BTBD9,XR
HSPA9,NEDD4L,POLH,ATN1,ACTB,CTSC,ODF2L,NUDC,USP22,MED18,ACTG1,DPYSL3,TUBA1A,C
NEDD4L,ACTB,CTSC,ODF2L,USP22,MED18,DPYSL3,CFL1,BTBD9,MAVS,ACTR2,VIM,UBC,KIDINS2
HTT,UBC,EIF4G2,ZNF316,DBN1,FLNA,FADS1,PRPF8,FSTL1,TUBB2B,COL5A1,COL4A1,H3-3B,CL
YSL3,TUBA1A,CFL1,MAVS,HTT,ZNF124,HSP90AA1,FTH1,KIDINS220,EIF4G2,ZNF316,DBN1,ENO1
FL,RPL8,IGFBP2,HSPA1B,ENO2,DVL3,GPI,SPTBN1,GNB1,GDI1,PTMS,SPEN,EIF1,CHSY1,TMSB10
ODF2L,NUDC,MED18,ACTG1,EEF2,CFL1,BTBD9,MAVS,ZNF124,HSP90AA1,ACTR2,UBC,KIDINS22
BD9,XRCC2,HSP90AA1,TUBB,ENO1,TUBA1B,RPL8,TMF1,PSAP,RPL13A,GPI,BSG,P4HB,RPL5,TME
,NRBP2,USP22,DPYSL3,CFL1,HTT,VIM,ZNF316,DBN1,ENO1,MAP1B,SLC9A7,ALDOA,COL6A1,NEF
C,USP22,MED18,ACTG1,EEF2,DPYSL3,TUBA1A,BTBD9,XRCC2,MAVS,HSP90AB1,ZNF124,HSP90A
,COPA,HNRNPA1,ARSD,MS4A10,HSPA9,NEDD4L,POLH,ATN1,ACTB,CTSC,ODF2L,NUDC,NRBP2,U
D4L,CTSC,ODF2L,NRBP2,USP22,MED18,ACTG1,EEF2,TUBA1A,CFL1,HTT,HSP90AB1,ZNF124,HSP
EDD4L,POLH,NRBP2,DPYSL3,BTBD9,HTT,HSP90AA1,VIM,NUMA1,ZNF316,DBN1,FLNA,NCL,RPL7A
CTB,NUDC,USP22,CFL1,BTBD9,MAVS,HSP90AA1,VIM,ZNF316,DBN1,TUBA1B,MAP1B,SLC9A7,PK
ED18,ACTG1,CFL1,BTBD9,MAVS,HTT,HSP90AA1,ACTR2,FTH1,UBC,KIDINS220,NUMA1,RPL3,RPS
NS220,TMSB4X,NUMA1,ZNF316,DBN1,ENO1,TUBA1B,MAP1B,PKM,COL6A1,FBXL4,UBA1,ABHD2
RCC2,MAVS,HTT,HSP90AB1,KIDINS220,NUMA1,EIF4G2,ZNF316,DBN1,TUBA1B,MAP1B,COL6A1,H
,POLH,ATN1,ACTB,CTSC,NUDC,NRBP2,DPYSL3,TUBA1A,CFL1,BTBD9,ZNF124,HSP90AA1,ACTR2
B,CTSC,ODF2L,NUDC,USP22,MED18,ACTG1,DPYSL3,CFL1,BTBD9,MAVS,HSP90AB1,ZNF124,HSP
2,USP22,MED18,ACTG1,DPYSL3,TUBA1A,CFL1,MAVS,HTT,HSP90AB1,ZNF124,ACTR2,KIDINS220
0,HSPA9,NEDD4L,POLH,ATN1,ACTB,CTSC,ODF2L,NUDC,NRBP2,USP22,MED18,ACTG1,DPYSL3,T
D9,MAVS,HTT,HSP90AA1,FTH1,KIDINS220,EIF4G2,DBN1,ENO1,PIK3R2,MAP1B,NCL,RPL7A,SLC9
NPA1,ARSD,MS4A10,HSPA9,NEDD4L,POLH,ACTB,NUDC,USP22,EEF2,TUBA1A,CFL1,BTBD9,XRCC
1S4A10,HSPA9,NEDD4L,POLH,ACTB,CTSC,ODF2L,NUDC,NRBP2,USP22,MED18,ACTG1,EEF2,DPY
3,ODF2L,NRBP2,ACTG1,EEF2,DPYSL3,TUBA1A,BTBD9,MAVS,HTT,HSP90AB1,ZNF124,ACTR2,TUE
BN1,ALDOA,YBX1,ATP2B4,TUBB2B,YWHAE,GNAS,GPI,GDI1,FSCN1,PTMS,GSTP1,CALM1,PLXNB2
,COPA,HNRNPA1,ARSD,MS4A10,HSPA9,NEDD4L,POLH,ATN1,ACTB,CTSC,ODF2L,NUDC,NRBP2,U
4L,POLH,ACTB,ODF2L,NUDC,USP22,MED18,ACTG1,EEF2,TUBA1A,BTBD9,XRCC2,MAVS,HTT,HSI
,ACTG1,DPYSL3,TUBA1A,XRCC2,MAVS,HTT,HSP90AB1,HSP90AA1,UBC,KIDINS220,EIF4G2,RPL3
/S,HTT,HSP90AB1,ZNF124,FTH1,EIF4G2,RPL3,ZNF316,ENO1,TUBA1B,PIK3R2,NCL,ALDOA,PKM,
IK3R2,FLNA,SLC9A7,PKM,COL6A1,FTL,HSP90B1,FBXL4,UBA1,FSTL1,HSPA1B,COL4A1,PGK1,PT
CTG1,TUBA1A,CFL1,HTT,HSP90AA1,FTH1,KIDINS220,NUMA1,ENO1,PIK3R2,FLNA,NCL,COL6A1,

LNA,PKM,RPL8,FBXL4,ABHD2,TUBB2B,PGK1,CANX,DDX5,KAT6A,FASN,ALDH6A1,GNB1,GDI1,CA
K3R2,FLNA,PKM,COL6A1,FBXL4,PTMA,BSG,RPL7,GDI1,MARCKS,CALM1,USP11,IGF2,EIF4B,TRI
16,RPL4,TUBA1B,NCL,RPL7A,ALDOA,FADS1,RPL8,YBX1,CALR,TMF1,TUBB2B,COL5A1,H3-3B,PT
D,HSPA9,NEDD4L,POLH,ACTB,NUDC,NRBP2,USP22,MTRNR2L2,MED18,ACTG1,EEF2,CFL1,BTBD
RSD,MS4A10,HSPA9,NEDD4L,POLH,ACTB,CTSC,ODF2L,NUDC,USP22,MED18,ACTG1,DPYSL3,CF
3N1,FLNA,MAP1B,RPL8,ABHD2,TUBB2B,BSG,ALDH6A1,STMN2,RPL7,MARCKS,NPM1,PLXNB2,US
P90AB1,FTH1,KIDINS220,NUMA1,NCL,PKM,COL6A1,RPL8,FBXL4,UBA1,ABHD2,DDX5,ENO2,FKE
ACTB,CTSC,ODF2L,NUDC,USP22,MED18,ACTG1,DPYSL3,CFL1,MAVS,HSP90AB1,ZNF124,HSP90
,UBC,TMSB4X,DBN1,TUBA1B,MAP1B,SLC9A7,PKM,GANAB,FTL,YBX1,HSPA1B,COL4A1,DDX5,KA
,UBC,TMSB4X,DBN1,TUBA1B,MAP1B,SLC9A7,PKM,GANAB,FTL,YBX1,HSPA1B,COL4A1,DDX5,KA
,UBC,TMSB4X,DBN1,TUBA1B,MAP1B,SLC9A7,PKM,GANAB,FTL,YBX1,HSPA1B,COL4A1,DDX5,KA
SC,NUDC,NRBP2,USP22,CFL1,XRCC2,ZNF124,HSP90AA1,ACTR2,VIM,FTH1,NUMA1,RPL3,ZNF31
ZNF316,DBN1,PIK3R2,RPL7A,YBX1,FSTL1,PTMA,DDX5,RPL13A,KAT6A,KIF1A,RACK1,FASN,CALM
RSD,NEDD4L,POLH,ACTB,CTSC,ODF2L,NRBP2,USP22,MTRNR2L2,ACTG1,DPYSL3,TUBA1A,BTB
RSD,MS4A10,HSPA9,NEDD4L,POLH,ACTB,CTSC,ODF2L,USP22,MED18,ACTG1,EEF2,DPYSL3,CFL
LH,ATN1,ACTB,CTSC,ODF2L,NUDC,USP22,MED18,ACTG1,EEF2,DPYSL3,CFL1,BTBD9,MAVS,HSP
T1,COPA,HNRNPA1,ARSD,HSPA9,NEDD4L,POLH,ACTB,CTSC,NUDC,USP22,MED18,ACTG1,EEF2,
A1,FBXL4,KAT6A,GPI,BSG,SPTBN1,RPL7,P4HB,GDI1,USP11,IGF2,TRIM28,LRRC28,YWHAQ,DCH
NRBP2,MED18,DPYSL3,BTBD9,MAVS,VIM,TUBB,UBC,KIDINS220,TMSB4X,NUMA1,EIF4G2,ZNF31
,HTT,ZNF124,UBC,ZNF316,DBN1,ENO1,PIK3R2,MAP1B,PKM,COL6A1,FTL,NEFM,ATP2B4,CALR,
G1,EEF2,CFL1,HTT,ZNF124,ACTR2,VIM,UBC,TMSB4X,ZNF316,DBN1,ENO1,TUBA1B,MAP1B,NCL
UBC,KIDINS220,NUMA1,EIF4G2,RPL3,RPL4,ENO1,TUBA1B,PIK3R2,FLNA,NCL,RPL7A,ALDOA,PK
FL1,HTT,VIM,UBC,KIDINS220,TMSB4X,ZNF316,DBN1,ENO1,TUBA1B,PIK3R2,MAP1B,RPL7A,PKM
RSD,MS4A10,HSPA9,NEDD4L,POLH,ACTB,CTSC,ODF2L,USP22,MED18,ACTG1,DPYSL3,CFL1,BTB
124,HSP90AA1,DBN1,MAP1B,NCL,YBX1,COL5A1,PTMA,YWHAE,GNAS,SPTBN1,GNB1,SHISA9,PT
D,NEDD4L,POLH,ACTB,NUDC,ACTG1,EEF2,DPYSL3,CFL1,BTBD9,XRCC2,MAVS,HTT,HSP90AB1,Z
FL1,XRCC2,HTT,HSP90AA1,ACTR2,TUBB,FTH1,UBC,NUMA1,RPS6,ZNF316,DBN1,TUBA1B,FLNA,
USP22,ACTG1,CFL1,BTBD9,HTT,HSP90AB1,ZNF124,NUMA1,EIF4G2,ZNF316,MAP1B,NCL,PKM,G
D18,DPYSL3,BTBD9,VIM,UBC,KIDINS220,NUMA1,EIF4G2,RPL3,RPL4,TUBA1B,PIK3R2,FLNA,NCL
DD4L,POLH,ACTB,ODF2L,NUDC,USP22,MED18,ACTG1,DPYSL3,CFL1,XRCC2,MAVS,HSP90AB1,Z
PIK3R2,MAP1B,RPL7A,PKM,HSP90B1,FBXL4,TUBB2B,COL5A1,PTMA,KAT6A,GNAS,BSG,FASN,SP
G1,MAVS,HSP90AB1,ZNF124,HSP90AA1,ACTR2,KIDINS220,RPL3,RPS6,ZNF316,DBN1,RPL4,PIK3
D,NEDD4L,POLH,NUDC,NRBP2,USP22,DPYSL3,TUBA1A,CFL1,BTBD9,MAVS,SPARC,TUBB,UBC,Z
ED18,DPYSL3,BTBD9,MAVS,VIM,TUBB,UBC,KIDINS220,NUMA1,EIF4G2,ZNF316,RPL4,ENO1,TUB
,KIDINS220,TMSB4X,EIF4G2,RPL3,RPL4,GANAB,DDX5,RPL13A,KAT6A,YWHAE,DDX17,RACK1,AL
9,NEDD4L,POLH,ODF2L,MTRNR2L2,MED18,DPYSL3,TUBA1A,CFL1,BTBD9,XRCC2,HTT,HSP90A
LH,ACTB,ODF2L,NUDC,USP22,MED18,ACTG1,CFL1,BTBD9,XRCC2,MAVS,HSP90AB1,ZNF124,HS
HSPA9,NEDD4L,POLH,ACTB,ODF2L,NUDC,USP22,MED18,ACTG1,EEF2,DPYSL3,TUBA1A,CFL1,BT
IM,SPARC,FTH1,UBC,NUMA1,RPL3,ALDOA,NEFL,COL5A1,COL4A1,RPLP0,CLU,PTMA,GNAS,HTR
RSD,HSPA9,NEDD4L,ACTB,CTSC,ODF2L,USP22,MED18,ACTG1,DPYSL3,CFL1,BTBD9,XRCC2,MA
9,NEDD4L,POLH,ATN1,ACTB,CTSC,ODF2L,NUDC,ACTG1,DPYSL3,TUBA1A,CFL1,BTBD9,XRCC2,H
TBD9,XRCC2,HSP90AB1,HSP90AA1,TUBB,FTH1,UBC,KIDINS220,RPL3,DBN1,ENO1,TUBA1B,FLN
RSD,MS4A10,HSPA9,NEDD4L,POLH,ACTB,CTSC,ODF2L,USP22,MED18,ACTG1,EEF2,DPYSL3,CFL
L3,BTBD9,VIM,UBC,KIDINS220,NUMA1,EIF4G2,TUBA1B,FLNA,NCL,RPL7A,ALDOA,PKM,NEFL,RP
SD,NEDD4L,ACTB,USP22,EEF2,DPYSL3,CFL1,BTBD9,MAVS,HTT,HSP90AB1,ACTR2,VIM,TUBB,FT

P90AB1,ZNF124,HSP90AA1,ACTR2,UBC,RPL3,RPS6,ZNF316,DBN1,PIK3R2,NCL,GANAB,FADS1,YB
L3,SPARC,ZNF316,DBN1,MAP1B,COL6A1,NEFM,UBA1,FSTL1,COL5A1,CES3,PTMA,ENO2,YWHAE,
MAVS,ACTR2,TUBA1B,PIK3R2,FLNA,NCL,SLC9A7,ALDOA,PKM,NEFM,RPL8,FBXL4,TUBB2B,COL
PA9,NEDD4L,POLH,ACTB,ODF2L,NUDC,USP22,MED18,ACTG1,DPYSL3,CFL1,XRCC2,MAVS,HSP9
3,CTSC,ODF2L,NUDC,USP22,MED18,ACTG1,EEF2,DPYSL3,CFL1,BTBD9,MAVS,HSP90AB1,ZNF12
SC,NUDC,NRBP2,USP22,CFL1,ZNF124,VIM,FTH1,RPL3,ZNF316,DBN1,ENO1,TUBA1B,PIK3R2,FL
L4,YBX1,COL4A1,ENO2,GNAS,SPTBN1,SHISA9,MYH9,P4HB,IDS,PTMS,CALM1,USP11,IGF2,TRIM
B,NUDC,NRBP2,USP22,MTRNR2L2,ACTG1,EEF2,DPYSL3,TUBA1A,CFL1,BTBD9,HTT,HSP90AB1,H
A1,MS4A10,NEDD4L,ACTB,CTSC,ODF2L,NUDC,NRBP2,USP22,ACTG1,EEF2,TUBA1A,CFL1,BTBD9
RCC2,HSP90AB1,HSP90AA1,TUBB,FTH1,UBC,KIDINS220,RPL3,DBN1,ENO1,TUBA1B,FLNA,NCL,M
KM,FBXL4,ABHD2,TUBB2B,CANX,DDX5,KAT6A,FASN,ALDH6A1,GNB1,PLXNB2,USP11,IGF2,EIF4E
,HTT,HSP90AB1,ZNF124,FTH1,NUMA1,DBN1,ENO1,NCL,RPL7A,SLC9A7,PKM,GANAB,HSP90B1,F
TUBB2B,KAT6A,FASN,ALDH6A1,MYH9,USP11,IGF2,TRIM28,YWHAG,LRRC28,DCHS1,SET,CHSY1,
,HTT,ZNF124,UBC,KIDINS220,NUMA1,MAP1B,NCL,PKM,GANAB,ABHD2,TUBB2B,ENO2,DVL3,GP
A9,NEDD4L,POLH,ACTB,NUDC,USP22,ACTG1,MAVS,HTT,HSP90AB1,HSP90AA1,FTH1,EIF4G2,ZNF
NF124,HSP90AA1,ACTR2,SPARC,TUBB,FTH1,UBC,EIF4G2,RPL3,ZNF316,ENO1,TUBA1B,PIK3R2,H
G1,TUBA1A,XRCC2,HTT,ZNF124,HSP90AA1,FTH1,KIDINS220,RPL3,DBN1,PIK3R2,FLNA,MAP1B,N
B,CTSC,ODF2L,NUDC,MED18,ACTG1,DPYSL3,TUBA1A,CFL1,BTBD9,XRCC2,MAVS,HSP90AB1,ZN
UBA1B,FLNA,RPL7A,ALDOA,NEFL,RPL8,FBXL4,TUBB2B,CANX,PTMA,DDX5,KIF1A,GNAS,DDX17,M
C,ODF2L,NUDC,USP22,MED18,ACTG1,DPYSL3,CFL1,BTBD9,MAVS,HSP90AB1,ZNF124,HSP90AA
2,ACTG1,TUBA1A,BTBD9,MAVS,HTT,ACTR2,TUBB,UBC,KIDINS220,RPL3,ZNF316,RPL4,ENO1,FL
,MED18,EEF2,HSP90AB1,FTH1,UBC,KIDINS220,TMSB4X,NUMA1,EIF4G2,RPL3,RPS6,DBN1,RPL4
B,CTSC,ODF2L,NUDC,USP22,MED18,ACTG1,CFL1,XRCC2,MAVS,HSP90AB1,ZNF124,HSP90AA1,
A,BTBD9,HTT,HSP90AB1,ZNF124,KIDINS220,NUMA1,EIF4G2,ZNF316,NCL,COL6A1,HSP90B1,NE
UMA1,EIF4G2,TUBA1B,RPL7A,PKM,GANAB,COL6A1,CANX,PTMA,YWHAE,GNAS,BSG,FASN,SPTB
VS,HSP90AB1,ZNF124,HSP90AA1,ACTR2,KIDINS220,NUMA1,RPL3,RPS6,ZNF316,DBN1,RPL4,PIK
S,ZNF124,HSP90AA1,ACTR2,KIDINS220,NUMA1,RPS6,ZNF316,DBN1,RPL4,ENO1,TUBA1B,PIK3R
SL3,CFL1,BTBD9,ZNF124,HSP90AA1,ACTR2,TMSB4X,RPS6,ZNF316,PIK3R2,NCL,RPL7A,PKM,GA
A1,MS4A10,NEDD4L,ATN1,ACTB,ODF2L,NRBP2,USP22,MTRNR2L2,MED18,ACTG1,EEF2,DPYSL3
C,ODF2L,NUDC,USP22,MED18,ACTG1,CFL1,MAVS,HSP90AB1,ZNF124,HSP90AA1,ACTR2,FTH1,K
90AA1,ENO1,TUBA1B,RPL8,TMF1,PSAP,RPL13A,BSG,SEPTIN2,RPL5,TMBIM6,NPM1,PLD3,HSPD
CTB,CTSC,ODF2L,NRBP2,USP22,ACTG1,TUBA1A,CFL1,BTBD9,XRCC2,HTT,HSP90AB1,ZNF124,A
,KAT6A,ALDH6A1,GNB1,STMN2,MYH9,P4HB,GDI1,CALM1,TRIM28,YWHAG,LRRC28,HACD3,SET,
22,MED18,ACTG1,EEF2,DPYSL3,CFL1,BTBD9,MAVS,HSP90AB1,ZNF124,HSP90AA1,VIM,FTH1,UB
,NEDD4L,POLH,ACTB,ODF2L,NUDC,MED18,ACTG1,EEF2,TUBA1A,CFL1,BTBD9,MAVS,HSP90AB1
,USP22,MED18,ACTG1,CFL1,XRCC2,MAVS,HSP90AB1,ZNF124,HSP90AA1,ACTR2,FTH1,KIDINS22
SPA9,NEDD4L,POLH,ACTB,CTSC,ODF2L,MTRNR2L2,MED18,DPYSL3,TUBA1A,BTBD9,XRCC2,HTT
90AA1,VIM,NUMA1,DBN1,MAP1B,NCL,RPL7A,COL6A1,HSP90B1,NEFM,YBX1,TUBB2B,PTMA,YWH
,ZNF124,HSP90AA1,ACTR2,NUMA1,RPL3,RPS6,ZNF316,PIK3R2,NCL,GANAB,HSP90B1,NEFL,H3-
F124,FTH1,KIDINS220,NUMA1,NCL,PKM,GANAB,HSP90B1,PRPF8,HSPA1B,TUBB2B,DVL3,GPI,BS
D4L,POLH,ACTB,CTSC,ODF2L,MED18,ACTG1,DPYSL3,CFL1,BTBD9,XRCC2,MAVS,HSP90AB1,ZN
1,FTH1,FLNA,NCL,SLC9A7,GANAB,FADS1,NEFL,RPL8,YBX1,UBA1,COL5A1,CES3,RPLP0,H3-3B,C
A9,NEDD4L,ACTB,CTSC,NUDC,NRBP2,USP22,MTRNR2L2,ACTG1,EEF2,DPYSL3,BTBD9,MAVS,HT
P22,MTRNR2L2,ACTG1,DPYSL3,CFL1,BTBD9,XRCC2,MAVS,HTT,HSP90AB1,HSP90AA1,ACTR2,VIM
S,TUBA1B,RPL8,TMF1,PSAP,CANX,SEPTIN2,HSPD1,SMIM14,TRIM28,SPEN,EIF1,SERINC1,POLR2

EDD4L,POLH,ACTB,ACTG1,DPYSL3,MAVS,HSP90AB1,FTH1,UBC,DCX,EIF4G2,ZNF316,DBN1,ENO1
124,HSP90AA1,ACTR2,KIDINS220,NUMA1,RPS6,ZNF316,PIK3R2,NCL,HSP90B1,NEFL,NEFM,RPL8
11,ENO1,TUBA1B,FADS1,FBXL4,YBX1,TUBB2B,PSAP,SEC62,YWHAE,KIF1A,GNAS,SLC2A3,HTRA4,
KAT6A,DVL3,GNAS,IDS,SMG1,PLXNB2,HSPD1,USP11,VCP,DCHS1,SET,INA,ITGB1,TUBB4B,AGRN,
L,NUDC,USP22,MED18,ACTG1,CFL1,BTBD9,MAVS,HSP90AB1,ZNF124,HSP90AA1,ACTR2,VIM,FT
2,MED18,ACTG1,CFL1,MAVS,HSP90AB1,ZNF124,HSP90AA1,ACTR2,FTH1,KIDINS220,NUMA1,RPL
1,ARSD,MS4A10,NEDD4L,ACTB,CTSC,ODF2L,NUDC,NRBP2,EEF2,TUBA1A,CFL1,BTBD9,HTT,HSF
ARSD,MS4A10,HSPA9,NEDD4L,POLH,ACTB,CTSC,ODF2L,NUDC,USP22,MED18,ACTG1,DPYSL3,C
3,DDX5,SPTBN1,GNB1,MYH9,RPL7,P4HB,IDS,PTMS,CALM1,IGF2,CKB,DDB1,PKD1,TPT1,ITGB1,E
ARSD,MS4A10,HSPA9,NEDD4L,POLH,ACTB,CTSC,ODF2L,NUDC,USP22,MED18,ACTG1,DPYSL3,C
IO1,TUBA1B,FLNA,NCL,RPL7A,ALDOA,RPL8,FBXL4,TUBB2B,RPLP0,PTMA,ENO2,DDX17,BSG,FAS
F2L,NUDC,USP22,MED18,ACTG1,CFL1,BTBD9,XRCC2,MAVS,HSP90AB1,ZNF124,HSP90AA1,ACTR
C,USP22,MED18,ACTG1,CFL1,XRCC2,MAVS,HSP90AB1,ZNF124,HSP90AA1,ACTR2,FTH1,KIDINS2
HNRNPA1,ARSD,MS4A10,NEDD4L,POLH,ACTB,CTSC,ODF2L,NUDC,NRBP2,USP22,ACTG1,EEF2,D
JDC,MED18,ACTG1,DPYSL3,CFL1,HSP90AA1,SPARC,FTH1,KIDINS220,EIF4G2,RPS6,PIK3R2,NCL,
D18,DPYSL3,TUBA1A,BTBD9,MAVS,VIM,TUBB,UBC,KIDINS220,NUMA1,EIF4G2,ZNF316,RPL4,ENC
,SEPTIN2,NPM1,ATP5F1A,EIF4B,PPP2R1A,VAPA,MORF4L1,WSB1,AP3D1,ARF1,NFE2L1,NACA,CIF
TL,YBX1,PTMA,DDX5,SOX4,SPTBN1,SHISA9,MYH9,IDS,PTMS,IGF2,AFF4,YWHAG,YWHAQ,DCHS1,
0,HSPA9,NEDD4L,POLH,ATN1,ACTB,NUDC,NRBP2,USP22,MTRNR2L2,MED18,ACTG1,EEF2,DPYS
DPYSL3,BTBD9,MAVS,ACTR2,VIM,TUBB,UBC,KIDINS220,NUMA1,EIF4G2,ZNF316,RPL4,ENO1,TU
IPA1,ARSD,HSPA9,NEDD4L,POLH,ACTB,CTSC,ODF2L,NUDC,NRBP2,USP22,MTRNR2L2,MED18,AC
S4A10,HSPA9,NEDD4L,POLH,ACTB,CTSC,ODF2L,NUDC,NRBP2,USP22,MED18,ACTG1,EEF2,DPYS
SP22,MED18,EEF2,DPYSL3,TUBA1A,CFL1,BTBD9,XRCC2,HSP90AB1,HSP90AA1,ACTR2,SPARC,TU
ARSD,HSPA9,NEDD4L,POLH,ACTB,CTSC,NUDC,NRBP2,MTRNR2L2,ACTG1,EEF2,CFL1,MAVS,HTT,
D18,EEF2,CFL1,BTBD9,ZNF124,HSP90AA1,ACTR2,SPARC,FTH1,UBC,EIF4G2,RPL3,ZNF316,DBN1
,USP22,MED18,CFL1,MAVS,HSP90AB1,ZNF124,HSP90AA1,ACTR2,FTH1,KIDINS220,NUMA1,RPL3
CFL1,XRCC2,MAVS,HTT,FTH1,UBC,TMSB4X,DCX,ZNF316,DBN1,ENO1,TUBA1B,PIK3R2,RPL7A,AL
1,ENO1,TUBA1B,RPL8,TMF1,PSAP,RPL13A,BSG,SEPTIN2,RPL5,TMBIM6,NPM1,PLD3,HSPD1,EIF4
A1,EIF4G2,MAP1B,PKM,COL6A1,NEFM,FBXL4,UBA1,IGFBP2,FSTL1,CES3,H3-3B,PTMA,YWHAE,G
CTG1,CFL1,BTBD9,ZNF124,HSP90AA1,ACTR2,ZNF316,DBN1,ENO1,FLNA,NCL,ALDOA,PKM,FTL,F
HNRNPA1,ARSD,MS4A10,HSPA9,NEDD4L,POLH,ATN1,ACTB,CTSC,ODF2L,NUDC,NRBP2,USP22,AC
RPL7A,ALDOA,FBXL4,YBX1,PTMA,SOX4,SEC62,DDX17,GNB1,RPL7,P4HB,RPL5,NPM1,PLXNB2,CS
3,ODF2L,NUDC,USP22,MED18,ACTG1,DPYSL3,CFL1,XRCC2,MAVS,HSP90AB1,ZNF124,HSP90AA1
,NEDD4L,ACTB,ODF2L,NUDC,NRBP2,ACTG1,EEF2,CFL1,HTT,ACTR2,VIM,TUBB,FTH1,UBC,KIDIN
S4A10,HSPA9,NEDD4L,POLH,ACTB,CTSC,ODF2L,USP22,MED18,ACTG1,DPYSL3,CFL1,BTBD9,XR
L,CTSC,ODF2L,NRBP2,USP22,MED18,ACTG1,EEF2,DPYSL3,CFL1,BTBD9,XRCC2,MAVS,HSP90A
PM1,HSPD1,USP11,VCP,TPI1,CHSY1,EIF4A1,GTF3C4,RPS3,AGRN,MARCHF6,SBF1,HNRNPU,TSPA
PA1,MS4A10,NEDD4L,ACTB,CTSC,ODF2L,NUDC,USP22,ACTG1,EEF2,DPYSL3,TUBA1A,CFL1,BTBD
IDC,USP22,MED18,ACTG1,DPYSL3,CFL1,MAVS,HSP90AB1,ZNF124,HSP90AA1,ACTR2,FTH1,KIDIN
FL1,MAVS,ZNF124,HSP90AA1,ACTR2,KIDINS220,NUMA1,RPL3,RPS6,ZNF316,DBN1,RPL4,PIK3R2
PTMA,YWHAE,SPTBN1,CALM3,GNB1,IDS,GDI1,PTMS,SMG1,PLXNB2,AFF4,YWHAG,YWHAQ,HACD
4,HSP90AA1,ACTR2,KIDINS220,NUMA1,RPS6,ZNF316,DBN1,TUBA1B,PIK3R2,NCL,GANAB,HSP90
L,POLH,ATN1,ACTB,ODF2L,NRBP2,USP22,MED18,TUBA1A,CFL1,BTBD9,XRCC2,MAVS,HTT,ZNF1
1,ARSD,ACTB,ODF2L,NUDC,USP22,MED18,ACTG1,DPYSL3,CFL1,MAVS,HSP90AB1,ZNF124,HSP9
,MED18,CFL1,MAVS,HSP90AB1,ZNF124,HSP90AA1,ACTR2,FTH1,UBC,KIDINS220,NUMA1,RPL3,R

22,MED18,ACTG1,EEF2,DPYSL3,TUBA1A,CFL1,XRCC2,MAVS,HTT,ZNF124,TUBB,TMSB4X,DCX,ZN
BA1B,RPL7A,PKM,GANAB,COL6A1,FADS1,CANX,PTMA,GNAS,BSG,FASN,SPTBN1,GNB1,MYH9,P4
L5A1,PTMA,GNAS,FASN,SHISA9,MYH9,PLXNB2,EIF4B,MARCKSL1,MATR3,DCHS1,SET,RPL15,SSE
L3,GNAS,FASN,GNB1,FSCN1,COL4A2,SPEN,SET,CHSY1,PKD1,HMGB1,AGRN,MIDN,CDC42BPB,HI
EDD4L,ACTB,CTSC,ODF2L,MTRNR2L2,MED18,ACTG1,EEF2,DPYSL3,BTBD9,XRCC2,MAVS,HSP90
TIN2,MARCKS,USP11,UBB,CKB,HACD3,DCHS1,EIF4A1,MRC2,LDHB,SERINC1,VAPA,POLR2A,MOF
TSC,ODF2L,NRBP2,USP22,ACTG1,EEF2,DPYSL3,TUBA1A,CFL1,BTBD9,XRCC2,HTT,HSP90AB1,Z
L2,ACTG1,EEF2,CFL1,BTBD9,HTT,ACTR2,UBC,TMSB4X,RPS6,DBN1,RPL4,MAP1B,NCL,SLC9A7,N
VS,HTT,ZNF124,HSP90AA1,EIF4G2,ZNF316,ENO1,TUBA1B,PIK3R2,NCL,ALDOA,PKM,RPL8,FBXL
P90AA1,DBN1,PRPF8,HSPA1B,PGK1,GNAS,BSG,FASN,ALDH6A1,SPTBN1,GNB1,APP,FSCN1,NPM
E,SPTBN1,GDI1,PTMS,SMG1,PLXNB2,IGF2,MARCKSL1,AFF4,YWHAG,YWHAQ,HACD3,DCHS1,PKD
,ACTB,CTSC,MED18,ACTG1,DPYSL3,CFL1,BTBD9,XRCC2,MAVS,HSP90AB1,ZNF124,ACTR2,FTH1
X1,CANX,RPL6,KIF1A,GNAS,FASN,GNB1,APP,GDI1,FSCN1,SPTAN1,HSPD1,EIF3C,ATP5F1A,YWHA
,COPA,HNRNPA1,ARSD,MS4A10,HSPA9,NEDD4L,POLH,ATN1,ACTB,CTSC,ODF2L,NUDC,NRBP2,U
TB,ODF2L,NUDC,USP22,MED18,ACTG1,CFL1,XRCC2,MAVS,HSP90AB1,ZNF124,HSP90AA1,ACTR
D4L,POLH,ACTB,MED18,ACTG1,MAVS,HSP90AB1,HSP90AA1,ACTR2,VIM,TUBB,FTH1,NUMA1,EI
ACTG1,EEF2,MAVS,HTT,VIM,SPARC,UBC,EIF4G2,ZNF316,ENO1,FLNA,MAP1B,ALDOA,PKM,FTL,F
,ACTB,NRBP2,USP22,HSP90AB1,VIM,ZNF316,DBN1,MAP1B,COL6A1,NEFM,ATP2B4,FSTL1,COL5
KIDINS220,RPL3,RPS6,ZNF316,PIK3R2,GANAB,HSP90B1,H3-3B,RPL13A,RPL6,GNAS,FASN,P4HE
4,HSP90AA1,ACTR2,KIDINS220,RPS6,ZNF316,DBN1,RPL4,PIK3R2,NCL,GANAB,HSP90B1,RPL8,H
TSC,ODF2L,NRBP2,USP22,ACTG1,EEF2,DPYSL3,TUBA1A,CFL1,BTBD9,XRCC2,HTT,HSP90AB1,Z
8,ACTG1,MAVS,HSP90AB1,ZNF124,HSP90AA1,ACTR2,FTH1,KIDINS220,NUMA1,RPL3,RPS6,ZNF3
BA1A,CFL1,BTBD9,XRCC2,HTT,ZNF124,HSP90AA1,FTH1,UBC,KIDINS220,EIF4G2,ZNF316,DBN1,
BA1A,CFL1,BTBD9,XRCC2,HTT,ZNF124,HSP90AA1,FTH1,UBC,KIDINS220,EIF4G2,ZNF316,DBN1,
,ODF2L,NUDC,NRBP2,EEF2,TUBA1A,CFL1,BTBD9,HTT,HSP90AB1,VIM,FTH1,KIDINS220,TMSB4
,MED18,ACTG1,CFL1,MAVS,HSP90AB1,ZNF124,HSP90AA1,ACTR2,FTH1,KIDINS220,NUMA1,RPS6
FL1,BTBD9,HTT,HSP90AB1,HSP90AA1,ACTR2,FTH1,DCX,EIF4G2,ZNF316,DBN1,FLNA,NCL,FADS
2L,NUDC,USP22,MED18,CFL1,MAVS,HSP90AB1,ZNF124,HSP90AA1,ACTR2,FTH1,UBC,KIDINS22
M,FTH1,RPL7A,FTL,FSTL1,H3-3B,DDX5,SOX4,GNAS,BSG,FASN,SPTBN1,GNB1,SHISA9,APP,IDS,C
8,DPYSL3,XRCC2,MAVS,HTT,VIM,SPARC,ZNF316,PIK3R2,MAP1B,PKM,PRPF8,FBXL4,YBX1,UBA1
INS220,RPL3,RPL4,ENO1,PIK3R2,FTL,NEFL,NEFM,RPL8,IGFBP2,COL4A1,H3-3B,CANX,PTMA,DD
,RPL7A,ALDOA,HSP90B1,NEFL,FBXL4,TMF1,TUBB2B,PTMA,ENO2,DDX17,FASN,APP,RPL5,IGF2,
CFL1,HSP90AB1,ZNF124,HSP90AA1,FTH1,UBC,TMSB4X,EIF4G2,ZNF316,DBN1,TUBA1B,PIK3R2,
DC,USP22,MED18,CFL1,MAVS,HSP90AB1,ZNF124,HSP90AA1,ACTR2,FTH1,KIDINS220,NUMA1,R
,MAVS,HTT,ACTR2,VIM,FTH1,EIF4G2,PIK3R2,GANAB,NEFM,RPL8,YBX1,ATP2B4,CALR,TMF1,CO
A,NCL,RPL7A,ALDOA,RPL8,FBXL4,TUBB2B,COL4A1,PTMA,ENO2,GNAS,BSG,CALM3,SEPTIN2,P4
4,ACTR2,KIDINS220,RPS6,ZNF316,PIK3R2,HSP90B1,NEFM,FSTL1,TMF1,H3-3B,RPL13A,MAGED
F2L,NUDC,USP22,MED18,CFL1,XRCC2,MAVS,HSP90AB1,ZNF124,HSP90AA1,ACTR2,FTH1,KIDIN
0AA1,ACTR2,KIDINS220,RPS6,ZNF316,DBN1,PIK3R2,NCL,GANAB,HSP90B1,RPL8,H3-3B,FKBP1
S,ZNF124,HSP90AA1,ACTR2,KIDINS220,NUMA1,RPL3,RPS6,ZNF316,DBN1,PIK3R2,NCL,GANAB,
4,HSP90AA1,ACTR2,KIDINS220,RPS6,ZNF316,DBN1,PIK3R2,NCL,ALDOA,GANAB,RPL8,YBX1,PG
1,UBC,ZNF316,DBN1,TUBA1B,NCL,RPL7A,HSP90B1,FADS1,ATP2B4,FSTL1,ABHD2,HSPA1B,PTM
1,UBC,ZNF316,DBN1,TUBA1B,NCL,RPL7A,HSP90B1,FADS1,ATP2B4,FSTL1,ABHD2,HSPA1B,PTM
L,NUDC,USP22,MED18,ACTG1,DPYSL3,CFL1,XRCC2,MAVS,HSP90AB1,ZNF124,HSP90AA1,ACTR
SC,ODF2L,MTRNR2L2,MED18,ACTG1,EEF2,DPYSL3,BTBD9,XRCC2,MAVS,HSP90AB1,ZNF124,AC

\1,COL4A1,PTMA,KAT6A,KIF1A,GNAS,BSG,FASN,SPTBN1,GNB1,GSTP1,IGF2,MARCKSL1,CKB,C
T,VIM,FTH1,TMSB4X,EIF4G2,ZNF316,DBN1,MAP1B,FTL,HSP90B1,NEFM,RPL8,YBX1,UBA1,CALR,
,HSPA9,NEDD4L,ACTB,CTSC,MED18,ACTG1,DPYSL3,CFL1,BTBD9,XRCC2,MAVS,HSP90AB1,ZNF1
,FASN,SEPTIN2,GSTP1,CSDE1,EIF3C,YWHAG,NONO,SET,POLR2A,AP2M1,SEC61A1,TUBB4B,MO
IDINS220,RPS6,ZNF316,PIK3R2,NCL,ALDOA,HSP90B1,NEFM,PGK1,H3-3B,PTMA,DDX5,FKBP14,
2,HSP90AB1,UBC,NUMA1,RPL3,RPS6,RPL4,ENO1,FLNA,NCL,ALDOA,PKM,GANAB,COL6A1,HSP9
P2,USP22,ACTG1,XRCC2,HTT,VIM,FTH1,TMSB4X,EIF4G2,ZNF316,DBN1,FLNA,NCL,RPL7A,ALDO
C,USP22,MED18,ACTG1,CFL1,XRCC2,MAVS,HSP90AB1,ZNF124,HSP90AA1,ACTR2,FTH1,KIDINS2
HSP90AB1,ZNF124,HSP90AA1,ACTR2,KIDINS220,NUMA1,RPL3,RPS6,ZNF316,RPL4,PIK3R2,NCL
22,MED18,CFL1,MAVS,HSP90AB1,ZNF124,HSP90AA1,ACTR2,FTH1,KIDINS220,NUMA1,RPL3,RP
A1B,ALDOA,TMF1,RPL13A,YWHAE,BSG,FASN,SEPTIN2,P4HB,TMBIM6,NPM1,CSDE1,EIF4B,TRIM
,CFL1,XRCC2,MAVS,FTH1,UBC,TMSB4X,DCX,DBN1,TUBA1B,PIK3R2,RPL7A,ALDOA,PKM,COL6A
90AA1,ACTR2,KIDINS220,RPS6,ZNF316,PIK3R2,NCL,GANAB,RPL8,YBX1,ATP2B4,H3-3B,FKBP14
,VIM,FTH1,UBC,NUMA1,ZNF316,DBN1,RPL4,ENO1,PIK3R2,FLNA,NCL,FTL,NEFL,UGT8,RPL8,FB
L,POLH,ODF2L,NUDC,ACTG1,EEF2,DPYSL3,XRCC2,HSP90AB1,ZNF124,HSP90AA1,VIM,FTH1,UB
TR2,NUMA1,RPS6,ZNF316,PIK3R2,NCL,GANAB,HSP90B1,NEFM,RPL8,TMF1,H3-3B,RPL13A,MA
TSC,ODF2L,ACTG1,DPYSL3,CFL1,BTBD9,XRCC2,ZNF124,HSP90AA1,KIDINS220,RPL3,ZNF316,D
IEDD4L,POLH,ATN1,CTSC,ODF2L,NUDC,MED18,ACTG1,TUBA1A,CFL1,BTBD9,XRCC2,MAVS,HSP
L,POLH,ATN1,ACTB,ODF2L,NUDC,ACTG1,CFL1,XRCC2,HSP90AB1,ZNF124,HSP90AA1,KIDINS22
,CFL1,XRCC2,MAVS,SPARC,FTH1,UBC,TMSB4X,NUMA1,DCX,DBN1,ENO1,TUBA1B,PIK3R2,FLNA
C,USP22,MED18,ACTG1,CFL1,XRCC2,MAVS,HSP90AB1,ZNF124,HSP90AA1,ACTR2,VIM,FTH1,UB
,USP22,MED18,CFL1,XRCC2,MAVS,HSP90AB1,ZNF124,HSP90AA1,ACTR2,NUMA1,RPS6,ZNF316,
4,HSP90AA1,ACTR2,KIDINS220,RPS6,ZNF316,DBN1,PIK3R2,NCL,GANAB,YBX1,ATP2B4,H3-3B,F
4,HSP90AA1,ACTR2,KIDINS220,RPS6,ZNF316,DBN1,RPL4,PIK3R2,NCL,GANAB,HSP90B1,RPL8,A
90AB1,ZNF124,HSP90AA1,SPARC,TUBB,EIF4G2,ENO1,TUBA1B,MTRNR2L8,FTL,HSP90B1,UGT8,
2L,NUDC,NRBP2,TUBA1A,XRCC2,MAVS,ZNF124,HSP90AA1,VIM,TUBB,FTH1,UBC,KIDINS220,DCX
90AA1,ACTR2,KIDINS220,RPS6,ZNF316,DBN1,PIK3R2,GANAB,H3-3B,FKBP14,RPL6,RACK1,SHIS
HSPA9,ACTB,CTSC,USP22,MTRNR2L2,MED18,ACTG1,EEF2,DPYSL3,TUBA1A,BTBD9,HTT,HSP90
NRBP2,MTRNR2L2,MED18,EEF2,MAVS,HSP90AB1,FTH1,UBC,KIDINS220,TMSB4X,NUMA1,EIF4G
DN,CDC42BPB,ITM2B,PXDN,LAMA5,BRD3,KPNB1,NCS1,ZNF865,CHD4,JUND,CD81,ATP2A2,CR
IF316,DBN1,ENO1,FLNA,RPL7A,PKM,COL6A1,NEFM,RPL8,CALR,FSTL1,TUBB2B,PSAP,PTMA,KAT
1,MAVS,HSP90AB1,ZNF124,HSP90AA1,ACTR2,FTH1,NUMA1,RPL3,RPS6,ZNF316,DBN1,PIK3R2,M
R2,KIDINS220,RPS6,ZNF316,PIK3R2,NCL,ALDOA,HSP90B1,H3-3B,PTMA,DDX5,FKBP14,MAGED1
NUMA1,COL6A1,FBXL4,UBA1,DVL3,GPI,FASN,ALDH6A1,GDI1,PTMS,MARCKSL1,TRIM28,LRRC2
PA,ARSD,MS4A10,HSPA9,NEDD4L,POLH,ACTB,NUDC,NRBP2,USP22,MED18,ACTG1,EEF2,DPYSL3
S4A10,NEDD4L,ATN1,ACTB,NRBP2,USP22,ACTG1,MAVS,HTT,HSP90AB1,HSP90AA1,UBC,NUMA1,
,MED18,ACTG1,DPYSL3,CFL1,HSP90AB1,ZNF124,HSP90AA1,ACTR2,FTH1,KIDINS220,TMSB4X,N
DOA,FADS1,NEFM,TUBB2B,SOX4,KAT6A,DVL3,DDX17,CALM3,SEPTIN2,FSCN1,SPTAN1,MARCKS
2,RPS6,ZNF316,RPL4,PIK3R2,GANAB,HSP90B1,H3-3B,FKBP14,RPL6,DVL3,RACK1,GNB1,SHISA
BA1B,ALDOA,TMF1,PSAP,RPL13A,BSG,RPL7,SEPTIN2,RPL5,TMBIM6,NPM1,RPL37A,SPEN,EIF1,H
316,DBN1,ENO1,PIK3R2,FADS1,COL5A1,ENO2,RPL13A,KIF1A,GNAS,SPTBN1,GNB1,GDI1,IGF2,
P0,DDX5,RPL13A,YWHAE,DDX17,RACK1,BSG,SEPTIN2,USP11,EIF3C,ATP5F1A,MATR3,NONO,SET
OL6A1,HSP90B1,TMF1,PTMA,DDX5,ENO2,SEC62,KAT6A,DVL3,YWHAE,GNAS,DDX17,FASN,CAL
MRC2,TUBB4B,SBF1,HNRNPU,CTNNA1,LPCAT1,BRD3,NCS1,CLSTN1,JPT2,PDIA6,APLP1,MYO5A
,ZNF124,HSP90AA1,ZNF316,TUBA1B,PKM,GANAB,FADS1,NEFM,FBXL4,FSTL1,COL5A1,COL4A1

TUBA1A,MAVS,HSP90AB1,HSP90AA1,FTH1,ZNF316,PIK3R2,PKM,FADS1,FBXL4,YBX1,IGFBP2,CO
,FTH1,TMSB4X,RPS6,ZNF316,DBN1,RPL4,TUBA1B,PIK3R2,MAP1B,NCL,RPL7A,GANAB,HSP90B1
),HSPA9,NEDD4L,POLH,ATN1,CTSC,NUDC,USP22,MED18,ACTG1,EEF2,DPYSL3,TUBA1A,CFL1,BT
ODF2L,NRBP2,USP22,MED18,DPYSL3,CFL1,BTBD9,XRCC2,MAVS,HTT,HSP90AA1,ACTR2,VIM,SP
DC,USP22,MED18,CFL1,MAVS,HSP90AB1,ZNF124,HSP90AA1,ACTR2,FTH1,KIDINS220,NUMA1,RI
31,HSP90AA1,ZNF316,DBN1,ENO1,NCL,ALDOA,COL6A1,FTL,RPL8,IGFBP2,CALR,ZNF99,COL4A1
BD9,HSP90AA1,ACTR2,KIDINS220,TMSB4X,EIF4G2,RPS6,RPL4,ALDOA,GANAB,HSP90B1,NEFM,I
90AA1,ACTR2,KIDINS220,NUMA1,RPL3,ZNF316,DBN1,PIK3R2,NCL,GANAB,HSP90B1,RPL8,UBA1
LH,ATN1,ACTB,ODF2L,NUDC,NRBP2,USP22,ACTG1,EEF2,TUBA1A,XRCC2,MAVS,ZNF124,HSP90A
KAT6A,KIF1A,RPL13,FASN,SPTBN1,GNB1,SHISA9,RPS8,USP11,COL18A1,IGF2,SET,TMSB10,MR
4,HSP90AA1,ACTR2,KIDINS220,RPS6,ZNF316,TUBA1B,PIK3R2,NCL,GANAB,RPL8,HSPA1B,DDX5,
UBA1B,FLNA,NCL,RPL7A,ALDOA,FBXL4,PTMA,DDX17,FASN,APP,IGF2,MARCKSL1,MATR3,LRRC
C2,HTT,SPARC,UBC,KIDINS220,NUMA1,TUBA1B,PIK3R2,FLNA,NCL,RPL7A,SLC9A7,ALDOA,PKM,
P90AA1,ACTR2,VIM,UBC,NUMA1,RPL3,ZNF316,ENO1,TUBA1B,PIK3R2,MAP1B,SLC9A7,PKM,FTL,
L,CTSC,ODF2L,NUDC,NRBP2,USP22,ACTG1,CFL1,BTBD9,ZNF124,HSP90AA1,VIM,NUMA1,DCX,E
A,CFL1,BTBD9,HTT,FTH1,UBC,EIF4G2,ENO1,TUBA1B,NCL,ALDOA,GANAB,COL6A1,FTL,NEFM,F
32B,DDX5,SOX4,SPTBN1,SHISA9,MYH9,IDS,GDI1,PTMS,IGF2,YWHAG,CKB,DCHS1,TMSB10,MRC
R2,RPL7A,SLC9A7,YBX1,FSTL1,GNAS,BSG,SHISA9,IDS,GDI1,IGF2,AFF4,YWHAQ,CHSY1,TMSB10,
,COPA,ARSD,MS4A10,ATN1,ACTB,CTSC,ODF2L,NRBP2,USP22,MTRNR2L2,ACTG1,DPYSL3,CFL1
ODF2L,MED18,ACTG1,XRCC2,MAVS,HSP90AB1,ZNF124,HSP90AA1,ACTR2,KIDINS220,DCX,EIF4
,NUDC,USP22,CFL1,XRCC2,HTT,ACTR2,VIM,TUBB,FTH1,UBC,NUMA1,DBN1,ENO1,TUBA1B,NCL,
A10,HSPA9,NEDD4L,ACTB,ACTG1,EEF2,DPYSL3,TUBA1A,BTBD9,HTT,HSP90AB1,ZNF124,HSP90
L4,DVL3,GDI1,MARCKSL1,TRIM28,LRRC28,TMSB10,PKD1,HLA-C,ANXA2,AGRN,MIDN,AP3D1,CY
,XRCC2,MAVS,HSP90AB1,HSP90AA1,ACTR2,RPS6,ZNF316,DBN1,PIK3R2,NCL,HSP90B1,YBX1,IG
)9,XRCC2,MAVS,HTT,UBC,RPL3,ENO1,PIK3R2,FLNA,NCL,ALDOA,PKM,GANAB,NEFL,PRPF8,TUBE
3,ACTG1,EEF2,DPYSL3,CFL1,BTBD9,XRCC2,MAVS,HSP90AB1,SPARC,UBC,NUMA1,EIF4G2,RPL3,
4,KAT6A,GPI,ALDH6A1,GDI1,SPTAN1,CALM1,EIF4B,MARCKSL1,TRIM28,YWHAG,CKB,LRRC28,FN
ISB4X,EIF4G2,MAP1B,NCL,RPL7A,GANAB,UBA1,RPLP0,DDX5,RPL13A,KAT6A,DVL3,YWHAE,DDX1
1A,HSP90AB1,HSP90AA1,ACTR2,VIM,TUBB,UBC,NUMA1,RPL3,RPS6,RPL4,TUBA1B,PKM,GANAB,
RSD,MS4A10,HSPA9,NEDD4L,POLH,CTSC,ODF2L,NRBP2,USP22,MTRNR2L2,MED18,ACTG1,EEF
3P2,USP22,DPYSL3,XRCC2,MAVS,HTT,SPARC,DBN1,ENO1,MAP1B,SLC9A7,ALDOA,NEFM,TUBB2E
)C,USP22,MED18,ACTG1,EEF2,CFL1,BTBD9,XRCC2,MAVS,HSP90AB1,UBC,KIDINS220,NUMA1,RI
IDC,USP22,MED18,CFL1,MAVS,HSP90AB1,ZNF124,HSP90AA1,ACTR2,FTH1,KIDINS220,NUMA1,RI
P90AA1,ACTR2,KIDINS220,RPS6,ZNF316,PIK3R2,NCL,GANAB,HSP90B1,RPL8,RPL13A,RPL6,RAC
4,HSP90AA1,ACTR2,NUMA1,RPS6,ZNF316,DBN1,PIK3R2,NCL,GANAB,RPL8,H3-3B,DDX5,RPL13
,UBC,DBN1,FLNA,PKM,HSP90B1,FADS1,UGT8,YBX1,COL5A1,COL4A1,CES3,PTMA,ENO2,KAT6A
,MED18,HSP90AB1,ACTR2,TUBB,FTH1,UBC,TMSB4X,NUMA1,EIF4G2,RPL3,RPS6,RPL4,ENO1,G
,HSPD1,TRIM28,SPEN,SERINC1,CDC42BPB,ZNF770,TMEM170A,GTF2F1,PGAM1,SMARCA5,RPS
)0AB1,FTH1,UBC,ZNF316,ENO1,NCL,RPL7A,PKM,GANAB,COL6A1,NEFL,NEFM,RPL8,UBA1,ABHD
P22,MTRNR2L2,MED18,EEF2,DPYSL3,TUBA1A,CFL1,BTBD9,XRCC2,MAVS,HTT,ACTR2,SPARC,TU
3,TUBA1A,CFL1,BTBD9,MAVS,HTT,HSP90AA1,VIM,TUBB,KIDINS220,NUMA1,ZNF316,TUBA1B,FL
DDF2L,MED18,CFL1,HSP90AB1,ZNF124,HSP90AA1,ACTR2,VIM,FTH1,RPL3,ZNF316,PIK3R2,NCL,
,ZNF316,PIK3R2,NCL,ALDOA,HSP90B1,YBX1,CANX,DDX5,FKBP14,RPL13A,MAGED1,RPL6,DVL3,
GNB1,SHISA9,IDS,PTMS,IGF2,YWHAG,DCHS1,TMSB10,MRC2,PKD1,VAPA,TPT1,ITGB1,ZMIZ1,TIM
,HSP90AB1,HSP90AA1,ACTR2,RPS6,ZNF316,PIK3R2,NCL,ALDOA,HSP90B1,YBX1,ATP2B4,CALR,

NCL,RPL7A,NEFL,FBXL4,IGFBP2,TUBB2B,COL4A1,PTMA,ENO2,DDX17,BSG,RPL7,SEPTIN2,APP,F
22,MED18,CFL1,XRCC2,HSP90AB1,ZNF124,HSP90AA1,ACTR2,FTH1,KIDINS220,NUMA1,RPL3,RF
,MARCHF6,SMARCA5,RPS9,RPL17,RAB6B,EIF5,RPL26,SCRN1,DDX3X,RPS15,RPL12,RPS5,RPS7,
FL1,BTBD9,HTT,HSP90AB1,TUBB,FTH1,UBC,KIDINS220,TMSB4X,DBN1,ENO1,TUBA1B,MAP1B,F
P90AA1,ACTR2,KIDINS220,RPS6,ZNF316,PIK3R2,NCL,GANAB,HSP90B1,RPL8,TMF1,RPL13A,RPL
,ZNF316,PIK3R2,HSP90B1,YBX1,H3-3B,FKBP14,RPL13A,MAGED1,RPL6,DVL3,RACK1,SHISA9,P4
TN1,ACTB,CTSC,ODF2L,NUDC,USP22,MED18,ACTG1,DPYSL3,TUBA1A,CFL1,ZNF124,HSP90AA1,
,RPL8,UBA1,IGFBP2,ABHD2,COL4A1,RPLP0,GNAS,CALM3,RPS8,APP,PLXNB2,IGF2,CKB,COL4A
SP90AA1,ACTR2,KIDINS220,RPS6,ZNF316,PIK3R2,GANAB,HSP90B1,FBXL4,UBA1,H3-3B,RPL13
AVS,HTT,ZNF124,HSP90AA1,UBC,NUMA1,ZNF316,DBN1,MAP1B,SLC9A7,FTL,HSP90B1,PRPF8,FE
D4L,POLH,ACTB,CTSC,ODF2L,NUDC,NRBP2,USP22,MTRNR2L2,MED18,ACTG1,EEF2,DPYSL3,T
ISA9,MYH9,SEPTIN2,GDI1,IGF2,UBB,SET,EIF4A1,MORF4L1,AGRN,MIDN,DNAJA1,CYB5R3,CDC4
,CALM3,GNB1,SHISA9,FSCN1,SPTAN1,GSTP1,CALM1,MARCKSL1,SPEN,NONO,PSMD5,YWHAQ,E
RSD,MS4A10,HSPA9,NEDD4L,POLH,ATN1,ACTB,ODF2L,NUDC,MED18,ACTG1,DPYSL3,CFL1,BTB
NUDC,USP22,MED18,CFL1,MAVS,HSP90AB1,ZNF124,HSP90AA1,ACTR2,FTH1,KIDINS220,NUMA1
ZNF124,ACTR2,VIM,FTH1,UBC,ZNF316,FLNA,NCL,RPL7A,HSP90B1,NEFL,TUBB2B,COL5A1,CLU,
TH1,TMSB4X,EIF4G2,RPL3,RPS6,RPL4,NCL,RPLP0,DDX5,RPL13A,KAT6A,DVL3,YWHAE,DDX17,F
TH1,KIDINS220,EIF4G2,ZNF316,DBN1,PIK3R2,FLNA,NCL,RPL7A,PKM,GANAB,HSP90B1,PRPF8,F
BTBD9,HSP90AB1,VIM,FTH1,UBC,EIF4G2,PIK3R2,FLNA,NCL,MTRNR2L8,NEFL,NEFM,PRPF8,FB
UDC,MED18,DPYSL3,CFL1,HSP90AB1,ZNF124,HSP90AA1,ACTR2,FTH1,RPL3,ZNF316,RPL4,PIK3
F2L,NUDC,USP22,MED18,ACTG1,EEF2,XRCC2,HSP90AB1,ZNF124,HSP90AA1,ACTR2,FTH1,KIDI
4A10,NEDD4L,POLH,ATN1,ACTB,CTSC,ODF2L,NUDC,USP22,MTRNR2L2,ACTG1,DPYSL3,TUBA1A
UDC,NRBP2,MTRNR2L2,MED18,EEF2,MAVS,HSP90AB1,FTH1,UBC,KIDINS220,TMSB4X,NUMA1,
IK3R2,H3-3B,FKBP14,RPL6,DVL3,RACK1,SHISA9,P4HB,PLD3,USP11,SNRNP200,INA,PPP2R1A,S
CTSC,ODF2L,NRBP2,USP22,ACTG1,DPYSL3,CFL1,BTBD9,MAVS,HTT,HSP90AB1,UBC,ZNF316,DE
A,HNRNPA1,ARSD,MS4A10,HSPA9,NEDD4L,POLH,ATN1,ACTB,CTSC,ODF2L,NUDC,NRBP2,USP22
4G2,RPS6,RPL7A,ALDOA,PKM,GANAB,NEFM,FBXL4,YBX1,CALR,FSTL1,TMF1,CES3,PGK1,H3-3E
,HSP90AB1,ZNF124,HSP90AA1,ACTR2,FTH1,KIDINS220,NUMA1,RPS6,ZNF316,DBN1,PIK3R2,MA
,SPEN,POLR2A,SBF1,DNAJA1,CDC42BPB,GTF2F1,OGA,PGAM1,SMARCA5,RPS9,CIPC,PAPOLA,C
N1,PIK3R2,RPL7A,ALDOA,PKM,FTL,YBX1,CALR,COL5A1,PTMA,SEC62,KAT6A,YWHAE,RPL13,GN
,PIK3R2,HSPA1B,RACK1,CALM3,P4HB,TMBIM6,USP11,SMIM14,ATP5F1A,UBB,AFF4,EIF4A1,ZNF
,EIF4B,TRIM28,YWHAG,PKD1,STMN1,AGRN,SARAF,MIDN,CDC42BPB,SMARCA5,RPL23,NACA,B
,ACTR2,KIDINS220,RPS6,ZNF316,PIK3R2,NCL,GANAB,RPL8,FKBP14,RPL6,RACK1,SHISA9,P4HB,
,NCL,RPL13A,RPL13,BSG,FASN,GNB1,MYH9,P4HB,TMBIM6,HSPD1,EIF4B,UBB,TRIM28,GTF3C4
TH1,KIDINS220,EIF4G2,ZNF316,PIK3R2,FLNA,NCL,PKM,GANAB,HSP90B1,PRPF8,FBXL4,YBX1,U
,RPL10,SMARCA5,RPL23,RPS9,RPL17,C1ORF43,ENC1,EIF5,RPL26,SCRN1,PEBP1,DDX3X,RPS15
FBXL4,IGFBP2,TUBB2B,COL4A1,PTMA,SEPTIN2,APP,IGF2,UBB,COL4A2,YWHAQ,SSBP2,BASP1,F
TG1,EEF2,TUBA1A,MAVS,ZNF124,FTH1,KIDINS220,TMSB4X,EIF4G2,ZNF316,RPL4,PIK3R2,MAP
ENO1,PIK3R2,MAP1B,ALDOA,GANAB,NEFL,RPL8,UBA1,PGK1,CANX,MAGED1,RPL6,DVL3,GNAS,C
,DVL3,FASN,SPTBN1,MYH9,RPL5,FSCN1,PTMS,HSPD1,COL18A1,IGF2,CKB,EIF1,SET,CDKL1,ITG
FL1,BTBD9,HTT,SPARC,TUBB,FTH1,UBC,NUMA1,EIF4G2,MAP1B,NCL,SLC9A7,MTRNR2L8,FTL,H
SL3,CFL1,MAVS,HSP90AB1,ZNF124,HSP90AA1,ACTR2,FTH1,NUMA1,RPL3,RPS6,ZNF316,DBN1,R
OL4A1,PTMA,SEPTIN2,APP,IGF2,UBB,COL4A2,YWHAQ,CHSY1,SSBP2,BASP1,POLR2A,UCHL1,MC
K1,RPS8,SEPTIN2,GDI1,EIF4B,MATR3,SET,EIF4A1,PPP2R1A,ZNF43,VAPA,MORF4L1,AP3D1,SBF
H1,UBC,NUMA1,ZNF316,SLC9A7,FBXL4,YBX1,COL4A1,DDX5,SOX4,GPI,SHISA9,IDS,GDI1,TMBIM

2,FTH1,RPL3,RPS6,ZNF316,PIK3R2,NCL,ALDOA,GANAB,HSP90B1,DDX5,RPL13A,MAGED1,RPL6,
ACTG1,CFL1,MAVS,HTT,HSP90AA1,ACTR2,DBN1,FLNA,RPL7A,ALDOA,COL6A1,FTL,FADS1,COL4
ISP90B1,NEFL,IGFBP2,FSTL1,TUBB2B,COL4A1,H3-3B,RPL13A,SEC62,ALDH6A1,CALM3,RPS8,M
AS,MARCKS,PLXNB2,HACD3,SERINC1,POLR2A,MORF4L1,MARCHF6,FADS2,TPM4,LAMA5,KPNB1
DS,GDI1,PTMS,SMG1,YWHAG,YWHAQ,DCHS1,PKD1,TPT1,ITGB1,AGRN,SARAF,CDC42BPB,ITM2B
220,NUMA1,ZNF316,MAP1B,COL6A1,NEFM,UBA1,IGFBP2,FSTL1,COL5A1,RPLP0,ENO2,GNAS,FA
A1,MS4A10,NEDD4L,POLH,ATN1,ACTB,ODF2L,NUDC,NRBP2,USP22,MED18,EEF2,TUBA1A,CFL1,
KAT6A,GNB1,GDI1,VCP,MAPT,PKD1,STMN1,TIMP2,MIDN,SBF1,SF3B2,CDC42BPB,PXDN,CTNNA
L2,MED18,XRCC2,HSP90AB1,FTH1,UBC,KIDINS220,TMSB4X,NUMA1,EIF4G2,RPL3,RPS6,RPL4,M
,NCL,GANAB,HSP90B1,CALR,FSTL1,COL5A1,PGK1,SEC62,ALDH6A1,RPS8,SEPTIN2,TMBIM6,US
2,KIDINS220,RPS6,ZNF316,PIK3R2,NCL,GANAB,RPL8,YBX1,TMF1,MAGED1,RPL6,RACK1,P4HB,
CL,RPL7A,RPL8,DDX5,SOX4,FKBP14,RPL13A,RACK1,RPL7,SEPTIN2,PLXNB2,PLD3,HSPD1,USP11,
NUMA1,ZNF316,DBN1,MAP1B,COL6A1,NEFM,RPL8,UBA1,IGFBP2,FSTL1,ABHD2,COL5A1,SOX4,K
,PIK3R2,H3-3B,RPL6,DVL3,RACK1,TMBIM6,PLD3,SNRNP200,EIF4A1,LDHB,PKD1,SF3B1,ZNF43,
TR2,KIDINS220,ZNF316,DBN1,ENO1,PIK3R2,NCL,ALDOA,PKM,FTL,FADS1,YBX1,IGFBP2,PSAP,P
DF2L,USP22,MED18,EEF2,DPYSL3,CFL1,BTBD9,XRCC2,MAVS,HSP90AB1,FTH1,UBC,NUMA1,EIF
P14,P4HB,TMBIM6,USP11,SMIM14,ATP5F1A,RPL15,POLR2A,RPS3A,MORF4L1,DNAJA1,SF3B2,Z
NR2L2,MED18,HTT,HSP90AB1,FTH1,UBC,KIDINS220,TMSB4X,NUMA1,EIF4G2,RPL3,RPS6,RPL4,
HTT,HSP90AA1,SPARC,NUMA1,RPL3,RPS6,ZNF316,PIK3R2,FLNA,SLC9A7,HSP90B1,RPL8,FBXL4
PIK3R2,NCL,ALDOA,GANAB,DDX5,RPL13A,MAGED1,RPL6,DVL3,GNAS,ATP5F1B,SHISA9,P4HB,PL
2,ACTG1,EEF2,DPYSL3,CFL1,MAVS,HSP90AB1,VIM,SPARC,FTH1,UBC,NUMA1,RPS6,ZNF316,ENO
RNR2L2,MED18,TUBA1A,HTT,HSP90AB1,ACTR2,FTH1,UBC,KIDINS220,TMSB4X,NUMA1,EIF4G2,
IPA1,ARSD,MS4A10,NEDD4L,ACTB,ODF2L,NRBP2,USP22,ACTG1,XRCC2,HSP90AB1,HSP90AA1,A
FBP2,BSG,FASN,GNB1,P4HB,HSPD1,EIF4B,TRIM28,CHSY1,ZNF770,FADS2,CTSD,EIF4H,CIPC,KP
,YWHAG,SPEN,YWHAQ,DCHS1,SET,CHSY1,CDKL1,ITGB1,CDC42BPB,CNN3,NCS1,JUND,VAT1,CC
MED18,XRCC2,MAVS,HSP90AB1,ZNF124,HSP90AA1,ACTR2,KIDINS220,NUMA1,RPL3,RPS6,ZNF31
A1,NEFM,FSTL1,CES3,KIF1A,SLC2A3,CALM3,GDI1,PLXNB2,MARCKSL1,DCHS1,TMSB10,MRC2,
SL3,CFL1,BTBD9,MAVS,UBC,NUMA1,RPL3,RPS6,DBN1,RPL4,ENO1,FLNA,SLC9A7,ALDOA,HSP90
RRC28,PKD1,HLA-C,ANXA2,AGRN,MIDN,AP3D1,CDC42BPB,CTSD,BRD3,ZNF865,CD81,ATP2A2,O
B1,SPTAN1,MARCKSL1,CKB,COL4A2,RPL37A,SF3B1,AGRN,TMEM170A,KPNB1,CLSTN1,EIF5,SR
ERINC1,CDC42BPB,GTF2F1,OGA,PGAM1,SMARCA5,CIPC,RHOA,CBX5,ZNF865,LENG8,VAT1,RT
ANX,DDX5,RPL13A,KAT6A,YWHAE,DDX17,RACK1,SEPTIN2,GDI1,MARCKS,CSDE1,ATP5F1A,MATR
9,NEDD4L,POLH,CTSC,ODF2L,NUDC,ACTG1,DPYSL3,CFL1,BTBD9,XRCC2,HSP90AB1,ZNF124,HS
,MS4A10,HSPA9,NEDD4L,POLH,ACTB,CTSC,ODF2L,NUDC,NRBP2,MED18,ACTG1,CFL1,BTBD9,
ENO2,KAT6A,DVL3,GNAS,MYH9,GDI1,TMBIM6,PTMS,EIF3C,EIF4B,CKB,SPEN,CHSY1,GTF3C4,SE
,ZNF316,PIK3R2,NCL,ALDOA,HSP90B1,DDX5,MAGED1,RPL6,GPI,RACK1,SHISA9,P4HB,TMBIM6
L2,MED18,HSP90AB1,FTH1,UBC,TMSB4X,NUMA1,EIF4G2,RPL3,RPS6,RPL4,GANAB,UBA1,FSTL1,
R,TUBB2B,SEC62,GNAS,SLC2A3,GNB1,PTMS,SMG1,NPM1,MARCKSL1,AFF4,NONO,TMSB10,CH
,IGFBP2,TUBB2B,PTMA,SEPTIN2,APP,IGF2,UBB,YWHAQ,SSBP2,BASP1,POLR2A,UCHL1,MORF4L
NPA1,ARSD,MS4A10,HSPA9,NEDD4L,ATN1,ACTB,NUDC,NRBP2,USP22,MED18,EEF2,DPYSL3,TUE
JSP22,MED18,EEF2,DPYSL3,CFL1,BTBD9,XRCC2,HSP90AB1,FTH1,UBC,KIDINS220,EIF4G2,RPL3
N2,FSCN1,SMG1,LDHB,PKD1,UCHL1,AGRN,MIDN,ARF1,CDC42BPB,GOLGA8A,HNRNPU,SCARB2,
16,PIK3R2,NCL,RPL8,YBX1,TMF1,H3-3B,RPL6,RACK1,SHISA9,P4HB,TMBIM6,PLD3,SNRNP200,
CTB,NRBP2,USP22,ACTG1,MAVS,HTT,HSP90AA1,UBC,NUMA1,ZNF316,PIK3R2,FLNA,NCL,MTRN
GNAS,FASN,MYH9,IDS,GDI1,IGF2,YWHAG,TMSB10,PPP2R1A,RPL15,PKD1,PDIA3,VAPA,TPT1,TUBE

NF124,HSP90AA1,RPS6,ZNF316,PIK3R2,NCL,SLC9A7,HSP90B1,IGFBP2,PTMA,DDX5,FKBP14,RPI
GK1,DDX5,RPL13A,KAT6A,KIF1A,GPI,ALDH6A1,SEPTIN2,P4HB,RPL5,FSCN1,PLD3,USP11,SNRNP2
DC,NRBP2,USP22,MTRNR2L2,EEF2,CFL1,VIM,TMSB4X,NUMA1,RPL3,ZNF316,DBN1,ENO1,PIK3R
UDC,USP22,MED18,MAVS,HSP90AB1,ZNF124,HSP90AA1,ACTR2,KIDINS220,NUMA1,RPL3,RPS6,Z
R2,RPL6,RACK1,SHISA9,P4HB,TMBIM6,PLD3,SNRNP200,EIF4A1,ZNF43,HMGB1,POLR2A,TPT1,E
AVS,VIM,SPARC,TMSB4X,NUMA1,RPL3,DBN1,MAP1B,COL6A1,FTL,FADS1,RPL8,ATP2B4,IGFBP2,
0,NEDD4L,ACTB,ODF2L,ACTG1,MAVS,HSP90AB1,VIM,UBC,KIDINS220,DBN1,TUBA1B,MAP1B,RF
16,PIK3R2,NCL,HSP90B1,CANX,PTMA,DDX5,MAGED1,RPL6,GPI,RACK1,P4HB,TMBIM6,USP11,EII
ALDOA,HSP90B1,YBX1,DDX5,RPL13A,MAGED1,RPL6,DVL3,RACK1,ATP5F1B,SHISA9,P4HB,SNRN
NO1,FSTL1,KAT6A,KIF1A,GNAS,P4HB,SMG1,IGF2,MARCKSL1,YWHAG,CKB,EIF1,TPI1,TMSB10,F
NS220,RPS6,ZNF316,PIK3R2,NCL,GANAB,HSP90B1,RPL8,TMF1,H3-3B,CANX,RPL13A,RPL6,RACI
ACTB,ODF2L,NUDC,USP22,MED18,ACTG1,EEF2,CFL1,BTBD9,XRCC2,MAVS,HSP90AB1,ZNF124,H
ACTB,ODF2L,NUDC,USP22,MED18,ACTG1,EEF2,CFL1,BTBD9,XRCC2,MAVS,HSP90AB1,ZNF124,H
9,NEDD4L,POLH,CTSC,ODF2L,NUDC,ACTG1,DPYSL3,CFL1,BTBD9,XRCC2,HSP90AB1,ZNF124,KII
L3,RPS6,ZNF316,PIK3R2,NCL,GANAB,HSP90B1,NEFL,DDX5,MAGED1,RPL6,GNAS,GPI,P4HB,USF
BTBD9,MAVS,HSP90AB1,ZNF124,TMSB4X,RPL3,ZNF316,RPL4,TUBA1B,PIK3R2,MAP1B,RPL7A,C
AA1,RPS6,ZNF316,TUBA1B,PIK3R2,NCL,HSP90B1,PRPF8,YBX1,ATP2B4,PTMA,DDX5,SOX4,MAGE
POLH,ACTB,CTSC,MTRNR2L2,EEF2,CFL1,BTBD9,MAVS,HTT,HSP90AA1,ACTR2,SPARC,NUMA1,RF
ED1,RPL6,SHISA9,P4HB,TMBIM6,RPL37A,RPL19,HMGB1,TUBB4B,BTF3,RPLP1,MARCHF6,SF3B2
BX1,RPL6,RACK1,P4HB,TMBIM6,PLD3,SNRNP200,SMIM14,EIF4B,ZNF43,HMGB1,POLR2A,TPT1,T
1,RPS6,ZNF316,PIK3R2,NCL,ALDOA,HSP90B1,DDX5,RPL13A,MAGED1,RPL6,DVL3,GPI,RACK1,AT
,SPTBN1,IGF2,VCP,MARCKSL1,AFF4,TRIM28,CKB,COL4A2,SET,TPI1,GTF3C4,PKD1,RPS3,VAPA,T
F1,RPL13A,RPL7,RPL5,TMBIM6,SPEN,SERINC1,POLR2A,DNAJA1,CDC42BPB,GTF2F1,PGAM1,SM
PA1,ARSD,HSPA9,NEDD4L,POLH,ATN1,ACTB,CTSC,ODF2L,USP22,MTRNR2L2,MED18,ACTG1,EEF
2,NUMA1,RPS6,ZNF316,PIK3R2,NCL,ALDOA,HSP90B1,CANX,DDX5,MAGED1,RPL6,GPI,SHISA9,P4
XNB2,UBB,DCHS1,SET,MRC2,SSBP2,TUBB4B,AGRN,PXDN,OGA,FADS2,SLC16A7,KPNB1,HK1,CC
L2,MED18,EEF2,TUBA1A,HSP90AB1,FTH1,UBC,KIDINS220,TMSB4X,NUMA1,EIF4G2,RPL3,RPS6,F
B1,HSP90AA1,ACTR2,FTH1,RPS6,ZNF316,ENO1,PIK3R2,NCL,ALDOA,GANAB,HSP90B1,CANX,CLU,
1,MARCKS,CALM1,IGF2,YWHAG,TMSB10,POLR2A,MORF4L1,NCAM1,AGRN,RPLP1,ZMIZ1,DNAJ
0,NEDD4L,NUDC,NRBP2,USP22,ACTG1,DPYSL3,TUBA1A,HSP90AB1,VIM,TUBB,KIDINS220,NUMA
VIM,UBC,KIDINS220,NUMA1,ENO1,MAP1B,ABHD2,TUBB2B,BSG,ALDH6A1,GNB1,SHISA9,STMN2
),POLH,ATN1,ACTB,NRBP2,MED18,ACTG1,DPYSL3,CFL1,MAVS,ZNF124,ACTR2,SPARC,RPL3,RPS
AB1,UBC,NUMA1,RPL3,RPL4,FLNA,HSP90B1,FADS1,RPL8,UBA1,TMF1,PSAP,DDX5,KIF1A,RPL13
L3,NCL,NEFL,PRPF8,YBX1,YWHAE,DDX17,BSG,FASN,EIF3C,AFF4,TRIM28,NONO,SET,RPL15,LDH
A1,RPL3,RPS6,ZNF316,PIK3R2,NCL,ALDOA,GANAB,NEFL,DDX5,MAGED1,RPL6,DVL3,GNAS,GPI,I
SD,MS4A10,NEDD4L,POLH,ACTB,CTSC,ODF2L,NUDC,NRBP2,USP22,MTRNR2L2,MED18,ACTG1,E
2,KIDINS220,RPS6,ZNF316,PIK3R2,NCL,GANAB,HSP90B1,RPL8,YBX1,CANX,MAGED1,RPL6,RACI
8,ACTG1,BTBD9,XRCC2,MAVS,HSP90AB1,ZNF124,HSP90AA1,ACTR2,KIDINS220,TMSB4X,EIF4G
DCHS1,PPP2R1A,SSBP2,STMN1,AP2B1,MAT2A,HNRNPA0,FADS2,TNS3,COL1A2,RHOA,CHD4,CRN
DCHS1,PPP2R1A,SSBP2,STMN1,AP2B1,MAT2A,HNRNPA0,FADS2,TNS3,COL1A2,RHOA,CHD4,CRN
RPL5,TMBIM6,PLD3,SNRNP200,PPP2R1A,SF3B1,HMGB1,POLR2A,TPT1,SF3B2,GTF2F1,RPL29,Z
F6,CDC42BPB,HNRNPU,LLGL1,CLSTN1,NR2F1,RAB6B,CXADR,CHD4,CRMP1,BEX3,MCL1,TLE5,D
S220,RPS6,ZNF316,PIK3R2,NCL,HSP90B1,RPL8,TMF1,RPL13A,RPL6,RACK1,P4HB,PLD3,SNRNP2
AP1B,ABHD2,ALDH6A1,GNB1,STMN2,P4HB,IDS,FSCN1,SPTAN1,CALM1,SMIM14,VCP,UBB,EGLN
KS,HSPD1,IGF2,SET,EGLN1,SSBP2,BASP1,STMN1,RPL23,CPE,TNS3,SLC22A17,KPNB1,C1ORF43

ISP90AB1,VIM,UBC,NUMA1,RPS6,ZNF316,DBN1,RPL7A,FADS1,NEFL,FSTL1,ABHD2,TUBB2B,DD
DC,USP22,MTRNR2L2,CFL1,VIM,NUMA1,RPL3,ZNF316,DBN1,ENO1,PIK3R2,MAP1B,ALDOA,MTR
GANAB,HSP90B1,IGFBP2,COL4A1,PTMA,SEC62,KAT6A,YWHAE,KIF1A,GNAS,HTRA4,CALM3,GNB
ZNF316,PIK3R2,ALDOA,PGK1,FKBP14,RPL13A,MAGED1,RPL6,GPI,RACK1,RPL5,GDI1,TMBIM6,F
RPL7,SPTAN1,ATP5F1A,CKB,LRRC28,NONO,TPI1,TMSB10,GTF3C4,PKD1,SSBP2,CHD3,VAPA,ITG
PL4,FADS1,RPL8,TMF1,TUBB2B,DDX5,KIF1A,DDX17,SEPTIN2,FSCN1,HSPD1,USP11,SNRNP200,A
,PIK3R2,NCL,GANAB,RPL13A,RPL6,P4HB,SNRNP200,EIF4B,PPP2R1A,HMGB1,POLR2A,TPT1,MA
NF124,HSP90AA1,ACTR2,RPS6,ZNF316,PIK3R2,MAP1B,NCL,ALDOA,HSP90B1,NEFL,UBA1,DDX5,
,FLNA,SLC9A7,GANAB,FTL,NEFL,PRPF8,YBX1,TMF1,TUBB2B,ENO2,RPL13A,SEC62,YWHAE,GN
,ZNF316,PIK3R2,HSP90B1,ATP2B4,TMF1,RPL13A,RACK1,CALM3,P4HB,RPL5,PLD3,TPI1,EIF4A1,
,ZNF316,PIK3R2,HSP90B1,ATP2B4,TMF1,RPL13A,RACK1,CALM3,P4HB,RPL5,PLD3,TPI1,EIF4A1,
4X,TUBA1B,MAP1B,ALDOA,HSP90B1,RPL8,YBX1,FSTL1,HSPA1B,TUBB2B,KAT6A,GNAS,STMN2,
K3R2,ALDOA,FKBP14,RPL13A,MAGED1,RPL6,GPI,RACK1,SHISA9,P4HB,GDI1,TMBIM6,HSPD1,SN
9,TUBB2B,H3-3B,CLU,PTMA,SEC62,RPL13,GNAS,DDX17,GDI1,FSCN1,PTMS,GSTP1,CSDE1,MAT
,ZNF124,FTH1,TMSB4X,NUMA1,ZNF316,ENO1,PIK3R2,MAP1B,NCL,RPL7A,COL6A1,FTL,HSP90E
,RPL3,RPL4,FLNA,FADS1,RPL8,TMF1,TUBB2B,DDX5,KIF1A,RPL13,DDX17,ALDH6A1,SEPTIN2,FS
MED18,EEF2,DPYSL3,CFL1,BTBD9,XRCC2,HTT,HSP90AB1,HSP90AA1,SPARC,KIDINS220,NUMA1
4L,ACTB,CTSC,NRBP2,USP22,EEF2,DPYSL3,TUBA1A,XRCC2,MAVS,HTT,ACTR2,TUBB,FTH1,TMS
B,ABHD2,SLC2A3,ALDH6A1,GNB1,STMN2,IDS,FSCN1,SPTAN1,CALM1,SMIM14,VCP,UBB,EGLN1
PKM,PGK1,PTMA,SEC62,YWHAE,FASN,GNB1,RPL7,SEPTIN2,SPTAN1,PLD3,UBB,YWHAG,YWHAQ,
,RPS6,ZNF316,PIK3R2,HSP90B1,TMF1,SOX4,RPL13A,MAGED1,RPL6,RACK1,P4HB,USP11,SNRN
2,MED18,TUBA1A,HSP90AB1,FTH1,UBC,KIDINS220,TMSB4X,NUMA1,EIF4G2,RPL3,RPS6,RPL4,
HD2,ALDH6A1,GNB1,STMN2,IDS,FSCN1,CALM1,SMIM14,VCP,UBB,EGLN1,CHSY1,PPP2R1A,PKD
PL13A,MAGED1,RPS8,TMBIM6,RPL37A,PPP2R1A,RPL15,SF3B1,RPS3,HMGB1,RPS3A,MAT2A,RP
LC9A7,HSP90B1,PTMA,DDX5,RPL13A,MAGED1,GPI,HTRA4,RPS8,TMBIM6,USP11,EIF3C,ATP5F1
,ZNF316,PIK3R2,NCL,ALDOA,HSP90B1,CANX,DDX5,RPL13A,MAGED1,RPL6,DVL3,GPI,RACK1,AT
7L3,CPSF2,SRSF5,U2AF2,CCNL2,BTBD2,PTBP1,ILF3,HNRNPUL1,SRSF3,MESD,ATRX,TRA2B,SMC
,ATN1,CTSC,ODF2L,MTRNR2L2,ACTG1,TUBA1A,CFL1,BTBD9,XRCC2,HTT,HSP90AB1,VIM,TUBE
2,KIDINS220,RPS6,ZNF316,PIK3R2,NCL,ALDOA,GANAB,HSP90B1,RPL8,H3-3B,PTMA,RPL13A,RP
BB2B,YWHAE,KIF1A,GNAS,RPS8,APP,CALM1,TMSB10,EIF4A1,HLA-C,RPL18,STMN1,AGRN,AP3D1
6,ZNF316,PIK3R2,NCL,ALDOA,HSP90B1,DDX5,MAGED1,RPL6,GPI,SHISA9,P4HB,USP11,EIF3C,AT
IDN,AP3D1,GOLGA8A,SCARB2,BRD3,ZNF865,CHD4,CD81,ATP2A2,OAZ1,SOX12,APLP1,MIF,APL
RBP2,CFL1,HTT,HSP90AA1,ACTR2,VIM,UBC,KIDINS220,ENO1,PIK3R2,FLNA,MAP1B,PRPF8,ABH
ANX,RPL13A,MAGED1,RPL6,GNAS,RACK1,TMBIM6,PLD3,SNRNP200,EIF4B,EIF1,PPP2R1A,SF3B1
,DPYSL3,MAVS,VIM,TMSB4X,EIF4G2,ZNF316,NCL,ALDOA,COL6A1,NEFM,FBXL4,UBA1,IGFBP2,
A,FADS1,TMF1,DDX5,KIF1A,RPL13,DDX17,ALDH6A1,SEPTIN2,FSCN1,HSPD1,USP11,SNRNP200,U
AE,SPTBN1,SHISA9,PLXNB2,TMSB10,MRC2,RPL15,PKD1,CHD3,TPT1,RPS17,MAT2A,HNRNPU,LA
1,HSP90AA1,NUMA1,ENO1,PKM,FTL,HSP90B1,RPL8,CLU,ENO2,DDX17,BSG,P4HB,RPL5,TMBIM
,HSPA9,NEDD4L,ATN1,ACTB,ODF2L,USP22,MTRNR2L2,ACTG1,EEF2,DPYSL3,TUBA1A,CFL1,BTB
SOX4,KAT6A,BSG,GNB1,MYH9,APP,NPM1,CSDE1,EIF4B,UBB,COL4A2,TMSB10,CHD3,VAPA,MOR
16,PIK3R2,NCL,ALDOA,HSP90B1,CANX,DDX5,MAGED1,RPL6,GPI,RACK1,SHISA9,P4HB,TMBIM6,
1A,GPI,BSG,FASN,GNB1,NPM1,HSPD1,COL18A1,EIF4B,AFF4,TRIM28,EIF4A1,GTF3C4,SSBP2,VA
M3,IDS,SMG1,PLXNB2,SET,GTF3C4,POLR2A,AGRN,PXDN,HNRNPU,FADS2,SLC16A7,KPNB1,HK
RPL13A,KIF1A,ALDH6A1,RPS8,SEPTIN2,P4HB,RPL5,FSCN1,GSTP1,PLD3,USP11,SNRNP200,SMI
,COL6A1,HSP90B1,FADS1,COL5A1,PTMA,DDX5,YWHAE,KIF1A,GNAS,RPS8,GDI1,GSTP1,HSPD1,

:FM,FBXL4,UBA1,HSPA1B,DVL3,GPI,ALDH6A1,SPTBN1,CALM3,GNB1,SEPTIN2,GDI1,MARCKS,TT(
RF1,ITM2B,GOLGA8A,SCARB2,ZNF865,CHD4,WDR1,ATP2A2,OAZ1,KHDRBS1,APLP1,TARDBP,AR(
NUMA1,RPL3,RPL4,FLNA,FADS1,TMF1,DDX5,KIF1A,RPL13,DDX17,ALDH6A1,SEPTIN2,FSCN1,USP
,HSP90B1,DVL3,EIF4B,PPP2R1A,HMGB1,POLR2A,GTF2F1,RHOA,ZNF865,RPS14,WDR1,RBBP4,F
M28,SET,ITGB1,TIMP2,CYB5R3,CDC42BPB,NCS1,ZNF865,JUND,LASP1,OAZ1,CCNI,MEGF8,RAB1
RC,TUBB,FTH1,RPS6,ZNF316,PIK3R2,PKM,ZNF99,DDX5,RPL6,KAT6A,YWHAE,KIF1A,FASN,ALDH6
PAPOLA,CBX5,ZNF865,KIF5B,LENG8,VAT1,RTN3,KHDRBS1,EIF5A,ACTN4,HNRNPH1,CTBP1,EID1
PA1,ARSD,HSPA9,NEDD4L,POLH,ATN1,ACTB,CTSC,NUDC,MED18,EEF2,DPYSL3,CFL1,BTBD9,AC
B,NPM1,CSDE1,MATR3,PSMD5,EIF4A1,RPS3A,SMARCA5,RPS9,RPL17,RAB6B,ENC1,EIF5,RPL26,
ZNF124,HSP90AA1,ACTR2,FTH1,KIDINS220,RPS6,ZNF316,PIK3R2,FLNA,NCL,ALDOA,HSP90B1,Y
,HNRNPA1,ARSD,NEDD4L,POLH,ATN1,ACTB,ODF2L,NRBP2,USP22,ACTG1,EEF2,TUBA1A,XRCC2,
I,GANAB,RPLP0,CANX,PTMA,DVL3,GNAS,FASN,GNB1,MYH9,NPM1,IGF2,EIF4B,MARCKSL1,CKB,
IBB2B,PTMA,SEPTIN2,APP,IGF2,UBB,YWHAQ,SSBP2,BASP1,POLR2A,MORF4L1,BTF3,VCL,DNAJA
.H,ODF2L,NUDC,ACTG1,XRCC2,HSP90AB1,ZNF124,HSP90AA1,KIDINS220,EIF4G2,RPL3,RPS6,ZN
JSP22,ACTG1,BTBD9,XRCC2,MAVS,HTT,HSP90AB1,ZNF124,HSP90AA1,ACTR2,VIM,FTH1,UBC,TN
VHAQ,AGRN,SARAF,ARF1,CDC42BPB,PJA2,CBX5,ZNF865,CHD4,CD81,HMGN2,ATP2A2,LASP1,PF
A,FADS1,TMF1,DDX5,KIF1A,RPL13,DDX17,ALDH6A1,SEPTIN2,FSCN1,USP11,SNRNP200,TRIM28,
1,ACTR2,NUMA1,ZNF316,PIK3R2,NCL,GANAB,RPL8,DDX5,RPL13A,RPL6,RACK1,SHISA9,P4HB,PL
Q,SET,CHSY1,UCHL1,MORF4L1,TIMP2,RPL10,AP3D1,CYB5R3,MAP4,LLGL1,CLSTN1,PSMD2,NL(
0AB1,ZNF124,HSP90AA1,KIDINS220,RPS6,ZNF316,RPL4,PIK3R2,NCL,SLC9A7,ALDOA,HSP90B1,
GANAB,ZNF99,RPLP0,DDX5,RPL13A,KAT6A,DVL3,YWHAE,DDX17,RACK1,STMN2,SEPTIN2,GDI1,T
F4G2,PKM,FTL,PRPF8,YBX1,UBA1,ATP2B4,CALR,TMF1,HSPA1B,PGK1,CANX,RPL6,KAT6A,FASN,
316,PIK3R2,RPL8,RPL13A,GNAS,P4HB,PPP2R1A,HMGB1,POLR2A,SF3B2,GTF2F1,RPL29,PAPOL
3B2B,RPLP0,KIF1A,GNB1,SEPTIN2,IDS,FSCN1,UBB,AFF4,MATR3,FN1,SPEN,GTF3C4,HMGB1,POI
NO1,MAP1B,ABHD2,TUBB2B,BSG,ALDH6A1,SHISA9,STMN2,RPL7,P4HB,IDS,FSCN1,SMIM14,VCP,
HNRNPA1,MS4A10,HSPA9,NEDD4L,POLH,ATN1,ACTB,CTSC,ODF2L,NUDC,NRBP2,USP22,MED18,
RCC2,HSP90AB1,UBC,RPL3,RPL4,ENO1,FLNA,COL6A1,HSP90B1,FADS1,FBXL4,TMF1,TUBB2B,R
,XRCC2,HSP90AB1,UBC,RPL3,RPL4,ENO1,FLNA,COL6A1,HSP90B1,FADS1,RPL8,FBXL4,UBA1,TN
,COPA,HNRNPA1,ARSD,MS4A10,HSPA9,NEDD4L,POLH,ATN1,ACTB,CTSC,ODF2L,NUDC,NRBP2,U
IUMA1,PKM,ATP2B4,FSTL1,PTMA,ENO2,SOX4,GNAS,SEPTIN2,PTMS,PLXNB2,IGF2,EIF4B,SPEN,N
FTH1,UBC,KIDINS220,PIK3R2,MAP1B,GANAB,PRPF8,ABHD2,TUBB2B,FKBP14,SLC2A3,ALDH6A1
3R2,NCL,HSP90B1,DDX5,MAGED1,RPL6,GPI,RACK1,P4HB,USP11,EIF3C,ATP5F1A,RPL19,SERINC
.DH6A1,STMN2,IDS,FSCN1,SMIM14,VCP,UBB,EGLN1,CHSY1,PPP2R1A,PKD1,SF3B1,CDKL1,STM
3R2,NCL,ALDOA,HSP90B1,DDX5,MAGED1,RPL6,GPI,SHISA9,P4HB,USP11,ATP5F1A,RPL19,SERIN
RNR2L2,MED18,EEF2,DPYSL3,TUBA1A,MAVS,HSP90AB1,FTH1,UBC,KIDINS220,TMSB4X,EIF4G2,
NPH1,ATP6V0C,SRSF5,U2AF2,ILF3,SRSF3,MESD,MAPK8IP1,CYP51A1,STARD7,HNRNPA2B1,KMT
2,EEF2,CFL1,BTBD9,MAVS,HTT,ZNF124,TUBB,NUMA1,EIF4G2,ZNF316,DBN1,RPL4,TUBA1B,SLC
.H,NUDC,NRBP2,MED18,CFL1,BTBD9,HTT,HSP90AB1,HSP90AA1,UBC,TMSB4X,NUMA1,ZNF316,F
USP22,ACTG1,EEF2,TUBA1A,BTBD9,HTT,ZNF124,HSP90AA1,FTH1,KIDINS220,NUMA1,ZNF316,D
NS220,RPS6,ZNF316,PIK3R2,ALDOA,HSP90B1,ABHD2,PGK1,H3-3B,CLU,FKBP14,RPL13A,MAGEL
AB1,ZNF124,HSP90AA1,FTH1,UBC,TMSB4X,NUMA1,RPL3,ZNF316,ENO1,PIK3R2,FLNA,NCL,PKM
SP90B1,HSPA1B,KAT6A,YWHAE,RPL13,DDX17,RACK1,GNB1,SHISA9,RPS8,P4HB,TMBIM6,SPTAN
HD4,JUND,CD81,ATP2A2,OAZ1,SOX12,MAGED2,APLP1,MIF,APLP2,CADM4,TRIB2,SQLE,CTDSP2,
),XRCC2,MAVS,HSP90AB1,UBC,NUMA1,RPL3,RPL4,ENO1,FLNA,COL6A1,HSP90B1,FADS1,RPL8,Y
6,ZNF316,PIK3R2,NCL,ALDOA,HSP90B1,DDX5,MAGED1,RPL6,GPI,RACK1,SHISA9,P4HB,GDI1,US

RPL4,ENO1,FLNA,HSP90B1,FADS1,TMF1,TUBB2B,DDX5,RPL13A,KIF1A,DDX17,ALDH6A1,SEPTIN
HSP90AB1,UBC,RPL3,RPL4,ENO1,FLNA,COL6A1,HSP90B1,FADS1,FBXL4,TMF1,TUBB2B,RPLP0,
RPL6,RACK1,ATP5F1B,SHISA9,SEPTIN2,P4HB,RPL5,PLD3,SNRNP200,EIF4B,EIF1,GTF3C4,HMGE
DDF2L,MED18,XRCC2,MAVS,HSP90AB1,ZNF124,HSP90AA1,ACTR2,KIDINS220,RPS6,ZNF316,RPL
AGRN,PXDN,SLC16A7,COL1A2,HK1,NR2F1,NLGN2,GLIPR1L2,ATP2A2,CCNI,TADA2B,RPS25,TKT,
B,NLGN2,GLIPR1L2,CHD4,ATP1A1,TKT,PIP4K2B,APLP1,CKAP4,COTL1,TMEM30A,PEA15,ACTR3,
.5A1,CES3,KIF1A,SLC2A3,IDS,SPTAN1,MARCKS,HSPD1,MATR3,RPL37A,DCHS1,SET,MRC2,SSBP2
HS1,SET,CHSY1,CYB5R3,RHOA,SLC7A5,NLGN2,CXADR,EIF5,LASP1,TLE5,NES,SOX12,CCN2,ARL4
L1,XRCC2,MAVS,HSP90AB1,ZNF124,HSP90AA1,ACTR2,FTH1,UBC,KIDINS220,DCX,RPS6,ZNF316
,BTBD9,HTT,ZNF124,HSP90AA1,ACTR2,FTH1,KIDINS220,NUMA1,DCX,ZNF316,DBN1,PIK3R2,FLI
.3,CFL1,BTBD9,HTT,TUBB,FTH1,EIF4G2,DBN1,PIK3R2,MAP1B,ALDOA,PKM,GANAB,NEFM,UBA1,
CKB,SERINC1,CHD3,TPT1,NCAM1,MARCHF6,SBF1,DNAJA1,AP2B1,CNN3,XRCC5,KPNB1,RPS20,
HSP90AB1,HSP90AA1,ACTR2,FTH1,KIDINS220,NUMA1,RPL3,RPS6,ZNF316,DBN1,ENO1,PIK3R2,N
I1,ODF2L,NRBP2,USP22,ACTG1,EEF2,DPYSL3,HSP90AB1,VIM,KIDINS220,RPL3,DBN1,ENO1,FLN
A1B,NCL,RPL7A,TUBB2B,PTMA,DVL3,GNAS,ATP5F1B,CALM3,SEPTIN2,P4HB,APP,UBB,PSMD5,EIF
B,USP11,SMIM14,SPEN,RPL15,HMGB1,POLR2A,RPL11,RPLP1,RPL10,SF3B2,RPS14,PFN1,RPL31
TB,ODF2L,USP22,MED18,ACTG1,TUBA1A,CFL1,XRCC2,MAVS,HSP90AB1,ZNF124,HSP90AA1,ACT
C2,HSP90AB1,TUBB,FTH1,UBC,NUMA1,EIF4G2,RPL3,RPL4,ENO1,FLNA,NCL,MTRNR2L8,FTL,HSP
2,DPYSL3,CFL1,BTBD9,XRCC2,MAVS,HSP90AB1,UBC,NUMA1,EIF4G2,RPL3,RPL4,ENO1,FLNA,R
NF124,FTH1,NUMA1,EIF4G2,ZNF316,PIK3R2,SLC9A7,ALDOA,PKM,CALR,TUBB2B,COL4A1,PSAP
AP1B,PKM,TUBB2B,CES3,PSAP,ENO2,YWHAE,GNAS,ALDH6A1,SPTBN1,GNB1,PTMS,PLXNB2,SMI
A,FADS1,TMF1,DDX5,KIF1A,RPL13,DDX17,ALDH6A1,SEPTIN2,FSCN1,USP11,SNRNP200,TRIM28,
A1,UBC,TMSB4X,NUMA1,ZNF316,DBN1,FLNA,PKM,FADS1,NEFM,FBXL4,YBX1,UBA1,TUBB2B,CC
USP11,SNRNP200,AFF4,CKB,FN1,EIF1,MAP9,POLR2A,TPT1,MORF4L1,MIDN,MARCHF6,WSB1,IT
LR2A,FADS2,HDLBP,HNRNPK,UBE2H,SEPTIN3,GPC1,NOVA2,HNRNPL,HNRNPA3,MDK,RAD23B,AS
D4L,ACTB,CTSC,NRBP2,CFL1,XRCC2,MAVS,HTT,HSP90AB1,HSP90AA1,ACTR2,SPARC,FTH1,TMS
D4L,POLH,ACTB,NRBP2,USP22,DPYSL3,BTBD9,HSP90AB1,VIM,NUMA1,RPS6,ZNF316,DBN1,ENC
3,RPL19,HMGB1,BTF3,RPLP1,MARCHF6,SF3B2,ZNF865,CXADR,LENG8,PRKCSH,EIF5A,PDIA6,ITN
FSCN1,COL4A2,RPL37A,TMSB10,STMN1,AGRN,TIMP2,CTSD,NACA,PAPOLA,NLGN2,CD81,TLE5,
KIDINS220,ZNF316,PIK3R2,FLNA,RPL7A,PKM,HSP90B1,FBXL4,YBX1,UBA1,TMF1,TUBB2B,COL5A
ACTR2,TUBB,FTH1,UBC,KIDINS220,TUBA1B,MAP1B,PKM,ABHD2,TUBB2B,PGK1,CANX,ENO2,KAT
PPP2R1A,PDIA3,VAPA,STMN1,UCHL1,ANXA2,AGRN,GOLGA8A,PGAM1,CTNNA1,RPL29,SCARB2,R
TN1,ACTB,NRBP2,EEF2,TUBA1A,HTT,HSP90AB1,ZNF124,HSP90AA1,VIM,TMSB4X,ZNF316,DBN1,F
A9,POLH,CTSC,NUDC,NRBP2,USP22,ACTG1,DPYSL3,TUBA1A,CFL1,BTBD9,HTT,HSP90AB1,HSP90
UBC,RPL3,RPS6,RPL4,ENO1,HSP90B1,RPL8,FBXL4,TMF1,TUBB2B,DVL3,KIF1A,GPI,BSG,GNB1,S
ADS1,NEFM,YBX1,UBA1,COL5A1,RPLP0,PTMA,RPL13A,KAT6A,APP,GDI1,FSCN1,IGF2,DDB1,EGLN
,RPS8,SPTAN1,GSTP1,SMG1,NPM1,EIF3C,TRIM28,YWHAG,TPI1,SF3B1,CHD3,POLR2A,TPT1,RPL
D18,EEF2,DPYSL3,CFL1,BTBD9,XRCC2,HSP90AB1,UBC,NUMA1,EIF4G2,RPL3,RPL4,ENO1,FLNA,
C2,MAVS,HSP90AB1,UBC,NUMA1,RPL3,RPL4,ENO1,FLNA,COL6A1,HSP90B1,FADS1,RPL8,YBX1,T
BB,FLNA,RPL7A,COL6A1,FTL,HSP90B1,PRPF8,UBA1,ATP2B4,HSPA1B,COL4A1,YWHAE,GPI,HTRA
3,CFL1,BTBD9,XRCC2,HSP90AB1,UBC,NUMA1,EIF4G2,RPL3,RPL4,ENO1,FLNA,RPL7A,ALDOA,CC
NB1,P4HB,GDI1,PTMS,PLXNB2,MARCKSL1,DCHS1,PPP2R1A,TPT1,ITGB1,MIDN,HNRNPU,TPM4,
,USP22,ACTG1,XRCC2,HSP90AA1,ACTR2,SPARC,FTH1,TMSB4X,EIF4G2,ZNF316,RPL4,NCL,SLC9
C9A7,PKM,FTL,HSP90B1,CALR,PSAP,ENO2,KAT6A,DVL3,SPTBN1,CALM3,RPL5,FSCN1,MARCKS,N
UBA1B,RPL7A,COL6A1,HSP90B1,RPL8,TMF1,CES3,PSAP,RPLP0,CANX,DDX5,ENO2,KAT6A,KIF1A,

AP1B,NCL,RPL7A,FTL,FADS1,NEFL,NEFM,PRPF8,UGT8,RPL8,ATP2B4,TMF1,ABHD2,COL4A1,PSA
ATN1,ACTB,CTSC,ODF2L,NUDC,USP22,MTRNR2L2,ACTG1,EEF2,DPYSL3,CFL1,BTBD9,XRCC2,H
LDH6A1,FSCN1,SMIM14,VCP,CHSY1,CDKL1,STMN1,UCHL1,AGRN,MIDN,AP3D1,AKAP12,GOLGA
3,CFL1,BTBD9,XRCC2,HSP90AB1,UBC,EIF4G2,RPL3,RPL4,ENO1,FLNA,RPL7A,ALDOA,COL6A1,H
SD,MS4A10,HSPA9,NEDD4L,POLH,ATN1,ACTB,CTSC,ODF2L,NRBP2,USP22,MED18,ACTG1,EEF2,D
RPS14,PFN1,MCL1,EIF5A,EIF3L,NGRN,RPLP2,DDOST,YWHAB,ARHGDIA,TM9SF3,MDH2,U2AF2,G
,PXDN,CTSD,MSN,CHD4,CD81,ATP2A2,BEX3,OAZ1,ASTN2,TADA2B,SPTBN2,DDX3X,RCN1,H1-0,JU
3,RPL19,HMGB1,BTF3,RPLP1,MARCHF6,SF3B2,ZNF865,CXADR,LENG8,PRKCSH,EIF5A,PDIA6,ITM
SC,ODF2L,USP22,MED18,ACTG1,DPYSL3,MAVS,HSP90AB1,HSP90AA1,VIM,FTH1,KIDINS220,NUM
28,CKB,FN1,EIF1,TPT1,RPL18,SARAF,MIDN,MARCHF6,WSB1,GTF2F1,HDLBP,PEBP1,TLN1,CHPF
L5,COL18A1,IGF2,SET,MRC2,RPL19,PPP2R1A,CDC42BPB,NFE2L1,CPE,EIF4H,LPCAT1,C1ORF43
R2L2,EEF2,CFL1,HSP90AA1,VIM,FTH1,NUMA1,RPL3,ENO1,MAP1B,NCL,MTRNR2L8,PKM,GANAB,
4L,ODF2L,NUDC,NRBP2,ACTG1,EEF2,CFL1,XRCC2,MAVS,ZNF124,HSP90AA1,VIM,KIDINS220,ZN
,MS4A10,HSPA9,NEDD4L,POLH,ATN1,ACTB,CTSC,ODF2L,NUDC,NRBP2,USP22,MED18,ACTG1,E
C,KIDINS220,NUMA1,EIF4G2,RPL3,RPS6,DBN1,ENO1,PIK3R2,NCL,SLC9A7,ALDOA,PKM,GANAB,N
PL13,GNB1,SEPTIN2,FSCN1,GSTP1,COL18A1,ATP5F1A,MAPT,DDB1,BASP1,VAPA,RPS3A,TUBB4B
RPL8,RPL6,GNAS,P4HB,SNRNP200,EIF4B,HMGB1,POLR2A,TPT1,SF3B2,GTF2F1,RPL29,PAPOLA
JMA1,RPL4,RPL7A,PKM,RPL8,CALR,CLU,ENO2,KIF1A,RPL13,GPI,RACK1,FASN,SPTBN1,APP,IDS,
1,ACTB,ODF2L,NUDC,NRBP2,USP22,ACTG1,EEF2,TUBA1A,MAVS,HTT,HSP90AB1,ZNF124,HSP90A
2,ABHD2,SLC2A3,ALDH6A1,FSCN1,SPTAN1,SNRNP200,SMIM14,CHSY1,VAPA,CDKL1,STMN1,UCH
NUMA1,RPL3,ENO1,MAP1B,NCL,MTRNR2L8,PKM,PRPF8,RPL8,YBX1,UBA1,PTMA,DDX5,SOX4,KA
L,POLH,ACTB,CTSC,ODF2L,NUDC,NRBP2,USP22,MTRNR2L2,ACTG1,EEF2,DPYSL3,ZNF124,HSP9
AB1,UBC,RPL3,RPL4,ENO1,FLNA,COL6A1,HSP90B1,FADS1,TMF1,TUBB2B,DDX5,KIF1A,RPL13,D
90B1,DDX5,MAGED1,RPL6,P4HB,USP11,RPL19,SERINC1,HMGB1,POLR2A,AP2M1,BTF3,RPLP1,M
4,HSP90AA1,ACTR2,FTH1,KIDINS220,NUMA1,ZNF316,DBN1,PIK3R2,FLNA,NCL,RPL7A,PKM,COL
RPL6,RACK1,SHISA9,SEPTIN2,P4HB,RPL5,TMBIM6,PLD3,SNRNP200,SMIM14,EIF4B,EIF1,GTF3C
S2,DDOST,COTL1,U2AF2,LMNB2,ACO2,RAPGEF1,ATG2B,ADAM9,RPL28,ARNT2,EXTL3,ECEL1,H
2,GSTP1,PLXNB2,SNRNP200,SMIM14,EIF3C,AFF4,MATR3,SET,EGLN1,SF3B1,RPS3A,MIDN,LAMA
HSP90B1,NEFM,YBX1,FSTL1,TUBB2B,CES3,RPL6,DVL3,YWHAE,RPL13,DDX17,BSG,ATP5F1B,MY
28,EXTL3,KIAA0895L,HNRNPA2B1,UBE2M,TLE1,EEF1B2,PRPF19,MTA2,FAF2,PCGF3,DEAF1,GSK
ICL,HSPA1B,DDX5,FKBP14,MAGED1,RPL6,DVL3,ATP5F1B,RPS8,MYH9,P4HB,RPL5,TMBIM6,SNR
P11,PPP2R1A,HNRNPA0,CNN3,RPL29,ZNF865,RPS14,MCL1,EIF5A,EIF3L,CHPF,NGRN,RPLP2,DD
IA1,MAP1B,ABHD2,ENO2,SLC2A3,ALDH6A1,STMN2,IDS,FSCN1,SPTAN1,SMIM14,VCP,UBB,EGLN
B,TUBB2B,RPLP0,H3-3B,PTMA,DDX5,SOX4,RPL13,ALDH6A1,P4HB,TMBIM6,MARCKS,PLD3,VCP,
A,FADS1,TMF1,KIF1A,ALDH6A1,SEPTIN2,FSCN1,USP11,SNRNP200,TRIM28,CKB,FN1,YWHAQ,TP
ATN1,ACTB,CTSC,ODF2L,NRBP2,USP22,ACTG1,EEF2,DPYSL3,HSP90AB1,SPARC,TUBB,NUMA1,D
,MAGED1,RPL6,P4HB,USP11,RPL19,SERINC1,HMGB1,POLR2A,AP2M1,BTF3,RPLP1,MARCHF6,D
PL8,YBX1,CALR,PSAP,CANX,DDX5,ENO2,RPL13A,KAT6A,HTRA4,BSG,FASN,ALDH6A1,SEPTIN2,GS
5,ENO2,FASN,SEPTIN2,EIF4A1,RPL15,LDHB,SERINC1,BASP1,POLR2A,TPT1,STMN1,MORF4L1,AG
,RPS7,RPS19,HNRNPK,LAMP1,GDI2,ARF3,PRDX1,ATP5PB,WDR6,RPS11,SUPT5H,CAND1,NKTR,M
2,PDIA3,AGRN,TIMP2,SBF1,MSN,IGFBP3,NCS1,ASTN2,TADA2B,CSNK1E,PFKL,ST3GAL2,CIRBP,S
A1,MS4A10,NEDD4L,POLH,ATN1,ACTB,CTSC,ODF2L,NUDC,NRBP2,USP22,ACTG1,EEF2,DPYSL3,T
,CHD3,WSB1,EIF4H,SLC16A7,PAPOLA,RHOA,ATF4,PFN1,PIP4K2B,MTSS2,CKAP5,RPS27,SMCHD
RPL3,RPL4,FLNA,FADS1,TMF1,DDX5,KIF1A,RPL13,GPI,ALDH6A1,SEPTIN2,FSCN1,GSTP1,USP11,
JSP22,MTRNR2L2,CFL1,XRCC2,UBC,TMSB4X,NUMA1,RPS6,RPL4,ENO1,TUBA1B,MTRNR2L8,FA

HSPA1B,TUBB2B,H3-3B,PTMA,DDX5,SOX4,RPL13,FASN,P4HB,VCP,UBB,PPP2R1A,HLA-C,PDIA3,Z
D4L,POLH,ACTB,ODF2L,ACTG1,EEF2,DPYSL3,MAVS,HTT,HSP90AB1,ZNF124,HSP90AA1,ACTR2,
AS,EIF4B,EIF1,HMGB1,SF3B2,RPL29,ZNF865,RPS14,PFN1,MCL1,PNN,RPS25,EIF5A,PDIA6,EIF3L
,ACTG1,EEF2,DPYSL3,TUBA1A,HTT,VIM,FTH1,UBC,EIF4G2,RPS6,ZNF316,TUBA1B,NCL,ALDOA,C
2,RPLP0,DDX5,RPL13A,KAT6A,DVL3,YWHAE,DDX17,RACK1,STMN2,SEPTIN2,GDI1,USP11,CSDE1
R2,HSP90B1,RPL6,SNRNP200,PPP2R1A,DNAH8,HMGB1,NCAM1,MARCHF6,RPS17,DNAJA1,SF3B
HD2,ALDH6A1,FSCN1,SMIM14,VCP,CHSY1,CDKL1,STMN1,UCHL1,AGRN,MIDN,AP3D1,AKAP12,G
F124,ACTR2,SPARC,NCL,COL6A1,HSP90B1,FADS1,YBX1,CALR,ABHD2,PTMA,DDX5,MAGED1,RPL
,CALM1,NPM1,IGF2,UBB,AFF4,EGLN1,MAP9,GTF3C4,TUBB4B,MAP4,HNRNPU,CTSD,MAGED2,T
16,MAP1B,RPL7A,RPL8,IGFBP2,FSTL1,TMF1,TUBB2B,COL5A1,COL4A1,H3-3B,PTMA,KAT6A,GN
IOA,VAT1,RTN3,KHDRBS1,HNRNPH1,MAGED4B,CTBP1,CDK16,BCLAF1,NNAT,HNRNPL,ABCA3,VI
,LENG8,PFN1,PRKCSH,KHDRBS1,ITM2C,RPS19,NGRN,RPL35A,RPLP2,DDOST,PTGES3,SND1,SR
3,CFL1,BTBD9,XRCC2,HSP90AB1,UBC,EIF4G2,RPL3,RPL4,ENO1,FLNA,RPL7A,ALDOA,COL6A1,H
,RPL19,HMGB1,AP2M1,BTF3,RPLP1,MARCHF6,DNAJA1,SF3B2,ZNF865,CXADR,LENG8,PNN,PRK
18,ACTG1,EEF2,CFL1,BTBD9,MAVS,HTT,HSP90AB1,VIM,SPARC,KIDINS220,TMSB4X,NUMA1,EIF
EF2,CFL1,HSP90AB1,ACTR2,VIM,TUBB,TUBA1B,NCL,SLC9A7,ALDOA,MTRNR2L8,GANAB,HSP90E
RN,TIMP2,NACA,KPNB1,IGFBP3,CHD4,CRMP1,BEX3,TLE5,KHDRBS1,MEGF8,DNAJC5,TAGLN,M
C,RPL3,FADS1,TMF1,DDX5,KIF1A,ALDH6A1,SEPTIN2,FSCN1,USP11,SNRNP200,EIF3C,AFF4,TRIM
ACTR2,EIF4G2,RPL3,ZNF316,DBN1,PIK3R2,MAP1B,ALDOA,GANAB,COL6A1,FTL,HSP90B1,RPL8,
ATN1,ODF2L,NUDC,MED18,BTBD9,XRCC2,ZNF124,HSP90AA1,UBC,EIF4G2,ZNF316,DBN1,TUBA
RPLP2,DDOST,MDH2,U2AF2,SUGP2,PRDX5,UBE4A,GCN1,PTBP1,CCAR2,BAG6,RAB11A,SERBP1,
HSP90B1,ENO2,RPL13A,KIF1A,GNAS,RACK1,CALM3,APP,CALM1,COL18A1,EIF4B,MARCKSL1,Y
RSD,MS4A10,HSPA9,POLH,ODF2L,NUDC,USP22,MTRNR2L2,ACTG1,DPYSL3,TUBA1A,CFL1,BTBD
A,FADS1,TMF1,DDX5,KIF1A,RPL13,ALDH6A1,SEPTIN2,FSCN1,USP11,SNRNP200,TRIM28,CKB,FN
,KIF1A,FSCN1,GSTP1,USP11,AFF4,TRIM28,CKB,FN1,EIF1,TPT1,RPL18,SARAF,MIDN,MARCHF6,
,GNAS,FASN,GNB1,MYH9,IGF2,MARCKSL1,CKB,SPEN,YWHAQ,RPL19,GTF3C4,PKD1,PDIA3,POL
L,POLH,ACTB,CTSC,NRBP2,USP22,MED18,ACTG1,DPYSL3,TUBA1A,HTT,HSP90AB1,ZNF124,HSP
HD2,PGK1,ENO2,KAT6A,ALDH6A1,MYH9,PTMS,EIF4B,MARCKSL1,YWHAG,FN1,EIF1,TMSB10,ST
DX3X,RPS15,RPL12,RPS7,RPS19,HNRNPK,LAMP1,U2AF2,GDI2,HSPA4,TNPO1,PSMB2,HDGF,TA
A,FADS1,TMF1,DDX5,KIF1A,RPL13,SEPTIN2,FSCN1,USP11,SNRNP200,TRIM28,CKB,FN1,TPT1,R
28,CKB,FN1,RPL18,WSB1,AP2B1,GTF2F1,LDHA,JUND,TMEM259,CCNI,GNB2,RPLP2,MYL6,MYH
DH,MLLT1,COPA,HNRNPA1,ARSD,MS4A10,HSPA9,NEDD4L,POLH,ATN1,ACTB,CTSC,ODF2L,NUD
IB,RPL19,HMGB1,POLR2A,SF3B2,XRCC5,RPL22,ZNF865,ATF4,LENG8,PRKCSH,KHDRBS1,EIF5A
D1,RPL6,RACK1,P4HB,TMBIM6,SMIM14,RPL19,PPP2R1A,HMGB1,POLR2A,MORF4L1,SF3B2,R
N2,FSCN1,USP11,SNRNP200,TRIM28,CKB,FN1,TPT1,RPL18,SARAF,MIDN,WSB1,AP2B1,GTF2F1,
HTT,VIM,NUMA1,MAP1B,RPL8,IGFBP2,TUBB2B,CLU,ENO2,KAT6A,KIF1A,SPTBN1,GNB1,SHISA9,
D1,PPP2R1A,POLR2A,DNAJA1,SF3B2,RPL22,ZNF865,RPS14,RBBP4,PFN1,RPS25,RAB7A,RPS19,
IO2,GNAS,ALDH6A1,CALM3,GDI1,CALM1,NPM1,MARCKSL1,COL4A2,EIF1,TIMP2,MIDN,CDC42B
TSC,USP22,ACTG1,EEF2,DPYSL3,CFL1,BTBD9,XRCC2,HSP90AB1,ZNF124,ACTR2,VIM,NUMA1,R
ZNF124,HSP90AA1,ACTR2,KIDINS220,RPS6,ZNF316,PIK3R2,NCL,ALDOA,HSP90B1,FBXL4,ATP2E
NGRN,RPLP2,DDOST,YWHAB,SRSF5,MDH2,U2AF2,PRDX5,ARPC2,SUPT6H,BAG6,RAB11A,CDC42
B,FN1,RPL18,WSB1,AP2B1,GTF2F1,LDHA,JUND,TMEM259,CCNI,RPLP2,MYL6,MYH10,UBE2H,S
A,FADS1,TMF1,DDX5,KIF1A,RPL13,ALDH6A1,SEPTIN2,FSCN1,USP11,SNRNP200,TRIM28,CKB,FN
0,NEDD4L,POLH,ATN1,CTSC,NUDC,NRBP2,USP22,ACTG1,EEF2,DPYSL3,CFL1,XRCC2,MAVS,HTT
,SEPTIN2,SPTAN1,SMG1,CALM1,AFF4,TPT1,CDC42BPB,CPE,LAMA5,KPNB1,RHOA,LASP1,BEX3,

2,RPS9,ZNF865,WDR1,PFN1,RPL31,EIF5A,EIF3L,RPLP2,DDOST,MDH2,U2AF2,TSR1,PRDX5,PRPF
PL7A,COL6A1,HSP90B1,NEFL,FSTL1,PTMA,ENO2,ALDH6A1,ATP5F1B,GNB1,RPL7,RPL5,PTMS,MA
3P2,RPL13A,SHISA9,RPS8,P4HB,USP11,SNRNP200,ATP5F1A,LRRC28,RPL19,GTF3C4,HMGB1,TPT
PDH,MLLT1,COPA,HNRNPA1,ARSD,MS4A10,NEDD4L,POLH,ATN1,ACTB,CTSC,ODF2L,NRBP2,USP2
UBC,TMSB4X,NUMA1,ENO1,ALDOA,GANAB,HSP90B1,NEFL,NEFM,RPL8,CALR,TMF1,HSPA1B,ZN
OA,ATP2B4,IGFBP2,TMF1,RPL13A,RPL6,RPL13,DDX17,P4HB,SNRNP200,EIF4A1,PPP2R1A,DNAH8
DH6A1,SEPTIN2,GSTP1,SMIM14,VCP,POLR2A,TPT1,MORF4L1,MIDN,MARCHF6,WSB1,ARF1,SF3I
FSTL1,RPLP0,DDX5,RPL13A,YWHAE,DDX17,RACK1,SEPTIN2,GDI1,USP11,CSDE1,ATP5F1A,MATR3
CFL1,BTBD9,XRCC2,HSP90AA1,TUBB,NUMA1,RPS6,RPL4,PIK3R2,FLNA,MAP1B,ALDOA,PKM,FTL
T8,IGFBP2,COL4A1,RPLP0,CANX,PTMA,DDX5,KAT6A,RPL13,GPI,BSG,FASN,ALDH6A1,GNB1,P4HE
ODF2L,MED18,ACTG1,XRCC2,MAVS,HSP90AB1,ZNF124,HSP90AA1,ACTR2,KIDINS220,NUMA1,R
FN1,EIF5A,EIF3L,RPLP2,DDOST,HSBP1,TM9SF3,MDH2,U2AF2,PRDX5,UBE4A,BAG6,RAB11A,CDC
PS14,RBBP4,PFN1,RAB7A,RPS19,NGRN,RPLP2,DDOST,YWHAB,RPS27,ARHGDIA,MDH2,U2AF2,G
HAG,COL4A2,DCHS1,EGLN1,PDIA3,TUBB4B,ANXA2,AGRN,ZMIZ1,MIDN,AP3D1,CYB5R3,CDC42BI
4,VCP,CHSY1,CDKL1,STMN1,UCHL1,MIDN,AKAP12,GOLGA8A,CTNNA1,SCARB2,RPS24,RPL22,ZN
NS220,TMSB4X,MAP1B,RPL7A,GANAB,PRPF8,HSPA1B,TUBB2B,FKBP14,ALDH6A1,IDS,ERVV-1,FS
3,RPL4,FADS1,TMF1,DDX5,KIF1A,ALDH6A1,SEPTIN2,FSCN1,USP11,SNRNP200,TRIM28,CKB,FN1,
,MARCHF6,WSB1,AP2B1,LDHA,SLC22A17,JUND,TMEM259,LENG8,CCNI,HDLBP,HNRNPDL,MYL6
A,TMSB10,SSBP2,CHD3,RPS3A,ZMIZ1,MAT2A,OGA,CNN3,TNS3,LPCAT1,RPS24,MSN,KPNB1,ZNF8
,RPLP0,DDX5,RPL13A,KAT6A,DVL3,YWHAE,DDX17,RACK1,SEPTIN2,GDI1,USP11,CSDE1,ATP5F1A
NUDC,ACTG1,EEF2,DPYSL3,CFL1,BTBD9,MAVS,HTT,HSP90AB1,ZNF124,HSP90AA1,TUBB,FTH1,T
CTR2,NUMA1,ZNF316,ENO1,PIK3R2,FADS1,UBA1,ATP2B4,TMF1,CANX,RPL13A,MAGED1,RPL6,RA
TD4,CTSB,MTRF1L,PRDX5,SKI,PDE7A,PSMD3,SDHA,ANP32A,H2AZ1,MGAT4B,FARP1,CUL3,ITGB
,BTBD9,XRCC2,MAVS,HSP90AB1,UBC,NUMA1,RPL3,RPL4,ENO1,FLNA,COL6A1,HSP90B1,FADS1,
PA1B,TUBB2B,RPLP0,H3-3B,PTMA,DDX5,SOX4,RPL13,FASN,ALDH6A1,P4HB,KLRD1,TMBIM6,PL
PS3,RPS3A,RPS16,LDHA,NACA,HK1,PFKL,RPS19,SQLE,RPL27,PDXK,NCKAP1,ACAP3,PTK7,RAB5B
CC2,HSP90AB1,UBC,TMSB4X,NUMA1,RPL3,RPL4,ENO1,HSP90B1,FADS1,TMF1,TUBB2B,COL5A
,FTH1,TMSB4X,NUMA1,EIF4G2,ENO1,TUBA1B,PIK3R2,MAP1B,GANAB,FADS1,YBX1,FSTL1,HSPA
1,DDX5,KIF1A,RPL13,SEPTIN2,FSCN1,USP11,TRIM28,CKB,FN1,SEC61A1,RPL18,SARAF,MIDN,WS
,COPA,HNRNPA1,ARSD,NEDD4L,POLH,ATN1,ACTB,CTSC,ODF2L,NRBP2,USP22,MED18,DPYSL3,
SOX4,AFF4,RPL15,ZMIZ1,XAF1,RPS16,LDHA,CNN3,RPL17,PAPOLA,LLGL1,RPL14,NCS1,CBX5,CL
RPL12,RPS7,RPS19,HNRNPK,LAMP1,GDI2,HSPA4,PSMB2,RPS15A,ARF3,PRDX1,ATP5PB,WDR6,RI
RPS6,ZNF316,PIK3R2,HSP90B1,FBXL4,TMF1,DDX5,FKBP14,RPL6,GPI,RACK1,P4HB,RPL5,TMBIM
32,RPS20,PFN1,RPL31,KHDRBS1,RPS25,EIF5A,ITM2C,EIF3L,RPLP2,DDOST,PTGES3,RPL23A,SRS
,ATP2B4,IGFBP2,CALR,TMF1,RPL13A,RPL6,RPL13,GNAS,RPS8,P4HB,SNRNP200,CSDE1,PPP2R1
I,CDC42BPB,AP2B1,PXDN,LAMA5,BRD3,NCS1,LENG8,HMGN2,TLE5,HDLBP,MYL6,DUX4,DNER,N
H2,U2AF2,PRDX5,BAG6,CDC42,EIF2AK1,PDAP1,YKT6,MCM3AP,TMEM167A,CHERP,CTTN,ERH,SI
RPS6,COL6A1,IGFBP2,SOX4,FKBP14,RPL13A,SEC62,YWHAE,RACK1,GNB1,STMN2,RPS8,MYH9,S
TSC,ODF2L,NUDC,USP22,MTRNR2L2,EEF2,CFL1,ZNF124,HSP90AA1,VIM,FTH1,KIDINS220,NUMA
A,NEDD4L,POLH,CTSC,ODF2L,NUDC,NRBP2,USP22,MED18,ACTG1,EEF2,DPYSL3,CFL1,BTBD9,X
,PFN1,EIF5A,EIF3L,RPLP2,DDOST,TM9SF3,MDH2,U2AF2,PRDX5,BAG6,CDC42,TRAF7,EIF2AK1,P
ZNF316,RPL4,ENO1,MAP1B,PKM,GANAB,COL6A1,HSP90B1,UGT8,UBA1,TMF1,COL5A1,COL4A1
11,TRIM28,CKB,FN1,RPL18,WSB1,AP2B1,GTF2F1,LDHA,JUND,TMEM259,LENG8,CCNI,GNB2,CH
N1,USP11,TRIM28,CKB,FN1,RPL18,WSB1,AP2B1,GTF2F1,LDHA,JUND,TMEM259,LENG8,CCNI,GN
N,AP3D1,CTSD,NCS1,MEGF8,ARL4C,RPS19,TLN1,COL6A2,B4GALNT4,H1-10,PLPPR3,HECTD4,S

)1,RPL6,GPI,GDI1,TMBIM6,SMIM14,PPP2R1A,POLR2A,RPS4X,SF3B2,RPL29,ZNF865,PFN1,MCL1
B4X,ZNF316,DBN1,RPL7A,HSPA1B,PSAP,MAGED1,SEC62,YWHAE,KIF1A,SLC2A3,FASN,RPL5,SM(
A,FSCN1,USP11,TRIM28,CKB,FN1,EIF1,RPL18,WSB1,AP2B1,GTF2F1,LDHA,RPL24,JUND,TMEM2
RIM28,CKB,FN1,RPL18,WSB1,AP2B1,GTF2F1,LDHA,JUND,TMEM259,LENG8,CCNI,GNB2,CHPF,R
RIM28,CKB,FN1,RPL18,WSB1,AP2B1,GTF2F1,LDHA,JUND,TMEM259,LENG8,CCNI,GNB2,CHPF,R
2,FSCN1,USP11,EIF3C,UBB,AFF4,FN1,POLR2A,TPT1,MIDN,MARCHF6,GTF2F1,GOLGA8A,LDHA,(
POLH,ATN1,ACTB,NRBP2,USP22,XRCC2,HTT,ACTR2,ZNF316,DBN1,RPL4,TUBA1B,MAP1B,ALDO
'1B,ABHD2,PGK1,CANX,KIF1A,SLC2A3,BSG,ALDH6A1,STMN2,IDS,FSCN1,TMBIM6,PLXNB2,TTC3,
2L,NUDC,USP22,MTRNR2L2,EEF2,CFL1,ZNF124,HSP90AA1,VIM,FTH1,NUMA1,RPL3,ENO1,PIK3R
FL1,BTBD9,XRCC2,HSP90AB1,ZNF124,ACTR2,VIM,NUMA1,RPL3,TUBA1B,ALDOA,HSP90B1,IGFB
FL1,BTBD9,XRCC2,HSP90AB1,ZNF124,ACTR2,VIM,NUMA1,RPL3,TUBA1B,ALDOA,HSP90B1,IGFB
2,HTT,HSP90AA1,UBC,ZNF316,FADS1,NEFM,UBA1,IGFBP2,CALR,COL5A1,RPLP0,H3-3B,PTMA,M
,EIF1,AP2B1,GTF2F1,LDHA,JUND,TMEM259,LENG8,CCNI,GNB2,MYL6,MYH10,UBE2H,LAPTM4B,
IS4A10,NEDD4L,ATN1,CTSC,ODF2L,NRBP2,USP22,MED18,DPYSL3,CFL1,XRCC2,HTT,HSP90AB1,
10,VAPA,MORF4L1,DNAJA1,FADS2,LDHA,CTSD,EIF4H,CIPC,C1ORF43,RHOA,CD81,PNN,SPTBN2,
M14,TPT1,SEC61A1,MORF4L1,MIDN,MARCHF6,ARF1,GOLGA8A,CPE,LPCAT1,SLC16A7,PEBP1,P
A10,HSPA9,ATN1,ACTB,CTSC,ODF2L,NUDC,MTRNR2L2,MED18,ACTG1,EEF2,DPYSL3,TUBA1A,CF
,DPYSL3,CFL1,BTBD9,XRCC2,HSP90AB1,UBC,TMSB4X,EIF4G2,RPL3,RPL4,ENO1,FLNA,RPL7A,
L2,ACTG1,DPYSL3,TUBA1A,BTBD9,ACTR2,VIM,KIDINS220,TMSB4X,NUMA1,RPS6,DBN1,RPL4,TU
HDLBP,RPS5,RHOB,MMP2,GLUL,CTDSP2,HNRNPL,ALKBH5,YEATS2,GRINA,MDK,TNPO1,ASXL1,S
F3C,SERINC1,HMGB1,BTF3,AGRN,RPLP1,MARCHF6,ZNF865,CXADR,WDR1,SRSF4,PFN1,PRKCS
MA1,RPS6,ALDOA,PKM,FADS1,NEFL,FBXL4,CALR,TUBB2B,COL4A1,DDX5,ENO2,SOX4,GPI,RACK
RPS27,ATP6V0C,NOP53,PABPN1,RBMX,PTBP1,ILF3,DIDO1,PHB2,SLC39A7,TRA2B,ZFR,SEPTIN7,
P90AA1,ZNF316,DBN1,ENO1,FLNA,SLC9A7,CALR,FSTL1,ABHD2,COL5A1,CES3,ENO2,SEC62,KIF
A,HNRNPA0,ZNF865,EIF5A,EIF3L,NGRN,RPLP2,DDOST,YWHAB,HSBP1,DDX39B,U2AF2,GNAI3,G
FL1,XRCC2,HSP90AB1,ACTR2,FTH1,UBC,NUMA1,RPL3,ZNF316,FLNA,MAP1B,NCL,ALDOA,FTL,N
1,TRIM28,CKB,FN1,RPL18,WSB1,AP2B1,GTF2F1,LDHA,JUND,TMEM259,LENG8,CCNI,GNB2,CH
RPLP0,DDX5,RPL13A,KAT6A,DVL3,YWHAE,DDX17,RACK1,SEPTIN2,GDI1,USP11,CSDE1,ATP5F1A,
POLH,ATN1,ACTB,CTSC,NUDC,NRBP2,EEF2,DPYSL3,TUBA1A,BTBD9,MAVS,HSP90AB1,ZNF124,H
CTG1,DPYSL3,TUBA1A,XRCC2,MAVS,HSP90AB1,ZNF124,HSP90AA1,VIM,KIDINS220,TMSB4X,EIF
A,EIF3L,RPLP2,DDOST,MDH2,U2AF2,PRDX5,UBE4A,CCAR2,BAG6,RAB11A,CDC42,TRAF7,EIF2AK
FL1,BTBD9,XRCC2,HSP90AB1,ZNF124,ACTR2,VIM,NUMA1,RPL3,TUBA1B,ALDOA,HSP90B1,IGFB
N1,USP11,TRIM28,CKB,FN1,RPL18,WSB1,AP2B1,GTF2F1,LDHA,RPL24,JUND,TMEM259,LENG8,(
RPL4,FLNA,FADS1,TMF1,TUBB2B,DDX5,KIF1A,RPL13,DDX17,ALDH6A1,SEPTIN2,FSCN1,USP11,
NN,RPS25,EIF5A,EIF3L,RPLP2,DDOST,TM9SF3,MDH2,U2AF2,CCT2,PRDX5,BAG6,CDC42,TRAF7,
,TMEM259,CHPF,RPLP2,MYL6,UBE2H,SMCHD1,ATP6V0C,LAPTM4B,MLXIP,HSPH1,AHCYL1,KLC
IF1A,FSCN1,TMBIM6,USP11,TRIM28,CKB,FN1,MAP9,RPL15,SEC61A1,RPL18,STMN1,SARAF,WSE
WHAG,PKD1,ZNF43,ZMIZ1,SBF1,NCS1,TMEM259,ATP2A2,LRPPRC,PDIA6,PLXNA1,ACTN4,RPS19,
ED1,RPL6,RACK1,SHISA9,P4HB,SNRNP200,EIF1,RPL19,HMGB1,TUBB4B,SF3B2,XRCC5,CIPC,RPL
2,RPL7A,TMF1,TUBB2B,DDX5,KIF1A,DDX17,SEPTIN2,P4HB,FSCN1,GSTP1,PLD3,USP11,TRIM28,
)6,CCT2,TBCD,PTOV1,MLEC,CORO1C,ISYNA1,RBBP7,WDR45B,LRRC4B,SLC39A6,NXN,PPP1CC,
N1,USP11,TRIM28,CKB,FN1,RPL18,WSB1,AP2B1,GTF2F1,LDHA,JUND,TMEM259,LENG8,CCNI,G
L4,FLNA,FADS1,TMF1,DDX5,KIF1A,RPL13,ALDH6A1,SEPTIN2,FSCN1,USP11,TRIM28,CKB,FN1,EI
A,FSCN1,USP11,TRIM28,CKB,FN1,EIF1,POLR2A,RPL18,MIDN,WSB1,AP2B1,GTF2F1,LDHA,JUND
9SF3,MDH2,U2AF2,PRDX5,BAG6,CDC42,EIF2AK1,PDAP1,YKT6,MCM3AP,TMEM167A,CHERP,CTT

A,FSCN1,USP11,TRIM28,CKB,FN1,RPL18,WSB1,AP2B1,GTF2F1,LDHA,RPL24,JUND,TMEM259,L
42BPB,AP2B1,PXDN,LAMA5,BRD3,NCS1,LENG8,HMGN2,TLE5,HDLBP,MYL6,DUX4,NOVA2,ANXA5,
,MYL6,UBE2H,SMCHD1,ATP6V0C,LAPTM4B,MLXIP,HSPH1,AHCYL1,KLC1,RAB11B,TUBB2A,PPP2
,MYL6,UBE2H,SMCHD1,ATP6V0C,LAPTM4B,MLXIP,HSPH1,AHCYL1,KLC1,RAB11B,TUBB2A,PPP2
B2B,KIF1A,FSCN1,USP11,TRIM28,CKB,FN1,EIF1,RPL18,WSB1,AP2B1,GTF2F1,LDHA,RPL24,JUN
8,CALR,FSTL1,PGK1,CANX,FKBP14,SEC62,DDX17,GPI,CALM3,RPL7,KLRD1,MARCKS,NPM1,PLD
TMF1,DDX5,KIF1A,FSCN1,USP11,TRIM28,CKB,FN1,EIF1,MAP9,RPL18,SARAF,WSB1,AP2B1,GTF
4,ENO1,NCL,ALDOA,YBX1,IGFBP2,PSAP,PGK1,CANX,DDX5,RPL13A,KAT6A,RPL13,FASN,ALDH6A
4,ENO1,NCL,ALDOA,YBX1,IGFBP2,PSAP,PGK1,CANX,DDX5,RPL13A,KAT6A,RPL13,FASN,ALDH6A
P200,MATR3,DDB1,TPI1,EIF4A1,MRC2,LDHB,HLA-C,SSBP2,TPT1,RPL18,BTF3,AGRN,CDC42BPB
MSB10,PKD1,VAPA,ITGB1,SARAF,AKAP12,CDC42BPB,ITM2B,SMARCA5,TPM4,ATP1A1,JUND,CC
OLH,NUDC,NRBP2,USP22,MTRNR2L2,ACTG1,EEF2,DPYSL3,CFL1,BTBD9,MAVS,HTT,HSP90AB1,
GF,BEX4,PGAP6,MED13,GNA13,THRAP3,EIF4G1,SC5D,EHMT2,RHEB,EIF3B,PRKAR2B,UBXN1,N
RF1,GOLGA8A,PRKAR1A,HNRNPDL,CKAP4,TM9SF3,ANXA5,PDXK,AHCYL1,PTOV1,ORAI2,KDELR
CX,RPS6,ZNF316,PIK3R2,NCL,ALDOA,HSP90B1,YBX1,CANX,CLU,DDX5,MAGED1,RPL6,GPI,RACK
1,TRIM28,CKB,FN1,RPL18,WSB1,AP2B1,GTF2F1,LDHA,JUND,TMEM259,LENG8,CCNI,GNB2,CH
,FADS1,TMF1,KIF1A,FSCN1,USP11,TRIM28,CKB,FN1,SEC61A1,RPL18,SARAF,WSB1,AP2B1,GTF2
1A,RPL19,PPP2R1A,ZNF865,MCL1,RPL31,LRPPRC,RPS5,DDOST,NARS1,ATP6V0C,DDX39B,U2AF
P2,DDOST,MDH2,U2AF2,PRDX5,UBE4A,BAG6,RAB11A,CDC42,TRAF7,EIF2AK1,PDAP1,YKT6,MC
TLE5,CD63,HNRNPDL,SON,KDELR1,RPL28,TUG1,KDM5B,FBRS,FLOT1,CUL3,PPP1R15A,BRK1,F
2,EEF2,HSP90AB1,SPARC,TUBB,UBC,TMSB4X,DCX,EIF4G2,RPS6,TUBA1B,MTRNR2L8,NEFL,ATP
SLC16A7,CBX5,KIF5B,RPS20,WDR1,HMGN2,PFN1,SPC25,PRKCSH,MAGED2,H1-10,RAB1A,CCT6
0AB1,ACTR2,ZNF316,TUBA1B,NCL,FTL,HSP90B1,PRPF8,CALR,HSPA1B,CES3,PTMA,RPL13A,MA
2,CAMSAP1,SNRNP70,ABCA2,TOP1,ODC1,ILF3,SSB,PLXNA3,H2AZ1,KIF21B,SRSF6,RABEP1,ACL
GK1,KIF1A,SLC2A3,ALDH6A1,FSCN1,TMBIM6,PLXNB2,TTC3,SMIM14,YWHAG,CHSY1,TMSB10,Z
,CHSY1,STMN1,UCHL1,ANXA2,AGRN,AP3D1,GOLGA8A,SCARB2,RPL22,CHD4,WDR1,OAZ1,NES,T
3,SMCHD1,RNF187,CAND1,MMP15,SRSF6,SMC3,IL6ST,LSM14B,YY1,SORT1,ATG4B,SNX2,STK3
MYH9,APP,TRIM28,SEC61A1,STMN1,MORF4L1,RPL10,RPL29,TNS3,EIF4H,C1ORF43,PAPOLA,R
RPL4,ENO1,PKM,COL6A1,HSP90B1,FADS1,TMF1,TUBB2B,PGK1,DDX5,KIF1A,RPL13,DDX17,GPI
3R2,FKBP14,RPL13A,RPL6,GPI,CALM3,SHISA9,TMBIM6,ATP5F1A,EIF1,EIF4A1,RPL19,RPS3,POL
,GOLGA8A,SCARB2,HK1,ZNF865,OAZ1,APLP1,SLC25A3,RAC1,MAP1A,CTDSP2,NOVA2,NUCB1,M
OP2B,SMCHD1,TBCD,PGD,VEGFA,BTBD2,TCF25,GIGYF1,ATXN2L,CYP51A1,YME1L1,VDAC1,TRI
S,HTT,HSP90AB1,VIM,UBC,KIDINS220,DBN1,NCL,HSP90B1,PRPF8,RPL8,YBX1,TMF1,COL5A1,C
RPL4,ENO1,PKM,COL6A1,HSP90B1,FADS1,TMF1,TUBB2B,PGK1,DDX5,KIF1A,RPL13,DDX17,GPI
A,FKBP14,RPL13A,YWHAE,FASN,RPS8,USP11,ATP5F1A,RPL19,PPP2R1A,HNRNPA0,RPS16,RPL22
0,PRKCSH,RPL35,ANXA6,HNRNPH1,RAB1A,CCT6A,MAGED4B,APPL1,PPP1CB,MTRF1L,SRSF1,P
WHAB,MDH2,U2AF2,PRDX5,TCP1,TAF7,ARPC2,BAG6,CDC42,EIF2AK1,PDAP1,PHB2,YKT6,TMEM
90AB1,ACTR2,VIM,TUBB,TUBA1B,NCL,SLC9A7,MTRNR2L8,HSP90B1,NEFL,RPL8,UBA1,CALR,FS
A,FSCN1,USP11,TRIM28,CKB,FN1,RPL18,WSB1,AP2B1,GTF2F1,LDHA,RPL24,JUND,TMEM259,L
EIF4B,FN1,STMN1,ANXA2,AP2B1,GOLGA8A,NCS1,OAZ1,APLP1,PTGES3,SEPTIN11,RAC1,ARHGDI
A,FSCN1,USP11,TRIM28,CKB,FN1,EIF1,POLR2A,RPL18,MIDN,WSB1,AP2B1,GTF2F1,LDHA,RPL2
1,USP11,TRIM28,CKB,FN1,RPL18,WSB1,AP2B1,GTF2F1,LDHA,JUND,TMEM259,LENG8,CCNI,G
2,DDOST,MDH2,U2AF2,TSR1,PRDX5,GCN1,PTBP1,CCAR2,BAG6,ARFGEF2,TRAF7,PRMT1,APMA
,EIF5A,PDIA6,EIF3L,DDOST,RPL23A,HSBP1,MAP1A,TM9SF3,U2AF2,PRDX5,UBE4A,BAG6,AKIRIN
A7,RAB1B,UBR5,TAX1BP1,MBD3,BNIP3,SCAP,GALNT10,RAN,SOBP,CACNA1H,TBKBP1,SS18L1,

[illegible]RPL13,GPI,FASN,ALDH6A1,SEPTIN2,NPM1,PLD3,HSPD1,COL18A1,CSDE1,AFF4,DDB1,LDHB,SF3B[illegible]
[illegible]24,ZNF865,CHD4,OAZ1,APLP1,RAC1,MAP1A,CTDSP2,NOVA2,PTOV1,SURF4,DDAH2,CORO1C,ST[illegible]

:D18,ACTG1,EEF2,DPYSL3,TUBA1A,CFL1,BTBD9,XRCC2,MAVS,HTT,HSP90AB1,ZNF124,HSP90AA
:F2,DPYSL3,TUBA1A,CFL1,BTBD9,XRCC2,MAVS,HTT,HSP90AB1,ZNF124,HSP90AA1,ACTR2,VIM,
EEF2,DPYSL3,TUBA1A,CFL1,BTBD9,XRCC2,MAVS,HTT,HSP90AB1,ZNF124,HSP90AA1,ACTR2,VIM
TG1,EEF2,DPYSL3,TUBA1A,CFL1,BTBD9,XRCC2,MAVS,HTT,ZNF124,HSP90AA1,ACTR2,VIM,SPAR
)AA1,ACTR2,VIM,FTH1,TMSB4X,EIF4G2,RPS6,ZNF316,DBN1,ENO1,TUBA1B,PIK3R2,FLNA,MAP1E
UBA1A,CFL1,BTBD9,XRCC2,MAVS,HTT,HSP90AB1,ZNF124,HSP90AA1,ACTR2,VIM,TUBB,FTH1,U
TG1,EEF2,DPYSL3,TUBA1A,CFL1,BTBD9,XRCC2,MAVS,HTT,HSP90AB1,ZNF124,HSP90AA1,ACTR2
I,TUBA1B,PIK3R2,FLNA,NCL,RPL7A,SLC9A7,ALDOA,PKM,GANAB,COL6A1,NEFL,RPL8,FBXL4,YB
3D9,XRCC2,MAVS,HTT,ZNF124,HSP90AA1,VIM,SPARC,TUBB,FTH1,UBC,KIDINS220,TMSB4X,NUM
CFL1,BTBD9,XRCC2,MAVS,HTT,ZNF124,HSP90AA1,ACTR2,VIM,TUBB,FTH1,UBC,KIDINS220,TMS
3,TUBA1A,CFL1,BTBD9,XRCC2,MAVS,HTT,HSP90AB1,ZNF124,HSP90AA1,ACTR2,VIM,SPARC,TUE
ITT,HSP90AB1,ZNF124,HSP90AA1,ACTR2,VIM,FTH1,KIDINS220,TMSB4X,NUMA1,EIF4G2,ZNF316
T,HSP90AB1,ZNF124,HSP90AA1,VIM,SPARC,TUBB,FTH1,KIDINS220,NUMA1,RPL3,ZNF316,DBN1
MAVS,HTT,HSP90AB1,ZNF124,HSP90AA1,VIM,SPARC,TUBB,FTH1,KIDINS220,NUMA1,RPL3,ZNF3
,HTT,ZNF124,HSP90AA1,ACTR2,SPARC,FTH1,UBC,KIDINS220,NUMA1,EIF4G2,RPL3,ZNF316,DBN
SL3,TUBA1A,CFL1,BTBD9,XRCC2,MAVS,HTT,HSP90AB1,ZNF124,HSP90AA1,ACTR2,VIM,TUBB,UB
TT,HSP90AB1,ZNF124,HSP90AA1,VIM,SPARC,TUBB,FTH1,UBC,KIDINS220,TMSB4X,NUMA1,RPL3
,HTT,HSP90AB1,ZNF124,HSP90AA1,ACTR2,VIM,KIDINS220,TMSB4X,NUMA1,RPL3,ZNF316,DBN
NF124,ACTR2,VIM,FTH1,KIDINS220,TMSB4X,NUMA1,ZNF316,DBN1,ENO1,PIK3R2,FLNA,MAP1B,
VS,HTT,HSP90AB1,ZNF124,HSP90AA1,ACTR2,VIM,SPARC,TUBB,FTH1,KIDINS220,TMSB4X,NUM
VS,HTT,ZNF124,HSP90AA1,VIM,TUBB,FTH1,UBC,TMSB4X,NUMA1,RPL3,RPS6,ZNF316,DBN1,RPL
EF2,DPYSL3,CFL1,BTBD9,XRCC2,MAVS,HTT,ZNF124,HSP90AA1,ACTR2,VIM,TUBB,FTH1,UBC,KID
TBD9,XRCC2,MAVS,HTT,ZNF124,HSP90AA1,ACTR2,VIM,TUBB,FTH1,UBC,TMSB4X,NUMA1,EIF4G
:D18,ACTG1,EEF2,DPYSL3,TUBA1A,CFL1,BTBD9,XRCC2,MAVS,HTT,HSP90AB1,ZNF124,HSP90AA
XRCC2,MAVS,HTT,HSP90AB1,ZNF124,HSP90AA1,VIM,SPARC,TUBB,FTH1,UBC,NUMA1,EIF4G2,Z
PYSL3,TUBA1A,CFL1,BTBD9,XRCC2,MAVS,HTT,HSP90AB1,ZNF124,HSP90AA1,ACTR2,VIM,SPAR
T,ZNF124,HSP90AA1,ACTR2,VIM,FTH1,UBC,TMSB4X,NUMA1,EIF4G2,RPS6,ZNF316,DBN1,RPL4,E
TUBA1A,CFL1,BTBD9,XRCC2,MAVS,HTT,HSP90AB1,ZNF124,HSP90AA1,ACTR2,VIM,TUBB,FTH1,
CTG1,EEF2,DPYSL3,TUBA1A,CFL1,BTBD9,XRCC2,MAVS,HTT,HSP90AB1,ZNF124,HSP90AA1,ACT
SP22,MED18,ACTG1,EEF2,DPYSL3,TUBA1A,CFL1,BTBD9,XRCC2,MAVS,HTT,HSP90AB1,ZNF124,
:D18,ACTG1,EEF2,DPYSL3,TUBA1A,CFL1,BTBD9,XRCC2,MAVS,HTT,HSP90AB1,ZNF124,HSP90A
2,MAVS,HTT,HSP90AB1,ZNF124,HSP90AA1,VIM,FTH1,KIDINS220,NUMA1,DCX,RPL3,ZNF316,DBN
CC2,MAVS,HTT,HSP90AB1,ZNF124,HSP90AA1,ACTR2,SPARC,FTH1,UBC,TMSB4X,EIF4G2,RPL3,Z
:F2,DPYSL3,TUBA1A,CFL1,BTBD9,XRCC2,MAVS,HTT,HSP90AB1,ZNF124,HSP90AA1,ACTR2,VIM,
TRNR2L2,MED18,ACTG1,EEF2,DPYSL3,TUBA1A,CFL1,BTBD9,XRCC2,MAVS,HTT,HSP90AB1,ZNF1
90AA1,VIM,KIDINS220,NUMA1,RPL3,ZNF316,DBN1,RPL4,ENO1,TUBA1B,PIK3R2,FLNA,MAP1B,N
CFL1,BTBD9,XRCC2,MAVS,HTT,HSP90AB1,ZNF124,HSP90AA1,ACTR2,VIM,SPARC,FTH1,UBC,TMS
AVS,HTT,HSP90AB1,ZNF124,HSP90AA1,ACTR2,VIM,TUBB,UBC,KIDINS220,TMSB4X,EIF4G2,ZNF
A1,VIM,FTH1,KIDINS220,NUMA1,EIF4G2,RPL3,ZNF316,DBN1,ENO1,TUBA1B,PIK3R2,FLNA,MAP1
CTG1,DPYSL3,TUBA1A,CFL1,XRCC2,MAVS,HTT,HSP90AB1,ZNF124,HSP90AA1,ACTR2,VIM,SPAR
EEF2,DPYSL3,TUBA1A,CFL1,BTBD9,XRCC2,MAVS,HTT,HSP90AB1,ZNF124,HSP90AA1,ACTR2,VIM
CTG1,EEF2,DPYSL3,TUBA1A,CFL1,BTBD9,XRCC2,MAVS,HTT,HSP90AB1,ZNF124,HSP90AA1,ACT
90AA1,ACTR2,VIM,FTH1,UBC,KIDINS220,TMSB4X,NUMA1,EIF4G2,ZNF316,DBN1,RPL4,ENO1,TU
A,CFL1,BTBD9,XRCC2,MAVS,HTT,HSP90AB1,ZNF124,HSP90AA1,ACTR2,VIM,TUBB,FTH1,KIDINS
PYSL3,TUBA1A,CFL1,BTBD9,XRCC2,MAVS,HTT,HSP90AB1,ZNF124,HSP90AA1,ACTR2,VIM,SPARC

|8,ACTG1,EEF2,DPYSL3,TUBA1A,CFL1,BTBD9,XRCC2,MAVS,HTT,HSP90AB1,ZNF124,HSP90AA1,
,HSP90AB1,ZNF124,HSP90AA1,VIM,FTH1,UBC,KIDINS220,NUMA1,DCX,EIF4G2,RPL3,ZNF316,DE
NF316,DBN1,ENO1,PIK3R2,FLNA,MAP1B,NCL,SLC9A7,PKM,COL6A1,HSP90B1,NEFM,UGT8,RPL
D18,ACTG1,EEF2,DPYSL3,TUBA1A,CFL1,BTBD9,XRCC2,MAVS,HTT,HSP90AB1,ZNF124,HSP90AA
MTRNR2L2,MED18,ACTG1,EEF2,DPYSL3,TUBA1A,CFL1,BTBD9,XRCC2,MAVS,HTT,HSP90AB1,ZN
,HTT,HSP90AB1,ZNF124,HSP90AA1,ACTR2,VIM,FTH1,UBC,KIDINS220,NUMA1,RPL3,RPS6,ZNF31
NF124,TUBB,FTH1,KIDINS220,NUMA1,RPL3,ZNF316,DBN1,RPL4,ENO1,TUBA1B,PIK3R2,FLNA,M
,MAVS,HTT,HSP90AB1,ZNF124,HSP90AA1,ACTR2,VIM,TUBB,FTH1,UBC,TMSB4X,NUMA1,EIF4G2
O1,TUBA1B,FLNA,MAP1B,NCL,SLC9A7,PKM,GANAB,COL6A1,FTL,HSP90B1,FADS1,NEFM,RPL8,F
,DPYSL3,CFL1,BTBD9,XRCC2,MAVS,HTT,HSP90AB1,ZNF124,HSP90AA1,VIM,TUBB,FTH1,UBC,TM
2,FLNA,MAP1B,NCL,PKM,COL6A1,HSP90B1,NEFM,RPL8,FBXL4,YBX1,IGFBP2,FSTL1,ABHD2,TUE
16,DBN1,ENO1,TUBA1B,PIK3R2,FLNA,MAP1B,NCL,SLC9A7,PKM,GANAB,COL6A1,HSP90B1,NEF
,ACTR2,VIM,UBC,NUMA1,ZNF316,DBN1,PIK3R2,FLNA,MAP1B,NCL,SLC9A7,PKM,GANAB,HSP90
20,NUMA1,RPL3,ZNF316,DBN1,RPL4,ENO1,TUBA1B,PIK3R2,FLNA,MAP1B,NCL,PKM,GANAB,CO
PYSL3,TUBA1A,CFL1,BTBD9,MAVS,HTT,HSP90AB1,ZNF124,HSP90AA1,VIM,FTH1,UBC,NUMA1,EI
A,SLC9A7,HSP90B1,RPL8,FBXL4,YBX1,FSTL1,TUBB2B,COL5A1,PSAP,PGK1,H3-3B,PTMA,DDX5,
,MED18,ACTG1,EEF2,DPYSL3,TUBA1A,CFL1,BTBD9,MAVS,HTT,HSP90AB1,ZNF124,HSP90AA1,AC
4,TUBB,KIDINS220,NUMA1,RPL3,ZNF316,DBN1,RPL4,ENO1,TUBA1B,PIK3R2,FLNA,MAP1B,NCL
UMA1,RPL3,DBN1,ENO1,FLNA,MAP1B,NCL,SLC9A7,PKM,GANAB,COL6A1,FTL,NEFM,RPL8,FBXL
0AB1,ZNF124,HSP90AA1,VIM,FTH1,KIDINS220,TMSB4X,NUMA1,DCX,RPL3,ZNF316,DBN1,RPL4,E
CFL1,BTBD9,MAVS,HTT,ZNF124,HSP90AA1,ACTR2,VIM,TUBB,FTH1,UBC,TMSB4X,NUMA1,EIF4G
316,DBN1,RPL4,ENO1,TUBA1B,FLNA,MAP1B,NCL,SLC9A7,PKM,GANAB,COL6A1,HSP90B1,FADS
SL3,TUBA1A,CFL1,BTBD9,XRCC2,MAVS,HTT,HSP90AB1,ZNF124,HSP90AA1,ACTR2,VIM,TUBB,FT
B1,ZNF124,HSP90AA1,ACTR2,VIM,FTH1,UBC,NUMA1,RPL3,ZNF316,DBN1,ENO1,PIK3R2,FLNA,
220,NUMA1,ZNF316,DBN1,ENO1,TUBA1B,PIK3R2,FLNA,MAP1B,NCL,SLC9A7,PKM,GANAB,COL6
TMSB4X,NUMA1,ZNF316,DBN1,ENO1,TUBA1B,FLNA,MAP1B,NCL,RPL7A,SLC9A7,PKM,GANAB,C
P90AB1,ZNF124,VIM,FTH1,UBC,KIDINS220,TMSB4X,NUMA1,EIF4G2,ZNF316,DBN1,RPL4,ENO1,
IM,FTH1,TMSB4X,NUMA1,DCX,RPL3,DBN1,ENO1,FLNA,MAP1B,NCL,RPL7A,SLC9A7,PKM,FTL,H
ACTG1,EEF2,DPYSL3,TUBA1A,CFL1,BTBD9,XRCC2,MAVS,HTT,HSP90AB1,ZNF124,HSP90AA1,ACT
TG1,DPYSL3,CFL1,XRCC2,MAVS,HTT,HSP90AB1,ZNF124,HSP90AA1,VIM,SPARC,TUBB,FTH1,UE
FL1,BTBD9,XRCC2,MAVS,HTT,ZNF124,HSP90AA1,VIM,SPARC,TUBB,FTH1,UBC,KIDINS220,TMSE
B1,ZNF124,HSP90AA1,ACTR2,VIM,TUBB,FTH1,UBC,KIDINS220,NUMA1,RPL3,ZNF316,DBN1,ENO
,TUBA1A,CFL1,BTBD9,XRCC2,MAVS,HTT,HSP90AB1,ZNF124,HSP90AA1,ACTR2,VIM,TUBB,FTH1,
UBA1A,CFL1,BTBD9,XRCC2,MAVS,HTT,HSP90AB1,ZNF124,HSP90AA1,VIM,FTH1,UBC,TMSB4X,N
C2,MAVS,HTT,HSP90AB1,ZNF124,HSP90AA1,ACTR2,VIM,TUBB,UBC,KIDINS220,TMSB4X,NUMA1
1,ACTR2,VIM,FTH1,KIDINS220,TMSB4X,NUMA1,ZNF316,DBN1,RPL4,ENO1,FLNA,MAP1B,NCL,RF
NF316,DBN1,ENO1,TUBA1B,PIK3R2,FLNA,MAP1B,NCL,SLC9A7,PKM,GANAB,COL6A1,HSP90B1,F
RNR2L2,ACTG1,EEF2,DPYSL3,TUBA1A,CFL1,BTBD9,XRCC2,MAVS,HTT,HSP90AB1,ZNF124,HSP
R2,VIM,FTH1,KIDINS220,TMSB4X,NUMA1,EIF4G2,ZNF316,DBN1,ENO1,TUBA1B,PIK3R2,FLNA,M
IM,FTH1,KIDINS220,NUMA1,RPL3,ZNF316,DBN1,RPL4,ENO1,TUBA1B,PIK3R2,FLNA,MAP1B,NCL
IM,FTH1,KIDINS220,NUMA1,RPL3,ZNF316,DBN1,RPL4,ENO1,TUBA1B,PIK3R2,FLNA,MAP1B,NCL
HSP90AA1,ACTR2,VIM,FTH1,UBC,NUMA1,RPL3,ZNF316,DBN1,ENO1,PIK3R2,FLNA,MAP1B,NCL,H
L2,MED18,ACTG1,EEF2,DPYSL3,TUBA1A,CFL1,BTBD9,XRCC2,MAVS,HTT,HSP90AB1,ZNF124,HS
6,DBN1,ENO1,PIK3R2,FLNA,MAP1B,NCL,RPL7A,SLC9A7,ALDOA,PKM,GANAB,COL6A1,FTL,HSP9
B1,ZNF124,VIM,TUBB,FTH1,UBC,NUMA1,ZNF316,DBN1,ENO1,TUBA1B,PIK3R2,FLNA,MAP1B,N

,UBC,KIDINS220,NUMA1,RPL3,ZNF316,DBN1,RPL4,ENO1,TUBA1B,PIK3R2,FLNA,MAP1B,NCL,SL
NUMA1,RPL3,ZNF316,DBN1,RPL4,ENO1,TUBA1B,PIK3R2,FLNA,MAP1B,NCL,MTRNR2L8,PKM,GA
A1,VIM,FTH1,NUMA1,DCX,RPL3,DBN1,ENO1,FLNA,MAP1B,NCL,SLC9A7,PKM,GANAB,FTL,HSP90
4G2,ZNF316,DBN1,TUBA1B,PIK3R2,FLNA,MAP1B,NCL,SLC9A7,PKM,COL6A1,HSP90B1,FADS1,N
BTBD9,XRCC2,MAVS,HTT,HSP90AB1,ZNF124,HSP90AA1,VIM,TMSB4X,NUMA1,DCX,EIF4G2,RPL3
SP22,MED18,ACTG1,EEF2,DPYSL3,TUBA1A,CFL1,BTBD9,XRCC2,MAVS,HTT,HSP90AB1,ZNF124,
A,MAP1B,NCL,PKM,GANAB,HSP90B1,FADS1,RPL8,FBXL4,YBX1,CALR,FSTL1,TUBB2B,CLU,PTM
0AA1,VIM,KIDINS220,NUMA1,RPL3,ZNF316,DBN1,RPL4,ENO1,TUBA1B,PIK3R2,FLNA,MAP1B,NC
ACTG1,EEF2,DPYSL3,TUBA1A,CFL1,BTBD9,XRCC2,MAVS,HTT,HSP90AB1,ZNF124,HSP90AA1,AC
VS,HTT,HSP90AB1,HSP90AA1,ACTR2,VIM,TUBB,FTH1,UBC,NUMA1,EIF4G2,ZNF316,RPL4,ENO1,
3,CFL1,XRCC2,MAVS,HTT,HSP90AB1,ZNF124,HSP90AA1,VIM,SPARC,TUBB,FTH1,UBC,NUMA1,E
124,VIM,FTH1,KIDINS220,NUMA1,RPL3,ZNF316,DBN1,RPL4,ENO1,TUBA1B,PIK3R2,FLNA,MAP1
NF124,HSP90AA1,ACTR2,VIM,FTH1,UBC,NUMA1,RPL3,ZNF316,DBN1,ENO1,PIK3R2,MAP1B,NCL,
HSP90AA1,VIM,FTH1,TMSB4X,NUMA1,DCX,RPL3,DBN1,ENO1,FLNA,MAP1B,NCL,SLC9A7,PKM,G
L2,ACTG1,EEF2,DPYSL3,TUBA1A,CFL1,BTBD9,XRCC2,MAVS,HTT,HSP90AB1,ZNF124,HSP90AA1,
HTT,HSP90AA1,VIM,TUBB,FTH1,UBC,NUMA1,EIF4G2,ZNF316,ENO1,TUBA1B,PIK3R2,MAP1B,NCL
KM,COL6A1,NEFM,RPL8,FBXL4,YBX1,UBA1,IGFBP2,TUBB2B,PTMA,ENO2,KAT6A,DVL3,GNAS,B
1A,CFL1,XRCC2,MAVS,HTT,HSP90AB1,ZNF124,HSP90AA1,ACTR2,VIM,FTH1,KIDINS220,TMSB4X
A,MAP1B,NCL,RPL7A,SLC9A7,ALDOA,PKM,NEFL,NEFM,PRPF8,RPL8,FBXL4,YBX1,IGFBP2,CALR,
3R2,FLNA,MAP1B,RPL7A,SLC9A7,PKM,GANAB,HSP90B1,NEFM,PRPF8,YBX1,IGFBP2,CALR,TUBE
UMA1,ZNF316,DBN1,PIK3R2,FLNA,NCL,RPL7A,SLC9A7,ALDOA,PKM,COL6A1,FTL,HSP90B1,NEF
FBXL4,YBX1,IGFBP2,FSTL1,TUBB2B,COL5A1,COL4A1,PSAP,RPLP0,H3-3B,PTMA,ENO2,SOX4,KA
LDOA,PKM,COL6A1,RPL8,FBXL4,YBX1,ATP2B4,ABHD2,TUBB2B,COL5A1,CES3,PTMA,ENO2,YWH
,FBXL4,IGFBP2,COL5A1,PSAP,PTMA,KAT6A,YWHAE,KIF1A,RPL13,GNAS,RACK1,BSG,FASN,SPTB
2,MAVS,HTT,HSP90AB1,ZNF124,HSP90AA1,ACTR2,VIM,TUBB,FTH1,UBC,KIDINS220,TMSB4X,NU
,COL4A1,H3-3B,PTMA,ENO2,KAT6A,DVL3,RPL13,GNAS,DDX17,FASN,GNB1,RPS8,MYH9,SEPTIN2
A1,ACTR2,VIM,TUBB,FTH1,UBC,KIDINS220,TMSB4X,NUMA1,DCX,ZNF316,DBN1,ENO1,TUBA1B,
4,HSP90AA1,ACTR2,VIM,SPARC,TUBB,FTH1,KIDINS220,EIF4G2,RPS6,ZNF316,DBN1,RPL4,ENO1
SP90AB1,VIM,SPARC,FTH1,TMSB4X,DCX,EIF4G2,RPL3,ZNF316,DBN1,ENO1,TUBA1B,PIK3R2,M
SL3,CFL1,BTBD9,XRCC2,MAVS,HTT,HSP90AB1,HSP90AA1,VIM,SPARC,FTH1,UBC,KIDINS220,TM
18,ACTG1,EEF2,DPYSL3,TUBA1A,CFL1,BTBD9,XRCC2,MAVS,HTT,HSP90AB1,ZNF124,HSP90AA1,
NF316,DBN1,ENO1,TUBA1B,PIK3R2,FLNA,MAP1B,NCL,SLC9A7,PKM,GANAB,COL6A1,FTL,HSP90
TL,NEFM,PRPF8,YBX1,ATP2B4,FSTL1,PTMA,ENO2,KAT6A,DVL3,YWHAE,GNAS,GPI,BSG,FASN,ALD
MSB4X,NUMA1,RPL3,ZNF316,DBN1,RPL4,ENO1,TUBA1B,PIK3R2,FLNA,NCL,PKM,GANAB,COL6A
1,BTBD9,MAVS,HTT,ZNF124,HSP90AA1,ACTR2,VIM,SPARC,FTH1,UBC,KIDINS220,NUMA1,DCX,Z
EF2,DPYSL3,TUBA1A,CFL1,XRCC2,MAVS,HTT,HSP90AB1,HSP90AA1,ACTR2,VIM,TUBB,FTH1,UB
BX1,TUBB2B,COL5A1,COL4A1,CANX,PTMA,ENO2,DVL3,GNAS,BSG,FASN,ATP5F1B,GNB1,GDI1,F
,ZNF316,DBN1,FLNA,MAP1B,SLC9A7,FTL,HSP90B1,NEFM,UGT8,YBX1,FSTL1,TUBB2B,COL4A1,
ABHD2,TUBB2B,PTMA,DDX5,SEC62,DVL3,YWHAE,GNAS,BSG,FASN,SPTBN1,GNB1,SHISA9,MYH
,VIM,FTH1,NUMA1,DCX,EIF4G2,RPL3,DBN1,ENO1,PIK3R2,FLNA,NCL,RPL7A,SLC9A7,ALDOA,PK
,GANAB,FADS1,NEFL,FBXL4,YBX1,CALR,ABHD2,TUBB2B,COL5A1,PSAP,PGK1,RPLP0,H3-3B,CL
PL7A,ALDOA,PKM,GANAB,FTL,NEFL,RPL8,FBXL4,YBX1,ATP2B4,IGFBP2,FSTL1,TUBB2B,COL5A
JBB,KIDINS220,NUMA1,RPL3,RPS6,ZNF316,DBN1,RPL4,ENO1,TUBA1B,PIK3R2,NCL,ALDOA,PK
1,RPL3,ZNF316,DBN1,ENO1,TUBA1B,PIK3R2,FLNA,MAP1B,NCL,PKM,GANAB,COL6A1,HSP90B1,
MED18,ACTG1,EEF2,DPYSL3,TUBA1A,CFL1,BTBD9,XRCC2,MAVS,HTT,HSP90AB1,ZNF124,HSP9

NS220,DCX,EIF4G2,ZNF316,DBN1,ENO1,PIK3R2,FLNA,MAP1B,RPL7A,SLC9A7,ALDOA,PKM,GANA
PL8,YBX1,IGFBP2,ABHD2,HSPA1B,CLU,PTMA,ENO2,RPL13A,DVL3,KIF1A,GNAS,GPI,BSG,FASN,SI
B1,FADS1,NEFM,FSTL1,TUBB2B,COL5A1,COL4A1,PSAP,RPLP0,CLU,PTMA,DDX5,KAT6A,DVL3,Y
,ACTR2,VIM,UBC,KIDINS220,TMSB4X,NUMA1,EIF4G2,RPL3,RPS6,ZNF316,ENO1,TUBA1B,PIK3R
TT,HSP90AA1,VIM,SPARC,FTH1,TMSB4X,NUMA1,RPL3,RPS6,ZNF316,DBN1,ENO1,TUBA1B,PIK3F
JBA1A,CFL1,BTBD9,MAVS,HTT,ZNF124,HSP90AA1,ACTR2,VIM,FTH1,TMSB4X,NUMA1,DCX,EIF4G
ZNF124,HSP90AA1,VIM,FTH1,UBC,KIDINS220,NUMA1,DBN1,RPL4,FLNA,MAP1B,NCL,PKM,GANA
24,VIM,TUBB,FTH1,NUMA1,EIF4G2,ZNF316,DBN1,RPL4,ENO1,TUBA1B,PIK3R2,NCL,RPL7A,ALD
A1A,CFL1,BTBD9,XRCC2,MAVS,HTT,HSP90AB1,ZNF124,HSP90AA1,ACTR2,TUBB,FTH1,UBC,TMS
,ACTG1,EEF2,DPYSL3,TUBA1A,CFL1,BTBD9,MAVS,HTT,HSP90AB1,ZNF124,HSP90AA1,ACTR2,VI
PF8,RPL8,YBX1,UBA1,CALR,HSPA1B,CANX,DDX5,RPL13A,SEC62,RPL6,KAT6A,DVL3,YWHAE,GNA
ABHD2,TUBB2B,DDX5,KAT6A,DVL3,YWHAE,GNAS,GPI,SPTBN1,CALM3,GNB1,STMN2,MYH9,RPL
,FBXL4,YBX1,UBA1,PGK1,DDX5,DVL3,BSG,FASN,ALDH6A1,SPTBN1,CALM3,MYH9,SEPTIN2,GDI1
P90AA1,ACTR2,TUBB,FTH1,KIDINS220,EIF4G2,RPS6,ZNF316,RPL4,PIK3R2,FLNA,MAP1B,NCL,SL
IDINS220,TMSB4X,DCX,EIF4G2,RPS6,ZNF316,DBN1,ENO1,TUBA1B,PIK3R2,FLNA,MAP1B,SLC9A
HSP90AB1,ZNF124,HSP90AA1,VIM,FTH1,UBC,NUMA1,EIF4G2,RPL3,RPS6,ZNF316,DBN1,ENO1,T
,FSTL1,H3-3B,ENO2,YWHAE,KIF1A,GNAS,SLC2A3,FASN,SPTBN1,GNB1,SHISA9,MYH9,SEPTIN2,P
BX1,UBA1,ATP2B4,IGFBP2,FSTL1,PGK1,PTMA,ENO2,KAT6A,DVL3,YWHAE,KIF1A,GNAS,DDX17,G
CTG1,DPYSL3,TUBA1A,CFL1,BTBD9,XRCC2,MAVS,HTT,HSP90AB1,ZNF124,HSP90AA1,VIM,TUBE
,CFL1,BTBD9,MAVS,HTT,HSP90AB1,ZNF124,HSP90AA1,VIM,FTH1,UBC,NUMA1,EIF4G2,RPL3,RF
AVS,HTT,ZNF124,HSP90AA1,ACTR2,VIM,SPARC,FTH1,UBC,TMSB4X,NUMA1,EIF4G2,RPS6,ZNF31
90AA1,VIM,FTH1,KIDINS220,TMSB4X,NUMA1,EIF4G2,RPL3,ZNF316,DBN1,RPL4,ENO1,TUBA1B,
1,ACTR2,VIM,SPARC,TUBB,FTH1,TMSB4X,RPS6,ZNF316,DBN1,RPL4,ENO1,TUBA1B,PIK3R2,FLN
MTRNR2L2,MED18,ACTG1,EEF2,DPYSL3,TUBA1A,CFL1,BTBD9,XRCC2,MAVS,HTT,HSP90AB1,ZN
BXL4,YBX1,CALR,HSPA1B,TUBB2B,PSAP,CANX,CLU,KAT6A,DVL3,RPL13,GNAS,DDX17,BSG,FAS
6,DBN1,ENO1,PIK3R2,RPL7A,SLC9A7,FTL,HSP90B1,RPL8,FBXL4,YBX1,ATP2B4,FSTL1,HSPA1B,T
GFBP2,CALR,HSPA1B,TUBB2B,COL5A1,PTMA,DDX5,SEC62,DVL3,YWHAE,GNAS,GPI,RACK1,FAS
SL3,TUBA1A,CFL1,MAVS,HTT,HSP90AB1,ZNF124,HSP90AA1,ACTR2,VIM,TUBB,FTH1,UBC,KIDIN
FLNA,MAP1B,NCL,PKM,GANAB,COL6A1,FTL,HSP90B1,NEFM,PRPF8,RPL8,FBXL4,YBX1,IGFBP2,F
A1A,CFL1,BTBD9,XRCC2,MAVS,HSP90AA1,ACTR2,VIM,SPARC,TUBB,UBC,KIDINS220,TMSB4X,N
X1,COL4A1,PTMA,KAT6A,DVL3,YWHAE,KIF1A,GNAS,SPTBN1,CALM3,SHISA9,STMN2,SEPTIN2,P4
NCL,PKM,GANAB,COL6A1,FADS1,RPL8,FBXL4,IGFBP2,ABHD2,TUBB2B,COL5A1,PGK1,DDX5,KA
4,HSP90AA1,ACTR2,FTH1,KIDINS220,NUMA1,EIF4G2,RPS6,RPL4,ENO1,TUBA1B,FLNA,RPL7A,SL
EIF4G2,ZNF316,DBN1,FLNA,MAP1B,NCL,RPL7A,SLC9A7,ALDOA,PKM,FTL,HSP90B1,FADS1,NEF
T,HSP90AB1,ZNF124,HSP90AA1,ACTR2,VIM,TUBB,FTH1,UBC,TMSB4X,NUMA1,EIF4G2,RPL3,RPS
DVL3,KIF1A,GNAS,BSG,FASN,ALDH6A1,SPTBN1,GNB1,STMN2,RPL7,SEPTIN2,MARCKS,SMG1,PL
K1,DDX5,KAT6A,DVL3,KIF1A,GNAS,SLC2A3,BSG,FASN,SPTBN1,GNB1,MYH9,RPL7,SEPTIN2,GDI1
C,KIDINS220,TMSB4X,NUMA1,EIF4G2,ZNF316,DBN1,TUBA1B,PIK3R2,FLNA,MAP1B,NCL,RPL7A,
A1,ACTR2,UBC,TMSB4X,NUMA1,DCX,DBN1,ENO1,TUBA1B,PIK3R2,MAP1B,NCL,SLC9A7,HSP90
RPL3,DBN1,RPL4,FLNA,MAP1B,NCL,PKM,GANAB,COL6A1,FADS1,RPL8,FBXL4,IGFBP2,FSTL1,A
90AA1,VIM,SPARC,TUBB,UBC,KIDINS220,TMSB4X,NUMA1,ENO1,TUBA1B,PIK3R2,FLNA,NCL,RPL
BN1,GNB1,STMN2,RPL7,SEPTIN2,MARCKS,SMG1,PLXNB2,USP11,EIF4B,MARCKSL1,AFF4,TRIM
C,TMSB4X,DBN1,RPL4,ENO1,TUBA1B,PIK3R2,MAP1B,RPL7A,SLC9A7,ALDOA,PKM,GANAB,COL6
C,TMSB4X,DBN1,RPL4,ENO1,TUBA1B,PIK3R2,MAP1B,RPL7A,SLC9A7,ALDOA,PKM,GANAB,COL6
C,TMSB4X,DBN1,RPL4,ENO1,TUBA1B,PIK3R2,MAP1B,RPL7A,SLC9A7,ALDOA,PKM,GANAB,COL6

RC,TUBB,UBC,KIDINS220,TMSB4X,NUMA1,EIF4G2,RPL3,ZNF316,DBN1,RPL4,ENO1,TUBA1B,PIK
IUMA1,RPL3,ZNF316,DBN1,PIK3R2,FLNA,NCL,SLC9A7,HSP90B1,FADS1,NEFM,RPL8,FBXL4,YBX1
A1,ACTR2,VIM,SPARC,TUBB,UBC,KIDINS220,TMSB4X,NUMA1,EIF4G2,RPL3,RPS6,ZNF316,DBN1
UBA1B,PIK3R2,NCL,SLC9A7,PKM,FTL,HSP90B1,FADS1,NEFM,UGT8,YBX1,IGFBP2,CALR,ABHD2,
AT6A,DVL3,GNAS,BSG,FASN,SPTBN1,STMN2,RPL7,SEPTIN2,FSCN1,MARCKS,CALM1,PLXNB2,US
SP22,ACTG1,EEF2,DPYSL3,TUBA1A,CFL1,BTBD9,XRCC2,MAVS,HTT,HSP90AB1,ZNF124,HSP90A
H1,TMSB4X,NUMA1,DBN1,FLNA,MAP1B,RPL7A,SLC9A7,PKM,GANAB,HSP90B1,UBA1,ABHD2,TU
F4G2,ZNF316,DBN1,RPL4,ENO1,TUBA1B,PIK3R2,NCL,RPL7A,ALDOA,PKM,GANAB,COL6A1,FTL,F
ACTR2,UBC,NUMA1,EIF4G2,RPL3,ZNF316,DBN1,ENO1,FLNA,MAP1B,NCL,SLC9A7,MTRNR2L8,CC
GANAB,COL6A1,FTL,NEFM,FBXL4,TUBB2B,PGK1,KAT6A,DVL3,GNAS,SLC2A3,BSG,FASN,SPTBN1
DI1,SPTAN1,GSTP1,CALM1,PLXNB2,PLD3,COL18A1,IGF2,MARCKSL1,TRIM28,CKB,FN1,DCHS1,S
P90AB1,ZNF124,HSP90AA1,VIM,TUBB,FTH1,UBC,KIDINS220,NUMA1,EIF4G2,RPL3,RPS6,ZNF316
BTBD9,MAVS,HTT,HSP90AB1,HSP90AA1,ACTR2,VIM,TUBB,FTH1,KIDINS220,TMSB4X,NUMA1,RP
CALR,TUBB2B,PTMA,SEC62,KAT6A,YWHAE,GNAS,FASN,SPTBN1,GNB1,SHISA9,MYH9,RPL7,P4HB,
SB4X,RPL3,ZNF316,DBN1,ENO1,TUBA1B,PIK3R2,FLNA,NCL,SLC9A7,PKM,GANAB,COL6A1,FADS
,XRCC2,MAVS,HSP90AB1,ZNF124,HSP90AA1,ACTR2,SPARC,TUBB,FTH1,KIDINS220,TMSB4X,NU
MED18,ACTG1,EEF2,DPYSL3,TUBA1A,CFL1,BTBD9,XRCC2,MAVS,HTT,HSP90AB1,ZNF124,HSP90A
CL,RPL7A,PKM,GANAB,COL6A1,FTL,YBX1,ATP2B4,CALR,TMF1,ABHD2,COL4A1,CANX,PTMA,ENO
EF2,CFL1,BTBD9,MAVS,HTT,HSP90AB1,HSP90AA1,ACTR2,VIM,TUBB,FTH1,UBC,KIDINS220,TMS
G2,RPL3,ZNF316,DBN1,RPL4,MAP1B,NCL,PKM,GANAB,COL6A1,FADS1,NEFM,FBXL4,UBA1,ABHD
AS,FASN,SPTBN1,CALM3,GNB1,MARCKS,SMG1,CALM1,IGF2,EIF4B,AFF4,YWHAQ,DCHS1,SET,EIF
L1,BTBD9,XRCC2,MAVS,HSP90AB1,ZNF124,HSP90AA1,VIM,FTH1,UBC,NUMA1,EIF4G2,RPL3,RPS
DCX,ZNF316,DBN1,ENO1,FLNA,MAP1B,SLC9A7,FTL,HSP90B1,FADS1,NEFM,UGT8,RPL8,YBX1,FST
C9A7,PKM,COL6A1,FBXL4,ABHD2,PGK1,CANX,DDX5,ENO2,SOX4,KAT6A,DVL3,KIF1A,GNAS,BSG,
K3R2,FLNA,MAP1B,NCL,PKM,COL6A1,HSP90B1,NEFM,UGT8,FBXL4,YBX1,UBA1,ATP2B4,FSTL1,
AA1,ACTR2,VIM,FTH1,UBC,KIDINS220,NUMA1,RPL3,ZNF316,RPL4,ENO1,TUBA1B,PIK3R2,FLNA
COL5A1,PTMA,KAT6A,RPL13,GNAS,HTRA4,BSG,FASN,GNB1,MYH9,RPL7,GDI1,FSCN1,SMG1,COL
,TUBA1A,CFL1,BTBD9,XRCC2,MAVS,ZNF124,HSP90AA1,ACTR2,TUBB,FTH1,KIDINS220,TMSB4X,
,UBC,KIDINS220,NUMA1,EIF4G2,RPL3,ZNF316,DBN1,ENO1,TUBA1B,PIK3R2,FLNA,MAP1B,SLC9
EIF4G2,RPL3,ZNF316,DBN1,RPL4,ENO1,PIK3R2,FLNA,NCL,MTRNR2L8,GANAB,COL6A1,FADS1,
RPL7A,SLC9A7,ALDOA,PKM,GANAB,HSP90B1,NEFL,RPL8,FBXL4,YBX1,TMF1,TUBB2B,COL5A1,P
4G2,ZNF316,DBN1,ENO1,PIK3R2,MAP1B,NCL,SLC9A7,PKM,COL6A1,NEFM,UGT8,RPL8,FBXL4,Y
COL6A1,FADS1,YBX1,FSTL1,TUBB2B,COL5A1,COL4A1,PTMA,SOX4,KAT6A,YWHAE,KIF1A,RPL13,
9,XRCC2,HTT,HSP90AA1,ACTR2,VIM,TUBB,UBC,KIDINS220,TMSB4X,NUMA1,EIF4G2,RPL4,ENO1
C,KIDINS220,TMSB4X,DCX,EIF4G2,RPL3,ZNF316,DBN1,ENO1,TUBA1B,MAP1B,NCL,SLC9A7,ALD
C,NRBP2,USP22,MTRNR2L2,MED18,ACTG1,EEF2,DPYSL3,TUBA1A,CFL1,BTBD9,XRCC2,MAVS,H
L13A,DVL3,YWHAE,KIF1A,GNAS,DDX17,BSG,FASN,ALDH6A1,SPTBN1,CALM3,SHISA9,MYH9,APP,F
,KIF1A,GNAS,GPI,FASN,SPTBN1,GNB1,SHISA9,MYH9,APP,GDI1,PTMS,SPTAN1,CALM1,IGF2,MAR
SEC62,YWHAE,GNAS,FASN,SPTBN1,GNB1,SHISA9,MYH9,APP,IDS,GDI1,PTMS,NPM1,USP11,IGF2,
TRNR2L2,MED18,ACTG1,EEF2,DPYSL3,TUBA1A,CFL1,BTBD9,XRCC2,MAVS,HTT,HSP90AB1,ZNF
,UBC,TMSB4X,ZNF316,ENO1,TUBA1B,MAP1B,NCL,RPL7A,ALDOA,PKM,GANAB,COL6A1,FADS1,N
B,FTH1,UBC,NUMA1,EIF4G2,ZNF316,DBN1,ENO1,TUBA1B,FLNA,NCL,RPL7A,ALDOA,GANAB,FTL,
IO1,TUBA1B,FLNA,NCL,RPL7A,SLC9A7,ALDOA,PKM,GANAB,FTL,HSP90B1,NEFL,RPL8,FBXL4,YB
BN1,FLNA,MAP1B,NCL,PKM,GANAB,FADS1,RPL8,ABHD2,TUBB2B,DDX5,ENO2,FKBP14,KAT6A,D
HSP90AB1,ZNF124,ACTR2,VIM,TUBB,FTH1,UBC,KIDINS220,TMSB4X,EIF4G2,ZNF316,DBN1,ENC

DINS220,TMSB4X,RPL3,RPS6,ZNF316,DBN1,ENO1,TUBA1B,PIK3R2,MAP1B,NCL,RPL7A,SLC9A7,A
,TUBB,FTH1,KIDINS220,TMSB4X,RPL3,RPS6,ZNF316,DBN1,ENO1,TUBA1B,PIK3R2,MAP1B,NCL,I
O1,FLNA,SLC9A7,PKM,HSP90B1,PRPF8,RPL8,FBXL4,YBX1,CALR,FSTL1,ABHD2,HSPA1B,COL5A1
,DBN1,RPL4,ENO1,PIK3R2,MAP1B,NCL,RPL7A,MTRNR2L8,COL6A1,NEFM,RPL8,FBXL4,UBA1,CA
NS220,TMSB4X,NUMA1,EIF4G2,DBN1,ENO1,TUBA1B,NCL,RPL7A,ALDOA,PKM,GANAB,FADS1,NE
.6A1,FADS1,PRPF8,YBX1,CALR,ABHD2,COL4A1,CANX,PTMA,YWHAE,GNAS,BSG,FASN,ALDH6A1,S
IF4G2,RPS6,ZNF316,TUBA1B,PIK3R2,FLNA,RPL7A,SLC9A7,ALDOA,MTRNR2L8,PKM,COL6A1,FAI
,RPL4,ENO1,TUBA1B,PIK3R2,MAP1B,NCL,PKM,GANAB,COL6A1,HSP90B1,NEFM,RPL8,FBXL4,YE
A1,RPL8,FBXL4,TUBB2B,DVL3,GNAS,SLC2A3,BSG,SPTBN1,GNB1,STMN2,RPL7,SEPTIN2,GDI1,SF
A1,RPL8,FBXL4,TUBB2B,DVL3,GNAS,SLC2A3,BSG,SPTBN1,GNB1,STMN2,RPL7,SEPTIN2,GDI1,SF
ACD3,DCHS1,EIF4A1,MRC2,LDHB,SSBP2,MORF4L1,AGRN,MARCHF6,PXDN,OGA,CTNNA1,FADS2,
B,PSAP,CANX,RPL13A,GPI,BSG,SEPTIN2,P4HB,APP,RPL5,FSCN1,TMBIM6,NPM1,PLXNB2,PLD3,H
R2L2,MED18,ACTG1,EEF2,DPYSL3,TUBA1A,CFL1,BTBD9,XRCC2,MAVS,HTT,HSP90AB1,ZNF124,H
SP90AA1,ACTR2,VIM,UBC,NUMA1,EIF4G2,RPL3,RPS6,ZNF316,DBN1,TUBA1B,PIK3R2,FLNA,RPL7
,FBXL4,TUBB2B,PGK1,PTMA,DVL3,YWHAE,GNAS,SLC2A3,GPI,BSG,FASN,SPTBN1,GNB1,STMN2,
BSG,MYH9,RPL7,GDI1,SPTAN1,MARCKS,USP11,EIF4B,MARCKSL1,AFF4,TRIM28,MATR3,YWHAG
HD2,TUBB2B,PGK1,DDX5,ENO2,KAT6A,DVL3,SLC2A3,BSG,SPTBN1,GNB1,MYH9,SEPTIN2,GDI1,S
TUBB2B,COL5A1,H3-3B,PTMA,KAT6A,DVL3,YWHAE,KIF1A,GNAS,GPI,BSG,FASN,CALM3,GNB1,M
ACTR2,FTH1,UBC,KIDINS220,NUMA1,DCX,EIF4G2,RPS6,ZNF316,DBN1,RPL4,ENO1,PIK3R2,FLNA,
,SPARC,FTH1,KIDINS220,EIF4G2,RPS6,DBN1,RPL4,ENO1,PIK3R2,NCL,RPL7A,ALDOA,PKM,GANA
2,MAVS,HTT,ACTR2,VIM,FTH1,UBC,KIDINS220,TMSB4X,NUMA1,EIF4G2,RPL3,RPS6,DBN1,RPL4,E
C2,MAVS,HTT,ZNF124,HSP90AA1,ACTR2,VIM,FTH1,UBC,TMSB4X,NUMA1,DCX,EIF4G2,RPL3,ZNF
BTBD9,XRCC2,HTT,HSP90AA1,ACTR2,VIM,SPARC,TUBB,UBC,KIDINS220,TMSB4X,NUMA1,EIF4G2
,DVL3,YWHAE,KIF1A,GNAS,SLC2A3,GPI,FASN,SPTBN1,GNB1,STMN2,MYH9,GDI1,PTMS,SPTAN1,C
N1,ENO1,PIK3R2,MAP1B,NCL,RPL7A,HSP90B1,YBX1,ABHD2,TUBB2B,COL5A1,PTMA,DDX5,SEC6
NS220,NUMA1,EIF4G2,ZNF316,DBN1,ENO1,MAP1B,NCL,RPL7A,ALDOA,PKM,GANAB,COL6A1,HSI
CC2,MAVS,HTT,ZNF124,ACTR2,VIM,FTH1,UBC,TMSB4X,NUMA1,DCX,ZNF316,DBN1,ENO1,PIK3R
1,TUBA1B,PIK3R2,FLNA,NCL,RPL7A,PKM,FTL,NEFL,NEFM,RPL8,FBXL4,YBX1,IGFBP2,TMF1,TUE
GNAS,FASN,SPTBN1,STMN2,MYH9,GDI1,PTMS,GSTP1,CALM1,IGF2,EIF4B,AFF4,TRIM28,MATR3,C
BA1B,PIK3R2,NCL,RPL7A,ALDOA,PKM,GANAB,COL6A1,FTL,FADS1,NEFM,YBX1,COL4A1,RPLP0,H
3,TUBA1A,CFL1,BTBD9,XRCC2,MAVS,HTT,ZNF124,HSP90AA1,ACTR2,VIM,SPARC,TUBB,FTH1,UB
,SLC2A3,BSG,FASN,SPTBN1,CALM3,GNB1,MYH9,SEPTIN2,GDI1,PTMS,SPTAN1,SMG1,CALM1,IG
COL6A1,NEFM,RPL8,FBXL4,TUBB2B,CANX,DDX5,FKBP14,KAT6A,DVL3,GPI,BSG,FASN,ALDH6A1,
1,FSTL1,CANX,DDX5,SEC62,KAT6A,DVL3,RPL13,GNAS,DDX17,GPI,BSG,ALDH6A1,ATP5F1B,MYH
NEFL,FBXL4,IGFBP2,TUBB2B,COL5A1,COL4A1,PTMA,ENO2,SEC62,KAT6A,GNAS,GPI,BSG,GNB1
B4X,EIF4G2,RPL3,RPS6,DBN1,RPL4,FLNA,MAP1B,NCL,RPL7A,GANAB,HSP90B1,NEFL,NEFM,PRF
,ALDOA,RPL8,FBXL4,YBX1,ATP2B4,ABHD2,TUBB2B,COL4A1,SOX4,SEC62,KAT6A,DVL3,YWHAE,K
1,NEFM,YBX1,FSTL1,ABHD2,TUBB2B,PTMA,DDX5,YWHAE,GNAS,SLC2A3,DDX17,FASN,SPTBN1,C
D18,ACTG1,DPYSL3,TUBA1A,CFL1,BTBD9,XRCC2,MAVS,HTT,HSP90AB1,ZNF124,ACTR2,FTH1,KI
FTH1,TMSB4X,NUMA1,RPS6,ZNF316,DBN1,ENO1,TUBA1B,PIK3R2,FLNA,RPL7A,PKM,PRPF8,RPL8
31,HSP90AA1,ACTR2,VIM,TUBB,FTH1,UBC,NUMA1,RPL3,RPS6,ZNF316,DBN1,PIK3R2,FLNA,RPL7
2,FTH1,UBC,EIF4G2,ZNF316,DBN1,PIK3R2,FLNA,MAP1B,NCL,RPL7A,SLC9A7,ALDOA,PKM,COL6
SN,MYH9,APP,FSCN1,SPTAN1,SMG1,NPM1,VCP,MARCKSL1,YWHAG,CKB,FN1,SPEN,SET,TPI1,CH
,DVL3,KIF1A,SLC2A3,GPI,FASN,ALDH6A1,CALM3,GNB1,STMN2,RPS8,SEPTIN2,P4HB,GDI1,FSCN
R2,VIM,NUMA1,ZNF316,DBN1,MAP1B,NCL,RPL7A,SLC9A7,PKM,COL6A1,FTL,HSP90B1,FADS1,NI

SP90AA1,ACTR2,FTH1,UBC,KIDINS220,NUMA1,RPL3,ZNF316,RPL4,ENO1,PIK3R2,FLNA,NCL,SLC
1A,CFL1,BTBD9,XRCC2,MAVS,HSP90AB1,ZNF124,HSP90AA1,ACTR2,SPARC,FTH1,UBC,KIDINS22
A1B,PIK3R2,RPL7A,SLC9A7,ALDOA,MTRNR2L8,PKM,COL6A1,FADS1,NEFL,NEFM,PRPF8,RPL8,F
MAVS,HTT,HSP90AB1,HSP90AA1,SPARC,FTH1,UBC,KIDINS220,EIF4G2,RPL3,RPS6,ZNF316,RPL4,
BB,FTH1,UBC,KIDINS220,TMSB4X,NUMA1,EIF4G2,RPL3,ZNF316,DBN1,RPL4,ENO1,TUBA1B,PIK
,FADS1,PRPF8,CALR,TUBB2B,COL5A1,CANX,DDX5,KAT6A,YWHAE,KIF1A,SLC2A3,BSG,FASN,ALD
S,FASN,SPTBN1,GNB1,SHISA9,MYH9,P4HB,IDS,PTMS,USP11,IGF2,EIF4B,TRIM28,YWHAG,CKB,Y
TL,FADS1,NEFL,NEFM,PRPF8,RPL8,YBX1,ATP2B4,IGFBP2,FSTL1,ABHD2,HSPA1B,COL4A1,PGK1
D9,XRCC2,HTT,ZNF124,HSP90AA1,ACTR2,SPARC,TUBB,FTH1,UBC,KIDINS220,TMSB4X,NUMA1,
R2,NCL,SLC9A7,PKM,GANAB,FADS1,PRPF8,RPL8,UBA1,IGFBP2,CALR,TUBB2B,COL5A1,COL4A
2,FLNA,NCL,RPL7A,ALDOA,MTRNR2L8,PKM,FTL,NEFL,RPL8,FBXL4,ATP2B4,IGFBP2,TUBB2B,CC
A,SLC9A7,ALDOA,PKM,COL6A1,FTL,HSP90B1,YBX1,FSTL1,TUBB2B,COL5A1,COL4A1,PTMA,KA
A7,ALDOA,PKM,HSP90B1,NEFL,RPL8,FBXL4,IGFBP2,TMF1,TUBB2B,COL5A1,PSAP,RPLP0,PTM
1,UBC,KIDINS220,NUMA1,EIF4G2,RPL3,ZNF316,DBN1,TUBA1B,PIK3R2,MAP1B,RPL7A,SLC9A7,
1A,CFL1,BTBD9,XRCC2,MAVS,HSP90AB1,ZNF124,HSP90AA1,ACTR2,SPARC,FTH1,UBC,KIDINS22
B,CLU,PTMA,KAT6A,DVL3,GNAS,BSG,FASN,SPTBN1,CALM3,GNB1,RPL7,APP,FSCN1,PLXNB2,PL
KIDINS220,TMSB4X,EIF4G2,RPL3,RPS6,ZNF316,DBN1,RPL4,ENO1,PIK3R2,MAP1B,NCL,RPL7A,
A1B,FLNA,NCL,RPL7A,SLC9A7,ALDOA,PKM,COL6A1,FADS1,NEFL,RPL8,FBXL4,UBA1,TMF1,TUBE
HSP90AB1,ZNF124,HSP90AA1,ACTR2,VIM,TUBB,FTH1,UBC,KIDINS220,TMSB4X,NUMA1,EIF4G2,
,UBC,NUMA1,RPS6,ZNF316,PIK3R2,FLNA,RPL7A,SLC9A7,MTRNR2L8,PKM,COL6A1,FTL,HSP90
L8,FBXL4,YBX1,UBA1,FSTL1,TMF1,HSPA1B,COL5A1,COL4A1,PGK1,CANX,ENO2,RPL13A,KAT6A,
YSL3,TUBA1A,CFL1,BTBD9,MAVS,HSP90AB1,ZNF124,HSP90AA1,ACTR2,VIM,SPARC,FTH1,UBC,K
,TUBB2B,DVL3,GNAS,SLC2A3,BSG,SPTBN1,GNB1,GDI1,SPTAN1,SMG1,USP11,IGF2,EIF4B,MAR
YSL3,TUBA1A,CFL1,XRCC2,HTT,HSP90AB1,ZNF124,HSP90AA1,ACTR2,SPARC,FTH1,UBC,KIDIN
DX5,ENO2,GNAS,GPI,FASN,SPTBN1,GNB1,SEPTIN2,APP,IDS,GDI1,FSCN1,PTMS,PLXNB2,USP11,
,FTH1,UBC,NUMA1,RPS6,ZNF316,DBN1,TUBA1B,NCL,RPL7A,PKM,COL6A1,FTL,HSP90B1,FADS
,FTH1,UBC,NUMA1,RPS6,ZNF316,DBN1,TUBA1B,NCL,RPL7A,PKM,COL6A1,FTL,HSP90B1,FADS
,NUMA1,EIF4G2,RPL3,RPS6,ZNF316,DBN1,RPL4,PIK3R2,NCL,GANAB,HSP90B1,NEFL,NEFM,RPL
,TUBB2B,COL4A1,CANX,DDX5,KAT6A,DVL3,GPI,BSG,ALDH6A1,SPTBN1,CALM3,GNB1,MYH9,SE
GFBP2,CES3,PTMA,ENO2,KAT6A,YWHAE,KIF1A,GNAS,FASN,CALM3,MYH9,SEPTIN2,PTMS,GSTP1,
,CES3,PTMA,YWHAE,GNAS,FASN,SPTBN1,CALM3,SHISA9,STMN2,SEPTIN2,APP,GDI1,SMG1,NPM1
2,CFL1,BTBD9,MAVS,HTT,HSP90AB1,HSP90AA1,ACTR2,VIM,TUBB,FTH1,UBC,KIDINS220,TMSB
0,ZNF316,DBN1,PIK3R2,FLNA,PKM,FTL,RPL8,FBXL4,YBX1,TUBB2B,PTMA,DDX5,KAT6A,DVL3,SL
6A1,FTL,HSP90B1,FADS1,NEFM,UGT8,RPL8,YBX1,IGFBP2,COL4A1,CES3,H3-3B,PTMA,ENO2,K
C9A7,HSP90B1,NEFM,PRPF8,RPL8,YBX1,HSPA1B,RPLP0,DDX5,RPL13A,RPL6,YWHAE,RPL13,GN
NF316,DBN1,ENO1,TUBA1B,FLNA,MAP1B,SLC9A7,PKM,GANAB,COL6A1,FADS1,NEFM,RPL8,UBA
,YBX1,IGFBP2,TUBB2B,COL4A1,DDX5,SOX4,RPL13A,MAGED1,YWHAE,KIF1A,RACK1,BSG,FASN,
VIM,TUBB,FTH1,UBC,KIDINS220,TMSB4X,NUMA1,DCX,EIF4G2,RPL3,ZNF316,DBN1,ENO1,TUBA1
2B,SEC62,DVL3,GNAS,SLC2A3,SPTBN1,GNB1,RPL7,SEPTIN2,GDI1,PTMS,GSTP1,SMG1,PLXNB2
9,XRCC2,MAVS,HTT,HSP90AB1,ZNF124,ACTR2,FTH1,UBC,TMSB4X,NUMA1,EIF4G2,RPL3,ZNF3
A1,VIM,TUBB,UBC,KIDINS220,NUMA1,EIF4G2,RPL3,RPS6,DBN1,RPL4,ENO1,TUBA1B,NCL,RPL7
HSP90B1,RPL8,ATP2B4,CALR,TMF1,TUBB2B,PGK1,RPLP0,CANX,PTMA,DDX5,ENO2,SOX4,SEC62,
7,GPI,RACK1,FASN,ALDH6A1,SEPTIN2,P4HB,GDI1,NPM1,USP11,SNRNP200,EIF3C,ATP5F1A,MAT
NUMA1,RPL3,RPS6,ZNF316,DBN1,PIK3R2,FLNA,RPL7A,COL6A1,FADS1,FBXL4,YBX1,CALR,FSTL1
,ZNF124,HSP90AA1,VIM,TUBB,UBC,NUMA1,EIF4G2,RPL3,DBN1,ENO1,FLNA,MAP1B,NCL,SLC9A

'BD9,XRCC2,MAVS,HSP90AB1,ZNF124,HSP90AA1,ACTR2,VIM,FTH1,UBC,KIDINS220,TMSB4X,NUI
COL5A1,H3-3B,PTMA,KAT6A,DVL3,KIF1A,GNAS,SLC2A3,BSG,ALDH6A1,SPTBN1,GNB1,MYH9,RPL
NA,MAP1B,NCL,RPL7A,SLC9A7,ALDOA,PKM,GANAB,HSP90B1,NEFL,NEFM,PRPF8,RPL8,FBXL4,Y
M28,YWHAG,RPL37A,SPEN,HACD3,SET,SF3B1,SERINC1,CHD3,VAPA,POLR2A,TUBB4B,BTF3,SBF
NUMA1,EIF4G2,RPS6,ZNF316,ENO1,TUBA1B,PIK3R2,FLNA,MAP1B,NCL,SLC9A7,MTRNR2L8,FAD
TBD9,XRCC2,MAVS,HTT,HSP90AB1,ZNF124,HSP90AA1,TUBB,UBC,KIDINS220,EIF4G2,RPL3,RPS
A1,EIF4G2,RPL3,RPS6,ZNF316,PIK3R2,NCL,GANAB,HSP90B1,NEFL,NEFM,RPL8,FBXL4,FSTL1,T
AA1,ACTR2,FTH1,UBC,KIDINS220,NUMA1,EIF4G2,RPL3,RPS6,ZNF316,DBN1,RPL4,PIK3R2,MAP
TAN1,MARCKS,SMG1,CALM1,NPM1,PLXNB2,USP11,IGF2,MARCKSL1,AFF4,MATR3,CKB,NONO,Y
CTR2,VIM,SPARC,FTH1,UBC,KIDINS220,DCX,EIF4G2,RPL3,ZNF316,ENO1,TUBA1B,PIK3R2,FLNA,
UBC,NUMA1,RPL3,RPS6,ZNF316,DBN1,PIK3R2,FLNA,RPL7A,SLC9A7,COL6A1,FTL,FBXL4,YBX1,
,SPTBN1,SHISA9,STMN2,MARCKS,CALM1,IGF2,EIF4B,TRIM28,YWHAG,CKB,DCHS1,SET,TMSB1
B,CANX,KAT6A,DVL3,GPI,BSG,FASN,GNB1,STMN2,SEPTIN2,GDI1,SPTAN1,SMG1,PLXNB2,SNRNF
CC2,MAVS,HTT,HSP90AB1,ZNF124,HSP90AA1,TUBB,KIDINS220,EIF4G2,RPL3,RPS6,ZNF316,DBN
FL1,BTBD9,MAVS,HSP90AB1,ZNF124,HSP90AA1,ACTR2,VIM,SPARC,FTH1,KIDINS220,NUMA1,DC
20,TMSB4X,NUMA1,EIF4G2,RPL4,ENO1,TUBA1B,FLNA,NCL,RPL7A,HSP90B1,NEFL,RPL8,FBXL4
U,KAT6A,YWHAE,GNAS,SPTBN1,CALM3,GNB1,MYH9,RPL7,APP,RPL5,PTMS,IGF2,TRIM28,COL4A
,PIK3R2,FLNA,MAP1B,RPL7A,SLC9A7,ALDOA,PKM,GANAB,COL6A1,HSP90B1,NEFL,FBXL4,YBX1
,PKD1,SSBP2,ITGB1,RPLP1,SARAF,SBF1,CYB5R3,HNRNPU,HNRNPA0,BRD3,PJA2,NCS1,CLSTN1,
20,NUMA1,RPL3,RPS6,ZNF316,DBN1,RPL4,TUBA1B,PIK3R2,NCL,ALDOA,GANAB,HSP90B1,NEFM,
IM6,PLD3,HSPD1,EIF4B,UBB,TRIM28,YWHAG,RPL37A,SPEN,HACD3,SET,SF3B1,SERINC1,CHD3,
M,FBXL4,UBA1,ATP2B4,HSPA1B,H3-3B,CANX,PTMA,ENO2,SOX4,YWHAE,GNAS,SPTBN1,STMN2,
A1,ACTR2,FTH1,KIDINS220,NUMA1,EIF4G2,RPL3,RPS6,ZNF316,PIK3R2,MAP1B,ALDOA,GANAB,H
SP22,MED18,ACTG1,EEF2,DPYSL3,TUBA1A,CFL1,BTBD9,XRCC2,MAVS,HTT,HSP90AB1,ZNF124,
90AA1,VIM,SPARC,TUBB,FTH1,UBC,TMSB4X,NUMA1,DCX,EIF4G2,RPL3,RPS6,ZNF316,DBN1,TUE
,COL6A1,FADS1,NEFL,YBX1,FSTL1,ABHD2,TUBB2B,PTMA,DDX5,RPL6,YWHAE,SLC2A3,FASN,SP
M,COL6A1,FTL,HSP90B1,UBA1,COL4A1,H3-3B,DDX5,FKBP14,KIF1A,GNAS,HTRA4,BSG,FASN,G
6,ZNF316,TUBA1B,PIK3R2,FLNA,RPL7A,SLC9A7,ALDOA,MTRNR2L8,PKM,COL6A1,FADS1,NEFL,
,TUBB2B,PGK1,CANX,DDX5,ENO2,FKBP14,KAT6A,DVL3,KIF1A,GPI,FASN,ALDH6A1,CALM3,GNB
HSP90B1,NEFM,RPL8,FBXL4,YBX1,CES3,SEC62,KAT6A,DVL3,GNAS,SLC2A3,FASN,SPTBN1,SHISA
,RPL3,ZNF316,ENO1,TUBA1B,NCL,RPL7A,ALDOA,COL6A1,FTL,HSP90B1,NEFL,RPL8,YBX1,IGFB
P90AA1,ACTR2,VIM,FTH1,UBC,KIDINS220,NUMA1,EIF4G2,RPL3,RPS6,ZNF316,DBN1,RPL4,ENO1
,TMSB4X,NUMA1,EIF4G2,ZNF316,RPL4,TUBA1B,ALDOA,HSP90B1,NEFL,PRPF8,UBA1,ATP2B4,FS
TUBA1A,CFL1,MAVS,ZNF124,HSP90AA1,ACTR2,SPARC,TUBB,FTH1,KIDINS220,DCX,EIF4G2,RPL3
A7,PKM,GANAB,HSP90B1,NEFM,PRPF8,RPL8,FBXL4,YBX1,UBA1,IGFBP2,ABHD2,COL5A1,COL4
2,MAVS,HTT,HSP90AB1,ZNF124,HSP90AA1,ACTR2,VIM,TUBB,UBC,KIDINS220,TMSB4X,ZNF316,D
SL3,TUBA1A,CFL1,XRCC2,MAVS,HTT,HSP90AB1,ZNF124,HSP90AA1,VIM,TUBB,FTH1,UBC,KIDIN
B,UBC,KIDINS220,TMSB4X,NUMA1,ZNF316,DBN1,ENO1,PIK3R2,FLNA,MAP1B,NCL,PKM,GANAE
,IGF2,MARCKSL1,TRIM28,CKB,DCHS1,SET,TPI1,MRC2,PKD1,PDIA3,BASP1,ITGB1,AGRN,TIMP2,
SP22,MTRNR2L2,MED18,ACTG1,EEF2,DPYSL3,TUBA1A,CFL1,BTBD9,XRCC2,MAVS,HTT,HSP90A
P90AB1,HSP90AA1,ACTR2,SPARC,FTH1,UBC,KIDINS220,NUMA1,DCX,EIF4G2,RPL3,ZNF316,DBN
,ZNF316,RPL4,ENO1,PIK3R2,PKM,GANAB,HSP90B1,FADS1,NEFM,FBXL4,YBX1,ATP2B4,CALR,TM
NEFM,FBXL4,CALR,ABHD2,HSPA1B,TUBB2B,COL5A1,COL4A1,PSAP,RPLP0,CANX,PTMA,DDX5,KI
MA,DDX5,RPL6,KAT6A,DVL3,KIF1A,GNAS,DDX17,FASN,SPTBN1,SEPTIN2,APP,GDI1,FSCN1,PTMS
PRPF8,RPL8,FBXL4,ABHD2,PGK1,CANX,SOX4,DVL3,BSG,ALDH6A1,STMN2,SEPTIN2,GDI1,SPTAN

LM1,USP11,IGF2,EIF4B,TRIM28,YWHAG,LRRC28,DCHS1,SET,CHSY1,TMSB10,PKD1,HLA-C,STM
M28,MATR3,LRRC28,YWHAQ,DCHS1,TMSB10,PKD1,HLA-C,TPT1,ITGB1,AGRN,TIMP2,MIDN,VCL
MA,SEC62,KAT6A,DDX17,GNB1,RPS8,MYH9,P4HB,NPM1,HSPD1,CSDE1,MARCKSL1,MATR3,LRR
)9,MAVS,HTT,HSP90AB1,HSP90AA1,ACTR2,VIM,TUBB,FTH1,UBC,KIDINS220,TMSB4X,NUMA1,EIF
L1,BTBD9,XRCC2,MAVS,HSP90AB1,ZNF124,HSP90AA1,ACTR2,FTH1,UBC,KIDINS220,TMSB4X,NU
SP11,IGF2,AFF4,TRIM28,YWHAG,DDB1,HACD3,DCHS1,SET,TMSB10,PKD1,VAPA,TPT1,STMN1,ITG
P14,KAT6A,DVL3,SLC2A3,FASN,ALDH6A1,GNB1,MYH9,SEPTIN2,GDI1,SMG1,MARCKSL1,TRIM28
A1,ACTR2,VIM,FTH1,UBC,KIDINS220,NUMA1,RPL3,RPS6,ZNF316,DBN1,RPL4,ENO1,TUBA1B,PI
T6A,GNAS,DDX17,ALDH6A1,SPTBN1,GNB1,MYH9,P4HB,PTMS,SPTAN1,CALM1,PLXNB2,IGF2,CO
T6A,GNAS,DDX17,ALDH6A1,SPTBN1,GNB1,MYH9,P4HB,PTMS,SPTAN1,CALM1,PLXNB2,IGF2,CO
T6A,GNAS,DDX17,ALDH6A1,SPTBN1,GNB1,MYH9,P4HB,PTMS,SPTAN1,CALM1,PLXNB2,IGF2,CO
,DBN1,ENO1,TUBA1B,PIK3R2,FLNA,MAP1B,SLC9A7,ALDOA,MTRNR2L8,PKM,COL6A1,PRPF8,R
3,GNB1,SEPTIN2,APP,IDS,TMBIM6,SPTAN1,SMG1,CALM1,PLXNB2,PLD3,HSPD1,IGF2,AFF4,TRIM
D9,MAVS,HTT,HSP90AB1,ZNF124,HSP90AA1,ACTR2,SPARC,TUBB,FTH1,ZNF316,DBN1,TUBA1B,
1,BTBD9,XRCC2,MAVS,HSP90AB1,ZNF124,HSP90AA1,ACTR2,FTH1,UBC,KIDINS220,TMSB4X,NU
90AB1,ZNF124,HSP90AA1,ACTR2,VIM,SPARC,FTH1,KIDINS220,NUMA1,RPL3,RPS6,ZNF316,DBN
DPYSL3,CFL1,BTBD9,MAVS,HTT,HSP90AB1,HSP90AA1,ACTR2,VIM,SPARC,TUBB,FTH1,NUMA1,EI
S1,PKD1,HLA-C,TPT1,ITGB1,AGRN,TIMP2,SARAF,MIDN,CYB5R3,MAP4,CDC42BPB,ITM2B,GOLG
6,RPL4,ENO1,TUBA1B,FLNA,NCL,RPL7A,HSP90B1,NEFL,RPL8,FBXL4,YBX1,ATP2B4,IGFBP2,TM
TUBB2B,KAT6A,KIF1A,SLC2A3,FASN,SPTBN1,GNB1,RPL7,SPTAN1,GSTP1,SMG1,CALM1,PLXNB2
,SLC9A7,ALDOA,PKM,COL6A1,HSP90B1,NEFM,RPL8,YBX1,UBA1,TUBB2B,COL5A1,PSAP,H3-3B,
M,NEFL,RPL8,FBXL4,IGFBP2,TUBB2B,COL4A1,PTMA,ENO2,RPL6,DVL3,GNAS,DDX17,BSG,ATP5
,COL6A1,NEFM,PRPF8,RPL8,FBXL4,YBX1,UBA1,TUBB2B,ENO2,SEC62,KAT6A,KIF1A,SLC2A3,DD
BD9,XRCC2,MAVS,HSP90AB1,ZNF124,HSP90AA1,ACTR2,FTH1,UBC,KIDINS220,TMSB4X,NUMA1,
MS,SMG1,PLXNB2,USP11,IGF2,AFF4,MATR3,YWHAG,YWHAQ,HACD3,DCHS1,SET,MRC2,PKD1,TP
NF124,HSP90AA1,FTH1,KIDINS220,NUMA1,EIF4G2,RPL3,ZNF316,DBN1,RPL4,ENO1,TUBA1B,PIK
MAP1B,RPL7A,ALDOA,PKM,FADS1,YBX1,FSTL1,TUBB2B,COL5A1,COL4A1,PGK1,PTMA,DDX5,KA
ANAB,COL6A1,FTL,FADS1,NEFM,YBX1,ATP2B4,IGFBP2,COL5A1,COL4A1,H3-3B,CLU,PTMA,GNA
,RPL7A,PKM,NEFL,NEFM,RPL8,FBXL4,IGFBP2,TUBB2B,COL4A1,PSAP,RPLP0,PTMA,ENO2,RPL1
NF124,HSP90AA1,ACTR2,FTH1,KIDINS220,TMSB4X,NUMA1,EIF4G2,RPL3,RPS6,ZNF316,DBN1,R
TBN1,GNB1,MYH9,SMG1,USP11,IGF2,MARCKSL1,LRRC28,SPEN,DDB1,DCHS1,SET,EGLN1,CHS
R2,NCL,GANAB,FTL,HSP90B1,RPL8,YBX1,UBA1,TMF1,HSPA1B,PGK1,H3-3B,CANX,FKBP14,RPL1
IF316,DBN1,PIK3R2,MAP1B,NCL,RPL7A,COL6A1,NEFL,NEFM,UGT8,UBA1,ATP2B4,IGFBP2,FSTL
A1B,FLNA,NCL,RPL7A,HSP90B1,NEFL,RPL8,FBXL4,YBX1,ATP2B4,IGFBP2,TMF1,TUBB2B,COL4A
DH6A1,RPS8,SEPTIN2,GDI1,NPM1,USP11,ATP5F1A,EIF4B,MATR3,PSMD5,SET,EIF4A1,PPP2R1A,D
1,SPARC,TUBB,FTH1,UBC,KIDINS220,NUMA1,DCX,TUBA1B,PIK3R2,FLNA,NCL,RPL7A,SLC9A7,A
SP90AA1,ACTR2,FTH1,KIDINS220,NUMA1,RPL3,RPS6,ZNF316,DBN1,PIK3R2,MAP1B,NCL,ALDOA,
BD9,XRCC2,MAVS,HSP90AB1,ZNF124,HSP90AA1,ACTR2,FTH1,UBC,KIDINS220,NUMA1,EIF4G2,F
A4,SPTBN1,CALM3,GNB1,SHISA9,P4HB,PTMS,SPTAN1,PLXNB2,HSPD1,MARCKSL1,CKB,COL4A2
VS,HSP90AB1,HSP90AA1,VIM,FTH1,UBC,KIDINS220,TMSB4X,NUMA1,DCX,EIF4G2,ZNF316,DBN1
SP90AB1,ZNF124,HSP90AA1,TUBB,KIDINS220,EIF4G2,RPL3,RPS6,ZNF316,DBN1,RPL4,ENO1,TU
A,NCL,MTRNR2L8,PKM,GANAB,COL6A1,FTL,FBXL4,YBX1,IGFBP2,CALR,FSTL1,TUBB2B,COL4A1
1,BTBD9,XRCC2,MAVS,HSP90AB1,ZNF124,HSP90AA1,ACTR2,FTH1,UBC,KIDINS220,TMSB4X,NU
8,FBXL4,TUBB2B,COL4A1,RPLP0,PTMA,ENO2,DVL3,KIF1A,GNAS,DDX17,BSG,ATP5F1B,CALM3,
H1,EIF4G2,DBN1,ENO1,TUBA1B,PIK3R2,MAP1B,SLC9A7,ALDOA,PKM,GANAB,FTL,NEFL,ATP2B4,

3X1,UBA1,ATP2B4,HSPA1B,PGK1,H3-3B,CANX,FKBP14,RPL13A,MAGED1,RPL6,DVL3,RACK1,GN
GNAS,SPTBN1,CALM3,SHISA9,IDS,GDI1,PTMS,SMG1,NPM1,PLXNB2,UBB,SPEN,NONO,YWHAQ,H
4A1,PSAP,RPLP0,H3-3B,PTMA,DDX5,ENO2,KAT6A,KIF1A,DDX17,HTRA4,BSG,GNB1,RPS8,SEPTIN
0AB1,ZNF124,HSP90AA1,ACTR2,FTH1,KIDINS220,TMSB4X,NUMA1,RPL3,RPS6,ZNF316,DBN1,RP
4,HSP90AA1,ACTR2,VIM,FTH1,KIDINS220,NUMA1,RPL3,RPS6,ZNF316,DBN1,RPL4,ENO1,TUBA1
NA,MAP1B,SLC9A7,ALDOA,PKM,COL6A1,PRPF8,RPL8,YBX1,UBA1,FSTL1,TUBB2B,H3-3B,PTMA,
28,CKB,COL4A2,YWHAQ,SET,CHSY1,MRC2,PKD1,SSBP2,VAPA,TPT1,ITGB1,AGRN,TIMP2,MARCH
HSP90AA1,VIM,SPARC,FTH1,UBC,KIDINS220,EIF4G2,ZNF316,DBN1,ENO1,TUBA1B,PIK3R2,FLNA,
),XRCC2,HTT,HSP90AB1,ZNF124,HSP90AA1,ACTR2,VIM,SPARC,FTH1,KIDINS220,TMSB4X,EIF4G
MTRNR2L8,PKM,GANAB,COL6A1,FTL,FBXL4,YBX1,IGFBP2,CALR,FSTL1,TUBB2B,COL4A1,PTMA,
3,TRIM28,YWHAG,LRRC28,HACD3,DCHS1,CHSY1,PKD1,HLA-C,SSBP2,STMN1,TUBB4B,UCHL1,A
FADS1,PRPF8,IGFBP2,ABHD2,TUBB2B,COL5A1,CANX,DDX5,DVL3,KIF1A,SLC2A3,HTRA4,BSG,SP
TMSB10,PKD1,STMN1,UCHL1,AGRN,SARAF,MIDN,AP3D1,CDC42BPB,GOLGA8A,HNRNPU,CTNN
1,BSG,FASN,ALDH6A1,GNB1,STMN2,IDS,FSCN1,SPTAN1,SMG1,PLXNB2,SMIM14,IGF2,VCP,UBB,
316,DBN1,ENO1,TUBA1B,PIK3R2,FLNA,MAP1B,NCL,ALDOA,COL6A1,FTL,NEFM,RPL8,FBXL4,YB
HSP90B1,NEFL,PRPF8,CALR,FSTL1,TMF1,ABHD2,TUBB2B,CANX,PTMA,DDX5,SOX4,RPL13A,MAG
CL,RPL7A,GANAB,HSP90B1,RPL8,FBXL4,YBX1,FSTL1,ABHD2,TUBB2B,COL5A1,COL4A1,RPLP0,
IF124,HSP90AA1,ACTR2,UBC,KIDINS220,NUMA1,EIF4G2,RPL3,RPS6,ZNF316,DBN1,RPL4,TUBA1
MYH9,P4HB,GDI1,FSCN1,CALM1,PLD3,IGF2,EIF4B,VCP,MARCKSL1,UBB,MATR3,HACD3,EGLN1,
1,ACTR2,VIM,FTH1,UBC,KIDINS220,NUMA1,RPL3,RPS6,ZNF316,DBN1,RPL4,ENO1,TUBA1B,PIK3
NA,MAP1B,NCL,RPL7A,PKM,GANAB,COL6A1,FTL,HSP90B1,RPL8,YBX1,ABHD2,HSPA1B,TUBB2B
,TUBA1B,FLNA,MAP1B,NCL,GANAB,PRPF8,RPL8,UBA1,CALR,RPLP0,DDX5,SOX4,RPL13A,SEC62,
ACTR2,FTH1,KIDINS220,NUMA1,RPL3,RPS6,ZNF316,DBN1,RPL4,TUBA1B,PIK3R2,MAP1B,NCL,S
FM,YBX1,IGFBP2,FSTL1,ABHD2,CES3,SEC62,YWHAE,KIF1A,GNAS,SLC2A3,SPTBN1,CALM3,GNB
N1,GNB1,MYH9,NPM1,HSPD1,MATR3,SPEN,YWHAQ,EIF1,TMSB10,SSBP2,VAPA,ITGB1,AGRN,MA
3R2,NCL,GANAB,FTL,HSP90B1,RPL8,YBX1,UBA1,TMF1,PGK1,H3-3B,CANX,FKBP14,RPL13A,RPL
2,NCL,GANAB,HSP90B1,RPL8,YBX1,ATP2B4,TMF1,HSPA1B,PGK1,H3-3B,DDX5,FKBP14,RPL6,KA
NAB,FTL,HSP90B1,RPL8,FBXL4,TMF1,COL4A1,H3-3B,PTMA,DDX5,SEC62,DVL3,GNAS,DDX17,G
,TUBA1A,CFL1,BTBD9,XRCC2,MAVS,HTT,HSP90AA1,ACTR2,VIM,SPARC,TUBB,FTH1,TMSB4X,NU
IDINS220,NUMA1,RPL3,RPS6,ZNF316,DBN1,RPL4,ENO1,TUBA1B,PIK3R2,MAP1B,NCL,ALDOA,G
1,EIF4B,TRIM28,YWHAG,RPL37A,SPEN,EIF1,HACD3,SF3B1,SERINC1,VAPA,POLR2A,BTF3,RPLP1,
CTR2,VIM,FTH1,UBC,KIDINS220,TMSB4X,EIF4G2,ZNF316,DBN1,ENO1,TUBA1B,PIK3R2,MAP1B,
CHSY1,TMSB10,PKD1,POLR2A,STMN1,UCHL1,AGRN,SARAF,MIDN,AP3D1,CDC42BPB,GOLGA8A
C,NUMA1,RPL3,RPS6,ZNF316,ENO1,TUBA1B,PIK3R2,FLNA,MAP1B,NCL,RPL7A,SLC9A7,NEFL,N
,ZNF124,HSP90AA1,ACTR2,SPARC,FTH1,UBC,TMSB4X,EIF4G2,RPL3,RPS6,ZNF316,DBN1,RPL4,
20,NUMA1,RPL3,RPS6,ZNF316,DBN1,RPL4,ENO1,PIK3R2,MAP1B,NCL,SLC9A7,ALDOA,PKM,GAN
,HSP90AA1,VIM,TUBB,UBC,KIDINS220,NUMA1,EIF4G2,RPL4,ENO1,TUBA1B,FLNA,NCL,RPL7A,SL
AE,GNAS,FASN,SPTBN1,CALM3,SHISA9,SEPTIN2,GDI1,SMG1,PLXNB2,IGF2,MARCKSL1,AFF4,Y
3B,RPL13A,MAGED1,RPL6,DVL3,GNAS,RACK1,P4HB,PLD3,USP11,SNRNP200,EIF4B,EIF1,PPP2R
G,FASN,ALDH6A1,GNB1,STMN2,FSCN1,SMG1,NPM1,PLXNB2,USP11,IGF2,AFF4,TRIM28,LRRC2
F124,HSP90AA1,ACTR2,FTH1,UBC,KIDINS220,TMSB4X,NUMA1,DCX,EIF4G2,RPS6,ZNF316,DBN1
LU,SEC62,KAT6A,BSG,ATP5F1B,GNB1,MYH9,RPL7,APP,RPL5,FSCN1,PTMS,ATP5F1A,VCP,MARC
T,HSP90AB1,VIM,KIDINS220,DCX,EIF4G2,RPS6,ZNF316,DBN1,ENO1,TUBA1B,MAP1B,RPL7A,ALD
M,FTH1,UBC,NUMA1,RPL3,RPS6,ZNF316,DBN1,PIK3R2,FLNA,RPL7A,SLC9A7,MTRNR2L8,GANAB
A,CDC42BPB,ZNF770,TMEM170A,GTF2F1,OGA,PGAM1,SMARCA5,FADS2,EIF4H,RPS9,XRCC5,CI

1,NCL,RPL7A,ALDOA,PKM,GANAB,COL6A1,FTL,HSP90B1,NEFL,NEFM,RPL8,FBXL4,CALR,ABHD2
3,ATP2B4,FSTL1,TMF1,H3-3B,PTMA,RPL13A,MAGED1,RPL6,DVL3,GNAS,RACK1,FASN,CALM3,SH
SPTBN1,P4HB,FSCN1,PTMS,IGF2,TRIM28,DCHS1,SET,CHSY1,PKD1,TPT1,ITGB1,MORF4L1,AGR
MARCHF6,HNRNPA0,RPSA,TSPAN3,LAMA5,BRD3,NCS1,NLGN2,JUND,LENG8,ATP2A2,CRMP1,CC
H1,KIDINS220,NUMA1,RPL3,RPS6,ZNF316,DBN1,RPL4,ENO1,TUBA1B,PIK3R2,FLNA,MAP1B,NCL
L3,RPS6,ZNF316,DBN1,RPL4,PIK3R2,MAP1B,NCL,ALDOA,GANAB,HSP90B1,NEFL,NEFM,FBXL4,Y
P90AB1,ZNF124,ACTR2,VIM,TUBB,FTH1,UBC,KIDINS220,TMSB4X,EIF4G2,ZNF316,DBN1,ENO1,T
CFL1,BTBD9,XRCC2,MAVS,HSP90AB1,ZNF124,HSP90AA1,ACTR2,VIM,FTH1,UBC,KIDINS220,TMSE
BTF3,RPS17,CDC42BPB,ITM2B,CTNNA1,CNN3,LAMA5,SLC22A17,NCS1,ATP1A1,JUND,EIF5,HMG
CFL1,BTBD9,XRCC2,MAVS,HSP90AB1,ZNF124,HSP90AA1,ACTR2,VIM,FTH1,UBC,KIDINS220,TMSE
SN,CALM3,APP,RPL5,IGF2,MARCKSL1,LRRC28,DDB1,YWHAQ,CHSY1,EIF4A1,RPL19,SF3B1,BASF
2,FTH1,KIDINS220,NUMA1,RPL3,RPS6,ZNF316,DBN1,RPL4,PIK3R2,MAP1B,NCL,SLC9A7,ALDOA
20,NUMA1,RPL3,RPS6,ZNF316,DBN1,RPL4,PIK3R2,MAP1B,NCL,ALDOA,GANAB,HSP90B1,NEFL,
DPYSL3,TUBA1A,CFL1,BTBD9,XRCC2,MAVS,HTT,HSP90AB1,ZNF124,HSP90AA1,ACTR2,VIM,TUBB,
RPL7A,PKM,NEFM,RPL8,FBXL4,YBX1,CALR,FSTL1,TMF1,CES3,PGK1,H3-3B,PTMA,DDX5,RPL13
D1,TUBA1B,NCL,RPL7A,HSP90B1,RPL8,FBXL4,YBX1,ATP2B4,IGFBP2,TMF1,TUBB2B,COL5A1,H3
PC,CBX5,ATF4,EIF5,OAZ1,NISCH,HDLBP,GLG1,SFPQ,PIP4K2B,ARCN1,RAC1,ATP6V0C,CPSF2,SRF
TMSB10,MRC2,PPP2R1A,PKD1,PDIA3,VAPA,TPT1,RPS3A,ITGB1,TUBB4B,ZMIZ1,TIMP2,MIDN,CD
L3,CFL1,BTBD9,XRCC2,MAVS,HSP90AB1,HSP90AA1,ACTR2,SPARC,FTH1,UBC,KIDINS220,TMSB
BA1B,FLNA,NCL,RPL7A,HSP90B1,NEFL,RPL8,FBXL4,YBX1,ATP2B4,IGFBP2,TMF1,TUBB2B,COL4
CTG1,EEF2,TUBA1A,BTBD9,XRCC2,MAVS,HTT,HSP90AB1,HSP90AA1,ACTR2,SPARC,TUBB,FTH1,U
SL3,CFL1,BTBD9,XRCC2,MAVS,HSP90AB1,ZNF124,HSP90AA1,ACTR2,VIM,FTH1,UBC,KIDINS220,
JBB,FTH1,UBC,TMSB4X,NUMA1,EIF4G2,RPL3,RPS6,RPL4,ENO1,TUBA1B,PIK3R2,MAP1B,RPL7A,
HSP90AB1,ZNF124,HSP90AA1,ACTR2,VIM,TUBB,FTH1,UBC,EIF4G2,RPL3,ZNF316,DBN1,RPL4,E
1,RPL4,PIK3R2,MAP1B,NCL,SLC9A7,GANAB,FTL,HSP90B1,RPL8,ATP2B4,CALR,FSTL1,ABHD2,PG
,RPS6,ZNF316,DBN1,RPL4,TUBA1B,PIK3R2,MAP1B,NCL,ALDOA,GANAB,HSP90B1,NEFL,RPL8,FE
LDOA,PKM,COL6A1,FTL,HSP90B1,FADS1,NEFM,YBX1,IGFBP2,CALR,COL5A1,COL4A1,PSAP,CLU,
B,TRIM28,YWHAG,RPL37A,SPEN,EIF1,HACD3,SF3B1,SERINC1,VAPA,POLR2A,MORF4L1,BTF3,RF
SNAS,SPTBN1,SHISA9,STMN2,PTMS,MARCKS,CALM1,PLXNB2,MARCKSL1,UBB,CKB,NONO,YWH
FADS1,PRPF8,FBXL4,YBX1,IGFBP2,ZNF99,COL4A1,PSAP,RPLP0,CLU,PTMA,DDX5,RPL6,KAT6A,Y
CTG1,EEF2,DPYSL3,TUBA1A,CFL1,BTBD9,XRCC2,MAVS,HTT,HSP90AB1,ZNF124,HSP90AA1,ACTF
SDE1,AFF4,MATR3,LRRC28,PSMD5,EIF4A1,GTF3C4,SSBP2,BASP1,SEC61A1,RPS3A,RPL18,UCHL
1,ACTR2,FTH1,KIDINS220,TMSB4X,NUMA1,RPL3,RPS6,ZNF316,DBN1,RPL4,PIK3R2,MAP1B,NCL,
S220,TMSB4X,EIF4G2,RPL3,ZNF316,DBN1,ENO1,TUBA1B,FLNA,MAP1B,RPL7A,SLC9A7,FTL,HSF
CC2,MAVS,HSP90AB1,ZNF124,HSP90AA1,ACTR2,TUBB,FTH1,UBC,KIDINS220,TMSB4X,NUMA1,D
B1,UBC,KIDINS220,TMSB4X,NUMA1,RPL3,RPS6,RPL4,ENO1,FLNA,NCL,SLC9A7,ALDOA,MTRNR2
N3,C1ORF43,SLC7A5,EIF5,CD81,ATP2A2,CCT7,OAZ1,PFN1,ACTN4,B4GALNT4,DNAJC5,MXD4,M
D9,XRCC2,MAVS,HTT,HSP90AB1,ZNF124,HSP90AA1,ACTR2,TUBB,FTH1,UBC,KIDINS220,TMSB4
NS220,NUMA1,RPL3,RPS6,ZNF316,DBN1,RPL4,TUBA1B,PIK3R2,MAP1B,NCL,ALDOA,GANAB,HSP
,NCL,GANAB,HSP90B1,RPL8,H3-3B,FKBP14,RPL13A,RPL6,DVL3,RACK1,ATP5F1B,GNB1,SHISA9
D3,DCHS1,MRC2,LDHB,PKD1,VAPA,TPT1,ITGB1,TUBB4B,AGRN,SARAF,COL11A1,VCL,CDC42BPB
0B1,RPL8,YBX1,ATP2B4,TMF1,HSPA1B,H3-3B,DDX5,FKBP14,RPL6,KAT6A,RACK1,SHISA9,RPL7,
24,HSP90AA1,ACTR2,VIM,SPARC,TUBB,FTH1,UBC,TMSB4X,NUMA1,EIF4G2,RPL3,RPS6,ZNF316,
90AA1,ACTR2,FTH1,UBC,KIDINS220,TMSB4X,NUMA1,DCX,EIF4G2,RPL3,RPS6,ZNF316,DBN1,RPL
PS6,ZNF316,DBN1,RPL4,TUBA1B,PIK3R2,MAP1B,NCL,ALDOA,GANAB,HSP90B1,NEFL,FBXL4,YB

NF316,DBN1,ENO1,FLNA,NCL,RPL7A,ALDOA,PKM,COL6A1,HSP90B1,FADS1,NEFM,PRPF8,RPL8,
4HB,GSTP1,CALM1,NPM1,MATR3,SPEN,YWHAQ,EIF1,PDIA3,SSBP2,VAPA,ITGB1,AGRN,COL11A1,
3P2,BASP1,HMGB1,AGRN,MIDN,MARCHF6,VCL,CYB5R3,CDC42BPB,CTNNA1,HK1,CBX5,ZNF865,
NRNPA0,CTSD,LPCAT1,BRD3,PJA2,LLGL1,CLSTN1,JUND,ATP2A2,OAZ1,JPT2,TADA2B,CDH2,ACT
AB1,ZNF124,ACTR2,TUBB,FTH1,UBC,KIDINS220,TMSB4X,NUMA1,DCX,EIF4G2,ZNF316,DBN1,R
RF4L1,VCL,FADS2,TPM4,PJA2,KPNB1,CXADR,CHD4,HMGN2,ATP2A2,CRMP1,TLE5,HNRNPC,CCNI
NF124,HSP90AA1,ACTR2,FTH1,UBC,KIDINS220,TMSB4X,NUMA1,EIF4G2,RPL3,RPS6,ZNF316,DB
MTRNR2L8,PKM,FTL,HSP90B1,FADS1,YBX1,FSTL1,HSPA1B,PGK1,H3-3B,PTMA,DDX5,ENO2,RPL
4,IGFBP2,FSTL1,TUBB2B,COL5A1,PTMA,SEC62,KAT6A,GNAS,BSG,FASN,ALDH6A1,CALM3,GNB
1,ATP5F1A,EIF4B,VCP,MARCKSL1,MATR3,CKB,COL4A2,SPEN,DCHS1,SET,TMSB10,INA,MRC2,RF
1,TPT1,ITGB1,AGRN,SARAF,COL11A1,CDC42BPB,ITM2B,PXDN,HNRNPU,TPM4,SLC22A17,KPNE
,UBC,NUMA1,EIF4G2,ZNF316,DBN1,ENO1,TUBA1B,NCL,RPL7A,SLC9A7,PKM,GANAB,NEFL,NEFN
G,CKB,SPEN,YWHAQ,DCHS1,EGLN1,TPI1,EIF4A1,RPL19,PDIA3,VAPA,HMGB1,CDKL1,RPS3A,NCA
SP22,MED18,ACTG1,EEF2,DPYSL3,TUBA1A,CFL1,BTBD9,XRCC2,MAVS,HTT,HSP90AB1,ZNF124,
2,FTH1,KIDINS220,NUMA1,RPL3,RPS6,ZNF316,DBN1,RPL4,TUBA1B,PIK3R2,MAP1B,NCL,SLC9A
F4G2,RPL3,DBN1,ENO1,TUBA1B,PIK3R2,MAP1B,SLC9A7,ALDOA,PKM,GANAB,HSP90B1,FADS1,N
ADS1,UBA1,IGFBP2,CALR,FSTL1,COL5A1,COL4A1,PSAP,RPLP0,CANX,PTMA,RPL13A,MAGED1,K
5A1,H3-3B,ENO2,MAGED1,GNAS,BSG,SHISA9,STMN2,APP,IDS,PTMS,GSTP1,SMG1,CALM1,PLXN
3,SNRNP200,EIF4B,EIF1,PPP2R1A,HMGB1,POLR2A,TPT1,RPS3A,RPLP1,MIDN,MARCHF6,RPS17,
ISPA1B,PGK1,H3-3B,DDX5,FKBP14,RPL13A,RPL6,KAT6A,RPL13,GPI,RACK1,SHISA9,RPS8,RPL7,
NF124,HSP90AA1,ACTR2,FTH1,UBC,KIDINS220,TMSB4X,NUMA1,EIF4G2,RPL3,RPS6,ZNF316,DB
16,DBN1,PIK3R2,MAP1B,NCL,ALDOA,GANAB,HSP90B1,NEFL,RPL8,FBXL4,UBA1,IGFBP2,H3-3B,
ENO1,PIK3R2,FLNA,NCL,RPL7A,SLC9A7,ALDOA,PKM,GANAB,FTL,HSP90B1,PRPF8,RPL8,FBXL4,
ENO1,PIK3R2,FLNA,NCL,RPL7A,SLC9A7,ALDOA,PKM,GANAB,FTL,HSP90B1,PRPF8,RPL8,FBXL4,
X,EIF4G2,DBN1,TUBA1B,FLNA,MAP1B,RPL7A,SLC9A7,ALDOA,GANAB,FTL,HSP90B1,NEFM,UGT8
,ZNF316,DBN1,PIK3R2,MAP1B,NCL,ALDOA,HSP90B1,NEFL,NEFM,UBA1,HSPA1B,PGK1,H3-3B,P
1,NEFL,NEFM,TUBB2B,COL4A1,CES3,PSAP,RPLP0,PTMA,SOX4,SEC62,KAT6A,GNAS,SPTBN1,MY
,NUMA1,RPL3,RPS6,ZNF316,DBN1,RPL4,TUBA1B,PIK3R2,MAP1B,NCL,ALDOA,GANAB,HSP90B1
ALM1,SMIM14,IGF2,TRIM28,YWHAG,TMSB10,INA,MRC2,PPP2R1A,RPL15,PKD1,PDIA3,SSBP2,V
,IGFBP2,FSTL1,TUBB2B,PTMA,DDX5,ENO2,KAT6A,KIF1A,GNAS,FASN,ALDH6A1,SPTBN1,RPS8,P
X5,MAGED1,RPL6,SLC2A3,DDX17,RPS8,P4HB,APP,NPM1,MARCKSL1,UBB,AFF4,TRIM28,COL4A
MARCKSL1,LRRC28,YWHAQ,CHSY1,SF3B1,BASP1,TPT1,SEC61A1,RPS3A,STMN1,MORF4L1,AGF
NCL,SLC9A7,ALDOA,PKM,COL6A1,HSP90B1,NEFL,NEFM,FBXL4,CALR,FSTL1,COL4A1,PSAP,PGK
PL3,RPS6,ZNF316,DBN1,PIK3R2,MAP1B,NCL,ALDOA,GANAB,HSP90B1,NEFL,RPL8,FBXL4,YBX1,
L5A1,CES3,RPLP0,PTMA,KAT6A,DVL3,KIF1A,BSG,APP,ERVV-1,RPL5,GSTP1,MARCKS,SMG1,PLXN
HB,APP,IGF2,MARCKSL1,UBB,CKB,COL4A2,HACD3,SET,CHSY1,RPL15,POLR2A,TPT1,STMN1,UC
1,RPL6,DVL3,GNAS,FASN,P4HB,USP11,SNRNP200,INA,PPP2R1A,DNAH8,HMGB1,SEC61A1,RPLP
S220,NUMA1,RPL3,RPS6,ZNF316,DBN1,RPL4,ENO1,PIK3R2,MAP1B,NCL,SLC9A7,ALDOA,GANAB
4,RPL6,RACK1,SHISA9,RPL7,P4HB,TMBIM6,SPTAN1,PLD3,USP11,SNRNP200,SMIM14,ATP5F1A,
HSP90B1,RPL8,YBX1,TMF1,HSPA1B,H3-3B,CANX,DDX5,RPL6,KAT6A,GPI,RACK1,SHISA9,RPS8,R
K1,H3-3B,DDX5,FKBP14,RPL13A,MAGED1,RPL6,DVL3,RACK1,GNB1,SHISA9,RPS8,RPL7,P4HB,R
1A,KAT6A,GNAS,FASN,SPTBN1,CALM3,GNB1,MYH9,SEPTIN2,GDI1,SMG1,CALM1,PLD3,SMIM14,
1A,KAT6A,GNAS,FASN,SPTBN1,CALM3,GNB1,MYH9,SEPTIN2,GDI1,SMG1,CALM1,PLD3,SMIM14,
2,FTH1,KIDINS220,NUMA1,RPL3,RPS6,ZNF316,DBN1,RPL4,PIK3R2,MAP1B,NCL,ALDOA,GANAB,
TR2,FTH1,UBC,KIDINS220,TMSB4X,NUMA1,DCX,EIF4G2,ZNF316,DBN1,ENO1,TUBA1B,PIK3R2,

OL4A2,SET,VAPA,SEC61A1,MORF4L1,AGRN,MIDN,RPL10,ARF1,CYB5R3,MAP4,CDC42BPB,ITM2E
,TUBB2B,COL5A1,ENO2,SEC62,KAT6A,YWHAE,KIF1A,GNAS,SLC2A3,DDX17,ALDH6A1,SPTBN1,C
24,ACTR2,FTH1,UBC,NUMA1,EIF4G2,ZNF316,DBN1,ENO1,TUBA1B,NCL,RPL7A,SLC9A7,PKM,G
RF4L1,RPS17,AP3D1,PGAM1,EIF4H,KPNB1,PTCD3,KIF5B,TMEM259,EIF5,BEX3,RBBP4,PFN1,PN
RPL13A,MAGED1,RPL6,DVL3,GPI,RACK1,ATP5F1B,GNB1,SHISA9,RPS8,P4HB,TMBIM6,USP11,SN
B1,FADS1,NEFL,YBX1,UBA1,TMF1,TUBB2B,COL5A1,PGK1,RPLP0,PTMA,DDX5,RPL13A,KAT6A,K
A,GANAB,HSP90B1,ATP2B4,CALR,FSTL1,TUBB2B,SEC62,YWHAE,GNAS,SLC2A3,FASN,SPTBN1,G
220,NUMA1,RPS6,ZNF316,DBN1,RPL4,PIK3R2,MAP1B,NCL,ALDOA,HSP90B1,NEFL,RPL8,FBXL4,
,ALDOA,GANAB,HSP90B1,NEFL,IGFBP2,H3-3B,PTMA,DDX5,FKBP14,RPL13A,MAGED1,RPL6,DVL
S6,ZNF316,DBN1,PIK3R2,MAP1B,NCL,ALDOA,GANAB,HSP90B1,NEFL,RPL8,FBXL4,YBX1,UBA1,A
28,EIF1,HACD3,SERINC1,CHD3,MORF4L1,BTF3,DNAJA1,CDC42BPB,GTF2F1,PGAM1,SMARCA5,
1,FTL,HSP90B1,FADS1,NEFM,FBXL4,YBX1,CALR,COL5A1,PSAP,PTMA,SEC62,KAT6A,YWHAE,KIF1
,RPL6,DVL3,RACK1,ATP5F1B,SHISA9,P4HB,TMBIM6,PLD3,USP11,SNRNP200,SMIM14,EIF4B,EIF
XL4,YBX1,UBA1,CALR,FSTL1,TUBB2B,COL4A1,PSAP,DDX5,YWHAE,FASN,SPTBN1,GNB1,RPS8,M
C,KIDINS220,EIF4G2,RPL3,RPS6,ZNF316,DBN1,RPL4,ENO1,FLNA,MAP1B,NCL,ALDOA,GANAB,C
GED1,RPL6,DVL3,GNAS,RACK1,P4HB,TMBIM6,PLD3,USP11,SNRNP200,EIF4B,EIF1,EIF4A1,PPP2
BN1,ENO1,NCL,ALDOA,PKM,FTL,HSP90B1,FADS1,PRPF8,UGT8,RPL8,FBXL4,IGFBP2,ZNF99,CO
90AB1,ZNF124,HSP90AA1,ACTR2,VIM,SPARC,TUBB,FTH1,KIDINS220,TMSB4X,NUMA1,EIF4G2,R
0,EIF4G2,RPL3,RPS6,ZNF316,DBN1,RPL4,ENO1,MAP1B,NCL,ALDOA,GANAB,COL6A1,FTL,HSP90
,RPL7A,ALDOA,PKM,GANAB,COL6A1,FTL,HSP90B1,FADS1,NEFM,FBXL4,YBX1,IGFBP2,CALR,TUE
C,KIDINS220,NUMA1,RPL3,RPS6,ZNF316,DBN1,RPL4,TUBA1B,PIK3R2,FLNA,MAP1B,NCL,ALDOA
DBN1,PIK3R2,MAP1B,NCL,ALDOA,HSP90B1,NEFL,PRPF8,YBX1,UBA1,ATP2B4,IGFBP2,CALR,TM
KBP14,RPL6,RACK1,SHISA9,P4HB,RPL5,TMBIM6,PLD3,USP11,SNRNP200,SMIM14,ATP5F1A,EIF
TP2B4,H3-3B,FKBP14,RPL13A,MAGED1,RPL6,DVL3,RACK1,GNB1,SHISA9,P4HB,RPL5,TMBIM6,
RPL8,YBX1,TUBB2B,COL4A1,PSAP,RPLP0,CLU,PTMA,SOX4,MAGED1,KAT6A,DVL3,YWHAE,KIF1A
,EIF4G2,ZNF316,ENO1,FLNA,MAP1B,NCL,MTRNR2L8,PKM,GANAB,COL6A1,FADS1,NEFL,RPL8,
A9,P4HB,RPL5,TMBIM6,PLD3,USP11,SNRNP200,SMIM14,ATP5F1A,EIF4B,SPEN,PPP2R1A,RPL15,
B1,ZNF124,VIM,FTH1,UBC,KIDINS220,TMSB4X,NUMA1,EIF4G2,RPL3,RPS6,ZNF316,DBN1,TUBA
2,RPL3,RPS6,RPL4,MAP1B,NCL,SLC9A7,GANAB,RPL8,YBX1,UBA1,IGFBP2,CALR,ZNF99,RPLP0,
IP1,LASP1,OAZ1,CCNI,SOX12,MIF,B4GALNT4,APLP2,TARDBP,CADM4,PLPPR3,STIP1,ATP6V0C,S
6A,DVL3,KIF1A,RPL13,GNB1,SHISA9,RPS8,SEPTIN2,IDS,RPL5,FSCN1,TMBIM6,GSTP1,PLXNB2,
AP1B,NCL,ALDOA,PKM,GANAB,FTL,HSP90B1,NEFL,YBX1,UBA1,ATP2B4,TMF1,HSPA1B,CANX,D
,RPL6,DVL3,RACK1,SHISA9,RPS8,P4HB,TMBIM6,USP11,SMIM14,RPL37A,RPL19,PPP2R1A,SF3B
,FN1,TMSB10,PKD1,HLA-C,VAPA,HMGB1,TUBB4B,ANXA2,AGRN,MIDN,AP3D1,ARF1,CYB5R3,MA
,CFL1,MAVS,HTT,HSP90AB1,HSP90AA1,ACTR2,VIM,SPARC,FTH1,UBC,KIDINS220,EIF4G2,RPL3,
ZNF316,DBN1,PIK3R2,FLNA,NCL,MTRNR2L8,GANAB,FADS1,NEFM,YBX1,UBA1,IGFBP2,CALR,AB
UMA1,RPL3,RPS6,ZNF316,DBN1,PIK3R2,MAP1B,NCL,ALDOA,GANAB,HSP90B1,NEFL,NEFM,YBX
,SMG1,NPM1,USP11,UBB,CKB,NONO,HACD3,DCHS1,TMSB10,EIF4A1,MRC2,RPL15,LDHB,SF3B
,P4HB,TMBIM6,PLD3,SNRNP200,SMIM14,EIF4B,PPP2R1A,RPL15,GTF3C4,SF3B1,ZNF43,HMGB
ACD3,SERINC1,POLR2A,BTF3,RPLP1,DNAJA1,CDC42BPB,TMEM170A,GTF2F1,OGA,PGAM1,SMA
MARCKSL1,TRIM28,CKB,SET,PKD1,VAPA,TUBB4B,UCHL1,MORF4L1,AGRN,MIDN,WSB1,SBF1,C
,SF3B1,AP2M1,TPT1,RPS3A,RPL18,TUBB4B,MORF4L1,XAF1,RPS17,AP3D1,CYB5R3,SF3B2,RPS
I3,SEPTIN2,P4HB,ERVV-1,GDI1,PTMS,SPTAN1,MARCKS,IGF2,UBB,MATR3,FN1,SPEN,NONO,PSM
,COL6A2,MXD4,RAC1,SLC35E2B,IPO5,CDKN1A,FAM199X,GDI2,PTPRA,PTOV1,ACAP3,SURF4,RA
,CANX,CLU,PTMA,FKBP14,KAT6A,YWHAE,RPL13,GPI,HTRA4,FASN,SPTBN1,GNB1,SHISA9,MYH9,

L4A1,CES3,CANX,PTMA,ENO2,RPL13A,DVL3,RPL13,GPI,HTRA4,RACK1,SPTBN1,RPL7,SEPTIN2,S
,FADS1,NEFL,PRPF8,IGFBP2,CALR,FSTL1,TUBB2B,COL4A1,PSAP,H3-3B,DDX5,ENO2,RPL13A,SE
BD9,XRCC2,HTT,HSP90AB1,ACTR2,VIM,FTH1,UBC,TMSB4X,DCX,EIF4G2,RPS6,ZNF316,DBN1,RI
ARC,FTH1,UBC,TMSB4X,NUMA1,ZNF316,DBN1,ENO1,MAP1B,ALDOA,PKM,GANAB,HSP90B1,FAD
PL3,RPS6,ZNF316,DBN1,RPL4,PIK3R2,MAP1B,NCL,ALDOA,GANAB,HSP90B1,NEFL,RPL8,FBXL4,Y
,DDX5,SOX4,KAT6A,RPL13,GPI,FASN,CALM3,GNB1,SHISA9,MYH9,APP,GDI1,NPM1,COL18A1,CSI
RPL8,FBXL4,TMF1,H3-3B,PTMA,SOX4,RPL13A,RPL6,DVL3,RACK1,FASN,ATP5F1B,RPS8,P4HB,SP
,HSPA1B,H3-3B,DDX5,RPL13A,RPL6,KAT6A,RPL13,GPI,RACK1,SHISA9,MYH9,RPL7,P4HB,SPTAN
A1,ACTR2,TUBB,FTH1,KIDINS220,TMSB4X,NUMA1,RPS6,ZNF316,DBN1,RPL4,TUBA1B,PIK3R2,N
C2,SSBP2,VAPA,POLR2A,TPT1,TUBB4B,MORF4L1,ANXA2,AGRN,TIMP2,MIDN,MARCHF6,DNAJA1,
,FKBP14,RPL6,RACK1,SHISA9,RPL7,P4HB,TMBIM6,SPTAN1,PLD3,USP11,SNRNP200,SMIM14,EIF
28,DDB1,EIF1,TMSB10,EIF4A1,LDHB,BASP1,POLR2A,TPT1,SEC61A1,RPS3A,STMN1,ITGB1,MORI
GANAB,COL6A1,FBXL4,UBA1,ATP2B4,TMF1,CANX,PTMA,DDX5,ENO2,SOX4,SEC62,KAT6A,DVL3,
HSP90B1,NEFL,NEFM,PRPF8,RPL8,FBXL4,IGFBP2,CALR,TUBB2B,COL4A1,PGK1,RPLP0,H3-3B,C
EIF4G2,RPS6,ZNF316,PIK3R2,MAP1B,NCL,GANAB,COL6A1,FADS1,PRPF8,RPL8,IGFBP2,CALR,HS
BXL4,UBA1,IGFBP2,HSPA1B,COL5A1,PGK1,DDX5,FKBP14,SEC62,KAT6A,YWHAE,RPL13,FASN,AL
2,PKD1,VAPA,TPT1,ITGB1,ZMIZ1,TIMP2,SARAF,COL11A1,MIDN,CDC42BPB,ITM2B,HNRNPU,TPM
MRC2,GTF3C4,PKD1,PDIA3,VAPA,ITGB1,TUBB4B,ZMIZ1,MIDN,CDC42BPB,TPM4,SCARB2,RHOA
,BTBD9,XRCC2,MAVS,HTT,HSP90AB1,ZNF124,ACTR2,VIM,SPARC,FTH1,UBC,TMSB4X,NUMA1,EII
G2,RPS6,ZNF316,RPL4,PIK3R2,NCL,SLC9A7,GANAB,HSP90B1,RPL8,FBXL4,YBX1,IGFBP2,TMF1,
RPL7A,SLC9A7,ALDOA,PKM,GANAB,FTL,FADS1,ATP2B4,FSTL1,HSPA1B,PSAP,RPLP0,CANX,PTMA
A1,ACTR2,VIM,UBC,KIDINS220,ZNF316,DBN1,ENO1,MAP1B,RPL7A,PKM,HSP90B1,RPL8,ATP2B
B5R3,MAP4,CDC42BPB,CTSD,BRD3,LLGL1,ZNF865,JUND,CD81,HMGN2,ATP2A2,OAZ1,PFKL,TK
FBP2,CALR,HSPA1B,H3-3B,PTMA,DDX5,FKBP14,RPL13A,MAGED1,RPL6,KAT6A,YWHAE,GPI,RAC
32B,COL4A1,CES3,RPLP0,PTMA,DDX5,SOX4,RPL6,RPL13,GNAS,BSG,FASN,MYH9,RPL7,ERVV-1,F
DBN1,RPL4,ENO1,FLNA,RPL7A,SLC9A7,HSP90B1,FADS1,NEFL,RPL8,FBXL4,YBX1,TMF1,TUBB2I
N1,TMSB10,HLA-C,UCHL1,ANXA2,AGRN,SARAF,MIDN,AP3D1,CYB5R3,MAP4,CDC42BPB,GOLGA8
7,RACK1,BSG,ATP5F1B,SEPTIN2,GDI1,MARCKS,USP11,COL18A1,CSDE1,EIF3C,ATP5F1A,EIF4B,
FTL,NEFM,YBX1,UBA1,ATP2B4,TMF1,ABHD2,H3-3B,SOX4,SEC62,KAT6A,YWHAE,RPL13,DDX17,C
2,DPYSL3,CFL1,BTBD9,XRCC2,MAVS,HTT,HSP90AB1,SPARC,TUBB,FTH1,UBC,NUMA1,EIF4G2,RI
3,COL4A1,ENO2,DVL3,YWHAE,GNAS,DDX17,GPI,ALDH6A1,SHISA9,RPS8,SEPTIN2,APP,GDI1,PTMS
PL3,ZNF316,RPL4,ENO1,FLNA,SLC9A7,ALDOA,PKM,GANAB,HSP90B1,NEFL,YBX1,CALR,TMF1,TU
PL3,RPS6,ZNF316,DBN1,PIK3R2,MAP1B,NCL,ALDOA,GANAB,HSP90B1,NEFL,RPL8,FBXL4,YBX1,I
K1,SHISA9,P4HB,TMBIM6,PLD3,SNRNP200,EIF4B,EIF1,EIF4A1,PPP2R1A,DNAH8,SF3B1,ZNF43,H
A,RPL6,GPI,RACK1,SHISA9,P4HB,TMBIM6,PLD3,USP11,SNRNP200,SMIM14,ATP5F1A,EIF4B,UBB,
,KIF1A,SLC2A3,STMN2,RPL7,IDS,SPTAN1,MARCKS,SMG1,NPM1,PLD3,HSPD1,IGF2,EIF4B,CKB,C
ANAB,COL6A1,UBA1,FSTL1,ZNF99,RPLP0,H3-3B,CANX,PTMA,DDX5,RPL13A,SEC62,KAT6A,YWHA
9,CIPC,RHOA,CBX5,LENG8,VAT1,RTN3,PFN1,CCNI,TADA2B,KHDRBS1,ACTN4,RPS5,DNAJC5,HNF
2,COL4A1,RPLP0,CANX,CLU,PTMA,DDX5,ENO2,MAGED1,KAT6A,HTRA4,FASN,SPTBN1,SHISA9,RI
BB,UBC,NUMA1,RPL3,DBN1,RPL4,ENO1,TUBA1B,PIK3R2,FLNA,MAP1B,NCL,SLC9A7,ALDOA,MT
NA,NCL,PKM,GANAB,FTL,NEFL,NEFM,YBX1,ATP2B4,ABHD2,ZNF99,COL4A1,RPLP0,PTMA,ENO2,I
ALDOA,GANAB,RPL8,YBX1,TMF1,HSPA1B,TUBB2B,CES3,H3-3B,CANX,CLU,DDX5,RPL13A,MAGE
,GPI,ATP5F1B,SHISA9,RPS8,P4HB,TMBIM6,USP11,SNRNP200,EIF3C,ATP5F1A,EIF4B,MATR3,RPL
IP2,SARAF,MIDN,CDC42BPB,ITM2B,HNRNPU,TPM4,SCARB2,RPS24,SLC22A17,CIPC,NCS1,RHOA
HSPA1B,DDX5,FKBP14,MAGED1,RPL6,GPI,RACK1,GNB1,SHISA9,P4HB,GDI1,TMBIM6,PLD3,HSPI

PLD3,IGF2,EIF3C,TRIM28,COL4A2,YWHAQ,HACD3,CHSY1,SSBP2,BASP1,VAPA,POLR2A,MORF4L1
PS6,ZNF316,DBN1,ENO1,PIK3R2,FLNA,MAP1B,NCL,ALDOA,PKM,GANAB,FTL,HSP90B1,NEFL,YBX
RPS19,HNRNPDL,WLS,HNRNPK,LAMP1,UBE2H,BCLAF1,HSBP1,C5ORF24,MDH2,SQSTM1,GDI2,S
RPL7A,PKM,HSP90B1,FADS1,YBX1,UBA1,ABHD2,TUBB2B,PGK1,CANX,DDX5,ENO2,KAT6A,RPL13
L6,GNAS,RACK1,GNB1,SHISA9,P4HB,PLD3,USP11,SNRNP200,EIF4B,EIF1,PPP2R1A,SF3B1,ZNF43
4HB,RPL5,TMBIM6,PLD3,USP11,SNRNP200,SMIM14,EIF4B,AFF4,EIF4A1,PPP2R1A,SF3B1,ZNF43,
ACTR2,SPARC,KIDINS220,DCX,EIF4G2,RPS6,ZNF316,RPL4,PIK3R2,RPL7A,ALDOA,PKM,GANAB,H
2,SPEN,DDB1,TMSB10,SF3B1,RPS3A,STMN1,AGRN,TIMP2,VCL,CTNNA1,CNN3,TPM4,TNS3,CTSD,
A,RPL6,DVL3,RACK1,FASN,P4HB,TMBIM6,PLD3,SNRNP200,EIF4B,EIF4A1,PPP2R1A,LDHB,PKD1,
BXL4,IGFBP2,FSTL1,ZNF99,TUBB2B,COL5A1,PTMA,ENO2,RPL13A,KAT6A,KIF1A,GNAS,SLC2A3,D
UBA1A,CFL1,BTBD9,HSP90AA1,ACTR2,VIM,SPARC,TUBB,FTH1,UBC,KIDINS220,NUMA1,DCX,RPL
2BPB,NACA,LAMA5,LLGL1,RAB6B,ATP1A1,JUND,CRMP1,PFN1,TLE5,PNN,NES,KLHL9,XPO1,SPTE
EIF4A1,LDHB,PDIA3,HMGB1,NCAM1,ZMIZ1,CDC42BPB,NFE2L1,HNRNPA0,TSPAN3,BRD3,IGFBP3
D9,XRCC2,HSP90AB1,ZNF124,HSP90AA1,FTH1,UBC,KIDINS220,TMSB4X,NUMA1,EIF4G2,RPL3,R
1,RPL3,RPS6,ZNF316,DBN1,ENO1,PIK3R2,MAP1B,NCL,ALDOA,GANAB,HSP90B1,NEFL,RPL8,FBX
DDX5,KAT6A,YWHAE,KIF1A,BSG,FASN,CALM3,GDI1,PTMS,MARCKS,CALM1,NPM1,PLXNB2,CSDE
RACK1,BSG,SEPTIN2,GDI1,GSTP1,NPM1,USP11,ATP5F1A,MATR3,NONO,SET,CHSY1,RPS3,AP2M1
FBXL4,YBX1,UBA1,IGFBP2,TMF1,TUBB2B,COL5A1,COL4A1,RPLP0,DDX5,KAT6A,DVL3,YWHAE,RF
XL4,ATP2B4,IGFBP2,FSTL1,ZNF99,TUBB2B,COL5A1,COL4A1,RPLP0,CLU,PTMA,YWHAE,DDX17,A
R2,NCL,ALDOA,GANAB,COL6A1,RPL8,YBX1,TMF1,HSPA1B,TUBB2B,CES3,H3-3B,CANX,CLU,DDX
NS220,NUMA1,RPS6,ZNF316,DBN1,RPL4,PIK3R2,FLNA,NCL,SLC9A7,ALDOA,GANAB,COL6A1,HSF
A,CFL1,BTBD9,MAVS,HSP90AB1,HSP90AA1,ACTR2,VIM,SPARC,TUBB,FTH1,UBC,KIDINS220,NUMA
DCX,EIF4G2,RPL3,RPS6,RPL4,MAP1B,NCL,RPL7A,GANAB,RPL8,UBA1,FSTL1,PGK1,RPLP0,H3-3E
F3B1,ZNF43,HMGB1,POLR2A,TPT1,SEC61A1,ITGB1,RPL11,MIDN,RPL10,SF3B2,GTF2F1,HNRNP
BN1,ENO1,TUBA1B,PIK3R2,MAP1B,ALDOA,COL6A1,FTL,HSP90B1,FADS1,NEFL,NEFM,UBA1,ATP2
,MTRNR2L2,MED18,ACTG1,EEF2,DPYSL3,CFL1,BTBD9,XRCC2,MAVS,HTT,HSP90AB1,HSP90AA1,
3,PTMA,DDX5,RPL13A,DVL3,DDX17,SPTBN1,ATP5F1B,RPS8,P4HB,SNRNP200,SMIM14,LRRC28,S
P1B,NCL,ALDOA,HSP90B1,NEFL,YBX1,H3-3B,CANX,CLU,PTMA,DDX5,FKBP14,RPL13A,MAGED1,F
CBX5,ZNF865,KIF5B,LENG8,VAT1,RTN3,PFN1,CCNI,KHDRBS1,EIF5A,ACTN4,RPS5,DNAJC5,HNRN
AS,DDX17,HTRA4,FASN,RPL7,IDS,RPL5,FSCN1,PTMS,SMG1,PLXNB2,HSPD1,IGF2,VCP,MARCKSL
43,POLR2A,TPT1,MORF4L1,AP3D1,SF3B2,RPS9,XRCC5,ZNF865,RPS14,RPS20,LENG8,WDR1,RP
RD3,PJA2,ZNF865,SLC7A5,CHD4,CD81,HMGN2,ATP2A2,LASP1,PFN1,PIP4K2B,MIF,TLN1,RPL36,C
,TMBIM6,PLD3,SNRNP200,SMIM14,EIF4B,EIF1,PPP2R1A,SF3B1,ZNF43,HMGB1,POLR2A,TPT1,TU
,HLA-C,SF3B1,CHD3,VAPA,SEC61A1,MORF4L1,BTF3,DNAJA1,MAT2A,FADS2,RPSA,TNS3,CTSD,EI
BA1,IGFBP2,TMF1,TUBB2B,COL5A1,COL4A1,RPLP0,DDX5,KAT6A,DVL3,YWHAE,RPL13,DDX17,G
5,RPS7,RPS19,HNRNPK,H1-10,LAMP1,MDH2,SQSTM1,GDI2,KLC1,ASXL1,ARF3,SRSF9,PRDX1,ATI
POLR2A,UCHL1,MORF4L1,BTF3,MIDN,DNAJA1,MAT2A,GTF2F1,PXDN,HNRNPA0,CTNNA1,FADS2,C
1B,RPL7A,SLC9A7,MTRNR2L8,PKM,GANAB,COL6A1,FTL,HSP90B1,FADS1,NEFL,NEFM,RPL8,ATP
GPI,P4HB,IGF2,EIF3C,YWHAG,LRRC28,EIF1,CHSY1,EIF4A1,GTF3C4,PDIA3,SERINC1,BASP1,HMGI
B1,TUBB4B,AGRN,ZMIZ1,XAF1,VCL,AP3D1,ARF1,CYB5R3,MAP4,ITM2B,HNRNPA0,RPS9,SLC16A
SP90B1,FADS1,NEFM,RPL8,TUBB2B,PSAP,RPLP0,H3-3B,CANX,DDX5,FKBP14,MAGED1,RPL6,KA
RPL4,PIK3R2,MAP1B,NCL,ALDOA,HSP90B1,NEFL,YBX1,UBA1,TUBB2B,H3-3B,CANX,DDX5,FKBP1
ORF4L1,BTF3,MIDN,DNAJA1,MAT2A,GTF2F1,PXDN,CTNNA1,FADS2,TPM4,EIF4H,BRD3,KPNB1,CE
1,ARF1,CLSTN1,ATF4,TMEM259,EIF5,RBBP4,NISCH,HDLBP,GLG1,SFPQ,PIP4K2B,NARS1,ARCN1
M6,PTMS,USP11,IGF2,MARCKSL1,MATR3,YWHAG,COL4A2,MRC2,PKD1,HLA-C,VAPA,TUBB4B,ZM

DVL3,GNAS,GPI,ATP5F1B,SHISA9,P4HB,PLD3,USP11,SNRNP200,IGF2,EIF3C,ATP5F1A,EIF4B,MA
A1,CES3,H3-3B,CLU,PTMA,RPL13A,KAT6A,KIF1A,RPL13,GNAS,GPI,HTRA4,BSG,SPTBN1,CALM3,
YH9,SEPTIN2,APP,GDI1,FSCN1,SPTAN1,HSPD1,COL4A2,SPEN,NONO,SET,TMSB10,EIF4A1,GTF3C
,CRMP1,EIF5A,CCND2,RPS5,HNRNPK,CAPRIN1,TOP2B,FAM199X,CTDSP2,NOVA2,HNRNPA3,GRI
,SMARCA5,SLC22A17,KPNB1,NCS1,RPS14,CXADR,CHD4,ATP1A1,JUND,HMGN2,ATP2A2,CCNI,DN
SN,SPTBN1,SHISA9,APP,FSCN1,SMG1,PLXNB2,SET,MRC2,GTF3C4,LDHB,PKD1,SSBP2,BASP1,P
MAVS,HTT,HSP90AB1,ZNF124,VIM,KIDINS220,TMSB4X,NUMA1,RPL3,RPL4,MAP1B,NCL,COL6A1
1,CNN3,RPSA,BRD3,EIF5,CD81,HMGN2,SRSF4,PFN1,JPT2,GPC3,PLXNA1,SOX12,PRDX2,DDX3X,
AP1B,NCL,RPL7A,GANAB,FBXL4,UBA1,FSTL1,ZNF99,PSAP,RPLP0,H3-3B,PTMA,DDX5,RPL13A,S
P11,CSDE1,EIF4B,MARCKSL1,TRIM28,YWHAG,SPEN,DCHS1,INA,MRC2,GTF3C4,DNAH8,PKD1,R
RPL5,TMBIM6,PLD3,SNRNP200,CSDE1,EIF4B,EIF1,EIF4A1,PPP2R1A,SF3B1,ZNF43,HMGB1,POL
,MATR3,MAP9,INA,EIF4A1,RPL19,POLR2A,SARAF,MARCHF6,ITM2B,SLC4A8,HNRNPA0,RPS16,RP
KAT6A,GNAS,ALDH6A1,CALM3,SHISA9,IDS,PTMS,SMG1,PLXNB2,YWHAQ,SET,TPI1,GTF3C4,POLR
HMGB1,TPT1,MIDN,SF3B2,GTF2F1,HNRNPA0,RPL29,RPS9,ZNF865,RPS14,RPS20,WDR1,PFN1,
GK1,CANX,PTMA,DDX5,KAT6A,KIF1A,RPL13,GPI,BSG,FASN,ALDH6A1,SEPTIN2,SMG1,NPM1,PLD
4G2,RPL3,DBN1,RPL4,ENO1,FLNA,RPL7A,ALDOA,COL6A1,HSP90B1,FADS1,NEFL,RPL8,FBXL4,Y
NF865,LENG8,SRSF4,RBBP4,PFN1,RPL31,PRKCSH,KHDRBS1,RAB7A,ITM2C,RPS5,RPS19,NGRN,
GANAB,UBA1,FSTL1,ZNF99,PGK1,RPLP0,H3-3B,CANX,CLU,PTMA,DDX5,RPL13A,SEC62,KAT6A,D
,TMF1,COL5A1,PSAP,RPLP0,PTMA,DDX5,RPL13A,YWHAE,RPL13,GPI,BSG,ALDH6A1,RPS8,MYH9,
LD3,SNRNP200,IGF2,EIF4B,VCP,MATR3,LRRC28,EIF1,RPL19,PDIA3,HMGB1,AP2M1,TPT1,RPS4X,
1,PIK3R2,FLNA,MAP1B,SLC9A7,MTRNR2L8,FTL,RPL8,YBX1,UBA1,FSTL1,ABHD2,TUBB2B,PSAP,
RPL3,RPS6,RPL4,ENO1,GANAB,COL6A1,UBA1,FSTL1,ZNF99,RPLP0,H3-3B,CANX,PTMA,DDX5,R
CTR2,VIM,TUBB,FTH1,KIDINS220,NUMA1,EIF4G2,DBN1,ENO1,RPL7A,SLC9A7,PKM,COL6A1,FTL
NB1,C1ORF43,PAPOLA,RPL14,RHOA,SLC7A5,ATP2A2,PFKL,PDIA6,MRFAP1,MIF,JUN,COL6A2,HN
NI,TADA2B,RPS25,SFPQ,CDH2,SOX12,H1-0,RPS5,RPS10,APLP2,RAB1A,SEPTIN3,CAPRIN1,CDK1
6,PIK3R2,MAP1B,NCL,GANAB,HSP90B1,NEFL,RPL8,FBXL4,ATP2B4,IGFBP2,FSTL1,H3-3B,CANX,
SSBP2,ITGB1,TUBB4B,AGRN,CDC42BPB,PXDN,CTSD,KPNB1,HK1,NCS1,NLGN2,JUND,VAT1,TLE
B1,NEFM,RPL8,FBXL4,YBX1,UBA1,TMF1,TUBB2B,PSAP,PGK1,PTMA,DDX5,DVL3,KIF1A,DDX17,G
AZ1,PFKL,TKT,PNMA1,SOX12,ARL4C,MIF,GPAT4,MYO5A,B4GALNT4,APLP2,TARDBP,CADM4,HN
SF4,PIP4K2B,HNRNPDL,HNRNPK,COTL1,CTBP1,SMCHD1,BCLAF1,U2AF2,PPP1CB,GAS7,PKN1,C
I3,PFN1,CCNI,KHDRBS1,EIF5A,ACTN4,HNRNPH1,ZNF454,MAGED4B,CTBP1,ARHGDIA,CDK16,BC
3,NONO,EIF4A1,POLR2A,AP2M1,MORF4L1,BTF3,XAF1,MAT2A,NACA,RPS24,RPL24,CBX5,CLSTN
SP90AA1,KIDINS220,EIF4G2,RPL3,RPS6,ZNF316,DBN1,RPL4,ENO1,NCL,ALDOA,PKM,GANAB,FTL
XRCC2,MAVS,HTT,HSP90AB1,ZNF124,HSP90AA1,ACTR2,VIM,TUBB,FTH1,UBC,KIDINS220,NUMA1
C61A1,RPLP1,MIDN,GTF2F1,CTNNA1,EIF4H,RPS9,SLC16A7,PAPOLA,RPL14,RHOA,ZNF865,NLGN
,HSPD1,USP11,EIF3C,ATP5F1A,UBB,RPL37A,RPL19,SERINC1,HMGB1,POLR2A,AP2M1,SEC61A1,
,RPLP0,H3-3B,PTMA,DDX5,RPL13A,SEC62,KAT6A,DVL3,YWHAE,KIF1A,DDX17,RACK1,BSG,ALDH
ID3,VAPA,MORF4L1,TIMP2,CYB5R3,CDC42BPB,AP2B1,NFE2L1,TNS3,LAMA5,CIPC,KPNB1,C1OR
1,MIDN,MAT2A,GTF2F1,PXDN,CTNNA1,FADS2,CNN3,EIF4H,BRD3,KPNB1,CBX5,CXADR,CD81,HM
A1A,CFL1,BTBD9,XRCC2,MAVS,HSP90AB1,HSP90AA1,ACTR2,VIM,SPARC,TUBB,FTH1,UBC,TMS
,DBN1,RPL4,ENO1,FLNA,RPL7A,ALDOA,MTRNR2L8,COL6A1,HSP90B1,FADS1,NEFM,RPL8,FBXL
BRD3,ZNF865,SLC7A5,CHD4,WDR1,HMGN2,ATP2A2,LASP1,OAZ1,SOX12,APLP1,MIF,TARDBP,C
SMIM14,ATP5F1A,EIF4B,AFF4,EIF1,EIF4A1,GTF3C4,ZNF43,HMGB1,POLR2A,TPT1,TUBB4B,RPL1
2L8,GANAB,FADS1,NEFM,YBX1,UBA1,CALR,COL5A1,CES3,RPLP0,H3-3B,PTMA,SOX4,MAGED1,
34B,ZMIZ1,TIMP2,SARAF,MIDN,SMARCA5,TPM4,RPS24,CIPC,C1ORF43,RHOA,CXADR,ATP1A1,LE

L13A,MAGED1,RPL6,YWHAE,GPI,HTRA4,FASN,RPS8,P4HB,TMBIM6,USP11,EIF3C,ATP5F1A,RPL37
200,EIF4B,UBB,AFF4,TRIM28,CKB,FN1,EIF1,MAP9,RPL19,PPP2R1A,SERINC1,BASP1,POLR2A,TPT
2,FLNA,MAP1B,RPL7A,SLC9A7,ALDOA,MTRNR2L8,PKM,GANAB,COL6A1,FADS1,PRPF8,RPL8,UB
ZNF316,DBN1,PIK3R2,MAP1B,NCL,ALDOA,GANAB,HSP90B1,NEFL,RPL8,FBXL4,UBA1,ATP2B4,PC
BTF3,RPL10,SF3B2,GTF2F1,RPL29,RPS9,ZNF865,RPS14,RPS20,WDR1,PFN1,RPL31,HNRNPC,KH
CALR,FSTL1,PSAP,H3-3B,PTMA,ENO2,SEC62,KAT6A,YWHAE,KIF1A,GNAS,DDX17,SPTBN1,CALM3
PL7A,SLC9A7,PKM,FTL,FADS1,YBX1,ATP2B4,TUBB2B,COL4A1,RPLP0,CLU,DDX5,ENO2,KIF1A,GN
F3C,ATP5F1A,RPL37A,RPL19,PPP2R1A,SERINC1,HMGB1,POLR2A,AP2M1,BTF3,AGRN,RPLP1,MA
IP200,SMIM14,ATP5F1A,MATR3,EIF1,RPL19,HMGB1,POLR2A,RPS4X,TUBB4B,BTF3,RPLP1,MAR
PKD1,PDIA3,SSBP2,RPS3A,STMN1,ANXA2,ZMIZ1,TIMP2,AP3D1,DNAJA1,MAP4,GOLGA8A,OGA,RPL
K1,P4HB,RPL5,TMBIM6,PLD3,SNRNP200,EIF4B,VCP,UBB,INA,EIF4A1,PPP2R1A,DNAH8,SF3B1,ZN
HSP90AA1,ACTR2,KIDINS220,NUMA1,RPS6,ZNF316,DBN1,RPL4,PIK3R2,MAP1B,NCL,GANAB,HSP
HSP90AA1,ACTR2,KIDINS220,NUMA1,RPS6,ZNF316,DBN1,RPL4,PIK3R2,MAP1B,NCL,GANAB,HSP
DINS220,EIF4G2,RPL3,RPS6,ZNF316,DBN1,RPL4,ENO1,NCL,ALDOA,PKM,GANAB,FTL,HSP90B1,
P11,SNRNP200,IGF2,EIF3C,ATP5F1A,EIF4B,LRRC28,RPL19,GTF3C4,PDIA3,SERINC1,HMGB1,POL
OL6A1,HSP90B1,FADS1,NEFL,RPL8,UBA1,IGFBP2,ABHD2,HSPA1B,COL4A1,RPLP0,CANX,PTMA,S
D1,RPL6,KAT6A,GPI,RACK1,CALM3,SHISA9,P4HB,TMBIM6,HSPD1,USP11,SNRNP200,SMIM14,EI
PL3,DBN1,PIK3R2,SLC9A7,ALDOA,MTRNR2L8,PKM,FTL,NEFM,RPL8,YBX1,UBA1,ABHD2,TUBB2B
2,XRCC5,CIPC,ZNF865,ATF4,CXADR,LENG8,RPL26,MCL1,PRKCSH,EIF5A,PDIA6,ITM2C,RPL30,EIF
TUBB4B,RPLP1,RPL10,RPS17,SF3B2,GTF2F1,RPL29,RPS9,XRCC5,CIPC,HK1,RHOA,ZNF865,RPS1
TP5F1B,SHISA9,P4HB,USP11,SNRNP200,EIF3C,ATP5F1A,MATR3,LRRC28,EIF1,RPL19,SERINC1,H
TPT1,ZMIZ1,TIMP2,MIDN,MARCHF6,ARF1,CYB5R3,SF3B2,ITM2B,FADS2,CPE,CNN3,TNS3,NCS1,A
MARCA5,CIPC,PAPOLA,CBX5,ZNF865,KIF5B,LENG8,VAT1,RTN3,CCNI,KHDRBS1,EIF5A,ACTN4,RPS
2,CFL1,BTBD9,XRCC2,MAVS,HTT,HSP90AB1,ZNF124,HSP90AA1,ACTR2,SPARC,TUBB,FTH1,TMS
4HB,TMBIM6,PLD3,USP11,EIF3C,ATP5F1A,EIF4B,LRRC28,RPL19,SERINC1,HMGB1,POLR2A,AP2
CNI,SLC3A2,TPM1,CDH2,DDX3X,CADM4,RAB1A,COTL1,L1CAM,NOVA2,LAMC1,BTBD2,PBX2,SCAF
RPL4,NCL,GANAB,RPL8,UBA1,FSTL1,ZNF99,RPLP0,H3-3B,PTMA,DDX5,RPL13A,SEC62,KAT6A,DV
,DDX5,FKBP14,RPL13A,MAGED1,RPL6,DVL3,GNAS,GPI,ATP5F1B,SHISA9,RPS8,P4HB,RPL5,TMBI
A1,MAT2A,PXDN,LDHA,NACA,TSPAN3,LAMA5,BRD3,C1ORF43,PAPOLA,NCS1,NR2F1,NLGN2,CHD4
1,EIF4G2,RPS6,DBN1,ENO1,FLNA,MAP1B,PKM,COL6A1,FTL,HSP90B1,NEFM,UGT8,RPL8,UBA1,A
2,RPL7,P4HB,IDS,FSCN1,SPTAN1,CALM1,SMIM14,VCP,UBB,EGLN1,CHSY1,MRC2,PPP2R1A,PKD1
6,ZNF316,DBN1,MAP1B,RPL7A,GANAB,COL6A1,FTL,HSP90B1,FADS1,NEFL,RPL8,FBXL4,FSTL1,
,DDX17,GPI,ALDH6A1,ATP5F1B,RPS8,SEPTIN2,P4HB,FSCN1,GSTP1,HSPD1,USP11,SNRNP200,EI
B,POLR2A,AP2M1,TPT1,TUBB4B,RPL11,MARCHF6,WSB1,AP3D1,ZNF770,CPE,EIF4H,TSPAN3,SL
P4HB,USP11,SNRNP200,IGF2,EIF3C,EIF4B,PDIA3,SERINC1,HMGB1,POLR2A,TPT1,TUBB4B,RPL1
EEF2,DPYSL3,TUBA1A,CFL1,BTBD9,MAVS,HSP90AB1,ZNF124,HSP90AA1,ACTR2,VIM,SPARC,TUB
K1,P4HB,RPL5,TMBIM6,PLD3,SNRNP200,CSDE1,EIF4B,EIF1,PPP2R1A,SF3B1,ZNF43,HMGB1,POI
2,RPS6,ZNF316,RPL4,TUBA1B,PIK3R2,NCL,SLC9A7,ALDOA,HSP90B1,FBXL4,YBX1,IGFBP2,TMF
MP1,SRSF4,RTN3,GLG1,EIF5A,APLP1,GPAT4,TLN1,HNRNPDL,CKAP5,LAMP1,PLPPR3,ARCN1,SKF
MP1,SRSF4,RTN3,GLG1,EIF5A,APLP1,GPAT4,TLN1,HNRNPDL,CKAP5,LAMP1,PLPPR3,ARCN1,SKF
ZNF865,RPS14,RPS20,WDR1,PFN1,MCL1,TLE5,PNN,KHDRBS1,RPS25,EIF5A,ITM2C,EIF3L,CHPF,N
DX3X,MIF,RHOB,SEPTIN3,MMP2,SKP1,MLXIP,PPP1CB,HNRNPL,ALKBH5,GRINA,COL2A1,SURF4,
00,EIF4B,EIF1,PPP2R1A,SF3B1,ZNF43,HMGB1,POLR2A,TPT1,RPS3A,MIDN,RPS17,DNAJA1,SF3B
1,CHSY1,PPP2R1A,PKD1,SF3B1,CDKL1,STMN1,UCHL1,AGRN,MIDN,AP3D1,AKAP12,CDC42BPB,C
3,GAP43,NR2F1,NLGN2,JUND,ATP2A2,CCNI,EIF5A,CCND2,DDX3X,RPL12,SHOX,WLS,NARS1,DUX4

K5,SOX4,RPL6,KAT6A,GNAS,SLC2A3,GPI,BSG,FASN,ALDH6A1,SHISA9,MYH9,IDS,GDI1,SMG1,NPM
NR2L8,PKM,COL6A1,PRPF8,RPL8,UBA1,FSTL1,ABHD2,TUBB2B,H3-3B,PTMA,DDX5,SOX4,MAGE
1,RPL7,SEPTIN2,APP,GDI1,GSTP1,MARCKS,CSDE1,COL4A2,RPL37A,LRRC28,EIF1,TMSB10,EIF4
PLD3,SNRNP200,SMIM14,EIF4B,VCP,EIF4A1,PPP2R1A,SF3B1,PDIA3,ZNF43,HMGB1,POLR2A,TPT
GB1,TUBB4B,UCHL1,NCAM1,RPLP1,MIDN,MARCHF6,RPS17,AP3D1,CDC42BPB,HNRNPU,LDHA,T
FF4,TRIM28,CKB,FN1,EIF1,TPT1,RPL18,SARAF,MIDN,WSB1,AP2B1,GTF2F1,GOLGA8A,HNRNPU,
RCHF6,SF3B2,GTF2F1,HNRNPA0,RPL23,RPL29,XRCC5,PAPOLA,RHOA,ZNF865,GLIPR1L2,RPS14
,MAGED1,RPL6,GPI,CALM3,SHISA9,P4HB,RPL5,PLD3,HSPD1,USP11,SNRNP200,EIF3C,ATP5F1A,
AS,BSG,SPTBN1,SEPTIN2,P4HB,TMBIM6,GSTP1,MARCKS,NPM1,COL18A1,EIF4B,VCP,AFF4,YWH
,SF3B1,ZNF43,HMGB1,POLR2A,RPL10,DNAJA1,SF3B2,GTF2F1,RPS9,XRCC5,ZNF865,RPS14,RP
,SF3B1,ZNF43,HMGB1,POLR2A,RPL10,DNAJA1,SF3B2,GTF2F1,RPS9,XRCC5,ZNF865,RPS14,RP
PTMS,SMG1,CALM1,IGF2,NONO,DDB1,EIF1,TMSB10,TUBB4B,ANXA2,AGRN,TIMP2,XAF1,SBF1,D
IRNP200,SMIM14,ATP5F1A,EGLN1,PPP2R1A,RPL15,GTF3C4,HMGB1,POLR2A,RPS4X,RPLP1,RPL
R3,CKB,EIF1,DCHS1,SET,MAP9,CHSY1,INA,RPL15,DNAH8,HLA-C,SEC61A1,RPS3A,MORF4L1,RPL
31,FADS1,RPL8,FBXL4,YBX1,UBA1,ATP2B4,IGFBP2,ABHD2,HSPA1B,COL4A1,PSAP,CLU,PTMA,SE
CN1,USP11,SNRNP200,TRIM28,CKB,FN1,YWHAQ,EIF1,SERINC1,POLR2A,TPT1,SEC61A1,RPS3A,
,DCX,EIF4G2,RPL3,RPS6,ZNF316,PIK3R2,FLNA,NCL,SLC9A7,HSP90B1,PRPF8,RPL8,FBXL4,TMF
B4X,NUMA1,DCX,RPS6,ENO1,TUBA1B,NCL,RPL7A,SLC9A7,ALDOA,GANAB,COL6A1,FTL,HSP90B
,CHSY1,PPP2R1A,PKD1,SF3B1,CDKL1,STMN1,UCHL1,AGRN,MIDN,AP3D1,AKAP12,CDC42BPB,G
CHSY1,TMSB10,GTF3C4,RPL18,TUBB4B,MORF4L1,DNAJA1,MAP4,LDHA,CNN3,TPM4,PAPOLA,N
P200,PPP2R1A,DNAH8,HMGB1,MIDN,MARCHF6,DNAJA1,SF3B2,MAT2A,GTF2F1,HNRNPA0,RPL2
GANAB,UBA1,FSTL1,ZNF99,RPLP0,H3-3B,PTMA,DDX5,RPL13A,SEC62,KAT6A,DVL3,YWHAE,KIF1
D1,SF3B1,CDKL1,STMN1,UCHL1,AGRN,MIDN,AP3D1,AKAP12,CDC42BPB,GOLGA8A,CTNNA1,SC
PS16,SLC16A7,ZNF865,CCT7,RBBP4,PFN1,MCL1,PRKCSH,LRPPRC,EIF5A,RAB7A,NGRN,DDOST,
A,RPL37A,INA,EIF4A1,PPP2R1A,RPL15,DNAH8,SERINC1,RPS3,HMGB1,POLR2A,RPS3A,BTF3,AGR
TP5F1B,SHISA9,P4HB,TMBIM6,USP11,SNRNP200,EIF3C,ATP5F1A,MATR3,LRRC28,EIF1,RPL19,SE
C3,PRRC2A,PPP1CC,ATG4B,DEK,HNRNPH3,KDM3A,ADNP,SSU72,FAF2,TASOR,DDX6,EXOC7,CRK
3,FTH1,NUMA1,RPL3,ZNF316,DBN1,RPL4,ENO1,PIK3R2,MTRNR2L8,GANAB,COL6A1,FTL,HSP90
L6,RACK1,ATP5F1B,P4HB,RPL5,TMBIM6,PLD3,HSPD1,SNRNP200,EIF4B,EIF4A1,PPP2R1A,DNAH
I,AKAP12,CTSD,NACA,NLGN2,CHD4,CRMP1,BEX3,TLE5,CSNK1E,PFKL,CDH6,PRDX2,ARL4C,APL
TP5F1A,EIF4B,LRRC28,RPL19,SERINC1,HMGB1,POLR2A,AP2M1,TPT1,TUBB4B,RPL11,BTF3,AG
P2,CADM4,SQLE,MAP1A,TIMP3,CTDSP2,NOVA2,CHD6,EEF1A2,CCT2,TBCD,VAMP2,GAS7,HSPA4,
D2,PGK1,CANX,SOX4,RPL6,DVL3,KIF1A,SLC2A3,BSG,ALDH6A1,GNB1,STMN2,IDS,FSCN1,TMBIM
I,HMGB1,POLR2A,TPT1,RPS4X,TUBB4B,MIDN,SF3B2,GTF2F1,RPL29,CIPC,RHOA,ZNF865,RPS1
FSTL1,ABHD2,H3-3B,ENO2,SOX4,KAT6A,GNAS,GPI,HTRA4,SPTBN1,SHISA9,APP,FSCN1,TRIM28,L
UBB,TRIM28,CKB,FN1,YWHAQ,EIF1,SERINC1,TPT1,SEC61A1,RPS3A,RPL18,RPLP1,SARAF,MIDN,
RP1,HNRNPA0,CTNNA1,LDHA,LPCAT1,CLSTN1,RAB6B,SLC7A5,TMEM259,PFN1,ASTN2,CCNI,TAD
S,NPM1,COL18A1,EIF4B,TRIM28,SET,EGLN1,EIF4A1,RPL19,PDIA3,ZNF43,TPT1,TUBB4B,MARCH
D9,MAVS,HTT,ZNF124,HSP90AA1,ACTR2,SPARC,TUBB,FTH1,KIDINS220,TMSB4X,EIF4G2,RPS6,Z
F4L1,DNAJA1,FADS2,LDHA,CTSD,EIF4H,CIPC,C1ORF43,RPL14,RHOA,CD81,LENG8,PNN,NISCH,
,USP11,EIF3C,ATP5F1A,UBB,RPL19,SERINC1,HMGB1,POLR2A,AP2M1,TUBB4B,BTF3,AGRN,RPL
PA,ZMIZ1,IGFBP3,RPL14,PTCD3,SLC7A5,ASTN2,NES,TADA2B,KHDRBS1,PFKL,CDH2,ARL4C,MRF
1,NR2F1,RAB6B,ENC1,NLGN2,CCNI,TADA2B,RPS25,TPM1,CDH2,B4GALNT4,SHOX,ATP1A2,CKA
114,UBB,AFF4,TRIM28,CKB,FN1,YWHAQ,EIF1,HACD3,MAP9,CHSY1,MRC2,PPP2R1A,SERINC1,PO
COL18A1,CSDE1,LRRC28,SPEN,DCHS1,HLA-C,SSBP2,CHD3,ANXA2,AGRN,XAF1,MIDN,VCL,MAP4,

C3,HSPD1,USP11,SNRNP200,EIF3C,VCP,TRIM28,TMSB10,GTF3C4,HLA-C,SERINC1,RPS3,VAPA,S
CN1,LAPTM4B,MAP1A,CTDSP2,CNN2,EEF1A2,CCT2,MORF4L2,PKN1,GRINA,PTOV1,MLEC,ACAP3
'11,SNRNP200,TRIM28,CKB,FN1,YWHAQ,EIF1,SERINC1,TPT1,SEC61A1,RPS3A,RPL18,RPLP1,SAF
'FN1,EIF5A,RAB7A,EIF3L,RPS19,CHPF,NGRN,RPLP2,DDOST,YWHAB,ARHGDIA,HSBP1,SRSF5,MDI
1A,TRIB2,CNN2,ZC3H7B,VAMP2,PTOV1,MDK,SRSF9,PLEC,UBE2R2,FKBP8,DLK1,COL27A1,SERB
A1,SPTBN1,CALM3,RPS8,RPL7,TMBIM6,CALM1,HSPD1,USP11,AFF4,RPL37A,HACD3,SET,TPI1,CI
,TOMM20,U2AF2,SUGP2,CPSF1,HNRNPL,ABCA3,ALKBH5,VPS35,SNRNP70,TOP1,VHL,TCF25,LU
TR2,VIM,TUBB,FTH1,UBC,EIF4G2,RPL3,RPS6,ZNF316,RPL4,ENO1,TUBA1B,FLNA,MAP1B,NCL,RF
DDX3X,RPS15,RPL12,RPS7,RPS19,TARDBP,HNRNPK,LAMP1,TTR,MDH2,GDI2,TSPYL4,HSPH1,AB
'BX1,TUBB2B,COL5A1,PGK1,CANX,CLU,PTMA,DDX5,FKBP14,RPL13A,MAGED1,RPL6,DVL3,GPI,F
HTT,HSP90AB1,ZNF124,HSP90AA1,VIM,SPARC,TUBB,FTH1,KIDINS220,TMSB4X,NUMA1,DCX,ZNI
SPEN,YWHAQ,HACD3,RPL19,GTF3C4,PKD1,PDIA3,POLR2A,BTF3,AGRN,MARCHF6,SBF1,DNAJA1,
1,MAT2A,GTF2F1,PXDN,CTNNA1,FADS2,EIF4H,BRD3,KPNB1,RPL14,CBX5,CXADR,CD81,WDR1,H
IF316,DBN1,RPL4,ENO1,MAP1B,NCL,ALDOA,GANAB,FTL,HSP90B1,FADS1,NEFM,UGT8,RPL8,YB
MSB4X,EIF4G2,ZNF316,DBN1,ENO1,PIK3R2,MAP1B,ALDOA,MTRNR2L8,PKM,GANAB,FTL,HSP90I
N1,PDIA6,APLP1,RPS19,MIF,SESN3,RPL36,MMP2,SQLE,SERF2,TIMP3,CHD6,TBCD,GAS7,GRINA,
CKB,FN1,YWHAQ,EIF1,SERINC1,TPT1,SEC61A1,RPS3A,RPL18,SARAF,MIDN,MARCHF6,WSB1,ARI
LD3,SNRNP200,ATP5F1A,EIF4B,EIF1,ZNF43,HMGB1,POLR2A,TPT1,RPS4X,RPS3A,RPL18,TUBB4E
GN2,JUND,ASTN2,JPT2,TLE5,TADA2B,SFPQ,PLXNA1,SOX12,PRDX2,TLN1,MYO5A,CDK16,APPL1,
IGFBP2,CANX,CLU,PTMA,DDX5,FKBP14,RPL13A,MAGED1,RPL6,YWHAE,GPI,HTRA4,RACK1,FASN
TC3,USP11,CSDE1,ATP5F1A,EIF4B,MATR3,NONO,CHSY1,RPS3,AP2M1,TPT1,RPL18,TUBB4B,MC
ALDH6A1,SHISA9,FSCN1,TMBIM6,PTMS,NPM1,COL18A1,CSDE1,SMIM14,MARCKSL1,UBB,AFF4,I
A,RHOA,ZNF865,RPS14,LENG8,WDR1,PFN1,TLE5,RPS25,RPL30,EIF3L,CD63,RPLP2,DDOST,CIRI
LR2A,TPT1,RPS4X,ITGB1,MARCHF6,SBF1,ARF1,GOLGA8A,HNRNPA0,CTSD,CIPC,PJA2,IGFBP3,NI
,UBB,EGLN1,CHSY1,MRC2,SSBP2,CDKL1,STMN1,TUBB4B,UCHL1,AGRN,ZMIZ1,MIDN,AP3D1,AK
ACTG1,DPYSL3,CFL1,XRCC2,MAVS,HTT,HSP90AB1,ZNF124,HSP90AA1,ACTR2,SPARC,TUBB,KIDI
PLP0,DDX5,FKBP14,SEC62,KIF1A,RPL13,DDX17,ALDH6A1,SEPTIN2,RPL5,FSCN1,GSTP1,PLD3,U
MF1,TUBB2B,RPLP0,DDX5,FKBP14,SEC62,KIF1A,RPL13,DDX17,ALDH6A1,SEPTIN2,RPL5,FSCN1,
ISP22,MED18,ACTG1,EEF2,DPYSL3,TUBA1A,CFL1,BTBD9,XRCC2,MAVS,HTT,HSP90AB1,ZNF124,I
ONO,PSMD5,HACD3,SET,EIF4A1,MRC2,PKD1,SSBP2,BASP1,UCHL1,AGRN,WSB1,AP2B1,MAT2A,I
,GNB1,STMN2,IDS,ERVV-1,FSCN1,SPTAN1,CALM1,NPM1,PLXNB2,USP11,SMIM14,VCP,UBB,AFF
:1,HMGB1,POLR2A,AP2M1,BTF3,AGRN,RPLP1,MARCHF6,DNAJA1,ARF1,SF3B2,XRCC5,RPL22,ZN
N1,UCHL1,ANXA2,AGRN,MIDN,AP3D1,AKAP12,CDC42BPB,GOLGA8A,CTNNA1,SCARB2,RPS24,G
C1,HMGB1,POLR2A,AP2M1,TUBB4B,BTF3,AGRN,RPLP1,MARCHF6,DNAJA1,ARF1,SF3B2,XRCC5
RPL3,RPS6,ZNF316,RPL4,ENO1,MAP1B,NCL,RPL7A,ALDOA,GANAB,FTL,HSP90B1,NEFL,NEFM,R
2E,MACROH2A2,MED13,TCEAL4,CAD,RBM4,PWWP3A,SRSF7,MEX3C,ASH1L,KDM2A,FARSA
:9A7,ALDOA,MTRNR2L8,PKM,GANAB,FADS1,NEFM,FBXL4,YBX1,UBA1,ATP2B4,CALR,TUBB2B,C
PIK3R2,FLNA,MAP1B,NCL,SLC9A7,ALDOA,PKM,GANAB,COL6A1,NEFL,NEFM,PRPF8,RPL8,FBXL4
BN1,RPL4,TUBA1B,PIK3R2,MAP1B,NCL,RPL7A,ALDOA,PKM,GANAB,FTL,HSP90B1,FADS1,RPL8,
D1,RPL6,DVL3,GPI,RACK1,GDI1,TMBIM6,HSPD1,USP11,SNRNP200,CSDE1,SMIM14,PPP2R1A,HL
,GANAB,FADS1,PRPF8,ATP2B4,TUBB2B,COL5A1,PSAP,PGK1,PTMA,DDX5,FKBP14,RPL13A,SEC6
I1,UBB,AFF4,RPL37A,LRRC28,SPEN,DDB1,TPI1,INA,SF3B1,VAPA,POLR2A,CDKL1,TPT1,RPL29,LF
CHD6,EEF1A2,CCT2,TBCD,VAMP2,HSPA4,PTOV1,MLEC,MDK,H3-3A,AZIN1,DDAH2,AEBP1,GRB2,
'BX1,UBA1,TMF1,TUBB2B,COL5A1,RPLP0,DDX5,FKBP14,RPL13A,KIF1A,RPL13,DDX17,HTRA4,B
P11,SNRNP200,EIF3C,ATP5F1A,EIF1,RPL19,SERINC1,HMGB1,POLR2A,AP2M1,TUBB4B,BTF3,AG

I2,P4HB,RPL5,FSCN1,PLD3,USP11,SNRNP200,CSDE1,SMIM14,UBB,AFF4,TRIM28,CKB,FN1,YWH
DDX5,FKBP14,SEC62,KIF1A,RPL13,DDX17,ALDH6A1,SEPTIN2,RPL5,FSCN1,GSTP1,PLD3,USP11,
31,TUBB4B,SF3B2,RPL23,RPL29,CIPC,HK1,ZNF865,ATF4,RPS14,LENG8,WDR1,MCL1,TLE5,PNN,
.4,PIK3R2,NCL,ALDOA,HSP90B1,RPL8,FBXL4,UBA1,ATP2B4,IGFBP2,CALR,FSTL1,TMF1,HSPA1B,
SHOX,COTL1,MAGED4B,CNTNAP2,HDGF,VEGFA,DSTN,BTBD2,PBX2,AEBP1,WDR6,UBE2R2,ATXN
HDGF,VEGFA,BTBD2,PBX2,NYNRIN,SCAF11,KDELR1,ZFAND5,STXBP1,UBQLN1,ZFAND3,COL27A
2,CHD3,BASP1,HMGB1,STMN1,ANXA2,AGRN,TMEM170A,CTNNA1,RPL23,CPE,TNS3,SLC22A17,K
4C,GNB2,MTSS2,APLP2,MXD4,SMCHD1,DNER,TIMP3,CTDSP2,NUCB1,CDH11,TBCD,SNX22,SQST
,DBN1,RPL4,TUBA1B,PIK3R2,FLNA,NCL,ALDOA,HSP90B1,PRPF8,YBX1,ATP2B4,CALR,TUBB2B,C
NA,NCL,RPL7A,PKM,GANAB,COL6A1,HSP90B1,RPL8,FBXL4,YBX1,UBA1,TMF1,ABHD2,TUBB2B,C
IGFBP2,CALR,FSTL1,COL5A1,COL4A1,PGK1,ENO2,SOX4,KIF1A,RPL13,GPI,HTRA4,BSG,FASN,G
COL5A2,HMGN2,VAT1,SRSF4,PFN1,PNN,NES,KHDRBS1,GPC3,ANXA6,NGRN,HNRNPDL,MYL6,WL
NCL,ALDOA,GANAB,HSP90B1,NEFL,CES3,H3-3B,CANX,CLU,PTMA,DDX5,FKBP14,RPL13A,MAGED
A,MAP1B,RPL7A,ALDOA,PKM,NEFL,PRPF8,UGT8,RPL8,UBA1,IGFBP2,TMF1,ABHD2,HSPA1B,TUB
F4A1,LDHB,SSBP2,BASP1,POLR2A,TPT1,STMN1,ITGB1,MORF4L1,BTF3,NCAM1,ANXA2,SARAF,M
I,PRKCSH,KHDRBS1,EIF5A,ITM2C,RPL30,EIF3L,NGRN,RPLP2,DDOST,NARS1,PTGES3,RPS12,ATF
R2,SPARC,UBC,ZNF316,RPL4,PIK3R2,NCL,SLC9A7,ALDOA,PKM,GANAB,COL6A1,HSP90B1,FADS
P90B1,FADS1,NEFL,PRPF8,TMF1,TUBB2B,RPLP0,H3-3B,PTMA,DDX5,FKBP14,RPL13A,KIF1A,RPL
PL7A,ALDOA,COL6A1,HSP90B1,FADS1,NEFL,RPL8,FBXL4,YBX1,UBA1,TMF1,TUBB2B,PSAP,RPLF
,RPLP0,CANX,DDX5,GPI,CALM3,SHISA9,SEPTIN2,P4HB,FSCN1,TMBIM6,GSTP1,NPM1,TTC3,USP1
IM14,IGF2,EIF1,DCHS1,SET,MRC2,PKD1,BASP1,TPT1,ANXA2,AGRN,TIMP2,XAF1,MARCHF6,SBF1
CKB,FN1,YWHAQ,SERINC1,TPT1,SEC61A1,RPS3A,RPL18,RPLP1,SARAF,MIDN,MARCHF6,WSB1,A
OL5A1,CES3,PTMA,KAT6A,KIF1A,GNAS,SLC2A3,FASN,SPTBN1,MYH9,RPL7,SEPTIN2,IDS,PTMS,SPT
TM2B,AP2B1,GOLGA8A,LPCAT1,RPL24,NCS1,ATP1A1,JUND,MCL1,CCNI,XPR1,EIF5A,DDR1,PRKAI
SXL1,ARF3,DSTN,DYNLL2,LUZP6,PCSK1N,CSNK1D,COL27A1,HNRNPD,AKIRIN1,DHX15,CIC,KIRRI
SB4X,EIF4G2,RPS6,ZNF316,DBN1,MAP1B,SLC9A7,ALDOA,PKM,GANAB,FTL,HSP90B1,FADS1,NEF
O1,MAP1B,RPL7A,PKM,FTL,HSP90B1,FADS1,RPL8,CALR,FSTL1,ABHD2,TUBB2B,COL5A1,PTMA,D
M2C,MAGED2,RPS19,NGRN,RPLP2,DDOST,ATP6V0C,RPL23A,DDX39B,TM9SF3,MDH2,TOMM20,C
GPC3,RPS15,COL6A2,B4GALNT4,DNAJC5,NDFIP1,SHOX,CIRBP,DUX4,PTGES3,LAMP1,RHOB,CKA
A1,COL4A1,RPLP0,DDX5,KAT6A,DVL3,YWHAE,RPL13,DDX17,HTRA4,BSG,SPTBN1,GNB1,MYH9,G
T6A,RPL13,ALDH6A1,MYH9,FSCN1,SPTAN1,SMIM14,EIF4B,AFF4,LRRC28,FN1,EIF1,HACD3,CHSY
RPS24,HK1,ZNF865,CHD4,OAZ1,PEBP1,PDIA6,PIP4K2B,ITM2C,APLP1,SLC25A3,RPLP2,MYH10,RA
RPL7A,SLC9A7,ALDOA,GANAB,COL6A1,FTL,FADS1,ATP2B4,IGFBP2,CALR,FSTL1,RPLP0,DDX5,EN
OAA1,SPARC,TUBB,FTH1,UBC,KIDINS220,TMSB4X,EIF4G2,RPL3,RPS6,ZNF316,RPL4,ENO1,TUBA
EPTIN2,P4HB,FSCN1,PLD3,HSPD1,USP11,SNRNP200,EIF3C,AFF4,CKB,LRRC28,FN1,SPEN,PSMD
N1,TPI1,EIF4A1,GTF3C4,PKD1,ZNF43,VAPA,TPT1,AGRN,ZMIZ1,MIDN,ARF1,ITM2B,TMEM170A,HN
.18,TIMP2,SF3B2,ITM2B,HNRNPA0,CTNNA1,SCARB2,RPL14,NCS1,SLC7A5,RPL26,OAZ1,NES,XPF
RPL7A,ALDOA,COL6A1,HSP90B1,FADS1,RPL8,FBXL4,YBX1,UBA1,TMF1,TUBB2B,COL5A1,PSAP,F
TMF1,TUBB2B,COL5A1,RPLP0,DDX5,FKBP14,RPL13A,KIF1A,RPL13,DDX17,GPI,ALDH6A1,ATP5F1
A4,ALDH6A1,CALM3,GNB1,P4HB,APP,RPL5,FSCN1,MARCKS,NPM1,HSPD1,MARCKSL1,YWHAG,C
OL6A1,HSP90B1,FADS1,RPL8,FBXL4,UBA1,TMF1,TUBB2B,PSAP,DDX5,RPL13A,KIF1A,RPL13,DDX
CTSD,KPNB1,RHOA,KIF5B,OAZ1,B4GALNT4,DNAJC5,APLP2,CALU,HECTD4,GPC1,TMEM132A,CD
A7,ALDOA,PKM,GANAB,COL6A1,HSP90B1,PRPF8,RPL8,ATP2B4,FSTL1,HSPA1B,YWHAE,KIF1A,G
NPM1,SMIM14,IGF2,EIF4B,UBB,YWHAG,RPL37A,FN1,EIF1,TMSB10,EIF4A1,RPL19,HLA-C,POLR2A
RPL13,SLC2A3,DDX17,BSG,GNB1,SHISA9,MYH9,SEPTIN2,ERVV-1,GSTP1,SMG1,PLXNB2,HSPD1,

\P,RPLP0,H3-3B,DDX5,MAGED1,SEC62,RPL6,RPL13,RACK1,BSG,SPTBN1,ATP5F1B,CALM3,SEPT
SP90AB1,ZNF124,ACTR2,VIM,TUBB,FTH1,NUMA1,RPL3,ENO1,TUBA1B,PIK3R2,NCL,SLC9A7,ALD
3A,CTNNA1,SCARB2,RPS24,GAP43,HK1,RPL22,ZNF865,CHD4,LENG8,WDR1,OAZ1,NES,TADA2B,I
ISP90B1,FADS1,RPL8,FBXL4,UBA1,TMF1,TUBB2B,PSAP,RPLP0,DDX5,FKBP14,SEC62,KIF1A,RPL
DPYSL3,TUBA1A,CFL1,BTBD9,XRCC2,MAVS,HTT,HSP90AB1,ACTR2,VIM,SPARC,FTH1,UBC,KIDIN
NAI3,PRDX5,UBE4A,SRP14,RPL37,ARPC2,CCAR2,BAG6,RAB11A,CDC42,TRAF7,EIF2AK1,PDAP1,I
JN,ANXA6,B4GALNT4,MTSS2,HNRNPH1,SQSTM1,TAF15,SKI,TMED2,SELENON,PRDX1,CAPZB,DU
/I2C,MAGED2,RPS19,NGRN,RPLP2,DDOST,ATP6V0C,RPL23A,DDX39B,TM9SF3,MDH2,TOMM20,I
/IA1,EIF4G2,RPS6,ZNF316,DBN1,TUBA1B,PIK3R2,NCL,SLC9A7,FTL,HSP90B1,NEFM,RPL8,FBXL4
,RPLP2,UBE2H,SMCHD1,ATP6V0C,U2AF2,MLXIP,AHCYL1,PPP2CA,PDE7A,CDC37,WDR6,SRSF3,L
,PAPOLA,PTCD3,KIF5B,ATP1A1,WDR1,ATP2A2,CCT7,MCL1,KHDRBS1,KLHL9,PIP4K2B,MMP14,D
,PRPF8,RPL8,YBX1,UBA1,PTMA,DDX5,SOX4,KAT6A,DVL3,KIF1A,ALDH6A1,SHISA9,RPL7,IDS,SPTA
F316,DBN1,FLNA,NCL,GANAB,NEFM,RPL8,FBXL4,CALR,FSTL1,HSPA1B,COL5A1,COL4A1,RPLP0
EF2,DPYSL3,CFL1,BTBD9,MAVS,HTT,ZNF124,ACTR2,TUBB,UBC,KIDINS220,TMSB4X,RPS6,ZNF3
IEFL,NEFM,YBX1,CALR,ABHD2,HSPA1B,COL5A1,PSAP,PGK1,RPLP0,CLU,PTMA,MAGED1,SEC62,I
3,SARAF,MARCHF6,AP2B1,PGAM1,CPE,TPM4,RPS24,BRD3,RPS9,SLC16A7,COL1A2,RPL9,CLSTN
\,RHOA,ZNF865,RPS14,RPS20,LENG8,WDR1,PFN1,MCL1,TLE5,PNN,PFKL,RPS25,EIF5A,PIP4K2B,
RPL5,TMBIM6,GSTP1,IGF2,EIF4B,HACD3,SET,EGLN1,EIF4A1,SERINC1,TPT1,TUBB4B,ANXA2,TIM
\A1,ACTR2,DCX,EIF4G2,RPL3,ZNF316,DBN1,RPL4,ENO1,PIK3R2,FLNA,MAP1B,ALDOA,MTRNR2L
HL1,ANXA2,AGRN,AP3D1,AKAP12,GOLGA8A,CTNNA1,RPL29,SCARB2,RPS24,CIPC,IGFBP3,GAP4
AT6A,DVL3,KIF1A,SHISA9,RPL7,IDS,SPTAN1,SMG1,NPM1,HSPD1,EIF3C,AFF4,RPL37A,MAPT,YWH
0AA1,ACTR2,VIM,TUBB,FTH1,NUMA1,EIF4G2,RPS6,ZNF316,DBN1,RPL4,FLNA,MAP1B,NCL,RPL
DX17,ALDH6A1,SEPTIN2,RPL5,FSCN1,GSTP1,PLD3,USP11,COL18A1,SNRNP200,SMIM14,TRIM2
/IARCHF6,DNAJA1,SF3B2,ZNF865,CXADR,LENG8,PRKCSH,SCRN1,EIF5A,PDIA6,ITM2C,EIF3L,MAG
5A1,HSP90B1,RPL8,FBXL4,YBX1,UBA1,TMF1,ABHD2,TUBB2B,COL5A1,COL4A1,PSAP,RPLP0,PTN
I,HMGB1,TUBB4B,MARCHF6,AP3D1,SF3B2,RPL23,RPL29,CIPC,PAPOLA,HK1,ZNF865,ATF4,RPS1
INRNPA2B1,CUL3,UBE2M,RALY,SRSF2,EEF1B2,RIC8A,PRPF19,SSBP4,ATG4B,FGFR1,CDC34,MT
5,RPL17,CBX5,RBBP4,HNRNPC,HDLBP,RPS25,PDIA6,ITM2C,RPL35,RPL36,YWHAB,HNRNPH1,RF
H9,IDS,ERVV-1,GDI1,FSCN1,PTMS,CALM1,CSDE1,SPEN,YWHAQ,RPL15,GTF3C4,SSBP2,CHD3,VA
3B,NDUFS1,NOLC1,ACIN1,ZC3H4
NP200,CSDE1,MATR3,RPL37A,EIF1,RPL19,PPP2R1A,SF3B1,HMGB1,VCL,SF3B2,GTF2F1,RPL23,I
DST,YWHAB,ARHGDIA,HSBP1,TM9SF3,MDH2,U2AF2,GNAI3,GDI2,PRDX5,UBE4A,TCP1,SRP14,RPI
1,CHSY1,PPP2R1A,PKD1,CDKL1,STMN1,UCHL1,AGRN,MIDN,AP3D1,AKAP12,CDC42BPB,GOLGA
UBB,FN1,EIF4A1,PPP2R1A,HLA-C,PDIA3,TUBB4B,AGRN,MIDN,SLC4A8,NFE2L1,LARP1,HNRNPA0
T1,RPS3A,RPL18,SARAF,MIDN,MARCHF6,WSB1,AP2B1,GTF2F1,GOLGA8A,LDHA,RPS9,JUND,TMI
DCX,RPS6,ZNF316,RPL4,ENO1,TUBA1B,PIK3R2,FLNA,MAP1B,NCL,RPL7A,SLC9A7,HSP90B1,NEFI
NAJA1,SF3B2,ZNF865,CXADR,LENG8,PRKCSH,EIF5A,PDIA6,ITM2C,EIF3L,MAGED2,RPS19,NGRN
TP1,PLD3,HSPD1,COL18A1,ATP5F1A,EIF4B,AFF4,SET,CHSY1,RPL19,LDHB,SF3B1,VAPA,HMGB1,
GRN,MARCHF6,MAT2A,LDHA,CNN3,CTSD,XRCC5,PAPOLA,KIF5B,PFN1,PNN,HDLBP,RPS25,RPS1C
IBTPS1,CUL3,SEPTIN7,TAOK1,UBE2D2,IREB2,LSM14A,TGOLN2,UBE2D3,HNRNPH3,USP47,SCAN
KP1,TRIB2,PEA15,CTDSP2,PGRMC1,TM9SF3,RTN1,PKN1,ASXL1,RAB11B,SRP14,PRPF6,SELENC
UBA1A,CFL1,BTBD9,HTT,ZNF124,HSP90AA1,ACTR2,VIM,TUBB,FTH1,UBC,KIDINS220,NUMA1,DC
1,BCLAF1,CDKN1A,TIMP3,TBCD,CPSF1,MEST,SSRP1,PDXK,SNRNP70,SMARCA4,SRSF1,AEBP1,A
SNRNP200,UBB,TRIM28,CKB,FN1,YWHAQ,EIF1,SERINC1,TPT1,SEC61A1,RPL18,SARAF,MIDN,MA
DS1,NEFL,PRPF8,RPL8,FSTL1,TUBB2B,RPLP0,H3-3B,ENO2,FKBP14,DDX17,RACK1,SHISA9,STM

NF43,AGRN,MAP4,SLC4A8,NFE2L1,LARP1,RTN4,CBX5,ATF4,ATP2A2,CCT7,CDH6,PEBP1,GPC3,A
TUBB,FTH1,TMSB4X,NUMA1,EIF4G2,ZNF316,RPL4,ENO1,TUBA1B,PIK3R2,MAP1B,ALDOA,GANA
,IK,NGRN,RPLP2,DDOST,RPL36,YWHAB,ARHGDIA,HSBP1,TM9SF3,MDH2,U2AF2,CCT2,GNAI3,NC
OL6A1,FTL,NEFL,UGT8,FBXL4,YBX1,UBA1,IGFBP2,ABHD2,TUBB2B,PSAP,H3-3B,CANX,CLU,DD
,ATP5F1A,EIF4B,MATR3,NONO,TMSB10,EIF4A1,RPS3,AP2M1,STMN1,MORF4L1,BTF3,XAF1,WSE
2,MAT2A,GTF2F1,HNRNPA0,ZNF865,RPS14,LENG8,WDR1,RPL26,SRSF4,RBBP4,PFN1,RPS25,RA
LGA8A,CTNNA1,SCARB2,RPS24,HK1,RPL22,ZNF865,CHD4,WDR1,VAT1,OAZ1,NES,RPL31,TADA
13,FASN,SHISA9,P4HB,PLXNB2,TTC3,PLD3,EIF4B,MARCKSL1,NONO,HACD3,EGLN1,CHSY1,RPL
LN1,MYO5A,CD63,MYL6,LAMP1,HECTD4,PPIB,HSBP1,DLG5,HS6ST1,PGRMC1,ANXA5,GNAI3,ML
AS,BSG,FASN,SPTBN1,GNB1,SHISA9,SEPTIN2,FSCN1,GSTP1,CALM1,EIF4B,VCP,MARCKSL1,COL
S35,SNRNP70,RABL6,LUZP6,SCAF11,LONP1,TXLNA,KDM5C,SF1,UPF1,SAFB2,ZNF664,VPS26B,
SF5,TM9SF3,MDH2,U2AF2,MORF4L2,MTRF1L,PRDX5,SRP14,NRDC,ARPC2,RABL6,SUPT5H,RPL
SP90B1,FADS1,RPL8,FBXL4,UBA1,TMF1,TUBB2B,RPLP0,DDX5,FKBP14,SEC62,KIF1A,RPL13,DD
CSH,RPS25,EIF5A,PDIA6,ACSL6,ITM2C,RPL30,EIF3L,MAGED2,RPS19,NGRN,RPLP2,DDOST,CIRE
4G2,RPL4,ENO1,PIK3R2,MAP1B,SLC9A7,ALDOA,MTRNR2L8,GANAB,COL6A1,HSP90B1,FADS1,N
1,NEFL,RPL8,UBA1,CALR,FSTL1,PSAP,PGK1,CANX,DDX5,FKBP14,SEC62,YWHAE,KIF1A,RPL13,D
YL6,WLS,DUX4,ARCN1,MMP2,TRIB2,EEF1A2,ACO2,ACAP3,MDK,TSPAN6,ARF3,CCND1,EIF3E,FKE
28,CKB,FN1,YWHAQ,EIF1,SERINC1,TPT1,RPL18,MORF4L1,SARAF,MIDN,WSB1,AP2B1,GTF2F1,
YBX1,UBA1,CALR,TUBB2B,COL5A1,PSAP,RPLP0,H3-3B,CANX,MAGED1,DDX17,ATP5F1B,P4HB,ID
1B,SLC9A7,COL6A1,FTL,NEFM,CALR,TMF1,TUBB2B,PGK1,CANX,CLU,KAT6A,HTRA4,BSG,ALDH
RFGEF2,CDC42,TRAF7,EIF2AK1,PRMT1,APMAP,YKT6,TMEM167A,CHERP,CTTN,LRP10,ERH,SEC
HAG,DCHS1,TMSB10,BTF3,ANXA2,AGRN,TIMP2,TNS3,LPCAT1,LLGL1,NLGN2,CHD4,CD81,HMG
9,XRCC2,MAVS,HSP90AB1,ACTR2,FTH1,UBC,KIDINS220,TMSB4X,NUMA1,EIF4G2,RPL3,RPS6,D
1,TPT1,SEC61A1,RPS3A,RPL18,SARAF,MIDN,MARCHF6,WSB1,AP2B1,GTF2F1,GOLGA8A,LDHA,
WSB1,AP2B1,GTF2F1,GOLGA8A,LDHA,RTN4,SLC22A17,JUND,TMEM259,LENG8,CCNI,XPR1,RPS
R2A,BTF3,AGRN,MARCHF6,DNAJA1,TMEM170A,BRD3,XRCC5,KPNB1,TMEM259,EIF5,HMGN2,JP
90AA1,TUBB,KIDINS220,NUMA1,RPS6,ZNF316,DBN1,ENO1,FLNA,SLC9A7,COL6A1,UBA1,ATP2B
MN1,ANXA2,AGRN,SARAF,AP3D1,AP2B1,GOLGA8A,CTSD,MSN,NCS1,SLC7A5,OAZ1,CCND2,GPC3
7,RPS15A,ARF3,PRDX1,ATP5PB,WDR6,RPS11,SUPT5H,CAND1,MAP2K2,DIDO1,RPS28,NKTR,H2
L18,SARAF,MIDN,WSB1,AP2B1,GTF2F1,LDHA,JUND,TMEM259,LENG8,CCNI,XPR1,HDLBP,GNB2
10,UBE2H,SMCHD1,ATP6V0C,LAPTM4B,MLXIP,HSPH1,AHCYL1,KLC1,CLSTN3,DDX42,RAB11B,T
C,NRBP2,USP22,MTRNR2L2,MED18,ACTG1,EEF2,DPYSL3,TUBA1A,CFL1,BTBD9,XRCC2,MAVS,H
,ITM2C,RPL30,NGRN,DDOST,CALU,ARCN1,SLC35E2B,ATP6V0C,FAM199X,DDX39B,MDH2,TOMM
S24,XRCC5,RPL22,ZNF865,ATF4,KIF5B,LENG8,RPL26,PRKCSH,KHDRBS1,EIF5A,ITM2C,RPL30,
DHA,JUND,TMEM259,LENG8,CCNI,HDLBP,GNB2,CHPF,RPLP2,MYL6,MYH10,SND1,UBE2H,ARC
RPS8,APP,PTMS,GSTP1,MARCKS,PLXNB2,VCP,MARCKSL1,TRIM28,MATR3,YWHAG,NONO,YWHA
CHPF,NGRN,RPLP2,DDOST,YWHAB,SND1,LAMP1,RPS27,ARHGDIA,SLC35E2B,SRSF5,MDH2,U2A
B,NFE2L1,LAMA5,ATP1A1,ATP2A2,LASP1,SFPQ,CDH2,PIP4K2B,ACTN4,GJA1,RPLP2,MXD4,CIRB
L3,NCL,SLC9A7,MTRNR2L8,PKM,GANAB,HSP90B1,NEFL,NEFM,RPL8,FBXL4,IGFBP2,CALR,FST
4,IGFBP2,FSTL1,RPLP0,H3-3B,DDX5,FKBP14,RPL13A,MAGED1,RPL6,GNAS,GPI,RACK1,BSG,AT
,EIF2AK1,PDAP1,YKT6,MCM3AP,TMEM167A,CHERP,CTTN,ERH,EIF3H,SEC14L1,NUDCD3,TUBGC
CHD1,ATP6V0C,LAPTM4B,MLXIP,HSPH1,AHCYL1,KLC1,CLSTN3,RAB11B,TUBB2A,PPP2CA,PDE
1,TPT1,RPL18,SARAF,MIDN,MARCHF6,WSB1,AP2B1,GTF2F1,GOLGA8A,LDHA,JUND,TMEM259,L
,HSP90AB1,ZNF124,HSP90AA1,ACTR2,VIM,SPARC,FTH1,TMSB4X,DCX,EIF4G2,RPL3,ZNF316,DB
PT2,NISCH,SPTBN2,SLC3A2,HDLBP,RPS25,PDIA6,GPC3,SOX12,CLTC,ARL6IP1,ANKRD52,HNRN

6,CCAR2,BAG6,CDC42,PPP1R9B,EIF2AK1,PDAP1,YKT6,MCM3AP,TMEM167A,CHERP,CTTN,ERH,
RCKS,SMG1,HSPD1,CSDE1,MATR3,LRRC28,NONO,DDB1,EIF1,MAP9,TPI1,SERINC1,BASP1,POL
T1,RPL10,SF3B2,GTF2F1,RPL23,RPL29,CIPC,ZNF865,RPS14,LENG8,WDR1,RPL26,MCL1,RPL31
22,MED18,EEF2,DPYSL3,TUBA1A,CFL1,XRCC2,MAVS,HTT,HSP90AB1,ZNF124,HSP90AA1,ACTR2,
F99,PSAP,PGK1,CANX,DDX5,MAGED1,SEC62,YWHAE,KIF1A,BSG,FASN,ATP5F1B,CALM3,KLRD1,
3,HMGB1,AP2M1,DNAJA1,ARF1,SF3B2,GTF2F1,CTNNA1,RPL29,RPS9,PJA2,ZNF865,RPL9,RPS14
B2,GOLGA8A,HNRNPA0,CPE,LPCAT1,RPS24,SLC22A17,CIPC,PJA2,RPL17,COL1A2,HDLBP,PEBP1
3,NONO,DNAH8,RPS3,AP2M1,MORF4L1,BTF3,XAF1,AP3D1,AKAP12,MAT2A,RPSA,RPL14,CBX5,A
,HSP90B1,NEFL,FBXL4,YBX1,UBA1,TMF1,HSPA1B,COL5A1,COL4A1,H3-3B,DDX5,SOX4,FKBP14
3,APP,PLD3,HSPD1,COL18A1,CSDE1,TRIM28,COL4A2,RPL19,GTF3C4,LDHB,HLA-C,SF3B1,VAPA,
PS6,ZNF316,RPL4,PIK3R2,NCL,ALDOA,FTL,HSP90B1,RPL8,FBXL4,YBX1,ATP2B4,IGFBP2,CALR,
42,TRAF7,EIF2AK1,PDAP1,PHB2,APMAP,PLXNB1,FARP1,YKT6,MCM3AP,TMEM167A,CHERP,CTT
NAI3,PRDX5,UBE4A,HSPA4,TCP1,MLEC,SRP14,PRPF6,TAF7,ARPC2,FAM168B,ATXN10,GNG2,DH
PB,ZNF770,OGA,FADS2,TNS3,LPCAT1,BRD3,KPNB1,LLGL1,JUND,RPL26,CRMP1,TLE5,CCND2,P
NF865,CHD4,WDR1,OAZ1,NES,RPL31,TADA2B,KHDRBS1,PIP4K2B,ITM2C,APLP1,SESN3,APLP2,D
SCN1,MARCKS,SMG1,VCP,AFF4,HACD3,CHSY1,SF3B1,SSBP2,CDKL1,AP2M1,STMN1,UCHL1,MO
,YWHAQ,EIF1,SERINC1,TPT1,RPL18,SARAF,MIDN,WSB1,AP2B1,GTF2F1,LDHA,CBX5,RPL9,ATP1A
,MYH10,SND1,UBE2H,LAPTM4B,MLXIP,ALKBH5,PDXK,AHCYL1,KLC1,CLSTN3,SCAF1,RAB11B,T
365,CLSTN1,NR2F1,LENG8,HMGN2,LASP1,RBBP4,TADA2B,XPO1,SLC3A2,RPS15,RPL12,MAGED
,MATR3,NONO,CHSY1,RPS3,TUBB4B,MORF4L1,BTF3,XAF1,MAT2A,RPSA,RPL24,RPL14,CBX5,T
MSB4X,NUMA1,EIF4G2,RPL3,ZNF316,RPL4,ENO1,TUBA1B,PIK3R2,NCL,RPL7A,SLC9A7,ALDOA,
CK1,RPL5,NPM1,PLD3,EIF4B,AFF4,EIF4A1,PKD1,ZNF43,HMGB1,TPT1,RPS4X,RPL10,RPS17,AP
5,CARM1,STRN4,ABI2,SLC6A8,PKDCC,PGAP6,SH3PXD2A,SLC20A1,TEAD1,LMAN2,KLF13,ANP32
,YBX1,TMF1,TUBB2B,DDX5,RPL13A,KIF1A,RPL13,DDX17,GPI,ALDH6A1,ATP5F1B,GNB1,SEPTIN2
D3,VCP,UBB,EIF4A1,PPP2R1A,HLA-C,PDIA3,TUBB4B,AGRN,RPLP1,MIDN,AP3D1,SLC4A8,NFE2L1
,DPYSL2,TOP1,TAF7,EIF3E,ARF4,TSC22D1,SDF4,LOC728392,VGF,FBRSL1,SEC23A,TP53BP1,DD
1,DDX5,RPL13A,KIF1A,DDX17,GPI,ALDH6A1,ATP5F1B,SEPTIN2,P4HB,KLRD1,FSCN1,TMBIM6,G
1B,COL4A1,RPLP0,H3-3B,ENO2,KAT6A,RPL13,GPI,SPTBN1,GNB1,SHISA9,STMN2,P4HB,APP,PT
B1,ARF1,AP2B1,GTF2F1,LDHA,RPS9,JUND,TMEM259,LENG8,CCNI,HDLBP,GNB2,RPS5,IK,CHPF
TUBA1A,CFL1,BTBD9,XRCC2,MAVS,HTT,HSP90AB1,ZNF124,HSP90AA1,ACTR2,SPARC,FTH1,TMS
STN1,RPS20,CCT7,CCN2,CLTC,MAGED2,SESN3,SHOX,CAPRIN1,SKP1,FAM199X,EID1,MDH2,TL
S11,SUPT5H,CAND1,MAP2K2,DIDO1,RPS28,MBTPS1,CUL3,SEPTIN7,FUS,PPP1R10,TAOK1,UBE
I6,CALM1,SNRNP200,EIF4A1,RPL15,DNAH8,SF3B1,HMGB1,POLR2A,TPT1,SEC61A1,RPS3A,RPL
F5,TM9SF3,MDH2,U2AF2,CCT2,GNAI3,NOP53,VCAN,PRDX5,BAG6,RAB11A,AKIRIN1,CCT5,CDC
A,HMGB1,AP2M1,ARF1,SF3B2,GTF2F1,CTNNA1,RPL29,PJA2,RPL24,ZNF865,RPL9,RPS14,RPS2
OVA2,ANXA5,MORF4L2,ASXL1,G3BP2,ID4,VGF,MEX3A,P3H3,CD99L2,YLPM1,PBX1,NOL4L,LRR
C14L1,NUDCD3,GNL1,TUBGCP6,INTS11,MTMR4,CAP1,PRPF19,AP1M1,TOMM70,ATP13A1,PAF
EPTIN2,APP,IDS,KLRD1,GDI1,SPTAN1,UBB,AFF4,MATR3,CKB,TPI1,PPP2R1A,LDHB,DNAH8,PDIA
1,RPL3,ZNF316,DBN1,ENO1,PIK3R2,FLNA,MAP1B,NCL,RPL7A,ALDOA,MTRNR2L8,PKM,GANAB,
RCC2,MAVS,ZNF124,HSP90AA1,ACTR2,FTH1,UBC,KIDINS220,TMSB4X,NUMA1,DCX,EIF4G2,RPL
DAP1,YKT6,MCM3AP,TMEM167A,CHERP,CTTN,ERH,SEC14L1,NUDCD3,TUBGCP6,RIC8A,INTS11,
,CES3,PSAP,RPLP0,CANX,DDX5,ENO2,RPL13A,MAGED1,RPL6,YWHAE,BSG,CALM3,SHISA9,SEPT
PF,RPLP2,MYL6,MYH10,UBE2H,SMCHD1,ATP6V0C,LAPTM4B,MLXIP,HSPH1,AHCYL1,KLC1,CLST
NB2,CHPF,RPLP2,MYL6,MYH10,UBE2H,SMCHD1,ATP6V0C,LAPTM4B,MLXIP,HSPH1,AHCYL1,KLC
LC35E2B,TMEM30A,SKP1,TRIB2,PEA15,NOVA2,SNX22,PABPN1,COL2A1,HDGF,RPL38,LAPTM4A,

,PNN,KHDRBS1,RPS25,EIF5A,ITM2C,EIF3L,CHPF,NGRN,RPLP2,DDOST,YWHAB,LAMP1,ARHGDIA
G1,NPM1,HSPD1,IGF2,MARCKSL1,AFF4,YWHAG,DDB1,EIF1,SET,RPL15,PKD1,PDIA3,SSBP2,VAPA
259,LENG8,CCNI,GNB2,CHPF,RPLP2,MYL6,MYH10,UBE2H,SMCHD1,ATP6V0C,LAPTM4B,MLXIP,H
RPLP2,MYL6,MYH10,UBE2H,SMCHD1,ATP6V0C,LAPTM4B,MLXIP,HSPH1,AHCYL1,KLC1,CLSTN3,D
RPLP2,MYL6,MYH10,UBE2H,SMCHD1,ATP6V0C,LAPTM4B,MLXIP,HSPH1,AHCYL1,KLC1,CLSTN3,D
CTSD,SLC22A17,CIPC,CHD4,JUND,ATP2A2,MCL1,CCNI,HDLBP,ARL4C,MRFAP1,HNRNPDL,MYL6,
A,PKM,GANAB,COL6A1,FTL,FADS1,PRPF8,RPL8,FBXL4,YBX1,UBA1,ATP2B4,CALR,FSTL1,COL5A
,SMIM14,VCP,UBB,YWHAG,EGLN1,CHSY1,TMSB10,ZNF43,STMN1,UCHL1,MORF4L1,ANXA2,AGR
2,FLNA,MAP1B,NCL,RPL7A,ALDOA,MTRNR2L8,PKM,GANAB,COL6A1,PRPF8,RPL8,YBX1,UBA1,IG
P2,CALR,FSTL1,ZNF99,TUBB2B,PSAP,PGK1,H3-3B,CANX,DDX5,FKBP14,SEC62,YWHAE,RPL13,SI
P2,CALR,FSTL1,ZNF99,TUBB2B,PSAP,PGK1,H3-3B,CANX,DDX5,FKBP14,SEC62,YWHAE,RPL13,SI
AGED1,RPL13,RPL5,FSCN1,PTMS,SMG1,IGF2,EIF3C,VCP,MARCKSL1,TRIM28,DDB1,PSMD5,HAC
PEA15,MLXIP,NOX4,AHCYL1,KLC1,CLSTN3,RAB11B,TUBB2A,PPP2CA,PDE7A,PSMD3,BNIP3L,CA
ZNF124,ACTR2,SPARC,UBC,TMSB4X,NUMA1,ZNF316,DBN1,ENO1,TUBA1B,PIK3R2,ALDOA,PKM,
PFKL,JUN,ARL6IP1,HNRNPK,SEPTIN11,CTBP1,GPC1,BCLAF1,TRIB2,EID1,EEF1A2,MEST,GDI2,HI
LXNA1,RAB7A,PRKAR1A,HNRNPDL,ST3GAL2,SND1,CKAP4,SMCHD1,TM9SF3,ANXA5,TM9SF2,MO
FL1,BTBD9,XRCC2,MAVS,ZNF124,ACTR2,VIM,SPARC,FTH1,KIDINS220,TMSB4X,DCX,RPL4,ENO1,
LDOA,PKM,COL6A1,HSP90B1,FADS1,RPL8,FBXL4,UBA1,TMF1,TUBB2B,PSAP,PGK1,RPLP0,DDX
BA1B,MTRNR2L8,FADS1,NEFL,RPL8,YBX1,UBA1,CALR,FSTL1,ZNF99,CES3,PSAP,PGK1,RPLP0,H
RSF1,CAPZB,HNRNPUL1,CASC3,ARF4,STMN3,CSNK1D,ZFAND3,TRIO,MTCH1,PCBP4,ACLY,YARS
H,EIF5A,PDIA6,MAGED2,RPS19,CHPF,JUN,NGRN,RPLP2,DDOST,ARHGDIA,RPL23A,DDX39B,SRSF
1,GNB1,MYH9,SMG1,PLD3,COL18A1,EIF3C,EIF4B,VCP,MARCKSL1,TRIM28,MATR3,YWHAG,CKB
PSMD8,SAFB2,EIF3H,KMT2E,PNISR,YY1,XPO6,ATG4B,ADNP,SART3,PPP4R3A,GID8,NSD1,CELSR
1A,RPL13,RACK1,FASN,GNB1,FSCN1,GSTP1,GTF3C4,SSBP2,VAPA,POLR2A,BTF3,ZMIZ1,TIMP2,
DI2,PRDX5,SRP14,ARPC2,CCAR2,BAG6,TXLNA,PDAP1,PHB2,UBA52,IARS1,MCM3AP,TMEM167A,
EFL,NEFM,PRPF8,RPL8,YBX1,UBA1,ATP2B4,FSTL1,ABHD2,HSPA1B,PSAP,PGK1,CANX,PTMA,DDX
PF,RPLP2,MYL6,MYH10,UBE2H,SMCHD1,ATP6V0C,LAPTM4B,MLXIP,HSPH1,MTRF1L,AHCYL1,KL
EIF4B,MATR3,NONO,TMSB10,RPS3,MORF4L1,BTF3,XAF1,WSB1,SBF1,SF3B2,MAT2A,RPSA,RPL
SP90AA1,UBC,NUMA1,RPL3,ENO1,PIK3R2,FLNA,NCL,RPL7A,SLC9A7,ALDOA,MTRNR2L8,PKM,G
4G2,ZNF316,DBN1,ENO1,PIK3R2,FLNA,RPL7A,SLC9A7,COL6A1,HSP90B1,RPL8,IGFBP2,CALR,F
1,PRMT1,PDAP1,FARP1,YKT6,MCM3AP,TMEM167A,CHERP,CTTN,ERH,SEC14L1,NUDCD3,GNL1,
P2,CALR,FSTL1,ZNF99,TUBB2B,PSAP,PGK1,H3-3B,CANX,DDX5,FKBP14,SEC62,YWHAE,RPL13,SI
CCNI,GNB2,CHPF,RPLP2,MYL6,MYH10,UBE2H,SMCHD1,ATP6V0C,LAPTM4B,MLXIP,HSPH1,UBE4
SNRNP200,UBB,TRIM28,CKB,FN1,YWHAQ,EIF1,MAP9,CHSY1,EIF4A1,SERINC1,BASP1,POLR2A,T
EIF2AK1,PDAP1,PHB2,YKT6,MCM3AP,TMEM167A,CHERP,CTTN,LRP10,ERH,SEC14L1,NUDCD3,G
1,RAB11B,TUBB2A,PPP2CA,PDE7A,PSMD3,CDC37,XPNPEP3,LARS1,YWHAH,UBQLN1,CSNK1D,IF
31,SBF1,AP2B1,GTF2F1,LDHA,JUND,TMEM259,LENG8,CCNI,CSNK1E,GNB2,CHPF,JUN,RPLP2,TA
GPAT4,APLP2,HNRNPH1,ATP1A2,CKAP4,ZC3H7B,RNF187,SUGP2,SQSTM1,NOP53,GDI2,AHCYL1
L22,ZNF865,ATF4,LENG8,RPL26,PRKCSH,KHDRBS1,EIF5A,ITM2C,RPL30,EIF3L,NGRN,DDOST,C
CKB,FN1,YWHAQ,EIF1,PPP2R1A,SERINC1,RPL18,ANXA2,SARAF,MARCHF6,WSB1,ARF1,AP2B1,G
MAP1LC3B,MTCH1,GPC2,ERGIC3,ANKRD12,POR,CNDP2,EHMT2,DRAP1,TBC1D14,DHFR,MRPS6
NB2,CHPF,RPLP2,MYL6,MYH10,UBE2H,SMCHD1,ATP6V0C,LAPTM4B,MLXIP,HSPH1,AHCYL1,KLC
F1,SERINC1,TPT1,RPL18,SARAF,MIDN,MARCHF6,WSB1,AP2B1,GTF2F1,GOLGA8A,SMARCA5,LD
,TMEM259,LENG8,CCNI,GNB2,CHPF,RPLP2,MYL6,MYH10,UBE2H,SMCHD1,ATP6V0C,LAPTM4B,
N,ERH,SEC14L1,NUDCD3,INTS11,MTMR4,CAP1,AP1M1,TOMM70,LYPLA2,ATP13A1,PAFAH1B2,P

ENG8,CCNI,GNB2,CHPF,RPLP2,MYL6,MYH10,UBE2H,SMCHD1,ATP6V0C,LAPTM4B,MLXIP,HSPH
,MORF4L2,ASXL1,G3BP2,ID4,VGF,MEX3A,P3H3,CD99L2,YLPM1,PBX1,NOL4L,LRRC4B,ATRX,ADI
CA,PDE7A,PSMD3,CDC37,XPNPEP3,LARS1,YWHAH,UBQLN1,CSNK1D,IFITM3,MLF2,AKIRIN1,YME
CA,PDE7A,PSMD3,CDC37,XPNPEP3,LARS1,YWHAH,UBQLN1,CSNK1D,IFITM3,MLF2,AKIRIN1,YME
D,TMEM259,LENG8,CCNI,GNB2,CHPF,RPLP2,MYL6,MYH10,UBE2H,SMCHD1,ATP6V0C,LAPTM4E
3,SNRNP200,CSDE1,ATP5F1A,EIF4B,MATR3,RPL37A,LRRC28,POLR2A,CDKL1,SEC61A1,NCAM1,
2F1,LDHA,PAPOLA,CBX5,RPL9,JUND,TMEM259,LENG8,MCL1,CCNI,CLTC,GNB2,CHPF,RPLP2,M
1,SEPTIN2,NPM1,PLD3,CSDE1,ATP5F1A,EIF4B,AFF4,DDB1,SET,CHSY1,RPL19,GTF3C4,LDHB,SF
1,SEPTIN2,NPM1,PLD3,CSDE1,ATP5F1A,EIF4B,AFF4,DDB1,SET,CHSY1,RPL19,GTF3C4,LDHB,SF
,RPL23,LDHA,CPE,EIF4H,C1ORF43,COL1A2,HK1,CBX5,ATP1A1,RPS20,LASP1,VAT1,PFKL,CCND
NI,PEBP1,PIP4K2B,MXD4,MAGED4B,TBCD,PGD,CAMSAP1,KLC1,VPS35,TUBB2A,TCF25,ACTN1,C
ZNF124,ACTR2,VIM,SPARC,FTH1,UBC,KIDINS220,TMSB4X,NUMA1,EIF4G2,RPS6,ZNF316,TUBA1
APA,BRI3,CNTN2,MEIS3,EIF1B,MCM7,NCOA6,RBBP6
2,ARF4,SEC23A,CTXN1,TXLNA,TXNDC5,SEC16A,BRD2,SLC7A6,GPSM1,POM121C,SPG7,SSR3,G
1,SHISA9,P4HB,RPL5,KLRD1,TMBIM6,HSPD1,USP11,CSDE1,SMIM14,EIF3C,ATP5F1A,UBB,RPL1
PF,RPLP2,MYL6,MYH10,UBE2H,SMCHD1,ATP6V0C,LAPTM4B,MLXIP,HSPH1,AHCYL1,KLC1,CLST
F1,LDHA,GLIPR1L2,JUND,TMEM259,LENG8,CCNI,GNB2,CHPF,JUN,B4GALNT4,RPLP2,MYL6,MY
2,CCT2,GNAI3,TM9SF2,HSPH1,MTRF1L,PRDX5,UBE4A,TCP1,MLEC,SRP14,TAF7,LUZP6,GNG2,Z
3AP,TMEM167A,CHERP,CTTN,ERH,SEC14L1,NUDCD3,RIC8A,INTS11,MOAP1,MTMR4,CAP1,AP1
US,RAB1B,GABARAPL1,MOAP1,KRT8,MTOR,TRIM8,FGFR1,UBE2K,HTATSF1,SCAMP1,TAOK2,R
2B4,FSTL1,COL5A1,RPLP0,CANX,PTMA,DDX5,YWHAE,DDX17,ATP5F1B,CALM3,RPS8,RPL7,SEPTI
A,MAGED4B,SRP9,TLN2,MTRF1L,PTOV1,SRSF1,DYNC1LI2,PRKDC,GABARAP,PFKM,YME1L1,DH
GED1,HTRA4,RACK1,P4HB,KLRD1,PLXNB2,EIF3C,EIF4B,AFF4,MATR3,NONO,TPI1,INA,LDHB,TPT
Y,BCL7A,PGAP6,MAZ,KIAA0232,KRT8,MKRN1,FOXK2,CREBBP,PHIP,TEAD1,MGAT3,CMPK1,CRIP
NF43,VAPA,CDKL1,STMN1,UCHL1,MORF4L1,ANXA2,AGRN,MIDN,AP3D1,GOLGA8A,CTNNA1,RPL
ADA2B,KHDRBS1,GLG1,PNMA1,H1-0,APLP1,RPLP2,INIP,SEPTIN11,RAC1,ARHGDIA,CDK16,BCLA
9,ANO8,DAAM1,SLK,TOB2,MED25,STC2,DUSP8,DPP7,ZNRF1,ARHGAP33,ZNF146,PIN1,YBX3,VA
OA,SLC7A5,EIF5,LENG8,LASP1,PFKL,DDX3X,RPS5,RPL35A,RPLP2,CTBP1,ARHGDIA,BCLAF1,PP
,ALDH6A1,SEPTIN2,FSCN1,NPM1,PLD3,USP11,SNRNP200,TRIM28,CKB,FN1,YWHAQ,EIF1,CHSY1
R2A,RPS3A,BTF3,RPL10,RPS16,XRCC5,COL1A2,GAP43,RPL22,ZNF865,RPL9,KIF5B,RPS20,WD
EST,PTOV1,MLEC,SURF4,DDAH2,TCF25,CORO1C,CD99,UBA52,NXN,EP400,TNKS,PPP1R15A,MA
,FLOT1,NXN,EVL,MASTL,KCTD12,HNRNPA2B1,NCOA4,RPS2,ZMYM4,HNRNPAB,CARM1,MAZ,NU
ES3,CANX,RPL13,GNAS,ALDH6A1,MYH9,IDS,GDI1,FSCN1,SPTAN1,NPM1,PLXNB2,HSPD1,IGF2,EI
,ALDH6A1,SEPTIN2,FSCN1,NPM1,PLD3,USP11,SNRNP200,TRIM28,CKB,FN1,EIF1,CHSY1,EIF4A1,
,ZNF865,RPL9,RPS14,PFN1,MCL1,RPL31,LRPPRC,RPS5,DDOST,NARS1,RPS27,ATP6V0C,HSBP1,
KDC,GABARAP,PFKM,DHX15,BAZ1B,GTPBP1,PI4KA,ADH5,DKK3,FUS,ERH,TPP1,LTBP4,TLE1,GT
167A,CHERP,CTTN,SERP1,ERH,SEC14L1,NUDCD3,GNL1,INTS11,MTMR4,CAP1,AP1M1,ATP13A1,
TL1,PSAP,PGK1,CANX,DDX5,FKBP14,SEC62,KIF1A,DDX17,GPI,CALM3,RPL7,RPL5,KLRD1,MARC
ENG8,CCNI,GNB2,CHPF,RPLP2,MYL6,MYH10,UBE2H,SMCHD1,ATP6V0C,LAPTM4B,MLXIP,HSPH
A,CDK16,L1CAM,CTDSP2,NOVA2,MEST,HNRNPL,TM9SF2,PTOV1,MLEC,RAB11B,PYGB,DPYSL2,
4,JUND,TMEM259,LENG8,CCNI,GNB2,CHPF,RPLP2,ARL6IP1,MYL6,MYH10,UBE2H,SMCHD1,ATP
NB2,CHPF,RPLP2,MYL6,MYH10,UBE2H,SMCHD1,ATP6V0C,LAPTM4B,MLXIP,HSPH1,AHCYL1,KLC
,PLXNB1,YKT6,TMEM167A,CHERP,CTTN,LRP10,ERH,SEC14L1,NUDCD3,GNL1,RIC8A,NCSTN,INT
1,TRAF7,PDAP1,PHB2,PLXNB1,DECR1,FARP1,MCM3AP,RPS27A,KLHDC3,ERH,RALY,SEC14L1,NU
CCSER2,PDHB,ASAP3,ARFGAP3,USP36

[illegible]1,VAPA,STMN1,UCHL1,RPL10,WSB1,HNRNPA0,RPL29,TNS3,CTSD,LAMA5,CIPC,C1ORF43,CHD4,
T3A,IDH1,NXN,CUL3,NDST1,TNKS,DKK3,PPP1R15A,MAP1LC3B,CARM1,GPC2,ELAVL1,ERGIC3,C[illegible]

1,ACTR2,VIM,SPARC,TUBB,FTH1,UBC,KIDINS220,TMSB4X,NUMA1,EIF4G2,RPL3,RPS6,ZNF316,D
SPARC,TUBB,FTH1,UBC,KIDINS220,TMSB4X,NUMA1,EIF4G2,RPL3,ZNF316,DBN1,RPL4,ENO1,TU
M,TUBB,FTH1,UBC,KIDINS220,TMSB4X,NUMA1,RPL3,RPS6,ZNF316,DBN1,RPL4,ENO1,TUBA1B,P
C,TUBB,FTH1,UBC,KIDINS220,TMSB4X,NUMA1,RPL3,RPS6,ZNF316,DBN1,ENO1,TUBA1B,PIK3R2
B,NCL,RPL7A,SLC9A7,ALDOA,PKM,GANAB,COL6A1,HSP90B1,NEFL,NEFM,PRPF8,RPL8,FBXL4,YB
BC,KIDINS220,TMSB4X,NUMA1,EIF4G2,RPL3,RPS6,ZNF316,DBN1,RPL4,ENO1,TUBA1B,PIK3R2,
2,VIM,SPARC,TUBB,FTH1,UBC,KIDINS220,TMSB4X,NUMA1,EIF4G2,RPL3,RPS6,ZNF316,DBN1,RP
X1,UBA1,IGFBP2,CALR,FSTL1,ABHD2,HSPA1B,COL5A1,COL4A1,H3-3B,PTMA,ENO2,SOX4,SEC62
MA1,RPL3,ZNF316,DBN1,ENO1,TUBA1B,PIK3R2,FLNA,MAP1B,NCL,RPL7A,SLC9A7,ALDOA,PKM,G
B4X,NUMA1,EIF4G2,RPL3,RPS6,ZNF316,DBN1,RPL4,ENO1,TUBA1B,PIK3R2,FLNA,MAP1B,NCL,F
3B,FTH1,KIDINS220,TMSB4X,NUMA1,EIF4G2,RPL3,ZNF316,DBN1,RPL4,ENO1,TUBA1B,PIK3R2,F
,DBN1,RPL4,ENO1,TUBA1B,PIK3R2,FLNA,MAP1B,NCL,RPL7A,SLC9A7,MTRNR2L8,PKM,GANAB,
,RPL4,ENO1,TUBA1B,PIK3R2,FLNA,MAP1B,NCL,SLC9A7,MTRNR2L8,PKM,GANAB,COL6A1,FTL,H
16,DBN1,RPL4,ENO1,TUBA1B,PIK3R2,FLNA,MAP1B,NCL,SLC9A7,MTRNR2L8,PKM,GANAB,COL6
N1,ENO1,TUBA1B,PIK3R2,FLNA,NCL,RPL7A,SLC9A7,PKM,GANAB,COL6A1,HSP90B1,FADS1,PRPF
C,KIDINS220,TMSB4X,NUMA1,EIF4G2,RPL3,RPS6,ZNF316,DBN1,RPL4,ENO1,TUBA1B,PIK3R2,FI
,ZNF316,DBN1,RPL4,ENO1,TUBA1B,PIK3R2,FLNA,MAP1B,NCL,RPL7A,SLC9A7,MTRNR2L8,PKM,
1,RPL4,ENO1,TUBA1B,PIK3R2,FLNA,MAP1B,NCL,SLC9A7,PKM,GANAB,COL6A1,FTL,HSP90B1,FA
NCL,SLC9A7,PKM,GANAB,COL6A1,HSP90B1,NEFM,RPL8,FBXL4,YBX1,UBA1,IGFBP2,ABHD2,TUE
A1,RPL3,ZNF316,DBN1,RPL4,ENO1,TUBA1B,PIK3R2,FLNA,MAP1B,NCL,RPL7A,SLC9A7,MTRNR2L
L4,ENO1,TUBA1B,PIK3R2,FLNA,MAP1B,NCL,RPL7A,SLC9A7,ALDOA,PKM,GANAB,FTL,FADS1,NEF
DINS220,TMSB4X,NUMA1,EIF4G2,RPL3,RPS6,ZNF316,ENO1,TUBA1B,PIK3R2,FLNA,MAP1B,NCL,
2,RPS6,ZNF316,ENO1,TUBA1B,PIK3R2,FLNA,MAP1B,NCL,SLC9A7,ALDOA,PKM,GANAB,FTL,HSP
1,ACTR2,VIM,SPARC,TUBB,FTH1,UBC,KIDINS220,TMSB4X,NUMA1,EIF4G2,RPL3,RPS6,ZNF316,D
NF316,DBN1,RPL4,ENO1,TUBA1B,PIK3R2,FLNA,MAP1B,NCL,RPL7A,SLC9A7,ALDOA,PKM,GANAE
C,TUBB,FTH1,UBC,KIDINS220,TMSB4X,NUMA1,EIF4G2,RPL3,ZNF316,DBN1,ENO1,TUBA1B,PIK3
ENO1,TUBA1B,PIK3R2,RPL7A,SLC9A7,ALDOA,PKM,GANAB,COL6A1,HSP90B1,FADS1,NEFM,PRPF
UBC,KIDINS220,TMSB4X,NUMA1,EIF4G2,ZNF316,DBN1,ENO1,TUBA1B,PIK3R2,FLNA,MAP1B,NCI
R2,VIM,TUBB,FTH1,UBC,KIDINS220,TMSB4X,NUMA1,EIF4G2,RPL3,RPS6,ZNF316,DBN1,RPL4,EN
HSP90AA1,ACTR2,VIM,SPARC,TUBB,FTH1,UBC,KIDINS220,TMSB4X,NUMA1,EIF4G2,RPL3,RPS6,
A1,ACTR2,VIM,SPARC,TUBB,FTH1,UBC,TMSB4X,NUMA1,EIF4G2,ZNF316,DBN1,RPL4,ENO1,TUB
N1,RPL4,ENO1,TUBA1B,PIK3R2,FLNA,MAP1B,NCL,SLC9A7,MTRNR2L8,PKM,GANAB,COL6A1,FTI
NF316,DBN1,ENO1,TUBA1B,PIK3R2,FLNA,MAP1B,NCL,SLC9A7,ALDOA,PKM,GANAB,COL6A1,FT
SPARC,TUBB,FTH1,UBC,KIDINS220,TMSB4X,NUMA1,EIF4G2,RPL3,RPS6,ZNF316,DBN1,RPL4,EN
24,HSP90AA1,ACTR2,VIM,SPARC,TUBB,FTH1,UBC,KIDINS220,TMSB4X,NUMA1,EIF4G2,RPL3,RF
ICL,SLC9A7,PKM,GANAB,COL6A1,FTL,HSP90B1,FADS1,NEFM,RPL8,FBXL4,YBX1,UBA1,IGFBP2,F
SB4X,NUMA1,EIF4G2,RPL3,RPS6,ZNF316,DBN1,RPL4,ENO1,TUBA1B,PIK3R2,FLNA,MAP1B,NCL,F
316,DBN1,RPL4,ENO1,TUBA1B,PIK3R2,FLNA,MAP1B,NCL,RPL7A,SLC9A7,ALDOA,PKM,HSP90B1,
B,NCL,RPL7A,SLC9A7,PKM,GANAB,COL6A1,HSP90B1,FADS1,NEFM,UGT8,RPL8,FBXL4,YBX1,IG
C,TUBB,FTH1,UBC,KIDINS220,NUMA1,EIF4G2,RPL3,ZNF316,DBN1,RPL4,ENO1,TUBA1B,PIK3R2,
,SPARC,TUBB,FTH1,UBC,KIDINS220,TMSB4X,NUMA1,EIF4G2,RPL3,ZNF316,DBN1,ENO1,TUBA1
R2,VIM,TUBB,FTH1,UBC,KIDINS220,TMSB4X,NUMA1,EIF4G2,RPS6,ZNF316,DBN1,RPL4,ENO1,TU
BA1B,PIK3R2,FLNA,NCL,RPL7A,SLC9A7,ALDOA,PKM,GANAB,FTL,HSP90B1,FADS1,NEFL,NEFM,U
220,TMSB4X,NUMA1,RPL3,ZNF316,DBN1,RPL4,ENO1,TUBA1B,PIK3R2,FLNA,MAP1B,NCL,RPL7A
C,TUBB,FTH1,UBC,KIDINS220,TMSB4X,NUMA1,EIF4G2,RPL3,ZNF316,DBN1,RPL4,ENO1,TUBA1B

ACTR2,VIM,SPARC,FTH1,UBC,KIDINS220,TMSB4X,NUMA1,EIF4G2,RPL3,ZNF316,DBN1,RPL4,EN(
3N1,ENO1,TUBA1B,PIK3R2,FLNA,MAP1B,NCL,RPL7A,SLC9A7,ALDOA,PKM,GANAB,COL6A1,HSP9
3,FBXL4,YBX1,IGFBP2,FSTL1,ABHD2,TUBB2B,COL5A1,CES3,PGK1,PTMA,ENO2,SOX4,SEC62,KA
1,VIM,TUBB,FTH1,UBC,TMSB4X,NUMA1,EIF4G2,RPL3,RPS6,ZNF316,DBN1,ENO1,TUBA1B,PIK3F
F124,HSP90AA1,ACTR2,VIM,SPARC,FTH1,UBC,KIDINS220,TMSB4X,NUMA1,EIF4G2,RPL3,RPS6,Z
6,DBN1,ENO1,PIK3R2,FLNA,MAP1B,NCL,SLC9A7,PKM,GANAB,HSP90B1,FADS1,RPL8,FBXL4,YB
AP1B,NCL,SLC9A7,PKM,GANAB,COL6A1,HSP90B1,FADS1,NEFM,RPL8,FBXL4,YBX1,IGFBP2,CALI
,RPL3,RPS6,ZNF316,DBN1,RPL4,ENO1,TUBA1B,PIK3R2,FLNA,NCL,RPL7A,SLC9A7,ALDOA,MTRN
FBXL4,YBX1,UBA1,ABHD2,HSPA1B,TUBB2B,PGK1,DDX5,ENO2,KAT6A,DVL3,KIF1A,GNAS,SLC2A3
ISB4X,NUMA1,EIF4G2,RPL3,RPS6,ZNF316,DBN1,ENO1,TUBA1B,PIK3R2,FLNA,MAP1B,NCL,RPL7
3B2B,CES3,PTMA,ENO2,KAT6A,DVL3,YWHAE,KIF1A,GNAS,SLC2A3,GPI,BSG,FASN,ALDH6A1,SPTE
1,PRPF8,RPL8,FBXL4,YBX1,IGFBP2,ABHD2,HSPA1B,TUBB2B,COL5A1,PGK1,DDX5,ENO2,SEC62,
31,FADS1,RPL8,FBXL4,YBX1,IGFBP2,CALR,FSTL1,ABHD2,TUBB2B,COL5A1,PSAP,PGK1,PTMA,DI
L6A1,HSP90B1,FADS1,NEFM,PRPF8,FBXL4,UBA1,CALR,ABHD2,TUBB2B,PGK1,CANX,DDX5,ENO
F4G2,RPL3,RPS6,ZNF316,DBN1,ENO1,TUBA1B,PIK3R2,FLNA,MAP1B,NCL,RPL7A,SLC9A7,ALDO
KAT6A,YWHAE,KIF1A,GNAS,SLC2A3,GPI,BSG,FASN,SPTBN1,GNB1,SHISA9,STMN2,RPS8,MYH9,SI
CTR2,VIM,FTH1,UBC,TMSB4X,NUMA1,EIF4G2,RPL3,RPS6,ZNF316,DBN1,ENO1,TUBA1B,PIK3R2,F
,PKM,GANAB,COL6A1,HSP90B1,FADS1,NEFM,RPL8,FBXL4,UBA1,IGFBP2,FSTL1,ABHD2,TUBB2E
L4,YBX1,IGFBP2,ABHD2,TUBB2B,PTMA,DDX5,ENO2,SOX4,KAT6A,DVL3,YWHAE,KIF1A,GNAS,SLC
ENO1,TUBA1B,PIK3R2,FLNA,MAP1B,NCL,SLC9A7,PKM,GANAB,COL6A1,FTL,HSP90B1,NEFM,UG
2,RPL3,ZNF316,DBN1,ENO1,TUBA1B,PIK3R2,FLNA,MAP1B,NCL,RPL7A,SLC9A7,ALDOA,PKM,GAI
1,RPL8,FBXL4,ABHD2,HSPA1B,TUBB2B,PGK1,DDX5,ENO2,KAT6A,DVL3,KIF1A,GNAS,SLC2A3,R
TH1,UBC,KIDINS220,TMSB4X,NUMA1,EIF4G2,RPL3,RPS6,ZNF316,DBN1,RPL4,ENO1,TUBA1B,PIK
MAP1B,NCL,SLC9A7,PKM,HSP90B1,FADS1,FBXL4,YBX1,CALR,TUBB2B,COL5A1,COL4A1,PTMA,D
A1,HSP90B1,RPL8,FBXL4,UBA1,CALR,ABHD2,HSPA1B,TUBB2B,PGK1,CANX,PTMA,DDX5,ENO2,K
OL6A1,HSP90B1,FADS1,PRPF8,RPL8,FBXL4,YBX1,UBA1,IGFBP2,FSTL1,ABHD2,TUBB2B,COL5A1
TUBA1B,PIK3R2,FLNA,MAP1B,NCL,RPL7A,SLC9A7,ALDOA,PKM,COL6A1,FTL,HSP90B1,FADS1,NI
SP90B1,NEFM,UGT8,YBX1,IGFBP2,ABHD2,TUBB2B,COL5A1,COL4A1,PGK1,CLU,PTMA,DDX5,EN
TR2,VIM,SPARC,FTH1,UBC,KIDINS220,TMSB4X,NUMA1,EIF4G2,RPL3,RPS6,ZNF316,DBN1,RPL4,
3C,NUMA1,EIF4G2,RPL3,ZNF316,DBN1,RPL4,ENO1,TUBA1B,PIK3R2,MAP1B,NCL,RPL7A,SLC9A7
34X,NUMA1,DCX,RPL3,RPS6,ZNF316,DBN1,RPL4,ENO1,TUBA1B,PIK3R2,FLNA,MAP1B,NCL,RPL7
1,PIK3R2,FLNA,MAP1B,NCL,RPL7A,SLC9A7,PKM,HSP90B1,FADS1,NEFM,RPL8,FBXL4,YBX1,CAL
UBC,KIDINS220,TMSB4X,NUMA1,DCX,EIF4G2,RPL3,RPS6,ZNF316,DBN1,RPL4,ENO1,TUBA1B,PI
IUMA1,EIF4G2,RPL3,RPS6,ZNF316,DBN1,ENO1,TUBA1B,PIK3R2,FLNA,MAP1B,NCL,RPL7A,SLC9
,RPL3,ZNF316,DBN1,RPL4,ENO1,TUBA1B,PIK3R2,FLNA,MAP1B,NCL,RPL7A,SLC9A7,PKM,GANA
L7A,SLC9A7,PKM,GANAB,COL6A1,HSP90B1,NEFM,RPL8,FBXL4,YBX1,UBA1,IGFBP2,FSTL1,ABH
RPL8,FBXL4,CALR,ABHD2,HSPA1B,TUBB2B,PGK1,DDX5,ENO2,KAT6A,DVL3,YWHAE,KIF1A,GNAS,
0AA1,ACTR2,VIM,SPARC,TUBB,FTH1,UBC,KIDINS220,TMSB4X,NUMA1,EIF4G2,RPL3,RPS6,ZNF3
IAP1B,NCL,RPL7A,SLC9A7,ALDOA,PKM,COL6A1,FADS1,NEFM,RPL8,FBXL4,YBX1,UBA1,ATP2B4,I
,MTRNR2L8,PKM,GANAB,COL6A1,FTL,HSP90B1,NEFM,RPL8,FBXL4,UBA1,IGFBP2,FSTL1,ABHD
,MTRNR2L8,PKM,GANAB,COL6A1,FTL,HSP90B1,NEFM,RPL8,FBXL4,UBA1,IGFBP2,FSTL1,ABHD
ISP90B1,FADS1,FBXL4,YBX1,CALR,TUBB2B,COL5A1,PTMA,DDX5,ENO2,SOX4,SEC62,KAT6A,DVL
P90AA1,ACTR2,VIM,SPARC,TUBB,FTH1,UBC,KIDINS220,TMSB4X,NUMA1,EIF4G2,RPL3,RPS6,ZNI
0B1,NEFM,PRPF8,FBXL4,YBX1,ATP2B4,IGFBP2,FSTL1,TUBB2B,COL5A1,COL4A1,PSAP,PGK1,RI
CL,RPL7A,SLC9A7,PKM,GANAB,COL6A1,FADS1,NEFL,PRPF8,RPL8,FBXL4,YBX1,IGFBP2,CALR,AE

C9A7,MTRNR2L8,PKM,GANAB,COL6A1,FTL,HSP90B1,NEFM,RPL8,FBXL4,YBX1,UBA1,IGFBP2,FS
NAB,COL6A1,FTL,HSP90B1,NEFM,RPL8,FBXL4,UBA1,FSTL1,TUBB2B,PGK1,CANX,DDX5,FKBP1
1,NEFM,UGT8,RPL8,YBX1,IGFBP2,FSTL1,ABHD2,TUBB2B,COL4A1,PGK1,CLU,PTMA,ENO2,SO
EFM,UGT8,RPL8,FBXL4,UBA1,TUBB2B,CES3,PTMA,ENO2,SOX4,SEC62,KAT6A,YWHAE,KIF1A,GN
,ZNF316,DBN1,ENO1,TUBA1B,FLNA,MAP1B,NCL,RPL7A,SLC9A7,PKM,GANAB,FTL,HSP90B1,NE
HSP90AA1,ACTR2,VIM,SPARC,TUBB,FTH1,UBC,KIDINS220,TMSB4X,NUMA1,EIF4G2,RPL3,RPS6,
,SEC62,KAT6A,DVL3,YWHAE,KIF1A,GNAS,HTRA4,FASN,SPTBN1,GNB1,SHISA9,MYH9,RPL7,P4HB
L,PKM,GANAB,COL6A1,HSP90B1,FADS1,NEFM,RPL8,FBXL4,YBX1,UBA1,IGFBP2,FSTL1,ABHD2,
TR2,VIM,SPARC,TUBB,FTH1,UBC,KIDINS220,TMSB4X,NUMA1,DCX,EIF4G2,RPL3,ZNF316,DBN1,
TUBA1B,PIK3R2,MAP1B,NCL,RPL7A,SLC9A7,PKM,GANAB,COL6A1,FTL,HSP90B1,FADS1,PRPF8,
IF4G2,ZNF316,DBN1,RPL4,ENO1,TUBA1B,PIK3R2,MAP1B,NCL,RPL7A,SLC9A7,ALDOA,PKM,GAN
B,NCL,SLC9A7,PKM,GANAB,COL6A1,FTL,HSP90B1,FADS1,NEFM,PRPF8,RPL8,FBXL4,YBX1,UBA
SLC9A7,HSP90B1,FBXL4,YBX1,CALR,TUBB2B,COL5A1,COL4A1,PTMA,DDX5,ENO2,SEC62,KAT6
ANAB,FTL,HSP90B1,NEFM,UGT8,RPL8,YBX1,IGFBP2,FSTL1,ABHD2,TUBB2B,COL5A1,COL4A1,P
ACTR2,VIM,SPARC,TUBB,FTH1,UBC,KIDINS220,TMSB4X,NUMA1,DCX,EIF4G2,RPL3,ZNF316,DB
,RPL7A,SLC9A7,PKM,GANAB,COL6A1,FTL,FADS1,NEFM,PRPF8,RPL8,YBX1,ATP2B4,CALR,FSTL
SG,FASN,ALDH6A1,SPTBN1,GNB1,SHISA9,STMN2,RPL7,SEPTIN2,GDI1,SPTAN1,MARCKS,SMG1,
,NUMA1,RPL3,ZNF316,DBN1,RPL4,ENO1,TUBA1B,PIK3R2,FLNA,MAP1B,NCL,RPL7A,MTRNR2L8,
FSTL1,ZNF99,COL5A1,COL4A1,CES3,PSAP,RPLP0,PTMA,DDX5,ENO2,SOX4,RPL13A,SEC62,KAT6
2B,COL5A1,PSAP,PGK1,PTMA,ENO2,SEC62,KAT6A,DVL3,YWHAE,KIF1A,GNAS,SLC2A3,HTRA4,B
L,NEFM,PRPF8,YBX1,ATP2B4,IGFBP2,CALR,FSTL1,TUBB2B,COL5A1,COL4A1,PSAP,PGK1,H3-3B
T6A,DVL3,YWHAE,KIF1A,GNAS,BSG,FASN,SPTBN1,GNB1,STMN2,RPL7,SEPTIN2,APP,GDI1,FSCN
IAE,KIF1A,GNAS,FASN,SPTBN1,CALM3,GNB1,SHISA9,STMN2,GDI1,PTMS,SMG1,CALM1,PLXNB2,
N1,GNB1,SHISA9,RPL7,P4HB,APP,SMG1,CALM1,HSPD1,IGF2,MARCKSL1,TRIM28,CKB,DDB1,YW
MA1,EIF4G2,ZNF316,DBN1,ENO1,FLNA,MAP1B,NCL,RPL7A,SLC9A7,PKM,GANAB,COL6A1,HSP90
,PTMS,SPTAN1,SMG1,PLXNB2,IGF2,MARCKSL1,AFF4,TRIM28,MATR3,YWHAG,CKB,COL4A2,SP
PIK3R2,FLNA,MAP1B,NCL,SLC9A7,ALDOA,PKM,GANAB,HSP90B1,FADS1,NEFM,FBXL4,YBX1,IGF
,PIK3R2,MAP1B,NCL,RPL7A,SLC9A7,ALDOA,PKM,GANAB,COL6A1,HSP90B1,NEFL,NEFM,PRPF8,
P1B,NCL,RPL7A,SLC9A7,ALDOA,PKM,GANAB,COL6A1,FTL,HSP90B1,NEFM,UGT8,RPL8,YBX1,U
SB4X,EIF4G2,RPS6,ZNF316,DBN1,ENO1,FLNA,MAP1B,NCL,SLC9A7,ALDOA,GANAB,COL6A1,HSP
ACTR2,VIM,TUBB,FTH1,UBC,KIDINS220,TMSB4X,NUMA1,EIF4G2,RPL3,RPS6,ZNF316,DBN1,RPL
B1,PRPF8,RPL8,YBX1,UBA1,ATP2B4,IGFBP2,FSTL1,TUBB2B,COL5A1,COL4A1,CES3,PSAP,PGK1
H6A1,SPTBN1,GNB1,STMN2,MYH9,RPL7,SEPTIN2,FSCN1,MARCKS,SMG1,CALM1,NPM1,PLXNB
1,FADS1,NEFM,PRPF8,RPL8,FBXL4,YBX1,UBA1,CALR,ABHD2,TUBB2B,PSAP,CANX,DDX5,ENO2,S
NF316,DBN1,ENO1,TUBA1B,PIK3R2,FLNA,MAP1B,NCL,SLC9A7,PKM,GANAB,COL6A1,FTL,HSP90
,TMSB4X,NUMA1,EIF4G2,ZNF316,DBN1,ENO1,TUBA1B,PIK3R2,MAP1B,NCL,RPL7A,SLC9A7,AL
SCN1,PTMS,SPTAN1,GSTP1,SMG1,PLXNB2,PLD3,USP11,EIF4B,MARCKSL1,COL4A2,RPL37A,SP
PSAP,PGK1,H3-3B,CLU,PTMA,ENO2,RPL13A,SEC62,KAT6A,DVL3,YWHAE,KIF1A,RPL13,GNAS,FA
,RPL7,P4HB,GDI1,PTMS,MARCKS,CALM1,PLXNB2,USP11,IGF2,EIF3C,EIF4B,AFF4,TRIM28,MAT
,COL6A1,FTL,HSP90B1,FADS1,NEFM,UGT8,RPL8,YBX1,IGFBP2,TUBB2B,COL5A1,COL4A1,PGK
,DDX5,RPL13A,SEC62,KAT6A,KIF1A,RPL13,GNAS,SLC2A3,BSG,FASN,SPTBN1,GNB1,SHISA9,ST
1,COL4A1,RPLP0,H3-3B,CANX,PTMA,ENO2,GNAS,DDX17,RACK1,BSG,CALM3,GNB1,RPL7,SEPTI
1,GANAB,COL6A1,FADS1,NEFM,PRPF8,RPL8,FBXL4,YBX1,UBA1,ATP2B4,CALR,FSTL1,ABHD2,HS
FBXL4,IGFBP2,ABHD2,TUBB2B,PGK1,CANX,PTMA,DDX5,ENO2,FKBP14,KAT6A,DVL3,YWHAE,SL
AA1,VIM,SPARC,TUBB,FTH1,UBC,KIDINS220,TMSB4X,NUMA1,DCX,RPL3,RPS6,ZNF316,DBN1,R

AB,COL6A1,FTL,HSP90B1,FADS1,NEFL,RPL8,FBXL4,YBX1,UBA1,IGFBP2,CALR,ABHD2,TUBB2B,C
PTBN1,GNB1,SHISA9,P4HB,ERVV-1,GDI1,PTMS,GSTP1,SMG1,USP11,SMIM14,IGF2,TRIM28,SPEN
WHAE,KIF1A,GNAS,HTRA4,BSG,FASN,SPTBN1,CALM3,GNB1,MYH9,RPL7,P4HB,IDS,RPL5,PTMS,S
2,FLNA,MAP1B,NCL,SLC9A7,ALDOA,PKM,HSP90B1,FADS1,NEFL,NEFM,PRPF8,RPL8,FBXL4,UBA
R2,FLNA,MAP1B,NCL,RPL7A,SLC9A7,ALDOA,PKM,COL6A1,HSP90B1,NEFL,NEFM,PRPF8,RPL8,FE
G2,RPS6,ZNF316,DBN1,ENO1,TUBA1B,FLNA,MAP1B,NCL,RPL7A,SLC9A7,ALDOA,PKM,GANAB,CO
B,HSP90B1,FADS1,PRPF8,RPL8,IGFBP2,FSTL1,ABHD2,TUBB2B,PGK1,DDX5,ENO2,FKBP14,DVL3
OA,PKM,GANAB,COL6A1,FTL,FADS1,NEFM,PRPF8,RPL8,YBX1,IGFBP2,CALR,ABHD2,COL5A1,CO
B4X,NUMA1,DCX,RPL3,RPS6,ZNF316,DBN1,RPL4,ENO1,PIK3R2,FLNA,MAP1B,NCL,RPL7A,SLC9A
M,TUBB,FTH1,UBC,TMSB4X,NUMA1,EIF4G2,RPL3,RPS6,ZNF316,DBN1,RPL4,ENO1,TUBA1B,PIK3
S,DDX17,GPI,RACK1,BSG,FASN,ALDH6A1,CALM3,SEPTIN2,P4HB,IDS,GDI1,GSTP1,NPM1,COL18A
7,SEPTIN2,P4HB,GDI1,FSCN1,PTMS,SPTAN1,GSTP1,MARCKS,SMG1,CALM1,USP11,CSDE1,IGF2,I
I,SPTAN1,GSTP1,MARCKS,SMG1,USP11,IGF2,EIF4B,MARCKSL1,TRIM28,MATR3,YWHAG,CKB,LF
C9A7,MTRNR2L8,COL6A1,FADS1,NEFL,NEFM,UGT8,RPL8,FBXL4,CALR,FSTL1,TMF1,ABHD2,TUE
A7,ALDOA,PKM,GANAB,FTL,HSP90B1,NEFL,RPL8,FBXL4,YBX1,UBA1,IGFBP2,ABHD2,TUBB2B,CC
TUBA1B,PIK3R2,FLNA,MAP1B,NCL,RPL7A,SLC9A7,ALDOA,COL6A1,FADS1,NEFL,NEFM,PRPF8,RF
4HB,APP,GDI1,PTMS,SPTAN1,GSTP1,MARCKS,CALM1,PLXNB2,HSPD1,USP11,IGF2,EIF4B,MARCK
PI,ALDH6A1,SPTBN1,CALM3,GNB1,SHISA9,STMN2,SEPTIN2,IDS,MARCKS,SMG1,NPM1,PLXNB2,U
3,FTH1,UBC,KIDINS220,TMSB4X,NUMA1,EIF4G2,RPL3,RPS6,ZNF316,DBN1,RPL4,ENO1,TUBA1B,
PS6,ZNF316,DBN1,ENO1,TUBA1B,PIK3R2,FLNA,MAP1B,NCL,RPL7A,SLC9A7,ALDOA,COL6A1,FAD
6,DBN1,ENO1,TUBA1B,PIK3R2,FLNA,MAP1B,NCL,SLC9A7,PKM,GANAB,COL6A1,FTL,HSP90B1,F
FLNA,MAP1B,NCL,PKM,COL6A1,FTL,HSP90B1,NEFM,UGT8,RPL8,FBXL4,UBA1,IGFBP2,CALR,FST
A,MAP1B,NCL,RPL7A,SLC9A7,PKM,GANAB,COL6A1,FTL,HSP90B1,NEFL,UGT8,RPL8,FBXL4,YBX
F124,HSP90AA1,ACTR2,VIM,SPARC,TUBB,FTH1,UBC,KIDINS220,TMSB4X,NUMA1,EIF4G2,RPL3,F
N,ATP5F1B,GNB1,RPS8,MYH9,SEPTIN2,P4HB,IDS,FSCN1,SPTAN1,GSTP1,SMG1,PLXNB2,HSPD1,U
TUBB2B,COL4A1,RPLP0,DDX5,ENO2,SOX4,GNAS,SLC2A3,GPI,HTRA4,BSG,FASN,SPTBN1,GNB1,S
N,ALDH6A1,SPTBN1,CALM3,GNB1,MYH9,SEPTIN2,GDI1,PTMS,SPTAN1,SMG1,CALM1,SMIM14,IG
S220,TMSB4X,NUMA1,EIF4G2,RPL3,RPS6,ZNF316,DBN1,RPL4,ENO1,TUBA1B,PIK3R2,MAP1B,NC
FSTL1,TUBB2B,PTMA,FKBP14,KAT6A,DVL3,YWHAE,KIF1A,GNAS,SLC2A3,GPI,BSG,FASN,SPTBN1,
UMA1,EIF4G2,RPL3,ZNF316,DBN1,RPL4,ENO1,TUBA1B,PIK3R2,FLNA,NCL,RPL7A,SLC9A7,ALDO
HB,FSCN1,MARCKS,CALM1,PLXNB2,HSPD1,COL18A1,IGF2,EIF4B,MARCKSL1,MATR3,YWHAG,CI
AT6A,DVL3,KIF1A,GNAS,GPI,BSG,FASN,ALDH6A1,GNB1,STMN2,RPS8,MYH9,P4HB,APP,GDI1,FSCI
C9A7,ALDOA,PKM,FADS1,NEFL,PRPF8,RPL8,FBXL4,TMF1,ZNF99,TUBB2B,CES3,PSAP,CANX,PTM
M,UGT8,RPL8,FBXL4,YBX1,ATP2B4,FSTL1,ABHD2,HSPA1B,TUBB2B,ENO2,SOX4,KAT6A,DVL3,YW
6,ZNF316,DBN1,RPL4,ENO1,TUBA1B,PIK3R2,MAP1B,NCL,RPL7A,MTRNR2L8,GANAB,COL6A1,F
XNB2,USP11,IGF2,ATP5F1A,EIF4B,MARCKSL1,AFF4,TRIM28,MATR3,YWHAG,CKB,NONO,YWHAQ
,SPTAN1,MARCKS,SMG1,CALM1,USP11,IGF2,EIF4B,MARCKSL1,AFF4,TRIM28,MATR3,YWHAG,C
ALDOA,PKM,COL6A1,HSP90B1,FADS1,NEFM,UGT8,RPL8,FBXL4,YBX1,UBA1,ATP2B4,IGFBP2,CA
B1,FBXL4,YBX1,TUBB2B,COL5A1,H3-3B,PTMA,DDX5,SEC62,YWHAE,KIF1A,GNAS,SLC2A3,DDX17
BHD2,TUBB2B,PGK1,CANX,DDX5,ENO2,KAT6A,DVL3,KIF1A,GNAS,SLC2A3,GPI,BSG,FASN,ALDH6
L7A,SLC9A7,ALDOA,PKM,GANAB,COL6A1,FTL,HSP90B1,NEFL,RPL8,FBXL4,UBA1,ATP2B4,CALR,
28,MATR3,YWHAG,CKB,NONO,YWHAQ,HACD3,DCHS1,TMSB10,MRC2,LDHB,PKD1,HLA-C,SSBP2
A1,FTL,FADS1,FBXL4,YBX1,ATP2B4,IGFBP2,CALR,FSTL1,HSPA1B,TUBB2B,COL4A1,PGK1,CANX,
A1,FTL,FADS1,FBXL4,YBX1,ATP2B4,IGFBP2,CALR,FSTL1,HSPA1B,TUBB2B,COL4A1,PGK1,CANX,
A1,FTL,FADS1,FBXL4,YBX1,ATP2B4,IGFBP2,CALR,FSTL1,HSPA1B,TUBB2B,COL4A1,PGK1,CANX,

:3R2,FLNA,NCL,RPL7A,SLC9A7,ALDOA,PKM,GANAB,FTL,HSP90B1,NEFL,NEFM,RPL8,FBXL4,YBX
I,CALR,FSTL1,TUBB2B,COL5A1,COL4A1,H3-3B,CLU,PTMA,SOX4,SEC62,KAT6A,YWHAE,KIF1A,GI
,RPL4,ENO1,TUBA1B,PIK3R2,FLNA,MAP1B,NCL,RPL7A,ALDOA,PKM,GANAB,FTL,HSP90B1,FADS
,TUBB2B,COL4A1,CLU,PTMA,DDX5,ENO2,SOX4,KAT6A,DVL3,YWHAE,KIF1A,GNAS,FASN,ALDH6A
P11,IGF2,ATP5F1A,EIF4B,MARCKSL1,TRIM28,MATR3,YWHAG,CKB,NONO,YWHAQ,HACD3,DCHS
A1,ACTR2,VIM,SPARC,TUBB,FTH1,UBC,KIDINS220,TMSB4X,NUMA1,DCX,EIF4G2,RPL3,RPS6,ZNF
BB2B,PSAP,PTMA,DVL3,KIF1A,GNAS,SLC2A3,BSG,FASN,ALDH6A1,SPTBN1,GNB1,SHISA9,STMN2
FADS1,RPL8,COL5A1,COL4A1,RPLP0,H3-3B,CANX,PTMA,ENO2,RPL6,KAT6A,DVL3,YWHAE,GNAS,
OL6A1,FADS1,NEFL,YBX1,UBA1,IGFBP2,CALR,FSTL1,ABHD2,TUBB2B,COL5A1,COL4A1,CES3,PS
,CALM3,GNB1,SHISA9,STMN2,MYH9,SEPTIN2,GDI1,PTMS,SPTAN1,GSTP1,PLXNB2,USP11,IGF2,I
SET,EGLN1,CHSY1,TMSB10,MRC2,PKD1,PDIA3,VAPA,SEC61A1,STMN1,TIMP2,SARAF,WSB1,VCL,
S,DBN1,RPL4,ENO1,TUBA1B,MAP1B,NCL,SLC9A7,ALDOA,PKM,GANAB,COL6A1,FTL,HSP90B1,FA
L3,RPS6,ZNF316,DBN1,ENO1,TUBA1B,PIK3R2,FLNA,MAP1B,NCL,RPL7A,SLC9A7,ALDOA,PKM,F
,APP,IDS,PTMS,SPTAN1,CALM1,USP11,IGF2,EIF4B,MARCKSL1,TRIM28,MATR3,YWHAG,CKB,DDE
S1,NEFM,PRPF8,RPL8,YBX1,UBA1,IGFBP2,CALR,FSTL1,TUBB2B,COL5A1,COL4A1,PSAP,PGK1,C
MA1,RPS6,ZNF316,ENO1,TUBA1B,PIK3R2,MAP1B,NCL,RPL7A,PKM,GANAB,FTL,HSP90B1,FADS1
AA1,ACTR2,VIM,SPARC,TUBB,FTH1,UBC,KIDINS220,TMSB4X,NUMA1,EIF4G2,RPL3,RPS6,ZNF31
2,YWHAE,GNAS,BSG,FASN,SPTBN1,GNB1,MYH9,SEPTIN2,GDI1,FSCN1,PTMS,SPTAN1,GSTP1,CA
B4X,NUMA1,EIF4G2,RPL3,RPS6,ZNF316,TUBA1B,PIK3R2,FLNA,RPL7A,SLC9A7,ALDOA,MTRNR2I
2,TUBB2B,CANX,DDX5,ENO2,KAT6A,DVL3,KIF1A,GNAS,GPI,BSG,FASN,ALDH6A1,CALM3,GNB1,S
F4A1,PKD1,TPT1,STMN1,ITGB1,MORF4L1,NCAM1,AGRN,ZMIZ1,MIDN,VCL,DNAJA1,ARF1,CDC42I
6,ZNF316,ENO1,TUBA1B,PIK3R2,FLNA,MAP1B,NCL,RPL7A,SLC9A7,ALDOA,MTRNR2L8,COL6A1,
TL1,TUBB2B,COL4A1,PGK1,PTMA,DDX5,ENO2,RPL13A,KAT6A,YWHAE,KIF1A,GNAS,SLC2A3,DDX
ALDH6A1,SPTBN1,GNB1,SHISA9,STMN2,MYH9,SEPTIN2,GDI1,TMBIM6,SPTAN1,CALM1,PLXNB2,
TUBB2B,COL5A1,CES3,H3-3B,PTMA,SEC62,DVL3,YWHAE,GNAS,SLC2A3,DDX17,GPI,FASN,SPTBN
,MAP1B,NCL,RPL7A,SLC9A7,ALDOA,PKM,GANAB,COL6A1,HSP90B1,NEFM,PRPF8,RPL8,FBXL4,U
L18A1,SMIM14,IGF2,MARCKSL1,TRIM28,CKB,LRRC28,SPEN,YWHAQ,HACD3,DCHS1,SET,EGLN1
,NUMA1,EIF4G2,RPL3,ZNF316,ENO1,TUBA1B,PIK3R2,FLNA,NCL,RPL7A,SLC9A7,ALDOA,PKM,CC
A7,ALDOA,PKM,GANAB,COL6A1,HSP90B1,FADS1,PRPF8,FBXL4,YBX1,UBA1,IGFBP2,TMF1,ABHD
PRPF8,YBX1,UBA1,IGFBP2,CALR,FSTL1,ABHD2,TUBB2B,COL5A1,COL4A1,PSAP,RPLP0,H3-3B,C
SAP,PGK1,RPLP0,H3-3B,CANX,PTMA,DDX5,ENO2,SOX4,RPL6,KAT6A,DVL3,YWHAE,KIF1A,DDX17
BX1,UBA1,ATP2B4,IGFBP2,FSTL1,COL5A1,COL4A1,CES3,H3-3B,PTMA,SOX4,YWHAE,GNAS,DDX
GNAS,DDX17,HTRA4,BSG,FASN,GNB1,MYH9,RPL7,P4HB,APP,FSCN1,SPTAN1,GSTP1,SMG1,NPM
,TUBA1B,PIK3R2,FLNA,NCL,RPL7A,SLC9A7,ALDOA,MTRNR2L8,PKM,COL6A1,HSP90B1,NEFL,RF
OA,PKM,COL6A1,FTL,HSP90B1,FADS1,NEFM,UGT8,RPL8,FBXL4,YBX1,ATP2B4,IGFBP2,ABHD2,
TT,HSP90AB1,ZNF124,HSP90AA1,ACTR2,VIM,SPARC,TUBB,FTH1,UBC,KIDINS220,TMSB4X,NUM
PTMS,MARCKS,HSPD1,USP11,SMIM14,IGF2,MARCKSL1,MATR3,YWHAG,CKB,PSMD5,DCHS1,TPI
CKSL1,AFF4,TRIM28,MATR3,YWHAG,CKB,YWHAQ,DCHS1,SET,CHSY1,PKD1,VAPA,TPT1,ITGB1,M
EIF4B,MARCKSL1,MATR3,YWHAG,CKB,YWHAQ,DCHS1,SET,CHSY1,PKD1,VAPA,TPT1,ITGB1,AGR
124,HSP90AA1,ACTR2,VIM,SPARC,TUBB,FTH1,UBC,KIDINS220,TMSB4X,NUMA1,EIF4G2,ZNF316
NEFM,PRPF8,RPL8,YBX1,UBA1,ATP2B4,IGFBP2,CALR,FSTL1,TMF1,ABHD2,COL5A1,COL4A1,PSA
HSP90B1,FADS1,NEFL,NEFM,RPL8,FBXL4,YBX1,UBA1,FSTL1,TMF1,ABHD2,HSPA1B,COL5A1,CC
X1,ATP2B4,TMF1,TUBB2B,COL4A1,PSAP,PGK1,H3-3B,CANX,PTMA,ENO2,MAGED1,DVL3,GNAS,I
VL3,KIF1A,GNAS,GPI,BSG,FASN,ALDH6A1,GNB1,STMN2,MYH9,RPL7,P4HB,IDS,ERVV-1,GDI1,FS
)1,TUBA1B,PIK3R2,FLNA,MAP1B,NCL,RPL7A,SLC9A7,ALDOA,PKM,COL6A1,FTL,HSP90B1,NEFM,

ALDOA,PKM,GANAB,COL6A1,HSP90B1,RPL8,FBXL4,YBX1,UBA1,IGFBP2,CALR,FSTL1,TUBB2B,C
RPL7A,SLC9A7,ALDOA,PKM,GANAB,COL6A1,HSP90B1,RPL8,FBXL4,YBX1,UBA1,IGFBP2,CALR,FS
,PGK1,H3-3B,PTMA,DDX5,ENO2,KAT6A,DVL3,KIF1A,RPL13,RACK1,BSG,FASN,SPTBN1,CALM3,G
ALR,ABHD2,HSPA1B,TUBB2B,PSAP,DDX5,ENO2,DVL3,GNAS,SPTBN1,CALM3,GNB1,STMN2,MYH9
EFM,PRPF8,RPL8,UBA1,IGFBP2,FSTL1,ABHD2,COL5A1,COL4A1,RPLP0,CANX,CLU,PTMA,DDX5,KA
SPTBN1,CALM3,GNB1,MYH9,P4HB,PTMS,SPTAN1,GSTP1,CALM1,NPM1,HSPD1,SMIM14,MARCKS
DS1,NEFL,NEFM,PRPF8,FBXL4,YBX1,ATP2B4,CALR,TMF1,HSPA1B,ZNF99,TUBB2B,COL4A1,RPLI
BX1,UBA1,TUBB2B,CANX,DDX5,FKBP14,KAT6A,DVL3,SLC2A3,GPI,BSG,FASN,ALDH6A1,CALM3,G
PTAN1,GSTP1,MARCKS,SMG1,USP11,IGF2,EIF4B,MARCKSL1,AFF4,TRIM28,YWHAG,CKB,LRRC28
PTAN1,GSTP1,MARCKS,SMG1,USP11,IGF2,EIF4B,MARCKSL1,AFF4,TRIM28,YWHAG,CKB,LRRC28
RPL23,CPE,LAMA5,PJA2,KPNB1,CXADR,CHD4,JUND,WDR1,CRMP1,BEX3,RBBP4,TLE5,CCNI,HDL
HSPD1,SMIM14,EIF4B,UBB,TRIM28,YWHAG,RPL37A,SPEN,EIF1,HACD3,DCHS1,SET,EGLN1,SF3B1
HSP90AA1,ACTR2,VIM,FTH1,UBC,KIDINS220,TMSB4X,NUMA1,EIF4G2,RPL3,RPS6,ZNF316,DBN1,
7A,SLC9A7,MTRNR2L8,GANAB,COL6A1,FTL,NEFL,FBXL4,YBX1,CALR,FSTL1,HSPA1B,TUBB2B,CC
SMG1,PLXNB2,USP11,IGF2,ATP5F1A,MARCKSL1,TRIM28,CKB,LRRC28,NONO,DDB1,YWHAQ,HA
G,CKB,LRRC28,YWHAQ,DCHS1,TMSB10,PKD1,HLA-C,VAPA,POLR2A,TPT1,STMN1,ITGB1,ANXA2,A
SPTAN1,MARCKS,SMG1,PLXNB2,IGF2,EIF4B,MARCKSL1,TRIM28,MATR3,YWHAG,CKB,LRRC28,F
MYH9,RPL7,APP,IDS,GDI1,GSTP1,SMG1,NPM1,HSPD1,IGF2,MARCKSL1,TRIM28,CKB,FN1,DCHS1
,NCL,RPL7A,PKM,GANAB,NEFL,NEFM,RPL8,FBXL4,YBX1,UBA1,CALR,FSTL1,ABHD2,HSPA1B,COL
AB,HSP90B1,NEFM,RPL8,FBXL4,YBX1,UBA1,CALR,FSTL1,TMF1,HSPA1B,TUBB2B,CES3,PSAP,PGK
ENO1,PIK3R2,MAP1B,NCL,SLC9A7,PKM,GANAB,HSP90B1,NEFL,NEFM,PRPF8,RPL8,FBXL4,YBX1,
316,DBN1,ENO1,MAP1B,NCL,RPL7A,SLC9A7,ALDOA,PKM,COL6A1,FTL,FADS1,NEFM,PRPF8,RPL
2,RPL3,RPS6,RPL4,ENO1,TUBA1B,PIK3R2,FLNA,MAP1B,NCL,RPL7A,SLC9A7,ALDOA,MTRNR2L8,
CALM1,PLXNB2,IGF2,EIF4B,MARCKSL1,AFF4,YWHAG,CKB,FN1,HACD3,DCHS1,HLA-C,SSBP2,TPT
62,YWHAE,GNAS,SLC2A3,DDX17,FASN,SPTBN1,GNB1,SHISA9,STMN2,APP,IDS,GDI1,PTMS,SMG1,
P90B1,FADS1,NEFM,UGT8,FBXL4,YBX1,UBA1,ATP2B4,IGFBP2,FSTL1,TUBB2B,COL5A1,COL4A1
2,FLNA,MAP1B,RPL7A,SLC9A7,ALDOA,PKM,COL6A1,FTL,HSP90B1,FADS1,FBXL4,YBX1,FSTL1,TU
BB2B,COL4A1,PSAP,PGK1,RPLP0,H3-3B,PTMA,ENO2,RPL13A,RPL6,RPL13,GNAS,DDX17,BSG,AT
CKB,YWHAQ,EGLN1,STMN1,ITGB1,UCHL1,MORF4L1,AGRN,ZMIZ1,TIMP2,MIDN,CYB5R3,CDC42
H3-3B,CANX,PTMA,ENO2,DVL3,YWHAE,GNAS,BSG,FASN,ALDH6A1,SPTBN1,CALM3,GNB1,MYH9,S
C,KIDINS220,TMSB4X,NUMA1,EIF4G2,RPS6,ZNF316,DBN1,RPL4,ENO1,TUBA1B,PIK3R2,FLNA,M
F2,EIF4B,MARCKSL1,TRIM28,YWHAG,CKB,LRRC28,FN1,YWHAQ,HACD3,DCHS1,SET,TMSB10,PK
,GNB1,STMN2,RPS8,SEPTIN2,GDI1,SPTAN1,SMG1,PLXNB2,HSPD1,USP11,SNRNP200,IGF2,EIF4E
9,SEPTIN2,FSCN1,GSTP1,SMG1,NPM1,PLXNB2,HSPD1,USP11,EIF4B,VCP,AFF4,YWHAG,SPEN,DD
,SEPTIN2,GDI1,FSCN1,SMG1,NPM1,COL18A1,IGF2,MARCKSL1,AFF4,TRIM28,CKB,COL4A2,LRR
PF8,RPL8,YBX1,UBA1,IGFBP2,CALR,HSPA1B,CES3,PGK1,RPLP0,H3-3B,CLU,DDX5,ENO2,FKBP14
KIF1A,GNAS,SLC2A3,GPI,HTRA4,BSG,SPTBN1,CALM3,GNB1,STMN2,RPS8,RPL7,P4HB,GDI1,FSC
CALM3,GNB1,SHISA9,SEPTIN2,IDS,GDI1,PTMS,MARCKS,SMG1,PLXNB2,USP11,IGF2,ATP5F1A,EI
DINS220,TMSB4X,NUMA1,DCX,EIF4G2,RPL3,ZNF316,DBN1,RPL4,TUBA1B,FLNA,MAP1B,NCL,RP
B,FBXL4,YBX1,FSTL1,ABHD2,ZNF99,CES3,PGK1,RPLP0,PTMA,DDX5,SEC62,RPL6,KAT6A,DVL3,Y
A,SLC9A7,MTRNR2L8,GANAB,COL6A1,FTL,HSP90B1,FADS1,NEFL,FBXL4,YBX1,ATP2B4,CALR,FS
A1,FTL,FADS1,NEFL,NEFM,PRPF8,RPL8,FBXL4,YBX1,ATP2B4,IGFBP2,CALR,ABHD2,HSPA1B,COL
SY1,MRC2,GTF3C4,PKD1,HLA-C,PDIA3,SSBP2,ZNF43,TPT1,ITGB1,TUBB4B,ANXA2,ZMIZ1,MIDN,
1,PTMS,SPTAN1,CALM1,PLXNB2,CSDE1,SMIM14,IGF2,EIF4B,MARCKSL1,AFF4,TRIM28,YWHAG,
EFM,YBX1,FSTL1,TUBB2B,COL5A1,CES3,H3-3B,PTMA,DDX5,SOX4,YWHAE,GNAS,DDX17,FASN,SI

C9A7,ALDOA,PKM,GANAB,COL6A1,FADS1,NEFL,NEFM,PRPF8,RPL8,YBX1,UBA1,CALR,TMF1,HSP
0,TMSB4X,NUMA1,EIF4G2,RPL3,RPS6,ZNF316,DBN1,RPL4,TUBA1B,PIK3R2,MAP1B,NCL,SLC9A
BXL4,YBX1,ATP2B4,CALR,HSPA1B,TUBB2B,COL4A1,H3-3B,CANX,CLU,PTMA,RPL13A,SEC62,GP
,ENO1,TUBA1B,PIK3R2,NCL,RPL7A,PKM,GANAB,COL6A1,HSP90B1,FADS1,NEFL,NEFM,FBXL4,Y
K3R2,FLNA,MAP1B,NCL,PKM,GANAB,COL6A1,HSP90B1,FADS1,NEFM,PRPF8,RPL8,FBXL4,YBX1,U
DH6A1,GNB1,STMN2,MYH9,P4HB,GDI1,SMG1,PLXNB2,USP11,IGF2,EIF4B,AFF4,TRIM28,YWHAG,
WHAQ,DCHS1,SET,EGLN1,CHSY1,INA,MRC2,PKD1,SSBP2,VAPA,TPT1,ITGB1,AGRN,ZMIZ1,TIMP2,
,CANX,PTMA,DDX5,ENO2,RPL13A,SEC62,GNAS,GPI,FASN,SPTBN1,GNB1,RPS8,SEPTIN2,APP,IDS,
EIF4G2,RPL3,RPS6,ZNF316,DBN1,ENO1,TUBA1B,FLNA,MAP1B,NCL,RPL7A,ALDOA,PKM,GANAB,
1,PSAP,PGK1,CANX,ENO2,RPL13A,KAT6A,DVL3,KIF1A,RPL13,GNAS,HTRA4,BSG,FASN,SPTBN1,C
OL4A1,PSAP,H3-3B,CANX,PTMA,ENO2,RPL13A,RPL6,KAT6A,DVL3,GNAS,DDX17,BSG,ATP5F1B,CA
T6A,YWHAE,RPL13,GNAS,DDX17,HTRA4,BSG,FASN,SPTBN1,GNB1,MYH9,APP,FSCN1,PTMS,SPTA
A,DDX5,ENO2,RPL6,DVL3,KIF1A,DDX17,HTRA4,BSG,FASN,ALDH6A1,GNB1,MYH9,SEPTIN2,APP,ID
PKM,COL6A1,HSP90B1,NEFM,PRPF8,UGT8,RPL8,FBXL4,YBX1,UBA1,IGFBP2,HSPA1B,TUBB2B,C
0,TMSB4X,NUMA1,EIF4G2,RPL3,RPS6,ZNF316,DBN1,RPL4,TUBA1B,PIK3R2,MAP1B,NCL,SLC9A
D3,COL18A1,SMIM14,IGF2,MARCKSL1,AFF4,TRIM28,COL4A2,YWHAQ,EIF1,DCHS1,SET,CHSY1,I
LDOA,MTRNR2L8,PKM,GANAB,FTL,HSP90B1,FADS1,NEFL,NEFM,UGT8,RPL8,FBXL4,YBX1,UBA1
B2B,PSAP,PTMA,DDX5,SOX4,KAT6A,DVL3,DDX17,FASN,ALDH6A1,ATP5F1B,CALM3,GNB1,MYH9,S
RPL3,ZNF316,DBN1,ENO1,TUBA1B,PIK3R2,FLNA,MAP1B,NCL,RPL7A,SLC9A7,ALDOA,PKM,GANA
B1,FBXL4,YBX1,CALR,FSTL1,HSPA1B,COL4A1,CANX,PTMA,DDX5,SOX4,SEC62,RPL6,YWHAE,RPL
KIF1A,RPL13,GNAS,DDX17,ALDH6A1,CALM3,GNB1,STMN2,RPS8,SEPTIN2,P4HB,RPL5,KLRD1,G
KIDINS220,NUMA1,DCX,EIF4G2,RPL3,RPS6,ZNF316,DBN1,RPL4,ENO1,TUBA1B,PIK3R2,FLNA,NC
CKSL1,TRIM28,YWHAG,CKB,LRRC28,YWHAQ,DCHS1,TMSB10,MRC2,PKD1,HLA-C,VAPA,POLR2A
S220,TMSB4X,EIF4G2,ZNF316,DBN1,ENO1,TUBA1B,FLNA,MAP1B,NCL,SLC9A7,ALDOA,PKM,GAN
,CSDE1,SMIM14,IGF2,EIF3C,UBB,TRIM28,YWHAG,CKB,SPEN,DDB1,CHSY1,GTF3C4,HLA-C,TUBE
1,YBX1,ATP2B4,FSTL1,ABHD2,HSPA1B,TUBB2B,PSAP,H3-3B,CANX,CLU,PTMA,DDX5,ENO2,SOX4
1,YBX1,ATP2B4,FSTL1,ABHD2,HSPA1B,TUBB2B,PSAP,H3-3B,CANX,CLU,PTMA,DDX5,ENO2,SOX4
.8,YBX1,FSTL1,TMF1,H3-3B,RPL13A,MAGED1,RPL6,DVL3,GNAS,RACK1,FASN,ATP5F1B,GNB1,SH
PTIN2,P4HB,GDI1,FSCN1,SPTAN1,CALM1,HSPD1,USP11,IGF2,EIF4B,MARCKSL1,TRIM28,YWHAG
,PLXNB2,SNRNP200,EIF4B,MARCKSL1,UBB,CKB,HACD3,DCHS1,SET,EIF4A1,MRC2,LDHB,SSBP2
I,PLXNB2,EIF4B,TRIM28,CKB,SPEN,YWHAQ,HACD3,DCHS1,SET,MRC2,LDHB,PKD1,SSBP2,VAPA,
4X,NUMA1,RPL3,RPS6,ZNF316,TUBA1B,PIK3R2,FLNA,RPL7A,SLC9A7,ALDOA,MTRNR2L8,PKM,G
LC2A3,GPI,HTRA4,BSG,FASN,SPTBN1,GNB1,SHISA9,MYH9,RPL7,P4HB,APP,IDS,PTMS,SPTAN1,GS
AT6A,YWHAE,KIF1A,GNAS,SLC2A3,FASN,CALM3,MYH9,SEPTIN2,PTMS,SPTAN1,GSTP1,PLXNB2,C
AS,DDX17,GPI,RACK1,BSG,FASN,ALDH6A1,SPTBN1,ATP5F1B,CALM3,SEPTIN2,RPL5,GDI1,GSTP1
1,IGFBP2,CALR,FSTL1,ABHD2,HSPA1B,TUBB2B,COL5A1,CES3,PSAP,RPLP0,H3-3B,CLU,PTMA,D
SPTBN1,GNB1,SHISA9,RPS8,P4HB,APP,FSCN1,TMBIM6,MARCKS,NPM1,HSPD1,COL18A1,CSDE1
B,FLNA,MAP1B,NCL,SLC9A7,ALDOA,PKM,COL6A1,FTL,HSP90B1,FADS1,NEFM,PRPF8,UGT8,RP
2,IGF2,EIF4B,MARCKSL1,TRIM28,MATR3,YWHAG,CKB,RPL37A,FN1,DDB1,HACD3,DCHS1,TMSB1
16,DBN1,PIK3R2,FLNA,MAP1B,NCL,RPL7A,SLC9A7,ALDOA,MTRNR2L8,PKM,COL6A1,FTL,HSP90
A,ALDOA,GANAB,FTL,HSP90B1,NEFM,PRPF8,UGT8,RPL8,FBXL4,YBX1,ATP2B4,IGFBP2,CALR,TM
,KAT6A,DVL3,YWHAE,GNAS,DDX17,HTRA4,BSG,FASN,ATP5F1B,SEPTIN2,P4HB,APP,GDI1,FSCN1,
R3,YWHAG,NONO,SET,EGLN1,TPI1,CHSY1,RPL15,GTF3C4,SF3B1,RPS3,POLR2A,TPT1,SEC61A1
,HSPA1B,TUBB2B,PGK1,DDX5,SOX4,RPL6,KAT6A,GNAS,SLC2A3,DDX17,GPI,BSG,FASN,SPTBN1,
7,ALDOA,GANAB,FTL,HSP90B1,FADS1,NEFL,NEFM,PRPF8,RPL8,YBX1,IGFBP2,FSTL1,HSPA1B,C

MA1,EIF4G2,RPL3,RPS6,ZNF316,DBN1,RPL4,ENO1,TUBA1B,PIK3R2,FLNA,MAP1B,NCL,RPL7A,SI
.7,P4HB,GDI1,PTMS,SPTAN1,GSTP1,CALM1,USP11,COL18A1,IGF2,EIF3C,MARCKSL1,UBB,CKB,I
'BX1,UBA1,IGFBP2,CALR,ABHD2,HSPA1B,COL5A1,COL4A1,CES3,PSAP,RPLP0,PTMA,ENO2,KAT6
'1,DNAJA1,ARF1,CDC42BPB,ZNF770,TMEM170A,GTF2F1,PXDN,OGA,PGAM1,SMARCA5,CTSD,EII
S1,NEFL,NEFM,PRPF8,RPL8,FBXL4,YBX1,ATP2B4,IGFBP2,FSTL1,TMF1,ZNF99,TUBB2B,COL5A1
6,ZNF316,DBN1,RPL4,ENO1,TUBA1B,FLNA,MAP1B,NCL,ALDOA,PKM,GANAB,COL6A1,FTL,HSP9(
'MF1,H3-3B,PTMA,RPL13A,MAGED1,RPL6,DVL3,GNAS,RACK1,FASN,ATP5F1B,MYH9,P4HB,TMBII
1B,NCL,ALDOA,PKM,GANAB,HSP90B1,NEFL,NEFM,RPL8,FBXL4,UBA1,IGFBP2,FSTL1,TMF1,PGK
WHAQ,DCHS1,SET,TMSB10,MRC2,LDHB,SSBP2,VAPA,AP2M1,TPT1,STMN1,ITGB1,UCHL1,MORF
MAP1B,NCL,SLC9A7,ALDOA,PKM,GANAB,FTL,HSP90B1,FADS1,NEFL,NEFM,RPL8,FBXL4,YBX1,I(
CALR,FSTL1,HSPA1B,TUBB2B,COL4A1,PGK1,DDX5,SOX4,RPL6,SLC2A3,DDX17,GPI,BSG,SPTBN
),BASP1,VAPA,STMN1,MORF4L1,ANXA2,AGRN,ZMIZ1,TIMP2,MIDN,MARCHF6,DNAJA1,CYB5R3,C
'200,IGF2,MARCKSL1,TRIM28,LRRC28,NONO,DDB1,HACD3,TMSB10,LDHB,PKD1,HLA-C,PDIA3,\
N1,RPL4,ENO1,TUBA1B,FLNA,MAP1B,NCL,ALDOA,PKM,GANAB,COL6A1,FTL,HSP90B1,FADS1,NE
X,EIF4G2,RPL3,RPS6,ZNF316,DBN1,RPL4,ENO1,PIK3R2,FLNA,NCL,RPL7A,ALDOA,PKM,GANAB,I
,YBX1,ATP2B4,IGFBP2,TMF1,TUBB2B,COL5A1,COL4A1,PSAP,H3-3B,CANX,PTMA,ENO2,MAGED1
\2,YWHAQ,EIF1,DCHS1,EIF4A1,RPL15,PKD1,PDIA3,VAPA,POLR2A,TPT1,STMN1,AGRN,ZMIZ1,MID
,UBA1,CALR,TMF1,TUBB2B,COL5A1,COL4A1,RPLP0,PTMA,DDX5,KAT6A,DVL3,YWHAE,RPL13,GI
,CD81,HMGN2,ATP2A2,ASTN2,MCL1,TLE5,TADA2B,CSNK1E,PFKL,SFPQ,EIF5A,CDH2,PIP4K2B,RI
,RPL8,YBX1,ATP2B4,FSTL1,HSPA1B,PGK1,H3-3B,PTMA,DDX5,SOX4,FKBP14,RPL13A,MAGED1,R
VAPA,POLR2A,TUBB4B,MORF4L1,BTF3,RPLP1,SBF1,DNAJA1,ARF1,CDC42BPB,ZNF770,TMEM1
RPL7,SEPTIN2,APP,FSCN1,PLXNB2,MARCKSL1,MATR3,YWHAG,FN1,YWHAQ,HACD3,DCHS1,SET,
ISP90B1,NEFL,NEFM,RPL8,FBXL4,YBX1,UBA1,IGFBP2,FSTL1,TMF1,H3-3B,PTMA,RPL13A,MAGE
HSP90AA1,ACTR2,VIM,SPARC,TUBB,FTH1,UBC,KIDINS220,TMSB4X,NUMA1,DCX,EIF4G2,RPL3,R
3A1B,PIK3R2,FLNA,NCL,SLC9A7,ALDOA,PKM,GANAB,COL6A1,FTL,HSP90B1,NEFL,NEFM,PRPF8,
TBN1,GNB1,MYH9,APP,IDS,GDI1,PTMS,SMG1,PLXNB2,USP11,ATP5F1A,EIF4B,MARCKSL1,MATR
NB1,SHISA9,RPS8,SEPTIN2,APP,GDI1,FSCN1,SPTAN1,GSTP1,USP11,SNRNP200,SMIM14,IGF2,EII
NEFM,PRPF8,RPL8,FBXL4,YBX1,UBA1,ATP2B4,IGFBP2,CALR,TMF1,HSPA1B,ZNF99,TUBB2B,CO
1,STMN2,MYH9,SEPTIN2,GDI1,TMBIM6,PTMS,SPTAN1,CALM1,IGF2,EIF4B,MARCKSL1,TRIM28,Y
9,SEPTIN2,GDI1,GSTP1,MARCKS,CALM1,NPM1,PLXNB2,EIF4B,MARCKSL1,CKB,RPL37A,FN1,YW
P2,FSTL1,TUBB2B,COL4A1,PGK1,CANX,PTMA,DDX5,SOX4,YWHAE,GNAS,DDX17,HTRA4,RACK1,E
,TUBA1B,PIK3R2,FLNA,MAP1B,NCL,RPL7A,ALDOA,PKM,GANAB,HSP90B1,FADS1,NEFL,PRPF8,RI
STL1,ABHD2,HSPA1B,COL4A1,PSAP,RPLP0,H3-3B,CANX,DDX5,ENO2,RPL13A,MAGED1,SEC62,KA
,RPS6,ZNF316,DBN1,RPL4,PIK3R2,FLNA,NCL,RPL7A,ALDOA,MTRNR2L8,PKM,GANAB,HSP90B1,
\1,PSAP,RPLP0,PTMA,ENO2,KAT6A,KIF1A,RPL13,GNAS,DDX17,GPI,HTRA4,BSG,FASN,SPTBN1,GI
)BN1,RPL4,ENO1,TUBA1B,MAP1B,NCL,RPL7A,SLC9A7,PKM,GANAB,COL6A1,FTL,HSP90B1,FADS
S220,TMSB4X,NUMA1,DCX,EIF4G2,RPL3,RPS6,ZNF316,DBN1,RPL4,ENO1,TUBA1B,PIK3R2,FLNA
3,COL6A1,FTL,HSP90B1,NEFM,RPL8,FBXL4,UBA1,ATP2B4,CALR,FSTL1,TUBB2B,PGK1,CANX,DD
XAF1,MARCHF6,AP3D1,SBF1,ARF1,CYB5R3,AKAP12,MAP4,CDC42BPB,PXDN,CTNNA1,FADS2,TN
B1,ZNF124,HSP90AA1,ACTR2,VIM,SPARC,TUBB,FTH1,UBC,KIDINS220,TMSB4X,NUMA1,DCX,EIF
1,ENO1,PIK3R2,RPL7A,ALDOA,GANAB,FTL,HSP90B1,NEFL,NEFM,RPL8,YBX1,CALR,FSTL1,TMF1
/IF1,TUBB2B,H3-3B,PTMA,DDX5,ENO2,SEC62,DVL3,YWHAE,KIF1A,GNAS,SLC2A3,DDX17,RACK1,
IF1A,RPL13,GNAS,DDX17,GPI,SPTBN1,CALM3,GNB1,SEPTIN2,P4HB,FSCN1,TMBIM6,GSTP1,SMC
,SPTAN1,CALM1,HSPD1,YWHAG,COL4A2,SPEN,DDB1,YWHAQ,DCHS1,SET,CHSY1,TMSB10,MRC
1,SMG1,PLXNB2,SNRNP200,IGF2,MARCKSL1,TRIM28,YWHAG,LRRC28,DDB1,CHSY1,TMSB10,PI

N1,TUBB4B,UCHL1,ANXA2,AGRN,SARAF,MIDN,VCL,AP3D1,CDC42BPB,GOLGA8A,HNRNPU,CTNN
,CYB5R3,MAP4,CDC42BPB,ITM2B,MAT2A,PXDN,HNRNPU,CTNNA1,CTSD,BRD3,CIPC,KPNB1,LLG
C28,SPEN,PSMD5,EIF4A1,GTF3C4,HLA-C,SERINC1,SSBP2,SEC61A1,RPS3A,ITGB1,UCHL1,BTF3,
4G2,RPL3,RPS6,ZNF316,TUBA1B,PIK3R2,FLNA,RPL7A,SLC9A7,ALDOA,MTRNR2L8,PKM,GANAB,
JMA1,DCX,EIF4G2,ZNF316,DBN1,ENO1,TUBA1B,PIK3R2,NCL,RPL7A,SLC9A7,PKM,GANAB,COL6
GB1,UCHL1,BTF3,AGRN,ZMIZ1,TIMP2,SARAF,MIDN,VCL,AP3D1,CDC42BPB,ITM2B,GOLGA8A,PXI
,YWHAG,LRRC28,TMSB10,PKD1,HLA-C,HMGB1,UCHL1,ANXA2,AGRN,SARAF,MIDN,VCL,AP3D1,
K3R2,FLNA,MAP1B,NCL,RPL7A,ALDOA,PKM,GANAB,FTL,HSP90B1,NEFL,NEFM,PRPF8,RPL8,FBX
L4A2,EIF1,DCHS1,TPI1,CHSY1,MRC2,RPL19,PPP2R1A,PKD1,SSBP2,VAPA,POLR2A,RPL18,STMN1
L4A2,EIF1,DCHS1,TPI1,CHSY1,MRC2,RPL19,PPP2R1A,PKD1,SSBP2,VAPA,POLR2A,RPL18,STMN1
L4A2,EIF1,DCHS1,TPI1,CHSY1,MRC2,RPL19,PPP2R1A,PKD1,SSBP2,VAPA,POLR2A,RPL18,STMN1
PL8,YBX1,UBA1,ATP2B4,FSTL1,ABHD2,TUBB2B,H3-3B,PTMA,DDX5,SOX4,KAT6A,DVL3,YWHAE,K
128,CKB,FN1,YWHAQ,HACD3,DCHS1,EGLN1,CHSY1,MRC2,HLA-C,CDKL1,ITGB1,TUBB4B,AGRN,
LNA,MAP1B,ALDOA,PKM,GANAB,FTL,HSP90B1,FADS1,NEFM,RPL8,FBXL4,YBX1,IGFBP2,FSTL1,
MA1,DCX,EIF4G2,ZNF316,DBN1,RPL4,ENO1,TUBA1B,PIK3R2,NCL,RPL7A,SLC9A7,PKM,GANAB,
1,RPL4,ENO1,TUBA1B,PIK3R2,FLNA,MAP1B,NCL,RPL7A,ALDOA,PKM,GANAB,HSP90B1,NEFL,RF
F4G2,RPL3,ZNF316,DBN1,ENO1,TUBA1B,PIK3R2,MAP1B,NCL,SLC9A7,ALDOA,PKM,GANAB,COL
A8A,CTNNA1,CTSD,BRD3,CIPC,LLGL1,NCS1,ZNF865,SLC7A5,JUND,CD81,HMGN2,ATP2A2,CRM
F1,TUBB2B,COL5A1,COL4A1,H3-3B,CANX,PTMA,DDX5,ENO2,MAGED1,SEC62,GNAS,DDX17,RAC
,HSPD1,IGF2,EIF4B,MARCKSL1,TRIM28,YWHAG,YWHAQ,HACD3,DCHS1,TPI1,TMSB10,PKD1,HL
,PTMA,DDX5,ENO2,RPL13A,SEC62,KAT6A,KIF1A,SLC2A3,GPI,FASN,SPTBN1,CALM3,GNB1,MYH9
1B,CALM3,RPS8,RPL7,SEPTIN2,P4HB,APP,PTMS,PLXNB2,IGF2,MARCKSL1,UBB,CKB,COL4A2,Y
X17,GPI,FASN,ALDH6A1,CALM3,GNB1,SHISA9,STMN2,MYH9,SEPTIN2,GDI1,TMBIM6,PTMS,SPTA
DCX,EIF4G2,ZNF316,DBN1,ENO1,TUBA1B,PIK3R2,NCL,RPL7A,SLC9A7,PKM,GANAB,COL6A1,FTL
T1,ITGB1,MORF4L1,AGRN,TIMP2,MIDN,MARCHF6,VCL,CYB5R3,CDC42BPB,ITM2B,PXDN,HNRN
3R2,FLNA,MAP1B,NCL,RPL7A,SLC9A7,ALDOA,PKM,GANAB,FTL,HSP90B1,NEFM,PRPF8,FBXL4,Y
T6A,DVL3,YWHAE,KIF1A,GNAS,FASN,SPTBN1,GNB1,SHISA9,MYH9,SEPTIN2,P4HB,GDI1,GSTP1,
S,BSG,FASN,ALDH6A1,GNB1,SHISA9,MYH9,RPL5,GSTP1,MARCKS,CALM1,NPM1,PLXNB2,COL18
3A,RPL6,DVL3,RPL13,GNAS,DDX17,BSG,ATP5F1B,CALM3,RPS8,RPL7,SEPTIN2,P4HB,APP,GDI1,
PL4,ENO1,PIK3R2,MAP1B,NCL,SLC9A7,ALDOA,PKM,GANAB,FTL,HSP90B1,FADS1,NEFL,NEFM,R
Y1,EIF4A1,PKD1,HLA-C,SSBP2,AP2M1,ITGB1,TUBB4B,ZMIZ1,MIDN,WSB1,VCL,AP3D1,ARF1,MAP
3A,RPL6,DVL3,RACK1,FASN,ATP5F1B,GNB1,SHISA9,P4HB,RPL5,TMBIM6,SPTAN1,NPM1,PLXNB
1,COL5A1,COL4A1,CES3,H3-3B,PTMA,ENO2,YWHAE,KIF1A,GNAS,SLC2A3,SPTBN1,CALM3,SHIS
1,H3-3B,CANX,PTMA,DDX5,ENO2,MAGED1,SEC62,GNAS,DDX17,RACK1,BSG,CALM3,RPS8,RPL7
NAH8,ZNF43,VAPA,SEC61A1,OR8K1,TUBB4B,MORF4L1,BTF3,XAF1,SARAF,AP3D1,SBF1,ARF1,C
LDOA,MTRNR2L8,PKM,GANAB,COL6A1,FTL,HSP90B1,NEFL,FBXL4,UBA1,ATP2B4,TMF1,HSPA1E
GANAB,HSP90B1,NEFL,NEFM,RPL8,FBXL4,YBX1,UBA1,ATP2B4,IGFBP2,TMF1,HSPA1B,H3-3B,C
RPL3,RPS6,ZNF316,DBN1,RPL4,TUBA1B,PIK3R2,MAP1B,NCL,SLC9A7,ALDOA,PKM,GANAB,FTL,H
,RPL37A,SPEN,NONO,YWHAQ,EIF1,EGLN1,INA,MRC2,SF3B1,SERINC1,SSBP2,VAPA,SEC61A1,ITG
,RPL4,ENO1,TUBA1B,PIK3R2,NCL,RPL7A,SLC9A7,ALDOA,PKM,GANAB,COL6A1,HSP90B1,NEFL,
JBA1B,MAP1B,NCL,ALDOA,PKM,GANAB,COL6A1,FTL,HSP90B1,FADS1,NEFL,NEFM,PRPF8,UGT8,
,PTMA,DDX5,SOX4,RPL13,DDX17,HTRA4,FASN,ATP5F1B,CALM3,SHISA9,MYH9,SEPTIN2,P4HB,A
MA1,DCX,EIF4G2,ZNF316,DBN1,RPL4,ENO1,TUBA1B,PIK3R2,NCL,RPL7A,SLC9A7,PKM,GANAB,
,RPL7,SEPTIN2,P4HB,APP,RPL5,GDI1,PTMS,PLD3,IGF2,EIF3C,EIF4B,UBB,CKB,COL4A2,LRRC28,
IGFBP2,CALR,FSTL1,PSAP,PGK1,CANX,PTMA,DDX5,ENO2,SOX4,SEC62,KAT6A,DVL3,YWHAE,RPL

B1,SHISA9,P4HB,RPL5,TMBIM6,NPM1,PLD3,HSPD1,USP11,COL18A1,SNRNP200,CSDE1,SMIM14
ACD3,DCHS1,SET,MRC2,GTF3C4,LDHB,SSBP2,BASP1,POLR2A,STMN1,MORF4L1,AGRN,TIMP2,N
I2,APP,IDS,FSCN1,PTMS,CALM1,SNRNP200,IGF2,EIF4B,UBB,TRIM28,COL4A2,HACD3,CHSY1,INA
L4,ENO1,PIK3R2,MAP1B,NCL,SLC9A7,ALDOA,PKM,GANAB,FTL,HSP90B1,FADS1,NEFL,NEFM,RP
3,PIK3R2,FLNA,MAP1B,NCL,RPL7A,SLC9A7,ALDOA,PKM,GANAB,HSP90B1,NEFL,RPL8,FBXL4,YB
DDX5,SOX4,KAT6A,DVL3,YWHAE,KIF1A,RPL13,GPI,BSG,FASN,GNB1,MYH9,RPL7,P4HB,TMBIM6,S
HF6,WSB1,DNAJA1,CDC42BPB,ITM2B,HNRNPU,CNN3,SCARB2,SLC22A17,BRD3,CIPC,LLGL1,NCS
NCL,RPL7A,SLC9A7,GANAB,HSP90B1,FADS1,RPL8,FBXL4,UBA1,IGFBP2,FSTL1,COL5A1,RPLP0,
2,ZNF316,DBN1,RPL4,TUBA1B,FLNA,MAP1B,NCL,RPL7A,SLC9A7,ALDOA,PKM,GANAB,COL6A1,F
DDX5,SOX4,RPL13,DDX17,HTRA4,FASN,ATP5F1B,SHISA9,MYH9,SEPTIN2,P4HB,APP,IDS,GDI1,FS
NXA2,AGRN,SARAF,MIDN,AP3D1,CDC42BPB,GOLGA8A,HNRNPU,CTNNA1,LPCAT1,LAMA5,BRD3,L
TBN1,GNB1,STMN2,MYH9,P4HB,CALM1,NPM1,USP11,IGF2,EIF3C,AFF4,TRIM28,MATR3,LRRC28
A1,BRD3,MSN,LLGL1,ZNF865,SLC7A5,CHD4,JUND,CD81,HMGN2,ATP2A2,OAZ1,PFN1,NISCH,TAD
TRIM28,LRRC28,NONO,DDB1,HACD3,CHSY1,LDHB,PKD1,HLA-C,CDKL1,TPT1,RPS3A,STMN1,UC
X1,IGFBP2,CALR,ZNF99,COL4A1,PSAP,PGK1,RPLP0,PTMA,DDX5,SOX4,KAT6A,DVL3,RPL13,GNA
ED1,SEC62,KAT6A,YWHAE,KIF1A,RPL13,BSG,FASN,ALDH6A1,CALM3,GNB1,SEPTIN2,P4HB,IDS,F
PTMA,ENO2,SEC62,KAT6A,DVL3,KIF1A,RPL13,GNAS,GPI,HTRA4,BSG,FASN,SPTBN1,GNB1,MYH9
B,PIK3R2,NCL,SLC9A7,GANAB,FTL,HSP90B1,NEFM,RPL8,FBXL4,YBX1,IGFBP2,FSTL1,TMF1,PGK
CHSY1,TMSB10,PKD1,BASP1,CDKL1,WSB1,VCL,ZNF770,FADS2,RPS16,LDHA,SLC22A17,BRD3,KI
R2,FLNA,MAP1B,NCL,RPL7A,ALDOA,GANAB,HSP90B1,NEFL,PRPF8,RPL8,FBXL4,YBX1,UBA1,ATP
,COL4A1,PSAP,PGK1,RPLP0,H3-3B,CLU,PTMA,DDX5,ENO2,SOX4,FKBP14,RPL13A,RPL6,YWHAE,
,KAT6A,DVL3,YWHAE,KIF1A,DDX17,RACK1,BSG,FASN,ALDH6A1,ATP5F1B,CALM3,STMN2,RPS8,S
LC9A7,ALDOA,GANAB,HSP90B1,NEFL,NEFM,RPL8,FBXL4,YBX1,UBA1,ATP2B4,IGFBP2,CALR,TMF
1,GDI1,PTMS,SMG1,PLXNB2,MARCKSL1,FN1,YWHAQ,HACD3,DCHS1,SET,EGLN1,TMSB10,MRC2
RCHF6,SBF1,ARF1,CDC42BPB,HNRNPU,HNRNPA0,CNN3,RPSA,EIF4H,LPCAT1,LAMA5,BRD3,CIPC
.6,DVL3,RACK1,FASN,ATP5F1B,GNB1,SHISA9,P4HB,RPL5,TMBIM6,NPM1,PLD3,USP11,SNRNP20
T6A,DVL3,RACK1,CALM3,GNB1,SHISA9,RPL7,P4HB,TMBIM6,SPTAN1,NPM1,PLD3,USP11,SNRNP
PI,RACK1,FASN,CALM3,RPS8,MYH9,P4HB,APP,TMBIM6,SMG1,USP11,IGF2,EIF3C,MARCKSL1,UE
MA1,EIF4G2,RPS6,ZNF316,DBN1,ENO1,PIK3R2,MAP1B,SLC9A7,ALDOA,PKM,GANAB,FTL,FADS1,
ANAB,HSP90B1,NEFL,RPL8,FBXL4,YBX1,UBA1,CALR,TMF1,HSPA1B,H3-3B,PTMA,DDX5,FKBP14,F
SBF1,DNAJA1,ARF1,CDC42BPB,TMEM170A,GTF2F1,OGA,PGAM1,SMARCA5,RPS16,CTSD,RPS9,
NCL,RPL7A,ALDOA,PKM,GANAB,COL6A1,FTL,HSP90B1,FADS1,NEFM,PRPF8,RPL8,FBXL4,YBX1,U
,CTNNA1,SCARB2,BRD3,HK1,ZNF865,ATF4,SLC7A5,CHD4,ATP1A1,CD81,HMGN2,ATP2A2,OAZ1,N
EFM,PRPF8,FBXL4,IGFBP2,FSTL1,TMF1,ZNF99,TUBB2B,COL5A1,COL4A1,CES3,RPLP0,PTMA,KA
ENO1,PIK3R2,FLNA,MAP1B,NCL,RPL7A,SLC9A7,PKM,GANAB,FTL,HSP90B1,FADS1,NEFL,RPL8,U
AB,FTL,HSP90B1,NEFL,RPL8,FBXL4,UBA1,ATP2B4,IGFBP2,CALR,TMF1,PGK1,H3-3B,CANX,PTMA
LC9A7,ALDOA,MTRNR2L8,PKM,COL6A1,HSP90B1,NEFL,FBXL4,YBX1,UBA1,TMF1,HSPA1B,TUBB
/HAG,FN1,YWHAQ,HACD3,DCHS1,SET,MRC2,LDHB,PKD1,VAPA,AP2M1,TPT1,ITGB1,MORF4L1,AG
1A,SF3B1,ZNF43,HMGB1,POLR2A,TPT1,RPS3A,TUBB4B,AGRN,RPLP1,MIDN,MARCHF6,DNAJA1,A
8,NONO,DDB1,HACD3,DCHS1,CHSY1,LDHB,PKD1,HLA-C,SSBP2,CDKL1,TPT1,RPS3A,STMN1,UCH
,ENO1,TUBA1B,PIK3R2,NCL,RPL7A,SLC9A7,PKM,GANAB,COL6A1,HSP90B1,NEFL,NEFM,RPL8,F
KSL1,TRIM28,MATR3,EIF1,DCHS1,CHSY1,PPP2R1A,PKD1,PDIA3,BASP1,VAPA,TPT1,UCHL1,AGRN
OA,PKM,GANAB,COL6A1,HSP90B1,NEFM,RPL8,FBXL4,UBA1,ATP2B4,IGFBP2,CALR,FSTL1,ABHD
,COL6A1,FTL,HSP90B1,FADS1,NEFL,RPL8,FBXL4,YBX1,UBA1,ATP2B4,CALR,FSTL1,ABHD2,HSPA
PC,IGFBP3,RHOA,CBX5,ZNF865,LENG8,VAT1,RTN3,PFN1,CCNI,TADA2B,KHDRBS1,HDLBP,EIF5A,

,HSPA1B,COL4A1,PGK1,RPLP0,CANX,CLU,PTMA,DDX5,ENO2,RPL13A,MAGED1,KAT6A,YWHAE,D
ISA9,P4HB,PLD3,HSPD1,USP11,SNRNP200,EIF4B,EIF1,TPI1,INA,EIF4A1,PPP2R1A,PKD1,SF3B1,Z
N,ZMIZ1,TIMP2,RPL10,WSB1,CYB5R3,AKAP12,MAP4,CDC42BPB,ITM2B,PXDN,OGA,PGAM1,CTN
T7,OAZ1,PFN1,PFKL,ARL4C,MRFAP1,RCN1,MTSS2,DNAJC5,APLP2,DUX4,SEPTIN11,HNRNPH1,C
,ALDOA,GANAB,HSP90B1,NEFL,RPL8,FBXL4,YBX1,UBA1,ATP2B4,CALR,TMF1,HSPA1B,COL4A1,
BX1,UBA1,CALR,TMF1,HSPA1B,H3-3B,CANX,PTMA,DDX5,FKBP14,RPL13A,MAGED1,RPL6,DVL3,
UBA1B,FLNA,MAP1B,RPL7A,SLC9A7,ALDOA,GANAB,FTL,HSP90B1,NEFM,PRPF8,UGT8,RPL8,YB
34X,NUMA1,DCX,EIF4G2,RPS6,ZNF316,DBN1,RPL4,ENO1,TUBA1B,PIK3R2,NCL,RPL7A,SLC9A7,
N2,OAZ1,TLE5,NISCH,CCNI,CSNK1E,PFKL,SFPQ,CDH2,PDIA6,MEGF8,ARL4C,RPL35,CHPF,B4GA
34X,NUMA1,DCX,EIF4G2,RPS6,ZNF316,DBN1,ENO1,TUBA1B,PIK3R2,NCL,RPL7A,SLC9A7,ALDOA
1,TPT1,RPS3A,STMN1,MORF4L1,BTF3,AGRN,RPLP1,MARCHF6,DNAJA1,PXDN,HNRNPA0,RPS16,
,GANAB,HSP90B1,NEFL,RPL8,FBXL4,YBX1,UBA1,IGFBP2,TMF1,H3-3B,PTMA,DDX5,FKBP14,RPL
RPL8,FBXL4,YBX1,UBA1,IGFBP2,TMF1,H3-3B,PTMA,DDX5,SOX4,FKBP14,RPL13A,MAGED1,RPL
,FTH1,UBC,KIDINS220,TMSB4X,NUMA1,EIF4G2,RPL3,RPS6,ZNF316,DBN1,ENO1,TUBA1B,FLNA,
A,SEC62,RPL6,KAT6A,DVL3,DDX17,RACK1,SPTBN1,ATP5F1B,SHISA9,RPS8,P4HB,APP,ERVV-1,G
-3B,CANX,PTMA,ENO2,MAGED1,SEC62,GNAS,DDX17,BSG,CALM3,RPS8,RPL7,SEPTIN2,P4HB,AP
9,SRSF5,U2AF2,NOP53,PABPN1,GDI2,ACO2,RBMX,CCNL2,BTBD2,RABL6,SRSF1,PTBP1,ILF3,AT
C42BPB,ITM2B,GOLGA8A,SMARCA5,TPM4,SCARB2,RPS24,SLC22A17,RPS9,CIPC,C1ORF43,NCS
4X,NUMA1,EIF4G2,RPL3,RPS6,ZNF316,DBN1,RPL4,ENO1,TUBA1B,PIK3R2,FLNA,MAP1B,NCL,SL
A1,CANX,PTMA,ENO2,MAGED1,GNAS,DDX17,BSG,CALM3,RPL7,SEPTIN2,APP,IDS,PTMS,SMG1,C
UBC,KIDINS220,TMSB4X,NUMA1,DCX,EIF4G2,RPL3,RPS6,ZNF316,DBN1,RPL4,ENO1,TUBA1B,PI
TMSB4X,NUMA1,DCX,EIF4G2,ZNF316,DBN1,RPL4,ENO1,TUBA1B,NCL,RPL7A,SLC9A7,ALDOA,PK
ALDOA,PKM,GANAB,HSP90B1,FADS1,NEFL,NEFM,PRPF8,RPL8,UBA1,IGFBP2,TMF1,ABHD2,HSP
NO1,PIK3R2,FLNA,NCL,MTRNR2L8,GANAB,COL6A1,FTL,NEFL,NEFM,RPL8,YBX1,UBA1,ATP2B4,
K1,H3-3B,DDX5,RPL13A,MAGED1,KAT6A,RPL13,GNAS,CALM3,GNB1,RPL7,P4HB,PTMS,SPTAN1
3XL4,YBX1,UBA1,ATP2B4,CALR,TMF1,HSPA1B,H3-3B,CANX,PTMA,DDX5,FKBP14,RPL13A,MAGE
PTMA,SEC62,KAT6A,YWHAE,KIF1A,RPL13,GNAS,DDX17,HTRA4,FASN,SPTBN1,GNB1,RPL7,IDS,R
LP1,SBF1,DNAJA1,ARF1,CDC42BPB,TMEM170A,GTF2F1,OGA,PGAM1,SMARCA5,RPS16,CTSD,F
AQ,HACD3,DCHS1,SET,EGLN1,MRC2,GTF3C4,SSBP2,BASP1,POLR2A,TPT1,TUBB4B,AGRN,TIMP
WHAE,KIF1A,RPL13,GPI,RACK1,FASN,GNB1,RPS8,MYH9,P4HB,APP,GDI1,FSCN1,SPTAN1,CSDE1,
R2,VIM,TUBB,FTH1,UBC,KIDINS220,TMSB4X,NUMA1,DCX,EIF4G2,RPL3,ZNF316,DBN1,RPL4,EN
1,RPLP1,RPL10,PXDN,SMARCA5,CTNNA1,RPL23,RPS9,KPNB1,RPL17,C1ORF43,LLGL1,GAP43,R
SLC9A7,ALDOA,GANAB,HSP90B1,NEFL,NEFM,FBXL4,YBX1,UBA1,ATP2B4,IGFBP2,TMF1,HSPA1
90B1,NEFM,RPL8,YBX1,UBA1,IGFBP2,CALR,FSTL1,TUBB2B,COL5A1,COL4A1,CLU,PTMA,DDX5,
CX,EIF4G2,RPS6,ZNF316,DBN1,RPL4,ENO1,TUBA1B,PIK3R2,NCL,RPL7A,SLC9A7,PKM,GANAB,C
L8,PKM,GANAB,COL6A1,HSP90B1,FADS1,NEFL,RPL8,FBXL4,YBX1,UBA1,TMF1,TUBB2B,COL5A
L6,CIRBP,DUX4,HECTD4,CTBP1,ARHGDIA,SLC35E2B,SMCHD1,IPO5,ATP6V0C,DNER,HSBP1,FA
,NUMA1,EIF4G2,ZNF316,RPL4,ENO1,PIK3R2,FLNA,MAP1B,NCL,RPL7A,SLC9A7,ALDOA,MTRNR
90B1,NEFL,NEFM,RPL8,FBXL4,YBX1,UBA1,ATP2B4,TMF1,HSPA1B,TUBB2B,PGK1,H3-3B,CANX,
,P4HB,TMBIM6,PLD3,USP11,SNRNP200,SMIM14,ATP5F1A,EIF4B,VCP,EIF1,INA,EIF4A1,PPP2R1A,
,ITM2B,SMARCA5,TPM4,KPNB1,NCS1,CXADR,CHD4,ATP1A1,JUND,HMGN2,ATP2A2,LASP1,PFN1,
4HB,TMBIM6,SPTAN1,PLD3,USP11,SNRNP200,SMIM14,ATP5F1A,EIF4B,UBB,AFF4,RPL37A,LRR
DBN1,RPL4,ENO1,TUBA1B,PIK3R2,NCL,RPL7A,SLC9A7,ALDOA,PKM,COL6A1,HSP90B1,NEFL,NE
4,ENO1,TUBA1B,PIK3R2,MAP1B,NCL,ALDOA,PKM,GANAB,FTL,HSP90B1,FADS1,NEFL,NEFM,PR
X1,UBA1,ATP2B4,CALR,TMF1,HSPA1B,H3-3B,CANX,PTMA,DDX5,FKBP14,RPL13A,MAGED1,RPL6

YBX1,IGFBP2,CALR,FSTL1,ABHD2,HSPA1B,TUBB2B,COL5A1,PSAP,PTMA,DDX5,SEC62,KAT6A,DV
MARCHF6,SBF1,DNAJA1,MAT2A,NFE2L1,SMARCA5,HNRNPA0,CNN3,RPSA,LPCAT1,LAMA5,XRCC
,CHD4,HMGN2,ATP2A2,LASP1,KHDRBS1,CDH2,PRDX2,RPS15,ITM2C,ARL4C,CHPF,APLP2,PLPPR
N4,GNB2,RPL35,GOLM1,B4GALNT4,MTSS2,NDFIP1,ANKRD52,H1-10,DUX4,YWHAB,HNRNPH1,C
PL4,ENO1,TUBA1B,PIK3R2,NCL,RPL7A,GANAB,COL6A1,HSP90B1,FADS1,NEFL,NEFM,RPL8,UBA
,HDLBP,EIF5A,RCN1,RPS5,APLP1,PRKAR1A,HNRNPK,RHOB,HNRNPH1,UBE2H,PLPPR3,HECTD4,
N1,TUBA1B,PIK3R2,FLNA,MAP1B,NCL,RPL7A,SLC9A7,ALDOA,PKM,GANAB,COL6A1,HSP90B1,FA
13A,SEC62,YWHAE,KIF1A,DDX17,RACK1,FASN,ATP5F1B,CALM3,GNB1,RPS8,RPL7,SEPTIN2,GDI1
1,RPS8,RPL7,SEPTIN2,APP,IDS,RPL5,GSTP1,NPM1,HSPD1,USP11,COL18A1,IGF2,TRIM28,CKB,S
PL19,CHD3,VAPA,TPT1,BTF3,AGRN,RPLP1,TIMP2,SBF1,AKAP12,SF3B2,CDC42BPB,MAT2A,GTF2
31,NCS1,ZNF865,CHD4,ATP1A1,JUND,HMGN2,ATP2A2,LASP1,OAZ1,CCNI,CSNK1E,EIF5A,CCND2,
M,RPL8,IGFBP2,TMF1,ABHD2,HSPA1B,COL4A1,PSAP,PGK1,RPLP0,CANX,PTMA,DDX5,ENO2,MAG
M1,AGRN,SARAF,VCL,SF3B2,CDC42BPB,MAT2A,GTF2F1,PXDN,HNRNPU,OGA,CTNNA1,FADS2,L
HSP90AA1,ACTR2,VIM,SPARC,TUBB,FTH1,UBC,KIDINS220,TMSB4X,NUMA1,DCX,EIF4G2,RPL3,R
7,ALDOA,GANAB,HSP90B1,NEFL,NEFM,RPL8,FBXL4,YBX1,UBA1,ATP2B4,IGFBP2,CALR,TMF1,HS
NEFL,NEFM,PRPF8,RPL8,IGFBP2,FSTL1,TMF1,HSPA1B,COL4A1,PGK1,H3-3B,CANX,CLU,PTMA,D
AT6A,DVL3,KIF1A,RPL13,SLC2A3,DDX17,BSG,FASN,ATP5F1B,CALM3,GNB1,SHISA9,RPS8,MYH9,
32,USP11,TRIM28,YWHAG,DDB1,DCHS1,SET,MAP9,TPI1,MRC2,PKD1,SSBP2,BASP1,TPT1,SEC61
GTF2F1,HNRNPA0,RPS16,RPL23,RPL29,PAPOLA,RHOA,PTCD3,ZNF865,TMEM259,LENG8,WDR1,
P4HB,TMBIM6,SPTAN1,PLD3,USP11,SNRNP200,SMIM14,ATP5F1A,EIF4B,UBB,RPL37A,EIF1,INA,
N1,TUBA1B,PIK3R2,FLNA,MAP1B,NCL,RPL7A,SLC9A7,ALDOA,PKM,GANAB,COL6A1,HSP90B1,FA
DDX5,RPL13A,MAGED1,RPL6,DVL3,GNAS,GPI,FASN,ATP5F1B,SHISA9,MYH9,P4HB,TMBIM6,SPT
YBX1,UBA1,IGFBP2,CALR,TMF1,HSPA1B,TUBB2B,COL5A1,COL4A1,CES3,RPLP0,PTMA,DDX5,EN
YBX1,UBA1,IGFBP2,CALR,TMF1,HSPA1B,TUBB2B,COL5A1,COL4A1,CES3,RPLP0,PTMA,DDX5,EN
,RPL8,YBX1,UBA1,CALR,FSTL1,TUBB2B,COL5A1,SEC62,GNAS,SLC2A3,DDX17,GPI,FASN,ALDH6
TMA,DDX5,FKBP14,RPL13A,MAGED1,RPL6,DVL3,GPI,RACK1,ATP5F1B,CALM3,GNB1,SHISA9,RP
H9,SEPTIN2,RPL5,PTMS,SMG1,CALM1,PLD3,SNRNP200,IGF2,EIF3C,EIF4B,VCP,MARCKSL1,AFF
,NEFL,RPL8,FBXL4,YBX1,UBA1,ATP2B4,CALR,TMF1,HSPA1B,H3-3B,CANX,PTMA,DDX5,FKBP14,
APA,TPT1,RPS3A,AGRN,ZMIZ1,TIMP2,SARAF,MIDN,RPL10,GTF2F1,NFE2L1,SMARCA5,TPM4,RPS
TMS,GSTP1,CALM1,PLXNB2,USP11,IGF2,EIF4B,MARCKSL1,MATR3,YWHAG,MAPT,SPEN,NONO,
2,YWHAQ,MAP9,RPL19,RPL15,SSBP2,BASP1,VAPA,UCHL1,MORF4L1,BTF3,AGRN,RPLP1,ZMIZ1,
RN,RPLP1,HNRNPA0,CNN3,RPSA,TNS3,MSN,KPNB1,C1ORF43,IGFBP3,RHOA,PTCD3,PSMD2,CXA
K1,CLU,FKBP14,RPL13A,SEC62,RPL6,KAT6A,YWHAE,KIF1A,GNAS,DDX17,GPI,FASN,GNB1,SHISA
UBA1,ATP2B4,TMF1,H3-3B,CANX,PTMA,DDX5,FKBP14,RPL13A,MAGED1,RPL6,DVL3,GNAS,GPI,F
NB2,HSPD1,COL18A1,SMIM14,IGF2,ATP5F1A,EIF4B,CKB,DDB1,YWHAQ,DCHS1,CHSY1,TMSB10,
CHL1,MORF4L1,AGRN,MIDN,DNAJA1,MAT2A,PXDN,HNRNPU,FADS2,CNN3,RPSA,EIF4H,LAMA5,BF
1,MIDN,MARCHF6,RPS17,DNAJA1,ARF1,SF3B2,MAT2A,GTF2F1,HNRNPA0,RPS16,RPL23,RPL29,H
,HSP90B1,NEFL,RPL8,FBXL4,YBX1,UBA1,IGFBP2,CALR,TMF1,H3-3B,CANX,PTMA,DDX5,ENO2,F
EIF4B,SPEN,EIF4A1,PPP2R1A,RPL15,GTF3C4,SF3B1,ZNF43,HMGB1,POLR2A,TPT1,RPS3A,TUBB
PL7,P4HB,TMBIM6,PTMS,SPTAN1,PLD3,USP11,SNRNP200,SMIM14,ATP5F1A,EIF4B,UBB,AFF4,R
PL5,TMBIM6,SPTAN1,PLD3,HSPD1,USP11,SNRNP200,CSDE1,SMIM14,ATP5F1A,EIF4B,SPEN,EIF1
MARCKSL1,YWHAG,CKB,SPEN,SET,MAP9,TMSB10,BASP1,VAPA,TUBB4B,ANXA2,TIMP2,SARAF,R
MARCKSL1,YWHAG,CKB,SPEN,SET,MAP9,TMSB10,BASP1,VAPA,TUBB4B,ANXA2,TIMP2,SARAF,R
HSP90B1,NEFL,NEFM,FBXL4,YBX1,UBA1,ATP2B4,IGFBP2,TMF1,HSPA1B,TUBB2B,PGK1,H3-3B,C
NCL,RPL7A,GANAB,COL6A1,HSP90B1,FADS1,NEFL,NEFM,RPL8,UBA1,TMF1,ABHD2,HSPA1B,ZNF

3,PXDN,OGA,HNRNPA0,CTNNA1,LPCAT1,CXADR,LASP1,OAZ1,SRSF4,PFN1,TADA2B,XPO1,PFKL,T
ALM3,GNB1,STMN2,MYH9,GDI1,PTMS,SPTAN1,SMG1,CALM1,NPM1,MARCKSL1,AFF4,FN1,NON(
ANAB,COL6A1,NEFL,NEFM,RPL8,IGFBP2,TMF1,ABHD2,HSPA1B,COL4A1,PSAP,PGK1,RPLP0,CAN
N,HDLBP,SFPQ,HNRNPDL,HNRNPK,H1-10,NARS1,HNRNPH1,CKAP4,RPS27,CCT6A,SMCHD1,ATP6
RNP200,EIF3C,ATP5F1A,EIF4B,MATR3,RPL37A,LRRC28,EIF1,INA,RPL19,PPP2R1A,LDHB,SF3B1,
KIF1A,RPL13,GPI,BSG,FASN,ALDH6A1,SPTBN1,RPS8,SEPTIN2,APP,RPL5,KLRD1,FSCN1,SMG1,NP
NB1,SHISA9,STMN2,MYH9,APP,IDS,GDI1,TMBIM6,PTMS,MARCKS,SMG1,NPM1,CSDE1,MARCKSL
YBX1,UBA1,IGFBP2,TMF1,H3-3B,PTMA,DDX5,SOX4,FKBP14,RPL13A,MAGED1,RPL6,DVL3,YWHA
3,YWHAE,GNAS,GPI,HTRA4,FASN,SHISA9,RPS8,MYH9,P4HB,TMBIM6,CALM1,USP11,SNRNP200,I
TP2B4,TMF1,H3-3B,CANX,CLU,PTMA,DDX5,FKBP14,RPL13A,MAGED1,RPL6,DVL3,GNAS,GPI,RAC
FADS2,EIF4H,LAMA5,CIPC,RPL14,RHOA,CBX5,ZNF865,GLIPR1L2,VAT1,RTN3,CCNI,KHDRBS1,HD
1A,RPL13,GNAS,DDX17,HTRA4,FASN,SPTBN1,RPL7,RPL5,FSCN1,PTMS,SMG1,NPM1,PLXNB2,HS
1,PPP2R1A,GTF3C4,SF3B1,ZNF43,HMGB1,POLR2A,TPT1,SEC61A1,TUBB4B,RPLP1,MIDN,RPL10
YH9,P4HB,APP,GDI1,PTMS,SPTAN1,SMG1,NPM1,PLXNB2,PLD3,SMIM14,IGF2,VCP,MARCKSL1,TF
OL6A1,FTL,HSP90B1,FADS1,NEFM,PRPF8,UGT8,RPL8,FBXL4,YBX1,IGFBP2,CALR,TMF1,ABHD2,
R1A,LDHB,SF3B1,ZNF43,HMGB1,POLR2A,TPT1,RPS3A,TUBB4B,MIDN,MARCHF6,RPS17,DNAJA1
L4A1,PSAP,RPLP0,CANX,PTMA,DDX5,RPL6,KAT6A,KIF1A,RPL13,GPI,BSG,FASN,ALDH6A1,SPTBN
PS6,ZNF316,DBN1,RPL4,TUBA1B,PIK3R2,SLC9A7,ALDOA,GANAB,FTL,HSP90B1,FADS1,NEFL,NE
0B1,FADS1,NEFM,UGT8,RPL8,YBX1,IGFBP2,CALR,TMF1,ABHD2,ZNF99,TUBB2B,COL4A1,PSAP,P
3B2B,COL5A1,COL4A1,PSAP,H3-3B,PTMA,SEC62,KAT6A,YWHAE,KIF1A,RPL13,GNAS,DDX17,HTR
A,GANAB,HSP90B1,NEFL,NEFM,RPL8,FBXL4,YBX1,UBA1,ATP2B4,CALR,TMF1,HSPA1B,PGK1,H3-
F1,HSPA1B,H3-3B,CANX,DDX5,FKBP14,RPL13A,MAGED1,RPL6,GPI,RACK1,CALM3,GNB1,SHISA
4B,LRRC28,SPEN,EIF1,PPP2R1A,RPL15,GTF3C4,SF3B1,ZNF43,HMGB1,POLR2A,TPT1,RPS3A,TU
PLD3,USP11,SNRNP200,CSDE1,SMIM14,ATP5F1A,EIF4B,SPEN,EIF1,INA,EIF4A1,PPP2R1A,RPL15,
A,GPI,BSG,FASN,SHISA9,MYH9,P4HB,TMBIM6,GSTP1,CALM1,NPM1,SNRNP200,CSDE1,VCP,UBB,
UBA1,IGFBP2,CALR,FSTL1,ABHD2,TUBB2B,COL5A1,COL4A1,RPLP0,CLU,SOX4,RPL13A,KAT6A,D
GTF3C4,SF3B1,ZNF43,HMGB1,POLR2A,TPT1,RPS3A,TUBB4B,RPL11,RPLP1,SARAF,RPL10,AP3D
A1B,PIK3R2,FLNA,MAP1B,NCL,SLC9A7,MTRNR2L8,COL6A1,FTL,HSP90B1,NEFL,NEFM,PRPF8,RP
H3-3B,PTMA,DDX5,RPL13A,MAGED1,SEC62,RPL6,KAT6A,DVL3,YWHAE,KIF1A,DDX17,RACK1,BSG
QLE,NOVA2,CHD6,EEF1A2,TBCD,VAMP2,GAS7,HSPA4,GRINA,PTOV1,MDK,LAMC1,ASXL1,H3-3A,F
GF2,ATP5F1A,MARCKSL1,TRIM28,DDB1,EIF1,DCHS1,SET,CHSY1,INA,EIF4A1,MRC2,RPL15,PKD1
DX5,RPL13A,MAGED1,RPL6,DVL3,GNAS,GPI,RACK1,ATP5F1B,CALM3,SHISA9,P4HB,RPL5,FSCN1,
1,SERINC1,HMGB1,POLR2A,AP2M1,SEC61A1,TUBB4B,BTF3,RPLP1,MIDN,MARCHF6,DNAJA1,SF
P4,CDC42BPB,MAT2A,CTSD,LAMA5,BRD3,CIPC,LLGL1,ZNF865,JUND,CD81,HMGN2,ATP2A2,CR
ZNF316,DBN1,ENO1,TUBA1B,FLNA,MAP1B,NCL,RPL7A,SLC9A7,ALDOA,MTRNR2L8,PKM,COL6A1
HD2,COL5A1,CES3,RPLP0,H3-3B,CLU,PTMA,SOX4,RPL13A,MAGED1,SEC62,KAT6A,KIF1A,RPL1
1,UBA1,ATP2B4,TMF1,HSPA1B,TUBB2B,H3-3B,CANX,DDX5,FKBP14,RPL13A,MAGED1,RPL6,DVL
1,PDIA3,SERINC1,VAPA,HMGB1,POLR2A,MORF4L1,VCL,CDC42BPB,PXDN,FADS2,RPS16,RPL23,
1,POLR2A,TPT1,SEC61A1,RPS3A,TUBB4B,MIDN,RPL10,SF3B2,GTF2F1,HNRNPA0,RPL29,RPS9,C
RCA5,RPS16,RPS9,CIPC,PAPOLA,CBX5,ZNF865,KIF5B,GLIPR1L2,LENG8,ATXN7L3B,VAT1,RTN3,
YB5R3,PXDN,HNRNPA0,CTNNA1,FADS2,CNN3,TNS3,CTSD,LPCAT1,LAMA5,SLC22A17,BRD3,RPL1
A,NACA,RPS24,RPS9,PJA2,RPL24,CBX5,CLSTN1,ATF4,KIF5B,CXADR,TMEM259,HMGN2,PNN,NIS
D5,EIF4A1,GTF3C4,LDHB,SERINC1,BASP1,VAPA,POLR2A,TPT1,STMN1,ITGB1,MORF4L1,NCAM1,
D23B,SKI,RPS15A,RAPGEF1,MAPK1,DYNLL2,DIDO1,STMN3,MPRIP,PSMA7,BCR,NOL4L,BAZ1B,G
P4HB,FSCN1,SPTAN1,GSTP1,PLD3,COL18A1,MARCKSL1,TRIM28,YWHAG,CKB,COL4A2,SPEN,NC

;PTAN1,PLD3,SNRNP200,IGF2,MARCKSL1,YWHAG,CKB,COL4A2,SPEN,NONO,YWHAQ,DCHS1,SET
EC62,GPI,HTRA4,FASN,ALDH6A1,SPTBN1,CALM3,GNB1,SHISA9,RPS8,MYH9,RPL7,SEPTIN2,APP,(
PL4,ENO1,TUBA1B,MAP1B,NCL,RPL7A,SLC9A7,ALDOA,PKM,GANAB,COL6A1,FTL,HSP90B1,FADS
)S1,PRPF8,UBA1,ATP2B4,IGFBP2,CALR,FSTL1,HSPA1B,TUBB2B,COL5A1,COL4A1,CES3,PSAP,PT
/BX1,UBA1,ATP2B4,TMF1,H3-3B,CANX,CLU,PTMA,DDX5,FKBP14,RPL13A,MAGED1,RPL6,DVL3,Y
DE1,ATP5F1A,MARCKSL1,TRIM28,COL4A2,DDB1,CHSY1,GTF3C4,PKD1,SEC61A1,ITGB1,RPLP1,
TAN1,CALM1,USP11,SNRNP200,CKB,RPL37A,INA,EIF4A1,RPL19,PPP2R1A,RPL15,LDHB,DNAH8,
1,SMG1,PLD3,USP11,SNRNP200,SMIM14,ATP5F1A,EIF4B,UBB,EIF1,EIF4A1,RPL15,GTF3C4,SF3
/IAP1B,NCL,RPL7A,SLC9A7,PKM,GANAB,COL6A1,FTL,HSP90B1,FADS1,NEFL,PRPF8,RPL8,ATP2E
PXDN,HNRNPU,NFE2L1,LLGL1,HK1,CLSTN1,NLGN2,CHD4,JUND,CD81,WDR1,ATP2A2,PFN1,AST
-4B,UBB,RPL37A,SPEN,EIF1,RPL15,GTF3C4,SF3B1,ZNF43,HMGB1,POLR2A,TPT1,RPL18,TUBB4
F4L1,RPLP1,HNRNPA0,CNN3,RPSA,RPS9,MSN,C1ORF43,RHOA,CXADR,RPS20,BEX3,CCT7,VAT1,
YWHAE,DDX17,BSG,FASN,ALDH6A1,ATP5F1B,CALM3,SEPTIN2,P4HB,APP,IDS,ERVV-1,GDI1,PTMS
CANX,DDX5,MAGED1,SEC62,KIF1A,DDX17,HTRA4,BSG,ALDH6A1,SPTBN1,CALM3,MYH9,SEPTIN2
PA1B,TUBB2B,COL5A1,COL4A1,CES3,PSAP,PGK1,CLU,ENO2,RPL13A,YWHAE,KIF1A,GNAS,BSG,
LDH6A1,SPTBN1,CALM3,GNB1,SEPTIN2,P4HB,GDI1,FSCN1,SMG1,NPM1,PLXNB2,PLD3,USP11,C
4,SCARB2,RPS24,SLC22A17,CIPC,NCS1,RHOA,CXADR,ATP1A1,JUND,CCNI,CSNK1E,SFPQ,PIP4K
,CBX5,CLSTN1,CXADR,ATP1A1,TADA2B,KHDRBS1,SFPQ,EIF5A,PEBP1,SOX12,MIF,MYO5A,SESN3
-4G2,ZNF316,DBN1,ENO1,PIK3R2,FLNA,MAP1B,SLC9A7,ALDOA,PKM,FADS1,NEFM,PRPF8,UGT8
PGK1,H3-3B,PTMA,DDX5,SOX4,FKBP14,RPL13A,MAGED1,RPL6,YWHAE,GPI,HTRA4,RACK1,FASN
,ENO2,KAT6A,DVL3,YWHAE,GNAS,BSG,FASN,ALDH6A1,GNB1,RPS8,MYH9,GDI1,TMBIM6,GSTP1
4,IGFBP2,CALR,FSTL1,TMF1,TUBB2B,COL5A1,COL4A1,CES3,PSAP,PGK1,ENO2,RPL13A,KAT6A,
(T,PNMA1,SOX12,ARL4C,MIF,GPAT4,MYO5A,B4GALNT4,APLP2,TARDBP,CADM4,HNRNPH1,HLA-
K1,FASN,CALM3,GNB1,SHISA9,RPS8,P4HB,TMBIM6,PLXNB2,HSPD1,USP11,SMIM14,EIF3C,ATP5
RPL5,TMBIM6,PTMS,GSTP1,MARCKS,CALM1,HSPD1,EIF3C,ATP5F1A,EIF4B,UBB,COL4A2,LRRC2
3,COL5A1,PSAP,PGK1,DDX5,YWHAE,KIF1A,RPL13,DDX17,ATP5F1B,GNB1,RPL7,SEPTIN2,P4HB,F
A,CTNNA1,CTSD,BRD3,ZNF865,SLC7A5,JUND,VAT1,PFN1,TLE5,XPR1,TKT,GPC3,SOX12,ARL4C,
MATR3,NONO,PSMD5,CHSY1,TMSB10,EIF4A1,DNAH8,RPS3,AP2M1,TPT1,SEC61A1,RPL18,TUBB
GPI,HTRA4,CALM3,STMN2,RPS8,MYH9,RPL7,P4HB,MARCKS,CALM1,NPM1,USP11,COL18A1,VCP
PL3,RPS6,ZNF316,DBN1,ENO1,TUBA1B,PIK3R2,FLNA,MAP1B,NCL,RPL7A,SLC9A7,MTRNR2L8,P
S,MARCKS,CALM1,PLXNB2,USP11,COL18A1,IGF2,ATP5F1A,VCP,MARCKSL1,MATR3,YWHAG,CKE
JBB2B,COL4A1,PSAP,PGK1,RPLP0,CANX,DDX5,FKBP14,RPL13A,MAGED1,KAT6A,DVL3,KIF1A,GF
UBA1,ATP2B4,CALR,TMF1,HSPA1B,H3-3B,CANX,DDX5,FKBP14,RPL13A,MAGED1,RPL6,DVL3,GN
MGB1,POLR2A,TPT1,RPS3A,TUBB4B,SARAF,RPL10,DNAJA1,SF3B2,GTF2F1,RPL23,RPL29,RPS9
,EIF1,EIF4A1,RPL15,GTF3C4,SF3B1,ZNF43,HMGB1,POLR2A,TPT1,RPS3A,TUBB4B,RPL11,SARAF
OL4A2,RPL37A,DCHS1,SET,EGLN1,MRC2,SSBP2,CHD3,BASP1,TPT1,STMN1,MORF4L1,ANXA2,A
E,KIF1A,DDX17,RACK1,BSG,ALDH6A1,STMN2,SEPTIN2,IDS,GDI1,MARCKS,NPM1,USP11,COL18A
RNPH1,MAGED4B,CTBP1,ARHGDIA,CDK16,BCLAF1,NNAT,SUGP2,PPP1CB,MEST,HNRNPL,ABCA3,
PL7,FSCN1,PTMS,SMG1,SNRNP200,SMIM14,IGF2,EIF3C,ATP5F1A,EIF4B,VCP,MARCKSL1,AFF4,
RNR2L8,PKM,GANAB,FADS1,NEFL,NEFM,PRPF8,RPL8,FBXL4,CALR,HSPA1B,TUBB2B,COL5A1,C
RPL6,YWHAE,KIF1A,RPL13,GNAS,GPI,HTRA4,RACK1,BSG,FASN,SPTBN1,ATP5F1B,CALM3,GNB1,
D1,RPL6,DVL3,RACK1,ATP5F1B,SHISA9,SEPTIN2,P4HB,RPL5,TMBIM6,MARCKS,SMG1,PLD3,HSF
.37A,LRRC28,EIF1,RPL19,PPP2R1A,SF3B1,SERINC1,HMGB1,POLR2A,AP2M1,TPT1,RPS4X,TUBB
,CXADR,ATP1A1,JUND,EIF5,HMGN2,CCNI,CSNK1E,PEBP1,B4GALNT4,DNAJC5,APLP2,MXD4,DUX
)1,USP11,SNRNP200,SMIM14,EIF3C,ATP5F1A,EIF4B,UBB,AFF4,LRRC28,SPEN,NONO,INA,RPL19,

I,BTF3,MIDN,DNAJA1,MAT2A,GTF2F1,PXDN,HNRNPA0,CTNNA1,FADS2,CNN3,RPSA,EIF4H,BRD3,I
K1,UBA1,ATP2B4,TMF1,HSPA1B,TUBB2B,COL5A1,PGK1,H3-3B,CANX,CLU,PTMA,DDX5,FKBP14,F
SSRP1,HSPH1,ACO2,HSPA4,KLC1,LAMC1,ASXL1,HDGF,TERF2IP,ARF3,VHL,SMARCA4,PRDX1,ATP
,SLC2A3,GPI,FASN,ALDH6A1,STMN2,MYH9,SEPTIN2,GDI1,FSCN1,TMBIM6,PTMS,SPTAN1,NPM1,
3,HMGB1,POLR2A,TPT1,RPS3A,TUBB4B,MIDN,RPL10,RPS17,DNAJA1,SF3B2,GTF2F1,HNRNPA0,I
,HMGB1,POLR2A,TPT1,SEC61A1,RPS3A,TUBB4B,MIDN,RPL10,DNAJA1,SF3B2,GTF2F1,HNRNPA(
HSP90B1,NEFM,RPL8,FBXL4,YBX1,UBA1,ATP2B4,CALR,FSTL1,TMF1,ABHD2,ZNF99,TUBB2B,COL
,LAMA5,SLC22A17,PJA2,HK1,ZNF865,CLSTN1,ATF4,RAB6B,SLC7A5,NLGN2,ATP1A1,CD81,WDR1
SF3B1,ZNF43,HMGB1,POLR2A,TPT1,SEC61A1,RPS4X,RPS3A,TUBB4B,MIDN,RPL10,DNAJA1,ARF
DX17,GPI,FASN,MYH9,GDI1,PTMS,SPTAN1,MARCKS,SMG1,PLXNB2,TTC3,USP11,SMIM14,IGF2,
L3,ZNF316,RPL4,TUBA1B,PIK3R2,MAP1B,NCL,SLC9A7,ALDOA,PKM,COL6A1,HSP90B1,NEFL,NEF
3N2,HDLBP,CDH6,CDH2,PIP4K2B,CCN2,RPL12,GNB2,B4GALNT4,APLP2,MXD4,WLS,MYH10,CAD
,RPL22,CLSTN1,ATF4,GLIPR1L2,HMGN2,ATP2A2,BEX3,CCNI,KHDRBS1,PFKL,GLG1,APLP1,HNRI
PS6,ZNF316,DBN1,ENO1,TUBA1B,NCL,RPL7A,SLC9A7,ALDOA,GANAB,COL6A1,FTL,HSP90B1,FA
L4,YBX1,UBA1,ATP2B4,CALR,FSTL1,TMF1,RPLP0,H3-3B,CANX,DDX5,RPL13A,MAGED1,RPL6,DV
E1,UBB,TRIM28,YWHAG,CKB,DDB1,EIF1,SET,TPI1,EIF4A1,ZNF43,CHD3,AP2M1,AGRN,TIMP2,MAI
I,TPT1,SEC61A1,RPS3A,TUBB4B,MORF4L1,BTF3,RPS17,AP3D1,CYB5R3,SF3B2,ITM2B,MAT2A,R
PL13,DDX17,GPI,HTRA4,BSG,SPTBN1,GNB1,RPS8,MYH9,GDI1,PTMS,MARCKS,SMG1,PLXNB2,US
TP5F1B,CALM3,RPS8,RPL7,SEPTIN2,PTMS,MARCKS,CALM1,NPM1,HSPD1,USP11,SNRNP200,IGI
X5,FKBP14,RPL13A,MAGED1,RPL6,DVL3,RACK1,ATP5F1B,SHISA9,SEPTIN2,P4HB,RPL5,GDI1,TM
P90B1,FADS1,NEFL,FBXL4,YBX1,ATP2B4,ABHD2,TUBB2B,COL5A1,CES3,PGK1,H3-3B,CANX,CLU
A1,EIF4G2,RPL3,RPS6,DBN1,RPL4,ENO1,TUBA1B,PIK3R2,MAP1B,NCL,SLC9A7,MTRNR2L8,PKM,
3,CANX,CLU,PTMA,DDX5,RPL13A,SEC62,KAT6A,DVL3,YWHAE,KIF1A,DDX17,RACK1,BSG,ALDH6A
A0,CNN3,RPL29,RPS9,HK1,ZNF865,RPS14,RPS20,WDR1,PFN1,MCL1,TLE5,RPL31,TKT,EIF5A,CD
2B4,FSTL1,TUBB2B,COL4A1,PSAP,PGK1,CANX,CLU,ENO2,SEC62,KIF1A,RPL13,GNAS,GPI,HTRA4
,ACTR2,SPARC,TUBB,FTH1,UBC,KIDINS220,TMSB4X,NUMA1,DCX,EIF4G2,RPL3,RPS6,ZNF316,DI
SPEN,DDB1,TPI1,PPP2R1A,RPL15,LDHB,CDKL1,ITGB1,TUBB4B,RPLP1,TIMP2,RPL10,WSB1,VCL,
RPL6,DVL3,GPI,RACK1,ATP5F1B,SHISA9,RPS8,P4HB,RPL5,TMBIM6,PLD3,USP11,SNRNP200,SMI
PH1,MAGED4B,CTBP1,ARHGDIA,CDK16,BCLAF1,NNAT,EID1,TOMM20,U2AF2,SUGP2,CPSF1,ME
1,TRIM28,CKB,SPEN,DDB1,YWHAQ,EIF1,SET,TPI1,TMSB10,GTF3C4,SSBP2,CHD3,AP2M1,TPT1,I
L26,SRSF4,RBBP4,PFN1,RPL31,PRKCSH,HNRNPC,KHDRBS1,ITM2C,RPL30,EIF3L,CHPF,NGRN,R
CADM4,HECTD4,L1CAM,SQLE,SERF2,TIMP3,CHD6,TBCD,LMNB2,GRINA,VPS35,MDK,JPT1,PTK7,
UBB4B,RPL10,SF3B2,GTF2F1,RPL23,RPL29,RPS9,XRCC5,CIPC,RPL24,PAPOLA,HK1,RHOA,ZNF8
F4H,SCARB2,RPL22,RHOA,GLIPR1L2,EIF5,LENG8,LASP1,PFN1,KHDRBS1,RPS19,HNRNPDL,PTG
GPI,HTRA4,BSG,SPTBN1,GNB1,MYH9,GDI1,PTMS,SMG1,PLXNB2,USP11,COL18A1,SMIM14,IGF2,
P5PB,GRB2,WDR6,RPS11,SUPT5H,CAND1,DIDO1,NKTR,LSS,HNRNPD,RAB11A,SSR1,SF1,PPM1G
CNN3,RPSA,TPM4,EIF4H,BRD3,RPL24,KPNB1,RPL14,CBX5,CXADR,RPS20,CD81,WDR1,HMGN2,F
2B4,IGFBP2,CALR,FSTL1,TUBB2B,COL4A1,CES3,H3-3B,PTMA,DDX5,SOX4,RPL13A,MAGED1,SE
B1,TPT1,SEC61A1,RPS3A,ITGB1,TUBB4B,BTF3,RPLP1,MIDN,RPL10,MARCHF6,CTNNA1,PAPOLA,
7,CLSTN1,NLGN2,JUND,TMEM259,SRSF4,PFN1,ASTN2,JPT2,TLE5,TADA2B,TPM1,EIF5A,CDH2,EI
T6A,YWHAE,RPL13,GNAS,DDX17,HTRA4,ATP5F1B,CALM3,GNB1,SEPTIN2,GDI1,FSCN1,TMBIM6,I
4,RPL13A,MAGED1,RPL6,DVL3,GPI,RACK1,ATP5F1B,CALM3,SHISA9,SEPTIN2,P4HB,RPL5,GDI1,T
3X5,CXADR,RPS20,CD81,WDR1,HMGN2,RBBP4,MCL1,PNN,RPL31,HNRNPC,HDLBP,EIF5A,CCND2
,RPS27,RAC1,ATP6V0C,SRP9,VAMP2,NOP53,PABPN1,HNRNPL,SCAF1,TERF2IP,RABL6,ILF3,ATXI
IZ1,MAP4,CDC42BPB,TMEM170A,CTNNA1,TPM4,SCARB2,RPS24,SLC22A17,COL1A2,RHOA,SLC

TR3,LRRC28,EIF1,RPL19,PDIA3,SERINC1,HMGB1,POLR2A,AP2M1,TPT1,RPS4X,TUBB4B,RPL11,I
GNB1,MYH9,ERVV-1,RPL5,PTMS,GSTP1,PLXNB2,USP11,COL18A1,IGF2,EIF3C,MARCKSL1,YWH/
C4,PDIA3,STMN1,TUBB4B,AGRN,ZMIZ1,ARF1,MAP4,CDC42BPB,MAT2A,HNRNPA0,CTSD,RPL24,H
NA,LAMC1,JPT1,CBX1,TNPO1,ASXL1,BTBD2,PBX2,SRSF1,PTBP1,CAPZB,UBE2R2,MAP7D1,ARF4
NAJC5,APLP2,RAB1A,CTBP1,ARHGDIA,NOVA2,CHD6,EEF1A2,TBCD,PGD,SQSTM1,VAMP2,ACTR3,
OLR2A,RPS4X,ITGB1,AGRN,TIMP2,VCL,GTF2F1,SLC4A8,PXDN,HNRNPU,OGA,CTNNA1,CNN3,CTS
,FTL,FADS1,UGT8,RPL8,FBXL4,UBA1,ATP2B4,IGFBP2,TUBB2B,CES3,CANX,CLU,PTMA,ENO2,FK
RL4C,APLP1,NDFIP1,SND1,COTL1,CTBP1,TOP2B,DLG5,CRPPA,CTDSP2,PGRMC1,CTSB,CHD6,P
EC62,KAT6A,DVL3,YWHAE,KIF1A,SLC2A3,DDX17,HTRA4,RACK1,BSG,ALDH6A1,ATP5F1B,STMN2,
PS3,POLR2A,AGRN,MARCHF6,AP3D1,ARF1,PXDN,PGAM1,RPS16,RPL23,LPCAT1,XRCC5,PJA2,C1
R2A,TPT1,RPS3A,TUBB4B,RPS17,SF3B2,GTF2F1,RPL23,RPL29,RPS9,XRCC5,CIPC,RPL24,PAPOL
L23,TNS3,RPL14,HK1,NCS1,NR2F1,KIF5B,RAB6B,CXADR,ATP1A1,HMGN2,BEX3,TADA2B,KHDRE
2A,ITGB1,AGRN,ZMIZ1,TIMP2,WSB1,VCL,GTF2F1,PXDN,HNRNPU,OGA,FADS2,BRD3,SLC16A7,K
MCL1,TLE5,TKT,EIF5A,PDIA6,RAB7A,EIF3L,CHPF,NGRN,HNRNPDL,RPLP2,DDOST,YWHAB,ATP1A2
3,HSPD1,COL18A1,CSDE1,IGF2,MARCKSL1,AFF4,DDB1,SET,CHSY1,LDHB,SF3B1,VAPA,HMGB1,C
BX1,UBA1,TMF1,HSPA1B,TUBB2B,COL5A1,PSAP,PGK1,RPLP0,H3-3B,PTMA,DDX5,FKBP14,RPL1
RPLP2,DDOST,NARS1,YWHAB,PTGES3,RPS12,RPS27,ARHGDIA,SLC35E2B,ATP6V0C,RPL23A,SEF
VL3,YWHAE,KIF1A,DDX17,RACK1,BSG,ALDH6A1,ATP5F1B,STMN2,RPS8,SEPTIN2,GDI1,MARCKS,
,SEPTIN2,GSTP1,HSPD1,SNRNP200,VCP,MARCKSL1,UBB,AFF4,MATR3,RPL37A,DDB1,HACD3,SE
RPS3A,TUBB4B,RPL11,BTF3,RPLP1,SARAF,MIDN,RPL10,MARCHF6,DNAJA1,CYB5R3,SF3B2,GTF
PGK1,RPLP0,RPL13A,KAT6A,DVL3,YWHAE,KIF1A,RPL13,GPI,RACK1,BSG,GNB1,RPL7,SEPTIN2,P
PL13A,SEC62,KAT6A,DVL3,YWHAE,DDX17,RACK1,BSG,ALDH6A1,ATP5F1B,STMN2,RPS8,SEPTIN2
,HSP90B1,PRPF8,RPL8,CALR,TUBB2B,COL4A1,PSAP,PGK1,CLU,PTMA,DDX5,ENO2,SOX4,FKBP1
RNPDL,PTGES3,CDK16,BCLAF1,C5ORF24,HNRNPL,MORF4L2,SSRP1,ABCA3,ALKBH5,PKN1,ACA
5,ATP6V0C,DLG5,CNN2,NOVA2,TM9SF3,CHD6,VAMP2,YEATS2,GRINA,PTOV1,MDK,DDX42,SNRNF
,PTMA,DDX5,RPL13A,MAGED1,RPL6,YWHAE,GNAS,GPI,FASN,SHISA9,MYH9,P4HB,TMBIM6,SPTAI
5,CCNI,TKT,TPM1,CDH2,PIP4K2B,MTSS2,APLP2,TARDBP,DUX4,RAB1A,GPC1,TOP2B,IPO5,NOVA2
GPI,BSG,GNB1,RPS8,SEPTIN2,P4HB,IDS,FSCN1,GSTP1,MARCKS,PLD3,HSPD1,USP11,SNRNP200,I
RNPH1,HLA-B,CTBP1,GPC1,SLC35E2B,IPO5,ATP6V0C,DNER,TRIB2,SQLE,TIMP3,CHD6,EEF1A2,T
HD2,ABCA2,DPYSL2,TOP1,VEGFA,VHL,SMARCA4,AZIN1,SRSF3,HECTD1,PFN2,P3H3,SERBP1,PRF
LAF1,NNAT,U2AF2,SUGP2,HNRNPL,ABCA3,ALKBH5,CAMSAP1,VPS35,DPYSL2,PPP2CA,SMARCA4
I1,ATF4,TMEM259,BEX3,SPC25,NISCH,PRKCSH,KLHL9,HDLBP,GLG1,SFPQ,PNMA1,RAB7A,NANC
,HSP90B1,FADS1,NEFM,PRPF8,UGT8,RPL8,FBXL4,YBX1,IGFBP2,TMF1,ABHD2,ZNF99,TUBB2B,C
,RPL3,RPS6,ZNF316,DBN1,ENO1,TUBA1B,PIK3R2,FLNA,MAP1B,NCL,SLC9A7,ALDOA,MTRNR2L8
I2,CXADR,LENG8,WDR1,LASP1,SRSF4,RBBP4,PFN1,TLE5,PNN,NES,NISCH,CDH6,PIP4K2B,SOX12
TUBB4B,BTF3,AGRN,RPLP1,RPL10,MARCHF6,DNAJA1,ARF1,SF3B2,TPM4,XRCC5,CIPC,RPL24,R
6A1,ATP5F1B,STMN2,RPS8,SEPTIN2,GDI1,NPM1,TTC3,USP11,COL18A1,EIF3C,ATP5F1A,EIF4B,U
F43,NLGN2,CHD4,HMGN2,CRMP1,BEX3,TLE5,SPTBN2,SLC3A2,CCND2,CDH2,GPC3,DDR1,ACSL6
IGN2,RBBP4,MCL1,PNN,HNRNPC,HDLBP,EIF5A,CCND2,SOX12,RPS15,ITM2C,ARL4C,RPL35,NGF
S4X,NUMA1,DCX,EIF4G2,RPL3,RPS6,ZNF316,DBN1,RPL4,ENO1,TUBA1B,PIK3R2,MAP1B,NCL,RP
4,UBA1,TMF1,ABHD2,TUBB2B,PSAP,RPLP0,DDX5,FKBP14,SEC62,KIF1A,RPL13,GNAS,DDX17,ALI
ADM4,ARCN1,COTL1,STIP1,SLC35E2B,IPO5,DNER,LAPTM4B,SQLE,MAP1A,TIMP3,CTDSP2,CNN2,
0,RPS17,SF3B2,GTF2F1,RPL23,RPL29,RPS9,XRCC5,CIPC,RPL24,HK1,RHOA,ZNF865,GLIPR1L2,
SEC62,KAT6A,HTRA4,ATP5F1B,CALM3,GNB1,SHISA9,APP,IDS,RPL5,GDI1,FSCN1,PTMS,SMG1,PL
ENG8,WDR1,RPL31,PRKCSH,TKT,PEBP1,PIP4K2B,SOX12,MYO5A,DNAJC5,RPLP2,NDFIP1,MXD4,F

7A,INA,EIF4A1,RPL19,PPP2R1A,RPL15,DNAH8,SF3B1,SERINC1,RPS3,HMGB1,POLR2A,AP2M1,RF
T1,RPS3A,RPL18,SARAF,MIDN,MARCHF6,WSB1,CYB5R3,AP2B1,GTF2F1,GOLGA8A,LDHA,CPE,CT
A1,FSTL1,ABHD2,HSPA1B,TUBB2B,PGK1,H3-3B,PTMA,DDX5,SOX4,MAGED1,KAT6A,DVL3,YWHA
GK1,H3-3B,CANX,DDX5,RPL13A,MAGED1,RPL6,DVL3,GNAS,GPI,SHISA9,MYH9,SEPTIN2,P4HB,RP
IDRBS1,EIF5A,RPL12,ITM2C,EIF3L,CHPF,RPLP2,DDOST,SLC35E2B,RPL23A,HSBP1,SRSF5,TM9S
3,SHISA9,APP,IDS,GDI1,PTMS,SPTAN1,SMG1,CALM1,NPM1,PLXNB2,USP11,CSDE1,IGF2,EIF4B,M
NAS,SLC2A3,GPI,FASN,ALDH6A1,SPTBN1,CALM3,APP,RPL5,GDI1,PTMS,SPTAN1,MARCKS,CALM1
RCHF6,DNAJA1,ARF1,SF3B2,HNRNPA0,RPL22,ZNF865,KIF5B,RPS14,CXADR,LENG8,WDR1,CCT7
CHF6,DNAJA1,SF3B2,GTF2F1,RPL23,RPL29,XRCC5,CIPC,RPL22,ZNF865,ATF4,CXADR,LENG8,R
L29,TNS3,LPCAT1,LAMA5,SLC22A17,CIPC,COL1A2,LLGL1,GAP43,NCS1,PSMD2,KIF5B,RAB6B,A
IF43,HMGB1,POLR2A,TPT1,RPS4X,RPS3A,TUBB4B,BTF3,DNAJA1,CYB5R3,SF3B2,GTF2F1,RPL2
90B1,NEFL,RPL8,FBXL4,UBA1,ATP2B4,IGFBP2,CALR,TMF1,CANX,CLU,PTMA,DDX5,FKBP14,RPL
90B1,NEFL,RPL8,FBXL4,UBA1,ATP2B4,IGFBP2,CALR,TMF1,CANX,CLU,PTMA,DDX5,FKBP14,RPL
FADS1,NEFM,PRPF8,UGT8,RPL8,FBXL4,YBX1,IGFBP2,TMF1,ABHD2,ZNF99,TUBB2B,COL4A1,PS
LR2A,AP2M1,TPT1,BTF3,AGRN,RPLP1,RPL10,MARCHF6,DNAJA1,ARF1,SF3B2,GTF2F1,RPL29,EIF
SEC62,YWHAE,GNAS,BSG,SPTBN1,RPS8,MYH9,P4HB,APP,RPL5,GDI1,MARCKS,SMG1,CALM1,NP
F3C,ATP5F1A,UBB,AFF4,RPL37A,SPEN,EGLN1,RPL19,RPL15,SERINC1,HMGB1,POLR2A,AP2M1,
,COL5A1,H3-3B,PTMA,DDX5,RPL13,GPI,RACK1,ALDH6A1,MYH9,P4HB,IDS,ERVV-1,PTMS,SPTAN
F3L,MAGED2,RPS19,NGRN,RPLP2,DDOST,CIRBP,YWHAB,CALU,ARCN1,ATP6V0C,LAPTM4B,TRIB
14,RPS20,WDR1,PFN1,MCL1,TLE5,PNN,KHDRBS1,EIF5A,MEGF8,ITM2C,EIF3L,RPS7,RPS19,CHP
HMGB1,POLR2A,AP2M1,TPT1,RPS4X,TUBB4B,RPL11,BTF3,AGRN,RPLP1,RPL10,MARCHF6,DNAJ
ATF4,JUND,EIF5,WDR1,HMGN2,ASTN2,TLE5,NES,TADA2B,SFPQ,PDIA6,GPC3,MEGF8,PRDX2,ARL
S5,DNAJC5,HNRNPH1,MAGED4B,CTBP1,ARHGDIA,CDK16,TMEM30A,BCLAF1,NNAT,CRPPA,EID1,
B4X,NUMA1,RPL3,RPS6,ZNF316,DBN1,TUBA1B,PIK3R2,MAP1B,NCL,RPL7A,SLC9A7,ALDOA,MT
M1,TPT1,TUBB4B,RPL11,BTF3,AGRN,RPLP1,RPL10,MARCHF6,DNAJA1,ARF1,SF3B2,XRCC5,CIP
F11,MSL1,STMN3,LSS,CSNK1D,IDH1,ZFAND3,HECTD1,SARS1,VANGL2,P3H3,KCNQ2,BCL9L,WD
L3,YWHAE,KIF1A,DDX17,RACK1,BSG,ALDH6A1,ATP5F1B,STMN2,RPS8,SEPTIN2,GDI1,MARCKS,
M6,USP11,SNRNP200,CSDE1,IGF2,EIF3C,ATP5F1A,EIF4B,MATR3,RPL37A,LRRC28,EIF1,RPL19,
4,CD81,LENG8,VAT1,PFN1,TLE5,XPO1,EIF5A,GNB2,WLS,DUX4,LGALS3BP,ATP1A2,ARCN1,MAGE
ATP2B4,IGFBP2,FSTL1,H3-3B,CLU,PTMA,DDX5,ENO2,SOX4,MAGED1,YWHAE,GNAS,GPI,CALM3,
,SF3B1,SSBP2,CDKL1,STMN1,TUBB4B,UCHL1,AGRN,ZMIZ1,MIDN,AP3D1,AKAP12,CDC42BPB,G
TMF1,TUBB2B,COL5A1,PSAP,PGK1,H3-3B,CANX,CLU,PTMA,DDX5,ENO2,SEC62,RPL6,KAT6A,RP
F3C,UBB,AFF4,TRIM28,CKB,FN1,YWHAQ,EIF1,HACD3,SERINC1,TPT1,SEC61A1,RPS3A,RPL18,RP
C22A17,CIPC,RHOA,PTCD3,CBX5,NLGN2,RBBP4,PFN1,TLE5,PRKCSH,XPO1,SFPQ,CLTC,B4GAL
1,BTF3,AGRN,RPLP1,RPL10,MARCHF6,DNAJA1,ARF1,SF3B2,GTF2F1,XRCC5,PAPOLA,RHOA,ZN
B,FTH1,UBC,KIDINS220,TMSB4X,NUMA1,DCX,EIF4G2,RPL3,RPS6,ZNF316,DBN1,RPL4,ENO1,TU
LR2A,TPT1,RPS3A,TUBB4B,RPS17,SF3B2,GTF2F1,RPL23,RPL29,RPS9,XRCC5,CIPC,PAPOLA,RH
1,H3-3B,PTMA,DDX5,FKBP14,RPL13A,MAGED1,RPL6,YWHAE,GPI,HTRA4,RACK1,FASN,GNB1,RP
1,SHISAL1,CDH11,ZC3H7B,ABCA3,PKN1,ACO2,PTOV1,MLEC,RBMX,H3-3A,HDGF,DPYSL2,ARF3,
1,SHISAL1,CDH11,ZC3H7B,ABCA3,PKN1,ACO2,PTOV1,MLEC,RBMX,H3-3A,HDGF,DPYSL2,ARF3,
NGRN,RPLP2,DDOST,YWHAB,ARHGDIA,SLC35E2B,RPL23A,HSBP1,TM9SF3,MDH2,U2AF2,CCT2,
SRSF9,DYNLL2,GIGYF1,STMN3,VGF,FLNC,ARNT2,DYNLL1,NAP1L4,TP53BP1,CTXN1,RBM15B,B
2,GTF2F1,HNRNPA0,RPL23,RPL29,EIF4H,RPS9,XRCC5,PAPOLA,RPL22,RHOA,ZNF865,RPS14,RF
GOLGA8A,CTNNA1,TPM4,SCARB2,RPS24,CIPC,GAP43,HK1,RPL22,ZNF865,CHD4,ATP1A1,WDR1,
4,CADM4,HECTD4,TMEM132A,EEF1A2,TGFBI,OPTN,SRSF9,BTBD2,PBX2,DYNLL2,ILF3,LUZP6,CA

/I1,PLD3,USP11,IGF2,MATR3,YWHAG,CKB,HACD3,DCHS1,TMSB10,PKD1,VAPA,RPS3A,RPL18,ITG
D1,KAT6A,DVL3,KIF1A,GPI,SPTBN1,GNB1,RPL7,IDS,SMG1,CALM1,NPM1,TTC3,HSPD1,USP11,SN
A1,MRC2,PPP2R1A,PKD1,AP2M1,UCHL1,AGRN,ZMIZ1,SARAF,MIDN,MAT2A,FADS2,TNS3,TSPAN3
1,RPS4X,TUBB4B,BTF3,CYB5R3,SF3B2,GTF2F1,RPL29,RPS9,CIPC,RPL24,RHOA,ZNF865,RPS14,
PM4,LAMA5,SLC22A17,LLGL1,RPL22,CLSTN1,NR2F1,RAB6B,SLC7A5,CXADR,CHD4,JUND,WDR1
LDHA,SLC22A17,PAPOLA,NCS1,ATP1A1,JUND,TMEM259,LENG8,ASTN2,MCL1,CCNI,TADA2B,XPF
,RPS20,LENG8,WDR1,RBBP4,PFN1,TLE5,PNN,PDIA6,PIP4K2B,RPL30,EIF3L,RPLP2,DDOST,YWH
EIF4A1,RPL19,SF3B1,SERINC1,ZNF43,HMGB1,POLR2A,AP2M1,TUBB4B,BTF3,AGRN,RPLP1,RPL
AG,CKB,RPL37A,SPEN,CHSY1,GTF3C4,LDHB,BASP1,POLR2A,TUBB4B,UCHL1,BTF3,RPLP1,RPL1
S20,WDR1,RPL26,RBBP4,PFN1,RPL31,SLC3A2,RPS25,EIF5A,RAB7A,RPL12,EIF3L,RPS5,RPS19,C
S20,WDR1,RPL26,RBBP4,PFN1,RPL31,SLC3A2,RPS25,EIF5A,RAB7A,RPL12,EIF3L,RPS5,RPS19,C
NAJA1,AKAP12,MAP4,CDC42BPB,SLC4A8,HNRNPU,NFE2L1,CTNNA1,TNS3,LAMA5,MSN,CBX5,RA
.10,SF3B2,GTF2F1,RPL23,RPL29,TPM4,XRCC5,HK1,ZNF865,RPS14,RPS20,LENG8,WDR1,SRSF4
.P1,SARAF,CYB5R3,MAP4,GOLGA8A,NFE2L1,HNRNPA0,RPL23,LDHA,LAMA5,RPS9,KPNB1,NCS1,
C62,KIF1A,FASN,ALDH6A1,SPTBN1,SHISA9,SEPTIN2,GDI1,FSCN1,PTMS,SPTAN1,MARCKS,SMG1,
RPL18,RPLP1,SARAF,MIDN,MARCHF6,WSB1,AP3D1,ARF1,ITM2B,AP2B1,GTF2F1,GOLGA8A,LDH
1,ABHD2,ZNF99,COL5A1,PSAP,RPLP0,PTMA,DDX5,RPL13A,KAT6A,DVL3,YWHAE,RPL13,GPI,BSG,
1,NEFL,PRPF8,UGT8,IGFBP2,FSTL1,TMF1,ABHD2,TUBB2B,PSAP,RPLP0,H3-3B,PTMA,DDX5,ENO
OLGA8A,CTNNA1,SCARB2,RPS24,CIPC,GAP43,HK1,RPL22,ZNF865,CHD4,ATP1A1,WDR1,OAZ1,P
CS1,CBX5,ATF4,PSMD2,CHD4,JUND,EIF5,ATP2A2,RPL26,ASTN2,TLE5,PRKCSH,KHDRBS1,PFKL,
3,RPL29,PTCD3,ZNF865,RPS14,LENG8,WDR1,RPL26,SRSF4,RBBP4,PFN1,MCL1,PRKCSH,RPS25
,DDX17,RACK1,BSG,ALDH6A1,ATP5F1B,STMN2,RPS8,SEPTIN2,GDI1,MARCKS,NPM1,TTC3,USP1
RB2,RPS24,GAP43,HK1,RPL22,ZNF865,CHD4,WDR1,OAZ1,NES,TADA2B,KHDRBS1,PFKL,PEBP1,
WHAB,RPS27,ARHGDIA,SLC35E2B,SMCHD1,ATP6V0C,DDX39B,NUCB1,TOMM20,U2AF2,RPN2,G
N,RPLP1,MARCHF6,MAT2A,HNRNPA0,RPS16,SLC16A7,ZNF865,RPL9,KIF5B,RPS14,CXADR,WDR1
ERINC1,HMGB1,POLR2A,AP2M1,TPT1,RPS4X,TUBB4B,BTF3,AGRN,RPLP1,RPL10,MARCHF6,DNA
.,PSMC5,WDR82,ASH1L,KDM2A,CDC42SE1,UBE2G1,TAB3,ABCE1,MSI2,NUP50,CCNL1,ZNF207,Z
B1,NEFM,UBA1,IGFBP2,HSPA1B,TUBB2B,COL4A1,PGK1,PTMA,ENO2,SOX4,SEC62,KAT6A,DVL3,
8,SF3B1,ZNF43,HMGB1,POLR2A,TPT1,RPS3A,RPL18,TUBB4B,DNAJA1,SF3B2,GTF2F1,RPL23,RF
P2,TAGLN,MYL6,SHOX,LGALS3BP,ATP1A2,CKAP4,PLPPR3,HECTD4,SLC35E2B,IPO5,ATP6V0C,NN
RN,RPLP1,RPL10,MARCHF6,DNAJA1,ARF1,SF3B2,XRCC5,CIPC,PAPOLA,ZNF865,ATF4,RPS14,CXA
/PS35,PTOV1,MLEC,CLSTN3,MDK,H3-3A,PYGB,DDAH2,PLCG1,CORO1C,LSS,STXBP1,ISYNA1,IDH
6,SPTAN1,CALM1,PLXNB2,TTC3,SMIM14,IGF2,VCP,UBB,TRIM28,YWHAG,EGLN1,CHSY1,TMSB1
,RPS20,WDR1,PFN1,MCL1,PNN,PRKCSH,RPS25,EIF5A,PDIA6,EIF3L,IK,CHPF,NGRN,HNRNPDL,R
.RRC28,SPEN,NONO,MRC2,GTF3C4,LDHB,PKD1,BASP1,POLR2A,TPT1,ITGB1,AGRN,TIMP2,MAR
MARCHF6,WSB1,ARF1,AP2B1,GTF2F1,GOLGA8A,LDHA,CPE,SLC22A17,RPS9,CIPC,PJA2,SLC16A7
DA2B,DDX3X,ARL4C,HNRNPDL,MYL6,SHOX,COTL1,MAGED4B,IPO5,L1CAM,TIMP3,SRP9,SRSF5,C
F6,RPS17,ZNF770,GTF2F1,HNRNPU,NFE2L1,CTNNA1,RPS16,CPE,CNN3,TPM4,RPS24,RPS9,SLC
NF316,DBN1,ENO1,PIK3R2,FLNA,MAP1B,NCL,ALDOA,MTRNR2L8,PKM,GANAB,COL6A1,HSP90B
KHDRBS1,SPTBN2,PFKL,SULF1,RPS19,JUN,ARL6IP1,HNRNPK,SEPTIN11,CALU,SEPTIN3,CTBP1,G
P1,MARCHF6,DNAJA1,ARF1,SF3B2,TPM4,XRCC5,CIPC,RPL22,ZNF865,ATF4,RPS14,CXADR,LENG
AP1,RPS19,MYO5A,DNAJC5,RPLP2,DDOST,DUX4,SEPTIN11,PLPPR3,GPC1,TRIB2,SQLE,NOVA2,C
P4,COTL1,MAGED4B,PTTG1IP,DLG5,VCAN,LAMC1,CNTNAP2,VEGFA,DSTN,BTBD2,PRDX1,AEBP1,
LR2A,TPT1,RPS3A,RPL18,RPLP1,SARAF,MIDN,MARCHF6,WSB1,ITM2B,AP2B1,GTF2F1,GOLGA8
ZNF770,GOLGA8A,PGAM1,SMARCA5,RPL23,CPE,CNN3,CTSD,EIF4H,LPCAT1,SLC22A17,BRD3,X

EC61A1,RPS3A,MORF4L1,RPLP1,MIDN,AP3D1,ARF1,CYB5R3,ITM2B,RPS16,CPE,EIF4H,TSPAN3,I
3,MDK,ASXL1,SMARCA4,AZIN1,NYNRIN,HSPB1,CASC3,QARS1,ZFAND5,CORO1C,KIF3C,SMARCC1
RAF,MIDN,MARCHF6,WSB1,ARF1,AP2B1,GTF2F1,GOLGA8A,LDHA,CPE,SLC22A17,RPS9,JUND,TM
H2,U2AF2,GNAI3,PRDX5,UBE4A,HSPA4,TCP1,MLEC,GCN1,SRP14,PRPF6,TAF7,RPL37,ARPC2,FAM
P1,KDM3B,RBM15B,DCTN1,MSMO1,SF1,SMC3,ZNF512B,CBX6,PRRC2A,PTPRF,MCM3AP,C1ORI
HSY1,TMSB10,MRC2,HLA-C,PDIA3,BASP1,HMGB1,STMN1,ITGB1,MORF4L1,AGRN,TIMP2,AP3D1
ZP6,SCAF11,POLR2B,SDF4,ATP6V1B2,HNRNPM,SDHA,LONP1,SARS1,CCT5,RBBP7,DFFB,UPF1,R
PL7A,ALDOA,MTRNR2L8,PKM,GANAB,FTL,HSP90B1,FADS1,NEFL,NEFM,PRPF8,UGT8,RPL8,FBXL
CA3,ACO2,HSPA4,ASXL1,PSMB2,RPS15A,ARF3,UBXN4,PRDX1,ATP5PB,GRB2,WDR6,RPS11,SUPT
RACK1,FASN,ATP5F1B,SHISA9,RPS8,P4HB,ERVV-1,RPL5,GDI1,TMBIM6,CALM1,HSPD1,USP11,SN
F316,DBN1,TUBA1B,RPL7A,SLC9A7,ALDOA,MTRNR2L8,GANAB,COL6A1,FTL,FADS1,NEFL,PRPF8
,TMEM170A,NFE2L1,RPS16,CNN3,CTSD,LPCAT1,BRD3,XRCC5,SLC16A7,KPNB1,RPL17,PAPOLA,
IMGN2,CRMP1,BEX3,RBBP4,MCL1,PNN,HNRNPC,HDLBP,CCND2,PNMA1,SOX12,ACTN4,RPS15,IT
X1,IGFBP2,CALR,TMF1,ABHD2,TUBB2B,COL4A1,PSAP,PGK1,CANX,CLU,PTMA,DDX5,RPL13A,KA
B1,NEFL,NEFM,PRPF8,YBX1,IGFBP2,FSTL1,TUBB2B,CES3,RPLP0,H3-3B,CANX,CLU,DDX5,RPL13
VPS35,MLEC,MDK,JPT1,H3-3A,SCAF1,CNTNAP2,RAB11B,TUBB2A,LAPTM4A,DDAH2,TCF25,ILF3,
F1,AP2B1,GTF2F1,GOLGA8A,LDHA,CPE,RPS9,JUND,TMEM259,LENG8,MCL1,CCNI,XPR1,HDLBP,
3,SARAF,RPL10,SF3B2,GTF2F1,RPL23,RPL29,RPS9,XRCC5,CIPC,PAPOLA,NCS1,RHOA,ZNF865,G
GLUL,EEF1A2,PABPN1,GDI2,NOX4,LMNB2,ALKBH5,PTPRA,CAMSAP1,ACAP3,LAMC1,RAD23B,NC
I,RPS8,P4HB,RPL5,TMBIM6,CALM1,HSPD1,USP11,CSDE1,EIF3C,ATP5F1A,RPL37A,PSMD5,INA,EI
RF4L1,BTF3,XAF1,AP3D1,SBF1,MAT2A,RPSA,NACA,RPS9,PJA2,RPL24,RPL17,RPL14,CBX5,CLST
LRRC28,TMSB10,EIF4A1,PPP2R1A,RPL15,GTF3C4,TPT1,SEC61A1,RPS4X,MORF4L1,RPL10,RPS
3P,YWHAB,SLC35E2B,HSBP1,TM9SF3,MDH2,U2AF2,FDFT1,CCT2,SUGP2,GNAI3,PRDX5,UBE4A,H
R2F1,GLIPR1L2,JUND,EIF5,LENG8,HMGN2,BEX3,ASTN2,PRKCSH,TADA2B,XPR1,HDLBP,EIF5A,PI
AP12,CDC42BPB,GOLGA8A,CTNNA1,TPM4,SCARB2,RPS24,IGFBP3,GAP43,HK1,RPL22,ZNF865,I
INS220,TMSB4X,EIF4G2,RPL3,RPS6,ZNF316,DBN1,PIK3R2,FLNA,MAP1B,RPL7A,PKM,GANAB,CC
SP11,COL18A1,SNRNP200,SMIM14,AFF4,TRIM28,CKB,FN1,YWHAQ,EIF1,HACD3,MAP9,CHSY1,IN
GSTP1,PLD3,USP11,COL18A1,SNRNP200,SMIM14,VCP,AFF4,TRIM28,CKB,RPL37A,FN1,YWHAQ,I
HSP90AA1,ACTR2,VIM,SPARC,TUBB,FTH1,UBC,KIDINS220,TMSB4X,NUMA1,DCX,EIF4G2,RPL3,R
PXDN,OGA,RPS16,SLC22A17,CIPC,KPNB1,ZNF865,NR2F1,NLGN2,GLIPR1L2,OAZ1,RTN3,HNRNP
4,YWHAG,DDB1,HACD3,DCHS1,EGLN1,CHSY1,TMSB10,PPP2R1A,PKD1,SF3B1,CHD3,VAPA,CDKL
NF865,ATF4,RPS14,CXADR,LENG8,WDR1,RPL26,SRSF4,PFN1,RPL31,PRKCSH,KHDRBS1,SCRN1,I
AP43,HK1,RPL22,ZNF865,CHD4,WDR1,OAZ1,NES,TADA2B,KHDRBS1,PFKL,GLG1,PEBP1,PNMA1,
,CIPC,ZNF865,ATF4,CXADR,LENG8,WDR1,RPL26,SRSF4,PFN1,RPL31,PRKCSH,SCRN1,RPS25,EII
PL8,YBX1,UBA1,CALR,FSTL1,CES3,PSAP,RPLP0,H3-3B,CANX,PTMA,DDX5,ENO2,RPL13A,MAGED

OL5A1,COL4A1,PSAP,RPLP0,CANX,DDX5,MAGED1,KIF1A,RPL13,DDX17,GPI,ALDH6A1,SPTBN1,C
,YBX1,UBA1,CALR,FSTL1,HSPA1B,ZNF99,TUBB2B,COL5A1,COL4A1,CES3,RPLP0,H3-3B,CLU,PT
ATP2B4,IGFBP2,CALR,FSTL1,HSPA1B,TUBB2B,COL4A1,DDX5,ENO2,RPL13A,YWHAE,KIF1A,GNAS
A-C,SF3B1,HMGB1,POLR2A,TPT1,SEC61A1,RPS4X,MIDN,DNAJA1,AKAP12,SF3B2,GTF2F1,HNRNI
2,KAT6A,RPL13,GNAS,DDX17,BSG,FASN,SPTBN1,GNB1,MYH9,P4HB,GDI1,FSCN1,TMBIM6,CALM
PCAT1,HK1,RPL22,CLSTN1,PSMD2,RAB6B,RPS20,MCL1,NES,CCNI,KHDRBS1,SLC3A2,EIF5A,CDH
PLCG1,CORO1C,LSS,STXBP1,ISYNA1,PLXNA3,IDH1,HYOU1,MAPK8IP1,SOD1,CTXN1,CD99L2,RB
SG,ALDH6A1,ATP5F1B,GNB1,RPS8,SEPTIN2,P4HB,RPL5,FSCN1,GSTP1,PLD3,HSPD1,USP11,SNR
GRN,RPLP1,MARCHF6,DNAJA1,ARF1,SF3B2,XRCC5,CIPC,RPL22,ZNF865,ATF4,RPS14,CXADR,LEN

AQ,EIF1,HACD3,MAP9,CHSY1,EIF4A1,PPP2R1A,SERINC1,POLR2A,TPT1,RPS3A,RPL18,NCAM1,RF
COL18A1,SNRNP200,SMIM14,VCP,AFF4,TRIM28,CKB,FN1,YWHAQ,EIF1,HACD3,MAP9,CHSY1,INA
KHDRBS1,HDLBP,TKT,EIF5A,DDR1,RPL12,ITM2C,EIF3L,CHPF,NGRN,DDOST,NARS1,DUX4,RPS1
,RPLP0,H3-3B,CANX,PTMA,DDX5,ENO2,SOX4,FKBP14,RPL13A,MAGED1,RPL6,YWHAE,RPL13,DD
N10,KDELR1,MAP2K2,HNRNPUL2-BSCL2,FBLN1,STT3B,STXBP1,UBQLN1,ARNT2,PRPF40A,WDR4
A1,MLF2,WDR45B,PRMT1,TXNDC5,FBRS,PBX1,NOL4L,BAZ1B,MMP15,CXXC5,GARS1,SFRP1,SV2
PNB1,C1ORF43,GAP43,ENC1,NLGN2,GLIPR1L2,JUND,CRMP1,LASP1,TLE5,CCNI,KHDRBS1,CSN
M1,VAMP2,ALKBH5,TCP1,AHCYL1,CAMSAP1,TMEM248,DDX42,SKI,ASXL1,KDELR2,PLEC,BTBD2
OL5A1,PGK1,H3-3B,CANX,CLU,PTMA,DDX5,FKBP14,RPL13A,MAGED1,RPL6,DVL3,GPI,RACK1,F
OL5A1,COL4A1,PSAP,RPLP0,PTMA,DDX5,SEC62,KAT6A,DVL3,YWHAE,RPL13,HTRA4,RACK1,BSG
NB1,SHISA9,MYH9,SEPTIN2,APP,IDS,RPL5,GDI1,FSCN1,TMBIM6,GSTP1,CALM1,PLXNB2,HSPD1,
S,HNRNPK,MYH10,DUX4,CADM4,CALU,CTBP1,STIP1,APPL1,SMCHD1,HDAC2,IPO5,TNC,DNER,SF
1,RPL6,DVL3,GNAS,GPI,RACK1,ATP5F1B,SHISA9,RPS8,P4HB,RPL5,TMBIM6,PLD3,USP11,SNRN
B2B,COL5A1,CES3,PSAP,CLU,DDX5,RPL6,YWHAE,DDX17,HTRA4,RACK1,FASN,ALDH6A1,SPTBN1
ARCHF6,MAT2A,CTNNA1,EIF4H,SLC22A17,RPS9,CBX5,TMEM259,RPS20,HMGN2,SRSF4,RBBP4,
P6V0C,RPL23A,DDX39B,SRSF5,MDH2,U2AF2,CCT2,GNAI3,ALKBH5,PRDX5,UBE4A,RAB11B,TSPA
1,NEFL,PRPF8,RPL8,YBX1,UBA1,CALR,ABHD2,TUBB2B,COL4A1,PSAP,RPLP0,PTMA,DDX5,FKBP1
13,DDX17,GPI,ALDH6A1,ATP5F1B,RPS8,SEPTIN2,P4HB,FSCN1,GSTP1,SMG1,PLD3,USP11,COL1
0,DDX5,FKBP14,RPL13A,SEC62,KIF1A,RPL13,DDX17,ALDH6A1,ATP5F1B,GNB1,RPS8,SEPTIN2,F
1,VCP,UBB,TRIM28,COL4A2,RPL37A,FN1,SPEN,NONO,DDB1,CHSY1,INA,PPP2R1A,SSBP2,ZNF43
,CDC42BPB,SLC4A8,PXDN,HNRNPU,EIF4H,LAMA5,CIPC,KPNB1,ZNF865,KIF5B,NLGN2,CXADR,A
RF1,AP2B1,GTF2F1,GOLGA8A,LDHA,CPE,RPS9,JUND,TMEM259,LENG8,CCNI,XPR1,HDLBP,PLX
TAN1,MARCKS,SMG1,NPM1,PLD3,HSPD1,COL18A1,SNRNP200,MARCKSL1,TRIM28,MATR3,RPL
R1A,B4GALNT4,HNRNPDL,MYL6,SHOX,MYH10,UBE2H,CKAP4,LAPTM4B,RPL23A,TM9SF3,RNF18
EL1,FLOT1,BRD2,SMC3,KMT2E,TMEM63B,REEP2,ITGAV,PCGF2,PAFAH1B2,TRIP12,HMGA1,ELN
L,FBXL4,YBX1,ATP2B4,CALR,ABHD2,HSPA1B,TUBB2B,COL4A1,CES3,PSAP,H3-3B,DDX5,ENO2,S
DX5,ENO2,SOX4,FKBP14,RPL13A,DVL3,GNAS,BSG,FASN,GNB1,MYH9,IDS,ERVV-1,GDI1,TMBIM
CCT2,GNAI3,MTRF1L,PRDX5,UBE4A,HSPA4,MLEC,MAPRE1,FAM168B,LUZP6,TNKS2,BAG6,HNRN
AP4,CALU,PLPPR3,IPO5,ATP6V0C,CHD6,EEF1A2,TBCD,MLXIP,KLHL24,LMNB2,PTPRA,ACTR3,CA
DI1,PTMS,MARCKS,SMG1,PLXNB2,USP11,COL18A1,SMIM14,IGF2,VCP,MARCKSL1,COL4A2,DD
1,PKD1,PDIA3,SEC61A1,STMN1,UCHL1,ANXA2,AGRN,AP3D1,SF3B2,AP2B1,GOLGA8A,PXDN,CTN
C1,PTTG1IP,MAP1A,CDKN1A,SERF2,CTDSP2,NOVA2,NUCB1,MEST,GAS7,ALKBH5,PDXK,PTOV1,
NO2,SEC62,GNAS,SLC2A3,DDX17,BSG,FASN,CALM3,GNB1,STMN2,SEPTIN2,TMBIM6,PTMS,SPTA
1B,MAP1B,SLC9A7,PKM,GANAB,COL6A1,HSP90B1,NEFL,NEFM,UGT8,YBX1,UBA1,ATP2B4,FSTL
5,EIF1,PPP2R1A,SERINC1,BASP1,POLR2A,TPT1,RPL18,TUBB4B,MIDN,MARCHF6,WSB1,ARF1,AP
IRNPU,PGAM1,CTNNA1,FADS2,CNN3,TPM4,TNS3,LAMA5,SLC22A17,BRD3,LLGL1,NCS1,ATF4,AT
1,DDX3X,RPS7,MIF,TLN1,JUN,MYO5A,H1-10,NARS1,SEPTIN11,UBE2H,SEPTIN3,CAPRIN1,APPL1,
RPLP0,DDX5,FKBP14,RPL13A,SEC62,KIF1A,RPL13,DDX17,ALDH6A1,GNB1,SEPTIN2,P4HB,APP,R
B,GNB1,RPS8,SEPTIN2,P4HB,RPL5,FSCN1,GSTP1,PLD3,HSPD1,USP11,SNRNP200,CSDE1,SMIM
OL4A2,FN1,SPEN,TMSB10,SSBP2,TPT1,BTF3,ZMIZ1,TIMP2,MIDN,RPL10,MARCHF6,AP3D1,DNAJ
17,GPI,ALDH6A1,ATP5F1B,SEPTIN2,P4HB,RPL5,FSCN1,GSTP1,SMG1,PLD3,USP11,COL18A1,SN
H11,GRINA,NCKAP1,ACAP3,PTK7,ASXL1,ABCA2,TUBB2A,ACTN1,CAND1,MAP7D1,DIDO1,CORO1
NAS,BSG,ALDH6A1,SHISA9,MYH9,APP,KLRD1,TMBIM6,MARCKS,NPM1,PLD3,COL18A1,CSDE1,U
A,STMN1,RPL11,ANXA2,TIMP2,SARAF,AP3D1,MAP4,SMARCA5,HNRNPA0,TNS3,KPNB1,C1ORF43,
COL18A1,EIF3C,VCP,UBB,YWHAG,NONO,YWHAQ,HACD3,DCHS1,SET,MAP9,TMSB10,MRC2,PPP

IN2,RPL5,GDI1,FSCN1,MARCKS,CALM1,NPM1,TTC3,PLD3,USP11,SNRNP200,IGF2,ATP5F1A,VCF
OA,MTRNR2L8,PKM,GANAB,HSP90B1,NEFL,NEFM,RPL8,FBXL4,YBX1,UBA1,IGFBP2,CALR,FSTL
KHDRBS1,PFKL,PEBP1,PNMA1,PIP4K2B,ITM2C,APLP1,SESN3,APLP2,INIP,DUX4,CKAP5,SEPTIN11
13,DDX17,ALDH6A1,SEPTIN2,APP,RPL5,FSCN1,GSTP1,PLD3,USP11,COL18A1,SNRNP200,SMIM1
S220,TMSB4X,NUMA1,DCX,EIF4G2,RPS6,RPL4,ENO1,TUBA1B,PIK3R2,FLNA,MAP1B,NCL,RPL7A,
PHB2,RER1,YKT6,MCM3AP,TMEM167A,CHERP,CTTN,ERH,EIF3H,SEC14L1,NUDCD3,GNL1,TUBG
SP4,MSL1,LONP1,PLXNA3,NBPF9,COL27A1,RAB11A,RPS13,SMS,DYNLL1,CTXN1,KIRREL1,MMP
J2AF2,CCT2,GNAI3,MTRF1L,PRDX5,UBE4A,HSPA4,MLEC,NRDC,MAPRE1,FAM168B,LUZP6,TNKS
,UBA1,ATP2B4,IGFBP2,TMF1,COL5A1,CLU,PTMA,DDX5,FKBP14,RPL13A,MAGED1,RPL6,DVL3,G
ARS1,YWHAH,CSNK1D,IFITM3,MLF2,AKIRIN1,YME1L1,TRAF4,ARIH1,CSNK1G2,GNG4,UBAP2L,M
DX3X,HNRNPDL,DDOST,MXD4,ANKRD52,RPL36,YWHAB,CALU,COTL1,CCT6A,SMCHD1,ATP6V0C
N1,SMG1,NPM1,PLD3,HSPD1,SNRNP200,EIF3C,AFF4,RPL37A,MAPT,YWHAQ,HACD3,TPI1,TMSB
,ENO2,KIF1A,GNAS,SLC2A3,GPI,BSG,FASN,ALDH6A1,CALM3,GNB1,RPS8,MYH9,P4HB,APP,GDI1
16,DBN1,TUBA1B,MAP1B,NCL,RPL7A,ALDOA,PKM,COL6A1,FTL,HSP90B1,NEFM,RPL8,YBX1,ATF
DVL3,YWHAE,KIF1A,SLC2A3,FASN,ALDH6A1,SPTBN1,GNB1,MYH9,SEPTIN2,APP,IDS,RPL5,GDI1,S
1,ATF4,LENG8,HDLBP,GLG1,CDH6,EIF5A,DDX3X,GPAT4,ANXA6,PTGES3,HMGCR,CAPRIN1,SLC3
,RPL30,EIF3L,IK,CHPF,CD63,RPLP2,DDOST,CIRBP,HSBP1,TM9SF3,MDH2,U2AF2,EEF1A2,FDFT1,
IP2,XAF1,COL11A1,RPS17,ZNF770,NFE2L1,CTNNA1,CPE,CNN3,TPM4,RPS24,SLC16A7,C1ORF43
8,GANAB,COL6A1,FTL,HSP90B1,NEFL,RPL8,FBXL4,YBX1,UBA1,IGFBP2,CALR,FSTL1,TUBB2B,C
3,NCS1,RPL22,CXADR,CHD4,ATP1A1,LENG8,WDR1,OAZ1,NES,TADA2B,KHDRBS1,GLG1,PNMA1,
AQ,HACD3,TPI1,TMSB10,SSBP2,RPS3,RPS3A,ITGB1,UCHL1,AGRN,MIDN,WSB1,DNAJA1,ARF1,G
A,SLC9A7,MTRNR2L8,PKM,GANAB,COL6A1,HSP90B1,FADS1,NEFL,ATP2B4,TMF1,ABHD2,HSPA
8,CKB,FN1,YWHAQ,EIF1,HACD3,MAP9,CHSY1,EIF4A1,PPP2R1A,BASP1,POLR2A,TPT1,SEC61A1,F
ED2,IK,RPS19,NGRN,RPLP2,DDOST,CIRBP,NARS1,ATP6V0C,RPL23A,DDX39B,TM9SF3,MDH2,TC
MA,DDX5,SEC62,KAT6A,DVL3,YWHAE,RPL13,RACK1,BSG,SPTBN1,ATP5F1B,GNB1,RPS8,MYH9,ID
14,LENG8,WDR1,MCL1,TLE5,PNN,KHDRBS1,HDLBP,TKT,EIF5A,CDH2,DDR1,DDX3X,RPL12,ITM2
A2,RUVBL2,EMC1,FAF2,PCGF3,CDK9,RTF2,WAC,GSK3B,PLXDC2,MBOAT7,NDUFS1,CTR9,RBM1
S12,ATP6V0C,HSBP1,KLHL24,MORF4L2,MTRF1L,PDXK,PKN1,GRINA,TMEM248,LAMC1,JPT1,RF
PA,HMGB1,ITGB1,ZMIZ1,TIMP2,DNAJA1,MAP4,SF3B2,OGA,NFE2L1,HNRNPA0,LAMA5,BRD3,PJA

RPL29,XRCC5,IGFBP3,COL1A2,RHOA,ZNF865,ATF4,LENG8,CCT7,RBBP4,ASTN2,MCL1,PNN,PRK
L37,ARPC2,CCAR2,SARS1,BAG6,RAB11A,TRAF7,EIF2AK1,PDAP1,PHB2,RER1,YKT6,MCM3AP,TM
A,CTNNA1,RPL29,CTSD,SCARB2,RPS24,CIPC,GAP43,HK1,RPL22,ZNF865,CHD4,ATP1A1,LENG8,
,RTN4,RPS24,RPS9,PJA2,KPNB1,CBX5,NLGN2,ATP2A2,CCT7,MCL1,SPC25,PRKCSH,HNRNPC,CD
EM259,LENG8,CCNI,XPR1,HDLBP,GNB2,PRKAR1A,CHPF,NGRN,RPLP2,MYL6,INIP,MYH10,SND1,
L,ATP2B4,FSTL1,ABHD2,TUBB2B,COL5A1,DDX5,ENO2,FKBP14,MAGED1,RPL6,YWHAE,KIF1A,GN
,RPLP2,DDOST,CIRBP,ATP6V0C,RPL23A,DDX39B,TM9SF3,MDH2,TOMM20,U2AF2,CCT2,GNAI3,
STMN1,WSB1,ARF1,AKAP12,HNRNPA0,RPL23,RPSA,CTSD,CIPC,MSN,SLC16A7,C1ORF43,RPL22,
,WLS,DUX4,PTGES3,SEPTIN3,APPL1,SMCHD1,HDAC2,TRIB2,TBCD,PPP1CB,GNAI3,GAS7,HSPH1,
IP5,CHMP1A,GGNBP2,NPTN,ADNP,ANKFY1,SRSF10,RPL27A,CCNG1,MARCHF7,EXOSC10,ZFP91,
N,ATG2B,AZIN1,KDELR2,RABL6,ADAM9,CAPZB,ACTN1,MAP7D1,YWHAH,PITPNM1,ZFAND3,PNM
X,EIF4G2,RPS6,ZNF316,DBN1,RPL4,TUBA1B,PIK3R2,FLNA,MAP1B,NCL,RPL7A,SLC9A7,ALDOA,
DAM9,NSG2,HNRNPUL2-BSCL2,DLK1,GIGYF1,DGKD,FBRSL1,BCL9L,LBH,SF1,CNTNAP1,KIF21E
RCHF6,WSB1,AP2B1,GTF2F1,GOLGA8A,LDHA,CPE,SLC22A17,RPS9,IGFBP3,JUND,TMEM259,LE
N2,RPL7,SEPTIN2,APP,GDI1,CALM1,NPM1,TTC3,PLD3,HSPD1,SNRNP200,CSDE1,SMIM14,EIF4B,

\RL6IP1,MYH10,SND1,RHOB,HLA-B,STIP1,SLC35E2B,IPO5,BCLAF1,RPL23A,GLUL,CNN2,CDH11,
B,FTL,HSP90B1,FADS1,NEFL,NEFM,PRPF8,RPL8,FBXL4,YBX1,UBA1,IGFBP2,FSTL1,HSPA1B,TUE
)P53,TM9SF2,MTRF1L,RPL27,PRDX5,UBE4A,TCP1,RPL36A,SRP14,TAF7,RPL37,ARPC2,ACADVL,(
K5,ENO2,RPL13A,SEC62,YWHAE,RPL13,RACK1,BSG,FASN,ALDH6A1,CALM3,SHISA9,STMN2,RPS8
31,MAT2A,RPSA,NACA,RPL14,CBX5,CLSTN1,ATF4,TMEM259,HMGN2,PNN,NISCH,PRKCSH,HDLBI
\B7A,EIF3L,RPS19,RPL35,NGRN,RPLP2,DDOST,YWHAB,SND1,ARCN1,RPS27,ARHGDIA,SLC35E2E
2B,KHDRBS1,PFKL,PEBP1,PIP4K2B,ITM2C,APLP1,SESN3,APLP2,INIP,DUX4,CKAP5,SEPTIN11,PTT
19,VAPA,HMGB1,POLR2A,CDKL1,TPT1,SEC61A1,RPS3A,TUBB4B,MIDN,RPL10,MARCHF6,SF3B2,
.EC,CBX1,MAPK1,PSMD3,UBE2R2,MAP7D1,CCT8,STMN3,LSS,STT3B,HECTD1,RAI1,NAP1L4,CCT5
.4A2,DDB1,SET,EIF4A1,GTF3C4,RPS4X,UCHL1,AGRN,MIDN,WSB1,VCL,CDC42BPB,GTF2F1,HNRI
=US,PPP1R10,SHC2,PLEKHB2,ELP1,ADAM10,BNIP3,SAE1,USO1,DIP2A,MZT2B,SPPL2B,FXR2,ARF
28,EIF3F,SF3A1,BAG6,RAB11A,WBP2,CCT5,CDC42,YME1L1,EIF2AK1,EXTL3,PDAP1,PHB2,APMAF
)X17,ALDH6A1,SEPTIN2,APP,RPL5,FSCN1,GSTP1,PLD3,USP11,COL18A1,SNRNP200,CSDE1,SMIM
3P,CALU,ARCN1,ATP6V0C,RPL23A,DDX39B,TM9SF3,MDH2,TOMM20,U2AF2,CCT2,GNAI3,TSR1,L
EFM,PRPF8,RPL8,FBXL4,YBX1,IGFBP2,FSTL1,TMF1,PSAP,CLU,DDX5,RPL6,DVL3,YWHAE,KIF1A,
)DX17,GPI,CALM3,GNB1,RPL7,RPL5,KLRD1,MARCKS,NPM1,PLD3,SNRNP200,CSDE1,ATP5F1A,E
3P8,FBLN1,HYOU1,MAPK8IP1,PFN2,HNRNPD,MEX3A,DYNLL1,ANP32A,MTA1,LBH,ZFP36L1,CXXC
LDHA,CTSD,CBX5,RPL9,JUND,TMEM259,LENG8,WDR1,CCNI,HDLBP,RAB7A,GNB2,PRKAR1A,CHI
)S,RPL5,FSCN1,COL18A1,RPL37A,EGLN1,CHSY1,INA,EIF4A1,PPP2R1A,SF3B1,PDIA3,SSBP2,ZNF
3A1,CALM3,GNB1,STMN2,RPS8,P4HB,APP,RPL5,TMBIM6,PTMS,SPTAN1,MARCKS,HSPD1,USP11,
C14L1,NUDCD3,GNL1,TUBGCP6,NCSTN,INTS11,MTMR4,CAP1,AP1M1,LYPLA2,SLC38A10,ATP13
3N2,BEX3,JPT2,PFKL,EIF5A,CCND2,PIP4K2B,ACSL6,ITM2C,RPS5,HMGCS1,ANXA6,MYL6,ATP1A2,
BN1,RPL4,ENO1,TUBA1B,NCL,RPL7A,MTRNR2L8,FTL,HSP90B1,FADS1,NEFL,NEFM,PRPF8,RPL8
CPE,JUND,TMEM259,LENG8,CCNI,XPR1,HDLBP,GNB2,PRKAR1A,CHPF,NGRN,RPLP2,MYL6,INIP,N
25,PEBP1,GNB2,TLN1,CHPF,JUN,HNRNPDL,TARDBP,RPLP2,MYL6,MYH10,UBE2H,RPS27,HMGCI
PT2,NES,KHDRBS1,GPC3,PRDX2,ITM2C,RPS19,HNRNPDL,MYL6,HNRNPK,INIP,CADM4,GPC1,PEA
4,IGFBP2,CALR,FSTL1,TMF1,ABHD2,TUBB2B,COL5A1,COL4A1,CES3,H3-3B,PTMA,ENO2,RPL13
,DDR1,MAGED2,APLP1,MTSS2,TARDBP,INIP,CIRBP,PTGES3,RAC1,ARHGDIA,CDK16,IPO5,ATP6V
AZ1,MBTPS1,CUL3,SEPTIN7,FUS,PPP1R10,TAOK1,UBE2D2,SMG5,IREB2,LSM14A,TCF3,YIPF3,M
2,CHPF,RPLP2,MYL6,MYH10,SND1,UBE2H,SMCHD1,ATP6V0C,LAPTM4B,SRSF5,MLXIP,HSPH1,ALI
JBB2A,PPP2CA,PDE7A,PSMD3,CDC37,PITPNA,XPNPEP3,LARS1,YWHAH,UBQLN1,CSNK1D,IFITM3
TT,HSP90AB1,ZNF124,HSP90AA1,ACTR2,VIM,SPARC,TUBB,FTH1,UBC,KIDINS220,TMSB4X,NUM
120,U2AF2,TSR1,MTRF1L,PRDX5,UBE4A,HSPA4,MLEC,LUZP6,RPL28,CCAR2,ARF4,NKTR,STMN3,
NGRN,APLP2,DDOST,CALU,ARCN1,SLC35E2B,ATP6V0C,BCLAF1,LAPTM4B,TRIB2,RPL23A,FAM1
N1,SMCHD1,ATP6V0C,LAPTM4B,MLXIP,HSPH1,ALKBH5,AHCYL1,ETNK1,KLC1,CLSTN3,DDX42,RA
Q,SET,TPI1,INA,PKD1,HLA-C,SSBP2,BASP1,HMGB1,AP2M1,TPT1,ITGB1,ANXA2,TIMP2,COL11A1,
=2,CCT2,GNAI3,NOP53,RPL27,PRDX5,UBE4A,HSPA4,TCP1,MLEC,ACAP3,SRP14,PRPF6,TAF7,RPL
P,MYH10,PLPPR3,COTL1,CCT6A,GPC1,BCLAF1,TIMP3,TMEM132A,TGFBI,PKN1,SURF4,TUBB2A
.1,ZNF99,TUBB2B,COL5A1,COL4A1,PSAP,PGK1,H3-3B,CANX,DDX5,FKBP14,SEC62,RPL6,YWHA
P5F1B,SHISA9,RPS8,P4HB,CALM1,USP11,SNRNP200,CSDE1,EIF3C,ATP5F1A,MATR3,LRRC28,TF
P6,NCSTN,INTS11,MTMR4,CAP1,PRPF19,AP1M1,TOMM70,ATP13A1,EIF4G1,PAFAH1B2,POLDIP
7A,PSMD3,CDC37,PITPNA,XPNPEP3,LARS1,YWHAH,UBQLN1,CSNK1D,IFITM3,MLF2,AKIRIN1,AM
.ENG8,CCNI,XPR1,HDLBP,GNB2,PRKAR1A,CHPF,NGRN,RPLP2,MYL6,MYH10,SND1,UBE2H,HMG
N1,RPL4,ENO1,PIK3R2,FLNA,MAP1B,NCL,RPL7A,SLC9A7,ALDOA,PKM,GANAB,COL6A1,FTL,FAD
PK,RAB1A,TTR,PLPPR3,TOP2B,CTSB,CDH11,ANXA5,PPP1CB,NOP53,PABPN1,PARP1,TSR1,ACTR

SEC14L1,NUDCD3,SSR3,GNL1,NCSTN,INTS11,MTMR4,NOC2L,AP1M1,UBE2I,C16ORF58,TOMM
R2A,RPS3A,MORF4L1,SARAF,COL11A1,RPL10,MARCHF6,CTNNA1,RPS16,CNN3,RPSA,BRD3,RPS
,LRPPRC,EIF3L,RPS5,MIF,DDOST,NARS1,CALU,ARCN1,SLC35E2B,ATP6V0C,DDX39B,U2AF2,FDF
SPARC,FTH1,UBC,KIDINS220,TMSB4X,NUMA1,EIF4G2,RPL3,RPS6,ZNF316,DBN1,ENO1,TUBA1B
PLD3,HSPD1,COL18A1,SNRNP200,VCP,UBB,MATR3,RPL37A,LRRC28,DDB1,PSMD5,EGLN1,TPI1,
,RPS20,LENG8,WDR1,RBBP4,PFN1,RPL31,HNRNPC,MEGF8,CLTC,EIF3L,H1-0,CHPF,RPLP2,DDO
,PLXNA1,RAB7A,DDX3X,CLTC,RCN1,PRKAR1A,TLN1,NGRN,DNAJC5,HNRNPDL,ST3GAL2,HNRNP
TF4,TMEM259,NISCH,PRKCSH,KLHL9,HDLBP,GLG1,SFPQ,PNMA1,RAB7A,NANOG,RPS7,CTNNB1
,SEC62,RPL13,DDX17,HTRA4,BSG,SPTBN1,ATP5F1B,GNB1,RPS8,APP,RPL5,KLRD1,FSCN1,MAR
SEC61A1,STMN1,MORF4L1,RPL10,WSB1,RPS17,AKAP12,HNRNPA0,FADS2,RPL29,RPSA,TNS3,L
STL1,TMF1,RPLP0,CANX,PTMA,DDX5,ENO2,SOX4,FKBP14,RPL13A,MAGED1,RPL6,YWHAE,RPL1
,LRP10,ERH,RALY,SEC14L1,NUDCD3,TUBGCP6,RIC8A,INTS11,MOAP1,MTMR4,CAP1,NASP,AP1
X30,BAG6,RAB11A,CDC42,YME1L1,TRAF7,EIF2AK1,PHB2,KLHL42,MASTL,YKT6,TMEM167A,TNK
P4K2B,GNB2,RCN1,RPL35,CHPF,CD63,HECTD4,TOP2B,HDAC2,IPO5,DNER,HS6ST1,TMEM132A,
UX4,CKAP5,SEPTIN11,PTTG1IP,MAP1A,TIMP3,CTDSP2,CNN2,NOVA2,NUCB1,ZC3H7B,U2AF2,CC
RF4L1,XAF1,MIDN,CYB5R3,AKAP12,MAP4,CDC42BPB,GOLGA8A,CTNNA1,SCARB2,RPS24,MSN,
1,JUND,TMEM259,LENG8,MCL1,CCNI,HDLBP,RAB7A,CLTC,GNB2,PRKAR1A,CHPF,RPLP2,MYL6,
JBB2A,PPP2CA,PDE7A,PSMD3,WDR6,GABARAP,ZFAND5,XPNPEP3,YWHAH,UBQLN1,AMOTL2,YM
2,ANXA6,NGRN,INIP,CIRBP,LGALS3BP,COTL1,ARHGDIA,CAPRIN1,GPC1,ATP6V0C,MMP2,DNER,
MEM259,HMGN2,NISCH,PRKCSH,KLHL9,HDLBP,GLG1,SFPQ,PNMA1,NANOG,RPS7,CTNNB1,RPL
PKM,GANAB,COL6A1,FTL,HSP90B1,FADS1,NEFM,PRPF8,FBXL4,YBX1,UBA1,ATP2B4,IGFBP2,CA
3D1,ARF1,SF3B2,GTF2F1,RPS9,PAPOLA,ZNF865,RPS14,RPS20,WDR1,PFN1,ASTN2,TLE5,PNN,PR
E,CDV3,TRMT112,CD44,DRAP1,STUB1,SRSF7,ZNF768,ZCCHC14,ATP6V0A1,SUDS3,MARK1,TRA
,P4HB,FSCN1,GSTP1,PLD3,USP11,COL18A1,SNRNP200,CSDE1,SMIM14,EIF3C,UBB,AFF4,TRIM2
,LARP1,HNRNPA0,RTN4,RPS9,RPL14,CBX5,ATF4,ATP2A2,CCT7,SPC25,PRKCSH,CDH6,PEBP1,G
IT4,PSMA7,ANP32A,VDAC1,EDC4,AP2A1,SLC39A6,SLC39A7,PPM1G,KLHL42,SLC35F5,RGMB,N
TP1,PLD3,USP11,SNRNP200,CSDE1,EIF3C,UBB,AFF4,TRIM28,CKB,FN1,PSMD5,YWHAQ,EIF1,HA
IS,SPTAN1,GSTP1,MARCKS,CALM1,PLXNB2,COL18A1,CSDE1,VCP,MARCKSL1,CKB,COL4A2,SP
,RPLP2,MYL6,MYH10,SND1,UBE2H,SMCHD1,ATP6V0C,LAPTM4B,MLXIP,MORF4L2,HSPH1,ALKB
B4X,DCX,EIF4G2,RPS6,ZNF316,DBN1,RPL4,ENO1,PIK3R2,MAP1B,NCL,RPL7A,SLC9A7,ALDOA,P
N2,MORF4L2,PKN1,NCKAP1,TMED2,NOTCH2,PDE7A,ARGLU1,PTBP1,TDRD5,RPL28,ZFAND5,STT
2D2,SMG5,IREB2,LSM14A,YIPF3,MSH6,TGOLN2,UBE2D3,HNRNPH3,NELL2,CASC4,USP47,SCAM
1,RPL29,XRCC5,SLC16A7,COL1A2,ZNF865,GLIPR1L2,RPS14,RPL26,RBBP4,PFN1,PNN,RPS25,E
2,PPP1R9B,EIF2AK1,RETREG2,PDAP1,PHB2,SLC39A7,TRA2B,YKT6,MCM3AP,TMEM167A,TNKS,
),LENG8,WDR1,PFN1,MCL1,RPL31,PRKCSH,HNRNPC,EIF5A,MEGF8,CLTC,EIF3L,RPS19,CHPF,CD
4B,ATRX,ADH5,PTPRF,GSE1,PCBP4,PPP1R10,TAOK1,PALM,RTL6,SHC2,RAB1B,UBE2M,TLE1,HD
AH1B2,POLDIP2,TRIP12,CHMP1A,TSNAX,EXOC5,TRMT112,CD2BP2,FAF2,KLC2,VPS51,FAM32A,E
3,STMN1,RPL11,MIDN,MARCHF6,RPS17,CDC42BPB,MAT2A,GTF2F1,HNRNPA0,FADS2,RPL23,SC
COL6A1,FADS1,PRPF8,RPL8,YBX1,UBA1,IGFBP2,ABHD2,HSPA1B,TUBB2B,PGK1,RPLP0,H3-3B,P
3,RPS6,ZNF316,ENO1,TUBA1B,PIK3R2,FLNA,MAP1B,NCL,SLC9A7,GANAB,FTL,HSP90B1,FADS1,
MOAP1,MTMR4,CAP1,AP1M1,TOMM70,XPO6,UBXN6,MYL12B,GAK,LYPLA2,ATP13A1,PAFAH1B2
N2,P4HB,IDS,ERVV-1,RPL5,GDI1,FSCN1,SPTAN1,ATP5F1A,VCP,TRIM28,CKB,COL4A2,EIF1,CHSY
N3,DDX42,RAB11B,TUBB2A,PPP2CA,G3BP2,PDE7A,PSMD3,CDC37,PITPNA,XPNPEP3,LARS1,YWH
1,CLSTN3,DDX42,RAB11B,TUBB2A,PPP2CA,G3BP2,PDE7A,PSMD3,CDC37,BNIP3L,PITPNA,XPNPE
,ARPC2,PBX2,AEBP1,GRB2,HSPB1,KDELR1,MAP2K2,MSL1,ISYNA1,ZFAND3,VANGL2,RAI1,FBRSL

,RPL23A,HSBP1,FAM199X,TM9SF3,MDH2,U2AF2,CCT2,GNAI3,NOP53,RPL27,PRDX5,TCP1,MLEC

,POLR2A,AP2M1,SEC61A1,RPL18,STMN1,NCAM1,ZMIZ1,TIMP2,SBF1,ARF1,AKAP12,CPE,TNS3,E

HSPH1,UBE4A,AHCYL1,KLC1,CLSTN3,DDX42,RAB11B,TUBB2A,PPP2CA,G3BP2,PDE7A,PSMD3,CD

DX42,RAB11B,TUBB2A,PPP2CA,G3BP2,PDE7A,PSMD3,CDC37,BNIP3L,PITPNA,XPNPEP3,LARS1,

DX42,RAB11B,TUBB2A,PPP2CA,G3BP2,PDE7A,PSMD3,CDC37,BNIP3L,PITPNA,XPNPEP3,LARS1,

UBE2H,RPS27,CTBP1,MORF4L2,PDXK,AHCYL1,CLSTN3,SCAF1,PDE7A,SRSF1,TCF25,WDR6,CAP

1,CES3,PSAP,H3-3B,CLU,PTMA,ENO2,SOX4,SEC62,KAT6A,YWHAE,KIF1A,GNAS,GPI,FASN,SPTBN

N,MIDN,AP3D1,GOLGA8A,RPL29,SCARB2,RPS24,GAP43,HK1,RPL22,RHOA,ZNF865,RPL9,CLST

FBP2,ABHD2,HSPA1B,TUBB2B,PGK1,RPLP0,H3-3B,PTMA,DDX5,SOX4,KAT6A,DVL3,YWHAE,KIF1

LC2A3,DDX17,GPI,RACK1,CALM3,STMN2,IDS,RPL5,GDI1,MARCKS,NPM1,TTC3,PLD3,HSPD1,USP

LC2A3,DDX17,GPI,RACK1,CALM3,STMN2,IDS,RPL5,GDI1,MARCKS,NPM1,TTC3,PLD3,HSPD1,USP

CD3,SET,GTF3C4,SF3B1,ZNF43,BASP1,VAPA,TPT1,RPS4X,ITGB1,AGRN,ZMIZ1,MIDN,RPL10,RPS1

SC3,PITPNA,XPNPEP3,YWHAH,UBQLN1,GHITM,SOD1,AMOTL2,YME1L1,TRAF4,STAT3,CSNK1G2,

,GANAB,COL6A1,FADS1,NEFM,RPL8,FBXL4,UBA1,IGFBP2,FSTL1,HSPA1B,TUBB2B,COL5A1,COL

NRNPL,ACO2,VPS35,ACAP3,TMEM248,SKI,ASXL1,ABCA2,VHL,PPP2CA,ODC1,SRSF1,UBE2R2,CAN

RF4L2,MTRF1L,PDXK,PTOV1,ORAI2,G3BP2,KDELR2,TCF25,GRB2,BNIP3L,KDELR1,ARF4,STT3A

PIK3R2,FLNA,MAP1B,NCL,RPL7A,GANAB,FTL,HSP90B1,NEFL,PRPF8,RPL8,FBXL4,UBA1,ATP2B4

5,FKBP14,RPL13A,SEC62,KIF1A,RPL13,DDX17,GPI,ALDH6A1,SEPTIN2,P4HB,APP,RPL5,KLRD1,F

3-3B,DDX5,FKBP14,SEC62,YWHAE,KIF1A,DDX17,GPI,RACK1,CALM3,STMN2,RPL7,SEPTIN2,APP,

1,STRN4,ABI2,KDM4A,UBE2I,SLC38A2,CNBP,ELAVL3,USP47,DAG1,IGSF3,OBSL1,TSPAN14,TNR

5,TM9SF3,MDH2,TOMM20,U2AF2,CCT2,GNAI3,LMNB2,PRDX5,MLEC,ACAP3,PRPF6,MAPRE1,FA

3,COL4A2,MAPT,FN1,MAP9,RPL19,SERINC1,SEC61A1,RPS3A,BTF3,RPLP1,ZMIZ1,SARAF,WSB1,F

3,MED25,PSMB5,SLC5A6,ZNF362,ELOB,STRIP1,FUBP1,RAD23A,CCDC124

MIDN,ARF1,PXDN,HNRNPA0,CTNNA1,FADS2,RPL23,CNN3,RPSA,EIF4H,SCARB2,LPCAT1,SLC22A1

CHERP,SERP1,ERH,PSMB4,EIF3H,SEC14L1,NUDCD3,INTS11,MTMR4,SPTY2D1,EIF3G,LYPLA2,PA

5,ENO2,FKBP14,SEC62,KAT6A,DVL3,KIF1A,GPI,FASN,ALDH6A1,ATP5F1B,SHISA9,RPS8,APP,ERV

LC1,CLSTN3,DDX42,RAB11B,TUBB2A,PPP2CA,G3BP2,PDE7A,PSMD3,CDC37,PITPNA,XPNPEP3,LA

24,RPL14,CBX5,ATF4,TMEM259,HMGN2,NISCH,PRKCSH,KLHL9,HDLBP,GLG1,SFPQ,PNMA1,RA

ANAB,FTL,HSP90B1,FADS1,NEFL,PRPF8,RPL8,YBX1,ATP2B4,ABHD2,HSPA1B,TUBB2B,COL5A1,F

STL1,HSPA1B,TUBB2B,COL5A1,CANX,ENO2,YWHAE,KIF1A,RPL13,DDX17,RACK1,BSG,FASN,RPL

RIC8A,NCSTN,INTS11,MOAP1,MTMR4,CAP1,AP1M1,TOMM70,GAK,LYPLA2,ATP13A1,PAFAH1B2,

LC2A3,DDX17,GPI,RACK1,CALM3,STMN2,IDS,RPL5,GDI1,MARCKS,NPM1,TTC3,PLD3,HSPD1,USP

A,AHCYL1,KLC1,CLSTN3,DDX42,RAB11B,TUBB2A,PPP2CA,G3BP2,PDE7A,PSMD3,CDC37,BNIP3L

PT1,SEC61A1,RPS3A,RPL18,NCAM1,RPLP1,SARAF,MIDN,MARCHF6,WSB1,ARF1,AP2B1,GTF2F1,

NL1,TUBGCP6,INTS11,MTMR4,CAP1,NASP,AP1M1,TOMM70,XPO6,UBXN6,MYL12B,GAK,LYPLA2

TM3,MLF2,AKIRIN1,YME1L1,TRAF4,ARIH1,SLC35F5,CSNK1G2,MASTL,EIF3H,GABARAPL1,TRAP

AGLN,MYL6,MYH10,UBE2H,RAB1A,SMCHD1,ATP6V0C,LAPTM4B,PEA15,MLXIP,TSPYL4,NOX4,HSP

,MDK,KDM6B,ABCA2,HNRNPUL2-BSCL2,PLTP,SDF4,GIGYF1,STMN3,LSS,STT3A,IDH1,MPRIP,SO

ALU,ARCN1,SLC35E2B,ATP6V0C,LAPTM4B,TRIB2,FAM199X,DDX39B,TM9SF3,MDH2,TOMM20,U

TF2F1,LDHA,RPSA,RTN4,RPS9,CIPC,RPL24,RPL9,JUND,TMEM259,LENG8,MCL1,CCNI,LRPPRC,H

,CACNA1H,MBOAT7,HIF3A,PRMT2,NECTIN2,PPP1R26,MIB2,SHB,PPFIA3,RAP1B,MYL6B,DTD1,EM

1,CLSTN3,DDX42,RAB11B,TUBB2A,PPP2CA,G3BP2,PDE7A,PSMD3,CDC37,BNIP3L,PITPNA,XPNP

HA,JUND,TMEM259,LENG8,MCL1,CCNI,HDLBP,CLTC,GNB2,CHPF,RPLP2,MYL6,MYH10,UBE2H,

MLXIP,HSPH1,AHCYL1,KLC1,CLSTN3,DDX42,RAB11B,TUBB2A,PPP2CA,G3BP2,PDE7A,PSMD3,C

OLDIP2,TRIP12,CHMP1A,TSNAX,EXOC5,TRMT112,CD2BP2,VPS51,FAM32A,EPN1,RTF2,UNC45A,

1,AHCYL1,KLC1,CLSTN3,DDX42,RAB11B,TUBB2A,PPP2CA,G3BP2,PDE7A,PSMD3,CDC37,BNIP3L
H5,PTPRF,GSE1,PCBP4,PPP1R10,TAOK1,PALM,RTL6,SHC2,RAB1B,UBE2M,TLE1,HDGFL2,NUCK
1L1,TRAF4,ARIH1,SLC35F5,MASTL,EIF3H,GABARAPL1,TRAP1,ELP1,NSD2,ELOVL5,SLC25A36,X
1L1,TRAF4,ARIH1,SLC35F5,MASTL,EIF3H,GABARAPL1,TRAP1,ELP1,NSD2,ELOVL5,SLC25A36,X
,MLXIP,HSPH1,UBE4A,AHCYL1,KLC1,CLSTN3,DDX42,RAB11B,TUBB2A,PPP2CA,G3BP2,PDE7A,P
RPLP1,GTF2F1,SMARCA5,CPE,TSPAN3,RPS9,RPL17,C1ORF43,RPL14,RHOA,PTCD3,CBX5,CXAD
L6,MYH10,SND1,UBE2H,ARCN1,SMCHD1,ATP6V0C,LAPTM4B,CCT2,MLXIP,HSPH1,ALKBH5,AHC
B1,CDKL1,STMN1,ITGB1,RPL11,RPL10,WSB1,AKAP12,MAT2A,HNRNPA0,RPL23,CNN3,CTSD,EI
B1,CDKL1,STMN1,ITGB1,RPL11,RPL10,WSB1,AKAP12,MAT2A,HNRNPA0,RPL23,CNN3,CTSD,EI
,PRDX2,ITM2C,MRFAP1,APLP1,CHPF,ANXA6,RPL35A,WLS,MYH10,YWHAB,RHOB,COTL1,HMGC
MIP,BNIP3L,DIDO1,LSS,SARS1,SREBF2,P3H3,TRAF4,MSMO1,LBH,ALDH3A2,HIP1,NOL4L,CD99,
B,PIK3R2,SLC9A7,ALDOA,MTRNR2L8,PKM,COL6A1,FTL,FADS1,NEFL,RPL8,FBXL4,UBA1,ATP2B

T2,PTPN23,YTHDF3,UBE2J1,GOLGA8B,ZBTB4,USP48,SNRPD2,METRNL,UNC13A,TSPAN13,ZN
,SERINC1,HMGB1,POLR2A,AP2M1,SEC61A1,RPS4X,TUBB4B,MORF4L1,BTF3,AGRN,RPLP1,RPL
N3,DDX42,RAB11B,TUBB2A,PPP2CA,G3BP2,PDE7A,PSMD3,CDC37,PITPNA,XPNPEP3,LARS1,YW
H10,UBE2H,RAB1A,SMCHD1,ATP6V0C,LAPTM4B,MLXIP,HSPH1,AHCYL1,KLC1,CLSTN3,DDX42,R
AND3,HGS,BAG6,HNRNPD,CDC42,EIF2AK1,MMP15,RAB2A,YKT6,MCM3AP,TMEM167A,CTTN,D
1,TOMM70,GAK,ATP13A1,PAFAH1B2,POLDIP2,TRIP12,CHMP1A,TSNAX,EXOC5,SF3A3,TRMT11
F1,ATP13A3,DNAJB1,NDUFA4,CD2BP2,FAF2,MTURN,KLC2,GOLPH3,GALNT17,DNAJC8,CELSR3,
N2,IDS,MARCKS,SMG1,NPM1,PLD3,USP11,SNRNP200,SMIM14,EIF4B,VCP,TRIM28,RPL37A,NON
X15,BAZ1B,GTPBP1,ZFP36L1,FARP1,PI4KA,UQCRC1,MASTL,TBC1D9B,HNRNPA2B1,ANP32B,ZM
1,ITGB1,NCAM1,XAF1,SARAF,COL11A1,RPS17,AP2B1,RPS16,LDHA,CNN3,RPL29,TPM4,CTSD,N
2,EWSR1,RNF157,HTRA1,DRAP1,TBC1D14,FAM8A1,ZCCHC14,CLPTM1L,SPOCK1,CACNA1H,CAS
29,SCARB2,RPS24,GAP43,NCS1,RPL22,ZNF865,RPL9,PSMD2,CXADR,CHD4,LENG8,WDR1,BEX3,
F1,MAP1A,CTDSP2,CNN2,NOVA2,CCT2,MEST,GAS7,TM9SF2,MORF4L2,PKN1,PTOV1,MLEC,SUR
/2,SPIN1,GIT1,DUS1L,EMD
B,TBCD,PGD,MEST,HNRNPL,PKN1,SKI,SNRNP70,ASXL1,HDGF,ABCA2,TMED2,MARS1,SELENON,
,EIF4A1,PPP2R1A,RPS3,BASP1,POLR2A,SEC61A1,RPS3A,RPL18,NCAM1,RPLP1,SARAF,MIDN,WS
1,ATXN7L3B,CCT7,ASTN2,PNN,PRKCSH,LRPPRC,KLHL9,RPS25,EIF5A,ITM2C,EIF3L,GPAT4,CD6
P1LC3B,TMOD2,CARM1,GPC2,IMPDH2,ERGIC3,CAP1,IST1,HSDL1,GAK,NELFB,ELAVL3,SNX3,PO
CKS1,LSM14B,ERGIC3,DCUN1D2,RNF145,SF3A2,UBE2I,TMCO3,CLIP2,GGNBP2,PFKP,OSBPL2,
F3C,EIF4B,MARCKSL1,TRIM28,MAPT,DDB1,PSMD5,DCHS1,SET,TMSB10,RPL19,PKD1,PDIA3,SE
,RPS3,BASP1,POLR2A,SEC61A1,RPS3A,RPL18,NCAM1,RPLP1,SARAF,MIDN,WSB1,RPS17,AP2B1,
,SERF2,DDX39B,SRSF5,U2AF2,CCT2,GNAI3,TM9SF2,HSPH1,MTRF1L,PRDX5,UBE4A,TCP1,MLEC,
F2I,MSH6,DEK,METRN,EIF4A3,DHX40,CNBP,AMFR,LGALS1,CHMP1A,ESYT1,LUC7L2,ELN,KIAA1
PAFAH1B2,POLDIP2,TRIP12,CHMP1A,TSNAX,EXOC5,TRMT112,CD2BP2,BANF1,FAF2,KLC2,VPS5
KS,NPM1,PLD3,SNRNP200,CSDE1,ATP5F1A,EIF4B,UBB,MATR3,RPL37A,LRRC28,MAPT,DDB1,SE
1,AHCYL1,KLC1,CLSTN3,DDX42,RAB11B,TUBB2A,PPP2CA,G3BP2,PDE7A,PSMD3,CDC37,BNIP3L
MAPK1,ODC1,TCF25,WDR6,CORO1C,SDHA,KIF5A,HIP1,NXN,EP400,RER1,PI4KA,PRRC2A,MAPRE
6V0C,LAPTM4B,MLXIP,HSPH1,UBE4A,AHCYL1,KLC1,CLSTN3,DDX42,RAB11B,TUBB2A,PPP2CA,G
1,CLSTN3,DDX42,RAB11B,TUBB2A,PPP2CA,G3BP2,PDE7A,PSMD3,CDC37,BNIP3L,HNRNPUL2-B
S11,MTMR4,CAP1,NASP,XPO6,GAK,LYPLA2,ATP13A1,POLDIP2,TRIP12,TSNAX,ZKSCAN1,SLC25A
DCD3,TUBGCP6,RIC8A,MOAP1,MTMR4,XPO6,UBXN6,MYL12B,GAK,LYPLA2,ATP13A1,POLDIP2,

ATP1A1,HMGN2,BEX3,RBBP4,JPT2,XPO1,HDLBP,RAB7A,DDX3X,RPS15,CD63,ANXA6,MTSS2,RPL
AP1,IST1,BAG5,USO1,TAPBP,POR,NOP56,NIBAN2,KLF13,FZR1,SUB1,FEZ1,CNDP2,RNF157,PITR

)BN1,RPL4,ENO1,TUBA1B,PIK3R2,FLNA,MAP1B,NCL,RPL7A,SLC9A7,ALDOA,PKM,GANAB,COL6A
JBA1B,PIK3R2,FLNA,MAP1B,NCL,RPL7A,SLC9A7,ALDOA,MTRNR2L8,PKM,GANAB,COL6A1,FTL,H
'IK3R2,FLNA,MAP1B,NCL,RPL7A,SLC9A7,ALDOA,PKM,GANAB,COL6A1,FTL,HSP90B1,FADS1,NEF
?,FLNA,MAP1B,NCL,RPL7A,SLC9A7,ALDOA,PKM,GANAB,FTL,HSP90B1,FADS1,NEFL,NEFM,PRPF8
3X1,UBA1,IGFBP2,CALR,FSTL1,TMF1,ABHD2,HSPA1B,COL5A1,COL4A1,PSAP,RPLP0,H3-3B,CLU,
FLNA,MAP1B,NCL,RPL7A,SLC9A7,ALDOA,PKM,GANAB,COL6A1,FTL,HSP90B1,FADS1,NEFL,NEFM
'L4,ENO1,TUBA1B,PIK3R2,FLNA,MAP1B,NCL,RPL7A,SLC9A7,ALDOA,MTRNR2L8,PKM,GANAB,CC
?,KAT6A,DVL3,YWHAE,KIF1A,RPL13,GNAS,DDX17,GPI,RACK1,BSG,FASN,ALDH6A1,SPTBN1,CALN
;ANAB,FTL,FADS1,NEFL,NEFM,PRPF8,RPL8,FBXL4,YBX1,IGFBP2,CALR,FSTL1,ABHD2,HSPA1B,ZI
RPL7A,SLC9A7,ALDOA,PKM,GANAB,FTL,HSP90B1,FADS1,NEFL,NEFM,PRPF8,RPL8,FBXL4,YBX1,U
LNA,MAP1B,NCL,RPL7A,SLC9A7,MTRNR2L8,PKM,GANAB,COL6A1,FTL,HSP90B1,FADS1,NEFL,N
COL6A1,FTL,HSP90B1,FADS1,NEFM,PRPF8,RPL8,FBXL4,YBX1,UBA1,IGFBP2,FSTL1,ABHD2,HSP.
HSP90B1,FADS1,NEFM,PRPF8,RPL8,FBXL4,YBX1,UBA1,IGFBP2,CALR,FSTL1,ABHD2,HSPA1B,TUI
A1,FTL,HSP90B1,FADS1,NEFM,PRPF8,RPL8,FBXL4,YBX1,UBA1,IGFBP2,CALR,FSTL1,ABHD2,HSF
F8,RPL8,FBXL4,YBX1,UBA1,IGFBP2,FSTL1,ABHD2,TUBB2B,COL5A1,COL4A1,PSAP,PGK1,H3-3B,
LNA,MAP1B,NCL,RPL7A,SLC9A7,ALDOA,PKM,GANAB,COL6A1,HSP90B1,FADS1,NEFL,NEFM,PRPI
GANAB,COL6A1,FTL,HSP90B1,FADS1,NEFM,PRPF8,RPL8,FBXL4,YBX1,UBA1,ATP2B4,IGFBP2,C/
ADS1,NEFM,PRPF8,RPL8,FBXL4,YBX1,UBA1,IGFBP2,CALR,FSTL1,ABHD2,HSPA1B,TUBB2B,COL5
3B2B,COL5A1,PGK1,PTMA,SOX4,SEC62,KAT6A,DVL3,YWHAE,KIF1A,GNAS,SLC2A3,GPI,BSG,FASI
_8,PKM,GANAB,COL6A1,FTL,HSP90B1,FADS1,NEFM,PRPF8,RPL8,FBXL4,YBX1,UBA1,ATP2B4,IGI
L,NEFM,PRPF8,RPL8,FBXL4,YBX1,UBA1,IGFBP2,FSTL1,ZNF99,TUBB2B,COL5A1,COL4A1,PSAP,I
SLC9A7,ALDOA,MTRNR2L8,PKM,GANAB,FTL,HSP90B1,FADS1,NEFL,NEFM,PRPF8,RPL8,FBXL4,Y
90B1,FADS1,NEFL,NEFM,PRPF8,RPL8,FBXL4,YBX1,IGFBP2,CALR,FSTL1,TMF1,TUBB2B,COL5A1
)BN1,RPL4,ENO1,TUBA1B,PIK3R2,FLNA,MAP1B,NCL,RPL7A,SLC9A7,ALDOA,MTRNR2L8,PKM,G/
3,COL6A1,FTL,FADS1,NEFL,PRPF8,RPL8,YBX1,ATP2B4,IGFBP2,CALR,FSTL1,TMF1,ABHD2,COL4
R2,FLNA,MAP1B,NCL,SLC9A7,ALDOA,PKM,GANAB,COL6A1,FTL,FADS1,NEFL,NEFM,PRPF8,RPL8
F8,RPL8,YBX1,UBA1,IGFBP2,FSTL1,TUBB2B,COL5A1,COL4A1,RPLP0,H3-3B,CANX,PTMA,DDX5,S
_,RPL7A,SLC9A7,ALDOA,PKM,GANAB,COL6A1,FTL,HSP90B1,FADS1,NEFM,PRPF8,UGT8,RPL8,FE
NO1,TUBA1B,PIK3R2,FLNA,MAP1B,NCL,RPL7A,SLC9A7,ALDOA,PKM,GANAB,COL6A1,FTL,HSP90
ZNF316,DBN1,RPL4,ENO1,TUBA1B,PIK3R2,FLNA,MAP1B,NCL,RPL7A,SLC9A7,ALDOA,PKM,GANA
A1B,PIK3R2,FLNA,MAP1B,NCL,RPL7A,SLC9A7,ALDOA,PKM,GANAB,COL6A1,FTL,HSP90B1,FADS1
_,HSP90B1,FADS1,NEFM,UGT8,RPL8,FBXL4,YBX1,UBA1,IGFBP2,CALR,FSTL1,ABHD2,HSPA1B,TU
L,HSP90B1,FADS1,PRPF8,RPL8,FBXL4,YBX1,IGFBP2,FSTL1,ABHD2,HSPA1B,TUBB2B,COL5A1,C
O1,TUBA1B,PIK3R2,FLNA,MAP1B,NCL,RPL7A,SLC9A7,ALDOA,MTRNR2L8,PKM,GANAB,COL6A1,
'S6,ZNF316,DBN1,RPL4,ENO1,TUBA1B,PIK3R2,FLNA,MAP1B,NCL,RPL7A,SLC9A7,ALDOA,MTRNF
FSTL1,ABHD2,TUBB2B,PSAP,PGK1,CANX,DDX5,ENO2,FKBP14,SEC62,KAT6A,DVL3,KIF1A,GNAS,S
RPL7A,SLC9A7,ALDOA,PKM,GANAB,COL6A1,FTL,HSP90B1,FADS1,NEFL,NEFM,PRPF8,RPL8,FBXI
,FADS1,NEFL,NEFM,PRPF8,RPL8,YBX1,UBA1,IGFBP2,CALR,FSTL1,ABHD2,HSPA1B,TUBB2B,COL
FBP2,CALR,FSTL1,ABHD2,HSPA1B,TUBB2B,COL5A1,CES3,PTMA,DDX5,ENO2,FKBP14,SEC62,KA
MAP1B,NCL,RPL7A,SLC9A7,ALDOA,PKM,GANAB,COL6A1,FTL,HSP90B1,FADS1,NEFM,PRPF8,RPI
B,PIK3R2,FLNA,MAP1B,NCL,RPL7A,SLC9A7,ALDOA,MTRNR2L8,PKM,GANAB,COL6A1,FTL,HSP9C
UBA1B,PIK3R2,FLNA,MAP1B,NCL,RPL7A,SLC9A7,ALDOA,MTRNR2L8,PKM,GANAB,COL6A1,FTL,I
JGT8,RPL8,FBXL4,YBX1,UBA1,ATP2B4,IGFBP2,FSTL1,TMF1,TUBB2B,COL5A1,COL4A1,PSAP,PG
A,SLC9A7,PKM,GANAB,COL6A1,FTL,HSP90B1,FADS1,NEFM,PRPF8,UGT8,RPL8,FBXL4,YBX1,UBA
,PIK3R2,FLNA,MAP1B,NCL,RPL7A,SLC9A7,ALDOA,PKM,GANAB,COL6A1,FTL,HSP90B1,FADS1,NE

D1,TUBA1B,PIK3R2,FLNA,MAP1B,NCL,RPL7A,SLC9A7,ALDOA,MTRNR2L8,PKM,GANAB,COL6A1,I
0B1,FADS1,NEFM,UGT8,RPL8,FBXL4,YBX1,UBA1,IGFBP2,FSTL1,ABHD2,HSPA1B,TUBB2B,COL5
.T6A,DVL3,YWHAE,KIF1A,GNAS,SLC2A3,GPI,BSG,FASN,ALDH6A1,SPTBN1,CALM3,GNB1,SHISA9,S
R2,FLNA,MAP1B,NCL,RPL7A,SLC9A7,ALDOA,MTRNR2L8,PKM,COL6A1,FTL,HSP90B1,FADS1,NEFI
:NF316,DBN1,RPL4,ENO1,TUBA1B,PIK3R2,FLNA,MAP1B,NCL,RPL7A,SLC9A7,MTRNR2L8,PKM,G
X1,IGFBP2,CALR,FSTL1,TUBB2B,COL5A1,PSAP,PGK1,CLU,PTMA,DDX5,SEC62,KAT6A,DVL3,YWH
R,FSTL1,ABHD2,HSPA1B,TUBB2B,PGK1,CANX,DDX5,ENO2,FKBP14,KAT6A,DVL3,YWHAE,KIF1A,C
IR2L8,PKM,GANAB,COL6A1,FTL,HSP90B1,FADS1,NEFL,NEFM,PRPF8,RPL8,FBXL4,YBX1,UBA1,AT
,RACK1,BSG,FASN,ALDH6A1,SPTBN1,GNB1,SHISA9,STMN2,MYH9,RPL7,SEPTIN2,P4HB,GDI1,TM
A,SLC9A7,ALDOA,PKM,GANAB,COL6A1,FTL,FADS1,NEFL,NEFM,PRPF8,RPL8,FBXL4,YBX1,UBA1,
3N1,CALM3,GNB1,SHISA9,STMN2,MYH9,RPL7,SEPTIN2,P4HB,GDI1,PTMS,SPTAN1,MARCKS,SMG
KAT6A,DVL3,KIF1A,GNAS,SLC2A3,GPI,BSG,FASN,SPTBN1,CALM3,GNB1,STMN2,MYH9,RPL7,SEF
DX5,ENO2,SEC62,KAT6A,DVL3,YWHAE,KIF1A,GNAS,SLC2A3,GPI,BSG,FASN,SPTBN1,GNB1,SHISA
2,SOX4,FKBP14,KAT6A,DVL3,YWHAE,KIF1A,SLC2A3,GPI,RACK1,BSG,FASN,ALDH6A1,SPTBN1,CA
A,MTRNR2L8,PKM,COL6A1,FADS1,NEFL,NEFM,PRPF8,RPL8,FBXL4,YBX1,ATP2B4,IGFBP2,FSTL
EPTIN2,P4HB,APP,GDI1,FSCN1,TMBIM6,PTMS,SPTAN1,MARCKS,SMG1,CALM1,PLXNB2,USP11,C
FLNA,MAP1B,NCL,RPL7A,SLC9A7,ALDOA,MTRNR2L8,PKM,COL6A1,FTL,FADS1,NEFL,NEFM,PRPF
3,PGK1,CANX,DDX5,ENO2,FKBP14,KAT6A,DVL3,KIF1A,GNAS,SLC2A3,GPI,BSG,FASN,ALDH6A1,SF
2A3,BSG,FASN,ALDH6A1,SPTBN1,GNB1,SHISA9,STMN2,MYH9,RPL7,SEPTIN2,GDI1,SPTAN1,GST
T8,RPL8,FBXL4,YBX1,UBA1,IGFBP2,FSTL1,ABHD2,TUBB2B,COL4A1,PGK1,CANX,CLU,PTMA,DDX
NAB,COL6A1,FTL,HSP90B1,FADS1,NEFL,NEFM,PRPF8,RPL8,FBXL4,YBX1,UBA1,IGFBP2,CALR,FS
ACK1,BSG,FASN,SPTBN1,CALM3,GNB1,SHISA9,STMN2,MYH9,RPL7,SEPTIN2,P4HB,GDI1,TMBIM6
3R2,MAP1B,NCL,SLC9A7,ALDOA,PKM,GANAB,FTL,HSP90B1,NEFL,NEFM,RPL8,FBXL4,YBX1,UBA
DX5,ENO2,SEC62,KAT6A,DVL3,YWHAE,GNAS,SLC2A3,HTRA4,FASN,SPTBN1,GNB1,SHISA9,STMN
KAT6A,DVL3,YWHAE,KIF1A,GNAS,SLC2A3,RACK1,BSG,FASN,SPTBN1,GNB1,SHISA9,STMN2,MYH9
,COL4A1,PSAP,H3-3B,PTMA,DDX5,ENO2,RPL13A,KAT6A,DVL3,KIF1A,RPL13,GNAS,DDX17,GPI,H
EFL,NEFM,PRPF8,UGT8,RPL8,FBXL4,YBX1,UBA1,ATP2B4,CALR,ABHD2,HSPA1B,TUBB2B,COL5A
O2,SOX4,KAT6A,DVL3,YWHAE,KIF1A,GNAS,SLC2A3,DDX17,GPI,BSG,FASN,ALDH6A1,SPTBN1,GN
ENO1,TUBA1B,PIK3R2,FLNA,MAP1B,NCL,RPL7A,SLC9A7,ALDOA,MTRNR2L8,PKM,GANAB,COL6
,ALDOA,PKM,GANAB,COL6A1,FTL,HSP90B1,FADS1,NEFM,PRPF8,RPL8,YBX1,ATP2B4,IGFBP2,CA
A,SLC9A7,ALDOA,PKM,COL6A1,HSP90B1,FADS1,NEFL,NEFM,PRPF8,RPL8,FBXL4,YBX1,UBA1,AT
R,TUBB2B,COL5A1,CANX,PTMA,DDX5,SEC62,KAT6A,DVL3,YWHAE,KIF1A,GNAS,SLC2A3,GPI,HTF
K3R2,FLNA,MAP1B,NCL,RPL7A,SLC9A7,ALDOA,PKM,GANAB,COL6A1,FTL,HSP90B1,FADS1,NEFN
A7,ALDOA,MTRNR2L8,PKM,COL6A1,FTL,FADS1,NEFL,NEFM,PRPF8,RPL8,FBXL4,ATP2B4,IGFBP2
3,COL6A1,FTL,HSP90B1,FADS1,NEFM,PRPF8,RPL8,FBXL4,UBA1,ABHD2,TUBB2B,PGK1,CANX,DI
D2,TUBB2B,COL5A1,CLU,PTMA,DDX5,ENO2,FKBP14,KAT6A,DVL3,YWHAE,KIF1A,GNAS,SLC2A3,I
,SLC2A3,RACK1,BSG,SPTBN1,GNB1,SHISA9,STMN2,MYH9,RPL7,SEPTIN2,P4HB,GDI1,TMBIM6,S
16,DBN1,RPL4,ENO1,TUBA1B,PIK3R2,FLNA,MAP1B,NCL,RPL7A,SLC9A7,ALDOA,MTRNR2L8,PKN
FSTL1,ABHD2,TUBB2B,COL5A1,CES3,PGK1,H3-3B,PTMA,DDX5,ENO2,SEC62,YWHAE,KIF1A,GNA
2,TUBB2B,PSAP,PGK1,CANX,DDX5,ENO2,SOX4,FKBP14,SEC62,KAT6A,DVL3,YWHAE,GNAS,SLC2
2,TUBB2B,PSAP,PGK1,CANX,DDX5,ENO2,SOX4,FKBP14,SEC62,KAT6A,DVL3,YWHAE,GNAS,SLC2
3,YWHAE,KIF1A,GNAS,SLC2A3,GPI,HTRA4,FASN,SPTBN1,CALM3,GNB1,SHISA9,STMN2,MYH9,RF
F316,DBN1,RPL4,ENO1,TUBA1B,PIK3R2,FLNA,MAP1B,NCL,RPL7A,SLC9A7,ALDOA,MTRNR2L8,PI
PLP0,H3-3B,CANX,CLU,PTMA,SEC62,KAT6A,DVL3,YWHAE,KIF1A,RPL13,GNAS,SLC2A3,GPI,RACK
3HD2,TUBB2B,COL4A1,RPLP0,H3-3B,CANX,PTMA,SOX4,RPL13A,DVL3,YWHAE,GNAS,DDX17,GPI,

STL1,TUBB2B,CES3,PGK1,CANX,DDX5,ENO2,FKBP14,SEC62,KAT6A,DVL3,YWHAE,GNAS,SLC2A3,
4,KAT6A,DVL3,YWHAE,GNAS,SLC2A3,GPI,BSG,FASN,ALDH6A1,SPTBN1,CALM3,GNB1,SHISA9,STN
X4,KAT6A,DVL3,YWHAE,KIF1A,GNAS,SLC2A3,GPI,BSG,FASN,ALDH6A1,SPTBN1,GNB1,SHISA9,STN
IAS,SLC2A3,GPI,FASN,ALDH6A1,SPTBN1,CALM3,GNB1,SHISA9,STMN2,MYH9,SEPTIN2,GDI1,PTM
M,PRPF8,UGT8,RPL8,FBXL4,YBX1,ATP2B4,IGFBP2,FSTL1,ABHD2,HSPA1B,TUBB2B,COL5A1,PG
ZNF316,DBN1,RPL4,ENO1,TUBA1B,PIK3R2,FLNA,MAP1B,NCL,RPL7A,SLC9A7,ALDOA,MTRNR2L8
,APP,RPL5,PTMS,SPTAN1,MARCKS,CALM1,PLXNB2,HSPD1,USP11,IGF2,EIF4B,MARCKSL1,AFF4,
HSPA1B,TUBB2B,PGK1,CANX,DDX5,ENO2,FKBP14,SEC62,KAT6A,DVL3,KIF1A,SLC2A3,GPI,BSG,I
RPL4,ENO1,TUBA1B,PIK3R2,FLNA,MAP1B,NCL,RPL7A,SLC9A7,ALDOA,MTRNR2L8,PKM,GANAB,C
RPL8,YBX1,ATP2B4,IGFBP2,CALR,FSTL1,TMF1,ABHD2,COL4A1,RPLP0,CANX,PTMA,DDX5,ENO2,
AB,COL6A1,FTL,HSP90B1,FADS1,NEFM,PRPF8,RPL8,YBX1,ATP2B4,IGFBP2,CALR,TMF1,ABHD2,
1,IGFBP2,FSTL1,ABHD2,TUBB2B,PGK1,RPLP0,CANX,DDX5,ENO2,FKBP14,KAT6A,DVL3,SLC2A3,C
A,YWHAE,GNAS,SLC2A3,HTRA4,FASN,SPTBN1,GNB1,SHISA9,STMN2,MYH9,SEPTIN2,P4HB,APP,ID
GK1,CLU,PTMA,ENO2,SOX4,KAT6A,DVL3,YWHAE,KIF1A,GNAS,SLC2A3,DDX17,GPI,BSG,FASN,ALI
N1,RPL4,ENO1,TUBA1B,PIK3R2,FLNA,MAP1B,NCL,RPL7A,SLC9A7,ALDOA,MTRNR2L8,PKM,GANA
1,TMF1,ABHD2,COL4A1,CANX,PTMA,ENO2,KAT6A,YWHAE,GNAS,BSG,FASN,ALDH6A1,SPTBN1,CA
PLXNB2,USP11,ATP5F1A,EIF4B,MARCKSL1,TRIM28,MATR3,YWHAG,CKB,LRRC28,NONO,YWHAQ
PKM,GANAB,COL6A1,HSP90B1,FADS1,NEFL,NEFM,RPL8,FBXL4,UBA1,ATP2B4,IGFBP2,CALR,AB
A,DVL3,YWHAE,KIF1A,RPL13,DDX17,GPI,HTRA4,FASN,SPTBN1,ATP5F1B,CALM3,SHISA9,STMN2,
SG,FASN,SPTBN1,CALM3,GNB1,SHISA9,MYH9,RPL7,RPL5,FSCN1,PTMS,MARCKS,SMG1,CALM1,
,CANX,ENO2,SOX4,KAT6A,DVL3,YWHAE,KIF1A,RPL13,GNAS,DDX17,BSG,FASN,SPTBN1,ATP5F1E
1,PTMS,SPTAN1,GSTP1,SMG1,CALM1,NPM1,COL18A1,SMIM14,IGF2,ATP5F1A,MARCKSL1,TRIM2
IGF2,ATP5F1A,MARCKSL1,UBB,YWHAG,CKB,FN1,SPEN,YWHAQ,HACD3,DCHS1,SET,TPI1,TMSB1
VHAQ,DCHS1,SET,EGLN1,CHSY1,TMSB10,PKD1,PDIA3,SSBP2,VAPA,SEC61A1,STMN1,ITGB1,TUBI
0B1,FADS1,PRPF8,RPL8,YBX1,UBA1,IGFBP2,FSTL1,ABHD2,HSPA1B,TUBB2B,COL5A1,PSAP,PGK1
EN,YWHAQ,EIF1,HACD3,DCHS1,SET,CHSY1,INA,EIF4A1,LDHB,PKD1,VAPA,TPT1,RPL18,STMN1,ITC
BP2,FSTL1,TUBB2B,COL5A1,COL4A1,PSAP,PGK1,RPLP0,H3-3B,CANX,CLU,PTMA,DDX5,SOX4,SE
RPL8,FBXL4,YBX1,IGFBP2,CALR,FSTL1,ABHD2,HSPA1B,TUBB2B,COL5A1,COL4A1,PSAP,RPLP0,
3A1,ATP2B4,IGFBP2,FSTL1,ABHD2,TUBB2B,COL4A1,CES3,PGK1,H3-3B,CLU,PTMA,DDX5,ENO2,
90B1,FADS1,NEFM,PRPF8,RPL8,YBX1,UBA1,IGFBP2,CALR,FSTL1,HSPA1B,TUBB2B,COL5A1,COI
4,ENO1,TUBA1B,PIK3R2,FLNA,NCL,RPL7A,SLC9A7,ALDOA,MTRNR2L8,PKM,GANAB,COL6A1,FT
,RPLP0,PTMA,ENO2,KAT6A,DVL3,YWHAE,KIF1A,RPL13,GNAS,DDX17,GPI,HTRA4,BSG,FASN,SPTI
2,IGF2,EIF4B,MARCKSL1,AFF4,TRIM28,MATR3,YWHAG,CKB,LRRC28,NONO,YWHAQ,HACD3,DCI
OX4,FKBP14,KAT6A,DVL3,SLC2A3,GPI,BSG,FASN,ALDH6A1,SPTBN1,CALM3,GNB1,STMN2,RPS8
B1,FADS1,NEFL,NEFM,PRPF8,UGT8,RPL8,FBXL4,YBX1,UBA1,ATP2B4,IGFBP2,FSTL1,HSPA1B,TU
DOA,PKM,GANAB,COL6A1,HSP90B1,FADS1,NEFM,PRPF8,RPL8,YBX1,UBA1,ATP2B4,IGFBP2,CALI
EN,YWHAQ,DCHS1,SET,CHSY1,INA,PKD1,SSBP2,HMGB1,POLR2A,RPS3A,ITGB1,BTF3,NCAM1,AN
SN,ALDH6A1,SPTBN1,CALM3,SHISA9,STMN2,MYH9,SEPTIN2,P4HB,APP,RPL5,FSCN1,TMBIM6,PTI
R3,YWHAG,CKB,LRRC28,YWHAQ,DCHS1,SET,EGLN1,CHSY1,INA,MRC2,PKD1,VAPA,TPT1,RPL18,S
1,H3-3B,CANX,CLU,PTMA,ENO2,SOX4,MAGED1,RPL6,KAT6A,DVL3,YWHAE,KIF1A,GNAS,SLC2A3,
MN2,RPS8,MYH9,RPL7,SEPTIN2,IDS,GDI1,FSCN1,PTMS,SPTAN1,GSTP1,SMG1,CALM1,PLXNB2,PI
N2,P4HB,APP,RPL5,FSCN1,SPTAN1,CALM1,PLXNB2,PLD3,HSPD1,IGF2,EIF3C,EIF4B,MARCKSL1,I
PA1B,TUBB2B,COL4A1,PGK1,CANX,DDX5,ENO2,FKBP14,RPL6,KAT6A,DVL3,KIF1A,GPI,RACK1,B
C2A3,GPI,BSG,FASN,ALDH6A1,SPTBN1,CALM3,GNB1,STMN2,RPS8,MYH9,RPL7,SEPTIN2,P4HB,II
PL4,ENO1,TUBA1B,PIK3R2,FLNA,MAP1B,NCL,RPL7A,SLC9A7,ALDOA,PKM,GANAB,COL6A1,FTL,I

COL5A1,COL4A1,PSAP,RPLP0,H3-3B,CANX,CLU,PTMA,SOX4,KAT6A,DVL3,YWHAE,KIF1A,RPL13,G
N,DDB1,EIF1,SET,EGLN1,CHSY1,TMSB10,INA,PKD1,HLA-C,SSBP2,POLR2A,TPT1,ITGB1,TUBB4B,
SPTAN1,GSTP1,SMG1,CALM1,PLXNB2,PLD3,HSPD1,USP11,SMIM14,IGF2,VCP,MARCKSL1,TRIM2
1,ATP2B4,FSTL1,TMF1,HSPA1B,TUBB2B,COL5A1,COL4A1,CES3,PSAP,RPLP0,H3-3B,CANX,PTMA
BXL4,YBX1,IGFBP2,CALR,FSTL1,TMF1,HSPA1B,ZNF99,TUBB2B,COL5A1,COL4A1,CES3,PSAP,PGI
L6A1,FTL,HSP90B1,FADS1,NEFM,UGT8,RPL8,YBX1,UBA1,ATP2B4,IGFBP2,CALR,TUBB2B,COL5
3,GNAS,SLC2A3,GPI,BSG,FASN,ALDH6A1,SPTBN1,GNB1,STMN2,RPL7,IDS,ERVV-1,GDI1,FSCN1,F
L4A1,RPLP0,H3-3B,CANX,PTMA,ENO2,SEC62,RPL6,DVL3,YWHAE,GNAS,DDX17,GPI,RACK1,BSG,
7,ALDOA,PKM,COL6A1,FTL,HSP90B1,FADS1,NEFL,NEFM,PRPF8,RPL8,FBXL4,YBX1,ATP2B4,IGFI
3R2,FLNA,MAP1B,NCL,RPL7A,SLC9A7,ALDOA,PKM,GANAB,COL6A1,FTL,HSP90B1,FADS1,NEFL,N
A1,SNRNP200,CSDE1,EIF3C,ATP5F1A,EIF4B,AFF4,TRIM28,MATR3,YWHAG,CKB,NONO,YWHAQ,H
EIF4B,MARCKSL1,AFF4,TRIM28,YWHAG,LRRC28,FN1,YWHAQ,DCHS1,SET,TPI1,TMSB10,PKD1,H
RRC28,YWHAQ,DCHS1,SET,TMSB10,INA,MRC2,LDHB,PKD1,HLA-C,VAPA,HMGB1,POLR2A,TPT1,S
BB2B,PSAP,PGK1,PTMA,DDX5,RPL13A,DVL3,YWHAE,KIF1A,RPL13,GNAS,DDX17,HTRA4,BSG,FAS
OL5A1,COL4A1,PGK1,RPLP0,PTMA,DDX5,ENO2,SEC62,KAT6A,DVL3,YWHAE,GNAS,GPI,HTRA4,BS
PL8,FBXL4,IGFBP2,FSTL1,TMF1,ABHD2,TUBB2B,COL5A1,COL4A1,PSAP,PGK1,RPLP0,H3-3B,CAN
KSL1,AFF4,MATR3,CKB,SPEN,NONO,YWHAQ,EIF1,HACD3,DCHS1,SET,TPI1,CHSY1,TMSB10,MRC
USP11,IGF2,ATP5F1A,EIF4B,MARCKSL1,UBB,MATR3,YWHAG,CKB,FN1,NONO,YWHAQ,HACD3,DC
,FLNA,MAP1B,NCL,SLC9A7,ALDOA,PKM,GANAB,COL6A1,FTL,HSP90B1,FADS1,NEFL,NEFM,PRPF
S1,NEFL,NEFM,PRPF8,RPL8,FBXL4,YBX1,ATP2B4,IGFBP2,FSTL1,TMF1,ABHD2,ZNF99,TUBB2B,C
ADS1,NEFL,NEFM,PRPF8,RPL8,FBXL4,YBX1,UBA1,ATP2B4,IGFBP2,CALR,FSTL1,TMF1,ABHD2,TU
TL1,ABHD2,TUBB2B,PGK1,DDX5,FKBP14,SEC62,DVL3,GNAS,SLC2A3,GPI,BSG,FASN,SPTBN1,CA
1,IGFBP2,FSTL1,TUBB2B,COL4A1,PSAP,PGK1,H3-3B,CANX,CLU,DDX5,SOX4,FKBP14,RPL13A,RF
RPS6,ZNF316,DBN1,RPL4,ENO1,TUBA1B,PIK3R2,FLNA,MAP1B,NCL,RPL7A,SLC9A7,ALDOA,MTRN
USP11,EIF4B,VCP,TRIM28,SPEN,DDB1,PSMD5,YWHAQ,HACD3,DCHS1,SET,CHSY1,INA,RPL15,GTF
SHISA9,MYH9,SEPTIN2,P4HB,APP,IDS,TMBIM6,PTMS,CALM1,PLXNB2,USP11,IGF2,ATP5F1A,MAR
F2,ATP5F1A,VCP,AFF4,TRIM28,MATR3,YWHAG,DDB1,YWHAQ,DCHS1,SET,EGLN1,GTF3C4,PKD1
CL,RPL7A,ALDOA,MTRNR2L8,GANAB,COL6A1,FTL,HSP90B1,FADS1,NEFL,NEFM,PRPF8,UGT8,RF
CALM3,GNB1,STMN2,RPL7,P4HB,GDI1,FSCN1,GSTP1,MARCKS,PLXNB2,USP11,IGF2,ATP5F1A,E
A,MTRNR2L8,PKM,COL6A1,FTL,HSP90B1,NEFL,NEFM,RPL8,FBXL4,YBX1,UBA1,ATP2B4,IGFBP2
KB,COL4A2,YWHAQ,HACD3,DCHS1,SET,TMSB10,EIF4A1,MRC2,LDHB,SSBP2,VAPA,POLR2A,STMI
N1,PTMS,MARCKS,SMG1,CALM1,PLXNB2,PLD3,HSPD1,USP11,CSDE1,SMIM14,IGF2,EIF4B,VCP,T
IA,DDX5,ENO2,YWHAE,KIF1A,RPL13,GNAS,DDX17,GPI,RACK1,BSG,FASN,ATP5F1B,CALM3,GNB1
VHAE,KIF1A,GNAS,SLC2A3,GPI,FASN,ALDH6A1,SPTBN1,GNB1,SHISA9,STMN2,MYH9,SEPTIN2,P4H
TL,HSP90B1,FADS1,NEFL,PRPF8,RPL8,FBXL4,YBX1,UBA1,IGFBP2,CALR,TMF1,TUBB2B,COL5A1
Q,HACD3,DCHS1,SET,TMSB10,MRC2,LDHB,PKD1,SSBP2,VAPA,POLR2A,TPT1,ITGB1,UCHL1,MOR
CKB,LRRC28,FN1,YWHAQ,DCHS1,TMSB10,MRC2,PKD1,HLA-C,PDIA3,VAPA,POLR2A,TPT1,STMN1
ALR,FSTL1,TUBB2B,COL5A1,COL4A1,CES3,H3-3B,PTMA,ENO2,SOX4,SEC62,KAT6A,DVL3,YWHAE
,FASN,SPTBN1,GNB1,SHISA9,STMN2,MYH9,SEPTIN2,APP,IDS,GDI1,TMBIM6,PTMS,MARCKS,SMC
A1,GNB1,STMN2,MYH9,P4HB,IDS,GDI1,FSCN1,PTMS,SPTAN1,MARCKS,SMG1,CALM1,PLXNB2,P
TMF1,ABHD2,TUBB2B,PSAP,PGK1,RPLP0,H3-3B,CANX,PTMA,DDX5,ENO2,SOX4,RPL13A,SEC62,K
2,VAPA,TPT1,ITGB1,UCHL1,MORF4L1,AGRN,TIMP2,MIDN,VCL,DNAJA1,ARF1,CDC42BPB,ITM2B,C
,PTMA,DDX5,KAT6A,KIF1A,RPL13,GNAS,DDX17,ALDH6A1,SPTBN1,CALM3,GNB1,SHISA9,RPS8,M
,PTMA,DDX5,KAT6A,KIF1A,RPL13,GNAS,DDX17,ALDH6A1,SPTBN1,CALM3,GNB1,SHISA9,RPS8,M
,PTMA,DDX5,KAT6A,KIF1A,RPL13,GNAS,DDX17,ALDH6A1,SPTBN1,CALM3,GNB1,SHISA9,RPS8,M

1,ATP2B4,IGFBP2,TMF1,TUBB2B,COL4A1,PSAP,PGK1,H3-3B,CANX,PTMA,ENO2,RPL13A,MAGED
NAS,SLC2A3,DDX17,BSG,FASN,SPTBN1,GNB1,STMN2,MYH9,RPL7,SEPTIN2,RPL5,GDI1,PTMS,SP
1,NEFL,NEFM,UGT8,RPL8,FBXL4,YBX1,UBA1,ATP2B4,IGFBP2,TMF1,HSPA1B,TUBB2B,COL4A1,F
1,CALM3,SHISA9,STMN2,SEPTIN2,P4HB,APP,RPL5,FSCN1,SPTAN1,MARCKS,PLXNB2,USP11,ATP
1,TMSB10,MRC2,LDHB,PKD1,SSBP2,VAPA,POLR2A,TPT1,ITGB1,UCHL1,MORF4L1,NCAM1,AGR
F316,DBN1,RPL4,ENO1,TUBA1B,PIK3R2,FLNA,MAP1B,NCL,RPL7A,SLC9A7,ALDOA,PKM,GANAB,
2,MYH9,RPL7,P4HB,IDS,GDI1,FSCN1,MARCKS,SMG1,CALM1,NPM1,PLXNB2,USP11,COL18A1,CS
DDX17,BSG,FASN,ALDH6A1,SPTBN1,CALM3,GNB1,MYH9,RPL7,SEPTIN2,P4HB,FSCN1,TMBIM6,
AP,RPLP0,H3-3B,CANX,CLU,PTMA,MAGED1,SEC62,KAT6A,DVL3,YWHAE,KIF1A,GNAS,BSG,FASN,
EIF4B,MARCKSL1,AFF4,TRIM28,MATR3,YWHAG,CKB,LRRC28,FN1,YWHAQ,DCHS1,TMSB10,MR
SBF1,DNAJA1,ARF1,CYB5R3,CDC42BPB,ITM2B,OGA,PGAM1,CTNNA1,EIF4H,LPCAT1,BRD3,COL
DS1,NEFL,NEFM,PRPF8,UGT8,RPL8,FBXL4,YBX1,IGFBP2,CALR,TMF1,ABHD2,ZNF99,TUBB2B,C
ADS1,NEFM,PRPF8,RPL8,FBXL4,YBX1,UBA1,FSTL1,ABHD2,ZNF99,TUBB2B,COL5A1,COL4A1,CE
31,YWHAQ,DCHS1,SET,CHSY1,INA,MRC2,PKD1,SSBP2,VAPA,TPT1,ITGB1,MORF4L1,AGRN,ZMIZ1
ANX,CLU,DDX5,ENO2,RPL13A,KAT6A,DVL3,KIF1A,RPL13,GNAS,GPI,HTRA4,BSG,FASN,SPTBN1,C
,NEFL,PRPF8,RPL8,FBXL4,YBX1,UBA1,ATP2B4,IGFBP2,CALR,TMF1,ABHD2,COL4A1,PGK1,H3-3
6,DBN1,RPL4,ENO1,TUBA1B,PIK3R2,FLNA,MAP1B,NCL,SLC9A7,ALDOA,PKM,GANAB,FTL,HSP90
LM1,NPM1,HSPD1,USP11,SMIM14,MARCKSL1,MATR3,CKB,COL4A2,FN1,SPEN,YWHAQ,EIF1,SET
L8,PKM,GANAB,COL6A1,FTL,FADS1,NEFL,NEFM,PRPF8,RPL8,FBXL4,YBX1,UBA1,ATP2B4,IGFBP
STMN2,MYH9,P4HB,GDI1,FSCN1,PTMS,SPTAN1,MARCKS,SMG1,CALM1,PLXNB2,HSPD1,USP11,C
BPB,ITM2B,MAT2A,PXDN,HNRNPU,HNRNPA0,CTNNA1,CNN3,LAMA5,BRD3,KPNB1,NCS1,RHOA,Z
FTL,HSP90B1,NEFL,NEFM,PRPF8,RPL8,FBXL4,ATP2B4,IGFBP2,FSTL1,TMF1,ABHD2,ZNF99,TUB
17,BSG,FASN,ALDH6A1,SPTBN1,CALM3,GNB1,SHISA9,MYH9,SEPTIN2,P4HB,GDI1,FSCN1,SPTAN
IGF2,EIF4B,MARCKSL1,TRIM28,YWHAG,LRRC28,FN1,SET,CHSY1,TMSB10,MRC2,PPP2R1A,PKD
N1,CALM3,GNB1,SHISA9,SEPTIN2,APP,GDI1,PTMS,GSTP1,MARCKS,SMG1,CALM1,NPM1,PLXNB2
JBA1,ABHD2,TUBB2B,COL5A1,PSAP,PGK1,CANX,DDX5,ENO2,SOX4,FKBP14,RPL6,KAT6A,DVL3,
,CHSY1,TMSB10,INA,MRC2,PKD1,HLA-C,PDIA3,SSBP2,VAPA,HMGB1,AP2M1,TPT1,SEC61A1,ITG
L6A1,FTL,HSP90B1,FADS1,NEFL,NEFM,RPL8,YBX1,UBA1,ATP2B4,IGFBP2,FSTL1,TUBB2B,COL4
2,TUBB2B,COL5A1,COL4A1,CES3,PSAP,PGK1,RPLP0,H3-3B,CANX,CLU,PTMA,SOX4,SEC62,RPL
ANX,CLU,PTMA,DDX5,ENO2,MAGED1,SEC62,KAT6A,DVL3,YWHAE,KIF1A,GNAS,BSG,FASN,SPTB
,HTRA4,BSG,FASN,ALDH6A1,ATP5F1B,CALM3,GNB1,RPS8,MYH9,SEPTIN2,P4HB,APP,IDS,RPL5,F
17,FASN,SPTBN1,CALM3,GNB1,SHISA9,STMN2,SEPTIN2,P4HB,APP,GDI1,SPTAN1,GSTP1,SMG1,
1,IGF2,VCP,MARCKSL1,TRIM28,YWHAG,CKB,COL4A2,FN1,SPEN,DDB1,YWHAQ,EIF1,DCHS1,SET
L8,FBXL4,YBX1,UBA1,ATP2B4,TMF1,HSPA1B,TUBB2B,COL5A1,COL4A1,PSAP,RPLP0,H3-3B,CA
TUBB2B,COL5A1,COL4A1,CES3,H3-3B,CLU,PTMA,DDX5,ENO2,SOX4,KAT6A,DVL3,YWHAE,KIF1A
A1,EIF4G2,RPL3,RPS6,ZNF316,DBN1,RPL4,ENO1,TUBA1B,PIK3R2,FLNA,MAP1B,NCL,RPL7A,SLC
1,TMSB10,EIF4A1,RPL19,PPP2R1A,RPL15,GTF3C4,PKD1,PDIA3,SSBP2,BASP1,VAPA,TUBB4B,BT
MORF4L1,AGRN,TIMP2,SARAF,COL11A1,MIDN,VCL,DNAJA1,CYB5R3,CDC42BPB,ITM2B,PXDN,H
N,ZMIZ1,TIMP2,COL11A1,MIDN,VCL,DNAJA1,CYB5R3,CDC42BPB,ITM2B,TMEM170A,HNRNPU,P
,DBN1,RPL4,ENO1,TUBA1B,PIK3R2,FLNA,MAP1B,NCL,RPL7A,SLC9A7,ALDOA,MTRNR2L8,PKM,
P,PGK1,H3-3B,PTMA,DDX5,ENO2,RPL13A,KAT6A,KIF1A,GNAS,BSG,FASN,ALDH6A1,CALM3,GNB
L4A1,CES3,PGK1,H3-3B,CANX,PTMA,DDX5,ENO2,SOX4,FKBP14,RPL13A,KAT6A,DVL3,KIF1A,RP
DX17,HTRA4,BSG,FASN,ALDH6A1,ATP5F1B,CALM3,RPS8,RPL7,SEPTIN2,P4HB,APP,IDS,RPL5,F
CN1,MARCKS,SMG1,CALM1,PLXNB2,USP11,CSDE1,SMIM14,IGF2,VCP,UBB,TRIM28,MATR3,YW
PRPF8,UGT8,RPL8,YBX1,UBA1,ATP2B4,IGFBP2,CALR,FSTL1,TMF1,ABHD2,TUBB2B,COL5A1,C

OL5A1,COL4A1,PSAP,CANX,PTMA,ENO2,KAT6A,DVL3,KIF1A,RPL13,GNAS,HTRA4,BSG,FASN,ATP
STL1,TUBB2B,COL5A1,COL4A1,PSAP,H3-3B,CANX,PTMA,ENO2,KAT6A,DVL3,KIF1A,RPL13,GNAS,
GNB1,SHISA9,RPS8,P4HB,GDI1,FSCN1,PTMS,MARCKS,PLXNB2,HSPD1,USP11,COL18A1,IGF2,MA
),RPL7,SEPTIN2,GDI1,GSTP1,SMG1,PLXNB2,USP11,EIF3C,EIF4B,MARCKSL1,AFF4,TRIM28,YWHA
AT6A,DVL3,YWHAE,KIF1A,RPL13,GNAS,SLC2A3,GPI,HTRA4,RACK1,FASN,ALDH6A1,GNB1,SHISA9
L1,TRIM28,MATR3,COL4A2,FN1,SPEN,YWHAQ,EIF1,HACD3,MRC2,RPL19,GTF3C4,PDIA3,SSBP2,
P0,H3-3B,CLU,PTMA,RPL13A,SEC62,YWHAE,KIF1A,RPL13,SLC2A3,GPI,HTRA4,RACK1,BSG,FASN
NB1,STMN2,RPS8,SEPTIN2,GDI1,TMBIM6,PTMS,SPTAN1,SMG1,NPM1,PLXNB2,HSPD1,USP11,SN
3,FN1,YWHAQ,DCHS1,TMSB10,MRC2,PKD1,HLA-C,VAPA,POLR2A,TPT1,ITGB1,TUBB4B,UCHL1,N
3,FN1,YWHAQ,DCHS1,TMSB10,MRC2,PKD1,HLA-C,VAPA,POLR2A,TPT1,ITGB1,TUBB4B,UCHL1,N
LBP,EIF5A,ITM2C,APLP1,SESN3,B4GALNT4,HNRNPK,INIP,DUX4,RAB1A,PLPPR3,SEPTIN3,CAPRIN
,SERINC1,CHD3,VAPA,POLR2A,AP2M1,TPT1,TUBB4B,MORF4L1,BTF3,RPLP1,RPL10,SBF1,DNAJ
ENO1,TUBA1B,PIK3R2,FLNA,MAP1B,NCL,SLC9A7,ALDOA,MTRNR2L8,PKM,COL6A1,FTL,FADS1,
L4A1,PGK1,RPLP0,CANX,PTMA,DDX5,SOX4,RPL6,GNAS,SLC2A3,DDX17,GPI,SPTBN1,GNB1,SHIS
ACD3,DCHS1,TMSB10,LDHB,PKD1,HLA-C,AP2M1,TPT1,RPS3A,STMN1,ITGB1,TUBB4B,UCHL1,MC
AGRN,TIMP2,SARAF,MIDN,MARCHF6,AP3D1,SBF1,CYB5R3,MAP4,CDC42BPB,ITM2B,MAT2A,PXD
FN1,YWHAQ,DCHS1,TMSB10,PKD1,HLA-C,PDIA3,POLR2A,STMN1,ITGB1,TUBB4B,NCAM1,ANXA2,
,SET,EGLN1,CHSY1,INA,EIF4A1,LDHB,PKD1,SSBP2,VAPA,AP2M1,ITGB1,TUBB4B,ANXA2,AGRN,R
L4A1,CES3,PGK1,RPLP0,H3-3B,CANX,PTMA,DDX5,FKBP14,RPL13A,SEC62,RPL6,KAT6A,DVL3,YW
K1,RPLP0,H3-3B,CANX,PTMA,DDX5,RPL13A,SEC62,RPL6,KAT6A,DVL3,YWHAE,RPL13,DDX17,GPI
,UBA1,IGFBP2,CALR,HSPA1B,CES3,RPLP0,CANX,DDX5,SOX4,RPL13A,MAGED1,SEC62,RPL6,KAT
8,FBXL4,YBX1,UBA1,ATP2B4,IGFBP2,FSTL1,HSPA1B,COL5A1,COL4A1,CES3,PSAP,PGK1,H3-3B,
PKM,COL6A1,FTL,HSP90B1,NEFL,NEFM,RPL8,FBXL4,YBX1,UBA1,ATP2B4,IGFBP2,TMF1,HSPA1
T1,STMN1,ITGB1,MORF4L1,AGRN,COL11A1,AP3D1,SBF1,DNAJA1,ARF1,CYB5R3,MAP4,CDC42B
,PLXNB2,USP11,IGF2,ATP5F1A,EIF4B,MARCKSL1,AFF4,TRIM28,MATR3,YWHAG,CKB,YWHAQ,HA
,CES3,H3-3B,CLU,PTMA,DDX5,ENO2,SEC62,KAT6A,DVL3,YWHAE,KIF1A,GNAS,SLC2A3,FASN,ALD
JBB2B,COL5A1,COL4A1,PGK1,H3-3B,PTMA,DDX5,SOX4,RPL13A,SEC62,KAT6A,DVL3,YWHAE,KI
TP5F1B,CALM3,RPS8,RPL7,SEPTIN2,P4HB,APP,RPL5,GDI1,FSCN1,MARCKS,PLXNB2,PLD3,SNRN
BPB,ITM2B,HNRNPU,CNN3,LAMA5,SLC22A17,PJA2,C1ORF43,LLGL1,HK1,NCS1,ZNF865,PSMD2,
EPTIN2,P4HB,GDI1,FSCN1,TMBIM6,PTMS,GSTP1,SMG1,CALM1,NPM1,HSPD1,USP11,COL18A1,
AP1B,NCL,RPL7A,SLC9A7,PKM,GANAB,COL6A1,FTL,HSP90B1,FADS1,NEFL,NEFM,PRPF8,UGT8,
KD1,HLA-C,PDIA3,POLR2A,TPT1,STMN1,ITGB1,TUBB4B,NCAM1,ANXA2,AGRN,TIMP2,SARAF,MID
3,MARCKSL1,TRIM28,LRRC28,NONO,DDB1,YWHAQ,HACD3,TMSB10,LDHB,PKD1,HLA-C,PDIA3,V
B1,DCHS1,SET,TPI1,CHSY1,INA,EIF4A1,GTF3C4,RPS3,RPS3A,NCAM1,ANXA2,AGRN,RPLP1,SARA
C28,FN1,YWHAQ,DCHS1,SET,EGLN1,CHSY1,TMSB10,EIF4A1,LDHB,PKD1,SSBP2,BASP1,AP2M1,
,RPL13A,SEC62,RPL6,KAT6A,DVL3,YWHAE,KIF1A,RPL13,GNAS,DDX17,GPI,RACK1,BSG,FASN,AL
N1,PTMS,GSTP1,CALM1,PLXNB2,USP11,COL18A1,IGF2,ATP5F1A,MARCKSL1,TRIM28,CKB,COL
F4B,MARCKSL1,AFF4,MATR3,YWHAG,CKB,YWHAQ,HACD3,DCHS1,SET,EGLN1,CHSY1,TMSB10,I
L7A,ALDOA,MTRNR2L8,COL6A1,HSP90B1,NEFL,NEFM,PRPF8,RPL8,YBX1,UBA1,ATP2B4,IGFBP2
WHAE,KIF1A,RPL13,GNAS,HTRA4,RACK1,BSG,FASN,SPTBN1,ATP5F1B,CALM3,GNB1,SHISA9,RPS
STL1,HSPA1B,TUBB2B,COL4A1,PGK1,RPLP0,H3-3B,CANX,DDX5,SOX4,RPL6,KAT6A,GNAS,DDX1
L4A1,PSAP,PGK1,H3-3B,PTMA,DDX5,ENO2,FKBP14,RPL13A,SEC62,KAT6A,DVL3,YWHAE,KIF1A,G
VCL,AP3D1,ARF1,ZNF770,TMEM170A,PXDN,HNRNPU,CTNNA1,FADS2,CPE,CNN3,LPCAT1,LAMA5
CKB,LRRC28,FN1,YWHAQ,HACD3,CHSY1,TMSB10,PKD1,HLA-C,PDIA3,HMGB1,SEC61A1,STMN1,
PTBN1,CALM3,GNB1,SHISA9,STMN2,MYH9,SEPTIN2,IDS,GDI1,FSCN1,TMBIM6,PTMS,MARCKS,S

A1B,COL5A1,COL4A1,CES3,PSAP,RPLP0,H3-3B,CANX,CLU,PTMA,DDX5,SOX4,SEC62,KAT6A,DVL3
7,ALDOA,PKM,GANAB,FTL,HSP90B1,NEFL,NEFM,RPL8,FBXL4,YBX1,UBA1,ATP2B4,IGFBP2,CALR
I,RACK1,BSG,FASN,SPTBN1,ATP5F1B,CALM3,GNB1,SHISA9,RPS8,MYH9,P4HB,IDS,GDI1,FSCN1,S
BX1,ATP2B4,CALR,FSTL1,TMF1,TUBB2B,RPLP0,H3-3B,CLU,PTMA,DDX5,ENO2,FKBP14,MAGED1
JBA1,ABHD2,TUBB2B,COL5A1,PGK1,RPLP0,CANX,DDX5,ENO2,FKBP14,KAT6A,DVL3,KIF1A,RPL1
LRRC28,NONO,DDB1,YWHAQ,HACD3,SET,EIF4A1,MRC2,PPP2R1A,RPL15,LDHB,PKD1,HLA-C,PDI
,MARCHF6,DNAJA1,CYB5R3,CDC42BPB,ITM2B,HNRNPU,CNN3,SLC22A17,BRD3,LLGL1,NCS1,RH
ERVV-1,GDI1,FSCN1,PTMS,SPTAN1,SMG1,PLXNB2,USP11,COL18A1,CSDE1,SMIM14,IGF2,EIF3C
COL6A1,FTL,HSP90B1,FADS1,NEFM,PRPF8,RPL8,YBX1,UBA1,ATP2B4,IGFBP2,FSTL1,TMF1,ABH
ALM3,GNB1,SHISA9,RPS8,RPL7,P4HB,GDI1,FSCN1,SPTAN1,SMG1,PLXNB2,USP11,SNRNP200,IG
ALM3,RPS8,RPL7,SEPTIN2,P4HB,APP,RPL5,GDI1,PTMS,SPTAN1,PLXNB2,IGF2,EIF4B,VCP,UBB,MA
N1,SMG1,NPM1,PLXNB2,IGF2,VCP,MARCKSL1,TRIM28,YWHAG,CKB,COL4A2,FN1,SPEN,DDB1,Y
DS,RPL5,FSCN1,SPTAN1,CALM1,USP11,IGF2,EIF3C,EIF4B,MARCKSL1,MATR3,CKB,COL4A2,LRRC
ES3,PGK1,CLU,PTMA,ENO2,SEC62,KAT6A,DVL3,GNAS,SLC2A3,FASN,SPTBN1,CALM3,GNB1,SHIS
7,ALDOA,PKM,GANAB,FTL,HSP90B1,NEFL,NEFM,RPL8,FBXL4,YBX1,UBA1,IGFBP2,FSTL1,TMF1,C
NA,EIF4A1,PPP2R1A,RPL15,GTF3C4,PKD1,PDIA3,VAPA,TPT1,STMN1,MORF4L1,AGRN,ZMIZ1,TIM
I,IGFBP2,FSTL1,ABHD2,TUBB2B,COL4A1,PGK1,H3-3B,CANX,PTMA,DDX5,ENO2,FKBP14,RPL13A
SEPTIN2,P4HB,APP,IDS,ERVV-1,RPL5,FSCN1,PTMS,SPTAN1,GSTP1,MARCKS,CALM1,USP11,SMIM
B,COL6A1,FTL,HSP90B1,FADS1,NEFL,NEFM,PRPF8,UGT8,FBXL4,YBX1,UBA1,ATP2B4,IGFBP2,CA
13,GNAS,DDX17,GPI,BSG,SHISA9,MYH9,P4HB,APP,IDS,RPL5,GDI1,TMBIM6,PTMS,CALM1,PLXNB
DI1,FSCN1,GSTP1,MARCKS,CALM1,NPM1,HSPD1,SNRNP200,SMIM14,EIF4B,AFF4,TRIM28,MATR
L,RPL7A,ALDOA,PKM,GANAB,COL6A1,FTL,HSP90B1,NEFL,NEFM,PRPF8,RPL8,FBXL4,YBX1,UBA1
,TPT1,ITGB1,TUBB4B,UCHL1,NCAM1,ANXA2,AGRN,SARAF,MIDN,AP3D1,CYB5R3,MAP4,CDC42BP
IAB,COL6A1,HSP90B1,FADS1,NEFM,PRPF8,RPL8,FBXL4,YBX1,UBA1,IGFBP2,CALR,FSTL1,ABHD2
B4B,NCAM1,ANXA2,ZMIZ1,SARAF,WSB1,SBF1,DNAJA1,ARF1,AKAP12,CDC42BPB,ITM2B,MAT2A,G
,KAT6A,DVL3,YWHAE,GNAS,BSG,FASN,ALDH6A1,SPTBN1,CALM3,GNB1,SHISA9,RPS8,MYH9,SEP
,KAT6A,DVL3,YWHAE,GNAS,BSG,FASN,ALDH6A1,SPTBN1,CALM3,GNB1,SHISA9,RPS8,MYH9,SEP
HISA9,MYH9,P4HB,FSCN1,TMBIM6,CALM1,PLD3,HSPD1,USP11,SNRNP200,SMIM14,IGF2,ATP5F1
,COL4A2,LRRC28,FN1,YWHAQ,CHSY1,TMSB10,INA,PKD1,HLA-C,PDIA3,BASP1,VAPA,POLR2A,UC
,BASP1,POLR2A,TUBB4B,MORF4L1,AGRN,MIDN,AKAP12,CDC42BPB,GOLGA8A,FADS2,LAMA5,P
POLR2A,TPT1,STMN1,ITGB1,MORF4L1,AGRN,TIMP2,COL11A1,VCL,CDC42BPB,AP2B1,SLC4A8,I
ANAB,COL6A1,FTL,FADS1,NEFL,NEFM,PRPF8,RPL8,FBXL4,YBX1,UBA1,ATP2B4,IGFBP2,CALR,FS
STP1,CALM1,PLXNB2,USP11,IGF2,ATP5F1A,AFF4,TRIM28,MATR3,YWHAG,CKB,LRRC28,NONO,D
OL18A1,SNRNP200,IGF2,EIF4B,MARCKSL1,UBB,AFF4,MATR3,YWHAG,CKB,COL4A2,HACD3,DCH
,NPM1,PLD3,USP11,SNRNP200,EIF3C,ATP5F1A,EIF4B,VCP,MATR3,CKB,NONO,YWHAQ,HACD3,S
DX5,ENO2,KAT6A,DVL3,YWHAE,KIF1A,GNAS,GPI,BSG,FASN,SPTBN1,GNB1,SHISA9,RPS8,MYH9,P
,IGF2,EIF4B,UBB,TRIM28,YWHAG,CKB,COL4A2,DDB1,HACD3,DCHS1,CHSY1,EIF4A1,MRC2,PPP2
L8,FBXL4,YBX1,ATP2B4,IGFBP2,CALR,ABHD2,HSPA1B,TUBB2B,COL5A1,COL4A1,CES3,H3-3B,C
0,MRC2,PKD1,HLA-C,CHD3,VAPA,HMGB1,POLR2A,TPT1,ITGB1,TUBB4B,UCHL1,ANXA2,AGRN,TI
B1,FADS1,NEFL,NEFM,PRPF8,RPL8,FBXL4,YBX1,ATP2B4,IGFBP2,CALR,FSTL1,ABHD2,HSPA1B,C
MF1,HSPA1B,TUBB2B,COL5A1,CES3,RPLP0,H3-3B,PTMA,DDX5,ENO2,SOX4,FKBP14,SEC62,RPL6
PTMS,MARCKS,NPM1,HSPD1,IGF2,EIF3C,UBB,MATR3,CKB,FN1,SPEN,PSMD5,YWHAQ,CHSY1,EIF
,TUBB4B,MORF4L1,SARAF,RPS17,AP3D1,CYB5R3,SF3B2,MAT2A,PGAM1,HNRNPA0,FADS2,RPSA
,GNB1,SHISA9,MYH9,APP,IDS,GDI1,PTMS,SPTAN1,PLXNB2,USP11,SMIM14,IGF2,ATP5F1A,UBB,A
OL5A1,COL4A1,PSAP,PGK1,RPLP0,CANX,ENO2,SOX4,SEC62,KAT6A,DVL3,YWHAE,KIF1A,RPL13,G

LC9A7,ALDOA,PKM,GANAB,HSP90B1,NEFL,NEFM,RPL8,FBXL4,YBX1,UBA1,ATP2B4,IGFBP2,CAL
LRRC28,SPEN,YWHAQ,DCHS1,SET,EGLN1,TPI1,CHSY1,EIF4A1,MRC2,PPP2R1A,PKD1,HLA-C,PDI
A,DVL3,KIF1A,RPL13,GNAS,DDX17,GPI,HTRA4,BSG,FASN,SPTBN1,GNB1,SHISA9,RPS8,MYH9,RP
F4H,LPCAT1,RPS9,CIPC,PAPOLA,RHOA,PTCD3,CBX5,ZNF865,KIF5B,LENG8,VAT1,SRSF4,RTN3,P
,COL4A1,PGK1,RPLP0,CANX,CLU,PTMA,DDX5,ENO2,SEC62,KAT6A,DVL3,YWHAE,KIF1A,RPL13,G
0B1,FADS1,NEFL,NEFM,PRPF8,UGT8,RPL8,FBXL4,YBX1,UBA1,IGFBP2,CALR,TMF1,ABHD2,ZNF9
M6,SPTAN1,CALM1,PLD3,USP11,SNRNP200,IGF2,EIF4B,VCP,RPL37A,NONO,EIF1,TPI1,INA,EIF4A
1,H3-3B,PTMA,DDX5,FKBP14,RPL13A,MAGED1,RPL6,DVL3,YWHAE,GNAS,GPI,HTRA4,RACK1,FA
4L1,AGRN,ZMIZ1,TIMP2,MIDN,DNAJA1,CDC42BPB,PXDN,CTNNA1,FADS2,TNS3,LAMA5,SLC22A1
GFBP2,FSTL1,HSPA1B,TUBB2B,COL5A1,PTMA,RPL13A,SEC62,KAT6A,YWHAE,KIF1A,RPL13,GNA
1,GNB1,SHISA9,MYH9,APP,IDS,GDI1,PTMS,SPTAN1,PLXNB2,USP11,SMIM14,IGF2,EIF3C,ATP5F1
DC42BPB,PXDN,CTSD,NACA,BRD3,KPNB1,C1ORF43,LLGL1,NCS1,NR2F1,NLGN2,CHD4,JUND,E
APA,POLR2A,TPT1,SEC61A1,RPS3A,TUBB4B,UCHL1,ANXA2,AGRN,MIDN,AP3D1,ARF1,CYB5R3,
FL,NEFM,PRPF8,UGT8,RPL8,FBXL4,YBX1,IGFBP2,CALR,TMF1,ABHD2,ZNF99,TUBB2B,COL4A1,
HSP90B1,NEFM,PRPF8,RPL8,FBXL4,YBX1,UBA1,ATP2B4,IGFBP2,CALR,FSTL1,TMF1,ABHD2,HSP
,KAT6A,KIF1A,GNAS,DDX17,HTRA4,BSG,ATP5F1B,CALM3,RPS8,RPL7,SEPTIN2,P4HB,APP,IDS,R
N,RPL10,MARCHF6,RPS17,ARF1,ITM2B,SLC4A8,PGAM1,HNRNPA0,FADS2,CPE,CNN3,TNS3,LPC
NAS,DDX17,GPI,HTRA4,BSG,FASN,SPTBN1,GNB1,SHISA9,RPS8,MYH9,APP,GDI1,FSCN1,PTMS,SP
PS15,ARL4C,RCN1,MIF,GPAT4,MYO5A,SESN3,MTSS2,APLP2,HNRNPDL,TARDBP,RPLP2,SHOX,DU
PL6,KAT6A,DVL3,GNAS,GPI,RACK1,SPTBN1,ATP5F1B,CALM3,GNB1,SHISA9,RPS8,MYH9,RPL7,S
70A,GTF2F1,PXDN,OGA,PGAM1,SMARCA5,FADS2,RPS16,CTSD,EIF4H,LPCAT1,RPS9,CIPC,PAPOL
TPI1,MRC2,LDHB,PKD1,SF3B1,SSBP2,BASP1,VAPA,HMGB1,POLR2A,STMN1,ITGB1,UCHL1,MOR
D1,RPL6,DVL3,YWHAE,GNAS,GPI,RACK1,FASN,GNB1,SHISA9,RPS8,MYH9,P4HB,TMBIM6,PTMS,
PS6,ZNF316,DBN1,RPL4,ENO1,TUBA1B,PIK3R2,FLNA,MAP1B,NCL,RPL7A,SLC9A7,ALDOA,MTRN
FBXL4,YBX1,ATP2B4,CALR,FSTL1,HSPA1B,COL5A1,COL4A1,PSAP,PGK1,RPLP0,H3-3B,CANX,CL
3,YWHAG,CKB,YWHAQ,HACD3,DCHS1,SET,EGLN1,CHSY1,INA,MRC2,LDHB,PKD1,HLA-C,VAPA,A
F4B,VCP,MARCKSL1,UBB,CKB,COL4A2,EGLN1,MAP9,TPI1,CHSY1,INA,MRC2,RPL19,PPP2R1A,R
L4A1,H3-3B,CANX,PTMA,DDX5,SOX4,RPL13A,SEC62,RPL13,SLC2A3,GPI,RACK1,BSG,FASN,SPT
WHAG,CKB,LRRC28,FN1,EIF1,HACD3,CHSY1,TMSB10,PKD1,HLA-C,PDIA3,VAPA,HMGB1,POLR2
HAQ,DCHS1,TMSB10,RPL19,PKD1,CHD3,BASP1,POLR2A,TPT1,TUBB4B,MORF4L1,ANXA2,AGR
BSG,FASN,SPTBN1,GNB1,SHISA9,RPS8,MYH9,P4HB,IDS,FSCN1,SPTAN1,SMG1,HSPD1,IGF2,EIF4
PL8,FBXL4,YBX1,UBA1,ATP2B4,CALR,TMF1,ABHD2,HSPA1B,TUBB2B,COL4A1,PGK1,H3-3B,CAN
T6A,DVL3,RPL13,DDX17,HTRA4,RACK1,BSG,ALDH6A1,SPTBN1,CALM3,GNB1,SHISA9,RPS8,MY
NEFM,RPL8,FBXL4,YBX1,UBA1,ATP2B4,CALR,FSTL1,TMF1,ABHD2,HSPA1B,TUBB2B,COL5A1,C
NB1,RPS8,MYH9,APP,GDI1,PTMS,SPTAN1,SMG1,USP11,COL18A1,SNRNP200,CSDE1,MARCKSL1
1,NEFL,NEFM,FBXL4,YBX1,UBA1,ATP2B4,IGFBP2,FSTL1,ABHD2,COL5A1,COL4A1,PSAP,PGK1,
,NCL,RPL7A,ALDOA,MTRNR2L8,GANAB,FTL,HSP90B1,FADS1,NEFL,NEFM,PRPF8,UGT8,RPL8,F
X5,SEC62,KAT6A,DVL3,KIF1A,GNAS,SLC2A3,DDX17,HTRA4,BSG,FASN,ALDH6A1,SPTBN1,GNB1,
S3,CTSD,EIF4H,LPCAT1,BRD3,IGFBP3,LLGL1,NCS1,ZNF865,CLSTN1,SLC7A5,JUND,CD81,ATP2
4G2,RPL3,RPS6,ZNF316,DBN1,RPL4,ENO1,TUBA1B,PIK3R2,FLNA,MAP1B,NCL,RPL7A,SLC9A7,A
,ABHD2,COL5A1,PSAP,PGK1,CLU,PTMA,DDX5,ENO2,FKBP14,RPL13A,SEC62,RPL6,DVL3,KIF1A,
ATP5F1B,CALM3,SHISA9,MYH9,RPL7,P4HB,RPL5,FSCN1,SPTAN1,GSTP1,SMG1,PLXNB2,PLD3,U
1,NPM1,SNRNP200,IGF2,EIF3C,EIF4B,VCP,UBB,AFF4,TRIM28,YWHAG,CKB,COL4A2,RPL37A,M
2,RPL15,GTF3C4,PKD1,HLA-C,TPT1,ITGB1,TUBB4B,UCHL1,MORF4L1,AGRN,ZMIZ1,TIMP2,RPL1
PP2R1A,PKD1,HLA-C,PDIA3,VAPA,STMN1,UCHL1,ANXA2,AGRN,MIDN,VCL,AP3D1,CYB5R3,MAP4,

A1,SCARB2,LPCAT1,BRD3,MSN,SLC16A7,LLGL1,ZNF865,SLC7A5,CHD4,ATP1A1,JUND,CD81,HM
GL1,NCS1,ZNF865,CHD4,JUND,CD81,HMGN2,ATP2A2,CRMP1,LASP1,OAZ1,TLE5,NISCH,CCNI,KH
AGRN,ZMIZ1,MARCHF6,DNAJA1,CYB5R3,SF3B2,SMARCA5,RPL23,RPS9,RPL17,C1ORF43,GAP43
,COL6A1,FTL,FADS1,NEFL,NEFM,PRPF8,RPL8,FBXL4,YBX1,UBA1,ATP2B4,IGFBP2,CALR,FSTL1,T
A1,FTL,HSP90B1,NEFL,NEFM,RPL8,FBXL4,UBA1,IGFBP2,TMF1,ABHD2,HSPA1B,COL4A1,PGK1,F
DN,HNRNPU,CTNNA1,CTSD,SCARB2,BRD3,KPNB1,LLGL1,NCS1,ZNF865,SLC7A5,CHD4,ATP1A1,J
ARF1,CYB5R3,MAP4,CDC42BPB,ITM2B,MAT2A,GOLGA8A,PXDN,HNRNPU,RPS16,CTSD,BRD3,CIF
L4,YBX1,UBA1,ATP2B4,CALR,TMF1,ABHD2,HSPA1B,TUBB2B,COL4A1,PGK1,H3-3B,CANX,PTMA,
,ITGB1,TUBB4B,MIDN,VCL,DNAJA1,MAP4,CDC42BPB,ITM2B,AP2B1,PXDN,PGAM1,SMARCA5,HN
,ITGB1,TUBB4B,MIDN,VCL,DNAJA1,MAP4,CDC42BPB,ITM2B,AP2B1,PXDN,PGAM1,SMARCA5,HN
,ITGB1,TUBB4B,MIDN,VCL,DNAJA1,MAP4,CDC42BPB,ITM2B,AP2B1,PXDN,PGAM1,SMARCA5,HN
IF1A,RPL13,GPI,BSG,FASN,ALDH6A1,GNB1,MYH9,RPL7,P4HB,TMBIM6,SPTAN1,SMG1,CALM1,N
XAF1,MIDN,SBF1,DNAJA1,ARF1,CYB5R3,AKAP12,MAT2A,TMEM170A,SLC4A8,HNRNPU,OGA,CTN
TMF1,TUBB2B,COL5A1,COL4A1,PGK1,RPLP0,H3-3B,CANX,PTMA,RPL13A,KAT6A,DVL3,YWHAE,K
COL6A1,FTL,HSP90B1,NEFL,NEFM,RPL8,FBXL4,UBA1,IGFBP2,TMF1,ABHD2,HSPA1B,ZNF99,COL
PL8,FBXL4,YBX1,UBA1,CALR,TMF1,ABHD2,HSPA1B,COL4A1,PGK1,H3-3B,PTMA,DDX5,FKBP14,R
6A1,FTL,HSP90B1,FADS1,NEFL,NEFM,PRPF8,RPL8,YBX1,ATP2B4,IGFBP2,CALR,FSTL1,TMF1,AE
P1,LASP1,CCNI,PFKL,PNMA1,PDIA6,MEGF8,SOX12,ARL4C,APLP1,MIF,GPAT4,CHPF,MYO5A,B4G
K1,BSG,CALM3,RPS8,RPL7,SEPTIN2,P4HB,APP,IDS,PTMS,SMG1,CALM1,NPM1,PLXNB2,PLD3,SN
A-C,PDIA3,SSBP2,AP2M1,STMN1,ITGB1,UCHL1,NCAM1,ANXA2,AGRN,ZMIZ1,TIMP2,XAF1,MIDN,S
,SEPTIN2,P4HB,GDI1,FSCN1,PTMS,SPTAN1,SMG1,NPM1,PLXNB2,COL18A1,EIF4B,MARCKSL1,AI
WHAQ,HACD3,SET,CHSY1,TMSB10,EIF4A1,RPL15,LDHB,SSBP2,BASP1,VAPA,POLR2A,TPT1,STMI
N1,SMG1,CALM1,NPM1,PLXNB2,MARCKSL1,AFF4,MATR3,YWHAG,CKB,LRRC28,FN1,EIF1,SET,T
,HSP90B1,NEFL,NEFM,RPL8,FBXL4,UBA1,IGFBP2,TMF1,ABHD2,HSPA1B,COL4A1,PGK1,RPLP0,
PU,TPM4,NACA,SLC22A17,PJA2,KPNB1,NCS1,RHOA,ZNF865,CXADR,CHD4,ATP1A1,JUND,EIF5,W
BX1,UBA1,IGFBP2,CALR,ABHD2,HSPA1B,ZNF99,TUBB2B,COL5A1,COL4A1,CES3,PSAP,RPLP0,H
MARCKS,NPM1,PLXNB2,TTC3,CSDE1,IGF2,EIF4B,UBB,YWHAG,CKB,COL4A2,SPEN,EIF1,SET,CHS
A1,SNRNP200,IGF2,EIF3C,EIF4B,MARCKSL1,AFF4,MATR3,YWHAG,COL4A2,RPL37A,NONO,YWH
PTMS,PLD3,SNRNP200,IGF2,EIF3C,VCP,UBB,CKB,COL4A2,PSMD5,YWHAQ,HACD3,CHSY1,TMSB
PL8,FBXL4,YBX1,UBA1,ATP2B4,IGFBP2,CALR,TMF1,ABHD2,HSPA1B,TUBB2B,PGK1,H3-3B,CANX
4,TMEM170A,HNRNPU,HNRNPA0,CTNNA1,CPE,TNS3,LPCAT1,LAMA5,BRD3,PAPOLA,IGFBP3,HK1
2,PLD3,USP11,SNRNP200,SMIM14,ATP5F1A,EIF4B,VCP,UBB,AFF4,EIF1,INA,EIF4A1,PPP2R1A,RPI
A9,STMN2,MYH9,IDS,GDI1,PTMS,GSTP1,SMG1,NPM1,PLXNB2,USP11,COL18A1,IGF2,EIF4B,UBB,
,SEPTIN2,P4HB,APP,IDS,PTMS,SMG1,NPM1,PLXNB2,PLD3,SNRNP200,SMIM14,IGF2,EIF3C,MAR(
YB5R3,ITM2B,MAT2A,NFE2L1,FADS2,RPSA,NACA,CIPC,PAPOLA,CBX5,CLSTN1,ATF4,CHD4,TMEN
3,COL4A1,PGK1,RPLP0,H3-3B,CANX,PTMA,DDX5,ENO2,SOX4,RPL13A,SEC62,KAT6A,DVL3,YWHA
ANX,DDX5,FKBP14,RPL13A,MAGED1,RPL6,DVL3,YWHAE,GNAS,GPI,RACK1,FASN,ATP5F1B,CALM
SP90B1,NEFL,NEFM,PRPF8,RPL8,FBXL4,YBX1,UBA1,ATP2B4,IGFBP2,CALR,FSTL1,TMF1,H3-3B,
B1,AGRN,MIDN,CDC42BPB,MAT2A,TMEM170A,PXDN,HNRNPU,OGA,PGAM1,SMARCA5,HNRNPA(
NEFM,RPL8,FBXL4,UBA1,IGFBP2,TMF1,ABHD2,HSPA1B,COL4A1,PGK1,RPLP0,CANX,CLU,PTMA,I
,RPL8,FBXL4,YBX1,IGFBP2,CALR,TMF1,ABHD2,ZNF99,TUBB2B,COL4A1,PSAP,PGK1,RPLP0,CAN
PP,IDS,GDI1,FSCN1,SPTAN1,MARCKS,CALM1,NPM1,PLD3,HSPD1,SNRNP200,VCP,TRIM28,MATR
COL6A1,FTL,HSP90B1,NEFL,NEFM,RPL8,FBXL4,UBA1,IGFBP2,TMF1,ABHD2,HSPA1B,ZNF99,COL
DDB1,PSMD5,HACD3,CHSY1,EIF4A1,RPL15,GTF3C4,LDHB,SF3B1,SSBP2,BASP1,POLR2A,TPT1,F
L13,GNAS,BSG,GNB1,SHISA9,MYH9,IDS,GDI1,PTMS,SPTAN1,SMG1,CALM1,PLXNB2,IGF2,EIF3C,

,ATP5F1A,EIF4B,VCP,UBB,AFF4,LRRC28,TMSB10,INA,EIF4A1,RPL19,PPP2R1A,RPL15,GTF3C4,L
MARCHF6,MAT2A,PXDN,OGA,HNRNPA0,FADS2,LAMA5,BRD3,SLC16A7,KPNB1,C1ORF43,PAPOLA
,GTF3C4,SF3B1,PDIA3,POLR2A,RPS3A,STMN1,UCHL1,MORF4L1,ANXA2,SARAF,MIDN,DNAJA1,S
L8,FBXL4,YBX1,UBA1,ATP2B4,IGFBP2,CALR,TMF1,HSPA1B,TUBB2B,PGK1,H3-3B,CANX,CLU,PT
X1,UBA1,CALR,TMF1,ABHD2,HSPA1B,COL4A1,PGK1,H3-3B,PTMA,DDX5,FKBP14,RPL13A,MAGE
SPTAN1,SMG1,CALM1,NPM1,PLXNB2,PLD3,USP11,SNRNP200,IGF2,EIF3C,VCP,MARCKSL1,AFF4
S1,RHOA,SLC7A5,ATP1A1,JUND,CD81,ATP2A2,CRMP1,NISCH,CCNI,TADA2B,CSNK1E,PFKL,SFPQ,
CLU,ENO2,RPL13A,MAGED1,DVL3,KIF1A,RPL13,GNAS,BSG,FASN,SPTBN1,GNB1,MYH9,SEPTIN2
TL,HSP90B1,FADS1,NEFM,UGT8,RPL8,YBX1,UBA1,ATP2B4,CALR,FSTL1,HSPA1B,TUBB2B,COL5
CN1,SPTAN1,CALM1,NPM1,PLD3,HSPD1,SNRNP200,VCP,TRIM28,MATR3,CKB,COL4A2,MAPT,SP
LGL1,ZNF865,SLC7A5,CHD4,ATP1A1,JUND,CD81,HMGN2,ATP2A2,CRMP1,LASP1,OAZ1,PFN1,N
,NONO,YWHAQ,HACD3,MRC2,PPP2R1A,RPL15,GTF3C4,PDIA3,VAPA,AP2M1,RPS3A,UCHL1,NCAM
A2B,KHDRBS1,PDIA6,GPC3,PIP4K2B,SOX12,ACTN4,ARL4C,APLP1,MIF,TLN1,SESN3,MTSS2,APL
HL1,AGRN,MIDN,AP3D1,ARF1,AKAP12,CDC42BPB,ITM2B,GOLGA8A,HNRNPU,CTNNA1,CNN3,CT
S,DDX17,GPI,HTRA4,FASN,SPTBN1,CALM3,GNB1,SHISA9,RPS8,MYH9,SEPTIN2,APP,ERVV-1,GDI1
RPL5,GDI1,PTMS,SMG1,NPM1,PLD3,HSPD1,IGF2,EIF4B,UBB,AFF4,YWHAG,MAPT,SPEN,NONO,DD
,RPL7,P4HB,GDI1,PTMS,GSTP1,MARCKS,SMG1,NPM1,TTC3,IGF2,VCP,MARCKSL1,AFF4,TRIM28
K1,H3-3B,PTMA,DDX5,SOX4,FKBP14,RPL13A,MAGED1,RPL6,DVL3,YWHAE,GNAS,GPI,HTRA4,RA
PNB1,C1ORF43,IGFBP3,PTCD3,PSMD2,JUND,EIF5,LENG8,WDR1,HMGN2,CRMP1,OAZ1,RBBP4,J
P2B4,CALR,TMF1,ABHD2,HSPA1B,TUBB2B,COL4A1,H3-3B,CANX,PTMA,DDX5,FKBP14,RPL13A,M
KIF1A,GNAS,SLC2A3,DDX17,GPI,HTRA4,RACK1,BSG,CALM3,GNB1,SHISA9,RPS8,RPL7,SEPTIN2,
EPTIN2,GDI1,GSTP1,MARCKS,NPM1,TTC3,USP11,COL18A1,SNRNP200,CSDE1,EIF3C,ATP5F1A,E
F1,HSPA1B,PGK1,H3-3B,CANX,PTMA,DDX5,FKBP14,RPL13A,MAGED1,RPL6,DVL3,YWHAE,GNAS,
,GTF3C4,SSBP2,POLR2A,SEC61A1,ITGB1,TUBB4B,AGRN,MARCHF6,AP3D1,CYB5R3,MAP4,CDC
,KPNB1,PAPOLA,LLGL1,NCS1,RHOA,CLSTN1,ATF4,ATP1A1,TMEM259,EIF5,HMGN2,CRMP1,LAS
0,SMIM14,ATP5F1A,EIF4B,VCP,UBB,AFF4,EIF1,INA,EIF4A1,PPP2R1A,RPL15,GTF3C4,LDHB,DNA
200,SMIM14,ATP5F1A,EIF4B,VCP,UBB,AFF4,RPL37A,LRRC28,SPEN,EIF1,INA,EIF4A1,PPP2R1A,R
B,YWHAG,COL4A2,SPEN,NONO,YWHAQ,EIF1,TMSB10,GTF3C4,SERINC1,ZNF43,HMGB1,POLR2
,NEFM,PRPF8,RPL8,FBXL4,YBX1,UBA1,ATP2B4,IGFBP2,FSTL1,TMF1,ABHD2,ZNF99,TUBB2B,CO
RPL13A,MAGED1,RPL6,DVL3,RPL13,GNAS,GPI,HTRA4,RACK1,ATP5F1B,CALM3,GNB1,SHISA9,M
CIPC,PAPOLA,RHOA,PTCD3,CBX5,ZNF865,KIF5B,GLIPR1L2,LENG8,ATXN7L3B,VAT1,SRSF4,RTN
UBA1,ABHD2,HSPA1B,TUBB2B,COL5A1,PGK1,CANX,DDX5,ENO2,SEC62,KAT6A,DVL3,YWHAE,KIF
NISCH,PFKL,GLG1,PEBP1,PIP4K2B,SOX12,ARL4C,APLP1,MIF,APLP2,HNRNPDL,TARDBP,MYH10,
AT6A,YWHAE,KIF1A,RPL13,GNAS,DDX17,HTRA4,RACK1,BSG,FASN,CALM3,RPS8,SEPTIN2,CALM1
BA1,ATP2B4,CALR,FSTL1,TMF1,ABHD2,HSPA1B,TUBB2B,COL4A1,PSAP,PGK1,H3-3B,CLU,DDX5
A,DDX5,FKBP14,RPL13A,MAGED1,RPL6,DVL3,YWHAE,GNAS,GPI,HTRA4,RACK1,FASN,ALDH6A1,A
2B,COL4A1,RPLP0,PTMA,DDX5,ENO2,SOX4,RPL13A,MAGED1,SEC62,RPL6,KAT6A,DVL3,YWHAE,
GRN,TIMP2,SARAF,COL11A1,MIDN,RPL10,DNAJA1,CDC42BPB,ITM2B,AP2B1,PXDN,TPM4,LAMA5
ARF1,SF3B2,GTF2F1,HNRNPA0,RPS16,RPL23,CNN3,RPL29,EIF4H,RPS9,CIPC,PAPOLA,HK1,RHO
HL1,BTF3,AGRN,ZMIZ1,TIMP2,MIDN,AP3D1,DNAJA1,ARF1,AKAP12,CDC42BPB,ITM2B,GOLGA8A
BXL4,UBA1,IGFBP2,FSTL1,TMF1,ABHD2,HSPA1B,ZNF99,COL4A1,CES3,PGK1,RPLP0,CANX,PTM
,ZMIZ1,MIDN,RPS17,VCL,AP3D1,SBF1,ARF1,CYB5R3,ITM2B,TMEM170A,SLC4A8,PGAM1,NFE2L
2,TUBB2B,COL5A1,COL4A1,CES3,PSAP,H3-3B,CLU,ENO2,MAGED1,SEC62,RPL6,DVL3,GNAS,G
A1B,TUBB2B,COL4A1,PGK1,H3-3B,DDX5,SOX4,RPL6,KAT6A,GNAS,DDX17,GPI,BSG,FASN,SPTBN
ACTN4,RPS5,DNAJC5,HNRNPH1,MAGED4B,CTBP1,ARHGDIA,CDK16,BCLAF1,NNAT,C5ORF24,SL

DX17,HTRA4,RACK1,FASN,ALDH6A1,SPTBN1,GNB1,SHISA9,STMN2,RPL7,P4HB,APP,FSCN1,PTMS
NF43,HMGB1,POLR2A,TPT1,SEC61A1,RPS3A,RPL18,TUBB4B,RPLP1,MIDN,RPL10,RPS17,VCL,DI
NA1,FADS2,LPCAT1,NCS1,ZNF865,SLC7A5,NLGN2,JUND,EIF5,CD81,CRMP1,LASP1,OAZ1,PFN1,N
TBP1,SLC35E2B,SMCHD1,IPO5,HSBP1,DLG5,ZC3H7B,SNX22,SQSTM1,LMNB2,ACTR3,GRINA,NC
H3-3B,CANX,DDX5,FKBP14,RPL13A,MAGED1,SEC62,RPL6,KAT6A,DVL3,GNAS,DDX17,GPI,RACK1
YWHAE,GNAS,GPI,RACK1,FASN,ATP5F1B,CALM3,GNB1,SHISA9,RPS8,MYH9,P4HB,RPL5,TMBIM6
K1,UBA1,ATP2B4,IGFBP2,CALR,FSTL1,HSPA1B,TUBB2B,COL5A1,DDX5,ENO2,RPL13A,SEC62,KA
ALDOA,PKM,GANAB,COL6A1,FTL,HSP90B1,NEFL,NEFM,RPL8,FBXL4,UBA1,ATP2B4,IGFBP2,CALF
LNT4,DNAJC5,APLP2,MXD4,SEPTIN11,PLPPR3,CTBP1,ARHGDIA,GPC1,TOP2B,TNC,ATP6V0C,SKF
A,PKM,GANAB,COL6A1,FTL,HSP90B1,NEFL,NEFM,RPL8,FBXL4,UBA1,IGFBP2,CALR,FSTL1,TMF1,
CNN3,RPSA,TNS3,CTSD,EIF4H,LAMA5,SLC22A17,BRD3,MSN,KPNB1,C1ORF43,PAPOLA,RPL14,F
13A,MAGED1,RPL6,DVL3,YWHAE,GNAS,GPI,HTRA4,RACK1,FASN,ATP5F1B,CALM3,SHISA9,RPS8,
6,DVL3,YWHAE,GNAS,GPI,RACK1,FASN,ATP5F1B,CALM3,GNB1,SHISA9,RPS8,MYH9,SEPTIN2,P4H
MAP1B,NCL,RPL7A,SLC9A7,ALDOA,PKM,GANAB,FTL,HSP90B1,FADS1,NEFM,PRPF8,UGT8,RPL8,
DI1,PTMS,SPTAN1,CALM1,PLXNB2,TTC3,HSPD1,COL18A1,SNRNP200,SMIM14,EIF3C,UBB,CKB,L
P,IDS,GDI1,PTMS,SMG1,NPM1,PLXNB2,PLD3,SNRNP200,IGF2,MARCKSL1,UBB,AFF4,TRIM28,M
XN10,SRSF3,SF3A1,MESD,RBM15B,KDM5C,ACTR1A,EP300,DGCR2,KDM5B,CSNK2A1,UBA52,SL
S1,RHOA,RAB6B,CXADR,ATP1A1,JUND,SRSF4,NISCH,CCNI,KHDRBS1,CSNK1E,SFPQ,PIP4K2B,SO
C9A7,ALDOA,MTRNR2L8,PKM,GANAB,COL6A1,FTL,HSP90B1,NEFM,PRPF8,RPL8,FBXL4,YBX1,U
CALM1,NPM1,PLXNB2,PLD3,IGF2,EIF3C,UBB,AFF4,TRIM28,CKB,COL4A2,SPEN,DDB1,PSMD5,YW
K3R2,NCL,RPL7A,ALDOA,PKM,GANAB,FTL,HSP90B1,FADS1,NEFL,NEFM,PRPF8,RPL8,YBX1,UBA1
M,GANAB,COL6A1,FTL,HSP90B1,NEFL,NEFM,PRPF8,RPL8,FBXL4,UBA1,ATP2B4,IGFBP2,CALR,T
A1B,TUBB2B,PGK1,RPLP0,H3-3B,PTMA,DDX5,ENO2,FKBP14,RPL13A,MAGED1,SEC62,RPL6,KAT
CALR,FSTL1,TMF1,ABHD2,HSPA1B,TUBB2B,COL5A1,COL4A1,CES3,PSAP,PGK1,RPLP0,H3-3B,C
,CALM1,NPM1,PLXNB2,HSPD1,USP11,SNRNP200,CSDE1,ATP5F1A,EIF4B,UBB,AFF4,CKB,NONO,
D1,RPL6,DVL3,YWHAE,GNAS,GPI,RACK1,FASN,ATP5F1B,CALM3,GNB1,SHISA9,RPS8,MYH9,RPL7
PL5,FSCN1,PTMS,SMG1,PLXNB2,HSPD1,USP11,COL18A1,IGF2,VCP,MARCKSL1,AFF4,TRIM28,C
RPS9,PAPOLA,CBX5,ZNF865,KIF5B,GLIPR1L2,LENG8,ATXN7L3B,VAT1,SRSF4,RTN3,PFN1,CCNI,T
2,MARCHF6,CDC42BPB,MAT2A,SLC4A8,PXDN,OGA,FADS2,RPS16,LAMA5,SLC16A7,KPNB1,C1OF
IGF2,MARCKSL1,TRIM28,COL4A2,SPEN,YWHAQ,TMSB10,RPL19,GTF3C4,ZNF43,VAPA,TPT1,SEC
D1,TUBA1B,PIK3R2,FLNA,MAP1B,NCL,RPL7A,SLC9A7,ALDOA,PKM,GANAB,COL6A1,FTL,HSP90B
HOA,RAB6B,ENC1,CXADR,EIF5,RPL26,CRMP1,OAZ1,SPTBN2,SCRN1,PEBP1,CDH2,DDX3X,RPS15
3,PGK1,H3-3B,CANX,PTMA,DDX5,FKBP14,RPL13A,MAGED1,RPL6,DVL3,YWHAE,GNAS,GPI,HTRA
ENO2,RPL13A,SEC62,KAT6A,YWHAE,KIF1A,GNAS,SLC2A3,DDX17,GPI,FASN,ALDH6A1,SPTBN1,C
COL6A1,FTL,HSP90B1,NEFL,NEFM,PRPF8,RPL8,FBXL4,UBA1,IGFBP2,FSTL1,TMF1,ABHD2,HSPA1
1,COL4A1,PSAP,PGK1,RPLP0,CANX,PTMA,DDX5,FKBP14,RPL13A,MAGED1,SEC62,KAT6A,KIF1A,
M199X,DLG5,CHD6,GDI2,TAF15,LMNB2,PDXK,TBC1D16,GRINA,ACAP3,ABCA2,PYGB,TERF2IP,TC
RL8,PKM,GANAB,COL6A1,FTL,HSP90B1,FADS1,NEFL,NEFM,RPL8,FBXL4,YBX1,UBA1,ATP2B4,IGF
PTMA,DDX5,FKBP14,RPL13A,MAGED1,RPL6,DVL3,GNAS,GPI,RACK1,ATP5F1B,CALM3,GNB1,SHIS
,RPL15,GTF3C4,DNAH8,SF3B1,ZNF43,HMGB1,POLR2A,TPT1,SEC61A1,RPS3A,TUBB4B,RPLP1,S
,TLE5,CCNI,EIF5A,H1-0,DNAJC5,APLP2,HNRNPDL,MYL6,ANKRD52,HNRNPK,INIP,MAGED4B,TOP2
C28,SPEN,EIF1,EIF4A1,PPP2R1A,RPL15,GTF3C4,SF3B1,ZNF43,HMGB1,POLR2A,TPT1,RPS3A,RP
EFM,PRPF8,UGT8,YBX1,CALR,FSTL1,TMF1,ABHD2,HSPA1B,TUBB2B,COL5A1,CES3,PGK1,RPLP0,
PF8,FBXL4,YBX1,UBA1,ATP2B4,CALR,TMF1,HSPA1B,TUBB2B,COL4A1,PGK1,H3-3B,CANX,CLU,F
,DVL3,GNAS,GPI,RACK1,ATP5F1B,CALM3,GNB1,SHISA9,RPS8,MYH9,SEPTIN2,P4HB,RPL5,GDI1,

L3,KIF1A,RPL13,GNAS,SLC2A3,DDX17,GPI,RACK1,FASN,ALDH6A1,SPTBN1,GNB1,SHISA9,RPS8,

5,CIPC,KPNB1,NCS1,RHOA,RPL9,ATP1A1,TMEM259,CD81,HMGN2,CRMP1,LASP1,VAT1,RTN3,PF

3,MAGED4B,CAPRIN1,STIP1,SLC35E2B,APPL1,PPIB,PEA15,TIMP3,NOVA2,CTSB,CHD6,U2AF2,EE

TBP1,ARHGDIA,IPO5,DLG5,CHD6,RNF187,ABCA3,RTN1,PDXK,CAMSAP1,GRINA,MDK,SKI,HDGF,

1,TMF1,ABHD2,HSPA1B,ZNF99,COL4A1,CES3,PGK1,RPLP0,CANX,PTMA,DDX5,ENO2,RPL13A,MA

CAPRIN1,HDAC2,MMP2,GLUL,NNAT,LUC7L3,FAM199X,CTDSP2,NOVA2,RPN2,HNRNPL,ALKBH5,

ADS1,NEFL,NEFM,PRPF8,RPL8,FBXL4,YBX1,ATP2B4,CALR,FSTL1,TMF1,HSPA1B,TUBB2B,COL4A

,PTMS,SPTAN1,PLD3,HSPD1,SNRNP200,EIF4B,UBB,AFF4,TRIM28,YWHAG,YWHAQ,EIF1,MAP9,C

PEN,DDB1,YWHAQ,SET,EGLN1,MAP9,CHSY1,TMSB10,EIF4A1,LDHB,PKD1,PDIA3,SEC61A1,STMN

F1,SLC4A8,HNRNPU,SMARCA5,FADS2,CPE,TPM4,TNS3,CTSD,LPCAT1,LAMA5,RPS24,SLC16A7,K

B4GALNT4,DNAJC5,APLP2,MYL6,ANKRD52,HNRNPK,RAB1A,PLPPR3,COTL1,TOP2B,TRIB2,CTD

ED1,SEC62,KAT6A,DVL3,YWHAE,KIF1A,DDX17,GPI,FASN,ALDH6A1,SPTBN1,GNB1,SHISA9,RPL7,

PCAT1,ZNF865,CLSTN1,JUND,EIF5,RTN3,PNN,RPL31,PFKL,TPM1,PEBP1,SOX12,RAB7A,CCN2,IT

PS6,ZNF316,DBN1,RPL4,ENO1,TUBA1B,PIK3R2,FLNA,MAP1B,NCL,RPL7A,SLC9A7,ALDOA,PKM,

PA1B,PGK1,H3-3B,CANX,CLU,PTMA,DDX5,FKBP14,RPL13A,MAGED1,RPL6,DVL3,YWHAE,GNAS,

DX5,ENO2,RPL13A,MAGED1,KAT6A,DVL3,GNAS,GPI,FASN,SPTBN1,ATP5F1B,GNB1,SHISA9,MYH

SEPTIN2,P4HB,RPL5,FSCN1,GSTP1,SMG1,CALM1,PLXNB2,PLD3,HSPD1,COL18A1,SNRNP200,IG

A1,MORF4L1,NCAM1,AGRN,ZMIZ1,TIMP2,COL11A1,MARCHF6,VCL,MAT2A,PXDN,HNRNPU,HNR

ATXN7L3B,PFN1,MCL1,PNN,PFKL,PDIA6,PIP4K2B,RAB7A,EIF3L,HNRNPDL,RPLP2,DDOST,YWHA

EIF4A1,PPP2R1A,RPL15,GTF3C4,SF3B1,ZNF43,HMGB1,POLR2A,TPT1,RPS3A,RPL18,ITGB1,TUB

ADS1,NEFL,NEFM,PRPF8,RPL8,FBXL4,YBX1,ATP2B4,CALR,FSTL1,TMF1,HSPA1B,TUBB2B,COL4A

AN1,CALM1,PLD3,USP11,SNRNP200,CSDE1,IGF2,EIF3C,ATP5F1A,EIF4B,MATR3,YWHAG,RPL37A

IO2,KAT6A,DVL3,YWHAE,RPL13,GNAS,DDX17,GPI,HTRA4,RACK1,BSG,FASN,SPTBN1,GNB1,RPS

IO2,KAT6A,DVL3,YWHAE,RPL13,GNAS,DDX17,GPI,HTRA4,RACK1,BSG,FASN,SPTBN1,GNB1,RPS

A1,SPTBN1,GNB1,STMN2,SEPTIN2,GDI1,TMBIM6,PTMS,SPTAN1,MARCKS,SMG1,CALM1,NPM1,

S8,P4HB,RPL5,TMBIM6,PLD3,HSPD1,USP11,SNRNP200,SMIM14,EIF3C,ATP5F1A,EIF4B,UBB,MA

,TRIM28,CKB,COL4A2,LRRC28,FN1,DDB1,HACD3,SET,TPI1,TMSB10,EIF4A1,MRC2,PPP2R1A,R

RPL13A,MAGED1,RPL6,DVL3,GNAS,GPI,RACK1,ATP5F1B,CALM3,GNB1,SHISA9,RPS8,MYH9,SEP

24,CIPC,C1ORF43,RHOA,ATF4,SLC7A5,CXADR,ATP1A1,JUND,LENG8,HMGN2,JPT2,RPL31,TKT,C

DDB1,YWHAQ,DCHS1,SET,TPI1,MRC2,PDIA3,SSBP2,BASP1,AP2M1,RPL18,STMN1,ITGB1,UCHL1,

ARF1,CYB5R3,ZNF770,GTF2F1,HNRNPU,NFE2L1,RPS16,TPM4,EIF4H,SCARB2,LAMA5,C1ORF43,

DR,RPS20,CD81,CCT7,SRSF4,RBBP4,PFN1,HNRNPC,RPL35,RPS10,ARL6IP1,WLS,HNRNPK,H1-10

,MYH9,RPL7,P4HB,PTMS,SPTAN1,CALM1,PLXNB2,USP11,CSDE1,SMIM14,ATP5F1A,EIF4B,MAR

RACK1,ATP5F1B,CALM3,SHISA9,RPS8,MYH9,P4HB,RPL5,TMBIM6,SPTAN1,PLD3,HSPD1,USP11,S

INA,EIF4A1,PKD1,PDIA3,SSBP2,BASP1,POLR2A,CDKL1,RPS4X,RPS3A,STMN1,TUBB4B,MORF4L1

RD3,KPNB1,PAPOLA,RPL14,TMEM259,EIF5,CD81,WDR1,HMGN2,RBBP4,PFN1,MCL1,JPT2,PNN,

HK1,RPL22,PTCD3,ZNF865,RPS14,COL5A2,LENG8,WDR1,ATXN7L3B,RPL26,CCT7,SRSF4,RBBP4

KBP14,RPL13A,MAGED1,RPL6,DVL3,YWHAE,GNAS,GPI,HTRA4,RACK1,FASN,ATP5F1B,CALM3,SH

4B,RPL11,RPLP1,SARAF,RPL10,AP3D1,SF3B2,GTF2F1,RPS16,RPL23,CNN3,RPL29,RPS9,XRCC5,

PL37A,LRRC28,EIF1,TMSB10,EIF4A1,RPL19,RPL15,GTF3C4,SF3B1,ZNF43,HMGB1,POLR2A,TPT

,INA,PPP2R1A,RPL15,GTF3C4,SF3B1,ZNF43,HMGB1,POLR2A,TPT1,SEC61A1,RPS4X,RPS3A,RPL

PL10,AP3D1,ARF1,HNRNPU,SMARCA5,LDHA,CIPC,KPNB1,RPL14,ATF4,RAB6B,SLC7A5,CXADR,AT

PL10,AP3D1,ARF1,HNRNPU,SMARCA5,LDHA,CIPC,KPNB1,RPL14,ATF4,RAB6B,SLC7A5,CXADR,AT

ANX,CLU,PTMA,DDX5,FKBP14,RPL13A,MAGED1,RPL6,DVL3,YWHAE,GNAS,GPI,RACK1,FASN,ATF

99,COL4A1,PGK1,RPLP0,CANX,PTMA,DDX5,ENO2,RPL13A,MAGED1,SEC62,RPL6,KAT6A,DVL3,Y

PM1,CHPF,COL6A2,B4GALNT4,APLP2,MXD4,CKAP5,SEPTIN11,COTL1,CTBP1,ARHGDIA,GPC1,C[illegible]
[illegible],EIF1,DCHS1,SET,TPI1,TMSB10,GTF3C4,SERINC1,CHD3,VAPA,AP2M1,TPT1,MORF4L1,ANXA2,T[illegible]
X,PTMA,DDX5,ENO2,MAGED1,SEC62,KAT6A,DVL3,YWHAE,KIF1A,DDX17,GPI,FASN,ALDH6A1,SPT[illegible]
[illegible]V0C,CPSF2,EID1,SRP9,SRSF5,U2AF2,VAMP2,PABPN1,HNRNPL,PARP1,HNRNPA3,PTPRA,ACTR3,[illegible]
SERINC1,HMGB1,POLR2A,AP2M1,TPT1,SEC61A1,RPS4X,ITGB1,TUBB4B,BTF3,AGRN,RPLP1,MID[illegible]
[illegible]M1,PLD3,HSPD1,USP11,COL18A1,SNRNP200,EIF3C,EIF4B,MARCKSL1,AFF4,TRIM28,CKB,FN1,Y[illegible]
[illegible]1,AFF4,YWHAG,CKB,FN1,NONO,YWHAQ,SET,TMSB10,GTF3C4,SSBP2,CHD3,BASP1,VAPA,POLR[illegible]
E,GPI,RACK1,FASN,ATP5F1B,CALM3,GNB1,SHISA9,RPS8,MYH9,P4HB,RPL5,TMBIM6,SPTAN1,PL[illegible]
GF2,EIF3C,ATP5F1A,EIF4B,RPL37A,LRRC28,EIF4A1,RPL19,PPP2R1A,RPL15,GTF3C4,DNAH8,SF[illegible]
[illegible]K1,ATP5F1B,CALM3,SHISA9,RPS8,MYH9,P4HB,RPL5,TMBIM6,SPTAN1,NPM1,PLD3,USP11,SNR[illegible]
[illegible]LBP,EIF5A,MEGF8,RPS5,DNAJC5,HNRNPDL,PTGES3,HNRNPH1,MAGED4B,CTBP1,ARHGDIA,CDK
[illegible]D1,IGF2,VCP,MARCKSL1,AFF4,TRIM28,CKB,SPEN,DDB1,YWHAQ,EIF1,SET,TPI1,TMSB10,EIF4A1
[illegible],RPS17,SF3B2,GTF2F1,HNRNPA0,RPL23,CNN3,RPL29,RPS9,XRCC5,CIPC,RPL24,PAPOLA,HK1,R[illegible]
[illegible]IM28,MATR3,YWHAG,COL4A2,YWHAQ,DCHS1,SET,INA,PPP2R1A,SSBP2,TPT1,RPL18,STMN1,IT[illegible]
ZNF99,TUBB2B,COL4A1,PSAP,PGK1,CANX,CLU,PTMA,DDX5,RPL13A,KAT6A,KIF1A,RPL13,GNAS,
,SF3B2,GTF2F1,HNRNPA0,RPS16,RPL23,RPL29,RPS9,XRCC5,CIPC,SLC16A7,PAPOLA,HK1,RHO[illegible]
1,CALM3,GNB1,MYH9,SEPTIN2,P4HB,APP,GDI1,FSCN1,SPTAN1,NPM1,PLD3,HSPD1,COL18A1,CS
FM,RPL8,FBXL4,YBX1,UBA1,IGFBP2,FSTL1,TMF1,ABHD2,COL4A1,PGK1,H3-3B,CANX,PTMA,DD[illegible]
GK1,RPLP0,CANX,CLU,PTMA,DDX5,SOX4,RPL13A,MAGED1,KAT6A,KIF1A,RPL13,GPI,HTRA4,RAC
[illegible]A4,FASN,SPTBN1,RPL7,RPL5,FSCN1,PTMS,SMG1,NPM1,PLXNB2,HSPD1,USP11,COL18A1,IGF2,[illegible]
[illegible]B,CANX,CLU,PTMA,DDX5,FKBP14,RPL13A,MAGED1,RPL6,KAT6A,DVL3,GNAS,DDX17,GPI,RACK[illegible]
[illegible],RPS8,P4HB,RPL5,FSCN1,TMBIM6,NPM1,PLXNB2,PLD3,HSPD1,USP11,SNRNP200,SMIM14,EIF3[illegible]
BB4B,RPL11,RPLP1,SARAF,RPL10,RPS17,AP3D1,SF3B2,GTF2F1,RPL23,RPL29,TPM4,RPS9,XRC[illegible]
GTF3C4,DNAH8,SF3B1,ZNF43,HMGB1,POLR2A,TPT1,SEC61A1,RPS3A,ITGB1,TUBB4B,RPL11,R[illegible]
COL4A2,YWHAQ,SET,EGLN1,RPL19,GTF3C4,CHD3,SEC61A1,RPS4X,MORF4L1,BTF3,AGRN,MAR
[illegible]VL3,KIF1A,GNAS,DDX17,GPI,FASN,ALDH6A1,CALM3,GNB1,RPS8,MYH9,P4HB,APP,FSCN1,MARC[illegible]
[illegible]1,SF3B2,GTF2F1,RPL23,RPL29,RPS9,XRCC5,CIPC,RPL24,PAPOLA,HK1,RHOA,ZNF865,GLIPR1L
[illegible]L8,FBXL4,YBX1,ATP2B4,IGFBP2,FSTL1,TMF1,ABHD2,HSPA1B,ZNF99,TUBB2B,COL5A1,COL4A1
,ALDH6A1,ATP5F1B,CALM3,STMN2,RPS8,SEPTIN2,GDI1,GSTP1,MARCKS,NPM1,TTC3,USP11,CC
[illegible]YGB,DSTN,ODC1,AEBP1,PTBP1,DYNLL2,ILF3,ACTN1,PLCG1,DIDO1,DUSP4,DLK1,CORO1C,LSS,
,VAPA,HMGB1,AP2M1,TUBB4B,AGRN,RPLP1,AP3D1,SBF1,ARF1,CYB5R3,CDC42BPB,MAT2A,TM[illegible]
,TMBIM6,NPM1,PLD3,HSPD1,USP11,SNRNP200,SMIM14,IGF2,EIF3C,ATP5F1A,EIF4B,VCP,UBB,A[illegible]
3B2,HNRNPA0,XRCC5,CIPC,HK1,RPL22,ZNF865,ATF4,PSMD2,KIF5B,CXADR,LENG8,RPL26,CCT[illegible]
MP1,OAZ1,TLE5,XPR1,PFKL,TKT,PNMA1,GPC3,DDR1,SOX12,ARL4C,MIF,GPAT4,MYO5A,B4GAL[illegible]
[illegible],FTL,HSP90B1,FADS1,NEFL,NEFM,PRPF8,RPL8,FBXL4,YBX1,UBA1,IGFBP2,CALR,FSTL1,ABHD2,
[illegible],HTRA4,FASN,ATP5F1B,CALM3,GNB1,SHISA9,MYH9,APP,IDS,RPL5,GDI1,FSCN1,PTMS,MARCKS,
3,GNAS,GPI,RACK1,ATP5F1B,CALM3,GNB1,SHISA9,RPS8,P4HB,RPL5,FSCN1,TMBIM6,NPM1,PL[illegible]
TPM4,LAMA5,PJA2,KPNB1,C1ORF43,CXADR,CHD4,ATP1A1,HMGN2,ATP2A2,CRMP1,SRSF4,PFN1
IPC,RPL24,PAPOLA,HK1,RHOA,ZNF865,GLIPR1L2,RPS14,RPS20,WDR1,RBBP4,PFN1,MCL1,TLE[illegible]
CCNI,KHDRBS1,HDLBP,EIF5A,MEGF8,ACTN4,RPS15,RPS5,MIF,DNAJC5,SLC25A3,PTGES3,HNRN
7,C1ORF43,ZNF865,RAB6B,CXADR,CHD4,JUND,WDR1,ATP2A2,CRMP1,VAT1,ASTN2,TLE5,CCNI,[illegible]
CH,PRKCSH,KLHL9,XPO1,HDLBP,GLG1,SFPQ,PNMA1,DDX3X,EIF3L,NANOG,MRFAP1,NGRN,RPL[illegible]
,AGRN,RPL10,MARCHF6,WSB1,AP3D1,DNAJA1,ARF1,MAT2A,SMARCA5,CTNNA1,RPS16,LDHA,CN
TPBP1,MGAT4B,KLHL42,PLXNB1,GLUD1,UPF1,RER1,ZNF512B,CEMIP2,CBX6,FMNL2,ANP32B,P
[illegible]NO,YWHAQ,EIF1,DCHS1,SET,CHSY1,TMSB10,PPP2R1A,SERINC1,SSBP2,TPT1,RPS4X,RPL18,ITG

T,MAP9,TMSB10,PPP2R1A,CHD3,VAPA,HMGB1,RPS4X,RPL18,ITGB1,ANXA2,SARAF,COL11A1,MID
GDI1,FSCN1,SPTAN1,MARCKS,PLD3,HSPD1,CSDE1,VCP,MARCKSL1,TRIM28,COL4A2,FN1,SPEN,I
S1,NEFL,NEFM,UGT8,RPL8,FBXL4,YBX1,UBA1,IGFBP2,CALR,FSTL1,ABHD2,HSPA1B,COL5A1,CO
MA,ENO2,DVL3,YWHAE,KIF1A,GNAS,SLC2A3,ALDH6A1,SPTBN1,GNB1,SHISA9,RPS8,APP,ERVV-1
WHAE,GNAS,GPI,RACK1,FASN,ATP5F1B,CALM3,SHISA9,RPS8,MYH9,P4HB,RPL5,TMBIM6,SPTAN
ZMIZ1,RPL10,VCL,DNAJA1,ARF1,HNRNPA0,FADS2,LDHA,CNN3,TNS3,EIF4H,TSPAN3,LPCAT1,LAM
PKD1,SF3B1,SSBP2,HMGB1,POLR2A,TPT1,ITGB1,RPL11,MORF4L1,MIDN,RPL10,MARCHF6,RPS
B1,SERINC1,ZNF43,HMGB1,POLR2A,TPT1,SEC61A1,RPS3A,RPL18,TUBB4B,RPL11,BTF3,RPLP1,
34,IGFBP2,CALR,FSTL1,HSPA1B,TUBB2B,COL5A1,COL4A1,PSAP,H3-3B,CANX,CLU,PTMA,DDX5,E
N2,JPT2,TLE5,CCNI,KHDRBS1,XPO1,ITM2C,ARL4C,CLTC,JUN,MYO5A,COL6A2,CD63,B4GALNT4
B,RPL11,RPLP1,RPL10,AP3D1,SF3B2,GTF2F1,RPL23,RPL29,RPS9,XRCC5,CIPC,RPL24,PAPOLA,I
SRSF4,PFN1,HNRNPC,KHDRBS1,CDH2,PDIA6,MMP14,ARL4C,MRFAP1,RPS19,NGRN,GOLM1,RPS
S,SPTAN1,MARCKS,NPM1,USP11,IGF2,MATR3,CKB,LRRC28,MAPT,FN1,SPEN,PSMD5,CHSY1,EIF4
,P4HB,KLRD1,FSCN1,GSTP1,PLXNB2,PLD3,SMIM14,ATP5F1A,EIF4B,UBB,AFF4,YWHAG,COL4A2,
CALM3,GNB1,RPS8,RPL7,P4HB,APP,FSCN1,PTMS,GSTP1,SMG1,CALM1,COL18A1,MARCKSL1,A
OL18A1,SNRNP200,IGF2,ATP5F1A,EIF4B,VCP,MARCKSL1,UBB,LRRC28,DDB1,YWHAQ,EIF1,MAP
2B,MIF,B4GALNT4,DNAJC5,APLP2,RPLP2,MXD4,DUX4,SEPTIN11,COTL1,CTBP1,ARHGDIA,TOP2E
3,DNAJC5,SLC25A3,RPLP2,NDFIP1,MXD4,SEPTIN11,CADM4,COTL1,CAPRIN1,STIP1,SLC35E2B,AT
3,RPL8,YBX1,UBA1,ATP2B4,TUBB2B,COL5A1,COL4A1,H3-3B,PTMA,DDX5,ENO2,RPL6,DVL3,YWH
N,GNB1,SHISA9,RPS8,P4HB,TMBIM6,CALM1,HSPD1,USP11,SNRNP200,CSDE1,EIF3C,ATP5F1A,AF
,NPM1,HSPD1,USP11,SNRNP200,IGF2,EIF4B,MARCKSL1,MATR3,CKB,LRRC28,MAPT,FN1,SPEN,
YWHAE,KIF1A,GNAS,SLC2A3,GPI,HTRA4,SPTBN1,CALM3,GNB1,SHISA9,SEPTIN2,P4HB,PTMS,GS
B,CTBP1,STIP1,GPC1,SLC35E2B,SMCHD1,IPO5,ATP6V0C,DNER,TRIB2,SQLE,TIMP3,CHD6,EEF1A
F1A,UBB,AFF4,RPL37A,SPEN,NONO,INA,RPL19,PPP2R1A,RPL15,GTF3C4,SF3B1,SERINC1,RPS3,
8,HACD3,DCHS1,TPI1,EIF4A1,MRC2,PPP2R1A,SSBP2,ZNF43,POLR2A,SEC61A1,RPL18,ITGB1,UC
RPL5,FSCN1,PLXNB2,PLD3,HSPD1,USP11,COL18A1,SNRNP200,CSDE1,SMIM14,EIF3C,AFF4,TRIM
APLP1,MIF,MYO5A,B4GALNT4,MTSS2,TARDBP,MXD4,INIP,CADM4,HNRNPH1,HLA-B,CTBP1,ARHG
4B,MORF4L1,BTF3,XAF1,WSB1,RPS17,AP3D1,SBF1,SF3B2,CDC42BPB,MAT2A,RPSA,TPM4,NACA
,MARCKSL1,UBB,RPL37A,LRRC28,MAPT,EIF1,HACD3,SET,MAP9,RPL19,GTF3C4,DNAH8,SERINC
KM,GANAB,FTL,HSP90B1,FADS1,NEFM,PRPF8,UGT8,RPL8,FBXL4,YBX1,UBA1,CALR,TMF1,ABHD
3,COL4A2,RPL37A,FN1,EIF1,DCHS1,TMSB10,INA,PPP2R1A,GTF3C4,BASP1,POLR2A,TPT1,NCAM1
PI,BSG,FASN,ALDH6A1,CALM3,GNB1,SEPTIN2,P4HB,RPL5,FSCN1,GSTP1,MARCKS,NPM1,TTC3,PI
NAS,GPI,RACK1,ATP5F1B,CALM3,GNB1,SHISA9,MYH9,P4HB,RPL5,FSCN1,TMBIM6,SPTAN1,NPM1
,XRCC5,CIPC,RPL24,PAPOLA,RPL22,RHOA,ZNF865,GLIPR1L2,RPS14,RPS20,CD81,LENG8,WDR1
F,RPL10,AP3D1,SF3B2,GTF2F1,RPS16,RPL23,CNN3,RPL29,RPS9,XRCC5,CIPC,RPL24,PAPOLA,HK
GRN,ZMIZ1,TIMP2,CDC42BPB,OGA,RPL23,CPE,TNS3,SLC22A17,PJA2,SLC16A7,KPNB1,C1ORF4
A1,CSDE1,ATP5F1A,EIF4B,UBB,MATR3,YWHAG,NONO,DDB1,PSMD5,SET,TMSB10,EIF4A1,PPP2R1
UBE4A,VPS35,ACAP3,DPYSL2,TOP1,PPP2CA,SMARCA4,RABL6,TCF25,LUZP6,GRB2,SCAF11,CLIF
MATR3,COL4A2,DDB1,SET,TMSB10,INA,RPL19,SF3B1,SSBP2,CHD3,HMGB1,STMN1,ITGB1,TUBE
OL4A1,CES3,PSAP,PGK1,RPLP0,H3-3B,CANX,CLU,PTMA,DDX5,SOX4,RPL13A,SEC62,RPL6,KAT6A
,MYH9,P4HB,RPL5,GSTP1,NPM1,TTC3,PLD3,HSPD1,COL18A1,IGF2,EIF3C,EIF4B,UBB,AFF4,TRIM
PD1,USP11,SNRNP200,CSDE1,SMIM14,EIF3C,EIF4B,UBB,AFF4,MATR3,RPL37A,NONO,DDB1,EIF1
4B,BTF3,AGRN,RPLP1,RPL10,MARCHF6,RPS17,DNAJA1,ARF1,SF3B2,GTF2F1,RPL23,RPL29,XRC
4,SEPTIN11,COTL1,CTBP1,ARHGDIA,TOP2B,SLC35E2B,TRIB2,GLUL,HS6ST1,CTDSP2,CNN2,NOV
,RPL15,GTF3C4,SERINC1,HMGB1,POLR2A,AP2M1,TPT1,SEC61A1,ITGB1,TUBB4B,RPL11,MORF

RPL24,KPNB1,RPL14,CBX5,RPS14,CXADR,RPS20,CD81,WDR1,HMGN2,RBBP4,MCL1,PNN,HNRN
RPL13A,MAGED1,RPL6,DVL3,GNAS,GPI,RACK1,FASN,ATP5F1B,CALM3,GNB1,SHISA9,RPS8,P4HB
5PB,PTBP1,GRB2,CDC37,WDR6,RPS11,SUPT5H,CAND1,DIDO1,DUSP4,NKTR,HNRNPD,NAP1L4,Y
COL18A1,SMIM14,EIF4B,MARCKSL1,AFF4,YWHAG,CKB,LRRC28,FN1,EIF1,HACD3,SET,CHSY1,T
RPL23,RPL29,EIF4H,RPS9,XRCC5,CIPC,PAPOLA,RPL22,RHOA,ZNF865,GLIPR1L2,RPS14,RPS20,C
,CNN3,RPL29,RPS9,CIPC,RPL24,HK1,RHOA,ZNF865,RPS14,RPS20,WDR1,RBBP4,PFN1,ASTN2,M
5A1,CES3,PSAP,PGK1,H3-3B,CANX,PTMA,DDX5,RPL13A,SEC62,RPL6,KAT6A,DVL3,KIF1A,DDX17
,BEX3,CCT7,ASTN2,MCL1,TLE5,PFKL,PDIA6,MEGF8,SOX12,ACTN4,MMP14,PRDX2,RPL12,ITM2
1,SF3B2,GTF2F1,HNRNPA0,RPL23,RPL29,RPS9,RPL24,SLC16A7,HK1,RHOA,ZNF865,RPS14,RPS
EIF3C,MARCKSL1,UBB,MATR3,YWHAG,CKB,MAPT,SPEN,DDB1,YWHAQ,DCHS1,SET,CHSY1,RPL1
M,PRPF8,UGT8,UBA1,ATP2B4,IGFBP2,CALR,TMF1,ABHD2,HSPA1B,TUBB2B,COL5A1,CES3,PGK
M4,RAB1A,ARCN1,SEPTIN3,MAGED4B,MAP1A,FAM199X,SHISAL1,TMEM132A,CTSB,KLHL24,CAN
NPDL,RPLP2,ARL6IP1,MXD4,SEPTIN11,RHOB,LGALS3BP,COTL1,ARHGDIA,STIP1,SMCHD1,IPO5,
DS1,NEFM,PRPF8,UGT8,YBX1,UBA1,ATP2B4,IGFBP2,CALR,TMF1,HSPA1B,TUBB2B,COL5A1,CO
L3,GNAS,GPI,RACK1,BSG,ATP5F1B,CALM3,SHISA9,MYH9,SEPTIN2,P4HB,RPL5,TMBIM6,SPTAN1
RCHF6,CYB5R3,CDC42BPB,TMEM170A,SLC4A8,GOLGA8A,PXDN,LDHA,CPE,TPM4,RPS9,RPL24,
PSA,EIF4H,RPS24,RPL24,RPL14,CBX5,ATF4,CXADR,TMEM259,EIF5,HMGN2,NISCH,PRKCSH,XP
P11,COL18A1,SMIM14,IGF2,VCP,MARCKSL1,COL4A2,DDB1,DCHS1,SET,EGLN1,TPI1,CHSY1,EI
F2,EIF3C,EIF4B,VCP,MARCKSL1,TRIM28,COL4A2,DDB1,PSMD5,HACD3,SET,CHSY1,TMSB10,GT
BIM6,MARCKS,SMG1,PLD3,HSPD1,USP11,SNRNP200,CSDE1,SMIM14,EIF3C,EIF4B,UBB,AFF4,M
,PTMA,DDX5,FKBP14,RPL13A,MAGED1,RPL6,KAT6A,DVL3,RPL13,GPI,HTRA4,RACK1,FASN,ATP5
,GANAB,FTL,HSP90B1,FADS1,NEFL,NEFM,RPL8,YBX1,UBA1,ATP2B4,TMF1,ABHD2,PSAP,PGK1,H
1,ATP5F1B,STMN2,RPS8,SEPTIN2,GDI1,MARCKS,NPM1,TTC3,USP11,COL18A1,CSDE1,EIF3C,AT
H2,EIF3L,CHPF,NGRN,RPLP2,DDOST,NARS1,YWHAB,RPS12,CTBP1,ARHGDIA,SLC35E2B,IPO5,A
,RACK1,FASN,ALDH6A1,SPTBN1,ATP5F1B,GNB1,SHISA9,RPS8,P4HB,APP,ERVV-1,GDI1,FSCN1,S
3N1,RPL4,ENO1,TUBA1B,PIK3R2,FLNA,MAP1B,NCL,RPL7A,SLC9A7,ALDOA,MTRNR2L8,PKM,GA
AP3D1,CYB5R3,GTF2F1,PXDN,HNRNPA0,RPL23,LDHA,RPSA,TPM4,TNS3,LAMA5,RPS24,RPL24,K
M14,EIF3C,ATP5F1A,EIF4B,MATR3,RPL37A,LRRC28,EIF1,RPL19,PPP2R1A,PKD1,SF3B1,SERINC1
ST,HNRNPL,ABCA3,ALKBH5,SNRPN,VPS35,ACAP3,CBX1,DPYSL2,TOP1,VHL,PPP2CA,SMARCA4,R
TGB1,MORF4L1,ANXA2,SARAF,MIDN,VCL,AP3D1,ARF1,MAP4,TMEM170A,HNRNPU,CTNNA1,CNN
PLP2,DDOST,NARS1,YWHAB,PTGES3,ATP1A2,RPS12,ARCN1,RPS27,CTBP1,SLC35E2B,ATP6V0C,
SKI,H3-3A,SCAF1,RAB11B,TUBB2A,DDAH2,TCF25,CDC37,CASC3,PLCG1,KDELR1,ZFAND5,FLNB
65,GLIPR1L2,RPS14,RPS20,CD81,LENG8,WDR1,PFN1,MCL1,TLE5,PNN,RPL31,KHDRBS1,EIF5A,
ES3,CKAP5,RHOB,HNRNPH1,CTBP1,BCLAF1,EID1,MDH2,MEST,HNRNPL,ABCA3,ALKBH5,PKN1,Y
,VCP,MARCKSL1,COL4A2,DDB1,DCHS1,SET,EGLN1,TPI1,CHSY1,EIF4A1,RPL19,DNAH8,PKD1,SS
,UBAP2L,MBTPS1,CUL3,RPS2,SEPTIN7,APEX1,PPP1R15A,TAOK1,CLPTM1,EIF3H,UBE2D2,SMG5,
BBP4,MCL1,PNN,RPL31,HNRNPC,HDLBP,EIF5A,CCND2,SOX12,ACTN4,RPS15,ITM2C,ARL4C,RPL
C62,DDX17,GPI,BSG,FASN,ALDH6A1,SPTBN1,CALM3,GNB1,SHISA9,STMN2,RPS8,MYH9,SEPTIN
,IGFBP3,RHOA,ZNF865,ATF4,GLIPR1L2,CXADR,CHD4,RPS20,LENG8,WDR1,LASP1,PFN1,TLE5,P
F3L,GNB2,APLP1,RPS7,RPS19,GPAT4,TLN1,COL6A2,CD63,GOLM1,MTSS2,RPLP2,RPL36,PTGE
MARCKS,SMG1,NPM1,USP11,CSDE1,SMIM14,ATP5F1A,UBB,YWHAG,SPEN,EIF1,DCHS1,TPI1,INA
TMBIM6,PLD3,USP11,SNRNP200,SMIM14,EIF3C,ATP5F1A,EIF4B,VCP,UBB,AFF4,MATR3,LRRC28
,SOX12,ACTN4,RPS15,ITM2C,ARL4C,RPL35,NGRN,MTSS2,APLP2,SLC25A3,RPLP2,WLS,H1-10,R
N10,DIDO1,GIGYF1,SF3A1,RBM15B,OS9,KDM5C,EP300,DGCR2,STARD7,SEC16A,SLC39A7,CD16
7A5,ATP1A1,TMEM259,ATP2A2,RPL26,LASP1,NISCH,HNRNPC,CCNI,TADA2B,PFKL,CDH6,SFPQ,P

3TF3,AGRN,RPLP1,MIDN,RPL10,MARCHF6,DNAJA1,ARF1,CYB5R3,SF3B2,GTF2F1,RPL23,RPL29
AG,COL4A2,DCHS1,SET,EGLN1,TPI1,TMSB10,MRC2,LDHB,PDIA3,SERINC1,RPL18,STMN1,TUBB4
IK1,RAB6B,SLC7A5,TMEM259,EIF5,CD81,OAZ1,TLE5,PNN,PFKL,HDLBP,SFPQ,PNMA1,PIP4K2B,M
,MSL1,STMN3,ZFAND3,HECTD1,VANGL2,PFN2,HNRNPD,CTXN1,PPP1R9B,ADAR,KCNQ2,WDR45E
HSPA4,AHCYL1,CAMSAP1,GRINA,VPS35,MLEC,ACAP3,MDK,H3-3A,VEGFA,CCND1,BTBD2,TCF25,
D,LAMA5,BRD3,RPS9,PJA2,SLC16A7,KPNB1,RPL17,COL1A2,HK1,NCS1,ZNF865,CLSTN1,NR2F1,
BP14,KAT6A,DVL3,YWHAE,KIF1A,GNAS,DDX17,RACK1,FASN,SPTBN1,CALM3,GNB1,SHISA9,RPS8
PP1CB,GAS7,TSR1,TAF15,RTN1,LAMC1,SKI,PYGB,SMARCA4,SRSF9,PDE7A,RABL6,CAPZB,SON,C
RPS8,RPL7,SEPTIN2,GDI1,MARCKS,NPM1,TTC3,USP11,COL18A1,CSDE1,EIF3C,ATP5F1A,EIF4B,
ORF43,IGFBP3,GAP43,HK1,ZNF865,CLSTN1,SLC7A5,ENC1,CHD4,EIF5,CD81,VAT1,OAZ1,RBBP4,
A,RHOA,ZNF865,GLIPR1L2,RPS14,RPS20,CD81,LENG8,WDR1,RBBP4,PFN1,ASTN2,MCL1,TLE5,P
3S1,KLHL9,CSNK1E,HDLBP,TKT,TPM1,PEBP1,SOX12,RCN1,IK,HMGCS1,ARL6IP1,NARS1,RPL36,P
PNB1,HK1,ZNF865,NR2F1,RAB6B,ENC1,NLGN2,GLIPR1L2,TLE5,HNRNPC,CCNI,TADA2B,PFKL,R
2,ARHGDIA,SLC35E2B,RPL23A,HSBP1,MAP1A,TM9SF3,MDH2,U2AF2,EEF1A2,GNAI3,GDI2,TSR1,M
CDKL1,STMN1,TUBB4B,UCHL1,MORF4L1,RPLP1,MIDN,RPL10,MARCHF6,WSB1,AKAP12,AP2B1,M
3A,MAGED1,SEC62,KIF1A,RPL13,DDX17,GPI,HTRA4,BSG,ALDH6A1,ATP5F1B,GNB1,RPS8,SEPTI
RF2,DDX39B,SRSF5,TM9SF3,MDH2,RAB27A,U2AF2,CCT2,GNAI3,GAS7,MORF4L2,MTRF1L,ALKB
,NPM1,TTC3,USP11,COL18A1,CSDE1,EIF3C,ATP5F1A,EIF4B,UBB,MATR3,NONO,PSMD5,YWHAQ,
T,EGLN1,MAP9,TPI1,PPP2R1A,GTF3C4,HLA-C,PDIA3,BASP1,VAPA,SEC61A1,ITGB1,TUBB4B,RPL1
F2F1,RPL23,RPL29,XRCC5,CIPC,RPL24,PAPOLA,RHOA,ZNF865,ATF4,GLIPR1L2,RPS14,CXADR,L
4HB,APP,IDS,RPL5,FSCN1,MARCKS,SMG1,NPM1,PLXNB2,PLD3,HSPD1,COL18A1,UBB,AFF4,TRI
,IDS,GDI1,MARCKS,NPM1,TTC3,USP11,COL18A1,CSDE1,EIF3C,ATP5F1A,EIF4B,UBB,MATR3,NO
4,KAT6A,DVL3,KIF1A,RPL13,GNAS,SLC2A3,DDX17,GPI,HTRA4,ALDH6A1,SPTBN1,CALM3,SHISA9
P3,CBX1,RAD23B,SNRNP70,PYGB,VEGFA,VHL,SMARCA4,ARPC2,PLEC,SRSF1,ARGLU1,PTBP1,IL
P70,ATP6AP1,NOTCH2,RNF10,SELENON,MVB12B,FAM171A2,RABL6,DYNLL2,WDR6,NPEPPS,FKE
N1,CALM1,USP11,SNRNP200,CSDE1,IGF2,EIF3C,ATP5F1A,EIF4B,MATR3,RPL37A,LRRC28,RPL19
2,TBCD,YEATS2,KLC1,GRINA,ACAP3,LAMC1,SELENON,BTBD2,PBX2,ACTN1,BNIP3L,FKBP8,DIDO
EIF3C,UBB,AFF4,CKB,FN1,SPEN,PSMD5,EIF1,HACD3,DCHS1,MAP9,RPL19,GTF3C4,PKD1,SERIN
TBCD,VAMP2,GAS7,YEATS2,HSPA4,GRINA,PTOV1,ACAP3,MDK,SKI,H3-3A,PYGB,SELENON,DDAH2
F40A,PCYOX1,PPP1R9B,SULF2,MMP15,CD164,SRSF6,MBTPS1,KCTD12,RGMB,CBX6,NPDC1,HI
,RABL6,TCF25,PTBP1,LUZP6,GRB2,SCAF11,CLIP3,SDF4,HNRNPM,SDHA,LONP1,ZFAND3,CCT5,F
G,MRFAP1,CTNNB1,MACF1,RPL35A,TSR2,TTR,ARCN1,RPS27,ATP6V0C,L1CAM,LUC7L3,CNN2,S
COL4A1,PSAP,PGK1,RPLP0,CANX,CLU,PTMA,DDX5,RPL13A,RPL6,KAT6A,KIF1A,RPL13,GPI,HTRA
3,PKM,FTL,HSP90B1,NEFL,NEFM,PRPF8,RPL8,FBXL4,YBX1,IGFBP2,CALR,FSTL1,HSPA1B,TUBB2
,DDX3X,GNB2,MIF,GPAT4,CHPF,COL6A2,CD63,RPLP2,DDOST,MXD4,H1-10,CIRBP,RPL36,SEPTI
PL22,ZNF865,ATF4,RPS14,CXADR,LENG8,WDR1,RPL26,CCT7,SRSF4,RBBP4,PFN1,MCL1,RPL31
BB,MATR3,NONO,PSMD5,SET,CHSY1,TMSB10,EIF4A1,PPP2R1A,RPL15,DNAH8,PKD1,SF3B1,ZNF
5,SOX12,ITM2C,TAGLN,ANKRD52,WLS,PTGES3,UBE2H,PLPPR3,MAGED4B,CDK16,IPO5,ATP6V0
N,MTSS2,APLP2,SLC25A3,RPLP2,WLS,RAC1,SMCHD1,NNAT,CPSF2,SRP9,U2AF2,TBCD,ANXA5,C
L7A,SLC9A7,ALDOA,PKM,GANAB,COL6A1,FTL,HSP90B1,FADS1,NEFL,NEFM,PRPF8,UGT8,RPL8,
DH6A1,SHISA9,SEPTIN2,P4HB,APP,RPL5,FSCN1,GSTP1,MARCKS,SMG1,PLD3,HSPD1,USP11,COL
,CHD6,EEF1A2,CCT2,TBCD,GAS7,MORF4L2,PKN1,GRINA,PTOV1,MLEC,MDK,PTK7,SKI,ASXL1,PY
RPS14,RPS20,LENG8,WDR1,SRSF4,PFN1,MCL1,TLE5,PNN,RPL31,HNRNPC,KHDRBS1,EIF5A,PDI
XNB2,TTC3,IGF2,EIF3C,VCP,MARCKSL1,AFF4,TRIM28,CKB,MAPT,DDB1,PSMD5,EIF1,HACD3,DC
RPL36,DUX4,COTL1,ATP6V0C,GLUL,SQLE,FAM199X,TIMP3,CTSB,ZC3H7B,RNF187,TBCD,PGD,M

'S3A,BTF3,AGRN,RPLP1,MARCHF6,DNAJA1,ARF1,SF3B2,MAT2A,HNRNPA0,RPS16,SLC16A7,ZNF8
TSD,SCARB2,SLC22A17,C1ORF43,CBX5,CHD4,JUND,TMEM259,BEX3,MCL1,PNN,CCNI,HDLBP,EI
E,KIF1A,RPL13,GPI,ALDH6A1,SPTBN1,GNB1,SHISA9,RPL7,P4HB,IDS,TMBIM6,SPTAN1,SMG1,CA
L5,SPTAN1,USP11,SNRNP200,CSDE1,IGF2,EIF3C,ATP5F1A,EIF4B,MATR3,YWHAG,LRRC28,EIF1,
F3,MDH2,U2AF2,EEF1A2,TSR1,PRDX5,UBE4A,GCN1,SKI,RAB11B,PRPF6,PTBP1,CAPZB,SUPT5H,
IATR3,SPEN,HACD3,EGLN1,CHSY1,TMSB10,INA,MRC2,PPP2R1A,RPL15,PKD1,SF3B1,CHD3,BASF
,NPM1,IGF2,MARCKSL1,COL4A2,RPL37A,YWHAQ,EIF1,TMSB10,LDHB,DNAH8,PKD1,AP2M1,MO
',SRSF4,PFN1,MCL1,RPL31,PRKCSH,KHDRBS1,SCRN1,RPS25,SFPQ,EIF5A,PDIA6,MEGF8,ITM2C
'L26,MCL1,PNN,PRKCSH,KHDRBS1,RPS25,EIF5A,PDIA6,ITM2C,RPL30,EIF3L,MAGED2,RPS19,NC
TP1A1,CRMP1,LASP1,RTN3,ASTN2,KHDRBS1,XPO1,CSNK1E,RPS25,TPM1,GPC3,MEGF8,ITM2C,
),RPS9,XRCC5,CIPC,RPL24,PAPOLA,RPL22,RHOA,ZNF865,GLIPR1L2,RPS14,RPS20,CD81,LENG8
13A,MAGED1,RPL6,KAT6A,YWHAE,KIF1A,GPI,HTRA4,RACK1,FASN,CALM3,SHISA9,RPS8,P4HB,R
13A,MAGED1,RPL6,KAT6A,YWHAE,KIF1A,GPI,HTRA4,RACK1,FASN,CALM3,SHISA9,RPS8,P4HB,R
AP,PGK1,RPLP0,CANX,PTMA,DDX5,RPL13A,RPL6,KAT6A,KIF1A,RPL13,GPI,HTRA4,BSG,FASN,ALD
'4H,XRCC5,PAPOLA,RHOA,ZNF865,ATF4,RPS14,CXADR,LENG8,WDR1,LASP1,SRSF4,PFN1,PNN,R
M1,PLD3,HSPD1,USP11,IGF2,EIF3C,MARCKSL1,UBB,TRIM28,COL4A2,RPL37A,DDB1,EIF1,SET,T
SEC61A1,RPL18,MORF4L1,BTF3,AGRN,RPLP1,RPL10,MARCHF6,DNAJA1,ARF1,SF3B2,TPM4,PJA
1,CALM1,NPM1,HSPD1,SNRNP200,MARCKSL1,UBB,MATR3,YWHAG,LRRC28,YWHAQ,EIF1,TPI1,T
2,RPL23A,DDX39B,TM9SF3,MDH2,TOMM20,U2AF2,CCT2,GNAI3,GDI2,TSR1,SSRP1,HSPH1,MTR
',NGRN,HNRNPDL,RPLP2,DDOST,SND1,RPS12,SLC35E2B,RPL23A,HSBP1,DDX39B,SRSF5,TM9S
A1,ARF1,SF3B2,GTF2F1,RPL23,RPL29,XRCC5,CIPC,RPL22,ZNF865,ATF4,RPS14,CXADR,LENG8,
4C,CHPF,JUN,CD63,NGRN,APLP2,HNRNPDL,MXD4,MYL6,H1-10,RPL36,SEPTIN11,CADM4,RAB1A
TOMM20,U2AF2,SUGP2,CPSF1,MEST,HNRNPL,TM9SF2,ABCA3,SNRPN,UBE4A,VPS35,SNRNP70,
RNR2L8,PKM,FTL,NEFL,NEFM,UGT8,RPL8,FBXL4,YBX1,UBA1,ATP2B4,CALR,TMF1,ABHD2,HSPA
C,PAPOLA,ZNF865,ATF4,RPS14,CXADR,LENG8,WDR1,RPL26,SRSF4,PFN1,RPL31,PRKCSH,SCRN
R45B,PBX1,AP2A1,LRRC4B,CST3,CXXC5,NXN,GLUD1,ZNF512B,ULK1,GPSM1,GSE1,BAP1,PALM,
NPM1,TTC3,USP11,COL18A1,CSDE1,EIF3C,ATP5F1A,EIF4B,UBB,MATR3,NONO,PSMD5,SET,CHS
PPP2R1A,SF3B1,PDIA3,SERINC1,HMGB1,POLR2A,AP2M1,TPT1,RPL18,ITGB1,TUBB4B,RPL11,BT
D4B,MMP2,LAPTM4B,TRIB2,RPL23A,CNN2,NOVA2,CDH11,MDH2,CHD6,MORF4L2,SSRP1,ABCA
GNB1,SHISA9,SEPTIN2,IDS,GDI1,FSCN1,PTMS,SPTAN1,MARCKS,CALM1,PLXNB2,SNRNP200,SM
OLGA8A,CTNNA1,TPM4,SCARB2,RPS24,CIPC,IGFBP3,GAP43,HK1,RPL22,ZNF865,RAB6B,CHD4,
L13,GNAS,DDX17,HTRA4,RACK1,FASN,SPTBN1,GNB1,MYH9,RPL7,SEPTIN2,IDS,ERVV-1,GDI1,T
L11,RPLP1,SARAF,MIDN,MARCHF6,WSB1,ARF1,AP2B1,GTF2F1,GOLGA8A,SMARCA5,FADS2,LD
IT4,DNAJC5,HNRNPDL,TSR2,HNRNPK,H1-10,DUX4,RHOB,HNRNPH1,CCT6A,IPO5,ATP6V0C,PEA1
'865,ATF4,RPS14,CXADR,LENG8,WDR1,LASP1,SRSF4,PFN1,PNN,PRKCSH,EIF5A,PDIA6,PIP4K2B,
IBA1B,PIK3R2,MAP1B,NCL,RPL7A,SLC9A7,ALDOA,MTRNR2L8,PKM,GANAB,COL6A1,HSP90B1,N
OA,ZNF865,GLIPR1L2,RPS14,RPS20,CD81,LENG8,WDR1,RBBP4,PFN1,ASTN2,MCL1,TLE5,PNN,K
S8,MYH9,P4HB,TMBIM6,HSPD1,USP11,SNRNP200,CSDE1,SMIM14,EIF3C,ATP5F1A,RPL37A,LRR
ATG2B,RABL6,AEBP1,NYNRIN,CAPZB,NKTR,LSS,TUG1,CSNK1D,ID4,STRAP,SREBF2,PFN2,HNRN
ATG2B,RABL6,AEBP1,NYNRIN,CAPZB,NKTR,LSS,TUG1,CSNK1D,ID4,STRAP,SREBF2,PFN2,HNRN
GNAI3,NOP53,TSR1,MTRF1L,RPL27,PRDX5,UBE4A,TCP1,GCN1,SRP14,PRPF6,TAF7,RPL37,ARPC
EX4,KIF5A,SGTA,MTA1,RERE,PBX1,H2AZ1,MMP15,GSPT1,CST3,GNG4,FARP1,RER1,ZNF512B,CE
'S20,CD81,LENG8,WDR1,LASP1,RBBP4,PFN1,MCL1,TLE5,RPL31,EIF5A,RAB7A,RPL30,EIF3L,IK,
OAZ1,NES,TADA2B,KHDRBS1,PFKL,GLG1,PEBP1,PNMA1,PIP4K2B,ITM2C,MAGED2,APLP1,SESN
PZB,UBE2R2,STMN3,STXBP1,STRAP,P3H3,PPP1R9B,STAT3,ARIH1,KCNQ2,BCL9L,NOL4L,LRRC4

3B1,TUBB4B,SARAF,COL11A1,MIDN,RPL10,VCL,AKAP12,SF3B2,CDC42BPB,ITM2B,TMEM170A,S
IRNP200,SMIM14,EIF3C,VCP,MARCKSL1,AFF4,CKB,RPL37A,MAPT,SPEN,DDB1,PSMD5,YWHAQ,T
,MSN,SLC16A7,KPNB1,PAPOLA,LLGL1,CXADR,CHD4,ATP1A1,EIF5,CRMP1,CCT7,OAZ1,NISCH,PI
,RPS20,WDR1,LASP1,PFN1,ASTN2,MCL1,TLE5,PNN,KHDRBS1,PFKL,RPS25,EIF5A,PDIA6,MEGF8,
,HMGN2,CRMP1,BEX3,MCL1,TLE5,SPC25,PFKL,GLG1,CCND2,PIP4K2B,SOX12,CCN2,DDX3X,RPI
R1,HDLBP,PEBP1,CCND2,RPL12,CLTC,GNB2,PRKAR1A,CHPF,NGRN,HNRNPDL,RPLP2,MYL6,ANK
AB,SLC35E2B,TM9SF3,MDH2,U2AF2,EEF1A2,FDFT1,SUGP2,GNAI3,TSR1,PRDX5,UBE4A,HSPA4,M
10,MARCHF6,AP3D1,DNAJA1,ARF1,SF3B2,GTF2F1,RPS9,XRCC5,CIPC,RHOA,ZNF865,ATF4,RPS1
0,DNAJA1,SF3B2,TMEM170A,GTF2F1,OGA,PGAM1,NFE2L1,SMARCA5,FADS2,BRD3,XRCC5,PJA2
HPF,NGRN,DNAJC5,RPLP2,DDOST,CIRBP,YWHAB,TTR,ARCN1,RPS27,CTBP1,ARHGDIA,SLC35E2I
HPF,NGRN,DNAJC5,RPLP2,DDOST,CIRBP,YWHAB,TTR,ARCN1,RPS27,CTBP1,ARHGDIA,SLC35E2I
AB6B,NLGN2,CD81,LENG8,CRMP1,BEX3,TLE5,PFKL,CDH6,SFPQ,TPM1,CCND2,CDH2,PDIA6,SOX
,RBBP4,PFN1,MCL1,PNN,RPL31,PRKCSH,KHDRBS1,RPS25,EIF5A,ACSL6,MMP14,RPL12,ITM2C,I
RPL9,NLGN2,GLIPR1L2,ATP1A1,HMGN2,RPL31,NISCH,SLC3A2,PFKL,TPM1,CDH2,PNMA1,PDIA6
,NPM1,PLXNB2,HSPD1,SNRNP200,CSDE1,IGF2,EIF4B,MARCKSL1,AFF4,TRIM28,CKB,COL4A2,M
A,CPE,SLC22A17,RPS9,NCS1,ATP1A1,JUND,TMEM259,LENG8,ASTN2,MCL1,CCNI,XPR1,HDLBP,PI
FASN,ALDH6A1,ATP5F1B,CALM3,SHISA9,RPS8,MYH9,RPL7,SEPTIN2,P4HB,GSTP1,HSPD1,SNRNI
2,SOX4,RPL13A,MAGED1,KAT6A,DVL3,YWHAE,RPL13,DDX17,RACK1,ALDH6A1,SPTBN1,ATP5F1E
FN1,NES,TADA2B,KHDRBS1,PFKL,GLG1,PEBP1,PNMA1,PIP4K2B,ITM2C,MAGED2,APLP1,SESN3,
HDLBP,PEBP1,CCN2,GNB2,RPS7,JUN,NDFIP1,MXD4,CKAP5,RHOB,HNRNPH1,SEPTIN3,HMGCR,A
,RAB7A,EIF3L,RPS19,NGRN,RPLP2,DDOST,YWHAB,SND1,ARCN1,RPS27,ARHGDIA,SLC35E2B,HS
I1,COL18A1,CSDE1,EIF3C,ATP5F1A,EIF4B,UBB,MATR3,NONO,PSMD5,YWHAQ,SET,CHSY1,TMSB
,PNMA1,PIP4K2B,ITM2C,APLP1,SESN3,APLP2,INIP,MYH10,DUX4,CKAP5,SEPTIN11,RAC1,BCLAF1
NAI3,HSPH1,RPL27,PRDX5,ACTR3,PKN1,HSPA4,MLEC,SRP14,RPL37,PPP2CA,ATG2B,ARPC2,PB
I,CCT7,SRSF4,RBBP4,PFN1,MCL1,NISCH,PRKCSH,LRPPRC,EIF5A,PDIA6,RAB7A,CLTC,MAGED2,F
JA1,ARF1,AKAP12,SF3B2,GTF2F1,RPL23,RPL29,XRCC5,CIPC,RPL22,ZNF865,ATF4,RPS14,CXADI
ZC3H4,CCDC124
YWHAE,KIF1A,GNAS,GPI,HTRA4,RACK1,CALM3,GNB1,SHISA9,STMN2,MYH9,RPL7,SEPTIN2,APP,
PL29,RPS9,XRCC5,CIPC,PAPOLA,RHOA,ZNF865,GLIPR1L2,RPS14,RPS20,CD81,LENG8,WDR1,RP
IAT,SHISAL1,NOVA2,ZC3H7B,RAB27A,EEF1A2,TBCD,SUGP2,SNX22,SQSTM1,VAMP2,NOP53,ABC
ADR,LENG8,WDR1,RPL26,SRSF4,PFN1,RPL31,PRKCSH,SCRN1,RPS25,EIF5A,PDIA6,MEGF8,ITM2
H1,HYOU1,MAPK8IP1,SOD1,CTXN1,CD99L2,RBBP7,WDR45B,EP300,LRRC4B,SLC39A6,SEC16A,G
0,PPP2R1A,PKD1,SF3B1,PDIA3,SSBP2,ZNF43,VAPA,CDKL1,STMN1,ITGB1,UCHL1,MORF4L1,ANX
PLP2,DDOST,RPL36,YWHAB,ARHGDIA,RPL23A,HSBP1,MAP1A,TM9SF3,MDH2,U2AF2,CCT2,GNAI
CHF6,VCL,ARF1,SF3B2,GTF2F1,PXDN,SMARCA5,CTNNA1,FADS2,RPS16,CNN3,SCARB2,LAMA5,E
,JUND,TMEM259,LENG8,MCL1,CCNI,XPR1,HDLBP,CCND2,PLXNA1,RAB7A,CLTC,GNB2,RPS5,IK,
CDH11,HNRNPL,CNTNAP2,HDGF,RPL38,MVB12B,AZIN1,BTBD2,PBX2,AEBP1,TCF25,PTBP1,DYNL
16A7,C1ORF43,PAPOLA,RHOA,PTCD3,CBX5,RPL9,ATF4,KIF5B,SLC7A5,TMEM259,LENG8,ATP2A
1,FADS1,NEFL,NEFM,RPL8,FBXL4,UBA1,ATP2B4,IGFBP2,CALR,FSTL1,TUBB2B,COL5A1,COL4A
PC1,APPL1,BCLAF1,TRIB2,EID1,EEF1A2,MEST,GDI2,HNRNPL,MORF4L2,PKN1,ACO2,VPS35,ACA
8,WDR1,RPL26,SRSF4,PFN1,RPL31,PRKCSH,KHDRBS1,SCRN1,RPS25,EIF5A,PDIA6,MEGF8,ITM
5ORF24,TOMM20,TBCD,SNX22,MEST,HNRNPL,ABCA3,PDXK,MLEC,ACAP3,TMEM248,LAMC1,PY
WDR6,UBE2R2,ATXN10,CLIP3,ZFAND5,HNRNPUL2-BSCL2,STMN3,STT3B,STXBP1,ZFAND3,ARNT
A,SMARCA5,HNRNPA0,CPE,SCARB2,LPCAT1,SLC22A17,RPS9,RPL24,C1ORF43,NCS1,ATP1A1,JU
RCC5,SLC16A7,RHOA,ZNF865,RAB6B,SLC7A5,ENC1,ATP1A1,EIF5,LENG8,WDR1,HMGN2,ATP2A2

RPS24,SLC22A17,SLC16A7,RPL14,RHOA,ATF4,CHD4,JUND,TMEM259,RPS20,EIF5,CD81,LENG8,

,HNRNPM,LSS,ISYNA1,HYOU1,MAPK8IP1,SOD1,DDIT4,RBBP7,WDR45B,EP300,RETREG2,SLC39

MEM259,LENG8,MCL1,CCNI,XPR1,HDLBP,PLXNA1,RAB7A,GNB2,RPS5,IK,PRKAR1A,CHPF,NGRN,

M168B,PTBP1,ATXN10,CCAR2,GNG2,SMARCC1,DHX30,BAG6,RAB11A,SERBP1,ARFGEF2,CDC42

F198,DAP,BAP1,FKBP4,PKDCC,PGAP6,SH3GL1,STK24,MACROH2A2,UBE2I,NMNAT2,SH3PXD2B,

,SBF1,DNAJA1,ARF1,CYB5R3,TPM4,TNS3,EIF4H,SCARB2,NACA,LAMA5,BRD3,RPS9,CIPC,MSN,K

AB3GAP2,EIF5B,SPINT2,FKBP1A,RABEP1,BRK1,SAFB2,VPS26B,CEP170,EIF2S2,EHMT1,UBE2M,

4,UBA1,ATP2B4,CALR,TMF1,ABHD2,ZNF99,COL4A1,PSAP,PGK1,RPLP0,H3-3B,CANX,CLU,PTMA,

5H,CAND1,MAP2K2,QARS1,DIDO1,RPS28,NKTR,CCT8,LSS,CSNK1D,SSR1,BAZ1B,KIF21B,SLC39

RNP200,CSDE1,SMIM14,EIF3C,ATP5F1A,MATR3,RPL37A,FN1,PSMD5,EIF1,EGLN1,RPL19,PPP2R

,ATP2B4,IGFBP2,CALR,FSTL1,TUBB2B,COL5A1,COL4A1,PSAP,PGK1,RPLP0,DDX5,ENO2,SEC62,

ATP1A1,JUND,TMEM259,EIF5,HMGN2,PFN1,JPT2,NES,KHDRBS1,GPC3,PRDX2,ITM2C,RPS7,RPS

M2C,ARL4C,RPL35,NGRN,MTSS2,APLP2,SLC25A3,RPLP2,DDOST,H1-10,ZNF454,RAC1,CDK16,S

T6A,KIF1A,RPL13,GNAS,GPI,HTRA4,BSG,FASN,ALDH6A1,RPS8,SEPTIN2,P4HB,APP,KLRD1,SMG1,

A,RPL6,KAT6A,DVL3,YWHAE,RPL13,GNAS,DDX17,RACK1,BSG,ALDH6A1,ATP5F1B,CALM3,SHISA

,CAPZB,PLCG1,QARS1,RPL28,ZFAND5,FLNB,LSS,ISYNA1,LOC728392,HYOU1,MAPK8IP1,MLF2,C

PLXNA1,CLTC,GNB2,RPS5,IK,PRKAR1A,CHPF,NGRN,RPLP2,MYL6,INIP,MYH10,SND1,UBE2H,HM

LIPR1L2,RPS14,RPS20,CD81,LENG8,WDR1,SRSF4,PFN1,MCL1,TLE5,PNN,RPL31,KHDRBS1,RPS

TCH2,RAPGEF1,SMARCA4,AZIN1,PDE7A,ODC1,ILF3,GRB2,PCSK1N,TNKS2,UBQLN1,CSNK1D,M

F4A1,RPL19,PPP2R1A,RPL15,DNAH8,SF3B1,SERINC1,RPS3,HMGB1,POLR2A,AP2M1,RPS4X,RPS

N1,ATF4,TMEM259,HMGN2,PNN,NISCH,PRKCSH,KLHL9,HDLBP,GLG1,SFPQ,PNMA1,RPS15,NAN

17,ARF1,CDC42BPB,PXDN,NFE2L1,HNRNPA0,FADS2,RPL23,LDHA,TPM4,XRCC5,COL1A2,RPL22,

ISPA4,MLEC,ZNF302,GCN1,SKI,PRPF6,TAF7,RPL38,RPL37,ARPC2,FAM168B,PTBP1,CCAR2,GNG

EBP1,CCND2,RPL12,MRFAP1,GNB2,RPS5,MYO5A,HNRNPDL,UBE2H,RAB1A,CTBP1,ARHGDIA,AT

RAB6B,CHD4,WDR1,OAZ1,PFN1,ASTN2,JPT2,NES,CCNI,TADA2B,KHDRBS1,PFKL,PEBP1,PIP4K2B

L6A1,FTL,HSP90B1,FADS1,NEFL,RPL8,FBXL4,UBA1,ATP2B4,FSTL1,TMF1,ABHD2,HSPA1B,TUBE

A,EIF4A1,PPP2R1A,SERINC1,BASP1,POLR2A,TPT1,SEC61A1,RPS3A,RPL18,MORF4L1,NCAM1,R

EIF1,HACD3,MAP9,CHSY1,EIF4A1,PPP2R1A,SERINC1,POLR2A,TPT1,SEC61A1,RPS3A,RPL18,MOF

PS6,ZNF316,DBN1,RPL4,ENO1,TUBA1B,PIK3R2,FLNA,MAP1B,NCL,RPL7A,SLC9A7,ALDOA,MTRN

C,CCNI,XPR1,HDLBP,CCND2,ACSL6,MMP14,DDX3X,MAGED2,GNB2,CTNNB1,RPS10,ARL6IP1,ND

1,SEC61A1,STMN1,UCHL1,AGRN,ZMIZ1,TIMP2,MIDN,AP3D1,DNAJA1,AKAP12,CDC42BPB,GOLG

RPS25,EIF5A,PDIA6,MEGF8,ITM2C,RPL30,EIF3L,MAGED2,RPS5,IK,RPS19,CHPF,JUN,NGRN,RPL

PIP4K2B,ITM2C,H1-0,APLP1,SESN3,APLP2,INIP,DUX4,CKAP5,SEPTIN11,RAC1,BCLAF1,PTTG1IP,

5A,PDIA6,ITM2C,RPL30,EIF3L,MAGED2,RPS5,IK,RPS19,CHPF,JUN,NGRN,RPLP2,DDOST,CIRBP,

1,SEC62,RPL6,KAT6A,DVL3,YWHAE,KIF1A,RPL13,DDX17,GPI,RACK1,BSG,FASN,ALDH6A1,SHISA

ALM3,STMN2,P4HB,GDI1,FSCN1,TMBIM6,PTMS,GSTP1,SMG1,NPM1,TTC3,PLD3,IGF2,EIF4B,VC

MA,DDX5,FKBP14,RPL13A,MAGED1,RPL6,DVL3,SLC2A3,DDX17,GPI,RACK1,BSG,ALDH6A1,SPTB

,BSG,FASN,ALDH6A1,SPTBN1,CALM3,GNB1,SHISA9,MYH9,RPL7,SEPTIN2,APP,GDI1,SPTAN1,GS

A0,RPL29,ZNF865,KIF5B,RPS14,WDR1,PFN1,MCL1,PNN,KHDRBS1,RPS25,EIF5A,MEGF8,ITM2C

1,PLD3,CSDE1,ATP5F1A,VCP,MARCKSL1,MATR3,YWHAG,RPL37A,LRRC28,NONO,DDB1,HACD3,

2,RPL12,MAGED2,RPS7,TLN1,NGRN,MTSS2,SLC25A3,ARL6IP1,DDOST,DUX4,RHOB,PLPPR3,RP

BP7,WDR45B,EP300,LRRC4B,SLC39A6,SLC35F5,GNG4,NXN,EP400,PPP1CC,GPSM1,TNKS,MAP1

NP200,CSDE1,SMIM14,EIF3C,UBB,AFF4,TRIM28,CKB,FN1,PSMD5,YWHAQ,EIF1,HACD3,MAP9,C

NG8,WDR1,RPL26,SRSF4,PFN1,RPL31,PRKCSH,KHDRBS1,SCRN1,RPS25,EIF5A,PDIA6,MEGF8,IT

PLP1,SARAF,MIDN,RPL10,MARCHF6,WSB1,ITM2B,AP2B1,GTF2F1,GOLGA8A,SMARCA5,LDHA,CPI
A,EIF4A1,PPP2R1A,SERINC1,BASP1,POLR2A,TPT1,SEC61A1,RPS3A,RPL18,MORF4L1,NCAM1,RPL
2,CALU,ARHGDIA,CAPRIN1,TOP2B,RPL23A,HSBP1,MAP1A,CTDSP2,DDX39B,TM9SF3,EEF1A2,PPI
X17,GPI,HTRA4,BSG,FASN,SPTBN1,ATP5F1B,GNB1,SHISA9,RPS8,P4HB,TMBIM6,SPTAN1,CALM1
45B,NAP1L1,PBX1,NOL4L,MMP15,CXXC5,CEMIP2,OAZ2,TNKS,OLFM1,INTS3,ABI2,SLC6A8,PGAP
A,CUL3,BRSK1,PPP1R15A,BAZ2A,CARM1,LINGO1,SOX2,RALY,MAZ,PLEKHA6,TBC1D9,LSM14B,
K1E,PFKL,TPM1,CCND2,DDX3X,RPL12,MAGED2,CD63,SHOX,WLS,NARS1,SEPTIN11,CADM4,CKA
,WDR6,DIDO1,PITPNA,CORO1C,MSL1,TNKS2,LOC728392,ATXN2L,HECTD1,FBRSL1,PSMA7,PPD
ASN,ATP5F1B,GNB1,SHISA9,RPS8,SEPTIN2,P4HB,ERVV-1,RPL5,GDI1,TMBIM6,CALM1,PLXNB2,H
,FASN,SPTBN1,ATP5F1B,GNB1,RPS8,MYH9,APP,IDS,RPL5,GDI1,PTMS,GSTP1,MARCKS,SMG1,N
COL18A1,SNRNP200,CSDE1,IGF2,MARCKSL1,YWHAG,CKB,COL4A2,FN1,SPEN,PSMD5,DCHS1,T
RP9,NUCB1,CHD6,HNRNPL,TAF15,TCP1,COL2A1,PTOV1,MLEC,CBX1,SNRNP70,H3-3A,SCAF1,MV
P200,CSDE1,IGF2,EIF3C,ATP5F1A,EIF4B,VCP,MATR3,RPL37A,LRRC28,EIF1,RPL19,PPP2R1A,PKD
,GNB1,SHISA9,STMN2,RPS8,RPL7,SEPTIN2,IDS,RPL5,GDI1,PTMS,SPTAN1,GSTP1,MARCKS,SMG
HNRNPC,HDLBP,CCND2,RPS15,RCN1,H1-0,ANXA6,MTSS2,RPLP2,ARL6IP1,WLS,HNRNPK,SLC35
N6,ATG2B,ARPC2,SUPT5H,HNRNPUL1,SON,RPL28,SUPT6H,LONP1,BAG6,RAB11A,AKIRIN1,CCT5
14,RPL13A,MAGED1,RPL6,DVL3,KIF1A,RPL13,DDX17,GPI,RACK1,BSG,FASN,SHISA9,STMN2,MYH
8A1,EIF4B,UBB,AFF4,TRIM28,CKB,FN1,PSMD5,YWHAQ,EIF1,HACD3,MAP9,CHSY1,INA,MRC2,PPI
P4HB,APP,RPL5,FSCN1,GSTP1,PLD3,HSPD1,USP11,COL18A1,SNRNP200,CSDE1,SMIM14,EIF3C,V
3,AP2M1,SEC61A1,TUBB4B,RPL11,ANXA2,RPLP1,ZMIZ1,MARCHF6,WSB1,SF3B2,AP2B1,PGAM1,
TP2A2,CRMP1,LASP1,BEX3,KHDRBS1,XPO1,CSNK1E,PFKL,TKT,TPM1,EIF5A,CCND2,CDH2,MEGF
NA1,GNB2,RPS5,IK,PRKAR1A,CHPF,NGRN,RPLP2,MYL6,INIP,MYH10,SND1,UBE2H,HMGCR,TOP2
37A,DCHS1,SET,EGLN1,TPI1,MRC2,LDHB,DNAH8,PKD1,SSBP2,CHD3,BASP1,HMGB1,RPS4X,STM
37,MLXIP,ANXA5,TM9SF2,MORF4L2,MTRF1L,PDXK,UBE4A,AHCYL1,ETNK1,KLC1,PTOV1,CLSTN3
,EWSR1,TSPAN14,FAM53C,TNRC6A,DRAP1,ATP6V0B,MAPRE3,ZBTB4,PLPPR2,RCOR2,STC2,ZNR
EC62,RPL6,YWHAE,GNAS,RACK1,FASN,ALDH6A1,CALM3,GNB1,SHISA9,STMN2,RPL7,SEPTIN2,A
6,SMG1,CALM1,PLXNB2,HSPD1,COL18A1,SMIM14,VCP,MARCKSL1,MATR3,YWHAG,EIF1,HACD3
IPD,CDC42,EDC4,WDR45B,PDAP1,PHB2,MCM3AP,RPL7L1,RPS27A,TMEM167A,KLHDC3,PSMD8,
MSAP1,SURF4,LAMC1,SNRNP70,HDGF,CCND1,AZIN1,SRSF1,WDR6,ATXN10,CLIP3,FKBP8,PLCG1
31,DCHS1,SET,EGLN1,TPI1,CHSY1,EIF4A1,RPL19,DNAH8,PKD1,SSBP2,AP2M1,TUBB4B,ANXA2,A
NA1,TPM4,SCARB2,LAMA5,RPS24,SLC22A17,NCS1,RPL22,ZNF865,PSMD2,RAB6B,CHD4,ATP1A1
MLEC,SURF4,SNRNP70,SRP14,PPP2CA,KDELR2,DDAH2,TCF25,PRKDC,PITPNA,CORO1C,STT3A,I
N1,CALM1,PLD3,CSDE1,IGF2,EIF4B,VCP,MARCKSL1,AFF4,TRIM28,YWHAG,CKB,DDB1,RPL19,P
1,TMF1,ABHD2,HSPA1B,TUBB2B,COL5A1,COL4A1,CES3,PSAP,PGK1,RPLP0,CANX,PTMA,DDX5,E
2B1,GTF2F1,GOLGA8A,FADS2,LDHA,CPE,CNN3,CTSD,RTN4,SLC22A17,CIPC,RPL24,SLC16A7,PA
P1A1,JUND,EIF5,WDR1,RPL26,ASTN2,MCL1,TLE5,PFKL,PDIA6,PLXNA1,MEGF8,MMP14,RPS15,R
SMCHD1,TRIB2,RPL23A,HSBP1,CPSF2,U2AF2,FDFT1,SQSTM1,KLHL24,PABPN1,MORF4L2,LMN
PL5,FSCN1,GSTP1,PLD3,HSPD1,USP11,COL18A1,SNRNP200,SMIM14,VCP,UBB,AFF4,TRIM28,CK
14,EIF3C,UBB,AFF4,TRIM28,CKB,FN1,PSMD5,YWHAQ,EIF1,HACD3,MAP9,CHSY1,EIF4A1,PPP2R1
A1,AKAP12,GOLGA8A,CTNNA1,RPS16,TSPAN3,PJA2,KPNB1,NCS1,ZNF865,PSMD2,SLC7A5,GLIP
RNP200,CSDE1,SMIM14,VCP,UBB,AFF4,TRIM28,CKB,RPL37A,FN1,DDB1,PSMD5,YWHAQ,EIF1,H
C,LSS,CSNK1D,LOC728392,MPRIP,RAI1,FBRSL1,MEX3A,DYNLL1,SYP,DCTN1,STARD7,HIP1,LRR
BB,AFF4,YWHAG,LRRC28,MAPT,FN1,NONO,DDB1,YWHAQ,DCHS1,TMSB10,PPP2R1A,GTF3C4,P
LLGL1,HK1,NCS1,PSMD2,NR2F1,CXADR,CHD4,ATP1A1,JUND,WDR1,HMGN2,ATP2A2,LASP1,BEX
2R1A,PKD1,ZNF43,CHD3,CDKL1,TPT1,SEC61A1,RPS3A,ITGB1,RPL11,MORF4L1,BTF3,ANXA2,RP

?,MARCKSL1,UBB,TRIM28,LRRC28,NONO,PSMD5,SET,CHSY1,PPP2R1A,RPL15,HLA-C,SF3B1,SEI
1,TMF1,ZNF99,TUBB2B,COL5A1,COL4A1,PSAP,PGK1,H3-3B,CANX,DDX5,FKBP14,SEC62,RPL6,Y
,RAC1,BCLAF1,PTTG1IP,MAP1A,TIMP3,CTDSP2,CNN2,NOVA2,NUCB1,C5ORF24,ZC3H7B,U2AF2,
4,VCP,UBB,AFF4,TRIM28,CKB,RPL37A,FN1,DDB1,YWHAQ,EIF1,HACD3,MAP9,CHSY1,INA,EIF4A1
SLC9A7,ALDOA,MTRNR2L8,PKM,GANAB,COL6A1,HSP90B1,FADS1,NEFM,PRPF8,UGT8,RPL8,FB)
CP6,INTS11,MTMR4,SPTY2D1,CAP1,COX6A1,AP1M1,TOMM70,XPO6,UBXN6,MYL12B,GAK,LYPL/
15,FLOT1,CXXC5,PLXNB1,ECEL1,ADH5,ULK1,OAZ2,GPSM1,DKK3,GSE1,GPC4,PRDX6,KIAA0232,
2,BAG6,HNRNPD,CDC42,EDC4,WDR45B,PDAP1,PHB2,MCM3AP,RPL7L1,RPS27A,TMEM167A,KLH
iPI,RACK1,BSG,FASN,ATP5F1B,CALM3,SHISA9,RPS8,MYH9,RPL7,P4HB,ERVV-1,RPL5,TMBIM6,SI
ASTL,APEX1,PPP1R15A,BRK1,BAZ2A,VAPB,EIF3H,GABARAPL1,TRAP1,RIC8A,ELP1,NSD2,ELOVL5
;,CDKN1A,PGRMC1,CTSB,CDH11,CHD6,GNAI3,HNRNPL,PARP1,SSRP1,PDXK,AHCYL1,PRPF6,DPY
10,EIF4A1,SSBP2,RPS3,RPS3A,ITGB1,UCHL1,AGRN,ZMIZ1,MIDN,RPL10,WSB1,DNAJA1,ARF1,TM
,FSCN1,HSPD1,USP11,MARCKSL1,TRIM28,CKB,COL4A2,RPL37A,MAPT,SPEN,DCHS1,SET,EGLN1
?2B4,CALR,FSTL1,TMF1,TUBB2B,CES3,RPLP0,CANX,PTMA,DDX5,ENO2,RPL13A,KAT6A,YWHAE,K
iPTAN1,GSTP1,NPM1,PLD3,USP11,COL18A1,SMIM14,IGF2,ATP5F1A,EIF4B,VCP,UBB,TRIM28,MA
5E2B,SMCHD1,ATP6V0C,BCLAF1,L1CAM,SERF2,TOMM20,GDI2,TBC1D16,AHCYL1,ZNF302,GCN
,CCT2,SUGP2,TSR1,PTPRA,PRDX5,UBE4A,HSPA4,ZNF302,GCN1,SKI,PRPF6,RPL37,PTBP1,CCAR2
,PAPOLA,RHOA,CBX5,KIF5B,RAB6B,SLC7A5,JUND,TMEM259,EIF5,LENG8,WDR1,ATP2A2,RPL26
OL5A1,COL4A1,CES3,PSAP,RPLP0,H3-3B,CANX,DDX5,RPL13A,MAGED1,SEC62,KIF1A,DDX17,ATI
?IP4K2B,ITM2C,H1-0,APLP1,SESN3,INIP,CKAP5,SEPTIN11,COTL1,RAC1,CDK16,BCLAF1,L1CAM,
TF2F1,OGA,NFE2L1,CNN3,TSPAN3,LPCAT1,LAMA5,XRCC5,MSN,KPNB1,C1ORF43,GAP43,RPL22,
1B,TUBB2B,COL5A1,COL4A1,CES3,PGK1,PTMA,DDX5,SOX4,RPL13A,RPL6,YWHAE,KIF1A,GNAS,I
RPS3A,RPL18,NCAM1,RPLP1,SARAF,MIDN,RPL10,MARCHF6,WSB1,RPS17,SF3B2,ITM2B,AP2B1,C
OMM20,U2AF2,CCT2,GNAI3,TM9SF2,LMNB2,MTRF1L,ALKBH5,PRDX5,UBE4A,HSPA4,MLEC,ACAI
)S,RPL5,GDI1,PTMS,GSTP1,MARCKS,SMG1,NPM1,PLXNB2,USP11,COL18A1,IGF2,EIF3C,VCP,MA
C,EIF3L,RPS7,CHPF,NGRN,DDOST,NARS1,DUX4,RPS12,CALU,ARHGDIA,CAPRIN1,TOP2B,PPIB,RP
7,ZNF362,SENP3,MIB2,SLC1A4,CACHD1,POMGNT1,GYS1,MSH2,LRP8,C8ORF33,ACIN1,CDC42E
?S15A,MARS1,RABL6,FAM168B,EIF3E,WDR6,SUPT5H,SON,CCAR2,ZFAND5,DIDO1,DLK1,DHX30,S
2,MSN,IGFBP3,HK1,NCS1,JUND,CD81,HMGN2,PFN1,ASTN2,MCL1,NISCH,PRKCSH,TADA2B,KHDI

(CSH,CCNI,LRPPRC,RPS25,EIF5A,DDR1,ACSL6,RAB7A,ITM2C,RPL30,RPL35,NGRN,DDOST,YWHA
EM167A,CHERP,CTTN,SERP1,ERH,PSMB4,EIF3H,SEC14L1,NUDCD3,TUBGCP6,NCSTN,MTMR4,SI
WDR1,CRMP1,OAZ1,NES,TADA2B,KHDRBS1,PFKL,PEBP1,PNMA1,PIP4K2B,ITM2C,MAGED2,APLI
)H6,PEBP1,CDH2,GPC3,PIP4K2B,ITM2C,APLP1,SESN3,HNRNPDL,DDOST,MXD4,MYH10,RHOB,HL
JBE2H,TOP2B,SMCHD1,ATP6V0C,LAPTM4B,MLXIP,MORF4L2,HSPH1,ALKBH5,UBE4A,TBC1D16,
NAS,RACK1,BSG,FASN,SPTBN1,CALM3,RPS8,APP,GDI1,FSCN1,TMBIM6,MARCKS,CALM1,PLXNB2
TM9SF2,LMNB2,MTRF1L,PRDX5,UBE4A,HSPA4,MLEC,ACAP3,SRP14,NRDC,PRPF6,MAPRE1,FAM
ATF4,CHD4,ATP1A1,HMGN2,BEX3,RBBP4,PRKCSH,LRPPRC,KHDRBS1,XPO1,HDLBP,EIF5A,PDIA6
PDXK,ACO2,TCP1,PTOV1,MLEC,CBX1,SNRNP70,DPYSL2,TAF7,RPL38,NOTCH2,G3BP2,AZIN1,SR
XPC,DDHD2,SRRT,QRICH1,TMCO1,MYEF2,PPP4R3B,RBM39
A8A,PCYOX1,PPDPF,BCL9L,NAP1L1,RERE,DGCR2,NOL4L,BAZ1B,MGAT4B,CXXC5,GLUD1,PI4KA,
PKM,GANAB,COL6A1,FTL,HSP90B1,FADS1,NEFL,RPL8,FBXL4,YBX1,ATP2B4,IGFBP2,CALR,FSTL
3,UBE2G2,FEM1B,CBX6,GPSM1,FUS,GPC2,GNL1,ZNF428,PRR12,EEF1B2,ELP1,LSM14B,PACS1,
ENG8,MCL1,CCNI,XPR1,HDLBP,PLXNA1,CLTC,GNB2,IK,PRKAR1A,TLN1,CHPF,NGRN,RPLP2,MYL6
VCP,UBB,MATR3,YWHAG,LRRC28,SPEN,DDB1,EIF1,SET,EGLN1,MAP9,EIF4A1,DNAH8,HLA-C,SF3

FDFT1,GDI2,HNRNPL,TAF15,VPS35,MLEC,RAB11B,DSTN,CAPZB,HNRNPUL1,LSS,HGS,RBM15B,(
3B2B,COL5A1,COL4A1,CES3,PSAP,RPLP0,H3-3B,CANX,PTMA,DDX5,ENO2,SOX4,SEC62,KAT6A,RI
CCAR2,ARF4,BAG6,RAB11A,CDC42,RBBP7,TRAF7,EIF2AK1,SF1,RETREG2,PDAP1,PHB2,APMAP,F
3,RPL7,SEPTIN2,IDS,FSCN1,TMBIM6,GSTP1,PLD3,USP11,CSDE1,EIF4B,MARCKSL1,UBB,TRIM28,
P,GLG1,SFPQ,PNMA1,NANOG,MRFAP1,RPS7,CTNNB1,RPL35A,HNRNPDL,TSR2,YWHAB,TTR,ARC
3,RPL23A,SRSF5,MDH2,U2AF2,FDFT1,GNAI3,HSPH1,PRDX5,UBE4A,HSPA4,TCP1,MLEC,RPL36A,S
TG1IP,MAP1A,TIMP3,CTDSP2,CNN2,NOVA2,NUCB1,ZC3H7B,U2AF2,CCT2,MEST,GAS7,MORF4L2,
,MAT2A,RPL23,TNS3,CTSD,RPS9,PJA2,SLC16A7,CBX5,ZNF865,KIF5B,NLGN2,RPS20,RPL26,MCL
5,RBBP7,ACTR1A,PRKAR2A,CSNK2A1,GRSF1,GSPT1,MAP6,SFRP1,CD164,GLUD1,CRTAP,ADH5,A
NPA0,RPL29,LAMA5,MSN,SLC16A7,LLGL1,ZNF865,RAB6B,CHD4,CD81,LENG8,MCL1,JPT2,PNN,
PC1A,RTF1,SIN3B,SLC25A6,PJA1,ATF5,RAN,DAD1,WRNIP1,RAB18,TMEM43,PGRMC2,TBCA,KDM
P,RAB2A,CD164,RER1,UBFD1,YKT6,TMEM167A,CHERP,CTTN,STK35,ERH,UBE2M,SEC14L1,NUD(
114,VCP,UBB,AFF4,TRIM28,CKB,RPL37A,FN1,DDB1,PSMD5,YWHAQ,EIF1,HACD3,MAP9,CHSY1,IN
LMNB2,MTRF1L,RPL27,PRDX5,UBE4A,HSPA4,MLEC,CHD2,SRP14,NRDC,PRPF6,MAPRE1,FAM168
SLC2A3,DDX17,HTRA4,RACK1,FASN,ATP5F1B,CALM3,GNB1,MYH9,SEPTIN2,RPL5,GSTP1,SMG1,
EIF4B,VCP,UBB,MATR3,RPL37A,LRRC28,MAPT,DDB1,PSMD5,HACD3,EGLN1,SERINC1,BASP1,VAF
5,HOXB5,NXN,TAGLN2,CBX6,BRSK1,GSE1,PCBP4,TAOK1,ERGIC1,SHC2,SLC6A8,GPC2,RAB14,F
PF,RPLP2,MYL6,MYH10,SND1,UBE2H,ARCN1,SMCHD1,ATP6V0C,LAPTM4B,CPSF2,SRSF5,CCT2,I
43,HMGB1,CDKL1,SEC61A1,RPL18,RPLP1,SARAF,MIDN,AP3D1,SBF1,MAP4,AP2B1,PGAM1,LARP
SNRNP200,EIF3C,VCP,UBB,AFF4,TRIM28,YWHAG,RPL37A,SPEN,NONO,YWHAQ,HACD3,SET,RPL
A1,PAFAH1B2,POLDIP2,TRIP12,CHMP1A,TSNAX,ZKSCAN1,SLC25A6,EXOC5,TRMT112,CD2BP2,RI
,PLPPR3,L1CAM,HS6ST1,MDH2,KLHL24,VCAN,RPL27,ACTR3,PKN1,TBC1D16,ACO2,PTOV1,SURF
3,YBX1,UBA1,CALR,FSTL1,TMF1,ABHD2,TUBB2B,COL5A1,COL4A1,CES3,PGK1,RPLP0,H3-3B,DD
MYH10,SND1,UBE2H,SMCHD1,ATP6V0C,LAPTM4B,SRSF5,MLXIP,MORF4L2,HSPH1,ALKBH5,UBE4
R,SMCHD1,ATP6V0C,LAPTM4B,SRSF5,U2AF2,MLXIP,SQSTM1,ALKBH5,UBE4A,ACO2,AHCYL1,ETI
15,NNAT,SRP9,RAB27A,RNF187,CPSF1,PPP1CB,VAMP2,HNRNPL,TAF15,COL2A1,MLEC,TSPAN6,D
A,SEC62,YWHAE,KIF1A,RPL13,GNAS,RACK1,BSG,FASN,ALDH6A1,GNB1,MYH9,FSCN1,GSTP1,MA
0C,BCLAF1,L1CAM,FAM199X,CTDSP2,NOVA2,EEF1A2,TBCD,MEST,TM9SF2,LMNB2,KLC1,VPS35
SH6,TGOLN2,UBE2D3,HNRNPH3,NELL2,CASC4,USP47,SCAMP5,CHMP1A,LUC7L2,CDV3,GGNBP2
KBH5,AHCYL1,KLC1,CLSTN3,DDX42,RAB11B,NRDC,TUBB2A,UBXN4,PPP2CA,G3BP2,PDE7A,PSM
,MLF2,AKIRIN1,AMOTL2,YME1L1,TRAF4,ARIH1,SF1,SLC35F5,UPF1,MASTL,PPP1R15A,GALNT11
A1,DCX,EIF4G2,RPL3,RPS6,ZNF316,DBN1,RPL4,ENO1,TUBA1B,PIK3R2,FLNA,MAP1B,NCL,RPL7A
,TNKS2,NBPF9,SARS1,BAG6,HNRNPD,CCT5,CDC42,PDAP1,PHB2,GSPT1,UBA52,MCM3AP,PREPL
99X,DDX39B,MDH2,TOMM20,U2AF2,TSR1,MTRF1L,PRDX5,UBE4A,HSPA4,MLEC,LUZP6,RPL28,C
AB11B,TUBB2A,PPP2CA,PDE7A,PSMD3,CDC37,WDR6,BNIP3L,ARF4,HNRNPUL2-BSCL2,PITPNA,XF
RPL10,MARCHF6,RPS17,ITM2B,OGA,PGAM1,RPS16,CTSD,NACA,LAMA5,BRD3,CIPC,MSN,KPNB1
L37,ATG2B,ARPC2,FAM168B,ATXN10,RPL28,GNG2,DHX30,BAG6,FBRSL1,RAB11A,TP53BP1,CDC
,MVB12B,SMARCA4,ARPC2,BTBD2,PBX2,FAM171A2,NYNRIN,HNRNPUL1,PLCG1,CORO1C,LSS,F
E,KIF1A,RPL13,SLC2A3,DDX17,GPI,RACK1,ATP5F1B,CALM3,STMN2,MYH9,RPL7,IDS,RPL5,KLRD
PI1,RPL19,PPP2R1A,GTF3C4,DNAH8,SERINC1,HMGB1,POLR2A,AP2M1,TPT1,BTF3,AGRN,RPLP1,I
2,TRIP12,CHMP1A,TSNAX,EXOC5,TRMT112,PSMD13,CD2BP2,BANF1,FAF2,VPS51,FAM32A,EPN1,
OTL2,YME1L1,TRAF4,ARIH1,SLC35F5,UPF1,MASTL,PPP1R15A,GALNT11,DAP,EIF3H,ELAVL1,GAB
CR,SMCHD1,ATP6V0C,LAPTM4B,SRSF5,MLXIP,MORF4L2,HSPH1,ALKBH5,UBE4A,AHCYL1,KLC1,
S1,NEFM,PRPF8,UGT8,RPL8,FBXL4,YBX1,UBA1,ATP2B4,CALR,FSTL1,TMF1,ABHD2,HSPA1B,ZNF
3,PTOV1,LAMC1,HDGF,SELENON,TUBB2A,ARF3,CCND1,PBX2,PCSK1N,SON,CLIP3,KDELR1,RPL2

70,LYPLA2,ATP13A1,IER2,PAFAH1B2,POLDIP2,TRIP12,CHMP1A,TSNAX,SLC25A6,EXOC5,TRMT11
9,SLC16A7,KPNB1,PAPOLA,CBX5,ATF4,RPS14,TMEM259,RPS20,CD81,HMGN2,BEX3,VAT1,OAZ1
T1,CCT2,GDI2,TM9SF2,HSPH1,MTRF1L,PRDX5,UBE4A,TCP1,ZNF302,GCN1,HDGF,SRP14,TAF7,F
,PIK3R2,FLNA,NCL,RPL7A,SLC9A7,ALDOA,PKM,GANAB,COL6A1,FTL,FADS1,NEFL,NEFM,RPL8,FI
INA,MRC2,PPP2R1A,RPL15,SF3B1,SERINC1,RPS3,BASP1,POLR2A,ITGB1,MORF4L1,ANXA2,AGRI
ST,CIRBP,LAMP1,HS6ST1,SRSF5,TM9SF3,MDH2,RAB27A,U2AF2,GNAI3,NOP53,LMNB2,PRDX5,U
K,CKAP4,RAB1A,CALU,CTBP1,STIP1,SMCHD1,TMEM132A,CNN2,TM9SF3,ANXA5,KLHL24,TM9SF
,RPL35A,TSR2,ARCN1,RPS27,HECTD4,ATP6V0C,BCLAF1,L1CAM,CNN2,SRP9,SRSF5,U2AF2,NOP
CKS,PLD3,USP11,UBB,AFF4,TRIM28,YWHAG,COL4A2,LRRC28,SPEN,DDB1,YWHAQ,EIF1,HACD3,
PCAT1,CIPC,C1ORF43,RHOA,ATF4,CHD4,ATP1A1,EIF5,LASP1,PFN1,PRKCSH,KHDRBS1,TKT,CCN
3,GNAS,DDX17,GPI,BSG,FASN,ALDH6A1,SPTBN1,GNB1,SHISA9,RPS8,P4HB,RPL5,SPTAN1,CALM
M1,TOMM70,XPO6,UBXN6,MYL12B,GAK,LYPLA2,ATP13A1,PAFAH1B2,POLDIP2,TRIP12,CHMP1A
S,PSMD8,CHERP,CTTN,SERP1,ERH,SEC14L1,NUDCD3,INTS11,MTMR4,SPTY2D1,NOC2L,AP1M1,
CTSB,CHD6,EEF1A2,VAMP2,PARP1,MTRF1L,PRDX5,HSPA4,KLC1,SURF4,SRP14,ABCA2,TOP1,VE
T2,MEST,GAS7,MORF4L2,SNRPN,PTOV1,MLEC,SURF4,SMARCA4,AZIN1,DDAH2,NYNRIN,WDR6,PI
HK1,NCS1,ZNF865,NLGN2,CHD4,JUND,WDR1,BEX3,OAZ1,SRSF4,NES,RPL31,HNRNPC,KHDRBS1
MYH10,SND1,UBE2H,ARCN1,SMCHD1,ATP6V0C,LAPTM4B,SRSF5,MLXIP,MORF4L2,HSPH1,ALKE
E1L1,CTNND1,TRAF4,MASTL,PRRC2A,CALD1,GALNT11,XRCC6,SSBP3,ELAVL1,GNL1,TRAP1,BLC
PTTG1IP,CRPPA,SRSF5,TM9SF3,SNX22,PGD,VAMP2,PABPN1,GDI2,ALKBH5,TCP1,PLOD1,MDK,TM
L35A,HNRNPDL,TSR2,TTR,RPS27,HECTD4,ATP6V0C,BCLAF1,L1CAM,CNN2,SRP9,MDH2,U2AF2,N
LR,ABHD2,HSPA1B,TUBB2B,COL5A1,COL4A1,PSAP,RPLP0,H3-3B,CANX,CLU,DDX5,MAGED1,SEC
KCSH,SLC3A2,RPS25,EIF5A,RPL12,EIF3L,IK,CHPF,HNRNPDL,RPLP2,DDOST,CIRBP,ATP1A2,PLPF
F3,SEPTIN9,CHPF2,SPIN1,C8ORF33,PTPN12,HADHB,SESN2,POGZ,AKIRIN2,CRAMP1
28,CKB,FN1,PSMD5,YWHAQ,EIF1,HACD3,MAP9,CHSY1,INA,EIF4A1,MRC2,PPP2R1A,SERINC1,CHD
PC3,PIP4K2B,ITM2C,APLP1,SESN3,HNRNPDL,MXD4,MYH10,RHOB,HLA-B,CCT6A,STIP1,SLC35E2I
AP1L3,ULK1,OAZ2,KLHDC3,ZNF664,VPS4A,TAX1BP1,PGAP6,KIFAP3,PNISR,AHSA1,DICER1,DHX38
CD3,MAP9,CHSY1,EIF4A1,PPP2R1A,RPL15,SERINC1,BASP1,POLR2A,TPT1,SEC61A1,RPS3A,RPL1
EN,PSMD5,EIF1,HACD3,DCHS1,EGLN1,RPL15,GTF3C4,HLA-C,SSBP2,VAPA,POLR2A,AP2M1,SEC6
H5,UBE4A,TBC1D16,AHCYL1,KLC1,CLSTN3,DDX42,RAB11B,NRDC,TUBB2A,UBXN4,PPP2CA,G3E
KM,GANAB,COL6A1,FTL,HSP90B1,FADS1,NEFL,NEFM,RPL8,FBXL4,YBX1,ATP2B4,CALR,FSTL1,A
3A,LOC728392,ATXN2L,ZSCAN18,ARNT2,PRPF40A,TXNIP,CIC,ST3GAL6,NAP1L1,ACTR1A,PRCC,I
P5,CHMP1A,LUC7L2,CDV3,GGNBP2,ADNP,ANKFY1,SRSF10,CCNG1,GID8,MARCHF7,ZCCHC14,P(
IF5A,PDIA6,ACTN4,RPL30,CLTC,EIF3L,H1-0,RPS19,RPL35,CHPF,GJA1,CD63,NGRN,RPLP2,DDOS
CHERP,CTTN,LRP10,RHBDD2,ERH,SEC14L1,NUDCD3,GNL1,HDGFL2,GALNT1,TUBGCP6,INTS11,
063,RPLP2,DDOST,CIRBP,LAMP1,ZNF454,SLC35E2B,DNER,RPL23A,HSBP1,HS6ST1,NUCB1,SRSI
DGFL2,NUCKS1,ELP1,AKT2,WNK1,SLC20A1,FGFR1,DHX40,FBXL19,AMFR,CELSR1,USF2,FXR2,SL
EPN1,RTF2,UNC45A,ZBTB4,HMGXB3,PDCD11,WARS1,SNRPB,KDM5A,SKIV2L,SHC1,SUDS3,MIB2,
ARB2,NACA,LAMA5,KPNB1,RPL17,NCS1,RPL22,PTCD3,ZNF865,CLSTN1,RPS20,LENG8,WDR1,AT
TMA,DDX5,SOX4,RPL13A,KAT6A,DVL3,YWHAE,KIF1A,RPL13,RACK1,ALDH6A1,SPTBN1,CALM3,SH
NEFL,UGT8,RPL8,FBXL4,YBX1,UBA1,ATP2B4,IGFBP2,CALR,TMF1,ZNF99,COL5A1,COL4A1,CES3
,POLDIP2,TRIP12,CHMP1A,TSNAX,SLC25A6,EXOC5,SF3A3,TRMT112,CD2BP2,VPS51,PPME1,FAN
Y1,INA,HLA-C,ZNF43,CHD3,VAPA,TPT1,STMN1,ITGB1,TUBB4B,BTF3,ANXA2,AGRN,ZMIZ1,COL11A
HAH,UBQLN1,CSNK1D,IFITM3,MLF2,AKIRIN1,AMOTL2,YME1L1,TRAF4,STAT3,ARIH1,SF1,SLC35F
EP3,LARS1,YWHAH,UBQLN1,CSNK1D,IFITM3,MLF2,AKIRIN1,AMOTL2,YME1L1,TRAF4,STAT3,ARII
L1,MEX3A,RAB11A,CIC,MTA1,VARS1,NOL4L,KIF21B,HOXB5,RAB2A,RAB3GAP2,GPSM1,RPS2,SLI

C,ACAP3,SRP14,TAF7,RPL37,ARPC2,CCAR2,SUPT6H,LARS1,PFKM,ID4,BAG6,RAB11A,AKIRIN1,CC

IF4H,XRCC5,PJA2,PAPOLA,ZNF865,RAB6B,JUND,WDR1,ATP2A2,SRSF4,RTN3,TLE5,NISCH,SLC3A

DC37,BNIP3L,HNRNPUL2-BSCL2,PITPNA,XPNPEP3,LARS1,YWHAH,UBQLN1,CSNK1D,IFITM3,SREE

YWHAH,UBQLN1,CSNK1D,IFITM3,MLF2,AKIRIN1,AMOTL2,YME1L1,TRAF4,STAT3,ARIH1,SF1,SLC

YWHAH,UBQLN1,CSNK1D,IFITM3,MLF2,AKIRIN1,AMOTL2,YME1L1,TRAF4,STAT3,ARIH1,SF1,SLC

ZB,GABARAP,ARF4,ZFAND5,KIF3C,ATP6V1B2,YWHAH,SREBF2,RPL39,ARNT2,AMOTL2,CTNND1,S

1,STMN2,RPS8,MYH9,APP,IDS,PTMS,SPTAN1,GSTP1,SMG1,CALM1,NPM1,PLXNB2,PLD3,HSPD1,

N1,PSMD2,CHD4,LENG8,WDR1,BEX3,OAZ1,PNN,NES,TADA2B,KHDRBS1,PFKL,GLG1,SFPQ,PEBP

A,RPL13,ALDH6A1,SPTBN1,SHISA9,RPL7,IDS,TMBIM6,SPTAN1,SMG1,CALM1,NPM1,PLXNB2,PLI

P11,COL18A1,SNRNP200,CSDE1,ATP5F1A,EIF4B,VCP,UBB,AFF4,MATR3,RPL37A,LRRC28,MAPT,F

P11,COL18A1,SNRNP200,CSDE1,ATP5F1A,EIF4B,VCP,UBB,AFF4,MATR3,RPL37A,LRRC28,MAPT,F

I7,AP3D1,ARF1,MAP4,SF3B2,ITM2B,TMEM170A,SLC4A8,HNRNPU,PGAM1,CTNNA1,CNN3,RPL29,

UPF1,MASTL,PPP1R15A,GALNT11,SPG7,ELAVL1,IMPDH2,GNL1,TRAP1,BLCAP,MOAP1,CLASP2,F

4A1,CES3,CLU,PTMA,DDX5,ENO2,RPL13A,MAGED1,SEC62,DVL3,YWHAE,KIF1A,GNAS,SLC2A3,G

ND1,POLR2B,SDF4,RPS28,YWHAH,LONP1,HGS,SREBF2,VGF,ARNT2,PSMA7,VDAC1,TRAF7,KIRRE

A,LOC728392,HYOU1,ZFAND3,ARNT2,ARFGEF2,CTXN1,SSR1,STAT3,TXLNA,XPOT,TXNDC5,SEC16

,CALR,FSTL1,TMF1,HSPA1B,COL4A1,CES3,PSAP,RPLP0,H3-3B,CANX,CLU,PTMA,DDX5,FKBP14,I

SCN1,TMBIM6,GSTP1,NPM1,PLD3,USP11,COL18A1,SNRNP200,CSDE1,SMIM14,VCP,UBB,AFF4,T

IDS,GDI1,MARCKS,CALM1,NPM1,TTC3,PLD3,HSPD1,COL18A1,SNRNP200,CSDE1,EIF4B,UBB,AF

C6A,NIPSNAP1,CDK9,ZNRF1,ELOB,SUDS3,DIPK1B,SMARCC2,PPM1A,SH3KBP1,ELOA

M168B,LUZP6,SON,CORO1C,SF3A1,BAG6,HNRNPD,RBBP7,TRAF7,EIF2AK1,EDC4,WDR45B,PDAI

PXDN,CTSD,NACA,LPCAT1,SLC16A7,C1ORF43,PAPOLA,HK1,RHOA,KIF5B,JUND,TMEM259,RPS20

I7,BRD3,MSN,KPNB1,PAPOLA,NCS1,SLC7A5,NLGN2,CHD4,COL5A2,LENG8,CRMP1,RBBP4,RTN3

FAH1B2,POLDIP2,TRIP12,TSNAX,EXOC5,TRMT112,CD2BP2,SUMO1,BANF1,EPN1,PSMC5,ZBTB4,

V-1,RPL5,GDI1,PTMS,MARCKS,SMG1,PLXNB2,USP11,COL18A1,CSDE1,SMIM14,EIF3C,VCP,UBB,

RS1,YWHAH,UBQLN1,CSNK1D,IFITM3,MLF2,AKIRIN1,AMOTL2,YME1L1,TRAF4,STAT3,ARIH1,SF1

37A,RPL30,NANOG,RPS7,CTNNB1,RPL35A,TSR2,TTR,RPS27,HECTD4,ATP6V0C,MMP2,L1CAM,LA

RPLP0,CANX,DDX5,SOX4,SEC62,RPL6,KAT6A,KIF1A,GNAS,HTRA4,BSG,ATP5F1B,CALM3,GNB1,SH

7,SEPTIN2,IDS,ERVV-1,TMBIM6,PTMS,GSTP1,SMG1,HSPD1,MARCKSL1,AFF4,CKB,LRRC28,SPEN

POLDIP2,TRIP12,CHMP1A,TSNAX,RGL2,EXOC5,SF3A3,TRMT112,COPE,CD2BP2,FAF2,KLC2,GOL

P11,COL18A1,SNRNP200,CSDE1,ATP5F1A,EIF4B,VCP,UBB,AFF4,MATR3,RPL37A,LRRC28,MAPT,F

,HNRNPUL2-BSCL2,PITPNA,XPNPEP3,LARS1,YWHAH,UBQLN1,CSNK1D,IFITM3,SREBF2,VGF,MLI

GOLGA8A,RPL23,LDHA,CPE,SCARB2,SLC22A17,RPS9,RPL24,CBX5,GLIPR1L2,JUND,TMEM259,L

,ATP13A1,PAFAH1B2,POLDIP2,TRIP12,CHMP1A,PSMB1,TSNAX,RGL2,SLC25A6,EXOC5,SF3A3,TR

1,ELP1,SPAG9,NSD2,ELOVL5,SLC25A36,XRN2,ZZEF1,ESYT1,GTF3C1,TNFAIP1,ALDH18A1,CNDP2

PH1,MTRF1L,AHCYL1,KLC1,CLSTN3,DDX42,RAB11B,TUBB2A,PPP2CA,G3BP2,AZIN1,PDE7A,PSMI

D1,FBRSL1,RBM15B,DHX15,BEX4,DCTN1,TXLNA,PDAP1,ALDH3A2,AP2A1,NOL4L,ANP32B,GPSM

2AF2,GDI2,TSR1,SSRP1,MTRF1L,PRDX5,UBE4A,ACO2,HSPA4,MLEC,NRDC,RPS15A,ARF3,LUZP6,

IDLBP,PLXNA1,GNB2,RPS5,IK,CHPF,JUN,NGRN,RPLP2,MYL6,MYH10,SND1,UBE2H,SMCHD1,ATP

MD

EP3,LARS1,YWHAH,UBQLN1,CSNK1D,IFITM3,MLF2,AKIRIN1,AMOTL2,YME1L1,TRAF4,STAT3,ARII

HMGCR,SMCHD1,ATP6V0C,LAPTM4B,SRSF5,MLXIP,HSPH1,HNRNPA3,UBE4A,ACO2,AHCYL1,ETNI

DC37,SCAF11,BNIP3L,HNRNPUL2-BSCL2,PITPNA,XPNPEP3,LARS1,YWHAH,FAU,UBQLN1,CSNK1D

ZBTB4,WARS1,SNRPB,KDM5A,SKIV2L,SHC1,SUDS3,MIB2,TNPO3,CUL4B,SZT2,SEC31A,AKAP17A

,HNRNPUL2-BSCL2,PITPNA,XPNPEP3,LARS1,YWHAH,UBQLN1,CSNK1D,IFITM3,VGF,MLF2,AKIRI
S1,AKT2,WNK1,SLC20A1,FGFR1,DHX40,FBXL19,AMFR,CELSR1,FXR2,SLC25A6,KIF26A,KAT2A,R
RN2,ZZEF1,ESYT1,GTF3C1,TNFAIP1,CNDP2,GLOD4,MTURN,DHX36,COLGALT1,TSPAN7,NUP98,C
RN2,ZZEF1,ESYT1,GTF3C1,TNFAIP1,CNDP2,GLOD4,MTURN,DHX36,COLGALT1,TSPAN7,NUP98,C
SMD3,CDC37,BNIP3L,HNRNPUL2-BSCL2,PITPNA,XPNPEP3,LARS1,YWHAH,UBQLN1,CSNK1D,IFIT
R,TMEM259,RPS20,RTN3,ASTN2,PNN,TKT,DDX3X,RPL30,CLTC,NANOG,RPS7,IK,HMGCS1,GJA1,
YL1,KLC1,CLSTN3,DDX42,RAB11B,NRDC,TUBB2A,OPTN,PPP2CA,G3BP2,AZIN1,PDE7A,PSMD3,S
4H,LAMA5,CIPC,C1ORF43,CBX5,ATF4,ENC1,CHD4,ATP1A1,CD81,HMGN2,BEX3,RBBP4,JPT2,PR
4H,LAMA5,CIPC,C1ORF43,CBX5,ATF4,ENC1,CHD4,ATP1A1,CD81,HMGN2,BEX3,RBBP4,JPT2,PR
R,SLC35E2B,TRIB2,PEA15,LUC7L3,DLG5,CNN2,RAB27A,U2AF2,SQSTM1,VCAN,RPL27,PRDX5,CE
ZFP36L1,GNG4,KCTD12,ZNF512B,FMNL2,ULK1,ZMYM4,DAP,TMOD2,C6ORF62,PKDCC,LTBP4,M
,CALR,FSTL1,TMF1,ZNF99,TUBB2B,COL5A1,COL4A1,CES3,PSAP,PGK1,CLU,PTMA,DDX5,ENO2,

598,CHPF2,BRPF3,PRNP,RNPS1,PRELID1,BRD1,VPS37B
10,MARCHF6,DNAJA1,ARF1,SF3B2,TMEM170A,TPM4,RPS24,XRCC5,CIPC,RPL24,SLC16A7,RPL2
AH,UBQLN1,CSNK1D,IFITM3,MLF2,AKIRIN1,AMOTL2,YME1L1,TRAF4,ARIH1,SF1,SLC35F5,CSN
AB11B,TOP1,TUBB2A,OPTN,PPP2CA,G3BP2,AZIN1,PDE7A,PSMD3,SRSF1,CDC37,BNIP3L,ZFAND
P,MTCH1,SERP1,RHBDD2,ERH,PSMB4,CLPTM1,EIF3H,SSR3,PNISR,RIC8A,AHSA1,PCNX3,ERGIC
2,COPE,CD2BP2,GOLPH3,VPS51,FAM32A,EPN1,ARPC3,RTF2,UNC45A,PSMF1,HMGXB3,WARS1,C
,MAP1S,RAVER1,MBOAT7,SLC12A7,SH3BP5,NKIRAS2,GUK1,NIPA1,NID1,SEMA6C,BSDC1,HMG2
O,SET,MRC2,RPL19,GTF3C4,SF3B1,SERINC1,BASP1,POLR2A,AP2M1,RPS4X,RPS3A,STMN1,TUE
YM4,FUS,YARS1,HSPG2,LTBP4,TLE1,MAZ,STK24,PRPF4B,MSH6,IGF2BP1,PCGF2,GNA13,METRI
CA,PAPOLA,LLGL1,HK1,ATP1A1,TMEM259,LENG8,LASP1,PFN1,ASTN2,NISCH,PRKCSH,XPR1,KH
KIN1,DPP7,RPL36AL,EPHB2,OBSCN,ANKRD13B,SKIL
OAZ1,PNN,NES,TADA2B,KHDRBS1,PFKL,GLG1,PEBP1,PNMA1,PIP4K2B,ITM2C,H1-0,APLP1,APLP
F4,ORAI2,SKI,SNRNP70,PYGB,DPYSL2,PPP2CA,MAPK1,SMARCA4,AZIN1,DDAH2,NYNRIN,TCF25,

LAPTM4A,PDE7A,PTBP1,ILF3,UBE2R2,KDM1A,LONP1,SOD1,NCOR2,TXNIP,CTNND1,TRAF7,KIRR
B1,RPS17,AP2B1,GTF2F1,RPL23,LDHA,CNN3,CTSD,SCARB2,NACA,PJA2,RPL24,CBX5,GLIPR1L2
3,TARDBP,RPLP2,DDOST,YWHAB,TTR,ARCN1,TOP2B,SLC35E2B,NNAT,MAP1A,DLG5,SRP9,NUCE
R,NIBAN2,ATP13A2,FZR1,SGSM3,SLC25A6,CNDP2,LIMK1,PITRM1,RAP2A,SOBP,TBXA2R,FAM32
RNF157,PCGF3,OTUD5,TMEM131,RTF2,PSIP1,DEGS1,P3H1,PIP5K1C,SETX,STAT1,ITPK1,COX5A
C61A1,ITGB1,TUBB4B,UCHL1,RPLP1,XAF1,MIDN,VCL,AP3D1,ARF1,MAP4,SLC4A8,HNRNPU,NFE2
GTF2F1,RPL23,LDHA,CNN3,CTSD,SCARB2,NACA,PJA2,RPL24,CBX5,GLIPR1L2,JUND,TMEM259,
,RPL36A,SRP14,TAF7,RPL38,PDE7A,FAM168B,LUZP6,QARS1,HNRNPUL2-BSCL2,POLR2B,GNG2
210,SLC38A1,CLUH,CD44,RASA4,NECTIN1,RNF41,SLC16A2,SRSF7,ZCCHC14,CLPTM1L,GPT2,RE
1,FAM32A,RTF2,UNC45A,ZBTB4,PDCD11,SNRPB,SKIV2L,SHC1,PCED1A,SUDS3,MIB2,CUL4B,VP
RINC1,BASP1,POLR2A,CDKL1,SEC61A1,NCAM1,RPLP1,ITM2B,ZNF770,GTF2F1,SMARCA5,CPE,T
,HNRNPUL2-BSCL2,PITPNA,XPNPEP3,LARS1,YWHAH,UBQLN1,CSNK1D,IFITM3,VGF,MLF2,AKIRI
2,PPP1R15A,MAP1LC3B,NDN,CARM1,ARHGEF17,HDGFL2,NCSTN,INTS11,SLC25A23,ERGIC3,CA
3BP2,PDE7A,PSMD3,CDC37,SCAF11,BNIP3L,HNRNPUL2-BSCL2,PITPNA,XPNPEP3,LARS1,YWHA
SCL2,PITPNA,XPNPEP3,LARS1,YWHAH,TNKS2,UBQLN1,CSNK1D,IFITM3,VGF,MLF2,AKIRIN1,AMO
6,SF3A3,TRMT112,CD2BP2,RPS21,BANF1,GOLPH3,PPP1CA,FAM32A,SYNGAP1,EPN1,ARPC3,RT
TRIP12,TSNAX,RGL2,PCM1,SLC25A6,SF3A3,TRMT112,COPE,EWSR1,GOLPH3,PPP1CA,PPME1,SY

.P2,RPL36,LAMP1,STIP1,SERF2,CTSB,U2AF2,HNRNPA3,PRDX5,PKN1,ACAP3,ZNF302,DDX42,ORA
M1,PCGF3,EXT2,SOBP,TBXA2R,OTUD5,GOLGA8B,MRPS6,CACNA1H,NAPB,TXN,DUSP8,EFNB3,R

A1,FTL,HSP90B1,FADS1,NEFL,NEFM,PRPF8,RPL8,FBXL4,YBX1,UBA1,IGFBP2,CALR,FSTL1,TMF1,
ISP90B1,FADS1,NEFL,NEFM,PRPF8,RPL8,FBXL4,YBX1,UBA1,ATP2B4,IGFBP2,CALR,FSTL1,TMF1
L,NEFM,PRPF8,RPL8,FBXL4,YBX1,UBA1,IGFBP2,CALR,FSTL1,ABHD2,HSPA1B,ZNF99,TUBB2B,C
,RPL8,FBXL4,YBX1,UBA1,IGFBP2,CALR,FSTL1,ABHD2,HSPA1B,ZNF99,TUBB2B,COL5A1,COL4A
,PTMA,DDX5,ENO2,SOX4,FKBP14,SEC62,KAT6A,DVL3,YWHAE,KIF1A,RPL13,GNAS,DDX17,GPI,HT
1,PRPF8,RPL8,FBXL4,YBX1,UBA1,ATP2B4,IGFBP2,CALR,FSTL1,ABHD2,HSPA1B,TUBB2B,COL5A
)L6A1,FTL,HSP90B1,FADS1,NEFL,NEFM,PRPF8,RPL8,FBXL4,YBX1,IGFBP2,CALR,FSTL1,TMF1,AE
13,GNB1,SHISA9,MYH9,SEPTIN2,P4HB,APP,IDS,GDI1,FSCN1,PTMS,SPTAN1,MARCKS,SMG1,CAL
NF99,TUBB2B,COL5A1,COL4A1,PSAP,PGK1,RPLP0,H3-3B,CANX,CLU,PTMA,ENO2,SOX4,RPL13A
JBA1,ATP2B4,IGFBP2,FSTL1,TMF1,HSPA1B,ZNF99,TUBB2B,COL5A1,COL4A1,CES3,PSAP,PGK1,
EFM,PRPF8,RPL8,FBXL4,YBX1,UBA1,ATP2B4,IGFBP2,CALR,FSTL1,ABHD2,HSPA1B,TUBB2B,COL
A1B,TUBB2B,COL5A1,PSAP,PGK1,H3-3B,PTMA,DDX5,ENO2,SEC62,KAT6A,DVL3,YWHAE,KIF1A,R
3B2B,COL4A1,PSAP,PGK1,CANX,PTMA,DDX5,ENO2,SOX4,FKBP14,SEC62,KAT6A,DVL3,YWHAE,K
PA1B,TUBB2B,COL4A1,PSAP,PGK1,CANX,PTMA,DDX5,ENO2,SOX4,FKBP14,SEC62,KAT6A,DVL3,Y
CANX,PTMA,DDX5,ENO2,RPL13A,KAT6A,DVL3,YWHAE,KIF1A,RPL13,GNAS,SLC2A3,GPI,HTRA4,R
F8,RPL8,FBXL4,YBX1,UBA1,IGFBP2,CALR,FSTL1,TMF1,ABHD2,HSPA1B,TUBB2B,COL5A1,COL4
LR,FSTL1,ABHD2,HSPA1B,TUBB2B,COL4A1,PSAP,PGK1,CANX,DDX5,ENO2,SOX4,FKBP14,SEC6
A1,PSAP,PGK1,CANX,PTMA,DDX5,ENO2,SOX4,FKBP14,SEC62,KAT6A,DVL3,YWHAE,KIF1A,GNAS,
,ALDH6A1,SPTBN1,CALM3,GNB1,SHISA9,STMN2,MYH9,RPL7,SEPTIN2,P4HB,APP,GDI1,PTMS,S
FBP2,CALR,FSTL1,ABHD2,HSPA1B,TUBB2B,PSAP,PGK1,H3-3B,CANX,DDX5,ENO2,SOX4,FKBP14,
PGK1,RPLP0,H3-3B,CANX,PTMA,DDX5,ENO2,RPL13A,SEC62,RPL6,KAT6A,DVL3,YWHAE,KIF1A,R
BX1,ATP2B4,IGFBP2,CALR,FSTL1,TMF1,ABHD2,HSPA1B,TUBB2B,COL5A1,COL4A1,CES3,PSAP,
,COL4A1,PSAP,RPLP0,H3-3B,CANX,PTMA,DDX5,ENO2,SOX4,RPL13A,MAGED1,KAT6A,DVL3,YWH
NAB,COL6A1,FTL,HSP90B1,FADS1,NEFL,NEFM,PRPF8,RPL8,FBXL4,YBX1,UBA1,ATP2B4,IGFBP2
A1,RPLP0,CANX,CLU,PTMA,DDX5,ENO2,SOX4,RPL13A,KAT6A,YWHAE,KIF1A,RPL13,GNAS,DDX17
,FBXL4,YBX1,UBA1,IGFBP2,FSTL1,ABHD2,HSPA1B,TUBB2B,COL5A1,COL4A1,PSAP,PGK1,RPLP
OX4,RPL13A,SEC62,RPL6,KAT6A,DVL3,YWHAE,KIF1A,RPL13,GNAS,DDX17,RACK1,BSG,FASN,AL
BXL4,YBX1,UBA1,ATP2B4,IGFBP2,CALR,FSTL1,ABHD2,HSPA1B,TUBB2B,COL5A1,COL4A1,CES3,
B1,FADS1,NEFL,NEFM,PRPF8,UGT8,RPL8,FBXL4,YBX1,UBA1,ATP2B4,IGFBP2,CALR,FSTL1,TMF
B,COL6A1,FTL,HSP90B1,FADS1,NEFL,NEFM,PRPF8,RPL8,FBXL4,YBX1,UBA1,ATP2B4,IGFBP2,C
,NEFL,NEFM,PRPF8,RPL8,YBX1,UBA1,ATP2B4,IGFBP2,CALR,FSTL1,TMF1,ABHD2,HSPA1B,TUBE
JBB2B,COL4A1,PSAP,PGK1,CANX,CLU,PTMA,DDX5,ENO2,SOX4,FKBP14,SEC62,KAT6A,DVL3,YW
OL4A1,PSAP,PGK1,H3-3B,CANX,PTMA,DDX5,ENO2,RPL13A,KAT6A,DVL3,YWHAE,KIF1A,RPL13,G
FTL,HSP90B1,FADS1,NEFL,NEFM,PRPF8,RPL8,FBXL4,YBX1,IGFBP2,CALR,FSTL1,TMF1,ABHD2,
2L8,PKM,GANAB,COL6A1,FTL,HSP90B1,FADS1,NEFL,NEFM,PRPF8,RPL8,FBXL4,YBX1,UBA1,AT
LC2A3,GPI,RACK1,BSG,FASN,ALDH6A1,SPTBN1,CALM3,GNB1,SHISA9,STMN2,RPS8,MYH9,RPL
4,YBX1,UBA1,ATP2B4,IGFBP2,FSTL1,ABHD2,TUBB2B,COL5A1,COL4A1,PSAP,PGK1,RPLP0,H3-
5A1,COL4A1,RPLP0,H3-3B,CANX,CLU,PTMA,DDX5,ENO2,SOX4,RPL13A,MAGED1,SEC62,RPL6,K
T6A,DVL3,YWHAE,KIF1A,GNAS,SLC2A3,GPI,BSG,FASN,ALDH6A1,SPTBN1,CALM3,GNB1,SHISA9,
8,FBXL4,YBX1,UBA1,ATP2B4,IGFBP2,CALR,FSTL1,TMF1,ABHD2,HSPA1B,TUBB2B,COL5A1,COL
B1,FADS1,NEFL,NEFM,PRPF8,RPL8,FBXL4,YBX1,UBA1,IGFBP2,CALR,FSTL1,ABHD2,HSPA1B,TU
HSP90B1,FADS1,NEFL,NEFM,PRPF8,UGT8,RPL8,FBXL4,YBX1,UBA1,IGFBP2,CALR,FSTL1,TMF1,
K1,RPLP0,H3-3B,CANX,PTMA,DDX5,ENO2,SOX4,MAGED1,RPL6,KAT6A,DVL3,KIF1A,RPL13,GNA
1,ATP2B4,IGFBP2,FSTL1,ABHD2,HSPA1B,TUBB2B,COL5A1,COL4A1,PSAP,PGK1,CANX,PTMA,DD
FM,PRPF8,UGT8,RPL8,FBXL4,YBX1,UBA1,ATP2B4,IGFBP2,CALR,FSTL1,ABHD2,HSPA1B,TUBB2

FTL,HSP90B1,FADS1,NEFM,PRPF8,RPL8,FBXL4,YBX1,UBA1,IGFBP2,CALR,FSTL1,ABHD2,HSPA1
A1,COL4A1,CES3,PGK1,H3-3B,CANX,PTMA,DDX5,ENO2,SOX4,SEC62,KAT6A,DVL3,YWHAE,KIF1A
STMN2,MYH9,RPL7,SEPTIN2,P4HB,GDI1,FSCN1,PTMS,SPTAN1,GSTP1,MARCKS,SMG1,CALM1,N
L,NEFM,PRPF8,RPL8,FBXL4,ATP2B4,IGFBP2,FSTL1,TMF1,ABHD2,HSPA1B,ZNF99,TUBB2B,COL
ANAB,COL6A1,FTL,HSP90B1,NEFL,NEFM,PRPF8,RPL8,FBXL4,YBX1,UBA1,ATP2B4,IGFBP2,CALR
HAE,KIF1A,RPL13,GNAS,SLC2A3,GPI,HTRA4,BSG,FASN,SPTBN1,CALM3,GNB1,SHISA9,STMN2,M
GNAS,SLC2A3,GPI,BSG,FASN,ALDH6A1,SPTBN1,CALM3,GNB1,SHISA9,STMN2,RPS8,MYH9,RPL7,
TP2B4,IGFBP2,CALR,TMF1,HSPA1B,ZNF99,TUBB2B,COL5A1,COL4A1,PSAP,RPLP0,H3-3B,CANX,
IBIM6,PTMS,SPTAN1,GSTP1,MARCKS,SMG1,CALM1,PLXNB2,USP11,IGF2,EIF3C,EIF4B,MARCKS
ATP2B4,IGFBP2,CALR,FSTL1,TMF1,ABHD2,ZNF99,TUBB2B,COL5A1,COL4A1,CES3,PSAP,RPLP0
i1,CALM1,PLXNB2,USP11,SMIM14,IGF2,EIF4B,MARCKSL1,AFF4,TRIM28,MATR3,YWHAG,CKB,L
PTIN2,P4HB,GDI1,PTMS,SPTAN1,GSTP1,MARCKS,SMG1,CALM1,PLXNB2,USP11,IGF2,EIF3C,EIF4
A9,STMN2,MYH9,RPL7,P4HB,APP,IDS,GDI1,PTMS,SPTAN1,GSTP1,MARCKS,SMG1,CALM1,PLXNB
LM3,GNB1,STMN2,RPS8,MYH9,SEPTIN2,P4HB,GDI1,TMBIM6,PTMS,SPTAN1,SMG1,CALM1,PLX
1,TMF1,ABHD2,ZNF99,TUBB2B,COL5A1,COL4A1,CES3,PSAP,PGK1,RPLP0,H3-3B,CANX,CLU,PTM
COL18A1,SMIM14,IGF2,EIF3C,ATP5F1A,EIF4B,MARCKSL1,AFF4,TRIM28,MATR3,YWHAG,CKB,YW
F8,RPL8,FBXL4,YBX1,ATP2B4,IGFBP2,FSTL1,TMF1,ABHD2,HSPA1B,ZNF99,TUBB2B,COL5A1,CO
PTBN1,CALM3,GNB1,SHISA9,STMN2,RPS8,MYH9,RPL7,SEPTIN2,P4HB,APP,GDI1,PTMS,SPTAN1,G
P1,MARCKS,SMG1,PLXNB2,USP11,IGF2,EIF3C,ATP5F1A,EIF4B,MARCKSL1,AFF4,TRIM28,MATR
X5,ENO2,SOX4,FKBP14,SEC62,KAT6A,DVL3,YWHAE,KIF1A,GNAS,SLC2A3,GPI,RACK1,BSG,FASN,
STL1,ABHD2,TUBB2B,COL5A1,COL4A1,PSAP,PGK1,RPLP0,H3-3B,CANX,PTMA,DDX5,ENO2,FKBP
,PTMS,SPTAN1,GSTP1,MARCKS,SMG1,CALM1,PLXNB2,USP11,IGF2,EIF3C,EIF4B,MARCKSL1,A
A1,IGFBP2,CALR,FSTL1,TMF1,ABHD2,TUBB2B,COL5A1,COL4A1,PGK1,RPLP0,H3-3B,CANX,PTM
N2,MYH9,RPL7,SEPTIN2,P4HB,APP,IDS,RPL5,GDI1,PTMS,SPTAN1,GSTP1,CALM1,NPM1,PLXNB2,U
),RPL7,SEPTIN2,P4HB,GDI1,PTMS,SPTAN1,GSTP1,MARCKS,SMG1,CALM1,PLXNB2,HSPD1,USP11
TRA4,BSG,FASN,SPTBN1,CALM3,GNB1,RPS8,MYH9,SEPTIN2,P4HB,APP,GDI1,FSCN1,TMBIM6,P
1,CES3,H3-3B,CLU,PTMA,DDX5,ENO2,SOX4,FKBP14,SEC62,KAT6A,DVL3,YWHAE,KIF1A,RPL13,G
IB1,SHISA9,STMN2,RPL7,SEPTIN2,RPL5,GDI1,TMBIM6,SPTAN1,GSTP1,MARCKS,SMG1,NPM1,PL
A1,FTL,HSP90B1,FADS1,NEFL,NEFM,PRPF8,UGT8,RPL8,FBXL4,YBX1,ATP2B4,IGFBP2,CALR,FST
LR,FSTL1,TMF1,ABHD2,HSPA1B,COL5A1,COL4A1,PGK1,RPLP0,H3-3B,CANX,CLU,PTMA,DDX5,E
TP2B4,IGFBP2,CALR,FSTL1,TMF1,HSPA1B,ZNF99,TUBB2B,COL5A1,COL4A1,CES3,PSAP,PGK1,F
RA4,BSG,FASN,ALDH6A1,SPTBN1,GNB1,SHISA9,STMN2,MYH9,RPL7,APP,IDS,GDI1,PTMS,SPTAN1
M,PRPF8,UGT8,RPL8,FBXL4,YBX1,UBA1,IGFBP2,FSTL1,ABHD2,HSPA1B,TUBB2B,COL5A1,COL4
2,FSTL1,TMF1,ABHD2,HSPA1B,ZNF99,TUBB2B,COL5A1,COL4A1,CES3,PSAP,PGK1,RPLP0,H3-3
DX5,ENO2,SOX4,FKBP14,RPL6,KAT6A,DVL3,KIF1A,RPL13,GNAS,SLC2A3,GPI,RACK1,BSG,FASN,
DDX17,BSG,FASN,ALDH6A1,SPTBN1,GNB1,SHISA9,STMN2,MYH9,RPL7,SEPTIN2,P4HB,IDS,GDI1,
PTAN1,GSTP1,MARCKS,SMG1,CALM1,PLXNB2,USP11,IGF2,EIF3C,EIF4B,MARCKSL1,AFF4,TRIM
M,GANAB,COL6A1,FTL,HSP90B1,FADS1,NEFL,NEFM,PRPF8,RPL8,FBXL4,YBX1,UBA1,ATP2B4,IG
S,SLC2A3,FASN,SPTBN1,CALM3,GNB1,SHISA9,STMN2,RPL7,SEPTIN2,GDI1,PTMS,SPTAN1,GSTP
A3,GPI,RACK1,BSG,FASN,ALDH6A1,SPTBN1,CALM3,GNB1,SHISA9,STMN2,RPS8,MYH9,RPL7,SE
A3,GPI,RACK1,BSG,FASN,ALDH6A1,SPTBN1,CALM3,GNB1,SHISA9,STMN2,RPS8,MYH9,RPL7,SE
PL7,SEPTIN2,P4HB,APP,IDS,RPL5,GDI1,PTMS,SPTAN1,CALM1,NPM1,PLXNB2,USP11,CSDE1,SMI
KM,GANAB,COL6A1,FTL,HSP90B1,FADS1,NEFL,NEFM,PRPF8,UGT8,RPL8,FBXL4,YBX1,UBA1,IG
1,BSG,FASN,SPTBN1,ATP5F1B,GNB1,SHISA9,RPS8,MYH9,RPL7,P4HB,APP,GDI1,FSCN1,PTMS,G
BSG,FASN,ALDH6A1,SPTBN1,ATP5F1B,CALM3,GNB1,MYH9,SEPTIN2,P4HB,APP,PTMS,SPTAN1,G

GPI,RACK1,BSG,FASN,ALDH6A1,SPTBN1,CALM3,GNB1,SHISA9,STMN2,RPS8,MYH9,RPL7,SEPTII
MN2,RPS8,RPL7,SEPTIN2,APP,GDI1,PTMS,SPTAN1,GSTP1,MARCKS,SMG1,PLXNB2,HSPD1,USP11
MN2,RPL7,SEPTIN2,RPL5,FSCN1,SPTAN1,GSTP1,MARCKS,SMG1,CALM1,PLXNB2,USP11,IGF2,EII
S,SPTAN1,GSTP1,MARCKS,SMG1,CALM1,NPM1,PLXNB2,HSPD1,USP11,IGF2,EIF4B,MARCKSL1,
K1,CLU,PTMA,DDX5,ENO2,SOX4,KAT6A,DVL3,YWHAE,KIF1A,GNAS,SLC2A3,DDX17,GPI,BSG,FAS
3,PKM,GANAB,COL6A1,FTL,HSP90B1,FADS1,NEFL,NEFM,PRPF8,RPL8,FBXL4,YBX1,UBA1,ATP2B
TRIM28,MATR3,YWHAG,CKB,DDB1,YWHAQ,DCHS1,SET,CHSY1,TMSB10,INA,MRC2,PKD1,HLA-C
FASN,ALDH6A1,SPTBN1,CALM3,GNB1,STMN2,RPS8,MYH9,RPL7,SEPTIN2,P4HB,GDI1,SPTAN1,G
COL6A1,FTL,HSP90B1,FADS1,NEFM,PRPF8,UGT8,RPL8,FBXL4,YBX1,UBA1,ATP2B4,IGFBP2,CAL
SEC62,KAT6A,YWHAE,GNAS,BSG,FASN,ALDH6A1,SPTBN1,CALM3,GNB1,SHISA9,MYH9,RPL7,SEF
COL5A1,COL4A1,RPLP0,H3-3B,CANX,PTMA,DDX5,ENO2,MAGED1,SEC62,RPL6,KAT6A,DVL3,YWI
GPI,BSG,FASN,ALDH6A1,SPTBN1,CALM3,GNB1,STMN2,RPS8,MYH9,RPL7,SEPTIN2,P4HB,GDI1,S
)S,PTMS,CALM1,PLXNB2,USP11,SMIM14,IGF2,ATP5F1A,EIF4B,MARCKSL1,UBB,AFF4,TRIM28,M
DH6A1,SPTBN1,GNB1,SHISA9,STMN2,RPL7,SEPTIN2,RPL5,GDI1,TMBIM6,SPTAN1,GSTP1,MARC
AB,COL6A1,FTL,HSP90B1,FADS1,NEFL,NEFM,PRPF8,UGT8,RPL8,FBXL4,YBX1,UBA1,ATP2B4,IGF
LM3,GNB1,SHISA9,MYH9,SEPTIN2,P4HB,GDI1,FSCN1,TMBIM6,PTMS,SPTAN1,GSTP1,CALM1,NF
,HACD3,DCHS1,TMSB10,MRC2,LDHB,PKD1,HLA-C,SSBP2,VAPA,HMGB1,POLR2A,TPT1,ITGB1,TU
HD2,HSPA1B,TUBB2B,COL5A1,COL4A1,PSAP,PGK1,CANX,PTMA,DDX5,ENO2,SOX4,FKBP14,SEC
RPS8,MYH9,P4HB,IDS,GDI1,FSCN1,PTMS,SPTAN1,SMG1,PLD3,HSPD1,IGF2,EIF3C,EIF4B,UBB,AI
PLXNB2,IGF2,ATP5F1A,EIF4B,AFF4,TRIM28,MATR3,YWHAG,CKB,YWHAQ,HACD3,DCHS1,SET,EG
3,CALM3,GNB1,SHISA9,MYH9,RPL7,SEPTIN2,P4HB,GDI1,FSCN1,TMBIM6,PTMS,SPTAN1,GSTP1,N
28,MATR3,YWHAG,CKB,COL4A2,LRRC28,NONO,PSMD5,YWHAQ,EIF1,HACD3,DCHS1,SET,CHSY1
0,PKD1,SSBP2,POLR2A,TPT1,SEC61A1,ITGB1,TUBB4B,AGRN,TIMP2,COL11A1,MIDN,MARCHF6,
34B,MORF4L1,ANXA2,AGRN,ZMIZ1,TIMP2,MIDN,MARCHF6,VCL,AP3D1,SBF1,ARF1,MAP4,CDC42
1,CLU,PTMA,DDX5,ENO2,FKBP14,KAT6A,DVL3,YWHAE,KIF1A,GNAS,SLC2A3,DDX17,GPI,HTRA4,E
GB1,TUBB4B,AGRN,TIMP2,MIDN,MARCHF6,AP3D1,DNAJA1,ARF1,SF3B2,CDC42BPB,ITM2B,MAT
C62,KAT6A,DVL3,YWHAE,KIF1A,RPL13,GNAS,HTRA4,RACK1,BSG,FASN,SPTBN1,ATP5F1B,CALM
H3-3B,CLU,PTMA,DDX5,ENO2,SOX4,RPL13A,MAGED1,SEC62,KAT6A,DVL3,YWHAE,KIF1A,GNAS,I
SOX4,RPL6,KAT6A,DVL3,YWHAE,KIF1A,GNAS,SLC2A3,DDX17,GPI,BSG,ALDH6A1,SPTBN1,CALM
4A1,CES3,PSAP,PTMA,DDX5,ENO2,RPL13A,MAGED1,SEC62,KAT6A,DVL3,YWHAE,GNAS,DDX17,C
L,HSP90B1,FADS1,NEFL,NEFM,PRPF8,RPL8,FBXL4,YBX1,UBA1,ATP2B4,IGFBP2,CALR,FSTL1,TM
3N1,ATP5F1B,GNB1,SHISA9,MYH9,RPL7,APP,RPL5,GDI1,FSCN1,PTMS,MARCKS,CALM1,PLD3,HS
HS1,TMSB10,MRC2,LDHB,PKD1,HLA-C,SSBP2,BASP1,VAPA,POLR2A,TPT1,RPL18,STMN1,ITGB1,
,MYH9,RPL7,SEPTIN2,ERVV-1,GDI1,FSCN1,TMBIM6,SPTAN1,MARCKS,SMG1,PLXNB2,HSPD1,US
JBB2B,COL5A1,COL4A1,CES3,PSAP,PGK1,H3-3B,PTMA,ENO2,SOX4,DVL3,YWHAE,KIF1A,RPL13,
R,FSTL1,TMF1,ABHD2,HSPA1B,TUBB2B,COL5A1,COL4A1,PSAP,PGK1,RPLP0,H3-3B,CANX,CLU,F
XA2,AGRN,ZMIZ1,TIMP2,MIDN,WSB1,AP3D1,SBF1,CYB5R3,CDC42BPB,MAT2A,TMEM170A,PXDN
MS,SPTAN1,MARCKS,SMG1,CALM1,NPM1,PLXNB2,HSPD1,IGF2,EIF4B,MARCKSL1,UBB,TRIM28,
STMN1,ITGB1,UCHL1,AGRN,TIMP2,SARAF,COL11A1,MIDN,RPL10,MARCHF6,VCL,SBF1,CYB5R3,
,FASN,ALDH6A1,SPTBN1,CALM3,SHISA9,STMN2,MYH9,RPL7,SEPTIN2,P4HB,RPL5,FSCN1,SPTAN
LD3,HSPD1,USP11,COL18A1,SMIM14,IGF2,VCP,MARCKSL1,AFF4,TRIM28,MATR3,CKB,LRRC28,I
JBB,TRIM28,YWHAG,CKB,COL4A2,LRRC28,SPEN,DDB1,HACD3,DCHS1,SET,CHSY1,TMSB10,INA
SG,FASN,ALDH6A1,SPTBN1,CALM3,GNB1,STMN2,RPS8,MYH9,SEPTIN2,P4HB,GDI1,FSCN1,TMBI
DS,GDI1,FSCN1,GSTP1,MARCKS,SMG1,CALM1,PLXNB2,HSPD1,USP11,IGF2,EIF3C,ATP5F1A,EIF
HSP90B1,FADS1,NEFL,NEFM,PRPF8,RPL8,FBXL4,YBX1,UBA1,ATP2B4,IGFBP2,CALR,FSTL1,TMF1

NAS,DDX17,GPI,HTRA4,RACK1,BSG,FASN,SPTBN1,GNB1,SHISA9,RPS8,MYH9,RPL7,APP,GDI1,FS
ANXA2,RPLP1,ZMIZ1,SARAF,RPS17,SBF1,DNAJA1,ARF1,CYB5R3,CDC42BPB,ITM2B,HNRNPU,OG
8,YWHAG,CKB,COL4A2,LRRC28,FN1,SPEN,YWHAQ,DCHS1,SET,EGLN1,TPI1,CHSY1,TMSB10,EIF
,DDX5,ENO2,RPL13A,MAGED1,SEC62,KAT6A,DVL3,YWHAE,KIF1A,GNAS,DDX17,HTRA4,RACK1,B
K1,RPLP0,H3-3B,PTMA,DDX5,ENO2,SOX4,FKBP14,RPL13A,SEC62,RPL6,KAT6A,DVL3,YWHAE,KIF
A1,COL4A1,PGK1,H3-3B,PTMA,ENO2,SOX4,RPL6,KAT6A,DVL3,YWHAE,KIF1A,GNAS,SLC2A3,DDX
TMS,GSTP1,MARCKS,SMG1,CALM1,NPM1,PLXNB2,PLD3,USP11,SMIM14,IGF2,EIF3C,ATP5F1A,
FASN,ALDH6A1,SPTBN1,CALM3,GNB1,STMN2,MYH9,RPL7,SEPTIN2,P4HB,GDI1,FSCN1,TMBIM6,
BP2,CALR,FSTL1,ABHD2,HSPA1B,TUBB2B,COL5A1,COL4A1,PSAP,PGK1,RPLP0,H3-3B,CANX,CL
NEFM,PRPF8,RPL8,FBXL4,YBX1,UBA1,ATP2B4,IGFBP2,CALR,FSTL1,ABHD2,TUBB2B,COL5A1,CC
ACD3,SET,EGLN1,TPI1,CHSY1,MRC2,PPP2R1A,RPL15,GTF3C4,LDHB,PKD1,PDIA3,ZNF43,RPS3,E
LA-C,PDIA3,SSBP2,VAPA,POLR2A,TPT1,STMN1,ITGB1,TUBB4B,NCAM1,ANXA2,AGRN,TIMP2,SAR
TMN1,ITGB1,TUBB4B,MORF4L1,ANXA2,AGRN,TIMP2,SARAF,MIDN,MARCHF6,VCL,AP3D1,ARF1,
N,ALDH6A1,SPTBN1,ATP5F1B,CALM3,GNB1,SHISA9,MYH9,RPL7,SEPTIN2,P4HB,APP,GDI1,TMBI
G,FASN,ALDH6A1,SPTBN1,GNB1,STMN2,RPS8,RPL7,SEPTIN2,IDS,RPL5,GDI1,FSCN1,PTMS,SPTA
X,CLU,PTMA,DDX5,ENO2,SOX4,SEC62,KAT6A,DVL3,YWHAE,KIF1A,RPL13,GNAS,SLC2A3,DDX17,
2,SSBP2,VAPA,AP2M1,STMN1,ITGB1,TUBB4B,UCHL1,MORF4L1,NCAM1,AGRN,TIMP2,COL11A1,
CHS1,SET,TMSB10,INA,MRC2,LDHB,PKD1,SSBP2,BASP1,VAPA,HMGB1,POLR2A,TPT1,ITGB1,UCH
8,UGT8,RPL8,FBXL4,YBX1,UBA1,IGFBP2,CALR,FSTL1,TMF1,ABHD2,HSPA1B,ZNF99,TUBB2B,CC
COL5A1,COL4A1,PSAP,PGK1,RPLP0,H3-3B,CANX,CLU,PTMA,DDX5,ENO2,SOX4,SEC62,KAT6A,DV
JBB2B,COL5A1,COL4A1,CES3,PSAP,PGK1,RPLP0,H3-3B,CANX,PTMA,ENO2,FKBP14,SEC62,RPL
LM3,GNB1,SHISA9,MYH9,RPL7,SEPTIN2,APP,GDI1,TMBIM6,PTMS,SPTAN1,GSTP1,MARCKS,SMG
PL6,KAT6A,KIF1A,RPL13,GNAS,GPI,HTRA4,RACK1,BSG,FASN,SPTBN1,ATP5F1B,GNB1,SHISA9,RF
NR2L8,PKM,GANAB,COL6A1,FTL,HSP90B1,FADS1,NEFL,NEFM,PRPF8,UGT8,RPL8,FBXL4,YBX1,U
F3C4,LDHB,DNAH8,PKD1,RPS3,HMGB1,RPS3A,ITGB1,TUBB4B,AGRN,ZMIZ1,MARCHF6,ARF1,CD
CKSL1,UBB,AFF4,TRIM28,MATR3,YWHAG,CKB,COL4A2,RPL37A,LRRC28,FN1,YWHAQ,EIF1,HAC
,ZNF43,VAPA,AP2M1,TPT1,STMN1,ITGB1,TUBB4B,MORF4L1,AGRN,AP3D1,ARF1,SLC4A8,PXDN,
L8,FBXL4,YBX1,UBA1,ATP2B4,IGFBP2,CALR,TMF1,HSPA1B,TUBB2B,COL5A1,COL4A1,CES3,PS
EIF4B,MARCKSL1,TRIM28,YWHAG,CKB,LRRC28,YWHAQ,HACD3,DCHS1,TMSB10,RPL15,LDHB,P
,FSTL1,TMF1,ABHD2,ZNF99,TUBB2B,COL4A1,PSAP,PGK1,H3-3B,CANX,CLU,PTMA,ENO2,RPL13
N1,MORF4L1,NCAM1,AGRN,RPLP1,ZMIZ1,VCL,DNAJA1,CDC42BPB,GOLGA8A,PXDN,HNRNPA0,F
TRIM28,YWHAG,CKB,LRRC28,NONO,DDB1,HACD3,DCHS1,SET,CHSY1,TMSB10,LDHB,PKD1,HLA
,STMN2,RPS8,MYH9,SEPTIN2,P4HB,RPL5,GDI1,FSCN1,PTMS,SPTAN1,GSTP1,SMG1,CALM1,PLD
B,GDI1,FSCN1,PTMS,SPTAN1,GSTP1,MARCKS,SMG1,CALM1,NPM1,PLXNB2,HSPD1,COL18A1,C
,RPLP0,H3-3B,CANX,CLU,PTMA,DDX5,ENO2,FKBP14,RPL13A,SEC62,RPL6,KAT6A,DVL3,YWHAE,
F4L1,AGRN,TIMP2,MIDN,MARCHF6,VCL,DNAJA1,ARF1,CDC42BPB,ITM2B,PXDN,HNRNPU,CTNNA
,ITGB1,TUBB4B,NCAM1,ANXA2,AGRN,TIMP2,SARAF,MIDN,MARCHF6,VCL,AP3D1,SBF1,CYB5R3,
,KIF1A,GNAS,SLC2A3,DDX17,GPI,BSG,FASN,ALDH6A1,SPTBN1,CALM3,GNB1,SHISA9,STMN2,M
1,PLXNB2,USP11,IGF2,ATP5F1A,EIF4B,MARCKSL1,AFF4,TRIM28,MATR3,YWHAG,CKB,FN1,DDE
LD3,HSPD1,USP11,SNRNP200,CSDE1,SMIM14,IGF2,EIF3C,EIF4B,VCP,TRIM28,YWHAG,CKB,LRR
KAT6A,DVL3,YWHAE,RPL13,GNAS,DDX17,HTRA4,RACK1,BSG,FASN,ALDH6A1,ATP5F1B,CALM3,G
GA,CTNNA1,FADS2,BRD3,CIPC,PJA2,KPNB1,NCS1,ZNF865,NLGN2,CXADR,CHD4,JUND,WDR1,H
YH9,RPL7,SEPTIN2,P4HB,TMBIM6,PTMS,SPTAN1,GSTP1,SMG1,CALM1,PLXNB2,PLD3,HSPD1,CC
YH9,RPL7,SEPTIN2,P4HB,TMBIM6,PTMS,SPTAN1,GSTP1,SMG1,CALM1,PLXNB2,PLD3,HSPD1,CC
YH9,RPL7,SEPTIN2,P4HB,TMBIM6,PTMS,SPTAN1,GSTP1,SMG1,CALM1,PLXNB2,PLD3,HSPD1,CC

)1,RPL6,DVL3,GNAS,DDX17,HTRA4,BSG,FASN,ALDH6A1,ATP5F1B,CALM3,RPS8,RPL7,SEPTIN2,P
'TAN1,GSTP1,CALM1,PLXNB2,HSPD1,USP11,CSDE1,IGF2,ATP5F1A,EIF4B,MARCKSL1,UBB,AFF4,
'SAP,PGK1,RPLP0,H3-3B,CANX,PTMA,DDX5,ENO2,RPL13A,MAGED1,RPL6,DVL3,RPL13,GNAS,DD
5F1A,EIF4B,MARCKSL1,UBB,MATR3,YWHAG,CKB,COL4A2,RPL37A,NONO,YWHAQ,HACD3,DCHS
I,ZMIZ1,TIMP2,MIDN,MARCHF6,VCL,DNAJA1,CDC42BPB,ITM2B,PXDN,HNRNPU,CTNNA1,NACA,B
COL6A1,FTL,HSP90B1,FADS1,NEFM,PRPF8,UGT8,RPL8,FBXL4,YBX1,UBA1,ATP2B4,IGFBP2,CAL
DE1,IGF2,UBB,AFF4,TRIM28,MATR3,YWHAG,CKB,LRRC28,DDB1,YWHAQ,HACD3,DCHS1,CHSY1
'TMS,SPTAN1,GSTP1,SMG1,CALM1,NPM1,HSPD1,COL18A1,SNRNP200,SMIM14,IGF2,EIF4B,VCF
SPTBN1,ATP5F1B,CALM3,GNB1,SHISA9,MYH9,RPL7,SEPTIN2,APP,IDS,ERVV-1,RPL5,FSCN1,PTM
C2,PKD1,HLA-C,VAPA,POLR2A,AP2M1,TPT1,ITGB1,TUBB4B,UCHL1,MORF4L1,NCAM1,ANXA2,AG
1A2,LLGL1,NCS1,ZNF865,CLSTN1,SLC7A5,ENC1,JUND,ATP2A2,CRMP1,LASP1,OAZ1,ASTN2,NIS
OL5A1,COL4A1,PSAP,PGK1,RPLP0,CANX,CLU,PTMA,DDX5,SOX4,RPL13A,MAGED1,RPL6,KAT6A,
S3,PSAP,PGK1,RPLP0,H3-3B,CLU,PTMA,DDX5,ENO2,RPL13A,SEC62,RPL6,KAT6A,DVL3,YWHAE,K
,TIMP2,MIDN,MARCHF6,VCL,DNAJA1,ARF1,CYB5R3,CDC42BPB,ITM2B,HNRNPU,PGAM1,CTNNA
ALM3,GNB1,SHISA9,RPS8,MYH9,RPL7,SEPTIN2,P4HB,APP,IDS,GDI1,FSCN1,PTMS,SPTAN1,SMG1
B,CLU,PTMA,DDX5,SOX4,FKBP14,RPL13A,MAGED1,SEC62,RPL6,KAT6A,DVL3,GNAS,SLC2A3,DD
B1,FADS1,NEFL,NEFM,PRPF8,RPL8,FBXL4,YBX1,UBA1,ATP2B4,IGFBP2,CALR,FSTL1,TMF1,ABHD
,TPI1,CHSY1,TMSB10,GTF3C4,LDHB,SSBP2,BASP1,VAPA,CDKL1,TPT1,STMN1,ITGB1,ANXA2,AG
2,CALR,FSTL1,TMF1,HSPA1B,ZNF99,TUBB2B,COL5A1,COL4A1,CES3,PSAP,RPLP0,H3-3B,CANX,
OL18A1,SNRNP200,CSDE1,SMIM14,IGF2,EIF4B,VCP,MARCKSL1,TRIM28,MATR3,YWHAG,CKB,L
NF865,CLSTN1,CHD4,ATP1A1,JUND,EIF5,LENG8,HMGN2,ATP2A2,CRMP1,LASP1,VAT1,OAZ1,PFN
B2B,COL5A1,COL4A1,CES3,PGK1,RPLP0,H3-3B,CANX,CLU,PTMA,DDX5,ENO2,MAGED1,SEC62,K
I1,MARCKS,SMG1,CALM1,NPM1,PLXNB2,USP11,IGF2,ATP5F1A,MARCKSL1,AFF4,MATR3,YWHAG
1,HLA-C,PDIA3,VAPA,POLR2A,STMN1,UCHL1,ANXA2,AGRN,TIMP2,SARAF,MIDN,AP3D1,SBF1,CYB
,USP11,IGF2,ATP5F1A,EIF4B,MARCKSL1,UBB,TRIM28,CKB,NONO,YWHAQ,HACD3,DCHS1,SET,T
KIF1A,SLC2A3,GPI,BSG,FASN,ALDH6A1,SPTBN1,CALM3,GNB1,STMN2,RPS8,MYH9,RPL7,SEPTIN
B1,TUBB4B,MORF4L1,NCAM1,ANXA2,AGRN,ZMIZ1,MIDN,WSB1,AP3D1,SBF1,ARF1,CYB5R3,MAF
A1,PSAP,PGK1,CANX,CLU,PTMA,DDX5,ENO2,SOX4,RPL13A,MAGED1,RPL6,YWHAE,KIF1A,RPL13
6,KAT6A,DVL3,YWHAE,KIF1A,GNAS,SLC2A3,DDX17,GPI,BSG,FASN,CALM3,GNB1,RPS8,MYH9,RF
I1,CALM3,GNB1,SHISA9,MYH9,RPL7,SEPTIN2,APP,IDS,RPL5,FSCN1,PTMS,SMG1,NPM1,PLXNB2,
SCN1,TMBIM6,PTMS,SMG1,CALM1,PLD3,USP11,IGF2,EIF3C,EIF4B,MARCKSL1,UBB,MATR3,CK
NPM1,PLXNB2,USP11,COL18A1,EIF4B,MARCKSL1,UBB,TRIM28,YWHAG,CKB,COL4A2,FN1,SPEN
,TPI1,CHSY1,MRC2,RPL15,GTF3C4,PKD1,HLA-C,PDIA3,SSBP2,ZNF43,TPT1,ITGB1,TUBB4B,MO
NX,PTMA,DDX5,ENO2,SOX4,RPL13A,MAGED1,SEC62,RPL6,KAT6A,DVL3,YWHAE,KIF1A,GNAS,DD
,GNAS,FASN,ALDH6A1,CALM3,SHISA9,STMN2,MYH9,SEPTIN2,RPL5,PTMS,SPTAN1,GSTP1,MARC
9A7,ALDOA,MTRNR2L8,PKM,GANAB,COL6A1,FTL,HSP90B1,NEFL,NEFM,PRPF8,UGT8,RPL8,FBX
F3,SARAF,RPS17,SBF1,DNAJA1,ARF1,CDC42BPB,ITM2B,GTF2F1,HNRNPU,PGAM1,FADS2,CNN3,
IRNPU,CTNNA1,CNN3,CTSD,RPS24,SLC22A17,BRD3,KPNB1,HK1,NCS1,RHOA,ZNF865,SLC7A5,C
GAM1,CNN3,CTSD,SCARB2,RPS24,SLC22A17,BRD3,LLGL1,NCS1,RHOA,ZNF865,KIF5B,SLC7A5,
GANAB,COL6A1,FTL,HSP90B1,FADS1,NEFL,NEFM,PRPF8,RPL8,FBXL4,YBX1,UBA1,IGFBP2,CALR,
1,SHISA9,STMN2,MYH9,SEPTIN2,GDI1,FSCN1,TMBIM6,PTMS,GSTP1,SMG1,PLXNB2,HSPD1,USP
L13,GNAS,DDX17,HTRA4,FASN,ALDH6A1,CALM3,GNB1,STMN2,RPS8,SEPTIN2,P4HB,APP,RPL5,K
SCN1,PTMS,SMG1,CALM1,PLXNB2,PLD3,USP11,SNRNP200,SMIM14,IGF2,EIF3C,MARCKSL1,UBE
HAG,CKB,LRRC28,NONO,DDB1,HACD3,SET,EGLN1,CHSY1,TMSB10,RPL19,PPP2R1A,LDHB,PKD1
)L4A1,PSAP,PGK1,H3-3B,CANX,CLU,PTMA,DDX5,ENO2,RPL13A,SEC62,KAT6A,YWHAE,KIF1A,GNA

5F1B,CALM3,GNB1,MYH9,RPL7,SEPTIN2,P4HB,IDS,GDI1,FSCN1,TMBIM6,SPTAN1,GSTP1,SMG1

,DDX17,HTRA4,BSG,FASN,ATP5F1B,CALM3,GNB1,MYH9,RPL7,SEPTIN2,P4HB,IDS,GDI1,FSCN1,T

RCKSL1,AFF4,SPEN,PSMD5,DCHS1,SET,TPI1,CHSY1,TMSB10,INA,MRC2,PPP2R1A,HLA-C,SSBP2

AG,LRRC28,FN1,YWHAQ,TMSB10,MRC2,GTF3C4,PKD1,HLA-C,SERINC1,HMGB1,POLR2A,TPT1,T

,MYH9,SEPTIN2,P4HB,APP,IDS,ERVV-1,TMBIM6,SPTAN1,CALM1,PLD3,HSPD1,COL18A1,SNRNP2

,VAPA,ITGB1,AGRN,COL11A1,MARCHF6,VCL,SBF1,DNAJA1,ARF1,CYB5R3,CDC42BPB,MAT2A,TM

,SPTBN1,ATP5F1B,CALM3,GNB1,SHISA9,RPS8,MYH9,P4HB,IDS,GDI1,FSCN1,SPTAN1,GSTP1,MA

RNP200,IGF2,EIF4B,MARCKSL1,TRIM28,YWHAG,CKB,LRRC28,FN1,NONO,DDB1,YWHAQ,HACD

ICAM1,ANXA2,AGRN,TIMP2,SARAF,MIDN,VCL,AP3D1,CYB5R3,MAP4,CDC42BPB,ITM2B,MAT2A,P

ICAM1,ANXA2,AGRN,TIMP2,SARAF,MIDN,VCL,AP3D1,CYB5R3,MAP4,CDC42BPB,ITM2B,MAT2A,P

1,STIP1,TOP2B,SLC35E2B,HDAC2,CTDSP2,NOVA2,CDH11,CCT2,HNRNPL,ABCA3,HNRNPA3,ACT

A1,ARF1,CDC42BPB,ZNF770,TMEM170A,GTF2F1,PXDN,OGA,PGAM1,SMARCA5,HNRNPA0,FADS

NEFL,NEFM,PRPF8,RPL8,FBXL4,YBX1,ATP2B4,IGFBP2,FSTL1,TMF1,ABHD2,HSPA1B,ZNF99,TUB

SA9,MYH9,APP,IDS,GDI1,TMBIM6,PTMS,SPTAN1,SMG1,PLXNB2,USP11,SMIM14,IGF2,MARCKSL

RF4L1,NCAM1,AGRN,COL11A1,MIDN,MARCHF6,AP3D1,ARF1,CDC42BPB,ITM2B,PXDN,HNRNPU

N,HNRNPU,CTNNA1,CTSD,LPCAT1,LAMA5,BRD3,CIPC,KPNB1,C1ORF43,LLGL1,NCS1,ZNF865,AT

AGRN,TIMP2,SARAF,MIDN,AP3D1,SBF1,ARF1,CYB5R3,MAP4,CDC42BPB,ITM2B,MAT2A,PXDN,HN

PLP1,ZMIZ1,SARAF,MIDN,VCL,AP3D1,SBF1,ARF1,CYB5R3,MAP4,SF3B2,CDC42BPB,ITM2B,ZNF7

VHAE,RPL13,DDX17,RACK1,FASN,ALDH6A1,SPTBN1,ATP5F1B,SHISA9,STMN2,RPS8,MYH9,RPL7,

,RACK1,FASN,ALDH6A1,SPTBN1,ATP5F1B,RPS8,SEPTIN2,P4HB,APP,GDI1,PTMS,SPTAN1,CALM1,

6A,DVL3,YWHAE,RPL13,GNAS,DDX17,GPI,HTRA4,RACK1,BSG,FASN,ALDH6A1,SPTBN1,ATP5F1B,

CANX,CLU,PTMA,DDX5,ENO2,SOX4,SEC62,KAT6A,DVL3,YWHAE,RPL13,GNAS,DDX17,HTRA4,FAS

B,TUBB2B,COL4A1,PSAP,PGK1,RPLP0,H3-3B,CANX,PTMA,DDX5,ENO2,SOX4,RPL13A,MAGED1,S

PB,ITM2B,PXDN,OGA,CTNNA1,CTSD,NACA,LAMA5,CIPC,KPNB1,C1ORF43,RHOA,CLSTN1,NLGN2

CD3,DCHS1,EGLN1,CHSY1,INA,LDHB,PKD1,HLA-C,VAPA,AP2M1,TPT1,RPL18,STMN1,ITGB1,TUB

H6A1,SPTBN1,CALM3,SHISA9,STMN2,RPL7,SEPTIN2,IDS,GDI1,FSCN1,PTMS,SPTAN1,GSTP1,MA

F1A,RPL13,GNAS,DDX17,HTRA4,BSG,FASN,SPTBN1,GNB1,MYH9,RPL7,SEPTIN2,P4HB,APP,FSCN

P200,SMIM14,IGF2,EIF3C,EIF4B,MARCKSL1,UBB,TRIM28,CKB,COL4A2,DDB1,YWHAQ,HACD3,SE

,NLGN2,CXADR,CHD4,JUND,EIF5,HMGN2,ATP2A2,CRMP1,LASP1,OAZ1,ASTN2,TLE5,NISCH,CCN

IGF2,EIF4B,MARCKSL1,TRIM28,MATR3,YWHAG,CKB,COL4A2,MAPT,FN1,SPEN,YWHAQ,EIF1,HA

RPL8,FBXL4,YBX1,UBA1,ATP2B4,IGFBP2,CALR,FSTL1,ABHD2,HSPA1B,ZNF99,TUBB2B,COL5A1,

N,AP3D1,SBF1,ARF1,CYB5R3,MAP4,CDC42BPB,ITM2B,MAT2A,PXDN,HNRNPU,CTNNA1,CNN3,CT

APA,HMGB1,POLR2A,TPT1,SEC61A1,RPS3A,TUBB4B,UCHL1,ANXA2,AGRN,MIDN,AP3D1,ARF1,C

F,MARCHF6,SBF1,MAP4,CDC42BPB,ITM2B,GOLGA8A,HNRNPU,HNRNPA0,CTNNA1,RPS16,RPSA,

TGB1,TUBB4B,RPL11,ANXA2,AGRN,ZMIZ1,SARAF,MIDN,WSB1,RPS17,ARF1,CYB5R3,MAP4,CDC

DH6A1,ATP5F1B,RPS8,SEPTIN2,P4HB,RPL5,GDI1,GSTP1,NPM1,PLXNB2,PLD3,USP11,COL18A1,

4A2,RPL37A,MAPT,FN1,YWHAQ,DCHS1,SET,TPI1,CHSY1,TMSB10,MRC2,RPL19,GTF3C4,PKD1,H

NA,MRC2,LDHB,PKD1,HLA-C,SERINC1,VAPA,AP2M1,TPT1,STMN1,ITGB1,TUBB4B,MORF4L1,BT

,FSTL1,TMF1,ABHD2,HSPA1B,ZNF99,TUBB2B,COL5A1,COL4A1,PSAP,PGK1,RPLP0,H3-3B,CANX

S8,MYH9,SEPTIN2,APP,GDI1,TMBIM6,PTMS,SPTAN1,GSTP1,MARCKS,SMG1,CALM1,PLD3,HSPD1

7,GPI,BSG,FASN,SPTBN1,GNB1,SHISA9,RPS8,MYH9,P4HB,APP,IDS,GDI1,PTMS,SPTAN1,GSTP1,C

GNAS,SLC2A3,GPI,HTRA4,BSG,FASN,ALDH6A1,SPTBN1,GNB1,SHISA9,RPS8,SEPTIN2,APP,IDS,GD

,BRD3,KPNB1,PAPOLA,IGFBP3,LLGL1,HK1,NCS1,RPL22,CLSTN1,ENC1,NLGN2,JUND,CD81,LEN

UCHL1,ANXA2,AGRN,XAF1,SARAF,MIDN,AP3D1,ARF1,CYB5R3,MAP4,SF3B2,CDC42BPB,GOLGA

MG1,NPM1,PLXNB2,USP11,CSDE1,IGF2,ATP5F1A,EIF4B,MARCKSL1,UBB,AFF4,YWHAG,CKB,FN1

3,GNAS,RACK1,BSG,FASN,SPTBN1,ATP5F1B,CALM3,GNB1,SHISA9,MYH9,RPL7,SEPTIN2,APP,IDS
R,FSTL1,TMF1,HSPA1B,TUBB2B,PGK1,H3-3B,CANX,PTMA,DDX5,SOX4,FKBP14,RPL13A,MAGED1
SPTAN1,GSTP1,MARCKS,SMG1,NPM1,PLD3,HSPD1,USP11,IGF2,EIF3C,VCP,MARCKSL1,UBB,TRI
,SEC62,DVL3,YWHAE,KIF1A,RPL13,GNAS,SLC2A3,DDX17,HTRA4,RACK1,FASN,ALDH6A1,SPTBN1
13,SLC2A3,GPI,RACK1,BSG,FASN,ALDH6A1,CALM3,GNB1,STMN2,RPS8,MYH9,SEPTIN2,P4HB,ER
IA3,HMGB1,AP2M1,STMN1,UCHL1,NCAM1,AGRN,TIMP2,SARAF,MIDN,RPS17,AP3D1,SBF1,DNAJ
HOA,ZNF865,SLC7A5,CHD4,ATP1A1,JUND,EIF5,CD81,HMGN2,ATP2A2,CRMP1,LASP1,OAZ1,PFN1
C,MARCKSL1,UBB,AFF4,TRIM28,MATR3,YWHAG,CKB,COL4A2,MAPT,SPEN,DDB1,EIF1,CHSY1,RP
HD2,HSPA1B,ZNF99,COL5A1,COL4A1,PSAP,RPLP0,CANX,CLU,PTMA,DDX5,ENO2,SOX4,RPL13A,M
GF2,EIF4B,VCP,MARCKSL1,UBB,TRIM28,YWHAG,COL4A2,DDB1,EIF1,DCHS1,SET,TPI1,CHSY1,TM
ATR3,YWHAG,CKB,COL4A2,SPEN,PSMD5,YWHAQ,HACD3,SET,CHSY1,TMSB10,EIF4A1,RPL15,LDH
YWHAQ,DCHS1,SET,TPI1,CHSY1,TMSB10,MRC2,GTF3C4,PKD1,HLA-C,PDIA3,SSBP2,ZNF43,CHD3
C28,FN1,SPEN,DDB1,PSMD5,YWHAQ,SET,CHSY1,TMSB10,EIF4A1,RPL19,RPL15,GTF3C4,LDHB,S
SA9,STMN2,MYH9,RPL7,SEPTIN2,P4HB,IDS,GDI1,TMBIM6,PTMS,GSTP1,MARCKS,SMG1,CALM1,
COL4A1,PGK1,H3-3B,PTMA,DDX5,SOX4,FKBP14,RPL13A,MAGED1,RPL6,DVL3,YWHAE,GNAS,GP
MP2,MIDN,MARCHF6,VCL,ARF1,ITM2B,TMEM170A,SLC4A8,HNRNPU,PGAM1,HNRNPA0,CTNNA1,
A,MAGED1,SEC62,RPL6,DVL3,KIF1A,GNAS,SLC2A3,DDX17,GPI,RACK1,BSG,ALDH6A1,CALM3,RP
14,IGF2,EIF3C,EIF4B,MARCKSL1,UBB,MATR3,CKB,LRRC28,FN1,DDB1,PSMD5,YWHAQ,EIF1,SE
ALR,FSTL1,HSPA1B,TUBB2B,COL5A1,COL4A1,CES3,PGK1,H3-3B,CANX,CLU,PTMA,DDX5,ENO2,
32,PLD3,USP11,COL18A1,SMIM14,IGF2,EIF3C,MARCKSL1,AFF4,MATR3,YWHAG,CKB,COL4A2,S
R3,YWHAG,COL4A2,LRRC28,SPEN,NONO,PSMD5,EIF1,DCHS1,SET,EGLN1,CHSY1,TMSB10,INA,E
1,ATP2B4,IGFBP2,CALR,FSTL1,TMF1,ABHD2,HSPA1B,TUBB2B,COL5A1,COL4A1,CES3,PSAP,PGK
PB,ITM2B,MAT2A,PXDN,HNRNPU,CTNNA1,RPS16,CNN3,CTSD,LPCAT1,BRD3,KPNB1,LLGL1,NCS
2,HSPA1B,ZNF99,TUBB2B,COL5A1,COL4A1,CES3,PSAP,PGK1,RPLP0,H3-3B,CANX,CLU,PTMA,DD
GTF2F1,HNRNPU,OGA,NFE2L1,SMARCA5,HNRNPA0,CNN3,LPCAT1,BRD3,PJA2,KPNB1,PAPOLA,R
TIN2,IDS,GDI1,SMG1,CALM1,PLXNB2,PLD3,HSPD1,USP11,COL18A1,SNRNP200,SMIM14,MARCK
TIN2,IDS,GDI1,SMG1,CALM1,PLXNB2,PLD3,HSPD1,USP11,COL18A1,SNRNP200,SMIM14,MARCK
1A,EIF4B,VCP,AFF4,RPL37A,NONO,EIF1,TPI1,INA,EIF4A1,RPL19,PPP2R1A,RPL15,GTF3C4,LDHB,
CHL1,ANXA2,AGRN,TIMP2,SARAF,COL11A1,MIDN,AP3D1,SBF1,CYB5R3,MAP4,SF3B2,CDC42BPB,
PJA2,KPNB1,COL1A2,NCS1,ZNF865,CLSTN1,ENC1,NLGN2,CHD4,ATP1A1,EIF5,WDR1,HMGN2,AT
PXDN,OGA,FADS2,LPCAT1,LAMA5,SLC22A17,BRD3,PJA2,SLC16A7,KPNB1,COL1A2,HK1,NCS1,R
STL1,TMF1,HSPA1B,ZNF99,TUBB2B,COL5A1,COL4A1,CES3,PSAP,RPLP0,H3-3B,CANX,CLU,PTMA
DDB1,YWHAQ,HACD3,DCHS1,CHSY1,EIF4A1,MRC2,PKD1,HLA-C,PDIA3,SSBP2,VAPA,CDKL1,TPT1,
HS1,SET,TMSB10,EIF4A1,MRC2,LDHB,SSBP2,CHD3,BASP1,POLR2A,TUBB4B,MORF4L1,ANXA2,A
SET,EGLN1,TPI1,CHSY1,PPP2R1A,GTF3C4,PKD1,SF3B1,PDIA3,ZNF43,VAPA,POLR2A,AP2M1,TPT1
RPL7,SEPTIN2,P4HB,APP,IDS,ERVV-1,GDI1,FSCN1,TMBIM6,PTMS,GSTP1,SMG1,CALM1,PLXNB2,
2R1A,LDHB,SF3B1,SSBP2,CHD3,BASP1,VAPA,TPT1,ITGB1,TUBB4B,MORF4L1,BTF3,AGRN,RPL1
LU,PTMA,DDX5,ENO2,SOX4,RPL6,KAT6A,DVL3,YWHAE,KIF1A,GNAS,SLC2A3,GPI,HTRA4,RACK1,
IMP2,SARAF,MIDN,VCL,CYB5R3,MAP4,CDC42BPB,ITM2B,AP2B1,MAT2A,PXDN,HNRNPU,CTNNA1
COL4A1,PSAP,PGK1,RPLP0,H3-3B,CANX,CLU,PTMA,DDX5,ENO2,FKBP14,RPL13A,SEC62,KAT6A,
5,KAT6A,DVL3,YWHAE,KIF1A,RPL13,GNAS,DDX17,BSG,SPTBN1,ATP5F1B,CALM3,GNB1,SHISA9,S
F4A1,RPL19,RPL15,GTF3C4,LDHB,PKD1,PDIA3,SERINC1,BASP1,VAPA,POLR2A,TPT1,SEC61A1,ST
A,EIF4H,NACA,RPL17,C1ORF43,RPL14,CBX5,ATF4,KIF5B,TMEM259,EIF5,HMGN2,RBBP4,PFN1,A
AFF4,MATR3,YWHAG,CKB,YWHAQ,HACD3,DCHS1,CHSY1,TMSB10,MRC2,PPP2R1A,GTF3C4,PKD1
GNAS,DDX17,GPI,BSG,FASN,ALDH6A1,SPTBN1,ATP5F1B,CALM3,GNB1,RPS8,APP,IDS,RPL5,FSCN

R,FSTL1,TMF1,ABHD2,HSPA1B,COL4A1,PGK1,H3-3B,CANX,PTMA,DDX5,SOX4,FKBP14,RPL13A,M
A3,CHD3,BASP1,VAPA,SEC61A1,STMN1,ITGB1,TUBB4B,UCHL1,AGRN,RPLP1,ZMIZ1,TIMP2,MIDN,
L7,APP,GDI1,FSCN1,PTMS,SPTAN1,MARCKS,SMG1,USP11,COL18A1,SNRNP200,CSDE1,IGF2,EIF
FN1,CCNI,TADA2B,KHDRBS1,EIF5A,MEGF8,ACTN4,PRDX2,RPS15,ITM2C,RPS5,DNAJC5,SLC25A3
GNAS,SLC2A3,DDX17,HTRA4,RACK1,FASN,SPTBN1,ATP5F1B,CALM3,STMN2,RPS8,RPL7,SEPTIN2
9,TUBB2B,COL5A1,COL4A1,PSAP,PGK1,RPLP0,CANX,CLU,PTMA,DDX5,ENO2,SOX4,RPL13A,MAG
A1,RPL19,PPP2R1A,DNAH8,PKD1,SF3B1,SSBP2,ZNF43,HMGB1,POLR2A,TPT1,SEC61A1,RPS3A,IT
SN,ATP5F1B,CALM3,GNB1,SHISA9,RPS8,MYH9,P4HB,RPL5,TMBIM6,PTMS,SPTAN1,CALM1,PLD3
7,BRD3,KPNB1,C1ORF43,LLGL1,NCS1,RHOA,CBX5,ZNF865,NR2F1,RAB6B,NLGN2,CHD4,JUND,
S,DDX17,GPI,RACK1,BSG,FASN,ALDH6A1,CALM3,GNB1,RPS8,RPL7,SEPTIN2,APP,IDS,RPL5,TMBI
A,UBB,AFF4,MATR3,YWHAG,CKB,COL4A2,NONO,YWHAQ,HACD3,DCHS1,CHSY1,TMSB10,MRC2,
F5,LENG8,HMGN2,CRMP1,BEX3,VAT1,PFN1,JPT2,TLE5,NISCH,CCNI,XPO1,EIF5A,PDIA6,SOX12,A
MAP4,CDC42BPB,ITM2B,PXDN,HNRNPU,CNN3,CTSD,LPCAT1,LAMA5,BRD3,CIPC,C1ORF43,LLGL
PSAP,PGK1,RPLP0,CANX,CLU,PTMA,DDX5,SOX4,RPL13A,MAGED1,RPL6,KAT6A,YWHAE,KIF1A,RF
A1B,TUBB2B,COL5A1,CES3,PSAP,PGK1,RPLP0,H3-3B,CANX,PTMA,DDX5,RPL13A,SEC62,RPL6,K
PL5,FSCN1,PTMS,SPTAN1,SMG1,CALM1,NPM1,PLXNB2,PLD3,SNRNP200,IGF2,EIF3C,EIF4B,VCP,
AT1,LAMA5,SLC22A17,BRD3,PAPOLA,COL1A2,LLGL1,NCS1,ZNF865,CLSTN1,RAB6B,CHD4,ATP1
TAN1,SMG1,NPM1,PLXNB2,USP11,COL18A1,SMIM14,IGF2,VCP,MARCKSL1,TRIM28,YWHAG,CO
JX4,YWHAB,CADM4,RHOB,CKAP4,PLPPR3,SEPTIN3,HECTD4,STIP1,CDK16,HDAC2,IPO5,TMEM30
EPTIN2,P4HB,TMBIM6,PTMS,SPTAN1,CALM1,PLD3,USP11,SNRNP200,SMIM14,EIF3C,ATP5F1A,E
A,RHOA,CBX5,ZNF865,KIF5B,RAB6B,GLIPR1L2,LENG8,ATXN7L3B,VAT1,SRSF4,RTN3,PFN1,CCN
F4L1,AGRN,TIMP2,XAF1,MIDN,MARCHF6,WSB1,AKAP12,MAT2A,SLC4A8,PXDN,OGA,CTNNA1,FAI
SPTAN1,CALM1,HSPD1,USP11,SNRNP200,CSDE1,IGF2,EIF3C,ATP5F1A,EIF4B,AFF4,MATR3,YWHA
IR2L8,PKM,GANAB,COL6A1,FTL,HSP90B1,FADS1,NEFL,NEFM,PRPF8,UGT8,RPL8,FBXL4,YBX1,U
LU,PTMA,DDX5,ENO2,SOX4,FKBP14,RPL13A,MAGED1,SEC62,RPL6,KAT6A,YWHAE,KIF1A,RPL13,
P2M1,TPT1,RPL18,STMN1,ITGB1,TUBB4B,BTF3,AGRN,TIMP2,SARAF,COL11A1,RPL10,RPS17,VC
PL15,GTF3C4,PKD1,RPS3,VAPA,POLR2A,AP2M1,SEC61A1,RPL18,STMN1,ITGB1,UCHL1,MORF4L
BN1,ATP5F1B,CALM3,GNB1,SHISA9,RPS8,MYH9,P4HB,IDS,GDI1,FSCN1,PTMS,GSTP1,MARCKS,S
A,STMN1,TUBB4B,UCHL1,ANXA2,AGRN,TIMP2,SARAF,MIDN,AP3D1,ARF1,CYB5R3,MAP4,CDC42E
I,ZMIZ1,TIMP2,MIDN,DNAJA1,CYB5R3,CDC42BPB,AP2B1,GOLGA8A,PXDN,NFE2L1,HNRNPA0,CT
B,UBB,TRIM28,YWHAG,CKB,COL4A2,FN1,SPEN,PSMD5,DCHS1,SET,TPI1,CHSY1,EIF4A1,MRC2,R
X,DDX5,FKBP14,RPL13A,MAGED1,SEC62,RPL6,KAT6A,DVL3,RPL13,GNAS,DDX17,GPI,RACK1,SP
H9,P4HB,APP,ERVV-1,GDI1,FSCN1,TMBIM6,GSTP1,SMG1,CALM1,NPM1,HSPD1,SNRNP200,IGF2,
ES3,PSAP,PGK1,RPLP0,H3-3B,CANX,CLU,PTMA,DDX5,RPL13A,SEC62,RPL6,KAT6A,DVL3,KIF1A,D
,AFF4,TRIM28,COL4A2,RPL37A,FN1,SPEN,NONO,DDB1,HACD3,SET,EGLN1,CHSY1,EIF4A1,MRC
H3-3B,DDX5,ENO2,SOX4,FKBP14,RPL6,KAT6A,KIF1A,GNAS,SLC2A3,DDX17,HTRA4,RACK1,BSG,F
BXL4,YBX1,UBA1,ATP2B4,IGFBP2,CALR,TMF1,ABHD2,HSPA1B,ZNF99,TUBB2B,COL5A1,COL4A1,
STMN2,RPL7,SEPTIN2,APP,SPTAN1,GSTP1,SMG1,CALM1,PLXNB2,PLD3,HSPD1,USP11,COL18A1,
A2,CRMP1,OAZ1,MCL1,TLE5,NISCH,CCNI,EIF5A,MEGF8,RPL12,GNB2,H1-0,RPS7,MIF,CHPF,COL
LDOA,MTRNR2L8,PKM,GANAB,COL6A1,FTL,HSP90B1,FADS1,NEFL,NEFM,PRPF8,RPL8,FBXL4,Y
RPL13,GNAS,DDX17,GPI,HTRA4,BSG,FASN,ALDH6A1,SPTBN1,CALM3,SHISA9,MYH9,RPL7,P4HB,
SP11,IGF2,EIF4B,VCP,MARCKSL1,UBB,MATR3,CKB,RPL37A,FN1,NONO,EIF1,MAP9,TPI1,CHSY1,
APT,FN1,DCHS1,SET,CHSY1,INA,MRC2,RPL19,PPP2R1A,GTF3C4,PDIA3,SSBP2,ZNF43,VAPA,HMG
0,WSB1,VCL,AP3D1,SBF1,CYB5R3,MAP4,ZNF770,SLC4A8,PGAM1,LDHA,TNS3,EIF4H,SCARB2,L
CDC42BPB,MAT2A,GOLGA8A,HNRNPU,PGAM1,CTNNA1,RPS16,RPL29,CTSD,SCARB2,RPS24,BR

1GN2,ATP2A2,OAZ1,PFN1,NISCH,TADA2B,GLG1,PDIA6,PIP4K2B,SOX12,ACTN4,ARL4C,APLP1,MI
DRBS1,PFKL,PNMA1,PDIA6,GPC3,MEGF8,SOX12,ARL4C,APLP1,MIF,GPAT4,CHPF,MYO5A,B4GA
,RHOA,ATF4,KIF5B,RAB6B,SLC7A5,ENC1,CXADR,EIF5,ATP2A2,RPL26,CRMP1,OAZ1,SRSF4,RTN
TMF1,HSPA1B,ZNF99,TUBB2B,COL5A1,COL4A1,PSAP,RPLP0,H3-3B,CANX,CLU,PTMA,DDX5,ENO
RPLP0,CANX,CLU,PTMA,DDX5,ENO2,MAGED1,SEC62,RPL6,KAT6A,DVL3,YWHAE,KIF1A,RPL13,DD
IUND,CD81,HMGN2,ATP2A2,CRMP1,LASP1,OAZ1,TLE5,NISCH,HNRNPC,CCNI,TADA2B,PDIA6,PIP4
C,PJA2,LLGL1,ZNF865,PSMD2,SLC7A5,CHD4,ATP1A1,JUND,CD81,WDR1,HMGN2,ATP2A2,CRMF
DDX5,FKBP14,RPL13A,MAGED1,RPL6,KAT6A,DVL3,YWHAE,RPL13,GNAS,DDX17,GPI,HTRA4,RAC
IRNPA0,RPSA,TPM4,CTSD,NACA,LAMA5,KPNB1,COL1A2,NCS1,RPL22,RHOA,ATF4,PSMD2,KIF5B,
IRNPA0,RPSA,TPM4,CTSD,NACA,LAMA5,KPNB1,COL1A2,NCS1,RPL22,RHOA,ATF4,PSMD2,KIF5B,
IRNPA0,RPSA,TPM4,CTSD,NACA,LAMA5,KPNB1,COL1A2,NCS1,RPL22,RHOA,ATF4,PSMD2,KIF5B,
PM1,PLXNB2,PLD3,USP11,SNRNP200,IGF2,EIF3C,VCP,MARCKSL1,AFF4,TRIM28,YWHAG,CKB,R
NA1,FADS2,CTSD,LPCAT1,BRD3,MSN,PAPOLA,HK1,NCS1,RHOA,CLSTN1,ENC1,NLGN2,CXADR,J
KIF1A,GNAS,SLC2A3,DDX17,GPI,HTRA4,BSG,FASN,ALDH6A1,SPTBN1,ATP5F1B,CALM3,GNB1,MY
.4A1,PGK1,RPLP0,CANX,CLU,PTMA,DDX5,ENO2,SOX4,RPL13A,MAGED1,SEC62,RPL6,KAT6A,DV
PL13A,MAGED1,SEC62,RPL6,KAT6A,DVL3,RPL13,GNAS,DDX17,GPI,RACK1,FASN,SPTBN1,ATP5F
HD2,HSPA1B,COL4A1,PSAP,PGK1,H3-3B,CANX,CLU,PTMA,DDX5,ENO2,SOX4,RPL13A,MAGED1,
ALNT4,DNAJC5,APLP2,TARDBP,MXD4,CADM4,HNRNPH1,PLPPR3,CTBP1,GPC1,SMCHD1,IPO5,M
IRNP200,SMIM14,IGF2,EIF3C,MARCKSL1,UBB,AFF4,TRIM28,MATR3,YWHAG,CKB,COL4A2,SPEN
BF1,ARF1,MAP4,CDC42BPB,ITM2B,GOLGA8A,PXDN,CNN3,RPL29,TNS3,EIF4H,SLC22A17,BRD3,
FF4,MATR3,FN1,EIF1,DCHS1,SET,TMSB10,MRC2,GTF3C4,PDIA3,SERINC1,SSBP2,BASP1,AP2M1,
N1,UCHL1,MORF4L1,BTF3,ANXA2,AGRN,RPLP1,ZMIZ1,SARAF,MIDN,MARCHF6,DNAJA1,ARF1,ZN
MSB10,GTF3C4,PDIA3,SERINC1,VAPA,HMGB1,TPT1,STMN1,TUBB4B,MORF4L1,ANXA2,AGRN,X
CANX,CLU,PTMA,DDX5,ENO2,MAGED1,SEC62,RPL6,KAT6A,DVL3,YWHAE,KIF1A,RPL13,DDX17,G
DR1,HMGN2,ATP2A2,CRMP1,LASP1,OAZ1,PFN1,TLE5,NISCH,CCNI,KHDRBS1,CSNK1E,EIF5A,CC
3-3B,CANX,PTMA,DDX5,MAGED1,SEC62,RPL6,KAT6A,YWHAE,KIF1A,RPL13,GNAS,DDX17,GPI,HT
Y1,TMSB10,EIF4A1,SSBP2,RPL18,MORF4L1,ANXA2,AGRN,XAF1,MIDN,WSB1,VCL,SBF1,DNAJA1,
IAQ,EIF1,HACD3,DCHS1,SET,EGLN1,TMSB10,MRC2,RPL15,LDHB,PKD1,SSBP2,BASP1,HMGB1,A
10,EIF4A1,RPL15,LDHB,SERINC1,SSBP2,BASP1,VAPA,POLR2A,TPT1,STMN1,ITGB1,RPL11,MORF
,PTMA,DDX5,FKBP14,RPL13A,MAGED1,RPL6,DVL3,YWHAE,KIF1A,RPL13,GNAS,GPI,HTRA4,RAC
,RPL22,CLSTN1,ATF4,RAB6B,ENC1,NLGN2,JUND,LENG8,PFN1,TLE5,TADA2B,SPTBN2,CSNK1E,
.15,GTF3C4,LDHB,DNAH8,PKD1,SF3B1,PDIA3,ZNF43,HMGB1,POLR2A,TPT1,SEC61A1,RPS4X,R
,TRIM28,COL4A2,LRRC28,FN1,SPEN,NONO,YWHAQ,EIF1,HACD3,DCHS1,SET,EGLN1,TPI1,TMSB
CKSL1,UBB,AFF4,TRIM28,MATR3,YWHAG,CKB,COL4A2,SPEN,DDB1,PSMD5,YWHAQ,EIF1,HACD3
1259,EIF5,HMGN2,OAZ1,RBBP4,NISCH,PRKCSH,HNRNPC,KLHL9,HDLBP,GLG1,SFPQ,PIP4K2B,
E,KIF1A,GNAS,DDX17,RACK1,FASN,ALDH6A1,ATP5F1B,CALM3,GNB1,RPL7,SEPTIN2,P4HB,APP,
3,GNB1,SHISA9,RPS8,MYH9,P4HB,RPL5,FSCN1,TMBIM6,PTMS,SPTAN1,CALM1,NPM1,PLD3,US
PTMA,DDX5,SOX4,FKBP14,RPL13A,MAGED1,SEC62,RPL6,KAT6A,DVL3,YWHAE,GNAS,GPI,HTRA
),FADS2,EIF4H,SCARB2,BRD3,KPNB1,GAP43,HK1,NCS1,CLSTN1,ATF4,KIF5B,RAB6B,SLC7A5,CX
DDX5,ENO2,SOX4,MAGED1,SEC62,RPL6,KAT6A,YWHAE,KIF1A,GNAS,DDX17,HTRA4,RACK1,FAS
X,CLU,PTMA,DDX5,SOX4,RPL13A,RPL6,KAT6A,KIF1A,RPL13,GPI,HTRA4,RACK1,BSG,FASN,ALD
3,CKB,COL4A2,MAPT,SPEN,NONO,EIF1,SET,EGLN1,TPI1,TMSB10,INA,MRC2,PKD1,PDIA3,RPS3,
.4A1,PGK1,RPLP0,CANX,CLU,PTMA,DDX5,ENO2,SOX4,RPL13A,MAGED1,SEC62,RPL6,KAT6A,DV
RPS3A,STMN1,ITGB1,RPL11,MORF4L1,BTF3,NCAM1,ANXA2,AGRN,SARAF,MIDN,MARCHF6,VCL,
EIF4B,VCP,MARCKSL1,CKB,FN1,SPEN,NONO,DDB1,EIF1,HACD3,DCHS1,MAP9,TPI1,CHSY1,TMS

DHB,PKD1,SF3B1,PDIA3,ZNF43,HMGB1,POLR2A,TPT1,SEC61A1,RPS3A,ITGB1,TUBB4B,BTF3,RI
,GAP43,HK1,NCS1,ZNF865,NR2F1,RAB6B,ENC1,NLGN2,GLIPR1L2,CXADR,CHD4,ATP1A1,JUND,I
F3B2,MAT2A,PXDN,HNRNPU,HNRNPA0,FADS2,RPSA,TNS3,EIF4H,SCARB2,BRD3,RPS9,SLC16A7,I
MA,DDX5,FKBP14,RPL13A,MAGED1,RPL6,DVL3,YWHAE,GNAS,GPI,HTRA4,RACK1,FASN,ALDH6A
D1,SEC62,RPL6,KAT6A,DVL3,YWHAE,RPL13,GNAS,DDX17,GPI,RACK1,FASN,ATP5F1B,CALM3,GN
,TRIM28,YWHAG,CKB,RPL37A,MAPT,SPEN,DDB1,YWHAQ,EIF1,HACD3,SET,TPI1,CHSY1,TMSB10,
,TPM1,CDH2,PDIA6,GPC3,DDX3X,GNB2,B4GALNT4,MTSS2,DNAJC5,APLP2,RPLP2,MXD4,DUX4,C
,P4HB,APP,RPL5,GDI1,SPTAN1,GSTP1,USP11,COL18A1,SNRNP200,CSDE1,SMIM14,IGF2,EIF3C,I
A1,COL4A1,CES3,RPLP0,PTMA,DDX5,ENO2,SEC62,YWHAE,KIF1A,GNAS,SLC2A3,DDX17,GPI,FASI
EN,EIF1,SET,EGLN1,TPI1,TMSB10,INA,MRC2,PKD1,PDIA3,RPS3,BASP1,VAPA,HMGB1,SEC61A1,R
ISCH,TADA2B,GLG1,PDIA6,DDR1,SOX12,ACTN4,ARL4C,APLP1,MIF,SESN3,MTSS2,APLP2,TARDBI
/I1,ANXA2,AGRN,SARAF,COL11A1,MIDN,RPS17,VCL,DNAJA1,ARF1,CDC42BPB,MAT2A,TMEM170
P2,HNRNPDL,TARDBP,INIP,RPL36,CADM4,COTL1,HECTD4,MAGED4B,CTBP1,RAC1,ARHGDIA,STI
SD,SCARB2,RPS24,BRD3,CIPC,HK1,RPL22,ZNF865,SLC7A5,CHD4,ATP1A1,CD81,WDR1,HMGN2,
I,FSCN1,SPTAN1,SMG1,CALM1,NPM1,PLD3,HSPD1,COL18A1,CSDE1,IGF2,EIF3C,ATP5F1A,MARC
DB1,EIF1,HACD3,SET,CHSY1,TMSB10,EIF4A1,MRC2,RPL19,RPL15,LDHB,SF3B1,PDIA3,VAPA,HM
,CKB,COL4A2,RPL37A,SPEN,DDB1,HACD3,DCHS1,SET,EGLN1,MAP9,CHSY1,EIF4A1,PDIA3,VAPA
CK1,FASN,ATP5F1B,GNB1,SHISA9,RPS8,RPL7,SEPTIN2,P4HB,FSCN1,TMBIM6,PTMS,CALM1,HSPI
IPT2,PNN,LRPPRC,TADA2B,KHDRBS1,PDIA6,GPC3,PIP4K2B,ACTN4,RPS15,RPL12,ITM2C,ARL4C,
IAGED1,SEC62,RPL6,KAT6A,DVL3,RPL13,GNAS,DDX17,GPI,RACK1,ATP5F1B,CALM3,GNB1,SHIS
APP,PTMS,SPTAN1,GSTP1,CALM1,PLXNB2,PLD3,HSPD1,USP11,COL18A1,IGF2,EIF3C,VCP,AFF4,
EIF4B,VCP,MATR3,YWHAG,RPL37A,NONO,PSMD5,SET,CHSY1,EIF4A1,PPP2R1A,DNAH8,PKD1,SF3
GPI,HTRA4,RACK1,FASN,ATP5F1B,CALM3,GNB1,SHISA9,RPS8,MYH9,SEPTIN2,P4HB,RPL5,GDI1,
42BPB,ITM2B,MAT2A,GTF2F1,PXDN,CTSD,LAMA5,XRCC5,MSN,KPNB1,LLGL1,HK1,NCS1,RHOA,
P1,VAT1,RTN3,PFN1,JPT2,SPC25,NES,PRKCSH,KHDRBS1,XPO1,PFKL,CDH2,PDIA6,GPC3,PIP4K2
I8,PKD1,SF3B1,PDIA3,ZNF43,HMGB1,POLR2A,TPT1,SEC61A1,RPS4X,RPS3A,ITGB1,TUBB4B,BTI
PL15,GTF3C4,LDHB,SF3B1,PDIA3,ZNF43,HMGB1,POLR2A,TPT1,SEC61A1,RPS3A,RPL18,ITGB1,
A,TPT1,SEC61A1,RPS3A,ITGB1,MORF4L1,RPLP1,SARAF,RPL10,AP3D1,ARF1,SF3B2,GTF2F1,NFE
L4A1,CES3,PSAP,PGK1,CLU,PTMA,DDX5,ENO2,SOX4,KAT6A,YWHAE,KIF1A,GNAS,SLC2A3,GPI,B
YH9,RPL7,P4HB,RPL5,TMBIM6,SPTAN1,PLXNB2,PLD3,HSPD1,USP11,SNRNP200,CSDE1,SMIM14
3,PFN1,CCNI,TADA2B,KHDRBS1,HDLBP,EIF5A,MEGF8,ACTN4,RPS15,RPS5,MIF,DNAJC5,SLC25A
1A,RPL13,SLC2A3,DDX17,GPI,HTRA4,RACK1,FASN,ALDH6A1,SPTBN1,CALM3,GNB1,SHISA9,STM
CADM4,HECTD4,RAC1,STIP1,SLC35E2B,DNER,BCLAF1,L1CAM,TRIB2,SQLE,MAP1A,FAM199X,TII
,HSPD1,USP11,COL18A1,SNRNP200,CSDE1,IGF2,EIF3C,EIF4B,VCP,MARCKSL1,UBB,TRIM28,MA
,SOX4,RPL13A,MAGED1,SEC62,KAT6A,YWHAE,RPL13,GNAS,HTRA4,RACK1,FASN,ATP5F1B,CALI
TP5F1B,CALM3,GNB1,SHISA9,RPS8,MYH9,SEPTIN2,P4HB,RPL5,TMBIM6,SPTAN1,NPM1,PLXNB
KIF1A,GNAS,DDX17,FASN,ALDH6A1,ATP5F1B,CALM3,RPL7,SEPTIN2,P4HB,APP,IDS,ERVV-1,GDI1
,SLC22A17,KPNB1,C1ORF43,NCS1,RHOA,ZNF865,NR2F1,RPS14,CXADR,CHD4,ATP1A1,JUND,WI
A,ZNF865,ATF4,GLIPR1L2,RPS14,TMEM259,RPS20,COL5A2,WDR1,ATXN7L3B,RPL26,LASP1,CC
,HNRNPU,CTNNA1,CNN3,SCARB2,RPS24,BRD3,NCS1,ZNF865,SLC7A5,CHD4,JUND,CD81,WDR1,I
A,DDX5,ENO2,MAGED1,SEC62,RPL6,KAT6A,DVL3,YWHAE,KIF1A,GNAS,DDX17,GPI,HTRA4,FASN,
1,SMARCA5,HNRNPA0,CTNNA1,CPE,CNN3,TNS3,LPCAT1,LAMA5,SLC22A17,BRD3,RPL24,LLGL1,
PI,BSG,ALDH6A1,SPTBN1,GNB1,SHISA9,STMN2,RPS8,MYH9,APP,IDS,GDI1,TMBIM6,PTMS,SPTAI
I1,SHISA9,MYH9,P4HB,IDS,GDI1,PTMS,SPTAN1,CALM1,PLXNB2,USP11,COL18A1,SMIM14,IGF2,
JGP2,PPP1CB,MEST,HNRNPL,ABCA3,ALKBH5,PRDX5,UBE4A,CCT3,VPS35,ACAP3,CBX1,SNRNP70

S,SPTAN1,SMG1,NPM1,HSPD1,COL18A1,SNRNP200,SMIM14,IGF2,EIF3C,ATP5F1A,EIF4B,VCP,M
NAJA1,ARF1,SF3B2,GTF2F1,HNRNPA0,RPL23,RPL29,EIF4H,RPS9,XRCC5,CIPC,RPL24,HK1,RPL2
MCL1,NISCH,CCNI,SPTBN2,MEGF8,RAB7A,GNB2,APLP1,IK,GPAT4,TLN1,CHPF,COL6A2,CD63,B4
KAP1,ACAP3,MDK,SURF4,TMEM248,SKI,PSMB2,RAB11B,ABCA2,PYGB,TERF2IP,VEGFA,RAPGEF
,ATP5F1B,CALM3,GNB1,SHISA9,RPS8,MYH9,RPL7,SEPTIN2,P4HB,RPL5,GDI1,TMBIM6,SPTAN1,
,SPTAN1,PLD3,HSPD1,USP11,SNRNP200,CSDE1,SMIM14,IGF2,EIF3C,ATP5F1A,EIF4B,VCP,UBB,
T6A,YWHAE,KIF1A,GNAS,SLC2A3,DDX17,GPI,FASN,ALDH6A1,SPTBN1,CALM3,GNB1,SHISA9,STM
R,FSTL1,TMF1,ABHD2,HSPA1B,COL5A1,COL4A1,CES3,PSAP,PGK1,RPLP0,CANX,CLU,PTMA,DDX
P1,TRIB2,HS6ST1,CTDSP2,CNN2,CHD6,EEF1A2,TBCD,PGD,SQSTM1,VAMP2,KLHL24,PDXK,PLOD
,ABHD2,HSPA1B,COL5A1,COL4A1,CES3,PSAP,PGK1,RPLP0,CANX,CLU,PTMA,DDX5,ENO2,SOX4,
PSMD2,TMEM259,RPS20,EIF5,CD81,LASP1,BEX3,OAZ1,SRSF4,RBBP4,PFN1,NES,HNRNPC,PRDX2
MYH9,P4HB,RPL5,TMBIM6,SPTAN1,PLXNB2,PLD3,HSPD1,USP11,SNRNP200,CSDE1,SMIM14,IG
HB,RPL5,TMBIM6,SPTAN1,PLXNB2,PLD3,HSPD1,USP11,SNRNP200,CSDE1,SMIM14,IGF2,EIF3C,
YBX1,UBA1,ATP2B4,IGFBP2,CALR,FSTL1,HSPA1B,TUBB2B,COL5A1,COL4A1,PSAP,PGK1,RPLP0,
RRC28,SPEN,DDB1,PSMD5,HACD3,SET,EGLN1,TPI1,CHSY1,EIF4A1,RPL19,PPP2R1A,RPL15,POL
ATR3,CKB,SPEN,DDB1,PSMD5,YWHAQ,EIF1,HACD3,CHSY1,TMSB10,RPL19,LDHB,SF3B1,SSBP2,
LC39A7,ATRX,CD164,TRA2B,ZFR,SMC3,ZNF512B,PRRC2A,TIMM50,PSMD8,FBXO21,SAFB2,FUS,
X12,DNAJC5,APLP2,SLC25A3,RPLP2,NDFIP1,MXD4,RPL36,DUX4,CADM4,COTL1,CTBP1,ARHGDI
BA1,ATP2B4,IGFBP2,CALR,FSTL1,TMF1,ABHD2,HSPA1B,TUBB2B,COL5A1,COL4A1,CES3,PSAP,
HAQ,EIF1,HACD3,SET,CHSY1,TMSB10,LDHB,SF3B1,SSBP2,BASP1,VAPA,POLR2A,STMN1,TUBB4
,IGFBP2,CALR,FSTL1,TMF1,ABHD2,HSPA1B,ZNF99,TUBB2B,COL5A1,PSAP,PGK1,RPLP0,H3-3B,
TMF1,ABHD2,HSPA1B,COL4A1,CES3,PGK1,RPLP0,CANX,CLU,PTMA,DDX5,ENO2,SOX4,FKBP14,R
6A,DVL3,YWHAE,RPL13,DDX17,RACK1,BSG,FASN,ALDH6A1,SPTBN1,ATP5F1B,CALM3,STMN2,R
ANX,PTMA,DDX5,ENO2,FKBP14,RPL13A,MAGED1,SEC62,KAT6A,DVL3,KIF1A,RPL13,SLC2A3,GPI,
EIF1,EGLN1,CHSY1,RPL15,SF3B1,VAPA,POLR2A,AP2M1,TPT1,RPS3A,RPL18,ITGB1,TUBB4B,AG
,P4HB,RPL5,GDI1,TMBIM6,SPTAN1,PLXNB2,PLD3,HSPD1,USP11,SNRNP200,CSDE1,SMIM14,IG
KB,COL4A2,SPEN,DDB1,YWHAQ,EIF1,SET,TPI1,CHSY1,TMSB10,EIF4A1,MRC2,GTF3C4,LDHB,DN
ADA2B,KHDRBS1,HDLBP,TPM1,EIF5A,ACTN4,RPS15,RPS5,APLP1,B4GALNT4,DNAJC5,SLC25A3,
F43,HK1,NCS1,ZNF865,CLSTN1,GLIPR1L2,EIF5,HMGN2,RBBP4,TLE5,HNRNPC,CCNI,XPR1,XPO
61A1,STMN1,RPL11,MORF4L1,AGRN,RPLP1,MIDN,RPL10,MARCHF6,AP2B1,MAT2A,TMEM170A,
1,FADS1,NEFM,PRPF8,RPL8,YBX1,UBA1,ATP2B4,IGFBP2,CALR,FSTL1,ABHD2,HSPA1B,TUBB2B,
,RPL12,EIF3L,GNB2,RCN1,RPS5,APLP1,RPS7,RPS19,HMGCS1,CHPF,COL6A2,RPS10,MTSS2,HN
,RACK1,FASN,ATP5F1B,CALM3,GNB1,SHISA9,RPS8,MYH9,P4HB,RPL5,FSCN1,TMBIM6,NPM1,PL
ALM3,GNB1,STMN2,MYH9,P4HB,RPL5,GDI1,PTMS,SPTAN1,SMG1,CALM1,NPM1,PLXNB2,CSDE1
B,ZNF99,COL4A1,CES3,PGK1,RPLP0,CANX,PTMA,DDX5,ENO2,SOX4,RPL13A,MAGED1,SEC62,R
RPL13,GPI,HTRA4,BSG,FASN,ALDH6A1,SPTBN1,GNB1,RPS8,SEPTIN2,P4HB,APP,ERVV-1,RPL5,K
P1,RAPGEF1,OPTN,MVB12B,KDELR2,LAPTM4A,BTBD2,RABL6,IPO7,ATXN10,CAND1,ZFAND5,HN
BP2,FSTL1,TMF1,ABHD2,HSPA1B,TUBB2B,COL5A1,COL4A1,PSAP,PGK1,CANX,CLU,PTMA,RPL1
SA9,RPS8,MYH9,SEPTIN2,P4HB,RPL5,GDI1,TMBIM6,SPTAN1,PLD3,HSPD1,USP11,SNRNP200,CS
ARAF,MIDN,RPL10,MARCHF6,AP3D1,DNAJA1,SF3B2,GTF2F1,HNRNPA0,RPS16,RPL23,CNN3,RPL
2B,TRIB2,CTDSP2,NOVA2,CHD6,EEF1A2,TBCD,PGD,VAMP2,ACTR3,PKN1,AHCYL1,CAMSAP1,GRI
L18,TUBB4B,RPL11,RPLP1,SARAF,RPL10,RPS17,AP3D1,SF3B2,GTF2F1,RPL23,CNN3,RPL29,TP
,CANX,PTMA,DDX5,SOX4,FKBP14,RPL13A,MAGED1,SEC62,KAT6A,YWHAE,KIF1A,RPL13,DDX17,
TMA,DDX5,FKBP14,RPL13A,MAGED1,RPL6,DVL3,GNAS,GPI,RACK1,FASN,ATP5F1B,CALM3,GNB
TMBIM6,SPTAN1,PLXNB2,PLD3,HSPD1,USP11,SNRNP200,CSDE1,SMIM14,IGF2,EIF3C,ATP5F1A,

RPL7,SEPTIN2,P4HB,APP,IDS,RPL5,GDI1,FSCN1,TMBIM6,PTMS,SPTAN1,GSTP1,SMG1,CALM1,PL
FN1,SPC25,NES,KHDRBS1,PFKL,CDH2,PDIA6,GPC3,PIP4K2B,MEGF8,MMP14,MAGED2,GNB2,APL
F1A2,HNRNPA3,PKN1,GRINA,COL2A1,MLEC,CLSTN3,MDK,JPT1,CBX1,PTK7,CHD2,ORAI2,RAB11
TOP1,MVB12B,MAPK1,RABL6,ODC1,ADAM9,PTBP1,HNRNPUL2-BSCL2,PHC2,STXBP1,ATXN2L,M
AGED1,SEC62,RPL6,KAT6A,DVL3,YWHAE,KIF1A,GNAS,DDX17,RACK1,FASN,ALDH6A1,SPTBN1,CAL
HNRNPA3,YEATS2,GRINA,COL2A1,MDK,LAMC1,JPT1,CBX1,RAD23B,TNPO1,ASXL1,NRDC,ARF3,SI
A1,PSAP,PGK1,RPLP0,H3-3B,CLU,PTMA,DDX5,ENO2,SOX4,FKBP14,RPL13A,MAGED1,SEC62,RPL
HSY1,TMSB10,INA,EIF4A1,MRC2,GTF3C4,POLR2A,RPL18,STMN1,TUBB4B,MORF4L1,RPLP1,MID
1,ITGB1,TUBB4B,RPL11,ANXA2,TIMP2,SARAF,MIDN,WSB1,AP3D1,MAP4,CDC42BPB,CTNNA1,CPI
PNB1,C1ORF43,PAPOLA,HK1,PTCD3,CLSTN1,ATF4,KIF5B,RPS14,CHD4,EIF5,CD81,WDR1,ATP2A
SP2,NOVA2,CHD6,EEF1A2,TBCD,PGD,SQSTM1,VAMP2,ACTR3,PDXK,PKN1,HSPA4,CAMSAP1,GRIN
SEPTIN2,APP,ERVV-1,FSCN1,PTMS,SMG1,CALM1,NPM1,HSPD1,SMIM14,IGF2,EIF3C,ATP5F1A,VC
M2C,MRFAP1,RPL35,JUN,CD63,HNRNPDL,SLC25A3,DDOST,MXD4,CIRBP,RPL36,YWHAB,SND1,R
GANAB,COL6A1,FTL,HSP90B1,FADS1,NEFL,NEFM,PRPF8,UGT8,RPL8,FBXL4,YBX1,UBA1,ATP2B4
GPI,HTRA4,RACK1,FASN,ATP5F1B,CALM3,GNB1,SHISA9,RPS8,MYH9,SEPTIN2,P4HB,RPL5,GDI1,T
9,RPL7,RPL5,GDI1,FSCN1,TMBIM6,PTMS,SPTAN1,GSTP1,CALM1,TTC3,HSPD1,USP11,COL18A1,
GF2,EIF3C,EIF4B,MARCKSL1,YWHAG,COL4A2,RPL37A,EGLN1,CHSY1,INA,MRC2,GTF3C4,PKD1,H
NPA0,FADS2,LDHA,CNN3,LAMA5,SLC22A17,XRCC5,SLC16A7,KPNB1,RPL17,C1ORF43,PAPOLA,H
3,ARCN1,CTBP1,ARHGDIA,SLC35E2B,LAPTM4B,NNAT,SQLE,TM9SF3,MDH2,U2AF2,FDFT1,SUGP
B4B,RPL11,RPLP1,SARAF,MIDN,RPL10,AP3D1,SF3B2,GTF2F1,RPS16,RPL23,CNN3,RPL29,RPS9,
A1,PSAP,PGK1,RPLP0,H3-3B,CLU,PTMA,DDX5,ENO2,SOX4,FKBP14,RPL13A,MAGED1,SEC62,RPL
,LRRC28,EIF1,RPL19,PPP2R1A,RPL15,GTF3C4,DNAH8,SF3B1,PDIA3,SERINC1,ZNF43,RPS3,HMG
3,MYH9,APP,IDS,RPL5,GDI1,PTMS,MARCKS,SMG1,PLXNB2,USP11,COL18A1,SMIM14,IGF2,VCP,M
3,MYH9,APP,IDS,RPL5,GDI1,PTMS,MARCKS,SMG1,PLXNB2,USP11,COL18A1,SMIM14,IGF2,VCP,M
PLXNB2,EIF4B,MARCKSL1,AFF4,MATR3,YWHAG,CKB,RPL37A,FN1,NONO,DDB1,HACD3,DCHS1,S
TR3,RPL37A,LRRC28,SPEN,NONO,EIF1,INA,EIF4A1,RPL19,PPP2R1A,RPL15,LDHB,PKD1,SF3B1,S
PL15,GTF3C4,LDHB,PKD1,SF3B1,RPS3,BASP1,VAPA,CDKL1,TPT1,RPL18,STMN1,ITGB1,TUBB4B
TIN2,P4HB,RPL5,GDI1,TMBIM6,SPTAN1,PLXNB2,PLD3,HSPD1,USP11,SNRNP200,CSDE1,SMIM14
DH6,PEBP1,PIP4K2B,SOX12,PRDX2,ARL4C,RPL35,MYO5A,DNAJC5,APLP2,HNRNPDL,TARDBP,RF
ANXA2,AGRN,MARCHF6,RPS17,DNAJA1,ARF1,MAP4,CDC42BPB,MAT2A,SLC4A8,GOLGA8A,PXDN
,PAPOLA,RPL14,PTCD3,CBX5,ATF4,SLC7A5,TMEM259,EIF5,CD81,WDR1,HMGN2,RPL26,CCT7,O
),MYH10,YWHAB,SEPTIN3,APPL1,SMCHD1,IPO5,ATP6V0C,LAPTM4B,TRIB2,NNAT,NUCB1,RNF187
CKSL1,UBB,AFF4,TRIM28,MATR3,COL4A2,RPL37A,FN1,SPEN,DCHS1,SET,EGLN1,TPI1,CHSY1,TM
NRNP200,CSDE1,SMIM14,IGF2,EIF3C,ATP5F1A,EIF4B,VCP,UBB,MATR3,YWHAG,RPL37A,LRRC28
,NCAM1,RPLP1,SARAF,MIDN,MARCHF6,ARF1,MAP4,TMEM170A,GTF2F1,GOLGA8A,PXDN,PGAM
RPL31,HDLBP,GLG1,EIF5A,PRDX2,RPS15,ITM2C,RPL30,ARL4C,IK,RPL35,NGRN,RPS10,MTSS2,AI
,PFN1,MCL1,PRKCSH,LRPPRC,RPS25,PDIA6,PLXNA1,RAB7A,RPS15,EIF3L,RPS19,CHPF,NGRN,RI
IISA9,RPS8,MYH9,SEPTIN2,P4HB,RPL5,TMBIM6,SPTAN1,CALM1,PLXNB2,PLD3,HSPD1,USP11,SN
CIPC,RPL24,PAPOLA,HK1,RHOA,ZNF865,GLIPR1L2,RPS14,RPS20,CD81,LENG8,WDR1,SRSF4,RI
1,SEC61A1,RPS3A,RPL18,ITGB1,TUBB4B,UCHL1,RPL11,RPLP1,SARAF,MIDN,RPL10,RPS17,AP3D
18,ITGB1,TUBB4B,RPL11,RPLP1,SARAF,MIDN,RPL10,RPS17,AP3D1,SF3B2,GTF2F1,HNRNPA0,R
TP1A1,JUND,EIF5,CD81,HMGN2,CCT7,VAT1,MCL1,JPT2,SPTBN2,PFKL,HDLBP,SOX12,ARL4C,APL
TP1A1,JUND,EIF5,CD81,HMGN2,CCT7,VAT1,MCL1,JPT2,SPTBN2,PFKL,HDLBP,SOX12,ARL4C,APL
P5F1B,CALM3,GNB1,SHISA9,RPS8,P4HB,RPL5,FSCN1,TMBIM6,NPM1,PLD3,HSPD1,USP11,SNRNI
WHAE,KIF1A,GNAS,DDX17,FASN,ALDH6A1,SPTBN1,CALM3,GNB1,SHISA9,RPS8,RPL7,P4HB,ERV\

DK16,TOP2B,SMCHD1,IPO5,SKP1,TRIB2,DLG5,HS6ST1,TIMP3,NOVA2,CTSB,CDH11,MLXIP,GAS7
IMP2,RPL10,SBF1,ARF1,CYB5R3,AKAP12,MAP4,CDC42BPB,AP2B1,MAT2A,GOLGA8A,NFE2L1,TN
BN1,CALM3,GNB1,SHISA9,RPL7,SEPTIN2,APP,ERVV-1,FSCN1,TMBIM6,PTMS,SMG1,CALM1,NPM
ACAP3,RAD23B,ATP6AP1,TERF2IP,RPS15A,TUBB2A,PPP2CA,ACADVL,PTBP1,ILF3,PLCG1,SRSF3,L
DN,RPL10,MARCHF6,DNAJA1,SF3B2,GTF2F1,HNRNPA0,RPL23,CNN3,RPL29,XRCC5,CIPC,PAPOLA
WHAQ,EIF1,HACD3,SET,CHSY1,INA,EIF4A1,MRC2,RPL19,LDHB,SF3B1,PDIA3,BASP1,HMGB1,TPT
2A,AP2M1,RPL18,MORF4L1,TIMP2,MIDN,SBF1,DNAJA1,ARF1,CYB5R3,MAP4,SF3B2,CDC42BPB,
D3,HSPD1,USP11,SNRNP200,CSDE1,SMIM14,EIF3C,ATP5F1A,EIF4B,MATR3,YWHAG,RPL37A,LRR
3B1,SERINC1,RPS3,HMGB1,POLR2A,AP2M1,TPT1,SEC61A1,RPS3A,BTF3,AGRN,RPLP1,MIDN,RPL
NP200,CSDE1,SMIM14,IGF2,EIF3C,ATP5F1A,EIF4B,VCP,UBB,AFF4,MATR3,YWHAG,RPL37A,LRRC
16,TMEM30A,BCLAF1,NNAT,CRPPA,EID1,U2AF2,SUGP2,MEST,HNRNPL,TM9SF2,SSRP1,ABCA3,A
I,MRC2,GTF3C4,DNAH8,HLA-C,SSBP2,CHD3,AP2M1,TPT1,ITGB1,MORF4L1,ANXA2,TIMP2,SARAF
RHOA,ZNF865,GLIPR1L2,RPS14,RPS20,CD81,LENG8,WDR1,PFN1,MCL1,TLE5,PNN,RPL31,PRKCS
GB1,ANXA2,ZMIZ1,TIMP2,MIDN,RPL10,WSB1,RPS17,VCL,CYB5R3,AKAP12,CDC42BPB,GTF2F1,H
GPI,HTRA4,RACK1,BSG,FASN,ALDH6A1,SPTBN1,ATP5F1B,CALM3,GNB1,SHISA9,RPS8,MYH9,SEI
A,ZNF865,GLIPR1L2,RPS14,RPS20,CD81,LENG8,WDR1,LASP1,RBBP4,PFN1,MCL1,TLE5,PNN,TKT
SDE1,IGF2,MARCKSL1,AFF4,TRIM28,COL4A2,NONO,DDB1,YWHAQ,SET,TPI1,RPL19,GTF3C4,LDH
X5,SOX4,FKBP14,RPL13A,MAGED1,RPL6,DVL3,YWHAE,GNAS,HTRA4,RACK1,BSG,FASN,ALDH6A1
CK1,BSG,FASN,ALDH6A1,GNB1,RPS8,MYH9,SEPTIN2,P4HB,APP,FSCN1,TMBIM6,SMG1,NPM1,PLI
VCP,MARCKSL1,AFF4,TRIM28,CKB,COL4A2,SPEN,DDB1,YWHAQ,EIF1,HACD3,DCHS1,SET,TPI1,C
1,ATP5F1B,CALM3,GNB1,SHISA9,RPS8,MYH9,RPL7,SEPTIN2,P4HB,RPL5,GDI1,TMBIM6,SPTAN1,
C,ATP5F1A,UBB,AFF4,RPL37A,LRRC28,SPEN,NONO,INA,EIF4A1,RPL19,RPL15,GTF3C4,SF3B1,SI
C5,CIPC,RPL24,PAPOLA,HK1,RHOA,ZNF865,GLIPR1L2,RPS14,RPS20,CD81,LENG8,WDR1,SRSF4
PLP1,SARAF,MIDN,RPL10,AP3D1,DNAJA1,SF3B2,GTF2F1,HNRNPA0,RPS16,RPL23,CNN3,RPL29,F
CHF6,WSB1,RPS17,MAP4,CDC42BPB,ITM2B,FADS2,RPS16,TNS3,EIF4H,LPCAT1,RPS9,SLC16A7,
KS,SMG1,CALM1,PLXNB2,HSPD1,IGF2,ATP5F1A,EIF4B,MARCKSL1,UBB,AFF4,YWHAG,CKB,COL
2,RPS14,RPS20,CD81,LENG8,WDR1,SRSF4,PFN1,MCL1,TLE5,PNN,RPL31,PRKCSH,KHDRBS1,KL
,PGK1,RPLP0,H3-3B,CANX,CLU,PTMA,DDX5,ENO2,FKBP14,MAGED1,SEC62,KAT6A,YWHAE,KIF1A
OL18A1,SNRNP200,EIF3C,ATP5F1A,EIF4B,UBB,AFF4,MATR3,CKB,RPL37A,NONO,PSMD5,SET,CHS
PITPNM1,STXBP1,ISYNA1,HYOU1,MAPK8IP1,MEX3A,FLNC,CTXN1,CD99L2,ANP32A,RBBP7,MSM
EM170A,PXDN,PGAM1,SMARCA5,HNRNPA0,FADS2,RPS16,CTSD,NACA,LPCAT1,LAMA5,RPS24,SL
FF4,MATR3,LRRC28,EIF1,HACD3,EIF4A1,RPL19,SF3B1,PDIA3,SERINC1,SSBP2,ZNF43,HMGB1,P
7,RBBP4,MCL1,PRKCSH,LRPPRC,TADA2B,XPR1,KHDRBS1,SCRN1,SFPQ,EIF5A,PDIA6,RAB7A,RPS
NT4,MTSS2,APLP2,TARDBP,CADM4,HNRNPH1,HLA-B,PLPPR3,CTBP1,STIP1,GPC1,SLC35E2B,SM
,TUBB2B,COL5A1,COL4A1,PSAP,PGK1,RPLP0,CANX,CLU,PTMA,DDX5,FKBP14,RPL13A,MAGED1,
SMG1,PLXNB2,TTC3,IGF2,EIF3C,VCP,MARCKSL1,AFF4,TRIM28,YWHAG,CKB,MAPT,SPEN,DDB1,
D3,USP11,COL18A1,SNRNP200,IGF2,EIF3C,ATP5F1A,EIF4B,AFF4,MATR3,RPL37A,LRRC28,EIF1,I
,TLE5,NES,HNRNPC,CCNI,XPO1,HDLBP,TPM1,EIF5A,SOX12,RPL12,RCN1,RPS5,APLP1,PRKAR1A,
5,PNN,RPL31,KHDRBS1,TKT,EIF5A,PDIA6,PIP4K2B,RAB7A,ITM2C,RPL30,EIF3L,RPS19,CHPF,NGF
IPH1,MAGED4B,CTBP1,ARHGDIA,CDK16,TMEM30A,BCLAF1,NNAT,CRPPA,EID1,TOMM20,U2AF2,
TADA2B,KHDRBS1,SFPQ,PEBP1,CCND2,SOX12,ITM2C,H1-0,APLP1,B4GALNT4,SHOX,H1-10,CIRE
35A,HNRNPDL,TSR2,WLS,HNRNPK,TTR,RPS27,RAC1,ATP6V0C,L1CAM,SQLE,DLG5,CNN2,SRSF5,
IN3,CTSD,SLC22A17,RPS9,XRCC5,PJA2,SLC16A7,PAPOLA,RPL9,ATF4,KIF5B,TMEM259,RPS20,BI
CBP4,ZNF664,CLCN7,SNN,SHC2,SSBP3,OLFM1,PKDCC,NUDCD3,MAZ,SH3GL1,PRR12,MSI1,NSD
GB1,TUBB4B,UCHL1,ZMIZ1,SARAF,MIDN,SBF1,CYB5R3,CDC42BPB,HNRNPA0,CNN3,SCARB2,LAM

DN,RPS17,ARF1,CYB5R3,MAP4,AP2B1,TMEM170A,GTF2F1,SMARCA5,HNRNPA0,CPE,CNN3,TPM4
NONO,SET,MAP9,TMSB10,EIF4A1,RPL15,GTF3C4,PDIA3,SERINC1,STMN1,TUBB4B,UCHL1,ANXA
L4A1,CES3,PGK1,RPLP0,CANX,PTMA,DDX5,ENO2,FKBP14,SEC62,RPL6,KAT6A,DVL3,YWHAE,RPL
,GDI1,FSCN1,PTMS,SPTAN1,GSTP1,MARCKS,CALM1,NPM1,PLXNB2,PLD3,USP11,COL18A1,CSD
I1,PLD3,HSPD1,USP11,SNRNP200,CSDE1,SMIM14,IGF2,EIF3C,ATP5F1A,EIF4B,VCP,MATR3,YWH
A5,CIPC,MSN,KPNB1,C1ORF43,RPL14,RHOA,CLSTN1,ATF4,SLC7A5,ENC1,ATP1A1,JUND,TMEM
17,AP3D1,CDC42BPB,MAT2A,GTF2F1,HNRNPA0,RPS16,RPL23,CNN3,RPL29,LAMA5,RPS24,RPS
SARAF,RPL10,AP3D1,SF3B2,MAT2A,GTF2F1,RPL23,CNN3,RPL29,NACA,LAMA5,BRD3,RPS9,XRC
ENO2,SOX4,RPL13A,MAGED1,SEC62,DVL3,YWHAE,DDX17,GPI,HTRA4,BSG,FASN,ALDH6A1,SPTB
,MTSS2,APLP2,MXD4,LGALS3BP,ATP1A2,RAB1A,SEPTIN3,MAGED4B,CTBP1,RAC1,GPC1,TOP2B
HK1,RHOA,ZNF865,ATF4,GLIPR1L2,RPS14,RPS20,CD81,LENG8,WDR1,PFN1,MCL1,TLE5,PNN,R
S10,APLP2,MYL6,WLS,HNRNPK,H1-10,PTGES3,SEPTIN3,GPC1,APPL1,SMCHD1,HDAC2,IPO5,LAP
A1,MRC2,RPL19,RPL15,LDHB,PKD1,SF3B1,PDIA3,SERINC1,BASP1,HMGB1,POLR2A,TPT1,RPS3
,LRRC28,MAPT,FN1,SPEN,NONO,DDB1,YWHAQ,EIF1,DCHS1,SET,EGLN1,MAP9,CHSY1,INA,PPP2
FF4,YWHAG,CKB,COL4A2,NONO,DCHS1,EGLN1,TMSB10,EIF4A1,MRC2,PKD1,HLA-C,SF3B1,BAS
9,CHSY1,INA,MRC2,GTF3C4,PDIA3,SSBP2,BASP1,HMGB1,CDKL1,TPT1,RPS4X,RPL18,STMN1,R
,SLC35E2B,TRIB2,GLUL,CTDSP2,NOVA2,ZC3H7B,CHD6,EEF1A2,TBCD,PGD,SQSTM1,VAMP2,AL
TP6V0C,SQLE,HS6ST1,CTDSP2,NOVA2,RAB27A,TBCD,PGD,HNRNPL,PDXK,HSPA4,VPS35,MLEC,
AE,GNAS,DDX17,GPI,RACK1,BSG,ALDH6A1,SPTBN1,CALM3,GNB1,SHISA9,STMN2,RPS8,SEPTIN
F4,MATR3,RPL37A,LRRC28,TPI1,INA,EIF4A1,RPL19,PPP2R1A,RPL15,GTF3C4,DNAH8,SF3B1,SE
YWHAQ,EIF1,HACD3,EGLN1,TPI1,CHSY1,TMSB10,RPL19,GTF3C4,PKD1,PDIA3,SERINC1,ZNF43,
TP1,SMG1,CALM1,COL18A1,SMIM14,IGF2,ATP5F1A,MARCKSL1,MATR3,YWHAG,CKB,COL4A2,
2,TBCD,VAMP2,GAS7,YEATS2,HSPA4,CAMSAP1,GRINA,PTOV1,ACAP3,MDK,SKI,H3-3A,RAB11B,
HMGB1,POLR2A,AP2M1,SEC61A1,RPS3A,ITGB1,RPL11,MORF4L1,BTF3,AGRN,RPLP1,SARAF,R
HL1,MORF4L1,ZMIZ1,COL11A1,WSB1,RPS17,VCL,AP3D1,SBF1,DNAJA1,ARF1,CYB5R3,PAFAH1
28,CKB,RPL37A,MAPT,FN1,SPEN,DDB1,PSMD5,YWHAQ,EIF1,HACD3,DCHS1,MAP9,CHSY1,PPP2
GDIA,GPC1,IPO5,ATP6V0C,DNER,L1CAM,TRIB2,SQLE,CTSB,CHD6,EEF1A2,TBCD,VAMP2,GAS7,L
,RPS9,RPL24,RPL14,CBX5,ZNF865,CLSTN1,ATF4,KIF5B,TMEM259,EIF5,HMGN2,CCT7,MCL1,N
1,BASP1,VAPA,HMGB1,POLR2A,SEC61A1,ITGB1,UCHL1,RPL11,ZMIZ1,SARAF,RPL10,WSB1,RPS1
2,HSPA1B,ZNF99,TUBB2B,COL5A1,PSAP,RPLP0,H3-3B,CANX,PTMA,DDX5,ENO2,SOX4,FKBP14,
,ANXA2,AGRN,ZMIZ1,TIMP2,XAF1,COL11A1,RPS17,AP3D1,DNAJA1,MAP4,AP2B1,PXDN,SMARCA
LD3,USP11,COL18A1,SNRNP200,SMIM14,EIF3C,ATP5F1A,EIF4B,MARCKSL1,UBB,AFF4,TRIM28,
,PLXNB2,PLD3,HSPD1,USP11,SNRNP200,SMIM14,IGF2,EIF3C,ATP5F1A,EIF4B,VCP,UBB,AFF4,M
,RPL26,SRSF4,RBBP4,PFN1,MCL1,TLE5,PNN,RPL31,KHDRBS1,EIF5A,PDIA6,PIP4K2B,RAB7A,IT
1,RHOA,ZNF865,GLIPR1L2,RPS14,RPS20,CD81,LENG8,WDR1,SRSF4,RBBP4,PFN1,MCL1,TLE5,
3,PAPOLA,COL1A2,GAP43,ZNF865,ATF4,NR2F1,NLGN2,CHD4,JUND,RPS20,EIF5,ATP2A2,LASP1,
1A,RPL15,DNAH8,PKD1,SF3B1,SERINC1,ZNF43,RPS3,VAPA,HMGB1,POLR2A,AP2M1,SEC61A1,O
3,SDF4,HNRNPM,SDHA,YWHAH,LONP1,ZFAND3,CCT5,RBBP7,DFFB,EIF2AK1,TXLNA,KDM5C,FB
4B,AGRN,RPLP1,ZMIZ1,TIMP2,SARAF,MIDN,RPL10,WSB1,RPS17,VCL,AP3D1,SBF1,DNAJA1,ARF
A,YWHAE,KIF1A,RPL13,GNAS,SLC2A3,DDX17,GPI,BSG,FASN,ALDH6A1,SPTBN1,ATP5F1B,CALM3
28,YWHAG,COL4A2,PSMD5,HACD3,SET,EGLN1,CHSY1,TMSB10,EIF4A1,RPL19,RPL15,GTF3C4,
,TMSB10,RPL19,RPL15,GTF3C4,PKD1,ZNF43,HMGB1,POLR2A,TPT1,RPS4X,RPS3A,ITGB1,TUB
C5,CIPC,PAPOLA,ZNF865,ATF4,RPS14,CXADR,LENG8,WDR1,RPL26,CCT7,SRSF4,RBBP4,PFN1,
A2,CHD6,EEF1A2,TBCD,PGD,SQSTM1,VAMP2,ALKBH5,PDXK,YEATS2,HSPA4,CAMSAP1,GRINA,A
4L1,BTF3,AGRN,RPLP1,RPL10,MARCHF6,DNAJA1,ARF1,SF3B2,SMARCA5,RPS16,TPM4,RPS24,B

PC,HDLBP,EIF5A,CCND2,RPS15,ITM2C,ARL4C,RPL35,NGRN,RPS10,MTSS2,APLP2,SLC25A3,RPL
,ERVV-1,RPL5,GDI1,TMBIM6,CALM1,NPM1,PLD3,HSPD1,USP11,SNRNP200,CSDE1,SMIM14,IGF
ME1L1,RBBP7,BAZ1B,KIF21B,EP400,MASTL,MBTPS1,HDAC6,KCTD12,TBC1D9B,CUL3,NPDC1,F
MSB10,PKD1,PDIA3,SERINC1,VAPA,HMGB1,CDKL1,STMN1,TUBB4B,UCHL1,ANXA2,AGRN,SARAF
D81,LENG8,WDR1,LASP1,RBBP4,PFN1,MCL1,TLE5,PNN,RPL31,EIF5A,PDIA6,PIP4K2B,RAB7A,R
CL1,TLE5,PNN,RPL31,HNRNPC,KHDRBS1,RPS25,TKT,EIF5A,PDIA6,RAB7A,RPL12,ITM2C,EIF3L,
,GPI,HTRA4,RACK1,BSG,FASN,ALDH6A1,SPTBN1,ATP5F1B,GNB1,STMN2,RPS8,SEPTIN2,P4HB,
,H1-0,RPS5,APLP1,RPS19,MYO5A,COL6A2,B4GALNT4,MTSS2,DNAJC5,ST3GAL2,SHOX,CIRBP,
20,CD81,WDR1,RPL26,LASP1,RBBP4,PFN1,MCL1,TLE5,PNN,HNRNPC,TKT,EIF5A,PDIA6,RAB7A,
9,BASP1,CDKL1,RPS3A,STMN1,ITGB1,RPL11,AGRN,TIMP2,MIDN,VCL,ARF1,CDC42BPB,ITM2B,
1,H3-3B,CLU,DDX5,ENO2,SOX4,FKBP14,RPL13A,MAGED1,SEC62,RPL6,KAT6A,DVL3,YWHAE,RP
SAP1,KLC1,ACAP3,LAMC1,CCT4,ASXL1,ABCA2,PRPF6,ATP6AP1,RPL38,TUBB2A,MVB12B,G3BP
SKP1,LUC7L3,PGRMC1,EEF1A2,MLXIP,SNX22,PABPN1,GDI2,HNRNPL,TLN2,HSPH1,ABCA3,ALKB
L4A1,CES3,PGK1,CANX,CLU,PTMA,DDX5,FKBP14,RPL13A,MAGED1,RPL6,KAT6A,DVL3,RPL13,D
,SMG1,CALM1,PLXNB2,TTC3,PLD3,HSPD1,USP11,SNRNP200,CSDE1,SMIM14,IGF2,EIF3C,ATP5
PNB1,COL1A2,ATF4,PSMD2,KIF5B,RAB6B,SLC7A5,CXADR,CHD4,ATP1A1,EIF5,WDR1,LASP1,CC
1,KLHL9,HDLBP,GLG1,SFPQ,PEBP1,PNMA1,PIP4K2B,EIF3L,NANOG,RPS7,CTNNB1,NGRN,RPL3
4A1,RPL19,DNAH8,PKD1,SSBP2,AP2M1,TUBB4B,ANXA2,AGRN,XAF1,MIDN,RPL10,WSB1,AP3D1,
3C4,VAPA,POLR2A,CDKL1,TPT1,RPS3A,TUBB4B,MORF4L1,BTF3,RPLP1,ZMIZ1,MIDN,RPL10,R
ATR3,RPL37A,DDB1,PSMD5,EIF1,TMSB10,RPL19,RPL15,GTF3C4,PKD1,ZNF43,HMGB1,POLR2A,
F1B,GNB1,SHISA9,RPS8,RPL7,P4HB,ERVV-1,RPL5,GDI1,TMBIM6,GSTP1,CALM1,PLXNB2,PLD3,
3-3B,DDX5,SOX4,RPL13A,MAGED1,SEC62,RPL6,KAT6A,YWHAE,RPL13,DDX17,GPI,HTRA4,RACK
5F1A,EIF4B,VCP,MARCKSL1,UBB,MATR3,YWHAG,NONO,PSMD5,YWHAQ,SET,CHSY1,TMSB10,E
TP6V0C,BCLAF1,HSBP1,DDX39B,SRSF5,TM9SF3,MDH2,U2AF2,EEF1A2,CCT2,GNAI3,GDI2,TSR1,
PTAN1,GSTP1,CALM1,NPM1,PLXNB2,COL18A1,IGF2,MARCKSL1,TRIM28,YWHAG,COL4A2,LRR
AB,COL6A1,FTL,HSP90B1,FADS1,NEFL,NEFM,PRPF8,RPL8,FBXL4,YBX1,UBA1,ATP2B4,IGFBP2,
PNB1,C1ORF43,NCS1,RHOA,PTCD3,ZNF865,ATF4,PSMD2,RPS14,RPL26,LASP1,TLE5,RPL31,CS
,HMGB1,POLR2A,AP2M1,TPT1,RPS4X,TUBB4B,RPL11,MORF4L1,BTF3,AGRN,RPLP1,RPL10,MA
BL6,TCF25,LUZP6,SCAF11,POLR2B,SDF4,ATP6V1B2,HNRNPM,SDHA,LONP1,ZFAND3,SARS1,VG
3,NACA,LPCAT1,LAMA5,LLGL1,HK1,NCS1,RPL22,ZNF865,CLSTN1,SLC7A5,NLGN2,CXADR,ATP1
RPL23A,DDX39B,SRSF5,TM9SF3,MDH2,RAB27A,U2AF2,CCT2,GNAI3,GAS7,TSR1,MORF4L2,MT
,PITPNA,SUPT6H,LSS,STXBP1,ISYNA1,LOC728392,HYOU1,MAPK8IP1,P3H3,MLF2,CD99L2,PPDP
PDIA6,PIP4K2B,ITM2C,RPL30,EIF3L,IK,CHPF,NGRN,DNAJC5,RPLP2,DDOST,YWHAB,CALU,ARHG
EATS2,CAMSAP1,SURF4,LAMC1,JPT1,SNRNP70,ASXL1,ABCA2,PRPF6,VHL,SMARCA4,SRSF9,PLE
P2,AP2M1,TUBB4B,ANXA2,AGRN,XAF1,MIDN,RPL10,WSB1,AP3D1,CYB5R3,MAP4,ITM2B,TMEM
MAZ,IREB2,LSM14A,ABCF1,YIPF3,NUDT19,RBPJ,EHD1,TGOLN2,YY1,SAP18,UBE2D3,SSBP4,MYL
35,NGRN,MTSS2,APLP2,SLC25A3,RPLP2,WLS,H1-10,CCT6A,RAC1,CDK16,SMCHD1,TMEM30A,D
,APP,RPL5,GDI1,FSCN1,TMBIM6,SPTAN1,GSTP1,SMG1,NPM1,HSPD1,SNRNP200,IGF2,VCP,MAF
N,PRKCSH,EIF5A,PDIA6,PIP4K2B,ACSL6,RPL30,CLTC,EIF3L,MAGED2,RPS19,MIF,TLN1,JUN,NG
3,SEPTIN11,LAMP1,HNRNPH1,CALU,COTL1,CTBP1,RAC1,ARHGDIA,CAPRIN1,GPC1,HDAC2,SERF
,EIF4A1,MRC2,RPL19,PPP2R1A,RPL15,HLA-C,SF3B1,SSBP2,CHD3,HMGB1,POLR2A,RPL11,MOR
,EIF1,TMSB10,EIF4A1,RPL19,RPL15,GTF3C4,PKD1,SF3B1,SERINC1,ZNF43,HMGB1,POLR2A,AP2
AC1,CDK16,SMCHD1,DNER,NNAT,CPSF2,SRP9,CTSB,RAB27A,U2AF2,TBCD,ANXA5,CPSF1,PABP
4,TRA2B,UQCRC1,HNRNPA2B1,TIMM50,TNKS,PSMD8,GSE1,SYMPK,FBXO21,AKAP9,PPP1R10,T
EBP1,GPC3,DNAJC5,APLP2,HNRNPDL,SLC25A3,RPLP2,HNRNPK,DUX4,CKAP5,SEPTIN11,RAB1A,

,XRCC5,CIPC,PAPOLA,ZNF865,ATF4,RPS14,CXADR,LENG8,WDR1,RPL26,SRSF4,PFN1,MCL1,PNN
4B,ANXA2,AGRN,RPLP1,ZMIZ1,SARAF,MIDN,MARCHF6,VCL,AP3D1,SBF1,DNAJA1,ARF1,CYB5R3,
MMP14,RPS15,RPL12,RPL30,RPS5,APLP1,GPAT4,CD63,SESN3,RPLP2,TAGLN,MYL6,INIP,CIRBP,S
3,PBX1,CST3,PRRC2A,GSE1,PCBP4,ZMYM3,SLC44A2,SHC2,SLC6A8,MAZ,STK24,PRR12,TMEM63
ACTN1,PLCG1,DIDO1,DLK1,CORO1C,GIGYF1,YWHAH,LSS,STXBP1,CSNK1D,HYOU1,ZFAND3,RA
,ENC1,NLGN2,GLIPR1L2,ATP2A2,CRMP1,PFN1,PNN,NES,CCNI,LRPPRC,TADA2B,XPO1,CSNK1E,F
3,SEPTIN2,APP,TMBIM6,PTMS,GSTP1,MARCKS,CALM1,NPM1,PLXNB2,HSPD1,CSDE1,SMIM14,AT
CLIP3,FKBP8,PITPNA,SSB,SRSF3,STT3B,LONP1,LOC728392,ATXN2L,UQCRC2,VANGL2,FBRSL1,
VCP,UBB,MATR3,FN1,NONO,PSMD5,SET,CHSY1,TMSB10,EIF4A1,PPP2R1A,RPL15,DNAH8,PKD1,
PFN1,MCL1,XPR1,KHDRBS1,XPO1,SPTBN2,SLC3A2,PTN,PFKL,HDLBP,SFPQ,PNMA1,PIP4K2B,RF
PNN,HNRNPC,KHDRBS1,RPS25,EIF5A,PDIA6,DDR1,PIP4K2B,ITM2C,RPL30,EIF3L,IK,CHPF,NGRN,
RHOB,ZNF454,RPS27,RAC1,STIP1,APPL1,SMCHD1,HDAC2,ATP6V0C,DNER,BCLAF1,TRIB2,PEA15
PS25,TPM1,CDH2,PDIA6,DDR1,PIP4K2B,DDX3X,ITM2C,CLTC,APLP1,COL6A2,RPS10,B4GALNT4,
MTRF1L,PRDX5,UBE4A,TCP1,GCN1,SKI,SRP14,PRPF6,TAF7,RPL37,CCNL2,ARPC2,PTBP1,CAPZB,
MAT2A,HNRNPA0,RPS16,RPL23,RPL29,TNS3,CTSD,EIF4H,LPCAT1,LAMA5,RPS24,RPS9,CIPC,MSN
N2,P4HB,APP,RPL5,FSCN1,GSTP1,MARCKS,PLD3,HSPD1,USP11,COL18A1,SNRNP200,CSDE1,SM
H5,RPL27,PRDX5,HSPA4,MLEC,ASXL1,SRP14,TSPAN6,PRPF6,ATG2B,ARPC2,FAM168B,SUPT5H,H
SET,CHSY1,TMSB10,EIF4A1,PPP2R1A,RPL15,DNAH8,PKD1,SF3B1,ZNF43,RPS3,VAPA,POLR2A,AF
11,MORF4L1,BTF3,ANXA2,AGRN,MIDN,MARCHF6,WSB1,AP3D1,SBF1,CYB5R3,SF3B2,CDC42BPE
ENG8,WDR1,RPL26,LASP1,MCL1,PNN,PRKCSH,TKT,EIF5A,PDIA6,PIP4K2B,ITM2C,RPL30,EIF3L,M
M28,MATR3,CKB,RPL37A,FN1,SPEN,DDB1,EIF1,HACD3,SET,MRC2,RPL19,RPL15,GTF3C4,LDHB,
NO,PSMD5,SET,CHSY1,TMSB10,EIF4A1,PPP2R1A,RPL15,DNAH8,PKD1,SF3B1,ZNF43,RPS3,CHD3
,APP,IDS,ERVV-1,RPL5,FSCN1,TMBIM6,PTMS,SPTAN1,MARCKS,NPM1,HSPD1,USP11,SMIM14,IG
F3,UBE2R2,KDM1A,HNRNPUL2-BSCL2,DLK1,CORO1C,MSL1,SMARCC1,CCT8,TUG1,ATXN2L,ZF
3P8,KDELR1,DIDO1,MSL1,SMARCC1,UBQLN1,TUG1,GPR107,MAPK8IP1,HNRNPD,MEX3A,BMPR2
),PPP2R1A,RPL15,GTF3C4,DNAH8,SF3B1,SERINC1,ZNF43,HMGB1,POLR2A,AP2M1,TPT1,RPS3A,
1,DUSP4,TSC22D1,MSL1,GIGYF1,STMN3,ISYNA1,CSNK1D,ATXN2L,IDH1,ZFAND3,MAPK8IP1,MEX
C1,ZNF43,BASP1,POLR2A,TPT1,RPS3A,RPL18,ITGB1,TUBB4B,MORF4L1,AGRN,MIDN,MARCHF6,
2,GRB2,CAPZB,BNIP3L,PLCG1,FLNB,CORO1C,LSS,STXBP1,ISYNA1,HYOU1,MAPK8IP1,FBRSL1,Z
F1A,MVD,ZNF664,VPS26B,PALM,SNN,MAZ,PRR12,PACS1,MSH6,UBE2D3,SSBP4,GNA13,METRN,E
RBBP7,DFFB,TXLNA,KDM5C,FBRS,SF1,RETREG2,SEC16A,UPF1,EIF5B,SPINT2,SAFB2,ZNF664,VP
SRSF5,MDH2,U2AF2,CPSF1,NOP53,PABPN1,GDI2,HNRNPA3,PTPRA,ACTR3,YEATS2,TBC1D16,ACC
4,BSG,FASN,ALDH6A1,GNB1,RPS8,MYH9,SEPTIN2,P4HB,APP,GDI1,SPTAN1,SMG1,CALM1,NPM1,
2B,COL5A1,COL4A1,CES3,PSAP,PGK1,RPLP0,H3-3B,CANX,CLU,DDX5,FKBP14,RPL13A,MAGED1,
N11,CADM4,MAGED4B,ARHGDIA,HDAC2,IPO5,DNER,GLUL,HSBP1,CDKN1A,DLG5,SHISAL1,HS6S
,PRKCSH,HNRNPC,KHDRBS1,SCRN1,RPS25,EIF5A,PDIA6,MEGF8,RAB7A,RPL12,ITM2C,RPL30,EI
F43,RPS3,VAPA,POLR2A,AP2M1,SEC61A1,STMN1,TUBB4B,MORF4L1,BTF3,TIMP2,XAF1,SARAF,R
C,SKP1,SHISAL1,CTDSP2,NOVA2,CTSB,RNF187,EEF1A2,ANXA5,VCAN,MORF4L2,SSRP1,ACTR3,R
PSF1,PABPN1,TSR1,ALKBH5,HNRNPA3,PDXK,CAMSAP1,CCT3,PTOV1,MDK,CCT4,SKI,SNRNP70,D
FBXL4,UBA1,IGFBP2,CALR,FSTL1,TMF1,ABHD2,HSPA1B,ZNF99,TUBB2B,COL5A1,CES3,PGK1,R
L18A1,SNRNP200,SMIM14,EIF3C,VCP,UBB,AFF4,TRIM28,YWHAG,CKB,RPL37A,FN1,SPEN,DDB1,Y
YGB,TERF2IP,SMARCA4,ATG2B,AZIN1,DDAH2,AEBP1,NYNRIN,ILF3,CASC3,QARS1,ZFAND5,DIDO1
A6,ITM2C,RPL30,EIF3L,CHPF,DNAJC5,HNRNPDL,RPLP2,DDOST,ATP1A2,SLC35E2B,RPL23A,HSB
HS1,SET,EGLN1,CHSY1,TMSB10,EIF4A1,RPL15,GTF3C4,LDHB,PKD1,SF3B1,PDIA3,ZNF43,BASP1
EST,ALKBH5,PDXK,CAMSAP1,KLC1,CCT3,VPS35,MLEC,RAD23B,SKI,NOTCH2,VEGFA,TUBB2A,AF

365,RPL9,KIF5B,RPS14,CXADR,LENG8,WDR1,CCT7,SRSF4,RBBP4,PFN1,MCL1,RPL31,PRKCSH,L
F5A,CCND2,RAB7A,PRKAR1A,CHPF,TARDBP,RPLP2,ARL6IP1,TSR2,MYL6,RPL36,SEPTIN11,LAMF
LM1,NPM1,TTC3,PLD3,HSPD1,USP11,SNRNP200,SMIM14,EIF3C,VCP,MARCKSL1,AFF4,CKB,RPL
EIF4A1,RPL19,GTF3C4,DNAH8,SF3B1,PDIA3,SERINC1,ZNF43,HMGB1,POLR2A,AP2M1,TPT1,RPS
SON,CCAR2,ARF4,SUPT6H,BAG6,RAB11A,SERBP1,ARFGEF2,AKIRIN1,CCT5,CDC42,PPP1R9B,TR
P1,HMGB1,TPT1,RPS3A,RPL18,STMN1,UCHL1,NCAM1,ZMIZ1,TIMP2,MIDN,MARCHF6,RPS17,SBF
RF4L1,ANXA2,ZMIZ1,TIMP2,XAF1,MIDN,AP3D1,AKAP12,CDC42BPB,MAT2A,GTF2F1,SLC4A8,HNF
,EIF3L,MAGED2,RPS5,IK,RPS19,CHPF,JUN,NGRN,RPLP2,DDOST,CIRBP,NARS1,RPL36,YWHAB,SI
GRN,APLP2,RPLP2,DDOST,CIRBP,CALU,ARCN1,COTL1,SLC35E2B,ATP6V0C,LAPTM4B,TRIB2,RPL
RPL30,ARL4C,CLTC,GNB2,APLP1,RPS19,TLN1,COL6A2,ANXA6,MTSS2,CIRBP,MYH10,ATP1A2,PL
,WDR1,ATXN7L3B,RPL26,SRSF4,RBBP4,PFN1,MCL1,TLE5,PNN,RPS25,EIF5A,PDIA6,PIP4K2B,RA
PL5,TMBIM6,SPTAN1,PLD3,HSPD1,USP11,SNRNP200,EIF3C,ATP5F1A,EIF4B,UBB,AFF4,RPL37A,L
PL5,TMBIM6,SPTAN1,PLD3,HSPD1,USP11,SNRNP200,EIF3C,ATP5F1A,EIF4B,UBB,AFF4,RPL37A,L
DH6A1,GNB1,RPS8,MYH9,SEPTIN2,P4HB,APP,GDI1,SPTAN1,SMG1,CALM1,NPM1,PLD3,HSPD1,CC
PL31,PRKCSH,SCRN1,RPS25,EIF5A,PDIA6,PIP4K2B,MEGF8,ITM2C,RPL30,EIF3L,MAGED2,GNB2
PI1,MRC2,RPL19,RPL15,GTF3C4,DNAH8,SF3B1,SERINC1,RPS3,CHD3,BASP1,TPT1,RPS4X,OR8K
2,RPL24,PAPOLA,HK1,RPL22,ZNF865,RPS14,CXADR,LENG8,WDR1,RPL26,SRSF4,PFN1,PNN,RPL
MSB10,INA,RPL19,PPP2R1A,RPL15,GTF3C4,SF3B1,BASP1,HMGB1,POLR2A,AP2M1,TPT1,SEC61
F1L,PRDX5,UBE4A,ACO2,HSPA4,MLEC,SRP14,NRDC,PRPF6,ARF3,MAPRE1,ARPC2,FAM168B,LUZ
F3,MDH2,U2AF2,EEF1A2,GNAI3,TSR1,MTRF1L,PRDX5,UBE4A,GCN1,PRPF6,MARS1,RPL37,CCNL
WDR1,RPL26,SRSF4,PFN1,MCL1,PNN,RPL31,PRKCSH,KHDRBS1,SCRN1,RPS25,EIF5A,PDIA6,ME
,CCT6A,SLC35E2B,CDKN1A,FAM199X,DLG5,TMEM132A,SRP9,ZC3H7B,RAB27A,GAS7,TSR1,TAF
DPYSL2,VHL,RABL6,TCF25,EIF3E,LUZP6,GRB2,SCAF11,CLIP3,POLR2B,MSL1,SDF4,SMARCC1,S
1B,TUBB2B,COL5A1,CES3,RPLP0,H3-3B,CANX,PTMA,DDX5,SOX4,FKBP14,MAGED1,RPL6,YWHA
1,RPS25,EIF5A,PDIA6,MEGF8,ITM2C,RPL30,EIF3L,MAGED2,RPS5,IK,RPS19,CHPF,JUN,NGRN,HN
TCAF1,SHC2,SSBP3,ABI2,UBR5,SLC6A8,PKDCC,MAZ,KIAA0232,PRR12,TMEM63B,DNAJB2,RAB6
Y1,TMSB10,EIF4A1,PPP2R1A,RPL15,DNAH8,PKD1,SF3B1,ZNF43,RPS3,VAPA,POLR2A,AP2M1,SEC
F3,AGRN,RPLP1,MIDN,RPL10,MARCHF6,DNAJA1,ARF1,SF3B2,GTF2F1,OGA,HNRNPA0,RPL29,TF
3,PRDX5,ACTR3,CLSTN3,ACAP3,ASXL1,PRPF6,RPL38,PBX2,EIF3E,PCSK1N,GABARAP,HNRNPUL1,
M14,IGF2,EIF4B,RPL37A,SPEN,NONO,PSMD5,HACD3,SET,TPI1,INA,EIF4A1,MRC2,PPP2R1A,GTF
ATP1A1,WDR1,OAZ1,PFN1,ASTN2,JPT2,NES,CCNI,TADA2B,KHDRBS1,PFKL,PEBP1,PIP4K2B,CCN
MBIM6,PTMS,SPTAN1,GSTP1,SMG1,PLXNB2,HSPD1,COL18A1,IGF2,EIF3C,EIF4B,VCP,MARCKSL1
HA,CPE,CTSD,SCARB2,RTN4,SLC22A17,RPS9,CIPC,SLC16A7,C1ORF43,PAPOLA,IGFBP3,CBX5,RF
5,CPSF2,PGRMC1,EID1,SRSF5,U2AF2,VAMP2,PARP1,PTPRA,HSPA4,ACAP3,RAD23B,ATP6AP1,TAF
ITM2C,RPL30,EIF3L,MAGED2,GNB2,RPS19,CHPF,JUN,NGRN,RPS10,RPLP2,DDOST,HNRNPK,CIF
EFL,NEFM,PRPF8,UGT8,RPL8,FBXL4,YBX1,UBA1,ATP2B4,IGFBP2,CALR,FSTL1,TMF1,ABHD2,HS
KHDRBS1,RPS25,EIF5A,PDIA6,DDR1,PIP4K2B,RAB7A,RPL30,EIF3L,IK,RPS19,CHPF,NGRN,RPLP2,
C28,INA,EIF4A1,RPL19,PPP2R1A,RPL15,GTF3C4,DNAH8,SF3B1,SERINC1,RPS3,HMGB1,POLR2A,
PD,VGF,CYP51A1,AKIRIN1,BMPR2,KCNQ2,EIF2AK1,BCL9L,BCR,CPD,TRIO,CNTNAP1,ALDH3A2,L
PD,VGF,CYP51A1,AKIRIN1,BMPR2,KCNQ2,EIF2AK1,BCL9L,BCR,CPD,TRIO,CNTNAP1,ALDH3A2,L
2,PTBP1,CCAR2,ARF4,BAG6,RAB11A,AKIRIN1,CCT5,CDC42,TRAF7,EIF2AK1,PRMT1,PDAP1,MMP
EMIP2,HNRNPA2B1,EML1,OAZ2,MCM3AP,BRSK1,GPSM1,HNRNPAB,GSE1,RABEP1,WSCD1,ZNF6
RPS19,CHPF,NGRN,DNAJC5,RPLP2,DDOST,YWHAB,ARCN1,ARHGDIA,SLC35E2B,IPO5,HSBP1,SRS
3,APLP2,INIP,MYH10,DUX4,CKAP5,SEPTIN11,RAC1,PTTG1IP,MAP1A,CDKN1A,TIMP3,CTDSP2,CN
4B,MGAT4B,CXXC5,SLC39A7,PPP2CB,PLXNB1,UBAP2L,ADCY6,ZNF512B,HNRNPA2B1,ULK1,GPS

;MARCA5,RPL23,RPSA,TPM4,SCARB2,RPS24,CIPC,C1ORF43,NCS1,PSMD2,RAB6B,ATP1A1,JUND
PI1,CHSY1,TMSB10,EIF4A1,DNAH8,SSBP2,RPS3,HMGB1,RPS3A,ITGB1,UCHL1,RPL11,AGRN,ZMI
RKCSH,XPO1,CSNK1E,HDLBP,TKT,PLXNA1,MEGF8,SOX12,MAGED2,CHPF,MYO5A,CD63,DNAJC5
ITM2C,EIF3L,CHPF,NGRN,RPLP2,DDOST,YWHAB,LAMP1,ARHGDIA,SLC35E2B,RPL23A,HSBP1,M/
L12,ITM2C,ARL4C,MIF,CHPF,MYO5A,HNRNPDL,ARL6IP1,SHOX,H1-10,RPL36,DUX4,PTGES3,LAM
RD52,INIP,MYH10,SND1,UBE2H,ARCN1,CTBP1,SMCHD1,ATP6V0C,SKP1,LAPTM4B,CPSF2,SRSF
1LEC,ZNF302,GCN1,SKI,PRPF6,RPL37,ARPC2,FAM168B,PTBP1,CDC37,CCAR2,ARF4,PITPNM1,A
4,CXADR,CHD4,LENG8,WDR1,RPL26,SRSF4,PFN1,ASTN2,TLE5,RPL31,PRKCSH,CSNK1E,PFKL,S(
,PAPOLA,IGFBP3,NCS1,RHOA,ZNF865,SLC7A5,NLGN2,GLIPR1L2,LENG8,ATXN7L3B,CRMP1,VA
B,ATP6V0C,LAPTM4B,HSBP1,SERF2,NUCB1,SRSF5,TM9SF3,MDH2,U2AF2,EEF1A2,CCT2,GNAI3,N
B,ATP6V0C,LAPTM4B,HSBP1,SERF2,NUCB1,SRSF5,TM9SF3,MDH2,U2AF2,EEF1A2,CCT2,GNAI3,N
12,ACTN4,TLN1,CHPF,MYO5A,COL6A2,TARDBP,DUX4,CAPRIN1,GPC1,TOP2B,SHISAL1,CRPPA,TI
RPL30,EIF3L,RPS7,RPS19,CHPF,NGRN,RPLP2,NDFIP1,DDOST,YWHAB,PTGES3,LAMP1,RPS12,RF
,GPC3,SOX12,RPL30,EIF3L,MRFAP1,CHPF,CD63,MACF1,HNRNPK,CADM4,RPS12,PLPPR3,ARCN
APT,FN1,SPEN,NONO,DDB1,YWHAQ,CHSY1,TMSB10,RPL15,GTF3C4,LDHB,PKD1,HLA-C,SSBP2,(
LXNA1,RAB7A,GNB2,RPS5,IK,PRKAR1A,CHPF,GJA1,NGRN,RPLP2,MYL6,INIP,MYH10,SND1,UBE2
200,VCP,MARCKSL1,UBB,AFF4,MATR3,RPL37A,SPEN,DDB1,PSMD5,HACD3,SET,EGLN1,MAP9,T
B,GNB1,MYH9,RPL7,SEPTIN2,APP,IDS,RPL5,TMBIM6,PTMS,MARCKS,SMG1,PLD3,HSPD1,USP11,
APLP2,INIP,MYH10,DUX4,CKAP5,SEPTIN11,RAC1,GLUL,PTTG1IP,MAP1A,CDKN1A,SERF2,TIMP3,(
PPL1,CDKN1A,CTDSP2,DDX39B,SQSTM1,CAMSAP1,KLC1,MLEC,CLSTN3,MDK,PTK7,RAB5B,OR/
BP1,SRSF5,TM9SF3,MDH2,U2AF2,FDFT1,GNAI3,MEST,HSPH1,PRDX5,UBE4A,HSPA4,TCP1,MLEC
10,EIF4A1,PPP2R1A,RPL15,DNAH8,PKD1,SF3B1,SERINC1,ZNF43,RPS3,VAPA,POLR2A,AP2M1,SE
,PTTG1IP,MAP1A,CDKN1A,TIMP3,CTDSP2,CNN2,NOVA2,NUCB1,ZC3H7B,U2AF2,CCT2,MEST,GA
2,FAM168B,ACADVL,LUZP6,CCAR2,GNG2,ISYNA1,ZFAND3,HGS,BAG6,HNRNPD,FBRSL1,RAB11/
RPS19,TLN1,CHPF,JUN,NGRN,RPLP2,ARL6IP1,DDOST,RPL36,YWHAB,ARCN1,RPS27,ARHGDIA,ST
R,LENG8,WDR1,RPL26,SRSF4,PFN1,MCL1,PNN,RPL31,PRKCSH,CCNI,KHDRBS1,SLC3A2,SCRN1,

GDI1,FSCN1,TMBIM6,PTMS,GSTP1,MARCKS,CALM1,CSDE1,ATP5F1A,VCP,MARCKSL1,COL4A2,F
L26,SRSF4,RBBP4,PFN1,MCL1,TLE5,PNN,RPS25,EIF5A,PDIA6,PIP4K2B,RAB7A,ITM2C,RPL30,EIF
A3,ALKBH5,ACTR3,CAMSAP1,GRINA,COL2A1,MDK,LAMC1,CHD2,ORAI2,HDGF,NRDC,VEGFA,RAF
C,RPL30,EIF3L,MAGED2,RPS5,IK,RPS19,CHPF,JUN,NGRN,RPLP2,DDOST,CIRBP,NARS1,SND1,CA
PX4,SLC35F5,GNG4,NXN,EP400,PPP1CC,TNKS,MAP1LC3B,MTCH1,ERGIC1,SLC44A2,HSPG2,C/
A2,AGRN,MIDN,AP3D1,AKAP12,MAP4,CDC42BPB,TMEM170A,SLC4A8,GOLGA8A,PGAM1,CTNNA
B,NOP53,TM9SF2,TSR1,MTRF1L,RPL27,PRDX5,UBE4A,TCP1,SURF4,GCN1,RPL36A,SRP14,PRPF
RD3,RPL17,HK1,ATF4,PSMD2,NR2F1,KIF5B,ENC1,NLGN2,GLIPR1L2,CHD4,ATP1A1,ATP2A2,CRI
PRKAR1A,CHPF,NGRN,RPLP2,MYL6,INIP,MYH10,SND1,UBE2H,ARCN1,HMGCR,TOP2B,SMCHD1,
L2,WDR6,SCAF11,BNIP3L,KIF3C,HNRNPM,STT3B,EIF3F,TUG1,ATXN2L,VANGL2,MAPK8IP1,RAB1
2,BEX3,CCT7,OAZ1,MCL1,LRPPRC,KHDRBS1,KLHL9,XPO1,CSNK1E,PFKL,CDH2,PIP4K2B,DDX3X
1,PGK1,RPLP0,PTMA,ENO2,SOX4,RPL13A,SEC62,DVL3,YWHAE,KIF1A,RPL13,GNAS,GPI,HTRA4,B
P3,TMEM248,LAMC1,SKI,ASXL1,ABCA2,VHL,PPP2CA,SMARCA4,ODC1,SRSF1,ARGLU1,ADAM9,P
2C,RPL30,EIF3L,MAGED2,RPS5,IK,RPS19,CHPF,JUN,NGRN,RPLP2,DDOST,CIRBP,NARS1,SND1,C
GB,DPYSL2,TMED2,DDAH2,PLEC,RABL6,ARGLU1,ADAM9,PTBP1,DYNLL2,ILF3,CDC37,WDR6,HN
2,PPDPF,ARIH1,PBX1,NOL4L,BAZ1B,CST3,CEMIP2,CUL3,NDST1,NPDC1,BRSK1,GPSM1,LRP10,S
ND,TMEM259,LENG8,WDR1,MCL1,CCNI,HDLBP,RPS25,EIF5A,PEBP1,CCND2,DDR1,PLXNA1,RAB
,LASP1,PNN,PRKCSH,CCNI,SLC3A2,PEBP1,CDH2,PDIA6,SOX12,ACTN4,ARL4C,MRFAP1,GNB2,AF

NISCH,CCNI,TADA2B,KHDRBS1,XPO1,HDLBP,RPS25,SFPQ,PEBP1,CLTC,MRFAP1,RPS5,IK,RPS19
A6,SEC16A,NXN,EP400,PPP1CC,CUL3,CHERP,CTTN,MAP1LC3B,C1ORF198,MTCH1,SAFB2,PRDX
CYFIP2,RPLP2,MYL6,INIP,MYH10,SND1,UBE2H,ARCN1,HMGCR,TOP2B,SMCHD1,ATP6V0C,LAPTM
,YME1L1,TRAF7,EIF2AK1,PHB2,APMAP,KLHL42,RER1,MASTL,YKT6,TMEM167A,TNKS,PSMD8,CH
SCAP,IER2,AUTS2,CLIP2,TSPYL2,WDR26,SCAMP4,PABPC4,LIMK1,MYCBP2,WIZ,DDX6,MAPKAPK2
PNB1,IGFBP3,RPL14,NCS1,RPL9,CLSTN1,RAB6B,CXADR,ATP1A1,JUND,CRMP1,CCT7,ASTN2,JPT
PNISR,INTS11,SPTY2D1,FBXW5,MBD3,SF3A2,AP1M1,BNIP3,SAE1,PRDX3,HSDL1,DENND5A,ACAT
,DDX5,ENO2,SOX4,MAGED1,SEC62,RPL6,YWHAE,KIF1A,RPL13,GNAS,SLC2A3,DDX17,GPI,HTRA4
A7,PPM1G,MBTPS1,HDAC6,CUL3,SLC7A6,SEPTIN7,ZMYM3,TRIR,ERH,TAOK1,CLPTM1,RAB1B,U
1A,GTF3C4,SF3B1,SERINC1,HMGB1,POLR2A,AP2M1,RPS4X,TUBB4B,MORF4L1,BTF3,AGRN,RP
,RPL6,KAT6A,YWHAE,KIF1A,GNAS,SLC2A3,DDX17,BSG,FASN,ALDH6A1,SPTBN1,ATP5F1B,CALM3
19,HNRNPDL,MYL6,WLS,HNRNPK,INIP,PTGES3,CADM4,RAB1A,CALU,CTBP1,STIP1,GPC1,SMCHD
SMCHD1,DNER,NNAT,U2AF2,TBCD,CPSF1,TSR1,TGFBI,ALKBH5,PDXK,CAMSAP1,CCT3,PTOV1,AC
,NPM1,PLD3,HSPD1,USP11,COL18A1,CSDE1,EIF3C,ATP5F1A,EIF4B,MARCKSL1,AFF4,TRIM28,CC
9,MYH9,APP,IDS,ERVV-1,GDI1,FSCN1,TMBIM6,PTMS,SPTAN1,SMG1,CALM1,NPM1,PLD3,USP11,
CTXN1,PCYOX1,CIC,RBBP7,PRMT1,RETREG2,LRRC4B,KIF21B,SEC16A,PHB2,SLC39A7,PRRC2A,
GCR,TOP2B,SMCHD1,ATP6V0C,LAPTM4B,SRSF5,TM9SF3,MLXIP,MORF4L2,HSPH1,ALKBH5,UBE
25,EIF5A,PDIA6,PIP4K2B,RPL12,RPL30,EIF3L,H1-0,IK,CHPF,CD63,DNAJC5,RPLP2,DDOST,CIRBF
PRIP,SOD1,FBRSL1,SMS,NAP1L4,CTXN1,PPDPF,BCL9L,XPOT,CAPN2,GRSF1,PI4KA,EIF5B,ANP32
3A,ITGB1,BTF3,AGRN,RPLP1,MARCHF6,DNAJA1,ARF1,SF3B2,MAT2A,HNRNPA0,RPS16,XRCC5,S
IOG,MRFAP1,RPS7,CTNNB1,MACF1,RPL35A,TSR2,NARS1,TTR,ARCN1,RPS27,HECTD4,ATP6V0C,
SLC7A5,ENC1,GLIPR1L2,TMEM259,RPS20,COL5A2,HMGN2,LASP1,RTN3,PFN1,TLE5,HNRNPC,R
2,YWHAH,BAG6,FBRSL1,RAB11A,SERBP1,CDC42,VDAC1,TRAF7,EIF2AK1,PRMT1,SF1,RETREG2,
P6V0C,MMP2,SKP1,LAPTM4B,RPL23A,CTDSP2,CNN2,TM9SF3,RNF187,VAMP2,GDI2,HNRNPL,RT
,CCN2,ITM2C,MAGED2,APLP1,SESN3,APLP2,INIP,DUX4,CKAP5,SEPTIN11,TNC,SKP1,TRIB2,PTTG
32B,COL5A1,PSAP,PGK1,RPLP0,H3-3B,CANX,CLU,PTMA,DDX5,ENO2,SOX4,RPL13A,MAGED1,SEC
PLP1,SARAF,MIDN,RPL10,MARCHF6,WSB1,RPS17,ARF1,SF3B2,ITM2B,AP2B1,GTF2F1,GOLGA8A
RF4L1,NCAM1,RPLP1,SARAF,MIDN,RPL10,MARCHF6,WSB1,RPS17,ARF1,SF3B2,ITM2B,AP2B1,GT
IR2L8,PKM,GANAB,COL6A1,FTL,HSP90B1,FADS1,NEFL,NEFM,PRPF8,UGT8,RPL8,FBXL4,YBX1,U
FIP1,TAGLN,RHOB,ATP1A2,PLPPR3,COTL1,PEA15,DLG5,NOVA2,RNF187,U2AF2,TBCD,MLXIP,SC
A8A,CTNNA1,RPL29,SCARB2,RPS24,CIPC,PJA2,GAP43,HK1,NCS1,RPL22,ZNF865,RPL9,CXADR,C
P2,DDOST,CIRBP,NARS1,SND1,CALU,ARCN1,ARHGDIA,SLC35E2B,ATP6V0C,RPL23A,FAM199X,DD
MAP1A,CDKN1A,TIMP3,CTDSP2,CNN2,NOVA2,NUCB1,ZC3H7B,U2AF2,CCT2,MEST,GAS7,TM9SF2
NARS1,CALU,ARCN1,ARHGDIA,ATP6V0C,LAPTM4B,TRIB2,RPL23A,DDX39B,SRSF5,TM9SF3,MDH
9,STMN2,RPS8,MYH9,RPL7,SEPTIN2,IDS,GDI1,SMG1,NPM1,PLXNB2,USP11,COL18A1,SNRNP200

P,AFF4,CKB,COL4A2,RPL37A,MAPT,FN1,SPEN,NONO,DDB1,EGLN1,CHSY1,INA,EIF4A1,RPL19,PF
N1,CALM3,GNB1,SHISA9,RPS8,MYH9,SEPTIN2,P4HB,IDS,RPL5,KLRD1,GDI1,SPTAN1,GSTP1,MA
STP1,PLXNB2,PLD3,HSPD1,COL18A1,CSDE1,SMIM14,IGF2,ATP5F1A,VCP,UBB,AFF4,YWHAG,CKB
,RPL30,EIF3L,MRFAP1,IK,CHPF,NGRN,HNRNPDL,RPLP2,DDOST,RPL36,YWHAB,PTGES3,LAMP1,
,EGLN1,INA,RPL19,PKD1,PDIA3,SSBP2,POLR2A,RPS4X,RPS3A,ANXA2,ZMIZ1,SARAF,RPL10,WSB1
S27,COTL1,CTBP1,RAC1,CAPRIN1,TOP2B,SLC35E2B,IPO5,ATP6V0C,TMEM30A,GLUL,HSBP1,HS
1LC3B,MTCH1,SLC44A2,HSPG2,CARM1,GPC2,TAX1BP1,LRATD2,CADM3,ACTR1B,MOAP1,PCNX3
HSY1,EIF4A1,PPP2R1A,SERINC1,POLR2A,TPT1,SEC61A1,RPS3A,RPL18,RPL11,MORF4L1,NCAM1
M2C,RPL30,EIF3L,MAGED2,RPS5,IK,RPS19,CHPF,JUN,NGRN,RPLP2,DDOST,CIRBP,NARS1,SND1

E,RPSA,SCARB2,LPCAT1,SLC22A17,RPS9,RPL24,NCS1,CBX5,GLIPR1L2,ATP1A1,JUND,TMEM259

LP1,SARAF,MIDN,RPL10,MARCHF6,WSB1,RPS17,ARF1,SF3B2,ITM2B,AP2B1,GTF2F1,GOLGA8A,I

P1CB,GDI2,PTPRA,PRDX5,UBE4A,TNPO1,ASXL1,PRPF6,PTBP1,SUPT5H,HNRNPUL1,SON,RPL28,C

,TTC3,USP11,SNRNP200,CSDE1,EIF3C,ATP5F1A,UBB,MATR3,LRRC28,NONO,MAP9,EIF4A1,RPL1

6,DCLK1,TBC1D9,DNAJB2,PHGDH,USP24,PHIP,UBE2D3,TRIM8,ADGRA3,COL16A1,TAOK2,SLC22

USP24,UBE2D3,WNK1,TRIM8,MYL12B,SH3PXD2B,IGLON5,LMO4,RSF1,AUTS2,SLC22A23,GGNBI

P4,COTL1,HECTD4,TOP2B,APPL1,ATP6V0C,TMEM30A,TMEM132A,PGRMC1,RNF187,EEF1A2,TG

PF,NAP1L1,TRIO,NOL4L,SEC16A,MAP6,NXN,MBTPS1,HNRNPA2B1,NCOA4,HIF1A,WSCD1,VPS26I

ISPD1,USP11,SNRNP200,CSDE1,SMIM14,EIF3C,ATP5F1A,UBB,AFF4,MATR3,RPL37A,LRRC28,FN1

PM1,PLXNB2,TTC3,USP11,COL18A1,IGF2,EIF3C,VCP,MARCKSL1,AFF4,MATR3,CKB,COL4A2,RPI

PI1,CHSY1,TMSB10,INA,LDHB,PKD1,PDIA3,SSBP2,ZNF43,HMGB1,AP2M1,TPT1,SEC61A1,STMN1

B12B,RABL6,SRSF1,AEBP1,NYNRIN,PTBP1,LUZP6,CDC37,PRKDC,HNRNPUL1,GIGYF1,YWHAH,PI

1,SF3B1,PDIA3,SERINC1,HMGB1,POLR2A,AP2M1,TPT1,RPS4X,RPS3A,ITGB1,TUBB4B,RPL11,BT

1,PLXNB2,USP11,EIF3C,CKB,MAPT,DDB1,HACD3,DCHS1,TPI1,TMSB10,INA,MRC2,RPL15,GTF3C

E2B,APPL1,SMCHD1,TMEM30A,TRIB2,DLG5,U2AF2,TBCD,PPP1CB,GDI2,TSR1,HSPH1,ALKBH5,H

,CDC42,PPP1R9B,TRAF7,DFFB,STAT3,EIF2AK1,TXLNA,PDAP1,AP2A1,GSPT1,UBA52,PLXNB1,CSN

49,RPL7,SEPTIN2,P4HB,ERVV-1,KLRD1,GDI1,TMBIM6,GSTP1,CALM1,NPM1,PLXNB2,TTC3,PLD3,

P2R1A,RPL15,PDIA3,SERINC1,HMGB1,POLR2A,TPT1,RPS3A,RPL18,RPL11,SARAF,MIDN,RPL10,M

VCP,UBB,AFF4,TRIM28,CKB,RPL37A,FN1,SPEN,DDB1,PSMD5,YWHAQ,EIF1,HACD3,MAP9,CHSY1,I

NFE2L1,SMARCA5,CTSD,NACA,RPS24,C1ORF43,PAPOLA,COL1A2,HK1,NCS1,ATF4,SLC7A5,ENC1

F8,SOX12,ITM2C,GNB2,CHPF,COL6A2,SESN3,APLP2,HNRNPDL,ARL6IP1,ANKRD52,H1-10,CIRBP,

B,SMCHD1,ATP6V0C,LAPTM4B,SRSF5,TM9SF3,MLXIP,MORF4L2,HSPH1,ALKBH5,UBE4A,TBC1D

N1,ANXA2,AGRN,ZMIZ1,TIMP2,XAF1,COL11A1,MARCHF6,VCL,AP3D1,DNAJA1,CDC42BPB,TMEM

,ORAI2,PSMB2,SCAF1,RAB11B,TUBB2A,OPTN,RPL37,PPP2CA,KDELR2,PSMD3,SRSF1,TCF25,GF

F1,TLK1,GNAZ,UBE2G1,SUDS3,PRR36,CTDNEP1,CRIM1,SYT1,SCAF8

PP,IDS,RPL5,FSCN1,PTMS,GSTP1,SMG1,PLXNB2,PLD3,CSDE1,SMIM14,IGF2,EIF4B,VCP,MARCKS

,TMSB10,PPP2R1A,RPL15,PKD1,CHD3,VAPA,SEC61A1,RPS3A,RPL18,STMN1,TUBB4B,MORF4L1

CHERP,CTTN,SERP1,VPS26B,ERH,CLPTM1,INTS3,ABI2,UBE2M,SEC14L1,ELAVL1,NUDCD3,FLOT2

I,RPS28,STT3B,STT3A,NBPF9,FBRSL1,CTXN1,PPP1R9B,PSMA7,KCNQ2,SGTA,PPP2CB,PLXNB1,S

GRN,XAF1,MIDN,WSB1,AP3D1,CYB5R3,MAP4,ITM2B,TMEM170A,GOLGA8A,HNRNPU,NFE2L1,SM

I,CD81,LENG8,WDR1,CRMP1,VAT1,OAZ1,RTN3,PFN1,NES,RPL31,TADA2B,KHDRBS1,PFKL,GLG1

DH1,SOD1,BEX4,BAZ1B,CD99,PHB2,UBA52,DIP2B,NXN,EP400,SELENOW,CUL3,ADH5,NDST1,ITG

PP2R1A,RPL15,PKD1,PDIA3,VAPA,POLR2A,TPT1,BTF3,ZMIZ1,TIMP2,WSB1,CYB5R3,MAP4,CDC42

NO2,FKBP14,RPL13A,MAGED1,RPL6,YWHAE,KIF1A,RPL13,DDX17,GPI,RACK1,BSG,FASN,ALDH6

APOLA,RPL9,RAB6B,ATP1A1,JUND,TMEM259,LENG8,WDR1,HMGN2,ATP2A2,RTN3,MCL1,CCNI,LI

PS19,MIF,JUN,CD63,DNAJC5,HNRNPDL,ARL6IP1,TSR2,H1-10,SEPTIN3,MAGED4B,ARHGDIA,STIP

B2,PRDX5,PKN1,ACO2,SURF4,TSPAN6,TAF7,RAPGEF1,CCNL2,PRDX1,ODC1,SRSF1,ADAM9,PTBP

KB,RPL37A,FN1,SPEN,DDB1,YWHAQ,EIF1,HACD3,MAP9,CHSY1,INA,EIF4A1,PPP2R1A,SERINC1,BA

A,SERINC1,POLR2A,TPT1,SEC61A1,RPS3A,RPL18,RPL11,MORF4L1,NCAM1,RPLP1,SARAF,MIDN

R1L2,CHD4,CD81,HMGN2,ATP2A2,CRMP1,LASP1,CCT7,VAT1,PFN1,RPL31,NISCH,LRPPRC,KLHL

ACD3,MAP9,CHSY1,INA,EIF4A1,MRC2,PPP2R1A,SERINC1,CHD3,BASP1,HMGB1,POLR2A,TPT1,SE

C4B,H2AZ1,RER1,ADH5,PTPRF,PODXL2,ZNF664,ZMYM3,VPS26B,BAZ2A,SEPTIN5,SLIT1,IARS2,YA

KD1,SF3B1,SSBP2,BASP1,VAPA,POLR2A,AP2M1,SEC61A1,STMN1,UCHL1,MORF4L1,NCAM1,ZMIZ

3,VAT1,CCNI,XPO1,SPTBN2,TPM1,EIF5A,CCND2,GPC3,PIP4K2B,MEGF8,ACTN4,DDX3X,ITM2C,RI

LP1,ZMIZ1,SBF1,SF3B2,CDC42BPB,TPM4,CTSD,BRD3,XRCC5,MSN,PAPOLA,LLGL1,NCS1,PTCD3

RINC1,ZNF43,RPS3,BASP1,VAPA,OR8K1,TUBB4B,UCHL1,MORF4L1,NCAM1,RPLP1,MIDN,WSB1,A
YWHAE,KIF1A,RPL13,SLC2A3,DDX17,GPI,RACK1,ATP5F1B,CALM3,GNB1,STMN2,MYH9,RPL7,SEF
CCT2,MEST,GAS7,TM9SF2,MORF4L2,SNRPN,PTOV1,MLEC,SURF4,PCDHGB6,PYGB,SMARCA4,A
,PPP2R1A,SERINC1,BASP1,POLR2A,TPT1,SEC61A1,RPS3A,RPL18,ITGB1,MORF4L1,NCAM1,ANXA
XL4,YBX1,ATP2B4,IGFBP2,CALR,FSTL1,TMF1,HSPA1B,ZNF99,TUBB2B,COL4A1,CES3,PSAP,PGK
A2,ATP13A1,PAFAH1B2,POLDIP2,TRIP12,CHMP1A,TSNAX,RGL2,SLC25A6,EXOC5,SF3A3,TRMT11
,EIF4G3,ATP1B2,AKT2,PCGF2,VASH1,CREBBP,SSBP4,GNA13,METRN,SH3PXD2B,ELAVL3,RNF11,
HDC3,PSMD8,CHERP,CTTN,SERP1,VPS26B,ERH,CLPTM1,INTS3,ABI2,UBE2M,SEC14L1,ELAVL1,N
PTAN1,CALM1,NPM1,PLD3,HSPD1,USP11,SNRNP200,CSDE1,SMIM14,EIF3C,ATP5F1A,EIF4B,VCP,
,SRPRA,SLC25A36,XRN2,ZZEF1,CDV3,FZR1,COPE,ATP1B3,CNDP2,GLOD4,MTURN,GOLPH3,PSM
YSL2,VEGFA,ARF3,MAPRE1,DDAH2,AEBP1,WDR6,NSG2,SMARCC1,PHC2,FAU,ZFAND3,STRAP,MAF
EM170A,GTF2F1,OGA,NFE2L1,CNN3,TSPAN3,LPCAT1,LAMA5,XRCC5,MSN,KPNB1,C1ORF43,GA
,TPI1,TMSB10,SSBP2,VAPA,SEC61A1,STMN1,UCHL1,MORF4L1,AGRN,RPLP1,TIMP2,XAF1,COL1
KIF1A,GNAS,SLC2A3,DDX17,GPI,HTRA4,FASN,SPTBN1,ATP5F1B,CALM3,GNB1,SHISA9,STMN2,RF
ATR3,YWHAG,RPL37A,FN1,PSMD5,HACD3,DCHS1,SET,CHSY1,INA,EIF4A1,MRC2,RPL19,PPP2R1A,
1,RPL36A,ABCA2,TAF7,TUBB2A,PPP2CA,MAPRE1,CCNL2,PDE7A,PSMD3,PTBP1,ILF3,CMIP,BNIP
2,YWHAH,PITPNM1,ZFAND3,BAG6,RAB11A,SERBP1,ARFGEF2,CDC42,TRAF7,EIF2AK1,PRMT1,SF
,CRMP1,CCT7,OAZ1,MCL1,NISCH,LRPPRC,KHDRBS1,KLHL9,XPO1,PFKL,TKT,PEBP1,CDH2,PIP4
P5F1B,CALM3,STMN2,RPS8,SEPTIN2,P4HB,IDS,ERVV-1,RPL5,GDI1,FSCN1,TMBIM6,GSTP1,MAR
GLUL,SQLE,MAP1A,SERF2,CTDSP2,CNN2,NOVA2,C5ORF24,ZC3H7B,U2AF2,CCT2,MEST,GAS7,T
,LENG8,WDR1,HMGN2,RBBP4,MCL1,PRKCSH,LRPPRC,XPO1,SLC3A2,TPM1,EIF5A,CCN2,DDX3X,
HTRA4,BSG,FASN,SPTBN1,CALM3,SHISA9,RPS8,MYH9,P4HB,APP,IDS,GDI1,FSCN1,TMBIM6,PTM
GTF2F1,LDHA,CPE,CNN3,RPSA,CTSD,RTN4,LPCAT1,LAMA5,PJA2,RPL24,CBX5,PSMD2,GLIPR1L2,
P3,SRP14,NRDC,PRPF6,MAPRE1,FAM168B,LUZP6,HSPB1,SON,CORO1C,DHX30,TNKS2,BAG6,HN
RCKSL1,MATR3,COL4A2,FN1,DDB1,DCHS1,SET,EGLN1,TPI1,CHSY1,EIF4A1,RPL19,GTF3C4,DNA
L23A,HSBP1,NNAT,MAP1A,DDX39B,TM9SF3,EEF1A2,PPP1CB,GDI2,MTRF1L,PTPRA,PRDX5,UBE4
BPA,PPFIA3,SCAF8,SLC39A1,USP36,ZC3H4
STT3B,ISYNA1,TMED9,TRAF7,EIF2AK1,PRMT1,MTA1,KDM5B,MMP15,PHB2,CST3,MAP6,KLHL42,
RBS1,PFKL,TKT,SFPQ,TPM1,CCND2,GPC3,PIP4K2B,DDX3X,ARL4C,MRFAP1,APLP1,MIF,MYO5A,

AB,LAMP1,MAP2,CALU,ARCN1,COTL1,ARHGDIA,ATP6V0C,BCLAF1,HSBP1,DDX39B,CTSB,TM9SF3
PTY2D1,CAP1,COX6A1,EIF3G,AP1M1,TOMM70,XPO6,UBXN6,MYL12B,GAK,LYPLA2,ATP13A1,PO
P1,SESN3,APLP2,INIP,DUX4,CKAP5,SEPTIN11,RAC1,BCLAF1,GLUL,PTTG1IP,MAP1A,CDKN1A,SEF
A-B,STIP1,SLC35E2B,IPO5,BCLAF1,GLUL,SQLE,CTDSP2,CNN2,CDH11,CHD6,FDFT1,RPN2,PABP
AHCYL1,KLC1,CLSTN3,RAB11B,PRPF6,TUBB2A,UBXN4,PPP2CA,G3BP2,KDELR2,PDE7A,PSMD3,
2,PLD3,COL18A1,CSDE1,IGF2,EIF4B,MARCKSL1,UBB,TRIM28,YWHAG,SPEN,DDB1,HACD3,SET,E
168B,LUZP6,SON,CORO1C,DHX30,TNKS2,BAG6,HNRNPD,CDC42,RBBP7,TRAF7,EIF2AK1,EDC4,
,RAB7A,DDX3X,RPS15,CLTC,CD63,ANXA6,MACF1,MTSS2,TSR2,H1-10,RPL36,DUX4,YWHAB,LAM
SF1,ARGLU1,RBP1,ILF3,CDC37,PRKDC,SCAF11,ACTN1,SON,SMARCC1,STT3B,PRPS1,HNRNPD,

UBFD1,HDAC6,PRDX6,ZNF664,PALM,BCL7A,PKDCC,TLE1,MAZ,DCLK1,PRR12,CADM3,RAB6A,SC
1,HSPA1B,TUBB2B,COL5A1,COL4A1,CES3,PSAP,PGK1,CANX,DDX5,ENO2,RPL13A,SEC62,RPL6,
ABCF1,RBPJ,NOC2L,NASP,AP2A2,RBM6,ATXN7L3,SSBP4,DPYSL4,DEK,CPXM1,ATP13A1,ADGRA3
,INIP,MYH10,SND1,UBE2H,ARCN1,HMGCR,TOP2B,SMCHD1,IPO5,ATP6V0C,LAPTM4B,SRSF5,TM
B1,SERINC1,RPS3,BASP1,POLR2A,RPS4X,UCHL1,RPL11,MORF4L1,RPLP1,COL11A1,MARCHF6,

CIC,TRAF7,DCTN1,EDC4,ACTR1A,STARD7,SF1,PBX1,AP2A1,GPX4,HJURP,SFRP1,IARS1,AGPAT1,I
PL13,GNAS,DDX17,GPI,HTRA4,FASN,SPTBN1,GNB1,SHISA9,STMN2,RPL7,SEPTIN2,P4HB,APP,IDS
LXNB1,DECR1,RER1,RAB3GAP2,YKT6,MCM3AP,TMEM167A,CHERP,CTTN,LRP10,ERH,EIF3H,SE
,CKB,RPL37A,SPEN,DDB1,YWHAQ,HACD3,DCHS1,SET,MAP9,TMSB10,RPL19,PPP2R1A,RPL15,GT
N1,RPS27,HECTD4,ATP6V0C,BCLAF1,L1CAM,CPSF2,SRP9,SRSF5,U2AF2,VAMP2,NOP53,PABPN1
RP14,PRPF6,TAF7,SELENON,RPL37,ATG2B,CCNL2,ARPC2,FAM168B,SRSF1,ATXN10,QARS1,GN
,SNRPN,PTOV1,MLEC,SURF4,SMARCA4,AZIN1,DDAH2,NYNRIN,WDR6,PRKDC,CASC3,RPL28,ZFA
1,PNN,SPC25,PRKCSH,CCNI,SPTBN2,PFKL,RPS25,EIF5A,CCND2,PDIA6,RAB7A,PRDX2,DDX3X,EI
NP32B,C1ORF198,BAP1,PPP1R10,SLIT1,TPP1,TRAM1,LSM14B,SLC7A1,PACS1,DICER1,SPTY2D
TADA2B,KLHL9,HDLBP,PEBP1,CCND2,PIP4K2B,CCN2,ACTN4,RCN1,GPAT4,GJA1,CD63,SESN3,S
2A,SRP54,DNAJB6,NARF,PCED1A,CAPZA2,TRAF3,CABIN1,MBD6,VPS16,PSMA4,NUP50,OBSCN,P
CD3,TUBGCP6,EEF1B2,NCSTN,INTS11,AHSA1,MTMR4,MKRN1,CAP1,LRRC41,PRPF19,USP24,AP
IA,EIF4A1,PPP2R1A,SERINC1,BASP1,POLR2A,TPT1,SEC61A1,RPS3A,RPL18,ITGB1,MORF4L1,NC
B,LUZP6,SON,DHX30,TNKS2,SARS1,BAG6,HNRNPD,AMOTL2,CDC42,TRAF7,EDC4,WDR45B,PDA
CALM1,PLXNB2,HSPD1,COL18A1,SMIM14,IGF2,MARCKSL1,AFF4,TRIM28,RPL37A,SPEN,NONO,
A,POLR2A,CDKL1,SEC61A1,MORF4L1,NCAM1,RPLP1,RPL10,AKAP12,ITM2B,ZNF770,GTF2F1,S
LEKHA6,SRSF2,CAPZA1,SOD2,DICER1,IST1,MN1,TUBB4A,IER2,CALCOCO1,IGLON5,LMO4,RNF
MLXIP,MORF4L2,HSPH1,ALKBH5,AHCYL1,ETNK1,KLC1,CLSTN3,DDX42,SCAF1,RAB11B,NRDC,T
1,CTNNA1,CTSD,SCARB2,RTN4,RPS24,XRCC5,C1ORF43,IGFBP3,HK1,PTCD3,ATF4,PSMD2,SLC7
19,GTF3C4,PKD1,HLA-C,SF3B1,ZNF43,BASP1,VAPA,POLR2A,CDKL1,TPT1,SEC61A1,RPS3A,OR8
PS21,BANF1,VPS51,FAM32A,SYNGAP1,EPN1,RTF2,CLPTM1L,GBA2,POLDIP3,UNC45A,RNMT,WA
4,TSPAN6,SELENON,PLEC,BTBD2,PRDX1,AEBP1,NYNRIN,NSG2,FKBP8,MSL1,SMARCE1,STMN3,
X5,ENO2,FKBP14,RPL13A,MAGED1,SEC62,YWHAE,KIF1A,RPL13,GNAS,DDX17,GPI,RACK1,FASN,
4A,AHCYL1,KLC1,CLSTN3,DDX42,RAB11B,TUBB2A,UBXN4,PPP2CA,G3BP2,KDELR2,PDE7A,PSM
NK1,KLC1,CLSTN3,RAB11B,SRP14,NRDC,TUBB2A,PPP2CA,PDE7A,PSMD3,SRSF1,ADAM9,EIF3E,
PYSL2,AZIN1,SRSF1,RPS11,HNRNPUL1,DLK1,STT3B,ISYNA1,SF3A1,GPR107,GHITM,MEX3A,ZS
RCKS,CALM1,COL18A1,SNRNP200,VCP,MARCKSL1,COL4A2,LRRC28,NONO,PSMD5,EIF1,DCHS
,PTOV1,MLEC,MDK,SURF4,ORAI2,SCAF1,RAB11B,PYGB,OPTN,DDAH2,DSTN,ODC1,NYNRIN,TCF
,ADNP,ANKFY1,SRSF10,CCNG1,GID8,MARCHF7,ZCCHC14,POLDIP3,EXOSC10,HMGXB3,ZFP91,
ID3,CDC37,WDR6,BNIP3L,HNRNPUL2-BSCL2,PITPNA,XPNPEP3,LARS1,YWHAH,UBQLN1,CSNK1D
,DAP,EIF3H,ELAVL1,IMPDH2,GABARAPL1,GNL1,TM9SF4,TRAP1,ELP1,MOAP1,CLASP2,NSD2,YY1
,SLC9A7,ALDOA,MTRNR2L8,PKM,GANAB,COL6A1,FTL,HSP90B1,FADS1,NEFL,NEFM,PRPF8,UGT
,TMEM167A,PSMD8,CTTN,SERP1,VPS26B,EIF3H,ELAVL1,NUDCD3,TAX1BP1,SH3GL1,NCSTN,INT
CAR2,ARF4,NKTR,CCT8,STMN3,TNKS2,NBPF9,SARS1,BAG6,HNRNPD,WBP2,AKIRIN1,CCT5,CDC
NPEP3,SRSF3,LARS1,YWHAH,UBQLN1,CSNK1D,IFITM3,VGF,MLF2,AKIRIN1,AMOTL2,YME1L1,T
,PAPOLA,NCS1,RPL22,RHOA,ENC1,NLGN2,JUND,EIF5,WDR1,ATXN7L3B,CRMP1,LASP1,PNN,PRK
42,YME1L1,TRAF7,EIF2AK1,SF1,RETREG2,PDAP1,PHB2,SLC39A7,KLHL42,IARS1,MASTL,YKT6,T
BLN1,STXBP1,ATXN2L,MAPK8IP1,RAI1,SREBF2,VGF,RAB11A,P3H3,WBP2,STARD7,KDM5B,HIP1,
1,GDI1,MARCKS,NPM1,TTC3,PLD3,HSPD1,USP11,COL18A1,SNRNP200,CSDE1,ATP5F1A,EIF4B,
MIDN,RPL10,MARCHF6,DNAJA1,ARF1,SF3B2,GTF2F1,PGAM1,HNRNPA0,RPS16,RPL23,RPL29,C
RTF2,UNC45A,ZBTB4,WARS1,SNRPB,KDM5A,ZDHHC5,SKIV2L,SHC1,SUDS3,MIB2,TNPO3,CUL4
ARAPL1,GNL1,TM9SF4,TRAP1,ELP1,MOAP1,CLASP2,NSD2,YY1,ELOVL5,SLC25A36,XRN2,SCAM
CLSTN3,DDX42,RAB11B,NRDC,TUBB2A,UBXN4,PPP2CA,G3BP2,KDELR2,PDE7A,PSMD3,CDC37,
99,TUBB2B,COL5A1,PSAP,PGK1,RPLP0,CLU,PTMA,DDX5,ENO2,FKBP14,RPL13A,MAGED1,SEC6
8,DUSP4,DHX30,LSS,FBLN1,CSNK1D,NBPF9,ZFAND3,SARS1,COL27A1,VANGL2,BAG6,RAI1,MEX

l2,CD2BP2,APEH,BANF1,VPS51,FAM32A,EPN1,RTF2,CLPTM1L,UNC45A,RNMT,EIF3M,WARS1,CTI
l,RBBP4,PFN1,MCL1,SPC25,HNRNPC,CCNI,RPL12,RPL35,CTNNB1,HNRNPDL,WLS,HNRNPK,MYH
RPL38,RPL37,CCNL2,FTO,LUZP6,UBE2R2,SUPT6H,MTRNR2L10,PITPNM1,ZFAND3,HGS,BAG6,HN
BXL4,YBX1,UBA1,ATP2B4,IGFBP2,CALR,FSTL1,TMF1,ABHD2,HSPA1B,TUBB2B,COL5A1,COL4A1
N,RPL10,WSB1,RPS17,VCL,ZNF770,MAT2A,GTF2F1,GOLGA8A,NFE2L1,RPS16,RPL23,EIF4H,TSP
IBE4A,HSPA4,KLC1,MLEC,TERF2IP,TAF7,RNF10,RPL37,G3BP2,ATG2B,FAM168B,LUZP6,CAPZB,S
'2,TSR1,MORF4L2,PDXK,AHCYL1,PTOV1,ORAI2,TNPO1,PYGB,G3BP2,KDELR2,SRSF9,TCF25,GRI
53,PABPN1,ACTR3,TBC1D16,ACO2,RBMX,CHD2,TERF2IP,TMED2,TUBB2A,PPP2CA,MAPK1,AZIN1
MAP9,RPL19,GTF3C4,PDIA3,HMGB1,POLR2A,TPT1,SEC61A1,RPS3A,RPL11,MORF4L1,RPL10,M
D2,RAB7A,CCN2,DDX3X,RPS15,RPS5,RPS19,CTNNB1,CD63,ANXA6,MTSS2,HNRNPDL,RPLP2,RPL
I1,NPM1,TTC3,USP11,SNRNP200,CSDE1,EIF3C,ATP5F1A,UBB,AFF4,MATR3,RPL37A,LRRC28,NOI
,TSNAX,RGL2,SLC25A6,EXOC5,SF3A3,TRMT112,COPE,CD2BP2,GOLPH3,PPP1CA,VPS51,PPME1,
NDUFA13,PSMC1,XPO6,SLC38A10,ATP13A1,NELFB,PAFAH1B2,ACAT2,POLDIP2,TRIP12,LMAN2,(
GFA,RAPGEF1,KDELR2,RABL6,ODC1,TCF25,PTBP1,FKBP8,PLCG1,DIDO1,PLTP,GIGYF1,PTP4A2
RKDC,CASC3,ZFAND5,CORO1C,SDF4,SMARCC1,HNRNPM,STT3A,PLXNA3,IDH1,SARS1,SOD1,CD
,KLHL9,SPTBN2,GLG1,PEBP1,CCND2,PIP4K2B,ITM2C,MAGED2,APLP1,SESN3,APLP2,ANKRD52,(
3H5,AHCYL1,ETNK1,KLC1,CLSTN3,DDX42,RAB11B,NRDC,PRPF6,TUBB2A,PPP2CA,G3BP2,PDE7A
AP,ELP1,CLASP2,PRPF19,PTPN23,CENPB,DHX40,RANGAP1,SLC25A36,ZZEF1,ESYT1,YTHDF3,MF
MEM248,ABCA2,ARPC2,DSTN,ODC1,ACADVL,ATP5PB,PCSK1N,MAP2K2,XPNPEP3,SUPT6H,HNRNF
OP53,PABPN1,HNRNPL,PTPRA,ACTR3,YEATS2,TBC1D16,ACO2,RBMX,CHD2,ASXL1,SCAF1,TMED
C62,KAT6A,KIF1A,RPL13,DDX17,GPI,HTRA4,RACK1,FASN,SPTBN1,CALM3,SHISA9,SEPTIN2,P4HB,
PR3,SLC35E2B,HSBP1,SRP9,NUCB1,MDH2,U2AF2,EEF1A2,CCT2,NOP53,PARP1,TM9SF2,TSR1,LI

)3,BASP1,HMGB1,POLR2A,TPT1,SEC61A1,RPS3A,RPL18,RPL11,NCAM1,SARAF,MIDN,MARCHF6,
B,IPO5,BCLAF1,GLUL,SQLE,CTDSP2,CNN2,NUCB1,CDH11,FDFT1,RPN2,HNRNPL,MORF4L2,NOX
3,CREBBP,DMWD,FGFR1,ARF5,IGLON5,CASC4,POLDIP2,WDTC1,HDGFL3,KIFC2,ANKHD1,GGNB
8,RPL11,NCAM1,SARAF,MIDN,RPL10,WSB1,CDC42BPB,ITM2B,AP2B1,GTF2F1,GOLGA8A,SMAR
1A1,TUBB4B,MIDN,RPS17,DNAJA1,MAP4,SF3B2,MAT2A,HNRNPU,HNRNPA0,FADS2,LDHA,CNN3,
3P2,PDE7A,PSMD3,CDC37,WDR6,BNIP3L,HNRNPUL2-BSCL2,PITPNA,XPNPEP3,LARS1,YWHAH,UB
BHD2,HSPA1B,ZNF99,TUBB2B,COL4A1,CES3,PSAP,H3-3B,CANX,CLU,PTMA,DDX5,ENO2,FKBP14
EIF3D,CST3,HOXB5,CD164,PTEN,SPINT2,BRSK1,CALD1,GPC4,C1ORF198,FBXO21,PCBP4,TCAF1
OLDIP3,EXOSC10,ZFP91,XPC,DDHD2,CNOT3,QRICH1,TMCO1,MYEF2,C6ORF89,RBM39
ST,YWHAB,RPS27,ARHGDIA,STIP1,TOP2B,LAPTM4B,TRIB2,PPIB,HSBP1,MAP1A,CRPPA,DDX39B,N
,MTMR4,MKRN1,ANKS1A,CAP1,PRPF19,AP1M1,TOMM70,DEK,ATP13A1,EIF4G1,PAFAH1B2,POLD
F5,TM9SF3,MDH2,RNF187,U2AF2,CCT2,GNAI3,NOP53,HSPH1,LMNB2,PRDX5,UBE4A,HSPA4,MLE
LC25A6,KIF26A,KAT2A,RNF157,RALGDS,GALNT10,SYT15,TASOR,WIZ,SOBP,LDB1,ZCCHC14,UBL
,TNPO3,CUL4B,SZT2,SEC31A,ACIN1,AKAP17A,MAPK8IP3,HARS1,MRPL3,DNMT1,CFAP298,DDX5
TP2A2,LASP1,BEX3,SRSF4,PFN1,PNN,HNRNPC,LRPPRC,SCRN1,HDLBP,TPM1,SOX12,MAGED2,IK,
HISA9,RPL7,IDS,TMBIM6,SPTAN1,SMG1,CALM1,NPM1,PLXNB2,PLD3,HSPD1,USP11,SNRNP200,C
3,H3-3B,CLU,PTMA,DDX5,SOX4,RPL13A,SEC62,KAT6A,DVL3,KIF1A,GNAS,GPI,HTRA4,BSG,FASN,
M32A,EPN1,ARPC3,RTF2,POLDIP3,UNC45A,ZBTB4,PSMF1,HMGXB3,WARS1,CLCN6,SNRPB,COX6
A1,MIDN,RPL10,WSB1,SBF1,DNAJA1,ARF1,CYB5R3,CDC42BPB,ITM2B,GOLGA8A,PXDN,OGA,CTN
5,CSNK1G2,UPF1,MASTL,MAPRE2,PPP1R15A,GALNT11,DAP,EIF3H,ELAVL1,IMPDH2,GABARAPL1
H1,SF1,SLC35F5,CSNK1G2,UPF1,MASTL,MAPRE2,PPP1R15A,GALNT11,DAP,EIF3H,ELAVL1,IMPD
T2,MTCH1,ZNF664,SHC2,SMG5,MAZ,SH3GL1,PLEKHA6,RIC8A,PLEKHB2,PHGDH,PSMD4,LRP3,T

:T5,CDC42,YME1L1,EIF2AK1,PDAP1,PHB2,SLC39A7,TRA2B,IARS1,YKT6,MCM3AP,TMEM167A,CH
A2,CSNK1E,TPM1,CCND2,PDIA6,PIP4K2B,MEGF8,CCN2,MMP14,PRDX2,GNB2,HMGCS1,JUN,COL
3F2,VGF,MLF2,AKIRIN1,AMOTL2,YME1L1,TRAF4,STAT3,ARIH1,SF1,SLC35F5,CSNK1G2,UPF1,MA
35F5,CSNK1G2,UPF1,MASTL,MAPRE2,PPP1R15A,GALNT11,DAP,EIF3H,ELAVL1,IMPDH2,GABARA
35F5,CSNK1G2,UPF1,MASTL,MAPRE2,PPP1R15A,GALNT11,DAP,EIF3H,ELAVL1,IMPDH2,GABARA
SSR1,XPOT,NAP1L1,CPD,SF1,HIP1,UBQLN2,SLC35F5,CD164,KIAA0895L,GLUD1,HDAC6,RPS2,SE
USP11,SNRNP200,SMIM14,IGF2,EIF4B,VCP,TRIM28,MATR3,YWHAG,RPL37A,LRRC28,NONO,DDI
1,PNMA1,MMP14,ITM2C,H1-0,APLP1,APLP2,SLC25A3,RPLP2,INIP,MYH10,DUX4,SEPTIN11,LGALS
D3,HSPD1,USP11,SNRNP200,EIF3C,MARCKSL1,AFF4,RPL37A,MAPT,SPEN,DDB1,PSMD5,YWHAQ,
PSMD5,SET,EGLN1,MAP9,CHSY1,INA,MRC2,LDHB,PKD1,PDIA3,RPS3,VAPA,POLR2A,SEC61A1,RPS
PSMD5,SET,EGLN1,MAP9,CHSY1,INA,MRC2,LDHB,PKD1,PDIA3,RPS3,VAPA,POLR2A,SEC61A1,RPS
,TNS3,LAMA5,RPL24,PAPOLA,COL1A2,LLGL1,NCS1,ATF4,JUND,EIF5,WDR1,ATP2A2,RPL26,MCL
RBM12,OGT,YY1,THRAP3,TUB,SLC25A36,ZZEF1,ESYT1,FAM50A,CNDP2,MTURN,GOLPH3,ECHS1,
PI,HTRA4,FASN,ALDH6A1,SPTBN1,SHISA9,STMN2,RPS8,RPL7,IDS,RPL5,GDI1,PTMS,SPTAN1,GST
L1,LBH,NDUFV1,GTPBP1,GRSF1,KIF21B,GPX4,DECR1,GNG4,PAPSS1,UBAP2L,UPF1,ZFR,NOMO
A,CST3,BRD2,CSNK1G2,RAB2A,SLC7A6,GPSM1,POM121C,TNKS,LRRC59,SERP1,SPG7,SSR3,ZN
RPL13A,SEC62,RPL6,KAT6A,DVL3,YWHAE,KIF1A,GNAS,DDX17,GPI,RACK1,ALDH6A1,SPTBN1,ATF
TRIM28,CKB,RPL37A,FN1,DDB1,PSMD5,YWHAQ,EIF1,HACD3,MAP9,CHSY1,INA,EIF4A1,PPP2R1A,I
F4,MATR3,YWHAG,LRRC28,PSMD5,EIF1,DCHS1,SET,EGLN1,MAP9,INA,EIF4A1,RPL19,PKD1,SF3I

P1,SEC16A,APMAP,UBFD1,YKT6,MCM3AP,RPL7L1,RPS27A,TMEM167A,KLHDC3,CHERP,CTTN,MT
,EIF5,LENG8,ATXN7L3B,JPT2,RPL31,NISCH,XPR1,KLHL9,EIF5A,PEBP1,CCND2,DDR1,MMP14,PR

3,ASTN2,MCL1,JPT2,PNN,PFKL,EIF5A,CDH2,MEGF8,GNB2,H1-0,RPL35,MIF,CHPF,MYO5A,COL6A
,PSMC4,PDCD11,WARS1,SNRPB,KDM5A,PMPCA,CFLAR,SHC1,PCED1A,CHMP5,MIB2,CUL4B,MCM
,TRIM28,CKB,MAPT,FN1,SPEN,LDHB,DNAH8,HLA-C,SF3B1,SERINC1,ZNF43,POLR2A,RPS4X,RPS
1,SLC35F5,CSNK1G2,UPF1,MASTL,MAPRE2,PPP1R15A,GALNT11,DAP,EIF3H,ELAVL1,IMPDH2,GA
APTM4B,CNN2,SRP9,MDH2,U2AF2,NOP53,PABPN1,PTPRA,ACTR3,YEATS2,TBC1D16,ACO2,RBMX
HISA9,P4HB,ERVV-1,RPL5,FSCN1,GSTP1,MARCKS,NPM1,PLD3,SMIM14,IGF2,EIF4B,AFF4,YWHAG
N,TPI1,CHSY1,TMSB10,GTF3C4,HLA-C,SF3B1,VAPA,POLR2A,CDKL1,RPS4X,AGRN,TIMP2,COL11A
PH3,VPS51,FAM32A,EPN1,ARPC3,RTF2,UNC45A,PSMF1,HMGXB3,MCFD2,WARS1,CLCN6,COX6B
PSMD5,SET,EGLN1,MAP9,CHSY1,INA,MRC2,LDHB,PKD1,PDIA3,RPS3,BASP1,VAPA,POLR2A,SEC61
F2,AKIRIN1,AMOTL2,YME1L1,TRAF4,STAT3,ARIH1,SF1,SLC35F5,CSNK1G2,UPF1,MASTL,PRRC2
ENG8,MCL1,PNN,CCNI,XPR1,HDLBP,PLXNA1,RAB7A,NANOG,GNB2,RPS5,IK,PRKAR1A,CHPF,GJA
MT112,CD2BP2,FAF2,KLC2,VPS51,PPME1,FAM32A,EPN1,RTF2,POLDIP3,UNC45A,ZBTB4,HMGXI
2,GLOD4,MTURN,DHX36,COLGALT1,TSPAN7,NUP98,OXSR1,EMC10,TMEM9,SYNGR1,RAB3A,SOX
D3,CDC37,HNRNPUL2-BSCL2,PITPNA,TPM3,XPNPEP3,LARS1,YWHAH,UBQLN1,CSNK1D,USP7,IFI
1,MTCH1,WSCD1,SEPTIN5,CEP170,EIF2S2,ELAVL1,HINT1,KIAA0232,TUBGCP6,SFXN1,PHGDH,S
RPL28,CCAR2,ARF4,SUPT6H,STMN3,TNKS2,SARS1,BAG6,HNRNPD,AKIRIN1,CCT5,CDC42,PPP1F
6V0C,MMP2,LAPTM4B,SRSF5,MLXIP,NOP53,HSPH1,MTRF1L,ALKBH5,PRDX5,PDXK,UBE4A,HSPA

H1,SF1,SLC35F5,CSNK1G2,UPF1,MASTL,MAPRE2,PPP1R15A,GALNT11,DAP,EIF3H,ELAVL1,IMPD
K1,KLC1,CLSTN3,DDX42,RAB11B,NRDC,TUBB2A,UBXN4,PPP2CA,G3BP2,PDE7A,PSMD3,SRSF1,T
),IFITM3,VGF,MLF2,AKIRIN1,TP53BP1,AMOTL2,YME1L1,TRAF4,STAT3,ARIH1,SF1,SLC35F5,CSNI
,MAPK8IP3,HARS1,MRPL3,DNMT1,CFAP298,DDX56,CPSF7,RAD50,JRK,TOMM40,COX5B

N1,AMOTL2,YME1L1,TRAF4,STAT3,ARIH1,SF1,SLC35F5,CSNK1G2,UPF1,MASTL,MAPRE2,PPP1R
NF157,RALGDS,GALNT10,SYT15,TASOR,WIZ,SOBP,LDB1,ZCCHC14,UBL3,ZSWIM6,BEX1,ILRUN,
XSR1,EMC10,TMEM9,SYNGR1,RPS29,OGDH,PLCD3,CTDNEP1,CMTR1,CLCN3,TRAPPC1,ABCE1,
XSR1,EMC10,TMEM9,SYNGR1,RPS29,OGDH,PLCD3,CTDNEP1,CMTR1,CLCN3,TRAPPC1,ABCE1,
M3,SREBF2,VGF,MLF2,AKIRIN1,AMOTL2,YME1L1,TRAF4,STAT3,ARIH1,SF1,SLC35F5,CSNK1G2
TARDBP,SLC25A3,HNRNPK,CIRBP,PTGES3,CKAP5,LAMP1,RHOB,RPS12,ZNF454,ARHGDIA,SMCH
RSF1,CDC37,WDR6,BNIP3L,HNRNPUL2-BSCL2,PITPNA,TPM3,XPNPEP3,SRSF3,LARS1,YWHAH,TN
KCSH,KHDRBS1,XPO1,HDLBP,EIF5A,CCND2,RAB7A,DDX3X,RPS15,GJA1,CD63,ANXA6,MTSS2,HN
KCSH,KHDRBS1,XPO1,HDLBP,EIF5A,CCND2,RAB7A,DDX3X,RPS15,GJA1,CD63,ANXA6,MTSS2,HN
X1,PTK7,CNTNAP2,ABCA2,RPL38,ARF3,AEBP1,GABARAP,HNRNPUL1,UBE2R2,FKBP8,SMARCE1,
AZ,LRATD2,TUBGCP6,KRT8,SCRIB,MBD3,YY1,TOMM70,MICAL1,DPYSL4,ATG4B,UNC119B,TUB
SOX4,FKBP14,MAGED1,GPI,HTRA4,RACK1,BSG,SPTBN1,CALM3,GNB1,SHISA9,RPS8,MYH9,RPL

2,ZNF865,ATF4,RPS14,CXADR,RPS20,LENG8,WDR1,RPL26,SRSF4,PFN1,ASTN2,PNN,RPL31,PRK
1G2,UPF1,MASTL,MAPRE2,PPP1R15A,GALNT11,DAP,EIF3H,ELAVL1,IMPDH2,GABARAPL1,GNL1
5,HNRNPUL2-BSCL2,PITPNA,XPNPEP3,RPS28,LARS1,YWHAH,TNKS2,UBQLN1,CSNK1D,IFITM3,V
,SPTY2D1,COX6A1,SART1,NASP,AP1M1,C16ORF58,TOMM70,PRDX3,DEK,GAK,ATP5MG,PAFAH1
LCN6,COX6B1,KDM5A,SKIV2L,SHC1,SUDS3,MIB2,TNPO3,NUP50,SZT2,SEC31A,AKAP17A,MAPK
B,SZT2,ACIN1,RAD50,RNF40,MMP24
B4B,UCHL1,MORF4L1,NCAM1,XAF1,VCL,AP3D1,DNAJA1,AKAP12,SF3B2,PXDN,HNRNPU,NFE2L1
,DHX40,USP5,IER2,CNBP,CAD,MDH1,LGALS1,CHMP1A,ESYT1,LUC7L2,ACVR1B,ELN,KIAA1210,
DRBS1,PTN,CSNK1E,PFKL,PDIA6,DDX3X,ITM2C,RPL30,MRFAP1,RCN1,HMGCS1,JUN,SESN3,RPL

2,INIP,DUX4,SEPTIN11,LGALS3BP,RAC1,GPC1,CDK16,BCLAF1,GLUL,PTTG1IP,SQLE,MAP1A,TIM
RKDC,TDRD5,CASC3,ZFAND5,PLTP,CORO1C,SDF4,SMARCC1,ATP6V1B2,HNRNPM,SDHA,YWHA

EL1,TRIO,CNTNAP1,GRSF1,CSNK1G2,GNG4,PAPSS1,UBAP2L,UBE2G2,EIF5B,ECEL1,CBX6,EML
,JUND,TMEM259,RPS20,EIF5,LENG8,CCT7,MCL1,PNN,CCNI,HDLBP,CLTC,NANOG,GNB2,RPS5,I
1,SRSF5,TM9SF3,MDH2,U2AF2,CCT2,NOP53,PARP1,TSR1,LMNB2,MTRF1L,SNRPN,PRDX5,UBE
A,RTF2,CLPX,GOLGA8B,CD300E,MRPS6,TXN,SLC5A6,GET1,RUSC1,PRMT2,ARHGAP33,SHC1,NE
,HEG1,CTDNEP1,FH
L1,LARP1,CTNNA1,FADS2,CNN3,CTSD,EIF4H,NACA,TSPAN3,BRD3,CIPC,PJA2,MSN,SLC16A7,PAP
EIF5,LENG8,CCT7,MCL1,PNN,CCNI,HDLBP,CLTC,NANOG,GNB2,RPS5,IK,HMGCS1,PRKAR1A,CH
,ATP6V1B2,ZFAND3,HGS,BAG6,HNRNPD,CDC42,EIF2AK1,MMP15,PHB2,ZFP36L1,RAB2A,YKT6,M
M10,WDR82,PGRMC2,VEZF1,MYL12A,NUP205,ERBB2,RNF4,PRMT2,RUSC2,ZNF598,SLC1A5,G
16,SZT2,SEC31A,AKAP17A,HARS1,MRPL3,DNMT1,CFAP298,CPSF7,RAD50,JRK,TOMM40,COX5
SPAN3,RPS24,RPS9,RPL17,C1ORF43,COL1A2,RPL14,RHOA,PTCD3,CBX5,GLIPR1L2,RPS14,CXA
N1,AMOTL2,YME1L1,TRAF4,STAT3,ARIH1,SF1,SLC35F5,CSNK1G2,UPF1,MASTL,MAPRE2,PPP1R
P1,IST1,DHCR7,BAG5,SLC20A1,FJX1,ELAVL3,PAFAH1B2,ABL1,HTATSF1,LGALS1,MTA2,POR,TAC
H,FAU,UBQLN1,CSNK1D,IFITM3,VGF,MLF2,AKIRIN1,TP53BP1,AMOTL2,YME1L1,TRAF4,STAT3,A
TL2,YME1L1,TRAF4,STAT3,ARIH1,SF1,SLC35F5,CSNK1G2,UPF1,MASTL,MAPRE2,PPP1R15A,GA
2,CLPTM1L,GBA2,POLDIP3,RNMT,HMGXB3,RPRM,WARS1,MBOAT7,OTUB1,HIRA,SHC1,CHMP5
NGAP1,EPN1,ARPC3,GBA2,POLDIP3,PSMF1,HMGXB3,MCFD2,RPRM,WARS1,CLCN6,STARD3,CO

I2,H3-3A,TOP1,MARS1,MVB12B,FAM168B,TCF25,PTBP1,ILF3,WDR6,GABARAP,TDRD5,BNIP3L,F
USC1,PRMT2,SHC1,NECTIN2,MAF1,HEG1,MAN1B1,PTPN12,PLXND1,MYL9,SH3KBP1,LDOC1,EPH

ABHD2,HSPA1B,ZNF99,TUBB2B,COL5A1,COL4A1,PSAP,PGK1,RPLP0,H3-3B,CANX,CLU,PTMA,DI
,ABHD2,HSPA1B,TUBB2B,COL5A1,COL4A1,PSAP,PGK1,RPLP0,H3-3B,CANX,CLU,PTMA,DDX5,EN
OL5A1,COL4A1,PSAP,PGK1,RPLP0,H3-3B,CANX,CLU,PTMA,DDX5,ENO2,SOX4,RPL13A,SEC62,K
1,PSAP,PGK1,RPLP0,H3-3B,CANX,CLU,PTMA,ENO2,SOX4,RPL13A,RPL6,KAT6A,DVL3,YWHAE,KIF
RA4,RACK1,BSG,FASN,ALDH6A1,SPTBN1,CALM3,GNB1,SHISA9,MYH9,SEPTIN2,P4HB,APP,IDS,G
1,COL4A1,PSAP,PGK1,RPLP0,H3-3B,CANX,CLU,PTMA,DDX5,ENO2,SOX4,RPL13A,SEC62,KAT6A,D
HD2,HSPA1B,TUBB2B,COL5A1,COL4A1,PSAP,PGK1,RPLP0,H3-3B,CANX,CLU,PTMA,DDX5,ENO2
I1,PLXNB2,HSPD1,USP11,COL18A1,SMIM14,IGF2,MARCKSL1,AFF4,TRIM28,MATR3,YWHAG,CK
,RPL6,KAT6A,DVL3,YWHAE,KIF1A,RPL13,GNAS,SLC2A3,GPI,HTRA4,RACK1,BSG,FASN,SPTBN1,C
RPLP0,H3-3B,CANX,PTMA,DDX5,ENO2,SOX4,RPL13A,SEC62,RPL6,KAT6A,DVL3,YWHAE,KIF1A,R
5A1,COL4A1,PSAP,PGK1,H3-3B,CANX,PTMA,DDX5,ENO2,SOX4,FKBP14,SEC62,KAT6A,DVL3,YW
PL13,GNAS,SLC2A3,GPI,RACK1,BSG,FASN,SPTBN1,CALM3,GNB1,SHISA9,STMN2,MYH9,RPL7,S
IF1A,GNAS,SLC2A3,GPI,RACK1,BSG,FASN,ALDH6A1,SPTBN1,CALM3,GNB1,SHISA9,STMN2,RPS8
WHAE,KIF1A,RPL13,GNAS,SLC2A3,GPI,RACK1,BSG,FASN,ALDH6A1,SPTBN1,CALM3,GNB1,SHIS
ACK1,BSG,FASN,SPTBN1,CALM3,GNB1,SHISA9,STMN2,RPS8,MYH9,RPL7,SEPTIN2,P4HB,APP,GD
1,CES3,PSAP,RPLP0,H3-3B,CANX,CLU,PTMA,DDX5,ENO2,SOX4,RPL13A,MAGED1,SEC62,RPL6,K
2,KAT6A,DVL3,YWHAE,KIF1A,RPL13,GNAS,SLC2A3,GPI,RACK1,BSG,FASN,ALDH6A1,SPTBN1,CAL
SLC2A3,GPI,RACK1,BSG,FASN,ALDH6A1,SPTBN1,CALM3,GNB1,SHISA9,STMN2,RPS8,MYH9,RPL
TAN1,GSTP1,MARCKS,SMG1,CALM1,PLXNB2,HSPD1,USP11,CSDE1,IGF2,ATP5F1A,EIF4B,MARC
SEC62,KAT6A,DVL3,YWHAE,KIF1A,RPL13,GNAS,SLC2A3,GPI,RACK1,BSG,FASN,ALDH6A1,SPTBN
L13,GNAS,DDX17,GPI,HTRA4,RACK1,BSG,FASN,SPTBN1,CALM3,GNB1,SHISA9,RPS8,MYH9,SEP
PGK1,RPLP0,H3-3B,CANX,CLU,PTMA,DDX5,ENO2,SOX4,RPL13A,MAGED1,KAT6A,DVL3,YWHAE,K
IAE,KIF1A,RPL13,GNAS,GPI,HTRA4,BSG,FASN,SPTBN1,ATP5F1B,CALM3,GNB1,SHISA9,MYH9,RP
,CALR,FSTL1,TMF1,ABHD2,HSPA1B,ZNF99,TUBB2B,COL5A1,COL4A1,PSAP,PGK1,RPLP0,H3-3B
,RACK1,BSG,FASN,ALDH6A1,SPTBN1,CALM3,GNB1,SHISA9,MYH9,RPL7,SEPTIN2,P4HB,APP,RPL
0,H3-3B,CANX,CLU,PTMA,DDX5,ENO2,SOX4,FKBP14,RPL13A,MAGED1,SEC62,KAT6A,DVL3,KIF1
DH6A1,SPTBN1,ATP5F1B,CALM3,GNB1,SHISA9,RPS8,MYH9,RPL7,SEPTIN2,P4HB,APP,IDS,GDI1,
PSAP,PGK1,H3-3B,CANX,CLU,PTMA,DDX5,ENO2,SOX4,SEC62,KAT6A,DVL3,YWHAE,KIF1A,RPL13
1,ABHD2,HSPA1B,ZNF99,TUBB2B,COL5A1,COL4A1,PSAP,PGK1,RPLP0,H3-3B,CANX,CLU,PTMA,D
LR,FSTL1,TMF1,ABHD2,HSPA1B,TUBB2B,COL5A1,COL4A1,PSAP,PGK1,RPLP0,H3-3B,CANX,CL
2B,COL5A1,COL4A1,PGK1,RPLP0,H3-3B,CANX,CLU,PTMA,DDX5,ENO2,SOX4,RPL13A,MAGED1,
HAE,KIF1A,GNAS,SLC2A3,GPI,RACK1,BSG,FASN,ALDH6A1,SPTBN1,CALM3,GNB1,SHISA9,STMN
NAS,SLC2A3,DDX17,GPI,HTRA4,RACK1,BSG,FASN,SPTBN1,GNB1,SHISA9,STMN2,RPS8,MYH9,SE
HSPA1B,ZNF99,TUBB2B,COL5A1,COL4A1,CES3,PSAP,PGK1,RPLP0,H3-3B,CANX,CLU,PTMA,DDX
P2B4,IGFBP2,CALR,FSTL1,TMF1,ABHD2,HSPA1B,TUBB2B,COL5A1,COL4A1,PSAP,PGK1,RPLP0,
7,SEPTIN2,P4HB,APP,GDI1,TMBIM6,PTMS,SPTAN1,GSTP1,MARCKS,SMG1,CALM1,PLXNB2,HSPD
B,CANX,CLU,PTMA,DDX5,SOX4,RPL13A,MAGED1,SEC62,RPL6,KAT6A,DVL3,YWHAE,KIF1A,RPL1
AT6A,DVL3,YWHAE,KIF1A,RPL13,GNAS,SLC2A3,DDX17,GPI,HTRA4,RACK1,BSG,FASN,SPTBN1,A
STMN2,MYH9,RPL7,SEPTIN2,P4HB,APP,GDI1,FSCN1,TMBIM6,PTMS,SPTAN1,GSTP1,MARCKS,SM
4A1,PGK1,RPLP0,H3-3B,CANX,CLU,PTMA,DDX5,ENO2,MAGED1,SEC62,RPL6,KAT6A,DVL3,YWH
BB2B,COL5A1,COL4A1,CES3,PSAP,PGK1,RPLP0,H3-3B,CANX,CLU,PTMA,DDX5,ENO2,SOX4,FK
BHD2,HSPA1B,TUBB2B,COL5A1,COL4A1,CES3,PSAP,PGK1,RPLP0,H3-3B,CANX,CLU,PTMA,DDX
,DDX17,HTRA4,RACK1,BSG,ALDH6A1,ATP5F1B,CALM3,GNB1,SHISA9,RPS8,MYH9,RPL7,SEPTIN
X5,ENO2,SOX4,FKBP14,SEC62,KAT6A,DVL3,YWHAE,KIF1A,GNAS,SLC2A3,GPI,HTRA4,BSG,FASN
B,COL5A1,COL4A1,CES3,PSAP,PGK1,H3-3B,CANX,PTMA,DDX5,ENO2,SOX4,SEC62,KAT6A,DVL3,

3,TUBB2B,COL5A1,COL4A1,PSAP,PGK1,H3-3B,CANX,CLU,PTMA,DDX5,ENO2,SOX4,RPL13A,RPL(
,GNAS,SLC2A3,DDX17,GPI,BSG,FASN,ALDH6A1,SPTBN1,CALM3,GNB1,SHISA9,STMN2,RPS8,MYI
PM1,PLXNB2,USP11,IGF2,EIF4B,MARCKSL1,AFF4,TRIM28,MATR3,YWHAG,CKB,RPL37A,LRRC28
5A1,COL4A1,CES3,PSAP,PGK1,RPLP0,H3-3B,CANX,CLU,PTMA,DDX5,ENO2,SOX4,RPL13A,MAGED
,FSTL1,TMF1,ABHD2,HSPA1B,TUBB2B,COL5A1,COL4A1,CES3,PGK1,RPLP0,H3-3B,CANX,CLU,PT
YH9,RPL7,P4HB,APP,IDS,RPL5,GDI1,PTMS,SPTAN1,GSTP1,MARCKS,CALM1,NPM1,PLXNB2,PLD3
SEPTIN2,P4HB,GDI1,FSCN1,PTMS,SPTAN1,GSTP1,MARCKS,SMG1,CALM1,PLXNB2,HSPD1,USP1
PTMA,DDX5,SOX4,RPL13A,SEC62,RPL6,KAT6A,DVL3,YWHAE,KIF1A,RPL13,GNAS,SLC2A3,DDX17
L1,AFF4,TRIM28,MATR3,YWHAG,CKB,LRRC28,FN1,YWHAQ,HACD3,DCHS1,SET,CHSY1,TMSB10,
,H3-3B,CANX,PTMA,ENO2,SOX4,RPL13A,RPL6,KAT6A,DVL3,YWHAE,RPL13,GNAS,DDX17,GPI,HT
RRC28,FN1,NONO,YWHAQ,HACD3,DCHS1,SET,TMSB10,MRC2,PKD1,HLA-C,SSBP2,VAPA,POLR2
B,MARCKSL1,AFF4,TRIM28,MATR3,YWHAG,CKB,LRRC28,FN1,YWHAQ,DCHS1,SET,EGLN1,TMSE
2,HSPD1,USP11,SMIM14,IGF2,EIF3C,ATP5F1A,EIF4B,MARCKSL1,AFF4,TRIM28,MATR3,YWHAG,(
NB2,HSPD1,USP11,COL18A1,SNRNP200,CSDE1,IGF2,EIF4B,MARCKSL1,AFF4,TRIM28,MATR3,YV
MA,DDX5,ENO2,SOX4,SEC62,KAT6A,DVL3,YWHAE,KIF1A,RPL13,GNAS,SLC2A3,DDX17,GPI,HTRA4
/HAQ,DCHS1,SET,CHSY1,TMSB10,MRC2,GTF3C4,PKD1,PDIA3,SSBP2,BASP1,VAPA,AP2M1,TPT1,
L4A1,CES3,PSAP,PGK1,RPLP0,H3-3B,CANX,CLU,PTMA,DDX5,ENO2,SOX4,SEC62,KAT6A,DVL3,YV
GSTP1,MARCKS,SMG1,CALM1,PLXNB2,HSPD1,USP11,COL18A1,SNRNP200,CSDE1,IGF2,EIF3C,A
3,YWHAG,CKB,LRRC28,FN1,NONO,YWHAQ,HACD3,DCHS1,SET,TMSB10,MRC2,LDHB,PKD1,HLA-
ALDH6A1,SPTBN1,CALM3,GNB1,SHISA9,STMN2,RPS8,RPL7,SEPTIN2,RPL5,GDI1,TMBIM6,SPTAN
14,RPL13A,SEC62,KAT6A,DVL3,YWHAE,KIF1A,RPL13,GNAS,SLC2A3,DDX17,GPI,HTRA4,RACK1,B
FF4,TRIM28,MATR3,YWHAG,CKB,LRRC28,FN1,YWHAQ,HACD3,DCHS1,SET,CHSY1,TMSB10,MRC
A,DDX5,ENO2,SOX4,RPL13A,MAGED1,SEC62,RPL6,KAT6A,DVL3,YWHAE,RPL13,GNAS,DDX17,HT
JSP11,CSDE1,SMIM14,IGF2,ATP5F1A,EIF4B,MARCKSL1,AFF4,TRIM28,MATR3,YWHAG,CKB,COL
,IGF2,EIF3C,EIF4B,MARCKSL1,AFF4,TRIM28,MATR3,YWHAG,CKB,LRRC28,FN1,YWHAQ,HACD3,
TMS,SPTAN1,SMG1,CALM1,PLXNB2,PLD3,USP11,COL18A1,IGF2,ATP5F1A,MARCKSL1,UBB,AFF4
GNAS,SLC2A3,DDX17,GPI,RACK1,BSG,FASN,ALDH6A1,SPTBN1,CALM3,GNB1,SHISA9,STMN2,RPS
XNB2,USP11,IGF2,ATP5F1A,EIF4B,MARCKSL1,UBB,AFF4,TRIM28,MATR3,YWHAG,CKB,COL4A2,
L1,TMF1,ABHD2,HSPA1B,ZNF99,TUBB2B,COL5A1,COL4A1,CES3,PSAP,PGK1,RPLP0,H3-3B,CAN
ENO2,MAGED1,SEC62,RPL6,KAT6A,DVL3,YWHAE,KIF1A,RPL13,GNAS,DDX17,GPI,BSG,FASN,ALDI
RPLP0,H3-3B,CANX,CLU,PTMA,DDX5,ENO2,SOX4,FKBP14,RPL13A,SEC62,RPL6,KAT6A,DVL3,YWI
,CALM1,NPM1,PLXNB2,USP11,CSDE1,IGF2,ATP5F1A,EIF4B,MARCKSL1,UBB,AFF4,TRIM28,MATI
A1,PSAP,PGK1,CLU,PTMA,DDX5,ENO2,SOX4,FKBP14,RPL13A,KAT6A,DVL3,YWHAE,KIF1A,GNAS,
3,CANX,CLU,PTMA,DDX5,ENO2,SOX4,RPL13A,MAGED1,SEC62,KAT6A,DVL3,YWHAE,KIF1A,RPL13
ALDH6A1,SPTBN1,CALM3,GNB1,SHISA9,STMN2,RPS8,MYH9,RPL7,SEPTIN2,P4HB,APP,IDS,GDI1,
FSCN1,TMBIM6,PTMS,SPTAN1,MARCKS,SMG1,CALM1,NPM1,PLXNB2,USP11,CSDE1,SMIM14,IG
28,MATR3,YWHAG,CKB,LRRC28,FN1,YWHAQ,DCHS1,SET,CHSY1,TMSB10,INA,MRC2,PKD1,HLA-
FBP2,CALR,FSTL1,TMF1,ABHD2,HSPA1B,ZNF99,TUBB2B,COL5A1,COL4A1,CES3,PSAP,PGK1,RPI
1,MARCKS,SMG1,CALM1,NPM1,PLXNB2,USP11,IGF2,EIF3C,ATP5F1A,EIF4B,MARCKSL1,UBB,TR
PTIN2,APP,GDI1,PTMS,SPTAN1,GSTP1,MARCKS,SMG1,PLXNB2,HSPD1,USP11,SNRNP200,SMIM1
PTIN2,APP,GDI1,PTMS,SPTAN1,GSTP1,MARCKS,SMG1,PLXNB2,HSPD1,USP11,SNRNP200,SMIM1
M14,IGF2,ATP5F1A,EIF4B,MARCKSL1,AFF4,TRIM28,MATR3,YWHAG,CKB,RPL37A,YWHAQ,HACD3
FBP2,CALR,FSTL1,TMF1,ABHD2,HSPA1B,ZNF99,TUBB2B,COL5A1,COL4A1,CES3,PSAP,PGK1,RPI
STP1,MARCKS,SMG1,CALM1,NPM1,HSPD1,USP11,COL18A1,SNRNP200,IGF2,MARCKSL1,TRIM2
STP1,SMG1,CALM1,NPM1,PLD3,HSPD1,USP11,COL18A1,SNRNP200,SMIM14,IGF2,EIF3C,EIF4B

N2,APP,GDI1,FSCN1,PTMS,SPTAN1,GSTP1,MARCKS,SMG1,CALM1,PLXNB2,HSPD1,USP11,SNRNI
I,SNRNP200,SMIM14,IGF2,ATP5F1A,EIF4B,MARCKSL1,AFF4,TRIM28,MATR3,YWHAG,CKB,RPL37
F3C,ATP5F1A,EIF4B,MARCKSL1,UBB,AFF4,TRIM28,MATR3,YWHAG,CKB,COL4A2,LRRC28,NONC
AFF4,TRIM28,MATR3,YWHAG,CKB,LRRC28,FN1,NONO,YWHAQ,EIF1,HACD3,DCHS1,SET,TPI1,TM
N,ALDH6A1,SPTBN1,GNB1,SHISA9,STMN2,RPS8,MYH9,RPL7,SEPTIN2,APP,GDI1,FSCN1,SPTAN1,
4,IGFBP2,CALR,FSTL1,TMF1,ABHD2,HSPA1B,ZNF99,TUBB2B,COL5A1,COL4A1,CES3,PSAP,PGK
,PDIA3,VAPA,TPT1,STMN1,ITGB1,TUBB4B,UCHL1,MORF4L1,BTF3,AGRN,ZMIZ1,TIMP2,MIDN,MA
STP1,MARCKS,SMG1,PLXNB2,HSPD1,USP11,SNRNP200,CSDE1,IGF2,EIF3C,ATP5F1A,EIF4B,MAR
R,FSTL1,ABHD2,HSPA1B,TUBB2B,COL5A1,COL4A1,CES3,PSAP,PGK1,H3-3B,CANX,CLU,PTMA,DI
PTIN2,P4HB,GDI1,FSCN1,TMBIM6,PTMS,SPTAN1,GSTP1,SMG1,CALM1,NPM1,PLXNB2,HSPD1,US
HAE,KIF1A,GNAS,DDX17,BSG,FASN,ALDH6A1,SPTBN1,CALM3,GNB1,SHISA9,MYH9,RPL7,SEPTIN
PTAN1,GSTP1,MARCKS,SMG1,PLXNB2,HSPD1,USP11,SNRNP200,IGF2,EIF3C,ATP5F1A,EIF4B,MA
ATR3,YWHAG,CKB,COL4A2,RPL37A,LRRC28,YWHAQ,HACD3,DCHS1,SET,EGLN1,CHSY1,INA,MRC
KS,SMG1,PLXNB2,USP11,IGF2,EIF3C,ATP5F1A,EIF4B,MARCKSL1,UBB,AFF4,TRIM28,MATR3,YWI
BP2,CALR,FSTL1,ABHD2,HSPA1B,TUBB2B,COL5A1,COL4A1,CES3,PSAP,PGK1,H3-3B,CANX,CLU
PM1,HSPD1,USP11,SMIM14,IGF2,MARCKSL1,AFF4,TRIM28,MATR3,CKB,COL4A2,LRRC28,FN1,SI
JBB4B,UCHL1,MORF4L1,ANXA2,AGRN,TIMP2,SARAF,MIDN,MARCHF6,VCL,AP3D1,ARF1,CYB5R3
62,KAT6A,DVL3,KIF1A,GNAS,SLC2A3,GPI,RACK1,BSG,FASN,ALDH6A1,SPTBN1,CALM3,GNB1,ST
FF4,TRIM28,MATR3,YWHAG,COL4A2,RPL37A,LRRC28,SPEN,NONO,YWHAQ,EIF1,HACD3,DCHS1,
LN1,CHSY1,TMSB10,EIF4A1,MRC2,RPL19,RPL15,GTF3C4,LDHB,PKD1,SSBP2,CDKL1,AP2M1,TF
MARCKS,SMG1,CALM1,NPM1,HSPD1,USP11,SMIM14,IGF2,EIF4B,MARCKSL1,UBB,MATR3,YWHA
,INA,RPL15,GTF3C4,PKD1,HLA-C,SSBP2,VAPA,STMN1,ITGB1,TUBB4B,UCHL1,MORF4L1,NCAM1
VCL,SBF1,CYB5R3,MAP4,CDC42BPB,ITM2B,AP2B1,MAT2A,GOLGA8A,PXDN,OGA,CTNNA1,FADS
2BPB,ITM2B,TMEM170A,PXDN,HNRNPU,HNRNPA0,CTNNA1,FADS2,CPE,CNN3,CTSD,LPCAT1,LAM
BSG,FASN,ALDH6A1,SPTBN1,GNB1,SHISA9,STMN2,MYH9,RPL7,P4HB,IDS,ERVV-1,GDI1,FSCN1,TI
2A,PXDN,HNRNPU,PGAM1,SMARCA5,CTNNA1,FADS2,CNN3,EIF4H,SCARB2,TSPAN3,LPCAT1,LAM
3,GNB1,SHISA9,RPS8,MYH9,RPL7,SEPTIN2,P4HB,APP,IDS,RPL5,GDI1,TMBIM6,PTMS,SPTAN1,GS
DDX17,GPI,HTRA4,RACK1,BSG,FASN,SPTBN1,ATP5F1B,CALM3,GNB1,SHISA9,STMN2,MYH9,RPL
3,GNB1,SHISA9,STMN2,RPS8,RPL7,SEPTIN2,APP,IDS,GDI1,PTMS,SPTAN1,MARCKS,SMG1,NPM1,
GPI,HTRA4,RACK1,BSG,FASN,ALDH6A1,SPTBN1,ATP5F1B,CALM3,GNB1,SHISA9,STMN2,RPS8,MY
MF1,ABHD2,HSPA1B,ZNF99,TUBB2B,COL5A1,COL4A1,CES3,PSAP,PGK1,RPLP0,H3-3B,CANX,CLU
SPD1,COL18A1,SNRNP200,IGF2,EIF4B,MARCKSL1,AFF4,TRIM28,MATR3,YWHAG,CKB,COL4A2,F
MORF4L1,NCAM1,AGRN,ZMIZ1,TIMP2,COL11A1,MIDN,MARCHF6,VCL,AP3D1,SBF1,DNAJA1,ARF
P11,SNRNP200,SMIM14,IGF2,EIF4B,MARCKSL1,TRIM28,YWHAG,CKB,LRRC28,FN1,NONO,DDB1
GNAS,SLC2A3,DDX17,HTRA4,BSG,FASN,ALDH6A1,SPTBN1,CALM3,GNB1,SHISA9,STMN2,MYH9,F
PTMA,DDX5,ENO2,SOX4,FKBP14,RPL13A,SEC62,RPL6,KAT6A,DVL3,KIF1A,GNAS,DDX17,GPI,HTR
,HNRNPU,HNRNPA0,FADS2,CNN3,RPL29,CTSD,EIF4H,SCARB2,TSPAN3,LPCAT1,BRD3,PJA2,C1OI
MATR3,YWHAG,CKB,COL4A2,YWHAQ,DCHS1,SET,EGLN1,TMSB10,RPL19,PPP2R1A,GTF3C4,PKD
CDC42BPB,ITM2B,PXDN,HNRNPU,CTNNA1,FADS2,CNN3,SLC22A17,BRD3,PJA2,KPNB1,LLGL1,N
1,MARCKS,CALM1,NPM1,PLXNB2,HSPD1,USP11,COL18A1,SNRNP200,CSDE1,IGF2,ATP5F1A,EIF
FN1,SPEN,YWHAQ,EIF1,DCHS1,SET,EGLN1,CHSY1,TMSB10,MRC2,LDHB,PKD1,HLA-C,PDIA3,SEF
,EIF4A1,PPP2R1A,RPL15,GTF3C4,SF3B1,PDIA3,SSBP2,BASP1,POLR2A,SEC61A1,RPL18,TUBB4B
M6,PTMS,SPTAN1,GSTP1,SMG1,CALM1,NPM1,PLXNB2,HSPD1,USP11,COL18A1,SNRNP200,CSD
4B,MARCKSL1,AFF4,TRIM28,MATR3,YWHAG,CKB,LRRC28,FN1,NONO,DDB1,YWHAQ,HACD3,DCI
I,ABHD2,HSPA1B,ZNF99,TUBB2B,COL5A1,COL4A1,CES3,PSAP,PGK1,RPLP0,H3-3B,CANX,CLU,P

;CN1,PTMS,SPTAN1,GSTP1,SMG1,NPM1,PLXNB2,USP11,COL18A1,SNRNP200,SMIM14,IGF2,VCF
;A,PGAM1,NFE2L1,SMARCA5,HNRNPA0,FADS2,RPL23,SCARB2,LAMA5,SLC22A17,BRD3,CIPC,PJ/
=4A1,MRC2,GTF3C4,LDHB,PKD1,SSBP2,VAPA,HMGB1,SEC61A1,ITGB1,TUBB4B,UCHL1,AGRN,ZI
SG,FASN,ATP5F1B,CALM3,GNB1,RPS8,RPL7,SEPTIN2,P4HB,APP,IDS,RPL5,FSCN1,PTMS,SPTAN1
=1A,RPL13,GNAS,SLC2A3,DDX17,GPI,HTRA4,RACK1,BSG,FASN,SPTBN1,ATP5F1B,CALM3,SHISA
(17,BSG,FASN,ALDH6A1,SPTBN1,CALM3,GNB1,SHISA9,STMN2,MYH9,SEPTIN2,IDS,GDI1,FSCN1,I
EIF4B,VCP,UBB,AFF4,TRIM28,MATR3,YWHAG,CKB,LRRC28,NONO,DDB1,YWHAQ,HACD3,DCHS1
,PTMS,SPTAN1,GSTP1,SMG1,CALM1,NPM1,HSPD1,COL18A1,SNRNP200,SMIM14,IGF2,EIF3C,EII
J,PTMA,ENO2,SOX4,RPL13A,KAT6A,DVL3,YWHAE,KIF1A,RPL13,GNAS,DDX17,GPI,HTRA4,RACK1,
)L4A1,PSAP,PGK1,RPLP0,H3-3B,CANX,CLU,PTMA,DDX5,ENO2,SOX4,FKBP14,RPL13A,SEC62,RPL
3ASP1,VAPA,HMGB1,POLR2A,AP2M1,TPT1,SEC61A1,RPS3A,TUBB4B,RPL11,MORF4L1,AGRN,TIM
AF,MIDN,MARCHF6,VCL,AP3D1,SBF1,ARF1,CYB5R3,MAP4,CDC42BPB,ITM2B,MAT2A,GOLGA8A,
CYB5R3,MAP4,CDC42BPB,ITM2B,MAT2A,SLC4A8,PXDN,HNRNPU,CTNNA1,CTSD,LAMA5,BRD3,C
M6,PTMS,MARCKS,SMG1,CALM1,NPM1,TTC3,HSPD1,USP11,CSDE1,SMIM14,IGF2,VCP,MARCKS
N1,SMG1,CALM1,NPM1,PLXNB2,PLD3,USP11,COL18A1,CSDE1,SMIM14,IGF2,MARCKSL1,AFF4,
,HTRA4,RACK1,FASN,SPTBN1,CALM3,GNB1,SHISA9,RPS8,RPL7,SEPTIN2,P4HB,APP,ERVV-1,FSCI
MIDN,MARCHF6,VCL,MAP4,CDC42BPB,ITM2B,MAT2A,PXDN,HNRNPA0,CTNNA1,FADS2,TNS3,CT
HL1,MORF4L1,AGRN,TIMP2,COL11A1,MIDN,MARCHF6,VCL,DNAJA1,CDC42BPB,ITM2B,MAT2A,F
)L5A1,COL4A1,PSAP,PGK1,RPLP0,H3-3B,CANX,CLU,PTMA,DDX5,ENO2,SOX4,RPL13A,MAGED1,F
/L3,YWHAE,KIF1A,RPL13,GNAS,SLC2A3,DDX17,GPI,HTRA4,RACK1,FASN,SPTBN1,CALM3,GNB1,
6,KAT6A,DVL3,YWHAE,KIF1A,RPL13,GNAS,SLC2A3,DDX17,GPI,HTRA4,RACK1,BSG,FASN,ALDH6/
G1,CALM1,NPM1,PLXNB2,USP11,SNRNP200,SMIM14,IGF2,EIF3C,ATP5F1A,EIF4B,MARCKSL1,AFI
PS8,MYH9,SEPTIN2,P4HB,APP,GDI1,FSCN1,GSTP1,MARCKS,SMG1,PLD3,HSPD1,USP11,COL18A1
JBA1,ATP2B4,IGFBP2,CALR,FSTL1,TMF1,ABHD2,HSPA1B,ZNF99,TUBB2B,COL5A1,COL4A1,CES
C42BPB,MAT2A,HNRNPU,PGAM1,NFE2L1,HNRNPA0,RPSA,CTSD,EIF4H,SCARB2,TSPAN3,LAMA5,
D3,DCHS1,SET,EGLN1,CHSY1,INA,MRC2,PKD1,HLA-C,PDIA3,SSBP2,VAPA,POLR2A,TPT1,RPS3A,
HNRNPU,OGA,NFE2L1,SMARCA5,HNRNPA0,CTNNA1,CNN3,TPM4,LAMA5,BRD3,PJA2,SLC16A7,KI
AP,RPLP0,H3-3B,CANX,CLU,PTMA,DDX5,ENO2,SOX4,FKBP14,RPL13A,SEC62,RPL6,KAT6A,DVL3,
KD1,HLA-C,PDIA3,HMGB1,TPT1,SEC61A1,RPS3A,RPL18,STMN1,ITGB1,UCHL1,ANXA2,AGRN,TIM
A,MAGED1,RPL6,KAT6A,DVL3,YWHAE,KIF1A,RPL13,GNAS,DDX17,HTRA4,RACK1,BSG,FASN,ALDI
ADS2,LDHA,TPM4,NACA,LAMA5,SLC22A17,BRD3,PJA2,KPNB1,C1ORF43,PAPOLA,NCS1,RHOA,CI
-C,VAPA,HMGB1,POLR2A,AP2M1,TPT1,RPS3A,STMN1,TUBB4B,UCHL1,BTF3,ANXA2,AGRN,TIMP2
3,HSPD1,USP11,SNRNP200,SMIM14,IGF2,EIF3C,EIF4B,VCP,MARCKSL1,UBB,TRIM28,MATR3,CK
SDE1,IGF2,EIF4B,MARCKSL1,UBB,AFF4,TRIM28,YWHAG,CKB,FN1,YWHAQ,HACD3,DCHS1,SET,E
KIF1A,RPL13,GNAS,SLC2A3,DDX17,GPI,RACK1,FASN,SPTBN1,ATP5F1B,CALM3,GNB1,SHISA9,RF
\1,FADS2,TPM4,SLC22A17,BRD3,PJA2,KPNB1,C1ORF43,NCS1,ZNF865,NLGN2,CXADR,CHD4,JUI
MAP4,CDC42BPB,ITM2B,MAT2A,PXDN,HNRNPU,CTNNA1,RPS16,CNN3,CTSD,LPCAT1,BRD3,CIPC
YH9,RPL7,SEPTIN2,APP,IDS,RPL5,GDI1,FSCN1,PTMS,SPTAN1,GSTP1,MARCKS,SMG1,CALM1,NPI
31,YWHAQ,HACD3,DCHS1,SET,EGLN1,CHSY1,INA,MRC2,RPL19,LDHB,PKD1,HLA-C,SERINC1,BAS
C28,NONO,DDB1,HACD3,DCHS1,SET,EGLN1,CHSY1,TMSB10,EIF4A1,LDHB,PKD1,HLA-C,SF3B1,
NB1,MYH9,SEPTIN2,P4HB,APP,IDS,ERVV-1,RPL5,GDI1,FSCN1,TMBIM6,PTMS,SPTAN1,MARCKS,
IMGN2,ATP2A2,CRMP1,LASP1,BEX3,VAT1,OAZ1,HNRNPC,CCNI,SPTBN2,SLC3A2,HDLBP,EIF5A,C
)L18A1,IGF2,EIF3C,EIF4B,UBB,AFF4,MATR3,CKB,COL4A2,LRRC28,FN1,NONO,DDB1,YWHAQ,EII
)L18A1,IGF2,EIF3C,EIF4B,UBB,AFF4,MATR3,CKB,COL4A2,LRRC28,FN1,NONO,DDB1,YWHAQ,EII
)L18A1,IGF2,EIF3C,EIF4B,UBB,AFF4,MATR3,CKB,COL4A2,LRRC28,FN1,NONO,DDB1,YWHAQ,EII

4HB,APP,IDS,RPL5,FSCN1,TMBIM6,PTMS,SPTAN1,MARCKS,SMG1,CALM1,NPM1,PLXNB2,PLD3,U
TRIM28,MATR3,YWHAG,CKB,COL4A2,FN1,SPEN,NONO,DDB1,YWHAQ,HACD3,DCHS1,SET,EGLN
DX17,HTRA4,RACK1,BSG,ALDH6A1,ATP5F1B,CALM3,RPS8,RPL7,SEPTIN2,P4HB,APP,IDS,RPL5,GI
1,SET,EGLN1,TPI1,TMSB10,EIF4A1,MRC2,LDHB,HLA-C,SERINC1,SSBP2,BASP1,VAPA,POLR2A,TF
RD3,PJA2,KPNB1,NCS1,ZNF865,NLGN2,CHD4,JUND,CD81,LENG8,WDR1,HMGN2,ATP2A2,CRMF
R,FSTL1,HSPA1B,TUBB2B,COL5A1,COL4A1,PSAP,PGK1,H3-3B,CANX,CLU,PTMA,DDX5,ENO2,SO
,TMSB10,MRC2,PPP2R1A,DNAH8,PKD1,SSBP2,BASP1,VAPA,POLR2A,AP2M1,TPT1,STMN1,ITGB1
P,MARCKSL1,UBB,TRIM28,MATR3,YWHAG,CKB,COL4A2,MAPT,FN1,SPEN,PSMD5,YWHAQ,EIF1,H
S,SMG1,CALM1,NPM1,PLXNB2,PLD3,USP11,COL18A1,SMIM14,IGF2,EIF3C,VCP,MARCKSL1,UBI
RN,TIMP2,SARAF,MIDN,MARCHF6,VCL,AP3D1,CYB5R3,MAP4,CDC42BPB,ITM2B,MAT2A,GOLGA
CH,TADA2B,SPTBN2,PFKL,HDLBP,TPM1,CCND2,SOX12,ARL4C,GNB2,H1-0,APLP1,RPS7,TLN1,CH
DVL3,YWHAE,KIF1A,RPL13,GNAS,GPI,HTRA4,RACK1,BSG,FASN,ALDH6A1,SPTBN1,CALM3,GNB1
IF1A,RPL13,GNAS,SLC2A3,GPI,HTRA4,RACK1,BSG,FASN,SPTBN1,ATP5F1B,CALM3,GNB1,SHISA
1,CNN3,LAMA5,SLC22A17,BRD3,LLGL1,NCS1,RHOA,ZNF865,KIF5B,SLC7A5,CHD4,ATP1A1,JUND
,CALM1,PLXNB2,PLD3,USP11,COL18A1,SNRNP200,IGF2,EIF3C,EIF4B,VCP,MARCKSL1,UBB,TRI
X17,GPI,RACK1,FASN,SPTBN1,CALM3,GNB1,SHISA9,RPS8,MYH9,SEPTIN2,P4HB,APP,RPL5,FSCN
D2,TUBB2B,COL5A1,COL4A1,PSAP,PGK1,RPLP0,H3-3B,CANX,CLU,PTMA,DDX5,ENO2,SOX4,FKBI
RN,TIMP2,MARCHF6,RPS17,VCL,AP3D1,SBF1,DNAJA1,ARF1,CDC42BPB,MAT2A,PXDN,HNRNPU,S
CLU,PTMA,DDX5,SOX4,RPL13A,SEC62,RPL6,KAT6A,DVL3,YWHAE,KIF1A,RPL13,GNAS,SLC2A3,DD
RRC28,NONO,DDB1,HACD3,DCHS1,SET,CHSY1,TMSB10,LDHB,PKD1,HLA-C,PDIA3,SSBP2,VAPA,I
I1,TLE5,CCNI,KHDRBS1,XPO1,CSNK1E,PFKL,TKT,SFPQ,CDH2,PDIA6,MEGF8,SOX12,CCN2,PRDX
KAT6A,DVL3,YWHAE,KIF1A,RPL13,GNAS,SLC2A3,DDX17,GPI,HTRA4,RACK1,BSG,FASN,ATP5F1B,
G,CKB,COL4A2,SPEN,NONO,YWHAQ,HACD3,DCHS1,SET,TPI1,CHSY1,TMSB10,EIF4A1,MRC2,PPP2
35R3,MAP4,CDC42BPB,MAT2A,TMEM170A,GOLGA8A,PXDN,HNRNPU,PGAM1,SMARCA5,CTNNA1
TMSB10,MRC2,GTF3C4,LDHB,SSBP2,BASP1,POLR2A,TPT1,STMN1,ITGB1,TUBB4B,UCHL1,MORI
2,IDS,GDI1,FSCN1,TMBIM6,SPTAN1,MARCKS,SMG1,CALM1,NPM1,PLXNB2,TTC3,HSPD1,USP11,
P4,CDC42BPB,ITM2B,TMEM170A,PXDN,HNRNPU,OGA,SMARCA5,HNRNPA0,CTNNA1,CPE,CNN3,L
,GNAS,DDX17,GPI,HTRA4,RACK1,BSG,FASN,ALDH6A1,SPTBN1,CALM3,GNB1,SHISA9,RPS8,MYH
PL7,SEPTIN2,P4HB,APP,IDS,RPL5,GDI1,FSCN1,GSTP1,MARCKS,SMG1,CALM1,NPM1,PLXNB2,HSI
PLD3,USP11,COL18A1,IGF2,EIF3C,VCP,MARCKSL1,UBB,AFF4,TRIM28,MATR3,YWHAG,CKB,COL
B,LRRC28,MAPT,FN1,SPEN,DDB1,PSMD5,YWHAQ,EIF1,HACD3,SET,TPI1,CHSY1,EIF4A1,RPL19,R
,NONO,YWHAQ,HACD3,DCHS1,SET,EGLN1,CHSY1,TMSB10,INA,MRC2,GTF3C4,LDHB,PKD1,HLA-
RF4L1,ANXA2,ZMIZ1,MIDN,RPL10,VCL,AP3D1,ARF1,CYB5R3,ZNF770,TMEM170A,SLC4A8,PXDN,I
X17,HTRA4,BSG,FASN,ALDH6A1,ATP5F1B,CALM3,GNB1,RPS8,MYH9,RPL7,SEPTIN2,P4HB,APP,ID
KS,CALM1,PLXNB2,USP11,COL18A1,SNRNP200,IGF2,ATP5F1A,EIF4B,MARCKSL1,UBB,AFF4,MA
L4,YBX1,UBA1,ATP2B4,IGFBP2,CALR,FSTL1,TMF1,ABHD2,HSPA1B,ZNF99,TUBB2B,COL5A1,CO
CTSD,LPCAT1,LAMA5,SLC22A17,CIPC,MSN,RPL24,IGFBP3,LLGL1,GAP43,HK1,NCS1,RPL22,CLS
HD4,ATP1A1,JUND,EIF5,CD81,WDR1,HMGN2,ATP2A2,CRMP1,LASP1,OAZ1,PFN1,TLE5,NISCH,CC
CHD4,ATP1A1,JUND,EIF5,CD81,WDR1,HMGN2,ATP2A2,CRMP1,LASP1,OAZ1,PFN1,ASTN2,TLE5,N
,FSTL1,TMF1,ABHD2,HSPA1B,TUBB2B,COL5A1,COL4A1,CES3,PSAP,PGK1,RPLP0,CANX,CLU,PTN
11,SMIM14,IGF2,VCP,MARCKSL1,UBB,TRIM28,MATR3,CKB,COL4A2,LRRC28,FN1,SPEN,YWHAQ
LRD1,GDI1,FSCN1,TMBIM6,GSTP1,MARCKS,CALM1,NPM1,HSPD1,SNRNP200,CSDE1,SMIM14,I
3,TRIM28,MATR3,YWHAG,CKB,COL4A2,LRRC28,FN1,PSMD5,YWHAQ,HACD3,SET,TPI1,CHSY1,TN
,HLA-C,SF3B1,SSBP2,POLR2A,CDKL1,AP2M1,TPT1,RPS3A,RPL18,STMN1,UCHL1,ANXA2,AGRN,T
AS,SLC2A3,DDX17,GPI,BSG,FASN,ALDH6A1,SPTBN1,ATP5F1B,CALM3,GNB1,SHISA9,MYH9,RPL7,

,PLD3,HSPD1,COL18A1,IGF2,EIF4B,VCP,MARCKSL1,AFF4,TRIM28,YWHAG,CKB,COL4A2,LRRC2
'MBIM6,SPTAN1,GSTP1,SMG1,PLD3,HSPD1,COL18A1,IGF2,EIF4B,VCP,MARCKSL1,AFF4,TRIM28
,BASP1,VAPA,POLR2A,CDKL1,TPT1,STMN1,ITGB1,TUBB4B,UCHL1,MORF4L1,NCAM1,ANXA2,AG
'UBB4B,ANXA2,AGRN,TIMP2,MIDN,MARCHF6,VCL,AP3D1,SBF1,ARF1,CYB5R3,MAP4,CDC42BPB,
00,SMIM14,IGF2,EIF4B,VCP,UBB,AFF4,TRIM28,MATR3,CKB,COL4A2,SPEN,PSMD5,DCHS1,CHSY
/IEM170A,HNRNPU,NFE2L1,SMARCA5,HNRNPA0,CNN3,RPSA,EIF4H,LPCAT1,LAMA5,BRD3,XRCC5
RCKS,SMG1,CALM1,NPM1,PLD3,HSPD1,USP11,IGF2,EIF3C,EIF4B,VCP,UBB,AFF4,TRIM28,MATR
3,TMSB10,LDHB,PKD1,HLA-C,PDIA3,VAPA,HMGB1,POLR2A,TPT1,SEC61A1,RPS3A,TUBB4B,UCHI
XDN,HNRNPU,CTNNA1,RPS16,CNN3,CTSD,LPCAT1,BRD3,CIPC,PJA2,KPNB1,COL1A2,LLGL1,NCS
XDN,HNRNPU,CTNNA1,RPS16,CNN3,CTSD,LPCAT1,BRD3,CIPC,PJA2,KPNB1,COL1A2,LLGL1,NCS
3,PKN1,GRINA,LAMC1,CBX1,RAD23B,CHD2,ORAI2,ASXL1,H3-3A,VEGFA,MVB12B,CCND1,G3BP2
2,RPS16,CTSD,EIF4H,LPCAT1,RPS9,XRCC5,CIPC,PAPOLA,IGFBP3,NCS1,RHOA,PTCD3,CBX5,ZNF
B2B,COL5A1,COL4A1,PGK1,RPLP0,H3-3B,CANX,CLU,PTMA,DDX5,ENO2,SOX4,FKBP14,MAGED1
1,UBB,AFF4,TRIM28,MATR3,YWHAG,CKB,COL4A2,LRRC28,NONO,PSMD5,YWHAQ,EIF1,HACD3,D
J,CTNNA1,CNN3,TNS3,NACA,LPCAT1,LAMA5,SLC22A17,BRD3,KPNB1,C1ORF43,LLGL1,NCS1,CB
F4,SLC7A5,CHD4,ATP1A1,JUND,CD81,HMGN2,ATP2A2,CRMP1,LASP1,OAZ1,PFN1,NISCH,CCNI,X
NRNPU,HNRNPA0,CTNNA1,CTSD,LPCAT1,BRD3,LLGL1,NCS1,ZNF865,CLSTN1,SLC7A5,CHD4,ATF
70,GTF2F1,PXDN,HNRNPU,OGA,HNRNPA0,CTNNA1,FADS2,CPE,CNN3,TSPAN3,LPCAT1,LAMA5,BI
SEPTIN2,P4HB,APP,ERVV-1,GDI1,FSCN1,TMBIM6,PTMS,SPTAN1,SMG1,NPM1,PLXNB2,HSPD1,SN
,PLXNB2,HSPD1,SNRNP200,CSDE1,SMIM14,ATP5F1A,UBB,CKB,LRRC28,SPEN,DDB1,PSMD5,YWI
CALM3,SHISA9,SEPTIN2,P4HB,RPL5,GDI1,FSCN1,SPTAN1,GSTP1,MARCKS,NPM1,PLXNB2,TTC3,
N,SPTBN1,CALM3,GNB1,SHISA9,STMN2,MYH9,RPL7,SEPTIN2,P4HB,APP,RPL5,GDI1,FSCN1,SPT
EC62,RPL6,KAT6A,DVL3,YWHAE,KIF1A,RPL13,GNAS,DDX17,GPI,HTRA4,BSG,FASN,ALDH6A1,ATF
,CHD4,JUND,ATP2A2,CRMP1,BEX3,VAT1,NISCH,HNRNPC,CCNI,KLHL9,SPTBN2,SLC3A2,PFKL,TK
B4B,UCHL1,AGRN,TIMP2,SARAF,COL11A1,MIDN,MARCHF6,WSB1,RPS17,VCL,CYB5R3,AKAP12,
RCKS,SMG1,CALM1,NPM1,PLXNB2,PLD3,HSPD1,USP11,COL18A1,SNRNP200,SMIM14,IGF2,ATP
1,PTMS,SPTAN1,GSTP1,SMG1,NPM1,PLXNB2,HSPD1,COL18A1,SMIM14,IGF2,VCP,MARCKSL1,TI
ET,CHSY1,TMSB10,EIF4A1,PPP2R1A,RPL15,SSBP2,BASP1,VAPA,POLR2A,TUBB4B,UCHL1,MORF4
,KHDRBS1,CSNK1E,EIF5A,PDIA6,SOX12,ARL4C,RCN1,APLP1,MIF,CHPF,JUN,SESN3,B4GALNT4,D
CD3,TPI1,CHSY1,TMSB10,EIF4A1,RPL19,PPP2R1A,RPL15,GTF3C4,PKD1,PDIA3,SERINC1,SSBP2,
COL4A1,PSAP,PGK1,RPLP0,H3-3B,CANX,CLU,PTMA,DDX5,ENO2,SOX4,FKBP14,RPL13A,MAGED
SD,LPCAT1,BRD3,COL1A2,LLGL1,NCS1,RHOA,ZNF865,CLSTN1,SLC7A5,CHD4,ATP1A1,JUND,CD
YB5R3,MAP4,CDC42BPB,ITM2B,MAT2A,PXDN,HNRNPU,RPS16,CNN3,CTSD,LPCAT1,LAMA5,BRD3
CTSD,EIF4H,SCARB2,TSPAN3,LPCAT1,MSN,KPNB1,C1ORF43,LLGL1,ZNF865,RAB6B,SLC7A5,JU
42BPB,ITM2B,PXDN,HNRNPU,PGAM1,CTNNA1,FADS2,CTSD,LPCAT1,LAMA5,BRD3,CIPC,PAPOLA
SNRNP200,CSDE1,EIF3C,ATP5F1A,EIF4B,UBB,AFF4,TRIM28,MATR3,YWHAG,CKB,RPL37A,NONO
LA-C,PDIA3,SSBP2,BASP1,POLR2A,AP2M1,TPT1,RPL18,ITGB1,MORF4L1,BTF3,ANXA2,AGRN,ZMI
3,AGRN,TIMP2,SARAF,COL11A1,RPS17,VCL,ARF1,CYB5R3,AKAP12,CDC42BPB,ITM2B,PXDN,HN
,PTMA,DDX5,ENO2,FKBP14,RPL13A,MAGED1,SEC62,RPL6,KAT6A,DVL3,YWHAE,RPL13,GNAS,DD
,USP11,CSDE1,EIF3C,EIF4B,MARCKSL1,UBB,AFF4,TRIM28,YWHAG,RPL37A,SPEN,DDB1,PSMD5,
ALM1,PLXNB2,USP11,SMIM14,IGF2,EIF3C,MARCKSL1,UBB,AFF4,TRIM28,YWHAG,COL4A2,LRR
I1,FSCN1,PTMS,SMG1,NPM1,PLXNB2,USP11,COL18A1,CSDE1,SMIM14,IGF2,EIF3C,VCP,MARCK
G8,WDR1,ASTN2,TLE5,RPL31,PRKCSH,HNRNPC,CSNK1E,PFKL,TKT,CDH2,PIP4K2B,SOX12,ARL4
3A,PXDN,HNRNPU,CTNNA1,RPS16,CTSD,LPCAT1,BRD3,MSN,C1ORF43,LLGL1,ZNF865,ATF4,SLC
I,SPEN,NONO,DDB1,YWHAQ,HACD3,DCHS1,SET,EGLN1,CHSY1,TMSB10,EIF4A1,MRC2,RPL19,RF

;,RPL5,FSCN1,PTMS,SMG1,CALM1,PLXNB2,PLD3,COL18A1,IGF2,EIF3C,ATP5F1A,VCP,MARCKSL
,RPL6,DVL3,YWHAE,GNAS,GPI,HTRA4,RACK1,BSG,FASN,ATP5F1B,CALM3,GNB1,SHISA9,RPS8,M
M28,MATR3,YWHAG,CKB,COL4A2,FN1,NONO,DDB1,PSMD5,YWHAQ,DCHS1,SET,EGLN1,INA,EIF
,ATP5F1B,CALM3,SHISA9,STMN2,MYH9,RPL7,SEPTIN2,P4HB,ERVV-1,RPL5,GDI1,FSCN1,PTMS,S
VV-1,GDI1,FSCN1,TMBIM6,PTMS,SPTAN1,SMG1,CALM1,NPM1,PLXNB2,TTC3,HSPD1,USP11,COI
A1,ARF1,SF3B2,CDC42BPB,GOLGA8A,HNRNPU,SMARCA5,RPS16,RPL23,NACA,LAMA5,BRD3,PJA
,ASTN2,TLE5,NISCH,CCNI,TADA2B,KHDRBS1,CSNK1E,PFKL,CDH6,SFPQ,CDH2,PDIA6,GPC3,SOX
L15,GTF3C4,LDHB,PKD1,HLA-C,SSBP2,BASP1,TPT1,TUBB4B,MORF4L1,NCAM1,ANXA2,RPLP1,T
IAGED1,RPL6,KAT6A,DVL3,YWHAE,KIF1A,RPL13,GNAS,SLC2A3,GPI,HTRA4,RACK1,BSG,FASN,AL
ISB10,MRC2,PPP2R1A,GTF3C4,PKD1,HLA-C,SSBP2,RPS3,VAPA,TPT1,RPS3A,STMN1,ITGB1,TUBE
HB,SF3B1,SSBP2,BASP1,VAPA,POLR2A,TPT1,RPS3A,STMN1,UCHL1,MORF4L1,BTF3,ANXA2,AGRI
,TPT1,ITGB1,TUBB4B,MORF4L1,ANXA2,ZMIZ1,SARAF,MIDN,WSB1,VCL,AP3D1,SBF1,ARF1,CYB5
SF3B1,PDIA3,SSBP2,BASP1,POLR2A,TPT1,SEC61A1,RPS3A,RPL18,STMN1,ITGB1,UCHL1,MORF4I
NPM1,PLXNB2,PLD3,HSPD1,USP11,SMIM14,IGF2,ATP5F1A,EIF4B,MARCKSL1,UBB,AFF4,TRIM28
I,HTRA4,RACK1,BSG,FASN,ATP5F1B,CALM3,GNB1,SHISA9,RPS8,MYH9,RPL7,SEPTIN2,P4HB,APF
FADS2,CPE,CNN3,TNS3,LPCAT1,LAMA5,SLC22A17,BRD3,IGFBP3,LLGL1,NCS1,CLSTN1,ATF4,RAI
S8,SEPTIN2,P4HB,APP,IDS,FSCN1,GSTP1,NPM1,PLXNB2,HSPD1,SMIM14,IGF2,ATP5F1A,MARCKS
T,CHSY1,TMSB10,EIF4A1,MRC2,RPL15,GTF3C4,LDHB,PKD1,SF3B1,PDIA3,SERINC1,SSBP2,CHD
SEC62,KAT6A,DVL3,YWHAE,KIF1A,RPL13,GNAS,SLC2A3,DDX17,GPI,BSG,FASN,ALDH6A1,SPTBN
PEN,NONO,PSMD5,YWHAQ,EIF1,CHSY1,TMSB10,EIF4A1,MRC2,RPL19,PPP2R1A,GTF3C4,PKD1,F
IF4A1,MRC2,GTF3C4,PKD1,HLA-C,SSBP2,VAPA,HMGB1,TPT1,RPS3A,STMN1,TUBB4B,UCHL1,MC
K1,RPLP0,H3-3B,CANX,CLU,PTMA,DDX5,ENO2,FKBP14,RPL13A,MAGED1,SEC62,RPL6,KAT6A,DV
1,ZNF865,CLSTN1,SLC7A5,CHD4,JUND,CD81,HMGN2,ATP2A2,CRMP1,LASP1,OAZ1,PFN1,TLE5,N
DX5,ENO2,FKBP14,MAGED1,SEC62,KAT6A,DVL3,YWHAE,KIF1A,RPL13,GNAS,GPI,BSG,FASN,SPTE
PL14,NCS1,ZNF865,ATF4,ATP1A1,HMGN2,LASP1,OAZ1,PFN1,ASTN2,JPT2,TADA2B,KHDRBS1,KL
KSL1,AFF4,MATR3,YWHAG,CKB,MAPT,SPEN,YWHAQ,EIF1,SET,EGLN1,MAP9,TPI1,CHSY1,TMSB1C
KSL1,AFF4,MATR3,YWHAG,CKB,MAPT,SPEN,YWHAQ,EIF1,SET,EGLN1,MAP9,TPI1,CHSY1,TMSB1C
DNAH8,PKD1,SF3B1,SSBP2,ZNF43,HMGB1,POLR2A,TPT1,SEC61A1,RPS3A,ITGB1,TUBB4B,RPL1
,MAT2A,PXDN,HNRNPU,SMARCA5,CTNNA1,RPS16,RPL23,CNN3,CTSD,LPCAT1,LAMA5,BRD3,CIPC
P2A2,CRMP1,TLE5,NISCH,CCNI,XPR1,XPO1,SLC3A2,HDLBP,EIF5A,CCND2,CDH2,PDIA6,DDX3X,RF
HOA,ZNF865,ATF4,NR2F1,RAB6B,ENC1,CXADR,CHD4,ATP1A1,JUND,ATP2A2,CRMP1,VAT1,PFN1,
,DDX5,ENO2,SOX4,RPL13A,SEC62,KAT6A,YWHAE,KIF1A,RPL13,SLC2A3,DDX17,GPI,HTRA4,RACI
RPS3A,STMN1,ITGB1,MORF4L1,BTF3,ZMIZ1,COL11A1,RPL10,MARCHF6,WSB1,RPS17,VCL,AP3
GRN,COL11A1,MIDN,VCL,AP3D1,AKAP12,MAP4,CDC42BPB,GOLGA8A,OGA,HNRNPA0,FADS2,RP
,SEC61A1,RPS3A,TUBB4B,RPL11,MORF4L1,SARAF,RPL10,WSB1,RPS17,AP3D1,SBF1,CYB5R3,S
PLD3,HSPD1,USP11,COL18A1,IGF2,EIF3C,ATP5F1A,VCP,MARCKSL1,AFF4,TRIM28,YWHAG,CKB,
),MARCHF6,DNAJA1,ARF1,ZNF770,GTF2F1,PGAM1,HNRNPA0,CTNNA1,FADS2,RPS16,CPE,CTSD,
BSG,FASN,ALDH6A1,SPTBN1,CALM3,SHISA9,STMN2,MYH9,SEPTIN2,IDS,RPL5,PTMS,SPTAN1,GS
,RPS16,CNN3,TNS3,CTSD,LAMA5,BRD3,CIPC,KPNB1,COL1A2,LLGL1,NCS1,ZNF865,RAB6B,CHD
DVL3,YWHAE,KIF1A,GNAS,SLC2A3,GPI,HTRA4,RACK1,BSG,FASN,ALDH6A1,SPTBN1,ATP5F1B,CAI
STMN2,RPS8,MYH9,RPL7,SEPTIN2,P4HB,APP,IDS,ERVV-1,RPL5,GDI1,TMBIM6,SPTAN1,MARCKS,(
TMN1,ITGB1,TUBB4B,MORF4L1,NCAM1,AGRN,MIDN,RPL10,MARCHF6,AP3D1,DNAJA1,MAT2A,G
STN2,MCL1,NISCH,PRKCSH,LRPPRC,KLHL9,PFKL,HDLBP,GLG1,SFPQ,DDX3X,ITM2C,RPL30,EIF
,HLA-C,VAPA,TPT1,RPL18,STMN1,ITGB1,TUBB4B,BTF3,ZMIZ1,TIMP2,SARAF,COL11A1,MIDN,WS
I1,SPTAN1,MARCKS,NPM1,HSPD1,SNRNP200,IGF2,EIF3C,ATP5F1A,EIF4B,VCP,MARCKSL1,MATR

MAGED1,SEC62,RPL6,KAT6A,DVL3,YWHAE,RPL13,GNAS,DDX17,GPI,HTRA4,RACK1,FASN,SPTBN1
,MARCHF6,VCL,AP3D1,SBF1,ARF1,CDC42BPB,MAT2A,TMEM170A,SLC4A8,OGA,LARP1,HNRNPA0
F3C,VCP,MARCKSL1,AFF4,TRIM28,CKB,COL4A2,RPL37A,LRRC28,FN1,SPEN,NONO,DDB1,HACD3
3,PTGES3,HNRNPH1,ZNF454,MAGED4B,CTBP1,ARHGDIA,CDK16,BCLAF1,NNAT,DDX39B,EID1,SR
,APP,ERVV-1,PTMS,SPTAN1,MARCKS,SMG1,CALM1,NPM1,PLXNB2,PLD3,HSPD1,USP11,COL18A
GED1,RPL6,KAT6A,YWHAE,KIF1A,RPL13,GNAS,GPI,HTRA4,RACK1,BSG,FASN,ALDH6A1,SPTBN1,A
GB1,TUBB4B,RPL11,AGRN,RPLP1,MIDN,RPL10,MARCHF6,RPS17,AP3D1,DNAJA1,ARF1,SF3B2,M
3,HSPD1,USP11,SNRNP200,CSDE1,IGF2,EIF3C,ATP5F1A,EIF4B,VCP,AFF4,MATR3,RPL37A,LRRC2
CD81,LENG8,HMGN2,ATP2A2,CRMP1,BEX3,VAT1,PFN1,TLE5,CCNI,XPO1,SPTBN2,CSNK1E,CCND
M6,GSTP1,SMG1,CALM1,NPM1,HSPD1,USP11,COL18A1,SMIM14,IGF2,EIF3C,VCP,MARCKSL1,U
PPP2R1A,GTF3C4,PKD1,HLA-C,PDIA3,VAPA,TPT1,RPS3A,STMN1,ITGB1,TUBB4B,BTF3,ZMIZ1,TIM
ARL4C,GNB2,H1-0,APLP1,MIF,HMGCS1,JUN,B4GALNT4,MTSS2,TARDBP,MYL6,ANKRD52,WLS,DU
1,ZNF865,SLC7A5,CHD4,JUND,CD81,WDR1,HMGN2,ATP2A2,CRMP1,LASP1,OAZ1,MCL1,NES,NIS
PL13,GNAS,GPI,HTRA4,RACK1,BSG,FASN,ALDH6A1,SPTBN1,ATP5F1B,CALM3,GNB1,SHISA9,RPS
AT6A,DVL3,YWHAE,KIF1A,RPL13,SLC2A3,DDX17,GPI,HTRA4,RACK1,BSG,FASN,ALDH6A1,SPTBN
,MARCKSL1,UBB,TRIM28,CKB,COL4A2,DDB1,PSMD5,YWHAQ,EIF1,HACD3,SET,CHSY1,TMSB10,I
A1,JUND,EIF5,CD81,ATP2A2,CRMP1,LASP1,OAZ1,SRSF4,PFN1,ASTN2,TLE5,NES,NISCH,TADA2B,
L4A2,LRRC28,FN1,SPEN,DDB1,YWHAQ,DCHS1,SET,EGLN1,MAP9,TPI1,CHSY1,EIF4A1,MRC2,RPL
A,DNER,SQLE,HS6ST1,NOVA2,CTSB,SRSF5,RAB27A,RPN2,TBCD,SUGP2,HNRNPL,PTPRA,CAMSAP
EIF4B,UBB,AFF4,RPL37A,SPEN,NONO,EIF1,EGLN1,TPI1,INA,EIF4A1,PPP2R1A,RPL15,GTF3C4,DNA
H,TADA2B,KHDRBS1,EIF5A,ACTN4,RPS15,ITM2C,RPS5,MIF,DNAJC5,SLC25A3,PTGES3,HNRNPH1
DS2,LPCAT1,LAMA5,SLC22A17,PJA2,SLC16A7,KPNB1,C1ORF43,LLGL1,HK1,NCS1,ZNF865,NLGN
AG,RPL37A,LRRC28,YWHAQ,EIF1,EGLN1,TPI1,TMSB10,INA,EIF4A1,RPL19,PPP2R1A,RPL15,GTF3
BA1,ATP2B4,IGFBP2,CALR,FSTL1,TMF1,ABHD2,HSPA1B,TUBB2B,COL5A1,COL4A1,CES3,PSAP,P
GNAS,SLC2A3,DDX17,GPI,HTRA4,FASN,ALDH6A1,SPTBN1,ATP5F1B,CALM3,GNB1,SHISA9,STMN2
L,ARF1,CYB5R3,AKAP12,CDC42BPB,ITM2B,AP2B1,PXDN,HNRNPU,SMARCA5,RPL23,RPSA,TPM4,
L1,BTF3,AGRN,TIMP2,SBF1,CYB5R3,PAFAH1B1,CDC42BPB,PXDN,PGAM1,NFE2L1,HNRNPA0,CTN
MG1,NPM1,PLD3,HSPD1,USP11,COL18A1,IGF2,EIF3C,EIF4B,VCP,UBB,TRIM28,MATR3,YWHAG,C
BPB,AP2B1,MAT2A,GOLGA8A,PXDN,HNRNPU,CTNNA1,CTSD,LPCAT1,LAMA5,BRD3,CIPC,MSN,LL
SD,LAMA5,BRD3,KPNB1,PAPOLA,COL1A2,NCS1,CLSTN1,ENC1,CXADR,CHD4,LENG8,WDR1,HMG
PL19,PPP2R1A,GTF3C4,LDHB,PKD1,HLA-C,PDIA3,SERINC1,SSBP2,CHD3,VAPA,TPT1,STMN1,TUB
TBN1,ATP5F1B,CALM3,GNB1,SHISA9,RPS8,MYH9,RPL7,SEPTIN2,P4HB,RPL5,GDI1,FSCN1,TMBIM
EIF3C,EIF4B,MARCKSL1,UBB,MATR3,YWHAG,COL4A2,LRRC28,SPEN,PSMD5,YWHAQ,EIF1,SET,M
DX17,GPI,RACK1,BSG,FASN,SPTBN1,ATP5F1B,GNB1,SHISA9,STMN2,RPS8,RPL7,SEPTIN2,P4HB,
2,DNAH8,PKD1,HLA-C,SSBP2,ZNF43,VAPA,HMGB1,AP2M1,TPT1,SEC61A1,RPS3A,STMN1,ITGB1,T
FASN,ALDH6A1,ATP5F1B,GNB1,SHISA9,RPS8,MYH9,SEPTIN2,APP,IDS,GDI1,FSCN1,TMBIM6,SPTA
,CES3,PSAP,PGK1,RPLP0,H3-3B,CLU,PTMA,DDX5,ENO2,SOX4,FKBP14,MAGED1,SEC62,RPL6,KA
,IGF2,EIF3C,ATP5F1A,EIF4B,MARCKSL1,AFF4,TRIM28,YWHAG,CKB,RPL37A,YWHAQ,HACD3,DCH
6A2,RPS10,B4GALNT4,DNAJC5,APLP2,RPLP2,MXD4,HNRNPK,H1-10,CIRBP,DUX4,CADM4,HLA-B,
BX1,UBA1,ATP2B4,IGFBP2,CALR,FSTL1,TMF1,ABHD2,HSPA1B,ZNF99,TUBB2B,COL5A1,COL4A1
RPL5,TMBIM6,PTMS,SPTAN1,SMG1,CALM1,NPM1,PLXNB2,PLD3,HSPD1,USP11,COL18A1,IGF2,E
INA,EIF4A1,RPL15,GTF3C4,DNAH8,PDIA3,SSBP2,HMGB1,CDKL1,AP2M1,TPT1,SEC61A1,ITGB1,T
B1,CDKL1,AP2M1,SEC61A1,TUBB4B,MORF4L1,ANXA2,RPLP1,WSB1,RPS17,AP3D1,DNAJA1,ARF
AMA5,BRD3,RPL17,C1ORF43,LLGL1,HK1,RPL22,RHOA,CLSTN1,PSMD2,NLGN2,GLIPR1L2,JUND
D3,CIPC,PJA2,LLGL1,GAP43,HK1,NCS1,ZNF865,CLSTN1,RAB6B,CHD4,JUND,CD81,WDR1,HMGN

=,SESN3,B4GALNT4,MTSS2,APLP2,HNRNPDL,TARDBP,MXD4,INIP,MYH10,RPL36,CADM4,HNRNPI
_NT4,DNAJC5,APLP2,TARDBP,INIP,CADM4,HNRNPH1,PLPPR3,MAGED4B,CTBP1,STIP1,GPC1,TOI
3,ASTN2,SPTBN2,SLC3A2,SCRN1,EIF5A,PEBP1,CDH2,DDX3X,RPS15,RPL12,EIF3L,MRFAP1,RPS5,
2,SOX4,FKBP14,RPL13A,SEC62,KAT6A,YWHAE,KIF1A,RPL13,GNAS,SLC2A3,DDX17,GPI,RACK1,E
)X17,GPI,HTRA4,RACK1,FASN,ALDH6A1,SPTBN1,CALM3,GNB1,SHISA9,RPS8,RPL7,SEPTIN2,P4HI
K2B,SOX12,APLP1,MIF,SESN3,B4GALNT4,MTSS2,DNAJC5,APLP2,MXD4,CADM4,PLPPR3,MAGEI
1,VAT1,OAZ1,NES,NISCH,TADA2B,SLC3A2,PFKL,HDLBP,TKT,PNMA1,GPC3,DDR1,PIP4K2B,SOX1
K1,FASN,ATP5F1B,CALM3,GNB1,SHISA9,RPS8,MYH9,RPL7,SEPTIN2,P4HB,RPL5,GDI1,FSCN1,TI
,RAB6B,ATP1A1,TMEM259,WDR1,HMGN2,ATP2A2,CRMP1,RTN3,PFN1,ASTN2,JPT2,TLE5,CCNI,XI
,RAB6B,ATP1A1,TMEM259,WDR1,HMGN2,ATP2A2,CRMP1,RTN3,PFN1,ASTN2,JPT2,TLE5,CCNI,XI
,RAB6B,ATP1A1,TMEM259,WDR1,HMGN2,ATP2A2,CRMP1,RTN3,PFN1,ASTN2,JPT2,TLE5,CCNI,XI
PL37A,MAPT,SPEN,DDB1,PSMD5,YWHAQ,EIF1,HACD3,SET,TPI1,CHSY1,TMSB10,EIF4A1,MRC2,DI
JND,ATP2A2,BEX3,VAT1,JPT2,NES,RPL31,TADA2B,SCRN1,HDLBP,TKT,CDH2,PDIA6,PIP4K2B,MEC
'H9,RPL7,P4HB,APP,RPL5,GDI1,FSCN1,TMBIM6,PTMS,SPTAN1,GSTP1,MARCKS,SMG1,CALM1,N
_3,YWHAE,KIF1A,RPL13,GNAS,DDX17,HTRA4,FASN,ALDH6A1,SPTBN1,CALM3,GNB1,SHISA9,RPS
1B,CALM3,GNB1,SHISA9,RPS8,MYH9,RPL7,SEPTIN2,P4HB,RPL5,GDI1,TMBIM6,PTMS,SPTAN1,S
SEC62,RPL6,KAT6A,DVL3,YWHAE,KIF1A,RPL13,GNAS,DDX17,GPI,HTRA4,BSG,FASN,ALDH6A1,SP
MP2,DNER,L1CAM,TRIB2,PEA15,SQLE,HS6ST1,NOVA2,CDH11,CHD6,EEF1A2,TBCD,ANXA5,SNX2
I,DDB1,PSMD5,YWHAQ,EIF1,HACD3,SET,CHSY1,TMSB10,RPL19,LDHB,SF3B1,SSBP2,BASP1,VAP
MSN,KPNB1,COL1A2,LLGL1,GAP43,NCS1,ZNF865,CLSTN1,KIF5B,RAB6B,SLC7A5,CHD4,ATP1A1
TPT1,SEC61A1,ITGB1,TUBB4B,UCHL1,MORF4L1,ANXA2,AGRN,ZMIZ1,TIMP2,COL11A1,MIDN,RI
IF770,MAT2A,GTF2F1,PXDN,HNRNPU,HNRNPA0,CTNNA1,FADS2,CNN3,RPSA,EIF4H,LAMA5,BRD3
AF1,SARAF,MIDN,ARF1,CYB5R3,MAP4,CDC42BPB,AP2B1,CTSD,LAMA5,MSN,KPNB1,LLGL1,NCS
PI,HTRA4,RACK1,FASN,ALDH6A1,SPTBN1,CALM3,GNB1,SHISA9,RPS8,RPL7,SEPTIN2,P4HB,APP,I
ND2,SOX12,GNB2,H1-0,MIF,CHPF,CD63,B4GALNT4,DNAJC5,APLP2,ANKRD52,WLS,HNRNPK,INIP
RA4,RACK1,FASN,ALDH6A1,SPTBN1,CALM3,GNB1,SHISA9,MYH9,SEPTIN2,P4HB,RPL5,GDI1,FSC
CYB5R3,MAP4,SF3B2,CDC42BPB,PXDN,LDHA,RPSA,SCARB2,NACA,TSPAN3,LAMA5,RPS24,BRD3
P2M1,STMN1,TUBB4B,UCHL1,NCAM1,AGRN,TIMP2,MIDN,MARCHF6,VCL,AP3D1,DNAJA1,ARF1,C
4L1,BTF3,NCAM1,ANXA2,AGRN,SARAF,MIDN,RPL10,MARCHF6,VCL,AP3D1,DNAJA1,SF3B2,MAT
K1,FASN,ALDH6A1,ATP5F1B,CALM3,GNB1,SHISA9,RPS8,MYH9,SEPTIN2,P4HB,RPL5,GDI1,FSCN1
PFKL,RPS25,TKT,TPM1,PEBP1,PDIA6,GPC3,PLXNA1,PIP4K2B,GNB2,TLN1,CHPF,MYO5A,CD63,N
S3A,ITGB1,TUBB4B,BTF3,RPLP1,SARAF,MIDN,RPL10,RPS17,AP3D1,DNAJA1,ARF1,CYB5R3,SF3I
10,MRC2,GTF3C4,LDHB,PKD1,SSBP2,CHD3,BASP1,VAPA,HMGB1,POLR2A,TPT1,RPL18,STMN1,T
,SET,CHSY1,TMSB10,RPL19,LDHB,SF3B1,SSBP2,BASP1,VAPA,POLR2A,STMN1,TUBB4B,UCHL1,I
IEGF8,DDX3X,HNRNPDL,NARS1,RAB1A,ARCN1,RPS27,RAC1,CAPRIN1,SMCHD1,ATP6V0C,BCLAF
DS,ERVV-1,GDI1,TMBIM6,PTMS,SPTAN1,MARCKS,NPM1,USP11,IGF2,EIF3C,VCP,UBB,MATR3,YV
11,SNRNP200,CSDE1,IGF2,EIF3C,ATP5F1A,EIF4B,VCP,AFF4,MATR3,YWHAG,CKB,RPL37A,LRRC
I,RACK1,FASN,CALM3,GNB1,SHISA9,RPS8,MYH9,P4HB,RPL5,FSCN1,TMBIM6,SPTAN1,CALM1,PL
ADR,ATP1A1,EIF5,CD81,ATP2A2,LASP1,SRSF4,RBBP4,RTN3,PFN1,ASTN2,CSNK1E,TPM1,GPC3,F
I,ALDH6A1,SPTBN1,CALM3,GNB1,SHISA9,RPS8,RPL7,APP,ERVV-1,GDI1,FSCN1,TMBIM6,PTMS,SI
6A1,GNB1,RPS8,MYH9,SEPTIN2,P4HB,APP,GDI1,FSCN1,TMBIM6,SPTAN1,SMG1,NPM1,PLD3,HS
BASP1,VAPA,HMGB1,TPT1,SEC61A1,RPS4X,RPS3A,STMN1,ITGB1,UCHL1,RPLP1,ZMIZ1,RPL10,W
_3,YWHAE,KIF1A,RPL13,GNAS,DDX17,HTRA4,RACK1,FASN,ALDH6A1,SPTBN1,CALM3,GNB1,SHIS
AP3D1,DNAJA1,MAT2A,PXDN,HNRNPA0,CTNNA1,FADS2,CNN3,RPSA,EIF4H,SLC22A17,BRD3,RPS
310,INA,MRC2,PPP2R1A,GTF3C4,PKD1,PDIA3,SERINC1,RPS3,BASP1,HMGB1,POLR2A,AP2M1,TP

PLP1,SARAF,MIDN,RPL10,RPS17,AP3D1,ARF1,CYB5R3,SF3B2,TMEM170A,GTF2F1,HNRNPA0,FAI
HMGN2,ATP2A2,CRMP1,PFN1,TLE5,CCNI,TADA2B,XPO1,SLC3A2,PFKL,RPS25,TKT,TPM1,EIF5A,C
KPNB1,IGFBP3,LLGL1,RPL14,NCS1,PSMD2,NR2F1,RPS14,TMEM259,EIF5,CD81,WDR1,HMGN2,C
1,ATP5F1B,CALM3,GNB1,SHISA9,RPS8,MYH9,P4HB,RPL5,FSCN1,TMBIM6,SPTAN1,NPM1,PLXNB
IB1,SHISA9,RPS8,MYH9,RPL7,SEPTIN2,P4HB,RPL5,GDI1,TMBIM6,PTMS,SPTAN1,SMG1,CALM1,F
EIF4A1,MRC2,DNAH8,SSBP2,CHD3,TPT1,RPS3A,ITGB1,UCHL1,BTF3,AGRN,ZMIZ1,SARAF,COL11
KAP5,CADM4,LAMP1,CTBP1,RAC1,ARHGDIA,STIP1,GPC1,TOP2B,ATP6V0C,TRIB2,HS6ST1,TIMP3
MARCKSL1,AFF4,TRIM28,MATR3,CKB,RPL37A,MAPT,NONO,DDB1,YWHAQ,HACD3,DCHS1,SET,M
N,ALDH6A1,SPTBN1,CALM3,GNB1,SHISA9,STMN2,MYH9,SEPTIN2,P4HB,APP,IDS,GDI1,TMBIM6,F
RPS4X,RPS3A,STMN1,ITGB1,UCHL1,RPLP1,ZMIZ1,WSB1,AP3D1,SF3B2,CDC42BPB,ITM2B,GOLG
P,MXD4,INIP,RPL36,CADM4,HNRNPH1,PLPPR3,HECTD4,CTBP1,RAC1,ARHGDIA,STIP1,SLC35E2B
A,SMARCA5,CPE,RPSA,NACA,LAMA5,BRD3,PJA2,KPNB1,LLGL1,GAP43,HK1,CBX5,ZNF865,NR2F1
P1,SLC35E2B,SMCHD1,IPO5,DNER,BCLAF1,L1CAM,SQLE,FAM199X,TIMP3,CTDSP2,NUCB1,CHD
ATP2A2,CRMP1,LASP1,OAZ1,NES,NISCH,TADA2B,KHDRBS1,PFKL,PEBP1,PNMA1,PDIA6,PIP4K2B
CKSL1,TRIM28,MATR3,CKB,COL4A2,SPEN,DDB1,HACD3,DCHS1,SET,TPI1,CHSY1,EIF4A1,RPL19,
GB1,CDKL1,AP2M1,SEC61A1,ITGB1,TUBB4B,BTF3,NCAM1,AGRN,RPL10,MARCHF6,WSB1,RPS1
,HMGB1,AP2M1,TPT1,SEC61A1,RPS3A,STMN1,ITGB1,TUBB4B,MORF4L1,NCAM1,ANXA2,AGRN,Z
D1,USP11,SNRNP200,CSDE1,IGF2,EIF3C,ATP5F1A,AFF4,MATR3,CKB,RPL37A,LRRC28,YWHAQ,E
EIF3L,MRFAP1,RCN1,RPL35,CD63,GOLM1,B4GALNT4,MTSS2,APLP2,TARDBP,SHOX,WLS,HNRN
A9,RPS8,MYH9,RPL7,SEPTIN2,P4HB,RPL5,GDI1,FSCN1,TMBIM6,PTMS,SPTAN1,SMG1,NPM1,PLX
TRIM28,MATR3,COL4A2,RPL37A,LRRC28,YWHAQ,HACD3,DCHS1,SET,MAP9,TMSB10,INA,EIF4A
B1,ZNF43,RPS3,BASP1,VAPA,POLR2A,AP2M1,TPT1,SEC61A1,RPS3A,OR8K1,RPL18,STMN1,TUB
TMBIM6,SPTAN1,PLXNB2,PLD3,HSPD1,USP11,SNRNP200,CSDE1,SMIM14,IGF2,EIF3C,ATP5F1A,
ZNF865,NLGN2,CHD4,JUND,CD81,CRMP1,VAT1,RBBP4,PFN1,MCL1,TLE5,HNRNPC,CCNI,TADA2
B,MEGF8,ACTN4,MMP14,PRDX2,GNB2,APLP1,RPS7,RPS19,GPAT4,JUN,CD63,SESN3,APLP2,MYL
F3,SARAF,MIDN,RPL10,MARCHF6,RPS17,AP3D1,DNAJA1,ARF1,CYB5R3,SF3B2,MAT2A,GTF2F1,H
TUBB4B,RPL11,MORF4L1,BTF3,RPLP1,SARAF,MIDN,RPL10,RPS17,AP3D1,CYB5R3,SF3B2,GTF2
E2L1,HNRNPA0,RPS16,RPL23,RPL29,RPSA,NACA,LAMA5,RPS24,BRD3,RPS9,XRCC5,HK1,RPL22,F
SG,FASN,ALDH6A1,SPTBN1,ATP5F1B,CALM3,GNB1,STMN2,RPS8,RPL7,RPL5,GDI1,FSCN1,PTMS
,IGF2,EIF3C,ATP5F1A,EIF4B,VCP,UBB,MATR3,YWHAG,RPL37A,LRRC28,SPEN,NONO,EIF1,TMSB
3,PTGES3,HNRNPH1,ZNF454,MAGED4B,CTBP1,ARHGDIA,CDK16,BCLAF1,NNAT,DDX39B,EID1,C
IN2,RPS8,MYH9,SEPTIN2,ERVV-1,GDI1,TMBIM6,PTMS,SPTAN1,MARCKS,SMG1,CALM1,NPM1,PL
MP3,CTDSP2,NOVA2,CHD6,EEF1A2,CCT2,TBCD,VAMP2,MEST,GAS7,TM9SF2,LMNB2,ALKBH5,PD
TR3,COL4A2,HACD3,SET,EGLN1,CHSY1,TMSB10,INA,PPP2R1A,RPL15,GTF3C4,LDHB,DNAH8,HL
M3,GNB1,SHISA9,MYH9,RPL7,P4HB,FSCN1,TMBIM6,PTMS,SPTAN1,GSTP1,SMG1,CALM1,NPM1,
2,PLD3,HSPD1,USP11,SNRNP200,CSDE1,IGF2,EIF3C,ATP5F1A,EIF4B,VCP,UBB,AFF4,MATR3,RPL
,TMBIM6,PTMS,SPTAN1,MARCKS,USP11,IGF2,EIF3C,EIF4B,VCP,UBB,MATR3,CKB,COL4A2,RPL
DR1,HMGN2,ATP2A2,CRMP1,LASP1,BEX3,VAT1,CCNI,SPTBN2,HDLBP,RPS25,CCND2,PIP4K2B,SC
T7,SRSF4,RBBP4,PFN1,MCL1,TLE5,PNN,PFKL,RPS25,TKT,EIF5A,PDIA6,PIP4K2B,RAB7A,EIF3L,H1
HMGN2,ATP2A2,CRMP1,LASP1,VAT1,OAZ1,PFN1,RPL31,NISCH,HNRNPC,PDIA6,PIP4K2B,MEGF8,
ALDH6A1,SPTBN1,CALM3,GNB1,SHISA9,RPS8,RPL7,SEPTIN2,P4HB,APP,ERVV-1,GDI1,FSCN1,TM
NCS1,RHOA,ZNF865,SLC7A5,ATP1A1,JUND,TMEM259,RPS20,EIF5,WDR1,HMGN2,RPL26,LASP1,
I1,GSTP1,SMG1,CALM1,NPM1,PLXNB2,USP11,EIF4B,VCP,TRIM28,MATR3,YWHAG,COL4A2,RPL
JBB,AFF4,TRIM28,YWHAG,CKB,COL4A2,NONO,PSMD5,EIF1,DCHS1,TMSB10,INA,MRC2,PPP2R1
,DPYSL2,TOP1,PPP2CA,SMARCA4,KDELR2,RABL6,TCF25,PTBP1,LUZP6,GRB2,SCAF11,CLIP3,CC

ARCKSL1,UBB,AFF4,MATR3,YWHAG,COL4A2,RPL37A,FN1,DDB1,HACD3,SET,MAP9,TPI1,CHSY1,
2,RHOA,PTCD3,ZNF865,ATF4,RPS14,RPS20,LENG8,WDR1,ATXN7L3B,RPL26,LASP1,CCT7,RBBP
GALNT4,RPLP2,CIRBP,RAB1A,PLPPR3,SEPTIN3,ZNF454,COTL1,GPC1,L1CAM,TRIB2,SQLE,HS6S
1,MVB12B,KDELR2,LAPTM4A,BTBD2,RABL6,ODC1,ATXN10,CASC3,ZFAND5,HNRNPUL2-BSCL2,
SMG1,PLD3,HSPD1,USP11,SNRNP200,CSDE1,SMIM14,IGF2,EIF3C,ATP5F1A,EIF4B,VCP,UBB,AFF
MATR3,RPL37A,LRRC28,NONO,EIF1,INA,EIF4A1,RPL19,PPP2R1A,RPL15,GTF3C4,LDHB,DNAH8,P
N2,MYH9,SEPTIN2,GDI1,TMBIM6,PTMS,SPTAN1,MARCKS,SMG1,CALM1,NPM1,PLXNB2,IGF2,EIF
,ENO2,SOX4,FKBP14,RPL13A,MAGED1,SEC62,RPL6,KAT6A,DVL3,YWHAE,KIF1A,RPL13,SLC2A3,
1,CAMSAP1,GRINA,ACAP3,MDK,LAMC1,H3-3A,ABCA2,VEGFA,TUBB2A,MAPRE1,BTBD2,ODC1,TC
FKBP14,RPL13A,MAGED1,SEC62,RPL6,KAT6A,DVL3,YWHAE,KIF1A,RPL13,SLC2A3,DDX17,GPI,HT
,RPS15,RPL30,CHPF,JUN,RPS10,WLS,HNRNPK,H1-10,MYH10,YWHAB,RAB1A,SEPTIN3,STIP1,GP
F2,EIF3C,ATP5F1A,EIF4B,VCP,UBB,MATR3,YWHAG,RPL37A,LRRC28,EIF1,INA,EIF4A1,RPL19,PPP
ATP5F1A,EIF4B,VCP,MATR3,YWHAG,RPL37A,LRRC28,EIF1,EGLN1,EIF4A1,RPL19,PPP2R1A,RPL1
,H3-3B,CLU,PTMA,DDX5,ENO2,RPL13A,SEC62,KAT6A,DVL3,YWHAE,KIF1A,GNAS,SLC2A3,DDX17,
R2A,CDKL1,AP2M1,RPS4X,RPS3A,STMN1,ITGB1,TUBB4B,RPLP1,TIMP2,RPL10,WSB1,VCL,AP3D1
BASP1,POLR2A,STMN1,TUBB4B,UCHL1,RPL11,MORF4L1,BTF3,ANXA2,ZMIZ1,MIDN,RPL10,VCL,
PPP1R10,STRN4,RAB14,KMT2E,SRSF2,EEF1B2,BLCAP,RBM5,LRRC41,IST1,NASP,EXOC4,CREBB
A,GPC1,ATP6V0C,TRIB2,GLUL,SQLE,CTDSP2,NOVA2,CTSB,RAB27A,EEF1A2,TBCD,ANXA5,PGD,A
RPLP0,H3-3B,PTMA,DDX5,ENO2,SOX4,RPL13A,MAGED1,SEC62,RPL6,KAT6A,DVL3,YWHAE,KIF1A
B,UCHL1,MORF4L1,BTF3,ZMIZ1,MIDN,RPL10,VCL,AP3D1,SBF1,DNAJA1,SF3B2,AP2B1,MAT2A,G
,CANX,CLU,PTMA,DDX5,ENO2,SOX4,FKBP14,RPL13A,SEC62,RPL6,KAT6A,DVL3,YWHAE,KIF1A,R
PL13A,MAGED1,SEC62,KAT6A,DVL3,YWHAE,KIF1A,GNAS,SLC2A3,DDX17,GPI,HTRA4,RACK1,BS
PL7,SEPTIN2,P4HB,APP,IDS,RPL5,FSCN1,TMBIM6,PTMS,SPTAN1,SMG1,NPM1,COL18A1,SNRNP
RACK1,BSG,FASN,ALDH6A1,RPS8,RPL7,P4HB,APP,IDS,ERVV-1,RPL5,GDI1,PTMS,SPTAN1,GSTP1
RN,RPLP1,SARAF,MIDN,RPL10,MARCHF6,AP3D1,ARF1,SF3B2,MAT2A,OGA,NFE2L1,SMARCA5,R
F2,EIF3C,ATP5F1A,EIF4B,VCP,UBB,MATR3,RPL37A,LRRC28,SPEN,NONO,EIF1,EGLN1,INA,EIF4A1
IAH8,HLA-C,SSBP2,CHD3,AP2M1,TPT1,ITGB1,MORF4L1,ANXA2,AGRN,RPLP1,TIMP2,SARAF,MID
PTGES3,HNRNPH1,ZNF454,MAGED4B,CTBP1,CDK16,BCLAF1,NNAT,TIMP3,DDX39B,EID1,C5ORF
1,SLC3A2,TKT,CDH6,TPM1,EIF5A,CCND2,CDH2,MEGF8,DDX3X,RPL12,GNB2,JUN,B4GALNT4,TA
,NFE2L1,HNRNPA0,FADS2,LDHA,RPL29,TNS3,CTSD,EIF4H,SCARB2,LPCAT1,RPS24,RPS9,CIPC,C
COL5A1,COL4A1,PGK1,RPLP0,H3-3B,CANX,CLU,PTMA,DDX5,ENO2,RPL13A,SEC62,RPL6,KAT6A,
RNPDL,TARDBP,WLS,HNRNPK,H1-10,PTGES3,CKAP5,LAMP1,TTR,CCT6A,RAC1,CAPRIN1,GPC1,C
LD3,USP11,COL18A1,SNRNP200,IGF2,EIF3C,ATP5F1A,EIF4B,VCP,UBB,AFF4,MATR3,RPL37A,LR
,IGF2,MARCKSL1,AFF4,YWHAG,CKB,COL4A2,RPL37A,FN1,NONO,YWHAQ,EIF1,DCHS1,SET,TPI1
PL6,KAT6A,DVL3,YWHAE,KIF1A,GNAS,DDX17,GPI,HTRA4,RACK1,FASN,ALDH6A1,SPTBN1,ATP5F1
LRD1,GDI1,FSCN1,TMBIM6,MARCKS,SMG1,NPM1,PLD3,HSPD1,USP11,COL18A1,SNRNP200,SM
IRNPUL2-BSCL2,SMARCE1,PHC2,LSS,FBLN1,TUG1,SREBF2,FBRSL1,KDM3B,BCL9L,NAP1L1,MT
3A,SEC62,YWHAE,KIF1A,RPL13,DDX17,GPI,RACK1,FASN,ALDH6A1,SPTBN1,ATP5F1B,CALM3,G
DE1,SMIM14,IGF2,EIF3C,ATP5F1A,EIF4B,VCP,UBB,AFF4,MATR3,YWHAG,RPL37A,LRRC28,SPEN,
29,EIF4H,RPS9,XRCC5,CIPC,RPL24,PAPOLA,COL1A2,HK1,RPL22,RHOA,ZNF865,GLIPR1L2,RPS1
NA,MLEC,CLSTN3,MDK,H3-3A,ABCA2,VEGFA,CCND1,BTBD2,NYNRIN,DYNLL2,PCSK1N,ACTN1,K
M4,RPS9,XRCC5,CIPC,RPL24,C1ORF43,PAPOLA,HK1,RHOA,ZNF865,ATF4,GLIPR1L2,RPS14,RPS
SG,FASN,ALDH6A1,ATP5F1B,CALM3,GNB1,SHISA9,STMN2,SEPTIN2,P4HB,APP,IDS,RPL5,KLRD1
1,SHISA9,RPS8,RPL7,SEPTIN2,P4HB,RPL5,KLRD1,GDI1,FSCN1,TMBIM6,PTMS,MARCKS,SMG1,
EIF4B,VCP,UBB,AFF4,MATR3,RPL37A,LRRC28,SPEN,NONO,EIF1,EGLN1,INA,EIF4A1,RPL19,PPP2

LXNB2,PLD3,HSPD1,COL18A1,SMIM14,IGF2,ATP5F1A,MARCKSL1,UBB,TRIM28,YWHAG,CKB,MA
LP1,RPS7,RPS19,HMGCS1,COL6A2,CD63,MTSS2,MYL6,INIP,PTGES3,CKAP4,MAGED4B,CTBP1,R
B,TERF2IP,VEGFA,BTBD2,PBX2,PTBP1,ILF3,WDR6,CAPZB,CLIP3,FKBP8,PLCG1,KDELR1,MAP2K
PRIP,HGS,SREBF2,SMS,RBM15B,TRAF4,KCNQ2,MTA1,RETREG2,AP2A1,BAZ1B,GPX4,GSPT1,CX
LM3,GNB1,SHISA9,RPS8,RPL7,P4HB,ERVV-1,GDI1,FSCN1,TMBIM6,PTMS,SPTAN1,SMG1,NPM1,F
RSF9,DSTN,PBX2,SRSF1,LUZP6,CAPZB,HNRNPUL1,UBE2R2,MAP7D1,CASC3,PLCG1,ARF4,MSL1,
6,KAT6A,DVL3,YWHAE,KIF1A,RPL13,GNAS,SLC2A3,DDX17,BSG,FASN,SPTBN1,CALM3,GNB1,SHI
DN,RPL10,RPS17,SBF1,DNAJA1,MAP4,ZNF770,AP2B1,MAT2A,GTF2F1,SLC4A8,OGA,PGAM1,SMA
E,CNN3,RPL29,LPCAT1,BRD3,KPNB1,LLGL1,NCS1,RPL22,ATF4,RAB6B,SLC7A5,CXADR,CHD4,AT
2,RPL26,OAZ1,JPT2,PNN,NISCH,XPO1,SPTBN2,PFKL,GLG1,TPM1,SOX12,RAB7A,CCN2,ITM2C,RI
NA,NCKAP1,MLEC,ACAP3,MDK,ORAI2,H3-3A,ABCA2,DPYSL2,TUBB2A,CCND1,SMARCA4,EIF3E,DY
CP,MARCKSL1,AFF4,MATR3,YWHAG,COL4A2,RPL37A,FN1,EIF1,HACD3,SET,EGLN1,TMSB10,INA,
AB1A,PLPPR3,TOP2B,SMCHD1,SKP1,BCLAF1,EID1,SRSF5,ZC3H7B,RNF187,U2AF2,MLXIP,PPP1C
4,IGFBP2,CALR,FSTL1,ABHD2,HSPA1B,TUBB2B,COL5A1,COL4A1,CES3,PSAP,PGK1,RPLP0,H3-3E
TMBIM6,SPTAN1,PLXNB2,PLD3,HSPD1,USP11,SNRNP200,CSDE1,SMIM14,IGF2,EIF3C,ATP5F1A,E
SNRNP200,IGF2,EIF3C,EIF4B,VCP,CKB,COL4A2,FN1,SPEN,PSMD5,HACD3,DCHS1,CHSY1,TMSB
HLA-C,PDIA3,SSBP2,ZNF43,VAPA,TPT1,SEC61A1,RPS3A,RPL18,STMN1,UCHL1,MORF4L1,NCAM1,
K1,ZNF865,NR2F1,KIF5B,RAB6B,ENC1,NLGN2,GLIPR1L2,ATP1A1,HMGN2,CCT7,NISCH,CCNI,TA
2,GDI2,TSR1,HSPH1,PTPRA,PRDX5,ACTR3,CAMSAP1,NCKAP1,MLEC,GCN1,SKI,RAB11B,PYGB,PF
XRCC5,CIPC,RPL24,C1ORF43,PAPOLA,IGFBP3,HK1,RPL22,RHOA,ZNF865,ATF4,GLIPR1L2,RPS1
6,KAT6A,DVL3,YWHAE,KIF1A,RPL13,GNAS,SLC2A3,DDX17,BSG,FASN,ALDH6A1,SPTBN1,ATP5F1
B1,POLR2A,AP2M1,TPT1,SEC61A1,RPS4X,RPS3A,BTF3,AGRN,RPLP1,MIDN,RPL10,MARCHF6,RF
MARCKSL1,TRIM28,COL4A2,LRRC28,SPEN,NONO,DDB1,HACD3,DCHS1,SET,EGLN1,TPI1,CHSY1,
MARCKSL1,TRIM28,COL4A2,LRRC28,SPEN,NONO,DDB1,HACD3,DCHS1,SET,EGLN1,TPI1,CHSY1,
SET,TMSB10,PDIA3,SERINC1,CHD3,BASP1,VAPA,HMGB1,POLR2A,AP2M1,TPT1,RPL18,ITGB1,TUE
SERINC1,ZNF43,HMGB1,POLR2A,AP2M1,TPT1,SEC61A1,RPS4X,ITGB1,TUBB4B,RPL11,BTF3,AGF
,UCHL1,ANXA2,AGRN,ZMIZ1,TIMP2,MIDN,MARCHF6,WSB1,RPS17,DNAJA1,ARF1,CYB5R3,SF3B2
,IGF2,EIF3C,ATP5F1A,EIF4B,VCP,UBB,AFF4,MATR3,YWHAG,RPL37A,LRRC28,SPEN,NONO,EIF1,E
PLP2,NDFIP1,MXD4,HNRNPK,DUX4,PTGES3,SEPTIN11,CADM4,RHOB,COTL1,CTBP1,RAC1,STIP1,
I,OGA,HNRNPA0,FADS2,CPE,CNN3,RPL29,EIF4H,LPCAT1,LAMA5,BRD3,CIPC,C1ORF43,LLGL1,NC
AZ1,RBBP4,MCL1,SPC25,RPL31,PRKCSH,RPS25,GLG1,PNMA1,PDIA6,SOX12,ACTN4,DDX3X,RPS
7,TBCD,ANXA5,PPP1CB,SQSTM1,GDI2,HNRNPL,TSR1,SSRP1,ALKBH5,HNRNPA3,PDXK,ACO2,TCF
MSB10,PPP2R1A,GTF3C4,PKD1,SF3B1,SSBP2,HMGB1,POLR2A,TPT1,STMN1,ITGB1,TUBB4B,NC
3,EIF1,EIF4A1,RPL19,PPP2R1A,RPL15,GTF3C4,DNAH8,PKD1,SF3B1,PDIA3,SERINC1,ZNF43,HMG
1,NFE2L1,RPS16,RPL23,SCARB2,MSN,KPNB1,PAPOLA,COL1A2,RPL14,NCS1,ZNF865,ATF4,PSMI
PLP2,SLC25A3,RPLP2,WLS,HNRNPK,H1-10,YWHAB,LAMP1,HNRNPH1,CCT6A,CDK16,TOP2B,APPI
PLP2,DDOST,YWHAB,SND1,TTR,ARCN1,RPS27,ARHGDIA,SLC35E2B,TMEM30A,BCLAF1,HSBP1,SI
RNP200,IGF2,EIF3C,ATP5F1A,EIF4B,VCP,MATR3,YWHAG,RPL37A,LRRC28,EIF1,INA,EIF4A1,RPL1
BBP4,PFN1,MCL1,TLE5,PNN,RPL31,PRKCSH,KHDRBS1,EIF5A,PDIA6,PIP4K2B,RAB7A,ITM2C,RPL
D1,SF3B2,GTF2F1,RPL23,CNN3,RPL29,TPM4,NACA,BRD3,RPS9,XRCC5,CIPC,RPL24,C1ORF43,P
PL23,CNN3,RPL29,RPS9,XRCC5,CIPC,RPL24,PAPOLA,IGFBP3,HK1,RHOA,ZNF865,ATF4,GLIPR1L
LP1,IK,RPL35,GPAT4,COL6A2,NGRN,B4GALNT4,MTSS2,DNAJC5,HNRNPDL,MXD4,MYL6,RPL36,C
LP1,IK,RPL35,GPAT4,COL6A2,NGRN,B4GALNT4,MTSS2,DNAJC5,HNRNPDL,MXD4,MYL6,RPL36,C
P200,CSDE1,SMIM14,IGF2,EIF3C,ATP5F1A,EIF4B,VCP,UBB,AFF4,MATR3,RPL37A,LRRC28,EIF1,II
/-1,GDI1,FSCN1,TMBIM6,PTMS,SPTAN1,SMG1,NPM1,SNRNP200,SMIM14,IGF2,EIF3C,ATP5F1A,\

,HNRNPL,ABCA3,PDXK,CAMSAP1,MLEC,MDK,ZNF302,CBX1,RAD23B,PTK7,TNPO1,SKI,KDM6B,
S3,LAMA5,CIPC,MSN,KPNB1,C1ORF43,RHOA,RAB6B,NLGN2,CHD4,HMGN2,CRMP1,LASP1,BEX
1,HSPD1,SMIM14,IGF2,EIF3C,ATP5F1A,VCP,MARCKSL1,AFF4,MATR3,YWHAG,COL4A2,RPL37A,
ONP1,CSNK1D,MESD,MAPK8IP1,CYP51A1,BEX4,PPDPF,TXNDC5,EP300,STARD7,AP2A1,CSNK2A
,ZNF865,ATF4,KIF5B,RPS14,CXADR,LENG8,WDR1,RPL26,CCT7,SRSF4,RBBP4,PFN1,MCL1,PNN
1,SEC61A1,RPS3A,RPL18,STMN1,ITGB1,AGRN,SARAF,MIDN,MARCHF6,WSB1,RPS17,ARF1,MAP
AP2B1,GTF2F1,NFE2L1,CTNNA1,TNS3,LAMA5,CIPC,MSN,SLC16A7,KPNB1,C1ORF43,IGFBP3,NC
C28,EIF1,EIF4A1,RPL19,PPP2R1A,RPL15,GTF3C4,DNAH8,PKD1,SF3B1,SERINC1,ZNF43,RPS3,H
10,MARCHF6,DNAJA1,ARF1,SF3B2,MAT2A,GTF2F1,HNRNPA0,RPS16,RPL29,XRCC5,CIPC,PAPO
28,EIF1,EIF4A1,RPL19,PPP2R1A,GTF3C4,DNAH8,PKD1,SF3B1,PDIA3,SERINC1,ZNF43,RPS3,HM
LKBH5,SNRPN,CAMSAP1,CCT3,VPS35,TNPO1,SNRNP70,PRPF6,VHL,SMARCA4,SRSF9,PDE7A,RA
,MIDN,WSB1,VCL,AP3D1,ARF1,PAFAH1B1,MAP4,SF3B2,CDC42BPB,ZNF770,TMEM170A,HNRNP
H,KHDRBS1,KLHL9,TKT,EIF5A,CDH2,PDIA6,PIP4K2B,RAB7A,RPL12,ITM2C,RPL30,EIF3L,RPS7,I
NRNPU,PGAM1,FADS2,CNN3,TPM4,CTSD,RPS24,SLC22A17,RPS9,KPNB1,RPL17,LLGL1,GAP43,
TIN2,P4HB,APP,KLRD1,GDI1,FSCN1,TMBIM6,MARCKS,SMG1,CALM1,NPM1,TTC3,PLD3,HSPD1,
,EIF5A,PDIA6,PIP4K2B,RAB7A,RPL30,EIF3L,IK,RPS19,CHPF,NGRN,HNRNPDL,RPLP2,DDOST,DU
B,HLA-C,SF3B1,ZNF43,VAPA,HMGB1,SEC61A1,STMN1,TUBB4B,UCHL1,MORF4L1,AGRN,RPLP1
,ATP5F1B,CALM3,GNB1,SHISA9,RPS8,MYH9,RPL7,SEPTIN2,P4HB,FSCN1,TMBIM6,SPTAN1,SMG
3,HSPD1,COL18A1,CSDE1,EIF3C,ATP5F1A,EIF4B,MARCKSL1,AFF4,TRIM28,MATR3,COL4A2,M
HSY1,TMSB10,EIF4A1,MRC2,GTF3C4,LDHB,DNAH8,HLA-C,SSBP2,CHD3,AP2M1,TPT1,STMN1,IT
CALM1,PLXNB2,PLD3,HSPD1,USP11,SNRNP200,CSDE1,SMIM14,IGF2,EIF3C,ATP5F1A,EIF4B,VC
RINC1,ZNF43,HMGB1,POLR2A,AP2M1,TPT1,SEC61A1,RPS3A,ITGB1,RPL11,MORF4L1,BTF3,AG
,PFN1,MCL1,TLE5,PNN,RPL31,PRKCSH,KHDRBS1,KLHL9,RPS25,EIF5A,PDIA6,PIP4K2B,MMP14,
PS9,XRCC5,CIPC,RPL24,PAPOLA,HK1,RPL22,RHOA,ZNF865,GLIPR1L2,RPS14,RPS20,CD81,LEN
PAPOLA,IGFBP3,COL1A2,RHOA,PTCD3,ATF4,KIF5B,GLIPR1L2,EIF5,LENG8,HMGN2,CRMP1,LAS
4A2,MAPT,SPEN,DDB1,EIF1,DCHS1,SET,TPI1,TMSB10,EIF4A1,MRC2,PKD1,HLA-C,SF3B1,BASP1,
HL9,RPS25,EIF5A,PDIA6,PIP4K2B,RPL12,ITM2C,RPL30,EIF3L,GNB2,RPS7,IK,RPS19,CHPF,NGR
A,RPL13,SLC2A3,DDX17,HTRA4,BSG,SPTBN1,ATP5F1B,CALM3,STMN2,RPS8,MYH9,RPL7,SEPTI
Y1,TMSB10,EIF4A1,PPP2R1A,RPL15,DNAH8,PKD1,SF3B1,PDIA3,ZNF43,RPS3,BASP1,VAPA,POL
1,WDR45B,NAP1L1,EP300,LRRC4B,SLC35F5,GNG4,ZNF512B,PPP1CC,OAZ2,GPSM1,KLHDC3,
C22A17,BRD3,XRCC5,MSN,KPNB1,C1ORF43,COL1A2,HK1,NCS1,CLSTN1,ATF4,ATP1A1,JUND,C
OLR2A,AP2M1,TPT1,SEC61A1,RPS4X,RPS3A,TUBB4B,RPL11,MORF4L1,BTF3,AGRN,RPLP1,MID
15,ITM2C,RPL30,EIF3L,MAGED2,IK,RPS19,CHPF,NGRN,RPLP2,DDOST,CIRBP,NARS1,RPL36,YW
HD1,IPO5,ATP6V0C,SKP1,DNER,TRIB2,HSBP1,SQLE,TIMP3,NUCB1,CHD6,EEF1A2,TBCD,SNX22,
,SEC62,KAT6A,DVL3,KIF1A,RPL13,GNAS,SLC2A3,DDX17,GPI,BSG,FASN,ALDH6A1,SPTBN1,ATP5
PSMD5,EIF1,HACD3,DCHS1,SET,EGLN1,TPI1,CHSY1,TMSB10,EIF4A1,RPL15,GTF3C4,LDHB,PKD
NA,EIF4A1,RPL19,PPP2R1A,LDHB,PKD1,SF3B1,PDIA3,SERINC1,ZNF43,HMGB1,POLR2A,AP2M1,
CD63,RPLP2,ARID1A,HNRNPK,DUX4,CKAP5,RHOB,HNRNPH1,UBE2H,PLPPR3,SEPTIN3,HECTD4,
N,DNAJC5,RPLP2,DDOST,DUX4,YWHAB,ARCN1,RPS27,CTBP1,ARHGDIA,SLC35E2B,BCLAF1,RPL
SUGP2,CPSF1,MEST,HNRNPL,TM9SF2,ABCA3,SNRPN,PRDX5,UBE4A,CCT3,VPS35,SNRNP70,DPY
P,SEPTIN11,CADM4,HNRNPH1,LGALS3BP,PLPPR3,HECTD4,MAGED4B,CTBP1,ARHGDIA,CAPRIN
MDH2,U2AF2,NOP53,PABPN1,HNRNPL,TSPYL4,LMNB2,HNRNPA3,ACTR3,CCT3,RBMX,CHD2,AS
EX3,CCT7,PFN1,PNN,PRKCSH,HNRNPC,KHDRBS1,HDLBP,RPS25,GPC3,RPS15,MRFAP1,RCN1,R
2,LRP3,AHCY,AP2A2,SMARCD1,CREBBP,BNIP3,XPO6,DPYSL4,REEP1,METRN,SGSM2,KDM3A,F
A5,SLC22A17,BRD3,SLC16A7,PAPOLA,IGFBP3,HK1,NCS1,RHOA,ZNF865,SLC7A5,NLGN2,CXAD

4,CTSD,EIF4H,LPCAT1,LAMA5,BRD3,KPNB1,COL1A2,HK1,NCS1,RHOA,CLSTN1,ATF4,NLGN2,CHD
2,AGRN,ZMIZ1,COL11A1,WSB1,ARF1,MAP4,CDC42BPB,MAT2A,OGA,HNRNPA0,TPM4,CTSD,PJA2
L13,SLC2A3,DDX17,RACK1,FASN,ALDH6A1,SPTBN1,ATP5F1B,CALM3,GNB1,SHISA9,RPS8,RPL7,S
E1,SMIM14,IGF2,ATP5F1A,MARCKSL1,AFF4,YWHAG,CKB,COL4A2,MAPT,DDB1,YWHAQ,EIF1,DCH
AG,RPL37A,LRRC28,EIF1,EIF4A1,RPL19,PPP2R1A,GTF3C4,DNAH8,PKD1,SF3B1,PDIA3,SERINC1,Z
259,CD81,LENG8,ATP2A2,CRMP1,LASP1,PFN1,JPT2,NES,LRPPRC,RPS25,TKT,EIF5A,CDH2,PIP4K
),XRCC5,PJA2,MSN,HK1,PTCD3,ZNF865,RPS14,RPS20,CD81,COL5A2,WDR1,ATXN7L3B,ATP2A2,I
C5,CIPC,RPL24,C1ORF43,PAPOLA,IGFBP3,HK1,RHOA,ZNF865,ATF4,GLIPR1L2,RPS14,RPS20,CD
N1,ATP5F1B,CALM3,GNB1,SHISA9,STMN2,RPS8,MYH9,RPL7,SEPTIN2,P4HB,APP,IDS,RPL5,GDI1,
,IPO5,FAM199X,CTDSP2,MDH2,CHD6,SNX22,SQSTM1,KLHL24,GAS7,LMNB2,CAMSAP1,PTOV1,C
PL31,PRKCSH,KHDRBS1,KLHL9,RPS25,EIF5A,PDIA6,PIP4K2B,RPL12,ITM2C,RPL30,EIF3L,GNB2,I
TM4B,TRIB2,CPSF2,SRP9,NUCB1,RNF187,U2AF2,TBCD,ANXA5,PPP1CB,SQSTM1,GNAI3,GAS7,GI
A,STMN1,RPL11,MORF4L1,NCAM1,ANXA2,AGRN,MARCHF6,AP3D1,SBF1,DNAJA1,PAFAH1B1,ITM
R1A,PKD1,HLA-C,SF3B1,PDIA3,SSBP2,ZNF43,BASP1,VAPA,HMGB1,POLR2A,AP2M1,TPT1,SEC61A
P1,HMGB1,POLR2A,SEC61A1,RPL18,STMN1,ANXA2,AGRN,ZMIZ1,XAF1,WSB1,AP3D1,SBF1,ARF1
PL11,MORF4L1,BTF3,ANXA2,RPLP1,MIDN,RPL10,WSB1,VCL,ARF1,MAP4,CDC42BPB,ITM2B,MAT
KBH5,PDXK,YEATS2,HSPA4,CAMSAP1,KLC1,GRINA,ACAP3,MDK,H3-3A,ABCA2,VEGFA,TUBB2A,M
IDK,SKI,RAPGEF1,ARF3,MVB12B,MAPK1,CCNL2,CAPZB,TDRD5,MAP7D1,PLCG1,DIDO1,DLK1,CC
2,APP,ERVV-1,RPL5,GDI1,FSCN1,PTMS,SPTAN1,MARCKS,CALM1,PLXNB2,TTC3,USP11,COL18A1
RINC1,RPS3,HMGB1,POLR2A,AP2M1,TPT1,SEC61A1,RPS3A,ITGB1,RPL11,BTF3,NCAM1,AGRN,R
CHD3,POLR2A,TPT1,STMN1,MORF4L1,BTF3,NCAM1,ANXA2,AGRN,TIMP2,COL11A1,MARCHF6,AI
YWHAQ,EIF1,HACD3,EGLN1,TPI1,CHSY1,TMSB10,INA,EIF4A1,PPP2R1A,RPL15,LDHB,DNAH8,PKD
PYGB,SELENON,DDAH2,AEBP1,GRB2,CAPZB,BNIP3L,PLCG1,FLNB,CORO1C,PHC2,LSS,STXBP1,IS
PL10,MARCHF6,AP3D1,DNAJA1,ARF1,SF3B2,MAT2A,SLC4A8,RPS16,RPL23,CNN3,TPM4,RPS24,XI
31,MAP4,CDC42BPB,MAT2A,SLC4A8,PGAM1,CPE,RPL29,TSPAN3,LPCAT1,CIPC,PAPOLA,IGFBP3,
R1A,SERINC1,CHD3,HMGB1,POLR2A,TPT1,SEC61A1,RPS3A,RPL18,TUBB4B,MORF4L1,ANXA2,AG
MNB2,YEATS2,KLC1,GRINA,VPS35,PTK7,ORAI2,SKI,H3-3A,SCAF1,RAB11B,PYGB,DPYSL2,OPTN,I
ISCH,PRKCSH,KLHL9,HDLBP,GLG1,SFPQ,PNMA1,RAB7A,RPL30,EIF3L,NANOG,MRFAP1,RPS5,RI
17,AKAP12,MAP4,AP2B1,HNRNPU,PGAM1,NFE2L1,SMARCA5,HNRNPA0,FADS2,RPSA,LPCAT1,BRI
MAGED1,RPL6,KAT6A,DVL3,YWHAE,KIF1A,RPL13,GNAS,SLC2A3,DDX17,HTRA4,RACK1,BSG,FASN
5,FADS2,RPS16,RPL23,CPE,RPL29,LPCAT1,LAMA5,SLC22A17,CIPC,PJA2,MSN,KPNB1,C1ORF43,
YWHAG,CKB,COL4A2,LRRC28,FN1,DDB1,YWHAQ,EIF1,HACD3,SET,EGLN1,MAP9,CHSY1,INA,RPL
ATR3,YWHAG,LRRC28,EIF1,EIF4A1,RPL19,RPL15,GTF3C4,PKD1,SF3B1,PDIA3,SERINC1,ZNF43,H
M2C,RPL30,EIF3L,IK,RPS19,CHPF,NGRN,DNAJC5,RPLP2,DDOST,DUX4,YWHAB,SND1,CALU,ARCN
PNN,RPL31,KHDRBS1,KLHL9,RPS25,EIF5A,PDIA6,PIP4K2B,ITM2C,RPL30,CLTC,EIF3L,GNB2,IK,I
BEX3,RTN3,PFN1,CCNI,XPO1,PFKL,EIF5A,CCND2,DDX3X,RPL12,MAGED2,GNB2,H1-0,GJA1,JUN,
R8K1,RPL18,STMN1,TUBB4B,MORF4L1,BTF3,NCAM1,XAF1,SARAF,RPL10,WSB1,AP3D1,SBF1,AI
RS,SF1,RETREG2,SEC16A,UPF1,SPINT2,BRK1,SAFB2,VPS26B,FUS,PPP1R10,SPG7,CEP170,EIF2S
1,SF3B2,CDC42BPB,MAT2A,TMEM170A,PXDN,HNRNPU,OGA,PGAM1,NFE2L1,FADS2,RPL23,CNN
,GNB1,RPS8,MYH9,RPL7,IDS,ERVV-1,RPL5,GDI1,TMBIM6,PTMS,SPTAN1,GSTP1,MARCKS,SMG1
LDHB,PKD1,HLA-C,SF3B1,PDIA3,SSBP2,ZNF43,RPS3,CHD3,VAPA,POLR2A,AP2M1,ITGB1,TUBB4I
34B,MORF4L1,AGRN,MARCHF6,RPS17,VCL,AP3D1,DNAJA1,CYB5R3,SF3B2,AP2B1,GTF2F1,OGA
MCL1,PNN,RPL31,PRKCSH,LRPPRC,SCRN1,RPS25,EIF5A,PDIA6,MEGF8,RAB7A,ITM2C,RPL30,EIF
CAP3,MDK,RAD23B,H3-3A,ABCA2,VEGFA,TCF25,CAPZB,ACTN1,HSPB1,PLCG1,CCAR2,DIDO1,CC
RD3,XRCC5,CIPC,RPL24,PAPOLA,IGFBP3,HK1,RPL22,ZNF865,ATF4,PSMD2,RPS14,CXADR,RPS2

P2,DDOST,WLS,HNRNPK,H1-10,HNRNPH1,RPS12,CCT6A,RAC1,CDK16,SMCHD1,TMEM30A,DNEF
2,EIF3C,ATP5F1A,EIF4B,VCP,UBB,AFF4,MATR3,RPL37A,LRRC28,FN1,PSMD5,EIF1,HACD3,EGLN1
RPS2,SEPTIN7,APEX1,ZMYM3,FUS,ERH,TAOK1,RAB1B,PKDCC,UBE2D2,SMG5,IREB2,LSM14A,YIP
F,MARCHF6,AP3D1,SBF1,ARF1,CYB5R3,MAP4,SF3B2,AP2B1,GOLGA8A,PXDN,HNRNPU,NFE2L1,S
PL30,EIF3L,IK,RPS19,CHPF,NGRN,DNAJC5,RPLP2,DDOST,YWHAB,CALU,ARCN1,CTBP1,ARHGDIA
RPS19,CHPF,NGRN,HNRNPDL,RPLP2,DDOST,YWHAB,ARCN1,RPS27,ARHGDIA,SLC35E2B,RPL23A
APP,ERVV-1,GDI1,TMBIM6,PTMS,SPTAN1,CALM1,NPM1,PLXNB2,SNRNP200,CSDE1,SMIM14,ATP5
CKAP4,PLPPR3,HECTD4,RAC1,STIP1,CDK16,SLC35E2B,MMP2,CDKN1A,CRPPA,NOVA2,NUCB1,CH
EIF3L,RPS5,RPS19,CHPF,NGRN,DNAJC5,HNRNPDL,RPLP2,DDOST,YWHAB,SND1,ATP1A2,CALU,PL
MAT2A,GOLGA8A,PXDN,HNRNPU,PGAM1,NFE2L1,HNRNPA0,CNN3,RPL29,TPM4,TNS3,CTSD,SCAF
L13,GNAS,DDX17,HTRA4,RACK1,BSG,ALDH6A1,ATP5F1B,GNB1,SHISA9,STMN2,RPS8,MYH9,APP
2,SMARCA4,KDELR2,DSTN,FAM171A2,ODC1,AEBP1,TCF25,UBE2R2,ATXN10,ACTN1,BNIP3L,FKE
H5,PDXK,PKN1,TBC1D16,TCP1,ETNK1,MLEC,ACAP3,TMEM248,LAMC1,PRPF6,DPYSL2,VEGFA,VI
DX17,GPI,HTRA4,RACK1,BSG,ALDH6A1,SPTBN1,CALM3,GNB1,SHISA9,STMN2,RPS8,MYH9,RPL7,
F1A,EIF4B,VCP,MATR3,YWHAG,LRRC28,DDB1,EIF1,DCHS1,TPI1,EIF4A1,RPL19,RPL15,GTF3C4,D
T7,TLE5,PNN,NISCH,PFKL,TPM1,EIF5A,PEBP1,PNMA1,PIP4K2B,MEGF8,RPS15,ITM2C,APLP1,B4
5A,HNRNPDL,ARL6IP1,TSR2,WLS,HNRNPK,PTGES3,RPS27,HECTD4,SMCHD1,ATP6V0C,BCLAF1,L
CYB5R3,MAP4,ITM2B,TMEM170A,GOLGA8A,HNRNPU,NFE2L1,SMARCA5,HNRNPA0,RPL23,RPL2
PS17,AP3D1,DNAJA1,SF3B2,GTF2F1,PXDN,HNRNPU,OGA,PGAM1,NFE2L1,SMARCA5,FADS2,LDHA
,TPT1,RPS4X,RPS3A,ITGB1,TUBB4B,MORF4L1,AGRN,MARCHF6,RPS17,VCL,AP3D1,DNAJA1,CYB
HSPD1,USP11,SNRNP200,CSDE1,SMIM14,EIF3C,ATP5F1A,VCP,MATR3,RPL37A,LRRC28,FN1,DDE
1,FASN,ALDH6A1,ATP5F1B,CALM3,GNB1,SHISA9,STMN2,RPS8,MYH9,RPL7,SEPTIN2,P4HB,APP,E
IF4A1,PPP2R1A,RPL15,GTF3C4,DNAH8,PKD1,SF3B1,SERINC1,ZNF43,RPS3,VAPA,POLR2A,AP2M
MTRF1L,ALKBH5,PRDX5,UBE4A,TCP1,TMEM248,GCN1,RAB11B,SRP14,TSPAN6,PRPF6,TAF7,RPL
C28,MAPT,FN1,EIF1,HACD3,DCHS1,EGLN1,TPI1,CHSY1,TMSB10,INA,MRC2,PPP2R1A,DNAH8,PKD
CALR,FSTL1,TMF1,ABHD2,HSPA1B,TUBB2B,COL5A1,COL4A1,CES3,PSAP,PGK1,RPLP0,H3-3B,C
IK1E,EIF5A,PEBP1,PDIA6,RPS15,RPL12,MAGED2,RPS19,RPL35,GJA1,GOLM1,SESN3,APLP2,TARI
RCHF6,DNAJA1,ARF1,AKAP12,SF3B2,GTF2F1,HNRNPA0,RPL23,RPL29,RPS24,XRCC5,CIPC,PAPC
F,CCT5,DFFB,EIF2AK1,TXLNA,SGTA,KDM5C,SF1,RETREG2,SEC16A,BRD2,UPF1,RAB3GAP2,EIF5
A1,JUND,COL5A2,LENG8,HMGN2,ATP2A2,CRMP1,VAT1,SRSF4,TLE5,PRKCSH,HNRNPC,PFKL,TPM
RF1L,ALKBH5,RPL27,HNRNPA3,PRDX5,UBE4A,HSPA4,TMEM248,GCN1,ASXL1,TSPAN6,PRPF6,AT
F,RBBP7,PRMT1,EP300,LBH,DGCR2,RETREG2,LRRC4B,KIF21B,SEC16A,GNG4,SRSF6,PPP1CC,C
DIA,SLC35E2B,IPO5,RPL23A,HSBP1,MAP1A,DDX39B,SRSF5,TM9SF3,MDH2,U2AF2,EEF1A2,FDFT
C,BTBD2,PDE7A,ODC1,SRSF1,PTBP1,ILF3,PRKDC,TDRD5,UBE2R2,SDF4,SMARCC1,ATP6V1B2,Y
170A,GOLGA8A,HNRNPU,NFE2L1,SMARCA5,HNRNPA0,RPL29,RPSA,TNS3,NACA,LPCAT1,LAMA5,
L12B,HNRNPH3,CENPB,NELFB,USP47,AGAP1,SCAMP5,CHMP1A,GGNBP2,NPTN,KIAA1191,ADNP,A
ONER,NNAT,CPSF2,SRP9,CTSB,RAB27A,U2AF2,TBCD,ANXA5,CPSF1,PABPN1,TSR1,ALKBH5,HNRN
RCKSL1,TRIM28,CKB,COL4A2,SPEN,NONO,DDB1,PSMD5,YWHAQ,HACD3,DCHS1,SET,TMSB10,EI
RN,RPS10,SESN3,HNRNPDL,RPLP2,DDOST,MYL6,CIRBP,RPL36,DUX4,CALU,PLPPR3,ZNF454,CTE
F2,HS6ST1,ZC3H7B,MDH2,CCT2,ANXA5,CPSF1,PPP1CB,KLHL24,MEST,ACTR3,PKN1,TBC1D16,AC
F4L1,BTF3,ANXA2,RPLP1,MIDN,MARCHF6,WSB1,RPS17,SBF1,MAP4,SF3B2,CDC42BPB,AP2B1,G
2M1,TPT1,SEC61A1,RPS4X,RPS3A,TUBB4B,RPL11,MORF4L1,BTF3,AGRN,RPLP1,SARAF,MIDN,RF
N1,TSR1,ALKBH5,HNRNPA3,PDXK,CAMSAP1,CCT3,PTOV1,MDK,CCT4,SKI,SNRNP70,DPYSL2,TAF
RIR,CLPTM1,STRN4,UBR5,RAB14,PRR12,EEF1B2,SFXN1,YIPF3,LRRC41,NASP,YY1,SMARCD1,CF
CALU,COTL1,CAPRIN1,STIP1,TOP2B,SLC35E2B,SMCHD1,IPO5,TNC,ATP6V0C,SKP1,CTDSP2,CNN

N,RPL31,PRKCSH,SCRN1,RPS25,TKT,EIF5A,PDIA6,PIP4K2B,MEGF8,ITM2C,RPL30,EIF3L,MAGED2
MAP4,CDC42BPB,ITM2B,ZNF770,AP2B1,GTF2F1,PXDN,OGA,FADS2,CNN3,TNS3,EIF4H,LPCAT1,S
EPTIN11,CADM4,RHOB,CKAP4,PLPPR3,ZNF454,MAGED4B,SLC35E2B,APPL1,IPO5,MMP2,DNER,
B,ATXN3,SLC4A2,QPRT,ATP1B2,REEP2,ITGAV,KDM4A,PHIP,UBE2D3,FJX1,DEK,UBE2K,CNBP,IGI
I1,FBRSL1,P3H3,AKIRIN1,TRAF4,RBBP7,MSMO1,TXLNA,TRIO,LBH,PBX1,HIP1,LRRC4B,CSNK1G
RPS25,TKT,TPM1,CCND2,DDR1,PIP4K2B,SOX12,DDX3X,RPS5,APLP1,CHPF,COL6A2,B4GALNT4,S
P5F1A,EIF4B,VCP,MARCKSL1,UBB,AFF4,TRIM28,MATR3,YWHAG,LRRC28,SPEN,NONO,DDB1,YW
MEX3A,IGFBP7,DFFB,BCR,TRIO,RERE,FBRS,ALDH3A2,HIP1,SEC16A,GNG4,SFRP1,RER1,MBTPS
SF3B1,SERINC1,ZNF43,RPS3,BASP1,VAPA,HMGB1,POLR2A,AP2M1,TPT1,SEC61A1,RPS3A,OR8K
PL30,MAGED2,GNB2,APLP1,COL6A2,NGRN,GOLM1,DNAJC5,HNRNPDL,NDFIP1,WLS,HNRNPK,CI
DNAJC5,HNRNPDL,RPLP2,DDOST,YWHAB,CALU,ARHGDIA,SLC35E2B,IPO5,RPL23A,HSBP1,MAP1
,DLG5,CRPPA,CTSB,CDH11,CCT2,ANXA5,PGD,TAF15,HSPH1,ABCA3,ALKBH5,HNRNPA3,PRDX5,U
MTSS2,SHOX,WLS,YWHAB,ATP1A2,CKAP4,COTL1,MAGED4B,CDK16,TMEM30A,MMP2,PTTG1IP,D
SON,CCAR2,ARF4,FBLN1,BAG6,RAB11A,ARNT2,SERBP1,ARFGEF2,CDC42,TRAF7,EIF2AK1,PRM
N,SLC16A7,C1ORF43,PAPOLA,COL1A2,RPL22,ATF4,SLC7A5,NLGN2,RPS14,CHD4,ATP1A1,JUND,
IM14,EIF3C,VCP,UBB,AFF4,TRIM28,CKB,RPL37A,FN1,SPEN,DDB1,PSMD5,YWHAQ,EIF1,HACD3,S
HNRNPUL1,ATXN10,SON,QARS1,RPL28,DHX30,EIF3F,SF3A1,BAG6,RAB11A,BMPR2,CCT5,CDC42
P2M1,SEC61A1,OR8K1,RPL18,STMN1,TUBB4B,MORF4L1,BTF3,TIMP2,XAF1,SARAF,RPL10,MAR
3,ITM2B,NFE2L1,TPM4,CTSD,SCARB2,RTN4,LPCAT1,LAMA5,BRD3,PJA2,MSN,SLC16A7,LLGL1,R
MAGED2,H1-0,RPS19,NGRN,HNRNPDL,RPLP2,DDOST,HNRNPK,CIRBP,CALU,ARCN1,COTL1,ATP6
PDIA3,SERINC1,SSBP2,ZNF43,HMGB1,POLR2A,TPT1,SEC61A1,RPS4X,RPS3A,RPL18,ITGB1,TUB
3,VAPA,POLR2A,AP2M1,SEC61A1,OR8K1,STMN1,TUBB4B,UCHL1,MORF4L1,BTF3,TIMP2,XAF1,S
F2,EIF4B,MARCKSL1,UBB,AFF4,TRIM28,YWHAG,COL4A2,RPL37A,FN1,NONO,YWHAQ,EIF1,DCH
AND3,NT5DC2,SOD1,BAG6,PFN2,IGFBP7,SMS,NAP1L4,DHX15,ANP32A,ST3GAL6,DFFB,WDR45B,
2,NAP1L4,SYP,RBM15B,BEX4,MSMO1,KIRREL1,TRIO,SF1,PRCC,KIF21B,SEC16A,MAP6,NXN,GLU
RPL18,ITGB1,BTF3,AGRN,RPLP1,MIDN,RPL10,MARCHF6,DNAJA1,ARF1,SF3B2,MAT2A,GTF2F1,
X3A,P3H3,AKIRIN1,BEX4,PPDPF,KCNQ2,BCL9L,WDR45B,PBX1,AP2A1,NOL4L,LRRC4B,CST3,ATR
WSB1,AP3D1,DNAJA1,ARF1,CYB5R3,ITM2B,AP2B1,TMEM170A,GTF2F1,GOLGA8A,SMARCA5,HN
SCAN18,CTXN1,CD99L2,PCYOX1,BEX4,TRAF4,RBBP7,WDR45B,EP300,DGCR2,STARD7,LRRC4B,
ELOVL5,ELAVL3,DIP2A,AGAP1,AUTS2,LRIG1,LUC7L2,NOP56,ATP13A3,RGPD5,IGSF3,MECP2,CRI
S26B,FUS,EIF2S2,EHMT1,SHC2,INTS3,RAB14,ZNF428,PNISR,PLEKHB2,PCNX3,ADAM10,FBXW5,
D2,RBMX,CHD2,SCAF1,TMED2,RAPGEF1,PPP2CA,MAPK1,DDAH2,SRSF1,PTBP1,ILF3,RPS11,KDM
,PLD3,HSPD1,COL18A1,CSDE1,IGF2,ATP5F1A,EIF4B,MARCKSL1,AFF4,TRIM28,COL4A2,MAPT,N
SEC62,RPL6,DVL3,YWHAE,KIF1A,RPL13,GNAS,SLC2A3,DDX17,GPI,HTRA4,BSG,ALDH6A1,SPTBN
T1,CTSB,MDH2,RNF187,U2AF2,FDFT1,CCT2,SUGP2,CPSF1,GDI2,PARP1,TSR1,PTPRA,PRDX5,CA
F3L,MAGED2,RPS5,IK,RPS19,CHPF,JUN,NGRN,DNAJC5,RPLP2,DDOST,CIRBP,NARS1,YWHAB,PT
RPL10,WSB1,AP3D1,SBF1,ARF1,CYB5R3,AKAP12,SF3B2,MAT2A,NFE2L1,FADS2,RPS16,RPL29,RP
TN1,KLC1,GRINA,NCKAP1,CLSTN3,MDK,LAMC1,PTK7,ORAI2,ASXL1,RAB11B,TUBB2A,ARF3,CCN
DPYSL2,TAF7,RNF10,LAPTM4A,SRSF9,PDE7A,PSMD3,RABL6,ODC1,IPO7,ARGLU1,TCF25,ADAM9,
PLP0,H3-3B,CLU,PTMA,DDX5,ENO2,SOX4,FKBP14,RPL13A,MAGED1,SEC62,RPL6,KAT6A,DVL3,Y
YWHAQ,EIF1,HACD3,MAP9,CHSY1,INA,EIF4A1,RPL19,PPP2R1A,SERINC1,ZNF43,BASP1,HMGB1,P
,CORO1C,KIF3C,SMARCC1,ATP6V1B2,HNRNPM,PFKM,PITPNM1,ISYNA1,HYOU1,SOD1,FBRSL1,
P1,MAP1A,DDX39B,SRSF5,TM9SF3,MDH2,U2AF2,EEF1A2,SUGP2,GNAI3,TSR1,MTRF1L,HNRNPA
I,VAPA,POLR2A,TPT1,RPS4X,RPL18,STMN1,ITGB1,BTF3,ANXA2,AGRN,ZMIZ1,MIDN,RPL10,RPS1
RF3,ARPC2,DDAH2,BTBD2,TCF25,CAPZB,RPS11,ACTN1,CMIP,PLCG1,CCAR2,PTP4A2,PLXNA3,AT

.RPPRC,SCRN1,RPS25,SFPQ,EIF5A,PDIA6,MEGF8,RAB7A,PRDX2,ITM2C,CLTC,EIF3L,MAGED2,RF
P1,UBE2H,CTBP1,APPL1,SMCHD1,ATP6V0C,LAPTM4B,SERF2,CPSF2,U2AF2,RPN2,MLXIP,ANXA5,S
37A,MAPT,SPEN,DDB1,PSMD5,YWHAQ,EIF1,HACD3,TPI1,CHSY1,TMSB10,EIF4A1,DNAH8,SSBP2,I
3A,RPL18,ITGB1,BTF3,AGRN,RPLP1,SARAF,MIDN,RPL10,MARCHF6,AP3D1,DNAJA1,ARF1,SF3B2
AF7,EIF2AK1,PRMT1,PDAP1,AP2A1,MMP15,SEC16A,PHB2,APMAP,SLC39A7,PLXNB1,FARP1,YKT6
1,MAP4,MAT2A,PXDN,HNRNPU,OGA,LARP1,HNRNPA0,CTNNA1,LDHA,TPM4,CTSD,LPCAT1,RPS24
RNPU,NFE2L1,RPS16,RPL23,RPL29,TNS3,LAMA5,PJA2,SLC16A7,KPNB1,LLGL1,RHOA,CBX5,PSM
ND1,LAMP1,ARHGDIA,SLC35E2B,IPO5,ATP6V0C,BCLAF1,RPL23A,HSBP1,FAM199X,CTDSP2,DDX
.23A,HSBP1,FAM199X,DDX39B,TM9SF3,MDH2,TOMM20,U2AF2,CCT2,GNAI3,GDI2,TM9SF2,TSR
PPR3,ARCN1,COTL1,MAGED4B,SMCHD1,MMP2,L1CAM,SHISAL1,PGRMC1,CDH11,CHD6,SNX22,
B7A,RPL12,ITM2C,RPL30,EIF3L,IK,RPS19,CHPF,CD63,NGRN,DNAJC5,RPLP2,DDOST,YWHAB,SNI
.RRC28,TPI1,EIF4A1,RPL19,PPP2R1A,RPL15,SF3B1,SERINC1,ZNF43,RPS3,HMGB1,POLR2A,AP2I
.RRC28,TPI1,EIF4A1,RPL19,PPP2R1A,RPL15,SF3B1,SERINC1,ZNF43,RPS3,HMGB1,POLR2A,AP2I
OL18A1,CSDE1,IGF2,ATP5F1A,EIF4B,MARCKSL1,AFF4,TRIM28,COL4A2,MAPT,DDB1,DCHS1,SET
,RPS5,IK,RPS19,CHPF,JUN,NGRN,RPLP2,DDOST,CIRBP,NARS1,SND1,CALU,ARCN1,ARHGDIA,TO
1,RPL18,RPL11,BTF3,NCAM1,AGRN,SARAF,MIDN,RPL10,VCL,AKAP12,CDC42BPB,AP2B1,SLC4A
.31,PRKCSH,HNRNPC,CCNI,KHDRBS1,RPS25,EIF5A,PDIA6,MEGF8,SOX12,RAB7A,DDX3X,RPL12,I
A1,RPS3A,RPL18,UCHL1,RPL11,MORF4L1,AGRN,RPLP1,ZMIZ1,SARAF,WSB1,VCL,MAP4,SLC4A8
ZP6,CCAR2,ARF4,SUPT6H,TNKS2,SARS1,BAG6,HNRNPD,AKIRIN1,CDC42,PPP1R9B,EDC4,TXLNA,V
2,PTBP1,HNRNPUL1,SON,CCAR2,ARF4,SUPT6H,PFKM,LONP1,SF3A1,BAG6,RAB11A,ARNT2,SERI
GF8,ITM2C,RPL30,EIF3L,MAGED2,RPS5,IK,RPS19,CHPF,JUN,NGRN,RPLP2,DDOST,CIRBP,NARS1
15,ABCA3,RPL27,PRDX5,AHCYL1,LAMC1,PTK7,SKI,SCAF1,ABCA2,PYGB,PRPF6,DPYSL2,RPL38,F
DHA,LONP1,ZFAND3,SARS1,WBP2,CCT5,EIF2AK1,TXLNA,SGTA,KDM5C,ACTR1A,SF1,RETREG2,S
E,KIF1A,RPL13,DDX17,GPI,HTRA4,RACK1,BSG,FASN,ALDH6A1,SPTBN1,CALM3,GNB1,MYH9,RPL
RNPDL,RPLP2,DDOST,CIRBP,NARS1,SND1,CALU,ARCN1,ARHGDIA,ATP6V0C,LAPTM4B,TRIB2,RF
A,MN1,TRIM8,SH3PXD2B,USP5,EIF4G1,IGLON5,ELAVL3,BGN,USF2,SLC22A23,CDV3,TCEA1,IGF1
61A1,OR8K1,STMN1,TUBB4B,MORF4L1,BTF3,TIMP2,XAF1,SARAF,RPL10,WSB1,RPS17,AP3D1,S
PM4,XRCC5,CIPC,PAPOLA,ZNF865,ATF4,GLIPR1L2,RPS14,CXADR,JUND,RPS20,LENG8,WDR1,RP
,UBE2R2,SCAF11,ACTN1,SON,PLCG1,GIGYF1,NKTR,YWHAH,SLIT3,TNKS2,ATXN2L,VGF,P3H3,PR
3C4,LDHB,PKD1,SSBP2,ZNF43,CHD3,BASP1,POLR2A,TPT1,RPS3A,UCHL1,NCAM1,ANXA2,AGRN,Z
2,ITM2C,MAGED2,APLP1,SESN3,APLP2,INIP,CIRBP,MYH10,DUX4,CKAP5,SEPTIN11,TNC,SKP1,TR
,YWHAG,CKB,MAPT,PSMD5,YWHAQ,EIF1,SET,EGLN1,TPI1,EIF4A1,PPP2R1A,GTF3C4,LDHB,SSBF
PL9,SLC7A5,CXADR,JUND,TMEM259,EIF5,LENG8,ATP2A2,MCL1,CCNI,XPR1,HDLBP,RPS25,PEBP
F7,RPS15A,PPP2CA,CCNL2,BTBD2,ARGLU1,PTBP1,ILF3,HNRNPUL1,QARS1,DUSP4,RPS28,FYN,H
RBP,CALU,ARCN1,ARHGDIA,CDK16,HDAC2,IPO5,ATP6V0C,SKP1,PPIB,RPL23A,HSBP1,DDX39B,SF
PA1B,TUBB2B,COL5A1,COL4A1,CES3,PGK1,RPLP0,H3-3B,CANX,CLU,PTMA,DDX5,ENO2,SOX4,FI
DDOST,YWHAB,LAMP1,CALU,RPS27,ARHGDIA,SLC35E2B,IPO5,RPL23A,HSBP1,MAP1A,SRSF5,TM
AP2M1,TPT1,SEC61A1,RPS3A,TUBB4B,BTF3,NCAM1,AGRN,RPLP1,MIDN,RPL10,MARCHF6,RPS1
RRC4B,MMP15,MGAT4B,PLXNB1,FARP1,NXN,IL6ST,SEPTIN7,CTTN,PCBP4,WSB2,SEPTIN5,PALM
RRC4B,MMP15,MGAT4B,PLXNB1,FARP1,NXN,IL6ST,SEPTIN7,CTTN,PCBP4,WSB2,SEPTIN5,PALM
15,PHB2,PLXNB1,FARP1,RER1,TRA2B,YKT6,MCM3AP,TMEM167A,CHERP,FKBP1A,CTTN,LRP10,
64,TRIR,CLCN7,SNN,SSBP3,ABI2,LSM14A,MACROH2A2,DNAJB2,CADM3,SPAG9,MOAP1,CLASP2,
F5,TM9SF3,MDH2,U2AF2,EEF1A2,FDFT1,GNAI3,PARP1,TSR1,MTRF1L,PRDX5,UBE4A,ACO2,HSP
N2,NOVA2,NUCB1,ZC3H7B,U2AF2,CCT2,MEST,GAS7,MORF4L2,ALKBH5,SNRPN,PDXK,PLOD1,GF
M1,SEPTIN7,LRRC59,KIAA0100,SLIT1,SHC2,FKBP4,SSBP3,UBR5,GABARAPL1,PGAP6,MAZ,KIAA

,WDR1,ATP2A2,RPL26,CRMP1,VAT1,MCL1,JPT2,TLE5,PNN,RPL31,CCNI,PEBP1,PIP4K2B,ACSL6,(
IZ1,COL11A1,MIDN,RPL10,WSB1,VCL,SBF1,ARF1,CYB5R3,ITM2B,MAT2A,TMEM170A,GTF2F1,O
,MYL6,ANKRD52,HNRNPK,CADM4,PLPPR3,COTL1,ARHGDIA,SMCHD1,IPO5,ATP6V0C,SKP1,LAPT
AP1A,FAM199X,TM9SF3,MDH2,RAB27A,U2AF2,EEF1A2,CCT2,GNAI3,NOP53,TSR1,MTRF1L,ALKE
IP1,RHOB,PLPPR3,SEPTIN3,ZNF454,CTBP1,CDK16,SLC35E2B,ATP6V0C,MMP2,SKP1,PPIB,HSBP
5,TM9SF3,CCT2,MLXIP,GDI2,MORF4L2,HSPH1,MTRF1L,ALKBH5,HNRNPA3,AHCYL1,ETNK1,KLC
TXN2L,HGS,BAG6,RAB11A,SERBP1,ARFGEF2,AKIRIN1,CDC42,YME1L1,VDAC1,TRAF7,EIF2AK1,PI
CRN1,RPS25,EIF5A,PDIA6,MEGF8,ITM2C,RPL30,EIF3L,MAGED2,RPS5,IK,RPS19,CHPF,JUN,NGRN
T1,RTN3,MCL1,PNN,HNRNPC,KHDRBS1,HDLBP,RPS25,CDH2,GPC3,ACTN4,RCN1,RPS5,APLP1,GI
NOP53,TSR1,MORF4L2,LMNB2,RPL27,PRDX5,UBE4A,HSPA4,TCP1,MLEC,RAB11B,SRP14,NRDC,I
NOP53,TSR1,MORF4L2,LMNB2,RPL27,PRDX5,UBE4A,HSPA4,TCP1,MLEC,RAB11B,SRP14,NRDC,I
MEM132A,NOVA2,CTSB,RNF187,EEF1A2,TBCD,ANXA5,MORF4L2,ABCA3,PKN1,GRINA,PTOV1,CL
PS27,ARHGDIA,STIP1,RPL23A,HSBP1,FAM199X,CTDSP2,PGRMC1,DDX39B,EID1,SRSF5,TM9SF3,I
1,SEPTIN3,HECTD4,SLC35E2B,PTTG1IP,CDKN1A,CRPPA,C5ORF24,ZC3H7B,PGD,VAMP2,GNAI3,S
CHD3,BASP1,TPT1,RPS3A,STMN1,TUBB4B,UCHL1,MORF4L1,NCAM1,AGRN,RPLP1,TIMP2,MIDN,
H,HMGCR,CAPRIN1,TOP2B,SMCHD1,IPO5,ATP6V0C,SKP1,LAPTM4B,SRSF5,TM9SF3,U2AF2,MLX
PI1,CHSY1,PPP2R1A,GTF3C4,HLA-C,PDIA3,CHD3,BASP1,VAPA,CDKL1,AP2M1,SEC61A1,RPL18,IT
SNRNP200,CSDE1,IGF2,EIF3C,ATP5F1A,VCP,UBB,TRIM28,MATR3,YWHAG,CKB,LRRC28,FN1,SPI
CTDSP2,CNN2,NOVA2,NUCB1,ZC3H7B,U2AF2,CCT2,MEST,GAS7,MORF4L2,ALKBH5,SNRPN,PDXI
AI2,ASXL1,VEGFA,MARS1,ARF3,SMARCA4,CCNL2,SRSF1,WDR6,ACTN1,SON,PLCG1,MSL1,SDF4,I
C,SRP14,PYGB,PRPF6,TAF7,SELENON,RPL37,ATG2B,CCNL2,ARPC2,FAM168B,SRSF1,ARGLU1,AT
C61A1,OR8K1,STMN1,TUBB4B,MORF4L1,BTF3,XAF1,SARAF,RPL10,MARCHF6,WSB1,AP3D1,SBI
S7,TM9SF2,MORF4L2,SNRPN,PDXK,PLOD1,GRINA,VPS35,PTOV1,MLEC,SURF4,PCDHGB6,SNRNI
A,MLF2,TP53BP1,YME1L1,DHX15,PSMA7,VDAC1,TRAF7,PDAP1,SLC39A6,PHB2,KLHL42,RER1,MA
TIP1,SLC35E2B,SMCHD1,ATP6V0C,BCLAF1,RPL23A,HSBP1,SERF2,DDX39B,SRSF5,TM9SF3,MDH
,RPS25,EIF5A,PDIA6,MEGF8,ACSL6,RPL12,ITM2C,RPL30,EIF3L,MAGED2,RPS5,IK,RPS19,CHPF,J

RPL37A,LRRC28,EIF1,DCHS1,TMSB10,EIF4A1,MRC2,RPL19,PPP2R1A,GTF3C4,LDHB,PKD1,PDIA
F3L,GNB2,IK,RPS19,CHPF,CD63,NGRN,DNAJC5,RPLP2,DDOST,CIRBP,YWHAB,SND1,LAMP1,CALU
PGEF1,MVB12B,CCND1,SRSF1,NYNRIN,MAP7D1,PLCG1,ZFAND5,DLK1,MSL1,RPS28,GIGYF1,STN
ALU,ARCN1,ARHGDIA,ATP6V0C,LAPTM4B,TRIB2,RPL23A,DDX39B,SRSF5,TM9SF3,MDH2,TOMM2
ARM1,GPC2,LRATD2,ACTR1B,MOAP1,ERGIC3,MSH6,IGF2BP1,IST1,MBD3,PSMC3,SMARCD1,CRE
1,RPS16,RPL29,EIF4H,SCARB2,RPS24,CIPC,GAP43,HK1,NCS1,RPL22,RHOA,ZNF865,RPL9,CLSTI
5,TAF7,RPL38,RPL37,ARPC2,AEBP1,ACADVL,PTBP1,CCAR2,ARF4,SMARCC1,HGS,BAG6,RAB11A,
MP1,LASP1,CCT7,RTN3,ASTN2,NES,CCNI,LRPPRC,KHDRBS1,XPO1,PFKL,RPS25,TKT,TPM1,CCND
ATP6V0C,LAPTM4B,CPSF2,SRSF5,TM9SF3,MLXIP,MORF4L2,HSPH1,ALKBH5,HNRNPA3,UBE4A,T
1A,ARFGEF2,IGFBP7,CD99L2,KIRREL1,LBH,PBX1,LRRC4B,PLXNB1,NXN,ZNF512B,CEMIP2,PRR
K,EIF3L,MRFAP1,RPS19,GPAT4,TLN1,JUN,RPS10,HNRNPDL,DDOST,ST3GAL2,MXD4,ANKRD52,IN
SG,FASN,SPTBN1,ATP5F1B,CALM3,GNB1,SHISA9,STMN2,MYH9,SEPTIN2,P4HB,APP,IDS,RPL5,KI
TBP1,CAPZB,UBE2R2,ACTN1,CAND1,POLR2B,SDF4,RPS28,PHC2,YWHAH,LONP1,ATXN2L,HGS,S
CALU,ARCN1,ARHGDIA,SLC35E2B,ATP6V0C,LAPTM4B,TRIB2,RPL23A,FAM199X,DDX39B,SRSF5,T
IRNPUL1,ACTN1,SON,PLCG1,SMARCC1,FAU,B2M,PITPNM1,HECTD1,PFN2,TCTN1,SERBP1,IGFBI
SPG7,OLFM1,INTS3,ABI2,SOX2,SLC6A8,PGAP6,MAZ,DCLK1,TBC1D9,PRR12,TMEM63B,MACROH
7A,CLTC,MRFAP1,RPS5,PRKAR1A,TLN1,CHPF,NGRN,HNRNPDL,TARDBP,SLC25A3,RPLP2,NDFIP1
PLP1,RPL35,GPAT4,CHPF,GJA1,MYO5A,COL6A2,ANXA6,APLP2,ARL6IP1,H1-10,CADM4,LGALS3BI

),RPL35,GJA1,HNRNPDL,RPLP2,ARL6IP1,CADM4,SND1,CKAP4,RPS12,HLA-B,RAB1A,HMGCR,APF
6,SNRPD3,GPC2,LTBP4,ELAVL1,TAX1BP1,DTX4,LRATD2,SLC4A2,FNBP1L,ACTR1B,MOAP1,KRT8
/I4B,SRSF5,TM9SF3,MLXIP,MORF4L2,HSPH1,ALKBH5,HNRNPA3,UBE4A,TBC1D16,AHCYL1,ETNK
HERP,FKBP1A,CTTN,LRP10,SERP1,ERH,EIF3H,RALY,SEC14L1,NUDCD3,FAM234A,GNL1,TUBGCP
2,MFAP4,SLC16A2,DHX36,UBE2Q1,ZNF532,CLPX,SLC43A2,PIP5K1C,SYNGR1,CCDC85C,CELSR3
'2,TLE5,PNN,NES,TADA2B,PTN,CSNK1E,PFKL,PEBP1,PDIA6,PIP4K2B,ACSL6,SOX12,RPL30,ARL4(
2,CDC34,FXR2,PINK1,USP10,KDM3A,NOP56,GTF3C1,TIA1,EWSR1,ATF5,DYNLRB1,ATF7IP,DAD1,
,RACK1,BSG,FASN,ALDH6A1,SPTBN1,ATP5F1B,CALM3,RPL7,SEPTIN2,RPL5,KLRD1,GDI1,FSCN1
BE2D2,IREB2,LSM14A,YIPF3,RBPJ,EHD1,TGOLN2,YY1,SAP18,UBE2D3,SLC20A1,HNRNPH3,USP4
LP1,MARCHF6,DNAJA1,ARF1,AKAP12,SF3B2,GTF2F1,PGAM1,HNRNPA0,RPL23,RPL29,RPS24,XR
,GNB1,STMN2,SEPTIN2,IDS,GDI1,FSCN1,TMBIM6,PTMS,SPTAN1,SMG1,CALM1,NPM1,PLXNB2,F
)1,HDAC2,DNER,PEA15,NNAT,FAM199X,SRP9,RAB27A,RNF187,CPSF1,PPP1CB,VAMP2,HNRNPL,
AP3,MDK,CCT4,SNRNP70,DPYSL2,TAF7,RNF10,CCNL2,DDAH2,PDE7A,PSMD3,RABL6,IPO7,ARGL
)L4A2,MAPT,NONO,DDB1,YWHAQ,DCHS1,SET,MAP9,CHSY1,EIF4A1,MRC2,RPL19,GTF3C4,LDHB
CSDE1,IGF2,EIF3C,UBB,AFF4,MATR3,LRRC28,SPEN,PSMD5,YWHAQ,EIF1,EGLN1,TMSB10,EIF4A
PPP1CC,CUL3,IL6ST,LMAN1,ZMYM3,EFR3B,TRIR,XRCC6,VAPB,RTL6,SHC2,FKBP4,SNRPD3,CARI
E4A,TBC1D16,AHCYL1,ETNK1,KLC1,CLSTN3,DDX42,RAB11B,NRDC,PRPF6,TUBB2A,UBXN4,PPP2
,CALU,ZNF454,SLC35E2B,RPL23A,HSBP1,MAP1A,TM9SF3,MDH2,U2AF2,EEF1A2,FDFT1,CCT2,S
B,NPLOC4,HNRNPAB,LOX,FUS,SPG7,HSPG2,UBE2D2,PGAP6,RAB14,GNA12,KIAA0232,PLEKHM2
LC16A7,RPL22,ZNF865,RPL9,KIF5B,RPS14,CXADR,LENG8,WDR1,RPL26,CCT7,SRSF4,RBBP4,PF
MMP2,BCLAF1,L1CAM,LUC7L3,DDX39B,CNN2,SRP9,SRSF5,MDH2,U2AF2,CPSF1,NOP53,PABPN
PS25,MMP14,MAGED2,GNB2,H1-0,RPS7,CD63,ANXA6,B4GALNT4,HNRNPDL,ARL6IP1,NDFIP1,SH
MMP15,PHB2,APMAP,RAB2A,RER1,RAB3GAP2,YKT6,TMEM167A,PSMD8,NPLOC4,CHERP,CTTN,
N1,PDXK,PKN1,PLOD1,AHCYL1,KLC1,PTOV1,CLSTN3,CCT4,PCDHGB6,SCAF1,TOP1,VEGFA,OPT
1IP,SQLE,MAP1A,CDKN1A,TIMP3,CTDSP2,CNN2,NOVA2,NUCB1,ZC3H7B,RAB27A,U2AF2,CCT2,C
62,RPL6,KAT6A,YWHAE,RPL13,GNAS,SLC2A3,DDX17,GPI,HTRA4,RACK1,BSG,FASN,ALDH6A1,SF
,LDHA,CPE,CNN3,RPSA,CTSD,SCARB2,RTN4,LPCAT1,LAMA5,RPS24,SLC22A17,PJA2,RPL24,NCS
TF2F1,GOLGA8A,LDHA,CPE,CNN3,RPSA,CTSD,SCARB2,RTN4,LPCAT1,LAMA5,RPS24,SLC22A17,F
BA1,ATP2B4,IGFBP2,CALR,FSTL1,TMF1,ABHD2,HSPA1B,ZNF99,TUBB2B,COL5A1,COL4A1,CES3
QSTM1,ACTR3,RTN1,AHCYL1,ETNK1,GRINA,ASXL1,SELENON,MVB12B,G3BP2,DSTN,BTBD2,DYN
HD4,ATP1A1,JUND,LENG8,WDR1,CRMP1,VAT1,OAZ1,NES,RPL31,PRKCSH,HNRNPC,TADA2B,KHI
)X39B,SRSF5,TM9SF3,MDH2,TOMM20,U2AF2,CCT2,GNAI3,NOP53,TM9SF2,TSR1,LMNB2,MTRF
2,MORF4L2,SNRPN,PLOD1,GRINA,VPS35,PTOV1,MLEC,SURF4,SKI,PCDHGB6,SNRNP70,PYGB,PF
2,TOMM20,U2AF2,CCT2,GNAI3,GDI2,TM9SF2,TSR1,SSRP1,LMNB2,MTRF1L,ALKBH5,PRDX5,UB
),CSDE1,IGF2,ATP5F1A,EIF4B,UBB,TRIM28,MATR3,YWHAG,RPL37A,MAPT,NONO,PSMD5,HACD3

P2R1A,SF3B1,PDIA3,SSBP2,ZNF43,CDKL1,AP2M1,TPT1,SEC61A1,TUBB4B,MORF4L1,ANXA2,ZM
RCKS,NPM1,TTC3,PLD3,HSPD1,USP11,SNRNP200,CSDE1,SMIM14,EIF3C,ATP5F1A,EIF4B,VCP,UE
,COL4A2,LRRC28,EIF1,DCHS1,SET,MAP9,CHSY1,TMSB10,MRC2,PKD1,SSBP2,ZNF43,CHD3,BASI
RPS12,HLA-B,ARHGDIA,SLC35E2B,RPL23A,HSBP1,FAM199X,DDX39B,TM9SF3,MDH2,U2AF2,FDI
1,RPS17,VCL,DNAJA1,CDC42BPB,ZNF770,MAT2A,SLC4A8,GOLGA8A,OGA,PGAM1,NFE2L1,RPL2
ST1,CDH11,U2AF2,EEF1A2,FDFT1,SQSTM1,LMNB2,MTRF1L,PKN1,YEATS2,ACO2,KLC1,DDX42,
,ERGIC3,MBD3,PSMC3,CREBBP,ANKRD12,BAG5,UBXN6,SLC20A1,DPYSL4,METRN,UNC119B,TA
,RPLP1,SARAF,MIDN,RPL10,MARCHF6,WSB1,RPS17,ITM2B,AP2B1,GTF2F1,GOLGA8A,SMARCA5
,CALU,ARCN1,ARHGDIA,SLC35E2B,ATP6V0C,LAPTM4B,TRIB2,RPL23A,FAM199X,DDX39B,SRSF5

),LENG8,WDR1,MCL1,PNN,CCNI,XPR1,HDLBP,EIF5A,PEBP1,CCND2,DDR1,PLXNA1,CLTC,NANOG,
LDHA,CPE,CNN3,RPSA,CTSD,SCARB2,RTN4,LPCAT1,LAMA5,RPS24,SLC22A17,RPS9,PJA2,RPL24,
CAR2,PHC2,SDHA,YWHAH,PFKM,TNKS2,LONP1,HYOU1,STRAP,COL27A1,BAG6,RAI1,ZSCAN18,S
19,PPP2R1A,GTF3C4,DNAH8,SERINC1,ZNF43,HMGB1,POLR2A,AP2M1,TPT1,RPS4X,RPS3A,RPL1
2A23,CDV3,GGNBP2,IGF1R,SLC25A6,FOXP4,STK39,MECP2,CUL1,WDR26,SHISA5,ZNF462,SALL2
P2,NOTCH1,SLC25A6,FOXP4,STK39,CUL1,DNAJB1,WDR26,ANO8,BNC2,PNRC2,SHISA5,ZNF462,
FBI,CLSTN3,PRPF6,RPL38,OPTN,BTBD2,PBX2,DYNLL2,LUZP6,CAPZB,UBE2R2,KDELR1,ARF4,ZF
3,SPG7,FKBP4,LINGO1,SOX2,TRAM1,ARHGEF17,SMG5,CNP,STK24,LSM14B,LTBP3,YY1,UBE2I,
1,SPEN,PSMD5,EIF1,EGLN1,EIF4A1,RPL19,PPP2R1A,RPL15,GTF3C4,HLA-C,SF3B1,SERINC1,HM
L37A,FN1,DDB1,DCHS1,SET,EGLN1,TPI1,CHSY1,EIF4A1,RPL19,GTF3C4,DNAH8,PKD1,SSBP2,VA
,ITGB1,TUBB4B,UCHL1,ANXA2,XAF1,SARAF,RPL10,MARCHF6,WSB1,RPS17,VCL,AP3D1,SBF1,DN
FKM,SF3A1,ATXN2L,PRPS1,GPR107,SREBF2,HNRNPD,MEX3A,P3H3,ZSCAN18,SERBP1,DHX15,ST
TF3,AGRN,RPLP1,SARAF,MIDN,RPL10,MARCHF6,RPS17,DNAJA1,ARF1,CYB5R3,SF3B2,GTF2F1,C
4,DNAH8,PKD1,SSBP2,RPS3,VAPA,POLR2A,TPT1,RPS4X,RPS3A,RPL18,STMN1,ITGB1,RPL11,MO
NRNPA3,TBC1D16,ACO2,TCP1,MLEC,ACAP3,RAD23B,CCT4,SNRNP70,ASXL1,H3-3A,DPYSL2,TME
NK1G2,FARP1,RER1,UBFD1,IARS1,COX7C,YKT6,MCM3AP,RPS27A,TMEM167A,CHERP,FKBP1A,C
USP11,SNRNP200,CSDE1,SMIM14,EIF4B,VCP,MARCKSL1,MATR3,YWHAG,CKB,COL4A2,RPL37A,
MARCHF6,WSB1,SBF1,AKAP12,SF3B2,ITM2B,AP2B1,GTF2F1,GOLGA8A,SMARCA5,HNRNPA0,RPL
NA,EIF4A1,PPP2R1A,SERINC1,BASP1,POLR2A,TPT1,SEC61A1,RPS3A,RPL18,ITGB1,RPL11,MOR
,JUND,TMEM259,EIF5,LENG8,ATXN7L3B,SRSF4,RPL31,NISCH,LRPPRC,HDLBP,EIF5A,CDH2,PDI
DUX4,LGALS3BP,PLPPR3,MAGED4B,ARHGDIA,APPL1,SMCHD1,MAP1A,FAM199X,DLG5,SERF2,T
16,AHCYL1,KLC1,CLSTN3,DDX42,RAB11B,NRDC,PRPF6,TUBB2A,UBXN4,PPP2CA,G3BP2,KDELR
170A,GTF2F1,SLC4A8,GOLGA8A,OGA,PGAM1,CTNNA1,FADS2,RPL23,CPE,TPM4,TNS3,SCARB2,
RB2,WDR6,KDELR1,ZFAND5,RPS28,YWHAH,UBQLN1,SF3A1,PRPS1,ZFAND3,IFITM3,HGS,SREBF2

SL1,UBB,YWHAG,CKB,COL4A2,LRRC28,FN1,DDB1,PSMD5,YWHAQ,DCHS1,SET,TMSB10,MRC2,F
,AGRN,TIMP2,SARAF,COL11A1,AP3D1,DNAJA1,CYB5R3,AKAP12,MAP4,SF3B2,CDC42BPB,ITM2B
2,EEF1B2,NCSTN,INTS11,QPRT,MTMR4,SPTY2D1,LRRC41,PRPF19,NOC2L,EIF3G,PACS2,AP1M1,
V2A,ECEL1,NDST1,ULK1,CHERP,GSE1,MAP1LC3B,WSCD1,ZNF664,SHC2,BCL7A,ARHGEF17,PGA
IARCA5,HNRNPA0,RPL29,TNS3,NACA,LPCAT1,LAMA5,PJA2,RPL22,RHOA,CLSTN1,ATF4,PSMD2,R
,PNMA1,GPC3,PIP4K2B,MMP14,RPL12,MAGED2,H1-0,APLP1,MIF,CD63,ANXA6,SESN3,APLP2,TA
B5,TNKS,DKK3,PPP1R15A,MAP1LC3B,TMOD2,PELP1,TCAF1,SLC44A2,CARM1,GPC2,IMPDH2,A
2BPB,AP2B1,PXDN,NFE2L1,RPS16,NACA,PJA2,MSN,KPNB1,C1ORF43,COL1A2,CLSTN1,ATF4,PSM
A1,ATP5F1B,CALM3,SHISA9,RPS8,MYH9,RPL7,SEPTIN2,P4HB,IDS,ERVV-1,RPL5,GDI1,FSCN1,SP
RPPRC,HDLBP,CCND2,DDR1,PLXNA1,DDX3X,ARL4C,GNB2,IK,CHPF,JUN,HNRNPDL,SLC25A3,RPL
P1,SLC35E2B,L1CAM,LAPTM4B,DLG5,HS6ST1,TMEM132A,CNN2,SRP9,RNF187,CCT2,MEST,PAR
P1,RPL28,CCAR2,HNRNPUL2-BSCL2,TSC22D1,DLK1,MSL1,YWHAH,FBLN1,COL27A1,TCTN1,RPL
SP1,POLR2A,TPT1,SEC61A1,RPS3A,RPL18,ITGB1,MORF4L1,NCAM1,ANXA2,RPLP1,SARAF,MIDN,
,RPL10,MARCHF6,WSB1,RPS17,CDC42BPB,ITM2B,AP2B1,GTF2F1,GOLGA8A,SMARCA5,LDHA,C
9,SLC3A2,SCRN1,SFPQ,EIF5A,ITM2C,CLTC,NANOG,H1-0,APLP1,RPS19,JUN,SESN3,MXD4,SHOX,
C61A1,RPS3A,RPL18,ITGB1,MORF4L1,NCAM1,ANXA2,RPLP1,SARAF,MIDN,RPL10,MARCHF6,WS
ARS1,SHC2,ANTXR1,RAB1B,UBE2M,ARHGEF17,MAZ,KIAA0232,DICER1,DHCR7,USP24,FXYD6,FB
Z1,TIMP2,MIDN,SBF1,AP2B1,GTF2F1,SMARCA5,HNRNPA0,LDHA,TPM4,SCARB2,NACA,LAMA5,SL
PL30,ARL4C,CLTC,GNB2,B4GALNT4,APLP2,MYL6,CIRBP,MYH10,NARS1,YWHAB,CADM4,HLA-B,
3,KIF5B,RAB6B,NLGN2,GLIPR1L2,ATP1A1,TMEM259,EIF5,CD81,ATP2A2,RPL26,CCT7,OAZ1,SRS

AP3D1,DNAJA1,PAFAH1B1,AKAP12,ITM2B,SLC4A8,GOLGA8A,PGAM1,FADS2,RPL23,RPSA,TNS3,
TIN2,IDS,RPL5,KLRD1,GDI1,FSCN1,GSTP1,MARCKS,SMG1,CALM1,NPM1,TTC3,PLD3,HSPD1,US
ZIN1,DDAH2,NYNRIN,PRKDC,TDRD5,CASC3,CLIP3,RPL28,ZFAND5,CORO1C,SDF4,SMARCC1,HN
2,RPLP1,SARAF,MIDN,RPL10,MARCHF6,WSB1,RPS17,ARF1,SF3B2,ITM2B,AP2B1,GTF2F1,GOL
1,RPLP0,CANX,CLU,DDX5,SOX4,FKBP14,SEC62,RPL6,KAT6A,DVL3,YWHAE,KIF1A,RPL13,GNAS,
2,COPE,CD2BP2,SUMO1,RPS21,BANF1,FAF2,KLC2,VPS51,PPME1,FAM8A1,FAM32A,EPN1,RTF2,
,AGAP1,USF2,PPP5C,NBPF15,NRXN2,KLF13,MKNK2,OBSL1,FZR1,MAPK8IP2,NSF,PPP1R14B,DC
UDCD3,FLOT2,TUBGCP6,EEF1B2,NCSTN,INTS11,QPRT,MTMR4,SPTY2D1,LRRC41,PRPF19,NOC
,AFF4,MATR3,RPL37A,LRRC28,PSMD5,EIF1,EGLN1,INA,EIF4A1,RPL19,PPP2R1A,RPL15,GTF3C4,
ME3,DHX36,HM13,COLGALT1,TSPAN7,NUP98,OXSR1,EMC10,PSMC4,BEX1,MEX3C,NSMF,DUSP8,
K8IP1,DGKD,SREBF2,PFN2,P3H3,CYP51A1,AKIRIN1,SMS,DYNLL1,ADAR,RBM15B,DHX15,BEX4,
P43,RPL22,RHOA,ZNF865,LENG8,WDR1,HMGN2,LASP1,OAZ1,RBBP4,MCL1,PRKCSH,LRPPRC,X
1A1,AP3D1,AKAP12,CDC42BPB,ZNF770,TMEM170A,PXDN,CTNNA1,FADS2,CNN3,CTSD,NACA,LP
L7,SEPTIN2,FSCN1,TMBIM6,SPTAN1,GSTP1,SMG1,CALM1,NPM1,PLXNB2,PLD3,USP11,COL18A
RPL15,DNAH8,HLA-C,SERINC1,SSBP2,POLR2A,TPT1,RPS3A,UCHL1,MORF4L1,ANXA2,AGRN,RPL
BL,FLNB,TSC22D1,PLTP,SSB,TNKS2,TUG1,PLXNA3,HYOU1,COL27A1,ARFGEF2,WBP2,DHX15,WD
1,PDAP1,MMP15,PHB2,APMAP,CST3,PLXNB1,GARS1,RAB2A,RER1,UBE2G2,RAB3GAP2,TAGLN2,
K2B,RPL12,ITM2C,EIF3L,MRFAP1,RPS19,GPAT4,CD63,MACF1,RPS10,CYFIP2,DDOST,MXD4,ANK
CKS,SMG1,CALM1,TTC3,COL18A1,SMIM14,VCP,UBB,AFF4,CKB,COL4A2,RPL37A,FN1,SPEN,NO
M9SF2,MORF4L2,ABCA3,PDXK,PTOV1,MLEC,SKI,PCDHGB6,PYGB,NOTCH2,PPP2CA,SMARCA4,A
RPS15,RPL12,MAGED2,H1-0,RPS19,TLN1,MYO5A,NGRN,RPL35A,DNAJC5,HNRNPDL,TARDBP,N
S,GSTP1,MARCKS,CALM1,PLD3,COL18A1,CSDE1,IGF2,EIF4B,VCP,MARCKSL1,UBB,AFF4,TRIM2
,JUND,TMEM259,EIF5,LENG8,WDR1,MCL1,PNN,CCNI,TADA2B,XPR1,HDLBP,PEBP1,DDR1,PLXNA
RNPD,CDC42,RBBP7,TRAF7,EIF2AK1,EDC4,WDR45B,PDAP1,MMP15,SEC16A,PHB2,CD164,IARS
AH8,PKD1,SSBP2,VAPA,AP2M1,ITGB1,TUBB4B,ANXA2,AGRN,XAF1,SARAF,MIDN,RPL10,WSB1,AP
A,TNPO1,ASXL1,PRPF6,G3BP2,CCNL2,PTBP1,HNRNPUL1,SON,RPL28,CCAR2,PHC2,SDHA,NKTR

HJURP,RAB2A,SELENOW,TBC1D9B,PRRC2A,ANP32B,SPINT2,PREPL,RPS27A,HNRNPAB,SYMPK,
COL6A2,GOLM1,SESN3,ARL6IP1,INIP,PTGES3,SEPTIN11,RHOB,SEPTIN3,COTL1,HECTD4,RAC1,

,MDH2,TOMM20,U2AF2,CCT2,RPN2,GNAI3,NOP53,TSR1,HSPH1,MTRF1L,RPL27,PRDX5,ACTR3,
LDIP2,TRIP12,XRN2,TSNAX,CRACD,SLC25A6,EXOC5,SF3A3,TRMT112,COPE,CD2BP2,SUMO1,RP
F2,TIMP3,CTDSP2,CNN2,NOVA2,NUCB1,C5ORF24,ZC3H7B,RAB27A,U2AF2,CCT2,ANXA5,MEST,
N1,GDI2,HNRNPL,MORF4L2,TAF15,VPS35,MLEC,CLSTN3,H3-3A,RAB11B,NRDC,UBXN4,G3BP2,
CDC37,WDR6,BNIP3L,PITPNA,XPNPEP3,SRSF3,LARS1,YWHAH,UBQLN1,CSNK1D,HYOU1,IFITM3,
GLN1,MAP9,CHSY1,TMSB10,INA,ZNF43,CHD3,AP2M1,RPS4X,STMN1,BTF3,NCAM1,ANXA2,AGRN
WDR45B,PDAP1,MMP15,SEC16A,PHB2,YKT6,MCM3AP,RPL7L1,RPS27A,TMEM167A,KLHDC3,TN
IP1,SEPTIN3,STIP1,ATP6V0C,LAPTM4B,SERF2,CTSB,MDH2,TOMM20,U2AF2,SUGP2,VAMP2,MES
PRPF40A,SMS,SYP,RBBP7,TRAF7,STAT3,EIF2AK1,WDR45B,SGTA,H2AZ1,SEC16A,ZFP36L1,PPM1

D2,RAB5C,CLASP2,PDPK1,SCRIB,LRRC41,LRP3,FXYD6,CIZ1,DENND4B,PACS2,APBB1,ANKRD1
KAT6A,DVL3,YWHAE,KIF1A,RPL13,GNAS,DDX17,HTRA4,RACK1,BSG,FASN,ALDH6A1,SPTBN1,ATP
,NTRK2,DIP2A,ACAT2,PSAT1,LUC7L2,ATP13A3,IGF1R,EFNB2,CLUH,EMC1,PNRC2,RASA4,RAN,M
9SF3,MLXIP,MORF4L2,HSPH1,ALKBH5,UBE4A,ACO2,AHCYL1,ETNK1,KLC1,CLSTN3,DDX42,RPL
VCL,AP3D1,ITM2B,HNRNPU,PGAM1,SMARCA5,HNRNPA0,CTNNA1,RPL23,RPSA,EIF4H,NACA,BRD

L6ST,MAPRE2,TMEM167A,LMAN1,HNRNPAB,PODXL2,FKBP1A,MVD,STK35,SEPTIN5,SLIT1,SOX2,
;,FSCN1,PTMS,SPTAN1,GSTP1,MARCKS,SMG1,CALM1,PLXNB2,TTC3,PLD3,HSPD1,USP11,COL18
C14L1,NUDCD3,GALNT1,TUBGCP6,RIC8A,INTS11,MOAP1,MTMR4,SPTY2D1,NUDT19,CAP1,COX6
TF3C4,RPS3,CHD3,VAPA,HMGB1,POLR2A,TPT1,SEC61A1,RPS3A,ITGB1,MORF4L1,NCAM1,ANXA2
,PTPRA,ACTR3,YEATS2,TBC1D16,ACO2,RBMX,ASXL1,SCAF1,TERF2IP,TMED2,PPP2CA,CCND1,M
G2,DHX30,EIF3F,IFITM3,HGS,BAG6,FBRSL1,RAB11A,KDM3B,CDC42,YME1L1,TRAF7,EIF2AK1,R
ND5,CORO1C,SDF4,SMARCC1,HNRNPM,STT3A,PLXNA3,IDH1,SARS1,SOD1,CYP51A1,CD99L2,DD
F3L,MRFAP1,HMGCS1,CHPF,RPLP2,ST3GAL2,TSR2,ANKRD52,RPL36,SEPTIN11,CADM4,LAMP1,H
1,NSD2,COPZ1,PSMC3,CHMP4B,CREBBP,UBE2I,DHX40,RAB11FIP3,EIF4G1,TRIP12,CELSR1,HMG
LC25A3,CIRBP,NARS1,SEPTIN11,SND1,RHOB,HNRNPH1,RAC1,APPL1,HDAC2,HSBP1,NNAT,PTTG
OXDC1,EIF2S1,KRI1,DNMT1,ATP6V1D,TBCB
IM1,TOMM70,LYPLA2,ATP13A1,ATP5MG,SRPRA,EIF4G1,PAFAH1B2,SLC30A9,POLDIP2,TRIP12,C
AM1,ANXA2,RPLP1,SARAF,MIDN,RPL10,MARCHF6,WSB1,RPS17,ARF1,SF3B2,CDC42BPB,ITM2B,
P1,MMP15,SEC16A,PHB2,GSPT1,UBA52,MCM3AP,RPL7L1,RPS27A,TMEM167A,KLHDC3,TNKS,PS
CHSY1,TMSB10,RPL19,GTF3C4,HLA-C,POLR2A,AP2M1,TPT1,SEC61A1,ITGB1,RPL11,ANXA2,RPL
MARCA5,RPS16,CPE,SCARB2,TSPAN3,RPS24,RPS9,RPL17,C1ORF43,COL1A2,RPL14,RHOA,PTCD
I1,SDCBP,ITFG1,SLC22A23,TSPYL2,ANKHD1,ATP13A3,ACVR1B,SMPD3,PFKP,FOXP4,DCAF5,ADN
UBB2A,OPTN,PPP2CA,G3BP2,PDE7A,PSMD3,SRSF1,TCF25,CDC37,WDR6,BNIP3L,ARF4,ZFAND5,
A5,ENC1,RPS20,EIF5,LENG8,ATXN7L3B,SRSF4,TLE5,NES,LRPPRC,XPR1,SPTBN2,TKT,GLG1,CDI
K1,ITGB1,TUBB4B,RPL11,MORF4L1,ANXA2,MIDN,RPL10,RPS17,VCL,AP3D1,DNAJA1,CYB5R3,ITI
RS1,OTUB1,HIRA,SKIV2L,SHC1,CHMP5,ASB6,SUDS3,GPS1,MIB2,TNPO3,THG1L,NUP50,SZT2,SEC
PLXNA3,ID4,NBPF9,VANGL2,HGS,RAI1,CD99L2,CDC42,BEX4,GRIPAP1,PSMA7,STAT3,WDR45B,E
,SHISA9,STMN2,RPS8,RPL7,SEPTIN2,P4HB,APP,IDS,RPL5,KLRD1,GDI1,TMBIM6,GSTP1,MARCKS
D3,CDC37,WDR6,BNIP3L,HNRNPUL2-BSCL2,PITPNA,XPNPEP3,ATP6V1B2,LARS1,YWHAH,UBQLN
CDC37,WDR6,ACTN1,BNIP3L,ZFAND5,PITPNA,SRSF3,LARS1,YWHAH,UBQLN1,CSNK1D,SARS1,IFI
CAN18,YLPM1,GRIPAP1,ARIH1,LBH,PBX1,ALDH3A2,CD99,SEC16A,UBA52,SLC39A7,PLXNB1,FARF
1,SET,EIF4A1,GTF3C4,DNAH8,HLA-C,SERINC1,SSBP2,CHD3,BASP1,VAPA,POLR2A,AP2M1,TPT1,I
25,WDR6,TDRD5,CASC3,PLCG1,CORO1C,SMARCC1,SDHA,YWHAH,SLIT3,LSS,PLXNA3,CSNK1D,N
XPC,DDHD2,CITED2,PNMA2,CNOT3,QRICH1,CLCN3,TMCO1,MYEF2,C6ORF89,RBM39
,IFITM3,VGF,MLF2,AKIRIN1,AMOTL2,YME1L1,TRAF4,STAT3,ARIH1,SF1,SLC35F5,CSNK1G2,RAB
,CCDC47,ELOVL5,SLC25A36,XRN2,SCAMP5,ZZEF1,AASDHPPT,ESYT1,GTF3C1,TNFAIP1,FAM50A
T8,RPL8,FBXL4,YBX1,UBA1,ATP2B4,IGFBP2,CALR,FSTL1,TMF1,ABHD2,HSPA1B,ZNF99,TUBB2B,
S11,MTMR4,CAP1,PRPF19,EIF3G,AP1M1,SSR2,NDUFA13,UBE2I,DAPK3,ATP13A1,IER2,EIF4G1,P
42,TXLNA,PDAP1,PHB2,GSPT1,UBA52,PLXNB1,MCM3AP,PREPL,TMEM167A,PSMD8,CTTN,SERP1
RAF4,ARIH1,SF1,SLC35F5,CSNK1G2,UPF1,MASTL,MAPRE2,APEX1,PPP1R15A,GALNT11,DAP,BRK
KCSH,HNRNPC,CCNI,XPR1,CSNK1E,PFKL,TKT,TPM1,EIF5A,PEBP1,CCND2,ACSL6,MMP14,CLTC,A
TMEM167A,TNKS,PSMD8,CHERP,CTTN,LRP10,SERP1,ACLY,ERH,SEC14L1,NUDCD3,TAX1BP1,FAM
NOL4L,LRRC4B,KIF21B,ZFP36L1,MGAT4B,BRD2,FARP1,KIAA0895L,NDST1,ULK1,GSE1,PCBP4,
VCP,UBB,AFF4,MATR3,CKB,COL4A2,RPL37A,LRRC28,MAPT,DDB1,PSMD5,HACD3,SET,EGLN1,M
PC,RPL24,NCS1,RPL22,ZNF865,RPS14,CXADR,RPS20,LENG8,WDR1,RPL26,SRSF4,PFN1,MCL1,F
3,MCM7,SZT2,SEC31A,ACIN1,AKAP17A,EIF2S1,MAPK8IP3,HARS1,MRPL3,DNMT1,CFAP298,DDX5
P5,ZZEF1,AASDHPPT,ESYT1,GTF3C1,TNFAIP1,FAM50A,CNDP2,GLOD4,MTURN,GOLPH3,DHX36,C
WDR6,BNIP3L,HNRNPUL2-BSCL2,PITPNA,XPNPEP3,ATP6V1B2,LARS1,YWHAH,UBQLN1,CSNK1D,I
2,KAT6A,DVL3,YWHAE,KIF1A,RPL13,GNAS,GPI,BSG,FASN,SPTBN1,CALM3,GNB1,SHISA9,STMN2,
3A,RAB11A,P3H3,AKIRIN1,GRIPAP1,KIF5A,BCL9L,XPOT,NAP1L1,KIRREL1,KDM5B,PBX1,CNTNAF

R9,OTUB1,KDM5A,ZDHHC5,HIRA,SKIV2L,RNF4,SHC1,SUDS3,GPS1,PSMA6,MIB2,TNPO3,SNX17,T
H10,HNRNPH1,HECTD4,CDK16,SLC35E2B,APPL1,SMCHD1,HDAC2,IPO5,SQLE,CTDSP2,CPSF2,NU
NRNPD,SEC23A,CDC42,TRAF7,EIF2AK1,TXLNA,MMP15,GSPT1,UBA52,GARS1,RAB2A,RAB3GAP2,
,CES3,PGK1,RPLP0,CLU,PTMA,DDX5,ENO2,SOX4,FKBP14,RPL13A,MAGED1,SEC62,KAT6A,DVL3
AN3,RPS24,KPNB1,RPL17,PAPOLA,IGFBP3,COL1A2,RPL14,NCS1,PTCD3,CBX5,KIF5B,SLC7A5,N
SUPT5H,KDELR1,HNRNPUL2-BSCL2,GNG2,SUPT6H,PTP4A2,NT5DC2,UQCRC2,BAG6,CDC42,YME
B2,GABARAP,HNRNPUL1,KDM1A,BNIP3L,KDELR1,MAP2K2,ARF4,SDF4,ATP6V1B2,ILF2,STT3A,L
I,DDAH2,PTBP1,FTO,ILF3,RPS11,CASC3,CCAR2,DIDO1,RPS28,HNRNPM,SRSF3,MESD,GPR107,G
ARCHF6,RPS17,VCL,AP3D1,SF3B2,AP2B1,GTF2F1,PXDN,PGAM1,NFE2L1,HNRNPA0,LDHA,CPE,T
.36,DUX4,PTGES3,SEPTIN11,LAMP1,RHOB,CTBP1,ARHGDIA,STIP1,HDAC2,RPL23A,SERF2,MDH2
NO,YWHAQ,EIF1,DCHS1,MAP9,TPI1,EIF4A1,RPL19,PPP2R1A,GTF3C4,DNAH8,SF3B1,PDIA3,SERI
FAM32A,SYNGAP1,EPN1,ARPC3,RTF2,GBA2,POLDIP3,UNC45A,PSMF1,HMGXB3,MCFD2,RPRM,
CHMP1A,TSNAX,PAIP2,EXOC5,TRMT112,PSMD13,CD2BP2,SUMO1,BANF1,COX8A,VPS51,DARS1,
,FAU,STMN3,B2M,ISYNA1,PLXNA3,IFITM3,PTPRS,DDIT4,BEX4,PPDPF,VDAC1,KCNQ2,EDC4,TXND
99L2,DDIT4,BEX4,SSR1,WDR45B,SLC39A6,CD99,GPX4,PPM1G,GNG4,NXN,EP400,RER1,UBFD1,
CIRBP,DUX4,PTGES3,CKAP5,SEPTIN11,UBE2H,MAGED4B,SKP1,L1CAM,TRIB2,PTTG1IP,MAP1A,S
,PSMD3,SRSF1,TCF25,CDC37,WDR6,BNIP3L,ARF4,HNRNPUL2-BSCL2,PITPNA,XPNPEP3,SRSF3,L
N2,TNFAIP1,COPE,CNDP2,MTURN,GOLPH3,RAP2A,DHX36,TSPAN7,NUP98,NACAD,OXSR1,EMC1
PM,PHC2,PFKM,EIF3F,LOC728392,VANGL2,P3H3,KDM3B,AKIRIN1,CD99L2,BEX4,PPDPF,VDAC1,
2,RAPGEF1,TUBB2A,PPP2CA,CCND1,MAPK1,AZIN1,DDAH2,PTBP1,FTO,ILF3,RPS11,CASC3,NSG
,ERVV-1,GDI1,FSCN1,TMBIM6,PTMS,GSTP1,SMG1,CALM1,NPM1,TTC3,PLD3,HSPD1,USP11,SMI
MNB2,MTRF1L,ALKBH5,HNRNPA3,PRDX5,UBE4A,SURF4,DDX42,GCN1,SRP14,PRPF6,TERF2IP,R

WSB1,RPS17,ARF1,SF3B2,CDC42BPB,ITM2B,AP2B1,GTF2F1,GOLGA8A,SMARCA5,LDHA,CPE,C
4,TAF15,LMNB2,VPS35,MLEC,CLSTN3,RAB11B,NRDC,UBXN4,MAPRE1,DSTN,CAPZB,HNRNPUL1
P2,TMX1,REEP5,PCM1,COPE,KIF26A,WDR26,CD2BP2,ACAP2,SALL2,ETF1,PPP1CA,RDH11,EPN1,
CA5,LDHA,RPSA,RTN4,SLC22A17,CIPC,RPL24,PAPOLA,CBX5,RPL9,NR2F1,GLIPR1L2,JUND,TME
TSPAN3,LPCAT1,LAMA5,RPS24,SLC22A17,BRD3,MSN,KPNB1,PAPOLA,LLGL1,NCS1,ZNF865,ATF
QLN1,CSNK1D,IFITM3,VGF,MLF2,AKIRIN1,AMOTL2,YME1L1,TRAF4,STAT3,ARIH1,SF1,SLC35F5,
,RPL13A,MAGED1,SEC62,RPL6,KAT6A,DVL3,YWHAE,KIF1A,GNAS,SLC2A3,DDX17,HTRA4,RACK1
I,C6ORF62,GABARAPL1,TLE1,AKT3,GTF2I,KIFAP3,DNAJB2,SH3PXD2A,SFXN1,SPAG9,RBM12,AB

IUCB1,TM9SF3,MDH2,RAB27A,U2AF2,FDFT1,GNAI3,NOP53,TSR1,HSPH1,MTRF1L,RPL27,PTPRA
DIP2,TRIP12,CHMP1A,PSMB1,TSNAX,SLC38A1,EXOC5,SF3A3,TRMT112,PSMD13,WDR26,CD2BP
C,ACAP3,HDGF,TERF2IP,TAF7,RPL38,UBXN4,CCNL2,FAM168B,EIF3E,LUZP6,CAPZB,SUPT5H,FK
L3,ZSWIM6,BEX1,ILRUN,CRTC1,ZFP91,RAVER1,CRABP2,SLC12A7,CASKIN1,COL7A1,NUMBL,GP
,CPSF7,RAD50,JRK,TOMM40,COX5B
RPL35,GJA1,TARDBP,NDFIP1,SHOX,ANKRD52,SEPTIN11,LAMP1,RPS12,ARCN1,RPS27,COTL1,H
CSDE1,EIF3C,VCP,MARCKSL1,AFF4,RPL37A,MAPT,SPEN,DDB1,PSMD5,YWHAQ,EIF1,HACD3,TPI1,
CALM3,SHISA9,STMN2,MYH9,RPL7,SEPTIN2,P4HB,APP,RPL5,GDI1,FSCN1,PTMS,SPTAN1,GSTP1,
B1,KDM5A,SKIV2L,SHC1,ASB6,SUDS3,MIB2,TNPO3,CUL4B,SZT2,SEC31A,AKAP17A,MAPK8IP3,
NNA1,TPM4,NACA,SLC22A17,BRD3,PJA2,MSN,COL1A2,HK1,NCS1,RHOA,SLC7A5,RPS14,CD81,C
I,GNL1,TM9SF4,TRAP1,PLEKHB2,ELP1,MOAP1,CLASP2,NSD2,YY1,CCDC47,ELOVL5,SLC25A36,
H2,GABARAPL1,GNL1,TM9SF4,TRAP1,PLEKHB2,ELP1,MOAP1,CLASP2,NSD2,YY1,CCDC47,ELOV
GOLN2,ATXN7L3,HSDL1,WBP1,UNC119B,FBXL19,OLFM2,SNX3,OGFR,SCAMP5,USF2,MTA2,HD

HERP,CTTN,LRP10,SERP1,ERH,EIF3H,SEC14L1,NUDCD3,TAX1BP1,GNL1,HDGFL2,SRSF2,CAPZA1
L6A2,ANXA6,RPS10,B4GALNT4,APLP2,HNRNPDL,ARL6IP1,SHOX,H1-10,RHOB,RPS12,PLPPR3,CO
ASTL,PRRC2A,MAPRE2,PPP1R15A,GALNT11,DAP,EIF3H,ELAVL1,IMPDH2,GABARAPL1,GNL1,TM9
APL1,GNL1,TM9SF4,TRAP1,PLEKHB2,ELP1,MOAP1,CLASP2,NSD2,YY1,CCDC47,ELOVL5,SLC25A
APL1,GNL1,TM9SF4,TRAP1,PLEKHB2,ELP1,MOAP1,CLASP2,NSD2,YY1,CCDC47,ELOVL5,SLC25A
PTIN7,PPP1R15A,GALNT11,FBXO21,SERP1,BAP1,SPG7,SSBP3,ELAVL1,IMPDH2,TLE1,LDLR,GNL
31,EIF1,DCHS1,SET,TPI1,CHSY1,EIF4A1,MRC2,PPP2R1A,GTF3C4,LDHB,PKD1,SSBP2,BASP1,VAP
S3BP,RAC1,ARHGDIA,GPC1,CDK16,SLC35E2B,BCLAF1,PTTG1IP,MAP1A,CDKN1A,TIMP3,CTDSP2
HACD3,TPI1,TMSB10,EIF4A1,PPP2R1A,SSBP2,RPS3,HMGB1,RPS3A,STMN1,ITGB1,UCHL1,RPL1
S4X,RPL11,MORF4L1,BTF3,NCAM1,ANXA2,RPLP1,ZMIZ1,RPS17,VCL,ITM2B,ZNF770,GTF2F1,SL
S4X,RPL11,MORF4L1,BTF3,NCAM1,ANXA2,RPLP1,ZMIZ1,RPS17,VCL,ITM2B,ZNF770,GTF2F1,SL
1,NES,PRKCSH,TADA2B,PFKL,CCND2,PDIA6,PLXNA1,ACSL6,PRDX2,CLTC,MIF,GPAT4,JUN,CD63,
,TRIB3,ADCY1,DHX36,HM13,TSPAN7,PSIP1,NUP98,NACAD,OXSR1,EMC10,SEMA4C,TMEM9,PSM
TP1,MARCKS,SMG1,CALM1,NPM1,PLD3,HSPD1,COL18A1,SNRNP200,EIF4B,MARCKSL1,MATR3,
1,EML1,BRSK1,PODXL2,FKBP1A,SAFB2,WSCD1,PPP1R10,ACLY,SSBP3,MAZ,IREB2,LSM14A,CN
NF428,SPAG9,CAP1,GOT2,ATG4B,PTPN23,DHX40,PAGR1,SRPRA,CALCOCO1,PLS3,SPPL2B,YTHD
P5F1B,CALM3,SHISA9,STMN2,RPS8,SEPTIN2,P4HB,RPL5,GDI1,SPTAN1,MARCKS,SMG1,CALM1,
RPL15,SERINC1,BASP1,POLR2A,TPT1,SEC61A1,RPS3A,RPL18,STMN1,ITGB1,MORF4L1,NCAM1,
31,PDIA3,SERINC1,RPS3,BASP1,VAPA,POLR2A,RPS3A,STMN1,RPL11,MORF4L1,BTF3,NCAM1,RF

TCH1,SERP1,RHBDD2,STK35,ERH,CLPTM1,RTL6,INTS3,ABI2,UBE2M,SEC14L1,NUDCD3,GNL1,F
DX2,RCN1,RPL35,MIF,PRKAR1A,NGRN,HNRNPDL,RPLP2,SHOX,HNRNPK,DUX4,PTGES3,LAMP1,

A2,CD63,GOLM1,CYFIP2,SLC25A3,SHOX,CIRBP,MYH10,DUX4,YWHAB,HNRNPH1,CKAP4,HLA-B,P
M7,VPS16,SZT2,EIF2S1,MAPK8IP3,DNMT1,CPSF7,TOMM40,ATP6V1D,M6PR,ATP5PD
3A,TUBB4B,BTF3,ANXA2,TIMP2,SBF1,CYB5R3,AKAP12,MAP4,ITM2B,ZNF770,SLC4A8,GOLGA8A,
ABARAPL1,GNL1,TM9SF4,TRAP1,PLEKHB2,ELP1,ACTR1B,MOAP1,CLASP2,NSD2,YY1,CCDC47,EL
,ASXL1,SCAF1,TERF2IP,TMED2,RAPGEF1,TUBB2A,PPP2CA,CCND1,MAPK1,DDAH2,PTBP1,FTO,IL
G,CKB,COL4A2,SPEN,NONO,DDB1,PSMD5,DCHS1,EGLN1,CHSY1,TMSB10,EIF4A1,MRC2,RPL15,
A1,MIDN,RPL10,WSB1,VCL,SBF1,ARF1,MAP4,PXDN,HNRNPU,FADS2,RPL23,CNN3,RPSA,LPCAT1,
31,KDM5A,SKIV2L,SHC1,CHMP5,SUDS3,GABARAPL2,MIB2,TNPO3,NUP50,SZT2,SEC31A,AKAP17
1A1,RPS4X,RPL11,MORF4L1,BTF3,NCAM1,ANXA2,RPLP1,ZMIZ1,RPS17,VCL,ITM2B,ZNF770,GTF
A,MAPRE2,PPP1R15A,GALNT11,DAP,EIF3H,ELAVL1,IMPDH2,GABARAPL1,GNL1,TM9SF4,TRAP1,F
A1,JUN,NGRN,RPLP2,ARL6IP1,MYL6,INIP,MYH10,SND1,UBE2H,ARCN1,HMGCR,TOP2B,SMCHD1,
B3,CRABP1,WARS1,SNRPB,KDM5A,SKIV2L,SHC1,ASB6,SUDS3,MIB2,TNPO3,CUL4B,SZT2,SEC31
X9,RPS29,NSMF,DUSP8,OGDH,ATP6V0A1,GUK1,ZNF146,PLCD3,CTDNEP1,CMTR1,CLCN3,TRAPP
TM3,MLF2,AKIRIN1,AMOTL2,YME1L1,CIC,TRAF4,STAT3,ARIH1,SF1,KPNA2,SLC35F5,CSNK1G2,
CRIB,LRP3,FKBP9,FBXW5,GSK3A,PRKACA,RBM6,CREBBP,NELFB,CALCOCO1,RIMS3,PSAT1,PIN
39B,TXLNA,PDAP1,PHB2,GSPT1,UBA52,RAB3GAP2,MCM3AP,PREPL,TMEM167A,PSMD8,FKBP1A,
A4,AHCYL1,KLC1,CLSTN3,DDX42,CHD2,RAB11B,NRDC,TUBB2A,OPTN,PPP2CA,G3BP2,PDE7A,PS

H2,GABARAPL1,GNL1,TM9SF4,TRAP1,PLEKHB2,ELP1,MOAP1,CLASP2,NSD2,YY1,CCDC47,ELOV
TCF25,ADAM9,EIF3E,CDC37,WDR6,BNIP3L,ARF4,HNRNPUL2-BSCL2,PITPNA,XPNPEP3,ATP6V1B2
K1G2,UPF1,TRA2B,MASTL,MAPRE2,PPP1R15A,GALNT11,DAP,EIF3H,ELAVL1,IMPDH2,GABARAPL

15A,GALNT11,DAP,EIF3H,ELAVL1,IMPDH2,GABARAPL1,GNL1,TM9SF4,TRAP1,PLEKHB2,ELP1,MC
CRTC1,ZFP91,CRABP2,SLC12A7,CASKIN1,COL7A1,NUMBL,GPR173,SGPL1,GPRIN1,MARVELD1,
MSI2,PAF1,PPM1A,RAD23A,EPHA2,STX12,VPS37B,ZC3H4,NDUFA10,CCDC124
MSI2,PAF1,PPM1A,RAD23A,EPHA2,STX12,VPS37B,ZC3H4,NDUFA10,CCDC124
,UPF1,MASTL,PRRC2A,MAPRE2,PPP1R15A,GALNT11,DAP,EIF3H,ELAVL1,IMPDH2,GABARAPL1,GI
HD1,TMEM30A,SKP1,CHD6,TOMM20,CCT2,CPSF1,GNAI3,SSRP1,HSPH1,HNRNPA3,C4ORF3,PKN1
NKS2,UBQLN1,CSNK1D,IFITM3,VGF,MLF2,AKIRIN1,AMOTL2,YME1L1,CTNND1,CIC,TRAF4,STAT3
IRNPDL,TSR2,MXD4,H1-10,CIRBP,RPL36,YWHAB,SEPTIN11,LAMP1,MAP2,SEPTIN3,CCT6A,STIP1,
IRNPDL,TSR2,MXD4,H1-10,CIRBP,RPL36,YWHAB,SEPTIN11,LAMP1,MAP2,SEPTIN3,CCT6A,STIP1,
SUPT6H,FBLN1,ISYNA1,LONP1,UBQLN1,TUG1,HYOU1,PTPRS,P3H3,FLNC,TCTN1,AKIRIN1,YME1
,TAPBP,OGFR,CGGBP1,CLIP2,KLF13,SIN3B,PFKP,REEP5,SLC25A6,OSBPL2,RNF157,RASA4,SAL
7,P4HB,IDS,FSCN1,PTMS,GSTP1,MARCKS,SMG1,NPM1,TTC3,PLD3,HSPD1,USP11,COL18A1,CSD

CSH,HNRNPC,KHDRBS1,SCRN1,RPS25,EIF5A,PDIA6,DDR1,MEGF8,ACSL6,RPL12,ITM2C,RPL30,
,TM9SF4,TRAP1,PLEKHB2,ELP1,MOAP1,CLASP2,NSD2,YY1,CCDC47,ELOVL5,SLC25A36,XRN2,S
GF,MLF2,AKIRIN1,DYNLL1,AMOTL2,YME1L1,CIC,TRAF4,STAT3,ARIH1,SF1,SLC35F5,CSNK1G2,C
B2,SEMA6B,CGGBP1,PSMB1,PCM1,TRMT112,CD2BP2,SUMO1,RPS21,BSCL2,SUB1,BANF1,GSI
BIP3,PRELID1,HARS1,MRPL3,DNMT1,CFAP298,DDX56,CPSF7,C2ORF68,RAD50,JRK,TOMM40,CC

,LARP1,FADS2,CNN3,NACA,BRD3,RPS9,XRCC5,CIPC,MSN,KPNB1,RPL14,CBX5,NR2F1,KIF5B,GL
RHGEF40,UBE4B,SLC38A1,CUL1,CRIP2,USP14,BSCL2,RASA4,EXT2,MAPKAPK2,SRSF7,GPT2,RE
.35A,DNAJC5,NDFIP1,TSR2,HNRNPK,H1-10,CIRBP,CADM4,HNRNPH1,UBE2H,SEPTIN3,ZNF454,C

P3,CTDSP2,DDX39B,CNN2,NOVA2,NUCB1,C5ORF24,ZC3H7B,CCT2,PGD,KLHL24,MEST,TM9SF2
H,SLIT3,IDH1,SOD1,BAG6,CYP51A1,SMS,CD99L2,DDIT4,SSR1,BCL9L,WDR45B,HIP1,BAZ1B,SLC

1,GPSM1,POM121C,HNRNPAB,GALNT11,SLC2A1,WSCD1,ACLY,RTL6,ARHGEF17,HINT1,RIC8A,E
K,HMGCS1,PRKAR1A,CHPF,GJA1,B4GALNT4,RPLP2,ARL6IP1,TSR2,MYL6,MYH10,RPL36,SEPTIN
4A,TCP1,PRPF6,TAF7,ATG2B,LUZP6,UBE2R2,ATXN10,NPEPPS,SON,RPL28,SUPT6H,NKTR,EIF3F,
CTIN2,MAF1,MAN1B1,ZDHHC17,PTPN12,SLC25A39,SMARCB1,PLXND1,LDOC1,SCCPDH,EIF3J,D

OLA,GAP43,RPL22,RHOA,CBX5,RPL9,PSMD2,RAB6B,ENC1,RPS20,EIF5,CD81,BEX3,CCT7,RBBP
F,GJA1,B4GALNT4,RPLP2,ARL6IP1,TSR2,MYL6,MYH10,RPL36,SEPTIN11,SND1,RHOB,UBE2H,S
ICM3AP,TMEM167A,TNKS,CTTN,LRRC59,DAP,MTCH1,SERP1,RHBDD2,ERH,PSMB4,IARS2,CLPTM
NL3,BCAR1,ZDHHC17,MFGE8,NF2,GPRC5B,CDK5RAP3,WDR33,TLNRD1,NOP58,ATP5PD
3,ATP5PD
DR,RPS20,CCT7,RTN3,ASTN2,PNN,HNRNPC,RPS25,TKT,CCND2,GPC3,SULF1,DDX3X,RPL30,CLT
15A,GALNT11,DAP,EIF3H,ELAVL1,IMPDH2,GABARAPL1,GNL1,TM9SF4,TRAP1,PLEKHB2,ELP1,MC
K2,RUVBL2,PFKP,ARHGEF40,MAPK8IP2,SLC25A6,MECP2,DNAJB1,KIF26A,CNDP2,RALGDS,PITF
RIH1,SF1,SLC35F5,CSNK1G2,UPF1,TRA2B,MASTL,MAPRE2,PPP1R15A,GALNT11,DAP,EIF3H,ELA
LNT11,DAP,EIF3H,ELAVL1,IMPDH2,GABARAPL1,GNL1,TM9SF4,TRAP1,PLEKHB2,ELP1,MOAP1,C
,ASB6,GPS1,THG1L,NUP50,SEC31A,FBXO7,MAPK8IP3,PRELID1,HARS1,MRPL3,DNMT1,CFAP298
X6B1,KDM5A,MYL12A,SHC1,ASB6,GABARAPL2,NUP50,SZT2,MAPK8IP3,PRELID1,DNMT1,DDX56

PLTP,SDF4,HNRNPM,GIGYF1,TUG1,PRPS1,IFITM3,UQCRC2,NCOR2,HNRNPD,VGF,FBRSL1,ARFG
HA2,EIF3J,DTD1

DX5,ENO2,SOX4,FKBP14,RPL13A,MAGED1,SEC62,RPL6,KAT6A,DVL3,YWHAE,KIF1A,RPL13,GNAS
O2,RPL13A,SEC62,KAT6A,DVL3,YWHAE,KIF1A,RPL13,GNAS,SLC2A3,GPI,HTRA4,RACK1,BSG,FAS
AT6A,DVL3,YWHAE,KIF1A,RPL13,GNAS,DDX17,GPI,HTRA4,RACK1,BSG,FASN,ALDH6A1,SPTBN1,A
1A,RPL13,GNAS,SLC2A3,DDX17,GPI,HTRA4,RACK1,BSG,FASN,ALDH6A1,SPTBN1,CALM3,GNB1,
DI1,FSCN1,PTMS,SPTAN1,GSTP1,MARCKS,SMG1,CALM1,PLXNB2,HSPD1,USP11,COL18A1,SNRN
DVL3,YWHAE,KIF1A,RPL13,GNAS,SLC2A3,DDX17,GPI,HTRA4,RACK1,BSG,FASN,ALDH6A1,SPTBN
,SOX4,RPL13A,SEC62,RPL6,KAT6A,DVL3,YWHAE,KIF1A,RPL13,GNAS,DDX17,GPI,HTRA4,RACK1,
B,COL4A2,SPEN,PSMD5,YWHAQ,EIF1,HACD3,DCHS1,SET,EGLN1,TPI1,CHSY1,INA,PPP2R1A,GTF
ALM3,GNB1,SHISA9,STMN2,MYH9,RPL7,SEPTIN2,P4HB,APP,ERVV-1,GDI1,FSCN1,TMBIM6,PTMS
PL13,GNAS,DDX17,GPI,HTRA4,RACK1,BSG,FASN,ALDH6A1,SPTBN1,ATP5F1B,CALM3,GNB1,SHIS
HAE,KIF1A,RPL13,GNAS,SLC2A3,GPI,HTRA4,RACK1,BSG,FASN,ALDH6A1,SPTBN1,CALM3,GNB1,
EPTIN2,P4HB,APP,GDI1,FSCN1,TMBIM6,PTMS,SPTAN1,GSTP1,MARCKS,SMG1,CALM1,PLXNB2,F
,MYH9,RPL7,SEPTIN2,P4HB,APP,IDS,GDI1,TMBIM6,PTMS,SPTAN1,GSTP1,MARCKS,SMG1,CALM
A9,STMN2,RPS8,MYH9,RPL7,SEPTIN2,P4HB,APP,IDS,GDI1,FSCN1,TMBIM6,PTMS,SPTAN1,GSTP1
DI1,FSCN1,TMBIM6,PTMS,SPTAN1,GSTP1,MARCKS,SMG1,CALM1,NPM1,PLXNB2,PLD3,HSPD1,U
KAT6A,DVL3,YWHAE,KIF1A,RPL13,GNAS,SLC2A3,DDX17,GPI,HTRA4,RACK1,BSG,FASN,ALDH6A1,
LM3,GNB1,SHISA9,STMN2,RPS8,MYH9,RPL7,SEPTIN2,P4HB,APP,GDI1,TMBIM6,PTMS,SPTAN1,GS
7,SEPTIN2,P4HB,APP,GDI1,FSCN1,TMBIM6,PTMS,SPTAN1,GSTP1,MARCKS,SMG1,CALM1,PLXNE
KSL1,AFF4,TRIM28,MATR3,YWHAG,CKB,RPL37A,LRRC28,SPEN,NONO,YWHAQ,HACD3,DCHS1,S
I1,CALM3,GNB1,SHISA9,STMN2,RPS8,MYH9,RPL7,SEPTIN2,P4HB,APP,GDI1,TMBIM6,PTMS,SPTA
TIN2,P4HB,APP,GDI1,FSCN1,PTMS,SPTAN1,GSTP1,MARCKS,SMG1,CALM1,NPM1,PLXNB2,PLD3,
KIF1A,RPL13,GNAS,GPI,HTRA4,BSG,FASN,SPTBN1,ATP5F1B,CALM3,GNB1,SHISA9,RPS8,MYH9,R
L7,SEPTIN2,P4HB,APP,RPL5,GDI1,TMBIM6,PTMS,SPTAN1,MARCKS,CALM1,NPM1,PLD3,HSPD1,U
,CANX,CLU,PTMA,DDX5,ENO2,SOX4,FKBP14,RPL13A,MAGED1,SEC62,RPL6,KAT6A,DVL3,YWHAE
5,GDI1,FSCN1,TMBIM6,PTMS,SPTAN1,GSTP1,SMG1,CALM1,NPM1,PLXNB2,HSPD1,USP11,COL1
A,RPL13,GNAS,DDX17,GPI,HTRA4,BSG,FASN,ALDH6A1,SPTBN1,CALM3,GNB1,SHISA9,STMN2,M
FSCN1,TMBIM6,PTMS,SPTAN1,GSTP1,CALM1,NPM1,PLXNB2,HSPD1,USP11,SNRNP200,CSDE1,IG
,GNAS,SLC2A3,DDX17,GPI,HTRA4,BSG,FASN,ALDH6A1,SPTBN1,CALM3,GNB1,SHISA9,STMN2,M
DDX5,ENO2,SOX4,FKBP14,RPL13A,SEC62,RPL6,KAT6A,DVL3,YWHAE,KIF1A,RPL13,GNAS,SLC2A
J,PTMA,DDX5,ENO2,SOX4,RPL13A,MAGED1,SEC62,RPL6,KAT6A,DVL3,YWHAE,KIF1A,RPL13,GN
SEC62,RPL6,KAT6A,DVL3,YWHAE,KIF1A,RPL13,GNAS,DDX17,HTRA4,RACK1,BSG,FASN,ALDH6A
,RPS8,MYH9,RPL7,SEPTIN2,APP,RPL5,GDI1,PTMS,SPTAN1,GSTP1,MARCKS,SMG1,PLXNB2,HSP
PTIN2,P4HB,APP,IDS,GDI1,FSCN1,TMBIM6,PTMS,SPTAN1,GSTP1,MARCKS,SMG1,CALM1,NPM1,
5,ENO2,SOX4,RPL13A,SEC62,RPL6,KAT6A,DVL3,YWHAE,KIF1A,RPL13,GNAS,DDX17,GPI,HTRA4,
H3-3B,CANX,CLU,PTMA,DDX5,ENO2,SOX4,FKBP14,RPL13A,MAGED1,SEC62,RPL6,KAT6A,DVL3,Y
1,USP11,COL18A1,SNRNP200,CSDE1,IGF2,EIF3C,ATP5F1A,EIF4B,MARCKSL1,AFF4,TRIM28,MA
3,GNAS,DDX17,GPI,HTRA4,RACK1,BSG,FASN,ALDH6A1,SPTBN1,ATP5F1B,CALM3,GNB1,SHISA9,
TP5F1B,CALM3,GNB1,SHISA9,MYH9,RPL7,SEPTIN2,P4HB,APP,GDI1,FSCN1,TMBIM6,PTMS,SPTA
1G1,CALM1,NPM1,PLXNB2,HSPD1,USP11,SNRNP200,CSDE1,SMIM14,IGF2,EIF3C,ATP5F1A,EIF4
AE,KIF1A,RPL13,GNAS,DDX17,GPI,RACK1,BSG,FASN,ALDH6A1,SPTBN1,CALM3,GNB1,SHISA9,ST
P14,RPL13A,MAGED1,SEC62,KAT6A,DVL3,YWHAE,KIF1A,RPL13,GNAS,DDX17,GPI,HTRA4,BSG,F
5,ENO2,SOX4,RPL13A,MAGED1,SEC62,RPL6,KAT6A,DVL3,YWHAE,KIF1A,RPL13,GNAS,SLC2A3,D
2,P4HB,APP,IDS,RPL5,FSCN1,PTMS,SPTAN1,SMG1,CALM1,NPM1,PLXNB2,PLD3,HSPD1,COL18A
,ALDH6A1,SPTBN1,CALM3,GNB1,SHISA9,STMN2,RPS8,MYH9,RPL7,SEPTIN2,P4HB,APP,IDS,RPL
YWHAE,KIF1A,GNAS,SLC2A3,DDX17,GPI,HTRA4,RACK1,BSG,FASN,ALDH6A1,SPTBN1,CALM3,GN

6,KAT6A,DVL3,YWHAE,KIF1A,RPL13,GNAS,SLC2A3,DDX17,GPI,HTRA4,RACK1,BSG,FASN,ALDH6/
H9,RPL7,SEPTIN2,P4HB,APP,RPL5,GDI1,FSCN1,TMBIM6,PTMS,SPTAN1,GSTP1,MARCKS,SMG1,C
,FN1,SPEN,NONO,YWHAQ,HACD3,DCHS1,SET,EGLN1,CHSY1,TMSB10,MRC2,LDHB,PKD1,HLA-C
)1,SEC62,KAT6A,DVL3,YWHAE,KIF1A,RPL13,GNAS,SLC2A3,DDX17,GPI,HTRA4,RACK1,BSG,FASN
TMA,DDX5,ENO2,SOX4,FKBP14,RPL13A,MAGED1,SEC62,RPL6,KAT6A,DVL3,YWHAE,KIF1A,RPL1
,HSPD1,USP11,CSDE1,SMIM14,IGF2,ATP5F1A,EIF4B,MARCKSL1,AFF4,TRIM28,MATR3,YWHAG,C
1,SNRNP200,CSDE1,IGF2,EIF3C,ATP5F1A,EIF4B,MARCKSL1,AFF4,TRIM28,MATR3,YWHAG,CKB,
,GPI,RACK1,BSG,FASN,SPTBN1,ATP5F1B,CALM3,GNB1,SHISA9,STMN2,RPS8,MYH9,RPL7,SEPTI
MRC2,PKD1,HLA-C,PDIA3,BASP1,VAPA,HMGB1,POLR2A,TPT1,STMN1,ITGB1,TUBB4B,UCHL1,M
RA4,RACK1,BSG,FASN,ALDH6A1,SPTBN1,ATP5F1B,CALM3,GNB1,SHISA9,RPS8,MYH9,RPL7,SEPT
,AP2M1,TPT1,STMN1,ITGB1,TUBB4B,UCHL1,MORF4L1,BTF3,NCAM1,ANXA2,AGRN,ZMIZ1,TIMF
310,INA,MRC2,LDHB,PKD1,HLA-C,PDIA3,BASP1,VAPA,POLR2A,TPT1,RPL18,STMN1,ITGB1,TUBB
CKB,LRRC28,SPEN,NONO,DDB1,YWHAQ,HACD3,DCHS1,SET,EGLN1,TPI1,CHSY1,TMSB10,INA,MF
VHAG,CKB,RPL37A,LRRC28,FN1,SPEN,NONO,YWHAQ,HACD3,DCHS1,SET,EGLN1,TMSB10,RPL1
,RACK1,FASN,SPTBN1,ATP5F1B,CALM3,GNB1,SHISA9,RPS8,MYH9,RPL7,SEPTIN2,P4HB,APP,ER
STMN1,ITGB1,TUBB4B,UCHL1,RPL11,MORF4L1,BTF3,NCAM1,AGRN,ZMIZ1,TIMP2,MIDN,WSB1,
VHAE,KIF1A,RPL13,GNAS,SLC2A3,DDX17,GPI,HTRA4,RACK1,BSG,FASN,SPTBN1,ATP5F1B,CALM
TP5F1A,EIF4B,MARCKSL1,AFF4,TRIM28,MATR3,YWHAG,CKB,RPL37A,LRRC28,FN1,NONO,DDB1
C,SSBP2,VAPA,POLR2A,TPT1,STMN1,ITGB1,TUBB4B,UCHL1,MORF4L1,NCAM1,ANXA2,AGRN,TI
I1,GSTP1,MARCKS,SMG1,PLXNB2,HSPD1,USP11,SNRNP200,IGF2,EIF3C,ATP5F1A,EIF4B,MARCK
SG,FASN,SPTBN1,ATP5F1B,CALM3,GNB1,SHISA9,STMN2,MYH9,SEPTIN2,P4HB,APP,IDS,ERVV-1,
2,PKD1,HLA-C,PDIA3,BASP1,VAPA,POLR2A,TPT1,STMN1,ITGB1,TUBB4B,UCHL1,NCAM1,ANXA2,
RA4,RACK1,BSG,FASN,SPTBN1,ATP5F1B,CALM3,GNB1,SHISA9,RPS8,MYH9,RPL7,SEPTIN2,P4HB
4A2,RPL37A,DDB1,YWHAQ,HACD3,DCHS1,SET,EGLN1,CHSY1,INA,MRC2,RPL19,PPP2R1A,LDHB,
,DCHS1,SET,CHSY1,TMSB10,EIF4A1,MRC2,PKD1,HLA-C,PDIA3,VAPA,POLR2A,TPT1,STMN1,ITGB
,TRIM28,MATR3,YWHAG,CKB,COL4A2,LRRC28,SPEN,DDB1,PSMD5,YWHAQ,EIF1,HACD3,DCHS1
S8,MYH9,SEPTIN2,P4HB,RPL5,GDI1,FSCN1,TMBIM6,PTMS,SPTAN1,GSTP1,MARCKS,SMG1,CALM
RPL37A,LRRC28,FN1,NONO,YWHAQ,HACD3,DCHS1,SET,TMSB10,INA,EIF4A1,MRC2,RPL19,LDH
X,CLU,PTMA,DDX5,ENO2,SOX4,RPL13A,MAGED1,SEC62,RPL6,KAT6A,DVL3,YWHAE,KIF1A,RPL13
H6A1,SPTBN1,CALM3,GNB1,SHISA9,MYH9,RPL7,SEPTIN2,P4HB,GDI1,FSCN1,TMBIM6,PTMS,SPT
HAE,KIF1A,RPL13,GNAS,SLC2A3,DDX17,GPI,HTRA4,RACK1,BSG,FASN,ALDH6A1,SPTBN1,ATP5F1
R3,YWHAG,CKB,RPL37A,LRRC28,YWHAQ,HACD3,DCHS1,SET,EGLN1,CHSY1,TMSB10,INA,EIF4A1
SLC2A3,DDX17,GPI,BSG,FASN,ALDH6A1,SPTBN1,GNB1,SHISA9,STMN2,MYH9,RPL7,SEPTIN2,P4H
,GNAS,SLC2A3,DDX17,GPI,HTRA4,RACK1,BSG,FASN,SPTBN1,ATP5F1B,CALM3,GNB1,SHISA9,RP
FSCN1,TMBIM6,SPTAN1,SMG1,CALM1,PLXNB2,TTC3,HSPD1,USP11,COL18A1,SNRNP200,CSDE1
F2,ATP5F1A,EIF4B,MARCKSL1,UBB,AFF4,TRIM28,MATR3,YWHAG,CKB,LRRC28,NONO,DDB1,Y
C,PDIA3,SSBP2,BASP1,VAPA,POLR2A,TPT1,STMN1,ITGB1,TUBB4B,UCHL1,NCAM1,ANXA2,AGRN
LP0,H3-3B,CANX,CLU,PTMA,DDX5,ENO2,SOX4,RPL13A,SEC62,RPL6,KAT6A,DVL3,YWHAE,KIF1A,
IM28,YWHAG,CKB,LRRC28,FN1,SPEN,NONO,DDB1,YWHAQ,HACD3,DCHS1,SET,EGLN1,TPI1,TM
4,IGF2,EIF3C,ATP5F1A,EIF4B,MARCKSL1,AFF4,TRIM28,MATR3,YWHAG,CKB,RPL37A,LRRC28,F
4,IGF2,EIF3C,ATP5F1A,EIF4B,MARCKSL1,AFF4,TRIM28,MATR3,YWHAG,CKB,RPL37A,LRRC28,F
,DCHS1,SET,EGLN1,CHSY1,INA,MRC2,RPL19,PPP2R1A,LDHB,PKD1,HLA-C,SSBP2,VAPA,AP2M1,
LP0,H3-3B,CANX,CLU,PTMA,DDX5,ENO2,SOX4,FKBP14,RPL13A,MAGED1,SEC62,RPL6,KAT6A,DV
8,YWHAG,CKB,COL4A2,RPL37A,FN1,NONO,DDB1,PSMD5,YWHAQ,HACD3,DCHS1,SET,EGLN1,M
,VCP,MARCKSL1,UBB,MATR3,YWHAG,COL4A2,FN1,SPEN,NONO,YWHAQ,EIF1,HACD3,SET,EGLN

P200,SMIM14,IGF2,EIF3C,ATP5F1A,EIF4B,MARCKSL1,AFF4,TRIM28,MATR3,YWHAG,CKB,RPL37
7A,LRRC28,FN1,NONO,DDB1,YWHAQ,HACD3,DCHS1,EGLN1,TMSB10,INA,MRC2,LDHB,PKD1,HLA
),DDB1,YWHAQ,HACD3,DCHS1,SET,TMSB10,INA,EIF4A1,MRC2,RPL19,LDHB,PKD1,HLA-C,SERIN
MSB10,MRC2,LDHB,PKD1,HLA-C,PDIA3,SSBP2,BASP1,POLR2A,TPT1,STMN1,ITGB1,TUBB4B,MOI
GSTP1,MARCKS,SMG1,CALM1,NPM1,PLXNB2,USP11,SMIM14,IGF2,EIF3C,ATP5F1A,EIF4B,MAR
1,RPLP0,H3-3B,CANX,CLU,PTMA,DDX5,ENO2,SOX4,RPL13A,SEC62,RPL6,KAT6A,DVL3,YWHAE,KI
RCHF6,RPS17,VCL,DNAJA1,ARF1,CYB5R3,CDC42BPB,ITM2B,PXDN,HNRNPU,PGAM1,HNRNPA0,
CKSL1,AFF4,TRIM28,MATR3,YWHAG,CKB,LRRC28,FN1,NONO,DDB1,YWHAQ,HACD3,DCHS1,SE
DX5,ENO2,SOX4,FKBP14,RPL13A,SEC62,RPL6,KAT6A,DVL3,YWHAE,KIF1A,RPL13,GNAS,SLC2A3,
P11,SMIM14,IGF2,EIF3C,EIF4B,MARCKSL1,AFF4,TRIM28,MATR3,YWHAG,CKB,COL4A2,LRRC28
2,P4HB,GDI1,FSCN1,TMBIM6,PTMS,SPTAN1,GSTP1,SMG1,CALM1,NPM1,PLXNB2,HSPD1,USP11,
RCKSL1,AFF4,TRIM28,MATR3,YWHAG,CKB,LRRC28,FN1,NONO,DDB1,YWHAQ,EIF1,HACD3,DCH
C2,RPL19,PPP2R1A,LDHB,PKD1,HLA-C,SSBP2,VAPA,AP2M1,TPT1,RPL18,STMN1,ITGB1,TUBB4B,
HAG,CKB,COL4A2,LRRC28,NONO,DDB1,YWHAQ,EIF1,HACD3,DCHS1,SET,TMSB10,INA,EIF4A1,M
,PTMA,DDX5,ENO2,SOX4,FKBP14,RPL13A,MAGED1,SEC62,KAT6A,DVL3,YWHAE,KIF1A,RPL13,G
PEN,PSMD5,YWHAQ,EIF1,HACD3,SET,EGLN1,TPI1,CHSY1,TMSB10,EIF4A1,GTF3C4,LDHB,PKD1,
,MAP4,CDC42BPB,ITM2B,MAT2A,PXDN,HNRNPU,CTNNA1,CTSD,LAMA5,BRD3,CIPC,PJA2,KPNB1,
MN2,RPS8,MYH9,RPL7,SEPTIN2,IDS,GDI1,TMBIM6,SPTAN1,GSTP1,MARCKS,SMG1,PLXNB2,PLD
SET,EGLN1,TPI1,CHSY1,INA,EIF4A1,MRC2,PPP2R1A,RPL15,GTF3C4,LDHB,PKD1,PDIA3,SSBP2,Z
T1,STMN1,ITGB1,MORF4L1,NCAM1,ANXA2,AGRN,ZMIZ1,TIMP2,COL11A1,MIDN,RPL10,MARCH
G,CKB,COL4A2,FN1,SPEN,DDB1,YWHAQ,EIF1,HACD3,DCHS1,SET,EGLN1,TPI1,CHSY1,TMSB10,I
,AGRN,RPLP1,ZMIZ1,TIMP2,MIDN,RPL10,MARCHF6,RPS17,VCL,SBF1,DNAJA1,ARF1,CYB5R3,M
2,RPS16,CPE,CNN3,CTSD,LPCAT1,LAMA5,SLC22A17,BRD3,CIPC,PJA2,SLC16A7,KPNB1,COL1A2
IA5,BRD3,KPNB1,PAPOLA,IGFBP3,COL1A2,LLGL1,HK1,NCS1,RPL22,RHOA,ZNF865,CLSTN1,CHD
MBIM6,PTMS,SPTAN1,GSTP1,MARCKS,SMG1,CALM1,NPM1,PLXNB2,USP11,COL18A1,CSDE1,SM
A5,BRD3,IGFBP3,LLGL1,NCS1,RPL22,RHOA,CLSTN1,ATF4,RAB6B,SLC7A5,ENC1,CXADR,ATP1A
STP1,SMG1,CALM1,PLXNB2,TTC3,PLD3,HSPD1,USP11,COL18A1,SNRNP200,SMIM14,IGF2,VCP,
.7,SEPTIN2,APP,RPL5,GDI1,FSCN1,TMBIM6,PTMS,SPTAN1,GSTP1,SMG1,CALM1,NPM1,USP11,C
PLXNB2,HSPD1,USP11,CSDE1,IGF2,ATP5F1A,EIF4B,VCP,MARCKSL1,UBB,AFF4,TRIM28,MATR3,
YH9,SEPTIN2,APP,IDS,GDI1,PTMS,SPTAN1,GSTP1,SMG1,CALM1,NPM1,PLXNB2,USP11,COL18A1
,PTMA,DDX5,SOX4,FKBP14,RPL13A,MAGED1,SEC62,RPL6,KAT6A,DVL3,YWHAE,KIF1A,RPL13,G
RPL37A,FN1,SPEN,YWHAQ,HACD3,DCHS1,SET,EGLN1,TPI1,CHSY1,TMSB10,INA,EIF4A1,MRC2,PP
F1,CDC42BPB,ITM2B,PXDN,HNRNPU,OGA,NFE2L1,CTNNA1,CNN3,TNS3,LAMA5,SLC22A17,BRD3,
,YWHAQ,EIF1,HACD3,SET,TMSB10,LDHB,PKD1,HLA-C,PDIA3,VAPA,HMGB1,POLR2A,AP2M1,TPT
RPL7,SEPTIN2,P4HB,APP,GDI1,FSCN1,PTMS,SPTAN1,GSTP1,MARCKS,SMG1,CALM1,NPM1,PLXN
A4,RACK1,BSG,FASN,ALDH6A1,SPTBN1,CALM3,GNB1,SHISA9,STMN2,RPS8,MYH9,SEPTIN2,P4H
RF43,LLGL1,NCS1,RHOA,CLSTN1,ATF4,JUND,EIF5,ATP2A2,RPL26,OAZ1,RBBP4,JPT2,CCNI,TAD
D1,SSBP2,BASP1,VAPA,POLR2A,TPT1,STMN1,ITGB1,MORF4L1,NCAM1,ANXA2,AGRN,ZMIZ1,TIM
ICS1,RHOA,ZNF865,SLC7A5,CHD4,ATP1A1,JUND,EIF5,CD81,WDR1,HMGN2,ATP2A2,CRMP1,LAS
4B,MARCKSL1,UBB,MATR3,YWHAG,CKB,COL4A2,FN1,NONO,PSMD5,YWHAQ,EIF1,HACD3,DCHS
RINC1,BASP1,VAPA,AP2M1,TPT1,SEC61A1,RPS4X,STMN1,ITGB1,TUBB4B,UCHL1,RPL11,NCAM1,
,UCHL1,MORF4L1,BTF3,AGRN,RPLP1,ZMIZ1,TIMP2,SARAF,MIDN,RPL10,VCL,DNAJA1,ARF1,CYB
E1,IGF2,ATP5F1A,EIF4B,MARCKSL1,AFF4,TRIM28,MATR3,YWHAG,CKB,COL4A2,LRRC28,FN1,N
HS1,TMSB10,INA,MRC2,RPL19,RPL15,LDHB,PKD1,HLA-C,VAPA,HMGB1,POLR2A,AP2M1,TPT1,SE
TMA,DDX5,ENO2,SOX4,FKBP14,RPL13A,SEC62,RPL6,KAT6A,DVL3,YWHAE,KIF1A,RPL13,GNAS,S

P,MARCKSL1,UBB,TRIM28,MATR3,YWHAG,CKB,COL4A2,LRRC28,FN1,SPEN,DDB1,YWHAQ,HACD
A2,C1ORF43,PAPOLA,RPL14,NCS1,RHOA,CLSTN1,PSMD2,NR2F1,EIF5,CD81,WDR1,ATXN7L3B,H
MIZ1,TIMP2,SARAF,MIDN,MARCHF6,WSB1,VCL,ARF1,CYB5R3,MAP4,SF3B2,CDC42BPB,ITM2B,M
,SMG1,CALM1,PLXNB2,PLD3,SNRNP200,CSDE1,IGF2,EIF3C,EIF4B,MARCKSL1,UBB,TRIM28,YW
9,STMN2,RPS8,MYH9,SEPTIN2,P4HB,APP,IDS,ERVV-1,RPL5,GDI1,FSCN1,PTMS,SPTAN1,GSTP1,SI
PTMS,SPTAN1,GSTP1,MARCKS,SMG1,CALM1,NPM1,PLXNB2,USP11,SNRNP200,CSDE1,IGF2,ATP
,SET,EGLN1,CHSY1,TMSB10,INA,EIF4A1,MRC2,RPL19,LDHB,PKD1,HLA-C,VAPA,POLR2A,CDKL1,
F4B,VCP,MARCKSL1,UBB,TRIM28,MATR3,YWHAG,CKB,COL4A2,MAPT,FN1,SPEN,PSMD5,YWHAQ
,BSG,FASN,ALDH6A1,SPTBN1,ATP5F1B,CALM3,GNB1,SHISA9,STMN2,RPS8,MYH9,RPL7,SEPTIN2
L6,KAT6A,DVL3,YWHAE,KIF1A,RPL13,GNAS,SLC2A3,DDX17,GPI,HTRA4,RACK1,BSG,FASN,SPTBN
MP2,MARCHF6,WSB1,RPS17,AP3D1,SBF1,ARF1,CYB5R3,ZNF770,AP2B1,GTF2F1,SLC4A8,HNRNP
,PXDN,HNRNPU,CTNNA1,CTSD,LPCAT1,BRD3,CIPC,PJA2,MSN,LLGL1,NCS1,RHOA,ZNF865,CLSTI
IPC,PJA2,KPNB1,LLGL1,NCS1,ZNF865,SLC7A5,CHD4,ATP1A1,JUND,CD81,WDR1,HMGN2,ATP2A
L1,UBB,MATR3,YWHAG,CKB,SPEN,NONO,DDB1,PSMD5,YWHAQ,DCHS1,TPI1,TMSB10,INA,EIF4A
TRIM28,CKB,COL4A2,LRRC28,FN1,SPEN,DDB1,YWHAQ,DCHS1,SET,EGLN1,MAP9,TPI1,CHSY1,T
N1,PTMS,SPTAN1,GSTP1,SMG1,CALM1,PLXNB2,PLD3,HSPD1,USP11,COL18A1,SNRNP200,CSDE
SD,EIF4H,LAMA5,SLC22A17,BRD3,PJA2,KPNB1,C1ORF43,LLGL1,HK1,NCS1,RHOA,ZNF865,CLST
PXDN,OGA,HNRNPA0,CTNNA1,FADS2,LDHA,CPE,CNN3,TPM4,LAMA5,SLC22A17,BRD3,CIPC,SLC1
RPL6,KAT6A,DVL3,YWHAE,KIF1A,RPL13,GNAS,GPI,HTRA4,RACK1,BSG,FASN,ALDH6A1,SPTBN1,A
SHISA9,RPS8,RPL7,SEPTIN2,P4HB,APP,ERVV-1,GDI1,FSCN1,PTMS,SPTAN1,GSTP1,MARCKS,SMG
A1,SPTBN1,ATP5F1B,CALM3,GNB1,SHISA9,STMN2,RPS8,MYH9,RPL7,P4HB,APP,RPL5,GDI1,FSCN
F4,TRIM28,MATR3,YWHAG,CKB,RPL37A,FN1,NONO,DDB1,YWHAQ,HACD3,DCHS1,SET,EGLN1,TI
I,SNRNP200,SMIM14,IGF2,EIF4B,VCP,MARCKSL1,UBB,TRIM28,MATR3,CKB,COL4A2,LRRC28,FI
3,PSAP,PGK1,RPLP0,H3-3B,CANX,CLU,PTMA,DDX5,ENO2,SOX4,RPL13A,MAGED1,SEC62,RPL6,K
BRD3,MSN,KPNB1,C1ORF43,IGFBP3,NCS1,RPL22,ZNF865,ATF4,NR2F1,NLGN2,ATP1A1,JUND,E
STMN1,ITGB1,TUBB4B,UCHL1,BTF3,AGRN,RPLP1,ZMIZ1,TIMP2,SARAF,COL11A1,MIDN,MARCHI
PNB1,C1ORF43,PAPOLA,HK1,RPL22,CBX5,ZNF865,ATF4,PSMD2,NR2F1,RAB6B,CHD4,ATP1A1,JU
YWHAE,KIF1A,RPL13,GNAS,SLC2A3,DDX17,GPI,RACK1,BSG,FASN,ALDH6A1,SPTBN1,ATP5F1B,C
P2,XAF1,SARAF,COL11A1,MIDN,MARCHF6,VCL,AP3D1,ARF1,CYB5R3,MAP4,CDC42BPB,ITM2B,M
H6A1,ATP5F1B,CALM3,RPS8,RPL7,SEPTIN2,P4HB,APP,IDS,RPL5,GDI1,FSCN1,TMBIM6,PTMS,SPT
LSTN1,NR2F1,ENC1,NLGN2,CHD4,ATP1A1,RPS20,LENG8,WDR1,ATP2A2,CRMP1,VAT1,RBBP4,PF
2,SARAF,MIDN,MARCHF6,VCL,AP3D1,SBF1,ARF1,AKAP12,MAP4,SF3B2,CDC42BPB,ITM2B,GOLG
B,FN1,NONO,DDB1,YWHAQ,HACD3,DCHS1,SET,EGLN1,TPI1,CHSY1,TMSB10,INA,EIF4A1,PPP2R1
EGLN1,TPI1,MRC2,HLA-C,SERINC1,SSBP2,BASP1,TPT1,SEC61A1,RPS4X,STMN1,ITGB1,TUBB4B,I
PS8,MYH9,RPL7,SEPTIN2,P4HB,APP,ERVV-1,RPL5,GDI1,FSCN1,TMBIM6,SPTAN1,GSTP1,MARCKS
ND,CD81,LENG8,WDR1,HMGN2,ATP2A2,CRMP1,LASP1,VAT1,OAZ1,PFN1,TLE5,NISCH,HNRNPC,C
C,PJA2,KPNB1,LLGL1,NCS1,ZNF865,CLSTN1,SLC7A5,CHD4,ATP1A1,JUND,CD81,HMGN2,ATP2A2
M1,PLXNB2,HSPD1,USP11,SNRNP200,CSDE1,SMIM14,IGF2,EIF3C,ATP5F1A,EIF4B,MARCKSL1,UI
P1,VAPA,AP2M1,TPT1,STMN1,ITGB1,TUBB4B,UCHL1,MORF4L1,BTF3,AGRN,TIMP2,COL11A1,M
VAPA,HMGB1,POLR2A,CDKL1,AP2M1,TPT1,RPS3A,STMN1,TUBB4B,UCHL1,BTF3,ANXA2,AGRN,Z
CALM1,NPM1,HSPD1,USP11,SNRNP200,CSDE1,IGF2,EIF3C,EIF4B,MARCKSL1,UBB,AFF4,MATR3,
CND2,CDH2,DDR1,PIP4K2B,SOX12,DDX3X,ITM2C,GNB2,H1-0,APLP1,MIF,SESN3,B4GALNT4,DNA
F1,DCHS1,EGLN1,MAP9,TPI1,CHSY1,EIF4A1,MRC2,RPL19,PPP2R1A,PKD1,SSBP2,ZNF43,RPS3,V
F1,DCHS1,EGLN1,MAP9,TPI1,CHSY1,EIF4A1,MRC2,RPL19,PPP2R1A,PKD1,SSBP2,ZNF43,RPS3,V
F1,DCHS1,EGLN1,MAP9,TPI1,CHSY1,EIF4A1,MRC2,RPL19,PPP2R1A,PKD1,SSBP2,ZNF43,RPS3,V

JSP11,SNRNP200,SMIM14,IGF2,EIF3C,EIF4B,MARCKSL1,UBB,AFF4,TRIM28,MATR3,YWHAG,CKI
1,CHSY1,EIF4A1,PPP2R1A,RPL15,LDHB,HLA-C,VAPA,HMGB1,AP2M1,TPT1,RPL18,STMN1,ITGB1,
DI1,FSCN1,TMBIM6,PTMS,MARCKS,SMG1,CALM1,NPM1,PLXNB2,TTC3,PLD3,HSPD1,SNRNP200,
PT1,STMN1,TUBB4B,UCHL1,MORF4L1,AGRN,TIMP2,COL11A1,MIDN,MARCHF6,VCL,SF3B2,CDC4
P1,LASP1,VAT1,OAZ1,TLE5,NISCH,HNRNPC,CCNI,XPO1,HDLBP,EIF5A,SOX12,DDX3X,H1-0,MIF,SE
X4,RPL13A,RPL6,KAT6A,DVL3,YWHAE,KIF1A,RPL13,GNAS,SLC2A3,DDX17,BSG,FASN,ALDH6A1,S
,UCHL1,MORF4L1,NCAM1,AGRN,RPLP1,ZMIZ1,TIMP2,SARAF,MIDN,AP3D1,DNAJA1,ARF1,CYB5F
IACD3,TPI1,CHSY1,TMSB10,EIF4A1,RPL19,RPL15,GTF3C4,LDHB,PKD1,SF3B1,PDIA3,SSBP2,CHD
3,AFF4,TRIM28,YWHAG,CKB,COL4A2,YWHAQ,EIF1,HACD3,DCHS1,SET,EGLN1,TPI1,CHSY1,TMSE
8A,PXDN,HNRNPU,CTNNA1,RPS16,CNN3,CTSD,NACA,LPCAT1,LAMA5,BRD3,CIPC,PJA2,KPNB1,LL
HPF,JUN,COL6A2,SESN3,B4GALNT4,MTSS2,DNAJC5,APLP2,MXD4,H1-10,MYH10,PTGES3,CKAP5
,SHISA9,RPS8,MYH9,SEPTIN2,P4HB,APP,GDI1,FSCN1,TMBIM6,SPTAN1,MARCKS,SMG1,CALM1,N
A9,STMN2,RPS8,MYH9,RPL7,SEPTIN2,P4HB,APP,IDS,GDI1,FSCN1,TMBIM6,PTMS,SPTAN1,GSTP1,
),EIF5,CD81,WDR1,HMGN2,ATP2A2,CRMP1,LASP1,OAZ1,PFN1,ASTN2,TLE5,NISCH,CCNI,TADA2E
M28,MATR3,YWHAG,COL4A2,RPL37A,DDB1,EIF1,DCHS1,SET,MAP9,TPI1,CHSY1,TMSB10,INA,MF
I1,TMBIM6,SPTAN1,MARCKS,SMG1,PLXNB2,PLD3,HSPD1,USP11,COL18A1,SNRNP200,SMIM14,I
P14,RPL13A,MAGED1,SEC62,RPL6,KAT6A,DVL3,YWHAE,KIF1A,RPL13,GNAS,DDX17,GPI,HTRA4,F
SMARCA5,HNRNPA0,FADS2,LDHA,CNN3,RPSA,TNS3,EIF4H,LPCAT1,LAMA5,BRD3,RPS9,XRCC5,CI
DX17,GPI,HTRA4,RACK1,BSG,FASN,ALDH6A1,SPTBN1,ATP5F1B,CALM3,GNB1,SHISA9,STMN2,RF
HMGB1,POLR2A,AP2M1,TPT1,SEC61A1,RPS3A,STMN1,TUBB4B,UCHL1,BTF3,ANXA2,AGRN,TIMF
2,DDX3X,ARL4C,GNB2,RCN1,APLP1,CHPF,SESN3,B4GALNT4,MTSS2,DNAJC5,APLP2,TARDBP,M
CALM3,GNB1,RPS8,RPL7,SEPTIN2,P4HB,APP,IDS,ERVV-1,FSCN1,PTMS,SPTAN1,GSTP1,SMG1,C
2R1A,LDHB,DNAH8,SF3B1,SSBP2,VAPA,AP2M1,TPT1,RPL18,STMN1,ITGB1,UCHL1,MORF4L1,NC
,RPS16,RPL29,CTSD,EIF4H,SCARB2,LPCAT1,RPS24,BRD3,CIPC,LLGL1,HK1,NCS1,ZNF865,CLST
F4L1,ANXA2,AGRN,ZMIZ1,TIMP2,COL11A1,MIDN,MARCHF6,VCL,AP3D1,DNAJA1,CYB5R3,MAP4,
SNRNP200,SMIM14,IGF2,EIF4B,VCP,MARCKSL1,UBB,AFF4,TRIM28,YWHAG,LRRC28,FN1,NONO
PCAT1,LAMA5,SLC22A17,BRD3,PAPOLA,IGFBP3,LLGL1,HK1,NCS1,RPL22,RHOA,ZNF865,CLSTN
9,RPL7,SEPTIN2,P4HB,APP,IDS,RPL5,GDI1,FSCN1,TMBIM6,SPTAN1,MARCKS,SMG1,CALM1,NPM
PD1,USP11,SNRNP200,SMIM14,IGF2,MARCKSL1,UBB,TRIM28,MATR3,CKB,COL4A2,LRRC28,FN
L4A2,MAPT,PSMD5,YWHAQ,EIF1,HACD3,DCHS1,SET,TPI1,CHSY1,TMSB10,INA,EIF4A1,PPP2R1A,F
PL15,GTF3C4,LDHB,PKD1,SF3B1,PDIA3,SERINC1,SSBP2,BASP1,VAPA,POLR2A,TPT1,SEC61A1,R
-C,SF3B1,SSBP2,BASP1,VAPA,HMGB1,POLR2A,TPT1,STMN1,ITGB1,TUBB4B,UCHL1,MORF4L1,A
HNRNPU,SMARCA5,CTNNA1,FADS2,CPE,CNN3,NACA,LPCAT1,LAMA5,BRD3,KPNB1,PAPOLA,IGFE
S,ERVV-1,RPL5,GDI1,FSCN1,TMBIM6,PTMS,SPTAN1,MARCKS,SMG1,CALM1,PLXNB2,PLD3,USP
TR3,YWHAG,CKB,COL4A2,NONO,YWHAQ,HACD3,DCHS1,SET,EGLN1,TMSB10,EIF4A1,MRC2,PPF
L4A1,CES3,PSAP,PGK1,RPLP0,H3-3B,CANX,CLU,PTMA,DDX5,ENO2,SOX4,FKBP14,RPL13A,MAGI
TN1,PSMD2,ENC1,RPS14,CHD4,EIF5,CD81,WDR1,CRMP1,CCT7,OAZ1,RTN3,NISCH,PRKCSH,CCI
CNI,KHDRBS1,XPO1,CSNK1E,PFKL,SFPQ,CDH2,PDIA6,MEGF8,SOX12,DDX3X,ARL4C,GNB2,APLP1
NISCH,CCNI,TADA2B,KHDRBS1,CSNK1E,PFKL,SFPQ,TPM1,CDH2,PDIA6,SOX12,DDX3X,ARL4C,GN
MA,DDX5,ENO2,SOX4,RPL13A,KAT6A,DVL3,YWHAE,KIF1A,RPL13,GNAS,SLC2A3,DDX17,GPI,RACK
,HACD3,SET,TPI1,TMSB10,RPL19,RPL15,GTF3C4,PKD1,PDIA3,SSBP2,BASP1,VAPA,POLR2A,CDK
GF2,EIF3C,ATP5F1A,EIF4B,VCP,UBB,AFF4,TRIM28,MATR3,YWHAG,CKB,COL4A2,LRRC28,FN1,SF
MSB10,EIF4A1,RPL19,RPL15,LDHB,SF3B1,PDIA3,SSBP2,BASP1,VAPA,POLR2A,TPT1,SEC61A1,STI
TIMP2,SARAF,MIDN,MARCHF6,VCL,AP3D1,ARF1,AKAP12,SF3B2,CDC42BPB,ITM2B,GOLGA8A,PXI
SEPTIN2,P4HB,APP,RPL5,GDI1,FSCN1,TMBIM6,PTMS,SPTAN1,GSTP1,MARCKS,SMG1,CALM1,NI

8,SPEN,DDB1,YWHAQ,EIF1,DCHS1,SET,EGLN1,TPI1,CHSY1,INA,EIF4A1,MRC2,PPP2R1A,RPL15,L
,YWHAG,CKB,COL4A2,LRRC28,SPEN,DDB1,YWHAQ,EIF1,HACD3,DCHS1,SET,EGLN1,TPI1,CHSY1
RN,RPLP1,TIMP2,MIDN,RPL10,MARCHF6,WSB1,VCL,AP3D1,DNAJA1,CYB5R3,ITM2B,GOLGA8A,F
,ITM2B,MAT2A,TMEM170A,GTF2F1,HNRNPU,NFE2L1,HNRNPA0,RPS16,CTSD,LPCAT1,LAMA5,BR
/1,TMSB10,MRC2,PPP2R1A,LDHB,DNAH8,PKD1,HLA-C,PDIA3,SERINC1,SSBP2,RPS3,VAPA,POLR2
,CIPC,KPNB1,PAPOLA,COL1A2,LLGL1,NCS1,RHOA,CBX5,CLSTN1,ATF4,ATP1A1,TMEM259,EIF5,
3,YWHAG,CKB,COL4A2,SPEN,NONO,YWHAQ,EIF1,HACD3,DCHS1,SET,EGLN1,CHSY1,TMSB10,IN
L1,ANXA2,AGRN,XAF1,SARAF,MIDN,AP3D1,ARF1,CYB5R3,MAP4,CDC42BPB,ITM2B,MAT2A,PXDN,
1,ZNF865,CLSTN1,SLC7A5,CHD4,JUND,CD81,HMGN2,ATP2A2,CRMP1,LASP1,OAZ1,PFN1,TLE5,N
1,ZNF865,CLSTN1,SLC7A5,CHD4,JUND,CD81,HMGN2,ATP2A2,CRMP1,LASP1,OAZ1,PFN1,TLE5,N
2,MAPK1,SMARCA4,SRSF9,DSTN,PBX2,DYNC1LI2,PTBP1,DYNLL2,ILF3,LUZP6,CAPZB,UBE2R2,AT
865,KIF5B,RAB6B,GLIPR1L2,LENG8,WDR1,ATXN7L3B,ATP2A2,VAT1,OAZ1,SRSF4,RTN3,PFN1,PN
,SEC62,KAT6A,DVL3,YWHAE,KIF1A,RPL13,GNAS,SLC2A3,DDX17,HTRA4,RACK1,BSG,FASN,SPTB
DCHS1,TMSB10,INA,MRC2,PPP2R1A,RPL15,GTF3C4,PKD1,PDIA3,SSBP2,CHD3,VAPA,TPT1,RPS3A
X5,ZNF865,SLC7A5,NLGN2,CHD4,ATP1A1,JUND,CD81,HMGN2,ATP2A2,CRMP1,LASP1,OAZ1,SRS
PR1,PFKL,TKT,SFPQ,PNMA1,GPC3,PIP4K2B,SOX12,ARL4C,APLP1,MIF,GPAT4,CHPF,MYO5A,CD6
P1A1,JUND,CD81,HMGN2,ATP2A2,CRMP1,LASP1,VAT1,OAZ1,PFN1,TLE5,NES,NISCH,CCNI,PFKL,H
RD3,KPNB1,PAPOLA,COL1A2,HK1,NCS1,RHOA,ZNF865,CLSTN1,ATF4,SLC7A5,NLGN2,CHD4,ATP
IRNP200,CSDE1,SMIM14,VCP,UBB,AFF4,CKB,COL4A2,LRRC28,SPEN,PSMD5,EIF1,HACD3,SET,EG
HAQ,HACD3,SET,EGLN1,TPI1,CHSY1,EIF4A1,RPL19,PPP2R1A,RPL15,GTF3C4,LDHB,SF3B1,PDIA3
,PLD3,USP11,COL18A1,SNRNP200,CSDE1,EIF3C,ATP5F1A,EIF4B,VCP,UBB,AFF4,TRIM28,MATR3
AN1,MARCKS,SMG1,NPM1,PLXNB2,PLD3,HSPD1,USP11,COL18A1,SNRNP200,IGF2,EIF4B,MARC
P5F1B,CALM3,RPS8,RPL7,SEPTIN2,P4HB,APP,IDS,ERVV-1,RPL5,GDI1,FSCN1,TMBIM6,PTMS,SPT
T,CCND2,MEGF8,ARL4C,H1-0,APLP1,MIF,CHPF,COL6A2,SESN3,B4GALNT4,MTSS2,APLP2,MXD4
CDC42BPB,ITM2B,AP2B1,GTF2F1,PXDN,HNRNPU,NFE2L1,SMARCA5,HNRNPA0,TPM4,SCARB2,N
5F1A,EIF4B,MARCKSL1,UBB,AFF4,MATR3,YWHAG,CKB,COL4A2,SPEN,NONO,YWHAQ,EIF1,HACD
RIM28,YWHAG,CKB,COL4A2,FN1,SPEN,DDB1,PSMD5,YWHAQ,EIF1,DCHS1,SET,EGLN1,TPI1,CHS
4L1,BTF3,AGRN,RPLP1,ZMIZ1,SARAF,MIDN,MARCHF6,VCL,DNAJA1,ARF1,SF3B2,ZNF770,MAT2A
DNAJC5,APLP2,MXD4,HNRNPK,RAB1A,PLPPR3,SEPTIN3,MAGED4B,CTBP1,ARHGDIA,CAPRIN1,STI
CHD3,VAPA,POLR2A,STMN1,ITGB1,UCHL1,MORF4L1,BTF3,AGRN,SARAF,COL11A1,MARCHF6,RF
1,SEC62,RPL6,KAT6A,DVL3,YWHAE,KIF1A,RPL13,GNAS,SLC2A3,DDX17,GPI,HTRA4,RACK1,BSG,
D81,WDR1,HMGN2,ATP2A2,CRMP1,LASP1,OAZ1,PFN1,TLE5,NES,CCNI,SPTBN2,PFKL,HDLBP,TKT
,CIPC,C1ORF43,LLGL1,ZNF865,SLC7A5,CHD4,JUND,CD81,WDR1,HMGN2,ATP2A2,CRMP1,LASP
ND,EIF5,CD81,LENG8,HMGN2,ATP2A2,RPL26,CRMP1,CCT7,VAT1,OAZ1,SRSF4,RBBP4,PFN1,AST
,LLGL1,HK1,NCS1,RHOA,ATF4,SLC7A5,CXADR,CHD4,LENG8,WDR1,ATXN7L3B,HMGN2,CRMP1,I
,PSMD5,HACD3,SET,EGLN1,TPI1,CHSY1,TMSB10,EIF4A1,MRC2,RPL19,PPP2R1A,RPL15,GTF3C4
IZ1,TIMP2,XAF1,MIDN,MARCHF6,WSB1,AP3D1,SBF1,ARF1,CYB5R3,AKAP12,MAP4,SF3B2,CDC42
RNPU,SMARCA5,FADS2,RPL23,TPM4,SCARB2,NACA,SLC22A17,BRD3,CIPC,KPNB1,C1ORF43,CC
DX17,HTRA4,RACK1,BSG,FASN,ALDH6A1,SPTBN1,ATP5F1B,CALM3,GNB1,SHISA9,STMN2,RPS8,N
,DCHS1,SET,TMSB10,INA,MRC2,GTF3C4,LDHB,PKD1,HLA-C,PDIA3,SSBP2,ZNF43,RPS3,BASP1,C
C28,FN1,NONO,PSMD5,EIF1,DCHS1,TPI1,CHSY1,TMSB10,INA,MRC2,PPP2R1A,RPL15,GTF3C4,PI
SL1,UBB,AFF4,TRIM28,MATR3,YWHAG,CKB,COL4A2,MAPT,SPEN,DDB1,EIF1,HACD3,DCHS1,CHS
C,APLP1,TLN1,CHPF,MYO5A,CD63,NGRN,B4GALNT4,DNAJC5,APLP2,HNRNPDL,NDFIP1,TSR2,M
7A5,JUND,ATP2A2,CRMP1,LASP1,CCT7,VAT1,OAZ1,PFN1,MCL1,TLE5,NES,NISCH,XPR1,KHDRBS
PL15,LDHB,PKD1,SF3B1,SERINC1,BASP1,VAPA,HMGB1,POLR2A,AP2M1,TPT1,RPL18,ITGB1,UCH

.1,AFF4,TRIM28,MATR3,CKB,COL4A2,MAPT,PSMD5,EIF1,DCHS1,SET,EGLN1,TPI1,CHSY1,TMSB1
IYH9,RPL7,SEPTIN2,P4HB,RPL5,FSCN1,TMBIM6,PTMS,SPTAN1,CALM1,NPM1,PLXNB2,PLD3,HSF
4A1,MRC2,RPL19,PPP2R1A,GTF3C4,SF3B1,PDIA3,SSBP2,VAPA,CDKL1,SEC61A1,RPS3A,STMN1,I
SPTAN1,GSTP1,SMG1,NPM1,PLXNB2,PLD3,USP11,COL18A1,SNRNP200,SMIM14,IGF2,ATP5F1A,I
L18A1,SNRNP200,CSDE1,SMIM14,IGF2,ATP5F1A,EIF4B,MARCKSL1,AFF4,TRIM28,YWHAG,CKB,L
2,KPNB1,C1ORF43,LLGL1,RPL14,HK1,CBX5,ZNF865,ATF4,PSMD2,SLC7A5,CHD4,ATP1A1,TMEN
12,DDX3X,ARL4C,GNB2,APLP1,CHPF,MYO5A,B4GALNT4,DNAJC5,APLP2,TARDBP,MXD4,CADM4,
IMP2,SARAF,MIDN,WSB1,SBF1,DNAJA1,ARF1,AKAP12,CDC42BPB,ITM2B,MAT2A,GTF2F1,SLC4A
DH6A1,SPTBN1,ATP5F1B,CALM3,GNB1,SHISA9,STMN2,RPS8,MYH9,SEPTIN2,P4HB,APP,IDS,ERV
34B,UCHL1,MORF4L1,ANXA2,AGRN,RPLP1,ZMIZ1,TIMP2,SARAF,MIDN,MARCHF6,WSB1,RPS17,V
N,ZMIZ1,SARAF,MIDN,RPL10,MARCHF6,VCL,AP3D1,DNAJA1,ARF1,ZNF770,MAT2A,GTF2F1,PXDN
R3,MAP4,ZNF770,TMEM170A,PXDN,HNRNPU,CTNNA1,FADS2,CPE,CNN3,LPCAT1,LAMA5,BRD3,K
L1,BTF3,NCAM1,ANXA2,AGRN,RPLP1,XAF1,SARAF,MIDN,RPL10,MARCHF6,HNRNPU,OGA,HNRNF
3,MATR3,YWHAG,CKB,RPL37A,FN1,YWHAQ,HACD3,DCHS1,SET,TMSB10,INA,MRC2,RPL19,GTF3
?,RPL5,FSCN1,TMBIM6,PTMS,SPTAN1,CALM1,PLD3,HSPD1,USP11,SNRNP200,CSDE1,IGF2,EIF3C
36B,CHD4,ATP1A1,JUND,RPS20,EIF5,CD81,WDR1,HMGN2,ATP2A2,CRMP1,LASP1,OAZ1,SRSF4,F
SL1,UBB,AFF4,TRIM28,YWHAG,CKB,COL4A2,FN1,SPEN,DDB1,PSMD5,YWHAQ,SET,MAP9,TPI1,TM
3,BASP1,HMGB1,POLR2A,TPT1,SEC61A1,RPS3A,RPL18,STMN1,ITGB1,UCHL1,MORF4L1,BTF3,N
1,CALM3,GNB1,SHISA9,STMN2,MYH9,RPL7,SEPTIN2,IDS,GDI1,FSCN1,PTMS,SPTAN1,GSTP1,MAF
'DIA3,SSBP2,VAPA,POLR2A,TPT1,RPS3A,ITGB1,TUBB4B,MORF4L1,NCAM1,AGRN,RPLP1,ZMIZ1,S
ORF4L1,AGRN,ZMIZ1,TIMP2,COL11A1,MIDN,RPL10,SBF1,DNAJA1,ARF1,MAP4,CDC42BPB,ITM2I
'L3,YWHAE,KIF1A,RPL13,SLC2A3,DDX17,GPI,RACK1,BSG,FASN,ALDH6A1,SPTBN1,ATP5F1B,CALI
VES,CCNI,PFKL,TKT,PNMA1,GPC3,PIP4K2B,SOX12,ITM2C,ARL4C,GNB2,MIF,GPAT4,MYO5A,COL
3N1,CALM3,GNB1,SHISA9,RPS8,MYH9,RPL7,SEPTIN2,P4HB,APP,IDS,ERVV-1,RPL5,GDI1,FSCN1,T
.HL9,HDLBP,TKT,GLG1,SFPQ,TPM1,EIF5A,CDH2,PNMA1,PLXNA1,SOX12,MMP14,PRDX2,DDX3X,I
),PPP2R1A,PKD1,HLA-C,SSBP2,BASP1,VAPA,CDKL1,TPT1,RPS4X,RPL18,STMN1,TUBB4B,UCHL1,
),PPP2R1A,PKD1,HLA-C,SSBP2,BASP1,VAPA,CDKL1,TPT1,RPS4X,RPL18,STMN1,TUBB4B,UCHL1,
1,AGRN,RPLP1,MIDN,RPL10,MARCHF6,RPS17,AP3D1,DNAJA1,ARF1,SF3B2,MAT2A,GTF2F1,HNF
C,PJA2,LLGL1,NCS1,ZNF865,ATF4,SLC7A5,CHD4,JUND,CD81,WDR1,HMGN2,ATP2A2,CRMP1,LAS
PL12,ITM2C,MAGED2,GNB2,RCN1,H1-0,APLP1,RPS19,JUN,MYO5A,COL6A2,CD63,ANXA6,B4GAL
CCNI,TADA2B,XPR1,XPO1,HDLBP,RPS25,TKT,GLG1,CCND2,CDH2,PIP4K2B,SOX12,DDX3X,APLP
K1,BSG,FASN,ALDH6A1,SPTBN1,ATP5F1B,CALM3,GNB1,SHISA9,STMN2,RPS8,MYH9,RPL7,SEPTI
D1,DNAJA1,ARF1,CYB5R3,MAP4,CDC42BPB,ITM2B,GTF2F1,HNRNPU,CTNNA1,CNN3,SCARB2,LF
S16,TPM4,LPCAT1,LAMA5,SLC22A17,PJA2,KPNB1,PAPOLA,COL1A2,NCS1,ZNF865,CLSTN1,ENC
F3B2,ITM2B,ZNF770,GTF2F1,SLC4A8,PGAM1,NFE2L1,FADS2,CPE,TPM4,EIF4H,TSPAN3,RPS24,S
COL4A2,SPEN,DDB1,PSMD5,YWHAQ,EIF1,DCHS1,EGLN1,TPI1,CHSY1,INA,PPP2R1A,RPL15,GTF3
EIF4H,LPCAT1,LAMA5,CIPC,SLC16A7,KPNB1,C1ORF43,PAPOLA,IGFBP3,RPL14,NCS1,RHOA,RPL
TP1,MARCKS,CALM1,NPM1,PLXNB2,HSPD1,USP11,COL18A1,SNRNP200,CSDE1,SMIM14,IGF2,A
4,JUND,CD81,WDR1,HMGN2,ATP2A2,CRMP1,LASP1,BEX3,OAZ1,PFN1,TLE5,NES,NISCH,CCNI,KH
LM3,GNB1,SHISA9,RPS8,SEPTIN2,P4HB,APP,IDS,ERVV-1,RPL5,GDI1,FSCN1,PTMS,SPTAN1,MAR(
CALM1,NPM1,PLXNB2,PLD3,HSPD1,USP11,SNRNP200,CSDE1,SMIM14,IGF2,EIF3C,EIF4B,UBB,AI
TF2F1,PXDN,CTNNA1,RPS16,LDHA,CNN3,TNS3,CTSD,EIF4H,LAMA5,SLC22A17,XRCC5,SLC16A7,
3L,H1-0,RPS5,MYO5A,HNRNPDL,ARID1A,NDFIP1,TSR2,WLS,HNRNPK,H1-10,SEPTIN11,RHOB,HN
SB1,RPS17,CYB5R3,AKAP12,MAP4,SF3B2,CDC42BPB,ITM2B,TMEM170A,GTF2F1,HNRNPU,NFE2
3,CKB,COL4A2,LRRC28,SPEN,NONO,DDB1,HACD3,DCHS1,SET,EGLN1,TPI1,TMSB10,INA,EIF4A1

|,ATP5F1B,CALM3,GNB1,SHISA9,RPS8,MYH9,RPL7,SEPTIN2,P4HB,RPL5,GDI1,FSCN1,TMBIM6,P
),CTNNA1,CNN3,EIF4H,TSPAN3,LAMA5,CIPC,COL1A2,LLGL1,HK1,NCS1,RHOA,ZNF865,CLSTN1,/
3,SET,EGLN1,MAP9,CHSY1,EIF4A1,MRC2,DNAH8,PKD1,HLA-C,SF3B1,PDIA3,SSBP2,ZNF43,VAPA,I
SF5,C5ORF24,TOMM20,U2AF2,SUGP2,CPSF1,PPP1CB,MEST,HNRNPL,TM9SF2,ABCA3,ALKBH5,
1,SNRNP200,CSDE1,IGF2,EIF3C,EIF4B,VCP,MARCKSL1,UBB,TRIM28,MATR3,CKB,COL4A2,DDB
TP5F1B,CALM3,GNB1,SHISA9,RPS8,MYH9,SEPTIN2,P4HB,APP,GDI1,FSCN1,TMBIM6,SPTAN1,M/
AT2A,GTF2F1,HNRNPU,HNRNPA0,RPS16,RPL23,CNN3,RPL29,EIF4H,RPS24,RPS9,XRCC5,CIPC,F
8,DDB1,EIF1,TPI1,INA,EIF4A1,RPL19,PPP2R1A,RPL15,GTF3C4,LDHB,DNAH8,PKD1,SF3B1,PDIA3
2,PDIA6,ITM2C,GNB2,APLP1,TLN1,B4GALNT4,MTSS2,DNAJC5,APLP2,MXD4,MYL6,ANKRD52,WI
BB,TRIM28,YWHAG,CKB,LRRC28,SPEN,DDB1,YWHAQ,DCHS1,SET,EGLN1,MAP9,CHSY1,TMSB1(
IP2,SARAF,COL11A1,MIDN,RPL10,WSB1,RPS17,DNAJA1,CYB5R3,AKAP12,MAP4,CDC42BPB,ITN
JX4,CADM4,UBE2H,ATP1A2,ARCN1,MAGED4B,ARHGDIA,SLC35E2B,MMP2,DNER,TRIB2,CNN2,N(
CH,SLC3A2,PFKL,HDLBP,TKT,GLG1,PNMA1,PDIA6,GPC3,PIP4K2B,MEGF8,SOX12,ARL4C,APLP1
8,MYH9,RPL7,SEPTIN2,P4HB,APP,GDI1,FSCN1,TMBIM6,SPTAN1,SMG1,CALM1,NPM1,PLXNB2,PI
1,ATP5F1B,GNB1,SHISA9,STMN2,RPS8,RPL7,SEPTIN2,P4HB,APP,IDS,ERVV-1,KLRD1,GDI1,FSCN
EIF4A1,MRC2,RPL19,RPL15,LDHB,SF3B1,SSBP2,BASP1,VAPA,POLR2A,SEC61A1,RPS3A,STMN1,
XPR1,KHDRBS1,SLC3A2,TPM1,CDH2,PDIA6,PIP4K2B,MEGF8,SOX12,CCN2,MMP14,PRDX2,DDX3
19,LDHB,DNAH8,PKD1,HLA-C,PDIA3,SSBP2,ZNF43,VAPA,HMGB1,AP2M1,TPT1,SEC61A1,RPS3A,I
P1,KLC1,GRINA,MLEC,ACAP3,MDK,DDX42,LAMC1,CBX1,SKI,SNRNP70,HDGF,ABCA2,RNF10,MVE
H8,SF3B1,ZNF43,RPS3,HMGB1,POLR2A,TPT1,SEC61A1,RPS3A,RPL18,ITGB1,TUBB4B,RPL11,M
,MAGED4B,CTBP1,ARHGDIA,CDK16,SKP1,BCLAF1,NNAT,DDX39B,EID1,SRSF5,TOMM20,U2AF2,
N2,GLIPR1L2,CXADR,CHD4,ATP1A1,CD81,WDR1,HMGN2,ATP2A2,RPL26,CRMP1,LASP1,OAZ1,SF
C4,LDHB,DNAH8,SF3B1,PDIA3,SERINC1,SSBP2,ZNF43,RPS3,HMGB1,POLR2A,CDKL1,AP2M1,TP
GK1,RPLP0,H3-3B,CANX,CLU,PTMA,DDX5,ENO2,SOX4,FKBP14,RPL13A,MAGED1,SEC62,RPL6,I
,RPS8,MYH9,RPL7,SEPTIN2,P4HB,RPL5,GDI1,FSCN1,PTMS,SPTAN1,GSTP1,MARCKS,CALM1,PL
,SCARB2,SLC22A17,BRD3,CIPC,KPNB1,C1ORF43,NCS1,RHOA,CBX5,ZNF865,ENC1,RPS14,CXAD
NNA1,CNN3,RPL29,TPM4,LPCAT1,MSN,KPNB1,IGFBP3,LLGL1,NCS1,CBX5,ATF4,ENC1,NLGN2,CI
KB,COL4A2,FN1,NONO,YWHAQ,HACD3,SET,EGLN1,CHSY1,TMSB10,INA,EIF4A1,MRC2,PPP2R1/
GL1,NCS1,RHOA,ZNF865,PSMD2,SLC7A5,CHD4,JUND,CD81,WDR1,HMGN2,ATP2A2,CRMP1,VAT
N2,ATP2A2,CRMP1,LASP1,BEX3,VAT1,OAZ1,RBBP4,PFN1,TLE5,NES,CCNI,XPR1,XPO1,PFKL,HDL
B4B,UCHL1,BTF3,NCAM1,ANXA2,AGRN,SARAF,RPL10,MARCHF6,RPS17,VCL,DNAJA1,ZNF770,P
I6,PTMS,SPTAN1,SMG1,NPM1,PLXNB2,PLD3,HSPD1,USP11,COL18A1,SNRNP200,CSDE1,SMIM1
MAP9,TPI1,INA,MRC2,RPL19,RPL15,PKD1,HLA-C,SF3B1,PDIA3,ZNF43,BASP1,VAPA,HMGB1,SEC6
APP,IDS,ERVV-1,GDI1,PTMS,SPTAN1,SMG1,CALM1,NPM1,PLXNB2,TTC3,PLD3,HSPD1,COL18A1,
TUBB4B,MORF4L1,NCAM1,ANXA2,AGRN,ZMIZ1,TIMP2,SARAF,MIDN,RPL10,WSB1,AP3D1,ARF1,(
N1,GSTP1,MARCKS,PLD3,HSPD1,USP11,COL18A1,SNRNP200,SMIM14,IGF2,EIF3C,ATP5F1A,EIF
T6A,DVL3,YWHAE,KIF1A,RPL13,GNAS,DDX17,GPI,RACK1,BSG,ALDH6A1,SPTBN1,ATP5F1B,CALN
IS1,EGLN1,TPI1,TMSB10,INA,MRC2,RPL19,PPP2R1A,LDHB,PKD1,HLA-C,PDIA3,SERINC1,SSBP2,
,RAB1A,PLPPR3,HECTD4,MAGED4B,ARHGDIA,GPC1,TRIB2,HS6ST1,NOVA2,CTSB,CDH11,RAB27/
,CES3,PSAP,PGK1,RPLP0,H3-3B,CANX,CLU,PTMA,DDX5,ENO2,SOX4,FKBP14,RPL13A,SEC62,RPI
EIF3C,MARCKSL1,AFF4,MATR3,RPL37A,FN1,SPEN,DDB1,YWHAQ,EIF1,SET,EGLN1,MAP9,TPI1,CH
UBB4B,UCHL1,RPL11,AGRN,RPLP1,TIMP2,MIDN,MARCHF6,RPS17,AP3D1,DNAJA1,ARF1,CYB5R
1,CYB5R3,MAP4,ZNF770,AP2B1,MAT2A,SLC4A8,PGAM1,SMARCA5,LDHA,TPM4,TNS3,CTSD,SCAI
,TMEM259,CD81,WDR1,HMGN2,CRMP1,BEX3,VAT1,PFN1,ASTN2,JPT2,TLE5,PNN,RPL31,TADA2E
I2,ATP2A2,CRMP1,LASP1,BEX3,VAT1,OAZ1,PFN1,JPT2,NES,NISCH,TADA2B,XPR1,PFKL,HDLBP,TI

H1,COTL1,HECTD4,MAGED4B,CTBP1,RAC1,ARHGDIA,STIP1,SLC35E2B,SMCHD1,IPO5,ATP6V0C,
P2B,SMCHD1,IPO5,DNER,TRIB2,PEA15,SQLE,SERF2,HS6ST1,TIMP3,NOVA2,CHD6,EEF1A2,TBCD,
,RPS7,RPS19,MIF,HMGCS1,JUN,COL6A2,HNRNPDL,TARDBP,SHOX,ANKRD52,WLS,HNRNPK,H1-1
SG,FASN,ALDH6A1,SPTBN1,ATP5F1B,CALM3,GNB1,SHISA9,STMN2,RPS8,MYH9,RPL7,SEPTIN2,
B,APP,ERVV-1,GDI1,FSCN1,TMBIM6,PTMS,SPTAN1,SMG1,CALM1,NPM1,PLD3,HSPD1,COL18A1,
04B,CTBP1,ARHGDIA,CAPRIN1,STIP1,GPC1,BCLAF1,PEA15,SQLE,MAP1A,TIMP3,CTDSP2,NOVA2,
2,ARL4C,APLP1,MIF,GPAT4,MYO5A,B4GALNT4,MTSS2,APLP2,TARDBP,INIP,RPL36,CADM4,HNRN
MBIM6,PTMS,SPTAN1,SMG1,NPM1,PLXNB2,PLD3,HSPD1,USP11,SNRNP200,CSDE1,SMIM14,IGF2
PR1,KHDRBS1,XPO1,PFKL,TKT,GLG1,TPM1,EIF5A,CDH2,GPC3,PIP4K2B,ACTN4,RPS15,ITM2C,A
PR1,KHDRBS1,XPO1,PFKL,TKT,GLG1,TPM1,EIF5A,CDH2,GPC3,PIP4K2B,ACTN4,RPS15,ITM2C,A
PR1,KHDRBS1,XPO1,PFKL,TKT,GLG1,TPM1,EIF5A,CDH2,GPC3,PIP4K2B,ACTN4,RPS15,ITM2C,A
NAH8,SSBP2,CHD3,TPT1,RPS3A,ITGB1,UCHL1,BTF3,AGRN,RPLP1,ZMIZ1,SARAF,COL11A1,MIDN
GF8,RPL12,ARL4C,CLTC,EIF3L,MRFAP1,RPS5,APLP1,RPS7,HMGCS1,TLN1,MYO5A,COL6A2,GOL
PM1,PLXNB2,USP11,COL18A1,SNRNP200,SMIM14,IGF2,EIF3C,EIF4B,VCP,MARCKSL1,UBB,MAT
8,RPL7,P4HB,APP,ERVV-1,GDI1,FSCN1,PTMS,SPTAN1,SMG1,NPM1,PLD3,HSPD1,COL18A1,SNRN
MG1,CALM1,PLXNB2,PLD3,HSPD1,USP11,COL18A1,SNRNP200,CSDE1,SMIM14,IGF2,EIF3C,ATP
TBN1,ATP5F1B,CALM3,GNB1,SHISA9,STMN2,RPS8,MYH9,RPL7,SEPTIN2,P4HB,APP,IDS,RPL5,GD
2,VAMP2,GAS7,PARP1,PKN1,YEATS2,CAMSAP1,GRINA,PTOV1,MDK,JPT1,ORAI2,SKI,H3-3A,ABCA
A,POLR2A,SEC61A1,STMN1,TUBB4B,UCHL1,MORF4L1,BTF3,ZMIZ1,MIDN,RPL10,RPS17,VCL,AP
,JUND,ATP2A2,CRMP1,LASP1,BEX3,ASTN2,JPT2,TLE5,NISCH,CCNI,TKT,GPC3,DDR1,MEGF8,SO
PL10,WSB1,VCL,AP3D1,DNAJA1,ARF1,CYB5R3,AKAP12,MAP4,SF3B2,CDC42BPB,ITM2B,MAT2A,T
,RPL24,KPNB1,PAPOLA,RPL14,CBX5,ATF4,SLC7A5,CXADR,JUND,TMEM259,RPS20,EIF5,CD81,W
1,RHOA,CLSTN1,RAB6B,SLC7A5,NLGN2,CHD4,LENG8,ATP2A2,CRMP1,LASP1,BEX3,VAT1,RBBP4
ERVV-1,GDI1,FSCN1,TMBIM6,PTMS,SPTAN1,SMG1,CALM1,NPM1,PLD3,HSPD1,COL18A1,SNRNP
,RAB1A,PLPPR3,ARCN1,SEPTIN3,COTL1,MAGED4B,CTBP1,ARHGDIA,CAPRIN1,STIP1,GPC1,TOP
N1,TMBIM6,GSTP1,SMG1,CALM1,NPM1,TTC3,PLD3,HSPD1,SNRNP200,SMIM14,IGF2,EIF3C,EIF
,KPNB1,IGFBP3,COL1A2,LLGL1,HK1,RPL22,CLSTN1,NR2F1,RAB6B,CXADR,CHD4,ATP1A1,JUND
YB5R3,MAP4,CDC42BPB,MAT2A,SLC4A8,HNRNPU,OGA,NFE2L1,HNRNPA0,CTNNA1,RPL29,NAC
A,GTF2F1,PXDN,PGAM1,HNRNPA0,CTNNA1,FADS2,CNN3,RPSA,TPM4,EIF4H,SLC22A17,BRD3,R
,TMBIM6,SPTAN1,CALM1,NPM1,PLXNB2,PLD3,HSPD1,USP11,COL18A1,SNRNP200,CSDE1,SMIM
GRN,GOLM1,B4GALNT4,APLP2,TARDBP,ARL6IP1,NDFIP1,TSR2,MXD4,ANKRD52,HNRNPK,H1-10
B2,MAT2A,GTF2F1,HNRNPA0,RPS16,RPL23,CNN3,RPL29,RPS9,XRCC5,CIPC,RPL24,SLC16A7,PA
UBB4B,MORF4L1,AGRN,TIMP2,XAF1,COL11A1,MARCHF6,VCL,MAP4,CDC42BPB,MAT2A,SLC4A
MORF4L1,BTF3,ZMIZ1,SARAF,MIDN,RPL10,RPS17,VCL,AP3D1,SBF1,DNAJA1,SF3B2,MAT2A,GTF
1,L1CAM,LAPTM4B,RPL23A,LUC7L3,MAP1A,DLG5,CPSF2,SRP9,SRSF5,MDH2,CHD6,EEF1A2,VAN
VHAG,CKB,COL4A2,RPL37A,LRRC28,MAPT,FN1,SPEN,NONO,DDB1,PSMD5,EIF1,MAP9,CHSY1,EI
28,EIF1,TPI1,INA,EIF4A1,RPL19,PPP2R1A,RPL15,GTF3C4,LDHB,DNAH8,PKD1,SF3B1,PDIA3,SER
LD3,HSPD1,USP11,SNRNP200,CSDE1,IGF2,EIF3C,ATP5F1A,EIF4B,VCP,UBB,AFF4,YWHAG,CKB,R
PIP4K2B,SOX12,DDX3X,RPL30,RPL35,MIF,DNAJC5,HNRNPDL,SHOX,HNRNPK,CKAP5,LAMP1,HNR
PTAN1,SMG1,NPM1,TTC3,PLD3,HSPD1,SNRNP200,CSDE1,SMIM14,IGF2,EIF3C,ATP5F1A,EIF4B,
PD1,COL18A1,SNRNP200,CSDE1,EIF3C,ATP5F1A,EIF4B,MARCKSL1,UBB,AFF4,TRIM28,MATR3,
/SB1,AP3D1,SF3B2,CDC42BPB,ITM2B,GOLGA8A,PXDN,RPS16,SCARB2,TSPAN3,LPCAT1,LAMA5,
A9,RPS8,RPL7,P4HB,APP,ERVV-1,GDI1,FSCN1,TMBIM6,PTMS,SPTAN1,SMG1,NPM1,PLD3,HSPD
,MSN,KPNB1,C1ORF43,LLGL1,RPL14,RHOA,CBX5,ATF4,ENC1,RPS14,CXADR,JUND,TMEM259,
T1,SEC61A1,RPS3A,RPL18,STMN1,UCHL1,BTF3,NCAM1,AGRN,RPLP1,MIDN,RPL10,MARCHF6,W

DS2,RPL23,CNN3,RPL29,TPM4,RPS9,XRCC5,CIPC,RPL24,SLC16A7,KPNB1,PAPOLA,HK1,RHOA,Z
CND2,CDH2,PIP4K2B,DDX3X,CLTC,APLP1,CHPF,COL6A2,ANXA6,SHOX,WLS,HNRNPK,NARS1,DU
OAZ1,RBBP4,ASTN2,MCL1,JPT2,TLE5,PNN,NES,LRPPRC,TADA2B,HDLBP,RPS25,TKT,EIF5A,CDH2
2,PLD3,HSPD1,USP11,COL18A1,SNRNP200,CSDE1,SMIM14,IGF2,EIF3C,ATP5F1A,EIF4B,VCP,UB
PLXNB2,PLD3,HSPD1,USP11,SNRNP200,CSDE1,SMIM14,IGF2,EIF3C,ATP5F1A,EIF4B,VCP,UBB,AF
1A1,MIDN,RPL10,AP3D1,SBF1,ARF1,CYB5R3,ITM2B,MAT2A,GTF2F1,OGA,NFE2L1,CNN3,EIF4H,T
,CTDSP2,NOVA2,CTSB,C5ORF24,CHD6,RNF187,EEF1A2,TBCD,SNX22,PGD,SQSTM1,VAMP2,LM
MAP9,PPP2R1A,PKD1,PDIA3,SSBP2,CHD3,VAPA,HMGB1,POLR2A,AP2M1,RPL18,STMN1,TUBB4B,
PTMS,SPTAN1,GSTP1,MARCKS,SMG1,CALM1,NPM1,PLXNB2,PLD3,CSDE1,EIF4B,MARCKSL1,AFF
A8A,PXDN,RPS16,SCARB2,TSPAN3,LPCAT1,LAMA5,RPS24,CIPC,C1ORF43,COL1A2,LLGL1,HK1,
,SMCHD1,IPO5,ATP6V0C,DNER,BCLAF1,L1CAM,TRIB2,SQLE,FAM199X,SERF2,TIMP3,CTDSP2,N
1,SLC7A5,CHD4,TMEM259,RPS20,EIF5,CD81,HMGN2,ATP2A2,LASP1,CCT7,OAZ1,PFN1,SPC25,N
06,EEF1A2,CCT2,TBCD,VAMP2,GAS7,TM9SF2,LMNB2,HSPA4,CAMSAP1,GRINA,VPS35,PTOV1,ML
,MEGF8,SOX12,ITM2C,APLP1,MIF,GPAT4,CHPF,GOLM1,SESN3,APLP2,TARDBP,INIP,DUX4,CKAP
GTF3C4,LDHB,PKD1,HLA-C,PDIA3,ZNF43,VAPA,CDKL1,TPT1,SEC61A1,STMN1,ITGB1,TUBB4B,U
7,MAP4,SF3B2,CDC42BPB,AP2B1,GTF2F1,OGA,NFE2L1,SMARCA5,LDHA,SLC22A17,BRD3,CIPC,
ZMIZ1,MIDN,RPL10,WSB1,AP3D1,ARF1,CYB5R3,MAP4,CDC42BPB,ITM2B,TMEM170A,GTF2F1,P
GLN1,TPI1,TMSB10,INA,EIF4A1,RPL19,PPP2R1A,RPL15,GTF3C4,LDHB,DNAH8,PKD1,SF3B1,SER
PK,RPL36,PTGES3,CADM4,LAMP1,RPS12,CALU,COTL1,HECTD4,RAC1,SMCHD1,HDAC2,ATP6V0
XNB2,PLD3,HSPD1,USP11,SNRNP200,CSDE1,SMIM14,IGF2,EIF3C,ATP5F1A,EIF4B,VCP,UBB,AFF4
1,RPL19,RPL15,GTF3C4,LDHB,HLA-C,SERINC1,HMGB1,POLR2A,RPS3A,STMN1,TIMP2,SARAF,C
B4B,RPL11,MORF4L1,BTF3,ANXA2,XAF1,SARAF,RPL10,WSB1,RPS17,AP3D1,SBF1,ARF1,CYB5R3
EIF4B,VCP,UBB,MATR3,YWHAG,RPL37A,LRRC28,SPEN,NONO,DDB1,EIF1,EGLN1,INA,EIF4A1,RP
B,XPR1,KLHL9,SLC3A2,RPS25,TKT,GLG1,CDH6,TPM1,CDH2,GPC3,PIP4K2B,ARL4C,CLTC,GNB2,
L6,ANKRD52,HNRNPK,H1-10,INIP,PTGES3,SEPTIN11,SND1,CKAP4,ZNF454,CTBP1,RAC1,STIP1,G
HNRNPA0,RPL23,CNN3,RPL29,RPS9,XRCC5,CIPC,RPL24,SLC16A7,PAPOLA,COL1A2,HK1,RPL22,
F1,HNRNPA0,FADS2,RPS16,RPL23,CNN3,RPL29,TPM4,RPS9,XRCC5,CIPC,RPL24,C1ORF43,PAP
RHOA,PTCD3,ZNF865,PSMD2,GLIPR1L2,RPS14,ATP1A1,RPS20,EIF5,LENG8,WDR1,RPL26,BEX3,
,SPTAN1,GSTP1,MARCKS,SMG1,CALM1,PLXNB2,TTC3,USP11,SNRNP200,CSDE1,SMIM14,IGF2,
10,EIF4A1,RPL19,PPP2R1A,RPL15,GTF3C4,DNAH8,PKD1,SF3B1,PDIA3,SERINC1,ZNF43,RPS3,H
5ORF24,TOMM20,U2AF2,SUGP2,CPSF1,PPP1CB,MEST,HNRNPL,TM9SF2,ABCA3,ALKBH5,SNRP
XNB2,HSPD1,IGF2,EIF3C,ATP5F1A,MARCKSL1,AFF4,TRIM28,MATR3,YWHAG,CKB,RPL37A,LRR
XK,ACO2,HSPA4,CAMSAP1,GRINA,VPS35,PTOV1,MLEC,CLSTN3,MDK,SURF4,CBX1,SKI,H3-3A,S
A-C,SF3B1,PDIA3,ZNF43,BASP1,HMGB1,POLR2A,CDKL1,TPT1,RPL18,TUBB4B,UCHL1,RPL11,M
PLXNB2,PLD3,HSPD1,USP11,COL18A1,SNRNP200,CSDE1,IGF2,EIF3C,ATP5F1A,EIF4B,VCP,UBB,
37A,LRRC28,EIF1,HACD3,INA,EIF4A1,RPL19,PPP2R1A,RPL15,GTF3C4,DNAH8,PKD1,SF3B1,PDIA
37A,LRRC28,FN1,NONO,DDB1,PSMD5,YWHAQ,EIF1,MAP9,CHSY1,EIF4A1,MRC2,RPL19,RPL15,G
X12,RAB7A,DDX3X,ARL4C,H1-0,COL6A2,DNAJC5,APLP2,MYL6,WLS,HNRNPK,H1-10,DUX4,RAB1
1-0,RPS19,CHPF,NGRN,HNRNPDL,RPLP2,DDOST,YWHAB,SND1,LAMP1,CALU,ARCN1,RPS27,ARH
,SOX12,ITM2C,MAGED2,APLP1,MIF,GPAT4,CHPF,JUN,GOLM1,SESN3,MTSS2,APLP2,TARDBP,CK
BIM6,PTMS,SPTAN1,SMG1,NPM1,PLD3,HSPD1,SNRNP200,SMIM14,IGF2,EIF3C,ATP5F1A,VCP,M
,VAT1,SRSF4,PFN1,ASTN2,MCL1,JPT2,TLE5,NES,NISCH,TADA2B,XPR1,SLC3A2,PFKL,PDIA6,GPC
7A,LRRC28,MAPT,FN1,NONO,DDB1,YWHAQ,HACD3,DCHS1,SET,MAP9,TPI1,TMSB10,MRC2,RPL
A,RPL15,GTF3C4,PKD1,PDIA3,CHD3,VAPA,TPT1,RPS3A,RPL18,ITGB1,TUBB4B,AGRN,RPLP1,ZMI
ORO1C,SDF4,HNRNPM,SDHA,YWHAH,LONP1,UBQLN1,ZFAND3,PFN2,CCT5,PSMA7,RBBP7,DFFB,

TMSB10,INA,EIF4A1,RPL19,GTF3C4,DNAH8,SF3B1,PDIA3,SSBP2,CHD3,HMGB1,CDKL1,RPS4X,R
4,PFN1,MCL1,TLE5,PNN,RPL31,PRKCSH,SLC3A2,CSNK1E,RPS25,TKT,EIF5A,PDIA6,SOX12,RAB7
T1,CTDSP2,CNN2,NOVA2,CTSB,NUCB1,ZC3H7B,EEF1A2,ANXA5,VAMP2,GAS7,PTPRA,PDXK,UBE
MSL1,SMARCE1,SDF4,HNRNPM,PHC2,STMN3,LSS,FBLN1,ISYNA1,TUG1,VANGL2,SREBF2,FBRSL
4,MATR3,YWHAG,COL4A2,RPL37A,LRRC28,SPEN,NONO,EIF1,EGLN1,TMSB10,INA,EIF4A1,RPL1
KD1,SF3B1,PDIA3,SERINC1,ZNF43,RPS3,HMGB1,POLR2A,AP2M1,TPT1,SEC61A1,RPS4X,RPS3A,
3C,ATP5F1A,EIF4B,MARCKSL1,AFF4,MATR3,YWHAG,CKB,RPL37A,FN1,NONO,DDB1,PSMD5,EIF
,DDX17,GPI,HTRA4,RACK1,BSG,FASN,ALDH6A1,SPTBN1,CALM3,GNB1,SHISA9,STMN2,RPS8,MY
F25,DYNLL2,ILF3,GRB2,ACTN1,CAND1,HSPB1,PLCG1,ZFAND5,DIDO1,FLNB,DLK1,CORO1C,MS
RA4,RACK1,BSG,FASN,ALDH6A1,SPTBN1,CALM3,GNB1,SHISA9,STMN2,RPS8,RPL7,SEPTIN2,P4
C1,APPL1,SMCHD1,IPO5,ATP6V0C,DNER,LAPTM4B,TRIB2,NNAT,FAM199X,SRP9,NUCB1,TBCD,S
2R1A,RPL15,GTF3C4,DNAH8,PKD1,SF3B1,PDIA3,SERINC1,ZNF43,RPS3,HMGB1,POLR2A,AP2M1
,GTF3C4,DNAH8,PKD1,SF3B1,PDIA3,SERINC1,ZNF43,RPS3,HMGB1,POLR2A,AP2M1,TPT1,SEC6
GPI,HTRA4,FASN,ALDH6A1,SPTBN1,CALM3,GNB1,SHISA9,STMN2,MYH9,SEPTIN2,P4HB,APP,IDS
,ARF1,CYB5R3,MAP4,SF3B2,ITM2B,TMEM170A,GTF2F1,PXDN,OGA,HNRNPA0,RPL23,CPE,CNN
,AP3D1,SBF1,DNAJA1,SF3B2,MAT2A,GTF2F1,PXDN,HNRNPU,SMARCA5,HNRNPA0,CTNNA1,FADS
P,ATG4B,DEK,DMWD,HNRNPH3,UBE2K,UBE2L3,RSF1,PUM1,KDM3A,TIA1,NSF,ADNP,SSU72,FAF
LKBH5,HSPA4,CAMSAP1,KLC1,CCT3,GRINA,VPS35,MDK,RAD23B,SKI,H3-3A,ABCA2,VEGFA,ARF
,RPL13,SLC2A3,DDX17,HTRA4,RACK1,BSG,FASN,ALDH6A1,SPTBN1,ATP5F1B,CALM3,GNB1,SHI
TF2F1,PXDN,SMARCA5,CTNNA1,FADS2,RPL23,CNN3,RPSA,TPM4,TNS3,CTSD,EIF4H,TSPAN3,LA
PL13,GNAS,DDX17,GPI,HTRA4,RACK1,BSG,FASN,ALDH6A1,SPTBN1,ATP5F1B,CALM3,GNB1,SHIS
,FASN,ALDH6A1,SPTBN1,CALM3,GNB1,SHISA9,STMN2,RPS8,MYH9,RPL7,P4HB,APP,ERVV-1,KL
200,IGF2,EIF4B,UBB,MATR3,YWHAG,CKB,RPL37A,SPEN,NONO,DDB1,PSMD5,YWHAQ,EIF1,HACD
,MARCKS,SMG1,PLXNB2,PLD3,HSPD1,COL18A1,SNRNP200,SMIM14,IGF2,ATP5F1A,EIF4B,VCP,
PS16,RPL23,LDHA,TPM4,NACA,LPCAT1,BRD3,RPS9,CIPC,PJA2,MSN,RPL24,C1ORF43,PAPOLA,H
,RPL19,PPP2R1A,RPL15,GTF3C4,DNAH8,PKD1,SF3B1,PDIA3,SERINC1,ZNF43,RPS3,HMGB1,POL
,MARCHF6,WSB1,VCL,AP3D1,ARF1,PAFAH1B1,MAP4,SF3B2,CDC42BPB,ZNF770,TMEM170A,SL
24,TOMM20,U2AF2,SUGP2,CPSF1,MEST,HNRNPL,ABCA3,ALKBH5,SNRPN,PRDX5,CCT3,GRINA,
RDBP,MYL6,HNRNPK,DUX4,CADM4,RAB1A,PLPPR3,COTL1,MAGED4B,CAPRIN1,TOP2B,TMEM30
1ORF43,PAPOLA,HK1,NCS1,RHOA,PTCD3,ATF4,NR2F1,SLC7A5,NLGN2,TMEM259,EIF5,CD81,LE
DVL3,YWHAE,KIF1A,RPL13,GNAS,SLC2A3,DDX17,GPI,HTRA4,BSG,FASN,ALDH6A1,SPTBN1,ATP5
DK16,HDAC2,TMEM30A,MMP2,DNER,BCLAF1,HSBP1,DLG5,CRPPA,HS6ST1,SRSF5,C5ORF24,M
RC28,EIF1,INA,EIF4A1,RPL19,PPP2R1A,RPL15,LDHB,PKD1,SF3B1,PDIA3,SERINC1,SSBP2,ZNF43,
,TMSB10,EIF4A1,GTF3C4,PKD1,HLA-C,SERINC1,SSBP2,CHD3,VAPA,HMGB1,POLR2A,AP2M1,TP
B,CALM3,GNB1,SHISA9,RPS8,RPL7,SEPTIN2,P4HB,APP,ERVV-1,GDI1,FSCN1,TMBIM6,PTMS,SP
IM14,IGF2,EIF3C,ATP5F1A,EIF4B,VCP,MARCKSL1,AFF4,TRIM28,MATR3,YWHAG,CKB,COL4A2,R
A1,SF1,HIP1,BAZ1B,KPNA2,UBA52,SRSF6,PAPSS1,UBAP2L,CRTAP,ZFR,CBX6,EML1,RPS2,GALN
B1,SHISA9,STMN2,RPS8,MYH9,RPL7,SEPTIN2,APP,IDS,KLRD1,GDI1,FSCN1,TMBIM6,PTMS,SPT
NONO,EIF1,EGLN1,INA,EIF4A1,RPL19,PPP2R1A,RPL15,GTF3C4,LDHB,DNAH8,PKD1,SF3B1,PDIA
14,RPS20,CD81,LENG8,WDR1,ATXN7L3B,RPL26,CRMP1,SRSF4,RBBP4,PFN1,MCL1,TLE5,PNN,R
DM1A,MAP7D1,PLCG1,DIDO1,DLK1,CORO1C,GIGYF1,YWHAH,LSS,STXBP1,ISYNA1,TUG1,CSNK1
20,CD81,LENG8,WDR1,ATXN7L3B,HMGN2,SRSF4,PFN1,MCL1,TLE5,PNN,RPL31,PRKCSH,HNRN
,GDI1,TMBIM6,PTMS,SPTAN1,GSTP1,SMG1,NPM1,PLXNB2,PLD3,HSPD1,SNRNP200,CSDE1,IGF2
NPM1,PLXNB2,PLD3,HSPD1,USP11,COL18A1,SNRNP200,CSDE1,SMIM14,IGF2,EIF3C,ATP5F1A,E
R1A,RPL15,GTF3C4,DNAH8,PKD1,SF3B1,PDIA3,SERINC1,ZNF43,HMGB1,POLR2A,AP2M1,TPT1,

PT,FN1,DDB1,YWHAQ,EIF1,HACD3,DCHS1,SET,EGLN1,CHSY1,TMSB10,INA,EIF4A1,MRC2,RPL15
AC1,STIP1,GPC1,TOP2B,SMCHD1,HDAC2,PEA15,DLG5,CPSF2,PGRMC1,SRP9,CTSB,CDH11,RAB
2,ZFAND5,DUSP4,MSL1,ATP6V1B2,STT3B,STXBP1,ISYNA1,ATXN2L,HYOU1,ZFAND3,VANGL2,SOI
XC5,BRD2,PPP2CB,RAB2A,GNG4,ZFR,ZNF512B,PTPRF,EML1,NPDC1,ANP32B,B4GAT1,GSE1,GP
PLD3,SNRNP200,SMIM14,IGF2,EIF3C,ATP5F1A,EIF4B,VCP,MARCKSL1,UBB,AFF4,MATR3,YWHAG
,STMN3,PFKM,FBLN1,CSNK1D,ZFAND3,PFN2,HNRNPD,VGF,FLNC,AKIRIN1,CTXN1,PPP1R9B,CIC,
SA9,STMN2,RPS8,MYH9,RPL7,SEPTIN2,P4HB,IDS,RPL5,KLRD1,GDI1,FSCN1,PTMS,SPTAN1,GSTI
RCA5,HNRNPA0,FADS2,LDHA,CNN3,TPM4,LPCAT1,LAMA5,SLC16A7,PAPOLA,RPL14,HK1,NCS1,C
P1A1,JUND,CD81,ATXN7L3B,HMGN2,ATP2A2,CRMP1,VAT1,OAZ1,ASTN2,PNN,NES,TADA2B,PFKL
PL30,MRFAP1,GPAT4,TLN1,JUN,COL6A2,CD63,NGRN,MTSS2,CYFIP2,ARL6IP1,NDFIP1,MXD4,HN
NLL2,ATXN10,ACTN1,MAP7D1,CLIP3,PLCG1,DIDO1,DLK1,CORO1C,GIGYF1,LSS,STXBP1,ISYNA
EIF4A1,RPL19,LDHB,DNAH8,PDIA3,SSBP2,CHD3,HMGB1,CDKL1,TPT1,SEC61A1,RPS4X,RPS3A,S
CB,PGD,MEST,GAS7,MORF4L2,TGFBI,HNRNPA3,YEATS2,HSPA4,AHCYL1,CCT3,NCKAP1,PTOV1,M
3,CANX,CLU,PTMA,DDX5,ENO2,RPL13A,MAGED1,SEC62,RPL6,KAT6A,DVL3,YWHAE,KIF1A,RPL13
EIF4B,VCP,UBB,AFF4,MATR3,YWHAG,RPL37A,LRRC28,SPEN,NONO,EIF1,EGLN1,INA,EIF4A1,RPL
10,INA,GTF3C4,PKD1,SF3B1,PDIA3,SERINC1,BASP1,VAPA,HMGB1,POLR2A,TPT1,SEC61A1,RPS3
,ANXA2,MIDN,VCL,AP3D1,CYB5R3,AKAP12,SF3B2,AP2B1,MAT2A,GTF2F1,SLC4A8,GOLGA8A,PG
DA2B,KLHL9,RPS25,TKT,SFPQ,CCND2,CDH2,PLXNA1,PIP4K2B,CCN2,DDX3X,CLTC,GNB2,H1-0,
RPF6,DPYSL2,TAF7,SELENON,RPL37,SMARCA4,ATG2B,CCNL2,ARPC2,FAM168B,SRSF1,ARGLU1
4,RPS20,CD81,LENG8,WDR1,ATXN7L3B,RPL26,SRSF4,RBBP4,PFN1,MCL1,TLE5,PNN,RPL31,PR
B,CALM3,GNB1,SHISA9,STMN2,RPS8,MYH9,RPL7,SEPTIN2,P4HB,IDS,RPL5,KLRD1,GDI1,FSCN1,
S17,DNAJA1,ARF1,CYB5R3,SF3B2,MAT2A,GTF2F1,HNRNPA0,RPS16,RPL23,RPL29,EIF4H,LAMA
,EIF4A1,MRC2,RPL19,DNAH8,PKD1,SSBP2,ZNF43,VAPA,AP2M1,ITGB1,TUBB4B,NCAM1,ANXA2,A
,EIF4A1,MRC2,RPL19,DNAH8,PKD1,SSBP2,ZNF43,VAPA,AP2M1,ITGB1,TUBB4B,NCAM1,ANXA2,A
B4B,MORF4L1,BTF3,ANXA2,AGRN,TIMP2,SBF1,DNAJA1,ARF1,CYB5R3,MAP4,SF3B2,CDC42BPE
N,RPLP1,MIDN,RPL10,MARCHF6,AP3D1,DNAJA1,ARF1,CYB5R3,SF3B2,GTF2F1,HNRNPA0,RPL2
,ITM2B,TMEM170A,SLC4A8,HNRNPA0,CTNNA1,FADS2,CPE,CNN3,RPL29,RPSA,TPM4,TNS3,LPCA
EGLN1,INA,EIF4A1,RPL19,PPP2R1A,RPL15,GTF3C4,DNAH8,PKD1,SF3B1,PDIA3,SERINC1,ZNF43,
TOP2B,SLC35E2B,APPL1,GLUL,SQLE,HS6ST1,CTDSP2,CNN2,NOVA2,CTSB,C5ORF24,RNF187,T
CS1,CBX5,PSMD2,KIF5B,RAB6B,NLGN2,RPS14,CHD4,JUND,HMGN2,CRMP1,BEX3,PFN1,PRKCSH
15,RPL12,RPL30,H1-0,RPL35,MIF,TLN1,GJA1,RPS10,SESN3,MTSS2,HNRNPDL,ARL6IP1,HNRNP
1,CAMSAP1,PTOV1,MLEC,SKI,SNRNP70,ASXL1,H3-3A,SCAF1,PYGB,DPYSL2,RPL38,NOTCH2,TUE
AM1,MIDN,VCL,DNAJA1,ITM2B,TMEM170A,GTF2F1,NFE2L1,FADS2,RPS16,RPL23,LDHA,CNN3,R
B1,POLR2A,AP2M1,TPT1,SEC61A1,RPS4X,RPS3A,RPL18,TUBB4B,RPL11,MORF4L1,BTF3,AGRN,
D2,KIF5B,SLC7A5,NLGN2,CXADR,CHD4,ATP1A1,JUND,TMEM259,CD81,HMGN2,ATP2A2,RPL26,C
L1,SMCHD1,IPO5,TMEM30A,DNER,CDKN1A,CPSF2,NUCB1,SRSF5,RAB27A,FDFT1,TBCD,SUGP2,
RSF5,TM9SF3,MDH2,U2AF2,FDFT1,GNAI3,MEST,GDI2,HSPH1,RPL27,PRDX5,ACTR3,UBE4A,HSP
19,PPP2R1A,RPL15,GTF3C4,DNAH8,PKD1,SF3B1,PDIA3,SERINC1,ZNF43,RPS3,HMGB1,POLR2A,
30,EIF3L,GNB2,IK,RPS19,CHPF,NGRN,DNAJC5,RPLP2,DDOST,NARS1,DUX4,YWHAB,PTGES3,RP
APOLA,HK1,RHOA,ZNF865,ATF4,GLIPR1L2,RPS14,RPS20,CD81,LENG8,WDR1,ATXN7L3B,HMGN
2,RPS14,RPS20,CD81,LENG8,WDR1,SRSF4,PFN1,ASTN2,MCL1,TLE5,PNN,RPL31,PRKCSH,KHD
KAP5,SEPTIN11,CALU,CTBP1,CDK16,APPL1,HDAC2,SKP1,GLUL,PEA15,SQLE,HS6ST1,TIMP3,CT
KAP5,SEPTIN11,CALU,CTBP1,CDK16,APPL1,HDAC2,SKP1,GLUL,PEA15,SQLE,HS6ST1,TIMP3,CT
NA,EIF4A1,RPL19,PPP2R1A,RPL15,LDHB,PKD1,SF3B1,PDIA3,SERINC1,ZNF43,RPS3,HMGB1,POL
/CP,MARCKSL1,UBB,AFF4,MATR3,YWHAG,CKB,COL4A2,RPL37A,LRRC28,SPEN,NONO,DDB1,EIF

RPF6,SELENON,MVB12B,AZIN1,SRSF9,DDAH2,FAM168B,NYNRIN,DYNLL2,WDR6,ATXN10,KDM1
3,TLE5,XPR1,KLHL9,SPTBN2,SLC3A2,PFKL,CCND2,CDH2,PDIA6,GPC3,DDR1,PIP4K2B,ACSL6,SC
LRRC28,FN1,DDB1,EIF1,HACD3,SET,EGLN1,TPI1,TMSB10,INA,EIF4A1,RPL19,LDHB,DNAH8,SF3B
1,PPM1G,HOXB5,DIP2B,UBFD1,PRRC2A,HNRNPA2B1,SEPTIN7,NPLOC4,FKBP1A,C1ORF198,MT
,PRKCSH,LRPPRC,TADA2B,RPS25,TKT,SFPQ,EIF5A,CDH2,PDIA6,RAB7A,RPS15,ITM2C,RPL30,EI
4,AP2B1,MAT2A,GTF2F1,GOLGA8A,PGAM1,RPL23,LDHA,CNN3,SCARB2,LPCAT1,RPS9,PJA2,RPL
S1,CBX5,CLSTN1,KIF5B,RAB6B,NLGN2,CXADR,CHD4,JUND,HMGN2,CRMP1,LASP1,BEX3,JPT2,
MGB1,POLR2A,AP2M1,TPT1,SEC61A1,RPS4X,RPS3A,RPL18,TUBB4B,RPL11,BTF3,NCAM1,AGRN
LA,RHOA,ZNF865,RPL9,ATF4,KIF5B,RPS14,CXADR,LENG8,WDR1,RPL26,LASP1,CCT7,SRSF4,RB
GB1,POLR2A,AP2M1,TPT1,SEC61A1,RPS4X,RPS3A,RPL18,TUBB4B,RPL11,MORF4L1,BTF3,AGR
BL6,ODC1,AEBP1,PTBP1,EIF3E,LUZP6,UBE2R2,SCAF11,CLIP3,MSL1,SDF4,SMARCC1,ATP6V1B2
U,CTNNA1,CNN3,CTSD,NACA,TSPAN3,LPCAT1,LAMA5,SLC22A17,KPNB1,LLGL1,HK1,NCS1,RPL2
,RPS19,CHPF,NGRN,DNAJC5,HNRNPDL,RPLP2,DDOST,YWHAB,PTGES3,RPS12,CALU,MAGED4B
RHOA,CLSTN1,KIF5B,SLC7A5,ENC1,NLGN2,CHD4,ATP1A1,EIF5,LENG8,HMGN2,RPL26,BEX3,OA
USP11,COL18A1,SNRNP200,CSDE1,IGF2,EIF3C,ATP5F1A,EIF4B,MARCKSL1,AFF4,TRIM28,MAT
X4,YWHAB,ATP1A2,CALU,ARCN1,CTBP1,ARHGDIA,TOP2B,SLC35E2B,LAPTM4B,RPL23A,HSBP1,
,ZMIZ1,COL11A1,RPL10,WSB1,RPS17,DNAJA1,ARF1,AKAP12,AP2B1,MAT2A,HNRNPA0,FADS2,L
1,CALM1,PLD3,HSPD1,USP11,COL18A1,SNRNP200,CSDE1,EIF3C,ATP5F1A,EIF4B,UBB,AFF4,MA
PT,DDB1,HACD3,DCHS1,SET,MAP9,CHSY1,EIF4A1,RPL19,GTF3C4,LDHB,HLA-C,SF3B1,CHD3,VA
GB1,TUBB4B,MORF4L1,BTF3,ANXA2,AGRN,RPLP1,TIMP2,XAF1,SARAF,MIDN,MARCHF6,WSB1,V
,UBB,AFF4,MATR3,YWHAG,RPL37A,LRRC28,SPEN,NONO,PSMD5,EIF1,EGLN1,INA,EIF4A1,RPL1
RN,RPLP1,SARAF,RPL10,MARCHF6,AP3D1,DNAJA1,ARF1,SF3B2,CDC42BPB,GTF2F1,SLC4A8,R
RPL12,ITM2C,RPL30,EIF3L,GNB2,RPS7,IK,RPS19,CHPF,NGRN,DNAJC5,HNRNPDL,RPLP2,DDOST
IG8,WDR1,RPL26,SRSF4,RBBP4,PFN1,ASTN2,MCL1,TLE5,PNN,RPL31,PRKCSH,KHDRBS1,RPS25
P1,RBBP4,PFN1,JPT2,RPL31,NISCH,KHDRBS1,XPO1,SPTBN2,PFKL,TKT,GLG1,CDH6,EIF5A,PEB
TPT1,RPS3A,RPL18,STMN1,UCHL1,BTF3,AGRN,TIMP2,VCL,PAFAH1B1,MAP4,CDC42BPB,ZNF77
,DNAJC5,RPLP2,DDOST,NARS1,DUX4,YWHAB,PTGES3,RPS12,CALU,MAGED4B,CTBP1,ARHGDIA
2,P4HB,APP,IDS,ERVV-1,RPL5,GDI1,PTMS,MARCKS,SMG1,CALM1,NPM1,PLXNB2,PLD3,HSPD1,
2A,AP2M1,TPT1,SEC61A1,RPS4X,RPS3A,OR8K1,RPL18,STMN1,TUBB4B,RPL11,MORF4L1,BTF3
CHERP,DAP,VPS26B,XRCC6,HSPG2,SHC2,CARM1,UBR5,PKDCC,GPC2,LTBP4,GABARAPL1,MAZ,S
D81,ATP2A2,LASP1,JPT2,NISCH,PRKCSH,TADA2B,PFKL,HDLBP,GLG1,SFPQ,PLXNA1,PIP4K2B,M
,RPL10,MARCHF6,AP3D1,DNAJA1,ARF1,CYB5R3,AKAP12,SF3B2,GTF2F1,RPL23,RPL29,TPM4,R
HAB,LAMP1,HNRNPH1,RPS12,CALU,ARCN1,ARHGDIA,SLC35E2B,IPO5,ATP6V0C,BCLAF1,LAPTM
,VAMP2,GAS7,ABCA3,YEATS2,HSPA4,CAMSAP1,KLC1,GRINA,VPS35,PTOV1,ACAP3,MDK,PTK7,O
1B,CALM3,GNB1,SHISA9,RPS8,MYH9,SEPTIN2,P4HB,APP,IDS,ERVV-1,RPL5,GDI1,FSCN1,TMBI
1,SF3B1,PDIA3,ZNF43,CHD3,BASP1,VAPA,TPT1,RPS4X,RPL18,STMN1,ITGB1,TUBB4B,BTF3,ANX
TPT1,SEC61A1,RPS4X,TUBB4B,RPL11,BTF3,AGRN,RPLP1,MIDN,RPL10,MARCHF6,RPS17,AP3D1,
CAPRIN1,GPC1,HDAC2,MMP2,GLUL,PEA15,NNAT,LUC7L3,CDKN1A,FAM199X,CTDSP2,PGRMC1,
23A,HSBP1,MAP1A,SERF2,DDX39B,SRSF5,TM9SF3,MDH2,U2AF2,EEF1A2,CCT2,SUGP2,GNAI3,G
SL2,TOP1,VHL,SMARCA4,LAPTM4A,PDE7A,RABL6,TCF25,EIF3E,LUZP6,GRB2,TDRD5,SCAF11,CL
1,HDAC2,ATP6V0C,TRIB2,DLG5,HS6ST1,CNN2,NOVA2,CDH11,CHD6,U2AF2,SUGP2,PABPN1,HN
L1,SCAF1,TMED2,RPS15A,RNF10,TUBB2A,PPP2CA,MAPK1,ARPC2,PLEC,SRSF1,PTBP1,ILF3,PCS
S7,RPS19,JUN,COL6A2,RPS10,NDFIP1,WLS,HNRNPK,H1-10,DUX4,PTGES3,SEPTIN3,APPL1,SMC
M171A1,GGNBP2,NOTCH1,IGF1R,APEH,PABPC4,HMGB3,MTURN,TSPAN14,TMEM98,NME2,GID
R,EIF5,HMGN2,RPL26,CRMP1,LASP1,SRSF4,PFN1,MCL1,JPT2,TLE5,NES,NISCH,PRKCSH,TADA2

04,ATP1A1,CD81,HMGN2,LASP1,TLE5,NES,NISCH,SLC3A2,PFKL,HDLBP,PEBP1,CDH2,PDIA6,RAB
2,RPL14,HK1,ZNF865,CLSTN1,ATF4,KIF5B,RAB6B,SLC7A5,NLGN2,ATP1A1,TMEM259,RPS20,EIF
SEPTIN2,P4HB,APP,IDS,RPL5,GDI1,FSCN1,TMBIM6,GSTP1,SMG1,NPM1,PLXNB2,PLD3,HSPD1,US
HS1,SET,EGLN1,TPI1,CHSY1,MRC2,RPL19,PPP2R1A,GTF3C4,DNAH8,PKD1,HLA-C,SSBP2,CHD3,E
ZNF43,RPS3,HMGB1,POLR2A,AP2M1,TPT1,RPS4X,RPS3A,RPL18,ITGB1,TUBB4B,RPL11,BTF3,AG
2B,PRDX2,CLTC,RCN1,RPS5,RPS19,CHPF,CD63,B4GALNT4,APLP2,HNRNPDL,RPLP2,MXD4,RPL
RPL26,SRSF4,PFN1,MCL1,TLE5,LRPPRC,TADA2B,SLC3A2,CDH2,PLXNA1,PIP4K2B,RAB7A,PRDX2
081,LENG8,WDR1,ATXN7L3B,CRMP1,SRSF4,PFN1,MCL1,TLE5,PNN,RPL31,PRKCSH,KHDRBS1,KI
FSCN1,TMBIM6,PTMS,SPTAN1,GSTP1,MARCKS,SMG1,NPM1,PLD3,HSPD1,COL18A1,CSDE1,IGF
LSTN3,MDK,SURF4,RAD23B,ASXL1,VEGFA,OPTN,MAPK1,AZIN1,SRSF9,DSTN,PBX2,AEBP1,NYNR
H1-0,RPS7,IK,RPS19,CHPF,NGRN,DNAJC5,HNRNPDL,RPLP2,DDOST,TSR2,NARS1,DUX4,YWHAB,F
DI2,SSRP1,HSPH1,HNRNPA3,PDXK,ACO2,TCP1,CAMSAP1,KLC1,PTOV1,MLEC,DDX42,CBX1,CCT4
12B,MAT2A,TMEM170A,GTF2F1,PXDN,NFE2L1,SMARCA5,CTNNA1,FADS2,LDHA,CNN3,TNS3,CTS
1,RPS4X,RPS3A,RPL18,STMN1,ITGB1,RPL11,MORF4L1,BTF3,AGRN,RPLP1,SARAF,MIDN,WSB1,
,AKAP12,MAT2A,TMEM170A,NFE2L1,HNRNPA0,CTNNA1,FADS2,RPS16,RPL23,CNN3,TPM4,TNS3
2A,GOLGA8A,PXDN,SMARCA5,HNRNPA0,CTNNA1,RPL23,CPE,TPM4,TNS3,SCARB2,LPCAT1,RPS2
IAPRE1,TCF25,CAPZB,ACTN1,BNIP3L,PLCG1,CCAR2,DIDO1,CORO1C,SMARCC1,NKTR,PTP4A2,L
ORO1C,SMARCC1,GIGYF1,NKTR,STMN3,TNKS2,PLXNA3,ATXN2L,SREBF2,VGF,P3H3,CYP51A1,AF
,SNRNP200,IGF2,ATP5F1A,EIF4B,VCP,MARCKSL1,UBB,AFF4,TRIM28,MATR3,YWHAG,CKB,COL4
PLP1,MIDN,RPL10,MARCHF6,WSB1,RPS17,DNAJA1,ARF1,SF3B2,MAT2A,GTF2F1,PXDN,HNRNPU
P3D1,SBF1,DNAJA1,AP2B1,MAT2A,TMEM170A,NFE2L1,SMARCA5,HNRNPA0,RPS16,CNN3,RPSA,T
1,SF3B1,PDIA3,SERINC1,SSBP2,ZNF43,VAPA,POLR2A,CDKL1,AP2M1,TPT1,SEC61A1,RPS3A,STM
SYNA1,CSNK1D,MESD,HYOU1,MAPK8IP1,FBRSL1,ZSCAN18,CTXN1,CD99L2,PCYOX1,RBM15B,BE
RCC5,RPL24,SLC16A7,PAPOLA,IGFBP3,COL1A2,HK1,RPL22,ZNF865,RPL9,PSMD2,RPS14,CXADF
LLGL1,RHOA,CBX5,CLSTN1,KIF5B,NLGN2,CHD4,TMEM259,EIF5,LENG8,WDR1,RPL26,SRSF4,R
GRN,RPLP1,SARAF,COL11A1,MIDN,MARCHF6,WSB1,VCL,AP3D1,ARF1,SF3B2,CDC42BPB,ITM2B,
DSTN,ODC1,TCF25,GRB2,CAPZB,TDRD5,ACTN1,BNIP3L,CASC3,FKBP8,KIF3C,SMARCC1,PHC2,L
PS7,HMGCS1,GJA1,CTNNB1,RPL35A,HNRNPDL,DDOST,TSR2,RAB1A,TTR,RPS27,HECTD4,RAC1,A
D3,PJA2,PAPOLA,IGFBP3,LLGL1,RPL14,HK1,RPL22,RHOA,PTCD3,CBX5,NR2F1,CHD4,ATP1A1,EI
N,ALDH6A1,SPTBN1,ATP5F1B,CALM3,GNB1,STMN2,SEPTIN2,P4HB,APP,KLRD1,GDI1,FSCN1,TME
LLGL1,NCS1,PSMD2,NLGN2,GLIPR1L2,CXADR,CHD4,WDR1,HMGN2,CRMP1,LASP1,BEX3,PFN1,
L19,PPP2R1A,GTF3C4,SF3B1,SERINC1,BASP1,VAPA,POLR2A,TPT1,RPS3A,OR8K1,RPL18,STMN1,
HMGB1,POLR2A,AP2M1,TPT1,SEC61A1,RPS4X,RPS3A,RPL18,TUBB4B,RPL11,MORF4L1,BTF3,AG
N1,RPS27,CTBP1,ARHGDIA,STIP1,SLC35E2B,IPO5,LAPTM4B,RPL23A,HSBP1,MAP1A,SERF2,SRSF
RPS19,MIF,CHPF,NGRN,DNAJC5,RPLP2,DDOST,NARS1,DUX4,YWHAB,CALU,CTBP1,ARHGDIA,CAF
,B4GALNT4,MTSS2,SHOX,WLS,NARS1,RPL36,DUX4,CADM4,CKAP4,HECTD4,MAGED4B,TOP2B,A
RF1,CYB5R3,AKAP12,SF3B2,ITM2B,MAT2A,NFE2L1,HNRNPA0,FADS2,CNN3,RPL29,RPSA,TPM4,N
S2,EHMT1,SHC2,INTS3,ZNF428,TTC37,PLEKHB2,PCNX3,ADAM10,FBXW5,MBD3,SF3A2,VASH1,SA
N3,TPM4,LPCAT1,LAMA5,SLC22A17,XRCC5,RPL24,RPL17,C1ORF43,RPL22,RHOA,CLSTN1,SLC7
,CALM1,PLD3,HSPD1,SNRNP200,SMIM14,IGF2,EIF3C,ATP5F1A,EIF4B,VCP,MARCKSL1,UBB,TRIM
3,MORF4L1,ANXA2,AGRN,ZMIZ1,RPL10,SBF1,ARF1,CYB5R3,AKAP12,GTF2F1,HNRNPU,NFE2L1,
,RPL23,RPL29,TNS3,EIF4H,LPCAT1,SLC22A17,BRD3,XRCC5,CIPC,RPL24,SLC16A7,KPNB1,PAPC
3L,MAGED2,RPS5,IK,RPS19,CHPF,JUN,NGRN,HNRNPDL,RPLP2,DDOST,CIRBP,NARS1,YWHAB,SN
ORO1C,SMARCC1,NKTR,PTP4A2,LSS,TUG1,PLXNA3,ATXN2L,RAI1,SREBF2,VGF,P3H3,WBP2,AKIR
0,LENG8,WDR1,RPL26,CRMP1,SRSF4,PFN1,MCL1,RPL31,PRKCSH,HNRNPC,XPR1,KHDRBS1,SC

R,NNAT,CPSF2,SRP9,SRSF5,U2AF2,FDFT1,TBCD,ANXA5,CPSF1,GDI2,TSR1,HSPH1,ALKBH5,PDXK
I,INA,EIF4A1,RPL19,PPP2R1A,GTF3C4,PKD1,SF3B1,PDIA3,SERINC1,SSBP2,ZNF43,HMGB1,POLR
F3,NUDT19,NSD2,RBPJ,EHD1,TGOLN2,YY1,TMEFF1,SAP18,UBE2I,UBE2D3,SSBP4,MYL12B,HNR
MARCA5,LARP1,CTNNA1,RPL23,CTSD,SCARB2,LPCAT1,LAMA5,RPS24,CIPC,MSN,NCS1,RPL22,F
,SLC35E2B,IPO5,RPL23A,HSBP1,MAP1A,SERF2,SRSF5,TM9SF3,MDH2,U2AF2,EEF1A2,FDFT1,SU
A,HSBP1,MAP1A,SERF2,DDX39B,NUCB1,SRSF5,TM9SF3,MDH2,U2AF2,EEF1A2,CCT2,SUGP2,GN
5F1A,UBB,CKB,RPL37A,LRRC28,SPEN,DDB1,PSMD5,HACD3,SET,EGLN1,MAP9,TPI1,CHSY1,INA,E
HD6,EEF1A2,MLXIP,CPSF1,HSPH1,LMNB2,ABCA3,PRDX5,ACTR3,HSPA4,KLC1,GRINA,VPS35,PTO
PPR3,ARCN1,RPS27,ARHGDIA,SLC35E2B,IPO5,RPL23A,HSBP1,MAP1A,SRSF5,TM9SF3,MDH2,U
RB2,LPCAT1,SLC22A17,C1ORF43,RPL14,NCS1,ZNF865,PSMD2,KIF5B,CHD4,JUND,TMEM259,EI
,ERVV-1,RPL5,KLRD1,GDI1,GSTP1,SMG1,NPM1,PLXNB2,PLD3,HSPD1,USP11,COL18A1,SNRNP2
3P8,DUSP4,STMN3,TUG1,MEX3A,P3H3,FLNC,BMPR2,CTXN1,YLPM1,DDIT4,BEX4,ANP32A,TRAF4,
HL,G3BP2,MAPK1,SMARCA4,BTBD2,PDE7A,ARGLU1,ILF3,WDR6,TDRD5,ATXN10,HSPB1,KDELR1
,SEPTIN2,P4HB,APP,IDS,RPL5,GDI1,FSCN1,TMBIM6,PTMS,SPTAN1,GSTP1,MARCKS,SMG1,TTC3
NAH8,PKD1,SF3B1,PDIA3,SERINC1,ZNF43,HMGB1,POLR2A,AP2M1,TPT1,RPS4X,RPS3A,RPL18,T
GALNT4,DNAJC5,NDFIP1,MXD4,MYL6,MYH10,YWHAB,SEPTIN11,CADM4,HNRNPH1,PLPPR3,COT
1CAM,LAPTM4B,RPL23A,SQLE,CNN2,SRP9,SRSF5,MDH2,U2AF2,NOP53,PABPN1,HNRNPL,PARP
9,RPSA,TNS3,NACA,LPCAT1,LAMA5,BRD3,PJA2,PAPOLA,RPL14,RPL22,RHOA,CLSTN1,ATF4,PSM
,CPE,LPCAT1,LAMA5,BRD3,RPS9,CIPC,MSN,RPL24,PAPOLA,LLGL1,RPL14,NCS1,PTCD3,KIF5B,
5R3,SF3B2,AP2B1,GTF2F1,OGA,RPS16,RPL23,RPL29,EIF4H,LPCAT1,SLC22A17,BRD3,XRCC5,C
31,PSMD5,YWHAQ,EIF1,EGLN1,TMSB10,EIF4A1,RPL19,PPP2R1A,GTF3C4,PKD1,HLA-C,SF3B1,SE
RVV-1,RPL5,GDI1,FSCN1,GSTP1,MARCKS,SMG1,CALM1,NPM1,PLXNB2,PLD3,HSPD1,USP11,CO
1,TPT1,SEC61A1,OR8K1,RPL18,STMN1,TUBB4B,MORF4L1,BTF3,TIMP2,XAF1,SARAF,RPL10,MA
L38,RPL37,ATG2B,CCNL2,ARPC2,PTBP1,SUPT5H,HNRNPUL1,SON,RPL28,CCAR2,ARF4,SUPT6H,
01,HLA-C,SF3B1,PDIA3,SERINC1,SSBP2,RPS3,CHD3,VAPA,AP2M1,SEC61A1,RPL18,STMN1,ITGB1
ANX,CLU,PTMA,DDX5,ENO2,SOX4,FKBP14,RPL13A,MAGED1,SEC62,RPL6,KAT6A,DVL3,YWHAE,K
DBP,DDOST,H1-10,MYH10,NARS1,RPL36,YWHAB,CADM4,CKAP4,RPS12,SEPTIN3,ZNF454,CCT6A
LA,HK1,RPL22,RHOA,ZNF865,ATF4,KIF5B,RPS14,CXADR,RPS20,LENG8,WDR1,RPL26,CCT7,SRS
5B,SPINT2,ZMYM4,FKBP1A,RABEP1,BRK1,SAFB2,VPS26B,PPP1R10,CEP170,EIF2S2,EHMT1,SHC
I1,PIP4K2B,SOX12,ACTN4,PRDX2,ARL4C,GNB2,RCN1,APLP1,RPS19,TLN1,COL6A2,B4GALNT4,M
G2B,ARPC2,CAPZB,SUPT5H,HNRNPUL1,SON,RPL28,CCAR2,ARF4,DIDO1,SUPT6H,EIF3F,SF3A1,E
CUL3,ADH5,IL6ST,LMAN1,ZMYM3,EFR3B,BAP1,TRIR,XRCC6,VAPB,RTL6,HSPG2,SHC2,SNRPD3,C
1,SUGP2,GNAI3,PARP1,TSR1,MTRF1L,PTPRA,PRDX5,UBE4A,ACO2,HSPA4,TCP1,ZNF302,DDX42,
WHAH,TNKS2,LOC728392,ATXN2L,HGS,BAG6,SREBF2,NCOR2,MEX3A,ZSCAN18,ARNT2,CCT5,P
BRD3,PAPOLA,RPL22,RHOA,CLSTN1,ATF4,PSMD2,RAB6B,SLC7A5,ENC1,NLGN2,RPS14,CHD4,J
ANKFY1,SRSF10,WRNIP1,ECHS1,RPL27A,CCNG1,GID8,TMEM131,MARCHF7,EXOSC10,RANBP9,
NPA3,PDXK,CAMSAP1,CCT3,PTOV1,MDK,CCT4,SKI,SNRNP70,DPYSL2,TAF7,NOTCH2,RNF10,LAP
F4A1,MRC2,GTF3C4,DNAH8,PDIA3,ZNF43,RPS4X,STMN1,TUBB4B,UCHL1,RPL11,BTF3,NCAM1,
3P1,ARHGDIA,CAPRIN1,CDK16,TOP2B,HDAC2,SKP1,PPIB,RPL23A,GLUL,HSBP1,SQLE,DDX39B,S
CO2,CAMSAP1,ACAP3,LAMC1,SKI,SNRNP70,ABCA2,TOP1,MVB12B,AZIN1,MAPRE1,SRSF9,ARPC2
TF2F1,HNRNPU,PGAM1,SMARCA5,LARP1,FADS2,TNS3,EIF4H,LPCAT1,SLC22A17,RPS9,PJA2,C1
L10,MARCHF6,RPS17,AP3D1,DNAJA1,ARF1,CYB5R3,AKAP12,SF3B2,GTF2F1,SMARCA5,RPS16,
7,NOTCH2,RNF10,PDE7A,PSMD3,RABL6,ODC1,SRSF1,IPO7,ARGLU1,TCF25,ADAM9,FTO,ILF3,W
REBBP,XPO6,PRDX3,ATG4B,DMWD,SNX2,UBE2L3,PUM1,SGSM2,USP10,TIA1,ADNP,SYNCRIP,SSU
I2,CTSB,TBCD,ANXA5,CPSF1,PGD,LMNB2,PDXK,ACAP3,MDK,H3-3A,VEGFA,RAPGEF1,MAPK1,MA

2,H1-0,RPS5,IK,RPS19,CHPF,JUN,NGRN,HNRNPDL,RPLP2,DDOST,HNRNPK,CIRBP,NARS1,SND1,C
LC22A17,BRD3,RPS9,PJA2,KPNB1,RPL17,LLGL1,NCS1,RHOA,ZNF865,CLSTN1,RAB6B,ENC1,GL
RPL23A,HSBP1,NNAT,SERF2,HS6ST1,NOVA2,CTSB,MDH2,EEF1A2,TBCD,MLXIP,SUGP2,ANXA5,P
LON5,ELAVL3,PAFAH1B2,AMFR,USP47,OLFM2,HMGA1,LUC7L2,RGL2,RBM25,MAPK8IP2,MGAT3
2,GNG4,EP400,UBFD1,KCTD12,ZNF512B,CUL3,ULK1,LMAN1,ZMYM4,PEG10,CHERP,MAP1LC3B
LC25A3,SHOX,HNRNPK,YWHAB,SEPTIN3,COTL1,MAGED4B,GPC1,TMEM30A,PEA15,PTTG1IP,DL
VHAQ,DCHS1,SET,MAP9,TPI1,INA,MRC2,RPL15,GTF3C4,DNAH8,PKD1,HLA-C,SSBP2,CHD3,BASP1
1,PTEN,SELENOW,C1ORF198,DAP,ZNF664,PELP1,IARS2,SNN,YARS1,HSPG2,OLFM1,PKDCC,ZMI
1,RPL18,STMN1,TUBB4B,MORF4L1,BTF3,NCAM1,RPLP1,TIMP2,XAF1,SARAF,RPL10,WSB1,AP3D
RBP,PTGES3,ATP1A2,SEPTIN3,MAGED4B,CAPRIN1,STIP1,SMCHD1,HDAC2,IPO5,PPIB,GLUL,SERF
A,SRSF5,TM9SF3,MDH2,U2AF2,EEF1A2,FDFT1,CCT2,SUGP2,GNAI3,NOP53,PARP1,TSR1,MTRF1
BE4A,TCP1,CAMSAP1,KLC1,ACAP3,RBMX,CBX1,CCT4,SKI,RPL36A,TSPAN6,TMED2,RPL38,VEGF
DLG5,TIMP3,PGRMC1,NOVA2,TM9SF3,VCAN,HSPH1,HNRNPA3,KLC1,LAMC1,RAB5B,CCT4,H3-3A
T1,XPOT,PDAP1,AP2A1,MMP15,FLOT1,SEC16A,PHB2,APMAP,MGAT4B,PLXNB1,DECR1,FARP1,RE
TMEM259,EIF5,LENG8,HMGN2,BEX3,RBBP4,JPT2,PRKCSH,HNRNPC,LRPPRC,KHDRBS1,XPO1,SI
SET,MAP9,CHSY1,INA,EIF4A1,MRC2,PPP2R1A,SERINC1,CHD3,BASP1,HMGB1,POLR2A,TPT1,SEC6
,YME1L1,TRAF7,DFFB,STAT3,EIF2AK1,EDC4,PDAP1,EIF3D,PHB2,APMAP,GSPT1,KLHL42,CSNK1C
CHF6,WSB1,AP3D1,SBF1,ARF1,CYB5R3,SF3B2,CDC42BPB,ITM2B,MAT2A,NFE2L1,SMARCA5,FAD
PL14,HK1,NCS1,RPL22,RHOA,SLC7A5,GLIPR1L2,TMEM259,CD81,WDR1,ATP2A2,CRMP1,LASP1,
V0C,SKP1,LAPTM4B,TRIB2,PPIB,RPL23A,HSBP1,SQLE,MAP1A,DDX39B,TM9SF3,MDH2,TOMM20
B4B,AGRN,SARAF,MIDN,MARCHF6,WSB1,SBF1,DNAJA1,ARF1,CYB5R3,AKAP12,TMEM170A,GTF2
ARAF,RPL10,WSB1,AP3D1,SBF1,ARF1,CYB5R3,AKAP12,SF3B2,ITM2B,MAT2A,NFE2L1,HNRNPA0,
S1,EGLN1,TMSB10,EIF4A1,MRC2,RPL19,PPP2R1A,LDHB,PKD1,HLA-C,SERINC1,SSBP2,BASP1,PC
,PRKAR2A,SF1,EIF3D,UBAP2L,GLUD1,UPF1,MBTPS1,TBC1D9B,NPDC1,IL6ST,FKBP1A,MAP1LC3
D1,PI4KA,ZFR,ZNF512B,CEMIP2,CBX6,NPDC1,IL6ST,ANP32B,BRSK1,SEPTIN7,GALNT11,SLIT2,D
HNRNPA0,RPS16,RPL23,RPL29,EIF4H,LAMA5,RPS9,CIPC,RPL24,PAPOLA,RHOA,ZNF865,RPL9,AT
X,GNG4,GLUD1,ZNF512B,CEMIP2,ULK1,BRSK1,GPSM1,DKK3,CTTN,DAP,PCBP4,PRDX6,RHBDD
IRNPA0,FADS2,RPS16,LDHA,CPE,CNN3,TPM4,CTSD,LPCAT1,SLC22A17,BRD3,RPS9,CIPC,RPL24,
SLC35F5,RAB2A,GNG4,EP400,ZFR,TBC1D9B,PPP1CC,IL6ST,SPINT2,C1ORF198,PRDX6,EFR3B,S
P2,IDH2,SYNCRIP,NTN1,ZNF462,CUL4A,ARHGAP5,CCDC88A,LDB1,STUB1,ZNF503,TNFRSF10B,S
MBD3,SF3A2,AP1M1,SAP18,BNIP3,SAE1,UBE2K,USO1,ATP5MG,NELFB,CDC34,USF2,SPPL2B,USI
1A,CASC3,CCAR2,DIDO1,HNRNPM,SRSF3,UBQLN1,MESD,GHITM,PFN2,RPL39,ARNT2,ADAR,BEX
ONO,DDB1,DCHS1,SET,MAP9,CHSY1,EIF4A1,RPL19,GTF3C4,LDHB,HLA-C,SF3B1,ZNF43,VAPA,HI
I1,ATP5F1B,CALM3,GNB1,SHISA9,STMN2,RPS8,MYH9,SEPTIN2,P4HB,ERVV-1,RPL5,KLRD1,FSCN
MSAP1,NCKAP1,COL2A1,RAB5B,GCN1,SKI,HDGF,PYGB,TAF7,RPL38,SELENON,RPL37,CCND1,G
GES3,SND1,CALU,ARCN1,ARHGDIA,STIP1,SLC35E2B,ATP6V0C,LAPTM4B,TRIB2,RPL23A,FAM19
PSA,TPM4,NACA,LAMA5,RPS24,BRD3,RPS9,CIPC,RPL24,C1ORF43,RPL14,PTCD3,CBX5,ZNF865,C
ID1,NYNRIN,TCF25,ATP5PB,PCSK1N,CAPZB,ACTN1,MAP7D1,FKBP8,PLTP,MSL1,HNRNPM,PHC2,
,FTO,ILF3,CAND1,SON,ZFAND5,SLIT3,STT3B,TUG1,ATXN2L,HECTD1,HGS,BAG6,SERBP1,SMS,RE
WHAE,RPL13,GNAS,DDX17,GPI,HTRA4,RACK1,BSG,FASN,ALDH6A1,SPTBN1,ATP5F1B,CALM3,GI
OLR2A,TPT1,SEC61A1,RPS3A,RPL18,ITGB1,MORF4L1,NCAM1,ANXA2,AGRN,RPLP1,ZMIZ1,SARA
MLF2,CTXN1,CD99L2,DDIT4,CIC,RBBP7,TRAF7,WDR45B,NAP1L1,CPD,EP300,DGCR2,RETREG2,
3,PRDX5,UBE4A,ACO2,HSPA4,ZNF302,DDX42,GCN1,SKI,RAB11B,PRPF6,TERF2IP,RPL37,ARPC2,
7,AP3D1,SBF1,ARF1,MAP4,SF3B2,ITM2B,TMEM170A,GTF2F1,SLC4A8,HNRNPU,OGA,PGAM1,NF
XN2L,ID4,VGF,P3H3,ARNT2,WBP2,AKIRIN1,SMS,NAP1L4,YME1L1,ADAR,TRAF4,RBBP7,TXLNA,TF

PS5,IK,RPS19,TLN1,CHPF,JUN,NGRN,RPLP2,DDOST,CIRBP,NARS1,RPL36,YWHAB,SND1,TTR,ARC
SQSTM1,VAMP2,GDI2,TM9SF2,MORF4L2,HSPH1,ALKBH5,HNRNPA3,PRDX5,PDXK,UBE4A,PKN1,H
RPS3,CHD3,HMGB1,RPS3A,ITGB1,UCHL1,RPL11,BTF3,AGRN,ZMIZ1,COL11A1,MIDN,RPL10,WSB
,GTF2F1,OGA,HNRNPA0,RPL23,RPL29,EIF4H,LAMA5,RPS9,XRCC5,CIPC,RPL24,PAPOLA,NCS1,RI
,ADH5,MCM3AP,RPS27A,TMEM167A,KLHDC3,CHERP,CTTN,LRP10,MTCH1,ERH,CYTH2,RAB1B,F
,RPL17,C1ORF43,GAP43,HK1,NCS1,PTCD3,CLSTN1,KIF5B,RAB6B,SLC7A5,ENC1,CHD4,JUND,T
D2,NLGN2,ATP1A1,RPS20,CD81,COL5A2,WDR1,ATP2A2,CRMP1,LASP1,BEX3,VAT1,NES,CCNI,KL
39B,SRSF5,TM9SF3,MDH2,TOMM20,U2AF2,CCT2,GNAI3,NOP53,TM9SF2,HSPH1,LMNB2,MTRF1
1,SSRP1,MTRF1L,PRDX5,UBE4A,ACO2,HSPA4,TCP1,AHCYL1,MLEC,NRDC,PRPF6,RPS15A,RPL37
VAMP2,PRDX5,PDXK,PKN1,GRINA,NCKAP1,COL2A1,PTOV1,CLSTN3,MDK,JPT1,RAD23B,ORAI2,T
D1,CALU,ARCN1,RPS27,ARHGDIA,STIP1,SLC35E2B,IPO5,RPL23A,HSBP1,MAP1A,SRSF5,TM9SF3,
M1,TPT1,SEC61A1,RPS3A,RPL18,ITGB1,TUBB4B,RPL11,BTF3,AGRN,RPLP1,MIDN,RPL10,MARCH
M1,TPT1,SEC61A1,RPS3A,RPL18,ITGB1,TUBB4B,RPL11,BTF3,AGRN,RPLP1,MIDN,RPL10,MARCH
,MAP9,CHSY1,EIF4A1,RPL19,GTF3C4,LDHB,HLA-C,SF3B1,ZNF43,VAPA,HMGB1,CDKL1,SEC61A1
P2B,HDAC2,ATP6V0C,SKP1,RPL23A,HSBP1,DDX39B,SRSF5,TM9SF3,MDH2,TOMM20,U2AF2,FDF
8,HNRNPU,OGA,NFE2L1,SMARCA5,RPL23,LDHA,CNN3,RPSA,TPM4,EIF4H,NACA,TSPAN3,RPS24,
ITM2C,RPL30,EIF3L,MAGED2,RPS5,RPS19,CHPF,JUN,NGRN,SESN3,RPL35A,DNAJC5,APLP2,RPL
,GOLGA8A,HNRNPU,OGA,PGAM1,FADS2,RPL23,LDHA,CNN3,RPSA,SCARB2,LPCAT1,RPS24,XRC
WDR45B,PDAP1,MMP15,PHB2,GSPT1,UBA52,RER1,RAB3GAP2,MCM3AP,RPL7L1,RPS27A,TMEM
BP1,WBP2,AKIRIN1,CCT5,CDC42,TRAF7,EIF2AK1,TXLNA,PRMT1,STARD7,PDAP1,MMP15,PHB2,U
,SND1,CALU,ARCN1,COTL1,ARHGDIA,SLC35E2B,ATP6V0C,LAPTM4B,TRIB2,RPL23A,HSBP1,FAM
RAPGEF1,TUBB2A,PBX2,PDE7A,FAM171A2,NYNRIN,PTBP1,DYNLL2,WDR6,PRKDC,GABARAP,ACT
EC16A,SLC39A7,RAB3GAP2,EIF5B,ZMYM4,NPLOC4,FKBP1A,RABEP1,SAFB2,ZNF664,VPS26B,PF
7,SEPTIN2,P4HB,IDS,ERVV-1,KLRD1,FSCN1,TMBIM6,PTMS,SPTAN1,MARCKS,CALM1,NPM1,PLX
L23A,HSBP1,DDX39B,SRSF5,TM9SF3,MDH2,TOMM20,U2AF2,EEF1A2,FDFT1,CCT2,GNAI3,NOP5
R,NPTN,MGAT3,DCAF5,WDR26,FXR1,SYNCRIP,ZNF462,FAM53C,SALL2,RBFOX2,MAPRE3,TNFR
BF1,ARF1,CYB5R3,AKAP12,SF3B2,CDC42BPB,ITM2B,MAT2A,NFE2L1,FADS2,RPS16,RPL23,RPL
L26,CCT7,SRSF4,RBBP4,PFN1,ASTN2,MCL1,PNN,RPL31,PRKCSH,LRPPRC,SLC3A2,SCRN1,RPS2
PF40A,CD99L2,YLPM1,ADAR,CTNND1,GRIPAP1,ANP32A,TRAF4,RETREG2,NOL4L,LRRC4B,BRD2
ZMIZ1,TIMP2,COL11A1,MIDN,RPL10,WSB1,VCL,DNAJA1,PAFAH1B1,MAP4,SF3B2,AP2B1,MAT2A,
IB2,GLUL,PTTG1IP,SQLE,MAP1A,CDKN1A,SERF2,TIMP3,CTDSP2,CNN2,NOVA2,NUCB1,ZC3H7B,
2,BASP1,VAPA,POLR2A,CDKL1,SEC61A1,RPL18,ITGB1,TUBB4B,BTF3,NCAM1,ANXA2,AGRN,ZMI
1,PLXNA1,RAB7A,CLTC,GNB2,RPS5,IK,GPAT4,PRKAR1A,TLN1,CHPF,NGRN,CYFIP2,HNRNPDL,SL
NRNPM,SRSF3,LARS1,YWHAH,LONP1,TUG1,MESD,MAPK8IP1,PFN2,HNRNPD,CYP51A1,RPS13,D
RSF5,TM9SF3,MDH2,TOMM20,U2AF2,FDFT1,CCT2,SUGP2,GNAI3,TSR1,LMNB2,PRDX5,UBE4A,A
KBP14,RPL13A,MAGED1,SEC62,RPL6,KAT6A,DVL3,YWHAE,KIF1A,RPL13,GNAS,SLC2A3,DDX17,G
M9SF3,MDH2,U2AF2,EEF1A2,FDFT1,CCT2,SUGP2,GNAI3,NOP53,PARP1,TSR1,MTRF1L,RPL27,PT
7,DNAJA1,ARF1,SF3B2,MAT2A,GTF2F1,HNRNPU,HNRNPA0,RPS16,RPL23,RPL29,EIF4H,XRCC5,S
,INTS3,MAZ,DCLK1,PRR12,CAPZA1,ATXN3,ELP1,SLC4A2,SPAG9,PLEKHM2,PDPK1,DCUN1D2,RE
,INTS3,MAZ,DCLK1,PRR12,CAPZA1,ATXN3,ELP1,SLC4A2,SPAG9,PLEKHM2,PDPK1,DCUN1D2,RE
SERP1,ERH,EIF3H,RALY,SEC14L1,NUDCD3,GNL1,HDGFL2,TUBGCP6,RIC8A,NCSTN,INTS11,MOA
ATP1B2,FOXK2,NSD2,AKT2,LRP3,MED13,VASH1,PACS2,YY1,TMEFF1,CREBBP,ATXN7L3,BNIP3,S
A4,TCP1,MLEC,ZNF302,DDX42,GCN1,SRP14,PRPF6,TAF7,RPL37,ATG2B,CCNL2,ARPC2,FAM168
RINA,VPS35,PTOV1,MLEC,ACAP3,SURF4,PCDHGB6,SNRNP70,PYGB,PRPF6,DPYSL2,RAPGEF1,PP
232,TBC1D9,PRR12,TMEM63B,EEF1B2,QPRT,SLC25A23,DCUN1D2,PTGFRN,YY1,GNA13,TRO,N

CCN2,MIF,COL6A2,GOLM1,B4GALNT4,DNAJC5,HNRNPDL,RPLP2,MXD4,MYL6,H1-10,CKAP5,SEF
GA,NFE2L1,CNN3,RPL29,TSPAN3,LPCAT1,XRCC5,MSN,KPNB1,C1ORF43,PAPOLA,RPL22,ZNF86
M4B,PEA15,DLG5,TIMP3,TMEM132A,PGRMC1,DDX39B,CNN2,TM9SF3,CHD6,RAB27A,EEF1A2,A
H5,RPL27,PRDX5,UBE4A,HSPA4,TCP1,MLEC,ACAP3,GCN1,TNPO1,SKI,SRP14,PRPF6,TAF7,RPL3
1,DLG5,HS6ST1,TIMP3,NOVA2,SRSF5,TBCD,SNX22,SQSTM1,MEST,GAS7,HNRNPL,ALKBH5,PRD
1,CLSTN3,DDX42,RAB11B,NRDC,PRPF6,TUBB2A,UBXN4,PPP2CA,PLEC,PDE7A,PSMD3,SRSF1,GF
RMT1,PDAP1,MMP15,PHB2,APMAP,KLHL42,PLXNB1,DECR1,RER1,UBE2G2,RAB3GAP2,MASTL,Y
N,TARDBP,RPLP2,DDOST,CIRBP,NARS1,SND1,CALU,PLPPR3,ARCN1,CTBP1,ARHGDIA,SLC35E2B,
PAT4,JUN,MXD4,MYH10,RPL36,PTGES3,RHOB,HNRNPH1,RPS12,PLPPR3,SEPTIN3,MAGED4B,CT
PYGB,PRPF6,TAF7,RPL38,OPTN,RPL37,ATG2B,CCNL2,ARPC2,FAM168B,ARGLU1,ACADVL,PTBP
PYGB,PRPF6,TAF7,RPL38,OPTN,RPL37,ATG2B,CCNL2,ARPC2,FAM168B,ARGLU1,ACADVL,PTBP
STN3,ACAP3,MDK,SURF4,ASXL1,RAPGEF1,TUBB2A,MAPRE1,ARPC2,FAM171A2,SRSF1,ACADVL,
MDH2,RAB27A,U2AF2,CCT2,GNAI3,NOP53,VCAN,TM9SF2,MTRF1L,RPL27,PRDX5,C4ORF3,TCP1,
SRP1,ALKBH5,UBE4A,AHCYL1,ETNK1,CCT3,NCKAP1,MLEC,CLSTN3,CBX1,PTK7,SKI,RPL36A,SR
SBF1,ARF1,AKAP12,MAP4,CDC42BPB,PXDN,HNRNPU,HNRNPA0,CPE,RPSA,TSPAN3,LPCAT1,RPS
IP,SQSTM1,MORF4L2,HSPH1,MTRF1L,ALKBH5,HNRNPA3,UBE4A,TBC1D16,AHCYL1,ETNK1,KLC
GB1,TUBB4B,UCHL1,RPL11,MORF4L1,BTF3,ANXA2,AGRN,MIDN,MARCHF6,WSB1,AP3D1,SBF1,
EN,NONO,DDB1,PSMD5,YWHAQ,EIF1,EGLN1,MAP9,TPI1,TMSB10,EIF4A1,MRC2,RPL15,GTF3C4,
K,PLOD1,GRINA,VPS35,PTOV1,MLEC,ACAP3,SURF4,PCDHGB6,SNRNP70,PYGB,PRPF6,DPYSL2,N
KIF3C,ID4,NBPF9,COL27A1,SREBF2,PFN2,MEX3A,CYP51A1,KDM3B,RPS13,WBP2,NAP1L4,CTXN
TXN10,QARS1,RPL28,GNG2,SMARCC1,SUPT6H,DHX30,EIF3F,HGS,BAG6,RAB11A,CCT5,CDC42,Y
F1,ARF1,CYB5R3,AKAP12,SF3B2,ITM2B,MAT2A,NFE2L1,FADS2,RPS16,RPL23,CPE,CNN3,RPL29,
P70,PYGB,PRPF6,RAPGEF1,SMARCA4,AZIN1,DDAH2,ARGLU1,NYNRIN,PRKDC,TDRD5,CASC3,RPL
ASTL,ECEL1,TMEM167A,PSMD8,CTTN,DAP,SERP1,AKAP9,VPS26B,PPP1R10,PSMB4,EIF3H,SEC14
2,TOMM20,U2AF2,CCT2,GNAI3,TM9SF2,HSPH1,RPL27,PRDX5,ACTR3,UBE4A,PKN1,HSPA4,TCP1
UN,ANXA6,NGRN,RPLP2,DDOST,CIRBP,NARS1,SND1,CALU,ARCN1,COTL1,ARHGDIA,SLC35E2B,

3,VAPA,POLR2A,CDKL1,AP2M1,SEC61A1,ITGB1,UCHL1,MORF4L1,AGRN,ZMIZ1,TIMP2,XAF1,SAF
J,ARCN1,ZNF454,RPS27,ARHGDIA,STIP1,SLC35E2B,IPO5,RPL23A,HSBP1,MAP1A,SRSF5,TM9SF3
MN3,STT3B,PITPNM1,ISYNA1,PLXNA3,LOC728392,ID4,COL27A1,HGS,SOD1,FBRSL1,MEX3A,SER
0,U2AF2,CCT2,GNAI3,NOP53,GDI2,TM9SF2,TSR1,SSRP1,LMNB2,MTRF1L,ALKBH5,SNRPN,PRD
EBBP,ANKRD12,XPO6,BAG5,UBXN6,SLC20A1,MICAL1,DPYSL4,METRN,UNC119B,SGSM2,CGGBF
N1,PSMD2,CXADR,CHD4,ATP1A1,LENG8,WDR1,BEX3,OAZ1,SRSF4,JPT2,PNN,NES,LRPPRC,TADA
SERBP1,ARFGEF2,AKIRIN1,CDC42,RBBP7,TRAF7,EIF2AK1,PRMT1,XPOT,SF1,RETREG2,PDAP1,
D2,PIP4K2B,SOX12,CCN2,PRDX2,DDX3X,ITM2C,MAGED2,APLP1,MIF,MYO5A,COL6A2,SESN3,TAF
BC1D16,AHCYL1,ETNK1,KLC1,CLSTN3,DDX42,RAB11B,NRDC,PRPF6,TUBB2A,UBXN4,PPP2CA,G
C2A,PTPRF,PSMD8,HIF1A,GPC4,C1ORF198,WSCD1,TRIR,CEP170,FKBP4,UBR5,SOX2,RALY,ZMI
IP,CIRBP,DUX4,TTR,CALU,SEPTIN3,RPS27,CCT6A,SMCHD1,HDAC2,ATP6V0C,TMEM30A,SKP1,L1
RD1,GDI1,FSCN1,TMBIM6,PTMS,SPTAN1,GSTP1,CALM1,NPM1,PLXNB2,HSPD1,COL18A1,SNRN
OD1,BAG6,SREBF2,PFN2,VGF,ARNT2,PPP1R9B,PSMA7,VDAC1,TRAF7,KIRREL1,LBH,NDUFV1,SF
M9SF3,MDH2,TOMM20,U2AF2,CCT2,GNAI3,NOP53,GDI2,VCAN,TM9SF2,TSR1,SSRP1,LMNB2,M
P7,SMS,TXNIP,CCT5,PSMA7,RBBP7,BCL9L,CAPN2,EXTL3,NDUFV1,SF1,GPX4,ZFP36L1,GARS1,N
2A2,DNAJB2,ITGAV,USP24,NOC2L,EIF3G,DENND4B,PTGFRN,YY1,PHIP,GNA13,FJX1,DEK,UNC11
,TSR2,MYL6,SHOX,INIP,MYH10,PTGES3,SND1,HNRNPH1,UBE2H,ARCN1,RPS27,HMGCR,SMCHD
P,CKAP4,TTR,RPS27,COTL1,GPC1,TRIB2,MAP1A,CDKN1A,CTDSP2,PGRMC1,EID1,C5ORF24,CDH

PL1,IPO5,LAPTM4B,PTTG1IP,DLG5,NOVA2,NUCB1,TM9SF3,RNF187,U2AF2,FDFT1,ANXA5,PPP1C
,PCNX3,ABCF1,ERGIC3,IST1,ATP6V0D1,CSE1L,PCGF2,MBD3,NASP,CREBBP,ANKRD12,BAG5,SL
1,KLC1,CLSTN3,DDX42,RAB11B,NRDC,PRPF6,TUBB2A,UBXN4,PPP2CA,G3BP2,KDELR2,PDE7A,
6,RIC8A,NCSTN,INTS11,AHSA1,MOAP1,MTMR4,PCNX3,ERGIC3,SPTY2D1,CAP1,NOC2L,COX6A1,
,ANKRD50,ATRN,ZBED4,ZFP91,DLG4,SLC12A7,DUSP8,METTL9,TBKBP1,ERBB2,CDC42EP4,NR2
C,CLTC,MAGED2,RPS7,CHPF,COL6A2,CD63,MTSS2,APLP2,TARDBP,SLC25A3,ST3GAL2,TSR2,INI
,MICAL3,RAB18,OTUD5,ZBTB4,CRTC1,TMEM43,SRP54,ACP1,RBM17,PRMT2,NADK,ZNF146,SEN
,GSTP1,MARCKS,SMG1,CALM1,NPM1,TTC3,PLD3,HSPD1,USP11,COL18A1,SNRNP200,SMIM14,
7,SCAMP5,CHMP1A,HDGFL3,LUC7L2,PAIP2,CLIC1,GGNBP2,NPTN,ADNP,SCAMP4,ADGRB2,HTR
CC5,CIPC,IGFBP3,RPL22,ZNF865,RPL9,ATF4,KIF5B,RPS14,CXADR,LENG8,WDR1,RPL26,LASP1,
PLD3,SNRNP200,CSDE1,IGF2,EIF4B,VCP,MARCKSL1,UBB,AFF4,TRIM28,YWHAG,CKB,COL4A2,M
TAF15,PDXK,TCP1,CAMSAP1,COL2A1,PTOV1,MLEC,ACAP3,SCAF1,TSPAN6,DPYSL2,AZIN1,PDE7A
LU1,TCF25,ADAM9,ILF3,SON,SLIT3,LONP1,ATXN2L,HECTD1,BAG6,SERBP1,RPL34,RBM15B,TXL
,HLA-C,SF3B1,VAPA,HMGB1,POLR2A,CDKL1,SEC61A1,STMN1,ITGB1,TUBB4B,UCHL1,RPL11,N
1,RPL15,GTF3C4,SSBP2,CHD3,VAPA,HMGB1,POLR2A,AP2M1,STMN1,ITGB1,UCHL1,MORF4L1,
M1,RALY,GPC2,FAM234A,LDLR,LRATD2,PRR12,LSM14B,SPAG9,PCNX3,DAZAP2,ERGIC3,SCRIB,
CA,G3BP2,KDELR2,PDE7A,PSMD3,TCF25,ADAM9,CDC37,WDR6,GABARAP,BNIP3L,HNRNPUL2-B
UGP2,PARP1,TSR1,LMNB2,MTRF1L,PTPRA,PRDX5,UBE4A,ACO2,HSPA4,ZNF302,DDX42,GCN1,S
,PHGDH,FOXK2,NUDT19,FBXW5,SF3A2,PACS2,PHIP,WNK1,FJX1,MYL12B,RCN2,EIF4G1,MZT2B,
N1,ASTN2,MCL1,PNN,RPL31,PRKCSH,LRPPRC,KHDRBS1,SCRN1,RPS25,SFPQ,EIF5A,PDIA6,DDR
1,HNRNPL,LMNB2,PTPRA,ACTR3,YEATS2,TBC1D16,ACO2,RBMX,CHD2,ASXL1,H3-3A,SCAF1,TER
IOX,HNRNPK,RPL36,DUX4,CADM4,SND1,ATP1A2,PLPPR3,MAGED4B,RAC1,TOP2B,SLC35E2B,PP
LRP10,SERP1,ERH,EIF2S2,EIF3H,RALY,SEC14L1,NUDCD3,GNL1,PNISR,STK24,TUBGCP6,RIC8A,
N,PPP2CA,PSMD3,ODC1,SRSF1,TCF25,ADAM9,DYNLL2,WDR6,CAPZB,PRKDC,SUPT5H,SON,MAP
CPSF1,MEST,GAS7,MORF4L2,SNRPN,PLOD1,VPS35,PTOV1,MLEC,ACAP3,SURF4,PCDHGB6,DPYS
PTBN1,GNB1,SHISA9,RPS8,MYH9,RPL7,SEPTIN2,P4HB,IDS,ERVV-1,RPL5,GDI1,FSCN1,TMBIM6,P
1,CBX5,PSMD2,GLIPR1L2,ATP1A1,JUND,TMEM259,EIF5,LENG8,WDR1,MCL1,PNN,CCNI,TADA2B
RPS9,PJA2,RPL24,SLC16A7,IGFBP3,NCS1,CBX5,GLIPR1L2,ATP1A1,JUND,TMEM259,LENG8,WD
,PSAP,PGK1,RPLP0,H3-3B,CANX,CLU,PTMA,DDX5,ENO2,SOX4,FKBP14,RPL13A,MAGED1,SEC62
C1LI2,NYNRIN,SUPT5H,ATXN10,MAP7D1,KDELR1,MAP2K2,HNRNPUL2-BSCL2,DLK1,MSL1,STM
DRBS1,PFKL,GLG1,PEBP1,PNMA1,DDR1,PIP4K2B,ITM2C,H1-0,APLP1,SESN3,MTSS2,APLP2,TSR
1L,ALKBH5,SNRPN,PRDX5,UBE4A,HSPA4,TCP1,MLEC,ACAP3,SRP14,NRDC,PRPF6,TAF7,MAPRE1
RPF6,PPP2CA,SMARCA4,AZIN1,DDAH2,ARGLU1,NYNRIN,TCF25,PRKDC,TDRD5,CASC3,RPL28,ZF
E4A,ACO2,HSPA4,TCP1,MLEC,ACAP3,SRP14,NRDC,PRPF6,TAF7,ARF3,MAPRE1,FAM168B,LUZP6
,DCHS1,SET,CHSY1,INA,EIF4A1,MRC2,PPP2R1A,RPL15,LDHB,DNAH8,PKD1,ZNF43,RPS3,BASP1,

IZ1,WSB1,RPS17,VCL,CYB5R3,MAP4,SF3B2,AP2B1,GTF2F1,PXDN,PGAM1,NFE2L1,SMARCA5,C
B,AFF4,TRIM28,MATR3,YWHAG,COL4A2,RPL37A,FN1,SPEN,PSMD5,YWHAQ,HACD3,DCHS1,SET,
P1,POLR2A,SEC61A1,STMN1,ITGB1,TUBB4B,UCHL1,MORF4L1,NCAM1,ANXA2,AGRN,TIMP2,MI
T1,CCT2,GNAI3,NOP53,TSR1,HSPH1,MTRF1L,RPL27,PRDX5,HSPA4,TCP1,CCT3,MLEC,ACAP3,G
3,LDHA,CPE,RPSA,TNS3,CTSD,EIF4H,NACA,RPS9,PJA2,RPL24,SLC16A7,KPNB1,C1ORF43,PAPOL
CBX1,RAB5B,ORAI2,SNRNP70,TSPAN6,PRPF6,TAF7,RAPGEF1,G3BP2,ATG2B,ARPC2,DDAH2,SR
PBP,CGGBP1,POR,RTF1,TOX4,NIBAN2,HIP1R,FZR1,PSMD13,PPP1R14B,CRIP2,PHPT1,CNDP2,E
,FADS2,LDHA,CPE,CNN3,RPSA,CTSD,SCARB2,LPCAT1,SLC22A17,RPS9,CIPC,PJA2,RPL24,C1OR
,TM9SF3,MDH2,TOMM20,U2AF2,CCT2,GNAI3,NOP53,GDI2,TM9SF2,TSR1,SSRP1,LMNB2,MTRF

,GNB2,RPS5,MIF,PRKAR1A,CHPF,NGRN,B4GALNT4,HNRNPDL,TARDBP,RPLP2,ARL6IP1,NDFIP1,
,SLC16A7,IGFBP3,CBX5,PSMD2,GLIPR1L2,JUND,TMEM259,EIF5,LENG8,WDR1,MCL1,PNN,CCNI,
ERBP1,AKIRIN1,CCT5,PPP1R9B,YME1L1,TXLNA,BCR,XPOT,CAPN2,PDAP1,GTPBP1,SEC16A,PHB
8,TUBB4B,BTF3,AGRN,RPLP1,ZMIZ1,MIDN,RPL10,MARCHF6,DNAJA1,ARF1,CYB5R3,AKAP12,MA
2,CAPN1,MAGEF1,CDK4,FAM168A,JAK1,SLC16A2,APPBP2,UBE2Q1,BCL9,RBFOX2,ZNF768,RABA
SALL2,FAM168A,RBFOX2,ZNF768,RABAC1,NSD1,WAC,IGF2BP2,ALCAM,PLXDC2,MED25,VEZF1,
AND5,MSL1,RPS28,LARS1,STMN3,FBLN1,STXBP1,STT3A,ATXN2L,STRAP,P3H3,RTL8C,AMOTL2,
TOMM70,TRIM8,SSBP4,GNA13,FJX1,SMO,TEAD1,CENPB,SCAP,AMFR,CELSR1,AGAP1,SGSM2,FA
GB1,POLR2A,AP2M1,SEC61A1,RPS4X,RPS3A,TUBB4B,MORF4L1,BTF3,AGRN,RPLP1,SARAF,RPL
PA,AP2M1,STMN1,ITGB1,TUBB4B,MORF4L1,ANXA2,AGRN,XAF1,SARAF,MIDN,RPL10,WSB1,AP3
JAJA1,ARF1,CYB5R3,SF3B2,CDC42BPB,ITM2B,MAT2A,SLC4A8,GOLGA8A,HNRNPU,SMARCA5,HN
3GAL6,EIF2AK1,SGTA,LBH,NDUFV1,ALDH3A2,BAZ1B,H2AZ1,SEC16A,GPX4,PPP2CB,PLXNB1,FA
GA,HNRNPA0,RPL23,RPL29,TPM4,XRCC5,CIPC,RPL24,KPNB1,PAPOLA,HK1,RPL22,RHOA,ZNF8
RF4L1,NCAM1,ZMIZ1,XAF1,SARAF,COL11A1,MARCHF6,RPS17,ARF1,CYB5R3,ITM2B,MAT2A,OG
ED2,TOP1,TAF7,RPL38,NOTCH2,RNF10,SMARCA4,AZIN1,CCNL2,PDE7A,PSMD3,PRDX1,RABL6,S
TTN,MAP1LC3B,SERP1,RHBDD2,STK35,ERH,EIF2S2,CYTH2,SNRPD3,UBE2M,SEC14L1,ELAVL1,N
,LRRC28,SPEN,NONO,PSMD5,EIF1,HACD3,SET,EGLN1,CHSY1,MRC2,RPL19,LDHB,PKD1,SF3B1,
23,LDHA,CPE,SCARB2,RTN4,LPCAT1,LAMA5,SLC22A17,RPS9,RPL24,C1ORF43,IGFBP3,NCS1,P
4L1,NCAM1,ANXA2,AGRN,RPLP1,SARAF,MIDN,RPL10,MARCHF6,WSB1,RPS17,ARF1,SF3B2,CDC
A6,SULF1,RAB7A,CCN2,RPL12,ITM2C,NANOG,RPS5,MIF,ANXA6,ST3GAL2,DUX4,CKAP5,LAMP1,F
MEM132A,CPSF2,CTSB,CDH11,TM9SF3,U2AF2,EEF1A2,VAMP2,NOP53,NOX4,PRDX5,PDXK,PKN1
2,PDE7A,PSMD3,TCF25,ADAM9,CDC37,WDR6,BNIP3L,HNRNPUL2-BSCL2,PITPNA,XPNPEP3,ATP6
LPCAT1,SLC22A17,PJA2,SLC16A7,KPNB1,RPL17,C1ORF43,PAPOLA,COL1A2,GAP43,CBX5,ZNF8
,NCOR2,SEC23A,ARNT2,RTL8C,ARFGEF2,WBP2,DYNLL1,NAP1L4,AMOTL2,YME1L1,CTNND1,TR

PP2R1A,RPL15,GTF3C4,PKD1,SF3B1,SSBP2,BASP1,POLR2A,TPT1,BTF3,NCAM1,ANXA2,AGRN,Z
,GOLGA8A,SMARCA5,CTNNA1,RPS16,RPL23,CPE,RPL29,TPM4,SCARB2,LPCAT1,CIPC,PJA2,MSN
NDUFA13,C16ORF58,XPO6,DAPK3,TRO,ATP13A1,PAFAH1B2,POLDIP2,TRIP12,CHMP1A,ITFG1,P
P6,LDLR,SSR3,TBC1D9,FLOT2,TMEM63B,TUBGCP6,BLCAP,MSI1,ATP1B2,SCRIB,NSD2,RBPJ,D
AB6B,SLC7A5,ENC1,NLGN2,RPS14,EIF5,LENG8,HMGN2,LASP1,OAZ1,PFN1,TLE5,NISCH,CCNI,T
RDBP,RPLP2,MXD4,INIP,CIRBP,PTGES3,SEPTIN11,RHOB,ARCN1,RAC1,ARHGDIA,CAPRIN1,CDK1
TXN3,ERGIC3,CAP1,IST1,NOC2L,KDM4A,PSMC3,BNIP3,ATP6V1G1,BAG5,ATG4B,HSDL1,GAK,AT
ID2,SLC7A5,NLGN2,CHD4,ATP1A1,TMEM259,HMGN2,ATP2A2,LASP1,BEX3,OAZ1,ASTN2,MCL1,J
TAN1,GSTP1,CALM1,PLXNB2,HSPD1,COL18A1,SNRNP200,ATP5F1A,VCP,MARCKSL1,AFF4,TRIM
P2,MYL6,SHOX,MYH10,PTGES3,SND1,LGALS3BP,UBE2H,CKAP4,HRNR,TOP2B,ATP6V0C,MMP2,
1,TAF15,ABCA3,RTN1,PDXK,PKN1,TBC1D16,AHCYL1,NCKAP1,PTOV1,CLSTN3,MDK,LAMC1,H3-
34,YME1L1,WDR45B,TRIO,EXTL3,VARS1,PPP2CB,KLHL42,CD164,KIAA0895L,IARS1,MASTL,NOM
,RPL10,MARCHF6,WSB1,RPS17,ARF1,CYB5R3,SF3B2,CDC42BPB,ITM2B,AP2B1,GTF2F1,GOLG
PE,CNN3,RPSA,CTSD,SCARB2,LPCAT1,SLC22A17,RPS9,CIPC,PJA2,RPL24,C1ORF43,PAPOLA,NC
H1-10,CIRBP,MYH10,DUX4,CADM4,UBE2H,HLA-B,HECTD4,MAGED4B,HDAC2,PEA15,SERF2,TIM
B1,RPS17,ARF1,SF3B2,CDC42BPB,ITM2B,AP2B1,GTF2F1,GOLGA8A,SMARCA5,LDHA,CPE,CNN3
XW5,CREBBP,SSBP4,ATG4B,FJX1,LYPLA2,USP5,DENND5A,MZT2B,LRIG1,HDGFL3,POR,SLC22A
C22A17,RPS9,CIPC,PJA2,MSN,SLC16A7,C1ORF43,IGFBP3,LLGL1,NCS1,CLSTN1,KIF5B,NLGN2,
RAB1A,CALU,PLPPR3,ARCN1,SEPTIN3,COTL1,RAC1,ARHGDIA,GPC1,GLUL,NNAT,TMEM132A,CP
F4,TLE5,NES,PRKCSH,CCNI,LRPPRC,TADA2B,PFKL,HDLBP,CDH6,PNMA1,PDIA6,PLXNA1,PIP4K2

CTSD,EIF4H,TSPAN3,RPS24,BRD3,XRCC5,PJA2,KPNB1,PAPOLA,PTCD3,PSMD2,NR2F1,KIF5B,RAE
P11,COL18A1,SNRNP200,CSDE1,ATP5F1A,EIF4B,VCP,UBB,AFF4,MATR3,CKB,COL4A2,RPL37A,L
RNPM,SLIT3,ISYNA1,STT3A,IDH1,SARS1,SOD1,BAG6,FBRSL1,CYP51A1,PRPF40A,CD99L2,DDIT4
GA8A,LDHA,CPE,CNN3,RPSA,CTSD,SCARB2,RTN4,LPCAT1,LAMA5,RPS24,SLC22A17,RPS9,CIPC,F
LC2A3,DDX17,GPI,HTRA4,RACK1,BSG,FASN,ALDH6A1,SPTBN1,ATP5F1B,CALM3,GNB1,STMN2,M
POLDIP3,UNC45A,HMGXB3,MAP1S,WARS1,KDM5A,SKIV2L,SHC1,PCED1A,ASB6,SUDS3,MIB2,TN
AF5,CD2BP2,LIMK1,MYCBP2,ZCCHC3,SLC25A1,SEPHS1,ZNF703,NUS1,SERPINE2,UBQLN4,C20C
2L,EIF3G,PACS2,AP1M1,NDUFA13,C16ORF58,XPO6,DAPK3,TRO,LYPLA2,ATP13A1,PAFAH1B2,PC
LDHB,DNAH8,SF3B1,SERINC1,RPS3,HMGB1,POLR2A,AP2M1,TPT1,SEC61A1,RPS4X,RPS3A,RPL1
,OGDH,GUK1,CTDNEP1,ZNF598,QRICH1,TFDP1,CLCN3,BRPF3,MSI2,SLC25A39,NBR1,RNPS1,PA
PSMA7,ST3GAL6,BCL9L,WDR45B,FARP1,NXN,EP400,UBAP2L,GLUD1,TAGLN2,TRA2B,IARS1,MB
PO1,SPTBN2,SLC3A2,TPM1,EIF5A,CCN2,DDX3X,RPS15,RPL12,ITM2C,NANOG,MAGED2,RCN1,H1
CAT1,LAMA5,SLC22A17,PAPOLA,IGFBP3,RPL14,NCS1,CLSTN1,PSMD2,RAB6B,SLC7A5,ENC1,NL
1,SMIM14,IGF2,EIF3C,EIF4B,MARCKSL1,TRIM28,YWHAG,LRRC28,NONO,HACD3,SET,TPI1,TMSE
P1,XAF1,COL11A1,RPS17,AP3D1,SBF1,ARF1,CYB5R3,SF3B2,CDC42BPB,MAT2A,GTF2F1,SLC4A
R45B,NAP1L1,KLHL42,SFRP1,ARID1B,KIAA0895L,RAB3GAP2,HDAC6,PRRC2A,HNRNPA2B1,PTP
YKT6,TMEM167A,SEPTIN7,CHERP,FKBP1A,CTTN,SLC25A5,LRP10,ERH,EIF2S2,RALY,SEC14L1,NU
RD52,INIP,CIRBP,DUX4,RHOB,HNRNPH1,CKAP4,TTR,CALU,SEPTIN3,COTL1,CCT6A,MAGED4B,A
NO,DDB1,PSMD5,YWHAQ,EIF1,HACD3,SET,EGLN1,MAP9,CHSY1,TMSB10,INA,EIF4A1,RPL19,PPP
ZIN1,DDAH2,NYNRIN,TCF25,PRKDC,TDRD5,CASC3,CLIP3,ZFAND5,PITPNA,CORO1C,SMARCC1,H
FIP1,MYL6,ANKRD52,HNRNPK,INIP,SEPTIN11,LAMP1,HNRNPH1,LGALS3BP,RPS12,COTL1,CAPF
8,YWHAG,CKB,COL4A2,FN1,SPEN,HACD3,DCHS1,SET,MAP9,CHSY1,TMSB10,EIF4A1,MRC2,GTF
1,DDX3X,CLTC,NANOG,GNB2,RPS5,IK,HMGCS1,PRKAR1A,CHPF,GJA1,JUN,NGRN,B4GALNT4,HN
1,YKT6,MCM3AP,RPL7L1,RPS27A,TMEM167A,KLHDC3,TNKS,PSMD8,CHERP,CTTN,SERP1,VPS26
3D1,DNAJA1,ARF1,CYB5R3,PAFAH1B1,MAP4,SF3B2,ITM2B,TMEM170A,GOLGA8A,HNRNPU,NFE2
,YWHAH,PFKM,TNKS2,LONP1,HYOU1,STRAP,BAG6,RAI1,ZSCAN18,SERBP1,AKIRIN1,CCT5,PPP1F

ERP1,NDN,TRIR,KIAA0100,PSMB4,FKBP4,BCL7A,C6ORF62,GPC2,NUDCD3,EFTUD2,NCSTN,SFX
APRIN1,GPC1,TOP2B,APPL1,HDAC2,LAPTM4B,MAP1A,SERF2,CRPPA,TMEM132A,PGRMC1,CTSE

UBE4A,HSPA4,AHCYL1,MLEC,SURF4,CHD2,ASXL1,SRP14,ABCA2,RPS15A,OPTN,RPL37,G3BP2,S
S21,BANF1,FAF2,PPME1,CSNK2B,FAM8A1,FAM32A,EPN1,RTF2,PSMC5,POLDIP3,PSMC4,HMGXE
GAS7,TM9SF2,MORF4L2,ABCA3,SNRPN,PDXK,PLOD1,VPS35,PTOV1,MLEC,CLSTN3,ACAP3,SURF
MAPRE1,DSTN,CAPZB,HNRNPUL1,CAND1,SON,RPL28,HNRNPUL2-BSCL2,TSC22D1,SDF4,LSS,EIF
VGF,MLF2,SEC23A,RTL8C,WBP2,AKIRIN1,AMOTL2,YME1L1,TRAF4,STAT3,ARIH1,HIP1,SLC35F5,
,TIMP2,COL11A1,MIDN,MARCHF6,WSB1,SBF1,CDC42BPB,MAT2A,TMEM170A,GTF2F1,GOLGA8
S,PSMD8,CHERP,CTTN,SERP1,VPS26B,ERH,CLPTM1,INTS3,ABI2,UBE2M,SEC14L1,ELAVL1,NUD
T,PABPN1,GDI2,MORF4L2,SNRPN,HNRNPA3,C4ORF3,PKN1,YEATS2,VPS35,ACAP3,MDK,DDX42,C
G,BRD2,KLHL42,GNG4,MASTL,HNRNPA2B1,PPP1CC,EML1,ANP32B,B4GAT1,KLHDC3,HNRNPAB

2,SLC20A1,SLC38A2,REEP1,PTPN23,IGLON5,PTPN11,RSF1,LGALS1,CHMP1A,AUTS2,CGGBP1,KL
5F1B,CALM3,GNB1,SHISA9,RPS8,MYH9,RPL7,SEPTIN2,P4HB,APP,IDS,GDI1,FSCN1,TMBIM6,PTM
YCBP2,TRIB3,HM13,CRKL,COLGALT1,LOXL2,NDRG1,SLK,RNMT,ZEB1,IGF2BP2,MRPS6,NUS1,M
36A,RAB11B,NRDC,PRPF6,TUBB2A,UBXN4,PPP2CA,G3BP2,KDELR2,PDE7A,PSMD3,SRSF1,ADAM
3,XRCC5,RPL24,RPL17,PAPOLA,PTCD3,RPL9,NR2F1,KIF5B,SLC7A5,ENC1,CXADR,TMEM259,RP

,FAM234A,MAZ,HDGFL2,INTS11,PRPF4B,WASF2,NSD2,CSE1L,COPZ1,COX6A1,MBD3,PRKACA,AT
3A1,SNRNP200,CSDE1,IGF2,ATP5F1A,EIF4B,VCP,MARCKSL1,MATR3,CKB,COL4A2,MAPT,FN1,SPI
5A1,NASP,AP1M1,C16ORF58,TOMM70,XPO6,UBXN6,MYL12B,GAK,LYPLA2,CENPB,ATP13A1,PAF/
2,RPLP1,ZMIZ1,TIMP2,SARAF,RPL10,MARCHF6,WSB1,SBF1,DNAJA1,ARF1,AKAP12,MAP4,SF3B2,
APK1,DDAH2,SRSF1,PTBP1,FTO,ILF3,RPS11,CASC3,NSG2,DIDO1,RPS28,HNRNPM,GIGYF1,MESI
ETREG2,PHB2,KLHL42,MASTL,YKT6,MCM3AP,RPL7L1,TMEM167A,TNKS,PSMD8,CHERP,CTTN,L
IT4,BEX4,SSR1,WDR45B,SLC39A6,CD99,GPX4,PPM1G,GNG4,NXN,EP400,RER1,UBFD1,MBTPS1,
HNRNPH1,ATP1A2,RPS12,SEPTIN3,ZNF454,CTBP1,STIP1,TMEM30A,SKP1,DNER,L1CAM,LAPTM4I
GA1,MTA2,KDM3A,ATP13A3,IGSF3,GGNBP2,CUL1,CLUH,DCAF5,SCAMP4,EWSR1,MAP4K4,PHPT1
1IP,TIMP3,CTDSP2,CTSB,SRSF5,RNF187,FDFT1,TBCD,SUGP2,GNAI3,KLHL24,GDI2,HNRNPL,SSF

CHMP1A,MTA2,TSNAX,EXOC5,TRMT112,PSMD13,CD2BP2,RPS21,CNDP2,FAF2,COX8A,DARS1,FAI
AP2B1,GTF2F1,GOLGA8A,SMARCA5,LDHA,CPE,CNN3,RPSA,CTSD,SCARB2,RTN4,LPCAT1,LAMA5
SMD8,CHERP,CTTN,SERP1,VPS26B,ERH,CLPTM1,INTS3,ABI2,UBE2M,SEC14L1,ELAVL1,NUDCD3,
LP1,ZMIZ1,TIMP2,XAF1,SARAF,RPL10,VCL,AP3D1,ARF1,CYB5R3,AKAP12,MAP4,TMEM170A,GTF2
D3,CBX5,GLIPR1L2,RPS14,CXADR,ATP1A1,TMEM259,RPS20,CCT7,RBBP4,RTN3,PFN1,ASTN2,PN
IP,BNC2,HNRNPR,EXOC7,RIMKLB,IGF2BP2,BEX1,PHKB,MCFD2,RALBP1,MAN2B1,DUSP8,IRS1,TI
HNRNPUL2-BSCL2,PITPNA,XPNPEP3,SRSF3,LARS1,YWHAH,TNKS2,UBQLN1,CSNK1D,HYOU1,IFIT
H6,PNMA1,PDIA6,PLXNA1,MEGF8,ACSL6,RAB7A,CCN2,RPL12,ARL4C,CLTC,ANXA6,DNAJC5,HNRI
M2B,MAT2A,GTF2F1,PGAM1,HNRNPA0,LDHA,CPE,TNS3,EIF4H,NACA,LPCAT1,RPS24,SLC22A17,C
C31A,AKAP17A,MAPK8IP3,PSMC6,PRELID1,HARS1,MRPL3,DNMT1,CFAP298,DDX56,CPSF7,RAD5
3CR,TXNDC5,MTA1,DGCR2,HIP1,LRRC4B,MMP15,ZFP36L1,CXXC5,PLXNB1,GNG4,EVL,SELENOW
,CALM1,NPM1,TTC3,PLD3,HSPD1,USP11,COL18A1,SNRNP200,CSDE1,SMIM14,EIF3C,ATP5F1A,E
1,CSNK1D,IFITM3,VGF,MLF2,SEC23A,RTL8C,AKIRIN1,AMOTL2,YME1L1,TRAF4,STAT3,ARIH1,SF
TM3,GHITM,DGKD,MLF2,ARNT2,AKIRIN1,AMOTL2,YME1L1,TRAF4,TRAF7,ARIH1,PRCC,GPX4,UE
P1,CD164,SMC3,PPP1CC,DKK3,HIF1A,LRRC59,ZMYM3,FUS,VAPB,SNN,RAB1B,UBE2M,RALY,NUD
TGB1,MORF4L1,BTF3,ANXA2,AGRN,ZMIZ1,TIMP2,MIDN,RPL10,MARCHF6,DNAJA1,ARF1,CYB5R
MAPK8IP1,BAG6,CYP51A1,KIF5A,STAT3,BCL9L,HIP1,CSNK2A1,GRSF1,MGAT4B,GNG4,NXN,KIAA

2A,UPF1,MASTL,MBTPS1,MAPRE2,APEX1,PPP1R15A,GALNT11,MVD,DAP,BAZ2A,XRCC6,VAPB,EII
,CNDP2,GLOD4,MTURN,GOLPH3,DHX36,COLGALT1,TSPAN7,NUP98,NACAD,OXSR1,EMC10,TME
COL5A1,COL4A1,CES3,PSAP,PGK1,RPLP0,H3-3B,CANX,CLU,PTMA,DDX5,ENO2,SOX4,FKBP14,RI
AFAH1B2,PLS3,POLDIP2,CHMP1A,NBPF15,GTF3C1,NDUFA4,TRMT112,WDR26,GPAA1,SUB1,ECH
,AKAP9,VPS26B,EIF3H,ELAVL1,NUDCD3,TAX1BP1,SH3GL1,NCSTN,INTS11,MTMR4,MKRN1,CAP1
K1,BAZ2A,VAPB,EIF3H,ELAVL1,IMPDH2,GABARAPL1,GNL1,TM9SF4,TRAP1,PLEKHB2,ELP1,MOAP
LPLP1,GPAT4,TLN1,CHPF,COL6A2,CD63,ANXA6,NGRN,RPS10,HNRNPDL,TARDBP,RPLP2,ST3GAL
M234A,HDGFL2,CNP,INTS11,AHSA1,MTMR4,ITGAV,SPTY2D1,CAP1,NOC2L,TGOLN2,AP1M1,NDUI
PRDX6,ZMYM3,SEPTIN5,PALM,RTL6,CCN1,CARM1,INTS3,C6ORF62,LTBP4,TLE1,MAZ,NUCKS1,A
AP9,CHSY1,INA,EIF4A1,LDHB,PKD1,PDIA3,SERINC1,SSBP2,ZNF43,RPS3,BASP1,VAPA,POLR2A,CD
RPL31,PRKCSH,LRPPRC,SCRN1,RPS25,EIF5A,PDIA6,PIP4K2B,MEGF8,SOX12,ITM2C,CLTC,EIF3L,
6,CPSF7,RAD50,JRK,TOMM40,ATP6V1D,COX5B
COLGALT1,TSPAN7,NUP98,NACAD,OXSR1,EMC10,TMEM9,SCG2,WDR82,SYNGR1,USP48,RPS29,[illegible]
FITM3,VGF,MLF2,RTL8C,AKIRIN1,AMOTL2,YME1L1,TRAF4,STAT3,ARIH1,SF1,SLC35F5,CSNK1G
,RPS8,MYH9,RPL7,SEPTIN2,P4HB,APP,IDS,ERVV-1,RPL5,GDI1,FSCN1,TMBIM6,PTMS,SPTAN1,GS
P1,LRRC4B,GRSF1,KIF21B,GSPT1,ZFP36L1,CXXC5,BRD2,HOXB5,SRSF6,PI4KA,RAB3GAP2,ZNF5

THG1L,SZT2,SEC31A,AKAP17A,MAPK8IP3,HARS1,MRPL3,DNMT1,CFAP298,DDX56,CPSF7,RAD50
JCB1,CHD6,TOMM20,TBCD,SUGP2,PPP1CB,VAMP2,GDI2,VCAN,TLN2,HNRNPA3,PRDX5,ACO2,TC
COX7C,YKT6,MCM3AP,TMEM167A,PPP1R15A,CTTN,LRRC59,DAP,MTCH1,SERP1,RHBDD2,ERH,
,YWHAE,KIF1A,RPL13,GNAS,SLC2A3,GPI,HTRA4,FASN,ALDH6A1,SPTBN1,CALM3,SHISA9,STMN2
LGN2,GLIPR1L2,CXADR,ATP1A1,LENG8,ATXN7L3B,ATP2A2,RPL26,LASP1,CCT7,RBBP4,RTN3,NE
E1L1,SSR1,TRAF7,EIF2AK1,TXLNA,PRMT1,SF1,PDAP1,EIF3D,SEC16A,PHB2,PAPSS1,IARS1,MASTL
OC728392,TMED9,HYOU1,FLNC,SEC23A,ARNT2,ARFGEF2,TP53BP1,CTXN1,RBM15B,SSR1,STAT
GHITM,PFN2,ARNT2,MTA1,CPD,ACTR1A,EP300,ALDH3A2,CSNK2A1,PHB2,SLC39A7,RAB2A,RER1,
PM4,SLC22A17,XRCC5,CIPC,PJA2,RPL24,RPL17,PAPOLA,IGFBP3,GAP43,RPL14,CBX5,NR2F1,CX
,TOMM20,TBCD,MEST,GDI2,HNRNPL,TSR1,HNRNPA3,PKN1,YEATS2,ZNF302,RAD23B,ORAI2,TN
NC1,SSBP2,HMGB1,POLR2A,AP2M1,TPT1,RPS4X,OR8K1,RPL18,ITGB1,BTF3,AGRN,RPLP1,ZMIZ
VARS1,CLCN6,COX6B1,KDM5A,MYL12A,SKIV2L,SHC1,ASB6,SUDS3,GABARAPL2,MIB2,TNPO3,N
FAM32A,EPN1,CDK9,RTF2,ZRANB2,UNC45A,ABHD17A,SNRPB,STARD3,RNF20,SKIV2L,RNF4,SHC
OC5,DGCR2,STARD7,KDM5B,ALDH3A2,VARS1,KPNA2,KLHL42,CSNK1G2,FARP1,RER1,ADCY6,PTE
MBTPS1,CUL3,NDST1,ITGB5,TIMM50,TNKS,APEX1,MAP1LC3B,MTCH1,PELP1,SLC44A2,CARM1,
SHISAL1,SERF2,TIMP3,CTDSP2,CNN2,NOVA2,NUCB1,ZC3H7B,U2AF2,EEF1A2,CCT2,SUGP2,MEST
LARS1,YWHAH,UBQLN1,CSNK1D,HYOU1,IFITM3,GHITM,NCOR2,HNRNPD,VGF,MLF2,WBP2,AKIR
0,TMEM9,WDR82,SYNGR1,USP48,RAB3A,RPS29,NSMF,DUSP8,NUP133,RRBP1,CACYBP,OGDH,D
KIF5A,STAT3,WDR45B,NAP1L1,KDM5C,EXTL3,LRRC4B,EIF3D,KLHL42,HOXB5,ARID1B,GLUD1,U
2,DIDO1,RPS28,MESD,GPR107,PFN2,ADAR,CPD,EP300,CSNK2A1,PHB2,SLC39A7,RAB2A,RER1,U
M14,EIF3C,EIF4B,VCP,UBB,AFF4,TRIM28,YWHAG,CKB,COL4A2,RPL37A,MAPT,FN1,SPEN,NONO,
PL38,RPL37,AEBP1,ACADVL,TCF25,PTBP1,LUZP6,SON,PLCG1,RPL28,CCAR2,ARF4,PLXNA3,TME

IN3,CTSD,SCARB2,RTN4,LPCAT1,LAMA5,SLC22A17,RPS9,CIPC,PJA2,RPL24,SLC16A7,C1ORF43,
,CAND1,SON,BNIP3L,QARS1,RPL28,TSC22D1,SDF4,LSS,EIF3F,B2M,TUG1,ID4,HGS,GHITM,PNM
ZNF503,PSIP1,MAPRE3,CHST3,NDRG4,RBM10,IGF2BP2,ILRUN,CRABP2,VEZF1,ARL8A,STAT1,DU
M259,LENG8,WDR1,HMGN2,MCL1,PNN,HNRNPC,CCNI,LRPPRC,XPR1,HDLBP,CCND2,DDR1,PIP4
4,RAB6B,SLC7A5,ENC1,CXADR,CHD4,ATP1A1,EIF5,CRMP1,BEX3,VAT1,OAZ1,JPT2,TLE5,NES,TAD
CSNK1G2,CD164,UPF1,UBFD1,MASTL,MBTPS1,ANP32B,MAPRE2,APEX1,PPP1R15A,GALNT11,D
,BSG,FASN,ALDH6A1,SPTBN1,ATP5F1B,CALM3,GNB1,SHISA9,STMN2,MYH9,RPL7,SEPTIN2,P4HE
CF1,ANKS1A,ITGAV,PSMC1,DPYSL4,AMBRA1,SLC38A2,CPXM1,TRO,POLDIP2,SDCBP,PINK1,CLT

,PRDX5,UBE4A,HSPA4,TCP1,NCKAP1,MLEC,GCN1,TNPO1,PCDHGB6,SNRNP70,ASXL1,SRP14,PY
2,CAB39,SUB1,FAF2,KLC2,VPS51,FAM32A,EPN1,RTF2,UNC45A,ZBTB4,STAT2,HMGXB3,ZEB1,PH
BP8,HNRNPUL2-BSCL2,SUPT6H,HNRNPM,ZFAND3,UQCRC2,HGS,BAG6,SERBP1,CDC42,YME1L1
R173,SGPL1,GPRIN1,MARVELD1,RFX7,PNMA2,TMEM151B,CUL4B,MBD6,FURIN,SCARB1,KDM2

RNR,ARHGDIA,MMP2,GLUL,HSBP1,NNAT,MAP1A,FAM199X,CRPPA,ZC3H7B,TM9SF3,MDH2,EEF1
TMSB10,EIF4A1,PPP2R1A,LDHB,PKD1,SSBP2,RPS3,BASP1,HMGB1,RPS3A,STMN1,ITGB1,UCHL1
,SMG1,CALM1,PLXNB2,PLD3,HSPD1,USP11,SNRNP200,SMIM14,IGF2,EIF3C,ATP5F1A,VCP,UBB,
PSMC6,HARS1,MRPL3,DNMT1,CFAP298,DDX56,CPSF7,C2ORF68,RAD50,JRK,DUS1L,TOMM40,AL
RMP1,OAZ1,RBBP4,JPT2,LRPPRC,SLC3A2,PFKL,HDLBP,SFPQ,CCND2,PLXNA1,MEGF8,ACSL6,R
XRN2,SCAMP5,ZZEF1,AASDHPPT,ESYT1,GTF3C1,TNFAIP1,FAM50A,CD2BP2,CNDP2,GLOD4,MTU
YL5,SLC25A36,XRN2,SCAMP5,ZZEF1,AASDHPPT,ESYT1,GTF3C1,TNFAIP1,FAM50A,CD2BP2,CNDP
GFL3,TAOK2,H6PD,SC5D,SIN3B,FZR1,CRIP2,ANO8,GALNT10,TM7SF2,CELF3,UBE2Q1,EXOC7,E

,TUBGCP6,NCSTN,INTS11,MTMR4,MKRN1,SPTY2D1,CAP1,NOC2L,COX6A1,NASP,AP1M1,ATXN7

TL1,MAGED4B,STIP1,GPC1,TOP2B,ATP6V0C,MMP2,L1CAM,FAM199X,SHISAL1,CTDSP2,NOVA2,

SF4,TRAP1,PLEKHB2,ELP1,MOAP1,CLASP2,RBM12,NSD2,YY1,CCDC47,ELOVL5,SRPRA,SLC30A

36,XRN2,SCAMP5,ZZEF1,AASDHPPT,ESYT1,GTF3C1,TNFAIP1,FAM50A,CD2BP2,CNDP2,GLOD4,M

36,XRN2,SCAMP5,ZZEF1,AASDHPPT,ESYT1,GTF3C1,TNFAIP1,FAM50A,CD2BP2,CNDP2,GLOD4,M

1,TRAP1,DNAJB2,BLCAP,ELP1,SPAG9,CLASP2,ATP6V0D1,FXYD6,CREBBP,PTPN23,CENPB,DHX4

A,HMGB1,POLR2A,AP2M1,SEC61A1,STMN1,ITGB1,UCHL1,RPL11,MORF4L1,BTF3,ANXA2,AGRN,

,DDX39B,CNN2,NOVA2,NUCB1,C5ORF24,CCT2,PGD,KLHL24,MEST,GAS7,TM9SF2,MORF4L2,AL

1,MORF4L1,AGRN,ZMIZ1,MIDN,RPL10,WSB1,SBF1,DNAJA1,ARF1,ITM2B,TMEM170A,GTF2F1,SL

C4A8,RPL23,CPE,RPSA,EIF4H,SCARB2,TSPAN3,RPS24,RPS9,CIPC,RPL24,C1ORF43,COL1A2,RPL

C4A8,RPL23,CPE,RPSA,EIF4H,SCARB2,TSPAN3,RPS24,RPS9,CIPC,RPL24,C1ORF43,COL1A2,RPL

APLP2,HNRNPDL,TARDBP,ARL6IP1,DDOST,MYL6,H1-10,SEPTIN11,TTR,PLPPR3,SEPTIN3,CCT6A,

F1,WDR82,SYNGR1,SHMT2,USP48,RPS29,STMN4,NSMF,DUSP8,NUP133,CHGB,COX4I1,OGDH,D

COL4A2,RPL37A,DDB1,PSMD5,DCHS1,SET,EGLN1,TPI1,TMSB10,EIF4A1,MRC2,RPL15,LDHB,PKD

P,TRAP1,LSM14B,PRPF4B,CSE1L,NOC2L,MBD3,UBE3A,AP2A2,YY1,TMEFF1,UBE2D3,SLC20A1,C

DF3,COPE,FEZ1,LIMK1,RHEB,UBE2J1,GOLGA8B,MEX3C,ALCAM,SNRPD2,MAN2B1,METRNL,SLC

IPM1,PLXNB2,PLD3,USP11,COL18A1,SNRNP200,CSDE1,SMIM14,IGF2,EIF3C,ATP5F1A,EIF4B,VC

ANXA2,RPLP1,SARAF,MIDN,RPL10,MARCHF6,WSB1,RPS17,ARF1,MAP4,SF3B2,ITM2B,AP2B1,GT

PLP1,XAF1,MARCHF6,ITM2B,MAT2A,TMEM170A,SLC4A8,HNRNPU,SMARCA5,HNRNPA0,RPS16,R

LOT2,EEF1B2,QPRT,MTMR4,ANKS1A,SPTY2D1,CAP1,LRRC41,PRPF19,NOC2L,SF3A2,PACS2,NA

HNRNPH1,UBE2H,HMGCR,RAC1,SMCHD1,TMEM30A,SKP1,BCLAF1,L1CAM,LAPTM4B,TRIB2,PPIE

LPPR3,CTBP1,GPC1,SMCHD1,IPO5,PEA15,TMEM132A,DDX39B,NOVA2,CTSB,ZC3H7B,RAB27A,E

HNRNPU,OGA,PGAM1,NFE2L1,LARP1,RPL29,SCARB2,NACA,RTN4,LPCAT1,BRD3,XRCC5,PJA2,M

OVL5,SLC25A36,XRN2,SCAMP5,ZZEF1,AASDHPPT,ESYT1,GTF3C1,TNFAIP1,FAM50A,CD2BP2,CN

F3,RPS11,CASC3,NSG2,DIDO1,RPS28,HNRNPM,GIGYF1,MESD,GPR107,GHITM,PFN2,ADAR,CPD

GTF3C4,LDHB,PKD1,HLA-C,SF3B1,PDIA3,SERINC1,SSBP2,RPS3,CHD3,VAPA,RPL18,RPL11,NCAM

LAMA5,BRD3,MSN,KPNB1,PAPOLA,LLGL1,NCS1,ZNF865,ATF4,SLC7A5,NLGN2,CHD4,JUND,ATP

A,MAPK8IP3,PRELID1,HARS1,MRPL3,DNMT1,CFAP298,DDX56,CPSF7,C2ORF68,RAD50,JRK,TOM

2F1,SLC4A8,RPL23,CPE,RPSA,EIF4H,SCARB2,TSPAN3,RPS24,RPS9,CIPC,RPL24,RPL17,C1ORF4

PLEKHB2,ELP1,MOAP1,CLASP2,RBM12,NSD2,YY1,CCDC47,ELOVL5,SRPRA,SLC25A36,XRN2,SCA

ATP6V0C,LAPTM4B,PEA15,SRSF5,TM9SF3,MLXIP,MORF4L2,HSPH1,MTRF1L,ALKBH5,PDXK,UBE

A,UBTD2,CCNK,AKAP17A,MAPK8IP3,PSMC6,HARS1,MRPL3,DNMT1,CFAP298,DDX56,NEDD8,CPS

C1,ABCE1,NOLC1,MSI2,PAF1,PPM1A,GNAO1,RAD23A,EPHA2,STX12,VPS37B,ZC3H4,NDUFA10,C

SRSF6,UPF1,TRA2B,MASTL,MAPRE2,BRSK1,GPSM1,NPLOC4,PPP1R15A,GALNT11,DAP,SHC2,EIF

K1,MTA2,MGAT5,NBPF15,HIP1R,MKNK2,PFKP,ARHGEF40,FZR1,SLC38A1,SAMD11,FAM50A,APE

CTTN,SLIT2,SERP1,WSCD1,VPS26B,EIF3H,ELAVL1,NUDCD3,SH3GL1,TUBGCP6,NCSTN,INTS11,M

MD3,TCF25,CDC37,CAPZB,PITPNA,GNG2,XPNPEP3,ATP6V1B2,LARS1,NKTR,YWHAH,TNKS2,UBQ

L5,SLC25A36,XRN2,SCAMP5,ZZEF1,AASDHPPT,ESYT1,GTF3C1,TNFAIP1,FAM50A,CD2BP2,CNDP

,SRSF3,LARS1,YWHAH,UBQLN1,CSNK1D,IFITM3,DGKD,VGF,MLF2,AKIRIN1,AMOTL2,YME1L1,C

1,GNL1,TM9SF4,TRAP1,PLEKHB2,ELP1,MOAP1,CLASP2,RBM12,NSD2,IST1,YY1,CCDC47,ELOVL

AP1,CLASP2,RBM12,NSD2,YY1,CCDC47,ELOVL5,SLC25A36,XRN2,SCAMP5,ZZEF1,AASDHPPT,E
RFX7,PNMA2,TMEM151B,CUL4B,MBD6,FURIN,SCARB1,KDM2B,ID3,DNMT1,NEDD8,SETD1B,ZC

NL1,TM9SF4,TRAP1,PLEKHB2,ELP1,MOAP1,CLASP2,RBM12,NSD2,YY1,CCDC47,ELOVL5,SRPRA
,ETNK1,CCT3,JPT1,GCN1,RPL36A,SRP14,TERF2IP,RPL38,RAPGEF1,RPL37,UBXN4,MAPRE1,CC
,ARIH1,NDUFV1,SF1,SLC35F5,CSNK1G2,UPF1,TRA2B,MASTL,MAPRE2,CALD1,APEX1,NPLOC4,P
,HDAC2,LAPTM4B,LUC7L3,SERF2,CTSB,U2AF2,VAMP2,MEST,GDI2,TSR1,MORF4L2,SNRPN,HNR
,HDAC2,LAPTM4B,LUC7L3,SERF2,CTSB,U2AF2,VAMP2,MEST,GDI2,TSR1,MORF4L2,SNRPN,HNR
L1,TRAF7,ST3GAL6,KIF5A,BCR,ACTR1A,LBH,RERE,AP2A1,NOL4L,BAZ1B,KIF21B,CD99,UBAP2L
L2,WIZ,OTUD5,GID8,TMEM131,RTF2,COLGALT1,SENP6,P3H1,PIP5K1C,DDAH1,IGF2BP2,MEX3C
E1,SMIM14,IGF2,EIF3C,EIF4B,VCP,MARCKSL1,AFF4,TRIM28,MATR3,YWHAG,CKB,COL4A2,LRR

IF3L,MAGED2,RPS5,IK,RPS19,CHPF,JUN,NGRN,GOLM1,DNAJC5,APLP2,RPLP2,DDOST,HNRNP
AMP5,ZZEF1,AASDHPPT,ESYT1,GTF3C1,TNFAIP1,FAM50A,CD2BP2,CNDP2,GLOD4,MTURN,GO
D164,UPF1,TRA2B,MASTL,MAPRE2,GPSM1,PPP1R15A,GALNT11,DAP,SHC2,EIF3H,ELAVL1,IMPD
,VPS51,FAM8A1,SLC25A1,FAM32A,JAK1,ZRANB2,PSMC5,NCBP2,PSMF1,RAVER1,STARD3,NKIR
X5B

IPR1L2,JUND,TMEM259,EIF5,COL5A2,HMGN2,LASP1,OAZ1,PFN1,MCL1,PRKCSH,HNRNPC,CCN
M10,RAVER1,VEZF1,ATP6V0E2,PSMA1,GPRIN1,NADK,NARF,NEURL4,PNMA2,ZNF598,SLC1A5,B
TL1,ARHGDIA,STIP1,CDK16,SLC35E2B,BCLAF1,HSBP1,FAM199X,PGRMC1,DDX39B,NUCB1,RA

,MORF4L2,ABCA3,ALKBH5,SNRPN,PRDX5,PTOV1,MLEC,ZNF302,SKI,PCDHGB6,PSMB2,PYGB,N
39A6,CD99,GPX4,HJURP,NXN,EP400,PI4KA,UBE2G2,MBTPS1,CUL3,ADH5,NCOA4,NDST1,PPP1R

P1,SLC7A1,ABCF1,FOXK2,YY1,TMEFF1,BNIP3,XPO6,SCD,TUBB4A,UBE2K,PTPN23,PLS3,SCAM
11,SND1,RHOB,UBE2H,ARCN1,SMCHD1,ATP6V0C,LAPTM4B,SRSF5,C5ORF24,ZC3H7B,U2AF2,M
2M,ISYNA1,MESD,DGKD,BAG6,SREBF2,HNRNPD,RAB11A,ARNT2,WBP2,AKIRIN1,RPL34,AMOTL
TD1

4,PFN1,MCL1,PRKCSH,CCNI,XPR1,KHDRBS1,SLC3A2,RPS25,CCND2,CDH2,GPC3,PLXNA1,PIP4K
CHD1,ATP6V0C,LAPTM4B,SRSF5,C5ORF24,ZC3H7B,U2AF2,MLXIP,PPP1CB,TSPYL4,HSPH1,AL
1,EIF3H,SSR3,PNISR,RIC8A,AHSA1,RAB5C,MTMR4,PCNX3,DAZAP2,ERGIC3,SPTY2D1,BORCS7,

C,NANOG,RPS7,IK,HMGCS1,GPAT4,GJA1,MTSS2,TARDBP,SLC25A3,HNRNPK,CIRBP,NARS1,PTG
AP1,CLASP2,RBM12,NSD2,YY1,CCDC47,ELOVL5,SLC25A36,XRN2,SCAMP5,ZZEF1,AASDHPPT,E
M1,FAM53C,VPS51,HNRNPR,SOBP,TBXA2R,LSM4,GOLGA8B,GSK3B,MRPS6,OAT,DDX41,TMEM
VL1,IMPDH2,GABARAPL1,GNL1,TM9SF4,TRAP1,PLEKHB2,ELP1,MOAP1,CLASP2,RBM12,NSD2,I
LASP2,RBM12,NSD2,YY1,CCDC47,ELOVL5,SLC25A36,XRN2,SCAMP5,ZZEF1,AASDHPPT,ESYT1,G
,DDX56,CPSF7,C2ORF68,TOMM40
,CPSF7,C2ORF68,JRK,SLC39A1,DUS1L,TOMM40,ALYREF

EF2,RBM15B,BEX4,VDAC1,SSR1,DFFB,SGTA,RETREG2,ALDH3A2,FLCN,LRRC4B,GTPBP1,BRD2,

;,SLC2A3,DDX17,GPI,HTRA4,RACK1,BSG,FASN,ALDH6A1,SPTBN1,ATP5F1B,CALM3,GNB1,SHISA9
SN,ALDH6A1,SPTBN1,CALM3,GNB1,SHISA9,STMN2,RPS8,MYH9,RPL7,SEPTIN2,P4HB,APP,IDS,GD
ATP5F1B,CALM3,GNB1,SHISA9,STMN2,RPS8,MYH9,RPL7,SEPTIN2,P4HB,APP,IDS,RPL5,GDI1,FSC
SHISA9,STMN2,RPS8,MYH9,RPL7,SEPTIN2,P4HB,APP,ERVV-1,GDI1,FSCN1,TMBIM6,PTMS,SPTAN
NP200,SMIM14,IGF2,ATP5F1A,VCP,MARCKSL1,UBB,AFF4,TRIM28,MATR3,YWHAG,CKB,COL4A2,
1,CALM3,GNB1,SHISA9,STMN2,RPS8,MYH9,RPL7,SEPTIN2,P4HB,APP,IDS,RPL5,GDI1,FSCN1,TMI
BSG,FASN,ALDH6A1,SPTBN1,ATP5F1B,CALM3,GNB1,SHISA9,STMN2,RPS8,MYH9,RPL7,SEPTIN2
3C4,PKD1,HLA-C,PDIA3,SSBP2,BASP1,VAPA,CDKL1,TPT1,SEC61A1,STMN1,ITGB1,TUBB4B,MORI
;,SPTAN1,GSTP1,MARCKS,SMG1,CALM1,NPM1,PLXNB2,PLD3,HSPD1,COL18A1,SNRNP200,CSDI
A9,STMN2,RPS8,MYH9,RPL7,SEPTIN2,P4HB,APP,GDI1,FSCN1,PTMS,SPTAN1,GSTP1,MARCKS,SN
,SHISA9,STMN2,RPS8,MYH9,RPL7,SEPTIN2,P4HB,APP,IDS,GDI1,FSCN1,TMBIM6,PTMS,SPTAN1,G
PLD3,USP11,COL18A1,SMIM14,IGF2,EIF3C,ATP5F1A,EIF4B,VCP,MARCKSL1,AFF4,TRIM28,MATR
1,PLXNB2,HSPD1,USP11,COL18A1,SNRNP200,CSDE1,SMIM14,IGF2,EIF3C,ATP5F1A,EIF4B,MAR
,MARCKS,SMG1,CALM1,PLXNB2,HSPD1,USP11,COL18A1,SNRNP200,CSDE1,SMIM14,IGF2,EIF3
SP11,COL18A1,SNRNP200,SMIM14,IGF2,EIF3C,ATP5F1A,EIF4B,MARCKSL1,UBB,AFF4,TRIM28,N
SPTBN1,ATP5F1B,CALM3,GNB1,SHISA9,STMN2,MYH9,RPL7,SEPTIN2,P4HB,APP,RPL5,GDI1,FSC
STP1,MARCKS,SMG1,CALM1,PLXNB2,HSPD1,USP11,COL18A1,SNRNP200,CSDE1,SMIM14,IGF2,I
32,HSPD1,USP11,SNRNP200,CSDE1,IGF2,EIF3C,ATP5F1A,EIF4B,MARCKSL1,AFF4,TRIM28,MATR
SET,EGLN1,TPI1,CHSY1,TMSB10,INA,MRC2,RPL15,GTF3C4,LDHB,PKD1,HLA-C,PDIA3,SSBP2,VAF
N1,GSTP1,MARCKS,SMG1,CALM1,PLXNB2,HSPD1,USP11,COL18A1,SNRNP200,CSDE1,SMIM14,
,HSPD1,USP11,SNRNP200,CSDE1,SMIM14,IGF2,EIF3C,EIF4B,VCP,MARCKSL1,UBB,TRIM28,MATI
PL7,SEPTIN2,P4HB,APP,RPL5,GDI1,TMBIM6,PTMS,SPTAN1,GSTP1,MARCKS,SMG1,CALM1,NPM1
JSP11,COL18A1,CSDE1,SMIM14,IGF2,ATP5F1A,EIF4B,VCP,MARCKSL1,UBB,AFF4,TRIM28,MATR
E,KIF1A,RPL13,GNAS,SLC2A3,DDX17,GPI,HTRA4,RACK1,BSG,FASN,ALDH6A1,SPTBN1,ATP5F1B,
18A1,SNRNP200,SMIM14,IGF2,EIF3C,EIF4B,MARCKSL1,AFF4,TRIM28,MATR3,YWHAG,CKB,COL4
YH9,RPL7,SEPTIN2,P4HB,APP,IDS,RPL5,GDI1,FSCN1,TMBIM6,PTMS,SPTAN1,GSTP1,MARCKS,SN
GF2,EIF3C,EIF4B,VCP,MARCKSL1,AFF4,TRIM28,MATR3,YWHAG,CKB,COL4A2,LRRC28,FN1,SPEN
YH9,RPL7,SEPTIN2,P4HB,APP,GDI1,FSCN1,TMBIM6,PTMS,SPTAN1,GSTP1,MARCKS,SMG1,CALM
3,DDX17,GPI,HTRA4,RACK1,BSG,FASN,ALDH6A1,SPTBN1,ATP5F1B,CALM3,GNB1,SHISA9,STMN2
AS,SLC2A3,DDX17,GPI,HTRA4,RACK1,BSG,FASN,ALDH6A1,SPTBN1,ATP5F1B,CALM3,GNB1,SHIS
1,SPTBN1,ATP5F1B,CALM3,GNB1,SHISA9,STMN2,RPS8,MYH9,RPL7,SEPTIN2,P4HB,APP,RPL5,GD
D1,USP11,SNRNP200,SMIM14,IGF2,EIF3C,ATP5F1A,EIF4B,MARCKSL1,UBB,AFF4,TRIM28,MATR
,PLXNB2,PLD3,HSPD1,USP11,COL18A1,SNRNP200,SMIM14,IGF2,EIF3C,ATP5F1A,EIF4B,MARCK
RACK1,BSG,FASN,ALDH6A1,SPTBN1,ATP5F1B,CALM3,GNB1,SHISA9,STMN2,RPS8,MYH9,RPL7,S
WHAE,KIF1A,RPL13,GNAS,SLC2A3,DDX17,GPI,HTRA4,RACK1,BSG,FASN,ALDH6A1,SPTBN1,CALI
TR3,YWHAG,CKB,RPL37A,LRRC28,FN1,SPEN,NONO,DDB1,YWHAQ,HACD3,DCHS1,SET,EGLN1,CI
,RPS8,MYH9,RPL7,SEPTIN2,P4HB,APP,IDS,GDI1,FSCN1,TMBIM6,PTMS,SPTAN1,GSTP1,SMG1,CA
N1,GSTP1,MARCKS,SMG1,CALM1,NPM1,PLXNB2,HSPD1,COL18A1,SMIM14,IGF2,EIF3C,EIF4B,V
3,MARCKSL1,UBB,AFF4,TRIM28,MATR3,YWHAG,CKB,LRRC28,FN1,SPEN,NONO,DDB1,YWHAQ,H
MN2,MYH9,RPL7,SEPTIN2,P4HB,GDI1,FSCN1,TMBIM6,PTMS,SPTAN1,GSTP1,SMG1,CALM1,NPM
ASN,ALDH6A1,SPTBN1,ATP5F1B,CALM3,GNB1,SHISA9,STMN2,RPS8,MYH9,RPL7,SEPTIN2,P4HB
DX17,GPI,HTRA4,RACK1,BSG,FASN,ALDH6A1,SPTBN1,ATP5F1B,CALM3,GNB1,SHISA9,STMN2,R
1,SNRNP200,SMIM14,IGF2,EIF3C,EIF4B,MARCKSL1,UBB,TRIM28,MATR3,YWHAG,CKB,COL4A2,
,GDI1,FSCN1,TMBIM6,PTMS,SPTAN1,GSTP1,MARCKS,SMG1,CALM1,NPM1,PLXNB2,HSPD1,USP
IB1,SHISA9,STMN2,MYH9,RPL7,SEPTIN2,P4HB,APP,IDS,RPL5,GDI1,FSCN1,TMBIM6,PTMS,SPTAN

A1,SPTBN1,CALM3,GNB1,SHISA9,STMN2,RPS8,MYH9,RPL7,SEPTIN2,P4HB,APP,IDS,ERVV-1,GDI1
CALM1,NPM1,PLXNB2,HSPD1,USP11,SNRNP200,CSDE1,SMIM14,IGF2,EIF3C,ATP5F1A,EIF4B,MAI
,SSBP2,BASP1,VAPA,HMGB1,POLR2A,AP2M1,TPT1,STMN1,ITGB1,TUBB4B,UCHL1,MORF4L1,NC
,ALDH6A1,SPTBN1,ATP5F1B,CALM3,GNB1,SHISA9,RPS8,MYH9,RPL7,SEPTIN2,P4HB,APP,IDS,ER
3,SLC2A3,DDX17,GPI,HTRA4,RACK1,BSG,FASN,ALDH6A1,SPTBN1,ATP5F1B,CALM3,GNB1,SHISA
CKB,RPL37A,LRRC28,NONO,DDB1,YWHAQ,HACD3,DCHS1,SET,EGLN1,CHSY1,TMSB10,INA,EIF4A
LRRC28,FN1,NONO,DDB1,YWHAQ,HACD3,DCHS1,SET,EGLN1,TPI1,CHSY1,TMSB10,INA,MRC2,RI
N2,P4HB,IDS,RPL5,GDI1,FSCN1,TMBIM6,PTMS,SPTAN1,GSTP1,MARCKS,SMG1,NPM1,PLD3,HSF
ORF4L1,NCAM1,ANXA2,AGRN,TIMP2,SARAF,MIDN,MARCHF6,VCL,AP3D1,SBF1,ARF1,CYB5R3,M
TIN2,P4HB,APP,FSCN1,TMBIM6,PTMS,SPTAN1,GSTP1,MARCKS,SMG1,CALM1,NPM1,PLXNB2,PLI
P2,SARAF,MIDN,MARCHF6,VCL,AP3D1,DNAJA1,ARF1,CYB5R3,MAP4,CDC42BPB,ITM2B,AP2B1,M
4B,MORF4L1,NCAM1,ANXA2,AGRN,TIMP2,SARAF,COL11A1,MIDN,MARCHF6,VCL,AP3D1,SBF1,AI
RC2,RPL19,LDHB,PKD1,HLA-C,VAPA,POLR2A,AP2M1,TPT1,RPS3A,RPL18,STMN1,ITGB1,TUBB4B
9,LDHB,PKD1,HLA-C,PDIA3,BASP1,VAPA,HMGB1,POLR2A,TPT1,SEC61A1,RPS3A,STMN1,ITGB1,T
VV-1,GDI1,FSCN1,PTMS,SPTAN1,GSTP1,MARCKS,SMG1,CALM1,PLXNB2,PLD3,HSPD1,USP11,CC
VCL,DNAJA1,ARF1,MAP4,CDC42BPB,ITM2B,MAT2A,PXDN,OGA,NFE2L1,HNRNPA0,CTNNA1,FADS
3,GNB1,SHISA9,STMN2,RPS8,MYH9,RPL7,SEPTIN2,P4HB,APP,IDS,ERVV-1,GDI1,FSCN1,PTMS,SP
1,YWHAQ,HACD3,DCHS1,SET,EGLN1,TMSB10,INA,MRC2,RPL19,PPP2R1A,LDHB,PKD1,HLA-C,PDI
MP2,SARAF,MIDN,MARCHF6,VCL,AP3D1,CYB5R3,MAP4,CDC42BPB,ITM2B,MAT2A,PXDN,HNRNPU
SL1,UBB,AFF4,TRIM28,MATR3,YWHAG,CKB,COL4A2,RPL37A,LRRC28,FN1,NONO,DDB1,YWHAQ
RPL5,GDI1,FSCN1,TMBIM6,PTMS,SPTAN1,GSTP1,MARCKS,SMG1,CALM1,NPM1,PLXNB2,PLD3,I
AGRN,TIMP2,SARAF,MIDN,RPL10,MARCHF6,VCL,AP3D1,SBF1,ARF1,CYB5R3,MAP4,CDC42BPB,I
,APP,IDS,RPL5,GDI1,FSCN1,PTMS,SPTAN1,GSTP1,MARCKS,SMG1,CALM1,NPM1,PLXNB2,PLD3,U
PKD1,HLA-C,SSBP2,VAPA,AP2M1,TPT1,RPL18,STMN1,ITGB1,TUBB4B,UCHL1,MORF4L1,BTF3,A
1,TUBB4B,UCHL1,NCAM1,ANXA2,AGRN,TIMP2,SARAF,MIDN,MARCHF6,VCL,AP3D1,SBF1,ARF1,C
,SET,MAP9,CHSY1,INA,EIF4A1,MRC2,RPL19,RPL15,GTF3C4,LDHB,PKD1,HLA-C,PDIA3,SERINC1,
I1,NPM1,PLXNB2,HSPD1,USP11,CSDE1,SMIM14,IGF2,ATP5F1A,EIF4B,MARCKSL1,UBB,AFF4,TRI
B,PKD1,HLA-C,SERINC1,SSBP2,VAPA,POLR2A,AP2M1,TPT1,RPL18,STMN1,ITGB1,TUBB4B,UCHL
,GNAS,SLC2A3,DDX17,GPI,HTRA4,RACK1,BSG,FASN,ALDH6A1,SPTBN1,ATP5F1B,CALM3,GNB1,
AN1,GSTP1,SMG1,CALM1,NPM1,PLXNB2,HSPD1,USP11,COL18A1,SNRNP200,CSDE1,SMIM14,IG
B,CALM3,GNB1,SHISA9,STMN2,RPS8,MYH9,RPL7,SEPTIN2,P4HB,APP,IDS,ERVV-1,RPL5,GDI1,FS
,MRC2,RPL19,PPP2R1A,RPL15,LDHB,PKD1,HLA-C,PDIA3,SSBP2,VAPA,AP2M1,TPT1,STMN1,ITGE
HB,IDS,ERVV-1,RPL5,GDI1,FSCN1,TMBIM6,PTMS,SPTAN1,GSTP1,MARCKS,SMG1,CALM1,NPM1,I
PS8,MYH9,RPL7,SEPTIN2,P4HB,APP,IDS,ERVV-1,GDI1,FSCN1,PTMS,SPTAN1,GSTP1,MARCKS,SM
1,SMIM14,IGF2,ATP5F1A,EIF4B,VCP,MARCKSL1,UBB,AFF4,TRIM28,YWHAG,CKB,LRRC28,FN1,N
VHAQ,EIF1,HACD3,DCHS1,SET,CHSY1,TMSB10,EIF4A1,MRC2,PPP2R1A,LDHB,DNAH8,PKD1,HLA-
,TIMP2,SARAF,MIDN,MARCHF6,VCL,AP3D1,SBF1,ARF1,CYB5R3,MAP4,CDC42BPB,ITM2B,MAT2A
RPL13,GNAS,DDX17,GPI,HTRA4,RACK1,BSG,FASN,ALDH6A1,SPTBN1,ATP5F1B,CALM3,GNB1,SH
SB10,INA,EIF4A1,MRC2,GTF3C4,PKD1,HLA-C,SSBP2,ZNF43,BASP1,VAPA,HMGB1,POLR2A,TPT1,
N1,NONO,DDB1,YWHAQ,HACD3,DCHS1,EGLN1,TMSB10,INA,MRC2,RPL19,GTF3C4,LDHB,PKD1,
N1,NONO,DDB1,YWHAQ,HACD3,DCHS1,EGLN1,TMSB10,INA,MRC2,RPL19,GTF3C4,LDHB,PKD1,
TPT1,RPL18,STMN1,ITGB1,TUBB4B,UCHL1,MORF4L1,BTF3,AGRN,ZMIZ1,TIMP2,XAF1,COL11A1
VL3,YWHAE,KIF1A,RPL13,GNAS,SLC2A3,DDX17,GPI,HTRA4,RACK1,BSG,FASN,ALDH6A1,SPTBN1,
AP9,TPI1,CHSY1,TMSB10,INA,EIF4A1,MRC2,RPL19,RPL15,DNAH8,PKD1,HLA-C,SF3B1,PDIA3,SSE
1,CHSY1,TMSB10,PPP2R1A,RPL15,GTF3C4,LDHB,SF3B1,PDIA3,SSBP2,RPS3,CHD3,VAPA,POLR2

A,LRRC28,FN1,NONO,DDB1,YWHAQ,HACD3,DCHS1,EGLN1,TMSB10,INA,MRC2,RPL19,GTF3C4,L
A-C,PDIA3,VAPA,HMGB1,POLR2A,AP2M1,TPT1,SEC61A1,RPS3A,STMN1,ITGB1,TUBB4B,UCHL1,N
C1,SSBP2,VAPA,POLR2A,AP2M1,TPT1,RPS3A,STMN1,ITGB1,TUBB4B,UCHL1,MORF4L1,NCAM1,A
RF4L1,NCAM1,ANXA2,AGRN,TIMP2,SARAF,MIDN,MARCHF6,VCL,SBF1,DNAJA1,ARF1,CYB5R3,AK
CKSL1,UBB,AFF4,TRIM28,MATR3,YWHAG,CKB,RPL37A,LRRC28,FN1,NONO,DDB1,YWHAQ,EIF1,I
F1A,RPL13,GNAS,SLC2A3,DDX17,GPI,HTRA4,RACK1,BSG,FASN,ALDH6A1,SPTBN1,ATP5F1B,CAL
CTNNA1,FADS2,CNN3,CTSD,LPCAT1,LAMA5,SLC22A17,BRD3,PJA2,KPNB1,COL1A2,LLGL1,NCS1
T,CHSY1,TMSB10,INA,MRC2,RPL19,LDHB,PKD1,HLA-C,PDIA3,SERINC1,VAPA,HMGB1,POLR2A,AI
DDX17,GPI,HTRA4,RACK1,BSG,FASN,ALDH6A1,SPTBN1,ATP5F1B,CALM3,GNB1,SHISA9,STMN2,
,FN1,SPEN,PSMD5,YWHAQ,EIF1,HACD3,SET,EGLN1,TPI1,CHSY1,TMSB10,EIF4A1,MRC2,RPL19,C
,COL18A1,SNRNP200,CSDE1,SMIM14,IGF2,EIF4B,VCP,MARCKSL1,UBB,AFF4,TRIM28,MATR3,YV
HS1,SET,TMSB10,INA,MRC2,RPL19,LDHB,PKD1,HLA-C,PDIA3,SERINC1,SSBP2,VAPA,HMGB1,POL
UCHL1,BTF3,AGRN,ZMIZ1,TIMP2,COL11A1,MIDN,RPL10,MARCHF6,WSB1,RPS17,VCL,SBF1,DNA
IRC2,RPL19,LDHB,PKD1,HLA-C,SERINC1,SSBP2,VAPA,POLR2A,AP2M1,TPT1,RPS3A,STMN1,ITGE
NAS,SLC2A3,DDX17,GPI,HTRA4,RACK1,BSG,FASN,ALDH6A1,SPTBN1,ATP5F1B,CALM3,GNB1,SHI
PDIA3,SSBP2,CHD3,BASP1,VAPA,CDKL1,TPT1,STMN1,ITGB1,UCHL1,ANXA2,AGRN,ZMIZ1,TIMP2,
,COL1A2,LLGL1,NCS1,ZNF865,CLSTN1,NLGN2,CXADR,CHD4,JUND,CD81,WDR1,HMGN2,ATP2A2
3,HSPD1,USP11,SNRNP200,SMIM14,IGF2,EIF3C,EIF4B,MARCKSL1,AFF4,TRIM28,MATR3,YWHAC
NF43,CHD3,BASP1,VAPA,CDKL1,TPT1,RPS4X,RPS3A,RPL18,ITGB1,TUBB4B,UCHL1,RPL11,MORI
F6,WSB1,VCL,AP3D1,DNAJA1,ARF1,CYB5R3,SF3B2,CDC42BPB,ITM2B,MAT2A,SLC4A8,PXDN,HNI
NA,EIF4A1,MRC2,RPL19,GTF3C4,LDHB,PKD1,SF3B1,SSBP2,RPS3,BASP1,VAPA,POLR2A,TPT1,RF
AP4,CDC42BPB,MAT2A,TMEM170A,PXDN,HNRNPU,PGAM1,HNRNPA0,CTNNA1,FADS2,RPS16,LDH
,LLGL1,HK1,NCS1,RHOA,CLSTN1,RAB6B,NLGN2,CHD4,ATP1A1,JUND,CD81,WDR1,HMGN2,ATP2
D4,JUND,EIF5,CD81,LENG8,WDR1,HMGN2,OAZ1,RBBP4,PFN1,TLE5,NES,HNRNPC,TADA2B,CSNK
IIM14,IGF2,EIF3C,ATP5F1A,EIF4B,VCP,UBB,AFF4,TRIM28,MATR3,YWHAG,CKB,LRRC28,SPEN,NC
1,JUND,RPS20,CD81,HMGN2,ATP2A2,CRMP1,CCT7,VAT1,SRSF4,JPT2,SPC25,NES,CCNI,TADA2B,
MARCKSL1,UBB,TRIM28,YWHAG,CKB,COL4A2,RPL37A,LRRC28,FN1,SPEN,DDB1,YWHAQ,DCHS1
OL18A1,CSDE1,SMIM14,IGF2,ATP5F1A,EIF4B,VCP,MARCKSL1,UBB,AFF4,TRIM28,MATR3,YWHAC
YWHAG,CKB,COL4A2,RPL37A,FN1,SPEN,NONO,YWHAQ,EIF1,HACD3,DCHS1,SET,MAP9,TMSB10,
,CSDE1,SMIM14,IGF2,EIF3C,ATP5F1A,EIF4B,VCP,MARCKSL1,AFF4,TRIM28,MATR3,YWHAG,COL
NAS,SLC2A3,DDX17,GPI,HTRA4,RACK1,BSG,FASN,ALDH6A1,SPTBN1,ATP5F1B,CALM3,GNB1,SHI
P2R1A,LDHB,PKD1,HLA-C,PDIA3,SSBP2,ZNF43,BASP1,VAPA,HMGB1,POLR2A,CDKL1,TPT1,SEC6
,CIPC,KPNB1,C1ORF43,LLGL1,NCS1,CBX5,ZNF865,CLSTN1,NLGN2,CXADR,CHD4,JUND,CD81,W
1,SEC61A1,RPS3A,STMN1,TUBB4B,UCHL1,MORF4L1,ANXA2,AGRN,ZMIZ1,TIMP2,XAF1,SARAF,M
B2,HSPD1,USP11,COL18A1,SNRNP200,SMIM14,IGF2,ATP5F1A,EIF4B,MARCKSL1,AFF4,MATR3,Y
B,APP,ERVV-1,GDI1,FSCN1,TMBIM6,PTMS,GSTP1,SMG1,CALM1,NPM1,PLXNB2,HSPD1,USP11,C
A2B,KLHL9,XPO1,TKT,EIF5A,CDH2,PDIA6,GPC3,PIP4K2B,SOX12,ACTN4,DDX3X,ARL4C,GNB2,RC
P2,XAF1,COL11A1,MIDN,MARCHF6,DNAJA1,CYB5R3,CDC42BPB,PXDN,CTNNA1,CNN3,TNS3,CTSI
P1,OAZ1,PFN1,ASTN2,NISCH,CCNI,KHDRBS1,CSNK1E,PFKL,CDH6,SFPQ,CDH2,PDIA6,GPC3,PIP4I
1,SET,EGLN1,TPI1,TMSB10,EIF4A1,MRC2,LDHB,PDIA3,SERINC1,SSBP2,BASP1,VAPA,POLR2A,CI
ANXA2,AGRN,ZMIZ1,TIMP2,XAF1,SARAF,COL11A1,MIDN,MARCHF6,WSB1,VCL,SBF1,DNAJA1,ARI
5R3,SF3B2,ITM2B,ZNF770,MAT2A,TMEM170A,PXDN,HNRNPU,OGA,NFE2L1,HNRNPA0,CTNNA1,I
ONO,DDB1,YWHAQ,HACD3,SET,CHSY1,TMSB10,INA,MRC2,PPP2R1A,RPL15,LDHB,PKD1,HLA-C,I
EC61A1,RPS3A,STMN1,ITGB1,UCHL1,NCAM1,ANXA2,AGRN,TIMP2,SARAF,MIDN,MARCHF6,RPS17
LC2A3,DDX17,GPI,HTRA4,RACK1,BSG,FASN,ALDH6A1,SPTBN1,ATP5F1B,CALM3,GNB1,SHISA9,S

)3,DCHS1,SET,EGLN1,MAP9,TPI1,CHSY1,TMSB10,INA,EIF4A1,MRC2,RPL19,LDHB,PKD1,HLA-C,S
IMGN2,ATP2A2,LASP1,BEX3,PFN1,ASTN2,MCL1,TLE5,NISCH,CCNI,TADA2B,XPR1,KHDRBS1,CSNI
IAT2A,SLC4A8,PXDN,HNRNPU,HNRNPA0,CTNNA1,FADS2,CPE,CNN3,EIF4H,NACA,TSPAN3,LPCAT1
/HAG,CKB,COL4A2,NONO,DDB1,PSMD5,YWHAQ,EIF1,HACD3,DCHS1,SET,CHSY1,TMSB10,INA,EII
MG1,CALM1,NPM1,PLD3,HSPD1,SNRNP200,CSDE1,SMIM14,IGF2,EIF3C,ATP5F1A,EIF4B,UBB,AF
'5F1A,EIF4B,MARCKSL1,UBB,AFF4,TRIM28,MATR3,YWHAG,CKB,COL4A2,RPL37A,SPEN,NONO,Y
AP2M1,TPT1,RPS3A,STMN1,ITGB1,UCHL1,BTF3,NCAM1,AGRN,ZMIZ1,TIMP2,XAF1,SARAF,MIDN,
),EIF1,HACD3,DCHS1,SET,TPI1,CHSY1,TMSB10,EIF4A1,MRC2,RPL19,PPP2R1A,RPL15,GTF3C4,LI
',P4HB,APP,IDS,GDI1,FSCN1,TMBIM6,PTMS,SPTAN1,GSTP1,MARCKS,SMG1,CALM1,NPM1,PLXNI
I1,ATP5F1B,CALM3,GNB1,SHISA9,STMN2,RPS8,MYH9,RPL7,SEPTIN2,P4HB,APP,IDS,ERVV-1,RPL
U,PGAM1,NFE2L1,HNRNPA0,CPE,CNN3,CTSD,EIF4H,NACA,TSPAN3,SLC22A17,BRD3,CIPC,MSN,K
N1,SLC7A5,CHD4,ATP1A1,JUND,EIF5,CD81,WDR1,HMGN2,ATP2A2,CRMP1,LASP1,OAZ1,PFN1,M(
2,CRMP1,LASP1,OAZ1,RBBP4,PFN1,NISCH,CCNI,PFKL,HDLBP,TKT,PNMA1,GPC3,DDR1,PIP4K2E
.1,MRC2,RPL19,PPP2R1A,RPL15,GTF3C4,PKD1,HLA-C,SF3B1,PDIA3,SERINC1,SSBP2,RPS3,BASF
MSB10,INA,EIF4A1,GTF3C4,LDHB,PKD1,HLA-C,SERINC1,SSBP2,ZNF43,CHD3,BASP1,VAPA,AP2M
1,IGF2,EIF4B,VCP,MARCKSL1,TRIM28,YWHAG,COL4A2,RPL37A,SPEN,YWHAQ,HACD3,DCHS1,SE
TN1,NR2F1,RAB6B,NLGN2,CHD4,ATP1A1,JUND,EIF5,CD81,HMGN2,ATP2A2,CRMP1,LASP1,BEX3,
6A7,KPNB1,C1ORF43,PAPOLA,COL1A2,NCS1,RHOA,ZNF865,CLSTN1,NR2F1,RAB6B,NLGN2,GLII
TP5F1B,CALM3,GNB1,SHISA9,RPS8,MYH9,SEPTIN2,P4HB,APP,GDI1,FSCN1,TMBIM6,PTMS,SPTA
1,CALM1,NPM1,PLXNB2,PLD3,HSPD1,USP11,COL18A1,SNRNP200,CSDE1,IGF2,EIF3C,ATP5F1A,
N1,PTMS,SPTAN1,GSTP1,MARCKS,SMG1,CALM1,PLXNB2,PLD3,HSPD1,USP11,COL18A1,SNRNP2
MSB10,INA,MRC2,RPL19,GTF3C4,LDHB,PKD1,HLA-C,PDIA3,CHD3,BASP1,VAPA,HMGB1,POLR2A
N1,DDB1,YWHAQ,EIF1,HACD3,SET,EGLN1,MAP9,TPI1,CHSY1,TMSB10,INA,EIF4A1,MRC2,RPL19,I
AT6A,DVL3,YWHAE,KIF1A,RPL13,GNAS,SLC2A3,DDX17,GPI,HTRA4,RACK1,BSG,FASN,ALDH6A1,S
IF5,CD81,LENG8,ATXN7L3B,ATP2A2,CRMP1,CCT7,OAZ1,SRSF4,RBBP4,PFN1,MCL1,NES,NISCH,
=6,WSB1,RPS17,SBF1,DNAJA1,ARF1,CYB5R3,MAP4,CDC42BPB,ITM2B,AP2B1,MAT2A,TMEM170
JND,TMEM259,EIF5,LENG8,WDR1,HMGN2,ATP2A2,LASP1,VAT1,PFN1,ASTN2,MCL1,NISCH,CCNI,
:ALM3,GNB1,SHISA9,RPS8,MYH9,RPL7,SEPTIN2,P4HB,APP,IDS,ERVV-1,RPL5,GDI1,FSCN1,TMBIN
MAT2A,GOLGA8A,PXDN,HNRNPU,NFE2L1,CTNNA1,RPS16,CNN3,RPL29,TNS3,CTSD,EIF4H,SCARB
AN1,MARCKS,SMG1,CALM1,NPM1,PLXNB2,TTC3,PLD3,USP11,SNRNP200,SMIM14,IGF2,EIF3C,I
N1,ASTN2,TLE5,CCNI,KLHL9,XPO1,HDLBP,EIF5A,PIP4K2B,SOX12,RPL12,ITM2C,ARL4C,GNB2,H
A8A,PXDN,HNRNPU,SMARCA5,HNRNPA0,CTNNA1,RPS16,RPL23,CNN3,CTSD,SCARB2,LPCAT1,LA
IA,RPL15,LDHB,PKD1,SF3B1,SSBP2,ZNF43,BASP1,HMGB1,POLR2A,CDKL1,TPT1,RPS4X,ITGB1,
MORF4L1,AGRN,ZMIZ1,TIMP2,XAF1,SARAF,COL11A1,MIDN,MARCHF6,VCL,AP3D1,SBF1,DNAJA1
;,NPM1,PLXNB2,PLD3,HSPD1,USP11,SNRNP200,CSDE1,SMIM14,IGF2,EIF3C,EIF4B,VCP,MARCKS
CNI,XPO1,HDLBP,EIF5A,CCND2,PIP4K2B,SOX12,DDX3X,H1-0,MIF,SESN3,B4GALNT4,DNAJC5,AP
,CRMP1,LASP1,VAT1,OAZ1,RBBP4,PFN1,TLE5,NES,NISCH,CCNI,PFKL,HDLBP,TKT,SFPQ,PNMA1,
3B,AFF4,TRIM28,MATR3,YWHAG,CKB,COL4A2,LRRC28,FN1,SPEN,NONO,DDB1,YWHAQ,EIF1,HA
IDN,MARCHF6,WSB1,RPS17,VCL,CYB5R3,MAP4,CDC42BPB,ITM2B,AP2B1,GTF2F1,GOLGA8A,P
MIZ1,TIMP2,SARAF,MIDN,MARCHF6,WSB1,VCL,AP3D1,SBF1,ARF1,CYB5R3,AKAP12,MAP4,SF3B
,YWHAG,CKB,LRRC28,MAPT,FN1,SPEN,NONO,PSMD5,YWHAQ,HACD3,SET,EGLN1,TPI1,CHSY1,T
JC5,APLP2,TARDBP,MXD4,ANKRD52,WLS,HNRNPK,DUX4,CADM4,UBE2H,RAB1A,PLPPR3,ARCN1
APA,HMGB1,POLR2A,TPT1,RPS4X,RPS3A,RPL18,STMN1,ITGB1,TUBB4B,UCHL1,NCAM1,AGRN,R
APA,HMGB1,POLR2A,TPT1,RPS4X,RPS3A,RPL18,STMN1,ITGB1,TUBB4B,UCHL1,NCAM1,AGRN,R
APA,HMGB1,POLR2A,TPT1,RPS4X,RPS3A,RPL18,STMN1,ITGB1,TUBB4B,UCHL1,NCAM1,AGRN,R

3,COL4A2,LRRC28,FN1,SPEN,DDB1,PSMD5,YWHAQ,EIF1,HACD3,SET,MAP9,TPI1,CHSY1,TMSB1(
,UCHL1,MORF4L1,AGRN,ZMIZ1,TIMP2,COL11A1,MIDN,WSB1,VCL,AP3D1,ARF1,CYB5R3,AKAP12
SMIM14,IGF2,EIF3C,EIF4B,VCP,MARCKSL1,UBB,AFF4,TRIM28,MATR3,YWHAG,CKB,COL4A2,SP
42BPB,GOLGA8A,PXDN,OGA,CTNNA1,FADS2,RPS16,RPL23,LDHA,CPE,TPM4,EIF4H,LAMA5,PJA2,
SN3,B4GALNT4,DNAJC5,APLP2,TARDBP,MXD4,WLS,HNRNPK,DUX4,CADM4,RAB1A,PLPPR3,ARCI
PTBN1,CALM3,GNB1,SHISA9,STMN2,RPS8,MYH9,RPL7,SEPTIN2,P4HB,APP,IDS,RPL5,GDI1,FSCN
3,CDC42BPB,ITM2B,GOLGA8A,PXDN,HNRNPU,NFE2L1,SMARCA5,HNRNPA0,CTNNA1,SCARB2,N
)3,VAPA,POLR2A,CDKL1,RPL18,STMN1,ITGB1,MORF4L1,BTF3,AGRN,SARAF,COL11A1,MARCHF(
310,INA,EIF4A1,PPP2R1A,RPL15,GTF3C4,LDHB,DNAH8,PKD1,HLA-C,PDIA3,BASP1,VAPA,HMGB1,
LGL1,NCS1,ZNF865,CLSTN1,SLC7A5,CHD4,JUND,EIF5,CD81,HMGN2,ATP2A2,CRMP1,LASP1,OA
,SEPTIN11,CKAP4,RAB1A,PLPPR3,MAGED4B,CTBP1,ARHGDIA,CAPRIN1,STIP1,GPC1,SMCHD1,IP
NPM1,PLD3,HSPD1,COL18A1,CSDE1,ATP5F1A,EIF4B,VCP,MARCKSL1,UBB,AFF4,TRIM28,MATR3,
MARCKS,SMG1,CALM1,PLXNB2,TTC3,PLD3,HSPD1,USP11,COL18A1,CSDE1,SMIM14,IGF2,EIF3(
,KHDRBS1,CSNK1E,PFKL,CDH6,SFPQ,CDH2,PDIA6,PIP4K2B,MEGF8,SOX12,ACTN4,DDX3X,ARL4
C2,PPP2R1A,GTF3C4,PKD1,HLA-C,SF3B1,SSBP2,RPS3,VAPA,HMGB1,TPT1,SEC61A1,RPS4X,RP
GF2,EIF3C,ATP5F1A,EIF4B,MARCKSL1,UBB,AFF4,YWHAG,COL4A2,RPL37A,LRRC28,FN1,SPEN,
ACK1,BSG,FASN,ALDH6A1,SPTBN1,ATP5F1B,CALM3,GNB1,SHISA9,RPS8,MYH9,RPL7,SEPTIN2,
PC,SLC16A7,KPNB1,PAPOLA,LLGL1,NCS1,RHOA,CBX5,RPL9,CLSTN1,ATP1A1,TMEM259,RPS20,
S8,MYH9,RPL7,SEPTIN2,P4HB,IDS,RPL5,KLRD1,GDI1,FSCN1,TMBIM6,PTMS,SPTAN1,GSTP1,MA
2,SARAF,MIDN,MARCHF6,AP3D1,SBF1,ARF1,CYB5R3,AKAP12,MAP4,SF3B2,CDC42BPB,ITM2B,
(D4,WLS,CADM4,RHOB,UBE2H,PLPPR3,ARCN1,MAGED4B,ARHGDIA,STIP1,GPC1,CDK16,SLC35E
LM1,NPM1,PLXNB2,PLD3,HSPD1,USP11,COL18A1,SNRNP200,CSDE1,SMIM14,IGF2,EIF3C,EIF4
AM1,AGRN,RPLP1,ZMIZ1,TIMP2,MIDN,RPL10,AP3D1,DNAJA1,AKAP12,MAP4,CDC42BPB,ITM2B,
N1,PSMD2,SLC7A5,CHD4,ATP1A1,JUND,CD81,WDR1,HMGN2,ATP2A2,CRMP1,LASP1,OAZ1,PFN1
DC42BPB,ITM2B,AP2B1,MAT2A,SLC4A8,GOLGA8A,PXDN,OGA,HNRNPA0,CTNNA1,FADS2,TPM4,
,DDB1,YWHAQ,HACD3,DCHS1,EGLN1,CHSY1,TMSB10,PPP2R1A,LDHB,PKD1,HLA-C,SF3B1,PDIA
1,ATF4,PSMD2,NR2F1,SLC7A5,ENC1,CXADR,CHD4,ATP1A1,JUND,EIF5,CD81,LENG8,PFN1,ASTN
11,PLXNB2,HSPD1,COL18A1,SNRNP200,CSDE1,IGF2,EIF3C,ATP5F1A,EIF4B,VCP,UBB,TRIM28,YV
1,YWHAQ,EIF1,DCHS1,SET,EGLN1,MAP9,CHSY1,TMSB10,INA,EIF4A1,MRC2,GTF3C4,LDHB,DNA
PL15,PKD1,SF3B1,PDIA3,BASP1,VAPA,HMGB1,POLR2A,TPT1,RPS3A,STMN1,ITGB1,TUBB4B,BT
PS3A,STMN1,ITGB1,UCHL1,RPL11,MORF4L1,BTF3,NCAM1,ANXA2,AGRN,RPLP1,XAF1,SARAF,MI
NXA2,AGRN,TIMP2,COL11A1,MIDN,MARCHF6,RPS17,VCL,AP3D1,DNAJA1,ARF1,CDC42BPB,ITM
P3,LLGL1,HK1,NCS1,RPL22,ZNF865,CLSTN1,ENC1,NLGN2,CXADR,JUND,CD81,LENG8,WDR1,S
11,SNRNP200,IGF2,EIF3C,EIF4B,VCP,UBB,TRIM28,MATR3,CKB,COL4A2,RPL37A,LRRC28,FN1,N
2R1A,LDHB,DNAH8,HLA-C,PDIA3,SERINC1,SSBP2,BASP1,VAPA,POLR2A,CDKL1,AP2M1,TPT1,TU
ED1,SEC62,RPL6,KAT6A,DVL3,YWHAE,KIF1A,RPL13,SLC2A3,DDX17,GPI,HTRA4,RACK1,BSG,FAS
I,KHDRBS1,SPTBN2,GLG1,TPM1,EIF5A,CCND2,CDH2,GPC3,MEGF8,SOX12,MMP14,DDX3X,RPS
,MIF,CHPF,JUN,MYO5A,B4GALNT4,DNAJC5,APLP2,RPLP2,MXD4,MYL6,MYH10,DUX4,CKAP5,CA
B2,APLP1,CHPF,JUN,MYO5A,COL6A2,B4GALNT4,DNAJC5,APLP2,RPLP2,MXD4,WLS,MYH10,RPL
(1,BSG,FASN,ALDH6A1,SPTBN1,ATP5F1B,CALM3,GNB1,SHISA9,STMN2,RPS8,MYH9,RPL7,SEPT
L1,TPT1,RPL18,ITGB1,TUBB4B,MORF4L1,BTF3,AGRN,ZMIZ1,COL11A1,MIDN,RPL10,MARCHF6,
PEN,NONO,PSMD5,EIF1,DCHS1,SET,EGLN1,CHSY1,TMSB10,INA,EIF4A1,MRC2,RPL19,GTF3C4,P
MN1,TUBB4B,UCHL1,MORF4L1,BTF3,NCAM1,ANXA2,AGRN,RPLP1,SARAF,MIDN,RPL10,MARCH
DN,HNRNPU,SMARCA5,CTNNA1,RPL23,CNN3,TPM4,CTSD,SCARB2,LPCAT1,RPS24,BRD3,KPNB1,
M1,PLXNB2,TTC3,PLD3,USP11,COL18A1,CSDE1,EIF3C,EIF4B,VCP,MARCKSL1,AFF4,MATR3,CK

DHB,PKD1,HLA-C,PDIA3,RPS3,BASP1,VAPA,HMGB1,TPT1,SEC61A1,RPS3A,STMN1,ITGB1,TUBB4
,INA,EIF4A1,MRC2,PPP2R1A,RPL15,LDHB,PKD1,HLA-C,PDIA3,RPS3,BASP1,VAPA,HMGB1,TPT1,S
PXDN,HNRNPU,NFE2L1,HNRNPA0,CTNNA1,FADS2,RPL23,LDHA,CNN3,EIF4H,LAMA5,SLC22A17,B
D3,CIPC,PJA2,KPNB1,C1ORF43,COL1A2,LLGL1,NCS1,RHOA,CBX5,ZNF865,CLSTN1,ATF4,SLC7
2A,SEC61A1,RPS4X,RPS3A,ITGB1,MORF4L1,BTF3,AGRN,RPLP1,ZMIZ1,TIMP2,MIDN,RPL10,RPS1
CD81,WDR1,HMGN2,CRMP1,LASP1,VAT1,SRSF4,RTN3,PFN1,ASTN2,JPT2,PNN,SPC25,NES,PRKC
A,EIF4A1,MRC2,RPL19,PPP2R1A,GTF3C4,SF3B1,PDIA3,SERINC1,SSBP2,ZNF43,VAPA,HMGB1,PC
,HNRNPU,NFE2L1,RPS16,CNN3,CTSD,LPCAT1,LAMA5,BRD3,CIPC,MSN,C1ORF43,LLGL1,ZNF865
NES,NISCH,HNRNPC,CCNI,LRPPRC,PFKL,HDLBP,TKT,PNMA1,GPC3,PIP4K2B,SOX12,ITM2C,ARL4
NES,NISCH,HNRNPC,CCNI,LRPPRC,PFKL,HDLBP,TKT,PNMA1,GPC3,PIP4K2B,SOX12,ITM2C,ARL4
TXN10,KDM1A,MAP7D1,ARF4,ZFAND5,HNRNPUL2-BSCL2,MSL1,ATP6V1B2,GIGYF1,PHC2,YWHA
NN,CCNI,TADA2B,KHDRBS1,PFKL,HDLBP,EIF5A,ACTN4,RPS15,ITM2C,RPS5,MIF,RPS10,MTSS2,D
N1,ATP5F1B,CALM3,GNB1,SHISA9,STMN2,RPS8,MYH9,RPL7,SEPTIN2,P4HB,APP,IDS,ERVV-1,GD
,STMN1,ITGB1,TUBB4B,MORF4L1,BTF3,AGRN,RPLP1,ZMIZ1,TIMP2,SARAF,COL11A1,MIDN,RPL
F4,RBBP4,MCL1,NISCH,CCNI,LRPPRC,SLC3A2,SFPQ,CDH2,PDIA6,MEGF8,SOX12,DDX3X,MAGE
63,B4GALNT4,DNAJC5,APLP2,HNRNPDL,TARDBP,MXD4,MYL6,RPL36,PTGES3,CADM4,HNRNPH1
HDLBP,TKT,SFPQ,PNMA1,GPC3,PIP4K2B,SOX12,ARL4C,GNB2,APLP1,MIF,GPAT4,MYO5A,COL6A
1A1,JUND,CD81,LENG8,WDR1,ATP2A2,CRMP1,TLE5,TADA2B,SPTBN2,PFKL,TPM1,EIF5A,PEBP1,
GLN1,MAP9,TPI1,CHSY1,INA,EIF4A1,RPL19,PPP2R1A,RPL15,GTF3C4,LDHB,SF3B1,PDIA3,SSBP2,
3,SSBP2,POLR2A,CDKL1,TPT1,RPS4X,RPS3A,STMN1,ITGB1,TUBB4B,RPL11,MORF4L1,AGRN,RF
,YWHAG,CKB,RPL37A,NONO,YWHAQ,EIF1,HACD3,SET,EGLN1,TPI1,CHSY1,TMSB10,MRC2,PPP2
KSL1,UBB,TRIM28,MATR3,YWHAG,CKB,COL4A2,LRRC28,FN1,SPEN,NONO,DDB1,YWHAQ,HACD
AN1,MARCKS,SMG1,CALM1,PLXNB2,PLD3,USP11,SNRNP200,IGF2,EIF3C,EIF4B,VCP,UBB,AFF4,
4,WLS,HNRNPK,H1-10,CIRBP,DUX4,CADM4,ATP1A2,RAB1A,PLPPR3,ARCN1,MAGED4B,CAPRIN1,
ACA,SLC22A17,BRD3,CIPC,PJA2,KPNB1,LLGL1,NCS1,RHOA,CBX5,ZNF865,RAB6B,SLC7A5,NLG
D3,DCHS1,SET,EGLN1,TPI1,TMSB10,EIF4A1,MRC2,RPL19,PPP2R1A,GTF3C4,LDHB,PKD1,HLA-C,
Y1,TMSB10,MRC2,PPP2R1A,RPL15,GTF3C4,PKD1,HLA-C,PDIA3,SSBP2,ZNF43,CHD3,VAPA,AP2M
,GTF2F1,PXDN,HNRNPU,PGAM1,NFE2L1,SMARCA5,HNRNPA0,CTNNA1,FADS2,RPL23,LDHA,CNN
P1,SMCHD1,IPO5,ATP6V0C,MMP2,TRIB2,CTDSP2,NOVA2,CTSB,CDH11,CHD6,EEF1A2,TBCD,SN
PS17,VCL,SBF1,DNAJA1,ARF1,MAP4,SF3B2,CDC42BPB,MAT2A,TMEM170A,PXDN,HNRNPU,NFE2
FASN,ALDH6A1,SPTBN1,ATP5F1B,CALM3,GNB1,SHISA9,STMN2,RPS8,MYH9,SEPTIN2,P4HB,APP,
,CDH2,PNMA1,GPC3,PIP4K2B,SOX12,ARL4C,APLP1,MIF,GPAT4,TLN1,MYO5A,COL6A2,CD63,B4
1,VAT1,OAZ1,PFN1,MCL1,NES,NISCH,XPR1,SLC3A2,PFKL,HDLBP,TKT,GLG1,PNMA1,PDIA6,GPC
TN2,TLE5,PNN,LRPPRC,TADA2B,PFKL,HDLBP,PDIA6,PIP4K2B,CCN2,ACTN4,DDX3X,ARL4C,GNB2,
PFN1,TLE5,NISCH,CCNI,TADA2B,CSNK1E,PFKL,HDLBP,TKT,CDH6,SFPQ,CCND2,CDH2,PNMA1,PD
,LDHB,DNAH8,PKD1,SF3B1,PDIA3,ZNF43,RPS3,BASP1,VAPA,HMGB1,POLR2A,CDKL1,AP2M1,TP
BPB,ITM2B,GTF2F1,PXDN,NFE2L1,HNRNPA0,CTNNA1,FADS2,LDHA,CNN3,RPL29,TNS3,CTSD,EI
L1A2,LLGL1,NCS1,RHOA,CBX5,ZNF865,RPS14,CXADR,CHD4,ATP1A1,JUND,WDR1,HMGN2,ATP2
MYH9,RPL7,SEPTIN2,P4HB,APP,ERVV-1,RPL5,GDI1,FSCN1,TMBIM6,SPTAN1,GSTP1,SMG1,CALM
DKL1,RPL18,ITGB1,TUBB4B,UCHL1,MORF4L1,AGRN,ZMIZ1,TIMP2,SARAF,MIDN,RPL10,RPS17,
KD1,PDIA3,SSBP2,CHD3,VAPA,HMGB1,TPT1,RPS3A,ITGB1,TUBB4B,UCHL1,MORF4L1,BTF3,NCA
Y1,INA,RPL15,GTF3C4,LDHB,PKD1,HLA-C,SERINC1,SSBP2,ZNF43,BASP1,VAPA,POLR2A,TPT1,R
XD4,TAGLN,ANKRD52,HNRNPK,H1-10,INIP,SEPTIN11,LGALS3BP,UBE2H,HLA-B,PLPPR3,CTBP1,
1,SLC3A2,PFKL,HDLBP,TKT,GLG1,SFPQ,CCND2,PNMA1,PDIA6,GPC3,PIP4K2B,SOX12,RPL12,AR
L1,MORF4L1,BTF3,AGRN,ZMIZ1,TIMP2,SARAF,COL11A1,MIDN,RPL10,DNAJA1,ARF1,CYB5R3,A

0,INA,EIF4A1,PPP2R1A,RPL15,GTF3C4,LDHB,DNAH8,PKD1,HLA-C,SF3B1,PDIA3,ZNF43,BASP1,V
D1,USP11,SNRNP200,CSDE1,SMIM14,IGF2,EIF3C,ATP5F1A,EIF4B,VCP,UBB,AFF4,MATR3,YWHA
TGB1,TUBB4B,UCHL1,RPL11,MORF4L1,BTF3,ANXA2,RPLP1,SARAF,MIDN,MARCHF6,WSB1,VCL
EIF4B,VCP,MARCKSL1,UBB,MATR3,YWHAG,CKB,RPL37A,FN1,SPEN,NONO,YWHAQ,EIF1,DCHS1,
RRC28,FN1,NONO,DDB1,YWHAQ,EIF1,HACD3,SET,CHSY1,TMSB10,GTF3C4,LDHB,PKD1,HLA-C,
1259,CD81,WDR1,HMGN2,ATP2A2,LASP1,BEX3,CCT7,OAZ1,RTN3,PFN1,NISCH,XPR1,XPO1,PFKL
,LAMP1,RHOB,RAB1A,PLPPR3,COTL1,CTBP1,RAC1,ARHGDIA,GPC1,TOP2B,SMCHD1,ATP6V0C,T
8,GOLGA8A,HNRNPU,OGA,NFE2L1,SMARCA5,LARP1,HNRNPA0,CNN3,RPL29,TPM4,LPCAT1,BRD
V-1,RPL5,FSCN1,TMBIM6,SPTAN1,MARCKS,SMG1,CALM1,NPM1,PLXNB2,PLD3,HSPD1,COL18A1
CL,AP3D1,SBF1,DNAJA1,ARF1,CYB5R3,MAP4,CDC42BPB,ZNF770,MAT2A,TMEM170A,GTF2F1,H
,HNRNPU,OGA,SMARCA5,HNRNPA0,CTNNA1,FADS2,RPS16,RPL23,CNN3,RPL29,RPSA,TNS3,EIF
PNB1,PAPOLA,IGFBP3,LLGL1,HK1,NCS1,RPL22,ZNF865,CLSTN1,RAB6B,ENC1,NLGN2,CXADR,
A0,FADS2,CNN3,RPSA,TNS3,CTSD,EIF4H,TSPAN3,SLC22A17,BRD3,RPS9,MSN,SLC16A7,KPNB1,
C4,LDHB,PKD1,HLA-C,PDIA3,SSBP2,CHD3,BASP1,HMGB1,POLR2A,TPT1,TUBB4B,MORF4L1,NC
,ATP5F1A,EIF4B,VCP,AFF4,MATR3,YWHAG,CKB,COL4A2,RPL37A,LRRC28,NONO,DDB1,YWHAQ,
FN1,ASTN2,JPT2,TLE5,NES,NISCH,TADA2B,XPR1,KHDRBS1,TKT,CDH2,PDIA6,GPC3,PLXNA1,PIP
ISB10,INA,MRC2,RPL19,PPP2R1A,RPL15,GTF3C4,LDHB,SF3B1,SERINC1,SSBP2,ZNF43,BASP1,V
ICAM1,ANXA2,AGRN,RPLP1,XAF1,COL11A1,MIDN,MARCHF6,SBF1,DNAJA1,CYB5R3,PAFAH1B1,I
CKS,SMG1,CALM1,NPM1,PLXNB2,PLD3,HSPD1,USP11,COL18A1,SNRNP200,CSDE1,SMIM14,IG
ARAF,COL11A1,MIDN,RPL10,MARCHF6,VCL,AKAP12,MAP4,CDC42BPB,MAT2A,TMEM170A,GTF
,ZNF770,MAT2A,HNRNPU,PGAM1,CTNNA1,FADS2,LDHA,TPM4,TNS3,CTSD,SCARB2,TSPAN3,RP
M3,GNB1,SHISA9,STMN2,RPS8,MYH9,RPL7,SEPTIN2,P4HB,APP,IDS,ERVV-1,GDI1,FSCN1,TMBIM
6A2,CD63,B4GALNT4,MTSS2,DNAJC5,APLP2,HNRNPDL,TARDBP,MXD4,INIP,YWHAB,CADM4,HN
MBIM6,PTMS,SPTAN1,GSTP1,MARCKS,SMG1,CALM1,PLXNB2,PLD3,HSPD1,USP11,COL18A1,SN
TM2C,RPL30,ARL4C,GNB2,RCN1,APLP1,RPL35,MIF,GPAT4,MYO5A,CD63,SESN3,MTSS2,APLP2,
ANXA2,AGRN,RPLP1,TIMP2,SARAF,COL11A1,RPL10,MARCHF6,WSB1,RPS17,AP3D1,SBF1,DNAJ
ANXA2,AGRN,RPLP1,TIMP2,SARAF,COL11A1,RPL10,MARCHF6,WSB1,RPS17,AP3D1,SBF1,DNAJ
NPA0,RPS16,RPL23,CNN3,RPL29,TPM4,EIF4H,RPS24,RPS9,XRCC5,CIPC,PJA2,MSN,RPL24,SLC1
P1,OAZ1,PFN1,TLE5,NES,TADA2B,SPTBN2,PFKL,HDLBP,TKT,PNMA1,PDIA6,GPC3,PIP4K2B,MEG
NT4,APLP2,HNRNPDL,HNRNPK,DUX4,CADM4,CKAP4,RAB1A,PLPPR3,SEPTIN3,CAPRIN1,STIP1,SL
1,MYO5A,ANXA6,B4GALNT4,APLP2,TARDBP,SHOX,WLS,HNRNPK,DUX4,SEPTIN11,CADM4,CKAP4
N2,P4HB,IDS,RPL5,KLRD1,GDI1,FSCN1,PTMS,SPTAN1,GSTP1,MARCKS,SMG1,CALM1,NPM1,TT
CAT1,LAMA5,SLC22A17,BRD3,CIPC,KPNB1,IGFBP3,HK1,NCS1,RHOA,ZNF865,CLSTN1,CXADR,A
1,NLGN2,CHD4,ATP1A1,RPS20,EIF5,WDR1,HMGN2,ATP2A2,CRMP1,PFN1,TLE5,PNN,NISCH,HNR
LC22A17,RPL24,KPNB1,C1ORF43,PTCD3,CBX5,ATF4,NR2F1,KIF5B,SLC7A5,ENC1,NLGN2,CXAD
C4,DNAH8,PKD1,HLA-C,SSBP2,BASP1,VAPA,HMGB1,POLR2A,AP2M1,TPT1,SEC61A1,RPS3A,ITG
9,CLSTN1,KIF5B,RAB6B,SLC7A5,NLGN2,GLIPR1L2,RPS20,LENG8,ATP2A2,BEX3,CCT7,OAZ1,M
TP5F1A,EIF4B,MARCKSL1,UBB,AFF4,MATR3,YWHAG,CKB,COL4A2,RPL37A,NONO,DDB1,YWHA
DRBS1,PFKL,HDLBP,TKT,CCND2,CDH2,PNMA1,GPC3,PIP4K2B,ACSL6,SOX12,MMP14,ITM2C,A
KS,SMG1,NPM1,PLXNB2,HSPD1,USP11,COL18A1,SNRNP200,CSDE1,SMIM14,IGF2,EIF3C,VCP,
FF4,MATR3,YWHAG,RPL37A,LRRC28,SPEN,NONO,PSMD5,YWHAQ,HACD3,DCHS1,SET,MAP9,TPI
KPNB1,PAPOLA,IGFBP3,RPL9,PSMD2,KIF5B,ATP1A1,JUND,TMEM259,RPS20,CD81,LASP1,CCT7
RNPH1,RPS27,CCT6A,HMGCR,RAC1,SMCHD1,ATP6V0C,BCLAF1,L1CAM,LAPTM4B,TRIB2,SQLE,
L1,SMARCA5,CTNNA1,RPL23,CNN3,RPSA,TPM4,TNS3,SCARB2,RPS24,SLC22A17,BRD3,CIPC,C1
,MRC2,RPL19,PPP2R1A,GTF3C4,PKD1,SF3B1,PDIA3,SSBP2,CHD3,VAPA,CDKL1,TPT1,SEC61A1,

TMS,SPTAN1,CALM1,PLXNB2,PLD3,HSPD1,USP11,COL18A1,SNRNP200,CSDE1,SMIM14,IGF2,EII
ATF4,PSMD2,NR2F1,KIF5B,RAB6B,SLC7A5,ENC1,NLGN2,CHD4,ATP1A1,JUND,EIF5,CD81,WDR1,I
HMGB1,AP2M1,TPT1,SEC61A1,RPS3A,STMN1,ITGB1,TUBB4B,MORF4L1,NCAM1,ANXA2,AGRN,Z
SNRPN,PRDX5,PKN1,CAMSAP1,GRINA,VPS35,ACAP3,CBX1,SNRNP70,PYGB,DPYSL2,TOP1,TAF7,\
1,PSMD5,HACD3,SET,EGLN1,TPI1,CHSY1,TMSB10,INA,MRC2,PPP2R1A,RPL15,GTF3C4,LDHB,DN
ARCKS,SMG1,CALM1,NPM1,PLXNB2,PLD3,HSPD1,COL18A1,SNRNP200,CSDE1,IGF2,EIF3C,ATP5
PJA2,MSN,PAPOLA,HK1,RPL22,RHOA,PTCD3,ZNF865,ATF4,GLIPR1L2,RPS14,TMEM259,RPS20,C
,SERINC1,SSBP2,ZNF43,RPS3,HMGB1,POLR2A,CDKL1,AP2M1,TPT1,SEC61A1,RPS4X,RPS3A,STN
LS,HNRNPK,H1-10,DUX4,CADM4,PLPPR3,ARCN1,MAGED4B,CTBP1,ARHGDIA,CAPRIN1,ATP6V0C
),INA,EIF4A1,PPP2R1A,GTF3C4,LDHB,PKD1,HLA-C,PDIA3,ZNF43,RPS3,CHD3,AP2M1,TPT1,SEC61
I2B,TMEM170A,GTF2F1,HNRNPU,NFE2L1,CTNNA1,CNN3,TPM4,TNS3,SCARB2,LPCAT1,RPS24,SL
OVA2,CTSB,CDH11,ZC3H7B,MDH2,RAB27A,U2AF2,TBCD,PABPN1,HNRNPL,MORF4L2,TAF15,PRI
,MIF,GPAT4,CHPF,MYO5A,CD63,GOLM1,B4GALNT4,APLP2,TARDBP,TAGLN,CADM4,HNRNPH1,H
LD3,HSPD1,COL18A1,SNRNP200,CSDE1,IGF2,EIF3C,ATP5F1A,EIF4B,MARCKSL1,UBB,AFF4,TRIM
1,TMBIM6,PTMS,SPTAN1,GSTP1,MARCKS,SMG1,CALM1,NPM1,PLXNB2,HSPD1,COL18A1,SNRNI
TUBB4B,UCHL1,MORF4L1,BTF3,ANXA2,AGRN,ZMIZ1,SARAF,MIDN,RPL10,VCL,AP3D1,DNAJA1,C
X,RPS15,ARL4C,MAGED2,RCN1,MIF,GPAT4,JUN,COL6A2,B4GALNT4,DNAJC5,APLP2,RPLP2,MXI
ITGB1,TUBB4B,NCAM1,ANXA2,AGRN,ZMIZ1,XAF1,MIDN,RPL10,WSB1,VCL,AP3D1,ARF1,CYB5R3
312B,MAPK1,SMARCA4,KDELR2,RABL6,PTBP1,ILF3,GRB2,CAPZB,UBE2R2,ZFAND5,PITPNA,MSL
IORF4L1,RPLP1,SARAF,MIDN,RPL10,MARCHF6,RPS17,VCL,AP3D1,DNAJA1,SF3B2,MAT2A,GTF2I
SUGP2,CPSF1,PPP1CB,MEST,HNRNPL,ABCA3,ALKBH5,PRDX5,C4ORF3,UBE4A,PKN1,CAMSAP1,C
RSF4,PFN1,ASTN2,TLE5,NES,CCNI,XPR1,XPO1,PFKL,HDLBP,TKT,TPM1,EIF5A,CCND2,PDIA6,PIP4I
T1,SEC61A1,RPS3A,BTF3,NCAM1,AGRN,RPLP1,MIDN,RPL10,MARCHF6,RPS17,AP3D1,DNAJA1,SI
KAT6A,DVL3,YWHAE,KIF1A,RPL13,GNAS,SLC2A3,DDX17,GPI,HTRA4,RACK1,BSG,FASN,ALDH6A1,
XNB2,USP11,COL18A1,CSDE1,SMIM14,IGF2,ATP5F1A,EIF4B,MARCKSL1,UBB,AFF4,TRIM28,MA
R,CHD4,ATP1A1,JUND,LENG8,HMGN2,ATP2A2,CRMP1,OAZ1,RBBP4,TLE5,NISCH,HNRNPC,CCNI,
HD4,TMEM259,CD81,WDR1,ATP2A2,BEX3,CCT7,OAZ1,ASTN2,JPT2,PRKCSH,CCNI,HDLBP,GLG1,
A,GTF3C4,SF3B1,PDIA3,SERINC1,SSBP2,ZNF43,VAPA,POLR2A,CDKL1,SEC61A1,RPS3A,STMN1,U
1,OAZ1,PFN1,TLE5,NES,TADA2B,XPR1,PFKL,HDLBP,TKT,GLG1,SFPQ,CCND2,PNMA1,GPC3,DDR
LBP,TKT,EIF5A,CCND2,CDH2,PNMA1,PDIA6,PIP4K2B,RPL12,ITM2C,GNB2,MIF,GJA1,MYO5A,COL
XDN,PGAM1,HNRNPA0,CTNNA1,FADS2,LDHA,RPSA,TPM4,CTSD,EIF4H,SCARB2,NACA,LPCAT1,RI
4,IGF2,EIF3C,ATP5F1A,EIF4B,VCP,UBB,AFF4,MATR3,YWHAG,COL4A2,RPL37A,LRRC28,SPEN,NC
61A1,RPL18,RPL11,MORF4L1,AGRN,ZMIZ1,TIMP2,XAF1,SARAF,MIDN,RPL10,WSB1,RPS17,VCL,A
SNRNP200,CSDE1,SMIM14,EIF3C,ATP5F1A,EIF4B,MARCKSL1,UBB,YWHAG,CKB,RPL37A,LRRC2
CYB5R3,MAP4,ITM2B,ZNF770,TMEM170A,GTF2F1,PXDN,HNRNPU,OGA,NFE2L1,SMARCA5,HNRN
4B,VCP,MARCKSL1,UBB,AFF4,TRIM28,MATR3,YWHAG,CKB,COL4A2,LRRC28,FN1,DDB1,DCHS1
M3,GNB1,SHISA9,STMN2,RPS8,MYH9,RPL7,SEPTIN2,P4HB,APP,IDS,ERVV-1,RPL5,GDI1,FSCN1,TN
ZNF43,CHD3,VAPA,HMGB1,AP2M1,TPT1,SEC61A1,STMN1,ITGB1,TUBB4B,UCHL1,NCAM1,ANXA2
A,RNF187,EEF1A2,FDFT1,TBCD,ANXA5,SNX22,GAS7,SSRP1,ALKBH5,PDXK,CAMSAP1,GRINA,PTO
L6,KAT6A,DVL3,YWHAE,KIF1A,RPL13,GNAS,SLC2A3,DDX17,GPI,HTRA4,RACK1,BSG,FASN,ALDH6
ISY1,INA,RPL19,PPP2R1A,GTF3C4,LDHB,PKD1,SF3B1,PDIA3,SSBP2,RPS3,BASP1,VAPA,HMGB1,F
3,AKAP12,SF3B2,CDC42BPB,ITM2B,ZNF770,MAT2A,SLC4A8,OGA,SMARCA5,CTNNA1,RPS16,LDI
RB2,NACA,LAMA5,RPS24,CIPC,C1ORF43,PAPOLA,IGFBP3,RPL14,HK1,ZNF865,PSMD2,SLC7A5,E
3,PFKL,HDLBP,RPS25,GLG1,SFPQ,TPM1,PEBP1,CCND2,PNMA1,PLXNA1,SOX12,PRDX2,DDX3X,IT
KT,GLG1,EIF5A,PEBP1,PNMA1,PDIA6,GPC3,DDR1,PIP4K2B,SOX12,ITM2C,ARL4C,APLP1,MIF,GPA

DNER,BCLAF1,L1CAM,SQLE,MAP1A,FAM199X,SERF2,TIMP3,CTDSP2,NOVA2,NUCB1,ZC3H7B,CH
,ANXA5,VAMP2,GAS7,YEATS2,HSPA4,CAMSAP1,GRINA,PTOV1,MLEC,CLSTN3,MDK,JPT1,ORAI2,S
0,MYH10,PTGES3,CADM4,LAMP1,UBE2H,TTR,CTBP1,APPL1,HDAC2,DNER,BCLAF1,HSBP1,CDKN
P4HB,IDS,GDI1,FSCN1,TMBIM6,PTMS,SPTAN1,GSTP1,MARCKS,SMG1,CALM1,NPM1,TTC3,PLD3,
SNRNP200,SMIM14,IGF2,EIF3C,ATP5F1A,EIF4B,VCP,MARCKSL1,UBB,AFF4,MATR3,YWHAG,CKB,
ZC3H7B,CHD6,EEF1A2,CCT2,TBCD,VAMP2,MEST,GAS7,TM9SF2,LMNB2,ACO2,HSPA4,CAMSAP1
NPH1,HLA-B,PLPPR3,MAGED4B,CTBP1,RAC1,STIP1,GPC1,SLC35E2B,SMCHD1,IPO5,ATP6V0C,SK
2,EIF3C,ATP5F1A,EIF4B,VCP,UBB,AFF4,MATR3,YWHAG,COL4A2,RPL37A,LRRC28,SPEN,NONO,EI
RL4C,GNB2,H1-0,RPS5,RPS7,RPS19,GPAT4,TLN1,MYO5A,COL6A2,CD63,MTSS2,APLP2,HNRNPD
RL4C,GNB2,H1-0,RPS5,RPS7,RPS19,GPAT4,TLN1,MYO5A,COL6A2,CD63,MTSS2,APLP2,HNRNPD
RL4C,GNB2,H1-0,RPS5,RPS7,RPS19,GPAT4,TLN1,MYO5A,COL6A2,CD63,MTSS2,APLP2,HNRNPD
I,RPL10,WSB1,AP3D1,SBF1,ARF1,CYB5R3,ITM2B,MAT2A,GTF2F1,OGA,NFE2L1,CTNNA1,CNN3,E
M1,DNAJC5,APLP2,DDOST,MXD4,HNRNPK,H1-10,INIP,MYH10,RPL36,SEPTIN11,CKAP4,RAB1A,S
R3,YWHAG,CKB,COL4A2,LRRC28,SPEN,DDB1,PSMD5,YWHAQ,HACD3,DCHS1,SET,EGLN1,MAP9,
NP200,SMIM14,IGF2,EIF3C,ATP5F1A,EIF4B,VCP,MARCKSL1,UBB,AFF4,MATR3,YWHAG,CKB,COL
5F1A,EIF4B,VCP,UBB,AFF4,MATR3,YWHAG,CKB,COL4A2,RPL37A,LRRC28,SPEN,NONO,EIF1,EG
DI1,FSCN1,TMBIM6,PTMS,SPTAN1,GSTP1,MARCKS,SMG1,CALM1,NPM1,PLXNB2,TTC3,HSPD1,U
2,PYGB,SELENON,TUBB2A,CCND1,AEBP1,DYNLL2,GRB2,CAPZB,UBE2R2,ACTN1,BNIP3L,FKBP8
P3D1,SBF1,DNAJA1,SF3B2,AP2B1,MAT2A,GTF2F1,PXDN,HNRNPU,SMARCA5,HNRNPA0,CTNNA1,F
X12,DDX3X,RPS15,ARL4C,CLTC,GNB2,H1-0,MIF,COL6A2,B4GALNT4,MTSS2,DNAJC5,APLP2,SHC
TMEM170A,SLC4A8,GOLGA8A,PXDN,HNRNPU,NFE2L1,CTNNA1,CPE,TPM4,TNS3,LPCAT1,LAMA5,
VDR1,HMGN2,RPL26,LASP1,OAZ1,SRSF4,RBBP4,PFN1,MCL1,JPT2,PNN,RPL31,HNRNPC,HDLBP,
,PFN1,TLE5,NES,XPR1,KLHL9,SLC3A2,PFKL,HDLBP,TKT,CCND2,CDH2,PDIA6,GPC3,DDR1,PIP4K
200,SMIM14,IGF2,EIF3C,ATP5F1A,EIF4B,VCP,MARCKSL1,UBB,AFF4,MATR3,YWHAG,CKB,COL4
2B,SKP1,TRIB2,CTDSP2,NOVA2,CTSB,CDH11,CHD6,EEF1A2,TBCD,SQSTM1,VAMP2,MORF4L2,A
4B,VCP,UBB,AFF4,TRIM28,YWHAG,CKB,COL4A2,RPL37A,MAPT,FN1,DDB1,PSMD5,HACD3,DCHS
,CRMP1,LASP1,OAZ1,SRSF4,PFN1,MCL1,JPT2,TLE5,PNN,SPC25,NES,TADA2B,XPR1,KHDRBS1,K
A,LAMA5,RPS24,SLC22A17,PJA2,KPNB1,C1ORF43,PAPOLA,LLGL1,HK1,NCS1,RHOA,CBX5,ZNF8
PS9,PJA2,RPL24,KPNB1,RPL14,CBX5,ATF4,ENC1,CXADR,JUND,TMEM259,RPS20,EIF5,CD81,WD
M14,IGF2,EIF3C,ATP5F1A,EIF4B,VCP,UBB,AFF4,MATR3,YWHAG,RPL37A,LRRC28,SPEN,NONO,EI
,MYH10,DUX4,HNRNPH1,LGALS3BP,UBE2H,CKAP4,HLA-B,PLPPR3,CTBP1,RAC1,ARHGDIA,GPC1
POLA,HK1,RPL22,RHOA,ZNF865,ATF4,KIF5B,SLC7A5,GLIPR1L2,RPS14,RPS20,CD81,LENG8,WD
8,PXDN,OGA,NFE2L1,SMARCA5,HNRNPA0,CTNNA1,FADS2,CNN3,TPM4,EIF4H,LPCAT1,LAMA5,SL
F2F1,PXDN,HNRNPU,SMARCA5,CTNNA1,FADS2,CNN3,RPSA,TPM4,TNS3,CTSD,EIF4H,TSPAN3,LAM
MP2,GAS7,NOP53,PABPN1,GDI2,HNRNPL,TSPYL4,TLN2,TAF15,LMNB2,ACTR3,ACO2,KLC1,RBMX
F4A1,MRC2,RPL19,RPL15,GTF3C4,LDHB,PKD1,SF3B1,PDIA3,SERINC1,SSBP2,CHD3,BASP1,VAP
NC1,SSBP2,ZNF43,RPS3,HMGB1,POLR2A,AP2M1,TPT1,SEC61A1,RPS4X,RPS3A,RPL18,TUBB4B,
PL37A,LRRC28,YWHAQ,EIF1,TPI1,TMSB10,INA,EIF4A1,RPL19,PPP2R1A,RPL15,GTF3C4,LDHB,DN
NPH1,CALU,COTL1,CTBP1,GPC1,SLC35E2B,SMCHD1,IPO5,DNER,BCLAF1,LAPTM4B,CDKN1A,S
VCP,MARCKSL1,UBB,AFF4,MATR3,YWHAG,COL4A2,RPL37A,LRRC28,FN1,SPEN,DDB1,PSMD5,EI
COL4A2,MAPT,FN1,DDB1,EIF1,HACD3,DCHS1,SET,MAP9,CHSY1,TMSB10,EIF4A1,RPL19,PPP2R1
RPS24,SLC22A17,CIPC,C1ORF43,COL1A2,LLGL1,HK1,NCS1,RPL22,RPL9,CLSTN1,ATF4,PSMD2,
1,COL18A1,SNRNP200,CSDE1,SMIM14,IGF2,EIF3C,ATP5F1A,EIF4B,VCP,MARCKSL1,UBB,AFF4,M
RPS20,CD81,WDR1,HMGN2,LASP1,CCT7,SRSF4,RBBP4,PFN1,MCL1,PNN,HNRNPC,KHDRBS1,HD
VSB1,RPS17,DNAJA1,MAP4,ITM2B,MAT2A,TMEM170A,GTF2F1,GOLGA8A,PXDN,HNRNPU,OGA,H

NF865,RPS14,RPS20,LENG8,WDR1,HMGN2,LASP1,SRSF4,RBBP4,PFN1,ASTN2,MCL1,TLE5,PNN,
X4,CADM4,CKAP4,RAB1A,PLPPR3,COTL1,MAGED4B,IPO5,TMEM30A,PEA15,DLG5,NOVA2,MLXIP
,PIP4K2B,CCN2,PRDX2,RPS15,ITM2C,RPL30,ARL4C,RPS19,RPL35,NGRN,GOLM1,RPS10,APLP2
B,AFF4,MATR3,YWHAG,RPL37A,LRRC28,EIF1,HACD3,MAP9,TMSB10,INA,EIF4A1,RPL19,PPP2R1
F4,MATR3,YWHAG,COL4A2,RPL37A,LRRC28,SPEN,NONO,EIF1,TMSB10,INA,EIF4A1,RPL19,PPP2
SPAN3,LPCAT1,RPS24,BRD3,XRCC5,MSN,KPNB1,C1ORF43,PAPOLA,LLGL1,GAP43,HK1,RPL22,
NB2,PDXK,CAMSAP1,GRINA,ACAP3,MDK,LAMC1,RAD23B,SKI,H3-3A,ABCA2,VEGFA,RAPGEF1,MA
UCHL1,RPL11,MORF4L1,ANXA2,AGRN,TIMP2,MIDN,RPL10,MARCHF6,WSB1,RPS17,VCL,AP3D1,
4,MATR3,YWHAG,CKB,COL4A2,RPL37A,LRRC28,MAPT,FN1,NONO,DDB1,YWHAQ,EIF1,HACD3,D
ICS1,RPL22,RPL9,CLSTN1,ATF4,PSMD2,KIF5B,RAB6B,SLC7A5,ENC1,NLGN2,CXADR,CHD4,ATP1
OVA2,NUCB1,CHD6,EEF1A2,CCT2,TBCD,VAMP2,MEST,GAS7,HNRNPL,TM9SF2,LMNB2,SNRPN,YE
ES,NISCH,KHDRBS1,TKT,CDH6,EIF5A,PNMA1,PDIA6,PLXNA1,PIP4K2B,MEGF8,MMP14,PRDX2,RP
EC,MDK,SURF4,CBX1,PTK7,ORAI2,SKI,H3-3A,SCAF1,RAB11B,ABCA2,PYGB,PPP2CA,MAPK1,DDA
5,SEPTIN11,CADM4,ARCN1,COTL1,HECTD4,CTBP1,RAC1,STIP1,SLC35E2B,IPO5,SKP1,DNER,L1
CHL1,MORF4L1,BTF3,ANXA2,RPLP1,ZMIZ1,MIDN,RPL10,WSB1,VCL,AP3D1,DNAJA1,ARF1,MAT2
PJA2,MSN,RPL24,C1ORF43,COL1A2,RPL14,HK1,RPL22,PTCD3,CLSTN1,ATF4,NR2F1,JUND,LEN
KDN,HNRNPU,OGA,HNRNPA0,CTNNA1,RPL23,CPE,CNN3,TNS3,SCARB2,LPCAT1,BRD3,SLC16A7,
INC1,SSBP2,RPS3,HMGB1,POLR2A,CDKL1,AP2M1,TPT1,SEC61A1,RPS3A,ITGB1,RPL11,BTF3,N
,LAPTM4B,RPL23A,NNAT,LUC7L3,SQLE,SERF2,CRPPA,HS6ST1,PGRMC1,EID1,RNF187,U2AF2,R
,MATR3,YWHAG,COL4A2,RPL37A,LRRC28,SPEN,NONO,EIF1,EGLN1,TMSB10,INA,EIF4A1,RPL19,
OL11A1,RPL10,MARCHF6,ARF1,CYB5R3,AKAP12,MAP4,CDC42BPB,ITM2B,MAT2A,GTF2F1,SLC4
,SF3B2,ITM2B,MAT2A,NFE2L1,FADS2,RPS16,RPL23,CPE,RPSA,TPM4,NACA,LAMA5,RPS24,BRD
L19,PPP2R1A,RPL15,GTF3C4,LDHB,DNAH8,PKD1,SF3B1,PDIA3,SERINC1,ZNF43,RPS3,HMGB1,P
,MIF,CHPF,MYO5A,COL6A2,MTSS2,APLP2,TARDBP,MXD4,MYL6,HNRNPK,INIP,DUX4,YWHAB,PTG
PC1,SMCHD1,HDAC2,DNER,PEA15,FAM199X,CPSF2,PGRMC1,DDX39B,SRP9,CTSB,CDH11,RAB2
RHOA,ZNF865,KIF5B,GLIPR1L2,RPS14,RPS20,CD81,LENG8,WDR1,ATXN7L3B,RPL26,LASP1,SRS
OLA,HK1,RHOA,ZNF865,ATF4,GLIPR1L2,RPS14,RPS20,CD81,LENG8,WDR1,ATXN7L3B,HMGN2,R
CCT7,SRSF4,RBBP4,PFN1,TLE5,PNN,RPL31,KHDRBS1,SPTBN2,CSNK1E,RPS25,EIF5A,PDIA6,ME
EIF3C,EIF4B,VCP,MARCKSL1,AFF4,TRIM28,MATR3,YWHAG,COL4A2,MAPT,SPEN,NONO,DDB1,PS
IGB1,POLR2A,AP2M1,TPT1,SEC61A1,RPS4X,RPS3A,RPL18,ITGB1,TUBB4B,UCHL1,RPL11,MORF
N,PRDX5,UBE4A,CCT3,VPS35,ACAP3,CBX1,SNRNP70,DPYSL2,TOP1,TAF7,VHL,PPP2CA,SMARCA4,
C28,FN1,SPEN,NONO,YWHAQ,EIF1,DCHS1,SET,TPI1,CHSY1,TMSB10,RPL15,GTF3C4,LDHB,DNAH
CAF1,RAB11B,ABCA2,PYGB,PRPF6,DDAH2,DYNC1LI2,TCF25,ADAM9,GRB2,WDR6,ACTN1,CASC3,
ORF4L1,RPLP1,TIMP2,SARAF,MIDN,RPL10,RPS17,AP3D1,DNAJA1,CYB5R3,MAP4,SF3B2,ITM2B,
AFF4,CKB,COL4A2,RPL37A,LRRC28,SPEN,NONO,DDB1,YWHAQ,EIF1,HACD3,EGLN1,TPI1,CHSY1,
3,SERINC1,SSBP2,ZNF43,RPS3,HMGB1,POLR2A,CDKL1,AP2M1,TPT1,SEC61A1,RPS4X,RPS3A,R
TF3C4,LDHB,PKD1,SF3B1,PDIA3,SERINC1,SSBP2,BASP1,VAPA,HMGB1,POLR2A,TPT1,RPS4X,RP
A,PLPPR3,SEPTIN3,COTL1,MAGED4B,CTBP1,CAPRIN1,TOP2B,TMEM30A,TRIB2,PEA15,NOVA2,C
GDIA,CAPRIN1,SLC35E2B,TMEM30A,BCLAF1,PPIB,RPL23A,HSBP1,SQLE,MAP1A,DDX39B,CTSB,
AP5,CADM4,RAB1A,ARCN1,COTL1,HECTD4,CTBP1,STIP1,SLC35E2B,IPO5,SKP1,DNER,L1CAM,L
ARCKSL1,UBB,AFF4,MATR3,YWHAG,CKB,COL4A2,RPL37A,LRRC28,FN1,SPEN,DDB1,YWHAQ,EIF
3,MEGF8,SOX12,PRDX2,DDX3X,ITM2C,RPS19,MIF,GPAT4,JUN,MYO5A,CD63,MTSS2,DNAJC5,HN
15,LDHB,PKD1,SF3B1,SSBP2,RPS3,BASP1,VAPA,POLR2A,TPT1,SEC61A1,RPS4X,RPS3A,RPL18,IT
Z1,TIMP2,SARAF,COL11A1,MIDN,RPL10,MARCHF6,WSB1,SBF1,AKAP12,SF3B2,CDC42BPB,TME
EIF2AK1,TXLNA,KDM5C,FBRS,SF1,RETREG2,SEC16A,GPX4,GLUD1,UPF1,EIF5B,NOMO1,NPDC1,

PS3A,STMN1,ITGB1,TUBB4B,UCHL1,MORF4L1,AGRN,RPLP1,ZMIZ1,TIMP2,XAF1,SARAF,MIDN,RI
A,DDX3X,RPS15,RPL30,CLTC,EIF3L,GNB2,H1-0,RPS5,IK,RPS19,CHPF,NGRN,DNAJC5,HNRNPDL,I
4A,YEATS2,CAMSAP1,KLC1,GRINA,VPS35,PTOV1,MDK,ORAI2,SKI,RAB11B,HDGF,ATP6AP1,RNF1(
.1,KDM3B,NAP1L4,CTXN1,PPP1R9B,PPDPF,BCL9L,TXNDC5,EP300,SF1,HIP1,KIF21B,UBA52,SRSI
9,PPP2R1A,RPL15,GTF3C4,DNAH8,PKD1,SF3B1,PDIA3,SERINC1,ZNF43,HMGB1,POLR2A,AP2M1,
TUBB4B,RPL11,MORF4L1,BTF3,AGRN,RPLP1,SARAF,MIDN,RPL10,MARCHF6,AP3D1,DNAJA1,AR
1,HACD3,DCHS1,SET,TPI1,TMSB10,EIF4A1,RPL15,GTF3C4,LDHB,DNAH8,SF3B1,PDIA3,SERINC1,
H9,RPL7,SEPTIN2,P4HB,APP,ERVV-1,GDI1,FSCN1,TMBIM6,PTMS,SPTAN1,SMG1,CALM1,NPM1,T
L1,GIGYF1,NKTR,STXBP1,TUG1,PLXNA3,ATXN2L,HYOU1,ID4,MPRIP,HGS,SOD1,RAI1,SREBF2,PF
HB,APP,ERVV-1,GDI1,FSCN1,TMBIM6,PTMS,SPTAN1,SMG1,CALM1,NPM1,TTC3,PLD3,HSPD1,CO
UGP2,ANXA5,CPSF1,PPP1CB,SQSTM1,GDI2,SSRP1,ALKBH5,HNRNPA3,PDXK,ACO2,TCP1,CAMSA
,TPT1,SEC61A1,RPS4X,RPS3A,RPL18,TUBB4B,RPL11,MORF4L1,BTF3,AGRN,RPLP1,SARAF,MIDN
1A1,RPS4X,RPS3A,RPL18,TUBB4B,RPL11,BTF3,NCAM1,AGRN,RPLP1,SARAF,MIDN,RPL10,MARC
,RPL5,GDI1,TMBIM6,PTMS,SPTAN1,GSTP1,MARCKS,SMG1,CALM1,NPM1,PLXNB2,TTC3,HSPD1,(
3,RPSA,TPM4,TNS3,NACA,TSPAN3,LPCAT1,LAMA5,RPS24,PJA2,RPL24,KPNB1,RPL17,NCS1,RHO/
2,CNN3,RPSA,TPM4,TNS3,CTSD,EIF4H,LAMA5,SLC22A17,BRD3,RPL24,SLC16A7,KPNB1,RPL14,(
2,LMF2,ECH1,IP6K1,GOLPH3,TASOR,PPP4R3A,CSNK2B,DDX6,AKAP8L,CRKL,PSMC5,RBM10,HS
3,CAPZB,TDRD5,ACTN1,CMIP,MAP7D1,PLCG1,CCAR2,DIDO1,DLK1,CORO1C,GIGYF1,PTP4A2,LS
SA9,STMN2,RPS8,MYH9,RPL7,SEPTIN2,P4HB,APP,ERVV-1,RPL5,GDI1,TMBIM6,PTMS,SPTAN1,GS
MA5,BRD3,PJA2,RPL24,SLC16A7,KPNB1,IGFBP3,RPL14,RHOA,CBX5,PSMD2,RAB6B,GLIPR1L2,R
SA9,RPS8,MYH9,RPL7,SEPTIN2,P4HB,APP,RPL5,FSCN1,TMBIM6,PTMS,SPTAN1,MARCKS,SMG1,C
RD1,GDI1,FSCN1,TMBIM6,PTMS,SPTAN1,GSTP1,SMG1,NPM1,TTC3,PLD3,HSPD1,COL18A1,SNRI
D3,DCHS1,SET,EGLN1,MAP9,CHSY1,INA,EIF4A1,MRC2,RPL19,PPP2R1A,RPL15,LDHB,PKD1,HLA-(
JBB,TRIM28,YWHAG,CKB,COL4A2,RPL37A,NONO,DDB1,PSMD5,YWHAQ,DCHS1,SET,CHSY1,TMS
K1,RHOA,PTCD3,ATF4,GLIPR1L2,RPS14,ATP1A1,RPS20,EIF5,CD81,COL5A2,WDR1,RPL26,LASP1,
R2A,AP2M1,TPT1,SEC61A1,RPS4X,RPS3A,RPL18,ITGB1,TUBB4B,RPL11,MORF4L1,BTF3,AGRN,
C4A8,HNRNPU,CTNNA1,RPS16,RPL23,CNN3,CTSD,SCARB2,NACA,TSPAN3,LPCAT1,LAMA5,SLC2
/PS35,ACAP3,CBX1,SNRNP70,TOP1,TAF7,VHL,PPP2CA,SMARCA4,LAPTM4A,SRSF9,PDE7A,RABL6
A,L1CAM,PEA15,LUC7L3,DLG5,NOVA2,CTSB,EEF1A2,PPP1CB,GDI2,TGFBI,SSRP1,HNRNPA3,PKN
NG8,HMGN2,ATP2A2,LASP1,PFN1,ASTN2,TLE5,PNN,NES,HNRNPC,SPTBN2,PFKL,SFPQ,PEBP1,C(
F1B,CALM3,GNB1,SHISA9,STMN2,RPS8,MYH9,SEPTIN2,P4HB,APP,IDS,ERVV-1,RPL5,GDI1,FSCN
DH2,RNF187,ANXA5,PGD,SQSTM1,GDI2,TSPYL4,TLN2,TSR1,SSRP1,HSPH1,ABCA3,HNRNPA3,YEA
RPS3,HMGB1,POLR2A,AP2M1,TPT1,SEC61A1,RPS4X,RPS3A,RPL18,TUBB4B,RPL11,BTF3,AGRN,
T1,RPS3A,RPL18,TUBB4B,MORF4L1,ANXA2,AGRN,ZMIZ1,TIMP2,XAF1,MIDN,RPL10,AP3D1,SBF1,
TAN1,SMG1,NPM1,PLD3,HSPD1,SNRNP200,SMIM14,IGF2,EIF3C,ATP5F1A,EIF4B,VCP,MARCKSL1
PL37A,LRRC28,FN1,DDB1,YWHAQ,EIF1,HACD3,DCHS1,SET,EGLN1,MAP9,CHSY1,INA,EIF4A1,MR(
T11,FKBP1A,RABEP1,MTCH1,SAFB2,NDN,ZNF664,ACLY,VAPB,CARM1,ARHGEF17,ELAVL1,VPS4A,
AN1,GSTP1,MARCKS,SMG1,CALM1,NPM1,PLXNB2,PLD3,HSPD1,USP11,COL18A1,SNRNP200,CSD
3,SERINC1,ZNF43,HMGB1,POLR2A,AP2M1,TPT1,SEC61A1,RPS4X,RPS3A,RPL18,ITGB1,TUBB4B,
PL31,PRKCSH,XPR1,KHDRBS1,TKT,EIF5A,CDH2,PDIA6,PIP4K2B,RAB7A,ITM2C,RPL30,CLTC,EIF3
D,HYOU1,ZFAND3,SARS1,MAPK8IP1,FBRSL1,MEX3A,P3H3,FLNC,SEC23A,AKIRIN1,DYNLL1,CTXN
PC,KHDRBS1,KLHL9,RPS25,SFPQ,EIF5A,PDIA6,PIP4K2B,RAB7A,MMP14,RPL12,ITM2C,RPL30,EIF
2,EIF4B,VCP,MARCKSL1,UBB,AFF4,TRIM28,YWHAG,CKB,RPL37A,LRRC28,MAPT,FN1,SPEN,NON(
F4B,VCP,UBB,AFF4,MATR3,COL4A2,RPL37A,LRRC28,SPEN,NONO,PSMD5,YWHAQ,EIF1,HACD3,I
SEC61A1,RPS4X,RPS3A,RPL18,ITGB1,TUBB4B,RPL11,MORF4L1,BTF3,AGRN,RPLP1,SARAF,MIDI

,PKD1,HLA-C,ZNF43,CHD3,BASP1,VAPA,HMGB1,AP2M1,OR8K1,RPL18,STMN1,ITGB1,TUBB4B,U
27A,GAS7,ACTR3,CAMSAP1,CCT3,NCKAP1,COL2A1,PTOV1,MLEC,ACAP3,MDK,SURF4,DDX42,RP
D1,FBRSL1,MLF2,PRPF40A,TP53BP1,CTXN1,RBBP7,KCNQ2,BCL9L,WDR45B,PRMT1,NAP1L1,KI
C4,MAP1LC3B,NDN,GRK2,BAP1,CLCN7,SNN,HSPG2,MAZ,DCLK1,LSM14A,DTX4,STK24,MACROH
,CKB,COL4A2,RPL37A,LRRC28,SPEN,NONO,DDB1,EIF1,HACD3,SET,EGLN1,MAP9,TMSB10,INA,
,DCTN1,KCNQ2,KIRREL1,TRIO,SF1,CNTNAP1,UBQLN2,FLOT1,SLC39A7,BRD2,TRA2B,COX7C,SM
1,SMG1,CALM1,NPM1,PLXNB2,USP11,COL18A1,SNRNP200,CSDE1,SMIM14,ATP5F1A,EIF4B,MA
BX5,ZNF865,RPL9,ATF4,PSMD2,KIF5B,RAB6B,SLC7A5,CHD4,ATP1A1,JUND,EIF5,WDR1,ATP2A2,
,HDLBP,PNMA1,DDR1,PIP4K2B,ACTN4,PRDX2,ARL4C,GNB2,RPS5,APLP1,TLN1,CHPF,MYO5A,CC
RNPK,CIRBP,MYH10,RPL36,SND1,RHOB,HNRNPH1,RPS12,HLA-B,PLPPR3,SEPTIN3,COTL1,MAG
1,TUG1,CSNK1D,HYOU1,ZFAND3,VANGL2,RAI1,PFN2,FBRSL1,MEX3A,P3H3,FLNC,KDM3B,AKIRI
TMN1,ITGB1,TUBB4B,UCHL1,MORF4L1,AGRN,RPLP1,ZMIZ1,TIMP2,XAF1,SARAF,MIDN,RPL10,W
LEC,ACAP3,TMEM248,RAD23B,SKI,ABCA2,TMED2,VEGFA,MAPK1,CCNL2,DYNC1LI2,ATP5PB,PT
,GNAS,SLC2A3,DDX17,GPI,HTRA4,RACK1,BSG,FASN,ALDH6A1,SPTBN1,ATP5F1B,CALM3,GNB1,
19,PPP2R1A,RPL15,GTF3C4,LDHB,DNAH8,PKD1,SF3B1,PDIA3,SERINC1,ZNF43,RPS3,HMGB1,PC
A,RPL18,STMN1,ITGB1,TUBB4B,UCHL1,RPL11,BTF3,NCAM1,AGRN,RPLP1,XAF1,MIDN,RPL10,M
AM1,SMARCA5,LARP1,CTNNA1,TPM4,TNS3,CTSD,EIF4H,SCARB2,TSPAN3,RPS24,KPNB1,C1ORF
PLP1,CHPF,GOLM1,B4GALNT4,MYL6,SHOX,ANKRD52,WLS,HNRNPK,RPL36,DUX4,YWHAB,CAD
,PTBP1,CDC37,UBE2R2,ACTN1,QARS1,CCAR2,ARF4,SDF4,EIF3F,PITPNM1,ISYNA1,LONP1,ATXN
KCSH,KHDRBS1,KLHL9,RPS25,SFPQ,EIF5A,PDIA6,PIP4K2B,RAB7A,DDX3X,RPL12,ITM2C,RPL30,
PTMS,SPTAN1,GSTP1,SMG1,CALM1,NPM1,PLXNB2,TTC3,USP11,COL18A1,SNRNP200,CSDE1,SM
,RPS9,XRCC5,CIPC,RPL24,PAPOLA,RHOA,ZNF865,ATF4,GLIPR1L2,RPS14,CXADR,CHD4,RPS20,
GRN,ZMIZ1,XAF1,MIDN,RPL10,WSB1,RPS17,VCL,AP3D1,SBF1,DNAJA1,ARF1,CYB5R3,MAP4,ITM
GRN,ZMIZ1,XAF1,MIDN,RPL10,WSB1,RPS17,VCL,AP3D1,SBF1,DNAJA1,ARF1,CYB5R3,MAP4,ITM
3,ITM2B,AP2B1,MAT2A,SLC4A8,GOLGA8A,HNRNPU,NFE2L1,CTNNA1,TNS3,LAMA5,CIPC,MSN,KP
3,CNN3,RPL29,TPM4,RPS9,XRCC5,CIPC,RPL24,PAPOLA,IGFBP3,HK1,RPL22,RHOA,ZNF865,ATF
T1,SLC22A17,KPNB1,COL1A2,LLGL1,NCS1,RPL22,CLSTN1,ATF4,RAB6B,CXADR,CHD4,ATP1A1,
HMGB1,POLR2A,AP2M1,TPT1,SEC61A1,RPS4X,RPS3A,RPL18,ITGB1,TUBB4B,RPL11,MORF4L1,
BCD,CPSF1,PGD,MEST,MORF4L2,ALKBH5,PDXK,HSPA4,CCT3,VPS35,MLEC,CLSTN3,ACAP3,MDK
,CCNI,TADA2B,KHDRBS1,CSNK1E,PFKL,TKT,SFPQ,EIF5A,CDH2,PDIA6,DDR1,PIP4K2B,SOX12,AC
,PTGES3,CKAP4,CCT6A,CAPRIN1,TOP2B,SMCHD1,ATP6V0C,TMEM30A,BCLAF1,HSBP1,CDKN1
B2A,AZIN1,LAPTM4A,SRSF9,PSMD3,RABL6,SRSF1,ARGLU1,TCF25,ADAM9,FTO,ILF3,LUZP6,PR
SA,TNS3,EIF4H,SCARB2,TSPAN3,LPCAT1,LAMA5,BRD3,RPS9,C1ORF43,PAPOLA,LLGL1,NCS1,R
RPLP1,SARAF,MIDN,RPL10,MARCHF6,RPS17,AP3D1,DNAJA1,ARF1,CYB5R3,SF3B2,GTF2F1,HN
RMP1,LASP1,BEX3,OAZ1,RBBP4,PFN1,MCL1,PNN,HNRNPC,CCNI,TADA2B,SLC3A2,PTN,EIF5A,P
ANXA5,CPSF1,PPP1CB,PABPN1,GDI2,ALKBH5,PTPRA,TCP1,CAMSAP1,CCT3,PTOV1,MLEC,CCT4,
4,TCP1,CAMSAP1,NCKAP1,MLEC,GCN1,RAB11B,SRP14,PYGB,PRPF6,TAF7,SELENON,RPL37,SM
P2M1,TPT1,RPS4X,RPS3A,RPL18,ITGB1,TUBB4B,RPL11,MORF4L1,BTF3,AGRN,RPLP1,SARAF,
12,CALU,ARCN1,RPS27,CTBP1,ARHGDIA,CAPRIN1,SLC35E2B,IPO5,ATP6V0C,BCLAF1,RPL23A,
2,SRSF4,PFN1,MCL1,TLE5,PNN,RPL31,PRKCSH,HNRNPC,KHDRBS1,KLHL9,RPS25,SFPQ,EIF5A,
BS1,KLHL9,RPS25,TKT,EIF5A,CDH2,PDIA6,DDR1,PIP4K2B,ACSL6,MMP14,RPL12,ITM2C,RPL30,
SP2,DDX39B,EEF1A2,TBCD,CPSF1,PGD,VAMP2,GNAI3,HSPH1,LMNB2,PTPRA,ACO2,KLC1,VPS3
SP2,DDX39B,EEF1A2,TBCD,CPSF1,PGD,VAMP2,GNAI3,HSPH1,LMNB2,PTPRA,ACO2,KLC1,VPS3
R2A,AP2M1,TPT1,SEC61A1,RPS4X,RPS3A,ITGB1,TUBB4B,RPL11,MORF4L1,BTF3,AGRN,RPLP1,
1,HACD3,SET,EGLN1,MAP9,TMSB10,INA,EIF4A1,RPL19,HLA-C,SF3B1,SSBP2,ZNF43,CHD3,VAPA

A,CASC3,CLIP3,FKBP8,PITPNA,MSL1,SDF4,GIGYF1,YWHAH,STMN3,LSS,PLXNA3,CSNK1D,ATXN2
X12,ACTN4,ITM2C,MAGED2,GNB2,RPS7,RPS19,TLN1,COL6A2,MTSS2,DNAJC5,ARL6IP1,TAGLN,
1,PDIA3,SSBP2,ZNF43,CHD3,HMGB1,CDKL1,TPT1,SEC61A1,RPS4X,RPS3A,STMN1,ITGB1,TUBB
CH1,SAFB2,ZNF664,PPP1R10,PELP1,ABI2,RAB1B,SSR3,GNL1,KMT2E,MACROH2A2,BLCAP,RBM
F3L,MAGED2,RPS19,CHPF,JUN,NGRN,RPLP2,DDOST,CIRBP,RPL36,YWHAB,LAMP1,HNRNPH1,CA
24,C1ORF43,IGFBP3,RPL14,CBX5,RPL9,ATF4,SLC7A5,GLIPR1L2,JUND,TMEM259,EIF5,LENG8,
TLE5,NISCH,XPR1,KHDRBS1,KLHL9,SPTBN2,SLC3A2,PFKL,EIF5A,CCND2,CDH2,GPC3,DDR1,PLX
,RPLP1,MIDN,RPL10,MARCHF6,RPS17,AP3D1,DNAJA1,ARF1,SF3B2,MAT2A,GTF2F1,HNRNPA0,R
BP4,PFN1,MCL1,PNN,RPL31,PRKCSH,LRPPRC,SCRN1,RPS25,SFPQ,EIF5A,PDIA6,DDR1,PIP4K2B,
,RPLP1,SARAF,MIDN,RPL10,MARCHF6,RPS17,AP3D1,DNAJA1,ARF1,CYB5R3,AKAP12,SF3B2,GT
,YWHAH,TNKS2,LONP1,ATXN2L,HGS,BAG6,CCT5,PPDPF,DFFB,TXLNA,SGTA,KDM5C,ACTR1A,FB
2,CBX5,ZNF865,CLSTN1,ATF4,SLC7A5,NLGN2,CXADR,ATP1A1,JUND,COL5A2,LENG8,HMGN2,AT
,CTBP1,ARHGDIA,SLC35E2B,IPO5,BCLAF1,RPL23A,HSBP1,MAP1A,CTDSP2,DDX39B,SRSF5,TM9
Z1,SRSF4,PFN1,ASTN2,MCL1,JPT2,TLE5,PNN,SPC25,NES,NISCH,PRKCSH,KLHL9,SPTBN2,CSNK1
3,YWHAG,COL4A2,MAPT,NONO,DDB1,YWHAQ,DCHS1,SET,MAP9,TPI1,CHSY1,EIF4A1,MRC2,RPL
QLE,MAP1A,DDX39B,TM9SF3,MDH2,U2AF2,EEF1A2,FDFT1,SUGP2,GNAI3,GDI2,PARP1,TSR1,SS
HA,RPL29,RPSA,TNS3,CTSD,EIF4H,SCARB2,TSPAN3,LPCAT1,CIPC,MSN,C1ORF43,NCS1,RPL22,
TR3,CKB,COL4A2,RPL37A,LRRC28,DDB1,PSMD5,YWHAQ,EIF1,HACD3,SET,EGLN1,TPI1,INA,EIF4
PA,HMGB1,CDKL1,TPT1,SEC61A1,STMN1,ITGB1,TUBB4B,UCHL1,RPL11,MORF4L1,BTF3,RPLP1
CL,AP3D1,SBF1,ARF1,CYB5R3,PAFAH1B1,MAP4,SF3B2,CDC42BPB,ZNF770,MAT2A,TMEM170A,
,PPP2R1A,RPL15,GTF3C4,LDHB,DNAH8,PKD1,SF3B1,PDIA3,SERINC1,ZNF43,RPS3,HMGB1,POL
L23,CNN3,TPM4,RPS24,RPS9,XRCC5,RPL24,SLC16A7,PAPOLA,IGFBP3,COL1A2,HK1,RPL22,RH
,NARS1,DUX4,YWHAB,PTGES3,RPS12,CALU,MAGED4B,CTBP1,ARHGDIA,SLC35E2B,IPO5,ATP6V
,TKT,EIF5A,CDH2,PDIA6,DDR1,PIP4K2B,RAB7A,RPL12,ITM2C,RPL30,CLTC,EIF3L,IK,RPS19,CHP
1,CDH2,PDIA6,PIP4K2B,DDX3X,RPS15,RPL30,NANOG,MRFAP1,RPS19,HMGCS1,MTSS2,HNRNP
,TMEM170A,GTF2F1,SLC4A8,CTNNA1,CNN3,RPSA,TPM4,TNS3,CTSD,EIF4H,LPCAT1,LAMA5,SLC
,SLC35E2B,IPO5,ATP6V0C,BCLAF1,RPL23A,HSBP1,MAP1A,CTDSP2,DDX39B,EID1,SRSF5,TM9SF
JSP11,COL18A1,SNRNP200,IGF2,EIF3C,ATP5F1A,EIF4B,VCP,MARCKSL1,TRIM28,MATR3,CKB,C
,ANXA2,TIMP2,XAF1,SARAF,RPL10,WSB1,RPS17,AP3D1,SBF1,ARF1,CYB5R3,SF3B2,ITM2B,MAT
H3PXD2A,FNBP1L,ACTR1B,MOAP1,PDPK1,PCNX3,LTBP3,PSMC3,PRKACA,YY1,SMARCD1,CREBE
EGF8,ACSL6,RAB7A,RPS15,ARL4C,GNB2,RCN1,H1-0,RPS5,APLP1,MIF,TLN1,CD63,ANXA6,NGRN,
PS24,RPS9,XRCC5,CIPC,MSN,RPL24,PAPOLA,RPL22,RHOA,ZNF865,ATF4,RPS14,CXADR,CHD4,R
4B,TRIB2,RPL23A,HSBP1,FAM199X,CTDSP2,DDX39B,ZC3H7B,TM9SF3,MDH2,TOMM20,U2AF2,
RAI2,SKI,H3-3A,RAB11B,PYGB,SELENON,OPTN,DDAH2,AEBP1,TCF25,GRB2,CAPZB,ACTN1,BNIP
I6,PTMS,GSTP1,SMG1,CALM1,PLXNB2,PLD3,HSPD1,COL18A1,SNRNP200,IGF2,EIF3C,EIF4B,VC
A2,AGRN,ZMIZ1,MIDN,RPL10,RPS17,AP3D1,SBF1,ARF1,MAP4,SF3B2,ITM2B,TMEM170A,GTF2F1
,DNAJA1,ARF1,CYB5R3,SF3B2,GTF2F1,HNRNPA0,RPL23,CNN3,RPL29,RPS9,XRCC5,CIPC,SLC16
NOVA2,NUCB1,CHD6,EEF1A2,RPN2,HNRNPL,PARP1,ALKBH5,HNRNPA3,YEATS2,GRINA,COL2A1,
DI2,PARP1,TSR1,MTRF1L,RPL27,PTPRA,PRDX5,UBE4A,ACO2,HSPA4,TCP1,CAMSAP1,MLEC,DDX
IP3,MAP2K2,POLR2B,MSL1,SDF4,SMARCC1,SDHA,TNKS2,LONP1,ZFAND3,SARS1,UQCRC2,PN
RNPL,LMNB2,ABCA3,ALKBH5,PRDX5,PDXK,PKN1,ETNK1,PTOV1,ACAP3,MDK,SURF4,CHD2,ORAI
K1N,RPS11,UBE2R2,CASC3,DIDO1,RPS28,SRSF3,MESD,GPR107,PFN2,ARFGEF2,ANP32A,EP300
HD1,HDAC2,TMEM30A,TRIB2,LUC7L3,SQLE,DLG5,NOVA2,SRP9,TOMM20,U2AF2,TBCD,SUGP2,
8,ZNF768,ZNF532,ZCCHC14,DLD,SEPHS1,IGF2BP2,UBE3C,ME2,CUEDC1,UBQLN4,SLC12A7,PO
B,XPR1,SLC3A2,CDH6,PEBP1,CDH2,GPC3,SOX12,RAB7A,CCN2,ITM2C,ARL4C,MRFAP1,GNB2,G

7A,ITM2C,MRFAP1,RPS7,MIF,GPAT4,GOLM1,MTSS2,APLP2,RPLP2,MYL6,H1-10,NARS1,SEPTIN1
5,CD81,HMGN2,CRMP1,BEX3,CCT7,OAZ1,SRSF4,JPT2,TLE5,PNN,TADA2B,SCRN1,HDLBP,SFPQ,
P11,COL18A1,SNRNP200,CSDE1,SMIM14,IGF2,ATP5F1A,EIF4B,VCP,MARCKSL1,UBB,TRIM28,M
BASP1,CDKL1,TPT1,RPL18,STMN1,ITGB1,TUBB4B,RPL11,BTF3,NCAM1,ANXA2,AGRN,ZMIZ1,TIM
RN,RPLP1,SARAF,MIDN,RPL10,MARCHF6,RPS17,AP3D1,DNAJA1,ARF1,CYB5R3,SF3B2,GTF2F1,
36,DUX4,YWHAB,PTGES3,SEPTIN11,CKAP4,PLPPR3,CTBP1,ARHGDIA,GPC1,SMCHD1,BCLAF1,RF
,RPS15,RPL30,EIF3L,H1-0,RPS5,IK,CHPF,CD63,NGRN,RPLP2,DDOST,H1-10,INIP,CIRBP,RPL36,Y
HL9,RPS25,EIF5A,PDIA6,PIP4K2B,MEGF8,RPL12,ITM2C,RPL30,EIF3L,GNB2,H1-0,RPS7,IK,RPS1
2,ATP5F1A,VCP,MARCKSL1,AFF4,TRIM28,CKB,COL4A2,FN1,SPEN,NONO,DDB1,PSMD5,YWHAQ,
IN,ILF3,CAPZB,HNRNPUL1,ATXN10,ACTN1,BNIP3L,MAP7D1,PLCG1,PITPNA,KIF3C,HNRNPM,SDH
PTGES3,RPS12,CALU,RPS27,MAGED4B,CTBP1,ARHGDIA,CAPRIN1,SLC35E2B,IPO5,ATP6V0C,PPI
,ORAI2,SNRNP70,ASXL1,H3-3A,SCAF1,PYGB,DPYSL2,TAF7,RPS15A,RPL38,NOTCH2,RNF10,TUB
D,EIF4H,SLC22A17,BRD3,RPS9,XRCC5,MSN,RPL17,C1ORF43,PAPOLA,RPL14,RPL9,NR2F1,KIF5
RPS17,AP3D1,SBF1,MAP4,ITM2B,AP2B1,GTF2F1,GOLGA8A,HNRNPU,PGAM1,SMARCA5,HNRNPA
,CTSD,EIF4H,NACA,LPCAT1,LAMA5,SLC22A17,RPS9,CIPC,PAPOLA,IGFBP3,HK1,NCS1,PTCD3,CE
4,RPS9,MSN,RPL24,COL1A2,LLGL1,RPL22,RPL9,ATF4,NLGN2,GLIPR1L2,RPS14,ATP1A1,JUND,
SS,TUG1,PLXNA3,ATXN2L,ID4,RAI1,SREBF2,P3H3,WBP2,SMS,POLR2E,TRAF4,RBBP7,MSMO1,T
RNT2,YME1L1,RBBP7,EDC4,TXLNA,TRIO,HIP1,VARS1,H2AZ1,GRSF1,CD99,CSNK1G2,NXN,EP400
A2,RPL37A,FN1,SPEN,NONO,EIF1,DCHS1,SET,TMSB10,INA,MRC2,RPL19,PPP2R1A,GTF3C4,PKD
,HNRNPA0,RPS16,RPL23,LDHA,RPL29,EIF4H,SCARB2,LAMA5,RPS9,XRCC5,CIPC,RPL24,SLC16A
NS3,CTSD,TSPAN3,LPCAT1,BRD3,XRCC5,SLC16A7,KPNB1,RPL17,PAPOLA,RPL9,KIF5B,ATP1A1,
N1,ITGB1,TUBB4B,RPL11,ANXA2,AGRN,RPLP1,ZMIZ1,TIMP2,XAF1,WSB1,VCL,AP3D1,SBF1,DNA
X4,PPDPF,CIC,TRAF4,RBBP7,WDR45B,EP300,DGCR2,STARD7,LRRC4B,CD99,SEC16A,SLC35F5,
R,LENG8,WDR1,ATXN7L3B,RPL26,CRMP1,CCT7,SRSF4,RBBP4,PFN1,MCL1,PNN,RPL31,PRKCSH
TN3,ASTN2,JPT2,NES,PRKCSH,HNRNPC,TADA2B,SPTBN2,CSNK1E,TKT,TPM1,EIF5A,PNMA1,PDIA
,AP2B1,MAT2A,TMEM170A,GTF2F1,GOLGA8A,PXDN,HNRNPU,SMARCA5,FADS2,RPS16,LDHA,CP
SS,ISYNA1,CSNK1D,HYOU1,MAPK8IP1,FBRSL1,ZSCAN18,BMPR2,NAP1L4,CD99L2,PPDPF,VDAC1
TP6V0C,MMP2,L1CAM,LAPTM4B,LUC7L3,SQLE,DLG5,SHISAL1,CNN2,SRP9,SRSF5,MDH2,U2AF
F5,ATXN7L3B,HMGN2,LASP1,CCT7,SRSF4,SPC25,PRKCSH,XPR1,XPO1,TPM1,EIF5A,CCND2,PNM
IM6,GSTP1,MARCKS,SMG1,CALM1,NPM1,PLD3,USP11,SNRNP200,CSDE1,SMIM14,IGF2,EIF3C,
ASTN2,PNN,SPC25,PRKCSH,CCNI,XPR1,XPO1,SLC3A2,HDLBP,TKT,TPM1,EIF5A,CCND2,CDH2,PIF
ITGB1,MORF4L1,NCAM1,AGRN,RPLP1,SARAF,MIDN,MARCHF6,WSB1,DNAJA1,ARF1,CYB5R3,AK
RN,RPLP1,SARAF,MIDN,RPL10,MARCHF6,RPS17,AP3D1,DNAJA1,ARF1,CYB5R3,SF3B2,GTF2F1,
5,TM9SF3,MDH2,U2AF2,EEF1A2,FDFT1,SUGP2,GNAI3,PARP1,TSR1,MTRF1L,PTPRA,PRDX5,UBE
RIN1,SLC35E2B,IPO5,ATP6V0C,BCLAF1,RPL23A,HSBP1,MAP1A,DDX39B,SRSF5,TM9SF3,MDH2,
PPL1,IPO5,PEA15,SQLE,FAM199X,TMEM132A,DDX39B,NOVA2,C5ORF24,RNF187,U2AF2,EEF1A2
ACA,LAMA5,RPS24,BRD3,XRCC5,CIPC,PJA2,RPL24,C1ORF43,PAPOLA,RPL14,RHOA,PTCD3,CBX
RT1,AP1M1,BNIP3,SAE1,GAK,UBE2K,USO1,ATP5MG,NELFB,DENND5A,DIP2A,CDC34,TMEM184
A5,ENC1,GLIPR1L2,CHD4,TMEM259,COL5A2,LENG8,ATXN7L3B,HMGN2,CRMP1,SRSF4,RBBP4,
28,YWHAG,COL4A2,LRRC28,FN1,SPEN,YWHAQ,EIF1,HACD3,DCHS1,TPI1,CHSY1,INA,EIF4A1,MF
HNRNPA0,CTNNA1,NACA,LPCAT1,CIPC,PJA2,KPNB1,C1ORF43,IGFBP3,COL1A2,RPL14,PTCD3,C
LA,COL1A2,LLGL1,RPL14,HK1,RHOA,ZNF865,ATF4,PSMD2,SLC7A5,GLIPR1L2,RPS14,ATP1A1,F
D1,CALU,ARCN1,COTL1,ARHGDIA,ATP6V0C,LAPTM4B,TRIB2,RPL23A,HSBP1,DDX39B,SRSF5,T
IN1,SMS,YME1L1,MSMO1,TXLNA,TRIO,LBH,SF1,HIP1,H2AZ1,CD99,KPNA2,ZFP36L1,KLHL42,CS
RN1,RPS25,EIF5A,CDH2,PDIA6,MEGF8,MMP14,RPL12,ITM2C,RPL30,CLTC,EIF3L,MAGED2,GNB

[illegible],UBE4A,TCP1,CAMSAP1,CCT3,VPS35,PTOV1,MDK,CCT4,SKI,SCAF1,DPYSL2,TAF7,RPL38,RNF10,
[illegible]2A,AP2M1,TPT1,RPS4X,RPS3A,TUBB4B,RPL11,MORF4L1,BTF3,AGRN,RPLP1,SARAF,MIDN,RPL1
NPH3,CENPB,PAFAH1B2,CASC4,USP47,SCAMP5,CHMP1A,LUC7L2,PAIP2,GGNBP2,NPTN,KIAA11[illegible]
[illegible]HOA,ZNF865,ATF4,PSMD2,RAB6B,SLC7A5,NLGN2,CXADR,CHD4,ATP1A1,LENG8,WDR1,ATP2A2,
[illegible]GP2,GNAI3,PARP1,TSR1,HSPH1,MTRF1L,PTPRA,PRDX5,UBE4A,ACO2,HSPA4,TCP1,MLEC,ZNF30
[illegible]I3,NOP53,GDI2,TSR1,MTRF1L,RPL27,PRDX5,UBE4A,ACO2,HSPA4,TCP1,MLEC,GCN1,SKI,RAB1[illegible]
[illegible]IF4A1,RPL19,PPP2R1A,RPL15,LDHB,SF3B1,ZNF43,BASP1,HMGB1,POLR2A,CDKL1,AP2M1,RPS4
V1,CLSTN3,ACAP3,SURF4,TMEM248,RAD23B,PTK7,ORAI2,ASXL1,SCAF1,HDGF,ABCA2,NRDC,PY[illegible]
2AF2,EEF1A2,GNAI3,PARP1,TSR1,MTRF1L,PRDX5,UBE4A,ACO2,HSPA4,TCP1,MLEC,ACAP3,GCN1
[illegible]5,ATP2A2,CRMP1,LASP1,BEX3,VAT1,CLEC7A,CCNI,TADA2B,KHDRBS1,SPTBN2,PFKL,RPS25,TK[illegible]
00,SMIM14,IGF2,EIF3C,ATP5F1A,VCP,UBB,AFF4,TRIM28,MATR3,YWHAG,CKB,RPL37A,MAPT,FN
TRAF7,MTA1,KIRREL1,EP300,PBX1,HIP1,AP2A1,NOL4L,LRRC4B,CSNK1G2,ATRX,FARP1,ULK1,N
,MAP2K2,DIDO1,XPNPEP3,RPS28,ATP6V1B2,HNRNPM,SSB,PHC2,SLIT3,CCT8,MESD,MPRIP,RAI1
,PLD3,HSPD1,USP11,COL18A1,SNRNP200,CSDE1,EIF3C,ATP5F1A,EIF4B,VCP,UBB,AFF4,TRIM28,
UBB4B,RPL11,BTF3,AGRN,RPLP1,SARAF,MIDN,RPL10,MARCHF6,RPS17,AP3D1,DNAJA1,ARF1,C
[illegible]1,MAGED4B,CTBP1,RAC1,TMEM30A,MMP2,SKP1,GLUL,SQLE,FAM199X,DLG5,TIMP3,CTDSP2,[illegible]
1,TAF15,PTPRA,ACTR3,YEATS2,TBC1D16,ACO2,CCT3,RBMX,TMEM248,RAD23B,CHD2,ASXL1,SC
D2,RAB6B,SLC7A5,ENC1,NLGN2,RPS14,CHD4,JUND,EIF5,CD81,LENG8,WDR1,HMGN2,LASP1,O[illegible]
[illegible]NC1,GLIPR1L2,CXADR,JUND,TMEM259,EIF5,CD81,HMGN2,CRMP1,LASP1,BEX3,OAZ1,RBBP4,[illegible]
IPC,SLC16A7,KPNB1,PAPOLA,COL1A2,LLGL1,HK1,RPL22,RHOA,ZNF865,ATF4,PSMD2,SLC7A5,[illegible]
[illegible]RINC1,HMGB1,POLR2A,AP2M1,TPT1,SEC61A1,RPS4X,ITGB1,TUBB4B,MORF4L1,BTF3,NCAM1,[illegible]
[illegible]L18A1,EIF3C,ATP5F1A,EIF4B,VCP,MARCKSL1,UBB,RPL37A,LRRC28,MAPT,FN1,SPEN,NONO,DD[illegible]
RCHF6,WSB1,RPS17,VCL,AP3D1,SBF1,ARF1,CYB5R3,SF3B2,CDC42BPB,ITM2B,AP2B1,MAT2A,N
PFKM,HYOU1,BAG6,RAB11A,CDC42,PPP1R9B,RBM15B,TRAF7,DFFB,STAT3,EIF2AK1,TXLNA,PR[illegible]
,TUBB4B,UCHL1,BTF3,NCAM1,ANXA2,AGRN,ZMIZ1,XAF1,SARAF,COL11A1,MIDN,MARCHF6,RPS
[illegible]IF1A,RPL13,GNAS,SLC2A3,DDX17,GPI,HTRA4,RACK1,BSG,FASN,ALDH6A1,SPTBN1,ATP5F1B,CA
[illegible],CTBP1,RAC1,GPC1,ATP6V0C,MMP2,DNER,BCLAF1,TRIB2,PEA15,CDKN1A,U2AF2,EEF1A2,CCT2
[illegible]F4,RBBP4,PFN1,MCL1,TLE5,PNN,RPL31,PRKCSH,CCNI,LRPPRC,KHDRBS1,SLC3A2,CSNK1E,SC
2,UBE2M,RAB14,ZNF428,PNISR,PLEKHB2,INTS11,PCNX3,ADAM10,SPTY2D1,FBXW5,MBD3,SF3A
[illegible]TSS2,APLP2,ARL6IP1,ST3GAL2,TSR2,HNRNPK,DUX4,SEPTIN11,CADM4,SND1,LAMP1,RHOB,HN[illegible]
[illegible]AG6,RAB11A,SERBP1,AKIRIN1,BMPR2,CCT5,CDC42,PPP1R9B,YME1L1,DFFB,STAT3,EIF2AK1,E[illegible]
ARM1,RAB1B,RALY,GPC2,FAM234A,HDGFL2,LRATD2,CNP,LSM14B,SPAG9,MOAP1,PCNX3,ERGI
GCN1,SKI,RAB11B,SRP14,PRPF6,TAF7,RPL37,ARPC2,ARGLU1,PTBP1,SUPT5H,HNRNPUL1,SON,[illegible]
PDPF,DFFB,BCL9L,TXNDC5,MTA1,LBH,FBRS,SF1,MMP15,SLC39A6,CSNK1G2,SRSF6,UPF1,UBE[illegible]
[illegible]ND,EIF5,CD81,LENG8,WDR1,HMGN2,LASP1,OAZ1,PFN1,TLE5,NISCH,CCNI,LRPPRC,TADA2B,SP[illegible]
ZFP91,RALBP1,ATP5F1C,ZDHHC5,XPC,CACYBP,SOS1,DDHD2,ZNF362,SRRT,ADIPOR1,QRICH1,F[illegible]
[illegible]M4A,SRSF9,PDE7A,PSMD3,RABL6,ODC1,SRSF1,IPO7,ARGLU1,TCF25,ADAM9,FTO,ILF3,WDR6,[illegible]
[illegible]NXA2,AGRN,ZMIZ1,SARAF,MIDN,RPL10,WSB1,SBF1,ARF1,MAP4,CDC42BPB,ITM2B,ZNF770,AP2
RSF5,TM9SF3,TOMM20,U2AF2,CCT2,SUGP2,GNAI3,TM9SF2,TSR1,PTPRA,PRDX5,UBE4A,ACO2,[illegible]
[illegible],RABL6,ODC1,DYNC1LI2,ILF3,UBE2R2,POLR2B,SMARCC1,SSB,GIGYF1,SDHA,SLIT3,STT3B,TNK
ORF43,PAPOLA,NCS1,CBX5,NR2F1,RAB6B,SLC7A5,RPS14,CHD4,JUND,TMEM259,EIF5,LENG8,H
[illegible]PL23,RPL29,TPM4,RPS24,BRD3,RPS9,XRCC5,CIPC,RPL24,KPNB1,PAPOLA,HK1,RPL22,RHOA,Z[illegible]
DR6,CAND1,SON,PLCG1,ZFAND5,DHX30,TUG1,ATXN2L,HECTD1,BAG6,SREBF2,SERBP1,SMS,RB
[illegible]72,COX8A,ECH1,IP6K1,GOLPH3,NBAS,DRAP1,PPP4R3A,TBC1D14,GID8,APPBP2,DEAF1,ATP5PO,
[illegible]PRE1,CCNL2,BTBD2,TCF25,CAPZB,CMIP,PLCG1,FLNB,SMARCC1,HNRNPM,GIGYF1,NKTR,PTP4[illegible]

CALU,ARCN1,COTL1,ARHGDIA,ATP6V0C,LAPTM4B,TRIB2,RPL23A,HSBP1,DDX39B,SRSF5,TM9SF
IPR1L2,JUND,LENG8,RPL26,CRMP1,CCT7,TLE5,NES,NISCH,PRKCSH,TPM1,EIF5A,CCND2,CDH2,
GD,SQSTM1,VAMP2,KLHL24,NOP53,GDI2,HNRNPL,LMNB2,ABCA3,RPL27,ACTR3,TCP1,PLOD1,G
,ADNP,EWSR1,EHMT2,TSPAN14,FAM53C,SALL2,CUL4A,PRPH,SYT15,TNRC6A,SRSF10,ASAH1,EF
,DAP,PCBP4,ZNF664,SEPTIN5,PALM,HSPG2,CARM1,LINGO1,INTS3,PKDCC,GPC2,LTBP4,GABAR
G5,PGRMC1,TBCD,VCAN,HNRNPA3,PRDX5,ACTR3,ETNK1,KLC1,LAMC1,CCT4,H3-3A,CNTNAP2,H
,HMGB1,AP2M1,TPT1,RPS3A,ITGB1,UCHL1,BTF3,NCAM1,ANXA2,AGRN,ZMIZ1,TIMP2,COL11A1,
Z2,TLE1,IREB2,LSM14A,KIAA0232,LSM14B,RAB6A,NMT1,SOD2,SLC7A1,NSD2,FBXW5,DENND4B
01,SBF1,ARF1,CYB5R3,SF3B2,CDC42BPB,ITM2B,MAT2A,NFE2L1,FADS2,RPS16,RPL23,CPE,RPL
F2,CTDSP2,CNN2,NUCB1,ZC3H7B,FDFT1,SUGP2,GAS7,PDXK,PKN1,TCP1,CAMSAP1,GRINA,SNRN
L,RPL27,PTPRA,PRDX5,UBE4A,ACO2,HSPA4,TCP1,ZNF302,DDX42,GCN1,SKI,SRP14,PRPF6,TAF7
A,MVB12B,AZIN1,KDELR2,SRSF9,PLEC,ODC1,SRSF1,IPO7,ILF3,GRB2,WDR6,PRKDC,HNRNPUL1
,CNTNAP2,HDGF,RPL38,NOTCH2,VEGFA,ARF3,CCND1,DSTN,BTBD2,PBX2,PRDX1,ARGLU1,AEBF
ER1,PI4KA,COX7C,YKT6,ADH5,MCM3AP,SPINT2,TMEM167A,CHERP,FKBP1A,CTTN,ATP6AP2,LRP
PTBN2,PFKL,HDLBP,RPS25,TKT,EIF5A,PNMA1,PLXNA1,RAB7A,PRDX2,DDX3X,RPS15,ARL4C,RPS
61A1,RPS3A,RPL18,ITGB1,RPL11,MORF4L1,NCAM1,ANXA2,RPLP1,SARAF,COL11A1,MIDN,RPL1
G2,RAB2A,RER1,UBFD1,IARS1,MASTL,COX7C,YKT6,RPL7L1,RPS27A,TMEM167A,PSMD8,CHERP,
DS2,RPS16,CNN3,RPL29,RPSA,TPM4,NACA,LAMA5,RPS24,BRD3,RPS9,CIPC,PJA2,RPL24,C1ORF4
,CCT7,VAT1,SRSF4,RTN3,PFN1,TLE5,KHDRBS1,KLHL9,SLC3A2,PFKL,HDLBP,GLG1,EIF5A,PEBP1
,U2AF2,EEF1A2,CCT2,SUGP2,GNAI3,GDI2,HNRNPL,TSR1,SSRP1,LMNB2,MTRF1L,PRDX5,UBE4A
2F1,GOLGA8A,PXDN,HNRNPU,SMARCA5,HNRNPA0,RPS16,RPL23,CNN3,TPM4,TNS3,CTSD,LPCA
,FADS2,RPS16,RPL23,CNN3,RPL29,RPSA,TPM4,EIF4H,NACA,LAMA5,RPS24,BRD3,RPS9,CIPC,PJA
OLR2A,CDKL1,AP2M1,TPT1,RPL18,STMN1,RPL11,MORF4L1,BTF3,ANXA2,AGRN,ZMIZ1,TIMP2,X
B,DAP,SLC2A1,FBXO21,SAFB2,ZNF664,VPS26B,FUS,VAPB,SLIT1,FKBP4,TLE1,HINT1,SH3GL1,DC
AP,STRN4,SLC6A8,PKDCC,LTBP4,AKT3,RDX,PRR12,TCF3,RNF145,NSD2,LRP3,CIZ1,GSK3A,CHM
F4,RPS14,CXADR,CHD4,RPS20,CD81,LENG8,WDR1,RPL26,CRMP1,CCT7,SRSF4,PFN1,ASTN2,M
2,BAP1,PALM,ERGIC1,TCAF1,SHC2,PKDCC,LTBP4,ARHGEF17,UBE2D2,VPS4A,PGAP6,PRR12,TM
SLC16A7,PAPOLA,IGFBP3,NCS1,NR2F1,RAB6B,SLC7A5,NLGN2,CHD4,ATP1A1,JUND,TMEM259,L
SLC44A2,HSPG2,CARM1,GPC2,VPS4A,ZMIZ2,GABARAPL1,RIC8A,SOD2,SPAG9,SLC7A1,ACTR1B,N
SLK,RBM10,RCC2,ADIPOR2,ARL8A,CDKN1B,WASF3,DUSP8,ZDHHC5,RBM14,ERBB2,RRBP1,GDF
P10,ARPC1A,RTF1,NOP56,GTF3C1,CDV3,TIA1,SIN3B,SLC25A6,KIF26A,JAM3,PJA1,ATF5,RAN,B4
K4,MTA1,CPD,ACTR1A,EP300,SF1,CSNK2A1,PHB2,SLC39A7,RAB2A,RER1,TRA2B,ZFR,SV2A,AGPA
MGB1,CDKL1,SEC61A1,STMN1,ITGB1,TUBB4B,UCHL1,RPL11,MORF4L1,AGRN,RPLP1,RPL10,W
1,TMBIM6,GSTP1,MARCKS,SMG1,NPM1,PLXNB2,TTC3,PLD3,COL18A1,SMIM14,IGF2,EIF3C,ATI
3BP2,FAM168B,AEBP1,PTBP1,FTO,SUPT5H,FKBP8,CCAR2,DLK1,XPNPEP3,SMARCC1,FYN,HNRN
9X,DDX39B,SRSF5,TM9SF3,MDH2,TOMM20,U2AF2,CCT2,GNAI3,NOP53,GDI2,VCAN,TM9SF2,TSF
CLSTN1,ATF4,KIF5B,CHD4,ATP1A1,TMEM259,EIF5,HMGN2,ATP2A2,OAZ1,RBBP4,PNN,NISCH,PRK
STMN3,FBLN1,CSNK1D,SARS1,MAPK8IP1,FBRSL1,MEX3A,WBP2,AKIRIN1,DYNLL1,TP53BP1,BEX
3M15B,NAP1L1,TXNDC5,GSPT1,PPM1G,KLHL42,GARS1,GNG4,UPF1,MBTPS1,RGMB,SELENOW,
NB1,STMN2,RPL7,SEPTIN2,P4HB,APP,IDS,RPL5,GDI1,FSCN1,TMBIM6,PTMS,SPTAN1,GSTP1,MAF
F,MIDN,RPL10,MARCHF6,WSB1,RPS17,AP3D1,ARF1,SF3B2,ITM2B,AP2B1,GTF2F1,GOLGA8A,HN
LRRC4B,SLC39A6,SEC16A,GNG4,NXN,EP400,PPP1CC,CUL3,ADH5,IL6ST,SPINT2,GPSM1,ZMYM4
ARGLU1,PTBP1,CAPZB,SUPT5H,SON,CCAR2,ARF4,SUPT6H,PITPNM1,CSNK1D,SF3A1,BAG6,RAB
E2L1,CTNNA1,FADS2,CNN3,RPL29,TPM4,TNS3,TSPAN3,LAMA5,SLC22A17,BRD3,MSN,RPL24,PAI
RIO,LBH,HIP1,H2AZ1,GRSF1,SLC39A6,CD99,KPNA2,ZFP36L1,KLHL42,CSNK1G2,NXN,EP400,UBA

:N1,RPS27,ARHGDIA,STIP1,SLC35E2B,SMCHD1,IPO5,ATP6V0C,BCLAF1,RPL23A,HSBP1,SERF2,D
HSPA4,AHCYL1,KLC1,CLSTN3,ORAI2,RAB11B,ABCA2,NRDC,PRPF6,TUBB2A,PPP2CA,CCND1,G3E
31,VCL,SBF1,ARF1,CYB5R3,ITM2B,MAT2A,TMEM170A,GTF2F1,OGA,NFE2L1,SMARCA5,CNN3,RI
HOA,ZNF865,ATF4,GLIPR1L2,RPS14,CXADR,CHD4,RPS20,CD81,LENG8,WDR1,RPL26,CRMP1,LA
RALY,SEC14L1,NUDCD3,SSR3,GNL1,STK24,TUBGCP6,RIC8A,NCSTN,LSM14B,INTS11,MOAP1,MT
MEM259,LENG8,HMGN2,CRMP1,CCT7,VAT1,OAZ1,RBBP4,PFN1,ASTN2,SPC25,NES,CCNI,TADA2
.HL9,XPO1,SPTBN2,SLC3A2,PFKL,CDH6,SFPQ,CCND2,CDH2,PDIA6,GPC3,PIP4K2B,MEGF8,ACTN
L,ALKBH5,RPL27,SNRPN,PRDX5,UBE4A,HSPA4,TCP1,MLEC,ACAP3,SNRNP70,RPL36A,SRP14,NR
,ARF3,MAPRE1,ARPC2,FAM168B,FTO,LUZP6,CAPZB,SON,RPL28,CCAR2,ARF4,SUPT6H,DHX30,N
NPO1,SNRNP70,ASXL1,HDGF,RPS15A,RNF10,SELENON,PPP2CA,AZIN1,DDAH2,PBX2,FAM171A2,
,MDH2,U2AF2,EEF1A2,FDFT1,CCT2,SUGP2,GNAI3,NOP53,PARP1,TSR1,MTRF1L,RPL27,PTPRA,P
HF6,RPS17,AP3D1,DNAJA1,ARF1,SF3B2,MAT2A,GTF2F1,HNRNPA0,RPS16,RPL29,TPM4,RPS9,XR
HF6,RPS17,AP3D1,DNAJA1,ARF1,SF3B2,MAT2A,GTF2F1,HNRNPA0,RPS16,RPL29,TPM4,RPS9,XR
,STMN1,ITGB1,TUBB4B,UCHL1,RPL11,MORF4L1,AGRN,RPLP1,RPL10,WSB1,RPS17,AP3D1,DN
T1,CCT2,SUGP2,GNAI3,NOP53,TM9SF2,TSR1,LMNB2,MTRF1L,ALKBH5,SNRPN,PRDX5,UBE4A,H
BRD3,MSN,SLC16A7,RPL17,HK1,NCS1,RPL22,RHOA,PTCD3,KIF5B,ENC1,JUND,CD81,CRMP1,BE
P2,DDOST,CIRBP,NARS1,DUX4,YWHAB,PTGES3,SEPTIN11,SND1,LAMP1,ATP1A2,RPS27,CCT6A,H
C5,MSN,C1ORF43,PAPOLA,IGFBP3,COL1A2,RPL14,HK1,RHOA,PTCD3,CLSTN1,KIF5B,RPS14,CX
167A,KLHDC3,TNKS,PSMD8,CHERP,FKBP1A,CTTN,SERP1,WSCD1,VPS26B,ERH,PSMB4,CLPTM1,
BA52,PLXNB1,FARP1,RER1,UBFD1,IARS1,COX7C,YKT6,CUL3,MCM3AP,RPL7L1,RPS27A,TMEM1
1199X,DDX39B,SRSF5,TM9SF3,MDH2,TOMM20,U2AF2,CCT2,GNAI3,NOP53,GDI2,TM9SF2,TSR1,
N1,CAND1,RPL28,ZFAND5,DIDO1,PITPNA,KIF3C,DHX30,FBLN1,STT3B,PITPNM1,TNKS2,LONP1,L
P1R10,XRCC6,CEP170,EIF2S2,CLPTM1,SHC2,INTS3,UBE2M,GPC2,IMPDH2,HINT1,RAB14,ZNF42
NB2,PLD3,HSPD1,USP11,COL18A1,SNRNP200,CSDE1,SMIM14,IGF2,EIF3C,ATP5F1A,EIF4B,MAR
3,GDI2,TM9SF2,TSR1,SSRP1,LMNB2,MTRF1L,ALKBH5,SNRPN,PRDX5,UBE4A,ACO2,HSPA4,TCP
SF10B,UBL3,NSD1,PLPPR2,CACNA1H,UBQLN4,ADIPOR2,ARHGEF9,DLG4,SLC12A7,NACC1,ERBE
29,RPSA,TPM4,NACA,LAMA5,RPS24,BRD3,RPS9,CIPC,PJA2,RPL24,RPL17,C1ORF43,PAPOLA,RP
5,SFPQ,EIF5A,PDIA6,DDR1,PIP4K2B,MEGF8,ACSL6,RAB7A,RPL12,ITM2C,RPL30,EIF3L,MAGED2
',MAP6,HOXB5,GNG4,NXN,UBAP2L,PI4KA,BRSK1,GSE1,C1ORF198,PCBP4,RHBDD2,PPP1R10,TA
SLC4A8,PXDN,OGA,HNRNPA0,FADS2,RPS16,RPL23,NACA,SLC22A17,BRD3,CIPC,PJA2,KPNB1,HK
RAB27A,U2AF2,CCT2,CPSF1,MEST,GAS7,MORF4L2,ABCA3,ALKBH5,SNRPN,PDXK,PLOD1,GRIN
Z1,TIMP2,XAF1,SARAF,MIDN,RPL10,MARCHF6,WSB1,RPS17,VCL,SBF1,ARF1,CYB5R3,PAFAH1B1
C25A3,RPLP2,NDFIP1,MYL6,INIP,MYH10,SND1,UBE2H,ARCN1,RPS27,HMGCR,TOP2B,SMCHD1,I
HX15,ANP32A,EDC4,EP300,STARD7,SF1,AP2A1,BAZ1B,PHB2,ATRX,PI4KA,TRA2B,COX7C,SMC3,C
CO2,HSPA4,MLEC,ACAP3,TMEM248,GCN1,SKI,RAB11B,SRP14,NRDC,PRPF6,RPL37,VHL,MAPRE
GPI,HTRA4,RACK1,BSG,FASN,ALDH6A1,ATP5F1B,CALM3,GNB1,SHISA9,STMN2,RPS8,MYH9,RPL7
PRA,PRDX5,UBE4A,ACO2,HSPA4,TCP1,ZNF302,DDX42,GCN1,SKI,SRP14,PRPF6,TAF7,RPL37,ATC
SLC16A7,RPL22,PTCD3,ZNF865,RPL9,KIF5B,RPS14,CXADR,TMEM259,RPS20,LENG8,WDR1,HMC
EP2,LRRC41,OGT,FBXW5,MBD3,UBE2D3,MICAL1,DPYSL4,TEAD1,USO1,PAGR1,DIP2A,PLS3,TMC
EP2,LRRC41,OGT,FBXW5,MBD3,UBE2D3,MICAL1,DPYSL4,TEAD1,USO1,PAGR1,DIP2A,PLS3,TMC
P1,MTMR4,MKRN1,YIPF3,SPTY2D1,CAP1,USP24,COX6A1,UBE3A,NASP,AP1M1,SMARCD1,TOMM
SSBP4,SORT1,MICAL1,HSDL1,PTPN23,ELOVL5,NELFB,IGFBP4,CDC34,RNF11,HMGA1,AGAP1,AB
B,ARGLU1,PTBP1,CAPZB,ATXN10,QARS1,CCAR2,ARF4,POLR2B,SUPT6H,DHX30,EIF3F,PITPNM1,
P2CA,SMARCA4,AZIN1,DDAH2,FAM168B,ARGLU1,NYNRIN,TCF25,ADAM9,PRKDC,TDRD5,CASC3,
ELL2,EIF4G1,IGLON5,LHFPL6,ELAVL3,TMCO3,HS2ST1,TAPBP,OGFR,USF2,KIFC2,ACVR1B,FZR1

PTIN11,RAB1A,CALU,HECTD4,MAGED4B,CTBP1,TOP2B,SMCHD1,IPO5,ATP6V0C,SKP1,BCLAF1,TI
5,PSMD2,NR2F1,RPS14,TMEM259,EIF5,LENG8,WDR1,HMGN2,RPL26,LASP1,VAT1,OAZ1,RBBP4,
NXA5,CPSF1,PABPN1,GDI2,PKN1,LAMC1,JPT1,ORAI2,SKI,CNTNAP2,RPL38,VEGFA,SELENON,RAI
7,UBXN4,G3BP2,ARPC2,ACADVL,PTBP1,SON,CCAR2,ARF4,SMARCC1,SUPT6H,FYN,LARS1,PFKM
X5,PDXK,UBE4A,HSPA4,GRINA,COL2A1,MLEC,RAB11B,HDGF,RAPGEF1,MVB12B,AZIN1,SRSF9,PI
B2,CDC37,WDR6,CAPZB,GABARAP,SCAF11,BNIP3L,ARF4,HNRNPUL2-BSCL2,DUSP4,PITPNA,XPN
KT6,RPS27A,TMEM167A,PSMD8,CHERP,FKBP1A,CTTN,LRP10,SERP1,ERH,EIF2S2,EIF3H,RALY,SI
,ATP6V0C,LAPTM4B,TRIB2,RPL23A,CDKN1A,SERF2,CPSF2,DDX39B,SRSF5,TM9SF3,MDH2,TOMI
BP1,CDK16,DNER,BCLAF1,NNAT,CDKN1A,HS6ST1,EID1,ZC3H7B,U2AF2,SUGP2,PGD,HNRNPL,TS
1,EIF3E,LUZP6,SUPT5H,ATXN10,QARS1,RPL28,CCAR2,ARF4,DIDO1,POLR2B,GNG2,SMARCE1,SU
1,EIF3E,LUZP6,SUPT5H,ATXN10,QARS1,RPL28,CCAR2,ARF4,DIDO1,POLR2B,GNG2,SMARCE1,SU
CDC37,WDR6,CAPZB,MAP7D1,CASC3,CLIP3,PLCG1,DIDO1,FLNB,MSL1,PITPNM1,TUG1,CSNK1D
,MLEC,ACAP3,ORAI2,SRP14,TAF7,RPS15A,MARS1,RPL37,G3BP2,ARPC2,RBP1,CAPZB,PRKDC,SU
P14,ABCA2,PYGB,ATP6AP1,TMED2,VHL,G3BP2,ATG2B,CCNL2,DSTN,BTBD2,PDE7A,ODC1,TCF2
24,BRD3,XRCC5,MSN,KPNB1,RPL17,IGFBP3,LLGL1,HK1,NCS1,CBX5,RPL9,CLSTN1,NR2F1,KIF5
C1,CLSTN3,DDX42,RAB11B,NRDC,PRPF6,TAF7,TUBB2A,OPTN,UBXN4,PPP2CA,G3BP2,KDELR2,PI
CYB5R3,SF3B2,CDC42BPB,ITM2B,MAT2A,OGA,NFE2L1,RPL29,TPM4,CTSD,SCARB2,TSPAN3,RT
PKD1,SF3B1,SERINC1,BASP1,VAPA,HMGB1,POLR2A,RPS4X,RPS3A,OR8K1,STMN1,ITGB1,MORF
NOTCH2,RAPGEF1,SMARCA4,AZIN1,DDAH2,ARGLU1,NYNRIN,TCF25,PRKDC,TDRD5,CASC3,RPL2
1,PSMA7,PPDPF,ARIH1,BCR,LBH,FBRS,MMP15,SLC39A6,KPNA2,MAP6,PLXNB1,DECR1,CSNK1G
ME1L1,DHX15,TRAF7,EIF2AK1,BCR,MMP15,PHB2,APMAP,KLHL42,RER1,MASTL,COX7C,YKT6,E
RPSA,TPM4,NACA,LAMA5,RPS24,BRD3,RPS9,CIPC,PJA2,RPL24,C1ORF43,PAPOLA,RPL14,PTCD
L28,ZFAND5,CORO1C,SDF4,SMARCC1,HNRNPM,SLIT3,ISYNA1,STT3A,IDH1,SARS1,SOD1,BAG6,F
L1,TAX1BP1,PGAP6,TM9SF4,RAB5C,SPAG9,DICER1,MTMR4,SPTY2D1,PRPF19,NOC2L,COX6A1,
,NCKAP1,MLEC,ACAP3,NRDC,TAF7,RPL37,ATG2B,MAPRE1,ARPC2,PDE7A,FAM168B,ARGLU1,LU
ATP6V0C,LAPTM4B,TRIB2,RPL23A,HSBP1,FAM199X,DDX39B,NUCB1,SRSF5,TM9SF3,MDH2,TOM

RAF,MIDN,MARCHF6,RPS17,SBF1,CYB5R3,MAP4,CDC42BPB,AP2B1,MAT2A,TMEM170A,HNRNPU
3,MDH2,U2AF2,EEF1A2,FDFT1,CCT2,SUGP2,GNAI3,PARP1,TSR1,MTRF1L,RPL27,PTPRA,PRDX5,U
BP1,DYNLL1,CTXN1,CD99L2,GRIPAP1,PSMA7,VDAC1,STAT3,PRMT1,EXTL3,VARS1,CSNK2A1,PPI
X5,UBE4A,ACO2,HSPA4,TCP1,MLEC,ACAP3,RAB11B,SRP14,NRDC,PRPF6,TAF7,ARF3,MAPRE1,FAI
P1,POR,RTF1,TOX4,NOP56,NIBAN2,HIP1R,H6PD,ATP13A2,FZR1,PSMD13,PPP1R14B,CRIP2,PHPT
2B,KHDRBS1,PFKL,TKT,GLG1,SFPQ,PEBP1,PNMA1,PDIA6,PIP4K2B,MMP14,ITM2C,RPL30,H1-0,
MMP15,PHB2,APMAP,PLXNB1,DECR1,HJURP,FARP1,RER1,TRA2B,YKT6,MCM3AP,RPS27A,TMEM
RDBP,SLC25A3,MYL6,MYH10,YWHAB,UBE2H,CKAP4,PLPPR3,COTL1,MAGED4B,CAPRIN1,GPC1,A
G3BP2,KDELR2,PDE7A,PSMD3,TCF25,ADAM9,CDC37,WDR6,GABARAP,BNIP3L,ARF4,HNRNPUL2-
Z2,TBC1D9,SH3PXD2A,MSI1,LZTR1,APBB1,ATXN7L3,THRAP3,DEK,SMO,FGFR1,EIF4A3,PTPN23,S
CAM,GLUL,PEA15,CDKN1A,TIMP3,PGRMC1,NOVA2,C5ORF24,CDH11,ZC3H7B,CHD6,TOMM20,F
IP200,CSDE1,SMIM14,IGF2,EIF3C,MARCKSL1,TRIM28,YWHAG,CKB,COL4A2,FN1,SPEN,DDB1,PS
1,GTPBP1,GRSF1,KIF21B,GPX4,DECR1,GARS1,GNG4,SRSF6,PAPSS1,UBAP2L,UPF1,UBE2G2,ZFI
TRF1L,ALKBH5,SNRPN,PRDX5,UBE4A,ACO2,HSPA4,TCP1,MLEC,ACAP3,RPL36A,SRP14,NRDC,PF
XN,EP400,CD164,SRSF6,KIAA0895L,UPF1,UBE2G2,EIF5B,TRA2B,IARS1,MBTPS1,ECEL1,CBX6,MA
9B,ADGRA3,LMO4,WDTC1,COL16A1,TAOK2,SLC22A23,TSPYL2,RGL2,CLIC1,IGF1R,FOXP4,MEC
1,ATP6V0C,LAPTM4B,RPL23A,NOVA2,SRSF5,RNF187,U2AF2,RPN2,MLXIP,ANXA5,SQSTM1,NOP5
H11,MDH2,RAB27A,ANXA5,VAMP2,MEST,GAS7,PABPN1,HNRNPL,VCAN,TSR1,NOX4,TGFBI,ABCA3

B,SNX22,VAMP2,PARP1,TM9SF2,MORF4L2,SSRP1,MTRF1L,ALKBH5,HNRNPA3,C4ORF3,TBC1D1
C20A1,DPYSL4,ATG4B,METRN,UNC119B,FBXL19,USP5,CASC4,TECR,TAPBP,AGAP1,ITFG1,SGSM
PSMD3,SRSF1,TCF25,ADAM9,CDC37,WDR6,GABARAP,BNIP3L,ARF4,HNRNPUL2-BSCL2,PITPNA,X
EIF3G,AP1M1,SAP18,NDUFA13,PSMC1,TOMM70,BAG5,UBXN6,GAK,LYPLA2,SLC38A10,ATP13A1,
F6,NEO1,MAP2K4,MEIS3,WDR5,GALNT2,SLC2A4RG,ZNF358,PRKCI,FAM20C,AKAP17A,POGZ,AD
P,CIRBP,DUX4,SEPTIN11,LGALS3BP,CKAP4,HLA-B,RAC1,SLC35E2B,APPL1,IPO5,ATP6V0C,MMP2
IP3,CAPZA2,CTDNEP1,MIB2,PCNA,RAB35,DYNLT1,SPIN1,PDXDC1,NBEA,KRI1,CCSER2,SLC35A4,
IGF2,ATP5F1A,EIF4B,VCP,MARCKSL1,UBB,AFF4,TRIM28,CKB,COL4A2,LRRC28,FN1,NONO,DDB
A1,ANKFY1,ETF1,SRSF10,ECHS1,RPL27A,DARS1,H2AX,CCNG1,GID8,MARCHF7,ZCCHC14,POLDI
CCT7,SRSF4,RBBP4,PFN1,ASTN2,MCL1,PNN,RPL31,NISCH,PRKCSH,CCNI,LRPPRC,KHDRBS1,SL
APT,FN1,DDB1,EIF1,HACD3,DCHS1,SET,EGLN1,TPI1,TMSB10,EIF4A1,RPL19,PPP2R1A,RPL15,LD
,SRSF1,PTBP1,CDC37,RPS11,PRKDC,GABARAP,HNRNPUL1,ACTN1,DLK1,PFKM,FBLN1,STT3B,IS
NA,NAP1L1,STARD7,GSPT1,PPM1G,BRD2,KLHL42,GARS1,MBTPS1,RGMB,SELENOW,ECEL1,HN
CAM1,RPLP1,SARAF,MIDN,RPL10,WSB1,RPS17,AP3D1,DNAJA1,ARF1,AKAP12,SF3B2,AP2B1,MAT
ANXA2,AGRN,ZMIZ1,TIMP2,XAF1,WSB1,RPS17,AP3D1,DNAJA1,ARF1,CYB5R3,PAFAH1B1,MAP4,S
NSD2,PSMC3,NASP,TTC28,UBXN6,DPYSL4,SLC38A2,SCD,HSDL1,SNX2,PAFAH1B2,HS2ST1,OLFM
SCL2,PITPNA,XPNPEP3,ATP6V1B2,SRSF3,LARS1,YWHAH,UBQLN1,CSNK1D,HYOU1,IFITM3,DGKD
KI,KDM6B,RAB11B,HDGF,PRPF6,TERF2IP,TMED2,TAF7,RPL37,MAPRE1,DDAH2,ARGLU1,RBP1,P
LMO4,USF2,CLTA,ARPC1A,FAM171A1,POR,SLC22A23,NRXN2,KLF13,CUL1,CLUH,PNRC2,GALNT
1,MEGF8,ACSL6,RAB7A,PRDX2,DDX3X,ITM2C,RPL30,CLTC,EIF3L,MAGED2,RPS5,IK,RPS19,TLN
F2IP,TMED2,RPS15A,RAPGEF1,TUBB2A,PPP2CA,CCND1,MAPK1,AZIN1,DDAH2,PTBP1,FTO,ILF3,
IB,GLUL,SERF2,MDH2,U2AF2,FDFT1,ANXA5,VAMP2,GNAI3,TSPYL4,TSR1,MTRF1L,HNRNPA3,PD
NCSTN,INTS11,MOAP1,MTMR4,PCNX3,ERGIC3,SPTY2D1,NUDT19,CAP1,ATP6V0D1,NOC2L,COX6
2K2,ZFAND5,PLTP,CORO1C,KIF3C,STXBP1,SREBF2,VGF,RPL39,ARNT2,ARFGEF2,DYNLL1,YME1
L2,TOP1,SMARCA4,AZIN1,DDAH2,FAM168B,ARGLU1,NYNRIN,ADAM9,ILF3,PRKDC,HSPB1,CASC
TMS,SPTAN1,GSTP1,MARCKS,SMG1,CALM1,NPM1,PLXNB2,HSPD1,COL18A1,SNRNP200,SMIM1
,XPR1,HDLBP,EIF5A,PEBP1,DDR1,PLXNA1,RAB7A,DDX3X,CLTC,NANOG,GNB2,RPS5,IK,HMGCS1
R1,MCL1,PNN,CCNI,TADA2B,XPR1,HDLBP,EIF5A,PEBP1,DDR1,PLXNA1,RAB7A,DDX3X,CLTC,NANO
,RPL6,KAT6A,DVL3,YWHAE,KIF1A,RPL13,GNAS,SLC2A3,DDX17,GPI,HTRA4,RACK1,BSG,FASN,AL
N3,STXBP1,CSNK1D,ZFAND3,VANGL2,VGF,FBRSL1,P3H3,ARFGEF2,AMOTL2,PPP1R9B,DHX15,C
2,INIP,CIRBP,MYH10,DUX4,CKAP5,SEPTIN11,RAC1,CAPRIN1,GPC1,BCLAF1,GLUL,PEA15,PTTG1
,FAM168B,ARGLU1,LUZP6,HSPB1,SON,RPL28,CCAR2,ARF4,CORO1C,DHX30,NKTR,STMN3,TNK
AND5,CORO1C,SDF4,SMARCC1,HNRNPM,YWHAH,SLIT3,ISYNA1,STT3A,IDH1,SARS1,SOD1,BAG6,
,HSPB1,SON,CCAR2,ARF4,CORO1C,SUPT6H,DHX30,TNKS2,SARS1,BAG6,HNRNPD,RAB11A,SERE
VAPA,POLR2A,AP2M1,TPT1,SEC61A1,RPS4X,RPS3A,RPL18,TUBB4B,RPL11,MORF4L1,BTF3,SAR

N3,TPM4,CTSD,EIF4H,NACA,RPS24,RPS9,RPL17,C1ORF43,PAPOLA,IGFBP3,RPL14,HK1,NCS1,R
EGLN1,CHSY1,TMSB10,INA,EIF4A1,MRC2,RPL19,RPL15,GTF3C4,LDHB,DNAH8,PKD1,HLA-C,SF
N,WSB1,RPS17,VCL,SBF1,DNAJA1,ARF1,MAP4,SF3B2,CDC42BPB,ZNF770,GTF2F1,GOLGA8A,P
CN1,RPL36A,SRP14,TAF7,RPL37,G3BP2,CCNL2,ARPC2,ARGLU1,PTBP1,HNRNPUL1,CCAR2,ARF
A,COL1A2,GAP43,RPL14,ZNF865,CLSTN1,ATF4,PSMD2,GLIPR1L2,RPS14,CHD4,COL5A2,LENG8,
F1,ARGLU1,AEBP1,ADAM9,PTBP1,GRB2,PRKDC,SUPT5H,SCAF11,RPL28,MSL1,HNRNPM,SSB,P
HMT2,RASA4,LIMK1,PITRM1,DRAP1,IQSEC1,WIZ,EXT2,TBC1D14,SLC25A1,MFAP4,STUB1,ZNF76
F43,IGFBP3,NCS1,CBX5,RPL9,NR2F1,GLIPR1L2,CXADR,ATP1A1,JUND,TMEM259,EIF5,LENG8,W
1L,ALKBH5,SNRPN,PRDX5,UBE4A,ACO2,HSPA4,TCP1,MLEC,ACAP3,SRP14,NRDC,PRPF6,TAF7,A

TSR2,MYL6,SHOX,INIP,MYH10,SND1,RHOB,UBE2H,ARCN1,HMGCR,SMCHD1,ATP6V0C,SKP1,LAP
,TADA2B,XPR1,HDLBP,EIF5A,PEBP1,DDR1,PLXNA1,RAB7A,DDX3X,CLTC,NANOG,GNB2,RPS5,IK,H
2,UBA52,BRD2,DECR1,PI4KA,UBE2G2,RAB3GAP2,UBFD1,MASTL,CUL3,MCM3AP,SPINT2,RPL7L1
P4,SF3B2,CDC42BPB,GTF2F1,PGAM1,SMARCA5,HNRNPA0,CTNNA1,RPS16,RPL23,RPL29,RPS24
C1,CHST3,NSD1,WAC,CRTC1,TUBA1C,PLXDC2,UBQLN4,IRF2BP2,SH3BP5,POGK,IRS1,NR2F2,IT
RS1,DSTYK,NADK,ZNF395,CAMK2N1,PRPF31,VOPP1,RAB5A,MARK1,CMTR1,CHPF2,LRRN1,HEXI
PPP1R9B,STAT3,ARIH1,KCNQ2,PRMT1,NAP1L1,KIRREL1,CNTNAP1,ALDH3A2,LRRC4B,ZFP36L1,
M171A1,SLC22A23,KIFC2,ELN,IGF1R,MGAT3,CMPK1,CRIP2,MAP4K4,ACAP2,ATP1B3,RNF157,H
10,MARCHF6,AP3D1,DNAJA1,ARF1,AKAP12,SF3B2,ITM2B,TMEM170A,GTF2F1,SLC4A8,PGAM1,S
01,DNAJA1,ARF1,CYB5R3,PAFAH1B1,MAP4,SF3B2,ITM2B,TMEM170A,GOLGA8A,PXDN,HNRNPU,I
IRNPA0,FADS2,LDHA,RPL29,TNS3,CTSD,SCARB2,TSPAN3,LPCAT1,SLC22A17,RPL24,C1ORF43,R
RP1,UQCRC1,HDAC6,ZFR,SMC3,HNRNPA2B1,PPP1CC,ULK1,ANP32B,LMAN1,ZMYM4,DKK3,HIF1
55,ATF4,GLIPR1L2,RPS14,CXADR,JUND,RPS20,LENG8,WDR1,RPL26,LASP1,CCT7,SRSF4,RBBP4,
A,PGAM1,NFE2L1,SMARCA5,FADS2,RPS16,LDHA,CPE,RPL29,CTSD,EIF4H,SCARB2,NACA,LAMA5
RSF1,IPO7,ARGLU1,RBP1,ADAM9,FTO,ILF3,PRKDC,SCAF11,SON,CCAR2,PTP4A2,PFKM,STT3B,U
IUDCD3,TUBGCP6,EEF1B2,RIC8A,NCSTN,INTS11,MOAP1,MTMR4,MKRN1,ABCF1,CAP1,LRRC41,
PDIA3,SERINC1,RPS3,CHD3,BASP1,VAPA,HMGB1,POLR2A,CDKL1,AP2M1,TPT1,SEC61A1,RPS4X,I
TCD3,CBX5,RPL9,NR2F1,SLC7A5,ATP1A1,JUND,TMEM259,CD81,COL5A2,LENG8,WDR1,ATXN7L3
C42BPB,ITM2B,AP2B1,GTF2F1,GOLGA8A,SMARCA5,LDHA,CPE,CNN3,RPSA,CTSD,SCARB2,RTN4,
RPS12,RAB1A,TTR,ZNF454,CTBP1,HMGCR,SLC35E2B,DNER,PPIB,RPL23A,PEA15,CDKN1A,CPSF2
,CCT3,GRINA,COL2A1,PTOV1,MDK,LAMC1,RAD23B,KDM6B,HDGF,VEGFA,ARF3,MVB12B,PPP2C
V1B2,SRSF3,LARS1,YWHAH,UBQLN1,CSNK1D,HYOU1,IFITM3,DGKD,VGF,MLF2,SEC23A,RTL8C,
65,ATF4,PSMD2,NR2F1,ENC1,NLGN2,GLIPR1L2,CHD4,ATP1A1,JUND,CD81,HMGN2,ATP2A2,CRM
AF4,XPOT,LBH,SLC35F5,CSNK1G2,RAB2A,CD164,KIAA0895L,MASTL,HDAC6,AGPAT1,ECEL1,SL

MIZ1,TIMP2,XAF1,COL11A1,MIDN,RPL10,RPS17,VCL,SBF1,DNAJA1,ARF1,CYB5R3,MAP4,CDC42
I,C1ORF43,PAPOLA,LLGL1,HK1,NCS1,PTCD3,CBX5,ZNF865,ATF4,PSMD2,SLC7A5,CXADR,CHD4,
SMB1,SLC25A6,EXOC5,NDUFA4,SF3A3,TRMT112,PSMD13,WDR26,GPAA1,GGT7,SUB1,CFDP1,C
ENND4B,FBLN5,SART1,MTOR,PRKACA,SMARCD1,XPO6,SSBP4,DPYSL4,TRO,NSG1,SLC38A10,FE
ADA2B,SPTBN2,CSNK1E,RPS25,TKT,CDH6,TPM1,EIF5A,PEBP1,CCND2,PDIA6,GPC3,PLXNA1,PIP4
6,IPO5,ATP6V0C,BCLAF1,L1CAM,LUC7L3,MAP1A,FAM199X,CTDSP2,PGRMC1,DDX39B,CNN2,N
P13A1,NELFB,USP5,RANGAP1,ELAVL3,TECR,SNX3,SPPL2B,AUTS2,POR,CLIP2,FAM118A,NOP56,N
IPT2,TLE5,NISCH,LRPPRC,KHDRBS1,KLHL9,XPO1,SPTBN2,SLC3A2,HDLBP,RPS25,TPM1,ACSL6,I
28,CKB,COL4A2,LRRC28,MAPT,EIF1,DCHS1,SET,EGLN1,CHSY1,INA,EIF4A1,MRC2,PPP2R1A,GTF
BCLAF1,LAPTM4B,TMEM132A,CPSF2,SRSF5,TM9SF3,MLXIP,ANXA5,NOP53,TM9SF2,SSRP1,HSP
3A,SCAF1,HDGF,PRPF6,DPYSL2,TOP1,RAPGEF1,TUBB2A,OPTN,AZIN1,SRSF9,PLEC,PBX2,FAM17
MO1,CEMIP2,HNRNPA2B1,CUL3,EML1,LMAN1,TNKS,PSMD8,IARS2,UBE2M,RALY,SMG5,ZMIZ2,TL
A8A,LDHA,CPE,CNN3,RPSA,CTSD,SCARB2,RTN4,LPCAT1,LAMA5,RPS24,SLC22A17,RPS9,CIPC,PJ
S1,CBX5,RPL9,ATF4,NR2F1,GLIPR1L2,CXADR,ATP1A1,JUND,TMEM259,LENG8,WDR1,HMGN2,M
P3,CTDSP2,PGRMC1,NOVA2,ZC3H7B,U2AF2,EEF1A2,ANXA5,SNX22,VAMP2,GDI2,TGFBI,ABCA3,H
,RPSA,CTSD,SCARB2,RTN4,LPCAT1,LAMA5,RPS24,SLC22A17,RPS9,CIPC,PJA2,RPL24,SLC16A7,I
23,ANP32E,PFKP,REEP5,FZR1,MAPK8IP2,SLC25A6,FOXP4,NUDT4,CRIP2,RALGDS,NTN1,LIMK1,S
CXADR,CHD4,JUND,EIF5,CD81,ATP2A2,CRMP1,LASP1,VAT1,OAZ1,SRSF4,RBBP4,PFN1,JPT2,TLE
SF2,PGRMC1,CNN2,CTSB,EEF1A2,SUGP2,CPSF1,SNX22,VAMP2,PABPN1,HNRNPL,VCAN,TAF15,A
B,ACTN4,DDX3X,RPS15,ARL4C,RPS19,MIF,GPAT4,MYO5A,NGRN,B4GALNT4,DNAJC5,APLP2,HN

36B,ENC1,GLIPR1L2,JUND,WDR1,CCT7,VAT1,OAZ1,PFN1,MCL1,PRKCSH,HNRNPC,KHDRBS1,HD
.RRC28,MAPT,FN1,NONO,DDB1,PSMD5,YWHAQ,EIF1,HACD3,SET,EGLN1,MAP9,CHSY1,TMSB10,
,BEX4,SSR1,BCL9L,WDR45B,HIP1,SLC39A6,CD99,GPX4,PPM1G,HJURP,GNG4,NXN,EP400,UBFD
JA2,RPL24,SLC16A7,IGFBP3,NCS1,CBX5,PSMD2,GLIPR1L2,ATP1A1,JUND,TMEM259,EIF5,LENG
MYH9,RPL7,SEPTIN2,P4HB,APP,ERVV-1,RPL5,GDI1,FSCN1,TMBIM6,SPTAN1,GSTP1,MARCKS,SM
PO3,MCM7,VPS16,NUP50,SZT2,SEC31A,AKAP17A,EIF2S1,MAPK8IP3,PSMC6,PRELID1,HARS1,MF
)RF194,KDM2A,DUSP8,IRS1,PRKAR2B,DPP7,COL7A1,RXRA,DNAJB6,RUSC1,AFDN,ELOB,GALNT
)LDIP2,TRIP12,CHMP1A,ITFG1,CGGBP1,PSMB1,TSNAX,SLC25A6,EXOC5,NDUFA4,SF3A3,TRMT1
18,STMN1,TUBB4B,UCHL1,BTF3,AGRN,RPLP1,TIMP2,SARAF,MIDN,MARCHF6,RPS17,VCL,AP3D1
F1,RAD23A,CTIF,EPHA2,SKIL,VPS37B,USP36,ZC3H4,NDUFA10,EMD
TPS1,CBX6,CUL3,IL6ST,RPS2,APEX1,PODXL2,RABEP1,SLC2A1,FBXO21,ERH,FKBP4,CARM1,ANT
-0,RPS19,TLN1,MYO5A,NGRN,SESN3,RPL35A,DNAJC5,HNRNPDL,TARDBP,NDFIP1,MYL6,ANKRD
GN2,CHD4,CD81,WDR1,RPL26,LASP1,BEX3,CCT7,RBBP4,PFN1,MCL1,TLE5,RPL31,CCNI,LRPPR
310,INA,EIF4A1,PPP2R1A,RPL15,GTF3C4,PKD1,HLA-C,SF3B1,PDIA3,SERINC1,SSBP2,RPS3,CHD3
8,PXDN,HNRNPU,NFE2L1,LARP1,HNRNPA0,CTNNA1,CPE,RPSA,TPM4,TNS3,EIF4H,SCARB2,TSPA
RF,KLHDC3,PPP1R15A,GPC4,LRRC59,BRK1,AKAP9,XRCC6,CLCN7,SHC2,SEC14L1,IMPDH2,TM9
JDCD3,GNL1,PNISR,STK24,TUBGCP6,RIC8A,NCSTN,INTS11,PRPF4B,MOAP1,MTMR4,PCNX3,ER
PPL1,SMCHD1,HDAC2,ATP6V0C,SKP1,BCLAF1,L1CAM,GLUL,CDKN1A,HS6ST1,TIMP3,PGRMC1,
2R1A,GTF3C4,LDHB,SF3B1,PDIA3,SERINC1,SSBP2,ZNF43,BASP1,HMGB1,POLR2A,CDKL1,AP2M
INRNPM,GIGYF1,NKTR,YWHAH,SLIT3,ISYNA1,STT3A,PLXNA3,CSNK1D,IDH1,SOD1,BAG6,FBRSL1
RIN1,GPC1,TOP2B,ATP6V0C,MMP2,SKP1,DNER,LAPTM4B,NNAT,PTTG1IP,CRPPA,HS6ST1,CTDSP
3C4,HLA-C,SF3B1,SSBP2,ZNF43,CHD3,BASP1,HMGB1,POLR2A,CDKL1,AP2M1,TPT1,RPS4X,RPS
IRNPDL,RPLP2,ARL6IP1,TSR2,MYL6,INIP,MYH10,SND1,RHOB,UBE2H,ARCN1,HMGCR,TOP2B,SM
B,ERH,CLPTM1,INTS3,ABI2,UBE2M,SEC14L1,ELAVL1,NUDCD3,SSR3,GNL1,FLOT2,TUBGCP6,EE
2L1,SMARCA5,HNRNPA0,RPS16,RPL23,CNN3,RPL29,TNS3,CTSD,NACA,TSPAN3,LPCAT1,LAMA5,F
39B,YME1L1,TXLNA,BCR,CAPN2,MTA1,PDAP1,PHB2,UBA52,PLXNB1,DECR1,CSNK1G2,PI4KA,UB

N1,AHSA1,MSI1,ADAM10,USP24,EHD1,EIF3G,GSK3A,DHX38,PRKACA,ANKRD12,MICAL1,LYPLA2,
3,SRSF5,U2AF2,MLXIP,PPP1CB,SNX22,SQSTM1,VAMP2,GDI2,HNRNPL,SSRP1,PRDX5,PDXK,PKN1

MARCA4,ARPC2,DDAH2,SRSF1,IPO7,ACADVL,EIF3E,FTO,LUZP6,CDC37,CAPZB,RPS11,MAP2K2,F
33,ACSL4,PDCD11,WARS1,PMPCA,SKIV2L,CFLAR,SHC1,PCED1A,CHMP5,ASB6,MIB2,TNPO3,MCM
4,PCDHGB6,PYGB,PRPF6,NOTCH2,SMARCA4,AZIN1,DDAH2,ARGLU1,NYNRIN,PRKDC,TDRD5,CA
3F,B2M,TUG1,ID4,GHITM,WBP2,PRPF40A,TXNIP,CTNND1,GRIPAP1,PPDPF,CIC,TRAF7,TXLNA,O
CSNK1G2,RAB2A,UPF1,MASTL,MAPRE2,APEX1,PPP1R15A,GALNT11,MVD,DAP,BRK1,BAZ2A,XRC
A,PXDN,FADS2,RPS16,CPE,RPL29,TPM4,TNS3,EIF4H,LPCAT1,SLC22A17,RPS9,MSN,KPNB1,C1O
CD3,GNL1,FLOT2,TUBGCP6,EEF1B2,RIC8A,NCSTN,INTS11,AHSA1,QPRT,MTMR4,SPTY2D1,LRR
)RAI2,SNRNP70,TSPAN6,PRPF6,MVB12B,MAPK1,ATG2B,ODC1,FAM168B,TCF25,PTBP1,ILF3,WD
,HIF1A,LRRC59,STK35,FUS,XRCC6,PELP1,SLIT1,ABI2,TRAM1,GPC2,IREB2,CNP,PRPF4B,ACTR1

.F13,PLXNA2,SC5D,ARHGEF40,FOXP4,CAB39,NTN1,MTURN,KLC2,FLII,SYT15,DYNLRB1,DRAP1,
MS,SPTAN1,GSTP1,MARCKS,SMG1,CALM1,NPM1,PLXNB2,TTC3,PLD3,HSPD1,USP11,COL18A1,S
MCFD2,RAB3A,STC2,RRP1B,NDUFS1,OTUB1,NSMF,DUSP8,DPP7,DGCR8,NEO1,DSTYK,PPP3CA,T
19,CDC37,WDR6,GABARAP,BNIP3L,ZFAND5,HNRNPUL2-BSCL2,PITPNA,XPNPEP3,ATP6V1B2,SRS
S20,WDR1,VAT1,RBBP4,RTN3,MCL1,NISCH,PRKCSH,HNRNPC,PTN,HDLBP,RPS25,CDH6,CCND2,

TXN7L3,MN1,SLC20A1,SORT1,DEK,RCN2,SLC38A10,WBP1,USP5,CNBP,TCEAL4,ELAVL3,ACAT2,T
EN,PSMD5,YWHAQ,EIF1,HACD3,DCHS1,SET,EGLN1,TPI1,CHSY1,EIF4A1,PPP2R1A,RPL15,GTF3C4
AH1B2,POLDIP2,TRIP12,XRN2,CHMP1A,PSMB1,TSNAX,GTF3C1,SLC25A6,EXOC5,SF3A3,TRMT11
ITM2B,ZNF770,AP2B1,MAT2A,TMEM170A,SLC4A8,GOLGA8A,PXDN,HNRNPU,OGA,NFE2L1,SMAF
D,GPR107,GHITM,PFN2,ADAR,CPD,EP300,DGCR2,ALDH3A2,CSNK2A1,PHB2,SLC39A7,RAB2A,RE
RP10,SERP1,RHBDD2,PPP1R10,ERH,RALY,SEC14L1,SMG5,NUDCD3,TAX1BP1,FAM234A,LRATD2
,CUL3,NDST1,ITGB5,TIMM50,TNKS,APEX1,PODXL2,MAP1LC3B,MTCH1,TMOD2,KIAA0100,PELP1
B,RPL23A,DLG5,TMEM132A,CPSF2,DDX39B,MDH2,RNF187,U2AF2,ANXA5,CPSF1,PPP1CB,NOP5
I,RAN,RHEB,TRIM27,ADRM1,HM13,COLGALT1,SEPHS1,SEZ6L2,OAT,ACSL4,CACNA1H,CUEDC1,Z
RP1,LMNB2,SNRPN,PTPRA,AHCYL1,CAMSAP1,GRINA,ACAP3,LAMC1,JPT1,CCT4,RAB11B,ABCA2,I

M32A,EPN1,CDK9,RTF2,ZRANB2,STUB1,TNFRSF10B,UNC45A,RNMT,ZBTB4,ACBD3,SETX,NDUFS
5,RPS24,SLC22A17,RPS9,CIPC,PJA2,RPL24,SLC16A7,PAPOLA,IGFBP3,NCS1,CBX5,PSMD2,GLIPF
,FLOT2,TUBGCP6,EEF1B2,RIC8A,NCSTN,INTS11,AHSA1,QPRT,MTMR4,SPTY2D1,CAP1,LRRC41,A
2F1,SLC4A8,HNRNPU,OGA,PGAM1,LARP1,HNRNPA0,CTNNA1,RPS16,LDHA,CNN3,TPM4,NACA,LAI
N,HNRNPC,XPR1,KLHL9,RPS25,TKT,CCND2,GPC3,SULF1,DDX3X,RPL30,CLTC,NANOG,MRFAP1,I
LK1,NEO1,LZTS2,GPR173,MARVELD1,MAP2K4,RFX7,SUDS3,ZNF358,TET3,GATAD2B,ADGRL1,M
TM3,GHITM,NCOR2,VGF,MLF2,WBP2,AKIRIN1,AMOTL2,YME1L1,CTNND1,TRAF4,STAT3,ARIH1,N
NPDL,SLC25A3,MYL6,ANKRD52,CKAP5,CADM4,SND1,UBE2H,RPS12,HLA-B,RAB1A,ZNF454,APPL
CIPC,SLC16A7,RPL17,C1ORF43,GAP43,RPL14,NCS1,CBX5,RPL9,ATF4,PSMD2,NR2F1,SLC7A5,RI
0,JRK,TOMM40,COX5B
V,PTPRF,OAZ2,LMAN1,APEX1,HIF1A,GSE1,CTTN,MAP1LC3B,SLC2A1,PRDX6,AKAP9,ZMYM3,PSMI
EIF4B,VCP,UBB,MATR3,YWHAG,COL4A2,RPL37A,LRRC28,SPEN,DDB1,YWHAQ,EIF1,HACD3,DCHS
1,SLC35F5,CSNK1G2,RAB2A,UPF1,MASTL,MBTPS1,MAPRE2,APEX1,PPP1R15A,GALNT11,MVD,DA
3A52,SLC35F5,CSNK1G2,RAB2A,GNG4,UBAP2L,UPF1,MASTL,TBC1D9B,MAPRE2,APEX1,PPP1R15
DCD3,CNP,SLC25A23,PACS1,CLASP2,ATP1B2,CAP1,PSMC3,GOT2,AP2A2,SSR2,KLHDC10,DOP1B,
3,AP2B1,PXDN,PGAM1,HNRNPA0,CTNNA1,FADS2,RPL23,CPE,CNN3,RPSA,TPM4,EIF4H,SCARB2,T
0895L,RER1,PI4KA,CUL3,NDST1,ULK1,MAPRE2,LMAN1,PPP1R15A,MAP1LC3B,NDN,BAP1,BAZ2A

F3H,ELAVL1,IMPDH2,GABARAPL1,GNL1,TM9SF4,ZNF428,TRAP1,PLEKHB2,ELP1,MOAP1,CLASP2
EM9,SCG2,WDR82,SYNGR1,USP48,RPS29,DUSP8,RNF20,NUP133,COX4I1,OGDH,DSTYK,GUK1,UI
PL13A,MAGED1,SEC62,RPL6,KAT6A,DVL3,YWHAE,KIF1A,RPL13,GNAS,SLC2A3,DDX17,GPI,HTRA
I1,DYNLRB1,TNRC6A,PPME1,CSNK2B,RPL27A,COX7A2L,FAM32A,OTUD5,EPN1,ARPC3,CSNK1A1
,PRPF19,COPZ1,EIF3G,RBM6,AP1M1,SMARCD1,SSR2,NDUFA13,UBE2I,DAPK3,ATP13A1,IER2,EII
P1,CLASP2,NSD2,YY1,CCDC47,ELOVL5,DHX40,SRPRA,IER2,SLC25A36,XRN2,SCAMP5,ZZEF1,ESY
2,ANKRD52,H1-10,DUX4,CADM4,ATP1A2,CKAP4,RAB1A,SEPTIN3,ZNF454,RPS27,COTL1,CAPRIN
FA13,PSMC1,XPO6,DEK,LYPLA2,SLC38A10,ATP13A1,NELFB,PAFAH1B2,ACAT2,POLDIP2,TRIP12,I
ATXN3,RAB6A,WASF2,MSI1,SCRIB,IGF2BP1,MED13,PCGF2,AP2A2,APBB1,WNK1,GAK,USP5,IER2,
DKL1,SEC61A1,RPL11,MORF4L1,BTF3,NCAM1,ANXA2,RPLP1,ZMIZ1,XAF1,RPS17,VCL,ITM2B,ZNI
,MAGED2,GNB2,H1-0,RPS5,IK,RPS19,MIF,CHPF,JUN,CD63,NGRN,RPLP2,DDOST,CIRBP,NARS1,S

DUSP8,RNF20,NUP133,COX4I1,OGDH,DSTYK,GUK1,UBN1,NECTIN2,PLCD3,CTDNEP1,CMTR1,CL
G2,RAB2A,UPF1,MASTL,MBTPS1,MAPRE2,APEX1,PPP1R15A,GALNT11,MVD,DAP,BRK1,SERP1,BA
GTP1,CALM1,NPM1,PLXNB2,HSPD1,USP11,COL18A1,SNRNP200,SMIM14,IGF2,EIF3C,ATP5F1A,EI
12B,ADH5,OAZ2,BRSK1,GPSM1,ZMYM4,KLHDC3,TNKS,PEG10,CHERP,LRP10,LRRC59,PCBP4,W

,JRK,TOMM40,UQCR11,COX5B
P1,CAMSAP1,CCT3,NCKAP1,MLEC,RBMX,TMEM248,RAD23B,SNRNP70,H3-3A,RPL36A,NRDC,PY
PSMB4,CLPTM1,EIF3H,RALY,SMG5,FAM234A,SSR3,PNISR,TUBGCP6,RIC8A,NMT1,AHSA1,MOAP
,RPS8,MYH9,RPL7,SEPTIN2,IDS,ERVV-1,RPL5,GDI1,PTMS,SPTAN1,GSTP1,MARCKS,SMG1,CALM
S,RPL31,NISCH,PRKCSH,HNRNPC,CCNI,LRPPRC,XPR1,CDH6,TPM1,EIF5A,GPC3,PIP4K2B,SOX1
,COX7C,YKT6,CUL3,ADH5,MCM3AP,TMEM167A,KLHDC3,TNKS,NPLOC4,CHERP,CTTN,LRP10,SE
3,TXLNA,XPOT,TXNDC5,SEC16A,BRD2,CSNK1G2,RAB2A,CD164,PI4KA,PTPRF,EML1,SLC7A6,MA
TRA2B,ZFR,PRRC2A,PREPL,TIMM50,SEPTIN7,PSMD8,HIF1A,SYMPK,SAFB2,PPP1R10,TRIR,ERH,E
ADR,ATP1A1,EIF5,LENG8,WDR1,HMGN2,RPL26,CRMP1,BEX3,RBBP4,RTN3,PFN1,MCL1,PNN,SP
PO1,SKI,ASXL1,H3-3A,ABCA2,PRPF6,TMED2,MARS1,MAPK1,MAPRE1,BTBD2,PRDX1,ODC1,FAM1
1,MIDN,RPL10,MARCHF6,DNAJA1,ARF1,CYB5R3,AKAP12,MAP4,SF3B2,CDC42BPB,GTF2F1,PXD
JP50,SZT2,SEC31A,AKAP17A,MAPK8IP3,PSMC6,PRELID1,HARS1,MRPL3,DNMT1,CFAP298,DDX5
1,PCED1A,SUDS3,ZSWIM8,CTDNEP1,KCTD10,MIB2,TNPO3,MCM7,RPL32,VPS16,SZT2,SEC31A,F
N,ZNF512B,CEMIP2,TBC1D9B,PTPRF,EML1,BRSK1,GPSM1,MAP1LC3B,MVD,SLC2A1,ZNF664,S
GPC2,PGAP6,GNL1,LRATD2,STK24,PLEKHB2,CADM3,SLC4A2,SLC25A23,ABCF1,ERGIC3,SPTY2
,MORF4L2,SNRPN,RTN1,PKN1,GRINA,PTOV1,MLEC,CLSTN3,SURF4,DDX42,RAD23B,TNPO1,SN
IN1,AMOTL2,YME1L1,CTNND1,TRAF4,STAT3,ARIH1,NDUFV1,SF1,GPX4,SLC35F5,CSNK1G2,RAB
STYK,ZNF146,CMTR1,CLCN3,TRAPPC1,AGPAT4,BRPF3,MSI2,CBX4,CCNL1,PDXDC1,PAF1,GNAO
PF1,EIF5B,HDAC6,FMNL2,GPSM1,LOX,SAFB2,PRDX6,VPS26B,BAP1,KIAA0100,SLC44A2,SSBP3,
BFD1,TRA2B,ZFR,YKT6,PRRC2A,ANP32B,PREPL,TIMM50,SEPTIN7,PSMD8,HIF1A,SAFB2,PPP1R1
DDB1,PSMD5,YWHAQ,EGLN1,MAP9,CHSY1,TMSB10,INA,EIF4A1,RPL19,PPP2R1A,SF3B1,PDIA3,S
D9,BAG6,RAB11A,SERBP1,AKIRIN1,CDC42,PPP1R9B,RBBP7,TRAF7,EIF2AK1,WDR45B,PRMT1,X

PAPOLA,NCS1,CBX5,RPL9,NR2F1,GLIPR1L2,CXADR,ATP1A1,JUND,TMEM259,EIF5,LENG8,WDR1
A8A,WBP2,TXNIP,CTNND1,GRIPAP1,CIC,TRAF7,OS9,ACTR1A,STARD7,SF1,PBX1,AP2A1,H2AZ1,K
JSP8,CHD8,ZDHHC5,SLC5A6,RRBP1,UBXN1,RETREG3,SKIV2L,NARF,ROCK2,UBE2G1,ATF6,PCE
K2B,RAB7A,RPL12,CLTC,NANOG,MAGED2,GNB2,RPS5,MIF,PRKAR1A,TLN1,CHPF,JUN,NGRN,M
A2B,XPR1,KHDRBS1,GLG1,TPM1,EIF5A,CCND2,CDH2,RAB7A,ARL4C,GNB2,RPS7,MIF,CHPF,JU
AP,BAZ2A,VAPB,EIF3H,ELAVL1,IMPDH2,GABARAPL1,GNL1,TM9SF4,TRAP1,PLEKHB2,ELP1,SPAG
3,APP,IDS,ERVV-1,RPL5,FSCN1,TMBIM6,PTMS,GSTP1,MARCKS,SMG1,PLXNB2,PLD3,HSPD1,USP
A,MTA2,TMEM94,DAG1,PAIP2,ATP13A3,RBM25,NOTCH1,SLC38A1,PPP1R14B,FEZ1,PSMD7,NCK

GB,PRPF6,TAF7,RPL37,ATG2B,CCNL2,ARPC2,FAM168B,ARGLU1,ADAM9,CDC37,RPL28,CCAR2,
KB,WARS1,SNRPB,KDM5A,SKIV2L,SHC1,SUDS3,MIB2,TRAPPC1,TNPO3,CUL4B,SZT2,SEC31A,AC
,SSR1,TRAF7,EIF2AK1,TXLNA,PRMT1,TRIO,SF1,PDAP1,MMP15,SEC16A,KPNA2,PHB2,SLC39A7,
3,ID3,DNMT1,NEDD8,SETD1B,ZC3H4

A2,FDFT1,CCT2,PGD,SQSTM1,MEST,TLN2,TSR1,MORF4L2,TGFBI,SSRP1,HSPH1,YEATS2,GRINA
,RPL11,MORF4L1,AGRN,ZMIZ1,SARAF,MIDN,RPL10,WSB1,SBF1,DNAJA1,ARF1,ITM2B,TMEM17
YWHAG,CKB,COL4A2,RPL37A,MAPT,DDB1,YWHAQ,EIF1,TPI1,TMSB10,INA,EIF4A1,RPL19,GTF3C
YREF,COX5B
AB7A,PRDX2,RPS15,ARL4C,RCN1,H1-0,APLP1,RPS19,MIF,HMGCS1,TLN1,CD63,DDOST,ST3GAL2
RN,GOLPH3,DHX36,HM13,COLGALT1,TSPAN7,NUP98,NACAD,OXSR1,EMC10,TMEM9,SCG2,WDR
2,GLOD4,MTURN,GOLPH3,ECHS1,DHX36,HM13,COLGALT1,TSPAN7,NUP98,NACAD,OXSR1,EMC
MILIN1,ZNF768,ZCCHC14,NDRG4,NACAD,HMGXB3,WDR82,BEX1,GDF1,SEZ6L2,CACNA1H,RCOR

L3,TOMM70,XPO6,UBXN6,DEK,MYL12B,LYPLA2,ATP13A1,PAFAH1B2,POLDIP2,TRIP12,XRN2,CH
CDH11,SUGP2,VAMP2,MEST,GDI2,HNRNPL,PARP1,TSPYL4,TAF15,TGFBI,ABCA3,HNRNPA3,C4OF
),SLC25A36,XRN2,SCAMP5,ZZEF1,AASDHPPT,ESYT1,GTF3C1,TNFAIP1,FAM50A,CD2BP2,ALDH18
TURN,GOLPH3,ECHS1,DHX36,HM13,COLGALT1,TSPAN7,NUP98,NACAD,OXSR1,EMC10,TMEM9,
TURN,GOLPH3,ECHS1,DHX36,HM13,COLGALT1,TSPAN7,NUP98,NACAD,OXSR1,EMC10,TMEM9,
),NTRK2,RANGAP1,MZT2B,TECR,SCAMP1,ZKSCAN1,YTHDF3,KLF13,CDV3,ATP13A2,MFN2,MAST
,XAF1,SARAF,COL11A1,MIDN,RPL10,MARCHF6,RPS17,VCL,AP3D1,SBF1,DNAJA1,AKAP12,ITM2B,
KBH5,SNRPN,PRDX5,PKN1,PLOD1,VPS35,PTOV1,MLEC,SURF4,ZNF302,ORAI2,SKI,PCDHGB6,SN
.C4A8,OGA,NFE2L1,LDHA,CNN3,RPL29,TPM4,TNS3,TSPAN3,LPCAT1,LAMA5,XRCC5,PJA2,MSN,K
.14,HK1,RHOA,PTCD3,CBX5,ATF4,NR2F1,GLIPR1L2,RPS14,CXADR,TMEM259,RPS20,ATP2A2,RPI
.14,HK1,RHOA,PTCD3,CBX5,ATF4,NR2F1,GLIPR1L2,RPS14,CXADR,TMEM259,RPS20,ATP2A2,RPI
ARHGDIA,CAPRIN1,STIP1,TOP2B,SLC35E2B,SMCHD1,MMP2,MAP1A,CDKN1A,FAM199X,HS6ST1,
STYK,GSTA4,NECTIN2,CMTR1,PPP2R5B,TRAPPC1,NOLC1,MSI2,CCNL1,MAPK8IP3,PAF1,RAP1B,(
)1,SSBP2,ZNF43,CHD3,BASP1,HMGB1,POLR2A,SEC61A1,RPS4X,STMN1,TUBB4B,UCHL1,MORF4
PXM1,HSDL1,TEAD1,HS2ST1,UBE2L3,CELSR1,ABL1,ZZEF1,LUC7L2,H6PD,ATP13A3,CLIC1,IGF1R
16A1,RBM14,UNC13A,TSPAN13,CTDNEP1,QRICH1,RPL36AL,TFDP1,CHPF2,BRPF3,SLC25A39,PR
P,MARCKSL1,TRIM28,MATR3,YWHAG,CKB,COL4A2,RPL37A,NONO,PSMD5,YWHAQ,EIF1,SET,EG
F2F1,GOLGA8A,SMARCA5,RPL23,LDHA,CPE,CNN3,RPSA,CTSD,SCARB2,NACA,RTN4,LPCAT1,LAN
PL23,RPSA,SCARB2,NACA,RTN4,RPS24,CIPC,RPL24,RPL17,COL1A2,RPL14,HK1,RHOA,PTCD3,CE

SP,AP1M1,NDUFA13,C16ORF58,TOMM70,XPO6,DAPK3,TRO,LYPLA2,ZER1,SLC38A10,ATP13A1,P
3,RPL23A,GLUL,NNAT,CRPPA,TIMP3,DDX39B,TOMM20,RNF187,FDFT1,RPN2,TBCD,MLXIP,CPSF

EF1A2,MLXIP,SNX22,VAMP2,TSR1,LMNB2,PRDX5,ACTR3,PKN1,VPS35,COL2A1,MLEC,SURF4,TM

ISN,RPL24,KPNB1,RPL17,COL1A2,LLGL1,RPL14,NCS1,RHOA,ZNF865,RPL9,CLSTN1,NR2F1,GLII
DP2,GLOD4,MTURN,GOLPH3,DHX36,HM13,COLGALT1,TSPAN7,NUP98,NACAD,OXSR1,EMC10,T
,EP300,ALDH3A2,CSNK2A1,PHB2,SLC39A7,RAB2A,RER1,TRA2B,ZFR,YKT6,ANP32B,PREPL,TIMM
11,ANXA2,ZMIZ1,TIMP2,MIDN,RPS17,AP3D1,DNAJA1,ARF1,CYB5R3,AKAP12,ITM2B,ZNF770,PXDN
2A2,CRMP1,LASP1,BEX3,RBBP4,RTN3,ASTN2,XPR1,KHDRBS1,XPO1,HDLBP,RPS25,SFPQ,TPM1,E
IM40,COX5B
3,COL1A2,RPL14,HK1,RHOA,PTCD3,CBX5,ATF4,NR2F1,GLIPR1L2,RPS14,CXADR,TMEM259,RPS
MP5,ZZEF1,AASDHPPT,ESYT1,GTF3C1,TNFAIP1,FAM50A,CD2BP2,CNDP2,GLOD4,LMF2,MTURN,
4A,TBC1D16,AHCYL1,ETNK1,KLC1,CCT3,CLSTN3,DDX42,RAB11B,NRDC,PRPF6,TUBB2A,UBXN4
SF7,RAD50,JRK,DUS1L,TOMM40,ALYREF,COX5B
CDC124
F3H,SLC6A8,ELAVL1,IMPDH2,GABARAPL1,GNL1,TM9SF4,TRAP1,PLEKHB2,ELP1,ACTR1B,MOAP1
H,PDLIM1,FEZ1,RALGDS,GALNT10,NTN1,LMF2,NCKAP5L,PITRM1,RHEB,UBE2J1,TMEM98,ECHS
MTMR4,CAP1,PRPF19,COPZ1,EIF3G,DHX38,AP1M1,SSR2,NDUFA13,UBE2I,UBE2D3,TOMM70,DAF
LN1,CSNK1D,IFITM3,DGKD,SREBF2,RPL39,CYP51A1,MLF2,SEC23A,AKIRIN1,NAP1L4,AMOTL2,Y

2,GLOD4,MTURN,GOLPH3,ECHS1,DHX36,HM13,COLGALT1,TSPAN7,NUP98,NACAD,OXSR1,EMC
TNND1,TRAF4,STAT3,ARIH1,SF1,GPX4,SLC35F5,CSNK1G2,ATRX,RAB2A,UBAP2L,UPF1,UBFD1,M
.5,SLC25A36,XRN2,SCAMP5,ZZEF1,AASDHPPT,ESYT1,GTF3C1,TNFAIP1,FAM50A,CD2BP2,CNDP2

SYT1,GTF3C1,TNFAIP1,FAM50A,CD2BP2,CNDP2,GLOD4,MTURN,GOLPH3,ECHS1,DHX36,HM13,
3H4

,SLC30A9,SLC25A36,XRN2,SCAMP5,ZZEF1,AASDHPPT,ESYT1,GTF3C1,TNFAIP1,FAM50A,CD2BP2
NL2,SRSF1,ATP5PB,EIF3E,GABARAP,SUPT5H,ATXN10,CAND1,CLIP3,MAP2K2,RPL28,ZFAND5,DID
PP1R15A,GALNT11,DAP,SPG7,BAZ2A,VAPB,EIF3H,ELAVL1,IMPDH2,GABARAPL1,GNL1,TM9SF4,T
NPA3,PKN1,ACAP3,DDX42,ORAI2,SNRNP70,H3-3A,PRPF6,TOP1,MVB12B,ATG2B,AZIN1,MAPRE1,
NPA3,PKN1,ACAP3,DDX42,ORAI2,SNRNP70,H3-3A,PRPF6,TOP1,MVB12B,ATG2B,AZIN1,MAPRE1,
,TAGLN2,MASTL,HDAC6,PPP1CC,PTPRF,IL6ST,SEPTIN7,KLHDC3,DKK3,GALNT11,GSE1,MTCH1,
,CCDC85C,ANKRD50,SETX,DIRAS1,SLC12A7,SH3BP5,PRKAR2B,ITPK1,NEO1,SGPL1,GPRIN1,CH
C28,MAPT,FN1,NONO,PSMD5,YWHAQ,HACD3,DCHS1,MAP9,TMSB10,RPL19,PPP2R1A,GTF3C4,L

,CIRBP,NARS1,PTGES3,SND1,LAMP1,MAP2,CALU,ARCN1,RPS27,RAC1,ARHGDIA,STIP1,SLC35E2
PH3,DHX36,HM13,COLGALT1,TSPAN7,NUP98,NACAD,OXSR1,EMC10,TMEM9,SCG2,WDR82,SYN
H2,GABARAPL1,GNL1,TM9SF4,TRAP1,PLEKHB2,ELP1,MOAP1,CLASP2,RBM12,NSD2,AP2A2,YY1
AS2,ZDHHC5,SRP54,PMPCA,UBXN1,RNF4,LZTS2,PCED1A,SUDS3,GABARAPL2,CNOT3,TRAPPC1,

,XPR1,KHDRBS1,KLHL9,PFKL,HDLBP,RPS25,SFPQ,CDH2,PNMA1,GPC3,SULF1,RAB7A,ACTN4,M
RPF3,FH,ZDHHC17,NOLC1,NUP58,NF2,GPRC5B,CDK5RAP3,PRDX4,AP3M1,NOP58,ATP6V1D
B27A,SUGP2,GAS7,PABPN1,VCAN,TSPYL4,TM9SF2,C4ORF3,YEATS2,PLOD1,AHCYL1,NCKAP1,PT

OTCH2,PPP2CA,SMARCA4,AZIN1,DDAH2,PRDX1,NYNRIN,TCF25,CAPZB,PRKDC,TDRD5,HSPB1,C
15A,FKBP1A,MAP1LC3B,MTCH1,CARM1,GPC2,ARHGEF17,RAB14,RDX,LRATD2,PRR12,SLC4A2,

1,H6PD,KLF13,CLIC1,CDV3,PFKP,IGF1R,SLC38A1,EWSR1,RNF157,KCTD20,HTRA1,EFNB1,ASNS
LXIP,PPP1CB,TSPYL4,HSPH1,ALKBH5,HNRNPA3,PDXK,UBE4A,PKN1,TBC1D16,ACO2,AHCYL1,KL
2,CCT5,CDC42,YME1L1,EIF2AK1,RETREG2,PDAP1,GTPBP1,CSNK2A1,SEC16A,PHB2,ZFP36L1,K

2B,ACSL6,PRDX2,ARL4C,CLTC,GNB2,RCN1,RPS5,APLP1,IK,RPS19,RPL35,CD63,B4GALNT4,APL
KBH5,HNRNPA3,PDXK,UBE4A,PKN1,TBC1D16,ACO2,AHCYL1,KLC1,CCT3,NCKAP1,CLSTN3,DDX4
COX6A1,SART1,DHX38,NASP,AP1M1,C16ORF58,PSMC1,TOMM70,PRDX3,DEK,CYB5B,GAK,SLC3

ES3,CKAP5,LAMP1,RHOB,HNRNPH1,RPS12,ZNF454,ARHGDIA,SMCHD1,TMEM30A,SKP1,DDX39
SYT1,GTF3C1,TNFAIP1,FAM50A,CD2BP2,CNDP2,GLOD4,MTURN,GOLPH3,ECHS1,DHX36,HM13,
43,TXN,MBOAT7,MYL12A,NUMBL,PRMT2,GUK1,NECTIN2,TMEM50A,DYNLT1,SPIN1,GYS1,PTPN
T1,YY1,CCDC47,ELOVL5,SRPRA,SLC30A9,SLC25A36,XRN2,SCAMP5,ZZEF1,AASDHPPT,ESYT1,G
TF3C1,TNFAIP1,COPE,FAM50A,CD2BP2,CNDP2,GLOD4,MTURN,GOLPH3,ECHS1,DHX36,HM13,C

KLHL42,DIP2B,CD164,UBAP2L,RER1,PI4KA,RAB3GAP2,SMC3,CBX6,MAPRE2,TIMM50,APEX1,RA

9,STMN2,RPS8,MYH9,RPL7,SEPTIN2,P4HB,APP,IDS,ERVV-1,RPL5,GDI1,FSCN1,TMBIM6,PTMS,SF
DI1,FSCN1,TMBIM6,PTMS,SPTAN1,GSTP1,MARCKS,SMG1,CALM1,NPM1,PLXNB2,PLD3,HSPD1,US
CN1,TMBIM6,PTMS,SPTAN1,GSTP1,MARCKS,SMG1,CALM1,NPM1,PLXNB2,PLD3,HSPD1,USP11,C
N1,GSTP1,MARCKS,SMG1,CALM1,NPM1,PLXNB2,PLD3,HSPD1,USP11,COL18A1,SNRNP200,CSDE
FN1,SPEN,NONO,DDB1,PSMD5,YWHAQ,EIF1,HACD3,DCHS1,SET,EGLN1,TPI1,CHSY1,TMSB10,IN
BIM6,PTMS,SPTAN1,GSTP1,MARCKS,SMG1,CALM1,NPM1,PLXNB2,PLD3,HSPD1,USP11,COL18A
,P4HB,APP,IDS,RPL5,GDI1,FSCN1,TMBIM6,PTMS,SPTAN1,GSTP1,MARCKS,SMG1,CALM1,NPM1,
F4L1,NCAM1,AGRN,ZMIZ1,TIMP2,SARAF,RPL10,MARCHF6,AP3D1,SBF1,ARF1,CYB5R3,MAP4,SF
E1,IGF2,EIF4B,VCP,MARCKSL1,UBB,TRIM28,MATR3,YWHAG,CKB,COL4A2,RPL37A,FN1,SPEN,NO
MG1,CALM1,NPM1,PLXNB2,PLD3,HSPD1,USP11,COL18A1,SNRNP200,CSDE1,SMIM14,IGF2,EIF3
STP1,MARCKS,SMG1,CALM1,PLXNB2,PLD3,HSPD1,USP11,COL18A1,SNRNP200,CSDE1,SMIM1
3,YWHAG,CKB,RPL37A,LRRC28,FN1,SPEN,NONO,DDB1,YWHAQ,EIF1,HACD3,DCHS1,SET,EGLN1
CKSL1,AFF4,TRIM28,MATR3,YWHAG,CKB,COL4A2,RPL37A,LRRC28,FN1,SPEN,NONO,DDB1,YWH
C,ATP5F1A,EIF4B,MARCKSL1,AFF4,TRIM28,MATR3,YWHAG,CKB,COL4A2,RPL37A,LRRC28,FN1,
MATR3,YWHAG,CKB,COL4A2,RPL37A,LRRC28,FN1,SPEN,NONO,DDB1,PSMD5,YWHAQ,EIF1,HAC
N1,TMBIM6,PTMS,SPTAN1,GSTP1,MARCKS,SMG1,CALM1,NPM1,PLXNB2,PLD3,HSPD1,USP11,C
EIF3C,ATP5F1A,EIF4B,MARCKSL1,AFF4,TRIM28,MATR3,YWHAG,CKB,COL4A2,RPL37A,LRRC28,
3,YWHAG,CKB,RPL37A,LRRC28,FN1,NONO,DDB1,PSMD5,YWHAQ,EIF1,HACD3,DCHS1,SET,EGL
A,HMGB1,POLR2A,CDKL1,AP2M1,TPT1,RPL18,STMN1,ITGB1,TUBB4B,UCHL1,MORF4L1,NCAM
IGF2,EIF3C,ATP5F1A,EIF4B,MARCKSL1,AFF4,TRIM28,MATR3,YWHAG,CKB,RPL37A,LRRC28,FN1
R3,YWHAG,CKB,COL4A2,RPL37A,LRRC28,FN1,SPEN,DDB1,YWHAQ,EIF1,HACD3,DCHS1,SET,EGL
1,PLD3,HSPD1,USP11,COL18A1,CSDE1,SMIM14,IGF2,ATP5F1A,EIF4B,VCP,MARCKSL1,UBB,AFF
3,YWHAG,CKB,COL4A2,RPL37A,LRRC28,FN1,SPEN,DDB1,EIF1,HACD3,DCHS1,SET,EGLN1,MAP
CALM3,GNB1,SHISA9,STMN2,RPS8,MYH9,RPL7,SEPTIN2,P4HB,APP,IDS,ERVV-1,RPL5,GDI1,FSC
4A2,LRRC28,FN1,SPEN,PSMD5,YWHAQ,EIF1,HACD3,DCHS1,SET,EGLN1,TPI1,CHSY1,TMSB10,EIF
MG1,CALM1,NPM1,PLXNB2,PLD3,HSPD1,USP11,COL18A1,SNRNP200,CSDE1,IGF2,ATP5F1A,EIF4
,NONO,DDB1,YWHAQ,EIF1,HACD3,DCHS1,SET,EGLN1,MAP9,TPI1,CHSY1,TMSB10,INA,EIF4A1,M
1,NPM1,PLXNB2,PLD3,HSPD1,USP11,COL18A1,SNRNP200,CSDE1,SMIM14,IGF2,EIF3C,ATP5F1
,RPS8,MYH9,RPL7,SEPTIN2,P4HB,APP,IDS,ERVV-1,RPL5,GDI1,FSCN1,TMBIM6,PTMS,SPTAN1,G
A9,STMN2,RPS8,MYH9,RPL7,SEPTIN2,P4HB,APP,IDS,RPL5,GDI1,FSCN1,TMBIM6,PTMS,SPTAN1,
DI1,FSCN1,TMBIM6,PTMS,SPTAN1,GSTP1,SMG1,CALM1,NPM1,PLXNB2,HSPD1,USP11,COL18A1,
3,YWHAG,CKB,COL4A2,RPL37A,LRRC28,FN1,NONO,DDB1,YWHAQ,HACD3,DCHS1,SET,EGLN1,T
SL1,AFF4,TRIM28,MATR3,YWHAG,CKB,COL4A2,RPL37A,LRRC28,FN1,SPEN,NONO,DDB1,YWHA
EPTIN2,P4HB,APP,IDS,ERVV-1,RPL5,GDI1,FSCN1,TMBIM6,PTMS,SPTAN1,GSTP1,MARCKS,SMG1
M3,GNB1,SHISA9,STMN2,RPS8,MYH9,RPL7,SEPTIN2,P4HB,APP,IDS,GDI1,FSCN1,TMBIM6,PTMS,
HSY1,TMSB10,INA,MRC2,RPL19,PPP2R1A,LDHB,PKD1,HLA-C,PDIA3,SSBP2,BASP1,VAPA,HMGB1
LM1,NPM1,PLXNB2,PLD3,HSPD1,USP11,COL18A1,SNRNP200,CSDE1,SMIM14,IGF2,EIF3C,EIF4E
CP,MARCKSL1,UBB,TRIM28,MATR3,YWHAG,CKB,COL4A2,SPEN,NONO,DDB1,PSMD5,YWHAQ,EI
ACD3,DCHS1,SET,EGLN1,CHSY1,TMSB10,INA,MRC2,RPL19,GTF3C4,LDHB,PKD1,HLA-C,PDIA3,S
1,PLXNB2,HSPD1,USP11,COL18A1,SNRNP200,CSDE1,SMIM14,IGF2,EIF3C,EIF4B,VCP,MARCKS
,APP,IDS,RPL5,GDI1,FSCN1,TMBIM6,PTMS,SPTAN1,GSTP1,MARCKS,SMG1,CALM1,NPM1,PLXNE
PS8,MYH9,RPL7,SEPTIN2,P4HB,APP,IDS,RPL5,GDI1,FSCN1,TMBIM6,PTMS,SPTAN1,GSTP1,MAR
LRRC28,FN1,SPEN,NONO,DDB1,PSMD5,YWHAQ,EIF1,HACD3,DCHS1,SET,EGLN1,TPI1,CHSY1,TM
11,CSDE1,SMIM14,IGF2,EIF3C,ATP5F1A,EIF4B,MARCKSL1,UBB,AFF4,TRIM28,MATR3,YWHAG,C
1,GSTP1,MARCKS,SMG1,CALM1,NPM1,PLXNB2,PLD3,HSPD1,USP11,SNRNP200,CSDE1,IGF2,EI

,FSCN1,TMBIM6,PTMS,SPTAN1,GSTP1,MARCKS,SMG1,CALM1,NPM1,PLXNB2,PLD3,HSPD1,USP
RCKSL1,UBB,AFF4,TRIM28,MATR3,YWHAG,CKB,COL4A2,RPL37A,LRRC28,FN1,SPEN,NONO,DDE
CAM1,ANXA2,AGRN,ZMIZ1,TIMP2,SARAF,MIDN,MARCHF6,VCL,AP3D1,SBF1,DNAJA1,CYB5R3,MAF
VV-1,GDI1,FSCN1,PTMS,SPTAN1,GSTP1,MARCKS,SMG1,CALM1,NPM1,PLXNB2,PLD3,HSPD1,USI
A9,STMN2,RPS8,MYH9,RPL7,SEPTIN2,P4HB,APP,RPL5,GDI1,FSCN1,TMBIM6,PTMS,SPTAN1,GSTF
A1,MRC2,RPL19,PPP2R1A,RPL15,LDHB,PKD1,HLA-C,PDIA3,SSBP2,VAPA,CDKL1,AP2M1,TPT1,RPI
PL19,LDHB,PKD1,HLA-C,PDIA3,SSBP2,BASP1,VAPA,HMGB1,POLR2A,TPT1,SEC61A1,RPS3A,STMI
PD1,USP11,COL18A1,CSDE1,SMIM14,IGF2,EIF3C,ATP5F1A,EIF4B,VCP,MARCKSL1,UBB,AFF4,TRI
AP4,CDC42BPB,ITM2B,MAT2A,TMEM170A,GTF2F1,GOLGA8A,PXDN,HNRNPU,NFE2L1,SMARCA5
D3,HSPD1,USP11,COL18A1,SNRNP200,SMIM14,IGF2,EIF3C,EIF4B,VCP,MARCKSL1,UBB,AFF4,TF
AT2A,GOLGA8A,PXDN,HNRNPU,NFE2L1,CTNNA1,FADS2,CNN3,CTSD,SCARB2,NACA,LPCAT1,LAM
RF1,CYB5R3,MAP4,SF3B2,CDC42BPB,ITM2B,MAT2A,SLC4A8,GOLGA8A,PXDN,HNRNPU,PGAM1,
,UCHL1,MORF4L1,BTF3,NCAM1,AGRN,ZMIZ1,TIMP2,XAF1,SARAF,COL11A1,MIDN,RPL10,MARC
UBB4B,UCHL1,NCAM1,ANXA2,AGRN,TIMP2,XAF1,SARAF,MIDN,RPL10,MARCHF6,WSB1,VCL,AP3
L18A1,SNRNP200,CSDE1,IGF2,EIF3C,EIF4B,VCP,MARCKSL1,UBB,AFF4,TRIM28,MATR3,YWHAG
2,RPS16,CPE,CNN3,TNS3,CTSD,EIF4H,NACA,TSPAN3,SLC22A17,BRD3,PJA2,KPNB1,C1ORF43,C
TAN1,GSTP1,MARCKS,SMG1,CALM1,NPM1,PLXNB2,PLD3,HSPD1,USP11,COL18A1,SNRNP200,C
A3,VAPA,HMGB1,POLR2A,AP2M1,TPT1,SEC61A1,RPS3A,STMN1,ITGB1,TUBB4B,UCHL1,NCAM1,
J,NFE2L1,HNRNPA0,CTNNA1,FADS2,RPS16,CNN3,CTSD,LPCAT1,LAMA5,BRD3,CIPC,PJA2,KPNB1
,HACD3,DCHS1,SET,EGLN1,TMSB10,INA,EIF4A1,MRC2,RPL19,GTF3C4,LDHB,PKD1,HLA-C,PDIA
HSPD1,USP11,COL18A1,SNRNP200,SMIM14,IGF2,EIF3C,ATP5F1A,EIF4B,VCP,MARCKSL1,UBB,AI
TM2B,MAT2A,TMEM170A,GTF2F1,PXDN,HNRNPU,NFE2L1,SMARCA5,HNRNPA0,CTNNA1,RPS16,C
USP11,COL18A1,SNRNP200,SMIM14,IGF2,ATP5F1A,VCP,MARCKSL1,AFF4,TRIM28,MATR3,YWH
GRN,ZMIZ1,TIMP2,XAF1,COL11A1,MIDN,RPL10,MARCHF6,WSB1,RPS17,VCL,AP3D1,SBF1,DNAJ
CYB5R3,MAP4,CDC42BPB,ITM2B,MAT2A,PXDN,HNRNPU,NFE2L1,HNRNPA0,CTNNA1,RPS16,CNN
SSBP2,VAPA,POLR2A,AP2M1,TPT1,SEC61A1,RPL18,STMN1,ITGB1,TUBB4B,RPL11,MORF4L1,NC
M28,MATR3,YWHAG,CKB,LRRC28,FN1,SPEN,NONO,YWHAQ,EIF1,HACD3,DCHS1,SET,TPI1,CHSY
1,MORF4L1,NCAM1,AGRN,TIMP2,COL11A1,MIDN,MARCHF6,VCL,AP3D1,DNAJA1,ARF1,CYB5R3,
SHISA9,STMN2,RPS8,MYH9,RPL7,SEPTIN2,P4HB,APP,IDS,RPL5,KLRD1,GDI1,FSCN1,TMBIM6,PT
GF2,EIF4B,VCP,MARCKSL1,UBB,AFF4,TRIM28,MATR3,YWHAG,CKB,COL4A2,RPL37A,LRRC28,M
SCN1,TMBIM6,PTMS,SPTAN1,GSTP1,MARCKS,SMG1,CALM1,NPM1,PLXNB2,PLD3,HSPD1,USP11,
B1,TUBB4B,UCHL1,MORF4L1,BTF3,NCAM1,AGRN,ZMIZ1,TIMP2,XAF1,SARAF,COL11A1,MIDN,RF
PLXNB2,PLD3,HSPD1,USP11,CSDE1,SMIM14,IGF2,EIF3C,ATP5F1A,EIF4B,VCP,MARCKSL1,UBB,A
G1,CALM1,NPM1,PLXNB2,PLD3,HSPD1,USP11,COL18A1,SNRNP200,CSDE1,SMIM14,IGF2,EIF3C,
ONO,DDB1,YWHAQ,HACD3,DCHS1,SET,EGLN1,TPI1,CHSY1,TMSB10,MRC2,PPP2R1A,LDHB,PKD
-C,SF3B1,SSBP2,CHD3,BASP1,VAPA,HMGB1,POLR2A,AP2M1,TPT1,RPL18,STMN1,ITGB1,UCHL1,
,GTF2F1,PXDN,HNRNPU,NFE2L1,SMARCA5,HNRNPA0,CTNNA1,RPS16,CNN3,CTSD,EIF4H,LPCAT
HISA9,STMN2,RPS8,MYH9,RPL7,SEPTIN2,P4HB,APP,IDS,RPL5,KLRD1,GDI1,FSCN1,TMBIM6,PTMS
SEC61A1,RPL18,ITGB1,TUBB4B,UCHL1,MORF4L1,AGRN,TIMP2,COL11A1,MIDN,MARCHF6,WSE
HLA-C,PDIA3,VAPA,HMGB1,POLR2A,AP2M1,TPT1,SEC61A1,RPS3A,STMN1,ITGB1,TUBB4B,UCHL
HLA-C,PDIA3,VAPA,HMGB1,POLR2A,AP2M1,TPT1,SEC61A1,RPS3A,STMN1,ITGB1,TUBB4B,UCHL
,MIDN,RPL10,MARCHF6,WSB1,RPS17,VCL,SBF1,DNAJA1,CYB5R3,AKAP12,MAP4,SF3B2,CDC42
ATP5F1B,CALM3,GNB1,SHISA9,STMN2,RPS8,MYH9,RPL7,SEPTIN2,P4HB,APP,IDS,RPL5,GDI1,FS
3P2,ZNF43,VAPA,CDKL1,AP2M1,SEC61A1,STMN1,ITGB1,TUBB4B,UCHL1,MORF4L1,NCAM1,ANX
A,CDKL1,AP2M1,TPT1,SEC61A1,RPS4X,RPS3A,ITGB1,TUBB4B,UCHL1,MORF4L1,AGRN,RPLP1,T

LDHB,DNAH8,PKD1,HLA-C,PDIA3,SERINC1,VAPA,HMGB1,POLR2A,AP2M1,TPT1,SEC61A1,RPS3A,
ICAM1,ANXA2,AGRN,TIMP2,XAF1,SARAF,MIDN,AP3D1,SBF1,ARF1,CYB5R3,MAP4,CDC42BPB,ITM
AGRN,RPLP1,ZMIZ1,TIMP2,COL11A1,MIDN,MARCHF6,VCL,AP3D1,DNAJA1,ARF1,CYB5R3,CDC42
AP12,MAP4,SF3B2,CDC42BPB,ITM2B,AP2B1,MAT2A,GOLGA8A,PXDN,OGA,NFE2L1,HNRNPA0,CT
HACD3,DCHS1,SET,MAP9,TPI1,TMSB10,INA,EIF4A1,MRC2,RPL19,LDHB,PKD1,HLA-C,PDIA3,SERI
LM3,GNB1,SHISA9,STMN2,RPS8,MYH9,RPL7,SEPTIN2,P4HB,APP,IDS,ERVV-1,RPL5,GDI1,FSCN1,T
,RHOA,ZNF865,KIF5B,SLC7A5,CXADR,CHD4,ATP1A1,JUND,EIF5,CD81,WDR1,HMGN2,ATP2A2,CI
P2M1,TPT1,SEC61A1,RPS3A,RPL18,STMN1,ITGB1,TUBB4B,UCHL1,MORF4L1,NCAM1,ANXA2,AG
RPS8,MYH9,RPL7,SEPTIN2,P4HB,APP,IDS,RPL5,GDI1,FSCN1,TMBIM6,PTMS,SPTAN1,GSTP1,MAI
GTF3C4,LDHB,PKD1,PDIA3,SSBP2,CHD3,BASP1,VAPA,POLR2A,CDKL1,TPT1,RPL18,STMN1,ITGB
WHAG,CKB,COL4A2,LRRC28,MAPT,FN1,SPEN,PSMD5,YWHAQ,EIF1,HACD3,DCHS1,SET,EGLN1,TI
LR2A,AP2M1,TPT1,SEC61A1,RPS3A,RPL18,STMN1,ITGB1,TUBB4B,UCHL1,MORF4L1,NCAM1,ANX
AJA1,ARF1,CYB5R3,MAP4,CDC42BPB,ITM2B,AP2B1,GTF2F1,GOLGA8A,PXDN,HNRNPU,PGAM1,N
31,TUBB4B,UCHL1,MORF4L1,NCAM1,AGRN,RPLP1,ZMIZ1,TIMP2,COL11A1,MIDN,MARCHF6,VCL
ISA9,STMN2,RPS8,MYH9,RPL7,SEPTIN2,P4HB,APP,IDS,RPL5,GDI1,FSCN1,TMBIM6,PTMS,SPTAN
COL11A1,MIDN,MARCHF6,RPS17,VCL,AP3D1,SBF1,DNAJA1,ARF1,CYB5R3,AKAP12,MAP4,SF3B2
2,CRMP1,LASP1,OAZ1,RBBP4,PFN1,MCL1,TLE5,NISCH,HNRNPC,CCNI,XPR1,PFKL,HDLBP,TKT,EI
G,COL4A2,RPL37A,LRRC28,FN1,SPEN,NONO,DDB1,YWHAQ,EIF1,HACD3,TMSB10,INA,MRC2,RPL
F4L1,BTF3,AGRN,RPLP1,ZMIZ1,TIMP2,MARCHF6,WSB1,RPS17,VCL,AP3D1,SBF1,DNAJA1,CYB5F
RNPU,NFE2L1,SMARCA5,HNRNPA0,CTNNA1,CPE,CNN3,RPL29,TNS3,CTSD,SCARB2,NACA,LAMA5
PS3A,STMN1,ITGB1,TUBB4B,UCHL1,MORF4L1,ANXA2,AGRN,RPLP1,ZMIZ1,TIMP2,SARAF,COL11
IA,CPE,CNN3,RPSA,TPM4,EIF4H,LPCAT1,BRD3,RPS9,KPNB1,RPL17,C1ORF43,IGFBP3,LLGL1,NC
2A2,CRMP1,LASP1,BEX3,VAT1,OAZ1,SRSF4,PFN1,ASTN2,MCL1,TLE5,NISCH,HNRNPC,CCNI,TADA
K1E,PFKL,TKT,GLG1,TPM1,PDIA6,PIP4K2B,MEGF8,SOX12,RAB7A,PRDX2,ARL4C,GNB2,H1-0,RPS
ONO,DDB1,YWHAQ,EIF1,HACD3,DCHS1,SET,EGLN1,TPI1,CHSY1,TMSB10,INA,MRC2,RPL19,PPP2
XPR1,XPO1,SPTBN2,CSNK1E,PFKL,HDLBP,SFPQ,EIF5A,CDH2,PNMA1,PDIA6,GPC3,MEGF8,SOX1
,SET,EGLN1,TPI1,CHSY1,TMSB10,INA,EIF4A1,MRC2,PPP2R1A,GTF3C4,LDHB,PKD1,HLA-C,SSBP
G,CKB,COL4A2,LRRC28,FN1,SPEN,NONO,PSMD5,YWHAQ,EIF1,HACD3,DCHS1,SET,EGLN1,MAP9
,INA,EIF4A1,MRC2,RPL19,RPL15,LDHB,PKD1,HLA-C,SF3B1,PDIA3,SERINC1,SSBP2,ZNF43,CHD3,
.4A2,SPEN,DDB1,PSMD5,YWHAQ,EIF1,HACD3,DCHS1,SET,EGLN1,CHSY1,INA,EIF4A1,RPL19,PPP2
SA9,STMN2,RPS8,MYH9,RPL7,SEPTIN2,P4HB,APP,IDS,RPL5,KLRD1,GDI1,FSCN1,TMBIM6,PTMS,
1A1,STMN1,ITGB1,TUBB4B,MORF4L1,BTF3,NCAM1,ANXA2,AGRN,ZMIZ1,TIMP2,SARAF,COL11A
/DR1,HMGN2,ATP2A2,CRMP1,LASP1,BEX3,VAT1,OAZ1,RTN3,JPT2,TLE5,NISCH,CCNI,SLC3A2,HD
IIDN,VCL,AP3D1,SBF1,ARF1,CYB5R3,MAP4,SF3B2,CDC42BPB,ITM2B,MAT2A,GOLGA8A,PXDN,HI
YWHAG,CKB,COL4A2,RPL37A,SPEN,NONO,YWHAQ,EIF1,HACD3,DCHS1,SET,EGLN1,TPI1,CHSY1,
OL18A1,SNRNP200,CSDE1,SMIM14,IGF2,EIF3C,EIF4B,VCP,MARCKSL1,UBB,AFF4,TRIM28,MATI
N1,RPS5,APLP1,RPL35,MIF,GPAT4,MYO5A,CD63,NGRN,GOLM1,B4GALNT4,MTSS2,HNRNPDL,T
D,EIF4H,NACA,TSPAN3,BRD3,PJA2,KPNB1,C1ORF43,PAPOLA,IGFBP3,LLGL1,NCS1,CLSTN1,NR2I
K2B,MEGF8,SOX12,ACTN4,DDX3X,ARL4C,H1-0,APLP1,MIF,CHPF,B4GALNT4,DNAJC5,APLP2,TAF
DKL1,TPT1,STMN1,TUBB4B,UCHL1,MORF4L1,NCAM1,AGRN,RPLP1,ZMIZ1,TIMP2,COL11A1,MID
F1,CYB5R3,MAP4,SF3B2,CDC42BPB,ITM2B,GTF2F1,GOLGA8A,PXDN,OGA,PGAM1,CTNNA1,LDH
FADS2,CNN3,RPSA,TPM4,CTSD,EIF4H,TSPAN3,LAMA5,BRD3,MSN,KPNB1,C1ORF43,PAPOLA,IGFI
PDIA3,ZNF43,BASP1,VAPA,HMGB1,POLR2A,TPT1,SEC61A1,RPS3A,STMN1,TUBB4B,UCHL1,ANXA
7,VCL,AP3D1,ARF1,CYB5R3,MAP4,CDC42BPB,ITM2B,MAT2A,PXDN,HNRNPU,NFE2L1,CTNNA1,RF
STMN2,RPS8,MYH9,RPL7,SEPTIN2,P4HB,APP,IDS,ERVV-1,RPL5,GDI1,FSCN1,TMBIM6,PTMS,SPTA

F3B1,PDIA3,SSBP2,ZNF43,BASP1,VAPA,HMGB1,AP2M1,TPT1,SEC61A1,RPS3A,STMN1,ITGB1,TU
K1E,PFKL,GLG1,CDH6,SFPQ,EIF5A,PEBP1,CCND2,CDH2,PDIA6,GPC3,PLXNA1,PIP4K2B,MEGF8,S
,LAMA5,BRD3,KPNB1,RPL17,PAPOLA,IGFBP3,COL1A2,LLGL1,HK1,NCS1,RHOA,ZNF865,CLSTN1
F4A1,MRC2,RPL19,RPL15,GTF3C4,LDHB,PKD1,SF3B1,PDIA3,ZNF43,BASP1,VAPA,HMGB1,POLR2
F4,TRIM28,MATR3,YWHAG,CKB,COL4A2,RPL37A,LRRC28,SPEN,NONO,DDB1,PSMD5,YWHAQ,EI
WHAQ,EIF1,HACD3,DCHS1,SET,EGLN1,TPI1,CHSY1,TMSB10,INA,EIF4A1,MRC2,PPP2R1A,LDHB,
,MARCHF6,VCL,AP3D1,ARF1,CYB5R3,AKAP12,CDC42BPB,ITM2B,GOLGA8A,PXDN,HNRNPU,NFE2
DHB,PKD1,SF3B1,PDIA3,SERINC1,SSBP2,CHD3,VAPA,HMGB1,POLR2A,CDKL1,AP2M1,TPT1,RPL
B2,PLD3,HSPD1,USP11,COL18A1,SNRNP200,SMIM14,IGF2,EIF3C,ATP5F1A,EIF4B,MARCKSL1,U
5,GDI1,FSCN1,TMBIM6,PTMS,SPTAN1,GSTP1,MARCKS,SMG1,CALM1,NPM1,PLXNB2,PLD3,HSPD
KPNB1,RPL17,C1ORF43,RPL14,RPL22,RHOA,PTCD3,CBX5,CLSTN1,ATF4,NR2F1,KIF5B,SLC7A5,
CL1,TLE5,NES,CCNI,XPR1,SPTBN2,SLC3A2,PFKL,HDLBP,TKT,SFPQ,PNMA1,PDIA6,GPC3,PIP4K2
,MEGF8,SOX12,ARL4C,APLP1,MIF,GPAT4,CHPF,MYO5A,COL6A2,CD63,SESN3,B4GALNT4,MTSS
1,VAPA,HMGB1,TPT1,STMN1,ITGB1,TUBB4B,BTF3,NCAM1,ANXA2,AGRN,ZMIZ1,TIMP2,XAF1,SA
11,SEC61A1,RPS3A,STMN1,ITGB1,TUBB4B,UCHL1,RPL11,ANXA2,AGRN,ZMIZ1,TIMP2,SARAF,MI
T,EGLN1,TPI1,CHSY1,TMSB10,INA,MRC2,PPP2R1A,RPL15,GTF3C4,LDHB,DNAH8,HLA-C,PDIA3,S
VAT1,OAZ1,SRSF4,RBBP4,PFN1,TLE5,CCNI,XPO1,SLC3A2,PFKL,HDLBP,TKT,CDH6,SFPQ,EIF5A,
PR1L2,CXADR,CHD4,JUND,EIF5,WDR1,ATP2A2,CRMP1,LASP1,BEX3,VAT1,OAZ1,RTN3,PFN1,HNR
N1,MARCKS,SMG1,CALM1,NPM1,PLD3,HSPD1,COL18A1,SNRNP200,CSDE1,IGF2,EIF3C,ATP5F1
EIF4B,VCP,MARCKSL1,UBB,TRIM28,MATR3,YWHAG,CKB,COL4A2,RPL37A,SPEN,DDB1,PSMD5,
00,SMIM14,IGF2,EIF3C,ATP5F1A,EIF4B,VCP,MARCKSL1,UBB,AFF4,TRIM28,MATR3,YWHAG,CK
,TPT1,SEC61A1,RPS3A,RPL18,STMN1,ITGB1,TUBB4B,UCHL1,MORF4L1,BTF3,ANXA2,AGRN,TIM
PPP2R1A,RPL15,GTF3C4,PKD1,PDIA3,SERINC1,SSBP2,RPS3,VAPA,POLR2A,SEC61A1,RPL18,STM
SPTBN1,ATP5F1B,CALM3,GNB1,SHISA9,STMN2,RPS8,MYH9,RPL7,SEPTIN2,P4HB,APP,IDS,RPL5,
TADA2B,KHDRBS1,PFKL,HDLBP,PLXNA1,RAB7A,CCN2,ARL4C,MRFAP1,GNB2,RCN1,RPS5,RPS7,
A,GTF2F1,GOLGA8A,HNRNPU,OGA,CTNNA1,CNN3,TPM4,TNS3,EIF4H,SCARB2,LPCAT1,RPS24,SL
,TADA2B,XPR1,SLC3A2,CSNK1E,HDLBP,RPS25,TKT,GLG1,SFPQ,TPM1,EIF5A,CDH2,PDIA6,GPC3,
M6,SPTAN1,GSTP1,MARCKS,CALM1,NPM1,PLXNB2,PLD3,HSPD1,USP11,SNRNP200,CSDE1,SMIM
2,NACA,TSPAN3,LAMA5,BRD3,CIPC,KPNB1,COL1A2,LLGL1,NCS1,RHOA,ZNF865,CLSTN1,SLC7A
EIF4B,VCP,MARCKSL1,UBB,AFF4,TRIM28,MATR3,YWHAG,CKB,COL4A2,LRRC28,FN1,SPEN,DDB
1-0,APLP1,GPAT4,GJA1,JUN,ANXA6,SESN3,B4GALNT4,APLP2,WLS,HNRNPK,H1-10,DUX4,SEPTIN
MA5,BRD3,MSN,SLC16A7,KPNB1,C1ORF43,LLGL1,HK1,RPL22,ZNF865,ATF4,PSMD2,SLC7A5,C
TUBB4B,RPL11,MORF4L1,NCAM1,ANXA2,TIMP2,MIDN,WSB1,RPS17,VCL,AP3D1,CYB5R3,CDC42
,ARF1,CYB5R3,MAP4,SF3B2,CDC42BPB,ITM2B,AP2B1,GTF2F1,SLC4A8,GOLGA8A,PXDN,OGA,C
L1,UBB,AFF4,TRIM28,MATR3,YWHAG,CKB,COL4A2,RPL37A,LRRC28,SPEN,NONO,DDB1,PSMD5
LP2,TARDBP,ARL6IP1,MXD4,ANKRD52,WLS,HNRNPK,DUX4,CADM4,UBE2H,RAB1A,PLPPR3,ARC
GPC3,PIP4K2B,MEGF8,SOX12,ITM2C,ARL4C,APLP1,IK,MIF,GPAT4,MYO5A,COL6A2,CD63,B4GA
CD3,DCHS1,SET,EGLN1,TMSB10,EIF4A1,MRC2,GTF3C4,LDHB,PKD1,HLA-C,PDIA3,SERINC1,SSB
DN,HNRNPU,NFE2L1,SMARCA5,HNRNPA0,CNN3,TPM4,SCARB2,NACA,LAMA5,SLC22A17,BRD3,
2,CDC42BPB,ITM2B,MAT2A,GOLGA8A,HNRNPU,SMARCA5,HNRNPA0,CTNNA1,RPS16,RPL23,CN
MSB10,EIF4A1,MRC2,RPL19,RPL15,GTF3C4,LDHB,PKD1,SF3B1,PDIA3,SERINC1,SSBP2,ZNF43,
,HECTD4,MAGED4B,CTBP1,CAPRIN1,STIP1,GPC1,TOP2B,SLC35E2B,MMP2,SKP1,L1CAM,TRIB2
PLP1,TIMP2,MIDN,RPS17,VCL,AP3D1,SBF1,DNAJA1,CYB5R3,MAP4,SF3B2,CDC42BPB,ITM2B,AP
PLP1,TIMP2,MIDN,RPS17,VCL,AP3D1,SBF1,DNAJA1,CYB5R3,MAP4,SF3B2,CDC42BPB,ITM2B,AP
PLP1,TIMP2,MIDN,RPS17,VCL,AP3D1,SBF1,DNAJA1,CYB5R3,MAP4,SF3B2,CDC42BPB,ITM2B,AP

),EIF4A1,MRC2,RPL19,RPL15,LDHB,SF3B1,PDIA3,SERINC1,SSBP2,BASP1,VAPA,HMGB1,POLR2A
2,MAP4,CDC42BPB,ITM2B,GTF2F1,PXDN,HNRNPU,PGAM1,NFE2L1,HNRNPA0,RPS16,CNN3,TPM4
EN,DDB1,PSMD5,YWHAQ,EIF1,HACD3,SET,EGLN1,MAP9,TPI1,CHSY1,TMSB10,INA,EIF4A1,RPL19
SLC16A7,KPNB1,PAPOLA,COL1A2,RHOA,ZNF865,CLSTN1,NR2F1,RAB6B,NLGN2,CXADR,CHD4,A
N1,HECTD4,MAGED4B,CAPRIN1,STIP1,GPC1,TOP2B,SLC35E2B,MMP2,TRIB2,PEA15,LUC7L3,SQ
N1,TMBIM6,PTMS,SPTAN1,GSTP1,MARCKS,SMG1,CALM1,NPM1,PLXNB2,HSPD1,USP11,SNRNP2
IACA,LAMA5,RPS24,SLC22A17,BRD3,CIPC,MSN,KPNB1,C1ORF43,IGFBP3,LLGL1,HK1,NCS1,RPL
6,RPS17,VCL,SBF1,DNAJA1,ARF1,CYB5R3,MAP4,CDC42BPB,AP2B1,MAT2A,TMEM170A,PXDN,HN
POLR2A,TPT1,RPS4X,RPL18,STMN1,ITGB1,MORF4L1,ANXA2,AGRN,ZMIZ1,TIMP2,SARAF,MIDN,I
Z1,RBBP4,RTN3,PFN1,TLE5,HNRNPC,CCNI,LRPPRC,XPR1,XPO1,PFKL,HDLBP,TKT,PNMA1,PDIA6,
O5,MMP2,SKP1,DNER,L1CAM,GLUL,PEA15,DLG5,TIMP3,CTDSP2,PGRMC1,CNN2,NOVA2,MDH2,
CKB,COL4A2,MAPT,FN1,SPEN,NONO,DDB1,EIF1,HACD3,DCHS1,SET,EGLN1,MAP9,CHSY1,TMSB
C,EIF4B,VCP,MARCKSL1,UBB,AFF4,TRIM28,YWHAG,COL4A2,RPL37A,FN1,SPEN,DDB1,PSMD5,Y
4C,GNB2,APLP1,CHPF,JUN,MYO5A,B4GALNT4,DNAJC5,APLP2,RPLP2,MXD4,RPL36,SEPTIN11,CA
S3A,RPL18,STMN1,ITGB1,TUBB4B,UCHL1,RPL11,MORF4L1,ANXA2,AGRN,RPLP1,ZMIZ1,TIMP2,S
NONO,DDB1,YWHAQ,EIF1,EGLN1,TPI1,CHSY1,TMSB10,INA,EIF4A1,RPL19,PPP2R1A,RPL15,GTF3
P4HB,APP,IDS,RPL5,GDI1,FSCN1,TMBIM6,PTMS,SPTAN1,GSTP1,MARCKS,SMG1,CALM1,NPM1,P
,EIF5,CD81,HMGN2,ATP2A2,CRMP1,LASP1,VAT1,SRSF4,RTN3,PFN1,ASTN2,JPT2,PNN,SPC25,NES
RCKS,SMG1,CALM1,NPM1,PLD3,HSPD1,USP11,CSDE1,SMIM14,IGF2,EIF3C,EIF4B,VCP,MARCKS
MAT2A,GOLGA8A,HNRNPU,SMARCA5,CTNNA1,RPS16,RPL23,CNN3,TNS3,CTSD,SCARB2,LPCAT1,
E2B,SMCHD1,MMP2,DNER,TRIB2,PEA15,DLG5,PGRMC1,NOVA2,CTSB,CDH11,TM9SF3,MDH2,CH
B,VCP,MARCKSL1,UBB,TRIM28,MATR3,CKB,COL4A2,RPL37A,LRRC28,SPEN,DDB1,PSMD5,YWH
AP2B1,MAT2A,PXDN,OGA,CTNNA1,FADS2,TPM4,TNS3,EIF4H,SCARB2,NACA,LAMA5,SLC22A17,BF
,JPT2,NES,TADA2B,PFKL,HDLBP,TKT,GLG1,PEBP1,PNMA1,PDIA6,GPC3,PIP4K2B,SOX12,ITM2C,
,LPCAT1,LAMA5,SLC22A17,PJA2,SLC16A7,KPNB1,C1ORF43,PAPOLA,COL1A2,LLGL1,GAP43,HK
3,ZNF43,CHD3,VAPA,HMGB1,POLR2A,CDKL1,AP2M1,TPT1,SEC61A1,RPS3A,STMN1,ITGB1,TUBB
2,TLE5,NISCH,HNRNPC,TADA2B,SPTBN2,CSNK1E,PFKL,TKT,TPM1,PEBP1,CCND2,PDIA6,PLXNA1
VHAG,CKB,COL4A2,LRRC28,FN1,SPEN,DDB1,PSMD5,YWHAQ,HACD3,DCHS1,SET,EGLN1,TPI1,CH
H8,PKD1,PDIA3,SSBP2,VAPA,AP2M1,SEC61A1,ITGB1,TUBB4B,UCHL1,RPL11,NCAM1,ANXA2,AGR
F3,AGRN,ZMIZ1,TIMP2,XAF1,MIDN,RPL10,MARCHF6,WSB1,RPS17,VCL,AP3D1,SBF1,ARF1,SF3B2
DN,MARCHF6,VCL,SBF1,DNAJA1,SF3B2,TMEM170A,PXDN,OGA,NFE2L1,HNRNPA0,CTNNA1,FADS
2B,AP2B1,SLC4A8,GOLGA8A,PXDN,OGA,SMARCA5,HNRNPA0,CTNNA1,FADS2,RPS16,CNN3,LPCA
RSF4,ASTN2,TLE5,RPL31,PRKCSH,HNRNPC,CSNK1E,PFKL,TKT,TPM1,CDH2,PIP4K2B,SOX12,ACT
ONO,DDB1,PSMD5,YWHAQ,EIF1,HACD3,SET,MAP9,CHSY1,TMSB10,EIF4A1,MRC2,RPL19,RPL15,
3B4B,UCHL1,MORF4L1,ANXA2,AGRN,TIMP2,COL11A1,MIDN,MARCHF6,VCL,AP3D1,DNAJA1,CYE
N,ALDH6A1,SPTBN1,ATP5F1B,CALM3,GNB1,SHISA9,STMN2,RPS8,MYH9,RPL7,SEPTIN2,P4HB,AF
515,RPL12,RPS7,RPS19,JUN,COL6A2,DNAJC5,ARL6IP1,MXD4,MYL6,SHOX,HNRNPK,MYH10,LAMI
DM4,LAMP1,RHOB,RAB1A,PLPPR3,COTL1,MAGED4B,CTBP1,RAC1,ARHGDIA,GPC1,SMCHD1,HD
.36,CKAP5,SEPTIN11,CADM4,LAMP1,RHOB,RAB1A,PLPPR3,SEPTIN3,COTL1,CTBP1,ARHGDIA,GF
N2,P4HB,APP,RPL5,GDI1,FSCN1,TMBIM6,PTMS,SPTAN1,GSTP1,MARCKS,SMG1,CALM1,NPM1,P
,VCL,AP3D1,SBF1,DNAJA1,MAT2A,TMEM170A,PXDN,HNRNPU,OGA,SMARCA5,CTNNA1,FADS2,LD
KD1,HLA-C,SSBP2,CHD3,VAPA,HMGB1,CDKL1,TPT1,SEC61A1,RPS3A,OR8K1,RPL18,STMN1,ITGI
F6,VCL,AP3D1,DNAJA1,ARF1,SF3B2,ZNF770,MAT2A,GTF2F1,PXDN,HNRNPU,NFE2L1,HNRNPA0,C
,C1ORF43,COL1A2,GAP43,HK1,NCS1,RPL22,PTCD3,ZNF865,CLSTN1,ATF4,PSMD2,SLC7A5,CHD
B,COL4A2,RPL37A,MAPT,FN1,NONO,PSMD5,YWHAQ,EIF1,HACD3,DCHS1,SET,TMSB10,INA,MRC

B,RPL11,MORF4L1,BTF3,NCAM1,ANXA2,RPLP1,ZMIZ1,SARAF,COL11A1,MIDN,RPL10,MARCHF6
EC61A1,RPS3A,STMN1,ITGB1,TUBB4B,RPL11,MORF4L1,BTF3,NCAM1,ANXA2,RPLP1,ZMIZ1,TIM
RD3,KPNB1,C1ORF43,LLGL1,HK1,RPL22,RPL9,CLSTN1,ATF4,NR2F1,SLC7A5,NLGN2,CHD4,ATP
A5,CHD4,ATP1A1,JUND,CD81,LENG8,WDR1,HMGN2,ATP2A2,CRMP1,LASP1,OAZ1,RBBP4,RTN3,F
17,VCL,AP3D1,ARF1,CYB5R3,SF3B2,ZNF770,MAT2A,TMEM170A,GOLGA8A,PGAM1,CTNNA1,FAD
SH,KHDRBS1,XPO1,PFKL,TKT,SFPQ,CDH2,PDIA6,GPC3,PIP4K2B,MEGF8,SOX12,ACTN4,MMP14
LR2A,CDKL1,SEC61A1,RPS3A,STMN1,TUBB4B,UCHL1,MORF4L1,BTF3,ANXA2,RPLP1,ZMIZ1,CC
,SLC7A5,NLGN2,CXADR,CHD4,JUND,CD81,WDR1,HMGN2,ATP2A2,CRMP1,LASP1,CCT7,VAT1,O
C,GNB2,MIF,GPAT4,MYO5A,COL6A2,CD63,B4GALNT4,MTSS2,DNAJC5,APLP2,HNRNPDL,TARDB
C,GNB2,MIF,GPAT4,MYO5A,COL6A2,CD63,B4GALNT4,MTSS2,DNAJC5,APLP2,HNRNPDL,TARDB
H,STMN3,PFKM,LSS,STT3B,ISYNA1,CSNK1D,ID4,ZFAND3,VANGL2,BAG6,HNRNPD,FBRSL1,FLNC,
NAJC5,APLP2,SLC25A3,MXD4,SHOX,H1-10,PTGES3,SEPTIN11,HNRNPH1,CKAP4,ZNF454,MAGED
I1,FSCN1,PTMS,SPTAN1,GSTP1,MARCKS,SMG1,CALM1,NPM1,PLXNB2,PLD3,HSPD1,USP11,COL
10,MARCHF6,WSB1,RPS17,VCL,ARF1,CYB5R3,AKAP12,CDC42BPB,ITM2B,AP2B1,TMEM170A,G
D2,GNB2,RPL35,MIF,GPAT4,CHPF,GOLM1,SESN3,B4GALNT4,DNAJC5,APLP2,TARDBP,MXD4,TA
,HLA-B,PLPPR3,HECTD4,CTBP1,ARHGDIA,STIP1,GPC1,SLC35E2B,SMCHD1,IPO5,ATP6V0C,DNEF
2,B4GALNT4,MTSS2,DNAJC5,APLP2,HNRNPDL,TARDBP,MXD4,MYL6,RPL36,CADM4,HNRNPH1,H
,CCND2,PDIA6,GPC3,DDR1,PLXNA1,PIP4K2B,MEGF8,ACTN4,PRDX2,ARL4C,GNB2,APLP1,GPAT4,
BASP1,POLR2A,CDKL1,AP2M1,TPT1,SEC61A1,RPS4X,RPS3A,STMN1,ITGB1,TUBB4B,RPL11,MO
LP1,ZMIZ1,TIMP2,SARAF,COL11A1,MIDN,RPL10,WSB1,VCL,AP3D1,ARF1,CYB5R3,AKAP12,MAP4
1A,RPL15,GTF3C4,LDHB,DNAH8,PKD1,HLA-C,SF3B1,PDIA3,SERINC1,SSBP2,ZNF43,RPS3,CHD3
3,DCHS1,SET,TPI1,CHSY1,EIF4A1,MRC2,RPL19,RPL15,LDHB,DNAH8,PKD1,SF3B1,SERINC1,SSB
TRIM28,MATR3,CKB,COL4A2,RPL37A,LRRC28,FN1,NONO,DDB1,PSMD5,YWHAQ,EIF1,HACD3,SE
TOP2B,IPO5,ATP6V0C,SKP1,TRIB2,PEA15,LUC7L3,SHISAL1,TIMP3,CTDSP2,CNN2,NOVA2,CTSB,
2,RPS14,CXADR,CHD4,ATP1A1,JUND,EIF5,WDR1,HMGN2,ATP2A2,CRMP1,LASP1,BEX3,OAZ1,SF
SERINC1,SSBP2,CHD3,BASP1,VAPA,HMGB1,POLR2A,AP2M1,TPT1,RPS4X,STMN1,TUBB4B,UCHL
I1,TPT1,STMN1,ITGB1,TUBB4B,MORF4L1,BTF3,ANXA2,AGRN,ZMIZ1,SARAF,MIDN,RPL10,WSB1,
3,RPSA,TPM4,CTSD,EIF4H,TSPAN3,LAMA5,BRD3,RPS9,PJA2,RPL24,KPNB1,C1ORF43,PAPOLA,LL
22,SQSTM1,VAMP2,GAS7,NOX4,PDXK,PKN1,PLOD1,CAMSAP1,GRINA,MLEC,MDK,LAMC1,ORAI2
1,SMARCA5,HNRNPA0,CTNNA1,RPS16,CNN3,RPSA,TNS3,CTSD,EIF4H,TSPAN3,LPCAT1,LAMA5,F
IDS,ERVV-1,GDI1,FSCN1,TMBIM6,PTMS,SPTAN1,GSTP1,MARCKS,SMG1,CALM1,NPM1,PLD3,HS
GALNT4,MTSS2,DNAJC5,APLP2,HNRNPDL,TARDBP,ARL6IP1,MXD4,INIP,RPL36,PTGES3,CADM4,
3,PIP4K2B,MEGF8,SOX12,RPL12,ARL4C,APLP1,MIF,GPAT4,CHPF,MYO5A,CD63,GOLM1,B4GAL
,RCN1,RPS5,APLP1,RPS7,RPL35,MIF,GPAT4,TLN1,COL6A2,NGRN,GOLM1,B4GALNT4,MTSS2,RF
IA6,PLXNA1,MEGF8,SOX12,RAB7A,ACTN4,PRDX2,ARL4C,GNB2,RCN1,H1-0,TLN1,COL6A2,SESN
T1,SEC61A1,RPS4X,RPS3A,OR8K1,RPL18,TUBB4B,RPL11,MORF4L1,BTF3,XAF1,SARAF,RPL10,M
4H,TSPAN3,LPCAT1,BRD3,MSN,KPNB1,IGFBP3,LLGL1,GAP43,NCS1,RPL22,ZNF865,CLSTN1,AT
A2,CRMP1,LASP1,BEX3,OAZ1,SRSF4,PFN1,TLE5,NISCH,HNRNPC,CCNI,TADA2B,XPR1,KHDRBS1,
1,NPM1,PLD3,HSPD1,USP11,SNRNP200,CSDE1,IGF2,EIF3C,ATP5F1A,EIF4B,MARCKSL1,UBB,AFF
VCL,SBF1,DNAJA1,CYB5R3,MAP4,TMEM170A,SLC4A8,PXDN,PGAM1,NFE2L1,SMARCA5,HNRNPA
M1,AGRN,RPLP1,ZMIZ1,TIMP2,SARAF,MIDN,RPL10,MARCHF6,WSB1,RPS17,SBF1,DNAJA1,AKAP
PS4X,RPS3A,STMN1,TUBB4B,MORF4L1,BTF3,NCAM1,ANXA2,RPLP1,ZMIZ1,TIMP2,XAF1,SARAF,
RHGDIA,GPC1,SMCHD1,IPO5,ATP6V0C,MMP2,TRIB2,GLUL,HSBP1,MAP1A,CDKN1A,DLG5,CTDS
L4C,MAGED2,APLP1,MIF,GPAT4,TLN1,MYO5A,CD63,GOLM1,SESN3,B4GALNT4,MTSS2,APLP2,T
AP12,CDC42BPB,ITM2B,AP2B1,PXDN,OGA,NFE2L1,SMARCA5,CTNNA1,FADS2,RPL23,CNN3,TP

APA,HMGB1,AP2M1,TPT1,SEC61A1,RPS3A,RPL18,STMN1,ITGB1,TUBB4B,UCHL1,ANXA2,AGRN,Z
G,CKB,RPL37A,LRRC28,NONO,DDB1,YWHAQ,EIF1,EGLN1,TPI1,CHSY1,TMSB10,INA,EIF4A1,RPL
,DNAJA1,ARF1,CYB5R3,MAP4,SF3B2,CDC42BPB,ZNF770,TMEM170A,GTF2F1,PXDN,PGAM1,NFE
MAP9,TPI1,CHSY1,INA,EIF4A1,RPL19,PPP2R1A,RPL15,GTF3C4,LDHB,DNAH8,PKD1,SF3B1,PDIA3
PDIA3,SERINC1,ZNF43,BASP1,VAPA,HMGB1,POLR2A,CDKL1,TPT1,SEC61A1,RPS3A,STMN1,TUB
,TKT,SFPQ,EIF5A,PDIA6,PIP4K2B,MEGF8,SOX12,RPS15,ARL4C,GNB2,RCN1,APLP1,RPS19,RPL3
RIB2,HS6ST1,TIMP3,CTDSP2,NOVA2,CTSB,CHD6,EEF1A2,TBCD,SNX22,PGD,SQSTM1,VAMP2,GA
D3,PJA2,RPL24,KPNB1,PAPOLA,IGFBP3,LLGL1,RPL14,NCS1,RHOA,ZNF865,RPL9,ATF4,RAB6B,A
,SNRNP200,CSDE1,SMIM14,IGF2,EIF4B,VCP,MARCKSL1,UBB,AFF4,TRIM28,MATR3,CKB,COL4A
HNRNPU,NFE2L1,SMARCA5,HNRNPA0,CTNNA1,FADS2,LDHA,CNN3,RPSA,CTSD,EIF4H,SCARB2,NA
4H,NACA,LAMA5,BRD3,RPS9,RPL24,SLC16A7,KPNB1,PAPOLA,IGFBP3,LLGL1,RPL14,RPL22,CBX
JUND,CD81,COL5A2,LENG8,WDR1,HMGN2,CRMP1,OAZ1,RTN3,ASTN2,TLE5,RPL31,PRKCSH,HN
C1ORF43,PAPOLA,IGFBP3,LLGL1,GAP43,RPL14,RPL22,RHOA,PTCD3,PSMD2,NR2F1,RPS14,CXA
AM1,ANXA2,AGRN,ZMIZ1,TIMP2,SARAF,MIDN,MARCHF6,VCL,AP3D1,DNAJA1,CYB5R3,MAP4,SF3
,EIF1,EGLN1,TPI1,CHSY1,TMSB10,INA,EIF4A1,RPL19,PPP2R1A,RPL15,GTF3C4,LDHB,DNAH8,PK
4K2B,MEGF8,SOX12,MMP14,PRDX2,ARL4C,RCN1,MIF,JUN,COL6A2,CD63,NGRN,MTSS2,DNAJC
/APA,RPL18,STMN1,TUBB4B,UCHL1,RPL11,MORF4L1,BTF3,AGRN,RPLP1,ZMIZ1,SARAF,RPS17,V
TM2B,MAT2A,SMARCA5,HNRNPA0,CTNNA1,FADS2,CNN3,RPSA,TNS3,CTSD,EIF4H,TSPAN3,SLC22
F2,ATP5F1A,EIF4B,MARCKSL1,UBB,AFF4,MATR3,YWHAG,CKB,COL4A2,RPL37A,FN1,SPEN,NON
2F1,GOLGA8A,PXDN,NFE2L1,CTNNA1,CNN3,TPM4,TNS3,EIF4H,SCARB2,RPS24,SLC22A17,CIPC,
S24,BRD3,RPS9,C1ORF43,PAPOLA,IGFBP3,LLGL1,GAP43,HK1,RPL22,ZNF865,RPL9,CLSTN1,AT
6,PTMS,SPTAN1,SMG1,CALM1,NPM1,PLXNB2,PLD3,HSPD1,COL18A1,SNRNP200,CSDE1,SMIM1
RNPH1,HLA-B,PLPPR3,MAGED4B,CTBP1,ARHGDIA,STIP1,GPC1,SLC35E2B,SMCHD1,IPO5,ATP6V
IRNP200,IGF2,EIF3C,ATP5F1A,VCP,MARCKSL1,UBB,AFF4,TRIM28,MATR3,YWHAG,CKB,COL4A2,
RPLP2,NDFIP1,TSR2,ANKRD52,MYH10,NARS1,DUX4,YWHAB,CKAP5,SEPTIN11,RHOB,UBE2H,CK
A1,ARF1,CYB5R3,SF3B2,CDC42BPB,TMEM170A,GTF2F1,SLC4A8,GOLGA8A,HNRNPU,SMARCA5,
A1,ARF1,CYB5R3,SF3B2,CDC42BPB,TMEM170A,GTF2F1,SLC4A8,GOLGA8A,HNRNPU,SMARCA5,
6A7,PAPOLA,HK1,RPL22,RHOA,PTCD3,ZNF865,ATF4,GLIPR1L2,RPS14,TMEM259,RPS20,CD81,
GF8,SOX12,ACTN4,ARL4C,MAGED2,GNB2,APLP1,RPS19,MIF,GPAT4,CHPF,MYO5A,COL6A2,CD63
C35E2B,ATP6V0C,L1CAM,DLG5,TMEM132A,CTDSP2,CNN2,NOVA2,RNF187,U2AF2,EEF1A2,TBC
,RAB1A,PLPPR3,ZNF454,COTL1,HECTD4,MAGED4B,CAPRIN1,GPC1,CDK16,TOP2B,IPO5,ATP6V0
C3,PLD3,HSPD1,USP11,COL18A1,CSDE1,SMIM14,IGF2,EIF3C,EIF4B,VCP,MARCKSL1,UBB,AFF4,
TP1A1,JUND,EIF5,CD81,HMGN2,CRMP1,LASP1,OAZ1,RTN3,PFN1,ASTN2,TLE5,NISCH,HNRNPC,C
NPC,CCNI,XPR1,KHDRBS1,XPO1,SLC3A2,HDLBP,TPM1,EIF5A,CCND2,CDH2,PDIA6,DDX3X,RPL12
DR,TMEM259,EIF5,CRMP1,BEX3,CCT7,RBBP4,PFN1,MCL1,PNN,NISCH,PRKCSH,TADA2B,XPR1,H
31,TUBB4B,UCHL1,NCAM1,ANXA2,AGRN,RPLP1,TIMP2,XAF1,MIDN,WSB1,RPS17,VCL,AP3D1,SB
CL1,NISCH,HNRNPC,CCNI,KHDRBS1,PFKL,PDIA6,PIP4K2B,CCN2,PRDX2,DDX3X,RPL12,ARL4C,M
Q,EIF1,HACD3,DCHS1,SET,EGLN1,TMSB10,EIF4A1,MRC2,PPP2R1A,RPL15,LDHB,DNAH8,HLA-C,P
RL4C,MIF,MYO5A,COL6A2,B4GALNT4,MTSS2,DNAJC5,APLP2,HNRNPDL,TARDBP,TAGLN,ANKRD
MARCKSL1,UBB,AFF4,TRIM28,MATR3,YWHAG,CKB,COL4A2,MAPT,FN1,SPEN,DDB1,PSMD5,EIF1,
,CHSY1,TMSB10,EIF4A1,RPL15,GTF3C4,LDHB,DNAH8,PKD1,SF3B1,SERINC1,SSBP2,ZNF43,RP
,SRSF4,RBBP4,PFN1,MCL1,PNN,PRKCSH,KHDRBS1,XPO1,HDLBP,RPS25,EIF5A,GPC3,RPS15,IT
DLG5,SHISAL1,CPSF2,PGRMC1,DDX39B,SRSF5,MDH2,U2AF2,EEF1A2,PGD,VAMP2,NOP53,PABP
ORF43,PAPOLA,NCS1,RHOA,ZNF865,RAB6B,CXADR,ATP1A1,JUND,EIF5,CD81,LENG8,WDR1,HM
RPS4X,RPS3A,TUBB4B,RPL11,BTF3,NCAM1,ANXA2,AGRN,RPLP1,ZMIZ1,TIMP2,SARAF,MIDN,RPL

F3C,ATP5F1A,EIF4B,VCP,UBB,AFF4,MATR3,YWHAG,CKB,COL4A2,RPL37A,LRRC28,SPEN,NONO,I
HMGN2,ATP2A2,CRMP1,VAT1,PFN1,ASTN2,MCL1,TLE5,SPC25,LRPPRC,TADA2B,XPR1,SLC3A2,C
MIZ1,TIMP2,SARAF,MIDN,RPL10,WSB1,VCL,AP3D1,SBF1,DNAJA1,ARF1,CYB5R3,MAP4,CDC42BF
VHL,PPP2CA,SMARCA4,LAPTM4A,SRSF9,ARPC2,RABL6,ODC1,SRSF1,TCF25,PTBP1,ILF3,LUZP6,(
AH8,HLA-C,SF3B1,SSBP2,BASP1,HMGB1,POLR2A,CDKL1,TPT1,RPS3A,RPL18,ITGB1,TUBB4B,U(
F1A,EIF4B,MARCKSL1,UBB,AFF4,TRIM28,MATR3,COL4A2,MAPT,FN1,SPEN,NONO,DDB1,PSMD5
D81,COL5A2,LENG8,WDR1,ATXN7L3B,RPL26,LASP1,CCT7,SRSF4,RBBP4,PFN1,MCL1,TLE5,PNN
IN1,ITGB1,TUBB4B,RPL11,BTF3,AGRN,RPLP1,SARAF,MIDN,RPL10,MARCHF6,RPS17,VCL,AP3D1
,MMP2,FAM199X,DLG5,NOVA2,CTSB,MDH2,CHD6,EEF1A2,KLHL24,MORF4L2,PRDX5,GRINA,CL
A1,OR8K1,STMN1,ITGB1,TUBB4B,RPL11,MORF4L1,BTF3,ANXA2,AGRN,RPLP1,ZMIZ1,TIMP2,X
C22A17,BRD3,CIPC,C1ORF43,PAPOLA,NCS1,RHOA,ZNF865,SLC7A5,CXADR,ATP1A1,JUND,TME
X5,ACO2,ACAP3,MDK,SKI,ASXL1,HDGF,TSPAN6,PRPF6,RPL38,CCND1,AZIN1,MAPRE1,SRSF9,DS
LA-B,PLPPR3,ARCN1,COTL1,HECTD4,CTBP1,STIP1,GPC1,SLC35E2B,IPO5,ATP6V0C,SKP1,DNER
128,MATR3,COL4A2,MAPT,FN1,NONO,DDB1,YWHAQ,EIF1,HACD3,DCHS1,SET,MAP9,TPI1,CHSY1,
P200,CSDE1,SMIM14,EIF3C,ATP5F1A,EIF4B,MARCKSL1,UBB,TRIM28,MATR3,CKB,RPL37A,LRRC
YB5R3,SF3B2,ZNF770,AP2B1,MAT2A,GTF2F1,PXDN,HNRNPU,PGAM1,SMARCA5,HNRNPA0,CTNN
4,RPL36,RHOB,HNRNPH1,COTL1,HECTD4,CCT6A,MAGED4B,CTBP1,ARHGDIA,STIP1,GPC1,TOP
,MAP4,CDC42BPB,ITM2B,ZNF770,TMEM170A,GOLGA8A,PXDN,HNRNPU,NFE2L1,HNRNPA0,CTN
1,SMARCE1,GIGYF1,YWHAH,PTP4A2,SLIT3,UBQLN1,TUG1,SF3A1,ATXN2L,ID4,ZFAND3,MPRIP,S
F1,OGA,HNRNPA0,RPS16,RPL23,CNN3,RPL29,EIF4H,NACA,LPCAT1,BRD3,RPS9,XRCC5,CIPC,RPL
CT3,VPS35,ACAP3,DDX42,CBX1,SNRNP70,PYGB,PRPF6,DPYSL2,TOP1,TAF7,VHL,PPP2CA,SMAR
K2B,MEGF8,SOX12,CCN2,MMP14,RPL12,ITM2C,ARL4C,CLTC,MAGED2,APLP1,MIF,MYO5A,COL
F3B2,MAT2A,GTF2F1,PXDN,HNRNPU,HNRNPA0,RPS16,RPL23,LDHA,CNN3,RPL29,EIF4H,NACA,L
SPTBN1,ATP5F1B,CALM3,GNB1,SHISA9,STMN2,RPS8,MYH9,RPL7,SEPTIN2,P4HB,APP,IDS,ERVV
TR3,COL4A2,RPL37A,LRRC28,FN1,SPEN,DDB1,YWHAQ,EIF1,HACD3,DCHS1,SET,EGLN1,TPI1,CH
,TADA2B,CSNK1E,RPS25,SFPQ,EIF5A,CDH2,PIP4K2B,MEGF8,DDX3X,CLTC,MAGED2,H1-0,MYO5
TPM1,PNMA1,PDIA6,PLXNA1,PIP4K2B,ACSL6,SOX12,CCN2,RPL30,ARL4C,CLTC,MRFAP1,H1-0,A
CHL1,MORF4L1,BTF3,ANXA2,RPLP1,ZMIZ1,COL11A1,MIDN,MARCHF6,WSB1,DNAJA1,ARF1,CYB
1,PIP4K2B,SOX12,RPL12,ARL4C,MAGED2,APLP1,MIF,GPAT4,TLN1,MYO5A,CD63,B4GALNT4,MT
6A2,ANXA6,B4GALNT4,HNRNPDL,TARDBP,MYL6,WLS,HNRNPK,DUX4,CADM4,HNRNPH1,CKAP4,
S24,BRD3,CIPC,SLC16A7,KPNB1,C1ORF43,PAPOLA,IGFBP3,RPL14,HK1,NCS1,RHOA,CBX5,RPL
NO,EIF1,TMSB10,INA,EIF4A1,RPL19,RPL15,GTF3C4,DNAH8,PKD1,SF3B1,PDIA3,SERINC1,ZNF4
P3D1,SBF1,DNAJA1,CYB5R3,AKAP12,MAP4,CDC42BPB,ITM2B,AP2B1,MAT2A,PXDN,HNRNPU,PG
8,SPEN,DDB1,PSMD5,EIF1,HACD3,SET,EGLN1,TPI1,CHSY1,TMSB10,INA,EIF4A1,RPL19,PPP2R1A
PA0,CTNNA1,FADS2,RPL23,CPE,TPM4,TNS3,LPCAT1,LAMA5,RPS24,SLC22A17,BRD3,KPNB1,PA
,SET,EGLN1,MAP9,TPI1,CHSY1,TMSB10,INA,EIF4A1,MRC2,RPL19,PPP2R1A,RPL15,GTF3C4,LDH
BIM6,SPTAN1,GSTP1,MARCKS,CALM1,NPM1,PLXNB2,TTC3,PLD3,HSPD1,USP11,SNRNP200,CS
,AGRN,RPLP1,ZMIZ1,TIMP2,XAF1,MIDN,RPS17,VCL,SBF1,ARF1,MAP4,CDC42BPB,ITM2B,MAT2
V1,MDK,SURF4,PTK7,ORAI2,SKI,ASXL1,SCAF1,RAB11B,HDGF,PRPF6,TOP1,SELENON,PPP2CA,C
A1,SPTBN1,ATP5F1B,CALM3,GNB1,SHISA9,STMN2,RPS8,MYH9,RPL7,SEPTIN2,P4HB,APP,IDS,E
OLR2A,TPT1,RPS4X,RPS3A,TUBB4B,RPL11,BTF3,RPLP1,TIMP2,SARAF,MIDN,RPL10,WSB1,RPS1
A,CNN3,RPSA,NACA,LPCAT1,SLC22A17,BRD3,XRCC5,CIPC,RPL24,C1ORF43,PAPOLA,IGFBP3,L
NC1,RPS14,CXADR,JUND,TMEM259,EIF5,LENG8,WDR1,ATXN7L3B,LASP1,SRSF4,RBBP4,PFN1,A
M2C,ARL4C,GNB2,H1-0,TLN1,MYO5A,GOLM1,RPS10,DNAJC5,TARDBP,RPLP2,MXD4,WLS,RPL3
T4,MYO5A,B4GALNT4,MTSS2,APLP2,TARDBP,SLC25A3,RPLP2,ANKRD52,INIP,MYH10,DUX4,CA

ID6,EEF1A2,CCT2,TBCD,VAMP2,GAS7,HNRNPL,TM9SF2,LMNB2,PDXK,YEATS2,HSPA4,GRINA,VPS
KI,H3-3A,RAB11B,PYGB,SELENON,MVB12B,CCND1,MAPRE1,DDAH2,ODC1,AEBP1,TCF25,PTBP1
N1A,PGRMC1,CTSB,SRSF5,C5ORF24,ZC3H7B,MDH2,MLXIP,ANXA5,SQSTM1,VAMP2,GDI2,HNRNI
,HSPD1,USP11,COL18A1,CSDE1,IGF2,EIF3C,EIF4B,VCP,MARCKSL1,UBB,AFF4,TRIM28,MATR3,Y
,COL4A2,RPL37A,LRRC28,FN1,SPEN,DDB1,YWHAQ,EIF1,HACD3,SET,EGLN1,MAP9,TPI1,CHSY1,T
,GRINA,VPS35,PTOV1,MLEC,CLSTN3,MDK,SURF4,JPT1,H3-3A,SCAF1,PYGB,PRPF6,CCND1,AZIN
(P1,DNER,BCLAF1,L1CAM,TRIB2,SQLE,MAP1A,FAM199X,SERF2,TIMP3,CTDSP2,CTSB,NUCB1,CH
F1,HACD3,EGLN1,TMSB10,INA,EIF4A1,RPL19,PPP2R1A,RPL15,GTF3C4,LDHB,DNAH8,PKD1,SF3
)L,ARL6IP1,NDFIP1,MYL6,DUX4,YWHAB,PTGES3,CKAP5,CADM4,CKAP4,RAB1A,CALU,ARCN1,HEC
)L,ARL6IP1,NDFIP1,MYL6,DUX4,YWHAB,PTGES3,CKAP5,CADM4,CKAP4,RAB1A,CALU,ARCN1,HEC
)L,ARL6IP1,NDFIP1,MYL6,DUX4,YWHAB,PTGES3,CKAP5,CADM4,CKAP4,RAB1A,CALU,ARCN1,HEC
IF4H,TSPAN3,LPCAT1,RPS24,BRD3,XRCC5,MSN,KPNB1,C1ORF43,PAPOLA,LLGL1,GAP43,HK1,R
EPTIN3,CCT6A,MAGED4B,RAC1,GPC1,SLC35E2B,APPL1,IPO5,TRIB2,PPIB,GLUL,LUC7L3,CDKN1
TPI1,CHSY1,INA,EIF4A1,MRC2,PPP2R1A,GTF3C4,PKD1,HLA-C,PDIA3,SSBP2,ZNF43,CHD3,BASP1
.4A2,RPL37A,LRRC28,FN1,SPEN,DDB1,YWHAQ,EIF1,HACD3,SET,EGLN1,MAP9,TPI1,CHSY1,TMSE
LN1,TMSB10,INA,EIF4A1,RPL19,RPL15,GTF3C4,DNAH8,PKD1,SF3B1,PDIA3,SERINC1,ZNF43,HM
SP11,SNRNP200,IGF2,EIF3C,EIF4B,VCP,MARCKSL1,UBB,MATR3,YWHAG,CKB,COL4A2,FN1,SPE
,PLCG1,ZFAND5,DIDO1,DUSP4,FLNB,DLK1,PHC2,LSS,PITPNM1,STXBP1,ISYNA1,CSNK1D,MESD,
ADS2,CNN3,RPSA,TPM4,TNS3,CTSD,EIF4H,TSPAN3,LAMA5,SLC22A17,BRD3,PJA2,RPL24,SLC16
)X,H1-10,INIP,CADM4,ATP1A2,HLA-B,CALU,PLPPR3,SEPTIN3,COTL1,MAGED4B,CTBP1,ARHGDIA
CIPC,MSN,KPNB1,COL1A2,LLGL1,NCS1,RHOA,CLSTN1,RAB6B,NLGN2,CXADR,CHD4,CD81,HMG
GLG1,SFPQ,EIF5A,CCND2,PNMA1,SOX12,PRDX2,DDX3X,RPS15,ITM2C,RPL30,ARL4C,EIF3L,H1-0
2B,ACSL6,SOX12,RPL12,ITM2C,ARL4C,MAGED2,RCN1,APLP1,MIF,GPAT4,TLN1,MYO5A,COL6A2
A2,RPL37A,LRRC28,FN1,SPEN,DDB1,YWHAQ,EIF1,HACD3,SET,EGLN1,MAP9,TPI1,CHSY1,TMSB1
CTR3,PDXK,PKN1,HSPA4,CAMSAP1,GRINA,NCKAP1,MLEC,CLSTN3,MDK,RAD23B,ASXL1,H3-3A,A
1,SET,EGLN1,CHSY1,TMSB10,INA,EIF4A1,MRC2,RPL19,PPP2R1A,RPL15,GTF3C4,LDHB,PKD1,SF
LHL9,XPO1,SPTBN2,CSNK1E,PFKL,HDLBP,RPS25,CDH6,SFPQ,EIF5A,PEBP1,CDH2,PIP4K2B,CCN
55,CLSTN1,PSMD2,SLC7A5,NLGN2,RPS14,CXADR,CHD4,ATP1A1,TMEM259,EIF5,CD81,HMGN2,A
R1,HMGN2,RPL26,LASP1,BEX3,SRSF4,RBBP4,PFN1,MCL1,PNN,RPL31,HNRNPC,KHDRBS1,PFKL,
F1,HACD3,MAP9,TPI1,TMSB10,INA,EIF4A1,RPL19,PPP2R1A,RPL15,GTF3C4,LDHB,DNAH8,PKD1,S
,CDK16,TOP2B,SLC35E2B,SMCHD1,IPO5,TRIB2,MAP1A,CDKN1A,NOVA2,MDH2,CHD6,RPN2,MLX
R1,ATXN7L3B,RPL26,LASP1,SRSF4,RBBP4,PFN1,MCL1,TLE5,PNN,NES,RPL31,HNRNPC,CCNI,TA
LC22A17,BRD3,XRCC5,SLC16A7,KPNB1,RPL17,C1ORF43,PAPOLA,IGFBP3,COL1A2,LLGL1,GAP4
IA5,BRD3,PJA2,RPL24,SLC16A7,KPNB1,LLGL1,RPL14,CBX5,PSMD2,RAB6B,GLIPR1L2,RPS14,C
,ASXL1,SCAF1,TERF2IP,PPP2CA,MAPK1,AZIN1,CCNL2,BTBD2,RABL6,SRSF1,DYNC1LI2,ARGLU1
A,HMGB1,POLR2A,CDKL1,TPT1,RPS4X,RPS3A,STMN1,ITGB1,RPL11,MORF4L1,BTF3,NCAM1,AN
RPL11,BTF3,NCAM1,AGRN,RPLP1,SARAF,MIDN,RPL10,MARCHF6,RPS17,VCL,AP3D1,DNAJA1,AF
IAH8,PKD1,SF3B1,SERINC1,SSBP2,ZNF43,RPS3,HMGB1,POLR2A,CDKL1,AP2M1,TPT1,SEC61A1,
ERF2,CRPPA,CTSB,CDH11,CHD6,U2AF2,TBCD,MLXIP,PPP1CB,SNX22,GAS7,ABCA3,HNRNPA3,PKI
F1,HACD3,SET,EGLN1,MAP9,TPI1,CHSY1,TMSB10,INA,EIF4A1,MRC2,RPL19,PPP2R1A,LDHB,SF3E
A,GTF3C4,LDHB,HLA-C,SF3B1,SSBP2,ZNF43,CHD3,VAPA,HMGB1,CDKL1,TPT1,SEC61A1,STMN1,
KIF5B,RAB6B,SLC7A5,ENC1,NLGN2,CXADR,CHD4,ATP1A1,JUND,RPS20,EIF5,LENG8,WDR1,ATP2
IATR3,YWHAG,CKB,COL4A2,RPL37A,LRRC28,FN1,SPEN,DDB1,YWHAQ,EIF1,HACD3,SET,EGLN1,I
LBP,EIF5A,CCND2,RPS15,ITM2C,ARL4C,MRFAP1,RCN1,H1-0,RPS7,RPL35,CHPF,ANXA6,NGRN,RI
IRNPA0,CTNNA1,RPS16,LDHA,CNN3,RPL29,TPM4,TNS3,EIF4H,NACA,LPCAT1,XRCC5,SLC16A7,KI

RPL31,PRKCSH,HNRNPC,XPR1,KHDRBS1,RPS25,TKT,EIF5A,CDH2,PDIA6,MEGF8,ACSL6,RAB7A,I
P,VCAN,TGFBI,HNRNPA3,ACAP3,MDK,LAMC1,CBX1,RAB5B,CNTNAP2,HDGF,SRP14,VEGFA,SELEN
,HNRNPDL,SLC25A3,RPLP2,ARL6IP1,MXD4,HNRNPK,H1-10,MYH10,DUX4,SEPTIN11,LAMP1,RHO
A,RPL15,GTF3C4,LDHB,DNAH8,PKD1,SF3B1,PDIA3,SERINC1,SSBP2,ZNF43,RPS3,VAPA,HMGB1,F
R1A,RPL15,GTF3C4,DNAH8,PKD1,SF3B1,PDIA3,SERINC1,ZNF43,RPS3,HMGB1,POLR2A,AP2M1,
RHOA,ZNF865,CLSTN1,ATF4,PSMD2,NR2F1,SLC7A5,NLGN2,RPS14,TMEM259,EIF5,LENG8,WDR
APRE1,RABL6,ODC1,DYNLL2,ACTN1,MAP7D1,PLCG1,DIDO1,PITPNA,CORO1C,SDF4,NKTR,SLIT3,
BF1,ARF1,CYB5R3,MAP4,SF3B2,CDC42BPB,ZNF770,TMEM170A,SLC4A8,PXDN,NFE2L1,HNRNP
CHS1,SET,TMSB10,MRC2,RPL15,GTF3C4,PDIA3,SERINC1,SSBP2,CHD3,BASP1,VAPA,HMGB1,PO
A1,JUND,RPS20,EIF5,LENG8,WDR1,ATP2A2,RPL26,LASP1,OAZ1,SRSF4,RBBP4,PFN1,SPC25,NES
EATS2,HSPA4,GRINA,VPS35,PTOV1,MLEC,CLSTN3,MDK,SURF4,PTK7,ORAI2,SKI,H3-3A,SCAF1,RA
S15,CLTC,MAGED2,GNB2,RCN1,APLP1,RPS19,RPL35,MIF,CHPF,JUN,CTNNB1,COL6A2,SESN3,T
H2,RABL6,DYNC1LI2,NYNRIN,TCF25,ADAM9,GRB2,ATXN10,HSPB1,KDM1A,CASC3,FKBP8,PLCG
CAM,LAPTM4B,TRIB2,LUC7L3,PTTG1IP,SQLE,MAP1A,CDKN1A,TIMP3,CTDSP2,CNN2,NOVA2,NUC
A,PXDN,HNRNPA0,CTNNA1,FADS2,RPL23,LDHA,CNN3,RPSA,TPM4,TNS3,CTSD,EIF4H,SCARB2,TS
G8,WDR1,ATP2A2,SRSF4,PFN1,PNN,HNRNPC,TADA2B,KHDRBS1,SLC3A2,HDLBP,RPS25,TPM1,C
PAPOLA,CLSTN1,NR2F1,RAB6B,ENC1,CXADR,JUND,RPS20,CD81,CRMP1,LASP1,VAT1,OAZ1,SRS
CAM1,AGRN,RPLP1,MIDN,RPL10,MARCHF6,WSB1,RPS17,DNAJA1,ARF1,SF3B2,CDC42BPB,MAT2
PN2,TBCD,SUGP2,ANXA5,PPP1CB,SNX22,MEST,GAS7,HNRNPL,TSR1,LMNB2,HNRNPA3,PDXK,TC
PPP2R1A,RPL15,GTF3C4,DNAH8,PKD1,SF3B1,PDIA3,SERINC1,ZNF43,HMGB1,POLR2A,AP2M1,T
A8,PXDN,HNRNPU,PGAM1,NFE2L1,SMARCA5,HNRNPA0,CTNNA1,RPS16,RPL23,CNN3,TPM4,TNS
,RPS9,CIPC,PJA2,RPL24,RPL17,C1ORF43,PAPOLA,RPL14,PTCD3,CBX5,ZNF865,CLSTN1,ATF4,K
OLR2A,AP2M1,TPT1,SEC61A1,RPS4X,RPS3A,RPL18,ITGB1,TUBB4B,RPL11,MORF4L1,BTF3,AGF
ES3,CADM4,HNRNPH1,ATP1A2,CKAP4,RAB1A,PLPPR3,ARCN1,COTL1,ARHGDIA,GPC1,TOP2B,IP
7A,GAS7,PLOD1,CAMSAP1,KLC1,GRINA,COL2A1,PTOV1,MLEC,ACAP3,SURF4,DDX42,CCT4,RPL
F4,RBBP4,PFN1,MCL1,TLE5,PNN,RPL31,HNRNPC,CCNI,TADA2B,XPR1,KHDRBS1,PFKL,RPS25,TK
PL26,SRSF4,RBBP4,PFN1,MCL1,TLE5,PNN,RPL31,PRKCSH,HNRNPC,KHDRBS1,KLHL9,RPS25,T
GF8,RAB7A,RPS15,RPL12,ITM2C,RPL30,CLTC,EIF3L,GNB2,H1-0,RPS5,RPS7,IK,RPS19,MIF,NGR
MD5,YWHAQ,EIF1,DCHS1,SET,TPI1,CHSY1,TMSB10,INA,MRC2,RPL19,LDHB,PKD1,HLA-C,PDIA3,
4L1,BTF3,AGRN,RPLP1,SARAF,MIDN,RPL10,MARCHF6,RPS17,AP3D1,DNAJA1,ARF1,CYB5R3,SF
,LAPTM4A,SRSF9,PDE7A,RABL6,ODC1,SRSF1,TCF25,ILF3,LUZP6,GRB2,TDRD5,SCAF11,BNIP3L,
8,PKD1,HLA-C,SF3B1,PDIA3,SERINC1,CHD3,VAPA,HMGB1,POLR2A,CDKL1,AP2M1,TPT1,SEC61A
PLCG1,CORO1C,NKTR,LSS,STXBP1,ISYNA1,PLXNA3,IDH1,HYOU1,ID4,MAPK8IP1,SOD1,FBRSL1,
MAT2A,TMEM170A,GOLGA8A,PXDN,HNRNPU,OGA,PGAM1,HNRNPA0,FADS2,RPS16,CPE,CNN3,T
,TMSB10,EIF4A1,RPL19,PPP2R1A,RPL15,GTF3C4,DNAH8,SF3B1,PDIA3,SSBP2,ZNF43,RPS3,CHD
PL18,ITGB1,TUBB4B,RPL11,MORF4L1,BTF3,AGRN,RPLP1,SARAF,MIDN,RPL10,MARCHF6,RPS17
S3A,STMN1,ITGB1,RPL11,MORF4L1,BTF3,NCAM1,ANXA2,AGRN,SARAF,MARCHF6,VCL,AP3D1,D
TSB,MDH2,CHD6,U2AF2,EEF1A2,TBCD,ANXA5,PPP1CB,VAMP2,MORF4L2,TGFBI,HNRNPA3,PRD
SRSF5,TM9SF3,MDH2,U2AF2,EEF1A2,FDFT1,SUGP2,CPSF1,GNAI3,GDI2,PARP1,TSR1,MTRF1L,R
APTM4B,TRIB2,PEA15,LUC7L3,SQLE,MAP1A,SERF2,TIMP3,CTDSP2,CNN2,NOVA2,ZC3H7B,CHD6
1,HACD3,SET,EGLN1,MAP9,TPI1,CHSY1,TMSB10,INA,EIF4A1,RPL19,LDHB,DNAH8,HLA-C,SF3B1,
RNPDL,TARDBP,DDOST,MYL6,HNRNPK,H1-10,SEPTIN11,LAMP1,HNRNPH1,SEPTIN3,CCT6A,MAG
GB1,RPL11,MORF4L1,BTF3,NCAM1,AGRN,ZMIZ1,TIMP2,SARAF,COL11A1,MARCHF6,RPS17,VC
M170A,HNRNPU,NFE2L1,SMARCA5,RPL23,RPSA,TPM4,TNS3,SCARB2,RPS24,CIPC,RPL24,C1OR
,SPINT2,ZMYM4,RABEP1,DAP,BRK1,SAFB2,VPS26B,FUS,PPP1R10,SPG7,CEP170,EIF2S2,EHMT1,

PL10,WSB1,RPS17,VCL,AP3D1,SBF1,DNAJA1,ARF1,SF3B2,CDC42BPB,MAT2A,TMEM170A,PXDN,
RPLP2,DDOST,CIRBP,DUX4,YWHAB,PTGES3,SND1,ARCN1,ZNF454,RPS27,CTBP1,ARHGDIA,CAPF
),SELENON,TUBB2A,ARF3,UBXN4,PPP2CA,CCND1,SMARCA4,SRSF9,PLEC,PBX2,PRDX1,FAM168I
F6,UPF1,ZFR,ZNF512B,CBX6,FKBP1A,RABEP1,SLC2A1,FBXO21,MTCH1,NDN,ZNF664,STK35,SPC
TPT1,SEC61A1,RPS4X,RPS3A,RPL18,ITGB1,TUBB4B,RPL11,MORF4L1,BTF3,AGRN,RPLP1,SARA
F1,CYB5R3,SF3B2,MAT2A,GTF2F1,HNRNPA0,RPS16,RPL23,CNN3,RPL29,TPM4,EIF4H,LAMA5,RI
,SSBP2,CHD3,BASP1,VAPA,HMGB1,POLR2A,CDKL1,AP2M1,TPT1,RPL18,ITGB1,TUBB4B,UCHL1,I
TC3,PLD3,HSPD1,COL18A1,SNRNP200,CSDE1,SMIM14,IGF2,EIF3C,ATP5F1A,EIF4B,VCP,MARCK
N2,HNRNPD,CYP51A1,FLNC,CTXN1,CD99L2,YME1L1,MSMO1,PRMT1,MTA1,TRIO,EP300,SF1,HIF
L18A1,SNRNP200,SMIM14,IGF2,EIF3C,ATP5F1A,EIF4B,VCP,MARCKSL1,UBB,AFF4,MATR3,YWHA
P1,PTOV1,MLEC,ZNF302,RAD23B,CCT4,SKI,SNRNP70,ASXL1,H3-3A,SCAF1,DPYSL2,TOP1,TAF7,I
N,RPL10,MARCHF6,RPS17,AP3D1,DNAJA1,ARF1,CYB5R3,SF3B2,MAT2A,GTF2F1,SLC4A8,HNRNP
CHF6,RPS17,AP3D1,DNAJA1,ARF1,CYB5R3,SF3B2,MAT2A,GTF2F1,HNRNPA0,RPS16,RPL23,RPL2
COL18A1,CSDE1,IGF2,EIF3C,ATP5F1A,EIF4B,MARCKSL1,UBB,AFF4,MATR3,YWHAG,CKB,COL4A
A,PTCD3,ZNF865,ATF4,PSMD2,KIF5B,RPS14,RPS20,EIF5,LENG8,WDR1,RPL26,LASP1,OAZ1,PFN1
CBX5,PSMD2,RAB6B,GLIPR1L2,RPS14,CXADR,ATP1A1,JUND,RPS20,CD81,WDR1,ATXN7L3B,HMC
F1,WDR82,ASH1L,RAVER1,NDUFS1,KDM2A,SPCS2,PMPCA,CDC42SE1,CACYBP,MED12,TLK1,UBI
S,STXBP1,TUG1,ATXN2L,ID4,SOD1,RAI1,SREBF2,PFN2,PNMA8A,P3H3,WBP2,IGFBP7,PRPF40A,SI
TP1,MARCKS,SMG1,CALM1,NPM1,PLXNB2,PLD3,HSPD1,USP11,COL18A1,SNRNP200,CSDE1,SM
PS14,CXADR,ATP1A1,JUND,RPS20,CD81,WDR1,ATXN7L3B,HMGN2,RPL26,CRMP1,LASP1,BEX3,(
ALM1,NPM1,PLXNB2,TTC3,PLD3,HSPD1,USP11,COL18A1,CSDE1,IGF2,EIF3C,EIF4B,MARCKSL1,
NP200,CSDE1,SMIM14,IGF2,EIF3C,ATP5F1A,EIF4B,VCP,MARCKSL1,UBB,AFF4,MATR3,YWHAG,C
C,SF3B1,SERINC1,ZNF43,BASP1,VAPA,HMGB1,POLR2A,AP2M1,TPT1,SEC61A1,RPS3A,RPL18,TUI
SB10,INA,EIF4A1,MRC2,PPP2R1A,GTF3C4,DNAH8,PKD1,HLA-C,PDIA3,SSBP2,RPS3,CHD3,BASP1,
,CCT7,SRSF4,RBBP4,PFN1,ASTN2,PRKCSH,HNRNPC,KLHL9,SLC3A2,CSNK1E,PFKL,RPS25,TPM1
RPLP1,SARAF,MIDN,RPL10,MARCHF6,RPS17,AP3D1,DNAJA1,ARF1,CYB5R3,SF3B2,MAT2A,GTF2
2A17,KPNB1,LLGL1,HK1,NCS1,RPL22,CBX5,ZNF865,CLSTN1,ATF4,SLC7A5,NLGN2,CXADR,ATP1
,ODC1,SRSF1,TCF25,ILF3,LUZP6,TDRD5,SCAF11,BNIP3L,CLIP3,MAP2K2,ARF4,POLR2B,SDF4,A
N1,AHCYL1,ETNK1,GRINA,ACAP3,MDK,LAMC1,PTK7,TNPO1,ABCA2,TSPAN6,RPL38,OPTN,SRSF9,
CND2,PNMA1,PDIA6,RAB7A,DDX3X,RPL12,MRFAP1,RPS5,RPS19,MIF,HMGCS1,TLN1,JUN,MYO5A
1,TMBIM6,PTMS,SPTAN1,GSTP1,MARCKS,SMG1,CALM1,NPM1,PLXNB2,PLD3,HSPD1,COL18A1,(
TS2,ACO2,HSPA4,CAMSAP1,KLC1,VPS35,ACAP3,ZNF302,DDX42,LAMC1,JPT1,SNRNP70,ASXL1,F
,RPLP1,SARAF,MIDN,RPL10,MARCHF6,RPS17,AP3D1,DNAJA1,ARF1,CYB5R3,SF3B2,MAT2A,GTF2
,DNAJA1,ARF1,CYB5R3,AKAP12,MAP4,SF3B2,CDC42BPB,AP2B1,MAT2A,GTF2F1,SLC4A8,GOLG
,UBB,AFF4,MATR3,YWHAG,CKB,COL4A2,RPL37A,LRRC28,FN1,SPEN,NONO,DDB1,YWHAQ,EIF1,
C2,RPL19,PPP2R1A,GTF3C4,LDHB,PKD1,SF3B1,PDIA3,SERINC1,SSBP2,BASP1,VAPA,HMGB1,PO
,RAB14,MAZ,HDGFL2,IREB2,NUCKS1,LSM14B,SOD2,PLEKHM2,FOXK2,EDF1,MSH6,LRRC41,PRF
DE1,SMIM14,IGF2,EIF3C,EIF4B,VCP,MARCKSL1,AFF4,TRIM28,CKB,COL4A2,RPL37A,MAPT,FN1,S
RPL11,MORF4L1,BTF3,AGRN,RPLP1,SARAF,MIDN,RPL10,MARCHF6,RPS17,AP3D1,DNAJA1,ARF
L,IK,RPS19,CHPF,NGRN,DNAJC5,RPLP2,DDOST,DUX4,YWHAB,PTGES3,SND1,LAMP1,CALU,ARCI
N1,YLPM1,KIF5A,MSMO1,TXLNA,WDR45B,MTA1,TRIO,LBH,RETREG2,PBX1,HIP1,LRRC4B,FLOT1
F3L,GNB2,H1-0,RPS7,IK,RPS19,CHPF,NGRN,DNAJC5,HNRNPDL,RPLP2,DDOST,TSR2,NARS1,DUX
O,DDB1,PSMD5,EIF1,HACD3,DCHS1,SET,CHSY1,TMSB10,EIF4A1,MRC2,RPL19,PPP2R1A,RPL15,(
EGLN1,TMSB10,INA,EIF4A1,RPL19,PPP2R1A,RPL15,GTF3C4,LDHB,PKD1,SF3B1,PDIA3,SERINC1,
N,RPL10,MARCHF6,AP3D1,DNAJA1,ARF1,CYB5R3,AKAP12,SF3B2,GTF2F1,SLC4A8,SMARCA5,HN

JCHL1,BTF3,ANXA2,AGRN,RPLP1,TIMP2,SARAF,MIDN,MARCHF6,RPS17,VCL,AP3D1,SBF1,ARF1,C
'L36A,VEGFA,ARF3,DSTN,ODC1,SRSF1,ARGLU1,NYNRIN,PTBP1,ILF3,LUZP6,CDC37,HNRNPUL1,S
RREL1,PBX1,AP2A1,BAZ1B,KIF21B,SEC16A,ZFP36L1,MGAT4B,CXXC5,SLC39A7,CSNK1G2,NXN,L
H2A2,PLEKHM2,ACTR1B,DICER1,MTMR4,NSD2,LRP3,USP24,FKBP9,SF3A2,FBLN5,PTGFRN,PRKA
EIF4A1,RPL19,PPP2R1A,RPL15,LDHB,HLA-C,SF3B1,SSBP2,ZNF43,CHD3,VAPA,HMGB1,CDKL1,TF
MC3,SELENOW,HNRNPA2B1,CUL3,BRSK1,MTCH1,PCBP4,VPS26B,ACLY,SEPTIN5,ERGIC1,YARS1,
ARCKSL1,UBB,AFF4,TRIM28,MATR3,YWHAG,CKB,COL4A2,RPL37A,LRRC28,FN1,SPEN,DDB1,PSM
,RPL26,CRMP1,LASP1,VAT1,PFN1,ASTN2,JPT2,TLE5,NES,RPL31,NISCH,PRKCSH,XPR1,KHDRBS1
OL6A2,SESN3,B4GALNT4,MTSS2,APLP2,ARL6IP1,ST3GAL2,MYH10,DUX4,CKAP5,LAMP1,RHOB,H
ED4B,CTBP1,ARHGDIA,SMCHD1,TMEM30A,DNER,PTTG1IP,SQLE,SRSF5,MDH2,U2AF2,TBCD,ML
N1,PRPF40A,CTXN1,YLPM1,SYP,TRAF4,RBBP7,MSMO1,MTA1,TRIO,LBH,SF1,PBX1,HIP1,AP2A1,N
VSB1,VCL,AP3D1,SBF1,DNAJA1,ARF1,SF3B2,CDC42BPB,ITM2B,MAT2A,TMEM170A,PXDN,HNRNF
BP1,LUZP6,GRB2,CAPZB,SCAF11,ACTN1,CAND1,MAP7D1,MAP2K2,XPNPEP3,KIF3C,SMARCC1,Y
SHISA9,STMN2,RPS8,MYH9,SEPTIN2,P4HB,APP,IDS,ERVV-1,RPL5,GDI1,FSCN1,TMBIM6,PTMS,SF
OLR2A,AP2M1,TPT1,SEC61A1,RPS4X,RPS3A,RPL18,ITGB1,TUBB4B,RPL11,MORF4L1,BTF3,AGRN
ARCHF6,DNAJA1,MAP4,TMEM170A,GTF2F1,HNRNPU,OGA,PGAM1,NFE2L1,SMARCA5,HNRNPA0,
43,IGFBP3,HK1,PTCD3,SLC7A5,RPS14,CHD4,ATP1A1,JUND,TMEM259,EIF5,CD81,LENG8,WDR1,I
M4,UBE2H,CKAP4,COTL1,MAGED4B,STIP1,TOP2B,SMCHD1,HDAC2,TNC,PEA15,DLG5,NOVA2,MD
2L,HGS,BAG6,SEC23A,ARFGEF2,PPP1R9B,ANP32A,VDAC1,TRAF7,EIF2AK1,EDC4,PRMT1,BCR,EX
CLTC,EIF3L,GNB2,H1-0,RPS7,IK,RPS19,TLN1,CHPF,NGRN,DNAJC5,HNRNPDL,RPLP2,DDOST,TSI
MIM14,ATP5F1A,EIF4B,MARCKSL1,UBB,AFF4,TRIM28,MATR3,YWHAG,CKB,COL4A2,RPL37A,LRF
CD81,LENG8,WDR1,ATP2A2,RPL26,LASP1,CCT7,SRSF4,RBBP4,PFN1,ASTN2,MCL1,TLE5,PNN,RF
2B,MAT2A,TMEM170A,GTF2F1,GOLGA8A,PXDN,HNRNPU,NFE2L1,SMARCA5,LARP1,HNRNPA0,RI
2B,MAT2A,TMEM170A,GTF2F1,GOLGA8A,PXDN,HNRNPU,NFE2L1,SMARCA5,LARP1,HNRNPA0,RI
NB1,C1ORF43,GAP43,CBX5,RPL9,KIF5B,RAB6B,SLC7A5,NLGN2,CXADR,CHD4,CD81,WDR1,HM(
I,PSMD2,KIF5B,RPS14,CXADR,CHD4,RPS20,LENG8,WDR1,RPL26,CRMP1,CCT7,SRSF4,RBBP4,PI
JUND,EIF5,WDR1,HMGN2,RPL26,CRMP1,RTN3,PFN1,ASTN2,JPT2,TLE5,NES,CCNI,TADA2B,KLHL
BTF3,AGRN,RPLP1,SARAF,MIDN,RPL10,MARCHF6,RPS17,AP3D1,DNAJA1,ARF1,CYB5R3,AKAP12,
(,TMEM248,RAD23B,SKI,SNRNP70,PYGB,VEGFA,RAPGEF1,TUBB2A,ARF3,MAPK1,DDAH2,PLEC,B
CTN4,RPL12,ITM2C,ARL4C,CLTC,GNB2,H1-0,APLP1,RPL35,CHPF,MYO5A,COL6A2,SESN3,B4GAL
A,CTDSP2,CPSF2,CTSB,SRSF5,CDH11,TOMM20,RAB27A,TBCD,SUGP2,ANXA5,PPP1CB,PABPN1,A
KDC,HNRNPUL1,SCAF11,ZFAND5,DIDO1,DUSP4,HNRNPM,PFKM,LSS,TNKS2,PRPS1,HECTD1,BAG
HOA,CLSTN1,KIF5B,CHD4,JUND,TMEM259,RPS20,CD81,HMGN2,RPL26,CRMP1,LASP1,BEX3,CC
RNPA0,RPL23,RPL29,TPM4,EIF4H,LAMA5,RPS24,RPS9,XRCC5,CIPC,RPL24,PAPOLA,HK1,NCS1,RI
EBP1,PNMA1,GPC3,MEGF8,SOX12,RAB7A,CCN2,ITM2C,ARL4C,MRFAP1,RCN1,APLP1,RPL35,CD6
SKI,SNRNP70,DPYSL2,RPL38,NOTCH2,TUBB2A,MAPK1,SMARCA4,AZIN1,LAPTM4A,SRSF9,CCNL2
MARCA4,ATG2B,CCNL2,ARPC2,FAM168B,SRSF1,ARGLU1,ACADVL,CDC37,UBE2R2,ATXN10,ACTN
MIDN,RPL10,MARCHF6,RPS17,AP3D1,DNAJA1,ARF1,CYB5R3,MAP4,SF3B2,MAT2A,GTF2F1,PGAM
HSBP1,MAP1A,SERF2,DDX39B,SRSF5,TM9SF3,MDH2,RAB27A,U2AF2,EEF1A2,FDFT1,CCT2,SUGP
PDIA6,PIP4K2B,MEGF8,RAB7A,MMP14,RPL12,ITM2C,RPL30,EIF3L,MAGED2,GNB2,H1-0,RPS7,IK,
,EIF3L,GNB2,H1-0,RPS7,IK,RPS19,CHPF,NGRN,DNAJC5,HNRNPDL,RPLP2,DDOST,HNRNPK,NARS
5,PTOV1,MLEC,CLSTN3,MDK,RAD23B,PTK7,SCAF1,PRPF6,SELENON,TUBB2A,OPTN,MVB12B,PL
5,PTOV1,MLEC,CLSTN3,MDK,RAD23B,PTK7,SCAF1,PRPF6,SELENON,TUBB2A,OPTN,MVB12B,PL
SARAF,MIDN,RPL10,MARCHF6,RPS17,AP3D1,DNAJA1,ARF1,CYB5R3,AKAP12,SF3B2,MAT2A,GTF
,HMGB1,CDKL1,TPT1,SEC61A1,RPS4X,RPS3A,STMN1,ITGB1,TUBB4B,MORF4L1,AGRN,RPLP1,Z

L,ZFAND3,HECTD1,COL27A1,RAI1,PFN2,FBRSL1,MEX3A,FLNC,SMS,NAP1L4,AMOTL2,RBM15B,[illegible]
ANKRD52,WLS,H1-10,CIRBP,MYH10,RPL36,PTGES3,CKAP5,CADM4,UBE2H,PLPPR3,MAGED4B,[illegible]
4B,UCHL1,MORF4L1,AGRN,RPLP1,ZMIZ1,TIMP2,XAF1,SARAF,MIDN,RPL10,WSB1,RPS17,VCL,AP[illegible]
5,FOXK2,LRRC41,MED13,UBE3A,YY1,NDUFA13,TRIM8,DMWD,HNRNPH3,ATP13A1,SRPRA,EIF4G[illegible]
LU,ARCN1,COTL1,ARHGDIA,IPO5,ATP6V0C,BCLAF1,LAPTM4B,TRIB2,RPL23A,HSBP1,MAP1A,CT[illegible]
WDR1,HMGN2,BEX3,RBBP4,MCL1,PNN,CCNI,TADA2B,KHDRBS1,PFKL,HDLBP,GLG1,EIF5A,PEBP[illegible]
NA1,PIP4K2B,ACSL6,SOX12,DDX3X,RPL12,ITM2C,NANOG,H1-0,APLP1,RPS19,MIF,MYO5A,CD63,[illegible]
PS16,RPL23,RPL29,EIF4H,LAMA5,RPS9,XRCC5,CIPC,RPL24,SLC16A7,PAPOLA,HK1,NCS1,RPL22[illegible]
MEGF8,RAB7A,PRDX2,RPS15,ITM2C,RPL30,CLTC,EIF3L,MAGED2,RPS5,IK,RPS19,TLN1,CHPF,JU[illegible]
F2F1,HNRNPA0,RPS16,RPL23,RPL29,TPM4,EIF4H,LAMA5,RPS24,RPS9,XRCC5,CIPC,RPL24,PAPO[illegible]
RS,SF1,RETREG2,AP2A1,SEC16A,UBA52,SLC39A7,UPF1,UBE2G2,EIF5B,NOMO1,CBX6,NPDC1,S[illegible]
P2A2,CRMP1,VAT1,SRSF4,RTN3,ASTN2,TLE5,PRKCSH,HNRNPC,KHDRBS1,SPTBN2,PFKL,TPM1,[illegible]
SF3,MDH2,U2AF2,EEF1A2,FDFT1,CCT2,SUGP2,GNAI3,GDI2,PARP1,TSR1,MTRF1L,PTPRA,PRDX5[illegible]
E,GLG1,PNMA1,PDIA6,PLXNA1,PIP4K2B,RAB7A,CCN2,MMP14,ARL4C,RCN1,H1-0,APLP1,GPAT4,[illegible]
19,PPP2R1A,GTF3C4,LDHB,PKD1,HLA-C,SF3B1,ZNF43,BASP1,VAPA,HMGB1,POLR2A,CDKL1,TP[illegible]
RP1,MTRF1L,PTPRA,PRDX5,ACTR3,UBE4A,ACO2,HSPA4,CAMSAP1,MLEC,ZNF302,TMEM248,DD[illegible]
RHOA,PTCD3,ATF4,SLC7A5,CXADR,CHD4,ATP1A1,EIF5,CD81,COL5A2,LENG8,HMGN2,LASP1,BE[illegible]
A1,RPL19,PPP2R1A,RPL15,GTF3C4,LDHB,DNAH8,PKD1,SF3B1,SERINC1,SSBP2,ZNF43,HMGB1,[illegible]
,ZMIZ1,RPL10,WSB1,RPS17,AP3D1,DNAJA1,ARF1,AKAP12,MAP4,SF3B2,AP2B1,MAT2A,HNRNPA[illegible]
SLC4A8,PXDN,HNRNPU,CTNNA1,RPS16,RPL23,CNN3,RPL29,RPSA,CTSD,SCARB2,NACA,TSPAN3,[illegible]
R2A,AP2M1,TPT1,SEC61A1,RPS4X,RPS3A,RPL18,ITGB1,TUBB4B,RPL11,MORF4L1,BTF3,AGRN,[illegible]
OA,ZNF865,ATF4,PSMD2,RPS14,CXADR,CHD4,RPS20,LENG8,WDR1,ATXN7L3B,RPL26,CRMP1,C[illegible]
0C,BCLAF1,RPL23A,HSBP1,MAP1A,CTDSP2,DDX39B,EID1,SRSF5,TM9SF3,MDH2,RAB27A,U2AF2[illegible]
F,NGRN,DNAJC5,RPLP2,DDOST,DUX4,YWHAB,PTGES3,SND1,ATP1A2,RPS12,CALU,ARCN1,RPS27[illegible]
L,TARDBP,RPLP2,ANKRD52,PTGES3,CKAP5,RHOB,HNRNPH1,TTR,ARCN1,SEPTIN3,ZNF454,RPS[illegible]
22A17,BRD3,CIPC,PJA2,LLGL1,HK1,RHOA,ZNF865,CLSTN1,ATF4,PSMD2,KIF5B,RAB6B,SLC7A5[illegible]
3,MDH2,RAB27A,U2AF2,EEF1A2,FDFT1,CCT2,SUGP2,GNAI3,PARP1,TSR1,MTRF1L,ALKBH5,RPL[illegible]
OL4A2,RPL37A,DDB1,PSMD5,YWHAQ,HACD3,SET,CHSY1,TMSB10,INA,EIF4A1,RPL15,GTF3C4,L[illegible]
A,NFE2L1,SMARCA5,FADS2,RPS16,RPL23,RPL29,RPSA,TPM4,EIF4H,NACA,LPCAT1,LAMA5,RPS2[illegible]
P,PHIP,ANKRD12,UBXN6,SLC20A1,MICAL1,DPYSL4,METRN,UNC119B,IGLON5,BGN,OLFM2,CEL[illegible]
MACF1,B4GALNT4,CYFIP2,DNAJC5,SLC25A3,RPLP2,NDFIP1,TSR2,MXD4,H1-10,MYH10,DUX4,C[illegible]
PS20,LENG8,WDR1,RPL26,SRSF4,PFN1,ASTN2,MCL1,TLE5,PNN,RPL31,PRKCSH,HNRNPC,CCNI,[illegible]
CT2,RPN2,SUGP2,GNAI3,MEST,GAS7,GDI2,TM9SF2,TSR1,SSRP1,HSPH1,LMNB2,MTRF1L,ALKB[illegible]
L,PLCG1,DIDO1,FLNB,CORO1C,KIF3C,PHC2,LSS,STXBP1,ISYNA1,CSNK1D,MESD,HYOU1,MAPK[illegible]
,MARCKSL1,AFF4,MATR3,YWHAG,COL4A2,RPL37A,SPEN,NONO,DDB1,EIF1,HACD3,DCHS1,EGL[illegible]
,SLC4A8,HNRNPU,OGA,PGAM1,NFE2L1,CTNNA1,FADS2,CPE,CNN3,RPL29,TPM4,TNS3,TSPAN3,[illegible]
A7,PAPOLA,HK1,RHOA,ZNF865,CLSTN1,ATF4,GLIPR1L2,RPS14,CXADR,CHD4,RPS20,LENG8,WD[illegible]
MDK,LAMC1,JPT1,CBX1,RAD23B,ORAI2,TNPO1,ASXL1,NRDC,RNF10,ARF3,MVB12B,SRSF9,DST[illegible]
42,GCN1,SKI,RAB11B,SRP14,PRPF6,TAF7,RPL37,ATG2B,CCNL2,ARPC2,FAM168B,ARGLU1,PTB[illegible]
A8A,WBP2,CCT5,SYP,PPDPF,RBBP7,DFFB,EIF2AK1,TXLNA,SGTA,KDM5C,ACTR1A,SF1,RETREG2[illegible]
,TNPO1,SKI,SCAF1,TOP1,SELENON,RAPGEF1,TUBB2A,MVB12B,PPP2CA,CCND1,AZIN1,SRSF9,P[illegible]
,STARD7,SF1,CSNK2A1,PHB2,SLC39A7,SRSF6,RER1,UBFD1,TRA2B,COX7C,ZFR,NAP1L3,AGPAT[illegible]
PP1CB,GNAI3,GAS7,PARP1,TLN2,MORF4L2,HSPH1,HNRNPA3,PDXK,ACO2,TCP1,PTOV1,MLEC,[illegible]
K,NSMF,DPP7,NELFCD,SLC5A7,ITPK1,SOS1,MCRIP1,ZNF362,KPNA3,ROCK2,CAMK2N1,MAP2K[illegible]
AT4,MYO5A,COL6A2,NGRN,GOLM1,SESN3,B4GALNT4,MTSS2,RPL35A,DNAJC5,APLP2,MXD4,H[illegible]

1,RHOB,HNRNPH1,LGALS3BP,CKAP4,SEPTIN3,COTL1,CCT6A,CTBP1,GPC1,CDK16,SLC35E2B,H
TPM1,EIF5A,PEBP1,CCND2,PNMA1,GPC3,DDR1,PIP4K2B,ACSL6,SOX12,MMP14,DDX3X,RPS15,R
ATR3,YWHAG,CKB,COL4A2,RPL37A,LRRC28,DDB1,PSMD5,YWHAQ,EIF1,DCHS1,SET,EGLN1,MA
IP2,XAF1,COL11A1,MARCHF6,AP3D1,SBF1,DNAJA1,CDC42BPB,MAT2A,TMEM170A,SLC4A8,PXD
HNRNPA0,RPS16,RPL23,RPL29,EIF4H,LAMA5,RPS9,XRCC5,CIPC,RPL24,PAPOLA,HK1,NCS1,RPL
L23A,SERF2,CTDSP2,RNF187,U2AF2,TBCD,MLXIP,SUGP2,PGD,MEST,GDI2,TSR1,MORF4L2,LM
WHAB,SND1,RPS12,TTR,CALU,PLPPR3,ARCN1,RPS27,MAGED4B,ARHGDIA,CAPRIN1,SLC35E2B,
19,MIF,CHPF,CD63,NGRN,DNAJC5,HNRNPDL,RPLP2,DDOST,TSR2,NARS1,DUX4,YWHAB,PTGES3,
DCHS1,SET,MAP9,CHSY1,TMSB10,EIF4A1,RPL15,GTF3C4,LDHB,DNAH8,PDIA3,SERINC1,ZNF43,
A,PTP4A2,FAU,LSS,B2M,STXBP1,ISYNA1,PLXNA3,ATXN2L,HECTD1,HGS,SREBF2,FBRSL1,MEX3A
B,RPL23A,HSBP1,MAP1A,DDX39B,SRSF5,TM9SF3,MDH2,U2AF2,EEF1A2,FDFT1,CCT2,SUGP2,G
B2A,SRSF9,PDE7A,PSMD3,RABL6,ODC1,SRSF1,IPO7,ARGLU1,TCF25,ADAM9,FTO,ILF3,LUZP6,C
B,ATP1A1,TMEM259,RPS20,CD81,LASP1,CCT7,SRSF4,PFN1,JPT2,PNN,PRKCSH,HNRNPC,CCNI,K
0,CTNNA1,TPM4,EIF4H,SCARB2,NACA,RTN4,RPS9,CIPC,RPL24,SLC16A7,RPL17,C1ORF43,RPL1
X5,CLSTN1,RAB6B,NLGN2,RPS14,CHD4,ATP1A1,TMEM259,CD81,WDR1,CRMP1,LASP1,BEX3,R
EIF5,ATXN7L3B,LASP1,CCT7,PFN1,MCL1,NES,CCNI,XPO1,SLC3A2,PTN,PFKL,HDLBP,GLG1,SFP
LNA,TRIO,LBH,SF1,HIP1,H2AZ1,CD99,ZFP36L1,KLHL42,CSNK1G2,GNG4,NXN,EP400,KCTD12,
,CD164,MASTL,HDAC6,KCTD12,PTEN,HNRNPA2B1,ULK1,EML1,ITGB5,RPS2,ZMYM4,HNRNPAB,
1,HLA-C,SF3B1,ZNF43,BASP1,POLR2A,TPT1,RPL18,UCHL1,MORF4L1,NCAM1,ANXA2,AGRN,ZM
7,C1ORF43,RPL22,PTCD3,ZNF865,RPL9,KIF5B,RPS14,CXADR,ATP1A1,TMEM259,RPS20,COL5A
JUND,TMEM259,RPS20,EIF5,COL5A2,WDR1,HMGN2,CRMP1,LASP1,BEX3,VAT1,SRSF4,RTN3,PF
A1,ARF1,MAP4,CDC42BPB,AP2B1,TMEM170A,GTF2F1,SLC4A8,GOLGA8A,PXDN,OGA,NFE2L1,C
,RAB2A,GNG4,EP400,PAPSS1,CRTAP,ZFR,TBC1D9B,PPP1CC,IL6ST,SPINT2,ITGB5,MAP1LC3B,C1
,HNRNPC,LRPPRC,XPR1,KHDRBS1,SPTBN2,SCRN1,RPS25,EIF5A,PDIA6,MEGF8,SOX12,RAB7A,P
6,GPC3,PLXNA1,PIP4K2B,SOX12,CCN2,ACTN4,DDX3X,ITM2C,ARL4C,RCN1,GOLM1,MTSS2,HNR
E,CNN3,RPSA,CTSD,RTN4,LPCAT1,LAMA5,RPS24,SLC22A17,CIPC,PJA2,RPL24,SLC16A7,PAPOLA
,TRAF4,TRAF7,WDR45B,DGCR2,STARD7,KDM5B,KIF21B,SEC16A,GPX4,SLC35F5,RAB2A,GNG4,
2,VAMP2,MEST,NOP53,PABPN1,HNRNPL,LMNB2,HNRNPA3,PTPRA,ACTR3,YEATS2,TBC1D16,AC
A1,GPC3,MEGF8,CCN2,DDX3X,RPS15,CLTC,MRFAP1,RPS5,RPL35,MIF,JUN,COL6A2,SESN3,MT
ATP5F1A,EIF4B,MARCKSL1,UBB,AFF4,TRIM28,MATR3,YWHAG,CKB,LRRC28,MAPT,FN1,SPEN,N
4K2B,MEGF8,ACSL6,RAB7A,MMP14,ITM2C,MRFAP1,MAGED2,GNB2,CHPF,COL6A2,GOLM1,B4
AP12,ITM2B,AP2B1,GTF2F1,GOLGA8A,SMARCA5,CTNNA1,LDHA,CPE,CNN3,TNS3,CTSD,SCARB
RPL23,RPL29,TPM4,EIF4H,LAMA5,RPS24,RPS9,XRCC5,CIPC,RPL24,PAPOLA,COL1A2,HK1,NCS1
4A,ACO2,HSPA4,TCP1,MLEC,ZNF302,DDX42,GCN1,SKI,RAB11B,SRP14,PRPF6,TAF7,RAPGEF1,R
RAB27A,U2AF2,EEF1A2,FDFT1,CCT2,SUGP2,PARP1,TSR1,MTRF1L,RPL27,PTPRA,PRDX5,UBE4A
,ANXA5,HNRNPL,TGFBI,ALKBH5,PKN1,GRINA,CLSTN3,ACAP3,DDX42,LAMC1,CHD2,ORAI2,ASXL
5,CLSTN1,ATF4,KIF5B,CHD4,TMEM259,EIF5,ATXN7L3B,ATP2A2,BEX3,CCT7,OAZ1,RBBP4,RTN3
,USF2,SPPL2B,FXR2,CHMP1A,PINK1,USP10,KDM3A,ARPC1A,TAOK2,RTF1,NOP56,GTF3C1,TIA1
PFN1,ASTN2,PNN,RPL31,NISCH,HNRNPC,TADA2B,KHDRBS1,XPO1,PFKL,SCRN1,TKT,EIF5A,CDH
C2,PPP2R1A,RPL15,GTF3C4,HLA-C,PDIA3,SERINC1,SSBP2,ZNF43,RPS3,BASP1,VAPA,HMGB1,P
3X5,CLSTN1,PSMD2,RAB6B,SLC7A5,NLGN2,CXADR,ATP1A1,JUND,TMEM259,EIF5,CD81,LENG8,
RPS20,LENG8,WDR1,RPL26,SRSF4,MCL1,TLE5,PNN,PRKCSH,HNRNPC,CCNI,TADA2B,XPR1,KHD
M9SF3,MDH2,TOMM20,U2AF2,CCT2,RPN2,GNAI3,NOP53,GDI2,TM9SF2,TSR1,SSRP1,HSPH1,LM
NK1G2,GNG4,FARP1,NXN,MASTL,HDAC6,KCTD12,ZNF512B,CUL3,ULK1,IL6ST,MCM3AP,ITGB5,
,RPS5,IK,RPS19,CHPF,JUN,MYO5A,NGRN,DNAJC5,APLP2,HNRNPDL,RPLP2,DDOST,TSR2,CIRBP

,PPP2CA,AZIN1,LAPTM4A,SRSF9,PDE7A,FAM171A2,PSMD3,RABL6,ODC1,IPO7,ARGLU1,TCF25,A
0,MARCHF6,AP3D1,DNAJA1,ARF1,CYB5R3,AKAP12,SF3B2,GTF2F1,PGAM1,HNRNPA0,FADS2,RP
91,PJA1,ADNP,SCAMP4,ADGRB2,HTRA1,ANKFY1,ETF1,ARHGAP5,SRSF10,ECHS1,RPL27A,TBC1D
CRMP1,VAT1,OAZ1,RTN3,PFN1,TLE5,NES,RPL31,TADA2B,XPR1,KHDRBS1,SPTBN2,PFKL,HDLBP
2,TMEM248,DDX42,GCN1,SKI,RAB11B,SRP14,PRPF6,TERF2IP,TAF7,RPL37,ATG2B,CCNL2,ARPC
1B,SRP14,PRPF6,TAF7,RPL37,PPP2CA,CCNL2,ARPC2,FAM168B,PTBP1,CAPZB,SUPT5H,ATXN10,
4X,RPS3A,STMN1,ITGB1,TUBB4B,RPL11,AGRN,RPLP1,ZMIZ1,TIMP2,COL11A1,RPL10,WSB1,RPS
GB,TOP1,SELENON,OPTN,ATG2B,AZIN1,MAPRE1,KDELR2,SRSF9,DSTN,PLEC,BTBD2,PBX2,PSMI
I,SKI,SRP14,PRPF6,TAF7,RPL37,PPP2CA,CCNL2,ARPC2,FAM168B,ARGLU1,PTBP1,CAPZB,ATXN1
T,GLG1,SFPQ,EIF5A,PEBP1,CDH2,PNMA1,PIP4K2B,SOX12,CCN2,ACTN4,DDX3X,RPS15,GNB2,H1
1,NONO,PSMD5,YWHAQ,HACD3,DCHS1,SET,MAP9,TPI1,CHSY1,TMSB10,INA,EIF4A1,RPL19,PPP2
IPDC1,IL6ST,GPSM1,GPC4,PCBP4,TMOD2,ZNF664,ZMYM3,PALM,ERGIC1,SHC2,FKBP4,LINGO1,
,PFN2,HNRNPD,FBRSL1,TCTN1,SERBP1,BEX4,VDAC1,TRAF4,RBBP7,TRAF7,ST3GAL6,MSMO1,X
YWHAG,COL4A2,RPL37A,LRRC28,SPEN,NONO,PSMD5,YWHAQ,HACD3,SET,TMSB10,INA,EIF4A1,
YB5R3,SF3B2,GTF2F1,SLC4A8,PGAM1,RPL23,RPL29,TPM4,EIF4H,LAMA5,RPS9,XRCC5,CIPC,PJ
PGRMC1,NOVA2,CTSB,ZC3H7B,TBCD,SNX22,SQSTM1,VAMP2,VCAN,NOX4,TAF15,TGFBI,SSRP1,
CAF1,TERF2IP,TMED2,TOP1,RPS15A,RNF10,RAPGEF1,TUBB2A,PPP2CA,MAPK1,AZIN1,ARPC2,DD
AZ1,PFN1,TLE5,NISCH,CCNI,LRPPRC,TADA2B,SPTBN2,CSNK1E,RPS25,TKT,GLG1,CDH6,SFPQ,T
MCL1,JPT2,PNN,SPC25,HNRNPC,LRPPRC,TADA2B,SLC3A2,CSNK1E,PFKL,HDLBP,SFPQ,EIF5A,RA
GLIPR1L2,RPS14,ATP1A1,TMEM259,RPS20,LENG8,WDR1,RPL26,OAZ1,SRSF4,MCL1,TLE5,PNN,F
AGRN,RPLP1,XAF1,MIDN,RPL10,MARCHF6,RPS17,DNAJA1,ARF1,AKAP12,SF3B2,ITM2B,GTF2F1,
B1,PSMD5,YWHAQ,EIF1,HACD3,SET,EGLN1,MAP9,TMSB10,EIF4A1,RPL19,GTF3C4,DNAH8,PKD1,
FE2L1,FADS2,RPS16,RPL23,CPE,CNN3,RPL29,RPSA,TPM4,NACA,LAMA5,RPS24,SLC22A17,BRD3
MT1,PDAP1,MMP15,PHB2,GSPT1,UBA52,PLXNB1,CSNK1G2,FARP1,RER1,UBFD1,IARS1,COX7C,
17,VCL,SBF1,DNAJA1,ARF1,AKAP12,ITM2B,TMEM170A,SLC4A8,GOLGA8A,LARP1,FADS2,CPE,CN
LM3,GNB1,SHISA9,STMN2,RPS8,MYH9,RPL7,SEPTIN2,P4HB,APP,IDS,ERVV-1,RPL5,GDI1,TMBIM
,RPN2,PPP1CB,NOP53,NOX4,HSPH1,PTPRA,AHCYL1,ACAP3,DDX42,JPT1,CBX1,PTK7,CCT4,ASXL
RN1,RPS25,SFPQ,EIF5A,PDIA6,MEGF8,ACSL6,RAB7A,RPL12,ITM2C,RPL30,EIF3L,MAGED2,RPS5
2,VASH1,SART1,AP1M1,SAP18,BNIP3,SAE1,PRDX3,HSDL1,UBE2K,USO1,ATP5MG,NELFB,DENND
RNPH1,UBE2H,RPS12,CCT6A,CTBP1,CAPRIN1,GPC1,TOP2B,SMCHD1,IPO5,DNER,LAPTM4B,GLU
DC4,TXLNA,PRMT1,PDAP1,AP2A1,MMP15,EIF3D,SEC16A,PHB2,APMAP,GSPT1,UBA52,PLXNB1,RA
C3,SCRIB,MSH6,NOC2L,KDM4A,PSMC3,NASP,UBE2I,XPO6,UBXN6,DAPK3,DPYSL4,SLC38A2,HSD
CCAR2,ARF4,SMARCC1,SUPT6H,PITPNM1,CSNK1D,BAG6,RAB11A,SERBP1,ARFGEF2,WBP2,AKIF
2G2,EIF5B,NOMO1,FEM1B,ECEL1,CBX6,PTPRF,MCM3AP,SPINT2,LMAN1,KLHDC3,CHERP,GALN
TBN2,CSNK1E,RPS25,TKT,GLG1,CDH6,SFPQ,TPM1,EIF5A,PEBP1,CCND2,PDIA6,GPC3,PLXNA1,P
GF13,SETD1A,FUBP1,TMCO1,MYEF2,TLE4,LRP6,PRDX4,PPP4R3B,DUS1L,RBM39,EMD
CAND1,SON,ZFAND5,DUSP4,TUG1,ATXN2L,HECTD1,BAG6,SREBF2,SERBP1,SMS,RBM15B,TXLNA
2B1,MAT2A,GTF2F1,SLC4A8,GOLGA8A,HNRNPU,PGAM1,HNRNPA0,CTNNA1,FADS2,LDHA,CNN3,
MLEC,ACAP3,SKI,PRPF6,DPYSL2,SELENON,VHL,MAPRE1,FAM168B,AEBP1,PTBP1,CDC37,SUPT5
KS2,PLXNA3,ID4,BAG6,FBRSL1,ZSCAN18,WBP2,IGFBP7,SMS,NAP1L4,DDIT4,ADAR,RBM15B,VDA
IMGN2,ATP2A2,CCT7,VAT1,RTN3,RPL31,PRKCSH,HNRNPC,HDLBP,GLG1,PEBP1,CCND2,CDH2,P
NF865,ATF4,PSMD2,RPS14,CXADR,CHD4,RPS20,LENG8,WDR1,RPL26,LASP1,SRSF4,PFN1,ASTN
M15B,TXLNA,NAP1L1,TXNDC5,LBH,CSNK2A1,GSPT1,PPM1G,KLHL42,HOXB5,GARS1,GNG4,UPF
HSF1,WDFY1,NUS1,MAP1S,PRMT5,MED25,RAVER1,NDUFS1,OTUB1,KARS1,SPCS2,NKIRAS2,PM
A2,STMN3,LSS,TNKS2,TUG1,PLXNA3,ATXN2L,TMED9,IDH1,SARS1,MPRIP,UQCRC2,SOD1,SREBF

3,MDH2,TOMM20,U2AF2,EEF1A2,CCT2,SUGP2,GNAI3,NOP53,GDI2,TM9SF2,TSR1,SSRP1,LMNB
PDIA6,GPC3,PIP4K2B,MEGF8,PRDX2,RPL12,CLTC,GNB2,RCN1,H1-0,RPL35,CHPF,CD63,GOLM1
RINA,NCKAP1,PTOV1,RBMX,LAMC1,JPT1,KDM6B,PYGB,PRPF6,TOP1,VEGFA,VHL,G3BP2,AZIN1,
N1,PSIP1,MAPRE3,NSD1,ZBTB4,RIMKLB,ZEB1,PLPPR2,ASH1L,RAB3A,ANKRD50,STC2,DLG4,EG
APL1,RAB14,MAZ,IREB2,LRATD2,NUCKS1,LSM14B,FNBP1L,KRT8,DCUN1D2,LTBP3,ITGAV,SCRI
HDGF,RPL38,NOTCH2,VEGFA,MVB12B,KDELR2,DSTN,BTBD2,PBX2,AEBP1,NYNRIN,RBP1,WDR6,
,MIDN,RPL10,MARCHF6,RPS17,VCL,CYB5R3,AKAP12,CDC42BPB,ITM2B,ZNF770,MAT2A,GTF2F1
,VASH1,PACS2,TMEFF1,BNIP3,PHIP,WNK1,BAG5,UBXN6,SLC20A1,SORT1,CPXM1,DMWD,SCD,F
29,RPSA,TPM4,NACA,LAMA5,RPS24,BRD3,RPS9,XRCC5,CIPC,PJA2,RPL24,RPL17,C1ORF43,PAP
IP70,ASXL1,H3-3A,TMED2,NOTCH2,VEGFA,SELENON,ARF3,KDELR2,AEBP1,NYNRIN,EIF3E,CDC3
,RPL37,ARPC2,FAM168B,ARGLU1,PTBP1,SON,CCAR2,ARF4,SMARCC1,SUPT6H,PITPNM1,CSNK1
,CAND1,NPEPPS,KDM1A,FKBP8,ZFAND5,DIDO1,DUSP4,DLK1,SUPT6H,SRSF3,NKTR,STT3B,STXB
1,NYNRIN,ILF3,WDR6,UBE2R2,ATXN10,CLIP3,ZFAND5,HNRNPUL2-BSCL2,MSL1,SLIT3,ILF2,STM
10,MTCH1,VPS26B,ERH,CYTH2,EIF3H,SNRPD3,RAB1B,RALY,SEC14L1,NUDCD3,SSR3,GNL1,STK
5,CHPF,CD63,ANXA6,B4GALNT4,MTSS2,RPL35A,RPLP2,ARL6IP1,NDFIP1,RPL36,DUX4,YWHAB,S
,MARCHF6,WSB1,RPS17,SBF1,ARF1,CYB5R3,SF3B2,CDC42BPB,ITM2B,AP2B1,TMEM170A,GTF
FKBP1A,CTTN,DAP,SERP1,AKAP9,RHBDD2,STK35,ERH,EIF2S2,RAB1B,UBE2M,SEC14L1,NUDCD
3,PAPOLA,RPL14,PTCD3,CBX5,ZNF865,CLSTN1,ATF4,KIF5B,CHD4,ATP1A1,TMEM259,EIF5,COL
,CCND2,PDIA6,GPC3,PLXNA1,DDX3X,ARL4C,GNB2,H1-0,RPS19,MIF,HMGCS1,TLN1,MYO5A,COL
,ACO2,HSPA4,AHCYL1,MLEC,TNPO1,SKI,KDM6B,RAB11B,SRP14,NRDC,PRPF6,DPYSL2,RPS15A,
T1,LAMA5,RPS24,SLC22A17,BRD3,CIPC,PJA2,SLC16A7,IGFBP3,COL1A2,NCS1,RHOA,ZNF865,CL
2,RPL24,C1ORF43,PAPOLA,RPL14,PTCD3,CBX5,ZNF865,CLSTN1,ATF4,KIF5B,ATP1A1,TMEM25
AF1,SARAF,MIDN,RPL10,AP3D1,MAP4,CDC42BPB,SLC4A8,HNRNPU,OGA,PGAM1,SMARCA5,HNR
LK1,TMEM63B,ELP1,RAB6A,ERGIC3,FOXK2,AKT2,ARPP19,MBD3,AP2A2,YY1,TMEFF1,BNIP3,PH
IP4B,PACS2,RBM6,CREBBP,ANKRD12,CCDC47,MICAL1,ATG4B,DMWD,CROCC,TUBB4A,TECR,L
L1,TLE5,PNN,RPL31,PRKCSH,LRPPRC,CSNK1E,PFKL,SCRN1,RPS25,EIF5A,PDIA6,DDR1,PIP4K2
EM63B,ELP1,SLC7A1,PDPK1,MSH6,NOC2L,ANKRD12,WNK1,TRIM8,MICAL1,DPYSL4,CROCC,NS
ENG8,WDR1,HMGN2,ATP2A2,BEX3,VAT1,MCL1,CCNI,TADA2B,XPR1,HDLBP,GLG1,SFPQ,EIF5A,
MOAP1,PDPK1,LTBP3,PSMC3,PACS2,CREBBP,UBE2I,ANKRD12,UBXN6,DPYSL4,METRN,UNC119
10,CITED2,MCRIP1,ARPC5,PPP1R26,SLC25A29,HEG1,QRICH1,TFDP1,YBX3,FAM20C,EPHB2,LR
GALT5,ATF7IP,DAD1,RAB18,ATP5PO,CRTC1,TMEM43,ZBED4,ZFP91,FBL,NAPB,TBCA,HSPE1,ATP
T1,PREPL,TIMM50,ZMYM4,SEPTIN7,HIF1A,SYMPK,FBXO21,SAFB2,PPP1R10,TRIR,ERH,EIF3H,AE
SB1,RPS17,AP3D1,DNAJA1,ARF1,AKAP12,SF3B2,AP2B1,MAT2A,HNRNPA0,FADS2,RPL23,LDHA,C
5F1A,EIF4B,UBB,AFF4,MATR3,YWHAG,CKB,COL4A2,RPL37A,LRRC28,MAPT,FN1,SPEN,NONO,
PM,YWHAH,FBLN1,PITPNM1,CSNK1D,SF3A1,ATXN2L,NT5DC2,HECTD1,SARS1,MPRIP,VANGL2,
1,SSRP1,HSPH1,LMNB2,MTRF1L,ALKBH5,SNRPN,PRDX5,UBE4A,ACO2,HSPA4,TCP1,MLEC,ACA
CSH,HNRNPC,TADA2B,XPR1,KLHL9,SPTBN2,HDLBP,GLG1,SFPQ,PEBP1,PNMA1,PDIA6,SULF1,P
4,KCNQ2,BCL9L,WDR45B,MTA1,KIRREL1,LBH,NDUFV1,KDM5B,PBX1,NOL4L,BAZ1B,LRRC4B,K
ECEL1,TBC1D9B,HNRNPA2B1,EML1,HNRNPAB,GLO1,FUS,TRIR,PELP1,VAPB,CCN1,HSPG2,UBE2
CKS,SMG1,CALM1,NPM1,PLXNB2,PLD3,USP11,COL18A1,SNRNP200,CSDE1,IGF2,EIF3C,ATP5F
RNPU,HNRNPA0,LDHA,CPE,CNN3,RPSA,CTSD,SCARB2,RTN4,LPCAT1,LAMA5,RPS24,SLC22A17,
,CHERP,MAP1LC3B,C1ORF198,MTCH1,XRCC6,SHC2,SNRPD3,SOX2,GPC2,ARHGEF17,GABARA
11A,SERBP1,ARFGEF2,WBP2,AKIRIN1,BMPR2,SMS,CCT5,CDC42,PPP1R9B,YME1L1,TRAF7,EIF2
OLA,COL1A2,LLGL1,RPL14,HK1,NCS1,CBX5,CLSTN1,ATF4,RAB6B,GLIPR1L2,CXADR,ATP1A1,J
P2L,MASTL,HDAC6,KCTD12,SMC3,SELENOW,CUL3,FMNL2,ULK1,IL6ST,OAZ2,MCM3AP,POM12

DX39B,NUCB1,SRSF5,TM9SF3,MDH2,TOMM20,U2AF2,CCT2,RPN2,GNAI3,NOP53,TM9SF2,HSPH
P2,PDE7A,PSMD3,TCF25,PTBP1,ILF3,CDC37,WDR6,SCAF11,BNIP3L,ARF4,DUSP4,PITPNA,PLTP,
L29,TNS3,CTSD,EIF4H,TSPAN3,LPCAT1,LAMA5,RPS24,BRD3,XRCC5,MSN,KPNB1,C1ORF43,PAF
SP1,SRSF4,PFN1,ASTN2,MCL1,TLE5,PNN,RPL31,PRKCSH,CSNK1E,PFKL,SCRN1,HDLBP,RPS25,
MR4,MKRN1,PCNX3,YIPF3,CAP1,ATP6V0D1,PRPF19,USP24,NOC2L,NASP,AP1M1,SMARCD1,UBE
B,XPR1,KHDRBS1,SLC3A2,CSNK1E,PFKL,TKT,CDH6,CDH2,MEGF8,ACSL6,SOX12,PRDX2,DDX3X,
4,NANOG,GNB2,RPS19,GJA1,COL6A2,CD63,DNAJC5,RPLP2,MXD4,TAGLN,HNRNPK,H1-10,CIRB
DC,PRPF6,TAF7,RPL38,RPL37,MAPRE1,ARPC2,FAM168B,ARGLU1,LUZP6,HSPB1,SON,RPL28,C
KTR,STMN3,PFKM,TNKS2,STT3A,LONP1,HYOU1,ID4,NBPF9,SARS1,VANGL2,BAG6,HNRNPD,RAB
AEBP1,NYNRIN,PTBP1,ILF3,PCSK1N,SUPT5H,ACTN1,HSPB1,SON,BNIP3L,MAP7D1,CLIP3,FKBP8,
RDX5,UBE4A,ACO2,HSPA4,TCP1,MLEC,DDX42,GCN1,SKI,RAB11B,SRP14,PRPF6,TAF7,RAPGEF1,
CC5,PJA2,MSN,RPL24,SLC16A7,KPNB1,PAPOLA,RPL22,RHOA,PTCD3,ZNF865,RPL9,ATF4,GLIPR
CC5,PJA2,MSN,RPL24,SLC16A7,KPNB1,PAPOLA,RPL22,RHOA,PTCD3,ZNF865,RPL9,ATF4,GLIPR
JA1,ARF1,AKAP12,SF3B2,AP2B1,MAT2A,HNRNPA0,FADS2,RPL23,LDHA,CNN3,RPL29,RPSA,TNS3
SPA4,TCP1,MLEC,ACAP3,TMEM248,GCN1,SKI,RAB11B,SRP14,NRDC,PRPF6,TAF7,VHL,MAPRE1,
X3,OAZ1,RBBP4,RTN3,JPT2,TLE5,PNN,CLEC7A,NES,RPL31,PRKCSH,KLHL9,CSNK1E,RPS25,SFP
MGCR,ARHGDIA,STIP1,SLC35E2B,IPO5,ATP6V0C,RPL23A,FAM199X,SERF2,CPSF2,DDX39B,EID1
ADR,EIF5,LENG8,WDR1,ATXN7L3B,ATP2A2,CRMP1,LASP1,CCT7,SRSF4,ASTN2,MCL1,TLE5,PNN,
,EIF3H,INTS3,ABI2,UBE2M,SEC14L1,ELAVL1,NUDCD3,TM9SF4,SH3GL1,FLOT2,TUBGCP6,EEF1B
67A,KLHDC3,CHERP,FKBP1A,CTTN,STK35,ERH,CYTH2,SNRPD3,RALY,SEC14L1,ELAVL1,NUDCD
,SSRP1,LMNB2,MTRF1L,ALKBH5,SNRPN,PRDX5,UBE4A,ACO2,HSPA4,TCP1,AHCYL1,MLEC,ACAP
OC728392,ATXN2L,COL27A1,FBRSL1,KDM3B,ARFGEF2,BMPR2,SMS,CTXN1,RBM15B,PPDPF,S
8,KIFAP3,TTC37,PNISR,PLEKHB2,INTS11,PCNX3,ABCF1,ADAM10,SPTY2D1,BORCS7,OGT,COX6
KSL1,UBB,AFF4,MATR3,YWHAG,RPL37A,LRRC28,YWHAQ,EIF1,HACD3,DCHS1,SET,EGLN1,TPI1
1,MLEC,ACAP3,GCN1,TNPO1,RAB11B,SRP14,NRDC,PRPF6,TAF7,ARF3,MAPRE1,FAM168B,ARGL
2,KPNA3,AFDN,CAMK2N1,MEIS3,RFX7,NIPA1,GALNT2,PPP1R26,PPP3CA,CTDNEP1,SLC1A4,ZNF
14,PTCD3,CBX5,ZNF865,CLSTN1,ATF4,KIF5B,CHD4,ATP1A1,JUND,TMEM259,EIF5,HMGN2,ATP
,RPS5,IK,RPS19,CHPF,JUN,NGRN,RPL35A,HNRNPDL,RPLP2,DDOST,HNRNPK,CIRBP,NARS1,RPL
OK1,VAPB,PALM,ERGIC1,RTL6,CCN1,SHC2,ANTXR1,STRN4,RAB1B,ELAVL1,TLE1,PGAP6,MAZ,H
1,CBX5,ZNF865,CLSTN1,ATF4,NR2F1,KIF5B,RAB6B,NLGN2,GLIPR1L2,CXADR,CHD4,ATP1A1,R
,VPS35,PTOV1,MLEC,ACAP3,SURF4,PCDHGB6,SNRNP70,PRPF6,DPYSL2,TOP1,NOTCH2,RAPGE
,AKAP12,SF3B2,CDC42BPB,ITM2B,GTF2F1,PXDN,HNRNPU,OGA,PGAM1,FADS2,LDHA,CNN3,CT
PO5,ATP6V0C,SKP1,LAPTM4B,CPSF2,SRSF5,TM9SF3,U2AF2,CCT2,MLXIP,MORF4L2,HSPH1,MT
BX6,PRRC2A,HNRNPA2B1,ANP32B,MCM3AP,RPS2,FKBP1A,SAFB2,PPP1R10,ERH,PELP1,SNRPD
1,FAM168B,PTBP1,FTO,LUZP6,CDC37,SUPT5H,SON,CCAR2,ARF4,DIDO1,POLR2B,CORO1C,XPN
,SEPTIN2,P4HB,APP,IDS,ERVV-1,RPL5,KLRD1,GDI1,FSCN1,TMBIM6,GSTP1,MARCKS,SMG1,CAL
2B,ARPC2,FAM168B,ARGLU1,PTBP1,ATXN10,SON,CCAR2,ARF4,POLR2B,SMARCC1,DHX30,PIT
N2,ATP2A2,RPL26,LASP1,CCT7,SRSF4,RBBP4,RTN3,PFN1,MCL1,PNN,RPL31,NISCH,PRKCSH,L
O3,USP47,LMO4,PTPN11,RSF1,USP10,DAG1,GTF3C1,H6PD,MKNK2,GGNBP2,PPP1R14B,KIF26
O3,USP47,LMO4,PTPN11,RSF1,USP10,DAG1,GTF3C1,H6PD,MKNK2,GGNBP2,PPP1R14B,KIF26
70,XPO6,UBXN6,MYL12B,GAK,LYPLA2,ATP13A1,PAFAH1B2,POLDIP2,TRIP12,XRN2,CHMP1A,PS
L1,FXR2,SGSM2,ZZEF1,SLC22A23,H6PD,KLF13,RGL2,SMPD3,OBSL1,CLUH,PPP1R14B,DCAF5,U
HGS,BAG6,RAB11A,SERBP1,AKIRIN1,CDC42,PPP1R9B,YME1L1,VDAC1,TRAF7,EIF2AK1,PRMT1,
RPL28,ZFAND5,CORO1C,SDF4,SMARCC1,HNRNPM,NKTR,YWHAH,ISYNA1,STT3A,PLXNA3,IDH1,S
,NPTN,SLC25A6,NUDT4,DCAF5,EHMT2,NCKAP5L,AMPD2,ZNF462,FAM53C,ARHGAP5,MOGS,CA

RIB2,GLUL,SQLE,TIMP3,PGRMC1,SRP9,ZC3H7B,CHD6,U2AF2,EEF1A2,TBCD,MLXIP,ANXA5,CPSI
MCL1,JPT2,PRKCSH,LRPPRC,TADA2B,XPO1,SPTBN2,SLC3A2,GLG1,CDH6,TPM1,EIF5A,CDH2,GI
PGEF1,TUBB2A,RPL37,ARF3,VHL,MAPRE1,SRSF9,CCNL2,DDAH2,DSTN,PBX2,RABL6,AEBP1,NYNI
,ID4,NT5DC2,BAG6,RAB11A,SERBP1,ARFGEF2,WBP2,AKIRIN1,AMOTL2,CCT5,CDC42,YME1L1,D
BX2,AEBP1,DYNLL2,GRB2,CLIP3,DHX30,GIGYF1,STMN3,FBLN1,PITPNM1,LONP1,CSNK1D,HYOU
NPEP3,ATP6V1B2,SRSF3,LARS1,YWHAH,TNKS2,UBQLN1,CSNK1D,IFITM3,NCOR2,VGF,RPL39,ML
EC14L1,SMG5,NUDCD3,GNL1,PNISR,STK24,FLOT2,TUBGCP6,RIC8A,NCSTN,INTS11,MOAP1,MTI
M20,U2AF2,EEF1A2,CCT2,CPSF1,GNAI3,NOP53,GDI2,TM9SF2,TSR1,SSRP1,LMNB2,MTRF1L,ALK
R1,HSPH1,ABCA3,NCKAP1,COL2A1,PTOV1,ACAP3,TMEM248,SNRNP70,PRPF6,TOP1,VEGFA,VHL
JPT6H,DHX30,PTP4A2,TUG1,TMED9,STRAP,BAG6,RAB11A,KDM3B,WBP2,SMS,AMOTL2,CDC42,F
JPT6H,DHX30,PTP4A2,TUG1,TMED9,STRAP,BAG6,RAB11A,KDM3B,WBP2,SMS,AMOTL2,CDC42,F
,ATXN2L,NBPF9,PTP4A1,MAPK8IP1,VGF,FBRSL1,MEX3A,MLF2,WBP2,BMPR2,DYNLL1,CCT5,DH
JPT5H,HNRNPUL1,SON,RPL28,CCAR2,SUPT6H,LARS1,PFKM,EIF3F,LONP1,SF3A1,ID4,BAG6,RAB
,PTBP1,ILF3,GRB2,RPS11,SUPT5H,UBE2R2,SCAF11,ZFAND5,PITPNA,SUPT6H,HNRNPM,STT3A,
B,RAB6B,SLC7A5,CHD4,JUND,WDR1,CRMP1,LASP1,BEX3,CCT7,NES,CCNI,SPTBN2,CSNK1E,HDL
DE7A,PSMD3,SRSF1,TCF25,ADAM9,ILF3,GRB2,CDC37,PCSK1N,WDR6,SCAF11,BNIP3L,ARF4,ZF
N4,LPCAT1,LAMA5,BRD3,PJA2,MSN,SLC16A7,C1ORF43,LLGL1,RPL14,HK1,NCS1,RPL22,RHOA,
4L1,ANXA2,SARAF,COL11A1,RPL10,MARCHF6,RPS17,AP3D1,DNAJA1,PAFAH1B1,AKAP12,CDC42
8,ZFAND5,CORO1C,SDF4,SMARCC1,HNRNPM,YWHAH,ISYNA1,STT3A,IDH1,SARS1,SOD1,BAG6,N
,FARP1,MBTPS1,HDAC6,PTEN,SELENOW,SAFB2,PRDX6,ZNF664,TRIR,CYTH2,HSPG2,SSBP3,CA
CEL1,HNRNPA2B1,RPS27A,TMEM167A,TNKS,PSMD8,CHERP,CTTN,SERP1,RHBDD2,STK35,PPP1
,CBX5,ZNF865,CLSTN1,ATF4,KIF5B,CHD4,ATP1A1,TMEM259,EIF5,HMGN2,ATP2A2,BEX3,CCT7,
BRSL1,CYP51A1,AKIRIN1,CD99L2,DDIT4,BEX4,VDAC1,SSR1,BCL9L,WDR45B,SGTA,ALDH3A2,HI
TGOLN2,YY1,AP1M1,EXOC4,NDUFA13,SLC38A10,DHX40,ATP13A1,ATP5MG,SRPRA,NELFB,PAFA
JZP6,QARS1,CCAR2,GNG2,CORO1C,ATP6V1B2,ISYNA1,SF3A1,ZFAND3,HECTD1,BAG6,HNRNPD,F
IM20,U2AF2,CCT2,GNAI3,NOP53,GDI2,TM9SF2,TSR1,SSRP1,LMNB2,MTRF1L,ALKBH5,RPL27,SN

,FADS2,RPS16,RPL23,CPE,CNN3,RPL29,TNS3,TSPAN3,CIPC,PJA2,MSN,SLC16A7,KPNB1,C1ORF
JBE4A,ACO2,HSPA4,TCP1,MLEC,DDX42,GCN1,SKI,RAB11B,SRP14,PRPF6,TAF7,RPL38,RPL37,AT
P2CB,PLXNB1,GNG4,FARP1,UBFD1,PRRC2A,ULK1,PODXL2,CTTN,C1ORF198,NDN,VPS26B,SEPT
M168B,ARGLU1,PTBP1,LUZP6,CDC37,SUPT5H,HSPB1,SON,CCAR2,ARF4,POLR2B,CORO1C,SUP
1,CNDP2,RNF157,EHMT2,RASA4,LIMK1,PITRM1,DRAP1,RHEB,IQSEC1,WIZ,TBC1D14,TBXA2R,SL
APLP1,SESN3,APLP2,SLC25A3,RPLP2,INIP,MYH10,DUX4,CKAP5,SEPTIN11,LAMP1,LGALS3BP,TT
167A,KLHDC3,CHERP,FKBP1A,CTTN,LRP10,SERP1,ERH,EIF3H,RALY,SEC14L1,NUDCD3,FAM234
PPL1,IPO5,TMEM30A,PPIB,PEA15,CDKN1A,DLG5,HS6ST1,PGRMC1,DDX39B,NUCB1,CDH11,TM
BSCL2,PITPNA,XPNPEP3,ATP6V1B2,SRSF3,LARS1,YWHAH,FAU,UBQLN1,CSNK1D,HYOU1,IFITM3
SH3PXD2B,AMFR,TRIP12,CDC34,LUC7L2,KIFC2,YTHDF3,CDV3,TIA1,OBSL1,CLUH,DCAF5,CRIP2,
DFT1,MLXIP,PPP1CB,GAS7,NOP53,HNRNPL,PARP1,TLN2,SSRP1,PRDX5,PDXK,ACO2,AHCYL1,GF
MD5,DCHS1,SET,EGLN1,MAP9,TPI1,CHSY1,TMSB10,INA,EIF4A1,RPL15,GTF3C4,LDHB,DNAH8,P
R,NOMO1,PPP1CC,EML1,OAZ2,BRSK1,PODXL2,FKBP1A,FBXO21,SAFB2,WSCD1,ZNF664,FUS,PF
RPF6,TAF7,ARF3,MAPRE1,FAM168B,ARGLU1,LUZP6,HSPB1,SON,RPL28,CCAR2,ARF4,CORO1C,S
APRE2,SEPTIN7,PODXL2,FKBP1A,MVD,DAP,MTCH1,SAFB2,EFR3B,YARS1,CARM1,RAB1B,ELAVL1
P2,CUL1,WDR26,SHISA5,ZNF462,CUL4A,MYCBP2,ARHGAP5,CCDC88A,CAPN1,WRNIP1,MAGEF1,
3,MORF4L2,HSPH1,MTRF1L,ALKBH5,PRDX5,UBE4A,ACO2,HSPA4,AHCYL1,ETNK1,PTOV1,DDX4
,PTPRA,PKN1,HSPA4,PLOD1,MDK,PTK7,ABCA2,VEGFA,SELENON,RAPGEF1,OPTN,SMARCA4,AZI

6,AHCYL1,VPS35,RBMX,DDX42,RAB5B,ORAI2,ASXL1,SRP14,PRPF6,ATP6AP1,TMED2,RAPGEF1,
12,POR,FAM118A,RTF1,NOP56,NIBAN2,RUVBL2,SC5D,FZR1,FOXP4,DNAJB1,PPP1R14B,KAT2A,C
PNPEP3,ATP6V1B2,SRSF3,LARS1,YWHAH,UBQLN1,CSNK1D,HYOU1,IFITM3,DGKD,VGF,MLF2,SE
,NELFB,PAFAH1B2,ACAT2,POLDIP2,TRIP12,LMAN2,CHMP1A,CLTA,USP10,ZZEF1,TSNAX,PAIP2,Z
AM23,GPRC5B,SETD1B,MEX3D,MYL6B
,SKP1,DNER,TRIB2,GLUL,PEA15,FAM199X,HS6ST1,CTDSP2,PGRMC1,CTSB,CDH11,TM9SF3,CH
PCMTD2,ARRDC3,DPF2,TOLLIP
1,PSMD5,HACD3,SET,MAP9,CHSY1,TMSB10,EIF4A1,MRC2,RPL19,PPP2R1A,RPL15,GTF3C4,LDH
P3,EXOSC10,ZFP91,XPC,DDHD2,SRRT,STRIP1,QRICH1,BCAR1,FAM20C,FUBP1,TMCO1,MYEF2,
C3A2,PFKL,SCRN1,RPS25,SFPQ,EIF5A,PDIA6,DDR1,MEGF8,ACSL6,RAB7A,DDX3X,RPL12,ITM2C,
HB,DNAH8,PKD1,PDIA3,CHD3,VAPA,HMGB1,POLR2A,TPT1,SEC61A1,RPL18,UCHL1,MORF4L1,
YNA1,SF3A1,ATXN2L,PRPS1,GPR107,GHITM,HNRNPD,MEX3A,ZSCAN18,SERBP1,KDM3B,YLPM1,
RNPA2B1,HNRNPAB,FUS,PELP1,VAPB,CCN1,UBE2M,ARHGEF17,TLE1,RDX,MAZ,SRSF2,CADM3,S
2A,HNRNPA0,RPL23,LDHA,CNN3,RPL29,RPSA,TNS3,CTSD,EIF4H,SCARB2,RTN4,LPCAT1,LAMA5,
F3B2,AP2B1,TMEM170A,GTF2F1,SLC4A8,GOLGA8A,OGA,NFE2L1,SMARCA5,HNRNPA0,RPL23,L
2,FAM210B,RSF1,MTA2,CGGBP1,SLC22A23,TOX4,TSPYL2,NIBAN2,KIFC2,ANKHD1,RGL2,RUVBL
,VGF,MLF2,SEC23A,TCTN1,RTL8C,WBP2,AKIRIN1,AMOTL2,YME1L1,CTNND1,TRAF4,STAT3,ARI
TBP1,LUZP6,CDC37,CAPZB,SUPT5H,SON,CCAR2,ARF4,SUPT6H,FAU,PFKM,PITPNM1,CSNK1D,SF
T10,RAN,GOLPH3,PRPH,GALNT17,RHEB,PPP4R3A,MOGS,NECTIN1,SOBP,TMEM131,CDK9,CLPT
1,CHPF,JUN,NGRN,RPLP2,DDOST,CIRBP,NARS1,RPL36,YWHAB,PTGES3,SND1,LAMP1,MAP2,TTR
RPS11,CASC3,NSG2,DIDO1,RPS28,HNRNPM,SRSF3,TUG1,MESD,GPR107,GHITM,SREBF2,PFN2,
K,TCP1,PLOD1,PTOV1,RBMX,TMEM248,DDX42,SNRNP70,DPYSL2,TERF2IP,RNF10,ADAM9,CDC
A1,NASP,TGOLN2,AP1M1,UBE2I,PSMC1,GAK,LYPLA2,SLC38A10,ATP13A1,NELFB,PAFAH1B2,PO
L1,CTNND1,TRAF4,SSR1,KIF5A,PRMT1,XPOT,NAP1L1,CPD,TRIO,EXTL3,LBH,PRKAR2A,HIP1,GTP
,RPL28,ZFAND5,CORO1C,SDF4,SMARCC1,HNRNPM,NKTR,ISYNA1,STT3A,PLXNA3,IDH1,SARS1,
4,IGF2,EIF3C,ATP5F1A,EIF4B,VCP,MARCKSL1,TRIM28,MATR3,YWHAG,CKB,RPL37A,LRRC28,M
,PRKAR1A,CHPF,GJA1,JUN,NGRN,B4GALNT4,DNAJC5,HNRNPDL,RPLP2,ARL6IP1,ST3GAL2,TSR2
G,GNB2,RPS5,HMGCS1,PRKAR1A,TLN1,CHPF,GJA1,JUN,NGRN,B4GALNT4,DNAJC5,HNRNPDL,
DH6A1,SPTBN1,ATP5F1B,CALM3,GNB1,SHISA9,STMN2,RPS8,MYH9,RPL7,SEPTIN2,P4HB,APP,ID
C,KIF5A,KCNQ2,BCL9L,PRMT1,SGTA,FBRS,PBX1,ALDH3A2,AP2A1,KIF21B,ZFP36L1,CXXC5,BRD
P,SQLE,MAP1A,CDKN1A,SERF2,TIMP3,CTDSP2,CNN2,NOVA2,NUCB1,C5ORF24,ZC3H7B,U2AF2,
S2,SF3A1,NBPF9,SARS1,BAG6,HNRNPD,RAB11A,SERBP1,CCT5,CDC42,YME1L1,RBBP7,TRAF7,E
FBRSL1,CYP51A1,CD99L2,DDIT4,BEX4,VDAC1,SSR1,BCL9L,WDR45B,SGTA,ALDH3A2,HIP1,SLC
P1,AKIRIN1,CDC42,PPP1R9B,YME1L1,RBBP7,TRAF7,EIF2AK1,EDC4,TXLNA,WDR45B,PDAP1,M
AF,MIDN,RPL10,MARCHF6,WSB1,RPS17,AP3D1,SBF1,ARF1,CYB5R3,MAP4,SF3B2,CDC42BPB,IT

HOA,CBX5,ZNF865,CLSTN1,PSMD2,SLC7A5,CXADR,CHD4,JUND,TMEM259,EIF5,CD81,ATXN7L3
B1,SERINC1,RPS3,VAPA,POLR2A,TPT1,SEC61A1,RPS3A,ITGB1,TUBB4B,RPL11,MORF4L1,RPLP
DN,NFE2L1,RPS16,RPL23,CPE,RPL29,TPM4,TNS3,CTSD,NACA,TSPAN3,LPCAT1,LAMA5,SLC22A
4,SMARCC1,SUPT6H,FYN,LARS1,NKTR,YWHAH,PFKM,PRPS1,ID4,SARS1,BAG6,RAB11A,ARNT2,W
,HMGN2,VAT1,OAZ1,RBBP4,RTN3,PFN1,MCL1,CLEC7A,TADA2B,XPR1,KLHL9,SCRN1,GLG1,EIF5
HC2,ILF2,PRPS1,GPR107,UQCRC2,COL27A1,SOD1,BAG6,RAB11A,SEC23A,ARNT2,NAP1L4,RBM1
8,NDRG1,DHFR,MRPS6,CRTC1,CACNA1H,SETX,ZBED4,MBOAT7,MAN2B1,CASKIN1,CHKA,HIF3A,
DR1,SRSF4,MCL1,PNN,RPL31,HNRNPC,CCNI,TADA2B,XPR1,HDLBP,EIF5A,PEBP1,CCND2,DDR1,
RF3,MAPRE1,FAM168B,ARGLU1,LUZP6,HSPB1,SON,RPL28,CCAR2,ARF4,CORO1C,SUPT6H,DHX

'TM4B,RPL23A,CPSF2,NOVA2,SRSF5,RNF187,U2AF2,CCT2,RPN2,MLXIP,ANXA5,NOP53,MORF4L[illegible]
HMGCS1,PRKAR1A,CHPF,GJA1,JUN,NGRN,B4GALNT4,DNAJC5,HNRNPDL,RPLP2,ARL6IP1,ST3GA
I,TMEM167A,KLHDC3,CHERP,FKBP1A,MAP1LC3B,SERP1,WSCD1,AKAP9,TAOK1,CYTH2,SNRPD3
4,RPS9,CIPC,PJA2,RPL24,SLC16A7,PAPOLA,NCS1,RPL22,RHOA,PTCD3,ZNF865,RPL9,RPS14,CXA
'PK1,INF2,NEO1,MARVELD1,CAMK2N1,RFX7,RAB5A,PPP3CA,MARK1,COL6A3,PAIP1,LRRN1,HEXI
IM1,ZNF516,FAM20C,MSI2,LRP8,TUT4,CRIM1,PTPN12,CBX4,SAMD1,SESN2,KLF3,SYT4,PFKFB3,
MGAT4B,CXXC5,UBAP2L,CTNND2,ADCY6,ZFR,ZNF512B,CUL3,PTPRF,ULK1,ANP32B,BRSK1,GPS
MGB3,NCKAP5L,HTRA1,B4GALT5,CUL4A,GALNT17,UBE2J1,PPP4R3A,NECTIN1,ECHS1,SOBP,MA
SMARCA5,HNRNPA0,FADS2,RPS16,RPL23,CNN3,RPL29,TPM4,RPS24,BRD3,XRCC5,CIPC,RPL24,S
NFE2L1,SMARCA5,HNRNPA0,CTNNA1,RPS16,RPL23,CNN3,RPL29,TNS3,CTSD,SCARB2,NACA,TSF
RPL14,NCS1,RPL22,PTCD3,RAB6B,SLC7A5,GLIPR1L2,CHD4,JUND,CD81,LENG8,WDR1,RPL26,CF
A,RABEP1,LRP10,FUS,VAPB,YARS1,SHC2,RALY,C6ORF62,GPC2,LDLR,EFTUD2,STK24,ACTR1B,S
,PFN1,ASTN2,MCL1,TLE5,PNN,RPL31,PRKCSH,LRPPRC,KHDRBS1,SLC3A2,CSNK1E,SCRN1,RPS2
,RPS24,SLC22A17,XRCC5,CIPC,MSN,RPL24,SLC16A7,KPNB1,RPL17,COL1A2,NCS1,ZNF865,NR2
BQLN1,LOC728392,PRPS1,MAPK8IP1,BAG6,PFN2,SERBP1,PRPF40A,SMS,RPL34,TP53BP1,RBM
PRPF19,SART1,AP1M1,SSR2,TOMM70,ATG4B,GAK,LYPLA2,ATP13A1,EIF4G1,PAFAH1B2,POLDIP
RPS3A,OR8K1,ITGB1,TUBB4B,UCHL1,RPL11,AGRN,MIDN,RPL10,MARCHF6,WSB1,VCL,ARF1,MA
3B,ATP2A2,ASTN2,MCL1,PRKCSH,HNRNPC,CCNI,TADA2B,XPR1,KHDRBS1,SLC3A2,HDLBP,RPS25
LPCAT1,LAMA5,RPS24,SLC22A17,RPS9,CIPC,PJA2,RPL24,SLC16A7,C1ORF43,PAPOLA,IGFBP3,N
2,TOMM20,RNF187,RPN2,TBCD,MLXIP,PPP1CB,PGD,NOP53,PABPN1,TM9SF2,MORF4L2,ALKBH5
CA,CCND1,AZIN1,SRSF9,ARPC2,PLEC,PBX2,FAM171A2,FAM168B,AEBP1,DYNLL2,WDR6,CAPZB,U
WBP2,AKIRIN1,AMOTL2,YME1L1,TRAF4,STAT3,ARIH1,LBH,SF1,HIP1,SLC35F5,CSNK1G2,RAB2A
MP1,LASP1,BEX3,SRSF4,RTN3,PFN1,TLE5,RPL31,NISCH,CCNI,KHDRBS1,XPO1,SLC3A2,CSNK1E,I
C7A6,IL6ST,RPS2,CALD1,GALNT11,LRRC59,DAP,BRK1,SERP1,BAP1,BAZ2A,XRCC6,VAPB,RTL6,SI

BPB,ITM2B,AP2B1,GTF2F1,PXDN,HNRNPU,OGA,PGAM1,CTNNA1,FADS2,RPS16,RPL23,CPE,RPL2
ATP1A1,EIF5,WDR1,BEX3,CCT7,RTN3,JPT2,PRKCSH,KHDRBS1,SPTBN2,SLC3A2,PFKL,GLG1,CD
UL4A,TNRC6A,PPME1,MAGEF1,RPL27A,FAM32A,OTUD5,EPN1,CDK9,RTF2,CLPTM1L,CSNK1A1,F
3XL19,DIP2A,POLDIP2,IGFBP4,HS2ST1,USP47,OLFM2,OGFR,AGAP1,SCAMP5,HTATSF1,LGALS1,
K2B,ACSL6,RAB7A,CCN2,DDX3X,RPL12,ITM2C,GNB2,RCN1,RPL35,TLN1,MYO5A,COL6A2,CD63,
OVA2,CDH11,CCT2,TBCD,SUGP2,PGD,MEST,GAS7,GDI2,HNRNPL,TM9SF2,MORF4L2,ABCA3,SNF
IIBAN2,RUVBL2,OBSL1,ATP13A2,IGF1R,FZR1,SGSM3,SLC25A6,OSBPL2,COPE,SUMO1,NDUFC2,
TM2C,CLTC,GNB2,H1-0,RPS19,TLN1,CD63,MTSS2,DNAJC5,APLP2,MYL6,SHOX,ANKRD52,WLS,H
3C4,LDHB,HLA-C,PDIA3,ZNF43,CHD3,BASP1,VAPA,TPT1,RPL18,STMN1,ITGB1,TUBB4B,BTF3,NC
H1,MTRF1L,ALKBH5,PDXK,UBE4A,HSPA4,AHCYL1,KLC1,PTOV1,CLSTN3,ACAP3,DDX42,CHD2,OF
'1A2,PRDX1,NYNRIN,RBP1,PTBP1,WDR6,GABARAP,HNRNPUL1,UBE2R2,SCAF11,CAND1,KDM1A,
LE1,KIAA0232,KIFAP3,EEF1B2,RIC8A,PLEKHB2,LSM14B,INTS11,SLC4A2,SLC25A23,PDPK1,MSH6
A2,RPL24,SLC16A7,PAPOLA,IGFBP3,NCS1,CBX5,RPL9,PSMD2,GLIPR1L2,ATP1A1,JUND,TMEM25
CL1,PNN,HNRNPC,CCNI,LRPPRC,XPR1,HDLBP,RPS25,EIF5A,PEBP1,CCND2,DDR1,PLXNA1,RAB7
HNRNPA3,C4ORF3,PLOD1,CAMSAP1,GRINA,MLEC,CLSTN3,ORAI2,ASXL1,ABCA2,DPYSL2,MARS1,
PAPOLA,IGFBP3,NCS1,CBX5,PSMD2,NR2F1,SLC7A5,GLIPR1L2,ATP1A1,JUND,TMEM259,EIF5,LE
ALL2,WIZ,TBC1D14,DDX6,PRXL2A,JAK1,GID8,STUB1,NACAD,OXSR1,SLK,RANBP9,ZSWIM6,BEX1
5,RPL31,NISCH,HNRNPC,KHDRBS1,XPO1,SLC3A2,PFKL,GLG1,EIF5A,CCND2,CDH2,PDIA6,DDR1,
BCA3,HNRNPA3,PRDX5,ACO2,AHCYL1,COL2A1,CLSTN3,PTK7,RAB5B,HDGF,G3BP2,KDELR2,LAF
RNPDL,ARL6IP1,NDFIP1,TSR2,SHOX,CIRBP,DUX4,SEPTIN11,RHOB,HNRNPH1,ATP1A2,CKAP4,ARC

LBP,RPS25,TKT,CDH6,GPC3,RAB7A,ACTN4,DDX3X,RPS15,RPL12,EIF3L,MRFAP1,RCN1,IK,ANXA
INA,EIF4A1,MRC2,LDHB,PKD1,SF3B1,PDIA3,SERINC1,SSBP2,ZNF43,RPS3,BASP1,VAPA,HMGB1,
1,MBTPS1,SELENOW,CUL3,ADH5,NDST1,ITGB5,TIMM50,TNKS,APEX1,PPP1R15A,MAP1LC3B,M
8,WDR1,ATXN7L3B,MCL1,PNN,CCNI,TADA2B,XPR1,HDLBP,EIF5A,PEBP1,DDR1,PLXNA1,RAB7A,
G1,CALM1,NPM1,PLXNB2,PLD3,HSPD1,COL18A1,SMIM14,IGF2,EIF3C,EIF4B,MARCKSL1,UBB,A
RPL3,DNMT1,CFAP298,DDX56,CPSF7,RAD50,JRK,DUS1L,TOMM40,ALYREF,ATP6V1D,COX5B
2,SLC25A29,CTDNEP1,DYNLT1,AGPAT4,TUT4,HMG20B,UBE2E1,NBEA,GNAL,RAD23A,SETD1B,N
12,PSMD13,WDR26,CD2BP2,GPAA1,GGT7,SUB1,BANF1,CFDP1,CUL4A,TNRC6A,PPME1,MAGEF1
,DNAJA1,ARF1,CYB5R3,AKAP12,SF3B2,ITM2B,GTF2F1,SLC4A8,NFE2L1,HNRNPA0,CTNNA1,RPS

XR1,STRN4,BCL7A,RAB1B,UBR5,SLC6A8,TAX1BP1,GNL1,SH3GL1,DCLK1,TMEM63B,RAB6A,FO
52,HNRNPK,INIP,CIRBP,SEPTIN11,LAMP1,HNRNPH1,LGALS3BP,RPS12,RAB1A,ZNF454,COTL1,C
,PFKL,CDH6,SFPQ,PDIA6,GPC3,MEGF8,ACTN4,MMP14,PRDX2,RPS15,ARL4C,EIF3L,MIF,GPAT4
,BASP1,AP2M1,TPT1,SEC61A1,RPS4X,RPS3A,STMN1,ITGB1,TUBB4B,NCAM1,ANXA2,AGRN,RPL
N3,BRD3,PJA2,MSN,RPL24,SLC16A7,KPNB1,RPL17,C1ORF43,IGFBP3,LLGL1,GAP43,CBX5,ZNF
SF4,SH3GL1,TUBGCP6,PLEKHB2,NCSTN,ELP1,AHSA1,CLASP2,WASF2,MKRN1,WBP11,SPTY2D1,
GIC3,NUDT19,CAP1,ATP6V0D1,USP24,COX6A1,NASP,AP1M1,UBE2I,C16ORF58,TOMM70,UBXN6
NOVA2,CTSB,C5ORF24,CDH11,ZC3H7B,MDH2,CHD6,TOMM20,FDFT1,TBCD,HNRNPL,PARP1,TL
1,TPT1,SEC61A1,RPL18,TUBB4B,UCHL1,MORF4L1,AGRN,RPLP1,SARAF,MIDN,WSB1,RPS17,VC
,CYP51A1,SEC23A,ARNT2,PRPF40A,CD99L2,DDIT4,BEX4,VDAC1,RBBP7,SSR1,TXLNA,BCL9L,WD
2,SRSF5,TM9SF3,RAB27A,U2AF2,RPN2,SNX22,GDI2,TSR1,PKN1,HSPA4,CLSTN3,MDK,JPT1,PTK
A,OR8K1,RPL18,STMN1,UCHL1,MORF4L1,NCAM1,ANXA2,AGRN,ZMIZ1,TIMP2,XAF1,MIDN,MAR
CHD1,ATP6V0C,MMP2,LAPTM4B,RPL23A,NOVA2,SRSF5,RNF187,U2AF2,MLXIP,ANXA5,SNX22,N
F1B2,RIC8A,NCSTN,INTS11,AHSA1,QPRT,MTMR4,YIPF3,SPTY2D1,LRRC41,PRPF19,NOC2L,EIF3
JA2,MSN,KPNB1,RPL14,RPL22,RHOA,CLSTN1,ATF4,PSMD2,RAB6B,SLC7A5,ENC1,NLGN2,RPS1
E2G2,RAB3GAP2,UBFD1,IARS1,MASTL,YKT6,CUL3,MCM3AP,SPINT2,RPL7L1,TMEM167A,KLHD

SLC38A10,ATP13A1,HS2ST1,MZT2B,PSAT1,MTA2,TXNRD1,TMX1,MDM2,SLC38A1,CUL1,CD2BP
,TBC1D16,AHCYL1,CAMSAP1,KLC1,MLEC,ZNF302,RBMX,LAMC1,PTK7,ORAI2,TNPO1,ASXL1,HD

PL28,CCAR2,ARF4,GNG2,FYN,PFKM,TNKS2,STT3A,LONP1,ID4,SARS1,VANGL2,HGS,BAG6,HNR
17,VPS16,NUP50,SZT2,SEC31A,COG4,EIF2S1,MAPK8IP3,PSMC6,PRELID1,HARS1,DNMT1,CFAP2
SC3,CLIP3,RPL28,ZFAND5,CORO1C,SDF4,SMARCC1,HNRNPM,NKTR,SLIT3,STXBP1,ISYNA1,STT
9,ACTR1A,STARD7,SF1,PBX1,AP2A1,H2AZ1,KPNA2,BRD2,SFRP1,UBFD1,FBXO11,IARS1,RGMB,
C6,VAPB,EIF3H,ELAVL1,IMPDH2,GABARAPL1,SSR3,GNL1,TM9SF4,ZNF428,TRAP1,PLEKHB2,EL
RF43,PAPOLA,LLGL1,HK1,RHOA,PSMD2,RAB6B,SLC7A5,CXADR,CHD4,EIF5,CD81,LENG8,WDR1
41,PRPF19,NOC2L,EIF3G,PACS2,NASP,AP1M1,NDUFA13,C16ORF58,PSMC1,XPO6,UBXN6,DAP
R6,GABARAP,TDRD5,ACTN1,BNIP3L,MAP7D1,PLCG1,PITPNA,PLTP,CORO1C,SDF4,RPS28,HNRN
,MSI1,CAP1,MSH6,NSD2,FXYD6,MBD3,GSK3A,PACS2,NASP,TMEFF1,BNIP3,ATP5MC2,SLC20A1,

MICAL3,STUB1,NDRG1,CHST3,GPT2,OXSR1,NSD1,HSF1,RIMKLB,CLCN4,DNAJC8,TMEM43,SNR
NRNP200,CSDE1,SMIM14,IGF2,EIF3C,ATP5F1A,EIF4B,VCP,UBB,AFF4,TRIM28,MATR3,YWHAG,C
RAF3,TFDP1,FYTTD1,CABIN1,BCAR1,GIT1,TP53I11,SAMD1,UBE2E1,OBSCN,CDC42BPA,PRELID1
F3,LARS1,YWHAH,UBQLN1,CSNK1D,HYOU1,IFITM3,DGKD,VGF,MLF2,SEC23A,RTL8C,WBP2,AKI
PLXNA1,PIP4K2B,MEGF8,EIF3L,MRFAP1,MAGED2,RCN1,RPS7,MIF,GPAT4,GJA1,DNAJC5,TARDB

ECR,UBE2L3,PSAT1,FXR2,LGALS1,CLTA,KDM3A,PSMB1,LUC7L2,RBM4,CLIC1,HSPA1A,IGSF3,A
,PKD1,HLA-C,SF3B1,SERINC1,SSBP2,ZNF43,CHD3,VAPA,POLR2A,AP2M1,SEC61A1,TUBB4B,UC
2,COPE,CD2BP2,SUMO1,RPS21,SART3,BANF1,GOLPH3,PPP1CA,VPS51,PPME1,FAM8A1,FAM32
CA5,HNRNPA0,CTNNA1,RPS16,LDHA,CPE,TPM4,SCARB2,LAMA5,RPS24,SLC22A17,BRD3,CIPC,
R1,TRA2B,ZFR,SMC3,YKT6,PRRC2A,ANP32B,PREPL,TIMM50,SEPTIN7,PSMD8,HIF1A,SAFB2,SER
,TUBGCP6,RIC8A,INTS11,AHSA1,MTMR4,ITGAV,SPTY2D1,BORCS7,NOC2L,COX6A1,CHMP4B,T
,SLC44A2,CARM1,GPC2,IMPDH2,PGAP6,RAB14,LDLR,GNL1,EFTUD2,LRATD2,STK24,PRR12,PL
3,TSR1,MTRF1L,ABCA3,RPL27,PRDX5,C4ORF3,PKN1,ACO2,AHCYL1,GRINA,CLSTN3,LAMC1,JPT
ZYX,TUBA1C,RAB3A,ATRN,EI24,SLC12A7,WASF3,NSMF,ATP5F1E,INF2,SOS1,MGRN1,SGPL1,SET
PYGB,PRPF6,ATP6AP1,DPYSL2,RAPGEF1,OPTN,MVB12B,G3BP2,MAPK1,SMARCA4,MAPRE1,PLE

1,ATMIN,SNRPB,ZDHHC5,SKIV2L,SHC1,COX7A2,SUDS3,MIB2,CUL7,UQCRH,TRAPPC1,CUL4B,S
1L2,ATP1A1,JUND,TMEM259,EIF5,LENG8,WDR1,HMGN2,MCL1,PNN,CCNI,LRPPRC,TADA2B,XP
KT2,PRPF19,NOC2L,EIF3G,PACS2,AP1M1,NDUFA13,UBE2I,C16ORF58,PSMC1,XPO6,UBXN6,DA
IA5,BRD3,CIPC,MSN,KPNB1,PAPOLA,COL1A2,ATF4,NR2F1,RAB6B,ENC1,NLGN2,ATP1A1,JUND,
RPS7,IK,HMGCS1,GPAT4,GJA1,MTSS2,TARDBP,SLC25A3,HNRNPK,CIRBP,NARS1,PTGES3,CKAP
SI2,SAMD1,KLF3,DNM1,GNAO1,NFIB,CPNE2,SRSF8,CSTB,SKIL
DUFV1,SF1,GPX4,SLC35F5,CSNK1G2,RAB2A,UBAP2L,UPF1,UBFD1,MASTL,ZFR,MAPRE2,POM12
1,DNER,BCLAF1,RPL23A,PEA15,LUC7L3,CDKN1A,CRPPA,CPSF2,NOVA2,EEF1A2,TBCD,MLXIP,P
S14,CXADR,EIF5,CD81,WDR1,LASP1,SRSF4,RBBP4,PFN1,RPL31,CCNI,LRPPRC,TADA2B,KHDRB

34,VAPB,TCAF1,INTS3,RAB1B,UBE2M,SLC6A8,LRATD2,FLOT2,TMEM63B,NUCKS1,SFXN1,RBM1
1,SET,EGLN1,MAP9,INA,EIF4A1,MRC2,RPL19,RPL15,LDHB,DNAH8,HLA-C,SF3B1,PDIA3,SERINC
P,BRK1,SERP1,BAZ2A,XRCC6,VAPB,EIF3H,ELAVL1,IMPDH2,GABARAPL1,GNL1,TM9SF4,ZNF428
5A,GALNT11,DAP,BRK1,PPP1R10,SPG7,BAZ2A,XRCC6,VAPB,CLPTM1,SNN,EIF3H,ELAVL1,IMPDH
,FGFR1,HNRNPH3,LHFPL6,MZT2B,LMO4,SLC25A36,WDTC1,SUN1,MACROH2A1,ANP32E,CLIC1,
SPAN3,LPCAT1,SLC22A17,BRD3,XRCC5,PJA2,MSN,KPNB1,C1ORF43,PAPOLA,LLGL1,NCS1,CBX
,SEPTIN5,CARM1,RAB1B,RALY,ARHGEF17,ZMIZ2,HDGFL2,LRATD2,PRR12,FLOT2,TUBGCP6,NU

,RBM12,WBP11,NSD2,YY1,CCDC47,ELOVL5,SRPRA,SLC25A36,XRN2,SCAMP5,ZZEF1,AASDHPPT
BN1,NECTIN2,PLCD3,CTDNEP1,CMTR1,CLCN3,TRAPPC1,ABCE1,MSI2,CCNL1,PAF1,DDX19A,PP
4,RACK1,BSG,FASN,ALDH6A1,SPTBN1,ATP5F1B,CALM3,GNB1,SHISA9,STMN2,RPS8,MYH9,RPL7
,MAPRE3,ZBTB4,HMGXB3,FBL,WARS1,VEZF1,SNRPB,NKIRAS2,ZDHHC5,XPC,SKIV2L,CFLAR,SH
4G1,PAFAH1B2,PLS3,POLDIP2,CHMP1A,NBPF15,GTF3C1,MFN2,NDUFA4,SF3A3,TRMT112,WD
T1,GTF3C1,CDV3,FZR1,TNFAIP1,FAM50A,CD2BP2,ALDH18A1,ATP1B3,CNDP2,GLOD4,MTURN,G
1,GPC1,SLC35E2B,APPL1,SMCHD1,HDAC2,L1CAM,TRIB2,PPIB,PEA15,CDKN1A,CRPPA,CDH11,R
MAN2,CHMP1A,PSMB1,TSNAX,PAIP2,PCM1,CUL1,EXOC5,OSBPL2,TRMT112,PSMD13,CD2BP2,
EIF4G1,TUB,SLC25A36,OLFM2,FAM210B,COL16A1,FXR2,NBPF15,TSPYL2,NIBAN2,HSPA1A,MA
770,GTF2F1,SLC4A8,PXDN,PGAM1,NFE2L1,SMARCA5,RPS16,RPL23,CPE,EIF4H,SCARB2,TSPAN
ND1,LAMP1,MAP2,CALU,ARCN1,RPS27,HRNR,ARHGDIA,SLC35E2B,ATP6V0C,RPL23A,HS6ST1,D

CN3,TRAPPC1,ABCE1,MSI2,CCNL1,PAF1,DDX19A,PPM1A,GNAO1,RAD23A,EPHA2,SKIL,STX12,VP
Z2A,XRCC6,VAPB,EIF3H,ELAVL1,IMPDH2,GABARAPL1,GNL1,TM9SF4,ZNF428,TRAP1,PLEKHB2,E
F4B,VCP,MARCKSL1,UBB,AFF4,MATR3,YWHAG,CKB,MAPT,FN1,SPEN,DDB1,PSMD5,YWHAQ,EIF
SCD1,GRK2,VPS26B,BAZ2A,CLCN7,CEP170,TAOK1,PALM,SHC2,SSBP3,CARM1,STRN4,INTS3,C

GB,DPYSL2,TMED2,TOP1,TAF7,RPL37,VHL,AZIN1,LAPTM4A,SRSF9,ARGLU1,RBP1,ATP5PB,ADAM
1,PCNX3,DAZAP2,ERGIC3,WBP11,NUDT19,IGF2BP1,ATP6V0D1,USP24,COX6A1,SART1,NASP,AP1
1,NPM1,PLXNB2,PLD3,HSPD1,USP11,COL18A1,SNRNP200,CSDE1,IGF2,ATP5F1A,EIF4B,MARCKS
2,DDX3X,RPS15,ITM2C,RPL30,ARL4C,NANOG,MRFAP1,MAGED2,RPS5,RPS19,HMGCS1,TLN1,JUI
ERP1,ERH,EIF2S2,SLIT1,IARS2,ABI2,RAB1B,RALY,SEC14L1,NUDCD3,TAX1BP1,HINT1,CNP,PNISR,
PRE2,GPSM1,POM121C,LMAN1,TNKS,CTTN,LOX,LRRC59,DAP,SERP1,PRDX6,SPG7,CLCN7,TRAN
EIF3H,ABI2,RAB14,SRP72,TM9SF4,KMT2E,PNISR,EIF4G3,CAPZA1,NCSTN,QPRT,LRRC41,MED13,
C25,PRKCSH,HNRNPC,CCNI,XPR1,SPTBN2,CSNK1E,GLG1,EIF5A,PEBP1,CCND2,PLXNA1,PIP4K2
68B,IPO7,TCF25,PTBP1,EIF3E,ILF3,WDR6,GABARAP,TDRD5,UBE2R2,QARS1,POLR2B,SDF4,RPS
N,PGAM1,SMARCA5,HNRNPA0,CTNNA1,RPS16,RPL23,RPL29,RPS24,RPS9,XRCC5,CIPC,PJA2,RPL
6,CPSF7,C2ORF68,RAD50,JRK,DUS1L,TOMM40,ALYREF,COX5B
PIGT,RNPS1,AKAP17A,TMCO1,HARS1,MRPL3,DNMT1,CFAP298,CPSF7,RAD50,JRK,TOMM40,COX
TK35,CLCN7,TAOK1,OLFM1,CARM1,STRN4,INTS3,SLC6A8,TRAM1,GABARAPL1,PGAP6,HDGFL2,
D1,CAP1,IST1,ATP6V0D1,MBD3,GSK3A,SMARCD1,APBB1,ANKRD12,BAG5,SLC20A1,MICAL1,ATG
RNP70,ARF3,PPP2CA,CCND1,SMARCA4,AZIN1,DDAH2,PLEC,FAM171A2,ACADVL,NYNRIN,TCF25,A
2A,GNG4,UBAP2L,UPF1,UBFD1,MASTL,MBTPS1,MAPRE2,POM121C,CALD1,SEPTIN7,APEX1,PPP
1,RAD23A,EPHA2,STX12,VPS37B,USP36,NDUFA10,CCDC124
ABI2,SOX2,LTBP4,ARHGEF17,ELAVL1,GABARAPL1,TLE1,LDLR,LSM14A,MOAP1,DICER1,PCNX3,M
0,ERH,EIF3H,ABI2,RAB14,SRP72,TM9SF4,KMT2E,PNISR,EIF4G3,PRR12,CAPZA1,QPRT,RBM5,LR
SERINC1,SSBP2,ZNF43,VAPA,HMGB1,CDKL1,AP2M1,TPT1,SEC61A1,RPL18,STMN1,TUBB4B,RPL
POT,SF1,RETREG2,PDAP1,AP2A1,MMP15,FLOT1,SEC16A,GPX4,PHB2,MGAT4B,SLC39A7,DECR1,

,HMGN2,MCL1,PNN,CCNI,LRPPRC,TADA2B,XPR1,HDLBP,EIF5A,PEBP1,CCND2,DDR1,PLXNA1,RA
PNA2,GPX4,SFRP1,FBXO11,IARS1,HDAC6,RGMB,AGPAT1,PRRC2A,CUL3,IL6ST,MAPRE2,RPS2,TM
D1A,SS18L1,SUDS3,MAF1,SLC25A29,ZSWIM8,UFM1,TMEM59,FER,GATAD2B,MCM7,COPB1,VPS1
ACF1,B4GALNT4,CYFIP2,TARDBP,RPLP2,ARL6IP1,TSR2,TAGLN,MYL6,ANKRD52,INIP,MYH10,SN
N,MYO5A,COL6A2,ANXA6,GOLM1,B4GALNT4,DNAJC5,HNRNPDL,RPLP2,ARL6IP1,ST3GAL2,MXD
9,MOAP1,CLASP2,RBM12,WBP11,NSD2,YY1,CCDC47,ELOVL5,SRPRA,RIMS3,SLC30A9,ACAT2,SL
P11,CSDE1,SMIM14,IGF2,EIF4B,VCP,MARCKSL1,UBB,AFF4,TRIM28,YWHAG,CKB,COL4A2,RPL37
AP5L,PCGF3,TNRC6A,HNRNPR,CAPN1,WRNIP1,LSM4,KPNA6,CDK9,ZRANB2,SRSF7,SENP6,ZNF5

ARF4,HNRNPUL2-BSCL2,POLR2B,SUPT6H,LSS,EIF3F,PITPNM1,CSNK1D,TMED9,ID4,SARS1,HGS,
IN1,AKAP17A,EIF2S1,MAPK8IP3,HARS1,MRPL3,DNMT1,CFAP298,DDX56,NEDD8,CPSF7,RAD50,J
KLHL42,CSNK1G2,GARS1,PAPSS1,RER1,RAB3GAP2,IARS1,MASTL,COX7C,YKT6,CUL3,ADH5,MC

,NCKAP1,MLEC,RBMX,TMEM248,ASXL1,RPL36A,SCAF1,RAB11B,HDGF,PYGB,TERF2IP,TMED2,N
0A,GTF2F1,SLC4A8,OGA,NFE2L1,LDHA,CNN3,RPL29,TPM4,TNS3,TSPAN3,LPCAT1,LAMA5,XRCC
4,PKD1,SF3B1,ZNF43,RPS3,CHD3,POLR2A,AP2M1,TPT1,SEC61A1,RPS4X,RPL18,ITGB1,UCHL1,

,TAGLN,SHOX,H1-10,CIRBP,DUX4,YWHAB,CKAP5,SEPTIN11,SND1,LAMP1,HNRNPH1,CKAP4,HLA
82,SYNGR1,USP48,RPS29,NSMF,DUSP8,RNF20,NUP133,CHGB,COX4I1,OGDH,DSTYK,GUK1,UB
10,TMEM9,SCG2,WDR82,SYNGR1,USP48,RPS29,NSMF,DUSP8,RNF20,NUP133,CHGB,COX4I1,O
2,AGPAT3,DTX3,TXN,IRF2BP2,WDR47,RHOBTB2,NELFCD,EFNB3,ZNF362,SHANK1,MEIS3,GABA

MP1A,PSMB1,FAM118A,TSNAX,RGL2,CRACD,SLC25A6,EXOC5,SF3A3,TRMT112,CD2BP2,SUMO1
RF3,PDXK,PTOV1,MLEC,CLSTN3,SURF4,CHD2,ORAI2,SKI,ASXL1,RAB11B,HDGF,TSPAN6,TOP1,SE
A1,CNDP2,GLOD4,LMF2,MTURN,GOLPH3,ECHS1,DHX36,HM13,CRKL,COLGALT1,TSPAN7,NUP9
SCG2,WDR82,SYNGR1,USP48,RPS29,NSMF,DUSP8,RNF20,NUP133,CHGB,COX4I1,OGDH,DSTYK
SCG2,WDR82,SYNGR1,USP48,RPS29,NSMF,DUSP8,RNF20,NUP133,CHGB,COX4I1,OGDH,DSTYK
2,TNFAIP1,DNAJB1,FAM50A,ALDH18A1,PABPC4,CNDP2,LIMK1,GOLPH3,RHEB,RAP2A,RNF41,TR
,ZNF770,GOLGA8A,PXDN,HNRNPU,OGA,SMARCA5,HNRNPA0,CTNNA1,FADS2,RPS16,CNN3,TPM4
IRNP70,PSMB2,PYGB,DPYSL2,PPP2CA,MAPK1,SMARCA4,AZIN1,DDAH2,PRDX1,ARGLU1,NYNRIN,
PNB1,C1ORF43,PAPOLA,GAP43,RPL22,RHOA,ZNF865,NR2F1,NLGN2,RPS14,CXADR,LENG8,WDI
L26,LASP1,CCT7,RTN3,PFN1,ASTN2,NES,KLHL9,TKT,GLG1,CCND2,GPC3,SULF1,PRDX2,DDX3X,
L26,LASP1,CCT7,RTN3,PFN1,ASTN2,NES,KLHL9,TKT,GLG1,CCND2,GPC3,SULF1,PRDX2,DDX3X,
,TMEM132A,CTDSP2,SRP9,CHD6,RNF187,CCT2,ANXA5,MEST,HNRNPL,PARP1,TAF15,ABCA3,C4C
GNAO1,RAD23A,BTG1,EPHA2,STX12,VPS37B,RBBP6,ZC3H4,PPP2R5C,NDUFA10,CCDC124
4L1,BTF3,NCAM1,ANXA2,AGRN,XAF1,COL11A1,MARCHF6,WSB1,VCL,AP3D1,DNAJA1,ARF1,AKAP
,PJA1,SUB1,ANO8,GLOD4,PCGF3,CFDP1,FLII,WRNIP1,LDB1,ZNF503,TSPAN7,DEGS1,SLK,SYNG
NP,UBTD2,RNPS1,PRELID1,MARF1,CTIF,PIEZO1,KLHL7,BRD1,LEMD2,SKIL,VPS37B
LN1,MAP9,TPI1,CHSY1,TMSB10,INA,EIF4A1,PPP2R1A,RPL15,GTF3C4,LDHB,DNAH8,PKD1,HLA-C
MA5,RPS24,SLC22A17,RPS9,CIPC,PJA2,RPL24,SLC16A7,IGFBP3,NCS1,CBX5,RPL9,PSMD2,GLIPF
3X5,RPL9,ATF4,NR2F1,KIF5B,RAB6B,ENC1,GLIPR1L2,RPS14,CXADR,TMEM259,RPS20,WDR1,RF

AFAH1B2,TRIP12,CHMP1A,ITFG1,CGGBP1,ARPC1A,PSMB1,PAIP2,SLC25A6,EXOC5,SF3A3,TRMT
1,SQSTM1,VAMP2,GNAI3,TSPYL4,SSRP1,ALKBH5,PLOD1,CCT3,GRINA,ACAP3,SURF4,ZNF302,TN

MEM248,RAD23B,SKI,ASXL1,VEGFA,OPTN,RPL37,VHL,MVB12B,ARPC2,DSTN,BTBD2,PBX2,PDE7A

PR1L2,TMEM259,CD81,LENG8,HMGN2,ATP2A2,RPL26,BEX3,CCT7,ASTN2,JPT2,TLE5,NISCH,TAD
MEM9,SCG2,WDR82,SYNGR1,USP48,RPS29,NSMF,DUSP8,RNF20,NUP133,THBS2,CHGB,COX4I1
I50,SEPTIN7,PSMD8,HIF1A,SAFB2,SERP1,PPP1R10,ERH,EIF3H,ABI2,RAB14,SRP72,TM9SF4,KMT
N,HNRNPU,OGA,PGAM1,NFE2L1,HNRNPA0,CTNNA1,CPE,CNN3,TPM4,EIF4H,NACA,LPCAT1,BRD3
EIF5A,CDH2,GPC3,DDR1,SULF1,PIP4K2B,MEGF8,RPL12,ARL4C,GNB2,H1-0,IK,RPS19,RPL35,MIF

20,ATP2A2,RPL26,LASP1,CCT7,RTN3,PFN1,ASTN2,NES,KLHL9,TKT,GLG1,CCND2,GPC3,SULF1,
GOLPH3,ECHS1,DHX36,HM13,COLGALT1,TSPAN7,NUP98,NACAD,OXSR1,EMC10,SEMA4C,TMEM
,PPP2CA,CCND1,G3BP2,KDELR2,PDE7A,PSMD3,TCF25,ADAM9,CDC37,WDR6,SCAF11,BNIP3L,H

,CLASP2,RBM12,NSD2,DHCR7,EHD1,AP2A2,YY1,CCDC47,ELOVL5,SLC25A36,XRN2,PTPN11,SCA
1,TBC1D14,TBXA2R,PMEPA1,SLC25A1,STUB1,LOXL2,ZCCHC14,DEGS1,CHST3,PCOLCE,SEMA4
PK3,LYPLA2,ATP13A1,IER2,PAFAH1B2,PLS3,POLDIP2,CHMP1A,CGGBP1,TSNAX,GTF3C1,HIP1R,N
ME1L1,TRAF4,STAT3,ARIH1,LBH,NDUFV1,KDM5B,SF1,PDAP1,EIF3D,PLXNB1,DECR1,SLC35F5,C

10,TMEM9,SCG2,WDR82,SYNGR1,USP48,RPS29,NSMF,DUSP8,RNF20,NUP133,CHGB,COX4I1,O
ASTL,MBTPS1,MAPRE2,SEPTIN7,APEX1,PPP1R15A,GALNT11,MVD,DAP,BRK1,BAP1,PPP1R10,SP
,GLOD4,MTURN,GOLPH3,ECHS1,DHX36,HM13,COLGALT1,TSPAN7,NUP98,NACAD,OXSR1,EMC1

COLGALT1,TSPAN7,NUP98,NACAD,OXSR1,EMC10,SEMA4C,TMEM9,SCG2,WDR82,SYNGR1,USP4

,ALDH18A1,CNDP2,GLOD4,LMF2,MTURN,GOLPH3,ECHS1,APPBP2,DHX36,HM13,CRKL,COLGAL
O1,POLR2B,SMARCC1,SUPT6H,RPS28,HNRNPM,SDHA,NKTR,CCT8,PFKM,STXBP1,TMED9,HGS,
RAP1,PLEKHB2,ELP1,RAB6A,MOAP1,KRT8,CLASP2,RBM12,WBP11,SRSF11,NSD2,YY1,CCDC47,
PRDX1,ODC1,FAM168B,IPO7,TCF25,PTBP1,ILF3,CDC37,WDR6,GABARAP,TDRD5,BNIP3L,PLCG1
PRDX1,ODC1,FAM168B,IPO7,TCF25,PTBP1,ILF3,CDC37,WDR6,GABARAP,TDRD5,BNIP3L,PLCG1
BRK1,BAP1,TRIR,PALM,ANTXR1,LINGO1,SLC6A8,GABARAPL1,RAB14,PLEKHA6,KMT2E,TTC37,R
MP5,GPS1,MAF1,EPB41L1,HEG1,CNOT3,QRICH1,API5,MSH2,TUT4,TSN,SAMD1,HADHB,TRIM24,
LDHB,SF3B1,PDIA3,SERINC1,SSBP2,RPS3,CHD3,HMGB1,AP2M1,TPT1,SEC61A1,RPS3A,OR8K1,S

2B,ATP6V0C,TRIB2,RPL23A,FAM199X,SERF2,CPSF2,DDX39B,SRSF5,TM9SF3,MDH2,TOMM20,U2
IGR1,USP48,RPS29,STMN4,NSMF,DUSP8,RNF20,NUP133,CHGB,COX4I1,OGDH,DSTYK,GUK1,UE
,CCDC47,ELOVL5,SLC25A36,XRN2,PTPN11,SCAMP5,ZZEF1,AASDHPPT,ESYT1,GTF3C1,TNFAIP1,
RPL32,VPS16,PSMA4,SZT2,UBE2E1,SCOC,TMCO1,EIF2S1,HARS1,NEDD8,MARF1,CDK5RAP3,RAD

MP14,RPL12,H1-0,APLP1,RPS19,HMGCS1,TLN1,MYO5A,CD63,NGRN,MACF1,RPL35A,TARDBP,W

TOV1,MLEC,CLSTN3,ACAP3,PTK7,GCN1,SELENON,RPL37,CCND1,G3BP2,SMARCA4,SRSF1,IPO7,

ASC3,CLIP3,ZFAND5,CORO1C,SMARCC1,HNRNPM,YWHAH,SLIT3,ISYNA1,STT3A,CSNK1D,IDH1,S
ABCF1,ERGIC3,CAP1,IST1,ATP6V0D1,DHCR7,NOC2L,GSK3A,SMARCD1,ANKRD12,SSBP4,BAG5,S

,SRSF10,CAPN1,ASAH1,JAK1,CDK9,SENP6,CLPX,RBM10,P3H1,MRPS6,MEX3C,ACSL4,CRTC1,CC
C1,CCT3,NCKAP1,CLSTN3,DDX42,SCAF1,RAB11B,NRDC,TOP1,TUBB2A,OPTN,UBXN4,PPP2CA,C
LHL42,PLXNB1,HJURP,RER1,RAB3GAP2,UBFD1,EIF5B,MASTL,COX7C,YKT6,SV2A,NOMO1,AGPA

P2,HNRNPDL,ST3GAL2,TAGLN,HNRNPK,H1-10,NARS1,RPL36,PTGES3,LAMP1,RAB1A,TTR,PLPPR
2,SCAF1,RAB11B,NRDC,TOP1,TUBB2A,UBXN4,PPP2CA,CCND1,G3BP2,PDE7A,PSMD3,SRSF1,ILF
3A10,ATP5MG,PAFAH1B2,LMAN2,CLTA,SEMA6B,CGGBP1,PSMB1,TCEA1,PCM1,OSBPL2,TRMT1

,CDH11,CHD6,TOMM20,CCT2,CPSF1,GNAI3,SSRP1,HSPH1,HNRNPA3,C4ORF3,PKN1,ETNK1,CC
COLGALT1,TSPAN7,NUP98,NACAD,OXSR1,EMC10,SEMA4C,TMEM9,SCG2,WDR82,SYNGR1,USP4
12,SMARCB1,PLXND1,CCSER2,LDOC1,EIF3J,NAMPT,DTD1,EMD
TF3C1,TNFAIP1,FAM50A,CD2BP2,SYNCRIP,CNDP2,GLOD4,LMF2,MTURN,GOLPH3,ECHS1,DHX3
OLGALT1,TSPAN7,NUP98,NACAD,OXSR1,EMC10,SEMA4C,TMEM9,SCG2,WDR82,SYNGR1,USP4

ABEP1,C1ORF198,MTCH1,ZMYM3,FUS,CLCN7,SHC2,NUDCD3,PGAP6,SSR3,SH3GL1,PLEKHA6,EE

TAN1,GSTP1,MARCKS,SMG1,CALM1,NPM1,PLXNB2,PLD3,HSPD1,USP11,COL18A1,SNRNP200,C
P11,COL18A1,SNRNP200,CSDE1,SMIM14,IGF2,EIF3C,ATP5F1A,EIF4B,VCP,MARCKSL1,UBB,AFF
OL18A1,SNRNP200,CSDE1,SMIM14,IGF2,EIF3C,ATP5F1A,EIF4B,VCP,MARCKSL1,UBB,AFF4,TRIM
1,SMIM14,IGF2,EIF4B,VCP,MARCKSL1,UBB,TRIM28,MATR3,YWHAG,CKB,COL4A2,RPL37A,FN1
A,MRC2,PPP2R1A,GTF3C4,PKD1,HLA-C,SF3B1,PDIA3,SSBP2,RPS3,BASP1,VAPA,POLR2A,CDKL1,
1,SNRNP200,CSDE1,SMIM14,IGF2,EIF3C,ATP5F1A,EIF4B,VCP,MARCKSL1,UBB,AFF4,TRIM28,MA
PLXNB2,PLD3,HSPD1,COL18A1,SNRNP200,CSDE1,SMIM14,IGF2,EIF3C,ATP5F1A,EIF4B,VCP,MA
3B2,CDC42BPB,ITM2B,MAT2A,TMEM170A,GTF2F1,SLC4A8,HNRNPU,OGA,PGAM1,NFE2L1,HNR
NO,DDB1,PSMD5,YWHAQ,EIF1,HACD3,DCHS1,SET,EGLN1,MAP9,TPI1,CHSY1,TMSB10,INA,EIF4
C,ATP5F1A,EIF4B,VCP,MARCKSL1,UBB,TRIM28,MATR3,YWHAG,CKB,COL4A2,RPL37A,LRRC28,
,IGF2,EIF3C,ATP5F1A,EIF4B,VCP,MARCKSL1,AFF4,TRIM28,MATR3,YWHAG,CKB,COL4A2,RPL3
,CHSY1,TMSB10,INA,EIF4A1,MRC2,GTF3C4,PKD1,HLA-C,PDIA3,SSBP2,BASP1,VAPA,HMGB1,PO
AQ,HACD3,DCHS1,SET,EGLN1,TPI1,CHSY1,TMSB10,INA,MRC2,RPL19,PPP2R1A,RPL15,GTF3C4
SPEN,NONO,DDB1,YWHAQ,HACD3,DCHS1,SET,EGLN1,TPI1,CHSY1,TMSB10,INA,EIF4A1,MRC2,R
D3,DCHS1,SET,EGLN1,TPI1,CHSY1,TMSB10,EIF4A1,MRC2,PPP2R1A,RPL15,GTF3C4,PKD1,HLA-
OL18A1,SMIM14,IGF2,EIF3C,EIF4B,VCP,MARCKSL1,UBB,TRIM28,MATR3,YWHAG,CKB,COL4A2,
FN1,SPEN,NONO,DDB1,YWHAQ,HACD3,DCHS1,SET,EGLN1,TPI1,CHSY1,TMSB10,INA,MRC2,RPL1
1,CHSY1,TMSB10,INA,MRC2,RPL19,PPP2R1A,RPL15,GTF3C4,LDHB,PKD1,HLA-C,PDIA3,SSBP2,
1,ANXA2,AGRN,ZMIZ1,TIMP2,XAF1,SARAF,COL11A1,MIDN,MARCHF6,RPS17,VCL,AP3D1,SBF1,D
,SPEN,NONO,DDB1,YWHAQ,HACD3,DCHS1,SET,EGLN1,TPI1,CHSY1,TMSB10,INA,MRC2,RPL19,
N1,TPI1,CHSY1,TMSB10,INA,EIF4A1,MRC2,PPP2R1A,RPL15,GTF3C4,LDHB,PKD1,SF3B1,PDIA3,
,TRIM28,MATR3,YWHAG,CKB,COL4A2,RPL37A,LRRC28,FN1,SPEN,DDB1,EIF1,HACD3,DCHS1,S
,TPI1,CHSY1,INA,EIF4A1,MRC2,RPL19,PPP2R1A,PKD1,HLA-C,SF3B1,PDIA3,SERINC1,SSBP2,RPS
1,TMBIM6,PTMS,SPTAN1,GSTP1,MARCKS,SMG1,CALM1,NPM1,PLXNB2,PLD3,HSPD1,USP11,C
4A1,PPP2R1A,RPL15,GTF3C4,LDHB,PKD1,PDIA3,SSBP2,ZNF43,CHD3,BASP1,VAPA,CDKL1,AP2
B,VCP,MARCKSL1,UBB,AFF4,TRIM28,MATR3,YWHAG,CKB,COL4A2,RPL37A,FN1,SPEN,NONO,D
RC2,RPL15,GTF3C4,LDHB,PKD1,PDIA3,SSBP2,ZNF43,RPS3,BASP1,VAPA,HMGB1,POLR2A,TPT1,
,EIF4B,VCP,MARCKSL1,UBB,AFF4,TRIM28,MATR3,YWHAG,CKB,COL4A2,RPL37A,LRRC28,FN1,
STP1,MARCKS,SMG1,CALM1,NPM1,PLXNB2,PLD3,HSPD1,USP11,COL18A1,SNRNP200,CSDE1,S
GSTP1,MARCKS,SMG1,CALM1,NPM1,PLXNB2,TTC3,PLD3,HSPD1,USP11,COL18A1,SNRNP200,C
,SNRNP200,CSDE1,SMIM14,IGF2,EIF3C,EIF4B,VCP,MARCKSL1,UBB,AFF4,TRIM28,MATR3,YWHA
MSB10,INA,EIF4A1,MRC2,RPL19,PPP2R1A,GTF3C4,LDHB,PKD1,HLA-C,PDIA3,SERINC1,SSBP2,V
,HACD3,DCHS1,SET,EGLN1,TPI1,CHSY1,TMSB10,EIF4A1,MRC2,PPP2R1A,GTF3C4,PKD1,HLA-C,
,CALM1,NPM1,PLXNB2,PLD3,HSPD1,USP11,COL18A1,SNRNP200,CSDE1,SMIM14,IGF2,EIF3C,A
SPTAN1,GSTP1,MARCKS,SMG1,CALM1,NPM1,PLXNB2,PLD3,HSPD1,USP11,COL18A1,SNRNP20
,POLR2A,AP2M1,TPT1,SEC61A1,RPS3A,STMN1,ITGB1,TUBB4B,UCHL1,MORF4L1,NCAM1,ANXA
,VCP,MARCKSL1,UBB,AFF4,TRIM28,MATR3,YWHAG,CKB,COL4A2,RPL37A,LRRC28,FN1,SPEN,
F1,HACD3,DCHS1,SET,EGLN1,MAP9,TPI1,CHSY1,TMSB10,INA,EIF4A1,MRC2,PPP2R1A,RPL15,GT
SBP2,BASP1,VAPA,HMGB1,POLR2A,CDKL1,AP2M1,TPT1,SEC61A1,RPS3A,RPL18,STMN1,ITGB1,
1,UBB,AFF4,TRIM28,MATR3,YWHAG,CKB,COL4A2,RPL37A,LRRC28,MAPT,FN1,SPEN,PSMD5,Y
2,PLD3,HSPD1,USP11,COL18A1,SNRNP200,CSDE1,SMIM14,IGF2,ATP5F1A,EIF4B,VCP,MARCKS
KS,SMG1,CALM1,NPM1,PLXNB2,PLD3,HSPD1,USP11,COL18A1,SMIM14,IGF2,EIF3C,ATP5F1A,E
SB10,INA,EIF4A1,RPL19,PPP2R1A,RPL15,GTF3C4,LDHB,SF3B1,PDIA3,SSBP2,RPS3,BASP1,VAP
KB,COL4A2,RPL37A,LRRC28,FN1,SPEN,NONO,DDB1,YWHAQ,EIF1,HACD3,DCHS1,SET,EGLN1,T
3C,ATP5F1A,EIF4B,MARCKSL1,AFF4,TRIM28,MATR3,YWHAG,CKB,COL4A2,RPL37A,LRRC28,FN

11,COL18A1,SNRNP200,SMIM14,IGF2,EIF3C,ATP5F1A,EIF4B,MARCKSL1,UBB,AFF4,TRIM28,MA
31,YWHAQ,HACD3,DCHS1,SET,EGLN1,TPI1,CHSY1,TMSB10,INA,MRC2,RPL19,GTF3C4,LDHB,DN/
P4,CDC42BPB,ITM2B,AP2B1,SLC4A8,PXDN,HNRNPU,OGA,PGAM1,NFE2L1,SMARCA5,CTNNA1,FA
P11,COL18A1,SNRNP200,CSDE1,SMIM14,IGF2,EIF3C,ATP5F1A,EIF4B,VCP,MARCKSL1,UBB,AFF4
P1,MARCKS,SMG1,CALM1,NPM1,PLXNB2,PLD3,HSPD1,USP11,COL18A1,SNRNP200,SMIM14,IGF
L18,STMN1,ITGB1,TUBB4B,UCHL1,MORF4L1,BTF3,NCAM1,ANXA2,AGRN,ZMIZ1,TIMP2,XAF1,CC
N1,ITGB1,TUBB4B,UCHL1,NCAM1,ANXA2,AGRN,TIMP2,XAF1,SARAF,MIDN,MARCHF6,VCL,AP3D1
M28,MATR3,YWHAG,CKB,COL4A2,MAPT,FN1,SPEN,NONO,DDB1,PSMD5,YWHAQ,HACD3,DCHS1
,HNRNPA0,CTNNA1,RPS16,CNN3,TNS3,CTSD,EIF4H,LPCAT1,LAMA5,BRD3,CIPC,PJA2,MSN,KPNB
RIM28,MATR3,YWHAG,CKB,COL4A2,LRRC28,MAPT,FN1,SPEN,NONO,DDB1,PSMD5,YWHAQ,EIF1,
IA5,BRD3,CIPC,MSN,KPNB1,C1ORF43,COL1A2,LLGL1,NCS1,RHOA,ZNF865,CLSTN1,RAB6B,SLC
NFE2L1,CTNNA1,CNN3,RPL29,CTSD,SCARB2,LPCAT1,LAMA5,BRD3,CIPC,PJA2,MSN,SLC16A7,K
HF6,WSB1,RPS17,VCL,AP3D1,SBF1,ARF1,CYB5R3,AKAP12,CDC42BPB,ITM2B,AP2B1,GTF2F1,G
D1,SBF1,ARF1,CYB5R3,MAP4,SF3B2,CDC42BPB,ITM2B,AP2B1,MAT2A,GTF2F1,GOLGA8A,PXDN
,CKB,COL4A2,RPL37A,SPEN,DDB1,YWHAQ,EIF1,HACD3,DCHS1,SET,EGLN1,MAP9,TPI1,CHSY1,T
OL1A2,LLGL1,HK1,NCS1,RHOA,ZNF865,CLSTN1,KIF5B,RAB6B,NLGN2,CXADR,CHD4,ATP1A1,JU
SDE1,IGF2,EIF3C,ATP5F1A,EIF4B,VCP,MARCKSL1,UBB,AFF4,TRIM28,MATR3,YWHAG,CKB,COL4
ANXA2,AGRN,TIMP2,XAF1,SARAF,MIDN,MARCHF6,AP3D1,SBF1,ARF1,CYB5R3,MAP4,SF3B2,CDC
,C1ORF43,COL1A2,LLGL1,NCS1,RHOA,ZNF865,CLSTN1,ATF4,PSMD2,SLC7A5,NLGN2,CHD4,AT
3,SERINC1,SSBP2,VAPA,HMGB1,POLR2A,AP2M1,TPT1,SEC61A1,RPS3A,STMN1,ITGB1,TUBB4B,
FF4,TRIM28,MATR3,YWHAG,CKB,COL4A2,LRRC28,FN1,SPEN,NONO,DDB1,PSMD5,YWHAQ,EIF1,
CNN3,TNS3,CTSD,EIF4H,LPCAT1,BRD3,CIPC,PJA2,MSN,KPNB1,C1ORF43,COL1A2,LLGL1,NCS1,
AG,CKB,COL4A2,LRRC28,FN1,SPEN,NONO,DDB1,YWHAQ,EIF1,HACD3,DCHS1,SET,EGLN1,TPI1,
A1,ARF1,CYB5R3,AKAP12,MAP4,CDC42BPB,ITM2B,TMEM170A,GTF2F1,PXDN,HNRNPU,PGAM1,
3,TNS3,CTSD,EIF4H,NACA,TSPAN3,LPCAT1,LAMA5,BRD3,CIPC,PJA2,MSN,KPNB1,C1ORF43,COL1
CAM1,AGRN,ZMIZ1,TIMP2,XAF1,SARAF,MIDN,RPL10,MARCHF6,RPS17,AP3D1,SBF1,DNAJA1,ARF
1,TMSB10,INA,MRC2,RPL19,PPP2R1A,GTF3C4,LDHB,PKD1,HLA-C,PDIA3,SERINC1,SSBP2,CHD3,
MAP4,CDC42BPB,ITM2B,MAT2A,PXDN,HNRNPU,OGA,NFE2L1,CTNNA1,FADS2,TPM4,TNS3,CTSD
MS,SPTAN1,GSTP1,MARCKS,SMG1,CALM1,NPM1,PLXNB2,PLD3,HSPD1,USP11,COL18A1,SNRN
APT,FN1,SPEN,PSMD5,YWHAQ,EIF1,HACD3,DCHS1,SET,EGLN1,TPI1,CHSY1,TMSB10,EIF4A1,MR
COL18A1,SNRNP200,CSDE1,SMIM14,IGF2,EIF3C,ATP5F1A,EIF4B,VCP,MARCKSL1,UBB,AFF4,TR
PL10,MARCHF6,WSB1,RPS17,VCL,AP3D1,SBF1,DNAJA1,ARF1,CYB5R3,AKAP12,MAP4,CDC42BPB
FF4,TRIM28,MATR3,YWHAG,CKB,COL4A2,RPL37A,LRRC28,NONO,DDB1,YWHAQ,EIF1,HACD3,D
ATP5F1A,EIF4B,VCP,MARCKSL1,UBB,TRIM28,MATR3,YWHAG,CKB,COL4A2,RPL37A,LRRC28,SF
1,HLA-C,SF3B1,PDIA3,SSBP2,ZNF43,BASP1,VAPA,HMGB1,POLR2A,CDKL1,TPT1,SEC61A1,RPS3A
MORF4L1,BTF3,NCAM1,AGRN,ZMIZ1,TIMP2,XAF1,SARAF,MIDN,MARCHF6,VCL,AP3D1,DNAJA1,
1,BRD3,CIPC,PJA2,MSN,KPNB1,C1ORF43,COL1A2,LLGL1,NCS1,RHOA,CBX5,ZNF865,CLSTN1,A
S,SPTAN1,GSTP1,MARCKS,SMG1,CALM1,NPM1,PLXNB2,PLD3,HSPD1,COL18A1,SNRNP200,CSDE
31,RPS17,VCL,SBF1,DNAJA1,ARF1,CYB5R3,AKAP12,MAP4,CDC42BPB,ITM2B,AP2B1,MAT2A,SLC
1,NCAM1,ANXA2,AGRN,TIMP2,XAF1,SARAF,MIDN,MARCHF6,VCL,AP3D1,SBF1,ARF1,CYB5R3,MA
1,NCAM1,ANXA2,AGRN,TIMP2,XAF1,SARAF,MIDN,MARCHF6,VCL,AP3D1,SBF1,ARF1,CYB5R3,MA
3PB,ITM2B,TMEM170A,GTF2F1,PXDN,HNRNPU,PGAM1,NFE2L1,CTNNA1,FADS2,CNN3,TPM4,TN
CN1,TMBIM6,PTMS,SPTAN1,GSTP1,MARCKS,SMG1,CALM1,NPM1,PLXNB2,PLD3,HSPD1,USP11,
A2,AGRN,ZMIZ1,TIMP2,MIDN,RPL10,MARCHF6,WSB1,VCL,AP3D1,SBF1,ARF1,CYB5R3,MAP4,CD
IMP2,SARAF,COL11A1,RPL10,MARCHF6,WSB1,RPS17,VCL,SBF1,DNAJA1,ARF1,CYB5R3,MAP4,S

STMN1,ITGB1,TUBB4B,UCHL1,MORF4L1,NCAM1,ANXA2,AGRN,RPLP1,TIMP2,XAF1,SARAF,COL1
I2B,MAT2A,PXDN,HNRNPU,NFE2L1,HNRNPA0,CTNNA1,RPS16,CNN3,TNS3,CTSD,EIF4H,NACA,TSF
BPB,ITM2B,PXDN,HNRNPU,OGA,NFE2L1,CTNNA1,FADS2,CNN3,TPM4,TNS3,CTSD,NACA,LAMA5,
TNNA1,FADS2,RPS16,CTSD,LPCAT1,LAMA5,BRD3,CIPC,PJA2,MSN,KPNB1,C1ORF43,COL1A2,LLG
NC1,SSBP2,BASP1,VAPA,HMGB1,POLR2A,AP2M1,TPT1,RPS3A,RPL18,STMN1,ITGB1,TUBB4B,UC
TMBIM6,PTMS,SPTAN1,GSTP1,MARCKS,SMG1,CALM1,NPM1,PLXNB2,TTC3,PLD3,HSPD1,USP11,
RMP1,LASP1,OAZ1,SRSF4,PFN1,ASTN2,TLE5,NISCH,CCNI,TADA2B,KHDRBS1,XPO1,CSNK1E,PFK
RN,TIMP2,XAF1,SARAF,MIDN,RPL10,MARCHF6,VCL,AP3D1,SBF1,ARF1,CYB5R3,MAP4,SF3B2,CD
RCKS,SMG1,CALM1,NPM1,PLXNB2,PLD3,HSPD1,USP11,COL18A1,SNRNP200,CSDE1,SMIM14,IG
1,UCHL1,ANXA2,AGRN,ZMIZ1,TIMP2,COL11A1,MIDN,MARCHF6,RPS17,VCL,AP3D1,SBF1,DNAJA1
PI1,CHSY1,TMSB10,EIF4A1,RPL19,RPL15,GTF3C4,LDHB,PKD1,SF3B1,PDIA3,SERINC1,SSBP2,ZN
XA2,AGRN,TIMP2,XAF1,SARAF,COL11A1,MIDN,MARCHF6,VCL,AP3D1,ARF1,CYB5R3,MAP4,CDC4
NFE2L1,CTNNA1,CNN3,TNS3,SCARB2,NACA,LPCAT1,SLC22A17,BRD3,CIPC,PJA2,MSN,KPNB1,RP
.,AP3D1,DNAJA1,ARF1,CYB5R3,MAP4,CDC42BPB,ITM2B,GOLGA8A,PXDN,HNRNPU,OGA,NFE2L1
1,GSTP1,MARCKS,SMG1,CALM1,NPM1,PLXNB2,PLD3,HSPD1,USP11,COL18A1,SNRNP200,CSDE
,CDC42BPB,ITM2B,MAT2A,TMEM170A,SLC4A8,PXDN,HNRNPU,NFE2L1,SMARCA5,HNRNPA0,FAD
F5A,PNMA1,GPC3,DDR1,PIP4K2B,SOX12,DDX3X,ARL4C,MAGED2,H1-0,APLP1,MIF,GPAT4,MYO5
.19,RPL15,GTF3C4,LDHB,PKD1,HLA-C,PDIA3,SERINC1,BASP1,VAPA,HMGB1,POLR2A,AP2M1,TPT
R3,AKAP12,MAP4,CDC42BPB,TMEM170A,GTF2F1,OGA,PGAM1,NFE2L1,HNRNPA0,CTNNA1,FADS
,RPS24,SLC22A17,BRD3,CIPC,PJA2,KPNB1,PAPOLA,IGFBP3,LLGL1,NCS1,RPL22,RHOA,CBX5,ZN
A1,MIDN,WSB1,RPS17,AP3D1,SBF1,DNAJA1,ARF1,AKAP12,MAP4,SF3B2,CDC42BPB,ITM2B,PXDN
CS1,RHOA,PTCD3,CLSTN1,ATF4,KIF5B,RAB6B,SLC7A5,ENC1,NLGN2,CXADR,CHD4,ATP1A1,JUND
A2B,KLHL9,SLC3A2,CSNK1E,PFKL,RPS25,TKT,CDH6,SFPQ,CCND2,CDH2,PNMA1,PDIA6,GPC3,PIF
S5,APLP1,TLN1,CHPF,MYO5A,COL6A2,CD63,NGRN,RPS10,B4GALNT4,MTSS2,APLP2,HNRNPDL,M
R1A,RPL15,LDHB,DNAH8,PKD1,HLA-C,SF3B1,SSBP2,RPS3,CHD3,BASP1,VAPA,HMGB1,POLR2A,
2,PRDX2,RPS15,ARL4C,GNB2,RCN1,H1-0,RPS5,RPL35,TLN1,MYO5A,COL6A2,SESN3,B4GALNT4
2,RPS3,CHD3,BASP1,VAPA,HMGB1,CDKL1,AP2M1,TPT1,SEC61A1,ITGB1,TUBB4B,UCHL1,RPL11
,TPI1,CHSY1,TMSB10,INA,EIF4A1,MRC2,PPP2R1A,RPL15,GTF3C4,PKD1,HLA-C,SSBP2,ZNF43,CH
,BASP1,VAPA,HMGB1,POLR2A,AP2M1,TPT1,RPS4X,ITGB1,TUBB4B,UCHL1,MORF4L1,BTF3,AGR
R1A,GTF3C4,PKD1,PDIA3,SSBP2,ZNF43,RPS3,VAPA,HMGB1,POLR2A,CDKL1,AP2M1,TPT1,RPS3
,SPTAN1,GSTP1,MARCKS,SMG1,CALM1,NPM1,PLXNB2,TTC3,PLD3,HSPD1,USP11,COL18A1,SNR
1,MIDN,RPS17,VCL,AP3D1,SBF1,ARF1,CYB5R3,MAP4,SF3B2,CDC42BPB,ITM2B,MAT2A,PXDN,HN
LBP,EIF5A,CCND2,CDH2,PDIA6,DDR1,PIP4K2B,MEGF8,SOX12,DDX3X,ITM2C,ARL4C,MAGED2,G
NRNPU,NFE2L1,SMARCA5,RPS16,RPL23,CNN3,RPL29,CTSD,SCARB2,LPCAT1,LAMA5,RPS24,BRD
TMSB10,MRC2,PPP2R1A,LDHB,DNAH8,SERINC1,SSBP2,BASP1,VAPA,HMGB1,AP2M1,TPT1,STMN
R3,CKB,COL4A2,RPL37A,LRRC28,FN1,SPEN,DDB1,YWHAQ,EIF1,HACD3,DCHS1,SET,EGLN1,TPI1,
ARDBP,RPLP2,NDFIP1,TSR2,MXD4,ANKRD52,H1-10,MYH10,DUX4,YWHAB,CKAP5,SND1,LAMP1,F
F1,RAB6B,NLGN2,CXADR,CHD4,ATP1A1,JUND,RPS20,EIF5,CD81,LENG8,HMGN2,CRMP1,LASP1,
RDBP,MXD4,MYL6,HNRNPK,INIP,CADM4,RHOB,RAB1A,PLPPR3,SEPTIN3,COTL1,HECTD4,MAGED
N,MARCHF6,VCL,AP3D1,DNAJA1,AKAP12,MAP4,CDC42BPB,MAT2A,GOLGA8A,PXDN,OGA,HNRN
A,CNN3,TNS3,CTSD,EIF4H,LPCAT1,LAMA5,SLC22A17,BRD3,MSN,KPNB1,PAPOLA,COL1A2,LLGL
BP3,LLGL1,GAP43,RPL14,RHOA,CBX5,ATF4,PSMD2,NR2F1,SLC7A5,CXADR,JUND,TMEM259,RPS
2,AGRN,ZMIZ1,TIMP2,XAF1,SARAF,COL11A1,MIDN,RPL10,MARCHF6,RPS17,VCL,AP3D1,SBF1,DI
S16,CNN3,TNS3,CTSD,NACA,LPCAT1,LAMA5,SLC22A17,BRD3,CIPC,PJA2,KPNB1,IGFBP3,COL1A
N1,GSTP1,MARCKS,SMG1,CALM1,NPM1,PLXNB2,TTC3,PLD3,HSPD1,USP11,COL18A1,SNRNP20

BB4B,MORF4L1,BTF3,NCAM1,ANXA2,AGRN,RPLP1,ZMIZ1,TIMP2,MIDN,RPL10,WSB1,RPS17,VCL
SOX12,CCN2,ACTN4,RPS15,ARL4C,EIF3L,RCN1,APLP1,IK,RPS19,RPL35,MIF,GPAT4,MYO5A,COL
1,ATF4,KIF5B,RAB6B,SLC7A5,NLGN2,CXADR,ATP1A1,JUND,RPS20,CD81,LENG8,WDR1,HMGN2,A
2A,AP2M1,SEC61A1,RPS3A,RPL18,STMN1,TUBB4B,UCHL1,MORF4L1,BTF3,ANXA2,AGRN,SARAF,
F1,HACD3,DCHS1,SET,EGLN1,TPI1,CHSY1,TMSB10,INA,EIF4A1,MRC2,RPL19,PPP2R1A,RPL15,G
PKD1,SF3B1,SERINC1,SSBP2,CHD3,BASP1,VAPA,HMGB1,POLR2A,AP2M1,TPT1,SEC61A1,RPL18,
2L1,SMARCA5,HNRNPA0,CTNNA1,FADS2,RPL23,CNN3,TNS3,CTSD,SCARB2,NACA,RPS24,SLC22A
18,STMN1,ITGB1,TUBB4B,UCHL1,MORF4L1,BTF3,ANXA2,AGRN,TIMP2,SARAF,COL11A1,MARCH
3B,TRIM28,MATR3,YWHAG,CKB,COL4A2,LRRC28,FN1,SPEN,NONO,DDB1,YWHAQ,EIF1,HACD3,D
D1,USP11,COL18A1,SNRNP200,CSDE1,SMIM14,IGF2,EIF3C,ATP5F1A,EIF4B,VCP,MARCKSL1,UBE
NLGN2,TMEM259,EIF5,CD81,COL5A2,CRMP1,BEX3,SRSF4,RBBP4,RTN3,PFN1,ASTN2,TLE5,PNN
3,SOX12,ARL4C,APLP1,MIF,GPAT4,CHPF,MYO5A,COL6A2,NGRN,B4GALNT4,MTSS2,APLP2,HNRN
S2,DNAJC5,APLP2,TARDBP,MXD4,HNRNPK,INIP,MYH10,RPL36,CADM4,HNRNPH1,HLA-B,PLPPR3,
RAF,MIDN,MARCHF6,RPS17,VCL,AP3D1,SBF1,DNAJA1,ARF1,PAFAH1B1,MAP4,CDC42BPB,ITM2B
DN,RPL10,WSB1,RPS17,VCL,AP3D1,SBF1,ARF1,CYB5R3,MAP4,SF3B2,CDC42BPB,ITM2B,TMEM1
SSBP2,RPS3,CHD3,BASP1,VAPA,HMGB1,POLR2A,CDKL1,TPT1,SEC61A1,RPL18,STMN1,ITGB1,TU
CCND2,CDH2,PDIA6,MEGF8,ITM2C,CLTC,MAGED2,GNB2,RCN1,APLP1,RPL35,MIF,TLN1,COL6A2
NPC,CCNI,XPO1,SLC3A2,HDLBP,TKT,EIF5A,CCND2,CDH2,SOX12,DDX3X,ITM2C,CLTC,MAGED2,(
A,EIF4B,VCP,MARCKSL1,UBB,AFF4,TRIM28,MATR3,CKB,COL4A2,MAPT,FN1,SPEN,NONO,DDB1,
WHAQ,HACD3,DCHS1,SET,EGLN1,TPI1,CHSY1,TMSB10,INA,MRC2,PPP2R1A,RPL15,GTF3C4,LDH
3,COL4A2,RPL37A,FN1,SPEN,PSMD5,YWHAQ,EIF1,HACD3,DCHS1,SET,EGLN1,TPI1,CHSY1,TMSB
P2,SARAF,MIDN,MARCHF6,VCL,AP3D1,DNAJA1,ARF1,CYB5R3,MAP4,SF3B2,CDC42BPB,ITM2B,A
IN1,ITGB1,TUBB4B,UCHL1,MORF4L1,BTF3,AGRN,ZMIZ1,TIMP2,SARAF,MIDN,SBF1,DNAJA1,ARF
KLRD1,GDI1,FSCN1,TMBIM6,PTMS,SPTAN1,GSTP1,MARCKS,SMG1,CALM1,NPM1,PLXNB2,PLD3
RPL35,MIF,TLN1,COL6A2,ANXA6,NGRN,GOLM1,B4GALNT4,MTSS2,RPL35A,DNAJC5,APLP2,TAR
LC22A17,BRD3,CIPC,MSN,IGFBP3,COL1A2,LLGL1,HK1,NCS1,RHOA,ZNF865,ATF4,RAB6B,SLC7A
PIP4K2B,SOX12,RPL30,CLTC,MRFAP1,RCN1,H1-0,APLP1,MYO5A,COL6A2,SESN3,MTSS2,DNAJC
I14,IGF2,EIF3C,EIF4B,VCP,MARCKSL1,UBB,AFF4,TRIM28,MATR3,YWHAG,CKB,COL4A2,RPL37A,
A5,CHD4,JUND,CD81,WDR1,HMGN2,ATP2A2,CRMP1,LASP1,BEX3,VAT1,OAZ1,MCL1,TLE5,PNN,NI
1,PSMD5,YWHAQ,EIF1,HACD3,SET,EGLN1,MAP9,TPI1,CHSY1,TMSB10,EIF4A1,MRC2,RPL19,PPP2
I11,CADM4,RHOB,LGALS3BP,ATP1A2,RAB1A,PLPPR3,ARCN1,SEPTIN3,COTL1,MAGED4B,CTBP1,
HD4,ATP1A1,JUND,EIF5,CD81,WDR1,HMGN2,ATP2A2,RPL26,LASP1,OAZ1,PFN1,TLE5,NES,NISCH
BPB,ITM2B,ZNF770,MAT2A,TMEM170A,PXDN,HNRNPU,PGAM1,HNRNPA0,FADS2,RPS16,LDHA,C
TNNA1,FADS2,RPS16,CNN3,CTSD,SCARB2,NACA,LAMA5,BRD3,XRCC5,CIPC,PJA2,KPNB1,C1ORF
5,YWHAQ,EIF1,HACD3,DCHS1,SET,EGLN1,MAP9,TPI1,CHSY1,TMSB10,EIF4A1,RPL19,RPL15,GTF3
N1,COTL1,HECTD4,MAGED4B,CAPRIN1,STIP1,GPC1,TOP2B,SLC35E2B,ATP6V0C,MMP2,SKP1,L1
LNT4,MTSS2,DNAJC5,APLP2,HNRNPDL,TARDBP,MXD4,INIP,MYH10,RPL36,CADM4,HNRNPH1,HL
P2,BASP1,HMGB1,POLR2A,AP2M1,TPT1,SEC61A1,STMN1,ITGB1,TUBB4B,UCHL1,MORF4L1,NC
CIPC,PJA2,SLC16A7,KPNB1,C1ORF43,COL1A2,LLGL1,NCS1,RHOA,CBX5,ZNF865,KIF5B,RAB6B,S
N3,TNS3,CTSD,SCARB2,NACA,LPCAT1,LAMA5,RPS24,BRD3,CIPC,MSN,SLC16A7,KPNB1,C1ORF4
RPS3,BASP1,VAPA,HMGB1,POLR2A,CDKL1,TPT1,SEC61A1,RPS4X,RPS3A,STMN1,ITGB1,TUBB4B,
,GLUL,PEA15,LUC7L3,SQLE,FAM199X,CTDSP2,NOVA2,CTSB,CDH11,MDH2,CHD6,EEF1A2,TBCD,
2B1,MAT2A,TMEM170A,GTF2F1,GOLGA8A,PXDN,OGA,PGAM1,NFE2L1,SMARCA5,HNRNPA0,CTN
2B1,MAT2A,TMEM170A,GTF2F1,GOLGA8A,PXDN,OGA,PGAM1,NFE2L1,SMARCA5,HNRNPA0,CTN
2B1,MAT2A,TMEM170A,GTF2F1,GOLGA8A,PXDN,OGA,PGAM1,NFE2L1,SMARCA5,HNRNPA0,CTN

,TPT1,SEC61A1,RPS3A,STMN1,TUBB4B,UCHL1,RPL11,MORF4L1,BTF3,NCAM1,ANXA2,AGRN,RF
,SCARB2,TSPAN3,LPCAT1,LAMA5,SLC22A17,BRD3,CIPC,PJA2,KPNB1,C1ORF43,LLGL1,HK1,NC
,PPP2R1A,RPL15,LDHB,SF3B1,SERINC1,SSBP2,BASP1,VAPA,POLR2A,SEC61A1,STMN1,TUBB4B,
TP1A1,JUND,RPS20,EIF5,COL5A2,LENG8,WDR1,HMGN2,ATP2A2,CRMP1,BEX3,VAT1,OAZ1,SRS
LE,CTDSP2,NOVA2,MDH2,CHD6,EEF1A2,CCT2,TBCD,VAMP2,MORF4L2,HNRNPA3,PRDX5,HSPA4,
00,CSDE1,SMIM14,IGF2,ATP5F1A,EIF4B,VCP,MARCKSL1,UBB,AFF4,TRIM28,MATR3,YWHAG,CK
22,RHOA,ZNF865,ATF4,RAB6B,SLC7A5,NLGN2,CHD4,ATP1A1,JUND,CD81,LENG8,WDR1,HMGN
RNPU,NFE2L1,SMARCA5,HNRNPA0,CTNNA1,RPS16,CNN3,RPSA,TNS3,CTSD,EIF4H,LPCAT1,LAM
RPL10,MARCHF6,RPS17,VCL,AP3D1,ARF1,CYB5R3,SF3B2,CDC42BPB,ITM2B,ZNF770,TMEM170
GPC3,PIP4K2B,SOX12,RPL12,ITM2C,ARL4C,MAGED2,GNB2,MIF,GPAT4,MYO5A,COL6A2,CD63,
CHD6,EEF1A2,RPN2,TBCD,MLXIP,CPSF1,SNX22,TGFBI,ABCA3,PDXK,PKN1,CAMSAP1,MLEC,ACA
10,INA,EIF4A1,MRC2,RPL19,PPP2R1A,GTF3C4,LDHB,PKD1,HLA-C,SF3B1,SSBP2,ZNF43,CHD3,V
WHAQ,EIF1,HACD3,DCHS1,SET,EGLN1,MAP9,CHSY1,TMSB10,INA,EIF4A1,MRC2,PPP2R1A,RPL15
DM4,LAMP1,RHOB,RAB1A,PLPPR3,COTL1,CTBP1,RAC1,ARHGDIA,GPC1,TOP2B,SMCHD1,HDAC
SARAF,MIDN,MARCHF6,WSB1,RPS17,VCL,AP3D1,SBF1,DNAJA1,ARF1,CYB5R3,MAP4,CDC42BPB,
C4,PKD1,SF3B1,SERINC1,ZNF43,RPS3,BASP1,VAPA,HMGB1,POLR2A,TPT1,SEC61A1,RPS4X,RPS
LXNB2,PLD3,HSPD1,USP11,COL18A1,SNRNP200,CSDE1,SMIM14,IGF2,ATP5F1A,VCP,MARCKSL1
S,PRKCSH,KHDRBS1,XPO1,SPTBN2,PFKL,HDLBP,RPS25,TKT,EIF5A,CDH2,PNMA1,PDIA6,GPC3,P
L1,UBB,AFF4,TRIM28,MATR3,YWHAG,CKB,COL4A2,LRRC28,FN1,SPEN,NONO,DDB1,PSMD5,YW
LAMA5,RPS24,BRD3,CIPC,PJA2,MSN,SLC16A7,KPNB1,C1ORF43,LLGL1,HK1,RPL22,ZNF865,AT
D6,U2AF2,EEF1A2,TBCD,SUGP2,SQSTM1,KLHL24,HNRNPL,MORF4L2,PRDX5,PKN1,AHCYL1,CA
AQ,HACD3,DCHS1,SET,EGLN1,TPI1,CHSY1,TMSB10,INA,MRC2,RPL19,PPP2R1A,RPL15,GTF3C4,
RD3,CIPC,KPNB1,C1ORF43,PAPOLA,LLGL1,HK1,NCS1,RHOA,CBX5,ZNF865,NR2F1,RAB6B,SLC7
ARL4C,APLP1,MIF,GPAT4,MYO5A,COL6A2,CD63,B4GALNT4,MTSS2,APLP2,HNRNPDL,TARDBP,S
1,NCS1,ZNF865,CLSTN1,NR2F1,RAB6B,SLC7A5,ENC1,NLGN2,GLIPR1L2,CXADR,CHD4,JUND,EIF
4B,UCHL1,MORF4L1,ANXA2,AGRN,ZMIZ1,TIMP2,XAF1,SARAF,COL11A1,MIDN,VCL,AP3D1,ARF1,
,PIP4K2B,MEGF8,SOX12,DDX3X,GNB2,APLP1,GPAT4,TLN1,CHPF,MYO5A,CD63,ANXA6,GOLM1,
HSY1,INA,EIF4A1,MRC2,RPL19,PPP2R1A,RPL15,GTF3C4,LDHB,DNAH8,PKD1,HLA-C,SF3B1,PDIA
N,RPLP1,ZMIZ1,TIMP2,XAF1,SARAF,MIDN,RPS17,VCL,AP3D1,SBF1,DNAJA1,ARF1,CYB5R3,MAP4,
2,CDC42BPB,ITM2B,MAT2A,TMEM170A,GTF2F1,SLC4A8,PXDN,HNRNPU,OGA,PGAM1,NFE2L1,H
S2,RPS16,CNN3,RPSA,TNS3,CTSD,EIF4H,TSPAN3,LPCAT1,LAMA5,SLC22A17,BRD3,RPS9,XRCC5,
T1,LAMA5,SLC22A17,BRD3,PJA2,SLC16A7,KPNB1,RPL17,C1ORF43,PAPOLA,COL1A2,LLGL1,H
TN4,MMP14,ARL4C,APLP1,RPS19,GPAT4,TLN1,CHPF,MYO5A,CD63,NGRN,GOLM1,B4GALNT4,M
,GTF3C4,LDHB,PKD1,SF3B1,PDIA3,SERINC1,SSBP2,BASP1,VAPA,HMGB1,POLR2A,TPT1,SEC61A
5R3,AKAP12,MAP4,CDC42BPB,ITM2B,MAT2A,SLC4A8,GOLGA8A,PXDN,OGA,HNRNPA0,CTNNA1,
P,IDS,RPL5,GDI1,FSCN1,TMBIM6,PTMS,SPTAN1,GSTP1,MARCKS,SMG1,CALM1,NPM1,PLXNB2,
P1,CKAP4,RAB1A,PLPPR3,ZNF454,MAGED4B,CTBP1,STIP1,TOP2B,SMCHD1,HDAC2,MMP2,DNE
AC2,TNC,ATP6V0C,MMP2,DNER,TRIB2,PEA15,SQLE,HS6ST1,CTDSP2,NOVA2,ZC3H7B,CHD6,U2
C1,TOP2B,HDAC2,ATP6V0C,TRIB2,TIMP3,CTDSP2,NOVA2,CHD6,EEF1A2,TBCD,CPSF1,SNX22,P
LXNB2,TTC3,PLD3,USP11,COL18A1,SNRNP200,CSDE1,IGF2,EIF3C,ATP5F1A,EIF4B,VCP,MARCK
HA,CNN3,TNS3,EIF4H,LAMA5,BRD3,XRCC5,CIPC,KPNB1,C1ORF43,PAPOLA,NCS1,RPL22,RHOA,
31,TUBB4B,UCHL1,MORF4L1,ANXA2,AGRN,RPLP1,ZMIZ1,TIMP2,COL11A1,MIDN,RPL10,MARCH
TNNA1,FADS2,RPL23,CNN3,RPL29,RPSA,TPM4,TNS3,CTSD,EIF4H,SCARB2,NACA,TSPAN3,LAMA
4,ATP1A1,JUND,CD81,WDR1,HMGN2,ATP2A2,CRMP1,LASP1,VAT1,OAZ1,PFN1,JPT2,NES,RPL31,
2,RPL15,GTF3C4,LDHB,DNAH8,PKD1,SF3B1,PDIA3,SERINC1,SSBP2,BASP1,VAPA,HMGB1,POLR

,RPS17,VCL,SBF1,DNAJA1,ARF1,CYB5R3,MAP4,SF3B2,CDC42BPB,ZNF770,MAT2A,TMEM170A,G
MP2,SARAF,COL11A1,MIDN,RPL10,MARCHF6,RPS17,VCL,SBF1,DNAJA1,ARF1,CYB5R3,MAP4,SF3
1A1,JUND,RPS20,CD81,WDR1,ATP2A2,CRMP1,BEX3,VAT1,OAZ1,RBBP4,PFN1,ASTN2,JPT2,TLE5,
PFN1,JPT2,TLE5,PNN,NES,NISCH,PRKCSH,HNRNPC,XPR1,KHDRBS1,XPO1,SPTBN2,SLC3A2,PFKL
S2,RPS16,RPL23,RPL29,TPM4,TNS3,CTSD,TSPAN3,LPCAT1,CIPC,PJA2,KPNB1,C1ORF43,IGFBP3
,PRDX2,ITM2C,MAGED2,GNB2,RPS5,APLP1,RPS7,RPS19,HMGCS1,GPAT4,JUN,COL6A2,CD63,G
OL11A1,MIDN,MARCHF6,VCL,DNAJA1,ARF1,CYB5R3,MAP4,SF3B2,CDC42BPB,ZNF770,MAT2A,TM
AZ1,PFN1,MCL1,TLE5,NES,NISCH,TADA2B,XPR1,SLC3A2,PFKL,HDLBP,RPS25,TKT,GLG1,CCND2,
P,MXD4,HNRNPK,INIP,YWHAB,CADM4,HNRNPH1,HLA-B,PLPPR3,MAGED4B,CTBP1,ARHGDIA,STI
P,MXD4,HNRNPK,INIP,YWHAB,CADM4,HNRNPH1,HLA-B,PLPPR3,MAGED4B,CTBP1,ARHGDIA,STI
AKIRIN1,CTXN1,PPP1R9B,ANP32A,CIC,KCNQ2,BCL9L,PRMT1,KIRREL1,TRIO,SF1,PBX1,AP2A1,H
D4B,CTBP1,ARHGDIA,CDK16,HDAC2,SKP1,BCLAF1,NNAT,MAP1A,HS6ST1,CTDSP2,DDX39B,EID1,
L18A1,SNRNP200,CSDE1,IGF2,EIF3C,ATP5F1A,EIF4B,VCP,MARCKSL1,UBB,TRIM28,MATR3,CKB,
TF2F1,GOLGA8A,PXDN,HNRNPU,NFE2L1,SMARCA5,CNN3,TPM4,TNS3,SCARB2,RPS24,SLC22A1
GLN,MYL6,HNRNPK,INIP,CADM4,PLPPR3,ARCN1,COTL1,HECTD4,CTBP1,ARHGDIA,CAPRIN1,STIP
,L1CAM,TRIB2,PEA15,SQLE,FAM199X,SERF2,TIMP3,NOVA2,CTSB,NUCB1,CHD6,EEF1A2,TBCD,
HLA-B,PLPPR3,HECTD4,CTBP1,ARHGDIA,STIP1,GPC1,SLC35E2B,SMCHD1,IPO5,ATP6V0C,DNER,L
TLN1,CHPF,JUN,COL6A2,CD63,GOLM1,B4GALNT4,MTSS2,DNAJC5,APLP2,HNRNPDL,ARL6IP1,N
RF4L1,NCAM1,ANXA2,AGRN,RPLP1,ZMIZ1,TIMP2,MIDN,RPL10,WSB1,VCL,AP3D1,ARF1,CYB5R3,
,SF3B2,ITM2B,TMEM170A,GTF2F1,PXDN,PGAM1,NFE2L1,HNRNPA0,FADS2,RPL23,LDHA,CPE,C
,BASP1,VAPA,HMGB1,POLR2A,AP2M1,TPT1,SEC61A1,RPS4X,RPS3A,RPL18,TUBB4B,RPL11,MO
P2,CHD3,BASP1,VAPA,HMGB1,POLR2A,TPT1,SEC61A1,STMN1,ITGB1,TUBB4B,UCHL1,MORF4L1
ET,MAP9,CHSY1,TMSB10,EIF4A1,MRC2,RPL19,RPL15,GTF3C4,LDHB,PKD1,SF3B1,PDIA3,SERINC
CDH11,MDH2,CHD6,RNF187,EEF1A2,TBCD,PPP1CB,SQSTM1,VAMP2,SSRP1,PRDX5,ACTR3,PKN1
RSF4,PFN1,TLE5,NISCH,HNRNPC,CCNI,TADA2B,KHDRBS1,XPO1,SPTBN2,SLC3A2,CSNK1E,RPS25
1,MORF4L1,NCAM1,AGRN,ZMIZ1,TIMP2,XAF1,COL11A1,MIDN,MARCHF6,RPS17,VCL,DNAJA1,C
VCL,AP3D1,SBF1,ARF1,CYB5R3,MAP4,CDC42BPB,ZNF770,TMEM170A,SLC4A8,PXDN,HNRNPU,
LGL1,RPL14,RHOA,PTCD3,CBX5,ATF4,SLC7A5,ENC1,CXADR,JUND,TMEM259,RPS20,EIF5,CD81,
,H3-3A,DPYSL2,TOP1,TUBB2A,MVB12B,CCND1,ATG2B,SRSF9,CCNL2,PBX2,FAM171A2,AEBP1,
RPS24,BRD3,XRCC5,CIPC,PJA2,SLC16A7,KPNB1,RPL17,PAPOLA,COL1A2,LLGL1,NCS1,RHOA,CB
PD1,USP11,COL18A1,SNRNP200,SMIM14,IGF2,EIF3C,ATP5F1A,EIF4B,VCP,MARCKSL1,UBB,AFF
HNRNPH1,HLA-B,PLPPR3,ARCN1,HECTD4,CTBP1,RAC1,ARHGDIA,GPC1,CDK16,SLC35E2B,SMC
NT4,APLP2,TARDBP,TAGLN,INIP,MYH10,RPL36,CADM4,HNRNPH1,HLA-B,PLPPR3,ARCN1,COTL1,
PL35A,DNAJC5,APLP2,TARDBP,RPLP2,ARL6IP1,NDFIP1,DDOST,MXD4,MYL6,SHOX,CIRBP,DUX4,
I3,B4GALNT4,MTSS2,DNAJC5,APLP2,NDFIP1,MXD4,HNRNPK,H1-10,MYH10,NARS1,DUX4,CKAP5,
MARCHF6,WSB1,RPS17,VCL,AP3D1,SBF1,ARF1,CYB5R3,SF3B2,ITM2B,MAT2A,GTF2F1,SLC4A8,P
TF4,RAB6B,SLC7A5,NLGN2,CXADR,ATP1A1,JUND,CD81,WDR1,HMGN2,ATP2A2,CRMP1,LASP1,O
,KLHL9,SPTBN2,SLC3A2,CSNK1E,HDLBP,RPS25,CDH6,EIF5A,CCND2,CDH2,PIP4K2B,MEGF8,SO
F4,TRIM28,MATR3,YWHAG,CKB,COL4A2,LRRC28,FN1,SPEN,DDB1,PSMD5,YWHAQ,EIF1,HACD3,
0,CTNNA1,FADS2,RPS16,CNN3,RPL29,TPM4,TNS3,CTSD,TSPAN3,LPCAT1,LAMA5,RPS24,BRD3,
12,MAP4,CDC42BPB,ITM2B,TMEM170A,GTF2F1,PXDN,HNRNPU,NFE2L1,SMARCA5,LARP1,CTN
MARCHF6,WSB1,RPS17,VCL,SBF1,DNAJA1,ARF1,CYB5R3,AKAP12,CDC42BPB,ITM2B,ZNF770,M
P2,CPSF2,NOVA2,TM9SF3,MDH2,CHD6,EEF1A2,FDFT1,MLXIP,SUGP2,SNX22,VAMP2,KLHL24,GA
ARDBP,MXD4,H1-10,INIP,CIRBP,MYH10,RPL36,YWHAB,PTGES3,CADM4,RHOB,HNRNPH1,LGAL
M4,SCARB2,NACA,LAMA5,SLC22A17,BRD3,MSN,SLC16A7,KPNB1,C1ORF43,LLGL1,HK1,NCS1,R

ZMIZ1,MIDN,RPL10,RPS17,VCL,AP3D1,SBF1,DNAJA1,ARF1,CYB5R3,MAP4,SF3B2,CDC42BPB,ITM
19,PPP2R1A,RPL15,GTF3C4,LDHB,DNAH8,PKD1,SF3B1,PDIA3,SERINC1,SSBP2,ZNF43,RPS3,HMG
E2L1,SMARCA5,FADS2,RPS16,CPE,CNN3,RPL29,TPM4,CTSD,EIF4H,LPCAT1,LAMA5,RPS24,SLC22
,SERINC1,SSBP2,ZNF43,RPS3,BASP1,HMGB1,POLR2A,CDKL1,AP2M1,TPT1,SEC61A1,OR8K1,ITG
B4B,UCHL1,MORF4L1,ANXA2,AGRN,TIMP2,XAF1,SARAF,MIDN,MARCHF6,AP3D1,SBF1,ARF1,CYB
5,MIF,TLN1,CHPF,COL6A2,GOLM1,SESN3,TARDBP,ST3GAL2,MXD4,MYL6,H1-10,RPL36,PTGES3
S7,PRDX5,PDXK,PKN1,HSPA4,CAMSAP1,GRINA,NCKAP1,MLEC,ACAP3,MDK,LAMC1,SKI,H3-3A,A
TP1A1,TMEM259,EIF5,WDR1,HMGN2,ATP2A2,LASP1,BEX3,OAZ1,PFN1,ASTN2,MCL1,JPT2,NES,N
2,LRRC28,FN1,SPEN,DDB1,PSMD5,YWHAQ,EIF1,DCHS1,SET,TPI1,CHSY1,TMSB10,INA,MRC2,RP
CA,LAMA5,RPS24,BRD3,SLC16A7,KPNB1,C1ORF43,IGFBP3,HK1,NCS1,RPL22,RHOA,CLSTN1,AT
5,RPL9,ATF4,CXADR,JUND,TMEM259,RPS20,EIF5,CD81,WDR1,ATXN7L3B,HMGN2,RPL26,LASP1
RNPC,TADA2B,XPO1,CSNK1E,PFKL,TKT,TPM1,PEBP1,CDH2,PNMA1,DDR1,PIP4K2B,SOX12,ACTN
DR,ATP1A1,TMEM259,RPS20,EIF5,CD81,ATXN7L3B,BEX3,CCT7,OAZ1,SRSF4,RBBP4,PFN1,MCL
B2,CDC42BPB,ITM2B,AP2B1,GOLGA8A,PXDN,HNRNPU,NFE2L1,HNRNPA0,CTNNA1,FADS2,CNN3
D1,SF3B1,PDIA3,SERINC1,SSBP2,ZNF43,RPS3,HMGB1,POLR2A,CDKL1,AP2M1,TPT1,SEC61A1,R
5,APLP2,HNRNPDL,MXD4,ANKRD52,H1-10,SEPTIN11,CADM4,RHOB,HNRNPH1,PLPPR3,SEPTIN3,
VCL,SBF1,ARF1,CYB5R3,AKAP12,SF3B2,ZNF770,GTF2F1,GOLGA8A,PXDN,HNRNPU,NFE2L1,SMA
A17,RPS9,CIPC,MSN,KPNB1,C1ORF43,PAPOLA,IGFBP3,LLGL1,GAP43,RPL14,RHOA,PTCD3,PSM
O,YWHAQ,EIF1,HACD3,DCHS1,SET,EGLN1,TPI1,TMSB10,INA,EIF4A1,MRC2,RPL19,PPP2R1A,RPL
C1ORF43,COL1A2,LLGL1,RHOA,CBX5,CLSTN1,CXADR,CHD4,ATP1A1,JUND,TMEM259,LENG8,W
F4,PSMD2,SLC7A5,ENC1,CXADR,ATP1A1,JUND,EIF5,CD81,HMGN2,ATP2A2,RPL26,BEX3,CCT7,O
4,IGF2,EIF3C,ATP5F1A,EIF4B,VCP,UBB,AFF4,CKB,COL4A2,RPL37A,LRRC28,FN1,SPEN,DDB1,PS
OC,DNER,TRIB2,PEA15,SQLE,SERF2,HS6ST1,TIMP3,NOVA2,CTSB,NUCB1,CDH11,CHD6,RNF187,
,MAPT,FN1,SPEN,DDB1,PSMD5,YWHAQ,EIF1,DCHS1,SET,EGLN1,TPI1,CHSY1,TMSB10,INA,EIF4A
AP4,RAB1A,PLPPR3,HECTD4,ARHGDIA,CAPRIN1,STIP1,CDK16,TOP2B,APPL1,HDAC2,ATP6V0C,TM
HNRNPA0,CTNNA1,FADS2,RPS16,RPL23,LDHA,CNN3,RPL29,RPSA,TPM4,CTSD,TSPAN3,LPCAT1,S
HNRNPA0,CTNNA1,FADS2,RPS16,RPL23,LDHA,CNN3,RPL29,RPSA,TPM4,CTSD,TSPAN3,LPCAT1,S
COL5A2,LENG8,WDR1,ATXN7L3B,RPL26,LASP1,CCT7,SRSF4,RBBP4,PFN1,MCL1,TLE5,PNN,NES
3,B4GALNT4,MTSS2,APLP2,HNRNPDL,TARDBP,ST3GAL2,MXD4,INIP,RPL36,PTGES3,CADM4,RHO
O,MLXIP,ANXA5,PPP1CB,VAMP2,GDI2,PARP1,HNRNPA3,PTPRA,PRDX5,PKN1,YEATS2,COL2A1,AC
OC,PEA15,DLG5,CTDSP2,NOVA2,TM9SF3,EEF1A2,TBCD,PPP1CB,PABPN1,HNRNPL,HNRNPA3,ACT
TRIM28,MATR3,YWHAG,CKB,COL4A2,MAPT,FN1,NONO,DDB1,PSMD5,YWHAQ,HACD3,DCHS1,SE
CCNI,LRPPRC,TADA2B,SLC3A2,CSNK1E,PFKL,SFPQ,CDH2,PNMA1,PDIA6,PIP4K2B,MEGF8,SOX12
,ITM2C,MAGED2,GNB2,RCN1,H1-0,APLP1,RPS19,CHPF,JUN,MYO5A,COL6A2,CD63,ANXA6,B4GA
OLBP,GLG1,SFPQ,TPM1,PEBP1,PNMA1,PIP4K2B,SOX12,ACTN4,PRDX2,CLTC,EIF3L,H1-0,APLP1,
F1,DNAJA1,ARF1,MAP4,CDC42BPB,MAT2A,TMEM170A,GTF2F1,SLC4A8,HNRNPU,SMARCA5,LAR
RFAP1,MIF,GPAT4,JUN,COL6A2,GOLM1,RPS10,DNAJC5,APLP2,HNRNPDL,RPLP2,DDOST,MXD4,
DIA3,SERINC1,SSBP2,CHD3,BASP1,VAPA,POLR2A,CDKL1,AP2M1,TPT1,SEC61A1,RPS4X,TUBB4B
52,WLS,YWHAB,CADM4,HNRNPH1,PLPPR3,COTL1,MAGED4B,CTBP1,ARHGDIA,STIP1,GPC1,SLC3
HACD3,DCHS1,CHSY1,INA,RPL15,GTF3C4,LDHB,DNAH8,PKD1,HLA-C,PDIA3,SERINC1,SSBP2,ZN
S3,CDKL1,AP2M1,SEC61A1,RPS4X,RPS3A,RPL18,STMN1,ITGB1,TUBB4B,RPL11,BTF3,AGRN,MID
M2C,RCN1,RPS7,RPS19,GPAT4,JUN,COL6A2,GOLM1,RPS10,DNAJC5,APLP2,SLC25A3,RPLP2,ARL
N1,HNRNPL,PARP1,HSPH1,HNRNPA3,ACTR3,ACO2,HSPA4,CCT3,ZNF302,RBMX,RAD23B,ASXL1,S
GN2,ATP2A2,RPL26,CRMP1,LASP1,OAZ1,SRSF4,RBBP4,JPT2,TLE5,RPL31,NISCH,HNRNPC,CCNI
10,WSB1,RPS17,SBF1,DNAJA1,ARF1,CYB5R3,AKAP12,SF3B2,CDC42BPB,MAT2A,TMEM170A,GT

DDB1,EIF1,SET,EGLN1,TPI1,CHSY1,TMSB10,INA,EIF4A1,RPL19,PPP2R1A,RPL15,GTF3C4,LDHB,I
SNK1E,PFKL,GLG1,TPM1,EIF5A,CDH2,PDIA6,DDR1,PLXNA1,MEGF8,RAB7A,ITM2C,ARL4C,MIF,TI
'B,ITM2B,ZNF770,TMEM170A,GTF2F1,PXDN,HNRNPU,OGA,NFE2L1,SMARCA5,LARP1,HNRNPA0,
GRB2,PCSK1N,PRKDC,TDRD5,UBE2R2,SCAF11,HSPB1,BNIP3L,CLIP3,PLCG1,MAP2K2,ARF4,FLN
CHL1,RPL11,MORF4L1,BTF3,ANXA2,RPLP1,ZMIZ1,TIMP2,COL11A1,MIDN,RPL10,MARCHF6,RPS
,YWHAQ,EIF1,HACD3,DCHS1,SET,EGLN1,MAP9,TPI1,CHSY1,TMSB10,INA,EIF4A1,MRC2,RPL19,F
,LRPPRC,SPTBN2,SLC3A2,CSNK1E,PFKL,RPS25,TKT,EIF5A,CDH2,PDIA6,PLXNA1,PIP4K2B,RAB7
1,DNAJA1,ARF1,CYB5R3,SF3B2,MAT2A,GTF2F1,HNRNPU,OGA,HNRNPA0,RPS16,RPL23,CNN3,RF
STN3,ACAP3,MDK,JPT1,RAD23B,CHD2,ORAI2,ASXL1,HDGF,PRPF6,SELENON,OPTN,ARF3,CCND1
AF1,SARAF,MIDN,RPL10,WSB1,VCL,AP3D1,SBF1,ARF1,CYB5R3,MAP4,CDC42BPB,ITM2B,AP2B1,
M259,EIF5,LENG8,WDR1,HMGN2,ATP2A2,RPL26,CRMP1,LASP1,OAZ1,SRSF4,RBBP4,RTN3,TLE5
STN,DYNC1LI2,AEBP1,EIF3E,DYNLL2,PCSK1N,CAPZB,UBE2R2,SCAF11,ACTN1,BNIP3L,PLCG1,DU
,L1CAM,LAPTM4B,TRIB2,LUC7L3,SQLE,DLG5,HS6ST1,TIMP3,CHD6,EEF1A2,CCT2,TBCD,SNX22,
,TMSB10,EIF4A1,MRC2,RPL19,PPP2R1A,GTF3C4,LDHB,PKD1,HLA-C,SF3B1,SSBP2,ZNF43,CHD3
C28,SPEN,NONO,DDB1,PSMD5,YWHAQ,HACD3,SET,EGLN1,TPI1,CHSY1,TMSB10,INA,EIF4A1,RPL
IA1,FADS2,RPL23,CNN3,RPSA,TPM4,TNS3,CTSD,EIF4H,LAMA5,BRD3,PJA2,RPL24,SLC16A7,KPNE
2B,SLC35E2B,HDAC2,IPO5,PEA15,FAM199X,DLG5,CTDSP2,NOVA2,CTSB,CHD6,RAB27A,RNF187
NA1,FADS2,RPL23,CPE,CNN3,RPL29,RPSA,TNS3,NACA,LPCAT1,LAMA5,SLC22A17,BRD3,PJA2,SL
OD1,SREBF2,KDM3B,CTXN1,PPP1R9B,RBM15B,PSMA7,PPDPF,ARIH1,BCR,SGTA,XPOT,CAPN2,T
24,C1ORF43,PAPOLA,IGFBP3,COL1A2,RPL14,HK1,RPL22,RHOA,PTCD3,ZNF865,ATF4,GLIPR1L
CA4,AZIN1,LAPTM4A,SRSF9,ARPC2,PLEC,PDE7A,RABL6,ODC1,TCF25,PTBP1,ILF3,LUZP6,GRB2,I
6A2,MTSS2,TARDBP,ANKRD52,HNRNPK,INIP,CIRBP,SEPTIN11,UBE2H,ATP1A2,CKAP4,PLPPR3,SE
PCAT1,LAMA5,RPS9,XRCC5,CIPC,PJA2,MSN,RPL24,PAPOLA,IGFBP3,HK1,NCS1,RHOA,PTCD3,ZNI
-1,RPL5,GDI1,FSCN1,TMBIM6,PTMS,SPTAN1,GSTP1,MARCKS,SMG1,CALM1,NPM1,PLXNB2,TTC
SY1,TMSB10,INA,EIF4A1,PPP2R1A,RPL15,GTF3C4,DNAH8,PKD1,SF3B1,PDIA3,SSBP2,ZNF43,VAP
A,ANXA6,GOLM1,SESN3,B4GALNT4,DNAJC5,APLP2,HNRNPDL,MXD4,MYL6,HNRNPK,H1-10,INIP,
PLP1,RPS7,RPL35,COL6A2,B4GALNT4,MTSS2,APLP2,HNRNPDL,ARL6IP1,MXD4,HNRNPK,MYH10
5R3,MAP4,SF3B2,MAT2A,PXDN,PGAM1,NFE2L1,SMARCA5,RPS16,CNN3,RPL29,CTSD,EIF4H,LPC
SS2,APLP2,TARDBP,MXD4,ANKRD52,INIP,CIRBP,MYH10,RPL36,PTGES3,CADM4,RHOB,HNRNPH
RAB1A,PLPPR3,ARCN1,COTL1,HECTD4,MAGED4B,CTBP1,STIP1,GPC1,IPO5,MMP2,SKP1,DNER,L
9,CLSTN1,ATF4,KIF5B,SLC7A5,NLGN2,GLIPR1L2,RPS20,LENG8,HMGN2,ATP2A2,RPL26,OAZ1,M
3,VAPA,HMGB1,POLR2A,AP2M1,TPT1,SEC61A1,RPS4X,RPS3A,RPL18,ITGB1,TUBB4B,UCHL1,RPI
AM1,NFE2L1,HNRNPA0,CTNNA1,RPS16,LDHA,TNS3,CTSD,EIF4H,TSPAN3,LPCAT1,RPS24,BRD3,F
,RPL15,GTF3C4,LDHB,PKD1,SF3B1,PDIA3,SSBP2,ZNF43,BASP1,HMGB1,POLR2A,CDKL1,AP2M1
POLA,IGFBP3,LLGL1,RPL14,HK1,NCS1,CLSTN1,ATF4,NR2F1,RAB6B,ENC1,NLGN2,CXADR,CHD4,
B,DNAH8,PKD1,PDIA3,SERINC1,SSBP2,ZNF43,RPS3,VAPA,POLR2A,AP2M1,TPT1,SEC61A1,RPL18
DE1,SMIM14,IGF2,EIF3C,ATP5F1A,EIF4B,MARCKSL1,UBB,AFF4,TRIM28,MATR3,YWHAG,CKB,CC
,GTF2F1,GOLGA8A,PXDN,HNRNPU,OGA,NFE2L1,CTNNA1,FADS2,CNN3,RPL29,TPM4,TNS3,CTSD
CND1,SRSF9,DSTN,PLEC,AEBP1,TCF25,ATP5PB,ILF3,PCSK1N,WDR6,CAPZB,GABARAP,HNRNPUI
RVV-1,RPL5,KLRD1,GDI1,FSCN1,TMBIM6,PTMS,SPTAN1,GSTP1,MARCKS,SMG1,CALM1,NPM1,PI
7,VCL,AP3D1,SBF1,DNAJA1,ARF1,CYB5R3,AP2B1,MAT2A,PXDN,NFE2L1,HNRNPA0,CTNNA1,FAD
LGL1,RPL14,HK1,RHOA,ZNF865,ATF4,NR2F1,RAB6B,NLGN2,GLIPR1L2,RPS14,ATP1A1,RPS20,C
STN2,TLE5,NES,CCNI,LRPPRC,TADA2B,HDLBP,RPS25,TKT,GLG1,CDH6,EIF5A,PEBP1,CCND2,CD
6,DUX4,PTGES3,HNRNPH1,UBE2H,CKAP4,RPS12,ARCN1,COTL1,ARHGDIA,STIP1,GPC1,CDK16,A
DM4,HNRNPH1,HLA-B,PLPPR3,ARCN1,COTL1,CTBP1,RAC1,ARHGDIA,STIP1,GPC1,SLC35E2B,SM

S35,PTOV1,MLEC,CLSTN3,MDK,SURF4,CBX1,PTK7,ORAI2,SKI,SNRNP70,H3-3A,SCAF1,RAB11B,A
,DYNLL2,ILF3,GRB2,CAPZB,ACTN1,BNIP3L,CASC3,FKBP8,PLCG1,ZFAND5,DIDO1,DUSP4,FLNB,[
PL,PARP1,TSPYL4,TLN2,TSR1,MORF4L2,SSRP1,HSPH1,MTRF1L,ABCA3,RPL27,PTPRA,ACTR3,AC
WHAG,CKB,COL4A2,RPL37A,MAPT,FN1,NONO,DDB1,PSMD5,YWHAQ,HACD3,DCHS1,SET,EGLN1
TMSB10,INA,EIF4A1,MRC2,RPL19,PPP2R1A,GTF3C4,LDHB,DNAH8,HLA-C,SF3B1,PDIA3,SSBP2,ZN
1,MAPRE1,DDAH2,RABL6,DYNC1LI2,AEBP1,NYNRIN,TCF25,DYNLL2,ACTN1,CASC3,FKBP8,PLCG
HD6,RNF187,EEF1A2,CCT2,TBCD,SNX22,PGD,SQSTM1,VAMP2,GNAI3,MEST,GAS7,GDI2,TM9SF2
B1,PDIA3,SERINC1,SSBP2,ZNF43,RPS3,HMGB1,POLR2A,AP2M1,TPT1,SEC61A1,RPS4X,RPS3A,R
CTD4,RAC1,ARHGDIA,GPC1,CDK16,TOP2B,APPL1,BCLAF1,PPIB,GLUL,PTTG1IP,CDKN1A,FAM199
CTD4,RAC1,ARHGDIA,GPC1,CDK16,TOP2B,APPL1,BCLAF1,PPIB,GLUL,PTTG1IP,CDKN1A,FAM199
CTD4,RAC1,ARHGDIA,GPC1,CDK16,TOP2B,APPL1,BCLAF1,PPIB,GLUL,PTTG1IP,CDKN1A,FAM199
RPL22,RHOA,ZNF865,CLSTN1,ATF4,PSMD2,NR2F1,SLC7A5,NLGN2,RPS14,TMEM259,EIF5,LENG8
A,FAM199X,HS6ST1,CTDSP2,CTSB,TM9SF3,CHD6,TBCD,SUGP2,VAMP2,GDI2,HNRNPL,TGFBI,LM
,VAPA,SEC61A1,RPL18,STMN1,ITGB1,TUBB4B,UCHL1,AGRN,RPLP1,ZMIZ1,TIMP2,XAF1,SARAF,(
310,INA,EIF4A1,RPL19,PPP2R1A,GTF3C4,LDHB,DNAH8,HLA-C,SF3B1,PDIA3,SSBP2,ZNF43,CHD3,
GB1,POLR2A,AP2M1,TPT1,SEC61A1,RPS4X,RPS3A,RPL18,ITGB1,TUBB4B,UCHL1,RPL11,MORF4
N,NONO,DDB1,PSMD5,YWHAQ,EIF1,HACD3,DCHS1,EGLN1,MAP9,TPI1,CHSY1,TMSB10,INA,EIF4A
IDH1,HYOU1,ZFAND3,MAPK8IP1,SOD1,PFN2,FBRSL1,MEX3A,ZSCAN18,DYNLL1,CTXN1,CD99L2,
A7,KPNB1,IGFBP3,RPL14,RHOA,CBX5,PSMD2,RAB6B,GLIPR1L2,RPS14,CXADR,ATP1A1,JUND,RF
A,STIP1,HDAC2,IPO5,MMP2,L1CAM,TRIB2,PPIB,GLUL,PEA15,MAP1A,FAM199X,DLG5,TIMP3,PGR
GN2,ATP2A2,CRMP1,LASP1,BEX3,VAT1,SRSF4,RBBP4,PFN1,ASTN2,TLE5,PRKCSH,TADA2B,XPR1,
),IK,RPL35,ANXA6,NGRN,RPS10,MTSS2,APLP2,SLC25A3,RPLP2,DDOST,ANKRD52,WLS,HNRNPK,
2,CD63,ANXA6,MTSS2,APLP2,TARDBP,ANKRD52,WLS,INIP,CIRBP,MYH10,RPL36,PTGES3,CADM4
0,INA,EIF4A1,MRC2,RPL19,PPP2R1A,GTF3C4,LDHB,DNAH8,HLA-C,SF3B1,PDIA3,SSBP2,ZNF43,C
BCA2,DPYSL2,TOP1,VEGFA,TUBB2A,ARF3,CCND1,ATG2B,BTBD2,SRSF1,NYNRIN,PTBP1,DYNLL2
F3B1,PDIA3,SSBP2,ZNF43,CHD3,BASP1,VAPA,HMGB1,POLR2A,CDKL1,AP2M1,SEC61A1,RPL18,T
2,ACTN4,RPL12,ARL4C,GNB2,H1-0,MIF,HMGCS1,GPAT4,TLN1,CHPF,GJA1,CD63,B4GALNT4,API
ATP2A2,LASP1,PFN1,MCL1,PNN,NISCH,KHDRBS1,XPO1,SLC3A2,PFKL,TKT,CDH6,CCND2,CDH2,S
,HDLBP,EIF5A,CCND2,PIP4K2B,SOX12,RPS15,ITM2C,ARL4C,MRFAP1,RCN1,H1-0,RPS7,IK,RPL35
SF3B1,PDIA3,SERINC1,SSBP2,ZNF43,RPS3,VAPA,HMGB1,POLR2A,CDKL1,AP2M1,TPT1,SEC61A1,
KIP,SUGP2,SNX22,GAS7,HNRNPL,TAF15,AHCYL1,GRINA,NCKAP1,VPS35,PTOV1,ACAP3,ZNF302,T
DA2B,KHDRBS1,PFKL,TKT,EIF5A,CDH2,PDIA6,PIP4K2B,MEGF8,RAB7A,PRDX2,RPL12,ITM2C,RPL
3,HK1,NCS1,ZNF865,ATF4,NR2F1,RAB6B,SLC7A5,ENC1,NLGN2,GLIPR1L2,RPS14,CXADR,CHD4,
XADR,ATP1A1,JUND,RPS20,CD81,WDR1,ATXN7L3B,HMGN2,RPL26,CRMP1,LASP1,BEX3,OAZ1,SF
,ATP5PB,ADAM9,PTBP1,ILF3,RPS11,SUPT5H,HNRNPUL1,UBE2R2,ATXN10,KDM1A,FKBP8,DIDO1
XA2,AGRN,SARAF,MIDN,RPL10,MARCHF6,WSB1,AP3D1,SBF1,DNAJA1,ARF1,PAFAH1B1,ITM2B,AI
RF1,CYB5R3,SF3B2,MAT2A,GTF2F1,PXDN,HNRNPA0,RPS16,RPL23,CNN3,RPL29,EIF4H,LPCAT1,L
RPS4X,RPS3A,RPL18,TUBB4B,UCHL1,RPL11,MORF4L1,BTF3,NCAM1,AGRN,RPLP1,SARAF,MIDN
N1,YEATS2,ACO2,CAMSAP1,KLC1,NCKAP1,MLEC,ACAP3,CBX1,RAD23B,CHD2,SKI,H3-3A,CNTNA
31,PDIA3,SSBP2,CHD3,HMGB1,CDKL1,TPT1,SEC61A1,RPS4X,RPS3A,STMN1,ITGB1,TUBB4B,NCA
ITGB1,TUBB4B,UCHL1,RPL11,MORF4L1,BTF3,NCAM1,AGRN,RPLP1,ZMIZ1,RPL10,WSB1,RPS17
2A2,RPL26,LASP1,OAZ1,SRSF4,RBBP4,PFN1,SPC25,NES,RPL31,PRKCSH,CCNI,TADA2B,KHDRBS
MAP9,TPI1,CHSY1,TMSB10,INA,EIF4A1,RPL19,PPP2R1A,RPL15,GTF3C4,LDHB,DNAH8,HLA-C,SF3
PS10,MTSS2,APLP2,SLC25A3,RPLP2,ARL6IP1,DDOST,WLS,HNRNPK,H1-10,YWHAB,LAMP1,RPS12
PNB1,PAPOLA,GAP43,RPL14,RHOA,CBX5,ZNF865,ATF4,NR2F1,KIF5B,RAB6B,ENC1,NLGN2,ATP1

MMP14,RPL12,ITM2C,RPL30,EIF3L,MRFAP1,MAGED2,H1-0,RPS7,RPS19,CHPF,JUN,NGRN,B4GA
NON,OPTN,ARF3,CCND1,DSTN,BTBD2,PBX2,FAM171A2,PRDX1,AEBP1,NYNRIN,DYNLL2,WDR6,CA
B,RPS12,RAB1A,CALU,COTL1,CDK16,SMCHD1,IPO5,TMEM30A,DNER,LAPTM4B,NNAT,SERF2,CF
POLR2A,CDKL1,AP2M1,TPT1,SEC61A1,RPS4X,RPS3A,RPL18,ITGB1,TUBB4B,RPL11,MORF4L1,BT
TPT1,SEC61A1,RPS4X,RPS3A,RPL18,ITGB1,TUBB4B,UCHL1,RPL11,MORF4L1,BTF3,AGRN,RPLP
1,HMGN2,CRMP1,LASP1,OAZ1,PFN1,ASTN2,MCL1,TLE5,PRKCSH,LRPPRC,TADA2B,XPO1,SLC3A
STMN3,LSS,STXBP1,UBQLN1,TUG1,ID4,RAI1,SREBF2,PFN2,VGF,FBRSL1,P3H3,CYP51A1,FLNC,A
A0,CTNNA1,FADS2,CNN3,TNS3,CTSD,EIF4H,LPCAT1,LAMA5,SLC22A17,BRD3,RPS9,CIPC,MSN,SL
LR2A,AP2M1,TPT1,RPL18,STMN1,ITGB1,TUBB4B,MORF4L1,BTF3,NCAM1,ANXA2,AGRN,ZMIZ1,
,RPL31,PRKCSH,CCNI,TADA2B,KHDRBS1,SPTBN2,PFKL,HDLBP,GLG1,CDH6,SFPQ,EIF5A,PEBP1
AB11B,PYGB,SELENON,OPTN,PPP2CA,MAPK1,MAPRE1,DDAH2,NYNRIN,TCF25,ADAM9,GRB2,WD
ARDBP,SLC25A3,ST3GAL2,MXD4,H1-10,INIP,RPL36,CADM4,SND1,RHOB,PLPPR3,ARCN1,HECTD
1,ZFAND5,FLNB,PITPNA,CORO1C,PHC2,YWHAH,LSS,TNKS2,STXBP1,ISYNA1,MESD,IDH1,HYOU1,
CB1,ZC3H7B,CHD6,RAB27A,U2AF2,EEF1A2,CCT2,TBCD,ANXA5,SNX22,VAMP2,MEST,GAS7,TM9S
PAN3,LPCAT1,LAMA5,BRD3,RPS9,CIPC,MSN,KPNB1,C1ORF43,PAPOLA,RPL14,HK1,RPL22,RHOA
CND2,CDH2,PDIA6,GPC3,PIP4K2B,MEGF8,RAB7A,CCN2,ACTN4,DDX3X,RPL12,EIF3L,MAGED2,G
F4,RBBP4,PFN1,TLE5,NISCH,HNRNPC,TADA2B,XPR1,XPO1,SPTBN2,SLC3A2,CSNK1E,PFKL,RPS2
2A,GTF2F1,PXDN,HNRNPU,SMARCA5,HNRNPA0,RPS16,RPL23,LDHA,CNN3,RPL29,RPSA,TPM4,EI
CP1,KLC1,NCKAP1,MDK,RAD23B,ORAI2,SKI,ASXL1,H3-3A,SCAF1,PYGB,DPYSL2,TMED2,RNF10,V
TPT1,SEC61A1,RPS4X,RPS3A,RPL18,ITGB1,TUBB4B,UCHL1,RPL11,MORF4L1,BTF3,AGRN,RPLP1
3,EIF4H,SLC22A17,SLC16A7,KPNB1,PAPOLA,COL1A2,RPL14,HK1,NCS1,ZNF865,CLSTN1,NR2F1
EIF5B,ENC1,NLGN2,CXADR,CHD4,TMEM259,EIF5,HMGN2,CRMP1,OAZ1,SRSF4,RBBP4,PFN1,MC
RN,RPLP1,SARAF,MIDN,RPL10,MARCHF6,RPS17,VCL,AP3D1,DNAJA1,ARF1,CYB5R3,AKAP12,SF3
O5,TMEM30A,SKP1,DNER,PEA15,SHISAL1,NOVA2,TM9SF3,CHD6,U2AF2,TBCD,ANXA5,CPSF1,PP
36A,TSPAN6,PYGB,RPS15A,VEGFA,ARF3,MVB12B,DSTN,FAM171A2,ODC1,SRSF1,PTBP1,ILF3,LU
XT,EIF5A,CDH2,PDIA6,PIP4K2B,MEGF8,RAB7A,PRDX2,RPL12,ITM2C,RPL30,CLTC,EIF3L,MRFAP1
XT,SFPQ,EIF5A,CDH2,PDIA6,PIP4K2B,MEGF8,RAB7A,MMP14,RPL12,ITM2C,RPL30,EIF3L,GNB2,
N,RPS10,DNAJC5,RPLP2,DDOST,H1-10,NARS1,DUX4,YWHAB,SND1,HNRNPH1,PLPPR3,ARCN1,SE
SERINC1,SSBP2,ZNF43,CHD3,BASP1,AP2M1,RPS3A,RPL18,STMN1,ITGB1,TUBB4B,UCHL1,MOR
3B2,MAT2A,GTF2F1,SLC4A8,HNRNPA0,CTNNA1,RPS16,RPL23,LDHA,CNN3,RPL29,TPM4,EIF4H,N
CLIP3,MAP2K2,ARF4,FLNB,POLR2B,SDF4,SMARCC1,ATP6V1B2,HNRNPM,SDHA,YWHAH,STT3B,T
A1,STMN1,ITGB1,TUBB4B,UCHL1,MORF4L1,ANXA2,AGRN,TIMP2,XAF1,SARAF,MIDN,RPL10,MAR
MLF2,ZSCAN18,WBP2,CTXN1,CD99L2,PCYOX1,CIC,TRAF4,RBBP7,KIF5A,BCL9L,WDR45B,PRMT
NS3,SCARB2,TSPAN3,LPCAT1,BRD3,RPS9,CIPC,RPL24,SLC16A7,KPNB1,C1ORF43,PAPOLA,IGFB
3,VAPA,HMGB1,POLR2A,CDKL1,AP2M1,TPT1,SEC61A1,RPS3A,RPL18,ITGB1,TUBB4B,UCHL1,RP
,AP3D1,DNAJA1,ARF1,CYB5R3,AKAP12,SF3B2,MAT2A,GTF2F1,HNRNPA0,RPS16,RPL23,RPL29,T
NAJA1,ARF1,PAFAH1B1,ITM2B,AP2B1,MAT2A,GTF2F1,OGA,SMARCA5,HNRNPA0,CTNNA1,FADS2,
K5,GRINA,VPS35,MLEC,ACAP3,MDK,PTK7,ORAI2,ASXL1,H3-3A,PRPF6,OPTN,ARF3,CCND1,SMAR
PL27,PTPRA,PRDX5,ACTR3,UBE4A,ACO2,HSPA4,TCP1,CAMSAP1,NCKAP1,MLEC,TMEM248,PTK7
,U2AF2,EEF1A2,CCT2,TBCD,SNX22,VAMP2,MEST,GAS7,TM9SF2,MORF4L2,PDXK,PKN1,HSPA4,G
PDIA3,SSBP2,ZNF43,CHD3,VAPA,HMGB1,CDKL1,TPT1,SEC61A1,RPS4X,RPS3A,STMN1,ITGB1,TU
ED4B,CTBP1,ARHGDIA,STIP1,SLC35E2B,HDAC2,LAPTM4B,TRIB2,DLG5,HS6ST1,TMEM132A,NO
L,AP3D1,ARF1,CYB5R3,MAP4,ITM2B,ZNF770,MAT2A,GTF2F1,PXDN,HNRNPU,SMARCA5,LARP1,H
F43,PAPOLA,NCS1,RHOA,RPL9,RAB6B,CXADR,ATP1A1,JUND,TMEM259,LENG8,WDR1,ATP2A2,RF
SHC2,INTS3,RAB1B,SH3GL1,ZNF428,TTC37,PNISR,SRSF2,PLEKHB2,PCNX3,ADAM10,ATP6V0D1,

HNRNPU,OGA,PGAM1,NFE2L1,FADS2,RPL23,CNN3,RPL29,TPM4,CTSD,LPCAT1,LAMA5,RPS24,S[illegible]
RIN1,STIP1,TOP2B,SLC35E2B,IPO5,ATP6V0C,TMEM30A,LAPTM4B,TRIB2,HSBP1,NNAT,MAP1A,FA[illegible]
B,AEBP1,TCF25,ATP5PB,PTBP1,DYNLL2,ILF3,PCSK1N,WDR6,HNRNPUL1,UBE2R2,HSPB1,SON,C[illegible]
G7,ACLY,SEPTIN5,VAPB,SSBP3,CARM1,ARHGEF17,ELAVL1,VPS4A,NUDCD3,GNA12,MAZ,LSM14A,
F,MIDN,RPL10,MARCHF6,RPS17,AP3D1,DNAJA1,ARF1,CYB5R3,SF3B2,MAT2A,GTF2F1,OGA,SM[illegible]
PS24,RPS9,XRCC5,CIPC,RPL24,PAPOLA,COL1A2,HK1,RPL22,RHOA,ZNF865,RPL9,ATF4,PSMD2,[illegible]
MORF4L1,BTF3,ANXA2,AGRN,TIMP2,XAF1,MIDN,RPL10,AP3D1,SBF1,DNAJA1,ARF1,CYB5R3,AKA
SL1,UBB,AFF4,MATR3,YWHAG,CKB,COL4A2,RPL37A,LRRC28,FN1,SPEN,DDB1,PSMD5,YWHAQ,E[illegible]
P1,NOL4L,GTPBP1,FLOT1,KPNA2,PPP2CB,GNG4,EP400,MASTL,KCTD12,ZFR,ZNF512B,PTPRF,A[illegible]
AG,CKB,COL4A2,RPL37A,LRRC28,FN1,SPEN,DDB1,PSMD5,YWHAQ,EIF1,HACD3,SET,EGLN1,MAP
RPL38,NOTCH2,RNF10,SMARCA4,AZIN1,SRSF9,CCNL2,PSMD3,RABL6,SRSF1,ARGLU1,TCF25,A[illegible]
A0,RPS16,RPL23,RPL29,TPM4,EIF4H,LAMA5,RPS24,RPS9,XRCC5,CIPC,MSN,RPL24,SLC16A7,PA
9,EIF4H,LAMA5,RPS9,XRCC5,CIPC,RPL24,SLC16A7,PAPOLA,HK1,NCS1,RPL22,RHOA,PTCD3,ZN[illegible]
A2,RPL37A,FN1,SPEN,NONO,DDB1,PSMD5,YWHAQ,EIF1,HACD3,DCHS1,SET,MAP9,TPI1,TMSB10,[illegible]
I,MCL1,NISCH,HNRNPC,LRPPRC,XPO1,SLC3A2,PTN,RPS25,TKT,EIF5A,PEBP1,CCND2,CDH2,PDIA
GN2,RPL26,CRMP1,LASP1,BEX3,OAZ1,SRSF4,RBBP4,MCL1,JPT2,TLE5,PNN,RPL31,HNRNPC,TAD
E2G1,ELOB,CNOT3,UQCRH,MAN1B1,TMEM59,BRPF3,ABCE1,CCNL1,CDC42BPA,NUDT3,FAM19[illegible]
MS,CTXN1,YME1L1,KIF5A,MSMO1,KCNQ2,EDC4,TXLNA,TRIO,LBH,FBRS,HIP1,VARS1,NOL4L,H2[illegible]
IM14,IGF2,ATP5F1A,EIF4B,VCP,MARCKSL1,UBB,AFF4,TRIM28,MATR3,YWHAG,CKB,COL4A2,RF[illegible]
OAZ1,SRSF4,RBBP4,MCL1,JPT2,TLE5,PNN,RPL31,PRKCSH,HNRNPC,XPR1,SLC3A2,PTN,PFKL,HD
UBB,AFF4,TRIM28,MATR3,YWHAG,CKB,RPL37A,LRRC28,FN1,SPEN,NONO,DDB1,YWHAQ,EIF1,[illegible]
OL4A2,RPL37A,LRRC28,FN1,SPEN,DDB1,PSMD5,YWHAQ,EIF1,HACD3,SET,EGLN1,MAP9,TPI1,C[illegible]
B4B,RPL11,MORF4L1,AGRN,ZMIZ1,TIMP2,SARAF,COL11A1,MARCHF6,RPS17,VCL,AP3D1,SBF1
VAPA,HMGB1,POLR2A,CDKL1,AP2M1,RPS4X,RPS3A,STMN1,TUBB4B,UCHL1,RPL11,MORF4L1,[illegible]
,PDIA6,MEGF8,SOX12,RAB7A,DDX3X,RPS15,EIF3L,H1-0,RPS5,RPS7,IK,RPS19,MIF,MYO5A,GOL[illegible]
F1,SLC4A8,SMARCA5,HNRNPA0,RPS16,RPL23,CNN3,RPL29,TPM4,EIF4H,LAMA5,RPS24,BRD3,R
A1,JUND,COL5A2,LENG8,WDR1,ATXN7L3B,HMGN2,ATP2A2,CRMP1,VAT1,SRSF4,RTN3,ASTN2,T
TP6V1B2,HNRNPM,SDHA,NKTR,YWHAH,STT3B,TNKS2,LONP1,UBQLN1,NBPF9,SARS1,SREBF2,P[illegible]
BTBD2,PBX2,FAM171A2,AEBP1,NYNRIN,DYNLL2,ILF3,WDR6,UBE2R2,SCAF11,ATXN10,DUSP4,P[illegible]
,CTNNB1,ANXA6,NGRN,B4GALNT4,MTSS2,RPL35A,TARDBP,RPLP2,NDFIP1,DDOST,HNRNPK,MY[illegible]
CSDE1,SMIM14,ATP5F1A,EIF4B,VCP,MARCKSL1,UBB,AFF4,MATR3,YWHAG,CKB,COL4A2,RPL37[illegible]
PSMB2,H3-3A,DPYSL2,TAF7,RPS15A,VEGFA,RAPGEF1,OPTN,ARF3,UBXN4,ATG2B,SRSF9,PSMD3,
2F1,HNRNPA0,RPS16,RPL23,CNN3,RPL29,TPM4,EIF4H,LAMA5,RPS9,XRCC5,CIPC,RPL24,SLC16[illegible]
A8A,NFE2L1,SMARCA5,LDHA,CPE,RPL29,TPM4,TNS3,CTSD,LAMA5,XRCC5,CIPC,PJA2,MSN,SLC1
HACD3,SET,EGLN1,MAP9,TPI1,CHSY1,TMSB10,INA,EIF4A1,RPL19,RPL15,GTF3C4,LDHB,DNAH8,
LR2A,CDKL1,TPT1,SEC61A1,RPS4X,RPS3A,OR8K1,RPL18,STMN1,ITGB1,TUBB4B,UCHL1,BTF3,[illegible]
PF19,NOC2L,GSK3A,SF3A2,ATP5MC3,PRKACA,NASP,AP2A2,CERS2,DPYSL4,DMWD,METRN,RCN2
SPEN,NONO,DDB1,YWHAQ,EIF1,DCHS1,EGLN1,CHSY1,TMSB10,INA,RPL19,RPL15,GTF3C4,LDH[illegible]
1,CYB5R3,AKAP12,SF3B2,GTF2F1,SMARCA5,HNRNPA0,RPS16,RPL23,CNN3,RPL29,TPM4,EIF4H,
N1,RPS27,MAGED4B,CTBP1,ARHGDIA,CAPRIN1,STIP1,TOP2B,SLC35E2B,IPO5,BCLAF1,RPL23A,[illegible]
,ZFP36L1,CSNK1G2,ATRX,GNG4,EP400,ZNF512B,CUL3,ADH5,ANP32B,BRSK1,ZMYM4,PEG10,D
K4,YWHAB,PTGES3,SND1,ATP1A2,RPS12,CALU,RPS27,CCT6A,MAGED4B,CTBP1,RAC1,ARHGDIA,[illegible]
GTF3C4,LDHB,PKD1,HLA-C,SF3B1,PDIA3,SERINC1,ZNF43,RPS3,CHD3,VAPA,HMGB1,POLR2A,CD
,SSBP2,ZNF43,VAPA,HMGB1,POLR2A,AP2M1,TPT1,SEC61A1,RPS4X,RPS3A,ITGB1,TUBB4B,UCH[illegible]
IRNPA0,RPS16,RPL23,CNN3,RPL29,TPM4,EIF4H,LAMA5,RPS24,BRD3,RPS9,XRCC5,CIPC,RPL24,[illegible]

CYB5R3,SF3B2,CDC42BPB,MAT2A,TMEM170A,GTF2F1,PXDN,PGAM1,SMARCA5,LARP1,HNRNPA(
SCAF11,RPL28,ZFAND5,HNRNPUL2-BSCL2,KIF3C,DHX30,YWHAH,PTP4A2,FAU,STMN3,PFKM,LSS
JBFD1,HDAC6,ZNF512B,PRRC2A,HNRNPA2B1,PPP1CC,PTPRF,NPDC1,IL6ST,BRSK1,GPSM1,LMA
ACA,YY1,XPO6,TEAD1,METRN,ARF5,ELAVL3,DIP2A,AMFR,POLDIP2,UBE2L3,CDC34,CELSR1,HMG
PT1,SEC61A1,RPS4X,RPS3A,STMN1,ITGB1,TUBB4B,MORF4L1,ANXA2,AGRN,RPLP1,ZMIZ1,TIMP2
SHC2,STRN4,ABI2,UBR5,MAZ,PRR12,TMEM63B,ATXN3,RAB5C,QPRT,TCF3,ATP1B2,RNF145,REI
MD5,YWHAQ,EIF1,DCHS1,SET,EGLN1,TPI1,TMSB10,INA,EIF4A1,MRC2,RPL19,PPP2R1A,RPL15,G
,XPO1,SPTBN2,SLC3A2,PFKL,HDLBP,TKT,SFPQ,PEBP1,PDIA6,GPC3,PIP4K2B,ACSL6,SOX12,CCN
INRNPH1,SEPTIN3,COTL1,CTBP1,RAC1,STIP1,GPC1,TOP2B,SLC35E2B,APPL1,LAPTM4B,TRIB2,H
XIP,PPP1CB,PGD,MEST,GAS7,TSPYL4,MORF4L2,SSRP1,HNRNPA3,PTPRA,PDXK,AHCYL1,PTOV1,I
IOL4L,ZFP36L1,CSNK1G2,GNG4,EP400,UBFD1,KCTD12,ZNF512B,PPP1CC,ULK1,BRSK1,ZMYM4
PU,PGAM1,NFE2L1,FADS2,RPL23,CNN3,RPL29,TPM4,LPCAT1,LAMA5,RPS24,SLC22A17,XRCC5,C
WHAH,ATXN2L,MPRIP,BAG6,PNMA8A,FBRSL1,ARNT2,IGFBP7,TXNIP,RBM15B,CTNND1,TRAF7,BC
PTAN1,GSTP1,MARCKS,SMG1,CALM1,NPM1,PLXNB2,TTC3,PLD3,HSPD1,COL18A1,SNRNP200,CS
J,RPLP1,SARAF,MIDN,RPL10,MARCHF6,RPS17,AP3D1,DNAJA1,ARF1,CYB5R3,AKAP12,SF3B2,MA
CTNNA1,LDHA,CNN3,RPL29,TPM4,EIF4H,LAMA5,RPS24,RPS9,XRCC5,SLC16A7,KPNB1,PAPOLA,I
LASP1,OAZ1,SRSF4,JPT2,TLE5,PRKCSH,LRPPRC,TADA2B,KHDRBS1,PFKL,HDLBP,TKT,GLG1,TPM
DH2,U2AF2,MLXIP,MEST,GDI2,HSPH1,LMNB2,HNRNPA3,ACTR3,CLSTN3,LAMC1,ASXL1,CNTNAP2
TL3,STARD7,PDAP1,HIP1,PHB2,APMAP,KLHL42,PLXNB1,CSNK1G2,FARP1,EP400,UPF1,RER1,YK
R2,NARS1,DUX4,YWHAB,PTGES3,SND1,RPS12,CALU,PLPPR3,ARCN1,RPS27,MAGED4B,CTBP1,R
RC28,FN1,SPEN,DDB1,PSMD5,YWHAQ,EIF1,DCHS1,SET,EGLN1,TPI1,TMSB10,INA,EIF4A1,MRC2,F
PL31,PRKCSH,LRPPRC,CSNK1E,PFKL,SCRN1,RPS25,EIF5A,PDIA6,DDR1,PIP4K2B,MEGF8,RAB7A,
PL23,CPE,CNN3,RPL29,RPSA,TNS3,NACA,TSPAN3,LPCAT1,LAMA5,RPS24,SLC22A17,BRD3,PJA2,I
PL23,CPE,CNN3,RPL29,RPSA,TNS3,NACA,TSPAN3,LPCAT1,LAMA5,RPS24,SLC22A17,BRD3,PJA2,I
GN2,CRMP1,BEX3,VAT1,JPT2,TLE5,XPR1,KHDRBS1,KLHL9,SPTBN2,SLC3A2,CSNK1E,PFKL,HDLE
FN1,ASTN2,MCL1,TLE5,PNN,RPL31,PRKCSH,HNRNPC,LRPPRC,TADA2B,XPR1,KHDRBS1,CSNK1E
.9,XPO1,PFKL,HDLBP,SFPQ,EIF5A,PDIA6,GPC3,PLXNA1,PIP4K2B,MEGF8,ACSL6,MMP14,PRDX2,I
,SF3B2,GTF2F1,SLC4A8,SMARCA5,HNRNPA0,FADS2,RPS16,RPL23,CNN3,RPL29,TPM4,EIF4H,LA
TBD2,SRSF1,TCF25,CAPZB,RPS11,TDRD5,CAND1,HSPB1,PLCG1,CCAR2,ZFAND5,DIDO1,KIF3C,S
NT4,APLP2,HNRNPDL,ARL6IP1,WLS,CIRBP,MYH10,NARS1,YWHAB,CKAP5,SEPTIN11,CADM4,RHC
LKBH5,HNRNPA3,PTPRA,TBC1D16,CAMSAP1,MDK,RBMX,DDX42,PTK7,RAB5B,GCN1,TNPO1,SKI
G6,SREBF2,HNRNPD,MEX3A,SERBP1,PRPF40A,SMS,AMOTL2,RBBP7,TRAF7,MSMO1,EIF2AK1,PR
CT7,VAT1,OAZ1,SRSF4,PFN1,ASTN2,RPL31,NISCH,CCNI,TADA2B,XPR1,KHDRBS1,XPO1,SLC3A2,F
PL22,RHOA,ZNF865,ATF4,PSMD2,KIF5B,GLIPR1L2,RPS14,CXADR,CHD4,RPS20,CD81,LENG8,WD
G3,GOLM1,SESN3,DNAJC5,APLP2,HNRNPDL,ARL6IP1,TSR2,TAGLN,MYL6,HNRNPK,H1-10,RPL36
2,PDE7A,FAM171A2,RABL6,ODC1,FAM168B,SRSF1,IPO7,ARGLU1,TCF25,ADAM9,ILF3,LUZP6,PR
I1,HSPB1,QARS1,RPL28,CCAR2,GNG2,SDF4,SMARCC1,SUPT6H,DHX30,EIF3F,B2M,PITPNM1,TNK
11,HNRNPA0,RPS16,RPL23,RPL29,TPM4,EIF4H,LAMA5,RPS24,RPS9,XRCC5,CIPC,RPL24,SLC16A
2,GNAI3,PARP1,TSR1,MTRF1L,ALKBH5,RPL27,PTPRA,PRDX5,UBE4A,ACO2,HSPA4,TCP1,MLEC,T
,RPS19,TLN1,CHPF,JUN,MYO5A,NGRN,B4GALNT4,DNAJC5,HNRNPDL,RPLP2,DDOST,TSR2,NARS
S1,DUX4,YWHAB,PTGES3,LAMP1,RPS12,CALU,RPS27,MAGED4B,CTBP1,ARHGDIA,CAPRIN1,SLC3
LEC,FAM171A2,TCF25,LUZP6,CAPZB,RPS11,UBE2R2,ACTN1,BNIP3L,PLCG1,ZFAND5,DIDO1,POLI
LEC,FAM171A2,TCF25,LUZP6,CAPZB,RPS11,UBE2R2,ACTN1,BNIP3L,PLCG1,ZFAND5,DIDO1,POLI
2F1,HNRNPA0,RPS16,RPL23,CNN3,RPL29,TPM4,RPS24,RPS9,XRCC5,CIPC,RPL24,SLC16A7,PAP(
MIZ1,TIMP2,SARAF,MIDN,RPL10,VCL,AP3D1,SBF1,DNAJA1,ARF1,PAFAH1B1,AKAP12,MAP4,SF3E

DHX15,VDAC1,RBBP7,TRAF7,SULF2,DFFB,PRMT1,BCR,TXNDC5,MTA1,TRIO,EP300,DGCR2,SF1,F
CAPRIN1,GPC1,CDK16,IPO5,ATP6V0C,SKP1,L1CAM,PPIB,SQLE,SHISAL1,CTDSP2,NOVA2,CTSB,R
3D1,SBF1,DNAJA1,ARF1,AKAP12,SF3B2,CDC42BPB,ITM2B,MAT2A,TMEM170A,PXDN,HNRNPU,O
1,TCEAL4,CAD,MZT2B,SPPL2B,PINK1,ITFG1,USP10,KDM3A,RBM4,CLIC1,PFKP,CUL1,PPP1R14B,
DSP2,DDX39B,SRSF5,TM9SF3,MDH2,TOMM20,U2AF2,CCT2,RPN2,SUGP2,GNAI3,MEST,GDI2,TM
P1,RAB7A,RPL12,CLTC,MRFAP1,GNB2,RPS5,IK,CHPF,JUN,B4GALNT4,MTSS2,SLC25A3,RPLP2,AF
,ANXA6,RPS10,B4GALNT4,TAGLN,ANKRD52,WLS,CIRBP,MYH10,DUX4,YWHAB,PTGES3,UBE2H,A
,RHOA,PTCD3,ZNF865,RPL9,ATF4,GLIPR1L2,RPS14,CXADR,CHD4,TMEM259,RPS20,CD81,LENG
JN,NGRN,HNRNPDL,RPLP2,DDOST,INIP,CIRBP,NARS1,RPL36,YWHAB,SND1,LAMP1,CALU,ARCN1
OLA,HK1,NCS1,RPL22,RHOA,ZNF865,ATF4,KIF5B,GLIPR1L2,RPS14,CXADR,CHD4,RPS20,CD81,LE
PINT2,NPLOC4,CHERP,FKBP1A,SYMPK,SAFB2,SERP1,ZNF664,VPS26B,BAP1,FUS,PPP1R10,XRC
PEBP1,CCND2,PDIA6,PIP4K2B,SOX12,ACTN4,PRDX2,ARL4C,EIF3L,GNB2,RCN1,APLP1,RPS19,MI
,UBE4A,ACO2,HSPA4,TCP1,CAMSAP1,ZNF302,DDX42,GCN1,SKI,RAB11B,SRP14,PRPF6,TAF7,MA
,GJA1,JUN,MYO5A,COL6A2,B4GALNT4,MTSS2,DNAJC5,APLP2,RPLP2,ARL6IP1,RPL36,DUX4,SEP
T1,SEC61A1,STMN1,ITGB1,TUBB4B,UCHL1,RPL11,MORF4L1,BTF3,NCAM1,ANXA2,AGRN,RPLP1
X42,GCN1,SKI,ASXL1,SRP14,ABCA2,PYGB,PRPF6,TERF2IP,TMED2,TAF7,RPL37,SMARCA4,ATG2E
X3,OAZ1,RBBP4,PFN1,ASTN2,JPT2,TLE5,PNN,NES,RPL31,NISCH,PRKCSH,HNRNPC,KHDRBS1,SP
POLR2A,AP2M1,TPT1,SEC61A1,RPS4X,RPL18,STMN1,ITGB1,TUBB4B,RPL11,MORF4L1,NCAM1,
0,FADS2,RPL23,LDHA,CNN3,RPL29,RPSA,TNS3,CTSD,EIF4H,TSPAN3,RTN4,LPCAT1,LAMA5,RPS2
LPCAT1,LAMA5,SLC22A17,XRCC5,SLC16A7,KPNB1,LLGL1,HK1,NCS1,RPL22,CBX5,ZNF865,CLS
RPLP1,SARAF,MIDN,RPL10,MARCHF6,RPS17,AP3D1,DNAJA1,ARF1,CYB5R3,AKAP12,SF3B2,MAT
CT7,SRSF4,RBBP4,PFN1,ASTN2,MCL1,TLE5,PNN,RPL31,PRKCSH,HNRNPC,XPR1,KHDRBS1,SPTE
,EEF1A2,FDFT1,CCT2,SUGP2,GNAI3,PARP1,TSR1,MTRF1L,ALKBH5,RPL27,PTPRA,PRDX5,UBE4A
,MAGED4B,CTBP1,ARHGDIA,STIP1,SLC35E2B,IPO5,BCLAF1,RPL23A,HSBP1,MAP1A,SERF2,CTD
27,CCT6A,CTBP1,HMGCR,RAC1,SMCHD1,HDAC2,ATP6V0C,BCLAF1,LAPTM4B,TRIB2,CDKN1A,H
,NLGN2,RPS14,CHD4,ATP1A1,TMEM259,CD81,LENG8,WDR1,HMGN2,LASP1,BEX3,CCT7,VAT1,O
27,PTPRA,PRDX5,UBE4A,ACO2,HSPA4,TCP1,TMEM248,DDX42,GCN1,SKI,ASXL1,RAB11B,SRP14,
DHB,SF3B1,BASP1,VAPA,HMGB1,POLR2A,CDKL1,AP2M1,TPT1,RPS3A,STMN1,ITGB1,TUBB4B,UC
24,SLC22A17,BRD3,RPS9,CIPC,RPL24,RPL17,C1ORF43,PAPOLA,RPL14,PTCD3,CBX5,CLSTN1,AT
LSR1,HMGA1,USF2,ABL1,RSF1,LGALS1,AUTS2,CGGBP1,RTF1,TOX4,HIP1R,H6PD,RGL2,ACVR1B
ADM4,LAMP1,PLPPR3,HRNR,HECTD4,ARHGDIA,GPC1,TOP2B,ATP6V0C,MMP2,PPIB,MAP1A,CDK
KHDRBS1,SLC3A2,CSNK1E,PFKL,SCRN1,RPS25,TKT,EIF5A,PDIA6,PIP4K2B,MEGF8,RPL12,ITM2
H5,RPL27,PRDX5,C4ORF3,UBE4A,ACO2,HSPA4,TCP1,MLEC,ACAP3,CCT4,SNRNP70,RPL36A,SRF
8IP1,FBRSL1,ZSCAN18,PRPF40A,NAP1L4,CTXN1,CD99L2,PCYOX1,RBM15B,BEX4,PPDPF,CIC,TR
N1,MAP9,TPI1,CHSY1,TMSB10,INA,EIF4A1,MRC2,PPP2R1A,GTF3C4,LDHB,PKD1,HLA-C,PDIA3,S
LPCAT1,LAMA5,SLC22A17,BRD3,RPL24,SLC16A7,PAPOLA,IGFBP3,COL1A2,LLGL1,RPL14,HK1,N
R1,RPL26,LASP1,CCT7,SRSF4,RBBP4,PFN1,ASTN2,MCL1,TLE5,PNN,RPL31,PRKCSH,HNRNPC,LF
I,PBX2,SRSF1,DYNLL2,FTO,LUZP6,GRB2,PCSK1N,CAPZB,HNRNPUL1,UBE2R2,HSPB1,BNIP3L,M
P1,SUPT5H,ATXN10,QARS1,CCAR2,ARF4,POLR2B,SMARCC1,SUPT6H,DHX30,PFKM,ATXN2L,ID4,
,SEC16A,UBA52,SLC39A7,RAB3GAP2,EIF5B,ZFR,AGPAT1,ZMYM4,NPLOC4,FKBP1A,RABEP1,SYN
LEC,PBX2,FAM171A2,ODC1,SRSF1,AEBP1,PTBP1,ILF3,CAPZB,HNRNPUL1,TDRD5,ATXN10,MAP7
1,PRRC2A,NPDC1,ANP32B,PREPL,TIMM50,SEPTIN7,HIF1A,PPP1R15A,FBXO21,SAFB2,FUS,PPP1F
CCT4,SNRNP70,ASXL1,H3-3A,DPYSL2,TOP1,TAF7,RPL38,NOTCH2,RNF10,SMARCA4,AZIN1,PDE7
4,CTBP2,WDR5,SS18L1,SLC25A29,KLHL36,HEG1,MIB2,SP1,BTBD1,DYNLT1,SPIN1,FAM20C,MIC
1-10,RPL36,SND1,RHOB,HNRNPH1,COTL1,CCT6A,ARHGDIA,GPC1,APPL1,SMCHD1,TNC,TMEM3

IDAC2,IPO5,PPIB,HSBP1,DLG5,HS6ST1,TIMP3,TMEM132A,CTDSP2,CNN2,CTSB,TM9SF3,MDH2,F
PL12,RPL30,ARL4C,CLTC,EIF3L,MRFAP1,RCN1,RPS5,GPAT4,GJA1,CD63,SESN3,B4GALNT4,MTS
P9,CHSY1,TMSB10,INA,EIF4A1,MRC2,GTF3C4,PKD1,HLA-C,SF3B1,PDIA3,SERINC1,SSBP2,CHD3,
N,HNRNPU,OGA,NFE2L1,LDHA,CPE,CNN3,RPL29,RPSA,TNS3,EIF4H,NACA,LPCAT1,LAMA5,SLC22
22,RHOA,ZNF865,RPL9,ATF4,KIF5B,GLIPR1L2,RPS14,CXADR,CHD4,JUND,RPS20,CD81,LENG8,V
NB2,PRDX5,RTN1,YEATS2,TCP1,PLOD1,MLEC,ACAP3,SURF4,DDX42,PTK7,ASXL1,ABCA2,DPYSL2
L1CAM,LAPTM4B,PPIB,GLUL,HSBP1,LUC7L3,SERF2,CRPPA,CPSF2,TM9SF3,MDH2,U2AF2,EEF1A
RPS12,CALU,PLPPR3,ARCN1,ZNF454,COTL1,CCT6A,MAGED4B,CTBP1,RAC1,CAPRIN1,STIP1,TO
CHD3,HMGB1,POLR2A,AP2M1,RPS4X,STMN1,TUBB4B,UCHL1,RPL11,MORF4L1,BTF3,NCAM1,AN
,P3H3,PSMA7,ANP32A,TRAF4,EDC4,BCR,KIRREL1,EXTL3,RERE,KDM5B,VARS1,UBQLN2,MMP15,
IAI3,PARP1,VCAN,TSR1,NOX4,MTRF1L,ABCA3,ALKBH5,RPL27,PTPRA,PRDX5,UBE4A,ACO2,HSPA
DC37,HNRNPUL1,SCAF11,ACTN1,CAND1,BNIP3L,DIDO1,DUSP4,PITPNA,SUPT6H,GIGYF1,STMN3
HDRBS1,XPO1,SCRN1,HDLBP,RPS25,SFPQ,EIF5A,CCND2,GPC3,PIP4K2B,RPS15,ITM2C,ARL4C,N
4,NCS1,PTCD3,ZNF865,CLSTN1,PSMD2,NR2F1,RAB6B,SLC7A5,ENC1,ATP1A1,TMEM259,RPS20,
BBP4,PFN1,TLE5,NES,TADA2B,XPR1,KLHL9,SLC3A2,CSNK1E,PFKL,RPS25,CDH6,PDIA6,GPC3,DD
Q,EIF5A,PEBP1,GPC3,PLXNA1,CCN2,DDX3X,ARL4C,EIF3L,MRFAP1,GNB2,RCN1,RPS5,APLP1,IK,R
ZNF512B,CUL3,FMNL2,PTPRF,ULK1,IL6ST,MCM3AP,POM121C,RPS2,LMAN1,ZMYM4,PODXL2,GA
PODXL2,GSE1,MAP1LC3B,MTCH1,TMOD2,SEPTIN5,CLCN7,CEP170,HSPG2,FKBP4,CARM1,INTS3
IZ1,TIMP2,XAF1,COL11A1,MIDN,RPL10,MARCHF6,RPS17,AP3D1,SBF1,DNAJA1,ARF1,CYB5R3,AK
2,LENG8,WDR1,ATXN7L3B,ATP2A2,RPL26,LASP1,CCT7,SRSF4,RBBP4,PFN1,MCL1,RPL31,PRKCS
N1,JPT2,PNN,SPC25,NES,NISCH,PRKCSH,KHDRBS1,SPTBN2,GLG1,EIF5A,PDIA6,GPC3,DDR1,PIP4
TNNA1,FADS2,LDHA,CPE,RPL29,TNS3,CTSD,EIF4H,TSPAN3,LPCAT1,LAMA5,SLC22A17,BRD3,CIP
ORF198,PRDX6,EFR3B,BAP1,TRIR,XRCC6,EHMT1,ERGIC1,RTL6,SLC44A2,HSPG2,SNRPD3,CARI
RDX2,RPL12,ITM2C,RPL30,CLTC,EIF3L,MAGED2,GNB2,RCN1,RPS5,IK,RPS19,CHPF,JUN,MYO5A
NPDL,TARDBP,SLC25A3,TAGLN,WLS,HNRNPK,NARS1,DUX4,SND1,LAMP1,RPS12,RAB1A,SEPTIN3
,IGFBP3,NCS1,CBX5,NR2F1,RAB6B,SLC7A5,NLGN2,GLIPR1L2,CHD4,ATP1A1,JUND,TMEM259,E
NXN,EP400,RER1,PI4KA,CRTAP,ZFR,TBC1D9B,PPP1CC,RPS2,DKK3,MAP1LC3B,MVD,C1ORF198
D2,HSPA4,CAMSAP1,CCT3,RBMX,TMEM248,CHD2,ASXL1,H3-3A,SCAF1,ATP6AP1,TERF2IP,TMED2
S2,HNRNPDL,SLC25A3,ARL6IP1,DDOST,MYL6,HNRNPK,CIRBP,NARS1,PTGES3,CKAP5,HNRNPH1
ONO,PSMD5,YWHAQ,EIF1,DCHS1,TPI1,CHSY1,TMSB10,INA,EIF4A1,MRC2,RPL19,PPP2R1A,RPL15
GALNT4,MXD4,TAGLN,MYL6,CIRBP,RPL36,DUX4,CADM4,RHOB,HNRNPH1,UBE2H,ATP1A2,RPS1
2,LPCAT1,SLC22A17,RPS9,CIPC,PJA2,RPL24,RPL17,C1ORF43,NCS1,RHOA,CBX5,ATF4,NLGN2,G
,RPL22,RHOA,ZNF865,ATF4,PSMD2,GLIPR1L2,RPS14,CXADR,CHD4,RPS20,LENG8,WDR1,RPL26
PL37,ATG2B,CCNL2,ARPC2,FAM168B,ARGLU1,PTBP1,CAPZB,SUPT5H,UBE2R2,ATXN10,SON,RP
,ACO2,HSPA4,TCP1,MLEC,ZNF302,TMEM248,DDX42,ORAI2,GCN1,SKI,ASXL1,RAB11B,TSPAN6,P
1,SRP14,DPYSL2,RPL38,VEGFA,OPTN,CCND1,MAPK1,SMARCA4,ATG2B,SRSF9,BTBD2,PBX2,FAM
,PNN,SPC25,NES,NISCH,PRKCSH,HNRNPC,TADA2B,XPR1,KHDRBS1,KLHL9,XPO1,HDLBP,GLG1,C
,SIN3B,SLC25A6,KIF26A,JAM3,PJA1,ATF5,RAN,IP6K1,DAD1,RAB18,OTUD5,USP48,CRTC1,TMEM
2,MEGF8,SOX12,MMP14,PRDX2,DDX3X,RPL12,RPL30,GNB2,RPS5,RPS19,GPAT4,TLN1,JUN,CD63
OLR2A,AP2M1,TPT1,SEC61A1,RPL18,STMN1,ITGB1,UCHL1,RPL11,MORF4L1,NCAM1,ANXA2,AG
LASP1,OAZ1,ASTN2,MCL1,NES,RPL31,NISCH,LRPPRC,TADA2B,KHDRBS1,PFKL,TKT,GLG1,CDH6
RBS1,KLHL9,SCRN1,HDLBP,RPS25,TKT,SFPQ,EIF5A,CDH2,PDIA6,DDR1,PIP4K2B,MEGF8,ACSL6,C
NB2,MTRF1L,ALKBH5,RPL27,SNRPN,PRDX5,UBE4A,ACO2,HSPA4,TCP1,AHCYL1,MLEC,ACAP3,RA
RPS2,ZMYM4,PODXL2,GALNT11,GSE1,MAP1LC3B,DAP,MTCH1,TMOD2,ZNF664,SEPTIN5,CLCN7,
,NARS1,YWHAB,PTGES3,SND1,RPS12,CALU,ARCN1,RPS27,CTBP1,ARHGDIA,STIP1,SLC35E2B,IPC

DAM9,FTO,ILF3,LUZP6,PRKDC,GABARAP,SCAF11,CAND1,SON,PLCG1,ZFAND5,DIDO1,DUSP4,DH
L23,RPL29,RPS24,RPS9,XRCC5,CIPC,MSN,RPL24,PAPOLA,IGFBP3,HK1,RPL22,RHOA,ZNF865,RI
14,CCNG1,GID8,MARCHF7,ZCCHC14,DEGS1,EXOSC10,UBL3,SYNGR1,RNF216,ZFP91,EIF3M,TB
,RPS25,GLG1,EIF5A,CCND2,CDH2,PNMA1,GPC3,DDR1,PIP4K2B,SOX12,ACTN4,RPL12,ITM2C,MI
2,FAM168B,SRSF1,ARGLU1,PTBP1,CDC37,CAPZB,SUPT5H,UBE2R2,ATXN10,QARS1,CCAR2,ARF
SON,QARS1,CCAR2,ARF4,POLR2B,SMARCC1,SUPT6H,DHX30,BAG6,FBRSL1,RAB11A,ARNT2,SEF
17,VCL,AP3D1,ARF1,CYB5R3,AKAP12,SF3B2,CDC42BPB,GTF2F1,GOLGA8A,PXDN,OGA,NFE2L1,
D3,ADAM9,PTBP1,DYNLL2,ILF3,PCSK1N,HSPB1,MAP7D1,FKBP8,KDELR1,QARS1,DLK1,PLTP,MSI
0,SON,RPL28,CCAR2,ARF4,POLR2B,GNG2,SMARCC1,SUPT6H,DHX30,FBLN1,PITPNM1,BAG6,R
-0,RPS5,APLP1,RPS10,SESN3,MTSS2,DNAJC5,APLP2,RPLP2,TSR2,SHOX,MYH10,DUX4,CADM4,L
R1A,RPL15,HLA-C,SF3B1,SSBP2,ZNF43,RPS3,CHD3,BASP1,POLR2A,CDKL1,SEC61A1,RPS3A,UC
STRN4,ELAVL1,GNL1,MAZ,STK24,PRR12,TMEM63B,SLC25A23,PACS1,DAZAP2,MSI1,ANKS1A,SC
POT,OS9,TXNDC5,KDM5C,PRKAR2A,STARD7,NDUFV1,SF1,RETREG2,BAZ1B,GTPBP1,SLC39A6,C
RPL19,GTF3C4,LDHB,PKD1,HLA-C,SF3B1,SERINC1,ZNF43,RPS3,BASP1,VAPA,POLR2A,CDKL1,T
A2,RPL24,PAPOLA,COL1A2,HK1,NCS1,RPL22,RHOA,ZNF865,RPL9,ATF4,GLIPR1L2,RPS14,CXADI
HSPH1,TBC1D16,ACO2,TCP1,PLOD1,KLC1,CCT3,GRINA,NCKAP1,PTOV1,MLEC,SURF4,DDX42,R
AH2,SRSF1,ARGLU1,PTBP1,FTO,ILF3,RPS11,ATXN10,CASC3,DIDO1,RPS28,SRSF3,TUG1,MESD,I
PM1,EIF5A,PEBP1,CCND2,PDIA6,GPC3,PLXNA1,PIP4K2B,ACSL6,SOX12,RAB7A,CCN2,DDX3X,RPL
B7A,ACTN4,PRDX2,RPS15,ARL4C,GNB2,RPS7,IK,RPL35,CHPF,COL6A2,CD63,ANXA6,NGRN,GOL
PRKCSH,HNRNPC,CCNI,TADA2B,XPR1,KHDRBS1,KLHL9,SCRN1,HDLBP,RPS25,TKT,SFPQ,EIF5A,C
OGA,PGAM1,HNRNPA0,RPL23,RPL29,RPS24,SLC22A17,XRCC5,CIPC,SLC16A7,KPNB1,IGFBP3,H
,PDIA3,SERINC1,SSBP2,RPS3,BASP1,VAPA,HMGB1,POLR2A,SEC61A1,RPS4X,RPS3A,OR8K1,ITGE
,RPS9,CIPC,PJA2,RPL24,RPL17,C1ORF43,PAPOLA,RPL14,PTCD3,CBX5,ZNF865,CLSTN1,ATF4,P
YKT6,CUL3,MCM3AP,SPINT2,RPS27A,TMEM167A,CHERP,FKBP1A,CTTN,ATP6AP2,DAP,SERP1,RH
NN3,RPL29,TPM4,CTSD,EIF4H,TSPAN3,LPCAT1,LAMA5,RPS24,SLC22A17,BRD3,CIPC,MSN,C1OR
6,PTMS,SPTAN1,GSTP1,MARCKS,SMG1,CALM1,NPM1,PLXNB2,PLD3,HSPD1,USP11,COL18A1,SI
1,RPL36A,KDM6B,RAB11B,SRP14,TSPAN6,PYGB,TOP1,TAF7,RPS15A,MARS1,SELENON,TUBB2A
,IK,RPS19,CHPF,JUN,ANXA6,NGRN,APLP2,HNRNPDL,RPLP2,DDOST,CIRBP,NARS1,RPL36,YWHAE
5A,ACAT2,CDC34,TMEM184B,USF2,SPPL2B,FXR2,PINK1,USP10,KDM3A,ARPC1A,TAOK2,RTF1,N
JL,DLG5,SHISAL1,CTDSP2,DDX39B,SRSF5,RAB27A,U2AF2,EEF1A2,SNX22,VAMP2,GAS7,ALKBH5,
AB2A,RER1,UBFD1,IARS1,MASTL,COX7C,YKT6,RGMB,MAPRE2,RPL7L1,RPS27A,TMEM167A,CHE
DL1,FBXL19,HS2ST1,SLC25A36,OLFM2,FAM210B,ABL1,RSF1,PINK1,TOX4,NIBAN2,KIFC2,HIP1R,
RIN1,SMS,CCT5,CDC42,PPP1R9B,YME1L1,TRAF7,EIF2AK1,TXLNA,PRMT1,XPOT,EXTL3,STARD7,F
T11,FKBP1A,CTTN,SYMPK,SAFB2,SERP1,ZNF664,BAP1,FUS,KIAA0100,IARS2,CARM1,ABI2,NUDC
IP4K2B,ACSL6,SOX12,RAB7A,CCN2,DDX3X,RPL12,ITM2C,RPL30,GNB2,RCN1,RPL35,TLN1,MYO

,NAP1L1,TXNDC5,LBH,CSNK2A1,KPNA2,GSPT1,PPM1G,BRD2,KLHL42,HOXB5,GARS1,GNG4,UPI
RPL29,RPSA,TPM4,CTSD,LAMA5,SLC22A17,PJA2,RPL24,SLC16A7,KPNB1,PAPOLA,IGFBP3,COL1
H,CCAR2,ARF4,CORO1C,XPNPEP3,YWHAH,PTP4A2,PITPNM1,TNKS2,ISYNA1,STT3A,LONP1,CSNK
C1,RBBP7,EDC4,NOL4L,H2AZ1,SLC39A6,KIF21B,KLHL42,CSNK1G2,NXN,SRSF6,ARID1B,RER1,E
NMA1,GPC3,PIP4K2B,MEGF8,ACSL6,RAB7A,ACTN4,PRDX2,RPL12,MAGED2,RPS5,APLP1,RPS7,IK
2,MCL1,TLE5,PNN,RPL31,PRKCSH,CCNI,XPR1,KHDRBS1,KLHL9,SLC3A2,CSNK1E,PFKL,SCRN1,H
F1,RGMB,SELENOW,ECEL1,TBC1D9B,HNRNPA2B1,APEX1,HNRNPAB,GLO1,FUS,PSMB4,PELP1,V
PCA,THOC2,CACYBP,MED12,TLK1,CHGB,VKORC1L1,ELOB,UBN1,STRIP1,MED16,BTF3L4,CNOT
2,HNRNPD,VGF,FBRSL1,P3H3,CYP51A1,ARNT2,WBP2,AKIRIN1,YME1L1,CTNND1,DHX15,PSMA7,

2,MTRF1L,ALKBH5,SNRPN,PRDX5,UBE4A,ACO2,HSPA4,TCP1,AHCYL1,MLEC,ACAP3,TNPO1,SKI,
,SESN3,B4GALNT4,H1-10,INIP,CIRBP,MYH10,DUX4,SEPTIN11,LAMP1,UBE2H,ATP1A2,CKAP4,RA
,MAPRE1,SRSF9,CCNL2,DSTN,BTBD2,NYNRIN,ADAM9,DYNLL2,HNRNPUL1,KDM1A,FKBP8,PLCG
LN2,NACC1,THOC2,ZNRF1,SRRT,ARHGAP33,AFDN,UBE2G1,CAMK2N1,RFX7,NIPA1,PPP3CA,SLC
B,IGF2BP1,NSD2,AKT2,TEX2,FBXW5,MBD3,UBE2I,MICAL1,DPYSL4,UNC119B,EIF4G1,TMCO3,TA
JBE2R2,SCAF11,ATXN10,MAP7D1,KDELR1,MAP2K2,ZFAND5,HNRNPUL2-BSCL2,PITPNA,XPNPEP
,SLC4A8,PXDN,OGA,PGAM1,HNRNPA0,CTNNA1,RPS16,LDHA,CNN3,RPL29,CTSD,EIF4H,NACA,RT
GFR1,EIF4A3,CENPB,ADGRA3,SNX2,SLC25A36,CELSR1,HTATSF1,SGSM2,USP10,HDGFL3,CLIP2,
LA,GAP43,RPL14,RHOA,PTCD3,CBX5,ZNF865,RPL9,CLSTN1,ATF4,KIF5B,NLGN2,CHD4,ATP1A1,
7,HNRNPUL1,SCAF11,MAP2K2,DUSP4,MSL1,ATP6V1B2,HNRNPM,GIGYF1,FAU,ILF2,LSS,PITPNM
D,BAG6,FBRSL1,RAB11A,SERBP1,ARFGEF2,WBP2,AKIRIN1,RPL34,CCT5,CDC42,YME1L1,VDAC
P1,ATXN2L,HNRNPD,RAB11A,TCTN1,RTL8C,CTXN1,PPP1R9B,RBM15B,MSMO1,CAPN2,KDM5C,
N3,FBLN1,STT3B,PITPNM1,STXBP1,ID4,ZFAND3,VANGL2,BAG6,MEX3A,SEC23A,ARNT2,RTL8C,
24,TUBGCP6,RIC8A,NCSTN,LSM14B,INTS11,QPRT,MOAP1,MTMR4,YIPF3,SPTY2D1,CAP1,ATP6V
ND1,LAMP1,UBE2H,SEPTIN3,ZNF454,ARHGDIA,STIP1,MMP2,PPIB,PTTG1IP,SERF2,CTDSP2,SRP9
2F1,GOLGA8A,SMARCA5,HNRNPA0,FADS2,RPS16,LDHA,CPE,CNN3,RPSA,CTSD,SCARB2,RTN4,L
,TAX1BP1,TUBGCP6,NCSTN,INTS11,AHSA1,RAB5C,SPAG9,MTMR4,PDPK1,MKRN1,SPTY2D1,CA
5A2,HMGN2,ATP2A2,BEX3,CCT7,OAZ1,RBBP4,MCL1,JPT2,PNN,NISCH,PRKCSH,HNRNPC,TADA2
6A2,NGRN,MTSS2,DNAJC5,TARDBP,RPLP2,NDFIP1,DDOST,TSR2,ANKRD52,CIRBP,MYH10,YWH
RPL37,ARF3,MAPRE1,ARPC2,FAM168B,PTBP1,FTO,LUZP6,CDC37,CAPZB,SUPT5H,SON,CCAR2,
STN1,NR2F1,KIF5B,SLC7A5,NLGN2,GLIPR1L2,CHD4,ATP1A1,JUND,TMEM259,EIF5,COL5A2,LE
9,EIF5,ATXN7L3B,HMGN2,ATP2A2,BEX3,CCT7,OAZ1,RBBP4,PNN,NISCH,PRKCSH,HNRNPC,TADA
NPA0,CTNNA1,RPS16,RPL23,CPE,CNN3,TNS3,CTSD,EIF4H,SCARB2,NACA,LPCAT1,SLC22A17,BR
IP,TTC28,TOMM70,ANKRD12,SSBP4,GNA13,DPYSL4,SLC38A2,CPXM1,SCD,SCAP,DIP2A,MZT2B,
IO4,PTPN11,WDTC1,USF2,FXR2,AUTS2,ZZEF1,ARPC1A,CLIP2,SLC22A23,H6PD,CDV3,OBSL1,SG
B,MEGF8,RAB7A,RPL12,ITM2C,RPL30,EIF3L,MAGED2,GNB2,H1-0,RPS5,IK,RPS19,MIF,TLN1,CH
G1,UNC119B,SH3PXD2B,IGLON5,LMO4,CELSR1,FAM210B,USF2,COL16A1,AUTS2,CLIP2,LUC7L
EBP1,CCND2,DDR1,PLXNA1,PIP4K2B,SOX12,DDX3X,RPS15,RPL12,ITM2C,ARL4C,MRFAP1,GNB
,CDC34,CELSR1,AGAP1,RSF1,FAM118A,RTF1,TOX4,NIBAN2,HIP1R,H6PD,RGL2,SIN3B,PSMD13,
8,PDXP,ABCD3,KLF3,GNAL,RAP1B,SMARCB1,LRIG3,MAFB,NAMPT
5F1C,KDM2A,SRP54,RETREG3,NELFCD,DNAJB6,ACP1,RBM17,PRMT2,PPP1R12A,SHC1,NARF,PC
I2,C6ORF62,RAB14,TM9SF4,KMT2E,PNISR,EIF4G3,PRR12,CAPZA1,QPRT,WBP11,LRRC41,MED
NN3,RPL29,RPSA,TNS3,CTSD,EIF4H,RTN4,LPCAT1,LAMA5,RPS24,RPS9,CIPC,MSN,SLC16A7,C1C
DB1,PSMD5,YWHAQ,EIF1,DCHS1,SET,EGLN1,MAP9,CHSY1,TMSB10,INA,EIF4A1,MRC2,PPP2R1A
AI1,HNRNPD,FBRSL1,SEC23A,SERBP1,RPS13,ARFGEF2,AKIRIN1,BMPR2,DHX15,POLR2E,VDAC1
3,RAB11B,SRP14,NRDC,PRPF6,TAF7,TUBB2A,ARF3,MAPRE1,ARPC2,FAM168B,ARGLU1,RBP1,L
IP4K2B,RAB7A,DDX3X,RPS15,RPL12,RPL30,EIF3L,NANOG,MRFAP1,RPS7,RPL35,MIF,GJA1,CTN
IF21B,GPX4,CST3,ZFP36L1,MGAT4B,BRD2,ATRX,RAB2A,UBAP2L,TAGLN2,YKT6,SV2A,CBX6,UL
M,TRAM1,RDX,MAZ,ZNF428,SRSF2,CADM3,SLC4A2,PDPK1,MSI1,PSMD4,SCRIB,LRRC41,ATP6V
1A,EIF4B,VCP,MARCKSL1,UBB,AFF4,TRIM28,MATR3,YWHAG,CKB,RPL37A,LRRC28,MAPT,FN1,S
RPS9,CIPC,PJA2,MSN,RPL24,SLC16A7,PAPOLA,IGFBP3,NCS1,PTCD3,CBX5,CLSTN1,PSMD2,ENC
L1,FAM234A,LRATD2,SLC4A2,FNBP1L,SOD2,ACTR1B,MOAP1,PCNX3,ABCF1,ERGIC3,ANKS1A,S
K1,EDC4,TXLNA,PRMT1,XPOT,EXTL3,STARD7,PDAP1,AP2A1,MMP15,FLOT1,EIF3D,SEC16A,PHB
UND,TMEM259,EIF5,CD81,WDR1,ATXN7L3B,ATP2A2,RPL26,OAZ1,SRSF4,ASTN2,MCL1,PNN,NES
1C,RPS2,LMAN1,ZMYM4,PODXL2,GSE1,TMOD2,EFR3B,FUS,SPG7,CLCN7,IARS2,HSPG2,SSBP3,

11,LMNB2,MTRF1L,ALKBH5,RPL27,SNRPN,PRDX5,ACTR3,UBE4A,PKN1,HSPA4,TCP1,NCKAP1,ML
XPNPEP3,SUPT6H,RPS28,HNRNPM,GIGYF1,SRSF3,LARS1,YWHAH,FAU,PITPNM1,TNKS2,LONP1,L
OLA,GAP43,RPL22,ZNF865,PSMD2,NR2F1,SLC7A5,RPS14,TMEM259,EIF5,LENG8,WDR1,HMGN
EIF5A,PDIA6,DDR1,PIP4K2B,MEGF8,ITM2C,RPL30,EIF3L,MAGED2,GNB2,H1-0,RPS5,IK,RPS19,MI
E2I,C16ORF58,TOMM70,XPO6,UBXN6,MYL12B,GAK,LYPLA2,ATP13A1,IER2,EIF4G1,PAFAH1B2,P
,ARL4C,MAGED2,RPS5,RPL35,MIF,TLN1,JUN,CTNNB1,MTSS2,APLP2,HNRNPDL,SLC25A3,TSR2,T
P,MYH10,YWHAB,CKAP4,PLPPR3,SEPTIN3,COTL1,CCT6A,MAGED4B,GPC1,TOP2B,SLC35E2B,MI
CAR2,CORO1C,DHX30,LARS1,NKTR,STMN3,TNKS2,SF3A1,NBPF9,BAG6,HNRNPD,RAB11A,SERBF
11A,ARNT2,WBP2,AKIRIN1,KDM4B,CCT5,CDC42,PPP1R9B,YME1L1,TRAF7,EIF2AK1,EDC4,TXLN
,KDELR1,MAP2K2,DIDO1,PITPNA,TPM3,RPS28,PHC2,LARS1,YWHAH,PTP4A2,STMN3,PFKM,LSS,F
RPL37,G3BP2,ATG2B,ARPC2,FAM168B,ARGLU1,PTBP1,CAPZB,SUPT5H,ATXN10,SON,RPL28,CC
1L2,RPS14,CXADR,CHD4,ATP1A1,JUND,RPS20,COL5A2,LENG8,WDR1,RPL26,CCT7,SRSF4,RBBF
1L2,RPS14,CXADR,CHD4,ATP1A1,JUND,RPS20,COL5A2,LENG8,WDR1,RPL26,CCT7,SRSF4,RBBF
,CTSD,EIF4H,RTN4,LPCAT1,LAMA5,RPS24,RPS9,CIPC,MSN,SLC16A7,C1ORF43,IGFBP3,RPL22,F
FAM168B,ARGLU1,PTBP1,FTO,LUZP6,SUPT5H,SON,CCAR2,ARF4,DIDO1,CORO1C,XPNPEP3,SM
Q,CDH2,MEGF8,ACSL6,SOX12,ACTN4,RPL12,CLTC,EIF3L,MRFAP1,MAGED2,GNB2,IK,RPL35,HM
,CTSB,SRSF5,TM9SF3,MDH2,TOMM20,RAB27A,U2AF2,CCT2,GNAI3,KLHL24,GAS7,NOP53,GDI2
SPC25,NISCH,CCNI,TADA2B,XPR1,KHDRBS1,KLHL9,SPTBN2,CSNK1E,RPS25,SFPQ,EIF5A,CCND
2,RIC8A,NCSTN,INTS11,AHSA1,QPRT,MTMR4,SPTY2D1,LRRC41,PRPF19,COPZ1,NOC2L,COX6A
3,SSR3,GNL1,PNISR,TUBGCP6,EEF1B2,RIC8A,NCSTN,SFXN1,INTS11,MOAP1,MTMR4,MKRN1,PC
3,SRP14,NRDC,PRPF6,TAF7,RPS15A,RPL37,ARF3,MAPRE1,ARPC2,FAM168B,ARGLU1,PTBP1,FT
TAT3,ARIH1,BCR,CPD,ACTR1A,TRIO,EP300,STARD7,BAZ1B,MMP15,KIF21B,SEC16A,KPNA2,CST
A1,SF3A2,VASH1,SART1,AP1M1,SAP18,BNIP3,ATP5MC2,SAE1,PRDX3,HSDL1,UBE2K,USO1,ATP5
,TMSB10,INA,EIF4A1,MRC2,RPL19,PPP2R1A,RPL15,GTF3C4,LDHB,DNAH8,PKD1,SF3B1,PDIA3,S
U1,PTBP1,LUZP6,CDC37,SUPT5H,HSPB1,SON,CCAR2,ARF4,POLR2B,CORO1C,SUPT6H,DHX30,T
516,MSI2,MBD6,APOE,TUT4,NCOA6,SAMD1,SESN2,KLF3,PPM1A,FZD7,POU3F3,LRP6,CRB2,HM
2A2,CCT7,OAZ1,RBBP4,PNN,NISCH,PRKCSH,HNRNPC,TADA2B,XPR1,KLHL9,SPTBN2,HDLBP,GL
36,YWHAB,SND1,MAP2,CALU,ARCN1,COTL1,HRNR,ARHGDIA,ATP6V0C,LAPTM4B,TRIB2,RPL23A,
DGFL2,PLEKHA6,CAPZA1,SFXN1,ELP1,SOD2,DAZAP2,MSI1,AKT2,PRPF19,MED13,EHD1,FXYD6,M
PS20,COL5A2,CRMP1,OAZ1,RTN3,SPC25,NISCH,PRKCSH,HNRNPC,CCNI,TADA2B,XPR1,PTN,HDL
F1,SMARCA4,AZIN1,DDAH2,FAM168B,ARGLU1,ACADVL,NYNRIN,ADAM9,ILF3,PRKDC,HSPB1,CAS
SD,EIF4H,NACA,LAMA5,RPS9,XRCC5,RPL17,PAPOLA,RPL14,HK1,NCS1,RPL22,ZNF865,CLSTN1,K
RF1L,ALKBH5,HNRNPA3,UBE4A,TBC1D16,ACO2,AHCYL1,ETNK1,KLC1,VPS35,CLSTN3,ZNF302,D
3,IMPDH2,TAX1BP1,SSR3,GNL1,HDGFL2,DTX4,KMT2E,SRSF2,MACROH2A2,BLCAP,FNBP1L,DICE
PEP3,SMARCC1,DHX30,PFKM,TNKS2,STT3A,LONP1,SF3A1,ATXN2L,GPR107,SARS1,VANGL2,GHI
M1,NPM1,PLXNB2,PLD3,HSPD1,USP11,COL18A1,SNRNP200,SMIM14,IGF2,EIF3C,ATP5F1A,VCP,
PNM1,ISYNA1,CSNK1D,BAG6,FBRSL1,RAB11A,SERBP1,ARFGEF2,WBP2,AKIRIN1,RPL34,CCT5,C
RPPRC,XPR1,SCRN1,RPS25,SFPQ,EIF5A,CDH2,PDIA6,SULF1,PIP4K2B,MEGF8,RAB7A,PRDX2,RPS
A,JAM3,FXR1,CYFIP1,PDLIM1,RALGDS,GALNT10,SALL2,SYT15,GALNT17,PPP1CA,ARHGAP5,IQS
A,JAM3,FXR1,CYFIP1,PDLIM1,RALGDS,GALNT10,SALL2,SYT15,GALNT17,PPP1CA,ARHGAP5,IQS
MB1,TSNAX,ZKSCAN1,RGL2,CRACD,TCEA1,SLC25A6,EXOC5,SF3A3,TRMT112,COPE,CD2BP2,SU
SP14,TBC1D10B,MOGS,SRSF10,NECTIN1,SLC25A1,ADCY1,SLC16A2,TMEM131,LDB1,ZCCHC14
BCR,PDAP1,CSNK2A1,MMP15,EIF3D,PHB2,KLHL42,PLXNB1,FARP1,RER1,UBE2G2,RAB3GAP2,M
ARS1,SOD1,BAG6,FBRSL1,CYP51A1,SEC23A,AKIRIN1,CD99L2,DDIT4,BEX4,VDAC1,SSR1,BCL9L
PN1,MAPKAPK2,ZNF503,SRSF7,GSK3B,SEPHS1,SEZ6L2,PLPPR2,ZNF703,CRTC1,CACNA1H,ZYX,F

F1,PGD,VAMP2,HSPH1,LMNB2,PRDX5,ACTR3,YEATS2,HSPA4,CAMSAP1,ETNK1,KLC1,GRINA,VPS3
PC3,CCN2,DDX3X,RPS15,RPL12,ITM2C,EIF3L,NANOG,MAGED2,GNB2,RCN1,H1-0,RPS7,RPS19,H
RIN,PTBP1,GRB2,PCSK1N,HNRNPUL1,TDRD5,SCAF11,CLIP3,KDELR1,HNRNPUL2-BSCL2,FLNB,D
HX15,TRAF7,EIF2AK1,WDR45B,PRMT1,XPOT,STARD7,PDAP1,AP2A1,MMP15,SEC16A,PHB2,APMA
J1,ZFAND3,MPRIP,VGF,FBRSL1,MEX3A,FLNC,ARNT2,ARFGEF2,DYNLL1,NAP1L4,TP53BP1,CTXN1
LF2,ARNT2,ARFGEF2,AKIRIN1,PRPF40A,TP53BP1,AMOTL2,YME1L1,TRAF4,SSR1,STAT3,ARIH1,PI
MR4,PCNX3,ERGIC3,YIPF3,SPTY2D1,CAP1,ATP6V0D1,USP24,NOC2L,COX6A1,EIF3G,CHMP4B,NA
KBH5,SNRPN,PRDX5,UBE4A,ACO2,HSPA4,TCP1,MLEC,ACAP3,ZNF302,TMEM248,GCN1,SKI,RAB1
L,PPP2CA,SMARCA4,ATG2B,LAPTM4A,DSTN,PDE7A,PRDX1,RABL6,SRSF1,IPO7,AEBP1,RBP1,TCF2
PPP1R9B,YME1L1,TRAF7,EIF2AK1,PDAP1,AP2A1,MMP15,EIF3D,PHB2,APMAP,KLHL42,FARP1,REF
PPP1R9B,YME1L1,TRAF7,EIF2AK1,PDAP1,AP2A1,MMP15,EIF3D,PHB2,APMAP,KLHL42,FARP1,REF
X15,BEX4,KCNQ2,BCL9L,KDM5C,ACTR1A,STARD7,NDUFV1,RETREG2,HIP1,VARS1,NOL4L,LRRC4
11A,KDM3B,AKIRIN1,BMPR2,AMOTL2,CCT5,CDC42,PPP1R9B,YME1L1,TRAF7,ST3GAL6,EIF2AK1
ATXN2L,HECTD1,HGS,BAG6,VGF,P3H3,SERBP1,CTXN1,PPP1R9B,DHX15,BEX4,PSMA7,CIC,RBBP7
LBP,RPS25,TPM1,CCND2,PDIA6,GPC3,DDR1,PIP4K2B,ACSL6,CCN2,MMP14,PRDX2,DDX3X,RPS15
AND5,HNRNPUL2-BSCL2,PITPNA,GNG2,SDF4,XPNPEP3,ATP6V1B2,SRSF3,LARS1,YWHAH,CCT8,PI
PTCD3,SLC7A5,GLIPR1L2,CXADR,TMEM259,EIF5,CD81,WDR1,ATXN7L3B,ATP2A2,CRMP1,LASP1
BPB,ITM2B,AP2B1,PXDN,HNRNPU,OGA,NFE2L1,HNRNPA0,CTNNA1,RPS16,CNN3,RPL29,RPSA,CT
COR2,FBRSL1,CYP51A1,SEC23A,AKIRIN1,CD99L2,DDIT4,BEX4,VDAC1,RBBP7,SSR1,BCL9L,WDR
RM1,RALY,SRSF2,LSM14B,RAB6A,MTMR4,LZTR1,CAP1,MSH6,AKT2,MED13,TEX2,COX6A1,KDM4
R10,ERH,EIF3H,VPS41,RALY,SEC14L1,SMG5,NUDCD3,TAX1BP1,FAM234A,SRSF2,FLOT2,TUBGC
OAZ1,RBBP4,RTN3,PNN,NISCH,PRKCSH,HNRNPC,TADA2B,XPR1,KHDRBS1,KLHL9,SPTBN2,HDLB
P1,SLC39A6,CD99,SEC16A,GPX4,PPM1G,SLC35F5,HJURP,GNG4,NXN,EP400,UBFD1,MBTPS1,RG
H1B2,SLC30A9,ACAT2,SCAMP5,LMAN2,CHMP1A,CLTA,SEMA6B,HDGFL3,TOX4,ANKHD1,YTHDF3
FBRSL1,MLF2,TP53BP1,YME1L1,PSMA7,VDAC1,RBBP7,TRAF7,EIF2AK1,EDC4,WDR45B,PDAP1,EI
NRPN,PRDX5,UBE4A,ACO2,HSPA4,TCP1,AHCYL1,MLEC,ACAP3,CHD2,SRP14,NRDC,PRPF6,TAF7,F

43,PAPOLA,LLGL1,NCS1,RPL22,CBX5,KIF5B,RAB6B,ENC1,NLGN2,CXADR,CHD4,ATP1A1,TMEM2
G2B,ARPC2,FAM168B,ACADVL,PTBP1,CAPZB,SUPT5H,ATXN10,CCAR2,ARF4,POLR2B,GNG2,SMA
IN5,VAPB,SLIT1,SHC2,UBE2M,RALY,LTBP4,RDX,EFTUD2,FLOT2,TMEM63B,PLEKHB2,SH3PXD2A
T6H,DHX30,TNKS2,SF3A1,SARS1,BAG6,HNRNPD,RAB11A,SERBP1,AKIRIN1,CDC42,PPP1R9B,YM
LC25A1,MFAP4,STUB1,ZNF768,GOLGA8B,NDRG1,DHFR,MRPS6,OAT,ABHD17A,CRTC1,CACNA1H
R,COTL1,RAC1,ARHGDIA,GPC1,CDK16,SLC35E2B,BCLAF1,RPL23A,GLUL,PTTG1IP,SQLE,MAP1A
A,GNL1,HDGFL2,GALNT1,TUBGCP6,RIC8A,NCSTN,INTS11,ACTR1B,MOAP1,DICER1,MTMR4,PCN
9SF3,PGD,MEST,HSPH1,MTRF1L,HNRNPA3,ACTR3,ETNK1,KLC1,COL2A1,MLEC,TNPO1,H3-3A,HD
,DGKD,VGF,MLF2,SEC23A,RTL8C,WBP2,AKIRIN1,TP53BP1,AMOTL2,YME1L1,CTNND1,TRAF4,ST
CD2BP2,AAMP,FXR1,CYFIP1,SUB1,EHMT2,MTURN,ZNF462,SALL2,HNRNPR,CAPN1,RNF41,SH3B
RINA,VPS35,CLSTN3,SURF4,CCT4,SKI,RPL36A,ABCA2,PYGB,MARS1,PPP2CA,CCNL2,DDAH2,DSTN
KD1,PDIA3,SSBP2,ZNF43,CHD3,VAPA,HMGB1,AP2M1,TPT1,SEC61A1,RPS3A,RPL18,STMN1,ITGB
P1R10,ACLY,IARS2,SSBP3,ABI2,SRP72,MAZ,SH3GL1,IREB2,LSM14A,CNP,TRAP1,ELP1,LSM14B,
SUPT6H,DHX30,NKTR,STMN3,TNKS2,SF3A1,NBPF9,SARS1,BAG6,HNRNPD,RAB11A,SERBP1,AKIRI
,UBE2D2,TLE1,HINT1,RAB14,GNL1,DCLK1,DTX4,ZNF428,TMEM63B,TUBGCP6,LSM14B,RAB6A,I
,FAM168A,JAK1,SLC16A2,APPBP2,UBE2Q1,BCL9,MARCHF7,RBFOX2,CHST3,NSD1,ZBTB4,WAC,C
2,CCT4,ORAI2,PSMB2,SCAF1,NRDC,TUBB2A,OPTN,UBXN4,PPP2CA,CCND1,G3BP2,KDELR2,DSTI
N1,CCNL2,DSTN,BTBD2,PBX2,PDE7A,DYNC1LI2,CDC37,CAPZB,UBE2R2,CLIP3,KDELR1,MAP2K2,

ZIN1,ARPC2,DSTN,PRDX1,TCF25,DYNLL2,GRB2,PCSK1N,WDR6,SCAF11,BNIP3L,CLIP3,PLCG1,A
NDP2,EHMT2,LIMK1,AMPD2,TASOR,COPS6,DRAP1,PPP1CA,IQSEC1,VPS51,WIZ,TBXA2R,ASAH1,Z
EC23A,RTL8C,WBP2,AKIRIN1,AMOTL2,YME1L1,CTNND1,TRAF4,STAT3,ARIH1,LBH,SF1,HIP1,SLC
KSCAN1,RGL2,PCM1,EXOC5,TRMT112,PSMD13,CD2BP2,SUMO1,RPS21,BANF1,COX8A,TASOR,T

D6,EEF1A2,SNX22,GAS7,GDI2,HSPH1,RTN1,PKN1,CAMSAP1,KLC1,VPS35,ACAP3,TMEM248,CBX

B,DNAH8,PKD1,HLA-C,SF3B1,PDIA3,SERINC1,ZNF43,RPS3,CHD3,BASP1,VAPA,HMGB1,POLR2A,I
RDX4,PPP4R3B,C6ORF89,DUS1L,RBM39
,RPL30,EIF3L,NANOG,MAGED2,RPS5,IK,RPS19,CHPF,JUN,MYO5A,ANXA6,NGRN,APLP2,RPLP2,DI
TF3,NCAM1,RPLP1,ZMIZ1,TIMP2,COL11A1,RPL10,WSB1,RPS17,DNAJA1,CYB5R3,MAP4,SF3B2,C
,GRIPAP1,TRAF7,ARIH1,EIF2AK1,SGTA,NAP1L1,TXNDC5,LBH,DGCR2,NDUFV1,PBX1,ALDH3A2,G
LC4A2,PDPK1,MSI1,FOXK2,PSMD4,NSD2,ATP6V0D1,SH3BP4,ARPP19,SF3A2,NASP,AP2A2,UBE2D
RPS24,RPS9,CIPC,MSN,SLC16A7,C1ORF43,IGFBP3,HK1,RPL22,RHOA,PTCD3,CBX5,ATF4,PSMD
DHA,CPE,RPL29,TPM4,EIF4H,TSPAN3,RTN4,LAMA5,SLC22A17,BRD3,CIPC,PJA2,MSN,SLC16A7,K
L2,PDHA1,MECP2,DNAJB1,EFNB2,OSBPL2,PPP1R14B,CRIP2,BSCL2,SYNCRIP,ACAP2,PHPT1,KAT
H1,LBH,SF1,HIP1,SLC39A6,GPX4,SLC35F5,CSNK1G2,RAB2A,FARP1,CD164,UPF1,UBFD1,MASTL
3A1,GHITM,BAG6,FBRSL1,RAB11A,SERBP1,ARFGEF2,WBP2,AKIRIN1,BMPR2,SMS,CCT5,CDC42
M1L,SLK,RANBP9,BEX1,MEX3C,CACNA1H,RCC2,ZBED4,VEZF1,ARL8A,CASKIN1,POGK,SPCS2,KI
,ARCN1,RPS27,HRNR,ARHGDIA,STIP1,SLC35E2B,SMCHD1,IPO5,ATP6V0C,BCLAF1,RPL23A,HSB
RPL39,ADAR,CTNND1,RBBP7,EDC4,CPD,EP300,DGCR2,ALDH3A2,CSNK2A1,PHB2,SLC39A7,RAB
37,PCSK1N,CAPZB,ACTN1,HSPB1,SON,PLCG1,KDELR1,RPL28,CCAR2,ARF4,HNRNPUL2-BSCL2,
LDIP2,TRIP12,PTPN11,LMAN2,CHMP1A,CLTA,USP10,TSNAX,GTF3C1,ZKSCAN1,TCEA1,PFKP,PC
BP1,GRSF1,CST3,GARS1,NXN,EP400,ADCY6,TRA2B,MASTL,MBTPS1,KCTD12,RPS2,APEX1,PPP1
SOD1,BAG6,NCOR2,VGF,CYP51A1,SERBP1,SMS,CD99L2,DDIT4,BEX4,SSR1,WDR45B,KDM5C,PB
APT,FN1,NONO,PSMD5,YWHAQ,EIF1,SET,EGLN1,TPI1,TMSB10,EIF4A1,PPP2R1A,GTF3C4,LDHB,H
,MYL6,INIP,MYH10,SND1,RHOB,UBE2H,CKAP4,ARCN1,CTBP1,HMGCR,TOP2B,SMCHD1,ATP6V0
RPLP2,ARL6IP1,NDFIP1,ST3GAL2,TSR2,MYL6,SHOX,HNRNPK,INIP,MYH10,SND1,RHOB,UBE2H,C
S,ERVV-1,RPL5,KLRD1,GDI1,FSCN1,TMBIM6,PTMS,SPTAN1,GSTP1,MARCKS,SMG1,CALM1,NPM
2,HOXB5,DIP2B,RGMB,SELENOW,HNRNPA2B1,CUL3,ULK1,SLC7A6,OAZ2,BRSK1,GPSM1,POM1
CCT2,SUGP2,MEST,GAS7,TM9SF2,MORF4L2,ABCA3,ALKBH5,SNRPN,PDXK,PLOD1,CAMSAP1,GR
EIF2AK1,EDC4,TXLNA,WDR45B,PDAP1,MMP15,SEC16A,PHB2,APMAP,GSPT1,UBA52,RAB2A,CD16
9A6,CD99,SEC16A,GPX4,PPM1G,SLC35F5,HJURP,GNG4,NXN,EP400,UBE2G2,UBFD1,MBTPS1,F
P15,SEC16A,PHB2,APMAP,GSPT1,UBA52,RAB2A,CD164,RAB3GAP2,UBFD1,IARS1,MASTL,YKT6,
M2B,MAT2A,NFE2L1,SMARCA5,FADS2,RPL23,CNN3,RPSA,TPM4,EIF4H,NACA,TSPAN3,LAMA5,BR

B,SRSF4,RBBP4,ASTN2,CLEC7A,NISCH,PRKCSH,CCNI,LRPPRC,XPR1,TKT,EIF5A,PEBP1,PDIA6,GI
1,TIMP2,SARAF,MIDN,RPL10,MARCHF6,AP3D1,DNAJA1,ARF1,CYB5R3,AKAP12,SF3B2,ITM2B,GT
17,BRD3,RPS9,SLC16A7,KPNB1,IGFBP3,LLGL1,RPL14,HK1,RHOA,RPL9,ATF4,NR2F1,RAB6B,SL
BP2,AKIRIN1,BMPR2,CCT5,CDC42,YME1L1,SSR1,TRAF7,EIF2AK1,PRMT1,PDAP1,BAZ1B,MMP15
A,CCND2,GPC3,PLXNA1,PIP4K2B,SOX12,MMP14,PRDX2,RPS15,RPL30,EIF3L,NANOG,GNB2,RPS
15B,DHX15,BEX4,GRIPAP1,ANP32A,RBBP7,STAT3,CAPN2,KDM5C,TRIO,EP300,EXTL3,STARD7,SF
FAM20B,BRI3,EFNB3,PRMT2,SHC1,VKORC1L1,NECTIN2,GABARAPL2,PPP1R26,SLC2A4RG,TME
LXNA1,PIP4K2B,RAB7A,RPL12,ARL4C,CLTC,NANOG,GNB2,RPS5,IK,MIF,HMGCS1,PRKAR1A,TL
0,STMN3,LSS,TNKS2,SF3A1,SARS1,BAG6,HNRNPD,RAB11A,SERBP1,AKIRIN1,CCT5,CDC42,PPP

2,HSPH1,MTRF1L,ALKBH5,PRDX5,UBE4A,TBC1D16,ACO2,HSPA4,AHCYL1,ETNK1,KLC1,PTOV1,C
L2,TSR2,MYL6,SHOX,INIP,MYH10,SND1,RHOB,UBE2H,CKAP4,ARCN1,CTBP1,HMGCR,TOP2B,SM
,RAB1B,VPS41,SEC14L1,ELAVL1,NUDCD3,GNL1,TUBGCP6,RIC8A,INTS11,ACTR1B,MTMR4,PHGI
ADR,CHD4,RPS20,CD81,LENG8,WDR1,HMGN2,RPL26,CRMP1,SRSF4,RBBP4,PFN1,ASTN2,MCL1,
M1,HMGB2,MSI2,TUT4,CRIM1,UBE2E1,KDM2B,SMARCC2,KLF3,PLPP3,GNAO1,SETD1B,MEX3B,
,ASAP3,UBTF,CASK,COX5B,TCF20
SM1,RPL7L1,CTTN,DAP,AKAP9,ZNF664,BAZ2A,KIAA0100,SLIT1,ERGIC1,SHC2,FKBP4,SSBP3,UBF
APKAPK2,GID8,CDK9,NSD1,SPOCK1,SEMA4C,SYNGR1,ZNF703,ALCAM,RCOR2,MAP1S,SLC12A7,
LC16A7,KPNB1,RPL17,PAPOLA,IGFBP3,COL1A2,HK1,RPL22,ZNF865,RPL9,ATF4,PSMD2,KIF5B,I
PAN3,LPCAT1,LAMA5,PJA2,MSN,SLC16A7,KPNB1,PAPOLA,RPL14,NCS1,RPL22,RHOA,CLSTN1,AT
RMP1,LASP1,BEX3,VAT1,OAZ1,TLE5,PNN,RPL31,NISCH,CSNK1E,HDLBP,TKT,CCND2,CDH2,PIP4K
LC25A23,MSI1,CAP1,MSH6,NSD2,PRPF19,FXYD6,GSK3A,PACS2,GOT2,AP2A2,TGOLN2,BNIP3,CD
25,TKT,SFPQ,EIF5A,PDIA6,DDR1,PIP4K2B,MEGF8,ACSL6,RAB7A,ITM2C,RPL30,EIF3L,MAGED2,H
2F1,KIF5B,SLC7A5,CXADR,RPS20,RPL26,CRMP1,CCT7,OAZ1,RBBP4,PFN1,JPT2,CLEC7A,RPL31,
15B,VDAC1,EIF2AK1,PRMT1,NAP1L1,EP300,H2AZ1,GSPT1,ZFP36L1,BRD2,KLHL42,DIP2B,GARS
P2,TRIP12,CHMP1A,TSNAX,EXOC5,SF3A3,TRMT112,PSMD13,COPE,CD2BP2,GPAA1,BANF1,GLOD
AP4,SF3B2,ITM2B,MAT2A,TMEM170A,GTF2F1,GOLGA8A,HNRNPU,PGAM1,NFE2L1,LARP1,HNRNI
5,EIF5A,PEBP1,CCND2,DDR1,PLXNA1,MEGF8,RAB7A,DDX3X,CLTC,MRFAP1,GNB2,RPS5,RPS19,M
NCS1,CBX5,ATF4,PSMD2,NR2F1,SLC7A5,NLGN2,GLIPR1L2,CXADR,CHD4,ATP1A1,JUND,TMEM25
5,RPL27,PDXK,ACO2,HSPA4,TCP1,COL2A1,CLSTN3,ACAP3,SURF4,ZNF302,CBX1,RAB5B,ORAI2,S
UBE2R2,ATXN10,ACTN1,HSPB1,MAP7D1,KDELR1,MAP2K2,QARS1,CCAR2,HNRNPUL2-BSCL2,DLK
,FARP1,CD164,UPF1,UBFD1,MASTL,MBTPS1,ANP32B,MAPRE2,APEX1,PPP1R15A,GALNT11,MVD,
PFKL,HDLBP,TPM1,EIF5A,CCND2,CDH2,PDIA6,MEGF8,MMP14,PRDX2,DDX3X,RPL12,CLTC,MAGI
NN,ELAVL1,GABARAPL1,GNL1,TRAP1,ELP1,SOD2,SPAG9,SYT11,PRPF19,DAPK3,PTPN23,CENPB,

29,SCARB2,NACA,LAMA5,SLC22A17,XRCC5,MSN,SLC16A7,KPNB1,RPL17,C1ORF43,PAPOLA,GAF
H6,PEBP1,DDR1,PLXNA1,PIP4K2B,CCN2,RPS15,ARL4C,CLTC,EIF3L,APLP1,IK,RPS19,MIF,GPAT4
PHKB,USP19,NDUFS1,CTR9,VEZF1,ATMIN,SNRPB,NKIRAS2,ZDHHC5,SKIV2L,MCRIP1,GPRIN1,CFI
CHMP1A,SGSM2,NBPF15,RGPD5,MKNK2,NOTCH1,PDHA1,REEP5,OSBPL2,DCAF5,IDH2,GPAA1,S
,NGRN,GOLM1,B4GALNT4,APLP2,TARDBP,ARL6IP1,NDFIP1,ST3GAL2,TSR2,MXD4,MYL6,HNRNP
RPN,PTPRA,PKN1,YEATS2,AHCYL1,CAMSAP1,KLC1,VPS35,PTOV1,MLEC,SURF4,DDX42,CBX1,OR
BSCL2,SUB1,FEZ1,CNDP2,RNF157,BNC2,KCTD20,LIMK1,PITRM1,ANKFY1,PCGF3,DYNLRB1,PPI
H1-10,MYH10,YWHAB,PTGES3,CKAP5,RHOB,UBE2H,ATP1A2,RAB1A,PLPPR3,ZNF454,HECTD4,M
CAM1,ANXA2,AGRN,RPLP1,ZMIZ1,SARAF,COL11A1,MIDN,RPL10,MARCHF6,WSB1,RPS17,AP3D1,S
RAI2,PSMB2,SCAF1,RAB11B,TSPAN6,NRDC,PRPF6,TOP1,TUBB2A,OPTN,RPL37,PPP2CA,G3BP2,A
ZFAND5,HNRNPUL2-BSCL2,DUSP4,DLK1,MSL1,SMARCE1,XPNPEP3,GIGYF1,PTP4A2,FBLN1,STT
6,PRPF19,ARPP19,CREBBP,BNIP3,TTC28,SLC38A2,ATG4B,PTPN23,WBP1,TMEM184B,MTA2,FAM
59,EIF5,LENG8,WDR1,ATXN7L3B,MCL1,PNN,HNRNPC,CCNI,LRPPRC,TADA2B,XPR1,HDLBP,EIF5A
A,RPL12,CLTC,NANOG,MAGED2,GNB2,RPS5,IK,MIF,HMGCS1,PRKAR1A,TLN1,CHPF,GJA1,JUN,N
SELENON,SMARCA4,ARPC2,DDAH2,PLEC,FAM171A2,ACADVL,NYNRIN,TCF25,ATP5PB,FTO,PCSK
NG8,WDR1,ATXN7L3B,HMGN2,ATP2A2,MCL1,PNN,CCNI,TADA2B,XPR1,HDLBP,EIF5A,PEBP1,CCN
,ILRUN,CACNA1H,ZYX,ANKRD50,SOX9,ZFP91,SLC12A7,CASKIN1,OTUB1,EGLN2,NACC1,PMPCA,
PIP4K2B,ACSL6,DDX3X,RPL12,RPL30,CLTC,NANOG,MRFAP1,MAGED2,GNB2,RPS7,HMGCS1,JUI
PTM4A,ARPC2,PLEC,BTBD2,PBX2,IPO7,NYNRIN,TCF25,ATP5PB,LUZP6,WDR6,CAPZB,HNRNPUL1,
CN1,COTL1,MAGED4B,RAC1,ARHGDIA,IPO5,MMP2,SKP1,MAP1A,CRPPA,TIMP3,CPSF2,DDX39B,C

5,MTSS2,CYFIP2,HNRNPDL,TARDBP,DDOST,WLS,H1-10,PTGES3,CKAP5,SEPTIN11,HNRNPH1,UBI
POLR2A,CDKL1,SEC61A1,RPS4X,RPS3A,RPL18,STMN1,RPL11,MORF4L1,BTF3,NCAM1,ANXA2,RI
/D,MTCH1,TMOD2,KIAA0100,PELP1,CLPTM1,SLC44A2,OLFM1,CARM1,GPC2,IMPDH2,PGAP6,RA
DDX3X,CLTC,NANOG,GNB2,RPS5,RPS7,IK,HMGCS1,PRKAR1A,TLN1,CHPF,GJA1,JUN,NGRN,B4G
FF4,TRIM28,MATR3,CKB,COL4A2,RPL37A,LRRC28,MAPT,FN1,SPEN,NONO,DDB1,HACD3,EGLN1

NL,KLF10,VPS37B
,RPL27A,FAM32A,OTUD5,EPN1,CDK9,RTF2,CLPTM1L,CSNK1A1,POLDIP3,PSMF1,PHKB,USP19,N
16,RPL23,RPL29,EIF4H,BRD3,RPS9,XRCC5,PJA2,MSN,SLC16A7,C1ORF43,PAPOLA,IGFBP3,COL1

K2,YIPF3,IGF2BP1,AKT2,LRP3,RBPJ,IVD,FBXW5,MBD3,AP2A2,CREBBP,BNIP3,DAPK3,MICAL1,M
CT6A,CAPRIN1,GPC1,TOP2B,ATP6V0C,MMP2,SKP1,DNER,LAPTM4B,NNAT,PTTG1IP,CRPPA,HS6S
,CHPF,MYO5A,COL6A2,ANXA6,NGRN,B4GALNT4,MTSS2,DNAJC5,APLP2,NDFIP1,MXD4,SHOX,H1
1,ZMIZ1,TIMP2,XAF1,COL11A1,MIDN,MARCHF6,WSB1,VCL,AP3D1,SBF1,ARF1,CYB5R3,PAFAH1I
365,CLSTN1,RAB6B,SLC7A5,CHD4,JUND,TMEM259,EIF5,CD81,WDR1,HMGN2,CRMP1,BEX3,OAZ
IST1,FKBP9,NDUFA13,BNIP3,ATP6V1G1,TTC28,ANKRD12,CERS2,TRIM8,HSDL1,FGFR1,GAK,EIF
,GAK,LYPLA2,SLC38A10,ATP13A1,PAFAH1B2,POLDIP2,TRIP12,PTPN11,CHMP1A,CLTA,USP10,PS
N2,TSR1,MORF4L2,SSRP1,MTRF1L,ALKBH5,PRDX5,PDXK,AHCYL1,ETNK1,GRINA,VPS35,MDK,SU
L,AP3D1,SBF1,DNAJA1,MAP4,AP2B1,PGAM1,SMARCA5,LARP1,CTNNA1,LDHA,CPE,RPL29,RPSA,T
R45B,PRMT1,TXNDC5,HIP1,H2AZ1,SLC39A6,CD99,GPX4,HJURP,GNG4,NXN,EP400,PI4KA,UBE2C
7,PSMB2,RAB11B,TSPAN6,TMED2,TAF7,RPS15A,G3BP2,MAPK1,ATG2B,AZIN1,ARPC2,PSMD3,OD
CHF6,WSB1,VCL,SBF1,DNAJA1,ARF1,MAP4,SF3B2,CDC42BPB,MAT2A,TMEM170A,GTF2F1,PXDN
OP53,MORF4L2,HSPH1,MTRF1L,ALKBH5,HNRNPA3,UBE4A,TBC1D16,ACO2,HSPA4,AHCYL1,ETN
G,PACS2,UBE3A,NASP,AP1M1,NDUFA13,C16ORF58,PSMC1,XPO6,UBXN6,DAPK3,PRDX3,TRO,GA
4,ATP1A1,TMEM259,EIF5,LENG8,HMGN2,LASP1,OAZ1,SRSF4,PFN1,ASTN2,TLE5,NISCH,PRKCSH
C3,CHERP,FKBP1A,CTTN,LRP10,MAP1LC3B,BRK1,SERP1,WSCD1,AKAP9,TAOK1,CYTH2,SNRPD3,

2,NDUFC2,APEH,BANF1,ARL6IP5,RALGDS,KCTD20,MAEA,NCKAP5L,PCGF3,DAD1,TBC1D14,LSM4
GF,TMED2,SELENON,TUBB2A,RPL37,G3BP2,SMARCA4,AZIN1,KDELR2,PLEC,FAM168B,AEBP1,T

NPD,FBRSL1,RAB11A,ARNT2,KDM4B,RPL34,AMOTL2,CDC42,DHX15,POLR2E,KDM5C,RETREG2,
98,DDX56,CPSF7,JRK,DUS1L,TOMM40,ALYREF,ATP6V1D,M6PR,ATP5PD
3A,PLXNA3,CSNK1D,IDH1,SARS1,SOD1,BAG6,FBRSL1,CYP51A1,SEC23A,PRPF40A,CD99L2,DDIT
AGPAT1,PRRC2A,IL6ST,RPS2,LMAN1,HNRNPAB,HIF1A,PODXL2,MVD,DAP,PPP1R10,SEPTIN5,CYT
P1,MOAP1,CLASP2,NSD2,AKT2,YY1,APBB1,ELOVL5,DHX40,SRPRA,SLC30A9,SLC25A36,XRN2,SC
,RPL26,LASP1,BEX3,OAZ1,RTN3,PFN1,ASTN2,MCL1,JPT2,TLE5,PNN,RPL31,NISCH,LRPPRC,SPTE
K3,PRDX3,TRO,GAK,LYPLA2,PTPN23,ATP13A1,NELFB,PAFAH1B2,POLDIP2,TRIP12,CHMP1A,ITFG
PM,GIGYF1,FAU,LSS,STT3B,EIF3F,TNKS2,LONP1,TUG1,PLXNA3,PRPS1,IFITM3,UQCRC2,PTPRS,
DEK,SMO,EIF4A3,DHX40,NELFB,CNBP,RANGAP1,CHMP1A,CLTA,AUTS2,ESYT1,SLC22A23,LUC7L

PD2,AGPAT3,MAP1S,CRABP2,HSPE1,CHKA,DUSP3,CCDC6,KIAA2013,TSPAN11,NUMBL,ACP1,ARP
KB,COL4A2,LRRC28,FN1,SPEN,DDB1,YWHAQ,EIF1,HACD3,DCHS1,SET,MAP9,TPI1,CHSY1,TMSB1
,RAD23A,SCAF8,PLXND1,FZD7,POU3F3,PRCP,MEX3B,PCMTD2,UBTF,SKIL,ARFGAP3,USP36,CTS
RIN1,AMOTL2,YME1L1,CTNND1,TRAF4,STAT3,ARIH1,LBH,SF1,SLC39A6,GPX4,SLC35F5,CSNK1G
P,RPLP2,NDFIP1,TSR2,MYL6,INIP,CKAP5,LAMP1,RHOB,RPS12,HLA-B,CALU,ARCN1,RPS27,HMG

CVR1B,SC5D,GGNBP2,NPTN,SLC25A6,OSBPL2,TMA7,CD44,RNF157,RASA4,HLA-A,PTPN1,TBC1D
HL1,BTF3,NCAM1,ANXA2,AGRN,ZMIZ1,TIMP2,MIDN,RPL10,MARCHF6,WSB1,RPS17,DNAJA1,ARF
A,SYNGAP1,EPN1,ARPC3,RTF2,GBA2,NDRG1,POLDIP3,UNC45A,PSMF1,HMGXB3,ACBD3,PHKB,R
RPL24,KPNB1,PAPOLA,NCS1,RPL22,PTCD3,CBX5,ZNF865,ATF4,PSMD2,KIF5B,NLGN2,CHD4,RPS
P1,PPP1R10,ERH,EIF3H,ABI2,C6ORF62,RAB14,TM9SF4,KMT2E,PNISR,EIF4G3,PRR12,CAPZA1,N
GOLN2,AP1M1,SAP18,NDUFA13,PSMC1,XPO6,UBXN6,GAK,LYPLA2,PTPN23,SLC38A10,ATP13A1,
EKHB2,CADM3,SLC4A2,SLC25A23,ABCF1,ERGIC3,SPTY2D1,CAP1,IST1,ATP6V0D1,DHCR7,MBD3,
1,ORAI2,SKI,SNRNP70,TSPAN6,RPL38,SELENON,TUBB2A,MVB12B,G3BP2,ARPC2,DSTN,RABL6,
D3,VKORC1L1,UBE2G1,PRPF31,MED16,PCNA,TFDP1,GNL3,OPA1,ZC3H7A,PPT1,CCNK,MAML1,K
C,BTBD2,DYNC1LI2,AEBP1,TCF25,ADAM9,ILF3,CDC37,RPS11,KDELR1,ARF4,CORO1C,XPNPEP3,

NX17,THG1L,RPL32,SZT2,SEC31A,ACIN1,AKAP17A,HARS1,DNMT1,CFAP298,CEBPZ,CPSF7,JRK,T
R1,HDLBP,EIF5A,PEBP1,CCND2,DDR1,PLXNA1,RAB7A,DDX3X,CLTC,NANOG,GNB2,RPS5,IK,HMG
PK3,PRDX3,TRO,GAK,LYPLA2,PTPN23,ATP13A1,NELFB,IER2,PAFAH1B2,POLDIP2,TRIP12,CHMP1
TMEM259,CD81,LENG8,HMGN2,CRMP1,LASP1,CCT7,VAT1,OAZ1,SRSF4,RBBP4,RTN3,JPT2,TLE
,LAMP1,RHOB,HNRNPH1,RPS12,ZNF454,ARHGDIA,SMCHD1,TMEM30A,SKP1,BCLAF1,TIMP3,DD

1C,RPS2,CALD1,APEX1,NPLOC4,PPP1R15A,GALNT11,DAP,BRK1,BAP1,GRN,BAZ2A,XRCC6,PELF
P1CB,PGD,GAS7,PABPN1,TM9SF2,MORF4L2,ALKBH5,PDXK,ACO2,HSPA4,TCP1,PLOD1,GRINA,
S1,SPTBN2,PFKL,SFPQ,SOX12,CCN2,ITM2C,ARL4C,EIF3L,MRFAP1,IK,GPAT4,PRKAR1A,CHPF,G

2,MSI1,ATP1B2,IGF2BP1,USP24,MED13,PCGF2,FBXW5,CHMP4B,APBB1,C16ORF58,MN1,TTC28
1,ZNF43,RPS3,BASP1,VAPA,POLR2A,SEC61A1,RPS4X,RPS3A,STMN1,UCHL1,RPL11,MORF4L1,BT
,TRAP1,PLEKHB2,ELP1,MOAP1,CLASP2,RBM12,NSD2,YY1,APBB1,CCDC47,ELOVL5,DHX40,SRPR
2,GABARAPL1,GNL1,TRAP1,DNAJB2,RIC8A,PLEKHB2,ELP1,RAB6A,CLASP2,NSD2,ATP6V0D1,PRF
SGSM3,TNFAIP1,SF3A3,PHPT1,ATF5,ERP29,HMGB3,LMF2,EFNB1,RHEB,RAP2A,CDK4,NUP153,A
5,RAB6B,SLC7A5,NLGN2,CXADR,CHD4,JUND,RPS20,EIF5,CD81,COL5A2,LENG8,WDR1,HMGN2,
CKS1,ATXN3,INTS11,RAB6A,SLC4A2,KRT8,PDPK1,WASF2,ERGIC3,EDF1,SCRIB,SPTY2D1,CAP1,

,ESYT1,GTF3C1,YTHDF3,KLF13,CDV3,FZR1,TNFAIP1,FAM50A,CD2BP2,PIK3C2B,ALDH18A1,CND
11A,GNAO1,RAD23A,EPHA2,SKIL,STX12,VPS37B,RBBP6,ZC3H4,NDUFA10,CCDC124
,SEPTIN2,P4HB,APP,IDS,ERVV-1,RPL5,GDI1,FSCN1,TMBIM6,PTMS,SPTAN1,GSTP1,MARCKS,SM
C1,MEIS3,CTDNEP1,MIB2,TMEM59,TRAPPC1,SPG11,TNPO3,CUL4B,PSMD12,TPX2,NUP50,SZT2,
26,SCAMP4,GPAA1,SUB1,KCTD20,ECH1,DYNLRB1,TNRC6A,PPME1,CSNK2B,RPL27A,COX7A2L,
OLPH3,ECHS1,ALDH7A1,DHX36,HM13,COLGALT1,TSPAN7,NUP98,NACAD,OXSR1,EMC10,PSMC4
NF187,RPN2,MLXIP,PPP1CB,GNAI3,GAS7,HNRNPL,VCAN,TM9SF2,TSR1,ABCA3,PTPRA,AHCYL1,K
SUMO1,BANF1,COX8A,CUL4A,VPS51,DARS1,NME2,FAM32A,OTUD5,EPN1,CDK9,RTF2,ZRANB2,P
K8IP2,CRIP2,ADGRB2,ATP1B3,RALGDS,GALNT10,LMF2,SALL2,FLII,NBAS,VPS51,WIZ,NECTIN1,S
I3,RPS24,BRD3,RPS9,CIPC,MSN,RPL24,RPL17,C1ORF43,COL1A2,RPL14,HK1,RHOA,PTCD3,CBX
DX39B,SRSF5,TM9SF3,MDH2,TOMM20,U2AF2,FDFT1,CCT2,GNAI3,NOP53,GDI2,TM9SF2,NOX4,

S37B,RBBP6,ZC3H4,NDUFA10,CCDC124
LP1,MOAP1,CLASP2,RBM12,NSD2,YY1,CCDC47,ELOVL5,DHX40,SRPRA,SLC25A36,XRN2,SCAMP
1,HACD3,DCHS1,SET,EGLN1,TPI1,CHSY1,TMSB10,INA,EIF4A1,MRC2,RPL19,RPL15,GTF3C4,DNA
ORF62,LTBP4,VPS4A,PGAP6,MAZ,TMEM63B,QPRT,MSI1,LTBP3,YIPF3,MED13,MTOR,SMARCD1

/I9,PTBP1,ILF3,LUZP6,PRKDC,ACTN1,KDM1A,BNIP3L,KDELR1,ARF4,ZFAND5,MSL1,NKTR,FAU,C(
M1,C16ORF58,UBE2D3,TOMM70,PRDX3,DEK,GAK,LYPLA2,SLC38A10,IER2,PAFAH1B2,CGGBP1,
SL1,UBB,TRIM28,MATR3,YWHAG,COL4A2,RPL37A,MAPT,FN1,SPEN,DDB1,PSMD5,EIF1,DCHS1,S
N,NGRN,MACF1,B4GALNT4,MTSS2,CYFIP2,DNAJC5,HNRNPDL,TARDBP,RPLP2,DDOST,ST3GAL2,
,SRSF2,CAPZA1,TUBGCP6,RIC8A,NCSTN,INTS11,AHSA1,RAB5C,QPRT,CLASP2,MTMR4,PDPK1,YI
/I1,LDLR,SSR3,PLEKHA6,ZNF428,NCSTN,ELP1,PACS1,DICER1,CAP1,LRRC41,LRP3,DHCR7,EHD1
COX6A1,NASP,YY1,SMARCD1,CREBBP,ATXN7L3,XPO6,AMBRA1,ATG4B,UBE2L3,SPPL2B,ZZEF1,I
3,RAB7A,DDX3X,RPS15,RPL30,ARL4C,GNB2,RCN1,APLP1,RPS7,IK,MIF,PRKAR1A,CHPF,GJA1,NG
S28,SSB,GIGYF1,STT3B,TNKS2,LONP1,TUG1,PRPS1,UQCRC2,HGS,SOD1,PFN2,NCOR2,HNRNPD,
.24,SLC16A7,NCS1,RPL22,PTCD3,ZNF865,RPL9,KIF5B,GLIPR1L2,RPS14,CXADR,RPS20,CD81,LE

(5B,ATP5PD
,EFTUD2,STK24,TMEM63B,LSM14B,SLC4A2,FNBP1L,SOD2,SPAG9,PLEKHM2,TCF3,DCUN1D2,AT
4B,REEP1,DMWD,HSDL1,LYPLA2,SH3PXD2B,USP5,IER2,RANGAP1,SLC30A9,AMFR,TECR,TAPBP,
TP5PB,WDR6,PRKDC,HSPB1,MAP7D1,CASC3,KDELR1,RPL28,ZFAND5,TSC22D1,POLR2B,CORO
1R15A,GALNT11,DAP,BRK1,BAP1,SPG7,BAZ2A,XRCC6,VAPB,EIF3H,ELAVL1,IMPDH2,GABARAPL1

MSI1,LTBP3,SCRIB,USP24,MED13,TEX2,KDM4A,PRKACA,TMEFF1,MN1,UBE2D3,TTC28,TRIM8,TH
RC41,MED13,UBE3A,ATP5MC3,NASP,YY1,SMARCD1,CREBBP,XPO6,AMBRA1,ATG4B,UBE2L3,SP
11,MORF4L1,ANXA2,RPLP1,ZMIZ1,TIMP2,COL11A1,MIDN,MARCHF6,WSB1,RPS17,VCL,DNAJA1,(
,RAB2A,FARP1,SRSF6,RER1,PI4KA,UBE2G2,RAB3GAP2,TAGLN2,TRA2B,COX7C,YKT6,MCM3AP,T

AB7A,DDX3X,RPL12,CLTC,NANOG,MAGED2,GNB2,RPS5,IK,MIF,HMGCS1,PRKAR1A,TLN1,CHPF,G
MEM167A,LMAN1,ZMYM4,TNKS,HNRNPAB,HIF1A,PODXL2,FKBP1A,MVD,VPS26B,PPP1R10,SEPTI
I6,TRPC4AP,DYNC1I2,CCNL1,POGZ,PES1,ATP6V1F,KHDC4,SECISBP2L,QSOX1,CYTH3
D1,RHOB,HNRNPH1,UBE2H,RAB1A,CALU,ARCN1,RPS27,HRNR,CCT6A,HMGCR,GPC1,SMCHD1,A
4,SHOX,H1-10,CIRBP,MYH10,RPL36,YWHAB,CADM4,SND1,RHOB,HNRNPH1,UBE2H,COTL1,HEC
.C25A36,XRN2,SCAMP5,ZZEF1,AASDHPPT,ESYT1,GTF3C1,FZR1,TNFAIP1,COPE,FAM50A,CD2BP2
A,LRRC28,MAPT,FN1,NONO,DDB1,PSMD5,YWHAQ,EIF1,DCHS1,SET,EGLN1,MAP9,TPI1,TMSB10,
32,MAPRE3,CHST3,YWHAZ,NDUFB8,DDX41,ACSL4,CRTC1,ALCAM,ADIPOR2,KIF3A,TXN,HOXA5,S

BAG6,RAB11A,CYP51A1,ARNT2,SERBP1,KDM3B,AKIRIN1,KDM4B,TP53BP1,CCT5,CDC42,PPP1R
RK,TOMM40,ATP6V1D,COX5B
/I3AP,SPINT2,TMEM167A,KLHDC3,TNKS,CHERP,CTTN,SLC25A5,LRP10,SERP1,BAP1,FUS,ACLY,E

IOTCH2,MARS1,SELENON,RPL37,CCND1,G3BP2,CCNL2,ARPC2,SRSF1,ARGLU1,PTBP1,DYNLL2,
5,PJA2,MSN,KPNB1,C1ORF43,PAPOLA,GAP43,HK1,RPL22,RHOA,ZNF865,NR2F1,NLGN2,RPS14,(
RPL11,MORF4L1,BTF3,NCAM1,ANXA2,AGRN,RPLP1,ZMIZ1,TIMP2,XAF1,SARAF,MIDN,RPL10,RPS

-B,PLPPR3,MAGED4B,CTBP1,GPC1,APPL1,SMCHD1,IPO5,ATP6V0C,SKP1,BCLAF1,HSBP1,LUC7L
N1,NECTIN2,PLCD3,CTDNEP1,CMTR1,CLCN3,TRAPPC1,ABCE1,MSI2,CCNL1,MAPK8IP3,PAF1,DD
GDH,DSTYK,GSTA4,GUK1,UBN1,NECTIN2,PLCD3,CTDNEP1,CMTR1,CLCN3,TRAPPC1,ABCE1,NO
RAPL2,THRA,TRAF3,AMD1,LRP8,TRPC4AP,CBX4,HADHB,POGZ,MAPK8IP3,PLPP3,ITGA7,PLXND1,

I,PIK3C2B,BANF1,FAF2,KLC2,ATF7IP,VPS51,PPME1,CDK4,NME2,FAM32A,EPN1,RTF2,PSMC5,PC
ELENON,RAPGEF1,OPTN,MAPK1,AZIN1,MAPRE1,ARPC2,PLEC,PBX2,FAM171A2,AEBP1,PRKDC,GA
8,NACAD,OXSR1,EMC10,SEMA4C,TMEM9,SCG2,WDR82,SYNGR1,USP48,PDCD11,RPS29,NSMF,D
K,GSTA4,GUK1,UBN1,NECTIN2,PLCD3,CTDNEP1,CMTR1,CLCN3,TRAPPC1,ABCE1,MSI2,CCNL1,M
K,GSTA4,GUK1,UBN1,NECTIN2,PLCD3,CTDNEP1,CMTR1,CLCN3,TRAPPC1,ABCE1,MSI2,CCNL1,M
IB3,SLC25A1,POLD2,TSPAN7,GOLGA8B,UBL3,EMC10,ZBTB4,P3H1,WDR82,SOX9,RPS29,MBOAT
4,CTSD,EIF4H,RTN4,SLC22A17,XRCC5,PJA2,MSN,SLC16A7,KPNB1,C1ORF43,PAPOLA,IGFBP3,LL
TCF25,CAPZB,PRKDC,TDRD5,HSPB1,CASC3,CLIP3,RPL28,ZFAND5,PITPNA,PLTP,CORO1C,SDF4,
R1,HMGN2,RPL26,LASP1,OAZ1,SRSF4,RBBP4,MCL1,SPC25,PRKCSH,LRPPRC,XPO1,SPTBN2,SLC
RPS15,ITM2C,RPL30,CLTC,NANOG,RPS7,HMGCS1,GPAT4,GJA1,NGRN,MACF1,RPS10,MTSS2,DN
RPS15,ITM2C,RPL30,CLTC,NANOG,RPS7,HMGCS1,GPAT4,GJA1,NGRN,MACF1,RPS10,MTSS2,DN
ORF3,PDXK,TBC1D16,AHCYL1,ETNK1,PTOV1,MLEC,ZNF302,LAMC1,CBX1,CHD2,H3-3A,SCAF1,HI

12,MAP4,SF3B2,CDC42BPB,ITM2B,MAT2A,TMEM170A,SLC4A8,GOLGA8A,PXDN,OGA,NFE2L1,SI
R1,MRPS6,GDF1,ACSL4,CCDC85C,RAB3A,RCOR2,STC2,DTX3,NAPB,RAVER1,HOXA5,IRF2BP2,EC

C,SF3B1,PDIA3,SERINC1,SSBP2,ZNF43,RPS3,VAPA,HMGB1,POLR2A,TPT1,SEC61A1,RPS4X,OR8K
R1L2,ATP1A1,JUND,TMEM259,EIF5,LENG8,WDR1,ATXN7L3B,MCL1,PNN,CCNI,TADA2B,XPR1,HDL
PL26,LASP1,CCT7,VAT1,RBBP4,RTN3,ASTN2,MCL1,NES,KLHL9,HDLBP,RPS25,SFPQ,CCND2,PIP4

T112,PSMD13,WDR26,CD2BP2,GPAA1,GGT7,APEH,SUB1,FAF2,CFDP1,CUL4A,TNRC6A,PPP1CA,VI
IPO1,ASXL1,RPL36A,RAB11B,HDGF,DPYSL2,TMED2,TAF7,MVB12B,G3BP2,SMARCA4,KDELR2,SF

A,RABL6,ACADVL,PTBP1,DYNLL2,ILF3,CDC37,ATXN10,HSPB1,CMIP,KDELR1,MAP2K2,CCAR2,DH

DA2B,XPR1,KHDRBS1,KLHL9,PFKL,RPS25,TKT,GLG1,CDH6,SFPQ,TPM1,EIF5A,PEBP1,DDR1,PIP4
,OGDH,DSTYK,GUK1,UBN1,NECTIN2,PLCD3,CTDNEP1,CMTR1,CLCN3,TRAPPC1,ABCE1,MSI2,CC
2E,PNISR,EIF4G3,PRR12,CAPZA1,QPRT,LRRC41,MED13,UBE3A,ATP5MC3,NASP,YY1,SMARCD1,C
,RPS9,CIPC,PJA2,MSN,KPNB1,C1ORF43,PAPOLA,IGFBP3,LLGL1,RPL14,RPL22,RHOA,PTCD3,CB
F,HMGCS1,GPAT4,TLN1,JUN,MYO5A,COL6A2,ANXA6,GOLM1,SESN3,CYFIP2,DDOST,ST3GAL2,TA

PRDX2,DDX3X,RPS15,ITM2C,RPL30,CLTC,NANOG,RPS7,HMGCS1,GPAT4,GJA1,NGRN,MACF1,GC
M9,SCG2,WDR82,SYNGR1,USP48,RPS29,STMN4,NSMF,DUSP8,RNF20,NUP133,CHGB,COX4I1,OG
NRNPUL2-BSCL2,PITPNA,XPNPEP3,SUPT6H,ATP6V1B2,SRSF3,LARS1,YWHAH,FAU,UBQLN1,CSNK

AMP5,ZZEF1,AASDHPPT,ESYT1,GTF3C1,TNFAIP1,FAM50A,CD2BP2,CNDP2,RNF157,GLOD4,MTUF
C,DDX41,PGRMC2,ARHGEF9,TXN,IRF2BP2,HSPE1,ZNF592,POGK,KDM2A,EGLN2,CEBPG,DPP7,P
IDUFA4,TRMT112,WDR26,CD2BP2,SCAMP4,GPAA1,SUB1,BANF1,KCTD20,DYNLRB1,TNRC6A,PPI
SNK1G2,UPF1,MASTL,NAP1L3,PRRC2A,PPP1CC,IL6ST,ANP32B,MAPRE2,CALD1,NPLOC4,DKK3,F

GDH,DSTYK,GSTA4,GUK1,UBN1,NECTIN2,PLCD3,CTDNEP1,CMTR1,CLCN3,TRAPPC1,ABCE1,NO
G7,BAZ2A,XRCC6,VAPB,CLPTM1,EIF3H,ELAVL1,IMPDH2,GABARAPL1,FAM234A,GNL1,TM9SF4,ZI
0,ZBTB4,SEMA4C,TMEM9,SCG2,WDR82,SYNGR1,USP48,RPS29,NSMF,DUSP8,RNF20,NUP133,C

8,RPS29,NSMF,DUSP8,RNF20,NUP133,CHGB,COX4I1,OGDH,DSTYK,GSTA4,GUK1,UBN1,NECTIN

.T1,TSPAN7,NUP98,NACAD,OXSR1,EMC10,RNMT,SEMA4C,TMEM9,SCG2,WDR82,SYNGR1,USP48
MEX3A,ZSCAN18,ARFGEF2,WBP2,NAP1L4,TXNIP,RPL34,TP53BP1,CCT5,RBM15B,POLR2E,SSR1,
THRAP3,ELOVL5,USP5,IER2,DENND5A,SLC25A36,XRN2,SCAMP5,SGSM2,ZZEF1,AASDHPPT,ESYT
,ARF4,PLTP,SDF4,RPS28,HNRNPM,GIGYF1,STT3B,EIF3F,LONP1,TUG1,PLXNA3,PRPS1,IFITM3,U
,ARF4,PLTP,SDF4,RPS28,HNRNPM,GIGYF1,STT3B,EIF3F,LONP1,TUG1,PLXNA3,PRPS1,IFITM3,U
IC8A,CLASP2,ERGIC3,PCGF2,FBXW5,PSMC3,SART1,NASP,TGOLN2,MN1,FJX1,METRN,HNRNPH3
POGZ,BRAT1,TLE4,PDHB,GPRC5B,ZNF207,SH3KBP1,KLF10,JRK,SKIL,SCCPDH,DTD1
TMN1,ITGB1,UCHL1,MORF4L1,BTF3,ANXA2,AGRN,RPLP1,ZMIZ1,SARAF,COL11A1,MARCHF6,WS

AF2,CCT2,GNAI3,NOP53,GDI2,VCAN,TM9SF2,TSR1,SSRP1,LMNB2,MTRF1L,ALKBH5,RPL27,SNF
3N1,NECTIN2,PLCD3,CTDNEP1,CMTR1,CLCN3,TRAPPC1,ABCE1,MSI2,CCNL1,MAPK8IP3,PAF1,DD
COPE,FAM50A,CD2BP2,CNDP2,GLOD4,MTURN,GOLPH3,RAP2A,ECHS1,COX7A2L,DHX36,HM13,
)50,SEC11A,TOMM40,ATP6V1D,M6PR,USP36

VLS,H1-10,INIP,CIRBP,CALU,RPS27,HRNR,CCT6A,CAPRIN1,CDK16,APPL1,SMCHD1,HDAC2,L1CA

DYNC1LI2,ARGLU1,ACADVL,ILF3,ATXN10,CAND1,MAP7D1,NSG2,FKBP8,QARS1,RPL28,HNRNPU

ARS1,SOD1,BAG6,FBRSL1,RPL39,CYP51A1,ZSCAN18,PRPF40A,CD99L2,DDIT4,DHX15,RBBP7,SS
LC20A1,MICAL1,ATG4B,DMWD,NELFB,USP5,IER2,RANGAP1,ELAVL3,SLC30A9,MZT2B,TECR,TA

DC85C,PGRMC2,NAPB,IRF2BP2,CCDC93,NSMF,DUSP8,METRNL,SLC16A1,RBM14,KIAA2013,CA
CND1,G3BP2,PDE7A,PSMD3,ODC1,SRSF1,ILF3,CDC37,WDR6,RPS11,SCAF11,BNIP3L,ZFAND5,H
T1,IL6ST,MCM3AP,TMEM167A,KLHDC3,CHERP,PPP1R15A,CTTN,LRP10,DAP,AKAP9,ACLY,KIAA0

3,ARCN1,SEPTIN3,MAGED4B,HMGCR,CAPRIN1,CDK16,IPO5,SKP1,DNER,PEA15,HSBP1,DLG5,H
3,CDC37,WDR6,RPS11,SCAF11,BNIP3L,ZFAND5,HNRNPUL2-BSCL2,DUSP4,PITPNA,XPNPEP3,SU
12,CD2BP2,SUMO1,RPS21,BSCL2,SUB1,BANF1,COX8A,GSR,VPS51,FAM8A1,SLC25A1,FAM32A,J

T3,ZNF302,JPT1,CHD2,ORAI2,GCN1,H3-3A,RPL36A,CNTNAP2,SRP14,TERF2IP,RPL38,MARS1,RA
8,RPS29,NSMF,DUSP8,RNF20,NUP133,CHGB,COX4I1,OGDH,DSTYK,GSTA4,GUK1,UBN1,NECTIN

5,HM13,CRKL,COLGALT1,TSPAN7,NUP98,NACAD,OXSR1,EMC10,ZBTB4,SEMA4C,TMEM9,PLK2,
3,RPS29,NSMF,DUSP8,RNF20,NUP133,CHGB,COX4I1,OGDH,DSTYK,GSTA4,GUK1,UBN1,NECTIN2

:F1B2,ATXN3,SLC4A2,IGF2BP1,LRP3,ATP6V0D1,MED13,ATP6V1G1,ANKRD12,CCDC47,UBXN6,SL

SDE1,SMIM14,IGF2,EIF3C,ATP5F1A,EIF4B,VCP,MARCKSL1,UBB,AFF4,TRIM28,MATR3,YWHAG,C
F4,TRIM28,MATR3,YWHAG,CKB,COL4A2,RPL37A,LRRC28,MAPT,FN1,SPEN,NONO,DDB1,PSMD5,
M28,MATR3,YWHAG,CKB,COL4A2,RPL37A,LRRC28,FN1,SPEN,NONO,DDB1,YWHAQ,EIF1,HACD3,
,SPEN,NONO,DDB1,PSMD5,YWHAQ,EIF1,HACD3,DCHS1,SET,EGLN1,MAP9,TPI1,CHSY1,TMSB10,
,TPT1,SEC61A1,RPS4X,RPL18,STMN1,ITGB1,TUBB4B,MORF4L1,BTF3,NCAM1,AGRN,ZMIZ1,TIM
TR3,YWHAG,CKB,COL4A2,RPL37A,LRRC28,FN1,SPEN,NONO,DDB1,PSMD5,YWHAQ,EIF1,HACD3
RCKSL1,UBB,AFF4,TRIM28,MATR3,YWHAG,CKB,COL4A2,RPL37A,LRRC28,FN1,SPEN,NONO,DD
NPA0,CTNNA1,FADS2,CPE,CNN3,CTSD,EIF4H,SCARB2,TSPAN3,LPCAT1,LAMA5,SLC22A17,BRD3,
A1,MRC2,RPL19,PPP2R1A,RPL15,GTF3C4,LDHB,PKD1,HLA-C,SF3B1,PDIA3,SERINC1,SSBP2,ZNF
FN1,SPEN,NONO,DDB1,YWHAQ,EIF1,HACD3,DCHS1,SET,EGLN1,MAP9,TPI1,CHSY1,TMSB10,INA,
A,LRRC28,MAPT,FN1,SPEN,NONO,DDB1,PSMD5,YWHAQ,EIF1,HACD3,DCHS1,SET,EGLN1,TPI1,C
R2A,AP2M1,TPT1,RPL18,STMN1,ITGB1,TUBB4B,UCHL1,NCAM1,ANXA2,AGRN,TIMP2,SARAF,MI
,LDHB,PKD1,HLA-C,PDIA3,SSBP2,BASP1,VAPA,HMGB1,POLR2A,AP2M1,TPT1,SEC61A1,RPS3A,R
PL19,PPP2R1A,RPL15,GTF3C4,LDHB,PKD1,HLA-C,PDIA3,SSBP2,BASP1,VAPA,HMGB1,POLR2A,A
,PDIA3,SSBP2,ZNF43,BASP1,VAPA,POLR2A,AP2M1,TPT1,RPL18,STMN1,ITGB1,TUBB4B,UCHL1,
SPEN,NONO,DDB1,PSMD5,YWHAQ,EIF1,HACD3,DCHS1,SET,EGLN1,MAP9,TPI1,CHSY1,TMSB10,
9,PPP2R1A,GTF3C4,LDHB,PKD1,HLA-C,PDIA3,SERINC1,SSBP2,BASP1,VAPA,HMGB1,POLR2A,AF
BASP1,VAPA,HMGB1,POLR2A,AP2M1,TPT1,SEC61A1,RPS3A,RPL18,STMN1,ITGB1,TUBB4B,UCH
NAJA1,ARF1,CYB5R3,MAP4,CDC42BPB,ITM2B,SLC4A8,GOLGA8A,PXDN,HNRNPU,NFE2L1,SMAR
PPP2R1A,GTF3C4,LDHB,PKD1,HLA-C,PDIA3,SSBP2,BASP1,VAPA,HMGB1,POLR2A,AP2M1,TPT1,S
SSBP2,ZNF43,BASP1,VAPA,HMGB1,CDKL1,TPT1,SEC61A1,RPS4X,RPS3A,STMN1,ITGB1,TUBB4B
T,EGLN1,MAP9,TPI1,CHSY1,INA,EIF4A1,MRC2,RPL19,PPP2R1A,GTF3C4,PKD1,HLA-C,SF3B1,PD
3,CHD3,BASP1,VAPA,HMGB1,POLR2A,TPT1,STMN1,ITGB1,TUBB4B,UCHL1,MORF4L1,BTF3,NC
L18A1,SNRNP200,CSDE1,SMIM14,IGF2,EIF3C,ATP5F1A,EIF4B,VCP,MARCKSL1,UBB,AFF4,TRIM
1,TPT1,RPL18,STMN1,ITGB1,TUBB4B,UCHL1,ANXA2,AGRN,RPLP1,ZMIZ1,TIMP2,COL11A1,MID
B1,PSMD5,YWHAQ,EIF1,HACD3,DCHS1,SET,EGLN1,MAP9,TPI1,CHSY1,TMSB10,INA,EIF4A1,MR
,SEC61A1,RPL18,ITGB1,TUBB4B,UCHL1,RPL11,MORF4L1,AGRN,TIMP2,COL11A1,MIDN,RPL10,
SPEN,NONO,DDB1,YWHAQ,EIF1,HACD3,DCHS1,SET,EGLN1,TPI1,CHSY1,TMSB10,INA,EIF4A1,MR
MIM14,IGF2,EIF3C,ATP5F1A,EIF4B,VCP,MARCKSL1,UBB,AFF4,TRIM28,MATR3,YWHAG,CKB,COL
SDE1,SMIM14,IGF2,EIF3C,ATP5F1A,EIF4B,VCP,MARCKSL1,UBB,AFF4,TRIM28,MATR3,YWHAG,C
G,CKB,COL4A2,RPL37A,LRRC28,MAPT,FN1,SPEN,NONO,PSMD5,YWHAQ,EIF1,HACD3,DCHS1,SE
PA,HMGB1,POLR2A,AP2M1,TPT1,SEC61A1,RPS3A,RPL18,STMN1,ITGB1,TUBB4B,UCHL1,MORF
PDIA3,SSBP2,ZNF43,BASP1,VAPA,HMGB1,POLR2A,AP2M1,TPT1,STMN1,ITGB1,TUBB4B,UCHL1,
TP5F1A,EIF4B,VCP,MARCKSL1,UBB,AFF4,TRIM28,MATR3,YWHAG,CKB,COL4A2,RPL37A,LRRC2
,CSDE1,SMIM14,IGF2,EIF3C,ATP5F1A,EIF4B,VCP,MARCKSL1,UBB,AFF4,TRIM28,MATR3,YWHAG
,AGRN,TIMP2,XAF1,SARAF,MIDN,MARCHF6,VCL,AP3D1,SBF1,ARF1,CYB5R3,MAP4,SF3B2,CDC4
ONO,DDB1,PSMD5,YWHAQ,EIF1,HACD3,DCHS1,SET,EGLN1,MAP9,TPI1,CHSY1,TMSB10,INA,EIF
F3C4,LDHB,PKD1,HLA-C,SSBP2,ZNF43,RPS3,BASP1,VAPA,HMGB1,CDKL1,TPT1,SEC61A1,RPS3
TUBB4B,UCHL1,MORF4L1,BTF3,NCAM1,ANXA2,AGRN,ZMIZ1,TIMP2,XAF1,SARAF,MIDN,MARCH
WHAQ,EIF1,HACD3,DCHS1,SET,EGLN1,TPI1,CHSY1,TMSB10,INA,EIF4A1,MRC2,RPL19,PPP2R1A,
L1,UBB,AFF4,TRIM28,MATR3,YWHAG,CKB,COL4A2,RPL37A,FN1,SPEN,NONO,DDB1,PSMD5,YW
IF4B,VCP,MARCKSL1,UBB,TRIM28,MATR3,YWHAG,CKB,COL4A2,RPL37A,LRRC28,FN1,SPEN,NC
,HMGB1,POLR2A,TPT1,SEC61A1,RPS3A,RPL18,STMN1,TUBB4B,UCHL1,MORF4L1,BTF3,AGRN,
PI1,CHSY1,TMSB10,INA,MRC2,RPL19,PPP2R1A,RPL15,GTF3C4,LDHB,PKD1,HLA-C,PDIA3,SSBP2
1,SPEN,NONO,DDB1,YWHAQ,EIF1,HACD3,DCHS1,SET,EGLN1,TPI1,CHSY1,TMSB10,INA,EIF4A1,

.TR3,YWHAG,CKB,COL4A2,RPL37A,LRRC28,MAPT,FN1,SPEN,NONO,DDB1,PSMD5,YWHAQ,EIF1,H
AH8,PKD1,HLA-C,PDIA3,SSBP2,BASP1,VAPA,HMGB1,POLR2A,CDKL1,AP2M1,TPT1,SEC61A1,RPS
ADS2,RPS16,CNN3,TPM4,CTSD,EIF4H,LPCAT1,LAMA5,SLC22A17,BRD3,CIPC,PJA2,MSN,KPNB1,C
4,TRIM28,MATR3,YWHAG,CKB,COL4A2,RPL37A,LRRC28,SPEN,DDB1,PSMD5,YWHAQ,EIF1,HACD
2,EIF3C,ATP5F1A,EIF4B,VCP,MARCKSL1,UBB,AFF4,TRIM28,MATR3,CKB,COL4A2,RPL37A,LRRC
OL11A1,MIDN,RPL10,MARCHF6,WSB1,RPS17,VCL,AP3D1,SBF1,DNAJA1,ARF1,CYB5R3,AKAP12,M
,SBF1,ARF1,CYB5R3,MAP4,SF3B2,CDC42BPB,ITM2B,AP2B1,MAT2A,GOLGA8A,PXDN,HNRNPU,N
,SET,EGLN1,CHSY1,TMSB10,INA,MRC2,RPL19,PPP2R1A,RPL15,GTF3C4,PKD1,SF3B1,PDIA3,SER
1,C1ORF43,COL1A2,LLGL1,NCS1,RHOA,ZNF865,CLSTN1,ATF4,PSMD2,SLC7A5,NLGN2,CHD4,AT
,HACD3,DCHS1,SET,EGLN1,TPI1,CHSY1,TMSB10,INA,EIF4A1,RPL19,PPP2R1A,RPL15,GTF3C4,LD
7A5,NLGN2,CHD4,ATP1A1,JUND,EIF5,CD81,WDR1,HMGN2,ATP2A2,CRMP1,LASP1,BEX3,VAT1,O
PNB1,C1ORF43,COL1A2,LLGL1,NCS1,RHOA,CBX5,ZNF865,CLSTN1,ATF4,RAB6B,SLC7A5,CHD4,
OLGA8A,PXDN,HNRNPU,PGAM1,NFE2L1,SMARCA5,CTNNA1,FADS2,RPL23,CNN3,TNS3,CTSD,SCA
I,HNRNPU,NFE2L1,SMARCA5,HNRNPA0,CTNNA1,FADS2,RPS16,RPL23,CNN3,CTSD,EIF4H,SCARB
MSB10,INA,EIF4A1,MRC2,PPP2R1A,RPL15,GTF3C4,LDHB,DNAH8,PKD1,HLA-C,SF3B1,PDIA3,SSB
ND,TMEM259,EIF5,CD81,WDR1,HMGN2,RPL26,CRMP1,LASP1,BEX3,OAZ1,PFN1,ASTN2,TLE5,NIS
4A2,RPL37A,SPEN,DDB1,PSMD5,YWHAQ,EIF1,HACD3,DCHS1,SET,EGLN1,MAP9,TPI1,CHSY1,TMS
42BPB,ITM2B,MAT2A,GOLGA8A,PXDN,HNRNPU,NFE2L1,SMARCA5,HNRNPA0,CTNNA1,RPS16,RP
P1A1,JUND,EIF5,CD81,WDR1,HMGN2,ATP2A2,CRMP1,LASP1,OAZ1,RBBP4,RTN3,PFN1,MCL1,JPT
JCHL1,MORF4L1,NCAM1,ANXA2,AGRN,TIMP2,XAF1,SARAF,COL11A1,MIDN,MARCHF6,VCL,AP3D
HACD3,DCHS1,SET,EGLN1,MAP9,TPI1,CHSY1,TMSB10,INA,MRC2,RPL19,PPP2R1A,RPL15,GTF3C
RHOA,ZNF865,CLSTN1,ATF4,PSMD2,SLC7A5,CHD4,ATP1A1,JUND,CD81,WDR1,HMGN2,ATP2A2,C
CHSY1,TMSB10,INA,EIF4A1,MRC2,PPP2R1A,RPL15,GTF3C4,LDHB,PKD1,HLA-C,SSBP2,RPS3,CHD
NFE2L1,CTNNA1,FADS2,CNN3,TNS3,CTSD,SCARB2,NACA,LPCAT1,LAMA5,RPS24,SLC22A17,BRD3
1A2,LLGL1,HK1,NCS1,RHOA,ZNF865,CLSTN1,ATF4,PSMD2,SLC7A5,CHD4,ATP1A1,JUND,CD81,H
1,CYB5R3,SF3B2,CDC42BPB,ITM2B,AP2B1,MAT2A,PXDN,HNRNPU,PGAM1,SMARCA5,HNRNPA0,C
,BASP1,VAPA,HMGB1,POLR2A,AP2M1,TPT1,SEC61A1,STMN1,ITGB1,TUBB4B,UCHL1,MORF4L1,N
,NACA,LAMA5,SLC22A17,BRD3,CIPC,PJA2,MSN,KPNB1,C1ORF43,PAPOLA,IGFBP3,COL1A2,NCS
P200,CSDE1,SMIM14,IGF2,EIF3C,ATP5F1A,EIF4B,VCP,MARCKSL1,UBB,AFF4,TRIM28,MATR3,YWH
C2,RPL19,PPP2R1A,RPL15,GTF3C4,LDHB,PKD1,SF3B1,PDIA3,SERINC1,SSBP2,ZNF43,CHD3,BASP
RIM28,MATR3,YWHAG,CKB,COL4A2,RPL37A,LRRC28,FN1,SPEN,NONO,DDB1,PSMD5,YWHAQ,EIF
,ITM2B,GTF2F1,PXDN,HNRNPU,PGAM1,NFE2L1,SMARCA5,CTNNA1,FADS2,RPL23,CNN3,TNS3,C
CHS1,SET,EGLN1,CHSY1,TMSB10,INA,EIF4A1,MRC2,RPL19,PPP2R1A,LDHB,DNAH8,PKD1,HLA-C
PEN,DDB1,YWHAQ,EIF1,HACD3,DCHS1,SET,EGLN1,MAP9,TPI1,CHSY1,TMSB10,INA,MRC2,RPL19,
,STMN1,ITGB1,TUBB4B,UCHL1,MORF4L1,ANXA2,AGRN,TIMP2,XAF1,SARAF,MIDN,RPL10,MARC
ARF1,CYB5R3,MAP4,CDC42BPB,ITM2B,AP2B1,MAT2A,GOLGA8A,PXDN,HNRNPU,NFE2L1,SMARC
TF4,PSMD2,SLC7A5,CHD4,ATP1A1,JUND,CD81,HMGN2,ATP2A2,CRMP1,LASP1,OAZ1,RTN3,PFN1
E1,SMIM14,IGF2,EIF3C,ATP5F1A,EIF4B,VCP,MARCKSL1,UBB,AFF4,TRIM28,MATR3,YWHAG,CKB,
4A8,GOLGA8A,PXDN,OGA,HNRNPA0,CTNNA1,FADS2,RPS16,LDHA,CPE,CNN3,RPL29,TNS3,CTSD,
AP4,CDC42BPB,ITM2B,MAT2A,GTF2F1,PXDN,HNRNPU,NFE2L1,HNRNPA0,CTNNA1,RPS16,CNN3,T
AP4,CDC42BPB,ITM2B,MAT2A,GTF2F1,PXDN,HNRNPU,NFE2L1,HNRNPA0,CTNNA1,RPS16,CNN3,T
S3,CTSD,SCARB2,NACA,LAMA5,RPS24,SLC22A17,BRD3,CIPC,PJA2,SLC16A7,KPNB1,RPL17,COL
COL18A1,CSDE1,SMIM14,IGF2,EIF3C,ATP5F1A,EIF4B,VCP,MARCKSL1,UBB,AFF4,TRIM28,MATR
C42BPB,ITM2B,ZNF770,TMEM170A,GOLGA8A,PXDN,HNRNPU,OGA,HNRNPA0,CTNNA1,FADS2,CP
SF3B2,CDC42BPB,ITM2B,MAT2A,PXDN,HNRNPU,OGA,PGAM1,NFE2L1,SMARCA5,HNRNPA0,FADS

1A1,MIDN,MARCHF6,VCL,AP3D1,SBF1,DNAJA1,ARF1,CYB5R3,MAP4,SF3B2,CDC42BPB,ITM2B,M
PAN3,LPCAT1,LAMA5,SLC22A17,BRD3,CIPC,PJA2,KPNB1,C1ORF43,COL1A2,LLGL1,NCS1,RHOA,
SLC22A17,BRD3,CIPC,PJA2,KPNB1,PAPOLA,COL1A2,LLGL1,NCS1,RHOA,CBX5,ZNF865,CLSTN1,
GL1,HK1,NCS1,RHOA,ZNF865,CLSTN1,ATF4,RAB6B,SLC7A5,ENC1,CHD4,ATP1A1,JUND,EIF5,CD8
CHL1,MORF4L1,NCAM1,AGRN,ZMIZ1,TIMP2,COL11A1,MIDN,MARCHF6,WSB1,VCL,AP3D1,SBF1,
COL18A1,SNRNP200,CSDE1,SMIM14,IGF2,EIF3C,ATP5F1A,EIF4B,VCP,MARCKSL1,UBB,AFF4,TR
L,TKT,SFPQ,CDH2,PDIA6,GPC3,PIP4K2B,MEGF8,SOX12,ACTN4,PRDX2,DDX3X,ARL4C,GNB2,H1-
DC42BPB,ITM2B,MAT2A,GOLGA8A,PXDN,HNRNPU,NFE2L1,SMARCA5,CTNNA1,RPS16,RPL23,CNN
F2,EIF3C,ATP5F1A,EIF4B,MARCKSL1,UBB,AFF4,TRIM28,MATR3,YWHAG,CKB,COL4A2,RPL37A,L
,ARF1,CYB5R3,AKAP12,MAP4,SF3B2,CDC42BPB,ITM2B,MAT2A,TMEM170A,SLC4A8,PXDN,HNRN
F43,CHD3,BASP1,VAPA,POLR2A,CDKL1,AP2M1,TPT1,RPL18,STMN1,ITGB1,RPL11,MORF4L1,BT
2BPB,ITM2B,MAT2A,SLC4A8,GOLGA8A,PXDN,HNRNPU,NFE2L1,SMARCA5,CTNNA1,RPS16,RPL2
L17,IGFBP3,COL1A2,LLGL1,HK1,NCS1,RPL22,RHOA,CBX5,ZNF865,KIF5B,RAB6B,SLC7A5,RPS1
,CTNNA1,FADS2,RPL23,CNN3,TPM4,TNS3,CTSD,NACA,LAMA5,SLC22A17,BRD3,CIPC,PJA2,MSN,
1,SMIM14,IGF2,EIF3C,ATP5F1A,EIF4B,VCP,MARCKSL1,UBB,AFF4,TRIM28,MATR3,YWHAG,CKB,C
DS2,LDHA,CNN3,RPSA,TNS3,EIF4H,TSPAN3,LPCAT1,LAMA5,SLC22A17,BRD3,RPS9,XRCC5,CIPC,S
A,CD63,SESN3,B4GALNT4,MTSS2,DNAJC5,APLP2,TARDBP,MXD4,HNRNPK,INIP,DUX4,CADM4,HN
T1,SEC61A1,RPS3A,RPL18,STMN1,TUBB4B,UCHL1,ANXA2,AGRN,TIMP2,XAF1,SARAF,COL11A1,M
2,RPL23,CNN3,TPM4,EIF4H,TSPAN3,LPCAT1,LAMA5,RPS24,BRD3,RPS9,CIPC,KPNB1,PAPOLA,IG
NF865,CLSTN1,SLC7A5,RPS14,CXADR,CHD4,ATP1A1,JUND,EIF5,CD81,LENG8,HMGN2,ATP2A2,C
N,OGA,PGAM1,HNRNPA0,CTNNA1,FADS2,LDHA,CPE,CNN3,RPSA,TPM4,CTSD,NACA,LPCAT1,LAM
D,EIF5,CD81,LENG8,HMGN2,ATP2A2,RPL26,CRMP1,LASP1,BEX3,OAZ1,RBBP4,PFN1,ASTN2,TLE5
P4K2B,MEGF8,SOX12,DDX3X,ITM2C,ARL4C,GNB2,APLP1,MIF,GPAT4,CHPF,MYO5A,COL6A2,B4G
MXD4,HNRNPK,H1-10,MYH10,DUX4,YWHAB,CKAP5,CADM4,LGALS3BP,UBE2H,CKAP4,HLA-B,PLP
CDKL1,AP2M1,TPT1,SEC61A1,RPS3A,OR8K1,RPL18,STMN1,ITGB1,TUBB4B,UCHL1,MORF4L1,E
,DNAJC5,APLP2,HNRNPDL,RPLP2,MXD4,H1-10,MYH10,SEPTIN11,RHOB,UBE2H,CKAP4,HLA-B,P
,MORF4L1,ANXA2,AGRN,RPLP1,ZMIZ1,TIMP2,SARAF,MIDN,MARCHF6,WSB1,RPS17,VCL,ARF1,C
HD3,BASP1,VAPA,CDKL1,AP2M1,SEC61A1,RPL18,STMN1,ITGB1,TUBB4B,UCHL1,MORF4L1,NCAM
N,TIMP2,COL11A1,MIDN,MARCHF6,WSB1,VCL,DNAJA1,ARF1,CYB5R3,MAP4,SF3B2,CDC42BPB,I
A,RPL18,STMN1,ITGB1,TUBB4B,UCHL1,MORF4L1,ANXA2,AGRN,TIMP2,XAF1,SARAF,COL11A1,M
NP200,CSDE1,SMIM14,IGF2,EIF3C,ATP5F1A,EIF4B,VCP,MARCKSL1,UBB,AFF4,TRIM28,MATR3,Y
NRNPU,PGAM1,NFE2L1,SMARCA5,HNRNPA0,CTNNA1,LDHA,CPE,CNN3,RPSA,TPM4,TNS3,CTSD,E
NB2,APLP1,MIF,COL6A2,SESN3,B4GALNT4,DNAJC5,APLP2,RPLP2,ARL6IP1,MXD4,MYL6,ANKRD
D3,CIPC,PJA2,MSN,KPNB1,C1ORF43,COL1A2,LLGL1,GAP43,NCS1,RPL22,ZNF865,CLSTN1,ATF4
I1,ITGB1,TUBB4B,UCHL1,MORF4L1,NCAM1,ANXA2,AGRN,TIMP2,XAF1,SARAF,COL11A1,MIDN,M
,CHSY1,TMSB10,INA,EIF4A1,MRC2,RPL19,RPL15,GTF3C4,LDHB,PKD1,PDIA3,SERINC1,SSBP2,ZN
RHOB,HNRNPH1,UBE2H,CKAP4,RAB1A,CALU,CTBP1,RAC1,ARHGDIA,STIP1,TOP2B,APPL1,SMCHD
BEX3,VAT1,OAZ1,RBBP4,PFN1,ASTN2,JPT2,TLE5,NISCH,CCNI,TADA2B,KHDRBS1,XPO1,SPTBN2,
4B,CTBP1,ARHGDIA,CAPRIN1,STIP1,GPC1,TOP2B,APPL1,SMCHD1,IPO5,ATP6V0C,MMP2,DNER,T
PA0,CTNNA1,FADS2,RPS16,RPL23,LDHA,CNN3,TPM4,NACA,TSPAN3,LPCAT1,LAMA5,SLC22A17,B
1,GAP43,HK1,NCS1,ZNF865,CLSTN1,RAB6B,SLC7A5,ENC1,NLGN2,CXADR,JUND,EIF5,CD81,LEN
S20,EIF5,CD81,WDR1,HMGN2,CRMP1,CCT7,OAZ1,SRSF4,RBBP4,RTN3,MCL1,JPT2,PNN,NES,RP
NAJA1,ARF1,CYB5R3,AKAP12,MAP4,SF3B2,CDC42BPB,ITM2B,AP2B1,MAT2A,GOLGA8A,PXDN,HN
2,LLGL1,NCS1,RHOA,CBX5,ZNF865,CLSTN1,SLC7A5,CHD4,ATP1A1,JUND,TMEM259,CD81,HMG
0,CSDE1,SMIM14,IGF2,EIF3C,ATP5F1A,EIF4B,VCP,MARCKSL1,UBB,AFF4,TRIM28,MATR3,YWHA

.,AP3D1,SBF1,ARF1,CYB5R3,MAP4,SF3B2,CDC42BPB,ITM2B,ZNF770,TMEM170A,GOLGA8A,PXD
6A2,SESN3,B4GALNT4,MTSS2,APLP2,HNRNPDL,TARDBP,RPLP2,NDFIP1,SHOX,ANKRD52,HNRNF
ATP2A2,CRMP1,VAT1,OAZ1,SRSF4,RTN3,ASTN2,JPT2,TLE5,NES,PRKCSH,TADA2B,SPTBN2,CSNK
,MIDN,RPL10,MARCHF6,RPS17,AP3D1,DNAJA1,CYB5R3,SF3B2,CDC42BPB,MAT2A,GTF2F1,PXDN
TF3C4,LDHB,DNAH8,PKD1,PDIA3,SSBP2,ZNF43,CHD3,BASP1,VAPA,HMGB1,POLR2A,CDKL1,TPT
STMN1,ITGB1,TUBB4B,UCHL1,MORF4L1,BTF3,NCAM1,ANXA2,AGRN,RPLP1,ZMIZ1,TIMP2,XAF1
17,BRD3,CIPC,PJA2,KPNB1,COL1A2,LLGL1,GAP43,HK1,NCS1,RPL22,RHOA,CBX5,ZNF865,CLST
IF6,RPS17,VCL,SBF1,DNAJA1,ARF1,CYB5R3,AKAP12,MAP4,SF3B2,CDC42BPB,AP2B1,MAT2A,TM
DCHS1,SET,EGLN1,MAP9,TPI1,CHSY1,TMSB10,INA,EIF4A1,MRC2,RPL19,PPP2R1A,RPL15,GTF3C
3,AFF4,TRIM28,MATR3,YWHAG,CKB,COL4A2,LRRC28,MAPT,FN1,SPEN,NONO,DDB1,PSMD5,YWH
,NISCH,PRKCSH,TADA2B,XPR1,XPO1,PFKL,HDLBP,GLG1,SFPQ,PLXNA1,PIP4K2B,SOX12,ACTN4,
NPDL,TARDBP,MXD4,INIP,CIRBP,RPL36,YWHAB,CADM4,HNRNPH1,HLA-B,PLPPR3,ARCN1,COTL1
ARCN1,COTL1,HECTD4,MAGED4B,CTBP1,ARHGDIA,CAPRIN1,STIP1,GPC1,SLC35E2B,SMCHD1,I
,ZNF770,MAT2A,TMEM170A,GTF2F1,SLC4A8,PXDN,HNRNPU,PGAM1,NFE2L1,HNRNPA0,FADS2,F
70A,GTF2F1,SLC4A8,PXDN,HNRNPU,PGAM1,NFE2L1,SMARCA5,HNRNPA0,CTNNA1,FADS2,CPE,C
JBB4B,UCHL1,MORF4L1,ANXA2,AGRN,RPLP1,ZMIZ1,TIMP2,SARAF,MIDN,RPL10,MARCHF6,WSB
2,CD63,SESN3,B4GALNT4,MTSS2,DNAJC5,APLP2,HNRNPDL,TARDBP,MXD4,MYL6,HNRNPK,INIP,C
GNB2,RCN1,H1-0,APLP1,GJA1,COL6A2,B4GALNT4,APLP2,TARDBP,RPLP2,SHOX,WLS,HNRNPK,H
,PSMD5,YWHAQ,EIF1,HACD3,DCHS1,SET,EGLN1,MAP9,TPI1,CHSY1,TMSB10,INA,EIF4A1,MRC2,F
HB,DNAH8,HLA-C,SF3B1,PDIA3,SSBP2,RPS3,CHD3,BASP1,VAPA,HMGB1,POLR2A,CDKL1,TPT1,SE
10,INA,EIF4A1,MRC2,PPP2R1A,RPL15,GTF3C4,LDHB,DNAH8,PKD1,HLA-C,SF3B1,PDIA3,SSBP2,Z
P2B1,MAT2A,GTF2F1,GOLGA8A,PXDN,HNRNPU,NFE2L1,HNRNPA0,CTNNA1,RPS16,CNN3,TNS3,C
1,CYB5R3,AKAP12,MAP4,CDC42BPB,ITM2B,ZNF770,MAT2A,GOLGA8A,PXDN,HNRNPU,PGAM1,N
,HSPD1,USP11,COL18A1,SNRNP200,CSDE1,SMIM14,IGF2,EIF3C,ATP5F1A,EIF4B,VCP,MARCKSL
DBP,ARL6IP1,NDFIP1,MXD4,TAGLN,DUX4,CKAP5,SEPTIN11,CADM4,RHOB,HNRNPH1,RAB1A,PLF
A5,CXADR,ATP1A1,JUND,TMEM259,EIF5,CD81,LENG8,HMGN2,ATP2A2,RPL26,CRMP1,LASP1,OA
C5,APLP2,HNRNPDL,MYL6,SHOX,WLS,HNRNPK,H1-10,YWHAB,HNRNPH1,UBE2H,CKAP4,RAB1A,A
,LRRC28,SPEN,NONO,DDB1,PSMD5,YWHAQ,EIF1,HACD3,DCHS1,SET,EGLN1,MAP9,TPI1,CHSY1,
SCH,CCNI,TADA2B,KHDRBS1,SPTBN2,CSNK1E,PFKL,TKT,SFPQ,CDH2,PNMA1,PDIA6,PIP4K2B,M
2R1A,RPL15,GTF3C4,LDHB,SF3B1,PDIA3,SERINC1,SSBP2,CHD3,BASP1,VAPA,HMGB1,POLR2A,T
CAPRIN1,STIP1,GPC1,SLC35E2B,MMP2,TRIB2,FAM199X,CRPPA,TMEM132A,CTDSP2,CNN2,NOV
H,TADA2B,XPR1,KHDRBS1,XPO1,SFPQ,PNMA1,PDIA6,GPC3,PIP4K2B,MEGF8,SOX12,ACTN4,RPL1
NN3,TNS3,EIF4H,SCARB2,LAMA5,SLC22A17,BRD3,RPS9,XRCC5,PJA2,MSN,KPNB1,C1ORF43,IG
F43,IGFBP3,LLGL1,HK1,NCS1,RHOA,ZNF865,CLSTN1,ENC1,NLGN2,CHD4,ATP1A1,JUND,WDR1,H
3C4,LDHB,DNAH8,PKD1,HLA-C,PDIA3,SERINC1,SSBP2,RPS3,BASP1,VAPA,HMGB1,CDKL1,AP2M1
CAM,TRIB2,PEA15,LUC7L3,SQLE,CTDSP2,NOVA2,CTSB,MDH2,CHD6,EEF1A2,CCT2,TBCD,ANXA
A-B,PLPPR3,HECTD4,CTBP1,RAC1,ARHGDIA,STIP1,GPC1,SLC35E2B,SMCHD1,IPO5,ATP6V0C,DN
AM1,ANXA2,AGRN,ZMIZ1,TIMP2,XAF1,COL11A1,MIDN,MARCHF6,VCL,AP3D1,DNAJA1,CYB5R3,A
SLC7A5,NLGN2,RPS14,CXADR,CHD4,ATP1A1,JUND,EIF5,WDR1,HMGN2,ATP2A2,CRMP1,LASP1,E
3,LLGL1,GAP43,HK1,RPL22,ZNF865,ATF4,PSMD2,SLC7A5,CHD4,ATP1A1,JUND,EIF5,CD81,WDR
,RPL11,MORF4L1,BTF3,NCAM1,ANXA2,AGRN,TIMP2,SARAF,MIDN,RPL10,MARCHF6,WSB1,RPS1
SQSTM1,VAMP2,MORF4L2,HNRNPA3,PRDX5,ACTR3,HSPA4,CAMSAP1,GRINA,PTOV1,CLSTN3,AC
NNA1,FADS2,RPSA,TPM4,TNS3,CTSD,NACA,TSPAN3,LAMA5,PJA2,MSN,RPL24,SLC16A7,KPNB1,C
NNA1,FADS2,RPSA,TPM4,TNS3,CTSD,NACA,TSPAN3,LAMA5,PJA2,MSN,RPL24,SLC16A7,KPNB1,C
NNA1,FADS2,RPSA,TPM4,TNS3,CTSD,NACA,TSPAN3,LAMA5,PJA2,MSN,RPL24,SLC16A7,KPNB1,C

PLP1,ZMIZ1,SARAF,MIDN,RPL10,MARCHF6,VCL,AP3D1,SBF1,DNAJA1,ARF1,SF3B2,ZNF770,AP2B
S1,RPL22,RHOA,ZNF865,PSMD2,KIF5B,SLC7A5,ENC1,NLGN2,RPS14,CXADR,CHD4,ATP1A1,JUND
,UCHL1,RPL11,MORF4L1,BTF3,ANXA2,AGRN,RPLP1,ZMIZ1,TIMP2,SARAF,MIDN,RPL10,MARCHF
F4,RBBP4,ASTN2,MCL1,TLE5,PRKCSH,HNRNPC,CCNI,XPR1,XPO1,HDLBP,TKT,EIF5A,CCND2,PIP4
,CAMSAP1,GRINA,PTOV1,CLSTN3,ACAP3,MDK,LAMC1,JPT1,CBX1,RAD23B,PTK7,CHD2,ASXL1,H3
B,COL4A2,RPL37A,SPEN,NONO,DDB1,YWHAQ,EIF1,HACD3,DCHS1,SET,EGLN1,TPI1,CHSY1,TMSB
2,ATP2A2,CRMP1,LASP1,BEX3,VAT1,OAZ1,RTN3,PFN1,JPT2,TLE5,PNN,NES,NISCH,PRKCSH,HNRN
A5,RPS24,SLC22A17,BRD3,RPS9,XRCC5,CIPC,RPL24,SLC16A7,KPNB1,RPL17,PAPOLA,COL1A2,
A,SLC4A8,PXDN,HNRNPU,PGAM1,NFE2L1,SMARCA5,HNRNPA0,CTNNA1,FADS2,CPE,CNN3,RPL29
B4GALNT4,MTSS2,DNAJC5,APLP2,HNRNPDL,TARDBP,MXD4,HNRNPK,INIP,RPL36,YWHAB,PTGES
P3,SURF4,LAMC1,JPT1,RAD23B,SKI,H3-3A,ABCA2,DPYSL2,NOTCH2,VEGFA,SELENON,TUBB2A,M
APA,HMGB1,CDKL1,TPT1,SEC61A1,RPS4X,STMN1,ITGB1,TUBB4B,UCHL1,RPL11,MORF4L1,BTF
,GTF3C4,LDHB,DNAH8,PKD1,HLA-C,PDIA3,SSBP2,ZNF43,RPS3,BASP1,VAPA,CDKL1,TPT1,RPS3A
2,ATP6V0C,TRIB2,SQLE,HS6ST1,TIMP3,CTDSP2,NOVA2,CTSB,ZC3H7B,CHD6,EEF1A2,TBCD,CPSF
ITM2B,ZNF770,MAT2A,TMEM170A,GTF2F1,GOLGA8A,PXDN,HNRNPU,NFE2L1,SMARCA5,HNRNP
3A,ITGB1,TUBB4B,UCHL1,RPL11,MORF4L1,BTF3,NCAM1,AGRN,RPLP1,XAF1,SARAF,MIDN,RPL
1,UBB,AFF4,TRIM28,MATR3,YWHAG,CKB,COL4A2,LRRC28,FN1,SPEN,NONO,DDB1,PSMD5,YWHA
IP4K2B,MEGF8,SOX12,ACTN4,MMP14,PRDX2,DDX3X,ITM2C,ARL4C,GNB2,APLP1,RPS7,RPS19,G
WHAQ,EIF1,HACD3,DCHS1,SET,EGLN1,MAP9,TPI1,CHSY1,TMSB10,INA,EIF4A1,MRC2,RPL19,PPP2
F4,PSMD2,SLC7A5,CHD4,ATP1A1,JUND,EIF5,CD81,WDR1,HMGN2,ATP2A2,RPL26,CRMP1,LASP1,
MSAP1,GRINA,NCKAP1,MLEC,CLSTN3,ACAP3,MDK,DDX42,LAMC1,JPT1,SKI,ASXL1,CNTNAP2,DPY
LDHB,DNAH8,HLA-C,SF3B1,PDIA3,SSBP2,ZNF43,CHD3,BASP1,VAPA,HMGB1,POLR2A,CDKL1,TPT
A5,NLGN2,RPS14,CXADR,CHD4,ATP1A1,JUND,RPS20,CD81,LENG8,WDR1,HMGN2,ATP2A2,CRMP
LC25A3,RPLP2,INIP,MYH10,RPL36,YWHAB,PTGES3,CKAP5,CADM4,HNRNPH1,HLA-B,PLPPR3,AR
5,CD81,HMGN2,ATP2A2,CRMP1,LASP1,BEX3,VAT1,RBBP4,PFN1,MCL1,TLE5,CCNI,TADA2B,XPR
,CYB5R3,PAFAH1B1,AKAP12,MAP4,CDC42BPB,ITM2B,AP2B1,MAT2A,TMEM170A,SLC4A8,GOLG
B4GALNT4,APLP2,NDFIP1,MXD4,TAGLN,HNRNPK,H1-10,MYH10,NARS1,DUX4,YWHAB,CKAP5,SEF
3,SERINC1,SSBP2,RPS3,CHD3,BASP1,VAPA,TPT1,SEC61A1,STMN1,ITGB1,TUBB4B,UCHL1,MORF
,SF3B2,CDC42BPB,ITM2B,ZNF770,GTF2F1,SLC4A8,GOLGA8A,PXDN,HNRNPU,OGA,NFE2L1,HNF
NRNPA0,CTNNA1,FADS2,CPE,CNN3,RPL29,RPSA,TNS3,CTSD,EIF4H,SCARB2,LPCAT1,LAMA5,SLC
MSN,SLC16A7,KPNB1,C1ORF43,PAPOLA,IGFBP3,LLGL1,RPL14,RHOA,PTCD3,ATF4,PSMD2,NR2
K1,NCS1,RHOA,ZNF865,CLSTN1,ATF4,NR2F1,RAB6B,SLC7A5,ENC1,NLGN2,GLIPR1L2,RPS14,CX
TSS2,DNAJC5,APLP2,HNRNPDL,NDFIP1,TSR2,MXD4,TAGLN,ANKRD52,HNRNPK,H1-10,INIP,MYH
1,RPS4X,RPS3A,RPL18,STMN1,ITGB1,TUBB4B,UCHL1,RPL11,MORF4L1,BTF3,NCAM1,ANXA2,AG
FADS2,RPS16,TPM4,CTSD,SCARB2,TSPAN3,LPCAT1,LAMA5,SLC22A17,PJA2,KPNB1,PAPOLA,CO
PLD3,HSPD1,USP11,COL18A1,SNRNP200,CSDE1,SMIM14,IGF2,EIF3C,ATP5F1A,EIF4B,VCP,MARC
2,L1CAM,LAPTM4B,HSBP1,FAM199X,CRPPA,TMEM132A,CNN2,SRSF5,CHD6,EEF1A2,ANXA5,PPP1
F2,EEF1A2,TBCD,CPSF1,PGD,SQSTM1,VAMP2,GAS7,PDXK,PKN1,HSPA4,CAMSAP1,GRINA,NCKA
GD,SQSTM1,VAMP2,GAS7,PRDX5,PDXK,PKN1,HSPA4,CAMSAP1,GRINA,NCKAP1,PTOV1,MLEC,AC
SL1,UBB,AFF4,TRIM28,MATR3,YWHAG,CKB,COL4A2,RPL37A,MAPT,FN1,SPEN,NONO,DDB1,YWH
CBX5,ATF4,NR2F1,SLC7A5,CXADR,TMEM259,EIF5,HMGN2,RPL26,LASP1,VAT1,SRSF4,RBBP4,RT
IF6,SBF1,DNAJA1,ARF1,MAP4,SF3B2,CDC42BPB,ITM2B,ZNF770,MAT2A,TMEM170A,GTF2F1,SLC
5,BRD3,RPL24,SLC16A7,KPNB1,C1ORF43,PAPOLA,IGFBP3,LLGL1,RPL14,RHOA,PTCD3,CBX5,NF
NISCH,PRKCSH,TADA2B,KHDRBS1,PFKL,GLG1,SFPQ,PEBP1,PNMA1,PDIA6,PIP4K2B,MEGF8,SOX
2A,AP2M1,TPT1,SEC61A1,RPS4X,RPL18,ITGB1,TUBB4B,UCHL1,MORF4L1,ANXA2,AGRN,RPLP1,Z

GTF2F1,SLC4A8,PXDN,HNRNPU,OGA,PGAM1,SMARCA5,CTNNA1,FADS2,LDHA,CPE,CNN3,TPM4,C
B2,CDC42BPB,ZNF770,MAT2A,TMEM170A,GTF2F1,SLC4A8,PXDN,HNRNPU,OGA,PGAM1,SMAR(
,NISCH,PRKCSH,CCNI,KHDRBS1,XPO1,SPTBN2,CSNK1E,GLG1,SFPQ,TPM1,CDH2,PNMA1,GPC3,
.,HDLBP,TKT,SFPQ,CDH2,PNMA1,PDIA6,GPC3,PIP4K2B,MEGF8,SOX12,ARL4C,GNB2,MIF,HMGC
,RPL14,HK1,NCS1,RHOA,PTCD3,CLSTN1,ATF4,PSMD2,NR2F1,KIF5B,RAB6B,ENC1,CHD4,ATP1A
OLM1,SESN3,B4GALNT4,MTSS2,APLP2,HNRNPDL,MXD4,MYL6,ANKRD52,HNRNPK,H1-10,INIP,P
IEM170A,PXDN,PGAM1,NFE2L1,SMARCA5,HNRNPA0,FADS2,RPS16,CNN3,RPL29,TPM4,TNS3,CT
CDH2,PNMA1,PDIA6,GPC3,DDR1,PIP4K2B,MEGF8,SOX12,RPL12,ARL4C,MRFAP1,MAGED2,APLF
P1,GPC1,SLC35E2B,SMCHD1,IPO5,ATP6V0C,DNER,TRIB2,PEA15,SQLE,SERF2,HS6ST1,TIMP3,N(
P1,GPC1,SLC35E2B,SMCHD1,IPO5,ATP6V0C,DNER,TRIB2,PEA15,SQLE,SERF2,HS6ST1,TIMP3,N(
I2AZ1,UBQLN2,CST3,ZFP36L1,CXXC5,BRD2,GNG4,NXN,PI4KA,COX7C,ZNF512B,CEMIP2,HNRNP
SRSF5,C5ORF24,ZC3H7B,TOMM20,U2AF2,SUGP2,CPSF1,PPP1CB,MEST,HNRNPL,ABCA3,ALKBH
COL4A2,RPL37A,LRRC28,SPEN,DDB1,PSMD5,YWHAQ,HACD3,DCHS1,SET,EGLN1,TPI1,CHSY1,T
7,BRD3,RPS9,CIPC,SLC16A7,C1ORF43,PAPOLA,IGFBP3,LLGL1,NCS1,RHOA,RAB6B,CXADR,CHD
1,SLC35E2B,IPO5,ATP6V0C,SKP1,DNER,L1CAM,LAPTM4B,TRIB2,PEA15,LUC7L3,SQLE,TIMP3,D
SNX22,PGD,SQSTM1,VAMP2,GAS7,GDI2,HNRNPL,TM9SF2,LMNB2,YEATS2,HSPA4,CAMSAP1,GRI
L1CAM,TRIB2,PEA15,SQLE,FAM199X,SERF2,HS6ST1,TIMP3,NOVA2,CTSB,NUCB1,CHD6,RNF187,
IDFIP1,ST3GAL2,MYL6,HNRNPK,H1-10,MYH10,DUX4,PTGES3,CKAP5,CADM4,LAMP1,RHOB,HNR
AKAP12,SF3B2,ITM2B,ZNF770,MAT2A,TMEM170A,GTF2F1,PXDN,HNRNPU,PGAM1,NFE2L1,HNR
NN3,RPL29,RPSA,TPM4,TNS3,TSPAN3,LPCAT1,LAMA5,BRD3,RPS9,XRCC5,PJA2,RPL24,KPNB1,R
RF4L1,ANXA2,AGRN,RPLP1,TIMP2,SARAF,RPL10,MARCHF6,WSB1,RPS17,VCL,AP3D1,SBF1,DNA
,ANXA2,AGRN,TIMP2,XAF1,COL11A1,MIDN,RPL10,MARCHF6,WSB1,RPS17,VCL,DNAJA1,ARF1,C
1,SSBP2,BASP1,VAPA,HMGB1,POLR2A,TPT1,SEC61A1,RPS4X,RPS3A,STMN1,ITGB1,TUBB4B,UC
,GRINA,ACAP3,MDK,LAMC1,JPT1,CHD2,ORAI2,SKI,ASXL1,H3-3A,HDGF,SRP14,TSPAN6,SELENO
,CDH6,SFPQ,EIF5A,CCND2,CDH2,PNMA1,PDIA6,SULF1,MEGF8,SOX12,DDX3X,NANOG,GNB2,RC
YB5R3,MAP4,CDC42BPB,MAT2A,TMEM170A,GOLGA8A,PXDN,PGAM1,NFE2L1,HNRNPA0,FADS2,
SMARCA5,LARP1,HNRNPA0,CTNNA1,FADS2,CPE,CNN3,CTSD,EIF4H,SCARB2,NACA,LPCAT1,LAMA
WDR1,HMGN2,RPL26,BEX3,OAZ1,RBBP4,MCL1,JPT2,PNN,SPC25,RPL31,PRKCSH,HNRNPC,LRP
NYNRIN,TCF25,PTBP1,EIF3E,DYNLL2,LUZP6,GRB2,PCSK1N,UBE2R2,ATXN10,ACTN1,BNIP3L,MA
X5,RPL9,CLSTN1,ATF4,KIF5B,RAB6B,GLIPR1L2,ATP1A1,JUND,TMEM259,RPS20,EIF5,CD81,WD
,TRIM28,MATR3,YWHAG,CKB,COL4A2,LRRC28,FN1,NONO,DDB1,YWHAQ,EIF1,HACD3,DCHS1,S
D1,IPO5,ATP6V0C,SKP1,DNER,L1CAM,TRIB2,PEA15,HSBP1,SQLE,FAM199X,TIMP3,NOVA2,CTS
HECTD4,CTBP1,STIP1,GPC1,SLC35E2B,SMCHD1,IPO5,ATP6V0C,SKP1,DNER,L1CAM,LAPTM4B,
TGES3,CKAP5,SEPTIN11,LAMP1,HNRNPH1,PLPPR3,COTL1,HECTD4,CTBP1,ARHGDIA,STIP1,SLC
SEPTIN11,LAMP1,RHOB,HNRNPH1,CKAP4,HLA-B,COTL1,CTBP1,RAC1,STIP1,GPC1,TOP2B,SMC
GAM1,NFE2L1,HNRNPA0,FADS2,CPE,CNN3,RPSA,TPM4,TNS3,CTSD,EIF4H,NACA,TSPAN3,LPCAT
Z1,ASTN2,MCL1,TLE5,NISCH,HNRNPC,CCNI,XPR1,KLHL9,XPO1,SLC3A2,PFKL,TKT,SFPQ,TPM1,
12,PRDX2,DDX3X,CLTC,MAGED2,GNB2,H1-0,MIF,MYO5A,ANXA6,SESN3,B4GALNT4,DNAJC5,AP
CHS1,SET,MAP9,TPI1,CHSY1,INA,EIF4A1,MRC2,RPL19,PPP2R1A,RPL15,GTF3C4,PKD1,HLA-C,S
JA2,SLC16A7,KPNB1,COL1A2,HK1,NCS1,RHOA,ZNF865,CLSTN1,PSMD2,SLC7A5,NLGN2,ATP1A
IA1,RPS16,RPL23,CNN3,TPM4,TNS3,EIF4H,SCARB2,LPCAT1,RPS24,SLC22A17,RPS9,CIPC,RPL24
AT2A,TMEM170A,GTF2F1,SLC4A8,GOLGA8A,HNRNPU,OGA,PGAM1,NFE2L1,SMARCA5,HNRNPA0
S7,PABPN1,HNRNPL,ALKBH5,PTPRA,UBE4A,PKN1,YEATS2,KLC1,VPS35,PTOV1,ACAP3,SURF4,T
3BP,HLA-B,PLPPR3,CTBP1,RAC1,ARHGDIA,GPC1,SLC35E2B,SMCHD1,IPO5,ATP6V0C,DNER,BCL
HOA,CBX5,ZNF865,NR2F1,RAB6B,SLC7A5,ENC1,NLGN2,RPS14,CXADR,CHD4,ATP1A1,JUND,EIF

M2B,MAT2A,TMEM170A,SLC4A8,PXDN,HNRNPU,OGA,PGAM1,NFE2L1,SMARCA5,LARP1,HNRNPA
GB1,POLR2A,CDKL1,AP2M1,TPT1,SEC61A1,RPS4X,RPS3A,RPL18,STMN1,ITGB1,TUBB4B,UCHL1
2A17,BRD3,XRCC5,C1ORF43,PAPOLA,RPL14,HK1,RHOA,PTCD3,ZNF865,RPL9,ATF4,PSMD2,ENC
GB1,TUBB4B,UCHL1,RPL11,BTF3,AGRN,RPLP1,TIMP2,MIDN,RPL10,MARCHF6,WSB1,RPS17,VCL
35R3,MAP4,SF3B2,CDC42BPB,ITM2B,AP2B1,MAT2A,GOLGA8A,PXDN,HNRNPU,NFE2L1,SMARCA
,CADM4,SND1,HNRNPH1,PLPPR3,ARCN1,COTL1,HECTD4,CTBP1,ARHGDIA,STIP1,SLC35E2B,APP
3CA2,DPYSL2,VEGFA,TUBB2A,ARF3,CCND1,SMARCA4,ATG2B,MAPRE1,FAM171A2,RABL6,ODC1
IISCH,TADA2B,KHDRBS1,KLHL9,HDLBP,RPS25,TKT,GLG1,SFPQ,TPM1,EIF5A,CDH2,PNMA1,DDR
PL19,PPP2R1A,RPL15,LDHB,DNAH8,PKD1,HLA-C,SF3B1,PDIA3,SERINC1,SSBP2,RPS3,BASP1,VAP
TF4,RAB6B,SLC7A5,ENC1,NLGN2,ATP1A1,JUND,CD81,WDR1,HMGN2,RPL26,CRMP1,VAT1,OAZ1,
,BEX3,OAZ1,SRSF4,RBBP4,PFN1,MCL1,JPT2,PNN,RPL31,PRKCSH,HNRNPC,CCNI,LRPPRC,TADA
I4,MMP14,ITM2C,ARL4C,GNB2,RCN1,APLP1,TLN1,CHPF,MYO5A,COL6A2,CD63,NGRN,B4GALNT
1,PNN,NES,RPL31,HNRNPC,XPR1,HDLBP,RPS25,TPM1,EIF5A,PEBP1,CCND2,CDH2,PDIA6,PIP4K2
,CTSD,EIF4H,NACA,LAMA5,BRD3,KPNB1,PAPOLA,COL1A2,LLGL1,NCS1,PTCD3,ZNF865,CLSTN1,
PS4X,RPS3A,RPL18,STMN1,ITGB1,TUBB4B,UCHL1,RPL11,BTF3,NCAM1,AGRN,RPLP1,SARAF,MI
HECTD4,CCT6A,MAGED4B,CTBP1,ARHGDIA,STIP1,GPC1,TOP2B,SLC35E2B,HDAC2,IPO5,ATP6V0
RCA5,LARP1,HNRNPA0,CTNNA1,FADS2,RPS16,RPL23,CNN3,RPL29,RPSA,TPM4,TNS3,CTSD,EIF4
ID2,KIF5B,SLC7A5,CXADR,TMEM259,RPS20,EIF5,CD81,WDR1,ATXN7L3B,HMGN2,ATP2A2,CRMP
15,GTF3C4,LDHB,DNAH8,PKD1,HLA-C,SERINC1,SSBP2,CHD3,BASP1,VAPA,HMGB1,POLR2A,AP2
VDR1,ATP2A2,RPL26,CRMP1,SRSF4,RBBP4,SPC25,RPL31,PRKCSH,TADA2B,XPR1,KHDRBS1,CSN
AZ1,SRSF4,PFN1,MCL1,JPT2,TLE5,NES,TADA2B,KHDRBS1,SPTBN2,CSNK1E,PFKL,HDLBP,RPS2
MD5,YWHAQ,EIF1,HACD3,SET,EGLN1,MAP9,TPI1,CHSY1,TMSB10,INA,EIF4A1,RPL19,PPP2R1A,R
EEF1A2,TBCD,SNX22,SQSTM1,VAMP2,GAS7,PARP1,YEATS2,HSPA4,CAMSAP1,GRINA,PTOV1,ACA
1,MRC2,PPP2R1A,RPL15,GTF3C4,DNAH8,PKD1,HLA-C,SSBP2,ZNF43,BASP1,VAPA,HMGB1,POLR
MEM30A,DNER,BCLAF1,LAPTM4B,TRIB2,SQLE,MAP1A,NOVA2,SRSF5,MDH2,CHD6,U2AF2,ANXA5
SLC22A17,BRD3,CIPC,MSN,KPNB1,RPL17,C1ORF43,PAPOLA,IGFBP3,RPL14,HK1,PTCD3,CBX5,
SLC22A17,BRD3,CIPC,MSN,KPNB1,RPL17,C1ORF43,PAPOLA,IGFBP3,RPL14,HK1,PTCD3,CBX5,
,RPL31,HNRNPC,LRPPRC,XPR1,KHDRBS1,SPTBN2,SLC3A2,CSNK1E,PFKL,RPS25,TKT,EIF5A,CD
B,HNRNPH1,HLA-B,PLPPR3,ARCN1,COTL1,HECTD4,CTBP1,ARHGDIA,STIP1,GPC1,CDK16,SLC35
AP3,MDK,LAMC1,JPT1,CBX1,PTK7,CHD2,ASXL1,H3-3A,SRP14,ABCA2,DPYSL2,VEGFA,CCND1,SM
R3,AHCYL1,GRINA,COL2A1,ACAP3,LAMC1,CBX1,RAB5B,SKI,ASXL1,H3-3A,CNTNAP2,HDGF,RNF
T,EGLN1,MAP9,CHSY1,TMSB10,INA,EIF4A1,MRC2,RPL19,PPP2R1A,GTF3C4,LDHB,SF3B1,PDIA3
,ACTN4,DDX3X,ARL4C,CLTC,GNB2,APLP1,RPL35,MIF,GPAT4,CHPF,JUN,COL6A2,GOLM1,SESN3
LNT4,APLP2,HNRNPDL,RPLP2,MYL6,HNRNPK,H1-10,MYH10,DUX4,YWHAB,CADM4,CKAP4,RAB1
RPS7,RPL35,CHPF,MYO5A,CTNNB1,RPL35A,DNAJC5,HNRNPDL,SLC25A3,ARL6IP1,NDFIP1,TSR2
P1,HNRNPA0,FADS2,CNN3,RPL29,TPM4,TNS3,CTSD,EIF4H,TSPAN3,LPCAT1,LAMA5,RPS24,SLC2
H1-10,CIRBP,NARS1,PTGES3,CADM4,LAMP1,RHOB,HNRNPH1,CKAP4,RPS12,SEPTIN3,CTBP1,HM
,UCHL1,MORF4L1,ANXA2,AGRN,TIMP2,XAF1,COL11A1,MIDN,MARCHF6,VCL,AP3D1,DNAJA1,CY
35E2B,SMCHD1,IPO5,ATP6V0C,SKP1,DNER,TRIB2,HS6ST1,TIMP3,NOVA2,CTSB,NUCB1,CDH11,C
F43,CHD3,BASP1,POLR2A,AP2M1,TPT1,RPS4X,RPS3A,STMN1,ITGB1,TUBB4B,MORF4L1,BTF3,N
N,RPL10,MARCHF6,VCL,ARF1,CYB5R3,MAP4,SF3B2,CDC42BPB,ITM2B,ZNF770,AP2B1,MAT2A,
6IP1,MXD4,MYL6,WLS,HNRNPK,H1-10,INIP,DUX4,YWHAB,PTGES3,CADM4,HNRNPH1,CKAP4,RA
CAF1,PRPF6,ATP6AP1,TERF2IP,TOP1,RPS15A,RAPGEF1,TUBB2A,ARF3,UBXN4,PPP2CA,MAPK1,
,TADA2B,KHDRBS1,CSNK1E,PFKL,TKT,SFPQ,PEBP1,CDH2,PNMA1,PDIA6,GPC3,PIP4K2B,SOX12,
F2F1,SLC4A8,PXDN,HNRNPU,PGAM1,NFE2L1,SMARCA5,HNRNPA0,CTNNA1,FADS2,LDHA,CPE,C

DNAH8,PKD1,SF3B1,PDIA3,SERINC1,SSBP2,ZNF43,RPS3,HMGB1,POLR2A,AP2M1,TPT1,SEC61A1
LN1,CHPF,MYO5A,COL6A2,CD63,SESN3,B4GALNT4,MTSS2,DNAJC5,APLP2,HNRNPDL,ST3GAL2,
CTNNA1,FADS2,RPL23,CPE,RPL29,TPM4,TNS3,TSPAN3,LPCAT1,LAMA5,RPS24,SLC22A17,BRD3,
IB,POLR2B,SDF4,SMARCC1,ATP6V1B2,HNRNPM,SDHA,NKTR,YWHAH,LSS,STT3B,TNKS2,LONP1,
17,VCL,AP3D1,DNAJA1,CYB5R3,MAP4,SF3B2,ITM2B,MAT2A,TMEM170A,GTF2F1,GOLGA8A,PXD
PP2R1A,GTF3C4,LDHB,PKD1,HLA-C,SF3B1,SSBP2,ZNF43,CHD3,VAPA,HMGB1,CDKL1,TPT1,SEC
A,DDX3X,RPS15,RPL30,CLTC,EIF3L,H1-0,IK,RPS19,CHPF,NGRN,HNRNPDL,RPLP2,DDOST,INIP,N
L29,RPSA,EIF4H,NACA,LPCAT1,LAMA5,RPS9,XRCC5,CIPC,PJA2,RPL24,SLC16A7,PAPOLA,IGFBP
,SMARCA4,DSTN,AEBP1,ATP5PB,PTBP1,EIF3E,DYNLL2,PCSK1N,CAPZB,UBE2R2,SCAF11,ATXN1
SLC4A8,SMARCA5,LARP1,HNRNPA0,CTNNA1,CPE,CNN3,RPL29,RPSA,TNS3,CTSD,TSPAN3,LPCAT
,RPL31,NISCH,HNRNPC,CCNI,TADA2B,KHDRBS1,SLC3A2,CSNK1E,PFKL,CDH6,SFPQ,PEBP1,CDH
JSP4,DLK1,SDF4,PTP4A2,SLIT3,STMN3,LONP1,PLXNA3,ATXN2L,ID4,ZFAND3,VANGL2,RAI1,PFN2,
,SQSTM1,VAMP2,GAS7,GDI2,PARP1,TM9SF2,PDXK,PKN1,YEATS2,HSPA4,GRINA,VPS35,PTOV1,M
,VAPA,HMGB1,CDKL1,TPT1,SEC61A1,STMN1,ITGB1,TUBB4B,UCHL1,RPL11,MORF4L1,BTF3,NC
19,PPP2R1A,RPL15,GTF3C4,LDHB,PKD1,HLA-C,SF3B1,PDIA3,SSBP2,ZNF43,CHD3,BASP1,VAPA,H
31,IGFBP3,LLGL1,RPL14,RHOA,CBX5,ATF4,PSMD2,NR2F1,RAB6B,GLIPR1L2,RPS14,CXADR,ATP
,EEF1A2,CCT2,TBCD,SUGP2,ANXA5,CPSF1,VAMP2,PABPN1,HNRNPL,PARP1,TAF15,ABCA3,ALKB
C16A7,KPNB1,PAPOLA,IGFBP3,LLGL1,HK1,NCS1,RPL22,RHOA,CLSTN1,ATF4,PSMD2,NR2F1,RA
XNDC5,EP300,SF1,RETREG2,BAZ1B,SEC16A,MGAT4B,CXXC5,SLC39A7,MAP6,DIP2B,ATRX,GNG
2,RPS14,ATP1A1,RPS20,CD81,LENG8,WDR1,ATXN7L3B,RPL26,LASP1,SRSF4,RBBP4,PFN1,MCL
PRKDC,TDRD5,UBE2R2,SCAF11,BNIP3L,CLIP3,PLCG1,MAP2K2,HNRNPUL2-BSCL2,FLNB,POLR2
PTIN3,ZNF454,COTL1,MAGED4B,ARHGDIA,CAPRIN1,STIP1,GPC1,CDK16,HDAC2,IPO5,DNER,PE
F865,RPL9,ATF4,GLIPR1L2,RPS14,CXADR,CHD4,JUND,TMEM259,RPS20,LENG8,WDR1,ATXN7L3
3,PLD3,HSPD1,USP11,COL18A1,SNRNP200,CSDE1,SMIM14,IGF2,EIF3C,ATP5F1A,EIF4B,VCP,MA
A,HMGB1,POLR2A,CDKL1,AP2M1,TPT1,SEC61A1,STMN1,ITGB1,TUBB4B,UCHL1,RPL11,BTF3,N
CADM4,RHOB,RAB1A,PLPPR3,COTL1,MAGED4B,CTBP1,ARHGDIA,TOP2B,SMCHD1,IPO5,SKP1,B
,NARS1,CKAP5,CADM4,UBE2H,CKAP4,RPS12,PLPPR3,SEPTIN3,RAC1,ARHGDIA,GPC1,SMCHD1,A
AT1,LAMA5,RPS24,SLC22A17,BRD3,XRCC5,C1ORF43,PAPOLA,RPL14,HK1,RHOA,PTCD3,ZNF86
1,LGALS3BP,HLA-B,PLPPR3,ARCN1,HECTD4,CTBP1,RAC1,ARHGDIA,STIP1,GPC1,CDK16,SLC35
1CAM,TRIB2,PEA15,HSBP1,FAM199X,DLG5,TMEM132A,CTDSP2,CPSF2,NOVA2,CTSB,CDH11,M
ICL1,NES,RPL31,NISCH,CCNI,LRPPRC,KHDRBS1,SPTBN2,PFKL,TKT,TPM1,CCND2,CDH2,PDIA6,
L11,MORF4L1,BTF3,AGRN,RPLP1,SARAF,MIDN,RPL10,MARCHF6,WSB1,RPS17,VCL,AP3D1,DNA
RPS9,PJA2,KPNB1,C1ORF43,PAPOLA,HK1,RPL22,RPL9,CLSTN1,RAB6B,SLC7A5,ATP1A1,JUND,T
,TPT1,RPS4X,RPS3A,OR8K1,STMN1,ITGB1,TUBB4B,RPL11,MORF4L1,ANXA2,AGRN,RPLP1,ZMIZ
,JUND,TMEM259,EIF5,CD81,LENG8,WDR1,ATXN7L3B,HMGN2,RPL26,CRMP1,LASP1,PFN1,ASTN
,STMN1,ITGB1,TUBB4B,UCHL1,MORF4L1,BTF3,NCAM1,AGRN,TIMP2,MARCHF6,SBF1,CYB5R3,
L4A2,RPL37A,LRRC28,SPEN,NONO,DDB1,PSMD5,YWHAQ,EIF1,HACD3,DCHS1,SET,EGLN1,MAP
,EIF4H,SCARB2,NACA,SLC22A17,BRD3,PJA2,MSN,KPNB1,C1ORF43,COL1A2,LLGL1,GAP43,NC
1,UBE2R2,HSPB1,SON,CLIP3,FKBP8,DLK1,MSL1,GIGYF1,SDHA,NKTR,STMN3,LSS,FBLN1,STT3
XNB2,TTC3,PLD3,HSPD1,USP11,COL18A1,SNRNP200,CSDE1,SMIM14,IGF2,EIF3C,ATP5F1A,EIF
S2,RPS16,RPL23,LDHA,CNN3,RPSA,TPM4,TNS3,CTSD,EIF4H,SCARB2,NACA,LPCAT1,LAMA5,RPS
D81,COL5A2,ATXN7L3B,ATP2A2,RPL26,SRSF4,RBBP4,PFN1,MCL1,TLE5,PNN,NISCH,LRPPRC,X
H2,PNMA1,PDIA6,GPC3,PIP4K2B,RAB7A,CCN2,ACTN4,RPL12,ITM2C,EIF3L,H1-0,RPS5,APLP1,R
PPL1,ATP6V0C,TMEM30A,GLUL,PEA15,DLG5,HS6ST1,CPSF2,PGRMC1,C5ORF24,CDH11,TM9SF
CHD1,IPO5,ATP6V0C,DNER,L1CAM,TRIB2,PTTG1IP,SQLE,MAP1A,CDKN1A,SERF2,TIMP3,CTDSP

ABCA2,PYGB,PRPF6,OPTN,PPP2CA,MAPK1,MAPRE1,DDAH2,RABL6,DYNC1LI2,NYNRIN,TCF25,AD
DLK1,PHC2,YWHAH,STMN3,LSS,PITPNM1,STXBP1,ISYNA1,TUG1,MESD,HYOU1,ID4,ZFAND3,MAP
O2,HSPA4,ETNK1,KLC1,VPS35,ACAP3,LAMC1,JPT1,ORAI2,SKI,ASXL1,PSMB2,HDGF,PRPF6,ATP6
,MAP9,CHSY1,TMSB10,INA,EIF4A1,MRC2,RPL19,PPP2R1A,GTF3C4,LDHB,PKD1,SF3B1,PDIA3,SE
NF43,CHD3,HMGB1,CDKL1,TPT1,SEC61A1,RPS4X,RPS3A,STMN1,ITGB1,TUBB4B,UCHL1,MORF4
1,ZFAND5,DIDO1,DUSP4,DLK1,CORO1C,LSS,PITPNM1,STXBP1,ISYNA1,STT3A,PLXNA3,USP7,IDH
,LMNB2,PRDX5,YEATS2,ACO2,HSPA4,PLOD1,CAMSAP1,KLC1,GRINA,VPS35,PTOV1,MLEC,CLSTN
PL18,ITGB1,TUBB4B,UCHL1,RPL11,MORF4L1,BTF3,AGRN,RPLP1,SARAF,MIDN,RPL10,MARCHF
9X,SHISAL1,HS6ST1,NUCB1,SRSF5,C5ORF24,CDH11,ZC3H7B,RNF187,TBCD,SQSTM1,KLHL24,M
9X,SHISAL1,HS6ST1,NUCB1,SRSF5,C5ORF24,CDH11,ZC3H7B,RNF187,TBCD,SQSTM1,KLHL24,M
9X,SHISAL1,HS6ST1,NUCB1,SRSF5,C5ORF24,CDH11,ZC3H7B,RNF187,TBCD,SQSTM1,KLHL24,M
3,WDR1,HMGN2,CRMP1,LASP1,VAT1,OAZ1,SRSF4,RBBP4,PFN1,ASTN2,MCL1,TLE5,PRKCSH,LRP
MNB2,RPL27,PTPRA,PRDX5,UBE4A,CAMSAP1,KLC1,VPS35,COL2A1,PTOV1,MLEC,ACAP3,SURF4,
COL11A1,MIDN,RPL10,MARCHF6,VCL,AP3D1,SBF1,ARF1,AKAP12,MAP4,CDC42BPB,ITM2B,ZNF7
VAPA,HMGB1,CDKL1,TPT1,SEC61A1,RPS4X,RPS3A,STMN1,ITGB1,TUBB4B,UCHL1,MORF4L1,AN
4L1,BTF3,AGRN,RPLP1,SARAF,MIDN,RPL10,MARCHF6,WSB1,RPS17,VCL,AP3D1,DNAJA1,ARF1,C
A1,MRC2,PPP2R1A,RPL15,GTF3C4,PKD1,HLA-C,SF3B1,PDIA3,SERINC1,ZNF43,RPS3,BASP1,VAPA
SYP,PSMA7,PPDPF,CIC,TRAF4,RBBP7,MSMO1,WDR45B,NAP1L1,KIRREL1,EP300,DGCR2,KDM5B
PS20,CD81,WDR1,ATXN7L3B,HMGN2,RPL26,CRMP1,LASP1,BEX3,OAZ1,SRSF4,RBBP4,MCL1,JPT
MC1,NOVA2,CTSB,CDH11,ZC3H7B,CHD6,EEF1A2,TBCD,PGD,VAMP2,GDI2,ACTR3,PKN1,CAMSAP
KHDRBS1,KLHL9,SPTBN2,SLC3A2,HDLBP,GLG1,PEBP1,CCND2,CDH2,PDIA6,GPC3,DDR1,PLXNA
H1-10,YWHAB,LAMP1,HNRNPH1,CCT6A,RAC1,STIP1,CDK16,TOP2B,SLC35E2B,APPL1,SMCHD1,I
,RHOB,HNRNPH1,UBE2H,RAB1A,PLPPR3,MAGED4B,CTBP1,ARHGDIA,CAPRIN1,STIP1,GPC1,CDK
HD3,HMGB1,CDKL1,TPT1,SEC61A1,RPS4X,RPS3A,STMN1,ITGB1,TUBB4B,UCHL1,MORF4L1,NC
2,PCSK1N,CAPZB,ATXN10,ACTN1,KDM1A,MAP7D1,CLIP3,FKBP8,PLCG1,ZFAND5,DIDO1,DLK1,CC
UBB4B,UCHL1,RPL11,MORF4L1,ANXA2,AGRN,RPLP1,TIMP2,SARAF,MIDN,RPL10,MARCHF6,WSI
LP2,HNRNPDL,RPLP2,MXD4,MYL6,ANKRD52,WLS,HNRNPK,MYH10,DUX4,CADM4,HNRNPH1,CKA
ULF1,MEGF8,ACSL6,SOX12,CCN2,MMP14,PRDX2,RPS15,ITM2C,ARL4C,MAGED2,GNB2,RCN1,R
5,ANXA6,NGRN,RPS10,MTSS2,APLP2,SLC25A3,RPLP2,ARL6IP1,DDOST,ANKRD52,WLS,HNRNPK,H
,RPS4X,RPS3A,RPL18,ITGB1,TUBB4B,RPL11,MORF4L1,BTF3,AGRN,RPLP1,SARAF,MIDN,RPL10,
MEM248,LAMC1,CCT4,ORAI2,GCN1,SKI,PRPF6,TERF2IP,VEGFA,SELENON,TUBB2A,MVB12B,MA
L30,CLTC,EIF3L,MRFAP1,MAGED2,H1-0,IK,RPS19,GPAT4,CHPF,ANXA6,NGRN,DNAJC5,HNRNPDL
,ATP1A1,JUND,HMGN2,ATP2A2,CRMP1,LASP1,BEX3,VAT1,RTN3,PFN1,TLE5,CCNI,TADA2B,KLHL9
RSF4,RBBP4,MCL1,JPT2,TLE5,PNN,SPC25,RPL31,NISCH,PRKCSH,HNRNPC,KHDRBS1,SLC3A2,PT
,RPS28,HNRNPM,GIGYF1,SRSF3,FAU,TUG1,SF3A1,MESD,GPR107,SREBF2,PPP6R1,ARNT2,ARFG
P2B1,MAT2A,TMEM170A,GTF2F1,PXDN,PGAM1,NFE2L1,SMARCA5,HNRNPA0,CTNNA1,FADS2,RP
LAMA5,RPS9,XRCC5,CIPC,PJA2,MSN,RPL24,SLC16A7,PAPOLA,IGFBP3,COL1A2,HK1,NCS1,RHOA
N,RPL10,MARCHF6,WSB1,RPS17,VCL,AP3D1,DNAJA1,ARF1,CYB5R3,SF3B2,MAT2A,GTF2F1,PXDN
P2,HDGF,ABCA2,PYGB,DPYSL2,TOP1,VEGFA,VHL,MVB12B,SMARCA4,AZIN1,ARGLU1,CDC37,CA
AM1,AGRN,RPLP1,ZMIZ1,TIMP2,XAF1,SARAF,MIDN,RPL10,MARCHF6,WSB1,RPS17,VCL,AP3D1,S
7,VCL,AP3D1,DNAJA1,ARF1,AKAP12,SF3B2,ZNF770,AP2B1,MAT2A,PXDN,HNRNPA0,FADS2,RPL23
1,SPTBN2,PFKL,HDLBP,GLG1,CDH6,SFPQ,EIF5A,PEBP1,CDH2,SOX12,RAB7A,CCN2,PRDX2,DDX
3B1,PDIA3,SSBP2,ZNF43,CHD3,VAPA,HMGB1,CDKL1,TPT1,SEC61A1,RPS4X,RPS3A,STMN1,ITGB
2,SEPTIN3,HECTD4,CCT6A,RAC1,CDK16,SLC35E2B,APPL1,SMCHD1,IPO5,ATP6V0C,TMEM30A,M
1A1,JUND,EIF5,LENG8,WDR1,ATXN7L3B,HMGN2,ATP2A2,CRMP1,LASP1,BEX3,CCT7,SRSF4,RBB

LNT4,HNRNPDL,RPLP2,NDFIP1,DDOST,HNRNPK,DUX4,YWHAB,PTGES3,SND1,LAMP1,ATP1A2,RF
APZB,HNRNPUL1,UBE2R2,SCAF11,BNIP3L,MAP7D1,KDELR1,MAP2K2,ZFAND5,HNRNPUL2-BSCL2
RPPA,CPSF2,PGRMC1,MDH2,RAB27A,U2AF2,EEF1A2,FDFT1,TBCD,SUGP2,ANXA5,CPSF1,PPP1CB
TF3,AGRN,RPLP1,SARAF,MIDN,RPL10,MARCHF6,RPS17,VCL,AP3D1,DNAJA1,ARF1,CYB5R3,AKAP
P1,SARAF,MIDN,RPL10,MARCHF6,RPS17,VCL,AP3D1,DNAJA1,ARF1,CYB5R3,SF3B2,MAT2A,GTF2I
2,CSNK1E,TPM1,EIF5A,CDH2,PDIA6,GPC3,CCN2,MMP14,PRDX2,DDX3X,RPS15,RPL12,ITM2C,AR
ARNT2,SMS,CTXN1,YME1L1,RBM15B,POLR2E,PSMA7,VDAC1,TRAF4,RBBP7,MSMO1,KCNQ2,TXL
LC16A7,RPL17,C1ORF43,PAPOLA,IGFBP3,COL1A2,LLGL1,NCS1,ZNF865,RAB6B,NLGN2,GLIPR1
TIMP2,XAF1,MIDN,RPL10,VCL,SBF1,DNAJA1,ARF1,CYB5R3,MAP4,SF3B2,CDC42BPB,ITM2B,AP2I
,CDH2,SOX12,RAB7A,CCN2,PRDX2,RPS15,RPL12,ITM2C,RPL30,EIF3L,MRFAP1,RPS5,RPS7,GPAT
R6,ATXN10,ACTN1,HSPB1,KDM1A,CASC3,FKBP8,PLCG1,ZFAND5,FLNB,PITPNA,CORO1C,SMARC
4,MAGED4B,CTBP1,STIP1,GPC1,APPL1,HDAC2,IPO5,MMP2,BCLAF1,L1CAM,LAPTM4B,PPIB,SQL
MAPK8IP1,SOD1,FBRSL1,P3H3,ZSCAN18,CTXN1,CD99L2,PPDPF,CIC,TRAF4,RBBP7,TRAF7,BCL9
F2,MORF4L2,SNRPN,PDXK,PKN1,HSPA4,PLOD1,GRINA,VPS35,PTOV1,MLEC,CLSTN3,MDK,SURF4
A,PTCD3,CBX5,CLSTN1,ATF4,SLC7A5,ENC1,RPS14,CXADR,CHD4,ATP1A1,JUND,TMEM259,EIF5,C
NB2,RPS5,APLP1,RPS19,CHPF,JUN,CD63,NGRN,GOLM1,SESN3,DNAJC5,SLC25A3,RPLP2,ARID1A
5,TKT,SFPQ,TPM1,PEBP1,CCND2,CDH2,PNMA1,PDIA6,GPC3,MEGF8,ACSL6,ITM2C,EIF3L,GNB2,
F4H,SCARB2,NACA,RPS9,XRCC5,CIPC,PJA2,MSN,RPL24,SLC16A7,C1ORF43,IGFBP3,GAP43,HK1,
EGFA,SELENON,RAPGEF1,TUBB2A,VHL,MVB12B,MAPK1,AZIN1,SRSF9,CCNL2,PSMD3,RABL6,FA
,SARAF,MIDN,RPL10,MARCHF6,RPS17,VCL,AP3D1,DNAJA1,ARF1,CYB5R3,AKAP12,SF3B2,CDC42
,KIF5B,RAB6B,ENC1,GLIPR1L2,CXADR,ATP1A1,JUND,RPS20,COL5A2,ATXN7L3B,HMGN2,ATP2A2
L1,PNN,NISCH,PRKCSH,HNRNPC,TADA2B,XPR1,KLHL9,XPO1,SPTBN2,SCRN1,HDLBP,GLG1,SFPQ
B2,MAT2A,GTF2F1,SLC4A8,OGA,SMARCA5,HNRNPA0,RPS16,RPL23,CNN3,RPL29,RPSA,TPM4,EIF
P1CB,SNX22,PABPN1,GDI2,TM9SF2,MORF4L2,TGFBI,SSRP1,HNRNPA3,PRDX5,PKN1,YEATS2,CAI
ZP6,CDC37,WDR6,HNRNPUL1,SCAF11,QARS1,HNRNPUL2-BSCL2,PITPNA,DLK1,MSL1,KIF3C,PH(
,H1-0,IK,RPS19,GPAT4,CHPF,ANXA6,NGRN,DNAJC5,HNRNPDL,RPLP2,DDOST,DUX4,YWHAB,SND
H1-0,RPS7,IK,RPS19,CHPF,NGRN,DNAJC5,HNRNPDL,RPLP2,DDOST,TSR2,NARS1,DUX4,YWHAB,F
PTIN3,ZNF454,RPS27,CTBP1,RAC1,ARHGDIA,CDK16,TOP2B,SLC35E2B,HDAC2,ATP6V0C,BCLAF
F4L1,ANXA2,AGRN,TIMP2,COL11A1,MIDN,MARCHF6,WSB1,RPS17,VCL,SBF1,DNAJA1,ARF1,AKA
NACA,LAMA5,RPS24,RPS9,XRCC5,CIPC,RPL24,PAPOLA,COL1A2,LLGL1,HK1,NCS1,RPL22,RHOA,Z
TNKS2,LONP1,UBQLN1,NBPF9,ZFAND3,SARS1,HGS,SREBF2,PNMA8A,VGF,BMPR2,CCT5,SYP,PO
CHF6,VCL,DNAJA1,ARF1,CYB5R3,AKAP12,MAP4,SF3B2,CDC42BPB,AP2B1,GOLGA8A,PXDN,HNR
1,EP300,LBH,DGCR2,KDM5B,HIP1,LRRC4B,SLC39A6,SEC16A,GPX4,SLC35F5,GNG4,NXN,EP400
P3,LLGL1,RPL14,NCS1,PTCD3,RPL9,PSMD2,GLIPR1L2,RPS14,CXADR,JUND,TMEM259,EIF5,CD8
L11,BTF3,AGRN,RPLP1,TIMP2,SARAF,MIDN,RPL10,MARCHF6,WSB1,RPS17,AP3D1,DNAJA1,ARF
PM4,CTSD,EIF4H,LAMA5,RPS9,XRCC5,CIPC,MSN,RPL24,SLC16A7,PAPOLA,COL1A2,HK1,NCS1,F
,CNN3,TPM4,TNS3,CTSD,EIF4H,LAMA5,SLC22A17,RPS9,PJA2,MSN,SLC16A7,KPNB1,C1ORF43,P
CA4,PLEC,BTBD2,PBX2,NYNRIN,TCF25,DYNLL2,LUZP6,PCSK1N,UBE2R2,ACTN1,KDM1A,MAP7D1
,GCN1,SKI,RAB11B,SRP14,PRPF6,DPYSL2,TMED2,TAF7,SELENON,RPL37,MAPK1,SMARCA4,ATG
RINA,VPS35,PTOV1,MLEC,CLSTN3,MDK,CBX1,RAD23B,PTK7,SKI,ASXL1,H3-3A,ABCA2,PYGB,TE
JBB4B,UCHL1,MORF4L1,ANXA2,AGRN,RPLP1,ZMIZ1,TIMP2,XAF1,SARAF,MIDN,RPL10,MARCHF6
VA2,CTSB,RNF187,CCT2,TBCD,SUGP2,CPSF1,SQSTM1,KLHL24,MEST,GAS7,PABPN1,PARP1,TAF
INRNPA0,CTNNA1,FADS2,RPS16,LDHA,CNN3,NACA,LPCAT1,LAMA5,SLC22A17,RPS9,XRCC5,CIPC
PL26,SRSF4,RBBP4,JPT2,RPL31,PRKCSH,CCNI,KHDRBS1,CSNK1E,PFKL,TKT,CDH6,PEBP1,CDH2
FBXW5,MBD3,SF3A2,VASH1,SART1,AP1M1,SAP18,BNIP3,ATP5MC2,SAE1,GAK,UBE2K,PTPN23,U

LC22A17,XRCC5,RPL24,KPNB1,RPL17,C1ORF43,COL1A2,LLGL1,RPL22,RHOA,CLSTN1,SLC7A5,
M199X,SERF2,TMEM132A,DDX39B,NUCB1,SRSF5,TM9SF3,MDH2,U2AF2,EEF1A2,FDFT1,GNAI3,
ASC3,CLIP3,FKBP8,KDELR1,MAP2K2,DIDO1,FLNB,TSC22D1,DLK1,DHX30,SRSF3,PHC2,SDHA,NK
,STK24,PRR12,SFXN1,LSM14B,FOXK2,NUDT19,LRRC41,NSD2,IST1,NOC2L,GSK3A,PRKACA,MN1
ARCA5,HNRNPA0,RPS16,RPL23,CNN3,RPL29,TPM4,TNS3,EIF4H,LAMA5,RPS24,BRD3,RPS9,XRCC
KIF5B,GLIPR1L2,RPS14,CXADR,CHD4,RPS20,LENG8,WDR1,RPL26,CRMP1,LASP1,CCT7,SRSF4,F
AP12,MAP4,SF3B2,CDC42BPB,ITM2B,AP2B1,MAT2A,SLC4A8,GOLGA8A,HNRNPU,OGA,NFE2L1,C
EIF1,HACD3,SET,EGLN1,MAP9,TPI1,CHSY1,TMSB10,INA,EIF4A1,MRC2,RPL19,PPP2R1A,GTF3C4,
NP32B,RPS2,HNRNPAB,MAP1LC3B,DAP,SAFB2,ZNF664,EFR3B,BAP1,TRIR,SEPTIN5,EHMT1,PALM
9,TPI1,CHSY1,TMSB10,INA,EIF4A1,MRC2,RPL19,PPP2R1A,GTF3C4,LDHB,DNAH8,HLA-C,SF3B1,
DAM9,PTBP1,FTO,ILF3,LUZP6,PRKDC,HNRNPUL1,SCAF11,ACTN1,SON,ZFAND5,DIDO1,DUSP4,PI
POLA,HK1,NCS1,RPL22,RHOA,PTCD3,ZNF865,RPL9,ATF4,KIF5B,GLIPR1L2,RPS14,CXADR,CHD4
F865,RPL9,ATF4,GLIPR1L2,RPS14,CXADR,CHD4,ATP1A1,TMEM259,RPS20,CD81,LENG8,WDR1,
NA,EIF4A1,MRC2,PPP2R1A,RPL15,GTF3C4,LDHB,DNAH8,PKD1,HLA-C,SF3B1,PDIA3,SERINC1,SS
A6,PIP4K2B,SOX12,ACTN4,MMP14,PRDX2,DDX3X,RPS15,RPL12,MRFAP1,MAGED2,GNB2,H1-0,R
DA2B,SLC3A2,PTN,PFKL,HDLBP,RPS25,SFPQ,TPM1,EIF5A,CCND2,PNMA1,SOX12,CCN2,ACTN4,P
3B,PDHB,RNF40,ZC3H4,CNOT1,CCDC124
AZ1,CD99,GPX4,GSPT1,KLHL42,CSNK1G2,GNG4,NXN,UBAP2L,GLUD1,MASTL,HDAC6,KCTD12,S
PL37A,LRRC28,MAPT,FN1,SPEN,NONO,DDB1,PSMD5,YWHAQ,EIF1,DCHS1,SET,EGLN1,TPI1,CHSY
DLBP,RPS25,SFPQ,TPM1,EIF5A,CCND2,PNMA1,PDIA6,SOX12,RAB7A,CCN2,ACTN4,PRDX2,RPS15,
HACD3,DCHS1,SET,EGLN1,MAP9,TPI1,CHSY1,INA,MRC2,RPL19,PPP2R1A,GTF3C4,LDHB,PKD1,SF
HSY1,TMSB10,INA,EIF4A1,MRC2,RPL19,PPP2R1A,GTF3C4,LDHB,DNAH8,PKD1,HLA-C,SF3B1,PDI
1,ARF1,CYB5R3,AKAP12,MAP4,CDC42BPB,AP2B1,MAT2A,GTF2F1,SLC4A8,GOLGA8A,PXDN,HNR
BTF3,NCAM1,ANXA2,RPLP1,TIMP2,XAF1,SARAF,MIDN,RPL10,MARCHF6,RPS17,VCL,ARF1,MAP4,
M1,SESN3,DNAJC5,HNRNPDL,RPLP2,DDOST,INIP,DUX4,SEPTIN11,SND1,RPS12,TTR,ZNF454,RPS
PS9,XRCC5,CIPC,RPL24,C1ORF43,PAPOLA,IGFBP3,COL1A2,HK1,NCS1,RPL22,RHOA,ZNF865,R
LE5,PRKCSH,HNRNPC,KHDRBS1,SPTBN2,PFKL,TPM1,PEBP1,CCND2,PDIA6,PIP4K2B,SOX12,ACT
NMA8A,VGF,BMPR2,CCT5,SYP,POLR2E,PPDPF,RBBP7,DFFB,TXLNA,SGTA,KDM5C,PRKAR2A,FBF
TPNA,MSL1,ATP6V1B2,GIGYF1,STMN3,PFKM,LSS,STXBP1,CSNK1D,IDH1,ZFAND3,STRAP,HECTD
H10,DUX4,CKAP5,SEPTIN11,SND1,RHOB,HNRNPH1,LGALS3BP,ARCN1,CTBP1,RAC1,ARHGDIA,S
A,LRRC28,MAPT,FN1,NONO,YWHAQ,EIF1,HACD3,DCHS1,SET,TPI1,TMSB10,EIF4A1,MRC2,RPL1
PRDX1,ODC1,ATP5PB,GRB2,WDR6,CAPZB,RPS11,PRKDC,SUPT5H,HNRNPUL1,UBE2R2,SCAF11,
A7,PAPOLA,HK1,RPL22,RHOA,ZNF865,RPL9,CLSTN1,ATF4,KIF5B,SLC7A5,GLIPR1L2,RPS14,CXA
6A7,KPNB1,C1ORF43,LLGL1,RHOA,CLSTN1,PSMD2,KIF5B,RAB6B,NLGN2,CXADR,CHD4,ATP1A1
HLA-C,SF3B1,PDIA3,SSBP2,ZNF43,CHD3,VAPA,HMGB1,CDKL1,AP2M1,TPT1,SEC61A1,RPS4X,R
NCAM1,AGRN,RPLP1,XAF1,SARAF,MIDN,MARCHF6,WSB1,RPS17,VCL,ARF1,CYB5R3,MAP4,SF3B
2,SLC38A10,NELFB,RIMS3,AMFR,OGFR,AGAP1,CHMP1A,RTF1,KIFC2,H6PD,ACVR1B,SIN3B,SLC2
3,DNAH8,PKD1,HLA-C,SSBP2,ZNF43,CHD3,BASP1,AP2M1,TPT1,RPS3A,OR8K1,RPL18,STMN1,ITG
LAMA5,RPS24,BRD3,RPS9,XRCC5,CIPC,RPL24,PAPOLA,HK1,NCS1,RPL22,RHOA,ZNF865,ATF4,F
HSBP1,MAP1A,SERF2,DDX39B,SRSF5,TM9SF3,MDH2,U2AF2,EEF1A2,FDFT1,CCT2,SUGP2,CPSF1
KK3,GSE1,LRP10,GPC4,MAP1LC3B,DAP,PCBP4,WSCD1,ZNF664,VPS26B,TRIR,TAOK1,IARS2,PA
CAPRIN1,STIP1,SLC35E2B,IPO5,ATP6V0C,BCLAF1,PPIB,RPL23A,HSBP1,MAP1A,CTDSP2,DDX39B,
DKL1,AP2M1,TPT1,SEC61A1,RPS4X,RPS3A,OR8K1,ITGB1,TUBB4B,UCHL1,MORF4L1,BTF3,NCAM
L1,RPL11,MORF4L1,BTF3,AGRN,RPLP1,SARAF,MIDN,RPL10,MARCHF6,RPS17,VCL,AP3D1,DNAJ
PAPOLA,IGFBP3,COL1A2,HK1,RPL22,RHOA,ZNF865,ATF4,PSMD2,KIF5B,GLIPR1L2,RPS14,CXAD

),CTNNA1,FADS2,RPS16,CTSD,SCARB2,NACA,TSPAN3,LPCAT1,LAMA5,RPS24,SLC22A17,BRD3,X
,STT3B,ISYNA1,TUG1,LOC728392,SF3A1,ID4,GPR107,MPRIP,HNRNPD,FBRSL1,ZSCAN18,SERBF
N1,HIF1A,CHERP,SLIT2,GPC4,PCBP4,ZMYM3,FUS,XRCC6,ERGIC1,YARS1,FKBP4,SOX2,SLC6A8,I
A1,USF2,USP10,LRIG1,SCAMP1,SLC22A23,DAG1,KIFC2,KLF13,PLXNA2,CDV3,IGF1R,MGAT3,SL
,XAF1,SARAF,MIDN,RPL10,RPS17,VCL,AP3D1,SBF1,DNAJA1,ARF1,PAFAH1B1,AKAP12,MAP4,SF3
EP2,ITGAV,DHCR7,MED13,KDM4A,APBB1,UBE2I,SLC38A2,UBE2K,FBXL19,CNBP,ELAVL3,PAFAH1
TF3C4,DNAH8,PKD1,HLA-C,SF3B1,PDIA3,SSBP2,ZNF43,CHD3,BASP1,VAPA,HMGB1,POLR2A,AP2
N2,PRDX2,RPS15,ITM2C,MAGED2,GNB2,H1-0,RPS5,RPS7,IK,GPAT4,JUN,ANXA6,SESN3,B4GALNT
HSBP1,CRPPA,HS6ST1,CTDSP2,PGRMC1,CNN2,SRP9,CTSB,ZC3H7B,MDH2,CHD6,TOMM20,RAB2
MLEC,CLSTN3,ACAP3,TMEM248,JPT1,CHD2,ORAI2,SKI,ASXL1,CNTNAP2,ABCA2,PRPF6,VEGFA,S
,PEG10,CHERP,SLC25A5,GPC4,MAP1LC3B,DAP,WSCD1,ZNF664,VPS26B,TRIR,SEPTIN5,IARS2,P
CIPC,RPL24,RPL17,C1ORF43,PAPOLA,COL1A2,RPL22,RHOA,ZNF865,CLSTN1,SLC7A5,ENC1,GLI
CR,NAP1L1,LBH,PRKAR2A,BAZ1B,GTPBP1,MMP15,GPX4,SLC39A7,KLHL42,DECR1,RAB2A,GNG4
SDE1,SMIM14,IGF2,EIF3C,ATP5F1A,EIF4B,VCP,MARCKSL1,UBB,AFF4,MATR3,YWHAG,CKB,COL4
T2A,GTF2F1,SLC4A8,SMARCA5,HNRNPA0,RPS16,RPL23,CNN3,RPL29,TPM4,EIF4H,LAMA5,RPS2
IGFBP3,RPL14,HK1,NCS1,RPL22,RHOA,CBX5,ZNF865,ATF4,KIF5B,ENC1,NLGN2,CXADR,CHD4,J
M1,EIF5A,PEBP1,CCND2,PNMA1,PDIA6,PLXNA1,PIP4K2B,MEGF8,SOX12,RAB7A,CCN2,PRDX2,RP
,HDGF,ARF3,G3BP2,ARPC2,DSTN,BTBD2,PRDX1,SRSF1,AEBP1,NYNRIN,DYNLL2,WDR6,CAPZB,U
T6,RGMB,ECEL1,MCM3AP,RPS27A,TMEM167A,SEPTIN7,CHERP,FKBP1A,CTTN,LRP10,SERP1,RH
AC1,ARHGDIA,CAPRIN1,STIP1,SLC35E2B,IPO5,ATP6V0C,BCLAF1,PPIB,RPL23A,HSBP1,MAP1A,CD
RPL19,PPP2R1A,RPL15,GTF3C4,DNAH8,PKD1,HLA-C,SF3B1,PDIA3,SSBP2,ZNF43,CHD3,BASP1,V
RPS15,ITM2C,RPL30,EIF3L,MAGED2,GNB2,H1-0,RPS5,IK,RPS19,MIF,CHPF,JUN,CD63,NGRN,RF
RPL24,SLC16A7,PAPOLA,IGFBP3,RPL14,HK1,NCS1,RPL22,RHOA,CLSTN1,ATF4,PSMD2,NR2F1,F
RPL24,SLC16A7,PAPOLA,IGFBP3,RPL14,HK1,NCS1,RPL22,RHOA,CLSTN1,ATF4,PSMD2,NR2F1,F
3P,RPS25,TPM1,CCND2,CDH2,PDIA6,GPC3,DDR1,SULF1,PLXNA1,PIP4K2B,ACSL6,SOX12,CCN2,N
,PFKL,SCRN1,RPS25,TKT,SFPQ,EIF5A,CDH2,PDIA6,MEGF8,RAB7A,RPS15,RPL12,ITM2C,RPL30,
RPS15,RPL12,ARL4C,CLTC,GNB2,RCN1,RPS5,APLP1,RPS7,RPS19,MIF,TLN1,CHPF,JUN,COL6A2,
MA5,RPS24,BRD3,RPS9,XRCC5,CIPC,RPL24,PAPOLA,IGFBP3,COL1A2,HK1,RPL22,RHOA,ZNF86
SMARCC1,GIGYF1,NKTR,YWHAH,PTP4A2,SLIT3,STMN3,PLXNA3,ATXN2L,IDH1,ZFAND3,NT5DC2,S
DB,ATP1A2,CKAP4,RPS12,SEPTIN3,RAC1,ARHGDIA,STIP1,CDK16,APPL1,SMCHD1,HDAC2,ATP6V
,SNRNP70,PSMB2,ATP6AP1,TERF2IP,NOTCH2,CCND1,MAPK1,SMARCA4,AZIN1,CCNL2,FAM168E
KAR2A,DGCR2,SF1,BAZ1B,H2AZ1,KPNA2,PHB2,UBA52,CXXC5,KLHL42,DIP2B,GNG4,UBAP2L,PI
PFKL,RPS25,GLG1,EIF5A,PDIA6,SULF1,PIP4K2B,SOX12,MMP14,DDX3X,RPL12,RPL30,EIF3L,MRF
R1,RPL26,LASP1,CCT7,SRSF4,RBBP4,PFN1,ASTN2,MCL1,TLE5,PNN,RPL31,PRKCSH,HNRNPC,L
,YWHAB,CADM4,RHOB,HNRNPH1,LGALS3BP,UBE2H,HLA-B,RAB1A,CCT6A,MAGED4B,STIP1,SLC
KDC,GABARAP,SCAF11,SON,ZFAND5,DUSP4,DHX30,HNRNPM,STMN3,PFKM,TNKS2,ATXN2L,PRF
KS2,CSNK1D,ATXN2L,HGS,BAG6,RAB11A,RPL39,SEC23A,CDC42,YME1L1,DHX15,BEX4,GRIPAP1,
7,PAPOLA,COL1A2,HK1,NCS1,RPL22,RHOA,ZNF865,RPL9,ATF4,KIF5B,GLIPR1L2,RPS14,CXADR,
MEM248,DDX42,GCN1,SKI,ASXL1,RAB11B,SRP14,TSPAN6,PRPF6,TAF7,RPL38,MARS1,RPL37,AT
1,DUX4,YWHAB,PTGES3,SND1,ATP1A2,RPS12,CALU,PLPPR3,CCT6A,MAGED4B,CTBP1,RAC1,CA
5E2B,IPO5,ATP6V0C,BCLAF1,RPL23A,HSBP1,MAP1A,CTDSP2,DDX39B,EID1,SRSF5,TM9SF3,MD
R2B,MSL1,KIF3C,YWHAH,PTP4A2,LSS,FBLN1,STT3B,LOC728392,ATXN2L,IDH1,ID4,GPR107,RPL
R2B,MSL1,KIF3C,YWHAH,PTP4A2,LSS,FBLN1,STT3B,LOC728392,ATXN2L,IDH1,ID4,GPR107,RPL
OLA,HK1,NCS1,RPL22,RHOA,ZNF865,RPL9,ATF4,PSMD2,KIF5B,GLIPR1L2,RPS14,CXADR,CHD4,
2,CDC42BPB,ITM2B,MAT2A,TMEM170A,SLC4A8,GOLGA8A,PXDN,HNRNPU,PGAM1,NFE2L1,FAD

RETREG2,MMP15,SLC39A6,PPP2CB,DECR1,FARP1,UPF1,RER1,TAGLN2,TBC1D9B,PPP1CC,ULK1
NF187,EEF1A2,TBCD,ANXA5,PPP1CB,VCAN,TLN2,MORF4L2,TGFBI,SSRP1,ABCA3,ACTR3,RTN1,F
GA,PGAM1,NFE2L1,FADS2,RPL23,CNN3,RPL29,TPM4,LPCAT1,LAMA5,RPS24,SLC22A17,XRCC5,
,ADNP,SSU72,ECH1,RAN,DRAP1,PPP1CA,PPP4R3A,WIZ,SRSF10,DDX6,PWWP3A,APPBP2,DEAF1,SI
9SF2,TSR1,SSRP1,HSPH1,LMNB2,MTRF1L,RPL27,PRDX5,UBE4A,ACO2,HSPA4,TCP1,AHCYL1,CA
RL6IP1,TSR2,MYL6,MYH10,RPL36,SEPTIN11,SND1,LAMP1,RHOB,HNRNPH1,UBE2H,CKAP4,PLPPI
TP1A2,PLPPR3,SEPTIN3,ZNF454,MAGED4B,ARHGDIA,CDK16,IPO5,ATP6V0C,SKP1,TRIB2,SHISAL
8,WDR1,ATP2A2,RPL26,LASP1,CCT7,SRSF4,RBBP4,PFN1,ASTN2,MCL1,TLE5,PNN,RPL31,PRKCS
,RPS27,ARHGDIA,CDK16,SLC35E2B,SMCHD1,HDAC2,IPO5,ATP6V0C,SKP1,BCLAF1,LAPTM4B,TF
ENG8,WDR1,RPL26,LASP1,CCT7,SRSF4,RBBP4,PFN1,ASTN2,MCL1,TLE5,PNN,RPL31,PRKCSH,HI
C6,CLPTM1,SHC2,INTS3,ABI2,BCL7A,GPC2,TLE1,HINT1,RAB14,MAZ,SH3GL1,DCLK1,ZNF428,PN
F,TLN1,COL6A2,B4GALNT4,MTSS2,APLP2,SLC25A3,ARL6IP1,ST3GAL2,TSR2,HNRNPK,CIRBP,DL
RS1,SELENON,RPL37,CCNL2,ARPC2,ARGLU1,PTBP1,SUPT5H,HNRNPUL1,SON,RPL28,CCAR2,AF
TIN11,CADM4,RHOB,ATP1A2,CALU,SEPTIN3,COTL1,HECTD4,CTBP1,RAC1,ARHGDIA,GPC1,TOP2
,SARAF,COL11A1,MIDN,RPL10,MARCHF6,WSB1,RPS17,AP3D1,DNAJA1,ARF1,AKAP12,MAP4,SF3
3,CCNL2,ARPC2,DDAH2,FAM168B,ARGLU1,PTBP1,CDC37,CAPZB,UBE2R2,ATXN10,SON,QARS1,F
TBN2,CSNK1E,PFKL,HDLBP,TKT,EIF5A,PEBP1,CCND2,PNMA1,MEGF8,RAB7A,CCN2,DDX3X,RPS1
RPLP1,TIMP2,COL11A1,MIDN,RPL10,MARCHF6,WSB1,RPS17,VCL,AP3D1,DNAJA1,ARF1,SF3B2,C
4,RPS9,CIPC,MSN,SLC16A7,C1ORF43,PAPOLA,IGFBP3,RPL14,HK1,RPL22,RHOA,PTCD3,CBX5,A
TN1,ATF4,SLC7A5,NLGN2,RPS14,CXADR,ATP1A1,JUND,COL5A2,LENG8,HMGN2,ATP2A2,RPL26,C
2A,GTF2F1,SLC4A8,OGA,SMARCA5,HNRNPA0,FADS2,RPS16,RPL23,CNN3,RPL29,TPM4,TNS3,EIF
3N2,CSNK1E,PFKL,SCRN1,RPS25,EIF5A,PDIA6,MEGF8,RAB7A,RPL12,ITM2C,RPL30,CLTC,EIF3L,I
,ACO2,HSPA4,TCP1,ZNF302,TMEM248,DDX42,GCN1,SKI,ASXL1,RAB11B,SRP14,TSPAN6,PRPF6,
SP2,DDX39B,NUCB1,SRSF5,TM9SF3,MDH2,U2AF2,EEF1A2,FDFT1,CCT2,SUGP2,GNAI3,NOP53,G
S6ST1,TIMP3,NOVA2,SRP9,ZC3H7B,U2AF2,EEF1A2,TBCD,MLXIP,CPSF1,MEST,NOP53,PABPN1,H
AZ1,ASTN2,MCL1,TLE5,NISCH,TADA2B,PFKL,GLG1,EIF5A,PEBP1,PDIA6,GPC3,MEGF8,ACSL6,SC
TSPAN6,PRPF6,TAF7,RPL38,MARS1,RPL37,G3BP2,ATG2B,CCNL2,ARPC2,ARGLU1,PTBP1,CAPZE
CHL1,MORF4L1,BTF3,ANXA2,RPLP1,ZMIZ1,TIMP2,SARAF,COL11A1,MIDN,RPL10,RPS17,VCL,AP3
TF4,PSMD2,KIF5B,SLC7A5,NLGN2,CXADR,CHD4,ATP1A1,TMEM259,EIF5,HMGN2,ATP2A2,CRMP1
,PFKP,FOXP4,PSMD13,PPP1R14B,CRIP2,PHPT1,EHMT2,RASA4,NTN1,LIMK1,PITRM1,DRAP1,RHE
N1A,DLG5,SHISAL1,TIMP3,TMEM132A,CTDSP2,SRP9,RAB27A,EEF1A2,MLXIP,SUGP2,ANXA5,CPSI
C,RPL30,EIF3L,MAGED2,H1-0,RPS5,IK,RPS19,CHPF,JUN,ANXA6,NGRN,DNAJC5,APLP2,HNRNPDL
P14,NRDC,PYGB,PRPF6,TAF7,RPL38,SELENON,RPL37,ARF3,MAPRE1,SRSF9,CCNL2,ARPC2,FAM1
AF4,RBBP7,TRAF7,BCL9L,WDR45B,EP300,DGCR2,STARD7,SF1,LRRC4B,CD99,SEC16A,SLC35F5
SBP2,ZNF43,CHD3,BASP1,VAPA,HMGB1,AP2M1,TPT1,SEC61A1,RPS3A,RPL18,STMN1,UCHL1,MC
ICS1,CBX5,ZNF865,CLSTN1,ATF4,RAB6B,GLIPR1L2,CXADR,ATP1A1,JUND,TMEM259,EIF5,CD81,
RPPRC,TADA2B,CSNK1E,PFKL,SCRN1,RPS25,TKT,SFPQ,EIF5A,CDH2,PDIA6,PIP4K2B,MEGF8,RAE
AP7D1,CASC3,PLCG1,KDELR1,ARF4,ZFAND5,DLK1,TPM3,MSL1,GIGYF1,NKTR,PTP4A2,STMN3,P
BAG6,RAB11A,SERBP1,ARFGEF2,WBP2,AKIRIN1,SMS,CCT5,CDC42,PPP1R9B,RBM15B,VDAC1,T
MPK,MVD,BRK1,SAFB2,ZNF664,VPS26B,PPP1R10,XRCC6,CEP170,EIF2S2,EHMT1,VAPB,CLPTM1,
D1,PLCG1,MAP2K2,CCAR2,DIDO1,HNRNPUL2-BSCL2,PITPNA,TSC22D1,MSL1,GIGYF1,SRSF3,SD
R10,TAOK1,EIF3H,ARHGEF17,TM9SF4,KMT2E,GTF2I,PNISR,SRSF2,PRR12,CAPZA1,DICER1,RBM
A,PSMD3,PRDX1,RABL6,SRSF1,IPO7,ARGLU1,RBP1,ATP5PB,ADAM9,FTO,ILF3,GRB2,CDC37,PRK
CALL1,MCM7,ADGRL1,LRP8,NCOA6,SAMD1,SLC25A39,KLF3,PIEZO1,CPNE2,DUS1L,MAFB,GLB1,
DA,DNER,LAPTM4B,HSBP1,PTTG1IP,CDKN1A,SHISAL1,TIMP3,TMEM132A,CTDSP2,PGRMC1,CNN

RAB27A,RNF187,EEF1A2,MLXIP,PPP1CB,SQSTM1,VAMP2,GAS7,TSPYL4,LMNB2,ALKBH5,TCP1,M
SS2,APLP2,SLC25A3,RPLP2,NDFIP1,MYL6,HNRNPK,H1-10,INIP,CIRBP,DUX4,CKAP5,SEPTIN11,SN
,BASP1,VAPA,HMGB1,POLR2A,CDKL1,TPT1,SEC61A1,RPS4X,RPL18,STMN1,ITGB1,TUBB4B,UCH
A17,BRD3,XRCC5,CIPC,PJA2,KPNB1,C1ORF43,PAPOLA,RPL14,NCS1,RPL22,ZNF865,PSMD2,KIF
WDR1,RPL26,LASP1,CCT7,SRSF4,RBBP4,PFN1,ASTN2,MCL1,TLE5,PNN,RPL31,PRKCSH,LRPPRC,
,TOP1,VEGFA,SMARCA4,MAPRE1,PLEC,ODC1,IPO7,DYNC1LI2,AEBP1,TCF25,ATP5PB,PTBP1,DYN
A2,FDFT1,CCT2,SUGP2,CPSF1,MEST,NOP53,GDI2,PARP1,TM9SF2,TSR1,SSRP1,HSPH1,MTRF1L,
P2B,SLC35E2B,IPO5,ATP6V0C,BCLAF1,PPIB,RPL23A,HSBP1,MAP1A,CDKN1A,DDX39B,SRSF5,TM
NXA2,AGRN,ZMIZ1,SARAF,COL11A1,RPL10,MARCHF6,WSB1,SBF1,ARF1,MAP4,CDC42BPB,ITM2
,CXXC5,GNG4,RER1,PI4KA,PPP1CC,ULK1,EML1,BRSK1,GPSM1,POM121C,PODXL2,GSE1,DAP,G
4,TCP1,MDK,ZNF302,TMEM248,DDX42,GCN1,SKI,ASXL1,RPL36A,RAB11B,SRP14,TSPAN6,PRPF
,PFKM,LSS,STT3B,TNKS2,PRPS1,HGS,MAPK8IP1,HNRNPD,MEX3A,SERBP1,PRPF40A,SMS,AMOT
MRFAP1,MAGED2,RCN1,H1-0,RPS7,RPS19,PRKAR1A,JUN,CTNNB1,COL6A2,ANXA6,NGRN,RPS10
EIF5,LENG8,ATXN7L3B,HMGN2,RPL26,CRMP1,SRSF4,ASTN2,MCL1,PNN,RPL31,NISCH,PRKCSH
R1,PLXNA1,MEGF8,PRDX2,RPS15,RPL12,ITM2C,RPL30,ARL4C,EIF3L,NANOG,RPS5,GPAT4,TLN
PS19,HMGCS1,GPAT4,PRKAR1A,CHPF,GJA1,CD63,MACF1,DNAJC5,RPLP2,TSR2,DUX4,SEPTIN11
ALNT11,GSE1,MAP1LC3B,DAP,TMOD2,ZNF664,ZMYM3,SEPTIN5,PALM,HSPG2,SSBP3,CARM1,PK
,RAB1B,UBE2M,MAZ,DCLK1,IREB2,LRATD2,TBC1D9,NUCKS1,LSM14B,SLC7A1,MOAP1,ERGIC3,
KAP12,MAP4,CDC42BPB,ITM2B,AP2B1,MAT2A,GOLGA8A,PXDN,OGA,NFE2L1,SMARCA5,LARP1,F
SH,LRPPRC,XPR1,KLHL9,SCRN1,RPS25,SFPQ,EIF5A,PDIA6,PLXNA1,PIP4K2B,MEGF8,RAB7A,ACT
4K2B,SOX12,MMP14,PRDX2,RPL12,ITM2C,RPL30,NANOG,MRFAP1,MAGED2,GNB2,H1-0,RPS5,A
PC,C1ORF43,COL1A2,LLGL1,GAP43,HK1,NCS1,CBX5,PSMD2,KIF5B,RAB6B,SLC7A5,ENC1,CHD4
M1,GPC2,LTBP4,VPS4A,ZMIZ2,GABARAPL1,SRSF2,RIC8A,SOD2,SPAG9,SLC7A1,ACTR1B,MOAP1
A,NGRN,SESN3,DNAJC5,APLP2,RPLP2,DDOST,TSR2,CIRBP,NARS1,DUX4,YWHAB,PTGES3,SEPTIN
3,ZNF454,CTBP1,RAC1,ARHGDIA,GPC1,SLC35E2B,HDAC2,IPO5,ATP6V0C,DNER,BCLAF1,LAPTM4
IF5,LENG8,WDR1,HMGN2,ASTN2,MCL1,PNN,CCNI,LRPPRC,TADA2B,XPR1,KHDRBS1,HDLBP,RPS
,MTCH1,PRDX6,EFR3B,BAP1,EHMT1,RTL6,SLC44A2,HSPG2,CARM1,RALY,LTBP4,VPS4A,ZMIZ2,
2,RPS15A,RAPGEF1,TUBB2A,ARF3,PPP2CA,CCND1,MAPK1,ARPC2,DDAH2,PLEC,PTBP1,FTO,ILF3
1,RPS12,TTR,CALU,PLPPR3,ARCN1,ZNF454,RPS27,HRNR,CAPRIN1,STIP1,CDK16,SLC35E2B,APP
5,LDHB,PKD1,HLA-C,SF3B1,SERINC1,SSBP2,ZNF43,RPS3,CHD3,BASP1,VAPA,HMGB1,POLR2A,C
2,HLA-B,PLPPR3,COTL1,HECTD4,GPC1,SLC35E2B,SMCHD1,HDAC2,IPO5,TNC,SHISAL1,CRPPA,T
LIPR1L2,CXADR,CHD4,ATP1A1,JUND,TMEM259,EIF5,LENG8,WDR1,HMGN2,BEX3,CCT7,RBBP4,
5,LASP1,SRSF4,PFN1,ASTN2,MCL1,TLE5,PNN,RPL31,PRKCSH,HNRNPC,XPR1,KHDRBS1,CSNK1E,
L28,CCAR2,ARF4,POLR2B,GNG2,SMARCC1,SUPT6H,DHX30,PHC2,PITPNM1,ISYNA1,CSNK1D,SF
RPF6,TAF7,MARS1,RPL37,ATG2B,ARPC2,FAM168B,ARGLU1,PTBP1,CAPZB,SUPT5H,ACTN1,SON
M171A2,AEBP1,DYNLL2,ILF3,LUZP6,CAPZB,UBE2R2,ATXN10,CLIP3,RPL28,PITPNA,GNG2,MSL1,
CDH6,SFPQ,PNMA1,PDIA6,SULF1,PIP4K2B,RAB7A,DDX3X,RPL12,ITM2C,RPL30,EIF3L,NANOG,MF
43,PGRMC2,ZFP91,FBL,TBCA,RALBP1,HSPE1,ATP5F1C,KDM2A,SRP54,NELFCD,ATF6B,DNAJB6,
3,GOLM1,B4GALNT4,MTSS2,RPL35A,APLP2,SLC25A3,NDFIP1,TAGLN,SHOX,CIRBP,NARS1,DUX4
RN,ZMIZ1,XAF1,COL11A1,MIDN,RPL10,MARCHF6,WSB1,RPS17,VCL,AP3D1,SBF1,DNAJA1,ARF1,
5,PEBP1,CCND2,CDH2,PDIA6,PIP4K2B,ACTN4,PRDX2,DDX3X,RPS15,RPL12,ARL4C,MRFAP1,RCN
CCN2,MMP14,DDX3X,RPL12,ITM2C,RPL30,CLTC,EIF3L,MRFAP1,RPS19,MIF,GPAT4,CHPF,JUN,N
AB11B,SRP14,NRDC,PRPF6,TAF7,RPS15A,RPL37,ARF3,MAPRE1,ARPC2,FAM168B,ARGLU1,ACAD
,PALM,HSPG2,SSBP3,CARM1,PKDCC,LTBP4,ARHGEF17,IMPDH2,ZMIZ2,GABARAPL1,RAB14,MAZ
O5,ATP6V0C,BCLAF1,LAPTM4B,TRIB2,RPL23A,HSBP1,FAM199X,SERF2,CPSF2,DDX39B,EID1,SR

HX30,SLIT3,STT3B,TNKS2,ATXN2L,NBPF9,HECTD1,HGS,BAG6,SREBF2,SERBP1,WBP2,SMS,RBM
PL9,ATF4,KIF5B,GLIPR1L2,RPS14,CXADR,CHD4,RPS20,CD81,LENG8,WDR1,HMGN2,RPL26,LASP
CA,KARS1,NSMF,CEBPG,ZDHHC5,SLC5A6,XPC,CACYBP,SLC5A7,ITPK1,DDHD2,PPP1R12A,ZNF36
RFAP1,MAGED2,H1-0,APLP1,RPS19,MIF,TLN1,CD63,ANXA6,NGRN,SESN3,MTSS2,APLP2,TARDBP
4,POLR2B,GNG2,SMARCC1,SUPT6H,DHX30,PHC2,EIF3F,PITPNM1,ISYNA1,CSNK1D,SF3A1,GPR1
RBP1,ARFGEF2,WBP2,AKIRIN1,CCT5,CDC42,PPP1R9B,YME1L1,RBM15B,TRAF7,EIF2AK1,TXLNA,
,HNRNPA0,RPL23,LDHA,CPE,CNN3,RPL29,RPSA,TPM4,TNS3,CTSD,NACA,LPCAT1,LAMA5,RPS24,
L1,YWHAH,SLIT3,STMN3,CSNK1D,HECTD1,IFITM3,MAPK8IP1,GHITM,PNMA8A,FBRSL1,DYNLL1,
AB11A,SERBP1,ARFGEF2,AKIRIN1,CCT5,CDC42,YME1L1,RBM15B,VDAC1,TRAF7,EIF2AK1,PRMT
AMP1,RPS12,RAB1A,TTR,PLPPR3,ARCN1,SEPTIN3,CTBP1,RAC1,CAPRIN1,STIP1,CDK16,TOP2B,S
HL1,RPL11,BTF3,NCAM1,ANXA2,RPLP1,ZMIZ1,MIDN,WSB1,RPS17,VCL,SBF1,AKAP12,MAP4,SF3
RIB,AKT2,DHCR7,IVD,PCGF2,TEX2,MBD3,YY1,CREBBP,ANKRD12,CERS2,DAPK3,GNA13,DPYSL4
D99,GPX4,APMAP,KLHL42,CSNK1G2,RAB2A,SFRP1,KIAA0895L,GLUD1,PI4KA,RAB3GAP2,SELEN
PT1,SEC61A1,RPS4X,RPS3A,OR8K1,STMN1,ITGB1,TUBB4B,RPL11,MORF4L1,NCAM1,ANXA2,RP
R,CHD4,JUND,TMEM259,RPS20,CD81,LENG8,WDR1,RPL26,LASP1,SRSF4,PFN1,ASTN2,MCL1,TL
AB5B,ASXL1,TOP1,RPS15A,VEGFA,SELENON,TUBB2A,MVB12B,MAPK1,SMARCA4,MAPRE1,DDAH
PFN2,RPL39,TP53BP1,ADAR,EP300,SF1,CSNK2A1,PHB2,SLC39A7,PPM1G,RAB2A,SRSF6,RER1,U
12,ITM2C,RPL30,GNB2,RCN1,RPL35,TLN1,MYO5A,COL6A2,CD63,NGRN,GOLM1,B4GALNT4,MT
M1,RPS10,RPLP2,TSR2,H1-10,NARS1,SND1,HNRNPH1,UBE2H,RPS12,CALU,ZNF454,RPS27,HEC
DH2,PDIA6,DDR1,PIP4K2B,MEGF8,ACSL6,CCN2,MMP14,DDX3X,RPL12,ITM2C,RPL30,CLTC,EIF
IK1,RPL22,PTCD3,ZNF865,RPL9,ATF4,PSMD2,KIF5B,RPS14,CXADR,RPS20,LENG8,WDR1,RPL26,
31,UCHL1,RPL11,MORF4L1,BTF3,ANXA2,ZMIZ1,SARAF,MIDN,RPL10,MARCHF6,WSB1,RPS17,VC
SMD2,KIF5B,CHD4,ATP1A1,JUND,TMEM259,EIF5,HMGN2,ATP2A2,CRMP1,BEX3,CCT7,OAZ1,RB
BDD2,STK35,ERH,PSMB4,EIF2S2,CYTH2,EIF3H,SNRPD3,RALY,SEC14L1,NUDCD3,SSR3,GNL1,TU
F43,IGFBP3,COL1A2,LLGL1,GAP43,NCS1,RPL22,RHOA,CBX5,ZNF865,CLSTN1,ATF4,PSMD2,KIF
NRNP200,CSDE1,SMIM14,IGF2,EIF3C,ATP5F1A,EIF4B,VCP,MARCKSL1,UBB,AFF4,TRIM28,MATR
,RPL37,SMARCA4,MAPRE1,CCNL2,PDE7A,SRSF1,ARGLU1,ADAM9,PTBP1,DYNLL2,PRKDC,ATXN1
3,SND1,LAMP1,CALU,ARCN1,COTL1,ARHGDIA,CAPRIN1,SLC35E2B,IPO5,ATP6V0C,BCLAF1,LAPT
OP56,GTF3C1,TIA1,SIN3B,SLC25A6,KIF26A,JAM3,PJA1,EWSR1,ATF5,RAN,DYNLRB1,ATF7IP,DAD
RPL27,PTPRA,PDXK,YEATS2,TCP1,VPS35,MDK,DDX42,LAMC1,RAD23B,RAB5B,GCN1,SKI,H3-3A,
RP,FKBP1A,CTTN,MTCH1,SERP1,AKAP9,RHBDD2,STK35,ERH,EIF2S2,CYTH2,EIF3H,SNRPD3,RA
,RGL2,RUVBL2,PDHA1,DNAJB1,EFNB2,CRIP2,SYNCRIP,PHPT1,KAT2A,RALGDS,EHMT2,PSMD7,L
PDAP1,AP2A1,CSNK2A1,MMP15,FLOT1,EIF3D,SEC16A,PHB2,APMAP,CST3,UBA52,PLXNB1,DECR1
D3,HINT1,GNL1,MAZ,SH3GL1,PNISR,TRAP1,ATXN3,ELP1,LSM14B,SLC4A2,ABCF1,SPTY2D1,MB
5A,COL6A2,CD63,NGRN,GOLM1,B4GALNT4,MTSS2,APLP2,TARDBP,ARL6IP1,NDFIP1,ST3GAL2,T

F1,MBTPS1,SMC3,RGMB,SELENOW,ECEL1,TBC1D9B,HNRNPA2B1,APEX1,HNRNPAB,GLO1,FUS,
A2,RPL14,HK1,NCS1,RPL22,ZNF865,CLSTN1,KIF5B,RAB6B,SLC7A5,NLGN2,RPS14,CHD4,ATP1A
1D,ATXN2L,HYOU1,GPR107,SARS1,VANGL2,GHITM,BAG6,RAI1,HNRNPD,FBRSL1,RAB11A,SEC2
IF5B,PTEN,ECEL1,TBC1D9B,ATP6AP2,RABEP1,SYMPK,LOX,PCBP4,ZNF664,ACLY,CLCN7,PALM,
,RPS19,GPAT4,TLN1,GJA1,MYO5A,RPS10,MXD4,MYL6,WLS,H1-10,CKAP5,SEPTIN11,CADM4,CK
HDLBP,RPS25,TKT,EIF5A,CDH2,PDIA6,MEGF8,ACSL6,RAB7A,MMP14,RPL12,ITM2C,RPL30,EIF3L,
APB,CCN1,HSPG2,UBE2M,RDX,MAZ,HDGFL2,ZNF428,SRSF2,CADM3,SLC4A2,SLC7A1,PDPK1,MS
3,UQCRH,MAN1B1,TMEM59,SPG11,BRPF3,FUBP1,ZC3H7A,MAML1,CDC42BPA,NUDT3,PAF1,RA
VDAC1,TXLNA,MTA1,TRIO,LBH,SF1,ALDH3A2,HIP1,FLOT1,SLC39A6,CD99,GPX4,ZFP36L1,PLXN

RAB11B,SRP14,NRDC,PRPF6,TAF7,RPS15A,RPL37,ARF3,MAPRE1,ARPC2,FAM168B,ARGLU1,PTE
\B1A,ARCN1,SEPTIN3,COTL1,HECTD4,CTBP1,RAC1,ARHGDIA,TOP2B,SMCHD1,HDAC2,IPO5,ATP6
1,CCAR2,ARF4,ZFAND5,DIDO1,DHX30,SRSF3,SDHA,LARS1,STMN3,LSS,STT3A,TUG1,PLXNA3,CSI
25A29,CTDNEP1,TFDP1,AGPAT4,PHF3,ZNF516,MBD6,SAMD1,SYT1,UBE2E1,ARHGAP23,SMARCC
\PBP,OLFM2,WDTC1,USF2,PINK1,AUTS2,CGGBP1,POR,CLIP2,SLC22A23,TSPYL2,H6PD,REEP5,F
3,GIGYF1,SLIT3,STMN3,FBLN1,STT3B,STXBP1,UBQLN1,ATXN2L,NT5DC2,PTPRS,PNMA8A,P3H3,
TN4,LAMA5,SLC22A17,BRD3,XRCC5,CIPC,PJA2,MSN,SLC16A7,KPNB1,PAPOLA,GAP43,HK1,NCS1
NIBAN2,KLF13,RGPD5,MKNK2,SIN3B,MFN2,ARHGEF40,CUL1,NSF,SCAMP4,EWSR1,PABPC4,PDL
,JUND,TMEM259,EIF5,HMGN2,ATP2A2,BEX3,CCT7,OAZ1,SRSF4,RBBP4,MCL1,PNN,NISCH,PRKC
1,TMED9,IDH1,FBRSL1,MEX3A,CYP51A1,SEC23A,ARFGEF2,AKIRIN1,KDM4B,TP53BP1,AMOTL2,I
1,TRAF7,EIF2AK1,PRMT1,XPOT,EXTL3,STARD7,PDAP1,AP2A1,CSNK2A1,MMP15,FLOT1,SEC16A,I
TRIO,PRKAR2A,NDUFV1,FBRS,SF1,H2AZ1,SLC39A6,MGAT4B,DECR1,HOXB5,NXN,EP400,CD164,
ARFGEF2,PRPF40A,BMPR2,TXNIP,CTXN1,CD99L2,AMOTL2,YLPM1,CDC42,PPP1R9B,DHX15,PPDI
0D1,USP24,EHD1,NOC2L,COX6A1,EIF3G,UBE3A,NASP,TGOLN2,AP1M1,UBE2I,C16ORF58,TOMM
,CTSB,NUCB1,CDH11,RNF187,U2AF2,EEF1A2,FDFT1,SUGP2,ANXA5,PPP1CB,MEST,GAS7,GDI2,T
PCAT1,LAMA5,RPS24,SLC22A17,RPS9,CIPC,PJA2,RPL24,SLC16A7,C1ORF43,PAPOLA,IGFBP3,N
P1,LRRC41,PRPF19,NOC2L,SART1,AP1M1,SSR2,NDUFA13,PSMC1,TOMM70,THRAP3,LYPLA2,SL
3,XPR1,KLHL9,SPTBN2,HDLBP,GLG1,SFPQ,PEBP1,PNMA1,PDIA6,SULF1,PIP4K2B,MEGF8,RAB7
\B,CADM4,RHOB,HNRNPH1,LGALS3BP,ATP1A2,SEPTIN3,CTBP1,RAC1,ATP6V0C,SKP1,DNER,PPI
\RF4,POLR2B,XPNPEP3,SMARCC1,SUPT6H,DHX30,PFKM,TNKS2,STT3A,LONP1,CSNK1D,HYOU1,
NG8,WDR1,HMGN2,BEX3,CCT7,RBBP4,ASTN2,MCL1,PRKCSH,TADA2B,XPR1,CSNK1E,HDLBP,RP
2B,XPR1,KLHL9,SPTBN2,HDLBP,RPS25,GLG1,SFPQ,PEBP1,PNMA1,PDIA6,SULF1,PIP4K2B,MEG
D3,XRCC5,CIPC,PJA2,RPL24,KPNB1,C1ORF43,PAPOLA,LLGL1,GAP43,RPL14,HK1,NCS1,RHOA,
SLC25A36,CELSR1,USF2,FXR2,USP10,LUC7L2,RGPD5,ANP32E,PLXNA2,NOMO2,MGAT3,EFNB2,
SM3,FOXP4,TMA7,CMPK1,FAM50A,WDR26,ADNP,ADGRB2,MAP4K4,CCNY,RASA4,HMGB3,LMF2
PF,JUN,CD63,NGRN,RPS10,RPLP2,DDOST,CIRBP,NARS1,DUX4,YWHAB,SND1,LAMP1,MAP2,CAL
2,NIBAN2,HIP1R,CDV3,ARHGEF40,FOXP4,DCAF5,WDR26,SYNCRIP,EHMT2,RASA4,HMGB3,LMF2
2,RPS5,IK,PRKAR1A,CHPF,GJA1,JUN,MACF1,GOLM1,B4GALNT4,HNRNPDL,SLC25A3,MYL6,SHO
,FAM50A,CRIP2,PHPT1,EHMT2,RASA4,HMGB3,LIMK1,SHISA5,PITRM1,CUL4A,DRAP1,IQSEC1,WI

ED1A,CAPZA2,CTDNEP1,MAN1B1,TFDP1,SPIN1,CABIN1,MBD6,VPS16,PSMA4,NUP50,PIP5K1A,A
13,COX6A1,ATP5MC3,NASP,YY1,SMARCD1,CREBBP,ATXN7L3,XPO6,AMBRA1,ATG4B,UNC119B,T
RF43,PAPOLA,IGFBP3,RPL22,RHOA,PTCD3,CBX5,ATF4,SLC7A5,ENC1,GLIPR1L2,CHD4,ATP1A1,
,GTF3C4,LDHB,PKD1,HLA-C,SF3B1,PDIA3,SERINC1,SSBP2,ZNF43,BASP1,VAPA,HMGB1,POLR2A
,TRAF7,EIF2AK1,TXLNA,PRMT1,BCR,MTA1,EXTL3,LBH,DGCR2,RETREG2,ALDH3A2,BAZ1B,PHB
UZP6,PRKDC,HSPB1,SON,RPL28,CCAR2,ARF4,CORO1C,SUPT6H,DHX30,NKTR,STMN3,TNKS2,S
NB1,NGRN,RPL35A,CYFIP2,TARDBP,SLC25A3,DDOST,TSR2,MXD4,ANKRD52,WLS,INIP,NARS1,PT
1,BRSK1,GPSM1,LMAN1,GSE1,MTCH1,PCBP4,WSCD1,BAZ2A,TAOK1,SHC2,INTS3,SLC6A8,TRA
0D1,SH3BP4,ARPP19,SF3A2,VASH1,NASP,AP2A2,SSBP4,SLC20A1,AMBRA1,CPXM1,REEP1,METR
PEN,NONO,DDB1,PSMD5,YWHAQ,EIF1,HACD3,DCHS1,SET,EGLN1,MAP9,TPI1,CHSY1,TMSB10,IN
1,GLIPR1L2,ATP1A1,JUND,TMEM259,EIF5,CD81,LENG8,WDR1,ATXN7L3B,CCT7,ASTN2,MCL1,P
CRIB,IST1,ATP6V0D1,CSE1L,PCGF2,MBD3,PACS2,NASP,CREBBP,ANKRD12,XPO6,BAG5,UBXN6,
2,APMAP,GSPT1,UBA52,PLXNB1,DECR1,FARP1,RER1,UBE2G2,IARS1,MASTL,COX7C,YKT6,ADH5
,NISCH,PRKCSH,TADA2B,KHDRBS1,XPO1,PFKL,SFPQ,TPM1,EIF5A,CCND2,CDH2,PDIA6,PLXNA1,
ARM1,RAB1B,UBE2M,IMPDH2,RAB14,MAZ,DCLK1,IREB2,ZNF428,LRATD2,STK24,SRSF2,TUBG

.EC,ACAP3,SRP14,NRDC,PRPF6,TAF7,RPL38,RPL37,PPP2CA,ATG2B,MAPRE1,ARPC2,PBX2,PDE7/
JBQLN1,TUG1,CSNK1D,PRPS1,IFITM3,UQCRC2,HNRNPD,MLF2,ARFGEF2,WBP2,AKIRIN1,NAP1L
2,RPL26,LASP1,VAT1,OAZ1,RBBP4,MCL1,JPT2,PNN,NISCH,PRKCSH,LRPPRC,TADA2B,XPO1,SPTE
IF,GPAT4,CHPF,JUN,CD63,NGRN,RPS10,RPL35A,DNAJC5,HNRNPDL,TARDBP,RPLP2,DDOST,CIRI
OLDIP2,TRIP12,CHMP1A,PSMB1,TSNAX,ZKSCAN1,RGL2,TCEA1,PCM1,SLC25A6,EXOC5,SF3A3,T
TAGLN,MYL6,SHOX,WLS,H1-10,CIRBP,YWHAB,RHOB,UBE2H,TTR,ARCN1,COTL1,MAGED4B,SLC
MP2,BCLAF1,L1CAM,DCN,MAP1A,FAM199X,SHISAL1,TIMP3,TMEM132A,CTDSP2,CPSF2,PGRMC
P1,CCT5,CDC42,YME1L1,RBBP7,TRAF7,EIF2AK1,EDC4,WDR45B,PDAP1,MMP15,GRSF1,SEC16A,
A,WDR45B,KDM5C,PDAP1,MMP15,PHB2,GSPT1,UBA52,PLXNB1,RAB3GAP2,MASTL,YKT6,ADH5,
FBLN1,PITPNM1,ISYNA1,TUG1,IDH1,COL27A1,GHITM,NCOR2,HNRNPD,MEX3A,FLNC,KDM3B,IGF
AR2,ARF4,POLR2B,GNG2,SMARCC1,SUPT6H,DHX30,YWHAH,PITPNM1,ISYNA1,CSNK1D,SF3A1,G
P4,PFN1,ASTN2,MCL1,TLE5,PNN,RPL31,PRKCSH,HNRNPC,LRPPRC,KHDRBS1,KLHL9,SLC3A2,CS
P4,PFN1,ASTN2,MCL1,TLE5,PNN,RPL31,PRKCSH,HNRNPC,LRPPRC,KHDRBS1,KLHL9,SLC3A2,CS
RHOA,PTCD3,CBX5,ATF4,SLC7A5,ENC1,GLIPR1L2,CHD4,ATP1A1,TMEM259,EIF5,CD81,LENG8,H
ARCC1,DHX30,YWHAH,PFKM,TNKS2,STT3A,SF3A1,GPR107,SARS1,VANGL2,GHITM,DGKD,BAG6,
IGCS1,CHPF,MYO5A,NGRN,SESN3,MTSS2,NDFIP1,TSR2,WLS,HNRNPK,INIP,CIRBP,SND1,HNRNP
,VCAN,TM9SF2,MORF4L2,HSPH1,LMNB2,MTRF1L,RPL27,PRDX5,UBE4A,PKN1,HSPA4,TCP1,CCT
2,PLXNA1,PIP4K2B,RPS15,ITM2C,RPL30,CLTC,MRFAP1,MAGED2,GNB2,RPS5,IK,RPS19,MIF,HM
1,EIF3G,DHX38,PACS2,AP1M1,NDUFA13,UBE2I,C16ORF58,TOMM70,XPO6,DAPK3,PRDX3,TRO,G
NX3,ERGIC3,YIPF3,CAP1,IST1,USP24,COPZ1,NOC2L,UBE3A,NASP,AP1M1,SMARCD1,UBE2I,C16
D,LUZP6,CDC37,CAPZB,HSPB1,SON,RPL28,CCAR2,ARF4,POLR2B,CORO1C,SUPT6H,DHX30,NKT
3,KLHL42,CSNK1G2,EP400,RER1,UBFD1,TAGLN2,ZNF512B,ECEL1,PRRC2A,PPP1CC,CUL3,EML1
MG,NELFB,SCAP,DENND5A,ACAT2,TMEM184B,USF2,SPPL2B,FXR2,PINK1,ARPC1A,TAOK2,RTF1,
ERINC1,CHD3,BASP1,VAPA,HMGB1,POLR2A,CDKL1,AP2M1,TPT1,SEC61A1,RPS4X,RPS3A,RPL18
NKS2,SF3A1,SARS1,BAG6,HNRNPD,RAB11A,SERBP1,ARFGEF2,AKIRIN1,CDC42,PPP1R9B,YME1L
GA2,SKIL,MMP24
G1,SFPQ,TPM1,PEBP1,PNMA1,PDIA6,SULF1,PIP4K2B,MEGF8,RAB7A,DDX3X,RPS15,RPL12,RPL3
HSBP1,SQLE,MAP1A,DDX39B,SRSF5,TM9SF3,MDH2,TOMM20,U2AF2,CCT2,RPN2,SUGP2,GNAI3
IBD3,UBE3A,ANKRD12,WNK1,SLC20A1,FGFR1,NSG1,DHX40,FBXL19,SCAP,IER2,AMFR,MZT2B,B
BP,PEBP1,CCND2,CDH2,SULF1,ACSL6,SOX12,MMP14,DDX3X,RPL30,CLTC,MAGED2,GNB2,H1-0
C3,RPL28,ZFAND5,CORO1C,SDF4,SMARCC1,HNRNPM,NKTR,ISYNA1,STT3A,PLXNA3,IDH1,SARS1
KIF5B,ATP1A1,JUND,TMEM259,EIF5,CRMP1,CCT7,VAT1,OAZ1,SRSF4,RTN3,ASTN2,TLE5,NES,RPL
DX42,CCT4,PSMB2,RPL36A,RAB11B,NRDC,PRPF6,TUBB2A,OPTN,UBXN4,PPP2CA,G3BP2,KDELF
ER1,WASF2,MKRN1,HSD17B4,OGT,DHCR7,MED13,COX6A1,DHX38,NDUFA13,MN1,ATG4B,DEK,H
TM,DGKD,BAG6,SREBF2,HNRNPD,RAB11A,ARNT2,SERBP1,ARFGEF2,AKIRIN1,SMS,CDC42,PPP1
MARCKSL1,UBB,AFF4,TRIM28,MATR3,YWHAG,CKB,COL4A2,RPL37A,LRRC28,MAPT,FN1,SPEN,
DC42,YME1L1,VDAC1,RBBP7,TRAF7,EIF2AK1,PRMT1,XPOT,STARD7,SF1,RETREG2,PDAP1,CSNK
15,ITM2C,RPL30,CLTC,EIF3L,MAGED2,H1-0,RPS5,IK,RPS19,RPL35,MIF,TLN1,CHPF,JUN,ANXA6,
EC1,RNF41,TRIB3,PBXIP1,CDK9,SRSF7,HERPUD1,NDRG4,RANBP9,RNMT,ME2,ZNF703,CACNA1
EC1,RNF41,TRIB3,PBXIP1,CDK9,SRSF7,HERPUD1,NDRG4,RANBP9,RNMT,ME2,ZNF703,CACNA1
MO1,RPS21,BANF1,FAF2,ECH1,KLC2,GOLPH3,VPS51,PPME1,AREL1,FAM8A1,TRIB3,C1QBP,FAM
,TOB2,WAC,ZSWIM6,HSF1,RIMKLB,LGR4,PA2G4,UBQLN4,TFRC,DIRAS1,VEZF1,WASF3,STARD3,
ASTL,YKT6,HNRNPA2B1,MCM3AP,RPS27A,TMEM167A,SEPTIN7,KLHDC3,TNKS,PSMD8,CHERP,F
,WDR45B,SGTA,ALDH3A2,SLC39A6,CD99,SEC16A,GPX4,PPM1G,SLC35F5,GNG4,NXN,EP400,UB
RCOR2,DTX3,NDUFS1,DLG4,HSPE1,KDM2A,ZNRF1,NR2F6,ANOS1,ARPC5,AFDN,CAMK2N1,MEIS

35,CLSTN3,ACAP3,JPT1,H3-3A,SCAF1,PRPF6,ATP6AP1,VEGFA,RAPGEF1,TUBB2A,MVB12B,CCND
IMGCS1,MYO5A,CTNNB1,NGRN,MTSS2,DNAJC5,HNRNPDL,TARDBP,NDFIP1,MXD4,MYL6,ANKRD
DLK1,POLR2B,SMARCE1,FYN,NKTR,STMN3,PFKM,UBQLN1,TUG1,PLXNA3,ID4,COL27A1,VANGL
AP,SLC39A7,PLXNB1,DECR1,FARP1,EP400,RER1,PI4KA,TRA2B,IARS1,COX7C,YKT6,CUL3,MCM3
,RBM15B,BEX4,ANP32A,TRAF4,KIF5A,KCNQ2,SGTA,MTA1,KIRREL1,EP300,RERE,ALDH3A2,VAR
RMT1,XPOT,NAP1L1,CPD,SF1,SLC39A6,GPX4,SLC35F5,CSNK1G2,ATRX,RAB2A,GNG4,CD164,GL
ASP,TGOLN2,AP1M1,NDUFA13,UBE2I,PSMC1,XPO6,MYL12B,GAK,LYPLA2,SLC38A10,ATP13A1,N
1B,SRP14,NRDC,PRPF6,TAF7,RAPGEF1,ARF3,MAPRE1,CCNL2,FAM168B,ARGLU1,RBP1,PTBP1,
25,LUZP6,PCSK1N,TDRD5,SCAF11,KDM1A,BNIP3L,DLK1,SDF4,SDHA,CCT8,STMN3,B2M,TNKS2,
R1,UBFD1,TAGLN2,MASTL,YKT6,ECEL1,HNRNPA2B1,MAPRE2,TMEM167A,SEPTIN7,TNKS,PSMD8
R1,UBFD1,TAGLN2,MASTL,YKT6,ECEL1,HNRNPA2B1,MAPRE2,TMEM167A,SEPTIN7,TNKS,PSMD8
4B,CSNK2A1,GRSF1,CD99,GPX4,GSPT1,CSNK1G2,GNG4,FARP1,NXN,UBFD1,TAGLN2,CRTAP,YK
,EDC4,TXLNA,PDAP1,AP2A1,MMP15,GPX4,PHB2,GSPT1,UBA52,SLC39A7,HJURP,RER1,UBFD1,T
7,DFFB,BCR,XPOT,OS9,CAPN2,NAP1L1,KIRREL1,KDM5C,DGCR2,RETREG2,PBX1,VARS1,AP2A1,
,RPL12,CLTC,GNB2,APLP1,RPS7,RPL35,GPAT4,CHPF,GJA1,JUN,COL6A2,CD63,MACF1,B4GALN
TPNM1,UBQLN1,CSNK1D,SF3A1,HYOU1,IFITM3,GHITM,DGKD,NCOR2,VGF,MLF2,PPP6R1,SEC2
,CCT7,VAT1,SRSF4,RTN3,PFN1,JPT2,TLE5,NES,LRPPRC,KHDRBS1,KLHL9,SPTBN2,SLC3A2,PTN
TSD,SCARB2,RPS24,BRD3,RPS9,PJA2,RPL24,SLC16A7,KPNB1,RPL17,PAPOLA,IGFBP3,RPL14,H
R45B,SGTA,ALDH3A2,SLC39A6,CD99,SEC16A,GPX4,PPM1G,SLC35F5,GNG4,NXN,EP400,UBFD1,
4A,AP1M1,CREBBP,BNIP3,MN1,WNK1,GNA13,ATG4B,TRO,NELFB,SH3PXD2B,ADGRA3,TCEAL4,C
P6,RIC8A,NCSTN,INTS11,AHSA1,MTMR4,PDPK1,ITGAV,SPTY2D1,BORCS7,NUDT19,CAP1,ATP6V
BP,GLG1,SFPQ,PEBP1,PNMA1,PDIA6,SULF1,PIP4K2B,MEGF8,RAB7A,DDX3X,RPS15,RPL12,ITM2
GMB,SELENOW,CUL3,ADH5,NDST1,ITGB5,TIMM50,TNKS,PSMD8,APEX1,DKK3,PPP1R15A,GALNT
,TMX1,PCM1,OSBPL2,TRMT112,PSMD13,WDR26,CD2BP2,SUMO1,RPS21,PJA1,ADNP,GPAA1,SU
F3D,GRSF1,SLC39A6,PHB2,APMAP,KLHL42,RER1,UBFD1,MASTL,YKT6,RPL7L1,RPS27A,TMEM1
RPS15A,OPTN,RPL37,ARF3,G3BP2,MAPRE1,ARPC2,FAM168B,ARGLU1,PTBP1,FTO,LUZP6,CAPZE

59,EIF5,LENG8,WDR1,ATP2A2,CRMP1,LASP1,CCT7,OAZ1,SRSF4,RTN3,PFN1,ASTN2,TLE5,NISCH
ARCC1,SUPT6H,DHX30,YWHAH,PITPNM1,ISYNA1,CSNK1D,SF3A1,STRAP,MPRIP,BAG6,RAB11A,S
,NCSTN,FNBP1L,PLEKHM2,KRT8,TCF3,ABCF1,ATP1B2,FOXK2,ADAM10,IGF2BP1,LRP3,RBPJ,US
E1L1,RBBP7,TRAF7,EIF2AK1,EDC4,TXLNA,WDR45B,PRMT1,PDAP1,ALDH3A2,CSNK2A1,MMP15,
,SETX,ZBED4,MBOAT7,MAN2B1,CHKA,HIF3A,FAM20B,BRI3,EFNB3,RUSC1,PRMT2,SHC1,RPS6K
,CDKN1A,SERF2,TIMP3,CTDSP2,DDX39B,CNN2,NOVA2,NUCB1,C5ORF24,ZC3H7B,TOMM20,U2
IX3,ERGIC3,YIPF3,SPTY2D1,NUDT19,CAP1,USP24,EHD1,COX6A1,UBE3A,NASP,AP1M1,C16ORF5
DGF,NRDC,DPYSL2,NOTCH2,VEGFA,SELENON,MVB12B,ATG2B,KDELR2,LAPTM4A,DSTN,BTBD2,
TAT3,ARIH1,LBH,FBRS,SF1,HIP1,SLC39A6,GPX4,SLC35F5,CSNK1G2,RAB2A,GNG4,FARP1,CD164
GRL3,ADRM1,MFAP4,SLC16A2,SYNGAP1,CDK9,MARCHF7,RBFOX2,MTRNR2L4,NDRG4,IGF2BP
,SRSF1,AEBP1,RBP1,PTBP1,DYNLL2,FTO,ILF3,CDC37,CAPZB,GABARAP,SUPT5H,HNRNPUL1,AC
1,TUBB4B,UCHL1,MORF4L1,NCAM1,ANXA2,AGRN,RPLP1,ZMIZ1,TIMP2,XAF1,SARAF,COL11A1,
PRPF4B,CSE1L,NOC2L,MBD3,UBE3A,NASP,AP2A2,YY1,TMEFF1,UBE2I,UBE2D3,TTC28,ATP5MC2
IN1,CCT5,CDC42,PPP1R9B,YME1L1,RBBP7,TRAF7,EIF2AK1,EDC4,TXLNA,WDR45B,PDAP1,MMP1
MOAP1,ATP1B2,FOXK2,REEP2,AKT2,MBD3,PACS2,GOT2,AP2A2,XPO6,TRIM8,SSBP4,UBXN6,SLC
GSK3B,MEX3C,ASH1L,CRTC1,ZYX,TUBA1C,UBQLN4,DIRAS1,DTX3,IRF2BP2,SH3BP5,IRS1,ERBB
,PDE7A,PSMD3,SRSF1,TCF25,GRB2,CDC37,WDR6,RPS11,SCAF11,KDELR1,ARF4,ZFAND5,DUS
,RPL28,DUSP4,FLNB,CORO1C,MSL1,SDHA,LARS1,FBLN1,ATXN2L,IDH1,ZFAND3,PTP4A1,MEX3A,

ARF4,ZFAND5,HNRNPUL2-BSCL2,PITPNA,PLTP,DHX30,FAU,SLIT3,CCT8,MTRNR2L10,PITPNM1,ST
ZNF768,GOLGA8B,DEGS1,UBL3,FAM160B2,MRPS6,PHKB,NAPB,RAVER1,MBOAT7,RALBP1,WASI
39A6,GPX4,SLC35F5,CSNK1G2,RAB2A,FARP1,CD164,UPF1,UBFD1,MASTL,MBTPS1,ANP32B,MA
NRC6A,VPS51,FAM8A1,DARS1,TRIB3,C1QBP,FAM32A,SYNGAP1,EPN1,CDK9,RTF2,ZRANB2,PSM

1,RAB5B,ASXL1,RAB11B,SRP14,PRPF6,TERF2IP,VEGFA,SELENON,RAPGEF1,TUBB2A,ARF3,UBXN

RPS4X,OR8K1,ITGB1,TUBB4B,UCHL1,RPL11,MORF4L1,BTF3,NCAM1,ANXA2,AGRN,RPLP1,TIMP

DOST,CIRBP,NARS1,RPL36,YWHAB,PTGES3,SND1,LAMP1,MAP2,CALU,ARCN1,COTL1,ARHGDIA,S
CDC42BPB,ZNF770,AP2B1,GTF2F1,PXDN,NFE2L1,SMARCA5,HNRNPA0,FADS2,RPS16,RPL23,CNN
TPBP1,MMP15,CD99,SEC16A,CST3,UBA52,MGAT4B,SLC39A7,PPM1G,KLHL42,PLXNB1,FARP1,C
3,CERS2,SLC20A1,AMBRA1,METRN,EIF4A3,PTPN23,SLC38A10,CENPB,ATP5MG,SRPRA,NELFB,C
2,SLC7A5,ENC1,GLIPR1L2,CXADR,CHD4,ATP1A1,TMEM259,EIF5,CD81,LENG8,HMGN2,LASP1,BE
PNB1,C1ORF43,PAPOLA,IGFBP3,HK1,NCS1,PTCD3,ZNF865,ATF4,RAB6B,ENC1,NLGN2,JUND,TM
2A,EHMT2,PSMD7,LMF2,LIMK1,AMPD2,PITRM1,SUN2,DRAP1,IQSEC1,NECTIN1,RAP2A,TBC1D14
,MBTPS1,ANP32B,MAPRE2,SEPTIN7,APEX1,PPP1R15A,GALNT11,MVD,DAP,BRK1,SERP1,PPP1R1
,PPP1R9B,YME1L1,TRAF7,DFFB,EIF2AK1,EDC4,PRMT1,XPOT,STARD7,PDAP1,AP2A1,CSNK2A1,M
AA2013,RRBP1,NR2F2,GET1,DNAJB6,TNPO2,AFDN,PPP1R26,SLC25A29,KLHL36,CAPZA2,PLCD3,
P1,FAM199X,SERF2,CTDSP2,DDX39B,NUCB1,SRSF5,TM9SF3,MDH2,TOMM20,RAB27A,U2AF2,CC
2A,RER1,UBFD1,TRA2B,ZFR,YKT6,SV2A,NAP1L3,PRRC2A,ANP32B,PREPL,TIMM50,ZMYM4,SEPTI
SLIT3,ILF2,TNKS2,STT3A,LONP1,TUG1,SARS1,HGS,GHITM,ZSCAN18,SERBP1,KDM3B,AKIRIN1,A
M1,SLC38A1,SLC25A6,EXOC5,OSBPL2,TRMT112,PSMD13,CD2BP2,RPS21,BANF1,PSMD7,COX8A
R15A,FBXO21,SAFB2,SPG7,BAZ2A,PELP1,VAPB,SLC44A2,UBE2M,ELAVL1,GABARAPL1,TLE1,FAM
X1,FLOT1,SLC39A6,CD99,SEC16A,GPX4,PPM1G,PPP2CB,SLC35F5,GNG4,NXN,EP400,UBFD1,MI
HLA-C,PDIA3,SERINC1,SSBP2,RPS3,CHD3,BASP1,VAPA,HMGB1,POLR2A,CDKL1,SEC61A1,RPS3A,
C,MMP2,LAPTM4B,RPL23A,NOVA2,SRSF5,TM9SF3,MDH2,RNF187,U2AF2,MLXIP,ANXA5,SNX22,N
KAP4,ARCN1,CTBP1,HMGCR,TOP2B,SMCHD1,ATP6V0C,LAPTM4B,RPL23A,NOVA2,SRSF5,TM9SF
1,PLXNB2,TTC3,PLD3,HSPD1,USP11,COL18A1,SNRNP200,CSDE1,SMIM14,IGF2,EIF3C,ATP5F1A,
21C,TIMM50,PEG10,GSE1,SYMPK,SLIT2,MTCH1,RHBDD2,ZMYM3,STK35,SEPTIN5,KIAA0100,PA
INA,VPS35,PTOV1,MLEC,SURF4,CBX1,SKI,PCDHGB6,SNRNP70,PYGB,PRPF6,NOTCH2,RAPGEF1,
4,UBFD1,IARS1,MASTL,YKT6,MCM3AP,SPINT2,PREPL,RPL7L1,RPS27A,TMEM167A,KLHDC3,TNK
RGMB,SELENOW,CUL3,ADH5,NDST1,ITGB5,TIMM50,TNKS,PSMD8,APEX1,DKK3,PPP1R15A,FKBP
MCM3AP,RPL7L1,RPS27A,TMEM167A,KLHDC3,TNKS,PSMD8,CHERP,FKBP1A,CTTN,LRP10,MTC
D3,CIPC,PJA2,MSN,RPL24,SLC16A7,KPNB1,RPL17,RPL14,PTCD3,ZNF865,CLSTN1,ATF4,KIF5B,

PC3,SULF1,MEGF8,SOX12,RAB7A,CCN2,MMP14,RPL12,ITM2C,CLTC,NANOG,RPS5,RPS7,IK,RPS
F2F1,NFE2L1,RPS16,LDHA,CPE,CNN3,EIF4H,SCARB2,TSPAN3,LPCAT1,RPS24,SLC22A17,RPL24,
C7A5,NLGN2,CHD4,ATP1A1,JUND,EIF5,CD81,WDR1,ATP2A2,RPL26,CRMP1,SRSF4,RTN3,PFN1,A
5,PHB2,UBA52,SLC39A7,PLXNB1,GARS1,TAGLN2,TRA2B,IARS1,COX7C,YKT6,MCM3AP,PREPL,TN
7,IK,RPS19,MIF,TLN1,JUN,CTNNB1,ANXA6,NGRN,RPL35A,CYFIP2,DNAJC5,ARID1A,TAGLN,HNRN
1,FLCN,GRSF1,GPX4,UBA52,MGAT4B,CXXC5,BRD2,PPP2CB,GNG4,CD164,KIAA0895L,RER1,FEI
M151B,MED16,TRAF3,MIB2,SHB,HID1,TRIM24,AKAP17A,PPFIA3,RAP1B,PLXND1,MYL9,GPRC5B,
N1,CHPF,GJA1,JUN,NGRN,MACF1,RPS10,B4GALNT4,CYFIP2,HNRNPDL,TARDBP,SLC25A3,RPLP2
1R9B,YME1L1,RBBP7,TRAF7,EIF2AK1,EDC4,TXLNA,WDR45B,PDAP1,MMP15,SEC16A,PHB2,APMA

CLSTN3,DDX42,CCT4,ORAI2,PSMB2,SCAF1,RAB11B,TUBB2A,OPTN,UBXN4,PPP2CA,CCND1,G3BP
MCHD1,ATP6V0C,MMP2,LAPTM4B,RPL23A,NOVA2,SRSF5,TM9SF3,MDH2,RNF187,U2AF2,MLXIP,
DH,ADAM10,CAP1,LRRC41,EHD1,PCGF2,COPZ1,NOC2L,SMARCD1,SSR2,NDUFA13,UBE2I,UBE2D
PNN,RPL31,PRKCSH,HNRNPC,LRPPRC,SLC3A2,CSNK1E,PFKL,SCRN1,RPS25,EIF5A,PDIA6,PIP4K
,CRB2,RBBP6

R5,RALY,GABARAPL1,PGAP6,MAZ,DTX4,KIAA0232,PRR12,EEF1B2,CADM3,SLC4A2,QPRT,DCUN1D
CASKIN1,WASF3,SH3BP5,DPP7,SLC5A6,NR2F2,FAM20B,NR2F6,ANOS1,GNAZ,TNPO2,NADK,MEIS
RPS14,CXADR,RPS20,LENG8,WDR1,ATXN7L3B,HMGN2,RPL26,LASP1,CCT7,SRSF4,RBBP4,PFN1
F4,PSMD2,RAB6B,SLC7A5,ENC1,NLGN2,RPS14,CXADR,ATP1A1,JUND,TMEM259,EIF5,CD81,LEN
2B,MEGF8,PRDX2,RPS15,ITM2C,ARL4C,RCN1,APLP1,IK,RPS19,MIF,GPAT4,TLN1,CHPF,JUN,MY
D74,SAE1,SLC38A2,DEK,FGFR1,DHX40,IER2,CNBP,DIP2A,MZT2B,LMO4,PTPN11,SUN1,LMAN2,C
1-0,RPS5,IK,RPS19,CHPF,JUN,NGRN,RPL35A,HNRNPDL,RPLP2,DDOST,ST3GAL2,HNRNPK,CIRBP
CCNI,TADA2B,KHDRBS1,XPO1,SLC3A2,CSNK1E,SCRN1,HDLBP,TKT,SFPQ,EIF5A,CCND2,PNMA1,
1,GNG4,GLUD1,UQCRC1,MASTL,COX7C,MBTPS1,KCTD12,HNRNPA2B1,PPP1CC,EML1,B4GAT1,
D4,FAF2,METAP2,GOLPH3,VPS51,MAGEF1,RPL27A,TRIB3,FAM32A,JAK1,EPN1,ARPC3,RTF2,STUB
PA0,FADS2,RPS16,RPL23,LDHA,RPL29,TPM4,TNS3,CTSD,EIF4H,NACA,TSPAN3,SLC22A17,RPS9,C
MIF,HMGCS1,GPAT4,PRKAR1A,TLN1,CHPF,GJA1,JUN,NGRN,CYFIP2,HNRNPDL,TARDBP,SLC25A3
59,EIF5,LENG8,WDR1,ATXN7L3B,HMGN2,MCL1,PNN,CCNI,LRPPRC,TADA2B,XPR1,HDLBP,RPS25
SKI,SCAF1,HDGF,ABCA2,PYGB,ATP6AP1,DPYSL2,TMED2,OPTN,PPP2CA,SMARCA4,ATG2B,MAPRE
K1,TPM3,MSL1,KIF3C,SUPT6H,PFKM,TNKS2,STXBP1,UBQLN1,TUG1,PLXNA3,LOC728392,ZFAND
,DAP,BRK1,SERP1,PPP1R10,BAZ2A,XRCC6,VAPB,EIF3H,ELAVL1,IMPDH2,GABARAPL1,SSR3,GNL
ED2,GNB2,APLP1,RPS7,CHPF,CD63,ANXA6,GOLM1,B4GALNT4,MTSS2,DNAJC5,TARDBP,TAGLN,
DHX40,SRPRA,ARF5,DENND5A,USP47,AGAP1,SCAMP5,ZZEF1,YTHDF3,CDV3,SIN3B,MFN2,TNFAI

P43,NCS1,RPL22,RHOA,PTCD3,CBX5,KIF5B,RAB6B,ENC1,NLGN2,CXADR,CHD4,ATP1A1,JUND,RP
,CHPF,COL6A2,CD63,ANXA6,MACF1,GOLM1,B4GALNT4,CYFIP2,DNAJC5,HNRNPDL,SLC25A3,RP
LAR,SHC1,MEIS3,ASB6,CTDNEP1,MIB2,UQCRH,TMEM59,TRAPPC1,SPG11,CUL4B,PSMD12,RPL3
YNCRIP,ALDH18A1,FEZ1,BNC2,CUL4A,DRAP1,UBE2J1,PPP4R3A,TBXA2R,AKAP8L,MICAL3,MAPK
K,H1-10,MYH10,NARS1,DUX4,HNRNPH1,HLA-B,RAB1A,TTR,PLPPR3,ARCN1,SEPTIN3,HECTD4,CC
AI2,SKI,SNRNP70,RAB11B,HDGF,PYGB,DPYSL2,PPP2CA,MAPK1,SMARCA4,AZIN1,MAPRE1,DDAH
P4R3A,RAP2A,SOBP,TBXA2R,FAM32A,OTUD5,RTF2,HM13,CLPX,GOLGA8B,NDRG1,EXOSC10,CD
AGED4B,GPC1,TOP2B,ATP6V0C,SKP1,L1CAM,PEA15,SQLE,FAM199X,SHISAL1,SERF2,TMEM132
SBF1,DNAJA1,ARF1,CYB5R3,AKAP12,SF3B2,CDC42BPB,ITM2B,AP2B1,TMEM170A,GOLGA8A,PX
AZIN1,PDE7A,PSMD3,SRSF1,ATP5PB,ADAM9,GRB2,WDR6,CAPZB,GABARAP,MAP2K2,ARF4,ZFAND
3B,PITPNM1,ISYNA1,ATXN2L,USP7,ID4,COL27A1,FBRSL1,MLF2,KDM3B,WBP2,KDM4B,DYNLL1,
M171A1,POR,SLC22A23,RBM4,PKIA,NOMO2,ARRDC4,MDM2,TNFAIP1,PSMD13,WDR26,ACAP2,BN
,PEBP1,CCND2,DDR1,PLXNA1,RAB7A,DDX3X,CLTC,NANOG,GNB2,RPS5,RPS7,IK,MIF,HMGCS1,P
IGRN,MACF1,RPS10,B4GALNT4,CYFIP2,HNRNPDL,TARDBP,SLC25A3,RPLP2,ARL6IP1,NDFIP1,ST
K1N,WDR6,HNRNPUL1,CMIP,CASC3,NSG2,MAP2K2,PITPNA,GNG2,PLTP,NKTR,PTP4A2,EIF3F,ISYN
ND2,DDR1,PLXNA1,RAB7A,DDX3X,RPL12,CLTC,NANOG,GNB2,RPS5,IK,MIF,HMGCS1,GPAT4,PRK
TSPAN11,CSRNP2,FAM20B,NR2F6,ARSB,GPRIN1,VKORC1L1,WDR5,KLHL36,CAPZA2,B4GALT2,
N,MYO5A,COL6A2,CD63,NGRN,B4GALNT4,MTSS2,ARID1A,TSR2,ANKRD52,H1-10,MYH10,RPL36
UBE2R2,ATXN10,ACTN1,CCAR2,POLR2B,MSL1,SMARCC1,YWHAH,PTP4A2,ILF2,STMN3,PFKM,L
C5ORF24,ZC3H7B,MDH2,CHD6,RAB27A,RNF187,MLXIP,SUGP2,SQSTM1,NOP53,GDI2,PARP1,TSP

E2H,RPS12,RPS27,COTL1,HECTD4,CCT6A,RAC1,STIP1,TOP2B,SLC35E2B,SMCHD1,MMP2,BCLA
PLP1,ZMIZ1,XAF1,COL11A1,RPL10,RPS17,VCL,DNAJA1,AKAP12,ITM2B,ZNF770,GTF2F1,SLC4A8,
B14,LDLR,GNL1,EFTUD2,LRATD2,STK24,PRR12,RIC8A,PLEKHB2,CADM3,SLC4A2,SLC25A23,AB
ALNT4,CYFIP2,DNAJC5,HNRNPDL,RPLP2,ARL6IP1,NDFIP1,ST3GAL2,TSR2,MYL6,SHOX,HNRNPK,
,TPI1,CHSY1,TMSB10,RPL19,PPP2R1A,RPL15,GTF3C4,DNAH8,HLA-C,SERINC1,CHD3,BASP1,HM

NDUFS1,CTR9,VEZF1,ATMIN,SNRPB,NKIRAS2,ZDHHC5,SKIV2L,MCRIP1,GPRIN1,CFLAR,SHC1,ME
A2,GAP43,RPL14,ZNF865,RPL9,CLSTN1,ATF4,KIF5B,GLIPR1L2,RPS14,CXADR,JUND,RPS20,CD

MYL12B,EIF4A3,ARF5,SCAP,PLS3,UBE2L3,CELSR1,SNX3,ESYT1,PSMB1,PPP5C,TSPYL2,HIP1R,MT
ST1,CTDSP2,SRSF5,TM9SF3,RAB27A,U2AF2,RPN2,SNX22,GDI2,TSR1,HSPH1,PKN1,HSPA4,TCP1,
-10,CIRBP,MYH10,RPL36,DUX4,PTGES3,SEPTIN11,CADM4,SND1,LAMP1,RHOB,CKAP4,CALU,PL
31,AKAP12,MAP4,SF3B2,CDC42BPB,ITM2B,AP2B1,TMEM170A,GTF2F1,SLC4A8,GOLGA8A,PXDN
1,RBBP4,PFN1,ASTN2,JPT2,TLE5,PNN,XPR1,SCRN1,HDLBP,TKT,EIF5A,CDH2,PNMA1,ACSL6,SOX
4A3,ZER1,IER2,IGLON5,SNX2,RIMS3,BGN,SLC25A36,CELSR1,HTATSF1,KDM3A,ZZEF1,PAIP2,AN
MB1,TSNAX,GTF3C1,ZKSCAN1,RGL2,PFKP,SLC38A1,SLC25A6,EXOC5,TRMT112,PSMD13,COPE,
RF4,TMEM248,CCT4,SKI,RPL36A,CNTNAP2,HDGF,SRP14,PYGB,ATP6AP1,ARF3,PPP2CA,ATG2B,
NS3,CTSD,SCARB2,NACA,TSPAN3,RTN4,LPCAT1,RPS24,SLC22A17,RPS9,XRCC5,CIPC,SLC16A7,
G2,UBFD1,MBTPS1,FEM1B,PRRC2A,CUL3,ADH5,PTPRF,NDST1,ULK1,ITGB5,TIMM50,TNKS,DKK
C1,SRSF1,ACADVL,CDC37,GABARAP,HNRNPUL1,UBE2R2,HSPB1,CLIP3,MAP2K2,ARF4,FLNB,TS
I,OGA,HNRNPA0,FADS2,LDHA,CPE,RPL29,RPSA,TPM4,CTSD,NACA,LPCAT1,LAMA5,SLC22A17,BR
K1,KLC1,PTOV1,CLSTN3,DDX42,CCT4,ORAI2,PSMB2,SCAF1,RAB11B,NRDC,TOP1,TUBB2A,OPT
K,LYPLA2,PTPN23,ATP13A1,NELFB,PAFAH1B2,POLDIP2,TRIP12,CHMP1A,ITFG1,MTA2,CGGBP1,
I,CCNI,TADA2B,SPTBN2,SLC3A2,CSNK1E,PFKL,RPS25,TKT,CDH6,TPM1,EIF5A,PDIA6,GPC3,PLXN
,RAB1B,VPS41,SEC14L1,ELAVL1,NUDCD3,GNL1,TUBGCP6,EEF1B2,RIC8A,SLC4A2,ACTR1B,MTM

,TYK2,OTUD5,GID8,CDK9,CELF3,PWWP3A,ATP6V0B,ARPC3,ZNF503,DAAM1,CLPTM1L,WDR11,
CF25,CDC37,PCSK1N,HNRNPUL1,UBE2R2,ACTN1,CLIP3,FKBP8,KDELR1,CCAR2,ARF4,ZFAND5,

PDAP1,PHB2,GSPT1,UBA52,SLC39A7,CSNK1G2,ARID1B,RER1,TRA2B,ADH5,MCM3AP,GPSM1,IT

4,BEX4,RBBP7,SSR1,BCL9L,WDR45B,HIP1,FLOT1,SLC39A6,CD99,SEC16A,GPX4,PPM1G,SLC35
H2,TCAF1,YARS1,SSBP3,SOX2,LTBP4,ELAVL1,MAZ,GTF2I,SRSF2,TRAP1,INTS11,RAB6A,PRPF4B,
AMP5,ZZEF1,ESYT1,GTF3C1,YTHDF3,KLF13,CDV3,FZR1,TNFAIP1,COPE,FAM50A,CD2BP2,ALDH1
3N2,PTN,SFPQ,TPM1,EIF5A,PEBP1,CCND2,PIP4K2B,ACSL6,SOX12,ACTN4,ITM2C,CLTC,RPS5,APL
1,MTA2,CGGBP1,PSMB1,TSNAX,SLC25A6,EXOC5,NDUFA4,SF3A3,TRMT112,PSMD13,WDR26,CD
HNRNPD,VGF,FBRSL1,MEX3A,ARFGEF2,WBP2,KDM4B,RBM15B,BEX4,VDAC1,SSR1,ST3GAL6,PF
2,NIBAN2,MKNK2,ELN,TMX1,TIA1,ARHGEF40,SGSM3,SLC38A1,EXOC5,CRIP2,CYFIP1,EWSR1,S

C5,KPNA3,ROCK2,MAF1,SLC25A29,TRAF3,PIN1,SPATS2L,UQCRH,LRRN1,SPIN1,BCAR1,SEMA6C
0,EIF4A1,MRC2,PPP2R1A,RPL15,GTF3C4,LDHB,PKD1,HLA-C,SSBP2,ZNF43,RPS3,CHD3,BASP1,
A,ELOA,TOLLIP
G2,ATRX,RAB2A,GNG4,FARP1,CD164,UPF1,UBFD1,MASTL,MBTPS1,ZFR,MAPRE2,SEPTIN7,APEX1
CR,SLC35E2B,SMCHD1,HDAC2,TMEM30A,MMP2,RPL23A,NNAT,LUC7L3,FAM199X,SERF2,DDX3

)10B,COPS6,TM7SF2,HSD17B12,H2AX,PBXIP1,CCNG1,NUP98,ZCCHC14,NSD1,SLC43A2,RBM10
1,CYB5R3,MAP4,SF3B2,CDC42BPB,AP2B1,MAT2A,TMEM170A,GTF2F1,GOLGA8A,PXDN,HNRNPU
PRM,WARS1,CLCN6,OTUB1,COX6B1,KDM5A,TBKBP1,ADAMTS7,SKIV2L,COX4I1,LZTS2,SHC1,PC
320,COL5A2,ATXN7L3B,ATP2A2,RPL26,SRSF4,RBBP4,RTN3,ASTN2,PNN,SPC25,NES,RPL31,NISCH
UCKS1,QPRT,LRRC41,SYT11,MED13,COX6A1,UBE3A,ATP5MC3,NASP,YY1,SMARCD1,CREBBP,XI
NELFB,PAFAH1B2,ACAT2,POLDIP2,TRIP12,LMAN2,CHMP1A,CLTA,ZZEF1,TSNAX,PAIP2,PFKP,PCM
,GSK3A,VASH1,SMARCD1,APBB1,ANKRD12,XPO6,BAG5,SLC20A1,SORT1,MICAL1,ATG4B,REEP1
ATP5PB,PTBP1,FTO,ILF3,WDR6,HNRNPUL1,TDRD5,SON,BNIP3L,PLCG1,CCAR2,ARF4,DUSP4,CO
KLF3,CFAP298,CPSF7,PITHD1,CSTB,ANAPC2,DPF2,RBBP6
KIF3C,DHX30,PHC2,SDHA,FAU,ILF2,STMN3,LSS,STT3B,STXBP1,TUG1,LOC728392,SF3A1,MESD

OMM40,LEMD2,UQCR11,COX5B,MYBBP1A
CS1,PRKAR1A,TLN1,CHPF,GJA1,JUN,NGRN,B4GALNT4,CYFIP2,DNAJC5,HNRNPDL,SLC25A3,RPL
1A,ITFG1,CGGBP1,PSMB1,FAM118A,TSNAX,GTF3C1,MKNK2,ACTA2,SLC25A6,EXOC5,NDUFA4,SI
5,PNN,NISCH,LRPPRC,KHDRBS1,SPTBN2,PTN,CSNK1E,HDLBP,RPS25,GLG1,SFPQ,PNMA1,PDIA6
DX39B,CDH11,CHD6,TOMM20,CCT2,CPSF1,PPP1CB,GNAI3,KLHL24,MEST,TSR1,TAF15,SSRP1,HS

P1,VAPB,EIF3H,ELAVL1,IMPDH2,GABARAPL1,GNL1,TM9SF4,TRAP1,TUBGCP6,PLEKHB2,ELP1,MC
COL2A1,MLEC,ACAP3,SURF4,ZNF302,RBMX,TMEM248,LAMC1,GCN1,TNPO1,SKI,PSMB2,RAB11I
JA1,CD63,NGRN,RPL35A,HNRNPDL,SLC25A3,RPLP2,ARL6IP1,CIRBP,RPL36,PTGES3,CKAP5,CAD

,TANC2,GNA13,DPYSL4,METRN,CROCC,WBP1,FBXL19,EIF4G1,ELAVL3,TRIP12,PSAT1,SNX3,HM
TF3,ANXA2,RPLP1,ZMIZ1,XAF1,SARAF,COL11A1,RPL10,MARCHF6,WSB1,VCL,AP3D1,SBF1,CYB5I
RA,SLC25A36,XRN2,SCAMP5,ZZEF1,AASDHPPT,ESYT1,GTF3C1,YTHDF3,KLF13,CDV3,FZR1,TNFA
PF19,YY1,GAK,EIF4A3,ELOVL5,SRPRA,IER2,CNBP,NTRK2,SLC25A36,XRN2,SCAMP5,ZZEF1,ANKH
DCY1,MFAP4,CELF3,UBE2Q1,ZNF768,NUP98,HERPUD1,ZBTB4,WDR82,DHFR,SYNGR1,MEX3C,R
ATP2A2,CRMP1,LASP1,OAZ1,SRSF4,RBBP4,RTN3,PFN1,ASTN2,MCL1,JPT2,PNN,SPC25,PRKCSH,
MSH6,IST1,DHCR7,NASP,APBB1,ATXN7L3,SSBP4,SLC20A1,DMWD,SRPRA,RIMS3,ELAVL3,PAFAH1

DP2,GALNT10,GLOD4,MTURN,GOLPH3,ECHS1,ALDH7A1,DHX36,HM13,COLGALT1,TSPAN7,NUP9

G1,CALM1,NPM1,PLXNB2,TTC3,PLD3,HSPD1,USP11,COL18A1,SNRNP200,CSDE1,SMIM14,IGF2,I
COG4,SCOC,EIF2S1,MAPK8IP3,MRPL3,DDX19A,DNMT1,C2ORF68,TOMM40,ATP6V1D,VPS37B
FAM32A,OTUD5,EPN1,ARPC3,CSNK1A1,MAPRE3,TNFRSF10B,ZBTB4,STAT2,HMGXB3,ACBD3,FBI
4,TMEM9,SCG2,WDR82,SYNGR1,MEX3C,USP48,RAB3A,RPS29,NSMF,DUSP8,RNF20,KDM5A,RBM
LC1,MLEC,CLSTN3,MDK,LAMC1,PTK7,SNRNP70,KDM6B,CNTNAP2,RAB11B,RNF10,VEGFA,ARF3
SMC5,POLDIP3,UNC45A,ZBTB4,STAT2,ABHD17A,PHKB,PDCD11,WARS1,SNRPB,STARD3,RNF20,S
SH3BGRL3,AKAP8L,MFAP4,TOM1L2,DEAF1,ZNF768,RABAC1,ZSWIM6,ZEB1,SEZ6L2,MEX3C,CACI
5,ZNF865,ATF4,NR2F1,GLIPR1L2,RPS14,CXADR,ATP1A1,TMEM259,RPS20,ATXN7L3B,RPL26,LA
HSPH1,LMNB2,MTRF1L,ALKBH5,SNRPN,HNRNPA3,PRDX5,UBE4A,HSPA4,TCP1,MLEC,ACAP3,ZNI

P5,ZZEF1,AASDHPPT,ESYT1,GTF3C1,YTHDF3,KLF13,CDV3,FZR1,TNFAIP1,FAM50A,CD2BP2,PIK3C
H8,PKD1,HLA-C,SSBP2,ZNF43,RPS3,BASP1,VAPA,HMGB1,POLR2A,AP2M1,TPT1,SEC61A1,RPS4X
,CREBBP,ATXN7L3,TANC2,WNK1,CERS2,DAPK3,SLC20A1,THRAP3,EIF4A3,PTPN23,FBXL19,EIF4

CT8,STT3B,MESD,PTPRS,PCBP2,RAB11A,SEC23A,ARNT2,SERBP1,KDM3B,PRPF40A,SMS,TXNIP,F
,ZZEF1,PSMB1,GTF3C1,HIP1R,ZKSCAN1,MACROH2A1,TCEA1,TRMT112,COPE,TMA7,CD2BP2,SL
ET,EGLN1,TPI1,TMSB10,EIF4A1,MRC2,PPP2R1A,RPL15,GTF3C4,LDHB,DNAH8,PKD1,HLA-C,SF3
MYL6,SHOX,WLS,H1-10,YWHAB,CKAP5,SND1,LAMP1,RPS12,CALU,ARCN1,ZNF454,CCT6A,STIP1
PF3,SPTY2D1,BORCS7,CAP1,IST1,ATP6V0D1,PRPF19,SART1,NASP,TGOLN2,AP1M1,EXOC4,SAP1
,GOT2,APBB1,KLHDC10,UBXN6,SCD,CROCC,PTPN23,USO1,DHX40,PAGR1,SRPRA,CALCOCO1,S
KTN1,MGAT5,YTHDF3,MAPK8IP2,NSF,ADNP,SART3,PNRC2,HMGB3,METAP2,ECH1,CFDP1,ETF1,
RN,CYFIP2,SLC25A3,ARL6IP1,NDFIP1,MXD4,SHOX,WLS,H1-10,CIRBP,RPL36,YWHAB,SEPTIN11,
,VGF,FBRSL1,MEX3A,ARFGEF2,RBM15B,CTNND1,VDAC1,SSR1,TRAF7,DFFB,TXNDC5,MTA1,TRI
NG8,WDR1,HMGN2,RPL26,CRMP1,SRSF4,RBBP4,PFN1,MCL1,PNN,NES,RPL31,NISCH,PRKCSH,

TP1B2,FOXK2,NSD2,USP24,FBXW5,SF3A2,DENND4B,PHIP,TRIM8,DAPK3,SORT1,CD74,SAE1,PRD
SPPL2B,ITFG1,SGSM2,AUTS2,MTA2,CGGBP1,ESYT1,TMEM94,HDGFL3,POR,CLIP2,FAM118A,NC
1C,SDF4,SMARCC1,HNRNPM,ISYNA1,STT3A,PLXNA3,IDH1,SARS1,COL27A1,MAPK8IP1,SOD1,ME
,GNL1,TM9SF4,ZNF428,TRAP1,TUBGCP6,PLEKHB2,ELP1,RAB6A,SPAG9,MOAP1,KRT8,CLASP2,

IRAP3,TRO,UBE2K,ZER1,SRPRA,SH3PXD2B,RANGAP1,TCEAL4,CAD,MDH1,CDC34,CELSR1,SNX3,
PL2B,ZZEF1,MGAT5,YTHDF3,NSF,ADNP,SART3,PNRC2,HMGB3,METAP2,ECH1,ETF1,TNRC6A,PP
CYB5R3,AKAP12,MAP4,SF3B2,AP2B1,MAT2A,PGAM1,NFE2L1,SMARCA5,CTNNA1,RPL23,CNN3,R
TMEM167A,NPLOC4,CHERP,FKBP1A,CTTN,SLC25A5,ATP6AP2,LRP10,MTCH1,VPS26B,FUS,ERH,C

GJA1,JUN,NGRN,B4GALNT4,CYFIP2,DNAJC5,HNRNPDL,TARDBP,RPLP2,ARL6IP1,NDFIP1,TSR2,M
N5,TCAF1,SSBP3,SOX2,LTBP4,ELAVL1,MAZ,GTF2I,SRSF2,INTS11,RAB6A,PRPF4B,MTMR4,WASF

TP6V0C,SKP1,LAPTM4B,PEA15,CPSF2,NUCB1,SRSF5,U2AF2,CCT2,MLXIP,NOP53,TSPYL4,MORF
TD4,CCT6A,MAGED4B,CTBP1,GPC1,HDAC2,IPO5,ATP6V0C,MMP2,DNER,LAPTM4B,TRIB2,FAM19
,PIK3C2B,ALDH18A1,CNDP2,GALNT10,GLOD4,MTURN,GOLPH3,ECHS1,DHX36,HM13,COLGALT
,INA,EIF4A1,MRC2,RPL19,PPP2R1A,RPL15,GTF3C4,LDHB,PKD1,SF3B1,PDIA3,SSBP2,ZNF43,RPS
H3BP5,OTUB1,POGK,CHD8,NKIRAS2,LARGE1,RHOBTB2,PMPCA,CDC42EP4,NUP133,CNTN1,BA

9B,YME1L1,DHX15,KIF5A,EIF2AK1,WDR45B,PRMT1,PDAP1,MMP15,EIF3D,PHB2,KLHL42,PLXNB

RH,EIF2S2,SLIT1,IARS2,SLC44A2,INTS3,ABI2,RAB1B,UBE2M,RALY,SEC14L1,C6ORF62,ELAVL1,

TDRD5,UBE2R2,CAND1,QARS1,RPL28,ARF4,SDF4,KIF3C,SMARCC1,PTP4A2,FBLN1,EIF3F,ID4,Z
CXADR,LENG8,WDR1,HMGN2,RPL26,CRMP1,LASP1,OAZ1,SRSF4,RBBP4,RTN3,MCL1,SPC25,PR
17,VCL,AP3D1,ARF1,MAP4,SF3B2,CDC42BPB,ITM2B,AP2B1,MAT2A,TMEM170A,GTF2F1,SLC4A

3,FAM199X,DLG5,SERF2,TIMP3,TMEM132A,CTDSP2,CNN2,EID1,NOVA2,CTSB,NUCB1,C5ORF24
X19A,PPM1A,GNAO1,RAD23A,SLC39A1,EPHA2,SKIL,STX12,VPS37B,RBBP6,ZC3H4,NDUFA10,FK
LC1,MSI2,CCNL1,RNPS1,MAPK8IP3,PAF1,DDX19A,PPM1A,GNAO1,RAD23A,SLC39A1,EPHA2,SKIL
SECISBP2L,EPHA2,HES6,NAMPT,ABCA7

LDIP3,UNC45A,ZBTB4,STAT2,P3H1,HMGXB3,DHFR,ACBD3,CRABP1,ABHD17A,PHKB,PDCD11,W
ABARAP,SCAF11,HSPB1,SON,BNIP3L,CLIP3,FKBP8,PLCG1,KDELR1,CCAR2,XPNPEP3,GIGYF1,SD
USP8,RNF20,NUP133,CHGB,COX4I1,OGDH,DSTYK,GSTA4,GUK1,UBN1,NECTIN2,PLCD3,CTDNE
APK8IP3,PAF1,DDX19A,PPM1A,GNAO1,RAD23A,SLC39A1,EPHA2,SKIL,STX12,VPS37B,RBBP6,ZC
APK8IP3,PAF1,DDX19A,PPM1A,GNAO1,RAD23A,SLC39A1,EPHA2,SKIL,STX12,VPS37B,RBBP6,ZC
T7,IRF2BP2,DUSP8,RBM14,NUP133,CACYBP,DNAJB6,ADIPOR1,PLCD3,PCDHB2,ZNF598,QRICH1
GL1,GAP43,NCS1,RPL22,RHOA,PTCD3,ATF4,PSMD2,NR2F1,KIF5B,RAB6B,ENC1,NLGN2,CXADR,
XPNPEP3,SMARCC1,ATP6V1B2,HNRNPM,SDHA,YWHAH,SLIT3,PITPNM1,ISYNA1,STT3A,IDH1,SAR
3A2,CDH6,TPM1,EIF5A,DDR1,SULF1,CCN2,DDX3X,RPS15,RPL12,ITM2C,EIF3L,NANOG,MAGED2
NAJC5,TARDBP,SLC25A3,RPLP2,TAGLN,WLS,HNRNPK,INIP,CIRBP,NARS1,PTGES3,CKAP5,LAMP1
NAJC5,TARDBP,SLC25A3,RPLP2,TAGLN,WLS,HNRNPK,INIP,CIRBP,NARS1,PTGES3,CKAP5,LAMP1
GF,SRP14,ABCA2,PRPF6,DPYSL2,TMED2,RPL38,RAPGEF1,TUBB2A,OPTN,CCND1,KDELR2,SRS

MARCA5,LARP1,HNRNPA0,CTNNA1,FADS2,RPS16,RPL23,CPE,RPL29,TPM4,TNS3,SCARB2,NACA,
LN2,MYL12A,CCDC6,HIRA,ACP1,NEO1,PPP1R12A,LMTK3,DSTYK,SETD3,ZNF395,GUK1,WDR5,S

1,RPL18,TUBB4B,UCHL1,MORF4L1,BTF3,ANXA2,RPLP1,TIMP2,XAF1,SARAF,COL11A1,RPL10,M
BP,EIF5A,PEBP1,CCND2,DDR1,PLXNA1,RAB7A,DDX3X,CLTC,NANOG,GNB2,RPS5,RPS7,IK,MIF,H
K2B,MEGF8,DDX3X,RPS15,RPL30,CLTC,EIF3L,NANOG,MRFAP1,GNB2,RCN1,RPS7,MIF,HMGCS

S51,PPME1,MAGEF1,RPL27A,SLC25A1,FAM32A,OTUD5,EPN1,CDK9,RTF2,CLPTM1L,CSNK1A1,
SF9,ARPC2,PLEC,PRDX1,FAM168B,IPO7,DYNC1LI2,AEBP1,ILF3,PCSK1N,SCAF11,SON,BNIP3L,

X30,HNRNPM,GIGYF1,YWHAH,SLIT3,STMN3,ISYNA1,PLXNA3,CSNK1D,ATXN2L,IDH1,ZFAND3,PTF

K2B,PRDX2,DDX3X,ITM2C,RPL30,ARL4C,EIF3L,MRFAP1,GNB2,RCN1,APLP1,RPS7,MIF,GPAT4,P
NL1,MAPK8IP3,PAF1,DDX19A,PPM1A,GNAO1,RAD23A,MYEF2,SLC39A1,EPHA2,SKIL,STX12,VPS
REBBP,XPO6,AMBRA1,ATG4B,UBE2L3,SPPL2B,ZZEF1,MGAT5,YTHDF3,NSF,ADNP,SART3,PNRC
X5,ZNF865,CLSTN1,PSMD2,KIF5B,RAB6B,SLC7A5,ENC1,NLGN2,CXADR,ATP1A1,JUND,TMEM25
GLN,SHOX,ANKRD52,CIRBP,MYH10,NARS1,RPL36,DUX4,YWHAB,PTGES3,SEPTIN11,UBE2H,ATF

LM1,RPS10,MTSS2,DNAJC5,TARDBP,SLC25A3,RPLP2,TAGLN,WLS,HNRNPK,INIP,CIRBP,NARS1,
DH,DSTYK,GSTA4,GUK1,UBN1,NECTIN2,PLCD3,CTDNEP1,CMTR1,CLCN3,PPP2R5B,TRAPPC1,A
1D,HYOU1,IFITM3,GHITM,DGKD,SREBF2,VGF,MLF2,SEC23A,RTL8C,WBP2,AKIRIN1,NAP1L4,TP

N,GOLPH3,PDGFRB,NECTIN1,COX7A2L,SH3BGRL3,DHX36,HM13,COLGALT1,DAAM1,TSPAN7,N
MPCA,PER1,BACE1,NR2F6,ANOS1,DDHD2,GRHPR,GPRIN1,EPB41L1,SLC25A29,KLHL36,B4GALT
ME1,CSNK2B,RPL27A,FAM32A,OTUD5,EPN1,ARPC3,CSNK1A1,MAPRE3,POLDIP3,PSMF1,HMGXB
PP1R15A,GALNT11,DAP,BAZ2A,VAPB,RTL6,EIF3H,SNRPD3,ELAVL1,IMPDH2,VPS4A,GABARAPL1,

LC1,MSI2,CCNL1,RNPS1,MAPK8IP3,PAF1,DDX19A,PPM1A,GNAO1,RAD23A,SLC39A1,EPHA2,SKIL
NF428,TRAP1,TUBGCP6,PLEKHB2,ELP1,RAB6A,MOAP1,CLASP2,RBM12,WBP11,SRSF11,NSD2,G
HGB,COX4I1,OGDH,DSTYK,GSTA4,GUK1,UBN1,NECTIN2,PLCD3,CTDNEP1,RNASEK,CMTR1,CLC

I2,PLCD3,CTDNEP1,CMTR1,CLCN3,TRAPPC1,ABCE1,NOLC1,MSI2,CCNL1,RNPS1,MAPK8IP3,PAF

,PDCD11,NREP,RPS29,NSMF,DUSP8,RNF20,NUP133,CHGB,COX4I1,OGDH,DSTYK,GSTA4,GUK1,

KIF5A,XPOT,TXNDC5,ACTR1A,PDAP1,HIP1,APMAP,HOXB5,NXN,PAPSS1,UBAP2L,RER1,PI4KA,CRT

1,TSNAX,GTF3C1,ZKSCAN1,KLF13,TNFAIP1,FAM50A,CD2BP2,ALDH18A1,CNDP2,RNF157,GALNT

QCRC2,HNRNPD,VGF,FBRSL1,MEX3A,RPL39,ARFGEF2,WBP2,KDM4B,RBM15B,BEX4,VDAC1,SS

QCRC2,HNRNPD,VGF,FBRSL1,MEX3A,RPL39,ARFGEF2,WBP2,KDM4B,RBM15B,BEX4,VDAC1,SS

,CROCC,RPS26,CNBP,IGLON5,SNX2,MDH1,TRIP12,LMO4,RNF11,PSAT1,TSPYL2,TSNAX,HSPA1A

B1,VCL,SBF1,DNAJA1,ARF1,PAFAH1B1,MAP4,SF3B2,ITM2B,AP2B1,GTF2F1,SLC4A8,GOLGA8A,

PN,PRDX5,UBE4A,HSPA4,TCP1,MLEC,ACAP3,SURF4,CHD2,RAB11B,SRP14,NRDC,PRPF6,TAF7,V

X19A,PPM1A,GNAO1,RAD23A,SLC39A1,EPHA2,SKIL,STX12,VPS37B,RBBP6,ZC3H4,NDUFA10,CC

COLGALT1,DAAM1,TSPAN7,RABAC1,NUP98,NACAD,OXSR1,EMC10,SEMA4C,TMEM9,SCG2,WDR

M,PPIB,HSBP1,NNAT,LUC7L3,FAM199X,DDX39B,SRP9,NUCB1,CDH11,TM9SF3,CHD6,RPN2,SUG

L2-BSCL2,TSC22D1,PLTP,MSL1,SMARCE1,CSNK1D,IDH1,NBPF9,MPRIP,COL27A1,DGKD,BAG6,

R1,BCL9L,WDR45B,TXNDC5,HIP1,H2AZ1,SLC39A6,CD99,GPX4,UBA52,PPM1G,HJURP,NXN,EP40

BP,SNX3,LGALS1,ITFG1,MTA2,ESYT1,POR,FAM118A,NOP56,KIFC2,H6PD,RUVBL2,SC5D,OBSL1

CYBP,FAM20B,ZNF395,ELOB,SENP3,MAPK3,ARL2BP,TRAPPC11,TRAF3,SLC1A5,SEMA6C,ADGRL

NRNPUL2-BSCL2,DUSP4,PITPNA,XPNPEP3,SUPT6H,RPS28,GIGYF1,LARS1,YWHAH,FAU,PFKM,PIT

100,ERH,PSMB4,EIF3H,SNRPD3,ANTXR1,ABI2,TPP1,RAB1B,UBE2M,VPS41,RALY,SEC14L1,C6OR

6ST1,TMEM132A,SRP9,CDH11,RAB27A,U2AF2,CCT2,TBCD,ANXA5,SNX22,VAMP2,PABPN1,GDI2,

T6H,RPS28,GIGYF1,LARS1,YWHAH,FAU,PFKM,PITPNM1,TNKS2,UBQLN1,TUG1,CSNK1D,IFITM3

AK1,OTUD5,CDK9,DHX36,ZRANB2,PSMC5,POLDIP3,TNFRSF10B,NCBP2,GPT2,PSMF1,PIP5K1C,

PGEF1,RPL37,VHL,UBXN4,MAPRE1,CCNL2,DSTN,PBX2,SRSF1,ATP5PB,EIF3E,ILF3,LUZP6,GABA

I2,PLCD3,CTDNEP1,CMTR1,CLCN3,TRAPPC1,ABCE1,NOLC1,MSI2,CCNL1,RNPS1,MAPK8IP3,PAF

CG2,WDR82,SYNGR1,USP48,RPS29,NSMF,DUSP8,RNF20,KDM5A,NUP133,CHGB,COX4I1,OGD

2,PLCD3,CTDNEP1,CMTR1,CLCN3,TRAPPC1,ABCE1,NOLC1,MSI2,CCNL1,RNPS1,MAPK8IP3,PAF1

_C20A1,GNA13,THRAP3,DOP1B,SAE1,FJX1,HSDL1,CROCC,RCN2,SH3PXD2B,EIF4G1,SNX2,TCEAl

:KB,COL4A2,RPL37A,LRRC28,FN1,SPEN,NONO,DDB1,PSMD5,YWHAQ,EIF1,HACD3,DCHS1,SET,E
YWHAQ,EIF1,HACD3,DCHS1,SET,EGLN1,TPI1,CHSY1,TMSB10,INA,EIF4A1,MRC2,RPL19,PPP2R1/
,DCHS1,SET,EGLN1,TPI1,CHSY1,TMSB10,INA,EIF4A1,MRC2,PPP2R1A,RPL15,GTF3C4,LDHB,DNA
INA,EIF4A1,MRC2,RPL19,PPP2R1A,RPL15,GTF3C4,LDHB,DNAH8,PKD1,HLA-C,SF3B1,PDIA3,SER
P2,SARAF,MIDN,RPL10,MARCHF6,RPS17,VCL,AP3D1,SBF1,ARF1,CYB5R3,MAP4,SF3B2,CDC42B
,DCHS1,SET,EGLN1,TPI1,CHSY1,TMSB10,INA,EIF4A1,MRC2,RPL19,PPP2R1A,RPL15,GTF3C4,LD
31,PSMD5,YWHAQ,EIF1,HACD3,DCHS1,SET,EGLN1,MAP9,TPI1,CHSY1,TMSB10,INA,EIF4A1,MRC
PJA2,KPNB1,C1ORF43,PAPOLA,LLGL1,HK1,NCS1,RHOA,CBX5,CLSTN1,ATF4,PSMD2,KIF5B,RAB
:43,RPS3,CHD3,BASP1,VAPA,HMGB1,CDKL1,TPT1,SEC61A1,RPS4X,RPS3A,STMN1,ITGB1,TUBB4
EIF4A1,MRC2,RPL19,PPP2R1A,RPL15,GTF3C4,LDHB,DNAH8,PKD1,SF3B1,PDIA3,SSBP2,ZNF43,F
:HSY1,TMSB10,INA,EIF4A1,MRC2,RPL19,PPP2R1A,GTF3C4,LDHB,PKD1,HLA-C,PDIA3,SSBP2,ZNF
DN,RPL10,MARCHF6,WSB1,VCL,AP3D1,SBF1,ARF1,CYB5R3,MAP4,CDC42BPB,ITM2B,MAT2A,TM
PL18,STMN1,ITGB1,TUBB4B,UCHL1,NCAM1,ANXA2,AGRN,TIMP2,XAF1,SARAF,MIDN,RPL10,MAF
P2M1,TPT1,SEC61A1,RPS3A,RPL18,STMN1,ITGB1,TUBB4B,UCHL1,MORF4L1,NCAM1,ANXA2,AG
RPL11,MORF4L1,BTF3,NCAM1,ANXA2,AGRN,RPLP1,ZMIZ1,TIMP2,COL11A1,MIDN,MARCHF6,WS
NA,EIF4A1,MRC2,PPP2R1A,GTF3C4,LDHB,PKD1,HLA-C,SF3B1,PDIA3,SSBP2,ZNF43,RPS3,BASP1
?2M1,TPT1,SEC61A1,RPS3A,RPL18,STMN1,ITGB1,TUBB4B,UCHL1,MORF4L1,NCAM1,ANXA2,AGI
L1,MORF4L1,NCAM1,ANXA2,AGRN,TIMP2,XAF1,SARAF,COL11A1,MIDN,RPL10,MARCHF6,VCL,AI
CA5,CTNNA1,FADS2,CNN3,RPL29,CTSD,EIF4H,SCARB2,NACA,TSPAN3,LPCAT1,LAMA5,SLC22A17
EC61A1,RPS3A,RPL18,STMN1,ITGB1,TUBB4B,UCHL1,NCAM1,ANXA2,AGRN,TIMP2,XAF1,SARAF,I
,UCHL1,MORF4L1,ANXA2,AGRN,RPLP1,ZMIZ1,TIMP2,MIDN,MARCHF6,WSB1,RPS17,VCL,AP3D1,
IA3,SERINC1,SSBP2,RPS3,CHD3,BASP1,VAPA,HMGB1,POLR2A,TPT1,STMN1,ITGB1,TUBB4B,UCI
AM1,ANXA2,AGRN,RPLP1,ZMIZ1,TIMP2,SARAF,MIDN,RPL10,MARCHF6,WSB1,VCL,AP3D1,SBF1,D
28,MATR3,YWHAG,CKB,COL4A2,RPL37A,LRRC28,FN1,SPEN,NONO,DDB1,PSMD5,YWHAQ,EIF1,I
I,MARCHF6,WSB1,RPS17,VCL,AP3D1,SBF1,DNAJA1,ARF1,CYB5R3,MAP4,SF3B2,CDC42BPB,ITM
C2,RPL19,PPP2R1A,GTF3C4,PKD1,HLA-C,SF3B1,PDIA3,SERINC1,SSBP2,ZNF43,RPS3,CHD3,BASI
MARCHF6,WSB1,RPS17,VCL,AP3D1,DNAJA1,ARF1,CYB5R3,SF3B2,CDC42BPB,ZNF770,AP2B1,M/
C2,RPL19,PPP2R1A,GTF3C4,LDHB,PKD1,HLA-C,SF3B1,PDIA3,SSBP2,CHD3,BASP1,VAPA,HMGB1
.4A2,RPL37A,LRRC28,MAPT,FN1,SPEN,NONO,DDB1,PSMD5,YWHAQ,EIF1,HACD3,DCHS1,SET,EG
KB,COL4A2,RPL37A,LRRC28,FN1,SPEN,NONO,DDB1,PSMD5,YWHAQ,EIF1,HACD3,DCHS1,SET,E(
:T,EGLN1,TPI1,CHSY1,TMSB10,INA,EIF4A1,MRC2,RPL19,PPP2R1A,RPL15,GTF3C4,LDHB,PKD1,S
:4L1,NCAM1,ANXA2,AGRN,TIMP2,XAF1,SARAF,COL11A1,MIDN,MARCHF6,VCL,AP3D1,SBF1,ARF1
RPL11,MORF4L1,BTF3,NCAM1,ANXA2,AGRN,ZMIZ1,TIMP2,SARAF,COL11A1,MIDN,MARCHF6,WS
3,FN1,SPEN,NONO,DDB1,PSMD5,YWHAQ,EIF1,HACD3,DCHS1,SET,EGLN1,MAP9,TPI1,CHSY1,TMS
3,CKB,COL4A2,RPL37A,LRRC28,MAPT,FN1,SPEN,NONO,DDB1,PSMD5,YWHAQ,EIF1,HACD3,DCH
2BPB,ITM2B,MAT2A,GTF2F1,GOLGA8A,PXDN,HNRNPU,NFE2L1,SMARCA5,CTNNA1,RPS16,RPL2
4A1,MRC2,PPP2R1A,RPL15,GTF3C4,LDHB,PKD1,SF3B1,PDIA3,SSBP2,ZNF43,RPS3,BASP1,VAPA,I
\,RPL18,STMN1,ITGB1,TUBB4B,UCHL1,RPL11,BTF3,ANXA2,AGRN,RPLP1,ZMIZ1,TIMP2,SARAF,M
F6,VCL,AP3D1,SBF1,DNAJA1,ARF1,CYB5R3,AKAP12,MAP4,SF3B2,CDC42BPB,ITM2B,AP2B1,MAT
RPL15,GTF3C4,LDHB,PKD1,SF3B1,PDIA3,SERINC1,SSBP2,ZNF43,CHD3,BASP1,VAPA,HMGB1,POI
HAQ,EIF1,HACD3,DCHS1,SET,EGLN1,MAP9,TPI1,CHSY1,TMSB10,INA,EIF4A1,MRC2,RPL19,PPP2F
)NO,DDB1,PSMD5,YWHAQ,EIF1,HACD3,DCHS1,SET,EGLN1,MAP9,TPI1,CHSY1,TMSB10,INA,EIF4/
RPLP1,ZMIZ1,TIMP2,SARAF,MIDN,RPL10,MARCHF6,WSB1,RPS17,VCL,AP3D1,SBF1,DNAJA1,ARF
,BASP1,VAPA,HMGB1,POLR2A,CDKL1,AP2M1,TPT1,SEC61A1,RPS3A,RPL18,STMN1,ITGB1,TUBB
/IRC2,RPL19,PPP2R1A,RPL15,GTF3C4,LDHB,DNAH8,PKD1,HLA-C,SF3B1,PDIA3,SERINC1,SSBP2,

HACD3,DCHS1,SET,EGLN1,TPI1,CHSY1,TMSB10,EIF4A1,MRC2,RPL19,PPP2R1A,RPL15,GTF3C4,D
3A,STMN1,ITGB1,TUBB4B,UCHL1,MORF4L1,NCAM1,ANXA2,AGRN,ZMIZ1,TIMP2,XAF1,SARAF,CO
1ORF43,COL1A2,LLGL1,HK1,NCS1,RHOA,ZNF865,CLSTN1,ATF4,RAB6B,SLC7A5,ENC1,NLGN2,C
3,DCHS1,SET,EGLN1,MAP9,TPI1,CHSY1,TMSB10,INA,MRC2,RPL19,PPP2R1A,RPL15,GTF3C4,LDH
28,MAPT,FN1,SPEN,NONO,DDB1,PSMD5,YWHAQ,EIF1,HACD3,DCHS1,SET,EGLN1,MAP9,TPI1,CH
MAP4,CDC42BPB,ITM2B,TMEM170A,GTF2F1,SLC4A8,PXDN,HNRNPU,PGAM1,NFE2L1,HNRNPA0,C
NFE2L1,SMARCA5,CTNNA1,RPS16,CNN3,RPL29,TNS3,CTSD,EIF4H,TSPAN3,LPCAT1,LAMA5,SLC2
RINC1,SSBP2,ZNF43,RPS3,VAPA,HMGB1,POLR2A,CDKL1,TPT1,SEC61A1,RPS3A,RPL18,STMN1,IT
TP1A1,JUND,CD81,WDR1,HMGN2,ATP2A2,CRMP1,LASP1,VAT1,OAZ1,SRSF4,RBBP4,RTN3,PFN1,
DHB,PKD1,SF3B1,PDIA3,SERINC1,SSBP2,ZNF43,RPS3,CHD3,BASP1,VAPA,HMGB1,POLR2A,CDKL
AZ1,SRSF4,PFN1,MCL1,JPT2,TLE5,NES,HNRNPC,CCNI,XPR1,KLHL9,SPTBN2,SLC3A2,PFKL,HDLB
ATP1A1,JUND,CD81,HMGN2,ATP2A2,CRMP1,LASP1,OAZ1,RBBP4,RTN3,PFN1,ASTN2,JPT2,TLE5,
ARB2,NACA,RPS24,SLC22A17,BRD3,CIPC,PJA2,MSN,KPNB1,COL1A2,LLGL1,HK1,NCS1,RPL22,R
2,NACA,LPCAT1,LAMA5,BRD3,CIPC,MSN,KPNB1,C1ORF43,COL1A2,LLGL1,NCS1,RHOA,ZNF865,
BP2,ZNF43,RPS3,CHD3,BASP1,VAPA,HMGB1,POLR2A,CDKL1,TPT1,SEC61A1,RPS4X,RPS3A,RPL1
SCH,CCNI,LRPPRC,TADA2B,KHDRBS1,CSNK1E,PFKL,HDLBP,RPS25,TKT,GLG1,SFPQ,EIF5A,CCNI
B10,INA,EIF4A1,MRC2,PPP2R1A,RPL15,GTF3C4,LDHB,DNAH8,PKD1,HLA-C,SF3B1,PDIA3,SSBP2
PL23,CNN3,TNS3,CTSD,EIF4H,SCARB2,NACA,LPCAT1,LAMA5,SLC22A17,BRD3,CIPC,MSN,KPNB1
T2,TLE5,NES,NISCH,HNRNPC,CCNI,LRPPRC,XPR1,PFKL,HDLBP,TKT,EIF5A,PNMA1,GPC3,PIP4K2
D1,SBF1,ARF1,CYB5R3,MAP4,CDC42BPB,ITM2B,MAT2A,PXDN,HNRNPU,OGA,NFE2L1,CTNNA1,FA
C4,LDHB,PKD1,HLA-C,PDIA3,SERINC1,SSBP2,RPS3,CHD3,BASP1,VAPA,HMGB1,POLR2A,TPT1,SE
CRMP1,LASP1,VAT1,OAZ1,SRSF4,RBBP4,RTN3,PFN1,JPT2,TLE5,NES,NISCH,HNRNPC,CCNI,LRPP
D3,BASP1,VAPA,HMGB1,POLR2A,CDKL1,AP2M1,TPT1,SEC61A1,RPS4X,RPL18,STMN1,ITGB1,TUB
3,CIPC,PJA2,SLC16A7,KPNB1,RPL17,IGFBP3,COL1A2,LLGL1,HK1,NCS1,RPL22,RHOA,CBX5,ZNF
MGN2,ATP2A2,CRMP1,LASP1,BEX3,VAT1,OAZ1,SRSF4,RTN3,PFN1,JPT2,TLE5,NES,NISCH,HNRN
CTNNA1,FADS2,RPS16,CPE,CNN3,RPL29,RPSA,CTSD,EIF4H,SCARB2,NACA,TSPAN3,LPCAT1,LAM
NCAM1,ANXA2,AGRN,ZMIZ1,TIMP2,XAF1,SARAF,COL11A1,MIDN,MARCHF6,WSB1,VCL,AP3D1,SE
1,RHOA,CBX5,ZNF865,CLSTN1,NR2F1,RAB6B,SLC7A5,NLGN2,CXADR,CHD4,ATP1A1,JUND,RPS2
HAG,CKB,COL4A2,RPL37A,LRRC28,MAPT,FN1,SPEN,NONO,DDB1,PSMD5,YWHAQ,EIF1,HACD3,D
P1,VAPA,POLR2A,CDKL1,AP2M1,TPT1,RPL18,STMN1,ITGB1,UCHL1,RPL11,MORF4L1,BTF3,NCA
1,HACD3,DCHS1,SET,EGLN1,MAP9,TPI1,CHSY1,TMSB10,INA,EIF4A1,MRC2,RPL19,PPP2R1A,RPL
TSD,SCARB2,NACA,LAMA5,RPS24,SLC22A17,BRD3,CIPC,PJA2,MSN,KPNB1,IGFBP3,COL1A2,LLG
,SF3B1,SERINC1,SSBP2,CHD3,BASP1,VAPA,HMGB1,POLR2A,CDKL1,AP2M1,TPT1,SEC61A1,RPS
PPP2R1A,RPL15,GTF3C4,LDHB,DNAH8,HLA-C,SF3B1,PDIA3,SSBP2,ZNF43,RPS3,CHD3,BASP1,VA
HF6,AP3D1,SBF1,ARF1,CYB5R3,AKAP12,MAP4,SF3B2,CDC42BPB,ITM2B,MAT2A,TMEM170A,GT
A5,HNRNPA0,CTNNA1,FADS2,TPM4,SCARB2,NACA,LAMA5,RPS24,BRD3,CIPC,PJA2,MSN,KPNB1,
,JPT2,TLE5,NES,NISCH,HNRNPC,CCNI,LRPPRC,XPR1,KHDRBS1,PFKL,HDLBP,TKT,SFPQ,CDH2,P
,COL4A2,RPL37A,LRRC28,MAPT,FN1,SPEN,NONO,DDB1,PSMD5,YWHAQ,EIF1,HACD3,DCHS1,SE
EIF4H,SCARB2,NACA,TSPAN3,LPCAT1,LAMA5,SLC22A17,BRD3,CIPC,PJA2,MSN,SLC16A7,KPNB1
TNS3,CTSD,EIF4H,NACA,TSPAN3,LPCAT1,LAMA5,SLC22A17,BRD3,XRCC5,CIPC,PJA2,KPNB1,C1O
TNS3,CTSD,EIF4H,NACA,TSPAN3,LPCAT1,LAMA5,SLC22A17,BRD3,XRCC5,CIPC,PJA2,KPNB1,C1O
1A2,LLGL1,GAP43,NCS1,RPL22,RHOA,CBX5,ZNF865,KIF5B,SLC7A5,ENC1,RPS14,CXADR,CHD4,
3,YWHAG,CKB,COL4A2,LRRC28,FN1,SPEN,NONO,DDB1,PSMD5,YWHAQ,EIF1,HACD3,DCHS1,SET
PE,CNN3,TNS3,CTSD,LPCAT1,LAMA5,SLC22A17,BRD3,CIPC,KPNB1,PAPOLA,IGFBP3,COL1A2,LL
2,RPS16,CPE,CNN3,RPSA,CTSD,EIF4H,NACA,LPCAT1,LAMA5,RPS24,SLC22A17,BRD3,XRCC5,CIP

/IAT2A,GTF2F1,GOLGA8A,PXDN,HNRNPU,NFE2L1,HNRNPA0,CTNNA1,RPS16,CNN3,TNS3,CTSD,E
CBX5,ZNF865,CLSTN1,RAB6B,SLC7A5,NLGN2,CHD4,ATP1A1,JUND,EIF5,CD81,WDR1,HMGN2,A
,NR2F1,SLC7A5,NLGN2,CXADR,CHD4,ATP1A1,JUND,RPS20,EIF5,CD81,LENG8,WDR1,HMGN2,AT
31,WDR1,HMGN2,ATP2A2,CRMP1,LASP1,BEX3,VAT1,RBBP4,RTN3,PFN1,JPT2,TLE5,NES,HNRNPC
DNAJA1,ARF1,CYB5R3,MAP4,CDC42BPB,ITM2B,MAT2A,SLC4A8,GOLGA8A,PXDN,HNRNPU,OGA,
IM28,MATR3,YWHAG,CKB,COL4A2,RPL37A,LRRC28,FN1,SPEN,NONO,DDB1,PSMD5,YWHAQ,EIF
0,APLP1,CHPF,JUN,MYO5A,COL6A2,SESN3,B4GALNT4,MTSS2,DNAJC5,APLP2,RPLP2,MXD4,ANK
N3,TNS3,CTSD,NACA,LPCAT1,LAMA5,SLC22A17,BRD3,XRCC5,CIPC,KPNB1,C1ORF43,COL1A2,LL
LRRC28,MAPT,FN1,SPEN,NONO,DDB1,YWHAQ,EIF1,HACD3,DCHS1,SET,EGLN1,TPI1,CHSY1,TMS
NPU,NFE2L1,SMARCA5,HNRNPA0,CTNNA1,FADS2,RPS16,LDHA,CNN3,RPSA,TNS3,EIF4H,TSPAN3,
F3,NCAM1,ANXA2,AGRN,TIMP2,SARAF,COL11A1,MARCHF6,RPS17,VCL,AP3D1,SBF1,DNAJA1,AR
3,CNN3,RPL29,TNS3,CTSD,NACA,LPCAT1,LAMA5,SLC22A17,BRD3,CIPC,PJA2,MSN,KPNB1,C1OR
4,CXADR,CHD4,ATP1A1,JUND,TMEM259,EIF5,CD81,WDR1,HMGN2,ATP2A2,CRMP1,LASP1,OAZ1
KPNB1,C1ORF43,PAPOLA,COL1A2,LLGL1,NCS1,RHOA,CBX5,ZNF865,CLSTN1,NR2F1,RAB6B,SL
COL4A2,RPL37A,LRRC28,FN1,SPEN,NONO,DDB1,YWHAQ,EIF1,HACD3,DCHS1,SET,EGLN1,TPI1,C
SLC16A7,KPNB1,RPL17,PAPOLA,COL1A2,LLGL1,HK1,NCS1,RHOA,CBX5,RPL9,CLSTN1,ATF4,NR
NRNPH1,HLA-B,RAB1A,PLPPR3,SEPTIN3,HECTD4,MAGED4B,CTBP1,ARHGDIA,CAPRIN1,STIP1,GP
MIDN,RPL10,MARCHF6,RPS17,VCL,AP3D1,SBF1,ARF1,CYB5R3,PAFAH1B1,AKAP12,MAP4,CDC42E
FBP3,LLGL1,GAP43,HK1,NCS1,RPL22,RHOA,ZNF865,CLSTN1,ATF4,PSMD2,KIF5B,RAB6B,ENC1
RMP1,LASP1,BEX3,VAT1,OAZ1,PFN1,ASTN2,TLE5,NISCH,CCNI,TADA2B,KHDRBS1,XPO1,CSNK1E
A5,RPS24,SLC22A17,BRD3,XRCC5,CIPC,KPNB1,C1ORF43,PAPOLA,IGFBP3,COL1A2,LLGL1,HK1,
,PNN,NISCH,TADA2B,SPTBN2,CSNK1E,PFKL,HDLBP,GLG1,SFPQ,TPM1,CCND2,CDH2,PNMA1,PD
GALNT4,DNAJC5,APLP2,TARDBP,MXD4,MYL6,ANKRD52,WLS,HNRNPK,H1-10,INIP,CIRBP,DUX4,C
PR3,SEPTIN3,COTL1,HECTD4,MAGED4B,CTBP1,ARHGDIA,CAPRIN1,STIP1,GPC1,SMCHD1,IPO5,
3TF3,NCAM1,ANXA2,AGRN,RPLP1,ZMIZ1,TIMP2,XAF1,SARAF,MIDN,MARCHF6,VCL,AP3D1,SBF1,D
LPPR3,SEPTIN3,COTL1,HECTD4,CTBP1,RAC1,ARHGDIA,CAPRIN1,STIP1,GPC1,SLC35E2B,SMCHD
YB5R3,MAP4,SF3B2,CDC42BPB,ITM2B,ZNF770,MAT2A,TMEM170A,GTF2F1,SLC4A8,PXDN,HNR
M1,AGRN,RPLP1,ZMIZ1,TIMP2,SARAF,COL11A1,MIDN,RPL10,MARCHF6,RPS17,VCL,AP3D1,SBF1
TM2B,MAT2A,GTF2F1,SLC4A8,PXDN,HNRNPU,OGA,NFE2L1,SMARCA5,LARP1,HNRNPA0,CTNNA
MARCHF6,WSB1,RPS17,AP3D1,SBF1,DNAJA1,ARF1,SF3B2,ITM2B,MAT2A,SLC4A8,PXDN,HNRNPU
WHAG,CKB,COL4A2,LRRC28,MAPT,FN1,SPEN,NONO,DDB1,PSMD5,YWHAQ,HACD3,DCHS1,SET,
IF4H,LPCAT1,LAMA5,SLC22A17,BRD3,CIPC,MSN,KPNB1,C1ORF43,IGFBP3,COL1A2,LLGL1,HK1
52,WLS,HNRNPK,H1-10,INIP,DUX4,CADM4,UBE2H,ATP1A2,RAB1A,PLPPR3,ARCN1,COTL1,MAGE
,PSMD2,RAB6B,SLC7A5,NLGN2,CXADR,CHD4,ATP1A1,JUND,CD81,LENG8,WDR1,HMGN2,ATP2A
ARCHF6,WSB1,VCL,SBF1,DNAJA1,ARF1,CYB5R3,AKAP12,MAP4,CDC42BPB,ITM2B,AP2B1,MAT2
NF43,CHD3,BASP1,VAPA,HMGB1,POLR2A,CDKL1,TPT1,RPL18,STMN1,ITGB1,TUBB4B,UCHL1,M
D1,IPO5,DNER,BCLAF1,HSBP1,LUC7L3,SQLE,MAP1A,CDKN1A,DLG5,CRPPA,HS6ST1,PGRMC1,C
SLC3A2,PFKL,HDLBP,RPS25,TKT,EIF5A,CDH2,PDIA6,GPC3,PIP4K2B,MEGF8,SOX12,DDX3X,RPL
TRIB2,PEA15,SQLE,TIMP3,CTDSP2,NOVA2,CTSB,CHD6,EEF1A2,CCT2,TBCD,CPSF1,SNX22,PGD,S
RD3,PJA2,KPNB1,RPL17,C1ORF43,PAPOLA,COL1A2,LLGL1,NCS1,RHOA,ZNF865,CLSTN1,NR2F
G8,WDR1,ATP2A2,CRMP1,LASP1,BEX3,OAZ1,SRSF4,RTN3,ASTN2,JPT2,NES,NISCH,CCNI,TADA2
L31,PRKCSH,HNRNPC,LRPPRC,CSNK1E,HDLBP,GLG1,TPM1,EIF5A,CDH2,PNMA1,PDIA6,SOX12,
NRNPU,NFE2L1,SMARCA5,CTNNA1,RPS16,RPL23,CNN3,TPM4,CTSD,TSPAN3,LPCAT1,LAMA5,RP
N2,ATP2A2,CRMP1,LASP1,BEX3,VAT1,OAZ1,RTN3,PFN1,MCL1,TLE5,NES,NISCH,HNRNPC,CCNI,
G,CKB,COL4A2,RPL37A,LRRC28,MAPT,FN1,SPEN,NONO,DDB1,PSMD5,YWHAQ,EIF1,HACD3,DC

N,HNRNPU,OGA,SMARCA5,HNRNPA0,CTNNA1,FADS2,CPE,CNN3,RPSA,TNS3,EIF4H,SCARB2,NA
K,MYH10,NARS1,RPL36,DUX4,YWHAB,CADM4,SND1,RHOB,UBE2H,CKAP4,HLA-B,RAB1A,PLPPR
1E,PFKL,TKT,EIF5A,PEBP1,CCND2,CDH2,PNMA1,GPC3,PIP4K2B,SOX12,ACTN4,MMP14,PRDX2,
,HNRNPU,PGAM1,HNRNPA0,FADS2,RPS16,RPL23,LDHA,CNN3,RPSA,TNS3,CTSD,EIF4H,SCARB2
1,RPS4X,RPS3A,RPL18,ITGB1,TUBB4B,UCHL1,RPL11,MORF4L1,BTF3,ANXA2,AGRN,RPLP1,ZMI
,SARAF,COL11A1,MIDN,MARCHF6,VCL,AP3D1,SBF1,DNAJA1,ARF1,CYB5R3,AKAP12,MAP4,SF3B
N1,ATF4,SLC7A5,CHD4,ATP1A1,JUND,EIF5,CD81,WDR1,HMGN2,ATP2A2,RPL26,CRMP1,LASP1,
EM170A,PXDN,HNRNPU,NFE2L1,SMARCA5,HNRNPA0,CTNNA1,RPS16,CNN3,RPSA,TNS3,CTSD,EI
4,LDHB,PKD1,SF3B1,PDIA3,SSBP2,RPS3,BASP1,VAPA,HMGB1,POLR2A,AP2M1,TPT1,RPS4X,RPS
AQ,EIF1,HACD3,DCHS1,SET,EGLN1,MAP9,TPI1,CHSY1,TMSB10,INA,EIF4A1,MRC2,RPL19,PPP2
RPL30,CLTC,H1-0,RPS5,APLP1,IK,MYO5A,NGRN,B4GALNT4,MTSS2,RPL35A,DNAJC5,APLP2,HN
,HECTD4,MAGED4B,CTBP1,ARHGDIA,CAPRIN1,STIP1,GPC1,TOP2B,SLC35E2B,SMCHD1,IPO5,AT
PO5,ATP6V0C,MMP2,DNER,TRIB2,PEA15,SQLE,DLG5,HS6ST1,TIMP3,PGRMC1,CNN2,NOVA2,CT
PS16,LDHA,CPE,CNN3,CTSD,EIF4H,SCARB2,NACA,LPCAT1,LAMA5,SLC22A17,BRD3,CIPC,PJA2,
NN3,RPL29,CTSD,NACA,LPCAT1,LAMA5,BRD3,CIPC,KPNB1,C1ORF43,PAPOLA,IGFBP3,LLGL1,
1,RPS17,VCL,AP3D1,DNAJA1,ARF1,CYB5R3,MAP4,SF3B2,CDC42BPB,ITM2B,MAT2A,TMEM170A,
CIRBP,RPL36,DUX4,YWHAB,CADM4,LGALS3BP,RAB1A,PLPPR3,COTL1,MAGED4B,CTBP1,ARHG
1-10,DUX4,PTGES3,CADM4,RHOB,UBE2H,ATP1A2,RAB1A,PLPPR3,ARCN1,COTL1,MAGED4B,CT
PL19,PPP2R1A,GTF3C4,LDHB,PKD1,HLA-C,SF3B1,PDIA3,SSBP2,ZNF43,CHD3,VAPA,HMGB1,CD
C61A1,RPS4X,RPL18,STMN1,ITGB1,TUBB4B,UCHL1,RPL11,MORF4L1,ANXA2,AGRN,RPLP1,ZMI
NF43,RPS3,BASP1,VAPA,HMGB1,POLR2A,CDKL1,AP2M1,TPT1,SEC61A1,RPS4X,RPS3A,STMN1,I
TSD,EIF4H,LPCAT1,LAMA5,BRD3,CIPC,KPNB1,COL1A2,LLGL1,GAP43,NCS1,RHOA,CBX5,ZNF86
FE2L1,SMARCA5,HNRNPA0,CTNNA1,RPL23,CPE,CNN3,RPL29,TPM4,EIF4H,SCARB2,NACA,LPCA
1,UBB,AFF4,TRIM28,MATR3,YWHAG,CKB,COL4A2,RPL37A,LRRC28,MAPT,FN1,SPEN,NONO,DDE
PR3,COTL1,HECTD4,CCT6A,CTBP1,ARHGDIA,STIP1,TOP2B,SLC35E2B,APPL1,SMCHD1,IPO5,DN
1,PFN1,JPT2,TLE5,SPC25,RPL31,NISCH,HNRNPC,CCNI,TADA2B,KHDRBS1,SPTBN2,CSNK1E,PF
RCN1,COTL1,MAGED4B,CAPRIN1,STIP1,GPC1,CDK16,APPL1,TNC,ATP6V0C,TMEM30A,DNER,PE
TMSB10,EIF4A1,MRC2,RPL19,RPL15,GTF3C4,LDHB,DNAH8,PKD1,HLA-C,SF3B1,PDIA3,SERINC1
EGF8,SOX12,PRDX2,DDX3X,RPS15,RPL12,ARL4C,RCN1,APLP1,MIF,GPAT4,CHPF,MYO5A,COL6A
PT1,SEC61A1,RPS3A,STMN1,TUBB4B,UCHL1,RPL11,MORF4L1,BTF3,NCAM1,ANXA2,AGRN,RPL
A2,CTSB,CDH11,MDH2,U2AF2,EEF1A2,MLXIP,SNX22,HNRNPL,PARP1,MORF4L2,ABCA3,HNRNPA
2,ITM2C,ARL4C,APLP1,RPL35,MIF,GPAT4,TLN1,CHPF,NGRN,GOLM1,SESN3,B4GALNT4,MTSS2
BP3,GAP43,RPL14,RPL22,RHOA,PTCD3,ATF4,PSMD2,RAB6B,SLC7A5,JUND,TMEM259,RPS20,E
HMGN2,ATP2A2,CRMP1,BEX3,VAT1,OAZ1,RBBP4,RTN3,PFN1,MCL1,JPT2,TLE5,NISCH,HNRNPC,
,SEC61A1,RPS3A,RPL18,STMN1,ITGB1,TUBB4B,UCHL1,RPL11,BTF3,ANXA2,AGRN,ZMIZ1,XAF1,
,VAMP2,MORF4L2,HNRNPA3,PRDX5,ACTR3,HSPA4,CAMSAP1,GRINA,PTOV1,CLSTN3,ACAP3,MD
ER,L1CAM,TRIB2,PEA15,SQLE,FAM199X,HS6ST1,TIMP3,PGRMC1,NOVA2,CTSB,NUCB1,CHD6,R
KAP12,MAP4,CDC42BPB,ITM2B,AP2B1,MAT2A,SLC4A8,GOLGA8A,PXDN,OGA,NFE2L1,HNRNPA0
EX3,OAZ1,SRSF4,PFN1,ASTN2,TLE5,NISCH,PRKCSH,HNRNPC,CCNI,TADA2B,XPR1,KHDRBS1,KL
1,HMGN2,ATP2A2,RPL26,CRMP1,LASP1,CCT7,OAZ1,RTN3,PFN1,JPT2,NES,RPL31,NISCH,CCNI,T
7,AP3D1,SBF1,DNAJA1,ARF1,PAFAH1B1,ITM2B,ZNF770,MAT2A,TMEM170A,GTF2F1,GOLGA8A,
AP3,MDK,LAMC1,JPT1,CBX1,RAD23B,PTK7,CHD2,ORAI2,ASXL1,H3-3A,RAB11B,HDGF,DPYSL2,
1ORF43,PAPOLA,IGFBP3,COL1A2,NCS1,RPL22,RHOA,CBX5,ZNF865,CLSTN1,ATF4,PSMD2,KIF5
1ORF43,PAPOLA,IGFBP3,COL1A2,NCS1,RPL22,RHOA,CBX5,ZNF865,CLSTN1,ATF4,PSMD2,KIF5
1ORF43,PAPOLA,IGFBP3,COL1A2,NCS1,RPL22,RHOA,CBX5,ZNF865,CLSTN1,ATF4,PSMD2,KIF5

1,MAT2A,GTF2F1,PXDN,HNRNPU,OGA,NFE2L1,SMARCA5,HNRNPA0,CTNNA1,FADS2,RPL23,CNN3
D,TMEM259,RPS20,EIF5,CD81,WDR1,HMGN2,ATP2A2,CRMP1,LASP1,BEX3,OAZ1,SRSF4,PFN1,AS
6,RPS17,VCL,AP3D1,SBF1,DNAJA1,ARF1,AKAP12,MAP4,SF3B2,ITM2B,ZNF770,AP2B1,MAT2A,GT
4K2B,SOX12,RAB7A,DDX3X,RPL12,ITM2C,ARL4C,MAGED2,GNB2,RCN1,H1-0,RPS5,APLP1,MIF,PI
3-3A,PRPF6,DPYSL2,VEGFA,TUBB2A,CCND1,SMARCA4,MAPRE1,SRSF9,DSTN,PBX2,ODC1,AEBP1
310,INA,EIF4A1,MRC2,PPP2R1A,RPL15,GTF3C4,LDHB,DNAH8,PKD1,HLA-C,SF3B1,SERINC1,SSB
NPC,CCNI,KHDRBS1,XPO1,SPTBN2,SLC3A2,PFKL,GLG1,SFPQ,PEBP1,PDIA6,DDR1,SULF1,PLXNA
LLGL1,NCS1,RHOA,RPL9,CLSTN1,ATF4,KIF5B,RAB6B,RPS14,ATP1A1,JUND,TMEM259,RPS20,EI
9,TNS3,EIF4H,TSPAN3,LPCAT1,LAMA5,SLC22A17,BRD3,KPNB1,RPL17,IGFBP3,COL1A2,LLGL1,N
S3,CADM4,RHOB,HNRNPH1,LGALS3BP,HLA-B,PLPPR3,ARCN1,SEPTIN3,CTBP1,ARHGDIA,CAPRIN
MVB12B,SMARCA4,AZIN1,MAPRE1,SRSF9,DSTN,PBX2,RABL6,ODC1,FAM168B,AEBP1,NYNRIN,PT
F3,ANXA2,AGRN,RPLP1,ZMIZ1,SARAF,COL11A1,MIDN,RPL10,MARCHF6,WSB1,RPS17,AP3D1,DNA
,RPL18,STMN1,ITGB1,TUBB4B,UCHL1,MORF4L1,NCAM1,ANXA2,AGRN,ZMIZ1,TIMP2,XAF1,SARA
F1,SNX22,PGD,SQSTM1,VAMP2,GAS7,PRDX5,PDXK,PKN1,HSPA4,PLOD1,CAMSAP1,GRINA,NCKAI
PA0,CTNNA1,FADS2,RPS16,RPL23,LDHA,CNN3,RPSA,CTSD,EIF4H,SCARB2,NACA,LPCAT1,LAMA5,
10,MARCHF6,WSB1,RPS17,VCL,AP3D1,DNAJA1,ARF1,AKAP12,SF3B2,MAT2A,TMEM170A,GTF2F1
AQ,EIF1,HACD3,DCHS1,SET,EGLN1,MAP9,TPI1,CHSY1,TMSB10,INA,EIF4A1,MRC2,RPL19,PPP2R1
PAT4,JUN,MYO5A,COL6A2,CD63,NGRN,GOLM1,SESN3,MTSS2,RPL35A,DNAJC5,APLP2,HNRNPD
R1A,RPL15,GTF3C4,LDHB,HLA-C,SF3B1,PDIA3,SERINC1,SSBP2,ZNF43,CHD3,VAPA,HMGB1,POL
,CCT7,OAZ1,RTN3,PFN1,NES,RPL31,NISCH,PRKCSH,CCNI,TADA2B,XPR1,KHDRBS1,SPTBN2,SLC
YSL2,TOP1,SELENON,CCND1,G3BP2,ATG2B,LAPTM4A,BTBD2,PBX2,ODC1,FAM168B,AEBP1,PTE
T1,RPS4X,RPS3A,RPL18,STMN1,ITGB1,TUBB4B,UCHL1,RPL11,MORF4L1,ANXA2,RPLP1,ZMIZ1,T
P1,LASP1,BEX3,VAT1,OAZ1,SRSF4,PFN1,TLE5,CCNI,LRPPRC,KLHL9,XPO1,SPTBN2,SLC3A2,CSNI
CN1,COTL1,HECTD4,MAGED4B,CTBP1,RAC1,ARHGDIA,STIP1,GPC1,CDK16,SLC35E2B,SMCHD1
1,KHDRBS1,KLHL9,XPO1,SLC3A2,CSNK1E,PFKL,HDLBP,RPS25,TKT,TPM1,EIF5A,CCND2,CDH2,G
A8A,PXDN,HNRNPU,PGAM1,CTNNA1,RPS16,CNN3,RPL29,CTSD,EIF4H,SCARB2,LPCAT1,LAMA5,R
PTIN11,LAMP1,HNRNPH1,UBE2H,CKAP4,HLA-B,RAB1A,PLPPR3,ARCN1,COTL1,HECTD4,CTBP1,A
4L1,BTF3,NCAM1,ANXA2,AGRN,RPLP1,ZMIZ1,SARAF,RPL10,MARCHF6,WSB1,RPS17,VCL,SBF1,
RNPA0,CTNNA1,FADS2,CPE,CNN3,RPSA,TNS3,CTSD,EIF4H,SCARB2,TSPAN3,LPCAT1,LAMA5,SLC2
22A17,BRD3,PJA2,SLC16A7,RPL17,PAPOLA,COL1A2,LLGL1,NCS1,CBX5,ZNF865,CLSTN1,ATF4,
F1,SLC7A5,RPS14,CXADR,CHD4,ATP1A1,TMEM259,RPS20,EIF5,CD81,WDR1,HMGN2,LASP1,BEX
ADR,CHD4,ATP1A1,JUND,RPS20,WDR1,ATP2A2,RPL26,CRMP1,VAT1,PFN1,ASTN2,TLE5,HNRNPC
10,NARS1,SEPTIN11,CADM4,LGALS3BP,UBE2H,HLA-B,PLPPR3,CTBP1,ARHGDIA,CAPRIN1,GPC1,
GRN,RPLP1,SARAF,MIDN,RPL10,MARCHF6,VCL,AP3D1,DNAJA1,ARF1,PAFAH1B1,SF3B2,ITM2B,A
L1A2,HK1,NCS1,ZNF865,CLSTN1,NR2F1,KIF5B,RAB6B,ENC1,NLGN2,GLIPR1L2,CXADR,CHD4,A
CKSL1,UBB,AFF4,TRIM28,MATR3,CKB,COL4A2,RPL37A,LRRC28,MAPT,FN1,SPEN,NONO,DDB1,F
CB,PGD,SQSTM1,VAMP2,KLHL24,HNRNPL,SSRP1,HNRNPA3,PTPRA,PKN1,CAMSAP1,KLC1,GRIN
P1,COL2A1,PTOV1,MLEC,ACAP3,MDK,DDX42,JPT1,PTK7,SKI,ASXL1,H3-3A,HDGF,ABCA2,VEGFA,
AP3,MDK,PTK7,SKI,ASXL1,H3-3A,ABCA2,DPYSL2,VEGFA,TUBB2A,ARF3,CCND1,MAPK1,SMARCA4
HAQ,EIF1,HACD3,DCHS1,SET,EGLN1,TPI1,TMSB10,EIF4A1,MRC2,RPL19,PPP2R1A,RPL15,GTF3C
TN3,PFN1,JPT2,PNN,NES,PRKCSH,HNRNPC,CCNI,LRPPRC,TADA2B,KHDRBS1,SLC3A2,CSNK1E,PI
C4A8,GOLGA8A,PXDN,HNRNPU,PGAM1,SMARCA5,HNRNPA0,CTNNA1,FADS2,RPS16,LDHA,CPE,C
R2F1,RAB6B,SLC7A5,GLIPR1L2,CXADR,ATP1A1,JUND,TMEM259,RPS20,EIF5,CD81,WDR1,ATXN7
K12,ITM2C,ARL4C,MAGED2,H1-0,APLP1,MIF,GPAT4,TLN1,CHPF,CD63,GOLM1,SESN3,B4GALNT
ZMIZ1,TIMP2,SARAF,COL11A1,MIDN,RPL10,WSB1,VCL,AP3D1,SBF1,DNAJA1,ARF1,CYB5R3,AKAF

:TSD,EIF4H,SCARB2,LPCAT1,LAMA5,SLC22A17,BRD3,CIPC,PJA2,RPL24,KPNB1,RPL17,C1ORF43
CA5,CTNNA1,FADS2,LDHA,CPE,CNN3,TPM4,CTSD,EIF4H,SCARB2,TSPAN3,LPCAT1,LAMA5,SLC22
PIP4K2B,SOX12,RAB7A,CCN2,ACTN4,PRDX2,DDX3X,RPL12,ITM2C,ARL4C,CLTC,GNB2,RPS7,RPL
S1,GPAT4,CHPF,MYO5A,COL6A2,CD63,B4GALNT4,APLP2,HNRNPDL,TARDBP,RPLP2,MXD4,TAGL
1,JUND,EIF5,CD81,WDR1,HMGN2,ATP2A2,RPL26,CRMP1,BEX3,OAZ1,SRSF4,RBBP4,PFN1,JPT2,
TGES3,SEPTIN11,SND1,LAMP1,CKAP4,RAB1A,PLPPR3,ZNF454,MAGED4B,CTBP1,RAC1,ARHGDIA
SD,EIF4H,LPCAT1,LAMA5,RPS24,SLC22A17,BRD3,XRCC5,SLC16A7,C1ORF43,PAPOLA,RPL14,HK
P1,MIF,GPAT4,CHPF,MYO5A,CD63,GOLM1,SESN3,B4GALNT4,MTSS2,APLP2,TARDBP,NDFIP1,TAG
OVA2,CTSB,NUCB1,CDH11,CHD6,RNF187,EEF1A2,TBCD,SNX22,SQSTM1,VAMP2,GAS7,GDI2,PAR
OVA2,CTSB,NUCB1,CDH11,CHD6,RNF187,EEF1A2,TBCD,SNX22,SQSTM1,VAMP2,GAS7,GDI2,PAR
A2B1,CUL3,BRSK1,GPSM1,LMAN1,ZMYM4,SEPTIN7,PEG10,GSE1,LOX,GPC4,PCBP4,AKAP9,ZNF
I5,SNRPN,PRDX5,ACTR3,C4ORF3,UBE4A,PKN1,HSPA4,CAMSAP1,CCT3,GRINA,VPS35,ACAP3,SUR
MSB10,INA,EIF4A1,MRC2,PPP2R1A,RPL15,GTF3C4,LDHB,DNAH8,HLA-C,SF3B1,PDIA3,SSBP2,CH
4,ATP1A1,JUND,TMEM259,EIF5,LENG8,WDR1,HMGN2,ATP2A2,RPL26,CRMP1,LASP1,CCT7,OAZ
DX39B,CNN2,NOVA2,CDH11,TM9SF3,CHD6,U2AF2,EEF1A2,CCT2,TBCD,SNX22,VAMP2,KLHL24,C
NA,VPS35,PTOV1,CLSTN3,ACAP3,MDK,SURF4,JPT1,CBX1,PTK7,RAB5B,ORAI2,SKI,H3-3A,SCAF1,
EEF1A2,TBCD,SNX22,PGD,SQSTM1,VAMP2,GAS7,PARP1,TM9SF2,LMNB2,YEATS2,HSPA4,CAMSA
NPH1,PLPPR3,ARCN1,COTL1,CTBP1,ARHGDIA,CAPRIN1,STIP1,GPC1,TOP2B,IPO5,MMP2,SKP1,E
NPA0,FADS2,RPL23,LDHA,CPE,CNN3,RPL29,RPSA,TPM4,TNS3,NACA,LPCAT1,LAMA5,RPS24,BRD
PL17,C1ORF43,LLGL1,NCS1,RHOA,PTCD3,ZNF865,CLSTN1,ATF4,PSMD2,SLC7A5,RPS14,CHD4,
JA1,ARF1,CYB5R3,SF3B2,ITM2B,ZNF770,TMEM170A,GTF2F1,SLC4A8,HNRNPU,PGAM1,NFE2L1,
YB5R3,AKAP12,MAP4,MAT2A,TMEM170A,SLC4A8,GOLGA8A,PXDN,OGA,HNRNPA0,CTNNA1,FADS
HL1,RPL11,MORF4L1,BTF3,NCAM1,ANXA2,AGRN,RPLP1,ZMIZ1,TIMP2,SARAF,MIDN,RPL10,MAP
N,OPTN,ARF3,MVB12B,CCND1,BTBD2,PBX2,PRDX1,AEBP1,NYNRIN,PTBP1,EIF3E,DYNLL2,PCSK1
N1,H1-0,MIF,MYO5A,CD63,SESN3,B4GALNT4,DNAJC5,APLP2,HNRNPDL,TARDBP,RPLP2,MXD4,M
LDHA,CPE,RPL29,NACA,TSPAN3,LPCAT1,LAMA5,SLC22A17,BRD3,CIPC,PJA2,MSN,SLC16A7,KPN
5,SLC22A17,BRD3,KPNB1,PAPOLA,IGFBP3,LLGL1,HK1,NCS1,RPL22,ZNF865,RPL9,CLSTN1,ATF
PRC,KHDRBS1,CSNK1E,PFKL,HDLBP,GLG1,EIF5A,CCND2,PNMA1,SOX12,RAB7A,ACTN4,PRDX2,D
P7D1,CLIP3,FKBP8,PLCG1,MAP2K2,ZFAND5,DIDO1,DUSP4,FLNB,PLTP,CORO1C,LSS,FBLN1,ISY
R1,HMGN2,CRMP1,LASP1,BEX3,VAT1,SRSF4,RTN3,PFN1,JPT2,PNN,SPC25,NES,PRKCSH,KHDRBS
ET,EGLN1,MAP9,TPI1,CHSY1,TMSB10,INA,EIF4A1,MRC2,RPL19,PPP2R1A,RPL15,GTF3C4,LDHB,
,CHD6,RAB27A,RNF187,U2AF2,EEF1A2,TBCD,SNX22,PGD,VAMP2,GAS7,GDI2,HNRNPL,PARP1,TI
TRIB2,HSBP1,LUC7L3,SQLE,DLG5,HS6ST1,TIMP3,NUCB1,CHD6,EEF1A2,CCT2,TBCD,SNX22,SQS
35E2B,APPL1,SMCHD1,IPO5,ATP6V0C,DNER,LAPTM4B,GLUL,HSBP1,FAM199X,DLG5,HS6ST1,C
HD1,IPO5,ATP6V0C,SKP1,TRIB2,GLUL,CDKN1A,SHISAL1,CRPPA,TM9SF3,MDH2,CHD6,RNF187,E
1,RPS24,SLC22A17,BRD3,RPS9,CIPC,PJA2,MSN,KPNB1,RPL17,C1ORF43,PAPOLA,RPL14,RPL22,
EIF5A,CCND2,DDR1,MEGF8,RAB7A,ACTN4,RPL12,ITM2C,RPL30,ARL4C,CLTC,GNB2,RCN1,H1-0
LP2,HNRNPDL,MXD4,MYL6,ANKRD52,WLS,HNRNPK,H1-10,INIP,CADM4,RHOB,RAB1A,PLPPR3,A
F3B1,PDIA3,SSBP2,ZNF43,RPS3,BASP1,VAPA,HMGB1,POLR2A,CDKL1,AP2M1,TPT1,SEC61A1,RP
1,JUND,CD81,LENG8,WDR1,ATP2A2,RPL26,CRMP1,LASP1,BEX3,VAT1,OAZ1,SRSF4,PFN1,ASTN2,
,C1ORF43,PAPOLA,LLGL1,NCS1,RHOA,ZNF865,ATF4,SLC7A5,RPS14,CXADR,ATP1A1,JUND,TME
,FADS2,CPE,CNN3,TPM4,TNS3,EIF4H,NACA,TSPAN3,LPCAT1,LAMA5,BRD3,PJA2,MSN,KPNB1,RP
MEM248,LAMC1,PTK7,CCT4,GCN1,SKI,RAB11B,HDGF,TERF2IP,RPS15A,RPL38,NOTCH2,MARS1,
LAF1,L1CAM,TRIB2,PPIB,HSBP1,SQLE,DLG5,HS6ST1,TIMP3,CTDSP2,CTSB,CHD6,RNF187,U2AF2
,WDR1,HMGN2,ATP2A2,CRMP1,LASP1,BEX3,VAT1,OAZ1,SRSF4,RBBP4,RTN3,PFN1,ASTN2,TLE

,CTNNA1,RPS16,CPE,CNN3,RPL29,TNS3,CTSD,EIF4H,TSPAN3,LPCAT1,LAMA5,SLC22A17,BRD3,
,RPL11,MORF4L1,BTF3,NCAM1,AGRN,RPLP1,SARAF,MIDN,RPL10,MARCHF6,WSB1,RPS17,VCL,
1,CXADR,CHD4,JUND,TMEM259,RPS20,EIF5,LENG8,WDR1,ATXN7L3B,ATP2A2,CRMP1,LASP1,BE
,AP3D1,SBF1,DNAJA1,ARF1,CYB5R3,AKAP12,MAP4,SF3B2,CDC42BPB,ITM2B,ZNF770,AP2B1,M
5,CTNNA1,RPS16,RPL23,CNN3,CTSD,SCARB2,LPCAT1,LAMA5,RPS24,BRD3,CIPC,MSN,C1ORF43
PL1,SMCHD1,IPO5,MMP2,DNER,BCLAF1,L1CAM,LAPTM4B,HSBP1,SQLE,FAM199X,DLG5,SERF2,
,AEBP1,EIF3E,DYNLL2,ACTN1,CAND1,SON,MAP7D1,CLIP3,PLCG1,DIDO1,DUSP4,FLNB,PITPNA,C
1,PLXNA1,MEGF8,SOX12,RAB7A,CCN2,MMP14,PRDX2,DDX3X,RPS15,ITM2C,RPL30,ARL4C,GNB
A,POLR2A,CDKL1,AP2M1,TPT1,SEC61A1,RPS4X,RPS3A,STMN1,ITGB1,TUBB4B,UCHL1,RPL11,M
SRSF4,RBBP4,PFN1,ASTN2,JPT2,TLE5,PNN,SPC25,CCNI,KHDRBS1,XPO1,PFKL,EIF5A,PDIA6,PLX
2B,SLC3A2,CSNK1E,HDLBP,GLG1,SFPQ,TPM1,EIF5A,CCND2,PNMA1,GPC3,SOX12,CCN2,PRDX2
4,DNAJC5,APLP2,HNRNPDL,SLC25A3,NDFIP1,TSR2,MXD4,TAGLN,ANKRD52,HNRNPK,H1-10,INIF
B,MMP14,RPS15,ITM2C,ARL4C,H1-0,RPS19,RPL35,MIF,CHPF,COL6A2,ANXA6,NGRN,GOLM1,RI
,ATF4,RAB6B,SLC7A5,ENC1,NLGN2,CXADR,CHD4,RPS20,EIF5,LENG8,WDR1,HMGN2,ATP2A2,CR
DN,RPL10,MARCHF6,WSB1,RPS17,VCL,AP3D1,DNAJA1,ARF1,CYB5R3,AKAP12,SF3B2,CDC42BP
C,LAPTM4B,TRIB2,RPL23A,PEA15,FAM199X,DLG5,HS6ST1,CTSB,CHD6,RNF187,TBCD,MLXIP,S
H,SCARB2,NACA,TSPAN3,LAMA5,RPS24,RPL24,SLC16A7,KPNB1,C1ORF43,PAPOLA,RPL14,PTC
1,LASP1,BEX3,CCT7,VAT1,OAZ1,SRSF4,RBBP4,RTN3,PFN1,MCL1,PNN,NES,RPL31,HNRNPC,CC
M1,TPT1,RPS4X,STMN1,ITGB1,TUBB4B,UCHL1,RPL11,MORF4L1,BTF3,NCAM1,ANXA2,AGRN,R
K1E,RPS25,TKT,CDH6,SFPQ,EIF5A,PEBP1,CDH2,GPC3,SOX12,RAB7A,ITM2C,CLTC,RPS5,RPL35
,GLG1,CDH6,CDH2,PNMA1,GPC3,PIP4K2B,SOX12,ACTN4,MMP14,PRDX2,DDX3X,RPS15,RPL12,
PL15,GTF3C4,LDHB,PKD1,SF3B1,PDIA3,SSBP2,ZNF43,RPS3,BASP1,VAPA,HMGB1,POLR2A,CDKL
P3,MDK,JPT1,PTK7,ORAI2,SKI,ASXL1,H3-3A,RAB11B,PYGB,TMED2,SELENON,TUBB2A,OPTN,M
2A,CDKL1,AP2M1,TPT1,SEC61A1,RPS4X,RPS3A,STMN1,ITGB1,TUBB4B,UCHL1,BTF3,NCAM1,AN
,SQSTM1,VAMP2,KLHL24,HNRNPL,TM9SF2,ALKBH5,PTPRA,PRDX5,TCP1,KLC1,GRINA,PTOV1,M
TF4,PSMD2,KIF5B,RAB6B,SLC7A5,CXADR,CHD4,ATP1A1,JUND,EIF5,CD81,COL5A2,WDR1,HMG
TF4,PSMD2,KIF5B,RAB6B,SLC7A5,CXADR,CHD4,ATP1A1,JUND,EIF5,CD81,COL5A2,WDR1,HMG
H2,PDIA6,PLXNA1,PIP4K2B,RAB7A,DDX3X,RPS15,ITM2C,RPL30,CLTC,EIF3L,H1-0,RPS5,IK,RPS1
E2B,SMCHD1,IPO5,ATP6V0C,DNER,L1CAM,LAPTM4B,TRIB2,PPIB,SQLE,CDKN1A,FAM199X,DLG
ARCA4,AZIN1,SRSF9,DSTN,BTBD2,PBX2,ODC1,AEBP1,PTBP1,DYNLL2,CDC37,CAPZB,UBE2R2,A
10,VEGFA,OPTN,ARF3,MVB12B,PPP2CA,CCND1,DSTN,BTBD2,FAM171A2,AEBP1,NYNRIN,PTBP1,
,SERINC1,SSBP2,ZNF43,RPS3,CHD3,VAPA,HMGB1,POLR2A,CDKL1,TPT1,SEC61A1,RPS4X,RPS3
,B4GALNT4,DNAJC5,APLP2,HNRNPDL,RPLP2,MXD4,INIP,DUX4,CKAP5,SEPTIN11,CADM4,RHOB,
A,PLPPR3,SEPTIN3,COTL1,HECTD4,CAPRIN1,STIP1,GPC1,SLC35E2B,ATP6V0C,TMEM30A,L1CA
,MXD4,WLS,HNRNPK,H1-10,NARS1,PTGES3,LAMP1,RHOB,HNRNPH1,CKAP4,RPS12,PLPPR3,ARC
2A17,BRD3,RPS9,PJA2,KPNB1,C1ORF43,PAPOLA,HK1,NCS1,ZNF865,ATF4,RAB6B,ENC1,NLGN2,
GCR,STIP1,GPC1,CDK16,TOP2B,SLC35E2B,ATP6V0C,TMEM30A,BCLAF1,LAPTM4B,HSBP1,SQL
5R3,AKAP12,MAP4,CDC42BPB,ITM2B,MAT2A,SLC4A8,GOLGA8A,PXDN,OGA,HNRNPA0,CTNNA1
HD6,RNF187,EEF1A2,TBCD,ANXA5,SNX22,SQSTM1,VAMP2,GDI2,PARP1,TAF15,ABCA3,PDXK,PK
CAM1,ANXA2,AGRN,RPLP1,ZMIZ1,TIMP2,XAF1,SARAF,MIDN,RPL10,MARCHF6,WSB1,RPS17,VCL
TF2F1,SLC4A8,HNRNPU,OGA,PGAM1,SMARCA5,HNRNPA0,CTNNA1,RPS16,RPL23,LDHA,CNN3,
1A,CALU,SEPTIN3,CCT6A,CTBP1,ARHGDIA,STIP1,GPC1,TOP2B,APPL1,SMCHD1,HDAC2,IPO5,A
DELR2,CCNL2,ARPC2,PLEC,SRSF1,ARGLU1,AEBP1,ATP5PB,PTBP1,ILF3,RPS11,HNRNPUL1,UBE
CCN2,DDX3X,MIF,COL6A2,GOLM1,SESN3,B4GALNT4,DNAJC5,APLP2,HNRNPDL,RPLP2,MXD4,M
N3,RPSA,TPM4,TNS3,CTSD,EIF4H,TSPAN3,LPCAT1,LAMA5,RPS24,BRD3,RPS9,CIPC,RPL24,SLC

,RPS4X,RPS3A,RPL18,STMN1,ITGB1,TUBB4B,UCHL1,RPL11,MORF4L1,BTF3,NCAM1,AGRN,RPL
MXD4,TAGLN,ANKRD52,HNRNPK,H1-10,INIP,CIRBP,MYH10,NARS1,DUX4,CADM4,ATP1A2,HLA-B
,CIPC,PJA2,RPL24,SLC16A7,KPNB1,PAPOLA,IGFBP3,LLGL1,RPL14,HK1,NCS1,CLSTN1,ATF4,PSN
UBQLN1,SF3A1,ATXN2L,NBPF9,ZFAND3,SARS1,HGS,BAG6,SREBF2,PNMA8A,VGF,BMPR2,CCT5,
N,HNRNPU,OGA,PGAM1,NFE2L1,SMARCA5,HNRNPA0,FADS2,RPS16,LDHA,CPE,RPL29,RPSA,TNS
61A1,RPS4X,STMN1,ITGB1,TUBB4B,UCHL1,RPL11,MORF4L1,BTF3,NCAM1,ANXA2,AGRN,RPLP1
ARS1,DUX4,YWHAB,SND1,LAMP1,HNRNPH1,TTR,CALU,ARCN1,RPS27,CTBP1,ARHGDIA,CAPRIN1
3,COL1A2,HK1,RHOA,PTCD3,ZNF865,RPL9,ATF4,NR2F1,KIF5B,GLIPR1L2,RPS14,CXADR,CHD4,
),ACTN1,CAND1,MAP7D1,FKBP8,PLCG1,DIDO1,DUSP4,DLK1,CORO1C,GIGYF1,NKTR,STMN3,LS
1,SLC22A17,BRD3,XRCC5,MSN,KPNB1,RPL17,PAPOLA,IGFBP3,COL1A2,LLGL1,HK1,NCS1,RPL2
2,PNMA1,PDIA6,PIP4K2B,SOX12,DDX3X,ARL4C,CLTC,RPS19,RPL35,MIF,COL6A2,GOLM1,SESN
,VGF,FBRSL1,MEX3A,P3H3,FLNC,KDM3B,WBP2,AKIRIN1,PRPF40A,BMPR2,DYNLL1,CTXN1,CD9
LEC,CLSTN3,ACAP3,MDK,RAD23B,PTK7,SKI,ASXL1,H3-3A,SCAF1,RAB11B,ABCA2,PYGB,TERF2I
AM1,ANXA2,AGRN,RPLP1,ZMIZ1,COL11A1,MIDN,RPL10,MARCHF6,WSB1,RPS17,VCL,AP3D1,DN
HMGB1,POLR2A,CDKL1,AP2M1,TPT1,SEC61A1,RPS4X,RPS3A,STMN1,ITGB1,TUBB4B,UCHL1,RP
1A1,JUND,TMEM259,RPS20,CD81,WDR1,ATXN7L3B,HMGN2,RPL26,LASP1,BEX3,SRSF4,RBBP4,
H5,PTPRA,PRDX5,ACTR3,RTN1,TBC1D16,GRINA,NCKAP1,PTOV1,MLEC,CLSTN3,LAMC1,CBX1,SK
B6B,SLC7A5,ENC1,NLGN2,RPS14,CXADR,CHD4,JUND,EIF5,CD81,LENG8,WDR1,HMGN2,ATP2A2
,NXN,RER1,KCTD12,PTEN,ECEL1,PRRC2A,PPP1CC,PTPRF,B4GAT1,RPS2,PSMD8,GALNT11,C1O
1,TLE5,PNN,RPL31,PRKCSH,XPR1,KHDRBS1,KLHL9,HDLBP,RPS25,TKT,SFPQ,EIF5A,CDH2,PDIA6
B,SDF4,ATP6V1B2,HNRNPM,SDHA,NKTR,YWHAH,PFKM,STT3B,TNKS2,LONP1,UBQLN1,ATXN2L,
15,FAM199X,DLG5,CTDSP2,PGRMC1,DDX39B,NOVA2,CTSB,CDH11,RAB27A,EEF1A2,TBCD,SUG
B,ATP2A2,RPL26,LASP1,CCT7,SRSF4,RBBP4,PFN1,ASTN2,MCL1,TLE5,PNN,RPL31,PRKCSH,CCN
RCKSL1,UBB,AFF4,TRIM28,MATR3,YWHAG,CKB,COL4A2,RPL37A,LRRC28,MAPT,FN1,SPEN,NON
CAM1,ANXA2,AGRN,RPLP1,ZMIZ1,XAF1,SARAF,MIDN,RPL10,MARCHF6,RPS17,VCL,AP3D1,DNAJ
CLAF1,LAPTM4B,TRIB2,PEA15,SQLE,CTDSP2,NOVA2,SRP9,CHD6,RNF187,U2AF2,EEF1A2,TBCD,
TP6V0C,SKP1,DNER,NNAT,SQLE,DLG5,CRPPA,HS6ST1,CTSB,SRSF5,ZC3H7B,TBCD,SUGP2,ANX
5,ATF4,PSMD2,SLC7A5,ENC1,NLGN2,CXADR,CHD4,JUND,TMEM259,EIF5,LENG8,WDR1,ATXN7L
2B,SMCHD1,IPO5,ATP6V0C,SKP1,DNER,BCLAF1,L1CAM,TRIB2,HSBP1,SQLE,FAM199X,SERF2,T
DH2,CHD6,RAB27A,EEF1A2,TBCD,ANXA5,GDI2,MORF4L2,HNRNPA3,PRDX5,PKN1,YEATS2,CAMS
GPC3,SOX12,RAB7A,PRDX2,ARL4C,MRFAP1,RPS5,RPS19,MIF,GPAT4,CHPF,JUN,CTNNB1,COL6A
JA1,ARF1,CYB5R3,AKAP12,SF3B2,CDC42BPB,AP2B1,MAT2A,GTF2F1,SLC4A8,OGA,PGAM1,SMA
MEM259,EIF5,CD81,LENG8,WDR1,ATP2A2,LASP1,BEX3,CCT7,OAZ1,SRSF4,RBBP4,RTN3,ASTN2,
Z1,TIMP2,SARAF,COL11A1,MIDN,RPL10,WSB1,RPS17,VCL,AP3D1,ARF1,CYB5R3,AKAP12,MAP4,S
2,JPT2,TLE5,NISCH,PRKCSH,HNRNPC,LRPPRC,TADA2B,XPO1,SPTBN2,CSNK1E,PFKL,RPS25,TKT
PAFAH1B1,AKAP12,MAP4,CDC42BPB,ITM2B,ZNF770,AP2B1,GOLGA8A,PXDN,HNRNPU,PGAM1,N
9,TPI1,CHSY1,TMSB10,EIF4A1,MRC2,RPL19,PPP2R1A,RPL15,GTF3C4,LDHB,DNAH8,PKD1,HLA-
S1,RHOA,CBX5,ZNF865,CLSTN1,KIF5B,RAB6B,SLC7A5,ENC1,NLGN2,RPS14,CXADR,CHD4,ATP1
B,ISYNA1,LONP1,CSNK1D,IDH1,HYOU1,HECTD1,COL27A1,VANGL2,PNMA8A,VGF,MEX3A,CYP51
4B,VCP,MARCKSL1,UBB,AFF4,TRIM28,MATR3,YWHAG,CKB,COL4A2,RPL37A,LRRC28,MAPT,FN1
24,BRD3,RPS9,XRCC5,PJA2,MSN,RPL24,SLC16A7,RPL17,PAPOLA,IGFBP3,COL1A2,LLGL1,RPL1
O1,SPTBN2,CSNK1E,RPS25,TKT,TPM1,PDIA6,GPC3,MEGF8,SOX12,CCN2,MMP14,PRDX2,DDX3
S19,RPL35,TLN1,JUN,RPS10,MTSS2,RPL35A,DNAJC5,HNRNPDL,TARDBP,RPLP2,ANKRD52,DUX
3,RAB27A,EEF1A2,TBCD,MLXIP,PPP1CB,SNX22,PGD,SQSTM1,MEST,PABPN1,GDI2,NOX4,HSPH1
2,NOVA2,NUCB1,CHD6,RNF187,EEF1A2,CCT2,TBCD,SNX22,PGD,VAMP2,GNAI3,MEST,GAS7,GDI

AM9,DYNLL2,GRB2,WDR6,ATXN10,ACTN1,HSPB1,KDM1A,CASC3,FKBP8,PLCG1,ZFAND5,FLNB,
K8IP1,FBRSL1,MEX3A,RAB11A,MLF2,FLNC,ZSCAN18,WBP2,DYNLL1,CTXN1,CD99L2,SYP,RBM15
AP1,TERF2IP,TMED2,TAF7,RPS15A,RAPGEF1,OPTN,ARF3,VHL,UBXN4,SMARCA4,ATG2B,KDELR
RINC1,SSBP2,ZNF43,RPS3,CHD3,VAPA,HMGB1,POLR2A,CDKL1,SEC61A1,RPS3A,STMN1,ITGB1,
L1,NCAM1,ANXA2,AGRN,RPLP1,ZMIZ1,TIMP2,XAF1,SARAF,MIDN,RPL10,MARCHF6,WSB1,RPS17
1,HYOU1,ZFAND3,MAPK8IP1,SOD1,FBRSL1,MEX3A,P3H3,MLF2,FLNC,WBP2,SMS,DYNLL1,CTXN
N3,ACAP3,MDK,TMEM248,PTK7,ORAI2,SKI,H3-3A,SCAF1,RAB11B,PYGB,PRPF6,NOTCH2,SELENO
6,RPS17,VCL,AP3D1,DNAJA1,ARF1,CYB5R3,AKAP12,SF3B2,CDC42BPB,MAT2A,GTF2F1,SLC4A8,
IEST,GAS7,GDI2,HNRNPL,ABCA3,HNRNPA3,PTPRA,PRDX5,CAMSAP1,KLC1,PTOV1,MLEC,CLSTN3
IEST,GAS7,GDI2,HNRNPL,ABCA3,HNRNPA3,PTPRA,PRDX5,CAMSAP1,KLC1,PTOV1,MLEC,CLSTN3
IEST,GAS7,GDI2,HNRNPL,ABCA3,HNRNPA3,PTPRA,PRDX5,CAMSAP1,KLC1,PTOV1,MLEC,CLSTN3
PRC,TADA2B,XPO1,SPTBN2,SLC3A2,CSNK1E,GLG1,TPM1,EIF5A,CDH2,PDIA6,GPC3,CCN2,MMP
RAD23B,ORAI2,GCN1,SKI,H3-3A,RAB11B,RPS15A,VEGFA,SELENON,RAPGEF1,TUBB2A,ARF3,UB
70,MAT2A,TMEM170A,SLC4A8,GOLGA8A,HNRNPU,OGA,LARP1,HNRNPA0,CTNNA1,FADS2,CNN3
NXA2,AGRN,RPLP1,ZMIZ1,TIMP2,XAF1,SARAF,MIDN,RPL10,MARCHF6,WSB1,RPS17,VCL,AP3D1,S
YB5R3,SF3B2,MAT2A,GTF2F1,SLC4A8,OGA,PGAM1,SMARCA5,CTNNA1,RPS16,RPL23,LDHA,CNN
,HMGB1,POLR2A,AP2M1,TPT1,SEC61A1,RPS3A,RPL18,STMN1,ITGB1,TUBB4B,UCHL1,RPL11,B
3,SF1,HIP1,NOL4L,LRRC4B,SEC16A,GPX4,MGAT4B,SLC35F5,RAB2A,GNG4,FARP1,NXN,CRTAP,K
T2,TLE5,PNN,SPC25,RPL31,NISCH,PRKCSH,HNRNPC,KHDRBS1,SLC3A2,PTN,PFKL,HDLBP,RPS25
1,GRINA,COL2A1,PTOV1,MLEC,CLSTN3,MDK,JPT1,PTK7,ORAI2,TNPO1,ASXL1,H3-3A,KDM6B,H
1,PIP4K2B,MEGF8,ACSL6,RAB7A,CCN2,MMP14,ITM2C,ARL4C,CLTC,MAGED2,GNB2,RCN1,RPS
HDAC2,IPO5,TMEM30A,DNER,BCLAF1,NNAT,CDKN1A,TMEM132A,CPSF2,SRP9,NUCB1,SRSF5,ZC
16,IPO5,ATP6V0C,SKP1,DNER,L1CAM,TRIB2,PPIB,PEA15,FAM199X,SHISAL1,TIMP3,CTDSP2,NO
AM1,ANXA2,AGRN,RPLP1,ZMIZ1,TIMP2,XAF1,SARAF,MIDN,RPL10,MARCHF6,WSB1,RPS17,VCL,A
RO1C,MSL1,GIGYF1,LSS,FBLN1,STXBP1,ISYNA1,TUG1,PLXNA3,CSNK1D,HYOU1,ZFAND3,VANG
31,RPS17,VCL,AP3D1,SBF1,DNAJA1,ARF1,CYB5R3,MAP4,SF3B2,ITM2B,ZNF770,AP2B1,MAT2A,G
P4,RAB1A,PLPPR3,ARCN1,SEPTIN3,HECTD4,CCT6A,MAGED4B,CAPRIN1,STIP1,HDAC2,MMP2,DN
PS19,CHPF,JUN,MYO5A,ANXA6,SESN3,B4GALNT4,MTSS2,APLP2,HNRNPDL,TARDBP,RPLP2,ARL
H1-10,MYH10,YWHAB,LAMP1,RPS12,SEPTIN3,HECTD4,CCT6A,RAC1,STIP1,CDK16,SLC35E2B,AP
MARCHF6,RPS17,VCL,AP3D1,DNAJA1,ARF1,CYB5R3,AKAP12,SF3B2,AP2B1,MAT2A,GTF2F1,SLC
PK1,ATG2B,SRSF9,PLEC,BTBD2,RABL6,ODC1,FAM168B,DYNC1LI2,TCF25,PTBP1,DYNLL2,WDR
,RPLP2,DDOST,ANKRD52,DUX4,YWHAB,SND1,ATP1A2,RPS12,CALU,PLPPR3,ARCN1,RPS27,MAG
,XPO1,SLC3A2,CSNK1E,PFKL,RPS25,TKT,TPM1,EIF5A,CCND2,CDH2,GPC3,PIP4K2B,MEGF8,SO
N,PFKL,HDLBP,RPS25,SFPQ,TPM1,EIF5A,CCND2,CDH2,PNMA1,PDIA6,SOX12,CCN2,ACTN4,PRD
EF2,YLPM1,YME1L1,ADAR,RBM15B,DHX15,PSMA7,ANP32A,PRMT1,OS9,MTA1,KDM5C,ACTR1A
S16,LDHA,CPE,CNN3,RPL29,TNS3,CTSD,EIF4H,LAMA5,SLC22A17,BRD3,RPS9,XRCC5,PJA2,MSN
,PTCD3,ZNF865,RPL9,CLSTN1,ATF4,GLIPR1L2,RPS14,CXADR,CHD4,ATP1A1,RPS20,CD81,COL5
,OGA,SMARCA5,HNRNPA0,RPS16,RPL23,CNN3,RPL29,RPSA,TPM4,EIF4H,NACA,LAMA5,RPS24,
ZB,ACTN1,KDM1A,PLCG1,DUSP4,MSL1,SDF4,XPNPEP3,SRSF3,STMN3,LSS,FBLN1,STT3B,LOC7
BF1,DNAJA1,ARF1,AKAP12,MAP4,SF3B2,CDC42BPB,ITM2B,MAT2A,TMEM170A,SLC4A8,PXDN,H
,LDHA,CNN3,RPL29,RPSA,TPM4,TNS3,CTSD,EIF4H,SCARB2,TSPAN3,RTN4,LPCAT1,LAMA5,RPS2
3X,RPS15,RPL12,ITM2C,RPL30,EIF3L,MRFAP1,RPS5,RPS7,GPAT4,COL6A2,CD63,ANXA6,B4GAL
1,TUBB4B,UCHL1,MORF4L1,ANXA2,AGRN,RPLP1,ZMIZ1,TIMP2,XAF1,SARAF,MIDN,RPL10,MARC
MP2,DNER,LAPTM4B,TRIB2,NNAT,DLG5,CPSF2,SRP9,NUCB1,SRSF5,RAB27A,RNF187,U2AF2,FD
P4,PFN1,MCL1,TLE5,PNN,NES,NISCH,PRKCSH,HNRNPC,CCNI,KHDRBS1,XPO1,RPS25,CDH6,TPM

'S12,PLPPR3,RPS27,CCT6A,CTBP1,RAC1,ARHGDIA,STIP1,GPC1,SLC35E2B,TMEM30A,BCLAF1,PI
2,DUSP4,PITPNA,GIGYF1,PHC2,YWHAH,STMN3,LSS,FBLN1,STT3B,STXBP1,PLXNA3,CSNK1D,ATX
3,MEST,TSR1,ALKBH5,PDXK,TCP1,CAMSAP1,CCT3,PTOV1,MLEC,ACAP3,TMEM248,LAMC1,SKI,AS
P12,SF3B2,AP2B1,MAT2A,GTF2F1,SLC4A8,HNRNPA0,FADS2,RPS16,RPL23,CNN3,RPL29,RPSA,TF
F1,SLC4A8,OGA,SMARCA5,HNRNPA0,CTNNA1,RPS16,RPL23,CNN3,RPL29,TPM4,TNS3,EIF4H,NA
L4C,MAGED2,RCN1,H1-0,RPS5,APLP1,RPS7,RPS19,GPAT4,TLN1,MYO5A,CTNNB1,CD63,NGRN,E
LNA,XPOT,MTA1,TRIO,EP300,SF1,HIP1,VARS1,NOL4L,GRSF1,GPX4,PPP2CB,CSNK1G2,GNG4,NX
L2,CXADR,CHD4,ATP1A1,JUND,RPS20,CD81,WDR1,HMGN2,ATP2A2,RPL26,CRMP1,LASP1,BEX3,
B1,MAT2A,GTF2F1,SLC4A8,GOLGA8A,PXDN,HNRNPU,NFE2L1,SMARCA5,HNRNPA0,CTNNA1,FAD
T4,COL6A2,CD63,ANXA6,B4GALNT4,MTSS2,DNAJC5,APLP2,SLC25A3,RPLP2,MXD4,ANKRD52,H1
C1,PHC2,YWHAH,LSS,PITPNM1,STXBP1,ISYNA1,CSNK1D,MESD,IDH1,HYOU1,MAPK8IP1,SOD1,F
LE,DLG5,SERF2,TIMP3,CTSB,CDH11,CHD6,EEF1A2,TBCD,MLXIP,SNX22,PGD,KLHL24,GAS7,PABF
DL,WDR45B,TRIO,EP300,LBH,DGCR2,HIP1,LRRC4B,SLC39A6,CD99,SEC16A,GPX4,PPM1G,SLC3
4,RAD23B,PTK7,SKI,PCDHGB6,ASXL1,H3-3A,ABCA2,PYGB,TERF2IP,SELENON,MAPK1,SMARCA4,
D81,LENG8,WDR1,ATP2A2,RPL26,CRMP1,LASP1,VAT1,PFN1,ASTN2,MCL1,JPT2,PNN,NES,NISCH
A,TSR2,TAGLN,HNRNPK,INIP,DUX4,PTGES3,CKAP5,SEPTIN11,CADM4,SND1,LAMP1,UBE2H,CKAP
RPL35,MIF,HMGCS1,GPAT4,TLN1,CHPF,MYO5A,COL6A2,CD63,NGRN,B4GALNT4,MTSS2,APLP2
,RPL22,RHOA,PTCD3,ZNF865,RPL9,KIF5B,RPS14,CXADR,ATP1A1,TMEM259,RPS20,COL5A2,LEN
M168B,ARGLU1,ACADVL,TCF25,PCSK1N,PRKDC,MAP7D1,CCAR2,DLK1,SDF4,ATP6V1B2,HNRNF
2BPB,MAT2A,GTF2F1,SLC4A8,OGA,SMARCA5,HNRNPA0,FADS2,RPS16,RPL23,LDHA,CNN3,RPL2
2,CRMP1,BEX3,RBBP4,RTN3,PFN1,MCL1,PNN,HNRNPC,TADA2B,KHDRBS1,KLHL9,SLC3A2,CSNK
Q,TPM1,PEBP1,PNMA1,PDIA6,PLXNA1,PIP4K2B,RAB7A,PRDX2,DDX3X,RPS15,RPL12,ITM2C,EIF3L
F4H,LAMA5,RPS24,BRD3,RPS9,XRCC5,CIPC,RPL24,SLC16A7,PAPOLA,IGFBP3,COL1A2,RPL14,HI
MSAP1,KLC1,GRINA,VPS35,ACAP3,MDK,SURF4,LAMC1,JPT1,PTK7,ORAI2,SRP14,TSPAN6,TMED2
C2,YWHAH,PTP4A2,STMN3,PFKM,LSS,STT3B,ISYNA1,TUG1,PLXNA3,GPR107,PFN2,HNRNPD,FBR
1,LAMP1,ATP1A2,RPS12,CALU,PLPPR3,ARCN1,RPS27,MAGED4B,ARHGDIA,CAPRIN1,STIP1,TOP2
PTGES3,SND1,ATP1A2,RPS12,CALU,RPS27,CCT6A,MAGED4B,CTBP1,RAC1,ARHGDIA,CAPRIN1,ST
1,RPL23A,NNAT,CDKN1A,SERF2,DDX39B,NUCB1,TM9SF3,MDH2,RAB27A,U2AF2,FDFT1,CCT2,RF
P12,MAP4,CDC42BPB,ITM2B,MAT2A,TMEM170A,SLC4A8,GOLGA8A,PXDN,HNRNPU,OGA,PGAM1
ZNF865,ATF4,PSMD2,KIF5B,GLIPR1L2,RPS14,CXADR,CHD4,ATP1A1,RPS20,CD81,LENG8,WDR1,
LR2E,PPDPF,RBBP7,DFFB,EIF2AK1,TXLNA,SGTA,KDM5C,ACTR1A,PRKAR2A,FBRS,SF1,RETREG2
NPU,NFE2L1,LARP1,CTNNA1,FADS2,CTSD,EIF4H,NACA,LPCAT1,LAMA5,RPS9,CIPC,PJA2,MSN,KF
),RER1,PI4KA,PPP1CC,CUL3,ADH5,ULK1,ANP32B,MAPRE2,TIMM50,LMAN1,TNKS,DKK3,PPP1R15
31,WDR1,HMGN2,ATP2A2,CRMP1,LASP1,VAT1,OAZ1,SRSF4,RBBP4,MCL1,JPT2,PNN,SPC25,HNR
1,CYB5R3,SF3B2,CDC42BPB,ITM2B,MAT2A,PXDN,HNRNPU,OGA,NFE2L1,SMARCA5,CTNNA1,RPS
RPL22,RHOA,ZNF865,RPL9,ATF4,KIF5B,GLIPR1L2,RPS14,CXADR,CHD4,RPS20,CD81,LENG8,WDI
APOLA,RPL14,CBX5,ATF4,KIF5B,RPS14,CXADR,ATP1A1,JUND,TMEM259,RPS20,WDR1,ATXN7L3
,CLIP3,FKBP8,PLCG1,ZFAND5,DIDO1,TSC22D1,DLK1,CORO1C,MSL1,YWHAH,PTP4A2,STMN3,L
2B,CCNL2,ARPC2,DDAH2,FAM168B,ARGLU1,ACADVL,PTBP1,CDC37,SUPT5H,ATXN10,ACTN1,Q
RF2IP,SELENON,MAPK1,SMARCA4,ATG2B,AZIN1,LAPTM4A,DDAH2,AEBP1,ACADVL,NYNRIN,TCF2
,WSB1,RPS17,VCL,AP3D1,SBF1,DNAJA1,ARF1,PAFAH1B1,AKAP12,MAP4,SF3B2,CDC42BPB,ITM2
15,LMNB2,ABCA3,ACTR3,PDXK,PKN1,TBC1D16,ACO2,HSPA4,PLOD1,GRINA,NCKAP1,PTOV1,AC
,PJA2,SLC16A7,KPNB1,RPL17,C1ORF43,PAPOLA,COL1A2,GAP43,HK1,NCS1,RPL22,RHOA,PTCD
,PIP4K2B,SOX12,CCN2,PRDX2,ITM2C,CLTC,EIF3L,MRFAP1,RPL35,MIF,HMGCS1,TLN1,MYO5A,C
SO1,ATP5MG,NELFB,DENND5A,DIP2A,CDC34,TMEM184B,USF2,SPPL2B,FXR2,CHMP1A,PINK1,A

,ENC1,GLIPR1L2,CHD4,ATP1A1,TMEM259,COL5A2,LENG8,ATXN7L3B,HMGN2,ATP2A2,CRMP1,C
NOP53,GDI2,PARP1,TM9SF2,TSR1,MORF4L2,SSRP1,HSPH1,LMNB2,MTRF1L,RPL27,PTPRA,PRD
(TR,FBLN1,STT3B,LONP1,CSNK1D,ZFAND3,COL27A1,MAPK8IP1,HNRNPD,VGF,FBRSL1,MEX3A,F
I,CERS2,DPYSL4,HSDL1,METRN,RCN2,CENPB,NELFB,ADGRA3,HMGA1,OGFR,AGAP1,USF2,ARPC
C5,CIPC,RPL24,C1ORF43,PAPOLA,IGFBP3,COL1A2,RPL14,HK1,NCS1,RPL22,RHOA,ZNF865,ATF4
RBBP4,PFN1,ASTN2,MCL1,TLE5,PNN,RPL31,PRKCSH,HNRNPC,LRPPRC,TADA2B,XPR1,KHDRBS1,
TNNA1,LDHA,TPM4,TNS3,CTSD,LPCAT1,LAMA5,BRD3,RPS9,XRCC5,CIPC,PJA2,MSN,SLC16A7,KF
LDHB,DNAH8,HLA-C,SF3B1,PDIA3,SSBP2,ZNF43,CHD3,VAPA,HMGB1,POLR2A,CDKL1,AP2M1,TP
M,SNN,HSPG2,SSBP3,CARM1,ABI2,PKDCC,TRAM1,GPC2,LTBP4,GABARAPL1,FAM234A,PGAP6,RA
PDIA3,SSBP2,ZNF43,CHD3,VAPA,HMGB1,POLR2A,CDKL1,AP2M1,TPT1,SEC61A1,RPS4X,RPS3A,S
TPNA,HNRNPM,STMN3,PFKM,FBLN1,STT3B,TNKS2,ISYNA1,ATXN2L,PRPS1,GPR107,HECTD1,MA
,ATP1A1,RPS20,CD81,LENG8,WDR1,RPL26,LASP1,CCT7,SRSF4,RBBP4,PFN1,ASTN2,MCL1,TLE5
ATP2A2,RPL26,LASP1,CCT7,SRSF4,RBBP4,PFN1,ASTN2,MCL1,TLE5,PNN,RPL31,PRKCSH,HNRNP
SBP2,CHD3,BASP1,VAPA,HMGB1,POLR2A,CDKL1,AP2M1,TPT1,RPS3A,RPL18,ITGB1,TUBB4B,UCI
PS7,RPS19,RPL35,GPAT4,CHPF,GJA1,MYO5A,CD63,NGRN,GOLM1,SESN3,APLP2,TARDBP,SLC25
RDX2,RPS15,ITM2C,ARL4C,CLTC,MAGED2,RCN1,H1-0,RPS7,IK,RPL35,HMGCS1,TLN1,CHPF,CD6

SMC3,ZNF512B,CUL3,IL6ST,MCM3AP,RPS2,PSMD8,PODXL2,GALNT11,GSE1,SLC25A5,ATP6AP2,I
'1,TMSB10,INA,EIF4A1,MRC2,RPL19,RPL15,GTF3C4,LDHB,DNAH8,PKD1,HLA-C,SF3B1,PDIA3,SS
ITM2C,ARL4C,CLTC,RCN1,H1-0,RPS7,RPL35,TLN1,CHPF,NGRN,RPS10,SESN3,MTSS2,APLP2,HN
F3B1,PDIA3,SERINC1,SSBP2,ZNF43,RPS3,CHD3,BASP1,VAPA,HMGB1,POLR2A,CDKL1,AP2M1,TP
A3,SSBP2,CHD3,HMGB1,POLR2A,CDKL1,TPT1,SEC61A1,RPS4X,RPS3A,STMN1,ITGB1,TUBB4B,U
NPU,OGA,PGAM1,SMARCA5,FADS2,RPS16,RPL23,LDHA,CNN3,RPL29,RPSA,TPM4,CTSD,EIF4H,S
CDC42BPB,ITM2B,AP2B1,MAT2A,TMEM170A,GTF2F1,SLC4A8,GOLGA8A,PXDN,HNRNPU,PGAM1,
27,HECTD4,CCT6A,ARHGDIA,STIP1,ATP6V0C,DNER,BCLAF1,PPIB,RPL23A,GLUL,PEA15,LUC7L3,
PL9,ATF4,PSMD2,KIF5B,GLIPR1L2,RPS14,CXADR,CHD4,RPS20,CD81,LENG8,WDR1,ATXN7L3B,R
N4,PRDX2,ARL4C,EIF3L,GNB2,RCN1,RPS5,APLP1,RPS19,MIF,TLN1,COL6A2,B4GALNT4,MTSS2,
RS,SF1,SEC16A,UBA52,BRD2,FARP1,UPF1,RAB3GAP2,EIF5B,UQCRC1,ZFR,AGPAT1,NPDC1,OAZ2
D1,SARS1,VANGL2,SOD1,VGF,FBRSL1,MEX3A,P3H3,MLF2,FLNC,AKIRIN1,PRPF40A,PPP1R9B,ADA
TIP1,GPC1,BCLAF1,TRIB2,PPIB,RPL23A,HSBP1,DLG5,TIMP3,SRSF5,C5ORF24,TM9SF3,CHD6,U2
,GTF3C4,LDHB,PKD1,HLA-C,SF3B1,PDIA3,SERINC1,SSBP2,CHD3,BASP1,VAPA,HMGB1,POLR2A,
CAND1,MAP2K2,QARS1,ZFAND5,DIDO1,PITPNA,MSL1,RPS28,SSB,NKTR,FAU,CCT8,LSS,STXBP1,
DR,CHD4,RPS20,CD81,COL5A2,LENG8,WDR1,RPL26,LASP1,CCT7,SRSF4,RBBP4,PFN1,ASTN2,M
,RPS20,LENG8,WDR1,HMGN2,ATP2A2,CRMP1,LASP1,BEX3,VAT1,PFN1,ASTN2,MCL1,JPT2,TLE5
PS3A,STMN1,ITGB1,TUBB4B,UCHL1,MORF4L1,ANXA2,AGRN,RPLP1,ZMIZ1,TIMP2,XAF1,SARAF,M
2,ZNF770,AP2B1,MAT2A,GTF2F1,GOLGA8A,PXDN,HNRNPU,OGA,PGAM1,CTNNA1,RPS16,RPL23,
5A6,OSBPL2,EWSR1,KAT2A,EHMT2,KCTD20,LMF2,CUL4A,SYT15,RHEB,PPP1CA,HNRNPR,ECHS1
B1,TUBB4B,UCHL1,RPL11,MORF4L1,NCAM1,ANXA2,AGRN,RPLP1,ZMIZ1,TIMP2,XAF1,COL11A1
PSMD2,KIF5B,GLIPR1L2,RPS14,CXADR,CHD4,RPS20,CD81,LENG8,WDR1,RPL26,CRMP1,LASP1,C
,GNAI3,GDI2,PARP1,TSR1,MTRF1L,RPL27,PTPRA,PRDX5,ACTR3,UBE4A,ACO2,HSPA4,TCP1,CAM
LM,HSPG2,SHC2,INTS3,SLC6A8,PKDCC,GPC2,LTBP4,GABARAPL1,RAB14,MAZ,PLEKHA6,PRR12,
,EID1,SRSF5,TM9SF3,MDH2,RAB27A,U2AF2,EEF1A2,FDFT1,CCT2,SUGP2,CPSF1,GNAI3,PARP1,V
11,AGRN,RPLP1,TIMP2,RPL10,MARCHF6,WSB1,RPS17,AP3D1,SBF1,DNAJA1,ARF1,MAP4,SF3B2,
A1,ARF1,CYB5R3,AKAP12,SF3B2,CDC42BPB,AP2B1,TMEM170A,GTF2F1,SLC4A8,GOLGA8A,SMA
R,CHD4,RPS20,CD81,LENG8,WDR1,ATXN7L3B,RPL26,CRMP1,LASP1,CCT7,SRSF4,RBBP4,PFN1,

RCC5,CIPC,PJA2,MSN,SLC16A7,KPNB1,C1ORF43,COL1A2,LLGL1,RPL14,HK1,NCS1,RPL22,RHC
P1,KDM3B,SMS,YME1L1,DHX15,PRMT1,SGTA,RERE,DGCR2,PBX1,ALDH3A2,NOL4L,BAZ1B,GTPE
PKDCC,TRAM1,LTBP4,ARHGEF17,VPS4A,LDLR,DTX4,KMT2E,KIAA0232,TMEM63B,NUCKS1,ACTF
LC25A6,DNAJB1,EFNB2,ADNP,CYFIP1,PABPC4,MAP4K4,PNRC2,KCTD20,HTRA1,LIMK1,KLC2,GAL
3B2,CDC42BPB,ITM2B,MAT2A,TMEM170A,SLC4A8,GOLGA8A,PXDN,HNRNPU,OGA,PGAM1,NFE2I
1B2,DIP2A,PLS3,HS2ST1,TRIP12,USP47,OLFM2,HMGA1,LUC7L2,DAG1,RGL2,MKNK2,IGSF3,ELN
M1,TPT1,SEC61A1,RPS4X,RPS3A,RPL18,STMN1,ITGB1,TUBB4B,UCHL1,MORF4L1,NCAM1,ANXA
T4,HNRNPDL,TARDBP,NDFIP1,ST3GAL2,MXD4,MYL6,H1-10,INIP,DUX4,CKAP5,CADM4,SND1,RHO
7A,RNF187,EEF1A2,TBCD,SNX22,PGD,KLHL24,PABPN1,GDI2,PARP1,ABCA3,PTPRA,PDXK,CAMS
ELENON,UBXN4,MAPK1,AZIN1,ARPC2,DSTN,PDE7A,PSMD3,PRDX1,SRSF1,IPO7,AEBP1,RBP1,ILF
ALM,HSPG2,SHC2,CARM1,LINGO1,STRN4,INTS3,ABI2,SLC6A8,PKDCC,GPC2,LTBP4,GABARAPL1
PR1L2,ATP1A1,TMEM259,COL5A2,LENG8,ATXN7L3B,HMGN2,ATP2A2,CRMP1,SRSF4,RBBP4,PFN
4,EP400,SRSF6,KIAA0895L,UPF1,ADCY6,TRA2B,MBTPS1,HDAC6,KCTD12,SMC3,ZNF512B,FMNL
A2,RPL37A,LRRC28,MAPT,FN1,SPEN,NONO,DDB1,PSMD5,YWHAQ,EIF1,HACD3,DCHS1,SET,MAP
4,BRD3,RPS9,XRCC5,CIPC,RPL24,SLC16A7,PAPOLA,IGFBP3,COL1A2,HK1,NCS1,RPL22,RHOA,Z
UND,TMEM259,CD81,COL5A2,LENG8,WDR1,CRMP1,LASP1,VAT1,SRSF4,RBBP4,PFN1,MCL1,TLE
L12,ARL4C,RPS5,APLP1,RPS19,RPL35,CHPF,COL6A2,CD63,ANXA6,B4GALNT4,MTSS2,HNRNPDL
UBE2R2,SCAF11,ACTN1,BNIP3L,MAP7D1,FKBP8,HNRNPUL2-BSCL2,DUSP4,TSC22D1,PLTP,MSL1
HBDD2,PPP1R10,ERH,EIF2S2,IARS2,EIF3H,SNRPD3,RALY,SEC14L1,VPS4A,NUDCD3,TAX1BP1,PG
DKN1A,SERF2,CTDSP2,DDX39B,SRSF5,TM9SF3,MDH2,RAB27A,U2AF2,EEF1A2,FDFT1,CCT2,SUG
APA,HMGB1,POLR2A,AP2M1,TPT1,SEC61A1,RPS4X,RPS3A,RPL18,STMN1,ITGB1,TUBB4B,UCHL
PS10,RPLP2,DDOST,CIRBP,NARS1,DUX4,YWHAB,SND1,LGALS3BP,CALU,PLPPR3,ARCN1,ZNF454,
RAB6B,SLC7A5,ENC1,NLGN2,RPS14,CHD4,JUND,EIF5,CD81,LENG8,WDR1,HMGN2,RPL26,LASP1
RAB6B,SLC7A5,ENC1,NLGN2,RPS14,CHD4,JUND,EIF5,CD81,LENG8,WDR1,HMGN2,RPL26,LASP1
MMP14,RPL12,ITM2C,NANOG,MRFAP1,MAGED2,GNB2,H1-0,APLP1,RPS7,RPS19,MIF,TLN1,CHPF
CLTC,EIF3L,MAGED2,GNB2,H1-0,RPS5,IK,RPS19,CHPF,JUN,MYO5A,NGRN,DNAJC5,HNRNPDL,T
CD63,NGRN,SESN3,RPL35A,DNAJC5,APLP2,HNRNPDL,TARDBP,ARL6IP1,NDFIP1,ST3GAL2,TSR2
5,ATF4,PSMD2,KIF5B,GLIPR1L2,RPS14,CXADR,CHD4,RPS20,CD81,LENG8,WDR1,ATXN7L3B,HM
SARS1,VGF,P3H3,ARNT2,SERBP1,WBP2,AKIRIN1,IGFBP7,SMS,NAP1L4,CTXN1,YME1L1,ADAR,VD
0C,DNER,L1CAM,TRIB2,GLUL,DLG5,SHISAL1,CRPPA,TIMP3,TMEM132A,CPSF2,PGRMC1,NOVA2,
3,SRSF1,ARGLU1,ADAM9,ILF3,WDR6,GABARAP,SCAF11,SON,KDM1A,RPL28,DUSP4,PITPNA,MSL
4KA,UBFD1,MBTPS1,KCTD12,SMC3,ECEL1,HNRNPA2B1,POM121C,LMAN1,HIF1A,CTTN,GPC4,D
FAP1,MAGED2,GNB2,APLP1,RPL35,TLN1,CHPF,MYO5A,COL6A2,CD63,ANXA6,NGRN,SESN3,B4G
RPPRC,XPR1,KHDRBS1,KLHL9,CSNK1E,PFKL,SCRN1,RPS25,TKT,SFPQ,EIF5A,CDH2,PDIA6,DDR1
35E2B,SMCHD1,ATP6V0C,MMP2,HSBP1,NNAT,FAM199X,DLG5,SERF2,HS6ST1,TIMP3,TMEM132
PS1,HECTD1,BAG6,RAI1,SREBF2,HNRNPD,PRPF40A,SMS,AMOTL2,YLPM1,RBM15B,CTNND1,POL
,ANP32A,TRAF7,EIF2AK1,EDC4,BCR,PDAP1,BAZ1B,MMP15,EIF3D,PHB2,APMAP,KLHL42,EP400,F
,CHD4,RPS20,CD81,LENG8,WDR1,RPL26,LASP1,CCT7,SRSF4,RBBP4,PFN1,ASTN2,MCL1,TLE5,P
TG2B,ARPC2,FAM168B,ARGLU1,PTBP1,CAPZB,SUPT5H,HNRNPUL1,ATXN10,SON,QARS1,RPL28,
PRIN1,STIP1,SLC35E2B,IPO5,ATP6V0C,TMEM30A,BCLAF1,PPIB,RPL23A,HSBP1,MAP1A,CDKN1A,
H2,RAB27A,U2AF2,EEF1A2,FDFT1,CCT2,SUGP2,GNAI3,NOP53,GDI2,PARP1,VCAN,TSR1,MTRF1L,
L39,ZSCAN18,RTL8C,RPS13,PRPF40A,NAP1L4,PCYOX1,DDIT4,YME1L1,ADAR,PSMA7,PPDPF,ANP
L39,ZSCAN18,RTL8C,RPS13,PRPF40A,NAP1L4,PCYOX1,DDIT4,YME1L1,ADAR,PSMA7,PPDPF,ANP
RPS20,LENG8,WDR1,RPL26,LASP1,CCT7,SRSF4,RBBP4,PFN1,ASTN2,MCL1,TLE5,PNN,NES,RPL3
S2,RPL23,CNN3,RPL29,TPM4,TNS3,LPCAT1,LAMA5,RPS24,SLC22A17,XRCC5,CIPC,RPL24,KPNB

,ANP32B,GPSM1,TNKS,DKK3,HNRNPAB,SLC25A5,RABEP1,DAP,MTCH1,GLO1,WSCD1,ZNF664,G
PKN1,KLC1,GRINA,NCKAP1,CLSTN3,MDK,SURF4,LAMC1,JPT1,PTK7,ORAI2,ASXL1,KDM6B,RAB1
CIPC,RPL24,RPL17,C1ORF43,PAPOLA,COL1A2,LLGL1,RPL22,RHOA,ZNF865,CLSTN1,SLC7A5,E
RSF7,PSMC5,EMC10,ZBTB4,HSF1,WDR82,UBE3C,GDF1,MEX3C,ASH1L,PHKB,BCOR,PRMT5,ALD
MSAP1,COL2A1,MLEC,ACAP3,CCT4,SNRNP70,RPL36A,RAB11B,SRP14,NRDC,PYGB,PRPF6,TAF7
R3,ARCN1,SEPTIN3,CTBP1,HMGCR,TOP2B,SMCHD1,IPO5,ATP6V0C,MMP2,SKP1,LAPTM4B,RPL
1,CTDSP2,CPSF2,NOVA2,CTSB,CDH11,TM9SF3,CHD6,RNF187,EEF1A2,FDFT1,TBCD,ANXA5,SNX
SH,HNRNPC,LRPPRC,KLHL9,CSNK1E,PFKL,SCRN1,RPS25,EIF5A,PDIA6,DDR1,PIP4K2B,MEGF8,R
RIB2,PPIB,RPL23A,HSBP1,MAP1A,SERF2,DDX39B,NUCB1,SRSF5,TM9SF3,MDH2,TOMM20,U2AF2
NRNPC,LRPPRC,KHDRBS1,KLHL9,SLC3A2,CSNK1E,PFKL,SCRN1,RPS25,TKT,SFPQ,EIF5A,PDIA6,
ISR,PLEKHB2,ELP1,LSM14B,SLC7A1,PCNX3,ABCF1,ADAM10,OGT,COX6A1,MBD3,SF3A2,VASH1,
JX4,PTGES3,CKAP5,SEPTIN11,CADM4,SND1,LAMP1,RHOB,HNRNPH1,UBE2H,CKAP4,RPS12,RPS
RF4,SDF4,SMARCC1,SUPT6H,PFKM,PITPNM1,STXBP1,LONP1,CSNK1D,SF3A1,ATXN2L,SARS1,GH
2B,APPL1,HDAC2,ATP6V0C,TMEM30A,L1CAM,TRIB2,PPIB,GLUL,SQLE,FAM199X,TMEM132A,NOV
3B2,ZNF770,AP2B1,MAT2A,GTF2F1,GOLGA8A,HNRNPA0,FADS2,RPL23,LDHA,CNN3,RPL29,RPSA
RPL28,CCAR2,ARF4,SDF4,DHX30,PHC2,PFKM,FBLN1,PITPNM1,TNKS2,ISYNA1,LONP1,CSNK1D,A
15,MRFAP1,H1-0,RPS5,RPS19,MIF,JUN,MYO5A,CTNNB1,CD63,ANXA6,B4GALNT4,MTSS2,RPL35
CDC42BPB,AP2B1,MAT2A,GTF2F1,SLC4A8,PXDN,HNRNPU,HNRNPA0,RPS16,RPL23,LDHA,CNN3,
TF4,SLC7A5,ENC1,GLIPR1L2,CHD4,ATP1A1,JUND,TMEM259,EIF5,CD81,LENG8,HMGN2,LASP1,
CRMP1,VAT1,SRSF4,RTN3,ASTN2,TLE5,PRKCSH,HNRNPC,LRPPRC,KHDRBS1,SPTBN2,PFKL,TPM
F4H,LAMA5,RPS24,BRD3,RPS9,XRCC5,CIPC,RPL24,SLC16A7,C1ORF43,PAPOLA,IGFBP3,COL1A2
MAGED2,GNB2,RCN1,RPS5,IK,RPS19,CHPF,JUN,MYO5A,NGRN,SESN3,DNAJC5,APLP2,TARDBP,
TAF7,RPL38,MARS1,RPL37,G3BP2,ATG2B,CCNL2,ARPC2,ARGLU1,PTBP1,CAPZB,SUPT5H,HNRN
DI2,PARP1,TSR1,MTRF1L,ALKBH5,RPL27,PTPRA,PRDX5,UBE4A,ACO2,HSPA4,TCP1,CAMSAP1,M
NRNPL,PARP1,TSPYL4,TM9SF2,MORF4L2,SSRP1,MTRF1L,ABCA3,SNRPN,PRDX5,PDXK,PKN1,HS
DX12,ACTN4,MMP14,PRDX2,RPS15,RPL12,ITM2C,MRFAP1,MAGED2,RCN1,H1-0,RPS5,APLP1,RP
3,SUPT5H,HNRNPUL1,SON,RPL28,CCAR2,ARF4,SMARCC1,SUPT6H,PFKM,EIF3F,LONP1,SF3A1,A
3D1,DNAJA1,AKAP12,MAP4,SF3B2,ITM2B,MAT2A,TMEM170A,GTF2F1,PXDN,HNRNPU,OGA,PGAM
,CCT7,OAZ1,SRSF4,RBBP4,MCL1,PNN,NISCH,PRKCSH,HNRNPC,TADA2B,XPR1,KLHL9,XPO1,HD
EB,IQSEC1,WIZ,HNRNPR,WRNIP1,ECHS1,TBC1D14,SLC25A1,ALDH7A1,MAPKAPK2,GID8,STUB1,Z
F1,SNX22,PGD,VAMP2,GAS7,HNRNPL,TSR1,HSPH1,MTRF1L,ABCA3,ALKBH5,HNRNPA3,PDXK,PK
,TARDBP,RPLP2,DDOST,HNRNPK,CIRBP,NARS1,PTGES3,SND1,ATP1A2,CALU,PLPPR3,ARCN1,CC
168B,ACADVL,LUZP6,HNRNPUL1,HSPB1,SON,QARS1,RPL28,CCAR2,ARF4,GNG2,CORO1C,SDF4,
5,RAB2A,GNG4,FARP1,EP400,SRSF6,PAPSS1,CRTAP,ZFR,TBC1D9B,PPP1CC,IL6ST,OAZ2,SPINT2,
ORF4L1,BTF3,NCAM1,ANXA2,AGRN,RPLP1,TIMP2,XAF1,MIDN,RPL10,MARCHF6,WSB1,VCL,AP3D
WDR1,ATXN7L3B,ATP2A2,RPL26,CRMP1,OAZ1,SRSF4,ASTN2,MCL1,TLE5,PNN,NES,NISCH,PRKC
37A,RPS15,ITM2C,RPL30,EIF3L,MAGED2,H1-0,RPS5,IK,RPS19,CHPF,JUN,NGRN,HNRNPDL,TARD
FKM,FBLN1,B2M,UBQLN1,CSNK1D,ATXN2L,USP7,ZFAND3,SARS1,COL27A1,PFN2,HNRNPD,VGF
TRAF7,EIF2AK1,TXLNA,PRMT1,XPOT,EXTL3,STARD7,PDAP1,AP2A1,MMP15,EIF3D,PHB2,APMAP,G
SHC2,INTS3,BCL7A,UBE2M,GPC2,IMPDH2,HINT1,RAB14,SH3GL1,ZNF428,KIFAP3,TTC37,PNISR,
DHA,PTP4A2,SLIT3,STMN3,FBLN1,PITPNM1,ISYNA1,UBQLN1,HYOU1,ZFAND3,IFITM3,PTPRS,VAN
12,RBM5,ERGIC3,LRRC41,SYT11,OGT,MED13,SF3A2,DHX38,YY1,SMARCD1,APBB1,NDUFA13,T
XDC,SCAF11,ACTN1,CAND1,SON,BNIP3L,QARS1,DUSP4,XPNPEP3,KIF3C,SMARCC1,STT3B,PITPN
,HES6,MEX3D,MYBBP1A,EMD
2,NUCB1,CDH11,ZC3H7B,TM9SF3,TBCD,SNX22,PGD,SQSTM1,VAMP2,GDI2,HNRNPL,TAF15,LM

LEC,CLSTN3,TMEM248,DDX42,LAMC1,RAD23B,CHD2,ORAI2,GCN1,HDGF,NRDC,PRPF6,VEGFA,
D1,RHOB,CKAP4,RPS12,PLPPR3,ARCN1,ZNF454,HECTD4,MAGED4B,GPC1,SLC35E2B,APPL1,HD
L1,RPL11,MORF4L1,BTF3,NCAM1,ANXA2,AGRN,RPLP1,TIMP2,SARAF,COL11A1,MIDN,RPL10,M
5B,SLC7A5,ENC1,NLGN2,RPS14,CXADR,CHD4,ATP1A1,TMEM259,RPS20,EIF5,WDR1,ATP2A2,CR
,KHDRBS1,KLHL9,SLC3A2,CSNK1E,PFKL,SCRN1,RPS25,TKT,SFPQ,EIF5A,PDIA6,DDR1,PIP4K2B,
NLL2,ILF3,CDC37,WDR6,TDRD5,ACTN1,CAND1,PLCG1,RPL28,POLR2B,SDF4,SMARCC1,FYN,SSB
ALKBH5,PRDX5,ACTR3,PDXK,UBE4A,YEATS2,ACO2,HSPA4,TCP1,CAMSAP1,MLEC,ACAP3,ZNF302
M9SF3,MDH2,RAB27A,U2AF2,EEF1A2,FDFT1,CCT2,SUGP2,CPSF1,PPP1CB,GAS7,PARP1,VCAN,TM
B,ZNF770,AP2B1,MAT2A,GTF2F1,SLC4A8,GOLGA8A,HNRNPU,OGA,PGAM1,HNRNPA0,FADS2,RP
LO1,BAP1,SPG7,TAOK1,EHMT1,IARS2,PALM,CLPTM1,SLC44A2,SHC2,FKBP4,SSBP3,BCL7A,RAB
,TOP1,TAF7,RPL38,MARS1,RPL37,G3BP2,ATG2B,KDELR2,CCNL2,ARPC2,ARGLU1,PTBP1,CDC3
L2,RBBP7,TRAF7,EIF2AK1,PRMT1,EP300,SF1,PDAP1,ALDH3A2,H2AZ1,GTPBP1,GPX4,GSPT1,BR
,RPL35A,DNAJC5,APLP2,HNRNPDL,SLC25A3,RPLP2,NDFIP1,MYL6,ANKRD52,WLS,HNRNPK,H1-1
,CCNI,XPR1,HDLBP,RPS25,GLG1,EIF5A,PEBP1,PNMA1,PDIA6,DDR1,PLXNA1,MEGF8,RAB7A,CCN
,JUN,CD63,ANXA6,RPS10,SESN3,DNAJC5,APLP2,TARDBP,TAGLN,MYL6,SHOX,HNRNPK,H1-10,C
,SND1,CKAP4,HLA-B,CALU,ARCN1,ZNF454,HMGCR,GPC1,CDK16,TOP2B,DNER,BCLAF1,PPIB,M
DCC,TRAM1,LTBP4,ARHGEF17,IMPDH2,GABARAPL1,RAB14,MAZ,DCLK1,IREB2,LRATD2,STK24,
,ATP1B2,RNF145,MSH6,VASH1,TGOLN2,UBE2I,UBE2D3,WNK1,SAE1,DPYSL4,ATG4B,CPXM1,DM
ADS2,RPS16,RPL23,CPE,RPL29,TPM4,EIF4H,TSPAN3,LPCAT1,LAMA5,SLC22A17,BRD3,CIPC,PJA
N4,PRDX2,RPS15,ITM2C,RPL30,CLTC,EIF3L,MAGED2,H1-0,RPS5,RPS7,IK,RPS19,RPL35,MIF,TL
PLP1,RPS7,RPS19,RPL35,GJA1,JUN,COL6A2,ANXA6,NGRN,RPS10,B4GALNT4,DNAJC5,HNRNPD
,ATP1A1,CD81,WDR1,CRMP1,LASP1,BEX3,CCT7,SRSF4,RTN3,PFN1,ASTN2,SPC25,NES,NISCH,H
,PDPK1,LTBP3,MSH6,EHD1,FBLN5,PSMC3,PACS2,CREBBP,BNIP3,UBE2I,ANKRD12,UBXN6,PRDX
11,SND1,ATP1A2,RPS12,ARCN1,RPS27,CTBP1,ARHGDIA,STIP1,SLC35E2B,SMCHD1,IPO5,ATP6V
B,PPIB,RPL23A,GLUL,LUC7L3,PTTG1IP,MAP1A,SERF2,TIMP3,CPSF2,NOVA2,NUCB1,SRSF5,CHD
25,GLG1,SFPQ,EIF5A,PEBP1,CCND2,PLXNA1,PIP4K2B,SOX12,RAB7A,DDX3X,RPS15,RPL12,ITM
RDX,KIAA0232,LRATD2,TMEM63B,TUBGCP6,RIC8A,INTS11,SPAG9,SLC7A1,PDPK1,ERGIC3,LTB
,RPS11,PRKDC,UBE2R2,CASC3,NSG2,DIDO1,RPS28,HNRNPM,GIGYF1,SRSF3,FAU,CCT8,MESD,
L1,SMCHD1,IPO5,ATP6V0C,SKP1,BCLAF1,PPIB,LUC7L3,SQLE,CDKN1A,DLG5,TIMP3,CTDSP2,CP
DKL1,AP2M1,TPT1,RPS3A,RPL18,UCHL1,RPL11,MORF4L1,BTF3,NCAM1,ANXA2,AGRN,RPLP1,Z
IMP3,TMEM132A,CTDSP2,PGRMC1,CNN2,NOVA2,CDH11,MDH2,ANXA5,VAMP2,KLHL24,MEST,T
MCL1,PNN,PRKCSH,CCNI,KHDRBS1,XPO1,HDLBP,RPS25,GLG1,EIF5A,PEBP1,CCND2,DDR1,PLX
PFKL,SCRN1,RPS25,TKT,EIF5A,PDIA6,DDR1,PIP4K2B,MEGF8,RPL12,ITM2C,RPL30,EIF3L,MAGE
3A1,HECTD1,BAG6,RAB11A,SERBP1,ARFGEF2,WBP2,AKIRIN1,SMS,CCT5,CDC42,PPP1R9B,YME1
,QARS1,RPL28,CCAR2,ARF4,SUPT6H,PFKM,PITPNM1,ISYNA1,LONP1,CSNK1D,SF3A1,ATXN2L,H
SMARCC1,LARS1,STMN3,PFKM,FBLN1,STT3B,STXBP1,ISYNA1,STT3A,TUG1,ATXN2L,ZFAND3,ST
RFAP1,MAGED2,RPS7,RPL35,MIF,GJA1,CTNNB1,CD63,ANXA6,MACF1,MTSS2,RPL35A,CYFIP2,H
ACP1,RBM17,PRMT2,PPP1R12A,SHC1,NARF,PCED1A,CAPZA2,CTDNEP1,TFDP1,RAB35,SPIN1,CA
,YWHAB,PTGES3,CADM4,SND1,ATP1A2,CKAP4,CALU,ARCN1,ZNF454,RPS27,COTL1,CCT6A,STIP
CYB5R3,PAFAH1B1,MAP4,CDC42BPB,ZNF770,MAT2A,TMEM170A,GTF2F1,SLC4A8,GOLGA8A,P
1,RPS5,RPS19,GPAT4,PRKAR1A,TLN1,JUN,MYO5A,COL6A2,RPS10,B4GALNT4,DNAJC5,HNRNPD
GRN,GOLM1,B4GALNT4,DNAJC5,HNRNPDL,TARDBP,DDOST,INIP,NARS1,DUX4,SND1,LAMP1,RH
VL,PTBP1,FTO,LUZP6,CAPZB,SUPT5H,HSPB1,SON,CCAR2,ARF4,GNG2,CORO1C,SUPT6H,DHX3
,DCLK1,IREB2,LRATD2,STK24,NUCKS1,ATXN3,SLC7A1,DICER1,DCUN1D2,LTBP3,ITGAV,TEX2,F
SF5,ZC3H7B,TM9SF3,MDH2,TOMM20,RAB27A,U2AF2,EEF1A2,CCT2,GNAI3,GAS7,NOP53,GDI2,

15B,POLR2E,PSMA7,RBBP7,NAP1L1,TXNDC5,BAZ1B,H2AZ1,KPNA2,PHB2,GSPT1,UBA52,PPM1G
P1,CCT7,SRSF4,RBBP4,PFN1,ASTN2,MCL1,TLE5,PNN,RPL31,NISCH,PRKCSH,HNRNPC,CCNI,LRP
52,SRRT,CAMK2N1,NECTIN2,HEG1,STRIP1,QRICH1,CLCN3,TUT4,CBX4,SETD1A,TMCO1,DDX56,
,RPLP2,ARL6IP1,NDFIP1,ST3GAL2,MXD4,ANKRD52,H1-10,INIP,CIRBP,MYH10,RPL36,DUX4,PTG
07,HGS,BAG6,RAB11A,SERBP1,ARFGEF2,AKIRIN1,SMS,YLPM1,CCT5,CDC42,PPP1R9B,YME1L1,
PRMT1,XPOT,STARD7,RETREG2,PDAP1,AP2A1,MMP15,FLOT1,EIF3D,SEC16A,PHB2,APMAP,UBA
RPS9,XRCC5,PJA2,RPL24,SLC16A7,KPNB1,RPL17,C1ORF43,IGFBP3,COL1A2,HK1,NCS1,RHOA,
NAP1L4,CTXN1,GRIPAP1,PSMA7,PPDPF,RBBP7,DFFB,KCNQ2,EDC4,WDR45B,PRMT1,MTA1,DGC
1,XPOT,STARD7,PDAP1,AP2A1,MMP15,FLOT1,EIF3D,SEC16A,PHB2,APMAP,CST3,MGAT4B,KLHL
LC35E2B,SMCHD1,ATP6V0C,TMEM30A,MMP2,LAPTM4B,TRIB2,NNAT,FAM199X,DLG5,SERF2,C
B2,ITM2B,MAT2A,GTF2F1,SLC4A8,HNRNPU,PGAM1,NFE2L1,SMARCA5,LARP1,CTNNA1,RPL23,
,DEK,FGFR1,UNC119B,NELFB,USP47,LMO4,CDC34,RNF11,HMGA1,WDTC1,TMEM184B,OGFR,
NOW,ECEL1,CBX6,TBC1D9B,NPDC1,BRSK1,PSMD8,GALNT11,CTTN,SAFB2,ZNF664,KIAA0100,E
LP1,TIMP2,SARAF,COL11A1,MIDN,RPL10,RPS17,VCL,AP3D1,DNAJA1,CYB5R3,SF3B2,ITM2B,MA
E5,PNN,RPL31,PRKCSH,HNRNPC,KHDRBS1,KLHL9,SPTBN2,SLC3A2,CSNK1E,PFKL,SCRN1,HDL
2,PDE7A,PSMD3,TCF25,CDC37,CAPZB,RPS11,GABARAP,BNIP3L,MAP7D1,CASC3,NSG2,CLIP3,C
BFD1,TRA2B,COX7C,ZFR,AGPAT1,PRRC2A,HNRNPA2B1,ADH5,NPDC1,ANP32B,PREPL,TIMM50,S
TSS2,APLP2,TARDBP,ARL6IP1,NDFIP1,ST3GAL2,TSR2,MXD4,MYL6,HNRNPK,H1-10,INIP,MYH10,
TD4,CCT6A,RAC1,CDK16,APPL1,SMCHD1,HDAC2,IPO5,DNER,BCLAF1,L1CAM,LAPTM4B,RPL23A
3L,MRFAP1,RPS7,RPS19,MIF,GPAT4,CHPF,JUN,NGRN,B4GALNT4,DNAJC5,APLP2,HNRNPDL,TAR
,LASP1,CCT7,SRSF4,RBBP4,PFN1,ASTN2,MCL1,PNN,RPL31,NISCH,PRKCSH,CCNI,LRPPRC,TADA
L,AP3D1,SBF1,ARF1,CYB5R3,AKAP12,MAP4,SF3B2,CDC42BPB,ITM2B,AP2B1,MAT2A,GTF2F1,SL
3P4,RTN3,PFN1,MCL1,JPT2,PNN,NISCH,PRKCSH,HNRNPC,TADA2B,XPR1,KHDRBS1,KLHL9,XPO1
JBGCP6,RIC8A,NCSTN,INTS11,MOAP1,MTMR4,ABCF1,YIPF3,SPTY2D1,CAP1,PRPF19,USP24,NOC
5B,RAB6B,SLC7A5,NLGN2,CXADR,CHD4,TMEM259,WDR1,ATP2A2,CRMP1,LASP1,BEX3,CCT7,V
3,YWHAG,CKB,COL4A2,RPL37A,LRRC28,MAPT,FN1,SPEN,NONO,DDB1,PSMD5,YWHAQ,EIF1,HAC
0,FKBP8,RPL28,CCAR2,ARF4,DIDO1,HNRNPUL2-BSCL2,FLNB,CORO1C,ATP6V1B2,GIGYF1,SDH
M4B,TRIB2,RPL23A,HSBP1,CDKN1A,FAM199X,CTDSP2,DDX39B,NUCB1,SRSF5,TM9SF3,MDH2,T
1,PPME1,RAB18,ZBTB4,CRTC1,TMEM43,ZFP91,TBCA,ATP5F1C,SRP54,NELFCD,ATF6B,DNAJB6,
RPL36A,ABCA2,RPL38,RNF10,RAPGEF1,MVB12B,MAPRE1,KDELR2,SRSF9,BTBD2,PBX2,PDE7A,
1B,UBE2M,RALY,SEC14L1,NUDCD3,FAM234A,SSR3,GNL1,STK24,TUBGCP6,NCSTN,INTS11,PRF
MF2,PITRM1,DRAP1,IQSEC1,WIZ,RAP2A,TBC1D14,CDK4,RNF41,TBXA2R,SLC25A1,NIPSNAP1,MA
,FARP1,RER1,PI4KA,UBE2G2,RAB3GAP2,UBFD1,IARS1,MASTL,COX7C,YKT6,CUL3,MCM3AP,RP
D3,SF3A2,BNIP3,TTC28,ATP5MC2,SSBP4,DPYSL4,FJX1,CPXM1,HSDL1,GAK,ATP13A1,SCAP,RPS2
SR2,MXD4,MYL6,HNRNPK,H1-10,INIP,MYH10,NARS1,DUX4,CKAP5,RHOB,HNRNPH1,HLA-B,RAB

XRCC6,PELP1,VAPB,CCN1,HSPG2,UBE2M,TRAM1,RDX,MAZ,ZNF428,SRSF2,CADM3,SLC4A2,SLC
1,TMEM259,RPS20,EIF5,CD81,WDR1,HMGN2,CRMP1,BEX3,VAT1,OAZ1,PFN1,MCL1,JPT2,TLE5,
3A,ARNT2,SERBP1,ARFGEF2,AKIRIN1,SMS,CDC42,PPP1R9B,VDAC1,RBBP7,TRAF7,EDC4,WDR45
RTL6,SLC44A2,SSBP3,BCL7A,SOX2,RALY,SLC6A8,PKDCC,ELAVL1,UBE2D2,TLE1,PGAP6,HINT1,R
AP4,RPS12,HLA-B,RAB1A,ZNF454,RPS27,MAGED4B,STIP1,CDK16,TOP2B,APPL1,SKP1,DNER,BC
,MRFAP1,MAGED2,H1-0,RPS5,RPS7,IK,RPS19,CHPF,JUN,CD63,ANXA6,NGRN,DNAJC5,APLP2,HN
I1,PSMD4,LRRC41,ATP6V0D1,SH3BP4,ARPP19,SF3A2,VASH1,NASP,AP2A2,WNK1,CERS2,SSBP4,
D23A,PDHB,KHDC4,ZNF207,MRPS27,WDR33,RNF40,DPF2,FKBP3
31,NXN,EP400,CD164,EVL,MASTL,KCTD12,ZFR,HNRNPA2B1,PPP1CC,FMNL2,NCOA4,ULK1,EML1

3P1,FTO,LUZP6,CDC37,CAPZB,SUPT5H,HSPB1,SON,CCAR2,ARF4,POLR2B,CORO1C,XPNPEP3,SL
5V0C,DNER,L1CAM,RPL23A,GLUL,DLG5,SHISAL1,HS6ST1,TMEM132A,CTDSP2,CPSF2,DDX39B,C
NK1D,ATXN2L,GPR107,HECTD1,COL27A1,HGS,PFN2,PNMA8A,FBRSL1,MEX3A,SEC23A,RTL8C,SI
C2,PPM1A,CENPT,UBTF,SKIL,MXRA8,NUTF2,ABCA7
OXP4,OSBPL2,PSMD13,CRIP2,SYNCRIP,PHPT1,EHMT2,RASA4,SALL2,WIZ,WRNIP1,JAK1,OTUD5,
MLF2,SEC23A,ARNT2,RTL8C,PRPF40A,BMPR2,CD99L2,AMOTL2,CCT5,YME1L1,CTNND1,PPDPF
,RPL22,RHOA,CBX5,ZNF865,CLSTN1,ATF4,PSMD2,RAB6B,ENC1,NLGN2,GLIPR1L2,ATP1A1,JUN
LIM1,SSU72,LMF2,NCKAP5L,EXT2,ECHS1,MAGEF1,FAM8A1,SH3BGRL3,ADCY1,FAM32A,LDB1,TS
SH,HNRNPC,TADA2B,XPR1,KHDRBS1,KLHL9,SPTBN2,HDLBP,RPS25,GLG1,SFPQ,TPM1,PEBP1,P
PPDPF,ANP32A,DFFB,ARIH1,WDR45B,PRMT1,OS9,TXNDC5,MTA1,ACTR1A,NDUFV1,HIP1,GTPBP1
PHB2,APMAP,CST3,SLC39A7,PLXNB1,DECR1,FARP1,RER1,PI4KA,UBE2G2,RAB3GAP2,TRA2B,MA
CEMIP2,CBX6,ANP32B,PREPL,RPL7L1,ITGB5,POM121C,RPS2,HNRNPAB,CHERP,GALNT11,GSE1
PF,CIC,RBBP7,ARIH1,WDR45B,PBX1,NOL4L,BAZ1B,LRRC4B,GSPT1,CST3,SLC39A7,NXN,CEMIP2
I70,XPO6,UBXN6,MYL12B,GAK,LYPLA2,ATP13A1,ATP5MG,IER2,PAFAH1B2,POLDIP2,TRIP12,XRN
TSPYL4,TSR1,MORF4L2,HNRNPA3,PRDX5,C4ORF3,PDXK,PKN1,GRINA,NCKAP1,MLEC,ACAP3,ZNF
CS1,CBX5,RPL9,CLSTN1,PSMD2,NR2F1,SLC7A5,GLIPR1L2,CXADR,ATP1A1,JUND,TMEM259,EIF5
C38A10,ATP13A1,ATP5MG,SRPRA,NELFB,EIF4G1,PAFAH1B2,SLC30A9,ACAT2,POLDIP2,TRIP12,L
A,DDX3X,RPS15,RPL12,ITM2C,RPL30,CLTC,EIF3L,NANOG,MRFAP1,MAGED2,RPS7,RPL35,MIF,HI
B,GLUL,PEA15,LUC7L3,FAM199X,CRPPA,CTDSP2,CPSF2,CNN2,ZC3H7B,RAB27A,RNF187,TBCD,
ID4,GPR107,SARS1,VANGL2,DGKD,BAG6,SREBF2,HNRNPD,RAB11A,CYP51A1,ARNT2,SERBP1,AI
S25,SFPQ,EIF5A,PEBP1,CCND2,PNMA1,DDR1,PIP4K2B,SOX12,PRDX2,DDX3X,RPL12,ARL4C,MRF
F8,RAB7A,DDX3X,RPS15,RPL12,ITM2C,RPL30,EIF3L,NANOG,MRFAP1,MAGED2,RPS7,RPL35,HM
PTCD3,CBX5,CLSTN1,PSMD2,NR2F1,NLGN2,CXADR,CHD4,ATP1A1,JUND,LENG8,WDR1,HMGN2,
WDR26,PARK7,SYNCRIP,RALGDS,HTRA1,RAN,PCGF3,ASNS,WRNIP1,SOBP,TRIB3,JAK1,LDB1,EPN
,LIMK1,ECHS1,DDX6,FAM168A,SLC16A2,GID8,LDB1,CDK9,EXOC7,NSD1,RANBP9,MEX3C,ACBD
J,PLPPR3,ARCN1,RPS27,HRNR,CTBP1,ARHGDIA,CDK16,SLC35E2B,SMCHD1,HDAC2,IPO5,ATP6V
,NCKAP5L,LIMK1,TSPAN14,CUL4A,RHEB,IQSEC1,WIZ,CAPN1,WRNIP1,MAPKAPK2,GID8,RBFOX2
X,MYH10,PTGES3,SND1,RHOB,UBE2H,CKAP4,ARCN1,RPS27,HRNR,CCT6A,CTBP1,ARHGDIA,TOF
Z,RPL27A,TBC1D14,SLC25A1,MAPKAPK2,RTF2,STUB1,TSPAN7,ZNF768,PIP5K1C,WDR82,DHFR,N

CIN1,PDXDC1,KRI1,CCSER2,PCMTD2,PITHD1,UQCR11,CASK,MYBBP1A,TBCB
TCEAL4,UBE2L3,LGALS1,ZZEF1,KTN1,MGAT5,YTHDF3,RBM4,MTREX,SLC38A1,MAPK8IP2,NSF,A
TMEM259,EIF5,CD81,LENG8,HMGN2,LASP1,BEX3,RBBP4,PFN1,ASTN2,MCL1,JPT2,TLE5,PNN,N
,CDKL1,AP2M1,TPT1,SEC61A1,RPS4X,RPS3A,RPL18,STMN1,ITGB1,UCHL1,RPL11,MORF4L1,BT
2,APMAP,UBA52,PLXNB1,CSNK1G2,FARP1,NXN,EP400,RER1,UBE2G2,UBFD1,KCTD12,AGPAT1,C
F3A1,NBPF9,STRAP,SARS1,BAG6,HNRNPD,RAB11A,SERBP1,RTL8C,AKIRIN1,CCT5,CDC42,PPP1F
GES3,SEPTIN11,LGALS3BP,RAB1A,TTR,ARCN1,RPS27,HECTD4,RAC1,ARHGDIA,STIP1,SMCHD1,
M1,GPC2,GNA12,MAZ,DTX4,NUCKS1,SLC4A2,KRT8,PCNX3,MSI1,LTBP3,ATP1B2,MSH6,DENND4
N,EIF4A3,SLC38A10,CENPB,DHX40,ATP5MG,SRPRA,NELFB,CASC4,MDH1,BGN,CDC34,PSAT1,LG
A,EIF4A1,MRC2,RPL19,PPP2R1A,RPL15,LDHB,DNAH8,PKD1,HLA-C,SF3B1,PDIA3,SERINC1,SSBP
NN,PRKCSH,CCNI,LRPPRC,TADA2B,XPR1,PTN,HDLBP,RPS25,GLG1,EIF5A,PEBP1,CCND2,DDR1,F
SLC20A1,MICAL1,PRDX3,ATG4B,METRN,NSG1,UNC119B,FBXL19,ATP13A1,USP5,CASC4,TECR,T
,MCM3AP,RPS27A,TMEM167A,SEPTIN7,KLHDC3,CHERP,FKBP1A,CTTN,LRP10,MAP1LC3B,MTCH
PIP4K2B,MEGF8,SOX12,MMP14,PRDX2,DDX3X,RPS15,RPL12,ITM2C,ARL4C,CLTC,MAGED2,RPS
CP6,NUCKS1,ATXN3,SLC7A1,RNF145,SCRIB,MSH6,MBD3,VASH1,YY1,CREBBP,UBE2I,UBE2D3,T

A,FAM168B,ARGLU1,ACADVL,LUZP6,ATXN10,HSPB1,SON,QARS1,CCAR2,GNG2,CORO1C,DHX30
4,TP53BP1,AMOTL2,YME1L1,CTNND1,VDAC1,TRAF4,STAT3,ARIH1,LBH,KDM5B,RETREG2,FLCN,
3N2,SLC3A2,GLG1,CDH6,TPM1,EIF5A,CDH2,PDIA6,GPC3,SULF1,PLXNA1,CCN2,PRDX2,DDX3X,R
3P,NARS1,DUX4,SND1,LAMP1,LGALS3BP,CALU,PLPPR3,ARCN1,ZNF454,HRNR,MAGED4B,CTBP1
RMT112,PSMD13,COPE,CD2BP2,RPS21,PARK7,APEH,BANF1,FAF2,AMPD2,KLC2,GOLPH3,PPP1C
35E2B,SMCHD1,IPO5,BCLAF1,L1CAM,LUC7L3,CDKN1A,FAM199X,SHISAL1,HS6ST1,TIMP3,PGRM
1,CTSB,SRSF5,MDH2,CHD6,RNF187,EEF1A2,TBCD,ANXA5,SNX22,GNAI3,PABPN1,NOX4,TAF15,T
PHB2,APMAP,RAB2A,CD164,RER1,UBFD1,TRA2B,IARS1,MASTL,YKT6,MCM3AP,SPINT2,PREPL,R
MCM3AP,PREPL,GPSM1,RPL7L1,RPS27A,TMEM167A,KLHDC3,TNKS,PSMD8,NPLOC4,CHERP,FK
BP7,PRPF40A,BMPR2,DYNLL1,CTXN1,AMOTL2,DDIT4,CTNND1,POLR2E,PSMA7,CIC,SSR1,DCTN
PR107,STRAP,BAG6,RAB11A,SERBP1,ARFGEF2,AKIRIN1,CDC42,YME1L1,VDAC1,TRAF7,EIF2AK
NK1E,PFKL,SCRN1,RPS25,SFPQ,EIF5A,PDIA6,MEGF8,ACSL6,SOX12,RAB7A,PRDX2,RPL12,ITM2
NK1E,PFKL,SCRN1,RPS25,SFPQ,EIF5A,PDIA6,MEGF8,ACSL6,SOX12,RAB7A,PRDX2,RPL12,ITM2
MGN2,LASP1,BEX3,RBBP4,PFN1,ASTN2,MCL1,JPT2,TLE5,PNN,PRKCSH,HNRNPC,LRPPRC,KHDR
SREBF2,HNRNPD,RAB11A,ARNT2,SERBP1,ARFGEF2,SMS,CDC42,YME1L1,VDAC1,RBBP7,TRAF7
H1,LGALS3BP,RPS12,TTR,CALU,ARCN1,COTL1,HECTD4,STIP1,CDK16,TOP2B,SLC35E2B,TNC,T
3,MLEC,ACAP3,DDX42,CCT4,RAB11B,SRP14,TSPAN6,NRDC,ATP6AP1,TOP1,TAF7,RAPGEF1,TUB
GCS1,GPAT4,TLN1,CHPF,JUN,COL6A2,NGRN,GOLM1,SESN3,MTSS2,APLP2,HNRNPDL,SLC25A3,
AK,LYPLA2,ATP13A1,IER2,PAFAH1B2,POLDIP2,TRIP12,CHMP1A,ITFG1,CGGBP1,PSMB1,TSNAX,
ORF58,UBE2D3,TOMM70,XPO6,CCDC47,UBXN6,DPYSL4,DEK,TRO,MYL12B,GAK,LYPLA2,ATP13
R,STMN3,PFKM,TNKS2,STT3A,LONP1,SF3A1,HYOU1,ID4,NBPF9,SARS1,VANGL2,BAG6,HNRNPD,
1,NPDC1,IL6ST,MAPRE2,ZMYM4,HNRNPAB,ZNF664,RHBDD2,ZMYM3,FUS,BAZ2A,PALM,CCN1,HS
GTF3C1,YTHDF3,CDV3,TIA1,SIN3B,SLC25A6,KIF26A,JAM3,PJA1,EWSR1,ATF5,RAN,IP6K1,DYNLR
,ITGB1,TUBB4B,UCHL1,RPL11,MORF4L1,BTF3,NCAM1,ANXA2,AGRN,RPLP1,ZMIZ1,TIMP2,SAR
1,RBBP7,TRAF7,EIF2AK1,EDC4,TXLNA,WDR45B,PRMT1,BCR,PDAP1,ALDH3A2,AP2A1,CSNK2A1,

0,EIF3L,NANOG,MRFAP1,RPS5,RPS7,IK,RPL35,MIF,GJA1,CTNNB1,ANXA6,NGRN,RPL35A,CYFIP2
,NOP53,GDI2,TM9SF2,TSR1,SSRP1,HSPH1,LMNB2,MTRF1L,ALKBH5,RPL27,SNRPN,PRDX5,UBE
GN,LMO4,CELSR1,RNF11,HMGA1,WDTC1,OGFR,USF2,PUM1,FXR2,LGALS1,HDGFL3,ZKSCAN1,
,RPS5,CTNNB1,RPS10,RPLP2,ARL6IP1,ARID1A,NDFIP1,ST3GAL2,TAGLN,ANKRD52,WLS,H1-10,C
1,SOD1,BAG6,NCOR2,VGF,FBRSL1,CYP51A1,SEC23A,ARNT2,SERBP1,AKIRIN1,SMS,CD99L2,PCY
31,NISCH,PRKCSH,TADA2B,XPO1,SLC3A2,HDLBP,RPS25,TKT,CDH6,TPM1,CDH2,PDIA6,DDR1,P
R2,PLEC,PDE7A,PSMD3,SRSF1,TCF25,ADAM9,EIF3E,CDC37,PCSK1N,WDR6,CAPZB,GABARAP,TD
INRNPH3,PAGR1,SRPRA,USP5,ADGRA3,RANGAP1,TCEAL4,CAD,PLS3,MZT2B,USP47,SNX3,USF2,
R9B,VDAC1,RBBP7,TRAF7,EIF2AK1,EDC4,WDR45B,PRMT1,XPOT,KDM5C,EXTL3,PDAP1,ALDH3
NONO,DDB1,PSMD5,YWHAQ,EIF1,HACD3,DCHS1,SET,EGLN1,MAP9,TPI1,CHSY1,TMSB10,INA,EIF
2A1,MMP15,FLOT1,SEC16A,PHB2,APMAP,CST3,PLXNB1,DECR1,FARP1,RER1,PI4KA,UBE2G2,RA
NGRN,RPLP2,ARL6IP1,NDFIP1,DDOST,INIP,CIRBP,NARS1,RPL36,YWHAB,SND1,TTR,CALU,ARCN
H,UBQLN4,DUSP8,BRI3,NR2F6,ITPK1,DDHD2,CITED2,MCRIP1,RPS6KA2,SLC2A4RG,CTDNEP1,Z
H,UBQLN4,DUSP8,BRI3,NR2F6,ITPK1,DDHD2,CITED2,MCRIP1,RPS6KA2,SLC2A4RG,CTDNEP1,Z
M32A,EPN1,ARPC3,RTF2,ZRANB2,CLPTM1L,POLDIP3,UNC45A,RNMT,ZBTB4,STAT2,PSMF1,HMG
ATP6V0E2,KIAA2013,NAPA,RXRA,ZNRF1,FAM20B,ITPK1,RUSC1,NARF,AFDN,ROCK2,ZDHHC9,UB
KBP1A,CTTN,LRP10,SERP1,ERH,EIF2S2,EIF3H,SNRPD3,RALY,SEC14L1,SMG5,NUDCD3,TAX1BP1
FD1,MBTPS1,RGMB,SELENOW,CUL3,ADH5,NDST1,NPDC1,ITGB5,TIMM50,TNKS,PSMD8,APEX1,
,RFX7,CUL4B,ZDHHC17,ADGRL1,NCOA6,RYBP,CBX4,SAMD1,SELENOF,SESN2,MYADM,POU3F3

)1,SMARCA4,MAPRE1,DDAH2,BTBD2,TCF25,DYNLL2,ILF3,LUZP6,CAPZB,RPS11,HNRNPUL1,UBE2
52,HNRNPK,H1-10,INIP,NARS1,CKAP5,SEPTIN11,CADM4,LAMP1,HNRNPH1,LGALS3BP,CKAP4,RF
2,HGS,VGF,RPL39,FLNC,ARNT2,RTL8C,CTXN1,AMOTL2,POLR2E,ANP32A,KCNQ2,TRIO,EP300,NI
AP,RPS27A,TMEM167A,KLHDC3,CHERP,FKBP1A,CTTN,SLC25A5,LRP10,FBXO21,MTCH1,SERP1,I
S1,NOL4L,H2AZ1,MMP15,GRSF1,GPX4,GSPT1,CST3,ZFP36L1,SLC39A7,SLC35F5,GNG4,FARP1,I
LUD1,UPF1,MASTL,HDAC6,ZFR,ECEL1,MAPRE2,APEX1,PPP1R15A,GALNT11,MAP1LC3B,MVD,DAF
ELFB,RANGAP1,PAFAH1B2,ACAT2,POLDIP2,TRIP12,CHMP1A,CLTA,MTA2,USP10,TSNAX,ZKSCAN
LUZP6,PRKDC,SUPT5H,SCAF11,HSPB1,SON,CMIP,CCAR2,ARF4,DIDO1,CORO1C,SUPT6H,DHX30,
LONP1,HYOU1,RAI1,SREBF2,NCOR2,PNMA8A,P3H3,SEC23A,KDM3B,NAP1L4,CTXN1,CCT5,PCYO
3,CHERP,CTTN,SLC25A5,LRP10,DAP,SERP1,STK35,PPP1R10,ERH,CYTH2,EIF3H,ANTXR1,INTS3,UI
3,CHERP,CTTN,SLC25A5,LRP10,DAP,SERP1,STK35,PPP1R10,ERH,CYTH2,EIF3H,ANTXR1,INTS3,UI
T6,SV2A,ZNF512B,CEMIP2,SELENOW,ADH5,PTPRF,NDST1,ULK1,OAZ2,ZMYM4,FBXO21,PCBP4,V
RA2B,FBXO11,IARS1,COX7C,YKT6,RGMB,FEM1B,NDST1,IL6ST,MCM3AP,RPL7L1,TMEM167A,TN
GRSF1,SEC16A,SLC39A7,MAP6,SLC35F5,HJURP,EP400,CD164,GLUD1,TRA2B,RGMB,PRRC2A,O
IT4,MTSS2,DNAJC5,RPLP2,ARL6IP1,DDOST,TAGLN,ANKRD52,HNRNPK,CIRBP,MYH10,NARS1,DU
3A,TCTN1,RTL8C,WBP2,AKIRIN1,TP53BP1,AMOTL2,YME1L1,CTNND1,PSMA7,TRAF4,STAT3,ARIH
,CSNK1E,PFKL,HDLBP,GLG1,CDH6,EIF5A,PEBP1,CCND2,PDIA6,GPC3,DDR1,PLXNA1,SOX12,ACT
K1,RHOA,PTCD3,CBX5,RPL9,ATF4,NR2F1,KIF5B,ENC1,RPS14,CHD4,TMEM259,RPS20,EIF5,CD81
MBTPS1,RGMB,SELENOW,FEM1B,CUL3,ADH5,NDST1,ULK1,ITGB5,TIMM50,TNKS,PSMD8,APEX1
ELSR1,FAM210B,HMGA1,SDCBP,CHMP1A,PINK1,DAG1,TOX4,NBPF15,NIBAN2,KIFC2,IGSF3,KIA
D1,PRPF19,CSE1L,NOC2L,COX6A1,EIF3G,SF3A2,CHMP4B,SART1,DHX38,PACS2,TGOLN2,AP1M
C,RPL30,EIF3L,NANOG,MRFAP1,RPS7,RPL35,MIF,GJA1,CTNNB1,ANXA6,NGRN,RPL35A,CYFIP2,H
11,GPC4,MAP1LC3B,MTCH1,TMOD2,KIAA0100,CEP170,PELP1,CLPTM1,SLC44A2,CARM1,GPC2
B1,BANF1,GALNT10,PTPN1,COX8A,GSR,TASOR,TNRC6A,PPME1,RPL27A,FAM8A1,DARS1,TRIB3,
67A,KLHDC3,TNKS,PSMD8,CHERP,CTTN,GPC4,LRRC59,MTCH1,BRK1,SERP1,AKAP9,RHBDD2,V
3,HSPB1,SON,RPL28,CCAR2,ARF4,CORO1C,SUPT6H,DHX30,NKTR,STMN3,PFKM,TNKS2,STT3A,L

H,PRKCSH,HNRNPC,LRPPRC,XPO1,PTN,CSNK1E,HDLBP,RPS25,TKT,TPM1,PNMA1,PDIA6,DDR1,PI
ERBP1,RPS13,ARFGEF2,AKIRIN1,BMPR2,CCT5,CDC42,YME1L1,VDAC1,TRAF7,EIF2AK1,EDC4,PR
P24,FBXW5,AP2A2,CREBBP,DPYSL4,METRN,GAK,ZER1,FBXL19,ARF5,RIMS3,PAFAH1B2,DIP2A,F
SEC16A,PHB2,APMAP,GSPT1,UBA52,MGAT4B,PLXNB1,RAB2A,FARP1,CD164,RER1,RAB3GAP2,UI
A2,NECTIN2,PPP1R26,MAF1,SLC2A4RG,HEG1,TMEM151B,MED16,TRAF3,MIB2,SHB,HID1,SRCIN
AF2,CCT2,RPN2,PGD,GNAI3,KLHL24,MEST,GAS7,TM9SF2,MORF4L2,LMNB2,ABCA3,ALKBH5,SNI
3,TOMM70,XPO6,UBXN6,MICAL1,MYL12B,GAK,LYPLA2,CENPB,ATP13A1,ATP5MG,PAFAH1B2,PC
PBX2,AEBP1,NYNRIN,DYNLL2,WDR6,GABARAP,UBE2R2,SCAF11,ATXN10,BNIP3L,MAP7D1,CLIP3
4,UBAP2L,KIAA0895L,UPF1,UBFD1,MASTL,MBTPS1,ANP32B,MAPRE2,POM121C,SEPTIN7,APEX1,
2,ZYX,TUBA1C,RCOR2,SETX,PRMT5,ZFP91,ALDH9A1,SNRPB,SORCS2,ATP5F1C,POGK,METTL9,N
TN1,CAND1,KDM1A,BNIP3L,NSG2,CCAR2,DIDO1,TSC22D1,POLR2B,SMARCC1,GIGYF1,PHC2,CC
MIDN,RPL10,MARCHF6,WSB1,RPS17,VCL,AP3D1,SBF1,DNAJA1,ARF1,CYB5R3,AKAP12,MAP4,SF3
,SSBP4,SLC20A1,THRAP3,DPYSL4,CPXM1,HSDL1,TEAD1,ATP13A1,DIP2A,HS2ST1,UBE2L3,CELSI
5,SEC16A,PHB2,APMAP,GSPT1,UBA52,RAB2A,CD164,RAB3GAP2,UBFD1,TRA2B,IARS1,MASTL,Y
38A2,FJX1,TEAD1,ZER1,PAGR1,SRPRA,CNBP,LHFPL6,ELAVL3,PAFAH1B2,ACAT2,UBE2L3,HMGA
2,XPC,NR2F2,BRI3,RBM17,NEO1,WBP1L,MARVELD1,RFX7,RAB5A,PPP3CA,MARK1,CNOT3,COL6
P4,PITPNA,XPNPEP3,SUPT6H,RPS28,SRSF3,LARS1,YWHAH,FAU,TNKS2,STT3A,UBQLN1,CSNK1D,L
RPL39,MLF2,RPS13,WBP2,NAP1L4,CTXN1,CCT5,GRIPAP1,SSR1,ST3GAL6,MSMO1,SULF2,ARIH1

TXBP1,ISYNA1,STT3A,LONP1,LOC728392,ATXN2L,PRPS1,MPRIP,HGS,SREBF2,RTL8C,ARFGEF2,[illegible]
F3,ATMIN,METTL9,SLC5A6,NAPA,FAM20B,RUSC1,ACP1,PRMT2,SHC1,NEURL4,MAP2K4,RAB5A,[illegible]
PRE2,POM121C,CALD1,SEPTIN7,APEX1,PPP1R15A,GALNT11,MVD,DAP,BRK1,SERP1,PPP1R10,S[illegible]
C5,CLPTM1L,GBA2,POLDIP3,UNC45A,RNMT,ZBTB4,PSMF1,ABHD17A,PDCD11,RPRM,WARS1,CL[illegible]

N4,MVB12B,SMARCA4,ATG2B,AZIN1,MAPRE1,KDELR2,LAPTM4A,SRSF9,ARPC2,RABL6,ADAM9,F[illegible]

2,SARAF,MIDN,WSB1,VCL,AP3D1,SBF1,DNAJA1,ARF1,CYB5R3,PAFAH1B1,AKAP12,MAP4,ITM2B,[illegible]

SLC35E2B,IPO5,ATP6V0C,BCLAF1,LAPTM4B,TRIB2,RPL23A,HSBP1,FAM199X,HS6ST1,CTDSP2,P[illegible]
I3,TPM4,TNS3,CTSD,NACA,TSPAN3,LPCAT1,LAMA5,XRCC5,PJA2,MSN,RPL24,KPNB1,C1ORF43,I[illegible]
D164,SRSF6,UQCRC1,HDAC6,SMC3,SELENOW,PRRC2A,HNRNPA2B1,PPP1CC,EML1,ANP32B,DK[illegible]
CASC4,PSAT1,AUTS2,HDGFL3,RTF1,DAG1,NIBAN2,ANKHD1,RUVBL2,TMX1,MDM2,SAMD11,CLU[illegible]
EX3,RBBP4,RTN3,PFN1,MCL1,JPT2,PNN,NES,RPL31,NISCH,PRKCSH,LRPPRC,KHDRBS1,XPO1,SF[illegible]
MEM259,RPS20,CD81,WDR1,HMGN2,ATP2A2,RPL26,LASP1,CCT7,OAZ1,RBBP4,PFN1,ASTN2,MCL[illegible]
,CDK4,TBXA2R,SLC25A1,DAAM1,EMC10,DHFR,MRPS6,NUS1,ABHD17A,CCDC85C,RAB3A,PGRM[illegible]
0,SPG7,BAZ2A,XRCC6,VAPB,EIF3H,ELAVL1,IMPDH2,GABARAPL1,FAM234A,SSR3,GNL1,TM9SF4,
MMP15,EIF3D,PHB2,APMAP,CST3,PLXNB1,DECR1,CSNK1G2,RAB2A,FARP1,RER1,UBE2G2,RAB3[illegible]
,TRAF3,CNOT3,MIB2,FYTTD1,RAB35,VAV2,HEXIM1,FAM20C,GIT1,MSI2,C8ORF33,SETD1A,ZC3H[illegible]
CT2,RPN2,GNAI3,NOP53,TSPYL4,TM9SF2,HSPH1,LMNB2,MTRF1L,ALKBH5,RPL27,SNRPN,PRDX5
N7,PSMD8,NPLOC4,HIF1A,SAFB2,PPP1R10,WSB2,ERH,EIF3H,ABI2,SEC14L1,C6ORF62,RAB14,S[illegible]
MOTL2,DFFB,STAT3,ARIH1,PRMT1,MTA1,KDM5C,KDM5B,FBRS,PDAP1,PRCC,AP2A1,BAZ1B,FLC
,TNRC6A,VPS51,DARS1,FAM32A,SYNGAP1,EPN1,CDK9,RTF2,ZRANB2,EXOC7,PSMC5,CLPTM1L,[illegible]
M234A,PGAP6,LDLR,KMT2E,TRAP1,BLCAP,FNBP1L,SPAG9,DICER1,CLASP2,RBM12,DAZAP2,WBF
BTPS1,ZFR,SELENOW,CUL3,FMNL2,ADH5,NDST1,NPDC1,ITGB5,TIMM50,TNKS,APEX1,DKK3,PPF
RPL18,ITGB1,TUBB4B,BTF3,NCAM1,ANXA2,AGRN,ZMIZ1,TIMP2,XAF1,SARAF,COL11A1,MIDN,R[illegible]
NOP53,MORF4L2,HSPH1,MTRF1L,ALKBH5,HNRNPA3,PDXK,UBE4A,YEATS2,TBC1D16,ACO2,HSP[illegible]
F3,RNF187,U2AF2,RPN2,MLXIP,ANXA5,SNX22,NOP53,PARP1,TM9SF2,MORF4L2,HSPH1,MTRF1L
,EIF4B,VCP,MARCKSL1,UBB,AFF4,TRIM28,MATR3,YWHAG,CKB,COL4A2,RPL37A,LRRC28,MAPT,
LM,YARS1,SSBP3,ANTXR1,RAB1B,UBR5,SLC6A8,GABARAPL1,MAZ,IREB2,KMT2E,KIAA0232,LRA
,PPP2CA,SMARCA4,AZIN1,DDAH2,FAM171A2,ARGLU1,AEBP1,ACADVL,NYNRIN,TCF25,GRB2,WD[illegible]
S,PSMD8,CHERP,CTTN,LRP10,MTCH1,GLO1,SERP1,RHBDD2,VPS26B,STK35,ERH,CLPTM1,RTL[illegible]
1A,GPC4,MAP1LC3B,MTCH1,TMOD2,KIAA0100,CEP170,PELP1,CLPTM1,SLC44A2,OLFM1,CARM
H1,SERP1,WSCD1,RHBDD2,VPS26B,STK35,ERH,CLPTM1,RTL6,EIF3H,INTS3,ABI2,UBE2M,SEC14
NLGN2,RPS14,CXADR,ATP1A1,TMEM259,EIF5,CD81,COL5A2,HMGN2,ATP2A2,RPL26,CCT7,PFN1

19,MIF,HMGCS1,GPAT4,NGRN,DNAJC5,HNRNPDL,ANKRD52,HNRNPK,INIP,CIRBP,DUX4,PTGES3,
SLC16A7,C1ORF43,COL1A2,GAP43,RPL14,HK1,RHOA,RPL9,ATF4,PSMD2,KIF5B,SLC7A5,ENC1,[illegible]
STN2,TLE5,NES,RPL31,NISCH,PRKCSH,CCNI,LRPPRC,KHDRBS1,PFKL,HDLBP,SFPQ,TPM1,EIF5A
MEM167A,KLHDC3,NPLOC4,CHERP,CTTN,LRP10,BRK1,SERP1,AKAP9,STK35,ERH,EIF2S2,EIF3H,
PK,INIP,CIRBP,RPL36,DUX4,SEPTIN11,HNRNPH1,RPS12,ARCN1,HECTD4,RAC1,CAPRIN1,STIP1,[illegible]
M1B,ECEL1,TBC1D9B,HNRNPA2B1,CUL3,NDST1,POM121C,LMAN1,SEPTIN7,TNKS,VPS26B,ERH,
,FGD1,CPNE2,SH3KBP1,ASAP3,EPHA2,MYL6B,ARSA,DTD1,EMD
,ARL6IP1,NDFIP1,ST3GAL2,TSR2,MYL6,SHOX,ANKRD52,WLS,INIP,MYH10,SND1,RHOB,HNRNPH[illegible]
AP,GSPT1,UBA52,RAB2A,CD164,RAB3GAP2,UBFD1,IARS1,MASTL,YKT6,MCM3AP,SPINT2,PREPL,

P2,ATG2B,KDELR2,DSTN,PDE7A,PSMD3,SRSF1,TCF25,ATP5PB,LUZP6,GRB2,CDC37,WDR6,RPS1
ANXA5,SNX22,NOP53,MORF4L2,HSPH1,MTRF1L,ALKBH5,HNRNPA3,PDXK,UBE4A,YEATS2,TBC1D
D3,XPO6,CCDC47,UBXN6,THRAP3,MICAL1,FJX1,MYL12B,LYPLA2,FBXL19,ATP13A1,SRPRA,USP5,
2B,MEGF8,SOX12,MMP14,RPS15,ITM2C,RPL30,CLTC,EIF3L,MRFAP1,MAGED2,GNB2,H1-0,RPS

D2,RNF145,ADAM10,PCGF2,PTGFRN,YY1,CERS2,GNA13,SCD,TRO,CROCC,NELL2,UNC119B,SH3
S3,WDR5,SS18L1,GPS1,SLC2A4RG,MARK1,HEG1,TRAPPC11,TRAF3,CNOT3,ZNF629,PCNA,FYTT
,ASTN2,MCL1,PNN,RPL31,NISCH,PRKCSH,HNRNPC,CCNI,LRPPRC,XPR1,KHDRBS1,KLHL9,SPTBN
IG8,HMGN2,LASP1,OAZ1,SRSF4,PFN1,ASTN2,TLE5,NISCH,PRKCSH,CCNI,TADA2B,SPTBN2,SLC3
D5A,CTNNB1,COL6A2,CD63,ANXA6,NGRN,GOLM1,SESN3,B4GALNT4,MTSS2,APLP2,HNRNPDL,S
HMP1A,AUTS2,MTA2,ESYT1,LUC7L2,NIBAN2,ANP32E,CLIC1,ELN,IGF1R,SGSM3,SLC38A1,SLC25
P,NARS1,RPL36,YWHAB,SND1,MAP2,CALU,ARCN1,COTL1,HRNR,ARHGDIA,CAPRIN1,SLC35E2B,I
GPC3,DDR1,PIP4K2B,MEGF8,SOX12,RAB7A,CCN2,ACTN4,MMP14,PRDX2,RPS15,RPL12,ITM2C,
APEX1,PEG10,DKK3,HIF1A,SYMPK,VPS26B,STK35,FUS,XRCC6,PELP1,SLC44A2,UBE2M,SMG5,T
B1,GBA2,UNC45A,RNMT,ZBTB4,PSMF1,HMGXB3,MCFD2,EIF3M,WARS1,CLCN6,RAVER1,NDUFS1
CIPC,PJA2,MSN,RPL24,SLC16A7,KPNB1,C1ORF43,IGFBP3,GAP43,RPL14,NCS1,RHOA,CBX5,ZNF
,RPLP2,ARID1A,NDFIP1,TSR2,MYL6,SHOX,INIP,MYH10,PTGES3,SND1,HNRNPH1,UBE2H,ARCN1,
,EIF5A,PEBP1,CCND2,DDR1,PLXNA1,RAB7A,DDX3X,CLTC,NANOG,MAGED2,GNB2,RPS5,RPS7,IK
E1,KDELR2,DSTN,PSMD3,PRDX1,RABL6,FAM168B,SRSF1,IPO7,AEBP1,DYNLL2,CDC37,WDR6,AC
D3,IFITM3,COL27A1,VANGL2,HNRNPD,P3H3,PPP6R1,RPS13,DYNLL1,YLPM1,DDIT4,SYP,CTNND1,
1,TM9SF4,ZNF428,TRAP1,PLEKHB2,ELP1,SPAG9,MOAP1,CLASP2,RBM12,WBP11,NSD2,AKT2,GC
MYL6,SHOX,ANKRD52,WLS,CIRBP,MYH10,NARS1,RPL36,SEPTIN11,CADM4,LGALS3BP,CKAP4,P
P1,PIK3C2B,ALDH18A1,PABPC4,CNDP2,MTURN,LIMK1,GOLPH3,RHEB,DHX36,COLGALT1,TSPAN

PS20,EIF5,CD81,COL5A2,WDR1,HMGN2,ATP2A2,CRMP1,LASP1,BEX3,CCT7,RBBP4,TLE5,NISCH,
PLP2,ST3GAL2,MXD4,MYL6,H1-10,CIRBP,MYH10,NARS1,RPL36,YWHAB,PTGES3,SEPTIN11,CADM
2,TPX2,SZT2,ACIN1,COG4,SCOC,AKAP17A,MAPK8IP3,PAF1,PRELID1,MRPL3,DNMT1,CFAP298,N
APK2,JAK1,PPP2R5D,LOXL2,ZSWIM6,IGF2BP2,GDF1,OAT,LGR4,SNRPD2,RCOR2,MAP1S,RASSF
CT6A,RAC1,ARHGDIA,CAPRIN1,GPC1,CDK16,TOP2B,SLC35E2B,APPL1,SMCHD1,IPO5,TRIB2,CDK
2,DSTN,PLEC,ODC1,NYNRIN,TCF25,LUZP6,PCSK1N,WDR6,PRKDC,TDRD5,SON,CASC3,KDELR1,
300E,MRPS6,CRTC1,CACNA1H,SETX,MED25,TXN,C20ORF194,SLC5A6,GET1,NELFCD,RUSC1,PR
A,CPSF2,CNN2,CTSB,CDH11,RAB27A,EEF1A2,SUGP2,PPP1CB,SNX22,VAMP2,GNAI3,NOP53,VCAN
DN,OGA,PGAM1,NFE2L1,SMARCA5,HNRNPA0,CTNNA1,CNN3,RPL29,TPM4,CTSD,EIF4H,SCARB2,
D5,HNRNPUL2-BSCL2,PITPNA,XPNPEP3,SUPT6H,RPS28,ATP6V1B2,YWHAH,PTP4A2,FBLN1,TNKS
CTXN1,PPDPF,TRAF4,RBBP7,NAP1L1,TXNDC5,CPD,EP300,DGCR2,STARD7,NDUFV1,FBRS,RETRE
NC2,FAF2,METAP2,MAEA,PCGF3,GOLPH3,HIPK2,TNRC6A,VPS51,NUP153,DARS1,H2AX,RAB18,RT
RKAR1A,TLN1,CHPF,GJA1,JUN,NGRN,B4GALNT4,CYFIP2,DNAJC5,HNRNPDL,TARDBP,SLC25A3,
3GAL2,TSR2,MYL6,SHOX,ANKRD52,INIP,MYH10,SND1,RHOB,HNRNPH1,UBE2H,CKAP4,RAB1A,C
NA1,STT3A,LONP1,UBQLN1,PLXNA3,SARS1,COL27A1,SOD1,SREBF2,NCOR2,HNRNPD,VGF,MEX3
AR1A,TLN1,CHPF,GJA1,JUN,NGRN,B4GALNT4,CYFIP2,DNAJC5,HNRNPDL,TARDBP,RPLP2,ARL6I
TRAF3,CUL7,TET3,FYTTD1,CHPF2,VAV2,MXRA5,HEXIM1,ZNF516,ADGRL1,FURIN,SAMD1,SRCIN1
,DUX4,YWHAB,SEPTIN11,UBE2H,RPS12,ARCN1,MAGED4B,ARHGDIA,TOP2B,SLC35E2B,TMEM30
SS,B2M,UBQLN1,PLXNA3,LOC728392,COL27A1,VANGL2,BAG6,RAI1,SREBF2,VGF,MEX3A,RAB1
R1,TAF15,HSPH1,LMNB2,MTRF1L,ALKBH5,PDXK,TBC1D16,ACO2,HSPA4,PLOD1,AHCYL1,CAMSA

F1,LAPTM4B,TRIB2,RPL23A,NNAT,LUC7L3,CDKN1A,FAM199X,DLG5,CRPPA,PGRMC1,DDX39B,C
,PXDN,HNRNPU,OGA,PGAM1,NFE2L1,SMARCA5,HNRNPA0,RPS16,RPL23,CPE,RPSA,EIF4H,SCARE
CF1,ERGIC3,CAP1,IST1,ATP6V0D1,DHCR7,MBD3,GSK3A,VASH1,SART1,SMARCD1,APBB1,ANKRD
INIP,MYH10,SND1,RHOB,UBE2H,CKAP4,ARCN1,CTBP1,HMGCR,TOP2B,SMCHD1,IPO5,ATP6V0C,
IGB1,POLR2A,CDKL1,AP2M1,TPT1,SEC61A1,RPL18,STMN1,ITGB1,UCHL1,RPL11,BTF3,NCAM1,

IS3,ASB6,CTDNEP1,MIB2,UQCRH,TMEM59,TRAPPC1,SPG11,CUL4B,PSMD12,RPL32,TPX2,SZT2,
31,COL5A2,LENG8,WDR1,HMGN2,RPL26,LASP1,CCT7,OAZ1,SRSF4,RBBP4,PFN1,ASTN2,MCL1,T

REX,IGSF3,PFKP,TRIM2,WDR26,CYFIP1,SYNCRIP,ALDH18A1,PHPT1,GALNT10,GLOD4,SSU72,HT
GRINA,CLSTN3,MDK,JPT1,PTK7,PSMB2,RAB11B,TSPAN6,ATP6AP1,TMED2,TAF7,RPS15A,G3BP2
PPR3,SEPTIN3,HECTD4,SMCHD1,HDAC2,IPO5,ATP6V0C,MMP2,BCLAF1,LAPTM4B,NNAT,CDKN1A
I,OGA,SMARCA5,RPS16,CPE,TNS3,CTSD,SCARB2,TSPAN3,LAMA5,SLC22A17,RPS9,CIPC,MSN,RP
12,RAB7A,DDX3X,RPS15,ARL4C,EIF3L,NANOG,GNB2,APLP1,IK,RPL35,GPAT4,TLN1,CHPF,GJA1,
P32E,PDHA1,FZR1,MAPK8IP2,SLC25A6,STK39,SYNCRIP,SSU72,LIMK1,ECH1,GOLPH3,TASOR,PF
,CD2BP2,RPS21,APEH,BANF1,PSMD7,TNRC6A,PPP1CA,VPS51,PMEPA1,C1QBP,FAM32A,OTUD5,S
CCNL2,ARPC2,DSTN,FAM171A2,SRSF1,DYNC1LI2,AEBP1,ACADVL,DYNLL2,CDC37,WDR6,ACTN1,
C1ORF43,PAPOLA,IGFBP3,COL1A2,RPL14,HK1,NCS1,RHOA,PTCD3,ZNF865,CLSTN1,ATF4,PSMD
3,PPP1R15A,FKBP1A,GPC4,MAP1LC3B,MVD,MTCH1,PELP1,CLPTM1,SLC44A2,HSPG2,CARM1,AE
C22D1,SUPT6H,DHX30,RPS28,ATP6V1B2,HNRNPM,LARS1,STMN3,MTRNR2L10,CSNK1D,ID4,SAR
D3,RPS9,CIPC,SLC16A7,KPNB1,C1ORF43,IGFBP3,COL1A2,LLGL1,RPL14,RPL22,RHOA,ZNF865,
N,UBXN4,PPP2CA,CCND1,G3BP2,KDELR2,DSTN,PDE7A,PSMD3,ODC1,SRSF1,RBP1,TCF25,ADAM
PSMB1,TSNAX,SLC25A6,EXOC5,NDUFA4,SF3A3,TRMT112,PSMD13,WDR26,CD2BP2,GPAA1,GGT
A1,PIP4K2B,ACSL6,RAB7A,CCN2,RPL12,ITM2C,ARL4C,CLTC,EIF3L,GNB2,RCN1,H1-0,RPS19,RP
IR4,ADAM10,SPTY2D1,CAP1,LRRC41,IST1,EHD1,PCGF2,COPZ1,NOC2L,SMARCD1,SSR2,NDUFA1

NCBP2,NSD1,RNMT,ACBD3,TMEM43,SOX9,EIF3M,TXN,RAVER1,HOXA5,STARD3,LARGE1,SRP54,
KIF3C,SUPT6H,DHX30,HNRNPM,PTP4A2,PFKM,LSS,TMED9,SREBF2,RPL39,SERBP1,RPS13,IGFB

GB5,TMEM167A,PSMD8,CTTN,SYMPK,LRP10,SLIT2,A2M,DAP,SERP1,WSCD1,NDN,VPS26B,PSME

F5,HJURP,GNG4,NXN,EP400,UBE2G2,UBFD1,MBTPS1,SELENOW,FEM1B,CUL3,ADH5,NDST1,UL
ACTR1B,WASF2,NSD2,DHCR7,CSE1L,IVD,COPZ1,KDM4A,MBD3,PACS2,PRKACA,ATXN7L3,SLC2
8A1,ATP1B3,CNDP2,GLOD4,MTURN,GOLPH3,ECHS1,ALDH7A1,DHX36,HM13,COLGALT1,TSPAN
P1,IK,RPS19,MIF,HMGCS1,GJA1,JUN,COL6A2,ANXA6,B4GALNT4,MTSS2,SLC25A3,TSR2,MXD4,
2BP2,GPAA1,GGT7,BSCL2,SUB1,BANF1,FAF2,COX8A,CFDP1,CUL4A,TNRC6A,PPME1,MOGS,MA
MT1,SGTA,TXNDC5,MTA1,EXTL3,LBH,RETREG2,ALDH3A2,FLCN,H2AZ1,BRD2,KLHL42,DIP2B,CD
YNCRIP,ACAP2,PTPN1,LMF2,DRAP1,SRSF10,ADCY1,FAM32A,LSM4,MAPKAPK2,UBE2Q1,DEAF1,M

C,ARHGAP23,KLF3,DNMT1,SMARCB1,SCAF8,FZD8,SPSB3,SH3KBP1,ATP8A2,MEX3D
VAPA,HMGB1,POLR2A,AP2M1,SEC61A1,RPS4X,STMN1,ITGB1,TUBB4B,UCHL1,RPL11,MORF4L1,

,PPP1R15A,GALNT11,MVD,DAP,BRK1,BAP1,PPP1R10,SPG7,BAZ2A,XRCC6,VAPB,EIF3H,ELAVL1,
9B,CNN2,EID1,SRP9,C5ORF24,TM9SF3,MDH2,RNF187,CCT2,MLXIP,CPSF1,GNAI3,MEST,GDI2,H

,GSK3B,MEX3C,OAT,TUBA1C,RAB3A,ARHGEF9,WARS1,CHKA,DUSP8,ZDHHC5,ATP6V0E2,UNC13
J,SMARCA5,LARP1,HNRNPA0,CTNNA1,FADS2,RPS16,LDHA,CPE,CNN3,RPSA,CTSD,EIF4H,SCARB2
CED1A,ASB6,SUDS3,PNMA2,PCDHB2,MIB2,VAV2,TNPO3,MCM7,VPS16,NUP50,SZT2,SEC31A,AKAI
H,PRKCSH,LRPPRC,TADA2B,KHDRBS1,SPTBN2,HDLBP,TKT,SFPQ,EIF5A,PEBP1,PNMA1,GPC3,AC
PO6,AMBRA1,ATG4B,PAFAH1B2,SLC25A36,UBE2L3,SPPL2B,ZZEF1,MGAT5,YTHDF3,RBM4,MTRI
I1,EXOC5,TRMT112,PSMD13,CD2BP2,SUMO1,RPS21,CAB39,APEH,BANF1,GALNT10,FAF2,COX8
,DMWD,HSDL1,LYPLA2,SH3PXD2B,USP5,IER2,RANGAP1,LHFPL6,SLC30A9,AMFR,IGFBP4,TECR,
RO1C,SUPT6H,RPS28,HNRNPM,GIGYF1,SRSF3,NKTR,STMN3,PFKM,STT3A,LONP1,TUG1,MESD,

,SARS1,PTPRS,HGS,BAG6,SREBF2,RAB11A,CYP51A1,FLNC,ARFGEF2,IGFBP7,TP53BP1,CTXN1,C

P2,ARL6IP1,NDFIP1,ST3GAL2,TSR2,MYL6,SHOX,INIP,MYH10,SND1,RHOB,UBE2H,CKAP4,ARCN1
F3A3,TRMT112,PSMD13,WDR26,CD2BP2,GPAA1,GGT7,BSCL2,SUB1,BANF1,COX8A,CFDP1,CUL
,ACSL6,MMP14,RPS15,RPL12,ARL4C,CLTC,MAGED2,GNB2,RPS5,APLP1,RPS19,RPL35,MIF,GPA
SPH1,HNRNPA3,C4ORF3,PKN1,ETNK1,CCT3,ZNF302,RBMX,JPT1,CBX1,CHD2,CCT4,ORAI2,GCN1

DAP1,KRT8,CLASP2,RBM12,NSD2,AKT2,YY1,SAP18,CCDC47,PTPN23,ELOVL5,DHX40,SRPRA,IER2
B,SRP14,ABCA2,NRDC,PRPF6,TMED2,RNF10,OPTN,ATG2B,LAPTM4A,CCNL2,DSTN,PLEC,PDE7A,
DM4,SND1,LAMP1,RPS12,HLA-B,MAP2,RAB1A,TTR,CALU,PLPPR3,ZNF454,CCT6A,MAGED4B,CTBI

GA1,OGFR,AGAP1,LGALS1,AUTS2,PPP5C,NBPF15,GTF3C1,NRXN2,KIFC2,ZKSCAN1,KLF13,RBM
R3,PAFAH1B1,SF3B2,ITM2B,ZNF770,AP2B1,MAT2A,TMEM170A,PXDN,HNRNPU,PGAM1,SMARCA
IP1,COPE,FAM50A,CD2BP2,PIK3C2B,ALDH18A1,PABPC4,CNDP2,GLOD4,MTURN,GOLPH3,ECHS
D1,YTHDF3,KLF13,CDV3,TNFAIP1,COPE,FAM50A,CD2BP2,SART3,PABPC4,CNDP2,GLOD4,MTUR
AVER1,MBOAT7,SNRPB,STARD3,FARSA,SRP54,THOC2,UBXN1,ZNRF1,FAM20B,BRI3,DDHD2,RUS
CCNI,SPTBN2,SLC3A2,CSNK1E,PFKL,TPM1,EIF5A,CDH2,GPC3,DDR1,PIP4K2B,MEGF8,RAB7A,IT
B2,TAPBP,ABL1,LGALS1,CHMP1A,MTA2,POR,NIBAN2,KIFC2,ZKSCAN1,CLIC1,RUVBL2,ACVR1B,

8,NACAD,OXSR1,EMC10,SEMA4C,TMEM9,SCG2,WDR82,SYNGR1,MEX3C,USP48,ACSL4,RPS29,N

EIF3C,ATP5F1A,EIF4B,VCP,MARCKSL1,UBB,AFF4,TRIM28,MATR3,YWHAG,CKB,COL4A2,RPL37A

L,WARS1,VEZF1,SNRPB,KDM5A,NKIRAS2,ZDHHC5,XPC,SKIV2L,CFLAR,SHC1,MEIS3,COX7A2,CTD
I14,NUP133,CHGB,COX4I1,OGDH,GSTA4,GUK1,UBN1,NECTIN2,PLCD3,CTDNEP1,ZNF598,QRICH
3,UBXN4,G3BP2,SMARCA4,AZIN1,KDELR2,SRSF9,ARPC2,DDAH2,DSTN,PLEC,PBX2,PDE7A,FAM17
SKIV2L,RNF4,LZTS2,SHC1,PCED1A,SUDS3,ZSWIM8,CTDNEP1,PCDHB2,KCTD10,BTF3L4,MIB2,V
NA1H,SOX9,MAP1S,STC2,FBL,RALBP1,SLC12A7,GET1,SLC5A7,NUMBL,PHB,GPR173,MARVELD1
SP1,CCT7,RBBP4,RTN3,PFN1,ASTN2,PNN,CLEC7A,SPC25,NES,CCNI,LRPPRC,XPR1,KLHL9,TKT,C
F302,GCN1,HDGF,SRP14,NRDC,PRPF6,TERF2IP,TAF7,RPL38,VEGFA,RPL37,UBXN4,G3BP2,ATG2

C2B,ALDH18A1,PABPC4,CNDP2,GALNT10,GLOD4,MTURN,GOLPH3,ECHS1,ALDH7A1,DHX36,HM1
K,RPS3A,RPL18,ITGB1,TUBB4B,UCHL1,MORF4L1,BTF3,NCAM1,ANXA2,AGRN,RPLP1,TIMP2,XAF1
G1,CALCOCO1,RIMS3,ELAVL3,DENND5A,PLS3,SLC25A36,PTPN11,WDTC1,RSF1,SGSM2,NBPF15

POLR2E,TRAF7,EIF2AK1,NAP1L1,TXNDC5,KIRREL1,EP300,RERE,PBX1,APMAP,ZFP36L1,KLHL42,
JMO1,RPS21,BSCL2,SUB1,BANF1,PSMD7,KCTD20,GSR,VPS51,FAM8A1,SLC25A1,FAM32A,JAK1,
31,SERINC1,SSBP2,ZNF43,CHD3,BASP1,VAPA,HMGB1,POLR2A,CDKL1,AP2M1,TPT1,SEC61A1,RF
,SLC35E2B,SMCHD1,SKP1,DNER,BCLAF1,PTTG1IP,FAM199X,PGRMC1,SRSF5,CDH11,U2AF2,CC
8,PSMC1,DEK,GAK,LYPLA2,SLC38A10,ATP13A1,RANGAP1,RIMS3,PAFAH1B2,POLDIP2,TRIP12,X
NX2,CASC4,PLS3,PUM1,SGSM2,MTA2,CLIP2,FAM118A,YTHDF3,PLXNA2,HSPA1A,CDV3,UBE4B,M
TASOR,TNRC6A,PPP4R3A,SOBP,DDX6,AKAP8L,KPNA6,GID8,DEAF1,EXOC7,ZNF532,PSMC5,CSNK
CADM4,SND1,RPS12,ARCN1,RPS27,GPC1,TOP2B,IPO5,ATP6V0C,TMEM30A,MMP2,BCLAF1,L1CA
D,LBH,RETREG2,CNTNAP1,ALDH3A2,FLCN,LRRC4B,GTPBP1,MMP15,GRSF1,GPX4,BRD2,CSNK1
HNRNPC,LRPPRC,XPR1,KLHL9,SLC3A2,CSNK1E,SCRN1,RPS25,SFPQ,EIF5A,PEBP1,PDIA6,PIP4K2

X3,DPYSL4,SCD,HSDL1,METRN,TUBB4A,ZER1,SLC38A10,PAGR1,UNC119B,FBXL19,SRPRA,ARF
DP56,NIBAN2,H6PD,RUVBL2,SC5D,GRIK5,OBSL1,ATP13A2,FZR1,SLC25A6,DNAJB1,PPP1R14B,S
X3A,SEC23A,AKIRIN1,KDM4B,CTXN1,AMOTL2,DDIT4,BEX4,TRAF4,KCNQ2,EDC4,WDR45B,XPOT,
RBM12,NSD2,AKT2,YY1,SAP18,CCDC47,THRAP3,ATG4B,PTPN23,ELOVL5,DHX40,SRPRA,IER2,NT

USF2,PINK1,TMEM94,DAG1,NIBAN2,SGSM3,FOXP4,RPS21,CAB39,ANO8,KAT2A,RNF157,SSU72,
P4R3A,VPS51,WIZ,SOBP,DDX6,AKAP8L,KPNA6,GID8,DEAF1,EXOC7,PSMC5,EXOSC10,NSD1,UBE3
PSA,TPM4,TNS3,CTSD,NACA,RPS24,SLC22A17,BRD3,RPS9,SLC16A7,RPL17,C1ORF43,PAPOLA,I
CLPTM1,CYTH2,INTS3,RAB1B,RALY,SEC14L1,NUDCD3,FAM234A,SSR3,GNL1,SH3GL1,STK24,CA

YL6,SHOX,ANKRD52,INIP,MYH10,SND1,RHOB,HNRNPH1,UBE2H,CALU,ARCN1,RPS27,CTBP1,HM
2,NSD2,CSE1L,COPZ1,KDM4A,MBD3,PACS2,PRKACA,ATXN7L3,MN1,SLC20A1,SORT1,DEK,ELOV

4L2,NOX4,HSPH1,MTRF1L,ALKBH5,SNRPN,PRDX5,C4ORF3,UBE4A,TBC1D16,ACO2,HSPA4,TCP1
9X,DLG5,SHISAL1,CRPPA,TIMP3,CNN2,EID1,SRP9,ZC3H7B,MDH2,CHD6,RNF187,U2AF2,EEF1A2
1,TSPAN7,NUP98,NACAD,OXSR1,EMC10,SEMA4C,PSMC4,TMEM9,SCG2,WDR82,SYNGR1,MEX3C
3,BASP1,VAPA,HMGB1,POLR2A,AP2M1,TPT1,RPL18,STMN1,TUBB4B,UCHL1,BTF3,NCAM1,ANXA
CE1,MED12,SLC5A7,LONP2,DDHD2,NRP2,DSTYK,ELOB,COX7A2,BTF3L4,TRAF3,MIB2,PIGS,FER,

1,SLC35F5,CD164,UBAP2L,RER1,PI4KA,UBFD1,IARS1,MASTL,COX7C,YKT6,PPP1CC,MCM3AP,SF

NUDCD3,TAX1BP1,HINT1,CNP,SRSF2,CAPZA1,TUBGCP6,RIC8A,NCSTN,ATXN3,INTS11,AHSA1,RA

FAND3,HGS,MAPK8IP1,DGKD,PFN2,VGF,RPL39,P3H3,SEC23A,KDM3B,WBP2,IGFBP7,KDM4B,DY
KCSH,LRPPRC,XPO1,SPTBN2,SLC3A2,CSNK1E,CDH6,TPM1,EIF5A,GPC3,DDR1,SULF1,CCN2,MM
3,HNRNPU,SMARCA5,LARP1,CTNNA1,RPL23,CNN3,RPL29,RPSA,TNS3,EIF4H,RTN4,LAMA5,RPS2

,CHD6,TOMM20,RAB27A,RNF187,EEF1A2,MLXIP,CPSF1,SQSTM1,VAMP2,GAS7,HNRNPL,LMNB2
3P3,CCDC124
.,STX12,VPS37B,RBBP6,ZC3H4,NDUFA10,FKBP3,CCDC124

ARS1,VEZF1,SNRPB,KDM5A,RBM14,SRP54,LMNA,UQCRFS1,SKIV2L,SOS1,LZTS2,SHC1,PCED1A,
HA,PTP4A2,STMN3,PFKM,MTRNR2L10,FBLN1,STXBP1,ISYNA1,LOC728392,USP7,IDH1,COL27A1
P1,CMTR1,CLCN3,TRAPPC1,ABCE1,NOLC1,MSI2,CCNL1,NBR1,RNPS1,AKAP17A,MAPK8IP3,PAF
3H4,NDUFA10,FKBP3,CCDC124
3H4,NDUFA10,FKBP3,CCDC124
,CLCN3,HEXIM1,UFM1,TRAPPC1,AGPAT4,BRPF3,PSMD12,CBX4,SLC25A39,PDXDC1,GNAO1,SM
,CHD4,JUND,TMEM259,EIF5,COL5A2,HMGN2,ATP2A2,LASP1,RBBP4,RTN3,PFN1,MCL1,TLE5,CL
S1,SOD1,BAG6,VGF,FBRSL1,RPL39,CYP51A1,ZSCAN18,PRPF40A,SMS,NAP1L4,CD99L2,PCYOX1
,RCN1,H1-0,RPS5,RPS19,HMGCS1,TLN1,MYO5A,CTNNB1,NGRN,RPL35A,DNAJC5,HNRNPDL,TA
,RHOB,HNRNPH1,UBE2H,RPS12,ARCN1,ZNF454,HMGCR,ARHGDIA,CAPRIN1,STIP1,SLC35E2B,A
,RHOB,HNRNPH1,UBE2H,RPS12,ARCN1,ZNF454,HMGCR,ARHGDIA,CAPRIN1,STIP1,SLC35E2B,A
F9,PLEC,BTBD2,PBX2,FAM171A2,IPO7,NYNRIN,RBP1,WDR6,GABARAP,ACTN1,CAND1,PLCG1,M

TSPAN3,LPCAT1,RPS24,SLC22A17,BRD3,XRCC5,CIPC,PJA2,MSN,RPL24,SLC16A7,KPNB1,C1OR
S18L1,PNMA2,CAPZA2,MARK1,TRAF3,POTEC,QRICH1,TFDP1,ZC3H15,RAB35,SPIN1,CABIN1,ZN

ARCHF6,WSB1,RPS17,VCL,AP3D1,SBF1,DNAJA1,ARF1,CYB5R3,MAP4,SF3B2,CDC42BPB,ITM2B,
IMGCS1,PRKAR1A,TLN1,CHPF,GJA1,JUN,NGRN,B4GALNT4,CYFIP2,DNAJC5,HNRNPDL,TARDBP,
,GJA1,MACF1,RPS10,SESN3,MTSS2,DNAJC5,HNRNPDL,TARDBP,SLC25A3,MYL6,HNRNPK,INIP,C

POLDIP3,PSMC2,NCBP2,UNC45A,RNMT,PSMF1,CRABP1,ABHD17A,PHKB,PDCD11,USP19,WARS1
LCG1,ARF4,DIDO1,HNRNPUL2-BSCL2,DUSP4,TPM3,POLR2B,XPNPEP3,ATP6V1B2,SDHA,B2M,U

RS,VANGL2,VGF,MEX3A,P3H3,MLF2,FLNC,ARNT2,RTL8C,ARFGEF2,AKIRIN1,KDM4B,SMS,CTXN

RKAR1A,COL6A2,CD63,ANXA6,NGRN,GOLM1,SESN3,B4GALNT4,MTSS2,DNAJC5,APLP2,TARDB
37B,RBBP6,ZC3H4,NDUFA10,FKBP3,CCDC124
2,HMGB3,ECH1,ETF1,TNRC6A,PPP4R3A,SOBP,DDX6,AKAP8L,KPNA6,GID8,DEAF1,EXOC7,PSMC5
9,LENG8,WDR1,BEX3,CCT7,OAZ1,SRSF4,JPT2,SPC25,NES,RPL31,NISCH,HNRNPC,CCNI,LRPPRC
1A2,CKAP4,HLA-B,SEPTIN3,COTL1,CTBP1,GPC1,SLC35E2B,APPL1,L1CAM,RPL23A,PEA15,DLG5

PTGES3,CKAP5,LAMP1,RHOB,HNRNPH1,UBE2H,RPS12,ARCN1,ZNF454,HMGCR,ARHGDIA,CAPR
BCE1,NOLC1,MSI2,CCNL1,RNPS1,AKAP17A,MAPK8IP3,PAF1,DDX19A,RAP1B,PPM1A,GNAO1,RAD
53BP1,AMOTL2,YME1L1,CTNND1,TRAF4,STAT3,ARIH1,LBH,KDM5B,SF1,HIP1,SLC39A6,GPX4,G

UP98,NACAD,OXSR1,EMC10,TMEM9,SCG2,WDR82,SYNGR1,USP48,RPS29,STMN4,NSMF,DUSP8
T2,CUL7,SLC12A4,MXRA5,MCM7,SETD1A,SLC25A39,TCEA2,PRELID1,POU3F3,GET4,MAFB,TLNR
3,ADIPOR2,FBL,WARS1,CLCN6,VEZF1,SNRPB,KDM5A,NKIRAS2,ZDHHC5,XPC,SKIV2L,CFLAR,SH
GNL1,TM9SF4,TRAP1,DNAJB2,TUBGCP6,PLEKHB2,ELP1,SPAG9,PLEKHM2,MOAP1,KRT8,CLASP

.,STX12,VPS37B,RBBP6,ZC3H4,NDUFA10,FKBP3,CCDC124
OT2,YY1,CCDC47,THRAP3,ELOVL5,SRPRA,IER2,DENND5A,SLC25A36,XRN2,SCAMP5,HTATSF1,S
N3,TRAPPC1,ABCE1,NOLC1,MSI2,CCNL1,NBR1,RNPS1,MAPK8IP3,PAF1,DDX19A,RAP1B,PPM1A,

1,DDX19A,RAP1B,PPM1A,GNAO1,RAD23A,SLC39A1,EPHA2,SKIL,STX12,VPS37B,RBBP6,ZC3H4,N

,UBN1,NECTIN2,PLCD3,CTDNEP1,CMTR1,CLCN3,TRAPPC1,ABCE1,NOLC1,MSI2,CCNL1,NBR1,RI
TAP,TRA2B,IARS1,MBTPS1,HDAC6,PTEN,FEM1B,HNRNPA2B1,EML1,IL6ST,ANP32B,MAPRE2,RPL
10,GLOD4,MAEA,MTURN,GOLPH3,ECHS1,ADCY1,ALDH7A1,DHX36,HM13,COLGALT1,TSPAN7,N
R1,ST3GAL6,DFFB,STAT3,SGTA,TXNDC5,MTA1,EXTL3,RETREG2,ALDH3A2,FLCN,H2AZ1,BRD2,K
R1,ST3GAL6,DFFB,STAT3,SGTA,TXNDC5,MTA1,EXTL3,RETREG2,ALDH3A2,FLCN,H2AZ1,BRD2,K
,TIA1,PFKP,ARHGEF40,MAPK8IP2,TRMT112,CRIP2,ADNP,PNRC2,EHMT2,HLA-A,MTURN,TSPAN1

PXDN,PGAM1,SMARCA5,CPE,CNN3,RPL29,RPSA,TNS3,CTSD,EIF4H,NACA,TSPAN3,RTN4,LPCAT1,

EGFA,RAPGEF1,TUBB2A,OPTN,ARF3,G3BP2,MAPRE1,DDAH2,FAM168B,IPO7,ARGLU1,RBP1,LUZ
DC124
32,SYNGR1,USP48,RPS29,NSMF,DUSP8,RNF20,NUP133,CACYBP,CHGB,COX4I1,OGDH,DSTYK,P

P2,CPSF1,GNAI3,HNRNPL,TSPYL4,TM9SF2,MORF4L2,HSPH1,MTRF1L,HNRNPA3,PRDX5,C4ORF

HNRNPD,VGF,FBRSL1,P3H3,KDM3B,RTL8C,BEX4,POLR2E,PSMA7,CIC,TRAF4,EIF2AK1,EDC4,WD

0,PI4KA,UBE2G2,RAB3GAP2,MBTPS1,SELENOW,CUL3,ADH5,NDST1,RPL7L1,ITGB5,CALD1,RPS
,FZR1,SGSM3,UBE4B,SLC25A6,DNAJB1,OSBPL2,CRIP2,AAMP,SUB1,CNDP2,KCTD20,LMF2,LIM

1,TP53I11,RYBP,CRIM1,OBSCN,TCEA2,ATP6V1F,MEX3B,SRSF8,ALYREF,SKIL,ARFGAP3
PNM1,TNKS2,UBQLN1,TUG1,CSNK1D,IFITM3,UQCRC2,PCBP2,MAPK8IP1,DGKD,SREBF2,HNRN
F62,VPS4A,NUDCD3,GNL1,LRATD2,PNISR,FLOT2,TUBGCP6,RIC8A,NCSTN,ATXN3,INTS11,SEMA

,HNRNPL,TGFBI,LMNB2,PRDX5,TBC1D16,ACO2,TCP1,PLOD1,CAMSAP1,ETNK1,KLC1,CCT3,MLE
,UQCRC2,PCBP2,MAPK8IP1,DGKD,SREBF2,HNRNPD,VGF,MLF2,ARFGEF2,AKIRIN1,SMS,DYNLL1
DHFR,SHMT2,RAVER1,STARD3,NKIRAS2,ZDHHC5,SRP54,PMPCA,UBXN1,CACYBP,RNF4,LZTS2,C

AP,SUPT5H,ATXN10,CAND1,CLIP3,MAP2K2,RPL28,ZFAND5,DIDO1,TSC22D1,POLR2B,SMARCC1
1,DDX19A,RAP1B,PPM1A,GNAO1,RAD23A,MYEF2,SLC39A1,EPHA2,SKIL,STX12,VPS37B,RBBP6,Z

H,DSTYK,GSTA4,GUK1,UBN1,NECTIN2,PLCD3,CTDNEP1,RNASEK,CMTR1,CLCN3,TRAPPC1,ABCE
,DDX19A,RAP1B,PPM1A,GNAO1,RAD23A,SLC39A1,EPHA2,SKIL,STX12,VPS37B,RBBP6,ZC3H4,N

_4,OLFM2,OGFR,MTA2,TMEM94,NOP56,ANKHD1,CDV3,SC5D,MFN2,MDM2,UBE4B,NPTN,SLC25

GLN1,MAP9,TPI1,CHSY1,TMSB10,IN
A,RPL15,GTF3C4,LDHB,PKD1,HLA-C,
AH8,PKD1,HLA-C,PDIA3,SSBP2,ZNF
INC1,SSBP2,ZNF43,RPS3,CHD3,BAS
PB,ITM2B,AP2B1,MAT2A,TMEM170A,
HB,DNAH8,PKD1,HLA-C,SF3B1,PDI
2,RPL19,PPP2R1A,RPL15,GTF3C4,LD
6B,ENC1,NLGN2,CXADR,ATP1A1,JUND,
B,UCHL1,MORF4L1,BTF3,NCAM1,ANX
RPS3,BASP1,VAPA,HMGB1,POLR2A
F43,BASP1,VAPA,HMGB1,POLR2A,AP
IEM170A,GTF2F1,GOLGA8A,PXDN,HNRNP
RCHF6,WSB1,RPS17,VCL,AP3D1,SBF1,
GRN,TIMP2,XAF1,SARAF,MIDN,RPL10,
SB1,RPS17,VCL,AP3D1,SBF1,DNAJA
,VAPA,HMGB1,POLR2A,CDKL1,TPT1
RN,TIMP2,XAF1,SARAF,COL11A1,MIDN,
P3D1,SBF1,ARF1,CYB5R3,MAP4,SF3B2,
',BRD3,CIPC,PJA2,MSN,KPNB1,C1ORF43,
MIDN,RPL10,MARCHF6,WSB1,VCL,AP3
SBF1,DNAJA1,ARF1,CYB5R3,MAP4
HL1,MORF4L1,BTF3,NCAM1,ANXA2,
NAJA1,ARF1,CYB5R3,MAP4,CDC42
HACD3,DCHS1,SET,EGLN1,MAP9,TPI1
2B,MAT2A,TMEM170A,GTF2F1,SLC4A
P1,VAPA,HMGB1,POLR2A,TPT1,SEC
AT2A,TMEM170A,SLC4A8,PXDN,HNRN
,POLR2A,CDKL1,AP2M1,TPT1,SEC6
LN1,MAP9,TPI1,CHSY1,TMSB10,INA
GLN1,TPI1,CHSY1,TMSB10,INA,EIF4
SF3B1,PDIA3,SERINC1,SSBP2,ZNF43,R
,CYB5R3,MAP4,SF3B2,CDC42BPB,ITM2
SB1,RPS17,VCL,SBF1,DNAJA1,ARF1,CY
SB10,INA,EIF4A1,MRC2,RPL19,PPP
S1,SET,EGLN1,MAP9,TPI1,CHSY1,TMSB1
3,CNN3,RPL29,TNS3,CTSD,EIF4H,SCARB2
HMGB1,POLR2A,CDKL1,TPT1,SEC6
MIDN,MARCHF6,WSB1,RPS17,VCL,AP
T2A,GTF2F1,GOLGA8A,PXDN,HNRNPU,OGA
LR2A,CDKL1,AP2M1,TPT1,RPL18,STM
R1A,RPL15,GTF3C4,PKD1,HLA-C,SF3B
A1,MRC2,PPP2R1A,RPL15,GTF3C4,LD
1,CYB5R3,MAP4,SF3B2,ITM2B,ZNF
4B,UCHL1,MORF4L1,NCAM1,ANXA2,
CHD3,BASP1,VAPA,HMGB1,POLR2

DNAH8,PKD1,HLA-C,SF3B1,PDIA3,SSB
DL11A1,MIDN,RPL10,MARCHF6,RPS17,V
CXADR,CHD4,ATP1A1,JUND,EIF5,CD81,WDR1
HB,DNAH8,PKD1,HLA-C,SF3B1,PDIA3
SY1,TMSB10,INA,EIF4A1,MRC2,RPL19
CTNNA1,FADS2,LDHA,CPE,CNN3,TNS3,
2A17,BRD3,CIPC,PJA2,MSN,C1ORF43,C
TGB1,TUBB4B,UCHL1,RPL11,MORF4L
JPT2,TLE5,NES,NISCH,HNRNPC,CCNI,LRPPRC,X
1,AP2M1,TPT1,SEC61A1,RPS4X,RPS3
3P,TKT,SFPQ,CCND2,CDH2,PNMA1,PDIA6,GP
,NES,NISCH,HNRNPC,CCNI,TADA2B,XPR1,SP
HOA,CBX5,ZNF865,CLSTN1,KIF5B,RAB6
,CLSTN1,ATF4,PSMD2,RAB6B,SLC7A5,CHD4,AT
8,STMN1,ITGB1,TUBB4B,UCHL1,
D2,CDH2,PNMA1,PDIA6,GPC3,SOX12,P
,ZNF43,RPS3,CHD3,BASP1,VAPA,H
,COL1A2,LLGL1,NCS1,RHOA,CBX5,ZNF865,
B,SOX12,DDX3X,ITM2C,ARL4C,GNB2,H1-0
ADS2,RPS16,CNN3,TPM4,TNS3,CTSD,NACA
EC61A1,RPS4X,RPS3A,RPL18,STMN
RC,XPR1,SPTBN2,PFKL,HDLBP,TKT,GLG1,SF
3B4B,UCHL1,RPL11,NCAM1,ANXA2,AGR
F865,CLSTN1,KIF5B,RAB6B,SLC7A5,EN
PC,CCNI,LRPPRC,XPR1,KHDRBS1,SLC3A
A5,SLC22A17,BRD3,CIPC,PJA2,SLC1
3F1,DNAJA1,ARF1,CYB5R3,AKAP12,MA
0,EIF5,CD81,WDR1,HMGN2,ATP2A2,CRMP1,
DCHS1,SET,EGLN1,MAP9,TPI1,CHSY1,
M1,ANXA2,AGRN,ZMIZ1,TIMP2,SARAF,
L15,GTF3C4,LDHB,DNAH8,PKD1,H
GL1,HK1,NCS1,RHOA,CBX5,ZNF865,K
3A,RPL18,STMN1,ITGB1,TUBB4B,UC
APA,HMGB1,POLR2A,CDKL1,TPT1,S
F2F1,SLC4A8,GOLGA8A,PXDN,HNRNPU,P
C1ORF43,PAPOLA,LLGL1,GAP43,NCS1,ZNF8
NMA1,PDIA6,GPC3,PIP4K2B,MEGF8,SOX12
T,EGLN1,MAP9,TPI1,CHSY1,TMSB10,I
,IGFBP3,COL1A2,LLGL1,GAP43,HK1,N
RF43,COL1A2,LLGL1,NCS1,RHOA,CBX5,Z
RF43,COL1A2,LLGL1,NCS1,RHOA,CBX5,Z
ATP1A1,JUND,EIF5,CD81,WDR1,HMGN
T,EGLN1,MAP9,TPI1,CHSY1,TMSB10,IN
GL1,HK1,NCS1,RPL22,RHOA,ZNF865,CLST
PC,PJA2,MSN,SLC16A7,KPNB1,RPL17,PAP

IF4H,NACA,TSPAN3,LPCAT1,LAMA5,SLC22A1
TP2A2,RPL26,CRMP1,LASP1,BEX3,CCT7,VA
P2A2,CRMP1,LASP1,VAT1,OAZ1,SRSF4,RB
C,CCNI,XPR1,KLHL9,SPTBN2,SLC3A2,PFK
NFE2L1,SMARCA5,HNRNPA0,CTNNA1,FAD
1,HACD3,DCHS1,SET,EGLN1,MAP9,T
KRD52,WLS,HNRNPK,MYH10,RPL36,PTG
LGL1,NCS1,RHOA,PTCD3,CBX5,ZNF865,CLS
B10,INA,EIF4A1,MRC2,RPL19,PPP2R
LPCAT1,LAMA5,SLC22A17,BRD3,RPS9,
F1,CYB5R3,MAP4,CDC42BPB,ITM2B,AP2
F43,COL1A2,LLGL1,HK1,NCS1,RHOA,CBX
,SRSF4,PFN1,ASTN2,JPT2,TLE5,NISCH,
C7A5,NLGN2,CXADR,CHD4,ATP1A1,JUND,R
HSY1,TMSB10,INA,EIF4A1,MRC2,RPL
2F1,KIF5B,ATP1A1,TMEM259,RPS20,EI
C1,TOP2B,SLC35E2B,SMCHD1,IPO5,ATP6V0C,
BPB,ITM2B,MAT2A,TMEM170A,GTF2F1,PX
,JUND,EIF5,CD81,LENG8,WDR
,PFKL,TKT,GLG1,CDH6,SFPQ,TPM1,EIF
NCS1,RHOA,PTCD3,CBX5,ZNF865,CLS
IA6,GPC3,PIP4K2B,MEGF8,SOX12,RAB7A,
ADM4,LAMP1,HNRNPH1,CKAP4,RAB1A,CALU
MMP2,SKP1,TRIB2,HSBP1,NNAT,DLG5,HS6ST
NAJA1,ARF1,CYB5R3,AKAP12,MAP4,
D1,HDAC2,IPO5,ATP6V0C,SKP1,LAP
NPU,PGAM1,NFE2L1,SMARCA5,HNRNPA
I,DNAJA1,ARF1,CYB5R3,MAP4,CDC42B
1,FADS2,RPS16,RPL23,LDHA,CPE,CNN
,OGA,NFE2L1,SMARCA5,HNRNPA0,CTN
EGLN1,MAP9,TPI1,CHSY1,TMSB10,INA
,NCS1,RHOA,ZNF865,CLSTN1,ATF4,PS
D4B,CAPRIN1,STIP1,GPC1,TOP2B,SLC35E2B,
2,CRMP1,LASP1,CCT7,VAT1,OAZ1,PFN1,MCL
A,TMEM170A,SLC4A8,GOLGA8A,PXDN,OGA,
ORF4L1,BTF3,NCAM1,ANXA2,AGRN,Z
N2,EID1,C5ORF24,ZC3H7B,TM9SF3,CHD6,
12,ITM2C,ARL4C,NANOG,MAGED2,GNB2
QSTM1,VAMP2,KLHL24,GAS7,TAF15,P
1,RAB6B,SLC7A5,ENC1,NLGN2,CXADR,CHD4
3,KHDRBS1,KLHL9,SPTBN2,PFKL,H
ACTN4,PRDX2,RPS15,ITM2C,RPL30,AR
S24,BRD3,XRCC5,CIPC,PJA2,MSN,KPNB1,C
TADA2B,XPR1,KHDRBS1,SLC3A2,PFKL,HDL
HS1,SET,EGLN1,MAP9,TPI1,CHSY1,T

CA,TSPAN3,LPCAT1,LAMA5,RPS24,S
3,ARCN1,SEPTIN3,COTL1,HECTD4
ARL4C,GNB2,H1-0,RPS5,APLP1,RPS19,G
,TSPAN3,RPS24,BRD3,RPS9,CIPC,PJA2
Z1,TIMP2,XAF1,SARAF,COL11A1
2,CDC42BPB,ITM2B,AP2B1,MAT2A,SLC4
/AT1,OAZ1,RTN3,JPT2,TLE5,PNN,NES,
F4H,TSPAN3,LPCAT1,LAMA5,RPS24,BRD3,X
3A,RPL18,STMN1,ITGB1,TUBB4B,U
R1A,RPL15,GTF3C4,LDHB,PKD1,HLA
RNPDL,ARL6IP1,ARID1A,TSR2,MYL
TP6V0C,MMP2,DNER,L1CAM,TRIB2,PEA15,
SB,NUCB1,CHD6,U2AF2,EEF1A2,CCT2,TBCD,SN
MSN,RPL24,IGFBP3,LLGL1,GAP43,HK
HK1,NCS1,RPL22,RHOA,ZNF865,CLS
GTF2F1,GOLGA8A,PXDN,HNRNPU,OGA
DIA,CAPRIN1,STIP1,IPO5,ATP6V0C,MMP2,
BP1,CAPRIN1,STIP1,GPC1,TOP2B,IPO5,MMP2
KL1,TPT1,SEC61A1,RPS4X,RPS3A,OR
Z1,TIMP2,SARAF,MIDN,RPL10,MAR
TGB1,TUBB4B,MORF4L1,BTF3,NCA
65,RPL9,CLSTN1,KIF5B,RAB6B,SLC7A5,NL
T1,LAMA5,RPS24,SLC22A17,BRD3,
31,PSMD5,YWHAQ,EIF1,HACD3,DCHS1,S
IER,LAPTM4B,HSBP1,LUC7L3,FAM199X,
KL,HDLBP,TKT,CDH6,SFPQ,TPM1,CD
A15,PTTG1IP,SQLE,DLG5,HS6ST1,CPSF2,PGR
,SSBP2,ZNF43,RPS3,CHD3,BASP1
2,CD63,GOLM1,B4GALNT4,MTSS2,DNAJC
P1,ZMIZ1,SARAF,MIDN,RPL10,MARCHF6
A3,PTPRA,PRDX5,ACTR3,PLOD1,ETNK1,GRINA
,APLP2,HNRNPDL,TARDBP,ST3GAL2,MXD4,IN
IF5,CD81,LENG8,WDR1,HMGN2,ATP2A2,
CCNI,XPR1,KHDRBS1,KLHL9,XPO1,SPT
COL11A1,MIDN,RPL10,MARCHF6,
OK,LAMC1,JPT1,CBX1,RAD23B,PTK7,CHD2,ORAI2
NF187,EEF1A2,CCT2,TBCD,SNX22,PGD,S
,CTNNA1,FADS2,RPS16,RPL23,LDHA,CPE
HL9,XPO1,SPTBN2,SLC3A2,CSNK1E,PFKL,HD
ADA2B,XPR1,KHDRBS1,XPO1,SLC3A2,PF
PXDN,HNRNPU,NFE2L1,SMARCA5,HNRNPA0
VEGFA,TUBB2A,ARF3,CCND1,SMARCA4,MAPRE1,S
3,RAB6B,CXADR,CHD4,ATP1A1,TMEM259,E
3,RAB6B,CXADR,CHD4,ATP1A1,TMEM259,E
3,RAB6B,CXADR,CHD4,ATP1A1,TMEM259,E

3,RPL29,RPSA,TPM4,TNS3,CTSD,EIF4H
STN2,JPT2,TLE5,NISCH,HNRNPC,CCNI,
TF2F1,SLC4A8,PXDN,HNRNPU,OGA,
RKAR1A,GJA1,JUN,COL6A2,CD63,ANXA6
,PTBP1,EIF3E,DYNLL2,ILF3,PCSK1N,CAPZB,UBE
P2,CHD3,BASP1,VAPA,HMGB1,POLR
A1,PIP4K2B,MEGF8,ACSL6,SOX12,CCN2,DDX3X
F5,CD81,WDR1,HMGN2,RPL26,CRMP1,LASP
ICS1,RHOA,CBX5,ZNF865,CLSTN1,A
1,STIP1,GPC1,SLC35E2B,SMCHD1,IPO5,ATP6V
TBP1,DYNLL2,ILF3,PCSK1N,UBE2R2,AT
AJA1,ARF1,CYB5R3,AKAP12,SF3B2,ZNF
AF,MIDN,RPL10,WSB1,RPS17,VC
P1,PTOV1,MLEC,ACAP3,MDK,LAMC1,PTK7,
RPS24,BRD3,SLC16A7,KPNB1,RPL1
1,SLC4A8,PXDN,HNRNPU,NFE2L1,SM
1A,RPL15,GTF3C4,LDHB,DNAH8,PKD1,
OL,SLC25A3,MXD4,MYL6,ANKRD52,WLS,HN
LR2A,CDKL1,TPT1,SEC61A1,RPS4X
3A2,PFKL,TKT,GLG1,SFPQ,PNMA1,PDIA6,GP
3P1,EIF3E,PCSK1N,WDR6,SCAF11,ACTN1,CASC3,CLI
IMP2,SARAF,MIDN,RPL10,RPS17,AP
K1E,PFKL,HDLBP,CDH6,SFPQ,EIF5A,CCND2
,IPO5,ATP6V0C,DNER,BCLAF1,L1CAM,TRIB
GPC3,PIP4K2B,MEGF8,SOX12,DDX3X,ARL4C
RPS24,BRD3,CIPC,PJA2,KPNB1,C1ORF43,IG
RHGDIA,CAPRIN1,STIP1,GPC1,CDK16,TOP2B,
DNAJA1,ARF1,CYB5R3,AKAP12,MA
2A17,BRD3,XRCC5,KPNB1,RPL17,P
RAB6B,ENC1,GLIPR1L2,CXADR,CH
C3,OAZ1,SRSF4,RBBP4,PFN1,MCL1
,CCNI,LRPPRC,TADA2B,XPR1,KLHL9,XPO1,S
SMCHD1,IPO5,ATP6V0C,MMP2,SKP1,DNER
P2B1,MAT2A,GTF2F1,PXDN,HNRNPU,O
TP1A1,RPS20,EIF5,LENG8,WDR1,HMGN2,AT
PSMD5,YWHAQ,EIF1,HACD3,DCHS1,SET,
IA,VPS35,MLEC,CLSTN3,MDK,RBMX,LAMC1,J
TUBB2A,ARF3,CCND1,SMARCA4,ATG2B,MAP
4,ATG2B,MAPRE1,DSTN,PLEC,FAM171A2,
4,LDHB,DNAH8,PKD1,HLA-C,SF3B1,PD
FKL,HDLBP,GLG1,CDH2,PNMA1,PDIA6,GPC3
NN3,RPL29,RPSA,TPM4,TNS3,CTSD,EIF4H,
L3B,HMGN2,RPL26,LASP1,CCT7,OAZ1,
4,MTSS2,DNAJC5,APLP2,HNRNPDL,TARDBP,S
P12,MAP4,SF3B2,CDC42BPB,ITM2B,

3,PAPOLA,COL1A2,LLGL1,RPL14,HK
A17,BRD3,CIPC,PJA2,RPL24,KPNB
L35,MIF,HMGCS1,GPAT4,CHPF,GJA1,J
LN,MYL6,HNRNPK,INIP,NARS1,YWHAB,PTGES3,CAD
TLE5,PNN,NES,NISCH,CCNI,LRP
A,STIP1,GPC1,TOP2B,SMCHD1,HDAC2,DNER,PEA15,
K1,RHOA,PTCD3,ZNF865,ATF4,PSMD2,E
GLN,ANKRD52,INIP,CIRBP,MYH10,NARS1,RPL
RP1,PRDX5,PDXK,YEATS2,HSPA4,CAMSAP1,GR
RP1,PRDX5,PDXK,YEATS2,HSPA4,CAMSAP1,GR
664,ZMYM3,VPS26B,STK35,ACLY,SEPT
RF4,DDX42,CBX1,RAD23B,SNRNP70,CN
D3,BASP1,VAPA,HMGB1,POLR2A,CDKL
1,SRSF4,RBBP4,JPT2,TLE5,RPL31,NISCH,PR
GAS7,GDI2,TM9SF2,PRDX5,PDXK,PKN1,HSPA4,CAMS
RAB11B,ABCA2,PYGB,PRPF6,DPYSL2,SEL
AP1,GRINA,PTOV1,CLSTN3,ACAP3,MDK,JPT1,P
BCLAF1,TRIB2,DLG5,CRPPA,PGRMC1,
D3,XRCC5,PJA2,RPL24,KPNB1,RP
ATP1A1,TMEM259,RPS20,EIF5,L
SMARCA5,HNRNPA0,FADS2,RPS16,R
S2,RPL23,LDHA,TPM4,TNS3,EIF4H,SCARB2
RCHF6,VCL,AP3D1,DNAJA1,ARF1,P
N,CAPZB,SCAF11,ACTN1,MAP7D1,FKB
MYL6,ANKRD52,HNRNPK,H1-10,CADM4,RHOB,
B1,C1ORF43,PAPOLA,COL1A2,HK1,NCS1,RP
4,RAB6B,SLC7A5,ENC1,NLGN2,CXADR,
DDX3X,RPS15,ITM2C,RPL30,ARL4C,EI
NA1,TUG1,CSNK1D,ATXN2L,HYOU1,ID4,
S1,KLHL9,XPO1,PFKL,HDLBP,GLG1,SFPQ,
,DNAH8,PKD1,PDIA3,SERINC1,SSB
M9SF2,LMNB2,ACTR3,PDXK,YEATS2,CAMSA
STM1,VAMP2,GAS7,GDI2,PARP1,TM9SF2,PD
TDSP2,NOVA2,ZC3H7B,CHD6,RNF18
EEF1A2,TBCD,PPP1CB,SNX22,SQSTM1,KLH
PTCD3,CBX5,ZNF865,CLSTN1,
,APLP1,RPS7,MIF,GPAT4,CHPF,MYO
RCN1,SEPTIN3,COTL1,MAGED4B,CTBP1,RAC1
S4X,RPS3A,RPL18,STMN1,UCHL1,RPL
,MCL1,JPT2,TLE5,NISCH,PRKCSH,TA
EM259,EIF5,LENG8,WDR1,HMGN2,ATP
L17,PAPOLA,LLGL1,RPL14,NCS1,
SELENON,TUBB2A,UBXN4,MVB12B,MAPK1,
2,EEF1A2,CCT2,TBCD,SNX22,PGD,VAMP2
5,NES,PRKCSH,HNRNPC,CCNI,TADA2B,XPR1,

PJA2,RPL24,SLC16A7,KPNB1,R
AP3D1,DNAJA1,ARF1,CYB5R3,AKAP1
EX3,CCT7,OAZ1,SRSF4,RBBP4,ASTN2,JPT2
AT2A,GTF2F1,SLC4A8,GOLGA8A,H
,LLGL1,NCS1,RPL22,RHOA,ZNF865,CLSTN1
TIMP3,PGRMC1,DDX39B,CTSB,CHD6,EEF1A2
ORO1C,SDF4,NKTR,SLIT3,STMN3,LSS,S
2,RCN1,APLP1,RPS7,MIF,GPAT4,MYO
ORF4L1,BTF3,NCAM1,ANXA2,AGRN
NA1,PIP4K2B,MEGF8,SOX12,ACTN4
,DDX3X,RPS15,ITM2C,RPL30,ARL4C
P,SEPTIN11,LAMP1,HNRNPH1,LGALS3BP,UBE2H
PS10,SESN3,B4GALNT4,MTSS2,AP
MP1,LASP1,BEX3,VAT1,OAZ1,RBBP4,PFN1,
B,MAT2A,TMEM170A,GTF2F1,PXDN,
JGP2,ANXA5,KLHL24,GAS7,PABPN1
D3,CBX5,ATF4,PSMD2,KIF5B,SLC7A5
NI,XPR1,KHDRBS1,XPO1,SCRN1,HDLBP,
PLP1,ZMIZ1,TIMP2,XAF1,COL11A1,MI
,MIF,GPAT4,TLN1,JUN,MYO5A,CD63,GOLM1
ARL4C,EIF3L,MRFAP1,RCN1,H1-0,RPS5,
L1,AP2M1,TPT1,SEC61A1,RPS
APRE1,DDAH2,DSTN,ODC1,AEBP1,TCF25,DYNL
NXA2,AGRN,RPLP1,ZMIZ1,TIMP2,XA
LEC,ACAP3,MDK,DDX42,RAB5B,ORAI2,
N2,ATP2A2,RPL26,CRMP1,BEX3,CCT7,VAT
N2,ATP2A2,RPL26,CRMP1,BEX3,CCT7,VAT
),CHPF,NGRN,DNAJC5,HNR
5,SERF2,HS6ST1,TIMP3,PGRMC1,CNN2,CT
TXN10,KDM1A,CLIP3,MAP2K2,RPL28,
DYNLL2,ILF3,WDR6,UBE2R2,ATXN10,ACTN
A,STMN1,ITGB1,TUBB4B,UCHL1,RPL1
RPS12,RAB1A,PLPPR3,SEPTIN3,CO
M,PEA15,HSBP1,FAM199X,DLG5,SHISAL1,T
CN1,RPS27,COTL1,HRNR,HECTD4,
CXADR,CHD4,ATP1A1,JUND,TMEM2
E,HS6ST1,PGRMC1,CNN2,EID1,C5ORF24,RAB
,FADS2,RPS16,RPL23,CNN3,TPM4,CTSD,
N1,YEATS2,CAMSAP1,KLC1,GRINA,NCKAP1,PTO
L,SBF1,DNAJA1,ARF1,CYB5R3,AKA
RPL29,TNS3,CTSD,SCARB2,NACA,TS
TP6V0C,TMEM30A,DNER,BCLAF1,TRIB
E2R2,SON,KDM1A,MAP7D1,CASC3,PLC
YL6,H1-10,INIP,RPL36,DUX4,CKAP5,S
16A7,KPNB1,C1ORF43,PAPOLA,COL

.P1,SARAF,MIDN,RPL10,MARCHF6
,PLPPR3,ARCN1,CTBP1,STIP1,GPC1
/ID2,NR2F1,RAB6B,ENC1,NLGN2,C
SYP,POLR2E,PPDPF,RBBP7,DFFB,EIF2AK1,
S3,CTSD,EIF4H,SCARB2,TSPAN3,LPCAT1
I,ZMIZ1,XAF1,COL11A1,MIDN,RPL10,M
,CDK16,TOP2B,SLC35E2B,I
JUND,TMEM259,RPS20,CD81
S,FBLN1,PITPNM1,CSNK1D,ATXN2L,ID4,ZFAND3,SAR
?2,ZNF865,CLSTN1,ATF4,KIF5B,
3,B4GALNT4,DNAJC5,APLP2,HNRNPDL,
)L2,YLPM1,GRIPAP1,PSMA7,ANP32A,TR
P,TMED2,SELENON,MAPK1,ATG2B,LAPTM4
AJA1,ARF1,AKAP12,MAP4,SF3B2,ZNF7
L11,MORF4L1,NCAM1,ANXA2,AGR
PFN1,MCL1,JPT2,TLE5,PNN,NES,RPL
KI,ASXL1,H3-3A,RAB11B,ABCA2,PYGB
,RPL26,LASP1,OAZ1,PFN1,ASTN2,TLE
RF198,ZNF664,STK35,CLCN7,RTL6,CCN
-,PIP4K2B,RAB7A,DDX3X,RPL
NBPF9,ZFAND3,STRAP,SARS1,HGS,BAG6,
iP2,PGD,HNRNPL,HSPH1,ABCA3,ALKBH5,HN
NI,LRPPRC,XPR1,SLC3A2,PFKL,
NO,DDB1,PSMD5,YWHAQ,EIF1,HACD3,
A1,ARF1,CYB5R3,AKAP12,SF3B2,
,MLXIP,PGD,SQSTM1,VAMP2,NOX4,HSPH1,PR
A5,CPSF1,SNX22,VAMP2,GAS7,PABPN1,
3B,ATP2A2,RPL26,CRMP1,LASP1,BEX3,
IMP3,CTDSP2,DDX39B,NOVA2,CTSB,NUCB1,CH
AP1,ETNK1,GRINA,COL2A1,ACAP3,MDK,LAMC1,P
2,ANXA6,GOLM1,RPS10,MTSS2,DNA
RCA5,HNRNPA0,FADS2,RPS16,RPL23
MCL1,JPT2,NES,RPL31,PRKCSH,C
SF3B2,CDC42BPB,ITM2B,MAT2A
T,GLG1,SFPQ,TPM1,EIF5A,PEBP1,CCND2
IFE2L1,SMARCA5,HNRNPA0,CTNNA1,F
C,SF3B1,PDIA3,SERINC1,SSBP2,ZN
A1,JUND,WDR1,HMGN2,ATP2A2,CRMP1,
A1,FLNC,SERBP1,WBP2,AKIRIN1,C
I,SPEN,NONO,DDB1,PSMD5,YWHAQ,EI
4,HK1,RHOA,PTCD3,CBX5,ZNF
K,RPL12,ITM2C,RPL30,CLTC,EIF
4,CKAP5,SEPTIN11,SND1,LAMP1,R
,LMNB2,ALKBH5,PTPRA,PRDX5,ACTR3,C4OR
2,HNRNPL,TM9SF2,ABCA3,ALKBH5,PR

PITPNA,CORO1C,ATP6V1B2,PHC2,YWHAH,LSS
5B,PPDPF,CIC,TRAF4,RBBP7,KIF5A,MS
2,DSTN,PSMD3,PRDX1,ODC1,DYNC1LI2
TUBB4B,UCHL1,RPL11,MORF4L1,BTF3
,VCL,AP3D1,SBF1,DNAJA1,ARF1,AK
1,CD99L2,CIC,TRAF4,RBBP7,KIF5A,MS
N,OPTN,MAPRE1,DDAH2,ODC1,AEBP1,TCF25
OGA,SMARCA5,HNRNPA0,FADS2,RPS
3,DDX42,LAMC1,CBX1,ORAI2,H3-3A,RAB11B,
3,DDX42,LAMC1,CBX1,ORAI2,H3-3A,RAB11B,
3,DDX42,LAMC1,CBX1,ORAI2,H3-3A,RAB11B,
14,PRDX2,DDX3X,RPS15,RPL12,ITM2C,
XN4,MVB12B,MAPK1,ATG2B,DSTN,RAB
,TNS3,CTSD,EIF4H,TSPAN3,LPCAT1,LAMA5
SBF1,DNAJA1,ARF1,PAFAH1B1,AK
N3,RPL29,TPM4,TNS3,EIF4H,NACA,
TF3,NCAM1,AGRN,RPLP1,ZMIZ1,TIM
KCTD12,ZFR,ZNF512B,TBC1D9B,PPP1CC,
,SFPQ,TPM1,EIF5A,CCND2,CDH2,PNMA1,
DGF,SRP14,ABCA2,PRPF6,TMED2,T
5,APLP1,RPS7,RPL35,TLN1,MYO5A,COL6A2
C3H7B,TOMM20,RAB27A,U2AF2,FDFT1,T
VA2,CTSB,CHD6,EEF1A2,TBCD,SUGP2
AP3D1,SBF1,DNAJA1,ARF1,AKAP12,
L2,RAI1,HNRNPD,FBRSL1,MEX3A,RAB11A,ML
GTF2F1,SLC4A8,GOLGA8A,PXDN,PGA
IER,TRIB2,GLUL,PEA15,MAP1A,CDKN1A,
SIP1,DDOST,HNRNPK,H1-10,CIRBP,YWHAB,PTGES3,S
PL1,SMCHD1,HDAC2,IPO5,TMEM3
4A8,SMARCA5,HNRNPA0,FADS2,R
6,ATXN10,HSPB1,PLCG1,MAP2K2,ZFAND5,HN
ED4B,CTBP1,ARHGDIA,STI
X12,DDX3X,CLTC,GNB2,APLP1,MIF,HMGCS
X2,RPS15,ITM2C,ARL4C,CLTC,RCN1,H1
,EP300,EXTL3,DGCR2,STARD7,KDM5B,RET
,SLC16A7,RPL17,C1ORF43,PAPOLA,IGF
A2,LENG8,WDR1,ATXN7L3B,AT
RPS9,XRCC5,CIPC,PJA2,MSN,RPL2
28392,ATXN2L,TMED9,HYOU1,ID4,HECTD1,MP
NRNPU,OGA,PGAM1,NFE2L1,SMARCA5
4,RPS9,CIPC,MSN,SLC16A7,C1OR
NT4,MTSS2,DNAJC5,APLP2,SLC2
HF6,WSB1,RPS17,VCL,AP3D1,SBF
FT1,TBCD,ANXA5,CPSF1,PPP1CB,SQST
1,CCND2,CDH2,PIP4K2B,SOX12,ACTN4

PIB,RPL23A,HSBP1,NNAT,MAP1A,FAM
N2L,ZFAND3,STRAP,HECTD1,VANGL2,FBRSL1,MEX3A
SXL1,PSMB2,SCAF1,PYGB,DPYSL2,RPS
PM4,CTSD,EIF4H,LAMA5,RPS24,RP
CA,LPCAT1,LAMA5,RPS24,BRD3,R
34GALNT4,MTSS2,DNAJC5,HNRNPDL,T
N,EP400,CD164,GLUD1,RER1,MASTL,HDAC
VAT1,RBBP4,RTN3,ASTN2,TLE5,NES,H
DS2,LDHA,CPE,TPM4,TNS3,CTSD,SCARB2,NA
-10,CIRBP,MYH10,CKAP5,SEP
BRSL1,ZSCAN18,WBP2,SMS,DYNLL1,CTXN1,
PN1,HNRNPL,TM9SF2,TSR1,RPL27,PKN1,A
5F5,GNG4,NXN,EP400,PAPSS1,RER1,PI4KA
,ATG2B,AZIN1,LAPTM4A,DDAH2,AEBP1,NYN
I,HNRNPC,LRPPRC,XPO1,SPTB
4,CALU,COTL1,HECTD4,RAC1,ARHGDI
,TARDBP,MXD4,WLS,H1-10,CIRBP,MYH1
IG8,WDR1,ATXN7L3B,ATP2A2,RPL2
PM,GIGYF1,LARS1,FAU,LSS,FBLN1,STT3B,PITP
9,TPM4,TNS3,EIF4H,NACA,LAMA5,R
K1E,PFKL,HDLBP,TKT,GLG1,CDH6,TPM1
L,NANOG,MRFAP1,H1-0,RPS5,RPS
K1,NCS1,RPL22,RHOA,PTCD3,ZN
2,RNF10,SELENON,TUBB2A,MVB12B,BTBD2,PBX2
SL1,MEX3A,MLF2,PPP6R1,ZSCAN18,SERBP1
2B,SLC35E2B,IPO5,TMEM30
TIP1,SLC35E2B,IPO5,ATP6V0C,
PN2,TBCD,CPSF1,PPP1CB,GNAI3,G
I,NFE2L1,SMARCA5,HNRNPA0,FADS2,LDHA,C
ATXN7L3B,RPL26,CRMP1,LASP1,C
2,SEC16A,UBA52,BRD2,UPF1,RAB3GAP2,E
PNB1,C1ORF43,LLGL1,HK1,NCS1,RHOA
5A,GALNT11,CTTN,GPC4,MAP1LC3B,C1ORF198,
NPC,LRPPRC,TADA2B,KLHL9,SLC3A2,P
S16,RPL23,LDHA,CNN3,RPL29,TPM4,SC
R1,ATXN7L3B,HMGN2,RPL26,L
B,HMGN2,RPL26,LASP1,BEX3,CCT7,SR
SS,FBLN1,STXBP1,TUG1,PLXNA3,CSNK1D,AT
ARS1,RPL28,CCAR2,ARF4,POLR2B,SD
25,DYNLL2,ILF3,GRB2,WDR6,CAPZB,CASC
2B,MAT2A,TMEM170A,SLC4A8,GOL
AP3,DDX42,LAMC1,CBX1,ORAI2,H3-3
03,CBX5,ZNF865,ATF4,PSMD2,NR2F1,KIF
COL6A2,GOLM1,B4GALNT4,MTSS2,DNAJ
UTS2,USP10,KDM3A,ARPC1A,TAOK2,RTF1,

)AZ1,SRSF4,RBBP4,PFN1,ASTN2,PNN,RPL31
)X5,UBE4A,YEATS2,ACO2,HS
RAB11A,CYP51A1,PPP6R1,SERBP1,KDM
C1A,TAOK2,RTF1,NOP56,KIFC2,MACROH2A1,MKN
I,PSMD2,GLIPR1L2,RPS14,CXADR
KLHL9,SPTBN2,CSNK1E,PFKL,SCR
NB1,C1ORF43,LLGL1,GAP43,NCS1,R
T1,SEC61A1,RPS4X,RPS3A,STMN1,
AB14,MAZ,DCLK1,IREB2,LRATD2,MACRO
TMN1,ITGB1,TUBB4B,UCHL1,MORF
PK8IP1,BAG6,SREBF2,HNRNPD,MEX3A,
,PNN,RPL31,PRKCSH,HNRNPC
C,LRPPRC,XPR1,KHDRBS1,KLHL
HL1,MORF4L1,BTF3,ANXA2,AGRN,Z
A3,RPLP2,ARL6IP1,DDOST,M
3,ANXA6,NGRN,RPS10,MTSS2,RPL35A,A

MAP1LC3B,DAP,SLC2A1,MTCH1,PRDX6
BP2,RPS3,CHD3,BASP1,VAPA,HMGB1,P
IRNPDL,SLC25A3,RPLP2,DDOST,WLS
T1,SEC61A1,RPS4X,RPS3A,RPL18,I
JCHL1,MORF4L1,BTF3,NCAM1,ANXA2,
CARB2,NACA,TSPAN3,LAMA5,RPS24,XRCC5
,NFE2L1,SMARCA5,HNRNPA0,CTNNA1,RPS1
SQLE,MAP1A,CDKN1A,FAM19
PL26,CRMP1,LASP1,CCT7,SRSF
APLP2,TARDBP,SLC25A3,ARL6IP1,ST3
,SPINT2,ZMYM4,CHERP,FKBP1A,RABEP1,
AR,SYP,PPDPF,ANP32A,CIC,TRAF7,ARIH1,KCN
AF2,RPN2,TBCD,PPP1CB,PGD,MEST,G
AP2M1,TPT1,RPS3A,RPL18,ITGB1,
CSNK1D,ATXN2L,PRPS1,HECTD1,UQCRC2
CL1,TLE5,PNN,NES,RPL31,PR
,NES,CCNI,XPR1,KHDRBS1,KLHL9,XPO
IIDN,RPL10,MARCHF6,WSB1,RPS1
LDHA,CPE,CNN3,RPSA,TPM4,CTSD,EIF4H
I,FAM8A1,SLC25A1,POLD2,CDK9,CLPX,DEGS1,G
,MIDN,RPL10,MARCHF6,RPS17,VCL,S
CCT7,SRSF4,RBBP4,PFN1,ASTN2,
SAP1,MLEC,ZNF302,DDX42,G
TMEM63B,NUCKS1,ELP1,LSM14B,FNBP1L,KRT8
CAN,TSR1,NOX4,MTRF1L,AB
CDC42BPB,AP2B1,MAT2A,TMEM170A,GT
ARCA5,HNRNPA0,FADS2,RPS16,RPL23,C
ASTN2,MCL1,TLE5,PNN,RPL31

A,ZNF865,CLSTN1,ATF4,PSMD2,KIF5
P1,KIF21B,SEC16A,GPX4,PHB2,CST3,SLC39A7,G
1B,PACS1,RBM12,PCNX3,DAZAP2,ANKS1A,
NT17,RHEB,PPP4R3A,NECTIN1,RAP2A,EXT2,FAM16
1,SMARCA5,HNRNPA0,FADS2,RPL23,CN
,GGNBP2,OBSL1,RBM25,NOTCH1,MAST2,MAPK8I
2,AGRN,RPLP1,ZMIZ1,TIMP2,XAF1,
B,HNRNPH1,UBE2H,RAB1A,CALU,P
P1,VPS35,PTOV1,MDK,SURF4,RAD23B
3,CDC37,PCSK1N,GABARAP,HNRNPUL1
,RAB14,MAZ,PLEKHA6,LRATD2,PRR12,TMEM
N1,ASTN2,MCL1,PNN,RPL31,NISCH,HNR
2,NPDC1,IL6ST,POM121C,RPS2,LMAN1,TNK
9,TPI1,TMSB10,INA,EIF4A1,MRC2,R
NF865,RPL9,ATF4,PSMD2,KIF5B
5,PNN,SPC25,NES,RPL31,NISCH,PRKCSH,
,SLC25A3,ST3GAL2,TAGLN,MYL6,AN
,GIGYF1,SRSF3,STMN3,LSS,STT3B,EIF3F,STXBP
P6,IREB2,PNISR,STK24,TUBGCP6,R
P2,CPSF1,GNAI3,GDI2,
1,MORF4L1,NCAM1,ANXA2,AGRN,RP
RPS27,CTBP1,ARHGDIA,CDK
,CCT7,OAZ1,PFN1,ASTN2,TLE5,NIS
,CCT7,OAZ1,PFN1,ASTN2,TLE5,NIS
,MYO5A,COL6A2,ANXA6,MACF1,RPS10,SESN3
RDBP,RPLP2,DDOST,TSR2,HNRNP
,MXD4,MYL6,H1-10,CIRBP,RP
GN2,RPL26,CRMP1,LASP1,CCT7,
C1,TRAF4,KIF5A,KCNQ2,TXLNA,TR
CDH11,TM9SF3,TOMM20,EEF1A2,TBCD,VA
1,SMARCE1,HNRNPM,PFKM,LSS,MTRNR2L10,FB
P,ZMYM3,BAP1,STK35,FUS,XRCC6,ERH,PSMB4
LNT4,DNAJC5,SLC25A3,MYH10,YW
,PIP4K2B,MEGF8,RAB7A,RPL12
A,CTDSP2,CTSB,SRSF5,CDH11,FDFT1,TBCD,M
R2E,PSMA7,RBBP7,EIF2AK1,TXLNA,WDR45B,TX
ER1,CRTAP,MASTL,COX7C,YKT6,RGMB,EC
NN,SPC25,RPL31,PRKCSH,LRP
CCAR2,ARF4,POLR2B,GNG2,SM
DDX39B,EID1,SRSF5,ZC3H7B,T
,ABCA3,ALKBH5,RPL27,PTPR
32A,TRAF4,RBBP7,NAP1L1,MTA1,TRIO
32A,TRAF4,RBBP7,NAP1L1,MTA1,TRIO
1,PRKCSH,HNRNPC,CCNI,LRPP
1,RPL17,C1ORF43,COL1A2,LLGL1,HK1,

;RK2,EFR3B,GRN,SEPTIN5,ERGIC1,SHC2,FKBP4
1B,SELENON,RAPGEF1,TUBB2A,OPTN,ARF
NC1,GLIPR1L2,ATP1A1,TMEM259,COL
)H9A1,RRP1B,NAPB,RAVER1,KARS1,DUSP3,
,RPS15A,RPL38,SELENON,RPL37,AR
23A,GLUL,PTTG1IP,SERF2,HS6ST1,NU
22,PGD,SQSTM1,GDI2,VCAN,MORF4L2,TA
AB7A,PRDX2,RPS15,ITM2C,RPL3
,FDFT1,CCT2,RPN2,SUGP2,GNAI
DDR1,PIP4K2B,MEGF8,ACSL6,
SART1,ATP5MC3,SAP18,BNIP3,TTC28,AT
27,CCT6A,CTBP1,HMGCR,CAPRIN1,GPC1
ITM,BAG6,RAB11A,RPL39,ARNT2,
/A2,CTSB,NUCB1,CDH11,ZC3H7B,CHD
,TNS3,CTSD,EIF4H,SCARB2,TSPAN3,RT
TXN2L,HGS,BAG6,RAB11A,SERBP1,A
A,HNRNPDL,TARDBP,RPLP2,ARL6IP
RPL29,RPSA,TPM4,EIF4H,SCARB2,NAC
BEX3,RBBP4,PFN1,MCL1,JPT2,PNN
1,EIF5A,PEBP1,CCND2,PDIA6,PIP4K2B,SO
2,RPL14,HK1,NCS1,RPL22,RHOA,
RPLP2,DDOST,TSR2,CIRBP,NARS1,DU
IPUL1,SON,RPL28,CCAR2,ARF4,
LEC,ZNF302,DDX42,GCN1,S
SPA4,AHCYL1,CAMSAP1,VPS35,LAMC1,SKI,SN
S7,IK,RPS19,CHPF,MYO5A,COL6A2,
TXN2L,HYOU1,ID4,BAG6,RAB11
I1,NFE2L1,SMARCA5,HNRNPA0,FADS2,R
LBP,GLG1,SFPQ,PEBP1,PNMA1,PDI
ZNF768,WDR82,DHFR,MRPS6,NUS1,CRTC1,CACNA
N1,ACO2,TCP1,PLOD1,COL2A1,PTOV1
)TL1,CTBP1,RAC1,ARHGDIA,SL
SMARCC1,SUPT6H,DHX30,LARS1,NKTR,ILF2
,ITGB5,KLHDC3,DKK3,CTTN,MAP1LC3B
1,SBF1,ARF1,CYB5R3,PAFAH1B1,A
CSH,TADA2B,XPR1,KHDRBS1,XPO1
BP,RPLP2,DDOST,HNRNPK,CIR
,MLF2,FLNC,AKIRIN1,CTXN1,PPP1R9B,SYP,DHX1
SPT1,CST3,UBA52,BRD2,PLXNB1,D
SRSF2,PLEKHB2,ELP1,INTS11,PCNX3,AB
GL2,RAI1,PFN2,HNRNPD,VGF,MEX3A,P3H3
TC28,XPO6,BAG5,SAE1,ATG4B,DEK,EIF
M1,UBQLN1,LOC728392,PRPS1,NBPF9,NT5D

NB2,ABCA3,ALKBH5,PDXK,YEATS2,TCP1,AHCYL1,C

TUBB2A,OPTN,MVB12B,CCND1,AZIN1,KDELR
)AC2,IPO5,TMEM30A,SKP1,DNER,L
ARCHF6,WSB1,RPS17,VCL,AP3D1,DNA
MP1,LASP1,BEX3,CCT7,OAZ1,SRSF4,RT
MEGF8,ACSL6,RAB7A,ITM2C
;,SLIT3,STT3B,ISYNA1,LONP1,HGS
?,TMEM248,DDX42,CCT4,GCN1,RP
19SF2,TSR1,NOX4,SSRP1,MT
S16,LDHA,RPL29,TPM4,CTSD,LAMA5,RPS9,PJ
1B,SLC6A8,TRAM1,GPC2,ARHGEF17,VPS4
7,SUPT5H,HNRNPUL1,HSPB1,S
D2,KLHL42,CSNK1G2,DIP2B,GARS1,GN
0,MYH10,DUX4,YWHAB,PTGES3,CADM4,C
2,ACTN4,MMP14,PRDX2,DDX3X,RPS15,
IRBP,RPL36,YWHAB,CADM4,LAMP1,RHOB,
AP1A,HS6ST1,CTDSP2,CPSF2,NOVA2
TUBGCP6,NUCKS1,SLC7A1,DICER1,DCUN1D
WD,PTPN23,USO1,RCN2,SLC38A10,ARF5,PAF
2,MSN,SLC16A7,KPNB1,RPL17,C1ORF
N1,CHPF,JUN,CD63,NGRN,D
_,SLC25A3,RPLP2,NDFIP1,MYL6,ANKRD52
NRNPC,TADA2B,XPR1,KHDRBS1,XP
3,DPYSL4,METRN,PTPN23,UNC119B,SNX2,
)C,BCLAF1,RPL23A,FAM199X,SERF2
)6,RNF187,U2AF2,RPN2,TBCD,PPP1CB
2C,ARL4C,CLTC,MAGED2,GNB2,RPS5,IK
3,ANKS1A,MSH6,DHCR7,NOC2L,FBLN5,
GPR107,PTPRS,GHITM,SOD1,PFN2,ADAR,CTNN
SF2,EID1,CTSB,U2AF2,TBCD,MLXIP,SUG
IIZ1,TIMP2,SARAF,COL11A1,MIDN,RPL10,
SPYL4,NOX4,SSRP1,PRDX5,ACTR3,PKN1,TCP1,
IA1,SOX12,RAB7A,DDX3X,RPS15,CLTC,GNB
D2,GNB2,RCN1,H1-0,RPS5,IK,
L1,VDAC1,TRAF7,EIF2AK1,ED
/OU1,ID4,ZFAND3,HECTD1,BA
RAP,SARS1,IFITM3,VANGL2,BAG6,SR
IRNPDL,TARDBP,SLC25A3,DDOST,TSR2,
BIN1,OPA1,MBD6,VPS16,PSMA4,NUP50,
1,GPC1,TOP2B,SLC35E2B,ATP6V0C,DNER,LA
GAM1,NFE2L1,HNRNPA0,RPL23,CPE,CNN
_,TARDBP,RPLP2,ARL6IP1,DDOST,TA
OB,CKAP4,RPS12,HLA-B,CALU,ARCN
),PFKM,TNKS2,STT3A,LONP1,SF3A1,H
3XW5,VASH1,PRKACA,CREBBP,UBE2I,MI
/CAN,TM9SF2,TSR1,MORF4L2,NOX4,S

,BRD2,KLHL42,HOXB5,HJURP,GARS1,GN
PRC,KHDRBS1,SLC3A2,CSNK1E,P
MYEF2,TLE4,PRDX4,ID2,SH3KBP1,PPP4R3B,C
ES3,SEPTIN11,RHOB,LGALS3BP,UBE2H,
,VDAC1,TRAF7,EIF2AK1,PRMT1
52,SLC39A7,KLHL42,PLXNB1,DECR1
PTCD3,ZNF865,ATF4,PSMD2,NR2F
R2,NDUFV1,LRRC4B,H2AZ1,MMP15,FLOT1,
42,PLXNB1,DECR1,FARP1,RER1,P
NN2,NOVA2,NUCB1,TM9SF3,CHD6,EEF1A
RPL29,TNS3,EIF4H,SCARB2,NACA,TSPA
USF2,LGALS1,USP10,HDGFL3,CLIP2,LUC7L2,T
IMT1,VAPB,RTL6,CCN1,SNN,CYTH2,SHC2,FKBP
T2A,GTF2F1,GOLGA8A,OGA,PGAM1,SM
3P,RPS25,TKT,EIF5A,PDIA6,DDR
CAR2,DLK1,POLR2B,XPNPEP3,KIF3C,SSB,LARS
EPTIN7,PSMD8,HIF1A,SAFB2,FUS,PPP1
NARS1,DUX4,CKAP5,RHOB,HNRNPH1,H
,NNAT,MAP1A,FAM199X,DLG5,CRPPA,HS
DBP,DDOST,ST3GAL2,ANKRD52,H1-1
2B,XPR1,KHDRBS1,SLC3A2,CSNK1
LC4A8,GOLGA8A,HNRNPU,PGAM1,NFE2L
,SPTBN2,SLC3A2,SCRN1,HDLBP,RPS2
C2L,COX6A1,EIF3G,SART1,UBE3A,N
AT1,OAZ1,PFN1,JPT2,TLE5,NES,RP
CD3,DCHS1,SET,EGLN1,MAP9,TPI1,CHSY
IA,STT3B,PITPNM1,SF3A1,ATXN2L,TM
TOMM20,U2AF2,EEF1A2,FDFT1,CC
ACP1,RBM17,PRMT2,PPP1R12A,NADK,
FAM171A2,AEBP1,ACADVL,ADAM9,PTBP1,
PF4B,AHSA1,MTMR4,PDPK1,MKRN1,YIPF3,
APKAPK2,STUB1,DAAM1,EMC10,ZSWIM6,
S27A,TMEM167A,SEPTIN7,KLHDC
6,DIP2A,MZT2B,FXR2,PAIP2,H6PD,CLIC
1A,TTR,PLPPR3,ARCN1,ZNF454,HECTD

7A1,PDPK1,MSI1,PSMD4,LRRC41,ATP6V0D1,SH3BP
PNN,HNRNPC,LRPPRC,TADA2B,SPTBN2,SLC
B,PRMT1,XPOT,KDM5C,EXTL3,RERE
AB14,MAZ,KIAA0232,TUBGCP6
CLAF1,CDKN1A,FAM199X,SHISAL1,SERF2,C
RNPDL,TARDBP,RPLP2,DDOST,
SLC20A1,AMBRA1,REEP1,METRN,EIF4A3,SLC38A

,PREPL,GPSM1,RPS2,ZMYM4,APEX1,NPLOC

JPT6H,DHX30,PFKM,TNKS2,STT3A,
;TSB,MDH2,CHD6,RAB27A,U2AF2,E
MS,CDC42,TRAF4,KCNQ2,MTA1,PRKAR2A

GID8,TMEM131,RTF2,COLGALT1,SENP6,RANB
,ANP32A,CIC,RBBP7,WDR45B,NAP1L1,TXNDC5,
D,EIF5,CD81,LENG8,WDR1,ATXN7L3B,HM
SPAN7,SEMA4C,ZEB1,WDR82,SYNGR1,MRPS
NMA1,PDIA6,SULF1,PIP4K2B,MEGF
,UBQLN2,MMP15,FLOT1,CD99,SEC16A,
ASTL,YKT6,MCM3AP,RPS27A,TMEM
,GPC4,LRRC59,RHBDD2,ACLY,XRCC6,TAO
2,CUL3,NDST1,ULK1,NPDC1,BRSK1,GPSM1,I
I2,CHMP1A,TSNAX,H6PD,ZKSCAN1,AN
F302,DDX42,ORAI2,SKI,H3-3A,SCA
,LENG8,WDR1,ATXN7L3B,HMGN2,ATP
LMAN2,CHMP1A,PSMB1,TSNAX,RUVBL2,PCM1
MGCS1,GJA1,CTNNB1,ANXA6,NGRN,MA
MLXIP,PGD,VAMP2,GAS7,NOP53,GDI2
RFGEF2,WBP2,AKIRIN1,KDM4B,CDC42,
FAP1,GNB2,RPS5,IK,RPL35,CHPF,JUN
GCS1,GJA1,CTNNB1,ANXA6,RPL35A
ATP2A2,CRMP1,LASP1,BEX3,VAT1,SRSF4
N1,HM13,STUB1,DEGS1,NSD1,PIP5K1C,
3,ANKRD50,SOX9,ZFP91,DIRAS1,NAPB,IRF2BP2
0C,SKP1,PPIB,RPL23A,HSBP1,S
,TNFRSF10B,MRPS6,MEX3C,ILRUN,PLPPR2,Z
P2B,MMP2,SKP1,BCLAF1,LAPTM4B,RPL23A
MRPS6,CRTC1,CACNA1H,SETX,ZBED4,MB

DNP,SART3,EWSR1,SUB1,ARL6IP5,PNRC2,NT
ISCH,PRKCSH,HNRNPC,LRPPRC,K
F3,NCAM1,AGRN,RPLP1,SARAF,COL11A
CBX6,SPINT2,TMEM167A,LMAN1,HNRNPAB,
R9B,YME1L1,RBM15B,RBBP7,TRAF7,EIF2A
ATP6V0C,MMP2,BCLAF1,L1CAM,L
3,FBLN5,AP2A2,CREBBP,APBB1,ATXN7L3,TTC28,WN
ALS1,SGSM2,AUTS2,SCAMP1,LUC7L2,RTF1,DA
2,ZNF43,BASP1,VAPA,HMGB1,POLR2A
PLXNA1,PIP4K2B,RAB7A,DDX3X,RP
APBP,CDC34,HMGA1,RSF1,LGALS1,ITFG1,SGSM2,
H1,RHBDD2,ERH,EIF2S2,CYTH2,SN
5,RPS19,RPL35,MIF,GPAT4,JUN,C
TC28,TOMM70,SORT1,CPXM1,REEP1,DM

,ATP6V1B2,TNKS2,ISYNA1,SF3A1,ZFA
BRD2,KLHL42,SLC35F5,CSNK1G2,DIP
PS15,RPL12,ITM2C,ARL4C,EIF3L,NANO
,ARHGDIA,CDK16,TOP2B,SLC3
CA,VPS51,PPME1,AREL1,C1QBP,FAM32A,
MC1,DDX39B,NOVA2,SRP9,CTSB,SRSF5,CDH1
GFBI,LMNB2,ABCA3,SNRPN,HNRNPA3,PRDX5,
PL7L1,RPS27A,TMEM167A,KLHDC3,TNKS,PSM
BP1A,CTTN,SLIT2,SERP1,WSCD1,VPS26B,ERH
I1,KCNQ2,BCR,MTA1,KIRREL1,RERE,STA
1,EDC4,PRMT1,XPOT,EXTL3,
C,RPL30,CLTC,EIF3L,MAGED2,
C,RPL30,CLTC,EIF3L,MAGED2,
BS1,XPO1,SPTBN2,CSNK1E,PFKL,
,EIF2AK1,EDC4,WDR45B,PRMT1,XPOT,PDA
MEM30A,SKP1,DNER,TRIB2,PPIB,GLU
B2A,RPL37,VHL,MAPK1,SMARCA4,AT
RPLP2,DDOST,TAGLN,RPL36,RHOB,UBE2H,AT
GTF3C1,HIP1R,YTHDF3,SLC25A6,EXOC5,NDUF
A1,IER2,EIF4G1,PAFAH1B2,POLDIP2,TRI
RAB11A,ARNT2,SERBP1,AKIRIN1,KDM4
PG2,SHC2,FKBP4,OLFM1,CARM1,UBR5
B1,ATF7IP,DAD1,PPME1,SRSF10,EX
AF,RPL10,MARCHF6,WSB1,RPS17,VCL,DN
MMP15,SEC16A,PHB2,APMAP,GSPT1,UBA

,HNRNPDL,TARDBP,SLC25A3,DD
4A,TBC1D16,ACO2,HSPA4,TCP1,AHCY
TCEA1,IGF1R,SLC25A6,WDR26,SUMO1,ADNP,KAT2
IRBP,RPL36,YWHAB,CADM4,SND1,
OX1,DDIT4,BEX4,VDAC1,SSR1,WDR45B,
XNA1,PIP4K2B,SOX12,ACTN4,
RD5,SON,BNIP3L,ARF4,ZFAND5,HNR
ABL1,HTATSF1,LMAN2,USP10,CGGBP1,KDM3
A2,CSNK2A1,MMP15,SEC16A,APMAP,GSPT1
4A1,MRC2,RPL19,PPP2R1A,RPL15,GTF3
B3GAP2,TRA2B,MASTL,YKT6,MCM
1,RPS27,HRNR,ARHGDIA,STIP
IF358,BTF3L4,UQCRH,TFDP1,LRRN1,GMF
IF358,BTF3L4,UQCRH,TFDP1,LRRN1,GMF
XB3,DHFR,ACBD3,CRABP1,PDCD11,MCFD2,EI
N1,PCED1A,VOPP1,PPP1R26,TGFBR1,THRA,TMEM1
,SSR3,GNL1,PNISR,STK24,SRSF2
KK3,PPP1R15A,GALNT11,GPC4,MAP1LC3B,MTCH
,CTIF,BTG1

2R2,ACTN1,HSPB1,CMIP,BNIP3L,FKBP8
PS12,HLA-B,RAB1A,SEPTIN3,ZNF454,COTL1,H
DUFV1,KDM5B,SF1,RETREG2,CNTNAP1,FLCN,N
ERH,EIF2S2,CYTH2,EIF3H,SNRPD3,
RER1,UQCRC1,MBTPS1,RGMB,ZNF512B,CEMIP
P,FBXO21,BRK1,SERP1,BAP1,GRN,SPG7,BAZ
1,TCEA1,PFKP,PCM1,SLC38A1,SLC25A
,TNKS2,SF3A1,NT5DC2,SARS1,BAG6,
X1,SYP,PPDPF,VDAC1,TRAF7,DFFB,ARIH1,SGTA
BE2M,VPS41,RALY,SEC14L1,SMG5,NU
BE2M,VPS41,RALY,SEC14L1,SMG5,NU
VPS26B,BAP1,SEPTIN5,VAPB,INTS3,RAB1B,GPC2,
NKS,CHERP,CTTN,LRP10,DAP,FBXO21,BR
AZ2,POM121C,LMAN1,TNKS,DKK3,A2M,
X4,CADM4,LAMP1,HNRNPH1,UBE2H,CKAP4
H1,PRMT1,LBH,STARD7,SF1,HIP1,SLC39
N4,DDX3X,ARL4C,CLTC,GNB2,H1
1,COL5A2,WDR1,ATXN7L3B,HMGN2,BEX3
,DKK3,PPP1R15A,GALNT11,GPC4,MAP1LC3B,MT
A1210,PFKP,FAM50A,DCAF5,CRIP2,CD2BP2,JAM3,R
1,SAP18,NDUFA13,PSMC1,TOMM70,XPO6,ATP5M
HNRNPDL,TARDBP,SLC25A3,DDOST,
,IMPDH2,PGAP6,RAB14,LDLR,GNL1,EFTUD2,LR
FAM32A,LSM4,OTUD5,EPN1,CDK9,ZRANB2,EXOC7,P
PS26B,STK35,PPP1R10,ERH,PSMB4,IARS2,
LONP1,SF3A1,HYOU1,ID4,NBPF9,SARS

LXNA1,MEGF8,SOX12,RAB7A,PRDX2,RPL12
MT1,XPOT,EXTL3,STARD7,PD
POLDIP2,PTPN11,OLFM2,CELSR1,HMGA1,AG
BFD1,IARS1,MASTL,YKT6,MCM3AP,SPINT
H1,TRIM24,AKAP17A,PPFIA3,RAP1B,PLXND1,MYL9,
RPN,PRDX5,C4ORF3,PDXK,PKN1,PLOD1,
OLDIP2,TRIP12,XRN2,CHMP1A,PSMB1,TSN
,KDELR1,RPL28,CCAR2,ZFAND5,DUSP4,PITPN
PPP1R15A,GALNT11,MVD,DAP,BRK1,SERP
NACC1,ERBB2,CSRNP2,ARSB,CHGB,NEO1,BME
CT8,LSS,STT3B,ATXN2L,IDH1,STRAP,HECT
3B2,CDC42BPB,ITM2B,ZNF770,MAT2
R1,ABL1,ZZEF1,LUC7L2,H6PD,ATP1
KT6,MCM3AP,SPINT2,PREPL,RPL7L1,RPS
1,AGAP1,SEMA6B,LUC7L2,RTF1,TSPYL2
A3,PAIP1,RAB35,VAV2,LRRN1,HEXIM1,EIF1B,HMG
LOC728392,SF3A1,PRPS1,HYOU1,ZFAND3,HEC
1,MTA1,KIRREL1,TRIO,EP300,RETREG2,

PRPF40A,NAP1L4,TP53BP1,CDC4
NECTIN2,PPP1R26,TMEM50A,PLCD3,RNASEK,TMEM1
PG7,BAZ2A,XRCC6,VAPB,EIF3H,ELAVL1,I
CN6,RAVER1,SNRPB,OTUB1,POGK,STARD3,COX6

TO,ILF3,WDR6,CAPZB,GABARAP,

ZNF770,MAT2A,GTF2F1,SLC4A8,GOLGA

GRMC1,DDX39B,NUCB1,SRSF5,TM9
GFBP3,COL1A2,LLGL1,HK1,NCS1,RHO
K3,HIF1A,LRP10,LRRC59,ZMYM3,FUS,ERH,VAPB
,CRIP2,HMGB3,COX8A,IQSEC1,PPME1,ZCCHC3,H
TBN2,CSNK1E,PFKL,SCRN1,HDLBP,T
1,JPT2,SPC25,NISCH,PRKCSH,CCNI,TA
C2,SETX,ZBED4,RAVER1,MBOAT7,SLC12A7
ZNF428,TRAP1,PLEKHB2,ELP1,SPAG9,MO
GAP2,TAGLN2,MASTL,YKT6,MCM3A
7A,PPT1,ARHGAP23,ANKRD13B,PLXND1,BTG1,
,ACTR3,UBE4A,PKN1,ACO2,HSPA4,T
RP72,TM9SF4,KMT2E,PNISR,EIF4G3,PRR1
T1,EIF3D,SLC39A6,KIF21B,CD99,SEC16A,G
POLDIP3,UNC45A,RNMT,ZBTB4,PSMF1,EIF
11,ANKS1A,SYT11,NOC2L,CHMP4B,CREBB
1R15A,GALNT11,FKBP1A,GPC4,MAP1LC3B,L
L10,MARCHF6,WSB1,RPS17,VCL,A
4,AHCYL1,ETNK1,KLC1,PTOV1,CLSTN3,
,ALKBH5,HNRNPA3,PDXK,UBE4A,YEATS2
,FN1,SPEN,NONO,DDB1,PSMD5,YWHAQ
TD2,PRR12,FLOT2,TMEM63B,DNAJB2,RIC8A,SH
R6,PRKDC,TDRD5,CASC3,CLIP3,RPL28,ZF
,EIF3H,INTS3,ABI2,UBE2M,SEC14L1,ELAVL
1,GPC2,IMPDH2,PGAP6,RAB14,LDLR,GNL1,EF
L1,ELAVL1,NUDCD3,SSR3,GNL1,SH3GL1,
,MCL1,PNN,CLEC7A,NISCH,PR

,SND1,LAMP1,LGALS3BP,RPS12,ZNF4
GLIPR1L2,RPS14,CXADR,CHD4,JU
,PEBP1,CCND2,PLXNA1,ACSL6,SOX
SNRPD3,RALY,SEC14L1,NUDCD3,TAX1B
GPC1,TOP2B,SLC35E2B,IPO5,BCLAF
PSMB4,IARS2,FKBP4,CARM1,STRN4,UBR5,

1,UBE2H,CKAP4,CALU,ARCN1,RPS27
RPL7L1,RPS27A,TMEM167A,KLHDC3,TN

1,SCAF11,ACTN1,BNIP3L,KDELR1,ARF4,ZF
)16,ACO2,HSPA4,AHCYL1,ETNK1,KLC1,P
,SCAP,IER2,POLDIP2,XRN2,LMAN2,CHMP
5,IK,RPS19,RPL35,MIF,PRKAR1A,TLN1

3PXD2B,NTRK2,LHFPL6,ELAVL3,PLS3,IGFBP4,T
D1,BCAR1,BRPF3,GIT1,ADGRL1,EPHB2,CDR2L,CBX
N2,SLC3A2,PFKL,SCRN1,HDLBP,RPS
A2,CSNK1E,PFKL,RPS25,TKT,CDH6,T
LC25A3,RPLP2,ARID1A,NDFIP1,DDOST
A6,TMA7,KIF26A,CD2BP2,RPS21,USP14,IDH2
PO5,ATP6V0C,SKP1,LAPTM4B,
CLTC,MAGED2,APLP1,IK,RPL35,TL
LE1,RDX,IREB2,SRSF2,PRR12,CADM3,NMT1,P
I,CTR9,SNRPB,COX6B1,KDM5A,ZDHHC5,XPC,
865,ATF4,PSMD2,KIF5B,SLC7A5,ENC1,N
RPS27,HMGCR,TOP2B,SMCHD1,ATP6V0
,MIF,HMGCS1,PRKAR1A,TLN1,CHP
TN1,CAND1,CASC3,PLCG1,QARS1,RPL2
,PPDPF,VDAC1,DCTN1,KCNQ2,WDR45B,MTA1,KIRR
OT2,YY1,APBB1,CCDC47,ELOVL5,CENPB,DH
LPPR3,ZNF454,COTL1,HECTD4,MAGED4B,STIP1,
N7,RABAC1,GOLGA8B,NACAD,DLD,PCOLCE,E

PRKCSH,XPR1,KHDRBS1,XPO1,SLC3A2,CS
M4,HNRNPH1,ATP1A2,CKAP4,RPS12,TTR,P
IUP58,CPSF7,C2ORF68,JRK,TOMM40,LEMD2,ALYREF
2,NAPB,TXN,ARL8A,STARD3,DUSP8,DPP7,NUMBL
KN1A,CRPPA,HS6ST1,TIMP3,CPSF2,NOV
ZFAND5,HNRNPUL2-BSCL2,PLTP,CORO1C,SDF
MT2,LZTS2,ARHGAP33,SHC1,RPS6KA2,NEC
N,TGFBI,LMNB2,ABCA3,ACTR3,RTN1,P
NACA,LPCAT1,LAMA5,SLC22A17,BRD
2,STXBP1,UBQLN1,CSNK1D,SF3A1,IFITM3
EG2,NOL4L,MMP15,KIF21B,GSPT1,PPP2
TF2,EIF3B,DEAF1,GBA2,TNFRSF10B,ZBT
RPLP2,ARL6IP1,NDFIP1,ST3GAL2
ALU,ARCN1,RPS27,CCT6A,HMGCR,TOP2B
3A,P3H3,MLF2,SERBP1,KDM3B,AKIRIN1,D
P1,NDFIP1,TSR2,MYL6,SHOX,ANKRD
,KDM2B,POGZ,SCAF8,SLC18A3,ATP8A
A,MMP2,SKP1,L1CAM,TRIB2,PPIB,PEA15,NNA
1A,RPL39,P3H3,CYP51A1,MLF2,FLNC,ARNT2,
AP1,KLC1,NCKAP1,VPS35,ACAP3,MDK,ZNF

:NN2,CTSB,NUCB1,SRSF5,C5ORF24,MDH2,T
32,NACA,TSPAN3,RPS24,BRD3,RPS9,X
)12,XPO6,BAG5,SLC20A1,MICAL1,ATG4B,REE
MMP2,LAPTM4B,RPL23A,PTTG1I
ANXA2,AGRN,RPLP1,ZMIZ1,TIMP2,XAF

ACIN1,COG4,SCOC,AKAP17A,MAPK8IP3,PAF1,PRELI
TLE5,PNN,RPL31,PRKCSH,KLH

TRA1,RAN,DYNLRB1,GALNT17,PPP4R3A,PD
,MAPK1,ATG2B,AZIN1,SRSF9,ARPC2,PS
,DLG5,HS6ST1,CNN2,NOVA2,ZC3H7B,CHD
L24,KPNB1,COL1A2,LLGL1,GAP43,N
MYO5A,COL6A2,ANXA6,NGRN,
P4R3A,IQSEC1,FAM8A1,TRIM27,SLC25A1
SYNGAP1,EPN1,CDK9,UBE2Q1,ARPC3,
,CAND1,BNIP3L,NSG2,PLCG1,KDELR1,QA
)2,SLC7A5,ENC1,RPS14,ATP1A1,RPS
3I2,GPC2,ARHGEF17,ELAVL1,TAX1BP1,PG
S1,VANGL2,PFN2,TCTN1,AKIRIN1,BMPR2,C
,ATF4,PSMD2,NR2F1,RAB6B,SLC7A5,NLGN2,
9,LUZP6,GRB2,CDC37,WDR6,RPS11,SCAF11,
T7,BSCL2,SUB1,BANF1,FAF2,COX8A,CFDP1,CUL
L35,TLN1,JUN,MYO5A,COL6A2,C
I3,UBE2I,UBE2D3,XPO6,CCDC47,UBXN6,M

JBXN1,ATF6B,ZNRF1,NR2F6,PPP1R12A,MCRIP1,
P7,PRPF40A,KDM4B,TP53BP1,CD99L2,PCYO

34,TAOK1,IARS2,EIF3H,LTBP4,ELAVL1,NUDCD3

K1,ITGB5,TIMM50,TNKS,APEX1,DKK3,PPP1R
)A1,SORT1,DEK,CYB5B,ELOVL5,SLC
7,NUP98,GOLGA8B,NACAD,OXSR1,EMC10,SE
TAGLN,MYL6,SHOX,WLS,H1-10,CIRBP,M
GEF1,RPL27A,FAM32A,OTUD5,EPN1,CDK9,RTF2,STU
)164,ARID1B,UBAP2L,PI4KA,RAB3GAP2,
MARCHF7,COLGALT1,DAAM1,SRSF7,ZBTB

,BTF3,NCAM1,ANXA2,AGRN,ZMIZ1

MPDH2,GABARAPL1,FAM234A,GNL1,TM9
NRNPL,TSPYL4,MORF4L2,TAF15,HSPH1,HNR

A,CNTN1,LRCH4,TSPAN13,DNAJB11,NAR
2,NACA,TSPAN3,LPCAT1,LAMA5,RPS2
P17A,EIF2S1,MAPK8IP3,PSMC6,PRELID1,HAR
SL6,SOX12,CCN2,PRDX2,ITM2C,RPL
EX,NSF,ADNP,SART3,PNRC2,HMGB3,FAF2,ECH
A,VPS51,FAM8A1,DARS1,H2AX,FAM32A,LSM4,
TAPBP,SPPL2B,ITFG1,SGSM2,AUTS2,MTA2,C
HYOU1,SARS1,IFITM3,PTPRS,HGS,BAG6,HN

CD99L2,YLPM1,SYP,PPDPF,TRAF4,TRA

,CCT6A,CTBP1,HMGCR,TOP2B,SMC
4A,DYNLRB1,TNRC6A,PPME1,MOGS,MAGEF1,RPL27
T4,TLN1,CHPF,JUN,COL6A2,CD63,SESN3
1,SNRNP70,PSMB2,H3-3A,RPL36A,KDM6B,CNT

2,NTRK2,DENND5A,SLC30A9,SLC25A36,XRN
PSMD3,DYNC1LI2,AEBP1,ACADVL,TCF2
P1,ARHGDIA,CAPRIN1,STIP1,CDK16,

4,ANP32E,HSPA1A,MKNK2,GRIK5,IGF1R,MDM2,
5,HNRNPA0,CTNNA1,FADS2,RPS16,RPL
1,ALDH7A1,DHX36,HM13,COLGALT1,TSPAN7
N,GOLPH3,ETF1,ECHS1,LDB1,PSME3,DH
C2,MCRIP1,KPNA3,ZNF395,NEURL4,MAP2K4,RGMA,
M2C,ARL4C,CLTC,NANOG,MRFAP1,GNB2,
PFKP,ARHGEF40,UBE4B,MAPK8IP2,SLC25A6,

NSMF,DUSP8,RNF20,KDM5A,RBM14,NUP133,CHG

,LRRC28,MAPT,FN1,SPEN,NONO,DDB1,

ONEP1,MIB2,XAB2,TMEM59,TRAPPC1,SPG11,TNPO3
H1,CMTR1,TFDP1,CLCN3,TRAPPC1,ABCE1,NOLC1
71A2,PTBP1,DYNLL2,PCSK1N,WDR6,TDRD5,
AV2,SPG11,TNPO3,CUL4B,MCM7,RPL32,VPS
,NADK,SHANK1,ZDHHC9,ELOB,VOPP1,GAL
CDH6,CCND2,PNMA1,GPC3,PIP
B,MAPRE1,CCNL2,BTBD2,FAM168B,

13,COLGALT1,TSPAN7,NUP98,GOLGA8B,NACAD,O
1,SARAF,COL11A1,MIDN,RPL10,
5,MACROH2A1,HSPA1A,MKNK2,GRIK5,PFK

DECR1,DIP2B,UBE2G2,TRA2B,COX7C,CEMIP2,EC
ZRANB2,EXOC7,PSMC5,POLDIP3,NCBP2,RNMT,G
PS4X,OR8K1,STMN1,ITGB1,TUBB4B,
CT2,RPN2,SUGP2,GNAI3,GDI2,HNRNPL,TM9
RN2,LMAN2,CHMP1A,CLTA,ZZE
MECP2,CLUH,IDH2,ALDH18A1,FEZ1,ACAP2,KAT2A
K1A1,NSD1,PLK2,BEX1,NUS1,MED25,CLCN
AM,RPL23A,GLUL,LUC7L3,CTDSP2,DD
G2,SRSF6,UBAP2L,GLUD1,PI4KA,UBE2G2,R
2B,MEGF8,SOX12,MMP14,RPS15,IT

5,ADGRA3,DENND5A,DIP2A,TMCO3,HTATSF1,LGAL
JB1,FEZ1,CNDP2,RNF157,KCTD20,EFNB1,SALL
KDM5C,ACTR1A,EP300,LBH,KDM5B,PBX1,CNTNAP
RK2,DENND5A,SLC30A9,SLC25A36,XRN2,SC

PSMD7,DYNLRB1,GALNT17,ARHGAP5,DAD1
3C,NUS1,CELSR3,MED25,CLCN6,PSMB5,FARS
GFBP3,COL1A2,LLGL1,RPL14,HK1,NC
PZA1,GALNT1,TUBGCP6,RIC8A,PLEKH

GCR,CAPRIN1,TOP2B,SMCHD1,ATP6V0C,
L5,SLC38A10,WBP1,CNBP,IGLON5,TCE

,AHCYL1,ETNK1,KLC1,VPS35,CLSTN3,ACAP3,Z
,FDFT1,TBCD,MLXIP,SUGP2,CPSF1,PPP1CB
C,USP48,ACSL4,SOX9,RPS29,NSMF,DUSP8,RNF2
A2,AGRN,RPLP1,ZMIZ1,TIMP2,XAF1,
RELA,GNAL,PPP2R2A,MRPS26,KHDC4,NFIB

PINT2,CALD1,RPS27A,TMEM167A,ZM

B5C,QPRT,CLASP2,MTMR4,PDPK1,

NLL1,TXNIP,SYP,DHX15,BEX4,KIF5A,SULF2
P14,DDX3X,RPS15,RPL12,ITM2C,ARL4C,E
4,BRD3,RPS9,XRCC5,PJA2,MSN

2,MTRF1L,ABCA3,ACTR3,RTN1,PDXK,PKN1

ASB6,SUDS3,PCDHB2,COL6A3,MIB2,TRAPP
,VANGL2,HGS,SOD1,DGKD,SREBF2,P3H
1,DDX19A,RAP1B,PPM1A,GNAO1,RAD23A,SLC39A1

ARCB1,RAD23A,NF2,MAFB,ANAPC2,STX12,
EC7A,NES,RPL31,NISCH,PRKCSH,CCNI
,DDIT4,DHX15,RBBP7,SSR1,BCL9L,WDR45B,
RDBP,NDFIP1,TAGLN,MYL6,ANKRD52,HNRNPK
PPL1,SMCHD1,TMEM30A,MMP2,SKP
PPL1,SMCHD1,TMEM30A,MMP2,SKP
AP2K2,DIDO1,DUSP4,DLK1,PLTP,SMARC

F43,PAPOLA,COL1A2,GAP43,NCS1,RPL22,CBX5,
F516,ADGRL1,LRP8,SAMD1,PRNP,SE

ZNF770,AP2B1,MAT2A,TMEM170A,
RPLP2,ARL6IP1,NDFIP1,ST3GAL2
CIRBP,NARS1,PTGES3,CKAP5,SEPTI

,NDUFS1,CTR9,VEZF1,ATMIN,SNRPB,NKIRAS2,ZDHHC
BQLN1,MESD,GPR107,GHITM,DGKD,BAG6,HNR

1,CDC42,DDIT4,PPP1R9B,GRIPAP1,PO

P,RPLP2,DDOST,ST3GAL2,MYL6,A

5,CSNK1A1,EXOSC10,NSD1,BEX1,UBE3C,NU
,TADA2B,KHDRBS1,XPO1,CSNK1E,PFK
5,SHISAL1,SERF2,CRPPA,TMEM132

IN1,STIP1,SLC35E2B,APPL1,SMCHD
D23A,SLC39A1,EPHA2,SKIL,STX12,VPS37B,RBBP6,
SPT1,PLXNB1,SLC35F5,CSNK1G2,RAB2A,F

,RNF20,NUP133,PER1,THBS2,CHGB,COX4I1,OGDH,
D1,CYTH3,PPP1R16A,ELOA
C1,MEIS3,CTDNEP1,MIB2,XAB2,TMEM59,TRAPP
P2,WBP11,BORCS7,SYT11,NSD2,IST1,PR

GSM2,ZZEF1,AASDHPPT,ESYT1,TSNAX,GTF3
GNAO1,RAD23A,MRPL28,SLC39A1,CYTH3,EPHA2

DUFA10,FKBP3,CCDC124

NPS1,AKAP17A,MAPK8IP3,PAF1,DDX19A,RAP1B,PP
7L1,POM121C,APEX1,LOX,MAP1LC3B,MVD,C
UP98,NACAD,OXSR1,EMC10,SEMA4C,TMEM9,S
LHL42,HOXB5,DIP2B,CD164,ARID1
LHL42,HOXB5,DIP2B,CD164,ARID1
4,FAM53C,TNRC6A,GALNT17,CSNK2B,RAP

RPS24,BRD3,SLC16A7,RPL17,C1ORF43,PA

ZP6,PRKDC,SUPT5H,HSPB1,SON,MAP2K2,

SMD10,GSTA4,GUK1,UBN1,PRPF31,NECTIN2,PLC

3,UBE4A,ACO2,HSPA4,TCP1,AHCYL1,PTOV1,RB

R45B,SGTA,KIRREL1,ACTR1A,EXTL3,SF1

27A,TIMM50,TNKS,APEX1,NPLOC4,DKK3,PPP
K1,PITRM1,DYNLRB1,VPS51,SOBP,TBXA2R,AS

PD,VGF,MLF2,ARFGEF2,AKIRIN1,SMS,DYN
3C,AHSA1,SPAG9,ACTR1B,KRT8,MTM

EC,ACAP3,DDX42,LAMC1,CBX1,PTK
1,NAP1L4,TP53BP1,AMOTL2,YME1L1,TRA
DX5A,PCED1A,SUDS3,GABARAPL2,RGMA,ZNF598,C

,SUPT6H,RPS28,HNRNPM,SDHA,NKTR,YW
C3H4,NDUFA10,FKBP3,CCDC124

1,NOLC1,MSI2,CCNL1,NBR1,RNPS1,MAPK8IP3,
DUFA10,FKBP3,CCDC124

A6,TRMT112,KIF26A,CD2BP2,USP14,EWSR1,S

| | | | | | | |
|---|---|---|---|---|---|---|
| **MIRNA** | hsa-miR-615- | MIRNA:hsa-m | 4.29E-72 | 71.3670862 | 892 | 2593 |
| **MIRNA** | hsa-miR-484 | MIRNA:hsa-m | 8.21E-62 | 61.0858935 | 891 | 2593 |
| **MIRNA** | hsa-miR-92a- | MIRNA:hsa-m | 3.58E-46 | 45.446627 | 1405 | 2593 |
| **MIRNA** | hsa-miR-16-5 | MIRNA:hsa-m | 1.71E-39 | 38.7658086 | 1556 | 2593 |
| **MIRNA** | hsa-miR-331- | MIRNA:hsa-m | 5.30E-39 | 38.275587 | 404 | 2593 |
| **MIRNA** | hsa-miR-877- | MIRNA:hsa-m | 7.13E-33 | 32.1471071 | 603 | 2593 |
| **MIRNA** | hsa-miR-320a | MIRNA:hsa-m | 2.11E-31 | 30.6756626 | 584 | 2593 |
| **MIRNA** | hsa-miR-744- | MIRNA:hsa-m | 3.12E-31 | 30.5059598 | 435 | 2593 |
| **MIRNA** | hsa-miR-193b | MIRNA:hsa-m | 6.11E-28 | 27.2142114 | 848 | 2593 |
| **MIRNA** | hsa-let-7b-5p | MIRNA:hsa-le | 1.26E-26 | 25.9012169 | 1213 | 2593 |
| **MIRNA** | hsa-miR-15b- | MIRNA:hsa-m | 2.85E-25 | 24.5457166 | 760 | 2593 |
| **MIRNA** | hsa-miR-218- | MIRNA:hsa-m | 2.46E-21 | 20.6094322 | 815 | 2593 |
| **MIRNA** | hsa-miR-423- | MIRNA:hsa-m | 9.10E-21 | 20.0407572 | 343 | 2593 |
| **MIRNA** | hsa-miR-222- | MIRNA:hsa-m | 1.72E-20 | 19.7635038 | 393 | 2593 |
| **MIRNA** | hsa-miR-186- | MIRNA:hsa-m | 4.34E-20 | 19.3628058 | 749 | 2593 |
| **MIRNA** | hsa-miR-324- | MIRNA:hsa-m | 1.37E-19 | 18.8628644 | 297 | 2593 |
| **MIRNA** | hsa-miR-324- | MIRNA:hsa-m | 1.50E-19 | 18.824702 | 337 | 2593 |
| **MIRNA** | hsa-miR-149- | MIRNA:hsa-m | 2.14E-19 | 18.6686903 | 394 | 2593 |
| **MIRNA** | hsa-miR-23b- | MIRNA:hsa-m | 1.74E-18 | 17.7604339 | 319 | 2593 |
| **MIRNA** | hsa-miR-423- | MIRNA:hsa-m | 3.62E-18 | 17.4410211 | 229 | 2593 |
| **MIRNA** | hsa-miR-1-3p | MIRNA:hsa-m | 3.68E-18 | 17.4341276 | 915 | 2593 |
| **MIRNA** | hsa-miR-30a- | MIRNA:hsa-m | 4.42E-18 | 17.3548507 | 730 | 2593 |
| **MIRNA** | hsa-miR-155- | MIRNA:hsa-m | 2.32E-16 | 15.6338947 | 900 | 2593 |
| **MIRNA** | hsa-miR-877- | MIRNA:hsa-m | 2.92E-15 | 14.5350381 | 233 | 2593 |
| **MIRNA** | hsa-miR-27b- | MIRNA:hsa-m | 3.74E-15 | 14.427616 | 419 | 2593 |
| **MIRNA** | hsa-miR-1226 | MIRNA:hsa-m | 1.05E-14 | 13.9792926 | 226 | 2593 |
| **MIRNA** | hsa-miR-30c- | MIRNA:hsa-m | 1.67E-14 | 13.777475 | 519 | 2593 |
| **MIRNA** | hsa-miR-15a- | MIRNA:hsa-m | 2.59E-14 | 13.5865923 | 717 | 2593 |
| **MIRNA** | hsa-miR-34a- | MIRNA:hsa-m | 4.54E-14 | 13.3425494 | 729 | 2593 |
| **MIRNA** | hsa-miR-195- | MIRNA:hsa-m | 1.19E-13 | 12.9241705 | 639 | 2593 |
| **MIRNA** | hsa-miR-100- | MIRNA:hsa-m | 2.46E-13 | 12.6092959 | 250 | 2593 |
| **MIRNA** | hsa-miR-486- | MIRNA:hsa-m | 4.12E-13 | 12.3849573 | 201 | 2593 |
| **MIRNA** | hsa-miR-17-5 | MIRNA:hsa-m | 6.00E-13 | 12.2217448 | 1174 | 2593 |
| **MIRNA** | hsa-miR-26a- | MIRNA:hsa-m | 3.07E-12 | 11.5128411 | 457 | 2593 |
| **MIRNA** | hsa-miR-29b- | MIRNA:hsa-m | 4.90E-12 | 11.3101764 | 260 | 2593 |
| **MIRNA** | hsa-miR-92b- | MIRNA:hsa-m | 8.52E-12 | 11.069465 | 708 | 2593 |
| **MIRNA** | hsa-miR-6797 | MIRNA:hsa-m | 1.33E-11 | 10.8754229 | 275 | 2593 |
| **MIRNA** | hsa-miR-1249 | MIRNA:hsa-m | 1.67E-11 | 10.7767583 | 272 | 2593 |
| **MIRNA** | hsa-miR-6778 | MIRNA:hsa-m | 2.10E-11 | 10.6768322 | 191 | 2593 |
| **MIRNA** | hsa-miR-1233 | MIRNA:hsa-m | 3.36E-11 | 10.4740175 | 196 | 2593 |
| **MIRNA** | hsa-miR-125b | MIRNA:hsa-m | 6.24E-11 | 10.204781 | 426 | 2593 |
| **MIRNA** | hsa-miR-221- | MIRNA:hsa-m | 9.06E-11 | 10.0428282 | 366 | 2593 |
| **MIRNA** | hsa-miR-185- | MIRNA:hsa-m | 1.84E-10 | 9.73567453 | 357 | 2593 |
| **MIRNA** | hsa-miR-651- | MIRNA:hsa-m | 6.11E-10 | 9.21422049 | 224 | 2593 |
| **MIRNA** | hsa-miR-183- | MIRNA:hsa-m | 7.30E-10 | 9.13678319 | 347 | 2593 |

| | | | | | | |
|---|---|---|---|---|---|---|
| **MIRNA** | hsa-miR-18a- | MIRNA:hsa-m | 8.97E-10 | 9.04704839 | 260 | 2593 |
| **MIRNA** | hsa-miR-939- | MIRNA:hsa-m | 1.47E-09 | 8.83298984 | 197 | 2593 |
| **MIRNA** | hsa-let-7e-5p | MIRNA:hsa-le | 1.55E-09 | 8.81008762 | 608 | 2593 |
| **MIRNA** | hsa-miR-608 | MIRNA:hsa-m | 1.82E-09 | 8.73950596 | 194 | 2593 |
| **MIRNA** | hsa-miR-134 | MIRNA:hsa-m | 2.26E-09 | 8.64628327 | 191 | 2593 |
| **MIRNA** | hsa-miR-296- | MIRNA:hsa-m | 2.37E-09 | 8.62543959 | 256 | 2593 |
| **MIRNA** | hsa-miR-317 | MIRNA:hsa-m | 2.53E-09 | 8.59619052 | 272 | 2593 |
| **MIRNA** | hsa-miR-27a- | MIRNA:hsa-m | 4.47E-09 | 8.34974907 | 427 | 2593 |
| **MIRNA** | hsa-miR-424- | MIRNA:hsa-m | 4.58E-09 | 8.33883267 | 470 | 2593 |
| **MIRNA** | hsa-miR-607 | MIRNA:hsa-m | 5.24E-09 | 8.28076105 | 279 | 2593 |
| **MIRNA** | hsa-miR-93-3 | MIRNA:hsa-m | 6.55E-09 | 8.18397599 | 268 | 2593 |
| **MIRNA** | hsa-miR-465 | MIRNA:hsa-m | 6.88E-09 | 8.16248166 | 191 | 2593 |
| **MIRNA** | hsa-miR-497- | MIRNA:hsa-m | 7.66E-09 | 8.11551596 | 460 | 2593 |
| **MIRNA** | hsa-miR-18a- | MIRNA:hsa-m | 1.20E-08 | 7.91909219 | 348 | 2593 |
| **MIRNA** | hsa-miR-314 | MIRNA:hsa-m | 1.24E-08 | 7.90559433 | 243 | 2593 |
| **MIRNA** | hsa-miR-93-5 | MIRNA:hsa-m | 1.95E-08 | 7.71043117 | 1213 | 2593 |
| **MIRNA** | hsa-miR-683 | MIRNA:hsa-m | 2.00E-08 | 7.69959255 | 431 | 2593 |
| **MIRNA** | hsa-miR-682 | MIRNA:hsa-m | 2.04E-08 | 7.68997634 | 380 | 2593 |
| **MIRNA** | hsa-miR-129 | MIRNA:hsa-m | 2.22E-08 | 7.65308283 | 119 | 2593 |
| **MIRNA** | hsa-miR-92a- | MIRNA:hsa-m | 2.61E-08 | 7.58320298 | 246 | 2593 |
| **MIRNA** | hsa-miR-30d- | MIRNA:hsa-m | 4.27E-08 | 7.3691958 | 367 | 2593 |
| **MIRNA** | hsa-miR-17-3 | MIRNA:hsa-m | 5.68E-08 | 7.24537052 | 172 | 2593 |
| **MIRNA** | hsa-miR-673 | MIRNA:hsa-m | 6.09E-08 | 7.21566402 | 352 | 2593 |
| **MIRNA** | hsa-miR-808 | MIRNA:hsa-m | 6.09E-08 | 7.21566402 | 352 | 2593 |
| **MIRNA** | hsa-miR-477 | MIRNA:hsa-m | 6.39E-08 | 7.19422599 | 286 | 2593 |
| **MIRNA** | hsa-miR-378 | MIRNA:hsa-m | 6.47E-08 | 7.18921235 | 165 | 2593 |
| **MIRNA** | hsa-miR-687 | MIRNA:hsa-m | 8.60E-08 | 7.06548118 | 247 | 2593 |
| **MIRNA** | hsa-miR-196 | MIRNA:hsa-m | 1.40E-07 | 6.85377944 | 302 | 2593 |
| **MIRNA** | hsa-miR-7-5p | MIRNA:hsa-m | 1.50E-07 | 6.82424691 | 576 | 2593 |
| **MIRNA** | hsa-miR-126 | MIRNA:hsa-m | 1.65E-07 | 6.78170379 | 270 | 2593 |
| **MIRNA** | hsa-miR-675 | MIRNA:hsa-m | 1.73E-07 | 6.76132238 | 246 | 2593 |
| **MIRNA** | hsa-miR-769- | MIRNA:hsa-m | 1.97E-07 | 6.70564618 | 165 | 2593 |
| **MIRNA** | hsa-miR-190 | MIRNA:hsa-m | 2.54E-07 | 6.59458147 | 181 | 2593 |
| **MIRNA** | hsa-miR-29a- | MIRNA:hsa-m | 2.64E-07 | 6.57885668 | 264 | 2593 |
| **MIRNA** | hsa-miR-672 | MIRNA:hsa-m | 3.48E-07 | 6.45837107 | 233 | 2593 |
| **MIRNA** | hsa-miR-10a- | MIRNA:hsa-m | 3.99E-07 | 6.39892904 | 462 | 2593 |
| **MIRNA** | hsa-miR-676 | MIRNA:hsa-m | 4.34E-07 | 6.36213841 | 246 | 2593 |
| **MIRNA** | hsa-miR-122 | MIRNA:hsa-m | 5.27E-07 | 6.27853746 | 207 | 2593 |
| **MIRNA** | hsa-miR-493- | MIRNA:hsa-m | 7.03E-07 | 6.15327029 | 87 | 2593 |
| **MIRNA** | hsa-miR-652- | MIRNA:hsa-m | 7.29E-07 | 6.13727249 | 162 | 2593 |
| **MIRNA** | hsa-miR-29c- | MIRNA:hsa-m | 8.35E-07 | 6.07849715 | 253 | 2593 |
| **MIRNA** | hsa-miR-30b- | MIRNA:hsa-m | 8.49E-07 | 6.07087701 | 414 | 2593 |
| **MIRNA** | hsa-miR-505- | MIRNA:hsa-m | 8.90E-07 | 6.05059524 | 133 | 2593 |
| **MIRNA** | hsa-miR-99b- | MIRNA:hsa-m | 9.37E-07 | 6.02840288 | 53 | 2593 |
| **MIRNA** | hsa-miR-491- | MIRNA:hsa-m | 1.0801E-06 | 5.96654977 | 171 | 2593 |

| | | | | | | |
|---|---|---|---|---|---|---|
| **MIRNA** | hsa-miR-453: | MIRNA:hsa-m | 1.1098E-06 | 5.95476656 | 230 | 2593 |
| **MIRNA** | hsa-miR-612( | MIRNA:hsa-m | 1.148E-06 | 5.94005125 | 300 | 2593 |
| **MIRNA** | hsa-miR-672: | MIRNA:hsa-m | 1.1776E-06 | 5.92900613 | 179 | 2593 |
| **MIRNA** | hsa-miR-23a- | MIRNA:hsa-m | 1.3307E-06 | 5.87592078 | 247 | 2593 |
| **MIRNA** | hsa-miR-476: | MIRNA:hsa-m | 1.3591E-06 | 5.86675064 | 280 | 2593 |
| **MIRNA** | hsa-miR-451( | MIRNA:hsa-m | 1.3942E-06 | 5.85566625 | 301 | 2593 |
| **MIRNA** | hsa-miR-765 | MIRNA:hsa-m | 1.4836E-06 | 5.82867196 | 348 | 2593 |
| **MIRNA** | hsa-miR-612 | MIRNA:hsa-m | 2.047E-06 | 5.68887904 | 303 | 2593 |
| **MIRNA** | hsa-miR-441( | MIRNA:hsa-m | 2.1305E-06 | 5.67151078 | 299 | 2593 |
| **MIRNA** | hsa-miR-613( | MIRNA:hsa-m | 2.1305E-06 | 5.67151078 | 299 | 2593 |
| **MIRNA** | hsa-miR-96-5 | MIRNA:hsa-m | 2.1979E-06 | 5.65799202 | 201 | 2593 |
| **MIRNA** | hsa-miR-30e- | MIRNA:hsa-m | 2.2246E-06 | 5.65273858 | 346 | 2593 |
| **MIRNA** | hsa-miR-568 | MIRNA:hsa-m | 2.2847E-06 | 5.64117386 | 136 | 2593 |
| **MIRNA** | hsa-miR-425- | MIRNA:hsa-m | 2.7128E-06 | 5.56659009 | 144 | 2593 |
| **MIRNA** | hsa-miR-687( | MIRNA:hsa-m | 2.7698E-06 | 5.55755501 | 198 | 2593 |
| **MIRNA** | hsa-miR-447( | MIRNA:hsa-m | 2.7698E-06 | 5.55755501 | 198 | 2593 |
| **MIRNA** | hsa-miR-20a- | MIRNA:hsa-m | 2.8277E-06 | 5.5485701 | 1064 | 2593 |
| **MIRNA** | hsa-miR-711( | MIRNA:hsa-m | 2.9192E-06 | 5.53473295 | 148 | 2593 |
| **MIRNA** | hsa-miR-120 | MIRNA:hsa-m | 3.655E-06 | 5.4371177 | 285 | 2593 |
| **MIRNA** | hsa-miR-320: | MIRNA:hsa-m | 4.2073E-06 | 5.37599848 | 269 | 2593 |
| **MIRNA** | hsa-miR-613: | MIRNA:hsa-m | 4.5324E-06 | 5.3436735 | 303 | 2593 |
| **MIRNA** | hsa-miR-679 | MIRNA:hsa-m | 5.9822E-06 | 5.22314266 | 162 | 2593 |
| **MIRNA** | hsa-miR-674: | MIRNA:hsa-m | 6.4965E-06 | 5.18731797 | 96 | 2593 |
| **MIRNA** | hsa-miR-760 | MIRNA:hsa-m | 7.1123E-06 | 5.14798796 | 194 | 2593 |
| **MIRNA** | hsa-miR-315: | MIRNA:hsa-m | 7.1428E-06 | 5.14613282 | 198 | 2593 |
| **MIRNA** | hsa-miR-673: | MIRNA:hsa-m | 7.1428E-06 | 5.14613282 | 198 | 2593 |
| **MIRNA** | hsa-miR-684: | MIRNA:hsa-m | 8.5289E-06 | 5.06910746 | 148 | 2593 |
| **MIRNA** | hsa-miR-675: | MIRNA:hsa-m | 8.5289E-06 | 5.06910746 | 148 | 2593 |
| **MIRNA** | hsa-miR-689 | MIRNA:hsa-m | 9.3726E-06 | 5.02813887 | 137 | 2593 |
| **MIRNA** | hsa-miR-471( | MIRNA:hsa-m | 1.0126E-05 | 4.99455646 | 164 | 2593 |
| **MIRNA** | hsa-miR-130I | MIRNA:hsa-m | 1.0497E-05 | 4.9789263 | 571 | 2593 |
| **MIRNA** | hsa-miR-673( | MIRNA:hsa-m | 1.1314E-05 | 4.94636923 | 204 | 2593 |
| **MIRNA** | hsa-miR-451( | MIRNA:hsa-m | 1.1314E-05 | 4.94636923 | 204 | 2593 |
| **MIRNA** | hsa-miR-317: | MIRNA:hsa-m | 1.1662E-05 | 4.93323013 | 134 | 2593 |
| **MIRNA** | hsa-let-7a-5p | MIRNA:hsa-le | 1.183E-05 | 4.92702681 | 637 | 2593 |
| **MIRNA** | hsa-miR-448 | MIRNA:hsa-m | 1.2907E-05 | 4.88916695 | 105 | 2593 |
| **MIRNA** | hsa-miR-318 | MIRNA:hsa-m | 1.4252E-05 | 4.84612708 | 185 | 2593 |
| **MIRNA** | hsa-miR-130 | MIRNA:hsa-m | 1.4337E-05 | 4.84354601 | 197 | 2593 |
| **MIRNA** | hsa-miR-185- | MIRNA:hsa-m | 1.4785E-05 | 4.83018454 | 109 | 2593 |
| **MIRNA** | hsa-miR-679 | MIRNA:hsa-m | 1.5083E-05 | 4.82150329 | 120 | 2593 |
| **MIRNA** | hsa-miR-539- | MIRNA:hsa-m | 1.7513E-05 | 4.75662805 | 210 | 2593 |
| **MIRNA** | hsa-miR-444 | MIRNA:hsa-m | 1.7802E-05 | 4.74952043 | 178 | 2593 |
| **MIRNA** | hsa-miR-447: | MIRNA:hsa-m | 1.7802E-05 | 4.74952043 | 178 | 2593 |
| **MIRNA** | hsa-miR-950( | MIRNA:hsa-m | 1.7894E-05 | 4.74729283 | 202 | 2593 |
| **MIRNA** | hsa-miR-317( | MIRNA:hsa-m | 1.7966E-05 | 4.7455451 | 182 | 2593 |

| | | | | | | |
|---|---|---|---|---|---|---|
| **MIRNA** | hsa-miR-766- | MIRNA:hsa-m | 1.9632E-05 | 4.70704361 | 231 | 2593 |
| **MIRNA** | hsa-miR-4747 | MIRNA:hsa-m | 2.1213E-05 | 4.67339157 | 303 | 2593 |
| **MIRNA** | hsa-miR-5196 | MIRNA:hsa-m | 2.1213E-05 | 4.67339157 | 303 | 2593 |
| **MIRNA** | hsa-miR-5703 | MIRNA:hsa-m | 2.2371E-05 | 4.65031844 | 203 | 2593 |
| **MIRNA** | hsa-miR-4434 | MIRNA:hsa-m | 2.2371E-05 | 4.65031844 | 203 | 2593 |
| **MIRNA** | hsa-miR-3937 | MIRNA:hsa-m | 2.3808E-05 | 4.62328472 | 59 | 2593 |
| **MIRNA** | hsa-miR-711 | MIRNA:hsa-m | 2.412E-05 | 4.61762968 | 257 | 2593 |
| **MIRNA** | hsa-miR-148a | MIRNA:hsa-m | 2.7044E-05 | 4.56792288 | 212 | 2593 |
| **MIRNA** | hsa-miR-6081 | MIRNA:hsa-m | 2.7772E-05 | 4.55639768 | 122 | 2593 |
| **MIRNA** | hsa-miR-4652 | MIRNA:hsa-m | 3.0725E-05 | 4.51251063 | 66 | 2593 |
| **MIRNA** | hsa-miR-4697 | MIRNA:hsa-m | 3.0788E-05 | 4.51161187 | 104 | 2593 |
| **MIRNA** | hsa-miR-503- | MIRNA:hsa-m | 3.1548E-05 | 4.50102182 | 250 | 2593 |
| **MIRNA** | hsa-miR-19b- | MIRNA:hsa-m | 3.2862E-05 | 4.48330472 | 712 | 2593 |
| **MIRNA** | hsa-miR-3150 | MIRNA:hsa-m | 3.7287E-05 | 4.42843828 | 230 | 2593 |
| **MIRNA** | hsa-miR-301a | MIRNA:hsa-m | 4.2934E-05 | 4.36719612 | 395 | 2593 |
| **MIRNA** | hsa-miR-4784 | MIRNA:hsa-m | 4.5609E-05 | 4.34094775 | 231 | 2593 |
| **MIRNA** | hsa-miR-935 | MIRNA:hsa-m | 4.6466E-05 | 4.33286327 | 120 | 2593 |
| **MIRNA** | hsa-miR-181b | MIRNA:hsa-m | 5.5423E-05 | 4.25631128 | 370 | 2593 |
| **MIRNA** | hsa-miR-22-3 | MIRNA:hsa-m | 5.6515E-05 | 4.24783935 | 163 | 2593 |
| **MIRNA** | hsa-miR-3916 | MIRNA:hsa-m | 5.7273E-05 | 4.24204871 | 187 | 2593 |
| **MIRNA** | hsa-miR-206 | MIRNA:hsa-m | 6.238E-05 | 4.20495503 | 121 | 2593 |
| **MIRNA** | hsa-miR-106b | MIRNA:hsa-m | 6.4845E-05 | 4.18812576 | 1084 | 2593 |
| **MIRNA** | hsa-miR-1237 | MIRNA:hsa-m | 8.0673E-05 | 4.09327279 | 107 | 2593 |
| **MIRNA** | hsa-miR-1205 | MIRNA:hsa-m | 8.8997E-05 | 4.05062563 | 100 | 2593 |
| **MIRNA** | hsa-miR-6848 | MIRNA:hsa-m | 8.8997E-05 | 4.05062563 | 100 | 2593 |
| **MIRNA** | hsa-miR-6846 | MIRNA:hsa-m | 8.8997E-05 | 4.05062563 | 100 | 2593 |
| **MIRNA** | hsa-miR-4723 | MIRNA:hsa-m | 0.00010238 | 3.98979419 | 252 | 2593 |
| **MIRNA** | hsa-miR-6791 | MIRNA:hsa-m | 0.00011077 | 3.95559409 | 123 | 2593 |
| **MIRNA** | hsa-miR-1252 | MIRNA:hsa-m | 0.00011295 | 3.94709663 | 154 | 2593 |
| **MIRNA** | hsa-miR-6859 | MIRNA:hsa-m | 0.00011409 | 3.94275539 | 182 | 2593 |
| **MIRNA** | hsa-miR-99a- | MIRNA:hsa-m | 0.00012293 | 3.91034252 | 131 | 2593 |
| **MIRNA** | hsa-miR-4292 | MIRNA:hsa-m | 0.0001383 | 3.85919287 | 120 | 2593 |
| **MIRNA** | hsa-miR-3125 | MIRNA:hsa-m | 0.00014232 | 3.8467352 | 183 | 2593 |
| **MIRNA** | hsa-miR-6870 | MIRNA:hsa-m | 0.00014762 | 3.83085089 | 254 | 2593 |
| **MIRNA** | hsa-miR-6758 | MIRNA:hsa-m | 0.00016837 | 3.77374619 | 259 | 2593 |
| **MIRNA** | hsa-miR-6813 | MIRNA:hsa-m | 0.00020585 | 3.68645413 | 74 | 2593 |
| **MIRNA** | hsa-miR-1296 | MIRNA:hsa-m | 0.00020743 | 3.68312453 | 99 | 2593 |
| **MIRNA** | hsa-miR-152- | MIRNA:hsa-m | 0.00023262 | 3.63335958 | 157 | 2593 |
| **MIRNA** | hsa-miR-130a | MIRNA:hsa-m | 0.00023847 | 3.62256639 | 398 | 2593 |
| **MIRNA** | hsa-miR-6856 | MIRNA:hsa-m | 0.00026537 | 3.57614014 | 253 | 2593 |
| **MIRNA** | hsa-miR-6803 | MIRNA:hsa-m | 0.00026834 | 3.57131148 | 111 | 2593 |
| **MIRNA** | hsa-miR-106b | MIRNA:hsa-m | 0.0002992 | 3.52404099 | 75 | 2593 |
| **MIRNA** | hsa-miR-4756 | MIRNA:hsa-m | 0.00030279 | 3.51885464 | 284 | 2593 |
| **MIRNA** | hsa-miR-31-5 | MIRNA:hsa-m | 0.00030439 | 3.51657036 | 224 | 2593 |
| **MIRNA** | hsa-miR-4688 | MIRNA:hsa-m | 0.00030946 | 3.5093996 | 104 | 2593 |

| | | | | | | |
|---|---|---|---|---|---|---|
| **MIRNA** | hsa-miR-762 | MIRNA:hsa-m | 0.00031714 | 3.49874595 | 123 | 2593 |
| **MIRNA** | hsa-miR-132 | MIRNA:hsa-m | 0.0003569 | 3.44745401 | 285 | 2593 |
| **MIRNA** | hsa-miR-319 | MIRNA:hsa-m | 0.00035764 | 3.4465573 | 72 | 2593 |
| **MIRNA** | hsa-miR-125a | MIRNA:hsa-m | 0.00038049 | 3.41965308 | 268 | 2593 |
| **MIRNA** | hsa-miR-646 | MIRNA:hsa-m | 0.0003972 | 3.40099213 | 251 | 2593 |
| **MIRNA** | hsa-miR-318 | MIRNA:hsa-m | 0.00046119 | 3.33611697 | 164 | 2593 |
| **MIRNA** | hsa-miR-505- | MIRNA:hsa-m | 0.00046223 | 3.33514501 | 222 | 2593 |
| **MIRNA** | hsa-miR-583 | MIRNA:hsa-m | 0.00046735 | 3.33036127 | 156 | 2593 |
| **MIRNA** | hsa-miR-683 | MIRNA:hsa-m | 0.00047513 | 3.32319063 | 87 | 2593 |
| **MIRNA** | hsa-miR-679 | MIRNA:hsa-m | 0.00051812 | 3.28556741 | 73 | 2593 |
| **MIRNA** | hsa-miR-612 | MIRNA:hsa-m | 0.0005343 | 3.2722128 | 181 | 2593 |
| **MIRNA** | hsa-miR-473 | MIRNA:hsa-m | 0.0005794 | 3.23701932 | 288 | 2593 |
| **MIRNA** | hsa-miR-427 | MIRNA:hsa-m | 0.00057984 | 3.23668921 | 236 | 2593 |
| **MIRNA** | hsa-miR-315 | MIRNA:hsa-m | 0.00058206 | 3.23503009 | 84 | 2593 |
| **MIRNA** | hsa-miR-678 | MIRNA:hsa-m | 0.00060874 | 3.21556475 | 232 | 2593 |
| **MIRNA** | hsa-miR-472 | MIRNA:hsa-m | 0.00060874 | 3.21556475 | 232 | 2593 |
| **MIRNA** | hsa-miR-454- | MIRNA:hsa-m | 0.00062319 | 3.20537732 | 396 | 2593 |
| **MIRNA** | hsa-miR-682 | MIRNA:hsa-m | 0.00069404 | 3.15861318 | 103 | 2593 |
| **MIRNA** | hsa-miR-464 | MIRNA:hsa-m | 0.0007041 | 3.15236757 | 250 | 2593 |
| **MIRNA** | hsa-miR-684 | MIRNA:hsa-m | 0.00072648 | 3.13877748 | 162 | 2593 |
| **MIRNA** | hsa-miR-125 | MIRNA:hsa-m | 0.00072648 | 3.13877748 | 162 | 2593 |
| **MIRNA** | hsa-miR-429 | MIRNA:hsa-m | 0.00079632 | 3.09891485 | 366 | 2593 |
| **MIRNA** | hsa-miR-427 | MIRNA:hsa-m | 0.00080377 | 3.09486963 | 44 | 2593 |
| **MIRNA** | hsa-let-7c-5p | MIRNA:hsa-le | 0.00081574 | 3.08844677 | 514 | 2593 |
| **MIRNA** | hsa-miR-449 | MIRNA:hsa-m | 0.00085069 | 3.07022675 | 119 | 2593 |
| **MIRNA** | hsa-miR-449 | MIRNA:hsa-m | 0.00085069 | 3.07022675 | 119 | 2593 |
| **MIRNA** | hsa-miR-500 | MIRNA:hsa-m | 0.00085069 | 3.07022675 | 119 | 2593 |
| **MIRNA** | hsa-miR-608 | MIRNA:hsa-m | 0.00086802 | 3.06147233 | 78 | 2593 |
| **MIRNA** | hsa-miR-559 | MIRNA:hsa-m | 0.00090247 | 3.04456585 | 89 | 2593 |
| **MIRNA** | hsa-miR-672 | MIRNA:hsa-m | 0.00090247 | 3.04456585 | 89 | 2593 |
| **MIRNA** | hsa-miR-920 | MIRNA:hsa-m | 0.00090247 | 3.04456585 | 89 | 2593 |
| **MIRNA** | hsa-miR-685 | MIRNA:hsa-m | 0.00091642 | 3.03790459 | 131 | 2593 |
| **MIRNA** | hsa-miR-715 | MIRNA:hsa-m | 0.00092866 | 3.03214269 | 135 | 2593 |
| **MIRNA** | hsa-miR-479 | MIRNA:hsa-m | 0.00093518 | 3.0291033 | 151 | 2593 |
| **MIRNA** | hsa-miR-342- | MIRNA:hsa-m | 0.00095111 | 3.02176789 | 300 | 2593 |
| **MIRNA** | hsa-miR-432- | MIRNA:hsa-m | 0.00107712 | 2.96773527 | 68 | 2593 |
| **MIRNA** | hsa-miR-103a | MIRNA:hsa-m | 0.00108302 | 2.96536408 | 451 | 2593 |
| **MIRNA** | hsa-miR-366 | MIRNA:hsa-m | 0.00112587 | 2.94851197 | 364 | 2593 |
| **MIRNA** | hsa-miR-625- | MIRNA:hsa-m | 0.00114713 | 2.9403878 | 227 | 2593 |
| **MIRNA** | hsa-miR-315 | MIRNA:hsa-m | 0.00115481 | 2.93749053 | 101 | 2593 |
| **MIRNA** | hsa-miR-361 | MIRNA:hsa-m | 0.00115481 | 2.93749053 | 101 | 2593 |
| **MIRNA** | hsa-miR-446 | MIRNA:hsa-m | 0.00117112 | 2.9313982 | 152 | 2593 |
| **MIRNA** | hsa-miR-676 | MIRNA:hsa-m | 0.00118051 | 2.92792913 | 148 | 2593 |
| **MIRNA** | hsa-miR-687 | MIRNA:hsa-m | 0.00120868 | 2.91769008 | 206 | 2593 |
| **MIRNA** | hsa-miR-675 | MIRNA:hsa-m | 0.00122531 | 2.91175316 | 105 | 2593 |

| | | | | | | |
|---|---|---|---|---|---|---|
| **MIRNA** | hsa-miR-430( | MIRNA:hsa-m | 0.00123042 | 2.90994552 | 90 | 2593 |
| **MIRNA** | hsa-miR-364( | MIRNA:hsa-m | 0.0013134 | 2.88160294 | 198 | 2593 |
| **MIRNA** | hsa-miR-301b | MIRNA:hsa-m | 0.001451 | 2.83833244 | 375 | 2593 |
| **MIRNA** | hsa-miR-133a | MIRNA:hsa-m | 0.00147609 | 2.83088745 | 129 | 2593 |
| **MIRNA** | hsa-miR-678( | MIRNA:hsa-m | 0.0015089 | 2.82133876 | 30 | 2593 |
| **MIRNA** | hsa-miR-296- | MIRNA:hsa-m | 0.00181373 | 2.74142699 | 73 | 2593 |
| **MIRNA** | hsa-miR-200a | MIRNA:hsa-m | 0.00184569 | 2.73384032 | 150 | 2593 |
| **MIRNA** | hsa-miR-678 | MIRNA:hsa-m | 0.00186101 | 2.73025056 | 126 | 2593 |
| **MIRNA** | hsa-miR-680 | MIRNA:hsa-m | 0.00186101 | 2.73025056 | 126 | 2593 |
| **MIRNA** | hsa-miR-682 | MIRNA:hsa-m | 0.00186571 | 2.72915639 | 49 | 2593 |
| **MIRNA** | hsa-miR-19a- | MIRNA:hsa-m | 0.00187438 | 2.72714168 | 569 | 2593 |
| **MIRNA** | hsa-miR-190( | MIRNA:hsa-m | 0.00188842 | 2.7239019 | 134 | 2593 |
| **MIRNA** | hsa-miR-674( | MIRNA:hsa-m | 0.00202346 | 2.69390448 | 103 | 2593 |
| **MIRNA** | hsa-miR-708- | MIRNA:hsa-m | 0.00202346 | 2.69390448 | 103 | 2593 |
| **MIRNA** | hsa-miR-360( | MIRNA:hsa-m | 0.002116 | 2.67448471 | 107 | 2593 |
| **MIRNA** | hsa-miR-430( | MIRNA:hsa-m | 0.00215222 | 2.66711416 | 248 | 2593 |
| **MIRNA** | hsa-miR-314 | MIRNA:hsa-m | 0.00220628 | 2.65633878 | 70 | 2593 |
| **MIRNA** | hsa-miR-444 | MIRNA:hsa-m | 0.00226654 | 2.6446358 | 218 | 2593 |
| **MIRNA** | hsa-miR-683 | MIRNA:hsa-m | 0.00234636 | 2.62960567 | 123 | 2593 |
| **MIRNA** | hsa-miR-342- | MIRNA:hsa-m | 0.00237076 | 2.62511278 | 127 | 2593 |
| **MIRNA** | hsa-miR-651( | MIRNA:hsa-m | 0.00237076 | 2.62511278 | 127 | 2593 |
| **MIRNA** | hsa-miR-751( | MIRNA:hsa-m | 0.00284807 | 2.5454494 | 152 | 2593 |
| **MIRNA** | hsa-miR-656- | MIRNA:hsa-m | 0.00299082 | 2.52420906 | 124 | 2593 |
| **MIRNA** | hsa-miR-468( | MIRNA:hsa-m | 0.00299082 | 2.52420906 | 124 | 2593 |
| **MIRNA** | hsa-miR-328- | MIRNA:hsa-m | 0.00301028 | 2.52139336 | 132 | 2593 |
| **MIRNA** | hsa-miR-545- | MIRNA:hsa-m | 0.00325243 | 2.48779228 | 165 | 2593 |
| **MIRNA** | hsa-miR-427( | MIRNA:hsa-m | 0.00338283 | 2.4707194 | 216 | 2593 |
| **MIRNA** | hsa-miR-609( | MIRNA:hsa-m | 0.0034731 | 2.45928211 | 86 | 2593 |
| **MIRNA** | hsa-miR-471( | MIRNA:hsa-m | 0.00352102 | 2.45333171 | 153 | 2593 |
| **MIRNA** | hsa-miR-677( | MIRNA:hsa-m | 0.00359513 | 2.44428597 | 225 | 2593 |
| **MIRNA** | hsa-miR-676( | MIRNA:hsa-m | 0.00359513 | 2.44428597 | 225 | 2593 |
| **MIRNA** | hsa-miR-582- | MIRNA:hsa-m | 0.00359621 | 2.44415519 | 149 | 2593 |
| **MIRNA** | hsa-miR-676( | MIRNA:hsa-m | 0.00371001 | 2.43062438 | 174 | 2593 |
| **MIRNA** | hsa-miR-315( | MIRNA:hsa-m | 0.00371001 | 2.43062438 | 174 | 2593 |
| **MIRNA** | hsa-miR-191 | MIRNA:hsa-m | 0.00371686 | 2.42982401 | 141 | 2593 |
| **MIRNA** | hsa-miR-392( | MIRNA:hsa-m | 0.00377327 | 2.42328266 | 121 | 2593 |
| **MIRNA** | hsa-miR-216a | MIRNA:hsa-m | 0.00394547 | 2.40390135 | 243 | 2593 |
| **MIRNA** | hsa-miR-368 | MIRNA:hsa-m | 0.00394547 | 2.40390135 | 243 | 2593 |
| **MIRNA** | hsa-miR-675 | MIRNA:hsa-m | 0.00401686 | 2.39611361 | 217 | 2593 |
| **MIRNA** | hsa-miR-197- | MIRNA:hsa-m | 0.00447239 | 2.3494606 | 407 | 2593 |
| **MIRNA** | hsa-miR-468( | MIRNA:hsa-m | 0.00479894 | 2.31885447 | 126 | 2593 |
| **MIRNA** | hsa-miR-548r | MIRNA:hsa-m | 0.00524489 | 2.28026332 | 95 | 2593 |
| **MIRNA** | hsa-miR-25-3 | MIRNA:hsa-m | 0.00533586 | 2.27279537 | 517 | 2593 |
| **MIRNA** | hsa-miR-558 | MIRNA:hsa-m | 0.00573509 | 2.24145968 | 143 | 2593 |
| **MIRNA** | hsa-miR-663a | MIRNA:hsa-m | 0.00573509 | 2.24145968 | 143 | 2593 |

| | | | | | | |
|---|---|---|---|---|---|---|
| **MIRNA** | hsa-miR-493- | MIRNA:hsa-m | 0.00600686 | 2.22135223 | 115 | 2593 |
| **MIRNA** | hsa-miR-467! | MIRNA:hsa-m | 0.00601249 | 2.22094536 | 73 | 2593 |
| **MIRNA** | hsa-miR-101- | MIRNA:hsa-m | 0.00610883 | 2.21404221 | 391 | 2593 |
| **MIRNA** | hsa-miR-10b- | MIRNA:hsa-m | 0.00611365 | 2.2136997 | 322 | 2593 |
| **MIRNA** | hsa-miR-361- | MIRNA:hsa-m | 0.00632417 | 2.19899633 | 160 | 2593 |
| **MIRNA** | hsa-miR-125: | MIRNA:hsa-m | 0.00654858 | 2.18385295 | 194 | 2593 |
| **MIRNA** | hsa-miR-711: | MIRNA:hsa-m | 0.00691543 | 2.16018087 | 148 | 2593 |
| **MIRNA** | hsa-miR-375 | MIRNA:hsa-m | 0.00719153 | 2.14317865 | 477 | 2593 |
| **MIRNA** | hsa-miR-28-5 | MIRNA:hsa-m | 0.00732887 | 2.13496325 | 104 | 2593 |
| **MIRNA** | hsa-miR-312( | MIRNA:hsa-m | 0.00757688 | 2.12050984 | 128 | 2593 |
| **MIRNA** | hsa-miR-675: | MIRNA:hsa-m | 0.0077774 | 2.10916563 | 85 | 2593 |
| **MIRNA** | hsa-miR-128- | MIRNA:hsa-m | 0.00822882 | 2.08466256 | 526 | 2593 |
| **MIRNA** | hsa-miR-320I | MIRNA:hsa-m | 0.00873098 | 2.05893701 | 145 | 2593 |
| **MIRNA** | hsa-miR-676: | MIRNA:hsa-m | 0.00910958 | 2.04050161 | 67 | 2593 |
| **MIRNA** | hsa-miR-684! | MIRNA:hsa-m | 0.00910958 | 2.04050161 | 67 | 2593 |
| **MIRNA** | hsa-let-7c-3p | MIRNA:hsa-le | 0.00923271 | 2.03467085 | 101 | 2593 |
| **MIRNA** | hsa-miR-316: | MIRNA:hsa-m | 0.0093804 | 2.0277786 | 339 | 2593 |
| **MIRNA** | hsa-miR-681( | MIRNA:hsa-m | 0.00942991 | 2.02549236 | 105 | 2593 |
| **MIRNA** | hsa-miR-182- | MIRNA:hsa-m | 0.0095157 | 2.0215591 | 179 | 2593 |
| **MIRNA** | hsa-miR-312( | MIRNA:hsa-m | 0.0095653 | 2.0193014 | 109 | 2593 |
| **MIRNA** | hsa-miR-677 | MIRNA:hsa-m | 0.00971685 | 2.01247433 | 82 | 2593 |
| **MIRNA** | hsa-miR-613 | MIRNA:hsa-m | 0.01032809 | 1.98597993 | 86 | 2593 |
| **MIRNA** | hsa-miR-548I | MIRNA:hsa-m | 0.01041908 | 1.9821705 | 171 | 2593 |
| **MIRNA** | hsa-miR-211( | MIRNA:hsa-m | 0.01155859 | 1.93709531 | 134 | 2593 |
| **MIRNA** | hsa-miR-519! | MIRNA:hsa-m | 0.01178471 | 1.92868092 | 40 | 2593 |
| **MIRNA** | hsa-miR-680! | MIRNA:hsa-m | 0.01189176 | 1.92475385 | 102 | 2593 |
| **MIRNA** | hsa-miR-680! | MIRNA:hsa-m | 0.01189176 | 1.92475385 | 102 | 2593 |
| **MIRNA** | hsa-miR-474! | MIRNA:hsa-m | 0.0119602 | 1.9222617 | 126 | 2593 |
| **MIRNA** | hsa-miR-448 | MIRNA:hsa-m | 0.0119602 | 1.9222617 | 126 | 2593 |
| **MIRNA** | hsa-miR-678: | MIRNA:hsa-m | 0.01295434 | 1.88758478 | 185 | 2593 |
| **MIRNA** | hsa-miR-472! | MIRNA:hsa-m | 0.01360489 | 1.86630484 | 831 | 2593 |
| **MIRNA** | hsa-miR-313 | MIRNA:hsa-m | 0.01367528 | 1.86406376 | 61 | 2593 |
| **MIRNA** | hsa-miR-474 | MIRNA:hsa-m | 0.01393696 | 1.85583183 | 72 | 2593 |
| **MIRNA** | hsa-miR-474: | MIRNA:hsa-m | 0.01419501 | 1.84786415 | 91 | 2593 |
| **MIRNA** | hsa-miR-715 | MIRNA:hsa-m | 0.01428012 | 1.84526809 | 135 | 2593 |
| **MIRNA** | hsa-miR-519 | MIRNA:hsa-m | 0.01428012 | 1.84526809 | 135 | 2593 |
| **MIRNA** | hsa-miR-391! | MIRNA:hsa-m | 0.01428084 | 1.8452461 | 160 | 2593 |
| **MIRNA** | hsa-miR-23c | MIRNA:hsa-m | 0.01460363 | 1.83553904 | 131 | 2593 |
| **MIRNA** | hsa-miR-678: | MIRNA:hsa-m | 0.0146516 | 1.83411508 | 95 | 2593 |
| **MIRNA** | hsa-miR-873- | MIRNA:hsa-m | 0.01465205 | 1.83410163 | 265 | 2593 |
| **MIRNA** | hsa-miR-464! | MIRNA:hsa-m | 0.01513588 | 1.81999223 | 76 | 2593 |
| **MIRNA** | hsa-miR-679: | MIRNA:hsa-m | 0.01622658 | 1.78977296 | 230 | 2593 |
| **MIRNA** | hsa-let-7f-5p | MIRNA:hsa-le | 0.01683953 | 1.77367 | 395 | 2593 |
| **MIRNA** | hsa-miR-442! | MIRNA:hsa-m | 0.01758063 | 1.75496566 | 136 | 2593 |
| **MIRNA** | hsa-miR-421 | MIRNA:hsa-m | 0.01768311 | 1.75244144 | 235 | 2593 |

| | | | | | | |
|---|---|---|---|---|---|---|
| **MIRNA** | hsa-miR-193a | MIRNA:hsa-m | 0.01845335 | 1.73392467 | 128 | 2593 |
| **MIRNA** | hsa-miR-432- | MIRNA:hsa-m | 0.0184563 | 1.73385538 | 92 | 2593 |
| **MIRNA** | hsa-miR-1227 | MIRNA:hsa-m | 0.02020793 | 1.69447816 | 48 | 2593 |
| **MIRNA** | hsa-miR-149- | MIRNA:hsa-m | 0.02158384 | 1.6658713 | 827 | 2593 |
| **MIRNA** | hsa-miR-6735 | MIRNA:hsa-m | 0.02220499 | 1.65354951 | 133 | 2593 |
| **MIRNA** | hsa-miR-34c- | MIRNA:hsa-m | 0.0226123 | 1.64565526 | 171 | 2593 |
| **MIRNA** | hsa-miR-6883 | MIRNA:hsa-m | 0.023332 | 1.63204795 | 818 | 2593 |
| **MIRNA** | hsa-miR-6785 | MIRNA:hsa-m | 0.023332 | 1.63204795 | 818 | 2593 |
| **MIRNA** | hsa-miR-196b | MIRNA:hsa-m | 0.02361061 | 1.62689286 | 150 | 2593 |
| **MIRNA** | hsa-miR-30a- | MIRNA:hsa-m | 0.02423382 | 1.61557805 | 193 | 2593 |
| **MIRNA** | hsa-miR-4664 | MIRNA:hsa-m | 0.02428007 | 1.61475008 | 97 | 2593 |
| **MIRNA** | hsa-miR-6875 | MIRNA:hsa-m | 0.02459716 | 1.60911504 | 109 | 2593 |
| **MIRNA** | hsa-miR-106a | MIRNA:hsa-m | 0.02466867 | 1.60785429 | 709 | 2593 |
| **MIRNA** | hsa-miR-5197 | MIRNA:hsa-m | 0.02493435 | 1.60320188 | 163 | 2593 |
| **MIRNA** | hsa-miR-1197 | MIRNA:hsa-m | 0.02533194 | 1.59633159 | 74 | 2593 |
| **MIRNA** | hsa-miR-181c | MIRNA:hsa-m | 0.02687484 | 1.57065413 | 283 | 2593 |
| **MIRNA** | hsa-miR-191- | MIRNA:hsa-m | 0.02691038 | 1.57008023 | 78 | 2593 |
| **MIRNA** | hsa-miR-6864 | MIRNA:hsa-m | 0.02691038 | 1.57008023 | 78 | 2593 |
| **MIRNA** | hsa-miR-6738 | MIRNA:hsa-m | 0.0272357 | 1.56486143 | 134 | 2593 |
| **MIRNA** | hsa-miR-130a | MIRNA:hsa-m | 0.02804915 | 1.55208034 | 130 | 2593 |
| **MIRNA** | hsa-miR-429 | MIRNA:hsa-m | 0.02854578 | 1.54445809 | 151 | 2593 |
| **MIRNA** | hsa-miR-1180 | MIRNA:hsa-m | 0.02923036 | 1.53416578 | 67 | 2593 |
| **MIRNA** | hsa-miR-4650 | MIRNA:hsa-m | 0.0292829 | 1.53338591 | 86 | 2593 |
| **MIRNA** | hsa-miR-4496 | MIRNA:hsa-m | 0.03006352 | 1.52196021 | 118 | 2593 |
| **MIRNA** | hsa-miR-3688 | MIRNA:hsa-m | 0.03054101 | 1.51511654 | 114 | 2593 |
| **MIRNA** | hsa-let-7g-3p | MIRNA:hsa-le | 0.03109926 | 1.50724991 | 106 | 2593 |
| **MIRNA** | hsa-let-7a-2-3 | MIRNA:hsa-le | 0.03114164 | 1.50665857 | 102 | 2593 |
| **MIRNA** | hsa-miR-107 | MIRNA:hsa-m | 0.03168668 | 1.49912326 | 298 | 2593 |
| **MIRNA** | hsa-miR-4436 | MIRNA:hsa-m | 0.03329103 | 1.4776728 | 135 | 2593 |
| **MIRNA** | hsa-miR-6132 | MIRNA:hsa-m | 0.03543613 | 1.45055367 | 127 | 2593 |
| **MIRNA** | hsa-miR-6836 | MIRNA:hsa-m | 0.03543613 | 1.45055367 | 127 | 2593 |
| **MIRNA** | hsa-miR-1275 | MIRNA:hsa-m | 0.03753025 | 1.42561853 | 161 | 2593 |
| **MIRNA** | hsa-miR-6747 | MIRNA:hsa-m | 0.03801551 | 1.42003913 | 115 | 2593 |
| **MIRNA** | hsa-miR-4519 | MIRNA:hsa-m | 0.0387078 | 1.41220152 | 91 | 2593 |
| **MIRNA** | hsa-miR-6732 | MIRNA:hsa-m | 0.03937075 | 1.40482626 | 99 | 2593 |
| **MIRNA** | hsa-miR-4679 | MIRNA:hsa-m | 0.03951925 | 1.40319131 | 68 | 2593 |
| **MIRNA** | hsa-miR-548b | MIRNA:hsa-m | 0.04130239 | 1.38402483 | 268 | 2593 |
| **MIRNA** | hsa-miR-548a | MIRNA:hsa-m | 0.04130239 | 1.38402483 | 268 | 2593 |
| **MIRNA** | hsa-miR-4282 | MIRNA:hsa-m | 0.04189351 | 1.37785323 | 291 | 2593 |
| **MIRNA** | hsa-miR-3915 | MIRNA:hsa-m | 0.04343951 | 1.36211506 | 128 | 2593 |
| **MIRNA** | hsa-miR-4665 | MIRNA:hsa-m | 0.04480087 | 1.34871358 | 162 | 2593 |
| **MIRNA** | hsa-miR-519c | MIRNA:hsa-m | 0.04866784 | 1.31275792 | 348 | 2593 |
| **MIRNA** | hsa-miR-4729 | MIRNA:hsa-m | 0.04923982 | 1.30768353 | 137 | 2593 |
| **MIRNA** | hsa-miR-637 | MIRNA:hsa-m | 0.0494246 | 1.3060568 | 104 | 2593 |
| **MIRNA** | hsa-miR-4515 | MIRNA:hsa-m | 0.04962049 | 1.30433896 | 40 | 2593 |

| | | |
|---|---|---|
| 382 | 14823 | EEF1A1,HSPA8,GAPDH,HNRNPA1,HSPA9,ATN1,EEF2,TUBA1A,HSP90A |
| 364 | 14823 | EEF1A1,GAPDH,MLLT1,HNRNPA1,ATN1,ACTB,NUDC,EEF2,CFL1,HSP |
| 461 | 14823 | EEF1A1,HSPA8,GAPDH,ACTB,EEF2,MAVS,HSP90AB1,TUBB,EIF4G2,R |
| 479 | 14823 | EEF1A1,ORC4,HSPA8,MLLT1,HNRNPA1,HSPA9,ACTB,ODF2L,ACTG1, |
| 187 | 14823 | IGFBP5,HSPA8,GAPDH,ARSD,ACTG1,EIF4G2,PRRC2B,FLNA,RPL7A,U |
| 232 | 14823 | GAPDH,ACTG1,TUBA1A,CFL1,HSP90AB1,DBN1,PRRC2B,TUBA1B,NC |
| 224 | 14823 | HSPA8,GAPDH,HSPA9,ACTB,USP22,ACTG1,EEF2,HSP90AB1,VIM,TUB |
| 183 | 14823 | PABPC1,COPA,ACTB,ACTG1,TUBA1A,RPL3,RPS6,RPL4,ENO1,TUBA1 |
| 283 | 14823 | IGFBP5,PABPC1,COPA,ACTG1,EEF2,CFL1,HSP90AB1,TUBB,PRRC2B |
| 365 | 14823 | EEF1A1,ORC4,PABPC1,HSPA8,GAPDH,MLLT1,ACTB,USP22,ACTG1,E |
| 255 | 14823 | EEF1A1,ORC4,HSPA8,HNRNPA1,ODF2L,HSP90AB1,ACTR2,TUBB,RPS |
| 258 | 14823 | GAPDH,HNRNPA1,ACTB,USP22,DPYSL3,ACTR2,RPL3,FLNA,PKM,HSP |
| 138 | 14823 | PABPC1,GAPDH,NEDD4L,ATN1,ACTG1,TUBB,PRRC2B,PKM,ATP2B4, |
| 151 | 14823 | ACTB,ACTG1,EEF2,HSP90AA1,DBN1,ENO1,FLNA,MAP1B,PKM,GANA |
| 239 | 14823 | EEF1A1,IGFBP5,HNRNPA1,EEF2,XRCC2,MAVS,HSP90AB1,HSP90AA1, |
| 123 | 14823 | HNRNPA1,ACTB,TUBA1A,CFL1,HTT,TUBB,TUBA1B,FLNA,ALDOA,GAN |
| 134 | 14823 | EEF1A1,IGFBP5,HSPA8,GAPDH,HSPA9,EEF2,HSP90AA1,TUBB,UBC,R |
| 149 | 14823 | PABPC1,HSPA8,GAPDH,HNRNPA1,ATN1,ACTG1,BTBD9,DBN1,FLNA,Y |
| 127 | 14823 | EEF1A1,GAPDH,COPA,FLNA,MAP1B,ALDOA,GANAB,PSAP,RACK1,MA |
| 101 | 14823 | EEF1A1,PABPC1,COPA,DPYSL3,KIDINS220,RPL3,RPL4,PKM,CALR,H |
| 272 | 14823 | LRP1,HNRNPA1,ACTB,CTSC,TMSB4X,DBN1,FLNA,HSP90B1,FADS1,U |
| 229 | 14823 | EEF1A1,PABPC1,HNRNPA1,NEDD4L,EEF2,VIM,KIDINS220,TUBA1B,P |
| 263 | 14823 | EEF2,MAVS,ACTR2,EIF4G2,DBN1,FLNA,GANAB,FADS1,UGT8,FSTL1, |
| 97 | 14823 | EEF1A1,HSPA8,MED18,ACTG1,TUBB,UBC,TUBA1B,TUBB2B,RACK1, |
| 146 | 14823 | ATN1,MAP1B,ALDOA,PRPF8,UBA1,PSAP,H3-3B,DDX17,FASN,CALM3, |
| 94 | 14823 | EEF1A1,IGFBP5,HSPA9,ACTG1,HSP90AB1,HSP90AA1,HSPA1B,RACK |
| 169 | 14823 | EEF1A1,CTSC,HSP90AA1,VIM,PIK3R2,PKM,SOX4,FASN,CSDE1,MARC |
| 215 | 14823 | ORC4,HSPA8,HNRNPA1,ODF2L,ACTR2,TUBB,GANAB,HSP90B1,ABHD |
| 217 | 14823 | MLLT1,ACTG1,CFL1,PRRC2B,MAP1B,GANAB,YBX1,HSPA1B,H3-3B,S |
| 195 | 14823 | ORC4,HSPA8,HNRNPA1,ODF2L,ACTR2,TUBB,GANAB,ABHD2,HSPA1E |
| 98 | 14823 | EEF1A1,ACTB,RPL7A,COL4A1,YWHAE,RPS8,RPL7,RPL5,SMG1,IGF2, |
| 84 | 14823 | CFL1,TUBB,CALR,FASN,CALM3,MYH9,PTMS,COL18A1,UBB,PKD1,C |
| 311 | 14823 | EEF1A1,IGFBP5,HSPA8,GAPDH,MLLT1,MED18,MAVS,HTT,HSP90AA1 |
| 148 | 14823 | HSPA8,BTBD9,EIF4G2,MAP1B,ABHD2,RPL13A,YWHAE,RPL13,FASN, |
| 98 | 14823 | COL1A1,SPARC,COL5A1,COL4A1,RACK1,CALM3,FSCN1,COL4A2,TP |
| 205 | 14823 | COPA,RPL3,PRRC2B,FLNA,MAP1B,NCL,ALDOA,RPL8,ATP2B4,TMF1, |
| 101 | 14823 | COL1A1,MLLT1,MAVS,TUBB,PKM,CANX,PTMA,PTMS,SMG1,HLA-C,A |
| 100 | 14823 | COL1A1,MLLT1,MAVS,TUBB,PKM,CANX,PTMA,PTMS,SMG1,HLA-C,A |
| 78 | 14823 | MLLT1,HSP90AB1,ALDOA,COL5A1,CES3,YWHAE,MYH9,PLD3,ATP5F |
| 79 | 14823 | MLLT1,HSP90AB1,ALDOA,COL5A1,CES3,YWHAE,MYH9,PLD3,ATP5F |
| 137 | 14823 | EEF1A1,PABPC1,RPL3,PKM,GANAB,HSPA1B,RPLP0,ATP5F1B,NPM1, |
| 122 | 14823 | EEF1A1,ACTB,ACTG1,UBC,FLNA,NCL,PKM,HSPA1B,RPLP0,YWHAE,F |
| 119 | 14823 | EEF1A1,IGFBP5,COL1A1,EEF2,BTBD9,HSP90AA1,TUBB,PKM,CANX,F |
| 84 | 14823 | ORC4,FTH1,UBC,PRRC2B,H3-3B,PTMA,TMBIM6,CSDE1,YWHAQ,CH |
| 115 | 14823 | POLH,USP22,HSP90AA1,TMSB4X,HSPA1B,RPLP0,H3-3B,SLC2A3,RA |

| | | |
|---|---|---|
| 93 | 14823 | EEF1A1,EEF2,UBC,RPL7A,CANX,DDX5,GNAS,RACK1,VCP,AFF4,SPEN, |
| 76 | 14823 | EEF2,HSP90AB1,PSAP,IDS,MARCKSL1,LARP1,MSN,COL1A2,NCS1,CE |
| 175 | 14823 | MS4A10,EIF4G2,TUBA1B,COL6A1,PRPF8,DVL3,YWHAE,SPTBN1,PLD |
| 75 | 14823 | MLLT1,DCX,PRRC2B,PKM,CALR,GDI1,TMBIM6,PLD3,TRIM28,AP2M |
| 74 | 14823 | EEF2,HSP90AB1,PSAP,IDS,MARCKSL1,LARP1,MSN,COL1A2,NCS1,CE |
| 91 | 14823 | EEF1A1,GAPDH,NUDC,EEF2,HSP90AB1,UBC,PRRC2B,TUBA1B,PIK3F |
| 95 | 14823 | MLLT1,HSP90B1,CALR,CLU,IDS,PLD3,COL18A1,MARCKSL1,MSN,NC |
| 132 | 14823 | ATN1,MAP1B,ALDOA,FSTL1,PSAP,H3-3B,TMBIM6,TTC3,AFF4,TMSB1 |
| 142 | 14823 | ORC4,HSPA8,HNRNPA1,ODF2L,ACTR2,EIF4G2,TMF1,ABHD2,HSPA1E |
| 96 | 14823 | IGFBP5,PRRC2B,MAP1B,CANX,CALM3,P4HB,GDI1,PDIA3,MIDN,AP3D |
| 93 | 14823 | EEF1A1,NEDD4L,ACTB,HSP90AB1,GANAB,PRPF8,H3-3B,CANX,ENO2 |
| 73 | 14823 | MLLT1,DCX,PRRC2B,PKM,CALR,GDI1,TMBIM6,PLD3,TRIM28,AP2M |
| 139 | 14823 | EEF1A1,ORC4,HSPA8,HNRNPA1,ODF2L,ACTR2,ABHD2,HSPA1B,CAN |
| 112 | 14823 | IGFBP5,PABPC1,TUBB,DBN1,ENO1,ALDOA,PKM,ATP2B4,HSPA1B,CC |
| 86 | 14823 | EEF2,VIM,TUBA1B,MAP1B,NCL,CALR,PTMA,CALM3,CALM1,YWHAQ, |
| 300 | 14823 | EEF1A1,HSPA8,GAPDH,MLLT1,MED18,MAVS,ACTR2,UBC,EIF4G2,PIK |
| 131 | 14823 | ORC4,HSPA8,HNRNPA1,ODF2L,ACTR2,ABHD2,HSPA1B,CANX,CLU,SL |
| 119 | 14823 | COL1A1,USP22,MAVS,PRRC2B,CALR,COL5A1,PSAP,CANX,YWHAE,ID |
| 52 | 14823 | HNRNPA1,NEDD4L,RPL3,GANAB,YBX1,HSPA1B,GPI,MYH9,VCP,MAR |
| 86 | 14823 | MLLT1,CFL1,PRRC2B,PKM,CALR,PSAP,GDI1,PLD3,MARCKSL1,AP2M |
| 115 | 14823 | PIK3R2,MAP1B,HSPA1B,RPLP0,SOX4,MARCKSL1,AFF4,VAPA,MARCH |
| 66 | 14823 | PABPC1,GAPDH,HSP90AB1,VIM,PKM,UBA1,DVL3,FSCN1,SMG1,POL |
| 111 | 14823 | MLLT1,CFL1,BTBD9,TUBB,DCX,CALR,CLU,YWHAE,PLD3,COL18A1,T |
| 111 | 14823 | MLLT1,CFL1,BTBD9,TUBB,DCX,CALR,CLU,YWHAE,PLD3,COL18A1,T |
| 95 | 14823 | COL1A1,MLLT1,CALR,CANX,YWHAE,KLRD1,IGF2,AP2M1,STMN1,MIL |
| 64 | 14823 | ACTB,HSP90AA1,VIM,ENO1,FLNA,HSPA1B,PTMA,P4HB,CYB5R3,LDH |
| 85 | 14823 | MLLT1,HSP90B1,CALR,COL5A1,CANX,YWHAE,SEPTIN2,PLD3,TPI1,SE |
| 98 | 14823 | ACTB,EEF2,TUBB,TUBA1B,FLNA,SLC9A7,PRPF8,ABHD2,CANX,CALM |
| 161 | 14823 | IGFBP5,XRCC2,FLNA,MAP1B,CANX,HTRA4,CALM3,SEPTIN2,MARCKS |
| 90 | 14823 | EEF1A1,COPA,HNRNPA1,RPL3,RPS6,RPL4,ENO1,ALDOA,PKM,UGT8, |
| 84 | 14823 | MLLT1,DCX,PRRC2B,PKM,CALR,PTMA,GDI1,AP2M1,MIDN,LARP1,MS |
| 63 | 14823 | IGFBP5,GAPDH,ACTG1,EEF2,UBC,COL4A1,RPL13A,EIF4A1,STMN1,A |
| 67 | 14823 | HSP90AB1,HSP90B1,CALR,CANX,CALM3,FSCN1,PLD3,PKD1,MIDN,V |
| 88 | 14823 | SPARC,FSTL1,COL4A1,CALM3,FSCN1,COL4A2,RPS4X,ITGB1,MORF |
| 80 | 14823 | MLLT1,NUMA1,CALR,RPL13A,CALM3,FSCN1,PTMS,COL18A1,PKD1, |
| 134 | 14823 | PABPC1,HSPA8,COPA,ACTG1,EEF2,TUBA1A,MAVS,HSP90AA1,SPARC |
| 83 | 14823 | MLLT1,DCX,PRRC2B,PKM,CALR,PTMA,GDI1,AP2M1,MIDN,LARP1,MS |
| 73 | 14823 | IGFBP5,GAPDH,HNRNPA1,MED18,EEF2,EIF4G2,RPL3,TUBA1B,RPL7 |
| 40 | 14823 | ACTB,RPL3,ALDOA,ABHD2,KIF1A,GNB1,CALM1,PLXNB2,CSDE1,DDE |
| 61 | 14823 | EEF1A1,ACTG1,RPS6,RPL4,TUBA1B,HSPA1B,H3-3B,KIF1A,BSG,ATP |
| 84 | 14823 | COL1A1,GAPDH,SPARC,COL4A1,CALM3,FSCN1,COL4A2,ITGB1,MOF |
| 122 | 14823 | SOX4,SNRNP200,MARCKSL1,AFF4,VAPA,RPS4X,MARCHF6,SBF1,ARF |
| 53 | 14823 | MLLT1,ATN1,HSP90AB1,TUBA1B,ALDOA,GDI1,PLD3,TRIM28,NONO, |
| 29 | 14823 | ACTG1,EEF2,ENO2,SMARCA5,HK1,PIP4K2B,RPS10,TMEM30A,PPP1C |
| 63 | 14823 | CFL1,PKM,IDS,MARCKSL1,TRIM28,HLA-C,SBF1,MAP4,LARP1,HNRN |

| | | |
|---|---|---|
| 78 | 14823 | COL1A1,DBN1,PRRC2B,PKM,CALR,ABHD2,GNB1,FSCN1,TMBIM6,M |
| 95 | 14823 | BTBD9,ACTR2,TUBB,DCX,UBA1,AP2M1,NFE2L1,HNRNPA0,MSN,CLS |
| 65 | 14823 | HSP90AB1,HSP90B1,CALR,CANX,CALM3,FSCN1,PLD3,PKD1,MIDN,V |
| 82 | 14823 | HSP90AA1,FLNA,MAP1B,PKM,PSAP,DDX5,SEPTIN2,CSDE1,ATP5F1A, |
| 90 | 14823 | EEF1A1,MLLT1,ARSD,USP22,CFL1,CALR,CALM3,FSCN1,RPL37A,NO |
| 95 | 14823 | BTBD9,ACTR2,TUBB,DCX,UBA1,AP2M1,NFE2L1,HNRNPA0,MSN,CLS |
| 106 | 14823 | EEF1A1,ORC4,COL1A1,USP22,PRRC2B,PKM,CALR,CANX,RPL13A,YW |
| 95 | 14823 | BTBD9,ACTR2,TUBB,DCX,UBA1,AP2M1,NFE2L1,HNRNPA0,MSN,CLS |
| 94 | 14823 | BTBD9,ACTR2,TUBB,DCX,UBA1,AP2M1,NFE2L1,HNRNPA0,MSN,CLS |
| 94 | 14823 | BTBD9,ACTR2,TUBB,DCX,UBA1,AP2M1,NFE2L1,HNRNPA0,MSN,CLS |
| 70 | 14823 | EEF1A1,GAPDH,HNRNPA1,PGK1,YWHAG,MORF4L1,ZMIZ1,PGAM1,C |
| 105 | 14823 | EEF1A1,BTBD9,HSP90AA1,RPLP0,CANX,SOX4,RPS8,MARCKSL1,AFF |
| 53 | 14823 | MLLT1,ACTB,TUBB,UBC,CALR,SMG1,SET,CHD3,AP2M1,MIDN,ATXN |
| 55 | 14823 | HSP90AA1,CLU,KAT6A,SNRNP200,EIF3C,AFF4,STMN1,MIDN,AP3D1, |
| 69 | 14823 | USP22,CFL1,ACTR2,DBN1,PRRC2B,ALDOA,CALR,ABHD2,RPL13A,ID |
| 69 | 14823 | USP22,CFL1,ACTR2,DBN1,PRRC2B,ALDOA,CALR,ABHD2,RPL13A,ID |
| 259 | 14823 | HSPA8,MLLT1,MED18,MAVS,ACTR2,TUBB,UBC,EIF4G2,FLNA,YBX1, |
| 56 | 14823 | COL1A1,DCX,PKM,CALR,P4HB,PLD3,MIDN,ARL4C,TLN1,MYO5A,MT |
| 90 | 14823 | EEF1A1,MLLT1,ARSD,USP22,CFL1,CALR,CALM3,FSCN1,RPL37A,NO |
| 86 | 14823 | USP22,CFL1,PRRC2B,PKM,RPL13A,IDS,TMBIM6,SET,TPI1,EIF4H,NC |
| 94 | 14823 | BTBD9,ACTR2,TUBB,DCX,UBA1,AP2M1,NFE2L1,HNRNPA0,MSN,CLS |
| 59 | 14823 | HSP90AB1,HSP90B1,UBA1,LARP1,MSN,SLC7A5,NLGN2,HDLBP,ITM2 |
| 41 | 14823 | MLLT1,HLA-C,MIDN,MAP4,HNRNPA0,MSN,PFN1,CDH6,RHOB,PABPN |
| 67 | 14823 | HNRNPA1,HSP90AA1,TUBA1B,PTMA,MAGED1,GPI,FSCN1,UBB,AKAP |
| 68 | 14823 | MAVS,PRRC2B,NCL,TMBIM6,SMG1,CHD3,CPE,ENC1,JUND,ATXN7L3 |
| 68 | 14823 | MAVS,PRRC2B,NCL,TMBIM6,SMG1,CHD3,CPE,ENC1,JUND,ATXN7L3 |
| 55 | 14823 | COL1A1,DCX,PKM,CALR,P4HB,PLD3,MIDN,ARL4C,TLN1,MYO5A,MT |
| 55 | 14823 | COL1A1,DCX,PKM,CALR,P4HB,PLD3,MIDN,ARL4C,TLN1,MYO5A,MT |
| 52 | 14823 | CANX,YWHAE,NPM1,IGF2,MARCKSL1,SET,GTF3C4,MIDN,MARCHF6, |
| 59 | 14823 | HSP90AB1,HSP90B1,UBA1,LARP1,MSN,SLC7A5,NLGN2,HDLBP,ITM2 |
| 153 | 14823 | IGFBP5,HSPA8,MED18,MAVS,UBC,HSP90B1,PSAP,DDX5,SOX4,KLRD |
| 69 | 14823 | MAVS,PRRC2B,NCL,TMBIM6,SMG1,CHD3,CPE,ENC1,JUND,ATXN7L3 |
| 69 | 14823 | ORC4,IGFBP5,HSP90B1,COL5A1,CANX,SOX4,SHISA9,SET,HNRNPU, |
| 51 | 14823 | YWHAE,NPM1,IGF2,MARCKSL1,SET,GTF3C4,MIDN,MARCHF6,SLC7 |
| 167 | 14823 | ACTB,EEF2,EIF4G2,RPL4,PKM,NEFM,DVL3,SPTBN1,APP,PLD3,IGF2, |
| 43 | 14823 | CALR,CLU,CALM3,EIF1,MIDN,SBF1,MSN,SLC7A5,GNB2,GPAT4,CDK |
| 64 | 14823 | PRRC2B,YWHAE,CALM3,TMBIM6,EIF1,MIDN,PGAM1,LARP1,NCS1,L |
| 67 | 14823 | COL1A1,CFL1,VIM,RPL3,ALDOA,PKM,COL6A1,UBA1,SPTBN1,KLRD1 |
| 44 | 14823 | HSP90AB1,PRRC2B,PKM,SOX4,IDS,FSCN1,SBF1,SLC7A5,DLG5,DDX |
| 47 | 14823 | EEF2,HSP90AB1,COL18A1,HMGB1,ZMIZ1,MIDN,MSN,NLGN2,SOX12 |
| 70 | 14823 | COPA,ACTG1,SOX4,SLC2A3,SHISA9,FSCN1,SMG1,CALM1,SF3B1,BA |
| 62 | 14823 | PRRC2B,PKM,CALR,PSAP,GDI1,PLD3,HLA-C,AP2M1,MIDN,MAP4,HN |
| 62 | 14823 | PRRC2B,PKM,CALR,PSAP,GDI1,PLD3,HLA-C,AP2M1,MIDN,MAP4,HN |
| 68 | 14823 | ACTR2,CALM3,MARCKSL1,CHSY1,MSN,SLC7A5,GLG1,ANKRD52,HL |
| 63 | 14823 | USP22,CFL1,PKM,HSP90B1,CALR,IDS,NONO,MIDN,MAT2A,SLC7A5, |

| | | |
|---|---|---|
| 75 | 14823 | COL1A1,PKM,CALR,RPL13A,GNB1,YWHAG,SET,MIDN,CHD4,PFN1,AS |
| 92 | 14823 | COL1A1,PKM,ABHD2,MYH9,FSCN1,MARCKSL1,AP2M1,MIDN,AP3D1, |
| 92 | 14823 | COL1A1,PKM,ABHD2,MYH9,FSCN1,MARCKSL1,AP2M1,MIDN,AP3D1, |
| 68 | 14823 | ORC4,IGFBP5,HSP90B1,COL5A1,CANX,SOX4,SHISA9,SET,HNRNPU,S |
| 68 | 14823 | ORC4,IGFBP5,HSP90B1,COL5A1,CANX,SOX4,SHISA9,SET,HNRNPU,S |
| 29 | 14823 | ALDH6A1,EIF3C,CBX5,CLSTN1,SLC7A5,CD81,B4GALNT4,ARHGDIA,C |
| 81 | 14823 | ACTB,USP22,HSP90AB1,PRRC2B,PKM,CALR,PSAP,GDI1,AP2M1,LARI |
| 70 | 14823 | IGFBP5,HSP90AA1,HSP90B1,SLC2A3,NONO,HLA-C,PDIA3,HMGB1,RI |
| 47 | 14823 | PRRC2B,RPL37A,TPI1,LARP1,ZC3H7B,RNF187,TNPO1,HDGF,MAPRE |
| 31 | 14823 | USP22,HSP90AB1,PKM,MARCKSL1,HLA-C,PGAM1,LARP1,MSN,VAT1 |
| 42 | 14823 | CALR,CLU,CALM3,EIF1,MIDN,SBF1,MSN,SLC7A5,GNB2,GPAT4,CDKI |
| 79 | 14823 | ORC4,COL1A1,HSPA8,HNRNPA1,ODF2L,ACTR2,CANX,SLC2A3,RPL15 |
| 181 | 14823 | HNRNPA1,ACTB,MAVS,DBN1,PKM,PSAP,PGK1,SOX4,TMBIM6,SMG1, |
| 74 | 14823 | EEF1A1,IGFBP5,COL1A1,PRRC2B,RPL13A,PLD3,EIF1,BASP1,AP2M1, |
| 112 | 14823 | IGFBP5,HSPA8,MED18,MAVS,UBC,SOX4,KLRD1,UBB,TUBB4B,TIMP2 |
| 74 | 14823 | EEF1A1,IGFBP5,COL1A1,PRRC2B,RPL13A,PLD3,EIF1,BASP1,AP2M1, |
| 46 | 14823 | ACTG1,TUBB,ALDOA,PKM,YBX1,CANX,VCP,CKB,EIF4A1,LDHA,RPS9, |
| 106 | 14823 | TUBB,GANAB,HSP90B1,TMF1,HSPA1B,FKBP14,RPL13A,RPS8,SEPTII |
| 57 | 14823 | MALAT1,HSPA1B,H3-3B,BSG,PTMS,HMGB1,RPS4X,BTF3,RPSA,RPL2 |
| 63 | 14823 | IGFBP5,CFL1,HSP90AA1,TUBB,DBN1,SMG1,VCP,HNRNPA0,CPE,RPS |
| 46 | 14823 | HNRNPA1,HSP90B1,FSTL1,H3-3B,PTMA,ANXA2,TKT,CCND2,GJA1,PF |
| 256 | 14823 | EEF1A1,HSPA8,MLLT1,MED18,MAVS,HTT,ACTR2,UBC,EIF4G2,YBX1, |
| 42 | 14823 | CALR,CLU,CALM3,EIF1,MIDN,SBF1,MSN,SLC7A5,GNB2,GPAT4,CDKI |
| 40 | 14823 | PKM,LAMA5,UBE2H,RAB1A,COTL1,PEA15,LMNB2,MTRF1L,PRDX5,R |
| 40 | 14823 | CALR,CLU,CALM3,FSCN1,MIDN,SBF1,MSN,GNB2,GPAT4,HS6ST1,DD |
| 40 | 14823 | CALR,CLU,CALM3,FSCN1,MIDN,SBF1,MSN,GNB2,GPAT4,HS6ST1,DD |
| 78 | 14823 | ACTB,USP22,HSP90AB1,PRRC2B,PKM,CALR,PSAP,GDI1,AP2M1,LARI |
| 46 | 14823 | KAT6A,P4HB,PKD1,ITGB1,MAT2A,NFE2L1,EIF5,CALU,KDM6B,ATP6A |
| 54 | 14823 | IGFBP5,CALR,ABHD2,GNAS,TMBIM6,CSDE1,GOLGA8A,EIF4H,RHOA, |
| 61 | 14823 | IGFBP5,CFL1,HSP90AA1,TUBB,DBN1,SMG1,VCP,HNRNPA0,CPE,RPS |
| 48 | 14823 | RPS6,COL4A1,SNRNP200,UBB,RPL37A,ITM2B,AP2B1,NFE2L1,SMAR |
| 45 | 14823 | KAT6A,P4HB,PKD1,ITGB1,MAT2A,NFE2L1,EIF5,CALU,KDM6B,ATP6A |
| 61 | 14823 | IGFBP5,CFL1,HSP90AA1,TUBB,DBN1,SMG1,VCP,HNRNPA0,CPE,RPS |
| 78 | 14823 | ACTB,USP22,HSP90AB1,PRRC2B,PKM,CALR,PSAP,GDI1,AP2M1,LARI |
| 79 | 14823 | COL1A1,HSP90AB1,PKM,ABHD2,DVL3,YWHAE,P4HB,RPS3,BASP1,HN |
| 32 | 14823 | PRRC2B,PKM,SOX4,IDS,FSCN1,PKD1,SLC7A5,ANKRD52,STIP1,DLG5 |
| 39 | 14823 | PKM,IDS,TRIM28,MIDN,SBF1,EIF5A,GNB2,GPAT4,ZC3H7B,TM9SF3,I |
| 54 | 14823 | IGFBP5,HSP90AA1,HSP90B1,HLA-C,SESN3,APLP2,ARL6IP1,YWHAB,C |
| 110 | 14823 | IGFBP5,HSPA8,MED18,MAVS,UBC,SOX4,APP,KLRD1,UBB,NFE2L1,TS |
| 77 | 14823 | COL1A1,HSP90AB1,PKM,ABHD2,DVL3,P4HB,RPS3,BASP1,HNRNPA0, |
| 42 | 14823 | ACTB,CLU,GDI1,PLD3,COL1A2,ARL4C,TLN1,ARHGDIA,DDX39B,ZC3H |
| 32 | 14823 | GAPDH,RPL37A,RPS3,MIDN,TKT,HNRNPK,MYH10,ZC3H7B,GRINA,HN |
| 84 | 14823 | MLLT1,CFL1,ACTR2,CALR,SEPTIN2,AP2M1,MIDN,VCL,MAT2A,PGAM |
| 70 | 14823 | PRRC2B,FLNA,SOX4,YWHAE,SMG1,NPM1,ATP5F1A,RPL37A,STMN1,A |
| 40 | 14823 | ACTG1,HSP90AB1,TUBB,SOX4,CALM3,MYH9,PKD1,CHD3,MIDN,PGA |

| | | |
|---|---|---|
| 45 | 14823 | COL1A1,PRRC2B,PKM,FSCN1,TRIM28,LLGL1,SPTBN2,HLA-B,DLG5, |
| 84 | 14823 | MLLT1,CFL1,ACTR2,CALR,SEPTIN2,AP2M1,MIDN,VCL,MAT2A,PGAM |
| 31 | 14823 | HSP90AB1,PKM,MARCKSL1,HLA-C,SBF1,PGAM1,LARP1,MSN,VAT1,[illegible] |
| 80 | 14823 | EEF1A1,ENO1,COL4A1,NPM1,TPI1,MAT2A,GOLGA8A,TPM4,SCARB2, |
| 76 | 14823 | ORC4,HSPA8,MLLT1,HNRNPA1,ODF2L,ACTR2,SPARC,CANX,SLC2A3, |
| 55 | 14823 | EEF2,COL4A1,PGK1,CANX,CALM3,VCP,CPE,MSN,JUND,ATP2A2,CDH |
| 69 | 14823 | EEF1A1,TUBB,PRRC2B,HSPA1B,COL4A1,RPLP0,H3-3B,CANX,TRIM2[illegible] |
| 53 | 14823 | IGFBP5,MAVS,DBN1,DDX5,APP,FSCN1,CALM1,YWHAQ,SARAF,CBX5, |
| 35 | 14823 | PLD3,RAB7A,ARL4C,TLN1,DDX39B,ZC3H7B,TM9SF3,LAMC1,ASXL1,[illegible] |
| 31 | 14823 | HLA-C,MAP4,HNRNPU,HNRNPA0,MSN,PFN1,CDKN1A,PLEC,ID4,TXLN |
| 59 | 14823 | EEF2,CFL1,VIM,TUBA1B,MAP1B,NCL,CALR,CALM3,CSDE1,YWHAQ,K |
| 84 | 14823 | MLLT1,CFL1,ACTR2,CALR,SEPTIN2,AP2M1,MIDN,VCL,MAT2A,PGAM |
| 72 | 14823 | CFL1,PKM,ABHD2,YWHAE,GDI1,SERINC1,MIDN,SBF1,MAT2A,EIF4H,[illegible] |
| 34 | 14823 | HLA-C,MAP4,HNRNPA0,MSN,PFN1,MDK,SKI,KDM6B,PLEC,GIGYF1,C |
| 71 | 14823 | CFL1,PKM,ABHD2,YWHAE,GDI1,SERINC1,MIDN,SBF1,MAT2A,EIF4H,[illegible] |
| 71 | 14823 | CFL1,PKM,ABHD2,YWHAE,GDI1,SERINC1,MIDN,SBF1,MAT2A,EIF4H,[illegible] |
| 108 | 14823 | IGFBP5,HSPA8,MED18,MAVS,UBC,RPL4,SOX4,KLRD1,UBB,POLR2A, |
| 39 | 14823 | HSP90AB1,PSAP,PKD1,MARCHF6,RPL22,ATXN7L3B,HMGN2,TLN1,M |
| 75 | 14823 | IGFBP5,COL1A1,BTBD9,PKM,PTMA,CALM3,SHISA9,PLD3,PKD1,BAS[illegible] |
| 54 | 14823 | COL1A1,HSP90AA1,GDI1,SMG1,DDB1,HNRNPA0,LDHA,RHOA,CCND2 |
| 54 | 14823 | ORC4,FTH1,NUMA1,PRRC2B,ABHD2,CANX,SOX4,GNB1,P4HB,EIF1,A |
| 101 | 14823 | IGFBP5,HSPA8,MED18,MAVS,UBC,SOX4,KLRD1,UBB,NFE2L1,TSPAN |
| 22 | 14823 | HSP90B1,UBA1,SOX4,GDI1,PKD1,LARP1,TARDBP,ANKRD52,CDK16,[illegible] |
| 133 | 14823 | ACTB,EIF4G2,RPL3,PRPF8,DVL3,PLD3,EIF3C,MARCKSL1,MIDN,VCL, |
| 43 | 14823 | COL1A1,PRRC2B,PKM,FSCN1,TRIM28,LLGL1,SPTBN2,HLA-B,DLG5, |
| 43 | 14823 | COL1A1,PRRC2B,PKM,FSCN1,TRIM28,LLGL1,SPTBN2,HLA-B,DLG5, |
| 43 | 14823 | COL1A1,PRRC2B,PKM,FSCN1,TRIM28,LLGL1,SPTBN2,HLA-B,DLG5, |
| 32 | 14823 | PRRC2B,PKM,SOX4,IDS,FSCN1,PKD1,SLC7A5,ANKRD52,STIP1,DLG[illegible] |
| 35 | 14823 | HSP90AB1,PKD1,MARCHF6,MSN,RPL22,HMGN2,GLG1,TLN1,ANKR[illegible] |
| 35 | 14823 | HSP90AB1,PKD1,MARCHF6,MSN,RPL22,HMGN2,GLG1,TLN1,ANKR[illegible] |
| 35 | 14823 | HSP90AB1,PKD1,MARCHF6,MSN,RPL22,HMGN2,GLG1,TLN1,ANKR[illegible] |
| 46 | 14823 | IGFBP5,PRRC2B,ZNF99,RPL13A,PLD3,EIF1,NLGN2,DDX3X,HMGCS1 |
| 47 | 14823 | EEF1A1,CFL1,FSCN1,RPL37A,TPI1,CHSY1,MIDN,HNRNPA0,LDHA,CP[illegible] |
| 51 | 14823 | FBXL4,PTMA,SOX4,GNB1,CHSY1,ZMIZ1,LARP1,HNRNPA0,MSN,CCN[illegible] |
| 86 | 14823 | IGFBP5,ACTG1,EEF2,CALR,CANX,IDS,RPL37A,YWHAQ,RPL15,VAPA,[illegible] |
| 29 | 14823 | ACTB,PKM,H3-3B,SOX4,GNAS,DDX17,CALM3,TPI1,TUBB4B,H3-3A,R |
| 119 | 14823 | ORC4,ODF2L,ACTR2,SLC2A3,GNB1,DNAJA1,MAP4,TMEM170A,SMA[illegible] |
| 100 | 14823 | IGFBP5,HSPA8,MED18,MAVS,UBC,SOX4,KLRD1,UBB,NFE2L1,TSPAN |
| 69 | 14823 | PRRC2B,PKM,CALR,PSAP,GDI1,CALM1,NPM1,AP2M1,EIF4H,MSN,CC |
| 38 | 14823 | SOX4,LARP1,LGALS3BP,UBE2H,RAB1A,PEA15,MLEC,SELENON,PLC[illegible] |
| 38 | 14823 | MLLT1,DCX,CALM3,TPI1,MIDN,ARF1,NFE2L1,RPS24,CBX5,ATXN7L3 |
| 51 | 14823 | COL1A1,PRRC2B,ALDOA,HSP90B1,CALR,YWHAE,GDI1,NPM1,RPL37[illegible] |
| 50 | 14823 | MLLT1,H3-3B,HLA-C,HNRNPA0,EIF4H,CD81,PLXNA1,SESN3,ARHGDI |
| 64 | 14823 | BTBD9,HSP90AB1,DCX,CALM1,PGAM1,HNRNPA0,ATXN7L3B,LASP1,[illegible] |
| 39 | 14823 | ACTB,CLU,PLD3,COL1A2,ARL4C,TLN1,ARHGDIA,DDX39B,ZC3H7B,T[illegible] |

| | | |
|---|---|---|
| 35 | 14823 | HSP90AB1,PKD1,MARCHF6,MSN,RPL22,HMGN2,GLG1,TLN1,ANKRD |
| 62 | 14823 | IGFBP5,UGT8,CALR,ABHD2,PGK1,APP,CSDE1,SET,EGLN1,GTF3C4, |
| 102 | 14823 | IGFBP5,HSPA8,MED18,MAVS,UBC,SOX4,KLRD1,UBB,NFE2L1,TSPAN |
| 45 | 14823 | COL1A1,PIK3R2,PKM,FTL,PTMA,SOX4,FSCN1,GSTP1,SMIM14,ANXA |
| 17 | 14823 | HSP90B1,SOX4,GDI1,PKD1,LARP1,ANKRD52,STIP1,MDK,IPO7,APEX |
| 30 | 14823 | SLC7A5,SOX12,ARL4C,GNB2,MTSS2,VEGFA,GIGYF1,HGS,POLR2E, |
| 50 | 14823 | MALAT1,PGK1,HMGB1,MARCHF6,BRD3,PSMD2,MCL1,RPL12,GJA1, |
| 44 | 14823 | CALR,FSCN1,SBF1,SLC7A5,JUND,MTSS2,ANKRD52,RHOB,HRNR,AR |
| 44 | 14823 | COL1A1,PKM,CANX,PTMA,GNAS,PTMS,MIDN,PGAM1,HNRNPA0,RPS |
| 23 | 14823 | HSP90B1,SOX4,GDI1,PKD1,LARP1,MSN,ATXN7L3B,HNRNPC,MDK,K |
| 143 | 14823 | ACTB,MAVS,DBN1,PKM,PSAP,PGK1,SOX4,TMBIM6,SMG1,CALM1,SE |
| 46 | 14823 | CALR,FSCN1,SBF1,SLC7A5,JUND,MTSS2,ANKRD52,RHOB,HRNR,AR |
| 38 | 14823 | ACTG1,HSP90AB1,TUBB,SOX4,CALM3,MYH9,PKD1,CHD3,MIDN,PGA |
| 38 | 14823 | TUBB,PRRC2B,MARCKSL1,GTF3C4,MORF4L1,SLC7A5,HNRNPC,MM |
| 39 | 14823 | COL1A1,SHISA9,GDI1,LASP1,TPM1,DDX3X,CALU,LUC7L3,TOMM20, |
| 73 | 14823 | IGFBP5,COL1A1,BTBD9,PKM,PTMA,CALM3,SHISA9,PLD3,PKD1,BAS |
| 29 | 14823 | HSP90AB1,PKM,MARCKSL1,HLA-C,PGAM1,LARP1,MSN,VAT1,EIF5A, |
| 66 | 14823 | EEF1A1,MLLT1,CFL1,CALM3,MARCKSL1,TRIM28,PKD1,MIDN,VCL,G |
| 43 | 14823 | ORC4,RPL13A,FSCN1,NONO,RPL18,NCS1,NLGN2,SND1,LGALS3BP, |
| 44 | 14823 | GAPDH,MLLT1,PRRC2B,TRIM28,NFE2L1,PFN1,CDH6,RHOB,BCLAF1 |
| 44 | 14823 | COL1A1,TUBB,PKM,PTMA,PTMS,SMG1,ZMIZ1,RAB6B,SLC7A5,ATXN |
| 50 | 14823 | PRRC2B,NONO,SET,TPI1,MIDN,MAT2A,GTF2F1,CPE,SLC7A5,PFN1,G |
| 43 | 14823 | NEDD4L,HSP90AA1,H3-3B,TMBIM6,DNAJA1,CCND2,ARL6IP1,TMEM |
| 43 | 14823 | ORC4,RPL13A,FSCN1,NONO,RPL18,NCS1,NLGN2,SND1,LGALS3BP, |
| 45 | 14823 | MED18,PKM,HSPA1B,BSG,APP,RPL5,SPEN,YWHAQ,HMGB1,MAP4,R |
| 53 | 14823 | LRP1,TUBB,HSPA1B,PTMA,YWHAQ,RPL15,RPL18,DDX3X,HNRNPDL, |
| 65 | 14823 | MLLT1,CFL1,CALM3,MARCKSL1,TRIM28,PKD1,MIDN,VCL,GTF2F1, |
| 33 | 14823 | EEF2,HSP90AB1,PRRC2B,PKD1,AP2M1,MARCHF6,COL1A2,RPL22,H |
| 50 | 14823 | NEDD4L,SPARC,ABHD2,RPL13A,P4HB,RPL10,NFE2L1,NLGN2,LASP1 |
| 67 | 14823 | ORC4,COL1A1,TUBB,HSP90B1,SEC61A1,HNRNPU,MSN,NLGN2,AST |
| 67 | 14823 | CFL1,BTBD9,ACTR2,TUBB,DCX,COL5A1,CANX,SEPTIN2,GTF2F1,NF |
| 49 | 14823 | ZNF124,HSP90AA1,EIF4G2,HSP90B1,GNB1,SET,TPI1,DNAJA1,TPM4, |
| 55 | 14823 | IDS,MARCKSL1,SERINC1,NCS1,CBX5,CLSTN1,VAT1,PFN1,EIF5A,RAC |
| 55 | 14823 | IDS,MARCKSL1,SERINC1,NCS1,CBX5,CLSTN1,VAT1,PFN1,EIF5A,RAC |
| 47 | 14823 | CFL1,CALR,YWHAE,CALM3,TMBIM6,IGF2,EIF1,LARP1,MSN,NCS1,LA |
| 42 | 14823 | IGFBP5,CFL1,HSP90B1,RPL13A,GNB1,HNRNPA0,GNB2,H1-0,COL6A |
| 71 | 14823 | H3-3B,CANX,GPI,TMBIM6,ATP5F1A,PKD1,AP3D1,RPS24,EIF5,BEX3, |
| 71 | 14823 | H3-3B,CANX,GPI,TMBIM6,ATP5F1A,PKD1,AP3D1,RPS24,EIF5,BEX3, |
| 65 | 14823 | MLLT1,CFL1,CALM3,MARCKSL1,TRIM28,PKD1,MIDN,VCL,GTF2F1, |
| 107 | 14823 | EEF1A1,IGFBP5,HNRNPA1,ACTG1,DPYSL3,TMSB4X,NCL,RPL7A,ALD |
| 43 | 14823 | IGFBP5,ZNF99,RPL13A,PLD3,EIF1,NLGN2,DDX3X,HMGCS1,CDKN1A |
| 35 | 14823 | HNRNPA1,FTH1,SLC9A7,MAP9,RPS3,CDKN1A,HNRNPA3,IPO7,NPEP |
| 130 | 14823 | EEF1A1,HSPA8,GAPDH,HNRNPA1,ATN1,HSP90AA1,TUBB,EIF4G2,PR |
| 47 | 14823 | MAVS,ACTR2,NCL,APP,SMG1,STMN1,CPE,SLC7A5,ENC1,GLG1,ITM2 |
| 47 | 14823 | CALR,FSCN1,SBF1,SLC7A5,JUND,MTSS2,ANKRD52,RHOB,HRNR,AR |

| | | |
|---|---|---|
| 40 | 14823 | ACTB,MTRNR2L2,NCL,CES3,SF3B1,RPS3,ZMIZ1,SARAF,SMARCA5,C |
| 29 | 14823 | COL1A1,MLLT1,DBN1,PKM,TRIM28,SBF1,RHOA,SPTBN2,HLA-B,DD) |
| 103 | 14823 | ZNF124,HSP90AA1,EIF4G2,HSP90B1,FKBP14,ATP5F1B,GNB1,APP,A |
| 88 | 14823 | PABPC1,ACTG1,MAVS,TUBA1B,PRPF8,HSPA1B,H3-3B,DVL3,SLC2A3, |
| 51 | 14823 | USP22,ACTG1,HSP90AA1,PRPF8,HSPA1B,BSG,SMG1,EIF3C,VCP,EIF |
| 59 | 14823 | COL1A1,TUBB,CALR,CANX,SOX4,CALM3,SEC61A1,HMGN2,ASTN2,IT |
| 48 | 14823 | NUMA1,RPL13A,FSCN1,TRIM28,NONO,CHSY1,RPL18,MSN,NLGN2,E |
| 121 | 14823 | CTSC,DPYSL3,HSP90AA1,MALAT1,SET,CHSY1,LDHB,ITGB1,NCAM1,F |
| 37 | 14823 | PRRC2B,GTF3C4,MORF4L1,CNN3,TPM4,SLC7A5,HNRNPC,CDKN1A, |
| 43 | 14823 | DCX,GNB1,APP,SEC61A1,ARL6IP1,LAPTM4B,RNF187,TUBB2A,CCNL |
| 32 | 14823 | CFL1,NUMA1,PRRC2B,TRIM28,CHSY1,HMGB1,LARP1,ATP1A1,EIF5A |
| 131 | 14823 | DCX,HSP90B1,H3-3B,CANX,GPI,HTRA4,TMBIM6,SPTAN1,ATP5F1A,M |
| 47 | 14823 | ATN1,TUBB,MTRNR2L8,GNAS,ATP5F1B,IGF2,TPI1,VCL,ARF1,ITM2B, |
| 27 | 14823 | ALDH6A1,HNRNPA0,RHOA,TAGLN,ANKRD52,U2AF2,HDGF,SELENON |
| 27 | 14823 | ALDH6A1,HNRNPA0,RHOA,TAGLN,ANKRD52,U2AF2,HDGF,SELENON |
| 36 | 14823 | NCL,CES3,PTMA,CALM1,RPS3,ZMIZ1,HNRNPU,SMARCA5,DDX3X,LM |
| 91 | 14823 | ORC4,MLLT1,PRRC2B,HSPA1B,H3-3B,PTMA,SOX4,SEPTIN2,CALM1, |
| 37 | 14823 | HSP90AB1,PKM,MARCKSL1,HLA-C,SERINC1,PGAM1,LARP1,MSN,CB |
| 55 | 14823 | TUBA1B,UBA1,RPL5,NPM1,RPS3,RPS4X,MORF4L1,CPE,PFN1,CCND |
| 38 | 14823 | MLLT1,ACTR2,TUBB,UBA1,PGAM1,MSN,SLC7A5,MTSS2,ARL6IP1,TM |
| 31 | 14823 | MLLT1,CALR,CLU,TRIM28,HLA-C,ZNF43,SBF1,SLC7A5,GPAT4,KDM |
| 32 | 14823 | HNRNPA1,HSP90B1,H3-3B,PTMA,PPIB,TIMP3,DLK1,WBP2,PSMA7,AN |
| 53 | 14823 | IGFBP5,ACTB,HSP90AA1,YWHAE,SHISA9,MARCKS,GOLGA8A,SPC25, |
| 44 | 14823 | CFL1,FASN,MARCKSL1,SERINC1,MIDN,SBF1,EIF4H,CBX5,VAT1,RAC |
| 19 | 14823 | DBN1,MIDN,SLC22A17,SLC7A5,CCT7,ASTN2,MYO5A,ARF3,SF1,PPP1 |
| 36 | 14823 | CFL1,ALDOA,HSP90B1,UBA1,FSCN1,EIF1,NCS1,SOX12,TARDBP,CDK |
| 36 | 14823 | DCX,PRRC2B,NPM1,HMGB1,AP2B1,NFE2L1,RPL23A,PARP1,TMEM2 |
| 42 | 14823 | MLLT1,ALDOA,PKM,MYH9,IDS,TRIM28,MIDN,SBF1,SLC7A5,CCT7,EI |
| 42 | 14823 | MLLT1,ALDOA,PKM,MYH9,IDS,TRIM28,MIDN,SBF1,SLC7A5,CCT7,EI |
| 56 | 14823 | CFL1,CALR,ABHD2,YWHAE,MAT2A,MSN,LENG8,ATXN7L3B,RTN3,MC |
| 192 | 14823 | MLLT1,USP22,CFL1,MAVS,ZNF124,ALDOA,CALR,COL5A1,CES3,MYH |
| 25 | 14823 | TUBA1B,ALDOA,SOX4,KAT6A,SET,PDIA3,CCND2,PPP1CB,CTNND1,Y |
| 28 | 14823 | COL1A1,MLLT1,PKM,TRIM28,SBF1,RHOA,SPTBN2,HLA-B,DDX39B,S |
| 33 | 14823 | EIF4G2,RPL13A,PTMS,HMGB1,MIDN,HNRNPA0,EIF4H,HMGN2,FDFT |
| 44 | 14823 | COL1A1,MLLT1,PKM,CALR,RPL13A,YWHAG,MIDN,MARCHF6,GTF2F |
| 44 | 14823 | CFL1,CALR,YWHAE,CALM3,TMBIM6,IGF2,EIF1,LARP1,MSN,NCS1,LA |
| 50 | 14823 | HSP90B1,ABHD2,PTMA,DVL3,SHISA9,P4HB,FSCN1,TRIM28,SET,EIF4 |
| 43 | 14823 | MALAT1,MAP1B,GTF3C4,TMEM170A,ATXN7L3B,PDIA6,HMGCS1,ARI |
| 34 | 14823 | USP22,DCX,CALR,P4HB,GDI1,TMBIM6,AP2M1,NFE2L1,LARP1,PFN1, |
| 74 | 14823 | CFL1,ALDOA,PKM,CLU,SHISA9,TMBIM6,TPI1,GTF3C4,NCS1,SLC7A5 |
| 29 | 14823 | HSP90AB1,PKM,ATP5F1A,ZMIZ1,HNRNPU,RPL12,CDKN1A,NUCB1,TF |
| 66 | 14823 | BTBD9,FADS1,PKD1,HMGB1,MAT2A,NCS1,PTCD3,JPT2,MYO5A,SND |
| 102 | 14823 | EIF4G2,DVL3,ATP5F1B,MYH9,CALM1,PLD3,EIF3C,MARCKSL1,YWHA |
| 44 | 14823 | CALR,SOX4,CALM3,TMBIM6,NPM1,IGF2,GTF3C4,MIDN,EIF4H,CCND |
| 67 | 14823 | EEF1A1,COPA,HTT,HSP90AA1,EIF4G2,TUBA1B,COL4A1,SEPTIN2,NPI |

| | | |
|---|---|---|
| 42 | 14823 | IGFBP5,SHISA9,SMIM14,HMGB1,LPCAT1,SLC7A5,MCL1,ACTN4,MM |
| 33 | 14823 | SPARC,TUBB,GNAS,CSDE1,NES,LRPPRC,DLG5,SURF4,TUBB2A,PITP |
| 21 | 14823 | COL18A1,UBB,HNRNPA0,RHOA,SLC7A5,SKI,KDELR1,BAP1,DAZAP2, |
| 190 | 14823 | MLLT1,USP22,CFL1,MAVS,ZNF124,ALDOA,CALR,COL5A1,CES3,MY |
| 43 | 14823 | MLLT1,RPL13A,TPI1,SEC61A1,MIDN,NFE2L1,RPS24,CBX5,ATXN7L3 |
| 52 | 14823 | PABPC1,MLLT1,HSP90AA1,HSPA1B,MAPT,SSBP2,SEC61A1,MAP4,NF |
| 188 | 14823 | MLLT1,USP22,CFL1,MAVS,ZNF124,ALDOA,CALR,COL5A1,CES3,MY |
| 188 | 14823 | MLLT1,USP22,CFL1,MAVS,ZNF124,ALDOA,CALR,COL5A1,CES3,MY |
| 47 | 14823 | TUBB,SLC9A7,ALDOA,ABHD2,RACK1,CALM3,CALM1,CKB,SARAF,M |
| 57 | 14823 | TUBA1A,YWHAE,YWHAG,EIF1,STMN1,HNRNPU,CLSTN1,GLG1,ARL6I |
| 34 | 14823 | MLLT1,PRRC2B,TRIM28,PFN1,CDH6,RHOB,DDX39B,PABPN1,LMNB |
| 37 | 14823 | MLLT1,ACTR2,TUBB,UBA1,PGAM1,MSN,SLC7A5,MTSS2,ARL6IP1,T |
| 166 | 14823 | HSPA8,MLLT1,MED18,MAVS,ACTR2,UBC,EIF4G2,PRPF8,SOX4,FKBP |
| 50 | 14823 | CALR,CALM3,NPM1,DDB1,CPE,KIF5B,SLC7A5,HMGN2,CCT7,TOMM |
| 28 | 14823 | UGT8,PSAP,PTMA,NONO,LASP1,CAPRIN1,CDKN1A,LMNB2,RAI1,PS |
| 77 | 14823 | HSP90AB1,TUBB,ENO1,HSP90B1,TMF1,HSPA1B,FKBP14,RPL13A,SE |
| 29 | 14823 | PSAP,SOX4,BASP1,CCND2,MTSS2,CTDSP2,NOTCH2,RPL37,RABL6,H |
| 29 | 14823 | FTH1,NONO,CDKL1,MIDN,MAT2A,CD81,PFN1,PDIA6,LGALS3BP,RAP |
| 43 | 14823 | CFL1,CALR,YWHAE,CALM3,TMBIM6,IGF2,EIF1,LARP1,MSN,NCS1,L |
| 42 | 14823 | MAP1B,GTF3C4,TMEM170A,ATXN7L3B,PDIA6,HMGCS1,ARL6IP1,CN |
| 47 | 14823 | PABPC1,TUBB,MALAT1,FSCN1,IGF2,PKD1,TIMP2,RBBP4,JUN,NGRN |
| 26 | 14823 | ACTG1,TUBB,RPL4,GPI,RACK1,EIF1,RPL19,RPLP1,ZMIZ1,SRSF4,RP |
| 31 | 14823 | PKM,CALM3,PLD3,TRIM28,CBX5,SLC7A5,DDR1,RPL35,HRNR,LMNB |
| 39 | 14823 | FTH1,NCL,HSPA1B,H3-3B,SHISA9,YWHAQ,TPM4,RTN4,KPNB1,KIF5 |
| 38 | 14823 | EEF2,ACTR2,TUBA1B,MAP1B,NCL,APP,YWHAQ,HMGN2,SRP9,ANXA5 |
| 36 | 14823 | NCL,CES3,FKBP14,MAGED1,APP,RPS3,HMGB1,ZMIZ1,SMARCA5,AR |
| 35 | 14823 | NCL,CES3,FKBP14,APP,RPS3,HMGB1,ZMIZ1,SMARCA5,ARID1A,LMN |
| 80 | 14823 | ORC4,ODF2L,ACTR2,TUBB,TUBA1B,GANAB,SLC2A3,DNAJA1,TMEM |
| 43 | 14823 | MLLT1,RPL13A,TPI1,SEC61A1,MIDN,NFE2L1,RPS24,CBX5,ATXN7L3 |
| 41 | 14823 | MLLT1,HSP90B1,CANX,PKD1,VCL,MAP4,LARP1,LAMA5,LLGL1,GLG |
| 41 | 14823 | MLLT1,HSP90B1,CANX,PKD1,VCL,MAP4,LARP1,LAMA5,LLGL1,GLG |
| 49 | 14823 | ACTB,PRRC2B,PKM,CALR,PSAP,CLU,GDI1,AP2M1,COL1A2,TLN1,AR |
| 38 | 14823 | COL1A1,MLLT1,PRRC2B,TRIM28,CDH6,MYO5A,RHOB,ARHGDIA,DD |
| 32 | 14823 | GNAS,SMG1,NFE2L1,RHOA,JPT2,CSNK1E,GPAT4,SESN3,PEA15,CPS |
| 34 | 14823 | EEF1A1,DCX,PKM,HSP90B1,UBA1,CALR,MIDN,TARDBP,CDK16,CDK |
| 26 | 14823 | HSP90AA1,DBN1,FSTL1,SMG1,SMARCA5,NCS1,GLG1,SURF4,YME1L |
| 73 | 14823 | IGFBP5,EEF2,BTBD9,PRRC2B,ZNF99,PGK1,SOX4,YWHAE,SMG1,CS |
| 73 | 14823 | IGFBP5,EEF2,BTBD9,PRRC2B,ZNF99,PGK1,SOX4,YWHAE,SMG1,CS |
| 78 | 14823 | ACTB,EEF2,XRCC2,DCX,DBN1,MAP1B,PTMA,ALDH6A1,MARCKS,SM |
| 41 | 14823 | IGFBP5,HSP90B1,ZNF99,RPL13A,DDX3X,H1-0,HMGCS1,PTK7,DYNL |
| 49 | 14823 | ACTB,PRRC2B,PKM,CALR,PSAP,CLU,GDI1,AP2M1,COL1A2,TLN1,AR |
| 90 | 14823 | HSPA8,MLLT1,MAVS,UBC,MAP1B,SOX4,SEPTIN2,KLRD1,CSDE1,TIM |
| 43 | 14823 | DBN1,HSPA1B,H3-3B,CALM1,SET,SLC16A7,RPL9,CHD4,CYFIP2,HNR |
| 35 | 14823 | USP22,PKM,CALR,COL4A1,MIDN,SBF1,CLSTN1,TLN1,ARHGDIA,CDK |
| 18 | 14823 | SET,SLC7A5,NUCB1,LMNB2,KDM6B,TPM3,RAI1,CBX6,SLC38A2,SC |

AB1,VIM,TUBB,UBC,RPL3,DBN1,PRRC2B,ENO1,TUBA1B,FLNA,NCL,RPL7A,ALDOA,GANAB,COL6A
90AB1,HSP90AA1,TUBB,EIF4G2,PRRC2B,ENO1,TUBA1B,FLNA,PKM,GANAB,HSP90B1,PRPF8,YBX
PL3,PRRC2B,ENO1,FLNA,MAP1B,NCL,RPL7A,PKM,GANAB,HSP90B1,PRPF8,RPL8,ATP2B4,TMF1
EEF2,TUBA1A,HTT,HSP90AA1,ACTR2,VIM,TUBB,FTH1,KIDINS220,RPL3,RPS6,RPL4,FLNA,GANAE
JBA1,TMF1,HSPA1B,RPLP0,GNAS,DDX17,GPI,FASN,SPTBN1,ATP5F1B,MYH9,SPTAN1,PLXNB2,HSI
L,ALDOA,PKM,COL6A1,HSPA1B,ZNF99,CANX,RPL13A,KAT6A,YWHAE,FASN,SHISA9,MYH9,SEPTIN
B,TUBA1B,RPL7A,MTRNR2L8,PKM,RPL8,CALR,CANX,RPL13A,DVL3,YWHAE,GNAS,ATP5F1B,CAL
B,NCL,RPL7A,YBX1,UBA1,HSPA1B,COL5A1,PTMA,RPL13,DDX17,RACK1,FASN,ATP5F1B,SEPTIN2
,TUBA1B,FLNA,NCL,PRPF8,RPL8,HSPA1B,COL4A1,KAT6A,GPI,FASN,SPTBN1,ATP5F1B,SHISA9,M
EF2,BTBD9,HTT,HSP90AA1,TUBB,KIDINS220,EIF4G2,PRRC2B,TUBA1B,FLNA,PRPF8,UGT8,UBA1
S6,GANAB,ABHD2,HSPA1B,RPLP0,CANX,CLU,DDX5,SLC2A3,BSG,FASN,GNB1,SEPTIN2,APP,MARC
90B1,FADS1,PRPF8,RPL8,FBXL4,ATP2B4,ABHD2,COL4A1,GNAS,DDX17,ATP5F1B,GNB1,TMBIM
PTMA,YWHAE,KIF1A,ATP5F1B,CALM3,TMBIM6,CSDE1,TRIM28,EIF1,MIDN,RPL10,GTF2F1,PGAM
B,RPL8,UBA1,CANX,FASN,APP,GDI1,PLXNB2,VCP,YWHAG,PPP2R1A,STMN1,MORF4L1,MIDN,RPS
,TUBB,EIF4G2,DBN1,RPL4,PRPF8,ABHD2,TUBB2B,PGK1,YWHAE,RACK1,FASN,RPL5,KLRD1,TME
NAB,COL6A1,PRPF8,UGT8,UBA1,HSPA1B,PGK1,H3-3B,RACK1,ATP5F1B,GNB1,HSPD1,CKB,SPEN
PL4,PRRC2B,ALDOA,PKM,FADS1,HSPA1B,PSAP,PGK1,RPLP0,GNAS,RACK1,FASN,ATP5F1B,RPS8
YBX1,CALR,FSTL1,HSPA1B,PSAP,RPLP0,YWHAE,KIF1A,GPI,ALDH6A1,MYH9,NPM1,SNRNP200,EIF
RCKS,NPM1,CSDE1,ATP5F1A,EIF4A1,RPL19,GTF3C4,SBF1,MAP4,TMEM170A,HK1,RPL22,ATP1A
SPA1B,PTMA,KIF1A,FASN,SPTBN1,APP,PTMS,EIF3C,EIF4B,SPEN,YWHAQ,SET,HLA-C,SF3B1,AP2M
JGT8,ATP2B4,CALR,TUBB2B,H3-3B,PTMA,DDX5,SEC62,FASN,CALM3,GNB1,MARCKS,CALM1,PLX
K3R2,NCL,PKM,SOX4,SEC62,YWHAE,FSCN1,PTMS,MARCKSL1,AFF4,TRIM28,EIF1,TPI1,GTF3C4,
GNAS,DDX17,TMBIM6,PTMS,MARCKS,EIF3C,COL4A2,INA,EIF4A1,ANXA2,AGRN,SARAF,MIDN,CD
MARCKS,SMG1,PLD3,IGF2,UCHL1,RPLP1,LARP1,HNRNPA0,NACA,RPS24,PAPOLA,EIF5A,CCND2,
TMBIM6,TTC3,AFF4,TMSB10,RPL19,TPT1,AP3D1,HNRNPU,OGA,CNN3,LPCAT1,RPS24,EIF5,BEX
1,CSDE1,EIF3C,CHD3,AP2M1,TUBB4B,MORF4L1,C1ORF43,WDR1,ATXN7L3B,ATP2A2,PFKL,ACT
KSL1,AFF4,DDB1,RPS3,VAPA,MARCHF6,SBF1,ARF1,HNRNPU,OGA,NFE2L1,LARP1,RTN4,LPCAT1
2,HSPA1B,H3-3B,CANX,CLU,KIF1A,SLC2A3,BSG,FASN,GNB1,SEPTIN2,APP,CSDE1,AFF4,TRIM28
LC2A3,GPI,SPTBN1,GDI1,PTMS,SPTAN1,LRRC28,SSBP2,HMGB1,POLR2A,AP2M1,SEC61A1,STM
,CANX,CLU,SLC2A3,BSG,FASN,SPTBN1,GNB1,SEPTIN2,APP,CSDE1,AFF4,YWHAQ,TPI1,SEC61A1,
ATP5F1A,VCP,UBB,HACD3,SET,RPL19,RPL15,LDHB,HMGB1,RPLP1,RPL10,PAFAH1B1,MAP4,ITM
HD3,RPL10,MAP4,PGAM1,NCS1,ATXN7L3B,LASP1,ARL4C,B4GALNT4,TAGLN,STIP1,PEA15,CDKN
,ACTR2,FTH1,UBC,EIF4G2,GANAB,PRPF8,ABHD2,CANX,CLU,DDX5,SOX4,FKBP14,KIF1A,GNAS,G
LDH6A1,SPTBN1,HSPD1,ATP5F1A,DDB1,SET,SSBP2,ARF1,MAT2A,HNRNPU,LARP1,HNRNPA0,RP
T1,RPS4X,ITGB1,MORF4L1,RPL22,COL5A2,LASP1,MCL1,SFPQ,CCND2,SOX12,CALU,HMGCR,M
HSPA1B,RPLP0,H3-3B,SOX4,FKBP14,YWHAE,FASN,SMG1,NPM1,CSDE1,NONO,PSMD5,EIF1,TPT
P2M1,STMN1,ZMIZ1,NFE2L1,CLSTN1,RAB6B,SLC7A5,NLGN2,ATXN7L3B,RTN3,ASTN2,HDLBP,C
P2M1,STMN1,ZMIZ1,NFE2L1,CLSTN1,RAB6B,SLC7A5,NLGN2,ATXN7L3B,RTN3,ASTN2,HDLBP,C
1A,TRIM28,SEC61A1,ZMIZ1,HNRNPU,PGAM1,RHOA,CDH6,PLXNA1,RPL12,ANKRD52,CDKN1A,TO
1A,TRIM28,SEC61A1,ZMIZ1,HNRNPU,PGAM1,RHOA,CDH6,PLXNA1,RPL12,ANKRD52,CDKN1A,TO
HSPD1,IGF2,SPEN,SET,RPS3A,VCL,GOLGA8A,RPL29,SCARB2,KPNB1,EIF5,MCL1,NES,PRDX2,RP
SCN1,TRIM28,CHSY1,RPL15,LDHB,STMN1,MIDN,MAT2A,HNRNPA0,RPS24,RHOA,ATP2A2,RPS7,G
TMA,CALM3,SHISA9,PLD3,SET,LDHB,PKD1,BASP1,NFE2L1,HNRNPA0,EIF4H,RHOA,LENG8,LASP
SY1,AP2M1,NCAM1,TPM4,EIF4H,MSN,ATP2A2,TADA2B,ITM2C,SESN3,SEPTIN3,RAC1,C5ORF24,M
K1,P4HB,KLRD1,SMG1,TRIM28,ITGB1,MORF4L1,AKAP12,LARP1,RPL23,RTN4,LENG8,ARHGDI

,SEC61A1,PAFAH1B1,ZNF770,RPL9,ATP1A1,SOX12,CCN2,ITM2C,HMGCS1,CKAP5,CCT6A,SMCHD
3X5,CLSTN1,PFN1,EIF5A,GPAT4,LGALS3BP,RAC1,CDKN1A,HNRNPA3,GRINA,VPS35,MDK,RAB11
3,MARCKSL1,YWHAG,YWHAQ,EIF4A1,PPP2R1A,HLA-C,HMGB1,TUBB4B,MIDN,RPL10,VCL,ZNF77
1,LARP1,CDH6,ARL4C,TLN1,MTSS2,RHOB,ARHGDIA,DDX39B,PABPN1,LMNB2,MDK,HDGF,SELEN
3X5,CLSTN1,PFN1,EIF5A,GPAT4,LGALS3BP,RAC1,CDKN1A,HNRNPA3,GRINA,VPS35,MDK,RAB11
R2,PKM,PRPF8,RPL8,HSPA1B,ZNF99,PTMA,FKBP14,KAT6A,GPI,RACK1,PTMS,NPM1,VCP,NONO,S
CS1,CBX5,CLSTN1,LENG8,LASP1,PFN1,EIF5A,RAC1,CDKN1A,C5ORF24,HNRNPA3,VPS35,MDK,KD
10,RPL19,TPT1,AP3D1,OGA,CNN3,LPCAT1,RPS24,PJA2,EIF5,BEX3,CDH6,ARL4C,TARDBP,ARL6IP
3,CANX,CLU,MAGED1,SLC2A3,FASN,GNB1,APP,YWHAQ,SEC61A1,DNAJA1,AP2B1,SMARCA5,RTN
D1,LPCAT1,CLSTN1,LASP1,ASTN2,HDLBP,PLXNA1,MTSS2,RNF187,MLXIP,PARP1,LMNB2,ASXL1,C
,RPL5,SPTAN1,NPM1,TRIM28,EIF4A1,SF3B2,EIF4H,ATP1A1,GPAT4,MTSS2,DNAJC5,TARDBP,TOM
1,LARP1,CDH6,ARL4C,TLN1,MTSS2,RHOB,ARHGDIA,DDX39B,PABPN1,LMNB2,MDK,HDGF,SELEN
X,CLU,SLC2A3,FASN,GNB1,APP,YWHAQ,SEC61A1,DNAJA1,AP2B1,RTN4,RPL14,KIF5B,ATXN7L3B,
OL4A1,PGK1,CLU,GNAS,DDX17,P4HB,ATP5F1A,UBB,SET,HMGB1,AP3D1,SF3B2,NFE2L1,FADS2,F
SET,STMN1,NCAM1,TPM4,RTN4,MSN,SLC7A5,ITM2C,H1-0,SESN3,APPL1,TM9SF3,ANXA5,PPP1C
K3R2,RPL7A,PRPF8,ABHD2,DDX5,SOX4,FKBP14,DVL3,GNAS,FASN,SPTBN1,ATP5F1B,MYH9,SEPT
LC2A3,FASN,GNB1,APP,YWHAQ,SEC61A1,DNAJA1,AP2B1,NFE2L1,RTN4,RPL14,KIF5B,ATXN7L3B
S,MARCKSL1,HLA-C,ZNF43,AP2M1,NCS1,CBX5,CLSTN1,SLC7A5,NLGN2,LENG8,CRMP1,PFN1,G
CKSL1,CHD3,RPS16,RPS9,KPNB1,HK1,KIF5B,SFPQ,PRKAR1A,RPS10,CAPRIN1,APPL1,IPO5,ILF3,
M1,MIDN,LARP1,XRCC5,MSN,RPL14,RHOA,CDH6,ARL4C,TLN1,RHOB,PEA15,DDX39B,PABPN1,GF
HF6,RPS17,SBF1,LARP1,LPCAT1,KPNB1,KIF5B,SLC7A5,ATP1A1,ATP2A2,MCL1,XPO1,PNMA1,PLX
R2A,TUBB4B,RPS16,NACA,XPO1,LGALS3BP,UBE2H,RAB1A,PEA15,TIMP3,RPL36A,SELENON,OPT
PI1,SEC61A1,HNRNPU,HNRNPA0,MSN,NCS1,RAB6B,SLC7A5,NLGN2,LENG8,LASP1,RBBP4,ASTN
PI1,SEC61A1,HNRNPU,HNRNPA0,MSN,NCS1,RAB6B,SLC7A5,NLGN2,LENG8,LASP1,RBBP4,ASTN
ON,RPS24,MSN,NCS1,RHOA,SLC7A5,LENG8,ASTN2,HDLBP,EIF5A,CCND2,SOX12,RAB7A,ITM2C,H
A,EIF4H,NLGN2,RTN3,CCND2,RPS7,NDFIP1,MXD4,LAMP1,LMNB2,TCP1,VEGFA,MAPK1,PTBP1,G
EC61A1,GTF2F1,HNRNPU,HMGN2,RBBP4,CDH6,CCND2,MTSS2,ARL6IP1,YWHAB,SND1,GLUL,HN
3,APP,CALM1,VCP,SPEN,SARAF,VCL,SBF1,MAP4,PGAM1,TSPAN3,LLGL1,ATP1A1,CCND2,NDFIP1
SL1,RPL15,PDIA3,RPS3A,HNRNPU,CNN3,SCARB2,XRCC5,KPNB1,KIF5B,SLC7A5,XPR1,HDLBP,EIF
ABHD2,APP,CKB,SET,SF3B1,POLR2A,NFE2L1,SLC7A5,CD81,CCT7,SLC3A2,RPS15,RPS7,GPAT4,
SN,LLGL1,CSNK1E,CDH6,SOX12,ARL4C,TLN1,RHOB,HLA-B,DDX39B,NUCB1,PABPN1,LMNB2,MD
NXA2,CTNNA1,LDHA,RPSA,C1ORF43,CD81,ATXN7L3B,EIF3L,HMGCS1,GPAT4,TOP2B,RPL23A,T
CL,NFE2L1,LAMA5,LLGL1,MCL1,GLG1,PDIA6,PLXNA1,ITM2C,STIP1,CDKN1A,C5ORF24,ZC3H7B,
4L1,PXDN,COL1A2,RPL22,COL5A2,OAZ1,MCL1,CCND2,HMGCR,MMP2,GLUL,MORF4L2,CCT4,K
MIDN,MAP4,PGAM1,NFE2L1,LARP1,RPS24,CBX5,SLC7A5,HMGN2,LASP1,ITM2C,ARL4C,B4GALN
,PRPF8,HSPA1B,H3-3B,DVL3,SLC2A3,FASN,USP11,COL4A2,EIF1,TPI1,MRC2,RPL15,POLR2A,PAF
SN,LLGL1,CSNK1E,CDH6,SOX12,ARL4C,TLN1,RHOB,HLA-B,DDX39B,NUCB1,PABPN1,LMNB2,MD
A,GPI,CSDE1,ATP5F1A,AP2M1,KPNB1,CHD4,SFPQ,RPS15,TARDBP,ARL6IP1,HNRNPK,CDKN1A,H
31,EIF4A1,SSBP2,HNRNPU,SLC7A5,XPR1,LGALS3BP,MAP2,SKP1,PEA15,LMNB2,HSPA4,VPS35,CF
5F1B,SEPTIN2,HMGB1,POLR2A,BTF3,NFE2L1,RPS16,CNN3,RPL29,LLGL1,RPL26,ACTN4,RPS19,
RF4L1,COL1A2,RPL22,COL5A2,MCL1,CCND2,JUN,COL6A2,HMGCR,MMP2,TMEM132A,MORF4L
F1,ZNF770,LARP1,LPCAT1,KIF5B,SLC7A5,HMGN2,ATP2A2,PNN,XPO1,PNMA1,PLXNA1,SOX12,PR
SET,PPP2R1A,RPL11,RPL10,LDHA,RPL22,ENC1,ATXN7L3B,HMGN2,LASP1,HNRNPC,DDX3X,RPLP
CB,NOX4,CNTNAP2,FAM171A2,H2AZ1,NPDC1,RPS2,AHSA1,ACTR1B,EDF1,IGF2BP1,MTOR,TANC2
PA0,CHD4,PFN1,EIF5A,MMP2,PTK7,PLEC,KDM4B,PPP1R9B,TXLNA,GTPBP1,KIAA0895L,PEG10,S

ARCKSL1,TPI1,AP2M1,EIF4H,COL1A2,HNRNPC,CDH6,ITM2C,RPL35A,ARID1A,LGALS3BP,CTBP1,
TN1,RAB6B,SLC7A5,NLGN2,ATXN7L3B,LASP1,ASTN2,HDLBP,CCND2,DDR1,PLXNA1,RAB7A,H1-0
CL,NFE2L1,LAMA5,LLGL1,MCL1,GLG1,PLXNA1,ITM2C,STIP1,CDKN1A,C5ORF24,ZC3H7B,RNF1
GTF3C4,LDHB,RPS4X,TMEM170A,LDHA,ATXN7L3B,HMGN2,PDIA6,ACTN4,GNB2,RPS5,HMGCS1,
NO,TPI1,CHSY1,PKD1,MIDN,MAP4,PGAM1,NFE2L1,LARP1,HNRNPA0,CPE,CBX5,MCL1,HDLBP,GL
TN1,RAB6B,SLC7A5,NLGN2,ATXN7L3B,LASP1,ASTN2,HDLBP,CCND2,DDR1,PLXNA1,RAB7A,H1-0
VHAE,YWHAG,AP2M1,MIDN,MAT2A,GTF2F1,CPE,MSN,SLC7A5,NLGN2,CRMP1,ASTN2,EIF5A,CCN
TN1,RAB6B,SLC7A5,NLGN2,ATXN7L3B,LASP1,ASTN2,HDLBP,CCND2,DDR1,PLXNA1,RAB7A,H1-0
TN1,RAB6B,SLC7A5,NLGN2,ATXN7L3B,LASP1,ASTN2,HDLBP,CCND2,DDR1,PLXNA1,RAB7A,H1-0
TN1,RAB6B,SLC7A5,NLGN2,ATXN7L3B,LASP1,ASTN2,HDLBP,CCND2,DDR1,PLXNA1,RAB7A,H1-0
PE,PFN1,CCND2,ACTN4,HMGCS1,PRKAR1A,HNRNPDL,APPL1,CDKN1A,CTSB,ALKBH5,RAB5B,NC
4,VAPA,MARCHF6,SBF1,LARP1,LPCAT1,KIF5B,SLC7A5,ATP2A2,XPO1,PNMA1,PLXNA1,SOX12,RP
L3B,HMGN2,PFN1,RAB7A,ITM2C,ARID1A,CDKN1A,C5ORF24,ZC3H7B,PABPN1,ALKBH5,HDGF,T
LDHA,SLC25A3,HDGF,DPYSL2,CCND1,RABL6,WDR6,CAND1,RPS28,PLXNA3,PNMA8A,BEX4,ARIH1
S,DDB1,SET,TPI1,MIDN,CNN3,EIF4H,SLC7A5,RPL35A,LGALS3BP,NOVA2,MLXIP,RAB5B,HDGF,PT
S,DDB1,SET,TPI1,MIDN,CNN3,EIF4H,SLC7A5,RPL35A,LGALS3BP,NOVA2,MLXIP,RAB5B,HDGF,PT
ABHD2,PGK1,DDX5,SOX4,FKBP14,GNAS,MYH9,SEPTIN2,APP,KLRD1,TMBIM6,CSDE1,CHSY1,PPP
SS2,RHOB,ARHGDIA,CDKN1A,DDX39B,ZC3H7B,TM9SF3,LMNB2,LAMC1,PLEC,FKBP8,TPM3,GIG
NO,TPI1,CHSY1,PKD1,MIDN,MAP4,PGAM1,NFE2L1,LARP1,HNRNPA0,CPE,CBX5,MCL1,HDLBP,GL
S1,SLC7A5,LENG8,HMGN2,HDLBP,GLG1,ARID1A,CDKN1A,DLG5,C5ORF24,RNF187,PABPN1,ALK
TN1,RAB6B,SLC7A5,NLGN2,ATXN7L3B,LASP1,ASTN2,HDLBP,CCND2,DDR1,PLXNA1,RAB7A,H1-0
C,SESN3,TARDBP,CDK16,GLUL,PABPN1,TSPYL4,GRINA,RAB11B,HDGF,CCND1,PSMD3,SON,SAF
I1,LMNB2,SELENON,PLEC,SDF4,PPP1R9B,TXLNA,PHB2,YKT6,CBX6,PEG10,PPP1R10,CARM1,MA
12,RPL29,RTN3,TLN1,ST3GAL2,FDFT1,PDXK,RAB5B,ORAI2,RPL36A,PBX2,PSMD3,GRB2,WDR6,
B,LASP1,MCL1,HDLBP,RAB7A,SESN3,STIP1,RNF187,LMNB2,ALKBH5,VPS35,HDGF,CCND1,PSM
B,LASP1,MCL1,HDLBP,RAB7A,SESN3,STIP1,RNF187,LMNB2,ALKBH5,VPS35,HDGF,CCND1,PSM
SS2,RHOB,ARHGDIA,CDKN1A,DDX39B,ZC3H7B,TM9SF3,LMNB2,LAMC1,PLEC,FKBP8,TPM3,GIG
SS2,RHOB,ARHGDIA,CDKN1A,DDX39B,ZC3H7B,TM9SF3,LMNB2,LAMC1,PLEC,FKBP8,TPM3,GIG
SLC7A5,CHD4,JUND,LENG8,ASTN2,CCND2,RAB7A,ITM2C,RPL35A,ARID1A,NOVA2,TNPO1,RPS15
C,SESN3,TARDBP,CDK16,GLUL,PABPN1,TSPYL4,GRINA,RAB11B,HDGF,CCND1,PSMD3,SON,SAF
1,UBB,ITGB1,MIDN,NFE2L1,TSPAN3,PAPOLA,CCND2,ARL6IP1,ANKRD52,NARS1,CKAP5,ARCN1,
B,LASP1,MCL1,HDLBP,RAB7A,SESN3,STIP1,RNF187,LMNB2,ALKBH5,VPS35,HDGF,CCND1,PSM
LC7A5,NLGN2,HMGN2,CRMP1,ASTN2,CCND2,APLP2,ANKRD52,SND1,MAP2,CDKN1A,PABPN1,A
5,CHD4,JUND,LENG8,ASTN2,CCND2,RAB7A,ITM2C,RPL35A,ARID1A,NOVA2,TNPO1,RPS15A,GRE
EIF3C,MARCKSL1,RPS4X,RPS3A,MIDN,VCL,ITM2B,LARP1,RPSA,TSPAN3,RPS24,CBX5,RPL9,ATX
N1A,HS6ST1,DDX39B,ZC3H7B,RAB11B,HDGF,ATP6AP1,RPL28,STMN3,SARS1,VGF,CTXN1,TXLN
SP1,SOX12,B4GALNT4,TAGLN,ANKRD52,PPIB,PEA15,RNF187,U2AF2,LMNB2,MTRF1L,HDGF,PE
,ATP5F1A,VCP,NONO,ZMIZ1,LARP1,RPS24,ATXN7L3B,HDLBP,RPS5,HMGCS1,RPL35A,TSR2,YW
39B,ZC3H7B,U2AF2,MDK,RAB5B,SKI,KDM6B,RAPGEF1,PPP2CA,MAPK1,GIGYF1,WBP2,CTXN1,T
,ITM2C,SESN3,CDKN1A,DLG5,DDX39B,RNF187,GNAI3,PABPN1,RAPGEF1,PTBP1,GIGYF1,HYOU1
SP1,RPS4X,HNRNPU,WDR1,PEBP1,ACSL6,RPL12,SESN3,ARL6IP1,CCT2,HDGF,DPYSL2,MSL1,GI
RNPA0,MSN,PFN1,ARL4C,GNB2,GPAT4,TLN1,DDX39B,ZC3H7B,TM9SF3,GRINA,LAMC1,KDM6B,
RNPA0,MSN,PFN1,ARL4C,GNB2,GPAT4,TLN1,DDX39B,ZC3H7B,TM9SF3,GRINA,LAMC1,KDM6B,
A-B,ARHGDIA,RPL23A,C5ORF24,PPP1CB,ALKBH5,MLEC,RAB5B,SKI,RAPGEF1,TUBB2A,PPP2CA,
LGN2,LENG8,PFN1,HDLBP,PDIA6,PABPN1,LMNB2,GRINA,RAPGEF1,CCND1,GIGYF1,GPR107,T

STN2,GLG1,GNB2,MTSS2,ARHGDIA,NOVA2,LMNB2,ALKBH5,RAB5B,CAPZB,HNRNPUL1,PLCG1,P

,SBF1,MAT2A,COL1A2,NCS1,KIF5B,SLC7A5,LENG8,HNRNPC,HDLBP,GLG1,CDH6,RPL12,ITM2C,

,SBF1,MAT2A,COL1A2,NCS1,KIF5B,SLC7A5,LENG8,HNRNPC,HDLBP,GLG1,CDH6,RPL12,ITM2C,

LC7A5,NLGN2,HMGN2,CRMP1,ASTN2,CCND2,APLP2,ANKRD52,SND1,MAP2,CDKN1A,PABPN1,AI

LC7A5,NLGN2,HMGN2,CRMP1,ASTN2,CCND2,APLP2,ANKRD52,SND1,MAP2,CDKN1A,PABPN1,AI

DKN1A,CPSF2,LMNB2,RAI1,SREBF2,NCOR2,YKT6,NPLOC4,CYTH2,ABI2,LTBP4,TMEM184B,FAM

P1,COL1A2,SLC7A5,NLGN2,CRMP1,TLN1,TARDBP,ANKRD52,NOVA2,TSPYL4,GRINA,MAP7D1,FK

PS17,CCNI,SESN3,APLP2,SLC25A3,ARL6IP1,YWHAB,CCT6A,CDKN1A,HSPA4,MLEC,KDM6B,DYNL

1,GRB2,CAPZB,HNRNPUL1,TPM3,RPS28,GIGYF1,POLR2E,KIF5A,LRRC4B,UBFD1,KLHDC3,YARS

,CALU,PABPN1,KDM6B,SON,GIGYF1,PPP1R9B,FBRS,KLHDC3,PPP1R10,MAZ,BLCAP,TCF3,ESYT1

N1A,HS6ST1,DDX39B,ZC3H7B,RAB11B,HDGF,ATP6AP1,RPL28,STMN3,SARS1,VGF,CTXN1,TXLNA

,SEC61A1,SBF1,AP2B1,LARP1,RPL23,CTSD,EIF4H,RTN4,KIF5B,CCND2,DDX3X,CLTC,RPS5,MYO

CALM1,SEC61A1,AP2B1,HNRNPU,CBX5,WDR1,PFN1,CCND2,DDX3X,EIF3L,HMGCS1,MACF1,SES

,CPE,NCS1,NLGN2,CRMP1,ASTN2,HMGCS1,ARL6IP1,ARID1A,LUC7L3,CDKN1A,MLXIP,PABPN1,LI

,NFE2L1,TSPAN3,CCND2,ARL6IP1,NARS1,CCT6A,SLC35E2B,TMEM30A,PPIB,DLG5,PARP1,MLEC

,CPE,NCS1,NLGN2,CRMP1,ASTN2,HMGCS1,ARL6IP1,ARID1A,LUC7L3,CDKN1A,MLXIP,PABPN1,LI

RPL26,ACTN4,LMNB2,H3-3A,SRSF9,SUPT5H,RPS28,USP7,KDM3B,CCT5,SF1,TIMM50,APEX1,HIF

N2,EIF3C,UBB,AFF4,HMGB1,LPCAT1,KPNB1,NR2F1,MCL1,EIF5A,PEBP1,PDIA6,DDX3X,H1-0,RPS

4,SLC7A5,MMP14,RPL35A,CDKN1A,C5ORF24,HNRNPA3,RAB5B,GRB2,SERBP1,DDIT4,MTA1,CS

24,GAP43,HMGN2,SRSF4,CCND2,ITM2C,LGALS3BP,TM9SF3,RNF187,TCP1,HDGF,TUBB2A,SON,

IB,PGD,VAMP2,VEGFA,CCND1,DLK1,SOD1,WBP2,PSMA7,SFRP1,ANP32B,LRRC59,SERP1,STK35

ABHD2,HSPA1B,DDX5,SOX4,FKBP14,GNAS,DDX17,MYH9,RPL7,SEPTIN2,APP,KLRD1,TMBIM6,CS

N1A,HS6ST1,DDX39B,ZC3H7B,RAB11B,HDGF,ATP6AP1,RPL28,STMN3,SARS1,VGF,CTXN1,TXLNA

AB5B,SKI,SELENON,PTBP1,KDELR1,SMARCE1,POLR2E,MTA1,SF1,CSNK2A1,SLC35F5,SNRPD3,

X39B,ZC3H7B,HDGF,ATP6AP1,RAPGEF1,RPL28,STMN3,SARS1,WBP2,CTXN1,TXLNA,WDR45B,F

X39B,ZC3H7B,HDGF,ATP6AP1,RAPGEF1,RPL28,STMN3,SARS1,WBP2,CTXN1,TXLNA,WDR45B,F

P1,COL1A2,SLC7A5,NLGN2,CRMP1,TLN1,TARDBP,ANKRD52,TSPYL4,GRINA,MAP7D1,FKBP8,ST

P1,PSMD3,SFRP1,CLCN7,GABARAPL1,MAZ,PACS2,ATXN7L3,PRDX3,DIP2A,SEMA6B,TMEM94,IP6

,INIP,EID1,SQSTM1,VEGFA,TUBB2A,ADAM9,CAND1,ID4,WDR45B,XPOT,TAGLN2,TRA2B,TMEM16

24,GAP43,HMGN2,SRSF4,CCND2,ITM2C,LGALS3BP,TM9SF3,RNF187,TCP1,HDGF,TUBB2A,SON,

CA5,ATP1A1,EIF5A,DDX3X,RPS15,TMEM30A,TRIB2,RNF187,PPP1CB,NOX4,DDX42,SKI,RPL36A,E

P1,PSMD3,SFRP1,CLCN7,GABARAPL1,MAZ,PACS2,ATXN7L3,PRDX3,DIP2A,SEMA6B,TMEM94,IP6

24,GAP43,HMGN2,SRSF4,CCND2,ITM2C,LGALS3BP,TM9SF3,RNF187,TCP1,HDGF,TUBB2A,SON,

P1,COL1A2,SLC7A5,NLGN2,CRMP1,TLN1,TARDBP,ANKRD52,TSPYL4,GRINA,MAP7D1,FKBP8,ST

NRNPA0,GAP43,B4GALNT4,LGALS3BP,LAPTM4B,SHISAL1,RNF187,U2AF2,MLXIP,TLN2,RAB5B,HI

,ZC3H7B,LMNB2,MTRF1L,MAPK1,GIGYF1,NCOR2,TRAF7,SF1,PHB2,INTS3,TMEM184B,AGAP1,

LMNB2,LAMC1,KDM6B,SELENON,BCL9L,EP300,PHB2,KIAA0895L,PTPRF,CARM1,SH3GL1,SLC4

CT6A,CDKN1A,MLEC,KDM6B,CCND1,PPP6R1,TXNIP,CSNK2A1,PTEN,RAB1B,RALY,RAB14,LDLR,

PAN3,CCND2,GJA1,ARL6IP1,NARS1,CCT6A,SLC35E2B,TMEM30A,DLG5,PARP1,MLEC,RAB5B,DF

GAP43,B4GALNT4,LGALS3BP,LAPTM4B,SHISAL1,RNF187,U2AF2,MLXIP,TLN2,RAB5B,HDGF,SEL

7B,TM9SF3,LAMC1,GIGYF1,PPP6R1,SERBP1,POLR2E,TXLNA,FBRS,BCL7A,RAB1B,SH3PXD2A,T

NRNPUL1,SSB,STMN3,PTP4A1,KDM4B,PHB2,PTEN,HIF1A,ERGIC1,SMG5,KIAA0232,SH3BP4,WNK

1,MSN,CLSTN1,LASP1,HDLBP,DDR1,PLXNA1,RAB7A,ARL4C,JUN,TAGLN,ANKRD52,CTDSP2,MLXI

ARF1,AP2B1,NFE2L1,XRCC5,RHOA,ATP2A2,RPL12,RPS7,MTSS2,RPL35A,DNAJC5,ARID1A,ANKR

M1,NLGN2,LASP1,B4GALNT4,ANKRD52,ARHGDIA,STIP1,PEA15,RNF187,U2AF2,MTRF1L,TUBB2

ZC3H7B,U2AF2,MDK,SKI,KDM6B,CDC37,TRAF7,KIAA0895L,CBX6,POM121C,INTS3,SLC7A1,ABL
1,MSN,CLSTN1,LASP1,HDLBP,DDR1,PLXNA1,RAB7A,ARL4C,JUN,TAGLN,ANKRD52,CTDSP2,MLXI
CALU,RAB11B,GIGYF1,MEX3A,PPP1R9B,KLHDC3,PPP1R10,CLCN7,TPP1,BLCAP,TCF3,ESYT1,MK
,RPL9,MCL1,XPO1,GPAT4,MACF1,RPL35A,CDKN1A,VEGFA,PPP2CA,ADAM9,PTBP1,CORO1C,PFK
,NPM1,SEC61A1,AP2B1,GOLGA8A,RTN4,KIF5B,CSNK1E,CCND2,DDX3X,MYO5A,SLC35E2B,SQST
6,APLP2,ANKRD52,RAB1A,GLUL,HS6ST1,CPSF2,VCAN,HSPA4,MPRIP,CTNND1,KDM5B,SF1,ZFP3
8,PJA2,RPL9,CCT7,HDLBP,CCND2,DDX3X,RPS15,MYH10,PPP1CB,MORF4L2,HNRNPA3,TNPO1,TU
JUND,PNN,HNRNPC,HMGCS1,HNRNPDL,TARDBP,ARL6IP1,ARID1A,RAC1,CDKN1A,LMNB2,GRB2,
TPM3,GIGYF1,PPP6R1,POLR2E,TXLNA,KDM5C,EP300,RAB1B,MAZ,TGOLN2,C16ORF58,NMNAT2
IA,PEG10,PPP1R10,MAZ,RAB5C,SLC25A23,AMFR,POLDIP2,IGFBP4,TMEM184B,ELN,ANO8,B4GA
KIF5B,LENG8,TADA2B,COTL1,RAC1,CDKN1A,ANXA5,PDXK,H3-3A,TPM3,CORO1C,PTP4A2,MLF2,A
1,MSN,CLSTN1,LASP1,HDLBP,DDR1,PLXNA1,RAB7A,ARL4C,JUN,TAGLN,ANKRD52,CTDSP2,MLXI
COL1A2,SLC7A5,JUND,LENG8,RTN3,HDLBP,ITM2C,TLN1,ANKRD52,GLUL,PABPN1,GRINA,PTK7,
CTXN1,TXLNA,BCL9L,MTA1,KIAA0895L,TIMM50,PEG10,PPP1R10,MAZ,SLC25A23,COPZ1,IGFBP4,
COL1A2,SLC7A5,JUND,LENG8,RTN3,HDLBP,ITM2C,TLN1,ANKRD52,GLUL,PABPN1,GRINA,PTK7,
COL1A2,SLC7A5,JUND,LENG8,RTN3,HDLBP,ITM2C,TLN1,ANKRD52,GLUL,PABPN1,GRINA,PTK7,
NFE2L1,TSPAN3,PRKCSH,CCND2,ARL6IP1,NARS1,CCT6A,SLC35E2B,TMEM30A,DLG5,PARP1,ML
TSS2,ANKRD52,LGALS3BP,CDKN1A,GRINA,HDGF,FKBP8,EXTL3,CBX6,TIMM50,EFTUD2,NUCKS
P1,HNRNPA0,EIF4H,LASP1,CSNK1E,CCND2,ITM2C,ARID1A,DLG5,TSPYL4,PTK7,RAB5B,CNTNAP2
2,DDR1,MTSS2,LUC7L3,CDKN1A,TSR1,MORF4L2,LAMC1,SKI,RAB11B,PLEC,DYNLL2,TPM3,CYP5
AP2M1,MIDN,LDHA,EIF4H,RAB6B,CDH6,CCND2,ITM2C,LGALS3BP,U2AF2,PDXK,HDGF,PHC2,MPR
I3,CCND2,ARL6IP1,NARS1,CCT6A,SLC35E2B,TMEM30A,CDKN1A,DLG5,PARP1,MLEC,RAB5B,DPY
MDK,IPO7,SON,APEX1,VPS4A,SRSF2,NMNAT2,HMGA1,AGAP1,SHMT2,SETD5,THRA,RNF40
LDHA,RPS24,CBX5,ATXN7L3B,ATP2A2,ACTN4,GPAT4,ARL6IP1,RHOB,CALU,TRIB2,CDKN1A,PGR
ZC3H7B,U2AF2,MDK,SKI,KDM6B,CDC37,TRAF7,KIAA0895L,CBX6,POM121C,INTS3,SLC7A1,ABL
ZC3H7B,U2AF2,MDK,SKI,KDM6B,CDC37,TRAF7,KIAA0895L,CBX6,POM121C,INTS3,SLC7A1,ABL
ZC3H7B,U2AF2,MDK,SKI,KDM6B,CDC37,TRAF7,KIAA0895L,CBX6,POM121C,INTS3,SLC7A1,ABL
5,ZC3H7B,LMNB2,MTRF1L,MAPK1,GIGYF1,NCOR2,TRAF7,SF1,PHB2,INTS3,TMEM184B,AGAP1,N
052,HS6ST1,HDGF,EXTL3,CBX6,TIMM50,CHERP,EFTUD2,NUCKS1,WASF2,C16ORF58,UBE2D3,SC
052,HS6ST1,HDGF,EXTL3,CBX6,TIMM50,CHERP,EFTUD2,NUCKS1,WASF2,C16ORF58,UBE2D3,SC
052,HS6ST1,HDGF,EXTL3,CBX6,TIMM50,CHERP,EFTUD2,NUCKS1,WASF2,C16ORF58,UBE2D3,SC
,CDKN1A,MLXIP,LMNB2,MAPK1,TSC22D1,PPP1R9B,TXLNA,XPOT,SF1,SEC16A,HOXB5,PEG10,TF
E,PFN1,MCL1,HDLBP,EIF3L,RAC1,TOMM20,ALKBH5,ASXL1,DYNLL2,RAI1,MLF2,CD99L2,CRTAP,
D2,MYO5A,LMNB2,ALKBH5,RAD23B,MAPK1,PLCG1,PHC2,GPR107,DYNLL1,TXLNA,BCL9L,PTPRF
RPS3A,SF3B2,PXDN,TNS3,RPS9,ATF4,CXADR,RPL26,RBBP4,PRKCSH,HNRNPC,CCND2,DDX3X,GJ
APGEF1,CCND1,ARGLU1,KDELR1,BCL9L,PHB2,MAPRE2,KLHDC3,YARS1,ABI2,CAPZA1,MKNK2,E
RCA5,RPS24,KIF5B,LASP1,ITM2C,ARL4C,MYO5A,CKAP5,CALU,PEA15,TIMP3,GNAI3,VCAN,CAMSA
I3,CCND2,ARL6IP1,NARS1,CCT6A,SLC35E2B,TMEM30A,DLG5,PARP1,MLEC,RAB5B,DPYSL2,RPS
OL1A2,SLC7A5,JUND,LASP1,HDLBP,ITM2C,TLN1,SESN3,ANKRD52,ARHGDIA,GLUL,PABPN1,GRIN
G1,KDELR1,SMARCE1,AKIRIN1,CTXN1,MTA1,SF1,CSNK2A1,POM121C,LSM14A,SLC7A1,NOC2L,C
B,EIF5A,CCND2,SESN3,CDKN1A,MLEC,HDGF,SELENON,FKBP8,MTA1,NPLOC4,TM9SF4,OGT,NO
A,TPI1,SEC61A1,WSB1,GOLGA8A,RPL24,LUC7L3,CAMSAP1,MDK,TNPO1,G3BP2,MAPRE1,CAPZE
A,SMCHD1,TM9SF3,PPP1CB,LMNB2,HSPA4,ARF3,MVB12B,CCND1,CCAR2,TPM3,GIGYF1,TXNIP,
DDR1,ARL4C,NARS1,TMEM30A,CDKN1A,SHISAL1,ZC3H7B,MLXIP,LMNB2,MTRF1L,MDK,RAB5B,
M9SF3,LAMC1,GIGYF1,PPP6R1,SERBP1,POLR2E,TXLNA,FBRS,RAB1B,SH3PXD2A,TGOLN2,C16C

52,HS6ST1,HDGF,EXTL3,CBX6,TIMM50,CHERP,EFTUD2,NUCKS1,WASF2,C16ORF58,UBE2D3,SC
ERINC1,MARCHF6,RHOA,PDIA6,INIP,EID1,CCT2,VEGFA,RAPGEF1,PPP2CA,G3BP2,ADAM9,ID4,TF
3,CCND2,ARL6IP1,NARS1,CCT6A,SLC35E2B,TMEM30A,DLG5,PARP1,MLEC,RAB5B,DPYSL2,RPS
2,MSN,LASP1,MCL1,CCNI,TPM1,CCN2,MMP14,SESN3,ARL6IP1,CDKN1A,CNN2,MLEC,VEGFA,TPM
1,VPS4A,SRSF2,HMGA1,AGAP1,UBQLN4,SETD5,RNF40
XN,KCTD12,ELAVL1,SCRIB,KLHDC10,FGFR1,ELAVL3,HMGA1,NOTCH1,VPS51,SRSF10,ZCCHC3,
CTNNB1,MYH10,IPO5,HSPA4,G3BP2,GRB2,TNKS2,YME1L1,STAT3,XPOT,H2AZ1,PHB2,HOXB5,AT
HGDIA,CDKN1A,SKI,RAB11B,PBX2,GRB2,RPL28,SDF4,GIGYF1,STMN3,VGF,WBP2,CTTN,CLCN7,
4,MLXIP,TLN2,HSPA4,KDM6B,HDGF,SELENON,PLEC,MEX3A,YME1L1,FBRS,RETREG2,TMEM16
M6B,IPO7,SREBF2,APEX1,VPS4A,SRSF2,UBE2D3,HMGA1,AGAP1,RNF41,SETD5,TNPO2,GATAD2
C61A1,HNRNPU,CBX5,WDR1,PFN1,CCND2,DDX3X,MACF1,SESN3,RHOB,RAB1A,DLG5,C5ORF24,
HGDIA,CDKN1A,SKI,RAB11B,PBX2,GRB2,RPL28,SDF4,GIGYF1,STMN3,VGF,WBP2,CTTN,CLCN7,
M1,NLGN2,LASP1,B4GALNT4,ANKRD52,ARHGDIA,STIP1,PEA15,RNF187,U2AF2,MTRF1L,TUBB2
P2,NNAT,CDKN1A,NOP53,PARP1,TUBB2A,CCND1,KDM1A,TSC22D1,PCYOX1,YKT6,TMEM167A,C
TSR1,MORF4L2,MAPK1,IFITM3,CYP51A1,ADAR,KIF5A,TXLNA,CSNK2A1,PEG10,KIAA0100,LDLR,
P1,HNRNPA0,EIF4H,LASP1,CSNK1E,CCND2,ITM2C,ARID1A,DLG5,TSPYL4,PTK7,RAB5B,CNTNAP2
CALU,U2AF2,PPP1R9B,BRD2,ZNF512B,PRRC2A,KLHDC3,HNRNPAB,PPP1R10,BLCAP,TCF3,ESYT
TF2F1,PGAM1,SLC7A5,ATXN7L3B,HMGN2,MCL1,GLG1,ARL6IP1,RPL36,SND1,ARHGDIA,LMNB2
HRNR,SURF4,SRSF1,GIGYF1,STAT3,WDR45B,ZFP36L1,BRD2,TAGLN2,LRP10,BAZ2A,MAZ,SCD,N
,DDX39B,PABPN1,LMNB2,SKI,HDGF,SELENON,CCND1,SMARCA4,VANGL2,PPP1R9B,FBRS,EIF3D
7L3B,RTN3,SESN3,ANKRD52,RAB5B,TUBB2A,GIGYF1,WBP2,TXNIP,BCL9L,CSNK1G2,KIAA0895L
LG1,PDIA6,PRKAR1A,ANKRD52,ALKBH5,MLEC,SKI,RAPGEF1,CAPZB,MAP7D1,GIGYF1,WBP2,CT
0A,L1CAM,SRP9,TM9SF3,LMNB2,ALKBH5,ARGLU1,PTP4A2,ID4,RAI1,ZFP36L1,ARID1B,TAGLN2,
HRNR,SURF4,SRSF1,GIGYF1,STAT3,WDR45B,ZFP36L1,BRD2,TAGLN2,LRP10,BAZ2A,MAZ,SCD,N
S9,MRFAP1,ARL6IP1,HNRNPK,GDI2,HNRNPL,GCN1,RPL36A,SMARCE1,STT3A,SERBP1,DYNLL1,
CAN,MLEC,TUBB2A,PPP2CA,CCND1,GRB2,TPM3,SYP,ARIH1,ZFR,RPL7L1,NPLOC4,XRCC6,GAB
GAM1,SLC7A5,ATXN7L3B,HMGN2,MCL1,GLG1,ARL6IP1,RPL36,SND1,ARHGDIA,LMNB2,ASXL1,
MGN2,PFN1,TLN1,ANKRD52,KDM6B,HDGF,FKBP8,EXTL3,CBX6,TIMM50,EFTUD2,NUCKS1,C16C
,PLXNA1,SHISAL1,CTSB,ZC3H7B,RNF187,MLXIP,TSR1,MORF4L2,LMNB2,HSPA4,SELENON,MLF
2,CCND2,ITM2C,MTSS2,ANKRD52,SND1,MAP2,CTBP1,PARP1,ALKBH5,PTK7,SKI,TUBB2A,ARF3,
2L1,HNRNPA0,RAB6B,SLC7A5,NLGN2,HMGN2,ASTN2,CCND2,ARL6IP1,RPL36,SND1,TSPYL4,RA
CD81,ATXN7L3B,MCL1,RAC1,CDKN1A,RAB27A,NCKAP1,ORAI2,IPO7,DYNC1LI2,CAPZB,DIDO1,S
1,CDKN1A,HNRNPA3,VPS35,MDK,RAB11B,MAPK1,PSMD3,FKBP8,HNRNPD,GTPBP1,BRD2,SFRP
1,CDKN1A,HNRNPA3,VPS35,MDK,RAB11B,MAPK1,PSMD3,FKBP8,HNRNPD,GTPBP1,BRD2,SFRP
SP1,PPIB,RNF187,PABPN1,LMNB2,PLCG1,MEX3A,MLF2,TXNIP,PPP1R9B,WDR45B,MTA1,RPL7L
2,PTK7,DYNLL2,GRB2,STMN3,PPP1R9B,DFFB,FLCN,MMP15,CRTAP,NPLOC4,BAZ2A,ERGIC1,FK
BBP4,CDH6,RNF187,TNPO1,SRSF1,DYNLL2,MPRIP,AKIRIN1,TXNIP,SSR1,STARD7,RETREG2,FLC
BBP4,CDH6,RNF187,TNPO1,SRSF1,DYNLL2,MPRIP,AKIRIN1,TXNIP,SSR1,STARD7,RETREG2,FLC
GAM1,SLC7A5,ATXN7L3B,HMGN2,MCL1,GLG1,ARL6IP1,RPL36,SND1,ARHGDIA,LMNB2,ASXL1,
OA,TMF1,HSPA1B,RPL13A,FASN,SHISA9,MYH9,HSPD1,CHSY1,LDHB,CDKL1,EIF4H,TSPAN3,RPL1
,MLXIP,LMNB2,MAPK1,TSC22D1,PPP1R9B,TXLNA,XPOT,SF1,SEC16A,HOXB5,PEG10,TRIR,FLOT
S,MPRIP,MLF2,CTNND1,CRTAP,HDAC6,EEF1B2,LRRC41,TSNAX,SLC38A1,DDX6,TRIM27,UBE2V1
C2B,MAP1B,RPL7A,PRPF8,YBX1,TMF1,HSPA1B,H3-3B,SOX4,NPM1,EIF4B,EIF1,RPS3A,ITGB1,T
C,UBE2H,HLA-B,CDKN1A,RNF187,PPP1CB,ALKBH5,HNRNPA3,PPP2CA,NPEPPS,DUSP4,HYOU1,M
HGDIA,CDKN1A,EEF1A2,SKI,RAB11B,PBX2,GRB2,RPL28,SDF4,GIGYF1,STMN3,VGF,WBP2,CTTN,

CN2,GOLM1,ANKRD52,L1CAM,CDKN1A,LMNB2,RAD23B,TRA2B,SMC3,PTPRF,ANP32B,LSM14A,
K39B,SKI,MAPK1,FBRS,POM121C,ZMIZ2,SLC7A1,MBD3,HMGA1,MAP1S,WARS1,SLC12A7,COX6
FF4,INA,VAPA,STMN1,DNAJA1,PAFAH1B1,HNRNPU,SMARCA5,NACA,RTN4,RHOA,CD81,MCL1,JUN
,FASN,RPS8,EIF1,POLR2A,PAFAH1B1,TMEM170A,TPM4,RPS9,HK1,HMGN2,TPM1,COL6A2,CTBP
4A1,XRCC5,PTCD3,PSMD2,PRKCSH,SND1,ARCN1,RAC1,TSR1,RAD23B,VEGFA,G3BP2,SON,RPL2
M2C,CTBP1,FAM199X,PABPN1,PARP1,ALKBH5,TUBB2A,CCND1,MAPRE1,CAND1,GPR107,CSNK
IF5A,INIP,SND1,LGALS3BP,SRSF1,CDC37,FKBP8,STAT3,ZFP36L1,BRD2,LRP10,LRRC59,BAZ2A,
PAFAH1B1,TMEM170A,CNN3,TNS3,LPCAT1,IGFBP3,RHOA,CXADR,WDR1,XPR1,PEBP1,CCN2,CHP
NOP53,TUBB2A,MAPK1,TSC22D1,SREBF2,PCYOX1,RETREG2,TMEM167A,CHERP,NUDCD3,DENN
2,GRB2,GABARAP,MPRIP,NAP1L4,SYP,MSMO1,TXNDC5,ZFR,CUL3,RPL7L1,DKK3,XRCC6,RAB14
,HRNR,CDKN1A,RNF187,SRSF1,FKBP8,WDR45B,KDM5B,PRRC2A,BRSK1,BAZ2A,PALM,ANTXR1,
ARCKSL1,AFF4,HACD3,INA,PKD1,SEC61A1,TUBB4B,AP3D1,LARP1,RPS24,KPNB1,KIF5B,EIF5,BE
,PFN1,MCL1,PNN,XPR1,GLUL,CCT3,MDK,ABCA2,HNRNPUL1,MAP7D1,PTP4A2,TXLNA,BCL9L,UB
,FKBP8,KDELR1,TXNIP,CTXN1,BCL9L,FBXO21,BAP1,NOC2L,SEMA6B,NIBAN2,COLGALT1,PER1,
,FKBP8,KDELR1,TXNIP,CTXN1,BCL9L,FBXO21,BAP1,NOC2L,SEMA6B,NIBAN2,COLGALT1,PER1,
NB2,RAD23B,LUZP6,CMIP,ID4,H2AZ1,CD164,TRA2B,SMC3,HNRNPA2B1,PTPRF,ANP32B,SEC14L
RPL37A,STMN1,MORF4L1,HNRNPU,LARP1,HMGN2,PFN1,CCND2,DDX3X,HNRNPDL,INIP,RHOB,
X5,VAT1,CALU,RAC1,MAPK1,FKBP8,GIGYF1,HNRNPD,PPP1R9B,KLHDC3,NPLOC4,PPP1R10,TPP
2,ACTN4,CDKN1A,RAB5B,NOTCH2,CASC3,TP53BP1,RBM15B,EP300,FLOT1,ZFP36L1,ADCY6,PTE
MEM30A,RPL23A,HS6ST1,ZC3H7B,MDK,HDGF,VEGFA,TUBB2A,ARF3,CCND1,GRB2,HYOU1,PHB2
6B,PLCG1,SEC16A,CHERP,CARM1,LTBP4,SMARCD1,TMEM184B,MKNK2,NECTIN1,ZCCHC3,EPN
NP32B,LRRC59,SERP1,STK35,ABI2,CAPZA1,RAB5C,ARPP19,CERS2,TOX4,KLF13,ANP32E,BSCL2,
ACTN4,DDX3X,CTNNB1,UBE2H,TMEM30A,ETNK1,SKI,H3-3A,RPS15A,PTP4A1,SERBP1,XPOT,CD1
1,PTK7,SKI,MAPK1,AZIN1,HNRNPUL1,FKBP8,NCOR2,HNRNPD,CSNK2A1,BRD4,KIAA0895L,POM1
R10,SRSF11,IGF2BP1,CERS2,B4GALT5,UBE2Q1,SRSF7,WDFY1,SETD1A,PPM1A
K16,CDKN1A,RNF187,RAPGEF1,SON,APEX1,BAZ2A,PRR12,MSI1,NMNAT2,HMGA1,NTN1,NDRG1,
48,RAB5B,RAPGEF1,MAPRE1,LAPTM4A,PHC2,AKIRIN1,TXNIP,TRAF7,FLOT1,CBX6,RPL7L1,BLCA
F5A,PPIB,GLUL,ZC3H7B,LMNB2,LAMC1,KDM6B,SELENON,EP300,PHB2,KIAA0895L,NPLOC4,GN
F5A,PPIB,GLUL,ZC3H7B,LMNB2,LAMC1,KDM6B,SELENON,EP300,PHB2,KIAA0895L,NPLOC4,GN
L1,HNRNPC,RAB7A,STIP1,PABPN1,KDM6B,IPO7,GRB2,GPR107,MPRIP,SREBF2,MEX3A,KDM5C,
9,PPP2R1A,HLA-C,SSBP2,AP2M1,ZNF770,MAT2A,PGAM1,PAPOLA,LLGL1,CLSTN1,SLC7A5,NLG
ARS1,PACS1,GNA13,SLC25A36,H6PD,TNRC6A,SH3BGRL3,CRKL,ALDH9A1,KDM2A,OLA1,MAP2K4
KI,MAPK1,FBRS,POM121C,ZMIZ2,SLC7A1,MBD3,HMGA1,MAP1S,WARS1,SLC12A7,COX6B1,RN
1,HSPA4,TMEM248,SELENON,PTP4A1,TXLNA,CEMIP2,RPL7L1,GRK2,XRCC6,RAB1B,SRSF2,SOD
1,CLSTN1,ARHGDIA,LMNB2,RAB5B,RAB11B,HNRNPUL1,PLCG1,GPR107,MEX3A,PPP6R1,WBP2,
SP1,PPIB,RNF187,PABPN1,LMNB2,MEX3A,MLF2,TXNIP,PPP1R9B,WDR45B,MTA1,RPL7L1,APEX1
H,PDIA6,ACTN4,TARDBP,LAPTM4B,HDGF,CCND1,G3BP2,MAPRE1,SON,CYP51A1,SF1,KLHDC3,C
6IP1,CNN2,GNAI3,ETNK1,TNPO1,DSTN,DYNLL2,TPM3,TXNIP,SSR1,STARD7,RPL7L1,POM121C,
TLN1,RPL23A,DDX39B,PABPN1,MDK,PLEC,SRSF1,PPP1R9B,APEX1,LDLR,CYB5B,SUMO1,SCAM
,PNN,DDR1,EIF3L,RAB1A,COTL1,CPSF2,TM9SF3,MLXIP,MTRF1L,PRDX5,KDM6B,RAB11B,HDGF,
M3,PTP4A1,CRTAP,NPLOC4,ABI2,RAB1B,KLHDC10,AMFR,POLDIP2,UBE2Q1,COLGALT1,YWHAZ
1,HRNR,MMP2,PEA15,RNF187,TBC1D16,CCND1,MAPK1,ARPC2,DUSP4,SMARCE1,TXNIP,BCL9L,
Q,MIDN,VCL,LARP1,TSPAN3,CBX5,ATXN7L3B,ATP2A2,GPAT4,MTSS2,ARL6IP1,NARS1,CALU,HDA
2,KLHL24,PTK7,TNPO1,CNTNAP2,RPS15A,TPM3,MSL1,DFFB,TXLNA,EP300,SF1,GRSF1,CD164,
M1,AFF4,EIF4A1,VAPA,NFE2L1,DDX3X,RPS10,RHOB,SMCHD1,ATP6V0C,SRP9,ZC3H7B,MLXIP,TS

P14,TLN1,STIP1,TMEM30A,CPSF2,MDH2,ALKBH5,LAMC1,SELENON,CCND1,HYOU1,STARD7,KL
NM1,ID4,ADAR,CPD,CSNK1G2,GNA12,PNISR,BLCAP,OGT,MDM2,MECP2,NBAS,SOBP,RBFOX2,Z
XPO6,ELOVL5,FXR2,SEMA6B,NIBAN2,COLGALT1,CDC42EP4,PER1,ACIN1,RNF40,SLC39A1,EPHA
19,PPP2R1A,HLA-C,SSBP2,AP2M1,ZNF770,MAT2A,PGAM1,PAPOLA,LLGL1,CLSTN1,SLC7A5,NLG
3,ARL6IP1,CDKN1A,LMNB2,SRSF1,FKBP8,GIGYF1,MEX3A,TXNIP,AMOTL2,RGMB,CBX6,NPLOC4
E2L1,LDHA,NANOG,TM9SF3,PDXK,SKI,CCND1,FKBP8,GIGYF1,TXNIP,TMEM167A,ZMYM4,PEG10
19,PPP2R1A,HLA-C,SSBP2,AP2M1,ZNF770,MAT2A,PGAM1,PAPOLA,LLGL1,CLSTN1,SLC7A5,NLG
19,PPP2R1A,HLA-C,SSBP2,AP2M1,ZNF770,MAT2A,PGAM1,PAPOLA,LLGL1,CLSTN1,SLC7A5,NLG
AT2A,TSPAN3,ATF4,SLC35E2B,SRP9,CHD2,MAPK1,MAPRE1,RTL8C,TXLNA,NAP1L1,TXNDC5,PBX1
P1,HLA-B,NUCB1,CAMSAP1,TUBB2A,YWHAH,FBLN1,NAP1L1,EP300,KPNA2,ZFP36L1,CEMIP2,SL
2,SKI,HDGF,SELENON,CCND1,VANGL2,PPP1R9B,FBRS,YKT6,CBX6,CHERP,RAB5C,TCF3,NMNAT
MEM30A,RPL23A,HS6ST1,ZC3H7B,MDK,VEGFA,TUBB2A,ARF3,CCND1,GRB2,HYOU1,PHB2,RAB1
14,SLC2A3,GNB1,MYH9,SEPTIN2,APP,KLRD1,SMG1,CSDE1,CHSY1,PPP2R1A,SERINC1,HMGB1,I
20,PABPN1,VCAN,ETNK1,RAB5B,CCND1,MAPRE1,GRB2,KDELR1,PTP4A2,GPR107,CUL3,TMEM1
IA7,APEX1,SLIT1,DENND4B,AMFR,TNFAIP1,TRIM2,ATP1B3,GALNT10,ANKFY1,WIZ,HNRNPR,PIP5
PTIN2,AFF4,HMGB1,PGAM1,LPCAT1,RPS14,PEBP1,PDIA6,DDX3X,H1-0,HNRNPH1,IPO5,TMEM30
HNRNPUL1,HNRNPM,HNRNPA2B1,NDST1,NPLOC4,CYTH2,TLE1,SLC7A1,DCUN1D2,PDHA1,HSD17
GEF1,FBRS,GLUD1,TAGLN2,KCTD12,GLO1,MSH6,UBE2D3,CCDC80,MDM2,MECP2,SCAMP4,PS
SP1,PPIB,RNF187,PABPN1,LMNB2,MEX3A,MLF2,TXNIP,PPP1R9B,WDR45B,MTA1,RPL7L1,APEX1
N2,GNAI3,ETNK1,TNPO1,DSTN,DYNLL2,TPM3,TXNIP,SSR1,STARD7,PTEN,RPL7L1,POM121C,GR
,ARHGDIA,VEGFA,TUBB2A,PLCG1,SEC23A,RTL8C,DDIT4,EP300,RERE,KLHL42,HOXB5,PTEN,SE
L35,STIP1,PSMD1,PRRC2A,BAZ2A,C6ORF62,MAZ,TRAP1,USP5,MKNK2,RPL27A,ZCCHC14,ZSWI
2,NCKAP1,ARF3,GIGYF1,PHC2,MSMO1,XPOT,AHCY,LYPLA2,MKNK2,FOXP4,PPP1R14B,GGT7,R
3,DDX3X,MDH2,LAPTM4A,DYNLL2,PRKAR2A,PHB2,RER1,CRTAP,FKBP1A,GLO1,SRSF2,FNBP1L,
,PRDX5,H3-3A,G3BP2,CTNND1,PSMA7,MSMO1,TAGLN2,CBX6,SERP1,CARM1,SLC6A8,SLC7A1,T
ID1A,LMNB2,RAD23B,FYN,PTP4A1,KPNA2,TRA2B,SMC3,PTPRF,ANP32B,ABI2,RAB5C,DAZAP2,LH
IB2,RAD23B,FYN,PTP4A1,KPNA2,TRA2B,SMC3,PTPRF,ANP32B,ABI2,RAB5C,DAZAP2,LHFPL6,CA
170A,SMARCA5,RPSA,RPS24,LASP1,CALU,PEA15,GNAI3,VCAN,RPL27,NOTCH2,VEGFA,PPP2CA,G
3,ARL6IP1,CDKN1A,CPSF2,LMNB2,SRSF1,FKBP8,GIGYF1,MEX3A,TXNIP,AMOTL2,RGMB,CBX6,N
,LMNB2,HNRNPA3,MDK,RAB5B,SKI,FKBP8,IFITM3,RTL8C,POLR2E,MAP1LC3B,STK35,FLOT2,C
,LMNB2,HNRNPA3,MDK,RAB5B,SKI,FKBP8,IFITM3,RTL8C,POLR2E,MAP1LC3B,STK35,FLOT2,C
HGDIA,GRINA,MAP7D1,FKBP8,STMN3,MEX3A,SERBP1,ARIH1,FBRS,PHB2,HOXB5,PTPRF,HNRNP
X39B,PABPN1,LMNB2,HDGF,SELENON,CCND1,PPP1R9B,FBRS,YKT6,CBX6,CHERP,RAB5C,TCF3,
F2,GRB2,FBRSL1,POLR2E,FEM1B,GLO1,FKBP4,VPS4A,CAPZA1,PHGDH,UBE2D3,IP6K1,DAD1,S
N1A,DLG5,ASXL1,SON,FKBP8,TPM3,HYOU1,MLF2,TXNIP,KDM5C,ZFP36L1,RAB2A,MAZ,SH3PXD2
1,CXXC5,PTPRF,SPINT2,RPL7L1,ERGIC1,ZMIZ2,EFTUD2,SUMO1,RPS21,TBXA2R,LOXL2,SEPHS1
DE1,EGLN1,STMN1,ITGB1,MORF4L1,LASP1,ITM2C,ARL4C,PRKAR1A,SESN3,RHOB,BCLAF1,LAPT
DE1,EGLN1,STMN1,ITGB1,MORF4L1,LASP1,ITM2C,ARL4C,PRKAR1A,SESN3,RHOB,BCLAF1,LAPT
G1,SET,GTF3C4,RPS3,DNAJA1,GOLGA8A,HNRNPA0,LDHA,KIF5B,PEBP1,DDX3X,TLN1,TARDBP,R
L2,GRB2,PPP1R9B,DFFB,TXLNA,XPOT,FLCN,MMP15,CRTAP,BAZ2A,FKBP4,NMNAT2,ESYT1,SYN
HGDIA,GRINA,MAP7D1,FKBP8,STMN3,MEX3A,SERBP1,ARIH1,FBRS,PHB2,HOXB5,PTPRF,HNRNP
P2,MIDN,LASP1,CCND2,SESN3,TMEM30A,CDKN1A,DLG5,PARP1,ETNK1,RAB5B,KDM6B,DPYSL2,
NPDL,INIP,ARHGDIA,EID1,TSPAN6,ILF3,YWHAH,ARIH1,IL6ST,TMEM167A,PRPF4B,YY1,PHIP,DEK,
N1A,HS6ST1,ZC3H7B,UBE4A,FKBP8,RPL28,STMN3,PPP1R9B,WDR45B,FBRS,HOXB5,CBX6,HNR
,NIBAN2,H6PD,CAB39,UBE2Q1,WASF3,RRM2,KHDC4,SLC39A1

A1,PRPF8,YBX1,UBA1,FSTL1,HSPA1B,TUBB2B,PSAP,CANX,PTMA,MAGED1,DVL3,RPL13,GNAS,FA
X1,TMF1,ABHD2,HSPA1B,RPLP0,FKBP14,YWHAE,FASN,MYH9,P4HB,APP,RPL5,TMBIM6,PLD3,CO
,ABHD2,HSPA1B,TUBB2B,COL4A1,H3-3B,DDX5,SOX4,FKBP14,RPL13A,KIF1A,DDX17,BSG,FASN,
3,FTL,HSP90B1,PRPF8,UGT8,YBX1,CALR,ABHD2,HSPA1B,COL4A1,RPLP0,H3-3B,CANX,CLU,RPL6
PD1,MARCKSL1,SPEN,TPI1,GTF3C4,RPS3,TUBB4B,ZMIZ1,MAP4,SF3B2,GOLGA8A,PGAM1,LDHA
N2,TTC3,COL18A1,SNRNP200,EIF3C,ATP5F1A,TRIM28,SPEN,SET,RPL19,BASP1,HMGB1,SBF1,AK
LM3,NPM1,PLD3,CSDE1,IGF2,EIF3C,CKB,SPEN,TPI1,GTF3C4,HMGB1,POLR2A,RPS4X,RPLP1,RPS
,HSPD1,TRIM28,CKB,NONO,EIF4A1,POLR2A,RPL18,STMN1,RPLP1,RPL10,AP3D1,SBF1,CYB5R3,
IYH9,PTMS,NPM1,SNRNP200,SMIM14,EIF4B,TRIM28,SPEN,NONO,DDB1,YWHAQ,EIF1,HACD3,TPI
,FSTL1,HSPA1B,DVL3,YWHAE,GNAS,DDX17,GPI,GNB1,SMG1,PLXNB2,PLD3,EIF3C,MARCKSL1,AI
CKS,NPM1,CSDE1,AFF4,YWHAQ,EIF4A1,RPS3,AP2M1,SEC61A1,RPS3A,VCL,DNAJA1,PAFAH1B1,AF
5,MARCKS,FN1,RPL15,HMGB1,POLR2A,ITGB1,UCHL1,ANXA2,CYB5R3,NFE2L1,CTNNA1,FADS2,F
1,LARP1,RPS9,NCS1,CLSTN1,CD81,LASP1,CCNI,CCND2,SOX12,GPAT4,RPS10,B4GALNT4,ARL6II
17,VCL,ZNF770,MAT2A,SCARB2,KPNB1,COL5A2,PFN1,SPTBN2,HDLBP,PEBP1,RPL12,TLN1,GJA1
3IM6,PLXNB2,CSDE1,EIF3C,SPEN,DDB1,EIF1,RPS4X,ITGB1,TUBB4B,BTF3,RPLP1,AKAP12,ITM2B
,LDHB,PKD1,POLR2A,ARF1,SF3B2,PXDN,PGAM1,LDHA,PSMD2,RBBP4,ACTN4,ITM2C,GNB2,ANX
,MYH9,RPS3,HMGB1,RPLP1,RPL10,SBF1,MAP4,MAT2A,NFE2L1,HNRNPA0,FADS2,PSMD2,CHD4,
3C,ATP5F1A,EIF4B,MARCKSL1,TRIM28,NONO,DDB1,RPL15,SERINC1,RPL18,SBF1,HNRNPU,RPS
1,ATXN7L3B,LASP1,CCT7,PNN,XPO1,PDIA6,ACTN4,HMGCS1,GJA1,DNAJC5,ARL6IP1,HNRNPK,CN
I1,RPLP1,HNRNPA0,FADS2,RPL26,RBBP4,PFKL,HDLBP,EIF5A,MRFAP1,H1-0,CYFIP2,APPL1,CDKN
XNB2,HSPD1,SMIM14,FN1,YWHAQ,HACD3,CHSY1,MRC2,PDIA3,SEC61A1,ANXA2,AGRN,AP3D1,AK
,VAPA,STMN1,TUBB4B,NCAM1,MARCHF6,RPS17,SBF1,DNAJA1,MAT2A,LARP1,LPCAT1,HK1,KIF5
C42BPB,CTNNA1,TSPAN3,PAPOLA,RHOA,RPS20,CD81,RTN3,NES,TADA2B,XPR1,XPO1,PTN,GLG1
RPL35,SLC25A3,RPS27,ARHGDIA,SMCHD1,TMEM30A,RPL23A,GLUL,SERF2,PGRMC1,MLXIP,SQS
3,KHDRBS1,CDH6,TARDBP,ARL6IP1,ANKRD52,LAPTM4B,CTSB,CDH11,MDH2,TSR1,LMNB2,TNPC
N4,RPL35,GPAT4,CTNNB1,CKAP4,SHISAL1,TBC1D16,GCN1,PSMB2,RPL36A,PRPF6,RABL6,FAM1
I,KIF5B,SLC7A5,HMGN2,ATP2A2,PFN1,MCL1,HNRNPC,XPO1,CSNK1E,TKT,CCND2,PNMA1,PLXNA
,YWHAQ,TPI1,SEC61A1,VCL,DNAJA1,PAFAH1B1,AP2B1,CNN3,RTN4,RPL14,KIF5B,SLC7A5,CHD4,
N1,VCL,MAP4,AP2B1,NFE2L1,LARP1,LDHA,RTN4,PTCD3,KIF5B,MCL1,KHDRBS1,HDLBP,TPM1,A
,RPL10,VCL,DNAJA1,PAFAH1B1,AP2B1,RTN4,RPL14,KIF5B,SLC7A5,ATXN7L3B,JPT2,LRPPRC,TAI
2B,SMARCA5,XRCC5,RPL14,ATP2A2,RPL26,RPL31,EIF5A,RPS15,ATP1A2,RPS27,SMCHD1,TMEM
1A,DDX39B,ZC3H7B,RNF187,U2AF2,SKI,KDM6B,ATP6AP1,TUBB2A,HNRNPUL1,GPR107,SARS1,II
PI,FASN,ATP5F1B,GNB1,MYH9,RPL7,SEPTIN2,APP,KLRD1,TMBIM6,CSDE1,NONO,CHSY1,TMSB1
S24,ATP1A1,EIF5,MCL1,SCRN1,TKT,GLG1,SFPQ,CCND2,CCN2,ACTN4,DDX3X,ARL4C,ANKRD52,I
MP2,MORF4L2,COL2A1,LAMC1,KDM6B,HDGF,NOTCH2,VEGFA,TUBB2A,VHL,CAND1,SMARCC1,F
1,RPS4X,RPS3A,ARF1,PAFAH1B1,GOLGA8A,PGAM1,SMARCA5,RPL23,RPL24,RPL9,KIF5B,RPS14,
CND2,PLXNA1,RAB7A,H1-0,SESN3,MTSS2,TAGLN,ANKRD52,CTDSP2,NOVA2,RNF187,TSPYL4,ML
CND2,PLXNA1,RAB7A,H1-0,SESN3,MTSS2,TAGLN,ANKRD52,CTDSP2,NOVA2,RNF187,TSPYL4,ML
MM20,HDGF,SELENON,DUSP4,PITPNA,GIGYF1,PHC2,PTP4A1,POLR2E,CRTAP,CBX6,TMEM167A,
MM20,HDGF,SELENON,DUSP4,PITPNA,GIGYF1,PHC2,PTP4A1,POLR2E,CRTAP,CBX6,TMEM167A,
S7,GPAT4,RPL35A,TARDBP,HNRNPK,RPS12,MMP2,LUC7L3,FAM199X,SUGP2,PARP1,TAF15,MTR
GJA1,CTNNB1,MTSS2,ARL6IP1,ARID1A,NDFIP1,YWHAB,RAB1A,SMCHD1,MMP2,TIMP3,TOMM20,C
1,CSNK1E,CCND2,ACTN4,ITM2C,ARID1A,GLUL,DLG5,TSPYL4,PTK7,RAB5B,GCN1,CNTNAP2,ATP6
DH2,RNF187,ALKBH5,PDXK,SKI,TERF2IP,CCND1,LAPTM4A,DYNC1LI2,CAND1,CORO1C,MLF2,CT
A,STIP1,SLC35E2B,LAPTM4B,GLUL,CTSB,C5ORF24,KLHL24,HNRNPL,TSR1,CCT3,RAB5B,RPS15A

D1,TOMM20,PPP1CB,TSR1,LMNB2,TCP1,ACAP3,TNPO1,CCND1,SON,MAP7D1,ZFAND5,GIGYF1,I
B,MAPRE1,PSMD3,FKBP8,AMOTL2,KIF5A,GTPBP1,FLOT1,BRD2,SFRP1,UBFD1,NPLOC4,CHERP,
70,RPSA,CBX5,PSMD2,ATXN7L3B,RBBP4,HNRNPC,RPL12,CLTC,GPAT4,MACF1,RPLP2,ARID1A,U
NON,CCND1,PLEC,SRSF1,FKBP8,TPM3,GIGYF1,STMN3,PPP6R1,CTXN1,CDC42,PPP1R9B,TXLNA,
B,MAPRE1,PSMD3,FKBP8,AMOTL2,KIF5A,GTPBP1,FLOT1,BRD2,SFRP1,UBFD1,NPLOC4,CHERP,
SET,EGLN1,RPLP1,SBF1,ARF1,LARP1,CTNNA1,C1ORF43,SLC7A5,CXADR,EIF5A,ACTN4,ITM2C,MA
DM6B,RAB11B,RAPGEF1,PSMD3,MAP7D1,FKBP8,MEX3A,TXNIP,AMOTL2,ARIH1,WDR45B,GTPBP
1,HMGCR,TMEM30A,LMNB2,VPS35,TNPO1,TERF2IP,CCND1,CCNL2,SRSF1,DYNLL2,GRB2,NPEP
4,RPL14,KIF5B,ATXN7L3B,JPT2,CSNK1E,CCND2,PDIA6,DDX3X,MYO5A,HNRNPDL,UBE2H,CALU,A
CNTNAP2,HDGF,CCND1,GRB2,CAND1,TPM3,SDF4,GIGYF1,SLIT3,AMOTL2,ARIH1,BCL9L,FLCN,B
MM20,PARP1,SURF4,MAPRE1,SUPT5H,HNRNPUL1,UBE2R2,ACTN1,DIDO1,DHX30,FYN,HYOU1,SE
NON,CCND1,PLEC,SRSF1,FKBP8,TPM3,GIGYF1,STMN3,PPP6R1,CTXN1,PPP1R9B,TXLNA,PRKAR2
,JPT2,CSNK1E,TPM1,CCND2,PDIA6,DDX3X,MYO5A,HNRNPDL,UBE2H,CALU,ARHGDIA,SLC35E2B,
RPL24,RPL26,RBBP4,RPL31,HNRNPC,KHDRBS1,SPTBN2,RPS15,ITM2C,ARID1A,RPS12,CDK16,CD
B,ALKBH5,TCP1,SKI,H3-3A,HDGF,CCND1,DYNC1LI2,DUSP4,TPM3,PTP4A2,PTP4A1,CCT5,CTNND
TIN2,KLRD1,TMBIM6,NPM1,CSDE1,IGF2,NONO,YWHAQ,SET,CHSY1,SERINC1,HMGB1,POLR2A,IT
,JPT2,CSNK1E,CCND2,PDIA6,DDX3X,MYO5A,HNRNPDL,UBE2H,CALU,ARHGDIA,SLC35E2B,LUC7L
LG1,EIF5A,PLXNA1,ANKRD52,RAC1,CDKN1A,TSPYL4,HNRNPA3,GRINA,VPS35,MLEC,MDK,KDM6
CAND1,PTP4A2,HYOU1,MPRIP,PPP6R1,DCTN1,CRTAP,UQCRC1,YKT6,HNRNPA2B1,SLC2A1,FUS,
RINA,MDK,PTK7,ORAI2,KDM6B,HDGF,SDF4,GIGYF1,STMN3,MEX3A,CTXN1,PPP1R9B,POLR2E,TX
NA1,SOX12,PRKAR1A,MACF1,APLP2,HNRNPH1,RPN2,MLXIP,RAD23B,TMED2,UBXN4,ATG2B,AZI
N,PTBP1,KDELR1,RPL28,SMARCE1,TXNIP,MTA1,PRKAR2A,SF1,KPNA2,PTEN,GPC4,IMPDH2,LSM
2,HNRNPC,CDH6,MTSS2,RPL36,YWHAB,GLUL,TSPYL4,NCKAP1,KDM6B,RAB11B,HDGF,ATP6AP1
2,HNRNPC,CDH6,MTSS2,RPL36,YWHAB,GLUL,TSPYL4,NCKAP1,KDM6B,RAB11B,HDGF,ATP6AP1
H1-0,MTSS2,LGALS3BP,CTDSP2,NOVA2,RNF187,PABPN1,TSPYL4,RAB11B,SELENON,FTO,GRB2,
RB2,CASC3,STXBP1,SOD1,TXNIP,DHX15,KDM5C,STARD7,UBA52,TRA2B,NDST1,PNISR,EEF1B2,I
IRNPA3,NCKAP1,RAB5B,HDGF,ATP6AP1,FTO,MAP7D1,GIGYF1,CSNK1D,PTP4A1,GHITM,WBP2,TX
,SLC35E2B,IPO5,GLUL,CDKN1A,SRP9,U2AF2,TAF15,MTRF1L,RBMX,SKI,NOTCH2,CCND1,MAPK
3L,DNAJC5,APLP2,TAGLN,LGALS3BP,CKAP4,CALU,SQSTM1,PARP1,LMNB2,GRINA,PTK7,ASXL1,
APLP2,TARDBP,SLC25A3,HNRNPK,LGALS3BP,RPS27,CDKN1A,TOMM20,LMNB2,HNRNPA3,ORAI2
DK,PTK7,RAB5B,HDGF,VEGFA,SELENON,TPM3,GIGYF1,STMN3,PPP6R1,CTXN1,PPP1R9B,TXLNA,
MED2,PLEC,KDELR1,ZFAND5,RAI1,PRKAR2A,BRD2,MBTPS1,TBC1D9B,RPL7L1,PEG10,ACLY,CLC
,RNF187,U2AF2,HNRNPA3,RAB5B,SKI,PLEC,PSMD3,HNRNPUL1,SARS1,MPRIP,MEX3A,RTL8C,PO
DM6B,HDGF,VEGFA,TUBB2A,VHL,CCND1,EIF3E,CAND1,HNRNPM,SARS1,PTP4A1,CDC42,VDAC1,
T4,STIP1,PEA15,CDKN1A,CTDSP2,ZC3H7B,U2AF2,PABPN1,HNRNPL,LMNB2,ABCA2,PSMD3,FKB
AH1B1,TMEM170A,GOLGA8A,TPM4,EIF4H,RPS9,HK1,HMGN2,OAZ1,SLC3A2,SFPQ,MMP14,PRD
DK,PTK7,RAB5B,HDGF,VEGFA,SELENON,TPM3,GIGYF1,STMN3,PPP6R1,CTXN1,PPP1R9B,TXLNA,
NRNPA3,AHCYL1,CLSTN3,MDK,IPO7,SUPT5H,HNRNPUL1,HNRNPD,PPP6R1,CTNND1,TRAF7,KDM
RTAP,IARS1,PPP1CC,DAZAP2,SRSF11,UBE2D3,FOXP4,CNDP2,PSMD7,MYCBP2,ARHGAP5,TRIM27
RPL35A,ARCN1,RAC1,MORF4L2,RPL27,TCP1,CAPZB,SUPT6H,SF3A1,SERBP1,CSNK2A1,NXN,IAR
2,LAMC1,KDM6B,HDGF,VEGFA,TUBB2A,VHL,SCAF11,CAND1,PTP4A1,CDC42,CTNND1,KDM5B,F
KAR1A,APLP2,MLXIP,PARP1,RAD23B,TMED2,UBXN4,AZIN1,ADAM9,CAND1,ZFAND5,GIGYF1,MTR
P2,CDKN1A,TBC1D16,KDM6B,MARS1,NSG2,CCAR2,KDM4B,TXNDC5,EXTL3,BAZ1B,BRD2,FKBP1
2,EIF4A3,MFN2,IGF1R,FAM8A1,DHFR,OAT,DDHD2,SP1
TK35,PPP1R10,MAZ,SH3GL1,SH3PXD2A,SLC25A23,DAZAP2,IGF2BP1,SLC38A2,IGFBP4,RNF11,E

CDKN1A,DLG5,NOVA2,PDXK,HDGF,TUBB2A,CCND1,TPM3,PHC2,MPRIP,WBP2,PPP1R9B,DFFB,G
,ARL6IP1,TAGLN,ANKRD52,RPL23A,SHISAL1,HS6ST1,CTDSP2,NOVA2,ZC3H7B,MLXIP,TSPYL4,R
37,U2AF2,HNRNPA3,RAB5B,SKI,PLEC,PSMD3,HNRNPUL1,SARS1,MEX3A,RTL8C,POLR2E,TRAF7,S
GJA1,ARL6IP1,MYH10,LAMP1,CNN2,GNAI3,SSRP1,ETNK1,TNPO1,TOP1,DSTN,TCF25,ADAM9,DY
G1,PLXNA1,ITM2C,EIF3L,RAC1,STIP1,CDKN1A,CTDSP2,C5ORF24,ZC3H7B,TOMM20,U2AF2,ML
,ARL6IP1,TAGLN,ANKRD52,RPL23A,SHISAL1,HS6ST1,CTDSP2,NOVA2,ZC3H7B,MLXIP,TSPYL4,R
D2,ITM2C,ARID1A,LGALS3BP,ARHGDIA,CTDSP2,RNF187,TSPYL4,LMNB2,RAB5B,PSMD3,HNRNP
,ARL6IP1,TAGLN,ANKRD52,RPL23A,SHISAL1,HS6ST1,CTDSP2,NOVA2,ZC3H7B,MLXIP,TSPYL4,R
,ARL6IP1,TAGLN,ANKRD52,RPL23A,SHISAL1,HS6ST1,CTDSP2,NOVA2,ZC3H7B,MLXIP,TSPYL4,R
,ARL6IP1,TAGLN,ANKRD52,RPL23A,SHISAL1,HS6ST1,CTDSP2,NOVA2,ZC3H7B,MLXIP,TSPYL4,R
TCH2,CCND1,KDELR1,CSNK1D,PPP1R9B,ZFP36L1,ADCY6,IARS1,RPL7L1,LOX,FBXO21,TAOK1,
S7,PRKAR1A,APLP2,CCT6A,MLXIP,RAD23B,TMED2,UBXN4,AZIN1,ADAM9,CAND1,ZFAND5,GIGYF
UBB2A,TPM3,ZFAND3,MLF2,STAT3,RGMB,RPS27A,TMEM167A,KLHDC3,FKBP1A,LRRC59,RAB5
,PPP2CB,PTEN,RPL7L1,PSMD8,BAZ2A,TAOK1,YARS1,KIFAP3,SH3PXD2A,SOD2,DICER1,WASF2,F
BP1,SMARCE1,PHC2,WBP2,TXNIP,CTXN1,AMOTL2,PPP1R9B,DFFB,FBRS,MMP15,NXN,RGMB,AN
BP1,SMARCE1,PHC2,WBP2,TXNIP,CTXN1,AMOTL2,PPP1R9B,DFFB,FBRS,MMP15,NXN,RGMB,AN
2R1A,LDHB,PKD1,SERINC1,HMGB1,ITGB1,MORF4L1,TIMP2,SARAF,MIDN,MARCHF6,WSB1,AP3
YF1,STMN3,PPP6R1,CTXN1,POLR2E,TXLNA,RAB2A,PTPRF,FKBP4,RAB1B,TCF3,TGOLN2,C16OR
G1,PLXNA1,ITM2C,EIF3L,RAC1,STIP1,CDKN1A,CTDSP2,C5ORF24,ZC3H7B,TOMM20,U2AF2,ML
BH5,PDXK,GRINA,MLEC,HDGF,RAPGEF1,CCND1,CAPZB,GPR107,WBP2,TXNIP,CTXN1,PPP1R9B,
,ARL6IP1,TAGLN,ANKRD52,RPL23A,SHISAL1,HS6ST1,CTDSP2,NOVA2,ZC3H7B,MLXIP,TSPYL4,R
RS1,MEX3A,PPP1R9B,NXN,TAGLN2,CBX6,PRRC2A,STK35,BAZ2A,HSPG2,ATXN7L3,CERS2,NMNAT
Z,SLC25A23,TCF3,IGFBP4,TMEM184B,MGAT5,ELN,FZR1,IP6K1,B4GALT5,EFNB1,RDH11,EPN1,T
ON,CCAR2,SDHA,ATXN2L,HYOU1,NT5DC2,BAG6,RPL34,HIP1,VARS1,CSNK2A1,MMP15,UBAP2L,
D3,GRB2,GABARAP,NPEPPS,KDELR1,CCAR2,HYOU1,PTP4A1,MEX3A,MLF2,KIAA0895L,RGMB,GF
D3,GRB2,GABARAP,NPEPPS,KDELR1,CCAR2,HYOU1,PTP4A1,MEX3A,MLF2,KIAA0895L,RGMB,GF
YF1,STMN3,PPP6R1,CTXN1,POLR2E,TXLNA,RAB2A,PTPRF,FKBP4,RAB1B,TCF3,TGOLN2,C16OR
YF1,STMN3,PPP6R1,CTXN1,POLR2E,TXLNA,RAB2A,PTPRF,FKBP4,RAB1B,TCF3,TGOLN2,C16OR
A,GRB2,TPM3,EP300,CSNK2A1,NXN,KLHDC3,WASF2,ARPP19,PACS2,CERS2,SCD,UNC119B,ABL
RS1,MEX3A,PPP1R9B,NXN,TAGLN2,CBX6,PRRC2A,STK35,BAZ2A,HSPG2,ATXN7L3,CERS2,NMNAT
CT6A,SLC35E2B,TMEM30A,MMP2,DLG5,PGRMC1,PARP1,MLEC,RAB5B,DPYSL2,RPS15A,MAPK
D3,GRB2,GABARAP,NPEPPS,KDELR1,CCAR2,HYOU1,PTP4A1,MEX3A,MLF2,KIAA0895L,RGMB,GF
KBH5,TAF7,MAP7D1,KDELR1,ZFAND5,GIGYF1,PHC2,STAT3,BCL9L,KDM5C,NXN,TMEM167A,SL
2,TPM3,EP300,CSNK2A1,NXN,KLHDC3,WASF2,ARPP19,PACS2,CERS2,SCD,UNC119B,ABL1,MEC
N7L3B,HMGN2,ATP2A2,CCND2,RPL30,H1-0,GPAT4,RPS10,RPL35A,HNRNPDL,CALU,GLUL,CDKN
,WDR45B,FBRS,CBX6,WSCD1,BCL7A,MSI1,MAPK8IP2,NCKAP5L,EFNB1,VPS51,H2AX,NDRG1,MA
X2,MEX3A,MLF2,TXNIP,TRAF7,WDR45B,MTA1,UBFD1,CBX6,BRSK1,RPL7L1,APEX1,CLCN7,STRN
HAB,HLA-B,SKP1,GLUL,MDH2,PARP1,SSRP1,CCT3,RAD23B,HDGF,ARGLU1,CAND1,SON,FKBP8,
RAF7,FBRS,KIAA0895L,BAP1,INTS3,SLC7A1,CNBP,HMGA1,ABL1,MKNK2,NCKAP5L,NECTIN1,FA
,SARS1,NCOR2,KDM5C,TAGLN2,CBX6,STK35,BAZ2A,GABARAPL1,SAP18,ATXN7L3,FBXL19,EIF4
YF1,STT3A,UBQLN1,LOC728392,ARFGEF2,TXLNA,SF1,SLC39A6,PHB2,CRTAP,TRA2B,CBX6,HN
HDGF,PLEC,STMN3,MEX3A,CTXN1,TXLNA,EP300,PTPRF,PEG10,PPP1R10,BCL7A,MAZ,SLC4A2,SL
HDGF,PLEC,STMN3,MEX3A,CTXN1,TXLNA,EP300,PTPRF,PEG10,PPP1R10,BCL7A,MAZ,SLC4A2,SL
CCND1,GRB2,CAPZB,NPEPPS,TPM3,SDF4,GIGYF1,CYP51A1,MLF2,WBP2,BCL9L,LRP10,GLO1,G
NIP,PPP1R9B,KDM5C,FBRS,GLUD1,TAGLN2,KCTD12,PRRC2A,KLHDC3,GLO1,TRIR,YARS1,INTS

HC2,STMN3,GPR107,DGKD,MEX3A,WBP2,PPP1R9B,PHB2,PLXNB1,NPLOC4,PEG10,CHERP,LRR
ANKRD52,CTBP1,ARHGDIA,CDKN1A,C5ORF24,RNF187,PABPN1,SSRP1,LMNB2,ALKBH5,GRINA,
ANKRD52,CTBP1,ARHGDIA,CDKN1A,C5ORF24,RNF187,PABPN1,SSRP1,LMNB2,ALKBH5,GRINA,
LKBH5,TAF7,MAP7D1,KDELR1,ZFAND5,GIGYF1,PHC2,BCL9L,KDM5C,NXN,TMEM167A,SLC2A1,F
LKBH5,TAF7,MAP7D1,KDELR1,ZFAND5,GIGYF1,PHC2,BCL9L,KDM5C,NXN,TMEM167A,SLC2A1,F
118A,GTF3C1,ZNF532,CACNA1H,MAP1S,SRCIN1,EPHA4,FZD7,VPS37B
BP8,PHC2,STMN3,MEX3A,SERBP1,PPP1R9B,ARIH1,FBRS,PHB2,HOXB5,NXN,CBX6,PRRC2A,PTPF
L2,PPP6R1,TXNIP,ANP32A,STAT3,CSNK2A1,RAB1B,VPS41,RALY,RAB14,LDLR,DTX4,SH3PXD2A,D
1,PLEKHA6,PNISR,FLOT2,NUCKS1,PACS1,IST1,UBE2D3,HMGA1,USF2,SALL2,AREL1,MAPKAPK2
,MKNK2,H2AX,LDB1,UBE2V1,WASF3,SEPTIN9,ZNF516,PLXND1,KHDC4
,WDR45B,FBRS,CBX6,WSCD1,BCL7A,MSI1,MAPK8IP2,NCKAP5L,EFNB1,VPS51,H2AX,NDRG1,MA
5A,HNRNPK,CDKN1A,TOMM20,SKI,HDGF,VEGFA,CCND1,DYNLL2,LUZP6,ATP6V1B2,TXNDC5,PR
N3,ARL6IP1,NDFIP1,ANKRD52,RHOB,RAB1A,DLG5,C5ORF24,CAMSAP1,MLEC,RAB5B,ABCA2,G3
MNB2,TUBB2A,MAPK1,TSC22D1,GPR107,XPOT,SF1,SEC16A,HOXB5,RAB2A,NXN,PEG10,FKBP1A
,RAB5B,DPYSL2,RPS15A,MAPK1,KDELR2,DYNC1LI2,LUZP6,KDELR1,ATP6V1B2,CCT8,SF3A1,PP
MNB2,TUBB2A,MAPK1,TSC22D1,GPR107,XPOT,SF1,SEC16A,HOXB5,RAB2A,NXN,PEG10,FKBP1A
1A,GPC4,PACS1,DAZAP2,USP24,TGOLN2,CREBBP,SLC20A1,COL12A1,GTF3C1,NOTCH1,MDM2,
7,HNRNPH1,RPS12,IPO5,TMEM30A,LAPTM4B,TIMP3,RNF187,KLHL24,RBMX,TNPO1,NRDC,TUBE
NK2A1,HDAC6,KCTD12,PTEN,RPS2,HIF1A,SLC2A1,EFR3B,ACLY,VAPB,CCN1,UBR5,SFXN1,PRKAC
,HGS,CTXN1,AMOTL2,CTNND1,PRKAR2A,PPP1CC,POM121C,TMEM167A,NPLOC4,FKBP1A,TRIR,
,ABI2,CAPZA1,RAB5C,ARPP19,CERS2,TOX4,KLF13,ANP32E,IGF1R,BSCL2,SALL2,ZCCHC3,CDK4,
DE1,SPEN,CHSY1,SERINC1,HMGB1,RPL18,ITGB1,TUBB4B,MORF4L1,MIDN,MARCHF6,WSB1,PA
,WDR45B,FBRS,CBX6,WSCD1,BCL7A,MSI1,MAPK8IP2,NCKAP5L,EFNB1,VPS51,H2AX,NDRG1,MA
SM14A,SLC4A2,SLC7A1,MSI1,ANKS1A,NOC2L,C16ORF58,POLDIP2,RNF11,TNFAIP1,CAPN1,HSD
BRS,CBX6,APEX1,BCL7A,PRR12,CNBP,HMGA1,MAPK8IP2,NCKAP5L,EFNB1,COLGALT1,FARSA,N
BRS,CBX6,APEX1,BCL7A,PRR12,CNBP,HMGA1,MAPK8IP2,NCKAP5L,EFNB1,COLGALT1,FARSA,N
MN3,MEX3A,SERBP1,PPP1R9B,ARIH1,FBRS,PHB2,HOXB5,NXN,CBX6,PRRC2A,PTPRF,HNRNPAB,
K1,FAM168A,APPBP2,PSME3,EXOC7,PSMF1,NACC1,UNC13A,TNPO2,ASB6,NIPA1,KLHL36,SEPTI
7A,GTF2I,CAPZA1,BLCAP,SOD2,MOAP1,CSE1L,ARPP19,CREBBP,COL12A1,TSNAX,IGSF3,TNRC6
,HGS,CTXN1,AMOTL2,PRKAR2A,PPP1CC,POM121C,TMEM167A,NPLOC4,FKBP1A,TRIR,ERGIC1,
XTL3,GLUD1,CUL3,RPL7L1,APEX1,GTF2I,MTOR,RCN2,RPS26,PUM1,LMAN2,IGF1R,SALL2,SRSF
K1,FAM168A,APPBP2,PSME3,EXOC7,PSMF1,NACC1,UNC13A,TNPO2,NIPA1,KLHL36,SEPTIN9,SL
,HGS,CTXN1,AMOTL2,PRKAR2A,PPP1CC,POM121C,TMEM167A,NPLOC4,FKBP1A,TRIR,ERGIC1,
MN3,MEX3A,SERBP1,PPP1R9B,ARIH1,FBRS,PHB2,HOXB5,NXN,CBX6,PRRC2A,PTPRF,HNRNPAB,
DGF,SELENON,TUBB2A,TSC22D1,GIGYF1,CTXN1,PCYOX1,PALLD,BCL9L,PRKAR2A,GTPBP1,GRS
NIBAN2,MKNK2,NECTIN1,TOB2,UBQLN4,SLC5A6,MCM7,ANKRD13B,NF2,RNF40
2,DAZAP2,LRP3,COPZ1,C16ORF58,TTC28,SLC38A2,ESYT1,MKNK2,TNFAIP1,SUMO1,CCNY,IP6K
DTX4,KIAA0232,SH3PXD2A,DICER1,UBE2D3,SLC38A2,IGF1R,FXR1,HLA-A,B4GALT5,TNRC6A,EC
YSL2,RPS15A,MAPK1,ATG2B,DYNC1LI2,LUZP6,KDELR1,ATP6V1B2,SF3A1,PPP6R1,BMPR2,TXN
LENON,TUBB2A,TSC22D1,GIGYF1,CTXN1,PCYOX1,PALLD,BCL9L,PRKAR2A,GTPBP1,CBX6,TAOK
CF3,TGOLN2,C16ORF58,NMNAT2,BAG5,CNBP,FXR2,IGF1R,SCAMP4,GALNT10,CCNY,ARL8A,NA
1,CLIC1,COPE,SALL2,NUP153,ADRM1,TOB2,KDM2A,TET3,PRELID1,UQCR11
P,LMNB2,ASXL1,RAB11B,TUBB2A,CDC37,GABARAP,FKBP8,TPM3,GIGYF1,GPR107,HGS,NCOR2
52,RAB27A,PARP1,RAB5B,LAPTM4A,PCSK1N,GIGYF1,ILF2,MPRIP,GHITM,TXNDC5,RPL7L1,LRR
A,SREBF2,RPL7L1,STRN4,SLC4A2,OGT,AGAP1,MGAT5,NIBAN2,MKNK2,ATP1B3,NSD1,UBE2V1,

.1,NIBAN2,NCKAP5L,NECTIN1,MAPKAPK2,ZNF703,MAP1S,SLC12A7,COX6B1,FARSA,MYL12A,XPC
P,LMNB2,ASXL1,RAB11B,TUBB2A,CDC37,GABARAP,FKBP8,TPM3,GIGYF1,GPR107,HGS,NCOR2
NK2,SIN3B,LDB1,UBE2V1,TRAF3,SEPTIN9,ZNF516,SRCIN1,PLXND1,PIEZO1
.M,STAT3,CSNK2A1,UBE2G2,SELENOW,PREPL,RPS2,PEG10,CLCN7,ELAVL1,KIAA0232,PDPK1,OC
M1,PABPN1,SKI,SRP14,VEGFA,SELENON,CCND1,DYNLL2,SLIT3,MPRIP,AKIRIN1,PCYOX1,PRKAR
6L1,TAGLN2,HNRNPA2B1,NCOA4,GLO1,ERH,YARS1,RAB1B,FBLN5,PTGFRN,SMARCD1,SAP18,NI
JBB2A,ATG2B,SRSF1,ID4,SERBP1,RTL8C,TP53BP1,SPINT2,XRCC6,LTBP4,MAZ,NUCKS1,FNBP1L
SON,CCAR2,ATP6V1B2,YWHAH,ARIH1,BCL9L,SPINT2,SSR3,PNISR,SOD2,WASF2,SCD,DIP2A,ABL
,FXR2,SCAMP4,GALNT10,CCNY,ZFP91,ARL8A,NACC1,CCDC6,ATG9A,POMGNT1,SECISBP2L,SET
LT5,EFNB1,SYNGAP1,ZNF703,MCFD2,RNF4,TMEM50A,PIN1,SETD1B,SIN3A
AMOTL2,CCT5,CTNND1,KDM5B,SF1,TAGLN2,ABCC9,SLC6A8,MAZ,PNISR,SOD2,NUDT19,NASP,C
P,LMNB2,ASXL1,RAB11B,TUBB2A,CDC37,GABARAP,FKBP8,TPM3,GIGYF1,GPR107,HGS,NCOR2
SKI,TUBB2A,MAPK1,AZIN1,MPRIP,ARIH1,KDM5C,CSNK2A1,GRSF1,KIAA0895L,TAGLN2,PRRC2A,
,TMEM184B,ELN,B4GALT5,EFNB1,PCGF3,COLGALT1,NID1,PIN1,TET3,SETD1B,RNF40,SIN3A
SKI,TUBB2A,MAPK1,AZIN1,MPRIP,ARIH1,KDM5C,CSNK2A1,GRSF1,KIAA0895L,TAGLN2,PRRC2A,
SKI,TUBB2A,MAPK1,AZIN1,MPRIP,ARIH1,KDM5C,CSNK2A1,GRSF1,KIAA0895L,TAGLN2,PRRC2A,
.EC,RAB5B,DPYSL2,RPS15A,MAPK1,DYNC1LI2,LUZP6,KDELR1,ATP6V1B2,SF3A1,FLNC,PPP6R1,T
1,C16ORF58,HMGA1,TMEM184B,ABL1,KIFC2,CLUH,ANKFY1,NCK2,MAPKAPK2,NUP98,NUS1,MC
,ABCA2,ATP6AP1,SELENON,RAPGEF1,CDC37,CAPZB,GPR107,SREBF2,MEX3A,MLF2,AMOTL2,AF
1A1,TXNIP,AMOTL2,FLCN,KPNA2,POM121C,TMOD2,KIAA0100,ERGIC1,CAPZA1,DAZAP2,PHGDH
IP,MLF2,SEC23A,WBP2,LRP10,BCL7A,SOD2,DAZAP2,PACS2,LYPLA2,CNBP,CCDC80,MDM2,MEC
YSL2,RPS15A,MAPK1,DYNC1LI2,LUZP6,KDELR1,ATP6V1B2,SF3A1,PPP6R1,TXNIP,MSMO1,CSNK2

MC1,C5ORF24,HSPA4,SURF4,RAB5B,PSMB2,NOTCH2,TUBB2A,CCND1,PBX2,IPO7,FLNB,SMARC
.1,NCKAP5L,NECTIN1,MAPKAPK2,ZNF703,MAP1S,SLC12A7,COX6B1,FARSA,MYL12A,XPC,RNF4,A
.1,NCKAP5L,NECTIN1,MAPKAPK2,ZNF703,MAP1S,SLC12A7,COX6B1,FARSA,MYL12A,XPC,RNF4,A
.1,NCKAP5L,NECTIN1,MAPKAPK2,ZNF703,MAP1S,SLC12A7,COX6B1,FARSA,MYL12A,XPC,RNF4,A
NIBAN2,MKNK2,NECTIN1,TOB2,UBQLN4,SLC5A6,MCM7,ANKRD13B,NF2,RNF40
CD,HMGA1,ABL1,ANKFY1,MAPKAPK2,NUP98,MCFD2,RPRM,ARL8A,NACC1,VAV2,RELA,NFASC,GF
CD,HMGA1,ABL1,ANKFY1,MAPKAPK2,NUP98,MCFD2,RPRM,ARL8A,NACC1,VAV2,RELA,NFASC,GF
CD,HMGA1,ABL1,ANKFY1,MAPKAPK2,NUP98,MCFD2,RPRM,ARL8A,NACC1,VAV2,RELA,NFASC,GF
RIR,FLOT2,SLC7A1,C16ORF58,NMNAT2,UNC119B,WDTC1,ESYT1,BSCL2,SYNCRIP,HMGB3,NECT
ZMYM4,MAP1LC3B,GNL1,MOAP1,KRT8,TGOLN2,KLHDC10,TEAD1,ESYT1,GTF3C1,KLF13,SUMC
F,LRP10,GRK2,TAOK1,LDLR,MAZ,PLEKHB2,RAB5C,SLC25A23,C16ORF58,PHIP,UBE2D3,FXR1,PI
A1,JUN,NGRN,UBE2H,EID1,EEF1A2,MLXIP,MTRF1L,TCP1,VPS35,CLSTN3,CBX1,RAD23B,ARF3,TN
EFNB1,RBFOX2,CCDC85C,ZNF516,TSN,POGZ,SECISBP2L
AP1,CCT3,SCAF1,PPP2CA,G3BP2,SRSF1,ARGLU1,UBE2R2,SON,KDELR1,SMARCE1,YWHAH,SF3A
15A,MAPK1,DYNC1LI2,LUZP6,KDELR1,ATP6V1B2,SF3A1,PPP6R1,TXNIP,MSMO1,CSNK2A1,GRSF
IA,TUBB2A,CCND1,PSMD3,MAP7D1,FKBP8,STMN3,MEX3A,SERBP1,ADAR,ARIH1,FBRS,PHB2,HOX
C16ORF58,POLDIP2,RNF11,TNFAIP1,FOXP4,CAPN1,PPP2R5D,CRKL,NUP98,TFRC,RALBP1,DUSP
C2L,ARPP19,REEP5,EFNB1,IQSEC1,PMEPA1,LDB1,CACNA1H,CDKN1B,ATG9A,NUMBL,PPP1R12A
3,RPS28,PTP4A1,RTL8C,WBP2,SMS,LRRC4B,RAB2A,YARS1,PLEKHA6,PLEKHB2,RAB5C,RBPJ,CHI
NAP1L1,PHB2,CSNK1G2,CBX6,CHERP,MTCH1,SH3PXD2A,ATXN3,ATP5MC3,C16ORF58,KLHDC1
VEGFA,CCND1,TPM3,GIGYF1,GPR107,WBP2,TRAF7,KDM5C,CAPZA1,SOD2,WASF2,IST1,SMARCD
ORF58,NMNAT2,BAG5,CNBP,FXR2,IGF1R,SCAMP4,GALNT10,CCNY,ARL8A,NACC1,MYL12A,CCDC

CD,HMGA1,ABL1,ANKFY1,MAPKAPK2,NUP98,MCFD2,RPRM,ARL8A,NACC1,VAV2,RELA,NFASC,GP
RA2B,KCTD12,PPP1CC,TMEM167A,CEP170,GTF2I,PNISR,CAPZA1,SOD2,FOXK2,CSE1L,SLC20A1,
15A,MAPK1,DYNC1LI2,LUZP6,KDELR1,ATP6V1B2,SF3A1,PPP6R1,TXNIP,MSMO1,CSNK2A1,GRSF
M3,CDC42,TAGLN2,RAB5C,RBPJ,PACS2,CERS2,ATP13A3,IGF1R,IP6K1,KPNA6,PSMC4,ZEB1,NR2

PSMF1,ARL8A,SLC12A7,CDC42EP4,PIN1,SP1,SETD1B,RNF40,VPS37B
TRX,PTEN,SEPTIN7,HNRNPAB,BAP1,TRAM1,SPAG9,DICER1,OGT,MED13,C16ORF58,GNA13,DEK,
CYTH2,XPO6,ABL1,SCAMP4,PCGF3,ZCCHC3,H2AX,SYNGR1,ARL8A,OTUB1,NACC1,UNC13A,THBS
7A,TRIR,RAB5C,SLC7A1,HSDL1,AGAP1,PKIA,IGF1R,SLC38A1,SUMO1,GALNT10,SYT15,TNRC6A,A
2B,RNF40
,CAMSAP1,MLEC,RAB5B,DPYSL2,CCND1,G3BP2,MAPK1,ATG2B,AZIN1,KDELR2,DYNC1LI2,HNRN
CYTH2,XPO6,ABL1,SCAMP4,PCGF3,ZCCHC3,H2AX,SYNGR1,ARL8A,OTUB1,NACC1,NKIRAS2,UNC
A,SREBF2,RPL7L1,STRN4,SLC4A2,OGT,AGAP1,MGAT5,MKNK2,NSD1,UBE2V1,UBQLN4,NACC1,C
CHERP,SEC14L1,SMG5,NUDCD3,AKT2,TUBB4A,CD44,RNF41,FAM168A,SEMA4C,SRF,ZFP91,NKIR
PLEKHA6,SOD2,MED13,KLHDC10,SYNCRIP,TNFRSF10B,ILRUN,ANKRD50,CDKN1B,ATP5F1C,COX
,ATP6AP1,SELENON,RAPGEF1,CDC37,CAPZB,GPR107,SREBF2,MEX3A,MLF2,AMOTL2,ARIH1,SF
1,MKNK2,LDB1,UBE2V1,GPRIN1,SEPTIN9,ZNF516,PLXND1,LDOC1
,ASXL1,RAB11B,SELENON,RAPGEF1,CDC37,CAPZB,FKBP8,RPL28,HGS,SERBP1,STAT3,RETREG2
ISG1,RANGAP1,TAPBP,LUC7L2,KLF13,FZR1,COX8A,EXT2,COLGALT1,OXSR1,CCDC85C,ZDHHC5
),ATRX,YKT6,CBX6,CHERP,RAB5C,TCF3,PSMC1,ANKRD12,NMNAT2,ELN,SCAMP4,IP6K1,EPN1,TC
.,UBFD1,TMEM167A,LRP10,ERGIC1,PRR12,RAB5C,DENND4B,UBE2D3,AMBRA1,FGFR1,MKNK2,
XN1,KDM5C,FBRS,GLUD1,TAGLN2,KCTD12,POM121C,GLO1,FKBP4,MSH6,SCAMP5,NIBAN2,KL
,RAB5C,PDPK1,TRIM8,AASDHPPT,SCAMP1,CCDC80,MDM2,EXOC5,SYNCRIP,RNF41,ZNF703,NR2
ISG1,RANGAP1,TAPBP,LUC7L2,KLF13,FZR1,COX8A,EXT2,COLGALT1,OXSR1,CCDC85C,ZDHHC5
SFRP1,CBX6,RPS2,BAZ2A,FKBP4,SLC7A1,SCD,TAPBP,ADNP,EWSR1,CD44,COPS6,H2AX,OTUD5,E
ARAPL1,AKT3,PLEKHB2,FNBP1L,SOD2,RBPJ,ATP5MC3,PHIP,PSAT1,COL12A1,ATP13A3,CDV3,CC
RAB11B,SELENON,RAPGEF1,CDC37,CAPZB,FKBP8,RPL28,HGS,SERBP1,STAT3,RETREG2,PHB2,Z
RF58,HMGA1,ABL1,ANKFY1,H2AX,MAPKAPK2,NUP98,MCFD2,RPRM,NACC1,VAV2,GPRC5B,SETD
2,MTA1,SF1,SEC16A,FKBP4,ANKS1A,C16ORF58,PHACTR4,TMEM94,MKNK2,IGF1R,TNFAIP1,EMC
CAND1,MAP7D1,GIGYF1,PTP4A2,TXLNA,KDM5C,CSNK2A1,NXN,TAGLN2,PPP1CC,VPS26B,TRIR,C
B5B,HDGF,SELENON,TUBB2A,ARF3,CAPZB,GIGYF1,MLF2,WBP2,STAT3,KDM5C,CBX6,TMEM167
REBF2,MSMO1,NAP1L1,KPNA2,PPP1CC,RPL7L1,TTC37,DICER1,DAZAP2,ELAVL3,TNFAIP1,EFNB2
1,NPLOC4,TPP1,PRR12,DAZAP2,MSI1,IGFBP4,TRMT112,SUMO1,SCAMP4,SSU72,B4GALT5,FAM5
1,NPLOC4,TPP1,PRR12,DAZAP2,MSI1,IGFBP4,TRMT112,SUMO1,SCAMP4,SSU72,B4GALT5,FAM5
1,APEX1,PPP1R10,CLCN7,RAB5C,KRT8,WASF2,VASH1,IP6K1,ARPC3,ILRUN,UBE2V1,SLC16A1,NA
BP4,DICER1,CLUH,SYNCRIP,ZCCHC3,GBA2,UBL3,ZBTB4,SRF,YWHAZ,ILRUN,MCFD2,STC2,ALDH
CN,SEC16A,ZFP36L1,SLC39A7,TAGLN2,NDST1,TMEM167A,APEX1,PEG10,VPS26B,TAOK1,EIF2S2
CN,SEC16A,ZFP36L1,SLC39A7,TAGLN2,NDST1,TMEM167A,APEX1,PEG10,VPS26B,TAOK1,EIF2S2
RAB11B,SELENON,RAPGEF1,CDC37,CAPZB,FKBP8,RPL28,HGS,SERBP1,STAT3,RETREG2,PHB2,Z
4,RPL22,CBX5,CHD4,ATP1A1,ATP2A2,CSNK1E,MYH10,HNRNPH1,ATP1A2,SKP1,CPSF1,SNX22,HS
2,SLC7A1,C16ORF58,NMNAT2,UNC119B,WDTC1,ESYT1,BSCL2,SYNCRIP,HMGB3,ZCCHC3,ILRUN
,RCC2,RPRM,STC2,RRP1B,CDKN1B,SLC16A1,FYTTD1,CLCN3,TMEM59,RNPS1,SYT4,LEPROT,FZ
JBB4B,MARCHF6,ARF1,ITM2B,GOLGA8A,SMARCA5,RPL23,RPSA,RPL24,COL1A2,RPL9,KIF5B,RI
ILF2,CTNND1,ARIH1,TMEM167A,STK35,TM9SF4,OGT,SCD,PAFAH1B2,SLC25A36,AGAP1,SCAMP
,CLCN7,CYTH2,HSPG2,XPO6,ABL1,SCAMP4,PCGF3,ZCCHC3,H2AX,SYNGR1,ARL8A,OTUB1,NACC

RAB5C,PDPK1,PCGF2,SCD,LHFPL6,RSF1,HIP1R,TRIM2,PCGF3,ARHGAP5,SRSF10,WAC,NDUFS1,
31,RNF4,THRA,ARHGEF18,RAD54L2,ADGRL1,PLXND1,GDE1
J,CTNNB1,MTSS2,ARID1A,SEPTIN11,RAC1,CDKN1A,NCKAP1,MLEC,ORAI2,TNPO1,KDM6B,VEGFA
1,SMCHD1,CDKN1A,GRINA,LAMC1,ORAI2,RPS15A,MAPRE1,SRSF1,SON,XPNPEP3,LSS,MTRNR2L
8,ARF4,SRSF3,WBP2,TP53BP1,DCTN1,GSPT1,HNRNPA2B1,RPS2,SLC25A5,MTCH1,ACLY,ABI2,AK
2A1,TAGLN2,PPP1CC,TMEM167A,TRIR,ERGIC1,FKBP4,SLC4A2,DAZAP2,MED13,CERS2,MYL12B,
3CL7A,MAZ,SCD,TMEM184B,AGAP1,LUC7L2,KLF13,ACVR1B,SIN3B,COX8A,COLGALT1,OXSR1,H
F,CTNNB1,CALU,ARHGDIA,CTDSP2,RPN2,PARP1,AHCYL1,MLEC,JPT1,RAPGEF1,PRDX1,LUZP6,S
ND4B,TUBB4A,SNX1,ZCCHC3,FAM168A,SEMA4C,SRF,ZFP91,OTUB1,NKIRAS2,CCDC6,ATG9A,TLK
,FNBP1L,SOD2,WASF2,RBM12,FKBP9,FBLN5,SCD,SNX3,COL12A1,SCAMP4,ACAP2,HMGB3,RAN
MAZ,LRATD2,SOD2,RANGAP1,AGAP1,MKNK2,SCAMP4,EXT2,YWHAZ,METRNL,AMD1
X3,RBBP4,PFKL,CDH6,NANOG,TAGLN,SMCHD1,TMEM30A,RNF187,GAS7,NOP53,PARP1,TNPO1,
A52,UQCRC1,ULK1,ACLY,FAM234A,SH3GL1,RBM5,RBPJ,SLC38A2,SF3A3,SYNCRIP,EHMT2,GOL
THBS2,ACIN1,RAD50,RNF40,SLC39A1,TOMM40,EPHA2
THBS2,ACIN1,RAD50,RNF40,SLC39A1,TOMM40,EPHA2
1,RAB5C,IGF2BP1,YY1,LHFPL6,PCGF3,CRKL,WAC,NDUFS1,VEZF1,DNAJB6,CITED2,TFDP1,HMG
HSBP1,TM9SF3,U2AF2,LMNB2,NCKAP1,VEGFA,TUBB2A,PTBP1,HNRNPUL1,SON,KDELR1,TPM3,I
1,BLCAP,TCF3,MSI1,COPZ1,ESYT1,MKNK2,TRMT112,HLA-A,TNRC6A,FAM168A,LDB1,UBE2V1,N
EN,CBX6,RPL7L1,LOX,GNL1,LSM14A,PNISR,RBM12,TTC28,TRIM8,USP5,ATP13A3,ACVR1B,SIN3E
,RAB1B,EIF4G1,CNBP,MKNK2,LMF2,ZCCHC3,PMEPA1,ARPC3,SRF,YWHAZ,NACC1,TNPO2,G6PC
1,ARL8A,CASKIN1,WASF3,PER1,TNPO2,GPRIN1,ASB6,CTDNEP1,ZNF516,MAPK8IP3
SALL2,ZCCHC3,CDK4,NDUFS1,ATP5F1E,CLCN3,NUP50,MYEF2,MRPS27,STX12
64,IL6ST,LRRC59,XRCC6,ERH,CCN1,LRATD2,CAPZA1,NUCKS1,ATXN3,CSE1L,ATP5MC3,TGOLN
121C,NPLOC4,STK35,TPP1,RALY,TRAM1,SH3GL1,SH3PXD2A,MSI1,COPZ1,RNF11,TRMT112,HLA

SHMT2,WARS1,CRABP2,CASKIN1,OTUB1,THRA,TRAF3,MIB2,FURIN,PPP2R2A,FZD7,RNF40,CTSA,
P,RBPJ,KLHDC10,HMGA1,PUM1,EFNB1,CAPN1,RPL27A,NSD1,SETD5,ASB6,RNASEK,SESN2,PPP2
L1,SH3GL1,EFTUD2,DAZAP2,LRP3,LYPLA2,MECP2,SUMO1,LDB1,UBE2Q1,NSD1,TOB2,MAP1S,R
L1,SH3GL1,EFTUD2,DAZAP2,LRP3,LYPLA2,MECP2,SUMO1,LDB1,UBE2Q1,NSD1,TOB2,MAP1S,R
GRSF1,PRRC2A,KLHDC3,CHERP,FKBP1A,FBXO21,BAZ2A,VPS4A,EDF1,SMARCD1,UBE2D3,THRA
N2,LENG8,LASP1,CLEC7A,EIF5A,PLXNA1,RAB7A,JUN,TAGLN,ANKRD52,GLUL,CDKN1A,CRPPA,CT
4,TFDP1,RELA,POGZ,NFIB
F4,THRA,ARHGEF18,RAD54L2,ADGRL1,PLXND1,GDE1
2,RBPJ,SRPRA,PSAT1,MDM2,CAPN1,SETD5,CDKN1B,PER1,TSN,TMEM214,NNT,MXRA7
GTPBP1,PLXNB1,CSNK1G2,NPLOC4,PEG10,CHERP,HSPG2,UBE2D3,XPO6,SCD,MGAT5,SCAMP4,
,CLCN7,RAB5C,KRT8,WASF2,VASH1,IP6K1,ARPC3,ILRUN,UBE2V1,SLC16A1,NACC1,ATG9A,OGD
CAPZA1,SLC7A1,PDPK1,CERS2,TUBB4A,UNC119B,USP5,RPS26,CNBP,FXR2,IGF1R,FOXP4,EXOC5
GRK2,ELAVL1,SOD2,SPTY2D1,PTGFRN,USP5,HS2ST1,LMAN2,MGAT5,TSNAX,KIAA1210,PNRC2,T
P4,IP6K1,MAPKAPK2,OTUD5,ARPC3,SETD5,NACC1,ATG9A,OGDH,CTDNEP1,DNMT1,MYL9
,SELENON,PTBP1,NPEPPS,TXNIP,ADAR,CTNND1,KIF5A,TXLNA,GSPT1,RGMB,CEMIP2,CBX6,ADH5
Z,ZNF703,MCFD2,ATG9A,ASB6,TMEM50A,RNASEK,EXT1,ZNF207,MRPS27
EXTL3,GLO1,ZNF664,SLIT1,SH3PXD2A,WASF2,VASH1,ATXN7L3,SH3PXD2B,LRIG1,MKNK2,MFN2
C2,GLUL,CDKN1A,PGRMC1,SURF4,TUBB2A,CCND1,PBX2,DYNC1LI2,HNRNPUL1,SMARCC1,SF3
TAGLN2,RAB1B,LDLR,SH3PXD2A,ARPP19,MECP2,EFNB1,DDX6,FAM168A,MAPRE3,WAC,SHMT2,N
R1,TUBB2A,ATG2B,FAM168B,CDC37,FKBP8,ATXN2L,ID4,RBBP7,DFFB,PRCC,PPP2CB,FBXO11,R

HL42,UBFD1,PTEN,BAZ2A,ABI2,SRSF2,LSM14B,YY1,RPS21,ANKFY1,C1QBP,YWHAZ,DDAH1,WDR8
CCHC14,PDE4DIP,P3H1,SETX,IRF2BP2,CASKIN1,UNC13A,DNAJB6,DSTYK,GALNT2
2
N2,LASP1,CLEC7A,EIF5A,PLXNA1,RAB7A,JUN,TAGLN,ANKRD52,GPC1,GLUL,CDKN1A,CRPPA,CTD
,TM9SF4,SOD2,NOC2L,ARPP19,RANGAP1,YTHDF3,REEP5,FZR1,ATP1B3,IQSEC1,EXT2,PMEPA1,L
,BAZ2A,SOX2,RALY,MAZ,SRSF2,SLC4A2,MOAP1,ANKS1A,COPZ1,YY1,KLHDC10,AASDHPPT,MKN
N2,LASP1,CLEC7A,EIF5A,PLXNA1,RAB7A,JUN,TAGLN,ANKRD52,GLUL,CDKN1A,CRPPA,CTDSP2,C
N2,LASP1,CLEC7A,EIF5A,PLXNA1,RAB7A,JUN,TAGLN,ANKRD52,GLUL,CDKN1A,CRPPA,CTDSP2,C
,IARS1,GNL1,RDX,MSH6,IGF2BP1,MTOR,PRKACA,HMGA1,RTF1,RGL2,CDV3,REEP5,ZCCHC3,ME
C7A6,GLO1,CCN1,LSM14A,SRSF2,SH3PXD2A,ARPP19,ESYT1,LUC7L2,NBPF15,PAIP2,NIBAN2,NO
2,ELN,SCAMP4,IP6K1,EPN1,TOB2,DNAJC8,TMEM43,ASB6,GPS1,UQCRH,SETD1B,UBTF
3,EIF4G1,CNBP,MKNK2,LMF2,ZCCHC3,PMEPA1,ARPC3,SRF,YWHAZ,NACC1,TNPO2,G6PC3,SRC
TGB1,MORF4L1,TIMP2,MIDN,MARCHF6,WSB1,TPM4,EIF4H,RPL14,CBX5,ATXN7L3B,LASP1,RTN
67A,TRIR,ERGIC1,MED13,PTGFRN,SAP18,KLHDC10,CERS2,RIMS3,SLC25A36,SDCBP,PLXNA2,CD
K1C,SETD5,NACC1,SRRT,NFIB,UBTF,EPHA2
A,LAPTM4B,RNF187,KLHL24,TNPO1,NRDC,TUBB2A,G3BP2,ATG2B,PBX2,DYNC1LI2,XPNPEP3,G
B12,SLC16A2,CDK9,MCFD2,RCC2,BCOR,TNPO2,ATF6,YBX3,TARS1
MD7,SRF,ASH1L,CCDC6,CITED2,AKAP17A,PPP2R2A,MYL9
,CLCN7,RAB5C,KRT8,WASF2,VASH1,IP6K1,ARPC3,ILRUN,UBE2V1,SLC16A1,NACC1,ATG9A,OGD
K2,SOD2,SPTY2D1,PTGFRN,USP5,HS2ST1,LMAN2,MGAT5,TSNAX,KIAA1210,PNRC2,TNRC6A,RN
PTIN7,HIF1A,BAP1,ABI2,SOX2,PDPK1,ATP5MC3,UNC119B,MKNK2,CAB39,ARPC3,CRKL,ZEB1,AC
M6,COX6B1,THOC2,GRHPR
NF157,IP6K1,IQSEC1,HM13,RBFOX2,POGK,ZNF598,ADGRL1,ZC3H4
WBP11,YY1,ATP6V1G1,TTC28,DIP2A,RNF11,YTHDF3,TMEM43,KIF3A,NUP205,ZNF146,ADGRL1,N
TC28,CYB5B,SLC38A1,SUMO1,SYNCRIP,KPNA6,CLPTM1L,RCC2,STC2,NDUFS1,NACC1,CCDC6,
FPL6,CAD,SCAMP1,IGF1R,CLUH,PCGF3,WAC,NDUFS1,VEZF1,SNRPB,CITED2,HMGB2,HMGA2,SK
,SCAMP1,IGF1R,CLUH,PCGF3,WAC,NDUFS1,VEZF1,SNRPB,CITED2,HMGB2,HMGA2,SKIL
3BP2,SRSF1,ARGLU1,WDR6,KDELR1,SMARCE1,YWHAH,CDC42,ARIH1,CSNK2A1,DECR1,CSNK1
IPLOC4,RAB1B,TM9SF4,SOD2,NOC2L,RANGAP1,YTHDF3,REEP5,FZR1,ATP1B3,IQSEC1,EXT2,PM
APZA1,SLC4A2,MSI1,FBXL19,RANGAP1,IGFBP4,KLF13,TNFAIP1,CLUH,OTUD5,PPP2R5D,CRABP2
APZA1,SLC4A2,MSI1,FBXL19,RANGAP1,IGFBP4,KLF13,TNFAIP1,CLUH,OTUD5,PPP2R5D,CRABP2
AB,TAOK1,SH3PXD2A,SLC4A2,PRPF4B,ARPP19,NMNAT2,BAG5,CNBP,TMEM184B,CLIP2,IGF1R,S
TGOLN2,FXR2,ELN,SCAMP4,IP6K1,EFNB1,EPN1,TOB2,DNAJC8,TMEM43,SETD5,NACC1,ASB6,U
RF,UBE2V1,STC2,ALDH9A1,NACC1,MSI2,MBD6,LDOC1,SIN3A,CASK
A,NMNAT2,FBXL19,CNBP,EFNB1,SHMT2,ZFP91,OTUB1,NACC1,GPS1,THRA,RNF40
,DUSP3,CTBP2,PPP1R26,KLHL7,ZNF207
TM4B,MORF4L2,DPYSL2,PTBP1,SON,PITPNA,KDM5B,KPNA2,CEP170,ATXN3,FNBP1L,RAB5C,SRS
TM4B,MORF4L2,DPYSL2,PTBP1,SON,PITPNA,KDM5B,KPNA2,CEP170,ATXN3,FNBP1L,RAB5C,SRS
HOB,TMEM30A,SKP1,HSPA4,ETNK1,SKI,VHL,GRB2,SON,TPM3,MLF2,SEC23A,SMS,TRA2B,ADH5,
CRIP,ZCCHC3,GBA2,UBL3,ZBTB4,SRF,YWHAZ,UBE2V1,MCFD2,PGRMC2,STC2,KIF3A,NDUFS1,CA
AB,TAOK1,SH3PXD2A,SLC4A2,PRPF4B,ARPP19,NMNAT2,BAG5,CNBP,TMEM184B,CLIP2,IGF1R,S
MAPK1,LAPTM4A,DYNC1LI2,HNRNPUL1,PITPNA,ADAR,MSMO1,SEC16A,PTEN,CEMIP2,FEM1B,CL
RNF11,CRACD,MDM2,SYNCRIP,TSPAN14,RBFOX2,SENP6,LIMS1,DDAH1,ZNF703,ASH1L,CDKN1
NPAB,TAOK1,PRPF4B,TGOLN2,SCAMP4,GALNT10,RASA4,NCKAP5L,IP6K1,TBXA2R,RBFOX2,ARL

SN,SPTBN1,ATP5F1B,RPL7,SEPTIN2,P4HB,RPL5,SMG1,PLXNB2,EIF3C,MARCKSL1,UBB,COL4A2

L18A1,EIF4B,TRIM28,NONO,SET,EIF4A1,PKD1,HLA-C,POLR2A,AP2M1,STMN1,AP3D1,CYB5R3,M

ATP5F1B,RPS8,MYH9,P4HB,GSTP1,SMG1,NPM1,COL18A1,CSDE1,EIF3C,ATP5F1A,EIF4B,TRIM2

,RPL13,SLC2A3,BSG,FASN,GNB1,SEPTIN2,APP,IDS,RPL5,TMBIM6,PLD3,HSPD1,CSDE1,EIF3C,AT

,RPL29,RPS9,XRCC5,NCS1,PSMD2,RPS14,PFKL,TKT,SFPQ,GNB2,CTNNB1,ARID1A,ANKRD52,NA

AP12,SF3B2,AP2B1,SMARCA5,LARP1,NACA,RPS9,KPNB1,RPL22,CHD4,WDR1,BEX3,CCT7,SRSF4

17,VCL,SBF1,ARF1,ITM2B,RPS16,RTN4,LAMA5,KPNB1,PAPOLA,CBX5,RPL9,ATP1A1,ATXN7L3B,

,AP2B1,LARP1,CTNNA1,KPNB1,NCS1,RPS14,HNRNPC,ACTN4,GNB2,MIF,CTNNB1,MTSS2,SLC25

1,RPS3,BASP1,HMGB1,SEC61A1,STMN1,UCHL1,DNAJA1,SF3B2,MAT2A,PGAM1,LPCAT1,RPL22,

FF4,TRIM28,CKB,EIF4A1,GTF3C4,SF3B1,CHD3,HMGB1,POLR2A,TPT1,RPS4X,RPL18,MIDN,MAR

2B1,TSPAN3,RTN4,C1ORF43,RPL14,KIF5B,SLC7A5,ATXN7L3B,ATP2A2,JPT2,LRPPRC,TADA2B,C

HOA,CLSTN1,TMEM259,WDR1,LASP1,VAT1,SCRN1,RPS25,GLG1,SFPQ,CDH2,PDIA6,PIP4K2B,R

P1,TAGLN,ANKRD52,ARHGDIA,BCLAF1,PPIB,RPL23A,GLUL,PEA15,TIMP3,RNF187,U2AF2,SQST

,NGRN,MACF1,ARL6IP1,ARID1A,NARS1,SEPTIN11,HNRNPH1,LGALS3BP,CAPRIN1,IPO5,TIMP3,M

,ZNF770,MAT2A,HNRNPU,PGAM1,SMARCA5,HNRNPA0,LDHA,RPL29,LPCAT1,LLGL1,RPL14,RPL

A6,SLC25A3,ARID1A,IPO5,HNRNPL,RPL27,PTK7,RPL37,BTBD2,GIGYF1,YWHAH,ATXN2L,WBP2,A

RPL31,PRKCSH,CCND2,RAB7A,ACTN4,RPS5,PRKAR1A,RPL35A,ARID1A,DDOST,ST3GAL2,CCT6A,

16,EIF4H,NACA,RPS9,KPNB1,NR2F1,EIF5,CSNK1E,PFKL,TKT,GPAT4,JUN,NGRN,CYFIP2,HNRNP

N2,RPN2,PPP1CB,GNAI3,HNRNPL,VCAN,TSR1,SSRP1,TCP1,PLOD1,ETNK1,RAB5B,TNPO1,PRPF6

1A,GCN1,VEGFA,SELENON,SRSF1,HNRNPUL1,NPEPPS,KDELR1,NT5DC2,HGS,FBRSL1,RBM15B

AP12,CDC42BPB,PXDN,HNRNPU,CNN3,TPM4,NCS1,CBX5,KIF5B,LASP1,SRSF4,PNN,TKT,TPM1,C

B,SLC7A5,ATP2A2,XPO1,CDH2,PNMA1,PLXNA1,SOX12,PRKAR1A,JUN,CTNNB1,APLP2,NDFIP1,CI

,CDH6,PEBP1,CCND2,CDH2,CLTC,HMGCS1,PRKAR1A,JUN,CTNNB1,NARS1,CKAP5,SEPTIN11,UB

TM1,CAMSAP1,SURF4,MAPRE1,TCF25,HNRNPUL1,CASC3,SRSF3,PTP4A2,HGS,DGKD,SREBF2,R

1,RPS15A,NOTCH2,RAPGEF1,CCNL2,SRSF1,DYNLL2,NPEPPS,PHC2,ID4,MPRIP,AKIRIN1,BMPR2

68B,IPO7,IDH1,BAG6,ADAR,EDC4,WDR45B,ATP6AP2,GPX1,ANTXR1,TM9SF4,MAZ,INTS11,QPRT,

1,SOX12,CCN2,PRKAR1A,APLP2,HNRNPDL,DDOST,HNRNPK,CKAP4,RPS12,ARCN1,RAC1,TM9SF

ATXN7L3B,MCL1,JPT2,PNN,LRPPRC,TADA2B,CSNK1E,CCND2,PDIA6,DDX3X,RPS5,JUN,MYO5A,

CSL6,NANOG,RCN1,RPS7,IK,CTNNB1,DNAJC5,H1-10,CKAP5,CAPRIN1,BCLAF1,L1CAM,PEA15,T

A2B,CSNK1E,CCND2,PDIA6,DDX3X,RPS5,MYO5A,HNRNPDL,RPL36,UBE2H,CALU,ARCN1,ARHG

30A,TM9SF3,PPP1CB,MORF4L2,TCP1,H3-3A,RPL36A,TERF2IP,RPS15A,KDELR2,PRDX1,ODC1,G

FITM3,SREBF2,TXLNA,MTA1,PTPRF,TIMM50,BAZ2A,CYTH2,STRN4,BCL7A,RALY,SH3PXD2A,SLC

0,PPP2R1A,SERINC1,HMGB1,ITGB1,TUBB4B,MORF4L1,MIDN,MARCHF6,WSB1,PAFAH1B1,ZNF7

MYH10,UBE2H,ARCN1,CTBP1,TMEM30A,CPSF2,MTRF1L,TBC1D16,TCP1,RPL37,KDELR1,DUSP4,

TP4A1,CDC42,STAT3,FBRS,MMP15,PTEN,FEM1B,CBX6,KLHDC3,LOX,LRP10,GRN,RTL6,ELAVL1

,ATP2A2,PFN1,MCL1,PRKCSH,XPR1,SFPQ,ACTN4,DDX3X,GNB2,IK,TLN1,MYO5A,SESN3,ANKRD5

EC,TUBB2A,DUSP4,GIGYF1,GPR107,NCOR2,MLF2,WBP2,PPP1R9B,BCL9L,SEC16A,PHB2,HOXB

EC,TUBB2A,DUSP4,GIGYF1,GPR107,NCOR2,MLF2,WBP2,PPP1R9B,BCL9L,SEC16A,PHB2,HOXB

NPLOC4,GPC4,GRK2,XRCC6,ABI2,RAB1B,GNL1,EFTUD2,PRR12,C16ORF58,KLHDC10,EIF4A3,Z

NPLOC4,GPC4,GRK2,XRCC6,ABI2,RAB1B,GNL1,EFTUD2,PRR12,C16ORF58,KLHDC10,EIF4A3,Z

F1L,GRINA,PTOV1,HDGF,NRDC,ARF3,PPP2CA,PTBP1,CASC3,SUPT6H,RPS28,PFKM,LSS,PCBP2,

CT3,TMEM248,GCN1,SKI,BNIP3L,ARF4,PITPNM1,TNKS2,GPR107,NT5DC2,HECTD1,HNRNPD,D

AP1,VEGFA,MARS1,SELENON,RAPGEF1,PBX2,CDC37,CAPZB,DUSP4,GPR107,SREBF2,MEX3A,M

TNND1,ARIH1,PRKAR2A,KDM5B,SF1,POM121C,RPS27A,GLO1,SERP1,XRCC6,MAZ,BLCAP,SOD2,

A,NOTCH2,ARF3,CCND1,HNRNPM,LARS1,PTP4A2,CSNK1D,HYOU1,MPRIP,TXNIP,TRAF4,CSNK2A

D4,SERBP1,TXNIP,SRSF6,PTEN,CEMIP2,TIMM50,HIF1A,ZMYM3,KIAA0100,TAOK1,PELP1,GNL1,P
PRR12,WASF2,DAZAP2,MSI1,UBE2D3,IGFBP4,HMGA1,TMEM184B,USF2,SUMO1,SCAMP4,SSU72
BE2H,CALU,ARCN1,RPS27,APPL1,GLUL,SQLE,CDKN1A,SERF2,TIMP3,PGRMC1,RPN2,SUGP2,PG
PRKAR2A,FBRS,YKT6,CBX6,PTPRF,APEX1,CHERP,RAB1B,ZMIZ2,RAB5C,TCF3,FBXL19,CNBP,SG
PRR12,WASF2,DAZAP2,MSI1,UBE2D3,IGFBP4,HMGA1,TMEM184B,USF2,SUMO1,SCAMP4,SSU72
ACF1,ARID1A,ANKRD52,LGALS3BP,HECTD4,GLUL,ZC3H7B,GRINA,MLEC,MARS1,HNRNPUL1,MA
1,SEC16A,BRD2,SFRP1,PTPRF,NPLOC4,TAOK1,FAM234A,PRR12,PRPF4B,DAZAP2,MSI1,LRP3,S
PS,PHC2,ID4,MPRIP,SERBP1,AKIRIN1,BMPR2,TXNIP,AMOTL2,ADAR,CTNND1,TXNDC5,RETREG2,
RHGDIA,SLC35E2B,LUC7L3,SSRP1,CAMSAP1,NCKAP1,LAMC1,RAD23B,SKI,HDGF,NOTCH2,VEG
RSK1,POM121C,FKBP1A,CLCN7,YARS1,SNRPD3,LINGO1,VPS4A,ATXN3,SOD2,RAB5C,PLEKHM2,
RBP1,BMPR2,CDC42,RBM15B,EIF2AK1,TRIO,VARS1,UBAP2L,GLUD1,TAGLN2,PRRC2A,NCOA4,K
A,FBRS,YKT6,CBX6,PTPRF,APEX1,CHERP,RAB1B,ZMIZ2,RAB5C,TCF3,FBXL19,CNBP,SGSM2,EL
LUC7L3,SSRP1,CAMSAP1,NCKAP1,LAMC1,RAD23B,SKI,HDGF,NOTCH2,VEGFA,TUBB2A,CCND1,
KN1A,SRP9,RNF187,LMNB2,GCN1,KDM6B,PRDX1,GRB2,HNRNPUL1,MAP7D1,RPL28,HNRNPM,
1,ZFP36L1,TAGLN2,PPP1CC,ABCC9,LMAN1,BAZ2A,SLC6A8,MAZ,HDGFL2,PNISR,SOD2,SLC7A1,
GB1,MORF4L1,ZMIZ1,MIDN,MARCHF6,WSB1,PAFAH1B1,ZNF770,PGAM1,TPM4,EIF4H,RPL17,P
L3,SSRP1,CAMSAP1,NCKAP1,LAMC1,RAD23B,SKI,HDGF,NOTCH2,VEGFA,TUBB2A,CCND1,SRSF1
B,RAB11B,PSMD3,FKBP8,STMN3,SREBF2,MEX3A,PPP1R9B,TRAF7,WDR45B,GTPBP1,BRD2,CSN
XRCC6,AKT2,GTF3C1,TNFAIP1,NTN1,SHISA5,WIZ,DDX6,RRP1B,CHD8,RNF20,TMPO,ERBB2,NFA
LNA,MTA1,GTPBP1,YKT6,CBX6,TIMM50,STK35,SLIT1,BCL7A,TCF3,ZER1,CNBP,RNF11,MGAT5,
N1,SRSF9,ADAM9,CAND1,BNIP3L,ZFAND5,GIGYF1,MTRNR2L10,PTP4A1,AKIRIN1,SGTA,NAP1L1,
14A,SOD2,RBM5,NOC2L,GOT2,C16ORF58,POLDIP2,RNF11,ABL1,TNFAIP1,BNC2,IP6K1,GSR,CA
,SELENON,RAPGEF1,TUBB2A,FTO,CAPZB,GIGYF1,CSNK1D,PTP4A1,GHITM,MEX3A,MLF2,AMOT
,SELENON,RAPGEF1,TUBB2A,FTO,CAPZB,GIGYF1,CSNK1D,PTP4A1,GHITM,MEX3A,MLF2,AMOT
HNRNPUL1,FKBP8,TPM3,GIGYF1,PRPS1,GPR107,MLF2,AMOTL2,YME1L1,GRSF1,KIF21B,SEC16
GF2BP1,YY1,TTC28,WNK1,SLC22A23,FAM118A,IGF1R,CAB39,ACAP2,ECH1,KPNA6,RDH11,GOL
XNIP,POLR2E,STAT3,KDM5C,SEC16A,SFRP1,PTPRF,TMEM167A,NPLOC4,GABARAPL1,GNL1,SRS
1,ATP6V1B2,UQCRC2,RTL8C,NAP1L4,PALLD,NAP1L1,CPD,TRA2B,SMC3,CEMIP2,RPS2,PSMD8,S
CNTNAP2,TMED2,PLEC,PRDX1,SRSF1,IPO7,ILF3,GRB2,CDC37,CAPZB,UBE2R2,MAP2K2,ARF4,P
2,RPL36A,SMARCE1,DHX30,STMN3,MEX3A,PRKAR2A,CSNK2A1,SFRP1,HNRNPA2B1,PTPRF,TME
CBX6,ULK1,ANP32B,TMEM167A,PPP1R15A,TAOK1,ABI2,BCL7A,RAB1B,SLC7A1,TCF3,C16ORF5
CN7,TRAP1,SOD2,PACS1,WASF2,IVD,YY1,HNRNPH3,NDUFC2,GALNT10,TSPAN14,DARS1,ADRM1,
LR2E,TRAF7,STK35,CARM1,CAPZA1,SLC4A2,SLC25A23,MSI1,ATXN7L3,TUBB4A,FBXL19,IGFBP4
TRAF4,KDM5B,FBRS,PTEN,FEM1B,CBX6,KLHDC3,LOX,RTL6,BCL7A,AKT3,MAZ,DICER1,AKT2,NA
P8,GIGYF1,SRSF3,GPR107,SARS1,IFITM3,SREBF2,MEX3A,MLF2,TXNIP,TRAF7,TXLNA,RAB2A,RG
X2,COL6A2,ATP1A2,SMCHD1,TAF15,HSPA4,DDX42,LAMC1,ORAI2,DPYSL2,RPS15A,MARS1,ARF3,
CBX6,ULK1,ANP32B,TMEM167A,PPP1R15A,TAOK1,ABI2,BCL7A,RAB1B,SLC7A1,TCF3,C16ORF5
5C,SLC39A7,BRD2,FMNL2,PTPRF,EML1,BAP1,ACLY,RTL6,SRP72,TM9SF4,GTF2I,PHGDH,IGF2B
,TRIB3,ZCCHC14,PDE4DIP,SLC43A2,YWHAZ,CITED2
S1,MASTL,HNRNPAB,CYTH2,VPS41,SLC7A1,UBE2I,USP10,TNRC6A,ATF7IP,NUP98,CSNK1A1,ZBT
BRS,MMP15,PTEN,FEM1B,CBX6,KLHDC3,LOX,WSB2,RTL6,AKT3,MAZ,DICER1,AKT2,NASP,FJX1,
NR2L10,PTP4A1,SERBP1,AKIRIN1,RBBP7,NAP1L1,TXNDC5,AP2A1,BAZ1B,ZFP36L1,RGMB,FEM1
A,GPX1,LTBP4,NUCKS1,WASF2,LZTR1,CAD,TMEM184B,KIFC2,SCAMP4,NCK2,TMEM9,NDUFB8,

SYT1,ELN,TNFAIP1,SUMO1,SCAMP4,SSU72,B4GALT5,EFNB1,LDB1,YWHAZ,ZNF703,DIRAS1,CAS

RSF1,NXN,POM121C,TMEM167A,CHERP,LRP10,TAOK1,BCL7A,RALY,LTBP4,NOC2L,SLC20A1,A
AB5B,TUBB2A,ARF3,GRB2,TPM3,GIGYF1,HYOU1,GPR107,NCOR2,MLF2,WBP2,PPP1R9B,TRAF7,
STK35,CARM1,CAPZA1,SLC4A2,SLC25A23,MSI1,ATXN7L3,TUBB4A,FBXL19,IGFBP4,CLUH,RNF15
NLL2,HNRNPUL1,TPM3,FAU,STT3B,KDM3B,TXNIP,CCT5,SSR1,STAT3,STARD7,PTEN,RPL7L1,POM
XIP,ALKBH5,MDK,ASXL1,SELENON,PSMD3,HNRNPUL1,FKBP8,SDF4,SARS1,IFITM3,RAI1,SREBF2
AB5B,TUBB2A,ARF3,GRB2,TPM3,GIGYF1,HYOU1,GPR107,NCOR2,MLF2,WBP2,PPP1R9B,TRAF7,
UL1,FKBP8,PLCG1,PITPNA,TPM3,STMN3,GPR107,MEX3A,MLF2,WBP2,KIF5A,PHB2,PLXNB1,HOX
AB5B,TUBB2A,ARF3,GRB2,TPM3,GIGYF1,HYOU1,GPR107,NCOR2,MLF2,WBP2,PPP1R9B,TRAF7,
AB5B,TUBB2A,ARF3,GRB2,TPM3,GIGYF1,HYOU1,GPR107,NCOR2,MLF2,WBP2,PPP1R9B,TRAF7,
AB5B,TUBB2A,ARF3,GRB2,TPM3,GIGYF1,HYOU1,GPR107,NCOR2,MLF2,WBP2,PPP1R9B,TRAF7,
DEK,USP5,EIF4G1,CGGBP1,FAM171A1,SIN3B,IGF1R,EFNB2,TSPAN14,FLII,TNRC6A,SH3BGRL3,T
1,MTRNR2L10,TNKS2,PTP4A1,AKIRIN1,NAP1L1,TXNDC5,ZFP36L1,PREPL,PEG10,TAOK1,EHMT1,
C,FOXK2,LRRC41,AHCY,ARPP19,C16ORF58,CNBP,SCAMP5,MGAT5,IGF1R,SUMO1,RNF41,POLD
FOXK2,DHCR7,EIF3G,ATP5MC3,SMARCD1,MDM2,RPS21,SYNCRIP,PPP4R3A,DDX6,CRKL,TNFRSF
P32B,CHERP,TAOK1,FKBP4,VPS4A,SOD2,SLC7A1,SMARCD1,LYPLA2,UNC119B,IGFBP4,BGN,NIB
P32B,CHERP,TAOK1,FKBP4,VPS4A,SOD2,SLC7A1,SMARCD1,LYPLA2,UNC119B,IGFBP4,BGN,NIB
1,PAFAH1B1,ZNF770,TPM4,EIF4H,RPL17,RPL14,CBX5,PSMD2,WDR1,ATXN7L3B,LASP1,MCL1,R
F58,NMNAT2,FBXL19,IER2,CNBP,IGFBP4,FXR2,SGSM2,SCAMP4,GALNT10,CCNY,EFNB1,ARL8A,
XIP,ALKBH5,MDK,ASXL1,SELENON,PSMD3,HNRNPUL1,FKBP8,SDF4,SARS1,IFITM3,RAI1,SREBF2
,BCR,KDM5C,MMP15,PRRC2A,TMEM167A,TAOK1,FKBP4,INTS3,FAM234A,SH3GL1,SH3PXD2A,N
AB5B,TUBB2A,ARF3,GRB2,TPM3,GIGYF1,HYOU1,GPR107,NCOR2,MLF2,WBP2,PPP1R9B,TRAF7,
2,EIF4G1,WDTC1,UBE4B,SUMO1,RNF157,PSME3,SRF,WAC,SHMT2,SETD5,SORCS2,KIAA2013,T
OB2,PIN1,RNPS1,SETD1B,UBTF,SIN3A
PRRC2A,ANP32B,RPS2,CHERP,SYMPK,FKBP4,RIC8A,PHIP,XRN2,PUM1,GTF3C1,KIAA1191,TRRA
K2,OGT,EHD1,AP1M1,THRAP3,EIF4A3,FBXL19,SLC25A36,ACVR1B,PDHA1,SLC25A6,SUMO1,BS
K2,OGT,EHD1,AP1M1,THRAP3,EIF4A3,FBXL19,SLC25A36,ACVR1B,PDHA1,SLC25A6,SUMO1,BS
F58,NMNAT2,FBXL19,IER2,CNBP,FXR2,SGSM2,SCAMP4,GALNT10,CCNY,EFNB1,ARL8A,OTUB1,
F58,NMNAT2,FBXL19,IER2,CNBP,FXR2,SGSM2,SCAMP4,GALNT10,CCNY,EFNB1,ARL8A,OTUB1,
1,MECP2,CAPN1,DDX6,FAM168A,GBA2,TOB2,HMGXB3,SORCS2,CDC42SE1,UBE2G1,B4GALT2,
2,EIF4G1,WDTC1,UBE4B,SUMO1,RNF157,PSME3,SRF,WAC,SHMT2,SETD5,SORCS2,KIAA2013,T
1,DYNC1LI2,DYNLL2,LUZP6,CAPZB,HNRNPUL1,SON,KDELR1,ATP6V1B2,SF3A1,PTP4A1,PPP6R1,
K2,OGT,EHD1,AP1M1,THRAP3,EIF4A3,FBXL19,SLC25A36,ACVR1B,PDHA1,SLC25A6,SUMO1,BS
C2A1,FUS,YARS1,CAPZA1,SLC4A2,SOD2,DAZAP2,HS2ST1,SLC25A36,FXR2,NIBAN2,PLXNA2,CCD
P2,CAPN1,DDX6,FAM168A,GBA2,TOB2,HMGXB3,SORCS2,CDC42SE1,UBE2G1,B4GALT2,BSDC1
1A,PGRMC1,PARP1,PLOD1,SURF4,ORAI2,GCN1,KDM6B,RPS15A,TUBB2A,CCND1,PBX2,FAM171
AP1S,FARSA,NACC1,DGCR8,G6PC3,MIB2,NF2,GDE1
N4,WASF2,TMEM184B,ABL1,PHACTR4,NIBAN2,MKNK2,NECTIN1,NSD1,SYNGR1,ILRUN,ARL8A,R
CORO1C,ATXN2L,WBP2,VDAC1,RPS2,KMT2E,BLCAP,CREBBP,RANGAP1,HMGA1,ATP13A3,RUVBL
RSA,NACC1,ASB6,THRA,CNOT3,TET3,POMGNT1,ANKRD13B,NF2,RNF40
G1,SCAMP4,RNF157,SALL2,COLGALT1,NDRG1,WAC,ZFP91,OTUB1,THBS2,TNPO2,MIB2,SETD1
RNPA2B1,SPINT2,PPP1R15A,MTCH1,GLO1,TRAM1,LDLR,SOD2,UBE2D3,PSMC1,IER2,FXR2,YTHD
C25A23,TCF3,LRP3,C16ORF58,CNBP,IGFBP4,TMEM184B,ELN,MAPK8IP2,SUMO1,CCNY,B4GAL
C25A23,TCF3,LRP3,C16ORF58,CNBP,IGFBP4,TMEM184B,ELN,MAPK8IP2,SUMO1,CCNY,B4GAL
RK2,STK35,INTS3,ABI2,BCL7A,GABARAPL1,GNA12,TM9SF4,SOD2,WASF2,OGT,SMARCD1,CERS
3,FAM234A,MSH6,OGT,NOC2L,SMARCD1,CERS2,FBXL19,RNF11,NIBAN2,UBE4B,SUMO1,SCAM

C59,SNRPD3,LINGO1,VPS4A,PRR12,RAB5C,SAP18,C16ORF58,UBE2D3,XPO6,UNC119B,MGAT5,K
MLEC,SKI,PSMB2,SRP14,ATP6AP1,RAPGEF1,TUBB2A,CCND1,GRB2,CAPZB,TPM3,MPRIP,CYP51A
MLEC,SKI,PSMB2,SRP14,ATP6AP1,RAPGEF1,TUBB2A,CCND1,GRB2,CAPZB,TPM3,MPRIP,CYP51A
TUS,YARS1,CAPZA1,SLC4A2,SOD2,DAZAP2,HS2ST1,SLC25A36,FXR2,NIBAN2,PLXNA2,CCDC80,TN
TUS,YARS1,CAPZA1,SLC4A2,SOD2,DAZAP2,HS2ST1,SLC25A36,FXR2,NIBAN2,PLXNA2,CCDC80,TN

RF,HNRNPAB,STK35,TAOK1,HSPG2,SLC4A2,PRPF4B,LRP3,ARPP19,ATXN7L3,XPO6,UNC119B,EIF
DICER1,UBE2D3,SLC38A2,PTPN23,WDTC1,FXR1,HLA-A,TNRC6A,MYCBP2,DDX6,PBXIP1,APPBP2,N
,GID8,TOB2,RPRM,EI24,CASKIN1,NACC1,HIF3A,UQCRFS1,ATG9A,VAV2,GYS1,PPP2R2A,SETD1B,

AP1S,FARSA,NACC1,DGCR8,G6PC3,MIB2,GDE1
KAR2A,SEC16A,DECR1,KIAA0895L,UBFD1,CUL3,NPLOC4,TAOK1,AKT3,NUCKS1,AHSA1,IVD,ATP5
BP2,MAPK1,ATG2B,AZIN1,DYNC1LI2,DYNLL2,HNRNPUL1,SON,CCAR2,ZFAND5,ATP6V1B2,GIGYF
A,TRIR,ERGIC1,MAZ,FLOT2,SLC4A2,SLC7A1,ATXN7L3,C16ORF58,UNC119B,EIF4G1,WDTC1,BSC
P6R1,TXNIP,MSMO1,RERE,CSNK2A1,GRSF1,SEC16A,MASTL,PTEN,CEMIP2,CUL3,RPS27A,TIMM5
A,TRIR,ERGIC1,MAZ,FLOT2,SLC4A2,SLC7A1,ATXN7L3,C16ORF58,UNC119B,EIF4G1,WDTC1,BSC
CAB39,ATP1B3,DHFR,POGK,NR2F2,TALDO1,SHB,TFDP1,NBEA,PCMTD2
B2A,G3BP2,ATG2B,DYNC1LI2,DIDO1,XPNPEP3,GIGYF1,ILF2,ID4,BMPR2,DDIT4,EP300,KPNA2,ZF
CA,SCD,NTRK2,LGALS1,PHACTR4,PDHA1,DAD1,DDX6,SRSF7,CLPTM1L,YWHAZ,RCC2,TFRC,ERBB
ERGIC1,BCL7A,GNL1,SOD2,SLC7A1,PLEKHM2,DIP2A,BGN,HMGA1,HDGFL3,LUC7L2,SC5D,CCDC
SOX9,STC2,NDUFS1,ATP5F1E,CLCN3,NUP50,SMARCB1,MYEF2,MRPS27,STX12
FAH1B1,ZNF770,AP2B1,MAT2A,TPM4,EIF4H,RPL17,C1ORF43,RPL14,CBX5,RPS14,WDR1,ATXN7
AP1S,FARSA,NACC1,DGCR8,G6PC3,MIB2,GDE1
D17B12,PPP2R5D,CRKL,NUP98,UBE2G1,SESN2,SETD1B,HES6
ACC1,THRA,G6PC3,TRAF3,SRCIN1,GDE1
ACC1,THRA,G6PC3,TRAF3,SRCIN1,GDE1
STK35,TAOK1,HSPG2,SLC4A2,PRPF4B,LRP3,ARPP19,ATXN7L3,XPO6,EIF4G1,TMEM184B,ABL1,C
N9,SLC1A5,SHB,VAV2,SP1,GMFB,FURIN,ACIN1,FAM3C,ZNF207,ITPKB,HES6,ARSA
A,ZCCHC3,KPNA6,YWHAZ,NDUFS1,ARL8A,NKIRAS2,SLC5A6,CCDC6,TYW5,TFDP1,HEXIM1,SLC39
BCL7A,GNL1,SOD2,PLEKHM2,DIP2A,BGN,HMGA1,HDGFL3,LUC7L2,SC5D,CCDC80,IGF1R,MDM2
10,DLD,ZBTB4,RARS1,POGK,CCDC6,ATP5F1E,MED12,SUDS3,HADHB,GNAL,PPM1A,LRIG3,GDE1
C1A5,SHB,VAV2,SP1,GMFB,FURIN,ACIN1,FAM3C,ZNF207,ITPKB,HES6,ARSA
BCL7A,GNL1,SOD2,PLEKHM2,DIP2A,BGN,HMGA1,HDGFL3,LUC7L2,SC5D,CCDC80,IGF1R,MDM2
STK35,TAOK1,HSPG2,SLC4A2,PRPF4B,LRP3,ARPP19,ATXN7L3,XPO6,EIF4G1,TMEM184B,ABL1,C
SF1,CBX6,TAOK1,ERGIC1,SNRPD3,ABI2,BCL7A,ZMIZ2,AKT3,TM9SF4,MAZ,DAZAP2,ARPP19,AMBR

K1,LDB1,MAP1S,ZNF516,HMGA2,UBTF
HS1,DDX6,NDRG1,ALCAM,MCFD2,RCC2,CDKN1B,SLC12A7,IRS1,CEBPG,NEURL4,ASB6,FURIN,A
P,MSMO1,CSNK2A1,GRSF1,SEC16A,MASTL,PTEN,CEMIP2,CUL3,RPS27A,TIMM50,CHERP,CEP17
1,ERGIC1,SNRPD3,ABI2,BCL7A,ZMIZ2,AKT3,TM9SF4,MAZ,DAZAP2,ARPP19,AMBRA1,SCD,SH3PX
CC1,MYL12A,CCDC6,POMGNT1,SECISBP2L,SETD1B,CPSF7,RNF40,QSOX1

,PPP1R9B,TRAF7,RETREG2,SEC16A,PHB2,ZFP36L1,HOXB5,CBX6,LRRC59,STK35,TRIR,RALY,AT
C59,VPS26B,BAP1,RAB1B,RDX,SFXN1,YY1,KLHDC10,XPO6,GNA13,CNBP,RANGAP1,TAPBP,NOP5
UBQLN4,NACC1,GPRIN1,NAV1,CTDNEP1,MBD6,MFGE8

C,RNF4,ASB6,TET3,RAD54L2,AMD1,ADGRL1,ANKRD13B,PRELID1,PLXND1,NF2,RNF40,GDE1
,PPP1R9B,TRAF7,RETREG2,SEC16A,PHB2,ZFP36L1,HOXB5,CBX6,LRRC59,STK35,TRIR,RALY,AT

GT,EHD1,COPZ1,MTOR,ATXN7L3,ATP5MC2,THRAP3,CASC4,DIP2A,PLS3,UBE2L3,ZZEF1,NIBAN2,
2A,CSNK2A1,DECR1,FEM1B,CUL3,PTPRF,KLHDC3,TAOK1,AKT3,ATP5MC3,SCD,ELOVL5,SRPRA,
ELL2,RIMS3,NUDT4,FXR1,SALL2,DAD1,WIZ,EXT2,KPNA6,SRF,TFRC,OTUB1,HIF3A,MAP2K4,CLCN
,SOD2,CSE1L,KDM4A,UBE2D3,SCD,AMFR,OGFR,MTA2,TXNRD1,CLUH,SUB1,ACAP2,DDX6,NCK2
1,ATP13A3,SC5D,IGF1R,KIAA1191,CCDC88A,PSME3,WAC,OTUB1,FARSA,NKIRAS2,TNPO2,B4GAL
D1B,RNF40,QSOX1

YB5B,EIF4A3,DIP2A,KLF13,CCDC80,IGF1R,SLC38A1,SSU72,RAN,TNRC6A,SLC12A7,CASKIN1,SL
,PPP1R9B,TRAF7,RETREG2,SEC16A,PHB2,ZFP36L1,HOXB5,CBX6,LRRC59,STK35,TRIR,RALY,AT
,POM121C,KLHDC3,CHERP,FKBP1A,STK35,TPP1,RALY,TRAM1,SH3GL1,SH3PXD2A,OGT,COPZ1,

,POM121C,KLHDC3,CHERP,FKBP1A,STK35,TPP1,RALY,TRAM1,SH3GL1,SH3PXD2A,COPZ1,SMAF
,POM121C,KLHDC3,CHERP,FKBP1A,STK35,TPP1,RALY,TRAM1,SH3GL1,SH3PXD2A,COPZ1,SMAF
TXNIP,MSMO1,CSNK2A1,GRSF1,SEC16A,MASTL,KCTD12,CEMIP2,CUL3,RPS27A,TIMM50,CHERP,
FD2,RPRM,NACC1,RPS6KA2,EPB41L1,VAV2,GPRC5B,SETD1B
RIH1,SF1,TAGLN2,ZNF512B,CBX6,WSB2,TAOK1,RAB1B,TM9SF4,SOD2,LRP3,SCD,FGFR1,ELAVL
,TUBB4A,HMGA1,ESYT1,LUC7L2,NIBAN2,ATP1B3,GALNT10,FAM53C,WIZ,TRIB3,GBA2,ZBTB4,U
P2,CCNY,SSU72,SALL2,CRKL,ZBTB4,TOB2,TMEM43,UBQLN4,ALDH9A1,SETD5,HIF3A,CCDC6,UE
2A1,GRSF1,SEC16A,MASTL,CEMIP2,CUL3,RPS27A,TIMM50,CHERP,CEP170,TAOK1,TPP1,UBE2D2

C1,PTP4A2,RPS13,STAT3,ARIH1,TXLNA,NAP1L1,BAZ1B,CSNK2A1,UBA52,BRD2,GARS1,MBTPS1,
ASB6,TET3,RAD54L2,ADGRL1,ANKRD13B,PRELID1,PLXND1,NF2,RNF40,GDE1
ASB6,TET3,RAD54L2,ADGRL1,ANKRD13B,PRELID1,PLXND1,NF2,RNF40,GDE1
ASB6,TET3,RAD54L2,ADGRL1,ANKRD13B,PRELID1,PLXND1,NF2,RNF40,GDE1

PRC5B,SETD1B
PRC5B,SETD1B
PRC5B,SETD1B
IN1,ZCCHC3,ILRUN,UBE2V1,PGRMC2,STC2,TNPO2,CAMK2N1,TMEM50A,ANKRD13B,CTIF,RAD5
D1,TNRC6A,CAPN1,SH3BGRL3,TOB2,KDM5A,NACC1,PPP1R12A,NAV1,EPHB2,NF2,RNF40,NECAP
K3C2B,CCNY,PCGF3,ZCCHC3,TNFRSF10B,DNAJC8,DIRAS1,ARL8A,SLC12A7,BTBD1,ADGRL1,PL
MED9,ID4,SREBF2,YME1L1,XPOT,TXNDC5,KPNA2,VAPB,TM9SF4,IREB2,TRAP1,SH3PXD2A,SOD2,

1,KIF5A,ARIH1,PRKAR2A,FLCN,AP2A1,GTPBP1,CSNK2A1,DECR1,UBFD1,PTEN,HNRNPA2B1,FKB
1,SEC16A,MASTL,CEMIP2,CUL3,RPS27A,TIMM50,CHERP,CEP170,TAOK1,TPP1,UBE2D2,RAB14,
KB5,TAGLN2,PTPRF,HNRNPAB,BAZ2A,TAOK1,SOX2,SLC4A2,PRPF4B,IGF2BP1,ARPP19,CERS2,FE
3,UBE2G1,ADD1,SESN2,SETD1B,BTG1,ID2
A,TNPO2,NUDT3,TMEM214,LDOC1
MP4B,C16ORF58,SCD,FZR1,TNFAIP1,MECP2,TNRC6A,MAPKAPK2,GOLGA8B,P3H1,TFRC,PER1,C
D,RAB11FIP3,MGAT5,TXNRD1,KIAA1191,WDR26,PIK3C2B,SSU72,PSME3,ZNF532,WAC,ZNF703,F
D1,CERS2,EIF4G1,HMGA1,AGAP1,SLC22A23,NIBAN2,MKNK2,SUMO1,RNF157,NTN1,IP6K1,CCDC
C6,POMGNT1,SECISBP2L,SETD1B,CPSF7,RNF40,QSOX1

RC5B,SETD1B

PAFAH1B2,PSAT1,TSNAX,IGSF3,IGF1R,NPTN,FXR1,TNRC6A,ZCCHC3,KPNA6,UBE2Q1,ARL8A,NKI

1,SEC16A,MASTL,PTEN,CEMIP2,CUL3,RPS27A,TIMM50,CHERP,CEP170,TAOK1,TPP1,UBE2D2,R

C2,ERBB2,THBS2,ARPC5,SP1,TMEM59,SESN2,PRELID1,ITPKB

CDV3,TRMT112,TNFRSF10B,SRF,ZEB1,TFRC,RASSF2,WASF3,COX6B1,UQCRFS1,SHC1,TGFB2,

S2,MARVELD1,ASB6,SLC2A4RG,CTDNEP1,ZNF516,SRCIN1,MYADM,TOMM40

DRM1,MCRIP1,TNPO2,ASB6,TMEM151B,SEPTIN9,SESN2,SETD1B,CPSF7,RNF40

IPUL1,ZFAND5,ATP6V1B2,GIGYF1,MPRIP,PFN2,PPP6R1,SERBP1,WBP2,BMPR2,MSMO1,WDR45E

13A,THBS2,MARVELD1,ASB6,SLC2A4RG,CTDNEP1,ZNF516,APOE,SRCIN1,MYADM,TOMM40

PRIN1,NAV1,CTDNEP1,MBD6,MFGE8

AS2,CCDC6,ATG9A,THRA,MSI2,CTIF,VPS37B

6B1,SENP3,KLHL36,PIN1,C8ORF33,PPP2R2A,GET4,EPHA2

1,TAGLN2,ZNF512B,CBX6,WSB2,TAOK1,RAB1B,TM9SF4,SOD2,LRP3,SCD,FGFR1,ELAVL3,WDTC

2,PHB2,ZFP36L1,CBX6,LRRC59,GRK2,ABI2,RALY,GNA12,TM9SF4,SLC25A23,OGT,ATXN7L3,HM

,INF2,CTDNEP1,TMEM151B,CTNNBIP1,CDR2L,SRCIN1,SETD1B,C6ORF89,HES6

B2,DNAJC8,TMEM43,ASB6,GPS1,UQCRH,RNPS1,SETD1B,UBTF,USP36

PSMF1,UBE2V1,ZFP91,RASSF2,SETD5,CASKIN1,ZDHHC9,GALNT2,BTF3L4,PPP2R2A,ATP6V1F,R

F13,DCAF5,SCAMP4,H2AX,TOB2,SRF,DNAJC8,MCFD2,ZFP91,DUSP8,MARVELD1,RELA,ID3,PPP2

C2,ASH1L,TFRC,SH3BP5,SLC16A1,ZDHHC5,CRIM1,SCOC,CCSER2,NFIB,PRMT7,DNAJC21

,INF2,CTDNEP1,TMEM151B,CTNNBIP1,CDR2L,SRCIN1,SETD1B,C6ORF89,HES6

COR,BACE1,ADGRL1,CBX4,FAM193B,DDX56,VPS28,UBTF,TOMM40

DC80,FXR1,HMGB3,ZCCHC3,MAGEF1,CDK4,DDX6,POLDIP3,NDRG4,EMC10,USP48,MCFD2,ZFP

ZFP36L1,CBX6,LRRC59,GRK2,ABI2,RALY,GNA12,TM9SF4,SLC25A23,OGT,ATXN7L3,HMGA1,AG

1B

1,ANKFY1,GSR,AREL1,ADRM1,NME2,GBA2,CACNA1H,COX6B1,ATG9A,MCRIP1,TNPO2,GALNT2,

APZA1,DAZAP2,FBLN5,MYL12B,USP5,SCAMP5,NIBAN2,IGF1R,TNFAIP1,SUMO1,LIMK1,SRSF10,

A,STK35,TM9SF4,CAPZA1,SCD,FGFR1,IGFBP4,WDTC1,AGAP1,SLC22A23,PLXNA2,CLUH,IP6K1,

,TNRC6A,DDX6,LDB1,PDE4DIP,RRM2,NACC1,CDC42EP4,NR2F2,NR2F6,AMD1,SLC39A10,PHF3,

53C,TNRC6A,NECTIN1,FAM168A,LDB1,YWHAZ,SYNGR1,DIRAS1,CDKN1B,CASKIN1,NACC1,UNC1

53C,TNRC6A,NECTIN1,FAM168A,LDB1,YWHAZ,SYNGR1,DIRAS1,CDKN1B,CASKIN1,NACC1,UNC1

CC1,ATG9A,OGDH,MCRIP1,NAV1,CTDNEP1,SEPTIN9,SP1,SRCIN1,PLXND1,WDR33,GDE1

9A1,KIF3A,NDUFS1,ZNF146,GATAD2B,PIP5K1A,SESN2,VPS37B

,HINT1,LDLR,LRATD2,SFXN1,DAZAP2,FBLN5,TGOLN2,SRPRA,HMGA1,WDTC1,PAIP2,MKNK2,KIA

,HINT1,LDLR,LRATD2,SFXN1,DAZAP2,FBLN5,TGOLN2,SRPRA,HMGA1,WDTC1,PAIP2,MKNK2,KIA

ZFP36L1,CBX6,LRRC59,GRK2,ABI2,RALY,GNA12,TM9SF4,SLC25A23,OGT,ATXN7L3,HMGA1,AG

PA4,KLC1,ZNF302,RAD23B,ORAI2,PPP2CA,MAPK1,PBX2,WDR6,LARS1,STMN3,ISYNA1,SOD1,HN

,UBE2V1,PGRMC2,STC2,TNPO2,CAMK2N1,TMEM50A,CTIF,RAD50,QSOX1,ZC3H4

ZD7,SECISBP2L

PS14,ATP2A2,RPL26,EIF5A,DDX3X,PRKAR1A,TLN1,SESN3,PGRMC1,C5ORF24,SSRP1,MLEC,ORA

,RNF157,TNRC6A,DARS1,H2AX,WAC,YWHAZ,HMGXB3,MCFD2,TNPO2,TET3,RAD54L2,RAD50,M

1,UNC13A,THBS2,MARVELD1,ASB6,SLC2A4RG,CTDNEP1,ZNF516,SRCIN1,MYADM,MYL9,TOMM

VEZF1,CITED2,UBE2G1,NUDT3,SYT4,HMGA2

\,VHL,CCND1,IPO7,DYNC1LI2,CAPZB,DIDO1,SREBF2,KDM3B,DDIT4,CAPN2,NAP1L1,EXTL3,GRSF
10,STT3A,ID4,NCOR2,NAP1L1,EXTL3,CSNK2A1,SFRP1,TRA2B,PTEN,FEM1B,TIMM50,SLC25A5,LI
<T3,PHACTR4,TNRC6A,MAGEF1,RDH11,ASH1L,RAVER1,DUSP3,CEBPG,RNF4,OGDH,MDC1,NCOA
SLC25A36,RNF11,PLXNA2,IGF1R,MECP2,SRSF10,MAPKAPK2,KPNA6,CRKL,RBFOX2,SLC43A2,T
IMGXB3,CCDC85C,NACC1,ZDHHC5,TSPAN11,TMEM151B,AMD1,CTNNBIP1,CDR2L,NFASC,SRCIN
ON,CORO1C,PTP4A2,DGKD,SEC23A,CDC42,STAT3,KCNQ2,NDST1,SLC7A6,FKBP1A,SERP1,ABI2,
<1,THRA,MSI2,RAP1B,ZNF106,CTIF,VPS37B
,SRSF10,OTUD5,NDRG4,CRTC1,IRS1,NUP133,GATAD2B,COX5B

,TSPAN6,SRSF1,DYNLL2,CASC3,RPL28,FYN,MPRIP,SREBF2,KDM3B,AKIRIN1,BMPR2,TXNIP,SSR1
PH3,ATF7IP,SRSF7,TMEM43,PDCD11,COX6B1,ASB6,VOPP1,MAPK3,ADD1,TRPC4AP,ELOA

A2,ZNF486
D4,ANP32A,XPOT,CSNK2A1,SLC39A6,SLC39A7,KLHL42,SMC3,RGMB,ANP32B,POM121C,CTTN,G
ACC1,SEPTIN9,ZNF516,PLXND1
3,IGF1R,TNRC6A,VPS51,SH3BGRL3,ATP5PO,NDRG1,DDAH1,GSK3B,RCC2,CDKN1B,KDM5A,UQC
3,SRCIN1,GSN,NDUFA5

2,PAFAH1B2,ESYT1,ANP32E,WDR26,SUB1,SSU72,ATF7IP,SRSF10,RBFOX2,RCC2,KIF3A,ATMIN,S
-A,FAM168A,ZNF703,UBE2V1,NACC1,NKIRAS2,WBP1L,SGPL1,TET3,SP1,POGZ

,TOLLIP
R2A,LDOC1,UBTF
PRM,RALBP1,PNMA2,ZNF516,GPRC5B,SETD1B,UBTF
PRM,RALBP1,PNMA2,ZNF516,GPRC5B,SETD1B,UBTF
P3,RNF11,HMGA1,MKNK2,ACVR1B,UBE4B,NTN1,LIMK1,ZCCHC3,RNF41,GBA2,DNAJC8,RAB3A,D
TDSP2,CPSF2,NOVA2,TOMM20,RNF187,MLXIP,GNAI3,TSPYL4,LMNB2,MLEC,RAB5B,ASXL1,KDM

,KLC2,CCNG1,NDRG1,SRF,CASKIN1,KDM2A,COX6B1,NACC1,ZDHHC9,THRA,MSI2,HIPK1
H,MCRIP1,NAV1,CTDNEP1,SEPTIN9,SP1,SRCIN1,PLXND1,GDE1
,TNRC6A,LDB1,DHX36,DSTYK,UBE2G1,HMGB2,RELA,COPS8,MAPK8IP3,NUP58,BTG1,ATP8A2,R
NRC6A,RNMT,MCFD2,NACC1,UQCRFS1,HMGB2,BSDC1,TLR6,SESN2,KLF10,DNAJC21

5,ANP32B,TIMM50,LMAN1,SERP1,RAB1B,AKT3,GNL1,HDGFL2,PRPF4B,ABCF1,SPTY2D1,IST1,SM

2,AREL1,EXOC7,SRSF7,OXSR1,NSD1,SLC43A2,SYNGR1,CCDC85C,CELSR3,UBQLN4,SOX9,RRP1
A1,ATXN2L,ARIH1,TXLNA,NAP1L1,BAZ1B,FARP1,UBAP2L,HNRNPA2B1,ADH5,B4GAT1,PEG10,RAE
NDUFS1,RALBP1,SNRPB,UBE2G1,SP1,NOLC1,MRPS27,ITPKB,TMED5
RGMB,SPINT2,KIAA0100,SLIT1,MAZ,TTC37,SFXN1,SOD2,PDPK1,CAP1,CSE1L,UBE2D3,SCD,USO1

32,ILRUN,ERBB2,RGMA,TET3,TGFB2,ELMO2,MEX3D,PPP2R5C

OSP2,CPSF2,NOVA2,TOMM20,RNF187,MLXIP,GNAI3,TSPYL4,LMNB2,MLEC,RAB5B,ASXL1,KDM6I
LDB1,MCFD2,CDKN1B,CDC42SE1,NUMBL,CTDNEP1,CCNL1,NUDT3,CTIF,TMEM214
K2,NOTCH1,FZR1,EFNB1,PDGFRB,CDK4,RDH11,MCFD2,CRABP2,CDKN1B,GMFB,GIT1,MBD6,TS
PSF2,NOVA2,TOMM20,RNF187,MLXIP,GNAI3,TSPYL4,LMNB2,MLEC,RAB5B,ASXL1,KDM6B,RAB
PSF2,NOVA2,TOMM20,RNF187,MLXIP,GNAI3,TSPYL4,LMNB2,MLEC,RAB5B,ASXL1,KDM6B,RAB
EIS1,RCC2,RRP1B,HOXA5,CDKN1B,SRRT,GPS1,AJUBA,ATP6V1F,MRPS26,LRIG3,HMGA2
TCH1,NPTN,MECP2,SUMO1,GALNT10,RPL27A,HSD17B12,C1QBP,NUP98,YWHAZ,WDR82,ALDH9

N1,GSN,NDUFA5
3,PRKAR1A,SESN3,ANKRD52,NARS1,ZNF454,PEA15,CDKN1A,SERF2,TSR1,COL2A1,CBX1,RAD23I
OV3,MECP2,WDR26,COX8A,TNRC6A,SRSF10,BCOR,RPRM,ZFP91,GPR173,UBE2G1,C8ORF33,SC

IGYF1,ID4,BMPR2,DDIT4,EP300,ZFP36L1,FBXO11,ULK1,LMAN1,FKBP1A,ZNF664,GRK2,BAZ2A,C

H,MCRIP1,NAV1,CTDNEP1,SEPTIN9,SP1,SRCIN1,GDE1
MT,MCFD2,NACC1,UQCRFS1,HMGB2,BSDC1,TLR6,SESN2,KLF10,DNAJC21
SL4,RASSF2,CDKN1B,WASF3,IRS1,DNAJB6,SHC1,SP1,DNMT1,ELMO2,SECISBP2L,M6PR

UP50,ARF6,NFASC,RAP1B,RAD50
MAP2K4,MSI2,HMGA2,SKIL
KIL

G2,PTEN,HNRNPA2B1,HIF1A,GRN,BAZ2A,RAB1B,VPS4A,EFTUD2,PNISR,NUCKS1,DICER1,PHGDH,
EPA1,LDB1,MCFD2,CDKN1B,CDC42SE1,NUMBL,CTDNEP1,CCNL1,NUDT3,CTIF
2,OTUB1,POGK,VDAC2,NFASC,ID3,HES6,NECAP1
2,OTUB1,POGK,VDAC2,NFASC,ID3,HES6,NECAP1
SUMO1,TBXA2R,DDX6,UBE2V1,NACC1,MYL12A,CCDC6,PIN1,POMGNT1,ANKRD13B,SECISBP2L,S
QCRH,MYL9,SETD1B,UBTF

SF11,YY1,SLC25A36,KLF13,IGF1R,SSU72,RAN,TNRC6A,ATP6V0B,CSNK1A1,NDRG1,ZBTB4,WAC,
SF11,YY1,SLC25A36,KLF13,IGF1R,SSU72,RAN,TNRC6A,ATP6V0B,CSNK1A1,NDRG1,ZBTB4,WAC,
HNRNPAB,HIF1A,FKBP1A,SERP1,PELP1,HINT1,PNISR,SFXN1,SOD2,RAB5C,DICER1,SPTY2D1,FKB
AMK2N1,ZNF146,TMEM50A,GATAD2B,PIP5K1A,SESN2,ZC3H4
SUMO1,TBXA2R,DDX6,UBE2V1,NACC1,MYL12A,CCDC6,PIN1,POMGNT1,ANKRD13B,SECISBP2L,S
UL3,RPS27A,HIF1A,CHERP,TAOK1,ELAVL1,TAX1BP1,LDLR,SRSF2,PLEKHB2,BLCAP,FNBP1L,DICE
3,DUSP8,TLK1,MDC1,SENP3,ARF6,FGFBP3,FAM3C,MYADM
.8A,NACC1,GDE1

,RPL37A,FN1,SPEN,NONO,HACD3,TPI1,TMSB10,EIF4A1,RPL15,GTF3C4,LDHB,PKD1,SF3B1,RPS

AP4,SF3B2,MAT2A,PGAM1,NFE2L1,LARP1,HNRNPA0,CTSD,TSPAN3,RPS24,RPS9,XRCC5,KPNB1,

3,CKB,COL4A2,SPEN,PSMD5,YWHAQ,EIF1,PPP2R1A,RPL15,SERINC1,HMGB1,TPT1,RPS3A,TUBB

P5F1A,EIF4B,AFF4,COL4A2,SPEN,YWHAQ,EIF1,TPI1,MRC2,PPP2R1A,GTF3C4,RPS3,CHD3,POLR

RS1,HNRNPH1,RPS12,RPS27,COTL1,TM9SF3,TOMM20,TCP1,SNRNP70,ASXL1,RPL36A,KDM6B,

,JPT2,PNN,HNRNPC,LRPPRC,KHDRBS1,HDLBP,CCND2,PIP4K2B,ITM2C,CLTC,MAGED2,HMGCS1

OAZ1,PFN1,MCL1,PNN,CCNI,CCND2,PIP4K2B,ACTN4,RPL30,EIF3L,NANOG,MIF,CTNNB1,GOLM1

A3,ANKRD52,PTGES3,COTL1,HRNR,ARHGDIA,CDK16,CNN2,TM9SF3,EEF1A2,GDI2,TSR1,LMNB2,

RPL9,CLSTN1,SLC7A5,CHD4,ATP1A1,RPL26,LASP1,PFN1,MCL1,JPT2,SPC25,XPO1,SPTBN2,SLC

CHF6,VCL,MAP4,PXDN,LARP1,FADS2,RPSA,TPM4,NACA,TSPAN3,RPS24,MSN,PAPOLA,HK1,CBX5,

SNK1E,CCND2,PDIA6,DDX3X,RPS5,CHPF,MYO5A,DNAJC5,HNRNPDL,TARDBP,NDFIP1,HNRNPK,R

PL12,ITM2C,RPL35,TLN1,GJA1,ARL6IP1,ARID1A,ST3GAL2,ANKRD52,YWHAB,RHOB,UBE2H,MAG

11,LMNB2,MTRF1L,KDM6B,HDGF,RPS15A,SELENON,PBX2,HNRNPUL1,MEX3A,MLF2,RTL8C,WB

IDH2,GNAI3,GDI2,GCN1,HDGF,RNF10,MARS1,SMARCA4,PBX2,RABL6,WDR6,SON,CASC3,ZFAND

,PSMD2,KIF5B,SLC7A5,JUND,EIF5,ATP2A2,MCL1,JPT2,HNRNPC,PLXNA1,PIP4K2B,MRFAP1,H1-

RID1B,RER1,RPS2,TIMM50,KLHDC3,TNKS,PALM,CLPTM1,RAB1B,MAZ,TRAP1,TMEM63B,TUBGC

,APPL1,U2AF2,KDM6B,RAB11B,G3BP2,KDELR2,ODC1,GRB2,CAPZB,SUPT5H,SON,ATXN2L,TMEI

1,HECTD4,CCT6A,GPC1,SMCHD1,HDAC2,GLUL,FDFT1,PARP1,PLOD1,CCT4,ORAI2,GCN1,SKI,P

,NOTCH2,VHL,G3BP2,DSTN,PRDX1,FAM168B,TCF25,DYNLL2,HNRNPUL1,ATXN10,KDM1A,TPM3

,TXLNA,PSMD1,BAZ1B,UPF1,TAGLN2,RPS2,PSMD8,XRCC6,PELP1,TLE1,FLOT2,QPRT,REEP2,NS

DH2,ACTN4,PRDX2,CLTC,MRFAP1,GNB2,GPAT4,GJA1,CD63,ARID1A,MXD4,WLS,H1-10,HNRNPH

RBP,RHOB,UBE2H,SMCHD1,NUCB1,RPN2,MLXIP,CPSF1,TGFBI,LMNB2,NCKAP1,RAD23B,SCAF1,

E2H,CALU,RAC1,PEA15,LUC7L3,FAM199X,TOMM20,EEF1A2,FDFT1,CCT2,MEST,HNRNPA3,YEAT

TL8C,AMOTL2,ANP32A,VDAC1,PRCC,SEC16A,BRD2,ECEL1,TMEM167A,SLC25A5,TPP1,RAB1B,G

,TXNIP,ADAR,CTNND1,TXNDC5,RETREG2,SEC16A,ZFP36L1,KCTD12,RGMB,FEM1B,HNRNPA2B1

USP24,EHD1,CCDC47,CYB5B,FGFR1,TUBB4A,CNBP,SCAMP1,IGF1R,AAMP,SCAMP4,SUN2,FAM5

3,MLXIP,GDI2,HSPH1,HSPA4,RAD23B,DPYSL2,TMED2,ARF3,UBXN4,AZIN1,ADAM9,CAND1,ZFAN

INRNPDL,RPL36,UBE2H,CALU,ARCN1,ARHGDIA,SLC35E2B,LUC7L3,CDKN1A,MLXIP,PGD,GDI2,T

19SF3,VAMP2,PDXK,KLC1,CCT3,GRINA,SURF4,TNPO1,SKI,H3-3A,PYGB,NOTCH2,VEGFA,RPL37,

IA,SLC35E2B,LUC7L3,CDKN1A,MLXIP,PGD,TM9SF2,SSRP1,UBE4A,CAMSAP1,ETNK1,NCKAP1,L

RB2,MAP7D1,TSC22D1,UBQLN1,PTP4A1,PCBP2,PFN2,NCOR2,SERBP1,BMPR2,BAZ1B,GTPBP1,

A2,KRT8,DAZAP2,OGT,COPZ1,ATXN7L3,ELOVL5,UNC119B,CAD,HMGA1,AGAP1,FXR2,MKNK2,S

70,HNRNPU,OGA,PGAM1,LARP1,CPE,RPSA,TPM4,EIF4H,RPL17,IGFBP3,RPL14,CBX5,CHD4,WD

,USP7,SARS1,VANGL2,SERBP1,AKIRIN1,DDIT4,DHX15,POLR2E,SSR1,ST3GAL6,KDM5C,EP300,R

,AKT3,MAZ,AKT2,NASP,FJX1,CNBP,AMFR,AGAP1,MDM2,WDR26,B4GALT5,FAM53C,PDGFRB,CO

2,MYH10,HMGCR,CAPRIN1,CDK16,APPL1,LUC7L3,C5ORF24,U2AF2,PPP1CB,PABPN1,SSRP1,AB

5,CSNK1G2,KIAA0895L,UBFD1,TMEM167A,LRP10,STK35,TRIR,ERGIC1,CARM1,INTS3,RAB1B,M

5,CSNK1G2,KIAA0895L,UBFD1,TMEM167A,LRP10,STK35,TRIR,ERGIC1,CARM1,INTS3,RAB1B,M

ER1,AMFR,POLDIP2,AGAP1,ABL1,SART3,LIMK1,IP6K1,KLC2,ADRM1,MAPKAPK2,ARPC3,COLGA

ER1,AMFR,POLDIP2,AGAP1,ABL1,SART3,LIMK1,IP6K1,KLC2,ADRM1,MAPKAPK2,ARPC3,COLGA

NCOR2,VDAC1,STAT3,EXTL3,CSNK2A1,UBAP2L,COX7C,CEMIP2,SLC7A6,DKK3,FUS,ACLY,CEP17

IT4,DHX15,TRAF4,EIF2AK1,SGTA,SF1,AP2A1,KPNA2,HOXB5,EVL,PTEN,CEMIP2,PTPRF,PEG10,F

LF2,AMOTL2,CDC42,ARIH1,SF1,UBA52,GLUD1,TAGLN2,ZNF512B,CBX6,PPP1CC,APEX1,CTTN,S

SLC7A1,PDPK1,DAZAP2,OGT,ARPP19,TGOLN2,YY1,TTC28,CERS2,CNBP,HS2ST1,RSF1,PUM1,M

1,GSPT1,ZFP36L1,PPP2CB,UQCRC1,MASTL,RPL7L1,DKK3,DAP,GLO1,SNRPD3,HINT1,SRSF2,CA

NISR,RAB5C,DICER1,LRRC41,RBPJ,CIZ1,CAD,PSAT1,RNF157,CUL4A,HNRNPR,RABAC1,WAC,DD/
,AMPD2,B4GALT5,FAM53C,CAPN1,NECTIN1,FAM168A,LDB1,YWHAZ,SYNGR1,RPRM,DIRAS1,ARL
D,VAMP2,SURF4,TUBB2A,CCND1,PBX2,RABL6,HNRNPUL1,ZFAND5,SSB,ATXN2L,NT5DC2,PCBP2,
SM2,ELN,SCAMP4,CD44,LIMK1,IP6K1,ANKFY1,PCGF3,MAPKAPK2,OTUD5,EPN1,ARPC3,TOB2,DN
,B4GALT5,FAM53C,CAPN1,NECTIN1,FAM168A,LDB1,YWHAZ,SYNGR1,RPRM,DIRAS1,ARL8A,CDK
P2K2,PTP4A2,MLF2,KDM5B,VARS1,BRD2,CRTAP,PRRC2A,NDST1,SEPTIN7,MAZ,SH3GL1,TMEM6
MARCD1,ELAVL3,IGFBP4,IGF1R,UBE4B,CLUH,SUMO1,SCAMP4,ADGRB2,CD44,BNC2,HLA-A,SSU
SEC16A,ZFP36L1,SFRP1,KCTD12,FEM1B,TMEM167A,KLHDC3,APEX1,PEG10,HIF1A,GSE1,SERP1
FA,TUBB2A,CCND1,SRSF1,DYNLL2,GRB2,TPM3,MSL1,YWHAH,SERBP1,TXNIP,POLR2E,ARIH1,PR
SLC25A23,PACS1,FOXK2,SPTY2D1,IST1,RBPJ,FBLN5,SMARCD1,CERS2,TEAD1,IGFBP4,RNF11,T
KLHDC3,BAP1,STK35,ACLY,BLCAP,NCSTN,NUCKS1,ELP1,SLC25A23,SCRIB,LRRC41,LRP3,NASP,
N,SCAMP4,LIMK1,IP6K1,ANKFY1,PCGF3,MAPKAPK2,OTUD5,EPN1,ARPC3,TOB2,DNAJC8,TMEM4
SRSF1,DYNLL2,GRB2,TPM3,MSL1,YWHAH,SERBP1,TXNIP,POLR2E,ARIH1,PRKAR2A,FLCN,DECR
ATXN2L,COL27A1,GHITM,HNRNPD,TXNIP,CIC,FBRS,PRCC,PRRC2A,CUL3,RPL7L1,RPS2,LMAN1,
NUDT19,NASP,ATP6V1G1,CYB5B,SCD,UNC119B,CNBP,PAFAH1B2,SLC25A36,HMGA1,SCAMP1,PD
APOLA,RPL14,HK1,CBX5,RPL9,WDR1,ATXN7L3B,LASP1,VAT1,MCL1,TADA2B,PLXNA1,RPL30,H1-
,DYNLL2,GRB2,TPM3,MSL1,YWHAH,SERBP1,TXNIP,POLR2E,ARIH1,PRKAR2A,FLCN,DECR1,CBX
IK1G2,NXN,SFRP1,CBX6,PRRC2A,BRSK1,NPLOC4,LRP10,STK35,TRIR,STRN4,LTBP4,PRR12,SLC
C,RNPS1,PAF1,CTSA
KLF13,FZR1,MAPK8IP2,SUMO1,SCAMP4,SSU72,IP6K1,PCGF3,DDX6,RDH11,EPN1,TOB2,YWHAZ,
TXNDC5,ZFP36L1,KCTD12,YKT6,FEM1B,PREPL,PEG10,TAOK1,VPS41,LDLR,MAZ,SH3GL1,IREB2,
PN1,RPL27A,CCNG1,PPP2R5D,CRKL,NUP98,YWHAZ,PDCD11,TFRC,NACC1,HIRA,PRMT2,UBE2G
L2,POLR2E,ARIH1,BCL9L,CBX6,NPLOC4,TAOK1,GABARAPL1,FAM234A,GNL1,TM9SF4,PRPF4B,
L2,POLR2E,ARIH1,BCL9L,CBX6,NPLOC4,TAOK1,GABARAPL1,FAM234A,GNL1,TM9SF4,PRPF4B,
A,HOXB5,NXN,PRRC2A,BRSK1,TMEM167A,NPLOC4,TRIR,SLIT1,HSPG2,INTS3,KRT8,VASH1,SAE
GA8B,OXSR1,TOB2,YWHAZ,ACSL4,RCC2,NOLC1,TGFB2,SEC31A,RNPS1,RAP1B,ID2,CPSF7,WDR
F2,CAPZA1,SOD2,DAZAP2,COX6A1,YY1,SMARCD1,ZER1,SGSM2,NIBAN2,PLXNA2,SUMO1,LIMK1,
LC6A8,GNL1,RDX,LSM14A,TRAP1,AHSA1,REEP2,ITGAV,IGF2BP1,LRRC41,USP24,MED13,PSMC3,
HC2,PTP4A2,SLIT3,ILF2,PFN2,BMPR2,POLR2E,VDAC1,EIF2AK1,BCL9L,CAPN2,ALDH3A2,HOXB5,
M167A,LMAN1,MTCH1,FUS,ERGIC1,ABI2,VPS4A,SLC7A1,WBP11,DHCR7,ATP5MC3,C16ORF58,M
8,CNBP,RNF11,MGAT5,FZR1,SLC38A1,SUMO1,SCAMP4,PTPN1,LIMK1,IP6K1,PCGF3,TNRC6A,AD
LDB1,TNFRSF10B,NSD1,EMC10,GOLGA4,STC2,NUP205,CACYBP,RNF4,ZNF395,RNASEK,ZNF598
,CLUH,RNF157,LIMK1,SALL2,MAPKAPK2,OTUD5,LDB1,PPP2R5D,NDRG1,WAC,DNAJC8,ZFP91,C
SP,FJX1,CNBP,AMFR,AGAP1,ABL1,DAG1,ELN,MDM2,WDR26,SYNCRIP,RAN,FAM53C,PDGFRB,C
MB,CBX6,TIMM50,NPLOC4,SNRPD3,CARM1,RALY,TM9SF4,WASF2,FXYD6,ATXN7L3,UNC119B,R
,RABL6,SRSF1,ATP5PB,SON,ZFAND5,XPNPEP3,SUPT6H,LSS,MTRNR2L10,STT3A,ID4,MAPK8IP1,
8,CNBP,RNF11,MGAT5,FZR1,SLC38A1,SUMO1,SCAMP4,PTPN1,LIMK1,IP6K1,PCGF3,TNRC6A,AD
P1,SAE1,SCD,SCAMP5,SEMA6B,DAG1,RUVBL2,TIA1,REEP5,TRRAP,TRIM27,PMEPA1,NME2,MAPK

B4,ZEB1,RPS29,CEBPG,DGCR8,G6PC3,CNOT3,YBX3,MSI2,RPL32,KDM2B,MXRA7,QSOX1,VPS37
CNBP,AMFR,AGAP1,CDV3,MDM2,WDR26,FAM53C,PDGFRB,SRSF10,COX7A2L,DDX6,UBE2Q1,LIM
B,PREPL,RPS27A,PEG10,TAOK1,VPS41,IMPDH2,LDLR,SH3GL1,IREB2,GALNT1,SH3PXD2A,SFXN
ZFP91,XPC,UNC13A,LONP2,ARSB,PRKCI,GATAD2B

KIN1,NACC1,NKIRAS2,GPR173,GPS1,PIN1,SEPTIN9,ZNF516,GIT1,ADGRL1,SECISBP2L,SETD1B,

MBRA1,LYPLA2,UNC119B,CNBP,BGN,WDTC1,COL12A1,SSU72,EFNB1,TNRC6A,TRIB3,GBA2,NDF
SF1,SEC16A,PHB2,HOXB5,KIAA0895L,TMEM167A,TRIR,PRR12,CAPZA1,IST1,UBE2D3,FGFR1,CN
57,LIMK1,SALL2,MAPKAPK2,OTUD5,LDB1,PPP2R5D,NDRG1,WAC,DNAJC8,ZFP91,OTUB1,POGK,
M121C,GRK2,BAZ2A,FKBP4,IMPDH2,SOD2,SPTY2D1,PTGFRN,PSMC3,USP5,HS2ST1,PTPN11,LM
,MEX3A,MLF2,TRAF7,CRTAP,CBX6,MAP1LC3B,CARM1,GNL1,TM9SF4,ATXN7L3,KLHDC10,TEAD
SEC16A,PHB2,HOXB5,KIAA0895L,TMEM167A,TRIR,PRR12,CAPZA1,IST1,UBE2D3,FGFR1,CNBP,I
KB5,RAB2A,NXN,PRRC2A,BRSK1,TMEM167A,NPLOC4,PEG10,CHERP,TRIR,LINGO1,INTS3,SLC4A
SEC16A,PHB2,HOXB5,KIAA0895L,TMEM167A,TRIR,PRR12,CAPZA1,IST1,UBE2D3,FGFR1,CNBP,I
SEC16A,PHB2,HOXB5,KIAA0895L,TMEM167A,TRIR,PRR12,CAPZA1,IST1,UBE2D3,FGFR1,CNBP,I
SEC16A,PHB2,HOXB5,KIAA0895L,TMEM167A,TRIR,PRR12,CAPZA1,IST1,UBE2D3,FGFR1,CNBP,I
RIB3,CCNG1,DDAH1,ZEB1,GSK3B,ASH1L,ZYX,PRMT5,RPS29,HOXA5,KDM5A,NKIRAS2,TMPO,PP
VPS41,LDLR,SH3GL1,IREB2,GALNT1,SH3PXD2A,SFXN1,ARPP19,VASH1,UBE2I,KLHDC10,UBE2D
IP3,SLC16A1,NACC1,KIAA2013,FYTTD1,POGZ,NF2,ID2
F10B,GSK3B,BCOR,RRM2,COX6B1,SLC16A1,GET1,RUSC1,NADK,MAFB
BAN2,REEP5,UBE4B,SSU72,IP6K1,FAM168A,ZBTB4,TOB2,HMGXB3,DNAJC8,TMEM43,STARD3,F
BAN2,REEP5,UBE4B,SSU72,IP6K1,FAM168A,ZBTB4,TOB2,HMGXB3,DNAJC8,TMEM43,STARD3,F
PL31,TADA2B,CCND2,PLXNA1,RPL30,PRKAR1A,GJA1,RPS10,SESN3,ANKRD52,NARS1,PTGES3,A
OTUB1,NACC1,CCDC6,GPS1,POMGNT1,SECISBP2L,SETD1B,RNF40,QSOX1
,MEX3A,MLF2,TRAF7,CRTAP,CBX6,MAP1LC3B,CARM1,GNL1,TM9SF4,ATXN7L3,KLHDC10,TEAD
OC2L,SMARCD1,CERS2,SLC20A1,AMBRA1,RNF11,COL12A1,NIBAN2,PLXNA2,UBE4B,IP6K1,EFN
SEC16A,PHB2,HOXB5,KIAA0895L,TMEM167A,TRIR,PRR12,CAPZA1,IST1,UBE2D3,FGFR1,CNBP,I
NPO2,EPB41L1,THRA,SP1,ADGRL1,SETD1A,SRCIN1,PLXND1,CCSER2,LDOC1,RNF40,NDUFA5,SN

P,ATP1B3,ANKFY1,PCGF3,HIPK2,PMEPA1,HM13,EMC10,ZNF703,TUBA1C,WBP1L,PAIP1,RAB35,N
CL2,RNF41,EPN1,WAC,HMGXB3,UBE3C,DNAJC8,UBE2V1,RAB3A,ZFP91,FARSA,NACC1,KIAA2013
CL2,RNF41,EPN1,WAC,HMGXB3,UBE3C,DNAJC8,UBE2V1,RAB3A,ZFP91,FARSA,NACC1,KIAA2013
NACC1,CCDC6,GPS1,POMGNT1,SECISBP2L,SETD1B,RNF40,QSOX1
NACC1,CCDC6,GPS1,POMGNT1,SECISBP2L,SETD1B,RNF40,QSOX1
BSDC1,GATAD2B,SESN2,CACNG7,NDUFA5,TMED5
NPO2,EPB41L1,THRA,SP1,ADGRL1,SETD1A,SRCIN1,PLXND1,CCSER2,LDOC1,RNF40,NDUFA5,SN
TXNIP,ADAR,MSMO1,STAT3,CSNK2A1,GRSF1,SEC16A,CD164,MASTL,PTEN,RGMB,CEMIP2,CUL3
CL2,RNF41,EPN1,WAC,HMGXB3,UBE3C,DNAJC8,UBE2V1,RAB3A,ZFP91,FARSA,CEBPG,NACC1,K
C80,TNFAIP1,MECP2,SUMO1,LIMK1,TRIB3,KPNA6,PDE4DIP,YWHAZ,RPRM,ZFP91,ALDH9A1,NKI
,GATAD2B,SESN2,CACNG7,NDUFA5,TMED5
A2,IPO7,WDR6,HNRNPUL1,BNIP3L,FKBP8,ATP6V1B2,SF3A1,ATXN2L,PTP4A1,RPS13,TRAF7,STAT

RM2,NACC1,SLC5A6,ATG9A,MCRIP1,GPRIN1,NAV1,CTDNEP1,SEPTIN9,SP1,MCM7,NFASC,MAPK
L2,SC5D,COPE,CAB39,TRRAP,ZCCHC14,GOLGA8B,WDFY1,SYNGR1,TFRC,TXN,TMPO,NUP133,AS

A,NFASC,UBTF
DF3,KLF13,CCDC80,IGF1R,KIAA1191,TRMT112,CAB39,CNDP2,RNF157,LIMK1,CDK4,PSME3,PDE
T5,EFNB1,DDX6,UBE2V1,MAP1S,FARSA,NACC1,DGCR8,TMEM50A,PIN1,ANKRD13B,SETD1B,PIE
T5,EFNB1,DDX6,UBE2V1,MAP1S,FARSA,NACC1,DGCR8,TMEM50A,PIN1,ANKRD13B,SETD1B,PIE
2,TEAD1,UNC119B,IGFBP4,SLC25A36,KLF13,TNFAIP1,SSU72,CAPN1,LDB1,PSMF1,WDR82,RAB
P4,IP6K1,FAM168A,SRF,HMGXB3,ASH1L,UBE2V1,SETD5,OTUB1,ZNF516,ADGRL1,PPP2R2A,MYL

KLF13,RNF157,PNRC2,SALL2,WIZ,FAM168A,CCNG1,GBA2,NDRG1,ZBTB4,TOB2,YWHAZ,HMGXB

A1,WBP2,KDM5C,PRKAR2A,HIP1,GRSF1,PRRC2A,KLHDC3,LRP10,STK35,INTS3,BCL7A,RALY,MAZ

A1,WBP2,KDM5C,PRKAR2A,HIP1,GRSF1,PRRC2A,KLHDC3,LRP10,STK35,INTS3,BCL7A,RALY,MAZ

IFAIP1,MECP2,SUMO1,LIMK1,TRIB3,KPNA6,PDE4DIP,YWHAZ,RPRM,ZFP91,ALDH9A1,NKIRAS2,M

IFAIP1,MECP2,SUMO1,LIMK1,TRIB3,KPNA6,PDE4DIP,YWHAZ,RPRM,ZFP91,ALDH9A1,NKIRAS2,M

4G1,TMEM184B,ABL1,CLIP2,IGF1R,UBE4B,SUMO1,RNF157,PCGF3,SALL2,TBXA2R,DDX6,PSME

NDRG1,ALCAM,MCFD2,RCC2,CDKN1B,SLC12A7,IRS1,CEBPG,DSTYK,NEURL4,ASB6,VAV2,TGFB2

SIN3A

MC3,FGFR1,SRPRA,PSAT1,TRAK1,IGF1R,EFNB2,SCAMP4,PNRC2,B4GALT5,RNF41,DHFR,ASH1L

1,MPRIP,PTP4A1,PFN2,PPP6R1,SERBP1,WBP2,BMPR2,PALLD,MSMO1,WDR45B,CPD,CSNK2A1,

L2,RNF157,PTPN1,HMGB3,FAM53C,WIZ,NDRG1,YWHAZ,ILRUN,UBE2V1,PGRMC2,ZFP91,RALBP

0,CHERP,CEP170,TAOK1,TPP1,UBE2D2,RAB14,LDLR,DTX4,SRSF2,BLCAP,NUCKS1,DICER1,ATP

L2,RNF157,PTPN1,HMGB3,FAM53C,WIZ,NDRG1,YWHAZ,ILRUN,UBE2V1,PGRMC2,ZFP91,RALBP

P36L1,FBXO11,PTEN,ULK1,LMAN1,FKBP1A,ZNF664,GRK2,FUS,BAZ2A,XRCC6,CARM1,LDLR,KM

32,LONP2,ARPC5,CAMK2N1,KCTD10,SP1,SPG11,FUBP1,BTG1,RBM39

C80,IGF1R,MDM2,CLUH,PNRC2,IP6K1,KPNA6,TMEM43,ZFP91,CASKIN1,COX6B1,FARSA,HIF3A,B

L3B,LASP1,MCL1,TADA2B,CCND2,PLXNA1,CLTC,PRKAR1A,SESN3,APLP2,MYL6,ANKRD52,NARS

CLIP2,IGF1R,UBE4B,SUMO1,RNF157,PCGF3,SALL2,TBXA2R,DDX6,PSME3,SRF,WAC,UBE2V1,RP

A10,COPS8,CCSER2,LRP6,HMGA2,RAD50,SKIL,TPGS2,ELOA

2,CLUH,PNRC2,IP6K1,KPNA6,TMEM43,ZFP91,CASKIN1,COX6B1,FARSA,HIF3A,BTF3L4,PIN1,REL

2,CLUH,PNRC2,IP6K1,KPNA6,TMEM43,ZFP91,CASKIN1,COX6B1,FARSA,HIF3A,BTF3L4,PIN1,REL

CLIP2,IGF1R,UBE4B,SUMO1,RNF157,PCGF3,SALL2,TBXA2R,DDX6,PSME3,SRF,WAC,UBE2V1,RP

RA1,SCD,SH3PXD2B,BGN,WDTC1,HDGFL3,PLXNA2,REEP5,BNC2,IP6K1,TNRC6A,LDB1,PPP2R5D,

KAP17A,DNMT1,KHDC4,SECISBP2L,MRPS27,PIK3R3,MAFB,VPS37B,KLF6

0,TAOK1,TPP1,UBE2D2,RAB14,LDLR,DTX4,SRSF2,BLCAP,DICER1,ATP6V0D1,YY1,UBE2D3,SLC3

D2B,BGN,WDTC1,HDGFL3,PLXNA2,REEP5,BNC2,IP6K1,TNRC6A,LDB1,PPP2R5D,PSME3,TNFRS

XN7L3,UBE2D3,SCD,CNBP,IGFBP4,HMGA1,GTF3C1,NIBAN2,KLF13,ACVR1B,CLUH,SUMO1,CD4

56,KLF13,TRRAP,MAP4K4,RASA4,EFNB1,RPL27A,TBXA2R,AKAP8L,TRIB3,TMEM9,WASF3,CDC42

'XN7L3,UBE2D3,SCD,CNBP,IGFBP4,HMGA1,GTF3C1,NIBAN2,KLF13,ACVR1B,CLUH,SUMO1,CD4

KLF13,MKNK2,IP6K1,VPS51,NME2,LSM4,PWWP3A,GOLGA8B,TNFRSF10B,UNC45A,SEMA4C,STA
TAPBP,PSAT1,TMEM184B,TRAK1,TOX4,NPTN,EFNB2,SCAMP4,PNRC2,DDX6,GID8,GOLGA8B,ASH
N3,TLR6,ARF6,LRIG3,NFIB,PFKFB3
,LSM4,OTUD5,CSNK1A1,NDRG1,PDE4DIP,LIMS1,MCFD2,MAP1S,RRP1B,UQCRFS1,BACE1,TYW5
_T2,TET3,MCM7,CRIM1,ARF6,NEDD8

C16A1,SLC5A6,CCDC6,EXT1,VAV2,SP1,BCAR1,SESN2,UBTF,USP36,SNU13
'XN7L3,UBE2D3,SCD,CNBP,IGFBP4,HMGA1,GTF3C1,NIBAN2,KLF13,ACVR1B,CLUH,SUMO1,CD4
SMARCD1,CERS2,RNF11,WDTC1,MKNK2,NTN1,LIMK1,ZCCHC3,GBA2,SHMT2,ZNF703,UBE2V1,C

RCD1,CERS2,RNF11,WDTC1,MKNK2,NTN1,LIMK1,ZCCHC3,GBA2,SHMT2,ZNF703,UBE2V1,OTUB
RCD1,CERS2,RNF11,WDTC1,MKNK2,NTN1,LIMK1,ZCCHC3,GBA2,SHMT2,ZNF703,UBE2V1,OTUB
,CEP170,TAOK1,TPP1,UBE2D2,RAB14,LDLR,DTX4,SRSF2,BLCAP,DICER1,ATP6V0D1,YY1,UBE2D3

3,WDTC1,MDM2,SLC38A1,BNC2,IP6K1,B4GALT5,FAM53C,TNRC6A,CCDC88A,PPP2R5D,PSME3,T
BE2V1,DIRAS1,COX6B1,CTBP2,TMEM59,SESN2,HMGA2,M6PR,TCF20
BE2G1,TAB3,AKAP17A,TMEM214,RNF40
2,RAB14,LDLR,DTX4,SRSF2,BLCAP,DICER1,ATP6V0D1,YY1,UBE2D3,SLC38A2,PAFAH1B2,RNF11,

,CBX6,PRRC2A,ADH5,PEG10,GLO1,RDX,SOD2,DICER1,MKRN1,WBP11,IGF2BP1,GSK3A,MTOR,T

0,QSOX1,ZC3H4
1
XND1,ZNF106,NF2,TMEM214,CPSF7,UBTF
TCF3,NUDT19,GOT2,TOMM70,PAGR1,SLC30A9,PSAT1,ABL1,ACVR1B,IGF1R,GALNT10,SNX1,TM

P1A,BAZ2A,RAB1B,VPS4A,GABARAPL1,EFTUD2,PNISR,NUCKS1,DICER1,MTMR4,ADAM10,OGT,M
_DLR,DTX4,SRSF2,BLCAP,DICER1,ATP6V0D1,YY1,UBE2D3,SLC38A2,PAFAH1B2,RNF11,KIAA1191
3XL19,HMGA1,WDTC1,TMEM184B,CLIP2,IGF1R,SUMO1,TBXA2R,DDX6,SHMT2,ASH1L,UBE2V1,M

OGDH,G6PC3,GATAD2B,ZNF106,ATP8A2,NDUFA10
RASSF2,CDKN1B,RRM2,FAM3C,ID2,NNT,MXRA8
C88A,DARS1,PMEPA1,OTUD5,LDB1,YWHAZ,UBE2V1,PDCD11,RRM2,NACC1,NKIRAS2,OGDH,TNPC

RAS2,CCDC6,ATF6B,TYW5,DDHD2,TFDP1,DYNLT1,SLC39A10,ARF6,COPS8,LRP6,SKIL,TPGS2,El
AB14,LDLR,DTX4,SRSF2,BLCAP,DICER1,ATP6V0D1,YY1,UBE2D3,SLC38A2,PAFAH1B2,RNF11,KIA

NMT1,ELMO2,EPHA2

,CSNK2A1,GRSF1,KLHL42,CD164,PTEN,CEMIP2,TNKS,CHERP,CEP170,BCL7A,SLC6A8,RAB14,LI

1,MDM2,SLC38A1,BNC2,IP6K1,B4GALT5,FAM53C,TNRC6A,CCDC88A,PPP2R5D,PSME3,TMEM9,

GA1,AGAP1,KIFC2,PLXNA2,ACVR1B,CLUH,CD44,IP6K1,TOB2,CCDC85C,NACC1,ERBB2,TSPAN11

IF40
2A,MYL9,USP36

91,SETD5,ARL2BP,BRPF3,CRIM1,NUP50,PPM1A
AP1,KIFC2,PLXNA2,ACVR1B,CLUH,CD44,IP6K1,TOB2,CCDC85C,NACC1,ERBB2,TSPAN11,THRA,T

KLHL36,SEPTIN9,TET3,PLXND1,TOMM40
TRIB3,CRKL,RBFOX2,SLC43A2,TOB2,YWHAZ,UBQLN4,ZFP91,ALDH9A1,POGK,MYL12A,CDC42SI
ANKFY1,CCDC88A,PSME3,RNMT,ZNF703,CCDC85C,UBE2V1,ALDH9A1,NKIRAS2,GPR173,MARVEI
HMGB2,TSN,CFAP298,USP36
3A,NR2F6,GPR173,RPS6KA2,GPS1,ADGRL1,SECISBP2L,GPRC5B,CPNE2,UBTF,SIN3A
3A,NR2F6,GPR173,RPS6KA2,GPS1,ADGRL1,SECISBP2L,GPRC5B,CPNE2,UBTF,SIN3A

A1191,DDX6,DARS1,APPBP2,LOXL2,GOLGA8B,YWHAZ,DDAH1,NUS1,SLC16A1,NR2F2,TSPAN13,
A1191,DDX6,DARS1,APPBP2,LOXL2,GOLGA8B,YWHAZ,DDAH1,NUS1,SLC16A1,NR2F2,TSPAN13,
AP1,KIFC2,PLXNA2,ACVR1B,CLUH,CD44,IP6K1,TOB2,CCDC85C,NACC1,ERBB2,TSPAN11,THRA,T
IRNPD,ADAR,RBM15B,PPDPF,VDAC1,BCR,PRKAR2A,H2AZ1,HNRNPA2B1,FUS,TAOK1,IARS2,FKBI

I2,NOTCH2,RAPGEF1,UBXN4,DYNC1LI2,CAPZB,RPS28,STXBP1,GOLGA8IP,PLXNA3,PRPS1,DDIT4
PR
40

F1,SLC39A6,KPNA2,SLC35F5,FBXO11,PPP1CC,RPL7L1,HNRNPAB,LRATD2,TTC37,DICER1,DAZAP
RRC59,ACLY,KIAA0100,CLPTM1,SYT11,LRP3,AHCY,DMWD,FXR2,ESYT1,FAM118A,H6PD,ANP32E
A6,SETD1A,EPHA4,ZNF207
OB2,UBE2V1,RPRM,SETD5,SRP54,GPR173,TNPO2,MDC1,TET3,TARS1,C8ORF33,SCOC,NUDT3,C
I1,RAD50,C6ORF89,HES6
SOX2,EFTUD2,KIAA0232,EIF4G3,RAB6A,SEMA3C,SPAG9,SLC7A1,KRT8,MKRN1,DAZAP2,RBPJ,SH

I,STARD7,RETREG2,FLCN,SEC16A,ZFP36L1,SLC39A7,UBE2G2,TAGLN2,PTEN,NDST1,TMEM167A

RK2,SLC6A8,PLEKHA6,PNISR,DAZAP2,MED13,PCGF2,PHIP,SSBP4,PRDX3,RABGAP1,SNX2,PAFAH

RFS1,CITED2,PPP1R12A,NUP50,SESN2,ATP8A2,PFKFB3,ARRDC3,CRAMP1

ORCS2,PSMA1,HMGB2,RYBP,FAM3C,PRELID1,LEPROT,GDE1

DUSP8,CHD8,NACC1,TNPO2,BTBD1,GATAD2B,FURIN,SESN2,PPFIA3,UBTF
6B,RAB11B,PDE7A,CDC37,CAPZB,FKBP8,DUSP4,PITPNA,PHC2,GPR107,SREBF2,NCOR2,MEX3A,C

NF40,GPM6B,TMED5

IARCD1,SCD,TEAD1,UNC119B,RANGAP1,CAD,POLDIP2,REEP5,UBE4B,MECP2,TRMT112,SCAMP4

3,SLC12A7,UNC13A,MAP2K4,TMEM151B,ADD1,ZNF629,SP1,UFM1,FER,AGPAT4,PHF3,GATAD2B
3EP1,GLO1,RDX,SOD2,IGF2BP1,TGOLN2,ATXN7L3,ATP6V1G1,SLC20A1,TUBB4A,EIF4A3,USP47,I

,SRPRA,TXNRD1,SLC25A6,TRIM27,FAM32A,LSM4,GID8,SRSF7,CSNK1A1,LIMS1,MCFD2,DUSP3,I

3,RAB11B,PDE7A,CDC37,CAPZB,FKBP8,DUSP4,PITPNA,PHC2,GPR107,SREBF2,NCOR2,MEX3A,C

SN,SESN2,NUDT3,CTIF,CYTH3,VPS37B
11B,PDE7A,CDC37,CAPZB,FKBP8,DUSP4,PITPNA,PHC2,GPR107,SREBF2,NCOR2,MEX3A,CYP51A
11B,PDE7A,CDC37,CAPZB,FKBP8,DUSP4,PITPNA,PHC2,GPR107,SREBF2,NCOR2,MEX3A,CYP51A

A1,CDKN1B,SLC16A1,ATP6V0E2,NR2F6,DSTYK,ARL2BP,XBP1,GATAD2B,COPS8,MAML1,AKAP17

B,RAB5B,ORAI2,KDM6B,TSPAN6,DPYSL2,NOTCH2,VEGFA,CCND1,MAPK1,LAPTM4A,DYNC1LI2,S
OC,MAPK8IP3,PPP2R2A,SETD1B,KLF10

CARM1,LDLR,KMT2E,SLC7A1,DAZAP2,NSD2,SORT1,SLC38A2,SCD,RSF1,TMEM94,RBM25,PNRC2

OGT,C16ORF58,RIMS3,PAFAH1B2,POLDIP2,ATP13A3,IGSF3,CDV3,CAB39,SSU72,ANKFY1,SUN2

SETD1B,PIEZO1,CPSF7,UBTF

YWHAZ,NUS1,ACSL4,ANKRD50,STC2,DUSP3,CCDC6,NR2F6,CITED2,ARPC5,SENP3,GMFB,HMGB
YWHAZ,NUS1,ACSL4,ANKRD50,STC2,DUSP3,CCDC6,NR2F6,CITED2,ARPC5,SENP3,GMFB,HMGB
P9,TGOLN2,CNBP,ATP13A3,YTHDF3,MDM2,NPTN,WDR26,ALDH18A1,PNRC2,RAN,CDK9,ISLR2,G

SETD1B,PIEZO1,CPSF7,UBTF
R1,DAZAP2,SPTY2D1,KLHDC10,FJX1,KIAA1191,MECP2,CMPK1,HMGB3,NTN1,LIMK1,RAN,DAD1,

3,POLR2A,TPT1,SEC61A1,MORF4L1,SARAF,RPS17,AP3D1,ARF1,SF3B2,AP2B1,GTF2F1,HNRNPU,
NCS1,CBX5,SLC7A5,RPS14,CHD4,ATP1A1,ATXN7L3B,ATP2A2,RTN3,HNRNPC,PFKL,HDLBP,TKT,
4B,UCHL1,RPL11,BTF3,RPLP1,SARAF,AP3D1,ARF1,CYB5R3,PAFAH1B1,MAP4,AP2B1,GOLGA8A,
2A,AP2M1,TPT1,SEC61A1,RPS3A,TUBB4B,MORF4L1,BTF3,AGRN,RPLP1,RPL10,MARCHF6,RPS1
SELENON,FAM168B,LUZP6,KDELR1,PITPNA,HNRNPM,GIGYF1,LARS1,PTP4A2,ILF2,UBQLN1,BAG
,ANXA6,GOLM1,TARDBP,ARL6IP1,ARID1A,ANKRD52,HNRNPK,YWHAB,SND1,HNRNPH1,CKAP4,A
,ANKRD52,RPL36,RPS12,CALU,RPS27,COTL1,RAC1,GLUL,MDH2,TAF15,RPL27,HSPA4,TCP1,CC
PRDX5,PDXK,SURF4,SKI,PSMB2,PYGB,NOTCH2,RPL37,FAM171A2,SRSF1,QARS1,DIDO1,PHC2,S
3A2,SFPQ,MEGF8,ACTN4,RPL12,TLN1,MYO5A,RPS10,ARID1A,HNRNPH1,ARCN1,COTL1,HMGCR,
CHD4,ATP1A1,CD81,ATXN7L3B,ATP2A2,PFN1,SPTBN2,SCRN1,CCND2,ACTN4,RPL12,HMGCS1,G
PL36,CKAP5,UBE2H,RAB1A,CALU,ARCN1,ARHGDIA,SLC35E2B,LUC7L3,CDKN1A,TM9SF3,MLXIP
ED4B,STIP1,SLC35E2B,IPO5,MMP2,SHISAL1,CTSB,CPSF1,SQSTM1,VCAN,HNRNPA3,ETNK1,CBX
P2,TXNIP,TRAF7,WDR45B,MTA1,GTPBP1,ZFP36L1,UBFD1,CBX6,PRRC2A,ULK1,BRSK1,RPL7L1,
5,SDHA,MESD,GPR107,PCBP2,BAG6,HNRNPD,YLPM1,RBM15B,KPNA2,MGAT4B,IARS1,PTEN,SV
),RPS7,HMGCS1,TLN1,GJA1,MTSS2,TARDBP,WLS,RPL36,PTGES3,CKAP5,SEPTIN11,RHOB,CCT6
P6,RAB6A,PACS1,DAZAP2,PSMD4,HSD17B4,MSH6,NSD2,ARPP19,SAP18,UBE2I,XPO6,SMO,SCD,
09,ID4,MPRIP,PTP4A1,GHITM,TCTN1,SMS,CTNND1,KDM5C,FBRS,VARS1,AP2A1,CXXC5,COX7C,T
RPF6,SELENON,ARPC2,SRSF1,ILF3,SUPT5H,CCAR2,DHX30,GIGYF1,ILF2,EIF3F,P3H3,SMS,RPL34
,ILF2,UQCRC2,GHITM,TXNIP,SSR1,EP300,DGCR2,STARD7,PTEN,RPL7L1,POM121C,RPS27A,GR
D2,AKT2,ATP6V0D1,WNK1,LYPLA2,FBXL19,SRPRA,PUM1,MTA2,UBE4B,SLC25A6,EWSR1,NTN1,F
1,LGALS3BP,ARCN1,COTL1,CTBP1,CAPRIN1,GPC1,HDAC2,L1CAM,PPIB,TIMP3,SRSF5,TBCD,CP
,HDGF,TMED2,UBXN4,MAPK1,AZIN1,ADAM9,PTBP1,ATXN10,CAND1,SON,BNIP3L,ZFAND5,POLR2
S2,NCKAP1,RAD23B,TNPO1,SKI,NOTCH2,VHL,CCND1,SMARCA4,SRSF1,ARGLU1,EIF3E,CDC37,S
NL1,SH3GL1,NUCKS1,WBP11,PTGFRN,ATP5MC3,SAP18,TTC28,THRAP3,HMGA1,TMEM184B,FA
,EML1,TMEM167A,KLHDC3,APEX1,PEG10,GALNT11,GSE1,SLC25A5,SERP1,VPS26B,BAZ2A,TAO
3C,VPS51,UBE2Q1,NUP98,RBM10,ZBTB4,LIMS1,RCC2,IRF2BP2,SH3BP5,KARS1,KDM2A,PMPCA
05,SMARCE1,GIGYF1,MTRNR2L10,IDH1,PTP4A1,NCOR2,RPL39,AKIRIN1,CDC42,DDIT4,NAP1L1,
M9SF2,SSRP1,UBE4A,CAMSAP1,ETNK1,NCKAP1,LAMC1,RAD23B,SKI,ASXL1,HDGF,ATP6AP1,NO
CCND1,G3BP2,SMARCA4,DDAH2,GRB2,CAPZB,SUPT5H,HNRNPUL1,SON,FKBP8,PLCG1,SMARC
AMC1,RAD23B,SKI,ASXL1,HDGF,NOTCH2,VEGFA,TUBB2A,CCND1,SRSF1,DYNLL2,GRB2,GABAR
KPNA2,PAPSS1,IARS1,RPL7L1,APEX1,XRCC6,CCN1,RAB5C,ACTR1B,DAZAP2,LRRC41,ARPP19,MT
C5D,NTN1,IP6K1,RNF41,NSD1,WAC,SYNGR1,CRABP2,CASKIN1,NACC1,ERBB2,CDC42EP4,RXRA,
R1,ATXN7L3B,LASP1,RTN3,MCL1,TADA2B,CCND2,PLXNA1,PRKAR1A,SESN3,SLC25A3,ANKRD52,
ETREG2,KPNA2,KLHL42,EP400,FBXO11,PTEN,CEMIP2,PPP1CC,MCM3AP,RPS27A,SLC25A5,SERF
X7A2L,DDX6,UBE2Q1,LOXL2,LIMS1,P3H1,GSK3B,NUS1,NREP,ZFP91,KDM2A,SLC16A1,NKIRAS2
CA3,CAMSAP1,MDK,ORAI2,TNPO1,SKI,SNRNP70,SCAF1,UBXN4,G3BP2,MAPK1,PSMD3,DYNC1LI
AZ,PRR12,DENND4B,UBE2D3,SCD,FGFR1,CNBP,IGFBP4,TMEM184B,GTF3C1,MKNK2,SUMO1,R
AZ,PRR12,DENND4B,UBE2D3,SCD,FGFR1,CNBP,IGFBP4,TMEM184B,GTF3C1,MKNK2,SUMO1,R
LT1,TNFRSF10B,YWHAZ,ZNF703,DNAJC8,MCFD2,RPRM,NACC1,ASB6,TMEM50A,CTDNEP1,EXT1,
LT1,TNFRSF10B,YWHAZ,ZNF703,DNAJC8,MCFD2,RPRM,NACC1,ASB6,TMEM50A,CTDNEP1,EXT1,
),EHMT1,SSR3,LSM14B,SLC7A1,MTMR4,EHD1,COPZ1,DHX38,CREBBP,WNK1,SORT1,THRAP3,S
JS,XRCC6,AKT3,SRSF2,EIF4G3,RAB5C,DICER1,PSMD4,YY1,RBM6,BNIP3,KLHDC10,CERS2,PAFA
TK35,WSB2,TAOK1,RAB1B,TM9SF4,PLEKHB2,SOD2,PDPK1,LRP3,XPO6,ATP5MC2,SCD,FGFR1,E
KNK2,CCDC80,MDM2,UBE4B,TNFAIP1,CAB39,SSU72,RBFOX2,YWHAZ,NUS1,ASH1L,TMEM43,UB
PZA1,PLEKHB2,ABCF1,FOXK2,YY1,WNK1,DMWD,TRO,EIF4A3,RCN2,CNBP,PTPN11,SIN3B,IGF1R

AH1,ZSWIM6,IGF2BP2,ZNF703,UBQLN4,PSMB5,POGK,NACC1,LARGE1,RNF4,NEO1,UBE2G1,MD
8A,CDKN1B,CASKIN1,NACC1,UNC13A,THBS2,NR2F6,GPR173,RPS6KA2,GPS1,SHB,ADGRL1,NF
,ARNT2,SERBP1,DHX15,MSMO1,STAT3,ARIH1,TXLNA,NAP1L1,STARD7,VARS1,BAZ1B,UBAP2L,PF
NAJC8,TMEM43,NACC1,NKIRAS2,CCDC6,ATG9A,OGDH,ASB6,CTDNEP1,UQCRH,ANKRD13B,SECI
N1B,CASKIN1,NACC1,UNC13A,THBS2,NR2F6,GPR173,RPS6KA2,GPS1,ADGRL1,NFASC,PPP2R2
3B,ELP1,DICER1,PHGDH,EDF1,LRP3,PCGF2,FGFR1,LYPLA2,HMGA1,PUM1,ESYT1,FZR1,MAPK8I
72,B4GALT5,PCGF3,FAM53C,NECTIN1,DDX6,FAM168A,LDB1,PPP2R5D,PSME3,SRF,YWHAZ,SYN
,VPS26B,TAOK1,EIF2S2,CARM1,UBR5,SLC6A8,HINT1,RAB14,LDLR,LRATD2,TBC1D9,SFXN1,DA
KAR2A,KDM5B,FLCN,DECR1,IARS1,CBX6,HNRNPA2B1,CUL3,POM121C,HIF1A,FKBP1A,BAZ2A,T
MEM184B,AGAP1,MKNK2,FXR1,RNF157,TNRC6A,ZCCHC3,RNF41,KPNA6,LDB1,EPN1,ZCCHC14,
GOT2,EXOC4,WNK1,DMWD,SCAP,ELAVL3,SC5D,PSMD13,FAM50A,CCNY,PITRM1,RTCB,ADRM1,C
3,NACC1,NKIRAS2,CCDC6,ATG9A,OGDH,ASB6,CTDNEP1,UQCRH,ANKRD13B,SECISBP2L,SETD1
1,CBX6,HNRNPA2B1,CUL3,POM121C,FKBP1A,BAZ2A,TAOK1,CARM1,VPS4A,GABARAPL1,AKT3,E
ACLY,KIAA0100,FKBP4,STRN4,PLEKHA6,ZNF428,FOXK2,EDF1,CIZ1,PTGFRN,KLHDC10,WNK1,UE
DHA1,IGF1R,UBE4B,SLC38A1,TNFAIP1,SLC25A6,PNRC2,SSU72,RAN,TNRC6A,RBFOX2,ZFP91,CD
,PRKAR1A,JUN,SESN3,MTSS2,APLP2,ANKRD52,NARS1,PTGES3,SND1,ARCN1,ZNF454,IPO5,LAP
,HNRNPA2B1,CUL3,POM121C,FKBP1A,BAZ2A,TAOK1,CARM1,VPS4A,GABARAPL1,AKT3,EFTUD
4A2,PRPF4B,RAB5C,DAZAP2,MSI1,SAP18,ATXN7L3,XPO6,EIF4G1,IGFBP4,ABL1,MKNK2,SUMO1,

ZNF703,UBE2V1,DIRAS1,CASKIN1,FARSA,NACC1,NKIRAS2,DGCR8,GPR173,SGPL1,GPS1,TMEM
,GALNT1,SH3PXD2A,SFXN1,NMT1,ARPP19,VASH1,KLHDC10,UBE2D3,GNA13,SAE1,ELOVL5,DHX
1,MDC1,FYTTD1,HEXIM1,SESN2,SETD1B,HMGA2
SOD2,LRP3,COX6A1,YY1,SCD,FGFR1,ZER1,ELAVL3,WDTC1,SGSM2,SLC22A23,IGF1R,UBE4B,CL
SOD2,LRP3,COX6A1,YY1,SCD,FGFR1,ZER1,ELAVL3,WDTC1,SGSM2,SLC22A23,IGF1R,UBE4B,CL
1,NELL2,ZER1,HMGA1,TMEM184B,ABL1,ACVR1B,NTN1,SSU72,CAPN1,DARS1,FAM168A,LDB1,P
33,KLF10,VPS37B
TBC1D10B,IP6K1,PCGF3,ATF7IP,DDX6,PSME3,TNFRSF10B,SEPHS1,ZNF703,UBE2V1,SETD5,KD
GOT2,SAP18,CCDC47,SLC20A1,HMGA1,FXR2,RGL2,CDV3,IGF1R,EWSR1,PCGF3,MYCBP2,ZCC
CRTAP,AGPAT1,TIMM50,LRRC59,GLO1,SERP1,TRIR,YARS1,TPP1,VPS4A,NUDCD3,AKT3,GNA12,
N1,SCD,UNC119B,USP5,EIF4G1,ELAVL3,PSAT1,NBPF15,GTF3C1,MKNK2,MFN2,IDH2,RAN,IP6K1
RM1,MAPKAPK2,OTUD5,EPN1,ARPC3,TOB2,YWHAZ,ZNF703,SETD5,NACC1,NKIRAS2,PMPCA,AT
,RPL32,MFGE8,PIP5K1A,FAM3C,ATP6V1F
TUB1,POGK,TNPO2,MIB2,NFASC,SESN2,FZD7,POU3F3,HES6
X7A2L,CDK4,DDX6,UBE2Q1,CRKL,LIMS1,P3H1,GSK3B,ZFP91,SLC16A1,NKIRAS2,CDC42SE1,P
ANGAP1,HMGA1,MKNK2,REEP5,IP6K1,FAM53C,IQSEC1,PSMF1,WAC,WDR82,CASKIN1,CDC42SE
NCOR2,SERBP1,CTNND1,VDAC1,BCR,NAP1L1,EXTL3,CSNK2A1,PHB2,PPM1G,SFRP1,TRA2B,PTE
RM1,MAPKAPK2,OTUD5,EPN1,ARPC3,TOB2,YWHAZ,ZNF703,SETD5,NACC1,NKIRAS2,PMPCA,AT
APK2,PSMC5,SRF,YWHAZ,PSMB5,CDC42EP4,CTCF,CTDNEP1,PBRM1,HADHB,NF2,CDIPT,PRDX4,

B,TMED5
S1,P3H1,RCC2,ZFP91,IRF2BP2,SLC16A1,NKIRAS2,CDC42SE1,PER1,CSRNP2,BACE1,COL7A1,R
1,ARPP19,VASH1,KLHDC10,UBE2D3,GNA13,SAE1,CYB5B,ELOVL5,DHX40,SRPRA,KDM3A,H6PD,N

NFIB,CPNE2,HMGA2,CPSF7,UBTF,SIN3A

RG1,TMEM9,SHMT2,TMEM43,UBE2V1,SETD5,COX6B1,NUP205,KIAA2013,CDC42SE1,SP1,GATAD
IBP,IGFBP4,HMGA1,SLC22A23,GTF3C1,NIBAN2,MKNK2,SUMO1,RNF157,HLA-A,NTN1,SSU72,IP6
TNPO2,MIB2,NFASC,SESN2,FZD7,POU3F3,HES6
AN2,MGAT5,TSNAX,HIP1R,KIAA1210,TRRAP,PNRC2,TNRC6A,ASNS,NUP153,RNMT,MCFD2,NACC1
1,TUBB4A,RANGAP1,HMGA1,GTF3C1,KLF13,FZR1,SUMO1,RNF157,LIMK1,IP6K1,DARS1,MAPKA
GFBP4,HMGA1,SLC22A23,GTF3C1,NIBAN2,MKNK2,SUMO1,RNF157,HLA-A,NTN1,SSU72,IP6K1,(
2,VASH1,ATXN7L3,UBE2D3,XPO6,CERS2,UNC119B,EIF4G1,HMGA1,TMEM184B,ABL1,MGAT5,KL
GFBP4,HMGA1,SLC22A23,GTF3C1,NIBAN2,MKNK2,SUMO1,RNF157,HLA-A,NTN1,SSU72,IP6K1,(
GFBP4,HMGA1,SLC22A23,GTF3C1,NIBAN2,MKNK2,SUMO1,RNF157,HLA-A,NTN1,SSU72,IP6K1,(
GFBP4,HMGA1,SLC22A23,GTF3C1,NIBAN2,MKNK2,SUMO1,RNF157,HLA-A,NTN1,SSU72,IP6K1,(
P1R12A,NIPA1,ZC3H15,RAB35,RELA,NUP50,SCARB1,SESN2,WDR33,ATP8A2,PFKFB3,SLC39A1,C
3,GNA13,SAE1,ELOVL5,DHX40,SRPRA,KDM3A,H6PD,NOTCH1,WDR26,PIK3C2B,PSMD7,TBC1D10

ARSA,TSPAN11,CDC42SE1,GATAD2B,NOLC1,SESN2,ID3,UBTF,SNU13
ARSA,TSPAN11,CDC42SE1,GATAD2B,NOLC1,SESN2,ID3,UBTF,SNU13
RCN1,ZNF454,RPS27,CDK16,PEA15,CDKN1A,DLG5,SERF2,TOMM20,MLXIP,SQSTM1,GDI2,TSR1

1,TUBB4A,RANGAP1,HMGA1,GTF3C1,KLF13,FZR1,SUMO1,RNF157,LIMK1,IP6K1,DARS1,MAPKA
B1,TRIB3,FAM168A,NDRG1,TMEM9,TOB2,HMGXB3,DNAJC8,UBE2V1,SETD5,COX6B1,NUP205,K
GFBP4,HMGA1,SLC22A23,GTF3C1,NIBAN2,MKNK2,SUMO1,RNF157,HLA-A,NTN1,SSU72,IP6K1,(
IU13

NCOA6,SETD1A,HMGA2,CPSF7,UBTF,CTSA,GDE1
,WBP1L,TNPO2,TGFBR1,EXT1,TET3,SP1,PIGS,FURIN,POGZ
,WBP1L,TNPO2,TGFBR1,EXT1,TET3,SP1,PIGS,FURIN,POGZ

IU13
,RPS27A,TIMM50,CHERP,CEP170,TAOK1,TPP1,RAB1B,UBE2D2,TAX1BP1,RAB14,LDLR,DTX4,SR
IAA2013,WBP1L,TNPO2,TGFBR1,EXT1,TET3,SP1,PIGS,FURIN,POGZ
RAS2,MYL12A,CDC42SE1,GPR173,TNPO2,UBE2G1,NID1,GMFB,SCOC,NF2,M6PR,USP36

'3,ARIH1,TXLNA,NAP1L1,TRIO,BAZ1B,CSNK2A1,ZFP36L1,UBAP2L,IARS1,ADH5,MCM3AP,PEG10,(

3IP3,ANKRD13B,PLXND1,NF2,PFKFB3,GDE1
SB6,PAIP1,MICALL1,DYNC1I2,SESN2,SKIL,NUTF2

E4DIP,ZEB1,SETX,SETD5,WASF3,PPP1R37,KDM5A,UBE2G1,CLCN3,RNPS1,CTIF,RNF40
ZO1,RNF40,SIN3A
ZO1,RNF40,SIN3A
3A,ASB6,ARHGEF18,AMD1,ZNF516,ADGRL1,EPHB2,NUDT3,ATP6V1F,GSN,USP36
.9,RNF40,UBTF,CTSA,SNU13

3,ILRUN,DNAJC8,SORCS2,STARD3,KDM2A,HIF3A,ZDHHC9,BTF3L4,ADGRL1,MSI2,SESN2,PPP2R
Z,SH3GL1,SH3PXD2A,RAB5C,DHCR7,PTGFRN,CERS2,UNC119B,RNF11,WDTC1,KLF13,PIK3C2B,
Z,SH3GL1,SH3PXD2A,RAB5C,DHCR7,PTGFRN,CERS2,UNC119B,RNF11,WDTC1,KLF13,PIK3C2B,
YL12A,CDC42SE1,GPR173,TNPO2,UBE2G1,NID1,GMFB,SCOC,NF2,M6PR,USP36
YL12A,CDC42SE1,GPR173,TNPO2,UBE2G1,NID1,GMFB,SCOC,NF2,M6PR,USP36

3,SRF,WAC,UBE2V1,RPRM,SETD5,SORCS2,NACC1,MYL12A,SLC2A4RG,THRA,PIN1,ADGRL1,SET
,FURIN,PRNP,AKAP17A,DNMT1,KHDC4,SECISBP2L,MRPS27,ARRDC3,MAFB,WAPL,VPS37B,KLF6

,USP48,FARSA,TLK1,DDHD2,KPNA3,ATF6,VOPP1,CAPZA2,EXT1,HEXIM1,MCM7,CBX4,NUP50,LA
GRSF1,KLHL42,CD164,PTEN,CEMIP2,TNKS,CHERP,CEP170,BCL7A,SLC6A8,RAB14,LDLR,BLCA
1,CASKIN1,KDM2A,CDC42SE1,MCRIP1,TNPO2,CAMK2N1,EPB41L1,TMEM50A,MBD6,FZD7,CTIF,
V0D1,YY1,UBE2D3,SLC38A2,GAK,CAD,PAFAH1B2,RNF11,KIAA1191,FXR1,RAN,SUN2,TNRC6A,A
1,CASKIN1,KDM2A,CDC42SE1,MCRIP1,TNPO2,CAMK2N1,EPB41L1,TMEM50A,MBD6,FZD7,CTIF,

T2E,SLC7A1,DAZAP2,NSD2,USP24,MTOR,XPO6,SORT1,SLC38A2,SCD,FGFR1,SCAP,RSF1,USP10,

TF3L4,PIN1,RELA,ADGRL1,CPNE2

1,PTGES3,ARCN1,ZNF454,MMP2,TRIB2,PEA15,CDKN1A,FAM199X,SERF2,CTDSP2,MLXIP,SQSTM

M,SETD5,SORCS2,NACC1,MYL12A,SLC2A4RG,THRA,PIN1,ADGRL1,SETD1A,NFASC,ANKRD13B,

,ADGRL1,CPNE2

,ADGRL1,CPNE2
M,SETD5,SORCS2,NACC1,MYL12A,SLC2A4RG,THRA,PIN1,ADGRL1,SETD1A,NFASC,ANKRD13B,
PSME3,GBA2,TNFRSF10B,PDE4DIP,NSD1,YWHAZ,SEPHS1,TMEM43,RALBP1,CASKIN1,SORCS2,

8A2,PAFAH1B2,RNF11,KIAA1191,MECP2,FXR1,RAN,SUN2,TNRC6A,ATF7IP,DAD1,CDK4,DDX6,RD
10B,PDE4DIP,NSD1,YWHAZ,SEPHS1,TMEM43,RALBP1,CASKIN1,SORCS2,MCRIP1,MARVELD1,N

4,RNF157,NTN1,SSU72,IP6K1,ANKFY1,CAPN1,ZCCHC3,RNMT,TOB2,SRF,ALDH9A1,RRM2,SORC
SE1,RETREG3,MED12,ARPC5,SP1,SESN2,NUDT3,PPP2R2A,DDX19A,NF2,MXRA7,PPP4R3B

4,RNF157,NTN1,SSU72,IP6K1,ANKFY1,CAPN1,ZCCHC3,RNMT,TOB2,SRF,ALDH9A1,RRM2,SORCS

AT2,ZSWIM6,TFRC,RRP1B,RALBP1,DUSP3,PPP1R37,ERBB2,RBM17,TNPO2,ARPC5,SGPL1,ZDHHC
H1L,USP48,UBQLN4,SETD5,TLK1,KPNA3,VOPP1,CAPZA2,EXT1,CBX4,NUP50,LANCL1,PPM1A,ZNF

,TLK1,PSMD10,MDC1,PNMA2,RUFY3,EPHA4,ANKRD13B,SYT4,ALYREF

4,RNF157,NTN1,SSU72,IP6K1,ANKFY1,CAPN1,ZCCHC3,RNMT,TOB2,SRF,ALDH9A1,RRM2,SORCS
OTUB1,DUSP8,NACC1,NKIRAS2,WBP1L,TNPO2,SGPL1,MARVELD1,TET3,SP1,SESN2,POGZ,PLXND

1,DUSP8,NACC1,NKIRAS2,WBP1L,TNPO2,SGPL1,MARVELD1,TET3,SP1,SESN2,POGZ,PLXND1
1,DUSP8,NACC1,NKIRAS2,WBP1L,TNPO2,SGPL1,MARVELD1,TET3,SP1,SESN2,POGZ,PLXND1
3,SLC38A2,PAFAH1B2,RNF11,KIAA1191,FXR1,RAN,SUN2,TNRC6A,ATF7IP,DAD1,CDK4,DDX6,RDH

TMEM9,ILRUN,UBE2V1,ALDH9A1,RALBP1,CASKIN1,POGK,KDM2A,MCRIP1,MDC1,CTDNEP1,TFDP

KIAA1191,FXR1,RAN,SUN2,TNRC6A,ATF7IP,DAD1,CDK4,DDX6,RDH11,MAPRE3,TNFRSF10B,ZBTB

GOLN2,SMARCD1,ATXN7L3,ATP6V1G1,SLC20A1,TUBB4A,EIF4A3,USP47,HMGA1,TMEM184B,US

EM98,RPL27A,DARS1,GID8,NDRG1,WAC,SEPHS1,PA2G4,TFRC,KIF3A,RRM2,CCDC6,CAMK2N1,C

ED13,ATP5MC3,BNIP3,C16ORF58,PAGR1,RIMS3,PAFAH1B2,POLDIP2,SDCBP,FAM171A1,ATP13A
1,FXR1,RAN,SUN2,TNRC6A,ATF7IP,DAD1,CDK4,DDX6,RDH11,MAPRE3,TNFRSF10B,ZBTB4,NUS1,
MYL12A,KIAA2013,TNPO2,MARVELD1,PIN1,SP1,ANKRD13B,PIEZO1,CPSF7,UBTF

O2,MARVELD1,ASB6,NAV1,AMD1,SNX17,NUDT3,SNU13

.OA,CRAMP1
A1191,FXR1,RAN,SUN2,TNRC6A,ATF7IP,DAD1,CDK4,DDX6,RDH11,MAPRE3,TNFRSF10B,ZBTB4,

DLR,BLCAP,DICER1,EHD1,ARPP19,MBD3,TGOLN2,YY1,UBE2D3,ANKRD12,WNK1,SLC38A2,SCD,

ILRUN,UBE2V1,ALDH9A1,RALBP1,CASKIN1,POGK,KDM2A,MCRIP1,MDC1,TFDP1,PLPP3,FZD7,SE

,THRA,TFDP1,AMD1,ZNF516,ADGRL1,CDR2L,MXRA7,EPHA2

FDP1,AMD1,ZNF516,ADGRL1,CDR2L,MXRA7,EPHA2

E1,TNPO2,UBE2G1,NID1,TET3,TARS1,RELA,MAPK8IP3,NF2,USP36
LD1,NAV1,NID1,TFDP1,AMD1,CDR2L,PPFIA3,ATP6V1F,HMGA2,NDUFA5,ARSA

SOS1,SGPL1,SETD3,TGFBR1,ADD1,SLC1A4,SP1,CCNK,RAP1B,MRPS27,PITHD1,TLNRD1,TMED5
SOS1,SGPL1,SETD3,TGFBR1,ADD1,SLC1A4,SP1,CCNK,RAP1B,MRPS27,PITHD1,TLNRD1,TMED5
FDP1,AMD1,ZNF516,ADGRL1,CDR2L,MXRA7,EPHA2
P4,ANTXR1,VPS41,ZMIZ2,TRAP1,ATXN3,SOD2,WASF2,MKRN1,MSI1,BORCS7,WNK1,SCD,USO1,C

,CIC,DCTN1,TXLNA,EP300,FLCN,PHB2,ARID1B,UBAP2L,PTEN,PRRC2A,IL6ST,FKBP1A,PSMB4,PE

ʼ2,MTOR,TGOLN2,SMARCD1,UBE2D3,WNK1,CERS2,SLC38A2,ELAVL3,MKNK2,NOTCH1,REEP5,T
,MKNK2,NOTCH1,TRIM2,NUDT4,PSMD13,CMPK1,SART3,GLOD4,LMF2,GSR,TASOR,CCNG1,ZEB

GRB10,NF2,SETD1B

H3BP4,UBE3A,PRDX3,DEK,RCN2,RIMS3,LMAN2,USP10,COL12A1,REEP5,IGF1R,KIAA1191,EFNB2

A,APEX1,PEG10,A2M,VPS26B,STK35,XRCC6,TAOK1,EIF2S2,SOX2,HINT1,LDLR,LRATD2,SFXN1,PD

H1B2,RNF11,FAM118A,ATP13A3,KLF13,TMX1,PFKP,IGF1R,EFNB2,SART3,PNRC2,METAP2,TNRC6

CYP51A1,SMS,PPP1R9B,YME1L1,ADAR,SSR1,MSMO1,WDR45B,RETREG2,PHB2,ZFP36L1,GNG4,

,SSU72,CRKL,GBA2,MEIS1,DNAJC8,SORCS2,CCDC6,CDC42SE1,MGRN1,SGPL1,PIN1,VAV2,SRC

,MICALL1,TRPC4AP,ARF6,C1ORF210,PPM1A,CRB2,ITPKB,ARRDC3
HMGA1,PHACTR4,CDV3,IGF1R,NUDT4,SUMO1,SART3,PCGF3,COPS6,ZCCHC3,AREL1,ALDH7A1,

RHOBTB2,UQCRFS1,BACE1,TLK1,ITPK1,SYT4,CNOT1

YP51A1,SMS,PPP1R9B,YME1L1,ADAR,SSR1,MSMO1,WDR45B,RETREG2,PHB2,ZFP36L1,GNG4,C

.1,SMS,PPP1R9B,YME1L1,ADAR,SSR1,MSMO1,WDR45B,RETREG2,PHB2,ZFP36L1,GNG4,CRTAP,
.1,SMS,PPP1R9B,YME1L1,ADAR,SSR1,MSMO1,WDR45B,RETREG2,PHB2,ZFP36L1,GNG4,CRTAP,

A,GNAL,HMGA2

ON,PITPNA,GIGYF1,SDHA,B2M,TNKS2,PTP4A1,PCBP2,TXNIP,ADAR,VDAC1,MSMO1,STAT3,HIP1,I

2,RAN,CCNG1,CSNK1A1,GOLGA8B,RNMT,ZBTB4,TFRC,ALDH9A1,KDM5A,MGRN1,PBRM1,CBX4,

,JAK1,UBE2Q1,CRKL,ZCCHC14,GOLGA8B,NUS1,ASH1L,EI24,METRNL,NACC1,XPC,PER1,BACE1,

2,CRIM1,EPHA4,AKAP17A,KLF3,PPP2R2A,GRB10,LEPROT,SECISBP2L,FZD8,BTG1,ID2,SH3KBP1,
2,CRIM1,EPHA4,AKAP17A,KLF3,PPP2R2A,GRB10,LEPROT,SECISBP2L,FZD8,BTG1,ID2,SH3KBP1,
GOLGA8B,KIF3A,CDKN1B,DUSP3,CLCN3,AMD1,MSI2,NUP50,COPS8,SCOC,EIF2S1,NUDT3,LEPRC

,RDH11,CSNK1A1,MAPRE3,TNFRSF10B,ZBTB4,NR2C2,ACSL4,ANKRD50,SLC12A7,ATMIN,PER1,N

PGAM1,LARP1,HNRNPA0,CTNNA1,RPL23,RPL29,RPSA,NACA,MSN,KPNB1,HK1,RPL9,PSMD2,CHD
PLXNA1,MEGF8,RPS15,CLTC,GNB2,H1-0,APLP1,TLN1,MYO5A,ANXA6,GOLM1,RPS10,RPLP2,ARI
NFE2L1,SMARCA5,LARP1,RPL23,SCARB2,RPS24,RPL24,KPNB1,RPL22,CBX5,RPL9,PSMD2,KIF5
7,VCL,AP3D1,SBF1,DNAJA1,PAFAH1B1,MAP4,AP2B1,LARP1,CNN3,RPSA,CTSD,NACA,TSPAN3,RT
6,SREBF2,P3H3,AKIRIN1,RPL34,YME1L1,CTNND1,BCL9L,ACTR1A,PSMD1,VARS1,GTPBP1,GRSF
RHGDIA,APPL1,IPO5,BCLAF1,GLUL,PEA15,SRP9,NUCB1,ZC3H7B,PABPN1,HNRNPL,TAF15,LMNB
T3,MDK,ORAI2,SCAF1,ABCA2,DPYSL2,RNF10,VEGFA,ARF3,MAPK1,ILF3,HNRNPUL1,NPEPPS,KD
SF3A1,ATXN2L,BAG6,RAI1,MLF2,ZSCAN18,WBP2,PPDPF,VDAC1,WDR45B,BCR,SEC16A,SFRP1,PF
TMEM30A,RPL23A,CTDSP2,NUCB1,ZC3H7B,TM9SF3,MDH2,CCT2,GDI2,HNRNPL,PARP1,SSRP1,
PAT4,TLN1,MTSS2,HNRNPDL,ARL6IP1,ANKRD52,WLS,RHOB,CALU,ARCN1,STIP1,APPL1,CDKN1
,PPP1CB,PGD,TM9SF2,SSRP1,UBE4A,TCP1,CAMSAP1,ETNK1,NCKAP1,VPS35,LAMC1,RAD23B,S
1,PTK7,CHD2,CNTNAP2,HDGF,ATP6AP1,RNF10,SRSF9,CCNL2,BTBD2,ODC1,EIF3E,CAPZB,PRKD
RPS2,APEX1,NPLOC4,CLCN7,YARS1,FKBP4,STRN4,PDPK1,WASF2,SRSF11,COPZ1,TEX2,UBE2D3,
2A,CEMIP2,PRRC2A,OAZ2,MCM3AP,RPS2,TIMM50,LRP10,FBXO21,GLO1,AKT3,MAZ,SRSF2,TRAF
A,GLUL,TM9SF3,HNRNPL,TSPYL4,TSR1,RPL27,TCP1,DDX42,CBX1,CCT4,KDM6B,HDGF,TERF2IP
ELOVL5,CENPB,NELFB,AGAP1,PUM1,GTF3C1,ADGRB2,METAP2,RAN,TBC1D10B,VPS51,ARFGAP
NKS,GSE1,SLC2A1,PPP1R10,ACLY,XRCC6,IMPDH2,VPS4A,NUDCD3,ZMIZ2,PLEKHB2,DAZAP2,MS
,POLR2E,EP300,PSMD1,PBX1,CALD1,KLHDC3,PSMD8,HNRNPAB,SLC2A1,ACLY,ERGIC1,FKBP4,
K2,IMPDH2,GNA12,GTF2I,INTS11,NMT1,SOD2,ABCF1,SPTY2D1,CAP1,MSH6,PTGFRN,PRDX3,US
LII,ETF1,MOGS,WIZ,RTCB,ADCY1,LSM4,EIF3B,ZNF503,SHMT2,ILRUN,PA2G4,SETD5,PSMB5,RXF
SF1,PGD,GNAI3,NOP53,HNRNPA3,HSPA4,ETNK1,DDX42,PYGB,NOTCH2,VEGFA,ARF3,CCND1,G3
2B,GNG2,CORO1C,ATP6V1B2,HNRNPM,GIGYF1,LSS,MTRNR2L10,IDH1,HECTD1,SARS1,PTP4A1,
UPT5H,ATXN10,KDM1A,FLNB,SMARCE1,PHC2,LARS1,EIF3F,UBQLN1,GHITM,RPL39,P3H3,CYP51
M171A1,CAB39,PHPT1,PNRC2,HMGB3,SHISA5,RNF41,PRXL2A,SLC25A1,ATP5PO,NCBP2,TOB2,I
K1,EIF2S2,CARM1,UBR5,HINT1,RAB14,LDLR,LRATD2,TBC1D9,SFXN1,SPAG9,DAZAP2,MED13,CC
,SGPL1,UBE2G1,ZNF395,GALNT2,EPB41L1,CTDNEP1,PBRM1,SLC1A5,ZNF629,CMTR1,CABIN1,
TXNDC5,MTA1,EP300,STARD7,ZFP36L1,KLHL42,TRA2B,UQCRC1,FEM1B,HNRNPA2B1,PREPL,RP
TCH2,VEGFA,TUBB2A,CCND1,ODC1,SRSF1,DYNLL2,GRB2,GABARAP,DLK1,TPM3,MSL1,YWHAH,
C1,GIGYF1,ATXN2L,STRAP,PCBP2,BAG6,CYP51A1,PRPF40A,TXNIP,RBM15B,AP2A1,UBQLN2,RAB
P,CAND1,TPM3,MSL1,YWHAH,HYOU1,NCOR2,SERBP1,TXNIP,CDC42,POLR2E,ARIH1,PRKAR2A,F
OR,CREBBP,GNA13,RCN2,ATP5MG,TAPBP,MACROH2A1,IGF1R,CUL1,NSF,NDUFC2,EWSR1,ATP
NAV1,CTDNEP1,NID1,PIN1,MBD6,MFGE8,SRCIN1,MYADM,FZD7,EPHA2,CTSA,GDE1
NARS1,PTGES3,SEPTIN11,ARCN1,ZNF454,CCT6A,SLC35E2B,MMP2,PEA15,CDKN1A,SERF2,TIMP
P1,ZNF664,GRK2,SHC2,SSR3,LSM14A,TRAP1,CAPZA1,EEF1B2,NUCKS1,SFXN1,SLC7A1,NUDT19,
,CDC42SE1,PER1,CSRNP2,BACE1,COL7A1,THBS2,DNAJB11,NID1,COL6A3,TET3,SP1,DYNLT1,PI
2,PTBP1,ILF3,GRB2,CAPZB,SUPT5H,SCAF11,FLNB,PITPNA,RPS28,LARS1,GOLGA8IP,PLXNA3,PR
NF157,HLA-A,SSU72,IP6K1,SYT15,CAPN1,PSME3,ARPC3,PSMF1,WAC,UBE2V1,ZFP91,CASKIN1,S
NF157,HLA-A,SSU72,IP6K1,SYT15,CAPN1,PSME3,ARPC3,PSMF1,WAC,UBE2V1,ZFP91,CASKIN1,S
ADGRL1,EPHB2,FURIN,NUDT3,PLXND1,GPRC5B,SETD1B,ZNF207,MRPS27,PFKFB3
ADGRL1,EPHB2,FURIN,NUDT3,PLXND1,GPRC5B,SETD1B,ZNF207,MRPS27,PFKFB3,UBTF
MO,LYPLA2,PSAT1,HMGA1,ABL1,PSMB1,KLF13,MKNK2,IGF1R,NPTN,EMC1,FXR1,PABPC4,CD44,
H1B2,TUB,SLC25A36,AGAP1,PAIP2,CLIC1,MFN2,MDM2,PNRC2,UBE2J1,DDX6,NME2,KPNA6,GID
LAVL3,HMGA1,WDTC1,ANKHD1,IGF1R,MDM2,SLC38A1,TRRAP,CYFIP1,BNC2,IP6K1,B4GALT5,F
QLN4,CDKN1B,PSMB5,SLC12A7,SRP54,CDC42SE1,CTDNEP1,BTF3L4,RAP1B,RNF40,MEX3D,CR
,PSMD13,WDR26,AAMP,IDH2,KAT2A,RALGDS,HLA-A,HIPK2,GSR,MYCBP2,PPP4R3A,ASNS,ARFGA

C1,RAB5A,THRA,PCNA,YBX3,TRAPPC1,NCOA6,CRIM1,ARF6,CCNL1,UBTD2,ACIN1,NBEA,FAM3C,
ASC,PPP2R2A,SECISBP2L,GPRC5B,CPNE2,UBTF,SIN3A
RC2A,ADH5,NDST1,TIMM50,PEG10,GLO1,RHBDD2,PPP1R10,KIAA0100,CARM1,DZIP1,RDX,GTF
SBP2L,SETD1B,UBTF
,SECISBP2L,GPRC5B,CPNE2,UBTF,SIN3A
P2,TRMT112,HIPK2,RPL27A,SLC25A1,PBXIP1,GBA2,ZNF703,DNAJC8,TUBA1C,SNRPD2,MAP1S,P
GR1,ILRUN,DIRAS1,CDKN1B,CASKIN1,NACC1,MYL12A,PER1,UNC13A,NR2F6,MCRIP1,GPR173,TI
ZAP2,MED13,COPZ1,ATP5MC3,TGOLN2,BNIP3,WNK1,WDTC1,PINK1,PAIP2,MKNK2,FOXP4,CUL1,
AOK1,CARM1,VPS4A,GABARAPL1,AKT3,EFTUD2,PNISR,TRAP1,FLOT2,GALNT1,PLEKHB2,NUCKS
,NSD1,S1PR3,YWHAZ,PA2G4,RAB3A,NREP,ZFP91,STC2,POGK,HIF3A,CCDC6,TNPO2,PNMA2,KLH
C1QBP,UBE2Q1,UNC45A,DHFR,ZNF703,OTUB1,RNF4,PPP1R12A,TMEM59,FER,POMGNT1,MCM7
B,UBTF
FTUD2,PNISR,FLOT2,GALNT1,PLEKHB2,NUCKS1,DICER1,OGT,RBPJ,SH3BP4,CHMP4B,MTOR,AT
BXN6,GNA13,SLC38A2,EIF4A3,ARF5,IER2,RANGAP1,TRIP12,TAPBP,HMGA1,SUN1,DAG1,MAST2,E
KN1B,SLC16A1,NUP205,SLC5A6,CCDC6,CDC42SE1,CITED2,SGPL1,CTDNEP1,TET3,BCAR1,NUP5
TM4B,PEA15,CDKN1A,SERF2,DDX39B,MLXIP,SQSTM1,TSR1,RPL27,LAMC1,CBX1,RAB5B,ORAI2,
2,PNISR,FLOT2,GALNT1,PLEKHB2,NUCKS1,OGT,RBPJ,SH3BP4,CHMP4B,ATP5MC3,C16ORF58,T
SCAMP4,RNF157,HLA-A,SSU72,COX8A,B4GALT5,EFNB1,KLC2,FAM53C,SALL2,NECTIN1,ZCCHC

50A,NID1,ADGRL1,RNPS1,NUDT3,ANKRD13B,SECISBP2L,SETD1B,NFIB,CPNE2,CPSF7,RNF40
40,SRPRA,DENND5A,KDM3A,H6PD,NOTCH1,PIK3C2B,MAP4K4,PSMD7,TBC1D10B,B4GALT5,CFD

UH,SUMO1,ADGRB2,CD44,BNC2,LIMK1,TBC1D10B,B4GALT5,PCGF3,ATF7IP,CCDC88A,FAM168
UH,SUMO1,ADGRB2,CD44,BNC2,LIMK1,TBC1D10B,B4GALT5,PCGF3,ATF7IP,CCDC88A,FAM168
PP2R5D,PSME3,ARPC3,STAT2,ZSWIM6,UBE2V1,SORCS2,COX6B1,NACC1,OGDH,MCRIP1,TNPO2,

M2A,NKIRAS2,MYL12A,GPR173,TNPO2,BTF3L4,PIN1,MCM7,ADGRL1,CDR2L,SETD1A,NUDT3,CC
IC3,GID8,RDH11,RANBP9,OAT,LGR4,USP19,RCC2,HOXA5,CDKN1B,VDAC2,ATG9A,MED12,NR2F
MAZ,PNISR,CAPZA1,ATXN3,SOD2,SLC25A23,PACS1,WASF2,DAZAP2,FOXK2,SPTY2D1,DHCR7,RB
,TRIM27,OTUD5,NUP98,MCFD2,KIF3A,CDKN1B,UNC13A,ATF6B,SLC1A5,TMEM59,GATAD2B,MC
G9A,NAV1,TMEM50A,CTDNEP1,RNASEK,ADGRL1,FURIN,RNPS1,ANKRD13B,SETD1B,RNF40,UBT

ER1,CSRNP2,BACE1,ATG9A,TET3,DYNLT1,PIGS,ABCE1,CTNNBIP1,SEC31A,PPT1,COL3A1,ID3,DN
1,NUMBL,PIN1,ADGRL1,FURIN,MFGE8,SESN2,ID3,MYADM,FZD7,EPHA2
N,NOMO1,FEM1B,CUL3,TIMM50,SLC25A5,SYMPK,FUS,KIAA0100,UBE2D2,GALNT1,PLEKHB2,NC
G9A,TMEM50A,CTDNEP1,RNASEK,ADGRL1,FURIN,RNPS1,ANKRD13B,SETD1B,RNF40,UBTF
ZNF207,RNF40

FX7,TET3,SP1,DYNLT1,PIGS,ABCE1,CTNNBIP1,SEC31A,PPT1,COL3A1,ID3,LRP6,MMP24
IOTCH1,PIK3C2B,PSMD7,TBC1D10B,B4GALT5,CFDP1,TNRC6A,PDGFRB,SOBP,FAM8A1,C1QBP,

D2B,NOLC1,SESN2,MAPK8IP3,PPP2R2A,MRPL3,TMEM214,GSN,TOMM40
SK1,CCDC88A,CAPN1,ZCCHC3,PSME3,SRF,YWHAZ,UBE2V1,RRM2,SORCS2,STARD3,NKIRAS2,OG

I,UQCRFS1,HMGB2,BSDC1,TLR6,SESN2,KLF3,KLF10,LEMD2,DNAJC21,CNOT1
PK2,NDRG1,TOB2,WAC,WDR82,ZNF703,NACC1,CDC42SE1,VDAC2,PPP1R12A,NAV1,ADGRL1,SES
CCDC88A,CAPN1,ZCCHC3,PSME3,SRF,YWHAZ,UBE2V1,SETD5,RRM2,SORCS2,STARD3,NKIRAS2,
LF13,ACVR1B,SCAMP4,CD44,NTN1,SSU72,FAM53C,NECTIN1,FAM168A,CCNG1,ARPC3,NDRG1,T
CCDC88A,CAPN1,ZCCHC3,PSME3,TOB2,SRF,YWHAZ,UBE2V1,RRM2,SORCS2,STARD3,NKIRAS2,
CCDC88A,CAPN1,ZCCHC3,PSME3,SRF,YWHAZ,UBE2V1,RRM2,SORCS2,STARD3,NKIRAS2,OGDH,
CCDC88A,CAPN1,ZCCHC3,PSME3,SRF,YWHAZ,UBE2V1,RRM2,SORCS2,STARD3,NKIRAS2,OGDH,
CRAMP1
DB,B4GALT5,CFDP1,TNRC6A,SOBP,FAM8A1,JAK1,KPNA6,SRSF7,ATP5PO,GOLGA8B,POLDIP3,TN

,MORF4L2,SSRP1,ABCA3,LAMC1,CBX1,RAB5B,ORAI2,SKI,KDM6B,TSPAN6,PYGB,DPYSL2,VEGF

PK2,NDRG1,TOB2,WAC,WDR82,ZNF703,NACC1,CDC42SE1,VDAC2,PPP1R12A,NAV1,ADGRL1,SES
IAA2013,GPR173,MARVELD1,ARHGEF18,ZNF516,ADGRL1,SESN2,ID3,MAPK8IP3,PPP2R2A,MRPL
CCDC88A,CAPN1,ZCCHC3,PSME3,SRF,YWHAZ,UBE2V1,RRM2,SORCS2,STARD3,NKIRAS2,OGDH,

SF2,SH3PXD2A,BLCAP,DICER1,MTMR4,RNF145,ATP6V0D1,YY1,UBE2D3,SLC38A2,SCD,PAFAH1B

GLO1,ZMYM3,BAZ2A,RDX,GTF2I,SRSF2,SOD2,DICER1,IGF2BP1,IVD,COPZ1,TGOLN2,RBM6,ATX

2A,RAD50,CACNG7,HIPK1,SNU13
SSU72,TNRC6A,TMEM98,CAPN1,GBA2,CLPX,WDR82,SHMT2,SETD5,POGK,TNPO2,MARVELD1,AF
SSU72,TNRC6A,TMEM98,CAPN1,GBA2,CLPX,WDR82,SHMT2,SETD5,POGK,TNPO2,MARVELD1,AF

D1A,NFASC,ANKRD13B,PPFIA3,CCSER2,PIEZO1,CPSF7,LDOC1,UBTF,NDUFA5

NCL1,PPM1A,SETD1B,NNT,CASK,CTSA,PPP2R5C
,DICER1,RNF145,SPTY2D1,EHD1,ARPP19,MBD3,TGOLN2,YY1,UBE2D3,ANKRD12,WNK1,SLC38A
CPNE2,RAD50,PFKFB3,QSOX1,HES6
TF7IP,DAD1,CDK4,DDX6,C1QBP,RDH11,MAPRE3,TNFRSF10B,ZBTB4,NUS1,ACSL4,ANKRD50,HO
CPNE2,RAD50,PFKFB3,QSOX1,HES6

,TMEM94,ELN,RBM25,IGF1R,TRRAP,PNRC2,RAN,CCNG1,RTF2,ZNF503,CSNK1A1,GOLGA8B,RN

1,HNRNPL,TSR1,LAMC1,JPT1,CBX1,RAD23B,RAB5B,ORAI2,SKI,KDM6B,TSPAN6,DPYSL2,VEGFA,

PPFIA3,CCSER2,PIEZO1,CPSF7,LDOC1,UBTF,NDUFA5

PPFIA3,CCSER2,PIEZO1,CPSF7,LDOC1,UBTF,NDUFA5
MCRIP1,MARVELD1,NAV1,EPB41L1,CTDNEP1,NID1,KCTD10,FURIN,CBX4,CDC42BPA,PFKFB3

H11,MAPRE3,TNFRSF10B,ZBTB4,NUS1,ACSL4,ANKRD50,HOXA5,PSMB5,SLC12A7,ATMIN,BMER
AV1,EPB41L1,CTDNEP1,NID1,KCTD10,FURIN,CBX4,CDC42BPA,PFKFB3,SIN3A

S2,STARD3,NACC1,TSPAN11,OGDH,THRA,ADGRL1,SRCIN1,SESN2,MXRA7,HES6,ARSA

S2,STARD3,NACC1,TSPAN11,OGDH,THRA,ADGRL1,SRCIN1,SESN2,MXRA7,HES6,ARSA

C9,ELOB,SLC1A5,SP1,TMEM59,TNPO3,HID1,NF2,STOX2,NNT,PAM
106,SETD1B,NNT,QSOX1,CASK,PPP2R5C

S2,STARD3,NACC1,TSPAN11,OGDH,THRA,ADGRL1,SRCIN1,SESN2,MXRA7,HES6,ARSA
)1

H11,MAPRE3,TNFRSF10B,ZBTB4,HMGXB3,NUS1,ACSL4,ANKRD50,HOXA5,PSMB5,SLC12A7,ATMI

P1,PLPP3,FZD7,SETD1B,TMEM214,PFKFB3

B4,NUS1,ACSL4,ANKRD50,HOXA5,PSMB5,SLC12A7,ATMIN,BMERB1,DSTYK,RFX7,NIPA1,KLHL36

SF2,PHACTR4,GTF3C1,ACVR1B,CDV3,IGF1R,SUMO1,TRRAP,EWSR1,METAP2,IP6K1,PCGF3,COPS

CTBP2,RAD54L2,TAB3,CRIM1,NFASC,FAM3C,HARS1,DNMT1,CRB2,ZNF486,UBTF

A3,IGSF3,CDV3,EXOC5,EFNB2,NSF,SF3A3,CAB39,SSU72,ANKFY1,SUN2,CUL4A,UBE2Q1,CRKL,Z
ACSL4,ANKRD50,HOXA5,PSMB5,SLC12A7,ATMIN,BMERB1,DSTYK,RFX7,NIPA1,KLHL36,THRA,K

NUS1,ACSL4,ANKRD50,HOXA5,PSMB5,SLC12A7,ATMIN,BMERB1,DSTYK,RFX7,NIPA1,KLHL36,TH

GAK,ELOVL5,DHX40,PAFAH1B2,DIP2A,RNF11,ESYT1,KLF13,CCDC80,RBM25,NPTN,MECP2,WDR

ETD1B,TMEM214,PFKFB3

NBP,RANGAP1,SEMA6B,RBM4,CLIC1,ACVR1B,GGT7,PHPT1,RAN,TSPAN14,SNX1,HNRNPR,NME2,

LP1,CYTH2,PKDCC,TRAP1,NUCKS1,MOAP1,RBPJ,FKBP9,KLHDC10,SRPRA,CAD,DENND5A,SLC25

NFAIP1,FOXP4,PIK3C2B,SUB1,ANKFY1,LDB1,BCL9,RANBP9,MEIS1,SRF,PIP5K1C,ZEB1,SOX9,CT
1,ILRUN,NR2C2,RALBP1,RBM17,DDHD2,NRP2,API5,CHPF2,NCOA6,EPHA4,PIEZO1,C6ORF89,AP

,SAMD11,CAB39,SUB1,SYNCRIP,PTPN1,SYT15,CCDC88A,C1QBP,PBXIP1,RDH11,PSIP1,HERPUD1

PK1,DAZAP2,FBLN5,MTOR,TGOLN2,PAGR1,SRPRA,TUB,HMGA1,WDTC1,PAIP2,HIP1R,ANP32E,M

A,HSD17B12,RAB18,ISLR2,RBFOX2,ZCCHC14,YWHAZ,ACSL4,NRP2,SENP3,PNMA2,BTF3L4,NOL

CRTAP,CBX6,RPL7L1,TMEM167A,NPLOC4,GPC4,STK35,TRIR,XRCC6,PELP1,CYTH2,SNRPD3,CA

IN1,ITPKB

CCNG1,RBFOX2,HERPUD1,NSD1,GOLGA4,SEMA4C,STAT2,YWHAZ,BCOR,ADIPOR2,SLC12A7,RR

CRTAP,CBX6,RPL7L1,TMEM167A,NPLOC4,GPC4,STK35,TRIR,XRCC6,PELP1,CYTH2,SNRPD3,CAR

CBX6,RPL7L1,TMEM167A,NPLOC4,GPC4,STK35,TRIR,XRCC6,PELP1,CYTH2,SNRPD3,CARM1,ST
CBX6,RPL7L1,TMEM167A,NPLOC4,GPC4,STK35,TRIR,XRCC6,PELP1,CYTH2,SNRPD3,CARM1,ST

KPNA2,SLC35F5,MASTL,PTEN,RGMB,FEM1B,PRRC2A,FMNL2,ULK1,RPS27A,HIF1A,GLO1,CEP17

ARF6,CCNK,FAM3C,PLPP3,RAP1B,HMGA2,ARRDC3

CAPZA2,MCM7,FURIN,NUP50,SETD1B,HMGA2,NNT,RBBP6,DMPK,PPP2R5C

,PFKFB3,SKIL
,PFKFB3,SKIL
OT,LRIG3,BTG1,NUTF2

NR2F2,DNAJB6,ASB6,NIPA1,KLHL36,KCTD10,BTF3L4,CLCN3,MCM7,PRNP,HADHB,EPHA4,POGZ,

D4,LENG8,WDR1,CCT7,MCL1,RPL31,HNRNPC,KHDRBS1,HDLBP,SFPQ,CCND2,CDH2,PLXNA1,ME
D1A,ST3GAL2,HNRNPK,RPL36,CKAP5,LAMP1,ATP1A2,ARHGDIA,CAPRIN1,STIP1,IPO5,GLUL,LUC
3,RPS14,ATP1A1,WDR1,ATP2A2,LASP1,CCT7,OAZ1,PFN1,MCL1,CCNI,XPR1,KHDRBS1,XPO1,SLC
N4,RPS24,RPL14,PTCD3,RPL9,KIF5B,SLC7A5,EIF5,ATXN7L3B,BEX3,MCL1,JPT2,PNN,RPL31,NIS
1,SEC16A,ATRX,IARS1,COX7C,ZNF512B,CEMIP2,PRRC2A,NDST1,RPS2,GSE1,LRRC59,STK35,FU
2,TBC1D16,CLSTN3,CBX1,RAB5B,RPL36A,RNF10,RAPGEF1,MVB12B,ARGLU1,BNIP3L,KDELR1,
ELR1,ZFAND5,ATP6V1B2,PTP4A2,PFKM,EIF3F,ISYNA1,PCBP2,RAB11A,ANP32A,VDAC1,TRAF4,CP
RC2A,PTPRF,TNKS,CHERP,SYMPK,SERP1,ZMYM3,PELP1,RTL6,CYTH2,FKBP4,INTS3,C6ORF62,
HSPH1,LMNB2,ALKBH5,PDXK,SURF4,LAMC1,CBX1,PTK7,CCT4,SNRNP70,ASXL1,HDGF,TMED2,M
A,PGRMC1,C5ORF24,CPSF1,NOP53,HNRNPL,YEATS2,AHCYL1,SURF4,GCN1,TNPO1,SKI,NRDC,T
I,ASXL1,HDGF,NOTCH2,VEGFA,TUBB2A,CCND1,MAPK1,SRSF1,PTBP1,DYNLL2,GRB2,GABARAP
C,ACTN1,FLNB,TPM3,CORO1C,MSL1,SDF4,PHC2,LARS1,PTP4A2,SLIT3,MPRIP,PTP4A1,PCBP2,G
,DIP2A,UBE2L3,TMEM184B,ABL1,PHACTR4,ARPC1A,DAG1,NIBAN2,MKNK2,HIPK2,DRAP1,NECTI
1,AHSA1,SOD2,DICER1,SART1,PACS2,ATXN7L3,KLHDC10,XPO6,CERS2,CCDC47,SORT1,DMWD,
,TMED2,TOP1,VEGFA,AZIN1,LAPTM4A,PSMD3,ILF3,CAND1,SON,MAP7D1,KDELR1,ATP6V1B2,SR
2,WIZ,SOBP,MAPKAPK2,GID8,PSME3,DEAF1,PSMC5,HMGXB3,CRTC1,PA2G4,TUBA1C,RARS1,FE
H6,FKBP9,CIZ1,CREBBP,SAP18,SAE1,EIF4A3,RANGAP1,TAPBP,HMGA1,PUM1,FXR2,MGAT5,TRA
NUDCD3,LDLR,GNA12,TRAP1,EEF1B2,RIC8A,NUCKS1,RAB5C,LTBP3,EDF1,LRRC41,AKT2,NOC2L
P5,HS2ST1,LMAN2,ITFG1,CLTA,MGAT5,TSNAX,KIAA1210,NOTCH1,EMC1,EWSR1,PNRC2,ECH1,C
RA,NARF,GPS1,MAPK3,MIB2,TNPO3,MICALL1,SZT2,AKIRIN2,NUP58,DDX56,ZC3H4,CNOT1
BP2,SRSF9,ARGLU1,ACADVL,PTBP1,SCAF11,ACTN1,CAND1,KDELR1,ARF4,FLNB,DLK1,TPM3,CO
SEC23A,AKIRIN1,PRPF40A,CTNND1,PRMT1,XPOT,CAPN2,NAP1L1,TXNDC5,TRIO,AP2A1,BAZ1B,C
IA1,PCYOX1,PALLD,STAT3,CPD,TRIO,PRKAR2A,PDAP1,ALDH3A2,UBQLN2,PPM1G,KLHL42,CSNK
GF2BP2,ACBD3,ZFP91,CDKN1B,CHD8,NR2F2,THRA,RNASEK,BTF3L4,PCNA,RELA,CDR2L,FBXO7
PZ1,ATP5MC3,TGOLN2,BNIP3,WNK1,WDTC1,PINK1,PAIP2,HIP1R,MKNK2,NOTCH1,CRACD,CUL1
SLC25A39,PIP5K1A,ACIN1,CENPF,FAM3C,PAF1,MRPS26,ZNF106,RNF40,NDUFA5,STX12,NDUFA1
S27A,PEG10,TAOK1,ABI2,VPS41,UBE2D2,LDLR,SH3GL1,IREB2,CAPZA1,GALNT1,SH3PXD2A,SFX
HYOU1,NCOR2,SERBP1,TXNIP,POLR2E,ARIH1,ACTR1A,PRKAR2A,FLCN,NOL4L,DECR1,CBX6,HN
3GAP2,ZFR,PPP1CC,RPS2,CALD1,TMEM167A,ZMYM4,SEPTIN7,PEG10,CHERP,CTTN,TMOD2,PP
LCN,NOL4L,DECR1,CBX6,HNRNPA2B1,CUL3,POM121C,FKBP1A,BAZ2A,KIAA0100,TAOK1,CARM
1B3,CCNG1,DEAF1,DDAH1,AGPAT3,RARS1,ALDH9A1,RRM2,MED12,COX5A,YBX3,RAP1B,DNMT1,

3,MLXIP,CPSF1,SQSTM1,TSR1,MORF4L2,MTRF1L,UBE4A,MDK,SURF4,LAMC1,JPT1,CBX1,RAB5
MSH6,OGT,EHD1,ARPP19,KLHDC10,WNK1,PRDX3,SLC38A2,RPS26,CNBP,DIP2A,TUB,SLC25A36,
GS,ABCE1,CTNNBIP1,TGFB2,SEC31A,COL3A1,ID3,DNMT1,SCAF8,LRP6,HMGA2,NNT,MMP24
PS1,HECTD1,BAG6,ZSCAN18,ARFGEF2,BMPR2,DDIT4,PSMA7,CIC,EIF2AK1,TXLNA,TRIO,RERE,FL
ORCS2,STARD3,GPR173,TNPO2,EPB41L1,BTF3L4,TET3,ZNF516,GATAD2B,ADGRL1,EPHB2,SRC
ORCS2,STARD3,GPR173,TNPO2,EPB41L1,BTF3L4,TET3,GATAD2B,ADGRL1,EPHB2,SRCIN1,PPP

,IP6K1,KLC2,ETF1,TASOR,PPP1CA,VPS51,ADRM1,SLC25A1,NME2,LSM4,PWWP3A,EXOC7,GOLG
8,LIMS1,TOB2,DDAH1,SEPHS1,ZYX,TUBA1C,NDUFS1,CDKN1B,PSMB5,RNF20,FARSA,NUP205,TS
AM53C,TNRC6A,CCDC88A,PPP2R5D,PSME3,EMC10,TMEM9,ILRUN,DNAJC8,ASH1L,UBE2V1,ALD
AMP1
P2,SRSF10,RNF41,RDH11,DDAH1,ZEB1,GSK3B,ASH1L,USP19,HOXA5,PSMB5,NACC1,THOC2,PR

TCEA2,DNMT1,KHDC4,LRIG3,ABR,STX12

2I,SRSF2,SOD2,SPAG9,IGF2BP1,LRRC41,IVD,PACS2,TGOLN2,CREBBP,ATXN7L3,ATP6V1G1,SLC

OGK,KDM2A,COX6B1,LMNA,SENP3,B4GALT2,CCNK
NPO2,RPS6KA2,GPS1,ADGRL1,SETD1A,CCSER2,SECISBP2L,GPRC5B,CPNE2,LDOC1,UBTF,NDUF
,CAB39,PNRC2,SUN2,CFDP1,HIPK2,TRIM27,NME2,APPBP2,NCBP2,GPT2,NSD1,YWHAZ,GSK3B,W
1,OGT,RBPJ,SH3BP4,CHMP4B,ATP5MC3,C16ORF58,THRAP3,FGFR1,PAGR1,SRPRA,IER2,RIMS3
L36,EXT1,RAD54L2,RELA,EPHB2,CRIM1,ATP6V1F,ZNF106,SECISBP2L,SETD1B,USP36,ZC3H4
,ACIN1,CENPF,FAM193B,LRIG3,SLC35A4,ABR,ARFGAP3,SIN3A,CNOT1,FKBP3

TP5MC3,C16ORF58,THRAP3,PAGR1,SRPRA,IER2,RIMS3,PAFAH1B2,PSAT1,TRAK1,CCDC80,TCEA
3SCL2,HIPK2,RHEB,LSM4,HM13,EXOC7,NUP98,LIMS1,SRF,DDAH1,AGPAT3,SETX,UBN1,NEURL4,
i0,NF2,MEX3D
SKI,KDM6B,HDGF,TSPAN6,DPYSL2,VEGFA,CCND1,MAPK1,ATG2B,LAPTM4A,ARPC2,DYNC1LI2,G
HRAP3,PAGR1,SRPRA,IER2,RIMS3,PAFAH1B2,PSAT1,TRAK1,CCDC80,EFNB2,CLUH,SCAMP4,PNF
3,FAM168A,LDB1,EPN1,WAC,YWHAZ,SYNGR1,RCC2,ARL8A,CDKN1B,CASKIN1,SORCS2,NACC1,U

P1,TNRC6A,PPP4R3A,SOBP,FAM8A1,JAK1,KPNA6,BCL9,SRSF7,GOLGA8B,TNFRSF10B,ZBTB4,LI

A,PPP2R5D,PSME3,TNFRSF10B,SRF,ILRUN,ZNF703,UBE2V1,SETD5,KDM2A,NACC1,MCRIP1,BTF
A,PPP2R5D,PSME3,TNFRSF10B,SRF,ILRUN,ZNF703,UBE2V1,SETD5,KDM2A,NACC1,MCRIP1,BTF
,EPB41L1,B4GALT2,GATAD2B,EPHB2,FURIN,SRCIN1,SESN2,TOMM40,NDUFA5,HES6,TCF20

SER2,SETD1B,HMGA2,UBTF,NDUFA5
3,NRP2,SRRT,ANXA1,GMFB,TUT4,NUP50,ATP6V1F,LRIG3,KLHL7,HMGA2
PJ,EHD1,SH3BP4,FBLN5,TGOLN2,YY1,AP1M1,SMARCD1,TRIM8,LYPLA2,IGFBP4,TAPBP,RNF11,L
M7,PRNP,DNMT1,PDHB,ZNF207,NNT,HIPK1,TLNRD1
F

MT1

STN,PDPK1,LRP3,AHCY,SCD,FGFR1,SCAP,USF2,PSMB1,FAM118A,H6PD,ANP32E,MKNK2,TRIM2,

JAK1,RAB18,KPNA6,BCL9,SRSF7,GOLGA8B,TNFRSF10B,DDAH1,WDR82,UBE3C,MCFD2,CELSR3

GDH,GPR173,MARVELD1,NAV1,EPB41L1,G6PC3,SRCIN1,PPFIA3,ATP6V1F,GSN,NDUFA5,HES6,SN

SN2,ID3,FZD7,NF2,SETD1B,POU3F3,NECAP1
OGDH,GPR173,MARVELD1,NAV1,EPB41L1,G6PC3,SRCIN1,PPFIA3,ATP6V1F,GSN,NDUFA5,HES6,
OB2,YWHAZ,UBE2V1,ZFP91,CASKIN1,SORCS2,STARD3,KDM2A,CDC42SE1,ZDHHC9,EPB41L1,TH
OGDH,GPR173,MARVELD1,NAV1,EPB41L1,G6PC3,SRCIN1,PPFIA3,ATP6V1F,GSN,NDUFA5,HES6,S
GPR173,MARVELD1,NAV1,EPB41L1,G6PC3,SRCIN1,PPFIA3,ATP6V1F,GSN,NDUFA5,HES6,SNU13
GPR173,MARVELD1,NAV1,EPB41L1,G6PC3,SRCIN1,PPFIA3,ATP6V1F,GSN,NDUFA5,HES6,SNU13

IFRSF10B,DDAH1,WDR82,UBE3C,CELSR3,SETD5,PSMB5,RRM2,ZNRF1,PPP1R12A,SETD3,ROCK2

A,CCND1,MAPK1,ATG2B,LAPTM4A,ARPC2,BTBD2,DYNC1LI2,SON,PITPNA,GIGYF1,B2M,TNKS2,P

SN2,ID3,FZD7,NF2,SETD1B,POU3F3,NECAP1
.3,TMEM214,RNF40,UBTF,TOMM40,CTSA,SNU13
GPR173,MARVELD1,NAV1,EPB41L1,G6PC3,SRCIN1,PPFIA3,ATP6V1F,GSN,NDUFA5,HES6,SNU13

2,RNF11,KIAA1191,TRIM2,CMPK1,FXR1,KCTD20,RAN,ANKFY1,SUN2,TNRC6A,ATF7IP,DAD1,CDK

N7L3,ATP6V1G1,SLC20A1,TUBB4A,EIF4A3,CNBP,SNX2,USP47,CDC34,HMGA1,PHACTR4,GTF3C1

RHGEF18,EXT1,SP1,ZNF516,ADGRL1,SESN2,ZNF207,RNF40,GSN,CTSA
RHGEF18,EXT1,SP1,ZNF516,ADGRL1,SESN2,ZNF207,RNF40,GSN,CTSA

2,SCD,GAK,ZER1,ELOVL5,DHX40,PAFAH1B2,DIP2A,RNF11,SEMA6B,ESYT1,KLF13,CCDC80,RBM

XA5,PSMB5,SLC12A7,ATMIN,BMERB1,DSTYK,RFX7,NIPA1,KLHL36,THRA,KCTD10,BTF3L4,CLCN

MT,ZBTB4,TFRC,ALDH9A1,FBL,KDM5A,DNAJB11,THOC3,PBRM1,CBX4,ARF6,CCNK,FAM3C,KRI1

,CCND1,MAPK1,ATG2B,LAPTM4A,FAM171A2,IPO7,DYNC1LI2,SON,PITPNA,FYN,GIGYF1,LARS1,P

31,DSTYK,RFX7,RAB5A,NIPA1,TGFBR1,KLHL36,THRA,KCTD10,BTF3L4,CLCN3,GMFB,PRNP,HAD

N,MED12,BMERB1,DSTYK,RFX7,NIPA1,KLHL36,THRA,KCTD10,BTF3L4,CLCN3,GMFB,PRNP,HAD

,THRA,KCTD10,BTF3L4,CLCN3,GMFB,PRNP,HADHB,TMCO1,POGZ,GRB10,AKIRIN2,NF2,SECISB

36,ZCCHC3,AREL1,SLC25A1,CCNG1,RBFOX2,HERPUD1,TNFRSF10B,NSD1,SLK,GOLGA4,SEMA4

CCHC14,GOLGA8B,RNMT,MEX3C,NUS1,ASH1L,TMEM43,ZYX,EI24,IRF2BP2,ARL8A,SH3BP5,NAC

CTD10,BTF3L4,CLCN3,GMFB,PRNP,HADHB,TMCO1,POGZ,GRB10,AKIRIN2,NF2,SECISBP2L,NFIE

RA,KCTD10,BTF3L4,CLCN3,GMFB,PRNP,HADHB,TMCO1,POGZ,GRB10,AKIRIN2,DNMT1,NF2,SE

26,CAB39,FXR1,RAN,TNRC6A,DAD1,DDX6,ADRM1,PMEPA1,RAB18,KPNA6,EXOC7,NDRG1,TNFR

,CCNG1,RDH11,SYNGR1,NUS1,PMPCA,ATF6B,DGCR8,ADIPOR1,SUDS3,MED16,TRAPPC1,MCM7,

A36,TMEM184B,CLTA,ANP32E,CLIC1,MKNK2,MDM2,EXOC5,ATF7IP,CDK4,PMEPA1,ARPC3,DEA

R9,RRM2,NACC1,CDC42EP4,NR2F2,NR2F6,SGPL1,RAB5A,TGFBR1,AMD1,PHF3,MSH2,TSN,CBX
3M1,PRMT7,DPF2,TMED5

,GOLGA4,YWHAZ,DNAJC8,NUS1,ACSL4,TFRC,ADIPOR2,ZFP91,CRABP2,DLG4,ERBB2,NELFCD,C

KNK2,KIAA1191,FOXP4,TRMT112,NDUFC2,PIK3C2B,ANKFY1,ETF1,ARHGAP5,DDX6,NCK2,DARS

C1,MSI2,MSH2,NUP50,COPS8,RNPS1,TMCO1,LRIG3,HIPK1,ZC3H4

RM1,STRN4,ABI2,RAB1B,RALY,FAM234A,HINT1,SSR3,MAZ,PRR12,PLEKHB2,ATXN3,PRPF4B,KR

I2,COX6B1,SLC5A6,PMPCA,ATG9A,GPS1,TGFBR1,SP1,AMD1,SNX17,MSI2,SYT1,C1ORF210,CO

M1,STRN4,ABI2,RAB1B,RALY,HINT1,SSR3,MAZ,PRR12,PLEKHB2,ATXN3,PRPF4B,KRT8,MSI1,NU[illegible]

RN4,ABI2,RAB1B,RALY,HINT1,SSR3,MAZ,PRR12,PLEKHB2,ATXN3,PRPF4B,KRT8,MSI1,NUDT19,[illegible]

RN4,ABI2,RAB1B,RALY,HINT1,SSR3,MAZ,PRR12,PLEKHB2,ATXN3,PRPF4B,KRT8,MSI1,NUDT19,[illegible]

0,ABI2,TAX1BP1,LDLR,KIAA0232,TBC1D9,SRSF2,FNBP1L,SOD2,PDPK1,FOXK2,PTGFRN,CCDC4[illegible]

GRB10,NFIB,BTG1,MFAP2,KLF10,UBTF,SKIL,RBBP6

:GF8,ACTN4,PRDX2,RPS15,CLTC,MRFAP1,H1-0,MACF1,MTSS2,ST3GAL2,HNRNPK,RPL36,LAMP1
7L3,DLG5,NUCB1,ZC3H7B,TM9SF3,MDH2,RPN2,SUGP2,PGD,SQSTM1,PARP1,ALKBH5,HNRNPA3
3A2,RPS25,TKT,PEBP1,DDX3X,RPS15,CLTC,RPS5,IK,HMGCS1,TLN1,MYO5A,MACF1,GOLM1,RPS
CH,LRPPRC,TADA2B,XPO1,SPTBN2,SLC3A2,CSNK1E,RPS25,CCND2,PDIA6,PIP4K2B,MEGF8,ACTN
IS,VAPB,ABI2,UBR5,SMG5,RDX,MAZ,IREB2,TRAP1,SH3PXD2A,PDPK1,PHGDH,RNF145,LRP3,PRK
CCAR2,GNG2,ATP6V1B2,GIGYF1,PFKM,USP7,HGS,KDM3B,STAT3,TXLNA,BCL9L,PRMT1,NAP1L1
D,KDM5C,RERE,SF1,FLOT1,BRD4,UBA52,RAB2A,EP400,UBAP2L,PTEN,ULK1,ANP32B,NPLOC4,FI
HINT1,LDLR,SH3GL1,PRR12,NMT1,PLEKHM2,PDPK1,PSMD4,AKT2,LRP3,GSK3A,MTOR,ATP5MC3
IARS1,SELENON,RAPGEF1,CCND1,ODC1,SRSF1,CDC37,HNRNPUL1,ACTN1,CASC3,DHX30,HNRNI
ERF2IP,MARS1,TUBB2A,CCND1,MAPK1,SMARCA4,PBX2,IPO7,FTO,LUZP6,HNRNPUL1,SON,BNIP3
,QARS1,ZFAND5,TPM3,MSL1,DHX30,HNRNPM,YWHAH,STT3B,STXBP1,PRPS1,MESD,HYOU1,NCO
HITM,BAG6,SREBF2,PFN2,RPS13,PALLD,SSR1,SULF2,KIF21B,CD99,SEC16A,GSPT1,PPP2CB,SLC
N1,NME2,EPN1,RABAC1,NDRG1,NSD1,YWHAZ,SYNGR1,ILRUN,PA2G4,ZYX,ARL8A,RRM2,CHD8,NA
PAFAH1B2,SLC25A36,HMGA1,AUTS2,MTA2,PHACTR4,LUC7L2,DAG1,PAIP2,MDM2,PARK7,GAS5,
SF3,YWHAH,B2M,MESD,ID4,PTP4A1,RAB11A,MLF2,KDM3B,SF1,RETREG2,PDAP1,CSNK2A1,RAB
3L,NDUFS1,NSMF,RUSC2,MDC1,COX5A,PIN1,RPL36AL,SP1,CACHD1,SLC39A10,RELA,SEC23IP,G
K1,MKNK2,RUVBL2,ARRDC4,BSCL2,NTN1,COX7A2L,SLC25A1,RAB18,HM13,EIF3B,CSNK1A1,EM
,GSK3A,PACS2,SSR2,WNK1,ATP5MC2,MICAL1,SCD,FGFR1,CENPB,NELFB,TRIP12,ABL1,LRIG1,C
UL4A,TNRC6A,ATF7IP,CCNG1,JAK1,CRKL,NSD1,UNC45A,RNMT,ZEB1,MCFD2,SORCS2,SLC16A1

RO1C,XPNPEP3,SMARCC1,ATP6V1B2,TNKS2,MPRIP,HNRNPD,P3H3,FLNC,WBP2,IGFBP7,CDC42,
D99,PHB2,ZFP36L1,CSNK1G2,ATRX,PPP1CC,PREPL,SEPTIN7,PEG10,LOX,MAP1LC3B,PPP1R10,T
1G2,RAB2A,MASTL,PTEN,FMNL2,ADH5,ITGB5,TMEM167A,HIF1A,LRRC59,ERGIC1,CCN1,YARS1,
,CENPF,GRB10,SLC39A1,CSTB
,CAB39,PNRC2,EHMT2,HMGB3,LMF2,CFDP1,CCNG1,APPBP2,NCBP2,NSD1,PLK2,WDFY1,NUS1,
0,SNU13
N1,MED13,ARPP19,VASH1,UBE2I,KLHDC10,UBE2D3,GNA13,SAE1,ELOVL5,DHX40,SRPRA,IER2,S
RNPA2B1,CUL3,POM121C,CALD1,FKBP1A,BAZ2A,TAOK1,CARM1,BCL7A,TRAM1,VPS4A,GABARA
P1R10,BAZ2A,CEP170,SOX2,RALY,IMPDH2,MAZ,PRR12,FLOT2,NUCKS1,SLC4A2,MOAP1,WASF2,
1,BCL7A,TRAM1,VPS4A,GABARAPL1,AKT3,EFTUD2,PNISR,FLOT2,GALNT1,PLEKHB2,NUCKS1,DI
,AP3M1,GDE1

B,ORAI2,SKI,KDM6B,TSPAN6,DPYSL2,RPS15A,NOTCH2,VEGFA,OPTN,RPL37,CCND1,MAPK1,ATG
RNF11,PSAT1,HMGA1,SDCBP,LUC7L2,RTF1,ZKSCAN1,CDV3,MDM2,METAP2,ETF1,TNRC6A,PPP

LCN,NOL4L,SLC39A6,KLHL42,ARID1B,IARS1,PTEN,PRRC2A,IL6ST,DKK3,FKBP1A,SLC25A5,SYMP
IN1,PPP2R2A,PPFIA3,ATP6V1F,RNF40,HES6,SNU13
2R2A,PPFIA3,ATP6V1F,RNF40,HES6,SNU13

A8B,TNFRSF10B,WDR11,SEMA4C,TMEM9,ZSWIM6,CRTC1,BCOR,STC2,SNRPB,ZNF592,PPP1R37
PAN13,RNF4,CTCF,ANXA1,B4GALT2,ADD1,ARHGEF18,GATAD2B,TSN,RNPS1,ACIN1,POGZ,PPP2F
H9A1,RALBP1,CASKIN1,POGK,KDM2A,NACC1,MCRIP1,GSTA4,MDC1,TFDP1,CACHD1,EPHB2,CO

RC1,DGCR8,NEO1,BMERB1,SFSWAP,API5,GNL3,MSH2,CRIM1,HARS1,ELAC2,MARF1,LRP6,WDR

20A1,SLC38A2,SCD,TUBB4A,EIF4A3,CNBP,USP47,HMGA1,PHACTR4,GTF3C1,RBM4,CDV3,TXNR

A5,SIN3A
DFY1,NUS1,AGPAT3,ALDH9A1,FARSA,SLC16A1,NUP133,NR2F2,RXRA,PHB,SOS1,SGPL1,UBE2G
,PAFAH1B2,PSAT1,TRAK1,ANKHD1,CCDC80,EFNB2,CLUH,SCAMP4,PNRC2,SSU72,ZCCHC3,UBE

1,IGF1R,EFNB2,CLUH,SCAMP4,PNRC2,SSU72,ZCCHC3,CDK4,UBE2Q1,CRKL,POLDIP3,TOB2,AS
MDC1,SENP3,CACHD1,SNX17,SETD1A,NBR1,COG4,CTIF,BRD1,MFAP2,SLC39A1,PIK3R3,UBTF

RB2,NPEPPS,SON,PITPNA,HNRNPM,GIGYF1,CCT8,B2M,TNKS2,PTP4A1,PCBP2,SEC23A,KDM3B,
C2,SSU72,ZCCHC3,UBE2Q1,CRKL,POLDIP3,TOB2,ASH1L,USP48,UBE2V1,MCFD2,RASSF2,CASK
NC13A,NR2F6,GPR173,GPRIN1,RPS6KA2,ASB6,GPS1,EPB41L1,SLC2A4RG,THRA,CTDNEP1,ZNF

MS1,DDAH1,WDR82,UBE3C,CELSR3,SETD5,RRM2,ZNRF1,NR2F6,PPP1R12A,SETD3,ROCK2,UBN

3L4,TRAF3,PIN1,TFDP1,AMD1,MCM7,ADGRL1,FURIN,SRCIN1,NUDT3,PPFIA3,GPRC5B,SETD1B,
3L4,TRAF3,PIN1,TFDP1,AMD1,MCM7,ADGRL1,FURIN,SRCIN1,NUDT3,PPFIA3,GPRC5B,SETD1B,

UC7L2,IGSF3,ACVR1B,IGF1R,NDUFA4,FXR1,GPAA1,ATP1B3,ATF5,SSU72,SHISA5,TNRC6A,TMEM

,DNAJB1,NUDT4,PSMD13,CMPK1,GLOD4,AMPD2,GSR,TASOR,RAP2A,FAM168A,CCNG1,RAB18,

,SETD5,RRM2,ATP5F1C,RBM14,RETREG3,ZNRF1,PPP1R12A,GNAZ,SHC1,SETD3,ROCK2,UBN1,

IU13

SNU13
IRA,GATAD2B,MSI2,MBD6,EPHB2,FZD7,CTIF,CPNE2,PFKFB3,CACNG7,HIPK1,VPS37B,HES6
SNU13

2,UBN1,PSMA6,PBRM1,STRIP1,TFDP1,FYTTD1,BTBD1,RELA,RAP1B,ZNF207,KLF10,PCMTD2,STX

TP4A1,PTPRS,PCBP2,NCOR2,SEC23A,ARFGEF2,BMPR2,TXNIP,ADAR,VDAC1,RBBP7,MSMO1,STA

K4,RNF41,DDX6,PBXIP1,RDH11,TOM1L2,SRSF7,ZCCHC14,MAPRE3,TNFRSF10B,ZBTB4,ZEB1,NU

1,ACVR1B,CDV3,IGF1R,NUDT4,SUMO1,USP14,TRRAP,EWSR1,MAP4K4,AMPD2,PCGF3,ZCCHC3,A

25,SGSM3,NPTN,TRIM2,MECP2,WDR26,CAB39,FXR1,KAT2A,KCTD20,RAN,TNRC6A,DAD1,RNF41

3,GMFB,PRNP,HADHB,TMCO1,POGZ,GRB10,AKIRIN2,NF2,SECISBP2L,NFIB,BTG1,QSOX1,VPS37

1,PLPP3,RAP1B,KLHL7,HMGA2,ARRDC3,MED14

TP4A2,EIF3F,B2M,TNKS2,PTP4A1,PCBP2,DGKD,SEC23A,BMPR2,TXNIP,CDC42,ADAR,VDAC1,RBE

HB,TMCO1,POGZ,GRB10,AKIRIN2,NF2,SECISBP2L,NFIB,BTG1,QSOX1,MAFB,VPS37B,KLF6,CTS

DHB,TMCO1,POGZ,GRB10,AKIRIN2,NF2,SECISBP2L,NFIB,BTG1,QSOX1,VPS37B,KLF6,CNOT1,AF

P2L,NFIB,BTG1,QSOX1,VPS37B,KLF6,ARSA

C,STAT2,YWHAZ,HSF1,BCOR,ADIPOR2,IRF2BP2,SLC12A7,RRM2,COX6B1,SLC5A6,PMPCA,ATG9/

C1,XPC,PER1,ATG9A,TLK1,CITED2,CAPZA2,SPATS2L,SLC39A10,POMGNT1,MCM7,ADGRL1,FURI
3,BTG1,QSOX1,VPS37B,KLF6,ARSA

:CISBP2L,NFIB,BTG1,QSOX1,VPS37B,KLF6,ARSA

;F10B,UBL3,ZBTB4,SEMA4C,WAC,NUS1,ACSL4,ANKRD50,RNF216,PRMT5,ADIPOR2,KIF3A,RASS[illegible]

MAML1,NUDT3,SETD1B,KLF10,C2ORF68,QSOX1,STX12,GDE1,SNU13

=1,RBFOX2,GOLGA8B,WDR11,SEMA4C,STAT2,PIP5K1C,DHFR,PRMT5,CASKIN1,PPP1R37,ZDHH[illegible]

4,PPP2R2A,RAP1B,ZNF207,AP3M1,USP36,KLF6,TMED5

CHGB,MAP2K4,MAPK3,THRA,ARL2BP,KCTD10,EXT1,FYTTD1,SP1,TNPO3,PHF3,OPA1,NOLC1,TGF

S1,JAK1,APPBP2,LOXL2,GOLGA8B,YWHAZ,DDAH1,DNAJC8,NUS1,ALDH9A1,SLC16A1,CCDC6,NR2

T8,MSI1,NUDT19,YY1,KLHDC10,SCD,EIF4A3,DHX40,CNBP,IGFBP4,PHACTR4,TOX4,GTF3C1,NIBA

PS8,EPHA4,PPP2R2A,POSTN,ATP6V1F,PLXND1,LRIG3,HMGA2,TMED5,ELOA

DT19,YY1,KLHDC10,SCD,FGFR1,EIF4A3,DHX40,CNBP,IGFBP4,PHACTR4,TOX4,GTF3C1,NIBAN2,[illegible]

[illegible]Y1,KLHDC10,SCD,EIF4A3,DHX40,CNBP,IGFBP4,PHACTR4,TOX4,GTF3C1,NIBAN2,MKNK2,CCDC

[illegible]Y1,KLHDC10,SCD,EIF4A3,DHX40,CNBP,IGFBP4,PHACTR4,TOX4,GTF3C1,NIBAN2,MKNK2,CCDC

[illegible],FJX1,SCD,RANGAP1,SLC22A23,MKNK2,ACVR1B,MFN2,PFKP,MDM2,KIAA1191,MECP2,EMC1,B[illegible]

,CKAP4,RPS12,HLA-B,HMGCR,BCLAF1,GLUL,MAP1A,HS6ST1,TMEM132A,CTDSP2,MDH2,U2AF2
3,PDXK,PLOD1,ORAI2,GCN1,SKI,ASXL1,TSPAN6,PYGB,TOP1,RNF10,MARS1,TUBB2A,G3BP2,SRS
10,SESN3,SLC25A3,ARID1A,MYL6,H1-10,UBE2H,ARCN1,HMGCR,ARHGDIA,CAPRIN1,CDK16,APF
N4,DDX3X,RPL12,RPL30,CLTC,MRFAP1,GNB2,RPS5,PRKAR1A,JUN,MYO5A,MACF1,HNRNPDL,NA
ACA,AP2A2,YY1,APBB1,WNK1,AMBRA1,SMO,SCD,NELL2,EIF4A3,ZER1,RANGAP1,CAD,HMGA1,TA
,FBRS,PBX1,PDAP1,CD99,BRD4,PPM1G,CSNK1G2,UQCRC1,CBX6,TBC1D9B,PRRC2A,HNRNPA2
KBP1A,SLC2A1,PRDX6,KIAA0100,INTS3,RALY,SEC14L1,SLC6A8,FAM234A,RAB14,SH3GL1,PRR1
3,PRKACA,AP2A2,SMARCD1,FGFR1,GAK,LYPLA2,PAGR1,FBXL19,EIF4G1,TUB,TECR,TAPBP,HMG
PM,GIGYF1,SF3A1,HYOU1,BAG6,SEC23A,PSMA7,CIC,SULF2,TXLNA,BCL9L,ACTR1A,EP300,STAR
3L,MAP7D1,KDELR1,MAP2K2,QARS1,SMARCC1,PTP4A2,PFKM,CSNK1D,ATXN2L,PRPS1,NT5DC2,
R2,CYP51A1,SERBP1,TXNIP,POLR2E,ARIH1,EDC4,TXLNA,PRKAR2A,FLCN,NOL4L,DECR1,EIF5B,
C35F5,HJURP,PI4KA,UBE2G2,PRRC2A,PPP1CC,CUL3,ADH5,NCOA4,MCM3AP,NPLOC4,FKBP1A,S
ACC1,SLC5A6,SRP54,PMPCA,ATG9A,LRCH4,MCRIP1,GPRIN1,NAV1,CTDNEP1,NID1,SEPTIN9,SP1,
PNRC2,ATF5,NCKAP5L,SUN2,FAM53C,HIPK2,DDX6,KPNA6,POLD2,EIF3B,EXOSC10,RBM10,PSM
2A,NXN,CD164,SRSF6,UPF1,KCTD12,ZNF512B,CEMIP2,FEM1B,ULK1,RPL7L1,RPS2,PSMD8,PEG
YS1,TRPC4AP,C8ORF33,PSMA4,PDXP,SCARB1,ACIN1,MYADM,KHDC4,TBCB,ATP5PD
IC10,RBM10,SRF,ZSWIM6,MED25,EIF3M,VDAC2,VOPP1,PNMA2,SLC1A5,AMD1,HEXIM1,NOLC1,
DBSL1,FXR1,EWSR1,KCTD20,FAF2,RAN,TBC1D10B,HIPK2,TBC1D14,SH3BGRL3,HM13,NUP98,N
,NACC1,UQCRFS1,SOS1,TAB3,HMGB2,BSDC1,TLR6,C8ORF33,SESN2,PPM1A,HMGA2,CPSF7,KL

,ADAR,DHX15,PSMA7,ANP32A,SSR1,XPOT,SEC16A,KLHL42,NXN,SRSF6,TAGLN2,TRA2B,FMNL2,
TAOK1,VPS41,RAB14,LDLR,SH3GL1,IREB2,CNP,GALNT1,SH3PXD2A,SFXN1,INTS11,PRPF4B,SOD
ANTXR1,ABI2,TRAM1,UBE2D2,GABARAPL1,RAB14,TTC37,STK24,SRSF2,SH3PXD2A,NUCKS1,RA

AGPAT3,FARSA,SLC16A1,SLC5A6,NUP133,UBXN1,NR2F2,THBS2,PHB,SOS1,SGPL1,SHC1,UBE2

NX2,CASC4,TMCO3,HMGA1,KDM3A,ESYT1,H6PD,NOTCH1,PIK3C2B,PSMD7,TBC1D10B,B4GALT
PL1,AKT3,EFTUD2,PNISR,FLOT2,GALNT1,PLEKHB2,NUCKS1,DICER1,MTMR4,TCF3,OGT,RBPJ,S
RBM12,WBP11,ANKS1A,DHCR7,CSE1L,EHD1,COPZ1,KDM4A,DHX38,NASP,AP2A2,YY1,BNIP3,KL
CER1,MTMR4,TCF3,OGT,RBPJ,SH3BP4,CHMP4B,ATP5MC3,C16ORF58,KLHDC10,THRAP3,PAG

G2B,AZIN1,LAPTM4A,ARPC2,DYNC1LI2,PTBP1,ILF3,SON,QARS1,PITPNA,GIGYF1,SDHA,B2M,TNK
4R3A,RAB18,KPNA6,CDK9,PPP2R5D,LOXL2,CSNK1A1,PDE4DIP,ZSWIM6,GSK3B,ZNF703,NUS1,R

K,FBXO21,PELP1,IARS2,GPX1,SNN,CYTH2,FKBP4,C6ORF62,LDLR,SH3PXD2A,NUCKS1,SFXN1,S

7,CEBPG,NKIRAS2,ZDHHC5,NUP205,ERBB2,CSRNP2,SGPL1,GPRIN1,ZNF395,MAN1B1,CHPF2,T
R2A,GRB10,CENPT,TLE4,LRP6,BRD1,NOP58,RBM39,CNOT1,UNC13B
PB1,SCARB1,PLPP3,DNMT1,FZD7,SETD1B,HMGA2,TMEM214,PFKFB3,AP3M1,IBTK

33,ZNF486,ITPKB,PFKFB3,ZC3H4,MYBBP1A

D1,IGF1R,SGSM3,TNFAIP1,SUMO1,TRRAP,SUB1,NCKAP5L,AMPD2,PCGF3,ZCCHC3,RPL27A,ARE

1,MAP2K4,THOC3,TGFBR1,ADD1,SP1,GYS1,ADGRL1,NOLC1,RYBP,CCNK,BRAT1,RAP1B,PLXND1
2Q1,CRKL,POLDIP3,TOB2,ASH1L,USP48,UBE2V1,MCFD2,RASSF2,CASKIN1,ATG9A,ZNRF1,TLK1,

H1L,USP48,UBE2V1,MCFD2,ALDH9A1,RASSF2,CASKIN1,ATG9A,ZNRF1,TLK1,KPNA3,VOPP1,CAPZ

BMPR2,TXNIP,ADAR,VDAC1,RBBP7,DCTN1,MSMO1,STAT3,TXLNA,HIP1,GRSF1,SEC16A,KPNA2,SI
IN1,ATG9A,ZNRF1,TLK1,KPNA3,VOPP1,CAPZA2,EXT1,VAV2,FURIN,CRIM1,CBX4,NUP50,RNPS1,L
516,ADGRL1,NFASC,MAPK8IP3,ANKRD13B,FZD7,SECISBP2L,GPRC5B,CPNE2,UBTF,NDUFA5,SIN

I1,PBRM1,STRIP1,TFDP1,FYTTD1,RAB35,BTBD1,RAP1B,KLF10,PCMTD2,STX12

CPNE2,LDOC1,VPS37B,SIN3A
CPNE2,LDOC1,VPS37B,SIN3A

I98,RNF41,PMEPA1,KPNA6,PSME3,CRKL,ZCCHC14,NSD1,UBE3C,TMEM43,PA2G4,ZYX,RAB3A,NF

UBE3C,NR2C2,RRP1B,RPS29,DLG4,RALBP1,ZNF592,DUSP3,NUP205,MED12,RBM17,PPP1R12A,A

RFX7,PBRM1,STRIP1,TFDP1,FYTTD1,BTBD1,ZDHHC17,LRP8,NCOA6,EIF2S1,RAP1B,DNMT1,KLF1

:12

.T3,TXLNA,NAP1L1,HIP1,SEC16A,KPNA2,SLC35F5,UBFD1,UQCRC1,MASTL,PTEN,RGMB,FEM1B,F

S1,ACSL4,ANKRD50,HOXA5,PSMB5,SLC12A7,ATMIN,SH3BP5,POGK,DUSP8,ERBB2,CCDC6,BMEI

AREL1,ALDH7A1,CCNG1,KPNA6,UBE2Q1,RBFOX2,DEGS1,HERPUD1,TNFRSF10B,PDE4DIP,NSD1,(

,DDX6,ADRM1,RAB18,KPNA6,RDH11,TOM1L2,EXOC7,SRSF7,ZCCHC14,NDRG1,MAPRE3,DEGS1

7B,KLF6,TMED5,ARSA

3P7,MSMO1,STAT3,TXLNA,HIP1,SEC16A,KPNA2,SLC35F5,UBFD1,MASTL,PTEN,RGMB,FEM1B,FM

SA,ARSA

RSA

A,GPS1,TGFBR1,MAF1,CNOT3,XBP1,AMD1,CACHD1,BSDC1,GATAD2B,SNX17,MSI2,SYT1,C1ORF

N,NUP50,ARF6,SLC25A39,TLE4,SETD1B,PIEZO1,ID2,CPSF7,SLC35A4,NNT,RBBP6,DMPK,PPP2R5

F2,NAPB,HOXA5,ARL8A,SLC12A7,ATMIN,NACC1,RNF4,TNPO2,NIPA1,KCTD10,BTF3L4,PIGS,GMF

C5,ERBB2,XPC,UQCRFS1,RNF4,ACP1,TNPO2,NARF,MAP2K4,ZNF598,PAIP1,SPATS2L,SLC12A4,AN

FB2,CRIM1,PTPN12,CENPF,FAM3C,EIF2S1,NUP58,FZD8,KLHL7,RB1CC1,TMED5,UNC13B

2F2,RXRA,TSPAN13,SOS1,SGPL1,SETD3,TGFBR1,EPB41L1,G6PC3,ADD1,AJUBA,SLC1A4,SP1,EPH

AN2,MKNK2,CCDC80,KIAA1210,MDM2,FOXP4,OSBPL2,SUMO1,SCAMP4,SART3,ADGRB2,CD44,A

MKNK2,CCDC80,KIAA1210,MDM2,FOXP4,OSBPL2,SUMO1,SCAMP4,SART3,CD44,ACAP2,RNF157

80,KIAA1210,MDM2,FOXP4,OSBPL2,SUMO1,SCAMP4,SART3,CD44,ACAP2,RNF157,RASA4,HLA-A
80,KIAA1210,MDM2,FOXP4,OSBPL2,SUMO1,SCAMP4,SART3,CD44,ACAP2,RNF157,RASA4,HLA-A

SCL2,ATP1B3,HMGB3,LIMK1,RAN,CUL4A,ARHGAP5,HNRNPR,PRXL2A,KPNA6,RDH11,ZNF532,CS

,RPN2,PPP1CB,PABPN1,HNRNPL,PARP1,TM9SF2,TAF15,ALKBH5,HSPA4,TCP1,DDX42,RAB5B,SN
F9,DSTN,PRDX1,RABL6,NYNRIN,ILF3,GRB2,CAPZB,SUPT5H,NPEPPS,MAP7D1,CASC3,RPL28,FLN
L1,HDAC2,IPO5,PPIB,CTSB,NUCB1,SRSF5,C5ORF24,U2AF2,PABPN1,TSR1,HSPH1,LMNB2,ABCA
RS1,RPL36,CKAP5,SND1,UBE2H,RPS12,RAB1A,CALU,ARCN1,RPS27,ARHGDIA,CAPRIN1,STIP1,SL
OK2,TIA1,MECP2,EFNB2,GGT7,KAT2A,PSMD7,KCTD20,KLC2,SALL2,FLII,DDX6,FAM168A,ADCY1
B1,ANP32B,RPL7L1,KLHDC3,DKK3,BAP1,ACLY,BAZ2A,XRCC6,VAPB,TCAF1,YARS1,FKBP4,ANTXR
2,INTS11,PACS1,TCF3,IGF2BP1,RBPJ,CSE1L,ARPP19,DENND4B,XPO6,WNK1,PRDX3,SLC38A2,TE
A1,OGFR,ABL1,FZR1,FOXP4,CLUH,ALDH18A1,CD44,ERP29,CCNY,NCKAP5L,RAN,IP6K1,FAM53C
D7,ALDH3A2,BAZ1B,UBQLN2,KPNA2,PHB2,KLHL42,GLUD1,UBFD1,COX7C,MBTPS1,PTEN,SMC3
HGS,PCBP2,BAG6,RAI1,SERBP1,YLPM1,SSR1,ARIH1,EIF2AK1,TXLNA,NAP1L1,EP300,PRKAR2A,B
CBX6,HNRNPA2B1,PPP1CC,CUL3,POM121C,FKBP1A,BAZ2A,TAOK1,CARM1,BCL7A,RAB1B,TRAM
LIT2,BAP1,HSPG2,SNRPD3,STRN4,LTBP4,ELAVL1,IMPDH2,GNA12,SH3GL1,EFTUD2,STK24,EIF4
,MICALL1,MCM7,NOLC1,NFASC,MFGE8,MAPK8IP3,ANKRD13B,PLXND1,NF2,PFKFB3,CSTB,AP3M
C4,TXN,IRF2BP2,CDKN1B,NSMF,SKIV2L,TSPAN13,RNF4,OLA1,SRRT,ELOB,FYTTD1,HEXIM1,POM
10,HIF1A,CHERP,SERP1,STK35,CLPTM1,YARS1,SHC2,STK24,SRSF2,TRAP1,NCSTN,RAB6A,NMT1

SETD1A,SLC25A39,FUBP1,KDM2B,POGZ,RAD23A,WDR33,SLC39A1,TOMM40,MEX3D
SD1,SNRPD2,AGPAT3,TXN,RRM2,ZNF146,NIPA1,ZSWIM8,TMEM50A,ZNF598,SP1,TMEM59,TNPO
F10,PCMTD2,DNAJC21,SNU13

PTPRF,ANP32B,POM121C,CTTN,SYMPK,LRRC59,SERP1,STK35,EHMT1,ABI2,UBR5,EFTUD2,TBC1
2,RBM5,PRPF19,ARPP19,VASH1,PTGFRN,KLHDC10,UBE2D3,GNA13,SAE1,SLC38A2,SCD,ELOVL
36A,RAB5C,SLC7A1,DCUN1D2,PHGDH,ADAM10,MSH6,RBPJ,CSE1L,EHD1,SH3BP4,EIF3G,NASP,U

G1,THOC3,TGFBR1,ADD1,SP1,CUL4B,GYS1,ADGRL1,NOLC1,RYBP,CCNK,POGZ,BRAT1,RAP1B,P

5,SUN2,KLC2,CFDP1,TNRC6A,SOBP,RPL27A,FAM8A1,C1QBP,JAK1,RAB18,KPNA6,BCL9,SRSF7,
H3BP4,CHMP4B,ATP5MC3,C16ORF58,KLHDC10,MN1,THRAP3,PRDX3,PAGR1,SRPRA,IER2,RIMS
HDC10,SORT1,ATG4B,CYB5B,EIF4G1,CAD,USP47,SNX3,MTA2,KTN1,AASDHPPT,GTF3C1,HSPA1A,
1,SRPRA,IER2,RIMS3,PAFAH1B2,PSAT1,HMGA1,SCAMP5,SUN1,TRAK1,ATP13A3,CDV3,ELN,CCD

S2,PTP4A1,PCBP2,SEC23A,BMPR2,TXNIP,ADAR,VDAC1,RBBP7,MSMO1,STAT3,ARIH1,TXLNA,NAP
NF216,KIF3A,HOXA5,CTR9,RRM2,NUP205,FAM20B,PHB,OLA1,CFLAR,COX5A,PNMA2,TET3,FER,

LC7A1,MOAP1,LTBP3,EDF1,ITGAV,ADAM10,RBPJ,FKBP9,DENND4B,ATP5MC3,AP2A2,KLHDC10,F

MEM59,TNPO3,RYBP,HID1,MYEF2,PLXND1,NF2,HMGA2,STOX2,SLC35A4,NNT,PAM,CCDC124,EM

L1,CCNG1,OTUD5,DAAM1,RBFOX2,HERPUD1,TNFRSF10B,NSD1,RNMT,GOLGA4,SEMA4C,STAT2

,ZNF106,SETD1B,MRPS27,PITHD1,TLNRD1,TMED5,NECAP1,MED14
KPNA3,ATF6,VOPP1,CAPZA2,EXT1,VAV2,FURIN,CRIM1,CBX4,NUP50,RNPS1,SCOC,LANCL1,GNAL

ZA2,EXT1,VAV2,FURIN,CRIM1,CBX4,NUP50,RNPS1,LANCL1,GNAL,PLPP3,PPM1A,SETD1B,BRD1,C

LC35F5,GARS1,CD164,UBAP2L,UPF1,PI4KA,UBFD1,UQCRC1,MASTL,PTEN,RGMB,FEM1B,FMNL2
ANCL1,GNAL,PLPP3,PPM1A,SETD1B,CPSF7,NNT,RBBP6,DMPK,PLRG1,CASK,PPP2R5C
I3A

REP,RCC2,UBQLN4,TFRC,POGK,IRS1,NACC1,CDC42EP4,ATG9A,RUSC1,RBM17,SGPL1,CFLAR,G

ARPC5,SEPTIN9,ZNF629,API5,AJUBA,EPHA4,GNAL,CFAP298,MYEF2,NF2,C6ORF89,UBTF,TMED5

0,PCMTD2,PRMT7,SKIL,STX12,TMED5

FMNL2,ULK1,RPS27A,TMEM167A,HIF1A,RABEP1,FBXO21,GLO1,KIAA0100,CEP170,INTS3,ABI2,U

RB1,DSTYK,RFX7,ASB6,RAB5A,NIPA1,KLHL36,THRA,HEG1,KCTD10,BTF3L4,CLCN3,GMFB,PHF3

GOLGA4,SEMA4C,STAT2,MEIS1,YWHAZ,UBE3C,SHMT2,BCOR,ADIPOR2,RAVER1,SLC12A7,RRM2

,TNFRSF10B,NCBP2,UBL3,ZBTB4,SEMA4C,WAC,NUS1,ACSL4,ANKRD50,RNF216,ADIPOR2,KIF3A

MNL2,ADH5,ULK1,RPS27A,TMEM167A,NPLOC4,HIF1A,SLC25A5,RABEP1,LRRC59,FBXO21,GLO1,

210,COPS8,EPHA4,CDC42BPA,PPP2R2A,DNMT1,NUP58,ATP6V1F,PLXND1,FZD8,LRIG3,HMGA2,S

5C

B,GATAD2B,SNX17,CCNL1,HADHB,POGZ,GRB10,RAP1B,DNMT1,CCSER2,KHDC4,SECISBP2L,NF

MD1,CABIN1,GATAD2B,PDXP,CCSER2,NF2,SETD1B,HMGA2,AP3M1,ELOA,EMD

HB2,ARF6,CCNK,NBEA,RAP1B,MRPS27,PITHD1,TLNRD1,CNOT1,TMED5

CAP2,RNF157,RASA4,HLA-A,SSU72,IP6K1,EFNB1,KLC2,SNX1,TASOR,WIZ,CAPN1,NECTIN1,ZCCH

7,RASA4,HLA-A,SSU72,IP6K1,EFNB1,KLC2,SNX1,TASOR,WIZ,CAPN1,NECTIN1,ZCCHC3,TBXA2R,H

A,SSU72,IP6K1,EFNB1,KLC2,SNX1,TASOR,WIZ,CAPN1,NECTIN1,ZCCHC3,TBXA2R,HSD17B12,ADF

A,SSU72,IP6K1,EFNB1,KLC2,SNX1,TASOR,WIZ,CAPN1,NECTIN1,ZCCHC3,TBXA2R,HSD17B12,ADF

SNK1A1,MAPRE3,DEGS1,TNFRSF10B,ZBTB4,WAC,YWHAZ,NR2C2,USP48,ACSL4,ANKRD50,EIF3M

RNP70,HDGF,NRDC,TMED2,MARS1,RAPGEF1,SMARCA4,PLEC,ACADVL,PTBP1,ILF3,CDC37,WDR
B,SDF4,DHX30,HNRNPM,GIGYF1,LARS1,FAU,ILF2,STT3A,UBQLN1,TMED9,PRPS1,HYOU1,NT5DC
3,ALKBH5,RPL27,PDXK,PKN1,AHCYL1,SURF4,LAMC1,RAD23B,PTK7,ORAI2,TNPO1,SKI,SNRNP7
LC35E2B,IPO5,L1CAM,LAPTM4B,LUC7L3,CDKN1A,TIMP3,MLXIP,PGD,SQSTM1,NOP53,HNRNPL,
,PSME3,EXOC7,ZNF503,POLDIP3,TNFRSF10B,RNMT,ZBTB4,PSMC4,SRF,WAC,IGF2BP2,RCC2,R
1,SLC6A8,ZMIZ2,MAZ,STK24,PLEKHB2,NUCKS1,SPAG9,WASF2,REEP2,SCRIB,MSH6,IGF2BP1,A
AD1,EIF4G1,PLS3,TUB,PTPN11,USF2,TSPYL2,ANKHD1,IGF1R,SF3A3,CLUH,USP14,EWSR1,SYNC
,GSR,RHEB,ATF7IP,PPP1CA,ARHGAP5,NECTIN1,SH3BGRL3,NUP153,ADRM1,SLC25A1,EPN1,HM
,ZNF512B,AGPAT1,TBC1D9B,PRRC2A,NDST1,SLC7A6,ANP32B,RPL7L1,TIMM50,PEG10,HNRNPA
AZ1B,CSNK2A1,EIF3D,CD99,SEC16A,GSPT1,PPM1G,BRD2,FARP1,NXN,PAPSS1,UBAP2L,RAB3G
1,VPS4A,GABARAPL1,AKT3,LSM14A,EFTUD2,PNISR,FLOT2,TMEM63B,GALNT1,PLEKHB2,NUCK
G3,BLCAP,SPAG9,PACS1,DICER1,MTMR4,RBM5,ITGAV,CAP1,SYT11,IST1,DHCR7,RBPJ,ARPP19,
1,TLNRD1,TPGS2,NECAP1,GDE1
GNT1,MCM7,NOLC1,C8ORF33,FUBP1,PIP5K1A,RNPS1,PDXDC1,EIF2S1,PPP2R2A,GRB10,SETD1
,RBM5,PSMD4,SRSF11,SCRIB,SPTY2D1,LRRC41,RBPJ,COX6A1,EIF3G,NASP,TGOLN2,YY1,SSR2

3,SEC23IP,GIT1,RNPS1,HARS1,DNMT1,ALYREF,M6PR,RBBP6,CRAMP1,TCF20

D9,TTC37,STK24,FLOT2,CAPZA1,PLEKHB2,BLCAP,SFXN1,ELP1,RAB5C,WASF2,EDF1,CAP1,MSH
,DHX40,SRPRA,USP5,PAFAH1B2,TRIP12,TAPBP,SNX3,ABL1,FXR2,SDCBP,KDM3A,DAG1,H6PD,N
BE2D3,ANKRD12,WNK1,BAG5,GNA13,DEK,SCD,HSDL1,RCN2,DHX40,ATP13A1,PSAT1,RSF1,SDC

LXND1,ZNF106,FZD7,SETD1B,LRIG3,MRPS27,PITHD1,TLNRD1,TMED5,NECAP1,MED14

GOLGA8B,TNFRSF10B,DDAH1,WDR82,UBE3C,CELSR3,SETD5,RRM2,RBM14,UBXN1,RETREG3,Z
3,PAFAH1B2,PSAT1,HMGA1,TMEM184B,SCAMP5,SUN1,TRAK1,TSPYL2,ATP13A3,HSPA1A,ACVR1
MKNK2,TXNRD1,NOTCH1,MFN2,IGF1R,FZR1,CMPK1,FXR1,ADNP,APEH,CD44,TSPAN14,B4GALT
C80,MECP2,EFNB2,CLUH,CAB39,SCAMP4,PNRC2,SSU72,KLC2,ZCCHC3,TBC1D14,CDK4,RNF41,

1L1,HIP1,SEC16A,KPNA2,SLC35F5,ATRX,UBFD1,TRA2B,MASTL,PTEN,RGMB,FEM1B,FMNL2,UL
COPB1,TUT4,SEC31A,PIGT,DNMT1,LRP6,ZNF207,HMGA2,SLC35A4,UBTF,UQCR11,RBM39,NAM

JX1,METRN,ATP13A1,SRPRA,CAD,SLC25A36,TMEM184B,OGFR,CLTA,KDM3A,ANP32E,MKNK2,

D

2,YWHAZ,PDCD11,PA2G4,ADIPOR2,SLC12A7,RRM2,OTUB1,COX6B1,SPCS2,RBM14,SLC5A6,LMN

.,PLPP3,PPM1A,SETD1B,CPSF7,NNT,RBBP6,DMPK,PLRG1,CASK,PPP2R5C

CPSF7,NNT,RBBP6,DMPK,PLRG1,CASK,PPP2R5C

2,ULK1,RPS27A,TMEM167A,HIF1A,RABEP1,FBXO21,GLO1,BAZ2A,CEP170,ABI2,UBR5,UBE2M,C

ALNT2,NAV1,PNMA2,RAD54L2,RELA,GATAD2B,MICALL1,GIT1,CRIM1,TSN,SLC25A39,LANCL1,G

BR5,TAX1BP1,LDLR,KIAA0232,SRSF2,FNBP1L,SOD2,PDPK1,MKRN1,FOXK2,NSD2,PTGFRN,TGC

,LRP8,PRNP,HADHB,EPHA4,TMCO1,POGZ,GRB10,HARS1,AKIRIN2,NF2,SECISBP2L,NFIB,BTG1,W

,COX6B1,FARSA,NKIRAS2,SLC5A6,LMNA,PMPCA,ATG9A,SFSWAP,ZNF629,VAV2,SP1,AMD1,BTBD

A,RASSF2,NAPB,HOXA5,MBOAT7,CTR9,ARL8A,SLC12A7,ATMIN,NACC1,BACE1,WBP1L,TNPO2,NI[illegible]

CEP170,ABI2,UBR5,TAX1BP1,LDLR,KIAA0232,TBC1D9,SRSF2,FNBP1L,SOD2,PDPK1,MKRN1,FO [illegible]LC35A4,TLNRD1,TMED5

IB,BTG1,STOX2,SH3KBP1,PIK3R3,ARRDC3,SKIL,STX12,VPS37B,CC2D1A

C3,TBXA2R,HSD17B12,ADRM1,ARPC3,ZNF503,CRKL,SRSF7,WAC,YWHAZ,SYNGR1,DNAJC8,WAR

ISD17B12,ADRM1,ARPC3,ZNF503,CRKL,SRSF7,WAC,YWHAZ,SYNGR1,DNAJC8,WARS1,ARL8A,CD

RM1,ARPC3,ZNF503,CRKL,SRSF7,WAC,YWHAZ,SYNGR1,DNAJC8,WARS1,ARL8A,CDKN1B,RRM2,
RM1,ARPC3,ZNF503,CRKL,SRSF7,WAC,YWHAZ,SYNGR1,DNAJC8,WARS1,ARL8A,CDKN1B,RRM2,

,SH3BP5,RRM2,SORCS2,COX6B1,CCDC6,DNAJB6,NR2F6,SGPL1,DSTYK,NIPA1,KLHL36,CAPZA2,

6,ACTN1,MAP7D1,KDELR1,CCAR2,DIDO1,PITPNA,TPM3,POLR2B,SUPT6H,HNRNPM,GIGYF1,PTP
2,GHITM,BAG6,ARFGEF2,CDC42,ADAR,DHX15,GRIPAP1,TRAF7,TXLNA,KDM5C,DGCR2,NDUFV1,
),KDM6B,HDGF,RPS15A,NOTCH2,MARS1,OPTN,UBXN4,CCND1,G3BP2,MAPRE1,ARPC2,PBX2,PS
TM9SF2,TSR1,TAF15,TGFBI,SSRP1,HSPH1,PDXK,UBE4A,TCP1,AHCYL1,CAMSAP1,ETNK1,CCT3,N
ARS1,RRP1B,ARL8A,NACC1,ZDHHC5,ERBB2,XPC,RRBP1,UNC13A,NRP2,TNPO2,RTRAF,TMEM15
KT2,MBD3,VASH1,PTGFRN,AP2A2,TGOLN2,AP1M1,EXOC4,UBE2D3,WNK1,THRAP3,EIF4A3,PAGF
RIP,CD44,PDLIM1,ECH1,RAN,ZNF462,TNRC6A,ATF7IP,TMEM98,TBC1D14,CDK4,RNF41,H2AX,AD
13,COLGALT1,TSPAN7,ZCCHC14,LIMS1,TOB2,GSK3B,MEX3C,PA2G4,AGPAT3,UBQLN4,EIF3M,A
B,SYMPK,GLO1,BAZ2A,XRCC6,PELP1,YARS1,SNRPD3,ABI2,SMG5,MAZ,SH3GL1,EFTUD2,TRAP1,
P2,KCTD12,CEMIP2,CBX6,PRRC2A,CUL3,ADH5,ITGB5,POM121C,TMEM167A,TIMM50,PEG10,HI
1,DICER1,MTMR4,TCF3,LZTR1,LRRC41,OGT,RBPJ,SH3BP4,CHMP4B,ATP5MC3,TGOLN2,C16O
TGFRN,MTOR,AP2A2,RBM6,CREBBP,PHIP,UBE2D3,SORT1,SMO,EIF4G1,PLS3,USP47,SLC25A36

B,PRDX4,STOX2
,PHIP,WNK1,UBXN6,AMBRA1,DEK,UBE2K,RPS26,CNBP,RIMS3,CAD,PAFAH1B2,PLS3,MZT2B,MD

6,IST1,COPZ1,NOC2L,ARPP19,PACS2,AP1M1,UBE2I,XPO6,CERS2,BAG5,GNA13,GAK,UNC119B,
TCH1,REEP5,IGF1R,SLC38A1,DNAJB1,FXR1,TRRAP,APEH,PIK3C2B,CD44,CNDP2,PSMD7,FAF2,
BP,CLTA,KDM3A,SCAMP1,PPP5C,TRAK1,LUC7L2,DAG1,ZKSCAN1,MACROH2A1,MTREX,CDV3,T

NRF1,PPP1R12A,SETD3,ROCK2,ZNF146,UBN1,WDR5,SENP3,PBRM1,STRIP1,TFDP1,FYTTD1,BT
B,CDV3,CCDC80,TIA1,TNFAIP1,EFNB2,CLUH,SCAMP4,PNRC2,SSU72,KLC2,ZCCHC3,TBC1D14,M
5,EFNB1,TASOR,PDGFRB,CAPN1,RAP2A,EXT2,CDK4,SH3BGRL3,RDH11,NUP98,NDRG1,DEGS1,T
MAPKAPK2,UBE2Q1,CRKL,POLDIP3,RNMT,TOB2,GSK3B,UBE3C,ILRUN,NR2C2,ASH1L,USP48,U

1,RPS27A,TMEM167A,APEX1,HIF1A,PPP1R15A,RABEP1,FBXO21,GLO1,BAZ2A,CEP170,CLPTM1,
PT,TPGS2

DM2,NPTN,EXOC5,NSF,FXR1,PARK7,GLOD4,RAN,EFNB1,ATF7IP,WRNIP1,PMEPA1,SYNGAP1,PS

NA,PMPCA,ATG9A,BRI3,DGCR8,SKIV2L,SHANK1,PSMA6,SLC12A4,AMD1,PIGS,PHF3,BSDC1,SEC2

6ORF62,TRAM1,ELAVL1,TAX1BP1,LDLR,GNA12,KIAA0232,SRSF2,SH3PXD2A,FNBP1L,SOD2,PDPI

JC1,PLPP3,ZNF106,SECISBP2L,SETD1B,LRP6,CPNE2,CACNG7,PIK3R3,DUS1L,AP3M1,ZC3H4,TP(

LN2,ANKRD12,CERS2,TRIM8,CCDC47,DAPK3,FJX1,SCD,GAK,PTPN23,RANGAP1,SCAMP5,USP10

/DR33,QSOX1,ARRDC3,VPS37B,KLF6,CNOT1,ARSA,TOLLIP

)1,TARS1,BSDC1,SNX17,NOLC1,MSI2,SYT1,C1ORF210,EPHA4,SESN2,AKAP17A,PPP2R2A,NUP58,

PA1,HEG1,KCTD10,BTF3L4,PIGS,GMFB,GATAD2B,SNX17,LRP8,SAMD1,CCNL1,HADHB,POGZ,GF

XK2,MSH6,NSD2,PTGFRN,TGOLN2,ANKRD12,TRIM8,CCDC47,FJX1,SCD,GAK,RANGAP1,TRIP12,

S1,ARL8A,CDKN1B,RRM2,STARD3,COX6B1,NACC1,PMPCA,CDC42EP4,OGDH,MCRIP1,GPR173,

KN1B,RRM2,STARD3,COX6B1,NACC1,PMPCA,CDC42EP4,OGDH,GPR173,ATF6,ASB6,NAV1,GPS

STARD3,COX6B1,NACC1,PMPCA,CDC42EP4,OGDH,GPR173,ATF6,ASB6,NAV1,GPS1,PNMA2,KLHI
STARD3,COX6B1,NACC1,PMPCA,CDC42EP4,OGDH,GPR173,ATF6,ASB6,NAV1,GPS1,PNMA2,KLHI

,MED16,ZNF598,BTF3L4,SLC1A5,HMGB2,PRNP,EPHA4,KLF3,EIF2S1,NUDT3,LEPROT,PDHB,NFIE

4A2,ILF2,CCT8,SF3A1,USP7,HYOU1,PCBP2,BAG6
,FBRS,VARS1,GTPBP1,UBQLN2,FLOT1,KIF
MD3,FAM168B,IPO7,DYNC1LI2,PTBP1,C
ICKAP1,LAMC1,RAD23B,ORAI
1B,ZNF598,SLC1A5,ZNF629,VAV2,PIGS,GA
R1,CNBP,IGLON5,ANK2,HMGA1,HTATSF1,PHACTR4,
DCY1,CRKL,SRSF7,GPT2,TOB2
TMIN,NACC1,UNC13A,TMEM14C,THBS2,MGRN1,
TUBGCP6,NUCKS1,NMT1,AHSA1,RAB5C,SPAG9,MT
F1A,GALNT11,GLO1,BAZ2A,IARS2,ANTXR1
RF58,KLHDC10,THRAP3,PRDX3,HNRNPH3,PAGR1,SR
6,PTPN11,HMGA1,ABL1,RSF1,AUTS2,LR

H1,XRN2,PSAT1,PUM1,MTA2,CGGBP1,A

ATP13A1,SH3PXD2B,EIF4G1,CAD,PLS3,HS2ST1
TBC1D10B,B4GALT5,CFDP1,TNRC6A,S
XNRD1,PKIA,TCEA1,MECP2,DNAJB1,CAB39,CYFI

3D1,ABCE1,RPL32,RNPS1,EIF2S1,RAP1B,DNMT1
APKAPK2,UBE2Q1,CRKL,POLDIP3,RNMT,PSMC4,T
TNFRSF10B,NCBP2,DDAH1,UBE3C,ACSL4,MCFD2,TF
BE2V1,MCFD2,RASSF2,PSMB5,CASKIN1,OTUB1,ATG9

ABI2,UBR5,TAX1BP1,LDLR,LSM14A,KIAA

ME3,RBFOX2,GOLGA8B,RBM10,GOLGA4,TO

3IP,SNX17,NOLC1,MSI2,SETD1A,SYT1,C1ORF210,E

K1,MKRN1,RNF145,FOXK2,PSMD4,CAP1,NSD2

GS2

,SLC22A23,MKNK2,ACVR1B,TCEA1,PFKP,REEP

ATP6V1F,PLXND1,NF2,LRIG3,HMGA2,SLC3

RB10,HARS1,RAP1B,DNMT1,CCSER2,KHDC4,SECISBP2L,L

SCAMP5,SLC22A23,MKNK2,ACVR1B,MFN2,PF

ATF6,ASB6,NAV1,GPS1,PNMA2,KLHL36,SLC2A4RG,

1,PNMA2,KLHL36,SLC2A4RG,PLCD3,THRA,CTDNEP

_36,SLC2A4RG,PLCD3,THRA,CTDNEP1,MED16,BTF

_36,SLC2A4RG,PLCD3,THRA,CTDNEP1,MED16,BTF

3,CDIPT,HMGA2,KLF10,QSOX1,SKIL,M6PR,PLRG

| | | | | | | |
|---|---|---|---|---|---|---|
| **HPA** | cerebral corte› | HPA:010000C | 1.80E-130 | 129.74542 | 8005 | 2014 |
| **HPA** | hippocampus | HPA:025000C | 8.87E-122 | 121.05231 | 6734 | 2014 |
| **HPA** | cerebellum | HPA:009000C | 5.91E-112 | 111.228219 | 7114 | 2014 |
| **HPA** | hippocampus | HPA:0250131 | 1.18E-111 | 110.92794 | 6143 | 2014 |
| **HPA** | colon | HPA:013000C | 1.94E-106 | 105.712668 | 8020 | 2014 |
| **HPA** | cerebral corte› | HPA:0100131 | 2.85E-105 | 104.545223 | 6624 | 2014 |
| **HPA** | caudate | HPA:008000C | 9.38E-103 | 102.027863 | 6874 | 2014 |
| **HPA** | pancreas | HPA:035000C | 1.12E-102 | 101.951274 | 7305 | 2014 |
| **HPA** | endometrium | HPA:016000C | 4.07E-102 | 101.39035 | 6954 | 2014 |
| **HPA** | thyroid gland | HPA:059000C | 2.55E-100 | 99.5930546 | 7022 | 2014 |
| **HPA** | thyroid gland; | HPA:0590051 | 2.55E-100 | 99.5930546 | 7022 | 2014 |
| **HPA** | endometrium | HPA:017000C | 3.68E-100 | 99.4335911 | 6814 | 2014 |
| **HPA** | pancreas; pan | HPA:0351181 | 1.36E-96 | 95.8677653 | 5540 | 2014 |
| **HPA** | epididymis; gl | HPA:0180051 | 9.89E-96 | 95.0046197 | 7232 | 2014 |
| **HPA** | epididymis | HPA:018000C | 9.89E-96 | 95.0046197 | 7232 | 2014 |
| **HPA** | endometrium | HPA:0160051 | 1.52E-93 | 92.8177957 | 6722 | 2014 |
| **HPA** | urinary bladde | HPA:061000C | 2.43E-93 | 92.6148004 | 7201 | 2014 |
| **HPA** | urinary bladde | HPA:0610831 | 2.43E-93 | 92.6148004 | 7201 | 2014 |
| **HPA** | breast | HPA:005000C | 4.65E-92 | 91.3322155 | 7084 | 2014 |
| **HPA** | endometrium | HPA:0170051 | 1.53E-91 | 90.814753 | 6627 | 2014 |
| **HPA** | ovary; ovarian | HPA:0340461 | 2.75E-91 | 90.5600091 | 3928 | 2014 |
| **HPA** | placenta | HPA:038000C | 1.93E-89 | 88.7147197 | 7385 | 2014 |
| **HPA** | adrenal gland | HPA:002000C | 1.04E-87 | 86.9839107 | 7319 | 2014 |
| **HPA** | adrenal gland; | HPA:0020051 | 1.18E-87 | 86.9269117 | 7313 | 2014 |
| **HPA** | cerebral corte› | HPA:0100132 | 2.08E-87 | 86.6827764 | 4923 | 2014 |
| **HPA** | hippocampus | HPA:0250132 | 3.24E-87 | 86.489136 | 4162 | 2014 |
| **HPA** | kidney | HPA:027000C | 5.04E-87 | 86.2973545 | 8002 | 2014 |
| **HPA** | cervix, uterine | HPA:011000C | 1.33E-85 | 84.8772199 | 6836 | 2014 |
| **HPA** | oral mucosa; : | HPA:0330221 | 2.39E-85 | 84.622447 | 6161 | 2014 |
| **HPA** | oral mucosa | HPA:033000C | 2.39E-85 | 84.622447 | 6161 | 2014 |
| **HPA** | fallopian tube | HPA:021000C | 4.04E-85 | 84.3935525 | 7359 | 2014 |
| **HPA** | seminal vesicl | HPA:043000C | 1.38E-84 | 83.8607715 | 6729 | 2014 |
| **HPA** | seminal vesicl | HPA:0430051 | 1.38E-84 | 83.8607715 | 6729 | 2014 |
| **HPA** | ovary | HPA:034000C | 2.79E-84 | 83.5539238 | 5348 | 2014 |
| **HPA** | urinary bladde | HPA:0610832 | 7.01E-84 | 83.1543865 | 5873 | 2014 |
| **HPA** | breast; glandu | HPA:0050051 | 4.13E-83 | 82.3841819 | 6717 | 2014 |
| **HPA** | cerebral corte› | HPA:0100121 | 2.59E-82 | 81.5859656 | 4857 | 2014 |
| **HPA** | prostate | HPA:039000C | 1.03E-81 | 80.9882913 | 6586 | 2014 |
| **HPA** | prostate; glan | HPA:0390051 | 1.03E-81 | 80.9882913 | 6586 | 2014 |
| **HPA** | epididymis; gl | HPA:0180052 | 6.84E-81 | 80.1650179 | 5601 | 2014 |
| **HPA** | endometrium | HPA:0160251 | 7.84E-81 | 80.1059135 | 4009 | 2014 |
| **HPA** | gallbladder; gl | HPA:0220051 | 1.37E-79 | 78.863476 | 7560 | 2014 |
| **HPA** | gallbladder | HPA:022000C | 1.37E-79 | 78.863476 | 7560 | 2014 |
| **HPA** | rectum; gland | HPA:0400051 | 1.88E-79 | 78.7264332 | 7555 | 2014 |
| **HPA** | rectum | HPA:040000C | 1.88E-79 | 78.7264332 | 7555 | 2014 |

| | | | | | | |
|---|---|---|---|---|---|---|
| **HPA** | kidney; cells ir | HPA:0270351 | 1.32E-78 | 77.8795336 | 5084 | 2014 |
| **HPA** | stomach 1 | HPA:054000C | 1.41E-78 | 77.8519719 | 7553 | 2014 |
| **HPA** | stomach 1; gl | HPA:0540051 | 1.41E-78 | 77.8519719 | 7553 | 2014 |
| **HPA** | lung | HPA:030000C | 1.52E-78 | 77.8182876 | 7220 | 2014 |
| **HPA** | pancreas; exo | HPA:0350471 | 1.53E-78 | 77.8143263 | 6841 | 2014 |
| **HPA** | colon; glandu | HPA:0130051 | 1.72E-78 | 77.7639682 | 7473 | 2014 |
| **HPA** | skin 1 | HPA:046000C | 2.35E-78 | 77.6281039 | 6996 | 2014 |
| **HPA** | caudate; neur | HPA:0080131 | 3.63E-78 | 77.4402756 | 6084 | 2014 |
| **HPA** | salivary gland; | HPA:0420051 | 1.05E-77 | 76.9790384 | 6802 | 2014 |
| **HPA** | salivary gland | HPA:042000C | 1.05E-77 | 76.9790384 | 6802 | 2014 |
| **HPA** | kidney; cells ir | HPA:0270361 | 1.35E-77 | 76.8697989 | 7710 | 2014 |
| **HPA** | skin 1; melan | HPA:0460671 | 2.82E-77 | 76.5496197 | 5423 | 2014 |
| **HPA** | esophagus | HPA:019000C | 6.52E-77 | 76.1854986 | 6583 | 2014 |
| **HPA** | esophagus; s | HPA:0190221 | 6.52E-77 | 76.1854986 | 6583 | 2014 |
| **HPA** | endometrium | HPA:0170251 | 4.95E-76 | 75.305484 | 3452 | 2014 |
| **HPA** | tonsil; squam | HPA:0600221 | 1.29E-75 | 74.8885824 | 6564 | 2014 |
| **HPA** | soft tissue 1 | HPA:050000C | 5.43E-75 | 74.2653516 | 5292 | 2014 |
| **HPA** | stomach 2; gl | HPA:0550051 | 1.33E-74 | 73.8774659 | 7512 | 2014 |
| **HPA** | stomach 2 | HPA:055000C | 1.33E-74 | 73.8774659 | 7512 | 2014 |
| **HPA** | skin 2; epider | HPA:0470681 | 2.14E-74 | 73.6699553 | 6667 | 2014 |
| **HPA** | skin 2 | HPA:047000C | 2.14E-74 | 73.6699553 | 6667 | 2014 |
| **HPA** | thyroid gland; | HPA:0590052 | 3.93E-74 | 73.4051468 | 5505 | 2014 |
| **HPA** | tonsil | HPA:060000C | 6.41E-73 | 72.1929768 | 7677 | 2014 |
| **HPA** | colon; periphe | HPA:0130241 | 7.00E-73 | 72.1548612 | 5073 | 2014 |
| **HPA** | nasopharynx | HPA:032000C | 3.09E-72 | 71.5098719 | 7093 | 2014 |
| **HPA** | endometrium | HPA:0170052 | 6.53E-72 | 71.1849597 | 4805 | 2014 |
| **HPA** | adrenal gland; | HPA:0020052 | 1.04E-71 | 70.9842557 | 5910 | 2014 |
| **HPA** | colon; endoth | HPA:0130201 | 1.09E-71 | 70.9637465 | 5709 | 2014 |
| **HPA** | smooth musc | HPA:049000C | 1.90E-71 | 70.7215525 | 5374 | 2014 |
| **HPA** | smooth musc | HPA:0490691 | 1.90E-71 | 70.7215525 | 5374 | 2014 |
| **HPA** | cervix, uterine | HPA:0110051 | 1.28E-70 | 69.8923863 | 5616 | 2014 |
| **HPA** | stomach 1; gl | HPA:0540052 | 1.76E-70 | 69.7537052 | 6256 | 2014 |
| **HPA** | caudate; glial | HPA:0080121 | 2.28E-70 | 69.6421875 | 4513 | 2014 |
| **HPA** | duodenum | HPA:015000C | 2.84E-70 | 69.5463382 | 7666 | 2014 |
| **HPA** | duodenum; g | HPA:0150051 | 2.84E-70 | 69.5463382 | 7666 | 2014 |
| **HPA** | bronchus | HPA:006000C | 2.91E-70 | 69.535676 | 7334 | 2014 |
| **HPA** | appendix; glar | HPA:0030051 | 5.74E-70 | 69.2413795 | 7202 | 2014 |
| **HPA** | gallbladder; gl | HPA:0220052 | 2.33E-69 | 68.6320536 | 6512 | 2014 |
| **HPA** | hippocampus | HPA:0250121 | 1.45E-68 | 67.8387576 | 4082 | 2014 |
| **HPA** | skin 1; keratin | HPA:0460661 | 1.60E-68 | 67.79696 | 6107 | 2014 |
| **HPA** | endometrium | HPA:0160052 | 2.93E-68 | 67.5332668 | 5066 | 2014 |
| **HPA** | seminal vesicl | HPA:0430052 | 8.50E-68 | 67.0707256 | 5149 | 2014 |
| **HPA** | fallopian tube; | HPA:0210052 | 1.08E-67 | 66.9651391 | 5074 | 2014 |
| **HPA** | lung; macropl | HPA:0300411 | 1.39E-67 | 66.8566617 | 6690 | 2014 |
| **HPA** | rectum; gland | HPA:0400052 | 2.16E-67 | 66.6655317 | 6468 | 2014 |

| | | | | | | |
|---|---|---|---|---|---|---|
| **HPA** | lung; alveolar | HPA:0301171 | 4.29E-67 | 66.3677643 | 5077 | 2014 |
| **HPA** | lymph node | HPA:0310000 | 4.81E-67 | 66.317541 | 6431 | 2014 |
| **HPA** | small intestine | HPA:0480051 | 6.86E-67 | 66.1634272 | 7653 | 2014 |
| **HPA** | small intestine | HPA:0480000 | 6.86E-67 | 66.1634272 | 7653 | 2014 |
| **HPA** | pancreas; pan | HPA:0351182 | 9.10E-67 | 66.0409447 | 3560 | 2014 |
| **HPA** | fallopian tube; | HPA:0210051 | 1.03E-66 | 65.9890594 | 6145 | 2014 |
| **HPA** | placenta; tropl | HPA:0380521 | 1.45E-66 | 65.8396923 | 6450 | 2014 |
| **HPA** | skin 2; epiderr | HPA:0470682 | 3.13E-66 | 65.5041713 | 4787 | 2014 |
| **HPA** | appendix | HPA:0030000 | 1.20E-65 | 64.9205909 | 7644 | 2014 |
| **HPA** | vagina | HPA:0620000 | 2.07E-64 | 63.6837079 | 5923 | 2014 |
| **HPA** | vagina; squan | HPA:0620221 | 2.07E-64 | 63.6837079 | 5923 | 2014 |
| **HPA** | salivary gland; | HPA:0420052 | 3.33E-64 | 63.4781852 | 4789 | 2014 |
| **HPA** | skin 1; fibrobl | HPA:0460651 | 1.21E-63 | 62.9168979 | 4765 | 2014 |
| **HPA** | parathyroid gl | HPA:0360000 | 2.41E-63 | 62.6173867 | 5824 | 2014 |
| **HPA** | parathyroid gl | HPA:0360051 | 2.41E-63 | 62.6173867 | 5824 | 2014 |
| **HPA** | soft tissue 1; | HPA:0500651 | 6.29E-63 | 62.2016393 | 4194 | 2014 |
| **HPA** | prostate; glan | HPA:0390052 | 2.26E-62 | 61.6467256 | 4730 | 2014 |
| **HPA** | tonsil; squam | HPA:0600222 | 3.05E-62 | 61.5152081 | 4887 | 2014 |
| **HPA** | placenta; deci | HPA:0380511 | 8.52E-62 | 61.0695415 | 4765 | 2014 |
| **HPA** | lymph node; ( | HPA:0310431 | 2.11E-61 | 60.6748665 | 4991 | 2014 |
| **HPA** | breast; glandu | HPA:0050052 | 2.29E-61 | 60.6406233 | 5024 | 2014 |
| **HPA** | testis | HPA:0570000 | 8.55E-61 | 60.0681263 | 8899 | 2014 |
| **HPA** | colon; periphe | HPA:0130242 | 1.13E-60 | 59.9488231 | 3433 | 2014 |
| **HPA** | tonsil; germin | HPA:0600431 | 2.82E-60 | 59.5504443 | 5899 | 2014 |
| **HPA** | endometrium | HPA:0160252 | 6.99E-60 | 59.1554369 | 2178 | 2014 |
| **HPA** | lymph node; r | HPA:0310441 | 7.76E-60 | 59.1100826 | 5969 | 2014 |
| **HPA** | kidney; cells ir | HPA:0270362 | 8.82E-60 | 59.0543511 | 6373 | 2014 |
| **HPA** | breast; myoep | HPA:0050091 | 1.10E-59 | 58.9593409 | 5607 | 2014 |
| **HPA** | stomach 2; gl | HPA:0550052 | 1.69E-59 | 58.7708616 | 6189 | 2014 |
| **HPA** | colon; glandu | HPA:0130052 | 2.63E-59 | 58.5798871 | 6333 | 2014 |
| **HPA** | tonsil; non-ge | HPA:0600441 | 5.09E-59 | 58.2930759 | 6022 | 2014 |
| **HPA** | cerebellum; ce | HPA:0090191 | 9.62E-59 | 58.0168374 | 4388 | 2014 |
| **HPA** | cerebral cortex | HPA:0100133 | 1.75E-58 | 57.7560894 | 1812 | 2014 |
| **HPA** | placenta; tropl | HPA:0380522 | 3.19E-58 | 57.4962956 | 5131 | 2014 |
| **HPA** | duodenum; g | HPA:0150052 | 4.86E-58 | 57.3137484 | 6537 | 2014 |
| **HPA** | liver | HPA:0290000 | 5.93E-58 | 57.2270466 | 6213 | 2014 |
| **HPA** | soft tissue 2 | HPA:0510000 | 8.44E-58 | 57.0736361 | 5132 | 2014 |
| **HPA** | skin 1; Langei | HPA:0460641 | 2.68E-57 | 56.5720445 | 4672 | 2014 |
| **HPA** | nasopharynx; | HPA:0320101 | 7.46E-57 | 56.1270314 | 6137 | 2014 |
| **HPA** | cervix, uterine | HPA:0110221 | 7.62E-57 | 56.1177907 | 5383 | 2014 |
| **HPA** | cerebellum; Pu | HPA:0090161 | 1.35E-56 | 55.868951 | 5498 | 2014 |
| **HPA** | caudate; neur | HPA:0080132 | 2.00E-56 | 55.6998563 | 4061 | 2014 |
| **HPA** | bronchus; res | HPA:0060101 | 3.54E-56 | 55.450665 | 6282 | 2014 |
| **HPA** | appendix; glar | HPA:0030052 | 4.12E-56 | 55.3847974 | 5821 | 2014 |
| **HPA** | esophagus; s | HPA:0190222 | 8.31E-56 | 55.0804945 | 4989 | 2014 |

| | | | | | | |
|---|---|---|---|---|---|---|
| **HPA** | cerebral cortex | HPA:0100201 | 1.04E-55 | 54.9834071 | 4585 | 2014 |
| **HPA** | cerebellum; Pu | HPA:0090162 | 1.55E-55 | 54.8083547 | 4074 | 2014 |
| **HPA** | endometrium | HPA:0170252 | 3.51E-55 | 54.4551155 | 1756 | 2014 |
| **HPA** | oral mucosa; s | HPA:0330222 | 5.25E-55 | 54.2799844 | 4303 | 2014 |
| **HPA** | spleen | HPA:0530000 | 1.34E-54 | 53.8712884 | 5811 | 2014 |
| **HPA** | ovary; ovarian | HPA:0340462 | 3.06E-54 | 53.5138823 | 2033 | 2014 |
| **HPA** | nasopharynx; | HPA:0320102 | 5.72E-54 | 53.2429005 | 5061 | 2014 |
| **HPA** | bone marrow | HPA:0040000 | 6.01E-54 | 53.2209203 | 6406 | 2014 |
| **HPA** | bone marrow; | HPA:0040081 | 6.01E-54 | 53.2209203 | 6406 | 2014 |
| **HPA** | pancreas; exo | HPA:0350472 | 1.01E-53 | 52.9943487 | 5212 | 2014 |
| **HPA** | spleen; cells ir | HPA:0530721 | 4.00E-53 | 52.3974384 | 4290 | 2014 |
| **HPA** | adipose tissue | HPA:0010000 | 5.45E-53 | 52.2639862 | 3893 | 2014 |
| **HPA** | adipose tissue | HPA:0010011 | 5.45E-53 | 52.2639862 | 3893 | 2014 |
| **HPA** | soft tissue 2; | HPA:0510651 | 7.57E-53 | 52.1206547 | 4073 | 2014 |
| **HPA** | liver; cholangi | HPA:0291161 | 2.55E-52 | 51.5936251 | 3908 | 2014 |
| **HPA** | bronchus; res | HPA:0060102 | 2.76E-52 | 51.5583269 | 5235 | 2014 |
| **HPA** | placenta; deci | HPA:0380512 | 2.79E-52 | 51.5549216 | 3499 | 2014 |
| **HPA** | cervix, uterine | HPA:0110052 | 4.84E-52 | 51.3153623 | 3973 | 2014 |
| **HPA** | appendix; lym | HPA:0030071 | 2.75E-51 | 50.5608839 | 5028 | 2014 |
| **HPA** | testis; Leydig | HPA:0570731 | 1.63E-50 | 49.7884717 | 7077 | 2014 |
| **HPA** | spleen; cells ir | HPA:0530711 | 9.45E-50 | 49.0246649 | 5249 | 2014 |
| **HPA** | vagina; squam | HPA:0620222 | 2.16E-49 | 48.6661284 | 3993 | 2014 |
| **HPA** | hippocampus | HPA:0250133 | 3.87E-49 | 48.4126435 | 1237 | 2014 |
| **HPA** | small intestine | HPA:0480052 | 9.82E-49 | 48.0080575 | 6516 | 2014 |
| **HPA** | skin 1; melano | HPA:0460672 | 1.09E-48 | 47.961778 | 3736 | 2014 |
| **HPA** | heart muscle; | HPA:0241101 | 1.20E-47 | 46.9201932 | 6242 | 2014 |
| **HPA** | heart muscle | HPA:0240000 | 1.20E-47 | 46.9201932 | 6242 | 2014 |
| **HPA** | skin 1; keratin | HPA:0460662 | 1.92E-47 | 46.715957 | 3942 | 2014 |
| **HPA** | cerebellum; ce | HPA:0090192 | 2.95E-47 | 46.5307295 | 2746 | 2014 |
| **HPA** | parathyroid gl | HPA:0360052 | 8.35E-47 | 46.0781083 | 4348 | 2014 |
| **HPA** | cerebral cortex | HPA:0100122 | 9.81E-47 | 46.0082238 | 2636 | 2014 |
| **HPA** | cervix, uterine | HPA:0110222 | 1.01E-46 | 45.9946619 | 3666 | 2014 |
| **HPA** | tonsil; germin | HPA:0600432 | 5.94E-46 | 45.2263902 | 4037 | 2014 |
| **HPA** | urinary bladde | HPA:0610833 | 9.94E-46 | 45.002743 | 1994 | 2014 |
| **HPA** | breast; adipoc | HPA:0050011 | 1.37E-44 | 43.863908 | 2750 | 2014 |
| **HPA** | caudate; glial | HPA:0080122 | 1.42E-44 | 43.8476542 | 2273 | 2014 |
| **HPA** | testis; Leydig | HPA:0570732 | 1.17E-42 | 41.9330713 | 5327 | 2014 |
| **HPA** | colon; endoth | HPA:0130202 | 3.11E-42 | 41.5077774 | 3687 | 2014 |
| **HPA** | lung; macroph | HPA:0300412 | 5.19E-42 | 41.2851901 | 4766 | 2014 |
| **HPA** | breast; myoep | HPA:0050092 | 5.28E-42 | 41.2771768 | 3736 | 2014 |
| **HPA** | fallopian tube; | HPA:0210053 | 5.91E-42 | 41.2284788 | 1771 | 2014 |
| **HPA** | liver; hepatocy | HPA:0290401 | 2.40E-41 | 40.6194073 | 5666 | 2014 |
| **HPA** | skin 1; fibrobl | HPA:0460652 | 2.97E-41 | 40.5267485 | 3340 | 2014 |
| **HPA** | placenta; tropl | HPA:0380523 | 2.58E-40 | 39.5887936 | 2030 | 2014 |
| **HPA** | cerebellum; ce | HPA:0090181 | 4.74E-40 | 39.3242978 | 4489 | 2014 |

| | | | | | | |
|---|---|---|---|---|---|---|
| **HPA** | lymph node; ( | HPA:0310432 | 5.46E-40 | 39.2630034 | 3257 | 2014 |
| **HPA** | appendix; lym | HPA:0030072 | 6.54E-40 | 39.1842943 | 3180 | 2014 |
| **HPA** | caudate; neur | HPA:0080133 | 9.59E-40 | 39.0179844 | 1083 | 2014 |
| **HPA** | lung; alveolar | HPA:0301172 | 1.42E-39 | 38.8489845 | 2895 | 2014 |
| **HPA** | lymph node; r | HPA:0310442 | 1.71E-39 | 38.7671279 | 3925 | 2014 |
| **HPA** | soft tissue 1; | HPA:0500652 | 1.99E-39 | 38.7001867 | 2683 | 2014 |
| **HPA** | skin 1; Langer | HPA:0460642 | 2.17E-39 | 38.6639748 | 3159 | 2014 |
| **HPA** | kidney; cells ir | HPA:0270352 | 1.51E-38 | 37.8214746 | 2959 | 2014 |
| **HPA** | hippocampus | HPA:0250122 | 2.22E-38 | 37.6529242 | 1970 | 2014 |
| **HPA** | skeletal muscl | HPA:0440341 | 4.17E-38 | 37.3800934 | 5816 | 2014 |
| **HPA** | skeletal muscl | HPA:0440000 | 4.17E-38 | 37.3800934 | 5816 | 2014 |
| **HPA** | soft tissue 2; | HPA:0510652 | 9.55E-38 | 37.0201192 | 2576 | 2014 |
| **HPA** | epididymis; gl | HPA:0180053 | 2.88E-37 | 36.5406752 | 1802 | 2014 |
| **HPA** | testis; sertoli c | HPA:0570801 | 1.01E-36 | 35.9946293 | 2279 | 2014 |
| **HPA** | pancreas; exo | HPA:0350473 | 1.64E-36 | 35.7856558 | 1914 | 2014 |
| **HPA** | cerebellum; Pu | HPA:0090163 | 3.78E-36 | 35.422244 | 1528 | 2014 |
| **HPA** | placenta; deci | HPA:0380513 | 4.43E-36 | 35.3540091 | 1151 | 2014 |
| **HPA** | endometrium | HPA:0170053 | 1.08E-35 | 34.9680473 | 1304 | 2014 |
| **HPA** | spleen; cells ir | HPA:0530722 | 2.49E-35 | 34.603888 | 2684 | 2014 |
| **HPA** | bone marrow; | HPA:0040082 | 3.91E-33 | 32.4079601 | 4179 | 2014 |
| **HPA** | stomach 2; gl | HPA:0550053 | 7.00E-33 | 32.1549014 | 2378 | 2014 |
| **HPA** | gallbladder; gl | HPA:0220053 | 8.29E-33 | 32.0812974 | 2811 | 2014 |
| **HPA** | nasopharynx; | HPA:0320103 | 8.29E-33 | 32.0812897 | 1870 | 2014 |
| **HPA** | tonsil; non-ge | HPA:0600442 | 1.15E-32 | 31.9380111 | 4126 | 2014 |
| **HPA** | esophagus; s | HPA:0190223 | 3.93E-32 | 31.4061406 | 1617 | 2014 |
| **HPA** | adipose tissue | HPA:0010012 | 5.44E-32 | 31.2641049 | 2201 | 2014 |
| **HPA** | testis; sertoli c | HPA:0570802 | 6.99E-32 | 31.1553706 | 1682 | 2014 |
| **HPA** | soft tissue 2; | HPA:0510701 | 9.06E-32 | 31.0428659 | 3145 | 2014 |
| **HPA** | endometrium | HPA:0160053 | 9.81E-32 | 31.0083482 | 1435 | 2014 |
| **HPA** | skin 2; epiderr | HPA:0470683 | 3.67E-31 | 30.4356142 | 1369 | 2014 |
| **HPA** | stomach 1; gl | HPA:0540053 | 4.83E-31 | 30.3162188 | 2407 | 2014 |
| **HPA** | bronchus; res | HPA:0060103 | 7.81E-31 | 30.1073314 | 1916 | 2014 |
| **HPA** | soft tissue 1; | HPA:0500701 | 1.06E-30 | 29.9746109 | 2760 | 2014 |
| **HPA** | breast; glandu | HPA:0050053 | 2.87E-30 | 29.5415981 | 1320 | 2014 |
| **HPA** | seminal vesicl | HPA:0430053 | 5.54E-30 | 29.256339 | 1431 | 2014 |
| **HPA** | rectum; gland | HPA:0400053 | 6.83E-29 | 28.1656442 | 2724 | 2014 |
| **HPA** | cerebral cortex | HPA:0100211 | 9.82E-29 | 28.0078386 | 4948 | 2014 |
| **HPA** | thyroid gland; | HPA:0590053 | 2.52E-28 | 27.5985482 | 1835 | 2014 |
| **HPA** | cerebellum; ce | HPA:0090182 | 2.78E-28 | 27.5552186 | 2885 | 2014 |
| **HPA** | oral mucosa; | HPA:0330223 | 3.32E-28 | 27.4785334 | 1133 | 2014 |
| **HPA** | liver; cholangi | HPA:0291162 | 4.28E-28 | 27.3681636 | 1934 | 2014 |
| **HPA** | adrenal gland; | HPA:0020053 | 1.03E-27 | 26.9874072 | 2222 | 2014 |
| **HPA** | tonsil; squam | HPA:0600223 | 9.99E-27 | 26.0002317 | 1452 | 2014 |
| **HPA** | cervix, uterine | HPA:0110053 | 9.24E-26 | 25.0342784 | 1158 | 2014 |
| **HPA** | salivary gland; | HPA:0420053 | 1.82E-25 | 24.7395955 | 1443 | 2014 |

| | | | | | | |
|---|---|---|---|---|---|---|
| **HPA** | cerebellum; ce | HPA:0090193 | 2.70E-25 | 24.568891 | 905 | 2014 |
| **HPA** | colon; periphe | HPA:0130243 | 3.44E-25 | 24.4630902 | 817 | 2014 |
| **HPA** | soft tissue 1; | HPA:0500111 | 2.37E-24 | 23.6256877 | 2157 | 2014 |
| **HPA** | heart muscle; | HPA:0241102 | 3.42E-24 | 23.4657076 | 4302 | 2014 |
| **HPA** | smooth musc | HPA:0490692 | 1.27E-23 | 22.8961214 | 3073 | 2014 |
| **HPA** | tonsil; germin | HPA:0600433 | 1.27E-23 | 22.8953223 | 1211 | 2014 |
| **HPA** | cerebral cortex | HPA:0100123 | 1.56E-23 | 22.8069886 | 726 | 2014 |
| **HPA** | soft tissue 1; | HPA:0500702 | 4.10E-23 | 22.3873096 | 1331 | 2014 |
| **HPA** | cervix, uterine | HPA:0110223 | 6.44E-23 | 22.1912473 | 1007 | 2014 |
| **HPA** | duodenum; g | HPA:0150053 | 7.33E-23 | 22.134911 | 2808 | 2014 |
| **HPA** | prostate; gland | HPA:0390053 | 7.46E-23 | 22.1270128 | 1304 | 2014 |
| **HPA** | appendix; glar | HPA:0030053 | 1.38E-22 | 21.8604278 | 2010 | 2014 |
| **HPA** | spleen; cells ir | HPA:0530712 | 2.79E-22 | 21.5541484 | 3086 | 2014 |
| **HPA** | breast; adipoc | HPA:0050012 | 7.46E-22 | 21.1272707 | 1469 | 2014 |
| **HPA** | cerebral cortex | HPA:0100202 | 1.35E-21 | 20.8681663 | 2450 | 2014 |
| **HPA** | colon; glandu | HPA:0130053 | 3.49E-21 | 20.4568724 | 2574 | 2014 |
| **HPA** | vagina; squan | HPA:0620223 | 1.22E-20 | 19.9147455 | 1113 | 2014 |
| **HPA** | testis; sperma | HPA:0570813 | 2.72E-20 | 19.5647538 | 1328 | 2014 |
| **HPA** | ovary; follicle | HPA:0340451 | 2.98E-20 | 19.5252088 | 3250 | 2014 |
| **HPA** | caudate; glial | HPA:0080123 | 3.80E-20 | 19.4196539 | 669 | 2014 |
| **HPA** | lung; alveolar | HPA:0301173 | 3.84E-20 | 19.4155695 | 807 | 2014 |
| **HPA** | pancreas; pan | HPA:0351183 | 6.81E-20 | 19.1666195 | 777 | 2014 |
| **HPA** | lymph node; | HPA:0310433 | 1.25E-19 | 18.9018703 | 930 | 2014 |
| **HPA** | skin 1; fibrobl | HPA:0460653 | 3.44E-19 | 18.4640232 | 839 | 2014 |
| **HPA** | testis; sperma | HPA:0570812 | 5.23E-19 | 18.2817175 | 2367 | 2014 |
| **HPA** | small intestine | HPA:0480053 | 5.70E-19 | 18.2443995 | 2762 | 2014 |
| **HPA** | endometrium | HPA:0160253 | 2.36E-18 | 17.6272698 | 415 | 2014 |
| **HPA** | ovary; follicle | HPA:0340452 | 2.41E-18 | 17.6186333 | 2203 | 2014 |
| **HPA** | liver; hepatocy | HPA:0290402 | 3.12E-18 | 17.5055929 | 3683 | 2014 |
| **HPA** | testis; sertoli c | HPA:0570803 | 9.75E-18 | 17.0108244 | 547 | 2014 |
| **HPA** | breast; myoep | HPA:0050093 | 4.21E-17 | 16.3753124 | 819 | 2014 |
| **HPA** | bone marrow; | HPA:0040083 | 6.81E-17 | 16.1669497 | 1423 | 2014 |
| **HPA** | skin 1; keratin | HPA:0460663 | 7.87E-17 | 16.1042963 | 869 | 2014 |
| **HPA** | kidney; cells ir | HPA:0270353 | 8.12E-17 | 16.0905033 | 764 | 2014 |
| **HPA** | testis; Leydig | HPA:0570733 | 5.33E-16 | 15.2735809 | 1739 | 2014 |
| **HPA** | lung; macroph | HPA:0300413 | 5.63E-16 | 15.2492797 | 1240 | 2014 |
| **HPA** | cerebral cortex | HPA:0100212 | 5.74E-16 | 15.2409369 | 2589 | 2014 |
| **HPA** | skin 1; melan | HPA:0460673 | 1.67E-15 | 14.7768861 | 870 | 2014 |
| **HPA** | skeletal muscl | HPA:0440342 | 5.35E-15 | 14.2716094 | 3726 | 2014 |
| **HPA** | testis; sperma | HPA:0570811 | 5.43E-15 | 14.2655688 | 2815 | 2014 |
| **HPA** | soft tissue 1; | HPA:0500112 | 7.75E-15 | 14.1104368 | 1423 | 2014 |
| **HPA** | ovary; ovarian | HPA:0340463 | 9.87E-15 | 14.0055133 | 371 | 2014 |
| **HPA** | parathyroid gl | HPA:0360053 | 1.01E-14 | 13.9938458 | 1508 | 2014 |
| **HPA** | tonsil; non-ge | HPA:0600443 | 1.17E-14 | 13.9318269 | 1294 | 2014 |
| **HPA** | appendix; lym | HPA:0030073 | 1.45E-14 | 13.8394342 | 836 | 2014 |

| | | | | | | |
|---|---|---|---|---|---|---|
| **HPA** | skin 1; Langer | HPA:0460643 | 1.67E-14 | 13.7761738 | 756 | 2014 |
| **HPA** | endometrium | HPA:0170253 | 1.81E-14 | 13.7419307 | 347 | 2014 |
| **HPA** | lymph node; r | HPA:0310443 | 2.08E-14 | 13.6824169 | 1190 | 2014 |
| **HPA** | testis; peritubu | HPA:0570772 | 1.04E-13 | 12.9822327 | 462 | 2014 |
| **HPA** | soft tissue 2; | HPA:0510702 | 2.73E-13 | 12.564337 | 1508 | 2014 |
| **HPA** | adipose tissue | HPA:0010013 | 5.69E-13 | 12.2446229 | 446 | 2014 |
| **HPA** | hippocampus | HPA:0250123 | 9.57E-13 | 12.0192612 | 537 | 2014 |
| **HPA** | testis; peritubu | HPA:0570771 | 3.25E-12 | 11.487955 | 639 | 2014 |
| **HPA** | testis; peritubu | HPA:0570773 | 4.41E-12 | 11.355562 | 199 | 2014 |
| **HPA** | colon; endoth | HPA:0130203 | 5.72E-12 | 11.2426204 | 792 | 2014 |
| **HPA** | soft tissue 1; | HPA:0500653 | 5.79E-12 | 11.2375244 | 530 | 2014 |
| **HPA** | testis; pachyte | HPA:0570763 | 1.25E-11 | 10.9022581 | 854 | 2014 |
| **HPA** | smooth musc | HPA:0490693 | 2.67E-11 | 10.5735023 | 605 | 2014 |
| **HPA** | testis; preleptc | HPA:0570782 | 3.59E-11 | 10.4443247 | 1328 | 2014 |
| **HPA** | cerebellum; ce | HPA:0090183 | 5.83E-11 | 10.2343808 | 738 | 2014 |
| **HPA** | kidney; cells ir | HPA:0270363 | 5.24E-10 | 9.28070669 | 2485 | 2014 |
| **HPA** | testis; preleptc | HPA:0570783 | 6.87E-10 | 9.16287737 | 642 | 2014 |
| **HPA** | soft tissue 2; | HPA:0510653 | 1.12E-09 | 8.94936458 | 501 | 2014 |
| **HPA** | heart muscle; | HPA:0241103 | 1.54E-09 | 8.81246703 | 1135 | 2014 |
| **HPA** | liver; cholangi | HPA:0291163 | 5.12E-09 | 8.29081901 | 314 | 2014 |
| **HPA** | spleen; cells ir | HPA:0530723 | 6.01E-09 | 8.22133433 | 833 | 2014 |
| **HPA** | skeletal muscl | HPA:0440343 | 9.42E-09 | 8.02595232 | 850 | 2014 |
| **HPA** | cerebellum; m | HPA:0091001 | 3.44E-08 | 7.46298555 | 343 | 2014 |
| **HPA** | cerebral cortex | HPA:0100213 | 8.62E-08 | 7.06443968 | 401 | 2014 |
| **HPA** | testis; preleptc | HPA:0570781 | 9.06E-08 | 7.04276733 | 1698 | 2014 |
| **HPA** | testis; cells in | HPA:0570742 | 1.08E-07 | 6.96854596 | 2601 | 2014 |
| **HPA** | testis; pachyte | HPA:0570762 | 1.41E-07 | 6.85075489 | 1713 | 2014 |
| **HPA** | cerebellum; m | HPA:0091002 | 2.67E-07 | 6.57429255 | 309 | 2014 |
| **HPA** | cerebellum; Pu | HPA:0090941 | 4.11E-07 | 6.386365 | 448 | 2014 |
| **HPA** | breast; adipoc | HPA:0050013 | 4.88E-07 | 6.31138854 | 304 | 2014 |
| **HPA** | cerebellum; m | HPA:0091012 | 5.49E-07 | 6.26027221 | 169 | 2014 |
| **HPA** | soft tissue 1; | HPA:0500703 | 6.42E-07 | 6.19235674 | 234 | 2014 |
| **HPA** | cerebellum; G | HPA:0090923 | 9.84E-07 | 6.0070996 | 107 | 2014 |
| **HPA** | testis; pachyte | HPA:0570761 | 1.8917E-06 | 5.72315715 | 2141 | 2014 |
| **HPA** | soft tissue 2; | HPA:0510703 | 2.0228E-06 | 5.6940543 | 283 | 2014 |
| **HPA** | cerebellum; m | HPA:0091011 | 2.5268E-06 | 5.59742214 | 186 | 2014 |
| **HPA** | cerebellum; G | HPA:0090922 | 2.9116E-06 | 5.53587001 | 179 | 2014 |
| **HPA** | cerebellum; G | HPA:0090921 | 5.6795E-06 | 5.24569257 | 197 | 2014 |
| **HPA** | cerebellum; Pu | HPA:0090942 | 6.5545E-06 | 5.18345739 | 368 | 2014 |
| **HPA** | testis; round c | HPA:0570793 | 9.9155E-06 | 5.00368696 | 917 | 2014 |
| **HPA** | ovary; follicle c | HPA:0340453 | 1.6786E-05 | 4.77506021 | 597 | 2014 |
| **HPA** | cerebral cortex | HPA:0100203 | 2.6934E-05 | 4.56969423 | 407 | 2014 |
| **HPA** | spleen; cells ir | HPA:0530713 | 3.7694E-05 | 4.42373119 | 805 | 2014 |
| **HPA** | testis; cells in | HPA:0570741 | 4.4267E-05 | 4.35392354 | 3355 | 2014 |
| **HPA** | cerebellum; gr | HPA:0090971 | 0.00011058 | 3.95633129 | 297 | 2014 |

| | | | | | | |
|---|---|---|---|---|---|---|
| **HPA** | fallopian tube; | HPA:0211092 | 0.00013856 | 3.85835446 | 328 | 2014 |
| **HPA** | cerebellum; m | HPA:0091003 | 0.00021105 | 3.6756159 | 182 | 2014 |
| **HPA** | cerebellum; Pu | HPA:0090943 | 0.00026252 | 3.58083209 | 191 | 2014 |
| **HPA** | cerebellum; m | HPA:0090991 | 0.00028337 | 3.54765236 | 265 | 2014 |
| **HPA** | cerebellum; sy | HPA:0091061 | 0.00036413 | 3.43874355 | 296 | 2014 |
| **HPA** | cerebellum; B | HPA:0090911 | 0.00054046 | 3.26723962 | 91 | 2014 |
| **HPA** | cerebellum; Pu | HPA:0090961 | 0.00063356 | 3.19821232 | 121 | 2014 |
| **HPA** | testis; cells in | HPA:0570743 | 0.00071816 | 3.14377877 | 827 | 2014 |
| **HPA** | cerebellum; w | HPA:0091071 | 0.00072924 | 3.13712995 | 292 | 2014 |
| **HPA** | testis; round c | HPA:0570792 | 0.00081728 | 3.0876294 | 1725 | 2014 |
| **HPA** | cerebellum; w | HPA:0091081 | 0.00099328 | 3.00292864 | 119 | 2014 |
| **HPA** | cerebellum; w | HPA:0091082 | 0.00113885 | 2.94353279 | 112 | 2014 |
| **HPA** | cerebellum; Pu | HPA:0090962 | 0.00147182 | 2.8321442 | 113 | 2014 |
| **HPA** | cerebellum; B | HPA:0090912 | 0.00169806 | 2.77004738 | 84 | 2014 |
| **HPA** | cerebellum; m | HPA:0090993 | 0.00281262 | 2.55088914 | 35 | 2014 |
| **HPA** | fallopian tube; | HPA:0211091 | 0.00295986 | 2.52872829 | 457 | 2014 |
| **HPA** | cerebellum; gr | HPA:0090982 | 0.00423556 | 2.37308873 | 129 | 2014 |
| **HPA** | cerebellum; gr | HPA:0090981 | 0.00442328 | 2.35425546 | 145 | 2014 |
| **HPA** | cerebellum; w | HPA:0091073 | 0.0050721 | 2.29481182 | 122 | 2014 |
| **HPA** | cerebellum; sy | HPA:0091063 | 0.00980065 | 2.00874528 | 125 | 2014 |
| **HPA** | cerebellum; w | HPA:0091072 | 0.01167482 | 1.93274978 | 225 | 2014 |
| **HPA** | cerebellum; Pu | HPA:0090963 | 0.01254751 | 1.90144232 | 80 | 2014 |
| **HPA** | cerebellum; m | HPA:0091013 | 0.01470025 | 1.83267524 | 119 | 2014 |
| **HPA** | cerebellum; sy | HPA:0091062 | 0.01507431 | 1.82176251 | 231 | 2014 |
| **HPA** | cerebellum; gr | HPA:0090972 | 0.01507431 | 1.82176251 | 231 | 2014 |
| **HPA** | cerebellum; Pu | HPA:0090953 | 0.01744071 | 1.75843595 | 108 | 2014 |
| **HPA** | cerebellum; m | HPA:0090992 | 0.01753416 | 1.75611499 | 148 | 2014 |
| **HPA** | soft tissue 1; | HPA:0500113 | 0.02421821 | 1.61585794 | 309 | 2014 |
| **HPA** | placenta; endo | HPA:0380202 | 0.02805964 | 1.55191786 | 163 | 2014 |
| **HPA** | cerebellum; Pu | HPA:0090952 | 0.03029758 | 1.51859205 | 210 | 2014 |
| **HPA** | fallopian tube; | HPA:0210851 | 0.03307962 | 1.48043954 | 612 | 2014 |
| **HPA** | cerebellum; sy | HPA:0091051 | 0.03825099 | 1.41735737 | 92 | 2014 |
| **HPA** | cerebellum; G | HPA:0090933 | 0.04737526 | 1.32444836 | 59 | 2014 |

| | | |
|---|---|---|
| 1877 | 10877 | EEF1A1,ORC4,PABPC1,COL1A1,LRP1,GAPDH,MLLT1,COPA,HNRNPA |
| 1690 | 10877 | EEF1A1,ORC4,PABPC1,COL1A1,LRP1,GAPDH,MLLT1,COPA,HNRNPA |
| 1730 | 10877 | EEF1A1,ORC4,PABPC1,COL1A1,LRP1,GAPDH,MLLT1,COPA,HNRNPA |
| 1579 | 10877 | EEF1A1,ORC4,PABPC1,COL1A1,LRP1,GAPDH,MLLT1,COPA,HNRNPA |
| 1846 | 10877 | EEF1A1,ORC4,PABPC1,COL1A1,GAPDH,MLLT1,COPA,HNRNPA1,ARS |
| 1644 | 10877 | EEF1A1,ORC4,PABPC1,COL1A1,LRP1,GAPDH,MLLT1,COPA,HNRNPA |
| 1678 | 10877 | EEF1A1,ORC4,PABPC1,COL1A1,LRP1,GAPDH,MLLT1,COPA,HNRNPA |
| 1742 | 10877 | EEF1A1,ORC4,PABPC1,COL1A1,GAPDH,MLLT1,COPA,HNRNPA1,ARS |
| 1689 | 10877 | EEF1A1,ORC4,PABPC1,COL1A1,LRP1,GAPDH,MLLT1,COPA,HNRNPA |
| 1696 | 10877 | EEF1A1,ORC4,PABPC1,COL1A1,GAPDH,MLLT1,COPA,HNRNPA1,ARS |
| 1696 | 10877 | EEF1A1,ORC4,PABPC1,COL1A1,GAPDH,MLLT1,COPA,HNRNPA1,ARS |
| 1664 | 10877 | EEF1A1,ORC4,PABPC1,COL1A1,LRP1,GAPDH,COPA,HNRNPA1,ARSD |
| 1447 | 10877 | EEF1A1,ORC4,PABPC1,COL1A1,GAPDH,MLLT1,COPA,HNRNPA1,ARS |
| 1719 | 10877 | EEF1A1,ORC4,PABPC1,COL1A1,GAPDH,MLLT1,COPA,HNRNPA1,ARS |
| 1719 | 10877 | EEF1A1,ORC4,PABPC1,COL1A1,GAPDH,MLLT1,COPA,HNRNPA1,ARS |
| 1637 | 10877 | EEF1A1,ORC4,COL1A1,GAPDH,MLLT1,COPA,HNRNPA1,ARSD,HSPA9 |
| 1710 | 10877 | EEF1A1,ORC4,PABPC1,COL1A1,GAPDH,MLLT1,COPA,HNRNPA1,ARS |
| 1710 | 10877 | EEF1A1,ORC4,PABPC1,COL1A1,GAPDH,MLLT1,COPA,HNRNPA1,ARS |
| 1690 | 10877 | EEF1A1,ORC4,PABPC1,COL1A1,LRP1,GAPDH,MLLT1,COPA,HNRNPA |
| 1618 | 10877 | EEF1A1,ORC4,PABPC1,COL1A1,GAPDH,COPA,HNRNPA1,ARSD,HSPA |
| 1134 | 10877 | EEF1A1,ORC4,COL1A1,LRP1,GAPDH,COPA,HNRNPA1,HSPA9,POLH,/ |
| 1730 | 10877 | EEF1A1,ORC4,PABPC1,COL1A1,LRP1,GAPDH,MLLT1,COPA,HNRNPA |
| 1717 | 10877 | EEF1A1,ORC4,PABPC1,COL1A1,GAPDH,MLLT1,COPA,HNRNPA1,ARS |
| 1716 | 10877 | EEF1A1,ORC4,PABPC1,COL1A1,GAPDH,MLLT1,COPA,HNRNPA1,ARS |
| 1316 | 10877 | EEF1A1,ORC4,PABPC1,COL1A1,LRP1,COPA,HNRNPA1,ARSD,HSPA9, |
| 1171 | 10877 | EEF1A1,ORC4,PABPC1,COL1A1,LRP1,MLLT1,COPA,HNRNPA1,ARSD, |
| 1813 | 10877 | EEF1A1,ORC4,PABPC1,COL1A1,GAPDH,MLLT1,COPA,HNRNPA1,ARS |
| 1639 | 10877 | EEF1A1,ORC4,PABPC1,COL1A1,GAPDH,MLLT1,COPA,HNRNPA1,ARS |
| 1529 | 10877 | EEF1A1,ORC4,PABPC1,COL1A1,GAPDH,MLLT1,COPA,HNRNPA1,ARS |
| 1529 | 10877 | EEF1A1,ORC4,PABPC1,COL1A1,GAPDH,MLLT1,COPA,HNRNPA1,ARS |
| 1718 | 10877 | EEF1A1,ORC4,PABPC1,COL1A1,GAPDH,MLLT1,COPA,HNRNPA1,ARS |
| 1620 | 10877 | EEF1A1,ORC4,PABPC1,COL1A1,GAPDH,MLLT1,COPA,HNRNPA1,ARS |
| 1620 | 10877 | EEF1A1,ORC4,PABPC1,COL1A1,GAPDH,MLLT1,COPA,HNRNPA1,ARS |
| 1386 | 10877 | EEF1A1,ORC4,COL1A1,LRP1,GAPDH,COPA,HNRNPA1,HSPA9,NEDD4 |
| 1477 | 10877 | EEF1A1,ORC4,PABPC1,COL1A1,GAPDH,MLLT1,COPA,HNRNPA1,ARS |
| 1615 | 10877 | EEF1A1,ORC4,PABPC1,COL1A1,GAPDH,MLLT1,COPA,HNRNPA1,ARS |
| 1292 | 10877 | EEF1A1,ORC4,COL1A1,LRP1,GAPDH,MLLT1,COPA,HNRNPA1,ARSD,I |
| 1591 | 10877 | EEF1A1,ORC4,PABPC1,GAPDH,MLLT1,COPA,HNRNPA1,ARSD,HSPA9 |
| 1591 | 10877 | EEF1A1,ORC4,PABPC1,GAPDH,MLLT1,COPA,HNRNPA1,ARSD,HSPA9 |
| 1423 | 10877 | EEF1A1,ORC4,PABPC1,COL1A1,GAPDH,MLLT1,COPA,HNRNPA1,ARS |
| 1126 | 10877 | EEF1A1,ORC4,PABPC1,COL1A1,LRP1,GAPDH,COPA,HNRNPA1,ARSD |
| 1737 | 10877 | EEF1A1,ORC4,PABPC1,COL1A1,LRP1,GAPDH,MLLT1,COPA,HNRNPA |
| 1737 | 10877 | EEF1A1,ORC4,PABPC1,COL1A1,LRP1,GAPDH,MLLT1,COPA,HNRNPA |
| 1736 | 10877 | EEF1A1,ORC4,PABPC1,COL1A1,GAPDH,MLLT1,COPA,HNRNPA1,ARS |
| 1736 | 10877 | EEF1A1,ORC4,PABPC1,COL1A1,GAPDH,MLLT1,COPA,HNRNPA1,ARS |

| | | |
|---|---|---|
| 1325 | 10877 | EEF1A1,ORC4,COL1A1,GAPDH,MLLT1,COPA,HNRNPA1,ARSD,HSPA9 |
| 1734 | 10877 | EEF1A1,ORC4,PABPC1,GAPDH,MLLT1,COPA,HNRNPA1,ARSD,HSPA9 |
| 1734 | 10877 | EEF1A1,ORC4,PABPC1,GAPDH,MLLT1,COPA,HNRNPA1,ARSD,HSPA9 |
| 1684 | 10877 | EEF1A1,ORC4,PABPC1,COL1A1,LRP1,GAPDH,MLLT1,COPA,HNRNPA |
| 1625 | 10877 | EEF1A1,ORC4,PABPC1,COL1A1,GAPDH,MLLT1,COPA,HNRNPA1,ARS |
| 1722 | 10877 | EEF1A1,ORC4,PABPC1,COL1A1,GAPDH,MLLT1,COPA,HNRNPA1,ARS |
| 1649 | 10877 | EEF1A1,ORC4,PABPC1,COL1A1,LRP1,GAPDH,MLLT1,COPA,HNRNPA |
| 1500 | 10877 | EEF1A1,ORC4,PABPC1,COL1A1,LRP1,GAPDH,MLLT1,COPA,HNRNPA |
| 1617 | 10877 | EEF1A1,ORC4,PABPC1,COL1A1,GAPDH,MLLT1,COPA,HNRNPA1,ARS |
| 1617 | 10877 | EEF1A1,ORC4,PABPC1,COL1A1,GAPDH,MLLT1,COPA,HNRNPA1,ARS |
| 1755 | 10877 | EEF1A1,ORC4,PABPC1,COL1A1,GAPDH,MLLT1,COPA,HNRNPA1,ARS |
| 1383 | 10877 | EEF1A1,ORC4,PABPC1,GAPDH,MLLT1,COPA,HNRNPA1,ARSD,HSPA9 |
| 1580 | 10877 | EEF1A1,ORC4,PABPC1,COL1A1,GAPDH,MLLT1,COPA,HNRNPA1,ARS |
| 1580 | 10877 | EEF1A1,ORC4,PABPC1,COL1A1,GAPDH,MLLT1,COPA,HNRNPA1,ARS |
| 1002 | 10877 | EEF1A1,ORC4,COL1A1,LRP1,GAPDH,HNRNPA1,ARSD,HSPA9,ATN1,A |
| 1574 | 10877 | EEF1A1,PABPC1,COL1A1,GAPDH,COPA,HNRNPA1,ARSD,HSPA9,NED |
| 1354 | 10877 | EEF1A1,COL1A1,LRP1,GAPDH,MLLT1,COPA,HNRNPA1,HSPA9,POLH, |
| 1720 | 10877 | EEF1A1,ORC4,PABPC1,COL1A1,GAPDH,MLLT1,COPA,HNRNPA1,ARS |
| 1720 | 10877 | EEF1A1,ORC4,PABPC1,COL1A1,GAPDH,MLLT1,COPA,HNRNPA1,ARS |
| 1588 | 10877 | EEF1A1,ORC4,PABPC1,COL1A1,GAPDH,MLLT1,COPA,HNRNPA1,ARS |
| 1588 | 10877 | EEF1A1,ORC4,PABPC1,COL1A1,GAPDH,MLLT1,COPA,HNRNPA1,ARS |
| 1390 | 10877 | EEF1A1,ORC4,COL1A1,GAPDH,MLLT1,COPA,HNRNPA1,ARSD,HSPA9 |
| 1741 | 10877 | EEF1A1,PABPC1,COL1A1,GAPDH,COPA,HNRNPA1,ARSD,HSPA9,NED |
| 1309 | 10877 | EEF1A1,ORC4,PABPC1,COL1A1,GAPDH,MLLT1,HNRNPA1,ARSD,HSP |
| 1651 | 10877 | EEF1A1,ORC4,PABPC1,COL1A1,LRP1,GAPDH,MLLT1,COPA,HNRNPA |
| 1257 | 10877 | EEF1A1,ORC4,PABPC1,COL1A1,COPA,HNRNPA1,ARSD,HSPA9,NEDD |
| 1455 | 10877 | EEF1A1,ORC4,COL1A1,GAPDH,MLLT1,COPA,HNRNPA1,ARSD,HSPA9 |
| 1420 | 10877 | EEF1A1,ORC4,COL1A1,GAPDH,MLLT1,COPA,HNRNPA1,HSPA9,POL |
| 1360 | 10877 | EEF1A1,ORC4,GAPDH,MLLT1,COPA,HNRNPA1,ARSD,HSPA9,POLH,A |
| 1360 | 10877 | EEF1A1,ORC4,GAPDH,MLLT1,COPA,HNRNPA1,ARSD,HSPA9,POLH,A |
| 1401 | 10877 | EEF1A1,MLLT1,COPA,HNRNPA1,ARSD,NEDD4L,POLH,ATN1,CTSC,N |
| 1511 | 10877 | EEF1A1,ORC4,PABPC1,GAPDH,MLLT1,COPA,HNRNPA1,ARSD,HSPA9 |
| 1198 | 10877 | EEF1A1,ORC4,COL1A1,LRP1,GAPDH,COPA,HNRNPA1,ARSD,HSPA9, |
| 1734 | 10877 | EEF1A1,ORC4,PABPC1,COL1A1,GAPDH,MLLT1,COPA,HNRNPA1,ARS |
| 1734 | 10877 | EEF1A1,ORC4,PABPC1,COL1A1,GAPDH,MLLT1,COPA,HNRNPA1,ARS |
| 1684 | 10877 | EEF1A1,PABPC1,COL1A1,LRP1,GAPDH,MLLT1,COPA,HNRNPA1,ARS |
| 1663 | 10877 | EEF1A1,ORC4,PABPC1,COL1A1,GAPDH,MLLT1,COPA,HNRNPA1,ARS |
| 1551 | 10877 | EEF1A1,ORC4,PABPC1,COL1A1,MLLT1,COPA,HNRNPA1,ARSD,HSPA |
| 1110 | 10877 | EEF1A1,ORC4,COL1A1,GAPDH,HNRNPA1,ARSD,HSPA9,POLH,ATN1, |
| 1481 | 10877 | EEF1A1,ORC4,PABPC1,COL1A1,GAPDH,MLLT1,HNRNPA1,ARSD,HSP |
| 1296 | 10877 | EEF1A1,ORC4,COL1A1,COPA,HNRNPA1,ARSD,HSPA9,POLH,ATN1,CT |
| 1310 | 10877 | EEF1A1,ORC4,PABPC1,GAPDH,MLLT1,COPA,HNRNPA1,ARSD,HSPA9 |
| 1296 | 10877 | EEF1A1,ORC4,COPA,HNRNPA1,ARSD,HSPA9,NEDD4L,POLH,ATN1,CT |
| 1576 | 10877 | EEF1A1,ORC4,PABPC1,COL1A1,LRP1,GAPDH,MLLT1,COPA,HNRNPA |
| 1539 | 10877 | EEF1A1,ORC4,PABPC1,COL1A1,GAPDH,MLLT1,COPA,HNRNPA1,ARS |

| | | |
|---|---|---|
| 1295 | 10877 | EEF1A1,ORC4,PABPC1,COL1A1,LRP1,GAPDH,MLLT1,COPA,HNRNPA |
| 1532 | 10877 | EEF1A1,ORC4,PABPC1,COL1A1,GAPDH,MLLT1,COPA,HNRNPA1,ARS |
| 1725 | 10877 | EEF1A1,ORC4,PABPC1,GAPDH,COPA,HNRNPA1,ARSD,HSPA9,NEDD4 |
| 1725 | 10877 | EEF1A1,ORC4,PABPC1,GAPDH,COPA,HNRNPA1,ARSD,HSPA9,NEDD4 |
| 1001 | 10877 | EEF1A1,MLLT1,COPA,HNRNPA1,ARSD,HSPA9,POLH,ATN1,NUDC,NRE |
| 1483 | 10877 | EEF1A1,ORC4,COL1A1,COPA,HNRNPA1,ARSD,HSPA9,NEDD4L,POLH |
| 1534 | 10877 | EEF1A1,ORC4,PABPC1,COL1A1,LRP1,GAPDH,MLLT1,COPA,HNRNPA |
| 1239 | 10877 | EEF1A1,ORC4,PABPC1,GAPDH,MLLT1,COPA,HNRNPA1,ARSD,HSPA9 |
| 1721 | 10877 | EEF1A1,ORC4,PABPC1,COL1A1,GAPDH,MLLT1,COPA,HNRNPA1,ARS |
| 1439 | 10877 | EEF1A1,ORC4,PABPC1,GAPDH,MLLT1,COPA,HNRNPA1,ARSD,HSPA9 |
| 1439 | 10877 | EEF1A1,ORC4,PABPC1,GAPDH,MLLT1,COPA,HNRNPA1,ARSD,HSPA9 |
| 1234 | 10877 | EEF1A1,ORC4,PABPC1,COL1A1,GAPDH,COPA,HNRNPA1,ARSD,HSPA |
| 1228 | 10877 | EEF1A1,COL1A1,LRP1,GAPDH,MLLT1,COPA,HNRNPA1,HSPA9,CTSC, |
| 1419 | 10877 | EEF1A1,ORC4,PABPC1,GAPDH,MLLT1,COPA,HNRNPA1,ARSD,HSPA9 |
| 1419 | 10877 | EEF1A1,ORC4,PABPC1,GAPDH,MLLT1,COPA,HNRNPA1,ARSD,HSPA9 |
| 1117 | 10877 | EEF1A1,COL1A1,LRP1,MLLT1,COPA,HNRNPA1,HSPA9,POLH,CTSC,N |
| 1218 | 10877 | EEF1A1,ORC4,PABPC1,GAPDH,COPA,HNRNPA1,ARSD,HSPA9,NEDD4 |
| 1247 | 10877 | EEF1A1,PABPC1,GAPDH,COPA,HNRNPA1,ARSD,HSPA9,POLH,ATN1,N |
| 1223 | 10877 | COL1A1,LRP1,GAPDH,MLLT1,COPA,HNRNPA1,ARSD,HSPA9,NEDD4L |
| 1264 | 10877 | EEF1A1,ORC4,PABPC1,COL1A1,GAPDH,COPA,HNRNPA1,ARSD,HSPA |
| 1270 | 10877 | EEF1A1,ORC4,PABPC1,GAPDH,MLLT1,COPA,HNRNPA1,ARSD,HSPA9 |
| 1886 | 10877 | EEF1A1,ORC4,PABPC1,COL1A1,LRP1,GAPDH,MLLT1,COPA,HNRNPA |
| 959 | 10877 | EEF1A1,ORC4,GAPDH,MLLT1,HNRNPA1,ARSD,HSPA9,NEDD4L,POLH |
| 1424 | 10877 | EEF1A1,PABPC1,COL1A1,GAPDH,COPA,HNRNPA1,ARSD,HSPA9,NEDI |
| 687 | 10877 | EEF1A1,ORC4,PABPC1,COL1A1,LRP1,GAPDH,COPA,HNRNPA1,ARSD |
| 1435 | 10877 | ORC4,PABPC1,COL1A1,GAPDH,MLLT1,COPA,HNRNPA1,ARSD,HSPA9 |
| 1504 | 10877 | EEF1A1,ORC4,COL1A1,GAPDH,MLLT1,COPA,HNRNPA1,ARSD,HSPA9 |
| 1371 | 10877 | EEF1A1,ORC4,COL1A1,MLLT1,HNRNPA1,ARSD,HSPA9,POLH,ACTB,C |
| 1472 | 10877 | EEF1A1,ORC4,PABPC1,COL1A1,GAPDH,MLLT1,COPA,HNRNPA1,ARS |
| 1496 | 10877 | EEF1A1,ORC4,PABPC1,MLLT1,COPA,HNRNPA1,ARSD,HSPA9,NEDD4I |
| 1442 | 10877 | EEF1A1,PABPC1,COL1A1,GAPDH,COPA,HNRNPA1,ARSD,HSPA9,POLI |
| 1143 | 10877 | EEF1A1,ORC4,PABPC1,COL1A1,LRP1,GAPDH,MLLT1,COPA,HNRNPA |
| 599 | 10877 | EEF1A1,HNRNPA1,ARSD,HSPA9,ATN1,EEF2,TUBA1A,NUMA1,EIF4G2, |
| 1281 | 10877 | EEF1A1,ORC4,PABPC1,LRP1,GAPDH,MLLT1,COPA,HNRNPA1,ARSD,I |
| 1527 | 10877 | EEF1A1,ORC4,PABPC1,COL1A1,MLLT1,COPA,HNRNPA1,ARSD,HSPA9 |
| 1472 | 10877 | EEF1A1,ORC4,LRP1,GAPDH,MLLT1,COPA,HNRNPA1,ARSD,HSPA9,PC |
| 1280 | 10877 | EEF1A1,ORC4,PABPC1,COL1A1,LRP1,GAPDH,MLLT1,COPA,HNRNPA |
| 1193 | 10877 | EEF1A1,ORC4,GAPDH,MLLT1,COPA,HNRNPA1,HSPA9,POLH,CTSC,NI |
| 1456 | 10877 | EEF1A1,ORC4,COL1A1,LRP1,COPA,HNRNPA1,ARSD,HSPA9,NEDD4L, |
| 1323 | 10877 | EEF1A1,ORC4,PABPC1,COL1A1,GAPDH,COPA,HNRNPA1,ARSD,POLH |
| 1343 | 10877 | EEF1A1,ORC4,PABPC1,COL1A1,LRP1,MLLT1,COPA,HNRNPA1,ARSD, |
| 1073 | 10877 | EEF1A1,PABPC1,LRP1,MLLT1,COPA,HNRNPA1,ARSD,HSPA9,ATN1,NI |
| 1479 | 10877 | EEF1A1,COL1A1,LRP1,COPA,HNRNPA1,ARSD,HSPA9,NEDD4L,POLH, |
| 1399 | 10877 | EEF1A1,ORC4,PABPC1,COL1A1,COPA,HNRNPA1,ARSD,HSPA9,NEDD4 |
| 1248 | 10877 | EEF1A1,ORC4,PABPC1,GAPDH,MLLT1,COPA,HNRNPA1,ARSD,HSPA9 |

| | | |
|---|---|---|
| 1172 | 10877 | EEF1A1,ORC4,PABPC1,LRP1,MLLT1,HNRNPA1,HSPA9,ACTB,CTSC,N |
| 1073 | 10877 | EEF1A1,ORC4,PABPC1,MLLT1,COPA,HNRNPA1,ARSD,HSPA9,POLH,A |
| 578 | 10877 | EEF1A1,ORC4,COL1A1,LRP1,HNRNPA1,ARSD,HSPA9,ACTB,USP22,E |
| 1116 | 10877 | EEF1A1,ORC4,PABPC1,COL1A1,GAPDH,MLLT1,COPA,HNRNPA1,ARS |
| 1393 | 10877 | EEF1A1,ORC4,PABPC1,GAPDH,COPA,HNRNPA1,ARSD,HSPA9,POLH,A |
| 640 | 10877 | COL1A1,LRP1,GAPDH,COPA,HNRNPA1,HSPA9,POLH,USP22,EEF2,DP |
| 1256 | 10877 | EEF1A1,ORC4,COL1A1,COPA,HNRNPA1,ARSD,HSPA9,POLH,ATN1,CT |
| 1494 | 10877 | EEF1A1,ORC4,PABPC1,COL1A1,GAPDH,MLLT1,COPA,HNRNPA1,ARS |
| 1494 | 10877 | EEF1A1,ORC4,PABPC1,COL1A1,GAPDH,MLLT1,COPA,HNRNPA1,ARS |
| 1283 | 10877 | EEF1A1,ORC4,COL1A1,GAPDH,COPA,HNRNPA1,ARSD,HSPA9,NEDD4 |
| 1108 | 10877 | EEF1A1,ORC4,PABPC1,COPA,HNRNPA1,ARSD,HSPA9,ACTB,USP22,A |
| 1030 | 10877 | ORC4,COL1A1,LRP1,HNRNPA1,HSPA9,ACTB,CTSC,NRBP2,USP22,AC |
| 1030 | 10877 | ORC4,COL1A1,LRP1,HNRNPA1,HSPA9,ACTB,CTSC,NRBP2,USP22,AC |
| 1065 | 10877 | EEF1A1,COL1A1,LRP1,GAPDH,MLLT1,COPA,HNRNPA1,HSPA9,CTSC, |
| 1031 | 10877 | EEF1A1,ORC4,COPA,HNRNPA1,ARSD,HSPA9,POLH,CTSC,NUDC,NRB |
| 1283 | 10877 | EEF1A1,COL1A1,COPA,HNRNPA1,ARSD,HSPA9,NEDD4L,POLH,ATN1, |
| 949 | 10877 | COL1A1,LRP1,GAPDH,MLLT1,COPA,HNRNPA1,ARSD,HSPA9,NEDD4L |
| 1043 | 10877 | EEF1A1,COPA,HNRNPA1,ARSD,NEDD4L,POLH,ATN1,CTSC,NUDC,NR |
| 1242 | 10877 | ORC4,PABPC1,COL1A1,COPA,HNRNPA1,ARSD,HSPA9,NEDD4L,POL |
| 1596 | 10877 | EEF1A1,ORC4,PABPC1,COL1A1,LRP1,GAPDH,MLLT1,COPA,HNRNPA |
| 1278 | 10877 | EEF1A1,ORC4,PABPC1,GAPDH,COPA,HNRNPA1,ARSD,HSPA9,POLH,A |
| 1039 | 10877 | EEF1A1,ORC4,PABPC1,GAPDH,MLLT1,COPA,HNRNPA1,ARSD,HSPA9 |
| 438 | 10877 | MLLT1,HNRNPA1,ARSD,HSPA9,TUBA1A,MAVS,NUMA1,TUBA1B,NCL, |
| 1498 | 10877 | EEF1A1,ORC4,PABPC1,COPA,HNRNPA1,ARSD,HSPA9,NEDD4L,POLH, |
| 986 | 10877 | EEF1A1,ORC4,PABPC1,GAPDH,MLLT1,COPA,HNRNPA1,ARSD,HSPA9 |
| 1448 | 10877 | ORC4,PABPC1,COL1A1,GAPDH,MLLT1,COPA,HNRNPA1,ARSD,HSPA |
| 1448 | 10877 | ORC4,PABPC1,COL1A1,GAPDH,MLLT1,COPA,HNRNPA1,ARSD,HSPA |
| 1023 | 10877 | EEF1A1,ORC4,PABPC1,COL1A1,GAPDH,MLLT1,HNRNPA1,ARSD,HSP |
| 778 | 10877 | EEF1A1,ORC4,COL1A1,COPA,HNRNPA1,ARSD,HSPA9,POLH,ATN1,NU |
| 1100 | 10877 | EEF1A1,ORC4,GAPDH,MLLT1,COPA,HNRNPA1,ARSD,HSPA9,POLH,A |
| 753 | 10877 | EEF1A1,ORC4,COL1A1,MLLT1,HNRNPA1,ARSD,HSPA9,POLH,ATN1,N |
| 966 | 10877 | EEF1A1,ORC4,PABPC1,HNRNPA1,ARSD,POLH,ATN1,NRBP2,EEF2,MA |
| 1037 | 10877 | EEF1A1,PABPC1,COL1A1,GAPDH,COPA,HNRNPA1,ARSD,HSPA9,POL |
| 609 | 10877 | EEF1A1,PABPC1,HNRNPA1,ARSD,HSPA9,ATN1,USP22,EEF2,TUBA1A, |
| 771 | 10877 | ORC4,COL1A1,LRP1,HNRNPA1,USP22,ACTG1,EEF2,TUBA1A,MAVS,H |
| 668 | 10877 | EEF1A1,ORC4,HNRNPA1,HSPA9,POLH,ATN1,EEF2,TUBA1A,MAVS,HS |
| 1270 | 10877 | EEF1A1,ORC4,COL1A1,LRP1,GAPDH,COPA,HNRNPA1,ARSD,HSPA9,P |
| 956 | 10877 | EEF1A1,ORC4,GAPDH,MLLT1,COPA,HNRNPA1,POLH,ACTB,NRBP2,U |
| 1164 | 10877 | ORC4,PABPC1,LRP1,GAPDH,COPA,HNRNPA1,ARSD,HSPA9,POLH,CT |
| 965 | 10877 | EEF1A1,ORC4,MLLT1,HNRNPA1,ARSD,HSPA9,POLH,ACTB,NUDC,US |
| 548 | 10877 | EEF1A1,COPA,HNRNPA1,ARSD,HSPA9,NUDC,USP22,EEF2,DPYSL3,TU |
| 1327 | 10877 | EEF1A1,ORC4,LRP1,GAPDH,MLLT1,COPA,HNRNPA1,ARSD,HSPA9,PO |
| 883 | 10877 | EEF1A1,COL1A1,LRP1,GAPDH,COPA,HNRNPA1,HSPA9,CTSC,USP22, |
| 602 | 10877 | ORC4,COPA,HNRNPA1,ARSD,HSPA9,ATN1,NRBP2,MAVS,HSP90AB1, |
| 1105 | 10877 | ORC4,COL1A1,LRP1,COPA,HNRNPA1,ARSD,HSPA9,POLH,ATN1,NUD |

| | | |
|---|---|---|
| 862 | 10877 | EEF1A1,PABPC1,HNRNPA1,ARSD,HSPA9,POLH,ACTB,NUDC,USP22,A |
| 846 | 10877 | PABPC1,COPA,HNRNPA1,ARSD,HSPA9,POLH,ACTB,NUDC,NRBP2,US |
| 378 | 10877 | HNRNPA1,ARSD,HSPA9,ATN1,TUBA1A,NUMA1,RPS6,TUBA1B,NCL,FT |
| 786 | 10877 | EEF1A1,ORC4,GAPDH,HNRNPA1,ARSD,HSPA9,POLH,NRBP2,USP22,E |
| 994 | 10877 | PABPC1,GAPDH,COPA,HNRNPA1,ARSD,HSPA9,POLH,ACTB,CTSC,NU |
| 741 | 10877 | EEF1A1,LRP1,COPA,HNRNPA1,HSPA9,POLH,USP22,EEF2,TUBA1A,CF |
| 840 | 10877 | ORC4,GAPDH,COPA,HNRNPA1,HSPA9,POLH,CTSC,NUDC,USP22,EEF |
| 796 | 10877 | ORC4,COPA,HNRNPA1,ARSD,HSPA9,POLH,ATN1,USP22,DPYSL3,TUE |
| 583 | 10877 | ORC4,HNRNPA1,ARSD,HSPA9,POLH,EEF2,TUBA1A,MAVS,HSP90AB1, |
| 1343 | 10877 | EEF1A1,ORC4,GAPDH,MLLT1,HNRNPA1,ARSD,POLH,ATN1,ACTB,CT |
| 1343 | 10877 | EEF1A1,ORC4,GAPDH,MLLT1,HNRNPA1,ARSD,POLH,ATN1,ACTB,CT |
| 713 | 10877 | EEF1A1,LRP1,GAPDH,COPA,HNRNPA1,HSPA9,USP22,EEF2,TUBA1A,C |
| 542 | 10877 | EEF1A1,COPA,HNRNPA1,ARSD,HSPA9,ACTB,USP22,ACTG1,EEF2,TUE |
| 646 | 10877 | COPA,HNRNPA1,ARSD,HSPA9,CTSC,NUDC,USP22,MAVS,HSP90AB1,V |
| 565 | 10877 | EEF1A1,GAPDH,COPA,HNRNPA1,ARSD,HSPA9,USP22,EEF2,TUBA1A,M |
| 476 | 10877 | EEF1A1,MLLT1,HNRNPA1,ARSD,HSPA9,ATN1,NUMA1,EIF4G2,RPS6,N |
| 386 | 10877 | GAPDH,COPA,HNRNPA1,HSPA9,POLH,ATN1,EEF2,MAVS,HSP90AB1,S |
| 422 | 10877 | EEF1A1,HNRNPA1,ARSD,HSPA9,EEF2,MAVS,NUMA1,NCL,HSP90B1,P |
| 728 | 10877 | HNRNPA1,ARSD,HSPA9,ACTB,USP22,ACTG1,EEF2,TUBA1A,CFL1,MA |
| 1021 | 10877 | EEF1A1,ORC4,PABPC1,COL1A1,GAPDH,MLLT1,COPA,HNRNPA1,ARS |
| 655 | 10877 | EEF1A1,PABPC1,COPA,HNRNPA1,ARSD,HSPA9,ATN1,USP22,TUBA1A, |
| 746 | 10877 | EEF1A1,COPA,HNRNPA1,ARSD,HSPA9,ACTB,CTSC,NRBP2,USP22,AC |
| 544 | 10877 | EEF1A1,HNRNPA1,ARSD,HSPA9,NRBP2,USP22,EEF2,TUBA1A,MAVS,H |
| 1009 | 10877 | EEF1A1,PABPC1,GAPDH,COPA,HNRNPA1,ARSD,HSPA9,POLH,ACTB,N |
| 485 | 10877 | EEF1A1,PABPC1,HNRNPA1,ARSD,HSPA9,USP22,EEF2,TUBA1A,MAVS, |
| 614 | 10877 | ORC4,LRP1,HNRNPA1,CTSC,USP22,TUBA1A,MAVS,VIM,UBC,KIDINS2 |
| 499 | 10877 | HNRNPA1,HSPA9,CTSC,USP22,MAVS,HSP90AB1,VIM,SPARC,TUBB,U |
| 811 | 10877 | ORC4,GAPDH,HNRNPA1,HSPA9,USP22,ACTG1,EEF2,DPYSL3,TUBA1 |
| 442 | 10877 | EEF1A1,ARSD,HSPA9,TUBA1A,MAVS,HSP90AB1,UBC,NUMA1,TUBA1 |
| 425 | 10877 | EEF1A1,GAPDH,MLLT1,HNRNPA1,ARSD,HSPA9,NRBP2,USP22,EEF2, |
| 655 | 10877 | EEF1A1,COPA,HNRNPA1,ARSD,HSPA9,ACTB,USP22,ACTG1,EEF2,TUE |
| 548 | 10877 | EEF1A1,HNRNPA1,ARSD,HSPA9,USP22,EEF2,TUBA1A,MAVS,HSP90A |
| 728 | 10877 | GAPDH,HNRNPA1,POLH,CTSC,NRBP2,DPYSL3,TUBA1A,MAVS,HSP90 |
| 411 | 10877 | EEF1A1,HNRNPA1,ARSD,HSPA9,EEF2,TUBA1A,MAVS,UBC,NUMA1,RF |
| 436 | 10877 | EEF1A1,COPA,HNRNPA1,ARSD,HSPA9,NRBP2,EEF2,TUBA1A,MAVS,V |
| 714 | 10877 | EEF1A1,GAPDH,COPA,HNRNPA1,ARSD,HSPA9,ATN1,NRBP2,USP22,E |
| 1150 | 10877 | EEF1A1,GAPDH,COPA,ARSD,HSPA9,POLH,NUDC,NRBP2,ACTG1,EEF |
| 522 | 10877 | EEF1A1,HNRNPA1,ARSD,HSPA9,POLH,NRBP2,USP22,EEF2,TUBA1A,M |
| 745 | 10877 | ORC4,COL1A1,COPA,HNRNPA1,ARSD,HSPA9,ATN1,NUDC,NRBP2,TU |
| 361 | 10877 | EEF1A1,GAPDH,HNRNPA1,ARSD,HSPA9,NRBP2,USP22,TUBA1A,MAV |
| 543 | 10877 | ORC4,COPA,HNRNPA1,HSPA9,CTSC,USP22,TUBA1A,MAVS,HSP90AB |
| 604 | 10877 | EEF1A1,COPA,HNRNPA1,ARSD,HSPA9,ATN1,NRBP2,TUBA1A,MAVS,H |
| 431 | 10877 | EEF1A1,PABPC1,HNRNPA1,ARSD,HSPA9,ATN1,EEF2,TUBA1A,MAVS,N |
| 360 | 10877 | EEF1A1,HNRNPA1,ARSD,TUBA1A,MAVS,UBC,NUMA1,TUBA1B,NCL,H |
| 425 | 10877 | EEF1A1,GAPDH,HNRNPA1,ARSD,HSPA9,ATN1,USP22,EEF2,TUBA1A,M |

| | | |
|---|---|---|
| 298 | 10877 | EEF1A1,ORC4,HNRNPA1,ATN1,HSP90AB1,UBC,NUMA1,NCL,HSP90B |
| 276 | 10877 | EEF1A1,HNRNPA1,HSPA9,ATN1,USP22,TUBA1A,HSP90AB1,NUMA1,D |
| 578 | 10877 | EEF1A1,GAPDH,HNRNPA1,HSPA9,ACTG1,EEF2,DPYSL3,TUBA1A,MAV |
| 1009 | 10877 | ORC4,MLLT1,COPA,HNRNPA1,ARSD,HSPA9,POLH,USP22,EEF2,MAV |
| 765 | 10877 | EEF1A1,ORC4,GAPDH,MLLT1,HNRNPA1,ARSD,HSPA9,POLH,ACTB,U |
| 366 | 10877 | PABPC1,HNRNPA1,ARSD,HSPA9,EEF2,TUBA1A,CFL1,MAVS,HSP90AB |
| 249 | 10877 | ORC4,HNRNPA1,ARSD,ATN1,EEF2,MAVS,FTH1,UBC,NUMA1,NCL,FTL |
| 392 | 10877 | GAPDH,HNRNPA1,CTSC,NRBP2,TUBA1A,HSP90AB1,TUBB,NUMA1,TU |
| 316 | 10877 | EEF1A1,HNRNPA1,ARSD,EEF2,MAVS,UBC,NUMA1,RPS6,NCL,UBA1,H |
| 708 | 10877 | EEF1A1,COPA,HNRNPA1,ARSD,HSPA9,ACTB,USP22,ACTG1,EEF2,TU |
| 385 | 10877 | HNRNPA1,ARSD,HSPA9,USP22,EEF2,TUBA1A,MAVS,HSP90AB1,UBC, |
| 540 | 10877 | COPA,HNRNPA1,ARSD,HSPA9,USP22,EEF2,TUBA1A,CFL1,MAVS,KIDI |
| 762 | 10877 | EEF1A1,ORC4,GAPDH,HNRNPA1,ARSD,HSPA9,ACTB,CTSC,USP22,AC |
| 419 | 10877 | ORC4,HNRNPA1,USP22,TUBA1A,MAVS,HSP90AB1,VIM,UBC,KIDINS2 |
| 629 | 10877 | PABPC1,HNRNPA1,HSPA9,CTSC,NUDC,EEF2,MAVS,VIM,SPARC,UBC, |
| 653 | 10877 | EEF1A1,COPA,HNRNPA1,ARSD,HSPA9,USP22,TUBA1A,CFL1,MAVS,N |
| 334 | 10877 | EEF1A1,GAPDH,HNRNPA1,ARSD,HSPA9,NRBP2,EEF2,MAVS,UBC,NU |
| 382 | 10877 | HNRNPA1,HSPA9,NUDC,HSP90AB1,VIM,SPARC,TUBB,NUMA1,DBN1, |
| 786 | 10877 | EEF1A1,ORC4,HSPA9,NEDD4L,CTSC,EEF2,TUBA1A,CFL1,MAVS,VIM, |
| 226 | 10877 | ORC4,HNRNPA1,HSPA9,MAVS,FTH1,UBC,NUMA1,NCL,FTL,HSP90B1, |
| 260 | 10877 | HNRNPA1,ARSD,HSPA9,USP22,EEF2,TUBA1A,MAVS,VIM,NUMA1,ENO |
| 252 | 10877 | HNRNPA1,HSPA9,POLH,TUBA1A,MAVS,NUMA1,TUBA1B,NCL,HSP90B |
| 288 | 10877 | HNRNPA1,HSPA9,ACTB,ACTG1,EEF2,TUBA1A,CFL1,MAVS,HSP90AB1 |
| 265 | 10877 | EEF1A1,COL1A1,GAPDH,HNRNPA1,HSPA9,EEF2,TUBA1A,VIM,SPARC, |
| 601 | 10877 | EEF1A1,COL1A1,COPA,HNRNPA1,ARSD,HSPA9,NUDC,HSP90AB1,VI |
| 682 | 10877 | EEF1A1,COPA,HNRNPA1,ARSD,HSPA9,ACTB,USP22,ACTG1,EEF2,TU |
| 156 | 10877 | HNRNPA1,ARSD,HSPA9,NUMA1,FLNA,NCL,HSP90B1,H3-3B,PTMS,N |
| 564 | 10877 | EEF1A1,ORC4,HSPA9,EEF2,TUBA1A,MAVS,RPS6,ENO1,TUBA1B,NCL, |
| 862 | 10877 | EEF1A1,ORC4,LRP1,GAPDH,MLLT1,COPA,HNRNPA1,ARSD,HSPA9,P |
| 189 | 10877 | HNRNPA1,HSPA9,VIM,SPARC,DCX,NCL,ATP2B4,PSAP,BSG,SEPTIN2, |
| 254 | 10877 | HNRNPA1,ARSD,HSPA9,ACTB,ACTG1,EEF2,TUBA1A,NUMA1,TUBA1B, |
| 391 | 10877 | ORC4,PABPC1,COPA,HNRNPA1,HSPA9,EEF2,TUBA1A,MAVS,HSP90A |
| 265 | 10877 | EEF1A1,GAPDH,HNRNPA1,ARSD,HSPA9,EEF2,TUBA1A,MAVS,UBC,NU |
| 240 | 10877 | HNRNPA1,VIM,SPARC,NUMA1,RPS6,FLNA,NCL,ATP2B4,TMF1,PSAP, |
| 456 | 10877 | COL1A1,HNRNPA1,ARSD,HSPA9,CTSC,USP22,EEF2,MAVS,VIM,SPAR |
| 347 | 10877 | HNRNPA1,ARSD,HSPA9,POLH,CTSC,USP22,EEF2,TUBA1A,MAVS,ACT |
| 633 | 10877 | COPA,ARSD,HSPA9,POLH,ACTG1,EEF2,DPYSL3,TUBA1A,SPARC,TUB |
| 261 | 10877 | EEF1A1,GAPDH,HNRNPA1,ARSD,HSPA9,EEF2,TUBA1A,MAVS,VIM,NU |
| 854 | 10877 | EEF1A1,ORC4,MLLT1,HNRNPA1,ARSD,POLH,ACTB,USP22,ACTG1,E |
| 674 | 10877 | EEF1A1,COL1A1,MLLT1,COPA,HNRNPA1,ARSD,HSPA9,NUDC,HSP90 |
| 383 | 10877 | EEF1A1,GAPDH,HNRNPA1,HSPA9,EEF2,DPYSL3,TUBA1A,HSP90AB1, |
| 136 | 10877 | COL1A1,HNRNPA1,HSPA9,DPYSL3,TUBA1A,MAVS,VIM,NUMA1,TUBA |
| 401 | 10877 | COPA,HNRNPA1,ARSD,USP22,UBC,NUMA1,PIK3R2,NCL,HSP90B1,U |
| 354 | 10877 | COPA,HNRNPA1,ARSD,HSPA9,ACTB,USP22,ACTG1,EEF2,TUBA1A,CF |
| 250 | 10877 | HNRNPA1,ARSD,HSPA9,ACTB,ACTG1,EEF2,TUBA1A,MAVS,ACTR2,VI |

| | | |
|---|---|---|
| 231 | 10877 | GAPDH,HNRNPA1,HSPA9,EEF2,TUBA1A,MAVS,VIM,UBC,NUMA1,TUB |
| 129 | 10877 | HNRNPA1,HSPA9,EEF2,MAVS,NUMA1,NCL,H3-3B,NPM1,ATP5F1A,M |
| 330 | 10877 | COPA,HNRNPA1,ARSD,HSPA9,ACTB,USP22,ACTG1,EEF2,CFL1,MAVS |
| 157 | 10877 | HNRNPA1,USP22,VIM,SPARC,NUMA1,FLNA,NCL,H3-3B,SPTBN1,SPT |
| 395 | 10877 | ORC4,GAPDH,HNRNPA1,HSPA9,USP22,TUBA1A,VIM,SPARC,UBC,NU |
| 151 | 10877 | HNRNPA1,VIM,NUMA1,NCL,COL6A1,PSAP,H3-3B,DDX5,DVL3,FASN, |
| 173 | 10877 | ORC4,HNRNPA1,ARSD,NUMA1,NCL,HSP90B1,UGT8,FBXL4,PSAP,H3- |
| 196 | 10877 | HNRNPA1,USP22,VIM,SPARC,NUMA1,FLNA,NCL,H3-3B,SPTBN1,GNB |
| 83 | 10877 | USP22,VIM,SPARC,NUMA1,FLNA,NCL,H3-3B,SPTBN1,SPTAN1,NPM1, |
| 231 | 10877 | ACTB,ACTG1,TUBA1A,VIM,NUMA1,TUBA1B,NCL,HSP90B1,TMF1,H3- |
| 169 | 10877 | HNRNPA1,EEF2,TUBA1A,VIM,NUMA1,TUBA1B,NCL,YBX1,H3-3B,DVL |
| 244 | 10877 | PABPC1,NUDC,USP22,VIM,SPARC,TUBB,NCL,PRPF8,ATP2B4,HSPA1 |
| 185 | 10877 | EEF1A1,HNRNPA1,ARSD,ACTB,ACTG1,UBC,NUMA1,FLNA,NCL,H3-3 |
| 347 | 10877 | COL1A1,HNRNPA1,NUDC,USP22,VIM,SPARC,NUMA1,NCL,RPL8,ATP2 |
| 215 | 10877 | HNRNPA1,ARSD,HSPA9,ATN1,UBC,NUMA1,NCL,H3-3B,PTMA,DDX5,E |
| 581 | 10877 | COL1A1,HNRNPA1,ARSD,HSPA9,CTSC,EEF2,MAVS,NUMA1,NCL,SLC |
| 189 | 10877 | HNRNPA1,NUDC,USP22,VIM,SPARC,NCL,ATP2B4,ATP5F1B,GDI1,NP |
| 155 | 10877 | HNRNPA1,EEF2,TUBA1A,VIM,NUMA1,TUBA1B,NCL,YBX1,H3-3B,CAN |
| 298 | 10877 | COPA,ARSD,HSPA9,NUMA1,NCL,ABHD2,HSPA1B,PSAP,H3-3B,DVL3, |
| 107 | 10877 | HSPA9,MAVS,NCL,PSAP,H3-3B,YWHAE,ATP5F1B,MYH9,GSTP1,NPM1 |
| 229 | 10877 | HNRNPA1,ARSD,HSPA9,ACTB,ACTG1,EEF2,CFL1,MAVS,ACTR2,NUM |
| 232 | 10877 | EEF1A1,HNRNPA1,ARSD,USP22,UBC,NUMA1,NCL,ALDOA,ABHD2,H3- |
| 112 | 10877 | SPARC,MAP1B,ALDOA,GANAB,FTL,PSAP,FASN,ALDH6A1,SEPTIN2,AP |
| 125 | 10877 | TUBA1A,SPARC,FTH1,TUBA1B,NEFL,ATP2B4,PTMA,ENO2,SPTBN1,G |
| 408 | 10877 | COL1A1,HNRNPA1,ARSD,NUDC,USP22,VIM,SPARC,NUMA1,NCL,PRP |
| 590 | 10877 | ORC4,GAPDH,POLH,ACTB,NRBP2,ACTG1,EEF2,TUBA1A,CFL1,HSP9 |
| 410 | 10877 | PABPC1,COPA,ARSD,HSPA9,NUDC,USP22,VIM,SPARC,TUBB,NCL,HS |
| 101 | 10877 | SPARC,MAP1B,ALDOA,GANAB,FTL,PSAP,FASN,ALDH6A1,SEPTIN2,AP |
| 134 | 10877 | MAP1B,ALDOA,GANAB,FTL,NEFM,PSAP,APP,CKB,MAPT,INA,UCHL1, |
| 99 | 10877 | VIM,NCL,H3-3B,FASN,NPM1,VCP,MATR3,NONO,SSBP2,CYB5R3,HN |
| 64 | 10877 | USP22,YBX1,DDX17,FSCN1,MARCKS,USP11,HMGB1,UCHL1,KPNB1, |
| 81 | 10877 | HNRNPA1,TUBA1A,TUBB,NUMA1,TUBA1B,NCL,H3-3B,PTMS,MARCK |
| 46 | 10877 | MAP1B,GANAB,PSAP,APP,PLD3,UCHL1,RPL10,CPE,RTN4,RPL9,EIF5, |
| 490 | 10877 | PABPC1,COPA,ARSD,HSPA9,NUDC,USP22,VIM,SPARC,TUBB,NUMA1, |
| 92 | 10877 | HNRNPA1,HSPA9,TUBA1A,VIM,UBC,NUMA1,RPS6,TUBA1B,MAP1B,N |
| 67 | 10877 | USP22,YBX1,DDX17,FSCN1,MARCKS,USP11,HMGB1,UCHL1,KPNB1, |
| 65 | 10877 | DBN1,MAP1B,ALDOA,GANAB,PSAP,SEPTIN2,APP,PLD3,UCHL1,RPL1 |
| 69 | 10877 | DBN1,MAP1B,ALDOA,GANAB,PSAP,SEPTIN2,APP,PLD3,UCHL1,RPL1 |
| 111 | 10877 | MAP1B,ALDOA,GANAB,FTL,PSAP,APP,CKB,INA,UCHL1,RPL10,AKAP1 |
| 233 | 10877 | PABPC1,HNRNPA1,ARSD,NUDC,SPARC,TUBB,NCL,ATP2B4,TMF1,HS |
| 162 | 10877 | HSPA9,EEF2,TUBA1A,TUBA1B,NCL,HSP90B1,RPLP0,DVL3,ALDH6A1, |
| 118 | 10877 | VIM,RPLP0,H3-3B,DVL3,SPTBN1,NPM1,SPEN,NONO,EGLN1,SSBP2, |
| 206 | 10877 | HNRNPA1,HSPA9,CFL1,MAVS,VIM,UBC,NUMA1,NCL,PSAP,H3-3B,CA |
| 718 | 10877 | ORC4,GAPDH,POLH,ACTB,NRBP2,ACTG1,EEF2,DPYSL3,TUBA1A,CF |
| 90 | 10877 | SPARC,FTL,UGT8,PSAP,ALDH6A1,FSCN1,PLD3,CKB,RPL10,MAP4,C |

| | | |
|---|---|---|
| 97 | 10877 | GAPDH,CFL1,ENO1,FLNA,GANAB,PRPF8,UGT8,YBX1,CLU,DDX5,SEP |
| 61 | 10877 | SPARC,MAP1B,FTL,PSAP,APP,PLD3,UCHL1,RPL10,MAP4,CPE,GAP43 |
| 63 | 10877 | MAP1B,ALDOA,PSAP,CKB,INA,UCHL1,RPL10,PJA2,RPL9,EIF5,ATP2A |
| 81 | 10877 | MAP1B,FTL,SEPTIN2,MARCKS,MAPT,BASP1,UCHL1,NCAM1,AKAP12, |
| 88 | 10877 | USP22,DBN1,ALDOA,ALDH6A1,APP,MARCKS,PLD3,UCHL1,CPE,RTN4 |
| 36 | 10877 | USP22,YBX1,DDX17,HMGB1,H1-10,PEA15,GAS7,TAF15,AEBP1,PTBP |
| 44 | 10877 | USP22,ALDOA,USP11,UCHL1,KPNB1,EIF5,TKT,CTNNB1,TAF15,IPO7, |
| 204 | 10877 | EEF2,TUBA1A,HSP90AA1,EIF4G2,RPS6,TUBA1B,YBX1,ABHD2,PTMA, |
| 86 | 10877 | SPARC,MAP1B,GANAB,FTL,ALDH6A1,SEPTIN2,APP,FSCN1,BASP1,NC |
| 388 | 10877 | PABPC1,COPA,HNRNPA1,ARSD,NUDC,SPARC,TUBB,NCL,HSP90B1,P |
| 43 | 10877 | USP22,DDX17,FSCN1,HMGB1,NLGN2,TKT,JUN,H1-10,PEA15,TAF15, |
| 41 | 10877 | USP22,DDX17,FSCN1,HMGB1,NLGN2,TKT,JUN,H1-10,PEA15,TAF15, |
| 41 | 10877 | USP22,ALDOA,USP11,UCHL1,KPNB1,EIF5,TKT,CTNNB1,TAF15,IPO7, |
| 33 | 10877 | USP22,YBX1,DDX17,HMGB1,H1-10,PEA15,TAF15,PTBP1,SOD1,ARNT |
| 18 | 10877 | MARCKS,BASP1,NCAM1,ATP1A2,L1CAM,ATP6V1B2,STXBP1,SV2A,T |
| 121 | 10877 | GAPDH,CFL1,ENO1,FLNA,GANAB,PRPF8,UGT8,YBX1,CLU,DDX5,SEP |
| 44 | 10877 | USP22,YBX1,DDX17,FSCN1,HMGB1,TKT,JUN,H1-10,PEA15,GAS7,TA |
| 48 | 10877 | USP22,YBX1,DDX17,FSCN1,HMGB1,TKT,JUN,H1-10,PEA15,GAS7,TA |
| 42 | 10877 | SPARC,FTL,ALDH6A1,APP,BASP1,MAP4,CPE,RTN4,MSN,ATP1A1,RTN |
| 42 | 10877 | DBN1,ALDH6A1,MARCKS,UCHL1,CPE,ATP1A1,RTN3,SPTBN2,CDH2,T |
| 66 | 10877 | SPARC,MAP1B,GANAB,FTL,ALDH6A1,SEPTIN2,APP,FSCN1,BASP1,NC |
| 30 | 10877 | USP22,ALDOA,USP11,UCHL1,KPNB1,EIF5,CTNNB1,TAF15,IPO7,KD |
| 40 | 10877 | USP22,MARCKS,USP11,HMGB1,UCHL1,KPNB1,GAP43,TKT,CTNNB1, |
| 67 | 10877 | DBN1,ALDH6A1,APP,MARCKS,UCHL1,CPE,GAP43,NLGN2,ATP1A1,RT |
| 67 | 10877 | SPARC,FTL,UGT8,PSAP,ALDH6A1,FSCN1,CKB,RPL10,MAP4,CPE,PJA |
| 37 | 10877 | MAP1B,ALDOA,CKB,UCHL1,PJA2,EIF5,ATP2A2,SPTBN2,MAGED2,MY |
| 47 | 10877 | MARCKS,MAPT,BASP1,NCAM1,CPE,GAP43,ATP1A1,ATP1A2,L1CAM,T |
| 84 | 10877 | GAPDH,HNRNPA1,TUBA1A,HSP90AB1,NUMA1,TUBA1B,NCL,H3-3B,D |
| 50 | 10877 | ACTB,USP22,TUBA1A,MAP1B,NEFM,HACD3,LDHB,STMN1,HK1,CBX5 |
| 61 | 10877 | MAP1B,ALDOA,FTL,NEFM,CKB,UCHL1,RPL10,AKAP12,RTN4,PJA2,EI |
| 149 | 10877 | PABPC1,GAPDH,MLLT1,MAP1B,GANAB,PRPF8,UGT8,YBX1,CLU,DDX |
| 32 | 10877 | ALDH6A1,MAPT,INA,BASP1,NCAM1,RTN4,PJA2,ATP1A2,L1CAM,PRD |
| 23 | 10877 | USP22,USP11,UCHL1,TKT,TAF15,FBLN1,ARNT2,ADAR,PRMT1,APEX1 |

1,ARSD,HSPA9,NEDD4L,POLH,ATN1,ACTB,CTSC,NUDC,NRBP2,USP22,ACTG1,EEF2,DPYSL3,TUB
1,ARSD,HSPA9,POLH,ATN1,CTSC,NUDC,NRBP2,USP22,ACTG1,EEF2,DPYSL3,TUBA1A,MAVS,HSP
1,ARSD,HSPA9,POLH,ATN1,CTSC,NUDC,NRBP2,USP22,ACTG1,EEF2,DPYSL3,TUBA1A,MAVS,HSP
1,ARSD,HSPA9,POLH,ATN1,CTSC,NUDC,NRBP2,USP22,ACTG1,EEF2,DPYSL3,TUBA1A,MAVS,HSP
D,HSPA9,NEDD4L,POLH,ATN1,ACTB,CTSC,NUDC,NRBP2,USP22,ACTG1,EEF2,DPYSL3,TUBA1A,C
1,ARSD,HSPA9,NEDD4L,POLH,ATN1,CTSC,NUDC,NRBP2,USP22,ACTG1,EEF2,DPYSL3,TUBA1A,M
1,ARSD,HSPA9,POLH,ATN1,CTSC,NUDC,NRBP2,USP22,ACTG1,EEF2,DPYSL3,TUBA1A,MAVS,HSP
D,HSPA9,NEDD4L,POLH,ATN1,ACTB,CTSC,NUDC,NRBP2,USP22,ACTG1,EEF2,DPYSL3,TUBA1A,M
1,ARSD,HSPA9,NEDD4L,POLH,ATN1,ACTB,CTSC,NUDC,NRBP2,USP22,ACTG1,EEF2,DPYSL3,TUB
D,HSPA9,NEDD4L,POLH,ATN1,CTSC,NUDC,NRBP2,USP22,ACTG1,EEF2,TUBA1A,CFL1,MAVS,HSP
D,HSPA9,NEDD4L,POLH,ATN1,CTSC,NUDC,NRBP2,USP22,ACTG1,EEF2,TUBA1A,CFL1,MAVS,HSP
,HSPA9,NEDD4L,POLH,ATN1,ACTB,CTSC,NUDC,NRBP2,USP22,ACTG1,EEF2,TUBA1A,MAVS,HSP9
D,HSPA9,POLH,ATN1,NUDC,NRBP2,USP22,ACTG1,EEF2,DPYSL3,TUBA1A,MAVS,HSP90AB1,ACT
D,HSPA9,NEDD4L,POLH,ATN1,ACTB,CTSC,NUDC,NRBP2,USP22,ACTG1,EEF2,DPYSL3,TUBA1A,C
D,HSPA9,NEDD4L,POLH,ATN1,ACTB,CTSC,NUDC,NRBP2,USP22,ACTG1,EEF2,DPYSL3,TUBA1A,C
,NEDD4L,POLH,ATN1,ACTB,CTSC,NUDC,NRBP2,USP22,ACTG1,EEF2,TUBA1A,CFL1,MAVS,HSP90
D,HSPA9,NEDD4L,POLH,ATN1,CTSC,NUDC,NRBP2,USP22,ACTG1,EEF2,TUBA1A,MAVS,HSP90AB
D,HSPA9,NEDD4L,POLH,ATN1,CTSC,NUDC,NRBP2,USP22,ACTG1,EEF2,TUBA1A,MAVS,HSP90AB
1,ARSD,HSPA9,NEDD4L,POLH,ATN1,ACTB,CTSC,NUDC,NRBP2,USP22,ACTG1,EEF2,DPYSL3,TUB
9,NEDD4L,POLH,ATN1,ACTB,CTSC,NUDC,NRBP2,USP22,ACTG1,EEF2,TUBA1A,MAVS,HSP90AB1,
TN1,NRBP2,USP22,ACTG1,EEF2,DPYSL3,TUBA1A,MAVS,HSP90AB1,VIM,TUBB,FTH1,UBC,NUMA
1,ARSD,HSPA9,NEDD4L,POLH,ATN1,ACTB,CTSC,NUDC,NRBP2,USP22,ACTG1,EEF2,TUBA1A,MA
D,HSPA9,NEDD4L,POLH,ATN1,CTSC,NUDC,NRBP2,USP22,ACTG1,EEF2,DPYSL3,TUBA1A,CFL1,M
D,HSPA9,NEDD4L,POLH,ATN1,CTSC,NUDC,NRBP2,USP22,ACTG1,EEF2,DPYSL3,TUBA1A,CFL1,M
NEDD4L,ATN1,NUDC,NRBP2,USP22,EEF2,DPYSL3,TUBA1A,HSP90AB1,ACTR2,TUBB,FTH1,UBC,T
,HSPA9,POLH,NUDC,NRBP2,USP22,EEF2,TUBA1A,MAVS,HSP90AB1,ACTR2,TUBB,UBC,NUMA1,D
D,HSPA9,NEDD4L,POLH,ATN1,ACTB,CTSC,NUDC,NRBP2,USP22,ACTG1,EEF2,DPYSL3,TUBA1A,C
D,NEDD4L,POLH,ATN1,CTSC,NUDC,NRBP2,USP22,ACTG1,EEF2,TUBA1A,CFL1,MAVS,HSP90AB1,
D,HSPA9,POLH,ATN1,CTSC,NUDC,NRBP2,USP22,ACTG1,EEF2,DPYSL3,TUBA1A,CFL1,MAVS,HSP
D,HSPA9,POLH,ATN1,CTSC,NUDC,NRBP2,USP22,ACTG1,EEF2,DPYSL3,TUBA1A,CFL1,MAVS,HSP
D,HSPA9,NEDD4L,POLH,ATN1,CTSC,NUDC,NRBP2,USP22,ACTG1,EEF2,DPYSL3,TUBA1A,CFL1,M
D,HSPA9,NEDD4L,POLH,ATN1,CTSC,NUDC,NRBP2,USP22,ACTG1,EEF2,TUBA1A,CFL1,MAVS,ACT
D,HSPA9,NEDD4L,POLH,ATN1,CTSC,NUDC,NRBP2,USP22,ACTG1,EEF2,TUBA1A,CFL1,MAVS,ACT
L,POLH,ATN1,CTSC,NRBP2,USP22,ACTG1,EEF2,DPYSL3,TUBA1A,CFL1,MAVS,HSP90AB1,VIM,TU
D,HSPA9,POLH,ATN1,CTSC,NUDC,NRBP2,USP22,EEF2,TUBA1A,MAVS,HSP90AB1,ACTR2,TUBB,U
D,HSPA9,NEDD4L,POLH,ATN1,ACTB,CTSC,NUDC,NRBP2,USP22,ACTG1,EEF2,DPYSL3,TUBA1A,C
HSPA9,POLH,ATN1,NUDC,NRBP2,USP22,ACTG1,EEF2,TUBA1A,MAVS,HSP90AB1,ACTR2,VIM,TUB
,NEDD4L,POLH,ATN1,NUDC,NRBP2,USP22,ACTG1,EEF2,TUBA1A,CFL1,MAVS,HSP90AB1,ACTR2,
,NEDD4L,POLH,ATN1,NUDC,NRBP2,USP22,ACTG1,EEF2,TUBA1A,CFL1,MAVS,HSP90AB1,ACTR2,
D,HSPA9,POLH,ATN1,ACTB,CTSC,NUDC,NRBP2,USP22,ACTG1,EEF2,DPYSL3,TUBA1A,MAVS,HSP
,HSPA9,POLH,ATN1,ACTB,CTSC,NRBP2,USP22,ACTG1,EEF2,DPYSL3,TUBA1A,MAVS,HSP90AB1,
1,ARSD,HSPA9,NEDD4L,POLH,ATN1,ACTB,CTSC,NUDC,NRBP2,USP22,ACTG1,EEF2,DPYSL3,TUB
1,ARSD,HSPA9,NEDD4L,POLH,ATN1,ACTB,CTSC,NUDC,NRBP2,USP22,ACTG1,EEF2,DPYSL3,TUB
D,HSPA9,NEDD4L,POLH,ATN1,ACTB,CTSC,NUDC,NRBP2,USP22,ACTG1,EEF2,TUBA1A,CFL1,MA
D,HSPA9,NEDD4L,POLH,ATN1,ACTB,CTSC,NUDC,NRBP2,USP22,ACTG1,EEF2,TUBA1A,CFL1,MA

,POLH,ATN1,ACTB,USP22,ACTG1,EEF2,DPYSL3,TUBA1A,CFL1,MAVS,HSP90AB1,ACTR2,VIM,SPA
,NEDD4L,POLH,ATN1,ACTB,CTSC,NUDC,NRBP2,USP22,ACTG1,EEF2,TUBA1A,CFL1,MAVS,HSP90
,NEDD4L,POLH,ATN1,ACTB,CTSC,NUDC,NRBP2,USP22,ACTG1,EEF2,TUBA1A,CFL1,MAVS,HSP90
1,ARSD,HSPA9,NEDD4L,POLH,ACTB,CTSC,NUDC,NRBP2,USP22,ACTG1,EEF2,DPYSL3,TUBA1A,C
D,HSPA9,NEDD4L,POLH,ACTB,CTSC,NUDC,NRBP2,USP22,ACTG1,EEF2,TUBA1A,MAVS,HSP90AB
D,HSPA9,NEDD4L,POLH,ATN1,ACTB,CTSC,NUDC,NRBP2,USP22,ACTG1,EEF2,TUBA1A,CFL1,MAV
1,ARSD,HSPA9,POLH,ATN1,CTSC,NUDC,NRBP2,USP22,ACTG1,EEF2,TUBA1A,CFL1,MAVS,HSP90
1,ARSD,HSPA9,POLH,ATN1,CTSC,NUDC,NRBP2,USP22,ACTG1,EEF2,DPYSL3,TUBA1A,MAVS,HSP
D,HSPA9,POLH,ATN1,ACTB,CTSC,NUDC,NRBP2,USP22,ACTG1,EEF2,TUBA1A,CFL1,MAVS,HSP90
D,HSPA9,POLH,ATN1,ACTB,CTSC,NUDC,NRBP2,USP22,ACTG1,EEF2,TUBA1A,CFL1,MAVS,HSP90
D,HSPA9,NEDD4L,POLH,ATN1,ACTB,CTSC,NUDC,NRBP2,USP22,ACTG1,EEF2,TUBA1A,CFL1,MAV
,POLH,ATN1,NUDC,NRBP2,USP22,ACTG1,EEF2,TUBA1A,CFL1,MAVS,HSP90AB1,ACTR2,VIM,TUB
D,HSPA9,POLH,ATN1,CTSC,NUDC,NRBP2,USP22,ACTG1,EEF2,TUBA1A,CFL1,MAVS,HSP90AB1,A
D,HSPA9,POLH,ATN1,CTSC,NUDC,NRBP2,USP22,ACTG1,EEF2,TUBA1A,CFL1,MAVS,HSP90AB1,A
CTB,USP22,EEF2,TUBA1A,MAVS,HSP90AB1,VIM,TUBB,FTH1,UBC,TMSB4X,NUMA1,EIF4G2,RPS6
D4L,POLH,ATN1,CTSC,NUDC,NRBP2,USP22,ACTG1,EEF2,DPYSL3,TUBA1A,CFL1,MAVS,HSP90AB
,CTSC,NRBP2,USP22,ACTG1,EEF2,DPYSL3,TUBA1A,CFL1,MAVS,HSP90AB1,ACTR2,VIM,SPARC,T
D,HSPA9,NEDD4L,POLH,ATN1,ACTB,NUDC,NRBP2,USP22,ACTG1,EEF2,TUBA1A,CFL1,MAVS,HSP
D,HSPA9,NEDD4L,POLH,ATN1,ACTB,NUDC,NRBP2,USP22,ACTG1,EEF2,TUBA1A,CFL1,MAVS,HSP
D,HSPA9,POLH,ATN1,CTSC,NUDC,NRBP2,USP22,ACTG1,EEF2,TUBA1A,CFL1,MAVS,HSP90AB1,A
D,HSPA9,POLH,ATN1,CTSC,NUDC,NRBP2,USP22,ACTG1,EEF2,TUBA1A,CFL1,MAVS,HSP90AB1,A
,NEDD4L,POLH,ATN1,CTSC,NUDC,NRBP2,USP22,EEF2,TUBA1A,MAVS,HSP90AB1,ACTR2,VIM,TU
D4L,POLH,ATN1,ACTB,CTSC,NUDC,NRBP2,USP22,ACTG1,EEF2,DPYSL3,TUBA1A,CFL1,MAVS,HSP
A9,NEDD4L,POLH,ATN1,NUDC,NRBP2,USP22,ACTG1,DPYSL3,TUBA1A,CFL1,MAVS,HSP90AB1,AC
1,ARSD,HSPA9,NEDD4L,POLH,ATN1,CTSC,NUDC,NRBP2,USP22,ACTG1,EEF2,DPYSL3,TUBA1A,C
4L,POLH,ATN1,CTSC,NUDC,NRBP2,USP22,EEF2,TUBA1A,MAVS,HSP90AB1,ACTR2,VIM,TUBB,UB
,POLH,ATN1,CTSC,NUDC,NRBP2,USP22,EEF2,TUBA1A,MAVS,HSP90AB1,ACTR2,SPARC,TUBB,UB
,ACTB,NRBP2,USP22,ACTG1,EEF2,DPYSL3,TUBA1A,CFL1,MAVS,HSP90AB1,ACTR2,VIM,TUBB,UB
TN1,ACTB,CTSC,NRBP2,USP22,ACTG1,EEF2,DPYSL3,TUBA1A,CFL1,MAVS,HSP90AB1,ACTR2,FTH
TN1,ACTB,CTSC,NRBP2,USP22,ACTG1,EEF2,DPYSL3,TUBA1A,CFL1,MAVS,HSP90AB1,ACTR2,FTH
UDC,NRBP2,USP22,ACTG1,TUBA1A,CFL1,MAVS,HSP90AB1,HSP90AA1,ACTR2,VIM,SPARC,TUBB,
,NEDD4L,POLH,ATN1,ACTB,CTSC,NUDC,NRBP2,USP22,ACTG1,EEF2,TUBA1A,MAVS,HSP90AB1,A
OLH,ATN1,NUDC,ACTG1,EEF2,TUBA1A,MAVS,HSP90AB1,ACTR2,VIM,SPARC,TUBB,FTH1,UBC,KI
D,HSPA9,NEDD4L,POLH,ATN1,ACTB,CTSC,NUDC,NRBP2,USP22,ACTG1,EEF2,TUBA1A,CFL1,MAV
D,HSPA9,NEDD4L,POLH,ATN1,ACTB,CTSC,NUDC,NRBP2,USP22,ACTG1,EEF2,TUBA1A,CFL1,MAV
D,HSPA9,NEDD4L,POLH,ATN1,CTSC,NUDC,NRBP2,USP22,ACTG1,EEF2,DPYSL3,TUBA1A,CFL1,M
D,HSPA9,NEDD4L,POLH,ATN1,ACTB,CTSC,NUDC,NRBP2,USP22,ACTG1,EEF2,TUBA1A,CFL1,MAV
9,NEDD4L,POLH,ATN1,ACTB,CTSC,NUDC,NRBP2,USP22,ACTG1,EEF2,TUBA1A,MAVS,HSP90AB1,
CTG1,EEF2,TUBA1A,MAVS,HSP90AB1,ACTR2,VIM,SPARC,FTH1,UBC,KIDINS220,NUMA1,EIF4G2,
A9,POLH,ATN1,CTSC,NUDC,NRBP2,USP22,ACTG1,EEF2,TUBA1A,MAVS,HSP90AB1,ACTR2,TUBB,
SC,NUDC,NRBP2,USP22,EEF2,TUBA1A,MAVS,HSP90AB1,ACTR2,VIM,FTH1,UBC,NUMA1,EIF4G2,
,NEDD4L,POLH,ATN1,CTSC,NUDC,NRBP2,USP22,EEF2,TUBA1A,MAVS,ACTR2,VIM,TUBB,UBC,KID
SC,NUDC,NRBP2,USP22,EEF2,DPYSL3,TUBA1A,MAVS,HSP90AB1,HSP90AA1,ACTR2,VIM,TUBB,U
1,ARSD,HSPA9,NEDD4L,POLH,ACTB,CTSC,NUDC,NRBP2,USP22,ACTG1,EEF2,DPYSL3,TUBA1A,C
D,HSPA9,NEDD4L,POLH,ATN1,ACTB,CTSC,NUDC,NRBP2,USP22,ACTG1,EEF2,TUBA1A,CFL1,MAV

1,ARSD,HSPA9,POLH,NRBP2,USP22,ACTG1,EEF2,DPYSL3,TUBA1A,CFL1,MAVS,HSP90AB1,ACTR
D,HSPA9,NEDD4L,POLH,ACTB,CTSC,NUDC,NRBP2,USP22,ACTG1,EEF2,TUBA1A,CFL1,MAVS,HSF
L,POLH,ATN1,ACTB,CTSC,NUDC,NRBP2,USP22,ACTG1,EEF2,TUBA1A,CFL1,MAVS,HSP90AB1,AC
L,POLH,ATN1,ACTB,CTSC,NUDC,NRBP2,USP22,ACTG1,EEF2,TUBA1A,CFL1,MAVS,HSP90AB1,AC
P2,USP22,EEF2,TUBA1A,MAVS,HSP90AB1,ACTR2,TUBB,FTH1,UBC,NUMA1,ENO1,TUBA1B,PIK3
,ATN1,CTSC,NUDC,NRBP2,USP22,ACTG1,EEF2,DPYSL3,TUBA1A,MAVS,HSP90AB1,HSP90AA1,AC
1,ARSD,HSPA9,NEDD4L,POLH,ATN1,CTSC,NUDC,NRBP2,ACTG1,EEF2,MAVS,HSP90AB1,HSP90A
,POLH,NUDC,NRBP2,USP22,EEF2,TUBA1A,CFL1,MAVS,HSP90AB1,SPARC,TUBB,UBC,KIDINS220,
D,HSPA9,NEDD4L,POLH,ATN1,ACTB,CTSC,NUDC,NRBP2,USP22,ACTG1,EEF2,TUBA1A,CFL1,MAV
,POLH,ATN1,NUDC,NRBP2,USP22,ACTG1,EEF2,TUBA1A,MAVS,HSP90AB1,ACTR2,VIM,TUBB,FTH
,POLH,ATN1,NUDC,NRBP2,USP22,ACTG1,EEF2,TUBA1A,MAVS,HSP90AB1,ACTR2,VIM,TUBB,FTH
9,ATN1,CTSC,NRBP2,USP22,EEF2,TUBA1A,MAVS,TUBB,UBC,KIDINS220,NUMA1,EIF4G2,RPS6,D
NUDC,NRBP2,USP22,ACTG1,EEF2,TUBA1A,CFL1,MAVS,HSP90AB1,ACTR2,VIM,SPARC,TUBB,FTH
,POLH,ATN1,NUDC,NRBP2,USP22,ACTG1,EEF2,DPYSL3,TUBA1A,CFL1,HSP90AB1,ACTR2,SPARC,
,POLH,ATN1,NUDC,NRBP2,USP22,ACTG1,EEF2,DPYSL3,TUBA1A,CFL1,HSP90AB1,ACTR2,SPARC,
RBP2,USP22,ACTG1,EEF2,DPYSL3,TUBA1A,CFL1,MAVS,HSP90AB1,ACTR2,VIM,SPARC,TMSB4X,
L,POLH,NRBP2,USP22,EEF2,TUBA1A,MAVS,HSP90AB1,ACTR2,TUBB,UBC,KIDINS220,NUMA1,EIF
UDC,NRBP2,USP22,EEF2,TUBA1A,MAVS,HSP90AB1,ACTR2,VIM,TUBB,UBC,KIDINS220,NUMA1,E
,POLH,ATN1,ACTB,ACTG1,EEF2,MAVS,HSP90AB1,ACTR2,VIM,SPARC,FTH1,UBC,NUMA1,EIF4G2,
9,NEDD4L,POLH,ACTB,NUDC,USP22,ACTG1,EEF2,TUBA1A,CFL1,MAVS,HSP90AB1,ACTR2,VIM,T
,POLH,ATN1,NUDC,NRBP2,USP22,EEF2,TUBA1A,MAVS,HSP90AB1,ACTR2,TUBB,FTH1,UBC,KIDI
1,ARSD,HSPA9,POLH,ATN1,ACTB,CTSC,NUDC,NRBP2,USP22,ACTG1,EEF2,DPYSL3,TUBA1A,CFL
,ATN1,NUDC,NRBP2,USP22,DPYSL3,TUBA1A,HSP90AB1,ACTR2,VIM,SPARC,TUBB,FTH1,UBC,KID
4L,POLH,ACTB,CTSC,NUDC,NRBP2,USP22,ACTG1,EEF2,TUBA1A,CFL1,MAVS,HSP90AB1,ACTR2
,HSPA9,ATN1,ACTB,CTSC,USP22,EEF2,TUBA1A,MAVS,SPARC,FTH1,UBC,NUMA1,RPS6,DBN1,TU
,POLH,ACTB,CTSC,NUDC,NRBP2,USP22,ACTG1,EEF2,TUBA1A,CFL1,MAVS,HSP90AB1,ACTR2,VI
,NEDD4L,POLH,ATN1,ACTB,CTSC,NUDC,NRBP2,USP22,ACTG1,EEF2,TUBA1A,MAVS,HSP90AB1,A
TSC,NUDC,NRBP2,USP22,ACTG1,EEF2,DPYSL3,TUBA1A,CFL1,HSP90AB1,ACTR2,VIM,TUBB,UB
D,HSPA9,NEDD4L,POLH,ATN1,ACTB,NUDC,NRBP2,USP22,ACTG1,EEF2,TUBA1A,CFL1,MAVS,HSF
L,POLH,ATN1,ACTB,CTSC,NUDC,NRBP2,USP22,ACTG1,EEF2,TUBA1A,CFL1,MAVS,HSP90AB1,AC
H,ACTB,CTSC,NUDC,NRBP2,USP22,ACTG1,EEF2,TUBA1A,CFL1,MAVS,HSP90AB1,ACTR2,VIM,TU
1,ARSD,HSPA9,POLH,ATN1,CTSC,NUDC,NRBP2,ACTG1,EEF2,DPYSL3,TUBA1A,MAVS,HSP90AB1,
RPS6,TUBA1B,MAP1B,NCL,GANAB,HSP90B1,PRPF8,RPL8,ABHD2,PSAP,RPLP0,H3-3B,CANX,DDX
SPA9,NEDD4L,POLH,ATN1,NUDC,NRBP2,EEF2,MAVS,HSP90AB1,ACTR2,SPARC,UBC,KIDINS220,
9,NEDD4L,POLH,ATN1,ACTB,CTSC,NUDC,NRBP2,USP22,ACTG1,EEF2,TUBA1A,CFL1,MAVS,HSP9
LH,ATN1,CTSC,NUDC,NRBP2,USP22,ACTG1,EEF2,TUBA1A,MAVS,HSP90AB1,ACTR2,VIM,TUBB,F
1,HSPA9,CTSC,NRBP2,USP22,ACTG1,EEF2,DPYSL3,TUBA1A,CFL1,MAVS,HSP90AB1,VIM,SPARC,
UDC,NRBP2,USP22,ACTG1,EEF2,TUBA1A,MAVS,HSP90AB1,VIM,TUBB,UBC,KIDINS220,TMSB4X,
POLH,ATN1,CTSC,NUDC,NRBP2,USP22,ACTG1,EEF2,DPYSL3,TUBA1A,MAVS,HSP90AB1,HSP90A
,ATN1,NUDC,NRBP2,ACTG1,EEF2,MAVS,HSP90AB1,ACTR2,VIM,TUBB,FTH1,UBC,KIDINS220,NU
,HSPA9,POLH,ATN1,NUDC,NRBP2,ACTG1,EEF2,TUBA1A,MAVS,HSP90AB1,HSP90AA1,ACTR2,UBC
BP2,USP22,EEF2,TUBA1A,HSP90AB1,FTH1,UBC,TMSB4X,NUMA1,DCX,EIF4G2,RPS6,TUBA1B,P
ATN1,CTSC,NUDC,NRBP2,USP22,ACTG1,EEF2,DPYSL3,TUBA1A,MAVS,HSP90AB1,HSP90AA1,ACT
4L,POLH,ATN1,NUDC,NRBP2,USP22,EEF2,TUBA1A,CFL1,MAVS,HSP90AB1,ACTR2,TUBB,UBC,KID
,POLH,ATN1,NUDC,NRBP2,USP22,EEF2,TUBA1A,MAVS,HSP90AB1,TUBB,UBC,KIDINS220,NUMA1

UDC,NRBP2,USP22,ACTG1,EEF2,DPYSL3,TUBA1A,MAVS,HSP90AB1,ACTR2,VIM,SPARC,FTH1,UB
TN1,NUDC,NRBP2,EEF2,TUBA1A,HSP90AB1,ACTR2,UBC,KIDINS220,NUMA1,EIF4G2,RPS6,TUBA
EF2,MAVS,UBC,NUMA1,EIF4G2,RPS6,DBN1,FLNA,NCL,HSP90B1,RPL8,YBX1,ATP2B4,HSPA1B,PS
D,HSPA9,POLH,ATN1,NUDC,NRBP2,USP22,EEF2,TUBA1A,MAVS,HSP90AB1,TUBB,UBC,KIDINS22
TN1,ACTB,CTSC,NRBP2,USP22,ACTG1,EEF2,TUBA1A,CFL1,MAVS,HSP90AB1,ACTR2,VIM,SPARC
YSL3,TUBA1A,MAVS,HSP90AB1,VIM,UBC,NUMA1,TUBA1B,FLNA,NCL,COL6A1,HSP90B1,RPL8,YE
SC,NUDC,NRBP2,USP22,EEF2,TUBA1A,MAVS,HSP90AB1,HSP90AA1,ACTR2,TUBB,FTH1,UBC,KID
D,HSPA9,POLH,ATN1,ACTB,CTSC,NUDC,NRBP2,USP22,ACTG1,EEF2,TUBA1A,CFL1,MAVS,HSP90
D,HSPA9,POLH,ATN1,ACTB,CTSC,NUDC,NRBP2,USP22,ACTG1,EEF2,TUBA1A,CFL1,MAVS,HSP90
L,POLH,NUDC,NRBP2,USP22,EEF2,TUBA1A,MAVS,HSP90AB1,ACTR2,VIM,TUBB,FTH1,UBC,KIDIN
CTG1,EEF2,TUBA1A,CFL1,MAVS,HSP90AB1,ACTR2,VIM,SPARC,TUBB,UBC,NUMA1,EIF4G2,ENO1
TG1,EEF2,DPYSL3,TUBA1A,MAVS,HSP90AB1,ACTR2,VIM,SPARC,TUBB,FTH1,UBC,KIDINS220,NU
TG1,EEF2,DPYSL3,TUBA1A,MAVS,HSP90AB1,ACTR2,VIM,SPARC,TUBB,FTH1,UBC,KIDINS220,NU
NRBP2,USP22,ACTG1,EEF2,TUBA1A,CFL1,HSP90AB1,VIM,SPARC,TUBB,UBC,TMSB4X,NUMA1,EI
P2,USP22,ACTG1,EEF2,TUBA1A,MAVS,HSP90AB1,ACTR2,VIM,TUBB,UBC,KIDINS220,NUMA1,EIF
CTSC,NUDC,NRBP2,USP22,EEF2,TUBA1A,MAVS,HSP90AB1,HSP90AA1,ACTR2,TUBB,UBC,NUMA1
,POLH,ATN1,EEF2,MAVS,HSP90AB1,ACTR2,SPARC,FTH1,NUMA1,EIF4G2,RPS6,PIK3R2,FLNA,MA
BP2,USP22,TUBA1A,MAVS,HSP90AB1,ACTR2,VIM,SPARC,UBC,KIDINS220,NUMA1,EIF4G2,RPS6,E
,ACTB,CTSC,NUDC,NRBP2,USP22,ACTG1,EEF2,TUBA1A,CFL1,MAVS,ACTR2,VIM,TUBB,KIDINS22
1,ARSD,HSPA9,POLH,ATN1,ACTB,CTSC,NUDC,NRBP2,USP22,ACTG1,EEF2,DPYSL3,TUBA1A,CFL
TN1,ACTB,CTSC,NRBP2,USP22,ACTG1,EEF2,TUBA1A,CFL1,MAVS,HSP90AB1,ACTR2,VIM,SPARC
,POLH,ATN1,NUDC,NRBP2,USP22,EEF2,TUBA1A,MAVS,UBC,KIDINS220,NUMA1,RPS6,ENO1,TUB
GANAB,HSP90B1,NEFL,RPL8,ABHD2,PSAP,RPLP0,H3-3B,CANX,DDX5,ENO2,SOX4,RPL13A,YWHA
,ACTB,NUDC,NRBP2,USP22,ACTG1,EEF2,TUBA1A,CFL1,MAVS,HSP90AB1,ACTR2,TUBB,FTH1,UB
,POLH,ATN1,NUDC,NRBP2,USP22,EEF2,TUBA1A,MAVS,HSP90AB1,ACTR2,VIM,TUBB,UBC,KIDINS
,NEDD4L,POLH,ATN1,ACTB,CTSC,NRBP2,USP22,ACTG1,EEF2,TUBA1A,MAVS,HSP90AB1,FTH1,U
,NEDD4L,POLH,ATN1,ACTB,CTSC,NRBP2,USP22,ACTG1,EEF2,TUBA1A,MAVS,HSP90AB1,FTH1,U
A9,POLH,ATN1,NRBP2,USP22,EEF2,TUBA1A,MAVS,HSP90AB1,FTH1,UBC,KIDINS220,NUMA1,EIF
DC,NRBP2,EEF2,HSP90AB1,ACTR2,UBC,NUMA1,EIF4G2,RPS6,ENO1,NCL,HSP90B1,PRPF8,RPL
TN1,NUDC,NRBP2,USP22,EEF2,TUBA1A,HSP90AB1,ACTR2,SPARC,UBC,KIDINS220,NUMA1,EIF4G
UDC,USP22,EEF2,TUBA1A,MAVS,HSP90AB1,FTH1,UBC,NUMA1,RPS6,ENO1,TUBA1B,NCL,FTL,HS
VS,HSP90AB1,VIM,UBC,NUMA1,EIF4G2,RPS6,PIK3R2,NCL,HSP90B1,FADS1,RPL8,FBXL4,YBX1,U
H,ACTB,CTSC,NUDC,NRBP2,USP22,ACTG1,EEF2,TUBA1A,CFL1,MAVS,HSP90AB1,ACTR2,VIM,TUE
MAVS,HSP90AB1,UBC,NUMA1,EIF4G2,RPS6,TUBA1B,PIK3R2,NCL,GANAB,HSP90B1,PRPF8,FBXL
SP90AB1,VIM,UBC,KIDINS220,NUMA1,EIF4G2,RPS6,ENO1,TUBA1B,PIK3R2,NCL,SLC9A7,HSP90
P90AB1,VIM,SPARC,FTH1,UBC,KIDINS220,NUMA1,EIF4G2,RPS6,ENO1,TUBA1B,NCL,FTL,HSP90E
OLH,ATN1,CTSC,NRBP2,USP22,EEF2,DPYSL3,TUBA1A,MAVS,HSP90AB1,ACTR2,VIM,SPARC,TUB
SP22,ACTG1,EEF2,DPYSL3,TUBA1A,MAVS,ACTR2,VIM,TUBB,UBC,KIDINS220,TMSB4X,NUMA1,R
SC,NRBP2,USP22,EEF2,TUBA1A,CFL1,MAVS,HSP90AB1,ACTR2,VIM,SPARC,FTH1,UBC,KIDINS22
22,ACTG1,EEF2,TUBA1A,HSP90AB1,ACTR2,VIM,TUBB,UBC,KIDINS220,NUMA1,EIF4G2,RPS6,EN
BA1A,MAVS,HSP90AB1,HSP90AA1,VIM,TUBB,UBC,NUMA1,RPS6,TUBA1B,PIK3R2,NCL,SLC9A7,H
LH,ATN1,CTSC,NUDC,NRBP2,USP22,ACTG1,EEF2,TUBA1A,MAVS,HSP90AB1,ACTR2,FTH1,UBC,N
EEF2,TUBA1A,CFL1,MAVS,HSP90AB1,VIM,SPARC,TUBB,FTH1,UBC,TMSB4X,NUMA1,EIF4G2,RPS
SPARC,NUMA1,EIF4G2,RPS6,TUBA1B,NCL,GANAB,HSP90B1,PRPF8,TMF1,ABHD2,PSAP,RPLP0,H
C,NRBP2,ACTG1,EEF2,TUBA1A,MAVS,HSP90AB1,FTH1,UBC,KIDINS220,NUMA1,EIF4G2,RPS6,EN

CTG1,EEF2,TUBA1A,CFL1,MAVS,HSP90AB1,ACTR2,VIM,TUBB,FTH1,UBC,NUMA1,EIF4G2,ENO1,T
P22,ACTG1,EEF2,TUBA1A,CFL1,MAVS,ACTR2,VIM,TUBB,TMSB4X,NUMA1,EIF4G2,RPS6,ENO1,TU
L,HSP90B1,NEFM,ABHD2,HSPA1B,PSAP,RPLP0,H3-3B,CANX,DDX5,ENO2,RPL13A,KAT6A,YWHAE
EF2,TUBA1A,MAVS,HSP90AB1,ACTR2,VIM,UBC,NUMA1,EIF4G2,ENO1,TUBA1B,PIK3R2,NCL,HSP
DC,NRBP2,USP22,ACTG1,EEF2,TUBA1A,CFL1,MAVS,HSP90AB1,ACTR2,VIM,TUBB,FTH1,UBC,TM
L1,HSP90AB1,ACTR2,VIM,SPARC,TMSB4X,NUMA1,RPS6,ENO1,TUBA1B,NCL,ALDOA,GANAB,CO
2,TUBA1A,MAVS,HSP90AB1,VIM,UBC,KIDINS220,TMSB4X,NUMA1,EIF4G2,RPS6,ENO1,TUBA1B,
A1A,MAVS,ACTR2,VIM,SPARC,UBC,KIDINS220,NUMA1,RPS6,TUBA1B,PIK3R2,FLNA,MAP1B,NCL,
SPARC,FTH1,KIDINS220,NUMA1,ENO1,TUBA1B,NCL,FTL,HSP90B1,FADS1,UGT8,FBXL4,CES3,P
C,NRBP2,USP22,ACTG1,EEF2,DPYSL3,MAVS,HSP90AB1,FTH1,UBC,KIDINS220,TMSB4X,NUMA1,
C,NRBP2,USP22,ACTG1,EEF2,DPYSL3,MAVS,HSP90AB1,FTH1,UBC,KIDINS220,TMSB4X,NUMA1,
FL1,HSP90AB1,VIM,SPARC,TUBB,UBC,TMSB4X,NUMA1,RPS6,ENO1,TUBA1B,NCL,SLC9A7,ALD
BA1A,MAVS,HSP90AB1,VIM,TUBB,NUMA1,EIF4G2,TUBA1B,NCL,GANAB,HSP90B1,FBXL4,ABHD2,
IM,SPARC,TUBB,UBC,DCX,ENO1,FLNA,NCL,GANAB,RPL8,UBA1,ATP2B4,CALR,TUBB2B,PSAP,H3
MAVS,NUMA1,EIF4G2,RPS6,TUBA1B,NCL,HSP90B1,RPL8,UBA1,PSAP,RPLP0,H3-3B,CANX,RPL13
CL,HSP90B1,RPL8,UBA1,ABHD2,RPLP0,PTMA,DDX5,RPL13A,SEC62,DVL3,RPL13,RPL7,RPL5,SF
PARC,NUMA1,FLNA,MAP1B,NCL,HSP90B1,UBA1,TMF1,PSAP,DDX5,DVL3,YWHAE,RPL13,P4HB,R
RPF8,RPL8,UBA1,TMF1,PSAP,H3-3B,CANX,DDX5,RPL13A,DVL3,YWHAE,ALDH6A1,SPTBN1,TMBI
VS,HSP90AB1,ACTR2,VIM,SPARC,TUBB,UBC,NUMA1,EIF4G2,ENO1,TUBA1B,NCL,ALDOA,GANAB,
D,HSPA9,POLH,ATN1,ACTB,NUDC,ACTG1,EEF2,TUBA1A,CFL1,MAVS,HSP90AB1,ACTR2,VIM,SPA
,CFL1,MAVS,UBC,KIDINS220,NUMA1,EIF4G2,RPS6,TUBA1B,NCL,HSP90B1,FBXL4,YBX1,ABHD2,
TG1,EEF2,TUBA1A,MAVS,UBC,KIDINS220,NUMA1,EIF4G2,RPS6,TUBA1B,NCL,HSP90B1,PRPF8,F
SP90AA1,TUBB,UBC,NUMA1,RPS6,TUBA1B,NCL,HSP90B1,FBXL4,ATP2B4,ABHD2,HSPA1B,PSAP
UDC,NRBP2,USP22,ACTG1,EEF2,TUBA1A,CFL1,MAVS,HSP90AB1,ACTR2,VIM,UBC,TMSB4X,NU
UBC,NUMA1,RPS6,TUBA1B,NCL,RPL8,YBX1,HSPA1B,H3-3B,DDX5,RPL13A,DVL3,RPL13,RPL7,ID
20,NUMA1,EIF4G2,RPS6,ENO1,TUBA1B,NCL,ALDOA,COL6A1,HSP90B1,FADS1,PRPF8,FBXL4,U
BC,DCX,FLNA,NCL,RPL8,ATP2B4,TUBB2B,PSAP,H3-3B,KAT6A,BSG,ATP5F1B,CALM3,SEPTIN2,A
,MAVS,VIM,SPARC,FTH1,UBC,KIDINS220,NUMA1,EIF4G2,RPS6,DBN1,ENO1,TUBA1B,MAP1B,NC
,PIK3R2,NCL,HSP90B1,UBA1,ATP2B4,PSAP,H3-3B,CANX,CLU,DDX5,DVL3,YWHAE,ALDH6A1,IDS,
TUBA1A,MAVS,UBC,NUMA1,RPS6,TUBA1B,NCL,PRPF8,RPL8,YBX1,UBA1,HSPA1B,H3-3B,DDX5,R
A1A,MAVS,KIDINS220,NUMA1,EIF4G2,RPS6,TUBA1B,NCL,HSP90B1,YBX1,ABHD2,PSAP,H3-3B,C
1,TUBB,UBC,NUMA1,EIF4G2,RPS6,TUBA1B,PIK3R2,NCL,SLC9A7,HSP90B1,UBA1,ATP2B4,ABH
AB1,ACTR2,SPARC,TUBB,FTH1,KIDINS220,NUMA1,EIF4G2,ENO1,TUBA1B,PIK3R2,MAP1B,NCL,
S6,TUBA1B,NCL,HSP90B1,RPL8,PSAP,PGK1,H3-3B,CANX,DDX5,DVL3,FASN,RPL5,FSCN1,PTMS,
M,NUMA1,TUBA1B,NCL,HSP90B1,UBA1,TMF1,HSPA1B,PSAP,H3-3B,CANX,DVL3,YWHAE,ALDH6A
EF2,TUBA1A,CFL1,MAVS,NUMA1,EIF4G2,RPS6,TUBA1B,PIK3R2,NCL,GANAB,HSP90B1,PRPF8,FE
,DPYSL3,TUBA1A,MAVS,HSP90AB1,SPARC,TUBB,FTH1,UBC,KIDINS220,TMSB4X,EIF4G2,RPS6,
AVS,VIM,UBC,NUMA1,TUBA1B,PIK3R2,NCL,GANAB,HSP90B1,PRPF8,UBA1,CALR,ABHD2,PSAP,
BA1A,MAVS,UBC,KIDINS220,NUMA1,EIF4G2,RPS6,ENO1,TUBA1B,NCL,SLC9A7,HSP90B1,FBXL4,
,UBC,NUMA1,RPS6,TUBA1B,NCL,RPL8,YBX1,UBA1,H3-3B,DDX5,RPL13,IDS,RPL5,NPM1,EIF3C,
1,ACTR2,UBC,NUMA1,ENO1,TUBA1B,FLNA,NCL,GANAB,FTL,HSP90B1,UBA1,HSPA1B,PSAP,H3-3
SP90AB1,SPARC,UBC,NUMA1,TUBA1B,PIK3R2,NCL,HSP90B1,PRPF8,UBA1,ATP2B4,ABHD2,PSA
UMA1,RPS6,TUBA1B,PIK3R2,NCL,RPL8,YBX1,PSAP,H3-3B,CLU,DVL3,YWHAE,RPL13,RPL5,GDI1,
SP90B1,PRPF8,UBA1,ABHD2,HSPA1B,PSAP,PGK1,H3-3B,CLU,PTMA,DDX5,SEC62,DVL3,ALDH6A1
AVS,UBC,NUMA1,TUBA1B,NCL,RPL8,UBA1,PSAP,H3-3B,CANX,SEC62,DVL3,SPTBN1,GNB1,P4HE

1,FBXL4,ABHD2,H3-3B,CANX,DDX5,KAT6A,YWHAE,RPL5,PTMS,NPM1,UBB,AFF4,TRIM28,MATR3
BN1,TUBA1B,NCL,UBA1,ABHD2,H3-3B,CANX,PTMA,DDX5,GNB1,PTMS,MARCKS,PLD3,ATP5F1A,
'S,HSP90AB1,SPARC,TUBB,FTH1,NUMA1,RPS6,TUBA1B,PIK3R2,NCL,SLC9A7,HSP90B1,RPL8,FB
S,HSP90AB1,KIDINS220,NUMA1,NCL,HSP90B1,FADS1,UGT8,FBXL4,TMF1,ABHD2,HSPA1B,COL4
SP22,ACTG1,DPYSL3,TUBA1A,MAVS,HSP90AB1,UBC,NUMA1,TUBA1B,FLNA,NCL,COL6A1,HSP90
1,ACTR2,VIM,TUBB,UBC,EIF4G2,TUBA1B,NCL,HSP90B1,RPL8,ABHD2,PSAP,H3-3B,DDX5,DVL3,
.,HSP90B1,UGT8,H3-3B,CANX,ALDH6A1,SEPTIN2,FSCN1,NPM1,EIF3C,VCP,UBB,AFF4,TRIM28,SF
JBA1B,MAP1B,NCL,SLC9A7,HSP90B1,NEFM,ATP2B4,ABHD2,PSAP,H3-3B,CANX,DVL3,SPTBN1,G
SPA1B,H3-3B,PTMA,DDX5,RPL13A,IDS,NPM1,EIF3C,ATP5F1A,EIF4B,VCP,UBB,TRIM28,MATR3,SF
3A1A,CFL1,MAVS,KIDINS220,NUMA1,EIF4G2,RPS6,TUBA1B,PIK3R2,NCL,GANAB,HSP90B1,RPL8,
NUMA1,TUBA1B,NCL,HSP90B1,FBXL4,UBA1,HSPA1B,PSAP,H3-3B,CANX,DDX5,DVL3,ALDH6A1,P4
NS220,NUMA1,EIF4G2,RPS6,TUBA1B,PIK3R2,NCL,GANAB,HSP90B1,CES3,PSAP,RPLP0,H3-3B,C
TG1,EEF2,CFL1,MAVS,ACTR2,VIM,SPARC,UBC,NUMA1,DBN1,ENO1,PIK3R2,NCL,ALDOA,GANAB,
20,NUMA1,ENO1,TUBA1B,NCL,HSP90B1,FADS1,H3-3B,CANX,PTMA,DDX5,DVL3,YWHAE,FASN,FS
KIDINS220,NUMA1,RPS6,FLNA,NCL,HSP90B1,ATP2B4,TMF1,COL4A1,RPLP0,H3-3B,CANX,DDX5,
JMA1,EIF4G2,TUBA1B,PIK3R2,NCL,HSP90B1,RPL8,YBX1,ABHD2,CES3,PSAP,RPLP0,H3-3B,CAN
IA1,RPS6,NCL,RPL8,H3-3B,DDX5,RPL13A,DVL3,RPL13,IDS,RPL5,PTMS,NPM1,EIF3C,ATP5F1A,E
NCL,HSP90B1,PRPF8,RPL8,ATP2B4,TUBB2B,RPLP0,H3-3B,DDX5,KAT6A,ATP5F1B,GDI1,NPM1,H
EIF4G2,RPS6,ENO1,TUBA1B,PIK3R2,NCL,HSP90B1,FADS1,ATP2B4,PGK1,RPLP0,DDX5,SEC62,K
,FBXL4,PSAP,H3-3B,CANX,DDX5,ALDH6A1,SEPTIN2,FSCN1,NPM1,ATP5F1A,UBB,AFF4,YWHAG,SF
1,TUBA1B,NCL,H3-3B,CANX,SPTBN1,MYH9,PTMS,SPTAN1,GSTP1,NPM1,COL18A1,ATP5F1A,VCF
31,UBA1,PSAP,H3-3B,DDX5,GDI1,NPM1,ATP5F1A,TRIM28,MATR3,SPEN,NONO,SET,UCHL1,RPS17
,ACTR2,VIM,TUBB,EIF4G2,TUBA1B,NCL,YBX1,ABHD2,PSAP,H3-3B,DDX5,DVL3,YWHAE,NPM1,EIF
,UBC,NUMA1,TUBA1B,NCL,COL6A1,HSP90B1,H3-3B,CANX,DDX5,DVL3,YWHAE,TMBIM6,MARCK
I,SPARC,TUBB,TMSB4X,NUMA1,DBN1,NCL,GANAB,HSP90B1,PRPF8,RPL8,UBA1,ATP2B4,TUBB2
3A1A,CFL1,MAVS,UBC,NUMA1,RPS6,TUBA1B,PIK3R2,NCL,GANAB,HSP90B1,YBX1,TMF1,ABHD2,
'M1,ATP5F1A,TRIM28,MATR3,YWHAG,SSBP2,HMGB1,DNAJA1,MAP4,CTNNA1,SCARB2,XRCC5,KI
,HSP90B1,ATP2B4,RPLP0,DDX5,KAT6A,DVL3,DDX17,ALDH6A1,ATP5F1B,MYH9,SEPTIN2,RPL5,FS
DLH,CTSC,NRBP2,USP22,EEF2,MAVS,HSP90AB1,FTH1,UBC,EIF4G2,ENO1,PIK3R2,NCL,GANAB,F
GDI1,FSCN1,SPTAN1,MARCKS,NPM1,ATP5F1A,MARCKSL1,UBB,TRIM28,MATR3,SSBP2,CHD3,VC
,NCL,H3-3B,CANX,DDX5,DVL3,PTMS,NPM1,ATP5F1A,MARCKSL1,TRIM28,MATR3,NONO,EGLN1,
31,VIM,FTH1,TMSB4X,NUMA1,RPS6,TUBA1B,FLNA,NCL,HSP90B1,FBXL4,YBX1,PSAP,H3-3B,CAN
IMA1,TUBA1B,NCL,HSPA1B,H3-3B,DDX5,DVL3,NPM1,ATP5F1A,VCP,MARCKSL1,UBB,TRIM28,MA
PGK1,H3-3B,CANX,DDX5,DVL3,SPTBN1,MYH9,SEPTIN2,FSCN1,SPTAN1,NPM1,COL18A1,VCP,TRI
C,NUMA1,NCL,FTL,RPL8,ATP2B4,TMF1,PSAP,PGK1,H3-3B,CANX,DVL3,P4HB,NPM1,HSPD1,COL1
R2,VIM,NUMA1,TUBA1B,FLNA,FTL,HSP90B1,UBA1,ABHD2,PSAP,CANX,RPL13A,DVL3,IDS,RPL5,N
B,FTH1,KIDINS220,TMSB4X,RPS6,DBN1,ENO1,TUBA1B,MAP1B,SLC9A7,ALDOA,FTL,NEFL,NEFM,
MA1,TUBA1B,NCL,GANAB,HSPA1B,H3-3B,CANX,DDX5,DVL3,TMBIM6,MARCKS,NPM1,ATP5F1A,V
EF2,MAVS,HSP90AB1,UBC,KIDINS220,NUMA1,NCL,ALDOA,FADS1,UGT8,ABHD2,HSPA1B,PSAP,PG
AB1,VIM,SPARC,TUBB,TMSB4X,NUMA1,DBN1,NCL,ALDOA,GANAB,HSP90B1,PRPF8,RPL8,UBA1,
TUBB,FTH1,NUMA1,RPS6,TUBA1B,PIK3R2,NCL,SLC9A7,HSP90B1,FBXL4,COL4A1,H3-3B,CANX,P
1B,FLNA,NCL,HSP90B1,H3-3B,DVL3,SPTBN1,SPTAN1,NPM1,COL18A1,MARCKSL1,TRIM28,MATF
3A1,ABHD2,PSAP,PGK1,RPLP0,H3-3B,DDX5,DVL3,YWHAE,ALDH6A1,SPTBN1,IDS,SPTAN1,NPM1,C
L1,MAVS,HSP90AB1,ACTR2,VIM,UBC,NUMA1,RPS6,TUBA1B,NCL,GANAB,TMF1,PSAP,PGK1,H3-3
I,EIF4G2,TUBA1B,NCL,YBX1,PSAP,H3-3B,DDX5,YWHAE,MYH9,SPTAN1,NPM1,EIF3C,ATP5F1A,VC

A1B,NCL,HSPA1B,H3-3B,DDX5,FSCN1,MARCKS,NPM1,ATP5F1A,VCP,MARCKSL1,UBB,TRIM28,M
RCKSL1,TRIM28,MRC2,SSBP2,HMGB1,XRCC5,CD81,RBBP4,HNRNPC,LRPPRC,KHDRBS1,SFPQ,
,HSP90AB1,ACTR2,VIM,NUMA1,NCL,GANAB,PSAP,PGK1,H3-3B,CANX,DDX5,DVL3,YWHAE,NPM1,
AN1,NPM1,TRIM28,MATR3,SPEN,NONO,SSBP2,HMGB1,POLR2A,ANXA2,TIMP2,HNRNPU,CNN3,XF
MA1,RPS6,TUBA1B,MAP1B,NCL,FBXL4,ATP2B4,PSAP,H3-3B,CANX,PTMA,SPTBN1,GNB1,FSCN1,F
TMS,NPM1,VCP,TRIM28,MATR3,SSBP2,CYB5R3,AP2B1,RPL29,C1ORF43,HNRNPC,SFPQ,DDX3X,
-3B,CANX,ALDH6A1,SEPTIN2,NPM1,AFF4,TRIM28,SET,BASP1,HMGB1,MARCHF6,MAP4,SF3B2,HI
1,SPTAN1,NPM1,MARCKSL1,TRIM28,MATR3,FN1,SPEN,NONO,SET,SSBP2,HMGB1,POLR2A,ANX
TRIM28,MATR3,NONO,SSBP2,POLR2A,HNRNPU,SLC16A7,RBBP4,KHDRBS1,SFPQ,H1-0,HNRNP
-3B,CANX,DDX5,SOX4,SPTBN1,PTMS,NPM1,COL18A1,ATP5F1A,VCP,TRIM28,MATR3,SPEN,NONO
3,NPM1,VCP,TRIM28,MATR3,RPL37A,EGLN1,MRC2,SSBP2,CHD3,HMGB1,MAP4,HNRNPU,RPL29,
3,TUBB2B,PSAP,CANX,SLC2A3,DDX17,ATP5F1B,GDI1,SPTAN1,NPM1,ATP5F1A,TRIM28,MATR3,S
,IDS,NPM1,COL18A1,UBB,TRIM28,MATR3,LRRC28,SPEN,NONO,EGLN1,HMGB1,ITGB1,BTF3,VC
B4,H3-3B,DDX5,ATP5F1B,GDI1,NPM1,VCP,TRIM28,MATR3,SPEN,NONO,SSBP2,POLR2A,VCL,SF
NO2,DVL3,NPM1,ATP5F1A,UBB,YWHAG,LRRC28,SPEN,EGLN1,SSBP2,SF3B2,XRCC5,HK1,PSMD
A7,FTL,HSP90B1,UBA1,CES3,PSAP,PGK1,RPLP0,H3-3B,CANX,DDX5,DVL3,RPL13,BSG,ALDH6A1
1,TRIM28,MATR3,NONO,SSBP2,POLR2A,SF3B2,GOLGA8A,HNRNPU,HNRNPA0,KPNB1,RBBP4,F
X,DDX5,DVL3,NPM1,VCP,TRIM28,SPEN,NONO,SSBP2,MAP4,HNRNPU,RPL17,C1ORF43,CLSTN1,J
TP5F1B,IDS,NPM1,COL18A1,ATP5F1A,TRIM28,MATR3,LRRC28,SPEN,NONO,DDB1,DCHS1,EGL
,ATP5F1A,MARCKSL1,LDHB,HMGB1,BTF3,ANXA2,CCT7,RBBP4,HNRNPC,SFPQ,CDH2,ACTN4,H1
1,NCL,PSAP,H3-3B,DVL3,YWHAE,MARCKS,NPM1,EIF3C,ATP5F1A,MARCKSL1,DCHS1,SET,EGLN
-3B,DVL3,TMBIM6,SMG1,NPM1,ATP5F1A,UBB,TRIM28,MATR3,NONO,EGLN1,SSBP2,HMGB1,HN
P,FSCN1,PLD3,MAPT,PDIA3,UCHL1,RPL10,MAP4,CPE,GAP43,RPL9,NLGN2,ATP1A1,EIF5,RTN3,P
NB1,SEPTIN2,GDI1,SPTAN1,ATP5F1A,CKB,BASP1,NCAM1,GAP43,WDR1,RTN3,PNN,SPTBN2,GLG
8,RPL8,ATP2B4,HSPA1B,H3-3B,DDX5,ATP5F1B,GDI1,NPM1,VCP,TRIM28,MATR3,SPEN,NONO,H
AA1,ACTR2,EIF4G2,RPS6,TUBA1B,PIK3R2,MAP1B,SLC9A7,FADS1,FBXL4,YBX1,ABHD2,CES3,PT
90B1,PRPF8,RPL8,ATP2B4,TMF1,HSPA1B,TUBB2B,PSAP,H3-3B,CANX,SLC2A3,DDX17,SPTBN1,
P,FSCN1,PLD3,PDIA3,UCHL1,RPL10,MAP4,CPE,GAP43,RPL9,NLGN2,ATP1A1,EIF5,RTN3,PFN1,T
RPL10,AKAP12,CPE,RTN4,PJA2,RPL9,EIF5,ATP2A2,RTN3,PFN1,SPTBN2,CDH2,RPS15,MAGED2,R
NPU,SRSF4,HNRNPC,H1-0,COL6A2,APLP2,TARDBP,HNRNPK,LUC7L3,DDX39B,HNRNPL,ALKBH5
GAP43,NLGN2,TKT,CTNNB1,H1-10,PEA15,TAF15,PTBP1,POLR2B,SOD1,ARNT2,TP53BP1,ADAR,
S,VCP,MATR3,SSBP2,HMGB1,CHD4,GLG1,SFPQ,DDX3X,RPS7,APLP2,HNRNPDL,HNRNPK,CKAP5,
RTN3,SPTBN2,TKT,RPS15,RPS5,APLP1,MTSS2,MAP2,MAP1A,PGRMC1,NUCB1,TOMM20,ACO2,G
NCL,HSP90B1,PRPF8,RPL8,ATP2B4,TMF1,HSPA1B,TUBB2B,PSAP,RPLP0,H3-3B,CANX,SLC2A3,D
CL,H3-3B,PTMS,MARCKS,NPM1,COL18A1,UBB,TRIM28,SPEN,NONO,SSBP2,ANXA2,MAP4,RBBP4
GAP43,NLGN2,TKT,CTNNB1,H1-10,PEA15,TAF15,AEBP1,PTBP1,POLR2B,SOD1,ARNT2,TP53BP1,
,CPE,RTN4,RPL9,EIF5,RTN3,SPTBN2,TKT,RPS15,RPS5,APLP1,MTSS2,DDOST,MAP2,MAP1A,PGR
,CPE,RTN4,RPL9,EIF5,RTN3,SPTBN2,TKT,RPS15,RPS5,APLP1,MTSS2,DDOST,MAP2,MAP1A,PGR
2,CPE,RTN4,PJA2,RPL9,EIF5,ATP2A2,RTN3,SPTBN2,RPS15,MAGED2,RPS5,CTNNB1,ANXA6,MTS
PA1B,TUBB2B,H3-3B,SLC2A3,DDX17,MARCKS,NPM1,VCP,AFF4,TRIM28,MATR3,SPEN,HMGB1,P
ATP5F1A,TRIM28,DDB1,DNAJA1,MAP4,NFE2L1,TPM4,BRD3,RPL24,RBBP4,PNN,KHDRBS1,XPO1,
CL,MAP4,HNRNPU,CNN3,XRCC5,GAP43,NES,HNRNPC,KHDRBS1,SLC3A2,SFPQ,H1-0,IK,COL6A2
NX,DVL3,SLC2A3,GDI1,NPM1,PLD3,ATP5F1A,UBB,TRIM28,EGLN1,HMGB1,STMN1,HNRNPU,SCA
1,HSP90AA1,ACTR2,FTH1,EIF4G2,RPS6,TUBA1B,PIK3R2,MAP1B,SLC9A7,FTL,FADS1,UGT8,FBX
E,PJA2,MSN,KPNB1,ATP1A1,PFN1,SPTBN2,TKT,RPS15,MAGED2,TLN1,GJA1,CTNNB1,ANXA6,AT

TIN2,COL18A1,CSDE1,MARCKSL1,VCL,CDC42BPB,HNRNPA0,LPCAT1,PJA2,CHD4,CDH6,PEBP1,J
,RPL9,ATP1A1,EIF5,RTN3,TKT,MAGED2,RPS5,TLN1,MAP2,COTL1,MAP1A,PGRMC1,NUCB1,TOM
2,SPTBN2,RPS15,MAGED2,RPS5,MYH10,MAP2,STIP1,MAP1A,TMEM132A,NUCB1,TOMM20,RPN2
CPE,RTN4,KPNB1,GAP43,NLGN2,ATP1A1,EIF5,RTN3,PFN1,CDH2,ANXA6,CADM4,ATP1A2,L1CAM
,GAP43,NLGN2,ATP1A1,EIF5,RTN3,SPTBN2,CDH2,TLN1,ANXA6,MYH10,MAP2,PGRMC1,TOMM2
1,SOD1,ARNT2,KDM3B,TP53BP1,ADAR,STAT3,CAPN2,BAZ1B,SNRPD3,SOX2,SAP18,PSAT1,HTAT
AEBP1,KDM3B,TP53BP1,ADAR,PRMT1,BAZ1B,PPM1G,PSMD8,SNRPD3,STRN4,PSMD4,CHMP4B,
DVL3,YWHAE,ALDH6A1,RPL5,SNRNP200,YWHAG,VAPA,RPS4X,DNAJA1,CYB5R3,PAFAH1B1,CTNN
AM1,RPL10,MAP4,CPE,RTN4,PJA2,MSN,KPNB1,ATP1A1,EIF5,RTN3,TKT,RPS15,MAGED2,APLP1,
RPF8,RPL8,ATP2B4,TMF1,HSPA1B,TUBB2B,PSAP,H3-3B,SLC2A3,DDX17,ATP5F1B,MARCKS,SMG
PTBP1,SOD1,ARNT2,KDM3B,TP53BP1,ADAR,PRMT1,EP300,BAZ1B,MAPRE2,APEX1,SNRPD3,SOX
PTBP1,SOD1,ARNT2,KDM3B,TP53BP1,ADAR,PRMT1,BAZ1B,MAPRE2,APEX1,SNRPD3,SOX2,SAP1
KDM3B,TP53BP1,ADAR,PRMT1,BAZ1B,PPM1G,PSMD8,SNRPD3,STRN4,PSMD4,CHMP4B,SART1,
2,KDM3B,TP53BP1,ADAR,STAT3,BAZ1B,SNRPD3,SOX2,SAP18,PSAT1,HTATSF1,MECP2,SUMO1,
IOD2,CADM3,IGLON5,RAB3A,NAPB,CNTN1,ATP6V0A1,PPP3CA,SLC1A3,GNAO1
TIN2,IDS,USP11,COL18A1,CSDE1,MARCKSL1,VCL,CDC42BPB,HNRNPA0,LPCAT1,PJA2,CHD4,SR
F15,PTBP1,ADAR,EP300,BAZ1B,MAPRE2,APEX1,SNRPD3,SOX2,SAP18,ELAVL3,PSAT1,HMGA1,HT
F15,PTBP1,SOD1,ADAR,PRMT1,EP300,SF1,BAZ1B,MAPRE2,APEX1,SNRPD3,SOX2,SAP18,ELAVL3
3,TKT,RPS15,APLP1,TLN1,CTNNB1,PEA15,PGRMC1,ACO2,CNTNAP2,MARS1,FKBP8,MAPRE2,BR
OMM20,PRDX5,ACO2,MARS1,ATP5PB,FKBP8,ATP6V1B2,STMN3,STXBP1,SYP,SV2A,TMOD2,AP2
AM1,RPL10,MAP4,CPE,RTN4,PJA2,MSN,KPNB1,ATP1A1,EIF5,RTN3,TKT,RPS15,MAGED2,APLP1,
I3B,TP53BP1,ADAR,PRMT1,BAZ1B,PSMD8,STRN4,PSMD4,CHMP4B,SAP18,HTATSF1,MECP2,ADN
,H1-10,TAF15,PTBP1,SOD1,TP53BP1,ADAR,BAZ1B,MAPRE2,APEX1,SAP18,ELAVL3,HMGA1,HTAT
N3,SPTBN2,CDH2,ANXA6,MAP2,TOMM20,PRDX5,ACO2,MARS1,ATP5PB,SCAF11,FKBP8,ATP6V1
2,MSN,ATP1A1,PFN1,SPTBN2,TKT,RPS15,MAGED2,TLN1,GJA1,CTNNB1,ANXA6,ATP1A2,MAP2,C
H10,MAP2,TMEM132A,RTN1,G3BP2,FKBP8,STMN3,FBRSL1,VDAC1,SF1,BRD4,MAP6,LINGO1,D
OMM20,GAS7,CNTNAP2,MARS1,ATP6V1B2,STXBP1,SYP,CD99,SV2A,SEPTIN7,TMOD2,SEPTIN5,
VL3,GDI1,TRIM28,SSBP2,DNAJA1,CYB5R3,SF3B2,HNRNPU,RPL29,RPL17,RPL14,KHDRBS1,SFP
,GJA1,ARID1A,TAGLN,NARS1,CKAP5,TOMM20,LAMC1,CBX1,MAP2K2,ATP6V1B2,LSS,LONP1,ZS
F5,ATP2A2,SPTBN2,MAGED2,MTSS2,DDOST,MYH10,MAP2,TMEM132A,RPN2,RTN1,G3BP2,HSPE
5,SPTBN1,SEPTIN2,IDS,COL18A1,CSDE1,AFF4,DNAJA1,HNRNPA0,CTSD,LPCAT1,PJA2,CHD4,ATP
5,CNTNAP2,MSMO1,CD99,SV2A,TMOD2,CADM3,MSH6,ANK2,USP47,ALDH7A1,NDRG4,ALCAM,N
,PSMD4,SAP18,HTATSF1,MECP2,ADNP,RTCB,AKAP8L,SOX9,FBL,RBM17,PNMA2,ACIN1,UBTF

A1A,MAVS,HSP90AB1,HSP90AA1,ACTR2,VIM,SPARC,TUBB,FTH1,UBC,KIDINS220,TMSB4X,NUMA
90AB1,ACTR2,VIM,SPARC,TUBB,FTH1,UBC,KIDINS220,NUMA1,DCX,EIF4G2,RPS6,DBN1,ENO1,TU
90AB1,HSP90AA1,ACTR2,SPARC,FTH1,UBC,KIDINS220,NUMA1,EIF4G2,RPS6,DBN1,ENO1,TUBA1
90AB1,ACTR2,SPARC,TUBB,FTH1,UBC,KIDINS220,NUMA1,DCX,EIF4G2,RPS6,DBN1,ENO1,TUBA1
FL1,MAVS,HSP90AB1,HSP90AA1,ACTR2,VIM,SPARC,TUBB,FTH1,UBC,KIDINS220,TMSB4X,NUMA
AVS,HSP90AB1,HSP90AA1,ACTR2,TUBB,FTH1,UBC,KIDINS220,TMSB4X,NUMA1,DCX,EIF4G2,RP
90AB1,ACTR2,VIM,SPARC,TUBB,FTH1,UBC,KIDINS220,TMSB4X,NUMA1,DCX,EIF4G2,RPS6,DBN1
IAVS,HSP90AB1,ACTR2,VIM,TUBB,FTH1,UBC,KIDINS220,TMSB4X,NUMA1,EIF4G2,RPS6,DBN1,EN
A1A,CFL1,MAVS,HSP90AB1,HSP90AA1,ACTR2,VIM,SPARC,TUBB,FTH1,UBC,KIDINS220,NUMA1,E
'90AB1,ACTR2,VIM,TUBB,FTH1,UBC,KIDINS220,NUMA1,EIF4G2,RPS6,ENO1,TUBA1B,PIK3R2,NC
'90AB1,ACTR2,VIM,TUBB,FTH1,UBC,KIDINS220,NUMA1,EIF4G2,RPS6,ENO1,TUBA1B,PIK3R2,NC
0AB1,ACTR2,VIM,TUBB,FTH1,UBC,KIDINS220,TMSB4X,NUMA1,EIF4G2,RPS6,DBN1,ENO1,TUBA
R2,TUBB,FTH1,UBC,KIDINS220,NUMA1,EIF4G2,RPS6,ENO1,TUBA1B,PIK3R2,MAP1B,NCL,ALDOA,
FL1,MAVS,HSP90AB1,HSP90AA1,ACTR2,VIM,TUBB,FTH1,UBC,KIDINS220,TMSB4X,NUMA1,DCX,E
FL1,MAVS,HSP90AB1,HSP90AA1,ACTR2,VIM,TUBB,FTH1,UBC,KIDINS220,TMSB4X,NUMA1,DCX,E
AB1,HSP90AA1,ACTR2,VIM,TUBB,FTH1,UBC,KIDINS220,NUMA1,EIF4G2,RPS6,DBN1,ENO1,TUBA
1,HSP90AA1,ACTR2,TUBB,FTH1,UBC,KIDINS220,TMSB4X,NUMA1,EIF4G2,RPS6,DBN1,ENO1,TUE
1,HSP90AA1,ACTR2,TUBB,FTH1,UBC,KIDINS220,TMSB4X,NUMA1,EIF4G2,RPS6,DBN1,ENO1,TUE
A1A,CFL1,MAVS,HSP90AB1,HSP90AA1,ACTR2,VIM,TUBB,FTH1,UBC,KIDINS220,NUMA1,EIF4G2,F
ACTR2,VIM,TUBB,FTH1,UBC,KIDINS220,NUMA1,EIF4G2,RPS6,DBN1,ENO1,TUBA1B,PIK3R2,NCL
1,EIF4G2,RPS6,DBN1,ENO1,TUBA1B,PIK3R2,FLNA,NCL,ALDOA,COL6A1,HSP90B1,FADS1,RPL8,F
'S,HSP90AB1,HSP90AA1,ACTR2,VIM,SPARC,FTH1,UBC,KIDINS220,NUMA1,EIF4G2,RPS6,ENO1,TU
AVS,HSP90AB1,HSP90AA1,ACTR2,VIM,SPARC,TUBB,FTH1,UBC,KIDINS220,TMSB4X,NUMA1,EIF4
AVS,HSP90AB1,HSP90AA1,ACTR2,VIM,SPARC,TUBB,FTH1,UBC,KIDINS220,TMSB4X,NUMA1,EIF4
MSB4X,NUMA1,DCX,EIF4G2,RPS6,TUBA1B,PIK3R2,MAP1B,NCL,GANAB,HSP90B1,PRPF8,RPL8,F
CX,EIF4G2,RPS6,TUBA1B,PIK3R2,NCL,SLC9A7,ALDOA,GANAB,HSP90B1,NEFL,RPL8,FBXL4,YBX1
FL1,MAVS,HSP90AB1,ACTR2,VIM,SPARC,TUBB,FTH1,UBC,KIDINS220,TMSB4X,NUMA1,DCX,EIF4
HSP90AA1,ACTR2,VIM,SPARC,TUBB,FTH1,UBC,KIDINS220,NUMA1,EIF4G2,RPS6,ENO1,TUBA1B,
90AB1,ACTR2,TUBB,FTH1,UBC,KIDINS220,NUMA1,DCX,EIF4G2,RPS6,DBN1,ENO1,TUBA1B,PIK3
90AB1,ACTR2,TUBB,FTH1,UBC,KIDINS220,NUMA1,DCX,EIF4G2,RPS6,DBN1,ENO1,TUBA1B,PIK3
AVS,HSP90AB1,HSP90AA1,ACTR2,VIM,TUBB,FTH1,UBC,KIDINS220,NUMA1,DCX,EIF4G2,RPS6,DI
R2,VIM,TUBB,FTH1,UBC,KIDINS220,TMSB4X,NUMA1,EIF4G2,RPS6,ENO1,TUBA1B,PIK3R2,NCL,
R2,VIM,TUBB,FTH1,UBC,KIDINS220,TMSB4X,NUMA1,EIF4G2,RPS6,ENO1,TUBA1B,PIK3R2,NCL,
BB,FTH1,UBC,NUMA1,EIF4G2,RPS6,DBN1,ENO1,TUBA1B,PIK3R2,FLNA,NCL,ALDOA,COL6A1,HS
BC,KIDINS220,TMSB4X,NUMA1,EIF4G2,RPS6,DBN1,ENO1,TUBA1B,PIK3R2,FLNA,NCL,SLC9A7,A
FL1,MAVS,HSP90AB1,HSP90AA1,ACTR2,TUBB,FTH1,UBC,KIDINS220,NUMA1,EIF4G2,RPS6,DBN1
B,FTH1,UBC,TMSB4X,NUMA1,EIF4G2,RPS6,ENO1,TUBA1B,MAP1B,NCL,GANAB,FTL,HSP90B1,FA
TUBB,FTH1,UBC,KIDINS220,NUMA1,EIF4G2,RPS6,DBN1,ENO1,TUBA1B,PIK3R2,MAP1B,NCL,ALD
TUBB,FTH1,UBC,KIDINS220,NUMA1,EIF4G2,RPS6,DBN1,ENO1,TUBA1B,PIK3R2,MAP1B,NCL,ALD
90AB1,HSP90AA1,ACTR2,VIM,TUBB,UBC,KIDINS220,NUMA1,DCX,EIF4G2,RPS6,DBN1,ENO1,TUE
IM,SPARC,FTH1,UBC,NUMA1,EIF4G2,RPS6,DBN1,ENO1,TUBA1B,PIK3R2,FLNA,NCL,GANAB,COL
A1A,CFL1,MAVS,HSP90AB1,HSP90AA1,ACTR2,TUBB,FTH1,UBC,KIDINS220,TMSB4X,NUMA1,DCX
A1A,CFL1,MAVS,HSP90AB1,HSP90AA1,ACTR2,TUBB,FTH1,UBC,KIDINS220,TMSB4X,NUMA1,DCX
'S,HSP90AB1,HSP90AA1,ACTR2,TUBB,FTH1,UBC,TMSB4X,NUMA1,DCX,EIF4G2,RPS6,DBN1,ENO
'S,HSP90AB1,HSP90AA1,ACTR2,TUBB,FTH1,UBC,TMSB4X,NUMA1,DCX,EIF4G2,RPS6,DBN1,ENO

RC,TUBB,UBC,KIDINS220,TMSB4X,NUMA1,EIF4G2,RPS6,DBN1,ENO1,TUBA1B,PIK3R2,FLNA,MA
AB1,ACTR2,TUBB,FTH1,UBC,KIDINS220,NUMA1,DCX,EIF4G2,RPS6,DBN1,ENO1,TUBA1B,PIK3R2
AB1,ACTR2,TUBB,FTH1,UBC,KIDINS220,NUMA1,DCX,EIF4G2,RPS6,DBN1,ENO1,TUBA1B,PIK3R2
FL1,MAVS,HSP90AB1,ACTR2,VIM,SPARC,TUBB,FTH1,UBC,KIDINS220,TMSB4X,NUMA1,EIF4G2,F
1,ACTR2,VIM,TUBB,FTH1,UBC,KIDINS220,TMSB4X,NUMA1,EIF4G2,RPS6,DBN1,ENO1,TUBA1B,P
/S,HSP90AB1,HSP90AA1,ACTR2,TUBB,FTH1,UBC,KIDINS220,TMSB4X,NUMA1,DCX,EIF4G2,RPS6,
AB1,ACTR2,VIM,SPARC,TUBB,FTH1,UBC,KIDINS220,TMSB4X,NUMA1,DCX,EIF4G2,RPS6,ENO1,T
90AB1,SPARC,TUBB,FTH1,UBC,TMSB4X,NUMA1,DCX,EIF4G2,RPS6,DBN1,ENO1,TUBA1B,PIK3R2
AB1,TUBB,FTH1,UBC,KIDINS220,NUMA1,EIF4G2,RPS6,DBN1,ENO1,TUBA1B,PIK3R2,NCL,SLC9A
AB1,TUBB,FTH1,UBC,KIDINS220,NUMA1,EIF4G2,RPS6,DBN1,ENO1,TUBA1B,PIK3R2,NCL,SLC9A
/S,HSP90AB1,ACTR2,SPARC,TUBB,FTH1,UBC,KIDINS220,TMSB4X,NUMA1,DCX,EIF4G2,RPS6,DB
3,FTH1,UBC,KIDINS220,NUMA1,EIF4G2,RPS6,ENO1,TUBA1B,PIK3R2,NCL,ALDOA,GANAB,FTL,H
CTR2,TUBB,FTH1,UBC,KIDINS220,NUMA1,EIF4G2,RPS6,DBN1,ENO1,TUBA1B,PIK3R2,FLNA,NCL
CTR2,TUBB,FTH1,UBC,KIDINS220,NUMA1,EIF4G2,RPS6,DBN1,ENO1,TUBA1B,PIK3R2,FLNA,NCL
,DBN1,ENO1,TUBA1B,FLNA,NCL,GANAB,FTL,HSP90B1,PRPF8,RPL8,YBX1,UBA1,ATP2B4,TMF1,A
1,ACTR2,VIM,TUBB,FTH1,UBC,KIDINS220,NUMA1,DCX,EIF4G2,RPS6,ENO1,TUBA1B,PIK3R2,FLN
JBB,FTH1,KIDINS220,TMSB4X,NUMA1,EIF4G2,RPS6,ENO1,TUBA1B,PIK3R2,MAP1B,NCL,SLC9A7
'90AB1,ACTR2,TUBB,UBC,KIDINS220,TMSB4X,NUMA1,DCX,EIF4G2,RPS6,DBN1,ENO1,TUBA1B,F
'90AB1,ACTR2,TUBB,UBC,KIDINS220,TMSB4X,NUMA1,DCX,EIF4G2,RPS6,DBN1,ENO1,TUBA1B,F
CTR2,SPARC,TUBB,FTH1,UBC,KIDINS220,TMSB4X,NUMA1,DCX,EIF4G2,RPS6,ENO1,TUBA1B,PIK
CTR2,SPARC,TUBB,FTH1,UBC,KIDINS220,TMSB4X,NUMA1,DCX,EIF4G2,RPS6,ENO1,TUBA1B,PIK
BB,UBC,KIDINS220,NUMA1,EIF4G2,RPS6,ENO1,TUBA1B,PIK3R2,NCL,SLC9A7,ALDOA,GANAB,HS
'90AB1,ACTR2,VIM,TUBB,FTH1,UBC,KIDINS220,TMSB4X,NUMA1,DCX,EIF4G2,RPS6,DBN1,ENO1
TR2,VIM,SPARC,TUBB,FTH1,UBC,KIDINS220,NUMA1,EIF4G2,RPS6,DBN1,ENO1,TUBA1B,PIK3R2
FL1,MAVS,HSP90AB1,HSP90AA1,ACTR2,VIM,TUBB,FTH1,UBC,KIDINS220,NUMA1,EIF4G2,RPS6,D
C,KIDINS220,NUMA1,EIF4G2,RPS6,DBN1,ENO1,TUBA1B,PIK3R2,NCL,SLC9A7,ALDOA,GANAB,HS
3C,NUMA1,EIF4G2,RPS6,ENO1,TUBA1B,PIK3R2,NCL,ALDOA,GANAB,HSP90B1,FADS1,PRPF8,UG
3C,KIDINS220,TMSB4X,NUMA1,EIF4G2,RPS6,DBN1,ENO1,TUBA1B,PIK3R2,FLNA,NCL,SLC9A7,A
11,UBC,NUMA1,EIF4G2,RPS6,ENO1,TUBA1B,FLNA,NCL,SLC9A7,ALDOA,GANAB,COL6A1,FTL,HSF
11,UBC,NUMA1,EIF4G2,RPS6,ENO1,TUBA1B,FLNA,NCL,SLC9A7,ALDOA,GANAB,COL6A1,FTL,HSF
JBC,KIDINS220,NUMA1,EIF4G2,RPS6,ENO1,TUBA1B,PIK3R2,NCL,SLC9A7,ALDOA,GANAB,HSP90
CTR2,TUBB,UBC,KIDINS220,NUMA1,EIF4G2,RPS6,DBN1,ENO1,TUBA1B,PIK3R2,NCL,SLC9A7,AL
DINS220,TMSB4X,NUMA1,EIF4G2,RPS6,ENO1,TUBA1B,NCL,GANAB,COL6A1,FTL,HSP90B1,FADS
/S,HSP90AB1,HSP90AA1,ACTR2,TUBB,FTH1,UBC,KIDINS220,NUMA1,EIF4G2,RPS6,ENO1,TUBA1
/S,HSP90AB1,HSP90AA1,ACTR2,TUBB,FTH1,UBC,KIDINS220,NUMA1,EIF4G2,RPS6,ENO1,TUBA1
AVS,HSP90AB1,HSP90AA1,ACTR2,VIM,TUBB,FTH1,UBC,NUMA1,DCX,EIF4G2,RPS6,ENO1,TUBA1E
/S,HSP90AB1,HSP90AA1,ACTR2,TUBB,FTH1,UBC,KIDINS220,TMSB4X,NUMA1,DCX,EIF4G2,RPS6,
HSP90AA1,ACTR2,TUBB,UBC,KIDINS220,TMSB4X,NUMA1,EIF4G2,RPS6,DBN1,ENO1,TUBA1B,PIK
RPS6,ENO1,TUBA1B,PIK3R2,MAP1B,NCL,GANAB,FTL,HSP90B1,FADS1,UGT8,RPL8,FBXL4,YBX1
FTH1,UBC,KIDINS220,TMSB4X,NUMA1,DCX,EIF4G2,RPS6,ENO1,TUBA1B,PIK3R2,FLNA,MAP1B,N
RPS6,ENO1,TUBA1B,PIK3R2,NCL,ALDOA,GANAB,HSP90B1,FADS1,RPL8,FBXL4,YBX1,UBA1,ATP2
INS220,TMSB4X,NUMA1,EIF4G2,RPS6,ENO1,TUBA1B,PIK3R2,NCL,ALDOA,GANAB,HSP90B1,PRI
BC,KIDINS220,NUMA1,EIF4G2,RPS6,TUBA1B,PIK3R2,NCL,SLC9A7,ALDOA,HSP90B1,RPL8,FBXL
FL1,MAVS,HSP90AB1,ACTR2,VIM,SPARC,TUBB,FTH1,UBC,KIDINS220,TMSB4X,NUMA1,EIF4G2,F
/S,HSP90AB1,ACTR2,TUBB,UBC,NUMA1,EIF4G2,RPS6,ENO1,TUBA1B,PIK3R2,NCL,SLC9A7,ALDC

2,VIM,SPARC,TUBB,UBC,NUMA1,EIF4G2,RPS6,ENO1,TUBA1B,PIK3R2,FLNA,NCL,SLC9A7,ALDOA
?90AB1,ACTR2,VIM,TUBB,FTH1,UBC,KIDINS220,TMSB4X,NUMA1,EIF4G2,RPS6,DBN1,ENO1,TUB
TR2,TUBB,FTH1,UBC,KIDINS220,NUMA1,DCX,EIF4G2,RPS6,ENO1,TUBA1B,PIK3R2,NCL,SLC9A7,
TR2,TUBB,FTH1,UBC,KIDINS220,NUMA1,DCX,EIF4G2,RPS6,ENO1,TUBA1B,PIK3R2,NCL,SLC9A7,
R2,MAP1B,NCL,ALDOA,FTL,HSP90B1,PRPF8,FBXL4,UBA1,ATP2B4,TMF1,ABHD2,PSAP,RPLP0,H3
TR2,VIM,TUBB,FTH1,UBC,KIDINS220,NUMA1,DCX,EIF4G2,RPS6,DBN1,TUBA1B,PIK3R2,NCL,SLC
A1,ACTR2,SPARC,UBC,KIDINS220,NUMA1,EIF4G2,RPS6,ENO1,TUBA1B,PIK3R2,NCL,ALDOA,GANA
NUMA1,RPS6,ENO1,TUBA1B,PIK3R2,FLNA,NCL,GANAB,FTL,HSP90B1,PRPF8,RPL8,FBXL4,YBX1,
VS,HSP90AB1,HSP90AA1,ACTR2,VIM,TUBB,FTH1,UBC,KIDINS220,TMSB4X,NUMA1,DCX,EIF4G2,F
1,UBC,KIDINS220,NUMA1,EIF4G2,RPS6,ENO1,TUBA1B,PIK3R2,MAP1B,NCL,SLC9A7,ALDOA,GAN
1,UBC,KIDINS220,NUMA1,EIF4G2,RPS6,ENO1,TUBA1B,PIK3R2,MAP1B,NCL,SLC9A7,ALDOA,GAN
BN1,ENO1,TUBA1B,PIK3R2,NCL,SLC9A7,GANAB,HSP90B1,PRPF8,RPL8,FBXL4,UBA1,TMF1,HSPA
H1,UBC,TMSB4X,NUMA1,EIF4G2,RPS6,ENO1,TUBA1B,PIK3R2,NCL,ALDOA,GANAB,COL6A1,FTL,H
,UBC,KIDINS220,NUMA1,EIF4G2,RPS6,DBN1,ENO1,TUBA1B,PIK3R2,NCL,SLC9A7,ALDOA,FTL,HS
,UBC,KIDINS220,NUMA1,EIF4G2,RPS6,DBN1,ENO1,TUBA1B,PIK3R2,NCL,SLC9A7,ALDOA,FTL,HS
NUMA1,EIF4G2,RPS6,ENO1,TUBA1B,PIK3R2,NCL,SLC9A7,ALDOA,GANAB,COL6A1,FTL,HSP90B1
4G2,RPS6,ENO1,TUBA1B,PIK3R2,NCL,ALDOA,GANAB,HSP90B1,FADS1,RPL8,FBXL4,YBX1,UBA1
IF4G2,RPS6,ENO1,TUBA1B,PIK3R2,FLNA,NCL,SLC9A7,ALDOA,GANAB,COL6A1,HSP90B1,FADS1,
RPS6,ENO1,PIK3R2,FLNA,MAP1B,NCL,SLC9A7,ALDOA,GANAB,HSP90B1,NEFM,RPL8,FBXL4,UBA
JBB,FTH1,UBC,KIDINS220,TMSB4X,NUMA1,EIF4G2,RPS6,ENO1,TUBA1B,PIK3R2,NCL,SLC9A7,AI
IS220,NUMA1,EIF4G2,RPS6,ENO1,TUBA1B,PIK3R2,NCL,SLC9A7,ALDOA,GANAB,HSP90B1,FADS1
1,MAVS,HSP90AB1,HSP90AA1,ACTR2,VIM,SPARC,TUBB,FTH1,UBC,KIDINS220,TMSB4X,NUMA1,D
DINS220,NUMA1,RPS6,DBN1,TUBA1B,PIK3R2,MAP1B,NCL,ALDOA,FADS1,NEFM,RPL8,FBXL4,UBA
,VIM,TUBB,FTH1,UBC,TMSB4X,NUMA1,EIF4G2,RPS6,ENO1,TUBA1B,PIK3R2,NCL,SLC9A7,ALDO
BA1B,PIK3R2,FLNA,NCL,GANAB,COL6A1,FTL,HSP90B1,RPL8,FBXL4,YBX1,UBA1,ATP2B4,CALR,P
M,TUBB,FTH1,UBC,KIDINS220,TMSB4X,NUMA1,EIF4G2,RPS6,DBN1,ENO1,TUBA1B,PIK3R2,FLNA
ACTR2,TUBB,FTH1,UBC,KIDINS220,NUMA1,EIF4G2,DBN1,ENO1,TUBA1B,PIK3R2,NCL,SLC9A7,AL
C,KIDINS220,NUMA1,EIF4G2,RPS6,DBN1,ENO1,TUBA1B,PIK3R2,FLNA,NCL,SLC9A7,ALDOA,GANA
?90AB1,ACTR2,TUBB,UBC,KIDINS220,NUMA1,DCX,EIF4G2,RPS6,DBN1,ENO1,TUBA1B,PIK3R2,NC
TR2,TUBB,UBC,KIDINS220,TMSB4X,NUMA1,EIF4G2,RPS6,DBN1,ENO1,TUBA1B,PIK3R2,NCL,ALD
BB,UBC,KIDINS220,TMSB4X,NUMA1,EIF4G2,RPS6,DBN1,ENO1,TUBA1B,PIK3R2,FLNA,NCL,ALDO
ACTR2,UBC,KIDINS220,NUMA1,EIF4G2,RPS6,ENO1,TUBA1B,NCL,SLC9A7,HSP90B1,PRPF8,RPL8
5,ENO2,SOX4,KAT6A,YWHAE,RPL5,TMBIM6,NPM1,PLD3,EIF3C,ATP5F1A,VCP,TRIM28,MATR3,YW
NUMA1,EIF4G2,RPS6,ENO1,TUBA1B,PIK3R2,NCL,ALDOA,GANAB,HSP90B1,FADS1,PRPF8,RPL8,
0AB1,HSP90AA1,ACTR2,UBC,KIDINS220,NUMA1,EIF4G2,RPS6,ENO1,TUBA1B,PIK3R2,NCL,SLC9
TH1,UBC,KIDINS220,NUMA1,EIF4G2,RPS6,ENO1,TUBA1B,PIK3R2,FLNA,NCL,SLC9A7,ALDOA,GA
TUBB,FTH1,UBC,KIDINS220,TMSB4X,NUMA1,EIF4G2,RPS6,DBN1,ENO1,TUBA1B,PIK3R2,MAP1B
NUMA1,EIF4G2,RPS6,ENO1,TUBA1B,PIK3R2,MAP1B,NCL,SLC9A7,ALDOA,GANAB,FTL,HSP90B1,F
A1,ACTR2,VIM,TUBB,FTH1,UBC,KIDINS220,NUMA1,EIF4G2,RPS6,DBN1,TUBA1B,PIK3R2,NCL,AL
MA1,EIF4G2,RPS6,PIK3R2,MAP1B,NCL,ALDOA,FTL,HSP90B1,FADS1,PRPF8,RPL8,FBXL4,YBX1,U
;,KIDINS220,NUMA1,EIF4G2,RPS6,TUBA1B,PIK3R2,NCL,HSP90B1,FADS1,NEFL,PRPF8,RPL8,FBX
K3R2,NCL,SLC9A7,GANAB,FTL,HSP90B1,NEFM,PRPF8,RPL8,FBXL4,YBX1,ABHD2,HSPA1B,PSAP,
R2,VIM,TUBB,FTH1,UBC,NUMA1,DCX,EIF4G2,RPS6,TUBA1B,PIK3R2,NCL,SLC9A7,ALDOA,FTL,H
DINS220,NUMA1,DCX,EIF4G2,RPS6,ENO1,TUBA1B,PIK3R2,NCL,ALDOA,GANAB,HSP90B1,FADS1,
,EIF4G2,RPS6,ENO1,TUBA1B,PIK3R2,FLNA,NCL,ALDOA,GANAB,FTL,HSP90B1,FADS1,PRPF8,RP

C,KIDINS220,NUMA1,EIF4G2,RPS6,DBN1,ENO1,TUBA1B,PIK3R2,FLNA,MAP1B,NCL,HSP90B1,FAI
1B,PIK3R2,NCL,HSP90B1,NEFL,PRPF8,RPL8,FBXL4,UBA1,CALR,ABHD2,RPLP0,H3-3B,CANX,PTM
AP,H3-3B,CANX,DDX5,RPL13A,SEC62,DVL3,RPL13,SPTBN1,ATP5F1B,SEPTIN2,FSCN1,PTMS,SPT
0,NUMA1,RPS6,ENO1,TUBA1B,PIK3R2,NCL,SLC9A7,ALDOA,GANAB,HSP90B1,FADS1,RPL8,YBX1,
,TUBB,FTH1,UBC,KIDINS220,TMSB4X,NUMA1,EIF4G2,RPS6,DBN1,ENO1,TUBA1B,PIK3R2,NCL,A
3X1,UBA1,ATP2B4,HSPA1B,PSAP,H3-3B,CANX,DDX5,RPL13A,DVL3,YWHAE,RPL13,DDX17,SPTBN1
INS220,NUMA1,EIF4G2,RPS6,TUBA1B,PIK3R2,NCL,ALDOA,HSP90B1,RPL8,FBXL4,UBA1,ATP2B4
AB1,ACTR2,VIM,SPARC,TUBB,FTH1,UBC,KIDINS220,TMSB4X,NUMA1,EIF4G2,RPS6,DBN1,ENO1,
AB1,ACTR2,VIM,SPARC,TUBB,FTH1,UBC,KIDINS220,TMSB4X,NUMA1,EIF4G2,RPS6,DBN1,ENO1,
IS220,NUMA1,EIF4G2,RPS6,ENO1,TUBA1B,PIK3R2,FLNA,NCL,SLC9A7,ALDOA,GANAB,HSP90B1,
,TUBA1B,PIK3R2,NCL,ALDOA,GANAB,HSP90B1,FADS1,PRPF8,RPL8,FBXL4,YBX1,UBA1,ATP2B4,
MA1,EIF4G2,RPS6,ENO1,TUBA1B,PIK3R2,NCL,ALDOA,COL6A1,HSP90B1,FADS1,PRPF8,RPL8,FE
MA1,EIF4G2,RPS6,ENO1,TUBA1B,PIK3R2,NCL,ALDOA,COL6A1,HSP90B1,FADS1,PRPF8,RPL8,FE
F4G2,RPS6,ENO1,TUBA1B,PIK3R2,NCL,SLC9A7,ALDOA,GANAB,COL6A1,FTL,HSP90B1,FADS1,R
4G2,ENO1,TUBA1B,PIK3R2,FLNA,NCL,ALDOA,GANAB,FTL,HSP90B1,FADS1,RPL8,YBX1,UBA1,ATI
,EIF4G2,RPS6,TUBA1B,PIK3R2,NCL,SLC9A7,ALDOA,FTL,HSP90B1,FADS1,RPL8,FBXL4,UBA1,AT
P1B,NCL,ALDOA,GANAB,HSP90B1,NEFM,RPL8,FBXL4,UBA1,ATP2B4,CALR,TMF1,ABHD2,PSAP,R
NO1,TUBA1B,PIK3R2,NCL,ALDOA,GANAB,HSP90B1,FADS1,PRPF8,RPL8,YBX1,UBA1,ATP2B4,TM
0,TMSB4X,NUMA1,EIF4G2,RPS6,ENO1,TUBA1B,PIK3R2,NCL,ALDOA,GANAB,FTL,HSP90B1,RPL8
1,MAVS,HSP90AB1,ACTR2,VIM,SPARC,TUBB,FTH1,UBC,KIDINS220,TMSB4X,NUMA1,DCX,EIF4G2
,FTH1,UBC,KIDINS220,TMSB4X,NUMA1,EIF4G2,RPS6,DBN1,ENO1,TUBA1B,PIK3R2,NCL,ALDOA,
A1B,PIK3R2,NCL,SLC9A7,FTL,HSP90B1,PRPF8,RPL8,YBX1,UBA1,ATP2B4,HSPA1B,PSAP,H3-3B,C
E,RPL13,RPL7,RPL5,TMBIM6,NPM1,EIF3C,ATP5F1A,VCP,TRIM28,MATR3,YWHAG,RPL37A,SPEN,
C,KIDINS220,NUMA1,EIF4G2,RPS6,ENO1,TUBA1B,PIK3R2,NCL,SLC9A7,GANAB,FTL,HSP90B1,FA
220,NUMA1,EIF4G2,RPS6,ENO1,TUBA1B,PIK3R2,NCL,ALDOA,GANAB,FTL,HSP90B1,FADS1,RPL
BC,KIDINS220,NUMA1,EIF4G2,RPS6,ENO1,TUBA1B,NCL,SLC9A7,FTL,HSP90B1,FADS1,UGT8,RF
BC,KIDINS220,NUMA1,EIF4G2,RPS6,ENO1,TUBA1B,NCL,SLC9A7,FTL,HSP90B1,FADS1,UGT8,RF
4G2,RPS6,ENO1,TUBA1B,FLNA,NCL,FTL,HSP90B1,FADS1,PRPF8,RPL8,FBXL4,YBX1,UBA1,ATP2E
,FBXL4,UBA1,TMF1,ABHD2,PGK1,RPLP0,H3-3B,CANX,DDX5,ENO2,SOX4,RPL13A,SEC62,KAT6A,
2,DBN1,ENO1,TUBA1B,PIK3R2,NCL,ALDOA,FTL,HSP90B1,FADS1,PRPF8,RPL8,FBXL4,YBX1,UBA
SP90B1,FADS1,PRPF8,UGT8,FBXL4,YBX1,TMF1,PSAP,H3-3B,CANX,DDX5,SOX4,SEC62,KAT6A,DV
JBA1,ATP2B4,HSPA1B,PSAP,RPLP0,H3-3B,CANX,PTMA,DDX5,RPL13A,SEC62,RPL6,KAT6A,DVL3,F
3B,UBC,TMSB4X,NUMA1,EIF4G2,RPS6,ENO1,TUBA1B,PIK3R2,NCL,SLC9A7,ALDOA,GANAB,FTL,H
4,ATP2B4,ABHD2,HSPA1B,PSAP,H3-3B,DDX5,DVL3,YWHAE,RPL13,DDX17,ATP5F1B,MYH9,P4HB,
B1,FADS1,RPL8,FBXL4,YBX1,UBA1,ATP2B4,COL4A1,CES3,PSAP,H3-3B,CANX,PTMA,DDX5,DVL3,
31,UGT8,RPL8,FBXL4,YBX1,TMF1,ABHD2,CES3,PSAP,H3-3B,CANX,DDX5,SOX4,RPL13A,SEC62,K
B,FTH1,UBC,NUMA1,EIF4G2,RPS6,ENO1,TUBA1B,PIK3R2,NCL,ALDOA,GANAB,FTL,FADS1,RPL8,
PS6,DBN1,TUBA1B,PIK3R2,FLNA,NCL,GANAB,COL6A1,HSP90B1,FADS1,FBXL4,ATP2B4,TMF1,C
),TMSB4X,NUMA1,EIF4G2,RPS6,TUBA1B,PIK3R2,FLNA,NCL,ALDOA,GANAB,FTL,HSP90B1,FADS1
IO1,TUBA1B,PIK3R2,FLNA,NCL,SLC9A7,ALDOA,HSP90B1,RPL8,YBX1,UBA1,TMF1,ABHD2,CES3,F
ISP90B1,RPL8,UBA1,ATP2B4,TMF1,ABHD2,PSAP,PGK1,RPLP0,H3-3B,CANX,PTMA,RPL13A,DVL3,
NUMA1,EIF4G2,RPS6,ENO1,TUBA1B,PIK3R2,NCL,SLC9A7,GANAB,FTL,HSP90B1,FADS1,UGT8,RF
6,TUBA1B,PIK3R2,NCL,ALDOA,COL6A1,HSP90B1,RPL8,FBXL4,YBX1,UBA1,ATP2B4,COL4A1,PSA
3-3B,CANX,DDX5,DVL3,YWHAE,BSG,MYH9,P4HB,RPL5,MARCKS,NPM1,SNRNP200,ATP5F1A,VCP,
O1,TUBA1B,PIK3R2,NCL,SLC9A7,HSP90B1,PRPF8,RPL8,FBXL4,UBA1,ATP2B4,TMF1,ABHD2,CES

TUBA1B,NCL,SLC9A7,ALDOA,FTL,HSP90B1,PRPF8,RPL8,YBX1,UBA1,TMF1,ABHD2,PSAP,PGK1,H
JBA1B,PIK3R2,NCL,ALDOA,FTL,HSP90B1,YBX1,UBA1,ATP2B4,PSAP,PGK1,H3-3B,CANX,PTMA,DD
,TMBIM6,NPM1,ATP5F1A,MATR3,YWHAG,RPL37A,LRRC28,SPEN,NONO,DDB1,SSBP2,BASP1,RPS
90B1,FADS1,UBA1,ATP2B4,CALR,PSAP,H3-3B,CANX,DDX5,KAT6A,DVL3,YWHAE,DDX17,SPTBN1,
SB4X,NUMA1,EIF4G2,RPS6,DBN1,ENO1,TUBA1B,PIK3R2,FLNA,NCL,ALDOA,GANAB,FTL,HSP90B
L6A1,HSP90B1,FBXL4,YBX1,UBA1,ATP2B4,TMF1,PSAP,H3-3B,CANX,DDX5,DVL3,RPL13,SPTBN1,
PIK3R2,NCL,GANAB,HSP90B1,FADS1,PRPF8,RPL8,FBXL4,YBX1,UBA1,ATP2B4,HSPA1B,PSAP,H3-
HSP90B1,FADS1,UBA1,ATP2B4,TMF1,HSPA1B,CES3,PSAP,PGK1,H3-3B,CANX,DDX5,SOX4,SEC62
SAP,RPLP0,H3-3B,CANX,PTMA,SOX4,RPL13A,DVL3,YWHAE,ALDH6A1,ATP5F1B,SEPTIN2,FSCN1,S
,EIF4G2,RPS6,ENO1,NCL,ALDOA,HSP90B1,FADS1,UGT8,RPL8,FBXL4,UBA1,ATP2B4,TMF1,ABHD
,EIF4G2,RPS6,ENO1,NCL,ALDOA,HSP90B1,FADS1,UGT8,RPL8,FBXL4,UBA1,ATP2B4,TMF1,ABHD
DA,GANAB,HSP90B1,FBXL4,YBX1,UBA1,ATP2B4,COL4A1,PSAP,H3-3B,CANX,PTMA,DDX5,DVL3,YW
HSPA1B,PSAP,H3-3B,CANX,CLU,DDX5,SEC62,DVL3,YWHAE,BSG,P4HB,RPL5,GDI1,GSTP1,NPM1,
-3B,CANX,DDX5,KAT6A,DDX17,BSG,ATP5F1B,CALM3,SEPTIN2,APP,GDI1,FSCN1,SPTAN1,MARCK
A,DVL3,RPL13,ALDH6A1,SPTBN1,ATP5F1B,RPL7,P4HB,RPL5,TMBIM6,SPTAN1,NPM1,EIF3C,ATP5
TAN1,SMG1,NPM1,EIF3C,ATP5F1A,VCP,TRIM28,MATR3,YWHAG,RPL37A,LRRC28,SPEN,NONO,D
PL5,PTMS,MARCKS,NPM1,IGF2,VCP,TRIM28,YWHAG,SPEN,NONO,SET,EGLN1,PDIA3,SSBP2,HMG
M6,NPM1,EIF3C,ATP5F1A,MARCKSL1,TRIM28,MATR3,YWHAG,NONO,SET,EGLN1,SSBP2,CHD3,R
HSP90B1,RPL8,YBX1,UBA1,PSAP,PGK1,H3-3B,CANX,PTMA,RPL13A,SEC62,DVL3,YWHAE,RPL13,
RC,TUBB,FTH1,UBC,TMSB4X,NUMA1,EIF4G2,RPS6,ENO1,TUBA1B,PIK3R2,FLNA,NCL,ALDOA,GA
PSAP,RPLP0,H3-3B,CANX,DDX5,YWHAE,RPL13,BSG,ALDH6A1,SPTBN1,ATP5F1B,MYH9,P4HB,IDS
BXL4,UBA1,TMF1,PSAP,PGK1,H3-3B,CANX,PTMA,DDX5,DVL3,RPL13,ALDH6A1,SPTBN1,ATP5F1E
,H3-3B,CANX,DVL3,RPL13,DDX17,PTMS,SPTAN1,GSTP1,MARCKS,NPM1,SNRNP200,ATP5F1A,VC
IA1,EIF4G2,RPS6,DBN1,ENO1,TUBA1B,PIK3R2,FLNA,NCL,ALDOA,GANAB,FTL,HSP90B1,PRPF8,R
S,RPL5,NPM1,EIF3C,ATP5F1A,EIF4B,VCP,UBB,TRIM28,MATR3,RPL37A,LRRC28,SPEN,NONO,DD
3A1,COL4A1,PSAP,H3-3B,CANX,PTMA,DDX5,DVL3,GPI,FASN,SPTBN1,FSCN1,PTMS,SPTAN1,SMG
PP,GDI1,FSCN1,SPTAN1,MARCKS,SMG1,CALM1,NPM1,HSPD1,EIF3C,ATP5F1A,MARCKSL1,UBB,T
L,ALDOA,GANAB,HSP90B1,NEFL,NEFM,FBXL4,ATP2B4,PSAP,H3-3B,CANX,PTMA,ENO2,SEC62,K
RPL5,GDI1,TMBIM6,SPTAN1,MARCKS,NPM1,COL18A1,EIF3C,ATP5F1A,VCP,MARCKSL1,UBB,TR
PL13A,SEC62,DVL3,RPL13,NPM1,EIF3C,ATP5F1A,EIF4B,UBB,TRIM28,MATR3,RPL37A,SPEN,NON
ANX,DDX5,RPL13A,DVL3,YWHAE,RPL13,DDX17,BSG,ALDH6A1,SPTBN1,ATP5F1B,P4HB,IDS,RPL5
D2,HSPA1B,PSAP,RPLP0,H3-3B,CANX,PTMA,RPL5,TMBIM6,SPTAN1,GSTP1,MARCKS,SMG1,NPM1
SLC9A7,COL6A1,HSP90B1,NEFM,RPL8,FBXL4,YBX1,ATP2B4,TMF1,ABHD2,PSAP,PGK1,H3-3B,CA
NPM1,EIF3C,ATP5F1A,VCP,MARCKSL1,UBB,TRIM28,MATR3,RPL37A,NONO,DDB1,SET,EGLN1,SS
1,SPTBN1,MYH9,P4HB,RPL5,GDI1,TMBIM6,SPTAN1,GSTP1,NPM1,ATP5F1A,VCP,MARCKSL1,TRI
3XL4,YBX1,UBA1,ABHD2,CES3,PSAP,RPLP0,H3-3B,CANX,DDX5,DVL3,YWHAE,RPL13,DDX17,BSG,
DBN1,ENO1,TUBA1B,PIK3R2,MAP1B,SLC9A7,ALDOA,FTL,NEFL,NEFM,FBXL4,UBA1,ATP2B4,ABHD
PGK1,H3-3B,CANX,DDX5,DVL3,ALDH6A1,SPTBN1,IDS,RPL5,TMBIM6,PTMS,GSTP1,NPM1,COL18
UBA1,CES3,PGK1,H3-3B,CANX,PTMA,DDX5,ENO2,SOX4,DVL3,SPTBN1,ATP5F1B,CALM3,GNB1,
ATP5F1A,EIF4B,VCP,UBB,TRIM28,MATR3,SPEN,NONO,SET,SSBP2,HMGB1,POLR2A,RPS4X,BTF3,
3,CANX,RPL13A,DVL3,YWHAE,ALDH6A1,SPTBN1,ATP5F1B,MYH9,RPL5,SPTAN1,GSTP1,NPM1,HS
,RPLP0,H3-3B,PTMA,DDX5,DVL3,YWHAE,SPTBN1,IDS,RPL5,GDI1,SPTAN1,SMG1,NPM1,HSPD1,A
,FSCN1,NPM1,EIF3C,ATP5F1A,VCP,TRIM28,MATR3,RPL37A,SPEN,SET,RPL19,SSBP2,VAPA,HMGE
,RPL5,TMBIM6,ATP5F1A,UBB,TRIM28,MATR3,SPEN,NONO,SET,SF3B1,SSBP2,VAPA,HMGB1,AN
3,SPTAN1,MARCKS,NPM1,EIF3C,ATP5F1A,UBB,TRIM28,MATR3,RPL37A,NONO,SET,RPL19,SF3B1

,YWHAG,RPL37A,SPEN,NONO,HACD3,SET,SSBP2,VAPA,RPS17,SF3B2,HNRNPU,RPL29,RPSA,CTSI
VCP,TRIM28,MATR3,RPL37A,MAPT,SPEN,YWHAQ,SSBP2,CHD3,STMN1,UCHL1,NCAM1,ANXA2,RF
XL4,YBX1,COL4A1,H3-3B,CANX,PTMA,DVL3,YWHAE,GPI,ATP5F1B,CALM3,GNB1,MYH9,GDI1,FSC
A1,PSAP,RPLP0,H3-3B,CANX,PTMA,DDX5,SOX4,KAT6A,DVL3,GPI,BSG,SPTBN1,ATP5F1B,IDS,GDI
31,FADS1,RPL8,FBXL4,UBA1,ATP2B4,HSPA1B,PSAP,H3-3B,CANX,DDX5,RPL13A,KAT6A,DVL3,DD
MYH9,MARCKS,NPM1,EIF3C,ATP5F1A,VCP,UBB,MATR3,SPEN,SET,EGLN1,RPL19,LDHB,SSBP2,HN
PEN,NONO,HACD3,SSBP2,BASP1,HMGB1,RPL10,DNAJA1,SF3B2,HNRNPU,RPL29,SCARB2,XRCC5
NB1,PTMS,SPTAN1,MARCKS,ATP5F1A,VCP,TRIM28,MATR3,YWHAG,MAPT,SPEN,SET,PDIA3,SSBP
PEN,NONO,SET,SSBP2,HMGB1,RPS3A,ANXA2,RPL10,DNAJA1,MAP4,SF3B2,GTF2F1,HNRNPU,RPS
YBX1,TMF1,CES3,PSAP,RPLP0,H3-3B,CANX,PTMA,DDX5,RPL13A,RPL13,ALDH6A1,SPTBN1,ATP5
HB,NPM1,ATP5F1A,VCP,MARCKSL1,UBB,TRIM28,MATR3,YWHAG,SPEN,NONO,EGLN1,PDIA3,SS
ANX,DDX5,DVL3,RPL13,SPTBN1,MYH9,IDS,RPL5,MARCKS,NPM1,EIF3C,ATP5F1A,VCP,MARCKSL1
FTL,HSP90B1,UBA1,ATP2B4,PSAP,H3-3B,CANX,PTMA,DDX5,SEC62,DVL3,YWHAE,SLC2A3,GPI,RF
CN1,PTMS,NPM1,ATP5F1A,VCP,UBB,AFF4,MATR3,YWHAG,RPL37A,LRRC28,SPEN,NONO,MRC2,S
SOX4,RPL13A,SEC62,DVL3,BSG,SPTBN1,GNB1,MYH9,SEPTIN2,FSCN1,PTMS,SPTAN1,GSTP1,NP
,DDX5,RPL13A,DVL3,BSG,SPTBN1,RPL7,P4HB,IDS,SPTAN1,MARCKS,SMG1,NPM1,EIF3C,ATP5F1
F4B,UBB,TRIM28,MATR3,SPEN,NONO,SET,SSBP2,HMGB1,BTF3,RPL10,SF3B2,HNRNPU,LARP1,
SPD1,USP11,ATP5F1A,TRIM28,MATR3,SPEN,NONO,SET,INA,SSBP2,HMGB1,POLR2A,SF3B2,ITM
AT6A,DVL3,YWHAE,DDX17,GPI,ALDH6A1,ATP5F1B,GNB1,MYH9,SEPTIN2,RPL5,GDI1,FSCN1,GST
PEN,NONO,SET,BASP1,HMGB1,RPS17,MAP4,SF3B2,GTF2F1,HNRNPU,NFE2L1,SCARB2,RTN4,XR
,TRIM28,MATR3,SPEN,NONO,EGLN1,HMGB1,RPL10,DNAJA1,CYB5R3,SF3B2,GTF2F1,HNRNPU,
,SF3B2,AP2B1,GTF2F1,HNRNPU,CPE,TPM4,SCARB2,HMGN2,RBBP4,HNRNPC,LRPPRC,KHDRBS
3C,ATP5F1A,VCP,MARCKSL1,MATR3,EGLN1,LDHB,SSBP2,HMGB1,RPS4X,STMN1,TUBB4B,RPL1
S,NPM1,ATP5F1A,VCP,UBB,TRIM28,MATR3,RPL37A,MRC2,SSBP2,CHD3,BASP1,HMGB1,ANXA2,
,RPLP0,H3-3B,DDX5,KAT6A,ATP5F1B,GDI1,NPM1,HSPD1,USP11,CSDE1,EIF3C,ATP5F1A,VCP,TR
CES3,PSAP,RPLP0,H3-3B,CANX,PTMA,DDX5,DVL3,RPL13,SPTBN1,RPL7,P4HB,IDS,RPL5,GDI1,TM
F5B,BEX3,RBBP4,RPL31,HNRNPC,KHDRBS1,RPS25,SFPQ,DDX3X,H1-0,GJA1,TARDBP,ARID1A,HN
CN1,MARCKS,CSDE1,ATP5F1A,EIF4B,TRIM28,MATR3,YWHAG,RPL37A,SPEN,NONO,DDB1,EGLN
TL,HSP90B1,FADS1,RPL8,UBA1,ATP2B4,ABHD2,CES3,PSAP,PGK1,RPLP0,H3-3B,CANX,PTMA,SO
L,AKAP12,HNRNPU,SCARB2,JUND,CD81,RBBP4,LRPPRC,KHDRBS1,SLC3A2,SFPQ,CDH2,MAGED
SSBP2,HMGB1,POLR2A,RPL10,SF3B2,HNRNPU,RPL23,RPSA,TPM4,EIF4H,XRCC5,RPL14,PSMD2,
,SEC62,DVL3,YWHAE,SLC2A3,TMBIM6,NPM1,ATP5F1A,VCP,TRIM28,MATR3,LRRC28,SPEN,NON
TR3,RPL37A,NONO,SET,SSBP2,VAPA,HMGB1,POLR2A,SF3B2,HNRNPU,RPS16,RPL23,RPSA,SCA
M28,MATR3,SPEN,NONO,SSBP2,HMGB1,AKAP12,GTF2F1,HNRNPU,TNS3,XRCC5,CLSTN1,CHD4,
8A1,EIF3C,ATP5F1A,TRIM28,MATR3,YWHAG,RPL37A,LRRC28,SPEN,NONO,HACD3,DCHS1,SET,E
PM1,PLD3,ATP5F1A,VCP,TRIM28,YWHAG,LRRC28,EGLN1,PDIA3,VAPA,HMGB1,POLR2A,RPS4X,A
,ATP2B4,TUBB2B,RPLP0,CANX,CLU,PTMA,ENO2,DVL3,YWHAE,SLC2A3,BSG,SPTBN1,ATP5F1B,G
CP,MARCKSL1,TRIM28,MATR3,RPL37A,NONO,SET,SSBP2,VAPA,HMGB1,ANXA2,DNAJA1,SF3B2,
K1,H3-3B,PTMA,DDX5,SOX4,KAT6A,DVL3,IDS,TMBIM6,PTMS,SMG1,NPM1,PLD3,ATP5F1A,EIF4E
ATP2B4,HSPA1B,TUBB2B,RPLP0,H3-3B,DDX5,KAT6A,ATP5F1B,RPL7,IDS,GDI1,NPM1,HSPD1,USP
TMA,DVL3,CALM3,MYH9,GDI1,FSCN1,PTMS,GSTP1,CALM1,ATP5F1A,TRIM28,RPL37A,YWHAQ,D
3,SSBP2,HMGB1,RPL10,XRCC5,CD81,BEX3,RBBP4,HNRNPC,KHDRBS1,XPO1,SFPQ,H1-0,TARD
OL18A1,ATP5F1A,VCP,UBB,MATR3,LRRC28,SPEN,NONO,DDB1,DCHS1,SET,SSBP2,BTF3,RPL10,
B,CANX,DDX5,DVL3,YWHAE,SLC2A3,MYH9,GDI1,NPM1,EIF3C,ATP5F1A,VCP,MARCKSL1,UBB,MA
P,MARCKSL1,MATR3,DDB1,DCHS1,SET,EGLN1,SSBP2,HMGB1,RPL10,SF3B2,HNRNPU,SMARCA

ATR3,RPL37A,NONO,SET,SSBP2,VAPA,SF3B2,HNRNPU,RPSA,XRCC5,RPS14,RBBP4,PFN1,NISCH,
CCN2,H1-0,GJA1,TARDBP,HNRNPK,H1-10,INIP,RPL36,LAMP1,CALU,HDAC2,LUC7L3,PGRMC1,H
ATP5F1A,VCP,MARCKSL1,MATR3,RPL37A,SET,EGLN1,RPL19,SF3B1,SSBP2,HMGB1,TPT1,STMN
RCC5,SLC16A7,PAPOLA,CD81,RBBP4,PNN,HNRNPC,KHDRBS1,SFPQ,DDX3X,H1-0,IK,COL6A2,TAR
PTMS,SPTAN1,MARCKS,NPM1,COL18A1,ATP5F1A,VCP,MARCKSL1,UBB,TRIM28,YWHAG,MAPT,S
H1-0,COL6A2,APLP2,TARDBP,HNRNPK,CKAP5,CAPRIN1,HDAC2,GLUL,LUC7L3,CPSF2,ALKBH5,F
NRNPU,SCARB2,XRCC5,GAP43,CBX5,RBBP4,RTN3,HNRNPC,SFPQ,H1-0,RPS7,HNRNPDL,TARDBF
A2,TIMP2,GTF2F1,HNRNPU,CNN3,XRCC5,SLC16A7,PAPOLA,JUND,CD81,RBBP4,PNN,HNRNPC,K
K,U2AF2,PABPN1,HNRNPL,MORF4L2,RAD23B,SNRNP70,HDGF,PTBP1,ILF3,HNRNPUL1,DIDO1,HN
,EGLN1,SSBP2,CHD3,HMGB1,RPS4X,MAP4,HNRNPU,CNN3,XRCC5,C1ORF43,CLSTN1,BEX3,RBE
,SLC16A7,C1ORF43,RPL14,RPS14,BEX3,RBBP4,HNRNPC,KHDRBS1,GLG1,SFPQ,COL6A2,APLP2
PEN,HACD3,HMGB1,POLR2A,RPL18,STMN1,SF3B2,HNRNPU,SMARCA5,HNRNPA0,RTN4,XRCC5,F
L,HNRNPU,XRCC5,SLC7A5,RBBP4,HNRNPC,KHDRBS1,KLHL9,SFPQ,TPM1,CCN2,DDX3X,H1-0,IK
3B2,GTF2F1,GOLGA8A,HNRNPU,SMARCA5,HNRNPA0,RPL29,XRCC5,KPNB1,PAPOLA,ATF4,RBBF
2,WDR1,SRSF4,RBBP4,PNN,HNRNPC,LRPPRC,XPR1,KHDRBS1,SFPQ,RAB7A,H1-0,RPS7,MYO5A,D
,SPTBN1,ATP5F1B,GNB1,P4HB,IDS,GDI1,TMBIM6,SPTAN1,MARCKS,SMG1,NPM1,HSPD1,COL18
PNN,PRKCSH,HNRNPC,KHDRBS1,XPO1,SFPQ,H1-0,IK,GOLM1,HNRNPDL,TARDBP,HNRNPK,MYH1
JUND,BEX3,RBBP4,HNRNPC,KHDRBS1,GLG1,SFPQ,DDX3X,CLTC,COL6A2,APLP2,HNRNPK,H1-10
N1,HMGB1,SF3B2,HNRNPU,CTNNA1,TPM4,SCARB2,BRD3,PJA2,PSMD2,SLC7A5,CHD4,WDR1,ATF
-0,CTNNB1,SEPTIN11,LAMP1,HLA-B,CALU,LUC7L3,PGRMC1,HNRNPL,HNRNPA3,ACO2,SNRNP70
1,HMGB1,RPS4X,RPL10,TPM4,XRCC5,RPL17,RPL14,CCT7,RBBP4,RPL31,HNRNPC,LRPPRC,KHD
RNPU,CTNNA1,TPM4,BRD3,CHD4,WDR1,ATP2A2,CCT7,HNRNPC,KHDRBS1,XPO1,SFPQ,TPM1,PF
FN1,SPTBN2,TKT,RPS15,MAGED2,RPS5,TLN1,ATP1A2,MAP2,COTL1,PPIB,MAP1A,PGRMC1,NUCE
1,GNB2,H1-0,GJA1,MYO5A,DNAJC5,ATP1A2,MAP2,SEPTIN3,CDK16,SKP1,L1CAM,GLUL,PABPN1
ACD3,SSBP2,POLR2A,TUBB4B,RPL11,VCL,SF3B2,GTF2F1,GOLGA8A,HNRNPU,SMARCA5,HNRNF
MA,SOX4,RPL13A,SEC62,DVL3,YWHAE,ALDH6A1,RPL5,COL18A1,SNRNP200,YWHAG,LRRC28,DD
ATP5F1B,GNB1,GDI1,SPTAN1,NPM1,HSPD1,EIF3C,ATP5F1A,VCP,TRIM28,MATR3,SPEN,HACD3,P
KT,RPS15,MAGED2,RPS5,TLN1,MAP2,COTL1,MAP1A,PGRMC1,NUCB1,TOMM20,GAS7,TM9SF2,F
PS5,CTNNB1,ANXA6,MTSS2,DDOST,MYH10,MAP2,STIP1,PPIB,MAP1A,TMEM132A,PGRMC1,NUCE
,TNPO1,SNRNP70,H3-3A,MAPK1,SMARCA4,HNRNPUL1,SUPT6H,HNRNPM,SRSF3,ILF2,MESD,KDN
PRMT1,SF1,BAZ1B,MAPRE2,PSMD8,APEX1,SNRPD3,SOX2,PSMD4,SAP18,ATXN7L3,ELAVL3,HMG
HDAC2,LUC7L3,TM9SF3,HNRNPL,CBX1,RAD23B,TNPO1,SNRNP70,H3-3A,SMARCA4,ILF3,PRKDC
G3BP2,FKBP8,SV2A,TPP1,SSR3,DCLK1,EEF1B2,REEP2,AP2A2,ELAVL3,FXR2,TRIM2,IDH2,NACAD,
DX17,SPTBN1,ATP5F1B,GNB1,GDI1,SPTAN1,NPM1,HSPD1,CSDE1,EIF3C,ATP5F1A,VCP,TRIM28,
,SFPQ,DDX3X,H1-0,TARDBP,HNRNPK,CAPRIN1,LUC7L3,DDX39B,HNRNPL,CBX1,SNRNP70,H3-3
,ADAR,PRMT1,EP300,SF1,BAZ1B,MAPRE2,PSMD8,APEX1,SNRPD3,SOX2,PSMD4,SAP18,ATXN7L3,
MC1,NUCB1,TOMM20,TM9SF2,PRDX5,ACO2,G3BP2,FKBP8,FAU,STMN3,MSMO1,SV2A,MAPRE2
MC1,NUCB1,TOMM20,TM9SF2,PRDX5,ACO2,G3BP2,FKBP8,FAU,STMN3,SYP,MSMO1,SV2A,MAF
S2,DDOST,MYH10,MAP2,STIP1,PPIB,MAP1A,TMEM132A,NUCB1,TOMM20,RPN2,TLN2,TM9SF2,R
OLR2A,RPL18,TUBB4B,ZMIZ1,SF3B2,GTF2F1,HNRNPU,SMARCA5,HNRNPA0,TPM4,RTN4,XRCC5,
,SFPQ,DDR1,MAGED2,HNRNPK,RAB1A,BCLAF1,LUC7L3,CTSB,MDH2,FDFT1,NOP53,GDI2,HNRNI
,TARDBP,HNRNPK,LAMP1,HLA-B,CAPRIN1,MMP2,LUC7L3,CPSF2,TM9SF3,HNRNPL,TSPYL4,TNP
RB2,XRCC5,RBBP4,PFN1,HNRNPC,KHDRBS1,GLG1,SFPQ,PRDX2,H1-0,TLN1,TARDBP,HNRNPK,H
L4,YBX1,ABHD2,CES3,PGK1,PTMA,SOX4,RPL13A,SEC62,DVL3,YWHAE,RPL13,GPI,ALDH6A1,RPL
P1A2,MAP2,COTL1,PPIB,PEA15,NUCB1,TOMM20,ACO2,G3BP2,IPO7,NSG2,POLR2B,SDF4,ATP6V

IUN,NARS1,CKAP4,HLA-B,L1CAM,RPL23A,CHD2,TERF2IP,VEGFA,MAPRE1,ACADVL,SUPT5H,KDM
M20,ACO2,G3BP2,FKBP8,SDF4,ATP6V1B2,KDM3B,MAP6,MAPRE2,AKAP9,SSR3,DCLK1,DTX4,EE
,RTN1,ACO2,CNTNAP2,G3BP2,FKBP8,ATP6V1B2,STMN3,FBRSL1,VDAC1,SF1,BRD4,MAP6,SV2A,
,PEA15,PGRMC1,TOMM20,GAS7,TLN2,PRDX5,ACO2,CNTNAP2,MARS1,ATP6V1B2,STMN3,STXBP
),PRDX5,RTN1,ACO2,MARS1,IPO7,ATP5PB,SCAF11,FKBP8,ATP6V1B2,STMN3,STXBP1,KDM3B,S
SF1,MECP2,SUMO1,ADNP,RTCB,AKAP8L,NDRG4,ZEB1,SOX9,FBL,MAN2B1,RBM17,ACIN1,UBTF
SART1,SAP18,ATXN7L3,HTATSF1,MECP2,CD2BP2,SUMO1,ADNP,RTCB,FBL,NACC1,RBM17,PNM
A1,PSMD2,KIF5B,CHD4,BEX3,CCT7,TADA2B,RPS25,RPS7,APLP2,ARCN1,CAPRIN1,SMCHD1,IPO5
TLN1,CTNNB1,ANXA6,PPIB,PEA15,TMEM132A,PGRMC1,ACO2,CNTNAP2,MARS1,G3BP2,AEBP1,A
i1,NPM1,HSPD1,VCP,MARCKSL1,AFF4,TRIM28,MATR3,SPEN,HACD3,HMGB1,POLR2A,RPL18,TU
2,SAP18,ELAVL3,PSAT1,HMGA1,HTATSF1,MECP2,ADNP,UBE2J1,RTCB,NDRG4,BACE1,RBM17,C
8,PSAT1,HMGA1,HTATSF1,MECP2,ADNP,UBE2J1,RTCB,NDRG4,BACE1,RBM17,CTCF,RFX7,APO
SAP18,ATXN7L3,HTATSF1,MECP2,CD2BP2,SUMO1,ADNP,RTCB,FBL,NACC1,RBM17,CMTR1,NFA
ADNP,RTCB,AKAP8L,NDRG4,ZEB1,SOX9,FBL,MAN2B1,RBM17,ACIN1,UBTF

SF4,CDH6,PEBP1,SULF1,JUN,NARS1,CKAP4,HLA-B,MMP2,L1CAM,RPL23A,CHD2,TERF2IP,VEGF
ATSF1,MECP2,ADNP,RTCB,AKAP8L,ZEB1,SEPHS1,SOX9,DLG4,NACC1,BACE1,RBM17,CTCF,RFX
,PSAT1,HMGA1,HTATSF1,MECP2,ADNP,RTCB,AKAP8L,ZEB1,SEPHS1,SOX9,DLG4,NACC1,BACE1,
SK1,AKAP9,TMOD2,TPP1,CNP,NMNAT2,SORT1,PSAT1,FXR2,TRIM2,MTURN,AKAP8L,ALDH7A1,N
A2,IGLON5,FXR2,CLTA,FAM168A,SEZ6L2,RAB3A,NAPB,CASKIN1,VDAC2,UQCRFS1,CHGB,GNAZ,
TLN1,CTNNB1,PEA15,TMEM132A,PGRMC1,ACO2,CNTNAP2,MARS1,G3BP2,ACADVL,FKBP8,POL
P,RTCB,FBL,RBM17,CMTR1,NFASC,ACIN1,ZNF326,UBTF
SF1,MECP2,ADNP,UBE2J1,RTCB,AKAP8L,ZEB1,FBL,MAN2B1,BACE1,RBM17,CTCF,RFX7,CMTR1,
32,STMN3,STXBP1,SYP,VDAC1,MSMO1,BRD4,SV2A,MAPRE2,SEPTIN7,TMOD2,SEPTIN5,SSR3,CA
)TL1,PEA15,NUCB1,TOMM20,ACO2,G3BP2,IPO7,POLR2B,SDF4,FBLN1,MAPRE2,AKAP9,TPP1,DT
CLK1,EEF1B2,SYT11,RCN2,ANK2,SCAMP5,TRIM2,SEZ6L2,ALCAM,VDAC2,UQCRFS1,ABCD3,PPT1
CADM3,REEP2,CHMP4B,AP2A2,IGLON5,USP47,CLTA,TRIM2,TSPAN7,NDRG4,MRPS6,RAB3A,NAP
Q,MAGED2,HNRNPK,CKAP5,CAPRIN1,MLXIP,PARP1,CBX1,H3-3A,PRPF6,VEGFA,SMARCA4,ILF3,P
CAN18,CD99L2,EDC4,CSNK2A1,SLC2A1,CLCN7,HDGFL2,RANGAP1,MKNK2,SUN2,HIPK2,WIZ,EC
31,NSG2,FKBP8,STMN3,FBRSL1,VDAC1,SF1,BRD4,MAP6,LINGO1,STRN4,DCLK1,EEF1B2,MSH6,
1A1,SRSF4,HDLBP,PEBP1,SULF1,PRKAR1A,JUN,GOLM1,WLS,NARS1,CKAP5,GLUL,CDKN1A,CTS
IAPB,MAN2B1,CNTN1,GNAZ,APOE,NFASC,ABCD3,SYT1,EPHA4,PPFIA3

1,DCX,EIF4G2,RPS6,DBN1,ENO1,TUBA1B,PIK3R2,FLNA,MAP1B,NCL,SLC9A7,ALDOA,GANAB,FTL

JBA1B,PIK3R2,MAP1B,NCL,SLC9A7,ALDOA,GANAB,FTL,HSP90B1,FADS1,NEFL,NEFM,PRPF8,UG

B,PIK3R2,MAP1B,NCL,SLC9A7,ALDOA,GANAB,FTL,HSP90B1,FADS1,NEFL,NEFM,PRPF8,UGT8,R

1B,PIK3R2,NCL,SLC9A7,ALDOA,GANAB,FTL,HSP90B1,NEFL,NEFM,PRPF8,RPL8,FBXL4,YBX1,UBA

1,DCX,EIF4G2,RPS6,DBN1,ENO1,TUBA1B,PIK3R2,FLNA,MAP1B,NCL,SLC9A7,ALDOA,GANAB,COI

S6,DBN1,ENO1,TUBA1B,PIK3R2,MAP1B,NCL,SLC9A7,ALDOA,GANAB,FTL,HSP90B1,FADS1,NEFM

,ENO1,TUBA1B,PIK3R2,NCL,SLC9A7,GANAB,COL6A1,FTL,HSP90B1,FADS1,NEFM,PRPF8,UGT8,

NO1,TUBA1B,PIK3R2,FLNA,MAP1B,NCL,SLC9A7,ALDOA,GANAB,FTL,HSP90B1,FADS1,PRPF8,RPL

IF4G2,RPS6,DBN1,ENO1,TUBA1B,PIK3R2,FLNA,NCL,ALDOA,GANAB,COL6A1,FTL,HSP90B1,FADS

L,SLC9A7,ALDOA,GANAB,COL6A1,FTL,HSP90B1,FADS1,PRPF8,UGT8,RPL8,FBXL4,YBX1,UBA1,A

L,SLC9A7,ALDOA,GANAB,COL6A1,FTL,HSP90B1,FADS1,PRPF8,UGT8,RPL8,FBXL4,YBX1,UBA1,A

1B,PIK3R2,FLNA,NCL,SLC9A7,ALDOA,GANAB,FTL,HSP90B1,FADS1,PRPF8,UGT8,RPL8,FBXL4,YE

,GANAB,FTL,HSP90B1,PRPF8,RPL8,FBXL4,UBA1,ATP2B4,TMF1,ABHD2,HSPA1B,CES3,PSAP,PGK

EIF4G2,RPS6,DBN1,ENO1,TUBA1B,PIK3R2,FLNA,NCL,SLC9A7,ALDOA,GANAB,FTL,HSP90B1,FADS

EIF4G2,RPS6,DBN1,ENO1,TUBA1B,PIK3R2,FLNA,NCL,SLC9A7,ALDOA,GANAB,FTL,HSP90B1,FADS

A1B,PIK3R2,FLNA,NCL,ALDOA,GANAB,HSP90B1,FADS1,PRPF8,UGT8,RPL8,FBXL4,YBX1,UBA1,AT

3A1B,PIK3R2,FLNA,NCL,SLC9A7,ALDOA,GANAB,FTL,HSP90B1,FADS1,PRPF8,RPL8,FBXL4,YBX1,U

3A1B,PIK3R2,FLNA,NCL,SLC9A7,ALDOA,GANAB,FTL,HSP90B1,FADS1,PRPF8,RPL8,FBXL4,YBX1,U

RPS6,DBN1,ENO1,TUBA1B,PIK3R2,FLNA,NCL,SLC9A7,ALDOA,GANAB,FTL,HSP90B1,FADS1,UGT8

,SLC9A7,ALDOA,GANAB,FTL,HSP90B1,FADS1,PRPF8,UGT8,RPL8,FBXL4,YBX1,UBA1,ATP2B4,CA

FBXL4,YBX1,UBA1,ATP2B4,TMF1,HSPA1B,COL4A1,PSAP,PGK1,RPLP0,H3-3B,CANX,DDX5,RPL13A

JBA1B,PIK3R2,FLNA,MAP1B,NCL,SLC9A7,ALDOA,GANAB,FTL,HSP90B1,FADS1,NEFM,PRPF8,UG

G2,RPS6,ENO1,TUBA1B,PIK3R2,MAP1B,NCL,SLC9A7,ALDOA,GANAB,HSP90B1,FADS1,PRPF8,UG

G2,RPS6,ENO1,TUBA1B,PIK3R2,MAP1B,NCL,SLC9A7,ALDOA,GANAB,HSP90B1,FADS1,PRPF8,UG

BXL4,UBA1,CALR,TMF1,ABHD2,HSPA1B,PSAP,PGK1,RPLP0,H3-3B,CANX,PTMA,DDX5,ENO2,SOX

,CALR,ABHD2,HSPA1B,PSAP,RPLP0,H3-3B,CANX,CLU,PTMA,DDX5,ENO2,SOX4,RPL13A,RPL6,KA

G2,RPS6,DBN1,ENO1,TUBA1B,PIK3R2,FLNA,MAP1B,NCL,SLC9A7,ALDOA,GANAB,FTL,HSP90B1,

PIK3R2,MAP1B,NCL,SLC9A7,ALDOA,GANAB,FTL,HSP90B1,FADS1,PRPF8,RPL8,FBXL4,YBX1,UBA

R2,FLNA,NCL,SLC9A7,ALDOA,GANAB,FTL,HSP90B1,FADS1,PRPF8,RPL8,FBXL4,YBX1,UBA1,ATP2

R2,FLNA,NCL,SLC9A7,ALDOA,GANAB,FTL,HSP90B1,FADS1,PRPF8,RPL8,FBXL4,YBX1,UBA1,ATP2

BN1,ENO1,TUBA1B,PIK3R2,FLNA,MAP1B,NCL,SLC9A7,ALDOA,GANAB,HSP90B1,FADS1,PRPF8,U

SLC9A7,ALDOA,GANAB,HSP90B1,FADS1,PRPF8,RPL8,FBXL4,YBX1,UBA1,ATP2B4,CALR,TMF1,AE

SLC9A7,ALDOA,GANAB,HSP90B1,FADS1,PRPF8,RPL8,FBXL4,YBX1,UBA1,ATP2B4,CALR,TMF1,AE

P90B1,FADS1,RPL8,FBXL4,YBX1,UBA1,ATP2B4,TMF1,HSPA1B,COL4A1,PSAP,PGK1,RPLP0,H3-3

LDOA,GANAB,HSP90B1,FADS1,PRPF8,RPL8,FBXL4,YBX1,UBA1,ATP2B4,TMF1,ABHD2,HSPA1B,C

,ENO1,TUBA1B,PIK3R2,NCL,SLC9A7,ALDOA,GANAB,FTL,HSP90B1,FADS1,UGT8,RPL8,FBXL4,YE

DS1,PRPF8,UGT8,RPL8,FBXL4,YBX1,UBA1,ATP2B4,TMF1,ABHD2,CES3,PSAP,PGK1,RPLP0,H3-3

OA,GANAB,FTL,HSP90B1,FADS1,RPL8,FBXL4,YBX1,UBA1,ATP2B4,CALR,TMF1,ABHD2,HSPA1B,

OA,GANAB,FTL,HSP90B1,FADS1,RPL8,FBXL4,YBX1,UBA1,ATP2B4,CALR,TMF1,ABHD2,HSPA1B,

3A1B,PIK3R2,NCL,SLC9A7,ALDOA,GANAB,HSP90B1,FADS1,PRPF8,RPL8,FBXL4,YBX1,UBA1,ATP2

6A1,FTL,HSP90B1,FADS1,RPL8,FBXL4,YBX1,UBA1,ATP2B4,CALR,COL4A1,PSAP,RPLP0,H3-3B,C

,EIF4G2,RPS6,DBN1,ENO1,TUBA1B,PIK3R2,NCL,SLC9A7,ALDOA,GANAB,FTL,HSP90B1,FADS1,P

,EIF4G2,RPS6,DBN1,ENO1,TUBA1B,PIK3R2,NCL,SLC9A7,ALDOA,GANAB,FTL,HSP90B1,FADS1,P

1,TUBA1B,PIK3R2,NCL,SLC9A7,ALDOA,GANAB,HSP90B1,FADS1,PRPF8,UGT8,RPL8,FBXL4,YBX1

1,TUBA1B,PIK3R2,NCL,SLC9A7,ALDOA,GANAB,HSP90B1,FADS1,PRPF8,UGT8,RPL8,FBXL4,YBX1

P1B,NCL,ALDOA,GANAB,HSP90B1,FADS1,PRPF8,RPL8,FBXL4,YBX1,UBA1,ATP2B4,TMF1,ABHD2,
2,FLNA,MAP1B,NCL,SLC9A7,ALDOA,GANAB,FTL,HSP90B1,FADS1,PRPF8,UGT8,RPL8,FBXL4,YBX
2,FLNA,MAP1B,NCL,SLC9A7,ALDOA,GANAB,FTL,HSP90B1,FADS1,PRPF8,UGT8,RPL8,FBXL4,YBX
RPS6,ENO1,TUBA1B,PIK3R2,FLNA,NCL,SLC9A7,ALDOA,GANAB,FTL,HSP90B1,FADS1,PRPF8,RPL8
IK3R2,FLNA,NCL,SLC9A7,ALDOA,GANAB,FTL,HSP90B1,FADS1,PRPF8,RPL8,FBXL4,YBX1,UBA1,A
,DBN1,ENO1,TUBA1B,PIK3R2,NCL,SLC9A7,ALDOA,GANAB,HSP90B1,FADS1,PRPF8,UGT8,RPL8,F
UBA1B,PIK3R2,FLNA,MAP1B,NCL,SLC9A7,ALDOA,GANAB,COL6A1,FTL,HSP90B1,FADS1,PRPF8,F
2,NCL,SLC9A7,GANAB,FTL,HSP90B1,FADS1,NEFM,PRPF8,RPL8,FBXL4,YBX1,ATP2B4,CALR,ABHD
A7,ALDOA,GANAB,FTL,HSP90B1,FADS1,PRPF8,RPL8,FBXL4,YBX1,UBA1,ATP2B4,TMF1,ABHD2,HS
A7,ALDOA,GANAB,FTL,HSP90B1,FADS1,PRPF8,RPL8,FBXL4,YBX1,UBA1,ATP2B4,TMF1,ABHD2,HS
N1,ENO1,TUBA1B,PIK3R2,FLNA,NCL,SLC9A7,ALDOA,GANAB,FTL,HSP90B1,FADS1,PRPF8,UGT8,
SP90B1,FADS1,PRPF8,RPL8,FBXL4,YBX1,UBA1,ATP2B4,CALR,HSPA1B,TUBB2B,PSAP,PGK1,RPLI
,SLC9A7,ALDOA,GANAB,FTL,HSP90B1,FADS1,PRPF8,RPL8,FBXL4,YBX1,UBA1,ATP2B4,TMF1,AB
,SLC9A7,ALDOA,GANAB,FTL,HSP90B1,FADS1,PRPF8,RPL8,FBXL4,YBX1,UBA1,ATP2B4,TMF1,AB
BHD2,HSPA1B,COL4A1,CES3,PSAP,RPLP0,H3-3B,CANX,DDX5,SOX4,RPL13A,SEC62,KAT6A,DVL3
IA,NCL,SLC9A7,ALDOA,GANAB,COL6A1,FTL,HSP90B1,FADS1,PRPF8,RPL8,FBXL4,YBX1,UBA1,AT
7,ALDOA,GANAB,COL6A1,FTL,HSP90B1,FADS1,NEFM,PRPF8,RPL8,FBXL4,YBX1,UBA1,ATP2B4,TI
PIK3R2,FLNA,NCL,SLC9A7,ALDOA,GANAB,FTL,HSP90B1,FADS1,PRPF8,UGT8,RPL8,FBXL4,YBX1,U
PIK3R2,FLNA,NCL,SLC9A7,ALDOA,GANAB,FTL,HSP90B1,FADS1,PRPF8,UGT8,RPL8,FBXL4,YBX1,U
K3R2,FLNA,MAP1B,NCL,SLC9A7,ALDOA,GANAB,FTL,HSP90B1,FADS1,PRPF8,RPL8,FBXL4,YBX1,U
K3R2,FLNA,MAP1B,NCL,SLC9A7,ALDOA,GANAB,FTL,HSP90B1,FADS1,PRPF8,RPL8,FBXL4,YBX1,U
SP90B1,PRPF8,YBX1,UBA1,ATP2B4,CALR,TMF1,ABHD2,HSPA1B,PSAP,PGK1,RPLP0,H3-3B,CANX,
,TUBA1B,PIK3R2,FLNA,NCL,SLC9A7,ALDOA,GANAB,COL6A1,FTL,HSP90B1,FADS1,PRPF8,RPL8,I
2,MAP1B,NCL,SLC9A7,ALDOA,GANAB,FADS1,NEFM,RPL8,FBXL4,YBX1,UBA1,ATP2B4,ABHD2,HSF
DBN1,ENO1,TUBA1B,PIK3R2,MAP1B,NCL,ALDOA,GANAB,FTL,HSP90B1,FADS1,PRPF8,RPL8,FBXL
P90B1,PRPF8,RPL8,FBXL4,YBX1,UBA1,ATP2B4,TMF1,ABHD2,HSPA1B,CES3,PSAP,PGK1,RPLP0,I
T8,FBXL4,UBA1,ATP2B4,ABHD2,HSPA1B,PSAP,PGK1,RPLP0,H3-3B,CANX,PTMA,DDX5,SOX4,SEC6
LDOA,GANAB,COL6A1,HSP90B1,FADS1,RPL8,FBXL4,YBX1,UBA1,ATP2B4,TMF1,ABHD2,HSPA1B,
P90B1,FADS1,RPL8,FBXL4,YBX1,UBA1,ATP2B4,TMF1,ABHD2,HSPA1B,COL4A1,CES3,PSAP,RPLP0
P90B1,FADS1,RPL8,FBXL4,YBX1,UBA1,ATP2B4,TMF1,ABHD2,HSPA1B,COL4A1,CES3,PSAP,RPLP0
B1,FADS1,PRPF8,RPL8,FBXL4,YBX1,UBA1,ATP2B4,CALR,TMF1,ABHD2,HSPA1B,CES3,PSAP,PGK
LDOA,GANAB,FTL,HSP90B1,FADS1,PRPF8,RPL8,FBXL4,YBX1,UBA1,ATP2B4,CALR,TMF1,ABHD2,F
S1,PRPF8,UGT8,RPL8,FBXL4,YBX1,ATP2B4,TMF1,ABHD2,HSPA1B,CES3,PSAP,RPLP0,H3-3B,CAN
B,PIK3R2,NCL,SLC9A7,ALDOA,GANAB,HSP90B1,FADS1,PRPF8,UGT8,RPL8,FBXL4,YBX1,UBA1,AT
B,PIK3R2,NCL,SLC9A7,ALDOA,GANAB,HSP90B1,FADS1,PRPF8,UGT8,RPL8,FBXL4,YBX1,UBA1,AT
B,PIK3R2,MAP1B,NCL,SLC9A7,ALDOA,GANAB,FTL,HSP90B1,FADS1,PRPF8,RPL8,FBXL4,YBX1,UE
,DBN1,ENO1,TUBA1B,PIK3R2,NCL,ALDOA,GANAB,HSP90B1,FADS1,PRPF8,UGT8,RPL8,FBXL4,YE
K3R2,NCL,SLC9A7,GANAB,FTL,HSP90B1,FADS1,PRPF8,UGT8,RPL8,FBXL4,YBX1,UBA1,ATP2B4,C
,ATP2B4,ABHD2,CES3,PSAP,PGK1,RPLP0,H3-3B,CANX,PTMA,SOX4,RPL13A,KAT6A,DVL3,YWHAE
NCL,SLC9A7,ALDOA,FTL,HSP90B1,FADS1,PRPF8,RPL8,FBXL4,YBX1,UBA1,ATP2B4,TMF1,HSPA1E
2B4,TMF1,ABHD2,CES3,PSAP,PGK1,RPLP0,H3-3B,CANX,CLU,PTMA,DDX5,RPL13A,SEC62,RPL6,K
PF8,RPL8,FBXL4,UBA1,TMF1,ABHD2,HSPA1B,PSAP,PGK1,RPLP0,H3-3B,CANX,PTMA,DDX5,RPL1
4,UBA1,ATP2B4,TMF1,ABHD2,HSPA1B,CES3,PSAP,PGK1,RPLP0,H3-3B,CANX,PTMA,RPL13A,SEC
RPS6,ENO1,TUBA1B,PIK3R2,FLNA,NCL,SLC9A7,ALDOA,GANAB,FTL,HSP90B1,FADS1,PRPF8,RPL8
DA,GANAB,HSP90B1,FADS1,PRPF8,UGT8,RPL8,FBXL4,YBX1,UBA1,ATP2B4,TMF1,ABHD2,CES3,PS

A,GANAB,HSP90B1,FADS1,PRPF8,RPL8,FBXL4,YBX1,UBA1,ATP2B4,CALR,TMF1,ABHD2,COL4A1,C
A1B,PIK3R2,FLNA,NCL,SLC9A7,ALDOA,GANAB,FTL,HSP90B1,FADS1,PRPF8,UGT8,RPL8,YBX1,UI
,ALDOA,GANAB,FTL,HSP90B1,FADS1,PRPF8,UGT8,RPL8,FBXL4,YBX1,UBA1,ATP2B4,CALR,TMF1
,ALDOA,GANAB,FTL,HSP90B1,FADS1,PRPF8,UGT8,RPL8,FBXL4,YBX1,UBA1,ATP2B4,CALR,TMF1
-3B,CANX,CLU,PTMA,DDX5,ENO2,DVL3,YWHAE,GPI,ATP5F1B,SEPTIN2,P4HB,RPL5,GDI1,SPTAN1
9A7,ALDOA,HSP90B1,FADS1,RPL8,FBXL4,UBA1,ATP2B4,CALR,TMF1,ABHD2,HSPA1B,TUBB2B,C
AB,FTL,HSP90B1,FADS1,PRPF8,UGT8,RPL8,FBXL4,YBX1,UBA1,ATP2B4,CALR,TMF1,ABHD2,HSP
,UBA1,ATP2B4,HSPA1B,PSAP,PGK1,RPLP0,H3-3B,CANX,PTMA,DDX5,RPL13A,SEC62,RPL6,KAT6
RPS6,DBN1,ENO1,TUBA1B,PIK3R2,NCL,ALDOA,GANAB,FTL,HSP90B1,FADS1,PRPF8,UGT8,RPL8,
IAB,FTL,HSP90B1,FADS1,PRPF8,RPL8,FBXL4,YBX1,UBA1,ATP2B4,HSPA1B,CES3,PSAP,PGK1,RPL
IAB,FTL,HSP90B1,FADS1,PRPF8,RPL8,FBXL4,YBX1,UBA1,ATP2B4,HSPA1B,CES3,PSAP,PGK1,RPL
A1B,CES3,PSAP,PGK1,RPLP0,H3-3B,CANX,DDX5,RPL13A,SEC62,KAT6A,DVL3,YWHAE,RPL13,GPI,
HSP90B1,FADS1,PRPF8,RPL8,FBXL4,YBX1,UBA1,ATP2B4,TMF1,COL4A1,PSAP,PGK1,RPLP0,H3-3
SP90B1,FADS1,PRPF8,UGT8,RPL8,FBXL4,YBX1,UBA1,ATP2B4,CALR,TMF1,ABHD2,HSPA1B,PSAP,
SP90B1,FADS1,PRPF8,UGT8,RPL8,FBXL4,YBX1,UBA1,ATP2B4,CALR,TMF1,ABHD2,HSPA1B,PSAP,
,FADS1,PRPF8,RPL8,FBXL4,YBX1,UBA1,ATP2B4,TMF1,ABHD2,COL4A1,PSAP,PGK1,H3-3B,CANX,
,CALR,TMF1,ABHD2,HSPA1B,PSAP,PGK1,RPLP0,H3-3B,CANX,PTMA,DDX5,SEC62,KAT6A,DVL3,Y
PRPF8,RPL8,FBXL4,YBX1,UBA1,ATP2B4,TMF1,ABHD2,HSPA1B,PSAP,PGK1,RPLP0,H3-3B,CANX,
A1,ATP2B4,CALR,TMF1,ABHD2,COL4A1,PSAP,RPLP0,H3-3B,CANX,DDX5,SOX4,RPL13A,SEC62,RF
LDOA,FTL,HSP90B1,FADS1,PRPF8,UGT8,RPL8,YBX1,UBA1,ATP2B4,TMF1,ABHD2,PSAP,PGK1,RP
,RPL8,YBX1,UBA1,TMF1,ABHD2,CES3,PSAP,PGK1,RPLP0,H3-3B,CANX,PTMA,DDX5,SOX4,RPL13
DCX,EIF4G2,RPS6,DBN1,ENO1,TUBA1B,PIK3R2,FLNA,MAP1B,NCL,SLC9A7,ALDOA,GANAB,COL6
A1,ABHD2,CES3,H3-3B,CANX,PTMA,DDX5,ENO2,SEC62,YWHAE,RPL13,SPTBN1,GNB1,APP,GDI1,
A,GANAB,FTL,HSP90B1,FADS1,PRPF8,RPL8,YBX1,UBA1,ATP2B4,TMF1,ABHD2,PSAP,PGK1,RPLP
SAP,H3-3B,CANX,DDX5,SOX4,RPL13A,DVL3,YWHAE,RPL13,SPTBN1,ATP5F1B,SEPTIN2,PTMS,SPT
A,NCL,SLC9A7,ALDOA,GANAB,FTL,HSP90B1,FADS1,PRPF8,UGT8,RPL8,YBX1,UBA1,ATP2B4,CAL
LDOA,GANAB,FTL,HSP90B1,FADS1,UGT8,RPL8,FBXL4,YBX1,UBA1,ATP2B4,CES3,PSAP,PGK1,RPL
AB,HSP90B1,FADS1,RPL8,FBXL4,YBX1,UBA1,ATP2B4,CALR,TMF1,ABHD2,CES3,PSAP,PGK1,RPL
CL,SLC9A7,ALDOA,GANAB,HSP90B1,FADS1,PRPF8,RPL8,FBXL4,YBX1,UBA1,ATP2B4,TMF1,ABHD
DOA,GANAB,HSP90B1,FADS1,UGT8,RPL8,FBXL4,YBX1,UBA1,ATP2B4,TMF1,ABHD2,CES3,PSAP,P
A,GANAB,FTL,HSP90B1,FADS1,PRPF8,RPL8,FBXL4,YBX1,UBA1,ATP2B4,CALR,TMF1,ABHD2,HSP
,FBXL4,UBA1,ATP2B4,CALR,TMF1,ABHD2,CES3,PGK1,RPLP0,H3-3B,CANX,DDX5,ENO2,SOX4,RF
VHAG,RPL37A,LRRC28,SPEN,NONO,DDB1,HACD3,RPL19,SF3B1,PDIA3,SSBP2,CHD3,BASP1,VAP
FBXL4,UBA1,CALR,TMF1,ABHD2,PSAP,PGK1,RPLP0,H3-3B,CANX,PTMA,DDX5,SOX4,RPL13A,SEC
A7,ALDOA,GANAB,HSP90B1,FADS1,UGT8,RPL8,YBX1,UBA1,ATP2B4,TMF1,ABHD2,CES3,PSAP,PG
ANAB,FTL,HSP90B1,FADS1,UGT8,RPL8,FBXL4,YBX1,UBA1,ATP2B4,CALR,TMF1,ABHD2,HSPA1B,C
,NCL,SLC9A7,ALDOA,GANAB,COL6A1,FTL,HSP90B1,FADS1,NEFL,NEFM,RPL8,FBXL4,YBX1,UBA
FADS1,PRPF8,RPL8,FBXL4,YBX1,UBA1,ATP2B4,CALR,HSPA1B,PSAP,PGK1,H3-3B,CANX,PTMA,DD
DOA,FTL,HSP90B1,FADS1,RPL8,FBXL4,UBA1,ATP2B4,CALR,TMF1,ABHD2,HSPA1B,TUBB2B,PSA
BA1,ATP2B4,CALR,HSPA1B,CES3,PSAP,RPLP0,H3-3B,CANX,PTMA,DDX5,SOX4,RPL13A,SEC62,RF
L4,UBA1,ATP2B4,CALR,ABHD2,HSPA1B,CES3,PGK1,RPLP0,H3-3B,CANX,PTMA,DDX5,ENO2,SOX
RPLP0,H3-3B,CANX,PTMA,DDX5,ENO2,SOX4,RPL13A,SEC62,RPL6,KAT6A,DVL3,YWHAE,RPL13,G
SP90B1,FADS1,RPL8,FBXL4,UBA1,ATP2B4,CALR,TMF1,ABHD2,HSPA1B,TUBB2B,PSAP,PGK1,RPL
UGT8,RPL8,FBXL4,YBX1,UBA1,ATP2B4,TMF1,ABHD2,CES3,PSAP,RPLP0,H3-3B,CANX,PTMA,DDX
L8,FBXL4,YBX1,UBA1,ATP2B4,HSPA1B,PSAP,RPLP0,H3-3B,CANX,PTMA,DDX5,SOX4,RPL13A,SEC

DS1,RPL8,FBXL4,UBA1,ATP2B4,TMF1,HSPA1B,TUBB2B,COL4A1,CES3,PSAP,PGK1,RPLP0,H3-3E
MA,DDX5,ENO2,SOX4,RPL13A,SEC62,RPL6,KAT6A,DVL3,YWHAE,RPL13,GPI,ATP5F1B,CALM3,GNI
AN1,MARCKS,NPM1,COL18A1,EIF3C,ATP5F1A,VCP,MARCKSL1,UBB,TRIM28,MATR3,RPL37A,LRI
,UBA1,ATP2B4,TMF1,HSPA1B,PSAP,H3-3B,CANX,PTMA,DDX5,RPL13A,KAT6A,YWHAE,RPL13,BSG
LDOA,GANAB,FTL,HSP90B1,FADS1,PRPF8,RPL8,FBXL4,YBX1,UBA1,ATP2B4,CALR,TMF1,ABHD2
,ATP5F1B,MYH9,SEPTIN2,FSCN1,PTMS,SPTAN1,MARCKS,NPM1,COL18A1,EIF3C,ATP5F1A,VCP,I
,TMF1,ABHD2,HSPA1B,PSAP,PGK1,H3-3B,CANX,PTMA,SOX4,RPL13A,SEC62,KAT6A,DVL3,YWHAI
TUBA1B,PIK3R2,FLNA,NCL,ALDOA,GANAB,FTL,HSP90B1,FADS1,PRPF8,RPL8,FBXL4,YBX1,UBA1
TUBA1B,PIK3R2,FLNA,NCL,ALDOA,GANAB,FTL,HSP90B1,FADS1,PRPF8,RPL8,FBXL4,YBX1,UBA1
FADS1,PRPF8,RPL8,FBXL4,YBX1,UBA1,ATP2B4,CALR,TMF1,CES3,PSAP,RPLP0,H3-3B,CANX,PTM
TMF1,ABHD2,COL4A1,PSAP,PGK1,RPLP0,H3-3B,CANX,PTMA,DDX5,SOX4,RPL13A,SEC62,KAT6A,
3XL4,YBX1,UBA1,ATP2B4,HSPA1B,COL4A1,PSAP,PGK1,H3-3B,CANX,PTMA,DDX5,KAT6A,DVL3,YV
3XL4,YBX1,UBA1,ATP2B4,HSPA1B,COL4A1,PSAP,PGK1,H3-3B,CANX,PTMA,DDX5,KAT6A,DVL3,YV
PL8,FBXL4,YBX1,UBA1,ATP2B4,ABHD2,COL4A1,PSAP,PGK1,H3-3B,CANX,PTMA,DDX5,DVL3,YWH
P2B4,CALR,ABHD2,HSPA1B,COL4A1,CES3,PSAP,RPLP0,H3-3B,CANX,PTMA,RPL13A,SEC62,DVL3
P2B4,TMF1,ABHD2,HSPA1B,PSAP,PGK1,RPLP0,H3-3B,CANX,PTMA,SOX4,RPL13A,SEC62,KAT6A,
RPLP0,H3-3B,CANX,DDX5,SOX4,RPL13A,SEC62,KAT6A,DVL3,YWHAE,RPL13,DDX17,SPTBN1,ATP5
IF1,ABHD2,HSPA1B,CES3,PSAP,PGK1,H3-3B,CANX,CLU,PTMA,DDX5,SOX4,SEC62,KAT6A,DVL3,Y
,YBX1,UBA1,ATP2B4,TMF1,ABHD2,CES3,PSAP,PGK1,H3-3B,CANX,PTMA,DDX5,RPL13A,SEC62,D
,RPS6,ENO1,TUBA1B,PIK3R2,MAP1B,NCL,SLC9A7,ALDOA,GANAB,COL6A1,FTL,FADS1,PRPF8,U
GANAB,FTL,HSP90B1,FADS1,PRPF8,RPL8,FBXL4,YBX1,UBA1,ATP2B4,CALR,TMF1,ABHD2,COL4
ANX,PTMA,DDX5,SOX4,RPL13A,KAT6A,DVL3,RPL13,ATP5F1B,CALM3,RPL7,IDS,RPL5,PTMS,MAF
NONO,DDB1,HACD3,RPL19,SF3B1,SSBP2,BASP1,RPS3A,UCHL1,BTF3,RPL10,RPS17,DNAJA1,ITM
DS1,UGT8,RPL8,FBXL4,YBX1,UBA1,ATP2B4,TMF1,ABHD2,CES3,PSAP,RPLP0,H3-3B,CANX,PTMA
,FBXL4,YBX1,ATP2B4,HSPA1B,PSAP,PGK1,H3-3B,CANX,PTMA,DDX5,RPL13A,SEC62,KAT6A,DVL
L8,FBXL4,YBX1,UBA1,ATP2B4,TMF1,ABHD2,HSPA1B,COL4A1,PSAP,PGK1,RPLP0,H3-3B,CANX,P
L8,FBXL4,YBX1,UBA1,ATP2B4,TMF1,ABHD2,HSPA1B,COL4A1,PSAP,PGK1,RPLP0,H3-3B,CANX,P
34,TMF1,HSPA1B,PSAP,H3-3B,CANX,PTMA,DDX5,RPL13A,SEC62,RPL6,DVL3,YWHAE,RPL13,DDX1
,DVL3,YWHAE,RPL13,GPI,ATP5F1B,RPL7,IDS,RPL5,GDI1,PTMS,SPTAN1,NPM1,EIF3C,ATP5F1A,V
1,ATP2B4,TMF1,ABHD2,PSAP,PGK1,RPLP0,H3-3B,CANX,PTMA,DDX5,RPL13A,SEC62,KAT6A,DVL
L3,YWHAE,RPL13,ALDH6A1,ATP5F1B,SEPTIN2,GDI1,FSCN1,MARCKS,NPM1,EIF3C,ATP5F1A,VCP
RPL13,DDX17,ATP5F1B,CALM3,RPL7,P4HB,IDS,RPL5,TMBIM6,PTMS,SPTAN1,GSTP1,MARCKS,C
HSP90B1,PRPF8,RPL8,YBX1,UBA1,ATP2B4,TMF1,ABHD2,PSAP,PGK1,RPLP0,H3-3B,CANX,PTMA,D
IDS,GDI1,TMBIM6,MARCKS,NPM1,SNRNP200,ATP5F1A,VCP,MARCKSL1,UBB,TRIM28,MATR3,YV
YWHAE,GPI,FASN,SPTBN1,ATP5F1B,SEPTIN2,FSCN1,PTMS,SPTAN1,GSTP1,NPM1,SNRNP200,CSI
AT6A,DVL3,YWHAE,ALDH6A1,ATP5F1B,SEPTIN2,GDI1,FSCN1,MARCKS,NPM1,EIF3C,ATP5F1A,VC
FBXL4,ATP2B4,CALR,TMF1,ABHD2,TUBB2B,CES3,PSAP,PGK1,RPLP0,H3-3B,CANX,CLU,PTMA,DI
OL4A1,PSAP,PGK1,RPLP0,H3-3B,CANX,PTMA,DDX5,SOX4,RPL13A,DVL3,YWHAE,RPL13,GPI,SPTE
,PRPF8,FBXL4,YBX1,UBA1,CALR,TMF1,ABHD2,PSAP,PGK1,H3-3B,CANX,PTMA,RPL13A,RPL6,DVI
PSAP,PGK1,H3-3B,CANX,PTMA,DDX5,SOX4,RPL13A,SEC62,DVL3,YWHAE,RPL13,ALDH6A1,ATP5F
YWHAE,RPL13,ALDH6A1,ATP5F1B,MYH9,RPL7,RPL5,SPTAN1,GSTP1,MARCKS,NPM1,HSPD1,SNF
L8,FBXL4,YBX1,UBA1,ATP2B4,CALR,TMF1,ABHD2,HSPA1B,CES3,PSAP,PGK1,RPLP0,H3-3B,CAN
P,H3-3B,CANX,PTMA,DDX5,DVL3,YWHAE,RPL13,DDX17,SPTBN1,P4HB,RPL5,FSCN1,TMBIM6,SPT
TRIM28,MATR3,YWHAG,LRRC28,SPEN,SET,EGLN1,PDIA3,SSBP2,VAPA,HMGB1,TPT1,RPS4X,RPL
S3,PGK1,H3-3B,CANX,PTMA,DDX5,ENO2,SOX4,KAT6A,DVL3,YWHAE,GPI,SPTBN1,ATP5F1B,CALM

3-3B,CANX,DDX5,RPL13A,SEC62,DVL3,YWHAE,SLC2A3,SPTBN1,ATP5F1B,MYH9,SEPTIN2,NPM1,
DX5,RPL13A,DVL3,YWHAE,RPL13,SLC2A3,SPTBN1,ATP5F1B,CALM3,MYH9,RPL7,SEPTIN2,RPL5,G
S3A,UCHL1,BTF3,RPL10,RPS17,DNAJA1,SF3B2,AP2B1,HNRNPU,HNRNPA0,RPS16,RPL23,CTSD,S
ATP5F1B,MYH9,SEPTIN2,P4HB,RPL5,PTMS,SPTAN1,GSTP1,MARCKS,NPM1,COL18A1,EIF3C,ATP
1,FADS1,RPL8,YBX1,UBA1,ATP2B4,ABHD2,PSAP,PGK1,RPLP0,H3-3B,CANX,PTMA,DDX5,RPL13A,
RPL5,GDI1,FSCN1,PTMS,SPTAN1,MARCKS,SMG1,NPM1,EIF3C,ATP5F1A,VCP,AFF4,TRIM28,MAT
3B,CANX,PTMA,DDX5,RPL13A,DVL3,YWHAE,RPL13,DDX17,SPTBN1,MYH9,RPL7,IDS,RPL5,GDI1,F
2,KAT6A,DVL3,YWHAE,SPTBN1,ATP5F1B,GNB1,MYH9,SEPTIN2,FSCN1,PTMS,SPTAN1,NPM1,COL
PTAN1,GSTP1,MARCKS,NPM1,EIF3C,ATP5F1A,VCP,AFF4,TRIM28,MATR3,YWHAG,CKB,SPEN,NON
2,HSPA1B,CES3,PSAP,PGK1,H3-3B,CANX,PTMA,DDX5,SOX4,KAT6A,DVL3,RPL13,DDX17,GPI,SPT
2,HSPA1B,CES3,PSAP,PGK1,H3-3B,CANX,PTMA,DDX5,SOX4,KAT6A,DVL3,RPL13,DDX17,GPI,SPT
WHAE,GPI,SPTBN1,P4HB,GDI1,FSCN1,SPTAN1,SMG1,NPM1,EIF3C,ATP5F1A,VCP,UBB,TRIM28,M
PLD3,COL18A1,EIF3C,ATP5F1A,VCP,TRIM28,MATR3,LRRC28,SPEN,NONO,SET,LDHB,PKD1,PDIA
S,SMG1,CALM1,NPM1,HSPD1,EIF3C,ATP5F1A,MARCKSL1,UBB,TRIM28,MATR3,RPL37A,SPEN,S
5F1A,TRIM28,MATR3,RPL37A,LRRC28,SPEN,NONO,SET,EGLN1,RPL19,PDIA3,VAPA,HMGB1,POL
DB1,HACD3,EGLN1,RPL19,SSBP2,VAPA,RPS4X,RPS3A,RPS17,SF3B2,ITM2B,AP2B1,GTF2F1,HN
GB1,POLR2A,ANXA2,RPL10,RPS17,MAP4,SF3B2,CDC42BPB,ITM2B,HNRNPU,CTNNA1,RPL23,EIF
PS4X,RPS3A,STMN1,BTF3,ANXA2,XAF1,RPL10,SF3B2,HNRNPU,CTNNA1,RPL23,RPL29,RPSA,TNS
SEPTIN2,RPL5,GDI1,MARCKS,NPM1,PLD3,COL18A1,SNRNP200,EIF3C,ATP5F1A,VCP,MARCKSL1
NAB,FTL,HSP90B1,FADS1,FBXL4,YBX1,UBA1,ATP2B4,CALR,TMF1,ABHD2,PSAP,PGK1,RPLP0,H3
,RPL5,GDI1,SPTAN1,SMG1,NPM1,COL18A1,SNRNP200,CSDE1,EIF3C,ATP5F1A,VCP,MARCKSL1,
,GNB1,MYH9,P4HB,IDS,TMBIM6,SPTAN1,SMG1,NPM1,EIF3C,ATP5F1A,VCP,MARCKSL1,UBB,TR
P,UBB,TRIM28,MATR3,RPL37A,LRRC28,SPEN,NONO,HACD3,SET,RPL19,PDIA3,SSBP2,VAPA,HM
PL8,YBX1,UBA1,ATP2B4,TMF1,ABHD2,PSAP,PGK1,RPLP0,H3-3B,CANX,PTMA,DDX5,RPL13A,SEC
B1,DCHS1,SET,SSBP2,VAPA,HMGB1,RPS4X,RPS3A,BTF3,ANXA2,RPL10,SF3B2,HNRNPU,SMARC
1,NPM1,ATP5F1A,VCP,UBB,TRIM28,MATR3,YWHAG,RPL37A,FN1,SPEN,NONO,SET,EGLN1,MRC2
RIM28,MATR3,RPL37A,SET,INA,LDHB,PDIA3,SSBP2,CHD3,POLR2A,UCHL1,ANXA2,VCL,AKAP12,
T6A,YWHAE,SPTBN1,ATP5F1B,CALM3,GNB1,P4HB,FSCN1,PTMS,SPTAN1,MARCKS,CALM1,NPM
IM28,MATR3,YWHAG,RPL37A,LRRC28,NONO,DCHS1,SET,EGLN1,LDHB,PKD1,PDIA3,HMGB1,RP
O,SET,EGLN1,SF3B1,SSBP2,HMGB1,POLR2A,RPS4X,RPS3A,BTF3,ANXA2,RPL10,DNAJA1,SF3B2
,GDI1,SPTAN1,NPM1,COL18A1,CSDE1,ATP5F1A,VCP,TRIM28,MATR3,LRRC28,SPEN,NONO,HAC
,HSPD1,SNRNP200,EIF3C,ATP5F1A,VCP,UBB,TRIM28,MATR3,RPL37A,LRRC28,SPEN,NONO,SET,
NX,ENO2,DVL3,GPI,SPTBN1,ATP5F1B,CALM3,GNB1,SEPTIN2,GDI1,PTMS,SPTAN1,MARCKS,CAL
BP2,HMGB1,POLR2A,BTF3,RPL10,DNAJA1,SF3B2,HNRNPU,RPS16,RPL23,RPSA,TPM4,EIF4H,XR
M28,MATR3,LRRC28,NONO,DDB1,SET,EGLN1,LDHB,PDIA3,SSBP2,BTF3,ANXA2,RPL10,HNRNPU
ALDH6A1,SPTBN1,ATP5F1B,MYH9,RPL7,P4HB,IDS,RPL5,SPTAN1,MARCKS,SMG1,NPM1,HSPD1,E
2,TUBB2B,CES3,PGK1,RPLP0,CANX,CLU,PTMA,ENO2,SOX4,SEC62,DVL3,YWHAE,SLC2A3,GPI,B
1,ATP5F1A,MARCKSL1,UBB,TRIM28,MATR3,RPL37A,SPEN,NONO,HACD3,SET,LDHB,PDIA3,SSB
GDI1,SPTAN1,SMG1,CALM1,NPM1,EIF3C,ATP5F1A,VCP,MARCKSL1,UBB,TRIM28,MATR3,YWHAG
ANXA2,RPL10,SF3B2,GTF2F1,HNRNPU,OGA,LARP1,CTNNA1,RPS16,RPL23,RPSA,TPM4,RPL24,H
PD1,ATP5F1A,VCP,MARCKSL1,UBB,TRIM28,YWHAG,SPEN,NONO,YWHAQ,DCHS1,SET,EGLN1,LD
TP5F1A,UBB,TRIM28,MATR3,LRRC28,SPEN,NONO,DDB1,HACD3,DCHS1,SET,EGLN1,SSBP2,VAP
1,POLR2A,RPS3A,BTF3,ANXA2,RPL10,SF3B2,GTF2F1,HNRNPU,NFE2L1,SMARCA5,LARP1,RPL2
A2,RPL10,DNAJA1,SF3B2,GTF2F1,HNRNPU,SMARCA5,RPL23,EIF4H,BRD3,RPS9,COL1A2,RPL14
,SSBP2,CHD3,HMGB1,RPS4X,BTF3,RPL10,SF3B2,GTF2F1,HNRNPU,LARP1,CTNNA1,RPS16,RPS

D,EIF4H,SCARB2,XRCC5,RPL24,RPL14,CBX5,CHD4,RPS20,WDR1,RBBP4,PNN,RPL31,HNRNPC,L
PL10,DNAJA1,PAFAH1B1,GTF2F1,CPE,CNN3,EIF4H,SCARB2,RTN4,PJA2,PAPOLA,GAP43,NCS1,SL
N1,PTMS,GSTP1,CALM1,SNRNP200,CSDE1,EIF3C,ATP5F1A,TRIM28,YWHAG,RPL37A,NONO,YW
1,TMBIM6,SMG1,NPM1,PLD3,COL18A1,CSDE1,ATP5F1A,EIF4B,VCP,AFF4,TRIM28,MATR3,YWHA
X17,ATP5F1B,SEPTIN2,IDS,RPL5,TMBIM6,PTMS,SMG1,NPM1,HSPD1,USP11,COL18A1,ATP5F1A,
MGB1,RPS4X,STMN1,TUBB4B,BTF3,RPL10,DNAJA1,SF3B2,HNRNPU,SMARCA5,LARP1,RPL23,RP
,GAP43,KIF5B,CHD4,WDR1,BEX3,SRSF4,RBBP4,RTN3,HNRNPC,KHDRBS1,RPS25,SFPQ,H1-0,RP
,HMGB1,TPT1,TUBB4B,NCAM1,VCL,SF3B2,HNRNPA0,EIF4H,RTN4,XRCC5,PJA2,C1ORF43,PAPC
16,RPL23,TPM4,XRCC5,RPL17,RPL14,RPS14,CHD4,RPS20,EIF5,RBBP4,RPL31,HNRNPC,TADA2
F1B,P4HB,IDS,RPL5,GDI1,SPTAN1,SMG1,NPM1,HSPD1,COL18A1,CSDE1,EIF3C,ATP5F1A,EIF4B,
BP2,BASP1,HMGB1,RPL10,WSB1,DNAJA1,SF3B2,HNRNPU,SCARB2,XRCC5,RPS14,CXADR,ATP1A
,TRIM28,MATR3,RPL37A,LRRC28,SPEN,NONO,DDB1,DCHS1,SET,EGLN1,PDIA3,SSBP2,VAPA,HM
PL7,IDS,GDI1,FSCN1,TMBIM6,MARCKS,NPM1,PLD3,COL18A1,EIF3C,ATP5F1A,VCP,MARCKSL1,U
SSBP2,HMGB1,POLR2A,ANXA2,CYB5R3,SF3B2,HNRNPU,NFE2L1,KPNB1,C1ORF43,PAPOLA,COL
M1,EIF3C,ATP5F1A,VCP,UBB,TRIM28,MATR3,CKB,RPL37A,LRRC28,FN1,SPEN,NONO,DDB1,SET,
A,VCP,MARCKSL1,TRIM28,MATR3,NONO,DDB1,HACD3,SET,EGLN1,RPL19,PDIA3,SSBP2,VAPA,H
CTNNA1,RPS16,RPL23,RPSA,HK1,RPS14,CHD4,JUND,RPS20,EIF5,RBBP4,PNN,HNRNPC,LRPPRC,
2B,GTF2F1,HNRNPU,OGA,SMARCA5,XRCC5,KPNB1,PTCD3,SRSF4,RBBP4,PNN,SPC25,PRKCSH,H
P1,MARCKS,PLD3,COL18A1,CSDE1,ATP5F1A,EIF4B,TRIM28,MATR3,YWHAG,RPL37A,SPEN,NONC
CC5,RPL24,GAP43,CLSTN1,HMGN2,CCT7,SRSF4,RBBP4,HNRNPC,RPS25,TKT,SFPQ,H1-0,HNRN
FADS2,LPCAT1,XRCC5,RPL14,HMGN2,RBBP4,PRKCSH,HNRNPC,KHDRBS1,XPO1,SFPQ,RAB7A,A
1,SLC3A2,RPS25,GLG1,SFPQ,PRDX2,MAGED2,H1-0,ANXA6,APLP2,TARDBP,HNRNPK,H1-10,HNR
10,HNRNPU,SMARCA5,LARP1,XRCC5,MSN,RPL24,RPL14,CHD4,RPS20,EIF5,RPL26,CCT7,RBBP4,
AP4,CDC42BPB,GTF2F1,HNRNPU,XRCC5,CLSTN1,BEX3,RBBP4,PFN1,HNRNPC,KHDRBS1,XPO1
IM28,MATR3,SPEN,NONO,HACD3,SET,INA,SSBP2,HMGB1,POLR2A,RPS3A,STMN1,TUBB4B,UCH
BIM6,SPTAN1,SMG1,NPM1,COL18A1,EIF3C,ATP5F1A,EIF4B,MARCKSL1,UBB,TRIM28,MATR3,R
NRNPK,H1-10,RPL36,LAMP1,CAPRIN1,HDAC2,LUC7L3,PGRMC1,CDH11,ANXA5,PABPN1,HNRNPL
1,MRC2,SF3B1,XAF1,RPL10,DNAJA1,PAFAH1B1,AKAP12,MAP4,ZNF770,GTF2F1,OGA,NFE2L1,CT
K4,RPL13A,DVL3,RPL13,GPI,ALDH6A1,SPTBN1,ATP5F1B,RPL7,P4HB,IDS,RPL5,GDI1,PTMS,SMG1
2,H1-0,GJA1,ANXA6,TARDBP,NDFIP1,HNRNPK,CKAP5,LAMP1,HNRNPH1,BCLAF1,CPSF2,MDH2,U
RPS20,EIF5,WDR1,RPL26,CCT7,RBBP4,NES,HNRNPC,LRPPRC,KHDRBS1,SFPQ,RAB7A,MAGED2,
O,SET,EGLN1,LDHB,SF3B1,SSBP2,HMGB1,TPT1,RPS4X,STMN1,RPL10,RPS17,VCL,DNAJA1,SF3
RB2,XRCC5,RPS14,CHD4,JUND,EIF5,RBBP4,RPL31,NISCH,HNRNPC,TADA2B,KHDRBS1,XPO1,SP
BEX3,RBBP4,NES,HNRNPC,KHDRBS1,CSNK1E,GLG1,SFPQ,MAGED2,H1-0,TLN1,HNRNPDL,TARD
GLN1,INA,MRC2,SSBP2,VAPA,POLR2A,RPS4X,XAF1,RPL10,CYB5R3,SF3B2,ITM2B,AP2B1,GTF2F
NXA2,AGRN,RPL10,CYB5R3,SF3B2,GTF2F1,HNRNPU,RPSA,CTSD,SCARB2,XRCC5,RPL24,C1OR
NB1,SEPTIN2,GDI1,TMBIM6,SPTAN1,MARCKS,COL18A1,EIF3C,ATP5F1A,EIF4B,VCP,MARCKSL1,
HNRNPU,RPSA,SCARB2,BRD3,XRCC5,C1ORF43,PAPOLA,CLSTN1,RPS14,EIF5,RBBP4,NISCH,HNR
,VCP,UBB,AFF4,TRIM28,MATR3,YWHAG,LRRC28,MAPT,SPEN,NONO,DDB1,SET,EGLN1,LDHB,SF
11,CSDE1,EIF3C,ATP5F1A,VCP,TRIM28,MATR3,SPEN,NONO,HACD3,SET,INA,LDHB,SSBP2,HMG
CHS1,SF3B1,PDIA3,SSBP2,TPT1,RPS3A,TUBB4B,MARCHF6,VCL,DNAJA1,CYB5R3,PAFAH1B1,SF
BP,HNRNPK,H1-10,LAMP1,HDAC2,BCLAF1,LUC7L3,DCN,CDH11,PABPN1,HNRNPL,ALKBH5,HNR
,SMARCA5,TPM4,BRD3,XRCC5,PTCD3,CBX5,PSMD2,KIF5B,SLC7A5,RPS14,CHD4,ATP1A1,EIF5,W
TR3,SET,EGLN1,SSBP2,BASP1,HMGB1,STMN1,RPL10,SF3B2,HNRNPU,XRCC5,MSN,RPL24,RPL1
5,XRCC5,RPL14,RPS20,RPL26,LASP1,CCT7,RBBP4,JPT2,HNRNPC,LRPPRC,XPO1,RPS25,GLG1,S

HNRNPC,TADA2B,KHDRBS1,XPO1,RPS25,SFPQ,MAGED2,H1-0,HNRNPDL,TARDBP,HNRNPK,RPL
NRNPL,SNRNP70,H3-3A,HDGF,SMARCA4,DSTN,SRSF1,PTBP1,ILF3,PRKDC,HNRNPUL1,KDM1A,D
1,RPL10,SF3B2,HNRNPU,LARP1,RPL23,RPSA,TPM4,XRCC5,RPL24,RPL14,CHD4,RPL26,RBBP4,H
RDBP,HNRNPK,TOP2B,CPSF2,TM9SF3,U2AF2,PABPN1,HNRNPL,MORF4L2,RBMX,LAMC1,CBX1,F
PEN,NONO,SET,EGLN1,SSBP2,VAPA,TPT1,UCHL1,NCAM1,ANXA2,DNAJA1,MAP4,SF3B2,HNRNPU,
RAD23B,TNPO1,SNRNP70,H3-3A,MAPK1,ILF3,PRKDC,CORO1C,SUPT6H,HNRNPM,SRSF3,ILF2,FB
P,HNRNPK,H1-10,RPL36,LAMP1,CALU,CAPRIN1,TOP2B,HDAC2,SKP1,BCLAF1,GLUL,LUC7L3,HNF
HDRBS1,SFPQ,DDX3X,MAGED2,H1-0,IK,COL6A2,TARDBP,HNRNPK,CKAP5,HNRNPH1,TOP2B,HD
NRNPM,SRSF3,ILF2,SMS,TXNIP,RBBP7,ZFR,ANP32B,CALD1,ZMYM3,FUS,XRCC6,PELP1,TCAF1,N
BP4,NES,HNRNPC,ACTN4,DDX3X,H1-0,TARDBP,NDFIP1,HNRNPK,H1-10,CKAP5,HNRNPH1,CAPRIN
,HNRNPDL,TARDBP,HNRNPK,CKAP5,CALU,CAPRIN1,LUC7L3,SQLE,CPSF2,DDX39B,TM9SF3,MLX
PJA2,KPNB1,EIF5,RBBP4,PNN,HNRNPC,KHDRBS1,PTN,SFPQ,EIF3L,IK,GOLM1,TARDBP,HNRNPK,
,ANXA6,HNRNPDL,TAGLN,HNRNPK,HNRNPH1,HDAC2,BCLAF1,LUC7L3,DDX39B,CDH11,PABPN1,
P4,PNN,PRKCSH,HNRNPC,KHDRBS1,XPO1,SFPQ,PDIA6,RPL12,MRFAP1,H1-0,IK,GOLM1,HNRNPD
DNAJC5,APLP2,TARDBP,ARID1A,HNRNPK,HNRNPH1,CAPRIN1,HDAC2,SKP1,BCLAF1,LUC7L3,MDH
A1,EIF3C,ATP5F1A,VCP,TRIM28,MATR3,LRRC28,FN1,SPEN,NONO,DDB1,HACD3,DCHS1,SET,EGL
0,HNRNPH1,HLA-B,CCT6A,BCLAF1,CPSF2,TOMM20,U2AF2,CCT2,SUGP2,PABPN1,GDI2,HNRNPL
,CALU,CAPRIN1,HDAC2,SKP1,LUC7L3,SQLE,CPSF2,SNRNP70,H3-3A,HDGF,VEGFA,MAPK1,IPO7,
P2A2,SRSF4,RBBP4,NISCH,HNRNPC,KHDRBS1,XPO1,SFPQ,TPM1,CDH2,RAB7A,DDX3X,H1-0,GJA
,H3-3A,SRSF1,ILF3,HNRNPUL1,DIDO1,FLNB,HNRNPM,SSB,SRSF3,LARS1,ILF2,SF3A1,VDAC1,RB
RBS1,RPS25,SFPQ,RAB7A,RPS15,MAGED2,RPS7,TLN1,APLP2,TARDBP,HNRNPK,H1-10,RPL36,Y
RDX2,DDX3X,H1-0,IK,TARDBP,MYL6,HNRNPK,HNRNPH1,HDAC2,BCLAF1,LUC7L3,DDX39B,RNF18
31,TOMM20,GAS7,TM9SF2,PRDX5,ACO2,CNTNAP2,MARS1,G3BP2,IPO7,ACADVL,ATP5PB,FKBP8
,CAMSAP1,RAB5B,CNTNAP2,DPYSL2,MAPK1,ATP6V1B2,STXBP1,PFN2,CD99L2,SYP,NAP1L1,RET
PA0,CNN3,RPL29,XRCC5,KPNB1,PAPOLA,ATF4,CRMP1,RBBP4,PNN,SPC25,PRKCSH,HNRNPC,KH
DB1,YWHAQ,DCHS1,EGLN1,SF3B1,VAPA,TPT1,RPS4X,MORF4L1,RPS17,DNAJA1,CYB5R3,PAFAH1
DIA3,HMGB1,POLR2A,RPL18,STMN1,VCL,SF3B2,GTF2F1,GOLGA8A,HNRNPU,SMARCA5,HNRNP
PRDX5,ACO2,CNTNAP2,MARS1,G3BP2,IPO7,ATP5PB,FKBP8,SDF4,ATP6V1B2,KDM3B,MAP6,MAP
31,TOMM20,RPN2,TLN2,TM9SF2,RTN1,ACO2,CNTNAP2,MARS1,G3BP2,ACADVL,ATP5PB,HSPB1,
M3B,TP53BP1,RBBP7,SF1,BRD4,ATRX,SRSF6,ZFR,HNRNPA2B1,CUL3,TNKS,APEX1,HNRNPAB,PR
A1,HTATSF1,MECP2,CD2BP2,SUMO1,ADNP,SSU72,UBE2J1,RTCB,AKAP8L,ZEB1,SEPHS1,FBL,MA
C,DIDO1,HNRNPM,SRSF3,ILF2,RAI1,KDM3B,NAP1L1,LBH,H2AZ1,ATRX,HNRNPAB,C1ORF198,FUS
NDUFB8,PRKAR2B,DPP7,BACE1,SYT1,PDHB,ARSA,FKBP3
MATR3,SPEN,HACD3,PDIA3,HMGB1,POLR2A,RPL18,STMN1,TUBB4B,VCL,SF3B2,GTF2F1,GOLG
A,HDGF,PRPF6,DPYSL2,PRKDC,SUPT6H,HNRNPM,SRSF3,ILF2,STXBP1,KDM3B,SYP,RBBP7,NAP1
,ELAVL3,HMGA1,HTATSF1,MECP2,CD2BP2,SUMO1,ADNP,SSU72,UBE2J1,RTCB,AKAP8L,ZEB1,SE
,AKAP9,TPP1,SSR3,DCLK1,EEF1B2,REEP2,AP2A2,ELAVL3,FXR2,TRIM2,IDH2,RPL27A,NACAD,ND
PRE2,AKAP9,TPP1,SSR3,DCLK1,EEF1B2,REEP2,AP2A2,NTRK2,ELAVL3,FXR2,TRIM2,COPE,IDH2,U
TN1,ACO2,CNTNAP2,MARS1,G3BP2,HSPB1,NSG2,FKBP8,POLR2B,ATP6V1B2,FAU,STMN3,FBRSL
,KPNB1,PNN,RPL31,HNRNPC,KHDRBS1,SFPQ,TPM1,IK,GOLM1,TARDBP,WLS,HNRNPK,CCT6A,B
PL,RPL27,ACO2,TCP1,RBMX,RAD23B,SNRNP70,TOP1,SRSF1,TCF25,ILF3,HSPB1,BNIP3L,PLCG1,
O1,SNRNP70,H3-3A,SRSF1,ARGLU1,NYNRIN,ILF3,PRKDC,HNRNPUL1,DIDO1,HNRNPM,SRSF3,PH
I1-10,RPL36,LAMP1,HLA-B,CALU,CAPRIN1,HDAC2,BCLAF1,TRIB2,LUC7L3,TM9SF3,MDH2,PGD,H
5,PLD3,COL18A1,SNRNP200,YWHAG,LRRC28,MAPT,DDB1,YWHAQ,DCHS1,EGLN1,MRC2,PKD1,S
1B2,FAU,FBLN1,CAPN2,SF1,BRD4,MAPRE2,AKAP9,LINGO1,TPP1,DTX4,EEF1B2,CADM3,PSMD4,

1A,SMARCC1,ATP6V1B2,ISYNA1,ATXN2L,SOD1,SSR1,PRMT1,TXNDC5,MTA1,TRIO,KDM5B,PRCC,
F1B2,REEP2,AP2A2,CD74,RCN2,FXR2,TRIM2,UBE2J1,NACAD,NDUFB8,SEZ6L2,DLG4,MAN2B1,DI
LINGO1,SSR3,DCLK1,DTX4,EEF1B2,REEP2,AP2A2,RCN2,ANK2,SCAMP5,FXR2,TRIM2,RPL27A,SE
'1,ARNT2,SYP,CAPN2,CD99,SV2A,SEPTIN7,TMOD2,SEPTIN5,IARS2,CADM3,REEP2,CHMP4B,AP2A
YP,VDAC1,MSMO1,BRD4,SV2A,MAPRE2,SEPTIN7,PSMD8,TMOD2,SEPTIN5,IARS2,SSR3,CADM3,R

A2,CMTR1,NCOA6,NFASC,ACIN1,KLF3,WDR33,ZNF326,UBTF,MYBBP1A
,RNF187,SSRP1,ALKBH5,TCP1,CCT3,PTOV1,DDX42,RAB5B,TNPO1,RNF10,OPTN,UBXN4,MAPK1
CADVL,FKBP8,POLR2B,MSMO1,PRMT1,CAPN2,CD99,BRD4,MAPRE2,BRSK1,AKAP9,TMOD2,TPP
3B4B,ZMIZ1,VCL,SF3B2,GTF2F1,HNRNPU,SMARCA5,HNRNPA0,TPM4,RTN4,XRCC5,KPNB1,PAPC
TCF,RFX7,APOE,ACIN1,AKAP17A,KLF3,GSN,UBTF
E,ACIN1,AKAP17A,KLF3,GSN,UBTF
SC,ACIN1,KLF3,WDR33,ZNF326,UBTF,MYBBP1A

,G3BP2,MAPRE1,ACADVL,SUPT5H,KDM1A,ZFAND5,SMARCC1,DHX30,ATP6V1B2,SLIT3,ISYNA1,A
7,APOE,NCOA6,ACIN1,AKAP17A,KLF3,GSN,UBTF
RBM17,CTCF,RFX7,SP1,APOE,NCOA6,ACIN1,AKAP17A,KLF3,GSN,UBTF
ACAD,VDAC2,UQCRFS1,ABCD3,GSN,ARSA
ATP6V0A1,SHC1,PPP3CA,FH,SYT1,DNM1,PPFIA3,PDHB
R2B,CD99,MAPRE2,BRSK1,AKAP9,TMOD2,TPP1,SSR3,CNP,CHMP4B,NMNAT2,SORT1,IGLON5,PS

,NCOA6,ACIN1,AKAP17A,GSN,UBTF
DM3,REEP2,AP2A2,SAE1,IGLON5,USP47,FXR2,CLTA,FAM168A,NDUFB8,MRPS6,SEZ6L2,RAB3A,N
TX4,EEF1B2,CHMP4B,ATP6V1G1,WNK1,NMNAT2,SORT1,CD74,RCN2,PSAT1,SCAMP5,FXR2,TRIM
,PDHB,FKBP3
3,MAN2B1,CNTN1,ATP6V0A1,GPRIN1,PPP3CA,SLC1A3,NFASC,EPHA4,PPFIA3,GNAO1,ATP6V1F,PI
RKDC,DIDO1,IDH1,STRAP,RBBP7,SGTA,LBH,ATRX,EP400,SRSF6,UBAP2L,TRA2B,HNRNPAB,FUS,
HS1,H2AX,SRF,CLCN6,MAN2B1,HSPE1,RXRA,PRPF31,TFDP1,MCM7,PPFIA3,CFAP298,CDK5RAP3
SYT11,ATP6V1G1,RCN2,IGLON5,ANK2,SCAMP5,FXR2,TRIM2,COPE,APEH,UBE2J1,NDUFB8,SEZ6I
B,ANXA5,PRDX5,RTN1,TERF2IP,VEGFA,G3BP2,CCNL2,ACADVL,RBP1,SUPT5H,DHX30,ATP6V1B2,

.,HSP90B1,FADS1,NEFL,NEFM,PRPF8,UGT8,RPL8,FBXL4,YBX1,UBA1,ATP2B4,CALR,TMF1,ABHD
T8,RPL8,FBXL4,YBX1,UBA1,ATP2B4,CALR,ABHD2,HSPA1B,TUBB2B,COL4A1,CES3,PSAP,PGK1,F
PL8,FBXL4,YBX1,UBA1,ATP2B4,CALR,TMF1,ABHD2,HSPA1B,CES3,PSAP,PGK1,RPLP0,H3-3B,CA
A1,ATP2B4,CALR,ABHD2,HSPA1B,TUBB2B,COL4A1,CES3,PSAP,PGK1,RPLP0,H3-3B,CANX,CLU,P
L6A1,HSP90B1,FADS1,NEFM,PRPF8,UGT8,RPL8,FBXL4,YBX1,UBA1,ATP2B4,TMF1,ABHD2,HSPA1
,PRPF8,RPL8,FBXL4,UBA1,ATP2B4,CALR,TMF1,ABHD2,HSPA1B,COL4A1,PSAP,PGK1,RPLP0,H3-
RPL8,FBXL4,YBX1,ATP2B4,CALR,TMF1,ABHD2,HSPA1B,COL4A1,CES3,PSAP,PGK1,RPLP0,H3-3B
.8,FBXL4,YBX1,UBA1,ATP2B4,CALR,TMF1,ABHD2,HSPA1B,CES3,PSAP,PGK1,RPLP0,H3-3B,CANX
S1,PRPF8,UGT8,RPL8,FBXL4,YBX1,UBA1,ATP2B4,CALR,TMF1,ABHD2,HSPA1B,COL4A1,CES3,PS
TP2B4,CALR,TMF1,ABHD2,HSPA1B,COL4A1,PSAP,PGK1,RPLP0,H3-3B,CANX,PTMA,DDX5,SOX4,
TP2B4,CALR,TMF1,ABHD2,HSPA1B,COL4A1,PSAP,PGK1,RPLP0,H3-3B,CANX,PTMA,DDX5,SOX4,
X1,UBA1,ATP2B4,CALR,TMF1,ABHD2,HSPA1B,COL4A1,CES3,PSAP,PGK1,RPLP0,H3-3B,CANX,CL
1,RPLP0,H3-3B,CANX,CLU,PTMA,DDX5,ENO2,RPL13A,SEC62,KAT6A,DVL3,YWHAE,DDX17,GPI,SF
S1,PRPF8,RPL8,FBXL4,YBX1,UBA1,ATP2B4,CALR,TMF1,ABHD2,HSPA1B,TUBB2B,CES3,PSAP,PG
S1,PRPF8,RPL8,FBXL4,YBX1,UBA1,ATP2B4,CALR,TMF1,ABHD2,HSPA1B,TUBB2B,CES3,PSAP,PG
P2B4,CALR,TMF1,ABHD2,HSPA1B,CES3,PSAP,PGK1,RPLP0,H3-3B,CANX,CLU,PTMA,DDX5,SOX4,
JBA1,ATP2B4,CALR,TMF1,ABHD2,HSPA1B,COL4A1,CES3,PSAP,PGK1,RPLP0,H3-3B,CANX,PTMA,
JBA1,ATP2B4,CALR,TMF1,ABHD2,HSPA1B,COL4A1,CES3,PSAP,PGK1,RPLP0,H3-3B,CANX,PTMA,
,RPL8,FBXL4,YBX1,UBA1,ATP2B4,CALR,TMF1,ABHD2,HSPA1B,TUBB2B,COL4A1,CES3,PSAP,PG
LR,TMF1,ABHD2,HSPA1B,CES3,PSAP,PGK1,RPLP0,H3-3B,CANX,CLU,PTMA,DDX5,SOX4,RPL13A,
A,SEC62,KAT6A,DVL3,YWHAE,RPL13,DDX17,GPI,ALDH6A1,SPTBN1,ATP5F1B,GNB1,MYH9,RPL7,
T8,RPL8,FBXL4,YBX1,UBA1,ATP2B4,CALR,TMF1,ABHD2,HSPA1B,COL4A1,PSAP,PGK1,RPLP0,H3
GT8,RPL8,FBXL4,YBX1,UBA1,ATP2B4,TMF1,ABHD2,HSPA1B,PSAP,PGK1,RPLP0,H3-3B,CANX,CLU
GT8,RPL8,FBXL4,YBX1,UBA1,ATP2B4,TMF1,ABHD2,HSPA1B,PSAP,PGK1,RPLP0,H3-3B,CANX,CLU
4,RPL13A,SEC62,RPL6,KAT6A,DVL3,YWHAE,RPL13,DDX17,ATP5F1B,CALM3,RPL7,P4HB,APP,IDS
T6A,DVL3,YWHAE,RPL13,DDX17,GPI,ATP5F1B,CALM3,GNB1,RPL7,P4HB,APP,IDS,RPL5,GDI1,TM
FADS1,PRPF8,UGT8,RPL8,FBXL4,YBX1,UBA1,ATP2B4,TMF1,ABHD2,HSPA1B,COL4A1,CES3,PSAF
A1,ATP2B4,CALR,TMF1,ABHD2,HSPA1B,CES3,PSAP,PGK1,RPLP0,H3-3B,CANX,CLU,PTMA,DDX5,S
2B4,TMF1,ABHD2,HSPA1B,TUBB2B,CES3,PSAP,PGK1,RPLP0,H3-3B,CANX,PTMA,DDX5,SOX4,RPL
2B4,TMF1,ABHD2,HSPA1B,TUBB2B,CES3,PSAP,PGK1,RPLP0,H3-3B,CANX,PTMA,DDX5,SOX4,RPL
GT8,RPL8,FBXL4,YBX1,UBA1,ATP2B4,CALR,TMF1,ABHD2,HSPA1B,TUBB2B,CES3,PSAP,PGK1,RF
HD2,HSPA1B,CES3,PSAP,PGK1,RPLP0,H3-3B,CANX,CLU,PTMA,DDX5,SOX4,RPL13A,SEC62,RPL6
HD2,HSPA1B,CES3,PSAP,PGK1,RPLP0,H3-3B,CANX,CLU,PTMA,DDX5,SOX4,RPL13A,SEC62,RPL6
B,CANX,DDX5,RPL13A,SEC62,KAT6A,DVL3,YWHAE,RPL13,DDX17,GPI,ALDH6A1,SPTBN1,ATP5F1
ES3,PSAP,PGK1,RPLP0,H3-3B,CANX,PTMA,DDX5,SOX4,RPL13A,KAT6A,DVL3,YWHAE,RPL13,DDX
X1,UBA1,ATP2B4,CALR,TMF1,ABHD2,HSPA1B,TUBB2B,CES3,PSAP,PGK1,RPLP0,H3-3B,CANX,P
B,CANX,PTMA,DDX5,ENO2,SOX4,RPL13A,SEC62,KAT6A,DVL3,YWHAE,RPL13,DDX17,ALDH6A1,SF
PSAP,PGK1,RPLP0,H3-3B,CANX,PTMA,DDX5,SOX4,RPL13A,SEC62,RPL6,KAT6A,DVL3,YWHAE,RP
PSAP,PGK1,RPLP0,H3-3B,CANX,PTMA,DDX5,SOX4,RPL13A,SEC62,RPL6,KAT6A,DVL3,YWHAE,RP
B4,CALR,TMF1,ABHD2,HSPA1B,PSAP,PGK1,RPLP0,H3-3B,CANX,CLU,PTMA,DDX5,RPL13A,SEC6
ANX,DDX5,SOX4,RPL13A,SEC62,KAT6A,DVL3,YWHAE,RPL13,DDX17,GPI,SPTBN1,ATP5F1B,RPL7,
RPF8,UGT8,RPL8,FBXL4,YBX1,UBA1,ATP2B4,CALR,TMF1,ABHD2,HSPA1B,CES3,PSAP,PGK1,RPL
RPF8,UGT8,RPL8,FBXL4,YBX1,UBA1,ATP2B4,CALR,TMF1,ABHD2,HSPA1B,CES3,PSAP,PGK1,RPL
,UBA1,ATP2B4,CALR,TMF1,ABHD2,HSPA1B,TUBB2B,CES3,PSAP,PGK1,RPLP0,H3-3B,CANX,PTM
,UBA1,ATP2B4,CALR,TMF1,ABHD2,HSPA1B,TUBB2B,CES3,PSAP,PGK1,RPLP0,H3-3B,CANX,PTM

,HSPA1B,COL4A1,CES3,PSAP,PGK1,H3-3B,CANX,DDX5,SOX4,SEC62,KAT6A,DVL3,YWHAE,SLC2A
1,UBA1,ATP2B4,CALR,TMF1,ABHD2,HSPA1B,CES3,PSAP,PGK1,RPLP0,H3-3B,CANX,CLU,PTMA,D
1,UBA1,ATP2B4,CALR,TMF1,ABHD2,HSPA1B,CES3,PSAP,PGK1,RPLP0,H3-3B,CANX,CLU,PTMA,D
3,FBXL4,YBX1,UBA1,ATP2B4,CALR,TMF1,ABHD2,HSPA1B,COL4A1,CES3,PSAP,PGK1,RPLP0,H3-3
ATP2B4,CALR,TMF1,ABHD2,CES3,PSAP,PGK1,RPLP0,H3-3B,CANX,PTMA,DDX5,RPL13A,SEC62,R
FBXL4,YBX1,UBA1,ATP2B4,TMF1,ABHD2,HSPA1B,TUBB2B,CES3,PSAP,PGK1,RPLP0,H3-3B,CANX
RPL8,FBXL4,YBX1,UBA1,ATP2B4,CALR,TMF1,HSPA1B,TUBB2B,COL4A1,PSAP,PGK1,RPLP0,H3-3
D2,HSPA1B,COL4A1,CES3,PSAP,PGK1,RPLP0,H3-3B,CANX,PTMA,DDX5,ENO2,SOX4,RPL13A,SEC
SPA1B,CES3,PSAP,PGK1,RPLP0,H3-3B,CANX,PTMA,DDX5,SOX4,RPL13A,SEC62,RPL6,KAT6A,DVL
SPA1B,CES3,PSAP,PGK1,RPLP0,H3-3B,CANX,PTMA,DDX5,SOX4,RPL13A,SEC62,RPL6,KAT6A,DVL
RPL8,FBXL4,YBX1,UBA1,ATP2B4,TMF1,ABHD2,COL4A1,CES3,PSAP,PGK1,RPLP0,H3-3B,CANX,P
P0,H3-3B,CANX,PTMA,DDX5,RPL13A,SEC62,KAT6A,DVL3,YWHAE,RPL13,DDX17,GPI,ALDH6A1,SP
HD2,HSPA1B,CES3,PSAP,PGK1,RPLP0,H3-3B,CANX,PTMA,DDX5,SOX4,RPL13A,SEC62,RPL6,KAT
HD2,HSPA1B,CES3,PSAP,PGK1,RPLP0,H3-3B,CANX,PTMA,DDX5,SOX4,RPL13A,SEC62,RPL6,KAT
,YWHAE,RPL13,DDX17,SPTBN1,ATP5F1B,CALM3,RPL7,SEPTIN2,GDI1,FSCN1,PTMS,SPTAN1,MA
P2B4,CALR,TMF1,ABHD2,HSPA1B,PSAP,PGK1,RPLP0,H3-3B,CANX,CLU,PTMA,DDX5,SOX4,SEC6
MF1,ABHD2,COL4A1,PSAP,PGK1,H3-3B,CANX,PTMA,DDX5,ENO2,KAT6A,DVL3,YWHAE,RPL13,GP
UBA1,ATP2B4,CALR,TMF1,ABHD2,HSPA1B,TUBB2B,CES3,PSAP,PGK1,RPLP0,H3-3B,CANX,CLU,D
UBA1,ATP2B4,CALR,TMF1,ABHD2,HSPA1B,TUBB2B,CES3,PSAP,PGK1,RPLP0,H3-3B,CANX,CLU,D
UBA1,ATP2B4,CALR,TMF1,ABHD2,HSPA1B,PSAP,PGK1,RPLP0,H3-3B,CANX,PTMA,DDX5,SOX4,RP
UBA1,ATP2B4,CALR,TMF1,ABHD2,HSPA1B,PSAP,PGK1,RPLP0,H3-3B,CANX,PTMA,DDX5,SOX4,RP
PTMA,DDX5,SOX4,RPL13A,SEC62,KAT6A,DVL3,YWHAE,RPL13,DDX17,GPI,ALDH6A1,SPTBN1,RP
FBXL4,YBX1,UBA1,ATP2B4,CALR,TMF1,ABHD2,HSPA1B,PSAP,PGK1,RPLP0,H3-3B,CANX,CLU,PT
PA1B,TUBB2B,COL4A1,CES3,PGK1,RPLP0,H3-3B,CANX,PTMA,DDX5,ENO2,SOX4,SEC62,RPL6,KA
L4,YBX1,UBA1,ATP2B4,CALR,TMF1,ABHD2,HSPA1B,TUBB2B,PSAP,PGK1,H3-3B,CANX,PTMA,DD
H3-3B,CANX,CLU,DDX5,SOX4,RPL13A,SEC62,RPL6,DVL3,YWHAE,RPL13,GPI,BSG,ALDH6A1,SPT
62,KAT6A,DVL3,YWHAE,RPL13,GPI,SPTBN1,ATP5F1B,GNB1,SEPTIN2,P4HB,IDS,RPL5,GDI1,FSC
TUBB2B,COL4A1,CES3,PSAP,PGK1,RPLP0,H3-3B,CANX,PTMA,DDX5,SOX4,RPL13A,KAT6A,DVL3,
),H3-3B,CANX,PTMA,DDX5,SOX4,RPL13A,SEC62,KAT6A,DVL3,YWHAE,RPL13,DDX17,SPTBN1,AT
),H3-3B,CANX,PTMA,DDX5,SOX4,RPL13A,SEC62,KAT6A,DVL3,YWHAE,RPL13,DDX17,SPTBN1,AT
1,RPLP0,H3-3B,CANX,CLU,PTMA,DDX5,SOX4,SEC62,KAT6A,DVL3,YWHAE,RPL13,DDX17,GPI,AL
PSAP,PGK1,RPLP0,H3-3B,CANX,CLU,PTMA,DDX5,SOX4,RPL13A,SEC62,RPL6,KAT6A,DVL3,YWHA
X,PTMA,DDX5,SOX4,RPL13A,SEC62,KAT6A,DVL3,YWHAE,DDX17,GPI,ALDH6A1,SPTBN1,ATP5F1B
TP2B4,CALR,TMF1,ABHD2,HSPA1B,TUBB2B,COL4A1,CES3,PSAP,PGK1,RPLP0,H3-3B,CANX,PTM
TP2B4,CALR,TMF1,ABHD2,HSPA1B,TUBB2B,COL4A1,CES3,PSAP,PGK1,RPLP0,H3-3B,CANX,PTM
A1,ATP2B4,CALR,TMF1,ABHD2,HSPA1B,TUBB2B,PSAP,PGK1,RPLP0,H3-3B,CANX,CLU,PTMA,DD
BX1,UBA1,ATP2B4,CALR,TMF1,ABHD2,HSPA1B,COL4A1,CES3,PSAP,PGK1,RPLP0,H3-3B,CANX,P
CALR,TMF1,ABHD2,HSPA1B,CES3,PSAP,PGK1,RPLP0,H3-3B,CANX,PTMA,DDX5,RPL13A,SEC62,K
,GPI,ALDH6A1,SPTBN1,ATP5F1B,SEPTIN2,APP,RPL5,GDI1,FSCN1,TMBIM6,PTMS,SPTAN1,GSTP1
3,PSAP,PGK1,RPLP0,H3-3B,CANX,PTMA,DDX5,SOX4,RPL13A,SEC62,RPL6,DVL3,YWHAE,RPL13,D
AT6A,DVL3,YWHAE,RPL13,GPI,BSG,ALDH6A1,SPTBN1,ATP5F1B,MYH9,RPL7,SEPTIN2,P4HB,IDS,
3A,SEC62,KAT6A,DVL3,YWHAE,RPL13,GPI,ALDH6A1,SPTBN1,ATP5F1B,GNB1,MYH9,RPL7,SEPTI
62,RPL6,KAT6A,DVL3,YWHAE,RPL13,GPI,ALDH6A1,ATP5F1B,CALM3,GNB1,MYH9,RPL7,P4HB,R
3,FBXL4,YBX1,UBA1,ATP2B4,CALR,TMF1,ABHD2,HSPA1B,COL4A1,CES3,PSAP,PGK1,RPLP0,H3-3
SAP,PGK1,RPLP0,H3-3B,CANX,PTMA,DDX5,RPL13A,SEC62,RPL6,KAT6A,DVL3,YWHAE,RPL13,DD

ES3,PSAP,RPLP0,H3-3B,CANX,PTMA,DDX5,SEC62,RPL6,KAT6A,DVL3,YWHAE,DDX17,GPI,SPTBN
A1,ATP2B4,CALR,TMF1,ABHD2,HSPA1B,PSAP,PGK1,RPLP0,H3-3B,CANX,PTMA,DDX5,SOX4,RPL
,ABHD2,HSPA1B,CES3,PSAP,PGK1,RPLP0,H3-3B,CANX,PTMA,DDX5,SOX4,RPL13A,SEC62,RPL6,
,ABHD2,HSPA1B,CES3,PSAP,PGK1,RPLP0,H3-3B,CANX,PTMA,DDX5,SOX4,RPL13A,SEC62,RPL6,
,MARCKS,SMG1,NPM1,PLD3,EIF3C,ATP5F1A,EIF4B,UBB,TRIM28,MATR3,YWHAG,RPL37A,LRRC
ES3,PSAP,PGK1,RPLP0,H3-3B,CANX,PTMA,SOX4,RPL13A,SEC62,RPL6,KAT6A,DVL3,YWHAE,RPL
A1B,PSAP,PGK1,RPLP0,H3-3B,CANX,PTMA,DDX5,SOX4,RPL13A,SEC62,RPL6,KAT6A,DVL3,YWHA
,DVL3,RPL13,DDX17,ATP5F1B,CALM3,MYH9,RPL7,SEPTIN2,P4HB,IDS,RPL5,GDI1,FSCN1,PTMS,
BXL4,YBX1,UBA1,ATP2B4,CALR,TMF1,ABHD2,HSPA1B,COL4A1,CES3,PSAP,PGK1,RPLP0,H3-3B
P0,H3-3B,CANX,PTMA,DDX5,SOX4,RPL13A,SEC62,RPL6,KAT6A,DVL3,YWHAE,RPL13,GPI,BSG,S
P0,H3-3B,CANX,PTMA,DDX5,SOX4,RPL13A,SEC62,RPL6,KAT6A,DVL3,YWHAE,RPL13,GPI,BSG,S
ALDH6A1,SPTBN1,ATP5F1B,GNB1,MYH9,RPL7,SEPTIN2,P4HB,IDS,RPL5,GDI1,TMBIM6,PTMS,SP
B,CANX,PTMA,DDX5,RPL13A,SEC62,DVL3,YWHAE,RPL13,DDX17,GPI,SPTBN1,ATP5F1B,CALM3,
PGK1,RPLP0,H3-3B,CANX,PTMA,DDX5,SOX4,RPL13A,SEC62,KAT6A,DVL3,YWHAE,RPL13,DDX17,
PGK1,RPLP0,H3-3B,CANX,PTMA,DDX5,SOX4,RPL13A,SEC62,KAT6A,DVL3,YWHAE,RPL13,DDX17,
,DDX5,KAT6A,DVL3,YWHAE,RPL13,GPI,SPTBN1,ATP5F1B,CALM3,GNB1,IDS,RPL5,GDI1,FSCN1,T
WHAE,RPL13,GPI,BSG,ALDH6A1,SPTBN1,ATP5F1B,CALM3,SEPTIN2,P4HB,RPL5,GDI1,PTMS,SPT
CLU,PTMA,DDX5,SOX4,SEC62,KAT6A,DVL3,YWHAE,RPL13,DDX17,GPI,CALM3,MYH9,RPL7,SEPTI
L6,KAT6A,DVL3,YWHAE,RPL13,DDX17,SPTBN1,ATP5F1B,MYH9,RPL7,SEPTIN2,P4HB,IDS,RPL5,F
LP0,H3-3B,CANX,PTMA,DDX5,SOX4,RPL13A,SEC62,RPL6,KAT6A,DVL3,YWHAE,RPL13,SLC2A3,D
A,SEC62,RPL6,KAT6A,DVL3,YWHAE,RPL13,FASN,ALDH6A1,SPTBN1,ATP5F1B,CALM3,MYH9,RPL
1,FTL,HSP90B1,FADS1,PRPF8,UGT8,RPL8,FBXL4,YBX1,UBA1,ATP2B4,CALR,TMF1,ABHD2,HSP
TMS,MARCKS,SMG1,PLD3,USP11,SNRNP200,EIF3C,ATP5F1A,VCP,MARCKSL1,UBB,TRIM28,MA
0,H3-3B,CANX,PTMA,DDX5,SOX4,RPL13A,SEC62,RPL6,DVL3,RPL13,SLC2A3,DDX17,GPI,SPTBN1
AN1,MARCKS,NPM1,ATP5F1A,VCP,MARCKSL1,UBB,TRIM28,MATR3,YWHAG,RPL37A,FN1,SPEN,
,TMF1,ABHD2,HSPA1B,PSAP,PGK1,RPLP0,H3-3B,CANX,PTMA,DDX5,SOX4,RPL13A,SEC62,RPL6
LP0,H3-3B,CANX,PTMA,DDX5,SOX4,RPL13A,SEC62,KAT6A,DVL3,YWHAE,RPL13,GPI,BSG,ALDH6
0,H3-3B,CANX,PTMA,DDX5,SOX4,RPL13A,SEC62,DVL3,YWHAE,RPL13,DDX17,GPI,ALDH6A1,SPT
2,HSPA1B,PSAP,PGK1,RPLP0,H3-3B,CANX,CLU,DDX5,RPL13A,SEC62,RPL6,KAT6A,DVL3,YWHAE
K1,RPLP0,H3-3B,CANX,PTMA,DDX5,RPL13A,RPL6,KAT6A,DVL3,YWHAE,RPL13,GPI,BSG,ALDH6
A1B,PSAP,PGK1,RPLP0,H3-3B,CANX,PTMA,DDX5,SOX4,RPL13A,SEC62,RPL6,KAT6A,DVL3,YWHA
L13A,SEC62,RPL6,KAT6A,DVL3,YWHAE,RPL13,GPI,SPTBN1,ATP5F1B,GNB1,RPL7,IDS,RPL5,GDI
,RPS4X,RPS3A,UCHL1,BTF3,RPL10,RPS17,DNAJA1,SF3B2,ITM2B,AP2B1,HNRNPU,LARP1,HNRN
62,KAT6A,DVL3,YWHAE,RPL13,SLC2A3,HTRA4,BSG,ATP5F1B,MYH9,SEPTIN2,P4HB,IDS,RPL5,T
K1,RPLP0,H3-3B,CANX,PTMA,DDX5,RPL13A,SEC62,KAT6A,DVL3,YWHAE,RPL13,GPI,BSG,ALDH6
OL4A1,CES3,PSAP,PGK1,RPLP0,H3-3B,CANX,PTMA,DDX5,SOX4,RPL13A,SEC62,RPL6,KAT6A,D
1,ATP2B4,ABHD2,COL4A1,PSAP,PGK1,H3-3B,CANX,PTMA,DDX5,ENO2,RPL13A,SEC62,KAT6A,DV
X5,SOX4,RPL13A,DVL3,YWHAE,RPL13,DDX17,GPI,SPTBN1,ATP5F1B,CALM3,MYH9,RPL7,P4HB,
,PGK1,H3-3B,CANX,PTMA,SOX4,RPL13A,SEC62,RPL6,KAT6A,DVL3,YWHAE,RPL13,DDX17,GPI,A
L6,KAT6A,DVL3,YWHAE,RPL13,DDX17,GPI,BSG,SPTBN1,ATP5F1B,CALM3,MYH9,RPL7,SEPTIN2,
,RPL13A,SEC62,RPL6,KAT6A,DVL3,YWHAE,RPL13,GPI,SPTBN1,ATP5F1B,CALM3,GNB1,RPL7,ID
PI,ATP5F1B,CALM3,RPL7,APP,IDS,RPL5,GDI1,TMBIM6,SPTAN1,MARCKS,SMG1,CALM1,NPM1,P
P0,H3-3B,CANX,PTMA,SOX4,RPL13A,SEC62,RPL6,KAT6A,YWHAE,RPL13,DDX17,GPI,ALDH6A1,A
5,RPL13A,SEC62,RPL6,DVL3,YWHAE,RPL13,DDX17,GPI,BSG,ALDH6A1,SPTBN1,ATP5F1B,CALM
62,KAT6A,DVL3,YWHAE,RPL13,DDX17,GPI,BSG,ALDH6A1,ATP5F1B,CALM3,MYH9,RPL7,SEPTIN2

3,CANX,PTMA,DDX5,SOX4,RPL13A,SEC62,DVL3,YWHAE,RPL13,GPI,BSG,SPTBN1,ATP5F1B,GNB1,
B1,RPL7,RPL5,GDI1,SPTAN1,SMG1,CALM1,NPM1,EIF3C,ATP5F1A,VCP,MARCKSL1,UBB,TRIM28,
RC28,NONO,SET,MRC2,PDIA3,SSBP2,HMGB1,RPS4X,RPL10,RPS17,VCL,DNAJA1,MAP4,SF3B2,HI
,ATP5F1B,CALM3,RPL7,P4HB,IDS,RPL5,GSTP1,MARCKS,CALM1,NPM1,SNRNP200,CSDE1,EIF3C
,COL4A1,PSAP,PGK1,RPLP0,H3-3B,CANX,PTMA,DDX5,SOX4,RPL13A,SEC62,KAT6A,DVL3,YWHAE
MARCKSL1,UBB,TRIM28,MATR3,YWHAG,SPEN,NONO,HACD3,SET,EGLN1,LDHB,PDIA3,SSBP2,CH
E,RPL13,DDX17,ATP5F1B,CALM3,GNB1,RPL7,P4HB,RPL5,GDI1,PTMS,SPTAN1,GSTP1,MARCKS,S
,ATP2B4,CALR,TMF1,ABHD2,PSAP,PGK1,RPLP0,H3-3B,CANX,CLU,PTMA,DDX5,SOX4,RPL13A,SE
,ATP2B4,CALR,TMF1,ABHD2,PSAP,PGK1,RPLP0,H3-3B,CANX,CLU,PTMA,DDX5,SOX4,RPL13A,SE
IA,DDX5,RPL13A,SEC62,RPL6,KAT6A,DVL3,YWHAE,RPL13,DDX17,ALDH6A1,SPTBN1,ATP5F1B,CA
DVL3,YWHAE,RPL13,SLC2A3,DDX17,GPI,SPTBN1,ATP5F1B,RPL7,SEPTIN2,RPL5,GDI1,SPTAN1,M
VHAE,RPL13,GPI,FASN,SPTBN1,ATP5F1B,GNB1,MYH9,P4HB,RPL5,FSCN1,PTMS,SPTAN1,SMG1,N
VHAE,RPL13,GPI,FASN,SPTBN1,ATP5F1B,GNB1,MYH9,P4HB,RPL5,FSCN1,PTMS,SPTAN1,SMG1,N
HAE,GPI,SPTBN1,ATP5F1B,CALM3,GNB1,SEPTIN2,P4HB,IDS,RPL5,GDI1,FSCN1,TMBIM6,PTMS,SI
,YWHAE,SLC2A3,ALDH6A1,SPTBN1,ATP5F1B,GNB1,MYH9,IDS,RPL5,GDI1,SPTAN1,GSTP1,NPM1,
YWHAE,RPL13,DDX17,GPI,ATP5F1B,RPL7,P4HB,RPL5,GDI1,TMBIM6,PTMS,SPTAN1,GSTP1,MAR
F1B,MYH9,RPL7,SEPTIN2,P4HB,RPL5,TMBIM6,PTMS,MARCKS,NPM1,USP11,COL18A1,SNRNP20
WHAE,RPL13,DDX17,ALDH6A1,ATP5F1B,P4HB,IDS,RPL5,GDI1,TMBIM6,SPTAN1,MARCKS,SMG1,
VL3,YWHAE,RPL13,SLC2A3,DDX17,SPTBN1,ATP5F1B,CALM3,MYH9,RPL7,SEPTIN2,P4HB,RPL5,G
GT8,RPL8,FBXL4,YBX1,ATP2B4,CALR,TMF1,ABHD2,TUBB2B,COL4A1,CES3,PSAP,PGK1,RPLP0,H
A1,PSAP,PGK1,RPLP0,H3-3B,CANX,PTMA,DDX5,SOX4,SEC62,KAT6A,DVL3,YWHAE,RPL13,SLC2A
RCKS,CALM1,NPM1,SNRNP200,EIF3C,ATP5F1A,EIF4B,VCP,UBB,AFF4,TRIM28,MATR3,RPL37A,SF
12B,AP2B1,HNRNPU,RPS16,RPL23,CPE,RPSA,CTSD,SCARB2,RPL24,RPL17,GAP43,RPL14,HK1,RI
A,DDX5,RPL13A,SEC62,RPL6,KAT6A,DVL3,YWHAE,RPL13,DDX17,GPI,BSG,ALDH6A1,SPTBN1,ATP
3,YWHAE,RPL13,ALDH6A1,SPTBN1,CALM3,MYH9,P4HB,RPL5,TMBIM6,SPTAN1,MARCKS,CALM1
TMA,DDX5,SOX4,SEC62,KAT6A,DVL3,RPL13,DDX17,GPI,BSG,ALDH6A1,SPTBN1,ATP5F1B,GNB1,
TMA,DDX5,SOX4,SEC62,KAT6A,DVL3,RPL13,DDX17,GPI,BSG,ALDH6A1,SPTBN1,ATP5F1B,GNB1,
17,SPTBN1,CALM3,MYH9,RPL7,P4HB,IDS,RPL5,PTMS,SPTAN1,MARCKS,CALM1,NPM1,COL18A1,
CP,UBB,AFF4,TRIM28,MATR3,YWHAG,RPL37A,SPEN,NONO,DDB1,HACD3,SET,EGLN1,RPL19,LDH
3,YWHAE,GPI,ALDH6A1,SPTBN1,ATP5F1B,GNB1,SEPTIN2,IDS,RPL5,PTMS,SPTAN1,NPM1,PLD3,H
,MARCKSL1,UBB,AFF4,TRIM28,MATR3,YWHAG,CKB,RPL37A,SPEN,NONO,YWHAQ,HACD3,SET,PI
ALM1,NPM1,SNRNP200,EIF3C,ATP5F1A,EIF4B,VCP,UBB,AFF4,TRIM28,MATR3,YWHAG,RPL37A,L
DX5,SOX4,RPL13A,SEC62,DVL3,RPL13,SLC2A3,DDX17,SPTBN1,ATP5F1B,MYH9,RPL7,SEPTIN2,I
VHAG,SPEN,NONO,HACD3,SET,EGLN1,RPL19,LDHB,CHD3,VAPA,HMGB1,POLR2A,RPS4X,BTF3,AI
DE1,EIF3C,ATP5F1A,VCP,UBB,AFF4,TRIM28,MATR3,YWHAG,RPL37A,LRRC28,FN1,SPEN,NONO,D
P,UBB,AFF4,TRIM28,MATR3,YWHAG,CKB,SPEN,NONO,HACD3,SET,EGLN1,LDHB,PDIA3,SSBP2,B
DX5,RPL13A,SEC62,KAT6A,DVL3,YWHAE,RPL13,GPI,ALDH6A1,SPTBN1,ATP5F1B,RPL7,P4HB,IDS,
3N1,CALM3,MYH9,RPL5,FSCN1,TMBIM6,PTMS,SPTAN1,CALM1,NPM1,USP11,COL18A1,EIF3C,AT
L3,SLC2A3,GPI,ATP5F1B,MYH9,P4HB,IDS,RPL5,TMBIM6,NPM1,PLD3,SNRNP200,EIF3C,ATP5F1A
1B,MYH9,P4HB,RPL5,GDI1,TMBIM6,PTMS,SPTAN1,NPM1,EIF3C,ATP5F1A,VCP,MARCKSL1,UBB,
RNP200,EIF3C,ATP5F1A,VCP,UBB,TRIM28,MATR3,RPL37A,NONO,DCHS1,SET,EGLN1,RPL19,PDI
X,PTMA,DDX5,SOX4,RPL13A,SEC62,RPL6,KAT6A,DVL3,RPL13,GPI,ALDH6A1,SPTBN1,ATP5F1B,G
TAN1,MARCKS,NPM1,SNRNP200,EIF3C,ATP5F1A,VCP,UBB,AFF4,TRIM28,MATR3,YWHAG,RPL37
.10,SF3B2,HNRNPU,SMARCA5,HNRNPA0,CTNNA1,FADS2,RPL23,RPSA,EIF4H,SCARB2,RPS9,XRC
3,GNB1,RPL7,IDS,GDI1,TMBIM6,PTMS,SPTAN1,SMG1,CALM1,NPM1,CSDE1,EIF3C,ATP5F1A,EIF

PLD3,SNRNP200,CSDE1,EIF3C,ATP5F1A,EIF4B,VCP,MARCKSL1,UBB,TRIM28,MATR3,YWHAG,RF
GDI1,SPTAN1,MARCKS,CALM1,NPM1,PLD3,HSPD1,SNRNP200,EIF3C,ATP5F1A,EIF4B,VCP,MARCK
CARB2,RPL24,KPNB1,RPL17,GAP43,RPL14,RPL9,PSMD2,CHD4,RPS20,WDR1,RPL26,BEX3,CCT7
5F1A,VCP,UBB,AFF4,TRIM28,MATR3,YWHAG,SPEN,NONO,HACD3,SET,EGLN1,SF3B1,PDIA3,SSBF
SEC62,DVL3,YWHAE,SLC2A3,SPTBN1,ATP5F1B,MYH9,SEPTIN2,P4HB,IDS,GDI1,FSCN1,SPTAN1,M
R3,YWHAG,RPL37A,FN1,SPEN,NONO,DDB1,SET,EGLN1,MRC2,SF3B1,PDIA3,SSBP2,CHD3,HMGB
SCN1,SPTAN1,MARCKS,NPM1,SNRNP200,EIF3C,ATP5F1A,VCP,MARCKSL1,UBB,TRIM28,MATR3,
18A1,CSDE1,ATP5F1A,VCP,MARCKSL1,UBB,AFF4,TRIM28,MATR3,YWHAG,RPL37A,MAPT,FN1,SF
NO,HACD3,SET,LDHB,SSBP2,BASP1,VAPA,HMGB1,RPS3A,STMN1,BTF3,RPL10,MARCHF6,RPS17,
BN1,ATP5F1B,IDS,RPL5,GDI1,TMBIM6,PTMS,SPTAN1,SMG1,NPM1,PLD3,CSDE1,EIF3C,ATP5F1A
BN1,ATP5F1B,IDS,RPL5,GDI1,TMBIM6,PTMS,SPTAN1,SMG1,NPM1,PLD3,CSDE1,EIF3C,ATP5F1A
ATR3,YWHAG,RPL37A,FN1,SPEN,NONO,DDB1,HACD3,SET,EGLN1,MRC2,PDIA3,SSBP2,CHD3,BAS
3,BASP1,HMGB1,TUBB4B,RPL10,DNAJA1,MAP4,SF3B2,ITM2B,GTF2F1,HNRNPU,NFE2L1,CTNNA
ET,INA,LDHB,PDIA3,SSBP2,CHD3,POLR2A,STMN1,UCHL1,ANXA2,RPL10,MARCHF6,VCL,AKAP12,
R2A,RPS4X,RPS3A,RPL10,RPS17,SF3B2,GTF2F1,HNRNPU,LARP1,CTNNA1,FADS2,RPS16,RPL23,
RNPU,LARP1,FADS2,RPS16,RPL23,RPL29,RPSA,SCARB2,XRCC5,RPL24,RPL17,C1ORF43,RPL14,N
4H,RPS9,XRCC5,RPL24,RPL14,CHD4,JUND,EIF5,BEX3,CCT7,RBBP4,MCL1,PRKCSH,HNRNPC,LRI
3,EIF4H,SCARB2,XRCC5,RPL24,RPL17,C1ORF43,RPL14,PSMD2,KIF5B,RPS14,WDR1,RPL26,BEX
,UBB,TRIM28,MATR3,YWHAG,RPL37A,LRRC28,NONO,DDB1,DCHS1,SET,EGLN1,RPL19,LDHB,SF
-3B,CANX,PTMA,DDX5,SOX4,SEC62,KAT6A,DVL3,YWHAE,RPL13,SLC2A3,DDX17,GPI,ATP5F1B,MY
UBB,TRIM28,MATR3,SPEN,NONO,HACD3,SET,EGLN1,RPL19,PKD1,PDIA3,SSBP2,VAPA,HMGB1,P
M28,MATR3,LRRC28,SPEN,NONO,DDB1,HACD3,DCHS1,SET,EGLN1,LDHB,SF3B1,SSBP2,CHD3,V
GB1,POLR2A,RPS4X,RPS3A,TUBB4B,ANXA2,RPL10,SF3B2,GTF2F1,HNRNPU,SMARCA5,CTNNA1,
62,DVL3,YWHAE,RPL13,SLC2A3,DDX17,SPTBN1,ATP5F1B,MYH9,RPL7,SEPTIN2,P4HB,GDI1,PTM
A5,LARP1,FADS2,RPS16,RPL23,RPSA,TPM4,EIF4H,XRCC5,RPL24,RPL17,RPL14,HK1,PTCD3,PSMI
,PDIA3,SSBP2,CHD3,HMGB1,RPS4X,RPS3A,ANXA2,RPL10,VCL,CYB5R3,PAFAH1B1,SF3B2,AP2B1
MAP4,GTF2F1,HNRNPU,HNRNPA0,CTSD,SCARB2,XRCC5,PJA2,KPNB1,RPL17,PAPOLA,NLGN2,CX
1,HSPD1,COL18A1,CSDE1,EIF3C,ATP5F1A,VCP,MARCKSL1,UBB,TRIM28,YWHAG,MAPT,SPEN,NC
S4X,STMN1,BTF3,ANXA2,RPL10,DNAJA1,SF3B2,ITM2B,HNRNPU,LARP1,RPL23,RPL29,RPSA,TPM
,HNRNPU,SMARCA5,LARP1,RPL23,RPSA,TPM4,XRCC5,RPL24,RPL17,KIF5B,CHD4,RPS20,CCT7,F
D3,SET,EGLN1,RPL19,PDIA3,VAPA,HMGB1,POLR2A,RPS4X,RPL10,SF3B2,ITM2B,AP2B1,GOLGA8
LDHB,BASP1,VAPA,HMGB1,RPS4X,TUBB4B,ANXA2,RPL10,SF3B2,GTF2F1,HNRNPU,CTNNA1,RPS
M1,CSDE1,EIF3C,ATP5F1A,VCP,AFF4,TRIM28,MATR3,YWHAG,MAPT,FN1,SPEN,NONO,DCHS1,SET
CC5,RPL24,RPL14,PSMD2,RPS14,ATP1A1,RPS20,EIF5,WDR1,RPL26,CCT7,RBBP4,PNN,RPL31,H
,CTNNA1,FADS2,RPL23,RPSA,TPM4,SCARB2,XRCC5,RPL24,RPL14,HK1,PTCD3,PSMD2,ATP1A1,E
EIF3C,ATP5F1A,EIF4B,VCP,MARCKSL1,TRIM28,MATR3,LRRC28,SPEN,NONO,DDB1,DCHS1,SET,E
SG,SPTBN1,ATP5F1B,CALM3,GNB1,SEPTIN2,IDS,GDI1,FSCN1,TMBIM6,SPTAN1,MARCKS,SMG1,
P2,HMGB1,XAF1,RPL10,RPS17,DNAJA1,SF3B2,GTF2F1,HNRNPU,NFE2L1,CTNNA1,RPL23,RPSA,E
,LRRC28,FN1,SPEN,NONO,HACD3,DCHS1,EGLN1,LDHB,SF3B1,SSBP2,VAPA,RPS3A,BTF3,PAFAH
K1,RPS14,CHD4,RPS20,EIF5,RPL26,CCT7,RBBP4,JPT2,HNRNPC,TADA2B,KHDRBS1,XPO1,RPS25
HB,PDIA3,HMGB1,BTF3,ANXA2,RPL10,CYB5R3,MAP4,SF3B2,HNRNPU,SCARB2,PAPOLA,RPL14,K
A,RPS4X,RPS3A,BTF3,RPL10,SF3B2,ITM2B,GTF2F1,HNRNPU,FADS2,RPSA,TPM4,EIF4H,SCARB2,
3,RPSA,TPM4,XRCC5,RPL24,RPL17,RPL14,CHD4,EIF5,RPL26,SRSF4,RBBP4,JPT2,PNN,RPL31,HN
4,SLC7A5,CHD4,JUND,RPL26,CCT7,SRSF4,RBBP4,HNRNPC,KHDRBS1,XPO1,SPTBN2,SFPQ,DDR1
A,TPM4,SCARB2,XRCC5,RPL24,RPL17,RPL14,SLC7A5,CHD4,ATP1A1,RPS20,EIF5,RPL26,CCT7,R

RPPRC,KHDRBS1,RPS25,GLG1,SFPQ,PRDX2,DDX3X,MRFAP1,H1-0,RPS7,HNRNPDL,TARDBP,ARI
C7A5,CHD4,RPL26,RBBP4,RTN3,HNRNPC,KHDRBS1,XPO1,CSNK1E,CDH2,RAB7A,ACTN4,DDX3X,
HAQ,DCHS1,MRC2,SF3B1,PDIA3,SSBP2,VAPA,HMGB1,POLR2A,TPT1,RPS3A,TUBB4B,AGRN,MAR
G,LRRC28,FN1,SPEN,NONO,DDB1,DCHS1,EGLN1,LDHB,SF3B1,SSBP2,HMGB1,POLR2A,BTF3,N
UBB,TRIM28,MATR3,LRRC28,FN1,SPEN,NONO,SET,EGLN1,LDHB,SF3B1,SSBP2,HMGB1,ITGB1,
L29,RPSA,TPM4,EIF4H,XRCC5,MSN,RPL24,RPL17,RPL14,KIF5B,CHD4,CD81,RPL26,BEX3,CCT7,
S7,IK,RPL35,HNRNPDL,TARDBP,HNRNPK,H1-10,RPL36,YWHAB,LAMP1,HNRNPH1,CALU,ARCN1,
LA,GAP43,RPL14,CHD4,HMGN2,BEX3,RTN3,NISCH,TADA2B,KHDRBS1,TKT,GLG1,SFPQ,PDIA6,M
,KHDRBS1,XPO1,RPS25,SFPQ,DDR1,RPS15,H1-0,RPS5,RPS7,IK,GJA1,CTNNB1,HNRNPDL,TARD
MARCKSL1,TRIM28,MATR3,LRRC28,NONO,HACD3,DCHS1,SET,EGLN1,RPL19,PDIA3,SSBP2,VAP
1,EIF5,CCT7,RBBP4,JPT2,HNRNPC,LRPPRC,KHDRBS1,KLHL9,SPTBN2,RPS25,GLG1,SFPQ,RAB7
IGB1,POLR2A,RPS4X,BTF3,RPL10,WSB1,DNAJA1,SF3B2,GTF2F1,HNRNPU,SMARCA5,LARP1,CTN
BB,TRIM28,MATR3,YWHAG,RPL37A,LRRC28,SPEN,NONO,DCHS1,SET,EGLN1,MRC2,SF3B1,SSB
1A2,LLGL1,RHOA,PSMD2,RPS14,CHD4,JUND,RPS20,HMGN2,SRSF4,PFN1,PNN,HNRNPC,LRPPRC
EGLN1,SF3B1,SSBP2,BASP1,HMGB1,RPS3A,BTF3,ANXA2,AGRN,RPL10,VCL,CYB5R3,MAP4,SF3E
HMGB1,POLR2A,RPS4X,BTF3,RPL10,WSB1,DNAJA1,SF3B2,GTF2F1,HNRNPU,SMARCA5,LARP1,C
TADA2B,KHDRBS1,XPO1,RPS25,SFPQ,RAB7A,RPS15,H1-0,RPS5,RPS7,IK,RPL35,GJA1,JUN,CTN
HNRNPC,LRPPRC,XPR1,KHDRBS1,XPO1,TKT,SFPQ,MRFAP1,H1-0,IK,CTNNB1,GOLM1,HNRNPDL,
,DDB1,DCHS1,EGLN1,MRC2,SF3B1,VAPA,RPS4X,ITGB1,XAF1,RPL10,RPS17,VCL,DNAJA1,CYB5
PDL,TARDBP,HNRNPK,H1-10,RPL36,YWHAB,LAMP1,CALU,CAPRIN1,TOP2B,HDAC2,SKP1,BCLAF1
CTN4,PRDX2,DDX3X,H1-0,IK,APLP2,HNRNPDL,TARDBP,HNRNPK,H1-10,LAMP1,HLA-B,CCT6A,CA
NPH1,HDAC2,BCLAF1,LUC7L3,CPSF2,CTSB,NUCB1,TM9SF3,MDH2,TOMM20,PABPN1,GDI2,HNF
PFN1,SPC25,HNRNPC,LRPPRC,KHDRBS1,RPS25,SFPQ,MAGED2,RPS5,TLN1,APLP2,TARDBP,ARI
,RPS25,GLG1,SFPQ,DDX3X,H1-0,APLP2,HNRNPDL,TARDBP,HNRNPK,H1-10,CKAP5,HNRNPH1,AF
1,RPL10,SF3B2,ITM2B,GTF2F1,GOLGA8A,HNRNPU,OGA,SMARCA5,RPS16,XRCC5,KPNB1,PAPC
PL37A,LRRC28,NONO,DCHS1,SET,EGLN1,SF3B1,PDIA3,SSBP2,VAPA,HMGB1,RPS4X,BTF3,XAF1,
,ACO2,RBMX,PTK7,SNRNP70,H3-3A,OPTN,SMARCA4,ARPC2,ILF3,PRKDC,HNRNPUL1,ACTN1,KD
NNA1,RPS16,TPM4,EIF4H,BRD3,PJA2,RPL24,LLGL1,HK1,KIF5B,SLC7A5,RPS14,CHD4,TMEM25
,NPM1,HSPD1,COL18A1,ATP5F1A,VCP,UBB,MATR3,YWHAG,RPL37A,LRRC28,SPEN,NONO,DDB1
2AF2,RPN2,PGD,GNAI3,PABPN1,GDI2,HNRNPL,LMNB2,ACO2,VPS35,RBMX,CHD2,SNRNP70,PYG
,H1-0,RPS7,RPL35,CTNNB1,APLP2,TARDBP,NDFIP1,TAGLN,WLS,HNRNPK,RPL36,HLA-B,CALU,R
32,HNRNPU,SMARCA5,RPL29,RPSA,RPS9,XRCC5,MSN,RPL24,C1ORF43,RPL14,RPL9,SLC7A5,RF
TBN2,RPS25,SFPQ,H1-0,GJA1,CTNNB1,HNRNPDL,TARDBP,ARID1A,HNRNPK,INIP,RPL36,CKAP5,
BP,HNRNPK,H1-10,LAMP1,HNRNPH1,HLA-B,CAPRIN1,TOP2B,HDAC2,BCLAF1,TRIB2,LUC7L3,CP
1,HNRNPU,NFE2L1,HNRNPA0,FADS2,RPL23,TPM4,SCARB2,LPCAT1,RPL24,RPL17,PTCD3,PSMD
43,RPL14,PSMD2,EIF5,BEX3,SRSF4,RBBP4,PFN1,PRKCSH,HNRNPC,KHDRBS1,GLG1,SFPQ,EIF5
CKB,LRRC28,MAPT,SPEN,YWHAQ,HACD3,DCHS1,EGLN1,INA,LDHB,SF3B1,BASP1,VAPA,STMN1,
NPC,KHDRBS1,XPO1,RPS25,SFPQ,MAGED2,H1-0,TARDBP,HNRNPK,RPL36,CKAP5,LAMP1,HNRN
3B1,SSBP2,VAPA,HMGB1,TPT1,RPS3A,RPL11,MORF4L1,BTF3,AGRN,RPL10,DNAJA1,CYB5R3,M
1,POLR2A,RPS3A,STMN1,TUBB4B,UCHL1,RPL11,XAF1,RPL10,MARCHF6,SF3B2,ITM2B,MAT2A,
3B2,HNRNPU,RPL29,EIF4H,LPCAT1,KPNB1,RPL17,RPL14,JUND,WDR1,BEX3,RBBP4,RTN3,TADA2
NPA3,CBX1,H3-3A,TOP1,PTBP1,ILF3,PRKDC,HNRNPUL1,KDM1A,DIDO1,HNRNPM,SSB,SRSF3,ILF
DR1,CCT7,RBBP4,HNRNPC,LRPPRC,XPR1,SLC3A2,PTN,CSNK1E,RPS25,SFPQ,PEBP1,DDR1,RAB
4,RPS14,CHD4,RPS20,WDR1,RPL26,BEX3,RBBP4,PFN1,MCL1,LRPPRC,KHDRBS1,RPS25,GLG1,
FPQ,CCN2,MAGED2,RPS7,TLN1,HNRNPK,H1-10,RPL36,SEPTIN11,LAMP1,HLA-B,COTL1,ARHGD

36,CKAP5,HNRNPH1,HLA-B,ARCN1,CAPRIN1,HDAC2,BCLAF1,LUC7L3,CPSF2,DDX39B,CTSB,TM9
IDO1,SMARCE1,SUPT6H,HNRNPM,SRSF3,ILF2,FBLN1,CSNK1D,P3H3,TP53BP1,RBBP7,LBH,SF1,
INRNPC,LRPPRC,KHDRBS1,RPS25,SFPQ,PDIA6,PRDX2,RPS15,MAGED2,H1-0,RPL35,TLN1,ANXA6
AD23B,SNRNP70,H3-3A,HDGF,ABCA2,CCNL2,SRSF1,TCF25,ATP5PB,PTBP1,ILF3,HNRNPUL1,ZF
SMARCA5,CNN3,EIF4H,XRCC5,C1ORF43,GAP43,ATF4,JUND,BEX3,SRSF4,RBBP4,PNN,NES,NISCH
LN1,KDM3B,RBBP7,ACTR1A,LBH,BRD4,ATRX,EP400,SRSF6,ZFR,HNRNPA2B1,CUL3,HNRNPAB,S
NPL,MORF4L2,LMNB2,ACO2,CCT3,CBX1,TNPO1,SNRNP70,H3-3A,DPYSL2,TOP1,MAPK1,SMAR
C2,CPSF2,TM9SF3,U2AF2,PPP1CB,PABPN1,HNRNPL,MORF4L2,TAF15,HNRNPA3,RBMX,LAMC1,
UCKS1,SRSF11,MSH6,USP24,YY1,CREBBP,THRAP3,SLC38A2,EIF4A3,CASC4,SUN1,MDM2,SF3A3
1,MMP2,BCLAF1,LUC7L3,CPSF2,TM9SF3,PABPN1,HNRNPL,TSPYL4,MORF4L2,TAF15,ALKBH5,F
IP,PARP1,MORF4L2,ALKBH5,RAD23B,TNPO1,SNRNP70,H3-3A,HDGF,VEGFA,MAPK1,SMARCA4,
LAMP1,HNRNPH1,HLA-B,CCT6A,HDAC2,BCLAF1,LUC7L3,SQLE,CPSF2,TM9SF3,TOMM20,U2AF2
HNRNPL,HNRNPA3,CBX1,SNRNP70,H3-3A,HDGF,DSTN,SRSF1,AEBP1,NYNRIN,ILF3,DIDO1,SMAR
L,TARDBP,DDOST,HNRNPK,MYH10,RPL36,HNRNPH1,HLA-B,MAP2,CCT6A,TOP2B,HDAC2,BCLAF
2,HNRNPL,MORF4L2,LMNB2,ALKBH5,HNRNPA3,C4ORF3,RBMX,CBX1,CHD2,SKI,SNRNP70,H3-
N1,LDHB,PKD1,PDIA3,SSBP2,HMGB1,UCHL1,BTF3,AGRN,XAF1,RPL10,WSB1,MAP4,SF3B2,GTF
,MORF4L2,TAF15,HNRNPA3,RBMX,CBX1,CCT4,SNRNP70,HDGF,ABCA2,TERF2IP,SMARCA4,MAP
FTO,ILF3,PRKDC,KDM1A,DIDO1,TPM3,CORO1C,SUPT6H,HNRNPM,SRSF3,ILF2,KDM3B,DDIT4,A
1,CTNNB1,APLP2,HNRNPDL,TARDBP,H1-10,LAMP1,TOP2B,HDAC2,BCLAF1,LUC7L3,CPSF2,DDX3
BP7,EXTL3,FLOT1,BRD4,TRA2B,UQCRC1,ZFR,HNRNPA2B1,HNRNPAB,PRDX6,FUS,VPS41,HDGFL
WHAB,SEPTIN11,LAMP1,HLA-B,COTL1,CAPRIN1,BCLAF1,TRIB2,LUC7L3,MDH2,TOMM20,HNRNF
7,MLXIP,SQSTM1,HNRNPL,TAF15,ALKBH5,PTPRA,TBC1D16,RBMX,CBX1,SNRNP70,PSMB2,H3-3
,SDF4,ATP6V1B2,STMN3,KDM3B,PRMT1,MAP6,MAPRE2,BRSK1,APEX1,AKAP9,TPP1,SSR3,DCLK
REG2,CNTNAP1,HIP1,AP2A1,MAP6,PPP2CB,GLUD1,EIF5B,SV2A,MCM3AP,MAPRE2,SEPTIN7,C1O
DRBS1,XPO1,SFPQ,PDIA6,RPL12,EIF3L,MRFAP1,H1-0,IK,RPL35,GOLM1,HNRNPDL,TARDBP,DDC
B1,ZNF770,AP2B1,PGAM1,NFE2L1,LARP1,CTNNA1,RPL23,RPSA,EIF4H,BRD3,RPL24,C1ORF43,C
A0,RPL29,TPM4,RTN4,XRCC5,PJA2,KPNB1,RPL17,PAPOLA,EIF5,RBBP4,PNN,SPC25,PRKCSH,HN
E2,BRSK1,APEX1,AKAP9,TPP1,SSR3,DCLK1,DTX4,EEF1B2,REEP2,PSMD4,AP2A2,SORT1,CD74,F
NSG2,FKBP8,POLR2B,ATP6V1B2,FAU,STMN3,SOD1,FBRSL1,SYP,VDAC1,CAPN2,SF1,BRD4,MAP6
DX6,ZMYM3,FUS,ERH,RALY,HDGFL2,EFTUD2,TRAP1,NUCKS1,RBM5,SPTY2D1,BORCS7,IST1,AN
N2B1,WASF3,NACC1,BACE1,RBM17,CTCF,RFX7,CMTR1,SP1,NCOA6,ACIN1,AKAP17A,KLF3,GSN,
,TCAF1,RALY,LTBP4,HDGFL2,EFTUD2,PRPF19,CALCOCO1,XRN2,LRIG1,MDM2,FAM50A,CFDP1,

A8A,HNRNPU,SMARCA5,HNRNPA0,RPL29,TPM4,RTN4,XRCC5,PJA2,KPNB1,RPL17,PAPOLA,ATF4,
L1,LBH,SF1,BRD4,UBA52,ATRX,SRSF6,HNRNPA2B1,MCM3AP,RPS27A,HNRNPAB,C1ORF198,FUS
PHS1,FBL,MAN2B1,WASF3,NACC1,BACE1,RBM17,CTCF,RFX7,PNMA2,CMTR1,SP1,NCOA6,ACIN
JFB8,ACSL4,DLG4,WASF3,PRKAR2B,DPP7,BACE1,FH,SYT1,PDHB,NNT,ZNF326,ARSA,FKBP3,MM
BE2J1,RPL27A,NACAD,NDUFB8,ACSL4,DLG4,WASF3,PRKAR2B,DPP7,BACE1,FH,SYT1,PDHB,NN
1,VDAC1,SF1,BRD4,MAP6,SV2A,MAPRE2,SEPTIN7,TMOD2,SEPTIN5,LINGO1,STRN4,SSR3,DCLK
CLAF1,LUC7L3,CPSF2,TM9SF3,U2AF2,CCT2,SUGP2,PABPN1,HNRNPL,TM9SF2,MORF4L2,HNRN
,MAP2K2,CORO1C,HNRNPM,GIGYF1,SRSF3,CCT8,LSS,RAI1,MLF2,KDM3B,SMS,RBBP7,SF1,PDA
IC2,ILF2,DDIT4,RBBP7,ACTR1A,PSMD1,HIP1,GTPBP1,BRD4,GLUD1,ZFR,AGPAT1,HNRNPAB,GSE
INRNPL,HNRNPA3,C4ORF3,ACO2,SNRNP70,H3-3A,OPTN,SMARCA4,ARPC2,PRDX1,SRSF1,ILF3,P
F3B1,VAPA,TPT1,RPS4X,MORF4L1,BTF3,AGRN,RPS17,DNAJA1,CYB5R3,PAFAH1B1,CDC42BPB,
CHMP4B,ATP6V1G1,WNK1,NMNAT2,SORT1,CD74,RCN2,ELAVL3,PSAT1,SCAMP5,FXR2,NOTCH1,

,BAZ1B,EP400,PI4KA,GLO1,ABI2,TPP1,C6ORF62,DICER1,NSD2,CSE1L,RBM6,RCN2,ANK2,CAD,P
PP7,CTCF,ATP6V0A1,FH,APOE,PPT1,EPHA4,PDHB,GSN,ARSA,FKBP3
Z6L2,ALCAM,WASF3,DPP7,VDAC2,UQCRFS1,BACE1,SLC1A3,NFASC,ABCD3,PPT1,PDHB,FKBP3,N
2,IGLON5,USP47,CLTA,NOTCH1,TRIM2,SSU72,UBE2J1,TSPAN7,NDRG4,MRPS6,RAB3A,SOX9,NA
EEP2,AP2A2,SAE1,RCN2,IGLON5,USP47,FXR2,CLTA,APEH,UBE2J1,FAM168A,NDRG4,NDUFB8,MI

,LAPTM4A,ARPC2,DSTN,PSMD3,NYNRIN,LUZP6,CAPZB,PLCG1,CORO1C,PFKM,EIF3F,TMED9,PCI
1,SSR3,CNP,PSMD4,CHMP4B,NMNAT2,SORT1,IGLON5,PSAT1,SCAMP5,FXR2,NOTCH1,TRIM2,ID
OLA,EIF5,RBBP4,PNN,SPC25,RPL31,PRKCSH,HNRNPC,KHDRBS1,XPO1,SFPQ,TPM1,ACTN4,DDX3

ATXN2L,SOD1,SSR1,PRMT1,TXNDC5,MTA1,TRIO,KDM5B,PRCC,BAZ1B,CD99,EP400,PI4KA,SEPTIN

SAT1,FXR2,NOTCH1,TRIM2,MTURN,AKAP8L,ALDH7A1,NACAD,NDUFB8,MAN2B1,VDAC2,UQCRFS

NAPB,DLG4,CASKIN1,VDAC2,UQCRFS1,BACE1,CHGB,GNAZ,ATP6V0A1,SHC1,PPP3CA,ADD1,FH,SY
2,IDH2,MTURN,FAM168A,SYNGAP1,NACAD,CNTN1,PPP3CA,APOE,ABCD3,PPT1,ATP6V1F,PDHB,W

DHB,VPS37B
PELP1,CARM1,SRP72,SSR3,PRR12,YIPF3,PRKACA,NDUFA13,CASC4,PLS3,FAM210B,CGGBP1,HS
,WAPL,SIN3A,CC2D1A
L2,CLCN4,ALCAM,VDAC2,UQCRFS1,FH,ABCD3,PPT1,PDHB,ANAPC2,DMPK,MYBBP1A,FKBP3
SLIT3,ATXN2L,HYOU1,SOD1,CD99L2,TRAF7,ST3GAL6,PRMT1,KDM5B,PRCC,BAZ1B,SEC16A,KPI

2,HSPA1B,TUBB2B,COL4A1,CES3,PSAP,PGK1,RPLP0,H3-3B,CANX,CLU,PTMA,DDX5,ENO2,SOX4,
RPLP0,H3-3B,CANX,CLU,PTMA,DDX5,ENO2,SOX4,RPL13A,SEC62,RPL6,KAT6A,DVL3,YWHAE,RPL1
NX,PTMA,DDX5,ENO2,SOX4,RPL13A,SEC62,RPL6,KAT6A,DVL3,YWHAE,RPL13,DDX17,GPI,FASN,A
TMA,DDX5,ENO2,SOX4,RPL13A,SEC62,RPL6,KAT6A,DVL3,YWHAE,RPL13,DDX17,GPI,SPTBN1,ATI
1B,TUBB2B,COL4A1,CES3,PSAP,PGK1,RPLP0,H3-3B,CANX,PTMA,DDX5,ENO2,SOX4,RPL13A,SEC
3B,CANX,PTMA,DDX5,ENO2,SOX4,RPL13A,SEC62,RPL6,KAT6A,DVL3,YWHAE,RPL13,SLC2A3,DDX
,CANX,PTMA,DDX5,ENO2,SOX4,RPL13A,SEC62,RPL6,KAT6A,DVL3,YWHAE,RPL13,DDX17,GPI,ALI
,CLU,PTMA,DDX5,ENO2,RPL13A,SEC62,RPL6,KAT6A,DVL3,YWHAE,RPL13,DDX17,GPI,BSG,ALDH
AP,PGK1,RPLP0,H3-3B,CANX,CLU,PTMA,DDX5,SOX4,RPL13A,SEC62,RPL6,KAT6A,DVL3,YWHAE,I
RPL13A,SEC62,RPL6,KAT6A,DVL3,YWHAE,RPL13,DDX17,GPI,ALDH6A1,SPTBN1,ATP5F1B,CALM3
RPL13A,SEC62,RPL6,KAT6A,DVL3,YWHAE,RPL13,DDX17,GPI,ALDH6A1,SPTBN1,ATP5F1B,CALM3
LU,PTMA,DDX5,SOX4,RPL13A,SEC62,RPL6,KAT6A,DVL3,YWHAE,RPL13,DDX17,GPI,BSG,ALDH6A
PTBN1,ATP5F1B,CALM3,SEPTIN2,P4HB,APP,RPL5,GDI1,PTMS,SPTAN1,MARCKS,SMG1,CALM1,NI
K1,RPLP0,H3-3B,CANX,CLU,PTMA,DDX5,SOX4,RPL13A,SEC62,RPL6,KAT6A,DVL3,YWHAE,RPL13
K1,RPLP0,H3-3B,CANX,CLU,PTMA,DDX5,SOX4,RPL13A,SEC62,RPL6,KAT6A,DVL3,YWHAE,RPL13
,RPL13A,SEC62,RPL6,KAT6A,DVL3,YWHAE,RPL13,DDX17,GPI,BSG,ALDH6A1,SPTBN1,ATP5F1B,M
DDX5,SOX4,RPL13A,RPL6,KAT6A,DVL3,YWHAE,RPL13,DDX17,GPI,ALDH6A1,SPTBN1,ATP5F1B,C
DDX5,SOX4,RPL13A,RPL6,KAT6A,DVL3,YWHAE,RPL13,DDX17,GPI,ALDH6A1,SPTBN1,ATP5F1B,C
K1,RPLP0,H3-3B,CANX,PTMA,DDX5,SOX4,RPL13A,SEC62,RPL6,KAT6A,DVL3,YWHAE,RPL13,SLC
SEC62,RPL6,KAT6A,DVL3,YWHAE,RPL13,DDX17,GPI,BSG,ALDH6A1,SPTBN1,ATP5F1B,CALM3,GI
SEPTIN2,P4HB,GDI1,FSCN1,PTMS,SPTAN1,GSTP1,MARCKS,NPM1,PLD3,HSPD1,COL18A1,CSDE1
-3B,CANX,PTMA,DDX5,SOX4,RPL13A,SEC62,RPL6,KAT6A,DVL3,YWHAE,RPL13,SLC2A3,DDX17,G
J,PTMA,DDX5,SOX4,RPL13A,SEC62,RPL6,KAT6A,DVL3,YWHAE,RPL13,DDX17,GPI,BSG,ALDH6A1,
J,PTMA,DDX5,SOX4,RPL13A,SEC62,RPL6,KAT6A,DVL3,YWHAE,RPL13,DDX17,GPI,BSG,ALDH6A1,
S,RPL5,GDI1,TMBIM6,PTMS,SPTAN1,SMG1,CALM1,NPM1,PLD3,USP11,CSDE1,EIF3C,ATP5F1A,VC
IBIM6,SPTAN1,SMG1,CALM1,NPM1,PLD3,USP11,EIF3C,ATP5F1A,VCP,UBB,TRIM28,MATR3,YWH
P,PGK1,RPLP0,H3-3B,CANX,PTMA,DDX5,SOX4,RPL13A,SEC62,RPL6,KAT6A,DVL3,YWHAE,RPL13,
SOX4,RPL13A,SEC62,RPL6,KAT6A,DVL3,YWHAE,RPL13,DDX17,GPI,BSG,ALDH6A1,SPTBN1,ATP5F
L13A,SEC62,RPL6,KAT6A,YWHAE,RPL13,SLC2A3,DDX17,GPI,BSG,SPTBN1,ATP5F1B,CALM3,RPL
L13A,SEC62,RPL6,KAT6A,YWHAE,RPL13,SLC2A3,DDX17,GPI,BSG,SPTBN1,ATP5F1B,CALM3,RPL
PLP0,H3-3B,CANX,CLU,PTMA,DDX5,SOX4,RPL13A,SEC62,RPL6,KAT6A,DVL3,YWHAE,RPL13,DDX1
S,KAT6A,DVL3,YWHAE,RPL13,DDX17,GPI,ALDH6A1,SPTBN1,ATP5F1B,CALM3,GNB1,MYH9,RPL7,
S,KAT6A,DVL3,YWHAE,RPL13,DDX17,GPI,ALDH6A1,SPTBN1,ATP5F1B,CALM3,GNB1,MYH9,RPL7,
3,GNB1,MYH9,RPL7,SEPTIN2,P4HB,RPL5,GDI1,FSCN1,PTMS,SPTAN1,GSTP1,MARCKS,NPM1,PLI
17,GPI,ALDH6A1,SPTBN1,ATP5F1B,CALM3,GNB1,MYH9,RPL7,SEPTIN2,P4HB,IDS,RPL5,GDI1,FS
TMA,DDX5,SOX4,RPL13A,SEC62,RPL6,KAT6A,DVL3,YWHAE,RPL13,SLC2A3,DDX17,GPI,FASN,ALD
PTBN1,ATP5F1B,SEPTIN2,IDS,GDI1,FSCN1,PTMS,SPTAN1,GSTP1,MARCKS,NPM1,PLD3,HSPD1,CS
L13,DDX17,GPI,BSG,ALDH6A1,SPTBN1,ATP5F1B,CALM3,MYH9,SEPTIN2,P4HB,APP,IDS,RPL5,GD
L13,DDX17,GPI,BSG,ALDH6A1,SPTBN1,ATP5F1B,CALM3,MYH9,SEPTIN2,P4HB,APP,IDS,RPL5,GD
2,KAT6A,DVL3,YWHAE,RPL13,SLC2A3,DDX17,GPI,BSG,ALDH6A1,SPTBN1,ATP5F1B,GNB1,MYH9,
,SEPTIN2,P4HB,RPL5,GDI1,PTMS,SPTAN1,MARCKS,NPM1,COL18A1,CSDE1,EIF3C,ATP5F1A,VCP,
LP0,H3-3B,CANX,CLU,PTMA,DDX5,SOX4,RPL13A,SEC62,RPL6,KAT6A,DVL3,YWHAE,RPL13,DDX17
LP0,H3-3B,CANX,CLU,PTMA,DDX5,SOX4,RPL13A,SEC62,RPL6,KAT6A,DVL3,YWHAE,RPL13,DDX17
A,DDX5,SOX4,RPL13A,SEC62,RPL6,KAT6A,DVL3,YWHAE,RPL13,DDX17,GPI,BSG,ALDH6A1,SPTBN
A,DDX5,SOX4,RPL13A,SEC62,RPL6,KAT6A,DVL3,YWHAE,RPL13,DDX17,GPI,BSG,ALDH6A1,SPTBN

3,SPTBN1,ATP5F1B,CALM3,GNB1,MYH9,RPL7,SEPTIN2,P4HB,RPL5,GDI1,FSCN1,TMBIM6,PTMS,
DX5,SOX4,RPL13A,SEC62,RPL6,KAT6A,DVL3,YWHAE,RPL13,SLC2A3,DDX17,GPI,BSG,ALDH6A1,S
DX5,SOX4,RPL13A,SEC62,RPL6,KAT6A,DVL3,YWHAE,RPL13,SLC2A3,DDX17,GPI,BSG,ALDH6A1,S
B,CANX,PTMA,DDX5,SOX4,RPL13A,SEC62,RPL6,KAT6A,DVL3,YWHAE,SLC2A3,DDX17,GPI,SPTBN
PL6,KAT6A,DVL3,YWHAE,RPL13,DDX17,GPI,BSG,ALDH6A1,SPTBN1,ATP5F1B,CALM3,GNB1,MYH
,PTMA,DDX5,SOX4,RPL13A,SEC62,RPL6,KAT6A,DVL3,YWHAE,RPL13,SLC2A3,GPI,BSG,ALDH6A1,
B,CANX,PTMA,DDX5,SOX4,RPL13A,SEC62,RPL6,KAT6A,DVL3,YWHAE,RPL13,DDX17,GPI,ALDH6A
62,RPL6,KAT6A,DVL3,YWHAE,RPL13,DDX17,GPI,SPTBN1,ATP5F1B,CALM3,GNB1,RPL7,P4HB,AP
3,YWHAE,RPL13,DDX17,GPI,ALDH6A1,SPTBN1,ATP5F1B,CALM3,GNB1,MYH9,RPL7,SEPTIN2,P4H
3,YWHAE,RPL13,DDX17,GPI,ALDH6A1,SPTBN1,ATP5F1B,CALM3,GNB1,MYH9,RPL7,SEPTIN2,P4H
TMA,DDX5,SOX4,RPL13A,SEC62,RPL6,KAT6A,DVL3,YWHAE,RPL13,DDX17,GPI,BSG,ALDH6A1,SP
TBN1,ATP5F1B,CALM3,MYH9,RPL7,SEPTIN2,P4HB,IDS,RPL5,TMBIM6,SPTAN1,GSTP1,MARCKS,
6A,DVL3,YWHAE,RPL13,DDX17,GPI,BSG,ALDH6A1,SPTBN1,ATP5F1B,CALM3,MYH9,RPL7,SEPTIN
6A,DVL3,YWHAE,RPL13,DDX17,GPI,BSG,ALDH6A1,SPTBN1,ATP5F1B,CALM3,MYH9,RPL7,SEPTIN
RCKS,CALM1,NPM1,COL18A1,CSDE1,EIF3C,ATP5F1A,VCP,MARCKSL1,UBB,TRIM28,MATR3,YWH
2,KAT6A,DVL3,YWHAE,RPL13,SLC2A3,DDX17,GPI,BSG,SPTBN1,ATP5F1B,CALM3,MYH9,RPL7,SE
I,SPTBN1,ATP5F1B,CALM3,GNB1,MYH9,SEPTIN2,IDS,RPL5,GDI1,FSCN1,TMBIM6,PTMS,SPTAN1,
DDX5,SOX4,RPL13A,SEC62,RPL6,KAT6A,DVL3,YWHAE,RPL13,DDX17,GPI,BSG,ALDH6A1,SPTBN1,
DDX5,SOX4,RPL13A,SEC62,RPL6,KAT6A,DVL3,YWHAE,RPL13,DDX17,GPI,BSG,ALDH6A1,SPTBN1,
L13A,SEC62,RPL6,KAT6A,DVL3,RPL13,DDX17,GPI,BSG,ALDH6A1,SPTBN1,ATP5F1B,CALM3,MYH
L13A,SEC62,RPL6,KAT6A,DVL3,RPL13,DDX17,GPI,BSG,ALDH6A1,SPTBN1,ATP5F1B,CALM3,MYH
L7,SEPTIN2,P4HB,IDS,RPL5,GDI1,TMBIM6,PTMS,SPTAN1,GSTP1,SMG1,NPM1,PLD3,COL18A1,SN
MA,DDX5,SOX4,RPL13A,SEC62,RPL6,KAT6A,DVL3,YWHAE,RPL13,SLC2A3,DDX17,GPI,BSG,SPTB
AT6A,YWHAE,RPL13,SLC2A3,GPI,SPTBN1,ATP5F1B,GNB1,SEPTIN2,P4HB,APP,IDS,GDI1,TMBIM6,
X5,SOX4,RPL13A,SEC62,RPL6,KAT6A,DVL3,YWHAE,RPL13,DDX17,GPI,SPTBN1,ATP5F1B,CALM3,
BN1,ATP5F1B,MYH9,RPL7,SEPTIN2,P4HB,IDS,RPL5,GDI1,TMBIM6,PTMS,SPTAN1,GSTP1,MARCK
1,SPTAN1,GSTP1,SMG1,NPM1,PLD3,HSPD1,USP11,CSDE1,EIF3C,ATP5F1A,VCP,UBB,AFF4,TRIM
YWHAE,RPL13,GPI,SPTBN1,ATP5F1B,CALM3,GNB1,MYH9,RPL7,SEPTIN2,P4HB,IDS,RPL5,FSCN1
P5F1B,CALM3,SEPTIN2,IDS,RPL5,GDI1,TMBIM6,PTMS,SPTAN1,GSTP1,SMG1,CALM1,NPM1,PLD3
P5F1B,CALM3,SEPTIN2,IDS,RPL5,GDI1,TMBIM6,PTMS,SPTAN1,GSTP1,SMG1,CALM1,NPM1,PLD3
DH6A1,SPTBN1,ATP5F1B,MYH9,SEPTIN2,P4HB,IDS,RPL5,GDI1,TMBIM6,PTMS,SPTAN1,MARCKS,
E,RPL13,DDX17,GPI,BSG,ALDH6A1,SPTBN1,ATP5F1B,CALM3,GNB1,RPL7,P4HB,IDS,RPL5,GDI1,
,SEPTIN2,APP,RPL5,GDI1,FSCN1,TMBIM6,PTMS,SPTAN1,GSTP1,MARCKS,NPM1,PLD3,HSPD1,SN
A,DDX5,SOX4,RPL13A,SEC62,RPL6,KAT6A,DVL3,YWHAE,RPL13,SLC2A3,DDX17,GPI,BSG,ALDH6
A,DDX5,SOX4,RPL13A,SEC62,RPL6,KAT6A,DVL3,YWHAE,RPL13,SLC2A3,DDX17,GPI,BSG,ALDH6
X5,SOX4,RPL13A,SEC62,RPL6,KAT6A,YWHAE,RPL13,DDX17,GPI,ALDH6A1,SPTBN1,ATP5F1B,CA
TMA,DDX5,RPL13A,SEC62,RPL6,KAT6A,DVL3,YWHAE,RPL13,SLC2A3,DDX17,GPI,BSG,ALDH6A1,S
AT6A,DVL3,YWHAE,RPL13,DDX17,GPI,BSG,ALDH6A1,SPTBN1,ATP5F1B,CALM3,GNB1,MYH9,RPL
,MARCKS,NPM1,HSPD1,CSDE1,EIF3C,ATP5F1A,VCP,MARCKSL1,UBB,AFF4,TRIM28,MATR3,YWH
DX17,GPI,SPTBN1,ATP5F1B,CALM3,GNB1,MYH9,RPL7,SEPTIN2,P4HB,IDS,RPL5,GDI1,FSCN1,TM
RPL5,GDI1,TMBIM6,SPTAN1,GSTP1,MARCKS,SMG1,NPM1,HSPD1,COL18A1,SNRNP200,CSDE1,
N2,P4HB,IDS,RPL5,GDI1,TMBIM6,PTMS,SPTAN1,GSTP1,MARCKS,NPM1,PLD3,CSDE1,EIF3C,ATP5
PL5,GDI1,TMBIM6,SPTAN1,GSTP1,MARCKS,SMG1,CALM1,NPM1,HSPD1,SNRNP200,EIF3C,ATP5F
B,CANX,PTMA,SOX4,RPL13A,RPL6,KAT6A,DVL3,YWHAE,SLC2A3,DDX17,GPI,SPTBN1,ATP5F1B,G
X17,GPI,BSG,ALDH6A1,SPTBN1,ATP5F1B,CALM3,GNB1,MYH9,RPL7,SEPTIN2,P4HB,IDS,RPL5,GD

1,ATP5F1B,CALM3,GNB1,MYH9,RPL7,SEPTIN2,P4HB,IDS,RPL5,FSCN1,PTMS,SPTAN1,GSTP1,MA
.13A,SEC62,RPL6,KAT6A,DVL3,YWHAE,RPL13,SLC2A3,DDX17,GPI,ALDH6A1,SPTBN1,ATP5F1B,M
KAT6A,DVL3,YWHAE,RPL13,DDX17,GPI,BSG,ALDH6A1,SPTBN1,ATP5F1B,CALM3,GNB1,MYH9,RP
KAT6A,DVL3,YWHAE,RPL13,DDX17,GPI,BSG,ALDH6A1,SPTBN1,ATP5F1B,CALM3,GNB1,MYH9,RP
28,SPEN,NONO,DDB1,YWHAQ,HACD3,DCHS1,SET,EGLN1,SF3B1,PDIA3,SSBP2,HMGB1,POLR2A,I
13,DDX17,GPI,ALDH6A1,ATP5F1B,CALM3,GNB1,MYH9,RPL7,P4HB,RPL5,GDI1,TMBIM6,SPTAN1,
E,RPL13,SLC2A3,DDX17,GPI,HTRA4,BSG,SPTBN1,ATP5F1B,MYH9,RPL7,SEPTIN2,P4HB,APP,IDS,I
SPTAN1,GSTP1,MARCKS,SMG1,CALM1,NPM1,HSPD1,SNRNP200,CSDE1,EIF3C,ATP5F1A,EIF4B,V
,CANX,PTMA,DDX5,RPL13A,SEC62,RPL6,KAT6A,DVL3,YWHAE,RPL13,SLC2A3,DDX17,GPI,BSG,AL
PTBN1,ATP5F1B,CALM3,MYH9,RPL7,SEPTIN2,P4HB,IDS,RPL5,FSCN1,TMBIM6,PTMS,SPTAN1,GS
PTBN1,ATP5F1B,CALM3,MYH9,RPL7,SEPTIN2,P4HB,IDS,RPL5,FSCN1,TMBIM6,PTMS,SPTAN1,GS
TAN1,MARCKS,SMG1,NPM1,COL18A1,CSDE1,EIF3C,ATP5F1A,EIF4B,VCP,MARCKSL1,UBB,AFF4,
/YH9,SEPTIN2,P4HB,RPL5,FSCN1,TMBIM6,PTMS,SPTAN1,MARCKS,SMG1,CALM1,NPM1,PLD3,SI
,GPI,BSG,ALDH6A1,SPTBN1,ATP5F1B,CALM3,GNB1,SEPTIN2,IDS,RPL5,PTMS,SPTAN1,GSTP1,CA
,GPI,BSG,ALDH6A1,SPTBN1,ATP5F1B,CALM3,GNB1,SEPTIN2,IDS,RPL5,PTMS,SPTAN1,GSTP1,CA
MBIM6,PTMS,SPTAN1,MARCKS,SMG1,CALM1,NPM1,HSPD1,SNRNP200,CSDE1,EIF3C,ATP5F1A,E
AN1,GSTP1,MARCKS,CALM1,NPM1,PLD3,HSPD1,SNRNP200,CSDE1,EIF3C,ATP5F1A,VCP,MARCK
N2,P4HB,RPL5,GDI1,FSCN1,TMBIM6,SPTAN1,CALM1,NPM1,PLD3,HSPD1,SNRNP200,CSDE1,EIF
SCN1,TMBIM6,PTMS,SPTAN1,GSTP1,MARCKS,NPM1,PLD3,HSPD1,USP11,COL18A1,SNRNP200,
DX17,GPI,SPTBN1,ATP5F1B,MYH9,SEPTIN2,P4HB,RPL5,FSCN1,SPTAN1,SMG1,NPM1,PLD3,SNRN
7,SEPTIN2,P4HB,IDS,RPL5,GDI1,FSCN1,PTMS,SPTAN1,MARCKS,CALM1,NPM1,COL18A1,CSDE1,
1B,TUBB2B,COL4A1,CES3,PSAP,PGK1,RPLP0,H3-3B,CANX,CLU,PTMA,DDX5,SOX4,RPL13A,SEC
TR3,YWHAG,RPL37A,MAPT,FN1,SPEN,NONO,YWHAQ,HACD3,DCHS1,SET,EGLN1,RPL19,SF3B1,P
,ATP5F1B,MYH9,RPL7,SEPTIN2,P4HB,IDS,RPL5,GDI1,FSCN1,SPTAN1,GSTP1,MARCKS,SMG1,NP
NONO,SET,EGLN1,MRC2,PDIA3,SSBP2,BASP1,HMGB1,RPS4X,RPL11,BTF3,RPL10,RPS17,VCL,D
,KAT6A,DVL3,YWHAE,RPL13,SLC2A3,GPI,ALDH6A1,SPTBN1,ATP5F1B,MYH9,RPL7,SEPTIN2,P4HE
1,SPTBN1,ATP5F1B,GNB1,RPL7,SEPTIN2,P4HB,IDS,RPL5,GDI1,TMBIM6,SPTAN1,GSTP1,MARCK
BN1,ATP5F1B,CALM3,MYH9,SEPTIN2,P4HB,APP,RPL5,GDI1,TMBIM6,PTMS,SPTAN1,CALM1,NPM
,RPL13,DDX17,GPI,BSG,ALDH6A1,SPTBN1,ATP5F1B,CALM3,GNB1,MYH9,RPL7,SEPTIN2,P4HB,ID
A1,SPTBN1,ATP5F1B,CALM3,GNB1,MYH9,RPL7,SEPTIN2,P4HB,IDS,RPL5,GDI1,TMBIM6,PTMS,SI
E,RPL13,SLC2A3,DDX17,GPI,SPTBN1,ATP5F1B,MYH9,RPL7,SEPTIN2,P4HB,IDS,GDI1,FSCN1,PTM
1,TMBIM6,PTMS,SPTAN1,NPM1,CSDE1,EIF3C,ATP5F1A,VCP,MARCKSL1,UBB,AFF4,TRIM28,MAT
PA0,RPS16,RPL23,RPL29,RPSA,CTSD,EIF4H,SCARB2,XRCC5,PJA2,RPL24,RPL17,C1ORF43,GAP4
/BIM6,PTMS,GSTP1,MARCKS,SMG1,NPM1,PLD3,HSPD1,COL18A1,SNRNP200,CSDE1,EIF3C,ATP
A1,SPTBN1,ATP5F1B,CALM3,GNB1,MYH9,RPL7,SEPTIN2,P4HB,IDS,RPL5,GDI1,TMBIM6,SPTAN1
/L3,YWHAE,RPL13,SLC2A3,GPI,ALDH6A1,SPTBN1,ATP5F1B,GNB1,MYH9,RPL7,P4HB,IDS,RPL5,G
L3,YWHAE,RPL13,GPI,ALDH6A1,SPTBN1,ATP5F1B,CALM3,GNB1,SEPTIN2,P4HB,IDS,RPL5,GDI1,
DS,RPL5,GDI1,FSCN1,TMBIM6,SPTAN1,GSTP1,MARCKS,CALM1,NPM1,PLD3,SNRNP200,CSDE1,
TP5F1B,CALM3,GNB1,RPL7,P4HB,RPL5,GDI1,TMBIM6,PTMS,SPTAN1,GSTP1,MARCKS,SMG1,CA
P4HB,IDS,RPL5,FSCN1,TMBIM6,PTMS,SPTAN1,GSTP1,MARCKS,CALM1,NPM1,PLD3,SNRNP200,
S,RPL5,GDI1,TMBIM6,PTMS,SPTAN1,SMG1,CALM1,NPM1,HSPD1,SNRNP200,CSDE1,EIF3C,ATP5
LD3,USP11,EIF3C,ATP5F1A,VCP,UBB,TRIM28,MATR3,YWHAG,RPL37A,LRRC28,SPEN,NONO,DDE
TP5F1B,CALM3,GNB1,MYH9,RPL7,P4HB,RPL5,GDI1,TMBIM6,PTMS,SPTAN1,GSTP1,MARCKS,SM
,GNB1,MYH9,RPL7,SEPTIN2,P4HB,IDS,RPL5,GDI1,SPTAN1,MARCKS,SMG1,CALM1,NPM1,HSPD1
,P4HB,IDS,RPL5,GDI1,PTMS,GSTP1,MARCKS,CALM1,NPM1,HSPD1,SNRNP200,CSDE1,EIF3C,ATI

,MYH9,SEPTIN2,P4HB,FSCN1,TMBIM6,PTMS,SPTAN1,GSTP1,MARCKS,SMG1,NPM1,PLD3,USP11

,MATR3,YWHAG,RPL37A,LRRC28,SPEN,NONO,DDB1,YWHAQ,HACD3,DCHS1,SET,EGLN1,RPL19,S

NRNPU,NFE2L1,SMARCA5,CTNNA1,RPS16,RPSA,SCARB2,XRCC5,RPL24,RPL17,PAPOLA,LLGL1,F

;,ATP5F1A,EIF4B,VCP,UBB,AFF4,TRIM28,MATR3,YWHAG,LRRC28,SPEN,NONO,DDB1,SET,EGLN1

:,RPL13,SLC2A3,DDX17,GPI,SPTBN1,ATP5F1B,RPL7,SEPTIN2,P4HB,IDS,RPL5,GDI1,FSCN1,TMBII

HD3,BASP1,VAPA,HMGB1,STMN1,ANXA2,RPL10,DNAJA1,CYB5R3,PAFAH1B1,AKAP12,MAP4,SF3B

SMG1,CALM1,NPM1,SNRNP200,EIF3C,ATP5F1A,EIF4B,VCP,UBB,TRIM28,MATR3,YWHAG,RPL37A

:C62,RPL6,KAT6A,DVL3,YWHAE,RPL13,SLC2A3,DDX17,GPI,ALDH6A1,SPTBN1,ATP5F1B,CALM3,M

:C62,RPL6,KAT6A,DVL3,YWHAE,RPL13,SLC2A3,DDX17,GPI,ALDH6A1,SPTBN1,ATP5F1B,CALM3,M

LM3,GNB1,RPL7,SEPTIN2,P4HB,IDS,RPL5,GDI1,TMBIM6,SPTAN1,GSTP1,MARCKS,SMG1,CALM1

IARCKS,NPM1,PLD3,HSPD1,USP11,COL18A1,SNRNP200,CSDE1,EIF3C,ATP5F1A,EIF4B,VCP,MAR(

NPM1,HSPD1,SNRNP200,CSDE1,EIF3C,ATP5F1A,VCP,UBB,AFF4,TRIM28,MATR3,YWHAG,CKB,RPI

NPM1,HSPD1,SNRNP200,CSDE1,EIF3C,ATP5F1A,VCP,UBB,AFF4,TRIM28,MATR3,YWHAG,CKB,RPI

'TAN1,MARCKS,SMG1,CALM1,NPM1,SNRNP200,CSDE1,EIF3C,ATP5F1A,VCP,UBB,AFF4,TRIM28,

HSPD1,CSDE1,EIF3C,ATP5F1A,VCP,MARCKSL1,UBB,TRIM28,MATR3,YWHAG,LRRC28,SPEN,NON

CKS,SMG1,NPM1,HSPD1,SNRNP200,EIF3C,ATP5F1A,EIF4B,VCP,UBB,TRIM28,MATR3,YWHAG,RF

)0,CSDE1,IGF2,EIF3C,ATP5F1A,VCP,AFF4,TRIM28,MATR3,YWHAG,RPL37A,LRRC28,FN1,SPEN,N(

COL18A1,SNRNP200,CSDE1,EIF3C,ATP5F1A,EIF4B,MARCKSL1,UBB,AFF4,TRIM28,MATR3,YWHA

DI1,FSCN1,PTMS,SPTAN1,GSTP1,MARCKS,CALM1,NPM1,PLD3,HSPD1,SNRNP200,CSDE1,EIF3C

I3-3B,CANX,CLU,PTMA,DDX5,SOX4,RPL13A,SEC62,RPL6,KAT6A,DVL3,YWHAE,RPL13,DDX17,GPI

3,DDX17,GPI,SPTBN1,ATP5F1B,RPL7,SEPTIN2,P4HB,IDS,RPL5,GDI1,FSCN1,TMBIM6,PTMS,SPTA

'EN,NONO,DDB1,DCHS1,SET,EGLN1,RPL19,SF3B1,PDIA3,SSBP2,CHD3,VAPA,HMGB1,POLR2A,TF

'L9,PSMD2,RPS14,CHD4,RPS20,WDR1,RPL26,BEX3,CCT7,SRSF4,RTN3,RPL31,PRKCSH,HNRNP(

5F1B,CALM3,GNB1,RPL7,P4HB,IDS,RPL5,GDI1,TMBIM6,SPTAN1,MARCKS,SMG1,CALM1,NPM1,I

,NPM1,SNRNP200,EIF3C,ATP5F1A,VCP,MARCKSL1,UBB,TRIM28,MATR3,YWHAG,RPL37A,LRRC2

SEPTIN2,P4HB,IDS,RPL5,GDI1,TMBIM6,SPTAN1,GSTP1,SMG1,NPM1,PLD3,HSPD1,USP11,COL1ε

SEPTIN2,P4HB,IDS,RPL5,GDI1,TMBIM6,SPTAN1,GSTP1,SMG1,NPM1,PLD3,HSPD1,USP11,COL1ε

EIF3C,ATP5F1A,EIF4B,VCP,MARCKSL1,UBB,AFF4,TRIM28,MATR3,YWHAG,RPL37A,LRRC28,NON

HB,SF3B1,SSBP2,VAPA,POLR2A,RPS4X,RPS3A,STMN1,RPL11,BTF3,RPS17,DNAJA1,PAFAH1B1,SF

ISPD1,COL18A1,CSDE1,EIF3C,ATP5F1A,VCP,UBB,TRIM28,MATR3,YWHAG,LRRC28,SPEN,NONO,I

DIA3,SSBP2,BASP1,VAPA,HMGB1,RPS3A,STMN1,BTF3,RPL10,DNAJA1,CYB5R3,PAFAH1B1,SF3B2

RRC28,SPEN,NONO,DDB1,SET,EGLN1,RPL19,SF3B1,SSBP2,CHD3,VAPA,HMGB1,POLR2A,RPS4X

RPL5,GDI1,MARCKS,SMG1,NPM1,PLD3,HSPD1,USP11,SNRNP200,CSDE1,EIF3C,ATP5F1A,EIF4B,

NXA2,RPL10,VCL,DNAJA1,MAP4,SF3B2,GTF2F1,HNRNPU,NFE2L1,SMARCA5,CTNNA1,FADS2,RPS

)DB1,DCHS1,SET,MRC2,SF3B1,PDIA3,SSBP2,HMGB1,POLR2A,RPS3A,BTF3,ANXA2,AGRN,RPL10,

ASP1,VAPA,HMGB1,RPL10,RPS17,PAFAH1B1,MAP4,SF3B2,GTF2F1,HNRNPU,NFE2L1,SMARCA5,

,RPL5,TMBIM6,PTMS,SPTAN1,GSTP1,SMG1,NPM1,PLD3,HSPD1,COL18A1,SNRNP200,CSDE1,EIF

P5F1A,VCP,UBB,AFF4,TRIM28,MATR3,YWHAG,FN1,SPEN,NONO,SET,EGLN1,MRC2,RPL19,SF3B1

,EIF4B,VCP,UBB,AFF4,TRIM28,MATR3,YWHAG,RPL37A,LRRC28,SPEN,NONO,DDB1,HACD3,SET,I

TRIM28,MATR3,YWHAG,LRRC28,SPEN,NONO,YWHAQ,HACD3,DCHS1,SET,EGLN1,MRC2,LDHB,PI

\3,SSBP2,CHD3,VAPA,HMGB1,TUBB4B,BTF3,RPL10,SF3B2,ITM2B,HNRNPU,SMARCA5,LARP1,CT

NB1,RPL7,P4HB,IDS,RPL5,GDI1,PTMS,SPTAN1,SMG1,NPM1,PLD3,HSPD1,COL18A1,CSDE1,EIF3

,LRRC28,FN1,NONO,DDB1,YWHAQ,HACD3,SET,EGLN1,MRC2,PDIA3,SSBP2,CHD3,BASP1,VAPA,H

C5,COL1A2,RPL14,RPS14,CHD4,JUND,RPS20,RPL26,BEX3,CCT7,RBBP4,PNN,RPL31,PRKCSH,HN

4B,VCP,MARCKSL1,UBB,AFF4,TRIM28,MATR3,YWHAG,LRRC28,FN1,SPEN,NONO,HACD3,DCHS1,

PL37A,SPEN,NONO,DDB1,YWHAQ,EGLN1,RPL19,LDHB,SSBP2,BASP1,VAPA,HMGB1,POLR2A,TPT
SL1,TRIM28,MATR3,YWHAG,LRRC28,SPEN,NONO,DDB1,HACD3,DCHS1,SET,EGLN1,LDHB,SF3B
,SRSF4,RPL31,HNRNPC,KHDRBS1,SPTBN2,RPS25,GLG1,SFPQ,RAB7A,PRDX2,MRFAP1,RCN1,H1
P2,HMGB1,TPT1,BTF3,ANXA2,RPL10,RPS17,DNAJA1,CYB5R3,SF3B2,CDC42BPB,GTF2F1,HNRNP
MARCKS,NPM1,PLD3,SNRNP200,EIF3C,ATP5F1A,VCP,MARCKSL1,UBB,TRIM28,MATR3,YWHAG,R
1,TPT1,RPS4X,MORF4L1,ANXA2,RPL10,RPS17,VCL,CYB5R3,MAP4,SF3B2,CDC42BPB,GTF2F1,H
YWHAG,RPL37A,LRRC28,NONO,DDB1,SET,EGLN1,PKD1,SF3B1,PDIA3,SSBP2,CHD3,VAPA,HMGB
PEN,NONO,DCHS1,SET,EGLN1,MRC2,LDHB,PDIA3,SSBP2,HMGB1,TPT1,ITGB1,ANXA2,RPL10,DNA
DNAJA1,PAFAH1B1,MAP4,SF3B2,GTF2F1,HNRNPU,NFE2L1,SMARCA5,RPS16,RPL29,RPSA,CTSD,
,EIF4B,VCP,UBB,AFF4,TRIM28,MATR3,YWHAG,CKB,LRRC28,MAPT,SPEN,NONO,DDB1,DCHS1,SE
,EIF4B,VCP,UBB,AFF4,TRIM28,MATR3,YWHAG,CKB,LRRC28,MAPT,SPEN,NONO,DDB1,DCHS1,SE
SP1,HMGB1,TPT1,TUBB4B,BTF3,ANXA2,RPL10,RPS17,CYB5R3,MAP4,SF3B2,CDC42BPB,GTF2F1
1,FADS2,SCARB2,BRD3,XRCC5,RPL24,PAPOLA,CLSTN1,PSMD2,JUND,EIF5,HMGN2,CCT7,RBBP
MAP4,ITM2B,GTF2F1,HNRNPU,HNRNPA0,CTSD,SCARB2,LPCAT1,XRCC5,PJA2,KPNB1,RPL17,PAF
RPL29,RPSA,TNS3,EIF4H,SCARB2,RPS9,RPL24,RPL17,PAPOLA,RPL14,RPL9,KIF5B,RPS14,CHD4,
NCS1,PTCD3,CLSTN1,PSMD2,CHD4,RPS20,WDR1,RPL26,CCT7,RBBP4,PNN,RPL31,NISCH,PRKCS
PPRC,KHDRBS1,XPO1,HDLBP,RPS25,GLG1,SFPQ,PDIA6,DDR1,DDX3X,RCN1,H1-0,RPL35,TLN1,H
3,CCT7,RBBP4,HNRNPC,LRPPRC,XPR1,KHDRBS1,XPO1,RPS25,GLG1,SFPQ,PDIA6,RAB7A,RPS1
F3B1,VAPA,HMGB1,RPS4X,RPS3A,STMN1,TUBB4B,RPL11,BTF3,RPL10,DNAJA1,MAP4,SF3B2,HN
YH9,RPL7,SEPTIN2,P4HB,RPL5,TMBIM6,PTMS,MARCKS,SMG1,NPM1,PLD3,HSPD1,COL18A1,SN
OLR2A,RPS4X,RPL10,DNAJA1,SF3B2,ZNF770,HNRNPU,SMARCA5,LARP1,HNRNPA0,CTNNA1,FAD
VAPA,HMGB1,POLR2A,BTF3,ANXA2,ZMIZ1,RPL10,WSB1,DNAJA1,SF3B2,CDC42BPB,ITM2B,GTF2
RPL23,RPSA,TPM4,EIF4H,SCARB2,RPL17,PAPOLA,RPL14,HK1,PSMD2,RPS14,RPS20,EIF5,RPL26
S,GSTP1,NPM1,PLD3,USP11,SNRNP200,CSDE1,EIF3C,ATP5F1A,EIF4B,VCP,MARCKSL1,UBB,TRI
D2,KIF5B,SLC7A5,RPS14,CHD4,JUND,RPL26,CCT7,SRSF4,RBBP4,RPL31,HNRNPC,LRPPRC,TADA
,HNRNPU,RPL23,RPL29,CTSD,SCARB2,KPNB1,RPL17,C1ORF43,PAPOLA,COL1A2,LLGL1,RPL14,
ADR,ATP1A1,JUND,EIF5,CD81,RBBP4,PNN,NES,PRKCSH,HNRNPC,LRPPRC,KHDRBS1,XPO1,SLC3
NO,SET,EGLN1,PDIA3,SSBP2,VAPA,POLR2A,TPT1,UCHL1,MORF4L1,BTF3,NCAM1,ANXA2,AGRN,
4,SCARB2,XRCC5,RPL24,RPL14,CLSTN1,PSMD2,KIF5B,JUND,RPS20,RPL26,CCT7,RBBP4,JPT2,
RBBP4,PNN,RPL31,HNRNPC,LRPPRC,TADA2B,KHDRBS1,XPO1,SPTBN2,RPS25,SFPQ,RPS15,H1-0
A,HNRNPU,SMARCA5,CTNNA1,FADS2,RPL23,RPSA,SCARB2,RPS9,XRCC5,RPL24,RPL17,RPL14,H
16,RPL23,RPSA,TPM4,EIF4H,SCARB2,BRD3,RPL24,RPL17,RPL14,HK1,PSMD2,KIF5B,SLC7A5,R
T,EGLN1,PDIA3,SSBP2,VAPA,HMGB1,POLR2A,TPT1,RPS4X,ITGB1,TUBB4B,MORF4L1,BTF3,NCA
NRNPC,LRPPRC,KHDRBS1,RPS25,SFPQ,RAB7A,RPS15,MAGED2,H1-0,RPS7,IK,RPL35,TLN1,CTN
EX3,RBBP4,PNN,RPL31,PRKCSH,HNRNPC,LRPPRC,TADA2B,KHDRBS1,KLHL9,RPS25,SFPQ,ACS
GLN1,RPL19,PDIA3,SSBP2,VAPA,HMGB1,POLR2A,BTF3,ANXA2,XAF1,RPL10,DNAJA1,SF3B2,GTF
CALM1,USP11,COL18A1,CSDE1,EIF3C,ATP5F1A,EIF4B,VCP,MARCKSL1,UBB,YWHAG,CKB,LRRC
EIF4H,SCARB2,BRD3,XRCC5,RPL24,RPL14,HK1,CLSTN1,PSMD2,CHD4,ATP1A1,EIF5,WDR1,HMG
1B1,SF3B2,HNRNPU,PGAM1,NFE2L1,SMARCA5,HNRNPA0,CNN3,RPSA,TPM4,TNS3,CTSD,SCARB
,TKT,SFPQ,DDR1,RAB7A,RPS15,H1-0,RPS5,RPS7,IK,JUN,CTNNB1,HNRNPDL,TARDBP,HNRNPK,
KIF5B,CXADR,ATP1A1,HMGN2,CCT7,SRSF4,RBBP4,JPT2,PRKCSH,HNRNPC,KHDRBS1,RPS25,TKT
BRD3,RPL24,RPL17,SLC7A5,CHD4,ATP1A1,JUND,EIF5,CCT7,RBBP4,MCL1,PNN,NISCH,HNRNPC,
RNPC,LRPPRC,TADA2B,XPR1,KHDRBS1,XPO1,RPS25,SFPQ,RAB7A,MAGED2,RPS5,RPS7,IK,GJA
,RAB7A,MAGED2,H1-0,CTNNB1,HNRNPDL,TARDBP,ARID1A,HNRNPK,H1-10,RPL36,YWHAB,CKA
BBP4,HNRNPC,LRPPRC,KHDRBS1,XPO1,RPS25,TKT,GLG1,SFPQ,RAB7A,DDX3X,RPS15,GNB2,R

D1A,HNRNPK,RPL36,YWHAB,SND1,LAMP1,HNRNPH1,UBE2H,CAPRIN1,TOP2B,HDAC2,SKP1,BCL/
GNB2,RPS7,IK,TARDBP,HNRNPK,RPL36,YWHAB,HNRNPH1,ARCN1,CAPRIN1,BCLAF1,L1CAM,LU(
CHF6,RPS17,VCL,DNAJA1,CYB5R3,PAFAH1B1,SF3B2,ITM2B,GTF2F1,HNRNPU,CTNNA1,RPL23,R
CAM1,AGRN,XAF1,RPL10,VCL,CYB5R3,MAP4,SF3B2,ITM2B,GTF2F1,HNRNPU,NFE2L1,SMARCA5
3TF3,RPL10,VCL,AKAP12,MAP4,SF3B2,ITM2B,GTF2F1,HNRNPU,NFE2L1,SMARCA5,LARP1,HNRN
RBBP4,MCL1,JPT2,PNN,SPC25,RPL31,LRPPRC,KHDRBS1,XPO1,RPS25,GLG1,SFPQ,RAB7A,RPS1
CAPRIN1,TOP2B,HDAC2,SKP1,BCLAF1,GLUL,LUC7L3,CPSF2,TM9SF3,HNRNPL,MORF4L2,LMNB
MMP14,PRDX2,DDX3X,MRFAP1,MAGED2,GNB2,H1-0,RPS7,CTNNB1,DNAJC5,APLP2,HNRNPDL,TA
BP,ARID1A,HNRNPK,INIP,RPL36,YWHAB,LAMP1,HNRNPH1,CKAP4,RPS12,CAPRIN1,TOP2B,HDAC
A,HMGB1,RPS4X,BTF3,XAF1,RPL10,DNAJA1,SF3B2,ITM2B,HNRNPU,SMARCA5,LARP1,CTNNA1,F
7A,PRDX2,DDX3X,MAGED2,H1-0,GJA1,CTNNB1,GOLM1,APLP2,HNRNPDL,TARDBP,HNRNPK,RPL3
NNA1,FADS2,RPL23,RPSA,TPM4,TNS3,EIF4H,SCARB2,XRCC5,RPL24,RPL17,RPL14,PTCD3,PSMD
P2,BASP1,VAPA,HMGB1,RPS4X,STMN1,BTF3,ANXA2,RPL10,VCL,DNAJA1,PAFAH1B1,SF3B2,CDC
C,XPR1,KHDRBS1,SFPQ,RAB7A,MMP14,DDX3X,MAGED2,H1-0,IK,COL6A2,ANXA6,APLP2,TARDBP,
32,HNRNPU,CTNNA1,CNN3,CTSD,EIF4H,SCARB2,LPCAT1,XRCC5,MSN,SLC16A7,C1ORF43,PAPOL
TNNA1,RPL23,RPL29,RPSA,TPM4,TNS3,EIF4H,SCARB2,XRCC5,PJA2,RPL24,RPL17,C1ORF43,RPI
NB1,HNRNPDL,TARDBP,HNRNPK,INIP,RPL36,CKAP5,LAMP1,ARCN1,RAC1,CAPRIN1,TOP2B,BCLA
TARDBP,HNRNPK,H1-10,MYH10,CKAP5,LAMP1,HNRNPH1,CCT6A,STIP1,HDAC2,BCLAF1,LUC7L3
R3,PAFAH1B1,AKAP12,MAP4,ZNF770,AP2B1,GTF2F1,OGA,PGAM1,NFE2L1,CTNNA1,RPS16,RPL2
1,GLUL,LUC7L3,CPSF2,TM9SF3,MDH2,HNRNPL,MORF4L2,TAF15,LMNB2,ALKBH5,HNRNPA3,AC
PRIN1,HDAC2,BCLAF1,LUC7L3,CPSF2,DDX39B,MDH2,HNRNPL,TSPYL4,MORF4L2,ABCA3,HNRN
RNPL,MORF4L2,ALKBH5,HNRNPA3,RAD23B,SNRNP70,H3-3A,HDGF,TERF2IP,G3BP2,SMARCA4,S
D1A,HNRNPK,H1-10,RPL36,YWHAB,SEPTIN11,LAMP1,RPS12,HLA-B,ARCN1,COTL1,CAPRIN1,TOF
RCN1,MAGED4B,CAPRIN1,HDAC2,BCLAF1,LUC7L3,CPSF2,DDX39B,NUCB1,TM9SF3,PABPN1,TSP
OLA,NCS1,PTCD3,RPL26,SRSF4,RBBP4,PFN1,PNN,SPC25,PRKCSH,HNRNPC,LRPPRC,XPR1,KHDR
RPL10,DNAJA1,MAP4,SF3B2,GOLGA8A,HNRNPU,CTNNA1,FADS2,RPL23,RPL29,RPSA,TPM4,TNS
M1A,DIDO1,SMARCE1,HNRNPM,SRSF3,ILF2,FBLN1,P3H3,TP53BP1,RBBP7,TXLNA,NAP1L1,LBH,S
9,RBBP4,MCL1,PNN,NISCH,PRKCSH,HNRNPC,KHDRBS1,XPO1,HDLBP,TKT,SFPQ,PDIA6,DDR1,CL
,HACD3,DCHS1,SET,EGLN1,RPL19,PKD1,SF3B1,VAPA,HMGB1,RPS3A,MORF4L1,BTF3,AGRN,RP
B,TERF2IP,MAPRE1,SRSF1,PTBP1,ILF3,PRKDC,HNRNPUL1,BNIP3L,CCAR2,FLNB,SMARCC1,HNF
AC1,CAPRIN1,BCLAF1,LUC7L3,DDX39B,CTSB,MDH2,TOMM20,HNRNPL,PARP1,MORF4L2,ALKBH
PS14,CHD4,RPS20,EIF5,HMGN2,CCT7,RBBP4,HNRNPC,LRPPRC,XPO1,RPS25,TKT,SFPQ,PDIA6,PI
HNRNPH1,CKAP4,HLA-B,RAC1,CAPRIN1,HDAC2,BCLAF1,LUC7L3,CPSF2,DDX39B,TM9SF3,TOMM
SF2,TM9SF3,HNRNPL,TSPYL4,MORF4L2,ALKBH5,SURF4,RBMX,RAD23B,TNPO1,SNRNP70,H3-3
2,SLC7A5,BEX3,SRSF4,RBBP4,MCL1,NES,HNRNPC,LRPPRC,KHDRBS1,XPO1,RPS25,TKT,GLG1,S
6A,DDR1,RAB7A,PRDX2,IK,TLN1,JUN,APLP2,TARDBP,NDFIP1,HNRNPK,CKAP5,LAMP1,UBE2H,CAL
TUBB4B,UCHL1,NCAM1,MARCHF6,PAFAH1B1,CDC42BPB,ITM2B,ZNF770,AP2B1,PGAM1,CTNNA
IPH1,HLA-B,RAC1,CAPRIN1,BCLAF1,LUC7L3,CPSF2,DDX39B,TM9SF3,TOMM20,MLXIP,PABPN1,H
AP4,SF3B2,GTF2F1,HNRNPU,NFE2L1,SMARCA5,CTNNA1,FADS2,TPM4,EIF4H,LPCAT1,BRD3,XRC
GTF2F1,GOLGA8A,HNRNPU,OGA,SMARCA5,RPS16,CNN3,LPCAT1,XRCC5,MSN,KPNB1,PAPOLA,N
2B,KHDRBS1,PFKL,RPS25,SFPQ,PEBP1,CCN2,RPL30,MAGED2,RCN1,RPS7,IK,GPAT4,TLN1,COL6
2,FBLN1,KDM3B,DDIT4,RBBP7,SF1,PBX1,CD99,BRD4,PPP2CB,ATRX,TRA2B,ZFR,AGPAT1,CALD1
7A,PRDX2,CLTC,MAGED2,H1-0,RPS7,GJA1,CTNNB1,APLP2,TARDBP,HNRNPK,INIP,YWHAB,LAMP
SFPQ,PRDX2,RPS15,MAGED2,H1-0,TLN1,ANXA6,APLP2,TARDBP,ARID1A,HNRNPK,H1-10,RPL36,
IA,TOP2B,HDAC2,BCLAF1,TRIB2,LUC7L3,CPSF2,TM9SF3,TOMM20,PABPN1,HNRNPL,ALKBH5,RI

SF3,TOMM20,MLXIP,PABPN1,MORF4L2,ALKBH5,RPL27,HNRNPA3,C4ORF3,RBMX,RAD23B,SNR
PBX1,CD99,BRD4,SLC39A7,TRA2B,UQCRC1,HNRNPA2B1,CALD1,APEX1,HNRNPAB,ZMYM3,FUS,
,APLP2,HNRNPDL,TARDBP,ARID1A,HNRNPK,H1-10,RPL36,YWHAB,SEPTIN11,LAMP1,HLA-B,ARC
AND5,DIDO1,HNRNPM,SSB,SRSF3,ILF2,STXBP1,NBPF9,ZSCAN18,KDM3B,SMS,TXNIP,TP53BP1,R
,TADA2B,KHDRBS1,CSNK1E,RPS25,TKT,GLG1,SFPQ,MMP14,DDX3X,GNB2,H1-0,IK,CTNNB1,HN
YMPK,MTCH1,PRDX6,FUS,ERH,TAOK1,RALY,SSR3,HDGFL2,EFTUD2,PNISR,SLC4A2,PLEKHM2,R
A4,PSMD3,SRSF1,ILF3,PRKDC,HNRNPUL1,DIDO1,SMARCE1,HNRNPM,SSB,GIGYF1,SRSF3,ILF2,
CBX1,RAD23B,SNRNP70,H3-3A,HDGF,ABCA2,PRPF6,CCNL2,SRSF1,TCF25,ATP5PB,PTBP1,ILF3,
,FAM50A,EWSR1,GALNT10,PDGFRB,WIZ,HNRNPR,PSME3,ZEB1,SEPHS1,ANKRD50,KDM2A,NACC
TPRA,RBMX,CBX1,TNPO1,SNRNP70,H3-3A,HDGF,MAPK1,SMARCA4,ATG2B,LAPTM4A,ARPC2,NY
PO7,ILF3,PRKDC,KDM1A,DIDO1,TPM3,CORO1C,SUPT6H,HNRNPM,SRSF3,ILF2,STRAP,KDM3B,D
,CCT2,SUGP2,PABPN1,GDI2,HNRNPL,TM9SF2,MORF4L2,HNRNPA3,ACO2,CBX1,SNRNP70,HDGF
CE1,HNRNPM,SRSF3,ILF2,FBLN1,SF3A1,STRAP,ZSCAN18,AKIRIN1,TP53BP1,DDIT4,PALLD,RBBP
1,LUC7L3,CRPPA,CPSF2,TOMM20,U2AF2,CCT2,SUGP2,PABPN1,GDI2,HNRNPL,TM9SF2,MORF4
A,HDGF,TERF2IP,TOP1,OPTN,SRSF9,ARPC2,SRSF1,ARGLU1,FTO,ILF3,PRKDC,HNRNPUL1,KDM
2F1,HNRNPU,PGAM1,NFE2L1,CTNNA1,TPM4,XRCC5,RPL24,PTCD3,PSMD2,KIF5B,SLC7A5,ATP1
RE1,SRSF1,PTBP1,ILF3,PRKDC,HNRNPUL1,CCAR2,DIDO1,SDF4,SMARCC1,HNRNPM,SSB,SRSF3
NP32A,RBBP7,TXLNA,NAP1L1,LBH,SF1,GTPBP1,BRD4,ATRX,EP400,SRSF6,ZFR,AGPAT1,HNRNPA
9B,MDH2,RNF187,MLXIP,SUGP2,HNRNPL,TM9SF2,MORF4L2,ALKBH5,PTPRA,TBC1D16,ACO2,C
2,TRAP1,FLOT2,NUCKS1,KRT8,PHGDH,IVD,THRAP3,SCD,HTATSF1,MTA2,TXNRD1,MDM2,MECP
L,PARP1,ABCA3,RPL27,HNRNPA3,ACO2,SNRNP70,H3-3A,OPTN,SMARCA4,ARPC2,PRDX1,SRSF1,
A,HDGF,RAPGEF1,SMARCA4,PDE7A,FAM168B,SRSF1,FTO,ILF3,PRKDC,KDM1A,DIDO1,TPM3,DH
1,DTX4,EEF1B2,REEP2,PSMD4,AP2A2,SORT1,CD74,RCN2,NTRK2,ELAVL3,PSAT1,FXR2,NOTCH1,
RF198,SNRPD3,SRP72,DCLK1,CNP,TRAP1,CADM3,PACS1,PHGDH,FXYD6,COX6A1,GSK3A,AP2A2,
ST,HNRNPK,MYH10,RPL36,HNRNPH1,HLA-B,MAP2,CCT6A,TOP2B,HDAC2,BCLAF1,LUC7L3,CDK
OL1A2,LLGL1,RPL14,RHOA,ZNF865,PSMD2,KIF5B,SLC7A5,GLIPR1L2,RPS14,CHD4,TMEM259,
RNPC,LRPPRC,KHDRBS1,XPO1,PTN,SFPQ,TPM1,ACTN4,DDX3X,EIF3L,MRFAP1,H1-0,IK,GOLM1,T
CN2,PSAT1,FXR2,NOTCH1,TRIM2,IDH2,UBE2J1,RTCB,RPL27A,FAM168A,ALDH7A1,NACAD,NDUF
,SV2A,MAPRE2,SEPTIN7,AKAP9,TMOD2,SEPTIN5,IARS2,LINGO1,STRN4,SSR3,DCLK1,DTX4,EEF1
KRD12,DAPK3,SCD,HTATSF1,SUN1,SGSM2,CGGBP1,MDM2,CUL1,FAM50A,AAMP,SUMO1,EWSR
,UBTF,KLF6,MYBBP1A
CDK9,PSIP1,FAM160B2,SEPHS1,ALCAM,TUBA1C,ZFP91,CDC42EP4,ZNRF1,TLK1,RBM17,CITED

EIF5,RBBP4,PNN,SPC25,PRKCSH,HNRNPC,LRPPRC,KHDRBS1,XPO1,PTN,SFPQ,TPM1,ACSL6,AC
,ERH,HDGFL2,EFTUD2,MOAP1,ATXN7L3,MN1,XRN2,HMGA1,LRIG1,REEP5,MDM2,SUN2,PRPH,T
1,AKAP17A,KLF3,GSN,UBTF,KLF6,MYBBP1A
P24
,ZNF326,ARSA,FKBP3,MMP24
1,DTX4,EEF1B2,CADM3,REEP2,MSH6,CHMP4B,AP2A2,ATP6V1G1,RCN2,IGLON5,ANK2,SCAMP5,
A3,VPS35,CBX1,SNRNP70,H3-3A,ABCA2,TERF2IP,TUBB2A,SRSF1,ARGLU1,ILF3,PRKDC,HNRNP
P1,GSPT1,BRD4,PPP2CB,DECR1,DIP2B,EP400,EIF5B,GPSM1,TIMM50,SYMPK,MTCH1,ZMYM3,F
1,A2M,SLC2A1,MTCH1,PRDX6,FUS,ERH,RALY,EFTUD2,TRAP1,PHGDH,SRSF11,SPTY2D1,ATXN7L
RKDC,HNRNPUL1,ACTN1,KDM1A,DIDO1,SMARCE1,HNRNPM,SSB,SRSF3,ILF2,SF3A1,RAI1,HNRN
ZNF770,AP2B1,PGAM1,NFE2L1,LARP1,CTNNA1,FADS2,RPL23,CPE,RPSA,EIF4H,BRD3,RPL24,C1
TRIM2,COPE,IDH2,SSU72,MTURN,FAM168A,SYNGAP1,TSPAN7,NACAD,NDUFB8,CRABP1,CNTN1,

LXNA2,TRIM2,EMC1,PABPC4,CD44,KCTD20,ECH1,ATF7IP,CCNG1,HM13,EXOSC10,TFRC,PMPCA

MMP24

PB,MAN2B1,CNTN1,CHGB,ATP6V0A1,GPRIN1,PPP3CA,SLC1A3,ADD1,FH,APOE,NFASC,ABCD3,SY

RPS6,SEZ6L2,ACSL4,RAB3A,NAPB,DLG4,CASKIN1,PRKAR2B,DPP7,VDAC2,UQCRFS1,BACE1,CHG

BP2,MAPK8IP1,RAI1,SREBF2,AKIRIN1,GRIPAP1,NAP1L1,LBH,PSMD1,GSPT1,COX7C,NOMO1,AGI

H2,SSU72,MTURN,AKAP8L,ALDH7A1,NACAD,NDUFB8,ACSL4,MAN2B1,PRKAR2B,VDAC2,UQCRFS

X,RPL12,H1-0,IK,NGRN,GOLM1,TARDBP,DDOST,WLS,HNRNPK,MYH10,NARS1,RPL36,CCT6A,HD

N7,GLO1,ABI2,TPP1,C6ORF62,LTBP4,SLC7A1,DICER1,NSD2,CSE1L,RBM6,RCN2,ANK2,CAD,PLXI

1,FH,APOE,ABCD3,GSN,ARSA,FKBP3

YT1,EPHA4,DNM1,GNAL,PPFIA3,PDHB,NNT

WDR33,GSN,VPS37B,ARSA,FKBP3

SPA1A,STK39,CUL1,CMPK1,CRIP2,PARK7,TRIB3,H2AX,EIF3B,CHST3,NSD1,TUBA1C,RBM17,OLA1

NA2,PPM1G,BRD2,MAP6,GLO1,INTS3,TPP1,C6ORF62,LTBP4,LRATD2,EIF4G3,QPRT,SLC7A1,DIC

RPL13A,SEC62,RPL6,KAT6A,DVL3,YWHAE,RPL13,SLC2A3,DDX17,GPI,BSG,ALDH6A1,SPTBN1,AT
13,DDX17,GPI,ALDH6A1,SPTBN1,ATP5F1B,CALM3,GNB1,RPL7,SEPTIN2,P4HB,APP,IDS,RPL5,GDI
LDH6A1,SPTBN1,ATP5F1B,CALM3,GNB1,RPL7,SEPTIN2,APP,IDS,RPL5,GDI1,FSCN1,TMBIM6,PT
P5F1B,CALM3,GNB1,RPL7,P4HB,APP,IDS,RPL5,GDI1,TMBIM6,PTMS,SPTAN1,SMG1,CALM1,NPM
62,RPL6,KAT6A,DVL3,YWHAE,RPL13,SLC2A3,GPI,BSG,ALDH6A1,SPTBN1,ATP5F1B,CALM3,GNB
X17,GPI,SPTBN1,ATP5F1B,CALM3,GNB1,RPL7,P4HB,APP,IDS,RPL5,GDI1,TMBIM6,PTMS,SPTAN1
DH6A1,SPTBN1,ATP5F1B,CALM3,GNB1,RPL7,SEPTIN2,P4HB,APP,IDS,RPL5,GDI1,FSCN1,TMBIM6
6A1,SPTBN1,ATP5F1B,CALM3,GNB1,MYH9,RPL7,SEPTIN2,P4HB,APP,IDS,RPL5,GDI1,TMBIM6,PT
RPL13,DDX17,GPI,BSG,ALDH6A1,SPTBN1,ATP5F1B,MYH9,RPL7,SEPTIN2,P4HB,IDS,RPL5,GDI1,T
,GNB1,RPL7,SEPTIN2,P4HB,IDS,RPL5,GDI1,TMBIM6,PTMS,SPTAN1,GSTP1,SMG1,CALM1,NPM1,
,GNB1,RPL7,SEPTIN2,P4HB,IDS,RPL5,GDI1,TMBIM6,PTMS,SPTAN1,GSTP1,SMG1,CALM1,NPM1,
1,SPTBN1,ATP5F1B,CALM3,GNB1,MYH9,RPL7,SEPTIN2,P4HB,APP,IDS,RPL5,GDI1,FSCN1,TMBIM
PM1,PLD3,HSPD1,CSDE1,EIF3C,ATP5F1A,EIF4B,VCP,UBB,AFF4,TRIM28,MATR3,YWHAG,RPL37A,
,SLC2A3,DDX17,GPI,BSG,ALDH6A1,SPTBN1,ATP5F1B,CALM3,GNB1,MYH9,RPL7,SEPTIN2,P4HB,
,SLC2A3,DDX17,GPI,BSG,ALDH6A1,SPTBN1,ATP5F1B,CALM3,GNB1,MYH9,RPL7,SEPTIN2,P4HB,
MYH9,RPL7,SEPTIN2,P4HB,IDS,RPL5,GDI1,TMBIM6,PTMS,SPTAN1,GSTP1,MARCKS,SMG1,NPM1,
ALM3,GNB1,MYH9,RPL7,SEPTIN2,P4HB,IDS,RPL5,GDI1,FSCN1,TMBIM6,PTMS,SPTAN1,GSTP1,M
ALM3,GNB1,MYH9,RPL7,SEPTIN2,P4HB,IDS,RPL5,GDI1,FSCN1,TMBIM6,PTMS,SPTAN1,GSTP1,M
2A3,DDX17,GPI,FASN,ALDH6A1,SPTBN1,ATP5F1B,CALM3,GNB1,MYH9,RPL7,SEPTIN2,P4HB,APP,
NB1,MYH9,RPL7,SEPTIN2,P4HB,APP,IDS,RPL5,GDI1,TMBIM6,PTMS,SPTAN1,GSTP1,MARCKS,SM
,EIF3C,ATP5F1A,VCP,MARCKSL1,UBB,TRIM28,MATR3,YWHAG,RPL37A,SPEN,NONO,YWHAQ,HA
PI,HTRA4,BSG,SPTBN1,ATP5F1B,MYH9,RPL7,SEPTIN2,P4HB,APP,IDS,RPL5,GDI1,FSCN1,TMBIM
SPTBN1,ATP5F1B,CALM3,GNB1,MYH9,SEPTIN2,P4HB,IDS,RPL5,GDI1,FSCN1,TMBIM6,PTMS,SPT
SPTBN1,ATP5F1B,CALM3,GNB1,MYH9,SEPTIN2,P4HB,IDS,RPL5,GDI1,FSCN1,TMBIM6,PTMS,SPT
CP,UBB,AFF4,TRIM28,MATR3,YWHAG,RPL37A,LRRC28,SPEN,NONO,DDB1,YWHAQ,HACD3,DCHS
AG,RPL37A,LRRC28,MAPT,SPEN,NONO,DDB1,YWHAQ,HACD3,DCHS1,SET,EGLN1,INA,RPL19,LD
SLC2A3,DDX17,GPI,BSG,ALDH6A1,SPTBN1,ATP5F1B,CALM3,GNB1,MYH9,RPL7,SEPTIN2,P4HB,I
1B,CALM3,MYH9,RPL7,SEPTIN2,P4HB,IDS,RPL5,GDI1,FSCN1,TMBIM6,PTMS,SPTAN1,GSTP1,M
7,SEPTIN2,P4HB,IDS,RPL5,GDI1,FSCN1,TMBIM6,PTMS,SPTAN1,GSTP1,MARCKS,CALM1,NPM1,P
7,SEPTIN2,P4HB,IDS,RPL5,GDI1,FSCN1,TMBIM6,PTMS,SPTAN1,GSTP1,MARCKS,CALM1,NPM1,P
17,GPI,ALDH6A1,SPTBN1,ATP5F1B,CALM3,GNB1,MYH9,RPL7,SEPTIN2,P4HB,IDS,RPL5,GDI1,TM
SEPTIN2,P4HB,APP,IDS,RPL5,GDI1,TMBIM6,PTMS,SPTAN1,GSTP1,MARCKS,SMG1,CALM1,NPM1
SEPTIN2,P4HB,APP,IDS,RPL5,GDI1,TMBIM6,PTMS,SPTAN1,GSTP1,MARCKS,SMG1,CALM1,NPM1
D3,HSPD1,COL18A1,CSDE1,EIF3C,ATP5F1A,EIF4B,VCP,MARCKSL1,UBB,TRIM28,MATR3,YWHAG
CN1,TMBIM6,SPTAN1,GSTP1,MARCKS,SMG1,CALM1,NPM1,PLD3,SNRNP200,CSDE1,EIF3C,ATP5
DH6A1,SPTBN1,ATP5F1B,CALM3,GNB1,MYH9,RPL7,SEPTIN2,P4HB,IDS,RPL5,GDI1,FSCN1,TMBIM
SDE1,EIF3C,ATP5F1A,VCP,MARCKSL1,UBB,AFF4,TRIM28,MATR3,YWHAG,CKB,RPL37A,LRRC28,
I1,PTMS,SPTAN1,GSTP1,MARCKS,SMG1,CALM1,NPM1,PLD3,HSPD1,SNRNP200,CSDE1,EIF3C,AT
I1,PTMS,SPTAN1,GSTP1,MARCKS,SMG1,CALM1,NPM1,PLD3,HSPD1,SNRNP200,CSDE1,EIF3C,AT
RPL7,SEPTIN2,P4HB,IDS,RPL5,GDI1,TMBIM6,SPTAN1,GSTP1,MARCKS,NPM1,PLD3,HSPD1,COL1
MARCKSL1,UBB,AFF4,TRIM28,MATR3,YWHAG,RPL37A,LRRC28,FN1,SPEN,NONO,YWHAQ,DCHS
,GPI,BSG,ALDH6A1,SPTBN1,ATP5F1B,CALM3,GNB1,MYH9,RPL7,SEPTIN2,P4HB,APP,IDS,RPL5,G
,GPI,BSG,ALDH6A1,SPTBN1,ATP5F1B,CALM3,GNB1,MYH9,RPL7,SEPTIN2,P4HB,APP,IDS,RPL5,G
N1,ATP5F1B,CALM3,GNB1,MYH9,RPL7,SEPTIN2,P4HB,APP,IDS,RPL5,GDI1,TMBIM6,PTMS,SPTAN
N1,ATP5F1B,CALM3,GNB1,MYH9,RPL7,SEPTIN2,P4HB,APP,IDS,RPL5,GDI1,TMBIM6,PTMS,SPTAN

,SPTAN1,CALM1,NPM1,COL18A1,SNRNP200,CSDE1,EIF3C,ATP5F1A,VCP,MARCKSL1,UBB,AFF4,

PTBN1,ATP5F1B,CALM3,GNB1,MYH9,RPL7,SEPTIN2,P4HB,IDS,RPL5,GDI1,TMBIM6,SPTAN1,GS

PTBN1,ATP5F1B,CALM3,GNB1,MYH9,RPL7,SEPTIN2,P4HB,IDS,RPL5,GDI1,TMBIM6,SPTAN1,GS

1,ATP5F1B,CALM3,GNB1,MYH9,RPL7,SEPTIN2,P4HB,IDS,RPL5,GDI1,FSCN1,TMBIM6,PTMS,SPT

9,RPL7,SEPTIN2,P4HB,IDS,RPL5,GDI1,TMBIM6,SPTAN1,GSTP1,MARCKS,SMG1,CALM1,NPM1,PL

,SPTBN1,ATP5F1B,CALM3,GNB1,MYH9,RPL7,SEPTIN2,P4HB,APP,IDS,RPL5,GDI1,TMBIM6,PTMS,

1,SPTBN1,ATP5F1B,CALM3,GNB1,MYH9,RPL7,SEPTIN2,P4HB,IDS,RPL5,GDI1,FSCN1,TMBIM6,P

,IDS,RPL5,GDI1,FSCN1,TMBIM6,PTMS,SPTAN1,MARCKS,SMG1,CALM1,NPM1,PLD3,HSPD1,US

B,IDS,RPL5,GDI1,TMBIM6,PTMS,SPTAN1,MARCKS,SMG1,CALM1,NPM1,PLD3,COL18A1,SNRNP

B,IDS,RPL5,GDI1,TMBIM6,PTMS,SPTAN1,MARCKS,SMG1,CALM1,NPM1,PLD3,COL18A1,SNRNP

TBN1,ATP5F1B,CALM3,GNB1,MYH9,RPL7,SEPTIN2,P4HB,IDS,RPL5,GDI1,TMBIM6,PTMS,SPTAN1

ALM1,NPM1,PLD3,HSPD1,USP11,SNRNP200,CSDE1,EIF3C,ATP5F1A,EIF4B,VCP,MARCKSL1,UB

2,P4HB,IDS,RPL5,GDI1,FSCN1,TMBIM6,PTMS,SPTAN1,GSTP1,MARCKS,SMG1,CALM1,NPM1,PL

2,P4HB,IDS,RPL5,GDI1,FSCN1,TMBIM6,PTMS,SPTAN1,GSTP1,MARCKS,SMG1,CALM1,NPM1,PL

AG,RPL37A,LRRC28,FN1,SPEN,NONO,DCHS1,SET,EGLN1,MRC2,SF3B1,PDIA3,SSBP2,BASP1,VA

PTIN2,P4HB,IDS,RPL5,GDI1,FSCN1,TMBIM6,SPTAN1,SMG1,CALM1,NPM1,PLD3,HSPD1,SNRNP2

,GSTP1,MARCKS,SMG1,CALM1,NPM1,HSPD1,SNRNP200,CSDE1,EIF3C,ATP5F1A,EIF4B,VCP,AFF

ATP5F1B,CALM3,GNB1,MYH9,RPL7,SEPTIN2,P4HB,IDS,RPL5,GDI1,TMBIM6,SPTAN1,GSTP1,MA

ATP5F1B,CALM3,GNB1,MYH9,RPL7,SEPTIN2,P4HB,IDS,RPL5,GDI1,TMBIM6,SPTAN1,GSTP1,MA

9,RPL7,SEPTIN2,P4HB,IDS,RPL5,GDI1,FSCN1,TMBIM6,PTMS,SPTAN1,GSTP1,MARCKS,SMG1,C

9,RPL7,SEPTIN2,P4HB,IDS,RPL5,GDI1,FSCN1,TMBIM6,PTMS,SPTAN1,GSTP1,MARCKS,SMG1,C

RNP200,CSDE1,EIF3C,ATP5F1A,EIF4B,VCP,MARCKSL1,UBB,AFF4,TRIM28,MATR3,YWHAG,RPL3

N1,ATP5F1B,CALM3,MYH9,RPL7,SEPTIN2,P4HB,IDS,RPL5,GDI1,FSCN1,TMBIM6,PTMS,SPTAN1,

TMS,MARCKS,SMG1,PLD3,USP11,SNRNP200,CSDE1,EIF3C,ATP5F1A,EIF4B,VCP,MARCKSL1,UB

GNB1,RPL7,SEPTIN2,P4HB,IDS,RPL5,GDI1,TMBIM6,PTMS,SPTAN1,GSTP1,MARCKS,SMG1,CAL

S,SMG1,NPM1,HSPD1,COL18A1,CSDE1,EIF3C,ATP5F1A,EIF4B,VCP,MARCKSL1,UBB,TRIM28,MA

28,MATR3,YWHAG,RPL37A,LRRC28,FN1,SPEN,NONO,DDB1,YWHAQ,HACD3,DCHS1,SET,EGLN1,

,TMBIM6,PTMS,SPTAN1,GSTP1,SMG1,CALM1,NPM1,PLD3,USP11,COL18A1,SNRNP200,CSDE1,

,HSPD1,USP11,COL18A1,SNRNP200,CSDE1,EIF3C,ATP5F1A,VCP,UBB,AFF4,TRIM28,MATR3,YW

,HSPD1,USP11,COL18A1,SNRNP200,CSDE1,EIF3C,ATP5F1A,VCP,UBB,AFF4,TRIM28,MATR3,YW

MG1,PLD3,HSPD1,COL18A1,SNRNP200,CSDE1,EIF3C,ATP5F1A,EIF4B,MARCKSL1,UBB,AFF4,T

SPTAN1,MARCKS,SMG1,CALM1,NPM1,HSPD1,COL18A1,CSDE1,EIF3C,ATP5F1A,EIF4B,VCP,MAR

RNP200,CSDE1,EIF3C,ATP5F1A,VCP,MARCKSL1,UBB,AFF4,TRIM28,MATR3,YWHAG,CKB,LRRC

1,SPTBN1,ATP5F1B,CALM3,GNB1,MYH9,RPL7,SEPTIN2,P4HB,APP,IDS,RPL5,GDI1,TMBIM6,SPT

1,SPTBN1,ATP5F1B,CALM3,GNB1,MYH9,RPL7,SEPTIN2,P4HB,APP,IDS,RPL5,GDI1,TMBIM6,SPT

LM3,GNB1,MYH9,RPL7,SEPTIN2,P4HB,IDS,RPL5,GDI1,TMBIM6,PTMS,SPTAN1,GSTP1,MARCKS,

PTBN1,ATP5F1B,CALM3,GNB1,MYH9,RPL7,SEPTIN2,P4HB,APP,IDS,RPL5,GDI1,TMBIM6,SPTAN1

7,SEPTIN2,P4HB,IDS,RPL5,GDI1,TMBIM6,SPTAN1,GSTP1,MARCKS,SMG1,CALM1,NPM1,PLD3,H

AG,CKB,RPL37A,LRRC28,SPEN,NONO,DDB1,HACD3,DCHS1,SET,EGLN1,LDHB,SF3B1,PDIA3,SS

BIM6,PTMS,SPTAN1,GSTP1,MARCKS,SMG1,CALM1,NPM1,PLD3,USP11,COL18A1,SNRNP200,C

IF3C,ATP5F1A,EIF4B,VCP,MARCKSL1,UBB,TRIM28,MATR3,YWHAG,RPL37A,LRRC28,SPEN,NON

F1A,EIF4B,VCP,MARCKSL1,UBB,TRIM28,MATR3,RPL37A,LRRC28,SPEN,NONO,DDB1,HACD3,D

1A,EIF4B,VCP,UBB,TRIM28,MATR3,YWHAG,RPL37A,LRRC28,SPEN,NONO,DDB1,YWHAQ,HACD3

NB1,MYH9,RPL7,SEPTIN2,P4HB,IDS,RPL5,GDI1,TMBIM6,SPTAN1,NPM1,PLD3,HSPD1,USP11,SN

I1,TMBIM6,PTMS,SPTAN1,MARCKS,SMG1,CALM1,NPM1,HSPD1,COL18A1,SNRNP200,CSDE1,EI

RCKS,CALM1,NPM1,PLD3,COL18A1,SNRNP200,CSDE1,EIF3C,ATP5F1A,VCP,MARCKSL1,UBB,AF
YH9,RPL7,SEPTIN2,P4HB,IDS,RPL5,GDI1,FSCN1,SPTAN1,GSTP1,MARCKS,SMG1,NPM1,PLD3,HS
L7,SEPTIN2,P4HB,IDS,RPL5,GDI1,TMBIM6,SPTAN1,MARCKS,SMG1,CALM1,NPM1,PLD3,HSPD1,
L7,SEPTIN2,P4HB,IDS,RPL5,GDI1,TMBIM6,SPTAN1,MARCKS,SMG1,CALM1,NPM1,PLD3,HSPD1,
RPS4X,TUBB4B,UCHL1,MORF4L1,BTF3,RPL10,RPS17,DNAJA1,PAFAH1B1,SF3B2,ITM2B,AP2B1,
GSTP1,MARCKS,SMG1,CALM1,NPM1,PLD3,HSPD1,SNRNP200,EIF3C,ATP5F1A,EIF4B,VCP,UBB,T
RPL5,GDI1,TMBIM6,PTMS,SPTAN1,GSTP1,MARCKS,SMG1,NPM1,PLD3,HSPD1,COL18A1,SNRNP
CP,UBB,AFF4,TRIM28,MATR3,YWHAG,RPL37A,LRRC28,SPEN,NONO,DDB1,DCHS1,SET,EGLN1,F
LDH6A1,SPTBN1,ATP5F1B,CALM3,GNB1,MYH9,RPL7,SEPTIN2,P4HB,APP,IDS,RPL5,GDI1,FSCN1,
TP1,MARCKS,CALM1,NPM1,PLD3,COL18A1,SNRNP200,CSDE1,EIF3C,ATP5F1A,EIF4B,VCP,MAR
TP1,MARCKS,CALM1,NPM1,PLD3,COL18A1,SNRNP200,CSDE1,EIF3C,ATP5F1A,EIF4B,VCP,MAR
,TRIM28,MATR3,YWHAG,RPL37A,LRRC28,SPEN,NONO,DDB1,YWHAQ,HACD3,DCHS1,SET,EGLN1
NRNP200,CSDE1,EIF3C,ATP5F1A,EIF4B,VCP,UBB,AFF4,TRIM28,MATR3,YWHAG,RPL37A,LRRC28
LM1,NPM1,PLD3,HSPD1,COL18A1,CSDE1,EIF3C,ATP5F1A,EIF4B,VCP,UBB,AFF4,TRIM28,MATR3
LM1,NPM1,PLD3,HSPD1,COL18A1,CSDE1,EIF3C,ATP5F1A,EIF4B,VCP,UBB,AFF4,TRIM28,MATR3
IF4B,VCP,AFF4,TRIM28,MATR3,YWHAG,RPL37A,FN1,SPEN,NONO,DDB1,YWHAQ,HACD3,DCHS1,
SL1,UBB,TRIM28,MATR3,YWHAG,CKB,RPL37A,LRRC28,SPEN,NONO,DDB1,YWHAQ,HACD3,DCH
3C,ATP5F1A,EIF4B,VCP,UBB,AFF4,TRIM28,MATR3,YWHAG,RPL37A,LRRC28,SPEN,NONO,DDB1,
CSDE1,IGF2,EIF3C,ATP5F1A,EIF4B,VCP,MARCKSL1,UBB,AFF4,TRIM28,MATR3,YWHAG,RPL37A,L
IP200,CSDE1,EIF3C,ATP5F1A,EIF4B,VCP,MARCKSL1,UBB,AFF4,TRIM28,MATR3,YWHAG,RPL37A,
EIF3C,ATP5F1A,EIF4B,VCP,MARCKSL1,UBB,TRIM28,MATR3,YWHAG,RPL37A,LRRC28,MAPT,SPE
62,RPL6,KAT6A,DVL3,YWHAE,RPL13,SLC2A3,DDX17,GPI,BSG,ALDH6A1,SPTBN1,ATP5F1B,CALM
DIA3,SSBP2,CHD3,VAPA,POLR2A,STMN1,TUBB4B,UCHL1,BTF3,NCAM1,ANXA2,AGRN,RPL10,RPS
M1,PLD3,HSPD1,USP11,SNRNP200,CSDE1,EIF3C,ATP5F1A,EIF4B,VCP,UBB,TRIM28,MATR3,YWH
NAJA1,CYB5R3,MAP4,SF3B2,GTF2F1,HNRNPU,NFE2L1,CTNNA1,CNN3,RPSA,TPM4,EIF4H,SCARE
3,IDS,RPL5,GDI1,FSCN1,SPTAN1,GSTP1,MARCKS,SMG1,NPM1,PLD3,HSPD1,SNRNP200,CSDE1,E
S,SMG1,NPM1,PLD3,HSPD1,COL18A1,SNRNP200,CSDE1,EIF3C,ATP5F1A,VCP,UBB,AFF4,TRIM2
1,PLD3,SNRNP200,CSDE1,EIF3C,ATP5F1A,VCP,MARCKSL1,UBB,AFF4,TRIM28,MATR3,YWHAG,L
DS,RPL5,GDI1,TMBIM6,SPTAN1,GSTP1,MARCKS,SMG1,CALM1,NPM1,HSPD1,COL18A1,SNRNP2
PTAN1,MARCKS,SMG1,CALM1,NPM1,HSPD1,COL18A1,SNRNP200,CSDE1,EIF3C,ATP5F1A,VCP,M
MS,SPTAN1,GSTP1,MARCKS,NPM1,PLD3,HSPD1,USP11,SNRNP200,CSDE1,EIF3C,ATP5F1A,EIF4B,
R3,YWHAG,RPL37A,LRRC28,FN1,SPEN,NONO,DDB1,HACD3,DCHS1,SET,EGLN1,RPL19,LDHB,SF
3,RPL14,HK1,NCS1,CBX5,RPL9,CLSTN1,RPS14,CHD4,RPS20,EIF5,RPL26,CCT7,RBBP4,RTN3,PN
5F1A,EIF4B,VCP,UBB,AFF4,TRIM28,MATR3,YWHAG,RPL37A,LRRC28,SPEN,NONO,YWHAQ,DCHS
,MARCKS,SMG1,CALM1,NPM1,HSPD1,COL18A1,SNRNP200,CSDE1,EIF3C,ATP5F1A,EIF4B,VCP,M
DI1,PTMS,SPTAN1,GSTP1,SMG1,NPM1,PLD3,HSPD1,COL18A1,CSDE1,EIF3C,ATP5F1A,VCP,MAR
FSCN1,TMBIM6,PTMS,SPTAN1,MARCKS,SMG1,CALM1,NPM1,HSPD1,COL18A1,SNRNP200,CSDE
EIF3C,ATP5F1A,VCP,MARCKSL1,UBB,TRIM28,MATR3,YWHAG,RPL37A,LRRC28,NONO,DDB1,HA
LM1,NPM1,PLD3,SNRNP200,EIF3C,ATP5F1A,EIF4B,VCP,UBB,TRIM28,MATR3,YWHAG,RPL37A,L
CSDE1,EIF3C,ATP5F1A,EIF4B,VCP,UBB,AFF4,TRIM28,MATR3,YWHAG,RPL37A,LRRC28,SPEN,NO
F1A,EIF4B,VCP,MARCKSL1,UBB,AFF4,TRIM28,MATR3,YWHAG,RPL37A,LRRC28,SPEN,NONO,DD
31,YWHAQ,HACD3,DCHS1,SET,EGLN1,INA,RPL19,LDHB,SF3B1,PDIA3,SSBP2,CHD3,BASP1,VAPA,
MG1,CALM1,NPM1,PLD3,HSPD1,SNRNP200,EIF3C,ATP5F1A,EIF4B,VCP,UBB,TRIM28,MATR3,YWH
1,SNRNP200,EIF3C,ATP5F1A,EIF4B,VCP,MARCKSL1,UBB,TRIM28,MATR3,YWHAG,RPL37A,LRRC
P5F1A,EIF4B,VCP,UBB,AFF4,TRIM28,MATR3,YWHAG,RPL37A,LRRC28,SPEN,NONO,DDB1,HACD3

,CSDE1,EIF3C,ATP5F1A,VCP,UBB,AFF4,TRIM28,MATR3,YWHAG,CKB,RPL37A,LRRC28,FN1,SPEN
SSBP2,VAPA,POLR2A,TPT1,RPS4X,RPS3A,RPL18,RPL11,MORF4L1,BTF3,AGRN,WSB1,RPS17,DN
RPL14,PSMD2,RPS14,JUND,TMEM259,RPS20,EIF5,CD81,HMGN2,RPL26,BEX3,CCT7,SRSF4,RBB
,RPL19,LDHB,SF3B1,PDIA3,SSBP2,VAPA,HMGB1,POLR2A,TPT1,RPS4X,RPS3A,RPL18,STMN1,TU
M6,PTMS,SPTAN1,GSTP1,MARCKS,NPM1,PLD3,HSPD1,USP11,COL18A1,SNRNP200,CSDE1,EIF3
2,CDC42BPB,HNRNPU,NFE2L1,SMARCA5,CNN3,RPSA,EIF4H,SCARB2,LPCAT1,XRCC5,PJA2,RPL2
,LRRC28,SPEN,NONO,DDB1,HACD3,DCHS1,SET,EGLN1,RPL19,LDHB,PKD1,SF3B1,PDIA3,SSBP2
MYH9,RPL7,SEPTIN2,P4HB,IDS,RPL5,GDI1,FSCN1,TMBIM6,PTMS,SPTAN1,MARCKS,SMG1,CALM
MYH9,RPL7,SEPTIN2,P4HB,IDS,RPL5,GDI1,FSCN1,TMBIM6,PTMS,SPTAN1,MARCKS,SMG1,CALM
1,NPM1,PLD3,HSPD1,COL18A1,EIF3C,ATP5F1A,EIF4B,MARCKSL1,UBB,TRIM28,MATR3,YWHAG,
CKSL1,UBB,AFF4,TRIM28,MATR3,YWHAG,RPL37A,LRRC28,FN1,SPEN,NONO,DDB1,DCHS1,SET,
L37A,LRRC28,MAPT,FN1,SPEN,NONO,DDB1,SET,EGLN1,MRC2,PDIA3,SSBP2,CHD3,HMGB1,POL
L37A,LRRC28,MAPT,FN1,SPEN,NONO,DDB1,SET,EGLN1,MRC2,PDIA3,SSBP2,CHD3,HMGB1,POL
MATR3,YWHAG,RPL37A,FN1,SPEN,NONO,DDB1,HACD3,DCHS1,SET,EGLN1,MRC2,LDHB,SF3B1,
O,DDB1,YWHAQ,HACD3,DCHS1,SET,EGLN1,LDHB,PDIA3,SSBP2,VAPA,HMGB1,RPS4X,TUBB4B,
L37A,LRRC28,SPEN,NONO,DDB1,HACD3,DCHS1,SET,EGLN1,RPL19,LDHB,PKD1,SF3B1,PDIA3,C
ONO,SET,EGLN1,RPL19,SF3B1,PDIA3,SSBP2,HMGB1,POLR2A,RPS3A,RPL18,RPL11,BTF3,ANXA2
G,SPEN,NONO,SET,EGLN1,PKD1,SF3B1,PDIA3,SSBP2,VAPA,HMGB1,POLR2A,TPT1,RPS4X,RPS3
,ATP5F1A,EIF4B,VCP,MARCKSL1,TRIM28,MATR3,YWHAG,RPL37A,LRRC28,SPEN,NONO,DDB1,H
,ALDH6A1,SPTBN1,ATP5F1B,MYH9,RPL7,SEPTIN2,P4HB,IDS,RPL5,TMBIM6,PTMS,SPTAN1,GSTP
N1,GSTP1,MARCKS,NPM1,PLD3,HSPD1,COL18A1,SNRNP200,CSDE1,EIF3C,ATP5F1A,VCP,MARC
T1,RPS4X,RPS3A,RPL18,STMN1,MORF4L1,BTF3,ANXA2,XAF1,RPL10,RPS17,DNAJA1,PAFAH1B1
,LRPPRC,RPS25,GLG1,SFPQ,PRDX2,RPL30,RCN1,H1-0,RPS7,IK,RPL35,APLP2,TARDBP,ST3GAL
HSPD1,COL18A1,SNRNP200,CSDE1,EIF3C,ATP5F1A,EIF4B,VCP,MARCKSL1,UBB,TRIM28,MATR3
8,NONO,DDB1,HACD3,SET,EGLN1,LDHB,SF3B1,PDIA3,SSBP2,CHD3,VAPA,HMGB1,POLR2A,TPT1
A1,CSDE1,EIF3C,ATP5F1A,EIF4B,VCP,UBB,AFF4,TRIM28,MATR3,YWHAG,CKB,LRRC28,MAPT,F
A1,CSDE1,EIF3C,ATP5F1A,EIF4B,VCP,UBB,AFF4,TRIM28,MATR3,YWHAG,CKB,LRRC28,MAPT,F
O,DDB1,HACD3,SET,EGLN1,SF3B1,PDIA3,SSBP2,CHD3,VAPA,HMGB1,POLR2A,TPT1,RPS4X,RPS
3B2,HNRNPU,PGAM1,NFE2L1,SMARCA5,LARP1,RPS16,RPL23,RPL29,RPSA,TPM4,CTSD,EIF4H,S
DDB1,YWHAQ,HACD3,DCHS1,SET,RPL19,LDHB,PKD1,SF3B1,PDIA3,SSBP2,VAPA,HMGB1,RPL11,
,CDC42BPB,ITM2B,ZNF770,AP2B1,HNRNPU,SMARCA5,LARP1,RPS16,RPL29,RPSA,CTSD,EIF4H,
,RPS3A,RPL18,STMN1,MORF4L1,BTF3,ANXA2,RPL10,RPS17,DNAJA1,PAFAH1B1,MAP4,SF3B2,C
VCP,UBB,TRIM28,MATR3,YWHAG,RPL37A,SPEN,NONO,DCHS1,SET,EGLN1,RPL19,LDHB,SF3B1,
16,RPSA,TPM4,EIF4H,SCARB2,BRD3,XRCC5,RPL24,PAPOLA,RPL14,HK1,PTCD3,PSMD2,KIF5B,S
RPS17,CYB5R3,SF3B2,GTF2F1,HNRNPU,NFE2L1,HNRNPA0,CTNNA1,RPL23,CTSD,KPNB1,C1OR
RPL29,RPSA,CTSD,EIF4H,SCARB2,RTN4,XRCC5,MSN,RPL24,PAPOLA,COL1A2,GAP43,RPL14,RH
3C,ATP5F1A,TRIM28,MATR3,YWHAG,RPL37A,LRRC28,FN1,SPEN,NONO,DDB1,HACD3,DCHS1,SE
1,PDIA3,SSBP2,CHD3,HMGB1,RPS4X,TUBB4B,BTF3,AGRN,RPL10,RPS17,VCL,CYB5R3,PAFAH1B
EGLN1,MRC2,LDHB,SF3B1,PDIA3,CHD3,VAPA,HMGB1,POLR2A,TPT1,RPS4X,RPS3A,RPL11,ANXA
DIA3,SSBP2,HMGB1,POLR2A,RPS4X,RPS3A,TUBB4B,RPL11,BTF3,ANXA2,RPL10,VCL,PAFAH1B1,
NNA1,RPS16,RPL29,TPM4,SCARB2,XRCC5,RPL24,RPL17,RPL14,KIF5B,SLC7A5,JUND,RPS20,EI
C,ATP5F1A,VCP,UBB,AFF4,TRIM28,MATR3,YWHAG,RPL37A,LRRC28,FN1,SPEN,NONO,DDB1,HA
HMGB1,RPS3A,TUBB4B,ANXA2,AGRN,RPL10,RPS17,VCL,CYB5R3,MAP4,SF3B2,CDC42BPB,GTF2
RNPC,LRPPRC,TADA2B,KHDRBS1,XPO1,SLC3A2,CSNK1E,HDLBP,RPS25,GLG1,SFPQ,DDR1,RAB
,SET,EGLN1,LDHB,SF3B1,SSBP2,VAPA,POLR2A,RPS4X,RPS3A,STMN1,BTF3,AGRN,RPS17,PAFAH

1,RPS4X,STMN1,TUBB4B,RPL11,BTF3,AGRN,RPL10,CYB5R3,PAFAH1B1,MAP4,HNRNPU,OGA,NF
1,PDIA3,SSBP2,BASP1,VAPA,HMGB1,POLR2A,RPS3A,STMN1,TUBB4B,BTF3,AGRN,RPL10,RPS17
I-0,RPS7,IK,APLP2,HNRNPDL,TARDBP,ARID1A,ST3GAL2,HNRNPK,RPL36,YWHAB,LAMP1,HNRNPI
U,NFE2L1,SMARCA5,FADS2,RPS16,CNN3,RPSA,TNS3,SCARB2,LPCAT1,BRD3,XRCC5,MSN,RPL24
PL37A,SPEN,DDB1,DCHS1,SET,EGLN1,RPL19,LDHB,SF3B1,SSBP2,BASP1,HMGB1,TPT1,RPS4X,S
NRNPU,NFE2L1,RPL29,EIF4H,SCARB2,BRD3,RPS9,XRCC5,PJA2,SLC16A7,RPL17,C1ORF43,PAPC
31,RPS4X,RPS3A,RPL18,MORF4L1,BTF3,ANXA2,RPL10,DNAJA1,MAP4,SF3B2,GTF2F1,HNRNPU,N
AJA1,CYB5R3,PAFAH1B1,AKAP12,MAP4,SF3B2,ITM2B,GTF2F1,HNRNPU,NFE2L1,SMARCA5,HNRN
,SCARB2,LPCAT1,XRCC5,PJA2,MSN,RPL24,PAPOLA,GAP43,RPL14,RHOA,CBX5,CLSTN1,ATF4,EIF
ET,EGLN1,MRC2,LDHB,SF3B1,SSBP2,CHD3,VAPA,HMGB1,POLR2A,TPT1,RPS3A,RPL18,RPL11,M
ET,EGLN1,MRC2,LDHB,SF3B1,SSBP2,CHD3,VAPA,HMGB1,POLR2A,TPT1,RPS3A,RPL18,RPL11,M
,HNRNPU,NFE2L1,LARP1,CNN3,RPL29,EIF4H,SCARB2,BRD3,XRCC5,PJA2,RPL17,C1ORF43,PAP
4,PNN,PRKCSH,HNRNPC,LRPPRC,TADA2B,KHDRBS1,XPO1,RPS25,GLG1,SFPQ,PDIA6,DDR1,RAB
POLA,NCS1,PTCD3,CLSTN1,NLGN2,CXADR,ATP1A1,JUND,EIF5,CD81,SRSF4,RBBP4,PNN,NES,PRI
ATP1A1,RPS20,EIF5,RPL26,CCT7,RBBP4,PNN,RPL31,PRKCSH,HNRNPC,LRPPRC,TADA2B,KHDRE
SH,HNRNPC,LRPPRC,KHDRBS1,RPS25,GLG1,SFPQ,RAB7A,PRDX2,RPL30,MRFAP1,RPS7,MYO5A,
HNRNPDL,TARDBP,ST3GAL2,HNRNPK,INIP,RPL36,SND1,LAMP1,HNRNPH1,UBE2H,CKAP4,RAB1A,
5,MAGED2,RCN1,H1-0,RPS7,RPL35,CTNNB1,GOLM1,APLP2,TARDBP,DDOST,HNRNPK,H1-10,INI
RNPU,OGA,SMARCA5,CTNNA1,RPS16,RPL23,RPL29,RPSA,TPM4,LPCAT1,XRCC5,MSN,RPL17,RPI
RNP200,CSDE1,ATP5F1A,VCP,UBB,AFF4,TRIM28,MATR3,YWHAG,RPL37A,LRRC28,SPEN,NONO,D
S2,RPL23,RPSA,TPM4,SCARB2,RPS9,XRCC5,RPL24,RPL17,RPL14,HK1,RPL9,PSMD2,KIF5B,SLC
F1,HNRNPU,CTNNA1,FADS2,RPL23,RPSA,TPM4,TNS3,CTSD,EIF4H,SCARB2,RPS9,XRCC5,RPL24,
,CCT7,RBBP4,MCL1,PNN,HNRNPC,LRPPRC,TADA2B,XPR1,KHDRBS1,XPO1,RPS25,TKT,GLG1,SF
M28,MATR3,YWHAG,RPL37A,LRRC28,SPEN,NONO,DCHS1,SET,EGLN1,LDHB,PDIA3,SSBP2,BASP
A2B,KHDRBS1,XPO1,CSNK1E,RPS25,TKT,SFPQ,DDR1,RAB7A,RPS15,MAGED2,H1-0,RPS5,RPS7,II
,RHOA,ATF4,RPS14,CHD4,HMGN2,BEX3,RBBP4,PNN,NISCH,HNRNPC,LRPPRC,XPR1,KHDRBS1,XI
A2,SFPQ,PEBP1,CDH2,PDIA6,RAB7A,RPL12,MAGED2,H1-0,IK,TLN1,GJA1,CTNNB1,ANXA6,NGRN
RPL10,RPS17,VCL,DNAJA1,AKAP12,MAP4,SF3B2,HNRNPU,SMARCA5,RPL23,CNN3,EIF4H,SCARE
RPL31,HNRNPC,LRPPRC,XPR1,KHDRBS1,RPS25,GLG1,SFPQ,PDIA6,RAB7A,RPS15,MAGED2,RCN
,RPS5,RPS7,RPL35,GJA1,JUN,CTNNB1,HNRNPDL,TARDBP,TSR2,HNRNPK,INIP,RPL36,YWHAB,CI
HK1,RPL9,KIF5B,SLC7A5,RPS14,CXADR,CHD4,ATP1A1,RPS20,EIF5,WDR1,RPL26,CCT7,SRSF4,M
PS14,EIF5,RPL26,CCT7,RBBP4,MCL1,JPT2,PNN,NISCH,HNRNPC,LRPPRC,XPR1,KHDRBS1,XPO1,
M1,AGRN,RPL10,VCL,PAFAH1B1,AKAP12,SF3B2,GTF2F1,OGA,HNRNPA0,CPE,EIF4H,RTN4,XRCC
NB1,RPS10,APLP2,TARDBP,HNRNPK,RPL36,LAMP1,HNRNPH1,HLA-B,CALU,ARCN1,RAC1,CAPRIN
L6,RAB7A,MAGED2,RCN1,H1-0,RPS7,CTNNB1,APLP2,HNRNPDL,DDOST,HNRNPK,RPL36,YWHAB,
2F1,GOLGA8A,HNRNPU,SMARCA5,LARP1,CTNNA1,FADS2,RPS16,RPL23,RPL29,RPSA,TPM4,TNS
28,MAPT,SPEN,DDB1,YWHAQ,HACD3,DCHS1,SET,EGLN1,INA,LDHB,SF3B1,SSBP2,BASP1,VAPA,P
N2,BEX3,CCT7,RBBP4,MCL1,PNN,RPL31,PRKCSH,HNRNPC,LRPPRC,KHDRBS1,KLHL9,XPO1,RPS
2,LPCAT1,BRD3,XRCC5,RPL24,C1ORF43,RPL14,HK1,NCS1,PTCD3,CBX5,PSMD2,SLC7A5,JUND,
RPL36,LAMP1,HNRNPH1,UBE2H,RPS12,CCT6A,CAPRIN1,TOP2B,HDAC2,BCLAF1,LUC7L3,CPSF2,
,GLG1,CDH6,SFPQ,PEBP1,CDH2,MEGF8,ACTN4,PRDX2,DDX3X,CLTC,MAGED2,H1-0,IK,MIF,TLN
LRPPRC,KHDRBS1,XPO1,RPS25,GLG1,SFPQ,PEBP1,CDH2,DDR1,RAB7A,PRDX2,DDX3X,MAGED2
1,CTNNB1,APLP2,HNRNPDL,TARDBP,ARID1A,HNRNPK,RPL36,LAMP1,UBE2H,RPS12,HLA-B,CCT6
P5,LAMP1,ARHGDIA,CAPRIN1,TOP2B,SKP1,BCLAF1,LUC7L3,CPSF2,DDX39B,CTSB,TM9SF3,MDH
CN1,H1-0,RPS7,IK,RPL35,CTNNB1,HNRNPDL,TARDBP,DDOST,HNRNPK,INIP,RPL36,SND1,LAMP1

AF1,LUC7L3,CPSF2,CTSB,TM9SF3,EEF1A2,SUGP2,HNRNPL,MORF4L2,LMNB2,ABCA3,ALKBH5,R
C7L3,PGRMC1,CTSB,TOMM20,RNF187,PABPN1,HNRNPL,PARP1,MORF4L2,C4ORF3,ACO2,RBM
PL29,EIF4H,LPCAT1,XRCC5,KPNB1,RPL17,RPL14,PTCD3,SLC7A5,CXADR,JUND,RPS20,WDR1,RP
,CTNNA1,FADS2,RPL23,RPSA,TPM4,EIF4H,SCARB2,LPCAT1,BRD3,XRCC5,PJA2,RPL24,RPL17,C1
IPA0,CNN3,RPSA,TPM4,SCARB2,LPCAT1,BRD3,XRCC5,KPNB1,PAPOLA,LLGL1,CLSTN1,PSMD2,S
I5,MAGED2,RPS7,TLN1,TARDBP,ARID1A,HNRNPK,RPL36,YWHAB,SEPTIN11,LAMP1,UBE2H,RPS1
2,ALKBH5,RPL27,HNRNPA3,PDXK,ACO2,CCT3,DDX42,CBX1,TNPO1,SNRNP70,H3-3A,SCAF1,TOP
RDBP,HNRNPK,YWHAB,CKAP5,LAMP1,HNRNPH1,CKAP4,CALU,ARHGDIA,GPC1,TOP2B,HDAC2,SI
2,BCLAF1,LUC7L3,CDKN1A,CPSF2,DDX39B,CTSB,TM9SF3,TOMM20,SQSTM1,PABPN1,HNRNPL,
ADS2,RPL23,RPL29,RPSA,TPM4,TNS3,EIF4H,SCARB2,RPS9,XRCC5,RPL24,RPL17,RPL14,PTCD3,
36,YWHAB,LAMP1,HNRNPH1,HLA-B,CAPRIN1,HDAC2,BCLAF1,TRIB2,LUC7L3,CPSF2,CTSB,NUCB
2,KIF5B,SLC7A5,RPS14,ATP1A1,RPS20,WDR1,RPL26,LASP1,CCT7,RBBP4,MCL1,JPT2,RPL31,PR
42BPB,GTF2F1,HNRNPU,NFE2L1,SMARCA5,RPSA,CTSD,SCARB2,LPCAT1,XRCC5,MSN,RPL24,RP
,HNRNPK,YWHAB,CKAP5,HNRNPH1,CALU,MAGED4B,ARHGDIA,CAPRIN1,BCLAF1,LUC7L3,DCN,C
LA,COL1A2,LLGL1,GAP43,RPL14,RHOA,PSMD2,KIF5B,SLC7A5,CHD4,JUND,WDR1,HMGN2,LASP1
L14,PTCD3,PSMD2,KIF5B,SLC7A5,RPS14,CHD4,ATP1A1,EIF5,WDR1,RPL26,LASP1,CCT7,RBBP4,
F1,LUC7L3,CDKN1A,CPSF2,DDX39B,TM9SF3,TOMM20,SQSTM1,NOP53,HNRNPL,MORF4L2,TAF
,CPSF2,DDX39B,TM9SF3,TOMM20,U2AF2,CCT2,SUGP2,PABPN1,GDI2,HNRNPL,MORF4L2,TAF1
9,TPM4,EIF4H,BRD3,PJA2,MSN,RPL24,LLGL1,HK1,PSMD2,KIF5B,SLC7A5,RPS14,CXADR,CHD4,T
O2,CCT3,CBX1,SNRNP70,H3-3A,TOP1,OPTN,MAPK1,SMARCA4,PSMD3,SRSF1,ARGLU1,ILF3,PRK
PA3,RBMX,TNPO1,SNRNP70,H3-3A,HDGF,OPTN,MAPK1,SMARCA4,NYNRIN,PTBP1,ILF3,PRKDC,H
RSF1,PTBP1,ILF3,PCSK1N,PRKDC,HNRNPUL1,ACTN1,FKBP8,MAP2K2,DIDO1,FLNB,SMARCE1,A
P2B,HDAC2,BCLAF1,LUC7L3,MDH2,TOMM20,U2AF2,PABPN1,HNRNPL,PARP1,MORF4L2,ALKBH
YL4,MORF4L2,ALKBH5,HNRNPA3,RBMX,CBX1,RAD23B,RAB5B,TNPO1,SNRNP70,H3-3A,HDGF,P
BS1,XPO1,SPTBN2,TKT,SFPQ,PNMA1,PDIA6,RAB7A,MMP14,DDX3X,RPL12,MRFAP1,H1-0,IK,RPL
3,SCARB2,RPS9,XRCC5,RPL24,RPL17,RPL14,CLSTN1,PSMD2,KIF5B,SLC7A5,RPS14,CHD4,ATP1
SF1,PBX1,CD99,BRD4,SLC39A7,GLUD1,TRA2B,AGPAT1,HNRNPA2B1,CALD1,HNRNPAB,SYMPK,Z
TC,MAGED2,GOLM1,HNRNPDL,DDOST,WLS,HNRNPK,MYH10,NARS1,SND1,HNRNPH1,UBE2H,RA
L10,RPS17,CYB5R3,SF3B2,AP2B1,HNRNPU,CTNNA1,FADS2,RPL23,RPL29,RPSA,TPM4,CTSD,EIF
RNPM,SSB,SRSF3,ILF2,ISYNA1,UBQLN1,SF3A1,MLF2,ZSCAN18,ADAR,CTNND1,ANP32A,RBBP7,K
H5,RPL27,HNRNPA3,ACTR3,ACO2,RAD23B,TNPO1,SNRNP70,H3-3A,HDGF,OPTN,MAPK1,PRKDC,
RDX2,H1-0,RPS7,TLN1,APLP2,TARDBP,ARID1A,HNRNPK,H1-10,RPL36,SEPTIN11,SND1,LAMP1,H
20,NOP53,HNRNPL,MORF4L2,ALKBH5,RPL27,HNRNPA3,C4ORF3,RBMX,RAD23B,SNRNP70,H3-
A,HDGF,G3BP2,SMARCA4,DSTN,SRSF1,ARGLU1,NYNRIN,PTBP1,ILF3,PRKDC,HNRNPUL1,ACTN1,
SFPQ,TPM1,PEBP1,DDR1,RAB7A,PRDX2,H1-0,IK,HMGCS1,TLN1,GJA1,ANXA6,GOLM1,APLP2,HN
LU,ARCN1,CCT6A,ARHGDIA,CAPRIN1,CDK16,HDAC2,EID1,CTSB,TM9SF3,MDH2,TOMM20,HNRN
1,CPE,TPM4,EIF4H,RTN4,LPCAT1,PJA2,RPL24,KPNB1,LLGL1,GAP43,HK1,NCS1,PSMD2,SLC7A5,
HNRNPL,MORF4L2,ALKBH5,RPL27,HNRNPA3,C4ORF3,RBMX,RAD23B,SNRNP70,H3-3A,HDGF,PR
C5,RPL24,C1ORF43,LLGL1,RPL14,HK1,ATF4,PSMD2,KIF5B,SLC7A5,CHD4,TMEM259,EIF5,WDR
NCS1,PTCD3,CBX5,ATF4,RPL26,SRSF4,RBBP4,PFN1,PNN,SPC25,PRKCSH,HNRNPC,LRPPRC,XPR
A2,CD63,ANXA6,NGRN,MACF1,TARDBP,WLS,HNRNPK,INIP,CKAP5,RAB1A,ARHGDIA,CAPRIN1,SM
,APEX1,HNRNPAB,C1ORF198,PRDX6,ZMYM3,FUS,XRCC6,ERH,HDGFL2,TRAP1,PHGDH,SRSF11,
1,HNRNPH1,UBE2H,CAPRIN1,CDK16,HDAC2,BCLAF1,LUC7L3,CPSF2,TM9SF3,MDH2,TOMM20,P
YWHAB,SEPTIN11,SND1,LAMP1,HLA-B,ARCN1,COTL1,ARHGDIA,CAPRIN1,CDK16,TOP2B,HDAC2,
PL27,SNRNP70,H3-3A,TERF2IP,TOP1,OPTN,G3BP2,SMARCA4,ARPC2,PTBP1,ILF3,GRB2,CAPZB,

:NP70,H3-3A,HDGF,PRPF6,DPYSL2,SMARCA4,SRSF1,PTBP1,ILF3,PRKDC,HNRNPUL1,HSPB1,DID(
ERH,PELP1,FKBP4,TLE1,HDGFL2,TRAP1,RAB6A,RBM5,SRSF11,MSH6,DHCR7,GSK3A,TGOLN2,C
N1,COTL1,CCT6A,CAPRIN1,CDK16,APPL1,HDAC2,BCLAF1,LUC7L3,MDH2,TOMM20,RNF187,HNI
BBP7,STAT3,KDM5B,H2AZ1,CD99,SLC39A7,SRSF6,TRA2B,ZFR,ANP32B,CALD1,APEX1,AKAP9,ZN
IRNPDL,TARDBP,HNRNPK,CKAP5,LAMP1,HNRNPH1,CKAP4,ARHGDIA,CAPRIN1,TOP2B,HDAC2,SK
BM5,SPTY2D1,BORCS7,IST1,PSMC3,MN1,ANKRD12,DAPK3,PRDX3,SCD,UBE2K,PTPN23,PTPN11
,SF3A1,HNRNPD,ZSCAN18,KDM3B,TP53BP1,RBBP7,KCNQ2,LBH,SF1,H2AZ1,UBQLN2,CSNK2A1,
PRKDC,HNRNPUL1,ZFAND5,DIDO1,SMARCE1,SMARCC1,HNRNPM,SSB,SRSF3,ILF2,STXBP1,NBPI
:1,LMNA,THOC2,RXRA,HIRA,DDHD2,ELOB,PRPF31,HEXIM1,FUBP1,ZNF207,ALYREF
'NRIN,TCF25,PTBP1,ILF3,PRKDC,HNRNPUL1,HSPB1,DIDO1,TPM3,CORO1C,SUPT6H,HNRNPM,SF
DDIT4,ANP32A,RBBP7,TRIO,H2AZ1,GTPBP1,BRD4,ATRX,SRSF6,ZFR,AGPAT1,HNRNPA2B1,ANP32I
',ABCA2,TERF2IP,TUBB2A,SMARCA4,SRSF1,ARGLU1,ILF3,PRKDC,HNRNPUL1,CASC3,CCAR2,DID
'7,TXLNA,PSMD1,SF1,PBX1,BRD4,UBA52,ATRX,TRA2B,ZFR,HNRNPA2B1,ADH5,CALD1,RPS27A,HI
L2,TAF15,HNRNPA3,RBMX,CBX1,CCT4,SNRNP70,KDM6B,HDGF,ABCA2,PRPF6,TERF2IP,TAF7,SN
1A,DIDO1,SMARCE1,HNRNPM,SSB,SRSF3,ILF2,PFKM,SF3A1,HNRNPD,MLF2,ZSCAN18,WBP2,RB
A1,WDR1,SRSF4,RBBP4,MCL1,HNRNPC,LRPPRC,TADA2B,KHDRBS1,SPTBN2,SLC3A2,TKT,GLG1,
,LARS1,ILF2,STT3B,ISYNA1,LONP1,UBQLN1,SF3A1,HNRNPD,SERBP1,KDM3B,NAP1L4,TP53BP1,
2B1,ANP32B,HNRNPAB,C1ORF198,FUS,ERH,HDGFL2,EFTUD2,SRSF2,PLEKHM2,RBM5,ABCF1,S
CT3,SURF4,TNPO1,PSMB2,H3-3A,HDGF,OPTN,PDE7A,PSMD3,FAM168B,SRSF1,ARGLU1,ACADVI
2,PIK3C2B,GLOD4,CCNY,KCTD20,NCKAP5L,SUN2,CAPN1,NECTIN1,ALDH7A1,MAPKAPK2,CDK9,I
ILF3,LUZP6,CAPZB,PRKDC,HNRNPUL1,KDM1A,DIDO1,SMARCE1,HNRNPM,SRSF3,ILF2,B2M,UB(
X30,HNRNPM,SRSF3,ILF2,PFKM,SF3A1,HNRNPD,FLNC,KDM3B,TP53BP1,DDIT4,VDAC1,RBBP7,T
TRIM2,IDH2,UBE2J1,RTCB,RPL27A,FAM168A,ALDH7A1,SYNGAP1,NACAD,NDUFB8,SEZ6L2,CRAE
,GAK,ANK2,SCAMP5,HTATSF1,LRIG1,AASDHPPT,TMEM94,SLC22A23,TSPYL2,IGF1R,NPTN,GPAA
N1A,CRPPA,CPSF2,CTSB,TM9SF3,TOMM20,U2AF2,CCT2,SUGP2,PABPN1,GDI2,HNRNPL,TM9SF
BEX3,CCT7,MCL1,JPT2,NISCH,TADA2B,CSNK1E,PFKL,RPS25,EIF5A,DDR1,CLTC,RPS7,MIF,HMG(
ARDBP,DDOST,HNRNPK,MYH10,NARS1,RPL36,LAMP1,HNRNPH1,HLA-B,CCT6A,TOP2B,HDAC2,E
FB8,SEZ6L2,ACSL4,DLG4,MAN2B1,WASF3,PRKAR2B,DPP7,BACE1,CTCF,ATP6V0A1,GPRIN1,SHC
B2,CADM3,REEP2,PSMD4,MSH6,SYT11,CHMP4B,AP2A2,ATP6V1G1,SORT1,RCN2,IGLON5,NTRK
1,KCTD20,NCKAP5L,CFDP1,PRXL2A,PSMC2,NSD1,SEPHS1,PPP1R37,PRKAR2B,LMNA,RXRA,TLK1

2,MGRN1,RTRAF,FUBP1,ARHGAP23,FAM193B,ZNF207,UBTF,ALYREF,NUTF2,UNC13B

TN4,DDX3X,RPL12,EIF3L,MRFAP1,H1-0,IK,RPL35,GOLM1,TARDBP,DDOST,HNRNPK,MYH10,NAR
NFRSF10B,RBM10,ALCAM,TUBA1C,SNRPD2,SOX9,KDM2A,CDC42EP4,ZNRF1,RBM17,MGRN1,D[illegible]

FXR2,CLTA,TRIM2,COPE,APEH,UBE2J1,RTCB,RPL27A,NACAD,NDUFB8,SEZ6L2,CLCN4,ALCAM,DI
JL1,FKBP8,CCAR2,DIDO1,TPM3,SDF4,SMARCC1,DHX30,HNRNPM,SSB,SRSF3,LARS1,ILF2,ISYNA
JS,PELP1,CLPTM1,RAB1B,HINT1,SRP72,HDGFL2,EFTUD2,GTF2I,TRAP1,SLC4A2,FNBP1L,MOAP1
3,ANKRD12,ATP5MC2,FGFR1,XRN2,TAPBP,SUN1,KTN1,OBSL1,RBM25,MDM2,CRIP2,AAMP,GGT
IPD,KDM3B,BMPR2,DDIT4,PSMA7,ANP32A,RBBP7,EDC4,TXLNA,LBH,NDUFV1,SF1,H2AZ1,FLOT1,
ORF43,COL1A2,LLGL1,RPL14,RHOA,ZNF865,PSMD2,KIF5B,SLC7A5,GLIPR1L2,RPS14,CHD4,TMI
PPP3CA,APOE,ABCD3,PRNP,PPT1,DNM1,GNAL,ATP6V1F,PDHB,WDR33,GSN,VPS37B,MYBBP1A,A

,LRCH4,LZTS2,SRRT,RFX7,B4GALT2,SEPTIN9,SLC1A5,PCNA,RAD54L2,TMEM59,BCAR1,OPA1,A

T1,PRNP,EPHA4,PPFIA3,GNAO1,ATP6V1F,PDHB,VPS37B
B,GNAZ,ATP6V0A1,SHC1,PPP3CA,ADD1,FH,TUT4,SYT1,EPHA4,DNM1,GNAL,PPFIA3,PDHB,NNT

PAT1,CUL3,ADH5,TNKS,HNRNPAB,SYMPK,C1ORF198,SAFB2,ERH,RTL6,FKBP4,NUDCD3,HINT1,L
1,FH,APOE,ABCD3,PRNP,GNAL,PPFIA3,ATP6V1F,NNT,ZNF326,GSN,MYBBP1A,ARSA,FKBP3
AC2,BCLAF1,LUC7L3,CRPPA,CPSF2,TM9SF3,TOMM20,U2AF2,CCT2,SUGP2,PABPN1,HNRNPL,P

NA2,TRIM2,EMC1,PABPC4,CD44,PDLIM1,ATP1B3,KCTD20,ECH1,ATF7IP,CCNG1,KPNA6,APPBP2,

,ELOB,SS18L1,RTRAF,PLCD3,COG4,LANCL1,FAM193B,CPSF7,ALYREF,GLB1,SCCPDH

ER1,NSD2,CSE1L,FBXW5,GOT2,RBM6,ATP6V1G1,BAG5,SORT1,CAD,OGFR,LRIG1,TSNAX,ATP1

P5F1B,CALM3,GNB1,MYH9,RPL7,SEPTIN2,P4HB,APP,IDS,RPL5,GDI1,FSCN1,TMBIM6,PTMS,SPT

1,FSCN1,TMBIM6,PTMS,SPTAN1,GSTP1,MARCKS,SMG1,CALM1,NPM1,PLD3,HSPD1,USP11,SNR

MS,SPTAN1,MARCKS,SMG1,CALM1,NPM1,PLD3,HSPD1,USP11,SNRNP200,CSDE1,EIF3C,ATP5F1

1,PLD3,HSPD1,USP11,SNRNP200,CSDE1,EIF3C,ATP5F1A,EIF4B,VCP,UBB,AFF4,TRIM28,MATR3,

1,MYH9,RPL7,SEPTIN2,P4HB,APP,IDS,RPL5,GDI1,FSCN1,TMBIM6,PTMS,SPTAN1,GSTP1,MARCK

,SMG1,CALM1,NPM1,PLD3,HSPD1,USP11,SNRNP200,CSDE1,EIF3C,ATP5F1A,EIF4B,VCP,UBB,AF

,PTMS,SPTAN1,GSTP1,MARCKS,SMG1,CALM1,NPM1,PLD3,HSPD1,USP11,SNRNP200,CSDE1,EIF

MS,SPTAN1,GSTP1,MARCKS,SMG1,CALM1,NPM1,PLD3,HSPD1,COL18A1,SNRNP200,CSDE1,EIF

MBIM6,PTMS,SPTAN1,GSTP1,MARCKS,SMG1,NPM1,PLD3,HSPD1,COL18A1,SNRNP200,CSDE1,E

,PLD3,HSPD1,USP11,COL18A1,SNRNP200,CSDE1,EIF3C,ATP5F1A,EIF4B,VCP,MARCKSL1,UBB,AF

,PLD3,HSPD1,USP11,COL18A1,SNRNP200,CSDE1,EIF3C,ATP5F1A,EIF4B,VCP,MARCKSL1,UBB,AF

I6,PTMS,SPTAN1,GSTP1,MARCKS,SMG1,CALM1,NPM1,PLD3,HSPD1,COL18A1,SNRNP200,CSDE1

LRRC28,MAPT,SPEN,NONO,DDB1,YWHAQ,HACD3,DCHS1,SET,EGLN1,RPL19,SF3B1,PDIA3,SSBF

APP,IDS,RPL5,GDI1,TMBIM6,PTMS,SPTAN1,GSTP1,MARCKS,SMG1,CALM1,NPM1,PLD3,HSPD1,U

APP,IDS,RPL5,GDI1,TMBIM6,PTMS,SPTAN1,GSTP1,MARCKS,SMG1,CALM1,NPM1,PLD3,HSPD1,U

PLD3,HSPD1,COL18A1,SNRNP200,CSDE1,EIF3C,ATP5F1A,EIF4B,VCP,MARCKSL1,UBB,AFF4,TRI

ARCKS,SMG1,CALM1,NPM1,PLD3,HSPD1,USP11,SNRNP200,CSDE1,EIF3C,ATP5F1A,EIF4B,VCP,

ARCKS,SMG1,CALM1,NPM1,PLD3,HSPD1,USP11,SNRNP200,CSDE1,EIF3C,ATP5F1A,EIF4B,VCP,

,IDS,RPL5,GDI1,FSCN1,TMBIM6,PTMS,SPTAN1,GSTP1,MARCKS,CALM1,NPM1,PLD3,HSPD1,COL

G1,CALM1,NPM1,PLD3,HSPD1,COL18A1,SNRNP200,CSDE1,EIF3C,ATP5F1A,EIF4B,VCP,MARCKS

CD3,DCHS1,SET,EGLN1,MRC2,RPL19,LDHB,PDIA3,SSBP2,CHD3,BASP1,VAPA,HMGB1,POLR2A,R

6,PTMS,SPTAN1,GSTP1,MARCKS,SMG1,NPM1,PLD3,HSPD1,USP11,COL18A1,SNRNP200,CSDE1,

AN1,GSTP1,SMG1,CALM1,NPM1,PLD3,HSPD1,USP11,COL18A1,CSDE1,EIF3C,ATP5F1A,EIF4B,V

AN1,GSTP1,SMG1,CALM1,NPM1,PLD3,HSPD1,USP11,COL18A1,CSDE1,EIF3C,ATP5F1A,EIF4B,V

1,SET,EGLN1,INA,MRC2,RPL19,LDHB,SF3B1,PDIA3,SSBP2,CHD3,BASP1,VAPA,POLR2A,TPT1,RP

B,SF3B1,PDIA3,SSBP2,CHD3,BASP1,VAPA,POLR2A,TPT1,RPS4X,RPS3A,RPL18,STMN1,TUBB4B,

DS,RPL5,GDI1,FSCN1,TMBIM6,PTMS,SPTAN1,GSTP1,MARCKS,SMG1,CALM1,NPM1,PLD3,HSPD

RCKS,SMG1,CALM1,NPM1,PLD3,HSPD1,COL18A1,SNRNP200,CSDE1,EIF3C,ATP5F1A,EIF4B,VC

LD3,HSPD1,SNRNP200,CSDE1,EIF3C,ATP5F1A,EIF4B,VCP,MARCKSL1,UBB,AFF4,TRIM28,MATR

LD3,HSPD1,SNRNP200,CSDE1,EIF3C,ATP5F1A,EIF4B,VCP,MARCKSL1,UBB,AFF4,TRIM28,MATR

BIM6,SPTAN1,GSTP1,MARCKS,SMG1,CALM1,NPM1,PLD3,HSPD1,USP11,COL18A1,SNRNP200,C

,PLD3,USP11,SNRNP200,CSDE1,EIF3C,ATP5F1A,EIF4B,VCP,MARCKSL1,UBB,AFF4,TRIM28,MAT

,PLD3,USP11,SNRNP200,CSDE1,EIF3C,ATP5F1A,EIF4B,VCP,MARCKSL1,UBB,AFF4,TRIM28,MAT

,RPL37A,SPEN,NONO,DDB1,YWHAQ,HACD3,DCHS1,SET,EGLN1,MRC2,RPL19,LDHB,SF3B1,PDIA

F1A,EIF4B,VCP,MARCKSL1,UBB,TRIM28,MATR3,YWHAG,RPL37A,LRRC28,SPEN,NONO,DDB1,H

I6,PTMS,SPTAN1,GSTP1,MARCKS,CALM1,NPM1,PLD3,HSPD1,COL18A1,SNRNP200,CSDE1,EIF3

SPEN,NONO,DDB1,YWHAQ,HACD3,DCHS1,SET,EGLN1,LDHB,SF3B1,PDIA3,SSBP2,BASP1,VAPA,H

TP5F1A,EIF4B,VCP,MARCKSL1,UBB,AFF4,TRIM28,MATR3,YWHAG,CKB,RPL37A,LRRC28,SPEN,N

TP5F1A,EIF4B,VCP,MARCKSL1,UBB,AFF4,TRIM28,MATR3,YWHAG,CKB,RPL37A,LRRC28,SPEN,N

8A1,SNRNP200,CSDE1,EIF3C,ATP5F1A,EIF4B,VCP,MARCKSL1,UBB,AFF4,TRIM28,MATR3,YWHA

1,SET,EGLN1,MRC2,SF3B1,PDIA3,SSBP2,BASP1,VAPA,HMGB1,RPS4X,RPS3A,RPL11,MORF4L1,

DI1,TMBIM6,SPTAN1,GSTP1,MARCKS,SMG1,CALM1,NPM1,PLD3,HSPD1,COL18A1,SNRNP200,C

DI1,TMBIM6,SPTAN1,GSTP1,MARCKS,SMG1,CALM1,NPM1,PLD3,HSPD1,COL18A1,SNRNP200,C

1,GSTP1,MARCKS,SMG1,CALM1,NPM1,PLD3,HSPD1,COL18A1,SNRNP200,CSDE1,EIF3C,ATP5F1

1,GSTP1,MARCKS,SMG1,CALM1,NPM1,PLD3,HSPD1,COL18A1,SNRNP200,CSDE1,EIF3C,ATP5F1

TRIM28,MATR3,YWHAG,RPL37A,LRRC28,MAPT,FN1,SPEN,NONO,YWHAQ,DCHS1,SET,EGLN1,MF
TP1,MARCKS,SMG1,CALM1,NPM1,PLD3,HSPD1,COL18A1,SNRNP200,CSDE1,EIF3C,ATP5F1A,EIF
TP1,MARCKS,SMG1,CALM1,NPM1,PLD3,HSPD1,COL18A1,SNRNP200,CSDE1,EIF3C,ATP5F1A,EIF
AN1,GSTP1,MARCKS,CALM1,NPM1,PLD3,HSPD1,USP11,COL18A1,SNRNP200,CSDE1,EIF3C,ATP
D3,HSPD1,COL18A1,SNRNP200,CSDE1,EIF3C,ATP5F1A,EIF4B,VCP,MARCKSL1,UBB,AFF4,TRIM
SPTAN1,MARCKS,SMG1,CALM1,NPM1,HSPD1,COL18A1,SNRNP200,CSDE1,EIF3C,ATP5F1A,EIF4
TMS,SPTAN1,GSTP1,MARCKS,SMG1,CALM1,NPM1,PLD3,HSPD1,USP11,COL18A1,SNRNP200,CS
11,SNRNP200,CSDE1,EIF3C,ATP5F1A,EIF4B,VCP,UBB,AFF4,TRIM28,MATR3,YWHAG,RPL37A,LR
200,CSDE1,EIF3C,ATP5F1A,EIF4B,VCP,MARCKSL1,UBB,AFF4,TRIM28,MATR3,YWHAG,RPL37A,L
200,CSDE1,EIF3C,ATP5F1A,EIF4B,VCP,MARCKSL1,UBB,AFF4,TRIM28,MATR3,YWHAG,RPL37A,L
,GSTP1,MARCKS,SMG1,CALM1,NPM1,PLD3,HSPD1,USP11,COL18A1,SNRNP200,CSDE1,EIF3C,A
3,AFF4,TRIM28,MATR3,YWHAG,RPL37A,LRRC28,NONO,DDB1,YWHAQ,HACD3,DCHS1,SET,EGLN
D3,HSPD1,SNRNP200,CSDE1,EIF3C,ATP5F1A,EIF4B,VCP,MARCKSL1,UBB,AFF4,TRIM28,MATR3,
D3,HSPD1,SNRNP200,CSDE1,EIF3C,ATP5F1A,EIF4B,VCP,MARCKSL1,UBB,AFF4,TRIM28,MATR3,
PA,HMGB1,RPS4X,RPS3A,TUBB4B,MORF4L1,BTF3,RPL10,RPS17,VCL,DNAJA1,CYB5R3,PAFAH1E
00,CSDE1,EIF3C,ATP5F1A,EIF4B,VCP,MARCKSL1,UBB,AFF4,TRIM28,MATR3,YWHAG,RPL37A,LR
4,TRIM28,MATR3,YWHAG,RPL37A,MAPT,FN1,SPEN,NONO,DDB1,YWHAQ,HACD3,DCHS1,SET,EGL
RCKS,SMG1,CALM1,NPM1,PLD3,HSPD1,COL18A1,SNRNP200,CSDE1,EIF3C,ATP5F1A,EIF4B,VCP,
RCKS,SMG1,CALM1,NPM1,PLD3,HSPD1,COL18A1,SNRNP200,CSDE1,EIF3C,ATP5F1A,EIF4B,VCP,
ALM1,NPM1,PLD3,HSPD1,SNRNP200,CSDE1,EIF3C,ATP5F1A,EIF4B,VCP,MARCKSL1,UBB,AFF4,TI
ALM1,NPM1,PLD3,HSPD1,SNRNP200,CSDE1,EIF3C,ATP5F1A,EIF4B,VCP,MARCKSL1,UBB,AFF4,TI
7A,LRRC28,SPEN,NONO,HACD3,DCHS1,SET,EGLN1,LDHB,PDIA3,SSBP2,CHD3,VAPA,HMGB1,TPT
GSTP1,MARCKS,SMG1,CALM1,NPM1,PLD3,HSPD1,USP11,SNRNP200,CSDE1,EIF3C,ATP5F1A,EIF
B,TRIM28,MATR3,YWHAG,RPL37A,LRRC28,MAPT,FN1,SPEN,NONO,YWHAQ,HACD3,DCHS1,SET,
1,NPM1,PLD3,COL18A1,SNRNP200,CSDE1,EIF3C,ATP5F1A,EIF4B,VCP,UBB,TRIM28,MATR3,YWH
TR3,YWHAG,RPL37A,LRRC28,SPEN,NONO,DDB1,YWHAQ,HACD3,DCHS1,SET,EGLN1,RPL19,LDH
NA,MRC2,RPL19,LDHB,PKD1,SF3B1,PDIA3,SSBP2,CHD3,VAPA,HMGB1,RPS4X,RPS3A,TUBB4B,R
IF3C,ATP5F1A,VCP,UBB,AFF4,TRIM28,MATR3,YWHAG,LRRC28,FN1,SPEN,NONO,DDB1,YWHAQ,
HAG,RPL37A,LRRC28,FN1,SPEN,NONO,HACD3,DCHS1,SET,EGLN1,MRC2,LDHB,SF3B1,PDIA3,SS
HAG,RPL37A,LRRC28,FN1,SPEN,NONO,HACD3,DCHS1,SET,EGLN1,MRC2,LDHB,SF3B1,PDIA3,SS
IM28,MATR3,YWHAG,RPL37A,LRRC28,SPEN,NONO,YWHAQ,DCHS1,SET,EGLN1,MRC2,PKD1,SF
CKSL1,UBB,AFF4,TRIM28,MATR3,YWHAG,CKB,RPL37A,LRRC28,SPEN,NONO,DDB1,HACD3,DCH
28,FN1,SPEN,NONO,YWHAQ,HACD3,DCHS1,SET,EGLN1,RPL19,LDHB,SF3B1,PDIA3,SSBP2,BASP1
AN1,MARCKS,SMG1,CALM1,NPM1,PLD3,HSPD1,COL18A1,SNRNP200,CSDE1,EIF3C,ATP5F1A,EIF
AN1,MARCKS,SMG1,CALM1,NPM1,PLD3,HSPD1,COL18A1,SNRNP200,CSDE1,EIF3C,ATP5F1A,EIF
SMG1,CALM1,NPM1,PLD3,HSPD1,USP11,COL18A1,SNRNP200,CSDE1,EIF3C,ATP5F1A,EIF4B,VCF
,GSTP1,MARCKS,SMG1,CALM1,NPM1,PLD3,HSPD1,COL18A1,SNRNP200,CSDE1,EIF3C,ATP5F1
SPD1,SNRNP200,CSDE1,EIF3C,ATP5F1A,EIF4B,VCP,MARCKSL1,UBB,AFF4,TRIM28,MATR3,YWHA
P2,BASP1,VAPA,HMGB1,POLR2A,RPS4X,RPS3A,STMN1,RPL11,BTF3,NCAM1,AGRN,XAF1,RPL10
SDE1,EIF3C,ATP5F1A,EIF4B,VCP,MARCKSL1,UBB,AFF4,TRIM28,MATR3,YWHAG,RPL37A,LRRC28
O,DDB1,YWHAQ,HACD3,DCHS1,SET,EGLN1,RPL19,LDHB,PKD1,SF3B1,PDIA3,VAPA,HMGB1,RPS
CHS1,SET,EGLN1,RPL19,LDHB,SF3B1,PDIA3,SSBP2,BASP1,VAPA,TPT1,RPS4X,RPS3A,TUBB4B,RF
,DCHS1,SET,EGLN1,RPL19,LDHB,PKD1,SF3B1,PDIA3,SSBP2,CHD3,VAPA,HMGB1,POLR2A,RPS4X
RNP200,CSDE1,EIF3C,ATP5F1A,EIF4B,VCP,UBB,AFF4,TRIM28,MATR3,YWHAG,RPL37A,LRRC28,
F3C,ATP5F1A,EIF4B,VCP,MARCKSL1,UBB,TRIM28,MATR3,YWHAG,CKB,RPL37A,LRRC28,SPEN,N

F4,TRIM28,MATR3,YWHAG,RPL37A,LRRC28,SPEN,NONO,YWHAQ,HACD3,DCHS1,SET,EGLN1,MF
PD1,SNRNP200,CSDE1,EIF3C,ATP5F1A,EIF4B,VCP,MARCKSL1,UBB,AFF4,TRIM28,MATR3,YWHA
COL18A1,SNRNP200,CSDE1,EIF3C,ATP5F1A,EIF4B,VCP,MARCKSL1,UBB,AFF4,TRIM28,MATR3,Y
COL18A1,SNRNP200,CSDE1,EIF3C,ATP5F1A,EIF4B,VCP,MARCKSL1,UBB,AFF4,TRIM28,MATR3,Y
GTF2F1,HNRNPU,PGAM1,NFE2L1,SMARCA5,LARP1,CTNNA1,RPL23,CPE,RPSA,TPM4,CTSD,EIF4
RIM28,MATR3,YWHAG,RPL37A,LRRC28,SPEN,NONO,DDB1,YWHAQ,HACD3,DCHS1,SET,EGLN1,
200,CSDE1,EIF3C,ATP5F1A,EIF4B,VCP,UBB,AFF4,TRIM28,MATR3,YWHAG,RPL37A,LRRC28,FN1,
RPL19,SF3B1,PDIA3,SSBP2,VAPA,HMGB1,POLR2A,TPT1,RPS4X,RPS3A,RPL18,STMN1,TUBB4B,R
TMBIM6,PTMS,SPTAN1,GSTP1,MARCKS,SMG1,CALM1,NPM1,PLD3,HSPD1,COL18A1,SNRNP200,
CKSL1,UBB,AFF4,TRIM28,MATR3,YWHAG,RPL37A,LRRC28,SPEN,NONO,DDB1,YWHAQ,HACD3,D
CKSL1,UBB,AFF4,TRIM28,MATR3,YWHAG,RPL37A,LRRC28,SPEN,NONO,DDB1,YWHAQ,HACD3,D
,RPL19,LDHB,SF3B1,PDIA3,SSBP2,CHD3,VAPA,HMGB1,POLR2A,TPT1,RPS4X,RPS3A,RPL18,TUB
,FN1,NONO,DDB1,YWHAQ,HACD3,DCHS1,SET,EGLN1,MRC2,LDHB,SF3B1,PDIA3,SSBP2,CHD3,B
,YWHAG,RPL37A,LRRC28,SPEN,NONO,DDB1,YWHAQ,HACD3,DCHS1,SET,RPL19,LDHB,PKD1,SF
,YWHAG,RPL37A,LRRC28,SPEN,NONO,DDB1,YWHAQ,HACD3,DCHS1,SET,RPL19,LDHB,PKD1,SF
SET,EGLN1,MRC2,LDHB,SF3B1,PDIA3,SSBP2,CHD3,VAPA,HMGB1,POLR2A,TPT1,RPS4X,RPS3A,
S1,SET,EGLN1,RPL19,LDHB,SF3B1,PDIA3,SSBP2,CHD3,BASP1,VAPA,HMGB1,RPS4X,RPS3A,RPL1
YWHAQ,HACD3,DCHS1,SET,EGLN1,RPL19,LDHB,PKD1,SF3B1,PDIA3,SSBP2,CHD3,VAPA,HMGB1,
LRRC28,FN1,SPEN,NONO,DDB1,SET,EGLN1,RPL19,SF3B1,PDIA3,SSBP2,BASP1,HMGB1,POLR2A
,SPEN,NONO,DDB1,YWHAQ,DCHS1,EGLN1,MRC2,RPL19,LDHB,SF3B1,PDIA3,SSBP2,BASP1,VAPA
N,NONO,DDB1,YWHAQ,HACD3,DCHS1,SET,EGLN1,LDHB,PDIA3,SSBP2,HMGB1,POLR2A,RPS4X,
3,GNB1,MYH9,RPL7,SEPTIN2,P4HB,APP,IDS,RPL5,GDI1,FSCN1,TMBIM6,PTMS,SPTAN1,GSTP1,M
S17,DNAJA1,CYB5R3,PAFAH1B1,SF3B2,CDC42BPB,ITM2B,GTF2F1,NFE2L1,SMARCA5,CTNNA1,F
AG,RPL37A,LRRC28,SPEN,NONO,DDB1,YWHAQ,HACD3,DCHS1,SET,EGLN1,MRC2,RPL19,LDHB,
32,XRCC5,PJA2,RPL24,RPL17,C1ORF43,PAPOLA,COL1A2,RPL14,CBX5,PSMD2,KIF5B,RPS14,CH
EIF3C,ATP5F1A,EIF4B,VCP,MARCKSL1,UBB,TRIM28,MATR3,YWHAG,RPL37A,LRRC28,SPEN,NON
8,MATR3,YWHAG,RPL37A,LRRC28,MAPT,FN1,SPEN,NONO,DDB1,HACD3,DCHS1,SET,EGLN1,LD
LRRC28,MAPT,SPEN,NONO,YWHAQ,HACD3,DCHS1,SET,EGLN1,MRC2,LDHB,SF3B1,PDIA3,SSBP2,
00,CSDE1,EIF3C,ATP5F1A,EIF4B,VCP,MARCKSL1,UBB,TRIM28,MATR3,YWHAG,CKB,RPL37A,LR
ARCKSL1,UBB,TRIM28,MATR3,YWHAG,CKB,RPL37A,LRRC28,FN1,SPEN,NONO,DDB1,YWHAQ,H
,VCP,MARCKSL1,UBB,TRIM28,MATR3,YWHAG,RPL37A,LRRC28,SPEN,NONO,DDB1,YWHAQ,HAC
3B1,SSBP2,VAPA,POLR2A,TPT1,RPS4X,RPS3A,STMN1,RPL11,MORF4L1,BTF3,AGRN,RPS17,DNA
NN,RPL31,PRKCSH,HNRNPC,LRPPRC,KHDRBS1,PTN,RPS25,GLG1,SFPQ,DDR1,RAB7A,PRDX2,DD
S1,SET,EGLN1,RPL19,SF3B1,PDIA3,SSBP2,CHD3,VAPA,HMGB1,TPT1,RPS4X,RPS3A,RPL11,MORF
MARCKSL1,UBB,TRIM28,MATR3,YWHAG,LRRC28,FN1,SPEN,NONO,DDB1,YWHAQ,HACD3,DCHS1
CKSL1,UBB,AFF4,TRIM28,MATR3,YWHAG,RPL37A,LRRC28,FN1,SPEN,NONO,DDB1,YWHAQ,HAC
1,EIF3C,ATP5F1A,VCP,MARCKSL1,UBB,AFF4,TRIM28,MATR3,YWHAG,RPL37A,MAPT,FN1,SPEN,N
CD3,DCHS1,SET,EGLN1,PKD1,SF3B1,PDIA3,SSBP2,CHD3,VAPA,HMGB1,POLR2A,TPT1,RPS4X,RP
RRC28,MAPT,FN1,SPEN,NONO,DDB1,YWHAQ,HACD3,DCHS1,SET,EGLN1,MRC2,RPL19,LDHB,PK
NO,DDB1,YWHAQ,HACD3,SET,EGLN1,MRC2,RPL19,PKD1,SF3B1,PDIA3,SSBP2,CHD3,BASP1,VAP
DB1,YWHAQ,HACD3,DCHS1,SET,EGLN1,RPL19,LDHB,SF3B1,SSBP2,VAPA,POLR2A,TPT1,RPS4X,R
POLR2A,RPS4X,RPS3A,UCHL1,RPL11,MORF4L1,BTF3,AGRN,RPL10,RPS17,DNAJA1,CYB5R3,PAF
HAG,RPL37A,LRRC28,FN1,SPEN,NONO,DDB1,HACD3,DCHS1,SET,EGLN1,MRC2,RPL19,LDHB,PK
28,SPEN,NONO,DDB1,YWHAQ,HACD3,DCHS1,SET,EGLN1,RPL19,PKD1,PDIA3,SSBP2,VAPA,HMG
,DCHS1,SET,EGLN1,MRC2,RPL19,SF3B1,PDIA3,SSBP2,CHD3,VAPA,HMGB1,POLR2A,TPT1,RPS4

l,NONO,DDB1,HACD3,DCHS1,SET,EGLN1,MRC2,SF3B1,PDIA3,SSBP2,CHD3,BASP1,VAPA,HMGB1
AJA1,CYB5R3,PAFAH1B1,SF3B2,CDC42BPB,ITM2B,ZNF770,AP2B1,MAT2A,GTF2F1,HNRNPU,PGA
P4,PNN,PRKCSH,HNRNPC,LRPPRC,KHDRBS1,XPO1,RPS25,TKT,GLG1,SFPQ,PDIA6,CCN2,PRDX2
BB4B,RPL11,MORF4L1,BTF3,ANXA2,RPL10,RPS17,VCL,DNAJA1,PAFAH1B1,SF3B2,CDC42BPB,I
C,ATP5F1A,EIF4B,VCP,MARCKSL1,UBB,AFF4,TRIM28,MATR3,YWHAG,RPL37A,LRRC28,FN1,SPEN
24,C1ORF43,PAPOLA,RPL14,CBX5,CHD4,JUND,EIF5,CD81,HMGN2,RPL26,BEX3,SRSF4,RBBP4,PI
,CHD3,VAPA,HMGB1,POLR2A,TPT1,RPS4X,RPS3A,TUBB4B,RPL11,BTF3,ANXA2,AGRN,RPL10,RP
1,NPM1,PLD3,HSPD1,USP11,COL18A1,SNRNP200,CSDE1,EIF3C,ATP5F1A,EIF4B,VCP,UBB,AFF4,T
1,NPM1,PLD3,HSPD1,USP11,COL18A1,SNRNP200,CSDE1,EIF3C,ATP5F1A,EIF4B,VCP,UBB,AFF4,T
RPL37A,LRRC28,SPEN,NONO,DDB1,YWHAQ,HACD3,DCHS1,SET,EGLN1,RPL19,PDIA3,SSBP2,VAF
EGLN1,RPL19,LDHB,SF3B1,SSBP2,BASP1,VAPA,HMGB1,RPS4X,RPS3A,RPL18,STMN1,TUBB4B,R
R2A,RPS4X,RPS3A,TUBB4B,BTF3,ANXA2,AGRN,RPL10,RPS17,VCL,CYB5R3,PAFAH1B1,AKAP12,S
R2A,RPS4X,RPS3A,TUBB4B,BTF3,ANXA2,AGRN,RPL10,RPS17,VCL,CYB5R3,PAFAH1B1,AKAP12,S
PDIA3,SSBP2,CHD3,BASP1,VAPA,HMGB1,POLR2A,TPT1,RPL18,TUBB4B,RPL11,BTF3,ANXA2,RPL
BTF3,ANXA2,AGRN,RPL10,WSB1,RPS17,VCL,CYB5R3,MAP4,SF3B2,CDC42BPB,GTF2F1,HNRNPU,
HD3,BASP1,VAPA,HMGB1,TPT1,RPS4X,RPL18,TUBB4B,RPL11,MORF4L1,BTF3,ANXA2,AGRN,XAF
,RPL10,RPS17,VCL,DNAJA1,PAFAH1B1,MAP4,SF3B2,CDC42BPB,ITM2B,GTF2F1,HNRNPU,NFE2L
A,RPL18,RPL11,BTF3,ANXA2,ZMIZ1,RPL10,RPS17,VCL,DNAJA1,PAFAH1B1,SF3B2,ITM2B,GTF2F
ACD3,DCHS1,SET,EGLN1,RPL19,LDHB,SF3B1,PDIA3,SSBP2,BASP1,VAPA,HMGB1,POLR2A,RPS4X
1,SMG1,NPM1,PLD3,HSPD1,COL18A1,SNRNP200,CSDE1,EIF3C,ATP5F1A,EIF4B,VCP,UBB,AFF4,T
KSL1,UBB,TRIM28,MATR3,YWHAG,RPL37A,LRRC28,FN1,SPEN,NONO,DDB1,YWHAQ,DCHS1,SET
,MAP4,SF3B2,ITM2B,MAT2A,GTF2F1,GOLGA8A,HNRNPU,NFE2L1,SMARCA5,LARP1,HNRNPA0,CT
2,HNRNPK,RPL36,YWHAB,LAMP1,HNRNPH1,MAP2,CALU,CCT6A,HMGCR,CAPRIN1,TOP2B,HDAC
,YWHAG,RPL37A,LRRC28,SPEN,NONO,DDB1,HACD3,DCHS1,SET,EGLN1,MRC2,RPL19,LDHB,PKI
l,RPS3A,TUBB4B,MORF4L1,BTF3,ANXA2,RPL10,VCL,DNAJA1,MAP4,SF3B2,MAT2A,GTF2F1,HNRI
N1,SPEN,NONO,DDB1,HACD3,DCHS1,SET,EGLN1,MRC2,LDHB,SF3B1,SSBP2,VAPA,HMGB1,POLR
N1,SPEN,NONO,DDB1,HACD3,DCHS1,SET,EGLN1,MRC2,LDHB,SF3B1,SSBP2,VAPA,HMGB1,POLR
3A,RPL18,MORF4L1,BTF3,ANXA2,RPL10,RPS17,VCL,PAFAH1B1,SF3B2,CDC42BPB,ITM2B,AP2B
CARB2,BRD3,XRCC5,RPL24,PAPOLA,COL1A2,RPL14,HK1,RHOA,CBX5,ZNF865,ATF4,RPS14,CHD
BTF3,XAF1,RPL10,RPS17,VCL,PAFAH1B1,SF3B2,ITM2B,AP2B1,GTF2F1,HNRNPU,SMARCA5,CTNI
SCARB2,LPCAT1,BRD3,XRCC5,PJA2,RPL24,C1ORF43,PAPOLA,COL1A2,LLGL1,GAP43,RPL14,PT
DC42BPB,AP2B1,MAT2A,GTF2F1,HNRNPU,NFE2L1,SMARCA5,HNRNPA0,CTNNA1,FADS2,RPS16,F
SSBP2,CHD3,BASP1,VAPA,HMGB1,POLR2A,RPS4X,RPS3A,RPL18,STMN1,TUBB4B,MORF4L1,BTF
SLC7A5,RPS14,CXADR,CHD4,JUND,WDR1,CCT7,RBBP4,JPT2,PNN,HNRNPC,LRPPRC,TADA2B,KHD
F43,PAPOLA,COL1A2,LLGL1,RHOA,RPL9,PSMD2,RPS14,CHD4,JUND,RPS20,HMGN2,RPL26,BEX
OA,CBX5,ZNF865,CLSTN1,ATF4,PSMD2,KIF5B,RPS14,CHD4,JUND,WDR1,HMGN2,BEX3,CCT7,SF
ET,EGLN1,INA,MRC2,LDHB,PKD1,SF3B1,PDIA3,SSBP2,CHD3,VAPA,HMGB1,POLR2A,RPS4X,RPS3
1,MAP4,SF3B2,CDC42BPB,ITM2B,GTF2F1,HNRNPU,NFE2L1,SMARCA5,HNRNPA0,CTNNA1,CPE,C
2,AGRN,RPL10,MARCHF6,RPS17,CYB5R3,PAFAH1B1,MAP4,SF3B2,CDC42BPB,ITM2B,GTF2F1,H
MAP4,SF3B2,CDC42BPB,ITM2B,MAT2A,GTF2F1,HNRNPU,NFE2L1,HNRNPA0,RPS16,RPL23,CNN
F5,RPL26,CCT7,RBBP4,JPT2,PNN,RPL31,PRKCSH,HNRNPC,LRPPRC,TADA2B,XPR1,KHDRBS1,XP
CD3,DCHS1,SET,EGLN1,RPL19,PKD1,SF3B1,PDIA3,VAPA,HMGB1,POLR2A,RPS4X,RPS3A,RPL18,I
2F1,HNRNPU,NFE2L1,HNRNPA0,CNN3,RPSA,TPM4,EIF4H,SCARB2,LPCAT1,BRD3,XRCC5,PJA2,RP
57A,ACTN4,PRDX2,RPS15,RPL12,RPL30,CLTC,EIF3L,MAGED2,RCN1,H1-0,RPL35,TLN1,CTNNB1,A
1B1,SF3B2,CDC42BPB,ITM2B,AP2B1,GTF2F1,HNRNPU,OGA,PGAM1,NFE2L1,SMARCA5,HNRNPA

E2L1,SMARCA5,LARP1,RPL23,RPSA,EIF4H,SCARB2,LPCAT1,XRCC5,PJA2,MSN,RPL24,KPNB1,LL
,VCL,DNAJA1,CYB5R3,PAFAH1B1,MAP4,SF3B2,GTF2F1,HNRNPU,OGA,NFE2L1,SMARCA5,LARP1
H1,MAP2,CALU,CCT6A,CAPRIN1,TOP2B,HDAC2,SKP1,BCLAF1,LUC7L3,CPSF2,CTSB,NUCB1,TM9
4,C1ORF43,PAPOLA,COL1A2,RPL14,KIF5B,RPS14,CHD4,JUND,RPS20,EIF5,HMGN2,BEX3,CCT7,S
STMN1,TUBB4B,BTF3,ANXA2,RPL10,VCL,DNAJA1,PAFAH1B1,MAP4,SF3B2,CDC42BPB,ITM2B,AP2
OLA,RPL14,RPL9,CLSTN1,RPS14,CHD4,JUND,RPS20,EIF5,CD81,HMGN2,BEX3,RBBP4,PFN1,PNN,
NFE2L1,SMARCA5,HNRNPA0,CTNNA1,FADS2,RPS16,RPL23,RPSA,TPM4,EIF4H,BRD3,XRCC5,PJA2
NPA0,CNN3,RPSA,TNS3,SCARB2,LPCAT1,BRD3,XRCC5,PJA2,MSN,C1ORF43,PAPOLA,RPL14,CBX
F5,WDR1,HMGN2,BEX3,CCT7,SRSF4,RBBP4,RTN3,MCL1,PNN,HNRNPC,XPO1,CSNK1E,RPS25,TK
ORF4L1,BTF3,AGRN,RPL10,VCL,DNAJA1,CYB5R3,PAFAH1B1,MAP4,SF3B2,AP2B1,GTF2F1,GOL
ORF4L1,BTF3,AGRN,RPL10,VCL,DNAJA1,CYB5R3,PAFAH1B1,MAP4,SF3B2,AP2B1,GTF2F1,GOL
OLA,COL1A2,LLGL1,RPL14,CLSTN1,ATF4,JUND,RPS20,CD81,HMGN2,BEX3,RBBP4,PFN1,PNN,N
7A,PRDX2,DDX3X,MAGED2,RCN1,RPS7,IK,TLN1,CTNNB1,APLP2,HNRNPDL,TARDBP,DDOST,HNR
KCSH,HNRNPC,LRPPRC,XPR1,KHDRBS1,XPO1,SLC3A2,SCRN1,SFPQ,PEBP1,CDH2,PNMA1,PDIA6
BS1,XPO1,SPTBN2,HDLBP,RPS25,TKT,SFPQ,EIF5A,PDIA6,DDR1,PRDX2,DDX3X,RPS15,RPL30,EIF
APLP2,TARDBP,ARID1A,HNRNPK,NARS1,RPL36,YWHAB,LAMP1,UBE2H,ARCN1,CAPRIN1,CDK16,
CALU,ARCN1,TOP2B,HDAC2,BCLAF1,LUC7L3,DCN,CPSF2,DDX39B,NUCB1,TM9SF3,PGD,NOP53
P,MYH10,RPL36,YWHAB,CKAP5,LAMP1,HNRNPH1,CKAP4,RPS12,CALU,CAPRIN1,HDAC2,BCLAF1
L14,HK1,PSMD2,SLC7A5,CHD4,EIF5,RPL26,LASP1,CCT7,SRSF4,RBBP4,PFN1,MCL1,JPT2,PNN,R
DDB1,HACD3,SET,EGLN1,LDHB,SF3B1,PDIA3,SSBP2,HMGB1,POLR2A,TPT1,RPS4X,RPS3A,STMN
7A5,RPS14,CHD4,ATP1A1,RPS20,RPL26,LASP1,BEX3,SRSF4,RBBP4,MCL1,JPT2,RPL31,HNRNPC,
C1ORF43,RPL14,PTCD3,CBX5,PSMD2,KIF5B,SLC7A5,RPS14,CXADR,CHD4,ATP1A1,JUND,RPS20
PQ,RAB7A,MRFAP1,MAGED2,H1-0,RPS7,IK,RPL35,TLN1,CTNNB1,APLP2,HNRNPDL,TARDBP,ARI
1,HMGB1,POLR2A,TPT1,RPS4X,STMN1,BTF3,RPL10,RPS17,VCL,DNAJA1,PAFAH1B1,MAP4,SF3B
K,HMGCS1,JUN,CTNNB1,RPS10,APLP2,HNRNPDL,TARDBP,HNRNPK,INIP,RPL36,YWHAB,CKAP5,L
PO1,RPS25,TKT,GLG1,SFPQ,RAB7A,MMP14,PRDX2,DDX3X,RPL30,MAGED2,H1-0,IK,MIF,GPAT4,
,TARDBP,RPLP2,ARID1A,NDFIP1,DDOST,ST3GAL2,WLS,HNRNPK,INIP,NARS1,YWHAB,CKAP5,SND
2,RTN4,XRCC5,KPNB1,C1ORF43,COL1A2,LLGL1,GAP43,RHOA,ATF4,RPS14,CXADR,JUND,BEX3
1,H1-0,RPS7,RPL35,TLN1,CTNNB1,GOLM1,APLP2,HNRNPDL,TARDBP,ARID1A,HNRNPK,RPL36,Y
KAP5,LAMP1,HNRNPH1,UBE2H,RPS12,HLA-B,RAC1,CAPRIN1,TOP2B,HDAC2,BCLAF1,LUC7L3,CP
CL1,JPT2,PRKCSH,HNRNPC,LRPPRC,KHDRBS1,XPO1,RPS25,GLG1,SFPQ,PDIA6,DDR1,RAB7A,PF
RPS25,TKT,GLG1,SFPQ,RAB7A,RPS15,MRFAP1,MAGED2,H1-0,RPS7,IK,TLN1,CTNNB1,APLP2,HN
5,PJA2,RPL17,C1ORF43,PAPOLA,LLGL1,GAP43,RPL14,RPS14,CHD4,HMGN2,BEX3,CCT7,RTN3,F
N1,SKP1,BCLAF1,LUC7L3,DDX39B,CTSB,TM9SF3,MDH2,TOMM20,HNRNPL,MORF4L2,TAF15,AL
LAMP1,HNRNPH1,CALU,ARCN1,CCT6A,HMGCR,CAPRIN1,BCLAF1,PPIB,LUC7L3,CPSF2,PGRMC1
3,EIF4H,SCARB2,XRCC5,PJA2,RPL24,RPL14,PTCD3,PSMD2,KIF5B,SLC7A5,RPS14,CHD4,ATP1A1
OLR2A,RPS3A,STMN1,TUBB4B,UCHL1,MORF4L1,BTF3,NCAM1,XAF1,RPL10,MARCHF6,DNAJA1,
S25,TKT,SFPQ,PEBP1,PDIA6,RAB7A,PRDX2,MAGED2,H1-0,IK,RPL35,JUN,CTNNB1,APLP2,HNRNP
WDR1,BEX3,CCT7,SRSF4,RBBP4,MCL1,PNN,NISCH,HNRNPC,LRPPRC,XPR1,KHDRBS1,GLG1,SFF
TM9SF3,TOMM20,U2AF2,SQSTM1,NOP53,PABPN1,HNRNPL,MORF4L2,TAF15,RPL27,HNRNPA3,
1,CTNNB1,GOLM1,HNRNPDL,TARDBP,HNRNPK,H1-10,NARS1,RPL36,YWHAB,SEPTIN11,LAMP1,H
,H1-0,RPS5,IK,GJA1,CTNNB1,ANXA6,APLP2,HNRNPDL,TARDBP,NDFIP1,WLS,HNRNPK,INIP,CKAF
A,RAC1,ARHGDIA,CAPRIN1,TOP2B,HDAC2,BCLAF1,LUC7L3,NUCB1,TM9SF3,TOMM20,RNF187,F
2,TOMM20,PABPN1,HNRNPL,MORF4L2,TAF15,RPL27,HNRNPA3,ACO2,PTOV1,RBMX,DDX42,CB
,HNRNPH1,RAB1A,ARCN1,HDAC2,BCLAF1,LUC7L3,CPSF2,TM9SF3,MDH2,RPN2,HNRNPL,ALKBH

PL27,HNRNPA3,C4ORF3,NCKAP1,RBMX,JPT1,CBX1,RAD23B,CHD2,SNRNP70,H3-3A,TERF2IP,TC
,CBX1,TNPO1,SNRNP70,H3-3A,HDGF,PRPF6,DPYSL2,MARS1,OPTN,G3BP2,SMARCA4,KDELR2,
L26,BEX3,RBBP4,RTN3,PFN1,MCL1,JPT2,PRKCSH,TADA2B,KHDRBS1,KLHL9,PFKL,RPS25,SFPC
ORF43,PAPOLA,COL1A2,LLGL1,RPL14,HK1,PSMD2,SLC7A5,CHD4,TMEM259,WDR1,HMGN2,AT
LC7A5,RPS14,CHD4,JUND,EIF5,CD81,WDR1,HMGN2,BEX3,SRSF4,RBBP4,PFN1,JPT2,PNN,RPL31
2,HLA-B,ARCN1,ARHGDIA,CAPRIN1,TOP2B,APPL1,HDAC2,BCLAF1,LUC7L3,MDH2,TOMM20,U2A
1,G3BP2,SMARCA4,SRSF1,ARGLU1,ILF3,PRKDC,HNRNPUL1,SON,DIDO1,SMARCE1,HNRNPM,SS
P1,L1CAM,LUC7L3,SRSF5,TM9SF3,U2AF2,MLXIP,HNRNPL,PARP1,TSPYL4,ABCA3,RPL27,HNRN
MORF4L2,TAF15,HNRNPA3,C4ORF3,CCT3,RBMX,RAD23B,SNRNP70,H3-3A,HDGF,TSPAN6,SELE
CLSTN1,PSMD2,KIF5B,SLC7A5,RPS14,CHD4,ATP1A1,JUND,RPS20,EIF5,RPL26,LASP1,CCT7,SRS
1,TM9SF3,MDH2,TOMM20,RAB27A,PABPN1,HNRNPL,TM9SF2,MORF4L2,ABCA3,ALKBH5,RPL27
KCSH,HNRNPC,LRPPRC,TADA2B,KHDRBS1,RPS25,GLG1,SFPQ,EIF5A,DDR1,RAB7A,MAGED2,RC
L14,HK1,ATF4,PSMD2,SLC7A5,EIF5,HMGN2,RPL26,LASP1,BEX3,CCT7,SRSF4,RBBP4,PFN1,JPT2
PSF2,PGRMC1,DDX39B,EID1,CTSB,MDH2,PGD,NOP53,PABPN1,GDI2,HNRNPL,PARP1,TSPYL4,M
,BEX3,RBBP4,MCL1,PNN,NES,NISCH,PRKCSH,HNRNPC,LRPPRC,XPR1,KHDRBS1,SLC3A2,RPS25
RPL31,PRKCSH,HNRNPC,LRPPRC,XPR1,KHDRBS1,XPO1,RPS25,GLG1,SFPQ,EIF5A,DDR1,RAB7A
15,RPL27,C4ORF3,ACO2,RBMX,DDX42,RAD23B,TNPO1,SNRNP70,H3-3A,HDGF,TSPAN6,TOP1,S
,HNRNPA3,ACO2,VPS35,RBMX,CBX1,RAD23B,CCT4,GCN1,SKI,SNRNP70,KDM6B,HDGF,PRPF6,
TMEM259,ATP2A2,RPL26,RBBP4,PFN1,MCL1,PNN,NISCH,PRKCSH,HNRNPC,TADA2B,KHDRBS1,X
DC,HNRNPUL1,DIDO1,SMARCE1,HNRNPM,SSB,GIGYF1,SRSF3,PHC2,ILF2,SF3A1,MAPK8IP1,HN
HNRNPUL1,KDM1A,DIDO1,FLNB,POLR2B,SMARCE1,SUPT6H,HNRNPM,SSB,SRSF3,ILF2,B2M,CS
TP6V1B2,HNRNPM,SSB,GIGYF1,SRSF3,ILF2,CSNK1D,SF3A1,ZSCAN18,TP53BP1,SYP,GRIPAP1,A
,RPL27,HNRNPA3,ACTR3,SNRNP70,PSMB2,H3-3A,TERF2IP,TAF7,G3BP2,SMARCA4,ARPC2,SRS
RPF6,DPYSL2,SRSF1,IPO7,ILF3,PRKDC,HNRNPUL1,KDM1A,DIDO1,TPM3,POLR2B,CORO1C,SUP
35,TLN1,CTNNB1,GOLM1,HNRNPDL,TARDBP,DDOST,HNRNPK,H1-10,MYH10,RPL36,CKAP5,SN
A1,JUND,EIF5,RPL26,LASP1,CCT7,RBBP4,MCL1,JPT2,RPL31,PRKCSH,HNRNPC,LRPPRC,KHDRB
MYM3,FUS,PELP1,FKBP4,STRN4,RALY,HDGFL2,GTF2I,TRAP1,TUBGCP6,RAB6A,MOAP1,RBM12,
B1A,STIP1,CDK16,APPL1,SKP1,BCLAF1,LUC7L3,CTSB,MDH2,RNF187,U2AF2,FDFT1,PPP1CB,PG
4H,SCARB2,RPS9,PJA2,RPL24,RPL17,COL1A2,LLGL1,RPL14,PTCD3,PSMD2,SLC7A5,RPS14,CXA
CNQ2,NOL4L,CD99,CST3,SLC39A7,DECR1,ZFR,ANP32B,RPS27A,NPLOC4,CTTN,ZMYM3,FUS,XR
HNRNPUL1,ACTN1,BNIP3L,FKBP8,DIDO1,CORO1C,SMARCE1,SUPT6H,HNRNPM,SRSF3,ILF2,SF3
NRNPH1,HLA-B,ARCN1,RAC1,CAPRIN1,TOP2B,SMCHD1,HDAC2,BCLAF1,LUC7L3,CPSF2,TOMM2
3A,HDGF,TSPAN6,TOP1,SMARCA4,SRSF1,PTBP1,ILF3,PRKDC,HNRNPUL1,HSPB1,SON,DIDO1,CO
,KDM1A,ZFAND5,DIDO1,FLNB,POLR2B,SMARCE1,SDF4,HNRNPM,SSB,GIGYF1,SRSF3,PHC2,ILF2
RNPDL,TARDBP,NDFIP1,ST3GAL2,RPL36,CKAP5,LAMP1,HNRNPH1,UBE2H,ARHGDIA,GLUL,LUC7
L,TSPYL4,TM9SF2,TAF15,ALKBH5,TBC1D16,ACO2,CCT3,VPS35,TNPO1,PRPF6,VEGFA,MARS1,S
NLGN2,CXADR,ATP1A1,CD81,WDR1,ATP2A2,CCT7,RTN3,MCL1,PNN,NISCH,LRPPRC,XPR1,XPO1,
PF6,DPYSL2,ARPC2,SRSF1,PTBP1,ILF3,PRKDC,HNRNPUL1,HSPB1,SON,ZFAND5,DIDO1,CORO1
1,HMGN2,ATP2A2,CCT7,SRSF4,RBBP4,RTN3,PFN1,PNN,NISCH,HNRNPC,LRPPRC,TADA2B,XPR1,
1,KHDRBS1,XPO1,SPTBN2,TKT,SFPQ,PNMA1,PDIA6,RAB7A,MMP14,DDX3X,RPL12,MRFAP1,H1-0
CHD1,HDAC2,IPO5,TOMM20,RNF187,MLXIP,PPP1CB,PABPN1,HNRNPL,PARP1,TSPYL4,TSR1,M
MSH6,TGOLN2,RBM6,THRAP3,XRN2,PTPN11,HTATSF1,SUN1,MTA2,CGGBP1,H6PD,REEP5,MDM
ABPN1,HNRNPL,MORF4L2,SSRP1,ALKBH5,HNRNPA3,PKN1,ACO2,VPS35,DDX42,CBX1,RAD23B,
BCLAF1,LUC7L3,MDH2,TOMM20,PABPN1,HNRNPL,PARP1,MORF4L2,ALKBH5,HNRNPA3,ACTR3,
RKDC,HNRNPUL1,ACTN1,MAP2K2,DIDO1,SMARCE1,XPNPEP3,SMARCC1,HNRNPM,SRSF3,ILF2,

D1,CORO1C,SUPT6H,HNRNPM,SRSF3,ILF2,SF3A1,SERBP1,KDM3B,BMPR2,PSMA7,RBBP7,TXLN
REBBP,XRN2,PTPN11,SUN1,MTA2,RTF1,MDM2,MECP2,CAB39,ARL6IP5,SUN2,CDK9,PSME3,RBF
RNPL,PARP1,MORF4L2,TAF15,ALKBH5,HNRNPA3,PTPRA,ACTR3,ACO2,SNRNP70,PSMB2,H3-3A,
MYM3,FUS,XRCC6,PELP1,ERGIC1,TCAF1,TPP1,GABARAPL1,HDGFL2,EFTUD2,SRSF2,TRAP1,NUC
P1,L1CAM,LUC7L3,CPSF2,PGRMC1,DDX39B,EID1,MDH2,TOMM20,U2AF2,ANXA5,PABPN1,HNRN
,HTATSF1,SUN1,LGALS1,SGSM2,MTA2,RTF1,REEP5,IGF1R,MDM2,FAM50A,AAMP,SUMO1,PSMD
BRD4,SLC39A7,GLUD1,TRA2B,UQCRC1,ZFR,AGPAT1,HNRNPA2B1,MAPRE2,C1ORF198,FUS,PEL
F9,GPR107,ZSCAN18,KDM3B,SMS,TXNIP,TP53BP1,POLR2E,RBBP7,STAT3,OS9,KDM5B,BAZ1B,

RSF3,ILF2,LSS,MESD,STRAP,MPRIP,KDM3B,TP53BP1,DDIT4,PSMA7,RBBP7,DCTN1,TXLNA,NAP1L
B,TNKS,HNRNPAB,C1ORF198,PRDX6,FUS,RAB14,HDGFL2,EFTUD2,SRSF2,RBM5,SPTY2D1,MSH
O1,SDF4,SMARCC1,DHX30,HNRNPM,SSB,SRSF3,SDHA,LARS1,ILF2,ISYNA1,UBQLN1,SF3A1,GPR
NRNPAB,ZMYM3,FUS,PELP1,ERGIC1,FKBP4,NUDCD3,TM9SF4,SRSF2,SLC7A1,MOAP1,RBM5,SR
MARCA4,MAPRE1,CCNL2,PBX2,SRSF1,ARGLU1,ATP5PB,PTBP1,ILF3,PRKDC,HNRNPUL1,SON,KD
BP7,LBH,CNTNAP1,H2AZ1,UBQLN2,UBA52,PPP2CB,SRSF6,TRA2B,COX7C,ZFR,CUL3,MCM3AP,F
CDH6,SFPQ,PEBP1,CDH2,DDR1,RAB7A,PRDX2,MAGED2,GNB2,H1-0,IK,JUN,CTNNB1,APLP2,HN
CCT5,RBBP7,MTA1,SF1,HIP1,PRCC,CD99,KPNA2,BRD2,UBE2G2,TRA2B,ZFR,HNRNPA2B1,ANP3
PTY2D1,MSH6,IST1,PSMC3,UBE3A,ATXN7L3,ANKRD12,DAPK3,PRDX3,EIF4A3,CASC4,XRN2,HTAT
L,ILF3,PRKDC,HNRNPUL1,HSPB1,KDM1A,CMIP,DIDO1,SMARCE1,HNRNPM,SRSF3,SDHA,ILF2,PF
PSME3,NDRG1,UNC45A,YWHAZ,ALCAM,SOX9,MED25,ZFP91,HSPE1,LMNA,RRBP1,CSRNP2,PHB,
QLN1,SF3A1,HNRNPD,SERBP1,KDM3B,DDIT4,PSMA7,ANP32A,RBBP7,EDC4,TXLNA,NDUFV1,SF1,
RAF7,ST3GAL6,SF1,PDAP1,GTPBP1,BRD4,UBA52,ATRX,TRA2B,MBTPS1,ZFR,NOMO1,AGPAT1,H
BP1,ACSL4,DLG4,MAN2B1,WASF3,PRKAR2B,DPP7,BACE1,CTCF,ATP6V0A1,GPRIN1,SHC1,FH,AP
1,CCNY,TBC1D14,TMEM131,RNMT,GSK3B,SYNGR1,TUBA1C,RAB3A,ANKRD50,CLCN6,NAPB,ST
2,MORF4L2,TAF15,ABCA3,HNRNPA3,RBMX,CBX1,CCT4,SNRNP70,H3-3A,KDM6B,HDGF,ABCA2,F
S1,GPAT4,MACF1,DNAJC5,APLP2,TSR2,PTGES3,UBE2H,RAB1A,ARCN1,CTBP1,ARHGDIA,CAPRI
BCLAF1,LUC7L3,SQLE,CPSF2,TM9SF3,TOMM20,U2AF2,CCT2,SUGP2,PABPN1,GDI2,HNRNPL,TM
1,FH,APOE,SYT1,PPT1,EPHA4,PDHB,WDR33,NNT,ZNF326,GSN,ARSA,FKBP3,MMP24
2,ANK2,ELAVL3,SCAMP5,FXR2,CLTA,TRIM2,COPE,APEH,UBE2J1,RTCB,RPL27A,FAM168A,NACA
,RBM17,NRP2,DSTYK,ZNF146,SEC31A,MAML1,EIF2S1,FAM193B,POSTN,SECISBP2L,ZNF207,C

S1,RPL36,LAMP1,HNRNPH1,ATP1A2,HLA-B,CCT6A,TOP2B,HDAC2,BCLAF1,LUC7L3,SQLE,CRPP
STYK,PRPF31,RTRAF,QRICH1,ABCD3,FAM193B,ZNF207,ALYREF

LG4,WASF3,DPP7,VDAC2,UQCRFS1,BACE1,ATP6V0A1,SLC1A3,FH,TUT4,NFASC,ABCD3,PRNP,PP
1,UBQLN1,SF3A1,GPR107,HNRNPD,SERBP1,ARFGEF2,TXNIP,CCT5,POLR2E,TRAF4,TRAF7,MTA
,KRT8,RBM5,YIPF3,IST1,DHCR7,GOT2,YY1,RBM6,NDUFA13,ACAT2,HTATSF1,CGGBP1,KTN1,PC
7,GLOD4,SUN2,WIZ,WDR11,MRPS6,TMEM43,ZFP91,ALDH9A1,CLCN6,KDM2A,SLC16A1,ZDHHC
CD99,BRD4,UBA52,ATRX,EP400,GLUD1,EVL,TRA2B,UQCRC1,ZFR,HNRNPA2B1,ANP32B,RPS27A
EM259,RPS20,WDR1,HMGN2,ATP2A2,BEX3,CCT7,MCL1,JPT2,NISCH,TADA2B,CSNK1E,PFKL,HD
RSA,FKBP3

3CD3,AKAP17A,GNAL,WDR33,RAD50,ZNF326,NOP58,PAM,NUTF2,CTSA,ARSA

SM14A,PNISR,RIC8A,NCSTN,PLEKHM2,MOAP1,RBM5,YIPF3,IST1,DHCR7,AHCY,GSK3A,PSMC3,U

ARP1,TM9SF2,MORF4L2,HNRNPA3,VPS35,CBX1,CCT4,SNRNP70,H3-3A,KDM6B,HDGF,ABCA2,PR

HM13,EXOSC10,CACNA1H,TFRC,CDKN1B,PMPCA,LRCH4,LZTS2,SRRT,MAP2K4,RFX7,B4GALT2,

3A3,RUVBL2,PCM1,EMC1,ALDH18A1,ATP1B3,PSMD7,KCTD20,HTRA1,ECH1,ATF7IP,FAM8A1,KPN

AN1,GSTP1,MARCKS,SMG1,CALM1,NPM1,PLD3,HSPD1,USP11,COL18A1,SNRNP200,CSDE1,EIF3
NP200,CSDE1,EIF3C,ATP5F1A,EIF4B,VCP,MARCKSL1,UBB,AFF4,TRIM28,MATR3,YWHAG,CKB,RI
A,EIF4B,VCP,MARCKSL1,UBB,AFF4,TRIM28,MATR3,YWHAG,CKB,RPL37A,LRRC28,MAPT,FN1,SP
YWHAG,RPL37A,LRRC28,MAPT,FN1,SPEN,NONO,DDB1,YWHAQ,HACD3,DCHS1,SET,EGLN1,INA,M
S,SMG1,CALM1,NPM1,PLD3,HSPD1,USP11,COL18A1,SNRNP200,CSDE1,EIF3C,ATP5F1A,EIF4B,V
F4,TRIM28,MATR3,YWHAG,RPL37A,LRRC28,SPEN,NONO,DDB1,YWHAQ,HACD3,DCHS1,SET,EGL
F3C,ATP5F1A,EIF4B,VCP,MARCKSL1,UBB,AFF4,TRIM28,MATR3,YWHAG,CKB,RPL37A,LRRC28,FN
3C,ATP5F1A,EIF4B,VCP,MARCKSL1,UBB,AFF4,TRIM28,MATR3,YWHAG,RPL37A,LRRC28,MAPT,S
EIF3C,ATP5F1A,EIF4B,VCP,MARCKSL1,UBB,AFF4,TRIM28,MATR3,YWHAG,RPL37A,LRRC28,FN1,S
FF4,TRIM28,MATR3,YWHAG,RPL37A,LRRC28,FN1,SPEN,NONO,DDB1,YWHAQ,HACD3,DCHS1,SET
FF4,TRIM28,MATR3,YWHAG,RPL37A,LRRC28,FN1,SPEN,NONO,DDB1,YWHAQ,HACD3,DCHS1,SET
1,EIF3C,ATP5F1A,EIF4B,VCP,MARCKSL1,UBB,AFF4,TRIM28,MATR3,YWHAG,RPL37A,LRRC28,FN
2,CHD3,VAPA,HMGB1,POLR2A,TPT1,RPS4X,RPS3A,TUBB4B,UCHL1,RPL11,MORF4L1,BTF3,XAF
USP11,COL18A1,SNRNP200,CSDE1,EIF3C,ATP5F1A,EIF4B,VCP,MARCKSL1,UBB,AFF4,TRIM28,M
USP11,COL18A1,SNRNP200,CSDE1,EIF3C,ATP5F1A,EIF4B,VCP,MARCKSL1,UBB,AFF4,TRIM28,M
M28,MATR3,YWHAG,RPL37A,LRRC28,SPEN,NONO,DDB1,YWHAQ,HACD3,DCHS1,SET,EGLN1,RPL
MARCKSL1,UBB,AFF4,TRIM28,MATR3,YWHAG,RPL37A,LRRC28,FN1,SPEN,NONO,DDB1,YWHAQ,
MARCKSL1,UBB,AFF4,TRIM28,MATR3,YWHAG,RPL37A,LRRC28,FN1,SPEN,NONO,DDB1,YWHAQ,
18A1,SNRNP200,CSDE1,EIF3C,ATP5F1A,EIF4B,VCP,MARCKSL1,UBB,AFF4,TRIM28,MATR3,YWH
L1,UBB,AFF4,TRIM28,MATR3,YWHAG,RPL37A,LRRC28,SPEN,NONO,DDB1,YWHAQ,HACD3,DCHS
PS4X,RPL18,STMN1,TUBB4B,BTF3,ANXA2,RPL10,RPS17,VCL,DNAJA1,CYB5R3,PAFAH1B1,AKAP
IGF2,EIF3C,ATP5F1A,EIF4B,VCP,MARCKSL1,UBB,AFF4,TRIM28,MATR3,YWHAG,RPL37A,LRRC2
CP,MARCKSL1,UBB,AFF4,TRIM28,MATR3,YWHAG,RPL37A,LRRC28,FN1,SPEN,NONO,DDB1,YWH
CP,MARCKSL1,UBB,AFF4,TRIM28,MATR3,YWHAG,RPL37A,LRRC28,FN1,SPEN,NONO,DDB1,YWH
S4X,RPS3A,RPL18,STMN1,TUBB4B,UCHL1,RPL11,MORF4L1,BTF3,AGRN,RPL10,RPS17,DNAJA1,
UCHL1,RPL11,MORF4L1,BTF3,AGRN,RPL10,WSB1,RPS17,DNAJA1,CYB5R3,PAFAH1B1,SF3B2,IT
1,USP11,COL18A1,SNRNP200,CSDE1,EIF3C,ATP5F1A,EIF4B,VCP,MARCKSL1,UBB,AFF4,TRIM28,
P,MARCKSL1,UBB,AFF4,TRIM28,MATR3,YWHAG,RPL37A,LRRC28,SPEN,NONO,DDB1,YWHAQ,HA
,YWHAG,LRRC28,FN1,SPEN,NONO,DDB1,YWHAQ,DCHS1,SET,EGLN1,MRC2,RPL19,LDHB,SF3B1
,YWHAG,LRRC28,FN1,SPEN,NONO,DDB1,YWHAQ,DCHS1,SET,EGLN1,MRC2,RPL19,LDHB,SF3B1
SDE1,EIF3C,ATP5F1A,EIF4B,VCP,MARCKSL1,UBB,AFF4,TRIM28,MATR3,YWHAG,RPL37A,LRRC2
R3,YWHAG,RPL37A,LRRC28,FN1,SPEN,NONO,DDB1,YWHAQ,HACD3,DCHS1,SET,EGLN1,MRC2,R
R3,YWHAG,RPL37A,LRRC28,FN1,SPEN,NONO,DDB1,YWHAQ,HACD3,DCHS1,SET,EGLN1,MRC2,R
3,SSBP2,CHD3,BASP1,VAPA,HMGB1,POLR2A,RPS4X,RPL18,STMN1,ITGB1,TUBB4B,BTF3,ANXA2
ACD3,DCHS1,SET,EGLN1,RPL19,LDHB,PKD1,SF3B1,PDIA3,SSBP2,CHD3,VAPA,HMGB1,POLR2A,T
C,ATP5F1A,EIF4B,VCP,MARCKSL1,UBB,AFF4,TRIM28,MATR3,YWHAG,RPL37A,LRRC28,MAPT,FN
MGB1,POLR2A,RPS3A,STMN1,TUBB4B,BTF3,AGRN,XAF1,RPL10,RPS17,DNAJA1,CYB5R3,PAFAH
ONO,DDB1,YWHAQ,HACD3,DCHS1,SET,EGLN1,MRC2,RPL19,LDHB,PKD1,SF3B1,PDIA3,SSBP2,C
ONO,DDB1,YWHAQ,HACD3,DCHS1,SET,EGLN1,MRC2,RPL19,LDHB,PKD1,SF3B1,PDIA3,SSBP2,C
G,RPL37A,LRRC28,SPEN,NONO,YWHAQ,HACD3,DCHS1,SET,EGLN1,LDHB,PKD1,SF3B1,PDIA3,S
BTF3,ANXA2,RPL10,RPS17,VCL,DNAJA1,CYB5R3,PAFAH1B1,AKAP12,MAP4,SF3B2,GTF2F1,HNRN
SDE1,EIF3C,ATP5F1A,EIF4B,VCP,MARCKSL1,UBB,AFF4,TRIM28,MATR3,YWHAG,RPL37A,LRRC2
SDE1,EIF3C,ATP5F1A,EIF4B,VCP,MARCKSL1,UBB,AFF4,TRIM28,MATR3,YWHAG,RPL37A,LRRC2
1A,EIF4B,VCP,MARCKSL1,UBB,AFF4,TRIM28,MATR3,YWHAG,CKB,RPL37A,LRRC28,FN1,SPEN,N
1A,EIF4B,VCP,MARCKSL1,UBB,AFF4,TRIM28,MATR3,YWHAG,CKB,RPL37A,LRRC28,FN1,SPEN,N

RC2,LDHB,SF3B1,PDIA3,SSBP2,CHD3,VAPA,HMGB1,TPT1,RPS4X,ITGB1,TUBB4B,BTF3,ANXA2,RP
4B,VCP,MARCKSL1,UBB,AFF4,TRIM28,MATR3,YWHAG,CKB,RPL37A,LRRC28,FN1,SPEN,NONO,D
4B,VCP,MARCKSL1,UBB,AFF4,TRIM28,MATR3,YWHAG,CKB,RPL37A,LRRC28,FN1,SPEN,NONO,D
5F1A,EIF4B,VCP,MARCKSL1,UBB,AFF4,TRIM28,MATR3,YWHAG,RPL37A,LRRC28,FN1,SPEN,NON
28,MATR3,YWHAG,RPL37A,LRRC28,SPEN,NONO,DDB1,YWHAQ,HACD3,DCHS1,SET,EGLN1,MRC2
B,VCP,MARCKSL1,UBB,AFF4,TRIM28,MATR3,YWHAG,CKB,RPL37A,LRRC28,FN1,SPEN,NONO,DD
DE1,EIF3C,ATP5F1A,EIF4B,VCP,MARCKSL1,UBB,AFF4,TRIM28,MATR3,YWHAG,RPL37A,LRRC28,
RC28,SPEN,NONO,DDB1,YWHAQ,HACD3,DCHS1,SET,EGLN1,INA,MRC2,RPL19,LDHB,SF3B1,PDIA
RRC28,MAPT,FN1,SPEN,NONO,DDB1,YWHAQ,HACD3,DCHS1,SET,EGLN1,MRC2,RPL19,LDHB,SF3
RRC28,MAPT,FN1,SPEN,NONO,DDB1,YWHAQ,HACD3,DCHS1,SET,EGLN1,MRC2,RPL19,LDHB,SF3
ATP5F1A,EIF4B,VCP,MARCKSL1,UBB,AFF4,TRIM28,MATR3,YWHAG,RPL37A,LRRC28,MAPT,FN1,S
1,LDHB,PKD1,SF3B1,PDIA3,SSBP2,CHD3,BASP1,VAPA,HMGB1,POLR2A,TPT1,RPS3A,RPL18,TUB
YWHAG,RPL37A,LRRC28,FN1,SPEN,NONO,DDB1,YWHAQ,HACD3,DCHS1,SET,EGLN1,MRC2,RPL1
YWHAG,RPL37A,LRRC28,FN1,SPEN,NONO,DDB1,YWHAQ,HACD3,DCHS1,SET,EGLN1,MRC2,RPL1
B1,MAP4,SF3B2,CDC42BPB,GTF2F1,HNRNPU,OGA,NFE2L1,SMARCA5,CTNNA1,RPS16,RPL23,CN
RC28,SPEN,NONO,DDB1,YWHAQ,HACD3,DCHS1,SET,EGLN1,RPL19,LDHB,PKD1,SF3B1,PDIA3,SS
LN1,MRC2,LDHB,SF3B1,PDIA3,SSBP2,CHD3,VAPA,HMGB1,POLR2A,TPT1,RPS4X,RPS3A,RPL18,IT
MARCKSL1,UBB,AFF4,TRIM28,MATR3,YWHAG,CKB,RPL37A,LRRC28,SPEN,NONO,DDB1,YWHAC
MARCKSL1,UBB,AFF4,TRIM28,MATR3,YWHAG,CKB,RPL37A,LRRC28,SPEN,NONO,DDB1,YWHAC
RIM28,MATR3,YWHAG,RPL37A,LRRC28,SPEN,NONO,DDB1,YWHAQ,HACD3,DCHS1,SET,EGLN1,M
RIM28,MATR3,YWHAG,RPL37A,LRRC28,SPEN,NONO,DDB1,YWHAQ,HACD3,DCHS1,SET,EGLN1,M
T1,RPS4X,RPS3A,TUBB4B,RPL11,MORF4L1,BTF3,NCAM1,AGRN,ZMIZ1,XAF1,RPL10,RPS17,VCL,
4B,VCP,MARCKSL1,UBB,AFF4,TRIM28,MATR3,YWHAG,RPL37A,LRRC28,SPEN,NONO,DDB1,YWH
EGLN1,RPL19,LDHB,SF3B1,PDIA3,SSBP2,CHD3,VAPA,POLR2A,STMN1,ITGB1,TUBB4B,UCHL1,M
HAG,RPL37A,LRRC28,MAPT,FN1,SPEN,NONO,DDB1,YWHAQ,HACD3,DCHS1,SET,EGLN1,MRC2,R
B,SF3B1,PDIA3,SSBP2,CHD3,BASP1,VAPA,HMGB1,POLR2A,RPS4X,RPS3A,STMN1,TUBB4B,RPL1
PL11,MORF4L1,BTF3,NCAM1,AGRN,XAF1,RPL10,RPS17,DNAJA1,CYB5R3,PAFAH1B1,SF3B2,ITM
HACD3,DCHS1,SET,EGLN1,MRC2,RPL19,PKD1,SF3B1,PDIA3,SSBP2,CHD3,VAPA,HMGB1,RPS4X,
BP2,CHD3,VAPA,HMGB1,POLR2A,RPS3A,RPL18,ITGB1,BTF3,NCAM1,AGRN,XAF1,RPL10,RPS17,
BP2,CHD3,VAPA,HMGB1,POLR2A,RPS3A,RPL18,ITGB1,BTF3,NCAM1,AGRN,XAF1,RPL10,RPS17,
3B1,PDIA3,SSBP2,BASP1,VAPA,HMGB1,POLR2A,TPT1,RPS4X,RPS3A,RPL18,TUBB4B,RPL11,MOF
S1,SET,EGLN1,RPL19,LDHB,PKD1,SF3B1,PDIA3,SSBP2,CHD3,BASP1,VAPA,HMGB1,POLR2A,TPT
,VAPA,HMGB1,POLR2A,STMN1,TUBB4B,RPL11,BTF3,AGRN,XAF1,RPL10,MARCHF6,RPS17,DNA
F4B,VCP,MARCKSL1,UBB,AFF4,TRIM28,MATR3,YWHAG,RPL37A,LRRC28,FN1,SPEN,NONO,DDB1
F4B,VCP,MARCKSL1,UBB,AFF4,TRIM28,MATR3,YWHAG,RPL37A,LRRC28,FN1,SPEN,NONO,DDB1
P,UBB,AFF4,TRIM28,MATR3,YWHAG,RPL37A,LRRC28,FN1,SPEN,NONO,DDB1,HACD3,DCHS1,SET
A,EIF4B,VCP,MARCKSL1,UBB,AFF4,TRIM28,MATR3,YWHAG,CKB,RPL37A,LRRC28,FN1,SPEN,NO
AG,RPL37A,LRRC28,FN1,SPEN,NONO,DDB1,HACD3,DCHS1,SET,EGLN1,MRC2,RPL19,LDHB,PKD
,MARCHF6,RPS17,DNAJA1,CYB5R3,PAFAH1B1,MAP4,SF3B2,ITM2B,GTF2F1,HNRNPU,NFE2L1,SN
,NONO,DDB1,YWHAQ,HACD3,DCHS1,SET,EGLN1,MRC2,PKD1,SF3B1,PDIA3,SSBP2,CHD3,VAPA,
4X,RPS3A,RPL18,STMN1,RPL11,MORF4L1,BTF3,ANXA2,XAF1,RPL10,RPS17,VCL,DNAJA1,PAFAH
PL11,BTF3,ANXA2,AGRN,XAF1,RPL10,MARCHF6,RPS17,VCL,DNAJA1,CYB5R3,PAFAH1B1,MAP4,S
X,RPS3A,RPL18,TUBB4B,RPL11,MORF4L1,BTF3,ANXA2,XAF1,RPL10,RPS17,CYB5R3,PAFAH1B1,
FN1,SPEN,NONO,DDB1,HACD3,DCHS1,SET,EGLN1,MRC2,RPL19,LDHB,PKD1,SF3B1,PDIA3,SSBP
NONO,DDB1,YWHAQ,HACD3,DCHS1,SET,EGLN1,RPL19,LDHB,PKD1,SF3B1,PDIA3,SSBP2,CHD3,VA

RC2,LDHB,SF3B1,PDIA3,SSBP2,CHD3,VAPA,HMGB1,POLR2A,TPT1,RPS4X,RPL18,TUBB4B,BTF3,A
G,RPL37A,LRRC28,SPEN,NONO,DDB1,YWHAQ,DCHS1,SET,EGLN1,MRC2,RPL19,LDHB,SF3B1,PD
WHAG,RPL37A,LRRC28,FN1,SPEN,NONO,DDB1,YWHAQ,HACD3,DCHS1,SET,EGLN1,MRC2,RPL19
WHAG,RPL37A,LRRC28,FN1,SPEN,NONO,DDB1,YWHAQ,HACD3,DCHS1,SET,EGLN1,MRC2,RPL19
H,SCARB2,LPCAT1,BRD3,PJA2,RPL24,KPNB1,PAPOLA,LLGL1,RPL14,RHOA,PTCD3,CBX5,ATF4,PS
MRC2,RPL19,LDHB,PKD1,SF3B1,PDIA3,SSBP2,CHD3,VAPA,HMGB1,POLR2A,TPT1,RPS4X,RPS3A,
SPEN,NONO,DDB1,YWHAQ,DCHS1,SET,EGLN1,MRC2,RPL19,SF3B1,PDIA3,SSBP2,CHD3,VAPA,HM
PL11,MORF4L1,BTF3,ANXA2,RPL10,RPS17,VCL,DNAJA1,PAFAH1B1,MAP4,SF3B2,CDC42BPB,ITM
,CSDE1,EIF3C,ATP5F1A,EIF4B,VCP,MARCKSL1,UBB,AFF4,TRIM28,MATR3,YWHAG,CKB,RPL37A,L
CHS1,SET,EGLN1,MRC2,RPL19,PKD1,SF3B1,PDIA3,SSBP2,CHD3,BASP1,VAPA,HMGB1,POLR2A,T
CHS1,SET,EGLN1,MRC2,RPL19,PKD1,SF3B1,PDIA3,SSBP2,CHD3,BASP1,VAPA,HMGB1,POLR2A,T
B4B,RPL11,MORF4L1,BTF3,AGRN,XAF1,RPL10,RPS17,VCL,DNAJA1,CYB5R3,PAFAH1B1,SF3B2,C
ASP1,VAPA,HMGB1,TPT1,RPS3A,TUBB4B,RPL11,BTF3,ANXA2,AGRN,XAF1,RPL10,WSB1,RPS17,V
3B1,PDIA3,SSBP2,BASP1,VAPA,HMGB1,POLR2A,RPS4X,RPS3A,RPL18,RPL11,MORF4L1,BTF3,XA
3B1,PDIA3,SSBP2,BASP1,VAPA,HMGB1,POLR2A,RPS4X,RPS3A,RPL18,RPL11,MORF4L1,BTF3,XA
RPL18,RPL11,MORF4L1,BTF3,ANXA2,AGRN,XAF1,RPL10,RPS17,VCL,CYB5R3,PAFAH1B1,AKAP12
8,TUBB4B,RPL11,ANXA2,RPL10,WSB1,RPS17,DNAJA1,PAFAH1B1,SF3B2,CDC42BPB,ITM2B,GTF
POLR2A,TPT1,RPS4X,RPS3A,RPL18,STMN1,TUBB4B,RPL11,BTF3,ANXA2,XAF1,RPL10,RPS17,VCL
,RPS3A,RPL18,RPL11,BTF3,ANXA2,XAF1,RPL10,WSB1,RPS17,VCL,DNAJA1,CYB5R3,PAFAH1B1,A
A,HMGB1,POLR2A,TPT1,RPS4X,RPL18,STMN1,TUBB4B,RPL11,MORF4L1,BTF3,ANXA2,AGRN,RPL
RPS3A,RPL18,STMN1,TUBB4B,RPL11,MORF4L1,BTF3,RPL10,MARCHF6,RPS17,VCL,DNAJA1,PAF
MARCKS,SMG1,CALM1,NPM1,PLD3,HSPD1,USP11,COL18A1,SNRNP200,CSDE1,EIF3C,ATP5F1A,E
ADS2,RPS16,CPE,CNN3,RPSA,TPM4,CTSD,EIF4H,SCARB2,RTN4,BRD3,PJA2,RPL24,KPNB1,PAPO
SF3B1,PDIA3,SSBP2,CHD3,BASP1,VAPA,HMGB1,POLR2A,TPT1,RPS4X,RPS3A,RPL18,STMN1,TUB
D4,JUND,TMEM259,RPS20,EIF5,CD81,BEX3,CCT7,RBBP4,PFN1,PNN,RPL31,PRKCSH,HNRNPC,LF
O,DDB1,YWHAQ,DCHS1,SET,EGLN1,MRC2,RPL19,LDHB,SF3B1,PDIA3,SSBP2,CHD3,BASP1,VAPA,
HB,PKD1,SF3B1,PDIA3,SSBP2,VAPA,HMGB1,RPS4X,RPL18,STMN1,TUBB4B,UCHL1,MORF4L1,BT
,HMGB1,POLR2A,TPT1,RPS4X,RPS3A,RPL18,ITGB1,TUBB4B,RPL11,BTF3,ANXA2,XAF1,RPL10,M
RC28,SPEN,NONO,DDB1,HACD3,SET,EGLN1,RPL19,PKD1,SF3B1,PDIA3,SSBP2,VAPA,HMGB1,POL
ACD3,DCHS1,SET,EGLN1,RPL19,LDHB,PKD1,SF3B1,PDIA3,SSBP2,CHD3,VAPA,HMGB1,POLR2A,T
D3,DCHS1,SET,EGLN1,MRC2,RPL19,LDHB,PDIA3,SSBP2,BASP1,VAPA,HMGB1,POLR2A,TPT1,RPS
JA1,PAFAH1B1,SF3B2,CDC42BPB,ITM2B,ZNF770,AP2B1,GTF2F1,HNRNPU,PGAM1,NFE2L1,SMA
X3X,MRFAP1,H1-0,RPS5,RPS7,IK,RPL35,HNRNPDL,TARDBP,ARID1A,NDFIP1,ST3GAL2,HNRNPK,H
4L1,ANXA2,RPL10,RPS17,VCL,CYB5R3,PAFAH1B1,MAP4,SF3B2,CDC42BPB,ITM2B,ZNF770,AP2
,SET,EGLN1,RPL19,LDHB,PKD1,SF3B1,PDIA3,SSBP2,VAPA,HMGB1,POLR2A,TPT1,RPS4X,RPS3A
D3,DCHS1,SET,EGLN1,RPL19,LDHB,PKD1,SF3B1,PDIA3,SSBP2,VAPA,HMGB1,POLR2A,RPS4X,RF
IONO,DDB1,HACD3,DCHS1,SET,EGLN1,MRC2,LDHB,SF3B1,PDIA3,SSBP2,CHD3,BASP1,VAPA,HM
S3A,RPL18,TUBB4B,MORF4L1,BTF3,ANXA2,RPL10,WSB1,RPS17,VCL,DNAJA1,MAP4,SF3B2,CDC
D1,SF3B1,PDIA3,SSBP2,CHD3,VAPA,HMGB1,POLR2A,TPT1,RPS4X,RPS3A,RPL18,TUBB4B,RPL11
A,HMGB1,POLR2A,TPT1,RPS4X,RPS3A,RPL18,STMN1,TUBB4B,MORF4L1,BTF3,ANXA2,RPL10,R
PS3A,RPL18,RPL11,MORF4L1,BTF3,AGRN,MARCHF6,WSB1,RPS17,DNAJA1,CYB5R3,PAFAH1B1,
AH1B1,SF3B2,ITM2B,ZNF770,AP2B1,HNRNPU,SMARCA5,LARP1,HNRNPA0,FADS2,RPS16,RPL23,
D1,SF3B1,PDIA3,CHD3,BASP1,VAPA,HMGB1,POLR2A,TPT1,RPS4X,RPL18,TUBB4B,RPL11,MORF
31,POLR2A,TPT1,RPS4X,RPS3A,RPL18,TUBB4B,RPL11,BTF3,ANXA2,XAF1,RPL10,WSB1,RPS17,V
X,RPS3A,RPL18,STMN1,TUBB4B,MORF4L1,BTF3,ANXA2,RPL10,RPS17,VCL,DNAJA1,PAFAH1B1,M

,POLR2A,RPS3A,ITGB1,RPL11,MORF4L1,BTF3,ANXA2,AGRN,RPL10,WSB1,RPS17,VCL,CYB5R3,
M1,SMARCA5,LARP1,CTNNA1,FADS2,RPS16,RPL23,RPL29,RPSA,TNS3,CTSD,EIF4H,SCARB2,LP
,DDX3X,RPS15,CLTC,MAGED2,RCN1,H1-0,RPS7,IK,GJA1,ANXA6,TARDBP,DDOST,HNRNPK,H1-1
TM2B,GTF2F1,GOLGA8A,HNRNPU,OGA,NFE2L1,SMARCA5,LARP1,CTNNA1,FADS2,RPS16,RPL23
N,NONO,DDB1,YWHAQ,DCHS1,SET,EGLN1,MRC2,RPL19,LDHB,SF3B1,PDIA3,SSBP2,BASP1,VAPA
NN,PRKCSH,HNRNPC,LRPPRC,KHDRBS1,XPO1,RPS25,TKT,GLG1,SFPQ,PEBP1,RAB7A,CCN2,PR
S17,PAFAH1B1,SF3B2,ITM2B,MAT2A,GTF2F1,HNRNPU,OGA,NFE2L1,SMARCA5,CTNNA1,FADS2,
TRIM28,MATR3,YWHAG,RPL37A,LRRC28,FN1,SPEN,NONO,DDB1,HACD3,DCHS1,SET,EGLN1,MR
TRIM28,MATR3,YWHAG,RPL37A,LRRC28,FN1,SPEN,NONO,DDB1,HACD3,DCHS1,SET,EGLN1,MR
A,HMGB1,POLR2A,TPT1,RPS4X,RPS3A,RPL18,TUBB4B,RPL11,MORF4L1,BTF3,ANXA2,RPL10,R
PL11,BTF3,RPL10,RPS17,DNAJA1,PAFAH1B1,MAP4,SF3B2,ITM2B,GTF2F1,HNRNPU,OGA,SMAR
F3B2,ZNF770,AP2B1,GTF2F1,HNRNPU,CTNNA1,RPL23,RPL29,RPSA,TPM4,CTSD,EIF4H,SCARB2
F3B2,ZNF770,AP2B1,GTF2F1,HNRNPU,CTNNA1,RPL23,RPL29,RPSA,TPM4,CTSD,EIF4H,SCARB2
10,RPS17,VCL,CYB5R3,MAP4,SF3B2,CDC42BPB,GTF2F1,HNRNPU,OGA,NFE2L1,LARP1,CTNNA1
,NFE2L1,LARP1,HNRNPA0,CTNNA1,FADS2,RPL23,CNN3,TPM4,CTSD,EIF4H,SCARB2,BRD3,C1OR
1,RPL10,RPS17,CYB5R3,PAFAH1B1,MAP4,SF3B2,ITM2B,MAT2A,GTF2F1,HNRNPU,OGA,PGAM1,
1,CTNNA1,RPL23,CNN3,RPL29,RPSA,EIF4H,LPCAT1,BRD3,RPS9,XRCC5,PJA2,RPL24,KPNB1,LL
1,HNRNPU,PGAM1,NFE2L1,SMARCA5,CTNNA1,FADS2,RPL23,RPL29,RPSA,TPM4,EIF4H,LPCAT1,
,RPS3A,RPL18,STMN1,TUBB4B,MORF4L1,BTF3,ANXA2,AGRN,RPL10,RPS17,VCL,DNAJA1,CYB5
RIM28,MATR3,YWHAG,RPL37A,LRRC28,FN1,SPEN,NONO,DDB1,YWHAQ,HACD3,DCHS1,SET,EG
,EGLN1,MRC2,RPL19,LDHB,SF3B1,PDIA3,SSBP2,BASP1,VAPA,HMGB1,TPT1,RPS4X,RPS3A,RPL1
TNNA1,RPS16,RPL23,RPL29,RPSA,TPM4,EIF4H,BRD3,XRCC5,RPL24,RPL17,C1ORF43,PAPOLA,C
2,SKP1,BCLAF1,LUC7L3,CPSF2,CTSB,NUCB1,TM9SF3,MDH2,TOMM20,EEF1A2,SUGP2,GAS7,NC
D1,SF3B1,PDIA3,SSBP2,CHD3,VAPA,HMGB1,POLR2A,TPT1,RPS4X,RPS3A,RPL18,STMN1,TUBB4
NPU,OGA,NFE2L1,SMARCA5,HNRNPA0,FADS2,RPS16,RPL23,RPSA,TPM4,CTSD,EIF4H,SCARB2,B
2A,RPS4X,RPS3A,RPL18,MORF4L1,BTF3,NCAM1,AGRN,XAF1,RPL10,RPS17,VCL,CYB5R3,PAFAH
2A,RPS4X,RPS3A,RPL18,MORF4L1,BTF3,NCAM1,AGRN,XAF1,RPL10,RPS17,VCL,CYB5R3,PAFAH
1,GTF2F1,HNRNPU,NFE2L1,SMARCA5,LARP1,HNRNPA0,CTNNA1,FADS2,RPS16,RPL23,RPSA,TP
4,TMEM259,RPS20,WDR1,HMGN2,RPL26,CCT7,SRSF4,RBBP4,MCL1,PNN,RPL31,NISCH,PRKCS
NA1,RPL23,CNN3,RPSA,TPM4,CTSD,EIF4H,LPCAT1,BRD3,XRCC5,PJA2,RPL24,SLC16A7,C1ORF4
CD3,ZNF865,CLSTN1,ATF4,PSMD2,KIF5B,CHD4,JUND,TMEM259,WDR1,HMGN2,BEX3,CCT7,SR
RPL23,RPL29,TPM4,EIF4H,LPCAT1,XRCC5,RPL24,RPL17,C1ORF43,PAPOLA,COL1A2,RPL14,HK1
3,RPL10,RPS17,DNAJA1,CYB5R3,PAFAH1B1,MAP4,SF3B2,GTF2F1,HNRNPU,OGA,PGAM1,NFE2
RBS1,XPO1,RPS25,TKT,SFPQ,RAB7A,ACTN4,PRDX2,RPS15,MAGED2,RCN1,H1-0,RPS5,RPS7,IK,
3,CCT7,SRSF4,RBBP4,PFN1,MCL1,PNN,HNRNPC,LRPPRC,TADA2B,XPR1,KHDRBS1,RPS25,TKT,S
SF4,RBBP4,RTN3,PNN,PRKCSH,HNRNPC,LRPPRC,TADA2B,XPO1,CSNK1E,RPS25,TKT,SFPQ,DD
A,RPL11,MORF4L1,BTF3,AGRN,XAF1,RPL10,RPS17,DNAJA1,CYB5R3,PAFAH1B1,MAP4,SF3B2,C
NN3,EIF4H,LPCAT1,BRD3,XRCC5,PJA2,MSN,RPL24,RPL17,C1ORF43,PAPOLA,COL1A2,RPL14,C
NRNPU,NFE2L1,SMARCA5,CTNNA1,FADS2,RPL23,RPL29,RPSA,TNS3,CTSD,EIF4H,SCARB2,RPS9
,RPSA,TPM4,CTSD,EIF4H,LPCAT1,RPS24,BRD3,XRCC5,PJA2,RPL24,RPL17,PAPOLA,COL1A2,LL
O1,RPS25,SFPQ,RAB7A,DDX3X,RPS15,MRFAP1,MAGED2,RCN1,H1-0,RPS7,IK,RPL35,TLN1,CTN
RPL11,MORF4L1,BTF3,AGRN,XAF1,RPL10,WSB1,RPS17,VCL,CYB5R3,SF3B2,CDC42BPB,AP2B1,
L24,C1ORF43,PAPOLA,COL1A2,RPL14,RPL9,CLSTN1,ATF4,KIF5B,RPS14,JUND,RPS20,EIF5,CD
NXA6,APLP2,TARDBP,DDOST,ST3GAL2,WLS,HNRNPK,INIP,MYH10,RPL36,YWHAB,SND1,LAMP1,
0,CTNNA1,CNN3,RPSA,TPM4,TNS3,CTSD,EIF4H,SCARB2,LPCAT1,BRD3,XRCC5,RPL24,C1ORF4

GL1,RPL14,PTCD3,RPL9,KIF5B,RPS14,CHD4,JUND,RPS20,EIF5,CD81,RPL26,LASP1,BEX3,CCT7,
,CTNNA1,RPL23,RPSA,TPM4,EIF4H,SCARB2,LPCAT1,XRCC5,MSN,RPL24,KPNB1,RPL17,PAPOLA,
SF3,TOMM20,EEF1A2,SUGP2,GAS7,NOP53,HNRNPL,PARP1,TM9SF2,MORF4L2,TAF15,LMNB2,F
SRSF4,RBBP4,PNN,PRKCSH,HNRNPC,LRPPRC,XPR1,KHDRBS1,XPO1,RPS25,SFPQ,RAB7A,CCN2,
2B1,GTF2F1,HNRNPU,OGA,NFE2L1,LARP1,CTNNA1,RPL23,RPSA,TPM4,EIF4H,SCARB2,LPCAT1,X
,RPL31,NISCH,HNRNPC,LRPPRC,TADA2B,XPR1,KHDRBS1,XPO1,CSNK1E,HDLBP,RPS25,TKT,GLG
,MSN,KPNB1,C1ORF43,PAPOLA,COL1A2,RPL14,HK1,ATF4,KIF5B,SLC7A5,RPS14,CXADR,CHD4,J
5,CLSTN1,KIF5B,RPS14,CXADR,CHD4,TMEM259,EIF5,HMGN2,BEX3,CCT7,SRSF4,RBBP4,PNN,NI
T,SFPQ,ACSL6,RAB7A,DDX3X,MAGED2,H1-0,APLP1,RPS7,IK,RPL35,GPAT4,TLN1,GJA1,GOLM1,I
GA8A,HNRNPU,OGA,NFE2L1,SMARCA5,HNRNPA0,CTNNA1,FADS2,RPL23,CNN3,RPL29,RPSA,TPM
GA8A,HNRNPU,OGA,NFE2L1,SMARCA5,HNRNPA0,CTNNA1,FADS2,RPL23,CNN3,RPL29,RPSA,TPM
ISCH,HNRNPC,LRPPRC,TADA2B,XPR1,KHDRBS1,HDLBP,RPS25,GLG1,SFPQ,PDIA6,MMP14,PRDX
NPK,INIP,RPL36,YWHAB,LAMP1,HNRNPH1,RPS12,RAB1A,ARCN1,HMGCR,CAPRIN1,TOP2B,HDAC
,RAB7A,DDX3X,RPL12,MAGED2,GNB2,RCN1,H1-0,IK,TLN1,CHPF,GJA1,MYO5A,CTNNB1,ANXA6,N
3L,MAGED2,H1-0,RPS5,RPS7,RPL35,CTNNB1,RPS10,APLP2,TARDBP,DDOST,HNRNPK,NARS1,RF
TOP2B,HDAC2,SKP1,DNER,BCLAF1,LUC7L3,CPSF2,TM9SF3,MDH2,EEF1A2,SUGP2,NOP53,HNRN
,HNRNPL,PARP1,MORF4L2,ALKBH5,ACTR3,DDX42,RAD23B,GCN1,TNPO1,SKI,SNRNP70,HDGF,T
,LUC7L3,CTSB,TM9SF3,MDH2,TOMM20,RPN2,MLXIP,HNRNPL,TM9SF2,MORF4L2,C4ORF3,TBC
PL31,PRKCSH,HNRNPC,LRPPRC,TADA2B,XPR1,KHDRBS1,XPO1,CSNK1E,RPS25,GLG1,SFPQ,DDI
1,TUBB4B,RPL11,MORF4L1,BTF3,ANXA2,RPL10,RPS17,VCL,DNAJA1,SF3B2,ITM2B,MAT2A,GTF2
LRPPRC,KHDRBS1,XPO1,RPS25,SFPQ,PDIA6,DDR1,RAB7A,RPS15,MAGED2,H1-0,RPS7,IK,RPL35
,EIF5,HMGN2,CCT7,RBBP4,MCL1,RPL31,NISCH,PRKCSH,HNRNPC,LRPPRC,TADA2B,KHDRBS1,F
D1A,NDFIP1,HNRNPK,H1-10,INIP,RPL36,YWHAB,LAMP1,HNRNPH1,ARCN1,CCT6A,CAPRIN1,TOP2
2,CDC42BPB,AP2B1,GTF2F1,GOLGA8A,HNRNPU,OGA,NFE2L1,CTNNA1,RPS16,RPL23,RPSA,TPM
LAMP1,RPS12,ARCN1,CCT6A,RAC1,CAPRIN1,TOP2B,HDAC2,BCLAF1,LUC7L3,CPSF2,DDX39B,TM
TLN1,COL6A2,ANXA6,MACF1,APLP2,TARDBP,WLS,HNRNPK,INIP,YWHAB,CKAP5,LAMP1,HNRNPH
D1,LAMP1,HNRNPH1,CALU,CCT6A,GPC1,TOP2B,HDAC2,BCLAF1,PPIB,SQLE,CPSF2,DDX39B,MDH
,CCT7,SRSF4,RBBP4,PFN1,PNN,NES,NISCH,HNRNPC,LRPPRC,TADA2B,XPR1,KHDRBS1,CSNK1E,
WHAB,SND1,LAMP1,UBE2H,CKAP4,RPS12,HLA-B,CALU,CAPRIN1,HDAC2,BCLAF1,LUC7L3,CRPP
SF2,DDX39B,TOMM20,HNRNPL,MORF4L2,TAF15,SSRP1,RPL27,HNRNPA3,C4ORF3,ACO2,NCKAI
RDX2,EIF3L,MAGED2,H1-0,RPS5,RPS7,IK,RPL35,HMGCS1,CTNNB1,GOLM1,RPS10,APLP2,TARDE
IRNPDL,TARDBP,ARID1A,DDOST,HNRNPK,H1-10,INIP,RPL36,SEPTIN11,LAMP1,HNRNPH1,UBE2H
PFN1,JPT2,NES,NISCH,LRPPRC,TADA2B,KHDRBS1,CSNK1E,SCRN1,HDLBP,RPS25,TKT,GLG1,SFF
KBH5,RPL27,HNRNPA3,ACTR3,ACO2,CCT3,RAD23B,TNPO1,SNRNP70,H3-3A,HDGF,TSPAN6,MAF
,CTSB,TM9SF3,MDH2,TOMM20,U2AF2,RPN2,PABPN1,HNRNPL,TM9SF2,MORF4L2,ABCA3,ALKB
,EIF5,WDR1,RPL26,LASP1,CCT7,RBBP4,MCL1,JPT2,PNN,RPL31,NISCH,PRKCSH,HNRNPC,LRPPF
PAFAH1B1,AKAP12,CDC42BPB,ITM2B,ZNF770,AP2B1,OGA,PGAM1,LARP1,CTNNA1,FADS2,CPE,C
DL,TARDBP,NDFIP1,ST3GAL2,HNRNPK,H1-10,INIP,RPL36,CKAP5,LAMP1,HNRNPH1,CKAP4,CALU
PQ,PEBP1,DDR1,RAB7A,PRDX2,DDX3X,CLTC,GNB2,H1-0,RPS7,IK,GPAT4,CHPF,MYO5A,GOLM1,D
ACO2,RAD23B,TNPO1,SNRNP70,SCAF1,HDGF,PRPF6,TOP1,SELENON,G3BP2,SMARCA4,AZIN1,A
HNRNPH1,UBE2H,HLA-B,CALU,CCT6A,CAPRIN1,BCLAF1,LUC7L3,SQLE,CDKN1A,PGRMC1,EID1,C
P5,LAMP1,HNRNPH1,UBE2H,CKAP4,CALU,CCT6A,CAPRIN1,HDAC2,BCLAF1,LUC7L3,CPSF2,DDX3
PGD,SQSTM1,NOP53,HNRNPL,PARP1,MORF4L2,TAF15,SSRP1,HNRNPA3,ACO2,NCKAP1,RBMX,R
X1,RAD23B,TNPO1,SNRNP70,H3-3A,HDGF,TSPAN6,UBXN4,MAPK1,SMARCA4,ARPC2,PLEC,PSMD
5,RPL27,ACO2,RBMX,CBX1,RAD23B,TNPO1,SNRNP70,H3-3A,HDGF,TMED2,VEGFA,MARS1,SELE

)P1,TAF7,OPTN,SMARCA4,KDELR2,SRSF9,PBX2,PSMD3,SRSF1,ARGLU1,ILF3,PRKDC,HNRNPUL1
ARPC2,PRKDC,HNRNPUL1,FKBP8,DIDO1,CORO1C,HNRNPM,GIGYF1,SRSF3,ILF2,LSS,STXBP1,ST
Q,PEBP1,CCN2,RPS15,RPL30,CLTC,MAGED2,GNB2,RCN1,RPS7,IK,GPAT4,TLN1,COL6A2,CD63,A
P2A2,BEX3,CCT7,SRSF4,RBBP4,MCL1,PNN,NES,NISCH,HNRNPC,LRPPRC,TADA2B,XPR1,KHDRB
I,NISCH,HNRNPC,TADA2B,XPR1,KHDRBS1,KLHL9,XPO1,RPS25,GLG1,SFPQ,TPM1,SULF1,RAB7A
F2,SQSTM1,HNRNPL,PARP1,MORF4L2,ALKBH5,RPL27,HNRNPA3,ACTR3,RBMX,TNPO1,SNRNP7
B,GIGYF1,SRSF3,ILF2,SF3A1,HNRNPD,ZSCAN18,KDM3B,TP53BP1,POLR2E,CIC,RBBP7,LBH,NDU
PA3,CAMSAP1,RBMX,CBX1,RAD23B,RAB5B,CHD2,TNPO1,SNRNP70,H3-3A,DPYSL2,TOP1,OPTN,
NON,G3BP2,SRSF1,ARGLU1,PTBP1,ILF3,PRKDC,HNRNPUL1,SON,ZFAND5,DIDO1,POLR2B,SMA
F4,RBBP4,MCL1,PNN,RPL31,PRKCSH,HNRNPC,LRPPRC,TADA2B,KHDRBS1,XPO1,CSNK1E,RPS2
,HNRNPA3,PKN1,ACO2,CCT3,RAD23B,SNRNP70,PSMB2,H3-3A,HDGF,PYGB,TMED2,VEGFA,OPT
N1,H1-0,RPS7,IK,RPL35,HMGCS1,CTNNB1,GOLM1,APLP2,TARDBP,DDOST,HNRNPK,NARS1,RPL
2,PNN,RPL31,PRKCSH,HNRNPC,LRPPRC,TADA2B,XPR1,KHDRBS1,XPO1,CSNK1E,RPS25,TKT,GL
ORF4L2,ALKBH5,PDXK,GRINA,PTOV1,RBMX,DDX42,LAMC1,JPT1,RAD23B,TNPO1,SNRNP70,H3-
5,GLG1,SFPQ,PEBP1,SULF1,RAB7A,ACTN4,MMP14,DDX3X,MAGED2,GNB2,H1-0,APLP1,IK,MIF,G
,ACTN4,RPS15,MAGED2,H1-0,RPS5,RPS7,IK,RPL35,HMGCS1,CTNNB1,APLP2,TARDBP,DDOST,H
ELENON,MAPK1,SRSF1,ARGLU1,ILF3,PRKDC,HNRNPUL1,HSPB1,SON,PLCG1,DIDO1,POLR2B,C
TERF2IP,RPS15A,MARS1,TUBB2A,SMARCA4,MAPRE1,SRSF9,SRSF1,ARGLU1,ATP5PB,PTBP1,ILF
PO1,SPTBN2,HDLBP,TKT,CDH6,SFPQ,PDIA6,DDR1,ACSL6,MMP14,RPS15,RPL12,CLTC,EIF3L,M
RNPD,ZSCAN18,TCTN1,KDM3B,TP53BP1,CCT5,RBBP7,KCNQ2,LBH,SF1,HIP1,H2AZ1,UBQLN2,B
NK1D,SF3A1,HNRNPD,KDM3B,TP53BP1,PSMA7,ANP32A,RBBP7,TRIO,LBH,SF1,HIP1,H2AZ1,BRD
NP32A,RBBP7,TXLNA,LBH,PDAP1,VARS1,UBQLN2,CD99,BRD4,SLC39A7,ATRX,TRA2B,UQCRC1,Z
1,PTBP1,EIF3E,ILF3,GRB2,CAPZB,PRKDC,HNRNPUL1,RPL28,DIDO1,XPNPEP3,SMARCC1,SUPT6
T6H,HNRNPM,SRSF3,ILF2,SF3A1,MESD,KDM3B,DDIT4,ANP32A,RBBP7,DCTN1,EDC4,LBH,SF1,H
)1,LAMP1,HNRNPH1,ATP1A2,MAP2,CCT6A,STIP1,HDAC2,BCLAF1,LUC7L3,CPSF2,DDX39B,TM9S
S1,XPO1,RPS25,SFPQ,EIF5A,PDIA6,DDR1,ACSL6,RAB7A,RPS15,MAGED2,H1-0,RPS7,IK,HMGCS1
RBM5,SRSF11,IST1,DHCR7,TGOLN2,RBM6,CREBBP,ATXN7L3,XRN2,PTPN11,LGALS1,MTA2,CG
D,SQSTM1,NOP53,GDI2,HNRNPL,PARP1,TSR1,SSRP1,LMNB2,RPL27,PRDX5,UBE4A,TBC1D16,A
DR,TMEM259,EIF5,HMGN2,RPL26,CCT7,SRSF4,RBBP4,PFN1,RPL31,PRKCSH,HNRNPC,LRPPRC
CC6,PELP1,CLPTM1,YARS1,TPP1,ZMIZ2,MAZ,HDGFL2,EFTUD2,SRSF2,EIF4G3,TRAP1,MACROH2
A1,STRAP,DGKD,KDM3B,AKIRIN1,TP53BP1,DDIT4,PSMA7,ANP32A,RBBP7,PRMT1,NAP1L1,LBH,
0,PGD,HNRNPL,PARP1,MORF4L2,TAF15,RPL27,HNRNPA3,ACTR3,C4ORF3,ACO2,CBX1,TNPO1,S
RO1C,SMARCE1,SUPT6H,HNRNPM,SRSF3,ILF2,LONP1,SF3A1,SERBP1,KDM3B,TP53BP1,DDIT4,
2,PFKM,CSNK1D,STRAP,HNRNPD,ZSCAN18,KDM3B,TP53BP1,ANP32A,RBBP7,ST3GAL6,TXLNA,N
L3,SQLE,CPSF2,PGRMC1,EID1,TM9SF3,MDH2,TOMM20,U2AF2,FDFT1,RPN2,SQSTM1,PABPN1,H
ELENON,OPTN,UBXN4,G3BP2,MAPK1,SMARCA4,ARPC2,PLEC,ACADVL,PTBP1,ILF3,PRKDC,HNRN
SPTBN2,CSNK1E,SCRN1,GLG1,CDH2,DDR1,MEGF8,RAB7A,ACTN4,PRDX2,CLTC,MRFAP1,MAGE
C,SMARCE1,SUPT6H,HNRNPM,SRSF3,ILF2,SF3A1,MESD,RAI1,KDM3B,DDIT4,ANP32A,RBBP7,ED
KHDRBS1,XPO1,CSNK1E,GLG1,CDH6,SFPQ,TPM1,DDR1,SULF1,RAB7A,PRDX2,DDX3X,MAGED2,
,IK,RPL35,TLN1,CTNNB1,GOLM1,HNRNPDL,TARDBP,NDFIP1,DDOST,HNRNPK,H1-10,MYH10,RP
RF4L2,ABCA3,PRDX5,PDXK,CCT3,VPS35,PTOV1,RBMX,DDX42,LAMC1,CBX1,CCT4,GCN1,SNRN
2,MECP2,AAMP,SART3,PCGF3,SUN2,CAPN1,CDK4,CDK9,RBFOX2,PSIP1,NDRG1,POLDIP3,SRF,T
RAB5B,TNPO1,SNRNP70,H3-3A,MARS1,OPTN,G3BP2,SRSF9,PSMD3,SRSF1,ACADVL,ATP5PB,PTE
,ACO2,RBMX,TNPO1,SNRNP70,PSMB2,H3-3A,TMED2,OPTN,G3BP2,MAPK1,SMARCA4,ARPC2,DS
LONP1,UBQLN1,SF3A1,SREBF2,HNRNPD,MLF2,SERBP1,KDM3B,RBBP7,EDC4,TXLNA,NAP1L1,LE

A,NAP1L1,LBH,PRCC,H2AZ1,BRD4,UBA52,EP400,SRSF6,PAPSS1,TRA2B,ZFR,AGPAT1,HNRNPA2E
OX2,SRF,ZEB1,SEPHS1,HSPE1,RBM14,LMNA,RRBP1,GET1,RXRA,HIRA,PHB,RBM17,CTCF,DSTYI
HDGF,TOP1,TAF7,OPTN,UBXN4,MAPK1,SMARCA4,ARPC2,SRSF1,PTBP1,ILF3,LUZP6,GRB2,CAPZI
KS1,PHGDH,SRSF11,MSH6,USP24,EHD1,YY1,CREBBP,THRAP3,SLC38A2,EIF4A3,SH3PXD2B,CAS
NPL,PARP1,MORF4L2,ABCA3,ACO2,RBMX,CBX1,RAD23B,RAB5B,CHD2,TNPO1,SNRNP70,H3-3A,
7,KCTD20,NCKAP5L,CFDP1,HIPK2,PPP4R3A,WIZ,ECHS1,PRXL2A,CCNG1,CDK9,EXOC7,PSMC2,T
P1,RALY,KIAA0232,CNP,TRAP1,NUCKS1,RBM5,PHGDH,SRSF11,SPTY2D1,SMARCD1,EIF4A3,NTR
2AZ1,CD99,SLC39A7,SRSF6,ARID1B,TRA2B,ZFR,HNRNPA2B1,ANP32B,CALD1,APEX1,PRDX6,AK

1,LBH,SF1,HIP1,NOL4L,H2AZ1,GTPBP1,BRD4,PPP2CB,ATRX,NXN,EP400,SRSF6,UQCRC1,MAST
,PRPF19,FKBP9,PSMC3,ANKRD12,DAPK3,PRDX3,EIF4A3,XRN2,HTATSF1,SUN1,SGSM2,CGGBP1
107,HNRNPD,MLF2,SERBP1,ARFGEF2,CCT5,TRAF4,RBBP7,OS9,KDM5B,RETREG2,FLCN,PRCC,
SF11,PRPF19,UBE3A,TGOLN2,RBM6,ATXN7L3,ATP5MC2,DAPK3,THRAP3,AMBRA1,EIF4A3,CALC
M1A,CASC3,FKBP8,CCAR2,DIDO1,SMARCE1,SDF4,SMARCC1,HNRNPM,SSB,SRSF3,PHC2,LARS1,
RPS27A,TIMM50,HNRNPAB,C1ORF198,MTCH1,ZMYM3,FUS,ERH,CLPTM1,RALY,NUDCD3,EFTUD2
RNPDL,TARDBP,RPLP2,WLS,HNRNPK,MYH10,RPL36,CKAP5,SEPTIN11,LAMP1,HNRNPH1,CAPRIN
2B,MCM3AP,CTTN,ZMYM3,FUS,XRCC6,PELP1,VAPB,SNN,STRN4,INTS3,RALY,EFTUD2,SRSF2,TR
SF1,SGSM2,ANP32E,REEP5,IGF1R,MDM2,FAM50A,SUMO1,CFDP1,WIZ,CDK9,PSMC2,TNFRSF10
KM,UBQLN1,SF3A1,NT5DC2,STRAP,UQCRC2,MLF2,AKIRIN1,PRPF40A,TP53BP1,PCYOX1,RBBP7
CITED2,LZTS2,PSMA6,FUBP1,PPT1,MYEF2,ZNF207,LEMD2,ALYREF,COX5B
FLOT1,PHB2,BRD4,DIP2B,ATRX,EP400,ARID1B,KIAA0895L,GLUD1,EVL,TRA2B,UQCRC1,ANP32B
INRNPA2B1,MCM3AP,RPS27A,HNRNPAB,SYMPK,MTCH1,FUS,TRIR,ERH,PELP1,ERGIC1,RALY,TLE
E,SYT1,PPT1,EPHA4,PDHB,WDR33,NNT,ZNF326,GSN,ARSA,FKBP3,MMP24
T1,UQCRFS1,THOC2,CNTN1,NAPA,GET1,RNF4,GNAZ,ATP6V0A1,GPRIN1,COX5A,EPB41L1,PPP3
RPF6,TERF2IP,TAF7,SMARCA4,MAPRE1,CCNL2,PBX2,ODC1,SRSF1,ARGLU1,ATP5PB,PTBP1,ILF
1,CDK16,SMCHD1,IPO5,SKP1,GLUL,EID1,SRSF5,RNF187,FDFT1,MLXIP,ANXA5,TSPYL4,SSRP1,
9SF2,MORF4L2,TAF15,HNRNPA3,ACO2,ETNK1,LAMC1,CBX1,SNRNP70,H3-3A,KDM6B,HDGF,AB

,NDUFB8,SEZ6L2,CLCN4,ACSL4,ALCAM,DLG4,CASKIN1,WASF3,DPP7,VDAC2,UQCRFS1,BACE1,
SF7,UBTF,ALYREF,M6PR

,CPSF2,DDX39B,TM9SF3,TOMM20,U2AF2,CCT2,SUGP2,PABPN1,GDI2,HNRNPL,TM9SF2,MORF

T1,ATP6V1F,PDHB,NNT,ZNF326,ANAPC2,VPS37B,MYBBP1A,FKBP3,MMP24
1,SF1,RETREG2,FLCN,PRCC,H2AZ1,CSNK2A1,KPNA2,GPX4,PPM1G,BRD2,EP400,SRSF6,TRA2B,
R,SLC22A23,SGSM3,DNAJB1,CMPK1,FAM50A,CAB39,FXR1,EWSR1,SYNCRIP,PABPC4,HTRA1,SL
,LMNA,CDC42EP4,UQCRFS1,GET1,TLK1,RBM17,NRP2,DSTYK,ZNF146,PCED1A,QRICH1,TARS1,
,HNRNPAB,ZMYM3,GRK2,FUS,CLCN7,CLPTM1,RALY,HDGFL2,TRAP1,MOAP1,PACS1,PDPK1,RBN
BP,RPS25,GLG1,EIF5A,DDR1,PRDX2,RPL30,CLTC,RPS7,MIF,HMGCS1,GPAT4,PRKAR1A,CD63,M

BE3A,PRKACA,KLHDC10,MN1,ANKRD12,DMWD,SCAP,CALCOCO1,SLC30A9,AMFR,LMAN2,SGS

PF6,TERF2IP,TAF7,TUBB2A,SMARCA4,MAPRE1,CCNL2,SRSF1,ARGLU1,ATP5PB,ILF3,PRKDC,SU

SEPTIN9,SLC1A5,PCNA,RAD54L2,TMEM59,BCAR1,OPA1,ABCD3,LANCL1,AKAP17A,GNAL,DDX19

A6,APPBP2,ZNF768,OXSR1,STAT2,SHMT2,SEZ6L2,TFRC,ADIPOR2,TXN,CDKN1B,PSMB5,SLC16A

C,ATP5F1A,EIF4B,VCP,MARCKSL1,UBB,AFF4,TRIM28,MATR3,YWHAG,CKB,RPL37A,LRRC28,MA
PL37A,LRRC28,MAPT,FN1,SPEN,NONO,DDB1,YWHAQ,HACD3,DCHS1,SET,EGLN1,INA,MRC2,RPL
EN,NONO,DDB1,YWHAQ,HACD3,DCHS1,SET,EGLN1,INA,RPL19,LDHB,SF3B1,PDIA3,SSBP2,BASP
RC2,RPL19,LDHB,PKD1,SF3B1,PDIA3,SSBP2,CHD3,BASP1,VAPA,HMGB1,POLR2A,TPT1,RPS4X,
CP,MARCKSL1,UBB,AFF4,TRIM28,MATR3,YWHAG,CKB,RPL37A,LRRC28,MAPT,FN1,SPEN,NONO
N1,INA,MRC2,RPL19,LDHB,PKD1,SF3B1,PDIA3,SSBP2,CHD3,BASP1,VAPA,HMGB1,POLR2A,TPT1
1,SPEN,NONO,DDB1,YWHAQ,HACD3,DCHS1,SET,EGLN1,INA,MRC2,RPL19,LDHB,SF3B1,PDIA3,
PEN,NONO,DDB1,YWHAQ,HACD3,DCHS1,SET,EGLN1,MRC2,RPL19,LDHB,PKD1,SF3B1,PDIA3,SS
SPEN,NONO,DDB1,YWHAQ,HACD3,DCHS1,SET,EGLN1,MRC2,RPL19,LDHB,PKD1,SF3B1,PDIA3,S
T,EGLN1,MRC2,RPL19,LDHB,SF3B1,PDIA3,SSBP2,CHD3,BASP1,VAPA,HMGB1,POLR2A,TPT1,RPS
T,EGLN1,MRC2,RPL19,LDHB,SF3B1,PDIA3,SSBP2,CHD3,BASP1,VAPA,HMGB1,POLR2A,TPT1,RPS
1,SPEN,NONO,DDB1,YWHAQ,HACD3,DCHS1,SET,EGLN1,MRC2,RPL19,LDHB,PKD1,SF3B1,PDIA3,
1,RPL10,RPS17,VCL,DNAJA1,CYB5R3,PAFAH1B1,SF3B2,ITM2B,ZNF770,AP2B1,GTF2F1,HNRNPU
ATR3,YWHAG,RPL37A,LRRC28,FN1,SPEN,NONO,DDB1,YWHAQ,HACD3,DCHS1,SET,EGLN1,MRC2
ATR3,YWHAG,RPL37A,LRRC28,FN1,SPEN,NONO,DDB1,YWHAQ,HACD3,DCHS1,SET,EGLN1,MRC2
L19,LDHB,PKD1,SF3B1,PDIA3,SSBP2,CHD3,BASP1,VAPA,HMGB1,TPT1,RPS4X,RPS3A,RPL18,STM
,HACD3,DCHS1,SET,EGLN1,RPL19,LDHB,PKD1,SF3B1,PDIA3,SSBP2,CHD3,BASP1,VAPA,HMGB1,
,HACD3,DCHS1,SET,EGLN1,RPL19,LDHB,PKD1,SF3B1,PDIA3,SSBP2,CHD3,BASP1,VAPA,HMGB1,
AG,RPL37A,LRRC28,MAPT,FN1,SPEN,NONO,DDB1,YWHAQ,HACD3,DCHS1,SET,EGLN1,MRC2,RPL
S1,SET,EGLN1,MRC2,RPL19,LDHB,PKD1,SF3B1,PDIA3,SSBP2,CHD3,BASP1,VAPA,HMGB1,POLR2
12,MAP4,SF3B2,CDC42BPB,ITM2B,GTF2F1,HNRNPU,NFE2L1,SMARCA5,LARP1,HNRNPA0,CTNN
,FN1,SPEN,NONO,DDB1,YWHAQ,HACD3,DCHS1,SET,EGLN1,MRC2,RPL19,LDHB,SF3B1,PDIA3,S
AQ,HACD3,DCHS1,SET,EGLN1,INA,MRC2,RPL19,LDHB,PKD1,SF3B1,PDIA3,SSBP2,CHD3,VAPA,H
AQ,HACD3,DCHS1,SET,EGLN1,INA,MRC2,RPL19,LDHB,PKD1,SF3B1,PDIA3,SSBP2,CHD3,VAPA,H
CYB5R3,PAFAH1B1,SF3B2,CDC42BPB,ITM2B,ZNF770,AP2B1,HNRNPU,OGA,NFE2L1,SMARCA5,L
M2B,ZNF770,AP2B1,HNRNPU,OGA,PGAM1,HNRNPA0,FADS2,RPS16,RPL23,CPE,CNN3,RPL29,R
MATR3,YWHAG,CKB,RPL37A,LRRC28,MAPT,FN1,SPEN,NONO,DDB1,YWHAQ,HACD3,DCHS1,SET,
CD3,DCHS1,SET,EGLN1,MRC2,RPL19,PKD1,SF3B1,PDIA3,SSBP2,CHD3,BASP1,VAPA,HMGB1,PO
,PDIA3,SSBP2,BASP1,VAPA,HMGB1,POLR2A,TPT1,RPS4X,RPS3A,RPL18,STMN1,TUBB4B,RPL11,
,PDIA3,SSBP2,BASP1,VAPA,HMGB1,POLR2A,TPT1,RPS4X,RPS3A,RPL18,STMN1,TUBB4B,RPL11,
,SPEN,NONO,DDB1,YWHAQ,HACD3,DCHS1,SET,EGLN1,MRC2,RPL19,LDHB,PKD1,SF3B1,PDIA3,
PL19,LDHB,PKD1,SF3B1,PDIA3,SSBP2,BASP1,VAPA,HMGB1,POLR2A,TPT1,RPS4X,RPS3A,RPL18
PL19,LDHB,PKD1,SF3B1,PDIA3,SSBP2,BASP1,VAPA,HMGB1,POLR2A,TPT1,RPS4X,RPS3A,RPL18
,XAF1,RPL10,RPS17,VCL,DNAJA1,CYB5R3,PAFAH1B1,AKAP12,MAP4,SF3B2,CDC42BPB,ITM2B,Z
TPT1,RPS4X,RPS3A,RPL18,TUBB4B,RPL11,MORF4L1,BTF3,ANXA2,AGRN,XAF1,RPL10,RPS17,VC
1,SPEN,NONO,DDB1,YWHAQ,HACD3,DCHS1,SET,EGLN1,RPL19,LDHB,SF3B1,PDIA3,SSBP2,CHD
1B1,SF3B2,CDC42BPB,ITM2B,ZNF770,AP2B1,MAT2A,GTF2F1,HNRNPU,NFE2L1,SMARCA5,LAR
HD3,BASP1,VAPA,HMGB1,POLR2A,TPT1,RPS4X,RPS3A,RPL18,TUBB4B,RPL11,MORF4L1,BTF3,A
HD3,BASP1,VAPA,HMGB1,POLR2A,TPT1,RPS4X,RPS3A,RPL18,TUBB4B,RPL11,MORF4L1,BTF3,A
SBP2,CHD3,BASP1,VAPA,HMGB1,POLR2A,RPS4X,RPS3A,RPL18,TUBB4B,RPL11,MORF4L1,BTF3,
PU,NFE2L1,SMARCA5,LARP1,HNRNPA0,CTNNA1,RPS16,RPL23,CNN3,RPSA,TPM4,CTSD,EIF4H,S
8,FN1,SPEN,NONO,DDB1,YWHAQ,HACD3,DCHS1,SET,EGLN1,MRC2,RPL19,LDHB,PKD1,SF3B1,P
8,FN1,SPEN,NONO,DDB1,YWHAQ,HACD3,DCHS1,SET,EGLN1,MRC2,RPL19,LDHB,PKD1,SF3B1,P
NO,DDB1,YWHAQ,HACD3,DCHS1,SET,EGLN1,MRC2,RPL19,LDHB,PKD1,SF3B1,PDIA3,SSBP2,C
NO,DDB1,YWHAQ,HACD3,DCHS1,SET,EGLN1,MRC2,RPL19,LDHB,PKD1,SF3B1,PDIA3,SSBP2,C

'L10,RPS17,VCL,DNAJA1,CYB5R3,PAFAH1B1,AKAP12,MAP4,SF3B2,CDC42BPB,ITM2B,GTF2F1,H
DDB1,YWHAQ,HACD3,DCHS1,SET,EGLN1,MRC2,RPL19,LDHB,PKD1,SF3B1,PDIA3,SSBP2,CHD3,B
DDB1,YWHAQ,HACD3,DCHS1,SET,EGLN1,MRC2,RPL19,LDHB,PKD1,SF3B1,PDIA3,SSBP2,CHD3,B
IO,DDB1,YWHAQ,HACD3,DCHS1,SET,EGLN1,MRC2,RPL19,LDHB,PKD1,SF3B1,PDIA3,SSBP2,CHD
2,RPL19,LDHB,PKD1,SF3B1,PDIA3,SSBP2,CHD3,VAPA,HMGB1,POLR2A,TPT1,RPS4X,RPS3A,RPL1
DB1,YWHAQ,HACD3,DCHS1,SET,EGLN1,MRC2,RPL19,LDHB,PKD1,SF3B1,PDIA3,SSBP2,CHD3,VA
FN1,NONO,DDB1,YWHAQ,HACD3,DCHS1,SET,EGLN1,MRC2,LDHB,PKD1,SF3B1,PDIA3,SSBP2,CH
A3,SSBP2,CHD3,BASP1,VAPA,POLR2A,RPS4X,RPS3A,RPL18,TUBB4B,UCHL1,RPL11,MORF4L1,BT
3B1,PDIA3,SSBP2,CHD3,VAPA,HMGB1,POLR2A,TPT1,RPS4X,RPS3A,RPL18,STMN1,TUBB4B,RPL1
3B1,PDIA3,SSBP2,CHD3,VAPA,HMGB1,POLR2A,TPT1,RPS4X,RPS3A,RPL18,STMN1,TUBB4B,RPL1
SPEN,NONO,DDB1,YWHAQ,HACD3,DCHS1,SET,EGLN1,MRC2,RPL19,LDHB,PKD1,SF3B1,PDIA3,SS
B4B,RPL11,MORF4L1,BTF3,ANXA2,XAF1,RPL10,WSB1,RPS17,VCL,DNAJA1,CYB5R3,PAFAH1B1,M
9,LDHB,PKD1,SF3B1,PDIA3,SSBP2,CHD3,VAPA,HMGB1,POLR2A,TPT1,RPS4X,RPS3A,RPL18,STM
9,LDHB,PKD1,SF3B1,PDIA3,SSBP2,CHD3,VAPA,HMGB1,POLR2A,TPT1,RPS4X,RPS3A,RPL18,STM
IN3,RPL29,RPSA,CTSD,EIF4H,SCARB2,BRD3,XRCC5,MSN,RPL24,RPL17,C1ORF43,PAPOLA,LLGL
SBP2,CHD3,VAPA,HMGB1,POLR2A,TPT1,RPS4X,RPS3A,RPL18,STMN1,TUBB4B,RPL11,BTF3,ANX
TGB1,TUBB4B,RPL11,MORF4L1,BTF3,NCAM1,ANXA2,AGRN,XAF1,RPL10,MARCHF6,RPS17,VCL,I
Q,HACD3,DCHS1,SET,EGLN1,MRC2,RPL19,LDHB,PKD1,SF3B1,PDIA3,SSBP2,BASP1,VAPA,HMGB1
Q,HACD3,DCHS1,SET,EGLN1,MRC2,RPL19,LDHB,PKD1,SF3B1,PDIA3,SSBP2,BASP1,VAPA,HMGB1
IRC2,RPL19,LDHB,PKD1,SF3B1,PDIA3,SSBP2,BASP1,VAPA,HMGB1,POLR2A,TPT1,RPS4X,RPS3A,
IRC2,RPL19,LDHB,PKD1,SF3B1,PDIA3,SSBP2,BASP1,VAPA,HMGB1,POLR2A,TPT1,RPS4X,RPS3A,
DNAJA1,CYB5R3,PAFAH1B1,SF3B2,CDC42BPB,ITM2B,MAT2A,GTF2F1,HNRNPU,OGA,PGAM1,NF
AQ,HACD3,DCHS1,SET,EGLN1,MRC2,RPL19,LDHB,PKD1,SF3B1,PDIA3,SSBP2,CHD3,BASP1,VAP
ORF4L1,BTF3,NCAM1,ANXA2,AGRN,RPL10,RPS17,VCL,DNAJA1,CYB5R3,PAFAH1B1,AKAP12,SF3
'L19,LDHB,PKD1,SF3B1,PDIA3,SSBP2,CHD3,VAPA,HMGB1,POLR2A,TPT1,RPS4X,RPS3A,RPL18,T
1,MORF4L1,BTF3,ANXA2,XAF1,RPL10,RPS17,DNAJA1,PAFAH1B1,MAP4,SF3B2,CDC42BPB,ITM2
I2B,AP2B1,GTF2F1,HNRNPU,PGAM1,NFE2L1,SMARCA5,LARP1,CTNNA1,FADS2,RPS16,RPL23,CN
RPS3A,RPL18,ITGB1,TUBB4B,MORF4L1,BTF3,ANXA2,AGRN,XAF1,RPL10,RPS17,VCL,CYB5R3,PA
VCL,DNAJA1,CYB5R3,PAFAH1B1,AKAP12,MAP4,SF3B2,ITM2B,GTF2F1,HNRNPU,PGAM1,NFE2L1,
VCL,DNAJA1,CYB5R3,PAFAH1B1,AKAP12,MAP4,SF3B2,ITM2B,GTF2F1,HNRNPU,PGAM1,NFE2L1,
RF4L1,BTF3,ANXA2,ZMIZ1,RPL10,MARCHF6,WSB1,RPS17,VCL,DNAJA1,CYB5R3,PAFAH1B1,SF3E
1,RPS4X,RPS3A,RPL18,STMN1,TUBB4B,RPL11,MORF4L1,BTF3,ANXA2,AGRN,XAF1,RPL10,MARC
JA1,CYB5R3,PAFAH1B1,MAP4,SF3B2,ITM2B,AP2B1,GTF2F1,HNRNPU,NFE2L1,SMARCA5,LARP1,I
,YWHAQ,HACD3,DCHS1,SET,EGLN1,MRC2,RPL19,LDHB,PKD1,SF3B1,PDIA3,SSBP2,CHD3,VAPA,I
,YWHAQ,HACD3,DCHS1,SET,EGLN1,MRC2,RPL19,LDHB,PKD1,SF3B1,PDIA3,SSBP2,CHD3,VAPA,I
',EGLN1,MRC2,RPL19,LDHB,PKD1,SF3B1,PDIA3,CHD3,BASP1,VAPA,HMGB1,POLR2A,TPT1,RPS4
NO,DDB1,YWHAQ,HACD3,DCHS1,SET,EGLN1,MRC2,RPL19,LDHB,PKD1,SF3B1,PDIA3,SSBP2,VAF
1,SF3B1,PDIA3,SSBP2,CHD3,VAPA,HMGB1,POLR2A,RPS4X,RPS3A,RPL18,TUBB4B,RPL11,MORF
IARCA5,LARP1,FADS2,RPS16,RPL23,RPL29,RPSA,TPM4,CTSD,EIF4H,SCARB2,RTN4,LPCAT1,BRI
HMGB1,POLR2A,TPT1,RPS4X,RPS3A,RPL18,STMN1,TUBB4B,RPL11,MORF4L1,BTF3,ANXA2,RPL
1B1,MAP4,SF3B2,CDC42BPB,ITM2B,GTF2F1,GOLGA8A,HNRNPU,NFE2L1,SMARCA5,LARP1,CTN
F3B2,CDC42BPB,ITM2B,MAT2A,GTF2F1,HNRNPU,OGA,PGAM1,NFE2L1,SMARCA5,LARP1,HNRN
MAP4,SF3B2,ITM2B,ZNF770,MAT2A,GTF2F1,GOLGA8A,HNRNPU,NFE2L1,SMARCA5,LARP1,CTNI
2,CHD3,VAPA,HMGB1,POLR2A,TPT1,RPS4X,RPS3A,RPL18,STMN1,TUBB4B,RPL11,MORF4L1,BT
APA,HMGB1,POLR2A,TPT1,RPS4X,RPS3A,RPL18,STMN1,TUBB4B,RPL11,MORF4L1,BTF3,ANXA2,

ANXA2,RPL10,MARCHF6,RPS17,VCL,DNAJA1,CYB5R3,SF3B2,CDC42BPB,GTF2F1,HNRNPU,NFE2
DIA3,SSBP2,CHD3,BASP1,VAPA,HMGB1,POLR2A,TPT1,RPS4X,RPS3A,RPL18,STMN1,TUBB4B,RPL
9,LDHB,PKD1,SF3B1,PDIA3,SSBP2,CHD3,VAPA,HMGB1,POLR2A,TPT1,RPS4X,RPS3A,RPL18,STMI
9,LDHB,PKD1,SF3B1,PDIA3,SSBP2,CHD3,VAPA,HMGB1,POLR2A,TPT1,RPS4X,RPS3A,RPL18,STMI
SMD2,KIF5B,SLC7A5,RPS14,CHD4,JUND,TMEM259,RPS20,EIF5,HMGN2,RPL26,CCT7,RBBP4,MC
RPL18,TUBB4B,RPL11,MORF4L1,BTF3,ANXA2,AGRN,XAF1,RPL10,MARCHF6,WSB1,RPS17,CYB
MGB1,TPT1,RPS4X,RPS3A,RPL18,RPL11,MORF4L1,BTF3,ANXA2,AGRN,XAF1,RPL10,RPS17,VCL,
M2B,AP2B1,MAT2A,GTF2F1,GOLGA8A,HNRNPU,PGAM1,NFE2L1,SMARCA5,LARP1,HNRNPA0,CTN
LRRC28,FN1,SPEN,NONO,DDB1,YWHAQ,HACD3,DCHS1,SET,EGLN1,MRC2,RPL19,LDHB,PKD1,SI
TPT1,RPS4X,RPS3A,RPL18,STMN1,TUBB4B,RPL11,MORF4L1,BTF3,ANXA2,AGRN,XAF1,RPL10,RF
TPT1,RPS4X,RPS3A,RPL18,STMN1,TUBB4B,RPL11,MORF4L1,BTF3,ANXA2,AGRN,XAF1,RPL10,RF
CDC42BPB,GTF2F1,GOLGA8A,HNRNPU,NFE2L1,LARP1,CTNNA1,RPS16,RPL23,CNN3,RPL29,RPS
VCL,CYB5R3,MAP4,SF3B2,CDC42BPB,GTF2F1,HNRNPU,OGA,NFE2L1,SMARCA5,HNRNPA0,CTNN
F1,RPL10,RPS17,VCL,PAFAH1B1,SF3B2,CDC42BPB,ITM2B,AP2B1,MAT2A,GTF2F1,HNRNPU,OG
F1,RPL10,RPS17,VCL,PAFAH1B1,SF3B2,CDC42BPB,ITM2B,AP2B1,MAT2A,GTF2F1,HNRNPU,OG
2,MAP4,SF3B2,CDC42BPB,GTF2F1,HNRNPU,OGA,NFE2L1,CTNNA1,RPL23,RPL29,RPSA,TPM4,TN
F2F1,HNRNPU,NFE2L1,SMARCA5,HNRNPA0,CTNNA1,RPL23,CPE,CNN3,RPL29,RPSA,TPM4,CTSD,
L,DNAJA1,PAFAH1B1,MAP4,SF3B2,CDC42BPB,ITM2B,MAT2A,GTF2F1,HNRNPU,NFE2L1,SMARC
AKAP12,MAP4,SF3B2,CDC42BPB,ITM2B,AP2B1,GTF2F1,HNRNPU,PGAM1,NFE2L1,LARP1,HNRNP
L10,RPS17,VCL,CYB5R3,PAFAH1B1,MAP4,ZNF770,MAT2A,GTF2F1,GOLGA8A,HNRNPU,OGA,PG
FAH1B1,MAP4,SF3B2,CDC42BPB,ITM2B,MAT2A,GTF2F1,GOLGA8A,HNRNPU,NFE2L1,LARP1,HNF
EIF4B,VCP,MARCKSL1,UBB,AFF4,TRIM28,MATR3,YWHAG,RPL37A,LRRC28,MAPT,FN1,SPEN,NON
LA,GAP43,RPL14,HK1,NCS1,CBX5,CLSTN1,PSMD2,SLC7A5,CHD4,JUND,RPL26,BEX3,CCT7,RBB
B4B,MORF4L1,BTF3,ANXA2,RPL10,RPS17,VCL,DNAJA1,CYB5R3,PAFAH1B1,MAP4,SF3B2,CDC4
RPPRC,TADA2B,XPR1,KHDRBS1,RPS25,TKT,GLG1,SFPQ,PDIA6,SULF1,RAB7A,CCN2,MMP14,DDX
,HMGB1,POLR2A,TPT1,RPS4X,RPS3A,STMN1,TUBB4B,RPL11,MORF4L1,BTF3,ANXA2,RPL10,VCI
TF3,ANXA2,AGRN,XAF1,RPL10,MARCHF6,WSB1,RPS17,VCL,DNAJA1,CYB5R3,PAFAH1B1,MAP4,SF
ARCHF6,RPS17,VCL,PAFAH1B1,MAP4,SF3B2,CDC42BPB,ITM2B,MAT2A,GTF2F1,HNRNPU,OGA,N
LR2A,TPT1,RPS4X,RPS3A,RPL18,TUBB4B,RPL11,MORF4L1,ANXA2,RPL10,WSB1,RPS17,VCL,DN
TPT1,RPS4X,RPS3A,RPL18,STMN1,TUBB4B,RPL11,BTF3,ANXA2,AGRN,XAF1,RPL10,WSB1,RPS17
S4X,STMN1,TUBB4B,RPL11,MORF4L1,BTF3,ANXA2,AGRN,RPL10,RPS17,VCL,DNAJA1,PAFAH1B1
RCA5,LARP1,HNRNPA0,RPS16,RPL23,CNN3,RPL29,RPSA,TPM4,CTSD,EIF4H,SCARB2,LPCAT1,BF
H1-10,INIP,RPL36,YWHAB,CKAP5,SND1,LAMP1,HNRNPH1,MAP2,CALU,CCT6A,HMGCR,CAPRIN1,
B1,GTF2F1,HNRNPU,NFE2L1,SMARCA5,LARP1,HNRNPA0,CTNNA1,FADS2,RPS16,RPL23,CNN3,RF
,RPL18,STMN1,RPL11,MORF4L1,BTF3,ANXA2,XAF1,RPL10,WSB1,RPS17,VCL,DNAJA1,PAFAH1B1
PS3A,RPL18,TUBB4B,RPL11,MORF4L1,BTF3,ANXA2,AGRN,XAF1,RPL10,WSB1,RPS17,VCL,CYB5F
GB1,POLR2A,TPT1,RPS3A,RPL18,TUBB4B,UCHL1,RPL11,MORF4L1,BTF3,NCAM1,ANXA2,AGRN,
42BPB,ZNF770,MAT2A,GTF2F1,HNRNPU,OGA,NFE2L1,SMARCA5,HNRNPA0,CTNNA1,FADS2,RPS
,MORF4L1,BTF3,ANXA2,AGRN,ZMIZ1,XAF1,RPL10,RPS17,CYB5R3,PAFAH1B1,SF3B2,ITM2B,ZNI
PS17,VCL,DNAJA1,CYB5R3,PAFAH1B1,MAP4,SF3B2,CDC42BPB,ITM2B,AP2B1,MAT2A,GTF2F1,G
SF3B2,CDC42BPB,ITM2B,ZNF770,AP2B1,MAT2A,GTF2F1,HNRNPU,OGA,PGAM1,NFE2L1,SMARC
,CNN3,RPSA,CTSD,EIF4H,SCARB2,BRD3,XRCC5,PJA2,RPL24,SLC16A7,KPNB1,RPL17,C1ORF43,
4L1,BTF3,ANXA2,AGRN,ZMIZ1,XAF1,RPL10,RPS17,CYB5R3,PAFAH1B1,MAP4,SF3B2,ITM2B,ZNF
CL,DNAJA1,PAFAH1B1,MAP4,SF3B2,CDC42BPB,ITM2B,GTF2F1,GOLGA8A,HNRNPU,OGA,NFE2L
MAP4,SF3B2,CDC42BPB,ITM2B,AP2B1,MAT2A,GTF2F1,GOLGA8A,HNRNPU,OGA,NFE2L1,SMARC

MAP4,SF3B2,CDC42BPB,ZNF770,HNRNPU,NFE2L1,SMARCA5,HNRNPA0,CTNNA1,FADS2,RPS16,F
CAT1,RPS24,BRD3,XRCC5,RPL24,RPL17,C1ORF43,COL1A2,LLGL1,RPL14,NCS1,PTCD3,CBX5,ZI
),INIP,RPL36,YWHAB,LAMP1,HNRNPH1,HLA-B,CALU,CTBP1,ARHGDIA,CAPRIN1,STIP1,TOP2B,SM
,RPSA,TPM4,EIF4H,BRD3,RPS9,XRCC5,PJA2,RPL24,KPNB1,RPL17,C1ORF43,PAPOLA,COL1A2,R
,HMGB1,TPT1,RPS4X,RPS3A,RPL18,STMN1,TUBB4B,RPL11,BTF3,ANXA2,AGRN,RPL10,RPS17,V
DX2,DDX3X,RPS15,MAGED2,RCN1,H1-0,RPS7,IK,RPL35,MIF,GPAT4,TLN1,COL6A2,ANXA6,HNRNF
RPS16,RPL23,CNN3,RPL29,RPSA,TPM4,EIF4H,SCARB2,BRD3,XRCC5,MSN,RPL24,KPNB1,RPL17,
C2,RPL19,LDHB,SF3B1,PDIA3,SSBP2,BASP1,VAPA,HMGB1,POLR2A,TPT1,RPS4X,RPS3A,RPL18,S
C2,RPL19,LDHB,SF3B1,PDIA3,SSBP2,BASP1,VAPA,HMGB1,POLR2A,TPT1,RPS4X,RPS3A,RPL18,S
PS17,VCL,DNAJA1,CYB5R3,PAFAH1B1,MAP4,SF3B2,CDC42BPB,ITM2B,GTF2F1,HNRNPU,NFE2L1
CA5,LARP1,HNRNPA0,CTNNA1,RPS16,RPL23,CNN3,RPL29,RPSA,TPM4,CTSD,EIF4H,LPCAT1,BRD
,XRCC5,SLC16A7,KPNB1,RPL17,C1ORF43,PAPOLA,COL1A2,LLGL1,RPL14,RHOA,CBX5,RPL9,AT
,XRCC5,SLC16A7,KPNB1,RPL17,C1ORF43,PAPOLA,COL1A2,LLGL1,RPL14,RHOA,CBX5,RPL9,AT
,FADS2,RPL23,CNN3,RPL29,RPSA,TNS3,EIF4H,SCARB2,BRD3,XRCC5,PJA2,RPL17,C1ORF43,PAI
F43,PAPOLA,COL1A2,LLGL1,RPL14,HK1,PTCD3,RPL9,ATF4,PSMD2,KIF5B,CXADR,ATP1A1,JUND
,NFE2L1,SMARCA5,LARP1,CTNNA1,FADS2,RPS16,RPL23,CNN3,RPL29,RPSA,TPM4,EIF4H,SCARB
GL1,RPL14,RPL9,CLSTN1,KIF5B,RPS14,CHD4,JUND,TMEM259,RPS20,EIF5,CD81,RPL26,LASP1,I
BRD3,RPS9,PJA2,RPL24,C1ORF43,PAPOLA,COL1A2,LLGL1,RPL14,HK1,CBX5,PSMD2,KIF5B,SLC
R3,PAFAH1B1,MAP4,SF3B2,CDC42BPB,ITM2B,ZNF770,GTF2F1,HNRNPU,OGA,NFE2L1,SMARCA
LN1,INA,MRC2,RPL19,LDHB,PKD1,SF3B1,PDIA3,SSBP2,CHD3,VAPA,HMGB1,POLR2A,TPT1,RPS4
8,STMN1,BTF3,ANXA2,AGRN,RPL10,RPS17,VCL,DNAJA1,CYB5R3,PAFAH1B1,SF3B2,CDC42BPB,
OL1A2,LLGL1,RPL14,HK1,ZNF865,PSMD2,KIF5B,RPS14,CXADR,CHD4,JUND,TMEM259,RPS20,E
OP53,HNRNPL,PARP1,TM9SF2,MORF4L2,TAF15,LMNB2,ALKBH5,RPL27,HNRNPA3,ACO2,NCKAP
3,RPL11,MORF4L1,BTF3,ANXA2,AGRN,XAF1,RPL10,WSB1,RPS17,VCL,DNAJA1,PAFAH1B1,MAP4,
RD3,XRCC5,PJA2,RPL24,KPNB1,RPL17,C1ORF43,PAPOLA,COL1A2,LLGL1,RPL14,HK1,RHOA,PT
I1B1,MAP4,SF3B2,CDC42BPB,ITM2B,ZNF770,GTF2F1,HNRNPU,PGAM1,NFE2L1,SMARCA5,LARP
I1B1,MAP4,SF3B2,CDC42BPB,ITM2B,ZNF770,GTF2F1,HNRNPU,PGAM1,NFE2L1,SMARCA5,LARP
M4,EIF4H,SCARB2,BRD3,XRCC5,RPL24,RPL17,C1ORF43,PAPOLA,COL1A2,LLGL1,RPL14,HK1,AT
SH,HNRNPC,LRPPRC,TADA2B,XPR1,KHDRBS1,XPO1,CSNK1E,RPS25,GLG1,SFPQ,PNMA1,PDIA6,R
3,COL1A2,LLGL1,RPL14,HK1,PTCD3,CBX5,PSMD2,KIF5B,SLC7A5,RPS14,CHD4,ATP1A1,TMEM2
SF4,RBBP4,RTN3,PFN1,MCL1,PNN,PRKCSH,HNRNPC,LRPPRC,KHDRBS1,CSNK1E,RPS25,TKT,SF
,ZNF865,KIF5B,RPS14,CXADR,CHD4,RPS20,EIF5,WDR1,RPL26,CCT7,SRSF4,RBBP4,MCL1,RPL3
L1,SMARCA5,LARP1,RPS16,RPL23,CNN3,RPL29,RPSA,TPM4,EIF4H,LPCAT1,XRCC5,PJA2,MSN,RF
RPL35,CTNNB1,APLP2,HNRNPDL,TARDBP,ARID1A,HNRNPK,RPL36,YWHAB,LAMP1,HNRNPH1,UB
SFPQ,PDIA6,RAB7A,MMP14,DDX3X,RPL12,RPL30,MAGED2,H1-0,IK,TLN1,JUN,COL6A2,ANXA6,API
R1,PRDX2,DDX3X,MAGED2,H1-0,APLP1,RPS7,GPAT4,TLN1,GJA1,GOLM1,RPS10,APLP2,HNRNPDI
DC42BPB,ITM2B,AP2B1,GTF2F1,GOLGA8A,HNRNPU,PGAM1,NFE2L1,LARP1,HNRNPA0,CTNNA1,
LSTN1,KIF5B,SLC7A5,RPS14,CHD4,JUND,TMEM259,WDR1,HMGN2,RPL26,BEX3,CCT7,SRSF4,RI
,XRCC5,PJA2,MSN,RPL24,KPNB1,RPL17,C1ORF43,PAPOLA,COL1A2,LLGL1,RPL14,HK1,RHOA,P
GL1,RPL14,HK1,PTCD3,RPL9,PSMD2,KIF5B,SLC7A5,CHD4,RPS20,EIF5,WDR1,HMGN2,RPL26,BE
NB1,APLP2,HNRNPDL,TARDBP,DDOST,HNRNPK,H1-10,MYH10,RPL36,YWHAB,LAMP1,ARCN1,CCT
GTF2F1,HNRNPU,PGAM1,NFE2L1,LARP1,CTNNA1,FADS2,RPS16,RPL23,CNN3,RPL29,RPSA,TPM
31,HMGN2,BEX3,SRSF4,RBBP4,PFN1,MCL1,PNN,RPL31,NISCH,PRKCSH,HNRNPC,TADA2B,XPR1,
HNRNPH1,UBE2H,CKAP4,CALU,ARCN1,CCT6A,HMGCR,CAPRIN1,CDK16,TOP2B,HDAC2,MMP2,S
3,PAPOLA,COL1A2,RPL14,HK1,NCS1,RHOA,PTCD3,CBX5,ATF4,PSMD2,SLC7A5,CHD4,JUND,TME

RBBP4,PFN1,MCL1,JPT2,PNN,SPC25,RPL31,NISCH,PRKCSH,HNRNPC,LRPPRC,TADA2B,KHDRBS
,RPL14,HK1,RPL9,KIF5B,RPS14,CHD4,JUND,RPS20,CD81,WDR1,RPL26,LASP1,BEX3,CCT7,RBBF
RPL27,SNRPN,ACO2,RBMX,TNPO1,SNRNP70,H3-3A,ABCA2,TOP1,RPL38,OPTN,UBXN4,G3BP2,MA
ACTN4,MMP14,PRDX2,DDX3X,CLTC,MAGED2,H1-0,IK,HMGCS1,GPAT4,TLN1,JUN,CTNNB1,CD63
XRCC5,MSN,RPL24,KPNB1,C1ORF43,PAPOLA,COL1A2,RPL14,HK1,RHOA,PTCD3,RPS14,CHD4,JU
G1,SFPQ,PDIA6,DDR1,MMP14,DDX3X,RPL12,RPL30,CLTC,MAGED2,RCN1,H1-0,RPS5,RPS7,IK,TL
JUND,TMEM259,RPS20,EIF5,CCT7,RBBP4,PFN1,MCL1,PNN,RPL31,NISCH,HNRNPC,LRPPRC,TAD
ES,RPL31,PRKCSH,HNRNPC,LRPPRC,TADA2B,XPR1,KHDRBS1,XPO1,CSNK1E,GLG1,SFPQ,RAB7A
RPS10,APLP2,HNRNPDL,TARDBP,ARID1A,ST3GAL2,HNRNPK,H1-10,RPL36,YWHAB,PTGES3,CKAF
14,EIF4H,SCARB2,LPCAT1,BRD3,XRCC5,PJA2,RPL24,SLC16A7,KPNB1,RPL17,C1ORF43,PAPOLA,
14,EIF4H,SCARB2,LPCAT1,BRD3,XRCC5,PJA2,RPL24,SLC16A7,KPNB1,RPL17,C1ORF43,PAPOLA,
2,DDX3X,RPL30,CLTC,MAGED2,RCN1,H1-0,IK,GPAT4,TLN1,CHPF,GJA1,COL6A2,APLP2,ST3GAL
C2,BCLAF1,PPIB,GLUL,LUC7L3,CPSF2,CTSB,NUCB1,TM9SF3,TOMM20,U2AF2,RPN2,PABPN1,HN
NGRN,MTSS2,HNRNPDL,TARDBP,RPLP2,ARID1A,NDFIP1,DDOST,ST3GAL2,WLS,HNRNPK,INIP,NA
PL36,SND1,LAMP1,HNRNPH1,CKAP4,RPS12,RAB1A,ARCN1,CCT6A,ARHGDIA,CDK16,HDAC2,BCLA
IPL,PARP1,MORF4L2,SSRP1,LMNB2,ABCA3,ALKBH5,RPL27,HNRNPA3,C4ORF3,TBC1D16,NCKAF
MED2,RPS15A,VEGFA,OPTN,G3BP2,SMARCA4,LAPTM4A,SRSF9,ARPC2,DSTN,PSMD3,ODC1,SRS
1D16,ACO2,CCT3,RBMX,CHD2,TNPO1,SNRNP70,H3-3A,HDGF,TSPAN6,PRPF6,TMED2,TOP1,OPT
R1,RAB7A,ACTN4,DDX3X,RPS15,MAGED2,H1-0,RPS5,RPS7,IK,GPAT4,PRKAR1A,TLN1,CHPF,COL
F1,HNRNPU,NFE2L1,SMARCA5,CTNNA1,FADS2,RPS16,RPL23,RPL29,RPSA,EIF4H,RPS9,XRCC5,M
,HMGCS1,CTNNB1,RPS10,APLP2,TARDBP,DDOST,HNRNPK,NARS1,RPL36,YWHAB,CKAP5,SEPTI
RPS25,TKT,GLG1,CDH6,SFPQ,PDIA6,DDR1,RAB7A,PRDX2,MAGED2,GNB2,H1-0,IK,JUN,CTNNB1,
B,HDAC2,BCLAF1,LUC7L3,CPSF2,DDX39B,SRSF5,TM9SF3,MDH2,TOMM20,RPN2,PABPN1,HNRN
14,EIF4H,SCARB2,LPCAT1,XRCC5,MSN,RPL24,RPL17,C1ORF43,PAPOLA,COL1A2,RPL14,HK1,RH
9SF3,MDH2,TOMM20,MLXIP,PGD,SQSTM1,NOP53,HNRNPL,MORF4L2,TAF15,HNRNPA3,ACO2,N
1,ARCN1,CCT6A,ARHGDIA,CAPRIN1,HDAC2,IPO5,SKP1,BCLAF1,PPIB,GLUL,LUC7L3,DCN,CDKN
H2,TOMM20,U2AF2,RPN2,PPP1CB,PGD,GNAI3,PABPN1,GDI2,HNRNPL,MORF4L2,TAF15,LMNB2,
PFKL,HDLBP,RPS25,TKT,GLG1,SFPQ,EIF5A,PDIA6,RAB7A,MMP14,DDX3X,RPL12,MAGED2,GNB2
A,CPSF2,CTSB,NUCB1,TM9SF3,MDH2,TOMM20,PABPN1,HNRNPL,RPL27,TBC1D16,ACO2,CCT3,
P1,RAD23B,SNRNP70,H3-3A,KDM6B,HDGF,PRPF6,TOP1,SELENON,OPTN,UBXN4,SMARCA4,LAPT
BP,DDOST,HNRNPK,NARS1,RPL36,YWHAB,CKAP5,SEPTIN11,SND1,LAMP1,HNRNPH1,CKAP4,RPS
,RPS12,ARCN1,CCT6A,CAPRIN1,HDAC2,SKP1,BCLAF1,LUC7L3,CPSF2,DDX39B,MDH2,TOMM20,
Q,PDIA6,RAB7A,MMP14,PRDX2,DDX3X,RPL30,MRFAP1,MAGED2,GNB2,H1-0,RPS7,PRKAR1A,TL
S1,OPTN,G3BP2,MAPK1,SMARCA4,SRSF1,ARGLU1,PRKDC,HNRNPUL1,ACTN1,FKBP8,DIDO1,FL
H5,RPL27,HNRNPA3,PKN1,ACO2,CBX1,RAD23B,RAB5B,CHD2,TNPO1,SNRNP70,PSMB2,H3-3A,H
C,TADA2B,XPR1,KHDRBS1,XPO1,RPS25,SFPQ,EIF5A,DDR1,RAB7A,RPS15,MAGED2,RCN1,H1-0,
CNN3,RPSA,TPM4,EIF4H,RTN4,LPCAT1,BRD3,PJA2,RPL24,KPNB1,C1ORF43,COL1A2,LLGL1,GAF
,ARCN1,CCT6A,ARHGDIA,CAPRIN1,CDK16,SKP1,BCLAF1,TRIB2,PPIB,LUC7L3,CPSF2,DDX39B,EI
DNAJC5,APLP2,HNRNPDL,TARDBP,ARID1A,HNRNPK,YWHAB,CKAP5,LAMP1,HNRNPH1,UBE2H,RAE
RPC2,SRSF1,ARGLU1,NYNRIN,PTBP1,ILF3,PRKDC,HNRNPUL1,HSPB1,SON,PLCG1,RPL28,DIDO1
CTSB,NUCB1,TM9SF3,MDH2,TOMM20,RNF187,CCT2,PABPN1,HNRNPL,PARP1,MORF4L2,LMNB2
9B,EID1,TM9SF3,MDH2,TOMM20,U2AF2,FDFT1,MLXIP,HNRNPL,TSPYL4,MORF4L2,TAF15,ABCA
AD23B,RAB5B,TNPO1,SNRNP70,KDM6B,HDGF,TSPAN6,TAF7,VEGFA,MARS1,OPTN,G3BP2,SMAF
D3,SRSF1,ARGLU1,FTO,ILF3,HNRNPUL1,ACTN1,SON,DIDO1,POLR2B,SMARCE1,SUPT6H,HNRNPM
ENON,OPTN,UBXN4,SMARCA4,DSTN,ODC1,SRSF1,ARGLU1,PTBP1,ILF3,PRKDC,HNRNPUL1,ACTN

,CCAR2,DIDO1,SMARCE1,HNRNPM,SSB,SRSF3,ILF2,PFKM,UBQLN1,SF3A1,RAI1,NCOR2,HNRNF
TRAP,PNMA8A,MLF2,KDM3B,TP53BP1,ANP32A,RBBP7,DCTN1,TXLNA,OS9,NAP1L1,PSMD1,HIP1,(
NXA6,NGRN,MACF1,GOLM1,HNRNPDL,TARDBP,WLS,HNRNPK,INIP,MYH10,CKAP5,CKAP4,RAB1A
S1,KLHL9,XPO1,CSNK1E,RPS25,GLG1,CDH6,SFPQ,TPM1,PEBP1,CCND2,CDH2,DDR1,MEGF8,RA
,CCN2,ACTN4,PRDX2,DDX3X,MAGED2,H1-0,IK,GPAT4,COL6A2,ANXA6,MACF1,APLP2,HNRNPDL,
0,H3-3A,TERF2IP,TOP1,RNF10,MARS1,OPTN,G3BP2,SMARCA4,ARPC2,DSTN,ARGLU1,PTBP1,EIF
JFV1,SF1,HIP1,VARS1,H2AZ1,UBQLN2,BRD4,UBA52,ATRX,SRSF6,GLUD1,TRA2B,ZFR,AGPAT1,M
MAPK1,SMARCA4,KDELR2,LAPTM4A,ARPC2,DSTN,SRSF1,IPO7,NYNRIN,PTBP1,FTO,ILF3,PRKDC,
RCE1,HNRNPM,SSB,SRSF3,EIF3F,SF3A1,HNRNPD,MLF2,SERBP1,KDM3B,TP53BP1,PSMA7,ANP32
5,GLG1,SFPQ,EIF5A,PDIA6,DDR1,RAB7A,RPS15,MAGED2,RCN1,H1-0,RPS7,IK,RPL35,HMGCS1,C
N,G3BP2,SMARCA4,TCF25,ILF3,PRKDC,HNRNPUL1,ACTN1,BNIP3L,MAP2K2,CCAR2,DIDO1,FLNB
36,LAMP1,HNRNPH1,CKAP4,HLA-B,ARCN1,HMGCR,ARHGDIA,CAPRIN1,TOP2B,HDAC2,BCLAF1,L
G1,SFPQ,PNMA1,RAB7A,ACTN4,PRDX2,DDX3X,CLTC,MAGED2,H1-0,IK,GPAT4,PRKAR1A,TLN1,CC
3A,HDGF,TOP1,TAF7,OPTN,MAPK1,SMARCA4,LAPTM4A,SRSF1,ARGLU1,PTBP1,FTO,PRKDC,HNF
PAT4,TLN1,COL6A2,MACF1,RPS10,DNAJC5,APLP2,TARDBP,HNRNPK,YWHAB,CKAP5,LAMP1,HNF
NRNPK,NARS1,RPL36,YWHAB,CKAP5,LAMP1,HNRNPH1,HLA-B,ARCN1,CCT6A,CAPRIN1,CDK16,S
ORO1C,SMARCE1,SMARCC1,SUPT6H,HNRNPM,SSB,SRSF3,ILF2,UBQLN1,CSNK1D,SF3A1,SREBF
3,PRKDC,HNRNPUL1,FKBP8,QARS1,CCAR2,DIDO1,SMARCE1,SMARCC1,HNRNPM,SSB,SRSF3,LA
AGED2,GNB2,RPS5,COL6A2,GOLM1,HNRNPDL,RPLP2,DDOST,WLS,HNRNPK,MYH10,NARS1,SND
RD4,UBA52,SLC39A7,SRSF6,GLUD1,TRA2B,UQCRC1,COX7C,ZFR,HNRNPA2B1,MAPRE2,RPS27A
4,SLC39A7,ATRX,EP400,SRSF6,TRA2B,UQCRC1,ZFR,AGPAT1,HNRNPA2B1,ANP32B,MCM3AP,HNI
FR,SV2A,HNRNPA2B1,ANP32B,GPSM1,HNRNPAB,SERP1,ZMYM3,FUS,ACLY,XRCC6,PELP1,VAPB,(
H,HNRNPM,SRSF3,ILF2,B2M,LONP1,UBQLN1,CSNK1D,SF3A1,ATXN2L,MPRIP,HNRNPD,SERBP1,I
2AZ1,GTPBP1,BRD4,UBA52,SLC39A7,PPP2CB,EP400,SRSF6,TRA2B,ZFR,AGPAT1,HNRNPA2B1,CU
F3,TOMM20,U2AF2,EEF1A2,CCT2,SUGP2,PPP1CB,NOP53,PABPN1,GDI2,HNRNPL,PARP1,TLN2,TI
,CTNNB1,RPS10,APLP2,TARDBP,DDOST,HNRNPK,RPL36,YWHAB,CKAP5,SEPTIN11,LAMP1,HNRN
GBP1,KTN1,RTF1,MTREX,MDM2,MECP2,ARL6IP5,ACAP2,SUN2,WIZ,PBXIP1,CDK9,PSME3,RBFOX
CO2,TCP1,PTOV1,SURF4,RBMX,DDX42,RAD23B,CHD2,CCT4,GCN1,SNRNP70,KDM6B,SCAF1,HD
,TADA2B,KHDRBS1,KLHL9,PFKL,RPS25,TKT,GLG1,SFPQ,PEBP1,CDH2,PDIA6,MEGF8,RAB7A,CCI
A2,NUCKS1,AHSA1,FOXK2,PSMD4,ITGAV,SRSF11,MSH6,PRPF19,CSE1L,NOC2L,SF3A2,AP2A2,YY
PSMD1,SF1,H2AZ1,BRD4,ATRX,SRSF6,GLUD1,TRA2B,HDAC6,ZFR,AGPAT1,HNRNPA2B1,ANP32B,
SKI,SNRNP70,H3-3A,HDGF,TOP1,G3BP2,SMARCA4,ARPC2,PSMD3,PRDX1,SRSF1,ARGLU1,TCF25
RBBP7,TXLNA,NAP1L1,LBH,NDUFV1,SF1,PRCC,H2AZ1,BRD4,UBA52,EP400,SRSF6,RER1,TRA2B,
AP1L1,KIRREL1,LBH,SF1,HIP1,FLOT1,BRD4,SLC39A7,MAP6,PPP2CB,ATRX,SRSF6,TRA2B,ZFR,HI
HNRNPL,MORF4L2,TAF15,SSRP1,ALKBH5,RPL27,PRDX5,ACO2,CCT3,PTOV1,ACAP3,RBMX,DDX4
NPUL1,ACTN1,MAP2K2,DIDO1,CORO1C,SMARCE1,DHX30,ATP6V1B2,HNRNPM,SSB,GIGYF1,SRSI
D2,GNB2,H1-0,APLP1,RPS7,GJA1,MYO5A,CTNNB1,RPS10,DNAJC5,APLP2,YWHAB,CKAP5,ATP1A
C4,TXLNA,NAP1L1,LBH,SF1,PRCC,H2AZ1,FLOT1,BRD4,EP400,SRSF6,RER1,TRA2B,COX7C,HDAC
H1-0,RPS5,IK,GPAT4,TLN1,NGRN,MACF1,APLP2,TARDBP,ARID1A,NDFIP1,DDOST,TSR2,MYL6,W
L36,YWHAB,CKAP5,SND1,LAMP1,HNRNPH1,ATP1A2,MAP2,CCT6A,RAC1,STIP1,HDAC2,BCLAF1,L
P70,H3-3A,SCAF1,HDGF,PRPF6,VEGFA,MARS1,SMARCA4,LAPTM4A,CCNL2,ARPC2,DSTN,SRSF1,
MEM43,TUBA1C,MED25,ZFP91,CDKN1B,RBM14,LMNA,THOC2,MED12,RXRA,TLK1,PHB,CTCF,DS
3P1,FTO,ILF3,PRKDC,HNRNPUL1,SON,KDM1A,KDELR1,ZFAND5,DIDO1,FLNB,POLR2B,CORO1C,
TN,SRSF1,PTBP1,ILF3,LUZP6,GRB2,CAPZB,PRKDC,HNRNPUL1,ACTN1,FKBP8,PLCG1,MAP2K2,D
3H,SF1,UBQLN2,KPNA2,BRD4,PPP2CB,KIAA0895L,GLUD1,EVL,TRA2B,UQCRC1,ZFR,CUL3,MCM

31,CUL3,NDST1,MCM3AP,POM121C,RPS27A,TNKS,APEX1,HNRNPAB,LRRC59,MTCH1,FUS,XRCC6
K,PRPF31,HEXIM1,MICALL1,SEC31A,FUBP1,POGZ,FAM193B,SMARCB1,SCAF8,FZD7,NF2,SECISE
B,PRKDC,HNRNPUL1,ACTN1,PLCG1,MAP2K2,DIDO1,POLR2B,CORO1C,SMARCE1,SUPT6H,FYN,H
C4,XRN2,RSF1,SUN1,POR,DAG1,RUVBL2,ELN,REEP5,MDM2,SF3A3,FAM50A,EWSR1,GALNT10,AT
SCAF1,HDGF,PRPF6,DPYSL2,TOP1,TAF7,OPTN,PPP2CA,ARPC2,PLEC,SRSF1,ARGLU1,NYNRIN,FT
NFRSF10B,NSD1,SEPHS1,TMEM43,SNRPD2,ZFP91,ALDH9A1,HSPE1,KDM2A,PPP1R37,PRKAR2B
K2,XRN2,PTPN11,HMGA1,SPPL2B,SUN1,MTA2,USP10,KTN1,TMEM94,MACROH2A1,IGF1R,MDM
AP9,ZMYM3,FUS,XRCC6,PELP1,ERGIC1,TCAF1,TPP1,GABARAPL1,HDGFL2,EFTUD2,SRSF2,TRAP

L,ZFR,AGPAT1,CUL3,APEX1,HNRNPAB,PODXL2,A2M,C1ORF198,MTCH1,ZMYM3,FUS,ERH,CARM
I,RTF1,H6PD,ANP32E,REEP5,IGF1R,MDM2,FAM50A,AAMP,ACAP2,HIPK2,WIZ,CDK9,TNFRSF10B,V
CSNK2A1,CD99,KPNA2,PPM1G,SFRP1,SRSF6,UPF1,TRA2B,HDAC6,ZFR,HNRNPA2B1,MCM3AP,GI
DCO1,USP47,XRN2,HTATSF1,SUN1,LGALS1,MTA2,CGGBP1,FAM171A1,PAIP2,ACTA2,ARRDC4,MI
ILF2,STT3B,ISYNA1,LONP1,UBQLN1,SF3A1,GPR107,HNRNPD,FBRSL1,SERBP1,KDM3B,ARFGEF
2,TRAP1,SFXN1,NMT1,PACS1,RBM5,SRSF11,FXYD6,TEX2,GSK3A,UBE3A,GOT2,TGOLN2,SMARCD
1,TOP2B,HDAC2,BCLAF1,L1CAM,TRIB2,LUC7L3,CPSF2,PGRMC1,EID1,CTSB,NUCB1,TM9SF3,MD
AP1,LSM14B,RAB6A,NMT1,AHSA1,PDPK1,PSMD4,SRSF11,MSH6,IGF2BP1,PRPF19,CSE1L,SF3A2,
B,WDR11,FAM160B2,PLK2,SEPHS1,DNAJC8,ALCAM,TUBA1C,SNRPD2,MED25,ZFP91,KARS1,KDI
,SULF2,NDUFV1,PSMD1,CSNK2A1,KPNA2,PHB2,GSPT1,BRD4,DECR1,ATRX,SRSF6,EIF5B,TRA2B

,HNRNPAB,RABEP1,ZMYM3,GRK2,FUS,XRCC6,CLPTM1,RALY,PGAP6,HDGFL2,TRAP1,FLOT2,EEI
E1,TM9SF4,HDGFL2,EFTUD2,KIAA0232,SRSF2,NUCKS1,FNBP1L,PLEKHM2,RBM5,SRSF11,PRPF1

CA,SLC1A3,SLC1A4,PSMD12,NFASC,COG4,LANCL1,EPHA4,DNM1,MAPK8IP3,PAF1,PPM1A,GNAO
3,PRKDC,HNRNPUL1,SON,KDM1A,CASC3,FKBP8,CCAR2,DIDO1,SMARCE1,SDF4,SMARCC1,HNRN
ALKBH5,RPL27,PTPRA,ACTR3,C4ORF3,UBE4A,PKN1,TCP1,CAMSAP1,CCT3,GRINA,PTOV1,DDX42
CA2,PRPF6,TERF2IP,TAF7,MARS1,TUBB2A,SMARCA4,MAPRE1,CCNL2,SRSF1,ARGLU1,ATP5PB,II

ATP6V0A1,PPP3CA,SLC1A3,FH,APOE,TUT4,NFASC,ABCD3,PRNP,PPT1,PPFIA3,ATP6V1F,PDHB,NN

4L2,TAF15,HNRNPA3,ACO2,ETNK1,LAMC1,CBX1,CCT4,SNRNP70,H3-3A,KDM6B,HDGF,ABCA2,PF

ZFR,HNRNPA2B1,MCM3AP,GPSM1,TMOD2,FUS,XRCC6,PELP1,BCL7A,RALY,PKDCC,GABARAPL1,
JN2,HIPK2,ETF1,HNRNPR,RTCB,RNF41,DDX6,AKAP8L,C1QBP,POLD2,STUB1,PSIP1,DLD,TUBA1C
,KLF3,FAM193B,SCAF8,ZNF207,CPSF7,DPF2,ALYREF
M5,SRSF11,SPTY2D1,CAP1,IST1,IVD,EIF3G,GSK3A,UBE3A,TGOLN2,SMARCD1,ATXN7L3,MN1,ATF
MACF1,DNAJC5,APLP2,TSR2,PTGES3,SEPTIN11,UBE2H,CKAP4,RPS12,RAB1A,ARCN1,CTBP1,HMG

/I2,ARPC1A,RTF1,TSNAX,PAIP2,GTF3C1,H6PD,GGNBP2,NOMO2,SGSM3,PPP1R14B,JAM3,FXR1,F

PT5H,HNRNPUL1,TDRD5,SCAF11,ACTN1,SON,FKBP8,CCAR2,DIDO1,TPM3,SDF4,SMARCC1,DHX3

A,WDR33,RAD50,ZNF326,NOP58,ANAPC2,PAM,ARMCX2,NUTF2,CTSA,ARSA

.1,PMPCA,ZNRF1,DNAJB6,GRHPR,LZTS2,SRRT,CTBP2,B4GALT2,SEPTIN9,SLC1A5,PCNA,CMTR1,

PT,FN1,SPEN,NONO,DDB1,YWHAQ,HACD3,DCHS1,SET,EGLN1,INA,MRC2,RPL19,LDHB,PKD1,SF3
19,LDHB,PKD1,SF3B1,PDIA3,SSBP2,CHD3,BASP1,VAPA,HMGB1,POLR2A,TPT1,RPS4X,RPS3A,RPL
1,VAPA,HMGB1,POLR2A,TPT1,RPS4X,RPS3A,RPL18,STMN1,UCHL1,RPL11,MORF4L1,BTF3,NCAM
RPS3A,RPL18,STMN1,TUBB4B,UCHL1,RPL11,MORF4L1,BTF3,AGRN,RPL10,MARCHF6,WSB1,RPS
,DDB1,YWHAQ,HACD3,DCHS1,SET,EGLN1,MRC2,RPL19,LDHB,PKD1,SF3B1,PDIA3,SSBP2,CHD3,
,RPS4X,RPS3A,RPL18,STMN1,TUBB4B,UCHL1,RPL11,MORF4L1,BTF3,AGRN,RPL10,WSB1,RPS1
SSBP2,CHD3,BASP1,VAPA,HMGB1,POLR2A,RPS4X,RPS3A,RPL18,STMN1,TUBB4B,UCHL1,RPL11,
BP2,CHD3,VAPA,HMGB1,POLR2A,TPT1,RPS4X,RPS3A,RPL18,STMN1,TUBB4B,UCHL1,RPL11,MC
SBP2,CHD3,BASP1,VAPA,HMGB1,TPT1,RPS4X,RPS3A,RPL18,STMN1,ITGB1,TUBB4B,RPL11,MOR
4X,RPS3A,RPL18,TUBB4B,RPL11,MORF4L1,BTF3,NCAM1,AGRN,ZMIZ1,XAF1,RPL10,MARCHF6,
4X,RPS3A,RPL18,TUBB4B,RPL11,MORF4L1,BTF3,NCAM1,AGRN,ZMIZ1,XAF1,RPL10,MARCHF6,
,SSBP2,CHD3,BASP1,VAPA,HMGB1,POLR2A,TPT1,RPS4X,RPS3A,RPL18,STMN1,TUBB4B,RPL11,
,PGAM1,NFE2L1,SMARCA5,LARP1,CTNNA1,FADS2,RPL23,CPE,CNN3,RPL29,RPSA,TPM4,CTSD,E
,RPL19,LDHB,PKD1,SF3B1,PDIA3,SSBP2,CHD3,BASP1,VAPA,HMGB1,POLR2A,TPT1,RPS4X,RPS3
,RPL19,LDHB,PKD1,SF3B1,PDIA3,SSBP2,CHD3,BASP1,VAPA,HMGB1,POLR2A,TPT1,RPS4X,RPS3
IN1,ITGB1,TUBB4B,RPL11,MORF4L1,BTF3,ANXA2,XAF1,RPL10,MARCHF6,RPS17,VCL,DNAJA1,C
POLR2A,TPT1,RPS4X,RPS3A,RPL18,STMN1,ITGB1,TUBB4B,RPL11,MORF4L1,BTF3,ANXA2,AGRN
POLR2A,TPT1,RPS4X,RPS3A,RPL18,STMN1,ITGB1,TUBB4B,RPL11,MORF4L1,BTF3,ANXA2,AGRN
_19,LDHB,SF3B1,PDIA3,SSBP2,CHD3,HMGB1,POLR2A,TPT1,RPS4X,RPS3A,RPL18,STMN1,ITGB1
A,TPT1,RPS4X,RPS3A,RPL18,STMN1,TUBB4B,RPL11,MORF4L1,BTF3,ANXA2,AGRN,XAF1,RPL10
1,RPL23,CNN3,RPL29,RPSA,TNS3,EIF4H,SCARB2,LPCAT1,BRD3,XRCC5,PJA2,MSN,RPL24,KPNE
SBP2,CHD3,BASP1,VAPA,HMGB1,POLR2A,TPT1,RPS4X,RPS3A,RPL18,STMN1,RPL11,MORF4L1,E
MGB1,POLR2A,TPT1,RPS4X,RPS3A,RPL18,TUBB4B,RPL11,MORF4L1,BTF3,NCAM1,AGRN,ZMIZ1
MGB1,POLR2A,TPT1,RPS4X,RPS3A,RPL18,TUBB4B,RPL11,MORF4L1,BTF3,NCAM1,AGRN,ZMIZ1
LARP1,HNRNPA0,FADS2,RPS16,RPL23,CPE,CNN3,RPL29,RPSA,TPM4,CTSD,EIF4H,SCARB2,LPCA
SA,TPM4,CTSD,EIF4H,SCARB2,RTN4,LPCAT1,RPS9,XRCC5,RPL24,SLC16A7,KPNB1,RPL17,C1O
,EGLN1,MRC2,RPL19,LDHB,PKD1,SF3B1,PDIA3,SSBP2,CHD3,VAPA,HMGB1,POLR2A,TPT1,RPS4
LR2A,TPT1,RPS4X,RPS3A,RPL18,STMN1,TUBB4B,RPL11,MORF4L1,BTF3,ANXA2,ZMIZ1,RPL10,
,MORF4L1,BTF3,ANXA2,XAF1,RPL10,RPS17,VCL,DNAJA1,CYB5R3,PAFAH1B1,MAP4,SF3B2,CDC4
,MORF4L1,BTF3,ANXA2,XAF1,RPL10,RPS17,VCL,DNAJA1,CYB5R3,PAFAH1B1,MAP4,SF3B2,CDC4
SSBP2,CHD3,VAPA,HMGB1,POLR2A,TPT1,RPS4X,RPS3A,RPL18,TUBB4B,RPL11,MORF4L1,BTF3
,ITGB1,TUBB4B,RPL11,BTF3,ANXA2,AGRN,XAF1,RPL10,MARCHF6,RPS17,VCL,DNAJA1,CYB5R3,
,ITGB1,TUBB4B,RPL11,BTF3,ANXA2,AGRN,XAF1,RPL10,MARCHF6,RPS17,VCL,DNAJA1,CYB5R3,
NF770,AP2B1,GTF2F1,HNRNPU,OGA,PGAM1,NFE2L1,SMARCA5,LARP1,HNRNPA0,CTNNA1,RPS1
L,DNAJA1,PAFAH1B1,MAP4,SF3B2,CDC42BPB,ITM2B,MAT2A,GTF2F1,HNRNPU,PGAM1,NFE2L1,
,HMGB1,POLR2A,RPS4X,RPS3A,RPL18,STMN1,TUBB4B,RPL11,MORF4L1,BTF3,ANXA2,XAF1,RF
1,HNRNPA0,FADS2,RPS16,RPL23,CNN3,RPL29,RPSA,TPM4,CTSD,EIF4H,SCARB2,RTN4,LPCAT1,
NXA2,RPL10,WSB1,RPS17,VCL,DNAJA1,CYB5R3,PAFAH1B1,MAP4,SF3B2,CDC42BPB,ITM2B,ZNF
NXA2,RPL10,WSB1,RPS17,VCL,DNAJA1,CYB5R3,PAFAH1B1,MAP4,SF3B2,CDC42BPB,ITM2B,ZNF
RPL10,RPS17,VCL,DNAJA1,PAFAH1B1,MAP4,SF3B2,CDC42BPB,ITM2B,ZNF770,MAT2A,GTF2F1,
CARB2,BRD3,XRCC5,PJA2,MSN,RPL24,RPL17,C1ORF43,PAPOLA,COL1A2,LLGL1,RPL14,HK1,RH
DIA3,SSBP2,CHD3,VAPA,HMGB1,POLR2A,TPT1,RPS4X,RPS3A,RPL18,TUBB4B,RPL11,MORF4L1,
DIA3,SSBP2,CHD3,VAPA,HMGB1,POLR2A,TPT1,RPS4X,RPS3A,RPL18,TUBB4B,RPL11,MORF4L1,
HD3,VAPA,HMGB1,POLR2A,TPT1,RPS4X,RPS3A,RPL18,STMN1,ITGB1,TUBB4B,RPL11,MORF4L1,
HD3,VAPA,HMGB1,POLR2A,TPT1,RPS4X,RPS3A,RPL18,STMN1,ITGB1,TUBB4B,RPL11,MORF4L1,

NRNPU,NFE2L1,SMARCA5,LARP1,HNRNPA0,CTNNA1,FADS2,RPS16,RPL23,CNN3,RPL29,RPSA,TF
ASP1,VAPA,HMGB1,POLR2A,TPT1,RPS4X,RPS3A,RPL18,STMN1,TUBB4B,RPL11,MORF4L1,BTF3,
ASP1,VAPA,HMGB1,POLR2A,TPT1,RPS4X,RPS3A,RPL18,STMN1,TUBB4B,RPL11,MORF4L1,BTF3,
)3,VAPA,HMGB1,POLR2A,TPT1,RPS4X,RPS3A,RPL18,STMN1,TUBB4B,RPL11,MORF4L1,BTF3,AN
8,STMN1,TUBB4B,RPL11,MORF4L1,BTF3,NCAM1,ANXA2,XAF1,RPL10,MARCHF6,WSB1,RPS17,
PA,HMGB1,POLR2A,TPT1,RPS4X,RPS3A,RPL18,STMN1,ITGB1,TUBB4B,RPL11,MORF4L1,BTF3,A
HD3,BASP1,VAPA,HMGB1,POLR2A,TPT1,RPS4X,RPS3A,RPL18,STMN1,TUBB4B,RPL11,MORF4L1,
TF3,AGRN,XAF1,RPL10,MARCHF6,WSB1,RPS17,DNAJA1,CYB5R3,PAFAH1B1,SF3B2,CDC42BPB,I
11,MORF4L1,BTF3,ANXA2,AGRN,XAF1,RPL10,MARCHF6,RPS17,VCL,DNAJA1,CYB5R3,PAFAH1B1
11,MORF4L1,BTF3,ANXA2,AGRN,XAF1,RPL10,MARCHF6,RPS17,VCL,DNAJA1,CYB5R3,PAFAH1B1
SBP2,CHD3,VAPA,HMGB1,POLR2A,TPT1,RPS4X,RPS3A,RPL18,STMN1,TUBB4B,UCHL1,MORF4L1
MAP4,SF3B2,CDC42BPB,MAT2A,GTF2F1,HNRNPU,OGA,NFE2L1,SMARCA5,LARP1,HNRNPA0,CTN
IN1,TUBB4B,RPL11,MORF4L1,BTF3,ANXA2,AGRN,ZMIZ1,XAF1,RPL10,RPS17,VCL,DNAJA1,CYB5
IN1,TUBB4B,RPL11,MORF4L1,BTF3,ANXA2,AGRN,ZMIZ1,XAF1,RPL10,RPS17,VCL,DNAJA1,CYB5
1,RPL14,HK1,RHOA,PTCD3,PSMD2,RPS14,CHD4,JUND,TMEM259,RPS20,EIF5,CD81,HMGN2,RP
A2,XAF1,RPL10,WSB1,RPS17,VCL,DNAJA1,CYB5R3,PAFAH1B1,MAP4,SF3B2,CDC42BPB,ITM2B,Z
DNAJA1,CYB5R3,PAFAH1B1,AKAP12,MAP4,SF3B2,CDC42BPB,ITM2B,GTF2F1,HNRNPU,OGA,NFE
,POLR2A,TPT1,RPS4X,RPS3A,RPL18,STMN1,TUBB4B,RPL11,MORF4L1,BTF3,ANXA2,AGRN,ZMIZ
,POLR2A,TPT1,RPS4X,RPS3A,RPL18,STMN1,TUBB4B,RPL11,MORF4L1,BTF3,ANXA2,AGRN,ZMIZ
RPL18,STMN1,TUBB4B,RPL11,MORF4L1,BTF3,ANXA2,AGRN,ZMIZ1,RPL10,MARCHF6,RPS17,VC
RPL18,STMN1,TUBB4B,RPL11,MORF4L1,BTF3,ANXA2,AGRN,ZMIZ1,RPL10,MARCHF6,RPS17,VC
FE2L1,SMARCA5,LARP1,HNRNPA0,CTNNA1,FADS2,RPS16,RPL23,CNN3,RPL29,RPSA,TPM4,TNS3,
A,HMGB1,POLR2A,TPT1,RPS4X,RPS3A,RPL18,STMN1,TUBB4B,RPL11,MORF4L1,BTF3,ANXA2,AG
B2,CDC42BPB,ITM2B,ZNF770,AP2B1,MAT2A,GTF2F1,OGA,PGAM1,NFE2L1,SMARCA5,CTNNA1,
TUBB4B,RPL11,MORF4L1,BTF3,ANXA2,AGRN,ZMIZ1,XAF1,RPL10,RPS17,VCL,CYB5R3,PAFAH1B
B,MAT2A,GTF2F1,HNRNPU,OGA,NFE2L1,SMARCA5,LARP1,HNRNPA0,CTNNA1,FADS2,RPS16,RPL
IN3,RPL29,RPSA,TPM4,EIF4H,SCARB2,LPCAT1,BRD3,XRCC5,PJA2,MSN,RPL24,SLC16A7,KPNB1,
AFAH1B1,MAP4,SF3B2,CDC42BPB,ITM2B,ZNF770,GTF2F1,HNRNPU,PGAM1,NFE2L1,SMARCA5,H
SMARCA5,LARP1,HNRNPA0,CTNNA1,FADS2,CNN3,RPSA,TPM4,CTSD,EIF4H,SCARB2,LPCAT1,LAN
SMARCA5,LARP1,HNRNPA0,CTNNA1,FADS2,CNN3,RPSA,TPM4,CTSD,EIF4H,SCARB2,LPCAT1,LAN
32,CDC42BPB,ITM2B,GTF2F1,HNRNPU,PGAM1,NFE2L1,SMARCA5,HNRNPA0,CTNNA1,FADS2,RPS
CHF6,RPS17,VCL,DNAJA1,CYB5R3,PAFAH1B1,SF3B2,CDC42BPB,ITM2B,ZNF770,AP2B1,GTF2F1,
FADS2,RPS16,RPL29,RPSA,CTSD,EIF4H,SCARB2,RTN4,LPCAT1,BRD3,XRCC5,PJA2,MSN,RPL24,K
HMGB1,POLR2A,TPT1,RPS4X,RPS3A,RPL18,STMN1,TUBB4B,RPL11,MORF4L1,BTF3,ANXA2,AGF
HMGB1,POLR2A,TPT1,RPS4X,RPS3A,RPL18,STMN1,TUBB4B,RPL11,MORF4L1,BTF3,ANXA2,AGF
X,RPL18,TUBB4B,RPL11,MORF4L1,BTF3,ANXA2,AGRN,ZMIZ1,XAF1,RPL10,RPS17,VCL,DNAJA1,
PA,HMGB1,POLR2A,TPT1,RPS4X,RPS3A,RPL18,STMN1,TUBB4B,RPL11,MORF4L1,BTF3,ANXA2,Z
4L1,BTF3,ANXA2,AGRN,ZMIZ1,XAF1,RPL10,MARCHF6,WSB1,RPS17,VCL,DNAJA1,PAFAH1B1,MA
D3,XRCC5,PJA2,MSN,RPL24,KPNB1,RPL17,PAPOLA,COL1A2,LLGL1,GAP43,RPL14,HK1,RHOA,PT
10,WSB1,RPS17,VCL,DNAJA1,CYB5R3,PAFAH1B1,MAP4,SF3B2,CDC42BPB,ITM2B,AP2B1,MAT2
NA1,FADS2,RPS16,RPL23,RPL29,RPSA,TPM4,TNS3,CTSD,EIF4H,SCARB2,LPCAT1,RPS24,BRD3,F
PA0,CTNNA1,FADS2,RPS16,RPL23,CNN3,RPL29,RPSA,TPM4,CTSD,EIF4H,SCARB2,LPCAT1,BRD3
NA1,FADS2,RPS16,RPL23,CNN3,RPL29,RPSA,TPM4,EIF4H,SCARB2,BRD3,RPS9,XRCC5,RPL24,KF
F3,ANXA2,AGRN,RPL10,MARCHF6,RPS17,DNAJA1,CYB5R3,PAFAH1B1,AKAP12,MAP4,SF3B2,CDC
,AGRN,XAF1,RPL10,RPS17,VCL,DNAJA1,PAFAH1B1,MAP4,SF3B2,CDC42BPB,ITM2B,GTF2F1,GO

L1,SMARCA5,HNRNPA0,CTNNA1,FADS2,RPS16,RPL23,CNN3,RPL29,RPSA,TPM4,TNS3,EIF4H,SC
11,MORF4L1,BTF3,ANXA2,AGRN,RPL10,RPS17,VCL,DNAJA1,CYB5R3,PAFAH1B1,MAP4,SF3B2,C
N1,TUBB4B,RPL11,MORF4L1,BTF3,ANXA2,AGRN,XAF1,RPL10,MARCHF6,WSB1,RPS17,VCL,DNA
N1,TUBB4B,RPL11,MORF4L1,BTF3,ANXA2,AGRN,XAF1,RPL10,MARCHF6,WSB1,RPS17,VCL,DNA
CL1,JPT2,PNN,RPL31,PRKCSH,HNRNPC,LRPPRC,TADA2B,XPR1,KHDRBS1,XPO1,SLC3A2,RPS25,T
5R3,PAFAH1B1,MAP4,SF3B2,ITM2B,ZNF770,MAT2A,GTF2F1,GOLGA8A,HNRNPU,OGA,PGAM1,N
CYB5R3,PAFAH1B1,MAP4,SF3B2,CDC42BPB,ITM2B,ZNF770,AP2B1,GTF2F1,HNRNPU,NFE2L1,S
NNA1,FADS2,RPL23,RPL29,RPSA,TPM4,EIF4H,SCARB2,RPS24,BRD3,RPS9,XRCC5,MSN,RPL24,K
F3B1,PDIA3,SSBP2,BASP1,VAPA,HMGB1,POLR2A,TPT1,RPS4X,RPS3A,RPL18,STMN1,TUBB4B,R
PS17,VCL,DNAJA1,CYB5R3,PAFAH1B1,MAP4,SF3B2,CDC42BPB,ITM2B,ZNF770,MAT2A,GTF2F1,
PS17,VCL,DNAJA1,CYB5R3,PAFAH1B1,MAP4,SF3B2,CDC42BPB,ITM2B,ZNF770,MAT2A,GTF2F1,
SA,TPM4,TNS3,EIF4H,SCARB2,LPCAT1,BRD3,RPS9,XRCC5,MSN,RPL24,SLC16A7,RPL17,C1ORF4
A1,RPS16,RPL23,CNN3,RPSA,TPM4,EIF4H,SCARB2,LPCAT1,BRD3,XRCC5,PJA2,MSN,RPL24,KP
A,PGAM1,SMARCA5,CTNNA1,RPL23,CNN3,RPL29,RPSA,TPM4,CTSD,EIF4H,LPCAT1,BRD3,XRCC5
A,PGAM1,SMARCA5,CTNNA1,RPL23,CNN3,RPL29,RPSA,TPM4,CTSD,EIF4H,LPCAT1,BRD3,XRCC5
NS3,EIF4H,SCARB2,LPCAT1,BRD3,RPS9,XRCC5,PJA2,SLC16A7,RPL17,C1ORF43,PAPOLA,LLGL1,
EIF4H,SCARB2,BRD3,RPS9,XRCC5,PJA2,RPL24,RPL17,PAPOLA,COL1A2,LLGL1,RPL14,HK1,PTC
A5,LARP1,HNRNPA0,CTNNA1,FADS2,RPL23,CNN3,RPSA,TPM4,CTSD,EIF4H,SCARB2,RPS24,BRD3
A0,CTNNA1,RPL23,CNN3,RPL29,RPSA,CTSD,EIF4H,LPCAT1,RPS24,BRD3,RPS9,XRCC5,PJA2,RPL
M1,NFE2L1,SMARCA5,LARP1,RPS16,RPL23,CNN3,RPL29,RPSA,TNS3,CTSD,EIF4H,SCARB2,LPC
RNPA0,CTNNA1,FADS2,RPS16,RPL23,CNN3,RPSA,TPM4,TNS3,CTSD,EIF4H,LPCAT1,RPS24,BRD3,
IO,DDB1,YWHAQ,HACD3,DCHS1,SET,EGLN1,INA,MRC2,RPL19,LDHB,PKD1,SF3B1,PDIA3,SSBP2,
P4,RTN3,PFN1,MCL1,RPL31,NISCH,PRKCSH,HNRNPC,TADA2B,XPR1,KHDRBS1,XPO1,CSNK1E,S
2BPB,ZNF770,GTF2F1,HNRNPU,OGA,PGAM1,NFE2L1,SMARCA5,LARP1,HNRNPA0,CTNNA1,RPS1
X3X,RPS15,MAGED2,RCN1,H1-0,RPL35,GPAT4,GJA1,COL6A2,ANXA6,GOLM1,TARDBP,ARID1A,W
L,DNAJA1,PAFAH1B1,MAP4,SF3B2,CDC42BPB,ITM2B,ZNF770,AP2B1,GTF2F1,GOLGA8A,HNRNP
F3B2,CDC42BPB,ITM2B,AP2B1,MAT2A,GTF2F1,HNRNPU,OGA,PGAM1,NFE2L1,SMARCA5,LARP1,
NFE2L1,LARP1,HNRNPA0,CTNNA1,FADS2,RPS16,RPL23,CNN3,RPSA,TPM4,CTSD,EIF4H,LPCAT1,
AJA1,PAFAH1B1,SF3B2,CDC42BPB,ITM2B,ZNF770,AP2B1,GTF2F1,GOLGA8A,HNRNPU,PGAM1,N
,VCL,DNAJA1,PAFAH1B1,MAP4,SF3B2,CDC42BPB,ITM2B,GTF2F1,GOLGA8A,HNRNPU,NFE2L1,S
,MAP4,SF3B2,CDC42BPB,ITM2B,AP2B1,GTF2F1,GOLGA8A,HNRNPU,OGA,PGAM1,NFE2L1,SMA
RD3,XRCC5,RPL24,RPL17,C1ORF43,PAPOLA,COL1A2,LLGL1,RPL14,HK1,NCS1,RHOA,PTCD3,CE
CDK16,TOP2B,HDAC2,SKP1,BCLAF1,LUC7L3,MAP1A,CPSF2,CTSB,NUCB1,TM9SF3,MDH2,TOM
PL29,RPSA,TPM4,CTSD,EIF4H,SCARB2,LPCAT1,BRD3,RPS9,XRCC5,MSN,RPL24,KPNB1,C1ORF4
1,MAP4,SF3B2,CDC42BPB,ITM2B,GTF2F1,HNRNPU,NFE2L1,SMARCA5,LARP1,CTNNA1,FADS2,R
R3,MAP4,SF3B2,CDC42BPB,AP2B1,GTF2F1,HNRNPU,PGAM1,NFE2L1,LARP1,HNRNPA0,CTNNA1,
RPL10,RPS17,VCL,DNAJA1,CYB5R3,AKAP12,MAP4,SF3B2,CDC42BPB,ITM2B,GTF2F1,HNRNPU,
16,RPL23,RPSA,TPM4,EIF4H,LPCAT1,RPS24,BRD3,RPS9,XRCC5,PJA2,MSN,RPL24,KPNB1,RPL1
F770,MAT2A,GTF2F1,HNRNPU,OGA,PGAM1,NFE2L1,SMARCA5,CTNNA1,FADS2,RPS16,RPL23,C
OLGA8A,HNRNPU,PGAM1,NFE2L1,SMARCA5,HNRNPA0,CTNNA1,FADS2,RPS16,RPL23,CNN3,RP
CA5,LARP1,HNRNPA0,CTNNA1,FADS2,RPS16,RPL23,CNN3,RPL29,RPSA,TPM4,TNS3,CTSD,EIF4H,
COL1A2,GAP43,RPL14,HK1,NCS1,PTCD3,CBX5,RPL9,CLSTN1,PSMD2,RPS14,CHD4,TMEM259,R
770,MAT2A,GTF2F1,GOLGA8A,HNRNPU,OGA,PGAM1,NFE2L1,SMARCA5,LARP1,CTNNA1,FADS2,
1,SMARCA5,LARP1,HNRNPA0,CTNNA1,FADS2,RPS16,RPL23,CPE,CNN3,RPL29,RPSA,TPM4,TNS3,
A5,LARP1,HNRNPA0,CTNNA1,FADS2,RPS16,RPL23,RPL29,RPSA,TPM4,EIF4H,LPCAT1,BRD3,RPS

RPL23,CNN3,RPL29,RPSA,TPM4,TNS3,CTSD,EIF4H,SCARB2,LPCAT1,RPS24,BRD3,XRCC5,PJA2,M
NF865,CLSTN1,PSMD2,SLC7A5,RPS14,CHD4,TMEM259,RPS20,WDR1,RPL26,CRMP1,CCT7,SRSF
CHD1,HDAC2,BCLAF1,PPIB,LUC7L3,SQLE,DCN,PGRMC1,DDX39B,CTSB,TM9SF3,MDH2,TOMM2
PL14,HK1,PTCD3,ZNF865,RPL9,ATF4,PSMD2,KIF5B,SLC7A5,RPS14,CHD4,JUND,TMEM259,RPS2
CL,DNAJA1,CYB5R3,PAFAH1B1,MAP4,SF3B2,CDC42BPB,ITM2B,GTF2F1,HNRNPU,OGA,NFE2L1,S
PDL,TARDBP,DDOST,ST3GAL2,HNRNPK,H1-10,MYH10,NARS1,YWHAB,PTGES3,CKAP5,SND1,LAM
,C1ORF43,PAPOLA,COL1A2,RPL14,HK1,RHOA,PTCD3,ZNF865,PSMD2,KIF5B,SLC7A5,RPS14,CX
TMN1,TUBB4B,RPL11,MORF4L1,BTF3,ANXA2,AGRN,XAF1,RPL10,RPS17,VCL,DNAJA1,PAFAH1B
TMN1,TUBB4B,RPL11,MORF4L1,BTF3,ANXA2,AGRN,XAF1,RPL10,RPS17,VCL,DNAJA1,PAFAH1B
1,SMARCA5,LARP1,HNRNPA0,CTNNA1,FADS2,RPS16,RPL23,RPL29,RPSA,TPM4,TNS3,CTSD,EIF4
3,XRCC5,MSN,KPNB1,RPL17,C1ORF43,PAPOLA,COL1A2,RPL14,HK1,RHOA,PTCD3,CBX5,RPL9,
F4,PSMD2,RPS14,CXADR,CHD4,RPS20,EIF5,CD81,WDR1,HMGN2,RPL26,BEX3,CCT7,SRSF4,RB
F4,PSMD2,RPS14,CXADR,CHD4,RPS20,EIF5,CD81,WDR1,HMGN2,RPL26,BEX3,CCT7,SRSF4,RB
POLA,COL1A2,LLGL1,RPL14,RPL9,CLSTN1,ATF4,RPS14,CXADR,JUND,RPS20,EIF5,CD81,HMGN2
,TMEM259,RPS20,EIF5,HMGN2,LASP1,CCT7,SRSF4,RBBP4,PFN1,MCL1,JPT2,PRKCSH,HNRNPC
2,BRD3,XRCC5,RPL24,KPNB1,RPL17,C1ORF43,PAPOLA,COL1A2,LLGL1,RPL14,HK1,PTCD3,CB
BEX3,CCT7,RBBP4,MCL1,PNN,RPL31,NISCH,PRKCSH,HNRNPC,LRPPRC,TADA2B,KHDRBS1,XPO1
7A5,RPS14,CXADR,CHD4,ATP1A1,JUND,RPS20,EIF5,HMGN2,RPL26,BEX3,CCT7,SRSF4,RBBP4,F
,LARP1,HNRNPA0,CTNNA1,RPS16,RPL23,CNN3,RPSA,TPM4,TNS3,CTSD,EIF4H,SCARB2,LPCAT1
X,RPS3A,RPL11,MORF4L1,BTF3,AGRN,XAF1,RPL10,MARCHF6,RPS17,DNAJA1,CYB5R3,PAFAH1B
GTF2F1,HNRNPU,NFE2L1,SMARCA5,CTNNA1,RPS16,RPL23,CNN3,RPSA,TPM4,CTSD,SCARB2,LF
IF5,WDR1,HMGN2,RPL26,CCT7,SRSF4,RBBP4,PFN1,MCL1,JPT2,PNN,RPL31,PRKCSH,HNRNPC,L
1,RBMX,DDX42,RAD23B,CHD2,TNPO1,SNRNP70,H3-3A,ABCA2,TERF2IP,TMED2,TOP1,MARS1,OF
SF3B2,CDC42BPB,ITM2B,GTF2F1,GOLGA8A,HNRNPU,NFE2L1,SMARCA5,LARP1,HNRNPA0,CTNI
CD3,CLSTN1,ATF4,PSMD2,RPS14,CHD4,JUND,TMEM259,RPS20,EIF5,HMGN2,CCT7,SRSF4,RBB
1,HNRNPA0,CTNNA1,FADS2,RPL23,CPE,CNN3,RPSA,TPM4,CTSD,EIF4H,SCARB2,LPCAT1,BRD3,X
1,HNRNPA0,CTNNA1,FADS2,RPL23,CPE,CNN3,RPSA,TPM4,CTSD,EIF4H,SCARB2,LPCAT1,BRD3,X
F4,KIF5B,SLC7A5,RPS14,CXADR,CHD4,JUND,RPS20,EIF5,RPL26,CCT7,SRSF4,RBBP4,MCL1,PNN
AB7A,PRDX2,DDX3X,RPL30,CLTC,MRFAP1,H1-0,RPS7,IK,RPL35,GPAT4,CHPF,MYO5A,GOLM1,A
59,RPS20,EIF5,WDR1,RPL26,BEX3,CCT7,SRSF4,RBBP4,MCL1,HNRNPC,LRPPRC,TADA2B,XPR1,K
PQ,PDIA6,DDR1,ACSL6,RAB7A,DDX3X,CLTC,MAGED2,H1-0,RPS7,IK,RPL35,GPAT4,TLN1,MACF1,
1,PRKCSH,HNRNPC,TADA2B,XPR1,KHDRBS1,XPO1,SPTBN2,SLC3A2,CSNK1E,RPS25,TKT,CDH6,S
L24,KPNB1,RPL17,C1ORF43,PAPOLA,LLGL1,RPL14,HK1,PTCD3,CBX5,RPL9,PSMD2,KIF5B,SLC
E2H,HLA-B,ARCN1,CCT6A,CAPRIN1,TOP2B,SMCHD1,HDAC2,BCLAF1,TRIB2,GLUL,LUC7L3,CPSF
LP2,TARDBP,WLS,HNRNPK,NARS1,RPL36,YWHAB,CKAP5,LAMP1,HNRNPH1,RAB1A,CALU,ARCN1
L,TARDBP,ARID1A,ST3GAL2,HNRNPK,H1-10,RPL36,YWHAB,CKAP5,LAMP1,HNRNPH1,ATP1A2,CA
FADS2,RPS16,RPL23,CNN3,RPL29,RPSA,TPM4,TNS3,CTSD,EIF4H,SCARB2,LPCAT1,BRD3,RPS9,X
BBP4,PFN1,MCL1,JPT2,PNN,NES,NISCH,PRKCSH,HNRNPC,LRPPRC,XPR1,KHDRBS1,XPO1,CSNK
TCD3,ATF4,PSMD2,KIF5B,SLC7A5,RPS14,CHD4,TMEM259,RPS20,EIF5,WDR1,HMGN2,RPL26,LA
EX3,CCT7,SRSF4,RBBP4,PFN1,MCL1,JPT2,PNN,NES,RPL31,NISCH,HNRNPC,LRPPRC,TADA2B,XP
T6A,CAPRIN1,TOP2B,HDAC2,BCLAF1,LUC7L3,CPSF2,DDX39B,TM9SF3,MDH2,TOMM20,U2AF2,R
4,CTSD,EIF4H,SCARB2,LPCAT1,BRD3,RPS9,PJA2,RPL24,RPL17,C1ORF43,COL1A2,LLGL1,RPL1
KHDRBS1,XPO1,CSNK1E,HDLBP,RPS25,GLG1,SFPQ,PDIA6,RAB7A,ACTN4,MMP14,DDX3X,RPL12
KP1,BCLAF1,PPIB,GLUL,LUC7L3,CPSF2,CTSB,TM9SF3,MDH2,RNF187,U2AF2,RPN2,SQSTM1,PA
M259,WDR1,HMGN2,RPL26,LASP1,BEX3,CCT7,SRSF4,RBBP4,MCL1,PNN,NISCH,PRKCSH,HNRN

S1,XPO1,CSNK1E,HDLBP,RPS25,GLG1,SFPQ,PDIA6,DDX3X,RPL30,CLTC,MAGED2,H1-0,RPS5,RPS
P4,PFN1,JPT2,PNN,RPL31,HNRNPC,LRPPRC,TADA2B,XPR1,KHDRBS1,XPO1,RPS25,GLG1,SFPQ,P
APK1,SMARCA4,SRSF9,ARPC2,SRSF1,ARGLU1,TCF25,ILF3,PRKDC,HNRNPUL1,NSG2,PLCG1,QAF
3,ANXA6,DNAJC5,APLP2,HNRNPDL,TARDBP,ARID1A,DDOST,HNRNPK,H1-10,INIP,NARS1,YWHAB,C
JND,RPS20,EIF5,WDR1,HMGN2,RPL26,LASP1,BEX3,CCT7,RBBP4,PFN1,MCL1,JPT2,PNN,RPL31,N
LN1,CHPF,COL6A2,APLP2,HNRNPDL,TARDBP,ST3GAL2,HNRNPK,MYH10,RPL36,YWHAB,CKAP5,LA
A2B,XPR1,KHDRBS1,XPO1,RPS25,TKT,GLG1,SFPQ,PDIA6,RAB7A,ACTN4,MMP14,PRDX2,DDX3X,
A,MMP14,DDX3X,MAGED2,GNB2,H1-0,RPS5,IK,MIF,TLN1,GJA1,COL6A2,ANXA6,MACF1,APLP2,HI
P5,LAMP1,HNRNPH1,ATP1A2,CALU,CTBP1,CAPRIN1,CDK16,TOP2B,HDAC2,SKP1,BCLAF1,PPIB,G
,COL1A2,LLGL1,RPL14,HK1,RHOA,PTCD3,RPL9,CLSTN1,ATF4,PSMD2,KIF5B,SLC7A5,RPS14,CX
,COL1A2,LLGL1,RPL14,HK1,RHOA,PTCD3,RPL9,CLSTN1,ATF4,PSMD2,KIF5B,SLC7A5,RPS14,CX
2,HNRNPK,H1-10,RPL36,YWHAB,CKAP5,LAMP1,HNRNPH1,CKAP4,CALU,ARCN1,ARHGDIA,CAPRI
RNPL,TM9SF2,MORF4L2,TAF15,LMNB2,ABCA3,ALKBH5,RPL27,HNRNPA3,PKN1,TBC1D16,ACO2
RS1,RPL36,YWHAB,CKAP5,SND1,LAMP1,HNRNPH1,CALU,SEPTIN3,CCT6A,GPC1,TOP2B,HDAC2,E
AF1,LUC7L3,CPSF2,DDX39B,EID1,TM9SF3,MDH2,TOMM20,RNF187,U2AF2,RPN2,SQSTM1,PABPI
P1,RBMX,DDX42,JPT1,CBX1,RAD23B,CHD2,TNPO1,SNRNP70,PSMB2,KDM6B,TAF7,OPTN,UBXN4
SF1,ARGLU1,NYNRIN,TCF25,PTBP1,ILF3,GRB2,CAPZB,PRKDC,HNRNPUL1,ACTN1,HSPB1,KDM1A
TN,UBXN4,G3BP2,SMARCA4,ARPC2,DSTN,SRSF1,ARGLU1,TCF25,PTBP1,ILF3,PRKDC,HNRNPUL1
6A2,ANXA6,RPS10,APLP2,TARDBP,ARID1A,TSR2,MYL6,HNRNPK,H1-10,NARS1,RPL36,YWHAB,CK
MSN,RPL24,SLC16A7,RPL17,C1ORF43,PAPOLA,COL1A2,RPL14,HK1,CBX5,RPL9,ATF4,KIF5B,SL
N11,LAMP1,HNRNPH1,CKAP4,RPS12,HLA-B,ARCN1,CCT6A,HMGCR,ARHGDIA,CAPRIN1,CDK16,T
GOLM1,APLP2,HNRNPDL,TARDBP,NDFIP1,DDOST,HNRNPK,H1-10,INIP,NARS1,RPL36,YWHAB,CK
NPL,PARP1,TSPYL4,MORF4L2,TAF15,ABCA3,RPL27,HNRNPA3,C4ORF3,PKN1,ACO2,VPS35,RBMX
HOA,PTCD3,PSMD2,SLC7A5,RPS14,CHD4,JUND,RPS20,EIF5,WDR1,HMGN2,RPL26,LASP1,BEX3,C
NCKAP1,RBMX,JPT1,RAD23B,TNPO1,SNRNP70,H3-3A,HDGF,TSPAN6,PRPF6,TOP1,SELENON,OPT
1A,CPSF2,PGRMC1,TM9SF3,MDH2,U2AF2,RPN2,MLXIP,ANXA5,PGD,PABPN1,HNRNPL,PARP1,TSF
PDXK,ACO2,VPS35,RBMX,RAD23B,CHD2,CCT4,GCN1,SNRNP70,PYGB,PRPF6,TERF2IP,TOP1,MAI
2,H1-0,RPS7,IK,PRKAR1A,TLN1,CHPF,GJA1,CTNNB1,COL6A2,NGRN,MACF1,HNRNPDL,TARDBP,N
VPS35,RBMX,RAD23B,CHD2,TNPO1,SNRNP70,PSMB2,H3-3A,HDGF,TSPAN6,TMED2,VEGFA,MAR
TM4A,SRSF9,CCNL2,ARPC2,PBX2,SRSF1,ARGLU1,ILF3,PRKDC,HNRNPUL1,ACTN1,HSPB1,SON,FI
12,HLA-B,CALU,ARCN1,HMGCR,CAPRIN1,CDK16,TOP2B,HDAC2,BCLAF1,LUC7L3,CPSF2,EID1,C
PABPN1,HNRNPL,PARP1,TSPYL4,MORF4L2,ALKBH5,RPL27,HNRNPA3,C4ORF3,PKN1,RBMX,DDX
N1,CHPF,GJA1,CTNNB1,ANXA6,NGRN,MACF1,DNAJC5,APLP2,HNRNPDL,TARDBP,NDFIP1,HNRNF
LNB,CORO1C,SMARCE1,SUPT6H,HNRNPM,SSB,SRSF3,LARS1,FAU,ILF2,EIF3F,SF3A1,DGKD,SERE
HDGF,TSPAN6,TMED2,VEGFA,MARS1,OPTN,UBXN4,ARPC2,DSTN,SRSF1,ARGLU1,TCF25,PTBP1,IL
RPS5,RPS7,IK,RPL35,HMGCS1,CTNNB1,GOLM1,APLP2,TARDBP,DDOST,HNRNPK,NARS1,RPL36,
P43,HK1,NCS1,RHOA,PTCD3,CBX5,CLSTN1,PSMD2,KIF5B,SLC7A5,NLGN2,RPS14,CXADR,ATP1A1
D1,CTSB,TM9SF3,MDH2,TOMM20,U2AF2,RPN2,MLXIP,HNRNPL,MORF4L2,ABCA3,ALKBH5,RPL2
31A,CALU,CTBP1,HMGCR,CAPRIN1,TOP2B,HDAC2,SKP1,BCLAF1,TRIB2,GLUL,LUC7L3,CPSF2,EI
,POLR2B,CORO1C,SMARCC1,SUPT6H,HNRNPM,SSB,SRSF3,ILF2,LSS,CSNK1D,SF3A1,PCBP2,SF
,ALKBH5,HNRNPA3,C4ORF3,ACO2,CCT3,RBMX,CBX1,RAD23B,GCN1,SNRNP70,PSMB2,H3-3A,H
3,ALKBH5,RPL27,HNRNPA3,TBC1D16,ACO2,CCT3,ACAP3,RBMX,DDX42,RAD23B,TNPO1,SNRNP7
RCA4,CCNL2,ARPC2,DSTN,PSMD3,ARGLU1,NYNRIN,TCF25,PTBP1,ILF3,PRKDC,HNRNPUL1,ACTN
M,SSB,SRSF3,ILF2,CSNK1D,SF3A1,NT5DC2,RAI1,PNMA8A,HNRNPD,MLF2,NAP1L4,TP53BP1,DDIT
I1,KDM1A,CMIP,KDELR1,ZFAND5,DIDO1,FLNB,POLR2B,CORO1C,SMARCE1,SUPT6H,HNRNPM,S

'D,MLF2,ZSCAN18,POLR2E,ANP32A,RBBP7,TRIO,LBH,NDUFV1,PBX1,AP2A1,H2AZ1,UBQLN2,CSN
JBQLN2,BRD4,PPM1G,ATRX,SRSF6,UBAP2L,UPF1,CRTAP,TRA2B,UQCRC1,ZFR,HNRNPA2B1,CUI
\,ARHGDIA,CAPRIN1,STIP1,TOP2B,SMCHD1,HDAC2,IPO5,RPL23A,LUC7L3,TOMM20,RNF187,U2/
\B7A,PRDX2,DDX3X,MAGED2,H1-0,RPS5,RPS7,IK,RPL35,GPAT4,PRKAR1A,CHPF,GJA1,CTNNB1,A
TARDBP,ST3GAL2,TAGLN,MYL6,WLS,HNRNPK,H1-10,RPL36,YWHAB,CKAP5,LAMP1,HNRNPH1,CI
'3E,ILF3,GRB2,PRKDC,HNRNPUL1,CASC3,FKBP8,PLCG1,MAP2K2,DIDO1,SMARCE1,XPNPEP3,SM
APRE2,RPS27A,HNRNPAB,PRDX6,FUS,CLPTM1,LINGO1,RALY,AKT3,HDGFL2,DCLK1,EFTUD2,SR
,HNRNPUL1,ATXN10,PLCG1,DIDO1,POLR2B,CORO1C,SMARCE1,SUPT6H,HNRNPM,SSB,GIGYF1,
'A,RBBP7,TXLNA,LBH,PSMD1,SF1,CSNK2A1,BRD4,UBA52,SRSF6,RER1,EIF5B,TRA2B,UQCRC1,Z
CTNNB1,GOLM1,RPS10,APLP2,TARDBP,DDOST,HNRNPK,INIP,RPL36,CKAP5,SEPTIN11,LAMP1,HI
,SMARCE1,SDF4,HNRNPM,SSB,SRSF3,PHC2,LARS1,ILF2,PFKM,FBLN1,STT3B,UBQLN1,SF3A1,TI
LUC7L3,CPSF2,CTSB,NUCB1,TM9SF3,MDH2,TOMM20,RAB27A,MLXIP,PABPN1,HNRNPL,TM9SF2
OL6A2,CD63,APLP2,TARDBP,ARID1A,ST3GAL2,MYL6,HNRNPK,H1-10,RPL36,YWHAB,CKAP5,SEPT
RNPUL1,KDM1A,DIDO1,CORO1C,SUPT6H,HNRNPM,SRSF3,ILF2,LSS,UBQLN1,CSNK1D,SF3A1,ME
RNPH1,CKAP4,HLA-B,RAB1A,CALU,MAGED4B,CAPRIN1,GPC1,HDAC2,TMEM30A,MMP2,BCLAF1,C
MCHD1,HDAC2,BCLAF1,LUC7L3,CPSF2,CTSB,NUCB1,TM9SF3,MDH2,TOMM20,RAB27A,U2AF2,I
'2,HNRNPD,SERBP1,PRPF40A,DDIT4,PSMA7,ANP32A,RBBP7,NAP1L1,LBH,SF1,PBX1,PRCC,BRD4,
\RS1,ILF2,CCT8,STT3B,ISYNA1,LONP1,UBQLN1,SF3A1,SOD1,BAG6,HNRNPD,VGF,SERBP1,KDM3
1,HNRNPH1,UBE2H,RPS12,RAB1A,CTBP1,CAPRIN1,STIP1,CDK16,TOP2B,APPL1,SKP1,BCLAF1,PI
,HNRNPAB,C1ORF198,FUS,VAPB,CLPTM1,SNRPD3,TPP1,RALY,TLE1,HDGFL2,EFTUD2,CNP,TRAF
RNPAB,C1ORF198,SAFB2,PRDX6,ZMYM3,FUS,TRIR,XRCC6,ERH,PELP1,SNRPD3,RALY,ELAVL1,TL
CLPTM1,SNRPD3,TPP1,RALY,SRP72,HDGFL2,EFTUD2,GTF2I,TRAP1,EEF1B2,NUCKS1,RBM5,YIPF
KDM3B,CCT5,POLR2E,PSMA7,ANP32A,VDAC1,RBBP7,SF1,UBQLN2,GRSF1,KPNA2,GSPT1,PPP2C
JL3,ANP32B,POM121C,RPS27A,HNRNPAB,CTTN,C1ORF198,MTCH1,GRK2,FUS,XRCC6,ERH,TAO
M9SF2,MORF4L2,TAF15,HNRNPA3,ACO2,VPS35,RBMX,CBX1,RAD23B,CCT4,GCN1,SKI,SNRNP7(
IPH1,RPS12,HLA-B,RAB1A,ARCN1,CCT6A,HMGCR,CAPRIN1,CDK16,TOP2B,BCLAF1,LUC7L3,CPS
K2,RBM10,SRF,ZEB1,MED25,ZFP91,HSPE1,KARS1,TMPO,LMNA,THOC2,RRBP1,MED12,RXRA,PHI
GF,ABCA2,PRPF6,TMED2,TOP1,NOTCH2,RNF10,VEGFA,MAPK1,MAPRE1,CCNL2,DSTN,PLEC,PDI
N2,MMP14,PRDX2,RPS15,RPL12,MAGED2,H1-0,RPS5,RPS7,IK,HMGCS1,CTNNB1,ANXA6,RPS10,I
'1,RBM6,CCDC47,THRAP3,REEP1,EIF4A3,RANGAP1,MTA2,USP10,LRIG1,PSMB1,TSPYL2,KIAA12
APEX1,HNRNPAB,RABEP1,PRDX6,ZMYM3,FUS,XRCC6,ERH,PELP1,HDGFL2,EFTUD2,GTF2I,TRAP
,PTBP1,ILF3,LUZP6,PRKDC,HNRNPUL1,KDM1A,MAP2K2,RPL28,DIDO1,POLR2B,CORO1C,SMAR
HDAC6,ZFR,HNRNPA2B1,CUL3,NDST1,MCM3AP,POM121C,RPS27A,APEX1,HNRNPAB,LRRC59,M
NRNPA2B1,ANP32B,MCM3AP,HNRNPAB,C1ORF198,PRDX6,FUS,XRCC6,ERH,PELP1,RALY,TLE1,H
2,RAB5B,TNPO1,SNRNP70,H3-3A,KDM6B,HDGF,RPS15A,RNF10,VEGFA,OPTN,UBXN4,ARPC2,PSI
F3,ILF2,PFKM,LSS,B2M,LONP1,PNMA8A,HNRNPD,MLF2,ZSCAN18,BMPR2,DDIT4,PSMA7,VDAC1,
2,MAP2,RAB1A,SEPTIN3,CTBP1,ARHGDIA,STIP1,CDK16,APPL1,HDAC2,IPO5,TMEM30A,SKP1,L1C
6,ZFR,HNRNPA2B1,CUL3,ANP32B,MCM3AP,APEX1,HNRNPAB,C1ORF198,LRRC59,MTCH1,FUS,X
LS,HNRNPK,H1-10,INIP,RPL36,CKAP5,HNRNPH1,CALU,CCT6A,CTBP1,HMGCR,CAPRIN1,CDK16,
UC7L3,CDKN1A,CPSF2,DDX39B,CTSB,TM9SF3,TOMM20,U2AF2,EEF1A2,CCT2,TBCD,SUGP2,PPF
ARGLU1,ILF3,CAPZB,PRKDC,HNRNPUL1,HSPB1,KDM1A,PLCG1,KDELR1,DIDO1,FLNB,SUPT6H,F
STYK,ZNF146,HEXIM1,FUBP1,FAM193B,SMARCB1,SCAF8,ZNF207,UBTF,LEMD2,ALYREF,EIF3J
SMARCE1,HNRNPM,SSB,GIGYF1,SRSF3,SDHA,PFKM,LSS,LONP1,UBQLN1,SF3A1,NBPF9,NT5DC
DIDO1,POLR2B,SMARCE1,SMARCC1,FYN,HNRNPM,SSB,SRSF3,ILF2,B2M,UBQLN1,SF3A1,TMED9
3AP,APEX1,PRDX6,ZMYM3,GRK2,FUS,XRCC6,ERH,EIF2S2,YARS1,RALY,HDGFL2,LSM14A,EFTUD

;,ERH,TAOK1,PELP1,TCAF1,RALY,TRAM1,TLE1,SH3GL1,HDGFL2,EFTUD2,SRSF2,TRAP1,EEF1B2
3P2L,ZNF207,LEMD2,ALYREF,PLRG1,COX5B
HNRNPM,SRSF3,ILF2,B2M,SF3A1,STRAP,HNRNPD,TCTN1,SERBP1,KDM3B,TP53BP1,POLR2E,PSI
TF7IP,PDGFRB,WIZ,HNRNPR,AKAP8L,ADRM1,PSME3,SRSF7,FAM160B2,ZEB1,SEPHS1,MRPS6,AI
O,CAPZB,PRKDC,HNRNPUL1,HSPB1,MAP2K2,DIDO1,SUPT6H,HNRNPM,GIGYF1,SRSF3,PHC2,ILF
,NACC1,LMNA,CDC42EP4,GET1,MED12,TLK1,RBM17,NRP2,DSTYK,ZNF146,SFSWAP,PRPF31,RT
2,MECP2,PARK7,ADNP,SYNCRIP,ERP29,PSMD7,ECH1,SUN2,HIPK2,WIZ,HNRNPR,H2AX,CDK9,PSI
1,NUCKS1,PHGDH,SRSF11,MSH6,USP24,EHD1,AP2A2,YY1,CREBBP,THRAP3,SLC38A2,EIF4A3,SH

1,TLE1,SRP72,TM9SF4,HDGFL2,EFTUD2,SLC7A1,PACS1,RBM5,FOXK2,PSMD4,SRSF11,SPTY2D1
WDR11,NSD1,FAM160B2,SEPHS1,TUBA1C,MCFD2,SNRPD2,MED25,ZFP91,KARS1,KDM2A,DUSP8
PSM1,SEPTIN7,HIF1A,TMOD2,FUS,XRCC6,PELP1,RALY,GABARAPL1,HDGFL2,EFTUD2,SRSF2,TR
DM2,NPTN,MECP2,PPP1R14B,AAMP,GGT7,EWSR1,KCTD20,PITRM1,SUN2,HIPK2,WIZ,SH3BGRL3
2,NAP1L4,TP53BP1,CCT5,PCYOX1,POLR2E,RBBP7,STAT3,MTA1,KDM5B,SF1,HIP1,FLCN,PRCC,C
D1,CREBBP,NDUFA13,BNIP3,TOMM70,THRAP3,DPYSL4,EIF4A3,CALCOCO1,CASC4,PTPN11,SUN
DH2,TOMM20,RNF187,U2AF2,SUGP2,PABPN1,HNRNPL,TSPYL4,MORF4L2,TAF15,ALKBH5,PRDX
SMARCD1,CREBBP,THRAP3,EIF4A3,HMGA1,HTATSF1,MTA2,KTN1,NOP56,MACROH2A1,RBM25,
M2A,NACC1,CDC42EP4,PRRC1,DGCR8,TLK1,RBM17,NRP2,MGRN1,DSTYK,SHC1,ELOB,ZNF146,
,UQCRC1,COX7C,MBTPS1,ZFR,NOMO1,HNRNPA2B1,GPSM1,HNRNPAB,SLC25A5,C1ORF198,MT

F1B2,MOAP1,PACS1,RBM5,SRSF11,SPTY2D1,CAP1,IST1,EIF3G,GSK3A,UBE3A,TGOLN2,SMARCD
9,TEX2,UBE3A,RBM6,NDUFA13,ATXN7L3,MN1,PSMC1,TRIM8,THRAP3,METRN,SCAP,USP47,XRN

1,LDOC1,RNF40,NDUFA5,GPM6B,COX5B
NPM,SSB,SRSF3,PHC2,LARS1,ILF2,STT3B,ISYNA1,LONP1,UBQLN1,SF3A1,GPR107,HNRNPD,VGF
,JPT1,RAB5B,TNPO1,PSMB2,SCAF1,DPYSL2,TMED2,RNF10,SELENON,OPTN,UBXN4,PPP2CA,MA
LF3,PRKDC,HNRNPUL1,ACTN1,SON,CASC3,FKBP8,CCAR2,DIDO1,TPM3,SMARCE1,SDF4,SMARC

IT,ZNF326,ANAPC2,VPS37B,MYBBP1A,FKBP3,MMP24

RPF6,TERF2IP,TAF7,MARS1,TUBB2A,SMARCA4,MAPRE1,CCNL2,ODC1,SRSF1,ARGLU1,ATP5PB,I

,HDGFL2,EFTUD2,SRSF2,TRAP1,RAB6A,AHSA1,RAB5C,SPAG9,SRSF11,MSH6,NSD2,PRPF19,CSE
;,SOX9,CDKN1B,RNF20,FARSA,PMPCA,CACYBP,TLK1,PHB,GRHPR,CTCF,OGDH,PPP1R12A,OLA1,

P6V1G1,WNK1,EIF4A3,SCAP,HS2ST1,XRN2,HMGA1,HTATSF1,RSF1,SPPL2B,MTA2,PAIP2,ANP32E,
GCR,ARHGDIA,CAPRIN1,CDK16,APPL1,SMCHD1,IPO5,SKP1,TRIB2,GLUL,FAM199X,PGRMC1,EID1

PJA1,SART3,PSMD7,FAF2,MAEA,IP6K1,KLC2,FAM53C,DYNLRB1,ARHGAP5,PPP4R3A,PRXL2A,TRII

30,HNRNPM,SSB,GIGYF1,SRSF3,LARS1,ILF2,CCT8,ISYNA1,LONP1,UBQLN1,SF3A1,GPR107,UQC

RAD54L2,TMEM59,BCAR1,HMGB2,MCM7,OPA1,ABCD3,PPT1,LANCL1,TRIM24,AKAP17A,DDX19A

3B1,PDIA3,SSBP2,CHD3,BASP1,VAPA,HMGB1,POLR2A,TPT1,RPS4X,RPS3A,RPL18,STMN1,ITGB1,
L18,STMN1,TUBB4B,UCHL1,RPL11,MORF4L1,BTF3,NCAM1,AGRN,XAF1,RPL10,MARCHF6,WSB1
M1,AGRN,RPL10,MARCHF6,WSB1,RPS17,DNAJA1,CYB5R3,PAFAH1B1,AKAP12,MAP4,SF3B2,CDC
S17,DNAJA1,CYB5R3,PAFAH1B1,SF3B2,CDC42BPB,ITM2B,ZNF770,AP2B1,GTF2F1,HNRNPU,OGA
VAPA,HMGB1,POLR2A,TPT1,RPS4X,RPS3A,RPL18,STMN1,ITGB1,TUBB4B,UCHL1,RPL11,MORF4
7,DNAJA1,CYB5R3,PAFAH1B1,SF3B2,CDC42BPB,ITM2B,ZNF770,AP2B1,GTF2F1,HNRNPU,OGA,
MORF4L1,BTF3,AGRN,XAF1,RPL10,MARCHF6,WSB1,RPS17,DNAJA1,CYB5R3,PAFAH1B1,MAP4,
ORF4L1,BTF3,NCAM1,ANXA2,XAF1,RPL10,MARCHF6,WSB1,RPS17,VCL,DNAJA1,CYB5R3,PAFAH1
F4L1,BTF3,ANXA2,XAF1,RPL10,MARCHF6,RPS17,VCL,DNAJA1,CYB5R3,PAFAH1B1,AKAP12,MAP
RPS17,VCL,DNAJA1,CYB5R3,PAFAH1B1,AKAP12,MAP4,SF3B2,CDC42BPB,ITM2B,ZNF770,MAT2A
RPS17,VCL,DNAJA1,CYB5R3,PAFAH1B1,AKAP12,MAP4,SF3B2,CDC42BPB,ITM2B,ZNF770,MAT2A
MORF4L1,BTF3,ANXA2,AGRN,XAF1,RPL10,MARCHF6,RPS17,VCL,DNAJA1,CYB5R3,PAFAH1B1,MA
EIF4H,SCARB2,LPCAT1,BRD3,RPS9,PJA2,RPL24,KPNB1,PAPOLA,COL1A2,LLGL1,RPL14,HK1,RHO
A,RPL18,TUBB4B,RPL11,MORF4L1,BTF3,AGRN,ZMIZ1,XAF1,RPL10,WSB1,RPS17,VCL,DNAJA1,
A,RPL18,TUBB4B,RPL11,MORF4L1,BTF3,AGRN,ZMIZ1,XAF1,RPL10,WSB1,RPS17,VCL,DNAJA1,
CYB5R3,PAFAH1B1,MAP4,SF3B2,CDC42BPB,ITM2B,ZNF770,MAT2A,GTF2F1,GOLGA8A,HNRNPU,
N,ZMIZ1,XAF1,RPL10,MARCHF6,RPS17,VCL,DNAJA1,CYB5R3,PAFAH1B1,MAP4,SF3B2,CDC42BPB
N,ZMIZ1,XAF1,RPL10,MARCHF6,RPS17,VCL,DNAJA1,CYB5R3,PAFAH1B1,MAP4,SF3B2,CDC42BPB
,TUBB4B,RPL11,MORF4L1,BTF3,ANXA2,AGRN,XAF1,RPL10,MARCHF6,RPS17,VCL,DNAJA1,CYB5
,MARCHF6,RPS17,VCL,DNAJA1,CYB5R3,PAFAH1B1,MAP4,SF3B2,CDC42BPB,ITM2B,ZNF770,AP2
B1,RPL17,C1ORF43,PAPOLA,LLGL1,RPL14,HK1,RHOA,PTCD3,CBX5,RPL9,CLSTN1,ATF4,KIF5B,F
BTF3,ANXA2,AGRN,XAF1,RPL10,WSB1,RPS17,VCL,DNAJA1,CYB5R3,PAFAH1B1,AKAP12,MAP4,SF
,XAF1,RPL10,WSB1,RPS17,VCL,DNAJA1,CYB5R3,PAFAH1B1,SF3B2,CDC42BPB,ITM2B,ZNF770,
,XAF1,RPL10,WSB1,RPS17,VCL,DNAJA1,CYB5R3,PAFAH1B1,SF3B2,CDC42BPB,ITM2B,ZNF770,
T1,BRD3,XRCC5,PJA2,RPL24,SLC16A7,KPNB1,RPL17,C1ORF43,PAPOLA,COL1A2,GAP43,RPL14,
RF43,PAPOLA,COL1A2,GAP43,RPL14,HK1,NCS1,RHOA,PTCD3,CBX5,RPL9,PSMD2,RPS14,CHD4,
X,RPS3A,RPL18,STMN1,ITGB1,TUBB4B,UCHL1,MORF4L1,BTF3,ANXA2,AGRN,ZMIZ1,XAF1,RPL1
MARCHF6,WSB1,RPS17,VCL,DNAJA1,CYB5R3,PAFAH1B1,MAP4,SF3B2,CDC42BPB,ITM2B,AP2B1
42BPB,ITM2B,ZNF770,AP2B1,GTF2F1,GOLGA8A,HNRNPU,OGA,PGAM1,NFE2L1,SMARCA5,LARP
42BPB,ITM2B,ZNF770,AP2B1,GTF2F1,GOLGA8A,HNRNPU,OGA,PGAM1,NFE2L1,SMARCA5,LARP
,ANXA2,AGRN,XAF1,RPL10,MARCHF6,WSB1,RPS17,VCL,DNAJA1,CYB5R3,PAFAH1B1,MAP4,SF3B
PAFAH1B1,MAP4,SF3B2,CDC42BPB,ITM2B,MAT2A,GTF2F1,GOLGA8A,HNRNPU,OGA,PGAM1,NF
PAFAH1B1,MAP4,SF3B2,CDC42BPB,ITM2B,MAT2A,GTF2F1,GOLGA8A,HNRNPU,OGA,PGAM1,NF
6,RPL23,CNN3,RPL29,RPSA,TPM4,TNS3,EIF4H,SCARB2,LPCAT1,BRD3,XRCC5,PJA2,MSN,RPL24
SMARCA5,LARP1,HNRNPA0,CTNNA1,FADS2,RPS16,RPL23,CNN3,RPL29,RPSA,TPM4,TNS3,CTSD,
PL10,MARCHF6,RPS17,VCL,DNAJA1,PAFAH1B1,MAP4,SF3B2,CDC42BPB,ITM2B,ZNF770,MAT2A,
BRD3,XRCC5,PJA2,MSN,RPL24,KPNB1,RPL17,C1ORF43,PAPOLA,COL1A2,LLGL1,GAP43,RPL14,
F770,GTF2F1,HNRNPU,OGA,PGAM1,NFE2L1,SMARCA5,LARP1,HNRNPA0,CTNNA1,FADS2,RPS16,
F770,GTF2F1,HNRNPU,OGA,PGAM1,NFE2L1,SMARCA5,LARP1,HNRNPA0,CTNNA1,FADS2,RPS16,
GOLGA8A,HNRNPU,PGAM1,NFE2L1,SMARCA5,HNRNPA0,CTNNA1,FADS2,RPS16,RPL23,CNN3,R
HOA,PTCD3,CBX5,CLSTN1,ATF4,PSMD2,KIF5B,SLC7A5,RPS14,CHD4,JUND,TMEM259,RPS20,EIF
BTF3,ANXA2,AGRN,ZMIZ1,XAF1,RPL10,MARCHF6,WSB1,RPS17,VCL,DNAJA1,CYB5R3,PAFAH1B1
BTF3,ANXA2,AGRN,ZMIZ1,XAF1,RPL10,MARCHF6,WSB1,RPS17,VCL,DNAJA1,CYB5R3,PAFAH1B1
BTF3,ANXA2,AGRN,ZMIZ1,XAF1,RPL10,MARCHF6,RPS17,VCL,DNAJA1,CYB5R3,PAFAH1B1,AKAP
BTF3,ANXA2,AGRN,ZMIZ1,XAF1,RPL10,MARCHF6,RPS17,VCL,DNAJA1,CYB5R3,PAFAH1B1,AKAP

PM4,TNS3,CTSD,EIF4H,SCARB2,LPCAT1,BRD3,XRCC5,PJA2,MSN,SLC16A7,KPNB1,RPL17,C1ORF
,ANXA2,AGRN,ZMIZ1,XAF1,RPL10,MARCHF6,RPS17,VCL,DNAJA1,CYB5R3,PAFAH1B1,SF3B2,CDC
,ANXA2,AGRN,ZMIZ1,XAF1,RPL10,MARCHF6,RPS17,VCL,DNAJA1,CYB5R3,PAFAH1B1,SF3B2,CDC
XA2,AGRN,RPL10,MARCHF6,RPS17,VCL,DNAJA1,CYB5R3,PAFAH1B1,AKAP12,MAP4,SF3B2,CDC4
VCL,DNAJA1,CYB5R3,PAFAH1B1,MAP4,SF3B2,CDC42BPB,ITM2B,AP2B1,MAT2A,GTF2F1,HNRNP
NXA2,AGRN,ZMIZ1,XAF1,RPL10,MARCHF6,WSB1,RPS17,VCL,DNAJA1,CYB5R3,PAFAH1B1,MAP4
BTF3,ANXA2,AGRN,XAF1,RPL10,WSB1,RPS17,VCL,DNAJA1,CYB5R3,PAFAH1B1,MAP4,SF3B2,CD
TM2B,ZNF770,AP2B1,GTF2F1,HNRNPU,OGA,PGAM1,NFE2L1,SMARCA5,LARP1,HNRNPA0,CTNN
,MAP4,SF3B2,CDC42BPB,ITM2B,AP2B1,GTF2F1,GOLGA8A,HNRNPU,PGAM1,NFE2L1,SMARCA5,
,MAP4,SF3B2,CDC42BPB,ITM2B,AP2B1,GTF2F1,GOLGA8A,HNRNPU,PGAM1,NFE2L1,SMARCA5,
,BTF3,ANXA2,AGRN,ZMIZ1,XAF1,RPL10,MARCHF6,WSB1,RPS17,VCL,DNAJA1,CYB5R3,PAFAH1B
NA1,FADS2,RPS16,RPL23,RPSA,TPM4,CTSD,EIF4H,SCARB2,LPCAT1,RPS24,BRD3,XRCC5,PJA2,M
R3,PAFAH1B1,MAP4,SF3B2,CDC42BPB,ITM2B,ZNF770,AP2B1,MAT2A,GTF2F1,GOLGA8A,HNRN
R3,PAFAH1B1,MAP4,SF3B2,CDC42BPB,ITM2B,ZNF770,AP2B1,MAT2A,GTF2F1,GOLGA8A,HNRN
L26,BEX3,CCT7,SRSF4,RBBP4,PFN1,MCL1,PNN,RPL31,NISCH,PRKCSH,HNRNPC,LRPPRC,TADA2
NF770,MAT2A,GTF2F1,GOLGA8A,HNRNPU,OGA,PGAM1,NFE2L1,SMARCA5,LARP1,HNRNPA0,CT
2L1,HNRNPA0,CTNNA1,RPL23,CPE,RPL29,RPSA,TPM4,TNS3,EIF4H,SCARB2,RTN4,LPCAT1,BRD3
Z1,XAF1,RPL10,MARCHF6,WSB1,RPS17,VCL,DNAJA1,CYB5R3,PAFAH1B1,MAP4,SF3B2,CDC42BP
Z1,XAF1,RPL10,MARCHF6,WSB1,RPS17,VCL,DNAJA1,CYB5R3,PAFAH1B1,MAP4,SF3B2,CDC42BP
CL,DNAJA1,CYB5R3,PAFAH1B1,MAP4,SF3B2,CDC42BPB,ITM2B,ZNF770,AP2B1,MAT2A,GTF2F1,G
CL,DNAJA1,CYB5R3,PAFAH1B1,MAP4,SF3B2,CDC42BPB,ITM2B,ZNF770,AP2B1,MAT2A,GTF2F1,G
CTSD,EIF4H,SCARB2,LPCAT1,BRD3,XRCC5,PJA2,RPL24,KPNB1,RPL17,C1ORF43,PAPOLA,COL1
RN,XAF1,RPL10,WSB1,RPS17,VCL,DNAJA1,CYB5R3,PAFAH1B1,MAP4,SF3B2,CDC42BPB,ITM2B,
FADS2,RPS16,CPE,CNN3,RPSA,TPM4,CTSD,EIF4H,SCARB2,RTN4,RPS24,BRD3,PJA2,RPL24,KPNB
1,SF3B2,CDC42BPB,ITM2B,ZNF770,AP2B1,MAT2A,GTF2F1,HNRNPU,OGA,PGAM1,NFE2L1,SMAR
23,RPL29,RPSA,TPM4,TNS3,CTSD,EIF4H,SCARB2,LPCAT1,BRD3,XRCC5,PJA2,MSN,RPL24,RPL1
RPL17,C1ORF43,PAPOLA,LLGL1,RPL14,HK1,RHOA,PTCD3,CBX5,RPL9,CLSTN1,PSMD2,SLC7A5
NRNPA0,CTNNA1,FADS2,RPS16,RPL23,CPE,CNN3,RPL29,RPSA,TPM4,CTSD,EIF4H,SCARB2,LPCA
MA5,BRD3,XRCC5,PJA2,MSN,RPL24,KPNB1,RPL17,C1ORF43,PAPOLA,COL1A2,LLGL1,RPL14,HK
MA5,BRD3,XRCC5,PJA2,MSN,RPL24,KPNB1,RPL17,C1ORF43,PAPOLA,COL1A2,LLGL1,RPL14,HK
S16,RPL23,CNN3,RPL29,RPSA,TPM4,TNS3,CTSD,EIF4H,SCARB2,LPCAT1,RPS24,BRD3,RPS9,PJA2
GOLGA8A,HNRNPU,PGAM1,NFE2L1,SMARCA5,LARP1,HNRNPA0,CTNNA1,FADS2,RPS16,RPL23,C
KPNB1,RPL17,PAPOLA,COL1A2,LLGL1,GAP43,RPL14,HK1,RHOA,PTCD3,CBX5,ZNF865,CLSTN1,A
RN,XAF1,RPL10,MARCHF6,WSB1,RPS17,VCL,DNAJA1,CYB5R3,PAFAH1B1,MAP4,SF3B2,CDC42BP
RN,XAF1,RPL10,MARCHF6,WSB1,RPS17,VCL,DNAJA1,CYB5R3,PAFAH1B1,MAP4,SF3B2,CDC42BP
CYB5R3,PAFAH1B1,MAP4,SF3B2,CDC42BPB,ITM2B,ZNF770,AP2B1,MAT2A,GTF2F1,GOLGA8A,H
MIZ1,XAF1,RPL10,WSB1,RPS17,VCL,DNAJA1,CYB5R3,PAFAH1B1,MAP4,SF3B2,CDC42BPB,ITM2B
P4,SF3B2,CDC42BPB,ITM2B,MAT2A,GTF2F1,GOLGA8A,HNRNPU,OGA,NFE2L1,SMARCA5,LARP1
TCD3,CBX5,CLSTN1,ATF4,PSMD2,KIF5B,RPS14,CXADR,CHD4,TMEM259,RPS20,EIF5,WDR1,HMG
A,GTF2F1,HNRNPU,PGAM1,NFE2L1,SMARCA5,LARP1,HNRNPA0,CTNNA1,FADS2,RPS16,RPL23,R
RPS9,XRCC5,RPL24,RPL17,C1ORF43,PAPOLA,COL1A2,RPL14,HK1,PTCD3,CBX5,RPL9,CLSTN1,P
,RPS9,XRCC5,PJA2,RPL24,KPNB1,RPL17,C1ORF43,PAPOLA,RPL14,HK1,PTCD3,CBX5,ATF4,PSM
PNB1,RPL17,C1ORF43,PAPOLA,COL1A2,LLGL1,RPL14,HK1,PTCD3,ZNF865,RPL9,PSMD2,KIF5B,
C42BPB,ITM2B,ZNF770,GTF2F1,HNRNPU,PGAM1,NFE2L1,SMARCA5,LARP1,CTNNA1,FADS2,RPS
LGA8A,HNRNPU,PGAM1,NFE2L1,SMARCA5,LARP1,HNRNPA0,CTNNA1,FADS2,RPS16,RPL23,CPE,

ARB2,LPCAT1,BRD3,XRCC5,PJA2,MSN,RPL24,SLC16A7,C1ORF43,PAPOLA,COL1A2,LLGL1,RPL1
DC42BPB,ITM2B,ZNF770,AP2B1,MAT2A,GTF2F1,GOLGA8A,HNRNPU,OGA,PGAM1,NFE2L1,SMAI
JA1,CYB5R3,PAFAH1B1,MAP4,SF3B2,CDC42BPB,ITM2B,ZNF770,AP2B1,GTF2F1,GOLGA8A,HNR
JA1,CYB5R3,PAFAH1B1,MAP4,SF3B2,CDC42BPB,ITM2B,ZNF770,AP2B1,GTF2F1,GOLGA8A,HNR
TKT,GLG1,CDH6,SFPQ,PEBP1,CDH2,PDIA6,DDR1,RAB7A,PRDX2,DDX3X,RPS15,RPL30,CLTC,MAC
FE2L1,SMARCA5,LARP1,CTNNA1,FADS2,RPS16,RPL23,CNN3,RPL29,RPSA,TPM4,EIF4H,SCARB2,
MARCA5,LARP1,HNRNPA0,CTNNA1,FADS2,RPS16,RPL23,CNN3,RPL29,RPSA,TPM4,CTSD,EIF4H,S
PNB1,RPL17,C1ORF43,PAPOLA,COL1A2,LLGL1,RPL14,HK1,PTCD3,CBX5,ZNF865,RPL9,ATF4,PS
L11,MORF4L1,BTF3,ANXA2,AGRN,ZMIZ1,XAF1,RPL10,WSB1,RPS17,VCL,DNAJA1,CYB5R3,PAFA
GOLGA8A,HNRNPU,OGA,NFE2L1,SMARCA5,LARP1,HNRNPA0,CTNNA1,FADS2,RPS16,RPL23,CNN
GOLGA8A,HNRNPU,OGA,NFE2L1,SMARCA5,LARP1,HNRNPA0,CTNNA1,FADS2,RPS16,RPL23,CNN
3,PAPOLA,COL1A2,RPL14,HK1,PTCD3,CBX5,RPL9,ATF4,PSMD2,KIF5B,SLC7A5,RPS14,CHD4,ATI
IB1,RPL17,C1ORF43,PAPOLA,COL1A2,LLGL1,RPL14,RPL9,CLSTN1,ATF4,KIF5B,SLC7A5,RPS14,
,PJA2,MSN,RPL24,SLC16A7,RPL17,C1ORF43,PAPOLA,COL1A2,LLGL1,RPL14,HK1,RHOA,PTCD3
,PJA2,MSN,RPL24,SLC16A7,RPL17,C1ORF43,PAPOLA,COL1A2,LLGL1,RPL14,HK1,RHOA,PTCD3
RPL14,RPL9,CLSTN1,SLC7A5,RPS14,CHD4,JUND,RPS20,EIF5,CD81,HMGN2,RPL26,BEX3,SRSF4
D3,RPL9,ATF4,PSMD2,SLC7A5,RPS14,CXADR,ATP1A1,JUND,TMEM259,RPS20,EIF5,WDR1,HMGI
,XRCC5,PJA2,RPL24,KPNB1,RPL17,PAPOLA,LLGL1,RPL14,HK1,PTCD3,CBX5,ZNF865,RPL9,PSM
24,SLC16A7,KPNB1,LLGL1,RPL14,RHOA,PTCD3,RPL9,CLSTN1,ATF4,KIF5B,RPS14,CXADR,CHD4
AT1,RPS24,XRCC5,PJA2,MSN,RPL24,KPNB1,RPL17,C1ORF43,COL1A2,LLGL1,RPL14,HK1,RHOA
RPS9,XRCC5,PJA2,RPL24,SLC16A7,RPL17,C1ORF43,PAPOLA,COL1A2,LLGL1,RPL14,HK1,PTCD
CHD3,VAPA,HMGB1,POLR2A,TPT1,RPS4X,RPS3A,RPL18,STMN1,TUBB4B,UCHL1,RPL11,MORF4L
CRN1,HDLBP,TKT,GLG1,CDH2,PDIA6,DDR1,RAB7A,ACTN4,PRDX2,DDX3X,RPL12,MAGED2,GNB2
6,RPL23,CNN3,RPL29,RPSA,TPM4,CTSD,EIF4H,SCARB2,LPCAT1,RPS9,XRCC5,PJA2,MSN,RPL24,
LS,HNRNPK,H1-10,MYH10,RPL36,YWHAB,LAMP1,HNRNPH1,UBE2H,CKAP4,HLA-B,CALU,MAGED
U,OGA,NFE2L1,SMARCA5,LARP1,HNRNPA0,CTNNA1,RPS16,RPL23,RPSA,TPM4,TNS3,CTSD,EIF4I
HNRNPA0,CTNNA1,FADS2,RPS16,RPL23,CNN3,RPL29,RPSA,TPM4,TNS3,CTSD,EIF4H,SCARB2,LI
RPS24,BRD3,XRCC5,PJA2,RPL24,SLC16A7,KPNB1,RPL17,C1ORF43,PAPOLA,COL1A2,LLGL1,RPL
FE2L1,SMARCA5,LARP1,HNRNPA0,CTNNA1,FADS2,RPS16,RPL23,CPE,RPL29,RPSA,TPM4,TNS3,C
MARCA5,LARP1,HNRNPA0,CTNNA1,FADS2,RPS16,RPL23,CPE,CNN3,RPL29,RPSA,TPM4,TNS3,CT
RCA5,LARP1,HNRNPA0,CTNNA1,RPS16,RPL23,CNN3,RPL29,RPSA,TPM4,TNS3,CTSD,EIF4H,SCARI
BX5,ZNF865,ATF4,PSMD2,SLC7A5,RPS14,CXADR,CHD4,JUND,TMEM259,RPS20,WDR1,HMGN2,R
M20,EEF1A2,MLXIP,SUGP2,GAS7,HNRNPL,PARP1,TM9SF2,MORF4L2,TAF15,LMNB2,ABCA3,ALKE
3,PAPOLA,COL1A2,LLGL1,RPL14,RHOA,ZNF865,RPL9,CLSTN1,ATF4,PSMD2,KIF5B,SLC7A5,RPS
PS16,RPL23,CPE,CNN3,RPL29,RPSA,TPM4,TNS3,CTSD,EIF4H,SCARB2,LPCAT1,RPS24,BRD3,RPS
FADS2,RPS16,RPL23,CNN3,RPL29,RPSA,TPM4,CTSD,EIF4H,SCARB2,LPCAT1,BRD3,RPS9,PJA2,F
OGA,NFE2L1,SMARCA5,LARP1,CTNNA1,FADS2,RPL23,CNN3,RPL29,RPSA,TNS3,EIF4H,SCARB2,R
7,C1ORF43,PAPOLA,COL1A2,LLGL1,RPL14,HK1,NCS1,ATF4,KIF5B,SLC7A5,RPS14,CXADR,CHD4
IN3,RPL29,RPSA,TPM4,EIF4H,SCARB2,BRD3,XRCC5,MSN,RPL24,SLC16A7,KPNB1,RPL17,C1ORI
L29,TPM4,EIF4H,LPCAT1,RPS24,XRCC5,PJA2,MSN,RPL24,KPNB1,RPL17,C1ORF43,PAPOLA,COL
SCARB2,LPCAT1,RPS24,BRD3,XRCC5,RPL24,SLC16A7,RPL17,C1ORF43,PAPOLA,COL1A2,LLGL
PS20,EIF5,WDR1,ATP2A2,RPL26,CRMP1,BEX3,CCT7,SRSF4,RBBP4,RTN3,MCL1,PNN,RPL31,PRK
RPS16,RPL23,CPE,CNN3,RPL29,RPSA,TPM4,EIF4H,SCARB2,BRD3,RPS9,XRCC5,RPL24,KPNB1,R
,EIF4H,SCARB2,BRD3,RPS9,XRCC5,PJA2,RPL24,RPL17,C1ORF43,PAPOLA,COL1A2,LLGL1,RPL1
9,XRCC5,PJA2,RPL24,KPNB1,RPL17,C1ORF43,PAPOLA,COL1A2,LLGL1,RPL14,HK1,PTCD3,ZNF

ISN,SLC16A7,C1ORF43,PAPOLA,COL1A2,LLGL1,GAP43,RPL14,RHOA,CBX5,CLSTN1,PSMD2,KIF

4,RBBP4,MCL1,PNN,RPL31,NISCH,PRKCSH,HNRNPC,LRPPRC,TADA2B,XPR1,KHDRBS1,XPO1,C

0,U2AF2,ANXA5,PABPN1,HNRNPL,PARP1,MORF4L2,TAF15,LMNB2,ABCA3,RPL27,ACO2,RBMX,J

0,EIF5,HMGN2,RPL26,CCT7,SRSF4,RBBP4,MCL1,JPT2,PNN,SPC25,RPL31,NISCH,HNRNPC,TAD

SMARCA5,LARP1,HNRNPA0,CTNNA1,RPS16,RPL23,CNN3,RPL29,RPSA,TPM4,CTSD,EIF4H,SCARB

IP1,HNRNPH1,UBE2H,CKAP4,ARCN1,CTBP1,ARHGDIA,CAPRIN1,STIP1,TOP2B,HDAC2,BCLAF1,P

DR,JUND,TMEM259,RPS20,EIF5,WDR1,HMGN2,RPL26,BEX3,CCT7,RBBP4,MCL1,JPT2,PNN,RPL

1,SF3B2,CDC42BPB,ITM2B,MAT2A,GTF2F1,HNRNPU,NFE2L1,SMARCA5,LARP1,HNRNPA0,CTNN

1,SF3B2,CDC42BPB,ITM2B,MAT2A,GTF2F1,HNRNPU,NFE2L1,SMARCA5,LARP1,HNRNPA0,CTNN

H,SCARB2,LPCAT1,RPS24,RPS9,XRCC5,PJA2,RPL24,SLC16A7,KPNB1,RPL17,C1ORF43,PAPOLA,

ATF4,PSMD2,SLC7A5,RPS14,CHD4,JUND,TMEM259,RPS20,EIF5,HMGN2,RPL26,LASP1,BEX3,CC

BP4,RTN3,PFN1,MCL1,PNN,NISCH,HNRNPC,LRPPRC,TADA2B,XPR1,KHDRBS1,XPO1,CSNK1E,RP

BP4,RTN3,PFN1,MCL1,PNN,NISCH,HNRNPC,LRPPRC,TADA2B,XPR1,KHDRBS1,XPO1,CSNK1E,RP

,RPL26,BEX3,RBBP4,PFN1,MCL1,PNN,RPL31,NISCH,PRKCSH,HNRNPC,LRPPRC,TADA2B,XPR1,K

,LRPPRC,TADA2B,XPR1,KHDRBS1,KLHL9,PFKL,SCRN1,RPS25,TKT,GLG1,CDH6,SFPQ,PEBP1,CD

X5,RPL9,PSMD2,KIF5B,SLC7A5,RPS14,CXADR,JUND,TMEM259,RPS20,EIF5,WDR1,HMGN2,ATP2

,CSNK1E,HDLBP,RPS25,GLG1,SFPQ,EIF5A,PDIA6,DDR1,ACTN4,DDX3X,RPS15,RPL12,RPL30,CL

FN1,MCL1,JPT2,PNN,RPL31,HNRNPC,LRPPRC,TADA2B,XPR1,KHDRBS1,KLHL9,XPO1,SPTBN2,C

,XRCC5,MSN,RPL24,KPNB1,RPL17,C1ORF43,PAPOLA,LLGL1,RPL14,HK1,RHOA,PTCD3,RPL9,PS

31,MAP4,SF3B2,CDC42BPB,ITM2B,AP2B1,GTF2F1,GOLGA8A,PXDN,HNRNPU,PGAM1,NFE2L1,LA

CAT1,BRD3,XRCC5,PJA2,MSN,RPL24,RPL17,C1ORF43,PAPOLA,COL1A2,RPL14,HK1,RHOA,PTC

LRPPRC,TADA2B,XPR1,KHDRBS1,XPO1,SPTBN2,CSNK1E,RPS25,TKT,GLG1,SFPQ,EIF5A,PDIA6,R

TN,G3BP2,MAPK1,SMARCA4,SRSF9,CCNL2,ARPC2,PSMD3,SRSF1,ARGLU1,TCF25,ILF3,PRKDC,

A1,FADS2,RPS16,RPL23,CPE,CNN3,RPL29,RPSA,TPM4,TNS3,CTSD,EIF4H,SCARB2,LPCAT1,RPS

P4,MCL1,PNN,RPL31,NISCH,PRKCSH,HNRNPC,LRPPRC,XPR1,KHDRBS1,XPO1,CSNK1E,RPS25,T

RCC5,PJA2,RPL24,KPNB1,RPL17,C1ORF43,PAPOLA,COL1A2,LLGL1,RPL14,HK1,RHOA,PTCD3,

RCC5,PJA2,RPL24,KPNB1,RPL17,C1ORF43,PAPOLA,COL1A2,LLGL1,RPL14,HK1,RHOA,PTCD3,

,RPL31,NISCH,PRKCSH,HNRNPC,LRPPRC,TADA2B,KHDRBS1,XPO1,SPTBN2,CSNK1E,RPS25,TK

PLP2,HNRNPDL,TARDBP,ARID1A,HNRNPK,INIP,NARS1,RPL36,YWHAB,SND1,LAMP1,HNRNPH1,U

KHDRBS1,SLC3A2,PTN,CSNK1E,RPS25,TKT,GLG1,SFPQ,EIF5A,PEBP1,PDIA6,DDR1,MEGF8,RAB7

GOLM1,RPS10,MTSS2,APLP2,HNRNPDL,TARDBP,ST3GAL2,HNRNPK,H1-10,NARS1,RPL36,YWHA

SFPQ,EIF5A,DDR1,ACSL6,RAB7A,CCN2,DDX3X,RPS15,RPL30,CLTC,MAGED2,H1-0,RPS5,RPS7,IK

7A5,RPS14,CHD4,JUND,EIF5,CD81,WDR1,RPL26,LASP1,BEX3,CCT7,RBBP4,PFN1,MCL1,JPT2,P

2,DDX39B,CTSB,NUCB1,TM9SF3,MDH2,TOMM20,U2AF2,RPN2,SUGP2,SQSTM1,PABPN1,HNRN

,CCT6A,MAGED4B,CTBP1,HMGCR,ARHGDIA,CAPRIN1,STIP1,GPC1,TOP2B,HDAC2,SKP1,BCLAF1

LU,CTBP1,CAPRIN1,STIP1,GPC1,TOP2B,HDAC2,TMEM30A,SKP1,BCLAF1,PPIB,GLUL,PEA15,LUC

RCC5,PJA2,RPL24,KPNB1,RPL17,C1ORF43,PAPOLA,COL1A2,LLGL1,RPL14,HK1,PTCD3,PSMD2

1E,RPS25,TKT,GLG1,PDIA6,RAB7A,ACTN4,MMP14,DDX3X,RPL30,CLTC,MAGED2,RCN1,H1-0,RP

SP1,BEX3,CCT7,SRSF4,RBBP4,PFN1,MCL1,JPT2,PNN,RPL31,PRKCSH,HNRNPC,LRPPRC,TADA2

1,KHDRBS1,XPO1,TKT,GLG1,SFPQ,PDIA6,DDR1,SULF1,RAB7A,ACTN4,DDX3X,RPS15,RPL30,CL

PN2,PABPN1,HNRNPL,MORF4L2,TAF15,LMNB2,ABCA3,ALKBH5,RPL27,HNRNPA3,PTPRA,C4ORF

4,HK1,PTCD3,ATF4,PSMD2,SLC7A5,RPS14,CXADR,CHD4,ATP1A1,TMEM259,RPS20,EIF5,HMGN2

,RPL30,CLTC,MAGED2,RCN1,H1-0,RPS5,RPS7,TLN1,CHPF,ANXA6,RPS10,APLP2,HNRNPDL,TAR

BPN1,HNRNPL,PARP1,TSPYL4,TM9SF2,MORF4L2,SSRP1,ALKBH5,RPL27,HNRNPA3,C4ORF3,PK

C,LRPPRC,TADA2B,XPR1,KHDRBS1,XPO1,CSNK1E,SCRN1,HDLBP,RPS25,GLG1,SFPQ,PEBP1,P

S7,IK,RPL35,GPAT4,PRKAR1A,TLN1,GJA1,COL6A2,ANXA6,RPS10,APLP2,TARDBP,RPLP2,ARID1A,
DIA6,DDR1,RAB7A,CCN2,ACTN4,PRDX2,DDX3X,RPS15,CLTC,MAGED2,H1-0,RPS5,RPS7,IK,RPL3
S1,RPL28,SMARCE1,DHX30,RPS28,HNRNPM,SRSF3,ILF2,PFKM,UBQLN1,CSNK1D,SF3A1,NT5D
KAP5,SND1,LAMP1,HNRNPH1,UBE2H,HLA-B,CALU,CCT6A,CTBP1,HMGCR,ARHGDIA,CAPRIN1,T
ISCH,PRKCSH,HNRNPC,LRPPRC,TADA2B,XPR1,KHDRBS1,XPO1,CSNK1E,HDLBP,RPS25,GLG1,S
AMP1,HNRNPH1,CKAP4,CALU,ARCN1,CCT6A,ARHGDIA,CAPRIN1,GPC1,SMCHD1,HDAC2,IPO5,SK
RPS15,RPL30,CLTC,MAGED2,H1-0,RPS5,RPS7,IK,RPL35,GJA1,CTNNB1,DNAJC5,APLP2,HNRNPD
NRNPDL,TARDBP,ARID1A,NDFIP1,MYL6,HNRNPK,H1-10,INIP,RPL36,YWHAB,CKAP5,LAMP1,HNRN
LUL,PEA15,LUC7L3,CPSF2,NUCB1,TM9SF3,MDH2,RPN2,SUGP2,GAS7,PABPN1,GDI2,HNRNPL,M
ADR,CHD4,JUND,TMEM259,EIF5,WDR1,HMGN2,ATP2A2,BEX3,CCT7,SRSF4,RBBP4,RTN3,PFN1,
ADR,CHD4,JUND,TMEM259,EIF5,WDR1,HMGN2,ATP2A2,BEX3,CCT7,SRSF4,RBBP4,RTN3,PFN1,
N1,GPC1,SMCHD1,HDAC2,SKP1,BCLAF1,PPIB,LUC7L3,SQLE,CPSF2,PGRMC1,DDX39B,EID1,MD
,CCT3,RBMX,RAD23B,RAB5B,CHD2,GCN1,TNPO1,SNRNP70,PSMB2,H3-3A,HDGF,TMED2,TOP1,
CLAF1,PPIB,SQLE,CRPPA,CPSF2,DDX39B,MDH2,TOMM20,U2AF2,RPN2,PPP1CB,PGD,SQSTM1,
1,HNRNPL,TM9SF2,MORF4L2,ALKBH5,RPL27,HNRNPA3,ACO2,CAMSAP1,VPS35,CBX1,RAD23B
,MAPK1,SMARCA4,ATG2B,KDELR2,SRSF9,ARPC2,DSTN,PSMD3,SRSF1,ARGLU1,ILF3,GRB2,PRK
,DIDO1,SDF4,SSB,SRSF3,ILF2,CSNK1D,SF3A1,RAI1,NCOR2,PNMA8A,MLF2,KDM3B,TP53BP1,PO
,ACTN1,HSPB1,BNIP3L,FKBP8,MAP2K2,DIDO1,FLNB,POLR2B,SMARCE1,SUPT6H,HNRNPM,SSB,
AP5,SEPTIN11,LAMP1,CKAP4,RPS12,HLA-B,ARCN1,COTL1,CCT6A,HMGCR,ARHGDIA,CAPRIN1,S
7A5,RPS14,CXADR,CHD4,TMEM259,RPS20,EIF5,HMGN2,RPL26,LASP1,BEX3,CCT7,SRSF4,RBB
OP2B,HDAC2,BCLAF1,LUC7L3,CPSF2,CTSB,SRSF5,TM9SF3,MDH2,TOMM20,RAB27A,U2AF2,FD
AP5,SEPTIN11,SND1,LAMP1,HNRNPH1,UBE2H,ARCN1,CCT6A,HMGCR,ARHGDIA,CAPRIN1,CDK1
,DDX42,CBX1,RAD23B,TNPO1,SKI,SNRNP70,H3-3A,HDGF,TSPAN6,PRPF6,TMED2,TOP1,MARS1,
CT7,RBBP4,PFN1,MCL1,JPT2,PNN,RPL31,PRKCSH,HNRNPC,LRPPRC,TADA2B,XPR1,KHDRBS1,X
N,G3BP2,SMARCA4,AZIN1,CCNL2,ARPC2,PSMD3,SRSF1,ARGLU1,NYNRIN,PTBP1,EIF3E,ILF3,PR
YL4,MORF4L2,LMNB2,ABCA3,ALKBH5,HNRNPA3,PDXK,PKN1,TBC1D16,CCT3,VPS35,PTOV1,SU
S1,TUBB2A,MAPRE1,CCNL2,PLEC,PBX2,SRSF1,ARGLU1,TCF25,PTBP1,FTO,ILF3,CDC37,PRKD
NDFIP1,TSR2,HNRNPK,H1-10,NARS1,YWHAB,CKAP5,LAMP1,HNRNPH1,ATP1A2,CKAP4,CTBP1,HM
S1,OPTN,UBXN4,G3BP2,SMARCA4,SRSF9,ARPC2,DSTN,SRSF1,ARGLU1,TCF25,PTBP1,ILF3,PRK
KBP8,PLCG1,DIDO1,FLNB,POLR2B,CORO1C,SMARCE1,HNRNPM,SSB,SRSF3,ILF2,UBQLN1,CSN
TSB,NUCB1,TM9SF3,MDH2,TOMM20,RAB27A,RNF187,U2AF2,RPN2,TBCD,SQSTM1,PABPN1,HN
42,CBX1,RAD23B,TNPO1,SNRNP70,PSMB2,H3-3A,HDGF,TSPAN6,PRPF6,TOP1,MARS1,OPTN,UB
K,H1-10,NARS1,YWHAB,PTGES3,CKAP5,LAMP1,HNRNPH1,ATP1A2,CKAP4,HLA-B,CALU,CTBP1,H
P1,KDM3B,TP53BP1,DDIT4,PSMA7,ANP32A,TRAF4,RBBP7,PRMT1,NAP1L1,LBH,NDUFV1,SF1,PD
F3,PRKDC,HNRNPUL1,ACTN1,BNIP3L,FKBP8,MAP2K2,DIDO1,FLNB,SMARCE1,XPNPEP3,HNRNP
YWHAB,CKAP5,LAMP1,HNRNPH1,RPS12,HLA-B,ARCN1,CCT6A,HMGCR,CAPRIN1,CDK16,TOP2B,
,EIF5,CD81,WDR1,ATP2A2,CRMP1,LASP1,BEX3,CCT7,RTN3,MCL1,PNN,NISCH,LRPPRC,TADA2B,
7,PRDX5,C4ORF3,TBC1D16,ACO2,CCT3,PTOV1,RBMX,DDX42,CBX1,RAD23B,TNPO1,SNRNP70,
D1,MDH2,U2AF2,MLXIP,SUGP2,GNAI3,HNRNPL,PARP1,TSPYL4,MORF4L2,LMNB2,ALKBH5,HNR
EBF2,HNRNPD,SERBP1,KDM3B,TP53BP1,RBBP7,TRAF7,TXLNA,NAP1L1,LBH,SF1,PRCC,GSPT1,
DGF,TSPAN6,TMED2,TOP1,VEGFA,MARS1,OPTN,UBXN4,SMARCA4,LAPTM4A,CCNL2,DSTN,PLEC
0,H3-3A,KDM6B,HDGF,TERF2IP,TMED2,TOP1,TAF7,RNF10,OPTN,UBXN4,G3BP2,MAPK1,SMAR
1,HSPB1,SON,CCAR2,DIDO1,FLNB,POLR2B,SMARCE1,SMARCC1,HNRNPM,SSB,SRSF3,FAU,ILF
4,PSMA7,ANP32A,RBBP7,NAP1L1,LBH,SF1,PBX1,H2AZ1,CSNK2A1,CD99,CST3,BRD4,UBA52,SL
SB,SRSF3,LARS1,ILF2,PFKM,CSNK1D,TMED9,MESD,HNRNPD,MLF2,SERBP1,KDM3B,WBP2,TP5

IK2A1,UBA52,SLC39A7,PPP2CB,ATRX,GARS1,EP400,SRSF6,ARID1B,GLUD1,TRA2B,UQCRC1,ZFF
L3,NPDC1,ANP32B,MAPRE2,DKK3,HNRNPAB,PODXL2,RABEP1,AKAP9,ZMYM3,FUS,XRCC6,PELP1
AF2,RPN2,MLXIP,PPP1CB,PABPN1,HNRNPL,PARP1,TSPYL4,TSR1,MORF4L2,ABCA3,PRDX5,PDXK,
NXA6,NGRN,MACF1,APLP2,HNRNPDL,TARDBP,RPLP2,ARID1A,TSR2,MYL6,HNRNPK,H1-10,INIP,F
KAP4,CALU,CTBP1,HMGCR,ARHGDIA,CAPRIN1,TOP2B,HDAC2,TMEM30A,BCLAF1,RPL23A,LUC7
MARCC1,HNRNPM,SRSF3,ILF2,EIF3F,B2M,LONP1,UBQLN1,SF3A1,MPRIP,HNRNPD,SERBP1,KDM
SF2,TRAP1,NUCKS1,MOAP1,RBM5,PHGDH,SRSF11,SPTY2D1,UBE3A,TGOLN2,RBM6,SMARCD1,
SRSF3,ILF2,STMN3,PFKM,STXBP1,MESD,STRAP,MPRIP,RAI1,NCOR2,P3H3,MLF2,ZSCAN18,TCT
FR,HNRNPA2B1,ANP32B,RPS27A,HNRNPAB,MTCH1,PRDX6,ZMYM3,FUS,PPP1R10,TRIR,XRCC6,E
NRNPH1,RPS12,RAB1A,ARCN1,CCT6A,CAPRIN1,CDK16,TOP2B,HDAC2,BCLAF1,LUC7L3,CRPPA,C
MED9,IDH1,GPR107,PNMA8A,HNRNPD,ZSCAN18,SERBP1,KDM3B,TP53BP1,ANP32A,RBBP7,NAP
,MORF4L2,ALKBH5,RPL27,HNRNPA3,PRDX5,ACO2,VPS35,ACAP3,SURF4,RBMX,RAD23B,RAB5B,
IN11,SND1,LAMP1,HNRNPH1,UBE2H,CKAP4,HLA-B,CALU,ARCN1,COTL1,RAC1,ARHGDIA,CAPRIN
ESD,SOD1,NCOR2,KDM3B,WBP2,TP53BP1,DDIT4,POLR2E,PSMA7,ANP32A,RBBP7,STAT3,ARIH1,
GLUL,LUC7L3,CPSF2,PGRMC1,CDH11,TM9SF3,MDH2,TOMM20,MLXIP,GNAI3,PABPN1,HNRNPL,
FDFT1,RPN2,MLXIP,HNRNPL,TSPYL4,TM9SF2,MORF4L2,TAF15,ALKBH5,RPL27,HNRNPA3,PRDX
,UBA52,ATRX,SRSF6,ARID1B,RER1,RAB3GAP2,TRA2B,ZFR,HNRNPA2B1,ANP32B,POM121C,RPS
B,NAP1L4,TP53BP1,CD99L2,CCT5,POLR2E,ANP32A,CIC,RBBP7,EDC4,TXLNA,OS9,MTA1,TRIO,S
PIB,LUC7L3,CTSB,SRSF5,MDH2,RNF187,U2AF2,FDFT1,CCT2,RPN2,SUGP2,PPP1CB,PGD,SQSTM
1,FLOT2,PHGDH,SRSF11,MSH6,PRPF19,PSMC3,AP2A2,SMARCD1,SAP18,THRAP3,EIF4A3,XRN2,
LE1,HDGFL2,SRSF2,TRAP1,NUCKS1,RBM5,WBP11,SRSF11,HSD17B4,MSH6,IST1,PSMC3,RBM6,
F3,SRSF11,MSH6,IST1,IVD,GSK3A,TGOLN2,NDUFA13,ATXN7L3,MN1,ATP6V1G1,THRAP3,SLC38A
B,DIP2B,GLUD1,EVL,TRA2B,UQCRC1,ZFR,ANP32B,HNRNPAB,RABEP1,ZMYM3,GRK2,FUS,XRCC
K1,TCAF1,CARM1,TLE1,HDGFL2,EFTUD2,TRAP1,SFXN1,PLEKHM2,RBM5,SPTY2D1,MSH6,IST1,F
),H3-3A,KDM6B,HDGF,PRPF6,TERF2IP,TAF7,RPS15A,MARS1,TUBB2A,G3BP2,SMARCA4,MAPRE1
F2,EID1,CTSB,NUCB1,TM9SF3,MDH2,TOMM20,RAB27A,RNF187,U2AF2,RPN2,MLXIP,PABPN1,HI
B,EFNB3,CTCF,DSTYK,ELOB,HEXIM1,GATAD2B,SEC31A,FUBP1,AKAP17A,SMARCC2,FAM193B,S
E7A,PSMD3,SRSF1,ARGLU1,TCF25,EIF3E,FTO,ILF3,CDC37,CAPZB,ATXN10,ACTN1,HSPB1,SON,B
DNAJC5,HNRNPDL,TARDBP,RPLP2,DDOST,WLS,HNRNPK,H1-10,RPL36,CKAP5,SEPTIN11,LAMP1
10,REEP5,DNAJB1,SF3A3,FAM50A,EWSR1,ERP29,MTURN,HNRNPR,ECHS1,AKAP8L,ADRM1,CDK
1,PACS1,WASF2,RBM5,PHGDH,SRSF11,SPTY2D1,BORCS7,MSH6,IST1,EIF3G,UBE3A,MN1,SCD,E
CE1,SMARCC1,SUPT6H,HNRNPM,SRSF3,ILF2,PFKM,B2M,UBQLN1,SF3A1,TMED9,SREBF2,HNRN
TCH1,ZMYM3,FUS,XRCC6,ERH,TAOK1,PELP1,TCAF1,EIF3H,TRAM1,TLE1,SH3GL1,HDGFL2,LSM
IDGFL2,SRSF2,NUCKS1,RBM5,WBP11,ITGAV,SPTY2D1,MSH6,PRPF19,EHD1,PSMC3,TGOLN2,RB
MD3,ARGLU1,RBP1,TCF25,ATP5PB,PTBP1,FTO,ILF3,HNRNPUL1,BNIP3L,FKBP8,DIDO1,FLNB,TP
RBBP7,TXLNA,OS9,TRIO,LBH,SF1,H2AZ1,GTPBP1,FLOT1,BRD4,DECR1,ATRX,RAB2A,GLUD1,EIF
AM,GLUL,CPSF2,PGRMC1,EID1,CDH11,MDH2,MLXIP,GNAI3,PABPN1,GDI2,TSPYL4,TLN2,MORF
RCC6,ERH,TCAF1,EIF3H,RALY,TLE1,SH3GL1,HDGFL2,LSM14A,EFTUD2,SRSF2,TRAP1,PLEKHM2
TOP2B,HDAC2,TMEM30A,BCLAF1,LUC7L3,SQLE,TMEM132A,CPSF2,DDX39B,EID1,MDH2,TOMM
1CB,NOP53,PABPN1,GDI2,HNRNPL,PARP1,TLN2,TM9SF2,MORF4L2,TAF15,HNRNPA3,ACO2,VP
HC2,PFKM,STT3B,EIF3F,IDH1,STRAP,UQCRC2,NCOR2,P3H3,ZSCAN18,SMS,DDIT4,POLR2E,PSM

2,RAI1,MLF2,ZSCAN18,SERBP1,KDM3B,TP53BP1,CCT5,VDAC1,RBBP7,EIF2AK1,EP300,LBH,PSM
,STRAP,HNRNPD,ZSCAN18,SERBP1,KDM3B,DDIT4,ANP32A,RBBP7,EDC4,TXLNA,TXNDC5,LBH,S
2,GTF2I,EEF1B2,MOAP1,RBM5,PHGDH,YIPF3,SRSF11,CAP1,MSH6,NSD2,DHCR7,EIF3G,RBM6,A

,PLEKHB2,RBM5,PHGDH,SRSF11,SPTY2D1,MSH6,UBE3A,RBM6,MN1,THRAP3,PRDX3,EIF4A3,CA

MA7,RBBP7,EDC4,TXLNA,SF1,PDAP1,VARS1,H2AZ1,BRD4,PPP2CB,DIP2B,RAB2A,EP400,GLUD1,E
NKRD50,KDM2A,NACC1,LMNA,THOC2,RXRA,HIRA,DDHD2,ELOB,ZNF146,PRPF31,HEXIM1,HMGB
2,PFKM,STXBP1,UBQLN1,CSNK1D,SF3A1,MESD,GPR107,MPRIP,PNMA8A,ZSCAN18,KDM3B,ARF
RAF,QRICH1,TNPO3,GATAD2B,SEC31A,NBR1,FUBP1,MAML1,EIF2S1,PSMC6,FAM193B,ELMO2,Z
ME3,PSIP1,NDRG1,RBM10,YWHAZ,DDAH1,SNRPD2,ZFP91,RRP1B,CLCN6,HSPE1,LMNA,CDC42EP
3PXD2B,RANGAP1,CASC4,XRN2,RSF1,SUN1,POR,DAG1,RUVBL2,ELN,REEP5,MDM2,SF3A3,FAM

,EHD1,ATXN7L3,MN1,ANKRD12,CCDC47,THRAP3,FGFR1,XRN2,HTATSF1,SUN1,MTA2,CGGBP1,
,NACC1,LMNA,CDC42EP4,MED12,ZNRF1,TLK1,PHB,RBM17,NRP2,MGRN1,DSTYK,ELOB,ZNF146
AP1,RAB6A,AHSA1,RAB5C,SPAG9,PDPK1,SRSF11,MSH6,NSD2,CSE1L,IVD,SF3A2,SMARCD1,WNK
,PBXIP1,CCNG1,PSME3,DHX36,RBFOX2,ZNF768,PSMC2,PLK2,MEIS1,SRF,HSF1,ZEB1,MRPS6,C
CSNK2A1,CD99,KPNA2,PPM1G,BRD2,EP400,SRSF6,KIAA0895L,UBE2G2,TRA2B,ZFR,FEM1B,HNR
1,MTA2,LRIG1,CGGBP1,ARPC1A,FAM171A1,RTF1,PAIP2,MACROH2A1,PDHA1,IGF1R,MDM2,STK
,ACTR3,C4ORF3,PKN1,TBC1D16,ACO2,CCT3,RBMX,RAD23B,RAB5B,GCN1,TNPO1,SNRNP70,PS
MDM2,FOXP4,DNAJB1,EFNB2,SF3A3,CAB39,ADNP,GPAA1,EWSR1,HNRNPR,WRNIP1,RNF41,AKA
PRPF31,NAV1,RTRAF,PLCD3,QRICH1,RAD54L2,ABCE1,ZDHHC17,NUP50,MAML1,PSMC6,FZD7,S
CH1,SAFB2,ZMYM3,FUS,XRCC6,PELP1,TPP1,RALY,TM9SF4,HDGFL2,EFTUD2,KIAA0232,SRSF2,

1,NDUFA13,SCAP,CAD,XRN2,TAPBP,HMGA1,HTATSF1,SGSM2,MTA2,USP10,CGGBP1,HDGFL3,S
2,HTATSF1,SUN1,MTA2,CGGBP1,CCDC80,NOMO2,ARRDC4,MDM2,MGAT3,MECP2,FAM50A,WD

,FBRSL1,MLF2,SERBP1,KDM3B,ARFGEF2,NAP1L4,TP53BP1,CCT5,PCYOX1,POLR2E,PSMA7,RB
PK1,KDELR2,LAPTM4A,ARPC2,DSTN,PSMD3,PRDX1,IPO7,NYNRIN,EIF3E,LUZP6,CAPZB,ATXN10,
C1,DHX30,HNRNPM,SSB,SRSF3,SDHA,LARS1,SLIT3,ILF2,CCT8,LSS,STXBP1,ISYNA1,LONP1,UBQ

F3,PRKDC,SUPT5H,HNRNPUL1,ACTN1,SON,CASC3,FKBP8,CCAR2,ZFAND5,DIDO1,TPM3,SMAR

1L,IVD,SF3A2,SMARCD1,TOMM70,WNK1,THRAP3,TUBB4A,EIF4A3,USO1,RCN2,SLC38A10,NELF
,PLCD3,KCTD10,HEXIM1,TRAPPC1,AGPAT4,RELA,ABCE1,OPA1,NUP50,HID1,AKAP17A,ID3,PAF1,

TXNRD1,MDM2,MECP2,FAM50A,EWSR1,SUB1,PDLIM1,ACAP2,ERP29,PCGF3,SUN2,HIPK2,WIZ,C
,SRSF5,RNF187,FDFT1,MLXIP,ANXA5,TSPYL4,SSRP1,ALKBH5,RPL27,PTPRA,ACTR3,C4ORF3,UE

M27,POLD2,GID8,DHX36,EIF3B,ZNF768,NDRG1,TNFRSF10B,RANBP9,RNMT,PLK2,YWHAZ,TUBA

RC2,HNRNPD,SERBP1,ARFGEF2,TXNIP,CCT5,POLR2E,TRAF4,RBBP7,TRAF7,MTA1,TRIO,KDM5B

A,POSTN,PDHB,CPNE2,MRPS27,WDR33,RAD50,ZNF326,NOP58,ANAPC2,GLB1,SCCPDH,NUTF2,C

TUBB4B,UCHL1,RPL11,MORF4L1,BTF3,NCAM1,ANXA2,AGRN,XAF1,RPL10,MARCHF6,WSB1,RPS
,RPS17,DNAJA1,CYB5R3,PAFAH1B1,MAP4,SF3B2,CDC42BPB,ITM2B,ZNF770,AP2B1,GTF2F1,HN
42BPB,ITM2B,ZNF770,AP2B1,MAT2A,GTF2F1,HNRNPU,OGA,PGAM1,NFE2L1,SMARCA5,LARP1,
A,PGAM1,NFE2L1,SMARCA5,LARP1,HNRNPA0,CTNNA1,FADS2,RPS16,RPL23,CPE,CNN3,RPL29,R
L1,BTF3,NCAM1,ANXA2,AGRN,ZMIZ1,XAF1,RPL10,MARCHF6,WSB1,RPS17,VCL,DNAJA1,CYB5R
PGAM1,NFE2L1,SMARCA5,LARP1,HNRNPA0,CTNNA1,FADS2,RPS16,RPL23,CPE,CNN3,RPL29,RPS
SF3B2,CDC42BPB,ITM2B,ZNF770,AP2B1,GTF2F1,HNRNPU,OGA,PGAM1,NFE2L1,SMARCA5,LAR
B1,MAP4,SF3B2,CDC42BPB,ITM2B,ZNF770,AP2B1,MAT2A,GTF2F1,HNRNPU,OGA,PGAM1,NFE2
4,SF3B2,CDC42BPB,ITM2B,ZNF770,MAT2A,GTF2F1,GOLGA8A,HNRNPU,OGA,PGAM1,NFE2L1,S
,GTF2F1,HNRNPU,OGA,PGAM1,NFE2L1,SMARCA5,LARP1,HNRNPA0,CTNNA1,FADS2,RPS16,RPL
,GTF2F1,HNRNPU,OGA,PGAM1,NFE2L1,SMARCA5,LARP1,HNRNPA0,CTNNA1,FADS2,RPS16,RPL
AP4,SF3B2,CDC42BPB,ITM2B,ZNF770,AP2B1,MAT2A,GTF2F1,GOLGA8A,HNRNPU,OGA,NFE2L1,
A,PTCD3,CBX5,ZNF865,RPL9,ATF4,PSMD2,KIF5B,SLC7A5,RPS14,CXADR,CHD4,JUND,TMEM25
YB5R3,PAFAH1B1,AKAP12,MAP4,SF3B2,CDC42BPB,ITM2B,ZNF770,AP2B1,MAT2A,GTF2F1,GO
YB5R3,PAFAH1B1,AKAP12,MAP4,SF3B2,CDC42BPB,ITM2B,ZNF770,AP2B1,MAT2A,GTF2F1,GO
OGA,PGAM1,NFE2L1,SMARCA5,LARP1,HNRNPA0,CTNNA1,FADS2,RPS16,RPL23,CPE,CNN3,RPL
,ITM2B,ZNF770,MAT2A,GTF2F1,GOLGA8A,HNRNPU,OGA,PGAM1,NFE2L1,SMARCA5,LARP1,HN
,ITM2B,ZNF770,MAT2A,GTF2F1,GOLGA8A,HNRNPU,OGA,PGAM1,NFE2L1,SMARCA5,LARP1,HN
5R3,PAFAH1B1,MAP4,SF3B2,CDC42BPB,ITM2B,ZNF770,MAT2A,GTF2F1,GOLGA8A,HNRNPU,OG
2B1,MAT2A,GTF2F1,GOLGA8A,HNRNPU,OGA,NFE2L1,SMARCA5,LARP1,HNRNPA0,CTNNA1,FADS
PS14,CHD4,JUND,TMEM259,RPS20,EIF5,CD81,HMGN2,RPL26,LASP1,BEX3,CCT7,SRSF4,RBBP4
3B2,CDC42BPB,ITM2B,ZNF770,AP2B1,GTF2F1,HNRNPU,PGAM1,NFE2L1,SMARCA5,LARP1,HNR
AP2B1,MAT2A,GTF2F1,HNRNPU,OGA,PGAM1,NFE2L1,SMARCA5,LARP1,HNRNPA0,CTNNA1,FADS
AP2B1,MAT2A,GTF2F1,HNRNPU,OGA,PGAM1,NFE2L1,SMARCA5,LARP1,HNRNPA0,CTNNA1,FADS
HK1,NCS1,RHOA,PTCD3,CBX5,ZNF865,RPL9,CLSTN1,PSMD2,SLC7A5,RPS14,CHD4,JUND,TME
TMEM259,RPS20,EIF5,WDR1,ATP2A2,RPL26,CRMP1,BEX3,CCT7,SRSF4,RBBP4,RTN3,PFN1,MC
,MARCHF6,WSB1,RPS17,VCL,DNAJA1,CYB5R3,PAFAH1B1,AKAP12,MAP4,SF3B2,CDC42BPB,IT
,MAT2A,GTF2F1,GOLGA8A,HNRNPU,PGAM1,NFE2L1,SMARCA5,HNRNPA0,CTNNA1,FADS2,RPS1
1,HNRNPA0,CTNNA1,FADS2,RPS16,RPL23,RPSA,TPM4,CTSD,EIF4H,SCARB2,RPS24,BRD3,RPS9,
1,HNRNPA0,CTNNA1,FADS2,RPS16,RPL23,RPSA,TPM4,CTSD,EIF4H,SCARB2,RPS24,BRD3,RPS9,
2,CDC42BPB,ITM2B,ZNF770,AP2B1,MAT2A,GTF2F1,GOLGA8A,PXDN,HNRNPU,OGA,PGAM1,NF
E2L1,SMARCA5,LARP1,HNRNPA0,CTNNA1,FADS2,RPS16,RPL23,CNN3,RPL29,RPSA,TPM4,CTSD,
E2L1,SMARCA5,LARP1,HNRNPA0,CTNNA1,FADS2,RPS16,RPL23,CNN3,RPL29,RPSA,TPM4,CTSD,
,KPNB1,RPL17,C1ORF43,PAPOLA,LLGL1,RPL14,HK1,RHOA,PTCD3,CBX5,RPL9,CLSTN1,ATF4,P
EIF4H,SCARB2,LPCAT1,BRD3,RPS9,XRCC5,PJA2,RPL24,KPNB1,RPL17,C1ORF43,PAPOLA,COL1
GTF2F1,GOLGA8A,HNRNPU,OGA,NFE2L1,SMARCA5,LARP1,HNRNPA0,CTNNA1,FADS2,RPS16,RP
,HK1,RHOA,PTCD3,CBX5,ZNF865,CLSTN1,ATF4,PSMD2,KIF5B,NLGN2,RPS14,CXADR,CHD4,JUN
RPL23,CPE,CNN3,RPL29,RPSA,TPM4,CTSD,EIF4H,SCARB2,LPCAT1,BRD3,RPS9,XRCC5,PJA2,MS
RPL23,CPE,CNN3,RPL29,RPSA,TPM4,CTSD,EIF4H,SCARB2,LPCAT1,BRD3,RPS9,XRCC5,PJA2,MS
PL29,RPSA,TPM4,CTSD,EIF4H,SCARB2,BRD3,RPS9,XRCC5,MSN,RPL24,KPNB1,RPL17,C1ORF43
5,CD81,HMGN2,RPL26,BEX3,CCT7,SRSF4,RBBP4,PFN1,MCL1,PNN,RPL31,PRKCSH,HNRNPC,LR
,MAP4,SF3B2,CDC42BPB,ITM2B,ZNF770,MAT2A,GTF2F1,GOLGA8A,HNRNPU,OGA,PGAM1,NFE
,MAP4,SF3B2,CDC42BPB,ITM2B,ZNF770,MAT2A,GTF2F1,GOLGA8A,HNRNPU,OGA,PGAM1,NFE
12,MAP4,SF3B2,CDC42BPB,ITM2B,ZNF770,MAT2A,GTF2F1,GOLGA8A,HNRNPU,OGA,PGAM1,NF
12,MAP4,SF3B2,CDC42BPB,ITM2B,ZNF770,MAT2A,GTF2F1,GOLGA8A,HNRNPU,OGA,PGAM1,NF

43,PAPOLA,COL1A2,RPL14,HK1,RHOA,PTCD3,CBX5,CLSTN1,ATF4,KIF5B,RPS14,CXADR,CHD4,
42BPB,ITM2B,ZNF770,AP2B1,GTF2F1,GOLGA8A,HNRNPU,PGAM1,NFE2L1,SMARCA5,LARP1,H
42BPB,ITM2B,ZNF770,AP2B1,GTF2F1,GOLGA8A,HNRNPU,PGAM1,NFE2L1,SMARCA5,LARP1,H
2BPB,ITM2B,ZNF770,GTF2F1,HNRNPU,PGAM1,NFE2L1,SMARCA5,LARP1,HNRNPA0,CTNNA1,FA
U,OGA,PGAM1,NFE2L1,SMARCA5,LARP1,HNRNPA0,CTNNA1,FADS2,RPS16,RPL23,CNN3,RPL29,
,SF3B2,CDC42BPB,ITM2B,ZNF770,MAT2A,GTF2F1,GOLGA8A,HNRNPU,OGA,PGAM1,NFE2L1,S
C42BPB,ITM2B,ZNF770,AP2B1,MAT2A,GTF2F1,HNRNPU,OGA,PGAM1,NFE2L1,SMARCA5,LARP1
1,FADS2,RPS16,RPL23,CNN3,RPSA,TPM4,CTSD,EIF4H,SCARB2,LPCAT1,RPS24,BRD3,RPS9,XRC
LARP1,HNRNPA0,CTNNA1,FADS2,RPS16,RPL23,CPE,CNN3,RPL29,RPSA,TPM4,TNS3,CTSD,EIF4
LARP1,HNRNPA0,CTNNA1,FADS2,RPS16,RPL23,CPE,CNN3,RPL29,RPSA,TPM4,TNS3,CTSD,EIF4
1,MAP4,SF3B2,CDC42BPB,ITM2B,ZNF770,AP2B1,MAT2A,GTF2F1,GOLGA8A,HNRNPU,OGA,PGA
SN,RPL24,KPNB1,RPL17,C1ORF43,PAPOLA,COL1A2,LLGL1,RPL14,HK1,NCS1,RHOA,PTCD3,R
U,OGA,PGAM1,NFE2L1,SMARCA5,LARP1,HNRNPA0,CTNNA1,FADS2,RPS16,RPL23,CNN3,RPL29
U,OGA,PGAM1,NFE2L1,SMARCA5,LARP1,HNRNPA0,CTNNA1,FADS2,RPS16,RPL23,CNN3,RPL29
B,XPR1,KHDRBS1,XPO1,CSNK1E,RPS25,TKT,GLG1,SFPQ,PDIA6,CCN2,ACTN4,MMP14,PRDX2,D
NNA1,FADS2,RPS16,RPL23,CNN3,RPSA,TPM4,TNS3,CTSD,EIF4H,SCARB2,LPCAT1,RPS24,BRD3,
,RPS9,XRCC5,PJA2,SLC16A7,KPNB1,RPL17,C1ORF43,PAPOLA,LLGL1,GAP43,RPL14,PTCD3,RP
B,ITM2B,ZNF770,AP2B1,GTF2F1,GOLGA8A,HNRNPU,OGA,PGAM1,NFE2L1,SMARCA5,LARP1,HN
B,ITM2B,ZNF770,AP2B1,GTF2F1,GOLGA8A,HNRNPU,OGA,PGAM1,NFE2L1,SMARCA5,LARP1,HN
OLGA8A,HNRNPU,PGAM1,NFE2L1,SMARCA5,LARP1,HNRNPA0,CTNNA1,FADS2,RPL23,CNN3,R
OLGA8A,HNRNPU,PGAM1,NFE2L1,SMARCA5,LARP1,HNRNPA0,CTNNA1,FADS2,RPL23,CNN3,R
A2,LLGL1,RPL14,HK1,RHOA,PTCD3,CBX5,RPL9,CLSTN1,ATF4,PSMD2,KIF5B,SLC7A5,RPS14,CX
,ZNF770,AP2B1,MAT2A,GTF2F1,GOLGA8A,HNRNPU,OGA,PGAM1,NFE2L1,SMARCA5,LARP1,HNR
1,PAPOLA,GAP43,RPL14,HK1,NCS1,CBX5,CLSTN1,PSMD2,KIF5B,SLC7A5,RPS14,CXADR,CHD4,
CA5,HNRNPA0,CTNNA1,FADS2,RPS16,RPL23,CNN3,RPL29,RPSA,TPM4,EIF4H,SCARB2,LPCAT1,
7,C1ORF43,LLGL1,RPL14,HK1,PTCD3,CBX5,PSMD2,KIF5B,SLC7A5,RPS14,CXADR,CHD4,ATP1A
,RPS14,CXADR,CHD4,ATP1A1,JUND,TMEM259,RPS20,EIF5,WDR1,HMGN2,RPL26,BEX3,CCT7,SR
T1,BRD3,XRCC5,PJA2,MSN,RPL24,SLC16A7,KPNB1,RPL17,C1ORF43,PAPOLA,COL1A2,LLGL1,
1,NCS1,RHOA,PTCD3,ZNF865,RPL9,CLSTN1,ATF4,PSMD2,KIF5B,SLC7A5,RPS14,CHD4,JUND,R
1,NCS1,RHOA,PTCD3,ZNF865,RPL9,CLSTN1,ATF4,PSMD2,KIF5B,SLC7A5,RPS14,CHD4,JUND,R
,MSN,RPL24,RPL17,C1ORF43,PAPOLA,COL1A2,LLGL1,RPL14,HK1,RHOA,PTCD3,CBX5,RPL9,C
PE,CNN3,RPL29,RPSA,TPM4,CTSD,EIF4H,SCARB2,LPCAT1,BRD3,RPS9,XRCC5,PJA2,RPL24,SLC
TF4,PSMD2,KIF5B,RPS14,CHD4,ATP1A1,JUND,TMEM259,RPS20,EIF5,WDR1,HMGN2,RPL26,LA
B,ITM2B,ZNF770,AP2B1,GTF2F1,GOLGA8A,HNRNPU,OGA,NFE2L1,SMARCA5,LARP1,HNRNPA0,
B,ITM2B,ZNF770,AP2B1,GTF2F1,GOLGA8A,HNRNPU,OGA,NFE2L1,SMARCA5,LARP1,HNRNPA0,
XDN,HNRNPU,OGA,PGAM1,NFE2L1,SMARCA5,LARP1,HNRNPA0,CTNNA1,FADS2,RPS16,RPL23,C
B,ZNF770,AP2B1,GTF2F1,GOLGA8A,HNRNPU,OGA,PGAM1,NFE2L1,SMARCA5,LARP1,HNRNPA0,
,HNRNPA0,CTNNA1,FADS2,RPS16,RPL23,CNN3,RPL29,RPSA,TPM4,TNS3,CTSD,EIF4H,SCARB2,L
N2,RPL26,LASP1,BEX3,CCT7,SRSF4,RBBP4,RTN3,PFN1,MCL1,PNN,RPL31,NISCH,PRKCSH,HNR
SA,TPM4,EIF4H,SCARB2,LPCAT1,RPS24,BRD3,XRCC5,PJA2,MSN,RPL24,KPNB1,RPL17,C1ORF4
SMD2,KIF5B,RPS14,CXADR,CHD4,ATP1A1,JUND,TMEM259,RPS20,HMGN2,RPL26,BEX3,CCT7,S
D2,KIF5B,SLC7A5,RPS14,CXADR,CHD4,ATP1A1,JUND,TMEM259,RPS20,EIF5,HMGN2,RPL26,BE
SLC7A5,RPS14,JUND,TMEM259,RPS20,EIF5,WDR1,HMGN2,RPL26,BEX3,CCT7,RBBP4,PFN1,MC
16,RPL23,CPE,CNN3,RPL29,RPSA,TPM4,TNS3,CTSD,EIF4H,SCARB2,BRD3,RPS9,XRCC5,PJA2,M
CNN3,RPL29,RPSA,TPM4,TNS3,EIF4H,SCARB2,LPCAT1,BRD3,RPS9,XRCC5,PJA2,RPL24,KPNB1,

4,HK1,RHOA,PTCD3,CBX5,ATF4,PSMD2,KIF5B,RPS14,CHD4,ATP1A1,JUND,RPS20,EIF5,HMGN2,
RCA5,LARP1,HNRNPA0,CTNNA1,RPS16,RPL23,CNN3,RPL29,RPSA,TPM4,TNS3,CTSD,EIF4H,SCAR
NPU,OGA,PGAM1,NFE2L1,SMARCA5,LARP1,HNRNPA0,CTNNA1,FADS2,RPS16,RPL23,CPE,CNN3,
NPU,OGA,PGAM1,NFE2L1,SMARCA5,LARP1,HNRNPA0,CTNNA1,FADS2,RPS16,RPL23,CPE,CNN3,
GED2,RCN1,H1-0,RPS5,RPS7,IK,MIF,TLN1,CHPF,CTNNB1,ANXA6,NGRN,GOLM1,RPS10,DNAJC5,
RPS24,BRD3,RPS9,XRCC5,MSN,RPL24,KPNB1,RPL17,C1ORF43,PAPOLA,COL1A2,LLGL1,RPL14,
SCARB2,LPCAT1,RPS24,BRD3,RPS9,XRCC5,PJA2,MSN,RPL24,SLC16A7,KPNB1,RPL17,C1ORF43,
MD2,KIF5B,SLC7A5,RPS14,CXADR,CHD4,JUND,RPS20,EIF5,HMGN2,RPL26,CCT7,SRSF4,RBBP4,
H1B1,MAP4,SF3B2,CDC42BPB,ITM2B,ZNF770,AP2B1,GTF2F1,GOLGA8A,HNRNPU,OGA,PGAM1,
3,RPL29,RPSA,TPM4,EIF4H,RPS24,BRD3,RPS9,XRCC5,PJA2,RPL24,RPL17,C1ORF43,PAPOLA,CC
3,RPL29,RPSA,TPM4,EIF4H,RPS24,BRD3,RPS9,XRCC5,PJA2,RPL24,RPL17,C1ORF43,PAPOLA,CC
P1A1,JUND,TMEM259,RPS20,EIF5,WDR1,HMGN2,RPL26,CCT7,SRSF4,RBBP4,PFN1,MCL1,JPT2,
CXADR,CHD4,JUND,RPS20,EIF5,CD81,HMGN2,RPL26,BEX3,SRSF4,RBBP4,RTN3,PFN1,MCL1,PN
,CBX5,RPL9,CLSTN1,ATF4,PSMD2,KIF5B,SLC7A5,RPS14,CXADR,CHD4,ATP1A1,JUND,TMEM259,
,CBX5,RPL9,CLSTN1,ATF4,PSMD2,KIF5B,SLC7A5,RPS14,CXADR,CHD4,ATP1A1,JUND,TMEM259,
,RBBP4,PFN1,MCL1,JPT2,PNN,RPL31,NISCH,PRKCSH,HNRNPC,LRPPRC,TADA2B,XPR1,KHDRBS
N2,RPL26,BEX3,CCT7,SRSF4,RBBP4,PFN1,MCL1,JPT2,PNN,RPL31,PRKCSH,HNRNPC,LRPPRC,TA
D2,SLC7A5,RPS14,CXADR,CHD4,ATP1A1,JUND,TMEM259,RPS20,EIF5,WDR1,HMGN2,RPL26,CC
,JUND,TMEM259,RPS20,EIF5,CD81,ATP2A2,RPL26,LASP1,BEX3,CCT7,RBBP4,RTN3,MCL1,PNN,
,PTCD3,CBX5,RPL9,KIF5B,SLC7A5,RPS14,CXADR,CHD4,JUND,RPS20,EIF5,CD81,WDR1,HMGN2,
3,CBX5,RPL9,PSMD2,KIF5B,RPS14,CHD4,ATP1A1,JUND,TMEM259,RPS20,EIF5,WDR1,HMGN2,R
1,BTF3,ANXA2,AGRN,ZMIZ1,TIMP2,XAF1,RPL10,MARCHF6,WSB1,RPS17,VCL,DNAJA1,CYB5R3,
,RPS7,IK,GPAT4,PRKAR1A,TLN1,CHPF,GJA1,JUN,CTNNB1,ANXA6,NGRN,MACF1,RPS10,DNAJC5,
,SLC16A7,KPNB1,RPL17,C1ORF43,PAPOLA,COL1A2,LLGL1,RPL14,HK1,RHOA,PTCD3,CBX5,RPL
4B,CTBP1,ARHGDIA,CAPRIN1,GPC1,TOP2B,HDAC2,MMP2,SKP1,BCLAF1,PPIB,LUC7L3,SQLE,DC
H,SCARB2,LPCAT1,BRD3,RPS9,XRCC5,PJA2,MSN,RPL24,KPNB1,C1ORF43,PAPOLA,COL1A2,RPL
PCAT1,BRD3,XRCC5,PJA2,MSN,RPL24,KPNB1,RPL17,PAPOLA,COL1A2,LLGL1,RPL14,HK1,PTCD
14,HK1,PTCD3,RPL9,PSMD2,KIF5B,SLC7A5,CXADR,CHD4,ATP1A1,JUND,RPS20,EIF5,WDR1,HM
CTSD,EIF4H,SCARB2,BRD3,RPS9,XRCC5,PJA2,RPL24,SLC16A7,KPNB1,RPL17,C1ORF43,PAPOLA
SD,EIF4H,SCARB2,LPCAT1,BRD3,RPS9,XRCC5,PJA2,MSN,RPL24,KPNB1,RPL17,C1ORF43,PAPO
B2,LPCAT1,BRD3,XRCC5,PJA2,MSN,RPL24,SLC16A7,KPNB1,RPL17,C1ORF43,PAPOLA,COL1A2,
PL26,CRMP1,LASP1,CCT7,SRSF4,RBBP4,MCL1,JPT2,PNN,RPL31,NISCH,PRKCSH,HNRNPC,LRPP
3H5,RPL27,HNRNPA3,C4ORF3,ACO2,NCKAP1,PTOV1,RBMX,DDX42,JPT1,CBX1,RAD23B,CHD2,G
14,CHD4,JUND,TMEM259,RPS20,EIF5,WDR1,HMGN2,RPL26,BEX3,CCT7,SRSF4,RBBP4,PFN1,M
9,XRCC5,PJA2,RPL24,KPNB1,RPL17,C1ORF43,PAPOLA,COL1A2,LLGL1,RPL14,RHOA,PTCD3,CB
RPL24,RPL17,C1ORF43,PAPOLA,COL1A2,LLGL1,RPL14,HK1,PTCD3,RPL9,ATF4,PSMD2,KIF5B,SL
TN4,BRD3,XRCC5,PJA2,KPNB1,RPL17,C1ORF43,PAPOLA,COL1A2,LLGL1,GAP43,RPL14,RHOA,
,JUND,TMEM259,RPS20,EIF5,HMGN2,RPL26,BEX3,CCT7,SRSF4,RBBP4,PFN1,MCL1,PNN,RPL31
F43,PAPOLA,COL1A2,LLGL1,RPL14,HK1,RHOA,PTCD3,CBX5,ZNF865,RPL9,PSMD2,KIF5B,SLC7A
1A2,RPL14,HK1,PTCD3,ZNF865,RPL9,ATF4,PSMD2,KIF5B,SLC7A5,RPS14,CXADR,CHD4,ATP1A1,
1,RPL14,HK1,NCS1,RHOA,PTCD3,CBX5,ZNF865,CLSTN1,PSMD2,KIF5B,SLC7A5,RPS14,CXADR,C
CSH,HNRNPC,LRPPRC,TADA2B,KHDRBS1,XPO1,SPTBN2,CSNK1E,RPS25,GLG1,SFPQ,PEBP1,PN
PL17,C1ORF43,PAPOLA,COL1A2,LLGL1,RPL14,HK1,RHOA,PTCD3,CBX5,ZNF865,RPL9,CLSTN1,
4,PTCD3,CBX5,RPL9,CLSTN1,ATF4,PSMD2,KIF5B,SLC7A5,RPS14,CXADR,CHD4,ATP1A1,JUND,T
365,PSMD2,KIF5B,SLC7A5,RPS14,CHD4,JUND,TMEM259,RPS20,EIF5,WDR1,RPL26,BEX3,CCT7,

5B,SLC7A5,RPS14,CHD4,JUND,EIF5,WDR1,HMGN2,RPL26,LASP1,BEX3,CCT7,SRSF4,RBBP4,PFN
SNK1E,PFKL,HDLBP,RPS25,GLG1,CDH6,SFPQ,PEBP1,PNMA1,DDR1,ACSL6,RAB7A,PRDX2,DDX3X
PT1,CBX1,RAD23B,PTK7,RAB5B,TNPO1,SNRNP70,H3-3A,SCAF1,HDGF,ABCA2,TAF7,NOTCH2,VE
A2B,KHDRBS1,XPO1,SPTBN2,RPS25,TKT,GLG1,SFPQ,EIF5A,DDR1,RAB7A,CCN2,MMP14,DDX3X,
2,LPCAT1,BRD3,XRCC5,PJA2,MSN,RPL24,KPNB1,RPL17,C1ORF43,PAPOLA,COL1A2,RPL14,HK1
IB,LUC7L3,DCN,CPSF2,PGRMC1,DDX39B,CDH11,TM9SF3,MDH2,TOMM20,U2AF2,RPN2,ANXA5,
31,NISCH,PRKCSH,HNRNPC,LRPPRC,TADA2B,XPR1,KHDRBS1,KLHL9,XPO1,CSNK1E,RPS25,TKT
A1,FADS2,RPS16,RPL23,CNN3,RPL29,RPSA,TPM4,TNS3,CTSD,EIF4H,SCARB2,LPCAT1,BRD3,RPS
A1,FADS2,RPS16,RPL23,CNN3,RPL29,RPSA,TPM4,TNS3,CTSD,EIF4H,SCARB2,LPCAT1,BRD3,RPS
COL1A2,LLGL1,RPL14,CBX5,RPL9,ATF4,PSMD2,KIF5B,SLC7A5,RPS14,CXADR,CHD4,ATP1A1,JU
T7,SRSF4,RBBP4,PFN1,MCL1,JPT2,PNN,SPC25,RPL31,PRKCSH,HNRNPC,LRPPRC,TADA2B,XPR1
S25,TKT,GLG1,SFPQ,PDIA6,DDR1,ACSL6,RAB7A,MMP14,PRDX2,DDX3X,RPL12,RPL30,MAGED2,
S25,TKT,GLG1,SFPQ,PDIA6,DDR1,ACSL6,RAB7A,MMP14,PRDX2,DDX3X,RPL12,RPL30,MAGED2,
KHDRBS1,HDLBP,RPS25,TKT,GLG1,SFPQ,EIF5A,PDIA6,RAB7A,MMP14,PRDX2,DDX3X,RPL12,RPL
H2,PDIA6,MEGF8,RAB7A,ACTN4,PRDX2,DDX3X,RPL12,CLTC,EIF3L,MAGED2,GNB2,RCN1,H1-0,I
A2,RPL26,BEX3,CCT7,RBBP4,PFN1,MCL1,JPT2,PNN,NISCH,PRKCSH,HNRNPC,LRPPRC,TADA2B,
TC,RCN1,H1-0,RPS5,RPS7,RPL35,HMGCS1,GPAT4,TLN1,CHPF,JUN,MYO5A,CTNNB1,RPS10,HN
SNK1E,HDLBP,RPS25,TKT,GLG1,SFPQ,EIF5A,DDR1,RAB7A,CCN2,MMP14,DDX3X,RPL12,RPL30,
MD2,KIF5B,SLC7A5,RPS14,CHD4,JUND,TMEM259,RPS20,CD81,WDR1,RPL26,LASP1,BEX3,CCT
RP1,HNRNPA0,CTNNA1,FADS2,RPS16,RPL23,CNN3,RPL29,RPSA,TPM4,TNS3,CTSD,EIF4H,SCAR
D3,CBX5,RPL9,ATF4,PSMD2,KIF5B,SLC7A5,RPS14,CXADR,CHD4,JUND,RPS20,EIF5,CD81,WDR1,
AB7A,CCN2,DDX3X,RPS15,RPL30,CLTC,MAGED2,H1-0,RPS5,RPS7,IK,RPL35,HMGCS1,GPAT4,C
HNRNPUL1,ACTN1,SON,NSG2,PLCG1,RPL28,CCAR2,CORO1C,SMARCE1,RPS28,ATP6V1B2,HNR
24,BRD3,RPS9,XRCC5,PJA2,RPL24,RPL17,C1ORF43,PAPOLA,COL1A2,LLGL1,RPL14,PTCD3,CB
KT,CDH6,SFPQ,PDIA6,DDR1,RAB7A,ACTN4,MMP14,PRDX2,DDX3X,RPS15,RPL30,MRFAP1,MAGE
ZNF865,RPL9,ATF4,PSMD2,KIF5B,SLC7A5,RPS14,CXADR,CHD4,JUND,TMEM259,RPS20,EIF5,CD
ZNF865,RPL9,ATF4,PSMD2,KIF5B,SLC7A5,RPS14,CXADR,CHD4,JUND,TMEM259,RPS20,EIF5,CD
T,GLG1,SFPQ,PDIA6,RAB7A,ACTN4,MMP14,PRDX2,DDX3X,RPS15,RPL30,CLTC,MRFAP1,MAGED
BE2H,CKAP4,CALU,CTBP1,HMGCR,CAPRIN1,TOP2B,HDAC2,TMEM30A,SKP1,BCLAF1,GLUL,LUC
A,PRDX2,DDX3X,RPS15,RPL30,CLTC,EIF3L,MAGED2,GNB2,H1-0,RPS7,IK,MIF,GPAT4,TLN1,GJA
AB,CKAP5,SND1,LAMP1,HNRNPH1,CALU,ARCN1,CTBP1,CAPRIN1,TOP2B,HDAC2,TMEM30A,SKP1
,RPL35,HMGCS1,GPAT4,CHPF,GJA1,JUN,CTNNB1,RPS10,DNAJC5,APLP2,HNRNPDL,TARDBP,RP
IN,SPC25,RPL31,NISCH,PRKCSH,HNRNPC,LRPPRC,TADA2B,XPR1,KHDRBS1,XPO1,HDLBP,RPS25
PL,TM9SF2,MORF4L2,TAF15,ALKBH5,RPL27,HNRNPA3,PTPRA,ACTR3,C4ORF3,ACO2,CAMSAP1,
,LUC7L3,DCN,CPSF2,PGRMC1,DDX39B,EID1,CTSB,TM9SF3,MDH2,U2AF2,PGD,NOP53,PABPN1,
C7L3,CPSF2,PGRMC1,NUCB1,TM9SF3,MDH2,CCT2,SUGP2,HNRNPL,MORF4L2,TAF15,LMNB2,AL
,KIF5B,SLC7A5,RPS14,CHD4,TMEM259,RPS20,RPL26,BEX3,CCT7,SRSF4,RBBP4,PFN1,MCL1,PN
S7,IK,GPAT4,PRKAR1A,TLN1,CHPF,CTNNB1,MACF1,DNAJC5,APLP2,HNRNPDL,TARDBP,NDFIP1,S
3,XPR1,KHDRBS1,XPO1,CSNK1E,RPS25,GLG1,CDH6,SFPQ,EIF5A,PDIA6,DDR1,RAB7A,CCN2,ACT
TC,EIF3L,MAGED2,H1-0,RPS5,RPS7,IK,RPL35,MIF,GPAT4,TLN1,CHPF,GJA1,CTNNB1,ANXA6,RPS
3,ACO2,PTOV1,SURF4,RBMX,RAD23B,TNPO1,SNRNP70,PSMB2,H3-3A,HDGF,TSPAN6,PRPF6,OF
,RPL26,LASP1,BEX3,CCT7,SRSF4,RBBP4,PFN1,MCL1,JPT2,PNN,RPL31,NISCH,PRKCSH,HNRNPC
BP,RPLP2,HNRNPK,H1-10,RPL36,YWHAB,CKAP5,SND1,LAMP1,HNRNPH1,ATP1A2,CKAP4,RAB1A
N1,TBC1D16,CCT3,VPS35,SURF4,RBMX,DDX42,RAD23B,GCN1,TNPO1,SNRNP70,PSMB2,H3-3A,
IA6,DDR1,MEGF8,RAB7A,PRDX2,DDX3X,CLTC,GNB2,RCN1,H1-0,RPS7,IK,MIF,GPAT4,CHPF,MY

HNRNPK,H1-10,RPL36,YWHAB,CKAP5,SEPTIN11,SND1,LAMP1,HNRNPH1,CKAP4,RPS12,HLA-B,C
5,GPAT4,TLN1,MYO5A,COL6A2,ANXA6,APLP2,ARID1A,MYL6,HNRNPK,H1-10,NARS1,RPL36,YWH
C2,MAPK8IP1,RAI1,PFN2,HNRNPD,MLF2,ZSCAN18,TCTN1,SERBP1,PRPF40A,TP53BP1,ADAR,RBE
OP2B,HDAC2,MMP2,BCLAF1,LUC7L3,CPSF2,PGRMC1,DDX39B,CNN2,EID1,NUCB1,CDH11,TM9S
SFPQ,PDIA6,DDR1,RAB7A,CCN2,PRDX2,DDX3X,RPS15,RPL12,MAGED2,H1-0,RPS7,IK,RPL35,GPA
P1,BCLAF1,PPIB,LUC7L3,SQLE,CPSF2,DDX39B,EID1,NUCB1,TM9SF3,MDH2,TOMM20,RNF187,
L,TARDBP,ARID1A,NDFIP1,HNRNPK,H1-10,INIP,RPL36,YWHAB,CKAP5,LAMP1,HNRNPH1,CKAP4,
NPH1,CKAP4,HLA-B,CALU,ARCN1,CTBP1,ARHGDIA,CAPRIN1,CDK16,TOP2B,APPL1,HDAC2,BCLAF
ORF4L2,LMNB2,ALKBH5,RPL27,ACO2,CAMSAP1,CCT3,RBMX,JPT1,CBX1,RAD23B,TNPO1,SNRN
MCL1,PNN,NES,RPL31,NISCH,HNRNPC,LRPPRC,TADA2B,XPR1,KHDRBS1,KLHL9,XPO1,CSNK1E,H
MCL1,PNN,NES,RPL31,NISCH,HNRNPC,LRPPRC,TADA2B,XPR1,KHDRBS1,KLHL9,XPO1,CSNK1E,H
H2,RNF187,U2AF2,MLXIP,ANXA5,PABPN1,HNRNPL,PARP1,TSPYL4,MORF4L2,TAF15,PTPRA,ACTF
RNF10,MARS1,UBXN4,G3BP2,SMARCA4,SRSF9,ARPC2,DSTN,PSMD3,SRSF1,ARGLU1,TCF25,PTE
GNAI3,NOP53,PABPN1,GDI2,HNRNPL,TLN2,MORF4L2,TAF15,LMNB2,HNRNPA3,PRDX5,PDXK,AC
,GCN1,TNPO1,SNRNP70,PSMB2,H3-3A,KDM6B,SCAF1,HDGF,TMED2,RPS15A,RNF10,VEGFA,MA
DC,HNRNPUL1,PLCG1,MAP2K2,QARS1,RPL28,CCAR2,ZFAND5,DIDO1,CORO1C,SMARCE1,HNRN
LR2E,PSMA7,ANP32A,RBBP7,EDC4,TXLNA,OS9,EP300,EXTL3,LBH,SF1,PBX1,PRCC,SEC16A,BRD
SRSF3,ILF2,FBLN1,EIF3F,UBQLN1,SF3A1,TMED9,MLF2,SERBP1,KDM3B,TP53BP1,PSMA7,ANP3
STIP1,TOP2B,APPL1,SMCHD1,HDAC2,IPO5,BCLAF1,TRIB2,LUC7L3,CPSF2,PGRMC1,CTSB,TM9SF
P4,PFN1,MCL1,JPT2,PNN,RPL31,NISCH,PRKCSH,HNRNPC,LRPPRC,TADA2B,XPR1,KHDRBS1,XPC
FT1,TBCD,SQSTM1,PABPN1,HNRNPL,TSPYL4,TM9SF2,MORF4L2,SSRP1,RPL27,HNRNPA3,PRDX
6,TOP2B,HDAC2,BCLAF1,LUC7L3,CPSF2,DDX39B,EID1,CTSB,NUCB1,TM9SF3,MDH2,TOMM20,F
SELENON,OPTN,UBXN4,SMARCA4,LAPTM4A,SRSF9,DSTN,PBX2,PSMD3,SRSF1,ARGLU1,NYNRIN
PO1,CSNK1E,SCRN1,HDLBP,RPS25,TKT,GLG1,SFPQ,EIF5A,PDIA6,DDR1,ACTN4,PRDX2,DDX3X,R
KDC,HNRNPUL1,HSPB1,SON,RPL28,CCAR2,DIDO1,CORO1C,SMARCE1,SMARCC1,SUPT6H,RPS2
RF4,CBX1,RAD23B,TNPO1,SNRNP70,H3-3A,SCAF1,HDGF,PRPF6,TAF7,MARS1,OPTN,MAPK1,SM
C,HNRNPUL1,SON,BNIP3L,CCAR2,DIDO1,FLNB,POLR2B,SMARCC1,RPS28,ATP6V1B2,HNRNPM,S
IGCR,ARHGDIA,CAPRIN1,STIP1,GPC1,TOP2B,HDAC2,IPO5,TMEM30A,SKP1,L1CAM,LUC7L3,CPS
DC,HNRNPUL1,ACTN1,HSPB1,BNIP3L,FKBP8,MAP2K2,DIDO1,FLNB,CORO1C,SMARCE1,SDF4,HN
IK1D,SF3A1,SREBF2,HNRNPD,SERBP1,KDM3B,PRPF40A,DDIT4,PSMA7,ANP32A,RBBP7,TXLNA,S
RNPL,TSPYL4,TM9SF2,MORF4L2,TAF15,RPL27,HNRNPA3,PRDX5,ACTR3,PKN1,ACO2,CCT3,VPS3
XN4,MAPK1,SMARCA4,SRSF9,SRSF1,IPO7,ARGLU1,NYNRIN,TCF25,PTBP1,PRKDC,HNRNPUL1,A
IMGCR,ARHGDIA,CAPRIN1,STIP1,GPC1,TOP2B,HDAC2,IPO5,SKP1,L1CAM,LUC7L3,DDX39B,SRS
AP1,VARS1,AP2A1,H2AZ1,UBQLN2,CSNK2A1,FLOT1,PHB2,GSPT1,BRD4,UBA52,SLC39A7,DIP2B
M,SSB,SRSF3,LARS1,ILF2,PFKM,LONP1,SF3A1,TMED9,IDH1,HNRNPD,ZSCAN18,KDM3B,WBP2,A
SMCHD1,BCLAF1,LUC7L3,CPSF2,CTSB,NUCB1,TM9SF3,MDH2,TOMM20,RAB27A,U2AF2,RPN2,
XPR1,XPO1,SPTBN2,SLC3A2,CSNK1E,SCRN1,HDLBP,RPS25,GLG1,CDH6,PEBP1,CCND2,CDH2,D
H3-3A,HDGF,TERF2IP,TOP1,RNF10,VEGFA,UBXN4,G3BP2,MAPK1,SMARCA4,SRSF9,PRDX1,SRSF
NPA3,ACTR3,C4ORF3,TBC1D16,CAMSAP1,CCT3,VPS35,RBMX,JPT1,CBX1,RAB5B,CHD2,GCN1,S
BRD4,UBA52,ATRX,EP400,SRSF6,ARID1B,PAPSS1,GLUD1,RER1,TRA2B,UQCRC1,COX7C,ZFR,HN
,SRSF1,ARGLU1,NYNRIN,TCF25,ILF3,WDR6,CAPZB,PRKDC,HNRNPUL1,ATXN10,ACTN1,KDM1A,
CA4,AZIN1,LAPTM4A,DSTN,PSMD3,SRSF1,ARGLU1,ACADVL,TCF25,PTBP1,FTO,ILF3,PRKDC,HNR
2,LSS,B2M,CSNK1D,DGKD,HNRNPD,ZSCAN18,SERBP1,KDM3B,PRPF40A,TP53BP1,DDIT4,ANP32
C39A7,EP400,SRSF6,GLUD1,UBE2G2,EIF5B,TRA2B,IARS1,UQCRC1,COX7C,ZFR,HNRNPA2B1,AN
BP1,PCYOX1,POLR2E,PSMA7,ANP32A,VDAC1,TRAF4,RBBP7,SSR1,SULF2,TXLNA,OS9,NAP1L1,C

R,AGPAT1,HNRNPA2B1,ANP32B,RPS27A,HNRNPAB,C1ORF198,SAFB2,PRDX6,ZMYM3,FUS,XRCC6
I,EIF2S2,VAPB,TPP1,TM9SF4,HDGFL2,EFTUD2,GTF2I,SRSF2,FLOT2,EEF1B2,SH3PXD2A,CADM3,
TCP1,CCT3,VPS35,PTOV1,RBMX,DDX42,LAMC1,CBX1,CCT4,GCN1,SNRNP70,H3-3A,SCAF1,HDG
RPL36,CKAP5,LAMP1,HNRNPH1,UBE2H,RAB1A,CALU,ARCN1,CCT6A,CTBP1,HMGCR,CAPRIN1,CD
L3,SQLE,CPSF2,DDX39B,CDH11,MLXIP,ANXA5,PPP1CB,SQSTM1,PABPN1,HNRNPL,PARP1,VCAN,
3B,PSMA7,VDAC1,RBBP7,EDC4,TXLNA,NAP1L1,LBH,SF1,UBQLN2,GRSF1,KPNA2,PHB2,GSPT1,B
ATXN7L3,ANKRD12,WNK1,CCDC47,THRAP3,EIF4A3,XRN2,PTPN11,HMGA1,SPPL2B,MTA2,KTN1,T
N1,KDM3B,BMPR2,NAP1L4,TP53BP1,CTNND1,POLR2E,ARIH1,BCR,SGTA,OS9,NAP1L1,LBH,HIP1
ERH,TAOK1,PELP1,TCAF1,FKBP4,EIF3H,TRAM1,LTBP4,NUDCD3,SH3GL1,LSM14A,EFTUD2,PNISF
CPSF2,CTSB,NUCB1,TM9SF3,MDH2,TOMM20,RAB27A,RNF187,U2AF2,RPN2,MLXIP,PABPN1,HNR
1L1,EXTL3,LBH,SF1,H2AZ1,UBQLN2,FLOT1,CD99,CST3,BRD4,UBA52,SLC39A7,RAB2A,SRSF6,G
GCN1,SNRNP70,PSMB2,H3-3A,HDGF,TOP1,RNF10,SELENON,OPTN,UBXN4,G3BP2,SMARCA4,C
N1,CDK16,TOP2B,SMCHD1,HDAC2,BCLAF1,TRIB2,PPIB,LUC7L3,CPSF2,EID1,CTSB,NUCB1,TM9S
EDC4,TXLNA,OS9,NAP1L1,ACTR1A,STARD7,SF1,PBX1,HIP1,FLCN,BRD4,UBA52,SLC39A7,ATRX,R
PARP1,TSPYL4,MORF4L2,LMNB2,ALKBH5,PTPRA,ACTR3,PDXK,TBC1D16,VPS35,LAMC1,CBX1,R
5,C4ORF3,ACO2,NCKAP1,VPS35,PTOV1,ACAP3,RBMX,CBX1,RAD23B,RAB5B,GCN1,TNPO1,SNRN
27A,HNRNPAB,MTCH1,SAFB2,ZMYM3,FUS,PPP1R10,TRIR,XRCC6,PELP1,EIF2S2,SNN,TCAF1,FKB
F1,HIP1,PRCC,BAZ1B,H2AZ1,CSNK2A1,CD99,KPNA2,PHB2,PPM1G,BRD2,DIP2B,EP400,SRSF6,K
I1,NOP53,GDI2,HNRNPL,PARP1,TSR1,SSRP1,LMNB2,RPL27,PTPRA,PRDX5,ACTR3,PDXK,UBE4A,
PTPN11,HMGA1,SPPL2B,SUN1,MTA2,USP10,KTN1,MACROH2A1,MDM2,FOXP4,MECP2,SF3A3,AD
ATXN7L3,MN1,ANKRD12,WNK1,CCDC47,THRAP3,SCD,XRN2,TAPBP,HMGA1,HTATSF1,RSF1,SUN
10,SCAP,RANGAP1,SLC30A9,XRN2,TMEM184B,HTATSF1,ITFG1,MTA2,USP10,CGGBP1,KTN1,AR
6,EIF2S2,RTL6,YARS1,RALY,IMPDH2,HDGFL2,GTF2I,FLOT2,EEF1B2,TCF3,RBM5,ABCF1,PHGDH
KBP9,AHCY,PSMC3,UBE3A,RBM6,EXOC4,ATXN7L3,ANKRD12,DAPK3,THRAP3,PRDX3,SCD,EIF4A
,SRSF9,CCNL2,PBX2,SRSF1,ARGLU1,ATP5PB,PTBP1,ILF3,GRB2,PRKDC,SUPT5H,HNRNPUL1,SC
NRNPL,TSPYL4,TM9SF2,MORF4L2,ALKBH5,RPL27,HNRNPA3,C4ORF3,ACO2,CCT3,NCKAP1,PTO
MARCB1,FZD7,NF2,SECISBP2L,ZNF207,LEMD2,ALYREF,M6PR,COX5B
NIP3L,PLCG1,MAP2K2,QARS1,ZFAND5,FLNB,DLK1,TPM3,CORO1C,XPNPEP3,SMARCC1,SUPT6H
,UBE2H,TTR,ARCN1,CCT6A,CAPRIN1,CDK16,TOP2B,BCLAF1,PPIB,GLUL,LUC7L3,CRPPA,PGRMC
),PSME3,ZRANB2,PSMC2,NSD1,HSF1,MCFD2,SNRPD2,SOX9,PSMB5,POGK,SLC16A1,NACC1,LMN
F4A3,SCAP,CAD,SLC30A9,XRN2,HTATSF1,SUN1,FXR2,PINK1,MTA2,USP10,CGGBP1,KTN1,PAIP2,
IPD,MLF2,SERBP1,KDM3B,DDIT4,POLR2E,PSMA7,ANP32A,RBBP7,TXLNA,NAP1L1,EP300,LBH,SF
14A,EFTUD2,GTF2I,SRSF2,TRAP1,EEF1B2,RBM5,PHGDH,SRSF11,SPTY2D1,MSH6,PRPF19,UBE3
M6,SMARCD1,ATXN7L3,ATP6V1G1,EIF4A3,CALCOCO1,RANGAP1,TAPBP,SUN1,MTA2,CGGBP1,K
M3,POLR2B,CORO1C,XPNPEP3,SMARCC1,DHX30,ATP6V1B2,HNRNPM,SSB,GIGYF1,SRSF3,SDHA
5B,TRA2B,IARS1,UQCRC1,MBTPS1,AGPAT1,HNRNPA2B1,HNRNPAB,SYMPK,C1ORF198,ZMYM3,
4L2,PTPRA,PRDX5,ACTR3,RTN1,PDXK,PKN1,ACO2,CAMSAP1,CCT3,VPS35,RAD23B,RAB5B,GCN1
,MOAP1,RBM5,WBP11,PHGDH,SRSF11,SPTY2D1,MSH6,PSMC3,SART1,UBE3A,RBM6,ATXN7L3,M
20,RNF187,EEF1A2,TBCD,MLXIP,SUGP2,SQSTM1,GNAI3,NOP53,HNRNPL,PARP1,TM9SF2,TSR1,
S35,RBMX,CBX1,RAD23B,PTK7,CHD2,CCT4,GCN1,SKI,SNRNP70,H3-3A,KDM6B,HDGF,ABCA2,PY
A7,RBBP7,TXLNA,BCR,SGTA,OS9,NAP1L1,LBH,GTPBP1,KIF21B,PHB2,PLXNB1,ATRX,GARS1,NXN

D1,SF1,PBX1,PRCC,UBQLN2,KIF21B,PHB2,BRD4,UBA52,SLC39A7,DECR1,DIP2B,EP400,ARID1B,
F1,EIF3D,BRD4,UBA52,PPP2CB,ATRX,GARS1,EP400,SRSF6,KIAA0895L,GLUD1,EVL,RAB3GAP2,T
TXN7L3,WNK1,THRAP3,CAD,PLS3,HS2ST1,PTPN11,TAPBP,HMGA1,HTATSF1,FXR2,MTA2,CGGBP

LCOCO1,RANGAP1,XRN2,TAPBP,HTATSF1,SUN1,SGSM2,MTA2,KTN1,PAIP2,H6PD,ANKHD1,MTR[illegible]

EVL,TRA2B,UQCRC1,ZFR,HNRNPA2B1,MCM3AP,LMAN1,HNRNPAB,RABEP1,SYMPK,ZMYM3,GRK[illegible]
2,NUP50,FUBP1,KLF3,SCAF8,ZNF207,SRSF8,CPSF7,ALYREF,CCDC124
GEF2,NAP1L4,TP53BP1,SYP,POLR2E,RBBP7,DCTN1,EIF2AK1,SGTA,OS9,NAP1L1,ACTR1A,LBH,F[illegible]
ZNF207,CPSF7,UBTF,ALYREF,M6PR
4,THOC2,MED12,TLK1,RBM17,GRHPR,ZNF146,PCED1A,PRPF31,HEXIM1,GATAD2B,PTPN12,NBF[illegible]
50A,EWSR1,BANF1,GALNT10,PRPH,ATF7IP,PDGFRB,WIZ,HNRNPR,AKAP8L,ADRM1,KPNA6,CDK9,

KTN1,MGAT5,RTF1,MKNK2,REEP5,MDM2,PPP1R14B,CMPK1,FAM50A,CRIP2,AAMP,PTPN1,MTU[illegible]
,SFSWAP,PRPF31,SS18L1,ANXA1,RTRAF,FYTTD1,HEXIM1,TNPO3,GATAD2B,ZDHHC17,FUBP1,C[illegible]
1,THRAP3,TUBB4A,EIF4A3,USO1,RCN2,SLC38A10,NELFB,XRN2,HMGA1,FXR2,MTA2,CGGBP1,K[illegible]
RTC1,ANKRD50,MED25,ZFP91,TBCA,SETD5,MYL12A,LMNA,THOC2,RRBP1,MED12,TLK1,PHB,R[illegible]
NPA2B1,ANP32B,MCM3AP,SEPTIN7,HIF1A,CTTN,LRRC59,PRDX6,AKAP9,ZMYM3,FUS,XRCC6,PE[illegible]
39,PPP1R14B,EWSR1,ERP29,SHISA5,SUN2,TNRC6A,WIZ,HNRNPR,ECHS1,TBC1D14,NUP153,H2A[illegible]
SMB2,H3-3A,SCAF1,HDGF,RPS15A,RNF10,OPTN,UBXN4,G3BP2,SMARCA4,SRSF9,DSTN,SRSF1,A[illegible]
P8L,H2AX,CCNG1,KPNA6,CDK9,PSME3,ARPC3,ZRANB2,MARCHF7,PSMC5,GOLGA8B,OXSR1,NS[illegible]
ECISBP2L,ZNF207,SRSF8,CPSF7,ALYREF,M6PR,COX5B
TRAP1,RIC8A,PLEKHM2,TCF3,SRSF11,PRPF19,FKBP9,TEX2,FBXW5,MTOR,UBE3A,GOT2,AP2A2,[illegible]

LC22A23,PAIP2,HIP1R,H6PD,ANKHD1,ANP32E,CDV3,IGF1R,MDM2,STK39,MECP2,FAM50A,SART[illegible]
R26,SUMO1,CAB39,SART3,BANF1,PSMD7,MAEA,SHISA5,PITRM1,IP6K1,PCGF3,SUN2,FAM53C,F[illegible]

BP7,STAT3,MTA1,KDM5B,SF1,HIP1,FLCN,PRCC,CSNK2A1,CD99,KPNA2,PPM1G,BRD2,GARS1,EP[illegible]
PLCG1,KDELR1,CORO1C,XPNPEP3,SUPT6H,PFKM,EIF3F,STT3A,ATXN2L,TMED9,MESD,USP7,HY[illegible]
LN1,SF3A1,GPR107,UQCRC2,BAG6,HNRNPD,MLF2,SERBP1,ARFGEF2,CCT5,TRAF4,RBBP7,STAT

E1,SDF4,SMARCC1,DHX30,HNRNPM,SSB,SRSF3,SDHA,LARS1,SLIT3,ILF2,CCT8,LSS,STT3B,STX[illegible]

,XRN2,HMGA1,MTA2,KTN1,AASDHPPT,PSMB1,MACROH2A1,HSPA1A,RUVBL2,KIAA1210,MDM2,[illegible]
HARS1,FAM193B,MRPS26,MYEF2,FZD7,NFIB,ZNF207,NOP58,RBM39,CNOT1

APN1,ASAH1,CDK9,PSME3,PSIP1,RBM10,YWHAZ,NDUFB8,MRPS6,SHMT2,SNRPD2,MED25,ZFP9[illegible]
E4A,PKN1,TBC1D16,TCP1,CAMSAP1,CCT3,GRINA,PTOV1,SURF4,DDX42,JPT1,RAB5B,TNPO1,PS

1C,PLXDC2,MED25,ZBED4,ZFP91,ALDH9A1,KIF3A,CLCN6,TBCA,SETD5,WDR47,HSPE1,RNF20,HI

,FBRS,SF1,RETREG2,HIP1,FLCN,PRCC,H2AZ1,CSNK2A1,SEC16A,KPNA2,GPX4,PPM1G,BRD2,PL[illegible]

[illegible]TSA,ARSA

317,VCL,DNAJA1,CYB5R3,PAFAH1B1,AKA
IRNPU,OGA,PGAM1,NFE2L1,SMARCA5,LAR
HNRNPA0,CTNNA1,FADS2,RPS16,RPL23,C
RPSA,TPM4,CTSD,EIF4H,SCARB2,RTN4,L
3,PAFAH1B1,AKAP12,MAP4,SF3B2,CDC42B
SA,TPM4,CTSD,EIF4H,SCARB2,RTN4,LPC
P1,HNRNPA0,CTNNA1,FADS2,RPS16,RPL23
L1,SMARCA5,LARP1,HNRNPA0,CTNNA1,FADS
MARCA5,LARP1,HNRNPA0,CTNNA1,FADS2
23,CNN3,RPL29,RPSA,TPM4,TNS3,CTSD,
23,CNN3,RPL29,RPSA,TPM4,TNS3,CTSD,
SMARCA5,LARP1,HNRNPA0,CTNNA1,FADS2
9,RPS20,EIF5,CD81,WDR1,HMGN2,AT
LGA8A,HNRNPU,PGAM1,NFE2L1,SMARCA
LGA8A,HNRNPU,PGAM1,NFE2L1,SMARCA
29,RPSA,TPM4,TNS3,CTSD,EIF4H,SCARB
RNPA0,CTNNA1,FADS2,RPS16,RPL23,CNN3
RNPA0,CTNNA1,FADS2,RPS16,RPL23,CNN3
A,NFE2L1,SMARCA5,LARP1,HNRNPA0,CTN
S2,RPS16,RPL23,CNN3,RPL29,RPSA,TPM4,
4,PFN1,MCL1,JPT2,PNN,RPL31,PRKCSH
NPA0,CTNNA1,FADS2,RPS16,RPL23,CN
2,RPS16,RPL23,CNN3,RPL29,RPSA,TPM4
2,RPS16,RPL23,CNN3,RPL29,RPSA,TPM4
I259,RPS20,EIF5,WDR1,ATP2A2,RPL26
L1,PNN,RPL31,PRKCSH,HNRNPC,LRPPR
I2B,ZNF770,AP2B1,MAT2A,GTF2F1,GOLG
6,RPL23,CNN3,RPL29,RPSA,TPM4,TNS3,CT
XRCC5,PJA2,RPL24,KPNB1,RPL17,C1O
XRCC5,PJA2,RPL24,KPNB1,RPL17,C1O
E2L1,SMARCA5,LARP1,HNRNPA0,CTNNA1,FA
EIF4H,SCARB2,LPCAT1,BRD3,RPS9,XRC
EIF4H,SCARB2,LPCAT1,BRD3,RPS9,XRC
SMD2,KIF5B,SLC7A5,RPS14,CXADR,CH
A2,LLGL1,RPL14,HK1,RHOA,PTCD3,CBX
L23,CNN3,RPSA,TPM4,TNS3,CTSD,EIF4H,
D,TMEM259,RPS20,EIF5,WDR1,HMGN2,RP
N,RPL24,KPNB1,RPL17,C1ORF43,PAPOL
N,RPL24,KPNB1,RPL17,C1ORF43,PAPOL
,PAPOLA,COL1A2,LLGL1,RPL14,HK1,CBX
PPRC,TADA2B,XPR1,KHDRBS1,XPO1,C
2L1,SMARCA5,LARP1,HNRNPA0,CTNNA1,FA
2L1,SMARCA5,LARP1,HNRNPA0,CTNNA1,FA
FE2L1,SMARCA5,LARP1,HNRNPA0,CTNNA1,FA
FE2L1,SMARCA5,LARP1,HNRNPA0,CTNNA1,FA

JUND,TMEM259,EIF5,WDR1,HMGN2,RPL2
NRNPA0,CTNNA1,FADS2,RPS16,RPL23,CPE
NRNPA0,CTNNA1,FADS2,RPS16,RPL23,CPE
ADS2,RPS16,RPL23,CPE,CNN3,RPL29,RPS
RPSA,TPM4,TNS3,CTSD,EIF4H,SCARB2,LPC
MARCA5,LARP1,HNRNPA0,CTNNA1,FADS2,RPS
,HNRNPA0,CTNNA1,FADS2,RPS16,RPL23,C
C5,PJA2,RPL24,SLC16A7,KPNB1,RPL17
H,SCARB2,LPCAT1,RPS24,BRD3,RPS9,XRC
H,SCARB2,LPCAT1,RPS24,BRD3,RPS9,XRC
M1,NFE2L1,SMARCA5,LARP1,HNRNPA0,CTNN
PL9,CLSTN1,ATF4,PSMD2,KIF5B,SLC7A
,RPSA,TPM4,EIF4H,SCARB2,LPCAT1,BRD
,RPSA,TPM4,EIF4H,SCARB2,LPCAT1,BRD
DX3X,RPS15,RPL12,CLTC,MAGED2,RCN1,H1
XRCC5,PJA2,RPL24,SLC16A7,KPNB1,
L9,CLSTN1,SLC7A5,RPS14,CXADR,CHD4
IRNPA0,CTNNA1,FADS2,RPS16,RPL23,CPE,C
IRNPA0,CTNNA1,FADS2,RPS16,RPL23,CPE,C
PL29,RPSA,TPM4,CTSD,EIF4H,SCARB2,LPC
PL29,RPSA,TPM4,CTSD,EIF4H,SCARB2,LPC
ADR,CHD4,ATP1A1,JUND,TMEM259,RP
NPA0,CTNNA1,FADS2,RPS16,RPL23,CNN3
ATP1A1,JUND,RPL26,LASP1,BEX3,CCT7
BRD3,XRCC5,MSN,RPL24,SLC16A7,KPNB
1,JUND,TMEM259,RPS20,EIF5,WDR1,H
SF4,RBBP4,PFN1,MCL1,PNN,RPL31,NI
RPL14,HK1,RHOA,RPL9,CLSTN1,ATF4,PS
PS20,EIF5,CD81,WDR1,HMGN2,RPL26,
PS20,EIF5,CD81,WDR1,HMGN2,RPL26,
LSTN1,ATF4,PSMD2,KIF5B,SLC7A5,RP
16A7,KPNB1,RPL17,C1ORF43,PAPOLA,C
SP1,BEX3,CCT7,SRSF4,RBBP4,RTN3,P
CTNNA1,FADS2,RPS16,RPL23,CPE,CNN3,RP
CTNNA1,FADS2,RPS16,RPL23,CPE,CNN3,RP
PE,CNN3,RPL29,RPSA,TPM4,TNS3,CTSD
CTNNA1,FADS2,RPS16,RPL23,CPE,CNN3,R
PCAT1,BRD3,RPS9,XRCC5,PJA2,MSN,RPL
NPC,LRPPRC,TADA2B,XPR1,KHDRBS1,
43,PAPOLA,COL1A2,LLGL1,RPL14,HK1,NC
RSF4,RBBP4,PFN1,MCL1,JPT2,PNN,RP
X3,CCT7,SRSF4,RBBP4,MCL1,JPT2,
L1,JPT2,PNN,SPC25,RPL31,NISCH,PRK
SN,RPL24,SLC16A7,KPNB1,RPL17,C
RPL17,PAPOLA,COL1A2,LLGL1,RPL14,HK1

RPL26,LASP1,BEX3,CCT7,SRSF4,RBBP4
B2,LPCAT1,RPS24,BRD3,RPS9,XRCC5,P
RPL29,RPSA,TPM4,TNS3,CTSD,EIF4H,SCA
RPL29,RPSA,TPM4,TNS3,CTSD,EIF4H,SCA
APLP2,HNRNPDL,TARDBP,ARID1A,TSR2,
,HK1,RHOA,PTCD3,CBX5,ZNF865,RPL9,CLS
PAPOLA,COL1A2,LLGL1,RPL14,RHO
,MCL1,JPT2,PNN,RPL31,NISCH,PRKC
NFE2L1,SMARCA5,LARP1,HNRNPA0,CTNNA
L1A2,LLGL1,RPL14,HK1,PTCD3,ZNF86
L1A2,LLGL1,RPL14,HK1,PTCD3,ZNF86
PNN,RPL31,NISCH,PRKCSH,HNRNPC,L
N,RPL31,NISCH,PRKCSH,HNRNPC,TADA2
,RPS20,EIF5,WDR1,HMGN2,RPL26,L
,RPS20,EIF5,WDR1,HMGN2,RPL26,L
1,XPO1,CSNK1E,HDLBP,RPS25,TKT,G
DA2B,XPR1,KHDRBS1,KLHL9,XPO1,SPTB
T7,SRSF4,RBBP4,MCL1,JPT2,PNN,R
RPL31,NISCH,PRKCSH,HNRNPC,L
RPL26,LASP1,BEX3,CCT7,SRSF4,RBBP
PL26,LASP1,BEX3,CCT7,SRSF4,RBBP4,P
PAFAH1B1,AKAP12,MAP4,SF3B2,CDC42
APLP2,TARDBP,RPLP2,ST3GAL2,HNRNPK,RP
9,PSMD2,KIF5B,SLC7A5,RPS14,CHD4,
N,CPSF2,PGRMC1,DDX39B,CTSB,CDH11,TM
14,HK1,RHOA,PTCD3,CBX5,RPL9,SLC7A
,CBX5,ZNF865,CLSTN1,PSMD2,KIF5B,S
GN2,RPL26,BEX3,CCT7,SRSF4,RBBP4
,COL1A2,LLGL1,RPL14,HK1,RHOA,PTCD
LA,COL1A2,LLGL1,RPL14,HK1,PTCD3,CBX
LLGL1,RPL14,HK1,RHOA,PTCD3,CBX5,P
RC,TADA2B,XPR1,KHDRBS1,XPO1,CSNK1E
CN1,TNPO1,SNRNP70,H3-3A,TERF2IP,T
CL1,JPT2,PNN,RPL31,NISCH,PRKCSH,H
X5,RPL9,CLSTN1,PSMD2,KIF5B,SLC7A
LC7A5,RPS14,CXADR,CHD4,ATP1A1,
RPL9,CLSTN1,ATF4,RPS14,CXADR,JUND,R
,NISCH,PRKCSH,HNRNPC,LRPPRC,TADA2
5,RPS14,CXADR,JUND,TMEM259,RPS20,
,TMEM259,RPS20,EIF5,WDR1,RPL26,
HD4,TMEM259,RPS20,WDR1,RPL26,CRMP
MA1,PDIA6,DDR1,RAB7A,PRDX2,DDX3
PSMD2,KIF5B,SLC7A5,RPS14,CXADR,JU
MEM259,RPS20,WDR1,HMGN2,RPL26,L
SRSF4,RBBP4,MCL1,JPT2,PNN,RPL3

1,MCL1,PNN,NES,NISCH,PRKCSH,HNRN
,RPL12,RPL30,CLTC,MRFAP1,GNB2,RPS7,I
GFA,UBXN4,G3BP2,MAPK1,SMARCA4,CCNL2
RPS15,RPL30,MAGED2,H1-0,RPS5,RPS
,RHOA,PTCD3,CBX5,RPL9,ATF4,PSM
PABPN1,HNRNPL,TSPYL4,MORF4L2,SSRP1,LMN
,GLG1,SFPQ,EIF5A,PDIA6,DDR1,MEGF8,
9,XRCC5,PJA2,MSN,RPL24,SLC16A7
9,XRCC5,PJA2,MSN,RPL24,SLC16A7
ND,TMEM259,RPS20,EIF5,HMGN2,RP
,KHDRBS1,XPO1,CSNK1E,SCRN1,RPS2
GNB2,H1-0,RPS5,IK,RPL35,MIF,GPAT4
GNB2,H1-0,RPS5,IK,RPL35,MIF,GPAT4
30,CLTC,EIF3L,MAGED2,GNB2,RCN1,
K,MIF,GPAT4,TLN1,CHPF,CTNNB1,NGRN
PR1,KHDRBS1,XPO1,CSNK1E,RPS25,TK
NPDL,TARDBP,RPLP2,DDOST,ST3G
MAGED2,RCN1,H1-0,RPS5,IK,RPL
,RBBP4,PFN1,MCL1,JPT2,PNN,RPL31,PR
2,RTN4,LPCAT1,RPS24,BRD3,RPS9,X
HMGN2,RPL26,LASP1,BEX3,CCT7,SR
PF,GJA1,JUN,CTNNB1,MACF1,RPS1
NPM,SRSF3,LARS1,ILF2,CCT8,PFKM,CSNK1D,TMED
X5,RPL9,CLSTN1,ATF4,PSMD2,KIF5B,S
D2,H1-0,RPS5,IK,TLN1,CHPF,CTNNB1,ANXA6
1,WDR1,HMGN2,ATP2A2,RPL26,BEX3,
1,WDR1,HMGN2,ATP2A2,RPL26,BEX3,
2,H1-0,RPS5,RPS7,IK,RPL35,TLN1,G
L3,CPSF2,NOVA2,CTSB,SRSF5,TM9SF3,M
1,CTNNB1,ANXA6,RPS10,DNAJC
,BCLAF1,TRIB2,PPIB,GLUL,LUC7L3,CPSF2,
P2,ARID1A,HNRNPK,INIP,NARS1,R
,TKT,GLG1,SFPQ,PDIA6,DDR1,RAB7A
CCT3,VPS35,RBMX,CBX1,RAD23B,RAB5B,GCN1,T
GDI2,HNRNPL,PARP1,TSPYL4,TM9SF2,MOR
KBH5,RPL27,HNRNPA3,ACTR3,C4ORF3,AC
N,NES,RPL31,NISCH,PRKCSH,HNRNPC
T3GAL2,MYL6,HNRNPK,H1-10,NARS1,RPL36
N4,MMP14,PRDX2,DDX3X,RPL30,CLT
10,APLP2,HNRNPDL,TARDBP,NDFI
TN,UBXN4,MAPK1,SMARCA4,ARPC2,DSTN,SRS
,LRPPRC,TADA2B,XPR1,KHDRBS1,
,CALU,ARCN1,CCT6A,MAGED4B,ARHGDIA,
KDM6B,HDGF,TSPAN6,TOP1,TAF7,RPS15A,RNF10,
5A,GOLM1,RPS10,DNAJC5,APLP2,HNRNPDL,

:ALU,ARCN1,COTL1,CTBP1,HMGCR,
AB,SEPTIN11,SND1,LAMP1,HNRNPH1,
3P7,TRIO,LBH,NDUFV1,VARS1,AP2A1,H2AZ1,UBQL
;F3,MDH2,U2AF2,CCT2,GNAI3,PABPN1,HNRNP
T4,PRKAR1A,TLN1,COL6A2,ANXA6,APL
J2AF2,RPN2,MLXIP,ANXA5,PABPN1,HNRNP
HLA-B,CALU,ARCN1,CCT6A,ARHGDIA
F1,TRIB2,LUC7L3,CPSF2,EID1,CTSB,NUC
IP70,H3-3A,HDGF,DPYSL2,TMED2,TOP1,RP
IDLBP,RPS25,TKT,GLG1,CDH6,SFPQ,TPM
IDLBP,RPS25,TKT,GLG1,CDH6,SFPQ,TPM
R3,C4ORF3,ETNK1,CCT3,VPS35,PTOV1,RB
3P1,FTO,ILF3,CAPZB,PRKDC,HNRNPUL1,ACTN1,HSP
O2,VPS35,RBMX,RAD23B,CHD2,CCT4,GCN1,
RS1,SELENON,UBXN4,G3BP2,MAPK1,SM
NPM,SSB,SRSF3,ILF2,PFKM,UBQLN1,CSNK1D,SF3A1,
)4,SLC39A7,ATRX,EP400,SRSF6,RER1,RAB3GAP
2A,RBBP7,TXLNA,PRMT1,NAP1L1,TXNDC5,TRIO,
F3,MDH2,TOMM20,RAB27A,U2AF2,SQSTM1
)1,PFKL,HDLBP,RPS25,TKT,GLG1,SFPQ
5,ACTR3,PKN1,CCT3,VPS35,PTOV1,RBMX,
RNF187,U2AF2,RPN2,MLXIP,PPP1CB,PGD,SQ
,TCF25,PTBP1,FTO,ILF3,PRKDC,HNRNPUL1,
RPS15,MAGED2,H1-0,RPS5,RPS7,IK,GP
8,HNRNPM,SSB,SRSF3,FAU,ILF2,LSS,EIF
ARCA4,AZIN1,LAPTM4A,ARPC2,SRSF1,ARGLU
SB,SRSF3,PHC2,LARS1,ILF2,CCT8,LSS,I
F2,PGRMC1,DDX39B,EID1,MDH2,TOMM20,U2AF
NRNPM,SSB,SRSF3,LARS1,ILF2,FBLN1,UBQLN1,
GTA,NAP1L1,EP300,LBH,SF1,PDAP1,PRCC,BRD
5,PTOV1,RBMX,CBX1,RAD23B,RAB5B,GCN
CTN1,HSPB1,CCAR2,DIDO1,FLNB,POLR2B,
F5,TM9SF3,U2AF2,MLXIP,PGD,HNRNPL,PA
,ATRX,SRSF6,KIAA0895L,GLUD1,CRTAP,TRA
KIRIN1,PRPF40A,TP53BP1,GRIPAP1,ANP32A,TRAF4,
MLXIP,NOP53,PABPN1,HNRNPL,TSPYL4,M
DR1,MEGF8,RAB7A,ACTN4,PRDX2,CLTC,M
1,ARGLU1,PTBP1,FTO,ILF3,PRKDC,HNRN
KI,SNRNP70,PSMB2,H3-3A,SCAF1,HDGF,PYGB,DPYS
RNPA2B1,NDST1,MCM3AP,POM121C,RPS27A,
FKBP8,MAP2K2,DIDO1,FLNB,CORO1C,SMARCE1,XPNPEP3,H
NPUL1,ACTN1,SON,KDM1A,DIDO1,FLNB,D
A,RBBP7,EDC4,TXLNA,NAP1L1,LBH,NDUFV1,S
P32B,MCM3AP,POM121C,RPS27A,HNRNPAB,SYMPK
PD,EXTL3,LBH,NDUFV1,SF1,PBX1,H2AZ1,CSNK2A

;,ERH,TAOK1,EIF3H,RALY,NUDCD3,TLE1,SRP72,HD
SLC7A1,MOAP1,RBM5,REEP2,SPTY2D1,PRPF19,MBD3,G
F,PRPF6,RNF10,VEGFA,MARS1,MAPK1,SMARC
)K16,TOP2B,SMCHD1,HDAC2,TMEM30A,BCLAF1
MORF4L2,TAF15,ALKBH5,HNRNPA3,PTPRA,PKN
RD4,UBA52,PPM1G,ATRX,GARS1,SRSF6,EVL,T
TMEM94,CLIP2,MACROH2A1,MDM2,FOXP4,MECP2,SF
,NOL4L,H2AZ1,CXXC5,MAP6,ATRX,GARS1,EP400,SR
R,SRSF2,TRAP1,EEF1B2,NUCKS1,RAB6A,NMT1
NPL,MORF4L2,ALKBH5,RPL27,HNR
LUD1,CRTAP,TRA2B,UQCRC1,ZFR,HNRNPA2B1,NPDC1,
CNL2,DSTN,SRSF1,NYNRIN,TCF25,PTBP1,ILF3,
F3,MDH2,RAB27A,RNF187,U2AF2,RPN2,
AB2A,EP400,SRSF6,EIF5B,CRTAP,TRA2B,ZFR,HNRNPA
AD23B,TNPO1,SKI,SNRNP70,H3-3A,HDGF,TMED2,
IP70,PSMB2,H3-3A,KDM6B,HDGF,PRPF6,TOP
P4,RALY,TRAM1,TLE1,HDGFL2,EFTUD2,GTF2
IAA0895L,UBE2G2,TRA2B,UQCRC1,ZFR,FEM1B,H
PKN1,TBC1D16,ACO2,TCP1,NCKAP1,PT
)NP,SYNCRIP,ERP29,GLOD4,PTPN1,PSMD7,ECH1,SUN2
1,MTA2,USP10,CGGBP1,KTN1,POR,RTF1,PAIP2,GTF3C
PC1A,CLIP2,PAIP2,HIP1R,ANP32E,MDM2,FOXP4,
I,SRSF11,SPTY2D1,MSH6,NSD2,FKBP9,IVD,EIF
3,AMFR,XRN2,HTATSF1,SUN1,MTA2,CGGBP1,LUC7L2,PA
AF11,SON,KDM1A,CASC3,FKBP8,MAP2K2,QARS1,
V1,ACAP3,SURF4,RBMX,RAD23B,GCN1,T

I,HNRNPM,GIGYF1,SRSF3,CCT8,PFKM,LSS,EIF3F
C1,EID1,CTSB,NUCB1,TM9SF3,MDH2,RNF
NA,PMPCA,UQCRFS1,RBM17,CITED2,ROCK2,ELOB
,ANP32E,CDV3,ACTA2,MDM2,SGSM3,FAM50A,FXR
1,H2AZ1,GTPBP1,CSNK2A1,FLOT1,CD99,SEC16
A,RBM6,ATXN7L3,MN1,THRAP3,DMWD,EIF4A3,NELF
KTN1,ARPC1A,ESYT1,ANP32E,MKNK2,MDM2,MECP2,FAM
A,LARS1,NKTR,ILF2,PFKM,LSS,IDH1,GPR107,S
FUS,ERH,PSMB4,PELP1,SNN,YARS1,SNRPD3,TP
,CNTNAP2,PYGB,DPYSL2,RPS15A,NOTCH2,M
MN1,ANKRD12,THRAP3,EIF4A3,NELFB,CALCOCO1,RANG
MORF4L2,TAF15,LMNB2,ABCA3,ALKBH5,HNRNP
GB,PRPF6,TERF2IP,TAF7,RPS15A,MARS1,TUBB2
N,EP400,SRSF6,PAPSS1,UBAP2L,RER1,TRA

,KIAA0895L,GLUD1,EIF5B,TRA2B,UQCRC1,COX
RA2B,UQCRC1,MASTL,ZFR,CUL3,ANP32B,MCM3AP,
1,KTN1,SLC22A23,RTF1,PAIP2,HIP1R,H6PD,CD

EX,MKNK2,RBM25,MDM2,SF3A3,FAM50A,SUMO1,ADNP,EW

2,FUS,XRCC6,IARS2,YARS1,RALY,PGAP6,SRP72,SS

PSMD1,SF1,NOL4L,BRD4,UBA52,MAP6,ATRX,GARS1,S

R1,FUBP1,PPT1,ACIN1,SMARCC2,POGZ,KLF3,MAPK8IP3,
PSME3,SRSF7,NCBP2,FAM160B2,ZEB1,SEPHS1,MR

RN,PITRM1,ANKFY1,SUN2,CFDP1,HIPK2,FLII,HSD17B1
DG4,MAML1,FAM193B,SCAF8,FZD7,SECISBP2L,ZNF207,
TN1,PSMB1,ANKHD1,MACROH2A1,HSPA1A,RUVBL2,KI
3M17,CTCF,NRP2,DSTYK,KPNA3,ELOB,MDC1,PCED1
LP1,VAPB,SNN,SNRPD3,STRN4,INTS3,ABI2,RALY,HDG
X,MICAL3,CDK9,PSME3,PSIP1,ATP5PO,DLD,TOB2,WAC,CR
RGLU1,NYNRIN,TCF25,ATP5PB,PTBP1,ILF3,LUZP6
D1,GOLGA4,MEIS1,SNRPD2,KARS1,FARSA,NACC1,THOC2,HI

SMARCD1,SAP18,NDUFA13,THRAP3,SCD,METRN,EIF4A3

3,SUB1,ACAP2,ERP29,GLOD4,PTPN1,NCKAP5L,SU
LII,WIZ,NUP153,PBXIP1,CCNG1,CDK9,PSME3,PSMC5,PSMC2

400,SRSF6,KIAA0895L,UPF1,RER1,UBE2G2,TRA2B,ZF
DU1,STRAP,PCBP2,MAPK8IP1,RAI1,SR
T3,OS9,MTA1,KDM5B,FBRS,SF1,RETREG2,HIP1,

3P1,ISYNA1,LONP1,UBQLN1,SF3A1,GPR107,U

PCM1,STK39,SF3A3,SUMO1,EWSR1,PDLIM1,ERP29,SSU7

1,CLCN6,HSPE1,KARS1,TMPO,THOC2,NAPA,RXRA
MB2,SCAF1,TSPAN6,DPYSL2,TMED2,RN

F3A,SRP54,CDC42EP4,UBXN1,NAPA,CACYBP,GE

‹NB1,EP400,SFRP1,SRSF6,UBE2G2,TRA2B,HDAC6,PT

| | | | | | | |
|---|---|---|---|---|---|---|
| **CORUM** | Ribosome, cy | CORUM:306 | 3.90E-31 | 30.4090632 | 80 | 1022 |
| **CORUM** | Nop56p-asso | CORUM:3055 | 1.66E-25 | 24.7793943 | 104 | 1022 |
| **CORUM** | 60S ribosoma | CORUM:308 | 6.03E-18 | 17.2195101 | 47 | 1022 |
| **CORUM** | 40S ribosoma | CORUM:338 | 2.10E-12 | 11.6767921 | 31 | 1022 |
| **CORUM** | 40S ribosoma | CORUM:305 | 5.00E-12 | 11.3014007 | 33 | 1022 |
| **CORUM** | Large Drosha | CORUM:1332 | 1.8616E-05 | 4.73011586 | 20 | 1022 |
| **CORUM** | TRBP contain | CORUM:5380 | 8.9083E-05 | 4.05020355 | 25 | 1022 |
| **CORUM** | PA700-20S-P | CORUM:193 | 0.00016086 | 3.79355109 | 36 | 1022 |
| **CORUM** | PA700 compl | CORUM:32 | 0.0002104 | 3.67695334 | 20 | 1022 |
| **CORUM** | TLE1 corepres | CORUM:1729 | 0.00030813 | 3.51126737 | 10 | 1022 |
| **CORUM** | LARC comple | CORUM:778 | 0.00060669 | 3.21703196 | 19 | 1022 |
| **CORUM** | Spliceosome | CORUM:351 | 0.00066543 | 3.17689488 | 141 | 1022 |
| **CORUM** | 26S proteasoı | CORUM:181 | 0.00176974 | 2.75209034 | 22 | 1022 |
| **CORUM** | BRG1-SIN3A- | CORUM:803 | 0.00252731 | 2.59734225 | 11 | 1022 |
| **CORUM** | TLE1 corepres | CORUM:3135 | 0.00392774 | 2.40585763 | 8 | 1022 |
| **CORUM** | CCT complex | CORUM:126 | 0.00392774 | 2.40585763 | 8 | 1022 |
| **CORUM** | Anti-HDAC2 c | CORUM:632 | 0.00484191 | 2.3149837 | 17 | 1022 |
| **CORUM** | ALL-1 superc | CORUM:1257 | 0.007079 | 2.1500281 | 28 | 1022 |
| **CORUM** | NUMAC comp | CORUM:86 | 0.00823188 | 2.08450088 | 10 | 1022 |
| **CORUM** | BRG1-SIN3A | CORUM:713 | 0.01293781 | 1.88813926 | 14 | 1022 |
| **CORUM** | HCF-1 compl | CORUM:2721 | 0.01294186 | 1.88800322 | 18 | 1022 |
| **CORUM** | p300-CBP-p2 | CORUM:570 | 0.01400828 | 1.8536151 | 7 | 1022 |
| **CORUM** | HEXIM1-DNA | CORUM:7285 | 0.0265456 | 1.57600751 | 9 | 1022 |
| **CORUM** | BAF complex | CORUM:566 | 0.0265456 | 1.57600751 | 9 | 1022 |
| **CORUM** | BRG1-associa | CORUM:807 | 0.0265456 | 1.57600751 | 9 | 1022 |
| **CORUM** | Brg1-associat | CORUM:710 | 0.0265456 | 1.57600751 | 9 | 1022 |
| **CORUM** | C complex sp | CORUM:1181 | 0.02810057 | 1.55128492 | 79 | 1022 |
| **CORUM** | SNW1 comple | CORUM:1335 | 0.03379324 | 1.4711702 | 17 | 1022 |
| **CORUM** | DGCR8 multi | CORUM:3082 | 0.03390247 | 1.46976861 | 11 | 1022 |
| **CORUM** | eIF3 complex | CORUM:1097 | 0.03663274 | 1.43613059 | 13 | 1022 |
| **CORUM** | BAF complex | CORUM:189 | 0.03663274 | 1.43613059 | 13 | 1022 |
| **CORUM** | ACTB-ANP32 | CORUM:7298 | 0.04992519 | 1.30168028 | 6 | 1022 |
| **CORUM** | Profilin 1 com | CORUM:2837 | 0.04992519 | 1.30168028 | 6 | 1022 |

| | | |
|---|---|---|
| 73 | 3627 | RPL3,RPS6,RPL4,RPL7A,RPL8,RPLP0,RPL13A,RPL6,RPL13,RPS8,RP |
| 82 | 3627 | EEF1A1,HNRNPA1,EEF2,TUBA1A,RPL3,RPS6,RPL4,NCL,RPL7A,RPL8, |
| 43 | 3627 | RPL3,RPL4,RPL7A,RPL8,RPLP0,RPL13A,RPL6,RPL13,RPL7,RPL5,RP |
| 29 | 3627 | RPS6,RPS8,RPS3,RPS4X,RPS3A,RPS17,RPS16,RPSA,RPS24,RPS9,RP |
| 30 | 3627 | RPS6,RPS8,RPS3,RPS4X,RPS3A,RPS17,RPS16,RPSA,RPS24,RPS9,RP |
| 17 | 3627 | DDX5,DDX17,HNRNPU,DDX3X,HNRNPDL,TARDBP,HNRNPH1,TAF15,I |
| 19 | 3627 | RPL7A,RPL13,RPL7,RPL37A,RPL19,RPL15,RPL11,RPLP1,RPL14,RPL |
| 24 | 3627 | PSMD5,PSMD2,PSMB2,PSMD3,PSMA7,PSMD1,PSMD8,PSMB4,PSMD |
| 16 | 3627 | PSMD5,PSMD2,PSMD3,PSMD1,PSMD8,PSMD4,PSMC3,PSMC1,PSMD |
| 10 | 3627 | ACTB,NCL,NPM1,NONO,MYH10,TOP2B,PARP1,HSPA4,TLE1,RAD50 |
| 15 | 3627 | ACTB,CHD4,RBBP4,HNRNPC,ARID1A,HDAC2,SMARCA4,SMARCE1,S |
| 64 | 3627 | PABPC1,PRPF8,DDX5,DDX17,SNRNP200,SPEN,SF3B1,SF3B2,SRSF4, |
| 16 | 3627 | PSMB2,PSMA7,PSMB4,PSMD4,PSMC3,PSMC1,PSMB1,PSMD13,PSM |
| 10 | 3627 | ARID1A,HDAC2,SMARCA4,SMARCE1,SMARCC1,SMARCD1,PRMT5,S |
| 8 | 3627 | NCL,NPM1,NONO,TOP2B,PARP1,HSPA4,TLE1,RAD50 |
| 8 | 3627 | CCT7,CCT6A,CCT2,TCP1,CCT3,CCT4,CCT8,CCT5 |
| 13 | 3627 | CHD3,CHD4,RBBP4,HDAC2,KDM1A,RBBP7,MTA1,GSE1,ZMYM3,GTF |
| 18 | 3627 | CHD3,SMARCA5,RBBP4,HDAC2,CPSF2,KDM1A,SMARCC1,RBBP7,SY |
| 9 | 3627 | ACTB,ARID1A,SMARCA4,SMARCE1,SMARCC1,CARM1,SMARCD1,SM |
| 11 | 3627 | RBBP4,ARID1A,HDAC2,SMARCA4,SMARCE1,SMARCC1,SMARCD1,PF |
| 13 | 3627 | HSPA8,HSP90AA1,RBBP4,HDAC2,HSPA4,RBBP7,OGT,SIN3B,WDR5,S |
| 7 | 3627 | ARID1A,SMARCA4,SMARCC1,EP300,CREBBP,SMARCC2,SMARCB1 |
| 8 | 3627 | MATR3,NONO,XRCC5,SFPQ,PRKDC,XRCC6,RBM14,HEXIM1 |
| 8 | 3627 | ACTB,ARID1A,SMARCA4,SMARCE1,SMARCC1,SMARCD1,SMARCC2, |
| 8 | 3627 | ARID1A,SMARCA4,SMARCE1,SMARCC1,SMARCD1,PRMT5,SMARCC2 |
| 8 | 3627 | ARID1A,SMARCA4,SMARCE1,SMARCC1,SMARCD1,PRMT5,SMARCC2 |
| 37 | 3627 | PABPC1,HNRNPA1,PRPF8,DDX5,SNRNP200,SF3B1,SF3B2,HNRNPU, |
| 12 | 3627 | EEF1A1,TUBB,PRPF8,SNRNP200,MATR3,DDB1,HNRNPU,PFKL,SFPQ, |
| 9 | 3627 | NCL,DDX5,DDX17,HNRNPU,HNRNPH1,ILF3,FUS,HNRNPR,DGCR8 |
| 10 | 3627 | EIF3C,EIF3L,EIF3E,EIF3F,EIF3D,EIF3H,EIF3G,EIF3B,EIF3M,EIF3J |
| 10 | 3627 | ACTB,ACTG1,ARID1A,SMARCA4,SMARCE1,SMARCC1,ARID1B,SMAR |
| 6 | 3627 | ACTB,PTMA,SLC25A3,ANP32A,C1QBP,PSMA1 |
| 6 | 3627 | HSPA8,ACTG1,TUBB2B,VCP,PFN1,CLTC |

L7,RPL5,RPL37A,RPL19,RPL15,RPS3,RPS4X,RPS3A,RPL18,RPL11,RPLP1,RPL10,RPS17,RPS16,F
,YBX1,RPLP0,RPL13A,RPL6,RPL13,RPS8,RPL7,RPL5,NPM1,RPL37A,RPL19,RPL15,RPS3A,RPL18,
L37A,RPL19,RPL15,RPL18,RPL11,RPLP1,RPL10,RPL23,RPL29,RPL24,RPL17,RPL14,RPL22,RPL
S14,RPS20,RPS25,RPS15,RPS5,RPS7,RPS19,RPS10,RPS12,RPS15A,RPS11,RPS28,FAU,RPS13,RF
S14,RPS20,RPS25,RPS15,RPS5,RPS7,RPS19,RPS10,RPS12,RPS27,RPS15A,RPS11,RPS28,FAU,RF
LF3,HNRNPUL1,HNRNPM,ILF2,DHX15,FUS,RALY,EWSR1,DGCR8
.12,RPL30,RPL35A,RPLP2,RPL36,RPL27,RPL28,RPL34,DICER1,RPL32
4,PSMC3,PSMC1,PSMB1,PSMD13,PSMD7,PSMC5,PSMC2,PSMC4,PSMB5,PSMA1,PSMD10,PSMA
13,PSMD7,PSMC5,PSMC2,PSMC4,PSMD10,PSMD12,PSMC6

ARCC1,MBD3,MTA2,GATAD2B,SMARCC2,SMARCB1,DPF2
DDX3X,IK,CIRBP,LUC7L3,DDX39B,SRSF5,U2AF2,SNRNP70,PRPF6,SRSF9,SRSF1,SRSF3,SF3A1,P
C5,PSMC2,PSMC4,PSMB5,PSMA1,PSMA6,PSMA4,PSMC6
ARCC2,SMARCB1,SIN3A

2I,MTA2,HMG20B,SIN3A
MPK,EFTUD2,MBD3,SAP18,MTA2,RAN,WDR5,SMARCC2,SMARCB1,SIN3A
ARCC2,SMARCB1
MT5,SMARCC2,SMARCB1,SIN3A
UDS3,SP1,SETD1A,SIN3A

SMARCB1
,SMARCB1
,SMARCB1
NN,HNRNPC,HNRNPK,HNRNPH1,HNRNPA3,RBMX,PRPF6,SRSF1,HNRNPM,SF3A1,HNRNPA2B1,SI
,UBR5,EFTUD2,MSH2

D1,SMARCC2,SMARCB1

RPL23,RPL29,RPSA,RPS24,RPS9,RPL24,RPL17,RPL14,RPL22,RPL9,RPS14,RPS20,RPL26,RPL31,
,RPL11,RPLP1,RPL10,RPS17,HNRNPU,RPS16,RPL23,RPL29,RPS24,RPS9,RPL24,RPL17,RPL14,R
9,RPL26,RPL31,RPL12,RPL30,RPL35,RPL35A,RPLP2,RPL36,RPL23A,RPL27,RPL36A,RPL38,RPL
PS2,RPS27A,RPS26,RPS21,RPS29
PS13,RPS2,RPS27A,RPS26,RPS21,RPS29

6,PSMD12,PSMA4,PSMC6

RPF40A,DHX15,SF1,PPM1G,SRSF6,TRA2B,TRIR,SNRPD3,EFTUD2,SRSF2,PRPF4B,WBP11,SRSF1

NRPD3,RALY,EFTUD2,PRPF4B,PRPF19,SF3A2,SART1,DHX38,EIF4A3,MTREX,SF3A3,SYNCRIP,HNF

RPS25,RPS15,RPL12,RPL30,RPS5,RPS7,RPS19,RPL35,RPS10,RPL35A,RPLP2,RPL36,RPS12,RPS[illegible]
PL22,RPL9,RPS14,RPL26,RPL31,RPS15,RPL12,RPL30,RPS7,RPL35,RPL35A,RPLP2,H1-10,RPL3[illegible]
37,RPL28,RPL39,RPL34,UBA52,RPL27A,RPL32

1,PRPF19,SF3A2,SART1,DHX38,EIF4A3,HTATSF1,MTREX,RBM25,SF3A3,CD2BP2,LSM4,SRSF7,N[illegible]

RNPR,DDX41,SNRPD2,SNRPB,XAB2,PLRG1

27,RPL23A,RPL27,RPL36A,RPS15A,RPL38,RPL37,RPS11,RPL28,RPS28,FAU,RPL39,RPS13,RPL3

;,RPS12,RPL23A,U2AF2,RPL27,RPL36A,SRP14,TOP1,RPS15A,ILF3,RPS11,RPL28,HNRNPM,ILF2,

CBP2,DNAJC8,DDX41,SNRPD2,SNRPB,THOC2,RBM17,SRRT,PRPF31,THOC3,XAB2,RNPS1,ACIN1

4,UBA52,RPS2,RPS27A,RPS26,RPS21,RPL27A,RPS29,RPL32

RPL39,RPS13,NAP1L1,RPS2,SLC25A5,IGF2BP1,LUC7L2,NOP56,ACTA2,RPL27A,FBL,RPS29,GNL3

,ZNF207,PLRG1,RBM39,SNU13

3,NOLC1,RPL32,NOP58,MYBBP1A,SNU13

| | | | | | | |
|---|---|---|---|---|---|---|
| **HP** | Abnormality o | HP:0003011 | 2.98E-23 | 22.5260964 | 2526 | 839 |
| **HP** | Abnormal mu: | HP:0011804 | 1.84E-20 | 19.7357088 | 2194 | 839 |
| **HP** | Abnormal mu: | HP:0003808 | 8.74E-18 | 17.058379 | 1838 | 839 |
| **HP** | Abnormality o | HP:0000925 | 1.12E-17 | 16.9504081 | 1313 | 839 |
| **HP** | Abnormal nen | HP:0012638 | 1.03E-16 | 15.9864437 | 3221 | 839 |
| **HP** | Abnormality o | HP:0000707 | 3.94E-16 | 15.4047925 | 3446 | 839 |
| **HP** | Hypoplasia of | HP:0002079 | 7.03E-16 | 15.1532089 | 475 | 839 |
| **HP** | Abnormal axia | HP:0009121 | 8.70E-16 | 15.0604877 | 2318 | 839 |
| **HP** | Abnormal nen | HP:0012639 | 2.50E-15 | 14.6025055 | 2677 | 839 |
| **HP** | Abnormality o | HP:0100886 | 3.30E-15 | 14.4814162 | 894 | 839 |
| **HP** | Abnormality o | HP:0100547 | 5.96E-15 | 14.2246711 | 1762 | 839 |
| **HP** | Abnormal cere | HP:0002060 | 7.64E-15 | 14.1170576 | 1733 | 839 |
| **HP** | Abnormality o | HP:0033127 | 1.16E-14 | 13.9338617 | 3353 | 839 |
| **HP** | Aplasia/Hypop | HP:0002977 | 1.20E-14 | 13.9224973 | 1619 | 839 |
| **HP** | Hypertelorism | HP:0000316 | 1.47E-14 | 13.8312324 | 668 | 839 |
| **HP** | Neurodevelop | HP:0012759 | 1.69E-14 | 13.7715242 | 2459 | 839 |
| **HP** | Abnormal sku | HP:0000929 | 2.96E-14 | 13.5279911 | 2053 | 839 |
| **HP** | Abnormality o | HP:0010674 | 4.44E-14 | 13.3528413 | 1020 | 839 |
| **HP** | Autosomal do | HP:0000006 | 8.74E-14 | 13.0585611 | 1783 | 839 |
| **HP** | Abnormality o | HP:0000177 | 2.23E-12 | 11.6515978 | 757 | 839 |
| **HP** | Abnormality o | HP:0000240 | 3.44E-12 | 11.4632539 | 1397 | 839 |
| **HP** | Aplasia/Hypop | HP:0007364 | 3.67E-12 | 11.4351084 | 1358 | 839 |
| **HP** | Abnormality o | HP:0012443 | 4.14E-12 | 11.3826974 | 2268 | 839 |
| **HP** | Abnormality o | HP:0011446 | 4.36E-12 | 11.360102 | 2434 | 839 |
| **HP** | Abnormality o | HP:0040064 | 4.49E-12 | 11.3474522 | 2010 | 839 |
| **HP** | Abnormality o | HP:0011842 | 5.08E-12 | 11.2939878 | 2730 | 839 |
| **HP** | Abnormal cere | HP:0002500 | 6.49E-12 | 11.1880507 | 918 | 839 |
| **HP** | Morphologica | HP:0002011 | 7.07E-12 | 11.1506861 | 2483 | 839 |
| **HP** | Neurodevelop | HP:0012758 | 8.46E-12 | 11.0725135 | 1989 | 839 |
| **HP** | Abnormality o | HP:0000377 | 1.00E-11 | 10.9996832 | 686 | 839 |
| **HP** | Abnormality o | HP:0000288 | 1.14E-11 | 10.9443008 | 561 | 839 |
| **HP** | Scoliosis | HP:0002650 | 1.16E-11 | 10.9373795 | 945 | 839 |
| **HP** | Neurological s | HP:0002167 | 1.22E-11 | 10.9124318 | 1407 | 839 |
| **HP** | Abnormal nas | HP:0005105 | 1.84E-11 | 10.7347082 | 1161 | 839 |
| **HP** | Inguinal herni | HP:0000023 | 2.88E-11 | 10.5401691 | 241 | 839 |
| **HP** | Abnormal cor | HP:0001273 | 3.08E-11 | 10.5120949 | 791 | 839 |
| **HP** | Abnormal app | HP:0011844 | 3.38E-11 | 10.4714519 | 1586 | 839 |
| **HP** | Abnormality o | HP:0010993 | 3.62E-11 | 10.4409146 | 1009 | 839 |
| **HP** | Abnormality o | HP:0001155 | 4.41E-11 | 10.3556643 | 1289 | 839 |
| **HP** | Abnormality o | HP:0000478 | 8.55E-11 | 10.0681697 | 2727 | 839 |
| **HP** | Abnormal thoı | HP:0000765 | 8.63E-11 | 10.0639786 | 784 | 839 |
| **HP** | Seizure | HP:0001250 | 9.11E-11 | 10.0403263 | 1707 | 839 |
| **HP** | Erythroid hyp | HP:0012133 | 1.33E-10 | 9.87653384 | 25 | 839 |
| **HP** | Microcephaly | HP:0000252 | 1.39E-10 | 9.85790955 | 1133 | 839 |
| **HP** | Normochromi | HP:0001895 | 1.44E-10 | 9.84263108 | 31 | 839 |

| | | | | | | |
|---|---|---|---|---|---|---|
| **HP** | Abnormality o | HP:0000924 | 1.59E-10 | 9.79762477 | 2831 | 839 |
| **HP** | Abnormality o | HP:0002817 | 2.00E-10 | 9.69826557 | 1405 | 839 |
| **HP** | Decreased hea | HP:0040195 | 2.15E-10 | 9.66723491 | 1146 | 839 |
| **HP** | Hypotonia | HP:0001252 | 2.88E-10 | 9.54074705 | 1574 | 839 |
| **HP** | Abnormality o | HP:0000315 | 3.87E-10 | 9.41261393 | 1346 | 839 |
| **HP** | Abnormality o | HP:0040068 | 4.06E-10 | 9.39187744 | 1476 | 839 |
| **HP** | Abnormality o | HP:0002813 | 4.06E-10 | 9.39187744 | 1476 | 839 |
| **HP** | Abnormal radi | HP:0031640 | 4.87E-10 | 9.31263179 | 22 | 839 |
| **HP** | Elevated red c | HP:0030270 | 4.87E-10 | 9.31263179 | 22 | 839 |
| **HP** | Radial artery a | HP:0020118 | 4.87E-10 | 9.31263179 | 22 | 839 |
| **HP** | Epicanthus | HP:0000286 | 5.00E-10 | 9.30109004 | 479 | 839 |
| **HP** | Abnormal skel | HP:0011805 | 5.84E-10 | 9.23334202 | 1621 | 839 |
| **HP** | Abnormality o | HP:0011337 | 6.71E-10 | 9.17358564 | 341 | 839 |
| **HP** | Malignant ger | HP:0006758 | 7.13E-10 | 9.14663295 | 26 | 839 |
| **HP** | Abnormality o | HP:0000152 | 7.67E-10 | 9.11544083 | 2875 | 839 |
| **HP** | Abnormal eyel | HP:0000492 | 8.45E-10 | 9.07307977 | 1123 | 839 |
| **HP** | Joint hypermo | HP:0001382 | 9.64E-10 | 9.01606259 | 307 | 839 |
| **HP** | Language imp | HP:0002463 | 1.05E-09 | 8.98019423 | 828 | 839 |
| **HP** | Abnormal digi | HP:0011297 | 1.16E-09 | 8.93483855 | 1296 | 839 |
| **HP** | Atrial septal de | HP:0001631 | 1.63E-09 | 8.78727369 | 365 | 839 |
| **HP** | Abnormality o | HP:0000153 | 1.67E-09 | 8.77715495 | 1937 | 839 |
| **HP** | Hernia | HP:0100790 | 1.71E-09 | 8.76600674 | 477 | 839 |
| **HP** | Abnormal atria | HP:0011994 | 2.52E-09 | 8.59800228 | 367 | 839 |
| **HP** | Abnormality o | HP:0000234 | 2.59E-09 | 8.58689386 | 2852 | 839 |
| **HP** | Abnormality o | HP:0032039 | 3.04E-09 | 8.51686023 | 1319 | 839 |
| **HP** | Macrocytic dy | HP:0005532 | 3.08E-09 | 8.5108277 | 23 | 839 |
| **HP** | Abnormal lip r | HP:0000159 | 4.06E-09 | 8.39153179 | 926 | 839 |
| **HP** | Thin corpus c | HP:0033725 | 4.21E-09 | 8.37565288 | 728 | 839 |
| **HP** | Sprengel anor | HP:0000912 | 4.93E-09 | 8.30727944 | 51 | 839 |
| **HP** | Wide nasal bri | HP:0000431 | 5.17E-09 | 8.28627524 | 500 | 839 |
| **HP** | Abnormality o | HP:0000290 | 5.23E-09 | 8.28171256 | 942 | 839 |
| **HP** | Abnormality o | HP:0000356 | 6.49E-09 | 8.18764561 | 1072 | 839 |
| **HP** | Generalized h | HP:0001290 | 7.17E-09 | 8.14463846 | 968 | 839 |
| **HP** | Abnormal ocu | HP:0030669 | 8.28E-09 | 8.08184057 | 1303 | 839 |
| **HP** | Delayed ability | HP:0031936 | 9.44E-09 | 8.0250803 | 155 | 839 |
| **HP** | Aplasia/Hypop | HP:0007370 | 1.04E-08 | 7.98185645 | 716 | 839 |
| **HP** | Abnormal eye | HP:0012373 | 1.14E-08 | 7.94429357 | 2218 | 839 |
| **HP** | High palate | HP:0000218 | 1.27E-08 | 7.89641096 | 632 | 839 |
| **HP** | Abnormal carc | HP:0001671 | 1.28E-08 | 7.89320155 | 579 | 839 |
| **HP** | Abnormal pala | HP:0000174 | 1.32E-08 | 7.88088953 | 968 | 839 |
| **HP** | Abnormal con | HP:0000549 | 1.43E-08 | 7.84573126 | 945 | 839 |
| **HP** | Abnormal aort | HP:0001679 | 1.61E-08 | 7.79428671 | 319 | 839 |
| **HP** | Abnormality o | HP:0001167 | 1.79E-08 | 7.74623107 | 1018 | 839 |
| **HP** | Abnormal foo | HP:0001760 | 2.45E-08 | 7.61038123 | 1219 | 839 |
| **HP** | Gait disturban | HP:0001288 | 2.62E-08 | 7.58091064 | 1002 | 839 |

| | | | | | | |
|---|---|---|---|---|---|---|
| **HP** | Abnormal oral | HP:0031816 | 3.00E-08 | 7.52358802 | 1855 | 839 |
| **HP** | Abnormal oral | HP:0000163 | 3.00E-08 | 7.52358802 | 1855 | 839 |
| **HP** | Abnormality o | HP:0002814 | 3.52E-08 | 7.45332388 | 1471 | 839 |
| **HP** | Abnormality o | HP:0002648 | 3.66E-08 | 7.43699254 | 775 | 839 |
| **HP** | Abnormal nas | HP:0000422 | 3.76E-08 | 7.42450397 | 929 | 839 |
| **HP** | Abnormality o | HP:0100022 | 3.91E-08 | 7.40736098 | 1842 | 839 |
| **HP** | Delayed gross | HP:0002194 | 4.61E-08 | 7.3365186 | 276 | 839 |
| **HP** | Involuntary m | HP:0004305 | 5.04E-08 | 7.29790555 | 993 | 839 |
| **HP** | Pure red cell a | HP:0012410 | 6.32E-08 | 7.19923254 | 25 | 839 |
| **HP** | Abnormal agg | HP:0006919 | 7.64E-08 | 7.11696434 | 310 | 839 |
| **HP** | Delayed speec | HP:0000750 | 1.26E-07 | 6.90065291 | 798 | 839 |
| **HP** | Persistence of | HP:0011904 | 1.30E-07 | 6.88575328 | 30 | 839 |
| **HP** | Abnormality o | HP:0000271 | 1.31E-07 | 6.883836 | 2602 | 839 |
| **HP** | Hypertonia | HP:0001276 | 1.57E-07 | 6.80334327 | 950 | 839 |
| **HP** | Depressed na | HP:0005280 | 1.70E-07 | 6.77018475 | 503 | 839 |
| **HP** | Abnormality o | HP:0009127 | 2.12E-07 | 6.67454542 | 474 | 839 |
| **HP** | Abnormal faci | HP:0001999 | 2.12E-07 | 6.67304842 | 1005 | 839 |
| **HP** | Abnormality o | HP:0002683 | 2.14E-07 | 6.66903209 | 891 | 839 |
| **HP** | Abnormality o | HP:0000002 | 2.21E-07 | 6.6551425 | 1343 | 839 |
| **HP** | Autistic behav | HP:0000729 | 2.35E-07 | 6.62942268 | 540 | 839 |
| **HP** | Osteosarcoma | HP:0002669 | 2.55E-07 | 6.59338142 | 33 | 839 |
| **HP** | Motor delay | HP:0001270 | 3.33E-07 | 6.47737874 | 663 | 839 |
| **HP** | Low-set ears | HP:0000369 | 3.68E-07 | 6.43417261 | 691 | 839 |
| **HP** | Global develo | HP:0001263 | 4.00E-07 | 6.3975267 | 1612 | 839 |
| **HP** | Strabismus | HP:0000486 | 4.92E-07 | 6.3078693 | 912 | 839 |
| **HP** | Abnormal eryt | HP:0030272 | 5.25E-07 | 6.28007818 | 29 | 839 |
| **HP** | Abnormality o | HP:0011729 | 5.43E-07 | 6.26500695 | 1105 | 839 |
| **HP** | Abnormal loca | HP:0000357 | 5.48E-07 | 6.26143631 | 726 | 839 |
| **HP** | Intellectual dis | HP:0001249 | 5.72E-07 | 6.24230529 | 1724 | 839 |
| **HP** | Abnormal cen | HP:0011442 | 5.82E-07 | 6.23487334 | 1719 | 839 |
| **HP** | Pectus excava | HP:0000767 | 6.76E-07 | 6.17015663 | 256 | 839 |
| **HP** | Adenocarcino | HP:0040276 | 7.22E-07 | 6.14155993 | 27 | 839 |
| **HP** | Wide mouth | HP:0000154 | 8.23E-07 | 6.08475919 | 158 | 839 |
| **HP** | Spasticity | HP:0001257 | 1.4199E-06 | 5.84773898 | 803 | 839 |
| **HP** | Abnormality o | HP:0000309 | 1.8729E-06 | 5.72749171 | 527 | 839 |
| **HP** | Abnormality o | HP:0025270 | 1.8984E-06 | 5.72161315 | 662 | 839 |
| **HP** | Abnormal ven | HP:0010438 | 1.9633E-06 | 5.707018 | 397 | 839 |
| **HP** | Abdominal wa | HP:0010866 | 2.0357E-06 | 5.69129404 | 410 | 839 |
| **HP** | Kyphosis | HP:0002808 | 2.0877E-06 | 5.68033748 | 423 | 839 |
| **HP** | Hernia of the a | HP:0004299 | 2.1329E-06 | 5.67103857 | 406 | 839 |
| **HP** | Ventricular sep | HP:0001629 | 2.1417E-06 | 5.66924539 | 389 | 839 |
| **HP** | Atrophy/Dege | HP:0007367 | 2.335E-06 | 5.63170812 | 779 | 839 |
| **HP** | Aplasia/hypop | HP:0009115 | 2.7902E-06 | 5.55436619 | 1052 | 839 |
| **HP** | Ptosis | HP:0000508 | 2.9775E-06 | 5.52615288 | 638 | 839 |
| **HP** | Hyperactivity | HP:0000752 | 3.6593E-06 | 5.43660662 | 448 | 839 |

| | | | | | | |
|---|---|---|---|---|---|---|
| **HP** | Pes planus | HP:0001763 | 3.7254E-06 | 5.42882819 | 292 | 839 |
| **HP** | Abnormality o | HP:0011339 | 3.789E-06 | 5.42147095 | 350 | 839 |
| **HP** | Abnormality o | HP:0008050 | 4.7505E-06 | 5.32325716 | 715 | 839 |
| **HP** | Abnormal loca | HP:0100542 | 4.7693E-06 | 5.32154149 | 164 | 839 |
| **HP** | Abnormality o | HP:0001227 | 5.5637E-06 | 5.25463361 | 29 | 839 |
| **HP** | Abnormality o | HP:0000464 | 5.7579E-06 | 5.23973365 | 548 | 839 |
| **HP** | Congenital ab | HP:0011361 | 6.1798E-06 | 5.20902782 | 254 | 839 |
| **HP** | Abnormal eye | HP:0012372 | 6.296E-06 | 5.20093833 | 2163 | 839 |
| **HP** | Abnormality o | HP:0000277 | 6.5481E-06 | 5.18388583 | 901 | 839 |
| **HP** | Short digit | HP:0011927 | 6.5866E-06 | 5.18134034 | 540 | 839 |
| **HP** | Upper motor r | HP:0002493 | 6.8759E-06 | 5.1626714 | 1188 | 839 |
| **HP** | Abnormality o | HP:0004298 | 6.9075E-06 | 5.16067858 | 452 | 839 |
| **HP** | Abnormal jaw | HP:0030791 | 7.2701E-06 | 5.13845834 | 945 | 839 |
| **HP** | Abnormal sys | HP:0011004 | 7.3662E-06 | 5.13275376 | 435 | 839 |
| **HP** | Thrombocytos | HP:0001894 | 8.7655E-06 | 5.05722417 | 51 | 839 |
| **HP** | Short attentior | HP:0000736 | 9.4979E-06 | 5.02237217 | 402 | 839 |
| **HP** | Abnormal mo | HP:0030962 | 1.0111E-05 | 4.9951869 | 579 | 839 |
| **HP** | Feeding difficu | HP:0011968 | 1.0296E-05 | 4.98734632 | 953 | 839 |
| **HP** | Abnormal cere | HP:0001317 | 1.0477E-05 | 4.979763 | 815 | 839 |
| **HP** | Aplasia/hypop | HP:0009815 | 1.0555E-05 | 4.97655612 | 643 | 839 |
| **HP** | Abnormality o | HP:0011843 | 1.1416E-05 | 4.94248259 | 1393 | 839 |
| **HP** | Abnormal ster | HP:0000766 | 1.1524E-05 | 4.93840943 | 386 | 839 |
| **HP** | Abnormality o | HP:0000496 | 1.2382E-05 | 4.90719407 | 1531 | 839 |
| **HP** | Adenocarcinoi | HP:0040275 | 1.38E-05 | 4.8601144 | 30 | 839 |
| **HP** | Abnormality o | HP:0100491 | 1.3846E-05 | 4.85866224 | 604 | 839 |
| **HP** | Abnormal carc | HP:0005120 | 1.3977E-05 | 4.85457281 | 426 | 839 |
| **HP** | Abnormality o | HP:0011138 | 1.4159E-05 | 4.84895469 | 1235 | 839 |
| **HP** | Abnormal nun | HP:0012131 | 1.6407E-05 | 4.78496722 | 38 | 839 |
| **HP** | Upper extremi | HP:0030310 | 1.7166E-05 | 4.76532656 | 90 | 839 |
| **HP** | Sarcoma | HP:0100242 | 1.7506E-05 | 4.7568189 | 165 | 839 |
| **HP** | Misalignment | HP:0000692 | 1.8832E-05 | 4.725107 | 292 | 839 |
| **HP** | Abnormality o | HP:0011282 | 2.0816E-05 | 4.68160374 | 821 | 839 |
| **HP** | Abnormality o | HP:0011283 | 2.0816E-05 | 4.68160374 | 821 | 839 |
| **HP** | Gastroesopha | HP:0002020 | 2.1916E-05 | 4.65923912 | 377 | 839 |
| **HP** | Vascular dilata | HP:0002617 | 2.2615E-05 | 4.64559661 | 185 | 839 |
| **HP** | Long philtrum | HP:0000343 | 2.2978E-05 | 4.6386967 | 339 | 839 |
| **HP** | Abnormal hair | HP:0010720 | 2.3377E-05 | 4.63121171 | 289 | 839 |
| **HP** | Abnormal hair | HP:0001595 | 2.8693E-05 | 4.54222613 | 1099 | 839 |
| **HP** | Increased mea | HP:0005518 | 3.3045E-05 | 4.48089268 | 39 | 839 |
| **HP** | Downslanted | HP:0000494 | 3.3505E-05 | 4.47488548 | 436 | 839 |
| **HP** | Abnormality o | HP:0000366 | 3.4115E-05 | 4.46705669 | 1522 | 839 |
| **HP** | Nonimmune h | HP:0001790 | 3.8333E-05 | 4.41643248 | 42 | 839 |
| **HP** | Abnormal cere | HP:0002538 | 5.7175E-05 | 4.24279327 | 638 | 839 |
| **HP** | Webbed neck | HP:0000465 | 6.5889E-05 | 4.18118622 | 111 | 839 |
| **HP** | Abnormality o | HP:0010936 | 7.0103E-05 | 4.15426266 | 714 | 839 |

| | | | | | | |
|---|---|---|---|---|---|---|
| **HP** | Macrocytic an | HP:0001972 | 7.2847E-05 | 4.13758868 | 61 | 839 |
| **HP** | Abnormality o | HP:0000539 | 7.3557E-05 | 4.13337816 | 535 | 839 |
| **HP** | Brain atrophy | HP:0012444 | 7.5871E-05 | 4.11992617 | 696 | 839 |
| **HP** | Abnormality o | HP:0002118 | 7.5976E-05 | 4.11932315 | 790 | 839 |
| **HP** | Aggressive be | HP:0000718 | 8.0354E-05 | 4.09498995 | 206 | 839 |
| **HP** | Adenocarcino | HP:0040273 | 9.0458E-05 | 4.0435531 | 35 | 839 |
| **HP** | Abnormal em | HP:0100851 | 9.6665E-05 | 4.01473172 | 510 | 839 |
| **HP** | Aortic aneurys | HP:0004942 | 0.00010647 | 3.97276311 | 138 | 839 |
| **HP** | Absent speech | HP:0001344 | 0.00012046 | 3.91915425 | 277 | 839 |
| **HP** | Abnormal sca | HP:0000782 | 0.00013134 | 3.88161222 | 150 | 839 |
| **HP** | Midface retrus | HP:0011800 | 0.00013636 | 3.86530043 | 257 | 839 |
| **HP** | Abnormality o | HP:0000436 | 0.00014199 | 3.84773656 | 303 | 839 |
| **HP** | Ventriculomeg | HP:0002119 | 0.00014565 | 3.83668869 | 618 | 839 |
| **HP** | Reticulocytop | HP:0001896 | 0.00014886 | 3.82721424 | 33 | 839 |
| **HP** | Obsessive-co | HP:0000722 | 0.00015432 | 3.81157244 | 103 | 839 |
| **HP** | Impairment in | HP:0031466 | 0.00015683 | 3.80457773 | 614 | 839 |
| **HP** | Abnormal han | HP:0005922 | 0.00015764 | 3.8023467 | 372 | 839 |
| **HP** | Short finger | HP:0009381 | 0.00016322 | 3.7872156 | 287 | 839 |
| **HP** | Joint dislocati | HP:0001373 | 0.00018143 | 3.74129895 | 347 | 839 |
| **HP** | Neoplasm of t | HP:0010622 | 0.00018225 | 3.7393416 | 93 | 839 |
| **HP** | Delayed myeli | HP:0012448 | 0.00018252 | 3.73868156 | 186 | 839 |
| **HP** | Aplasia/hypop | HP:0006496 | 0.00018872 | 3.7241927 | 506 | 839 |
| **HP** | Aplasia/hypop | HP:0005927 | 0.00020038 | 3.69814277 | 475 | 839 |
| **HP** | Abnormality o | HP:0000599 | 0.00020901 | 3.67982573 | 148 | 839 |
| **HP** | Aplasia/hypop | HP:0045060 | 0.0002104 | 3.67696248 | 598 | 839 |
| **HP** | Stereotypy | HP:0000733 | 0.00023475 | 3.62940201 | 219 | 839 |
| **HP** | Neoplasm by | HP:0011792 | 0.00028177 | 3.55010107 | 324 | 839 |
| **HP** | Attention defic | HP:0007018 | 0.00028286 | 3.54843192 | 307 | 839 |
| **HP** | Short stature | HP:0004322 | 0.00029057 | 3.53675025 | 1223 | 839 |
| **HP** | Short neck | HP:0000470 | 0.00030273 | 3.51894609 | 333 | 839 |
| **HP** | Cleft soft pala | HP:0000185 | 0.00031862 | 3.49673291 | 165 | 839 |
| **HP** | Elbow disloca | HP:0003042 | 0.00033887 | 3.46996395 | 81 | 839 |
| **HP** | Partial duplica | HP:0009944 | 0.0003432 | 3.46444774 | 37 | 839 |
| **HP** | Growth abnor | HP:0001507 | 0.00038171 | 3.41826574 | 2323 | 839 |
| **HP** | Acute myeloid | HP:0004808 | 0.0003913 | 3.40748892 | 68 | 839 |
| **HP** | Behavioral abr | HP:0000708 | 0.00043513 | 3.36137875 | 1661 | 839 |
| **HP** | Abnormality o | HP:0009553 | 0.0004406 | 3.3559528 | 214 | 839 |
| **HP** | Abnormality o | HP:0030047 | 0.00050017 | 3.30087909 | 96 | 839 |
| **HP** | Abnormal hea | HP:0001654 | 0.00052219 | 3.28216794 | 281 | 839 |
| **HP** | Patent forame | HP:0001655 | 0.00058662 | 3.23164365 | 69 | 839 |
| **HP** | Duplication of | HP:0009942 | 0.00061837 | 3.2087498 | 53 | 839 |
| **HP** | Abnormality o | HP:0004225 | 0.00063443 | 3.19761334 | 15 | 839 |
| **HP** | Hypermetropia | HP:0000540 | 0.00067316 | 3.17188188 | 208 | 839 |
| **HP** | Abnormality o | HP:0011329 | 0.00068043 | 3.16721858 | 291 | 839 |
| **HP** | Sleep disturba | HP:0002360 | 0.00070243 | 3.15339425 | 489 | 839 |

| | | | | | | |
|---|---|---|---|---|---|---|
| **HP** | Aplasia/Hypop | HP:0009239 | 0.00071334 | 3.14670136 | 13 | 839 |
| **HP** | Dilation of late | HP:0006956 | 0.00075817 | 3.1202354 | 73 | 839 |
| **HP** | Joint laxity | HP:0001388 | 0.00083176 | 3.08000384 | 205 | 839 |
| **HP** | Atrophy/Degei | HP:0007369 | 0.0008655 | 3.06273126 | 638 | 839 |
| **HP** | Abnormal nerv | HP:0001311 | 0.00086613 | 3.06241599 | 573 | 839 |
| **HP** | Aplasia/Hypop | HP:0009118 | 0.00088858 | 3.05130281 | 704 | 839 |
| **HP** | Aplasia/Hypop | HP:0009116 | 0.00094138 | 3.02623712 | 714 | 839 |
| **HP** | Abnormality o | HP:0011821 | 0.00094889 | 3.02278282 | 1122 | 839 |
| **HP** | Abnormality o | HP:0010938 | 0.00095306 | 3.02088123 | 819 | 839 |
| **HP** | Slanting of the | HP:0200006 | 0.00100291 | 2.99873943 | 616 | 839 |
| **HP** | Cleft palate | HP:0000175 | 0.00102564 | 2.98900671 | 556 | 839 |
| **HP** | Chronic const | HP:0012450 | 0.00105553 | 2.97652994 | 51 | 839 |
| **HP** | Micrognathia | HP:0000347 | 0.00106625 | 2.97214201 | 701 | 839 |
| **HP** | Aplasia/Hypop | HP:0006265 | 0.00111311 | 2.9534621 | 324 | 839 |
| **HP** | Developmenta | HP:0001087 | 0.00112137 | 2.95025069 | 64 | 839 |
| **HP** | Partial duplica | HP:0009999 | 0.00113664 | 2.94437633 | 39 | 839 |
| **HP** | Abnormal hard | HP:0100737 | 0.00115379 | 2.93787402 | 557 | 839 |
| **HP** | Mitral valve pr | HP:0001634 | 0.0011665 | 2.93311426 | 88 | 839 |
| **HP** | Abnormality o | HP:0003319 | 0.0013009 | 2.88575598 | 373 | 839 |
| **HP** | Abnormality o | HP:0001780 | 0.00130161 | 2.88552055 | 623 | 839 |
| **HP** | Coarctation of | HP:0001680 | 0.00144147 | 2.84119376 | 114 | 839 |
| **HP** | Abnormality o | HP:0001446 | 0.00151184 | 2.8204941 | 208 | 839 |
| **HP** | Thin vermilion | HP:0000233 | 0.00159032 | 2.79851492 | 335 | 839 |
| **HP** | Cerebral atrop | HP:0002059 | 0.00162197 | 2.7899583 | 625 | 839 |
| **HP** | Abnormal sup | HP:0031828 | 0.00176469 | 2.7533321 | 246 | 839 |
| **HP** | Babinski sign | HP:0003487 | 0.00176469 | 2.7533321 | 246 | 839 |
| **HP** | Synophrys | HP:0000664 | 0.00177794 | 2.75008259 | 126 | 839 |
| **HP** | Long fingers | HP:0100807 | 0.00187626 | 2.72670722 | 238 | 839 |
| **HP** | Functional abi | HP:0012719 | 0.00188513 | 2.72465759 | 923 | 839 |
| **HP** | Abnormality o | HP:0011338 | 0.00198469 | 2.70230705 | 230 | 839 |
| **HP** | Hypertrichosis | HP:0000998 | 0.00213629 | 2.67034041 | 218 | 839 |
| **HP** | Intellectual dis | HP:0001256 | 0.00230132 | 2.63802304 | 355 | 839 |
| **HP** | Abnormality o | HP:0009810 | 0.00241145 | 2.61772205 | 458 | 839 |
| **HP** | Abnormal mye | HP:0012447 | 0.00273406 | 2.56319141 | 500 | 839 |
| **HP** | Narrow mouth | HP:0000160 | 0.00279584 | 2.55348729 | 203 | 839 |
| **HP** | Lower limb m | HP:0007340 | 0.00296293 | 2.52827813 | 128 | 839 |
| **HP** | Focal-onset se | HP:0007359 | 0.00302234 | 2.51965654 | 266 | 839 |
| **HP** | Muscle weakn | HP:0001324 | 0.00303328 | 2.518087 | 899 | 839 |
| **HP** | Abnormality o | HP:0030178 | 0.00313616 | 2.50360143 | 492 | 839 |
| **HP** | Hypospadias | HP:0000047 | 0.00323691 | 2.48986964 | 305 | 839 |
| **HP** | Abnormality o | HP:0001421 | 0.00341791 | 2.46623876 | 95 | 839 |
| **HP** | Abnormal pate | HP:0003045 | 0.00341791 | 2.46623876 | 95 | 839 |
| **HP** | Prominent for | HP:0011220 | 0.00355494 | 2.44916763 | 192 | 839 |
| **HP** | Abnormal hem | HP:0011902 | 0.00404319 | 2.39327618 | 54 | 839 |
| **HP** | Aplasia/hypop | HP:0009122 | 0.00430133 | 2.36639749 | 801 | 839 |

| | | | | | | |
|---|---|---|---|---|---|---|
| HP | Deeply set eye | HP:0000490 | 0.00438427 | 2.35810243 | 189 | 839 |
| HP | Self-injurious l | HP:0100716 | 0.00449356 | 2.34740915 | 157 | 839 |
| HP | Tall stature | HP:0000098 | 0.00451543 | 2.34530111 | 177 | 839 |
| HP | Abnormal pos | HP:0000932 | 0.00454377 | 2.34258358 | 165 | 839 |
| HP | Pes cavus | HP:0001761 | 0.00492657 | 2.30745532 | 269 | 839 |
| HP | Facial hypertri | HP:0002219 | 0.00502126 | 2.29918753 | 134 | 839 |
| HP | Abnormality o | HP:0001574 | 0.00528185 | 2.27721377 | 2279 | 839 |
| HP | EEG abnorma | HP:0002353 | 0.00531295 | 2.27466443 | 469 | 839 |
| HP | Oral cleft | HP:0000202 | 0.00537566 | 2.26956811 | 594 | 839 |
| HP | Bladder diverti | HP:0000015 | 0.00587431 | 2.23104307 | 22 | 839 |
| HP | Bicuspid aorti | HP:0001647 | 0.00595727 | 2.22495289 | 86 | 839 |
| HP | Abnormality o | HP:0009602 | 0.00656826 | 2.18254939 | 187 | 839 |
| HP | Abnormal thu | HP:0001172 | 0.006729 | 2.17204952 | 345 | 839 |
| HP | Feeding difficu | HP:0008872 | 0.00675148 | 2.17060087 | 517 | 839 |
| HP | Joint hyperfle | HP:0005692 | 0.00681409 | 2.16659208 | 175 | 839 |
| HP | High forehead | HP:0000348 | 0.00686436 | 2.16339987 | 254 | 839 |
| HP | Acute leukemi | HP:0002488 | 0.00721343 | 2.1418579 | 105 | 839 |
| HP | Low anterior h | HP:0000294 | 0.00721343 | 2.1418579 | 105 | 839 |
| HP | Mental deterio | HP:0001268 | 0.007498 | 2.1250543 | 315 | 839 |
| HP | Abnormal mea | HP:0025065 | 0.0076099 | 2.11862119 | 49 | 839 |
| HP | Deviation of fi | HP:0004097 | 0.00768681 | 2.11425369 | 395 | 839 |
| HP | Abnormal atric | HP:0006705 | 0.00781659 | 2.10698254 | 132 | 839 |
| HP | Microtia | HP:0008551 | 0.00801491 | 2.09610157 | 136 | 839 |
| HP | Abnormality o | HP:0009811 | 0.0080435 | 2.09455471 | 238 | 839 |
| HP | Abnormal aort | HP:0031567 | 0.00807343 | 2.09294168 | 87 | 839 |
| HP | Pointed chin | HP:0000307 | 0.0085567 | 2.06769389 | 80 | 839 |
| HP | Morphologica | HP:0002492 | 0.00897591 | 2.04692144 | 43 | 839 |
| HP | Abnormal carc | HP:0001713 | 0.00920666 | 2.0358977 | 543 | 839 |
| HP | Abnormal 5th | HP:0004207 | 0.00930058 | 2.03148987 | 392 | 839 |
| HP | Polymicrogyria | HP:0002126 | 0.00950053 | 2.02225195 | 239 | 839 |
| HP | Agitation | HP:0000713 | 0.00986571 | 2.00587186 | 95 | 839 |
| HP | Abnormal soft | HP:0100736 | 0.01019787 | 1.99149035 | 210 | 839 |
| HP | Aplasia/Hypop | HP:0008772 | 0.01023215 | 1.99003307 | 141 | 839 |
| HP | Thick eyebrow | HP:0000574 | 0.01053533 | 1.97735172 | 114 | 839 |
| HP | Coarse facial f | HP:0000280 | 0.01078941 | 1.96700239 | 202 | 839 |
| HP | Thin upper lip | HP:0000219 | 0.01120118 | 1.95073617 | 240 | 839 |
| HP | Anteverted na | HP:0000463 | 0.01141475 | 1.94253347 | 545 | 839 |
| HP | Hypoplastic fi | HP:0011937 | 0.01186611 | 1.92569173 | 13 | 839 |
| HP | Abnormality o | HP:0000014 | 0.01230242 | 1.91000934 | 435 | 839 |
| HP | Absent thumb | HP:0009777 | 0.01295036 | 1.88771816 | 47 | 839 |
| HP | Coxa valga | HP:0002673 | 0.01311714 | 1.88216071 | 85 | 839 |
| HP | Lower limb sp | HP:0002061 | 0.01327212 | 1.87705959 | 199 | 839 |
| HP | Lower extremi | HP:0030311 | 0.01381565 | 1.85962867 | 267 | 839 |
| HP | Neoplasm of t | HP:0100273 | 0.01401347 | 1.85345433 | 78 | 839 |
| HP | Plagiocephaly | HP:0001357 | 0.01402889 | 1.85297669 | 100 | 839 |

| | | | | | | |
|---|---|---|---|---|---|---|
| **HP** | Hypoplastic fi | HP:0008398 | 0.0149127 | 1.82644362 | 11 | 839 |
| **HP** | Abnormality o | HP:0003028 | 0.0149168 | 1.82632428 | 127 | 839 |
| **HP** | Spastic parapl | HP:0001258 | 0.0149168 | 1.82632428 | 127 | 839 |
| **HP** | Non-motor se | HP:0033259 | 0.01512996 | 1.82016236 | 175 | 839 |
| **HP** | Enlarged post | HP:0005445 | 0.01584241 | 1.80017877 | 143 | 839 |
| **HP** | Cerebellar mal | HP:0002438 | 0.01605033 | 1.79451606 | 347 | 839 |
| **HP** | Clinodactyly | HP:0030084 | 0.01614024 | 1.79209012 | 474 | 839 |
| **HP** | Abnormality o | HP:0002644 | 0.01625595 | 1.78898763 | 539 | 839 |
| **HP** | Abnormal mitr | HP:0001633 | 0.0165698 | 1.78068272 | 112 | 839 |
| **HP** | Abnormality o | HP:0025031 | 0.01663772 | 1.77890627 | 2542 | 839 |
| **HP** | Triangular sha | HP:0009774 | 0.01675847 | 1.77576571 | 192 | 839 |
| **HP** | Abnormality o | HP:0005288 | 0.0167596 | 1.77573625 | 558 | 839 |
| **HP** | Delayed crania | HP:0000270 | 0.01681097 | 1.77440724 | 61 | 839 |
| **HP** | Abnormal mal | HP:0032076 | 0.01742548 | 1.75881514 | 321 | 839 |
| **HP** | Displacement | HP:0100627 | 0.01742548 | 1.75881514 | 321 | 839 |
| **HP** | Myoclonus | HP:0001336 | 0.01778604 | 1.74992079 | 330 | 839 |
| **HP** | Abnormal elas | HP:0010647 | 0.01795882 | 1.74572229 | 180 | 839 |
| **HP** | Deviation of tl | HP:0009484 | 0.01823575 | 1.73907647 | 411 | 839 |
| **HP** | Short thumb | HP:0009778 | 0.01827583 | 1.7381229 | 101 | 839 |
| **HP** | Myopia | HP:0000545 | 0.01830289 | 1.73748036 | 348 | 839 |
| **HP** | Abnormality o | HP:0030236 | 0.01891864 | 1.72311011 | 611 | 839 |
| **HP** | Neoplasm of t | HP:0100006 | 0.01960901 | 1.70754438 | 144 | 839 |
| **HP** | Abnormal fing | HP:0005918 | 0.020931 | 1.67920999 | 631 | 839 |
| **HP** | Abnormal hea | HP:0031653 | 0.02118217 | 1.67402949 | 296 | 839 |
| **HP** | Abnormality o | HP:0000032 | 0.02182069 | 1.66113143 | 1021 | 839 |
| **HP** | Abnormal ante | HP:0004328 | 0.02187693 | 1.66001361 | 1141 | 839 |
| **HP** | Sleep apnea | HP:0010535 | 0.02232217 | 1.6512636 | 98 | 839 |
| **HP** | Growth delay | HP:0001510 | 0.02250091 | 1.64779982 | 1617 | 839 |
| **HP** | Highly arched | HP:0002553 | 0.02253654 | 1.64711283 | 173 | 839 |
| **HP** | Congenital hip | HP:0001374 | 0.02290043 | 1.64015639 | 76 | 839 |
| **HP** | Osteolysis | HP:0002797 | 0.02290043 | 1.64015639 | 76 | 839 |
| **HP** | Shoulder dislo | HP:0003834 | 0.02300443 | 1.63818846 | 16 | 839 |
| **HP** | Myelodysplasi | HP:0002863 | 0.02367481 | 1.62571343 | 102 | 839 |
| **HP** | Bilateral tonic- | HP:0002069 | 0.02389662 | 1.62166348 | 275 | 839 |
| **HP** | Generalized-o | HP:0002197 | 0.02421289 | 1.6159534 | 319 | 839 |
| **HP** | Cerebellar hyp | HP:0001321 | 0.02535812 | 1.59588298 | 355 | 839 |
| **HP** | Abnormal hip | HP:0003272 | 0.02618533 | 1.58194202 | 483 | 839 |
| **HP** | Small for gest | HP:0001518 | 0.02628695 | 1.58025976 | 224 | 839 |
| **HP** | Thoracic aortic | HP:0012727 | 0.0289595 | 1.53820893 | 99 | 839 |
| **HP** | Abnormal join | HP:0001367 | 0.02956042 | 1.52928938 | 1010 | 839 |
| **HP** | Aplasia/Hypop | HP:0008771 | 0.02973523 | 1.52672876 | 146 | 839 |
| **HP** | Abnormality o | HP:0002693 | 0.03113986 | 1.50668336 | 183 | 839 |
| **HP** | Patent ductus | HP:0001643 | 0.03222621 | 1.49179078 | 264 | 839 |
| **HP** | Abnormality o | HP:0000795 | 0.03250124 | 1.48810001 | 375 | 839 |
| **HP** | Abnormal fear | HP:0100852 | 0.03254545 | 1.48750967 | 348 | 839 |

| | | | | | | |
|---|---|---|---|---|---|---|
| **HP** | Restlessness | HP:0000711 | 0.03305709 | 1.48073538 | 111 | 839 |
| **HP** | Abnormality o | HP:0000429 | 0.03381193 | 1.47093009 | 617 | 839 |
| **HP** | Abnormality o | HP:0010461 | 0.03475319 | 1.4590053 | 1052 | 839 |
| **HP** | Deviation of th | HP:0009179 | 0.03594989 | 1.44430247 | 291 | 839 |
| **HP** | Blepharophim | HP:0000581 | 0.0366186 | 1.4362983 | 135 | 839 |
| **HP** | Abnormality o | HP:0000164 | 0.0366912 | 1.43543807 | 840 | 839 |
| **HP** | Abnormality o | HP:0025032 | 0.03796801 | 1.4205822 | 1882 | 839 |
| **HP** | Gastrointestin | HP:0002672 | 0.03933243 | 1.40524927 | 67 | 839 |
| **HP** | Malignant gas | HP:0006749 | 0.03933243 | 1.40524927 | 67 | 839 |
| **HP** | Morphologica | HP:0002062 | 0.03977532 | 1.40038636 | 60 | 839 |
| **HP** | Abnormal eyel | HP:0000534 | 0.04157843 | 1.3811319 | 520 | 839 |
| **HP** | Abnormal hea | HP:0001627 | 0.04176434 | 1.37919435 | 1251 | 839 |
| **HP** | Slender finger | HP:0001238 | 0.04439488 | 1.35266708 | 124 | 839 |
| **HP** | Unsteady gait | HP:0002317 | 0.0450831 | 1.34598624 | 140 | 839 |
| **HP** | Abnormal uml | HP:0001551 | 0.04536943 | 1.34323665 | 262 | 839 |
| **HP** | Hyperlordosis | HP:0003307 | 0.04700442 | 1.32786131 | 177 | 839 |
| **HP** | Abnormality o | HP:0011121 | 0.04840374 | 1.31512104 | 1749 | 839 |
| **HP** | Abnormality o | HP:0000235 | 0.04971597 | 1.30350409 | 374 | 839 |
| **HP** | Aplasia/Hypop | HP:0100371 | 0.04978964 | 1.30286102 | 12 | 839 |
| **HP** | Abnormality o | HP:0010392 | 0.04978964 | 1.30286102 | 12 | 839 |
| **HP** | Aplasia/Hypop | HP:0010383 | 0.04978964 | 1.30286102 | 12 | 839 |
| **HP** | Abnormality o | HP:0010342 | 0.04978964 | 1.30286102 | 12 | 839 |
| **HP** | Aplasia/Hypop | HP:0010343 | 0.04978964 | 1.30286102 | 12 | 839 |

| | | |
|---|---|---|
| 605 | 4511 | ORC4,COL1A1,KCNQ1OT1,HNRNPA1,NEDD4L,ACTB,TUBA1A,HTT,S |
| 538 | 4511 | COL1A1,HNRNPA1,NEDD4L,ACTB,TUBA1A,HTT,SPARC,TUBB,KIDINS |
| 463 | 4511 | COL1A1,HNRNPA1,NEDD4L,ACTB,TUBA1A,HTT,SPARC,TUBB,KIDINS |
| 358 | 4511 | COL1A1,KCNQ1OT1,HNRNPA1,HSPA9,NEDD4L,ATN1,ACTB,ACTG1,X |
| 702 | 4511 | ORC4,COL1A1,KCNQ1OT1,HNRNPA1,HSPA9,NEDD4L,POLH,ATN1,A |
| 735 | 4511 | ORC4,COL1A1,KCNQ1OT1,HNRNPA1,HSPA9,NEDD4L,POLH,ATN1,A |
| 165 | 4511 | NEDD4L,ATN1,TUBA1A,TUBB,KIDINS220,DCX,PIK3R2,MAP1B,FBXL |
| 546 | 4511 | ORC4,COL1A1,KCNQ1OT1,HNRNPA1,HSPA9,NEDD4L,ATN1,ACTB,A |
| 608 | 4511 | ORC4,COL1A1,KCNQ1OT1,HNRNPA1,HSPA9,NEDD4L,ATN1,ACTB,C |
| 261 | 4511 | COL1A1,KCNQ1OT1,ATN1,ACTB,ACTG1,XRCC2,TUBB,KIDINS220,PI |
| 439 | 4511 | ORC4,HNRNPA1,HSPA9,NEDD4L,ATN1,ACTB,CTSC,ACTG1,TUBA1A, |
| 433 | 4511 | ORC4,HNRNPA1,HSPA9,NEDD4L,ATN1,ACTB,CTSC,ACTG1,TUBA1A, |
| 717 | 4511 | ORC4,COL1A1,KCNQ1OT1,COPA,HNRNPA1,HSPA9,NEDD4L,ATN1,A |
| 410 | 4511 | ORC4,KCNQ1OT1,HSPA9,NEDD4L,ATN1,ACTB,ACTG1,TUBA1A,XRC |
| 208 | 4511 | COL1A1,ACTB,ACTG1,XRCC2,TUBB,PIK3R2,FLNA,DVL3,YWHAE,ALD |
| 567 | 4511 | ORC4,COL1A1,KCNQ1OT1,HNRNPA1,HSPA9,NEDD4L,ATN1,ACTB,A |
| 492 | 4511 | ORC4,COL1A1,KCNQ1OT1,HNRNPA1,HSPA9,NEDD4L,ACTB,ACTG1, |
| 285 | 4511 | COL1A1,KCNQ1OT1,HNRNPA1,NEDD4L,ATN1,ACTB,ACTG1,XRCC2, |
| 439 | 4511 | COL1A1,KCNQ1OT1,COPA,HNRNPA1,HSPA9,NEDD4L,ATN1,ACTB,A |
| 222 | 4511 | ORC4,ATN1,ACTB,ACTG1,TUBB,FLNA,MAP1B,SLC9A7,UBA1,SOX4,K |
| 357 | 4511 | ORC4,ACTB,ACTG1,TUBA1A,XRCC2,TUBB,DCX,PIK3R2,MAP1B,SLC |
| 349 | 4511 | ORC4,HSPA9,NEDD4L,ATN1,ACTB,ACTG1,TUBA1A,XRCC2,TUBB,KID |
| 524 | 4511 | ORC4,COL1A1,KCNQ1OT1,HNRNPA1,HSPA9,NEDD4L,ATN1,ACTB,C |
| 554 | 4511 | ORC4,COL1A1,KCNQ1OT1,HNRNPA1,NEDD4L,ATN1,ACTB,ACTG1,E |
| 476 | 4511 | ORC4,COL1A1,KCNQ1OT1,HNRNPA1,HSPA9,NEDD4L,ATN1,ACTB,C |
| 606 | 4511 | ORC4,COL1A1,KCNQ1OT1,COPA,HNRNPA1,HSPA9,NEDD4L,ATN1,A |
| 256 | 4511 | HSPA9,NEDD4L,ATN1,ACTB,ACTG1,TUBA1A,HTT,TUBB,KIDINS220,D |
| 562 | 4511 | ORC4,COL1A1,KCNQ1OT1,HNRNPA1,HSPA9,NEDD4L,ATN1,ACTB,C |
| 471 | 4511 | ORC4,COL1A1,HSPA9,NEDD4L,ATN1,ACTB,ACTG1,TUBA1A,XRCC2, |
| 204 | 4511 | ORC4,KCNQ1OT1,HSPA9,ACTB,ACTG1,FLNA,MAP1B,COL6A1,FBXL |
| 175 | 4511 | ORC4,ATN1,ACTB,ACTG1,TUBB,FLNA,MAP1B,SLC9A7,UBA1,SOX4,K |
| 261 | 4511 | COL1A1,KCNQ1OT1,NEDD4L,ATN1,ACTB,ACTG1,XRCC2,HTT,SPARC |
| 357 | 4511 | COL1A1,HNRNPA1,NEDD4L,ATN1,ACTB,ACTG1,EEF2,HTT,SPARC,TU |
| 306 | 4511 | ORC4,COL1A1,HSPA9,ATN1,ACTB,ACTG1,TUBB,PIK3R2,FLNA,SLC9 |
| 94 | 4511 | COL1A1,KCNQ1OT1,TUBB,FLNA,UBA1,COL5A1,FKBP14,DVL3,YWH |
| 226 | 4511 | HSPA9,NEDD4L,ATN1,ACTB,ACTG1,TUBA1A,TUBB,KIDINS220,DCX,P |
| 391 | 4511 | ORC4,COL1A1,HNRNPA1,HSPA9,NEDD4L,ATN1,ACTB,CTSC,ACTG1, |
| 273 | 4511 | HSPA9,NEDD4L,ATN1,ACTB,ACTG1,TUBA1A,HTT,TUBB,KIDINS220,D |
| 331 | 4511 | ORC4,COL1A1,ATN1,ACTB,CTSC,ACTG1,XRCC2,HTT,SPARC,TUBB,F |
| 601 | 4511 | COL1A1,LRP1,KCNQ1OT1,HNRNPA1,NEDD4L,POLH,ATN1,ACTB,ACT |
| 223 | 4511 | ORC4,COL1A1,KCNQ1OT1,NEDD4L,ACTB,TUBB,PIK3R2,FLNA,MAP1 |
| 413 | 4511 | NEDD4L,ATN1,ACTB,ACTG1,TUBA1A,HTT,DCX,PIK3R2,FLNA,MAP1B, |
| 22 | 4511 | RPL5,RPL15,RPL18,RPL11,RPS17,RPS24,RPS14,RPL26,RPL31,RPS |
| 297 | 4511 | ORC4,ACTB,ACTG1,TUBA1A,XRCC2,TUBB,DCX,MAP1B,FBXL4,TUBB |
| 25 | 4511 | ALDOA,RPL5,TPI1,RPL15,SF3B1,RPL18,RPL11,RPS17,RPS24,HK1,R |

| | | |
|---|---|---|
| 618 | 4511 | ORC4,COL1A1,KCNQ1OT1,COPA,HNRNPA1,HSPA9,NEDD4L,ATN1,A |
| 352 | 4511 | ORC4,COL1A1,ATN1,ACTB,CTSC,ACTG1,XRCC2,HTT,SPARC,TUBB, |
| 299 | 4511 | ORC4,ACTB,ACTG1,TUBA1A,XRCC2,TUBB,DCX,MAP1B,FBXL4,TUBB |
| 385 | 4511 | COL1A1,NEDD4L,ACTB,TUBA1A,HTT,SPARC,TUBB,KIDINS220,DCX, |
| 339 | 4511 | COL1A1,LRP1,HSPA9,POLH,ATN1,ACTB,ACTG1,XRCC2,TUBB,PIK3R |
| 365 | 4511 | ORC4,COL1A1,HSPA9,NEDD4L,ATN1,ACTB,CTSC,ACTG1,XRCC2,HT |
| 365 | 4511 | ORC4,COL1A1,HSPA9,NEDD4L,ATN1,ACTB,CTSC,ACTG1,XRCC2,HT |
| 20 | 4511 | RPL5,RPL15,RPL18,RPL11,RPS17,RPS24,RPL26,RPL31,RPS7,RPS1 |
| 20 | 4511 | RPL5,RPL15,RPL18,RPL11,RPS17,RPS24,RPL26,RPL31,RPS7,RPS1 |
| 20 | 4511 | RPL5,RPL15,RPL18,RPL11,RPS17,RPS24,RPL26,RPL31,RPS7,RPS1 |
| 151 | 4511 | COL1A1,ATN1,ACTB,ACTG1,XRCC2,TUBB,FLNA,MAP1B,ALDOA,FBX |
| 393 | 4511 | ORC4,KCNQ1OT1,HNRNPA1,NEDD4L,ACTB,SPARC,KIDINS220,DCX, |
| 117 | 4511 | ORC4,ACTB,ACTG1,TUBB,PIK3R2,FLNA,SOX4,KAT6A,YWHAE,NONO, |
| 22 | 4511 | RPL5,RPL15,RPL18,RPL11,RPS17,RPS24,RPL26,RPL31,RPS7,RPS1 |
| 623 | 4511 | ORC4,COL1A1,LRP1,KCNQ1OT1,HNRNPA1,HSPA9,NEDD4L,POLH,A |
| 292 | 4511 | COL1A1,LRP1,POLH,ATN1,ACTB,ACTG1,XRCC2,TUBB,PIK3R2,FLNA, |
| 108 | 4511 | ORC4,COL1A1,NEDD4L,ATN1,FLNA,MAP1B,COL6A1,COL5A1,SOX4, |
| 229 | 4511 | HNRNPA1,NEDD4L,HTT,SPARC,TUBB,KIDINS220,DCX,FLNA,MAP1B, |
| 327 | 4511 | ORC4,COL1A1,NEDD4L,ATN1,ACTB,CTSC,ACTG1,XRCC2,TUBB,PIK |
| 122 | 4511 | COL1A1,HSPA9,XRCC2,PIK3R2,KAT6A,DVL3,RPL5,AFF4,NONO,RPL1 |
| 452 | 4511 | ORC4,COL1A1,KCNQ1OT1,HNRNPA1,HSPA9,NEDD4L,ATN1,ACTB,C |
| 149 | 4511 | COL1A1,KCNQ1OT1,NEDD4L,XRCC2,TUBB,FLNA,MAP1B,UBA1,COL |
| 122 | 4511 | COL1A1,HSPA9,XRCC2,PIK3R2,KAT6A,DVL3,RPL5,AFF4,NONO,RPL1 |
| 617 | 4511 | ORC4,COL1A1,LRP1,KCNQ1OT1,HNRNPA1,HSPA9,NEDD4L,POLH,A |
| 330 | 4511 | COL1A1,LRP1,HSPA9,POLH,ATN1,ACTB,ACTG1,XRCC2,TUBB,PIK3R |
| 20 | 4511 | RPL5,RPL15,RPL18,RPL11,RPS17,RPS24,RPL26,RPL31,RPS7,RPS1 |
| 248 | 4511 | ORC4,ATN1,ACTB,ACTG1,TUBB,FLNA,MAP1B,SLC9A7,FBXL4,UBA1, |
| 205 | 4511 | HSPA9,NEDD4L,ATN1,ACTB,ACTG1,TUBA1A,TUBB,KIDINS220,DCX,P |
| 32 | 4511 | FLNA,RPL5,RPL15,RPL18,RPL11,RPS17,RPS24,RPL26,RPL31,CDH2, |
| 153 | 4511 | ACTB,ACTG1,FLNA,PRPF8,UBA1,SOX4,DVL3,IDS,RPL5,SNRNP200,D |
| 251 | 4511 | COL1A1,KCNQ1OT1,ATN1,ACTB,ACTG1,XRCC2,KIDINS220,PIK3R2, |
| 278 | 4511 | ORC4,KCNQ1OT1,HSPA9,ATN1,ACTB,ACTG1,XRCC2,TUBB,FLNA,M |
| 256 | 4511 | COL1A1,ACTB,TUBA1A,TUBB,SLC9A7,COL6A1,FBXL4,PSAP,SOX4,K |
| 325 | 4511 | COL1A1,LRP1,HSPA9,POLH,ATN1,ACTB,ACTG1,XRCC2,TUBB,PIK3R |
| 65 | 4511 | SLC9A7,NONO,SET,CHD3,POLR2A,HK1,HNRNPK,NARS1,LMNB2,SN |
| 201 | 4511 | HSPA9,NEDD4L,ATN1,ACTB,ACTG1,TUBA1A,TUBB,KIDINS220,DCX,P |
| 501 | 4511 | LRP1,NEDD4L,POLH,ATN1,ACTB,ACTG1,EEF2,XRCC2,HTT,KIDINS22 |
| 182 | 4511 | ORC4,HSPA9,ATN1,XRCC2,TUBB,FLNA,MAP1B,COL6A1,NEFL,DVL3, |
| 170 | 4511 | COL1A1,HSPA9,ATN1,XRCC2,PIK3R2,FLNA,SOX4,KAT6A,DVL3,CAL |
| 255 | 4511 | ORC4,HSPA9,NEDD4L,ATN1,ACTB,XRCC2,TUBB,FLNA,MAP1B,COL6 |
| 250 | 4511 | NEDD4L,XRCC2,KIDINS220,FLNA,SLC9A7,COL6A1,FTL,UBA1,TUBB2 |
| 108 | 4511 | COL1A1,NEDD4L,XRCC2,PIK3R2,FLNA,MAP1B,GANAB,COL5A1,FKB |
| 265 | 4511 | ORC4,COL1A1,ATN1,ACTB,CTSC,ACTG1,XRCC2,TUBB,PIK3R2,FLNA |
| 306 | 4511 | COL1A1,NEDD4L,ATN1,ACTB,CTSC,XRCC2,HTT,FLNA,MAP1B,SLC9A |
| 261 | 4511 | HNRNPA1,ATN1,EEF2,HTT,FLNA,MAP1B,FTL,NEFL,UBA1,TUBB2B,P |

| | | |
|---|---|---|
| 431 | 4511 | ORC4,COL1A1,KCNQ1OT1,HSPA9,NEDD4L,ATN1,ACTB,CTSC,ACTG1 |
| 431 | 4511 | ORC4,COL1A1,KCNQ1OT1,HSPA9,NEDD4L,ATN1,ACTB,CTSC,ACTG1 |
| 356 | 4511 | ORC4,COL1A1,HSPA9,NEDD4L,ATN1,ACTB,CTSC,XRCC2,HTT,TUBB, |
| 212 | 4511 | ORC4,COL1A1,KCNQ1OT1,HNRNPA1,HSPA9,ACTB,ACTG1,XRCC2,TU |
| 245 | 4511 | COL1A1,ACTB,ACTG1,TUBB,PIK3R2,FLNA,SLC9A7,PRPF8,UBA1,SOX |
| 428 | 4511 | HNRNPA1,ATN1,ACTB,EEF2,XRCC2,HTT,KIDINS220,DCX,FLNA,MAP1 |
| 96 | 4511 | COL1A1,KIDINS220,SLC9A7,NONO,SET,PPP2R1A,CHD3,POLR2A,AG |
| 258 | 4511 | HNRNPA1,ATN1,HTT,FTL,NEFL,FBXL4,UBA1,PSAP,PGK1,KAT6A,KIF1 |
| 20 | 4511 | RPL5,RPL15,RPL18,RPL11,RPS17,RPS24,RPL26,RPL31,RPS7,RPS19 |
| 104 | 4511 | HTT,KIF1A,SLC2A3,VCP,MAPT,NONO,DCHS1,SET,AP2M1,ZMIZ1,RPL |
| 215 | 4511 | NEDD4L,HTT,SPARC,TUBB,KIDINS220,FLNA,MAP1B,SLC9A7,PGK1,K |
| 22 | 4511 | RPL5,RPL15,RPL18,RPL11,RPS17,RPS24,RPL26,RPL31,RPS7,RPS19 |
| 566 | 4511 | ORC4,COL1A1,LRP1,KCNQ1OT1,HNRNPA1,HSPA9,NEDD4L,POLH,AT |
| 247 | 4511 | HNRNPA1,TUBA1A,HTT,KIDINS220,DCX,FLNA,FTL,NEFL,COL4A1,PS |
| 149 | 4511 | COL1A1,TUBB,PIK3R2,FLNA,SLC9A7,SOX4,DVL3,GNAS,ALDH6A1,RP |
| 142 | 4511 | HNRNPA1,ACTB,KIDINS220,FLNA,COL6A1,NEFL,PSAP,KAT6A,KIF1A, |
| 258 | 4511 | COL1A1,KCNQ1OT1,NEDD4L,ACTB,ACTG1,XRCC2,TUBB,FLNA,SLC |
| 234 | 4511 | ORC4,COL1A1,KCNQ1OT1,HNRNPA1,HSPA9,ACTB,ACTG1,XRCC2,TU |
| 327 | 4511 | ORC4,COL1A1,KCNQ1OT1,HNRNPA1,HSPA9,ACTB,ACTG1,XRCC2,SP |
| 157 | 4511 | DCX,MAP1B,SLC9A7,KAT6A,KIF1A,GDI1,VCP,YWHAG,MAPT,NONO,S |
| 23 | 4511 | RPL5,RPL15,RPL18,RPL11,RPS17,RPS24,RPL26,RPL31,RPS7,RPS19 |
| 184 | 4511 | COL1A1,TUBA1A,SPARC,TUBB,KIDINS220,DCX,FLNA,SLC9A7,COL6 |
| 190 | 4511 | ORC4,ATN1,ACTB,TUBB,FLNA,COL4A1,SOX4,KAT6A,DVL3,YWHAE,R |
| 379 | 4511 | ORC4,HSPA9,NEDD4L,ATN1,ACTB,ACTG1,XRCC2,HTT,TUBB,KIDINS |
| 237 | 4511 | NEDD4L,XRCC2,KIDINS220,FLNA,SLC9A7,COL6A1,UBA1,TUBB2B,C |
| 21 | 4511 | RPL5,RPL15,RPL18,RPL11,RPS17,RPS24,RPL26,RPL31,RPS7,RPS19 |
| 277 | 4511 | ORC4,COL1A1,NEDD4L,ATN1,ACTB,ACTG1,DCX,PIK3R2,FLNA,MAP1 |
| 197 | 4511 | ORC4,ATN1,ACTB,TUBB,FLNA,MAP1B,COL4A1,SOX4,KAT6A,DVL3,Y |
| 400 | 4511 | ORC4,KCNQ1OT1,HNRNPA1,NEDD4L,ACTB,ACTG1,TUBA1A,XRCC2, |
| 399 | 4511 | HNRNPA1,ATN1,EEF2,TUBA1A,HTT,TUBB,KIDINS220,DCX,FLNA,SLC |
| 88 | 4511 | ACTB,TUBB,FLNA,COL5A1,KAT6A,DVL3,GNB1,PPP2R1A,POLR2A,ZM |
| 20 | 4511 | RPL5,RPL15,RPL18,RPL11,RPS17,RPS24,RPL26,RPL31,RPS7,RPS19 |
| 62 | 4511 | ACTB,ACTG1,SOX4,KAT6A,NONO,SET,ZMIZ1,GPC3,ARID1A,SNRPN,K |
| 212 | 4511 | HNRNPA1,TUBA1A,HTT,KIDINS220,DCX,FLNA,FTL,NEFL,COL4A1,PS |
| 151 | 4511 | COL1A1,LRP1,KCNQ1OT1,HSPA9,ACTB,ACTG1,KIDINS220,FLNA,SL |
| 181 | 4511 | ORC4,COL1A1,NEDD4L,ATN1,ACTB,HTT,DCX,FLNA,MAP1B,FTL,FBX |
| 121 | 4511 | PIK3R2,SOX4,KAT6A,DVL3,CALM3,RPL5,AFF4,NONO,RPL15,RPL18, |
| 124 | 4511 | COL1A1,KCNQ1OT1,XRCC2,TUBB,FLNA,UBA1,COL5A1,FKBP14,DVL |
| 127 | 4511 | COL1A1,ACTB,HTT,PIK3R2,FLNA,COL6A1,UBA1,SOX4,FKBP14,DVL3 |
| 123 | 4511 | COL1A1,KCNQ1OT1,XRCC2,TUBB,FLNA,UBA1,COL5A1,FKBP14,DVL |
| 119 | 4511 | PIK3R2,SOX4,KAT6A,DVL3,CALM3,RPL5,AFF4,NONO,RPL15,RPL18, |
| 206 | 4511 | HNRNPA1,ATN1,ACTB,ACTG1,HTT,KIDINS220,FTL,NEFL,FBXL4,UBA |
| 263 | 4511 | ORC4,COL1A1,NEDD4L,ACTB,ACTG1,XRCC2,HTT,TUBB,FLNA,SLC9 |
| 175 | 4511 | ACTB,ACTG1,XRCC2,PIK3R2,FLNA,ALDOA,NEFL,UBA1,SOX4,KAT6A, |
| 132 | 4511 | HTT,MAP1B,SLC9A7,TUBB2B,GDI1,IGF2,YWHAG,PPP2R1A,ZMIZ1,AP |

| | | |
|---|---|---|
| 95 | 4511 | COL1A1,CTSC,XRCC2,FLNA,SLC9A7,COL5A1,FKBP14,KAT6A,GDI1,F |
| 109 | 4511 | ATN1,ACTB,ACTG1,MAP1B,SLC9A7,KAT6A,ALDH6A1,GDI1,AFF4,NON |
| 191 | 4511 | ACTB,ACTG1,XRCC2,TUBB,PIK3R2,FLNA,SLC9A7,FBXL4,DVL3,YWH |
| 62 | 4511 | XRCC2,PIK3R2,SOX4,RPL5,AFF4,RPL15,RPL18,RPL11,RPS17,HNRN |
| 20 | 4511 | RPL5,RPL15,RPL18,RPL11,RPS17,RPS24,RPL26,RPL31,RPS7,RPS19 |
| 154 | 4511 | HSPA9,ATN1,ACTB,ACTG1,TUBB,FLNA,ALDOA,COL6A1,UBA1,DVL3, |
| 85 | 4511 | ACTB,ACTG1,FLNA,SLC9A7,ALDOA,RPL5,PPP2R1A,RPL15,RPL18,RP |
| 478 | 4511 | COL1A1,KCNQ1OT1,HNRNPA1,NEDD4L,POLH,ATN1,ACTB,ACTG1,X |
| 230 | 4511 | ORC4,COL1A1,NEDD4L,ACTB,ACTG1,XRCC2,TUBB,FLNA,COL6A1,U |
| 152 | 4511 | FLNA,SLC9A7,SOX4,KAT6A,DVL3,GNAS,RPL5,GDI1,IGF2,AFF4,CHSY |
| 289 | 4511 | HNRNPA1,ATN1,EEF2,TUBA1A,HTT,KIDINS220,DCX,FLNA,SLC9A7,FT |
| 132 | 4511 | COL1A1,KCNQ1OT1,HNRNPA1,XRCC2,TUBB,FLNA,UBA1,COL5A1,F |
| 239 | 4511 | ORC4,COL1A1,NEDD4L,ACTB,ACTG1,XRCC2,TUBB,FLNA,COL6A1,U |
| 128 | 4511 | COL1A1,NEDD4L,XRCC2,PIK3R2,FLNA,MAP1B,GANAB,UBA1,COL5A |
| 28 | 4511 | CALR,RPL5,RPL15,SF3B1,RPL18,RPL11,RPS17,RPSA,RPS24,RPS14, |
| 120 | 4511 | MAP1B,SLC9A7,TUBB2B,PSAP,IDS,GDI1,IGF2,YWHAG,UCHL1,ZMIZ1 |
| 160 | 4511 | COL1A1,NEDD4L,ACTB,XRCC2,PIK3R2,FLNA,MAP1B,GANAB,COL5A |
| 240 | 4511 | ORC4,COL1A1,ACTB,ACTG1,HTT,DCX,FLNA,COL6A1,PSAP,SOX4,KA |
| 211 | 4511 | KCNQ1OT1,ATN1,EEF2,TUBA1A,HTT,TUBB,FLNA,FTL,FBXL4,TUBB2 |
| 174 | 4511 | ORC4,ACTB,XRCC2,HTT,FLNA,SLC9A7,FBXL4,SOX4,KAT6A,DVL3,G |
| 329 | 4511 | ORC4,COL1A1,KCNQ1OT1,HNRNPA1,NEDD4L,ATN1,ACTB,ACTG1,N |
| 116 | 4511 | ACTB,TUBB,FLNA,COL5A1,KAT6A,DVL3,GNB1,FN1,PPP2R1A,POLR2 |
| 356 | 4511 | NEDD4L,ATN1,EEF2,XRCC2,HTT,KIDINS220,DCX,FLNA,SLC9A7,COL |
| 20 | 4511 | RPL5,RPL15,RPL18,RPL11,RPS17,RPS24,RPL26,RPL31,RPS7,RPS19 |
| 165 | 4511 | ORC4,COL1A1,NEDD4L,XRCC2,HTT,PIK3R2,FLNA,MAP1B,COL6A1,C |
| 125 | 4511 | COL1A1,HSPA9,XRCC2,PIK3R2,KAT6A,DVL3,RPL5,AFF4,NONO,RPL1 |
| 297 | 4511 | LRP1,HSPA9,ACTB,CTSC,ACTG1,TUBB,FLNA,MAP1B,SLC9A7,ALDOA |
| 23 | 4511 | RPL5,RPL15,SF3B1,RPL18,RPL11,RPS17,RPS24,RPS14,RPL26,RPL3 |
| 40 | 4511 | ORC4,COL1A1,NEDD4L,FLNA,MAP1B,COL5A1,SOX4,DVL3,ARF1,CO |
| 61 | 4511 | COL1A1,FLNA,GNAS,RPL5,RPL15,RPL18,RPL11,RPS17,RPS24,RPL2 |
| 93 | 4511 | FLNA,COL5A1,DVL3,GNAS,IDS,IGF2,NONO,DCHS1,SET,CHSY1,POLF |
| 211 | 4511 | KCNQ1OT1,ATN1,EEF2,TUBA1A,HTT,TUBB,FLNA,FTL,FBXL4,TUBB2 |
| 211 | 4511 | KCNQ1OT1,ATN1,EEF2,TUBA1A,HTT,TUBB,FLNA,FTL,FBXL4,TUBB2 |
| 113 | 4511 | ORC4,COL1A1,NEDD4L,ATN1,FLNA,MAP1B,FBXL4,COL5A1,PSAP,KA |
| 66 | 4511 | COL1A1,NEDD4L,FLNA,MAP1B,GANAB,COL5A1,COL4A1,AFF4,PKD1 |
| 104 | 4511 | ATN1,ACTB,ACTG1,TUBB,SLC9A7,UBA1,DVL3,ALDH6A1,GDI1,AFF4, |
| 92 | 4511 | ACTB,ACTG1,FLNA,SLC9A7,ALDOA,RPL5,PPP2R1A,RPL15,RPL18,RP |
| 268 | 4511 | LRP1,HSPA9,ACTB,CTSC,ACTG1,TUBB,FLNA,MAP1B,SLC9A7,ALDOA |
| 23 | 4511 | RPL5,RPL15,RPL18,RPL11,RPS17,RPS24,RPL26,RPL31,RPS7,RPS19 |
| 126 | 4511 | ACTB,ACTG1,FLNA,SLC9A7,FBXL4,DVL3,YWHAE,ALDH6A1,P4HB,GD |
| 352 | 4511 | ORC4,COL1A1,HSPA9,ATN1,ACTB,ACTG1,XRCC2,HTT,TUBB,NUMA1 |
| 24 | 4511 | COL1A1,RPL5,RPL15,RPL18,RPL11,RPS17,RPS24,COL1A2,RPL26,R |
| 170 | 4511 | NEDD4L,ATN1,ACTB,ACTG1,TUBA1A,TUBB,DCX,PIK3R2,MAP1B,TUB |
| 45 | 4511 | ACTB,ACTG1,DVL3,RPL5,RPL15,RPL18,RPL11,RPS17,RPS24,RPL26, |
| 186 | 4511 | ORC4,COL1A1,KCNQ1OT1,HNRNPA1,HSPA9,XRCC2,TUBB,FLNA,SL |

| | | |
|---|---|---|
| 30 | 4511 | UBA1,RPL5,RPL15,RPL18,RPL11,RPS17,RPS24,RPS14,RPL26,RPL3 |
| 147 | 4511 | XRCC2,HTT,KIDINS220,FLNA,COL5A1,COL4A1,FKBP14,KAT6A,COL1 |
| 182 | 4511 | HNRNPA1,ATN1,ACTB,ACTG1,HTT,KIDINS220,FTL,FBXL4,PSAP,YWH |
| 202 | 4511 | KCNQ1OT1,ACTB,CTSC,ACTG1,TUBA1A,XRCC2,HTT,TUBB,KIDINS22 |
| 70 | 4511 | HTT,SLC2A3,VCP,MAPT,NONO,AP2M1,ZMIZ1,DDX3X,CTNNB1,EEF1A |
| 21 | 4511 | RPL5,RPL15,RPL18,RPL11,RPS17,RPS24,RPL26,RPL31,RPS7,RPS1 |
| 141 | 4511 | HTT,DCX,PSAP,KIF1A,GNAS,SLC2A3,ATP5F1A,VCP,MAPT,NONO,DCH |
| 52 | 4511 | COL1A1,NEDD4L,FLNA,MAP1B,GANAB,COL5A1,AFF4,PKD1,ZMIZ1,A |
| 87 | 4511 | NEDD4L,HTT,KAT6A,KIF1A,PPP2R1A,RPL10,HNRNPU,NR2F1,TKT,CC |
| 55 | 4511 | ACTB,FLNA,NEFL,RPL5,VCP,RPL15,RPL18,RPL11,COL11A1,RPS17, |
| 82 | 4511 | COL1A1,KCNQ1OT1,HSPA9,ACTB,FLNA,SLC9A7,KAT6A,DVL3,KIF1A, |
| 93 | 4511 | HSPA9,ATN1,ACTB,ACTG1,FLNA,SLC9A7,KAT6A,ALDH6A1,GDI1,COL |
| 164 | 4511 | KCNQ1OT1,ACTB,ACTG1,TUBA1A,XRCC2,HTT,TUBB,KIDINS220,DC |
| 20 | 4511 | RPL5,RPL15,RPL18,RPL11,RPS17,RPS24,RPL26,RPL31,RPS7,RPS1 |
| 42 | 4511 | HTT,MAP1B,SLC2A3,VCP,YWHAG,MAPT,NR2F1,TKT,SQSTM1,SNRPN |
| 163 | 4511 | HNRNPA1,HTT,FTL,PSAP,PGK1,GNAS,SLC2A3,APP,VCP,YWHAG,MAP |
| 109 | 4511 | ATN1,FLNA,SLC9A7,SOX4,GNAS,RPL5,GDI1,IGF2,MAPT,RPL15,RPL1 |
| 89 | 4511 | FLNA,SLC9A7,SOX4,GNAS,RPL5,GDI1,IGF2,RPL15,RPL18,RPL11,ZM |
| 103 | 4511 | ORC4,COL1A1,NEDD4L,XRCC2,FLNA,MAP1B,COL6A1,COL5A1,SOX |
| 39 | 4511 | GNAS,RPL5,RPL15,RPL18,RPL11,RPS17,RPS24,RPS20,RPL26,RPL3 |
| 64 | 4511 | KIDINS220,DCX,FBXL4,ALDH6A1,GNB1,PPP2R1A,POLR2A,ZMIZ1,AP |
| 139 | 4511 | XRCC2,HTT,FLNA,SLC9A7,SOX4,DVL3,GNAS,RPL5,GDI1,IGF2,AFF4, |
| 132 | 4511 | XRCC2,HTT,FLNA,SLC9A7,SOX4,DVL3,GNAS,RPL5,GDI1,IGF2,AFF4, |
| 54 | 4511 | SLC9A7,RPL5,RPL15,RPL18,RPL11,RPS17,RPS24,RPL26,RPL31,LR |
| 159 | 4511 | ORC4,XRCC2,HTT,FLNA,SLC9A7,FBXL4,SOX4,KAT6A,DVL3,GNAS,R |
| 72 | 4511 | KAT6A,KIF1A,VCP,MAPT,SET,CHD3,NR2F1,TKT,CLTC,TARDBP,NARS |
| 97 | 4511 | COL1A1,KCNQ1OT1,POLH,ACTB,CTSC,TUBB,FLNA,GNAS,RPL5,COL |
| 93 | 4511 | MAP1B,SLC9A7,TUBB2B,GDI1,IGF2,YWHAG,ZMIZ1,ARF1,NR2F1,RP |
| 288 | 4511 | ORC4,COL1A1,HNRNPA1,HSPA9,ACTB,ACTG1,XRCC2,SPARC,TUBB, |
| 99 | 4511 | HSPA9,ACTB,ACTG1,TUBB,ALDOA,COL6A1,UBA1,DVL3,YWHAE,GNA |
| 58 | 4511 | ORC4,FLNA,COL4A1,FKBP14,RPL5,UBB,NONO,RPL15,RPL18,RPL11 |
| 35 | 4511 | ORC4,COL1A1,FLNA,COL5A1,SOX4,DVL3,COL5A2,PRKAR1A,ARID1 |
| 21 | 4511 | RPL5,CHSY1,RPL15,RPL18,RPL11,RPS17,RPS24,RPL26,RPL31,RPS |
| 498 | 4511 | ORC4,COL1A1,KCNQ1OT1,HNRNPA1,HSPA9,ACTB,ACTG1,XRCC2,H |
| 31 | 4511 | RPL5,NPM1,RPL15,SF3B1,RPL18,RPL11,RPS17,RPS24,RPS14,RPL2 |
| 373 | 4511 | COL1A1,HNRNPA1,POLH,ATN1,ACTB,ACTG1,HTT,NUMA1,DCX,MAP1 |
| 70 | 4511 | ACTB,ACTG1,SLC9A7,ALDOA,RPL5,RPL15,RPL18,RPL11,RPS17,RPS |
| 39 | 4511 | KIDINS220,DCX,TUBB2B,KIF1A,SET,CLTC,RAC1,GLUL,EEF1A2,SNR |
| 86 | 4511 | COL1A1,NEDD4L,ACTB,XRCC2,FLNA,MAP1B,GANAB,COL5A1,IDS,R |
| 31 | 4511 | HSPA9,DVL3,AFF4,NONO,COL11A1,TKT,RAC1,PPP1CB,SNRPN,ACAD |
| 26 | 4511 | ACTB,ACTG1,RPL5,CHSY1,RPL15,RPL18,RPL11,RPS17,RPS24,RPL2 |
| 12 | 4511 | SOX4,IGF2,GJA1,ARID1A,COL2A1,SMARCA4,SMARCE1,ARID1B,SMA |
| 68 | 4511 | KIDINS220,COL4A1,CHD3,TUBB4B,NR2F1,GJA1,CTNNB1,HNRNPK, |
| 88 | 4511 | ORC4,COL1A1,KCNQ1OT1,ACTB,ACTG1,FLNA,MAP1B,KAT6A,KIF1A |
| 133 | 4511 | ATN1,HTT,FTL,NEFL,KAT6A,KIF1A,SLC2A3,IDS,AFF4,MAPT,NONO,SE |

| | | |
|---|---|---|
| 11 | 4511 | SOX4,IGF2,ARID1A,COL2A1,SMARCA4,SMARCE1,ARID1B,SMARCD1, |
| 32 | 4511 | KIDINS220,DCX,TUBB2B,KIF1A,SET,CLTC,RAC1,GLUL,EEF1A2,SNRF |
| 67 | 4511 | COL1A1,FLNA,COL6A1,COL5A1,NONO,DCHS1,AGRN,COL1A2,COL5 |
| 165 | 4511 | ATN1,ACTB,ACTG1,HTT,KIDINS220,FTL,FBXL4,PSAP,YWHAE,KIF1A, |
| 151 | 4511 | DCX,MAP1B,NEFL,PSAP,YWHAE,KIF1A,GNB1,SPTAN1,VCP,YWHAG,M |
| 179 | 4511 | ORC4,COL1A1,NEDD4L,ACTB,ACTG1,XRCC2,TUBB,FLNA,COL6A1,U |
| 181 | 4511 | ORC4,COL1A1,NEDD4L,ACTB,ACTG1,XRCC2,TUBB,FLNA,COL6A1,U |
| 265 | 4511 | ORC4,COL1A1,NEDD4L,ACTB,ACTG1,XRCC2,TUBB,FLNA,COL6A1,U |
| 203 | 4511 | ORC4,COL1A1,HSPA9,ATN1,ACTB,ACTG1,FLNA,SLC9A7,PRPF8,FBXI |
| 160 | 4511 | ACTB,ACTG1,XRCC2,TUBB,FLNA,SLC9A7,FBXL4,DVL3,YWHAE,ALDH |
| 147 | 4511 | ORC4,NEDD4L,ATN1,ACTB,XRCC2,TUBB,FLNA,COL4A1,SOX4,FKBP |
| 25 | 4511 | COL1A1,COL5A1,COL5A2,TBCD,LMNB2,VPS35,ODC1,USP7,RAI1,TR |
| 178 | 4511 | ORC4,COL1A1,NEDD4L,ACTB,ACTG1,XRCC2,TUBB,FLNA,COL6A1,U |
| 95 | 4511 | XRCC2,FLNA,SLC9A7,SOX4,GNAS,RPL5,GDI1,IGF2,RPL15,RPL18,RF |
| 29 | 4511 | FLNA,RPL5,RPL15,RPL18,RPL11,RPS17,PXDN,RPS24,RPL26,RPL31, |
| 21 | 4511 | RPL5,CHSY1,RPL15,RPL18,RPL11,RPS17,RPS24,RPL26,RPL31,RPS |
| 147 | 4511 | ORC4,NEDD4L,ATN1,ACTB,XRCC2,TUBB,FLNA,COL4A1,SOX4,FKBP |
| 36 | 4511 | COL1A1,FLNA,GANAB,COL5A1,RPL5,DCHS1,PKD1,COL1A2,COL5A2 |
| 106 | 4511 | HSPA9,ACTB,ACTG1,TUBB,ALDOA,COL6A1,UBA1,DVL3,YWHAE,GNA |
| 161 | 4511 | NEDD4L,ATN1,ACTB,XRCC2,FLNA,SLC9A7,COL6A1,NEFL,SOX4,KAT |
| 43 | 4511 | RPL5,RPL15,RPL18,RPL11,RPS17,RPS24,CHD4,RPL26,RPL31,CDH2 |
| 67 | 4511 | FLNA,NEFL,PSAP,KAT6A,RPL5,VCP,RPL15,RPL18,RPL11,AGRN,RPS1 |
| 97 | 4511 | ATN1,ACTB,ACTG1,SLC9A7,KAT6A,GDI1,AFF4,NONO,AP2M1,ZMIZ1, |
| 161 | 4511 | ATN1,ACTB,ACTG1,HTT,KIDINS220,FBXL4,PSAP,YWHAE,KIF1A,SLC2 |
| 76 | 4511 | EEF2,HTT,SLC9A7,FTL,NEFL,COL4A1,PSAP,KIF1A,SLC2A3,GDI1,HSF |
| 76 | 4511 | EEF2,HTT,SLC9A7,FTL,NEFL,COL4A1,PSAP,KIF1A,SLC2A3,GDI1,HSF |
| 46 | 4511 | HSPA9,MAP1B,AFF4,CHSY1,HNRNPU,CDH2,SMCHD1,CDH11,ASXL1, |
| 74 | 4511 | CTSC,TUBB,FLNA,MAP1B,KIF1A,ZMIZ1,COL11A1,RPL10,MAT2A,COI |
| 223 | 4511 | ORC4,COL1A1,NEDD4L,ATN1,ACTB,HTT,DCX,FLNA,MAP1B,FTL,FBXI |
| 72 | 4511 | ATN1,UBA1,KAT6A,DVL3,KIF1A,IGF2,AFF4,NONO,DCHS1,PPP2R1A,C |
| 69 | 4511 | HSPA9,CTSC,MAP1B,IDS,AFF4,CHSY1,HNRNPU,CDH2,ARID1A,SMCHI |
| 101 | 4511 | KCNQ1OT1,ACTB,FLNA,KIF1A,IGF2,NONO,SET,ZMIZ1,PAFAH1B1,AT |
| 124 | 4511 | ORC4,COL1A1,XRCC2,FLNA,COL6A1,NEFL,UBA1,COL5A1,SOX4,DVL |
| 133 | 4511 | KIDINS220,DCX,NEFL,FBXL4,PSAP,KIF1A,ALDH6A1,GNB1,SPTAN1,H |
| 65 | 4511 | ORC4,TUBB,PIK3R2,FLNA,KAT6A,YWHAE,NONO,DCHS1,CHSY1,ZMIZ |
| 46 | 4511 | NEFL,KIF1A,HSPD1,VCP,MAPT,AGRN,HK1,L1CAM,VHL,PLEC,HSPB1, |
| 80 | 4511 | NEDD4L,DCX,FLNA,MAP1B,TUBB2B,KIF1A,GNB1,COL4A2,SET,AP2M |
| 217 | 4511 | COL1A1,HNRNPA1,HTT,SPARC,NUMA1,SLC9A7,COL6A1,NEFL,UBA1, |
| 131 | 4511 | DCX,MAP1B,NEFL,PSAP,YWHAE,KIF1A,GNB1,SPTAN1,VCP,YWHAG,M |
| 89 | 4511 | ORC4,XRCC2,TUBB,FLNA,FBXL4,UBA1,DVL3,RPL5,IGF2,AFF4,DCHS |
| 37 | 4511 | FLNA,NEFL,RPL5,VCP,RPL15,RPL18,RPL11,AGRN,RPS17,RPS24,RPL |
| 37 | 4511 | ORC4,COL1A1,NEDD4L,FLNA,MAP1B,COL5A1,SOX4,ARF1,COL5A2, |
| 62 | 4511 | KIDINS220,FLNA,DVL3,IGF2,CHD3,ZMIZ1,LRPPRC,CCND2,CDH2,PPF |
| 25 | 4511 | RPL5,RPL15,RPL18,RPL11,RPS17,CYB5R3,RPS24,RPL26,RPL31,RPS |
| 196 | 4511 | ORC4,COL1A1,NEDD4L,ACTB,ACTG1,XRCC2,TUBB,FLNA,COL6A1,U |

| | | |
|---|---|---|
| 61 | 4511 | ATN1,KIDINS220,SLC9A7,KAT6A,CDH2,GJA1,CTBP1,CRPPA,NOVA2,E |
| 53 | 4511 | KIF1A,DCHS1,SET,RPL10,ATP1A1,TKT,CDH2,CLTC,CTNNB1,EEF1A2, |
| 58 | 4511 | KCNQ1OT1,COL6A1,FKBP14,YWHAE,IGF2,NONO,ZMIZ1,PAFAH1B1, |
| 55 | 4511 | COL1A1,KCNQ1OT1,TUBB,FLNA,COL4A1,SOX4,IGF2,POLR2A,COL1 |
| 80 | 4511 | FLNA,MAP1B,NEFL,IDS,HSPD1,VCP,UCHL1,AGRN,ZMIZ1,HK1,ATP1A |
| 47 | 4511 | HSPA9,MAP1B,AFF4,CHSY1,HNRNPU,CDH2,SMCHD1,CDH11,ASXL1,I |
| 483 | 4511 | COL1A1,LRP1,KCNQ1OT1,HSPA9,NEDD4L,POLH,ACTB,CTSC,ACTG1 |
| 125 | 4511 | DCX,MAP1B,YWHAE,KIF1A,GNB1,SPTAN1,VCP,YWHAG,MAPT,AP2M1 |
| 152 | 4511 | ORC4,NEDD4L,ATN1,ACTB,ACTG1,XRCC2,TUBB,FLNA,COL4A1,SOX |
| 14 | 4511 | COL1A1,COL5A1,FKBP14,COL1A2,COL5A2,PLOD1,BAZ1B,LTBP4,G |
| 34 | 4511 | ACTB,FLNA,MAT2A,RPL26,MMP14,HNRNPK,RAC1,MMP2,FDFT1,RAI |
| 60 | 4511 | ACTB,ACTG1,XRCC2,FLNA,DVL3,GNAS,RPL5,CHSY1,RPL15,RPL18,F |
| 97 | 4511 | ACTB,ACTG1,XRCC2,FLNA,COL6A1,UBA1,DVL3,GNAS,RPL5,CHSY1,I |
| 135 | 4511 | COL1A1,DCX,FLNA,COL6A1,PSAP,SOX4,KIF1A,HSPD1,IGF2,UBB,AFF |
| 57 | 4511 | ORC4,COL1A1,FLNA,SOX4,P4HB,COL18A1,COL11A1,COL1A2,ARID1 |
| 76 | 4511 | ATN1,ACTB,PIK3R2,DVL3,YWHAE,ALDH6A1,YWHAG,POLR2A,ZMIZ1,I |
| 39 | 4511 | NUMA1,CALR,GNB1,RPL5,NPM1,RPL15,SF3B1,RPL18,RPL11,RPS17 |
| 39 | 4511 | RPL5,RPL15,RPL18,RPL11,RPS17,RPS24,RPL26,RPL31,LRPPRC,RPS |
| 90 | 4511 | HNRNPA1,ATN1,ACTB,HTT,FTL,COL4A1,PSAP,SLC2A3,APP,IDS,COL1 |
| 23 | 4511 | RPL5,RPL15,RPL18,RPL11,RPS17,RPS24,RPL26,RPL31,RPS7,RPS19 |
| 108 | 4511 | ORC4,ATN1,XRCC2,FLNA,MAP1B,SLC9A7,UBA1,DVL3,YWHAE,IGF2,N |
| 46 | 4511 | COL1A1,FLNA,GANAB,COL5A1,IDS,RPL5,NONO,DCHS1,PKD1,COL1 |
| 47 | 4511 | ORC4,HSPA9,MAP1B,RPL5,DCHS1,RPL15,RPL18,RPL11,RPS17,RPS |
| 72 | 4511 | ORC4,COL1A1,FLNA,COL6A1,COL5A1,SOX4,DVL3,COL5A2,PRKAR1 |
| 34 | 4511 | ACTB,FLNA,MAT2A,RPL26,MMP14,HNRNPK,RAC1,MMP2,FDFT1,RAI |
| 32 | 4511 | ACTB,ACTG1,FLNA,KAT6A,CDH2,CDH11,SKI,NOTCH2,MAPK1,MESD, |
| 21 | 4511 | VCP,TARDBP,L1CAM,SQSTM1,SOD1,RAI1,DCTN1,BAZ1B,FUS,SPG7, |
| 140 | 4511 | PIK3R2,FLNA,SOX4,KAT6A,DVL3,CALM3,RPL5,AFF4,NONO,RPL15,R |
| 107 | 4511 | ORC4,XRCC2,PIK3R2,FLNA,MAP1B,SLC9A7,SOX4,DVL3,YWHAE,GNA |
| 72 | 4511 | ATN1,ACTB,ACTG1,TUBA1A,TUBB,PIK3R2,MAP1B,TUBB2B,COL4A1, |
| 36 | 4511 | HNRNPA1,HTT,DCX,SLC2A3,APP,VCP,UCHL1,PFN1,PRKAR1A,TARDB |
| 65 | 4511 | ORC4,XRCC2,FLNA,COL4A1,FKBP14,RPL5,UBB,NONO,RPL15,RPL1 |
| 48 | 4511 | ORC4,HSPA9,MAP1B,RPL5,DCHS1,RPL15,RPL18,RPL11,RPS17,RPS |
| 41 | 4511 | FBXL4,SOX4,AFF4,COL1A2,ARID1A,CDH11,NOTCH2,SMARCA4,SMA |
| 63 | 4511 | KCNQ1OT1,ACTB,ACTG1,FLNA,SOX4,IDS,IGF2,AFF4,CHD4,GPC3,MI |
| 72 | 4511 | ATN1,ACTB,ACTG1,SLC9A7,KAT6A,GDI1,AFF4,NONO,AP2M1,RPL10, |
| 140 | 4511 | ACTB,SLC9A7,PRPF8,SOX4,KAT6A,DVL3,YWHAE,KIF1A,GDI1,SNRNP |
| 10 | 4511 | SOX4,ARID1A,SMARCA4,SMARCE1,ARID1B,SMARCD1,PPP2R5D,SMA |
| 116 | 4511 | COL1A1,KCNQ1OT1,HNRNPA1,HSPA9,FLNA,SLC9A7,NEFL,COL5A1,I |
| 22 | 4511 | RPL5,RPL15,RPL18,RPL11,RPS17,RPS24,RPL26,RPL31,RPS7,RPS19 |
| 33 | 4511 | COL1A1,FLNA,SOX4,DVL3,COL11A1,COL1A2,ARID1A,SHOX,ARCN1, |
| 62 | 4511 | KIDINS220,PSAP,KIF1A,HSPD1,VCP,UCHL1,GJA1,ARL6IP1,L1CAM,M |
| 78 | 4511 | COL1A1,NEDD4L,XRCC2,FLNA,MAP1B,COL6A1,COL5A1,DVL3,YWH |
| 31 | 4511 | RPL5,RPL15,RPL18,RPL11,RPS17,RPS24,RPS20,RPL26,RPL31,RPS7 |
| 37 | 4511 | KIDINS220,FBXL4,KAT6A,AFF4,NONO,SET,CHSY1,PPP2R1A,POLR2A |

| | | |
|---|---|---|
| 9 | 4511 | SOX4,ARID1A,SMARCA4,SMARCE1,ARID1B,SMARCD1,SMARCC2,SM |
| 44 | 4511 | HTT,FLNA,COL6A1,PSAP,KIF1A,NONO,UCHL1,RPL10,KLHL9,COL6A |
| 44 | 4511 | KIDINS220,KIF1A,HSPD1,VCP,UCHL1,ARL6IP1,L1CAM,MARS1,SDHA, |
| 56 | 4511 | SLC9A7,KIF1A,GDI1,SET,AP2M1,PAFAH1B1,CLTC,HNRNPK,ATP1A2, |
| 48 | 4511 | KCNQ1OT1,TUBB,COL4A1,SOX4,IGF2,POLR2A,CDH2,GPC3,ARID1A, |
| 96 | 4511 | KCNQ1OT1,ATN1,TUBA1A,HTT,TUBB,FLNA,COL4A1,SOX4,KIF1A,IG |
| 124 | 4511 | ORC4,XRCC2,FLNA,MAP1B,SLC9A7,SOX4,DVL3,YWHAE,GDI1,IGF2, |
| 138 | 4511 | COL1A1,ATN1,XRCC2,FLNA,COL6A1,COL5A1,SOX4,DVL3,YWHAE,KI |
| 40 | 4511 | COL1A1,FLNA,GANAB,COL5A1,IDS,RPL5,DCHS1,PKD1,COL1A2,COL |
| 528 | 4511 | ORC4,COL1A1,KCNQ1OT1,HNRNPA1,HSPA9,NEDD4L,ATN1,ACTB,C |
| 60 | 4511 | ACTB,ACTG1,XRCC2,FLNA,DVL3,GNAS,RPL5,CHSY1,RPL15,RPL18, |
| 142 | 4511 | ACTB,SLC9A7,PRPF8,SOX4,KAT6A,DVL3,YWHAE,KIF1A,GDI1,SNRNP |
| 26 | 4511 | ACTB,ACTG1,FLNA,IGF2,RPS19,MMP2,SKI,USP7,SEC23A,EP300,RE |
| 90 | 4511 | ORC4,XRCC2,TUBB,FLNA,FBXL4,UBA1,DVL3,RPL5,NPM1,IGF2,AFF4 |
| 90 | 4511 | ORC4,XRCC2,TUBB,FLNA,FBXL4,UBA1,DVL3,RPL5,NPM1,IGF2,AFF4 |
| 92 | 4511 | ATN1,HTT,FTL,PSAP,KIF1A,GNAS,SLC2A3,APP,SPTAN1,YWHAG,MAP |
| 57 | 4511 | COL1A1,SPARC,TUBB,FLNA,COL5A1,FKBP14,COL1A2,COL5A2,MEG |
| 110 | 4511 | ORC4,ATN1,XRCC2,FLNA,MAP1B,SLC9A7,UBA1,DVL3,YWHAE,IGF2, |
| 37 | 4511 | FLNA,GNAS,RPL5,RPL15,RPL18,RPL11,RPS17,RPS24,RPL26,RPL31, |
| 96 | 4511 | HTT,COL5A1,COL4A1,FKBP14,KAT6A,COL18A1,AFF4,NONO,UCHL1, |
| 153 | 4511 | KCNQ1OT1,HNRNPA1,ACTB,PIK3R2,FLNA,COL6A1,NEFL,FBXL4,UB |
| 48 | 4511 | TUBB,GNAS,RPS20,GPC3,CTNNB1,CDKN1A,MDH2,SKI,VHL,CCND1, |
| 157 | 4511 | ORC4,COL1A1,ACTB,CTSC,ACTG1,XRCC2,PIK3R2,FLNA,SLC9A7,CO |
| 84 | 4511 | COL1A1,NEDD4L,ACTB,PIK3R2,FLNA,MAP1B,COL5A1,KAT6A,DVL3, |
| 237 | 4511 | ORC4,KCNQ1OT1,NEDD4L,ATN1,ACTB,XRCC2,TUBB,DCX,FLNA,PRP |
| 261 | 4511 | COL1A1,HNRNPA1,POLH,ACTB,ACTG1,XRCC2,VIM,TUBB,KIDINS220 |
| 36 | 4511 | ATN1,IDS,AFF4,NONO,AGRN,CTNNB1,ARCN1,GNAI3,SNRPN,SKI,PRP |
| 354 | 4511 | ORC4,COL1A1,HNRNPA1,HSPA9,ACTB,ACTG1,XRCC2,SPARC,TUBB, |
| 55 | 4511 | HSPA9,ACTB,ACTG1,KAT6A,AFF4,CHSY1,ZMIZ1,LRPPRC,MEGF8,CY |
| 30 | 4511 | COL1A1,FLNA,YWHAE,AGRN,PAFAH1B1,COL5A2,GPC3,TBCD,PLOD1 |
| 30 | 4511 | HNRNPA1,CTSC,FLNA,KIF1A,VCP,SCARB2,CCN2,MMP14,CTNNB1,M |
| 11 | 4511 | COL1A1,NEDD4L,FLNA,MAP1B,COL5A1,ARF1,COL1A2,COL5A2,AEB |
| 37 | 4511 | HSPA9,XRCC2,UBA1,CALR,GNB1,RPL5,RPL15,SF3B1,RPL18,RPL11, |
| 79 | 4511 | SLC9A7,PSAP,KIF1A,GNB1,GDI1,VCP,COL4A2,MAPT,SET,AP2M1,RPL |
| 89 | 4511 | DCX,SLC9A7,FTL,PSAP,KIF1A,GDI1,SET,AP2M1,RPL10,MARCHF6,PA |
| 97 | 4511 | KCNQ1OT1,ATN1,TUBA1A,TUBB,FLNA,TUBB2B,COL4A1,SOX4,IGF2, |
| 125 | 4511 | COL1A1,ATN1,XRCC2,FLNA,COL6A1,COL5A1,SOX4,DVL3,YWHAE,KI |
| 67 | 4511 | COL1A1,ACTB,SLC9A7,FBXL4,RPL5,GDI1,IGF2,RPL15,RPL18,RPL11 |
| 36 | 4511 | COL1A1,GANAB,COL5A1,AFF4,PKD1,ZMIZ1,MAT2A,COL1A2,COL5A |
| 234 | 4511 | ORC4,COL1A1,COPA,NEDD4L,ACTB,ACTG1,XRCC2,HTT,PIK3R2,FL |
| 48 | 4511 | ORC4,HSPA9,MAP1B,RPL5,DCHS1,RPL15,RPL18,RPL11,RPS17,RPS |
| 57 | 4511 | COL1A1,KCNQ1OT1,TUBB,FLNA,COL4A1,SOX4,GNAS,IGF2,POLR2A |
| 76 | 4511 | NEDD4L,ACTB,XRCC2,FLNA,MAP1B,FKBP14,KAT6A,DVL3,CALM3,R |
| 101 | 4511 | ORC4,XRCC2,TUBB,FLNA,FBXL4,UBA1,DVL3,RPL5,NPM1,IGF2,AFF4 |
| 95 | 4511 | HNRNPA1,HTT,GNAS,SLC2A3,VCP,YWHAG,MAPT,NONO,UCHL1,ZMI |

| | | |
|---|---|---|
| 39 | 4511 | HNRNPA1,HTT,DCX,SLC2A3,APP,VCP,MAPT,UCHL1,PFN1,PRKAR1A,T |
| 153 | 4511 | ORC4,ACTB,SLC9A7,PRPF8,SOX4,KAT6A,DVL3,YWHAE,KIF1A,GDI1,S |
| 242 | 4511 | ORC4,KCNQ1OT1,NEDD4L,ATN1,ACTB,XRCC2,TUBB,DCX,FLNA,PRP |
| 82 | 4511 | ORC4,XRCC2,MAP1B,SLC9A7,DVL3,YWHAE,IGF2,NONO,CHSY1,PPP2 |
| 45 | 4511 | TUBB,DCHS1,ZMIZ1,RHOA,GJA1,RBMX,MAPK1,ODC1,DLK1,BCR,EP |
| 199 | 4511 | COL1A1,KCNQ1OT1,HSPA9,CTSC,FLNA,SLC9A7,COL5A1,SOX4,KAT6 |
| 403 | 4511 | ORC4,COL1A1,HNRNPA1,NEDD4L,ATN1,ACTB,ACTG1,HTT,NUMA1,D |
| 27 | 4511 | RPL5,RPL15,RPL18,RPL11,RPS17,RPS24,RPL26,RPL31,RPS7,RPS19 |
| 27 | 4511 | RPL5,RPL15,RPL18,RPL11,RPS17,RPS24,RPL26,RPL31,RPS7,RPS19 |
| 25 | 4511 | VCP,MAPT,TARDBP,L1CAM,SQSTM1,ATXN10,SOD1,RAI1,DCTN1,BAZ |
| 132 | 4511 | LRP1,HSPA9,ACTB,ACTG1,MAP1B,FBXL4,SOX4,KAT6A,NPM1,AFF4,C |
| 281 | 4511 | COL1A1,KCNQ1OT1,HNRNPA1,HSPA9,NEDD4L,ATN1,ACTB,XRCC2,F |
| 42 | 4511 | CTSC,FLNA,COL6A1,COL11A1,MAT2A,COL6A2,NARS1,PLOD1,COL2 |
| 46 | 4511 | HTT,NEFL,KIF1A,YWHAG,MAPT,TPI1,CHD3,CLTC,CYFIP2,EEF1A2,RA |
| 75 | 4511 | COL1A1,XRCC2,TUBB,FLNA,COL5A1,FKBP14,KAT6A,DVL3,IDS,IGF2 |
| 55 | 4511 | HNRNPA1,KIF1A,VCP,FN1,COL1A2,COL6A2,SHOX,DUX4,SMCHD1,L1 |
| 377 | 4511 | COL1A1,LRP1,KCNQ1OT1,HSPA9,NEDD4L,POLH,ACTB,CTSC,XRCC2 |
| 100 | 4511 | ORC4,COL1A1,KCNQ1OT1,ACTB,ACTG1,FLNA,MAP1B,KAT6A,KIF1A |
| 9 | 4511 | SOX4,ARID1A,SMARCA4,SMARCE1,ARID1B,SMARCD1,SMARCC2,SM |
| 9 | 4511 | SOX4,ARID1A,SMARCA4,SMARCE1,ARID1B,SMARCD1,SMARCC2,SM |
| 9 | 4511 | SOX4,ARID1A,SMARCA4,SMARCE1,ARID1B,SMARCD1,SMARCC2,SM |
| 9 | 4511 | SOX4,ARID1A,SMARCA4,SMARCE1,ARID1B,SMARCD1,SMARCC2,SM |
| 9 | 4511 | SOX4,ARID1A,SMARCA4,SMARCE1,ARID1B,SMARCD1,SMARCC2,SM |

PARC,TUBB,KIDINS220,NUMA1,DCX,PIK3R2,FLNA,SLC9A7,ALDOA,COL6A1,FTL,NEFL,FBXL4,UBA
220,NUMA1,DCX,PIK3R2,FLNA,SLC9A7,COL6A1,FTL,NEFL,FBXL4,UBA1,CALR,TUBB2B,COL5A1,
220,DCX,PIK3R2,FLNA,SLC9A7,COL6A1,FTL,NEFL,FBXL4,UBA1,TUBB2B,COL5A1,COL4A1,PSAP
RCC2,HTT,SPARC,TUBB,DCX,PIK3R2,FLNA,MAP1B,ALDOA,COL6A1,NEFL,FBXL4,UBA1,COL5A1,
CTB,ACTG1,EEF2,TUBA1A,XRCC2,HTT,SPARC,TUBB,KIDINS220,NUMA1,DCX,PIK3R2,FLNA,MAP1
CTB,CTSC,ACTG1,EEF2,TUBA1A,XRCC2,HTT,SPARC,TUBB,KIDINS220,NUMA1,DCX,PIK3R2,FLNA,
,TUBB2B,PSAP,SOX4,YWHAE,KIF1A,ALDH6A1,SPTAN1,NONO,DCHS1,SET,PPP2R1A,POLR2A,ZMI
CTG1,TUBA1A,XRCC2,HTT,SPARC,TUBB,KIDINS220,DCX,PIK3R2,FLNA,MAP1B,SLC9A7,ALDOA,C
TSC,ACTG1,EEF2,TUBA1A,XRCC2,HTT,SPARC,TUBB,KIDINS220,DCX,PIK3R2,FLNA,MAP1B,SLC9A
K3R2,FLNA,SLC9A7,KAT6A,DVL3,YWHAE,ALDH6A1,P4HB,RPL5,IGF2,AFF4,NONO,DCHS1,SET,CH
XRCC2,HTT,TUBB,KIDINS220,DCX,PIK3R2,MAP1B,FTL,FBXL4,TUBB2B,COL4A1,PSAP,SOX4,KAT6
XRCC2,HTT,TUBB,KIDINS220,DCX,PIK3R2,MAP1B,FTL,FBXL4,TUBB2B,COL4A1,PSAP,SOX4,KAT6
CTB,CTSC,ACTG1,TUBA1A,XRCC2,HTT,SPARC,TUBB,KIDINS220,NUMA1,DCX,PIK3R2,FLNA,MAP1
C2,HTT,TUBB,KIDINS220,DCX,PIK3R2,FLNA,MAP1B,FTL,NEFL,FBXL4,TUBB2B,COL4A1,PSAP,SO
H6A1,RPL5,AFF4,NONO,DCHS1,SET,CHSY1,PPP2R1A,RPL15,CHD3,POLR2A,RPL18,RPL11,ZMIZ1
CTG1,TUBA1A,XRCC2,HTT,SPARC,TUBB,KIDINS220,DCX,PIK3R2,FLNA,MAP1B,SLC9A7,ALDOA,C
TUBA1A,XRCC2,TUBB,KIDINS220,DCX,PIK3R2,FLNA,MAP1B,SLC9A7,COL6A1,FBXL4,UBA1,TUBE
HTT,SPARC,DCX,PIK3R2,FLNA,MAP1B,COL6A1,NEFL,FBXL4,UBA1,COL5A1,SOX4,FKBP14,DVL3,
CTG1,EEF2,TUBA1A,HTT,VIM,TUBB,FTH1,KIDINS220,PIK3R2,MAP1B,GANAB,COL6A1,FTL,NEFL,F
AT6A,DVL3,YWHAE,KIF1A,ALDH6A1,RPL5,GDI1,AFF4,NONO,DCHS1,CHSY1,PPP2R1A,AP2M1,ZMI
9A7,FBXL4,TUBB2B,COL4A1,PSAP,SOX4,KAT6A,DVL3,KIF1A,ALDH6A1,IDS,RPL5,GDI1,SPTAN1,H
INS220,DCX,PIK3R2,MAP1B,FBXL4,TUBB2B,COL4A1,PSAP,SOX4,KAT6A,YWHAE,KIF1A,ALDH6A1
TSC,ACTG1,EEF2,TUBA1A,XRCC2,HTT,SPARC,TUBB,KIDINS220,DCX,PIK3R2,FLNA,MAP1B,GANA
EF2,TUBA1A,XRCC2,HTT,SPARC,TUBB,KIDINS220,DCX,PIK3R2,FLNA,MAP1B,SLC9A7,ALDOA,FTL
TSC,ACTG1,XRCC2,HTT,SPARC,TUBB,KIDINS220,PIK3R2,FLNA,MAP1B,SLC9A7,COL6A1,NEFL,FE
CTB,CTSC,ACTG1,TUBA1A,XRCC2,HTT,SPARC,TUBB,KIDINS220,DCX,PIK3R2,FLNA,MAP1B,SLC9
CX,PIK3R2,MAP1B,FBXL4,TUBB2B,COL4A1,PSAP,SOX4,YWHAE,KIF1A,SLC2A3,ALDH6A1,IDS,SPT
TSC,ACTG1,EEF2,TUBA1A,XRCC2,HTT,SPARC,TUBB,KIDINS220,DCX,PIK3R2,FLNA,MAP1B,SLC9A
HTT,SPARC,TUBB,KIDINS220,DCX,PIK3R2,FLNA,MAP1B,SLC9A7,COL6A1,NEFL,FBXL4,TUBB2B,C
4,UBA1,COL5A1,COL4A1,KAT6A,KIF1A,GNB1,RPL5,IGF2,AFF4,DCHS1,SET,RPL15,RPL18,RPL11,
AT6A,DVL3,ALDH6A1,GDI1,AFF4,NONO,CHSY1,ZMIZ1,COL11A1,RPL10,AP3D1,HNRNPU,CDH2,C
C,DCX,PIK3R2,FLNA,MAP1B,COL6A1,NEFL,FBXL4,UBA1,COL5A1,SOX4,FKBP14,DVL3,KIF1A,RPL
BB,KIDINS220,DCX,FLNA,MAP1B,SLC9A7,FTL,NEFL,PSAP,PGK1,KAT6A,KIF1A,GNAS,SLC2A3,GN
A7,PRPF8,UBA1,SOX4,KAT6A,DVL3,YWHAE,KIF1A,GNAS,ALDH6A1,IDS,RPL5,GDI1,COL18A1,SNR
AE,IDS,IGF2,PPP2R1A,POLR2A,RPL10,PAFAH1B1,MAT2A,COL1A2,COL5A2,GPC3,SMCHD1,PLOD
IK3R2,MAP1B,FBXL4,TUBB2B,COL4A1,PSAP,SOX4,YWHAE,KIF1A,ALDH6A1,IDS,SPTAN1,YWHAG,
XRCC2,HTT,SPARC,TUBB,PIK3R2,FLNA,MAP1B,SLC9A7,COL6A1,NEFL,FBXL4,UBA1,COL5A1,COL
CX,PIK3R2,MAP1B,FTL,FBXL4,TUBB2B,COL4A1,PSAP,SOX4,YWHAE,KIF1A,GNAS,SLC2A3,ALDH6
IK3R2,FLNA,MAP1B,SLC9A7,COL6A1,NEFL,UBA1,COL5A1,SOX4,KAT6A,DVL3,YWHAE,KIF1A,GN
G1,EEF2,XRCC2,HTT,VIM,TUBB,KIDINS220,DCX,PIK3R2,FLNA,SLC9A7,ALDOA,COL6A1,FTL,NEF
B,NEFL,UBA1,COL5A1,SOX4,KAT6A,DVL3,GNB1,P4HB,IDS,RPL5,IGF2,VCP,FN1,DCHS1,PPP2R1
SLC9A7,FTL,FBXL4,UBA1,TUBB2B,COL4A1,PSAP,PGK1,SOX4,KAT6A,YWHAE,KIF1A,GNAS,SLC2
,RPS19,RPL35,RPS10,RPL35A,TSR2,RPS27,RPL27,RPS15A,RPS28,RPS26,RPS29,G6PC3
2B,COL4A1,SOX4,KAT6A,KIF1A,ALDH6A1,RPL5,SPTAN1,HSPD1,ATP5F1A,AFF4,YWHAG,DCHS1,S
PL26,RPL31,RPS7,RPS19,RPL35,RPS10,RPL35A,TSR2,RPS27,RPL27,RPS15A,RPS28,RPS26,RPS

TB,CTSC,ACTG1,TUBA1A,XRCC2,HTT,SPARC,TUBB,KIDINS220,NUMA1,DCX,PIK3R2,FLNA,MAP1
IK3R2,FLNA,MAP1B,SLC9A7,COL6A1,NEFL,UBA1,COL5A1,PSAP,SOX4,KAT6A,DVL3,YWHAE,KIF1
2B,COL4A1,PSAP,SOX4,KAT6A,KIF1A,ALDH6A1,RPL5,SPTAN1,HSPD1,ATP5F1A,AFF4,YWHAG,DC
IK3R2,FLNA,SLC9A7,COL6A1,NEFL,FBXL4,UBA1,TUBB2B,COL5A1,COL4A1,PSAP,SOX4,FKBP14
2,FLNA,MAP1B,SLC9A7,ALDOA,FTL,FBXL4,TUBB2B,COL5A1,SOX4,KAT6A,DVL3,YWHAE,KIF1A,G
,SPARC,TUBB,PIK3R2,FLNA,MAP1B,SLC9A7,COL6A1,NEFL,FBXL4,UBA1,COL5A1,COL4A1,PSAP
,SPARC,TUBB,PIK3R2,FLNA,MAP1B,SLC9A7,COL6A1,NEFL,FBXL4,UBA1,COL5A1,COL4A1,PSAP
,RPL35,RPS10,RPL35A,TSR2,RPS27,RPL27,RPS15A,RPS28,RPS26,RPS29
,RPL35,RPS10,RPL35A,TSR2,RPS27,RPL27,RPS15A,RPS28,RPS26,RPS29
,RPL35,RPS10,RPL35A,TSR2,RPS27,RPL27,RPS15A,RPS28,RPS26,RPS29
4,COL5A1,SOX4,KAT6A,DVL3,YWHAE,KIF1A,ALDH6A1,RPL5,COL18A1,NONO,DCHS1,RPL15,CH
IK3R2,FLNA,SLC9A7,ALDOA,COL6A1,FTL,NEFL,FBXL4,UBA1,TUBB2B,COL4A1,PSAP,PGK1,SOX
,DCHS1,SET,CHSY1,ZMIZ1,COL11A1,RPL10,PAFAH1B1,CCND2,GPC3,GJA1,ARID1A,GNAI3,SNRP
,RPL35,RPS10,RPL35A,TSR2,RPS27,RPL27,RPS15A,RPS28,MSH6,RPS26,RPS29,MSH2
N1,ACTB,CTSC,ACTG1,TUBA1A,XRCC2,HTT,TUBB,KIDINS220,NUMA1,DCX,PIK3R2,FLNA,MAP1
,MAP1B,SLC9A7,ALDOA,FBXL4,COL5A1,SOX4,KAT6A,DVL3,YWHAE,KIF1A,ALDH6A1,P4HB,RPL5,
KBP14,P4HB,COL18A1,SET,PPP2R1A,ZMIZ1,COL11A1,ARF1,COL1A2,COL5A2,COL6A2,ARID1A,
LC9A7,PGK1,KAT6A,KIF1A,GNAS,GNB1,APP,GDI1,IGF2,VCP,UBB,YWHAG,MAPT,NONO,SET,PPP
R2,FLNA,MAP1B,SLC9A7,COL6A1,NEFL,UBA1,COL5A1,SOX4,KAT6A,DVL3,YWHAE,KIF1A,GNAS,
5,RPL18,RPL11,COL11A1,RPL10,RPS17,RPS24,COL1A2,CHD4,RPL26,RPL31,TKT,GPC3,MEGF8
TSC,ACTG1,XRCC2,TUBB,NUMA1,PIK3R2,FLNA,MAP1B,SLC9A7,COL6A1,NEFL,FBXL4,UBA1,TUB
5A1,SOX4,FKBP14,DVL3,YWHAE,IDS,IGF2,PPP2R1A,POLR2A,COL11A1,RPL10,ARF1,PAFAH1B1,
5,RPL18,RPL11,COL11A1,RPL10,RPS17,RPS24,COL1A2,CHD4,RPL26,RPL31,TKT,GPC3,MEGF8
N1,ACTB,CTSC,ACTG1,TUBA1A,XRCC2,HTT,TUBB,KIDINS220,NUMA1,DCX,PIK3R2,FLNA,MAP1
2,FLNA,MAP1B,SLC9A7,ALDOA,FTL,FBXL4,TUBB2B,COL5A1,SOX4,KAT6A,DVL3,YWHAE,KIF1A,G
,RPL35,RPS10,RPL35A,TSR2,RPS27,RPL27,RPS15A,RPS28,RPS26,RPS29
SOX4,KAT6A,DVL3,YWHAE,KIF1A,ALDH6A1,IDS,RPL5,GDI1,AFF4,NONO,DCHS1,SET,CHSY1,PPP2
IK3R2,MAP1B,FBXL4,TUBB2B,COL4A1,PSAP,SOX4,YWHAE,KIF1A,ALDH6A1,SPTAN1,NONO,DCH
RPS7,RPS19,RPL35,RPS10,RPL35A,TSR2,RPS27,RPL27,RPS15A,RPS28,HSPG2,RPS26,PTPN11,
HS1,SET,RPL15,CHD3,RPL18,RPL11,ZMIZ1,COL11A1,RPL10,RPS17,CTSD,RPS24,RHOA,RPL26,
LNA,MAP1B,SLC9A7,KAT6A,DVL3,YWHAE,ALDH6A1,P4HB,RPL5,IGF2,AFF4,YWHAG,NONO,DCH
P1B,COL6A1,FBXL4,UBA1,COL5A1,COL4A1,SOX4,KAT6A,DVL3,YWHAE,KIF1A,GNB1,RPL5,IGF2,
T6A,KIF1A,ALDH6A1,GNB1,GDI1,SPTAN1,HSPD1,IGF2,ATP5F1A,YWHAG,NONO,DCHS1,SET,TPI1
2,FLNA,MAP1B,SLC9A7,ALDOA,FBXL4,TUBB2B,COL5A1,SOX4,KAT6A,DVL3,YWHAE,KIF1A,GNAS,
PN,RAB11B,ABCA2,PPP2CA,ODC1,KDM1A,DHX30,LSS,FBLN1,RAI1,KDM3B,TRIO,EP300,KDM5B
IK3R2,MAP1B,FBXL4,TUBB2B,COL4A1,PSAP,SOX4,YWHAE,KIF1A,ALDH6A1,SPTAN1,NONO,DCH
0,DCX,PIK3R2,FLNA,SLC9A7,ALDOA,COL6A1,FTL,NEFL,PRPF8,FBXL4,UBA1,CALR,TUBB2B,COL
YWHAE,ALDH6A1,GNB1,RPL5,IGF2,AFF4,NONO,DCHS1,SET,RPL15,CHD3,POLR2A,RPL18,RPL11
3,RPL5,AFF4,NONO,RPL15,RPL18,RPL11,COL11A1,RPL10,RPS17,HNRNPU,RPS24,COL1A2,CH
1,NEFL,COL4A1,SOX4,FKBP14,KAT6A,DVL3,YWHAE,KIF1A,ALDH6A1,GNB1,RPL5,IGF2,UBB,AF
B,COL4A1,SOX4,KAT6A,DVL3,GNAS,GNB1,RPL5,HSPD1,COL18A1,VCP,AFF4,COL4A2,MAPT,NO
P14,IDS,RPL5,AFF4,PPP2R1A,RPL15,PKD1,RPL18,RPL11,ZMIZ1,RPS17,ARF1,MAT2A,RPS24,COL
,MAP1B,SLC9A7,COL6A1,UBA1,COL5A1,SOX4,KAT6A,DVL3,YWHAE,KIF1A,GNAS,IDS,RPL5,GDI1,
7,COL6A1,NEFL,FBXL4,COL5A1,COL4A1,SOX4,FKBP14,KAT6A,KIF1A,GNAS,IDS,GDI1,HSPD1,IG
AP,KAT6A,KIF1A,SLC2A3,GNB1,APP,PLD3,HSPD1,VCP,YWHAG,MAPT,FN1,TPI1,PPP2R1A,CHD3,A

,XRCC2,TUBB,NUMA1,PIK3R2,FLNA,MAP1B,SLC9A7,COL6A1,NEFL,FBXL4,UBA1,COL5A1,COL4
,XRCC2,TUBB,NUMA1,PIK3R2,FLNA,MAP1B,SLC9A7,COL6A1,NEFL,FBXL4,UBA1,COL5A1,COL4
PIK3R2,FLNA,MAP1B,SLC9A7,COL6A1,NEFL,FBXL4,COL5A1,COL4A1,PSAP,SOX4,FKBP14,KAT6
JBB,KIDINS220,PIK3R2,FLNA,FBXL4,UBA1,KAT6A,DVL3,YWHAE,KIF1A,ALDH6A1,P4HB,IDS,COL1
(4,KAT6A,DVL3,GNAS,ALDH6A1,IDS,RPL5,GDI1,COL18A1,SNRNP200,NONO,DCHS1,SET,RPL15,
B,SLC9A7,COL6A1,FTL,NEFL,PRPF8,FBXL4,UBA1,TUBB2B,COL4A1,PSAP,PGK1,KAT6A,KIF1A,G
RN,ZMIZ1,PAFAH1B1,COL1A2,HK1,CDH2,HNRNPK,NARS1,LMNB2,SNRPN,PLOD1,CNTNAP2,RAB
A,GNAS,SLC2A3,APP,SPTAN1,HSPD1,VCP,YWHAG,MAPT,NONO,SET,TPI1,CHD3,AP2M1,UCHL1,M
,RPL35,RPS10,RPL35A,TSR2,RPS27,RPL27,RPS15A,RPS28,RPS26,RPS29
10,ATP1A1,TKT,CDH2,DDX3X,CLTC,CTNNB1,EEF1A2,SQSTM1,VAMP2,SNRPN,CHD2,SKI,CNTNAP
AT6A,KIF1A,GNAS,GNB1,GDI1,IGF2,UBB,YWHAG,NONO,SET,PPP2R1A,CHD3,ZMIZ1,RPL10,ARF
,RPL35,RPS10,RPL35A,TSR2,RPS27,RPL27,RPS15A,RPS28,RPS26,RPS29,SRP54,DNAJC21
N1,ACTB,CTSC,ACTG1,XRCC2,HTT,TUBB,KIDINS220,NUMA1,PIK3R2,FLNA,MAP1B,SLC9A7,ALD
P,SOX4,KAT6A,KIF1A,GNB1,APP,SPTAN1,HSPD1,VCP,YWHAG,COL4A2,MAPT,SET,TPI1,UCHL1,A
L5,GDI1,COL18A1,RPL15,RPL18,RPL11,ZMIZ1,COL11A1,RPS17,RPS24,COL1A2,RPL26,RPL31,
GNB1,RPL5,HSPD1,VCP,YWHAG,MAPT,RPL15,RPL18,RPL11,AGRN,RPS17,SBF1,RPS24,HK1,ATP
A7,COL6A1,FBXL4,SOX4,KAT6A,KIF1A,GNAS,ALDH6A1,IDS,IGF2,UBB,AFF4,NONO,DCHS1,SET,
JBB,KIDINS220,PIK3R2,FLNA,MAP1B,FBXL4,UBA1,KAT6A,DVL3,YWHAE,KIF1A,GNAS,ALDH6A1,P
ARC,TUBB,FLNA,ALDOA,COL6A1,TUBB2B,COL5A1,SOX4,FKBP14,KAT6A,DVL3,YWHAE,KIF1A,R
ET,POLR2A,AP2M1,UCHL1,ZMIZ1,RPL10,NR2F1,CDH2,DDX3X,CLTC,PRKAR1A,CTNNB1,CYFIP2,
,RPL35,RPS10,RPL35A,TSR2,RPS27,SQSTM1,RPL27,RPS15A,RPS28,RPS26,MDM2,RPS29,LMN
1,NEFL,TUBB2B,COL5A1,FKBP14,IGF2,NONO,SET,PPP2R1A,CHD3,POLR2A,AGRN,ZMIZ1,PAFA
PL5,IGF2,AFF4,CHSY1,RPL15,CHD3,RPL18,RPL11,AGRN,ZMIZ1,COL11A1,RPL10,RPS17,AP3D1,
220,PIK3R2,FLNA,MAP1B,SLC9A7,FBXL4,TUBB2B,COL4A1,PSAP,PGK1,SOX4,KAT6A,DVL3,YWH
DL4A1,SOX4,KAT6A,DVL3,GNAS,GNB1,RPL5,HSPD1,COL18A1,AFF4,COL4A2,NONO,RPL15,CHD
,RPL35,RPS10,RPL35A,TSR2,RPS27,RPL27,RPS15A,RPS28,PRPS1,RPS26,RPS29
B,COL6A1,NEFL,UBA1,COL5A1,SOX4,FKBP14,DVL3,KIF1A,GNAS,P4HB,IDS,HSPD1,COL18A1,VC
WHAE,RPL5,IGF2,AFF4,CHSY1,RPL15,CHD3,RPL18,RPL11,AGRN,ZMIZ1,COL11A1,RPL10,RPS17
HTT,TUBB,KIDINS220,DCX,PIK3R2,FLNA,MAP1B,SLC9A7,ALDOA,PRPF8,TUBB2B,COL4A1,PGK1,
A7,COL6A1,FTL,NEFL,FBXL4,TUBB2B,COL4A1,PSAP,PGK1,SOX4,KAT6A,YWHAE,KIF1A,SLC2A3
IZ1,COL1A2,GPC3,MEGF8,HNRNPK,NARS1,PPIB,CDH11,TBCD,PPP1CB,PLOD1,COL2A1,SKI,KD
,RPL35,RPS10,RPL35A,TSR2,RPS27,RPL27,RPS15A,RPS28,RPS26,RPS29
DM6B,MAPK1,SMARCA4,SMARCE1,ATP6V1B2,RAI1,SEC23A,KDM3B,CDC42,TRIO,FLCN,BAZ1B,
P,SOX4,KIF1A,SPTAN1,HSPD1,VCP,YWHAG,COL4A2,MAPT,SET,TPI1,UCHL1,ARF1,CYB5R3,PAFA
9A7,KAT6A,DVL3,KIF1A,GNAS,P4HB,IDS,GDI1,COL18A1,IGF2,AFF4,NONO,DCHS1,CHD3,ZMIZ1
L4,COL5A1,PSAP,KAT6A,KIF1A,GNAS,SLC2A3,APP,VCP,UBB,AFF4,YWHAG,MAPT,NONO,SET,POL
RPL11,RPL10,RPS17,RPS24,CHD4,RPL26,RPL31,TKT,CCND2,GPC3,MMP14,RPS7,RPS19,RPL35
3,YWHAE,IDS,IGF2,PPP2R1A,POLR2A,COL11A1,RPL10,PAFAH1B1,MAT2A,COL1A2,COL5A2,GP
,P4HB,IDS,MAPT,FN1,NONO,TPI1,AGRN,ZMIZ1,SCARB2,COL1A2,MEGF8,PRKAR1A,COL6A2,AR
3,YWHAE,IDS,IGF2,PPP2R1A,POLR2A,COL11A1,RPL10,PAFAH1B1,MAT2A,COL1A2,COL5A2,GP
RPL11,RPL10,RPS17,RPS24,CHD4,RPL26,RPL31,TKT,CCND2,GPC3,MMP14,RPS7,RPS19,RPL35
1,PSAP,YWHAE,KIF1A,SLC2A3,APP,IDS,SPTAN1,COL18A1,VCP,YWHAG,MAPT,SET,TPI1,UCHL1,Z
7,COL6A1,FBXL4,UBA1,SOX4,KAT6A,DVL3,GNAS,P4HB,RPL5,GDI1,IGF2,AFF4,FN1,DCHS1,CHS
DVL3,IDS,RPL5,VCP,AFF4,YWHAG,DCHS1,RPL15,RPL18,RPL11,AGRN,ZMIZ1,RPS17,RPS24,COL
F1,NR2F1,ATP1A1,RPS20,TKT,CDH2,DDX3X,CLTC,PRKAR1A,CYFIP2,EEF1A2,SNRPN,CHD2,KDM

N1,NONO,UCHL1,SBF1,MAT2A,COL1A2,HK1,COL5A2,RAB7A,HNRNPK,MMP2,HS6ST1,SNRPN,PL
NO,DCHS1,PPP2R1A,AP2M1,ZMIZ1,COL11A1,RPL10,HNRNPU,HK1,CDH2,CLTC,RPS7,CTNNB1,C
AE,ALDH6A1,P4HB,GDI1,AFF4,YWHAG,NONO,DCHS1,PPP2R1A,ZMIZ1,COL11A1,RPL10,PAFAH1B
PU,RPS24,RPL26,RPL31,CCND2,RPS7,RPS19,RPL35,RPS10,RPL35A,ARID1A,TSR2,RPS27,RPL27
,RPL35,RPS10,RPL35A,TSR2,RPS27,RPL27,RPS15A,RPS28,RPS26,RPS29
WHAE,GNAS,IDS,RPL5,VCP,MAPT,RPL15,RPL18,RPL11,COL11A1,RPL10,RPS17,PAFAH1B1,RPS
L11,ZMIZ1,RPS17,RPS24,RPL26,RPL31,LRPPRC,CDH2,RPS7,RPS19,RPL35,RPS10,RPL35A,ARID
RCC2,VIM,TUBB,KIDINS220,PIK3R2,FLNA,SLC9A7,FTL,NEFL,PRPF8,FBXL4,TUBB2B,COL5A1,CO
BA1,COL5A1,KAT6A,DVL3,P4HB,IDS,RPL5,IGF2,AFF4,DCHS1,CHSY1,RPL15,RPL18,RPL11,AGRN
1,RPL15,RPL18,RPL11,ZMIZ1,COL11A1,RPL10,RPS17,RPS24,RHOA,RPL26,RPL31,GPC3,MEGF
L,NEFL,TUBB2B,COL4A1,PSAP,SOX4,KAT6A,KIF1A,SLC2A3,GNB1,APP,GDI1,SPTAN1,HSPD1,VC
KBP14,KAT6A,DVL3,YWHAE,IDS,IGF2,PPP2R1A,POLR2A,COL11A1,RPL10,PAFAH1B1,MAT2A,COL
BA1,COL5A1,KAT6A,DVL3,P4HB,IDS,RPL5,NPM1,IGF2,UBB,AFF4,DCHS1,CHSY1,RPL15,AP2M1,
1,COL4A1,PSAP,FKBP14,APP,IDS,RPL5,AFF4,PPP2R1A,RPL15,PKD1,RPL18,RPL11,ZMIZ1,RPS17
RPL26,RPL31,RPS7,RPS19,RPL35,RPS10,RPL35A,TSR2,RPS27,RPL27,RPS15A,RPS28,BCR,TTC
,ARF1,NR2F1,RPS20,TKT,CDH2,CLTC,CYFIP2,CDKN1A,EEF1A2,SNRPN,MAPK1,ODC1,STXBP1,U
1,FKBP14,KAT6A,DVL3,CALM3,IDS,RPL5,COL18A1,IGF2,AFF4,FN1,NONO,PPP2R1A,RPL15,PKD1
T6A,KIF1A,ALDH6A1,GNB1,HSPD1,IGF2,UBB,AFF4,YWHAG,NONO,DCHS1,SET,PPP2R1A,POLR2A
B,COL4A1,SOX4,KIF1A,SPTAN1,PLD3,COL18A1,IGF2,ATP5F1A,NONO,DCHS1,PPP2R1A,UCHL1,Z
AS,RPL5,GDI1,IGF2,AFF4,FN1,DCHS1,CHSY1,RPL15,RPL18,RPL11,ZMIZ1,COL11A1,RPL10,RPS
UMA1,DCX,PIK3R2,FLNA,MAP1B,COL6A1,NEFL,UBA1,COL5A1,SOX4,FKBP14,DVL3,KIF1A,GNAS,
A,AGRN,ZMIZ1,MAT2A,COL1A2,GPC3,MEGF8,HNRNPK,NARS1,CTBP1,PPIB,CDH11,TBCD,PPP1C
A1,FTL,NEFL,PRPF8,FBXL4,UBA1,TUBB2B,COL4A1,PSAP,SOX4,KAT6A,DVL3,KIF1A,GNAS,SLC2
,RPL35,RPS10,RPL35A,TSR2,RPS27,RPL27,RPS15A,RPS28,RPS26,RPS29
OL5A1,PSAP,SOX4,KAT6A,DVL3,YWHAE,KIF1A,RPL13,VCP,AFF4,FN1,NONO,CHSY1,UCHL1,AGR
5,RPL18,RPL11,COL11A1,RPL10,RPS17,RPS24,COL1A2,CHD4,RPL26,RPL31,TKT,GPC3,MEGF8
,FTL,FBXL4,UBA1,SOX4,KAT6A,DVL3,KIF1A,GNAS,IDS,RPL5,NPM1,COL18A1,IGF2,AFF4,CHSY1,
1,RPS7,RPS19,RPL35,RPS10,RPL35A,TSR2,RPS27,RPL27,RPS15A,RPS28,RPS26,RPS29,G6PC3
L1A2,COL5A2,PRKAR1A,ARID1A,SHOX,SKI,ASXL1,NOTCH2,SMARCA4,AEBP1,FLNB,SMARCE1,C
6,RPL31,RPS7,RPS19,RPL35,PRKAR1A,CTNNB1,RPS10,RPL35A,TSR2,RPS27,CDKN1A,SQSTM1,
2A,RPL10,COL1A2,HK1,RHOA,GPC3,PRKAR1A,GJA1,HNRNPK,NARS1,SMCHD1,CDH11,TBCD,G
B,COL4A1,SOX4,KIF1A,SPTAN1,PLD3,COL18A1,IGF2,ATP5F1A,NONO,DCHS1,PPP2R1A,UCHL1,Z
B,COL4A1,SOX4,KIF1A,SPTAN1,PLD3,COL18A1,IGF2,ATP5F1A,NONO,DCHS1,PPP2R1A,UCHL1,Z
T6A,KIF1A,GNAS,AFF4,YWHAG,NONO,SET,POLR2A,AGRN,RPL10,ARF1,COL5A2,PRKCSH,CCN2,
,SEC61A1,ZMIZ1,RPL10,ARF1,MAT2A,COL1A2,COL5A2,PRKCSH,PRKAR1A,HNRNPK,HLA-B,PLO
ZMIZ1,COL11A1,RPL10,CDH2,CLTC,GJA1,CTNNB1,CDH11,COL2A1,SKI,NOTCH2,MAPK1,PRKDC
L11,ZMIZ1,RPS17,RPS24,RPL26,RPL31,LRPPRC,CDH2,RPS7,RPS19,RPL35,RPS10,RPL35A,ARID
,FTL,FBXL4,SOX4,KAT6A,DVL3,GNAS,IDS,RPL5,NPM1,COL18A1,AFF4,CHSY1,PPP2R1A,RPL15,R
,RPL35,RPS10,RPL35A,TSR2,RPS27,RPL27,RPS15A,RPS28,RPS26,RPS29,SRP54,PIEZO1,DNAJC
I1,AFF4,YWHAG,NONO,PPP2R1A,ZMIZ1,COL11A1,PAFAH1B1,CDH2,GPC3,CLTC,RPS19,GJA1,CY
,PIK3R2,FLNA,SLC9A7,PRPF8,FBXL4,UBA1,COL5A1,SOX4,KAT6A,DVL3,YWHAE,KIF1A,GNAS,SLC
PL31,RPS7,RPS19,RPL35,RPS10,RPL35A,TSR2,RPS27,RPL27,RPS15A,RPS28,RPS26,RPS29,PIEZ
B2B,COL4A1,YWHAE,KIF1A,GNB1,APP,COL18A1,VCP,COL4A2,MAPT,DCHS1,SET,ZMIZ1,CYB5R
,RPL31,GPC3,MEGF8,RPS7,RPS19,RPL35,RPS10,RPL35A,TSR2,RPS27,CTBP1,PPP1CB,RPL27,R
9A7,NEFL,FBXL4,UBA1,COL5A1,PSAP,FKBP14,DVL3,KIF1A,RPL5,GDI1,NPM1,HSPD1,COL18A1,

I,RPS7,RPS19,RPL35,RPS10,RPL35A,TSR2,RPS27,RPL27,RPS15A,PRDX1,RPS28,LARS1,PHGDH,I
18A1,AFF4,NONO,SF3B1,CHD3,TUBB4B,UCHL1,ZMIZ1,COL11A1,RPL10,COL1A2,HK1,RHOA,NR2
AE,KIF1A,SLC2A3,APP,SPTAN1,COL18A1,VCP,YWHAG,MAPT,SET,UCHL1,ZMIZ1,AP3D1,SBF1,CYE
20,DCX,PIK3R2,FLNA,TUBB2B,COL4A1,SOX4,YWHAE,KIF1A,GNAS,P4HB,IDS,COL18A1,IGF2,COL
2,SQSTM1,CHD2,CNTNAP2,ABCA2,MAPK1,ODC1,ATXN10,USP7,SARS1,KDM3B,KDM5C,TRIO,EP
),RPL35,RPS10,RPL35A,TSR2,RPS27,RPL27,RPS15A,RPS28,RPS26,RPS29,MSH2
IS1,SET,AP2M1,UCHL1,ZMIZ1,RPL10,ATP1A1,RPS20,TKT,CDH2,DDX3X,CLTC,PRKAR1A,CTNNB1,
RF1,MAT2A,COL1A2,COL5A2,HNRNPK,HLA-B,PLOD1,SKI,MAPK1,AEBP1,FLNB,TPM3,RAI1,ARFG
ND2,MACF1,CYFIP2,RAC1,MDH2,EEF1A2,TBCD,SNRPN,SKI,CNTNAP2,RAB11B,ODC1,SMARCE1,I
RPS24,RPL26,RPL31,CDH2,RPS7,RPS19,RPL35,RPS10,RPL35A,TSR2,RPS27,SMCHD1,CRPPA,SQ
P4HB,GDI1,COL18A1,IGF2,DCHS1,CHD3,ZMIZ1,COL11A1,LRPPRC,MEGF8,PRKAR1A,TSR2,SMC
1A2,HK1,DDX3X,PRKAR1A,CTNNB1,HNRNPK,MMP2,CDH11,SNRPN,RBMX,PPP2CA,ODC1,KDM1
(,TUBB2B,COL4A1,SOX4,YWHAE,KIF1A,COL18A1,IGF2,COL4A2,DCHS1,SET,PPP2R1A,CHD3,POL
),RPL35,RPS10,RPL35A,TSR2,RPS27,RPL27,RPS15A,RPS28,RPS26,RPS29
I,MAPK1,STXBP1,USP7,BCR,TRIO,EP300,BAZ1B,ARID1B,SMC3,NDN,GRN,EHMT1,GTF2I,UBE3A,C
T,NONO,AP2M1,UCHL1,ZMIZ1,RPS20,ATP2A2,PFN1,XPR1,CDH2,CLTC,PRKAR1A,CYFIP2,DNAJC5
8,RPL11,ZMIZ1,RPL10,RPS17,RPS24,RPL26,RPL31,GPC3,MMP14,RPS7,RPS19,RPL35,GJA1,RP
IIZ1,RPL10,RPS17,RPS24,RPL26,RPL31,GPC3,RPS7,RPS19,RPL35,GJA1,RPS10,RPL35A,ARID1A,
I,DVL3,YWHAE,AGRN,ARF1,PAFAH1B1,COL1A2,HK1,COL5A2,GPC3,PRKAR1A,GJA1,COL6A2,ARI
1,RPS7,RPS19,RPL35,PRKAR1A,RPS10,RPL35A,TSR2,SHOX,RPS27,SQSTM1,TAF15,RPL27,COL2
3D1,ARF1,CYB5R3,HNRNPU,NR2F1,CTNNB1,NARS1,MDH2,ASXL1,RAB11B,PPP2CA,KDM1A,DHX
DCHS1,CHSY1,RPL15,RPL18,RPL11,ZMIZ1,COL11A1,RPL10,RPS17,RPS24,RPL26,RPL31,GPC3,N
DCHS1,CHSY1,RPL15,RPL18,RPL11,ZMIZ1,COL11A1,RPL10,RPS17,RPS24,RPL26,RPL31,GPC3,N
PRC,RPS7,RPS19,RPL35,RPS10,RPL35A,ARID1A,TSR2,RPS27,NOVA2,RPL27,ASXL1,RPS15A,NO
PL5,GDI1,IGF2,AFF4,FN1,DCHS1,CHSY1,RPL15,RPL18,RPL11,ZMIZ1,COL11A1,RPL10,RPS17,RP
1,CTBP1,RAC1,NOVA2,EEF1A2,TBCD,SQSTM1,VAMP2,SNRPN,SKI,CNTNAP2,RAB11B,STXBP1,SA
18A1,IGF2,TRIM28,RPL15,SF3B1,RPL18,RPL11,RPS17,RPS24,RPS20,RPL26,RPL31,GPC3,RPS7
S20,TKT,CDH2,CLTC,CYFIP2,EEF1A2,SNRPN,MAPK1,ODC1,STXBP1,USP7,RAI1,KDM3B,SYP,CIC,I
FLNA,ALDOA,TUBB2B,COL5A1,SOX4,KAT6A,DVL3,KIF1A,RPL13,GNAS,P4HB,IDS,RPL5,NPM1,IGF
S,IDS,RPL5,RPL15,RPL18,RPL11,COL11A1,RPS17,PAFAH1B1,RPS24,RPL26,RPL31,GPC3,MEGF
I,COL11A1,RPS17,RPS24,RPL26,RPL31,DDX3X,RPS7,RPS19,RPL35,RPS10,RPL35A,TSR2,HNRNI
,SHOX,SKI,ASXL1,NOTCH2,SMARCA4,AEBP1,FLNB,SMARCE1,COL27A1,EP300,EXTL3,NXN,ARID
,RPS19,RPL35,RPS10,RPL35A,TSR2,RPS27,RPL27,RPS15A,RPS28,RPS26,RPS29
TT,SPARC,TUBB,KIDINS220,NUMA1,DCX,FLNA,SLC9A7,ALDOA,COL6A1,PRPF8,FBXL4,UBA1,CAL
6,RPL31,RPS7,RPS19,RPL35,RPS10,RPL35A,TSR2,RPS27,RPL27,ASXL1,RPS15A,RPS28,ADH5,SH
B,SLC9A7,FTL,NEFL,PRPF8,TUBB2B,PSAP,PGK1,KAT6A,KIF1A,GNAS,SLC2A3,APP,IDS,GDI1,NPN
24,RPL26,RPL31,LRPPRC,CDH2,RPS7,RPS19,RPL35,RPS10,RPL35A,ARID1A,TSR2,RPS27,CTBP1
N,ODC1,SON,KDM1A,MAP2K2,STXBP1,LONP1,RAB11A,CIC,EP300,KDM5B,ATP6AP2,GRN,TAOK
L5,NONO,DCHS1,PKD1,ARF1,MAT2A,COL1A2,COL5A2,RPL26,MMP14,DDX3X,PRKAR1A,GJA1,HI
VL,KDM1A,RAI1,EP300,RERE,FLCN,PKDCC,LTBP4,PACS1,PHGDH,YY1,CREBBP,TMEM94,NOTCH
6,RPL31,RPS7,RPS19,RPL35,RPS10,RPL35A,TSR2,RPS27,RPL27,RPS15A,RPS28,SMO,RPS26,SH
RCD1,SMARCC2,SMARCB1,DPF2
NRPN,PLOD1,SKI,RAB11B,ATP6AP1,SON,RAI1,YME1L1,KDM5C,EP300,RERE,FLCN,BAZ1B,FBXC
,P4HB,IGF2,AFF4,DCHS1,PPP2R1A,COL11A1,HNRNPU,COL1A2,CHD4,MEGF8,MMP14,RPS19,HN
T,POLR2A,UCHL1,AGRN,ZMIZ1,CLTC,CTNNB1,HNRNPK,ARCN1,EEF1A2,GNAI3,SNRPN,VPS35,SI

,SMARCC2,SMARCB1,DPF2
PN,ODC1,SON,STXBP1,LONP1,RAB11A,CIC,KDM5B,ATP6AP2,GRN,TAOK1,PTPN23,PSAT1,PDHA1,
A2,DDX3X,COL6A2,HNRNPK,RAC1,HS6ST1,PLOD1,COL2A1,SKI,NOTCH2,SELENON,MAPK1,FLNE
LC2A3,APP,SPTAN1,COL18A1,VCP,YWHAG,MAPT,SET,UCHL1,ZMIZ1,AP3D1,CYB5R3,PAFAH1B1,
APT,TPI1,AP2M1,AGRN,RPL10,MARCHF6,AP3D1,SBF1,PAFAH1B1,HNRNPU,SCARB2,HK1,RAB7A,
BA1,KAT6A,DVL3,P4HB,RPL5,IGF2,AFF4,DCHS1,CHSY1,RPL15,RPL18,RPL11,AGRN,ZMIZ1,COL1
BA1,KAT6A,DVL3,P4HB,RPL5,IGF2,AFF4,DCHS1,CHSY1,RPL15,RPL18,RPL11,AGRN,ZMIZ1,COL1
BA1,COL5A1,KAT6A,DVL3,KIF1A,GNAS,P4HB,IDS,RPL5,NPM1,IGF2,UBB,AFF4,NONO,DCHS1,CH
_4,SOX4,KAT6A,DVL3,YWHAE,KIF1A,ALDH6A1,GDI1,SNRNP200,AFF4,PPP2R1A,ZMIZ1,COL11A1,
I6A1,P4HB,GDI1,AFF4,YWHAG,NONO,PPP2R1A,ZMIZ1,COL11A1,PAFAH1B1,HNRNPU,NR2F1,CHD
14,KAT6A,DVL3,GNB1,RPL5,IGF2,UBB,NONO,CHSY1,RPL15,POLR2A,RPL18,RPL11,COL11A1,RF
IO,EP300,CNP,MED13,PCGF2,CREBBP,TANC2,SMO,EIF4G1,HS2ST1,ELN,ADNP,SETD5,GATAD2E
BA1,KAT6A,DVL3,P4HB,RPL5,IGF2,AFF4,DCHS1,CHSY1,RPL15,RPL18,RPL11,AGRN,ZMIZ1,COL1
PL11,ZMIZ1,RPL10,RPS17,RPS24,RPL26,RPL31,GPC3,MEGF8,RPS7,RPS19,RPL35,GJA1,RPS10,F
RPS7,RPS19,RPL35,RPS10,RPL35A,TSR2,RPS27,RPL27,RPS15A,RPS28,ADAR,PTEN,SH3PXD2B,F
7,RPS19,RPL35,RPS10,RPL35A,TSR2,RPS27,RPL27,RPS15A,RPS28,RPS26,RPS29
14,KAT6A,DVL3,GNB1,RPL5,IGF2,UBB,NONO,CHSY1,RPL15,POLR2A,RPL18,RPL11,COL11A1,RF
,MMP14,MMP2,PLOD1,COL2A1,SKI,SELENON,MAPK1,AEBP1,TPM3,EXTL3,BAZ1B,DZIP1,GTF2I,I
S,IDS,RPL5,RPL15,RPL18,RPL11,COL11A1,RPS17,PAFAH1B1,RPS24,RPL26,RPL31,GPC3,MEGF
5A,KIF1A,GNAS,GDI1,VCP,NONO,CHSY1,PPP2R1A,ZMIZ1,RPL10,COL1A2,RHOA,GPC3,MEGF8,RA
,MMP14,RPS7,RPS19,RPL35,GJA1,RPS10,RPL35A,TSR2,RPS27,MMP2,PPP1CB,RPL27,RPS15A,F
7,RPS24,HK1,RPL26,RPL31,KLHL9,RPS7,RPS19,RPL35,RPS10,RPL35A,HNRNPDL,TSR2,RPS27,S
RPL10,HNRNPU,HK1,CDH2,CLTC,GJA1,CTNNB1,GLUL,CDH11,PPP1CB,SNRPN,NOTCH2,MAPK1,
A3,APP,SPTAN1,COL18A1,VCP,YWHAG,MAPT,SET,UCHL1,ZMIZ1,AP3D1,CYB5R3,PAFAH1B1,CTSI
D1,VCP,UCHL1,TARDBP,MDH2,SQSTM1,ATXN10,SDHA,PRPS1,SOD1,CCT5,ADAR,SYP,KIF5A,KDM
D1,VCP,UCHL1,TARDBP,MDH2,SQSTM1,ATXN10,SDHA,PRPS1,SOD1,CCT5,ADAR,SYP,KIF5A,KDM
NOTCH2,KDM1A,DHX30,ATP6V1B2,RAI1,SMS,CDC42,TRIO,CSNK2A1,SMC3,ABCC9,EHMT1,PACS
_1A2,NR2F1,CHD4,GJA1,NARS1,LMNB2,PLOD1,COL2A1,SKI,ASXL1,RAB11B,MAPK1,ODC1,AEBF
L4,COL5A1,PSAP,KAT6A,KIF1A,GNAS,SLC2A3,APP,IDS,VCP,UBB,AFF4,YWHAG,MAPT,NONO,SET,
LTC,PRKAR1A,HNRNPK,CTBP1,RAC1,NOVA2,CDH11,EEF1A2,SNRPN,NOTCH2,SON,KDM1A,DLK
D1,CDH11,ASXL1,NOTCH2,MAPK1,SMARCA4,KDM1A,DHX30,ATP6V1B2,SDHA,RAI1,SMS,CDC42,
P2A2,SPTBN2,TKT,CLTC,L1CAM,EEF1A2,SNRPN,ABCA2,MARS1,DLK1,STXBP1,PRPS1,SARS1,RAI
_3,IDS,FN1,NONO,CHSY1,COL11A1,COL5A2,RPL26,GPC3,CCN2,MMP14,PRKAR1A,GJA1,COL6A
SPD1,YWHAG,PPP2R1A,POLR2A,ZMIZ1,AP3D1,SBF1,ARF1,CYB5R3,HNRNPU,HK1,NR2F1,LRPPR
Z1,COL11A1,RPL10,PAFAH1B1,CCND2,GJA1,GNAI3,COL2A1,SKI,NOTCH2,MAPK1,SON,DLK1,STX
TPM3,SMARCE1,FLNC,ADAR,TRAF7,DCTN1,KIF5A,EXTL3,GARS1,BAP1,SPG7,HINT1,REEP1,SMO,
I1,RPL10,MARCHF6,AP3D1,ARF1,PAFAH1B1,CDH2,CLTC,NARS1,ATP1A2,EEF1A2,VAMP2,YEATS2
,COL5A1,COL4A1,PSAP,PGK1,FKBP14,KIF1A,GNAS,SLC2A3,GPI,IDS,GDI1,NPM1,HSPD1,VCP,YWI
APT,AP2M1,AGRN,RPL10,MARCHF6,AP3D1,PAFAH1B1,HNRNPU,SCARB2,CLTC,CYFIP2,ATP1A2,
1,RPL15,RPL18,RPL11,ZMIZ1,RPL10,RPS17,RPS24,RPL26,RPL31,GPC3,RPS7,RPS19,RPL35,PR
_26,RPL31,KLHL9,RPS7,RPS19,RPL35,RPS10,RPL35A,TSR2,RPS27,SQSTM1,RPL27,RPS15A,RPS
ARID1A,SHOX,PLOD1,COL2A1,NOTCH2,SMARCA4,FLNB,SMARCE1,STXBP1,ARFGEF2,EP300,BAZ
P1CB,COL2A1,SKI,ASXL1,KDM6B,PRKDC,SON,KDM1A,FLNB,DLK1,COL27A1,ARNT2,CDC42,STA
S7,RPS19,RPL35,RPS10,RPL35A,TSR2,RPS27,APPL1,RPL27,RPS15A,RPS28,ATRX,RPS26,RPS29,
BA1,KAT6A,DVL3,P4HB,RPL5,IGF2,AFF4,FN1,DCHS1,CHSY1,RPL15,RPL18,RPL11,AGRN,ZMIZ1,

EEF1A2,SNRPN,SKI,MAPK1,PLEC,ODC1,PRKDC,SON,DLK1,PRPS1,RAI1,ARNT2,STAT3,BCR,KDM5
VAMP2,SNRPN,SKI,MAPK1,ODC1,STXBP1,USP7,RAI1,RAB11A,CIC,TRIO,EP300,RERE,KDM5B,ATI
MAT2A,COL1A2,GPC3,PRKAR1A,COL6A2,HNRNPK,CDKN1A,PLOD1,COL2A1,CCND1,DLK1,TPM3
A2,CDH2,GPC3,ARID1A,RAC1,CRPPA,PPP1CB,ASXL1,NOTCH2,SMARCA4,ACADVL,FTO,FLNB,SM
1,RAB7A,NARS1,MMP2,L1CAM,HS6ST1,PDXK,RAB11B,HSPB1,TPM3,ATP6V1B2,LONP1,PRPS1,KI
NOTCH2,KDM1A,DHX30,ATP6V1B2,RAI1,SMS,CDC42,TRIO,CSNK2A1,ARID1B,SMC3,ABCC9,EHM
,XRCC2,SPARC,TUBB,NUMA1,FLNA,MAP1B,SLC9A7,ALDOA,COL6A1,FTL,PRPF8,FBXL4,UBA1,C
,AGRN,RPL10,MARCHF6,AP3D1,PAFAH1B1,HNRNPU,SCARB2,CLTC,CYFIP2,ATP1A2,CTBP1,EEF1
4,FKBP14,KAT6A,DVL3,GNB1,RPL5,IGF2,UBB,NONO,CHSY1,RPL15,POLR2A,RPL18,RPL11,COL
TF2I,FBLN5,CLIP2,ELN,LIMK1,COL3A1
1,EP300,BAZ1B,NXN,SMC3,ABCC9,LOX,GTF2I,TTC37,PACS1,CREBBP,CLIP2,ELN,NOTCH1,ACTA
RPL11,ZMIZ1,RPS17,RPS24,RPL26,RPL31,GPC3,MEGF8,RPS7,RPS19,RPL35,RPS10,RPL35A,TSR
RPL15,RPL18,RPL11,ZMIZ1,RPS17,RPS24,RPL26,RPL31,GPC3,MEGF8,RPS7,RPS19,RPL35,GJA1
F4,NONO,DCHS1,SET,PPP2R1A,AGRN,ZMIZ1,CLEC7A,PRKCSH,CLTC,COL6A2,ARID1A,HNRNPK,R
1A,PLOD1,COL2A1,SKI,NOTCH2,MAPK1,SMARCA4,FLNB,SMARCE1,LONP1,SEC23A,STAT3,BCR,E
PAFAH1B1,HNRNPU,CCND2,CLTC,GJA1,CYFIP2,CTBP1,CDH11,EEF1A2,SKI,MAPK1,PRDX1,ODC1
,RPS24,RPS14,RPL26,RPL31,RPS7,RPS19,RPL35,RPS10,RPL35A,TSR2,RPS27,RPL27,ASXL1,RPS
S7,RPS19,RPL35,RPS10,RPL35A,ARID1A,TSR2,RPS27,RPL27,ASXL1,RPS15A,NOTCH2,SMARCE1,
8A1,VCP,YWHAG,MAPT,AP2M1,UCHL1,ITM2B,CTSD,SCARB2,XPR1,CLTC,PRKAR1A,CYFIP2,DNA
),RPL35,RPS10,RPL35A,TSR2,RPS27,RPL27,RPS15A,RPS28,RPS26,RPS29,SRP54,PIEZO1,DNAJC
NONO,CHSY1,PPP2R1A,COL11A1,RPL10,PAFAH1B1,GPC3,GJA1,SHOX,L1CAM,LMNB2,SNRPN,SK
A2,COL5A2,MMP14,GJA1,MMP2,PLOD1,COL2A1,SKI,SELENON,MAPK1,AEBP1,TPM3,EXTL3,BAZ
24,RPL26,RPL31,RPS7,RPS19,RPL35,RPS10,RPL35A,TSR2,HNRNPK,RPS27,CRPPA,RPL27,SKI,R
A,GJA1,COL6A2,ARID1A,SHOX,FDFT1,PPP1CB,PLOD1,COL2A1,SKI,ASXL1,NOTCH2,SELENON,M
1,EP300,BAZ1B,NXN,SMC3,ABCC9,LOX,GTF2I,TTC37,PACS1,CREBBP,CLIP2,ELN,NOTCH1,ACTA
KDM3B,BCR,RERE,BAZ1B,GTF2I,TUBGCP6,YY1,HS2ST1,ABL1,TMEM94,CLIP2,ELN,OBSL1,LIMK
GTF2I,CLIP2,ELN,NTN1,LIMK1,FLII,PRPH,DEAF1,NIPA1,SPG11,PDHB
PL18,RPL11,RPL10,RPS17,RPS24,COL1A2,CHD4,RPS20,RPL26,RPL31,TKT,CCND2,CDH2,GPC3,
AS,IGF2,NONO,CHSY1,PPP2R1A,COL11A1,RPL10,PAFAH1B1,CCND2,GPC3,MEGF8,GJA1,ARID1A
KIF1A,GNB1,COL18A1,COL4A2,SET,CCND2,CLTC,RAC1,CRPPA,EEF1A2,FDFT1,SNRPN,ODC1,SC
P,SQSTM1,TAF15,VPS35,OPTN,LSS,STXBP1,SOD1,DCTN1,UBQLN2,NDST1,FUS,GRN,VAPB,EIF4G
8,RPL11,COL11A1,RPS17,RPS24,RPL26,RPL31,DDX3X,RPS7,RPS19,RPL35,RPS10,RPL35A,TSR2
24,RPL26,RPL31,RPS7,RPS19,RPL35,RPS10,RPL35A,TSR2,HNRNPK,RPS27,CRPPA,GNAI3,RPL27
RCE1,ATP6V1B2,TRIO,EP300,ARID1B,SMC3,ABCC9,IARS2,SMARCD1,CREBBP,MN1,PHIP,AUTS2,
MP14,GJA1,ARID1A,HNRNPK,MMP2,L1CAM,COL2A1,RBMX,CNTNAP2,NOTCH2,MAPK1,SMARCA
HK1,CDH2,CLTC,CTNNB1,CDH11,SNRPN,MAPK1,ODC1,PRKDC,SON,KDM1A,SMARCE1,RAI1,SE
200,AFF4,PPP2R1A,COL11A1,PAFAH1B1,HK1,NR2F1,LRPPRC,GPC3,MEGF8,DDX3X,PRKAR1A,G
ARCC2,SMARCB1,DPF2
PSAP,FKBP14,DVL3,KIF1A,GDI1,HSPD1,COL18A1,IGF2,VCP,AFF4,MAPT,RPL11,ZMIZ1,SBF1,HNR
),RPL35,RPS10,RPL35A,TSR2,RPS27,RPL27,RPS15A,SON,RPS28,RPS26,RPS29,PSMD12
COL2A1,MAPK1,SMARCA4,KDELR2,DLK1,SMARCE1,STXBP1,BCR,EXTL3,ATRX,ARID1B,ABCC9,H
ARS1,ODC1,ATXN10,SDHA,STXBP1,SOD1,CCT5,ADAR,KIF5A,WDR45B,KDM5C,NDUFV1,ATRX,PI4
AE,AGRN,ARF1,PAFAH1B1,COL1A2,HK1,COL5A2,GPC3,GJA1,COL6A2,CTBP1,TBCD,PLOD1,COL2
,RPS19,RPL35,PRKAR1A,RPS10,RPL35A,TSR2,RPS27,RPL27,RPS15A,RPS28,SDHA,PALLD,FLCN
,COL11A1,CYFIP2,RAC1,TBCD,PPP2CA,MAPK1,DLK1,STXBP1,MESD,USP7,ADAR,ARID1B,FBXO1

ARCB1,DPF2
2,MMP2,SELENON,TPM3,SOD1,KIF5A,BAZ1B,RAB3GAP2,ATP6AP2,GTF2I,UBE3A,WNK1,REEP1,P
STXBP1,CCT5,KIF5A,WDR45B,KDM5C,NDUFV1,ATRX,RAB3GAP2,ZFR,SLC2A1,SPG7,REEP2,NDU
EEF1A2,CHD2,SON,STXBP1,RAB11A,SYP,CIC,KCNQ2,KDM5B,GLUD1,PI4KA,SLC2A1,TAOK1,EHM
,RAC1,CRPPA,PPP1CB,ASXL1,SMARCA4,ACADVL,FTO,FLNB,SMARCE1,CDC42,ARID1B,ABCC9,B4
F2,NONO,DCHS1,PAFAH1B1,CHD4,GPC3,MYO5A,ARID1A,RAC1,L1CAM,CRPPA,NOVA2,PPP1CB,S
NONO,DCHS1,CHSY1,PPP2R1A,ZMIZ1,COL11A1,RPL10,PAFAH1B1,RHOA,GPC3,GJA1,SHOX,NAR
F1A,IDS,VCP,AFF4,FN1,DCHS1,PPP2R1A,AGRN,ZMIZ1,COL11A1,PAFAH1B1,COL1A2,HK1,CHD4,
.5A2,MMP14,GJA1,MMP2,PLOD1,COL2A1,SKI,SELENON,MAPK1,AEBP1,TPM3,EXTL3,BAZ1B,DZI
TSC,ACTG1,XRCC2,HTT,TUBB,NUMA1,DCX,FLNA,MAP1B,SLC9A7,ALDOA,GANAB,COL6A1,FTL,FE
RPL11,ZMIZ1,RPS17,RPS24,RPL26,RPL31,GPC3,MEGF8,RPS7,RPS19,RPL35,RPS10,RPL35A,TSR
200,AFF4,PPP2R1A,COL11A1,PAFAH1B1,HK1,NR2F1,LRPPRC,GPC3,MEGF8,DDX3X,PRKAR1A,G
RE,CRTAP,SMC3,H19,ANTXR1,HSD17B4,CREBBP,SH3PXD2B,ALDH18A1,BANF1,PDGFRB,DDX6,P
,DCHS1,RPL15,RPL18,RPL11,ZMIZ1,RPL10,RPS17,RPS24,RPL26,RPL31,GPC3,RPS7,RPS19,RPL
,DCHS1,RPL15,RPL18,RPL11,ZMIZ1,RPL10,RPS17,RPS24,RPL26,RPL31,GPC3,RPS7,RPS19,RPL
T,SET,AP2M1,MARCHF6,SCARB2,PTCD3,CLTC,CYFIP2,DNAJC5,NARS1,EEF1A2,LMNB2,YEATS2,C
GF8,PRKAR1A,PLOD1,SKI,AEBP1,SON,MAP2K2,DLK1,ATP6V1B2,BAZ1B,GPX4,IARS1,PTEN,SPINT
NONO,CHSY1,PPP2R1A,COL11A1,RPL10,PAFAH1B1,GPC3,GJA1,SHOX,L1CAM,LMNB2,SNRPN,SK
RPS7,RPS19,RPL35,RPS10,RPL35A,TSR2,RPS27,CTBP1,LMNB2,RPL27,COL2A1,RPS15A,KDM1A
,COL11A1,RPL10,COL1A2,RHOA,NR2F1,GJA1,MYO5A,CTNNB1,HNRNPK,ARCN1,CPSF1,VCAN,TG
A1,TUBB2B,COL4A1,FKBP14,KAT6A,KIF1A,IDS,IGF2,VCP,AFF4,TPI1,POLR2A,AGRN,HK1,PFN1,KL
SMARCA4,SMARCE1,SDHA,IDH1,TRAF7,EP300,RERE,BRD4,PTEN,MAPRE2,GPC4,BAP1,SOX2,DIC
L6A1,UBA1,COL5A1,SOX4,KAT6A,DVL3,KIF1A,GNAS,IDS,RPL5,GDI1,IGF2,CHSY1,RPL15,RPL18,
RPL5,NONO,DCHS1,PKD1,ARF1,MAT2A,COL1A2,COL5A2,TPM1,CCND2,CDH2,GPC3,RPS19,PRK
PF8,FBXL4,UBA1,COL4A1,SOX4,KAT6A,DVL3,YWHAE,GNAS,RPL5,NPM1,SNRNP200,IGF2,AFF4,N
,FLNA,FTL,PRPF8,FBXL4,TUBB2B,COL5A1,COL4A1,SOX4,FKBP14,KAT6A,KIF1A,GNAS,ALDH6A1
PS1,USP7,SOD1,RAI1,DCTN1,EP300,RERE,FLCN,NDN,LTBP3,PCGF2,CREBBP,CCDC47,FXR1,PRP
DCX,FLNA,ALDOA,FBXL4,TUBB2B,COL5A1,SOX4,KAT6A,DVL3,YWHAE,KIF1A,RPL13,GNAS,GNB1
FIP2,CTBP1,RAC1,CDH11,MAPK1,KDM1A,STXBP1,MESD,TCTN1,CDC42,BCR,TRIO,EP300,CSNK2
,SELENON,TPM3,FBLN1,LONP1,GPC4,IARS2,HSPG2,COL12A1,OBSL1,ALDH18A1,SLC25A1,PPP
MP2,CDKN1A,SQSTM1,NOTCH2,IDH1,HNRNPA2B1,ELP1,WNK1,FGFR1,SH3PXD2B,IDH2,BANF1,
P1,ARFGEF2,CHST3
RPS17,RPS24,RPS14,RPL26,RPL31,RPS7,RPS19,RPL35,RPS10,RPL35A,TSR2,RPS27,RPL27,ASX
L10,MARCHF6,AP3D1,HNRNPU,CLTC,NARS1,ATP1A2,EEF1A2,TBCD,SQSTM1,VAMP2,YEATS2,CHD
FAH1B1,SCARB2,ATP1A1,CLTC,HNRNPK,NARS1,EEF1A2,TBCD,YEATS2,CHD2,CNTNAP2,MAPK1,
,ATP5F1A,NONO,DCHS1,PPP2R1A,RPL10,PAFAH1B1,GPC3,MYO5A,MACF1,ARID1A,RAC1,L1CAM
F1A,IDS,AFF4,PPP2R1A,AGRN,ZMIZ1,COL11A1,PAFAH1B1,COL1A2,HK1,CHD4,COL5A2,GPC3,C
,RPS17,CYB5R3,RPS24,COL1A2,PTCD3,RPL26,RPL31,RPS7,RPS19,RPL35,PRKAR1A,RPS10,RPL
2,HLA-B,SKI,AEBP1,TPM3,RERE,BAZ1B,LOX,KMT2E,GTF2I,MED13,FBLN5,TANC2,POR,CLIP2,EL
IA,MAP1B,COL6A1,NEFL,UBA1,COL5A1,PSAP,SOX4,KAT6A,DVL3,YWHAE,KIF1A,RPL13,GNAS,IDS
24,RPL26,RPL31,RPS7,RPS19,RPL35,RPS10,RPL35A,TSR2,HNRNPK,RPS27,CRPPA,GNAI3,RPL27
,COL1A2,CDH2,GPC3,ARID1A,RAC1,CRPPA,PPP1CB,ASXL1,NOTCH2,SMARCA4,ACADVL,FTO,FL
PL5,COL18A1,IGF2,AFF4,NONO,RPL11,ZMIZ1,ARF1,MAT2A,TKT,GPC3,MEGF8,GJA1,RAC1,PPP1
,DCHS1,RPL15,RPL18,RPL11,ZMIZ1,RPL10,RPS17,RPS24,RPL26,RPL31,GPC3,RPS7,RPS19,RPL
Z1,RPS20,PFN1,CDH2,PRKAR1A,TARDBP,HS6ST1,MDH2,SQSTM1,TAF15,SNRPN,ASXL1,OPTN,VH

ARDBP,SQSTM1,TAF15,VPS35,OPTN,LSS,STXBP1,SOD1,DCTN1,KDM5C,UBQLN2,NDST1,FUS,G
NRNP200,AFF4,PPP2R1A,COL11A1,PAFAH1B1,HK1,RHOA,NR2F1,LRPPRC,GPC3,MEGF8,DDX3X
F8,FBXL4,UBA1,COL4A1,SOX4,KAT6A,DVL3,YWHAE,GNAS,RPL5,NPM1,SNRNP200,IGF2,AFF4,N
R1A,COL11A1,RPL10,PAFAH1B1,GPC3,GJA1,SHOX,L1CAM,LMNB2,SNRPN,SKI,KDM6B,MAPK1,
300,RERE,DGCR2,BAZ1B,ABCC9,MAPRE2,HSPG2,KMT2E,GTF2I,RBPJ,MED13,CREBBP,TANC2,S
A,DVL3,GNAS,P4HB,IDS,GDI1,NPM1,IGF2,YWHAG,NONO,DCHS1,SET,CHSY1,CHD3,POLR2A,ZM
CX,FLNA,MAP1B,GANAB,COL6A1,FTL,FBXL4,CALR,COL5A1,PSAP,SOX4,KAT6A,KIF1A,GNAS,SL
,RPL35,CTNNB1,RPS10,RPL35A,TSR2,RPS27,RPL27,RPS15A,CCND1,RPS28,EP300,FLCN,MSH6,
,RPL35,CTNNB1,RPS10,RPL35A,TSR2,RPS27,RPL27,RPS15A,CCND1,RPS28,EP300,FLCN,MSH6,
1B,SLC2A1,FUS,SPG7,VAPB,GTF2I,CLIP2,ELN,NTN1,LIMK1,FLII,PRPH,DEAF1,NIPA1,SPG11,PDH
HSY1,ZMIZ1,COL11A1,HNRNPU,COL1A2,LRPPRC,CDH2,MEGF8,GJA1,CYFIP2,ARID1A,HNRNPK,
IK3R2,FLNA,MAP1B,GANAB,FBXL4,COL5A1,SOX4,KAT6A,DVL3,CALM3,IDS,RPL5,COL18A1,IGF2
A1,SKI,MAPK1,AEBP1,SON,SMARCE1,HYOU1,SMS,BCR,ATRX,LOX,EFTUD2,PACS1,NSD2,FBLN5,
B11B,ATXN10,STXBP1,SMS,EP300,KDM5B,GARS1,MCM3AP,ATP6AP2,PACS1,CIZ1,CREBBP,REE
,COL5A2,GPC3,MEGF8,GJA1,RAC1,CDH11,PLOD1,COL2A1,SKI,NOTCH2,PPP2CA,AEBP1,FTO,DL
CAM,COL2A1,SELENON,MAPK1,PLEC,SON,FLNB,TPM3,BCR,EXTL3,RERE,BAZ1B,GARS1,RAB3G
,SPARC,TUBB,NUMA1,FLNA,MAP1B,ALDOA,COL6A1,PRPF8,UBA1,CALR,COL5A1,PSAP,SOX4,FK
,P4HB,IGF2,AFF4,DCHS1,PPP2R1A,COL11A1,HNRNPU,CTSD,COL1A2,CHD4,MEGF8,MMP14,RPS
ARCB1,DPF2
ARCB1,DPF2
ARCB1,DPF2
ARCB1,DPF2
ARCB1,DPF2

\1,CALR,TUBB2B,COL5A1,COL4A1,PSAP,PGK1,SOX4,FKBP14,KAT6A,DVL3,YWHAE,KIF1A,GNAS,
COL4A1,PSAP,PGK1,SOX4,FKBP14,KAT6A,YWHAE,KIF1A,GNAS,SLC2A3,GPI,ALDH6A1,GNB1,P4H
,SOX4,FKBP14,KAT6A,YWHAE,KIF1A,ALDH6A1,GNB1,P4HB,APP,GDI1,SPTAN1,HSPD1,IGF2,ATP5
SOX4,FKBP14,DVL3,YWHAE,KIF1A,RPL13,GNAS,P4HB,IDS,RPL5,NPM1,HSPD1,IGF2,VCP,AFF4,Y\
B,SLC9A7,ALDOA,COL6A1,FTL,NEFL,PRPF8,FBXL4,UBA1,CALR,TUBB2B,COL5A1,COL4A1,PSAP,
,MAP1B,SLC9A7,ALDOA,GANAB,COL6A1,FTL,NEFL,PRPF8,FBXL4,UBA1,CALR,TUBB2B,COL5A1,(
Z1,PAFAH1B1,HK1,NR2F1,DDX3X,CLTC,CTNNB1,MACF1,ARID1A,HNRNPK,NARS1,RAC1,GLUL,CF
OL6A1,NEFL,FBXL4,UBA1,TUBB2B,COL5A1,COL4A1,PSAP,SOX4,FKBP14,KAT6A,DVL3,YWHAE,KI
A7,GANAB,COL6A1,FTL,NEFL,FBXL4,UBA1,CALR,TUBB2B,COL5A1,COL4A1,PSAP,SOX4,KAT6A,Y
ISY1,PPP2R1A,RPL15,CHD3,POLR2A,RPL18,RPL11,ZMIZ1,COL11A1,RPS17,AP3D1,PAFAH1B1,M/
6A,YWHAE,KIF1A,GNAS,SLC2A3,ALDH6A1,GNB1,APP,IDS,RPL5,SPTAN1,NPM1,HSPD1,COL18A1,A
6A,YWHAE,KIF1A,GNAS,SLC2A3,ALDH6A1,GNB1,APP,IDS,RPL5,SPTAN1,NPM1,HSPD1,COL18A1,A
B,SLC9A7,ALDOA,COL6A1,FTL,NEFL,PRPF8,FBXL4,UBA1,CALR,TUBB2B,COL5A1,COL4A1,PSAF
X4,KAT6A,YWHAE,KIF1A,SLC2A3,ALDH6A1,APP,RPL5,SPTAN1,HSPD1,COL18A1,IGF2,ATP5F1A,VC
I,COL11A1,RPS17,PAFAH1B1,MAT2A,HNRNPU,RPS24,COL1A2,CHD4,RPL26,RPL31,LRPPRC,CCN
OL6A1,NEFL,PRPF8,FBXL4,TUBB2B,COL5A1,COL4A1,PSAP,PGK1,SOX4,FKBP14,KAT6A,DVL3,YV
32B,COL5A1,COL4A1,PSAP,SOX4,KAT6A,DVL3,YWHAE,KIF1A,GNAS,ALDH6A1,P4HB,IDS,RPL5,GD
KIF1A,RPL13,GNAS,P4HB,IDS,NPM1,HSPD1,IGF2,VCP,YWHAG,MAPT,FN1,NONO,DCHS1,SET,TPI1
PRPF8,UBA1,CALR,TUBB2B,COL5A1,COL4A1,SOX4,KAT6A,DVL3,KIF1A,RPL13,GNAS,CALM3,GNI
IZ1,COL11A1,RPL10,AP3D1,PAFAH1B1,HNRNPU,HK1,CDH2,GPC3,DDX3X,CLTC,RPS7,RPS19,GJ/
SPD1,IGF2,ATP5F1A,AFF4,YWHAG,NONO,DCHS1,SET,PPP2R1A,RPL15,CHD3,POLR2A,RPL18,RPI
,RPL5,SPTAN1,HSPD1,ATP5F1A,AFF4,YWHAG,NONO,DCHS1,SET,PPP2R1A,RPL15,POLR2A,RPL1
3,FTL,FBXL4,CALR,TUBB2B,COL5A1,COL4A1,PSAP,SOX4,KAT6A,YWHAE,KIF1A,GNAS,SLC2A3,AI
.,NEFL,PRPF8,UBA1,TUBB2B,COL4A1,PSAP,PGK1,SOX4,KAT6A,DVL3,KIF1A,GNAS,SLC2A3,GPI,G
3XL4,UBA1,COL5A1,COL4A1,PSAP,SOX4,FKBP14,KAT6A,DVL3,YWHAE,KIF1A,RPL13,GNAS,GNB1
A7,ALDOA,COL6A1,NEFL,FBXL4,UBA1,TUBB2B,COL5A1,COL4A1,PSAP,SOX4,FKBP14,KAT6A,DVI
TAN1,VCP,YWHAG,MAPT,NONO,DCHS1,SET,PPP2R1A,POLR2A,AP2M1,ZMIZ1,PAFAH1B1,HNRNPU
\7,GANAB,FTL,NEFL,FBXL4,UBA1,CALR,TUBB2B,COL5A1,COL4A1,PSAP,SOX4,KAT6A,YWHAE,KII
COL5A1,COL4A1,PSAP,PGK1,SOX4,FKBP14,KAT6A,DVL3,YWHAE,KIF1A,GNAS,ALDH6A1,GNB1,P4
ZMIZ1,COL11A1,RPL10,RPS17,AP3D1,RPS24,NR2F1,CHD4,RPL26,RPL31,CDH2,GPC3,MEGF8,C
LTC,GJA1,CTNNB1,ARID1A,CTBP1,NOVA2,CDH11,PPP1CB,COL2A1,SKI,NOTCH2,PPP2CA,MAPK1
13,GNAS,P4HB,NPM1,HSPD1,IGF2,YWHAG,MAPT,FN1,NONO,DCHS1,SET,PPP2R1A,CHD3,RPL11,
B1,APP,GDI1,PLD3,IGF2,VCP,UBB,YWHAG,MAPT,NONO,SET,CHSY1,PPP2R1A,CHD3,AP2M1,AGR
NP200,AFF4,NONO,DCHS1,SET,CHSY1,PPP2R1A,RPL15,CHD3,RPL18,RPL11,ZMIZ1,COL11A1,RF
1,COL2A1,SKI,ASXL1,NOTCH2,PPP2CA,MAPK1,ODC1,AEBP1,FLNB,DLK1,KDM3B,CDC42,BCR,D(
,NONO,DCHS1,SET,PPP2R1A,POLR2A,ZMIZ1,PAFAH1B1,HNRNPU,HK1,NR2F1,CDH2,GPC3,DDX3
.4A1,PSAP,SOX4,KAT6A,DVL3,YWHAE,KIF1A,RPL13,GNAS,P4HB,IDS,RPL5,GDI1,NPM1,IGF2,VCP,
6A1,IDS,SPTAN1,VCP,YWHAG,MAPT,NONO,DCHS1,SET,PPP2R1A,POLR2A,AP2M1,ZMIZ1,CYB5R3,
AS,IDS,RPL5,GDI1,IGF2,VCP,AFF4,MAPT,NONO,DCHS1,CHSY1,PPP2R1A,RPL15,RPL18,RPL11,AG
L,PRPF8,FBXL4,UBA1,CALR,TUBB2B,COL5A1,COL4A1,PSAP,PGK1,SOX4,FKBP14,KAT6A,DVL3,Y
\,RPL15,POLR2A,RPL18,RPL11,AGRN,ZMIZ1,COL11A1,RPS17,ARF1,MAT2A,RPS24,COL1A2,CHD
A3,CALM3,GNB1,APP,IDS,GDI1,SPTAN1,CALM1,HSPD1,COL18A1,IGF2,ATP5F1A,VCP,YWHAG,CO

SET,PPP2R1A,RPL15,POLR2A,RPL18,RPL11,ZMIZ1,RPL10,RPS17,AP3D1,SBF1,ARF1,CYB5R3,PAF
29,UQCRFS1

B,SLC9A7,ALDOA,COL6A1,NEFL,FBXL4,UBA1,TUBB2B,COL5A1,COL4A1,PSAP,SOX4,FKBP14,K
A,RPL13,GNAS,IDS,RPL5,GDI1,IGF2,VCP,AFF4,MAPT,FN1,NONO,DCHS1,CHSY1,PPP2R1A,RPL15
HS1,SET,PPP2R1A,RPL15,POLR2A,RPL18,RPL11,ZMIZ1,RPL10,RPS17,AP3D1,SBF1,ARF1,CYB5
,KAT6A,YWHAE,KIF1A,ALDH6A1,GNB1,P4HB,GDI1,SPTAN1,HSPD1,IGF2,ATP5F1A,YWHAG,NONO,
NAS,ALDH6A1,P4HB,RPL5,GDI1,NPM1,COL18A1,AFF4,YWHAG,MAPT,NONO,DCHS1,CHSY1,PPP2
,SOX4,KAT6A,DVL3,YWHAE,KIF1A,RPL13,GNAS,P4HB,IDS,RPL5,GDI1,NPM1,IGF2,VCP,AFF4,FN1,
,SOX4,KAT6A,DVL3,YWHAE,KIF1A,RPL13,GNAS,P4HB,IDS,RPL5,GDI1,NPM1,IGF2,VCP,AFF4,FN1,

D3,RPL18,RPL11,ZMIZ1,COL11A1,RPL10,RPS17,PAFAH1B1,HNRNPU,RPS24,COL1A2,HK1,NR2F
4,FKBP14,KAT6A,DVL3,KIF1A,GNAS,GNB1,APP,IDS,RPL5,GDI1,HSPD1,IGF2,VCP,AFF4,YWHAG,M
N,COL2A1,SKI,KDM6B,NOTCH2,MAPK1,SMARCA4,SON,DLK1,SMARCE1,ATP6V1B2,STXBP1,RAI1

,SLC9A7,ALDOA,COL6A1,FTL,NEFL,PRPF8,FBXL4,UBA1,TUBB2B,COL5A1,COL4A1,PSAP,SOX4,
GDI1,NPM1,COL18A1,AFF4,YWHAG,NONO,DCHS1,PPP2R1A,RPL15,CHD3,RPL18,RPL11,ZMIZ1,
PPP1CB,LMNB2,PLOD1,COL2A1,SKI,KDM6B,NOTCH2,MAPK1,SMARCA4,KDELR2,AEBP1,SON,K
2R1A,CHD3,ZMIZ1,RPL10,ARF1,PAFAH1B1,HNRNPU,HK1,NR2F1,LRPPRC,SPTBN2,TKT,CCND2,C
DS,RPL5,GDI1,IGF2,VCP,AFF4,NONO,DCHS1,CHSY1,PPP2R1A,RPL15,RPL18,RPL11,ZMIZ1,COL
,MMP14,RPS7,RPS19,RPL35,GJA1,RPS10,RPL35A,TSR2,HNRNPK,RPS27,CTBP1,RAC1,MMP2,P
B2B,COL5A1,COL4A1,SOX4,FKBP14,KAT6A,DVL3,YWHAE,KIF1A,GNAS,ALDH6A1,GNB1,P4HB,ID
MAT2A,COL1A2,COL5A2,GPC3,MEGF8,MMP14,GJA1,ARID1A,RAC1,SMCHD1,MMP2,CDH11,PLO
,MMP14,RPS7,RPS19,RPL35,GJA1,RPS10,RPL35A,TSR2,HNRNPK,RPS27,CTBP1,RAC1,MMP2,P
,SLC9A7,ALDOA,COL6A1,FTL,NEFL,PRPF8,FBXL4,UBA1,TUBB2B,COL5A1,COL4A1,PSAP,SOX4,
NAS,ALDH6A1,P4HB,RPL5,GDI1,NPM1,COL18A1,AFF4,YWHAG,MAPT,NONO,DCHS1,CHSY1,PPP2

R1A,AP2M1,ZMIZ1,COL11A1,RPL10,AP3D1,CYB5R3,PAFAH1B1,HNRNPU,HK1,CLEC7A,CDH2,GP
S1,SET,PPP2R1A,POLR2A,ZMIZ1,PAFAH1B1,HNRNPU,HK1,NR2F1,CDH2,GPC3,DDX3X,CLTC,CTN
FNB1,TMEM43,RPS29,LMNA,SEPTIN9,FHL1,TMCO1,COL3A1,EMD
,RPL31,LRPPRC,GPC3,MEGF8,DDX3X,RPS7,RPS19,RPL35,PRKAR1A,GJA1,RPS10,RPL35A,ARID1
S1,PPP2R1A,RPL15,CHD3,POLR2A,RPL18,RPL11,AGRN,ZMIZ1,COL11A1,RPL10,RPS17,PAFAH1B
,AFF4,DCHS1,SET,CHSY1,RPL15,CHD3,RPL18,RPL11,AGRN,ZMIZ1,COL11A1,RPL10,RPS17,AP3D
,PPP2R1A,CHD3,POLR2A,AP2M1,RPL10,AP3D1,HNRNPU,COL1A2,CHD4,LRPPRC,SPTBN2,TKT,G
,ALDH6A1,P4HB,RPL5,GDI1,NPM1,COL18A1,AFF4,YWHAG,MAPT,NONO,DCHS1,CHSY1,PPP2R1A
,GARS1,FBXO11,TIMM50,FUS,KMT2E,PACS2,UBE3A,YY1,SMARCD1,CREBBP,SEMA6B,TMEM94,
S1,SET,PPP2R1A,POLR2A,ZMIZ1,PAFAH1B1,HNRNPU,HK1,NR2F1,CDH2,GPC3,DDX3X,CLTC,CTN
5A1,COL4A1,PSAP,PGK1,SOX4,FKBP14,KAT6A,DVL3,KIF1A,GNAS,SLC2A3,GNB1,APP,IDS,RPL5,
,AGRN,ZMIZ1,COL11A1,RPS17,PAFAH1B1,MAT2A,HNRNPU,RPS24,COL1A2,RPL26,RPL31,CDH2,
4,RPL26,RPL31,TKT,CCND2,CDH2,GPC3,MEGF8,MMP14,RPS7,RPS19,RPL35,GJA1,RPS10,RPL
4,NONO,DCHS1,SET,CHSY1,RPL15,CHD3,POLR2A,RPL18,RPL11,AGRN,ZMIZ1,COL11A1,RPS17,
O,RPL15,CHD3,POLR2A,RPL18,RPL11,AGRN,ZMIZ1,COL11A1,RPL10,RPS17,SBF1,CYB5R3,HN
1A2,CHD4,COL5A2,RPL26,RPL31,CDH2,MMP14,RPS7,RPS19,RPL35,GJA1,RPS10,RPL35A,TSR2
,IGF2,NONO,CHSY1,PPP2R1A,RPL15,RPL18,RPL11,ZMIZ1,COL11A1,RPL10,RPS17,PAFAH1B1,M
F2,VCP,FN1,NONO,DCHS1,CHSY1,PPP2R1A,CHD3,UCHL1,AGRN,ZMIZ1,COL11A1,RPL10,SBF1,
P2M1,UCHL1,AGRN,ZMIZ1,RPL10,MARCHF6,SBF1,SCARB2,HK1,CHD4,ATP1A1,RPS20,KLHL9,S

A1,SOX4,FKBP14,KAT6A,DVL3,YWHAE,KIF1A,GNAS,ALDH6A1,GNB1,P4HB,IDS,RPL5,GDI1,NPM1,
A1,SOX4,FKBP14,KAT6A,DVL3,YWHAE,KIF1A,GNAS,ALDH6A1,GNB1,P4HB,IDS,RPL5,GDI1,NPM1,
A,DVL3,YWHAE,KIF1A,RPL13,GNAS,IDS,GDI1,HSPD1,IGF2,VCP,AFF4,MAPT,FN1,NONO,DCHS1,CH
8A1,IGF2,VCP,AFF4,NONO,SET,CHSY1,PPP2R1A,CHD3,POLR2A,COL11A1,RPL10,PAFAH1B1,HN
CHD3,RPL18,RPL11,ZMIZ1,COL11A1,RPL10,RPS17,CTSD,RPS24,COL1A2,RHOA,NR2F1,RPL26,R
NAS,SLC2A3,ALDH6A1,GNB1,APP,GDI1,SPTAN1,PLD3,HSPD1,SNRNP200,VCP,YWHAG,MAPT,FN1,
11B,ABCA2,SELENON,PPP2CA,PLEC,ODC1,KDM1A,DLK1,DHX30,LARS1,LSS,FBLN1,STXBP1,MES
ARCHF6,CYB5R3,ITM2B,SCARB2,PTCD3,NR2F1,PFN1,LRPPRC,XPR1,SPTBN2,TKT,CLTC,MYO5A

2,ABCA2,MAPK1,ODC1,ATXN10,STXBP1,USP7,SARS1,RAI1,RAB11A,KDM3B,CIC,KDM5C,TRIO,EF
1,HNRNPU,HK1,NR2F1,LRPPRC,SPTBN2,TKT,CCND2,CLTC,PRKAR1A,MACF1,CYFIP2,ARID1A,NA

OA,COL6A1,FTL,NEFL,PRPF8,FBXL4,UBA1,TUBB2B,COL5A1,COL4A1,SOX4,FKBP14,KAT6A,DVL
RF1,CYB5R3,PAFAH1B1,ITM2B,CTSD,HK1,NR2F1,RPS20,PFN1,SPTBN2,DDX3X,CLTC,GJA1,MYO
CCND2,CDH2,GPC3,MEGF8,RPS7,RPS19,RPL35,PRKAR1A,GJA1,RPS10,RPL35A,ARID1A,TSR2,SH
1A1,RPL26,RPL31,KLHL9,RAB7A,CLTC,RPS7,RPS19,RPL35,COL6A2,RPS10,RPL35A,CYFIP2,HNF
CHSY1,PPP2R1A,AGRN,ZMIZ1,COL11A1,RPL10,RPS17,COL1A2,RHOA,NR2F1,CHD4,GPC3,MMP1
4HB,IDS,COL18A1,IGF2,VCP,AFF4,NONO,DCHS1,SET,CHSY1,PPP2R1A,CHD3,POLR2A,COL11A1,
PL13,GNAS,P4HB,IDS,RPL5,NPM1,IGF2,VCP,AFF4,YWHAG,FN1,NONO,DCHS1,SET,CHSY1,RPL15
NOVA2,EEF1A2,SQSTM1,VAMP2,SNRPN,CHD2,SKI,KDM6B,CNTNAP2,RAB11B,PPP2CA,SON,DLK1
A
H1B1,COL1A2,HK1,COL5A2,CDH2,CTNNB1,COL6A2,HNRNPK,NARS1,ARCN1,CTBP1,RAC1,CRPP
PAFAH1B1,CTSD,RPS24,CHD4,RPL26,RPL31,CDH2,GPC3,MEGF8,RPS7,RPS19,RPL35,GJA1,RPS
AE,KIF1A,ALDH6A1,GNB1,P4HB,APP,IDS,GDI1,NPM1,HSPD1,IGF2,AFF4,YWHAG,COL4A2,NONO,D
3,POLR2A,RPL18,RPL11,AGRN,ZMIZ1,COL11A1,RPL10,RPS17,SBF1,CYB5R3,HNRNPU,RPS24,C

P,AFF4,NONO,DCHS1,SET,CHSY1,PPP2R1A,UCHL1,AGRN,ZMIZ1,COL11A1,RPL10,ARF1,COL1A2
,AP3D1,PAFAH1B1,CTSD,RPS24,CHD4,RPL26,RPL31,CDH2,GPC3,MEGF8,RPS7,RPS19,RPL35,G
SOX4,KAT6A,DVL3,KIF1A,GNAS,GPI,GNB1,APP,IDS,GDI1,SPTAN1,HSPD1,SNRNP200,IGF2,VCP,A
,GPI,GNB1,APP,GDI1,SPTAN1,PLD3,HSPD1,COL18A1,VCP,YWHAG,COL4A2,MAPT,NONO,DCHS1,
M6B,SELENON,KDELR2,AEBP1,MAP2K2,FLNB,DLK1,TPM3,SMS,KDM5C,TRIO,EP300,EXTL3,BAZ

ATRX,NXN,ARID1B,HDAC6,ABCC9,GPC4,GTF2I,TTC37,PACS1,DHCR7,MED13,MTOR,PACS2,UBE
H1B1,ITM2B,CTSD,NR2F1,PFN1,SPTBN2,DDX3X,CLTC,GJA1,MYO5A,CTNNB1,MACF1,CYFIP2,TA
,COL11A1,HNRNPU,HK1,RHOA,LRPPRC,GPC3,MEGF8,PRKAR1A,TSR2,HNRNPK,CTBP1,SMCHD1
R2A,AGRN,RPL10,ARF1,SCARB2,COL5A2,PFN1,PRKCSH,CCN2,CLTC,MACF1,CYFIP2,TARDBP,D
,GJA1,RPS10,RPL35A,TSR2,HNRNPK,ARCN1,RPS27,CTBP1,RAC1,MMP2,PPP1CB,RPL27,ASXL1,
C3,MEGF8,MMP14,GJA1,RAC1,SMCHD1,MMP2,CDH11,PLOD1,COL2A1,SKI,ASXL1,NOTCH2,PPP2
D1A,NARS1,HLA-B,CTBP1,MMP2,L1CAM,PPIB,CRPPA,CDH11,SNRPN,PLOD1,COL2A1,SKI,NOTCH
C3,MEGF8,MMP14,GJA1,RAC1,SMCHD1,MMP2,CDH11,PLOD1,COL2A1,SKI,ASXL1,NOTCH2,PPP2
,GJA1,RPS10,RPL35A,TSR2,HNRNPK,ARCN1,RPS27,CTBP1,RAC1,MMP2,PPP1CB,RPL27,ASXL1,
IIZ1,AP3D1,SBF1,CYB5R3,PAFAH1B1,CTSD,PTCD3,PFN1,CLTC,MYO5A,CYFIP2,DNAJC5,TARDBP,
Y1,RPL15,RPL18,RPL11,AGRN,ZMIZ1,COL11A1,RPL10,RPS17,HNRNPU,RPS24,COL1A2,CHD4,R
1A2,CHD4,RPL26,RPL31,CLTC,RPS7,RPS19,RPL35,RPS10,RPL35A,CYFIP2,TARDBP,ARID1A,TSR
6B,CNTNAP2,MAPK1,ODC1,STXBP1,USP7,SARS1,RAI1,KDM3B,YME1L1,SYP,CIC,BCR,TRIO,EP30

.OD1,COL2A1,SKI,SELENON,MAPK1,KDELR2,AEBP1,FLNB,TPM3,DHX30,ATP6V1B2,STXBP1,SAR
/FIP2,HNRNPK,CDH11,EEF1A2,SNRPN,RAB11B,SELENON,MAPK1,ODC1,PRKDC,SON,KDM1A,DLI
1,HNRNPU,RHOA,NR2F1,CHD4,CDH2,GPC3,MEGF8,CLTC,RPS19,GJA1,CYFIP2,TSR2,HNRNPK,N
,RPS15A,SMARCA4,SON,SMARCE1,RPS28,RAI1,SMS,EXTL3,PBX1,BAZ1B,ARID1B,RAB3GAP2,AK

24,HK1,RPL26,RPL31,CDH2,GPC3,MEGF8,RPS7,RPS19,RPL35,COL6A2,RPS10,RPL35A,TSR2,SH
1A,TSR2,RPS27,CTBP1,NOVA2,PPP1CB,RPL27,SNRPN,ASXL1,RPS15A,NOTCH2,MAPK1,AEBP1,K
L4A1,PSAP,PGK1,SOX4,FKBP14,KAT6A,DVL3,YWHAE,KIF1A,GNAS,ALDH6A1,MYH9,P4HB,IDS,RPI
,ZMIZ1,COL11A1,RPL10,RPS17,AP3D1,MAT2A,HNRNPU,RPS24,COL1A2,COL5A2,RPL26,RPL31,
3,RPS7,RPS19,RPL35,PRKAR1A,GJA1,RPS10,RPL35A,ARID1A,TSR2,SHOX,RPS27,CTBP1,RAC1,CI
,YWHAG,COL4A2,MAPT,SET,TPI1,UCHL1,ARF1,CYB5R3,PAFAH1B1,ITM2B,CTSD,HK1,NR2F1,RP
.1A2,COL5A2,GPC3,MEGF8,MMP14,GJA1,CTNNB1,RAC1,SMCHD1,MMP2,CDH11,PLOD1,COL2A
RPL18,RPL11,AGRN,ZMIZ1,COL11A1,RPL10,RPS17,AP3D1,MAT2A,HNRNPU,RPS24,COL1A2,COL
,ARF1,MAT2A,RPS24,COL1A2,CHD4,COL5A2,RPL26,RPL31,TKT,CDH2,MMP14,RPS7,RPS19,RPL
37,RPS26,ABL1,RPS29,SKIV2L
SP7,SARS1,RAI1,KDM3B,SYP,CIC,BCR,TRIO,EP300,RERE,KDM5B,BAZ1B,CSNK2A1,GLUD1,FBXO
,RPL18,RPL11,ZMIZ1,RPL10,RPS17,ARF1,MAT2A,RPS24,COL1A2,CHD4,COL5A2,RPL26,RPL31,
,AGRN,ZMIZ1,AP3D1,PAFAH1B1,HK1,PTCD3,CLEC7A,PRKCSH,CLTC,COL6A2,CYFIP2,ARID1A,HN
MIZ1,RPL10,CYB5R3,PAFAH1B1,CTSD,SCARB2,HK1,CHD4,SPTBN2,GPC3,GJA1,MYO5A,MACF1,
17,RPS24,CHD4,RPL26,RPL31,GPC3,MEGF8,RPS7,RPS19,RPL35,PRKAR1A,GJA1,RPS10,RPL35
P4HB,IDS,NPM1,HSPD1,COL18A1,IGF2,VCP,AFF4,FN1,NONO,DCHS1,SET,CHSY1,PPP2R1A,UCHL
B,PLOD1,COL2A1,SKI,KDM6B,NOTCH2,SELENON,KDELR2,AEBP1,MAP2K2,FLNB,DLK1,TPM3,H
A3,GNB1,APP,RPL5,PLD3,HSPD1,COL18A1,SNRNP200,ATP5F1A,VCP,AFF4,YWHAG,COL4A2,MAP

N,COL11A1,RPL10,ARF1,PAFAH1B1,COL1A2,HK1,CHD4,COL5A2,KLHL9,GPC3,MEGF8,GJA1,COI
,MMP14,RPS7,RPS19,RPL35,GJA1,RPS10,RPL35A,TSR2,HNRNPK,RPS27,CTBP1,RAC1,MMP2,PF
PPP2R1A,RPL15,HLA-C,SF3B1,RPL18,RPL11,ZMIZ1,COL11A1,RPL10,RPS17,AP3D1,CYB5R3,HN

L27A1,ARFGEF2,EP300,EXTL3,NXN,ARID1B,SMC3,HSPG2,SMARCD1,CREBBP,MACROH2A1,EXT
TAF15,RPL27,ASXL1,RPS15A,CCND1,SMARCE1,RPS28,ATP6V1B2,SDHA,IDH1,TRAF7,EP300,FLCI
NAI3,SNRPN,PLOD1,SKI,NOTCH2,MAPK1,AEBP1,KDM1A,ATP6V1B2,RAI1,SMS,CDC42,KDM5C,TRI
MIZ1,RPL10,CYB5R3,PAFAH1B1,CTSD,SCARB2,HK1,CHD4,SPTBN2,GPC3,GJA1,MYO5A,MACF1,
MIZ1,RPL10,CYB5R3,PAFAH1B1,CTSD,SCARB2,HK1,CHD4,SPTBN2,GPC3,GJA1,MYO5A,MACF1,
CLTC,CYFIP2,DDOST,CTBP1,CDKN1A,EEF1A2,ABCA3,SKI,ASXL1,MAPK1,SON,STXBP1,LONP1,US
D1,SKI,MAPK1,AEBP1,FLNB,TPM3,RAI1,ARFGEF2,ADAR,STAT3,BCR,RERE,PBX1,BAZ1B,PTEN,LO
,MAP2K2,DLK1,TPM3,RAI1,SEC23A,CDC42,SYP,TRIO,EXTL3,RERE,BAZ1B,NXN,ARID1B,CRTAP,FI
1A,TSR2,RPS27,CTBP1,NOVA2,PPP1CB,RPL27,SNRPN,ASXL1,RPS15A,NOTCH2,MAPK1,AEBP1,K
PL18,RPL11,ZMIZ1,COL11A1,RPL10,RPS17,AP3D1,HNRNPU,RPS24,COL1A2,RHOA,ATP2A2,RPL
21
FIP2,TSR2,HNRNPK,NOVA2,CDH11,EEF1A2,PPP1CB,SKI,NOTCH2,MAPK1,ODC1,SON,KDM1A,MA
2A3,ALDH6A1,MYH9,IDS,RPL5,GDI1,NPM1,COL18A1,SNRNP200,AFF4,NONO,DCHS1,SET,CHSY1
O1,CTSA
,PAFAH1B1,CCND2,DDX3X,CLTC,MYO5A,MACF1,TARDBP,NARS1,RAC1,CRPPA,NOVA2,EEF1A2,F
PS15A,MAPK1,AEBP1,MAP2K2,RPS28,STXBP1,SMS,CDC42,GPC4,LZTR1,NSD2,RPS26,PTPN11,IG
IGF2,VCP,AFF4,MAPT,DCHS1,RPL15,RPL18,RPL11,ZMIZ1,RPL10,RPS17,SBF1,HNRNPU,RPS24,C

RPS26,DHFR,RPS29,SRP54,OPA1,PIEZO1,DNAJC21
2F1,CHD4,GJA1,MYO5A,CTNNB1,HNRNPK,ARCN1,CPSF1,VCAN,TGFBI,SNRPN,PLOD1,COL2A1,SI
35R3,PAFAH1B1,CTSD,PTCD3,CLTC,MYO5A,CYFIP2,TARDBP,NARS1,GLUL,NOVA2,EEF1A2,TBCD,
.4A2,DCHS1,SET,PPP2R1A,CHD3,POLR2A,ZMIZ1,PAFAH1B1,HNRNPU,HK1,CHD4,ATP1A1,XPR1,C
300,DGCR2,ARID1B,NDST1,TIMM50,SLC2A1,GRN,EHMT1,LINGO1,SLC6A8,KMT2E,PACS1,DHCR

TARDBP,HLA-B,EEF1A2,FDFT1,SQSTM1,VAMP2,SNRPN,VPS35,CHD2,SKI,CNTNAP2,RAB11B,ABC
EF2,ADAR,BCR,RERE,BAZ1B,PTEN,LOX,KMT2E,GTF2I,MED13,FBLN5,TANC2,POR,CLIP2,ELN,NC
DHX30,STXBP1,PRPS1,USP7,SOD1,SMS,CDC42,YME1L1,WDR45B,TRIO,EP300,EXTL3,RERE,CSN
STM1,RPL27,COL2A1,RPS15A,SELENON,FLNB,DLK1,TPM3,RPS28,GPX4,HSPG2,FGFR1,RPS26,F
HD1,CDH11,COL2A1,SKI,FLNB,RPS28,MESD,RAI1,SEC23A,CDC42,YME1L1,SYP,RERE,ATRX,NXN
A,FLNB,ATP6V1B2,STXBP1,MESD,KDM3B,SMS,CDC42,SYP,EXTL3,RERE,DGCR2,KDM5B,FLCN,AT
.R2A,ZMIZ1,PAFAH1B1,HNRNPU,HK1,CHD4,ATP1A1,XPR1,CCND2,GPC3,DDX3X,CLTC,CTNNB1,C

CREBBP,AUTS2,CLIP2,ELN,ADNP,TRRAP,LIMK1,SYNGAP1,CRKL,CLCN4,SETD5,MED12,GATAD2B
,TARDBP,CDKN1A,HS6ST1,NOVA2,MDH2,EEF1A2,SQSTM1,TAF15,SNRPN,VPS35,CHD2,ASXL1,OI
S10,RPL35A,ARID1A,TSR2,RPS27,CTBP1,RAC1,MMP2,L1CAM,LMNB2,RPL27,COL2A1,RPS15A,N
TSR2,RPS27,CTBP1,RAC1,LMNB2,RPL27,COL2A1,RPS15A,NOTCH2,MAPK1,SMARCA4,KDM1A,F
D1A,SHOX,CTBP1,TBCD,PLOD1,COL2A1,SKI,ASXL1,NOTCH2,SELENON,SMARCA4,AEBP1,FLNB,T
A1,RPS15A,RPS28,IDH1,ADAR,PTEN,MSH6,FGFR1,RPS26,PTPN11,MDM2,IDH2,EXT2,RPS29,LMN
30,STXBP1,PRPS1,ARNT2,KIF5A,EP300,RERE,FLCN,SLC2A1,SLC6A8,PKDCC,IREB2,KMT2E,PHGD
MEGF8,RPS7,RPS19,RPL35,PRKAR1A,GJA1,RPS10,RPL35A,ARID1A,TSR2,SHOX,RPS27,CTBP1,R
MEGF8,RPS7,RPS19,RPL35,PRKAR1A,GJA1,RPS10,RPL35A,ARID1A,TSR2,SHOX,RPS27,CTBP1,R
TCH2,KDM1A,SMARCE1,RPS28,ATP6V1B2,EP300,RERE,RAB3GAP2,SMC3,ABCC9,HSPG2,PACS1,
S24,CHD4,RPL26,RPL31,GPC3,MEGF8,RPS7,RPS19,RPL35,PRKAR1A,GJA1,RPS10,RPL35A,ARID
RS1,RAI1,RAB11A,CIC,TRIO,EP300,EXTL3,RERE,KDM5B,FLCN,NDST1,SLC2A1,GRN,TAOK1,EHM
,RPS19,RPL35,PRKAR1A,CTNNB1,RPS10,RPL35A,TSR2,RPS27,CDKN1A,MDH2,SQSTM1,TAF15,F
BCR,TRIO,RERE,BAZ1B,CSNK2A1,GLUD1,SMC3,ADH5,SLC2A1,NDN,SPG7,HSPG2,SLC6A8,KMT2
F2,VCP,AFF4,YWHAG,FN1,DCHS1,SET,CHSY1,RPL15,SEC61A1,RPL18,RPL11,COL11A1,RPL10,RF
3,RPS7,RPS19,RPL35,COL6A2,RPS10,RPL35A,TSR2,SHOX,RPS27,CDH11,PPP1CB,RPL27,SNRPN
PK,RPS27,SMCHD1,CRPPA,CDH11,GNAI3,RPL27,SNRPN,COL2A1,RPS15A,FTO,SON,DLK1,RPS28,
1B,SMC3,HSPG2,SMARCD1,CREBBP,MACROH2A1,EXT2,CHST3,SETD5,EXT1,SMARCC2,SMARCE

.R,TUBB2B,COL5A1,PSAP,SOX4,FKBP14,KAT6A,DVL3,YWHAE,KIF1A,RPL13,GNAS,SLC2A3,GNB1
3GL1,SRSF2,RPS26,MDM2,DDX41,RPS29,SRP54,DNAJC21
11,SNRNP200,IGF2,ATP5F1A,VCP,AFF4,YWHAG,MAPT,NONO,DCHS1,SET,PPP2R1A,POLR2A,AP2M
I,NOVA2,PPP1CB,RPL27,ASXL1,RPS15A,NOTCH2,MAPK1,AEBP1,KDM1A,MAP2K2,SMARCE1,RPS
1,CREBBP,PTPN23,PSAT1,PDHA1,ZNF462,VPS51,CSNK2B,SYNGAP1,PPP2R5D,DEAF1,ASH1L,MEI
NRNPK,HLA-B,RAC1,MMP2,FDFT1,PLOD1,COL2A1,SKI,SELENON,MAPK1,AEBP1,MAP2K2,TPM3,
1,IGF1R,TRRAP,DDX6,PPP2R5D,CHST3,STAT1,TALDO1,SETD1A
MT2,BCOR,RPS29

11,NDN,EHMT1,SLC6A8,GTF2I,OGT,PACS2,YY1,CREBBP,PHIP,CCDC47,CHMP1A,CLIP2,ELN,EM
RNPK,MMP2,PPIB,CDH11,PPP1CB,SKI,ASXL1,NOTCH2,MAPK1,KDELR2,SON,FLNB,DLK1,STXBP
I,ATXN10,SMARCE1,DHX30,STXBP1,CSNK1D,PRPS1,USP7,SARS1,SOD1,RAI1,RAB11A,ARNT2,CI

,VPS51,CSNK2B,SYNGAP1,PPP2R5D,DEAF1,ASH1L,MED25,EPB41L1,PPP3CA
3,DLK1,TPM3,LONP1,RAI1,SEC23A,EP300,BAZ1B,ARID1B,RAB3GAP2,IARS1,LTBP4,GTF2I,FBLN5
CTSD,PTCD3,CLTC,MYO5A,CYFIP2,NOVA2,EEF1A2,TBCD,SQSTM1,SNRPN,ACO2,VPS35,SKI,ASXL
,CLTC,CYFIP2,ATP1A2,CTBP1,EEF1A2,SQSTM1,VAMP2,SNRPN,YEATS2,CHD2,SKI,CNTNAP2,TUBI
I1A1,RPL10,RPS17,HNRNPU,RPS24,COL1A2,RPL26,RPL31,RPS7,RPS19,RPL35,GJA1,COL6A2,RI
I1A1,RPL10,RPS17,HNRNPU,RPS24,COL1A2,RPL26,RPL31,RPS7,RPS19,RPL35,PRKAR1A,GJA1,(
SY1,RPL15,AP2M1,RPL18,RPL11,AGRN,ZMIZ1,COL11A1,RPL10,RPS17,AP3D1,MAT2A,HNRNPU,F
RPL10,PAFAH1B1,COL1A2,HK1,RHOA,NR2F1,LRPPRC,GPC3,MEGF8,DDX3X,RPS19,PRKAR1A,G
)4,CDH2,GPC3,MEGF8,CLTC,RPS19,GJA1,CYFIP2,TSR2,HNRNPK,NARS1,NOVA2,CDH11,EEF1A2,
PS17,RPS24,RPL26,RPL31,GPC3,DDX3X,RPS7,RPS19,RPL35,GJA1,RPS10,RPL35A,ARID1A,TSR2,I
3,SETD1A,POGZ
I1A1,RPL10,RPS17,HNRNPU,RPS24,COL1A2,RPL26,RPL31,RPS7,RPS19,RPL35,GJA1,COL6A2,RI
RPL35A,ARID1A,TSR2,RPS27,CTBP1,RAC1,LMNB2,RPL27,COL2A1,RPS15A,NOTCH2,MAPK1,SMA
RPS26,DAG1,CHST3,RPS29,LARGE1,POMGNT1

PS17,RPS24,RPL26,RPL31,GPC3,DDX3X,RPS7,RPS19,RPL35,GJA1,RPS10,RPL35A,ARID1A,TSR2,I
LTBP3,PCGF2,SH3PXD2B,PTPN11,CLIP2,ELN,LIMK1,CHST3,BCOR,LMNA,MED12,TGFBR1,TGFB
3,RPS7,RPS19,RPL35,COL6A2,RPS10,RPL35A,TSR2,SHOX,RPS27,MMP2,CDH11,PPP1CB,RPL27,
AB7A,PRKAR1A,GJA1,COL6A2,ARID1A,SHOX,HNRNPK,ARCN1,CTBP1,MMP2,FDFT1,SNRPN,PDXK
RPS28,TRAF7,EP300,EHMT1,LZTR1,DHCR7,CREBBP,FGFR1,RPS26,PTPN11,ABL1,NOTCH1,TRRA
SMCHD1,CRPPA,SQSTM1,RPL27,RPS15A,SELENON,PLEC,HSPB1,FLNB,TPM3,RPS28,FLNC,DCTN
ODC1,PRKDC,SON,KDM1A,SMARCE1,STXBP1,RAI1,SEC23A,KDM3B,CDC42,SYP,BCR,KDM5C,EF
D,PTCD3,CLTC,MYO5A,CYFIP2,NOVA2,EEF1A2,TBCD,SQSTM1,SNRPN,ACO2,VPS35,SKI,ASXL1,PI
M5C,CNTNAP1,ZFR,ATP6AP2,SLC2A1,FUS,SPG7,TPP1,IREB2,CNP,ATXN3,REEP2,REEP1,NOP56,AT
M5C,CNTNAP1,ZFR,ATP6AP2,SLC2A1,FUS,SPG7,TPP1,IREB2,CNP,ATXN3,REEP2,REEP1,NOP56,AT
31,MED13,PACS2,PHIP,CCDC47,FGFR1,TMEM94,KLF13,IGF1R,FLII,NECTIN1,DEAF1,WAC,SETD5,
P1,SON,KDM1A,SMARCE1,LARS1,SMS,CDC42,BCR,KDM5C,TRIO,ATRX,FBXO11,ABCC9,MAPRE2,
POLR2A,RPL11,AGRN,RPL10,ARF1,SCARB2,RPS20,COL5A2,PFN1,PRKCSH,CCN2,CLTC,CTNNB1
1,RAI1,CDC42,STAT3,EXTL3,DGCR2,ATRX,NXN,FBXO11,SMC3,GPC4,NDN,EHMT1,HSPG2,SLC6A
TRIO,NDUFV1,CSNK2A1,ARID1B,SMC3,ABCC9,EHMT1,PACS1,LTBP3,MED13,PACS2,NDUFA13,PI
I1,RAB11A,SMS,ADAR,CIC,MSMO1,TRIO,EP300,KDM5B,FLCN,ARID1B,RAB3GAP2,HDAC6,PTEN,Z
2,ARID1A,SHOX,CTBP1,MMP2,L1CAM,FDFT1,PPP1CB,PLOD1,COL2A1,SKI,ASXL1,NOTCH2,SELEI
C,RAB7A,CLTC,CTNNB1,CYFIP2,DDOST,NARS1,GLUL,MDH2,EEF1A2,TBCD,ACO2,ASXL1,RAB11E
XBP1,SMS,BCR,EP300,EXTL3,RERE,FBXO11,PTEN,MAPRE2,IARS2,HSPG2,AKT3,KMT2E,TTC37,L
MFN2,BSCL2,ALDH18A1,CAPN1,SLC25A1,ASAH1,GBA2,NDRG1,LMNA,SLC5A7,DDHD2,DSTYK,NI
,CHD2,CNTNAP2,ATXN10,SMARCE1,SDHA,STXBP1,SARS1,RAB11A,ARFGEF2,SMS,CIC,TRAF7,KC
HAG,MAPT,NONO,TPI1,AGRN,AP3D1,SBF1,PAFAH1B1,COL1A2,HK1,ATP1A1,COL5A2,PFN1,KLHL
CTBP1,EEF1A2,SQSTM1,VAMP2,SNRPN,YEATS2,CHD2,SKI,CNTNAP2,TUBB2A,ATXN10,MAP2K2,L
KAR1A,RPS10,RPL35A,TSR2,ARCN1,RPS27,CTBP1,RAC1,SMCHD1,CDH11,FDFT1,RPL27,SKI,RPS
S28,FLNC,DCTN1,GARS1,HINT1,REEP1,RPS26,TIA1,MFN2,BSCL2,RPS29,TRAPPC11,SPG11,HARS
Z1B,ARID1B,PKDCC,GTF2I,YY1,SMARCD1,CREBBP,COL12A1,CLIP2,ELN,LIMK1,ZSWIM6,HIRA,SM
T3,EP300,MBTPS1,ABCC9,GPC4,H19,IARS2,PKDCC,KMT2E,TTC37,MED13,MTOR,CREBBP,MN1,
SRP54,DNAJC21
COL11A1,RPL10,RPS17,HNRNPU,RPS24,COL1A2,CHD4,RPL26,RPL31,RPS7,RPS19,RPL35,PRKAI

;C,EP300,EXTL3,RERE,FBXO11,IARS2,KMT2E,UBE3A,CREBBP,SH3PXD2B,ABL1,TMEM94,IGF1R,
RX,NDN,TAOK1,EHMT1,SLC6A8,KMT2E,DHCR7,CREBBP,TRIM8,FLII,CSNK2B,SYNGAP1,DEAF1,W
,SARS1,SMS,RERE,PTEN,TIMM50,LOX,GPC4,EHMT1,HSPG2,SLC6A8,KMT2E,DICER1,NSD2,MED
ARCE1,CDC42,EP300,ARID1B,ABCC9,B4GAT1,GPC4,PHGDH,DHCR7,PACS2,SMARCD1,CREBBP,
F5A,GARS1,SPG7,SLC6A8,HINT1,KMT2E,DNAJB2,REEP2,HSD17B4,MED13,PCGF2,COX6A1,FBL
T1,PACS1,MED13,PACS2,PHIP,CCDC47,FGFR1,TMEM94,KLF13,IGF1R,FLII,NECTIN1,DEAF1,WAC
ALR,COL5A1,PSAP,SOX4,FKBP14,KAT6A,DVL3,YWHAE,KIF1A,GNAS,GPI,MYH9,IDS,RPL5,SPTAN1,
A2,SQSTM1,VAMP2,SNRPN,YEATS2,CHD2,SKI,CNTNAP2,TUBB2A,ATXN10,MAP2K2,LSS,STXBP1,
11A1,RPS17,RPS24,RPL26,RPL31,GPC3,DDX3X,RPS7,RPS19,RPL35,GJA1,RPS10,RPL35A,ARID1

2,LIMK1,ZNF462,PPP2R5D,CHST3,WAC,BCOR,NR2F2,SKIV2L,TGFBR1,TGFB2
2,RPS27,RPL27,COL2A1,RPS15A,FLNB,DLK1,RPS28,ATP6V1B2,TRIO,EP300,NXN,PTEN,SMC3,G
I,COL6A2,RPS10,RPL35A,TSR2,RPS27,CTBP1,L1CAM,CRPPA,LMNB2,RPL27,COL2A1,RPS15A,S
AC1,EEF1A2,FDFT1,SNRPN,SKI,SELENON,SMARCA4,PRDX1,ODC1,SON,MAP2K2,DLK1,TPM3,SM
3AZ1B,ARID1B,CTNND2,PTEN,ABCC9,ANTXR1,SLC6A8,GTF2I,LZTR1,FBLN5,UBE3A,SMARCD1,P
,MAP2K2,ATP6V1B2,STXBP1,CTNND1,TRIO,BAZ1B,NXN,FBXO11,ANTXR1,AKT3,KMT2E,GTF2I,L
S15A,RPS28,BCR,ATRX,ADH5,SH3GL1,SRSF2,RPS26,ABL1,MDM2,NSD1,DDX41,RPS29,SRP54,M
RPS28,EP300,RAB3GAP2,SMC3,ABCC9,HSPG2,PACS1,CREBBP,RPS26,EMC1,BSCL2,RAB18,RP
JC5,TARDBP,ATP1A2,TTR,L1CAM,RAB27A,EEF1A2,SQSTM1,VPS35,CHD2,CNTNAP2,PRDX1,ATXN
21
(I,ASXL1,KDM6B,MAPK1,ODC1,PRKDC,KDM1A,FLNB,DLK1,LONP1,USP7,RAI1,CDC42,BCR,TRIO
1B,NXN,DZIP1,GTF2I,LTBP3,PCGF2,YY1,FGFR1,SH3PXD2B,PTPN11,CLIP2,ELN,NOTCH1,LIMK1,
PS15A,PLEC,DLK1,RPS28,RERE,PBX1,ATRX,FBXO11,MAPRE2,EFTUD2,TTC37,NSD2,PCGF2,RPS2
APK1,SMARCA4,AEBP1,MAP2K2,FLNB,TPM3,SMARCE1,STXBP1,COL27A1,EP300,EXTL3,NXN,ARI
2,LIMK1,ZNF462,PPP2R5D,CHST3,WAC,BCOR,NR2F2,SKIV2L,TGFBR1,TGFB2
(1,PDGFRB,CRKL,NSD1,CLCN4,CHD8,CUL7,MAN1B1,POGZ

MMP14,RPS7,RPS19,RPL35,GJA1,RPS10,RPL35A,TSR2,HNRNPK,ARCN1,RPS27,CTBP1,RAC1,M
,SHOX,RAC1,L1CAM,LMNB2,SNRPN,COL2A1,SKI,KDM6B,MAPK1,SMARCA4,ODC1,KDM1A,FLNB
)N,STXBP1,LONP1,SARS1,RAB11A,TCTN1,CIC,KDM5B,PI4KA,RAB3GAP2,PTEN,NDST1,EML1,B4G
G1,PINK1,IGF1R,MECP2,PARK7,PRPH,DEAF1,WAC,UNC13A,EPHA4,BRAT1,TOMM40,ABCA7
,HNRNPK,RPS27,SMCHD1,CRPPA,CDH11,GNAI3,RPL27,SNRPN,COL2A1,RPS15A,AEBP1,FTO,SO
,SKI,RPS15A,PLEC,DLK1,RPS28,RERE,PBX1,ATRX,FBXO11,MAPRE2,EFTUD2,TTC37,NSD2,PCGF
OBSL1,NBAS,CHST3,RBM10,WAC,ZSWIM6,BCOR,SETD5,MAN2B1,CTCF,RUSC2,CUL7,EXT1,PSM
I,FTO,MAP2K2,SMARCE1,STAT3,EXTL3,BAZ1B,ATRX,ARID1B,ABCC9,GPC4,EHMT1,GTF2I,CNP,P
C23A,KDM3B,CDC42,SYP,BCR,KDM5C,EP300,PBX1,CSNK2A1,ATRX,ARID1B,SMC3,NDN,PACS1,M
JA1,CTNNB1,ARID1A,HNRNPK,RAC1,GLUL,NOVA2,CDH11,PPP1CB,SNRPN,COL2A1,SKI,PRPF6,N

NPU,COL1A2,CHD4,COL5A2,GJA1,HNRNPK,HLA-B,TTR,CDH11,TBCD,SQSTM1,PLOD1,VPS35,AS

SPG2,SMARCD1,CCDC47,BGN,CRKL,MAN2B1,LMNA,COG4,SMARCC2,SMARCB1,DPF2,GLB1
KA,RAB3GAP2,ZFR,ATP6AP2,SLC2A1,FUS,SPG7,REEP2,GOT2,NDUFA13,REEP1,ATP13A2,PDHA1,
2A1,NOTCH2,SELENON,AEBP1,FLNB,TPM3,FBLN1,STXBP1,LONP1,ARNT2,ARFGEF2,EP300,BAZ1
,PTEN,MSH6,RPS26,MDM2,RPS29,MSH2,SH3KBP1,DMPK
1,SMC3,EHMT1,PKDCC,PCGF2,MN1,CCDC47,ADNP,EFNB1,PPP2R5D,MED12,PPP3CA,CNOT3,FA

SAT1,CHMP1A,COL12A1,CLIP2,DAG1,ELN,MFN2,LIMK1,CAPN1,RAB18,STUB1,GBA2,NSD1,MAN2
IFA13,REEP1,ATP13A2,PDHA1,NDUFA4,BSCL2,ALDH18A1,AMPD2,KLC2,CAPN1,ECHS1,SLC16A2,
T1,PACS1,PHGDH,UBE3A,TRIM8,MICAL1,MECP2,WDR26,CSNK2B,ASAH1,SYNGAP1,DEAF1,TSPA
GAT1,GPC4,PHGDH,DHCR7,PACS2,SMARCD1,FGFR1,DENND5A,PSAT1,DAG1,NDUFC2,ALDH18A
KI,ASXL1,RAB11B,NOTCH2,SMARCA4,ODC1,FTO,PRKDC,SON,QARS1,SMARCE1,STXBP1,LONP1
S1,L1CAM,LMNB2,SNRPN,SKI,KDM6B,RAB11B,MAPK1,ODC1,KDM1A,FLNB,DLK1,USP7,RAI1,CD
COL5A2,GPC3,CCN2,RPS19,GJA1,COL6A2,ARID1A,SHOX,HNRNPK,HLA-B,CTBP1,MMP2,TBCD,LM
P1,GTF2I,LTBP3,PCGF2,SH3PXD2B,PTPN11,CLIP2,ELN,NOTCH1,LIMK1,CHST3,BCOR,LMNA,MEI
3XL4,UBA1,CALR,COL5A1,PSAP,SOX4,FKBP14,KAT6A,DVL3,YWHAE,KIF1A,GNAS,SLC2A3,GPI,ALI
2,RPS27,RPL27,COL2A1,RPS15A,FLNB,DLK1,RPS28,ATP6V1B2,TRIO,EP300,NXN,PTEN,SMC3,GI
JA1,CTNNB1,ARID1A,HNRNPK,RAC1,SMCHD1,GLUL,NOVA2,CDH11,PPP1CB,SNRPN,COL2A1,SKI,
3H1,LMNA,MED12,FAM20C
.35,PRKAR1A,RPS10,RPL35A,TSR2,ARCN1,RPS27,CTBP1,RAC1,SMCHD1,CDH11,FDFT1,RPL27,S
.35,PRKAR1A,RPS10,RPL35A,TSR2,ARCN1,RPS27,CTBP1,RAC1,SMCHD1,CDH11,FDFT1,RPL27,S
HD2,SMARCE1,SDHA,STXBP1,RAB11A,SMS,ADAR,CIC,TRAF7,KIF5A,KCNQ2,STARD7,NDUFV1,KD
2,MAPRE2,ATP6AP2,EHMT1,ANTXR1,SLC6A8,LTBP4,AKT3,GTF2I,PHGDH,FBLN5,PRKACA,YY1,PT
KI,ASXL1,KDM6B,MAPK1,ODC1,PRKDC,KDM1A,FLNB,DLK1,LONP1,USP7,RAI1,CDC42,BCR,TRIO
,DLK1,RPS28,ATP6V1B2,SMC3,PKDCC,NSD2,DHCR7,EIF4A3,RPS26,BCOR,RPS29,SETD5,PSMD1
GFBI,SNRPN,PLOD1,COL2A1,SKI,ASXL1,NOTCH2,OPTN,AEBP1,SON,MAP2K2,PRPS1,USP7,RAI1,S
LHL9,GPC3,RAB7A,PRKAR1A,COL6A2,HNRNPDL,TARDBP,ARID1A,SHOX,HNRNPK,DUX4,SMCHD1
ER1,LZTR1,MSH6,YY1,CREBBP,MN1,SMO,FGFR1,PTPN11,MDM2,IDH2,EWSR1,PDGFRB,NSD1,Z
RPL11,ZMIZ1,COL11A1,RPS17,RPS24,COL5A2,RPL26,RPL31,CCND2,GPC3,MEGF8,RPS7,RPS19
AR1A,GJA1,HLA-B,PPP1CB,ABCA3,RPL27,SKI,NOTCH2,MAPK1,MAP2K2,FLNC,ARFGEF2,IGFBP7,
ONO,DCHS1,RPL15,RPL18,RPL11,ZMIZ1,RPL10,RPS17,PAFAH1B1,RPS24,CHD4,RPL26,RPL31,C
,MYH9,IDS,RPL5,NPM1,COL18A1,SNRNP200,VCP,AFF4,TRIM28,CHSY1,RPL15,SF3B1,CHD3,RPL
H,VPS51,SLC25A1,SLC5A7,SYT1,PIGT,POGZ,COL3A1,SLC18A3,GSN,PRMT7,DMPK
,P4HB,IDS,RPL5,NPM1,IGF2,ATP5F1A,VCP,AFF4,YWHAG,COL4A2,FN1,NONO,DCHS1,SET,CHSY1
A1,SMC3,ECEL1,ABCC9,EHMT1,KMT2E,PACS1,NSD2,MED13,CREBBP,MN1,TANC2,AUTS2,TMEM
2R5D,SNRPB,SLC5A7,THRA,CUL7,FHL1,COL3A1,ITGA7,SLC18A3,DNAJC21
PDGFRB,EXT2,CDKN1B,MAN2B1,LMNA,EXT1,COL3A1,ELMO2,MAFB

L1,RPS15A,RPS28,ATRX,ADH5,SRP72,SRSF2,RPS26,PTPN11,MDM2,DDX41,RPS29,SRP54,DNAJC
D2,STXBP1,LONP1,PRPS1,RAB11A,SYP,CIC,STAT3,KCNQ2,STARD7,KDM5B,ATP6AP2,SLC2A1,GR
ATXN10,SON,SDHA,STXBP1,SARS1,RAB11A,SMS,SYP,CIC,STAT3,KCNQ2,EXTL3,KDM5B,GLUD1,
,CRPPA,PPP1CB,ASXL1,RAB11B,SMARCA4,FTO,PRKDC,SON,MAP2K2,SMARCE1,TCTN1,CDC42,
CN2,RPS19,GJA1,COL6A2,ARID1A,SHOX,HNRNPK,HLA-B,CTBP1,MMP2,TBCD,LMNB2,SNRPN,PL
.35A,TSR2,RPS27,CTBP1,GPC1,RPL27,SNRPN,RPS15A,ACADVL,DLK1,RPS28,LONP1,SYP,FLCN,
N,NOTCH1,ACTA2,LIMK1,DDX6,CHST3,LMNA,MED12,TGFBR1,PSMD12,TGFB2,COL3A1,TCF20
,VCP,AFF4,FN1,NONO,CHSY1,HLA-C,SEC61A1,UCHL1,AGRN,COL11A1,RPL10,ARF1,PAFAH1B1,
,SKI,RPS15A,PLEC,DLK1,RPS28,RERE,PBX1,ATRX,FBXO11,MAPRE2,EFTUD2,TTC37,NSD2,PCGF
NB,DLK1,SMARCE1,CDC42,EP300,ARID1B,ABCC9,B4GAT1,GPC4,PHGDH,DHCR7,PACS2,SMARC
CB,SNRPN,SKI,NOTCH2,FTO,SON,STXBP1,ARFGEF2,CDC42,TRAF7,EP300,RERE,BAZ1B,ARID1B,
.35,PRKAR1A,RPS10,RPL35A,TSR2,HLA-B,ARCN1,RPS27,CTBP1,RAC1,SMCHD1,CDH11,FDFT1,R
L,MAPK1,SDHA,STXBP1,SOD1,RAI1,DCTN1,EP300,BAZ1B,UBQLN2,SMC3,SLC2A1,FUS,GRN,EHM

RN,VAPB,EIF4G1,PINK1,IGF1R,MECP2,PARK7,PRPH,DEAF1,WAC,UNC13A,PRNP,EPHA4,BRAT1,T

,PRKAR1A,GJA1,CTNNB1,ARID1A,HNRNPK,RAC1,GLUL,NOVA2,CDH11,PPP1CB,SNRPN,COL2A1,

NO,DCHS1,RPL15,RPL18,RPL11,ZMIZ1,RPL10,RPS17,PAFAH1B1,RPS24,CHD4,RPL26,RPL31,C

ODC1,KDM1A,FLNB,USP7,RAI1,BCR,TRIO,EP300,RERE,BAZ1B,ATRX,NXN,ARID1B,PTEN,SMC3,G

MO,CLIP2,ELN,MECP2,LIMK1,MOGS,DDX6,RAB18,CRKL,BCOR,SOX9,SETD5,LMNA,MED12,DG

IZ1,COL11A1,RPL10,COL1A2,HK1,RHOA,COL5A2,CLEC7A,GPC3,CCN2,CLTC,PRKAR1A,GJA1,CY

2A3,ALDH6A1,GNB1,APP,IDS,NPM1,HSPD1,IGF2,VCP,UBB,AFF4,TRIM28,YWHAG,MAPT,NONO,D

RPS26,RPS29,MSH2,PTPN12

RPS26,RPS29,MSH2,PTPN12

B

CTBP1,RAC1,SMCHD1,CDH11,TBCD,PPP1CB,SKI,ASXL1,NOTCH2,MAPK1,SMARCA4,ODC1,AEBP

,VCP,AFF4,FN1,NONO,DCHS1,TPI1,RPL15,PKD1,RPL18,RPL11,COL11A1,RPL10,RPS17,VCL,AR

ABL1,COL12A1,POR,ELN,ACTA2,STUB1,CRKL,DLG4,FARSA,CNTN1,MED12,HIRA,ZDHHC9,TGFB

1,NTRK2,TRAK1,DAG1,BSCL2,EXT2,SYNGAP1,PPP2R5D,STUB1,DEAF1,NDRG1,ZSWIM6,CLCN4,

K1,STXBP1,KDM3B,TRAF7,DGCR2,BAZ1B,ATRX,NXN,ARID1B,ABCC9,MAPRE2,GPC4,H19,EHMT

AP2,ECEL1,HNRNPA2B1,IARS2,HSPG2,GTF2I,PCGF2,WNK1,BGN,COL12A1,CLIP2,DAG1,ELN,OB

BP14,KAT6A,DVL3,YWHAE,KIF1A,GNAS,GPI,MYH9,IDS,RPL5,SPTAN1,NPM1,SNRNP200,IGF2,AFF

19,GJA1,HNRNPK,MMP2,PPIB,CDH11,PPP1CB,SKI,ASXL1,NOTCH2,MAPK1,KDELR2,SON,FLNB,

SLC2A3,GPI,ALDH6A1,GNB1,P4HB,APP,IDS,RPL5,GDI1,SPTAN1,NPM1,HSPD1,IGF2,ATP5F1A,VCF
HB,APP,IDS,GDI1,SPTAN1,NPM1,HSPD1,IGF2,ATP5F1A,VCP,YWHAG,COL4A2,MAPT,NONO,DCHS1
F1A,VCP,YWHAG,COL4A2,MAPT,NONO,DCHS1,SET,TPI1,PPP2R1A,CHD3,POLR2A,AP2M1,UCHL1
WHAG,MAPT,FN1,NONO,DCHS1,SET,TPI1,PPP2R1A,RPL15,CHD3,RPL18,RPL11,AGRN,ZMIZ1,COI
,PGK1,SOX4,FKBP14,KAT6A,DVL3,YWHAE,KIF1A,GNAS,SLC2A3,GPI,ALDH6A1,CALM3,GNB1,P4H
COL4A1,PSAP,PGK1,SOX4,FKBP14,KAT6A,DVL3,YWHAE,KIF1A,GNAS,SLC2A3,GPI,ALDH6A1,CALI
RPPA,NOVA2,MDH2,EEF1A2,FDFT1,TBCD,VAMP2,ACO2,ASXL1,TUBB2A,PPP2CA,SMARCA4,PRKD
IF1A,RPL13,GNAS,ALDH6A1,GNB1,P4HB,IDS,RPL5,GDI1,SPTAN1,NPM1,HSPD1,COL18A1,IGF2,A
WHAE,KIF1A,GNAS,SLC2A3,GPI,ALDH6A1,GNB1,P4HB,APP,IDS,RPL5,GDI1,SPTAN1,NPM1,PLD3,H
AT2A,HNRNPU,RPS24,COL1A2,CHD4,RPL26,RPL31,LRPPRC,CCND2,CDH2,GPC3,MEGF8,DDX3X,
TP5F1A,VCP,AFF4,YWHAG,COL4A2,MAPT,NONO,DCHS1,SET,PPP2R1A,RPL15,POLR2A,AP2M1,RI
TP5F1A,VCP,AFF4,YWHAG,COL4A2,MAPT,NONO,DCHS1,SET,PPP2R1A,RPL15,POLR2A,AP2M1,RI
,PGK1,SOX4,FKBP14,KAT6A,DVL3,YWHAE,KIF1A,RPL13,GNAS,SLC2A3,GPI,ALDH6A1,GNB1,P4H
CP,AFF4,YWHAG,MAPT,NONO,DCHS1,SET,PPP2R1A,RPL15,POLR2A,RPL18,UCHL1,RPL11,ZMIZ1,
ND2,CDH2,GPC3,MEGF8,DDX3X,RPS7,RPS19,RPL35,PRKAR1A,GJA1,RPS10,RPL35A,TSR2,NARS1
WHAE,KIF1A,GNAS,SLC2A3,GPI,ALDH6A1,GNB1,P4HB,APP,IDS,RPL5,GDI1,SPTAN1,NPM1,HSPD1,
I1,SPTAN1,NPM1,HSPD1,COL18A1,IGF2,ATP5F1A,VCP,UBB,AFF4,YWHAG,NONO,DCHS1,SET,CH
,PPP2R1A,CHD3,RPL11,AGRN,ZMIZ1,RPL10,SBF1,ARF1,PAFAH1B1,MAT2A,HNRNPU,SCARB2,CC
31,MYH9,P4HB,APP,RPL5,SPTAN1,CALM1,NPM1,PLD3,HSPD1,COL18A1,SNRNP200,IGF2,VCP,AF
A1,CTNNB1,CYFIP2,ARID1A,HNRNPK,CTBP1,NOVA2,CDH11,EEF1A2,PPP1CB,SNRPN,COL2A1,SKI
L11,ZMIZ1,RPL10,RPS17,AP3D1,SBF1,ARF1,CYB5R3,PAFAH1B1,HNRNPU,CTSD,RPS24,CHD4,RP
8,RPL11,ZMIZ1,RPL10,RPS17,AP3D1,SBF1,ARF1,CYB5R3,PAFAH1B1,HNRNPU,CTSD,RPS24,HK1
LDH6A1,GNB1,P4HB,APP,IDS,RPL5,SPTAN1,NPM1,PLD3,HSPD1,COL18A1,IGF2,ATP5F1A,VCP,AFI
GNB1,APP,IDS,RPL5,GDI1,SPTAN1,PLD3,HSPD1,COL18A1,SNRNP200,IGF2,VCP,UBB,AFF4,YWHA
,P4HB,IDS,RPL5,GDI1,NPM1,HSPD1,IGF2,VCP,AFF4,YWHAG,MAPT,FN1,NONO,DCHS1,CHSY1,PP
L3,YWHAE,KIF1A,RPL13,GNAS,ALDH6A1,GNB1,P4HB,IDS,RPL5,GDI1,SPTAN1,NPM1,HSPD1,COL
,HK1,NR2F1,CDH2,GPC3,DDX3X,CLTC,GJA1,CTNNB1,MACF1,CYFIP2,ARID1A,HNRNPK,NARS1,C
F1A,GNAS,SLC2A3,ALDH6A1,GNB1,P4HB,APP,IDS,RPL5,GDI1,SPTAN1,NPM1,PLD3,HSPD1,COL1
HB,APP,IDS,RPL5,GDI1,NPM1,HSPD1,IGF2,UBB,AFF4,YWHAG,COL4A2,NONO,DCHS1,SET,TPI1,
LTC,RPS7,RPS19,RPL35,GJA1,COL6A2,RPS10,RPL35A,ARID1A,TSR2,HNRNPK,NARS1,RPS27,CT
,SMARCA4,PRDX1,PRKDC,SON,MAP2K2,DLK1,TPM3,SMARCE1,ATP6V1B2,STXBP1,RAI1,SEC23
AGRN,ZMIZ1,RPL10,SBF1,ARF1,PAFAH1B1,MAT2A,HNRNPU,COL1A2,HK1,COL5A2,GPC3,MEGF
N,ZMIZ1,RPL10,ARF1,HNRNPU,SCARB2,COL1A2,HK1,NR2F1,RPS20,PFN1,LRPPRC,XPR1,SPTBN
L10,RPS17,PAFAH1B1,CTSD,RPS24,COL1A2,HK1,RHOA,NR2F1,RPL26,RPL31,LRPPRC,CCND2,C
GCR2,KDM5B,BAZ1B,NXN,ARID1B,CTNND2,MAPRE2,LOX,GPC4,H19,EHMT1,HSPG2,LTBP4,GTF2
X,CLTC,CTNNB1,MACF1,CYFIP2,ARID1A,HNRNPK,NARS1,CTBP1,RAC1,L1CAM,GLUL,CRPPA,NO
,AFF4,FN1,NONO,DCHS1,CHSY1,PPP2R1A,RPL15,RPL18,RPL11,AGRN,ZMIZ1,COL11A1,RPL10,R
PAFAH1B1,HNRNPU,HK1,NR2F1,XPR1,CDH2,GPC3,DDX3X,CLTC,GJA1,CTNNB1,MACF1,CYFIP2,
RN,ZMIZ1,COL11A1,RPL10,RPS17,PAFAH1B1,MAT2A,RPS24,COL1A2,HK1,NR2F1,CHD4,COL5A
WHAE,KIF1A,GNAS,SLC2A3,ALDH6A1,GNB1,MYH9,P4HB,APP,IDS,RPL5,NPM1,PLD3,HSPD1,COL
4,COL5A2,RPL26,RPL31,CDH2,GPC3,MEGF8,MMP14,RPS7,RPS19,RPL35,GJA1,RPS10,RPL35A,
L4A2,MAPT,NONO,SET,PPP2R1A,POLR2A,AP2M1,AGRN,ZMIZ1,RPL10,MARCHF6,AP3D1,ARF1,C

AH1B1,HNRNPU,CTSD,RPS24,RPL26,RPL31,XPR1,DDX3X,CLTC,RPS7,RPS19,RPL35,GJA1,CTNN

AT6A,DVL3,YWHAE,KIF1A,RPL13,GNAS,ALDH6A1,GNB1,P4HB,APP,IDS,RPL5,GDI1,SPTAN1,NPM1,
,RPL18,RPL11,AGRN,ZMIZ1,COL11A1,RPL10,RPS17,PAFAH1B1,MAT2A,RPS24,COL1A2,HK1,NR
3,PAFAH1B1,HNRNPU,CTSD,RPS24,RPL26,RPL31,XPR1,DDX3X,CLTC,RPS7,RPS19,RPL35,GJA1,
,DCHS1,SET,TPI1,PPP2R1A,CHD3,POLR2A,AP2M1,AGRN,ZMIZ1,RPL10,AP3D1,PAFAH1B1,HNRNF
R1A,RPL15,CHD3,RPL18,RPL11,ZMIZ1,COL11A1,RPL10,RPS17,PAFAH1B1,HNRNPU,RPS24,COL
,NONO,DCHS1,CHSY1,PPP2R1A,RPL15,RPL18,RPL11,ZMIZ1,COL11A1,RPL10,RPS17,SBF1,PAFA
,NONO,DCHS1,CHSY1,PPP2R1A,RPL15,RPL18,RPL11,ZMIZ1,COL11A1,RPL10,RPS17,SBF1,PAFA

1,CHD4,COL5A2,RPL26,RPL31,GPC3,MEGF8,RPS7,RPS19,RPL35,PRKAR1A,GJA1,RPS10,RPL35
IAPT,DCHS1,TPI1,PPP2R1A,RPL15,POLR2A,RPL18,UCHL1,RPL11,AGRN,ZMIZ1,COL11A1,RPL10,
,SEC23A,KDM3B,SMS,CDC42,BCR,TRIO,EP300,EXTL3,RERE,FLCN,BAZ1B,ATRX,NXN,ARID1B,FE

FKBP14,KAT6A,DVL3,YWHAE,KIF1A,GNAS,SLC2A3,ALDH6A1,GNB1,MYH9,P4HB,IDS,RPL5,GDI1,S
COL11A1,RPL10,RPS17,PAFAH1B1,HNRNPU,RPS24,COL1A2,HK1,RHOA,NR2F1,CHD4,COL5A2,RI
DM1A,FLNB,DLK1,SMARCE1,DHX30,ATP6V1B2,LONP1,SEC23A,KDM3B,ARFGEF2,SMS,STAT3,B
LTC,PRKAR1A,MACF1,CYFIP2,ARID1A,NARS1,CTBP1,RAC1,L1CAM,NOVA2,MDH2,EEF1A2,TBCD,
11A1,RPL10,RPS17,SBF1,PAFAH1B1,MAT2A,RPS24,COL1A2,RHOA,NR2F1,CHD4,COL5A2,RPL26
P1CB,RPL27,SNRPN,ASXL1,RPS15A,NOTCH2,MAPK1,ACADVL,SON,KDM1A,MAP2K2,FLNB,DLK1
S,RPL5,GDI1,NPM1,IGF2,VCP,UBB,AFF4,YWHAG,NONO,DCHS1,SET,CHSY1,PPP2R1A,RPL15,CHD
D1,COL2A1,SKI,ASXL1,NOTCH2,PPP2CA,MAPK1,SMARCA4,ODC1,AEBP1,FTO,FLNB,DLK1,SMAR
P1CB,RPL27,SNRPN,ASXL1,RPS15A,NOTCH2,MAPK1,ACADVL,SON,KDM1A,MAP2K2,FLNB,DLK1
FKBP14,KAT6A,DVL3,YWHAE,KIF1A,GNAS,SLC2A3,ALDH6A1,GNB1,MYH9,P4HB,IDS,RPL5,GDI1,S
R1A,RPL15,CHD3,RPL18,RPL11,ZMIZ1,COL11A1,RPL10,RPS17,PAFAH1B1,HNRNPU,RPS24,COL

C3,DDX3X,CLTC,RPS7,RPS19,GJA1,CTNNB1,CYFIP2,ARID1A,HNRNPK,CTBP1,GLUL,NOVA2,CDH
NB1,MACF1,ARID1A,HNRNPK,NARS1,CTBP1,RAC1,L1CAM,GLUL,CRPPA,NOVA2,MDH2,EEF1A2,F

A,TSR2,SHOX,HNRNPK,RPS27,CTBP1,GLUL,CDH11,RPL27,SKI,ASXL1,KDM6B,PRPF6,RPS15A,N
1,HNRNPU,CTSD,RPS24,COL1A2,HK1,RPL26,RPL31,LRPPRC,CCND2,CDH2,MEGF8,CLTC,RPS7,
D1,PAFAH1B1,PXDN,CTSD,RPS24,NR2F1,CHD4,RPL26,RPL31,CDH2,GPC3,MEGF8,CLTC,RPS7,RF
PC3,DDX3X,CLTC,MYO5A,CTNNB1,COL6A2,MACF1,CYFIP2,ARID1A,DDOST,HNRNPK,NARS1,CTE
,RPL15,CHD3,RPL18,RPL11,ZMIZ1,COL11A1,RPL10,RPS17,PAFAH1B1,HNRNPU,RPS24,COL1A2,
COL12A1,DAG1,TRIM2,WDR26,TRRAP,AMPD2,IQSEC1,CSNK2B,SLC25A1,PPP2R5D,ZSWIM6,SH
NB1,MACF1,ARID1A,HNRNPK,NARS1,CTBP1,RAC1,L1CAM,GLUL,CRPPA,NOVA2,MDH2,EEF1A2,F
NPM1,PLD3,HSPD1,COL18A1,SNRNP200,ATP5F1A,VCP,AFF4,YWHAG,COL4A2,MAPT,NONO,DCH
,GPC3,MEGF8,CLTC,RPS7,RPS19,RPL35,CTNNB1,COL6A2,RPS10,RPL35A,TSR2,SHOX,HNRNPK,
35A,TSR2,HNRNPK,ARCN1,RPS27,CTBP1,RAC1,MMP2,PPP1CB,RPL27,SNRPN,SKI,ASXL1,RPS15
,PAFAH1B1,MAT2A,HNRNPU,RPS24,COL1A2,RPL26,RPL31,CDH2,GPC3,MEGF8,DDX3X,CLTC,RP
RNPU,RPS24,COL1A2,HK1,RHOA,NR2F1,RPL26,RPL31,LRPPRC,SPTBN2,CDH2,DDX3X,RPS7,RP
,HNRNPK,HLA-B,RPS27,MMP2,PPP1CB,RPL27,PLOD1,SKI,RPS15A,MAPK1,AEBP1,FLNB,DLK1,T
AT2A,RPS24,COL1A2,NR2F1,CHD4,COL5A2,RPL26,RPL31,CCND2,GPC3,MEGF8,RPS7,RPS19,R
IAT2A,COL1A2,HK1,RHOA,ATP1A1,COL5A2,ATP2A2,GPC3,MEGF8,RAB7A,CCN2,MMP14,PRKAR1
PTBN2,RAB7A,DDX3X,CLTC,GJA1,CYFIP2,ARL6IP1,HNRNPK,NARS1,ATP1A2,HLA-B,ARCN1,CTBP1

,IGF2,VCP,UBB,AFF4,YWHAG,NONO,DCHS1,SET,CHSY1,PPP2R1A,RPL15,CHD3,POLR2A,AP2M1,
,IGF2,VCP,UBB,AFF4,YWHAG,NONO,DCHS1,SET,CHSY1,PPP2R1A,RPL15,CHD3,POLR2A,AP2M1,
SY1,PPP2R1A,CHD3,UCHL1,AGRN,ZMIZ1,COL11A1,RPL10,SBF1,ARF1,PAFAH1B1,MAT2A,SCAR
RNPU,COL1A2,HK1,CHD4,CD81,CDH2,MEGF8,MMP14,DDX3X,RPS19,PRKAR1A,GJA1,CYFIP2,SH
PL31,LRPPRC,CCND2,CDH2,GPC3,MEGF8,DDX3X,RPS7,RPS19,RPL35,PRKAR1A,GJA1,RPS10,R
NONO,SET,TPI1,PPP2R1A,CHD3,AP2M1,UCHL1,AGRN,ZMIZ1,RPL10,MARCHF6,AP3D1,SBF1,CY
D,RAI1,KDM3B,TRIO,EP300,EXTL3,KDM5B,GARS1,FBXO11,PTEN,TIMM50,FUS,CLCN7,EHMT1,
,MACF1,CYFIP2,DNAJC5,TARDBP,DDOST,NARS1,ATP1A2,TTR,CTBP1,RAC1,HS6ST1,NOVA2,MD

300,RERE,DGCR2,KDM5B,ATRX,ARID1B,NDST1,TIMM50,SLC2A1,NDN,GRN,TAOK1,EHMT1,LIN
S1,CTBP1,RAC1,L1CAM,NOVA2,MDH2,EEF1A2,TBCD,PPP1CB,LMNB2,SNRPN,SKI,CNTNAP2,RA

3,YWHAE,KIF1A,GNAS,SLC2A3,ALDH6A1,GNB1,MYH9,P4HB,IDS,RPL5,GDI1,NPM1,COL18A1,SN
5A,CTNNB1,MACF1,CYFIP2,TARDBP,ARL6IP1,NARS1,TTR,L1CAM,NOVA2,RAB27A,EEF1A2,TBCD,
OX,RPS27,GLUL,HS6ST1,CDH11,EEF1A2,FDFT1,RPL27,PLOD1,COL2A1,SKI,KDM6B,RPS15A,S
NPDL,TSR2,RPS27,SMCHD1,L1CAM,CRPPA,EEF1A2,SQSTM1,PABPN1,RPL27,PLOD1,RPS15A,M
4,DDX3X,CLTC,PRKAR1A,GJA1,CTNNB1,COL6A2,MACF1,ARID1A,HNRNPK,CTBP1,RAC1,GPC1,M
RPL10,PAFAH1B1,HNRNPU,CTSD,COL1A2,HK1,CHD4,CD81,CDH2,MEGF8,MMP14,DDX3X,RPS1
,SEC61A1,RPL18,RPL11,ZMIZ1,COL11A1,RPL10,RPS17,PAFAH1B1,MAT2A,HNRNPU,RPS24,COL
,LSS,STXBP1,USP7,SARS1,RAI1,RAB11A,ARNT2,KDM3B,ARFGEF2,SYP,CIC,KDM5C,TRIO,EP300,

,NOVA2,LMNB2,SNRPN,PLOD1,COL2A1,CNTNAP2,RAB11B,ABCA2,SELENON,PPP2CA,KDELR2,
10,RPL35A,TSR2,NARS1,RPS27,RAC1,GLUL,CRPPA,CDH11,EEF1A2,FDFT1,PPP1CB,GNAI3,RPL2
CHS1,SET,TPI1,CHSY1,PPP2R1A,CHD3,AP2M1,ZMIZ1,RPL10,AP3D1,CYB5R3,PAFAH1B1,HNRNP
L1A2,HK1,RHOA,NR2F1,RPL26,RPL31,LRPPRC,SPTBN2,CDH2,DDX3X,RPS7,RPS19,RPL35,PRK

,RPS20,COL5A2,RPL26,KLHL9,GPC3,MEGF8,CCN2,DDX3X,GJA1,CTNNB1,COL6A2,HNRNPDL,A
A1,RPS10,RPL35A,TSR2,NARS1,RPS27,RAC1,GLUL,CRPPA,CDH11,EEF1A2,FDFT1,PPP1CB,GNA
FF4,YWHAG,NONO,DCHS1,SET,CHSY1,PPP2R1A,AP2M1,AGRN,ZMIZ1,RPL10,MARCHF6,SBF1,CY
SET,TPI1,CHD3,AP2M1,UCHL1,AGRN,ZMIZ1,RPL10,ARF1,CYB5R3,PAFAH1B1,ITM2B,LDHA,CTSD
1B,NXN,PI4KA,RAB3GAP2,CRTAP,PTEN,SMC3,MAPRE2,LOX,GPC4,HSPG2,GTF2I,PACS1,LZTR1,H

A,SMARCD1,SH3PXD2B,HS2ST1,TRIP12,AUTS2,CLIP2,ELN,ADNP,TRRAP,LIMK1,SYNGAP1,WAC,
RDBP,ARL6IP1,TTR,L1CAM,NOVA2,RAB27A,EEF1A2,TBCD,SQSTM1,TAF15,CNTNAP2,RAB11B,M
,CDH11,GNAI3,SNRPN,COL2A1,SKI,ASXL1,KDM6B,NOTCH2,MAPK1,ODC1,AEBP1,SON,MAP2K2,
OST,HLA-B,CTBP1,CDKN1A,EEF1A2,TBCD,SQSTM1,PABPN1,ABCA3,VPS35,SKI,ASXL1,SELENO
PS15A,NOTCH2,MAPK1,ACADVL,FTO,MAP2K2,FLNB,DLK1,RPS28,SDHA,STXBP1,LONP1,RAI1,K
CA,MAPK1,ODC1,AEBP1,FTO,FLNB,DLK1,STXBP1,LONP1,VANGL2,KDM3B,CDC42,TRAF7,BCR,
2,SELENON,SMARCA4,AEBP1,SON,FLNB,DLK1,TPM3,SMARCE1,ATP6V1B2,STXBP1,PRPS1,MES
CA,MAPK1,ODC1,AEBP1,FTO,FLNB,DLK1,STXBP1,LONP1,VANGL2,KDM3B,CDC42,TRAF7,BCR,
PS15A,NOTCH2,MAPK1,ACADVL,FTO,FLNB,DLK1,RPS28,SDHA,STXBP1,LONP1,RAI1,KDM3B,BC
NARS1,GLUL,NOVA2,EEF1A2,TBCD,SQSTM1,TAF15,SNRPN,ACO2,VPS35,SKI,ASXL1,OPTN,PRDX
L26,RPL31,GPC3,MEGF8,RPS7,RPS19,RPL35,PRKAR1A,GJA1,COL6A2,RPS10,RPL35A,ARID1A,
2,HNRNPK,RPS27,CTBP1,HS6ST1,NOVA2,EEF1A2,PPP1CB,SQSTM1,GNAI3,PABPN1,RPL27,COL2
0,RERE,KDM5B,FLCN,BAZ1B,CSNK2A1,ATRX,GLUD1,FBXO11,SMC3,ADH5,NDST1,SLC2A1,NDN,

S1,RAI1,SEC23A,SYP,BCR,TRIO,EP300,BAZ1B,ATRX,GARS1,FBXO11,MAPRE2,ATP6AP2,LOX,IAR
K1,TPM3,SMARCE1,MESD,RAI1,SEC23A,KDM3B,CDC42,SYP,BCR,KDM5C,EP300,PBX1,CSNK2A1
ARS1,RAC1,NOVA2,CDH11,EEF1A2,TBCD,PPP1CB,SNRPN,RBMX,SKI,ASXL1,RAB11B,NOTCH2,M
T3,GTF2I,DHCR7,SMARCD1,FGFR1,RPS26,PTPN11,POR,CLIP2,SC5D,ELN,JAM3,LIMK1,FLII,DDX

OX,HNRNPK,ATP1A2,RPS27,CTBP1,CDH11,PPP1CB,SQSTM1,RPL27,SNRPN,COL2A1,SKI,RPS15
DM1A,MAP2K2,SMARCE1,RPS28,ATP6V1B2,EP300,RERE,RAB3GAP2,SMC3,ABCC9,NDN,HSPG2,
L5,NPM1,HSPD1,COL18A1,SNRNP200,IGF2,VCP,AFF4,TRIM28,YWHAG,MAPT,NONO,DCHS1,SET,
GPC3,MEGF8,RPS7,RPS19,RPL35,PRKAR1A,GJA1,COL6A2,RPS10,RPL35A,TSR2,SHOX,NARS1,A
DH11,PPP1CB,LMNB2,RPL27,SNRPN,COL2A1,SKI,ASXL1,RPS15A,NOTCH2,MAPK1,SMARCA4,FT
S20,PFN1,SPTBN2,DDX3X,CLTC,GJA1,MYO5A,CTNNB1,MACF1,CYFIP2,TARDBP,ARL6IP1,NARS1,
1,SKI,ASXL1,NOTCH2,PPP2CA,MAPK1,ODC1,AEBP1,FTO,FLNB,DLK1,STXBP1,LONP1,VANGL2,K
5A2,RPL26,RPL31,GPC3,MEGF8,RPS7,RPS19,RPL35,PRKAR1A,GJA1,COL6A2,RPS10,RPL35A,T
35,PRKAR1A,GJA1,RPS10,RPL35A,TSR2,HNRNPK,HLA-B,RPS27,MMP2,PPP1CB,RPL27,PLOD1,S

11,SMC3,ADH5,NDST1,SLC2A1,NDN,SPG7,HSPG2,SLC6A8,KMT2E,GTF2I,MSH6,DHCR7,MED13,
TKT,CDH2,GPC3,MEGF8,MMP14,RPS7,RPS19,RPL35,GJA1,RPS10,RPL35A,TSR2,HNRNPK,HLA-
IRNPK,ARCN1,CTBP1,RAC1,NOVA2,MDH2,EEF1A2,FDFT1,TBCD,GNAI3,SNRPN,SKI,ASXL1,RAB11
CYFIP2,DNAJC5,ARID1A,ATP1A2,CTBP1,RAC1,L1CAM,CRPPA,NOVA2,MDH2,TBCD,PPP1CB,SQST
A,ARID1A,TSR2,SHOX,ARCN1,RPS27,CTBP1,RAC1,PPIB,GLUL,CDH11,LMNB2,RPL27,SNRPN,COL
L1,AGRN,ZMIZ1,COL11A1,RPL10,ARF1,SCARB2,COL1A2,NR2F1,RPS20,COL5A2,RPL26,KLHL9,
YOU1,SMS,CDC42,KDM5C,TRIO,EP300,EXTL3,BAZ1B,NXN,ARID1B,PI4KA,RAB3GAP2,CRTAP,MB
T,NONO,RPL15,CHD3,POLR2A,RPL18,UCHL1,RPL11,AGRN,ZMIZ1,COL11A1,RPL10,RPS17,AP3

L6A2,ARL6IP1,ARID1A,SHOX,CTBP1,MMP2,HS6ST1,FDFT1,TBCD,LMNB2,SNRPN,PLOD1,COL2A1
P1CB,RPL27,SNRPN,ASXL1,RPS15A,NOTCH2,MAPK1,ACADVL,SON,KDM1A,MAP2K2,FLNB,DLK1
RNPU,RPS24,COL1A2,RHOA,ATP2A2,RPL26,CLEC7A,RPL31,LRPPRC,CDH2,GPC3,MEGF8,RAB7A

2,CHST3,SETD5,EXT1,SMARCC2,SMARCB1,KLHL7,DPF2
N,BRD4,PTEN,BAP1,SRSF2,DICER1,MSH6,SMO,RPS26,PTPN11,MDM2,IDH2,EWSR1,PDGFRB,EXT
IO,EP300,FLCN,BAZ1B,ATRX,NXN,RAB3GAP2,PTEN,SMC3,IL6ST,LOX,GPC4,H19,CLCN7,HSPG2,
CYFIP2,DNAJC5,ARID1A,ATP1A2,CTBP1,RAC1,L1CAM,CRPPA,NOVA2,MDH2,TBCD,PPP1CB,SQST
CYFIP2,DNAJC5,ARID1A,ATP1A2,CTBP1,RAC1,L1CAM,CRPPA,NOVA2,MDH2,TBCD,PPP1CB,SQST
P7,RAI1,RAB11A,SEC23A,ARNT2,ARFGEF2,ADAR,CIC,KCNQ2,EP300,RERE,KDM5B,CNTNAP1,FL
X,LTBP4,KMT2E,GTF2I,MED13,FBLN5,TANC2,BGN,PTPN11,POR,CLIP2,ELN,NOTCH1,ACTA2,AL
BXO11,PTEN,SMC3,ABCC9,MAPRE2,IARS2,HSPG2,ANTXR1,PKDCC,LTBP4,KMT2E,GTF2I,TTC37,
DM1A,MAP2K2,SMARCE1,RPS28,ATP6V1B2,STXBP1,KDM5C,EP300,RERE,RAB3GAP2,SMC3,ABC
26,RPL31,LRPPRC,CDH2,MEGF8,MMP14,RPS7,RPS19,RPL35,PRKAR1A,GJA1,MYO5A,RPS10,RP

P2K2,RPS28,ATP6V1B2,STXBP1,RAI1,CDC42,SYP,TRIO,EP300,RERE,KDM5B,FLCN,NXN,ARID1B,
,PPP2R1A,RPL15,CHD3,RPL18,UCHL1,RPL11,ZMIZ1,COL11A1,RPL10,RPS17,PAFAH1B1,CTSD,R

DFT1,TBCD,SQSTM1,LMNB2,SNRPN,ACO2,VPS35,SKI,ASXL1,CNTNAP2,TUBB2A,PRDX1,ODC1,F
F1R,CHST3,BCOR,RPS29,SNRPB,LMNA,SOS1,PIEZO1,MAFB
OL1A2,CHD4,COL5A2,RPL26,RPL31,GPC3,RPS7,RPS19,RPL35,PRKAR1A,GJA1,RPS10,RPL35A,

KI,ASXL1,RAB11B,ATP6AP1,NOTCH2,OPTN,AEBP1,SON,MAP2K2,STXBP1,PRPS1,USP7,RAI1,SMS,
,SQSTM1,SNRPN,ACO2,VPS35,SKI,ASXL1,PRDX1,PRKDC,SON,MAP2K2,QARS1,DHX30,FBLN1,ST
CND2,GPC3,DDX3X,CLTC,PRKAR1A,CTNNB1,CYFIP2,ARID1A,CTBP1,RAC1,L1CAM,GLUL,CRPPA,
7,CREBBP,PHIP,TRIP12,ATP13A2,MECP2,ADNP,IQSEC1,PPP2R5D,DEAF1,NSD1,WAC,SHMT2,CLC

A2,MAPK1,ODC1,ATXN10,SDHA,STXBP1,USP7,SARS1,RAI1,RAB11A,KDM3B,ADAR,CIC,DCTN1,KI
TCH1,ACTA2,ALDH18A1,LIMK1,DDX6,CRKL,CHST3,NSD1,LMNA,MED12,TGFBR1,PSMD12,TGFE
K2A1,RAB3GAP2,IARS1,TIMM50,SLC2A1,LINGO1,IREB2,PACS1,UBE3A,GOT2,YY1,CREBBP,CCD
PTPN11,COL12A1,OBSL1,EFNB1,EXT2,TMEM43,SOX9,RPS29,LMNA,SEPTIN9,CUL7,EXT1,TNPO3,
,RAB3GAP2,GPC4,H19,EHMT1,IARS2,ANTXR1,SLC6A8,LTBP4,IREB2,EFTUD2,LZTR1,MTOR,PRK
TRX,NXN,ARID1B,RAB3GAP2,SMC3,ECEL1,PTPRF,ABCC9,PACS1,LTBP3,PCGF2,PRKACA,YY1,SM
YFIP2,ARID1A,CTBP1,RAC1,L1CAM,GLUL,CRPPA,EEF1A2,TBCD,PPP1CB,LMNB2,SNRPN,SKI,ASX

,POGZ,DPF2,DMPK,SIN3A,TCF20
PTN,VHL,MAPK1,ATXN10,SMARCE1,SDHA,STXBP1,USP7,SARS1,SOD1,RAI1,TRAF7,DCTN1,BCR,E
OTCH2,MAPK1,SMARCA4,PRKDC,KDM1A,FLNB,DLK1,SMARCE1,RPS28,ATP6V1B2,STXBP1,SYP,
LNB,DLK1,SMARCE1,RPS28,ATP6V1B2,STXBP1,SYP,BCR,KDM5C,GPX4,NXN,ARID1B,SMC3,ABC
TPM3,SMARCE1,FBLN1,STXBP1,LONP1,COL27A1,ARNT2,ARFGEF2,EP300,EXTL3,BAZ1B,NXN,AR
A,EXT1,MSH2
H,AHCY,UBE3A,YY1,CREBBP,TRIM8,NTRK2,HS2ST1,PSAT1,TRRAP,ASNS,DDX6,SLC25A1,SLC16
AC1,CDH11,LMNB2,RPL27,SNRPN,COL2A1,RPS15A,NOTCH2,MAPK1,SMARCA4,SON,KDM1A,FL
AC1,LMNB2,RPL27,SNRPN,COL2A1,RPS15A,NOTCH2,MAPK1,SMARCA4,SON,KDM1A,FLNB,DLK1
NSD2,CREBBP,RPS26,HS2ST1,EMC1,ADNP,BSCL2,EFNB1,RAB18,NSD1,RPS29,SETD5,MAN2B1,
1A,TSR2,SHOX,RPS27,CTBP1,RAC1,PPIB,CDH11,LMNB2,RPL27,SNRPN,COL2A1,SKI,ASXL1,RPS
T1,SLC6A8,NSD2,UBE3A,CREBBP,TANC2,NTRK2,TECR,AUTS2,MECP2,WDR26,ADNP,FLII,IQSEC1
RPL27,SKI,ASXL1,TERF2IP,RPS15A,VHL,CCND1,SMARCA4,DLK1,SMARCE1,RPS28,ATP6V1B2,SD
E,GTF2I,MSH6,DHCR7,MED13,PCGF2,UBE3A,YY1,PHIP,TANC2,FGFR1,NTRK2,AUTS2,CLIP2,TRA
PS17,HNRNPU,RPS24,COL1A2,HK1,PTCD3,NR2F1,CHD4,RPL26,RPL31,TKT,CLTC,RPS7,RPS19,R
,COL2A1,RPS15A,NOTCH2,MAPK1,AEBP1,FTO,MAP2K2,FLNB,DLK1,RPS28,ATP6V1B2,STXBP1,G
,ATP6V1B2,SEC23A,SMS,B4GAT1,TTC37,PHGDH,SMARCD1,FGFR1,EIF4A3,RPS26,BGN,PSAT1,D
1,KLHL7,DPF2

,P4HB,IDS,RPL5,GDI1,NPM1,SNRNP200,IGF2,ATP5F1A,VCP,AFF4,TRIM28,YWHAG,COL4A2,FN1,

1,UCHL1,AGRN,ZMIZ1,RPL10,ARF1,ITM2B,COL1A2,HK1,NR2F1,ATP1A1,RPS20,ATP2A2,PFN1,L
28,ATP6V1B2,EP300,RERE,RAB3GAP2,SMC3,ABCC9,HSPG2,PACS1,LZTR1,NSD2,SMARCD1,CRE
25,PPP1R12A,EPB41L1,PPP3CA,SEC31A
RAI1,ARFGEF2,EP300,EXTL3,RERE,BAZ1B,ATRX,NXN,SMC3,ABCC9,LOX,CLCN7,DZIP1,GTF2I,TT

1,WDR26,ADNP,LIMK1,NBAS,PDGFRB,TMEM98,SOBP,DDX6,CHST3,NSD1,WAC,MED25,SETD5,
1,MESD,USP7,SEC23A,STAT3,EP300,EXTL3,RERE,KDM5B,CRTAP,PTEN,SMC3,IL6ST,H19,CLCN7,
C,TRAF7,DCTN1,TRIO,EP300,RERE,KDM5B,FLCN,BAZ1B,SMC3,NDST1,EML1,SLC2A1,NDN,BAP1

;,YY1,CREBBP,CCDC47,FGFR1,COL12A1,CLIP2,ELN,ADNP,LIMK1,EFNB1,SOBP,SLC25A1,WDR1
_1,PRDX1,MAP2K2,QARS1,DHX30,FBLN1,STXBP1,LONP1,SARS1,RAB11A,CDC42,ADAR,CIC,TRAF
32A,ATXN10,HSPB1,MAP2K2,LSS,STXBP1,LONP1,PRPS1,SARS1,RAI1,ARFGEF2,SMS,CCT5,KCNC
PS10,RPL35A,TSR2,SHOX,ARCN1,RPS27,CTBP1,MMP2,FDFT1,TBCD,GNAI3,RPL27,SNRPN,COL2
COL6A2,RPS10,RPL35A,TSR2,SHOX,ARCN1,RPS27,CTBP1,SMCHD1,MMP2,FDFT1,TBCD,GNAI3,F
RPS24,COL1A2,RHOA,COL5A2,RPL26,RPL31,LRPPRC,GPC3,MEGF8,RPS7,RPS19,RPL35,PRKAR
JA1,CTNNB1,ARID1A,HNRNPK,CTBP1,RAC1,MMP2,GLUL,NOVA2,CDH11,PPP1CB,SNRPN,COL2A1
TBCD,PPP1CB,SNRPN,SKI,ASXL1,RAB11B,NOTCH2,MAPK1,ODC1,SON,KDM1A,MAP2K2,RPS28,
HNRNPK,ARCN1,RPS27,CTBP1,SMCHD1,CRPPA,HS6ST1,CDH11,GNAI3,RPL27,SNRPN,COL2A1,A

PS10,RPL35A,TSR2,SHOX,ARCN1,RPS27,CTBP1,MMP2,FDFT1,TBCD,GNAI3,RPL27,SNRPN,COL2
RCA4,SON,KDM1A,FLNB,DLK1,SMARCE1,RPS28,ATP6V1B2,STXBP1,SYP,BCR,KDM5C,TRIO,GP

HNRNPK,ARCN1,RPS27,CTBP1,SMCHD1,CRPPA,HS6ST1,CDH11,GNAI3,RPL27,SNRPN,COL2A1,A
2,COL3A1
SNRPN,COL2A1,SKI,RPS15A,NOTCH2,MAPK1,AEBP1,FTO,MAP2K2,FLNB,DLK1,RPS28,ATP6V1B
,COL2A1,SKI,ASXL1,RAB11B,NOTCH2,MAPK1,SMARCA4,PLEC,ODC1,AEBP1,HSPB1,FLNB,DLK1,
P,WAC,GDF1,RPS29,NR2F2,MED12,CTCF,TALDO1
I1,KIF5A,GARS1,SPG7,HINT1,REEP1,RPS26,PTPN11,COL12A1,TIA1,OBSL1,MFN2,BSCL2,NDRG1
'300,KDM5B,PBX1,CSNK2A1,ATRX,ARID1B,SMC3,NDN,IARS2,KMT2E,PACS1,MED13,PCGF2,MTC
RDX1,MAP2K2,QARS1,DHX30,FBLN1,STXBP1,LONP1,SARS1,RAB11A,CDC42,CIC,TRAF7,KCNQ2,
TP13A2,MFN2,MECP2,NDUFA4,BSCL2,ALDH18A1,HTRA1,CAPN1,DARS1,SLC16A2,TSPAN7,GBA2,
TP13A2,MFN2,MECP2,NDUFA4,BSCL2,ALDH18A1,HTRA1,CAPN1,DARS1,SLC16A2,TSPAN7,GBA2,
CTCF,RUSC2,COX5A,TALDO1,SLC1A4,CUL4B,PSMD12,FHL1,TMCO1,KLHL7,HMGA2
LOX,EHMT1,IARS2,KMT2E,NSD2,PCGF2,FBLN5,YY1,PHIP,WNK1,ABL1,PUM1,TMEM94,COL12A1
,MACF1,CYFIP2,TARDBP,DDOST,HNRNPK,HLA-B,CTBP1,CDKN1A,EEF1A2,TBCD,SQSTM1,PABPN
8,PACS1,NSD2,OGT,MTOR,PACS2,SMARCD1,CCDC47,TUBB4A,DENND5A,TRIP12,SC5D,ELN,FLII,
HIP,CCDC47,FGFR1,CHMP1A,TMEM94,KLF13,PDHA1,IGF1R,NDUFA4,BSCL2,FLII,NECTIN1,ECHS
ZFR,NDST1,ABCC9,SLC2A1,NDN,H19,TAOK1,HINT1,KMT2E,TTC37,MED13,UBE3A,YY1,TANC2,TE
NON,MAPK1,SMARCA4,AEBP1,MAP2K2,FLNB,DLK1,TPM3,SMARCE1,FBLN1,STXBP1,LONP1,COL
,MARS1,PPP2CA,PRKDC,KDM1A,QARS1,DHX30,SDHA,STXBP1,PRPS1,RAI1,ARNT2,ADAR,KIF5A,E
TBP3,MED13,PCGF2,CREBBP,TANC2,EIF4A3,AUTS2,IGF1R,MDM2,MECP2,ALDH18A1,DDX6,SYNC
PA1,SPG11,FHL1,PRNP,HADHB,HARS1,SMARCB1,SLC18A3,NF2
NQ2,EXTL3,KDM5B,CTNND2,PI4KA,NDST1,BAP1,TAOK1,AKT3,PACS1,PHGDH,MTOR,TRIM8,MIC
.9,RAB7A,CCN2,CLTC,PRKAR1A,GJA1,COL6A2,CYFIP2,HNRNPDL,TARDBP,HNRNPK,ATP1A2,HLA-
SS,STXBP1,LONP1,PRPS1,SARS1,RAI1,ARFGEF2,SMS,KCNQ2,EP300,RERE,STARD7,KDM5B,FLC
15A,NOTCH2,KDM1A,RPS28,LSS,KDM3B,SMS,CDC42,EP300,RERE,KDM5B,ATRX,ARID1B,SMC3
1
MARCC2,SMARCB1,PRMT7,DPF2
TANC2,SH3PXD2B,FAM50A,ADNP,TRRAP,GPAA1,ALDH18A1,PDGFRB,EXT2,DDX6,PPP2R5D,CHST

R1A,GJA1,COL6A2,RPS10,RPL35A,TSR2,SHOX,ARCN1,RPS27,CTBP1,SMCHD1,MMP2,FDFT1,TB

EMC1,TRRAP,COX8A,FLII,SYNGAP1,PPP2R5D,DEAF1,CRKL,WAC,FARSA,CTCF,COX5A,SEPTIN9,C
VAC,CLCN4,ASH1L,BCOR,THOC2,EPB41L1,PPP3CA,SLC1A4,GATAD2B,SYT1,SMARCC2,TMCO1,P
13,FBLN5,PRKACA,TANC2,TUBB4A,POR,ELN,ACTA2,BSCL2,PDGFRB,DDX6,NSD1,DLG4,CDKN1B
FGFR1,DENND5A,PSAT1,DAG1,NDUFC2,ALDH18A1,VPS51,NSD1,ZSWIM6,BCOR,LARGE1,SLC5A6
N5,TANC2,REEP1,FGFR1,CHMP1A,ATP13A2,MFN2,PDHA1,TRIM2,MECP2,WDR26,BSCL2,ALDH18
C,SETD5,CTCF,RUSC2,COX5A,TALDO1,SLC1A4,CUL4B,PSMD12,FHL1,TMCO1,KLHL7,HMGA2
NPM1,COL18A1,SNRNP200,IGF2,AFF4,NONO,DCHS1,TPI1,CHSY1,PPP2R1A,RPL15,HLA-C,SF3B1
LONP1,PRPS1,SARS1,RAI1,ARFGEF2,SMS,KCNQ2,EP300,RERE,STARD7,KDM5B,FLCN,GLUD1,C
A,TSR2,HNRNPK,ARCN1,RPS27,CTBP1,SMCHD1,CRPPA,HS6ST1,CDH11,GNAI3,RPL27,SNRPN,CC

PC4,KMT2E,PACS1,MED13,CREBBP,TANC2,SMO,FGFR1,RPS26,ADNP,DDX6,CHST3,NSD1,SHMT2
DN,KDM1A,FLNB,DLK1,RPS28,ATP6V1B2,LONP1,CDC42,TRIO,EP300,EXTL3,BAZ1B,NXN,PTEN,SI
ARCE1,SDHA,STXBP1,LONP1,PRPS1,USP7,RAI1,RAB11A,TCTN1,KDM3B,CIC,TRIO,EP300,RERE,K
TPN11,CLIP2,ELN,OBSL1,ALDH18A1,LIMK1,EFNB1,CRKL,NSD1,LMNA,MED12,HIRA,SOS1,ZDHHC
ZTR1,HSD17B4,NSD2,MED13,MTOR,PHIP,TANC2,FGFR1,SH3PXD2B,NTRK2,MDH1,PTPN11,CLIP2
SH2,DNAJC21
S29,SETD5,MAN2B1,TMCO1,KLHL7
10,SDHA,STXBP1,DCTN1,EP300,STARD7,UBQLN2,CST3,HNRNPA2B1,A2M,FUS,GRN,ATXN3,CREE

,EP300,RERE,BAZ1B,ATRX,NXN,ARID1B,PI4KA,RAB3GAP2,PTEN,SMC3,GPC4,NDN,H19,PKDCC,E
CHST3,BCOR,LMNA,NR2F2,MED12,HIRA,CITED2,TGFBR1,TGFB2,COL3A1
6,ADNP,ZNF462,CHST3,RBM10,RPS29,LARGE1,MED12,PSMD12,SETD1A,CNOT1
D1B,SMC3,ECEL1,HSPG2,SMARCD1,CREBBP,FGFR1,HS2ST1,BGN,PTPN11,COL12A1,POR,MACI

MP2,PPP1CB,RPL27,ASXL1,RPS15A,NOTCH2,SELENON,MAPK1,ACADVL,FTO,MAP2K2,FLNB,DLK
,SMARCE1,USP7,RAI1,BCR,TRIO,EP300,RERE,BAZ1B,ATRX,NXN,ARID1B,PTEN,SMC3,GPC4,H19,
GAT1,NDN,TAOK1,AKT3,PHGDH,HSD17B4,MTOR,MN1,SMO,TECR,PSAT1,DAG1,PDHA1,MECP2,TF

N,DLK1,RPS28,ATP6V1B2,RAI1,SEC23A,SMS,DGCR2,NXN,B4GAT1,TTC37,PHGDH,SMARCD1,FGI
2,RPS26,ADNP,ZNF462,CHST3,RBM10,RPS29,LARGE1,MED12,PSMD12,SETD1A,CNOT1
D12,SMARCC2,TMCO1,SMARCB1,SETD1B,HMGA2,DPF2
ACS2,SMARCD1,CCDC47,SH3PXD2B,DENND5A,HS2ST1,CLIP2,ELN,WDR26,ADNP,BSCL2,LIMK1,
MED13,PCGF2,MTOR,PACS2,CREBBP,HS2ST1,TMEM94,IGF1R,MECP2,ADNP,TRRAP,PDGFRB,VPS
OTCH2,PPP2CA,MAPK1,SMARCA4,FTO,KDM1A,MAP2K2,FLNB,DLK1,STXBP1,LONP1,RAI1,SEC23

SXL1,CCND1,PLEC,ATXN10,SMARCE1,SDHA,VANGL2,ARNT2,SYP,TRAF7,KIF5A,TRIO,EP300,RERE

MECP2,NDUFA4,BSCL2,ALDH18A1,AMPD2,KLC2,CAPN1,ECHS1,SLC16A2,STUB1,GBA2,GPT2,CL
B,PTEN,SMC3,ECEL1,IL6ST,GPC4,IARS2,HSPG2,PKDCC,GTF2I,NSD2,DHCR7,FBLN5,CREBBP,CC

M20C,MAFB,TCF20

2B1,LMNA,NIPA1,COL6A3,SPG11,PSMD12,PIGT,ITGA7,MAFB
GBA2,GPT2,SETX,NDUFS1,MAN2B1,DDHD2,DSTYK,NIPA1,SPG11,OPA1,NDUFA10
N7,NSD1,WAC,DHFR,CLCN4,ASH1L,ACSL4,CACNA1H,MED12,CNTN2,EPB41L1,PPP3CA,GATAD2E
\1,VPS51,NSD1,BCOR,LARGE1,SLC5A6,PMPCA,NID1,CNOT3,POMGNT1,SMARCC2,POGZ,SMARC
,TCTN1,CDC42,EP300,RERE,BAZ1B,ARID1B,PI4KA,RAB3GAP2,B4GAT1,GPC4,GTF2I,PACS1,PHG
C42,SYP,TRAF7,BCR,KDM5C,TRIO,EP300,RERE,BAZ1B,CSNK2A1,ATRX,NXN,ARID1B,FBXO11,PT
MNB2,SNRPN,PLOD1,COL2A1,SKI,RAB11B,RPS15A,SELENON,SMARCA4,AEBP1,FLNB,DLK1,TPM
D12,CITED2,TGFBR1,TGFB2,COL3A1
DH6A1,GNB1,APP,IDS,RPL5,GDI1,NPM1,HSPD1,COL18A1,IGF2,VCP,UBB,AFF4,TRIM28,YWHAG,N
PC4,KMT2E,PACS1,MED13,CREBBP,TANC2,SMO,FGFR1,RPS26,ADNP,DDX6,CHST3,NSD1,SHMT2
,PRPF6,NOTCH2,PPP2CA,MAPK1,SMARCA4,FTO,KDM1A,MAP2K2,FLNB,DLK1,STXBP1,LONP1,R/

KI,RPS15A,NOTCH2,KDM1A,RPS28,LSS,KDM3B,SMS,CDC42,EP300,RERE,KDM5B,ATRX,ARID1B
KI,RPS15A,NOTCH2,KDM1A,RPS28,LSS,KDM3B,SMS,CDC42,EP300,RERE,KDM5B,ATRX,ARID1B
M5B,CTNND2,SLC2A1,BAP1,TAOK1,TPP1,AKT3,CIZ1,UBE3A,TRIM8,SMO,NTRK2,DENND5A,PSAT
PN11,PSAT1,ABL1,COL12A1,CLIP2,ELN,ALDH18A1,LIMK1,NBAS,PDGFRB,CHST3,NSD1,DLG4,M/
,EP300,RERE,BAZ1B,ATRX,NXN,ARID1B,PI4KA,RAB3GAP2,PTEN,SMC3,GPC4,NDN,H19,PKDCC,E
12,SIN3A
SMS,YME1L1,KDM5C,RERE,KDM5B,ALDH3A2,VARS1,BAZ1B,ATRX,ARID1B,PTEN,SMC3,LOX,NDN,
,CRPPA,MDH2,TBCD,SQSTM1,TAF15,LMNB2,SNRPN,COL2A1,SELENON,OPTN,SMARCA4,PLEC,F
SWIM6,CDKN1B,ERBB2,LMNA,RELA,FH,MSH2,EPHB2,SMARCB1,NF2,DMPK
,RPL35,PRKAR1A,GJA1,COL6A2,RPS10,RPL35A,ARID1A,TSR2,RPS27,CTBP1,MMP2,L1CAM,RPL
CDC42,TRAF7,BCR,EP300,BAZ1B,ATRX,SMC3,LOX,GPC4,EHMT1,AKT3,LDLR,DZIP1,GTF2I,TTC3
DH2,GPC3,MEGF8,RPS7,RPS19,RPL35,PRKAR1A,GJA1,RPS10,RPL35A,ARID1A,TSR2,HNRNPK,H
.18,RPL11,ZMIZ1,COL11A1,RPS17,AP3D1,ITM2B,MAT2A,PXDN,RPS24,COL1A2,HK1,RHOA,NR2F

,RPL15,SEC61A1,RPL18,RPL11,ZMIZ1,COL11A1,RPL10,RPS17,CYB5R3,PAFAH1B1,HNRNPU,SC/
194,IGF1R,TRRAP,ZNF462,NECTIN1,DDX6,CRKL,CHST3,SHMT2,BCOR,SETD5,MAN2B1,CTCF,RU

21
N,TAOK1,PHGDH,HSD17B4,UBE3A,TRIM8,MICAL1,PDHA1,MECP2,NDUFA4,WDR26,GPAA1,COX8
CTNND2,NDST1,SLC2A1,GRN,TAOK1,EHMT1,PHGDH,MED13,MTOR,UBE3A,TRIM8,TECR,MECP2,
YME1L1,RERE,GPX4,ARID1B,PI4KA,RAB3GAP2,HDAC6,B4GAT1,GPC4,EHMT1,TUBGCP6,PACS1,F
DD1,COL2A1,SKI,RAB11B,RPS15A,SELENON,SMARCA4,AEBP1,FLNB,DLK1,TPM3,SMARCE1,FBL
MBTPS1,NDN,H19,TTC37,NSD2,YY1,RPS26,AUTS2,POR,OBSL1,PDHA1,MECP2,NECTIN1,TSPAN7,

SCARB2,COL1A2,HK1,CHD4,COL5A2,RPL26,KLHL9,GPC3,MEGF8,CCN2,MMP14,MIF,PRKAR1A,G
2,RPS26,ADNP,ZNF462,CHST3,RBM10,RPS29,LARGE1,MED12,PSMD12,SETD1A,CNOT1
D1,CREBBP,FGFR1,DENND5A,PSAT1,DAG1,NDUFC2,ALDH18A1,VPS51,NSD1,ZSWIM6,BCOR,LAR
PTEN,SMC3,ABCC9,LOX,GPC4,SOX2,GTF2I,TTC37,PACS1,PHGDH,DHCR7,PCGF2,CREBBP,RPS2
PL27,SKI,RPS15A,NOTCH2,PLEC,KDM1A,RPS28,LSS,KDM3B,SMS,CDC42,EP300,RERE,KDM5B,E
IT1,VAPB,HSPG2,KMT2E,GTF2I,SRSF2,MSH6,MED13,CIZ1,YY1,CREBBP,PHIP,TANC2,FGFR1,PIN

OMM40,ABCA7

SKI,PRPF6,NOTCH2,PPP2CA,MAPK1,SMARCA4,PLEC,FTO,KDM1A,MAP2K2,FLNB,DLK1,SMARCE

DH2,GPC3,MEGF8,RPS7,RPS19,RPL35,PRKAR1A,GJA1,RPS10,RPL35A,ARID1A,TSR2,HNRNPK,HI

PC4,H19,GTF2I,PACS1,LZTR1,PACS2,UBE3A,CREBBP,EIF4A3,SH3PXD2B,PTPN11,CLIP2,MACRO

CR8,SEPTIN9,GATAD2B,CUL4B,PSMD12,NFIB,MAFB,TCF20

TFIP2,ARID1A,HNRNPK,NARS1,CTBP1,SMCHD1,MMP2,PPIB,HS6ST1,CDH11,EEF1A2,TBCD,PPP1C

CHS1,SET,PPP2R1A,POLR2A,UCHL1,RPL11,AGRN,ZMIZ1,COL11A1,RPL10,AP3D1,ARF1,PAFAH1I

1,SON,KDM1A,MAP2K2,SMARCE1,DHX30,RPS28,ATP6V1B2,LSS,STXBP1,MESD,RAI1,TCTN1,SMS

F1,MAT2A,HNRNPU,SCARB2,RPS24,COL1A2,CHD4,RPS20,COL5A2,RPL26,CLEC7A,RPL31,TKT,TI

R1,COL6A3,TGFB2,COL3A1,MAPK8IP3,SIN3A

NUS1,PMPCA,PPP3CA,QRICH1,PCNA,SZT2,PRNP,DNM1,MAPK8IP3,NF2,NECAP1

1,HSPG2,ANTXR1,LTBP4,GTF2I,PACS1,PHGDH,FBLN5,MTOR,CREBBP,TRIM8,SH3PXD2B,AUTS2,

SL1,MFN2,LIMK1,NECTIN1,CRKL,CHST3,TMEM43,BCOR,SETD5,LMNA,MED12,ARSB,TRAPPC11

4,NONO,DCHS1,TPI1,RPL15,SF3B1,RPL18,RPL11,AGRN,ZMIZ1,RPL10,RPS17,VCL,AP3D1,ARF1,

DLK1,STXBP1,MESD,USP7,SEC23A,STAT3,EP300,EXTL3,RERE,KDM5B,VARS1,GPX4,ATRX,CRTAP,

P,AFF4,YWHAG,COL4A2,MAPT,NONO,DCHS1,SET,TPI1,PPP2R1A,RPL15,CHD3,POLR2A,AP2M1,RI
,SET,TPI1,PPP2R1A,CHD3,POLR2A,AP2M1,UCHL1,AGRN,ZMIZ1,RPL10,MARCHF6,VCL,AP3D1,SE
,AGRN,ZMIZ1,RPL10,AP3D1,ARF1,CYB5R3,PAFAH1B1,ITM2B,HNRNPU,CTSD,COL1A2,HK1,NR2F
L11A1,RPL10,RPS17,SBF1,ARF1,PAFAH1B1,MAT2A,HNRNPU,SCARB2,RPS24,COL1A2,HK1,COL5
IB,APP,IDS,RPL5,GDI1,SPTAN1,CALM1,NPM1,PLD3,HSPD1,COL18A1,SNRNP200,IGF2,ATP5F1A,V
M3,GNB1,P4HB,APP,IDS,RPL5,GDI1,SPTAN1,CALM1,NPM1,PLD3,HSPD1,COL18A1,SNRNP200,IG
C,SON,KDM1A,MAP2K2,QARS1,SMARCE1,LSS,STXBP1,LONP1,PRPS1,USP7,SARS1,RAI1,RAB11A
TP5F1A,VCP,UBB,AFF4,YWHAG,MAPT,FN1,NONO,DCHS1,SET,TPI1,CHSY1,PPP2R1A,RPL15,CHD3
HSPD1,COL18A1,IGF2,ATP5F1A,VCP,AFF4,YWHAG,COL4A2,MAPT,FN1,NONO,DCHS1,SET,TPI1,PI
RPS7,RPS19,RPL35,PRKAR1A,GJA1,RPS10,RPL35A,TSR2,HNRNPK,NARS1,ARCN1,RPS27,CTBP1
PL18,UCHL1,RPL11,ZMIZ1,COL11A1,RPL10,RPS17,AP3D1,SBF1,ARF1,CYB5R3,PAFAH1B1,ITM2I
PL18,UCHL1,RPL11,ZMIZ1,COL11A1,RPL10,RPS17,AP3D1,SBF1,ARF1,CYB5R3,PAFAH1B1,ITM2I
B,APP,IDS,RPL5,GDI1,SPTAN1,NPM1,HSPD1,COL18A1,SNRNP200,IGF2,ATP5F1A,VCP,UBB,AFF4,
,RPL10,RPS17,AP3D1,SBF1,ARF1,CYB5R3,PAFAH1B1,HNRNPU,CTSD,RPS24,HK1,PTCD3,NR2F1,
I,RPS27,CTBP1,SMCHD1,MMP2,HS6ST1,CDH11,TBCD,PPP1CB,RPL27,SNRPN,COL2A1,SKI,ASXL
SNRNP200,IGF2,VCP,UBB,AFF4,YWHAG,COL4A2,MAPT,NONO,DCHS1,SET,TPI1,CHSY1,PPP2R1A
SY1,PPP2R1A,RPL15,CHD3,POLR2A,AP2M1,RPL18,RPL11,AGRN,ZMIZ1,COL11A1,RPL10,RPS17,
OL1A2,HK1,COL5A2,GPC3,MEGF8,DDX3X,CLTC,PRKAR1A,COL6A2,ARID1A,SHOX,HNRNPK,NARS
F4,YWHAG,COL4A2,MAPT,FN1,DCHS1,SET,EGLN1,PPP2R1A,RPL15,PKD1,CHD3,POLR2A,AP2M1
,ASXL1,RAB11B,NOTCH2,SELENON,PPP2CA,MAPK1,SMARCA4,PRDX1,ODC1,PRKDC,SON,KDM1
L26,RPL31,XPR1,CCND2,CDH2,GPC3,DDX3X,CLTC,RPS7,RPS19,RPL35,GJA1,CTNNB1,MACF1,R
,NR2F1,RPL26,RPL31,XPR1,CDH2,GPC3,DDX3X,CLTC,RPS7,RPS19,RPL35,GJA1,CTNNB1,MACF
F4,YWHAG,COL4A2,MAPT,FN1,NONO,DCHS1,SET,PPP2R1A,RPL15,PKD1,CHD3,POLR2A,AP2M1,I
G,MAPT,NONO,DCHS1,SET,CHSY1,PPP2R1A,RPL15,CHD3,AP2M1,SEC61A1,RPL18,UCHL1,RPL11
P2R1A,RPL15,CHD3,RPL18,UCHL1,RPL11,AGRN,ZMIZ1,COL11A1,RPL10,RPS17,SBF1,ARF1,PAF
18A1,IGF2,ATP5F1A,VCP,UBB,AFF4,YWHAG,MAPT,FN1,NONO,DCHS1,SET,TPI1,CHSY1,PPP2R1A,
TBP1,RAC1,L1CAM,GLUL,CRPPA,NOVA2,MDH2,EEF1A2,FDFT1,TBCD,SQSTM1,VAMP2,ACO2,SKI
3A1,IGF2,ATP5F1A,VCP,AFF4,YWHAG,COL4A2,MAPT,FN1,NONO,DCHS1,SET,TPI1,PPP2R1A,RPL1
CHSY1,PPP2R1A,RPL15,CHD3,POLR2A,AP2M1,RPL18,RPL11,AGRN,ZMIZ1,RPL10,RPS17,AP3D1,
BP1,RAC1,SMCHD1,CRPPA,CDH11,FDFT1,TBCD,PPP1CB,GNAI3,RPL27,PLOD1,RBMX,SKI,KDM6I
A,SMS,CDC42,SYP,BCR,KDM5C,TRIO,EP300,EXTL3,RERE,PBX1,FLCN,BAZ1B,ATRX,NXN,ARID1B,I
8,DDX3X,CLTC,PRKAR1A,COL6A2,ARID1A,SHOX,HNRNPK,NARS1,CTBP1,RAC1,MMP2,PPIB,CRPF
2,TKT,CCND2,GPC3,CLTC,PRKAR1A,GJA1,CTNNB1,MACF1,CYFIP2,TARDBP,ARID1A,DDOST,HNF
CDH2,GPC3,MEGF8,DDX3X,RPS7,RPS19,RPL35,PRKAR1A,GJA1,CTNNB1,RPS10,RPL35A,ARID1A,
2I,TTC37,PACS1,PHGDH,FBLN5,CREBBP,FGFR1,SH3PXD2B,AUTS2,CLIP2,ELN,NOTCH1,ACTA2,AI
/A2,MDH2,EEF1A2,FDFT1,TBCD,VAMP2,ACO2,SKI,ASXL1,TUBB2A,PPP2CA,SMARCA4,ODC1,PRKI
PS17,SBF1,ARF1,PAFAH1B1,MAT2A,RPS24,COL1A2,HK1,RHOA,NR2F1,CHD4,COL5A2,RPL26,RF
ARID1A,HNRNPK,NARS1,CTBP1,RAC1,L1CAM,GLUL,CRPPA,NOVA2,MDH2,EEF1A2,FDFT1,TBCD,S
2,ATP2A2,RPL26,RPL31,KLHL9,CCND2,GPC3,MEGF8,CCN2,MMP14,RPS7,RPS19,RPL35,PRKAR
.18A1,SNRNP200,IGF2,ATP5F1A,VCP,AFF4,TRIM28,YWHAG,COL4A2,MAPT,NONO,DCHS1,SET,TP
ARID1A,TSR2,HNRNPK,NARS1,RPS27,CTBP1,SMCHD1,MMP2,PPIB,CRPPA,CDH11,TBCD,PPP1CE
YB5R3,PAFAH1B1,HNRNPU,CTSD,SCARB2,COL1A2,HK1,NR2F1,ATP1A1,RPS20,ATP2A2,CLEC7A,I

B1,MACF1,RPS10,RPL35A,CYFIP2,ARID1A,TSR2,HNRNPK,NARS1,ARCN1,RPS27,CTBP1,RAC1,L1

,HSPD1,COL18A1,IGF2,ATP5F1A,VCP,UBB,AFF4,YWHAG,MAPT,FN1,NONO,DCHS1,SET,TPI1,CHS
2F1,CHD4,COL5A2,ATP2A2,RPL26,RPL31,KLHL9,CCND2,GPC3,MEGF8,CCN2,MMP14,RPS7,RPS
CTNNB1,MACF1,RPS10,RPL35A,CYFIP2,ARID1A,TSR2,HNRNPK,NARS1,ARCN1,RPS27,CTBP1,RA
U,COL1A2,HK1,NR2F1,CHD4,RPS20,COL5A2,LRPPRC,SPTBN2,TKT,CDH2,GPC3,DDX3X,CLTC,M
.1A2,HK1,RHOA,NR2F1,CHD4,COL5A2,RPL26,RPL31,LRPPRC,TKT,CCND2,CDH2,GPC3,MEGF8,M
H1B1,MAT2A,RPS24,COL1A2,RHOA,NR2F1,CHD4,COL5A2,RPL26,RPL31,CCND2,GPC3,MEGF8,F
H1B1,MAT2A,RPS24,COL1A2,RHOA,NR2F1,CHD4,COL5A2,RPL26,RPL31,CCND2,GPC3,MEGF8,F

A,ARID1A,TSR2,RPS27,CTBP1,EEF1A2,FDFT1,PPP1CB,RPL27,PLOD1,SKI,KDM6B,RPS15A,NOTCH
RPS17,VCL,SBF1,LDHA,RPS24,HK1,ATP1A1,RPS20,RPL26,PFN1,RPL31,KLHL9,SPTBN2,TPM1,G
XO11,HDAC6,PTEN,ABCC9,MAPRE2,GPC4,IARS2,HSPG2,AKT3,KMT2E,GTF2I,TTC37,PACS1,LTB

PTAN1,NPM1,HSPD1,COL18A1,SNRNP200,IGF2,ATP5F1A,VCP,UBB,AFF4,YWHAG,MAPT,NONO,D
PL26,RPL31,CCND2,CDH2,GPC3,MEGF8,CLTC,RPS7,RPS19,RPL35,PRKAR1A,GJA1,RPS10,RPL3
CR,EP300,BAZ1B,CSNK2A1,ARID1B,CTNND2,PTEN,ABCC9,LOX,ANTXR1,SLC6A8,PKDCC,KMT2E,
PPP1CB,SQSTM1,LMNB2,SNRPN,SKI,CNTNAP2,RAB11B,PPP2CA,MAPK1,ODC1,KDM1A,DLK1,SM
,RPL31,CCND2,GPC3,MEGF8,RAB7A,RPS7,RPS19,RPL35,PRKAR1A,GJA1,COL6A2,RPS10,RPL35
,SMARCE1,RPS28,LONP1,RAI1,BCR,EP300,RERE,KDM5B,FLCN,BAZ1B,GPX4,NXN,ARID1B,PTEN,
D3,POLR2A,AP2M1,RPL18,RPL11,AGRN,ZMIZ1,COL11A1,RPL10,RPS17,AP3D1,CYB5R3,PAFAH1E
CE1,RPS28,STXBP1,LONP1,VANGL2,KDM3B,ARFGEF2,CDC42,TRAF7,BCR,DGCR2,KDM5B,PBX1
,SMARCE1,RPS28,LONP1,RAI1,BCR,EP300,RERE,KDM5B,FLCN,BAZ1B,GPX4,NXN,ARID1B,PTEN,
PTAN1,NPM1,HSPD1,COL18A1,SNRNP200,IGF2,ATP5F1A,VCP,UBB,AFF4,YWHAG,MAPT,NONO,D
.1A2,HK1,RHOA,NR2F1,CHD4,COL5A2,RPL26,RPL31,LRPPRC,TKT,CCND2,CDH2,GPC3,MEGF8,C

11,EEF1A2,PPP1CB,SNRPN,COL2A1,RBMX,SKI,ASXL1,KDM6B,RAB11B,NOTCH2,SELENON,PPP2
DFT1,TBCD,VAMP2,ACO2,SKI,ASXL1,TUBB2A,PPP2CA,SMARCA4,PRKDC,SON,KDM1A,MAP2K2,C

OTCH2,MAPK1,SMARCA4,PRKDC,SON,KDM1A,FLNB,DLK1,SMARCE1,RPS28,ATP6V1B2,HYOU1,
RPS19,RPL35,GJA1,RPS10,RPL35A,CYFIP2,ARID1A,TSR2,HNRNPK,NARS1,RPS27,CTBP1,MMP2,
PS19,RPL35,GJA1,COL6A2,RPS10,RPL35A,ARID1A,TSR2,HNRNPK,NARS1,RPS27,CTBP1,RAC1,SM
BP1,RAC1,GLUL,CRPPA,NOVA2,EEF1A2,PPP1CB,GNAI3,SNRPN,ACO2,PLOD1,COL2A1,SKI,ASXL1,
,HK1,RHOA,NR2F1,CHD4,COL5A2,RPL26,RPL31,LRPPRC,TKT,CCND2,CDH2,GPC3,MEGF8,CLTC
MT2,SETD5,MBOAT7,PMPCA,SLC5A7,RUSC2,GALNT2,CNOT3,SPG11,TNPO3,NFASC,SYT1,DYNC1
DFT1,TBCD,VAMP2,ACO2,SKI,ASXL1,TUBB2A,PPP2CA,SMARCA4,PRKDC,SON,KDM1A,MAP2K2,C
S1,SET,PPP2R1A,RPL15,SF3B1,CHD3,POLR2A,RPL18,TUBB4B,UCHL1,RPL11,AGRN,ZMIZ1,COL1
ARCN1,RPS27,CTBP1,SMCHD1,CDH11,PPP1CB,RPL27,SNRPN,PLOD1,SKI,RPS15A,NOTCH2,SEL
A,NOTCH2,PPP2CA,MAPK1,ACADVL,FTO,SON,KDM1A,MAP2K2,FLNB,DLK1,SMARCE1,RPS28,SDH
S7,RPS19,RPL35,GJA1,CTNNB1,COL6A2,RPS10,RPL35A,ARID1A,TSR2,SHOX,HNRNPK,ARCN1,RF
S19,RPL35,PRKAR1A,GJA1,MYO5A,CTNNB1,COL6A2,MACF1,RPS10,RPL35A,TARDBP,ARID1A,DD
PM3,RPS28,RAI1,ARFGEF2,ADAR,TRAF7,BCR,EP300,RERE,DGCR2,BAZ1B,NXN,PTEN,LOX,EHMT1
PL35,PRKAR1A,GJA1,COL6A2,RPS10,RPL35A,ARID1A,TSR2,SHOX,NARS1,RPS27,CTBP1,RAC1,MI
A,GJA1,COL6A2,ARL6IP1,ARID1A,SHOX,HNRNPK,NARS1,ARCN1,CTBP1,MMP2,L1CAM,CRPPA,H
I,MMP2,L1CAM,CRPPA,HS6ST1,MDH2,EEF1A2,SQSTM1,VAMP2,PABPN1,LMNB2,SNRPN,PLOD1,

RPL18,RPL11,AGRN,ZMIZ1,COL11A1,RPL10,RPS17,AP3D1,CYB5R3,PAFAH1B1,MAT2A,HNRNPU,
RPL18,RPL11,AGRN,ZMIZ1,COL11A1,RPL10,RPS17,AP3D1,CYB5R3,PAFAH1B1,MAT2A,HNRNPU,
32,COL1A2,HK1,RHOA,CHD4,ATP1A1,COL5A2,ATP2A2,KLHL9,GPC3,MEGF8,RAB7A,CCN2,MMP1
OX,HNRNPK,ARCN1,CTBP1,RAC1,MMP2,CRPPA,NOVA2,CDH11,TBCD,PPP1CB,LMNB2,SNRPN,PL
PL35A,ARID1A,TSR2,SHOX,HNRNPK,RPS27,CTBP1,RAC1,MMP2,GLUL,HS6ST1,CDH11,EEF1A2,FD
35R3,ITM2B,SCARB2,HK1,PTCD3,NR2F1,CHD4,ATP1A1,RPS20,PFN1,LRPPRC,XPR1,KLHL9,SPTB
KMT2E,MED13,PACS2,UBE3A,YY1,SMARCD1,CREBBP,TANC2,SEMA6B,TMEM94,COL12A1,DAG1,
42,EEF1A2,TBCD,SQSTM1,VAMP2,LMNB2,SNRPN,YEATS2,ACO2,VPS35,CHD2,SKI,CNTNAP2,RAB

GO1,SLC6A8,KMT2E,PACS1,DHCR7,CREBBP,PHIP,TRIM8,TRIP12,ATP13A2,MECP2,ADNP,FLII,IQS
B11B,PPP2CA,MAPK1,ODC1,KDM1A,DLK1,SMARCE1,DHX30,SDHA,LSS,FBLN1,STXBP1,PRPS1,M

RNP200,IGF2,VCP,UBB,AFF4,YWHAG,MAPT,NONO,DCHS1,SET,CHSY1,PPP2R1A,RPL15,CHD3,PO
SQSTM1,TAF15,CNTNAP2,RAB11B,MARS1,OPTN,ODC1,FTO,ATXN10,SON,KDM1A,SDHA,FBLN1,
MARCA4,SON,MAP2K2,FLNB,DLK1,SMARCE1,RPS28,STXBP1,LONP1,VANGL2,RAI1,SYP,TRIO,EP3
ARS1,SELENON,VHL,PLEC,HSPB1,KDM1A,FLNB,TPM3,SMARCE1,RPS28,SDHA,STXBP1,PRPS1,RA
MP2,L1CAM,CDH11,EEF1A2,GNAI3,SNRPN,COL2A1,RBMX,ASXL1,KDM6B,CNTNAP2,RAB11B,NC
),PRKAR1A,GJA1,CYFIP2,SHOX,HNRNPK,ARCN1,CTBP1,RAC1,MMP2,PPIB,CRPPA,NOVA2,CDH11
1A2,HK1,PTCD3,NR2F1,CHD4,RPL26,RPL31,TKT,GPC3,CLTC,RPS7,RPS19,RPL35,PRKAR1A,GJA
EXTL3,RERE,NDUFV1,KDM5B,FLCN,BAZ1B,ATRX,ARID1B,IARS1,PTEN,SMC3,ADH5,NDST1,NDN,

PLEC,ODC1,AEBP1,KDM1A,DLK1,TPM3,DHX30,SDHA,LARS1,LSS,FBLN1,STXBP1,LONP1,PRPS1,M
7,COL2A1,SKI,ASXL1,RPS15A,NOTCH2,MAPK1,PRDX1,PRKDC,SON,MAP2K2,QARS1,FLNB,DLK1,
J,COL1A2,HK1,NR2F1,ATP1A1,LRPPRC,SPTBN2,TKT,CCND2,CDH2,GPC3,MEGF8,DDX3X,CLTC,F
AR1A,GJA1,MYO5A,CTNNB1,COL6A2,MACF1,RPS10,RPL35A,ARID1A,DDOST,TSR2,RPS27,CTBP1

RID1A,SHOX,HNRNPK,NARS1,CTBP1,RAC1,MMP2,L1CAM,GLUL,CRPPA,HS6ST1,FDFT1,TBCD,PPP
I3,RPL27,COL2A1,SKI,ASXL1,RPS15A,NOTCH2,MAPK1,PRDX1,PRKDC,SON,MAP2K2,QARS1,FLN
YB5R3,PAFAH1B1,HNRNPU,CTSD,SCARB2,COL1A2,HK1,NR2F1,CHD4,ATP2A2,SPTBN2,TKT,CDH2
,SCARB2,HK1,PTCD3,NR2F1,RPS20,PFN1,LRPPRC,XPR1,SPTBN2,DDX3X,CLTC,GJA1,MYO5A,CT
ISD17B4,PCGF2,FBLN5,CREBBP,CCDC47,SH3PXD2B,HS2ST1,PTPN11,ABL1,COL12A1,CLIP2,ELN

ZSWIM6,MED12,TALDO1,QRICH1,GATAD2B,CUL4B,FAM20C,SETD1A,SMARCC2,CENPF,TMCO1,S
ARS1,OPTN,ODC1,ATXN10,SON,SDHA,FBLN1,STXBP1,PRPS1,SARS1,SOD1,RAB11A,ARNT2,CCT5
FLNB,DLK1,RPS28,LARS1,STXBP1,MESD,RAI1,SEC23A,CDC42,YME1L1,SYP,BCR,EP300,EXTL3,F
N,OPTN,MAPK1,PLEC,ATXN10,SON,MAP2K2,TPM3,SDHA,STXBP1,LONP1,PRPS1,USP7,IDH1,SOD1
DM3B,BCR,EP300,RERE,DGCR2,NDUFV1,BAZ1B,ATRX,NXN,ARID1B,SMC3,GPC4,SOX2,AKT3,EFT
DGCR2,KDM5B,BAZ1B,ATRX,NXN,ARID1B,CTNND2,ABCC9,MAPRE2,LOX,GPC4,H19,EHMT1,HSP
D,USP7,SMS,WDR45B,TRIO,EXTL3,RERE,ALDH3A2,BAZ1B,ATRX,NXN,ARID1B,RAB3GAP2,MBTPS
DGCR2,KDM5B,BAZ1B,ATRX,NXN,ARID1B,CTNND2,ABCC9,MAPRE2,LOX,GPC4,H19,EHMT1,HSP
R,EP300,RERE,DGCR2,NDUFV1,BAZ1B,ATRX,NXN,ARID1B,SMC3,GPC4,SOX2,AKT3,EFTUD2,GTF
1,PRKDC,SON,MAP2K2,QARS1,DHX30,FBLN1,STXBP1,LONP1,SARS1,SOD1,RAB11A,CCT5,CDC4
TSR2,SHOX,ARCN1,RPS27,CTBP1,RAC1,SMCHD1,MMP2,PPIB,GLUL,CDH11,FDFT1,TBCD,GNAI3,
A1,SKI,RPS15A,SELENON,MAPK1,SMARCA4,PLEC,ODC1,AEBP1,KDM1A,MAP2K2,TPM3,SMARCE
SPG7,HSPG2,SLC6A8,KMT2E,GTF2I,MSH6,DHCR7,MED13,PCGF2,MTOR,UBE3A,YY1,CREBBP,P

S2,HSPG2,GTF2I,PACS1,AHCY,MTOR,CREBBP,FGFR1,BGN,ABL1,COL12A1,CLIP2,ELN,ACTA2,MI
,ATRX,NXN,ARID1B,SMC3,ABCC9,PREPL,GPC4,NDN,EHMT1,PACS1,MED13,PCGF2,MTOR,PACS2
APK1,ODC1,SON,KDM1A,MAP2K2,DLK1,RPS28,ATP6V1B2,STXBP1,RAI1,SMS,CDC42,SYP,BCR,K
5,RAB18,DEAF1,RBM10,WAC,RPS29,SNRPB,DSTYK,SMARCC2,SMARCB1,HMGA2,MAFB,DPF2

A,NOTCH2,SELENON,MAPK1,AEBP1,FTO,MAP2K2,FLNB,DLK1,RPS28,ATP6V1B2,STXBP1,SMS,CD
KMT2E,PACS1,LZTR1,NSD2,MED13,YY1,SMARCD1,CREBBP,TANC2,RPS26,HS2ST1,PTPN11,EMC
TPI1,CHSY1,PPP2R1A,RPL15,SF3B1,CHD3,POLR2A,RPL18,TUBB4B,UCHL1,RPL11,ZMIZ1,COL11
RCN1,RPS27,CTBP1,MMP2,CRPPA,CDH11,FDFT1,TBCD,GNAI3,RPL27,SNRPN,COL2A1,SKI,ASXL1
D,KDM1A,FLNB,DLK1,SMARCE1,RPS28,ATP6V1B2,LSS,STXBP1,LONP1,RAI1,SYP,BCR,KDM5C,TI
ATP1A2,HLA-B,TTR,L1CAM,HS6ST1,NOVA2,MDH2,RAB27A,EEF1A2,TBCD,SQSTM1,TAF15,SKI,CI
DM3B,CDC42,TRAF7,BCR,DGCR2,KDM5B,BAZ1B,ATRX,NXN,ARID1B,CTNND2,ABCC9,MAPRE2,L
SR2,SHOX,NARS1,ARCN1,RPS27,CTBP1,MMP2,CRPPA,CDH11,FDFT1,TBCD,GNAI3,RPL27,SNRPN
KI,RPS15A,VHL,MAPK1,AEBP1,SON,FLNB,DLK1,TPM3,RPS28,HYOU1,RAI1,ARFGEF2,IGFBP7,BM

PCGF2,AHCY,UBE3A,YY1,CREBBP,PHIP,TANC2,FGFR1,NTRK2,TECR,PINK1,AUTS2,CLIP2,TRAK1,
3,RPS27,RAC1,MMP2,PPP1CB,RPL27,SNRPN,PLOD1,COL2A1,SKI,RPS15A,NOTCH2,MAPK1,AEBP
B,SELENON,PPP2CA,SMARCA4,PRDX1,ODC1,ACADVL,SON,KDM1A,MAP2K2,DLK1,TPM3,SMARC
M1,ACO2,SKI,ASXL1,RAB11B,NOTCH2,VHL,CCND1,SMARCA4,ODC1,FTO,PRKDC,ATXN10,SON,K
2A1,SKI,ASXL1,RPS15A,NOTCH2,MAPK1,SMARCA4,SON,KDM1A,FLNB,DLK1,SMARCE1,RPS28,A
GPC3,MEGF8,RAB7A,CCN2,MMP14,DDX3X,PRKAR1A,GJA1,CTNNB1,COL6A2,HNRNPDL,ARID1A,D
TPS1,PTEN,SMC3,MAPRE2,LOX,GPC4,HSPG2,GTF2I,PACS1,LZTR1,HSD17B4,NSD2,PCGF2,FBLN
D1,SBF1,CYB5R3,PAFAH1B1,HNRNPU,RPS24,COL1A2,HK1,RHOA,PTCD3,NR2F1,RPL26,RPL31,LF

I,SKI,RPS15A,NOTCH2,SELENON,SMARCA4,AEBP1,MAP2K2,FLNB,TPM3,SMARCE1,SDHA,FBLN1
,SMARCE1,RPS28,LONP1,RAI1,FLNC,BCR,EP300,RERE,KDM5B,FLCN,BAZ1B,GPX4,NXN,ARID1B,
,MMP14,RPS7,RPS19,RPL35,PRKAR1A,GJA1,MYO5A,CTNNB1,RPS10,RPL35A,CYFIP2,ARID1A,TS

2,RPS29,CDKN1B,LMNA,SOS1,EXT1,FAM20C,FH,MSH2,PTPN12,SMARCB1,NF2,ELMO2
GTF2I,PACS1,LTBP3,DHCR7,PCGF2,UBE3A,PRKACA,CREBBP,TANC2,CCDC47,FGFR1,SH3PXD2B
M1,ACO2,SKI,ASXL1,RAB11B,NOTCH2,VHL,CCND1,SMARCA4,ODC1,FTO,PRKDC,ATXN10,SON,K
M1,ACO2,SKI,ASXL1,RAB11B,NOTCH2,VHL,CCND1,SMARCA4,ODC1,FTO,PRKDC,ATXN10,SON,K
CN,BAZ1B,ATRX,PI4KA,SMC3,TAOK1,EHMT1,LTBP4,GTF2I,ELP1,PACS1,PHGDH,NSD2,DHCR7,PC
DH18A1,LIMK1,EXT2,DDX6,CRKL,CHST3,NSD1,STAT1,LMNA,MED12,TGFBR1,EXT1,PSMD12,TGF
LTBP3,HSD17B4,DHCR7,MED13,MTOR,CREBBP,PHIP,TANC2,FGFR1,TRIP12,POR,CLIP2,SC5D,E
C9,NDN,HSPG2,ANTXR1,KMT2E,PACS1,LZTR1,NSD2,MED13,YY1,SMARCD1,CREBBP,TANC2,RPS
L35A,CYFIP2,ARID1A,TSR2,HNRNPK,DUX4,RPS27,CTBP1,RAC1,SMCHD1,MMP2,HS6ST1,NOVA2,(

CTNND2,RAB3GAP2,FBXO11,PTEN,MAPRE2,GPC4,PKDCC,KMT2E,EFTUD2,TTC37,PACS1,LZTR1
PS24,COL1A2,HK1,RHOA,NR2F1,CD81,COL5A2,RPL26,RPL31,LRPPRC,CCND2,CDH2,GPC3,MEC

TO,PRKDC,SON,MAP2K2,QARS1,DLK1,FBLN1,STXBP1,LONP1,SARS1,RAB11A,TCTN1,CDC42,CI(

TSR2,HNRNPK,HLA-B,TTR,ARCN1,RPS27,CTBP1,RAC1,SMCHD1,CDH11,FDFT1,TBCD,SQSTM1,RI

,YME1L1,KDM5C,EP300,RERE,KDM5B,ALDH3A2,VARS1,FLCN,BAZ1B,ATRX,ARID1B,FBXO11,PTE
XBP1,LONP1,SARS1,RAB11A,CDC42,YME1L1,ADAR,CIC,TRAF7,KCNQ2,RERE,NDUFV1,KDM5B,C
EEF1A2,TBCD,PPP1CB,LMNB2,SNRPN,SKI,ASXL1,NOTCH2,TUBB2A,PPP2CA,SMARCA4,PRDX1,O
N4,BCOR,MED25,MED12,DGCR8,RUSC2,MAN1B1,CUL4B,PSMD12,SETD1A,NFASC,DYNC1I2,PR

DM5C,TRIO,EP300,RERE,DGCR2,KDM5B,ATRX,ARID1B,NDST1,TIMM50,SLC2A1,NDN,FUS,GRN,T
32,COL3A1,TCF20
C47,FGFR1,NTRK2,DENND5A,CHMP1A,DAG1,MECP2,WDR26,TRRAP,ALDH18A1,IQSEC1,CCDC8
OPA1,FHL1,TMCO1,COL3A1,ITGA7,EMD
ACA,MN1,CCDC47,FGFR1,PTPN11,POR,ELN,OBSL1,MECP2,FAM50A,BANF1,AMPD2,FLII,PDGFR
ARCD1,CCDC47,SH3PXD2B,HS2ST1,AUTS2,SC5D,ELN,OBSL1,IGF1R,MECP2,WDR26,ADNP,TRRA
L1,TUBB2A,PPP2CA,SMARCA4,ODC1,FTO,SON,QARS1,SMARCE1,DHX30,STXBP1,LONP1,USP7,F

P300,NDUFV1,KDM5B,BAZ1B,UBQLN2,ATRX,SMC3,NDST1,SLC2A1,BAP1,FUS,GRN,EHMT1,VAP
BCR,KDM5C,CNTNAP1,GPX4,NXN,ARID1B,PI4KA,SMC3,ABCC9,GPC4,CLCN7,PKDCC,PACS1,PHG
C9,GPC4,CLCN7,PKDCC,NSD2,DHCR7,RBPJ,MTOR,SMARCD1,TRIM8,FGFR1,EIF4A3,RPS26,OB
ID1B,PTEN,SMC3,ECEL1,IL6ST,GPC4,CLCN7,IARS2,HSPG2,PKDCC,GTF2I,NSD2,DHCR7,FBLN5,

A2,DHFR,CLCN4,MAN2B1,NACC1,RHOBTB2,PPP3CA,CNOT3,GATAD2B,SETD1A,SYT1,POGZ,BRA
NB,DLK1,SMARCE1,RPS28,ATP6V1B2,B2M,STXBP1,LONP1,USP7,RAI1,SYP,BCR,KDM5C,TRIO,EX
I,SMARCE1,RPS28,ATP6V1B2,STXBP1,LONP1,USP7,RAI1,SYP,BCR,KDM5C,TRIO,EXTL3,GPX4,N
SOS1,SYT1,CCNK,TMCO1,FGD1,KLHL7,SIN3A
15A,NOTCH2,MAPK1,SMARCA4,SON,KDM1A,FLNB,DLK1,SMARCE1,RPS28,ATP6V1B2,B2M,STXB
,CSNK2B,EXT2,DDX6,SYNGAP1,PPP2R5D,DEAF1,SHMT2,ASH1L,CACNA1H,MED25,MBOAT7,NAC
HA,IDH1,PALLD,TRAF7,EP300,RERE,FLCN,BRD4,PTEN,MAPRE2,GPC4,BAP1,H19,SOX2,TTC37,SR
K1,NOP56,ELN,MECP2,ADNP,LIMK1,FLII,IQSEC1,DDX6,SYNGAP1,DEAF1,CRKL,TSPAN7,NSD1,W
PL35,PRKAR1A,GJA1,CTNNB1,COL6A2,MACF1,RPS10,RPL35A,CYFIP2,ARID1A,DDOST,TSR2,SH
GPX4,ATRX,NXN,CTNND2,SMC3,ECEL1,ABCC9,MAPRE2,GPC4,SLC2A1,HSPG2,PKDCC,LZTR1,PH
AG1,SHMT2,BCOR,RPS29,SETD5,SNRPB,LARGE1,TGFBR1,POMGNT1,TGFB2

NONO,DCHS1,SET,CHSY1,RPL15,POLR2A,SEC61A1,RPL18,RPL11,ZMIZ1,COL11A1,RPL10,RPS1

RPPRC,XPR1,TKT,CDH2,DDX3X,CLTC,PRKAR1A,CTNNB1,CYFIP2,DNAJC5,TARDBP,HNRNPK,HLA
BBP,RPS26,HS2ST1,PTPN11,EMC1,ADNP,BSCL2,EFNB1,VPS51,RAB18,CHST3,NSD1,RPS29,SET

C37,PACS1,LTBP3,LZTR1,NSD2,RBPJ,PCGF2,FBLN5,YY1,CREBBP,FGFR1,SH3PXD2B,PTPN11,C

MAN2B1,CTCF,PPP1R12A,ATF6,CNOT3,MAN1B1,GATAD2B,SYT1,PIGT,POGZ,COL3A1,HARS1,CE
HSPG2,ANTXR1,HSD17B4,YY1,SMARCD1,CREBBP,TANC2,WNK1,SMO,FGFR1,SH3PXD2B,PSAT1,
,TAOK1,EHMT1,SOX2,KMT2E,GTF2I,ATXN3,LTBP3,DHCR7,MED13,PCGF2,UBE3A,CREBBP,TAN

|,NSD1,P3H1,DLG4,MAN2B1,NSMF,THOC2,MED12,SLC5A7,ANOS1,TGFBR1,COL6A3,PIGS,GATA
F7,KCNQ2,RERE,KDM5B,CNTNAP1,VARS1,BAZ1B,ATRX,RAB3GAP2,SMC3,NDST1,TIMM50,ATP6A
Q2,EP300,RERE,STARD7,KDM5B,FLCN,GARS1,GLUD1,CTNND2,PI4KA,NDST1,TIMM50,SLC2A1,G
A1,SKI,ASXL1,RPS15A,NOTCH2,SELENON,MAPK1,AEBP1,FLNB,DLK1,TPM3,RPS28,ATP6V1B2,HY
RPL27,SNRPN,COL2A1,SKI,ASXL1,RPS15A,NOTCH2,SELENON,MAPK1,AEBP1,FLNB,DLK1,TPM3,F
1A,GJA1,CTNNB1,COL6A2,RPS10,RPL35A,TSR2,SHOX,NARS1,ARCN1,RPS27,CTBP1,SMCHD1,MN
,RBMX,SKI,ASXL1,PRPF6,NOTCH2,PPP2CA,MAPK1,SMARCA4,PLEC,ODC1,FTO,KDM1A,MAP2K2,
ATP6V1B2,STXBP1,RAI1,SMS,CDC42,SYP,KDM5C,TRIO,EP300,RERE,KDM5B,FLCN,ATRX,NXN,AR
SXL1,RPS15A,NOTCH2,MAPK1,SMARCA4,FTO,SON,MAP2K2,FLNB,DLK1,SMARCE1,RPS28,ATP6'

A1,SKI,ASXL1,RPS15A,NOTCH2,SELENON,MAPK1,AEBP1,FLNB,DLK1,TPM3,RPS28,ATP6V1B2,HY
K4,NXN,ARID1B,SMC3,ABCC9,GPC4,CLCN7,PKDCC,NSD2,DHCR7,RBPJ,MTOR,SMARCD1,TRIM8,

SXL1,RPS15A,NOTCH2,MAPK1,SMARCA4,FTO,SON,MAP2K2,FLNB,DLK1,SMARCE1,RPS28,ATP6'

2,STXBP1,EP300,EXTL3,GPX4,ATRX,NXN,CTNND2,SMC3,ECEL1,ABCC9,MAPRE2,GPC4,SLC2A1,H
SMARCE1,ATP6V1B2,LARS1,FBLN1,STXBP1,USP7,SOD1,RAI1,TCTN1,SMS,CDC42,SYP,BCR,TRIC

,TMEM43,RPS29,LMNA,DNAJB6,INF2,TRAPPC11,SEPTIN9,CUL7,SPG11,TNPO3,OPA1,FHL1,HARS
OR,PACS2,CREBBP,PHIP,TANC2,HS2ST1,TMEM94,SC5D,IGF1R,MECP2,ADNP,TRRAP,NBAS,PDGFI
RERE,KDM5B,CNTNAP1,VARS1,BAZ1B,ATRX,RAB3GAP2,SMC3,NDST1,TIMM50,ATP6AP2,SLC2A1
DEGS1,GPT2,SHMT2,CLCN4,ACSL4,SETX,NDUFS1,MAN2B1,LARGE1,MED12,DDHD2,RUSC2,DST
DEGS1,GPT2,SHMT2,CLCN4,ACSL4,SETX,NDUFS1,MAN2B1,LARGE1,MED12,DDHD2,RUSC2,DST

,POR,ELN,ACTA2,DDX6,STUB1,CRKL,ACSL4,DLG4,FARSA,CNTN1,MED12,HIRA,RUSC2,ZDHHC9,
I1,ABCA3,PLOD1,VPS35,SKI,ASXL1,SELENON,OPTN,CCND1,MAPK1,PLEC,ATXN10,SON,MAP2K2,
,CCDC88A,CSNK2B,EXT2,RAB18,SLC16A2,PPP2R5D,DEAF1,WAC,ZSWIM6,SHMT2,SETD5,MED12
1,RAB18,DEAF1,WAC,NDUFB8,NUS1,SETD5,NDUFS1,MAN2B1,CTCF,RUSC2,COX5A,TALDO1,SLC
CR,PTPN11,DAG1,KLF13,ATP13A2,MECP2,GPAA1,BSCL2,NTN1,ZNF462,IQSEC1,CSNK2B,EXT2,D
27A1,KIF5A,BCR,KDM5C,EP300,EXTL3,RERE,BAZ1B,NXN,ARID1B,MBTPS1,PTEN,SMC3,ECEL1,II
EP300,RERE,NDUFV1,CNTNAP1,ALDH3A2,FLCN,RAB3GAP2,ZFR,SLC2A1,FUS,SLC6A8,PKDCC,IRE
GAP1,CRKL,CHST3,LMNA,COL7A1,MED12,HIRA,CTCF,SEPTIN9,FAM20C,PSMD12,POGZ,COL3A1

AL1,SMO,PTPN23,TECR,IQSEC1,CSNK2B,SYNGAP1,DEAF1,EXOC7,NSD1,CLCN4,ASH1L,ARHGEF
-B,TTR,SMCHD1,L1CAM,CRPPA,HS6ST1,MDH2,EEF1A2,TBCD,SQSTM1,PABPN1,TAF15,PLOD1,SK
N,GLUD1,CTNND2,PI4KA,NDST1,TIMM50,SLC2A1,GRN,IARS2,ANTXR1,AKT3,IREB2,PHGDH,NSD2
,MAPRE2,ATP6AP2,GPC4,EHMT1,SOX2,TTC37,NSD2,OGT,DHCR7,MTOR,CREBBP,FGFR1,RPS26

3,NSD1,WAC,ZSWIM6,MAN2B1,MED12,SKIV2L,CNOT3,PSMD12,COG4,HMGA2,PRMT7,DPF2,CAS

CD,GNAI3,RPL27,SNRPN,COL2A1,SKI,ASXL1,RPS15A,NOTCH2,SELENON,MAPK1,AEBP1,FLNB,DL

:NOT3,GATAD2B,PSMD12,SETD1A,CENPF,COL3A1,MAPK8IP3,HMGA2,PRMT7,MAFB,SIN3A,TCF2
'OGZ,DNM1,GNAO1,CASK
;,MAN2B1,CHD8,MED12,ZDHHC9,TGFBR1,COL6A3,PSMD12,TGFB2,NNT,CNOT1,TCF20
;,PMPCA,NID1,CNOT3,POMGNT1,SMARCC2,POGZ,SMARCB1,SLC18A3,KLHL7,DPF2
A1,KLC2,CAPN1,DDX6,SLC25A1,GBA2,NDRG1,WDR11,NSD1,SETX,MED25,WARS1,KARS1,NSMF,

1,RPL18,RPL11,AGRN,ZMIZ1,COL11A1,RPL10,RPS17,VCL,AP3D1,ARF1,CYB5R3,PAFAH1B1,MAT
TNND2,PI4KA,NDST1,TIMM50,SLC2A1,GRN,IARS2,ANTXR1,AKT3,IREB2,PHGDH,NSD2,RBPJ,UBE
)L2A1,ASXL1,RPS15A,NOTCH2,MAPK1,SMARCA4,FTO,SON,MAP2K2,FLNB,DLK1,SMARCE1,RPS2

2,BCOR,RPS29,SETD5,MED12,G6PC3,PSMD12,MAFB,CNOT1,TCF20
MC3,ECEL1,ADH5,GPC4,PKDCC,KMT2E,EFTUD2,GTF2I,SEMA3C,PACS1,PHGDH,NSD2,DHCR7,MI
KDM5B,FLCN,ATRX,ARID1B,RAB3GAP2,SMC3,NDN,FUS,H19,TAOK1,HSPG2,SLC6A8,EFTUD2,ELF
C9,TGFBR1,CUL7,EXT1,CUL4B,TGFB2,SMARCC2,COL3A1,SMARCB1,FGD1,DPF2
2,TRAK1,ELN,LIMK1,EXT2,DDX6,SYNGAP1,NUS1,SOX9,MED25,SETD5,THOC2,MED12,SOS1,CTCI

3BP,TUBB4A,EIF4G1,NTRK2,PINK1,TRAK1,TIA1,ATP13A2,MFN2,MECP2,PARK7,BSCL2,ALDH18A

FTUD2,GTF2I,PACS1,LZTR1,PHGDH,DHCR7,PACS2,UBE3A,CREBBP,EIF4A3,SH3PXD2B,PTPN11,A

ROH2A1,OBSL1,ALDH18A1,EXT2,ASAH1,CHST3,TMEM43,BCOR,SETD5,SNRPB,LARGE1,LMNA,C

1,RPS28,SDHA,STXBP1,LONP1,RAI1,KDM3B,BMPR2,BCR,EP300,RERE,DGCR2,NDUFV1,BAZ1B,A
PKDCC,AKT3,GTF2I,PACS1,LZTR1,DHCR7,PACS2,UBE3A,SMARCD1,CREBBP,SMO,EIF4A3,SH3PX
RAP,IQSEC1,CCDC88A,CSNK2B,RAB18,SYNGAP1,DEAF1,SHMT2,ASH1L,MED25,MBOAT7,LARGI

FR1,EIF4A3,RPS26,BGN,PSAT1,DAG1,NECTIN1,SHMT2,BCOR,RPS29,SETD5,SNRPB,LARGE1,DG

EXT2,NSD1,WAC,CLCN4,MAN2B1,MED12,ARSB,PIGS,TARS1,CUL4B,FHL1,SMARCC2,SMARCB1,D
351,PPP2R5D,CRKL,TSPAN7,WAC,SHMT2,CLCN4,ACSL4,SETD5,RHOBTB2,MED12,CTCF,QRICH1
A,TCTN1,KDM3B,SMS,SYP,TRIO,EXTL3,RERE,PBX1,CSNK2A1,ATRX,NXN,ARID1B,RAB3GAP2,PTE

,PBX1,BAZ1B,ATRX,SMC3,HNRNPA2B1,TIMM50,SLC2A1,BAP1,FUS,SPG7,EHMT1,LTBP4,GTF2I,A

LCN4,BCOR,SETX,RARS1,NDUFS1,MAN2B1,DDHD2,DSTYK,NIPA1,SLC1A4,SPG11,GATAD2B,OPA1
CDC47,SMO,FGFR1,EIF4A3,COL12A1,CLIP2,ELN,OBSL1,ALDH18A1,LIMK1,COX8A,SLC25A1,PPP

3,SZT2,PIGT,NBEA,DNM1,GNAO1,SETD1B,JRK,CSTB,CASK
B1,SLC18A3,KLHL7,DPF2
DH,DHCR7,PCGF2,GOT2,SMARCD1,CREBBP,SMO,FGFR1,DENND5A,MDH1,PSAT1,PUM1,SEMA6
EN,SMC3,GPC4,NDN,H19,PKDCC,KMT2E,GTF2I,PACS1,LZTR1,PHGDH,MED13,PCGF2,PACS2,UE
3,SMARCE1,FBLN1,B2M,LONP1,USP7,SEC23A,ARNT2,KDM3B,EP300,EXTL3,RERE,BAZ1B,GPX4,

MAPT,NONO,DCHS1,SET,TPI1,PPP2R1A,RPL15,PKD1,SF3B1,POLR2A,RPL18,UCHL1,RPL11,AGRN
2,BCOR,RPS29,SETD5,MED12,G6PC3,PSMD12,MAFB,CNOT1,TCF20
AI1,SEC23A,TCTN1,KDM3B,SMS,SYP,TRIO,EXTL3,RERE,PBX1,CSNK2A1,ATRX,NXN,ARID1B,RAB3

,SMC3,MAPRE2,ATP6AP2,GPC4,EHMT1,SOX2,TTC37,NSD2,OGT,DHCR7,MTOR,CREBBP,FGFR1,
,SMC3,MAPRE2,ATP6AP2,GPC4,EHMT1,SOX2,TTC37,NSD2,OGT,DHCR7,MTOR,CREBBP,FGFR1,
T1,SEMA6B,TRAK1,NOP56,SC5D,ATP13A2,MECP2,NDUFA4,BSCL2,TNRC6A,CCDC88A,CSNK2B,A
AN2B1,LMNA,TGFBR1,MAN1B1,EXT1,CUL4B,TGFB2,COL3A1,FGD1,GSN
FTUD2,GTF2I,PACS1,LZTR1,PHGDH,DHCR7,PACS2,UBE3A,CREBBP,EIF4A3,SH3PXD2B,BGN,PTP

,HSPG2,ANTXR1,GTF2I,ELP1,PACS1,PACS2,UBE3A,TEAD1,TUB,PTPN11,CHMP1A,TMEM94,COL1
TO,HSPB1,DLK1,TPM3,ATP6V1B2,SDHA,PFKM,PRPS1,SOD1,RAI1,FLNC,CCT5,DCTN1,KIF5A,KD

27,COL2A1,SKI,ASXL1,RPS15A,NOTCH2,MAPK1,SMARCA4,AEBP1,FLNB,DLK1,SMARCE1,RPS28,
7,LZTR1,PCGF2,FBLN5,CREBBP,FGFR1,BGN,PTPN11,CLIP2,ELN,NOTCH1,ACTA2,ALDH18A1,LI
LA-B,ARCN1,RPS27,CTBP1,RAC1,SMCHD1,CRPPA,HS6ST1,CDH11,FDFT1,PPP1CB,LMNB2,RPL27
1,CHD4,COL5A2,RPL26,RPL31,XPR1,TKT,CDH2,GPC3,MEGF8,MMP14,RPS7,RPS19,RPL35,PRK

ARB2,RPS24,COL1A2,HK1,PTCD3,NR2F1,CHD4,RPL26,RPL31,XPR1,TKT,CLTC,RPS7,RPS19,RPL
SC2,PIGS,FAM20C,PSMD12,SZT2,TMCO1,TCF20

A,CSNK2B,ASAH1,ALDH7A1,SYNGAP1,DEAF1,TSPAN7,GPT2,NSD1,WAC,CLCN4,ASH1L,ACSL4,CA
WDR26,COX8A,IQSEC1,CSNK2B,ASAH1,ALDH7A1,SYNGAP1,DEAF1,TSPAN7,NSD1,WAC,DHFR,CL
PHGDH,DHCR7,PCGF2,AHCY,GOT2,SMARCD1,FGFR1,DENND5A,MDH1,PSAT1,CHMP1A,SEMA6B,
N1,B2M,LONP1,USP7,SEC23A,ARNT2,KDM3B,EP300,EXTL3,RERE,GPX4,NXN,ARID1B,CRTAP,HD
CLCN4,ACSL4,RPS29,SRP54,UQCRFS1,THOC2,CNTN1,MED12,SKIV2L,CTCF,TALDO1,CUL7,HAD

GJA1,COL6A2,ARL6IP1,ARID1A,SHOX,HLA-B,CTBP1,MMP2,L1CAM,HS6ST1,FDFT1,TBCD,PPP1CB

GE1,SLC5A6,PMPCA,NID1,CNOT3,POMGNT1,SMARCC2,POGZ,SMARCB1,SLC18A3,KLHL7,DPF2
6,PTPN11,TMEM94,CLIP2,ELN,ACTA2,TRRAP,LIMK1,EXT2,NSD1,WAC,ACSL4,BCOR,SNRPB,LMN
BAZ1B,ATRX,ARID1B,SMC3,MAPRE2,ATP6AP2,GPC4,EHMT1,SOX2,GTF2I,TTC37,NSD2,OGT,DHC
K1,CLIP2,ELN,IGF1R,MECP2,PARK7,ADNP,LIMK1,FLII,PRPH,PDGFRB,DDX6,STUB1,DEAF1,WDR1

1,ATP6V1B2,STXBP1,LONP1,PRPS1,RAI1,SEC23A,TCTN1,KDM3B,SMS,CDC42,SYP,BCR,TRIO,EX
LA-B,ARCN1,RPS27,CTBP1,RAC1,SMCHD1,CRPPA,HS6ST1,CDH11,FDFT1,PPP1CB,LMNB2,RPL27
H2A1,KLF13,ELN,OBSL1,TRRAP,LIMK1,ZNF462,EFNB1,FLII,EXT2,RAB18,DEAF1,CRKL,CHST3,G

B,GNAI3,SNRPN,PLOD1,COL2A1,SKI,KDM6B,NOTCH2,MAPK1,SMARCA4,PLEC,AEBP1,SON,KDM
31,SCARB2,HK1,PTCD3,RPS20,COL5A2,WDR1,PFN1,CLEC7A,PRKCSH,GPC3,CCN2,CLTC,MIF,PF

,CDC42,BCR,TRIO,EP300,RERE,CSNK2A1,ARID1B,SMC3,ECEL1,ABCC9,EHMT1,IARS2,HSPG2,A
M1,CCND2,CDH2,GPC3,MEGF8,MMP14,DDX3X,RPS7,RPS19,RPL35,MIF,PRKAR1A,GJA1,RPS10,

CLIP2,ELN,NOTCH1,ADNP,TRRAP,BSCL2,ALDH18A1,LIMK1,EFNB1,NECTIN1,NSD1,BCOR,MAN2
,CUL7,POMGNT1,FHL1,SYT1,ITGA7,GLB1,EMD
CYB5R3,PAFAH1B1,MAT2A,HNRNPU,SCARB2,RPS24,COL1A2,HK1,RHOA,CD81,COL5A2,WDR1,A
,PTEN,SMC3,IL6ST,H19,CLCN7,HSPG2,ANTXR1,HSD17B4,MTOR,YY1,SMARCD1,CREBBP,TANC2,

PL18,UCHL1,RPL11,AGRN,ZMIZ1,COL11A1,RPL10,MARCHF6,RPS17,VCL,AP3D1,SBF1,ARF1,CYE
3F1,ARF1,CYB5R3,PAFAH1B1,ITM2B,HNRNPU,LDHA,CTSD,COL1A2,HK1,NR2F1,CHD4,ATP1A1,RI
1,CHD4,RPS20,COL5A2,PFN1,LRPPRC,SPTBN2,TKT,CDH2,GPC3,DDX3X,CLTC,GJA1,MYO5A,CTN
A2,RPL26,RPL31,PRKCSH,GPC3,MEGF8,MMP14,DDX3X,CLTC,RPS7,RPS19,RPL35,PRKAR1A,GJ
CP,UBB,AFF4,YWHAG,COL4A2,MAPT,FN1,NONO,DCHS1,SET,TPI1,CHSY1,PPP2R1A,RPL15,CHD3
F2,ATP5F1A,VCP,UBB,AFF4,YWHAG,COL4A2,MAPT,FN1,NONO,DCHS1,SET,TPI1,CHSY1,PPP2R1
,ARNT2,ARFGEF2,CDC42,ADAR,CIC,KCNQ2,WDR45B,EXTL3,RERE,KDM5B,CNTNAP1,VARS1,FLC
3,POLR2A,AP2M1,RPL18,RPL11,AGRN,ZMIZ1,COL11A1,RPL10,RPS17,AP3D1,SBF1,ARF1,CYB5R
PP2R1A,RPL15,PKD1,CHD3,POLR2A,AP2M1,RPL18,UCHL1,RPL11,AGRN,ZMIZ1,COL11A1,RPL10
,SMCHD1,MMP2,CRPPA,HS6ST1,NOVA2,CDH11,EEF1A2,TBCD,PPP1CB,RPL27,SNRPN,COL2A1,S
B,HNRNPU,CTSD,RPS24,HK1,PTCD3,NR2F1,RPL26,RPL31,XPR1,CCND2,CDH2,GPC3,DDX3X,CLT
B,HNRNPU,CTSD,RPS24,HK1,PTCD3,NR2F1,RPL26,RPL31,XPR1,CCND2,CDH2,GPC3,DDX3X,CLT
,YWHAG,COL4A2,MAPT,FN1,NONO,DCHS1,SET,TPI1,CHSY1,PPP2R1A,RPL15,HLA-C,CHD3,POLR
RPL26,RPL31,XPR1,CDH2,GPC3,DDX3X,CLTC,RPS7,RPS19,RPL35,GJA1,MYO5A,CTNNB1,MACF
1,RPS15A,NOTCH2,PPP2CA,MAPK1,ODC1,KDM1A,MAP2K2,FLNB,TPM3,RPS28,ATP6V1B2,PRPS1
,RPL15,CHD3,POLR2A,AP2M1,RPL18,RPL11,AGRN,ZMIZ1,RPL10,MARCHF6,RPS17,AP3D1,SBF1
AP3D1,SBF1,ARF1,CYB5R3,PAFAH1B1,MAT2A,HNRNPU,CTSD,RPS24,COL1A2,HK1,RHOA,CHD4,
1,DUX4,HLA-B,CTBP1,RAC1,SMCHD1,MMP2,L1CAM,PPIB,CRPPA,CDH11,EEF1A2,TBCD,LMNB2,
,SEC61A1,RPL18,TUBB4B,UCHL1,RPL11,ZMIZ1,COL11A1,MARCHF6,RPS17,VCL,ARF1,PAFAH1E
A,MAP2K2,FLNB,DLK1,TPM3,SMARCE1,ATP6V1B2,STXBP1,MESD,RAI1,SEC23A,KDM3B,SMS,CD
PS10,RPL35A,CYFIP2,ARID1A,TSR2,HNRNPK,NARS1,ARCN1,RPS27,CTBP1,RAC1,L1CAM,CRPPA,
1,RPS10,RPL35A,CYFIP2,ARID1A,TSR2,HNRNPK,NARS1,ARCN1,RPS27,CTBP1,RAC1,L1CAM,GLU
RPL18,UCHL1,RPL11,ZMIZ1,COL11A1,RPL10,RPS17,AP3D1,SBF1,ARF1,CYB5R3,PAFAH1B1,ITM
,AGRN,ZMIZ1,RPL10,MARCHF6,RPS17,SBF1,ARF1,CYB5R3,PAFAH1B1,ITM2B,HNRNPU,CTSD,S
AH1B1,MAT2A,SCARB2,RPS24,COL1A2,HK1,RHOA,NR2F1,CHD4,ATP1A1,COL5A2,ATP2A2,RPL2
RPL15,HLA-C,CHD3,POLR2A,AP2M1,SEC61A1,RPL18,UCHL1,RPL11,AGRN,ZMIZ1,COL11A1,RPL
I,ASXL1,TUBB2A,PPP2CA,SMARCA4,ODC1,PRKDC,SON,KDM1A,MAP2K2,QARS1,SMARCE1,LSS,S
5,PKD1,CHD3,POLR2A,AP2M1,RPL18,UCHL1,RPL11,ZMIZ1,COL11A1,RPL10,RPS17,AP3D1,SBF
ARF1,CYB5R3,PAFAH1B1,HNRNPU,RPS24,COL1A2,HK1,NR2F1,ATP1A1,COL5A2,RPL26,RPL31,L
B,RAB11B,RPS15A,NOTCH2,MAPK1,PLEC,PRDX1,ODC1,AEBP1,SON,MAP2K2,DLK1,RPS28,ATP6
RAB3GAP2,CRTAP,FBXO11,HDAC6,PTEN,SMC3,PTPRF,ABCC9,MAPRE2,IARS2,HSPG2,ANTXR1,P
A,CDH11,EEF1A2,TBCD,LMNB2,SNRPN,PLOD1,COL2A1,SKI,NOTCH2,SELENON,MAPK1,SMARC
RNPK,NARS1,ATP1A2,HLA-B,TTR,CTBP1,RAC1,L1CAM,HS6ST1,NOVA2,MDH2,EEF1A2,TBCD,PPP1
TSR2,SHOX,HNRNPK,RPS27,CTBP1,RAC1,SMCHD1,MMP2,GLUL,HS6ST1,NOVA2,CDH11,EEF1A2
DNP,TRRAP,ALDH18A1,LIMK1,DDX6,CRKL,CHST3,NSD1,P3H1,ZEB1,GDF1,CLCN4,MED25,MAN2
DC,SON,KDM1A,MAP2K2,QARS1,SMARCE1,LSS,STXBP1,LONP1,PRPS1,USP7,SARS1,VANGL2,RA
PL31,CCND2,GPC3,MEGF8,RAB7A,CCN2,MMP14,RPS7,RPS19,RPL35,PRKAR1A,GJA1,COL6A2,R
SQSTM1,VAMP2,ACO2,SKI,ASXL1,TUBB2A,PPP2CA,SMARCA4,ODC1,PRKDC,SON,KDM1A,MAP2K
1A,GJA1,COL6A2,RPS10,RPL35A,HNRNPDL,ARID1A,TSR2,SHOX,HNRNPK,NARS1,RPS27,CTBP1,
I1,CHSY1,PPP2R1A,RPL15,SF3B1,CHD3,POLR2A,RPL18,TUBB4B,UCHL1,RPL11,AGRN,ZMIZ1,C
3,SQSTM1,ABCA3,RPL27,PLOD1,COL2A1,SKI,ASXL1,KDM6B,RPS15A,NOTCH2,SELENON,MAPK1,
LRPPRC,XPR1,CCND2,CDH2,GPC3,DDX3X,CLTC,GJA1,MYO5A,CTNNB1,MACF1,CYFIP2,DNAJC5,

CAM,CRPPA,CDH11,EEF1A2,TBCD,LMNB2,RPL27,SNRPN,ACO2,SKI,ASXL1,RAB11B,ABCA2,RPS1

Y1,PPP2R1A,RPL15,HLA-C,CHD3,POLR2A,AP2M1,SEC61A1,RPL18,UCHL1,RPL11,AGRN,ZMIZ1,C
19,RPL35,PRKAR1A,GJA1,COL6A2,RPS10,RPL35A,HNRNPDL,ARID1A,TSR2,SHOX,HNRNPK,NARS
C1,L1CAM,CRPPA,CDH11,EEF1A2,TBCD,LMNB2,RPL27,SNRPN,ACO2,SKI,ASXL1,RAB11B,ABCA2
YO5A,CTNNB1,COL6A2,MACF1,CYFIP2,SLC25A3,ARID1A,DDOST,HNRNPK,NARS1,CTBP1,RAC1,
MMP14,CLTC,RPS7,RPS19,RPL35,PRKAR1A,GJA1,RPS10,RPL35A,CYFIP2,ARID1A,TSR2,HNRNPK,
RAB7A,MMP14,RPS7,RPS19,RPL35,PRKAR1A,GJA1,COL6A2,RPS10,RPL35A,ARID1A,TSR2,SHOX,
RAB7A,MMP14,RPS7,RPS19,RPL35,PRKAR1A,GJA1,COL6A2,RPS10,RPL35A,ARID1A,TSR2,SHOX,

H2,PPP2CA,MAPK1,SMARCA4,SON,MAP2K2,QARS1,DLK1,SMARCE1,DHX30,RPS28,LONP1,TRAF
PC3,MEGF8,RAB7A,CCN2,CLTC,RPS7,RPS19,RPL35,PRKAR1A,GJA1,COL6A2,RPS10,RPL35A,CY
P3,DHCR7,MED13,PCGF2,MTOR,PACS2,UBE3A,SMARCD1,CREBBP,TANC2,EIF4A3,SH3PXD2B,H

CHS1,SET,CHSY1,PPP2R1A,RPL15,CHD3,POLR2A,AP2M1,RPL18,UCHL1,RPL11,AGRN,ZMIZ1,CC
5A,CYFIP2,ARID1A,TSR2,HNRNPK,NARS1,DUX4,HLA-B,RPS27,CTBP1,RAC1,GPC1,SMCHD1,NOV
GTF2I,LZTR1,MED13,FBLN5,UBE3A,YY1,SMARCD1,CREBBP,PHIP,TANC2,BGN,PTPN11,COL12A1
ARCE1,DHX30,SDHA,LSS,FBLN1,STXBP1,PRPS1,MESD,USP7,SARS1,SOD1,RAI1,SEC23A,KDM3E
A,ARID1A,TSR2,SHOX,HNRNPK,NARS1,ARCN1,RPS27,CTBP1,RAC1,MMP2,L1CAM,CRPPA,CDH11
SMC3,GPC4,HSPG2,PKDCC,LTBP4,EFTUD2,GTF2I,TTC37,PACS1,PHGDH,NSD2,DHCR7,PCGF2,Y
1,MAT2A,HNRNPU,SCARB2,RPS24,COL1A2,HK1,RHOA,COL5A2,WDR1,ATP2A2,RPL26,PFN1,CLE
,BAZ1B,ATRX,NXN,ARID1B,CTNND2,SMC3,ABCC9,MAPRE2,LOX,GPC4,H19,EHMT1,ERGIC1,HSP
SMC3,GPC4,HSPG2,PKDCC,LTBP4,EFTUD2,GTF2I,TTC37,PACS1,PHGDH,NSD2,DHCR7,PCGF2,Y
CHS1,SET,CHSY1,PPP2R1A,RPL15,CHD3,POLR2A,AP2M1,RPL18,UCHL1,RPL11,AGRN,ZMIZ1,CC
LTC,RPS7,RPS19,RPL35,PRKAR1A,GJA1,RPS10,RPL35A,CYFIP2,ARID1A,TSR2,HNRNPK,NARS1,

A,MAPK1,SMARCA4,PRDX1,ODC1,PRKDC,SON,KDM1A,MAP2K2,FLNB,DLK1,TPM3,SMARCE1,D
ARS1,SMARCE1,LSS,STXBP1,LONP1,PRPS1,USP7,SARS1,VANGL2,RAI1,RAB11A,TCTN1,ARNT2,

OL27A1,RAI1,SEC23A,CDC42,STAT3,EP300,RERE,PBX1,BAZ1B,CSNK2A1,ATRX,NXN,ARID1B,CT
CRPPA,NOVA2,CDH11,EEF1A2,PPP1CB,RPL27,SNRPN,COL2A1,SKI,ASXL1,KDM6B,RPS15A,NOTC
MCHD1,GLUL,CRPPA,CDH11,EEF1A2,FDFT1,TBCD,PPP1CB,GNAI3,RPL27,PLOD1,COL2A1,RBMX,
,KDM6B,CNTNAP2,RAB11B,SELENON,TUBB2A,PPP2CA,MAPK1,SMARCA4,PRDX1,ODC1,ACADVL,
,RPS7,RPS19,RPL35,PRKAR1A,GJA1,RPS10,RPL35A,CYFIP2,ARID1A,TSR2,HNRNPK,NARS1,DUX
I2,NBEA,BRAT1,MAPK8IP3,PRMT7,DPF2,TCF20
ARS1,SMARCE1,LSS,STXBP1,LONP1,PRPS1,USP7,SARS1,VANGL2,RAI1,RAB11A,TCTN1,ARNT2,
1A1,RPL10,MARCHF6,RPS17,AP3D1,SBF1,CYB5R3,PAFAH1B1,ITM2B,PXDN,HNRNPU,CTSD,RPS
ENON,PPP2CA,MAPK1,ODC1,AEBP1,SON,MAP2K2,FLNB,DLK1,TPM3,DHX30,RPS28,ATP6V1B2,M
A,STXBP1,LONP1,RAI1,KDM3B,CDC42,BCR,TRIO,EP300,RERE,DGCR2,NDUFV1,KDM5B,FLCN,B
S27,CTBP1,SMCHD1,CRPPA,HS6ST1,CDH11,PPP1CB,GNAI3,RPL27,SNRPN,PLOD1,COL2A1,SKI,
OST,TSR2,RPS27,CTBP1,L1CAM,DCN,CDH11,MDH2,SQSTM1,TGFBI,RPL27,SNRPN,ACO2,PLOD
,LDLR,KMT2E,GTF2I,LZTR1,DHCR7,MED13,FBLN5,CREBBP,TANC2,FGFR1,RPS26,BGN,PTPN11
MP2,L1CAM,CRPPA,LMNB2,ABCA3,RPL27,SNRPN,PLOD1,COL2A1,SKI,ASXL1,KDM6B,RAB11B,R
S6ST1,CTSB,FDFT1,LMNB2,SNRPN,PDXK,PLOD1,COL2A1,SKI,ASXL1,RAB11B,NOTCH2,SELENO
VPS35,COL2A1,CHD2,SKI,RAB11B,ABCA2,MARS1,SELENON,PLEC,ATXN10,HSPB1,TPM3,SMARC

SCARB2,RPS24,COL1A2,HK1,RHOA,COL5A2,WDR1,ATP2A2,RPL26,CLEC7A,RPL31,CCND2,CDH2
SCARB2,RPS24,COL1A2,HK1,RHOA,COL5A2,WDR1,ATP2A2,RPL26,CLEC7A,RPL31,CCND2,CDH2
I4,PRKAR1A,GJA1,COL6A2,HNRNPDL,ARL6IP1,ARID1A,SHOX,HNRNPK,NARS1,ARCN1,CTBP1,MM
LOD1,SKI,ASXL1,KDM6B,NOTCH2,PPP2CA,MAPK1,PLEC,FTO,SON,KDM1A,MAP2K2,FLNB,DLK1,
DFT1,RPL27,SNRPN,PLOD1,COL2A1,SKI,ASXL1,KDM6B,PRPF6,RPS15A,NOTCH2,MAPK1,SMARC
N2,TKT,CDH2,RAB7A,DDX3X,CLTC,GJA1,MYO5A,COL6A2,MACF1,CYFIP2,DNAJC5,HNRNPDL,TAF
MFN2,TRIM2,MECP2,WDR26,TRRAP,AMPD2,KLC2,IQSEC1,VPS51,CSNK2B,DDX6,SLC25A1,PPP2
11B,MARS1,OPTN,VHL,MAPK1,PRDX1,ATXN10,HSPB1,KDM1A,SMARCE1,XPNPEP3,DHX30,SDHA,

EC1,CSNK2B,SYNGAP1,PPP2R5D,DEAF1,NSD1,WAC,SHMT2,CLCN4,ASH1L,BCOR,MED25,THOC
ESD,USP7,SARS1,SOD1,RAI1,SEC23A,KDM3B,SMS,CDC42,YME1L1,ADAR,SYP,TRAF7,WDR45B,K

LR2A,AP2M1,RPL18,UCHL1,RPL11,AGRN,ZMIZ1,COL11A1,RPL10,RPS17,AP3D1,CYB5R3,PAFAH
STXBP1,PRPS1,SARS1,SOD1,RAB11A,ARNT2,CCT5,ADAR,CIC,TRAF7,DCTN1,KIF5A,KCNQ2,WDR
00,EXTL3,RERE,VARS1,GPX4,ATRX,NXN,ARID1B,RAB3GAP2,PTEN,SMC3,ABCC9,MAPRE2,PREPL
I1,FLNC,ADAR,TRAF7,DCTN1,KIF5A,KDM5C,EXTL3,ATRX,GARS1,PI4KA,MCM3AP,BAP1,SPG7,YA
TCH2,SELENON,MAPK1,SMARCA4,PRDX1,AEBP1,FTO,SON,MAP2K2,FLNB,DLK1,TPM3,SMARCE
,TBCD,PPP1CB,LMNB2,SNRPN,PLOD1,SKI,ASXL1,KDM6B,NOTCH2,PPP2CA,MAPK1,KDELR2,PLE
1,CTNNB1,COL6A2,MACF1,RPS10,RPL35A,CYFIP2,ARID1A,DDOST,TSR2,SHOX,HNRNPK,ARCN1,
RN,TAOK1,EHMT1,SOX2,SLC6A8,IREB2,KMT2E,GTF2I,PACS1,DHCR7,MED13,PCGF2,MTOR,PA

ESD,SARS1,RAI1,KDM3B,STAT3,KCNQ2,TRIO,EP300,EXTL3,KDM5B,PBX1,GARS1,FBXO11,PTEN
TPM3,DHX30,RPS28,STXBP1,PRPS1,MESD,USP7,VANGL2,SOD1,RAI1,TCTN1,SMS,CDC42,ADAR,
RKAR1A,GJA1,MYO5A,CTNNB1,MACF1,RPL35A,CYFIP2,ARID1A,DDOST,HNRNPK,NARS1,CTBP1,
,L1CAM,DCN,CDH11,MDH2,TGFBI,RPL27,SNRPN,ACO2,PLOD1,SKI,ASXL1,KDM6B,RAB11B,RPS

P1CB,GNAI3,LMNB2,PLOD1,COL2A1,SKI,ASXL1,KDM6B,NOTCH2,SELENON,MAPK1,SMARCA4,KI
B,DLK1,TPM3,DHX30,RPS28,STXBP1,PRPS1,MESD,USP7,VANGL2,SOD1,RAI1,TCTN1,SMS,CDC42
,GPC3,MEGF8,MMP14,DDX3X,CLTC,RPS19,PRKAR1A,GJA1,MYO5A,CTNNB1,CYFIP2,ARID1A,HN
NNB1,COL6A2,MACF1,CYFIP2,DNAJC5,TARDBP,ARL6IP1,DDOST,NARS1,ATP1A2,HLA-B,TTR,ARC
N,IGF1R,MECP2,ALDH18A1,LIMK1,EFNB1,SLC16A2,NSD1,RBM10,TMEM43,BCOR,SETD5,MAN2B

SMARCB1,DPF2,GLB1
,ADAR,CIC,DCTN1,KIF5A,KCNQ2,WDR45B,KDM5C,EXTL3,RERE,NDUFV1,KDM5B,CNTNAP1,ALDH
RERE,ATRX,NXN,ARID1B,CTNND2,RAB3GAP2,IARS1,GPC4,H19,EHMT1,IARS2,HSPG2,ANTXR1,SLC
I,RAI1,RAB11A,SEC23A,ARNT2,ARFGEF2,ADAR,CIC,DCTN1,KIF5A,KCNQ2,EP300,RERE,KDM5B,C
UD2,GTF2I,TTC37,PACS1,PHGDH,NSD2,DHCR7,CREBBP,NDUFA13,TRIM8,FGFR1,RPS26,PTPN1
G2,ANTXR1,LTBP4,GTF2I,TTC37,PACS1,PHGDH,DHCR7,FBLN5,MTOR,CREBBP,TRIM8,FGFR1,SH
1,PTEN,NDN,HSPG2,KMT2E,GTF2I,NSD2,DHCR7,MED13,SMARCD1,TANC2,TUBB4A,SH3PXD2B,H
G2,ANTXR1,LTBP4,GTF2I,TTC37,PACS1,PHGDH,FBLN5,MTOR,CREBBP,TRIM8,FGFR1,SH3PXD2B
2I,TTC37,PACS1,PHGDH,NSD2,DHCR7,CREBBP,NDUFA13,TRIM8,FGFR1,RPS26,PTPN11,ABL1,T
2,YME1L1,ADAR,CIC,TRAF7,DCTN1,KIF5A,KCNQ2,RERE,NDUFV1,KDM5B,CNTNAP1,VARS1,BAZ1
LMNB2,RPL27,SNRPN,COL2A1,SKI,ASXL1,RPS15A,NOTCH2,SELENON,MAPK1,SMARCA4,AEBP1,
1,RPS28,SDHA,STXBP1,LONP1,TCTN1,SMS,CDC42,ADAR,TRAF7,KIF5A,STAT3,TRIO,EP300,RERE
HIP,TANC2,TRIM8,FGFR1,NTRK2,TRIP12,TECR,AUTS2,CLIP2,TRAK1,NOP56,ELN,MECP2,ADNP,B

ECP2,BSCL2,LIMK1,FLII,EXT2,SLC16A2,DEAF1,CRKL,TSPAN7,NDRG1,CHST3,WDR11,NSD1,WAC
,YY1,CREBBP,FGFR1,DENND5A,HS2ST1,TMEM94,SC5D,IGF1R,MECP2,WDR26,ADNP,TRRAP,ZNI
DM5C,TRIO,EP300,RERE,DGCR2,KDM5B,FLCN,BAZ1B,ATRX,NXN,ARID1B,CTNND2,RAB3GAP2,F

C42,TRAF7,BCR,EP300,RERE,BAZ1B,GPX4,ATRX,NXN,CTNND2,RAB3GAP2,PTEN,SMC3,ECEL1,A
1,ADNP,BSCL2,EFNB1,VPS51,DDX6,RAB18,CHST3,NSD1,RPS29,SETD5,MAN2B1,LMNA,MED12,S
A1,RPL10,RPS17,AP3D1,CYB5R3,PAFAH1B1,ITM2B,MAT2A,PXDN,HNRNPU,CTNNA1,CTSD,RPS2
1,KDM6B,RPS15A,NOTCH2,SELENON,MAPK1,ODC1,AEBP1,FTO,FLNB,DLK1,TPM3,RPS28,ATP6V
RIO,EXTL3,RERE,CSNK2A1,GPX4,ATRX,NXN,ARID1B,RAB3GAP2,MBTPS1,PTEN,SMC3,ABCC9,GP
NTNAP2,RAB11B,MARS1,OPTN,ODC1,FTO,ATXN10,SON,KDM1A,SMARCE1,SDHA,FBLN1,STXBP1
OX,GPC4,H19,EHMT1,HSPG2,ANTXR1,LTBP4,GTF2I,TTC37,PACS1,LZTR1,PHGDH,DHCR7,FBLN5
,COL2A1,SKI,ASXL1,KDM6B,RPS15A,NOTCH2,SELENON,MAPK1,ODC1,AEBP1,FTO,SON,FLNB,D
PR2,ADAR,TRAF7,BCR,EP300,RERE,DGCR2,BAZ1B,NXN,PTEN,LOX,EHMT1,ANTXR1,LDLR,KMT2

NOP56,ELN,ATP13A2,MECP2,PARK7,ADNP,LIMK1,FLII,IQSEC1,SOBP,DDX6,SYNGAP1,DEAF1,CR
1,FTO,SON,FLNB,DLK1,TPM3,RPS28,STXBP1,RAI1,KDM3B,ARFGEF2,BMPR2,CDC42,ADAR,TRA
E1,DHX30,RPS28,SDHA,STT3B,STXBP1,STT3A,LONP1,PRPS1,USP7,SARS1,RAI1,RAB11A,TCTN1,
DM1A,MAP2K2,QARS1,SMARCE1,DHX30,STT3B,STXBP1,STT3A,LONP1,VANGL2,SOD1,TCTN1,A
TP6V1B2,B2M,STXBP1,LONP1,USP7,IDH1,RAI1,SYP,BCR,KDM5C,TRIO,EXTL3,RERE,GPX4,NXN,A
DOST,SHOX,HNRNPK,NARS1,HLA-B,ARCN1,CTBP1,RAC1,MMP2,L1CAM,PPIB,GLUL,CDKN1A,CF
5,CREBBP,CCDC47,SH3PXD2B,HS2ST1,BGN,PTPN11,ABL1,COL12A1,CLIP2,ELN,OBSL1,ACTA2,
PPRC,SPTBN2,CDH2,DDX3X,CLTC,RPS7,RPS19,RPL35,PRKAR1A,GJA1,MYO5A,CTNNB1,COL6A

,STXBP1,LONP1,SOD1,FLNC,ARNT2,ARFGEF2,KIF5A,EP300,EXTL3,CNTNAP1,BAZ1B,GPX4,ATRX
,PTEN,SMC3,GPC4,HSPG2,PKDCC,LTBP4,EFTUD2,GTF2I,TTC37,PACS1,PHGDH,NSD2,DHCR7,PC
R2,SHOX,HNRNPK,DUX4,RPS27,CTBP1,RAC1,SMCHD1,MMP2,HS6ST1,NOVA2,CDH11,RAB27A,F

,PTPN11,ABL1,CLIP2,ELN,OBSL1,WDR26,ADNP,BANF1,HMGB3,LIMK1,NECTIN1,SOBP,RDH11,C
DM1A,MAP2K2,QARS1,SMARCE1,DHX30,STT3B,STXBP1,STT3A,LONP1,VANGL2,SOD1,TCTN1,A
DM1A,MAP2K2,QARS1,SMARCE1,DHX30,STT3B,STXBP1,STT3A,LONP1,VANGL2,SOD1,TCTN1,A
GF2,CREBBP,WNK1,CCDC47,FGFR1,NTRK2,CHMP1A,CLIP2,TRAK1,ELN,PDHA1,MECP2,WDR26,
B2,COL3A1,CRB2,TCF20
LN,OBSL1,PDHA1,IGF1R,MECP2,ADNP,LIMK1,NBAS,EXT2,DDX6,PPP2R5D,EXOC7,TSPAN7,CHST
26,CNBP,HS2ST1,PTPN11,EMC1,ADNP,BSCL2,EFNB1,VPS51,DDX6,RAB18,CHST3,NSD1,RBM10
DH11,RAB27A,FDFT1,TBCD,PPP1CB,KLHL24,LMNB2,RPL27,SNRPN,SKI,ASXL1,ABCA2,TERF2IP

,NSD2,DHCR7,MED13,MTOR,PACS2,UBE3A,YY1,CREBBP,MN1,TANC2,FGFR1,SH3PXD2B,NTRK2
GF8,DDX3X,RPS7,RPS19,RPL35,PRKAR1A,GJA1,CTNNB1,RPS10,RPL35A,ARID1A,TSR2,SHOX,HN

,RERE,KDM5B,VARS1,BAZ1B,CSNK2A1,GPX4,ATRX,PI4KA,RAB3GAP2,PTEN,SMC3,NDST1,EML1

PL27,PLOD1,VPS35,SKI,ASXL1,RPS15A,NOTCH2,CCND1,PLEC,ATXN10,KDM1A,SMARCE1,RPS28

:N,SMC3,LOX,NDN,BAP1,EHMT1,HSPG2,ANTXR1,SLC6A8,KMT2E,GTF2I,ELP1,PACS1,OGT,MED1[illegible]
NTNAP1,VARS1,BAZ1B,ATRX,RAB3GAP2,SMC3,HNRNPA2B1,NDST1,TIMM50,ATP6AP2,SLC2A1,N[illegible]
DC1,FTO,SON,KDM1A,MAP2K2,QARS1,SMARCE1,DHX30,STXBP1,LONP1,USP7,VANGL2,RAI1,RA
NP,SMARCC2,NBEA,POGZ,DNM1,NFIB,JRK,UBTF,DMPK,SIN3A,TCF20

'AOK1,EHMT1,HSPG2,LINGO1,SLC6A8,KMT2E,CNP,PACS1,PHGDH,MSH6,DHCR7,UBE3A,CREBB

3A,SLC16A2,SYNGAP1,PPP2R5D,DEAF1,DEGS1,GPT2,WAC,ZSWIM6,CLCN4,ASH1L,MED25,MBOA

3,NECTIN1,PPP2R5D,DEAF1,TSPAN7,CHST3,WAC,CLCN4,ACSL4,MAN2B1,LMNA,MED12,SOS1,C[illegible]
P,GPAA1,EFNB1,EXT2,TSPAN7,WAC,ZSWIM6,CLCN4,ACSL4,BCOR,MED25,SETD5,NACC1,LMNA,[illegible]
RAI1,RAB11A,CDC42,ADAR,CIC,TRAF7,WDR45B,EXTL3,RERE,KDM5B,ARID1B,SMC3,EML1,B4GA[illegible]

3,HSPG2,KMT2E,GTF2I,SRSF2,ELP1,PHGDH,MSH6,MED13,CIZ1,UBE3A,PRKACA,YY1,CREBBP,N[illegible]
GDH,NSD2,DHCR7,RBPJ,MTOR,SMARCD1,TRIM8,FGFR1,EIF4A3,RPS26,PSAT1,TMEM94,OBSL1,[illegible]
SL1,NOTCH1,MECP2,ASAH1,CRKL,TSPAN7,CHST3,CLCN4,ACSL4,BCOR,RPS29,SETD5,LMNA,ME[illegible]
SMARCD1,CREBBP,CCDC47,SMO,FGFR1,EIF4A3,BGN,COL12A1,CLIP2,MACROH2A1,ELN,OBSL1

T1,MAPK8IP3,GNAO1,POU3F3,PRMT7,CASK
TL3,GPX4,NXN,ARID1B,CTNND2,RAB3GAP2,FBXO11,HDAC6,SMC3,ABCC9,MAPRE2,GPC4,NDN,[illegible]
XN,ARID1B,CTNND2,RAB3GAP2,FBXO11,HDAC6,SMC3,ABCC9,MAPRE2,GPC4,NDN,CLCN7,PKDC[illegible]

P1,LONP1,USP7,RAI1,SYP,BCR,KDM5C,TRIO,EXTL3,RERE,GPX4,NXN,ARID1B,CTNND2,RAB3GA[illegible]
C1,THOC2,EPB41L1,PPP3CA,MAN1B1,SLC1A4,SYT1,POGZ,JRK,CC2D1A
SF2,DICER1,MSH6,PCGF2,YY1,SMO,RPS26,PTPN11,MDM2,IDH2,EWSR1,PDGFRB,EXT2,CDK4,N[illegible]
AC,SHMT2,CLCN4,NUS1,ACSL4,CACNA1H,SETD5,MED12,HIRA,PPP1R12A,ZDHHC9,PPP3CA,TET3,
DX,ARCN1,RPS27,CTBP1,MMP2,L1CAM,PPIB,EEF1A2,PPP1CB,RPL27,SNRPN,COL2A1,SKI,ASXL1,
GDH,DHCR7,FGFR1,RPS26,HS2ST1,PTPN11,PSAT1,COL12A1,OBSL1,MECP2,EFNB1,NBAS,RDH1[illegible]

7,CYB5R3,PAFAH1B1,MAT2A,HNRNPU,SCARB2,RPS24,COL1A2,HK1,PTCD3,NR2F1,CHD4,RPS20

-B,ARCN1,RAC1,CDKN1A,HS6ST1,NOVA2,MDH2,EEF1A2,FDFT1,SQSTM1,VAMP2,GNAI3,TAF15,T
D5,MAN2B1,SOS1,CNOT3,SYT1,CCNK,TMCO1,FGD1,KLHL7,MAFB,SIN3A

LIP2,ELN,NOTCH1,ACTA2,ALDH18A1,LIMK1,ZNF462,PPP2R5D,CHST3,WAC,ACSL4,BCOR,LMNA,[illegible]

NPT,FGD1,RAD50,DPF2,DMPK,SIN3A,DNAJC21
,POR,SC5D,ALDH18A1,BANF1,ZNF462,EFNB1,PDGFRB,DDX6,CHST3,NSD1,RBM10,P3H1,MAN2B
C2,TRIM8,CCDC47,SMO,FGFR1,EIF4G1,TECR,PINK1,CLIP2,ELN,MFN2,MECP2,FXR1,PARK7,ADN[illegible]

.D2B,CUL4B,SYT1,POGZ,ITGA7,SLC18A3,GLB1,SIN3A
\P2,SLC2A1,NDN,GRN,SPG7,TAOK1,EHMT1,IARS2,TPP1,KMT2E,GTF2I,TUBGCP6,HSD17B4,MED
RN,IARS2,ANTXR1,TPP1,AKT3,IREB2,LTBP3,PHGDH,HSD17B4,NSD2,RBPJ,FBLN5,UBE3A,CREBE
YOU1,RAI1,KDM5C,TRIO,EP300,EXTL3,RERE,CNTNAP1,VARS1,FLCN,BAZ1B,CSNK2A1,ATRX,NXN
RPS28,ATP6V1B2,HYOU1,RAI1,KDM5C,TRIO,EP300,EXTL3,RERE,CNTNAP1,VARS1,FLCN,BAZ1B,C
MP2,CRPPA,CDH11,FDFT1,TBCD,GNAI3,RPL27,SNRPN,COL2A1,SKI,ASXL1,KDM6B,RPS15A,NOT
,FLNB,DLK1,SMARCE1,ATP6V1B2,STXBP1,LONP1,PRPS1,MESD,RAI1,SEC23A,TCTN1,KDM3B,SM
ID1B,CTNND2,RAB3GAP2,FBXO11,PTEN,MAPRE2,GPC4,NDN,EHMT1,PKDCC,KMT2E,EFTUD2,TT
V1B2,STXBP1,RAI1,SEC23A,ARNT2,SMS,CTNND1,STAT3,BCR,DGCR2,ARID1B,SMC3,ECEL1,MAP

YOU1,RAI1,KDM5C,TRIO,EP300,EXTL3,RERE,CNTNAP1,VARS1,FLCN,BAZ1B,CSNK2A1,ATRX,NXN
FGFR1,EIF4A3,RPS26,OBSL1,NOTCH1,MECP2,ASAH1,CRKL,TSPAN7,CHST3,CLCN4,ACSL4,BCO

V1B2,STXBP1,RAI1,SEC23A,ARNT2,SMS,CTNND1,STAT3,BCR,DGCR2,ARID1B,SMC3,ECEL1,MAP

HSPG2,PKDCC,LZTR1,PHGDH,DHCR7,CREBBP,FGFR1,RPS26,HS2ST1,BGN,PTPN11,PSAT1,COL1
),EP300,EXTL3,RERE,BAZ1B,GPX4,GARS1,NXN,ARID1B,RAB3GAP2,FBXO11,PTEN,SMC3,ABCC9,

S1,ITGA7,PLXND1,DMPK,EMD
RB,VPS51,DDX6,SYNGAP1,PPP2R5D,CRKL,TSPAN7,WAC,SHMT2,GDF1,CLCN4,ACSL4,SETD5,LM
I,NDN,GRN,SPG7,TAOK1,EHMT1,IARS2,TPP1,KMT2E,GTF2I,TUBGCP6,MED13,CIZ1,UBE3A,GOT
YK,NIPA1,SLC1A4,SPG11,OPA1,APOE,NFASC,PIGT,FBXO7,PRNP,HADHB,BRAT1,DNMT1,ARSA
YK,NIPA1,SLC1A4,SPG11,OPA1,APOE,NFASC,PIGT,FBXO7,PRNP,HADHB,BRAT1,DNMT1,ARSA

TGFBR1,CNOT3,GATAD2B,TGFB2,CCNK,COL3A1,SETD1B,SIN3A,CNOT1,TCF20
TPM3,SDHA,B2M,STXBP1,LONP1,PRPS1,USP7,IDH1,SOD1,RAI1,RAB11A,SEC23A,ARNT2,ARFGEF
,DGCR8,CNOT3,TET3,PIGT,POGZ,PLXND1,HMGA2,MAFB,SIN3A,TCF20
:1A4,CUL4B,PSMD12,FHL1,TMCO1,SMARCB1,KLHL7,HMGA2,GLB1,NDUFA10
DX6,DARS1,SYNGAP1,DEAF1,TSPAN7,GBA2,NSD1,WAC,CLCN4,ASH1L,BCOR,MED25,RARS1,MB
L6ST,GPC4,ERGIC1,HSPG2,HINT1,GTF2I,LZTR1,PHGDH,NSD2,SMARCD1,CREBBP,FGFR1,SH3PX
:B2,KMT2E,ELP1,PHGDH,HSD17B4,AHCY,COX6A1,UBE3A,YY1,CREBBP,NDUFA13,WNK1,TRIM8,T
,MAPK8IP3,CENPT,SIN3A,TCF20

9,MED25,RARS1,MBOAT7,CNTN2,GALNT2,EPB41L1,PPP3CA,TRAF3,MAN1B1,SZT2,SEC31A,NBE
KI,ABCA2,MARS1,SELENON,OPTN,VHL,CCND1,PLEC,ACADVL,HSPB1,MAP2K2,TPM3,SMARCE1,SI
2,RBPJ,UBE3A,CREBBP,TRIM8,CCDC47,PTPN23,NTRK2,DENND5A,MDH1,TECR,PSAT1,SEMA6B,T
,PTPN11,ABL1,TMEM94,POR,OBSL1,MECP2,EFNB1,SYNGAP1,NSD1,BCOR,SOX9,MED25,RPS29,

SK,TCF20

LK1,TPM3,RPS28,ATP6V1B2,LONP1,HYOU1,VANGL2,RAI1,KDM5C,TRIO,EP300,EXTL3,RERE,CNT

0

,LMNA,PMPCA,SLC5A7,ANOS1,INF2,RUSC2,NIPA1,TRAPPC11,PCNA,SPG11,CUL4B,PSMD12,OPA

2A,HNRNPU,SCARB2,RPS24,MSN,COL1A2,HK1,RHOA,RPS20,CD81,COL5A2,WDR1,ATP2A2,RPL2
3A,CREBBP,TRIM8,CCDC47,PTPN23,NTRK2,DENND5A,MDH1,TECR,PSAT1,TRAK1,DAG1,NOTCH
28,ATP6V1B2,STXBP1,VANGL2,RAI1,SEC23A,TCTN1,ARNT2,SMS,CTNND1,STAT3,BCR,RERE,DGC

ED13,PCGF2,CREBBP,TANC2,SMO,FGFR1,EIF4A3,RPS26,COL12A1,CLIP2,ELN,ADNP,ALDH18A1,
P1,LZTR1,HSD17B4,DHCR7,PCGF2,UBE3A,GOT2,YY1,SMARCD1,CREBBP,WNK1,FGFR1,PTPN23,

F,ZDHHC9,GALNT2,PPP3CA,TET3,CUL4B,PSMD12,SETD1A,SZT2,SYT1,PIGT,DNM1,NFIB,SIN3A,N

1,HTRA1,PDGFRB,ASAH1,SYNGAP1,GBA2,NDUFB8,NUS1,RNF216,COX6B1,PMPCA,SGPL1,PPP3

ABL1,TMEM94,CLIP2,MACROH2A1,KLF13,ELN,OBSL1,IGF1R,TRRAP,LIMK1,ZNF462,EFNB1,FLII,

OL7A1,MED12,SOS1,COL6A3,CUL7,EXT1,CUL4B,PSMD12,FHL1,SMARCC2,SMARCB1,ITGA7,KL

ATRX,NXN,ARID1B,SMC3,ABCC9,ATP6AP2,GPC4,SOX2,AKT3,LDLR,EFTUD2,GTF2I,TTC37,PACS1,
KD2B,PTPN11,AUTS2,CLIP2,MACROH2A1,KLF13,SC5D,ELN,OBSL1,TRRAP,LIMK1,ZNF462,EFNB1
E1,SLC5A6,PPP1R12A,EPB41L1,PPP3CA,MAN1B1,POMGNT1,CUL4B,FH,COL3A1,MAPK8IP3,CC2

CR8,TGFBR1,POMGNT1,TGFB2

DPF2,GLB1,CTSA
,MAN1B1,EXT1,GATAD2B,PSMD12,SYT1,SMARCC2,POGZ,MAPK8IP3,DPF2,TCF20
EN,SMC3,ECEL1,PTPRF,ABCC9,GPC4,EHMT1,ANTXR1,EFTUD2,TTC37,TUBGCP6,LTBP3,DHCR7,D

ATXN3,REEP2,FBLN5,CREBBP,REEP1,SMO,EIF4G1,PINK1,POR,CLIP2,ELN,ATP13A2,MECP2,ADN

I,FBXO7,NDUFA10
2R5D,CHST3,SOX9,SETD5,SNRPB,LARGE1,LMNA,SLC5A7,HIRA,DSTYK,THRA,COL6A3,CUL7,FHL

B,POR,CLIP2,DAG1,NOP56,SC5D,ELN,PDHA1,TRRAP,LIMK1,VPS51,RAB18,NSD1,RBM10,ZSWIM
E3A,CREBBP,PHIP,TANC2,FGFR1,EIF4A3,SH3PXD2B,TRIP12,PTPN11,ABL1,PUM1,POR,CLIP2,MA
,NXN,ARID1B,CRTAP,HDAC6,PTEN,SMC3,ECEL1,IL6ST,ABCC9,GPC4,NDN,CLCN7,IARS2,ERGIC1,

,ZMIZ1,COL11A1,RPL10,RPS17,AP3D1,ARF1,PAFAH1B1,MAT2A,HNRNPU,RPSA,SCARB2,RPS24,

GAP2,PTEN,SMC3,ECEL1,PTPRF,ABCC9,GPC4,EHMT1,ANTXR1,EFTUD2,TTC37,TUBGCP6,LTBP3

RPS26,PTPN11,ABL1,TMEM94,POR,OBSL1,MECP2,EFNB1,SYNGAP1,NSD1,BCOR,SOX9,MED25,
RPS26,PTPN11,ABL1,TMEM94,POR,OBSL1,MECP2,EFNB1,SYNGAP1,NSD1,BCOR,SOX9,MED25,
SAH1,SYNGAP1,DEAF1,DHFR,CLCN4,NUS1,ASH1L,CACNA1H,COX4I1,CNTN2,EPB41L1,PPP3CA,S

N11,ABL1,TMEM94,CLIP2,MACROH2A1,KLF13,ELN,OBSL1,IGF1R,TRRAP,LIMK1,ZNF462,EFNB1

2A1,CLIP2,DAG1,ELN,EMC1,WDR26,GPAA1,ALDH18A1,LIMK1,FLII,NBAS,PPP2R5D,DEAF1,NSD1,
M5C,CNTNAP1,BAZ1B,UBQLN2,ATRX,GARS1,PTEN,HNRNPA2B1,IL6ST,ABCC9,B4GAT1,GPC4,FU

ATP6V1B2,LONP1,RAI1,TCTN1,SYP,BCR,KDM5C,TRIO,EP300,EXTL3,RERE,GPX4,NXN,ARID1B,R
MK1,CRKL,CHST3,WAC,NDUFB8,GDF1,BCOR,MAN2B1,COX6B1,LMNA,NR2F2,SKIV2L,SOS1,CITE
,SNRPN,RBMX,SKI,KDM6B,PRPF6,RPS15A,NOTCH2,SELENON,MAPK1,SMARCA4,PLEC,ODC1,AE
R1A,GJA1,MYO5A,RPS10,RPL35A,HNRNPDL,ARID1A,TSR2,HLA-B,ARCN1,RPS27,CTBP1,SMCHD

35,PRKAR1A,GJA1,CTNNB1,COL6A2,MACF1,RPS10,RPL35A,CYFIP2,ARID1A,DDOST,TSR2,SHOX,

CNA1H,THOC2,MED12,CNTN2,NIPA1,EPB41L1,PPP3CA,TRAPPC11,SLC1A4,GYS1,SETD1A,PIGT,
CN4,NUS1,ASH1L,ACSL4,CACNA1H,ARHGEF9,MED25,MBOAT7,MED12,CNTN2,EPB41L1,PPP3CA
DAG1,ELN,WDR26,JAM3,TRRAP,GPAA1,ASNS,VPS51,SLC25A1,RAB18,STUB1,EXOC7,NSD1,RBM
AC6,PTEN,SMC3,ECEL1,IL6ST,ABCC9,GPC4,NDN,IARS2,ERGIC1,HSPG2,KMT2E,NSD2,DHCR7,ME
HB,HMGA2,SIN3A,DNAJC21,CNOT1

,LMNB2,SNRPN,PLOD1,COL2A1,SKI,ASXL1,RPS15A,NOTCH2,SELENON,MAPK1,SMARCA4,PLEC,

A,NR2F2,HIRA,SKIV2L,CTCF,TALDO1,TGFBR1,G6PC3,PSMD12,TGFB2,PIGT,TMCO1
R7,FBLN5,MTOR,CREBBP,FGFR1,RPS26,PTPN11,ABL1,TMEM94,POR,CLIP2,ELN,OBSL1,MECP2
1,NSD1,WAC,CLCN4,ASH1L,ACSL4,CACNA1H,SETD5,MAN2B1,NSMF,CHD8,THOC2,UNC13A,COL

TL3,RERE,DGCR2,PBX1,CSNK2A1,ATRX,NXN,ARID1B,RAB3GAP2,PTEN,SMC3,ECEL1,PTPRF,ABC
',SNRPN,RBMX,SKI,KDM6B,PRPF6,RPS15A,NOTCH2,SELENON,VHL,CCND1,MAPK1,SMARCA4,PL
DF1,BCOR,RNF216,SETD5,SNRPB,SOS1,CITED2,CTCF,CNOT3,CUL7,MAN1B1,PIGS,CUL4B,BRAT

1A,FLNB,SMARCE1,ATP6V1B2,STXBP1,LONP1,MESD,RAI1,SEC23A,SMS,CDC42,SYP,CTNND1,ST
KAR1A,CTNNB1,COL6A2,MACF1,CYFIP2,TARDBP,ARID1A,DDOST,HNRNPK,HLA-B,TTR,ARCN1,C

NTXR1,KMT2E,PACS1,LTBP3,LZTR1,NSD2,MED13,PACS2,YY1,SMARCD1,CREBBP,MN1,PHIP,TAN
,RPL35A,SLC25A3,ARID1A,TSR2,HNRNPK,HLA-B,TTR,ARCN1,RPS27,CTBP1,RAC1,MMP2,FDFT1,

31,LMNA,MED12,HIRA,DGCR8,ARSB,PIGS,TARS1,SEC31A,COL3A1,GNAO1,ELMO2,FGD1,CASK

TP2A2,RPL26,CLEC7A,RPL31,XPR1,TPM1,GPC3,MEGF8,CCN2,MMP14,DDX3X,RPS7,RPS19,RPL
,WNK1,SMO,FGFR1,SH3PXD2B,PSAT1,POR,SC5D,ALDH18A1,BANF1,ZNF462,EFNB1,PDGFRB,DD

35R3,PAFAH1B1,ITM2B,HNRNPU,LDHA,CTSD,RPS24,COL1A2,HK1,NR2F1,CHD4,ATP1A1,RPS20,C
PS20,COL5A2,PFN1,LRPPRC,KLHL9,SPTBN2,TKT,TPM1,CDH2,GPC3,RAB7A,CCN2,DDX3X,CLTC,
NNB1,COL6A2,MACF1,CYFIP2,TARDBP,SLC25A3,ARL6IP1,ARID1A,DDOST,HNRNPK,NARS1,TTR,C
A1,COL6A2,RPS10,RPL35A,ARID1A,TSR2,SHOX,HNRNPK,NARS1,DUX4,HLA-B,RPS27,CTBP1,RAC
,POLR2A,AP2M1,SEC61A1,RPL18,UCHL1,RPL11,AGRN,ZMIZ1,RPL10,MARCHF6,RPS17,AP3D1,S
A,RPL15,PKD1,CHD3,POLR2A,AP2M1,SEC61A1,RPL18,UCHL1,RPL11,AGRN,ZMIZ1,COL11A1,RPL
N,ARID1B,RAB3GAP2,ZFR,NDST1,MAPRE2,ATP6AP2,TAOK1,EHMT1,SOX2,SLC6A8,AKT3,CNP,PH
3,PAFAH1B1,MAT2A,HNRNPU,CTSD,SCARB2,RPS24,COL1A2,HK1,RHOA,CHD4,CD81,COL5A2,RF
,RPS17,AP3D1,SBF1,ARF1,CYB5R3,PAFAH1B1,ITM2B,MAT2A,HNRNPU,CTSD,SCARB2,RPS24,HK
KI,ASXL1,RPS15A,NOTCH2,PPP2CA,MAPK1,PLEC,ODC1,PRKDC,SON,KDM1A,MAP2K2,QARS1,FL
C,RPS7,RPS19,RPL35,GJA1,MYO5A,CTNNB1,MACF1,RPS10,RPL35A,CYFIP2,TARDBP,ARID1A,TS
C,RPS7,RPS19,RPL35,GJA1,MYO5A,CTNNB1,MACF1,RPS10,RPL35A,CYFIP2,TARDBP,ARID1A,TS
2A,AP2M1,SEC61A1,RPL18,UCHL1,RPL11,AGRN,ZMIZ1,COL11A1,RPL10,MARCHF6,RPS17,VCL,
1,RPS10,RPL35A,CYFIP2,TARDBP,ARID1A,TSR2,HNRNPK,NARS1,ARCN1,RPS27,CTBP1,RAC1,SM
,COL27A1,RAI1,SEC23A,SMS,CDC42,CTNND1,STAT3,TRIO,EP300,EXTL3,RERE,FLCN,CSNK2A1,
,ARF1,CYB5R3,PAFAH1B1,HNRNPU,CTSD,SCARB2,RPS24,COL1A2,HK1,PTCD3,NR2F1,CHD4,AT
CD81,COL5A2,RPL26,RPL31,LRPPRC,XPR1,CCND2,CDH2,GPC3,MEGF8,MMP14,DDX3X,CLTC,R
SNRPN,PLOD1,COL2A1,SKI,NOTCH2,SELENON,MAPK1,SMARCA4,KDELR2,PLEC,AEBP1,SON,MA
1,ITM2B,HNRNPU,CTNNA1,RPSA,RPS24,COL1A2,HK1,NR2F1,RPS14,CHD4,ATP1A1,COL5A2,ATF
C42,SYP,CTNND1,BCR,KDM5C,TRIO,EP300,EXTL3,RERE,PBX1,FLCN,BAZ1B,CSNK2A1,ATRX,NXN
CDH11,EEF1A2,TBCD,PPP1CB,GNAI3,LMNB2,RPL27,SNRPN,ACO2,COL2A1,SKI,ASXL1,CNTNAP2
L,CRPPA,NOVA2,CDH11,MDH2,EEF1A2,FDFT1,TBCD,VAMP2,LMNB2,RPL27,SNRPN,ACO2,SKI,AS
2B,MAT2A,HNRNPU,CTSD,SCARB2,RPS24,HK1,PTCD3,NR2F1,CHD4,ATP1A1,RPS20,COL5A2,RPL
ARB2,RPS24,COL1A2,HK1,NR2F1,CHD4,RPS20,ATP2A2,RPL26,PFN1,RPL31,LRPPRC,XPR1,SPT
6,RPL31,KLHL9,CCND2,GPC3,MEGF8,RAB7A,CCN2,MMP14,CLTC,RPS7,RPS19,RPL35,PRKAR1A
10,RPS17,AP3D1,SBF1,ARF1,CYB5R3,PAFAH1B1,MAT2A,HNRNPU,CTSD,SCARB2,RPS24,COL1A
TXBP1,LONP1,PRPS1,USP7,SARS1,VANGL2,RAI1,RAB11A,TCTN1,ARNT2,KDM3B,ARFGEF2,CDC
1,ARF1,CYB5R3,PAFAH1B1,ITM2B,MAT2A,HNRNPU,CTSD,SCARB2,RPS24,HK1,PTCD3,NR2F1,CH
RPPRC,SPTBN2,TKT,CCND2,CDH2,GPC3,MEGF8,DDX3X,CLTC,RPS7,RPS19,RPL35,PRKAR1A,GJ
V1B2,LSS,STXBP1,LONP1,MESD,RAI1,KDM3B,SMS,YME1L1,TRAF7,BCR,KDM5C,TRIO,EP300,RE
KDCC,LTBP4,IREB2,KMT2E,EFTUD2,GTF2I,TTC37,PACS1,LTBP3,PHGDH,HSD17B4,NSD2,DHCR7
A4,KDELR2,AEBP1,SON,MAP2K2,FLNB,DLK1,TPM3,SMARCE1,LSS,STXBP1,LONP1,PRPS1,MESD,
CB,SQSTM1,PABPN1,LMNB2,SNRPN,CHD2,SKI,CNTNAP2,RAB11B,ABCA2,OPTN,VHL,PPP2CA,M
,FDFT1,PPP1CB,RPL27,SNRPN,PLOD1,COL2A1,RBMX,SKI,ASXL1,KDM6B,PRPF6,RPS15A,NOTC
B1,MED12,HIRA,DGCR8,SKIV2L,ARSB,CTCF,TGFBR1,PIGS,CUL4B,TGFB2,SEC31A,TMCO1,POG
1,RAB11A,TCTN1,ARNT2,KDM3B,ARFGEF2,CDC42,ADAR,CIC,KCNQ2,WDR45B,EP300,EXTL3,RE
PS10,RPL35A,ARID1A,TSR2,SHOX,HNRNPK,NARS1,HLA-B,ARCN1,RPS27,CTBP1,RAC1,MMP2,L1
2,QARS1,SMARCE1,SDHA,LSS,STXBP1,LONP1,PRPS1,USP7,SARS1,VANGL2,RAI1,RAB11A,TCTN1
RAC1,MMP2,L1CAM,CRPPA,CTSB,SQSTM1,LMNB2,ABCA3,RPL27,SNRPN,PLOD1,COL2A1,SKI,AS
L11A1,RPL10,MARCHF6,RPS17,AP3D1,SBF1,CYB5R3,PAFAH1B1,ITM2B,MAT2A,PXDN,HNRNPU,
,SMARCA4,KDELR2,ODC1,AEBP1,SON,MAP2K2,FLNB,DLK1,TPM3,SMARCE1,RPS28,MESD,HYOU
ARID1A,DDOST,HNRNPK,NARS1,ATP1A2,HLA-B,TTR,ARCN1,CTBP1,RAC1,ARHGDIA,GPC1,L1CAM

5A,PPP2CA,MAPK1,SMARCA4,PRDX1,FTO,PRKDC,QARS1,DLK1,SMARCE1,DHX30,RPS28,LARS1,

COL11A1,RPL10,RPS17,AP3D1,SBF1,ARF1,CYB5R3,PAFAH1B1,MAT2A,HNRNPU,CTSD,SCARB2,RI
31,ATP1A2,RPS27,CTBP1,RAC1,SMCHD1,MMP2,L1CAM,CRPPA,CTSB,CDH11,FDFT1,PPP1CB,SQS
,RPS15A,PPP2CA,MAPK1,SMARCA4,PRDX1,FTO,PRKDC,QARS1,DLK1,SMARCE1,DHX30,RPS28,S
GLUL,CRPPA,HS6ST1,NOVA2,EEF1A2,TBCD,PPP1CB,VAMP2,GNAI3,SNRPN,ACO2,PLOD1,COL2A1
,NARS1,DUX4,HLA-B,RPS27,CTBP1,RAC1,GPC1,SMCHD1,MMP2,NOVA2,CDH11,MDH2,EEF1A2,FI
HNRNPK,NARS1,ARCN1,RPS27,CTBP1,RAC1,MMP2,L1CAM,PPIB,CRPPA,HS6ST1,CDH11,FDFT1,I
HNRNPK,NARS1,ARCN1,RPS27,CTBP1,RAC1,MMP2,L1CAM,PPIB,CRPPA,HS6ST1,CDH11,FDFT1,I

7,TRIO,EP300,RERE,PBX1,CNTNAP1,BAZ1B,CSNK2A1,ATRX,NXN,ARID1B,CTNND2,RAB3GAP2,AB
FIP2,HNRNPDL,TARDBP,SLC25A3,ARID1A,TSR2,SHOX,HNRNPK,NARS1,DUX4,ATP1A2,HLA-B,RPS
S2ST1,TRIP12,AUTS2,CLIP2,ELN,IGF1R,MDM2,MECP2,ADNP,TRRAP,ALDH18A1,LIMK1,DDX6,SYN

DL11A1,RPL10,RPS17,AP3D1,SBF1,ARF1,CYB5R3,PAFAH1B1,MAT2A,HNRNPU,CTSD,SCARB2,RPS
A2,CDH11,EEF1A2,FDFT1,TBCD,PPP1CB,RPL27,SNRPN,PLOD1,RBMX,SKI,ASXL1,KDM6B,RAB11I
,CLIP2,ELN,OBSL1,ALDH18A1,LIMK1,EFNB1,DDX6,CRKL,CHST3,NSD1,FARSA,LMNA,MED12,HIF
3,SMS,CDC42,YME1L1,ADAR,SYP,TRAF7,WDR45B,BCR,KDM5C,TRIO,EP300,EXTL3,RERE,KDM5E
,FDFT1,PPP1CB,LMNB2,ABCA3,RPL27,SNRPN,PDXK,PLOD1,COL2A1,SKI,ASXL1,KDM6B,RAB11I
Y1,CREBBP,FGFR1,RPS26,PTPN11,ABL1,AUTS2,TMEM94,POR,CLIP2,ELN,NOTCH1,IGF1R,TRRA
EC7A,RPL31,CCND2,CDH2,GPC3,MEGF8,CCN2,MMP14,DDX3X,CLTC,RPS7,RPS19,RPL35,PRKAR
G2,ANTXR1,LTBP4,GTF2I,TTC37,PACS1,PHGDH,NSD2,DHCR7,FBLN5,MTOR,SMARCD1,CREBBP,
Y1,CREBBP,FGFR1,RPS26,PTPN11,ABL1,AUTS2,TMEM94,POR,CLIP2,ELN,NOTCH1,IGF1R,TRRA
DL11A1,RPL10,RPS17,AP3D1,SBF1,ARF1,CYB5R3,PAFAH1B1,MAT2A,HNRNPU,CTSD,SCARB2,RPS
DUX4,HLA-B,RPS27,CTBP1,RAC1,GPC1,SMCHD1,NOVA2,CDH11,EEF1A2,FDFT1,TBCD,PPP1CB,R

HX30,ATP6V1B2,STXBP1,MESD,RAI1,SEC23A,KDM3B,SMS,CDC42,SYP,CTNND1,BCR,KDM5C,TRI
ARFGEF2,CDC42,ADAR,CIC,KCNQ2,WDR45B,EP300,EXTL3,RERE,KDM5B,CNTNAP1,VARS1,FLCN

NND2,RAB3GAP2,ABCC9,GPC4,HSPG2,PKDCC,LTBP4,KMT2E,GTF2I,TTC37,TUBGCP6,LTBP3,NS
CH2,PPP2CA,MAPK1,PRDX1,ODC1,PRKDC,SON,KDM1A,MAP2K2,QARS1,FLNB,DLK1,SMARCE1,RI
,SKI,ASXL1,KDM6B,RAB11B,RPS15A,NOTCH2,MAPK1,PLEC,PRDX1,ODC1,AEBP1,PRKDC,SON,M
ATXN10,SON,KDM1A,QARS1,TPM3,DHX30,ATP6V1B2,SDHA,LARS1,STT3B,STXBP1,STT3A,LONP
4,HLA-B,RPS27,CTBP1,RAC1,GPC1,SMCHD1,NOVA2,CDH11,EEF1A2,FDFT1,TBCD,PPP1CB,RPL27

ARFGEF2,CDC42,ADAR,CIC,KCNQ2,WDR45B,EP300,EXTL3,RERE,KDM5B,CNTNAP1,VARS1,FLCN
24,COL1A2,HK1,RHOA,PTCD3,NR2F1,CHD4,RPS20,RPL26,CLEC7A,RPL31,LRPPRC,SPTBN2,TKT
MESD,HYOU1,SEC23A,SMS,STAT3,BCR,KDM5C,TRIO,EP300,RERE,CNTNAP1,FLCN,CSNK2A1,ATR
AZ1B,GPX4,ATRX,NXN,ARID1B,PTEN,SMC3,GPC4,EHMT1,HSPG2,SOX2,PKDCC,LTBP4,AKT3,EFT
ASXL1,RPS15A,NOTCH2,SELENON,PPP2CA,MAPK1,SMARCA4,ODC1,AEBP1,FTO,SON,KDM1A,M
1,SKI,ASXL1,KDM6B,RAB11B,RPS15A,SMARCA4,AEBP1,SON,KDM1A,MAP2K2,DLK1,SMARCE1,D
,ABL1,POR,CLIP2,ELN,NOTCH1,ACTA2,WDR26,TRRAP,ALDH18A1,LIMK1,DDX6,CRKL,CHST3,NS
PS15A,NOTCH2,MAPK1,SMARCA4,ODC1,AEBP1,PRKDC,SON,KDM1A,FLNB,DLK1,SMARCE1,RPS
N,MAPK1,SMARCA4,KDELR2,PLEC,ODC1,AEBP1,HSPB1,SON,FLNB,DLK1,TPM3,SMARCE1,DHX3
E1,DHX30,SDHA,STXBP1,LONP1,PRPS1,USP7,RAI1,FLNC,TCTN1,SMS,CCT5,ADAR,TRAF7,DCTN1,

2,GPC3,MEGF8,CCN2,MMP14,DDX3X,CLTC,RPS7,RPS19,RPL35,PRKAR1A,GJA1,CTNNB1,COL6A
2,GPC3,MEGF8,CCN2,MMP14,DDX3X,CLTC,RPS7,RPS19,RPL35,PRKAR1A,GJA1,CTNNB1,COL6A
MP2,L1CAM,PPIB,CRPPA,HS6ST1,CTSB,CDH11,FDFT1,TBCD,SQSTM1,LMNB2,SNRPN,PDXK,PLO
ATP6V1B2,LARS1,STXBP1,LONP1,PRPS1,MESD,USP7,VANGL2,RAI1,SEC23A,TCTN1,SMS,ADAR,S
A4,PRKDC,SON,KDM1A,MAP2K2,FLNB,DLK1,SMARCE1,RPS28,ATP6V1B2,STXBP1,LONP1,HYOU
RDBP,ARL6IP1,DDOST,HNRNPK,NARS1,ATP1A2,HLA-B,TTR,ARCN1,CTBP1,RAC1,MMP2,L1CAM,
R5D,CHST3,ZSWIM6,SHMT2,SETD5,MBOAT7,PMPCA,SLC5A7,RUSC2,GALNT2,CNOT3,TET3,SPG
STXBP1,PRPS1,SARS1,SOD1,RAI1,RAB11A,TCTN1,SMS,CCT5,ADAR,CIC,TRAF7,DCTN1,KIF5A,KC

2,MED12,DGCR8,RUSC2,EPB41L1,PPP3CA,MAN1B1,SLC1A4,GATAD2B,CUL4B,PSMD12,SETD1A
DM5C,TRIO,EP300,EXTL3,RERE,KDM5B,PBX1,FLCN,CSNK2A1,NXN,ARID1B,PI4KA,RAB3GAP2,F

1B1,MAT2A,HNRNPU,CTSD,SCARB2,RPS24,COL1A2,HK1,RHOA,NR2F1,CHD4,CD81,COL5A2,WD
45B,KDM5C,EXTL3,RERE,NDUFV1,KDM5B,CNTNAP1,ALDH3A2,BAZ1B,UBQLN2,ATRX,PI4KA,RAB
,GPC4,IARS2,ANTXR1,AKT3,KMT2E,DICER1,LTBP3,HSD17B4,DHCR7,MED13,MTOR,SMARCD1,C
RS1,LTBP4,HINT1,DNAJB2,COX6A1,REEP1,SMO,RPS26,NTRK2,SLC30A9,PTPN11,COL12A1,TRA
1,DHX30,ATP6V1B2,STXBP1,LONP1,USP7,HYOU1,SARS1,RAI1,RAB11A,TCTN1,SMS,CDC42,CTN
C,FTO,SON,KDM1A,MAP2K2,FLNB,DLK1,ATP6V1B2,LARS1,STXBP1,LONP1,PRPS1,MESD,USP7,
RPS27,CTBP1,MMP2,L1CAM,PPIB,CDKN1A,HS6ST1,EEF1A2,PPP1CB,RPL27,SNRPN,PLOD1,COL
CS2,UBE3A,YY1,CREBBP,NDUFA13,TANC2,TRIM8,FGFR1,NTRK2,TECR,PINK1,AUTS2,CLIP2,TRA

,NDST1,MCM3AP,MAPRE2,PREPL,TIMM50,ATP6AP2,NDN,FUS,H19,CLCN7,EHMT1,IARS2,ANTXR
TRAF7,EP300,RERE,PBX1,CNTNAP1,VARS1,BAZ1B,CSNK2A1,ATRX,NXN,ARID1B,CTNND2,RAB3G
RAC1,GLUL,CRPPA,CDH11,MDH2,EEF1A2,FDFT1,TBCD,PPP1CB,VAMP2,GNAI3,LMNB2,SNRPN,A
15A,SMARCA4,AEBP1,SON,KDM1A,MAP2K2,DLK1,SMARCE1,DHX30,RPS28,SDHA,STXBP1,LONP

DELR2,PLEC,AEBP1,SON,KDM1A,FLNB,DLK1,TPM3,SMARCE1,DHX30,ATP6V1B2,SDHA,LSS,FBL
2,ADAR,TRAF7,EP300,RERE,PBX1,CNTNAP1,VARS1,BAZ1B,CSNK2A1,GPX4,ATRX,NXN,ARID1B,CT
RNPK,NARS1,ATP1A2,ARCN1,CTBP1,RAC1,ARHGDIA,SMCHD1,MMP2,L1CAM,CRPPA,NOVA2,CD
N1,CTBP1,L1CAM,CRPPA,HS6ST1,NOVA2,MDH2,RAB27A,EEF1A2,TBCD,SQSTM1,TAF15,LMNB2
1,LMNA,MED12,SOS1,ZDHHC9,TGFBR1,CUL7,FAM20C,FHL1,PIGT,COG4,COL3A1,ITGA7,PIEZO

3A2,BAZ1B,UBQLN2,ATRX,PI4KA,RAB3GAP2,IARS1,ZFR,NDST1,EML1,B4GAT1,TIMM50,ATP6A
6A8,LTBP4,IREB2,KMT2E,EFTUD2,LZTR1,NSD2,PCGF2,FBLN5,MTOR,PRKACA,YY1,CREBBP,M
NTNAP1,VARS1,FLCN,BAZ1B,UBQLN2,ATRX,PI4KA,ADCY6,SMC3,FUS,SPG7,TAOK1,EHMT1,VAP
1,ABL1,TMEM94,CLIP2,ELN,NOTCH1,PDHA1,IGF1R,NDUFA4,WDR26,NDUFC2,BSCL2,LIMK1,ZN
3PXD2B,AUTS2,CLIP2,ELN,NOTCH1,ACTA2,MECP2,ADNP,TRRAP,BSCL2,ALDH18A1,LIMK1,EFN
HS2ST1,BGN,PTPN11,AUTS2,COL12A1,CLIP2,ELN,OBSL1,MFN2,MECP2,ALDH18A1,HMGB3,LIM
3,AUTS2,CLIP2,ELN,NOTCH1,ACTA2,MECP2,ADNP,TRRAP,BSCL2,ALDH18A1,LIMK1,EFNB1,NEC
MEM94,CLIP2,ELN,NOTCH1,PDHA1,IGF1R,NDUFA4,WDR26,NDUFC2,LIMK1,ZNF462,EXT2,ECHS
B,UBQLN2,ATRX,RAB3GAP2,SMC3,HNRNPA2B1,NDST1,TIMM50,ATP6AP2,SLC2A1,NDN,FUS,GR
,SON,KDM1A,FLNB,DLK1,TPM3,SMARCE1,RPS28,ATP6V1B2,B2M,STXBP1,LONP1,USP7,IDH1,HY
,NDUFV1,KDM5B,CNTNAP1,CSNK2A1,ATRX,NXN,ARID1B,PTEN,SMC3,ECEL1,PREPL,GPC4,FUS,
SCL2,LIMK1,FLII,IQSEC1,DDX6,SYNGAP1,DEAF1,CRKL,TSPAN7,NSD1,WAC,ZSWIM6,SHMT2,CL

,CLCN4,ACSL4,DLG4,NSMF,CHD8,LMNA,PMPCA,MED12,ANOS1,ZDHHC9,TGFBR1,CNOT3,CUL7,
F462,FLII,PDGFRB,VPS51,PPP2R5D,DEAF1,CRKL,TSPAN7,WAC,ZSWIM6,SHMT2,CLCN4,ACSL4,M
BXO11,PTEN,ABCC9,MAPRE2,GPC4,NDN,EHMT1,HSPG2,PKDCC,KMT2E,EFTUD2,GTF2I,TTC37,F

BCC9,MAPRE2,GPC4,SLC2A1,FUS,GRN,ERGIC1,HSPG2,PKDCC,GTF2I,CNP,LZTR1,PHGDH,NSD2
OS1,CNOT3,MAN1B1,PSMD12,SYT1,CCNK,TMCO1,FGD1,KLHL7,MAFB,SIN3A,TCF20
4,COL1A2,HK1,RHOA,PTCD3,NR2F1,CHD4,COL5A2,RPL26,RPL31,LRPPRC,XPR1,TKT,CCND2,CD
1B2,MESD,HYOU1,RAI1,ARNT2,SMS,KDM5C,TRIO,EP300,EXTL3,RERE,DGCR2,CNTNAP1,VARS1,
C4,SLC2A1,CLCN7,EHMT1,IARS2,PKDCC,KMT2E,LTBP3,LZTR1,NSD2,DHCR7,RBPJ,MED13,MTO
,PRPS1,SARS1,SOD1,RAB11A,ARNT2,ARFGEF2,CCT5,ADAR,SYP,CIC,TRAF7,DCTN1,KIF5A,KCNQ2
,MTOR,CREBBP,TRIM8,FGFR1,SH3PXD2B,PTPN11,AUTS2,CLIP2,ELN,NOTCH1,ACTA2,MECP2,AD
LK1,TPM3,RPS28,ATP6V1B2,MESD,HYOU1,RAI1,SEC23A,ARNT2,SMS,KDM5C,TRIO,EP300,EXTL3
E,GTF2I,LZTR1,DHCR7,MED13,FBLN5,CREBBP,TANC2,FGFR1,RPS26,BGN,PTPN11,ABL1,POR,CI

KL,TSPAN7,NSD1,WAC,SHMT2,CLCN4,NUS1,ACSL4,CACNA1H,MED25,SETD5,MBOAT7,CDKN1B,I
F7,BCR,EP300,RERE,DGCR2,BAZ1B,NXN,ARID1B,CRTAP,PTEN,SMC3,ADH5,ABCC9,LOX,GPC4,CI
KDM3B,CIC,TRAF7,KIF5A,KCNQ2,TRIO,EP300,RERE,NDUFV1,KDM5B,PBX1,FLCN,CSNK2A1,ATR
RNT2,CDC42,YME1L1,TRAF7,EP300,RERE,CNTNAP1,VARS1,BAZ1B,GPX4,ARID1B,PI4KA,RAB3GA
RID1B,CTNND2,RAB3GAP2,CRTAP,FBXO11,HDAC6,SMC3,ABCC9,MAPRE2,GPC4,NDN,CLCN7,EF
RPPA,HS6ST1,FDFT1,TBCD,PPP1CB,SQSTM1,GNAI3,LMNB2,PLOD1,COL2A1,SKI,ASXL1,KDM6B,N
IGF1R,MECP2,ALDH18A1,LIMK1,EFNB1,SLC25A1,ASAH1,SLC16A2,CHST3,NSD1,RBM10,TMEM4
2,MACF1,RPS10,RPL35A,CYFIP2,TARDBP,ARID1A,DDOST,TSR2,ATP1A2,TTR,RPS27,CTBP1,GPC1

,ARID1B,RAB3GAP2,CRTAP,PTEN,SMC3,ECEL1,IL6ST,ATP6AP2,GPC4,NDN,CLCN7,IARS2,HSPG2,
GF2,YY1,CREBBP,FGFR1,RPS26,PTPN11,ABL1,AUTS2,TMEM94,POR,CLIP2,ELN,NOTCH1,IGF1R
DFT1,TBCD,PPP1CB,KLHL24,LMNB2,RPL27,SNRPN,COL2A1,SKI,ASXL1,ABCA2,TERF2IP,RPS15A

HST3,ZSWIM6,BCOR,SETD5,MAN2B1,LMNA,MED12,SOS1,ZDHHC9,PIGS,GATAD2B,CUL4B,PSMD
RNT2,CDC42,YME1L1,TRAF7,EP300,RERE,CNTNAP1,VARS1,BAZ1B,GPX4,ARID1B,PI4KA,RAB3GA
RNT2,CDC42,YME1L1,TRAF7,EP300,RERE,CNTNAP1,VARS1,BAZ1B,GPX4,ARID1B,PI4KA,RAB3GA
ADNP,ALDH18A1,LIMK1,FLII,CSNK2B,EXT2,SLC25A1,SYNGAP1,DEAF1,NSD1,WAC,ZSWIM6,CLCN

3,NSD1,SHMT2,CLCN4,ACSL4,BCOR,SETD5,SNRPB,MED12,HIRA,CTCF,PPP1R12A,CUL7,EXT1,TE
,RPS29,SETD5,MAN2B1,LMNA,MED12,SOS1,CNOT3,MAN1B1,CUL4B,PSMD12,SYT1,CCNK,TMC
,RPS15A,NOTCH2,MAPK1,SMARCA4,PLEC,ODC1,AEBP1,SON,KDM1A,MAP2K2,DLK1,SMARCE1,D

,BGN,PTPN11,AUTS2,TRAK1,SC5D,MECP2,ADNP,AMPD2,ZNF462,EFNB1,PDGFRB,NECTIN1,DDX
RNPK,RPS27,CTBP1,RAC1,SMCHD1,MMP2,GLUL,HS6ST1,NOVA2,CDH11,EEF1A2,FDFT1,PPP1CE

,B4GAT1,NDN,FUS,GRN,SPG7,TAOK1,EHMT1,IARS2,AKT3,KMT2E,GTF2I,TUBGCP6,PHGDH,HSD

,SDHA,LSS,VANGL2,ARNT2,KDM3B,SMS,CDC42,SYP,TRAF7,KIF5A,TRIO,EP300,RERE,KDM5B,PB

3,PACS2,UBE3A,YY1,CREBBP,PHIP,TANC2,CCDC47,TEAD1,TUB,PTPN11,CHMP1A,TMEM94,COL1
DN,FUS,GRN,SPG7,TAOK1,EHMT1,IARS2,TPP1,KMT2E,GTF2I,TUBGCP6,HSD17B4,MED13,CIZ1,
B11A,TCTN1,CDC42,ADAR,CIC,TRAF7,WDR45B,EP300,EXTL3,RERE,KDM5B,ARID1B,HDAC6,PTE

P,PHIP,TRIM8,FGFR1,EIF4G1,TRIP12,PINK1,ATP13A2,MECP2,WDR26,PARK7,ADNP,FLII,IQSEC1,

T7,NACC1,RHOBTB2,PPP3CA,TRAPPC11,SLC1A4,UFM1,CUL4B,SYT1,COG4,TMCO1,POGZ,BRA

CF,CUL7,POMGNT1,FAM20C,PSMD12,TMCO1,POGZ,TCF20
MED12,HIRA,DGCR8,CTCF,RUSC2,QRICH1,CUL7,EXT1,GATAD2B,CUL4B,SYT1,DYNC1I2,COG4,
T1,ATP6AP2,GPC4,NDN,GRN,TAOK1,EHMT1,ANTXR1,AKT3,ATXN3,PHGDH,HSD17B4,NSD2,DHCR

UFA13,PHIP,TANC2,SMO,FGFR1,EIF4G1,NTRK2,TECR,PSAT1,PINK1,CLIP2,TRAK1,ELN,ATP13A
NOTCH1,MECP2,FAM50A,BSCL2,ASNS,MOGS,ASAH1,CRKL,TSPAN7,CHST3,NSD1,CLCN4,ACSL4,
D12,COX4I1,CUL7,PIGS,PSMD12,COG4,SMARCC2,GNAO1,SMARCB1,FGD1,HMGA2,DPF2,SIN3A
,ALDH18A1,LIMK1,COX8A,EXT2,SLC25A1,PPP2R5D,CHST3,SOX9,SETD5,SNRPB,LARGE1,LMNA,

LCN7,ERGIC1,PKDCC,LTBP3,NSD2,DHCR7,RBPJ,PCGF2,MTOR,SMARCD1,TRIM8,FGFR1,EIF4A
C,LTBP3,NSD2,DHCR7,RBPJ,PCGF2,MTOR,SMARCD1,TRIM8,FGFR1,EIF4A3,RPS26,BGN,PUM1,

2,FBXO11,HDAC6,SMC3,ABCC9,MAPRE2,GPC4,NDN,CLCN7,ERGIC1,PKDCC,HINT1,KMT2E,LTB

D1,SOX9,RPS29,CDKN1B,ERBB2,LMNA,XPC,COL7A1,SKIV2L,SOS1,EXT1,RELA,FAM20C,FH,MS
,PSMD12,MSH2,SZT2,DYNC1I2,PRNP,TMCO1,NBEA,DNM1,FGD1,HMGA2,SH3KBP1,JRK,SIN3A,N
,RPS15A,NOTCH2,SELENON,MAPK1,SMARCA4,KDELR2,PLEC,SON,KDM1A,MAP2K2,FLNB,DLK1,
1,CHST3,WAC,SOX9,RPS29,SETD5,MAN2B1,MED12,HIRA,SOS1,ROBO3,TALDO1,COL6A3,CUL7,

,CD81,RPL26,RPL31,LRPPRC,XPR1,TKT,GPC3,MEGF8,DDX3X,CLTC,RPS7,RPS19,RPL35,PRKAR

GFBI,LMNB2,SNRPN,VPS35,CHD2,SKI,ASXL1,KDM6B,CNTNAP2,RAB11B,ABCA2,PRPF6,OPTN,V

NR2F2,MED12,HIRA,SKIV2L,ARSB,SOS1,CITED2,TGFBR1,TGFB2,COL3A1,GLB1

1,LMNA,MED12,HIRA,CTCF,TGFBR1,PPP3CA,THRA,TARS1,FAM20C,SETD1A,PIGT,TMCO1,KLHL
P,BSCL2,LIMK1,FLII,PRPH,IQSEC1,VPS51,CSNK2B,DDX6,SLC25A1,SYNGAP1,DEAF1,WAC,ASH1L

13,CIZ1,UBE3A,GOT2,TANC2,TRIM8,CCDC47,FGFR1,EIF4G1,NTRK2,TECR,CLIP2,TRAK1,SC5D,E
3P,WNK1,TRIM8,CCDC47,REEP1,PTPN23,NTRK2,DENND5A,MDH1,TECR,PSAT1,SEMA6B,TRAK1,D
,ARID1B,CTNND2,PI4KA,RAB3GAP2,HDAC6,PTEN,SMC3,ECEL1,ABCC9,MAPRE2,ATP6AP2,ERGIC
CSNK2A1,ATRX,NXN,ARID1B,CTNND2,PI4KA,RAB3GAP2,HDAC6,PTEN,SMC3,ECEL1,ABCC9,MAPR
CH2,SELENON,MAPK1,ODC1,AEBP1,FTO,SON,MAP2K2,FLNB,DLK1,TPM3,RPS28,ATP6V1B2,MES
IS,CDC42,ADAR,SYP,BCR,TRIO,EP300,EXTL3,RERE,DGCR2,KDM5B,PBX1,FLCN,CSNK2A1,ATRX,N
C37,PACS1,LZTR1,HSD17B4,NSD2,DHCR7,MED13,MTOR,PACS2,UBE3A,YY1,CREBBP,MN1,PHIP,
RE2,B4GAT1,GPC4,HSPG2,SOX2,EFTUD2,TTC37,PHGDH,NSD2,DHCR7,SMARCD1,CREBBP,TRIM

,ARID1B,CTNND2,PI4KA,RAB3GAP2,HDAC6,PTEN,SMC3,ECEL1,ABCC9,MAPRE2,ATP6AP2,ERGIC
R,RPS29,SETD5,LMNA,MED12,COX4I1,CUL7,PIGS,PSMD12,COG4,SMARCC2,GNAO1,SMARCB1,

RE2,B4GAT1,GPC4,HSPG2,SOX2,EFTUD2,TTC37,PHGDH,NSD2,DHCR7,SMARCD1,CREBBP,TRIM

2A1,OBSL1,MECP2,EFNB1,NBAS,RDH11,CHST3,WAC,SOX9,RPS29,SETD5,MAN2B1,LARGE1,MED
GPC4,HSPG2,LTBP4,KMT2E,GTF2I,PHGDH,HSD17B4,NSD2,DHCR7,RBPJ,MED13,PCGF2,YY1,SM

NA,RHOBTB2,MED12,CITED2,CTCF,TALDO1,QRICH1,MAN1B1,EXT1,GATAD2B,PSMD12,SYT1,SM
2,TANC2,TRIM8,CCDC47,FGFR1,EIF4G1,NTRK2,TECR,CLIP2,TRAK1,SC5D,ELN,ATP13A2,PDHA1,

F2,ADAR,PALLD,CIC,DCTN1,KIF5A,STAT3,KCNQ2,EP300,RERE,KDM5B,CNTNAP1,VARS1,FLCN,BA

OAT7,MAN2B1,SRP54,THOC2,HIRA,SKIV2L,EPB41L1,PPP3CA,QRICH1,PCNA,MAN1B1,POMGNT1
KD2B,HS2ST1,BGN,PTPN11,AUTS2,COL12A1,POR,CLIP2,MACROH2A1,ELN,OBSL1,ALDH18A1,LIN
TUBB4A,NTRK2,HS2ST1,PSAT1,TRAK1,DAG1,MFN2,PDHA1,TRIM2,NDUFA4,WDR26,TRRAP,HTRA

A,POGZ,BRAT1,DNM1,GNAO1,SMARCB1,NF2,CASK,CC2D1A
DHA,PFKM,STXBP1,PRPS1,SARS1,SOD1,RAI1,FLNC,YME1L1,ADAR,SYP,TRAF7,DCTN1,KIF5A,STA
TRAK1,DAG1,NOTCH1,PDHA1,MECP2,EMC1,GPAA1,FLII,IQSEC1,CCDC88A,ASNS,NECTIN1,SLC25
,SETD5,LMNA,MED12,HIRA,PPP1R12A,CUL7,GATAD2B,CUL4B,PSMD12,COL3A1,HMGA2,SIN3A

NAP1,VARS1,FLCN,BAZ1B,CSNK2A1,GPX4,ATRX,NXN,ARID1B,CTNND2,PI4KA,RAB3GAP2,HDAC6

1,FHL1,HADHB,HARS1,SLC18A3,MAFB,TCF20

!6,CLEC7A,RPL31,LRPPRC,XPR1,TKT,TPM1,CDH2,GPC3,MEGF8,RAB7A,CCN2,MMP14,DDX3X,RI
1,PDHA1,MECP2,EMC1,GPAA1,FLII,IQSEC1,CCDC88A,ASNS,NECTIN1,SLC25A1,ALDH7A1,SYNGAI
R2,NXN,ARID1B,SMC3,ECEL1,MAPRE2,B4GAT1,GPC4,HSPG2,SOX2,EFTUD2,TTC37,PHGDH,NSD

LIMK1,ZNF462,DDX6,STUB1,CHST3,NSD1,SHMT2,BCOR,MED25,RPS29,SETD5,NUP133,MED12,(
PTPN11,PSAT1,AUTS2,ELN,PDHA1,MECP2,WDR26,ADNP,COX8A,FLII,MOGS,CSNK2B,NECTIN1,SI

ECAP1,TCF20

:A,SPG11,OPA1,APOE,SZT2,FBXO7,PRNP,PPT1,DNM1,DNMT1,CSTB,UBTF,TOMM40,GLB1,DMPI

MOGS,EXT2,RAB18,DEAF1,CRKL,CHST3,SHMT2,GDF1,BCOR,RNF216,MED25,SETD5,SNRPB,LM

HL7,DPF2,EMD

PHGDH,MSH6,NSD2,DHCR7,FBLN5,CREBBP,NDUFA13,TRIM8,FGFR1,RPS26,PTPN11,ABL1,TME
,FLII,EXT2,RAB18,DEAF1,CRKL,CHST3,GDF1,BCOR,RNF216,SETD5,SNRPB,SOS1,CITED2,CTCF,
D1A

)HX38,CREBBP,MN1,PHIP,TRIM8,FGFR1,SH3PXD2B,HS2ST1,TUB,AUTS2,TMEM94,SC5D,ELN,OB

),TRRAP,ALDH18A1,BNC2,HTRA1,LIMK1,PDGFRB,CAPN1,DDX6,SLC16A2,STUB1,TSPAN7,GBA2,N

1,SEC31A,COL3A1,ITGA7,SLC18A3,DNAJC21

6,BCOR,SETX,DLG4,MAN2B1,LARGE1,PMPCA,THOC2,PPP1R12A,TGFBR1,NID1,POMGNT1,CUL4
ACROH2A1,KLF13,ELN,OBSL1,IGF1R,MECP2,TRRAP,LIMK1,ZNF462,EFNB1,FLII,EXT2,DDX6,RAB
HSPG2,ANTXR1,KMT2E,GTF2I,NSD2,DHCR7,MED13,FBLN5,SMARCD1,CREBBP,TANC2,WNK1,C

COL1A2,HK1,PTCD3,CHD4,RPS20,CD81,COL5A2,WDR1,ATP2A2,RPL26,PFN1,CLEC7A,RPL31,PR

,DHCR7,DHX38,CREBBP,MN1,PHIP,TRIM8,FGFR1,SH3PXD2B,HS2ST1,TUB,AUTS2,TMEM94,SC5

RPS29,SETD5,LMNA,MED12,HIRA,PPP1R12A,CUL7,GATAD2B,CUL4B,PSMD12,COL3A1,HMGA2,S
RPS29,SETD5,LMNA,MED12,HIRA,PPP1R12A,CUL7,GATAD2B,CUL4B,PSMD12,COL3A1,HMGA2,S
LC1A4,POMGNT1,APOE,NFASC,SZT2,FBXO7,PRNP,PPT1,BRAT1,DNM1,GNAO1,SMARCB1,NF2,J

,FLII,MOGS,EXT2,RAB18,DEAF1,CRKL,CHST3,SHMT2,GDF1,BCOR,RNF216,MED25,SETD5,SNR

RBM10,WAC,ZSWIM6,OAT,ACSL4,SETD5,MAN2B1,LARGE1,ATF6,TRAPPC11,TARS1,POMGNT1,P
S,EHMT1,VAPB,ERGIC1,YARS1,HSPG2,LTBP4,HINT1,GTF2I,DNAJB2,ATXN3,PHGDH,CIZ1,FBLN5,

AB3GAP2,IARS1,PTEN,SMC3,ABCC9,GPC4,CLCN7,ERGIC1,AKT3,HINT1,KMT2E,EFTUD2,PACS1,N
ED2,CTCF,TGFBR1,G6PC3,PSMD12,TGFB2,FHL1,HADHB,GLB1
EBP1,FTO,PRKDC,KDM1A,MAP2K2,FLNB,DLK1,TPM3,SMARCE1,RPS28,LSS,FBLN1,STT3B,STXB
1,MMP2,DCN,CRPPA,CDH11,MDH2,RAB27A,VCAN,TGFBI,RPL27,SNRPN,PLOD1,COL2A1,SKI,AS

HNRNPK,ARCN1,RPS27,CTBP1,APPL1,MMP2,L1CAM,PPIB,HS6ST1,EEF1A2,FDFT1,TBCD,PPP1C

NBEA,DNM1,GNAO1,SETD1B,JRK,CSTB,CASK
,MAN1B1,TARS1,GATAD2B,SZT2,SEC31A,PIGT,NBEA,BRAT1,DNM1,GNAO1,SETD1B,JRK,CSTB,
10,STAT2,WAC,DHFR,BCOR,LARGE1,RHOBTB2,PMPCA,THOC2,ROBO3,SLC1A3,NID1,MAN1B1,
ED13,FBLN5,SMARCD1,CREBBP,TANC2,WNK1,CCDC47,SMO,FGFR1,EIF4A3,SH3PXD2B,BGN,CO

AEBP1,MAP2K2,FLNB,DLK1,TPM3,SMARCE1,SDHA,PFKM,FBLN1,STXBP1,LONP1,PRPS1,COL27

,BNC2,LIMK1,EFNB1,SYNGAP1,NSD1,BCOR,SOX9,MED25,RPS29,SETD5,LMNA,NR2F2,COL7A1,
7A1,HIRA,ANOS1,TET3,FH,PSMD12,MSH2,SETD1A,NFASC,PRNP,EPHA4,TMCO1,HARS1,SETD1B,

C9,GPC4,EHMT1,ANTXR1,EFTUD2,TTC37,TUBGCP6,LTBP3,DHCR7,DHX38,CREBBP,MN1,PHIP,TI
LEC,ODC1,AEBP1,FTO,PRKDC,KDM1A,MAP2K2,FLNB,DLK1,TPM3,SMARCE1,RPS28,LSS,FBLN1,S
T1,MAPK8IP3,PLXND1,FGD1,HMGA2,TCF20

AT3,KDM5C,TRIO,EP300,ALDH3A2,FLCN,BAZ1B,ATRX,NXN,ARID1B,RAB3GAP2,PTEN,SMC3,IL6S
TBP1,RAC1,GPC1,CDKN1A,NOVA2,MDH2,RAB27A,EEF1A2,FDFT1,TBCD,SQSTM1,GNAI3,PABPN1

IC2,CCDC47,FGFR1,AUTS2,TMEM94,KLF13,ELN,OBSL1,IGF1R,WDR26,TRRAP,BANF1,ZNF462,FI
PPP1CB,RPL27,SNRPN,PLOD1,COL2A1,SKI,ASXL1,RPS15A,NOTCH2,SELENON,VHL,PPP2CA,MAP

35,PRKAR1A,GJA1,MYO5A,CTNNB1,COL6A2,RPS10,RPL35A,SLC25A3,ARID1A,DDOST,TSR2,HNR
OX6,PPP2R5D,CHST3,NSD1,RBM10,P3H1,ZSWIM6,SOX9,MAN2B1,LMNA,MED12,HIRA,CTCF,COX5

:OL5A2,RPL26,PFN1,RPL31,LRPPRC,KLHL9,SPTBN2,TKT,TPM1,CDH2,GPC3,MEGF8,RAB7A,CCN
PRKAR1A,GJA1,MYO5A,CTNNB1,COL6A2,MACF1,CYFIP2,HNRNPDL,TARDBP,SLC25A3,ARL6IP1,/
:TBP1,RAC1,L1CAM,GLUL,CRPPA,HS6ST1,NOVA2,RAB27A,EEF1A2,TBCD,PPP1CB,SQSTM1,VAMI
:1,SMCHD1,MMP2,L1CAM,PPIB,CRPPA,CDH11,EEF1A2,TBCD,PPP1CB,PABPN1,LMNB2,RPL27,SN
;BF1,ARF1,CYB5R3,PAFAH1B1,ITM2B,HNRNPU,LDHA,CTSD,SCARB2,RPS24,COL1A2,HK1,PTCD3
_10,MARCHF6,RPS17,AP3D1,SBF1,ARF1,CYB5R3,PAFAH1B1,ITM2B,MAT2A,HNRNPU,LDHA,CTSD
IGDH,HSD17B4,DHCR7,PCGF2,AHCY,UBE3A,GOT2,YY1,SMARCD1,TRIM8,FGFR1,PTPN23,HS2ST
PL26,RPL31,PRKCSH,LRPPRC,XPR1,CCND2,CDH2,GPC3,MEGF8,MMP14,DDX3X,CLTC,RPS7,RPS
:1,PTCD3,NR2F1,CHD4,ATP1A1,RPS20,COL5A2,RPL26,PFN1,RPL31,LRPPRC,XPR1,KLHL9,SPTBN
_NB,DLK1,TPM3,SMARCE1,RPS28,ATP6V1B2,PRPS1,COL27A1,VANGL2,RAI1,SEC23A,ARNT2,SMS
;R2,HNRNPK,NARS1,ATP1A2,ARCN1,RPS27,CTBP1,RAC1,SMCHD1,L1CAM,GLUL,CRPPA,HS6ST1,
;R2,HNRNPK,NARS1,ATP1A2,ARCN1,RPS27,CTBP1,RAC1,L1CAM,GLUL,CRPPA,NOVA2,CDH11,MD
AP3D1,SBF1,ARF1,CYB5R3,PAFAH1B1,ITM2B,MAT2A,HNRNPU,LDHA,CTSD,SCARB2,RPS24,COL1
CHD1,L1CAM,GLUL,CRPPA,NOVA2,CDH11,MDH2,EEF1A2,FDFT1,TBCD,PPP1CB,SQSTM1,VAMP2
ATRX,NXN,ARID1B,CTNND2,FBXO11,PTEN,ABCC9,SPINT2,MAPRE2,LOX,GPC4,EHMT1,HSPG2,AN
P1A1,RPS20,COL5A2,WDR1,ATP2A2,RPL26,RPL31,LRPPRC,SPTBN2,TKT,CCND2,CDH2,GPC3,ME
PS7,RPS19,RPL35,PRKAR1A,GJA1,CTNNB1,COL6A2,MACF1,RPS10,RPL35A,CYFIP2,ARID1A,TSR
P2K2,FLNB,DLK1,TPM3,SMARCE1,ATP6V1B2,LSS,STXBP1,LONP1,PRPS1,MESD,USP7,IDH1,COL
'2A2,RPL26,PFN1,PRKCSH,XPR1,SPTBN2,TPM1,CCND2,CDH2,GPC3,RAB7A,ACTN4,CLTC,RPS7,F
,ARID1B,RAB3GAP2,CRTAP,FBXO11,HDAC6,PTEN,SMC3,PTPRF,ABCC9,MAPRE2,PREPL,GPC4,N
,RAB11B,ABCA2,RPS15A,NOTCH2,PPP2CA,MAPK1,SMARCA4,PRDX1,ODC1,ACADVL,FTO,PRKDC
XL1,RAB11B,ABCA2,RPS15A,TUBB2A,PPP2CA,MAPK1,SMARCA4,PRDX1,FTO,PRKDC,SON,KDM1
_26,RPL31,XPR1,SPTBN2,CCND2,CDH2,GPC3,DDX3X,CLTC,RPS7,RPS19,RPL35,PRKAR1A,GJA1,
BN2,TKT,CCND2,CDH2,GPC3,MEGF8,MMP14,DDX3X,CLTC,RPS7,RPS19,RPL35,PRKAR1A,GJA1,
,GJA1,COL6A2,RPS10,RPL35A,CYFIP2,HNRNPDL,ARL6IP1,ARID1A,TSR2,SHOX,HNRNPK,NARS1,/
2,HK1,RHOA,NR2F1,CHD4,RPS20,CD81,COL5A2,RPL26,RPL31,PRKCSH,LRPPRC,XPR1,KLHL9,C
42,ADAR,CIC,KCNQ2,WDR45B,EP300,EXTL3,RERE,KDM5B,CNTNAP1,VARS1,FLCN,GPX4,ATRX,AI
ID4,ATP1A1,RPS20,COL5A2,RPL26,PFN1,RPL31,LRPPRC,XPR1,SPTBN2,CCND2,CDH2,GPC3,DDX
A1,MYO5A,CTNNB1,COL6A2,MACF1,RPS10,RPL35A,CYFIP2,ARID1A,DDOST,TSR2,HNRNPK,NARS
RE,KDM5B,PBX1,BAZ1B,ATRX,ARID1B,RAB3GAP2,FBXO11,MBTPS1,PTEN,SMC3,PTPRF,MAPRE2
,MED13,MTOR,PRKACA,SMARCD1,CREBBP,PHIP,TANC2,FGFR1,SH3PXD2B,DENND5A,HS2ST1,T
USP7,IDH1,COL27A1,RAI1,RAB11A,SEC23A,TCTN1,ARFGEF2,SMS,CDC42,ADAR,CIC,KIF5A,STAT
APK1,PLEC,ODC1,ATXN10,KDM1A,MAP2K2,DLK1,SMARCE1,DHX30,ATP6V1B2,SDHA,LSS,FBLN1,
H2,PPP2CA,MAPK1,SMARCA4,PLEC,ODC1,FTO,PRKDC,SON,KDM1A,MAP2K2,FLNB,DLK1,SMARC
Z,BRAT1,COL3A1,FGD1,KLHL7,HMGA2,GLB1,SIN3A
ERE,KDM5B,CNTNAP1,VARS1,FLCN,GPX4,ATRX,ARID1B,RAB3GAP2,ZFR,NDST1,EML1,MAPRE2,B
CAM,PPIB,CRPPA,HS6ST1,CDH11,FDFT1,TBCD,PPP1CB,SQSTM1,LMNB2,ABCA3,RPL27,SNRPN,F
,ARNT2,KDM3B,ARFGEF2,CDC42,ADAR,CIC,KCNQ2,WDR45B,EP300,EXTL3,RERE,NDUFV1,KDM
XL1,KDM6B,RAB11B,RPS15A,NOTCH2,SELENON,PPP2CA,MAPK1,SMARCA4,PLEC,ODC1,AEBP1,
CTNNA1,CTSD,RPS24,COL1A2,HK1,RHOA,PTCD3,NR2F1,CHD4,RPS20,COL5A2,RPL26,CLEC7A,I
J1,SEC23A,ARFGEF2,SMS,CDC42,KDM5C,TRIO,EP300,EXTL3,RERE,ALDH3A2,BAZ1B,GPX4,NXN,/
1,GLUL,CDKN1A,CRPPA,HS6ST1,NOVA2,CDH11,MDH2,RAB27A,EEF1A2,FDFT1,TBCD,SQSTM1,V/

,LSS,STT3B,STXBP1,STT3A,LONP1,MESD,USP7,SARS1,RAI1,RAB11A,ARNT2,ARFGEF2,CDC42,YN

PS24,COL1A2,HK1,RHOA,NR2F1,CHD4,RPS20,CD81,COL5A2,RPL26,CLEC7A,RPL31,PRKCSH,LR
STM1,LMNB2,ABCA3,RPL27,SNRPN,PLOD1,COL2A1,SKI,ASXL1,KDM6B,RAB11B,RPS15A,NOTCH
SDHA,LARS1,LSS,STT3B,STXBP1,STT3A,LONP1,MESD,USP7,SARS1,RAI1,RAB11A,ARNT2,ARFGEF
I,SKI,ASXL1,KDM6B,CNTNAP2,RAB11B,SELENON,TUBB2A,PPP2CA,MAPK1,SMARCA4,KDELR2,P
DFT1,TBCD,PPP1CB,RPL27,SNRPN,PLOD1,RBMX,SKI,ASXL1,KDM6B,RAB11B,RPS15A,NOTCH2,
PPP1CB,LMNB2,ABCA3,RPL27,SNRPN,PDXK,PLOD1,COL2A1,SKI,ASXL1,KDM6B,RAB11B,RPS15
PPP1CB,LMNB2,ABCA3,RPL27,SNRPN,PDXK,PLOD1,COL2A1,SKI,ASXL1,KDM6B,RAB11B,RPS15

CC9,MAPRE2,PREPL,GPC4,EFTUD2,GTF2I,TCF3,HSD17B4,NSD2,DHCR7,UBE3A,SMARCD1,CREE
27,CTBP1,SMCHD1,MMP2,L1CAM,GLUL,CRPPA,HS6ST1,MDH2,EEF1A2,FDFT1,TBCD,SQSTM1,F
NGAP1,CRKL,CHST3,WAC,ZSWIM6,LMNA,COL7A1,MED12,HIRA,CTCF,TALDO1,SEPTIN9,QRICH1,

S24,COL1A2,HK1,RHOA,NR2F1,CHD4,CD81,COL5A2,WDR1,ATP2A2,RPL26,PFN1,CLEC7A,RPL31
B,RPS15A,NOTCH2,PPP2CA,MAPK1,SMARCA4,PLEC,ODC1,SON,KDM1A,MAP2K2,QARS1,FLNB,D
RA,SOS1,ZDHHC9,TGFBR1,COL6A3,CUL7,MAN1B1,EXT1,TET3,CUL4B,PSMD12,TGFB2,SETD1A,N
,PBX1,FLCN,CSNK2A1,NXN,ARID1B,PI4KA,RAB3GAP2,FBXO11,IARS1,HNRNPA2B1,NDST1,MCM
B,RPS15A,NOTCH2,MARS1,MAPK1,SMARCA4,PLEC,ODC1,AEBP1,FTO,PRKDC,HSPB1,SON,KDM1
P,LIMK1,EXT2,DDX6,PPP2R5D,CRKL,CHST3,NSD1,RBM10,SHMT2,GDF1,BCOR,MED25,RPS29,S
1A,GJA1,CTNNB1,COL6A2,RPS10,RPL35A,CYFIP2,TARDBP,ARID1A,TSR2,SHOX,HNRNPK,NARS1,
MN1,TRIM8,FGFR1,SH3PXD2B,RPS26,ABL1,AUTS2,CLIP2,ELN,NOTCH1,ACTA2,MECP2,ADNP,TR
P,LIMK1,EXT2,DDX6,PPP2R5D,CRKL,CHST3,NSD1,RBM10,SHMT2,GDF1,BCOR,MED25,RPS29,S
S24,COL1A2,HK1,RHOA,NR2F1,CHD4,CD81,COL5A2,WDR1,ATP2A2,RPL26,PFN1,CLEC7A,RPL31
PL27,SNRPN,PLOD1,RBMX,SKI,ASXL1,KDM6B,RAB11B,RPS15A,NOTCH2,PPP2CA,MAPK1,SMAR

IO,EP300,EXTL3,RERE,KDM5B,PBX1,CNTNAP1,FLCN,BAZ1B,CSNK2A1,ATRX,NXN,ARID1B,RAB3
,GPX4,ATRX,ARID1B,RAB3GAP2,ZFR,NDST1,EML1,MAPRE2,B4GAT1,ATP6AP2,GPC4,TAOK1,EH

D2,DHCR7,MED13,DHX38,PACS2,SMARCD1,CREBBP,TANC2,FGFR1,SH3PXD2B,RPS26,HS2ST1,
PS28,ATP6V1B2,LARS1,STXBP1,COL27A1,RAI1,SEC23A,ARNT2,CDC42,CTNND1,STAT3,KDM5C,T
AP2K2,QARS1,FLNB,DLK1,TPM3,DHX30,RPS28,ATP6V1B2,LSS,STXBP1,LONP1,PRPS1,MESD,US
1,PRPS1,USP7,SARS1,RAI1,KDM3B,ARFGEF2,SMS,CDC42,YME1L1,CIC,TRAF7,KIF5A,KCNQ2,EP
7,SNRPN,PLOD1,RBMX,SKI,ASXL1,KDM6B,RAB11B,RPS15A,NOTCH2,PPP2CA,MAPK1,SMARCA4,

,GPX4,ATRX,ARID1B,RAB3GAP2,ZFR,NDST1,EML1,MAPRE2,B4GAT1,ATP6AP2,GPC4,TAOK1,EH
,CDH2,DDX3X,CLTC,RPS7,RPS19,RPL35,MIF,PRKAR1A,GJA1,MYO5A,CTNNB1,COL6A2,MACF1,F
X,ARID1B,CTNND2,RAB3GAP2,PTEN,SMC3,MCM3AP,LOX,GPC4,H19,HSPG2,ANTXR1,KMT2E,TC
UD2,GTF2I,TTC37,PACS1,LZTR1,PHGDH,NSD2,DHCR7,PCGF2,PACS2,PRKACA,YY1,CREBBP,ND
AP2K2,FLNB,DLK1,TPM3,SMARCE1,DHX30,RPS28,ATP6V1B2,STXBP1,MESD,HYOU1,RAI1,SEC23
HX30,RPS28,SDHA,STXBP1,LONP1,PRPS1,USP7,RAI1,SEC23A,TCTN1,ARNT2,CDC42,YME1L1,DC
D1,WAC,GDF1,BCOR,RPS29,LMNA,NR2F2,MED12,HIRA,DGCR8,CTCF,TALDO1,TGFBR1,PSMD1
28,ATP6V1B2,LARS1,STXBP1,LONP1,USP7,HYOU1,RAI1,TCTN1,SMS,CDC42,SYP,BCR,KDM5C,T
,ATP6V1B2,LARS1,FBLN1,STXBP1,LONP1,PRPS1,USP7,SARS1,SOD1,RAI1,SEC23A,TCTN1,SMS,
KIF5A,KCNQ2,WDR45B,KDM5C,TRIO,EP300,EXTL3,RERE,KDM5B,VARS1,BAZ1B,GARS1,MBTPS

2,RPS10,RPL35A,CYFIP2,ARID1A,TSR2,SHOX,HNRNPK,NARS1,ATP1A2,HLA-B,ARCN1,RPS27,CTE
2,RPS10,RPL35A,CYFIP2,ARID1A,TSR2,SHOX,HNRNPK,NARS1,ATP1A2,HLA-B,ARCN1,RPS27,CTE
)1,COL2A1,SKI,ASXL1,RAB11B,RPS15A,NOTCH2,SELENON,VHL,MAPK1,SMARCA4,KDELR2,PLEC
TAT3,TRIO,EP300,EXTL3,RERE,KDM5B,CNTNAP1,FLCN,GPX4,ATRX,NXN,ARID1B,PI4KA,RAB3GA
1,COL27A1,VANGL2,RAI1,SEC23A,TCTN1,SMS,CDC42,SYP,STAT3,BCR,KDM5C,TRIO,EP300,EXTL
GLUL,CRPPA,HS6ST1,NOVA2,MDH2,EEF1A2,TBCD,SQSTM1,VAMP2,PABPN1,LMNB2,SNRPN,PDXK
11,TNPO3,PSMD12,NFASC,SYT1,DYNC1I2,NBEA,BRAT1,MAPK8IP3,HARS1,PRMT7,DPF2,TCF20
NQ2,BCR,TRIO,EP300,EXTL3,RERE,STARD7,NDUFV1,KDM5B,FLCN,BAZ1B,UBQLN2,CTNND2,RA

,NFASC,SYT1,DYNC1I2,PRNP,SMARCC2,TMCO1,NBEA,POGZ,DNM1,GNAO1,NFIB,JRK,UBTF,DMF
3XO11,IARS1,NDST1,MCM3AP,MAPRE2,TIMM50,ATP6AP2,SLC2A1,NDN,EHMT1,LINGO1,TPP1,SL

R1,ATP2A2,RPL26,PFN1,CLEC7A,RPL31,LRPPRC,SPTBN2,TKT,CCND2,CDH2,GPC3,MEGF8,CCN2
3GAP2,IARS1,ZFR,SMC3,NDST1,EML1,B4GAT1,TIMM50,ATP6AP2,SLC2A1,FUS,SPG7,TAOK1,VAI
REBBP,MN1,TANC2,TRIM8,SMO,FGFR1,SH3PXD2B,RPS26,MDH1,TRIP12,PSAT1,TMEM94,POR,E
K1,DAG1,TIA1,OBSL1,ATP13A2,MFN2,EMC1,NDUFC2,BSCL2,ALDH18A1,CAPN1,SLC25A1,ASAH1
ND1,CIC,TRAF7,STAT3,BCR,TRIO,EP300,EXTL3,RERE,KDM5B,PBX1,FLCN,BAZ1B,ATRX,ARID1B,C
/ANGL2,RAI1,SEC23A,TCTN1,SMS,ADAR,STAT3,TRIO,EP300,EXTL3,RERE,KDM5B,CNTNAP1,VARS
2A1,SKI,ASXL1,RPS15A,NOTCH2,SELENON,CCND1,MAPK1,SMARCA4,KDELR2,PLEC,SON,KDM1
K1,ELN,MECP2,WDR26,PARK7,ADNP,TRRAP,NDUFC2,ALDH18A1,LIMK1,ZNF462,IQSEC1,CSNK2

1,SLC6A8,LTBP4,KMT2E,HSD17B4,MED13,AHCY,FBLN5,PACS2,UBE3A,YY1,SMARCD1,CREBBP,I
AP2,FBXO11,HDAC6,PTEN,SMC3,SPINT2,MAPRE2,PREPL,B4GAT1,ATP6AP2,GPC4,HSPG2,ANTXF
CO2,CHD2,SKI,ASXL1,KDM6B,CNTNAP2,RAB11B,ABCA2,MARS1,SELENON,TUBB2A,PPP2CA,MAI
1,PRPS1,USP7,RAI1,SEC23A,TCTN1,ARNT2,CDC42,YME1L1,KDM5C,TRIO,EP300,RERE,NDUFV1,I

N1,STXBP1,LONP1,USP7,IDH1,RAI1,FLNC,SEC23A,KDM3B,ARFGEF2,SMS,CDC42,ADAR,KIF5A,ST
TNND2,RAB3GAP2,FBXO11,MBTPS1,HDAC6,PTEN,SMC3,SPINT2,MAPRE2,PREPL,B4GAT1,ATP6AI
H11,EEF1A2,TBCD,PPP1CB,VAMP2,LMNB2,SNRPN,YEATS2,ACO2,COL2A1,RBMX,CHD2,SKI,ASXL
,SNRPN,ACO2,VPS35,CHD2,SKI,KDM6B,CNTNAP2,RAB11B,ABCA2,TERF2IP,MARS1,OPTN,VHL,M
1,FGD1,EMD

2,SLC2A1,FUS,SPG7,TAOK1,VAPB,LINGO1,SOX2,SLC6A8,IREB2,GTF2I,CNP,ATXN3,PHGDH,REEI
N1,CCDC47,FGFR1,TUBB4A,SH3PXD2B,RPS26,BGN,PTPN11,AUTS2,POR,SC5D,ELN,OBSL1,MECI
3,LINGO1,LTBP4,GTF2I,ATXN3,ELP1,PACS1,PHGDH,NSD2,DHCR7,PCGF2,UBE3A,YY1,CREBBP,W
=462,EXT2,ECHS1,PPP2R5D,CRKL,CHST3,NSD1,WAC,GDF1,BCOR,MED25,RPS29,SETD5,NDUFS1
31,NECTIN1,DDX6,CRKL,CHST3,NSD1,P3H1,ZEB1,GDF1,CLCN4,BCOR,MED25,MAN2B1,MED12,I
(1,KLC2,DDX6,SLC25A1,RAB18,GBA2,NDRG1,CHST3,NSD1,P3H1,TMEM43,BCOR,SOX9,MED25,
TIN1,DDX6,CRKL,CHST3,NSD1,P3H1,ZEB1,GDF1,CLCN4,BCOR,MED25,MAN2B1,MED12,HIRA,DG
1,PPP2R5D,CRKL,CHST3,NSD1,WAC,GDF1,BCOR,MED25,RPS29,SETD5,NDUFS1,SNRPB,UQCRF
N,SPG7,TAOK1,EHMT1,VAPB,IARS2,TPP1,KMT2E,GTF2I,TUBGCP6,ATXN3,HSD17B4,MED13,CIZ1
/OU1,VANGL2,RAI1,SYP,BCR,KDM5C,TRIO,EP300,EXTL3,RERE,CNTNAP1,VARS1,FLCN,BAZ1B,CS
ARS2,HSPG2,SLC6A8,IREB2,KMT2E,CNP,ATXN3,PACS1,LZTR1,NSD2,DHCR7,MED13,YY1,SMARC
N4,NUS1,ASH1L,ACSL4,CACNA1H,MED25,SETD5,MBOAT7,MED12,HIRA,RUSC2,PPP1R12A,ZDHH

,TET3,CUL4B,TGFB2,CCNK,TMCO1,FGD1
ED25,SETD5,RHOBTB2,MED12,CTCF,GALNT2,QRICH1,MAN1B1,EXT1,GATAD2B,PSMD12,SETD1
PACS1,LZTR1,HSD17B4,NSD2,DHCR7,RBPJ,MED13,PCGF2,MTOR,PACS2,UBE3A,YY1,CREBBP,M

2,DHCR7,PCGF2,CIZ1,SMARCD1,CREBBP,FGFR1,TUBB4A,RPS26,HS2ST1,PTPN11,PSAT1,COL12

)H2,GPC3,MEGF8,MMP14,DDX3X,CLTC,RPS7,RPS19,RPL35,MIF,PRKAR1A,GJA1,MYO5A,CTNNB
FLCN,BAZ1B,CSNK2A1,ATRX,NXN,ARID1B,CTNND2,PI4KA,RAB3GAP2,HDAC6,PTEN,SMC3,ECEL1
R,SMARCD1,TANC2,TRIM8,FGFR1,EIF4A3,SH3PXD2B,RPS26,HS2ST1,BGN,PTPN11,MACROH2A1
2,WDR45B,KDM5C,EXTL3,RERE,NDUFV1,KDM5B,CNTNAP1,ALDH3A2,BAZ1B,UBQLN2,ATRX,PI4K
ONP,TRRAP,BSCL2,ALDH18A1,LIMK1,EFNB1,NECTIN1,DDX6,CRKL,CHST3,NSD1,P3H1,ZEB1,GDF
3,RERE,DGCR2,CNTNAP1,VARS1,FLCN,BAZ1B,CSNK2A1,ATRX,NXN,ARID1B,CTNND2,PI4KA,RAB3
LIP2,SC5D,ELN,NOTCH1,ACTA2,WDR26,TRRAP,ALDH18A1,BANF1,HTRA1,LIMK1,PDGFRB,DDX6,

MAN2B1,SRP54,MED12,HIRA,PPP1R12A,ZDHHC9,PPP3CA,MAN1B1,TET3,PSMD12,MSH2,SETD1A
LCN7,EHMT1,SOX2,LTBP4,LDLR,KMT2E,GTF2I,TTC37,PACS1,LZTR1,PHGDH,DHCR7,RBPJ,MED1
X,ARID1B,PI4KA,RAB3GAP2,IARS1,SMC3,NDST1,PREPL,NDN,FUS,H19,TAOK1,EHMT1,HSPG2,LIN
AP2,HDAC6,PTEN,SMC3,B4GAT1,ATP6AP2,GPC4,BAP1,FUS,GRN,SPG7,CLCN7,EHMT1,TPP1,SOX2
RGIC1,HSPG2,PKDCC,HINT1,KMT2E,LTBP3,PHGDH,NSD2,DHCR7,RBPJ,MED13,PCGF2,MTOR,PF
NOTCH2,SELENON,VHL,CCND1,MAPK1,SMARCA4,KDELR2,PLEC,AEBP1,SON,KDM1A,FLNB,DLK1
3,BCOR,SOX9,MED25,SETD5,MAN2B1,LMNA,MED12,SLC5A7,ARSB,SOS1,ZDHHC9,TGFBR1,G6F
I,L1CAM,DCN,HS6ST1,CDH11,MDH2,EEF1A2,TBCD,SQSTM1,PABPN1,TGFBI,RPL27,SNRPN,ACO2

,PKDCC,GTF2I,NSD2,DHCR7,FBLN5,UBE3A,PRKACA,YY1,SMARCD1,CREBBP,WNK1,CCDC47,REE
,TRRAP,LIMK1,EXT2,DDX6,PPP2R5D,CRKL,CHST3,NSD1,RBM10,SHMT2,GDF1,BCOR,MED25,RF
A,NOTCH2,MAPK1,SMARCA4,PLEC,ODC1,AEBP1,FTO,SON,KDM1A,MAP2K2,FLNB,DLK1,SMARCE

D12,SETD1A,COL3A1,GLB1,SIN3A
AP2,HDAC6,PTEN,SMC3,B4GAT1,ATP6AP2,GPC4,BAP1,FUS,GRN,SPG7,CLCN7,EHMT1,TPP1,SOX2
AP2,HDAC6,PTEN,SMC3,B4GAT1,ATP6AP2,GPC4,BAP1,FUS,GRN,SPG7,CLCN7,EHMT1,TPP1,SOX2
N4,NUS1,ASH1L,SETD5,CDKN1B,SNRPB,COL7A1,MED12,SLC5A7,HIRA,CTCF,EPB41L1,PPP3CA,S

ET3,PSMD12,PIGT,CCNK,BRAT1,MAPK8IP3,NFIB,FGD1,PRMT7,GLB1,SIN3A,CASK,CNOT1,TCF20
D1,FGD1,KLHL7,MAFB,DMPK,SIN3A,TCF20
DHX30,RPS28,ATP6V1B2,SDHA,LSS,STXBP1,MESD,RAI1,SEC23A,TCTN1,SMS,CDC42,CTNND1,STA

6,SYNGAP1,RDH11,PPP2R5D,TSPAN7,CHST3,NSD1,WAC,ZSWIM6,CLCN4,NUS1,ACSL4,MED25,SE
3,RPL27,SNRPN,PLOD1,VPS35,COL2A1,RBMX,SKI,ASXL1,KDM6B,PRPF6,RPS15A,NOTCH2,PPP2C

17B4,DHCR7,MED13,CIZ1,MTOR,UBE3A,MN1,TANC2,TRIM8,SMO,FGFR1,EIF4G1,TECR,PSAT1,C

X1,BAZ1B,ATRX,ARID1B,SMC3,HNRNPA2B1,MAPRE2,TIMM50,ATP6AP2,GPC4,SLC2A1,BAP1,FUS

2A1,CLIP2,DAG1,ELN,EMC1,WDR26,ADNP,GPAA1,ALDH18A1,LIMK1,FLII,NBAS,PDGFRB,TMEM9
UBE3A,GOT2,TANC2,TRIM8,CCDC47,FGFR1,EIF4G1,NTRK2,CAD,TECR,CLIP2,TRAK1,SC5D,ELN,
N,SMC3,EML1,B4GAT1,ATP6AP2,GPC4,SLC2A1,NDN,BAP1,GRN,CLCN7,TAOK1,EHMT1,ANTXR1,

CSNK2B,SLC25A1,ASAH1,SLC16A2,SYNGAP1,PPP2R5D,DEAF1,NSD1,WAC,ZSWIM6,SHMT2,CLC

T1,DNM1,GNAO1,UBTF,CASK

CNK,POGZ,PRMT7,GLB1,CASK,TCF20
7,AHCY,MTOR,YY1,SMARCD1,SMO,FGFR1,PTPN23,DENND5A,BGN,PSAT1,DAG1,PDHA1,ACTA2,

2,PDHA1,IGF1R,MECP2,NDUFA4,PARK7,ADNP,LIMK1,FLII,PRPH,IQSEC1,PDGFRB,NECTIN1,ECHS
BCOR,MED25,RPS29,SETD5,LMNA,CNTN1,COL7A1,MED12,COX4I1,CUL7,PIGS,PSMD12,COG4,
,CASK
SLC5A7,HIRA,DSTYK,TGFBR1,THRA,COL6A3,CUL7,EXT1,TARS1,FHL1,SEC31A,SMARCC2,COL3

3,RPS26,BGN,PUM1,MACROH2A1,OBSL1,NOTCH1,IGF1R,MECP2,FAM50A,ADNP,ALDH18A1,EXT
OBSL1,NOTCH1,IGF1R,MECP2,FAM50A,ADNP,ALDH18A1,EXT2,ASAH1,CRKL,TSPAN7,CHST3,P3

P3,NSD2,DHCR7,RBPJ,MED13,PCGF2,MTOR,SMARCD1,TANC2,TRIM8,CCDC47,FGFR1,EIF4A3,

H2,PTPN12,SMARCB1,NF2,ELMO2,RAD50,DMPK
ECAP1,TCF20
TPM3,SMARCE1,RPS28,ATP6V1B2,SDHA,LSS,STXBP1,LONP1,PRPS1,MESD,USP7,IDH1,HYOU1,C
MAN1B1,CUL4B,FAM20C,PSMD12,FHL1,TMCO1,POGZ,FGD1,PRMT7,MAFB,GLB1,EMD

1A,GJA1,CTNNB1,COL6A2,MACF1,RPS10,RPL35A,CYFIP2,SLC25A3,ARID1A,DDOST,TSR2,SHOX,

L,PPP2CA,MAPK1,PLEC,ODC1,ADAM9,ATXN10,SON,DLK1,SMARCE1,DHX30,SDHA,LSS,FBLN1,

7,DPF2
,MED25,SETD5,MBOAT7,CHD8,NACC1,LARGE1,SLC5A7,CTCF,GALNT2,EPB41L1,PPP3CA,TRAF3

ELN,ATP13A2,PDHA1,MECP2,EMC1,ADNP,TRRAP,BSCL2,ALDH18A1,LIMK1,AMPD2,IQSEC1,MOG

AG1,NOTCH1,MFN2,PDHA1,MECP2,EMC1,GPAA1,BSCL2,FLII,IQSEC1,CCDC88A,ASNS,NECTIN1,

1,HSPG2,ANTXR1,PKDCC,LTBP4,KMT2E,EFTUD2,GTF2I,LZTR1,PHGDH,HSD17B4,NSD2,DHCR7,

E2,ATP6AP2,ERGIC1,HSPG2,ANTXR1,PKDCC,LTBP4,KMT2E,EFTUD2,GTF2I,LZTR1,PHGDH,HSD1

D,HYOU1,VANGL2,RAI1,SEC23A,ARNT2,SMS,BCR,KDM5C,TRIO,EP300,EXTL3,RERE,DGCR2,CN

XN,ARID1B,RAB3GAP2,HDAC6,PTEN,SMC3,ECEL1,PTPRF,ABCC9,GPC4,EHMT1,HSPG2,ANTXR1

,TANC2,FGFR1,SH3PXD2B,NTRK2,HS2ST1,BGN,TRIP12,PTPN11,AUTS2,TRAK1,SC5D,IGF1R,ME

8,SMO,FGFR1,EIF4A3,RPS26,HS2ST1,BGN,PTPN11,PSAT1,DAG1,FAM50A,WDR26,SUMO1,TRRA

1,HSPG2,ANTXR1,PKDCC,LTBP4,KMT2E,EFTUD2,GTF2I,LZTR1,PHGDH,HSD17B4,NSD2,DHCR7,

PLXND1,FGD1,HMGA2,MAFB,DPF2,SIN3A,CASK

8,SMO,FGFR1,EIF4A3,RPS26,HS2ST1,BGN,PTPN11,PSAT1,DAG1,FAM50A,WDR26,SUMO1,TRRA

12,HIRA,SOS1,ROBO3,TALDO1,COL6A3,CUL7,MAN1B1,CUL4B,FAM20C,PSMD12,FHL1,TMCO1

ARCD1,CREBBP,TANC2,WNK1,TRIM8,CCDC47,REEP1,SMO,FGFR1,EIF4A3,HS2ST1,TRIP12,TM

IARCC2,POGZ,BRAT1,COL3A1,MAPK8IP3,HMGA2,PRMT7,DPF2,SIN3A,TCF20

MECP2,EMC1,ADNP,TRRAP,BSCL2,ALDH18A1,LIMK1,AMPD2,IQSEC1,MOGS,CSNK2B,DDX6,ALD

Z1B,UBQLN2,ATRX,ARID1B,PI4KA,ADCY6,IARS1,PTEN,SMC3,LMAN1,FUS,SPG7,TAOK1,EHMT1,

,PSMD12,THG1L,POGZ,SETD1B,FGD1,HMGA2,PRMT7,GLB1,DMPK,SIN3A,DNAJC21,CC2D1A,T

K1,EFNB1,EXT2,ASAH1,SLC16A2,CRKL,CHST3,NSD1,TMEM43,BCOR,MED25,SETD5,SNRPB,LA

1,KLC2,ASNS,ECHS1,DDX6,DARS1,SLC25A1,SLC16A2,SYNGAP1,NDRG1,DEGS1,GPT2,DHFR,ND

T3,EXTL3,NDUFV1,CNTNAP1,UBQLN2,GARS1,PI4KA,ADCY6,PTEN,HNRNPA2B1,MCM3AP,PREPL,

A1,ALDH7A1,SYNGAP1,STUB1,DEAF1,NDRG1,CLCN4,NUS1,CACNA1H,ARHGEF9,MED25,MBOAT

,PTEN,SMC3,ECEL1,ABCC9,MAPRE2,ATP6AP2,ERGIC1,HSPG2,ANTXR1,SOX2,PKDCC,LTBP4,KM

PS7,RPS19,RPL35,MIF,PRKAR1A,GJA1,MYO5A,CTNNB1,COL6A2,RPS10,RPL35A,CYFIP2,SLC25A
P1,DEAF1,CLCN4,NUS1,CACNA1H,ARHGEF9,MED25,MBOAT7,NACC1,LARGE1,XPC,NUP133,SLC5
2,DHCR7,SMARCD1,CREBBP,TRIM8,SMO,FGFR1,EIF4A3,RPS26,HS2ST1,BGN,PTPN11,PSAT1,DA

G6PC3,COL6A3,PSMD12,PLXND1,MAFB,SIN3A,CNOT1,TCF20
LC25A1,SLC16A2,SYNGAP1,DEAF1,DEGS1,GPT2,NSD1,NDUFB8,ASH1L,BCOR,SETD5,FARSA,NAC

,NECAP1,ARSA,ABCA7

INA,CNTN1,MED12,SOS1,CITED2,CTCF,CNOT3,CUL7,MAN1B1,PIGS,CUL4B,BRAT1,MAPK8IP3,PL

M94,CLIP2,ELN,NOTCH1,PDHA1,IGF1R,NDUFA4,WDR26,NDUFC2,BSCL2,LIMK1,ZNF462,EXT2,E
COX4I1,ZDHHC9,TGFBR1,CNOT3,CUL7,MAN1B1,PIGS,CUL4B,SMARCC2,TMCO1,BRAT1,MAPK8

SL1,PDHA1,IGF1R,MECP2,WDR26,ADNP,TRRAP,GPAA1,ZNF462,FLII,VPS51,EXT2,RAB18,DEAF1,

NSD1,CLCN4,ACSL4,CACNA1H,SETX,SETD5,STAT1,COL7A1,MED12,HIRA,DDHD2,DSTYK,NIPA1,G

·B,THG1L,SMARCC2,CENPF,TMCO1,COL3A1,SMARCB1,SLC18A3,KLHL7,HMGA2,DPF2
·18,DEAF1,CRKL,TSPAN7,RBM10,GDF1,CLCN4,ACSL4,BCOR,RNF216,MED25,SETD5,SNRPB,NSN
CDC47,SMO,FGFR1,EIF4A3,SH3PXD2B,BGN,COL12A1,POR,CLIP2,ELN,OBSL1,WDR26,ALDH18A

KCSH,LRPPRC,XPR1,TKT,GPC3,MEGF8,CCN2,MMP14,CLTC,RPS7,RPS19,RPL35,MIF,PRKAR1A,(

D,ELN,OBSL1,PDHA1,IGF1R,MECP2,WDR26,ADNP,TRRAP,GPAA1,ZNF462,FLII,VPS51,EXT2,RAB1

SIN3A
SIN3A
RK,CSTB,TOMM40,CASK,NECAP1,ABCA7

PB,LMNA,CNTN1,MED12,SOS1,CITED2,CTCF,CNOT3,CUL7,MAN1B1,PIGS,CUL4B,BRAT1,MAPK8I

SMD12,THG1L,PIGT,POGZ,KLHL7,EPHA2,DNAJC21,CASK,TCF20
UBE3A,PRKACA,YY1,WNK1,CCDC47,REEP1,HS2ST1,PSAT1,COL12A1,CLIP2,DAG1,NOP56,ELN,TI

ISD2,DHCR7,RBPJ,MED13,PCGF2,MTOR,SMARCD1,CREBBP,TANC2,WNK1,SMO,FGFR1,EIF4A3,S

P1,STT3A,LONP1,USP7,SEC23A,ARNT2,KDM3B,SMS,CDC42,ADAR,KDM5C,EP300,RERE,DGCR2,K
XL1,ABCA2,PRPF6,RPS15A,NOTCH2,VHL,PPP2CA,MAPK1,SMARCA4,PLEC,AEBP1,MAP2K2,FLNB,S

3,RPL27,SNRPN,COL2A1,SKI,ASXL1,RPS15A,NOTCH2,SELENON,MAPK1,SMARCA4,KDELR2,PLEC

GLB1,CASK,CC2D1A
POMGNT1,THG1L,PIGT,SMARCC2,CENPF,TMCO1,COL3A1,MAPK8IP3,SMARCB1,SLC18A3,PDHB,
L12A1,OBSL1,WDR26,ALDH18A1,COX8A,DDX6,SLC25A1,PPP2R5D,CHST3,SOX9,SETD5,MAN2B1

A1,SOD1,FLNC,ARNT2,ARFGEF2,ADAR,KIF5A,STAT3,BCR,KDM5C,EP300,EXTL3,RERE,CNTNAP1,

MED12,HIRA,PPP1R12A,CUL7,GATAD2B,CUL4B,PSMD12,COL3A1,HMGA2,SIN3A
NFIB,JRK,DMPK,TCF20

RIM8,FGFR1,SH3PXD2B,HS2ST1,TUB,AUTS2,TMEM94,SC5D,ELN,OBSL1,PDHA1,IGF1R,MECP2,V
STT3B,STXBP1,STT3A,LONP1,USP7,SEC23A,ARNT2,KDM3B,SMS,CDC42,ADAR,KDM5C,EP300,RE

ST,MAPRE2,LOX,GPC4,NDN,H19,CLCN7,EHMT1,HSPG2,ANTXR1,KMT2E,GTF2I,PACS1,LTBP3,NSD
,TAF15,LMNB2,ABCA3,SNRPN,PLOD1,VPS35,COL2A1,SKI,ASXL1,RAB11B,MARS1,SELENON,OPT

LII,NBAS,NECTIN1,DDX6,DEAF1,CRKL,CHST3,RBM10,WAC,ZSWIM6,SHMT2,BCOR,MED25,SETD5
PK1,SMARCA4,AEBP1,ACADVL,FTO,SON,KDM1A,MAP2K2,FLNB,DLK1,TPM3,SMARCE1,RPS28,SD

NPK,HLA-B,RPS27,CTBP1,GPC1,MMP2,GLUL,CDKN1A,HS6ST1,CTSB,CDH11,MDH2,RAB27A,FDF
5A,TALDO1,TGFBR1,PPP3CA,THRA,TARS1,FAM20C,SETD1A,PIGT,COG4,TMCO1,KLHL7,DPF2

N2,DDX3X,CLTC,RPS7,RPS19,RPL35,PRKAR1A,GJA1,MYO5A,CTNNB1,COL6A2,MACF1,RPS10,RP
ARID1A,DDOST,HNRNPK,NARS1,DUX4,ATP1A2,HLA-B,TTR,CTBP1,RAC1,SMCHD1,L1CAM,GLUL,C
P2,GNAI3,TAF15,SNRPN,ACO2,PLOD1,COL2A1,SKI,ASXL1,KDM6B,CNTNAP2,RAB11B,MARS1,SEL
NRPN,PLOD1,COL2A1,SKI,ASXL1,RPS15A,NOTCH2,SELENON,VHL,CCND1,MAPK1,SMARCA4,KDE
,NR2F1,CHD4,ATP1A1,RPS20,COL5A2,WDR1,ATP2A2,RPL26,PFN1,CLEC7A,RPL31,LRPPRC,XPR1
),SCARB2,RPS24,COL1A2,HK1,PTCD3,NR2F1,CHD4,ATP1A1,RPS20,COL5A2,WDR1,ATP2A2,RPL2
1,TECR,PSAT1,CHMP1A,DAG1,ATP13A2,PDHA1,IGF1R,EMC1,WDR26,ADNP,TRRAP,AMPD2,ZNF4
19,RPL35,PRKAR1A,GJA1,CTNNB1,COL6A2,MACF1,RPS10,RPL35A,CYFIP2,ARID1A,TSR2,SHOX,
N2,CCND2,CDH2,GPC3,RAB7A,DDX3X,CLTC,RPS7,RPS19,RPL35,PRKAR1A,GJA1,MYO5A,CTNNB1
S,CDC42,CTNND1,TRAF7,STAT3,BCR,KDM5C,TRIO,EP300,EXTL3,RERE,FLCN,CSNK2A1,ATRX,NX
NOVA2,CDH11,MDH2,EEF1A2,FDFT1,TBCD,SQSTM1,VAMP2,LMNB2,RPL27,SNRPN,ACO2,VPS35
DH2,EEF1A2,FDFT1,TBCD,SQSTM1,VAMP2,LMNB2,RPL27,SNRPN,ACO2,VPS35,COL2A1,SKI,ASXL
1A2,HK1,RHOA,NR2F1,CHD4,ATP1A1,RPS20,CD81,COL5A2,RPL26,PFN1,CLEC7A,RPL31,PRKCSH
,LMNB2,RPL27,SNRPN,ACO2,VPS35,SKI,ASXL1,RAB11B,ABCA2,RPS15A,TUBB2A,PPP2CA,MAPK
NTXR1,LTBP4,AKT3,KMT2E,TTC37,PACS1,DICER1,TCF3,LZTR1,PHGDH,HSD17B4,NSD2,OGT,DH
EGF8,MMP14,DDX3X,CLTC,RPS7,RPS19,RPL35,PRKAR1A,GJA1,MYO5A,CTNNB1,COL6A2,MACF1
2,SHOX,HNRNPK,NARS1,ARCN1,RPS27,CTBP1,RAC1,SMCHD1,MMP2,L1CAM,PPIB,CRPPA,NOVA
27A1,RAI1,RAB11A,SEC23A,TCTN1,ARFGEF2,SMS,CDC42,ADAR,CIC,KIF5A,STAT3,WDR45B,BCR
RPS19,RPL35,PRKAR1A,GJA1,CTNNB1,COL6A2,MACF1,RPS10,RPL35A,CYFIP2,DNAJC5,HNRNPD
IDN,EHMT1,IARS2,HSPG2,ANTXR1,PKDCC,LTBP4,IREB2,KMT2E,EFTUD2,GTF2I,TTC37,PACS1,LT
C,SON,KDM1A,MAP2K2,QARS1,DLK1,SMARCE1,DHX30,RPS28,SDHA,LARS1,LSS,STT3B,STXBP1,S
1A,MAP2K2,QARS1,DLK1,SMARCE1,DHX30,RPS28,LARS1,LSS,STT3B,STXBP1,STT3A,LONP1,PRP
MYO5A,CTNNB1,MACF1,RPS10,RPL35A,CYFIP2,DNAJC5,TARDBP,ARID1A,TSR2,HNRNPK,NARS1,
MYO5A,CTNNB1,MACF1,RPS10,RPL35A,CYFIP2,DNAJC5,TARDBP,ARID1A,DDOST,TSR2,HNRNPK
ATP1A2,ARCN1,RPS27,CTBP1,RAC1,SMCHD1,MMP2,L1CAM,PPIB,GLUL,CRPPA,HS6ST1,CTSB,CD
CND2,CDH2,GPC3,MEGF8,RAB7A,CCN2,MMP14,DDX3X,CLTC,RPS7,RPS19,RPL35,MIF,PRKAR1A
RID1B,RAB3GAP2,ZFR,NDST1,EML1,MAPRE2,B4GAT1,TIMM50,ATP6AP2,GPC4,SLC2A1,GRN,SP
X3X,CLTC,RPS7,RPS19,RPL35,PRKAR1A,GJA1,MYO5A,CTNNB1,MACF1,RPS10,RPL35A,CYFIP2,DN
S1,ARCN1,RPS27,CTBP1,RAC1,L1CAM,GLUL,CRPPA,NOVA2,CDH11,MDH2,EEF1A2,FDFT1,TBCD,
2,B4GAT1,GPC4,SLC2A1,H19,EHMT1,HSPG2,ANTXR1,SLC6A8,KMT2E,EFTUD2,GTF2I,TTC37,TUE
RIP12,PSAT1,AUTS2,TMEM94,POR,CLIP2,SC5D,ELN,OBSL1,PDHA1,IGF1R,MECP2,EMC1,ADNP,T
3,WDR45B,TRIO,EP300,EXTL3,RERE,KDM5B,ALDH3A2,FLCN,BAZ1B,CSNK2A1,ATRX,GARS1,NXN
,STXBP1,PRPS1,MESD,USP7,SARS1,SOD1,RAI1,RAB11A,SEC23A,KDM3B,SMS,CDC42,YME1L1,AD
CE1,RPS28,ATP6V1B2,LSS,STXBP1,LONP1,PRPS1,MESD,HYOU1,COL27A1,VANGL2,RAI1,SEC23A

4GAT1,ATP6AP2,GPC4,TAOK1,EHMT1,SOX2,SLC6A8,AKT3,KMT2E,CNP,PHGDH,HSD17B4,NSD2,
PDXK,PLOD1,COL2A1,SKI,ASXL1,KDM6B,RAB11B,RPS15A,NOTCH2,MARS1,SELENON,MAPK1,SM
5B,CNTNAP1,VARS1,FLCN,GPX4,ATRX,ARID1B,RAB3GAP2,ZFR,NDST1,EML1,MAPRE2,B4GAT1,T
PRKDC,HSPB1,SON,KDM1A,MAP2K2,FLNB,DLK1,TPM3,SMARCE1,RPS28,ATP6V1B2,LARS1,LSS,
RPL31,LRPPRC,XPR1,SPTBN2,TKT,CCND2,CDH2,GPC3,MEGF8,MMP14,DDX3X,CLTC,RPS7,RPS1
ARID1B,PI4KA,RAB3GAP2,CRTAP,MBTPS1,HDAC6,PTEN,SMC3,ABCC9,MAPRE2,LOX,GPC4,SLC2
AMP2,LMNB2,SNRPN,YEATS2,ACO2,CHD2,SKI,ASXL1,KDM6B,CNTNAP2,RAB11B,ABCA2,ATP6AP1

ME1L1,ADAR,CIC,KIF5A,MSMO1,WDR45B,BCR,KDM5C,TRIO,EP300,EXTL3,RERE,DGCR2,NDUFV

:PPRC,XPR1,KLHL9,CCND2,CDH2,GPC3,MEGF8,RAB7A,CCN2,MMP14,DDX3X,CLTC,RPS7,RPS19
I2,SELENON,PPP2CA,MAPK1,SMARCA4,KDELR2,PLEC,ODC1,AEBP1,PRKDC,HSPB1,SON,KDM1A,
2,CDC42,YME1L1,ADAR,CIC,KIF5A,MSMO1,WDR45B,BCR,KDM5C,TRIO,EP300,EXTL3,RERE,DG(
RDX1,ODC1,AEBP1,ACADVL,ATXN10,SON,KDM1A,MAP2K2,QARS1,DLK1,TPM3,SMARCE1,DHX30
VHL,PPP2CA,CCND1,MAPK1,SMARCA4,PLEC,ODC1,AEBP1,SON,KDM1A,MAP2K2,QARS1,FLNB,DI
A,NOTCH2,MARS1,MAPK1,SMARCA4,KDELR2,PLEC,ODC1,AEBP1,FTO,PRKDC,HSPB1,SON,KDM1
A,NOTCH2,MARS1,MAPK1,SMARCA4,KDELR2,PLEC,ODC1,AEBP1,FTO,PRKDC,HSPB1,SON,KDM1

3BP,PHIP,FGFR1,RPS26,HS2ST1,TRIP12,PTPN11,AUTS2,CLIP2,SC5D,ELN,MECP2,TRRAP,LIMK1,Z
PABPN1,TAF15,LMNB2,RPL27,SNRPN,PLOD1,VPS35,COL2A1,SKI,ASXL1,RAB11B,RPS15A,MARS1
GATAD2B,CUL4B,FAM20C,PSMD12,SETD1A,SMARCC2,CENPF,TMCO1,POGZ,COL3A1,MAPK8IP3

,LRPPRC,XPR1,SPTBN2,TKT,CCND2,CDH2,GPC3,MEGF8,CCN2,MMP14,DDX3X,CLTC,RPS7,RPS1
LK1,SMARCE1,DHX30,RPS28,ATP6V1B2,LSS,STXBP1,LONP1,RAI1,SMS,CDC42,ADAR,SYP,CTNNI
NFASC,PIGT,SMARCC2,TMCO1,COL3A1,SMARCB1,FGD1,DPF2,SIN3A,DNAJC21,TCF20
3AP,MAPRE2,TIMM50,ATP6AP2,SLC2A1,NDN,GRN,EHMT1,LINGO1,TPP1,SLC6A8,LTBP4,IREB2,K
A,FLNB,DLK1,SMARCE1,RPS28,ATP6V1B2,LARS1,LSS,FBLN1,STXBP1,LONP1,USP7,HYOU1,SOD
ETD5,SNRPB,STAT1,LMNA,NR2F2,MED12,HIRA,SKIV2L,CITED2,CTCF,ZDHHC9,TALDO1,TGFBR1
,ATP1A2,HLA-B,ARCN1,RPS27,CTBP1,RAC1,SMCHD1,MMP2,PPIB,GLUL,CDKN1A,CRPPA,HS6ST1
RAP,BSCL2,ALDH18A1,LIMK1,EFNB1,NECTIN1,DDX6,CRKL,CHST3,NSD1,P3H1,ZEB1,GDF1,CLCI
ETD5,SNRPB,STAT1,LMNA,NR2F2,MED12,HIRA,SKIV2L,CITED2,CTCF,ZDHHC9,TALDO1,TGFBR1
,LRPPRC,XPR1,SPTBN2,TKT,CCND2,CDH2,GPC3,MEGF8,CCN2,MMP14,DDX3X,CLTC,RPS7,RPS1
CA4,PLEC,ODC1,AEBP1,SON,KDM1A,MAP2K2,QARS1,FLNB,DLK1,SMARCE1,DHX30,RPS28,ATP6

GAP2,CRTAP,FBXO11,HDAC6,PTEN,SMC3,PTPRF,ABCC9,MAPRE2,PREPL,GPC4,NDN,EHMT1,IARS
MT1,SOX2,SLC6A8,AKT3,CNP,PHGDH,HSD17B4,NSD2,DHCR7,PCGF2,AHCY,UBE3A,GOT2,YY1,SM

TUB,PTPN11,PUM1,AUTS2,TMEM94,CLIP2,SC5D,ELN,PDHA1,IGF1R,ADNP,TRRAP,GPAA1,LIMK1,
RIO,EP300,EXTL3,RERE,KDM5B,VARS1,FLCN,BAZ1B,ATRX,NXN,RAB3GAP2,FBXO11,MBTPS1,HI
P7,VANGL2,SOD1,RAI1,TCTN1,KDM3B,SMS,CDC42,YME1L1,ADAR,TRAF7,BCR,KDM5C,TRIO,EP3
300,EXTL3,RERE,NDUFV1,KDM5B,PBX1,CNTNAP1,VARS1,FLCN,CSNK2A1,GPX4,GARS1,ADCY6,FI
PLEC,ODC1,AEBP1,SON,KDM1A,MAP2K2,QARS1,FLNB,DLK1,SMARCE1,DHX30,RPS28,ATP6V1B2

MT1,SOX2,SLC6A8,AKT3,CNP,PHGDH,HSD17B4,NSD2,DHCR7,PCGF2,AHCY,UBE3A,GOT2,YY1,SM
RPS10,RPL35A,CYFIP2,TARDBP,ARID1A,DDOST,TSR2,HNRNPK,ATP1A2,HLA-B,TTR,ARCN1,RPS27
F3,LZTR1,HSD17B4,MED13,PCGF2,YY1,SMARCD1,CREBBP,MN1,PHIP,TANC2,CCDC47,FGFR1,EI
JFA13,TRIM8,FGFR1,RPS26,PTPN11,ABL1,AUTS2,TMEM94,POR,CLIP2,ELN,NOTCH1,PDHA1,IGF
A,ARNT2,SMS,CTNND1,STAT3,BCR,KDM5C,TRIO,EP300,RERE,DGCR2,CNTNAP1,FLCN,CSNK2A1
TN1,KDM5C,TRIO,EP300,RERE,NDUFV1,KDM5B,PBX1,BAZ1B,ATRX,NXN,ARID1B,FBXO11,PTEN,
2,APOE,TGFB2,COL3A1,TCF20
RIO,EP300,EXTL3,RERE,BAZ1B,GPX4,ATRX,NXN,ARID1B,PI4KA,RAB3GAP2,FBXO11,IARS1,PTEN,
CCT5,CDC42,SYP,KIF5A,BCR,KDM5C,TRIO,EP300,EXTL3,RERE,DGCR2,CNTNAP1,BAZ1B,GPX4,A
1,ZFR,HNRNPA2B1,MCM3AP,PREPL,ATP6AP2,SLC2A1,BAP1,FUS,GRN,SPG7,HSPG2,TPP1,SLC6A

3P1,RAC1,SMCHD1,MMP2,PPIB,GLUL,CDKN1A,CRPPA,HS6ST1,NOVA2,CDH11,EEF1A2,TBCD,PPF
3P1,RAC1,SMCHD1,MMP2,PPIB,GLUL,CDKN1A,CRPPA,HS6ST1,NOVA2,CDH11,EEF1A2,TBCD,PPF
;,ODC1,AEBP1,HSPB1,SON,MAP2K2,FLNB,DLK1,TPM3,SMARCE1,DHX30,ATP6V1B2,SDHA,LARS1
P2,FBXO11,HDAC6,PTEN,SMC3,HNRNPA2B1,ADH5,IL6ST,PREPL,NDN,CLCN7,EHMT1,ERGIC1,HS
L3,RERE,KDM5B,PBX1,VARS1,BAZ1B,CSNK2A1,GPX4,ATRX,NXN,ARID1B,CTNND2,RAB3GAP2,PT
K,YEATS2,ACO2,PLOD1,VPS35,COL2A1,CHD2,SKI,CNTNAP2,RAB11B,ABCA2,PRPF6,MARS1,SELE

AB3GAP2,PTEN,ZFR,HNRNPA2B1,NDST1,TIMM50,ATP6AP2,SLC2A1,BAP1,FUS,GRN,CLCN7,TAOK

PK,SIN3A,CASK,TCF20
LC6A8,LTBP4,IREB2,KMT2E,EFTUD2,ATXN3,PACS1,MED13,PACS2,UBE3A,GOT2,YY1,SMARCD1,C

2,MMP14,DDX3X,CLTC,RPS7,RPS19,RPL35,PRKAR1A,GJA1,CTNNB1,COL6A2,MACF1,RPS10,RPL
PB,HSPG2,LINGO1,SOX2,SLC6A8,IREB2,GTF2I,CNP,TUBGCP6,ATXN3,PHGDH,REEP2,HSD17B4,N
ELN,OBSL1,MECP2,WDR26,ADNP,TRRAP,FLII,DDX6,DEAF1,TSPAN7,CHST3,WDR11,WAC,ZSWIM6
,SYNGAP1,GBA2,NDRG1,NUS1,TMEM43,RARS1,RPS29,KARS1,LARGE1,LMNA,RHOBTB2,SLC5A7
CTNND2,CRTAP,FBXO11,IARS1,PTEN,SMC3,ECEL1,NDST1,ABCC9,MAPRE2,GPC4,NDN,H19,TAOK
S1,FLCN,GPX4,ATRX,NXN,ARID1B,PI4KA,RAB3GAP2,CRTAP,FBXO11,HDAC6,PTEN,SMC3,HNRNPA
A,MAP2K2,FLNB,DLK1,TPM3,SMARCE1,RPS28,ATP6V1B2,SDHA,LSS,STXBP1,LONP1,PRPS1,MES
B,EXT2,DDX6,SLC25A1,SYNGAP1,PPP2R5D,DEAF1,TSPAN7,NSD1,WAC,CLCN4,NUS1,ASH1L,ACSL

MN1,TANC2,TUBB4A,SH3PXD2B,HS2ST1,TECR,PUM1,SEMA6B,TMEM94,COL12A1,DAG1,MFN2,I
R1,IREB2,EFTUD2,GTF2I,TTC37,PACS1,LZTR1,HSD17B4,NSD2,DHCR7,PCGF2,YY1,CREBBP,MN1,
PK1,SMARCA4,PLEC,PRDX1,ODC1,AEBP1,FTO,SON,KDM1A,MAP2K2,QARS1,FLNB,DLK1,SMARC
KDM5B,PBX1,BAZ1B,ATRX,NXN,ARID1B,FBXO11,PTEN,SMC3,MCM3AP,SLC2A1,NDN,IARS2,HSPG

AT3,WDR45B,BCR,KDM5C,EP300,EXTL3,RERE,CNTNAP1,ALDH3A2,BAZ1B,CSNK2A1,ATRX,GARS
P2,GPC4,HSPG2,ANTXR1,IREB2,EFTUD2,GTF2I,TTC37,PACS1,LZTR1,HSD17B4,NSD2,DHCR7,PC
1,CNTNAP2,ABCA2,PRPF6,ATP6AP1,MARS1,SELENON,MAPK1,SMARCA4,PRDX1,SON,MAP2K2,FL
IAPK1,PLEC,PRDX1,ODC1,FTO,ATXN10,HSPB1,SON,KDM1A,DLK1,TPM3,SMARCE1,DHX30,SDHA,

P2,HSD17B4,GOT2,NDUFA13,TRIM8,REEP1,FGFR1,TUBB4A,PTPN23,NTRK2,DENND5A,TECR,PSA
P2,FAM50A,WDR26,ALDH18A1,BANF1,AMPD2,ZNF462,FLII,PDGFRB,CCDC88A,NECTIN1,EXT2,RD
VNK1,CCDC47,REEP1,FGFR1,TUBB4A,EIF4G1,NTRK2,CHMP1A,CLIP2,TRAK1,NOP56,ELN,ATP13A
I,SNRPB,UQCRFS1,NR2F2,MED12,HIRA,DGCR8,SKIV2L,SOS1,CITED2,TALDO1,TET3,TARS1,PSM
HIRA,DGCR8,SKIV2L,ARSB,CTCF,PPP1R12A,TGFBR1,THRA,PIGS,TARS1,CUL4B,TGFB2,SEC31A,T
SETD5,MAN2B1,SNRPB,LMNA,SLC5A7,PPP1R12A,DSTYK,ROBO3,COL6A3,CUL7,TARS1,SPG11,C
CR8,SKIV2L,ARSB,CTCF,PPP1R12A,TGFBR1,THRA,PIGS,TARS1,CUL4B,TGFB2,SEC31A,TMCO1,I
S1,NR2F2,MED12,HIRA,DGCR8,SKIV2L,SOS1,CITED2,TALDO1,TET3,TARS1,PSMD12,SEC31A,GN
I,UBE3A,GOT2,TANC2,TRIM8,CCDC47,FGFR1,EIF4G1,NTRK2,CAD,TECR,CLIP2,TRAK1,SC5D,ELI
SNK2A1,GPX4,ATRX,NXN,ARID1B,CTNND2,PI4KA,RAB3GAP2,CRTAP,FBXO11,HDAC6,PTEN,SMC3,
D1,CREBBP,NDUFA13,TANC2,CCDC47,FGFR1,RPS26,NTRK2,SLC30A9,PTPN11,PUM1,AUTS2,TR
HC9,PPP3CA,MAN1B1,TET3,SLC1A4,GATAD2B,CUL4B,PSMD12,MSH2,SZT2,DYNC1I2,PRNP,SMAF

A,SYT1,PIGT,SMARCC2,POGZ,BRAT1,MAPK8IP3,ITGA7,DPF2,DMPK,TCF20
N1,PHIP,TANC2,SMO,FGFR1,SH3PXD2B,NTRK2,HS2ST1,BGN,TRIP12,PTPN11,AUTS2,CLIP2,TRA

A1,CLIP2,ELN,OBSL1,ATP13A2,IGF1R,MECP2,BSCL2,LIMK1,EFNB1,NBAS,VPS51,SLC16A2,SYN

1,RPS10,RPL35A,CYFIP2,HNRNPDL,ARID1A,TSR2,HNRNPK,NARS1,DUX4,HLA-B,TTR,ARCN1,RPS
,ABCC9,MAPRE2,PREPL,ATP6AP2,LOX,GPC4,CLCN7,EHMT1,IARS2,ERGIC1,HSPG2,ANTXR1,SLC
,KLF13,OBSL1,NOTCH1,MECP2,ADNP,EFNB1,FLII,NBAS,PDGFRB,DDX6,ASAH1,RDH11,DEAF1,C
A,RAB3GAP2,IARS1,PTEN,ZFR,SMC3,HNRNPA2B1,NDST1,EML1,B4GAT1,TIMM50,ATP6AP2,SLC
1,CLCN4,BCOR,MED25,MAN2B1,LMNA,MED12,HIRA,DGCR8,SKIV2L,ARSB,SOS1,CTCF,PPP1R1
GAP2,HDAC6,PTEN,SMC3,ECEL1,ABCC9,MAPRE2,PREPL,ATP6AP2,LOX,GPC4,CLCN7,EHMT1,IA
ASAH1,CRKL,CHST3,NSD1,WAC,GDF1,BCOR,RPS29,STAT1,LMNA,NR2F2,MED12,HIRA,DGCR8,C

,SZT2,DYNC1I2,PRNP,TMCO1,NBEA,DNM1,FGD1,HMGA2,SH3KBP1,JRK,DMPK,SIN3A,DNAJC21,
13,PCGF2,FBLN5,CREBBP,TANC2,FGFR1,RPS26,BGN,PTPN11,ABL1,TMEM94,POR,CLIP2,ELN,N
GO1,SLC6A8,KMT2E,EFTUD2,CNP,ELP1,PACS1,LZTR1,PHGDH,HSD17B4,DHCR7,MED13,PCGF2
2,GTF2I,CNP,TUBGCP6,ATXN3,PACS1,PHGDH,HSD17B4,NSD2,DHCR7,PCGF2,AHCY,CIZ1,GOT2,
KACA,SMARCD1,TANC2,TRIM8,CCDC47,SMO,FGFR1,EIF4A3,RPS26,BGN,PSAT1,PUM1,MACRO
,TPM3,SMARCE1,DHX30,ATP6V1B2,SDHA,LSS,FBLN1,B2M,STXBP1,LONP1,MESD,USP7,IDH1,RA
C3,CUL7,MAN1B1,PIGS,FAM20C,TGFB2,FHL1,PIGT,COG4,COL3A1,ITGA7,SLC18A3,ELMO2,PIE
2,PLOD1,SKI,ASXL1,KDM6B,RAB11B,PRPF6,RPS15A,SELENON,MAPK1,SMARCA4,PLEC,PRDX1,A

P1,SMO,FGFR1,EIF4A3,SH3PXD2B,HS2ST1,BGN,PSAT1,CHMP1A,COL12A1,POR,CLIP2,DAG1,EL
S29,SETD5,SNRPB,STAT1,LMNA,NR2F2,MED12,HIRA,SKIV2L,CITED2,CTCF,ZDHHC9,TALDO1,T
1,DHX30,RPS28,ATP6V1B2,SDHA,LSS,STXBP1,MESD,RAI1,SEC23A,TCTN1,SMS,CDC42,CTNND1,

2,GTF2I,CNP,TUBGCP6,ATXN3,PACS1,PHGDH,HSD17B4,NSD2,DHCR7,PCGF2,AHCY,CIZ1,GOT2,
2,GTF2I,CNP,TUBGCP6,ATXN3,PACS1,PHGDH,HSD17B4,NSD2,DHCR7,PCGF2,AHCY,CIZ1,GOT2,
LC1A4,SZT2,SEC31A,SYT1,PIGT,POGZ,DNM1,CENPT,SLC18A3,PIEZO1,KLHL7,NECAP1

AT3,BCR,KDM5C,TRIO,EP300,RERE,NDUFV1,VARS1,CSNK2A1,NXN,ARID1B,RAB3GAP2,FBXO11,

TD5,CHD8,LMNA,MED12,HIRA,SOS1,GALNT2,PPP3CA,MAN1B1,CUL4B,FAM20C,PSMD12,TGFB
A,MAPK1,SMARCA4,PLEC,ODC1,FTO,PRKDC,ACTN1,SON,KDM1A,MAP2K2,FLNB,DLK1,SMARC

LIP2,TRAK1,DAG1,ELN,ATP13A2,PDHA1,MECP2,WDR26,TRRAP,ALDH18A1,LIMK1,AMPD2,IQSE

,SPG7,EHMT1,SOX2,LTBP4,GTF2I,TTC37,ATXN3,REEP2,NSD2,OGT,DHCR7,FBLN5,MTOR,CREB

8,SOBP,DDX6,C1QBP,PPP2R5D,DEAF1,CHST3,NSD1,RBM10,WAC,ZSWIM6,ZEB1,OAT,ACSL4,ME
ATP13A2,PDHA1,MECP2,EMC1,ADNP,TRRAP,BSCL2,ALDH18A1,LIMK1,COX8A,AMPD2,IQSEC1,M
SOX2,AKT3,ATXN3,PHGDH,HSD17B4,NSD2,DHCR7,RBPJ,AHCY,MTOR,YY1,SMARCD1,CREBBP,S

N4,ASH1L,BCOR,MED25,NDUFS1,NACC1,THOC2,MED12,DGCR8,RUSC2,EPB41L1,PPP3CA,MAN1

WDR26,JAM3,ADNP,COX8A,AMPD2,ZNF462,FLII,PDGFRB,CCDC88A,ASNS,VPS51,CSNK2B,SLC25

1,DDX6,SYNGAP1,STUB1,DEAF1,CRKL,WDR11,NSD1,WAC,CLCN4,NUS1,ASH1L,ACSL4,CACNA1H
SMARCC2,GNAO1,SMARCB1,FGD1,HMGA2,DPF2,SIN3A,CASK

A1,SMARCB1,ITGA7,SLC18A3,KLHL7,DPF2,DNAJC21

2,ASAH1,CRKL,TSPAN7,CHST3,RBM10,P3H1,WAC,CLCN4,ACSL4,BCOR,RPS29,SETD5,SNRPB,L
H1,WAC,CLCN4,ACSL4,BCOR,RPS29,SETD5,LMNA,MED12,COX4I1,GALNT2,CNOT3,CUL7,EXT1,P

RPS26,BGN,PUM1,MACROH2A1,OBSL1,NOTCH1,IGF1R,MECP2,FAM50A,ADNP,TRRAP,ALDH18A1

OL27A1,RAI1,RAB11A,SEC23A,ARNT2,KDM3B,SMS,YME1L1,ADAR,CIC,MSMO1,STAT3,BCR,KD

HNRNPK,HLA-B,TTR,ARCN1,RPS27,CTBP1,GPC1,APPL1,MMP2,L1CAM,PPIB,CDKN1A,HS6ST1,M

TXBP1,CSNK1D,PRPS1,USP7,SARS1,SOD1,RAI1,RAB11A,ARNT2,KDM3B,ARFGEF2,CDC42,YME1

,SEPTIN9,MAN1B1,TNPO3,GATAD2B,PSMD12,APOE,SETD1A,SYT1,PIGT,FBXO7,PRNP,PPT1,SMA

S,CSNK2B,DDX6,ALDH7A1,RAB18,SYNGAP1,STUB1,DEAF1,EXOC7,COLGALT1,GBA2,NSD1,DHFR
SLC25A1,ALDH7A1,SYNGAP1,STUB1,DEAF1,NDRG1,DEGS1,CLCN4,NUS1,CACNA1H,SETX,ARHG
MED13,YY1,CREBBP,PHIP,TANC2,SMO,FGFR1,EIF4A3,SH3PXD2B,RPS26,PTPN11,PSAT1,AUTS2,
17B4,NSD2,DHCR7,MED13,YY1,CREBBP,PHIP,TANC2,SMO,FGFR1,EIF4A3,SH3PXD2B,RPS26,PTP
TNAP1,VARS1,FLCN,BAZ1B,CSNK2A1,ATRX,NXN,ARID1B,CTNND2,PI4KA,RAB3GAP2,HDAC6,PTEN
,EFTUD2,TTC37,TUBGCP6,PACS1,LTBP3,PHGDH,NSD2,DHCR7,PCGF2,DHX38,PRKACA,YY1,SMA
CP2,ADNP,TRRAP,AMPD2,ZNF462,EFNB1,FLII,PDGFRB,NECTIN1,DDX6,SLC16A2,SYNGAP1,RDH1
P,EFNB1,FLII,NECTIN1,PPP2R5D,DEAF1,CRKL,CHST3,WDR11,RBM10,ZSWIM6,SHMT2,BCOR,SC

MED13,YY1,CREBBP,PHIP,TANC2,SMO,FGFR1,EIF4A3,SH3PXD2B,RPS26,PTPN11,PSAT1,AUTS2,

P,EFNB1,FLII,NECTIN1,PPP2R5D,DEAF1,CRKL,CHST3,WDR11,RBM10,ZSWIM6,SHMT2,BCOR,SC

,POGZ,FGD1,PRMT7,MAFB,GLB1,EMD
EM94,COL12A1,CLIP2,SC5D,ELN,NOTCH1,MFN2,IGF1R,MECP2,ADNP,BSCL2,LIMK1,EFNB1,FLII,

H7A1,RAB18,SYNGAP1,STUB1,DEAF1,EXOC7,COLGALT1,GBA2,NSD1,DHFR,CLCN4,NUS1,ASH1L

VAPB,IARS2,ERGIC1,LINGO1,SLC6A8,LTBP4,GTF2I,TTC37,ATXN3,ELP1,SEMA3C,PACS1,PHGDH,

CF20
RGE1,LMNA,NUP133,CNTN1,COL7A1,MED12,SOS1,CTCF,ZDHHC9,TGFBR1,COL6A3,CUL7,EXT1
JFB8,CLCN4,NUS1,RARS1,NDUFS1,MAN2B1,COX6B1,NACC1,LMNA,RHOBTB2,INF2,PPP3CA,TRA

B4GAT1,TIMM50,BAP1,FUS,SPG7,CLCN7,VAPB,YARS1,HSPG2,LTBP4,HINT1,IREB2,DNAJB2,ATX
7,NACC1,LARGE1,XPC,NUP133,SLC5A7,COX4I1,RUSC2,CNTN2,PPP3CA,TRAPPC11,TRAF3,MAN1

T2E,EFTUD2,GTF2I,LZTR1,PHGDH,HSD17B4,NSD2,DHCR7,MED13,PCGF2,YY1,CREBBP,PHIP,TA

3,ARID1A,DDOST,TSR2,SHOX,HNRNPK,DUX4,HLA-B,RPS27,CTBP1,RAC1,GPC1,SMCHD1,MMP2,
5A7,COX4I1,RUSC2,CNTN2,PPP3CA,TRAPPC11,TRAF3,MAN1B1,TET3,SLC1A4,POMGNT1,CUL4B,
AG1,FAM50A,WDR26,SUMO1,TRRAP,EFNB1,FLII,NECTIN1,PPP2R5D,DEAF1,CRKL,CHST3,WDR11

C1,SLC5A6,SRP54,UQCRFS1,CNTN1,COL7A1,SLC5A7,HIRA,SOS1,CTCF,TALDO1,EPB41L1,PPP3

XND1,FGD1,KLHL7,HMGA2,SIN3A,TCF20

CHS1,C1QBP,PPP2R5D,CRKL,CHST3,NSD1,WAC,GDF1,BCOR,MED25,RPS29,SETD5,NDUFS1,SN
IP3,SMARCB1,PLXND1,FGD1,HMGA2,DPF2,TCF20

TSPAN7,RBM10,WAC,ZSWIM6,CLCN4,ACSL4,BCOR,MED25,SETD5,NACC1,MED12,CTCF,PRPF31

ALNT2,PPP3CA,EXT1,SLC1A4,SPG11,PSMD12,FBXO7,PRNP,COL3A1,DNMT1,SMARCB1,NF2,JRK

/IF,LMNA,MED12,SOS1,CITED2,CTCF,G6PC3,CNOT3,CUL7,MAN1B1,PIGS,CUL4B,PSMD12,BRAT
1,LIMK1,COX8A,EXT2,DDX6,SLC25A1,PPP2R5D,CHST3,SOX9,SETD5,MAN2B1,SNRPB,LARGE1,SI

GJA1,CTNNB1,COL6A2,MACF1,RPS10,RPL35A,CYFIP2,TARDBP,ARID1A,DDOST,TSR2,HNRNPK,HI

8,DEAF1,TSPAN7,RBM10,WAC,ZSWIM6,CLCN4,ACSL4,BCOR,MED25,SETD5,NACC1,MED12,CTC

P3,PLXND1,FGD1,KLHL7,HMGA2,MAFB,SIN3A,TCF20

A1,MFN2,MECP2,BSCL2,ALDH18A1,LIMK1,AMPD2,KLC2,PRPH,CAPN1,SLC25A1,ASAH1,SLC16A2

SH3PXD2B,RPS26,HS2ST1,BGN,COL12A1,MACROH2A1,SC5D,NOTCH1,MECP2,ADNP,BANF1,EFN

KDM5B,PBX1,BAZ1B,ATRX,NXN,ARID1B,RAB3GAP2,PTEN,SMC3,MAPRE2,B4GAT1,ATP6AP2,GPC4
SMARCE1,RPS28,ATP6V1B2,SDHA,LSS,STXBP1,LONP1,RAI1,SEC23A,TCTN1,ADAR,MSMO1,EP30

C,FTO,PRKDC,SON,KDM1A,MAP2K2,FLNB,DLK1,TPM3,SMARCE1,RPS28,ATP6V1B2,SDHA,LSS,ST

KLHL7,DPF2,CASK
1,SNRPB,LARGE1,LMNA,THOC2,SLC5A7,ARSB,DSTYK,THRA,TRAPPC11,CNOT3,COL6A3,CUL7,EX

BAZ1B,GPX4,ATRX,NXN,ARID1B,RAB3GAP2,CRTAP,MBTPS1,PTEN,SMC3,ECEL1,IL6ST,LMAN1,A

VDR26,ADNP,TRRAP,GPAA1,ZNF462,FLII,VPS51,EXT2,RAB18,RDH11,DEAF1,CRKL,TSPAN7,RBM1
ERE,DGCR2,KDM5B,PBX1,BAZ1B,ATRX,NXN,ARID1B,RAB3GAP2,PTEN,SMC3,MAPRE2,B4GAT1,AT

D2,DHCR7,MED13,PCGF2,AHCY,UBE3A,PRKACA,YY1,SMARCD1,CREBBP,TANC2,CCDC47,FGFR1
TN,VHL,PPP2CA,CCND1,MAPK1,SMARCA4,PLEC,PRDX1,ODC1,ACADVL,ATXN10,SON,KDM1A,MAF

5,MAN2B1,LMNA,SOS1,CTCF,RUSC2,COX5A,TALDO1,CUL7,MAN1B1,EXT1,SLC1A4,PIGS,CUL4B,
HA,B2M,STXBP1,LONP1,RAI1,FLNC,TCTN1,KDM3B,ARFGEF2,BMPR2,CDC42,ADAR,TRAF7,STAT3

FT1,PPP1CB,GNAI3,KLHL24,LMNB2,ABCA3,RPL27,SNRPN,ACO2,PLOD1,COL2A1,CHD2,SKI,ASXL

L35A,CYFIP2,HNRNPDL,TARDBP,SLC25A3,ARL6IP1,ARID1A,DDOST,TSR2,SHOX,HNRNPK,NARS1,
RPPA,HS6ST1,NOVA2,MDH2,RAB27A,EEF1A2,TBCD,PPP1CB,SQSTM1,VAMP2,GNAI3,PABPN1,TA
ENON,TUBB2A,OPTN,PPP2CA,MAPK1,SMARCA4,KDELR2,PRDX1,ODC1,AEBP1,ACADVL,FTO,AT
LR2,PLEC,ODC1,AEBP1,FTO,SON,KDM1A,MAP2K2,FLNB,DLK1,TPM3,SMARCE1,RPS28,ATP6V1
,KLHL9,SPTBN2,TKT,CCND2,CDH2,GPC3,MEGF8,RAB7A,CCN2,MMP14,DDX3X,CLTC,RPS7,RPS
6,PFN1,CLEC7A,RPL31,PRKCSH,LRPPRC,XPR1,KLHL9,SPTBN2,TKT,CCND2,CDH2,GPC3,MEGF
62,EFNB1,IQSEC1,CCDC88A,ASNS,VPS51,MOGS,CSNK2B,DDX6,DARS1,RAB18,SYNGAP1,DEAF1,
HNRNPK,NARS1,DUX4,HLA-B,ARCN1,RPS27,CTBP1,RAC1,SMCHD1,MMP2,L1CAM,PPIB,CRPPA,N
,COL6A2,MACF1,RPS10,RPL35A,CYFIP2,DNAJC5,TARDBP,ARL6IP1,ARID1A,DDOST,TSR2,HNRN
N,ARID1B,CTNND2,RAB3GAP2,CRTAP,FBXO11,PTEN,ABCC9,SPINT2,MAPRE2,LOX,GPC4,BAP1,E
,COL2A1,SKI,ASXL1,CNTNAP2,RAB11B,ABCA2,RPS15A,TUBB2A,PPP2CA,MAPK1,SMARCA4,PRD
1,CNTNAP2,RAB11B,ABCA2,RPS15A,TUBB2A,PPP2CA,MAPK1,SMARCA4,PRDX1,ODC1,FTO,PRK
,LRPPRC,XPR1,KLHL9,SPTBN2,TKT,TPM1,CCND2,CDH2,GPC3,MEGF8,RAB7A,CCN2,MMP14,DD
1,SMARCA4,PRDX1,FTO,PRKDC,SON,KDM1A,MAP2K2,QARS1,DLK1,SMARCE1,DHX30,RPS28,LA
R7,MED13,FBLN5,MTOR,PACS2,SMARCD1,CREBBP,MN1,PHIP,TANC2,CCDC47,SMO,FGFR1,SH
,RPS10,RPL35A,CYFIP2,TARDBP,ARID1A,DDOST,TSR2,HNRNPK,NARS1,ATP1A2,HLA-B,ARCN1,R
,CDH11,EEF1A2,FDFT1,TBCD,PPP1CB,GNAI3,LMNB2,RPL27,SNRPN,ACO2,PLOD1,COL2A1,SKI,
,TRIO,EP300,EXTL3,RERE,KDM5B,ALDH3A2,FLCN,BAZ1B,CSNK2A1,ATRX,GARS1,NXN,ARID1B,C
L,TARDBP,ARID1A,SHOX,HNRNPK,NARS1,ATP1A2,TTR,ARCN1,RPS27,CTBP1,RAC1,APPL1,SMCH
BP3,PHGDH,HSD17B4,NSD2,DHCR7,MED13,PCGF2,MTOR,PACS2,PRKACA,YY1,SMARCD1,CREB
TT3A,LONP1,MESD,USP7,SARS1,RAI1,RAB11A,SEC23A,ARNT2,ARFGEF2,CDC42,YME1L1,ADAR,
S1,MESD,USP7,SARS1,VANGL2,RAI1,RAB11A,TCTN1,ARNT2,ARFGEF2,CDC42,YME1L1,ADAR,CI
ATP1A2,TTR,ARCN1,RPS27,CTBP1,RAC1,GPC1,SMCHD1,L1CAM,GLUL,CDKN1A,CRPPA,HS6ST1,
,NARS1,ATP1A2,HLA-B,TTR,ARCN1,RPS27,CTBP1,RAC1,ARHGDIA,SMCHD1,MMP2,L1CAM,CDKN
H11,EEF1A2,FDFT1,TBCD,PPP1CB,SQSTM1,PABPN1,LMNB2,ABCA3,RPL27,SNRPN,PDXK,PLOD
,GJA1,CTNNB1,COL6A2,MACF1,RPS10,RPL35A,CYFIP2,ARL6IP1,ARID1A,DDOST,TSR2,SHOX,H
G7,TAOK1,EHMT1,SOX2,SLC6A8,AKT3,KMT2E,CNP,PHGDH,HSD17B4,NSD2,DHCR7,RBPJ,MED1
AJC5,TARDBP,ARID1A,DDOST,TSR2,HNRNPK,NARS1,ATP1A2,HLA-B,TTR,ARCN1,RPS27,CTBP1,
PPP1CB,VAMP2,GNAI3,LMNB2,RPL27,SNRPN,ACO2,PLOD1,COL2A1,CHD2,SKI,ASXL1,KDM6B,C
GCP6,PACS1,LZTR1,PHGDH,NSD2,MED13,PCGF2,YY1,SMARCD1,CREBBP,MN1,PHIP,TANC2,CC
RRAP,LIMK1,ZNF462,FLII,NBAS,VPS51,CSNK2B,NECTIN1,EXT2,DDX6,RAB18,PPP2R5D,DEAF1,E
,ARID1B,CTNND2,RAB3GAP2,CRTAP,PTEN,ECEL1,IL6ST,MCM3AP,ATP6AP2,LOX,GPC4,NDN,FU
AR,SYP,CIC,TRAF7,DCTN1,WDR45B,KDM5C,TRIO,EP300,EXTL3,RERE,NDUFV1,KDM5B,PBX1,A
,TCTN1,KDM3B,SMS,CDC42,SYP,STAT3,BCR,KDM5C,TRIO,EP300,EXTL3,RERE,DGCR2,KDM5B,

DHCR7,MED13,PCGF2,AHCY,MTOR,UBE3A,GOT2,YY1,SMARCD1,CREBBP,TANC2,TRIM8,CCDC
ARCA4,KDELR2,PLEC,ODC1,AEBP1,FTO,PRKDC,HSPB1,SON,KDM1A,MAP2K2,FLNB,DLK1,TPM
IMM50,ATP6AP2,GPC4,SLC2A1,GRN,SPG7,TAOK1,EHMT1,SOX2,SLC6A8,AKT3,KMT2E,CNP,ATX
,FBLN1,STXBP1,LONP1,USP7,HYOU1,RAI1,FLNC,TCTN1,SMS,CCT5,CDC42,SYP,DCTN1,KIF5A,B
9,RPL35,MIF,PRKAR1A,GJA1,MYO5A,CTNNB1,COL6A2,MACF1,RPS10,RPL35A,CYFIP2,HNRNPD
A1,H19,CLCN7,IARS2,ERGIC1,HSPG2,ANTXR1,SOX2,HINT1,KMT2E,GTF2I,PACS1,LZTR1,HSD17
,TUBB2A,PPP2CA,MAPK1,SMARCA4,PRDX1,ODC1,FTO,PRKDC,ATXN10,SON,QARS1,DLK1,SMA

1,KDM5B,CNTNAP1,ALDH3A2,VARS1,FLCN,BAZ1B,CSNK2A1,ATRX,ARID1B,CTNND2,PI4KA,RAB3

,RPL35,MIF,PRKAR1A,GJA1,MYO5A,CTNNB1,COL6A2,MACF1,RPS10,RPL35A,CYFIP2,HNRNPDL,
MAP2K2,FLNB,DLK1,TPM3,SMARCE1,RPS28,ATP6V1B2,LARS1,LSS,FBLN1,B2M,STXBP1,LONP1
CR2,NDUFV1,KDM5B,CNTNAP1,ALDH3A2,VARS1,FLCN,BAZ1B,CSNK2A1,ATRX,ARID1B,CTNND2,
,ATP6V1B2,SDHA,LARS1,LSS,STT3B,STXBP1,STT3A,LONP1,PRPS1,USP7,SARS1,SOD1,RAI1,RAB
LK1,SMARCE1,DHX30,RPS28,ATP6V1B2,SDHA,LSS,STXBP1,LONP1,MESD,RAI1,TCTN1,SMS,CDC
1A,MAP2K2,FLNB,DLK1,SMARCE1,RPS28,ATP6V1B2,LARS1,LSS,FBLN1,B2M,STXBP1,LONP1,US
1A,MAP2K2,FLNB,DLK1,SMARCE1,RPS28,ATP6V1B2,LARS1,LSS,FBLN1,B2M,STXBP1,LONP1,US

ZNF462,NBAS,CCDC88A,VPS51,EXT2,DDX6,WAC,MED25,RPS29,MAN2B1,SNRPB,RHOBTB2,MED
,SELENON,OPTN,VHL,MAPK1,SMARCA4,PLEC,ACADVL,FTO,HSPB1,SON,KDM1A,FLNB,DLK1,TP
3,SMARCB1,CENPT,DPF2,GLB1,SIN3A,TCF20

9,RPL35,PRKAR1A,GJA1,CTNNB1,COL6A2,MACF1,RPS10,RPL35A,CYFIP2,TARDBP,ARID1A,TSR
D1,TRAF7,MSMO1,BCR,KDM5C,TRIO,EP300,RERE,DGCR2,KDM5B,PBX1,CNTNAP1,VARS1,FLCN,

MT2E,EFTUD2,ATXN3,PACS1,MED13,PACS2,UBE3A,GOT2,YY1,SMARCD1,CREBBP,MN1,TANC2,
)1,RAI1,TCTN1,SMS,CDC42,SYP,CTNND1,TRAF7,BCR,KDM5C,TRIO,EP300,EXTL3,RERE,BAZ1B,C
,G6PC3,SETD1A,SYT1,PIGT,TMCO1,PIEZO1
,NOVA2,CDH11,EEF1A2,TBCD,PPP1CB,SQSTM1,GNAI3,TAF15,LMNB2,RPL27,SNRPN,PLOD1,CO
N4,BCOR,MED25,SETD5,MAN2B1,NUP133,NR2F2,MED12,HIRA,DGCR8,SKIV2L,ARSB,CTCF,PPP1
,G6PC3,SETD1A,SYT1,PIGT,TMCO1,PIEZO1
9,RPL35,PRKAR1A,GJA1,CTNNB1,COL6A2,MACF1,RPS10,RPL35A,CYFIP2,TARDBP,ARID1A,TSR
V1B2,LSS,STXBP1,LONP1,MESD,RAI1,TCTN1,SMS,CDC42,ADAR,SYP,CTNND1,TRAF7,MSMO1,B

S2,HSPG2,ANTXR1,PKDCC,LTBP4,IREB2,KMT2E,EFTUD2,GTF2I,TTC37,PACS1,LTBP3,LZTR1,PH
IARCD1,CREBBP,TRIM8,SMO,FGFR1,PTPN23,HS2ST1,MDH1,TECR,PSAT1,CHMP1A,DAG1,ATP13

EFNB1,FLII,PDGFRB,NECTIN1,DDX6,RAB18,DEAF1,RBM10,WAC,ZSWIM6,RPS29,SETD5,LMNA,M
DAC6,PTEN,SMC3,ABCC9,PREPL,GPC4,NDN,H19,CLCN7,IARS2,HSPG2,ANTXR1,SOX2,SLC6A8,PK
300,RERE,KDM5B,PBX1,CNTNAP1,VARS1,BAZ1B,CSNK2A1,GPX4,ATRX,NXN,ARID1B,CTNND2,RA
BXO11,IARS1,PTEN,NDST1,MCM3AP,MAPRE2,PREPL,TIMM50,GPC4,NDN,CLCN7,EHMT1,IARS2,
2,LSS,STXBP1,LONP1,MESD,RAI1,TCTN1,SMS,CDC42,ADAR,SYP,CTNND1,TRAF7,MSMO1,BCR,K

IARCD1,CREBBP,TRIM8,SMO,FGFR1,PTPN23,HS2ST1,MDH1,TECR,PSAT1,CHMP1A,DAG1,ATP13
,CTBP1,RAC1,ARHGDIA,GPC1,SMCHD1,L1CAM,DCN,CRPPA,HS6ST1,TIMP3,NOVA2,CDH11,MDH
F4A3,SH3PXD2B,RPS26,TRIP12,PTPN11,PSAT1,ABL1,PUM1,COL12A1,SC5D,ELN,OBSL1,ACTA2,
1R,NDUFA4,WDR26,TRRAP,NDUFC2,BSCL2,LIMK1,ZNF462,EXT2,ECHS1,DDX6,PPP2R5D,CRKL,
,ATRX,NXN,ARID1B,CTNND2,RAB3GAP2,PTEN,SMC3,ECEL1,MCM3AP,MAPRE2,B4GAT1,LOX,GP
SMC3,MCM3AP,SLC2A1,NDN,FUS,SPG7,IARS2,HSPG2,ANTXR1,SOX2,SLC6A8,KMT2E,GTF2I,TU

SMC3,ECEL1,ADH5,ABCC9,MAPRE2,LOX,GPC4,NDN,H19,PSMB4,CLCN7,EHMT1,IARS2,ERGIC1,
ATRX,GARS1,NXN,ARID1B,PI4KA,RAB3GAP2,FBXO11,HDAC6,PTEN,SMC3,ECEL1,ABCC9,MCM3A
8,LTBP4,HINT1,KMT2E,GTF2I,DNAJB2,ATXN3,ELP1,PACS1,REEP2,HSD17B4,MSH6,MED13,COX

P1CB,GNAI3,LMNB2,RPL27,SNRPN,PLOD1,COL2A1,RBMX,SKI,ASXL1,KDM6B,RAB11B,RPS15A,N
P1CB,GNAI3,LMNB2,RPL27,SNRPN,PLOD1,COL2A1,RBMX,SKI,ASXL1,KDM6B,RAB11B,RPS15A,N
1,FBLN1,STXBP1,LONP1,PRPS1,USP7,SARS1,SOD1,RAI1,FLNC,SEC23A,TCTN1,ARNT2,ARFGEF2,
SPG2,ANTXR1,SOX2,PKDCC,AKT3,KMT2E,EFTUD2,TTC37,TUBGCP6,DICER1,PHGDH,HSD17B4,NS
EN,SMC3,ABCC9,MAPRE2,PREPL,GPC4,NDN,IARS2,HSPG2,ANTXR1,PKDCC,LTBP4,AKT3,KMT2E
NON,OPTN,VHL,MAPK1,PLEC,PRDX1,ATXN10,HSPB1,KDM1A,QARS1,TPM3,SMARCE1,XPNPEP3,D

1,EHMT1,VAPB,TPP1,SLC6A8,AKT3,HINT1,IREB2,GTF2I,ATXN3,PHGDH,REEP2,NSD2,CIZ1,UBE3A

CREBBP,MN1,TANC2,CCDC47,FGFR1,TUBB4A,NTRK2,DENND5A,HS2ST1,TRIP12,TECR,PUM1,CH

L35A,CYFIP2,TARDBP,ARID1A,TSR2,SHOX,HNRNPK,NARS1,DUX4,ATP1A2,HLA-B,ARCN1,RPS27,C
MSH6,DHCR7,RBPJ,GOT2,NDUFA13,TRIM8,REEP1,FGFR1,TUBB4A,PTPN23,NTRK2,DENND5A,SL
,CLCN4,ACSL4,SOX9,RPS29,SETD5,MAN2B1,NSMF,RHOBTB2,MED12,ARSB,ANOS1,SOS1,TALD
7,DNAJB6,INF2,DDHD2,DSTYK,NIPA1,PPP3CA,TRAPPC11,SEPTIN9,COL6A3,CUL7,SPG11,TNPO3,
K1,EHMT1,ERGIC1,HSPG2,SLC6A8,KMT2E,GTF2I,CNP,TTC37,PACS1,LTBP3,LZTR1,PHGDH,NSD2
A2B1,ADH5,IL6ST,ABCC9,PREPL,NDN,H19,CLCN7,EHMT1,ERGIC1,HSPG2,ANTXR1,SOX2,PKDCC,
D,USP7,IDH1,HYOU1,SARS1,COL27A1,RAI1,RAB11A,SEC23A,ARNT2,KDM3B,SMS,YME1L1,ADAR
L4,BCOR,MED25,RARS1,SETD5,MBOAT7,NDUFS1,DLG4,CHD8,SRP54,THOC2,MED12,HIRA,CTCF

GF1R,TRIM2,MECP2,NDUFA4,WDR26,FXR1,TRRAP,ALDH18A1,AMPD2,KLC2,IQSEC1,VPS51,CSN
,FGFR1,EIF4A3,SH3PXD2B,RPS26,HS2ST1,PTPN11,PSAT1,PUM1,AUTS2,TMEM94,POR,CLIP2,DA
E1,DHX30,RPS28,SDHA,LARS1,STT3B,STXBP1,STT3A,LONP1,PRPS1,MESD,USP7,SARS1,COL27A
G2,ANTXR1,SOX2,SLC6A8,KMT2E,GTF2I,TUBGCP6,PACS1,LZTR1,PHGDH,HSD17B4,NSD2,DHCR

S1,NXN,ARID1B,CTNND2,PI4KA,RAB3GAP2,ADCY6,IARS1,PTEN,SMC3,ECEL1,IL6ST,ABCC9,LOX,G
GF2,YY1,CREBBP,MN1,FGFR1,EIF4A3,SH3PXD2B,RPS26,HS2ST1,PTPN11,PSAT1,PUM1,AUTS2,T
LNB,DLK1,TPM3,SMARCE1,XPNPEP3,DHX30,ATP6V1B2,SDHA,LSS,FBLN1,STT3B,STXBP1,STT3A,
,FBLN1,STXBP1,LONP1,PRPS1,USP7,IDH1,SARS1,SOD1,RAI1,RAB11A,TCTN1,ARNT2,ARFGEF2,C

AT1,PUM1,CHMP1A,PINK1,AUTS2,CLIP2,TRAK1,ELN,ATP13A2,MFN2,PDHA1,MECP2,NDUFA4,WD
DH11,PPP2R5D,DEAF1,CRKL,TSPAN7,CHST3,WAC,GDF1,CLCN4,ACSL4,BCOR,MAN2B1,SNRPB,L
A2,PDHA1,MECP2,WDR26,ADNP,IDH2,ALDH18A1,LIMK1,FLII,PRPH,CSNK2B,NECTIN1,EXT2,SLC2
D12,SEC31A,GNAO1,CRB2,CASK,NDUFA10
TMCO1,POGZ,BRAT1,COL3A1,GNAO1,ELMO2,FGD1,KLHL7,HMGA2,GLB1,SIN3A,CASK
CUL4B,PSMD12,FHL1,SMARCC2,MAPK8IP3,SMARCB1,ITGA7,SLC18A3,NEMF,UBTF,DPF2,GLB1,S
POGZ,BRAT1,COL3A1,GNAO1,ELMO2,FGD1,KLHL7,HMGA2,GLB1,SIN3A,CASK
AO1,CRB2,CASK,NDUFA10
N,TIA1,ATP13A2,PDHA1,MECP2,EMC1,ADNP,TRRAP,BSCL2,ALDH18A1,LIMK1,COX8A,AMPD2,PR
,ECEL1,ABCC9,MAPRE2,ATP6AP2,GPC4,NDN,CLCN7,ERGIC1,HSPG2,ANTXR1,SOX2,PKDCC,LTB
AK1,NOP56,SC5D,PDHA1,MECP2,NDUFA4,ADNP,HMGB3,ZNF462,PDGFRB,ECHS1,DDX6,SLC25A
RCC2,TMCO1,NBEA,POGZ,DNM1,GNAO1,FGD1,HMGA2,SH3KBP1,JRK,UBTF,DPF2,SIN3A,DNAJC

K1,KLF13,SC5D,ELN,IGF1R,MECP2,ADNP,TRRAP,LIMK1,AMPD2,ZNF462,EFNB1,FLII,PDGFRB,M

GAP1,RDH11,PPP2R5D,CRKL,CHST3,WAC,BCOR,SOX9,RPS29,SETD5,MAN2B1,SNRPB,LMNA,MEI

27,CTBP1,APPL1,SMCHD1,MMP2,PPIB,DCN,CRPPA,HS6ST1,TIMP3,NOVA2,CDH11,MDH2,RAB27/
6A8,PKDCC,LTBP4,KMT2E,EFTUD2,GTF2I,LTBP3,LZTR1,PHGDH,HSD17B4,NSD2,DHCR7,MED13
RKL,TSPAN7,CHST3,WAC,GDF1,CLCN4,ACSL4,BCOR,SOX9,RPS29,SETD5,LMNA,MED12,SOS1,C
2A1,BAP1,FUS,GRN,SPG7,TAOK1,VAPB,ERGIC1,HSPG2,LINGO1,TPP1,SOX2,SLC6A8,AKT3,HINT
2A,TGFBR1,THRA,PIGS,TARS1,CUL4B,TGFB2,SEC31A,TMCO1,POGZ,BRAT1,COL3A1,GNAO1,ELI
RS2,ERGIC1,HSPG2,ANTXR1,SLC6A8,PKDCC,LTBP4,KMT2E,EFTUD2,GTF2I,LTBP3,LZTR1,PHGD
TCF,TALDO1,TGFBR1,PSMD12,APOE,TGFB2,COL3A1,GLB1,CTSA,TCF20

CC2D1A,NECAP1,ARSA,TCF20
OTCH1,ACTA2,WDR26,TRRAP,ALDH18A1,LIMK1,ZNF462,EXT2,DDX6,CRKL,CHST3,NSD1,RBM10,
,MTOR,UBE3A,GOT2,YY1,SMARCD1,CREBBP,MN1,TANC2,WNK1,CCDC47,FGFR1,EIF4A3,PTPN2
SMARCD1,CREBBP,TRIM8,SMO,FGFR1,TUBB4A,ELOVL5,DENND5A,MDH1,PSAT1,PUM1,CHMP1A
H2A1,OBSL1,NOTCH1,PDHA1,IGF1R,MECP2,FAM50A,ADNP,TRRAP,IDH2,ALDH18A1,EFNB1,NBAS
I1,FLNC,SEC23A,KDM3B,ARFGEF2,SMS,CCT5,CDC42,ADAR,KIF5A,MSMO1,STAT3,WDR45B,BCF
ZO1,FGD1,UBTF,GLB1,EMD
EBP1,ATXN10,SON,KDM1A,MAP2K2,DLK1,TPM3,SMARCE1,DHX30,RPS28,SDHA,STXBP1,STT3A,I

LN,OBSL1,MFN2,TRIM2,ALDH18A1,LIMK1,COX8A,CAPN1,EXT2,SLC25A1,ASAH1,RAB18,PPP2R5D
GFBR1,G6PC3,GYS1,SETD1A,SYT1,PIGT,TMCO1,CFAP298,PIEZO1
STAT3,BCR,KDM5C,TRIO,EP300,RERE,NDUFV1,VARS1,FLCN,BAZ1B,CSNK2A1,NXN,ARID1B,RAB

SMARCD1,CREBBP,TRIM8,SMO,FGFR1,TUBB4A,ELOVL5,DENND5A,MDH1,PSAT1,PUM1,CHMP1A
SMARCD1,CREBBP,TRIM8,SMO,FGFR1,TUBB4A,ELOVL5,DENND5A,MDH1,PSAT1,PUM1,CHMP1A

PTEN,SMC3,ECEL1,ABCC9,MAPRE2,PREPL,NDN,BAP1,CLCN7,EHMT1,IARS2,HSPG2,ANTXR1,KM

2,SETD1A,FHL1,SZT2,TMCO1,DNM1,CENPT,SETD1B,NFIB,FGD1,DPF2,SIN3A,DNAJC21,NECAP1,T
E1,RPS28,ATP6V1B2,LSS,STXBP1,LONP1,PRPS1,MESD,HYOU1,COL27A1,VANGL2,RAI1,SEC23A,T

C1,CCDC88A,ASNS,CSNK2B,DDX6,ALDH7A1,RAB18,SYNGAP1,STUB1,DEAF1,EXOC7,GBA2,NDUFI

BP,REEP1,SMO,FGFR1,EIF4G1,RPS26,PTPN11,ABL1,PINK1,TMEM94,POR,CLIP2,ELN,OBSL1,ATI

ED25,SETD5,MAN2B1,LARGE1,CTCF,PPP1R12A,ATF6,GALNT2,TRAPPC11,CNOT3,MAN1B1,TARS1
OGS,CSNK2B,DDX6,ASAH1,ALDH7A1,RAB18,SYNGAP1,STUB1,DEAF1,EXOC7,COLGALT1,GBA2,N
MO,FGFR1,PTPN23,DENND5A,BGN,PSAT1,POR,DAG1,NOTCH1,MFN2,PDHA1,ACTA2,MDM2,WDF

B1,SLC1A4,GATAD2B,CUL4B,PSMD12,MSH2,SETD1A,NFASC,SYT1,DYNC1I2,PRNP,PPT1,COG4,S

A1,ALDH7A1,RAB18,SYNGAP1,PPP2R5D,DEAF1,NSD1,WAC,ZSWIM6,NDUFB8,SHMT2,CLCN4,ASH

I,MED25,SETD5,MBOAT7,NDUFS1,CDKN1B,MAN2B1,NSMF,CHD8,THOC2,UNC13A,COL7A1,MED

MNA,MED12,COX4I1,GALNT2,CNOT3,CUL7,EXT1,PIGS,CUL4B,PSMD12,PIGT,COG4,SMARCC2,C
IGS,CUL4B,PSMD12,COG4,SMARCC2,COL3A1,MAPK8IP3,GNAO1,SMARCB1,PLXND1,FGD1,HMG

,EXT2,DDX6,ASAH1,CRKL,TSPAN7,CHST3,RBM10,P3H1,WAC,ZSWIM6,CLCN4,ACSL4,BCOR,SOX

I5C,EP300,EXTL3,RERE,DGCR2,KDM5B,ALDH3A2,FLCN,BAZ1B,GPX4,ATRX,GARS1,NXN,ARID1B,

DH2,EEF1A2,FDFT1,TBCD,PPP1CB,ABCA3,RPL27,SNRPN,ACO2,PLOD1,VPS35,COL2A1,RBMX,SK

L1,ADAR,SYP,CIC,TRAF7,DCTN1,STAT3,BCR,KDM5C,TRIO,EP300,EXTL3,RERE,DGCR2,NDUFV1,K

RCC2,POGZ,COL3A1,DNMT1,GNAO1,SMARCB1,SLC18A3,NF2,POU3F3,GSN,PRMT7,DMPK,SIN3A

:,CLCN4,NUS1,ASH1L,RNF216,MED25,RARS1,SETD5,MBOAT7,MAN2B1,NACC1,LARGE1,SLC5A6,
EF9,MED25,MBOAT7,NACC1,LARGE1,LMNA,XPC,NUP133,SLC5A7,COX4I1,RUSC2,CNTN2,PPP3C
COL12A1,CLIP2,SC5D,ELN,IGF1R,MECP2,EMC1,WDR26,SUMO1,TRRAP,BANF1,LIMK1,FLII,PDGF
'N11,PSAT1,AUTS2,COL12A1,CLIP2,SC5D,ELN,IGF1R,MECP2,EMC1,WDR26,SUMO1,TRRAP,BANI
I,SMC3,ECEL1,ABCC9,MAPRE2,PREPL,ATP6AP2,LOX,GPC4,PSMB4,CLCN7,EHMT1,IARS2,ERGIC
RCD1,CREBBP,MN1,PHIP,TRIM8,CCDC47,FGFR1,SH3PXD2B,HS2ST1,TUB,PTPN11,PSAT1,AUTS2
1,PPP2R5D,DEAF1,TSPAN7,CHST3,NSD1,WAC,ZSWIM6,CLCN4,NUS1,ACSL4,MED25,SETD5,CHD8
)X9,MED25,RPS29,SETD5,SNRPB,NSMF,LARGE1,LMNA,MED12,HIRA,DGCR8,ANOS1,CTCF,TGFB

COL12A1,CLIP2,SC5D,ELN,IGF1R,MECP2,EMC1,WDR26,SUMO1,TRRAP,BANF1,LIMK1,FLII,PDGF

)X9,MED25,RPS29,SETD5,SNRPB,NSMF,LARGE1,LMNA,MED12,HIRA,DGCR8,ANOS1,CTCF,TGFB

NBAS,NECTIN1,EXT2,DDX6,ASAH1,SLC16A2,PPP2R5D,DEAF1,CRKL,TSPAN7,NDRG1,CHST3,NSD

.,RNF216,MED25,RARS1,SETD5,MBOAT7,MAN2B1,NACC1,LARGE1,SLC5A6,RHOBTB2,PMPCA,XI

MSH6,NSD2,DHCR7,RBPJ,PCGF2,UBE3A,YY1,CREBBP,WNK1,CCDC47,REEP1,FGFR1,TUBB4A,EI

,CUL4B,PSMD12,FHL1,SMARCC2,SMARCB1,ITGA7,SLC18A3,FGD1,KLHL7,MAFB,DPF2,GLB1,EN
PPC11,CNOT3,SLC1A4,TARS1,SPG11,GATAD2B,SETD1A,SZT2,SYT1,POGZ,BRAT1,DNM1,MAPK8

N3,LZTR1,AHCY,COX6A1,FBLN5,PRKACA,NDUFA13,TRIM8,REEP1,SMO,FGFR1,ELOVL5,CNBP,NT
B1,TET3,SLC1A4,POMGNT1,CUL4B,OPA1,SZT2,SYT1,PIGT,PRNP,PPT1,NBEA,POGZ,DNM1,GNAC

NC2,SMO,FGFR1,EIF4A3,SH3PXD2B,RPS26,BGN,PTPN11,PSAT1,AUTS2,COL12A1,CLIP2,SC5D,EI

GLUL,CDKN1A,HS6ST1,NOVA2,CTSB,CDH11,MDH2,RAB27A,FDFT1,TBCD,PPP1CB,GNAI3,KLHL2
SZT2,SYT1,PIGT,PRNP,PPT1,NBEA,POGZ,DNM1,GNAO1,SLC18A3,SETD1B,JRK,CSTB,UBTF,CASI
,RBM10,ZSWIM6,SHMT2,BCOR,SOX9,MED25,RPS29,SETD5,SNRPB,NSMF,LARGE1,LMNA,MED1

CA,CNOT3,COL6A3,TET3,SLC1A4,SPG11,OPA1,PIGT,HADHB,SMARCC2,POGZ,SMARCB1,ITGA7,

RPB,COX6B1,LMNA,UQCRFS1,NR2F2,MED12,HIRA,DGCR8,SKIV2L,SOS1,CITED2,TALDO1,TRAPI

,ARL2BP,CNOT3,QRICH1,CUL7,ARHGEF18,EXT1,POMGNT1,GATAD2B,FAM20C,FH,SETD1A,DYN

,PRMT7,ARSA

1,MAPK8IP3,PLXND1,FGD1,HMGA2,DPF2,SIN3A,TCF20
RP54,LMNA,THOC2,SLC5A7,ARSB,DSTYK,THRA,TRAPPC11,CNOT3,COL6A3,CUL7,EXT1,TET3,PSI

LA-B,TTR,ARCN1,RPS27,CTBP1,RAC1,GPC1,APPL1,SMCHD1,MMP2,L1CAM,CDKN1A,NOVA2,CDI

F,PRPF31,ARL2BP,CNOT3,QRICH1,CUL7,ARHGEF18,EXT1,POMGNT1,GATAD2B,FAM20C,FH,SET

2,STUB1,GBA2,NDRG1,PIP5K1C,TMEM43,SETX,MED25,MAN2B1,LARGE1,LMNA,UNC13A,COL7A

IB1,FLII,PDGFRB,DDX6,SLC16A2,DEAF1,CRKL,TSPAN7,CHST3,NSD1,RBM10,SHMT2,CLCN4,ACS

4,NDN,H19,EHMT1,HSPG2,SOX2,KMT2E,GTF2I,TTC37,PACS1,DICER1,LZTR1,PHGDH,NSD2,OGT,
0,RERE,DGCR2,KDM5B,ALDH3A2,VARS1,BAZ1B,ARID1B,RAB3GAP2,PTEN,SMC3,HNRNPA2B1,IL

T3B,STXBP1,LONP1,PRPS1,MESD,USP7,IDH1,HYOU1,COL27A1,VANGL2,RAI1,RAB11A,FLNC,SE

XT1,TET3,PSMD12,FHL1,SEC31A,SMARCC2,COL3A1,SMARCB1,ITGA7,SLC18A3,DPF2,GLB1,DNA

TP6AP2,GPC4,NDN,PSMB4,CLCN7,IARS2,ERGIC1,HSPG2,SOX2,PKDCC,HINT1,GTF2I,ELP1,TCF3,

l0,WAC,ZSWIM6,CLCN4,ACSL4,BCOR,MED25,SETD5,NACC1,XPC,MED12,DGCR8,CTCF,PRPF31,
TP6AP2,GPC4,NDN,H19,EHMT1,HSPG2,SOX2,KMT2E,GTF2I,TTC37,PACS1,DICER1,LZTR1,PHGDH

,EIF4A3,SH3PXD2B,NTRK2,HS2ST1,PTPN11,ABL1,CLIP2,TRAK1,ELN,OBSL1,MECP2,WDR26,SUM
P2K2,DLK1,TPM3,SMARCE1,DHX30,RPS28,SDHA,LARS1,STT3B,B2M,STXBP1,STT3A,LONP1,PRPS

FAM20C,PSMD12,FHL1,SZT2,SYT1,CCNK,SMARCC2,TMCO1,COL3A1,SMARCB1,SETD1B,NFIB,K
3,BCR,TRIO,EP300,EXTL3,RERE,DGCR2,NDUFV1,KDM5B,PBX1,FLCN,BAZ1B,CSNK2A1,GPX4,ATI

.1,KDM6B,RAB11B,PRPF6,ATP6AP1,TERF2IP,RPS15A,NOTCH2,VHL,PPP2CA,MAPK1,SMARCA4,PI

,DUX4,ATP1A2,HLA-B,TTR,RPS27,CTBP1,RAC1,SMCHD1,MMP2,L1CAM,GLUL,CRPPA,HS6ST1,NC
\F15,SNRPN,ACO2,PLOD1,VPS35,COL2A1,SKI,ASXL1,KDM6B,CNTNAP2,RAB11B,ABCA2,MARS1,S
(N10,SON,KDM1A,MAP2K2,QARS1,DLK1,TPM3,SMARCE1,DHX30,ATP6V1B2,SDHA,LARS1,LSS,FE
32,LSS,STXBP1,LONP1,PRPS1,MESD,USP7,IDH1,COL27A1,VANGL2,RAI1,RAB11A,SEC23A,TCTN1
19,RPL35,PRKAR1A,GJA1,MYO5A,CTNNB1,COL6A2,MACF1,RPS10,RPL35A,CYFIP2,DNAJC5,HNF
3,RAB7A,CCN2,MMP14,DDX3X,CLTC,RPS7,RPS19,RPL35,PRKAR1A,GJA1,MYO5A,CTNNB1,COL6
EXOC7,GBA2,GPT2,WAC,ZSWIM6,SHMT2,CLCN4,ASH1L,BCOR,MED25,RARS1,MBOAT7,LARGE1
IOVA2,CDH11,EEF1A2,FDFT1,TBCD,PPP1CB,SQSTM1,GNAI3,PABPN1,LMNB2,ABCA3,RPL27,SNR
PK,NARS1,ATP1A2,HLA-B,TTR,ARCN1,RPS27,CTBP1,RAC1,GPC1,SMCHD1,L1CAM,GLUL,CDKN1
HMT1,IARS2,HSPG2,ANTXR1,PKDCC,LTBP4,AKT3,KMT2E,TTC37,ATXN3,PACS1,DICER1,TCF3,LZ
K1,ODC1,FTO,PRKDC,SON,KDM1A,MAP2K2,QARS1,DLK1,SMARCE1,DHX30,RPS28,SDHA,LARS1,
DC,SON,KDM1A,MAP2K2,QARS1,DLK1,SMARCE1,DHX30,RPS28,SDHA,LARS1,LSS,FBLN1,STT3E
X3X,CLTC,RPS7,RPS19,RPL35,MIF,PRKAR1A,GJA1,MYO5A,CTNNB1,COL6A2,MACF1,RPS10,RPL
RS1,LSS,FBLN1,STT3B,STXBP1,STT3A,LONP1,PRPS1,MESD,USP7,SARS1,VANGL2,RAI1,RAB11A,
I3PXD2B,RPS26,HS2ST1,BGN,TRIP12,PTPN11,PSAT1,AUTS2,TMEM94,POR,KLF13,ELN,ACTA2,EM
PS27,CTBP1,RAC1,ARHGDIA,SMCHD1,MMP2,L1CAM,GLUL,CRPPA,NOVA2,CDH11,MDH2,EEF1A
ASXL1,KDM6B,CNTNAP2,RAB11B,ABCA2,RPS15A,NOTCH2,SELENON,PPP2CA,MAPK1,SMARCA4,
TNND2,RAB3GAP2,CRTAP,MBTPS1,PTEN,ECEL1,HNRNPA2B1,IL6ST,MCM3AP,ATP6AP2,LOX,GP
D1,TNC,DCN,HS6ST1,TIMP3,NOVA2,CTSB,EEF1A2,ANXA5,CPSF1,PPP1CB,SQSTM1,VAMP2,GNAI
BP,PHIP,TANC2,SMO,FGFR1,SH3PXD2B,DENND5A,HS2ST1,TRIP12,PSAT1,AUTS2,TMEM94,POR,
SYP,CIC,KIF5A,MSMO1,WDR45B,BCR,KDM5C,TRIO,EP300,EXTL3,RERE,DGCR2,NDUFV1,KDM5E
C,KIF5A,MSMO1,KCNQ2,WDR45B,BCR,KDM5C,TRIO,EP300,EXTL3,RERE,DGCR2,NDUFV1,KDM5
NOVA2,CDH11,MDH2,EEF1A2,FDFT1,TBCD,PPP1CB,SQSTM1,VAMP2,GNAI3,LMNB2,RPL27,SNR
I1A,CRPPA,HS6ST1,NOVA2,CDH11,MDH2,RAB27A,EEF1A2,TBCD,PPP1CB,SQSTM1,VAMP2,PABPN
1,COL2A1,SKI,ASXL1,KDM6B,RAB11B,RPS15A,NOTCH2,MARS1,SELENON,VHL,PPP2CA,MAPK1,S
IRNPK,NARS1,DUX4,HLA-B,TTR,ARCN1,RPS27,CTBP1,RAC1,SMCHD1,MMP2,L1CAM,PPIB,GLUL,
3,PCGF2,AHCY,MTOR,UBE3A,GOT2,YY1,SMARCD1,CREBBP,TANC2,TRIM8,CCDC47,SMO,FGFR1
RAC1,GPC1,SMCHD1,L1CAM,GLUL,CDKN1A,CRPPA,HS6ST1,NOVA2,CDH11,MDH2,EEF1A2,FDFT
NTNAP2,RAB11B,ABCA2,RPS15A,MARS1,SELENON,TUBB2A,PPP2CA,MAPK1,SMARCA4,KDELR2,
DC47,FGFR1,EIF4A3,SH3PXD2B,RPS26,DENND5A,HS2ST1,PTPN11,PSAT1,POR,CLIP2,DAG1,KLF
XOC7,CRKL,TSPAN7,CHST3,NSD1,SHMT2,CLCN4,ACSL4,BCOR,MED25,SETD5,SNRPB,RHOBTB2
,SPG7,H19,TAOK1,IARS2,HSPG2,ANTXR1,LTBP4,HINT1,KMT2E,GTF2I,CNP,TUBGCP6,ELP1,PAC
DH3A2,FLCN,BAZ1B,UBQLN2,CSNK2A1,ATRX,NXN,ARID1B,PI4KA,RAB3GAP2,FBXO11,IARS1,PT
PBX1,VARS1,BAZ1B,CSNK2A1,GPX4,ATRX,NXN,ARID1B,CTNND2,RAB3GAP2,HDAC6,PTEN,SMC3,

17,SMO,FGFR1,PTPN23,NTRK2,HS2ST1,MDH1,TECR,PSAT1,CHMP1A,TRAK1,DAG1,ATP13A2,PDH
3,SMARCE1,RPS28,ATP6V1B2,LARS1,LSS,FBLN1,B2M,STXBP1,LONP1,USP7,IDH1,HYOU1,COL27
N3,PHGDH,HSD17B4,NSD2,DHCR7,RBPJ,MED13,PCGF2,AHCY,MTOR,UBE3A,GOT2,YY1,SMARCI
CR,KDM5C,TRIO,EP300,EXTL3,RERE,DGCR2,CNTNAP1,BAZ1B,GPX4,ATRX,GARS1,NXN,ARID1B,C
,TARDBP,ARID1A,DDOST,TSR2,HNRNPK,NARS1,DUX4,ATP1A2,HLA-B,TTR,ARCN1,RPS27,CTBP1
34,NSD2,DHCR7,MED13,PCGF2,FBLN5,MTOR,PRKACA,SMARCD1,CREBBP,TANC2,CCDC47,SMO
RCE1,XPNPEP3,DHX30,ATP6V1B2,SDHA,LARS1,LSS,FBLN1,STT3B,EIF3F,STXBP1,STT3A,LONP1,F

GAP2,FBXO11,IARS1,SMC3,ADH5,NDST1,MAPRE2,B4GAT1,SLC2A1,FUS,TAOK1,EHMT1,HSPG2,L

ARL6IP1,ARID1A,DDOST,TSR2,SHOX,HNRNPK,NARS1,DUX4,HLA-B,TTR,ARCN1,RPS27,CTBP1,RA
,USP7,HYOU1,COL27A1,RAI1,FLNC,TCTN1,SMS,CCT5,CDC42,SYP,DCTN1,KIF5A,BCR,KDM5C,TR
PI4KA,RAB3GAP2,FBXO11,IARS1,SMC3,ADH5,NDST1,MAPRE2,B4GAT1,SLC2A1,FUS,TAOK1,EHM
11A,TCTN1,KDM3B,ARFGEF2,SMS,CDC42,YME1L1,ADAR,CIC,TRAF7,KIF5A,STAT3,KCNQ2,WDR4
42,ADAR,SYP,CTNND1,TRAF7,MSMO1,BCR,KDM5C,TRIO,EP300,RERE,DGCR2,STARD7,KDM5B,P
P7,IDH1,HYOU1,COL27A1,SOD1,RAI1,TCTN1,SMS,CDC42,SYP,CTNND1,TRAF7,BCR,KDM5C,TRIO
P7,IDH1,HYOU1,COL27A1,SOD1,RAI1,TCTN1,SMS,CDC42,SYP,CTNND1,TRAF7,BCR,KDM5C,TRIO

12,HIRA,SOS1,CTCF,PPP1R12A,SEPTIN9,MAN1B1,TARS1,GATAD2B,CUL4B,SETD1A,SYT1,SMARC
M3,SMARCE1,RPS28,ATP6V1B2,SDHA,PFKM,LSS,B2M,STXBP1,LONP1,PRPS1,USP7,VANGL2,SOD

2,SHOX,HNRNPK,NARS1,DUX4,ATP1A2,HLA-B,ARCN1,RPS27,CTBP1,RAC1,GPC1,SMCHD1,MMP2
BAZ1B,CSNK2A1,ATRX,NXN,ARID1B,CTNND2,RAB3GAP2,FBXO11,PTEN,SMC3,ADH5,ABCC9,MA

CCDC47,FGFR1,TUBB4A,NTRK2,DENND5A,HS2ST1,TRIP12,TECR,PUM1,CHMP1A,SEMA6B,AUTS2
SNK2A1,GPX4,ATRX,GARS1,NXN,ARID1B,CTNND2,PI4KA,RAB3GAP2,FBXO11,IARS1,MBTPS1,PT

L2A1,RBMX,SKI,ASXL1,KDM6B,RAB11B,RPS15A,NOTCH2,SELENON,OPTN,PPP2CA,MAPK1,SMA
R12A,TGFBR1,THRA,PCNA,PIGS,TARS1,CUL4B,TGFB2,SEC31A,SMARCC2,TMCO1,POGZ,BRAT1

2,SHOX,HNRNPK,NARS1,DUX4,ATP1A2,HLA-B,ARCN1,RPS27,CTBP1,RAC1,GPC1,SMCHD1,MMP2
CR,KDM5C,TRIO,EP300,RERE,DGCR2,STARD7,KDM5B,PBX1,CNTNAP1,VARS1,FLCN,BAZ1B,CSN

GDH,HSD17B4,NSD2,DHCR7,MED13,PCGF2,MTOR,PACS2,PRKACA,YY1,SMARCD1,CREBBP,PHIP,
A2,PDHA1,IGF1R,EMC1,WDR26,ADNP,TRRAP,ALDH18A1,NTN1,AMPD2,ZNF462,EFNB1,FLII,IQSE

IED12,HIRA,SKIV2L,PRPF31,ARL2BP,G6PC3,ARHGEF18,MAN1B1,EXT1,SLC1A4,POMGNT1,GATA
KDCC,LTBP4,AKT3,KMT2E,GTF2I,TTC37,TUBGCP6,PACS1,DICER1,LZTR1,PHGDH,HSD17B4,NSD
B3GAP2,FBXO11,MBTPS1,HDAC6,PTEN,SMC3,PTPRF,SPINT2,MAPRE2,PREPL,B4GAT1,ATP6AP2,
SOX2,SLC6A8,LTBP4,KMT2E,ELP1,PACS1,NSD2,DHCR7,MTOR,PACS2,UBE3A,CREBBP,NDUFA13,
DM5C,TRIO,EP300,RERE,DGCR2,KDM5B,PBX1,CNTNAP1,VARS1,FLCN,BAZ1B,CSNK2A1,ATRX,N

A2,PDHA1,IGF1R,EMC1,WDR26,ADNP,TRRAP,NTN1,AMPD2,ZNF462,EFNB1,FLII,IQSEC1,CCDC8
2,EEF1A2,FDFT1,TBCD,CPSF1,PPP1CB,SQSTM1,VAMP2,GNAI3,PABPN1,VCAN,TGFBI,RPL27,SNR
IGF1R,MECP2,EFNB1,VPS51,MOGS,DDX6,SLC25A1,RAB18,SYNGAP1,PPP2R5D,EXOC7,CRKL,CH
CHST3,NSD1,RBM10,WAC,SHMT2,GDF1,BCOR,MED25,RPS29,SETD5,NDUFS1,SNRPB,STAT1,LM
C4,H19,HSPG2,ANTXR1,SOX2,KMT2E,EFTUD2,TTC37,TCF3,LZTR1,PHGDH,HSD17B4,NSD2,DHC
BGCP6,PACS1,LZTR1,PHGDH,HSD17B4,NSD2,DHCR7,RBPJ,MED13,AHCY,MTOR,PACS2,UBE3A,Y

PKDCC,AKT3,HINT1,KMT2E,EFTUD2,GTF2I,SEMA3C,PACS1,LZTR1,PHGDH,NSD2,DHCR7,RBPJ,M
P,MAPRE2,B4GAT1,ATP6AP2,LOX,GPC4,NDN,SPG7,EHMT1,IARS2,ERGIC1,YARS1,HSPG2,SLC6A8
6A1,CIZ1,MTOR,PACS2,UBE3A,GOT2,YY1,CREBBP,TANC2,REEP1,SMO,FGFR1,TUBB4A,ELOVL5,

OTCH2,SELENON,OPTN,PPP2CA,MAPK1,SMARCA4,PLEC,PRDX1,ODC1,AEBP1,FTO,PRKDC,SON,
OTCH2,SELENON,OPTN,PPP2CA,MAPK1,SMARCA4,PLEC,PRDX1,ODC1,AEBP1,FTO,PRKDC,SON,
SMS,CCT5,CDC42,SYP,KIF5A,BCR,KDM5C,TRIO,EP300,EXTL3,RERE,DGCR2,CNTNAP1,BAZ1B,G
SD2,DHCR7,RBPJ,PCGF2,MTOR,UBE3A,YY1,SMARCD1,CREBBP,MN1,TANC2,CCDC47,SMO,FGF
,GTF2I,TTC37,TUBGCP6,DICER1,LTBP3,HSD17B4,NSD2,DHCR7,MED13,MTOR,DHX38,PACS2,S
DHX30,SDHA,FBLN1,STXBP1,LONP1,PRPS1,USP7,IDH1,SARS1,SOD1,RAI1,RAB11A,FLNC,TCTN1,

,YY1,CREBBP,NDUFA13,TANC2,TRIM8,REEP1,SMO,FGFR1,TUBB4A,ELOVL5,EIF4G1,NTRK2,DE

IMP1A,SEMA6B,AUTS2,TMEM94,TRAK1,DAG1,IGF1R,MECP2,NDUFA4,EMC1,WDR26,ADNP,TRRA

TBP1,RAC1,GPC1,SMCHD1,MMP2,L1CAM,PPIB,GLUL,CDKN1A,CRPPA,HS6ST1,NOVA2,CDH11,M
C30A9,HS2ST1,MDH1,TECR,PSAT1,PUM1,CHMP1A,PINK1,AUTS2,CLIP2,TRAK1,NOP56,ELN,NOT
D1,SEPTIN9,CUL7,SLC1A4,GATAD2B,CUL4B,FAM20C,FH,PSMD12,PIGT,SMARCC2,POGZ,GNAO1
POMGNT1,OPA1,FHL1,SZT2,PRNP,HADHB,COG4,DNM1,HARS1,SMARCB1,ITGA7,PLXND1,SLC18
,MED13,PCGF2,AHCY,FBLN5,MTOR,PACS2,UBE3A,PRKACA,YY1,SMARCD1,CREBBP,MN1,PHIP,T
,AKT3,KMT2E,EFTUD2,TTC37,TUBGCP6,DICER1,PHGDH,HSD17B4,NSD2,DHCR7,RBPJ,PCGF2,M
,CIC,MSMO1,STAT3,BCR,KDM5C,EP300,EXTL3,RERE,DGCR2,KDM5B,ALDH3A2,FLCN,BAZ1B,G
,GALNT2,EPB41L1,PPP3CA,CNOT3,MAN1B1,TET3,GATAD2B,PSMD12,SETD1A,SZT2,SYT1,NBEA,

K2B,EXT2,DDX6,DARS1,SLC25A1,ASAH1,SYNGAP1,PPP2R5D,NDRG1,CHST3,WAC,ZSWIM6,SHMT
G1,ELN,IGF1R,MECP2,ADNP,TRRAP,ALDH18A1,LIMK1,ZNF462,DDX6,SLC25A1,RAB18,RDH11,P
1,RAI1,TCTN1,ARNT2,ARFGEF2,CDC42,YME1L1,ADAR,SYP,CIC,TRAF7,MSMO1,STAT3,KCNQ2,B
7,RBPJ,MED13,AHCY,MTOR,PACS2,UBE3A,YY1,SMARCD1,CREBBP,NDUFA13,MN1,PHIP,TANC2,T

GPC4,FUS,PSMB4,IARS2,ERGIC1,HSPG2,ANTXR1,SOX2,SLC6A8,PKDCC,LTBP4,HINT1,KMT2E,GT
MEM94,POR,CLIP2,DAG1,ELN,IGF1R,MECP2,ADNP,TRRAP,ALDH18A1,HMGB3,LIMK1,ZNF462,D
LONP1,PRPS1,USP7,SARS1,RAI1,RAB11A,TCTN1,ARNT2,KDM3B,ARFGEF2,SMS,CDC42,YME1L1,
CT5,CDC42,YME1L1,ADAR,SYP,CIC,TRAF7,DCTN1,KIF5A,STAT3,KCNQ2,WDR45B,BCR,KDM5C,T

R26,JAM3,PARK7,GPAA1,NDUFC2,BSCL2,ALDH18A1,HTRA1,LIMK1,COX8A,AMPD2,KLC2,PRPH,I
MNA,MED12,HIRA,SOS1,CITED2,CTCF,TGFBR1,CNOT3,CUL7,MAN1B1,PIGS,POMGNT1,FAM20C
5A1,ASAH1,SYNGAP1,STUB1,DEAF1,GBA2,NSD1,WAC,ZSWIM6,NDUFB8,CLCN4,NUS1,ASH1L,SE

SIN3A,EMD,TCF20

PH,IQSEC1,MOGS,CSNK2B,DDX6,ASAH1,ALDH7A1,RAB18,SYNGAP1,STUB1,DEAF1,EXOC7,COLG
P4,HINT1,KMT2E,EFTUD2,GTF2I,LTBP3,LZTR1,PHGDH,HSD17B4,NSD2,DHCR7,RBPJ,MED13,PC
1,C1QBP,SLC16A2,SYNGAP1,WDR11,ZSWIM6,NDUFB8,NUS1,TMEM43,BCOR,MED25,RPS29,SE
21,CASK,CC2D1A,NECAP1,TCF20

OGS,NECTIN1,DDX6,RAB18,SLC16A2,SYNGAP1,RDH11,PPP2R5D,DEAF1,CRKL,TSPAN7,CHST3,N

D12,HIRA,SOS1,ROBO3,TALDO1,TRAF3,CNOT3,COL6A3,CUL7,MAN1B1,PIGS,SPG11,CUL4B,FA

A,EEF1A2,FDFT1,TBCD,CPSF1,PPP1CB,VCAN,TGFBI,RPL27,SNRPN,ACO2,PLOD1,COL2A1,SKI,AS
3,PCGF2,UBE3A,YY1,CREBBP,PHIP,TANC2,SMO,FGFR1,EIF4A3,SH3PXD2B,RPS26,HS2ST1,PTPN1
ITED2,COX4I1,ZDHHC9,PPP3CA,CUL7,EXT1,PIGS,CUL4B,FAM20C,PSMD12,COG4,SMARCC2,PO
1,IREB2,GTF2I,CNP,TUBGCP6,ATXN3,PHGDH,REEP2,HSD17B4,MSH6,DHCR7,RBPJ,MTOR,GOT2
MO2,PIEZO1,FGD1,KLHL7,HMGA2,GLB1,SIN3A,CASK
H,HSD17B4,NSD2,DHCR7,MED13,PCGF2,UBE3A,PRKACA,YY1,CREBBP,PHIP,TANC2,SMO,FGFR1

,WAC,GDF1,ACSL4,BCOR,MED25,RPS29,SNRPB,LMNA,UQCRFS1,NR2F2,MED12,HIRA,DGCR8,S
3,NTRK2,HS2ST1,TECR,PTPN11,PSAT1,AUTS2,TRAK1,ELN,PDHA1,MECP2,WDR26,ADNP,TRRAP,
A,SEMA6B,POR,CLIP2,DAG1,NOP56,SC5D,ELN,ATP13A2,PDHA1,EMC1,WDR26,JAM3,TRRAP,GPA
S,EXT2,DDX6,ASAH1,CRKL,TSPAN7,CHST3,RBM10,P3H1,WAC,ZSWIM6,CLCN4,ACSL4,BCOR,SOX
R,KDM5C,EP300,EXTL3,RERE,CNTNAP1,ALDH3A2,BAZ1B,CSNK2A1,GPX4,ATRX,GARS1,NXN,ARID

LONP1,PRPS1,USP7,RAI1,SEC23A,TCTN1,ARNT2,CDC42,YME1L1,ADAR,TRAF7,DCTN1,KIF5A,BC

,STUB1,GBA2,CHST3,NSD1,ZSWIM6,BCOR,SOX9,SETD5,MAN2B1,SNRPB,LARGE1,LMNA,THOC

3GAP2,FBXO11,PTEN,SMC3,ECEL1,ABCC9,MAPRE2,PREPL,GPC4,NDN,BAP1,CLCN7,EHMT1,IAR

A,SEMA6B,POR,CLIP2,DAG1,NOP56,SC5D,ELN,ATP13A2,PDHA1,EMC1,WDR26,JAM3,TRRAP,GPA
A,SEMA6B,POR,CLIP2,DAG1,NOP56,SC5D,ELN,ATP13A2,PDHA1,EMC1,WDR26,JAM3,TRRAP,GPA

T2E,CNP,TTC37,TUBGCP6,PACS1,LTBP3,LZTR1,PHGDH,NSD2,DHCR7,RBPJ,MED13,PCGF2,AHC

TCF20
TCTN1,ARNT2,KDM3B,SMS,CDC42,ADAR,SYP,TRAF7,STAT3,BCR,KDM5C,TRIO,EP300,EXTL3,RE

38,SHMT2,CLCN4,NUS1,ASH1L,ARHGEF9,MED25,SETD5,MBOAT7,MAN2B1,LARGE1,SLC5A6,RH

P13A2,MECP2,ADNP,TRRAP,ALDH18A1,BNC2,HTRA1,LIMK1,EFNB1,PDGFRB,CAPN1,DDX6,SLC1

,POMGNT1,GATAD2B,PSMD12,THG1L,SYT1,PIGT,POGZ,COL3A1,HARS1,CENPT,FGD1,KLHL7,R
ISD1,DHFR,NDUFB8,CLCN4,NUS1,ASH1L,RNF216,MED25,RARS1,SETD5,MBOAT7,NDUFS1,MAN
26,JAM3,ADNP,COX8A,AMPD2,ZNF462,FLII,PDGFRB,CCDC88A,ASNS,VPS51,CSNK2B,SLC25A1,

MARCC2,TMCO1,NBEA,POGZ,DNM1,DNMT1,GNAO1,SMARCB1,NFIB,JRK,UBTF,DMPK,SIN3A,CA

1L,BCOR,SOX9,MED25,SETD5,COX6B1,LARGE1,RHOBTB2,PMPCA,THOC2,MED12,CTCF,EPB41

12,HIRA,ANOS1,PPP3CA,MAN1B1,TET3,GATAD2B,FH,PSMD12,MSH2,SETD1A,NFASC,SZT2,PRNP,

OL3A1,MAPK8IP3,GNAO1,SMARCB1,PLXND1,FGD1,HMGA2,PRMT7,MAFB,DPF2,SIN3A,CASK
A2,PRMT7,MAFB,DPF2,SIN3A,CASK

9,RPS29,SETD5,SNRPB,LMNA,MED12,COX4I1,GALNT2,CNOT3,CUL7,EXT1,PIGS,CUL4B,PSMD12

CTNND2,RAB3GAP2,CRTAP,FBXO11,MBTPS1,HDAC6,PTEN,SMC3,ECEL1,HNRNPA2B1,ADH5,MC

,ASXL1,PRPF6,RPS15A,NOTCH2,MARS1,SELENON,VHL,CCND1,MAPK1,SMARCA4,KDELR2,PLEC

DM5B,ALDH3A2,FLCN,BAZ1B,UBQLN2,CSNK2A1,ATRX,ARID1B,GLUD1,PI4KA,FBXO11,IARS1,PT

A,CASK,CNOT1,CC2D1A,TCF20

,RHOBTB2,PMPCA,XPC,COX4I1,EPB41L1,PPP3CA,TRAPPC11,MAN1B1,SLC1A4,UFM1,PIGS,TARS
A,TRAPPC11,TRAF3,MAN1B1,TET3,SLC1A4,POMGNT1,CUL4B,OPA1,NFASC,SZT2,SYT1,PIGT,PR
RB,ASNS,NECTIN1,EXT2,DDX6,SLC25A1,RAB18,RDH11,DEAF1,CHST3,NSD1,RBM10,SOX9,RPS2
1,LIMK1,FLII,PDGFRB,ASNS,NECTIN1,EXT2,DDX6,SLC25A1,RAB18,RDH11,DEAF1,CHST3,NSD1,
1,HSPG2,ANTXR1,SLC6A8,PKDCC,LTBP4,KMT2E,EFTUD2,GTF2I,TCF3,LTBP3,LZTR1,PHGDH,HSD
,TMEM94,SC5D,ELN,OBSL1,PDHA1,IGF1R,MECP2,WDR26,ADNP,TRRAP,GPAA1,ALDH18A1,BAN
,LMNA,MED12,HIRA,SOS1,PPP1R12A,GALNT2,PPP3CA,SEPTIN9,CNOT3,QRICH1,MAN1B1,GATA
R1,PPP3CA,NID1,G6PC3,SEPTIN9,POMGNT1,FAM20C,TGFB2,NFASC,SMARCC2,CENPF,TMCO1,

RB,ASNS,NECTIN1,EXT2,DDX6,SLC25A1,RAB18,RDH11,DEAF1,CHST3,NSD1,RBM10,SOX9,RPS2

R1,PPP3CA,NID1,G6PC3,SEPTIN9,POMGNT1,FAM20C,TGFB2,NFASC,SMARCC2,CENPF,TMCO1,

1,ZSWIM6,SHMT2,CLCN4,ACSL4,BCOR,SOX9,MED25,SETD5,LMNA,MED12,HIRA,INF2,CTCF,CO

C,COX4I1,EPB41L1,PPP3CA,TRAPPC11,MAN1B1,SLC1A4,UFM1,PIGS,TARS1,SPG11,FH,PSMD1

F4G1,NTRK2,CHMP1A,CLIP2,TRAK1,NOP56,ELN,NOTCH1,ATP13A2,PDHA1,ACTA2,MDM2,MECP

MD
IP3,HARS1,DNMT1,GNAO1,POU3F3,UBTF,PRMT7,CASK,NDUFA10,NECAP1,ARSA

TRK2,PTPN11,COL12A1,TRAK1,DAG1,NOP56,TIA1,ATP13A2,MFN2,TRIM2,MECP2,FXR1,BSCL2,A
1,SLC18A3,SETD1B,JRK,CSTB,UBTF,CASK,CC2D1A,NECAP1

N,IGF1R,MECP2,EMC1,WDR26,SUMO1,TRRAP,BANF1,LIMK1,EFNB1,FLII,PDGFRB,ASNS,NECTI

4,LMNB2,ABCA3,RPL27,SNRPN,ACO2,PLOD1,COL2A1,CHD2,SKI,ASXL1,KDM6B,RAB11B,ABCA2
K,CC2D1A,NECAP1
2,HIRA,DGCR8,ANOS1,CTCF,TGFBR1,PPP3CA,NID1,G6PC3,SEPTIN9,POMGNT1,FAM20C,TGFB2

PLXND1,SLC18A3,HMGA2,UBTF,DPF2,DMPK,SIN3A,CASK,ARSA

PC11,TET3,TARS1,GYS1,PSMD12,MSH2,FHL1,SEC31A,HADHB,GNAO1,CRB2,CASK,NDUFA10

C1I2,PIGT,CCNK,SMARCC2,POGZ,MAPK8IP3,GNAO1,SMARCB1,NFIB,FGD1,KLHL7,PRMT7,MAF

MD12,FHL1,SEC31A,SMARCC2,COL3A1,SMARCB1,ITGA7,SLC18A3,DPF2,GLB1,DNAJC21,TCF20

H11,MDH2,RAB27A,EEF1A2,FDFT1,TBCD,SQSTM1,GNAI3,PABPN1,TAF15,LMNB2,ABCA3,RPL27,S

TD1A,DYNC1I2,PIGT,CCNK,SMARCC2,POGZ,MAPK8IP3,GNAO1,SMARCB1,NFIB,FGD1,KLHL7,HM

1,SLC5A7,ARSB,INF2,OGDH,NIPA1,THRA,TRAPPC11,SEPTIN9,COL6A3,PIGS,SPG11,POMGNT1,C

L4,BCOR,SOX9,MED25,RPS29,SETD5,LMNA,NUP133,MED12,HIRA,ZDHHC9,TGFBR1,G6PC3,CO

,DHCR7,MED13,FBLN5,MTOR,DHX38,PACS2,YY1,SMARCD1,CREBBP,PHIP,TANC2,TRIM8,SMO,F
6ST,SPINT2,MAPRE2,B4GAT1,LOX,SLC2A1,NDN,BAP1,H19,EHMT1,IARS2,HSPG2,ANTXR1,SOX2,I

C23A,ARNT2,KDM3B,SMS,CDC42,YME1L1,ADAR,CIC,MSMO1,STAT3,BCR,KDM5C,EP300,EXTL3,I

JC21,TCF20

LZTR1,PHGDH,NSD2,DHCR7,FBLN5,UBE3A,PRKACA,YY1,SMARCD1,CREBBP,WNK1,CCDC47,RE

ARL2BP,CNOT3,QRICH1,CUL7,ARHGEF18,EXT1,POMGNT1,GATAD2B,FAM20C,FH,SETD1A,DYNC
H,NSD2,OGT,DHCR7,MED13,FBLN5,MTOR,DHX38,PACS2,YY1,SMARCD1,CREBBP,PHIP,TANC2,TI

MO1,ADNP,BANF1,HMGB3,LIMK1,EFNB1,FLII,PDGFRB,VPS51,NECTIN1,EXT2,SOBP,DDX6,SYNGA
S1,USP7,IDH1,HYOU1,SARS1,UQCRC2,SOD1,RAI1,RAB11A,SEC23A,TCTN1,ARNT2,KDM3B,ARFGI

LHL7,HMGA2,DPF2,SIN3A,DNAJC21,TCF20
RX,NXN,ARID1B,RAB3GAP2,PTEN,SMC3,HNRNPA2B1,ABCC9,ATP6AP2,LOX,GPC4,CLCN7,EHMT1

LEC,ODC1,AEBP1,ACADVL,FTO,SON,MAP2K2,FLNB,DLK1,SMARCE1,RPS28,ATP6V1B2,SDHA,PFK

)VA2,MDH2,RAB27A,EEF1A2,FDFT1,TBCD,PPP1CB,SQSTM1,VAMP2,GNAI3,PABPN1,TAF15,LMNE
SELENON,TUBB2A,OPTN,VHL,PPP2CA,CCND1,MAPK1,SMARCA4,KDELR2,PLEC,PRDX1,ODC1,AEE
3LN1,STT3B,STXBP1,STT3A,LONP1,PRPS1,USP7,SARS1,SOD1,RAI1,RAB11A,TCTN1,ARNT2,KDM
,ARFGEF2,SMS,CDC42,ADAR,PALLD,CIC,TRAF7,KIF5A,STAT3,WDR45B,BCR,TRIO,EP300,EXTL3,
RNPDL,TARDBP,ARL6IP1,ARID1A,DDOST,TSR2,HNRNPK,NARS1,ATP1A2,HLA-B,TTR,ARCN1,RPS2
A2,MACF1,RPS10,RPL35A,CYFIP2,DNAJC5,HNRNPDL,TARDBP,ARL6IP1,ARID1A,DDOST,TSR2,HN
I,SLC5A6,RHOBTB2,DDHD2,RUSC2,EPB41L1,PPP3CA,CNOT3,MAN1B1,SLC1A4,SPG11,CUL4B,D
PN,ACO2,PLOD1,COL2A1,SKI,ASXL1,KDM6B,CNTNAP2,RAB11B,ABCA2,RPS15A,NOTCH2,SELEN
A,CRPPA,HS6ST1,NOVA2,CDH11,MDH2,EEF1A2,FDFT1,TBCD,PPP1CB,SQSTM1,VAMP2,GNAI3,PA
ZTR1,PHGDH,HSD17B4,NSD2,OGT,DHCR7,MED13,FBLN5,MTOR,PACS2,UBE3A,SMARCD1,CREBI
LSS,FBLN1,STT3B,STXBP1,STT3A,LONP1,PRPS1,MESD,USP7,SARS1,VANGL2,RAI1,RAB11A,TCTI
3,STXBP1,STT3A,LONP1,PRPS1,MESD,USP7,SARS1,VANGL2,RAI1,RAB11A,TCTN1,ARNT2,KDM3B
L35A,CYFIP2,HNRNPDL,TARDBP,SLC25A3,ARL6IP1,ARID1A,DDOST,TSR2,SHOX,HNRNPK,NARS1,I
TCTN1,ARNT2,ARFGEF2,CDC42,YME1L1,ADAR,CIC,TRAF7,KIF5A,MSMO1,KCNQ2,WDR45B,BCR
MC1,TRRAP,GPAA1,ALDH18A1,BNC2,EFNB1,FLII,NBAS,VPS51,EXT2,DDX6,PPP2R5D,DEAF1,CHST
2,FDFT1,TBCD,PPP1CB,SQSTM1,VAMP2,GNAI3,LMNB2,RPL27,SNRPN,YEATS2,ACO2,PLOD1,COI
,KDELR2,PLEC,PRDX1,ODC1,AEBP1,ACADVL,FTO,PRKDC,SON,KDM1A,MAP2K2,QARS1,FLNB,DL
C4,NDN,FUS,SPG7,H19,TAOK1,IARS2,HSPG2,ANTXR1,LTBP4,HINT1,KMT2E,GTF2I,CNP,TUBGCP
3,KLHL24,PABPN1,VCAN,TGFBI,LMNB2,RPL27,SNRPN,YEATS2,VPS35,COL2A1,CHD2,SKI,ASXL1,I
CLIP2,SC5D,ELN,OBSL1,PDHA1,IGF1R,MECP2,EMC1,WDR26,SUMO1,ADNP,TRRAP,LIMK1,AMPD
3,CNTNAP1,ALDH3A2,VARS1,FLCN,BAZ1B,CSNK2A1,ATRX,NXN,ARID1B,CTNND2,PI4KA,RAB3GAF
B,CNTNAP1,ALDH3A2,VARS1,FLCN,BAZ1B,CSNK2A1,GPX4,ATRX,ARID1B,CTNND2,PI4KA,RAB3G
PN,ACO2,VPS35,COL2A1,CHD2,SKI,ASXL1,CNTNAP2,RAB11B,ABCA2,RPS15A,NOTCH2,TUBB2A,
N1,LMNB2,RPL27,SNRPN,YEATS2,ACO2,VPS35,COL2A1,RBMX,CHD2,SKI,ASXL1,KDM6B,CNTNAF
SMARCA4,KDELR2,PLEC,ODC1,AEBP1,FTO,PRKDC,HSPB1,SON,KDM1A,MAP2K2,FLNB,DLK1,TPN
CDKN1A,CRPPA,HS6ST1,NOVA2,CDH11,EEF1A2,FDFT1,TBCD,PPP1CB,SQSTM1,GNAI3,PABPN1,T
,PTPN23,NTRK2,HS2ST1,MDH1,TECR,PSAT1,CHMP1A,TRAK1,DAG1,NOTCH1,ATP13A2,PDHA1,A
T1,TBCD,PPP1CB,SQSTM1,VAMP2,GNAI3,TAF15,LMNB2,RPL27,SNRPN,ACO2,VPS35,COL2A1,CH
PLEC,PRDX1,ODC1,AEBP1,FTO,SON,KDM1A,MAP2K2,QARS1,FLNB,DLK1,TPM3,SMARCE1,DHX3
13,ELN,OBSL1,MECP2,EMC1,ADNP,BSCL2,ALDH18A1,HMGB3,LIMK1,AMPD2,ZNF462,ASNS,VPS
,MED12,HIRA,CTCF,PPP1R12A,ZDHHC9,TALDO1,GALNT2,CNOT3,QRICH1,CUL7,MAN1B1,EXT1,
S1,LZTR1,PHGDH,HSD17B4,NSD2,DHCR7,MED13,PCGF2,FBLN5,UBE3A,SMARCD1,CREBBP,TAN
EN,SMC3,HNRNPA2B1,NDST1,MCM3AP,MAPRE2,TIMM50,ATP6AP2,GPC4,SLC2A1,NDN,BAP1,FU
ECEL1,PTPRF,ABCC9,MAPRE2,PREPL,GPC4,NDN,EHMT1,IARS2,ERGIC1,HSPG2,ANTXR1,PKDCC,

HA1,IGF1R,EMC1,WDR26,ADNP,TRRAP,ALDH18A1,NTN1,AMPD2,ZNF462,EFNB1,FLII,IQSEC1,CCD
A1,SOD1,RAI1,SEC23A,TCTN1,ARNT2,KDM3B,ARFGEF2,SMS,CDC42,SYP,CTNND1,TRAF7,BCR,K
D1,CREBBP,NDUFA13,TANC2,TRIM8,CCDC47,SMO,FGFR1,PTPN23,NTRK2,HS2ST1,MDH1,TECR,
CTNND2,PI4KA,RAB3GAP2,FBXO11,IARS1,MBTPS1,HDAC6,PTEN,SMC3,ECEL1,ADH5,IL6ST,ABCC
,RAC1,ARHGDIA,GPC1,APPL1,SMCHD1,MMP2,L1CAM,PPIB,DCN,CRPPA,HS6ST1,TIMP3,NOVA2,C
,FGFR1,SH3PXD2B,RPS26,HS2ST1,BGN,PTPN11,ABL1,COL12A1,POR,CLIP2,SC5D,ELN,OBSL1,A
PRPS1,USP7,SARS1,RAI1,RAB11A,TCTN1,ARNT2,KDM3B,ARFGEF2,SMS,CDC42,ADAR,SYP,CIC,TF

LINGO1,SOX2,SLC6A8,KMT2E,EFTUD2,GTF2I,CNP,TUBGCP6,PACS1,PHGDH,NSD2,OGT,DHCR7,I

AC1,SMCHD1,MMP2,L1CAM,PPIB,GLUL,CDKN1A,CRPPA,HS6ST1,NOVA2,CDH11,EEF1A2,FDFT1,T
RIO,EP300,EXTL3,RERE,DGCR2,CNTNAP1,BAZ1B,GPX4,ATRX,GARS1,NXN,ARID1B,CTNND2,PI4K
IT1,HSPG2,LINGO1,SOX2,SLC6A8,KMT2E,EFTUD2,GTF2I,CNP,TUBGCP6,PACS1,PHGDH,NSD2,O
5B,KDM5C,EP300,EXTL3,RERE,DGCR2,NDUFV1,KDM5B,PBX1,CNTNAP1,ALDH3A2,VARS1,FLCN,
PBX1,CNTNAP1,VARS1,FLCN,BAZ1B,CSNK2A1,ATRX,NXN,ARID1B,CTNND2,RAB3GAP2,FBXO11,P
),EP300,EXTL3,RERE,BAZ1B,CSNK2A1,GPX4,ATRX,GARS1,NXN,ARID1B,CTNND2,PI4KA,RAB3GA
),EP300,EXTL3,RERE,BAZ1B,CSNK2A1,GPX4,ATRX,GARS1,NXN,ARID1B,CTNND2,PI4KA,RAB3GA

C2,TMCO1,BRAT1,COL3A1,SMARCB1,PLXND1,POU3F3,PIEZO1,FGD1,PRMT7,DPF2,SIN3A,DNA
1,RAI1,FLNC,KDM3B,SMS,CCT5,CDC42,YME1L1,ADAR,SYP,TRAF7,DCTN1,KIF5A,STAT3,WDR45

,L1CAM,PPIB,GLUL,CDKN1A,CRPPA,HS6ST1,NOVA2,CDH11,MDH2,EEF1A2,FDFT1,TBCD,PPP1C
PRE2,PREPL,GPC4,NDN,EHMT1,HSPG2,ANTXR1,PKDCC,AKT3,LDLR,KMT2E,EFTUD2,GTF2I,TTC

2,TMEM94,TRAK1,DAG1,IGF1R,MECP2,NDUFA4,EMC1,WDR26,ADNP,TRRAP,GPAA1,BSCL2,ALD
EN,SMC3,ECEL1,ADH5,IL6ST,ABCC9,MAPRE2,LOX,GPC4,SLC2A1,NDN,H19,PSMB4,CLCN7,EHM

RCA4,PLEC,PRDX1,ODC1,AEBP1,FTO,PRKDC,SON,KDM1A,MAP2K2,FLNB,DLK1,TPM3,SMARCE1
,COL3A1,GNAO1,SMARCB1,ELMO2,FGD1,KLHL7,HMGA2,DPF2,GLB1,SIN3A,DNAJC21,CASK

,L1CAM,PPIB,GLUL,CDKN1A,CRPPA,HS6ST1,NOVA2,CDH11,MDH2,EEF1A2,FDFT1,TBCD,PPP1C
K2A1,ATRX,NXN,ARID1B,CTNND2,RAB3GAP2,FBXO11,PTEN,SMC3,ECEL1,ADH5,ABCC9,MAPRE2

TANC2,SMO,FGFR1,SH3PXD2B,DENND5A,HS2ST1,MDH1,TRIP12,PTPN11,PSAT1,AUTS2,TMEM9
C1,CCDC88A,ASNS,VPS51,MOGS,CSNK2B,DDX6,DARS1,RAB18,SYNGAP1,STUB1,DEAF1,EXOC7,

D2B,PSMD12,NFASC,PIGT,CCNK,SMARCC2,CENPF,SMARCB1,FGD1,KLHL7,PRMT7,MAFB,DPF2,
2,DHCR7,MED13,PCGF2,MTOR,YY1,SMARCD1,CREBBP,MN1,PHIP,TANC2,TRIM8,CCDC47,FGFR
GPC4,SLC2A1,H19,EHMT1,HSPG2,ANTXR1,SLC6A8,IREB2,KMT2E,EFTUD2,GTF2I,TTC37,TUBG
,MN1,PHIP,TANC2,WNK1,CCDC47,NTRK2,CAD,ACAT2,TRIP12,TECR,PUM1,CHMP1A,AUTS2,TME
XN,ARID1B,CTNND2,RAB3GAP2,FBXO11,PTEN,SMC3,ECEL1,ADH5,ABCC9,MAPRE2,PREPL,GPC4

A,ASNS,VPS51,MOGS,CSNK2B,DDX6,DARS1,RAB18,SYNGAP1,DEAF1,EXOC7,GBA2,GPT2,NSD1,
PN,PDXK,YEATS2,ACO2,PLOD1,VPS35,COL2A1,SKI,ASXL1,KDM6B,RAB11B,PRPF6,ATP6AP1,RP
IST3,NSD1,RBM10,BCOR,MED25,RPS29,SETD5,SNRPB,LMNA,THOC2,CNTN1,MED12,SLC5A7,A
NA,UQCRFS1,NR2F2,MED12,HIRA,DGCR8,SKIV2L,SOS1,CITED2,CTCF,ZDHHC9,TALDO1,TGFBR
R7,MED13,PCGF2,YY1,SMARCD1,CREBBP,MN1,PHIP,TANC2,TRIM8,CCDC47,SMO,FGFR1,EIF4A
Y1,SMARCD1,CREBBP,NDUFA13,MN1,PHIP,TANC2,TRIM8,FGFR1,RPS26,TRIP12,PTPN11,CHMP

IED13,PCGF2,FBLN5,MTOR,PACS2,UBE3A,YY1,SMARCD1,CREBBP,PHIP,TANC2,WNK1,TRIM8,S
,LTBP4,HINT1,KMT2E,GTF2I,DNAJB2,PACS1,LTBP3,PHGDH,REEP2,HSD17B4,NSD2,DHCR7,RBP
EIF4G1,NTRK2,CAD,SLC30A9,PINK1,COL12A1,CLIP2,TRAK1,DAG1,NOP56,ELN,TIA1,ATP13A2,M

KDM1A,MAP2K2,FLNB,DLK1,TPM3,SMARCE1,DHX30,RPS28,ATP6V1B2,B2M,STXBP1,LONP1,ME
KDM1A,MAP2K2,FLNB,DLK1,TPM3,SMARCE1,DHX30,RPS28,ATP6V1B2,B2M,STXBP1,LONP1,ME
PX4,ATRX,GARS1,NXN,ARID1B,PI4KA,RAB3GAP2,CRTAP,FBXO11,HDAC6,PTEN,SMC3,ECEL1,IL6
R1,SH3PXD2B,DENND5A,BGN,PTPN11,PSAT1,AUTS2,POR,SC5D,OBSL1,NOTCH1,PDHA1,MECP2,
MARCD1,CREBBP,MN1,TANC2,TRIM8,SMO,FGFR1,SH3PXD2B,RPS26,HS2ST1,TUB,MDH1,TRIP12
SMS,CCT5,YME1L1,ADAR,SYP,CIC,TRAF7,DCTN1,KIF5A,KCNQ2,WDR45B,BCR,KDM5C,TRIO,EP3

NND5A,SLC30A9,ACAT2,TECR,PSAT1,PUM1,CHMP1A,PINK1,SEMA6B,AUTS2,CLIP2,TRAK1,NOP56

P,GPAA1,BSCL2,ALDH18A1,ZNF462,FLII,IQSEC1,CCDC88A,NECTIN1,EXT2,SOBP,DDX6,ALDH7A1

DH2,EEF1A2,FDFT1,TBCD,PPP1CB,SQSTM1,GNAI3,PABPN1,TAF15,LMNB2,RPL27,SNRPN,PLOD
CH1,ATP13A2,MFN2,PDHA1,MECP2,NDUFA4,EMC1,WDR26,JAM3,PARK7,ADNP,GPAA1,NDUFC2,E
,SMARCB1,SLC18A3,KLHL7,PRMT7,DPF2,GLB1,SIN3A,CASK,CNOT1,TCF20
A3,NF2,UBTF,DMPK,NECAP1,EMD
ANC2,CCDC47,SMO,TUBB4A,SH3PXD2B,NTRK2,DENND5A,HS2ST1,TRIP12,TECR,PTPN11,PUM1,
TOR,UBE3A,YY1,SMARCD1,CREBBP,MN1,TANC2,WNK1,CCDC47,SMO,FGFR1,SH3PXD2B,DENN
X4,ATRX,GARS1,NXN,ARID1B,CTNND2,RAB3GAP2,CRTAP,FBXO11,MBTPS1,HDAC6,PTEN,SMC3,
POGZ,DNM1,MAPK8IP3,GNAO1,PLXND1,SETD1B,POU3F3,NFIB,UBTF,DMPK,SIN3A,DNAJC21,CA

2,NUS1,BCOR,MED25,SETD5,MBOAT7,MAN2B1,ZNF592,LARGE1,LMNA,PMPCA,UQCRFS1,MED
P2R5D,EXOC7,CHST3,GPT2,NSD1,RBM10,BCOR,SOX9,MED25,RPS29,SETD5,SNRPB,LARGE1,LI
CR,TRIO,EP300,EXTL3,RERE,NDUFV1,KDM5B,PBX1,VARS1,FLCN,CSNK2A1,ATRX,NXN,ARID1B,G
RIM8,FGFR1,RPS26,TRIP12,PTPN11,CHMP1A,AUTS2,TMEM94,COL12A1,CLIP2,KLF13,ELN,NOT

F2I,LTBP3,LZTR1,PHGDH,MSH6,NSD2,MED13,FBLN5,UBE3A,YY1,SMARCD1,CREBBP,PHIP,TANC
DX6,SLC25A1,RAB18,SYNGAP1,RDH11,PPP2R5D,EXOC7,CHST3,GPT2,NSD1,RBM10,WAC,BCOR,
ADAR,SYP,CIC,KIF5A,MSMO1,STAT3,KCNQ2,WDR45B,BCR,KDM5C,TRIO,EP300,EXTL3,RERE,DC
RIO,EP300,EXTL3,RERE,STARD7,NDUFV1,KDM5B,CNTNAP1,ALDH3A2,BAZ1B,UBQLN2,ATRX,PI4

QSEC1,ASNS,CSNK2B,CAPN1,ECHS1,DARS1,ASAH1,RAB18,SLC16A2,SYNGAP1,STUB1,DEAF1,CC
,PSMD12,SETD1A,PIGT,COG4,TMCO1,POGZ,ELMO2,SETD1B,KLHL7,PRMT7,MAFB,TCF20
X,RARS1,SETD5,NDUFS1,CDKN1B,SNRPB,COL7A1,MED12,SLC5A7,HIRA,DNAJB6,DDHD2,CTCF,

ALT1,GBA2,DEGS1,NSD1,DHFR,NDUFB8,CLCN4,NUS1,ASH1L,SETX,RNF216,MED25,RARS1,SET
GF2,MTOR,PRKACA,YY1,SMARCD1,CREBBP,PHIP,TANC2,TRIM8,CCDC47,SMO,FGFR1,EIF4A3,SH
TD5,NDUFS1,NSMF,LMNA,MED12,SLC5A7,HIRA,ANOS1,SOS1,PPP1R12A,SGPL1,PPP3CA,SEPTIN

ISD1,RBM10,WAC,ZSWIM6,CLCN4,NUS1,ACSL4,BCOR,SOX9,MED25,SETD5,CHD8,LMNA,MED12,

M20C,PSMD12,TGFB2,FHL1,TMCO1,POGZ,GNAL,SLC18A3,PIEZO1,FGD1,PRMT7,MAFB,GLB1,EI

XL1,RAB11B,ABCA2,PRPF6,TERF2IP,RPS15A,NOTCH2,VHL,PPP2CA,CCND1,MAPK1,SMARCA4,PL
I1,PSAT1,AUTS2,COL12A1,CLIP2,SC5D,ELN,ACTA2,IGF1R,MECP2,EMC1,WDR26,SUMO1,TRRAP,
GZ,GNAO1,SMARCB1,PLXND1,FGD1,HMGA2,PRMT7,MAFB,DPF2,SIN3A,CASK,TCF20
,NDUFA13,TRIM8,REEP1,SMO,FGFR1,TUBB4A,PTPN23,NTRK2,DENND5A,SLC30A9,HS2ST1,MDH

,EIF4A3,SH3PXD2B,RPS26,HS2ST1,BGN,PTPN11,PSAT1,ABL1,AUTS2,COL12A1,CLIP2,SC5D,ELN

KIV2L,SOS1,CITED2,CTCF,TALDO1,TGFBR1,NID1,G6PC3,PSMD12,APOE,TGFB2,PIGT,TMCO1,CC
ALDH18A1,COX8A,ZNF462,FLII,IQSEC1,CCDC88A,ASNS,VPS51,MOGS,CSNK2B,NECTIN1,EXT2,DI
A1,ALDH18A1,LIMK1,PDGFRB,CCDC88A,ASNS,VPS51,SLC25A1,RAB18,STUB1,EXOC7,GBA2,DEG
9,RPS29,SETD5,SNRPB,LMNA,MED12,COX4I1,GALNT2,CNOT3,CUL7,EXT1,PIGS,CUL4B,FAM20C
)1B,CTNND2,PI4KA,RAB3GAP2,ADCY6,CRTAP,IARS1,PTEN,SMC3,ECEL1,HNRNPA2B1,IL6ST,ABC

R,KDM5C,TRIO,EP300,EXTL3,RERE,NDUFV1,KDM5B,PBX1,BAZ1B,ATRX,NXN,ARID1B,PI4KA,FBX

2,COL7A1,SLC5A7,HIRA,ARSB,DSTYK,NIPA1,THRA,COL6A3,CUL7,EXT1,SPG11,PSMD12,FHL1,SE

S2,HSPG2,ANTXR1,KMT2E,GTF2I,CNP,TTC37,TUBGCP6,PACS1,DICER1,LTBP3,LZTR1,PHGDH,N

A1,ALDH18A1,LIMK1,PDGFRB,CCDC88A,ASNS,VPS51,SLC25A1,RAB18,STUB1,EXOC7,GBA2,DEG
A1,ALDH18A1,LIMK1,PDGFRB,CCDC88A,ASNS,VPS51,SLC25A1,RAB18,STUB1,EXOC7,GBA2,DEG

Y,MTOR,PACS2,UBE3A,PRKACA,YY1,SMARCD1,CREBBP,NDUFA13,MN1,PHIP,TANC2,CCDC47,SM

RE,DGCR2,KDM5B,PBX1,VARS1,FLCN,BAZ1B,CSNK2A1,GPX4,ATRX,NXN,ARID1B,CTNND2,PI4KA,

OBTB2,XPC,NUP133,PPP1R12A,EPB41L1,PPP3CA,MAN1B1,PIGS,TARS1,SPG11,POMGNT1,CUL4

6A2,SYNGAP1,STUB1,TSPAN7,GBA2,NSD1,CLCN4,ACSL4,CACNA1H,BCOR,SETX,SOX9,MED25,RI

AD50,PRMT7,DPF2,EPHA2,DMPK,SIN3A,DNAJC21,CASK,TCF20
2B1,ZNF592,NACC1,LARGE1,SLC5A6,RHOBTB2,PMPCA,XPC,COX4I1,EPB41L1,PPP3CA,TRAPPC1
ALDH7A1,RAB18,SYNGAP1,PPP2R5D,DEAF1,CHST3,NSD1,WAC,ZSWIM6,NDUFB8,SHMT2,CLCN4,

ASK,TCF20

L1,PPP3CA,NID1,EXT1,TET3,TARS1,SPG11,POMGNT1,CUL4B,SEC31A,PRNP,SMARCC2,TMCO1,(

,PPT1,EPHA4,TMCO1,DNM1,HARS1,DNMT1,SMARCB1,NF2,SETD1B,NFIB,JRK,GSN,UBTF,TOMM

,PIGT,COG4,SMARCC2,COL3A1,MAPK8IP3,GNAO1,SMARCB1,PLXND1,FGD1,KLHL7,HMGA2,PR

M3AP,MAPRE2,PREPL,GPC4,SLC2A1,NDN,H19,CLCN7,TAOK1,EHMT1,IARS2,HSPG2,ANTXR1,SO

;,PRDX1,ODC1,ACADVL,FTO,PRKDC,SON,KDM1A,MAP2K2,FLNB,DLK1,TPM3,SMARCE1,RPS28,A

EN,SMC3,HNRNPA2B1,ADH5,NDST1,EML1,PREPL,TIMM50,ATP6AP2,GPC4,SLC2A1,NDN,BAP1,F

1,SPG11,FH,PSMD12,OPA1,APOE,SZT2,DYNC1I2,PIGT,PRNP,PPT1,COG4,TMCO1,POGZ,DNM1,M
NP,PPT1,NBEA,POGZ,DNM1,GNAO1,SLC18A3,SETD1B,JRK,CSTB,UBTF,CASK,CC2D1A,NECAP1,A
9,SETD5,SNRPB,LMNA,RHOBTB2,NUP133,MED12,SLC5A7,HIRA,SOS1,PPP1R12A,DSTYK,ZDHHC
RBM10,SOX9,RPS29,SETD5,SNRPB,LMNA,RHOBTB2,NUP133,MED12,SLC5A7,HIRA,SOS1,PPP1F
D17B4,NSD2,DHCR7,MED13,PCGF2,UBE3A,PRKACA,YY1,CREBBP,PHIP,TANC2,SMO,FGFR1,EIF4
F1,ZNF462,EFNB1,FLII,VPS51,EXT2,RAB18,RDH11,DEAF1,CRKL,TSPAN7,RBM10,WAC,ZSWIM6,C
D2B,CUL4B,FAM20C,PSMD12,TGFB2,SETD1A,FHL1,SZT2,DYNC1I2,PIGT,TMCO1,DNM1,MAPK8II
GNAO1,SMARCB1,PLXND1,SLC18A3,FGD1,KLHL7,HMGA2,MAFB,DPF2,CASK

9,SETD5,SNRPB,LMNA,RHOBTB2,NUP133,MED12,SLC5A7,HIRA,SOS1,PPP1R12A,DSTYK,ZDHHC

GNAO1,SMARCB1,PLXND1,SLC18A3,FGD1,KLHL7,HMGA2,MAFB,DPF2,CASK

K4I1,GALNT2,COL6A3,MAN1B1,GATAD2B,CUL4B,FAM20C,PSMD12,FHL1,SYT1,SMARCC2,TMCO

2,APOE,SZT2,DYNC1I2,PIGT,PRNP,PPT1,COG4,TMCO1,POGZ,DNM1,MAPK8IP3,DNMT1,GNAO1,K

2,WDR26,ADNP,IDH2,EWSR1,ALDH18A1,LIMK1,FLII,PRPH,PDGFRB,CSNK2B,NECTIN1,EXT2,CDK

LDH18A1,PRPH,CAPN1,EXT2,SLC25A1,ASAH1,SYNGAP1,TSPAN7,GBA2,NDRG1,WDR11,DHFR,ND

N1,EXT2,DDX6,SLC25A1,RAB18,RDH11,DEAF1,CHST3,NSD1,RBM10,BCOR,SOX9,RPS29,SETD5,S

,PRPF6,ATP6AP1,TERF2IP,RPS15A,NOTCH2,VHL,PPP2CA,MAPK1,SMARCA4,PLEC,ODC1,AEBP1,A

,NFASC,SMARCC2,CENPF,TMCO1,GNAO1,SMARCB1,PLXND1,SLC18A3,FGD1,KLHL7,HMGA2,MA

B,GLB1,SIN3A,CASK

SNRPN,PLOD1,VPS35,COL2A1,SKI,ASXL1,RAB11B,ATP6AP1,TERF2IP,RPS15A,NOTCH2,MARS1,SI

IGA2,PRMT7,MAFB,GLB1,SIN3A,CASK

UL4B,OPA1,FHL1,EPHA4,HARS1,SMARCB1,ITGA7,PLXND1,SLC18A3,HMGA2,NEMF,ATP8A2,MAFI

L6A3,EXT1,PIGS,GATAD2B,CUL4B,PSMD12,COG4,SMARCC2,TMCO1,COL3A1,SMARCB1,SLC18/

GFR1,RPS26,HS2ST1,TUB,PTPN11,ABL1,AUTS2,TMEM94,POR,CLIP2,DAG1,SC5D,ELN,OBSL1,AC
LDLR,KMT2E,GTF2I,TUBGCP6,ELP1,PHGDH,MSH6,NSD2,DHCR7,RBPJ,MED13,CHMP4B,DHX38,I

RERE,DGCR2,NDUFV1,KDM5B,PBX1,ALDH3A2,FLCN,BAZ1B,GPX4,ATRX,GARS1,NXN,ARID1B,CTI

EP1,SMO,FGFR1,EIF4A3,SH3PXD2B,HS2ST1,BGN,PTPN11,PSAT1,CHMP1A,AUTS2,COL12A1,POF

1I2,PIGT,CCNK,SMARCC2,POGZ,MAPK8IP3,GNAO1,SMARCB1,NFIB,FGD1,KLHL7,PRMT7,MAFB
RIM8,SMO,FGFR1,RPS26,HS2ST1,TUB,PTPN11,ABL1,AUTS2,TMEM94,POR,CLIP2,DAG1,SC5D,E

P1,RDH11,DEAF1,TSPAN7,CHST3,WDR11,NSD1,ZSWIM6,CLCN4,NUS1,ACSL4,BCOR,SETD5,MAN
EF2,ADAR,PALLD,CIC,TRAF7,DCTN1,KIF5A,STAT3,KCNQ2,BCR,TRIO,EP300,RERE,NDUFV1,KDM

,HSPG2,SOX2,PKDCC,LTBP4,AKT3,LDLR,DZIP1,EFTUD2,GTF2I,TTC37,PACS1,LTBP3,LZTR1,PH

M,LSS,STXBP1,LONP1,PRPS1,IDH1,VANGL2,SEC23A,ARNT2,ARFGEF2,BMPR2,SMS,CCT5,CDC4

32,RPL27,SNRPN,ACO2,PLOD1,VPS35,COL2A1,SKI,ASXL1,KDM6B,CNTNAP2,RAB11B,ABCA2,RPS
3P1,ACADVL,FTO,ATXN10,HSPB1,SON,KDM1A,MAP2K2,QARS1,DLK1,TPM3,SMARCE1,DHX30,AT
3B,ARFGEF2,SMS,CCT5,CDC42,YME1L1,ADAR,CIC,TRAF7,DCTN1,KIF5A,STAT3,KCNQ2,WDR45E
RERE,KDM5B,ALDH3A2,FLCN,BAZ1B,CSNK2A1,GPX4,ATRX,GARS1,NXN,ARID1B,CTNND2,RAB3(
7,CTBP1,RAC1,ARHGDIA,GPC1,SMCHD1,MMP2,L1CAM,GLUL,CDKN1A,CRPPA,HS6ST1,NOVA2,CI
RNPK,NARS1,ATP1A2,HLA-B,TTR,ARCN1,RPS27,CTBP1,RAC1,ARHGDIA,GPC1,SMCHD1,MMP2,L
YNC1I2,COG4,SMARCC2,TMCO1,POGZ,BRAT1,MAPK8IP3,GNAO1,SMARCB1,PDHB,POU3F3,KLI
ON,VHL,PPP2CA,CCND1,MAPK1,SMARCA4,KDELR2,PLEC,PRDX1,ODC1,AEBP1,ACADVL,FTO,PRK
BPN1,TAF15,LMNB2,RPL27,SNRPN,ACO2,PLOD1,VPS35,COL2A1,CHD2,SKI,ASXL1,CNTNAP2,RAI
3P,MN1,PHIP,TANC2,CCDC47,SMO,FGFR1,SH3PXD2B,RPS26,HS2ST1,BGN,TRIP12,PTPN11,PSAT
N1,ARNT2,KDM3B,ARFGEF2,SMS,CDC42,YME1L1,ADAR,CIC,TRAF7,KIF5A,MSMO1,KCNQ2,WDR
,ARFGEF2,SMS,CDC42,YME1L1,ADAR,CIC,TRAF7,KIF5A,MSMO1,KCNQ2,WDR45B,BCR,KDM5C,
DUX4,ATP1A2,HLA-B,TTR,ARCN1,RPS27,CTBP1,RAC1,SMCHD1,MMP2,L1CAM,PPIB,GLUL,CDKN
,KDM5C,TRIO,EP300,EXTL3,RERE,DGCR2,NDUFV1,KDM5B,CNTNAP1,ALDH3A2,VARS1,FLCN,BA
3,WDR11,NSD1,RBM10,WAC,ZSWIM6,SOX9,MED25,RPS29,MAN2B1,NSMF,CHD8,LMNA,NUP133,
L2A1,RBMX,CHD2,SKI,ASXL1,KDM6B,CNTNAP2,RAB11B,ABCA2,PRPF6,ATP6AP1,RPS15A,MARS1
K1,TPM3,SMARCE1,DHX30,RPS28,ATP6V1B2,SDHA,LARS1,LSS,STT3B,STXBP1,STT3A,LONP1,PF
S,ELP1,PACS1,LZTR1,PHGDH,HSD17B4,NSD2,DHCR7,MED13,PCGF2,FBLN5,UBE3A,SMARCD1,C
KDM6B,RAB11B,PRPF6,RPS15A,NOTCH2,MARS1,SELENON,TUBB2A,OPTN,VHL,PPP2CA,CCND1,
D2,ZNF462,EFNB1,FLII,NBAS,PDGFRB,VPS51,CSNK2B,NECTIN1,EXT2,DDX6,RAB18,PPP2R5D,DE
P2,FBXO11,IARS1,HDAC6,PTEN,SMC3,ADH5,NDST1,EML1,ABCC9,SPINT2,MAPRE2,B4GAT1,GPC
AP2,FBXO11,IARS1,ZFR,SMC3,ADH5,NDST1,EML1,MAPRE2,B4GAT1,ATP6AP2,GPC4,SLC2A1,FU
VHL,PPP2CA,CCND1,MAPK1,SMARCA4,PRDX1,ODC1,FTO,PRKDC,ATXN10,SON,KDM1A,MAP2K2,
P2,RAB11B,ABCA2,PRPF6,ATP6AP1,RPS15A,MARS1,SELENON,OPTN,VHL,PPP2CA,MAPK1,SMARC
M3,SMARCE1,DHX30,RPS28,ATP6V1B2,SDHA,LARS1,LSS,FBLN1,B2M,STXBP1,LONP1,PRPS1,USI
TAF15,LMNB2,ABCA3,RPL27,SNRPN,PDXK,ACO2,PLOD1,COL2A1,SKI,ASXL1,KDM6B,CNTNAP2,R
CTA2,IGF1R,EMC1,WDR26,ADNP,TRRAP,NDUFC2,ALDH18A1,NTN1,HTRA1,AMPD2,ZNF462,EFNB
D2,SKI,ASXL1,CNTNAP2,RAB11B,ABCA2,RPS15A,NOTCH2,TUBB2A,OPTN,VHL,PPP2CA,CCND1,M
0,RPS28,SDHA,LARS1,LSS,FBLN1,STT3B,STXBP1,STT3A,LONP1,PRPS1,MESD,USP7,SARS1,COL
51,NECTIN1,DDX6,RAB18,SLC16A2,SYNGAP1,RDH11,CRKL,CHST3,NSD1,RBM10,WAC,BCOR,RF
TET3,PIGS,GATAD2B,CUL4B,PSMD12,SYT1,PIGT,CCNK,SMARCC2,TMCO1,POGZ,BRAT1,COL3A1
IC2,CCDC47,FGFR1,TUBB4A,SH3PXD2B,HS2ST1,PTPN11,PSAT1,ABL1,AUTS2,TMEM94,COL12A1
S,GRN,SPG7,TAOK1,EHMT1,VAPB,HSPG2,LINGO1,TPP1,SLC6A8,LTBP4,IREB2,KMT2E,EFTUD2,(
,LTBP4,AKT3,KMT2E,EFTUD2,GTF2I,TTC37,TUBGCP6,PACS1,DICER1,LTBP3,HSD17B4,NSD2,DH

C88A,ASNS,VPS51,MOGS,CSNK2B,DDX6,DARS1,RAB18,SYNGAP1,PPP2R5D,STUB1,DEAF1,EXO(
KDM5C,TRIO,EP300,EXTL3,RERE,BAZ1B,CSNK2A1,GPX4,ATRX,GARS1,NXN,ARID1B,CTNND2,PI4I
PSAT1,CHMP1A,TRAK1,DAG1,NOTCH1,ATP13A2,PDHA1,ACTA2,IGF1R,NDUFA4,EMC1,WDR26,AD
9,MAPRE2,LOX,GPC4,NDN,H19,PSMB4,CLCN7,EHMT1,IARS2,ERGIC1,HSPG2,PKDCC,AKT3,HINT
CDH11,MDH2,RAB27A,EEF1A2,FDFT1,TBCD,CPSF1,PPP1CB,SQSTM1,VAMP2,GNAI3,PABPN1,VCA
ACTA2,IGF1R,MECP2,ALDH18A1,BANF1,LIMK1,EFNB1,PDGFRB,MOGS,EXT2,DDX6,SLC25A1,ASAI
RAF7,KIF5A,STAT3,KCNQ2,BCR,KDM5C,TRIO,EP300,EXTL3,RERE,STARD7,NDUFV1,KDM5B,CNTI

RBPJ,MED13,PCGF2,AHCY,FBLN5,UBE3A,GOT2,SMARCD1,CREBBP,TANC2,TRIM8,CCDC47,FGF

TBCD,PPP1CB,SQSTM1,GNAI3,PABPN1,TAF15,LMNB2,ABCA3,RPL27,SNRPN,PDXK,ACO2,PLOD1,
A,RAB3GAP2,FBXO11,IARS1,MBTPS1,HDAC6,PTEN,SMC3,ECEL1,ADH5,IL6ST,ABCC9,MAPRE2,AT
GT,DHCR7,RBPJ,MED13,PCGF2,AHCY,FBLN5,UBE3A,GOT2,SMARCD1,CREBBP,TANC2,TRIM8,C
,BAZ1B,CSNK2A1,GPX4,ATRX,GARS1,ARID1B,CTNND2,PI4KA,RAB3GAP2,ADCY6,FBXO11,IARS1,I
TEN,SMC3,ECEL1,ADH5,ABCC9,MAPRE2,PREPL,GPC4,NDN,PSMB4,EHMT1,IARS2,HSPG2,ANTXR
P2,CRTAP,FBXO11,IARS1,MBTPS1,HDAC6,PTEN,SMC3,ECEL1,ADH5,IL6ST,ABCC9,MAPRE2,B4G
P2,CRTAP,FBXO11,IARS1,MBTPS1,HDAC6,PTEN,SMC3,ECEL1,ADH5,IL6ST,ABCC9,MAPRE2,B4G

JC21,CASK,CNOT1,TCF20
B,BCR,KDM5C,EP300,EXTL3,RERE,NDUFV1,PBX1,CNTNAP1,BAZ1B,UBQLN2,ATRX,GARS1,NXN,A

3,SQSTM1,GNAI3,KLHL24,PABPN1,TAF15,LMNB2,RPL27,SNRPN,ACO2,PLOD1,VPS35,COL2A1,R
37,TUBGCP6,PACS1,TCF3,LTBP3,LZTR1,PHGDH,HSD17B4,NSD2,DHCR7,RBPJ,MED13,PCGF2,FE

H18A1,ZNF462,FLII,IQSEC1,CCDC88A,NECTIN1,EXT2,SOBP,DDX6,ALDH7A1,SLC16A2,SYNGAP1,F
T1,IARS2,ERGIC1,HSPG2,PKDCC,LTBP4,AKT3,HINT1,KMT2E,EFTUD2,GTF2I,SEMA3C,PACS1,LTB

,DHX30,RPS28,ATP6V1B2,FBLN1,B2M,STXBP1,LONP1,PRPS1,MESD,HYOU1,VANGL2,SOD1,RAI1

3,SQSTM1,GNAI3,KLHL24,PABPN1,TAF15,LMNB2,RPL27,SNRPN,ACO2,PLOD1,VPS35,COL2A1,R
,PREPL,GPC4,NDN,PSMB4,EHMT1,IARS2,HSPG2,ANTXR1,PKDCC,AKT3,LDLR,KMT2E,EFTUD2,G

4,POR,CLIP2,SC5D,ELN,OBSL1,PDHA1,IGF1R,MECP2,EMC1,WDR26,SUMO1,ADNP,TRRAP,LIMK1
GBA2,GPT2,NSD1,WAC,ZSWIM6,SHMT2,GDF1,CLCN4,ASH1L,BCOR,MED25,RARS1,MBOAT7,LA

SIN3A,CASK,TCF20
1,SH3PXD2B,RPS26,NTRK2,DENND5A,HS2ST1,MDH1,BGN,PTPN11,PSAT1,ABL1,PUM1,POR,CLIF
CP6,PACS1,TCF3,LZTR1,PHGDH,HSD17B4,NSD2,DHCR7,MED13,PCGF2,YY1,SMARCD1,CREBBP,I
M94,COL12A1,TRAK1,DAG1,SC5D,ELN,PDHA1,TRIM2,EMC1,WDR26,ADNP,TRRAP,GPAA1,ALDH1
4,NDN,PSMB4,EHMT1,IARS2,HSPG2,ANTXR1,PKDCC,AKT3,LDLR,KMT2E,EFTUD2,GTF2I,TTC37,T

WAC,ZSWIM6,SHMT2,GDF1,CLCN4,ASH1L,BCOR,MED25,RARS1,MBOAT7,LARGE1,SLC5A6,RHC
S15A,NOTCH2,SELENON,OPTN,VHL,PPP2CA,MAPK1,SMARCA4,PLEC,PRDX1,ODC1,AEBP1,ADAM
IOS1,SOS1,DDHD2,RUSC2,ZDHHC9,TGFBR1,G6PC3,COL6A3,TET3,TARS1,GATAD2B,CUL4B,FAM
1,G6PC3,TET3,TARS1,PSMD12,SETD1A,SEC31A,SYT1,PIGT,TMCO1,GNAO1,PIEZO1,CRB2,DPF2,C
3,SH3PXD2B,RPS26,HS2ST1,BGN,TRIP12,TECR,PTPN11,PSAT1,ABL1,PUM1,COL12A1,DAG1,SC5
1A,AUTS2,TMEM94,COL12A1,CLIP2,NOP56,KLF13,ELN,NOTCH1,ATP13A2,PDHA1,IGF1R,NDUFA4

MO,FGFR1,EIF4A3,SH3PXD2B,RPS26,HS2ST1,BGN,PTPN11,ABL1,PUM1,AUTS2,TMEM94,COL12A
J,MED13,PCGF2,AHCY,COX6A1,FBLN5,MTOR,YY1,SMARCD1,CREBBP,TANC2,WNK1,TRIM8,CCD
FN2,PDHA1,TRIM2,MECP2,EMC1,WDR26,PARK7,GPAA1,BSCL2,ALDH18A1,HTRA1,LIMK1,AMPD2

ESD,HYOU1,VANGL2,RAI1,SEC23A,TCTN1,ARNT2,KDM3B,SMS,CDC42,SYP,CTNND1,TRAF7,STAT3
ESD,HYOU1,VANGL2,RAI1,SEC23A,TCTN1,ARNT2,KDM3B,SMS,CDC42,SYP,CTNND1,TRAF7,STAT3
ST,ABCC9,MCM3AP,MAPRE2,B4GAT1,ATP6AP2,LOX,GPC4,NDN,SPG7,H19,CLCN7,EHMT1,IARS2
ADNP,ALDH18A1,EFNB1,FLII,NBAS,MOGS,EXT2,DDX6,RAB18,SLC16A2,PPP2R5D,DEAF1,CHST3,I
2,PTPN11,PSAT1,PUM1,AUTS2,TMEM94,POR,CLIP2,SC5D,ELN,OBSL1,PDHA1,IGF1R,MDM2,MECI
00,EXTL3,RERE,STARD7,NDUFV1,KDM5B,CNTNAP1,VARS1,FLCN,BAZ1B,UBQLN2,ATRX,GARS1,(

5,SC5D,ELN,ATP13A2,MFN2,PDHA1,MECP2,NDUFA4,WDR26,FXR1,PARK7,ADNP,GPAA1,BSCL2,AI

,SLC16A2,SYNGAP1,PPP2R5D,DEAF1,TSPAN7,DEGS1,GPT2,NSD1,WAC,ZSWIM6,SHMT2,CLCN4,N

1,VPS35,COL2A1,RBMX,SKI,ASXL1,KDM6B,CNTNAP2,RAB11B,PRPF6,RPS15A,NOTCH2,SELENON
BSCL2,ALDH18A1,HTRA1,LIMK1,COX8A,AMPD2,KLC2,PRPH,IQSEC1,ASNS,CSNK2B,CAPN1,ECHS

,AUTS2,TMEM94,CLIP2,KLF13,ELN,OBSL1,PDHA1,IGF1R,EMC1,WDR26,ADNP,GPAA1,BSCL2,ALD
D5A,BGN,PTPN11,PSAT1,AUTS2,POR,SC5D,OBSL1,NOTCH1,PDHA1,MECP2,ADNP,ALDH18A1,BAN
ECEL1,HNRNPA2B1,ADH5,MCM3AP,MAPRE2,PREPL,TIMM50,LOX,GPC4,SLC2A1,NDN,H19,CLCN
SK,CC2D1A,NECAP1,TCF20

12,SLC5A7,RUSC2,GALNT2,SLC1A3,CNOT3,COL6A3,MAN1B1,TET3,SLC1A4,SPG11,TNPO3,POM
MNA,MED12,SLC5A7,HIRA,SOS1,CTCF,RUSC2,PPP1R12A,ZDHHC9,TALDO1,GALNT2,SEPTIN9,CN
LUD1,CTNND2,PI4KA,RAB3GAP2,FBXO11,IARS1,PTEN,ZFR,SMC3,NDST1,EML1,MAPRE2,PREPL,
CH1,ATP13A2,PDHA1,IGF1R,NDUFA4,FAM50A,EMC1,WDR26,ADNP,ALDH18A1,HMGB3,LIMK1,EI

C2,CCDC47,FGFR1,SH3PXD2B,HS2ST1,BGN,PTPN11,PSAT1,ABL1,CHMP1A,AUTS2,COL12A1,POF
,SOX9,MED25,RPS29,SETD5,SNRPB,LARGE1,LMNA,MED12,SLC5A7,HIRA,SOS1,CTCF,RUSC2,PPI
GCR2,STARD7,NDUFV1,KDM5B,ALDH3A2,VARS1,FLCN,BAZ1B,CSNK2A1,DIP2B,ATRX,NXN,ARID1I
4KA,RAB3GAP2,IARS1,PTEN,ZFR,SMC3,HNRNPA2B1,NDST1,EML1,B4GAT1,TIMM50,ATP6AP2,A2I

LGALT1,GBA2,DEGS1,GPT2,NSD1,DLD,NDUFB8,SHMT2,CLCN4,NUS1,ASH1L,BCOR,SETX,MED2

EPB41L1,PPP3CA,PCNA,SLC1A4,SPG11,OPA1,APOE,SZT2,SEC31A,SYT1,PIGT,FBXO7,PRNP,POGZ

D5,MBOAT7,NDUFS1,MAN2B1,ZNF592,NACC1,LARGE1,SLC5A6,RHOBTB2,PMPCA,XPC,UNC13A,
H3PXD2B,RPS26,BGN,PTPN11,PSAT1,PUM1,AUTS2,COL12A1,CLIP2,MACROH2A1,SC5D,ELN,OBS
9,QRICH1,TET3,PSMD12,OPA1,TGFB2,FHL1,SZT2,SMARCC2,TMCO1,COL3A1,DNM1,SMARCB1,I

,HIRA,DGCR8,SOS1,PPP1R12A,GALNT2,PPP3CA,SEPTIN9,CNOT3,QRICH1,MAN1B1,GATAD2B,CL

MD

.EC,PRDX1,ODC1,AEBP1,ADAM9,PRKDC,SON,KDM1A,MAP2K2,QARS1,FLNB,DLK1,TPM3,SMARC
BSCL2,BANF1,BNC2,LIMK1,FLII,PDGFRB,CCDC88A,ASNS,MOGS,CSNK2B,NECTIN1,EXT2,DDX6,

H1,TECR,PSAT1,PUM1,CHMP1A,PINK1,AUTS2,CLIP2,TRAK1,NOP56,ELN,NOTCH1,ATP13A2,MFN2

I,ACTA2,IGF1R,MECP2,EMC1,WDR26,SUMO1,TRRAP,BSCL2,BANF1,BNC2,LIMK1,FLII,PDGFRB,C

DL3A1,CFAP298,TCF20
DX6,SLC25A1,ASAH1,SLC16A2,SYNGAP1,DEAF1,DEGS1,GPT2,NSD1,DLD,WAC,ZSWIM6,DHFR,ND
S1,NSD1,RBM10,STAT2,WAC,ZSWIM6,DHFR,BCOR,SETX,RNF216,SETD5,NDUFS1,DLG4,MAN2B
C,PSMD12,PIGT,COG4,SMARCC2,COL3A1,MAPK8IP3,GNAO1,SMARCB1,PLXND1,FGD1,KLHL7,HI
C9,LOX,GPC4,FUS,SPG7,H19,PSMB4,CLCN7,IARS2,ERGIC1,HSPG2,ANTXR1,SOX2,SLC6A8,PKDC

D11,PTEN,SMC3,MCM3AP,SLC2A1,NDN,BAP1,FUS,SPG7,CLCN7,IARS2,HSPG2,ANTXR1,TPP1,SO

C31A,PIGT,SMARCC2,COL3A1,SMARCB1,ITGA7,CENPT,SLC18A3,FGD1,PRMT7,MAFB,DPF2,GLE

SD2,DHCR7,RBPJ,MED13,PCGF2,AHCY,MTOR,PACS2,UBE3A,PRKACA,YY1,SMARCD1,CREBBP,NI

S1,NSD1,RBM10,STAT2,WAC,ZSWIM6,DHFR,BCOR,SETX,RNF216,SETD5,NDUFS1,DLG4,MAN2B
S1,NSD1,RBM10,STAT2,WAC,ZSWIM6,DHFR,BCOR,SETX,RNF216,SETD5,NDUFS1,DLG4,MAN2B

MO,FGFR1,RPS26,CNBP,DENND5A,HS2ST1,PTPN11,PSAT1,CHMP1A,AUTS2,TMEM94,POR,H6PD,I

,RAB3GAP2,HDAC6,PTEN,SMC3,ECEL1,PTPRF,ABCC9,SPINT2,MAPRE2,PREPL,LMAN1,GPC4,NDN

B,FH,PSMD12,APOE,DYNC1I2,PRNP,TMCO1,POGZ,COL3A1,MAPK8IP3,GNAO1,PDHB,KLHL7,UB

PS29,SETD5,STAT1,LMNA,NR2F2,COL7A1,MED12,HIRA,DDHD2,PPP1R12A,DSTYK,NIPA1,GALNT2

1,MAN1B1,SLC1A4,UFM1,PIGS,TARS1,SPG11,FH,PSMD12,OPA1,APOE,SZT2,DYNC1I2,PIGT,PRN
ASH1L,BCOR,SOX9,MED25,SETD5,MAN2B1,COX6B1,LARGE1,RHOBTB2,PMPCA,NUP133,THOC2

COL3A1,DNMT1,CFAP298,GNAO1,SMARCB1,SLC18A3,PDHB,KLHL7,CRB2,UBTF,DPF2,GLB1,DM

40,DMPK,CC2D1A,NDUFA10,NECAP1,ARSA,ABCA7,TCF20

MT7,MAFB,DPF2,SIN3A,CASK,TCF20

X2,SLC6A8,PKDCC,EFTUD2,GTF2I,TTC37,TUBGCP6,SEMA3C,LTBP3,LZTR1,PHGDH,HSD17B4,NS

TP6V1B2,SDHA,LARS1,LSS,STT3B,STXBP1,STT3A,LONP1,PRPS1,MESD,USP7,IDH1,HYOU1,SARS

US,GRN,SPG7,TAOK1,EHMT1,VAPB,HSPG2,ANTXR1,LINGO1,SOX2,SLC6A8,AKT3,IREB2,KMT2E,

/IAPK8IP3,DNMT1,GNAO1,KLHL7,ATP8A2,UBTF,TOMM40,GLB1,DMPK,CASK,CC2D1A,NECAP1,AI
\RSA
9,TGFBR1,PPP3CA,COL6A3,EXT1,TARS1,POMGNT1,GATAD2B,FAM20C,PSMD12,SEC31A,PIGT,C
R12A,DSTYK,ZDHHC9,TGFBR1,PPP3CA,COL6A3,EXT1,TARS1,POMGNT1,GATAD2B,FAM20C,PSMI
A3,SH3PXD2B,RPS26,HS2ST1,BGN,PTPN11,TAPBP,PSAT1,ABL1,AUTS2,COL12A1,POR,CLIP2,IGS
LCN4,ACSL4,BCOR,SOX9,MED25,SETD5,MAN2B1,NACC1,LMNA,XPC,MED12,HIRA,DGCR8,CTCF
P3,CENPT,SETD1B,NFIB,FGD1,KLHL7,DPF2,SIN3A,DNAJC21,NECAP1,TCF20

9,TGFBR1,PPP3CA,COL6A3,TARS1,POMGNT1,GATAD2B,FAM20C,PSMD12,SEC31A,PIGT,COG4,(

1,COL3A1,HARS1,GNAO1,SMARCB1,DPF2,SIN3A,CASK,TCF20

KLHL7,ATP8A2,UBTF,TOMM40,GLB1,DMPK,CASK,CC2D1A,NECAP1,ABCA7,TCF20

4,SLC25A1,ASAH1,JAK1,SYNGAP1,STUB1,DEAF1,GBA2,NSD1,WAC,ZSWIM6,NDUFB8,CLCN4,NUS

UFB8,SHMT2,OAT,CLCN4,NUS1,ACSL4,PHKB,TMEM43,BCOR,SETX,MED25,RARS1,IRF2BP2,NDI

SNRPB,LMNA,RHOBTB2,NUP133,MED12,SLC5A7,HIRA,ARSB,SOS1,PPP1R12A,DSTYK,ZDHHC9,T(

ACADVL,FTO,SON,KDM1A,MAP2K2,FLNB,DLK1,SMARCE1,DHX30,RPS28,ATP6V1B2,SDHA,PFKM,[illegible]

AFB,DPF2,CASK

ELENON,OPTN,VHL,PPP2CA,CCND1,MAPK1,SMARCA4,PLEC,PRDX1,ODC1,AEBP1,ACADVL,FTO,A

B,GLB1,DMPK,EMD

A3,FGD1,KLHL7,MAFB,DPF2,CNOT1,TCF20

CTA2,MDM2,MECP2,JAM3,FXR1,ADNP,TRRAP,EWSR1,ALDH18A1,BNC2,LIMK1,EFNB1,NECTIN1,E
PACS2,UBE3A,YY1,SMARCD1,CREBBP,TANC2,WNK1,SMO,FGFR1,TEAD1,SH3PXD2B,RPS26,CNBI

NND2,PI4KA,RAB3GAP2,CRTAP,FBXO11,IARS1,MBTPS1,HDAC6,PTEN,SMC3,ECEL1,HNRNPA2B1,

R,CLIP2,DAG1,MACROH2A1,ELN,OBSL1,MFN2,TRIM2,ALDH18A1,NTN1,LIMK1,COX8A,EFNB1,PD

;,GLB1,SIN3A,CASK

_N,OBSL1,ACTA2,MDM2,MECP2,JAM3,FXR1,ADNP,TRRAP,EWSR1,ALDH18A1,BNC2,LIMK1,EFNB

l2B1,SNRPB,NSMF,SRP54,LMNA,XPC,NUP133,COL7A1,MED12,HIRA,ANOS1,SOS1,CTCF,RUSC2,Z

5B,PBX1,CNTNAP1,VARS1,FLCN,BAZ1B,UBQLN2,CSNK2A1,ATRX,ARID1B,PI4KA,RAB3GAP2,ADC

GDH,MSH6,NSD2,DHCR7,RBPJ,PCGF2,AHCY,FBLN5,PACS2,PRKACA,YY1,CREBBP,NDUFA13,TRIM

2,ADAR,CTNND1,PALLD,KIF5A,MSMO1,STAT3,KCNQ2,KDM5C,EP300,EXTL3,RERE,ALDH3A2,FL

15A,MARS1,SELENON,TUBB2A,OPTN,VHL
P6V1B2,SDHA,LARS1,PFKM,LSS,FBLN1,STT3
,KDM5C,EP300,EXTL3,RERE,DGCR2,NDUFV
AP2,CRTAP,MBTPS1,HDAC6,PTEN,SMC3,E
DH11,MDH2,RAB27A,EEF1A2,FDFT1,TBCD,
1CAM,GLUL,CDKN1A,CRPPA,HS6ST1,NOVA2
L7,UBTF,PRMT7,DPF2,SIN3A,CASK,CC2D1A
DC,SON,KDM1A,MAP2K2,QARS1,FLNB,DLK1,T
B11B,ABCA2,RPS15A,NOTCH2,MARS1,
1,ABL1,AUTS2,TMEM94,POR,KLF13,ELN,ACT
45B,BCR,KDM5C,TRIO,EP300,EXTL3,RERE,
TRIO,EP300,EXTL3,RERE,DGCR2,NDUFV1,K
1A,CRPPA,HS6ST1,NOVA2,CDH11,MDH2,R
Z1B,CSNK2A1,GPX4,ATRX,ARID1B,CTNND2
,THOC2,CNTN1,MED12,HIRA,SKIV2L,ANOS1,S
,SELENON,TUBB2A,PPP2CA,MAPK1,SMARCA4,
PS1,MESD,USP7,HYOU1,SARS1,VANGL2,RAI1,
REBBP,TANC2,WNK1,CCDC47,FGFR1,TUBB4A,
MAPK1,SMARCA4,PLEC,ODC1,ATXN10,ACT
F1,EXOC7,CRKL,TSPAN7,CHST3,NSD1,WAC,Z
,SLC2A1,FUS,H19,CLCN7,TAOK1,EHMT
S,TAOK1,EHMT1,HSPG2,LINGO1,SOX2,SL
QARS1,FLNB,DLK1,SMARCE1,DHX30,RPS28,
A4,PLEC,PRDX1,ODC1,ACADVL,ATXN10,SON
7,IDH1,HYOU1,SARS1,COL27A1,SOD1,RAI1
AB11B,ABCA2,RPS15A,NOTCH2,MARS1,SE
1,FLII,IQSEC1,CCDC88A,ASNS,VPS51,MOGS,CSNK2
APK1,SMARCA4,PLEC,PRDX1,ODC1,FTO
27A1,SOD1,RAI1,SEC23A,TCTN1,ARNT2,KD
S29,SETD5,MAN2B1,LARGE1,LMNA,RHOBTB2,
,MAPK8IP3,SMARCB1,NFIB,FGD1,PRMT7,DPF
,POR,CLIP2,ELN,OBSL1,MFN2,ACTA2,MECP2,
TF2I,DNAJB2,ATXN3,PACS1,LZTR1,HSD1
CR7,MED13,MTOR,DHX38,PACS2,YY1,SMARC

7,GBA2,GPT2,NSD1,RBM10,WAC,ZSWIM6,SHMT2,G
A,RAB3GAP2,CRTAP,FBXO11,IARS1,MBTPS
NP,TRRAP,NDUFC2,BSCL2,ALDH18A1,NTN1,HTRA1,
1,KMT2E,EFTUD2,GTF2I,SEMA3C,PACS1,L
N,TGFBI,RPL27,SNRPN,PDXK,YEATS2,AC
H1,SLC16A2,CHST3,NSD1,RBM10,P3H1,TMEM43,
AP1,ALDH3A2,VARS1,FLCN,CSNK2A1,DIP

R1,TUBB4A,PTPN23,RPS26,NTRK2,DENND5A

COL2A1,SKI,ASXL1,KDM6B,CNTNAP2,RAB1
TP6AP2,LOX,GPC4,NDN,SPG7,H19,PSMB4,C
CDC47,FGFR1,TUBB4A,PTPN23,RPS26,NTR
PTEN,SMC3,NDST1,MCM3AP,MAPRE2,PREPL,
1,PKDCC,AKT3,LDLR,KMT2E,EFTUD2,GTF2I,TTC3
AT1,ATP6AP2,LOX,GPC4,SLC2A1,NDN,H1
AT1,ATP6AP2,LOX,GPC4,SLC2A1,NDN,H1

RID1B,PI4KA,RAB3GAP2,ADCY6,PTEN

BMX,SKI,ASXL1,KDM6B,CNTNAP2,RAB11B,AB
LN5,MTOR,PACS2,UBE3A,YY1,SMARCD1,CREBBP,

PP2R5D,DEAF1,CRKL,TSPAN7,DEGS1,GPT2,NS
P3,LZTR1,PHGDH,HSD17B4,NSD2,DHCR7

,SEC23A,TCTN1,ARNT2,KDM3B,SMS,CDC42,SYP,CT

BMX,SKI,ASXL1,KDM6B,CNTNAP2,RAB11B,AB
TF2I,TTC37,TUBGCP6,SLCO1B3,PACS1,TCF3,LT

,AMPD2,ZNF462,EFNB1,FLII,NBAS,PDGFRB,VPS
RGE1,SLC5A6,RHOBTB2,MED12,DDHD2,RUSC2,PPP1

2,TRAK1,SC5D,ELN,OBSL1,PDHA1,FAM50A,EM
MN1,PHIP,TANC2,CCDC47,FGFR1,EIF4A3,SH
A1,ZNF462,EFNB1,IQSEC1,VPS51,MOGS,CSNK
UBGCP6,SLCO1B3,PACS1,TCF3,LTBP3,LZTR1,P

BTB2,MED12,DDHD2,RUSC2,PPP1R12A,ZDHHC9,EPB
,PRKDC,ATXN10,SON,KDM1A,MAP2K2,D
I20C,FH,PSMD12,TGFB2,SEC31A,PIGT,TMCO1,POGZ,
CASK,NDUFA10
D,ELN,OBSL1,ACTA2,IGF1R,MECP2,FAM50A,WDR
,FAM50A,EMC1,WDR26,ADNP,ALDH18A1,HMGB3

1,POR,CLIP2,MACROH2A1,KLF13,SC5D,EL
C47,REEP1,SMO,FGFR1,EIF4A3,SH3PXD2B,H
,KLC2,FLII,IQSEC1,PDGFRB,VPS51,C

3,BCR,KDM5C,TRIO,EP300,EXTL3,RERE,DGCR2,KD
3,BCR,KDM5C,TRIO,EP300,EXTL3,RERE,DGCR2,KD
,ERGIC1,YARS1,HSPG2,SLC6A8,PKDCC,LT
NSD1,WAC,ZSWIM6,GDF1,RNF216,ARHGEF9,SETD
P2,WDR26,ADNP,TRRAP,GPAA1,LIMK1,EFNB1,
CTNND2,PI4KA,RAB3GAP2,ADCY6,MBTPS1

LDH18A1,LIMK1,AMPD2,FLII,PRPH,TN

NUS1,ASH1L,ACSL4,MED25,SETD5,MBOAT7,MA

N,OPTN,VHL,PPP2CA,CCND1,MAPK1,SMARCA4,PL
1,DARS1,ASAH1,RAB18,SLC16A2,SYNGAP1

DH18A1,LIMK1,EFNB1,FLII,NBAS,IQSEC1,CSNK2
NF1,ZNF462,EFNB1,FLII,NBAS,PDGFRB,MOGS,EXT
N7,TAOK1,EHMT1,IARS2,HSPG2,ANTXR1,SOX2,SL

GNT1,CUL4B,PSMD12,OPA1,THG1L,SETD1A,NFASC,SYT1,
IOT3,QRICH1,MAN1B1,EXT1,SLC1A4,
B4GAT1,TIMM50,GPC4,SLC2A1,NDN,FUS,E
FNB1,SALL2,FLII,NBAS,VPS51,TMEM98,EXT2,E

R,CLIP2,TRAK1,DAG1,MACROH2A1,ELN,OBSL
P1R12A,ZDHHC9,TALDO1,GALNT2,SEPT
B,GLUD1,CTNND2,PI4KA,RAB3GAP2,FBXO11,IA
M,SLC2A1,NDN,BAP1,FUS,GRN,SPG7,C

5,RARS1,MBOAT7,NDUFS1,MAN2B1,ZNF59

Z,DNM1,GNAO1,CENPT,PLXND1,SLC1

COX4I1,NIPA1,EPB41L1,PPP3CA,TRAPPC
SL1,NOTCH1,PDHA1,IGF1R,MECP2,FAM50A,
TGA7,PLXND1,SLC18A3,FGD1,GSN,MAFB,DP

JL4B,FAM20C,PSMD12,TGFB2,SETD1A,FHL1,SZT2

E1,RPS28,ATP6V1B2,SDHA,LSS,FBLN1,ST
SLC25A1,RAB18,RDH11,DEAF1,EXOC7,C

,PDHA1,MECP2,NDUFA4,EMC1,WDR26,J

CDC88A,ASNS,MOGS,CSNK2B,NECTIN1,E

UFB8,CLCN4,NUS1,ASH1L,BCOR,MED25,S
1,ZNF592,LARGE1,SLC5A6,RHOBTB2,PMPCA,NUP13
MGA2,PRMT7,MAFB,DPF2,SIN3A,CASK,TCF20
C,LTBP4,HINT1,KMT2E,GTF2I,SRSF2,ELP1,TCF

X2,SLC6A8,AKT3,KMT2E,GTF2I,TUBG

31,DNAJC21

DUFA13,MN1,PHIP,TANC2,WNK1,TRIM8,CC

1,ZNF592,LARGE1,SLC5A6,RHOBTB2,PMPCA,NUP13
1,ZNF592,LARGE1,SLC5A6,RHOBTB2,PMPCA,NUP13

KLF13,ELN,OBSL1,NOTCH1,PDHA1,IGF1R,MDM

N,BAP1,CLCN7,EHMT1,IARS2,ERGIC1,HSPG2

TF,TOMM40,DMPK,SIN3A,CASK,CC2D1A,ABCA7,TCF20

,PPP3CA,CUL7,EXT1,SLC1A4,SPG11,G

IP,PPT1,COG4,TMCO1,POGZ,DNM1,MAPK8IP3
2,MED12,HIRA,ARSB,CTCF,OGDH,PPP1R12A,TGFBR1,E

IPK,SIN3A,CASK

SD2,DHCR7,MTOR,UBE3A,PRKACA,GOT2,SMARC

1,COL27A1,VANGL2,MAPK8IP1,RAI1,RAB1

GTF2I,CNP,SRSF2,ATXN3,ELP1,PACS1

3CA7,TCF20

COG4,CENPF,TMCO1,BRAT1,COL3A1,ITG
D12,SEC31A,PIGT,COG4,CENPF,TMCO1,B
GF3,SC5D,ELN,OBSL1,ACTA2,IGF1R,MECP2
,RUSC2,PRPF31,ARL2BP,CNOT3,QRICH1,CUL7,A

CENPF,TMCO1,BRAT1,COL3A1,ITGA7,PL

S1,ASH1L,MCFD2,SETX,RARS1,SETD5,

JFS1,KARS1,NSMF,COX6B1,NACC1,LARGE1,

GFBR1,PPP3CA,COL6A3,CUL7,EXT1,TAR

LSS,B2M,STXBP1,LONP1,PRPS1,MESD,IDH1,
