## Supplemental Table 6 for "Downregulation of Neurodevelopmental Gene Expression in iPSC-Derived Cerebral Organoids Upon Infection by Human Cytomegalovirus"

| Gene Identifier | Forward 5'-3' | Reverse 5'-3' |
|---|---|---|
| GAPDH | GTGGACCTGACCTGCCGTCT | GGAGGAGTGGGTGTCGCTGT |
| CAMKV | AAGATGAGAGCAGGACACCC | CAAGCAGGCTTGCAGTCAGA |
| KCNF1 | GGGTTGGGTGTGGAGTTTTG | GGATTTAACCCAATCACCATGAC |
| CACNA1G | GCTGGGTCGACATCATGTACTTTG | CTGAGAACTGCGTGGCAAACC |
| FEZF2 | GTGGTGGAATTCGCCGCCGCCATGGCCAGCTCAGCT<br>TCCCTGGAGACCATGGTG | TGCTGGATATCAGCTCTGAACTGTCCTGGCTAG<br>GTCCTTTGCTGA |
| EMX1 | AGCCCCGTCTTAATGCAACA | CTAGGATTGCGGGGCTAGTG |
| CACNA1C | GCAACGGCTGGAACCTACTA | CGATGATGGCGTAGATGATG |
| FOXG1 | CGTTCAGCTACAACGCGCTCA | CAGATTGTGGCGGATGGAGTT |
| GJA1 | TACCAAACAGCAGCGGAGTT | TGGGCACCACTCTTTTGCTT |
| DMRTA2 | GCCTGCCTACGAAGTCTTTGGCTCGGTTT | CGTCTTGGGAAACAGATCAAACTTCTG |
| UL44 | | |
| UL99 | | |
| UL122 | | |
| UL123 | | |
