## Supplemental Figure 1 for "Downregulation of Neurodevelopmental Gene Expression in iPSC-Derived Cerebral Organoids Upon Infection by Human Cytomegalovirus"

O'Brien et al., 2021
Supplemental Fig 1

A

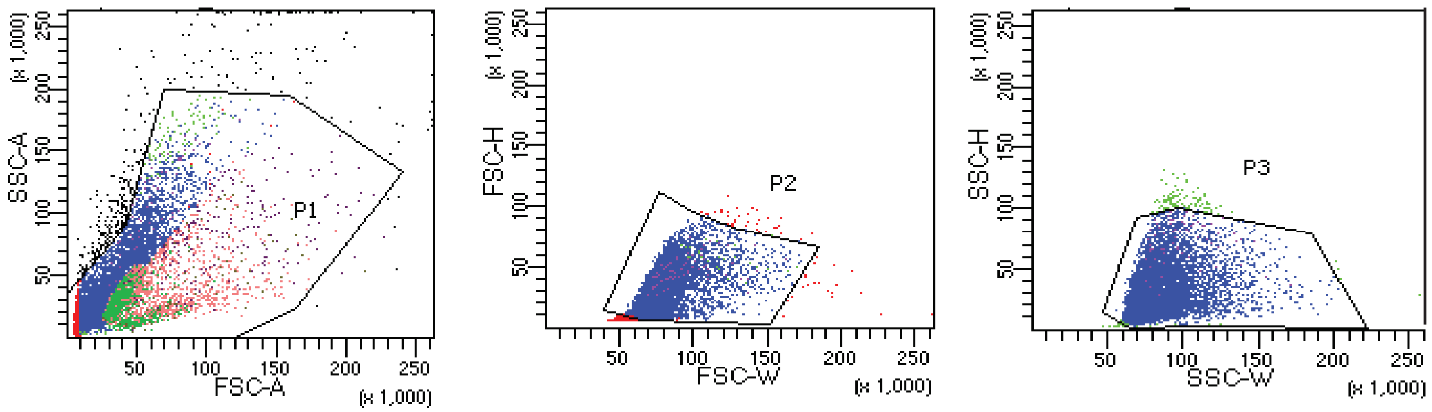


B

Tube: 141516

| Population | #Events | %Parent | %Total |
|---|---|---|---|
| All Events | 10,000 | #### | 100.0 |
| P1 | 9,628 | 96.3 | 96.3 |
| P2 | 8,195 | 85.1 | 82.0 |
| P3 | 8,053 | 98.3 | 80.5 |
| P4 | 67 | 0.8 | 0.7 |
| P5 | 3,019 | 30.2 | 30.2 |
| gfp int | 1,774 | 58.8 | 17.7 |
| gfp pos | 222 | 7.4 | 2.2 |
| gfp neg | 678 | 22.5 | 6.8 |

C

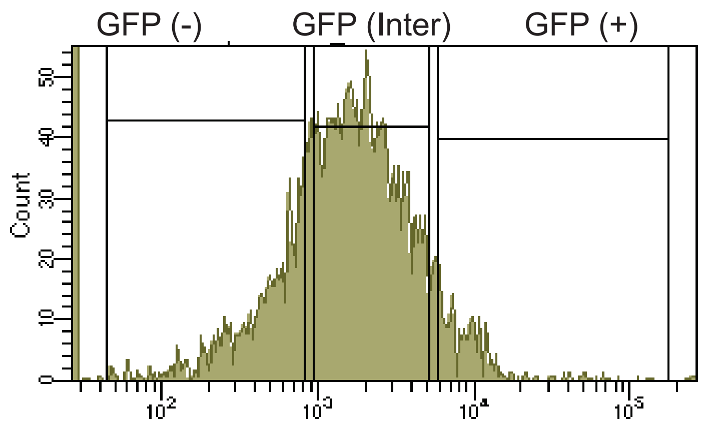


D

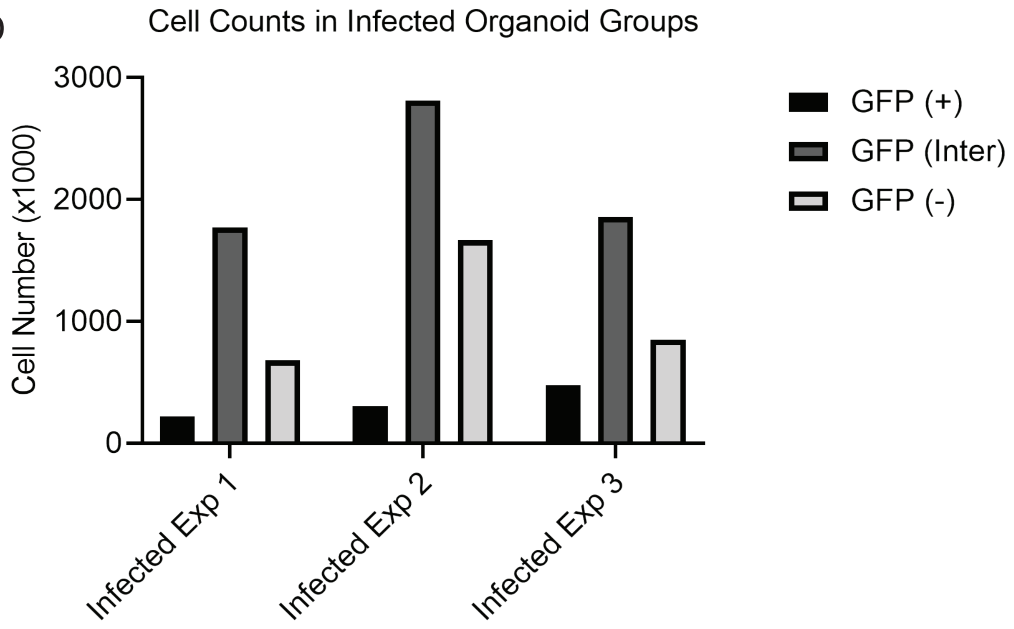
