## Supplementary figures and images for "Downregulation of Neurodevelopmental Gene Expression in iPSC-Derived Cerebral Organoids Upon Infection by Human Cytomegalovirus"

### Supplemental Figure 2

O'Brien et al., 2021
Supplemental Figure 2

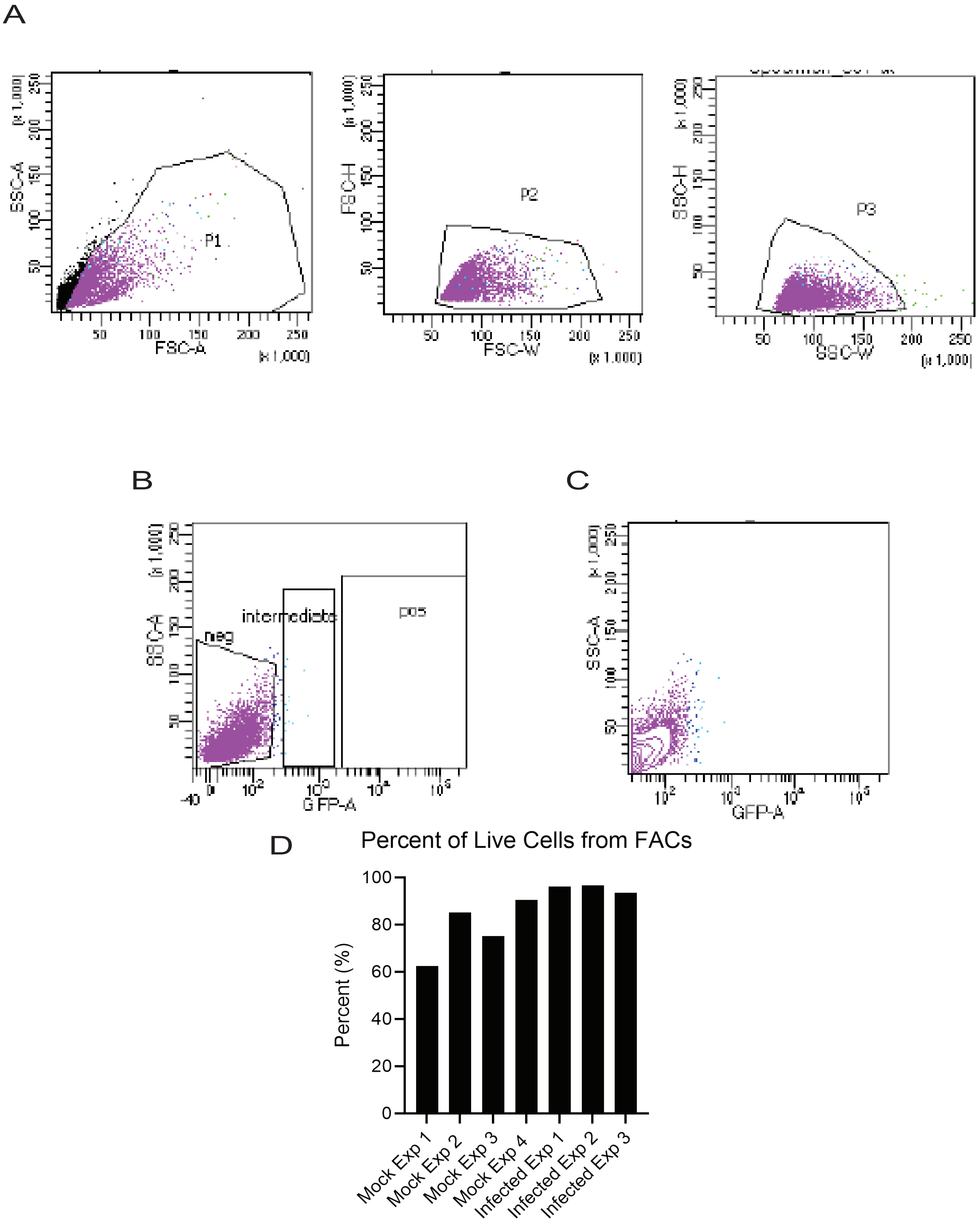

### Supplemental Figure 3

O’Brien et al., 2021
Supplemental Fig 3

A

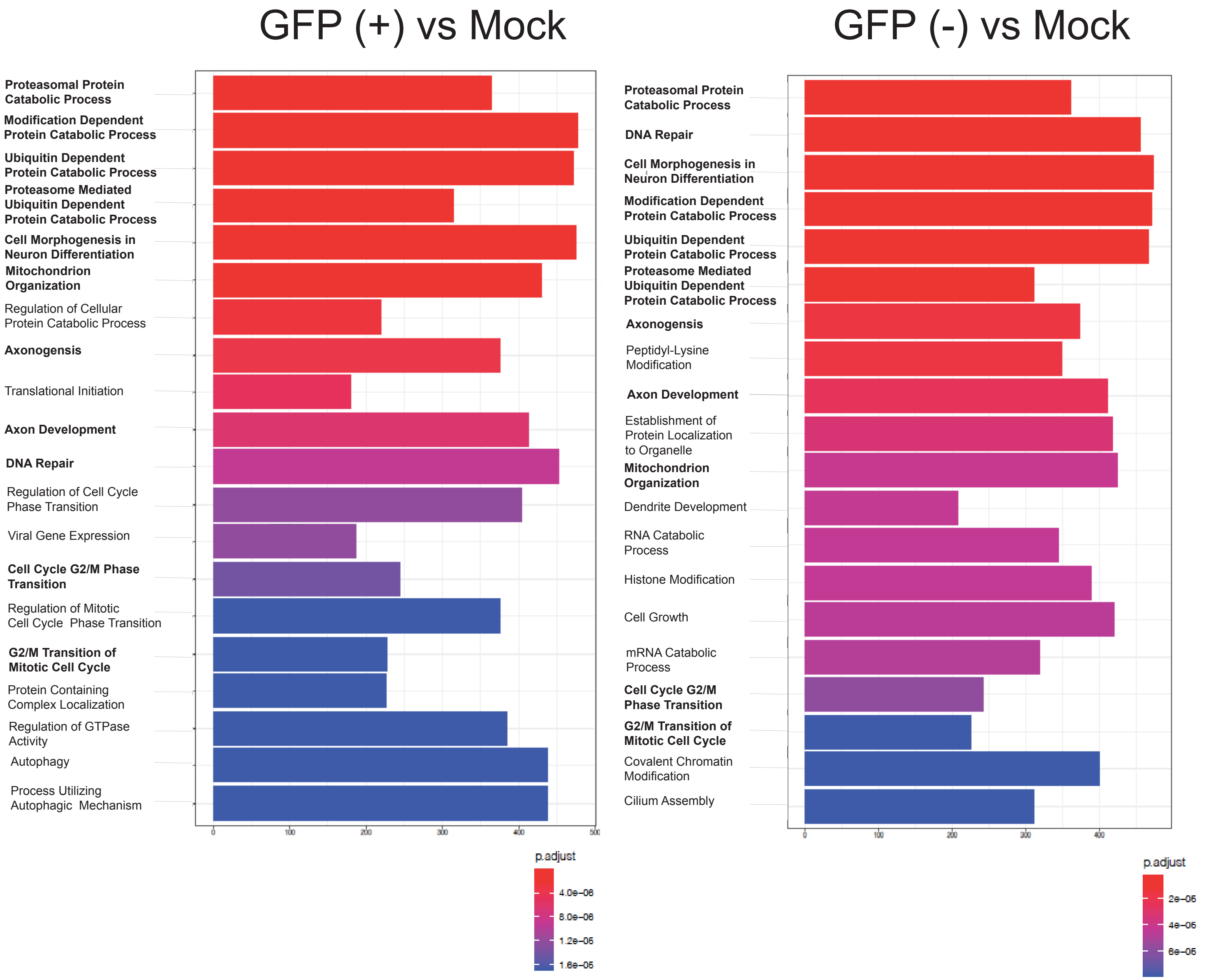

### Supplemental Figure 4

O’Brien et al., 2021
Supplemental Fig 4

C

RhoGDI Signaling

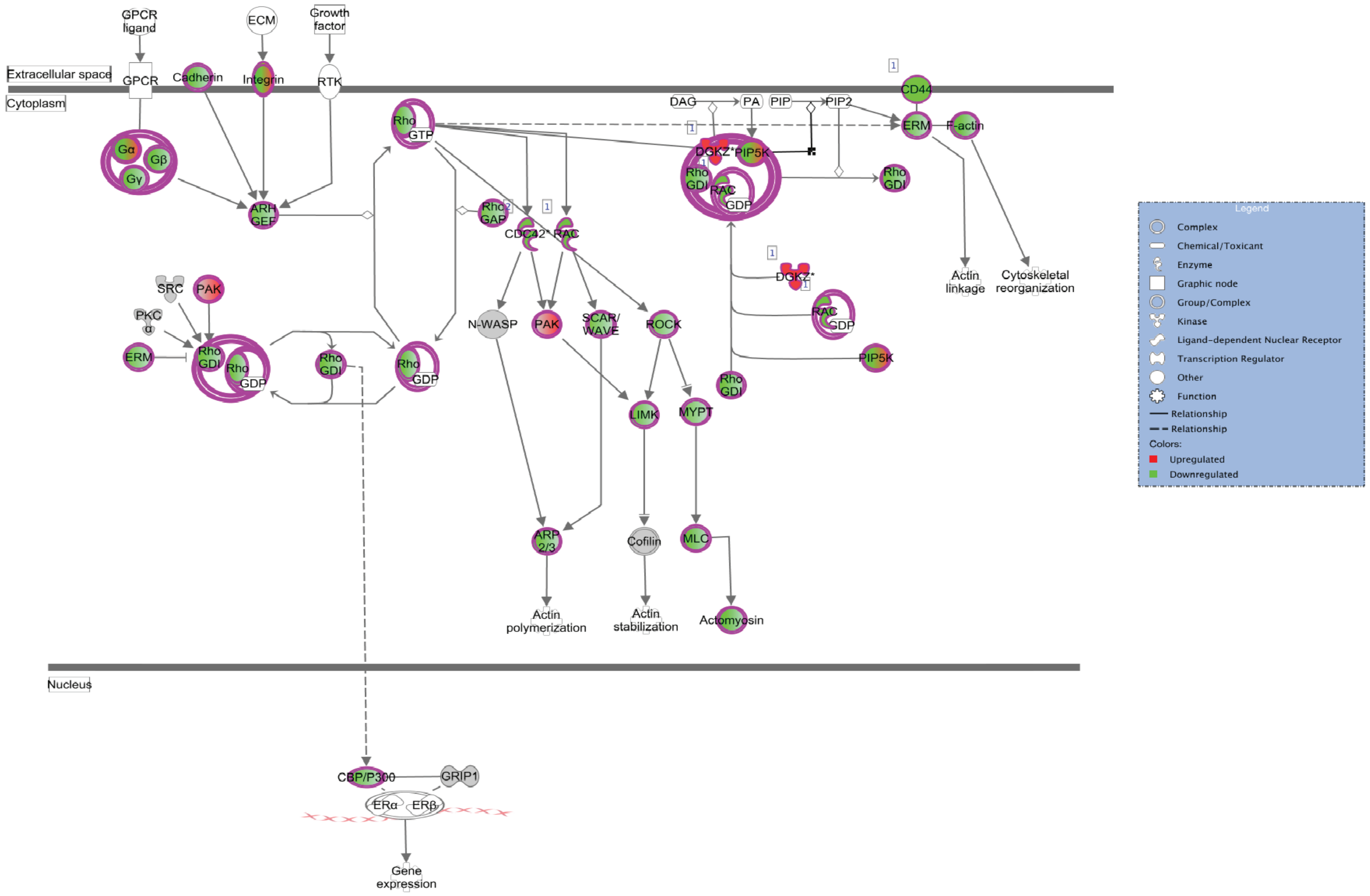

### Supplemental Figure 4

O'Brien et al., 2021
Supplemental Fig 4

A

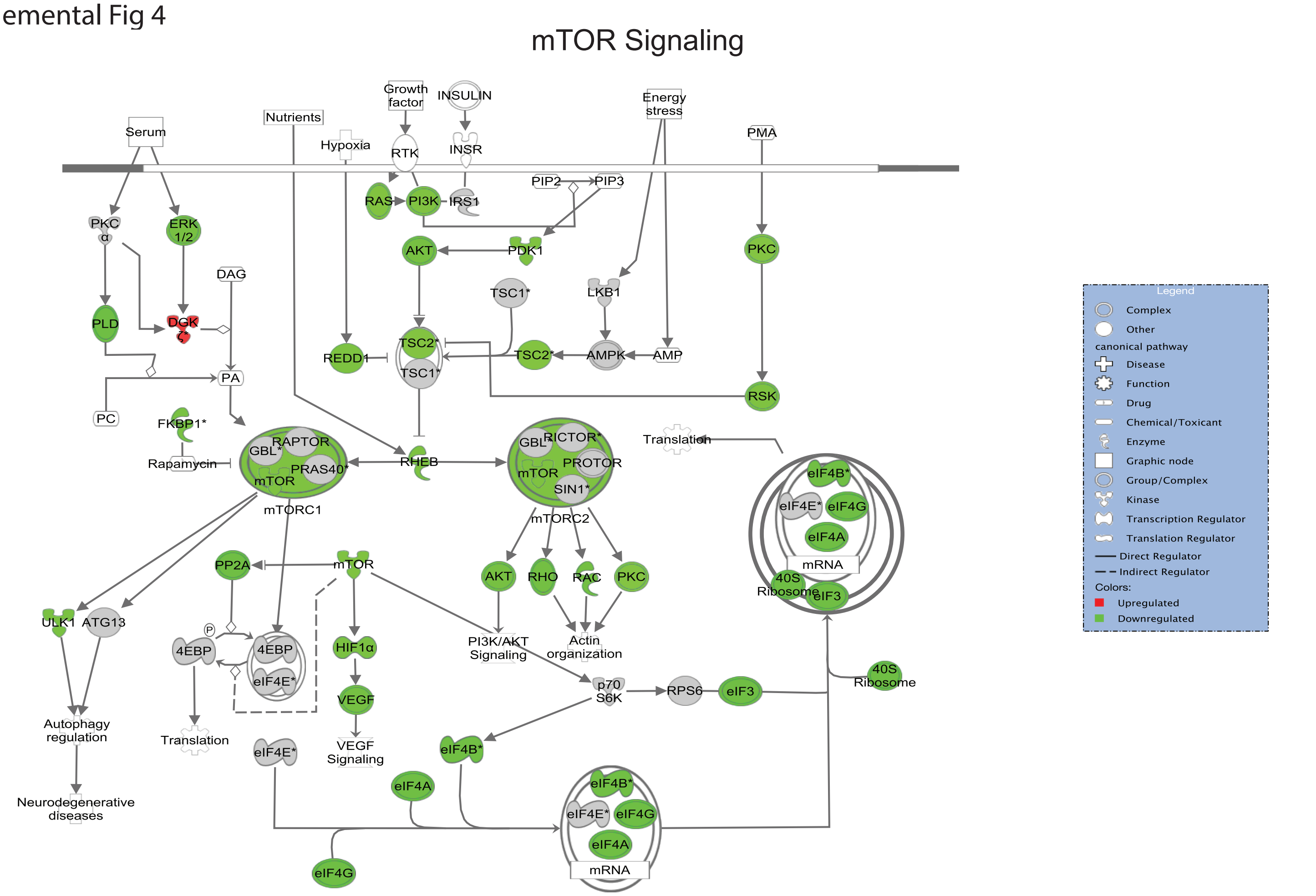


B

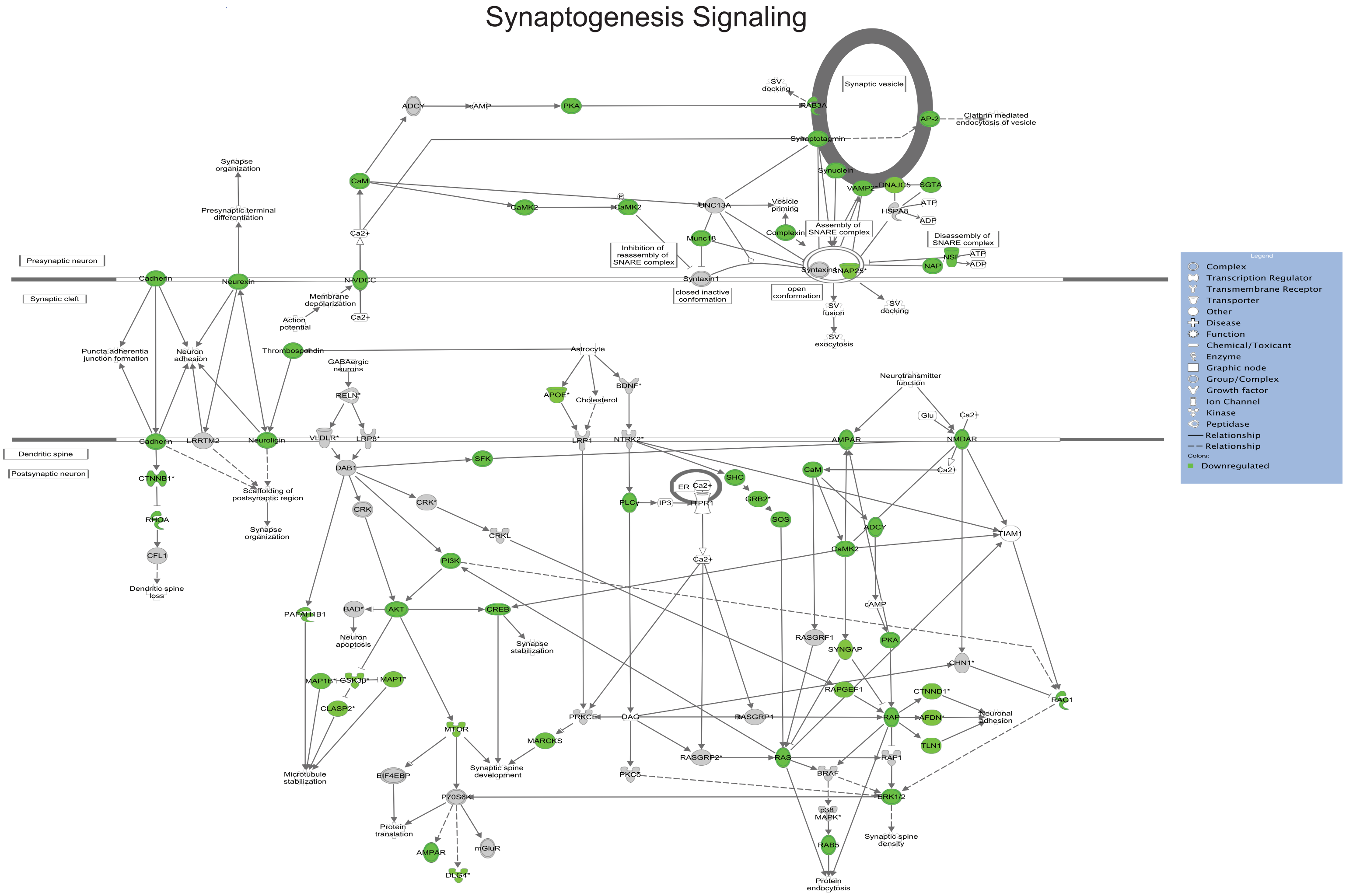


© 2000-2021 QIAGEN. All rights reserved.

### Supplemental Figure 5

O'Brien et al., 2021
Supplemental Fig S5

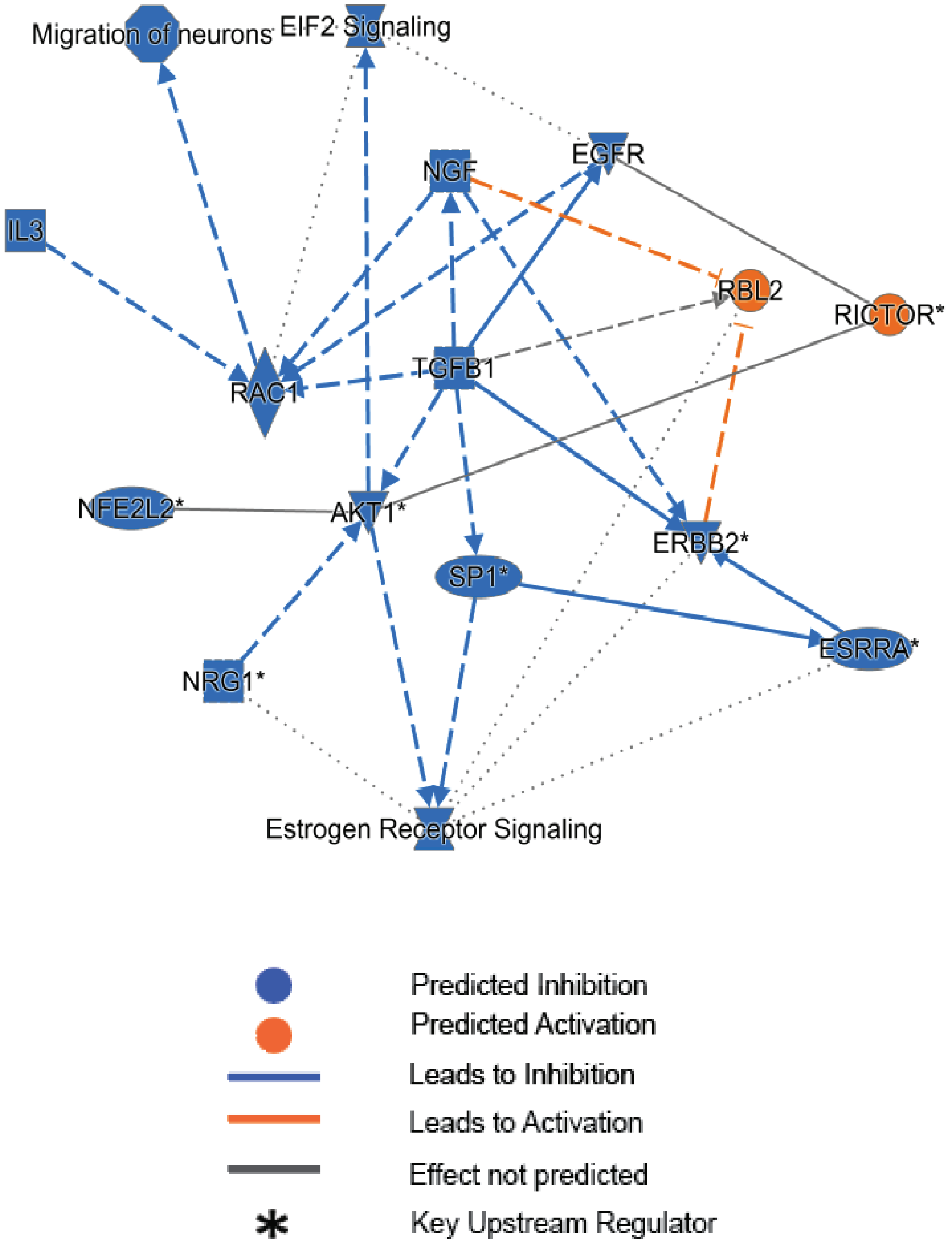
